# Detection and characterization of the SARS-CoV-2 lineage B.1.526 in New York

**DOI:** 10.1101/2021.02.14.431043

**Authors:** Anthony P. West, Joel O. Wertheim, Jade C. Wang, Tetyana I. Vasylyeva, Jennifer L. Havens, Moinuddin A. Chowdhury, Edimarlyn Gonzalez, Courtney E. Fang, Steve S. Di Lonardo, Scott Hughes, Jennifer L. Rakeman, Henry H. Lee, Christopher O. Barnes, Priyanthi N. P. Gnanapragasam, Zhi Yang, Christian Gaebler, Marina Caskey, Michel C. Nussenzweig, Jennifer R. Keeffe, Pamela J. Bjorkman

## Abstract

Wide-scale SARS-CoV-2 genome sequencing is critical to tracking viral evolution during the ongoing pandemic. Variants first detected in the United Kingdom, South Africa, and Brazil have spread to multiple countries. We developed the software tool, Variant Database (VDB), for quickly examining the changing landscape of spike mutations. Using VDB, we detected an emerging lineage of SARS-CoV-2 in the New York region that shares mutations with previously reported variants. The most common sets of spike mutations in this lineage (now designated as B.1.526) are L5F, T95I, D253G, E484K or S477N, D614G, and A701V. This lineage was first sequenced in late November 2020 when it represented <1% of sequenced coronavirus genomes that were collected in New York City (NYC). By February 2021, genomes from this lineage accounted for ^~^32% of 3288 sequenced genomes from NYC specimens. Phylodynamic inference confirmed the rapid growth of the B.1.526 lineage in NYC, notably the sub-clade defined by the spike mutation E484K, which has outpaced the growth of other variants in NYC. Pseudovirus neutralization experiments demonstrated that B.1.526 spike mutations adversely affect the neutralization titer of convalescent and vaccinee plasma, indicating the public health importance of this lineage.

## Introduction

After the early months of the SARS-CoV-2 pandemic in 2020, the vast majority of sequenced genomes contained the spike mutation D614G (along with 3 separate nucleotide changes)^1^. Following a period of gradual change, the fourth quarter of 2020 witnessed the emergence of several variants containing multiple mutations, many within the spike gene^2–5^. Multiple lines of evidence support escape from antibody selective pressure as a driving force for the development of these variants^6–9^.

Genomic surveillance of SARS-CoV-2 is now focused on monitoring the emergence of these variants and the functional impact that their mutations may have on the effectiveness of passive antibody therapies and the efficacy of vaccines to prevent mild or moderate COVID-19. While an increasing number of specimens are being sequenced, analysis of these genomes remains a challenge^10^. Here, we developed a simple and fast utility that permits rapid inspection of the mutational landscape revealed by genomic surveillance of SARS-CoV-2: Variant Database (**vdb**). With this tool, we uncovered several groups of recently sequenced genomes with mutations at critical antibody epitopes. Among this group is a new lineage emerging in NYC that has increased in frequency to now account for ^~^32% of sequenced genomes as of February 2021. We confirm the rapid spread of B.1.526 in NYC during early 2021 through phylodynamic inference. Furthermore, we evaluated the impact of the B.1.526 spike mutations on the neutralization titer of convalescent and vaccinee plasma.

## Results

### vdb

Phylogenetic analysis is critical to understand the relationships of viral genomes. However, other perspectives can be useful for detecting patterns in large numbers of sequences. We developed **vdb** as a utility to query the sets of spike mutations observed during genomic surveillance. Using the **vdb** tool to analyze SARS-CoV-2 sequences in the Global Initiative on Sharing Avian Influenza Data (GISAID) dataset^11,12^, we detected several clusters of sequences distinct from variants B.1.1.7, B.1.351, B.1.1.248, and B.1.429^2–5^ with spike mutations at sites known to be associated with resistance to antibodies against SARS-CoV-2^8,13^ (**Table 1**). The **vdb** program can find clusters of virus sharing identical sets of spike mutations, and then these patterns can be used to find potentially related sequences.

**Table 1.**
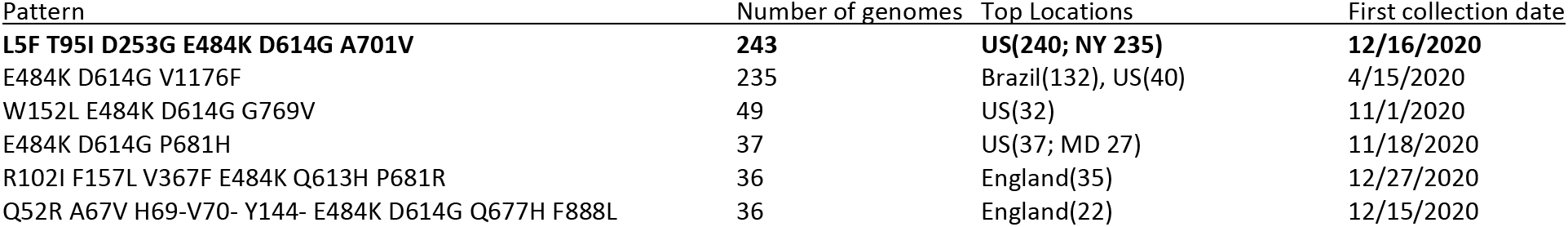
Mutation patterns of viruses with mutations at select Spike positions, excluding viruses related to variants B.1.1.7, B.1.351, B.1.1.248, and B.1.429. Mutations included in this analysis were E484K, N501Y, K417T, K417N, L452R, and A701V. In this table viruses are only included if their spike mutation pattern exactly matches the given pattern. Note about P681H/P681R: variant B.1.1.7 has P681H. Note about W152L: variant B.1.429 has W152C

### Defining mutations of B.1.526

One notable cluster of genome sequences was collected from the New York region and represents a distinct lineage, now designated as B.1.526 (**Figure 1, Supplementary Figure 1**). This variant is found within the 20.C clade and is distinguished by 3 defining spike mutations: L5F, T95I, and D253G. Within B.1.526, the largest sub-clade is defined by E484K and two distinct sub-clades are each defined by S477N; both of these mutations located within the receptor-binding domain (RBD) of spike (**Figure 2 and Supplementary Table 1**). We note that the evolutionary history at spike position 701 varies depending on whether the tree is rooted using a molecular clock (**Figure 1**) versus its sister clade (characterized by an L452R mutation; **Supplementary Figure 2**), the latter of which posits a substitution A701V followed by a reversion V701A. Among the nucleotide mutations in lineage B.1.526, the most characteristic include A16500C (NSP13 Q88H), A22320G (spike D253G), and T9867C (NSP4_L438P). Another notable feature of the B.1.526 lineage is the deletion of nucleotides 11288-11296 (NSP6 106-108), which also occurs in variants B.1.1.7, B.1.351, P.1, and B.1.525^14^.

**Figure 1.**
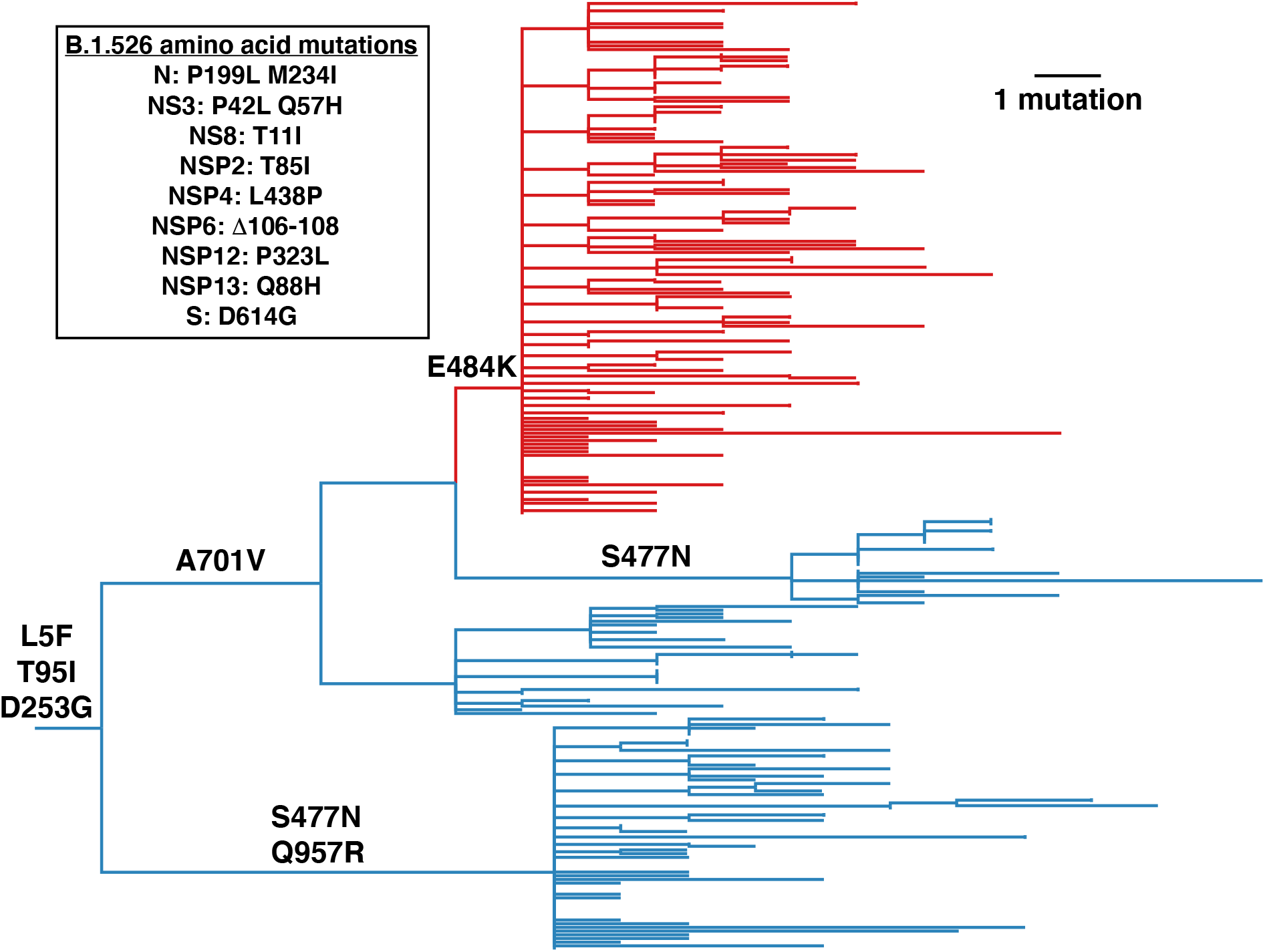
Phylogenetic tree of lineage B.1.526 indicating spike mutations. Maximum likelihood phylogeny of SARS-CoV-2 variant B.1.526 sampled by NYC PHL (n=258). Amino acid substitutions in the spike protein occurring on internal branches are labeled, including the three spike mutations characteristic of B.1.526. The B.1.526 clade defined by the E484K mutation is highlighted in red. Inset highlights non-spike amino acid substitutions and deletions differentiating the B.1.526 clade from the Hu-1 reference genome. For display purposes, only NYC PHL genomes are shown.

**Figure 2.**
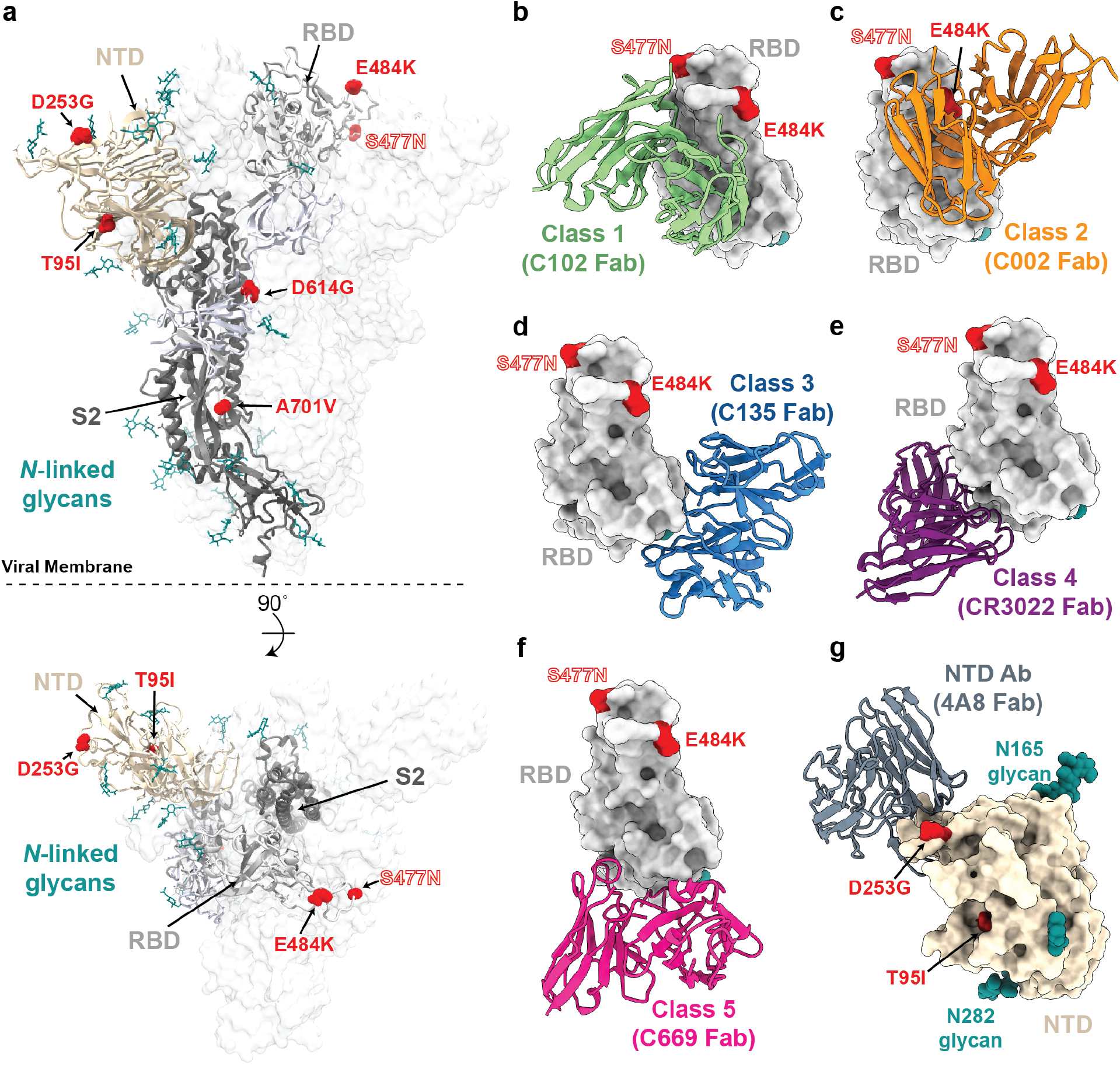
Structural locations of the spike mutations of lineage B.1.526. **a**, Side and top views of the SARS-CoV-2 spike trimer (PDB 7JJI) with mutations of lineage B.1.526 shown as spheres. **b-g**, Models of representative neutralizing antibodies (cartoon, VH-VL domain only) bound to RBD (**b-f**, gray surface) or NTD (**g**, wheat surface). Sites for B.1.526 lineage mutations are shown as red spheres. The S477N site is also shown for the branch containing this mutation instead of the E484K mutation (see **Figure 1**); **b**, Class 1 (PDB 7K8M); **c**, Class 2 (PDB 7K8S); **d**, Class 3 (PDB 7K8Z); **e**, Class 4 (PDB 6W41); **f**, Class 5 ^8^; **g**, NTD-specific antibody 4A8 (PDB 7C2L).

Regarding four of the spike mutations prevalent in this lineage: (1) E484K is known to attenuate neutralization of multiple anti-SARS-CoV-2 antibodies, particularly those found in class 2 anti-RBD neutralizing antibodies^13,15^, and is also present in variants B.1.351^4^ and P.1/B.1.1.248^2^, (2) D253G has been reported as an escape mutation from antibodies against the N-terminal domain^16^, (3) S477N has been identified in several earlier lineages^17^, is near the epitopes of multiple antibodies^18^, and has been implicated to increase viral infectivity through enhanced interactions with ACE2^19,20^, and (4) A701V sits adjacent to the S2’ cleavage site of the neighboring protomer and is shared with variant B.1.351^4^. The overall pattern of mutations in lineage B.1.526 (**Figure 2**) suggests that it arose in part in response to selective pressure from antibodies. Based on the dates of collection of these viruses, it appears that the frequency of this lineage has increased rapidly in New York (**Table 2**).

**Table 2.** Counts of virus genomes in lineage B.1.526 by month in New York State. The total number of sequenced genomes examined from GISAID from New York during these time periods is also listed. *Latest viral collection date was March 4, 2021. Note that geographic sampling may have varied over time as genome sequencing increased.

| Month | count | total sequences | fraction |
| --- | --- | --- | --- |
| Nov. 2020 | 2 | 524 | 0.4% |
| Dec. 2020 | 46 | 2209 | 2.1% |
| Jan. 2021 | 201 | 3148 | 6.4% |
| Feb. 2021 | 1207 | 3868 | 31.2% |
| March 2021* | 124 | 274 | 45.3% |

Viruses containing spike mutations L5F, T95I, D253G, E484K, D614G, and A701V (earliest collection date Dec. 16, 2020)
| Month | count | total sequences | fraction |
| --- | --- | --- | --- |
| Nov. 2020 | 0 |  |  |
| Dec. 2020 | 25 | 2209 | 1.1% |
| Jan. 2021 | 109 | 3148 | 3.5% |
| Feb. 2021 | 628 | 3868 | 16.2% |
| March 2021* | 61 | 274 | 22.3% |

### Trends in B.1.526 surveillance

As part of public health surveillance conducted by the New York City Public Health Laboratory (NYC PHL) and the Pandemic Response Lab (PRL) in New York, approximately 4.5 thousand SARS-CoV-2 genomes have been sequenced by NYC PHL and PRL from December 1, 2020 to February 28^th^, 2021. Of these genomes, approximately 25% are from lineage B.1.526. We separately analyzed these genomes, because viral genomic surveillance by PHL and PRL provides a less biased picture of viral diversity in NYC than genomes uploaded to GISAID. The proportion of B.1.526 genomes in NYC has steadily increased since this variant was first detected in NYC surveillance data in late 2020, and its weekly average exceeded 10% by 14 January 2021. From early January to early March, B.1.526 has been increasing by about 0.7% per day (segmented linear regression) and was at 43% the week prior to 03 March 2021 (**Figure 3A**). Around 54% (n=678) of the B.1.526 genomes contain the E484K mutation, which has also been rising in frequency since early 2021. The weekly average of B.1.526 genomes with E484K has been above 10% since 01 February 2021 and has been increasing around 0.4% per day (**Figure 3B**).

**Figure 3.**
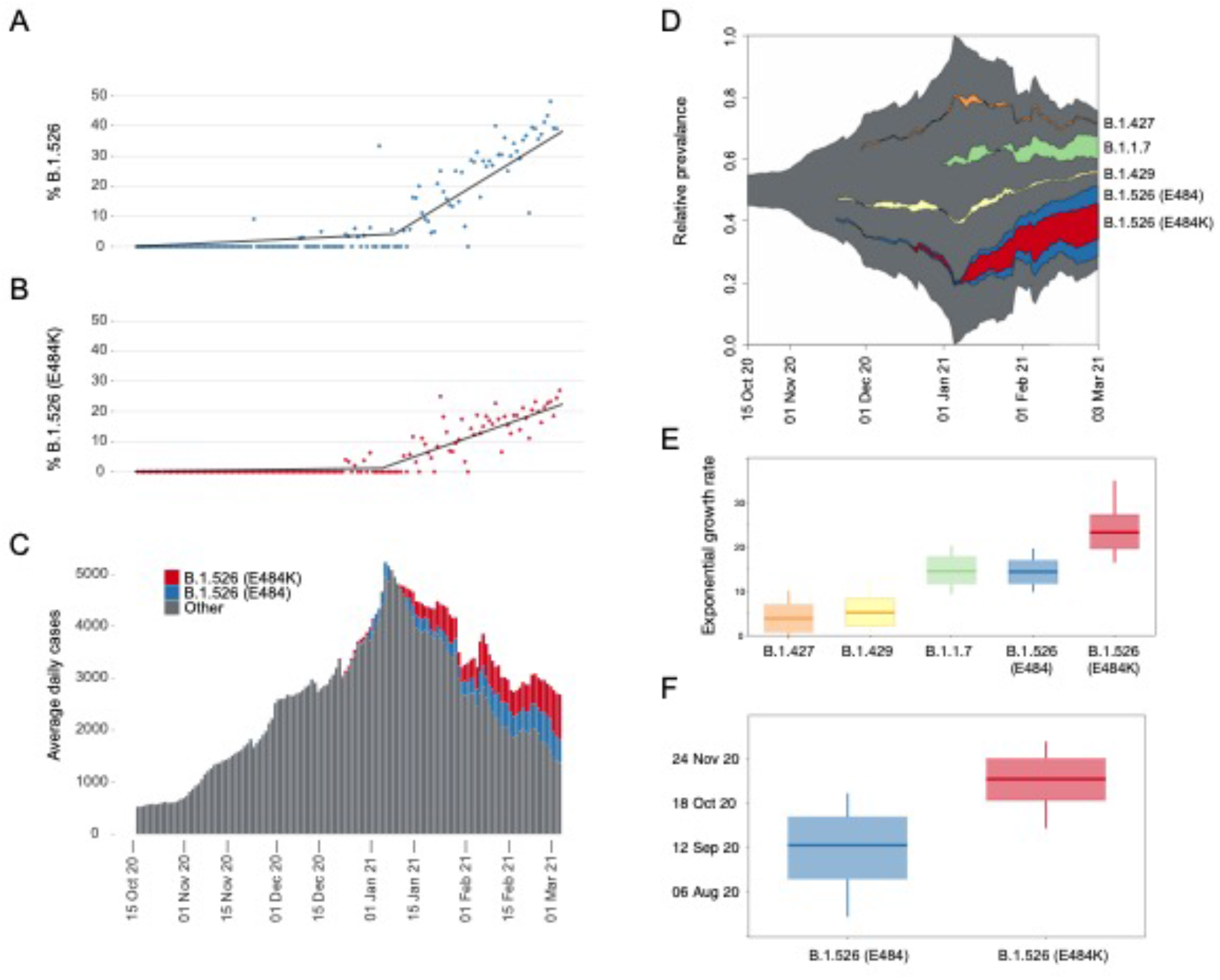
Rise of SARS-CoV-2 variants in New York City (NYC) in late-2020 and early 2021. (**A**) Relative frequency of B.1.526. Segmented linear regression is shown as a solid black line. (**B**) Relative frequency of B.1.526 with E484K mutation. Segmented linear regression is shown as a dashed gray line. (**C**) Rolling average number of total daily COVID-19 cases in NYC through time. Color indicates the estimated proportions of B.1.526 (blue) and B.1.526 E484K (red) extrapolated from a 7-day rolling average with an average of n=236 genomes sampled per week during this time period. (**D**) Muller plot depicting sampling, with pseudocounts, of SARS-CoV-2 variants scaled to the rolling average of total daily COVID-19 case counts. (**E**) Inferred exponential growth rates for SARS-CoV-2 variants in NYC; the horizontal line indicates the median growth rate estimate, the box outlines the interquartile range. (**F**) Inferred time of most recent common ancestor (TMRCA) estimates for B.1.526 (E484) and B.1.526 (E484K).

This increase in B.1.526 temporally coincides with the peak and subsequent decline of the second epidemic wave in NYC (**Figure 3C**). If we separate the approximated number of B.1.526 cases from the rest of second wave SARS-CoV-2, the non-B.1.526 virus has steadily declined since its peak in early January 2021. However, the increasing proportion of B.1.526 appears to have slowed the rate of decline in total COVID-19 case counts in NYC.

### Geographic distribution of B.1.526 in NYC

The New York City Public Health Laboratory and the PRL in New York have sequenced 4538 SARS-CoV-2 genomes from December 2020 thru February 2021 (**Figure 4A**). Geographic case distribution of specimens received at PHL and PRL for SARS-CoV-2 diagnostic nucleic acid amplification testing (NAAT) are representative of citywide testing efforts. Those SARS-CoV-2 positive specimens with NAAT cross-threshold values below 32 were selected at random to be sequenced. On a month-to-month basis using data generated by NYC PHL and PRL, we have observed an increasing number of B.1.526 genomes identified throughout NYC. The geographic distribution of over 600 B.1.526 E484K cases is similar (**Figure 4B**). While the B.1.526 lineage is not limited to NYC, almost 90% of genomes deposited to GISAID prior to March 2021, are from the New York region.

**Figure 4.**
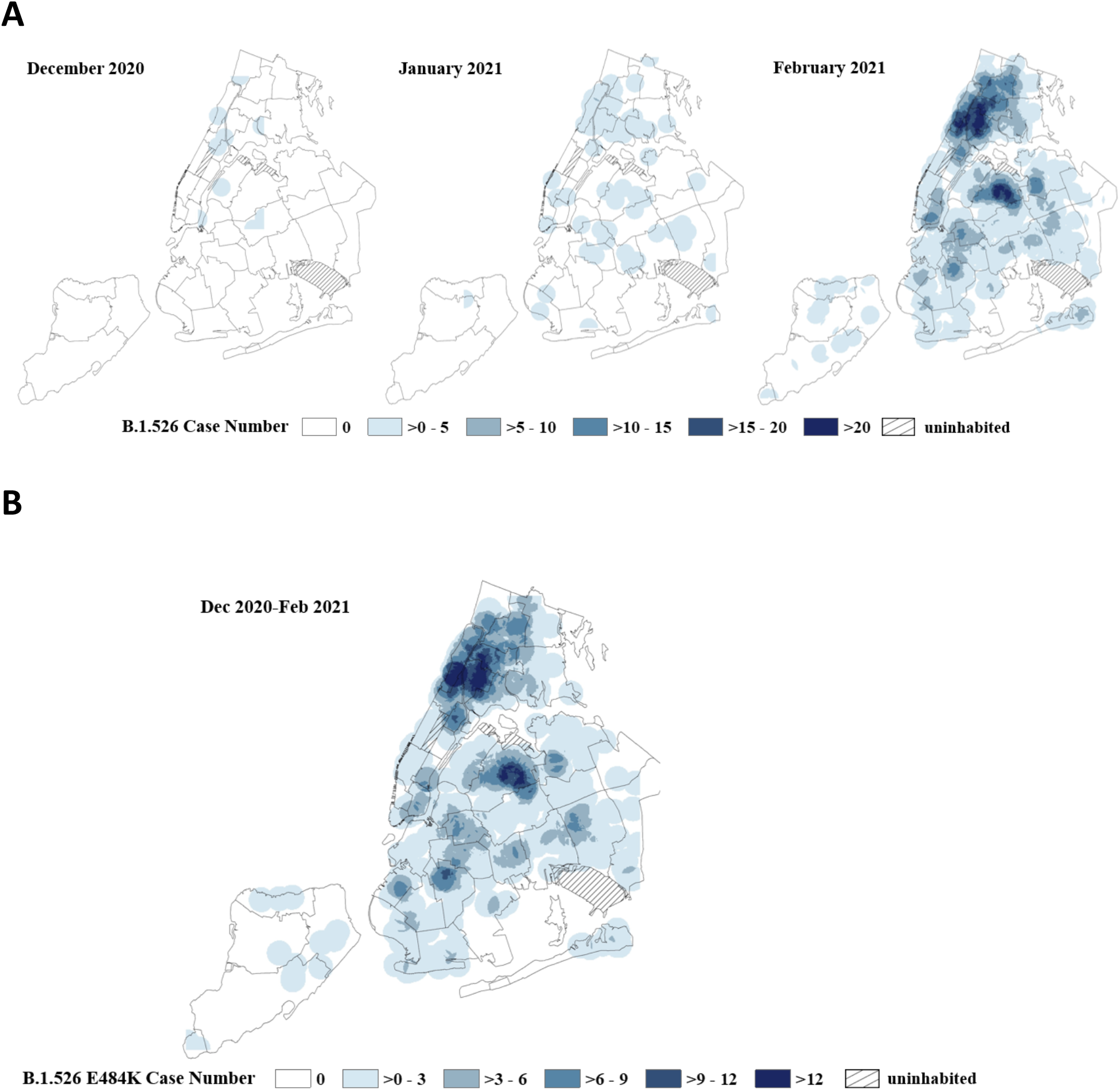
(**A**) Spaciotemporal increase of B.1.526 lineage in New York City (NYC). Point density of B.1.526 variants geo-located by case address overlayed on a map of NYC delineated by United Hospital Fund areas. Data for each month is based on specimen collection date. The NYC PHL and the PRL in New York have sequenced 4538 SARS-CoV-2 genomes from December 2020 thru February 2021. Data represents 11 B.1.526 variants out of 515 sequenced genomes in December 2020, 80 B.1.526 variants out of 735 sequenced genomes in January and 1063 B.1.526 variants identified out of a total of 3288 sequenced genomes in February 2021. (**B**) Distribution of B.1.526 E484K cases in NYC. Point density map of 608 B.1.526 E484K variant cases in NYC. Data is based on specimen collection period from December 1, 2020 through February 28th, 2021.

### Phylodynamic analysis

Other SARS-CoV-2 variants of concern or interest (B.1.1.7, B.1.427, and B.1.429) have also been circulating in NYC contemporaneously with the rise of B.1.526 and have all risen in relative frequency during the second wave of the NYC pandemic (**Figure 3D**). To compare the relative growth rates of these variants during this time-period, we fitted an exponential population growth model^21^ implemented in BEAST1.10^22^ to the sequences that correspond to these lineages of interest. Specifically, we estimated the growth rate for the B.1.1.7, B.1.427, and B.1.429 variants and for two subsets of the B.1.526 clade sequences (with and without the E484K mutation).

The B.1.526 E484K clade experienced more rapid exponential growth compared with other lineages: 23.2 (95% highest posterior density [HPD]: 19.6–27.1). B.1.526 with E484 and B.1.1.7 experienced similar growth rates: 14.3 (95% HPD: 11.7–16.9) and 14.5 (95% HPD 11.6 – 17.8), respectively. The B.1.427 and B.1.429 lineages experienced lower growth rates that were significantly greater than zero: 3.8 (95% HPD: 0.7–7.0) and 5.2 (95% HPD: 2.1–8.3), respectively. We caution that these lineage growth rates do not distinguish between per-contact transmissibility or per-virion infectiousness and speak only to the relative number of people detected with these variants in NYC during late 2020 and early 2021.

As part of the phylodynamic analysis, we inferred the time of most recent common ancestor (TMRCA) for the B.1.526 E484K clade to be 08 November 2020 (95% HPD: 22 October – 24 November). The TMRCA for the rest of the B.1.526 clade was estimated to be 15 September 2020 (95% HPD: 17 August – 08 October).

### Neutralization activity of convalescent and vaccinee plasma against B.1.526

The identification of several mutations associated with resistance to anti-SARS-CoV-2 antibodies in B.1.526 sequences raises the question of the impact on SARS-CoV-2 immunity. We generated HIV-based pseudoviruses expressing SARS-CoV-2 spike protein containing either the most common B.1.526 mutation pattern (v.1: L5F, T95I, D253G, E484K, D614G, and A701V), the 2^nd^ most common pattern (v.2: L5F, T95I, D253G, S477N, D614G, and Q957R), or only D614G. Pseudovirus neutralization titers were determined for human plasma samples from vaccinees [Moderna (mRNA-1273) or Pfizer-BioNTech(BNT162b2)]^8^ or convalescent plasma [at either 1.3^15^ or 6.2 months^13^ post-infection]. The E484K-containing B.1.526 pseudovirus had a statistically significant reduced neutralization titer compared to the D614G control: for vaccinee plasma, 4.5-fold reduced (*p* = 0.00005); for 1.3-month convalescent plasma, 6.0-fold reduced (*p* = 0.03); and for 6.2-month convalescent plasma, 4.8-fold reduced (*p* = 0.02) (**Figure 5a and Supplementary Table 2**). The smaller reduction of the titers in the 6.2-month convalescent plasma samples compared to the 1.3-month samples is consistent with the greater resistance of more matured anti-SARS-CoV-2 antibodies to viral escape mutations^23^. The S477N/Q957R-containing B.1.526 pseudovirus demonstrated a smaller effect on plasma neutralization (**Figure 5b**).

**Figure 5.**
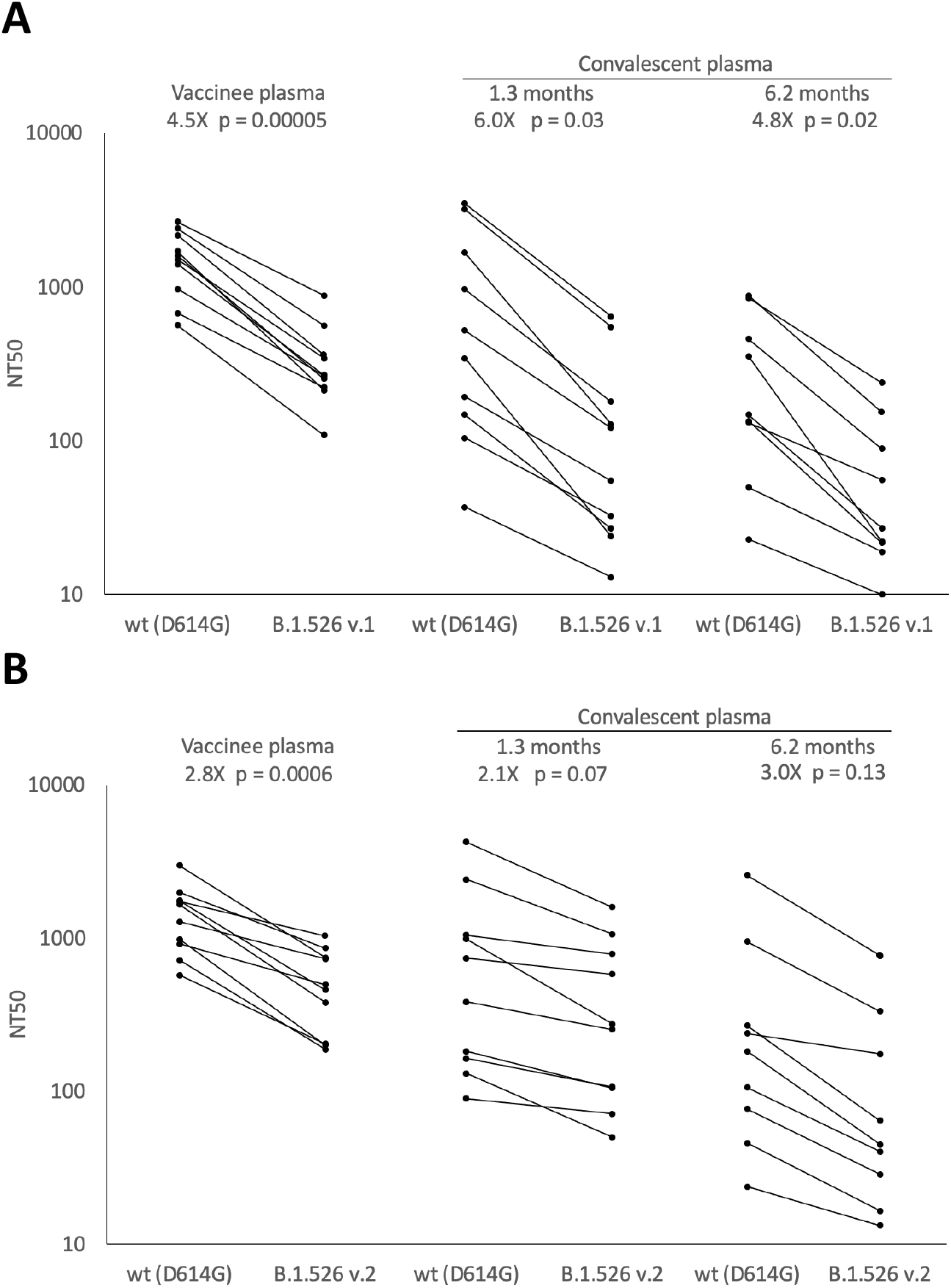
Plasma neutralizing activity against pseudoviruses with B.1.526 lineage spike mutations. SARS-CoV-2 pseudovirus neutralization assays were used to determine neutralization titer (NT50) for COVID-19 vaccinee (n=10) and convalescent plasma at 1.3 months (n=10) and 6.2 months (n=9) after infection. (**A**) Pseudovirus with spike mutations L5F, T95I, D253G, E484K, D614G (B.1.526 v.1), and A701V, (**B**) Pseudovirus with spike mutations L5F, T95I, D253G, S477N, D614G, and Q957R (B.1.526 v.2). Statistical significance was determined using paired two-tailed *t*-tests. Fold-differences of means are shown.

## Discussion

Genomic surveillance is a critical tool to monitor the progression of the COVID-19 pandemic and modelling suggests that sequencing at least 5% of specimens that test positive for SARS-Cov-2 in a geographic region is necessary to reliably detect the emergence of novel variants at a lower prevalence limit of between 0.1% to 1%^24^. Through the combination of increased sequencing efforts and the use of the software utility described here, we were able to identify the B.1.526 lineage and to begin to characterize its phylogenetic and phylodynamic patterns in NYC in early 2021. Based on sequences in GISAID as of March 2021, the majority of cases with sequence data are in the NYC region, but it is expected that the prevalence B.1.526 variants will continue to increase beyond the NYC region. The B.1.526 variant has also been described in other recent studies^25,26^.

Pseudovirus containing spike gene mutations associated with B.1.526 was significantly more resistant to neutralization by either convalescent or vaccinee plasma. The presence of E484K mutation likely plays a key role in facilitating increased viral transmission and reducing antibody neutralizing titers, as previously shown in other studies^7,27^. Continued monitoring for emerging variants with mutations such as E484K is important to maximize the impact of public health measures to mitigate the effects of the SARS-CoV-2 pandemic. For example, high frequencies of SARS-CoV-2 variants has potential impacts on selection of appropriate antibody therapeutics and vaccination strategies.

## Methods

### Variant Database Program

We developed a software tool named VDB (Variant Database). This tool consists of two Unix command line utilities: (1) **vdb**, a program for examining spike mutation patterns in a collection of sequenced viral genomes, and (2) **vdbCreate**, a program for generating a list of viral spike mutations from a multiple sequence alignment for use by **vdb**. The design goal for the query program **vdb** is to provide a fast, lightweight, and natural means to examine the landscape of SARS-CoV-2 spike mutations. These programs are written in Swift and are available for MacOS and Linux from the authors or from the Github repository: https://github.com/variant-database/vdb.

The **vdb** program implements a mutation pattern query language (see Supplemental Method) as a command shell. The first-class objects in this environment are a collection of viruses (a “cluster”) and a group of spike mutations (a “pattern”). These objects can be assigned to variables and are the return types of various commands. Generally, clusters can be obtained from searches for patterns, and patterns can be found by examining a given cluster. Clusters can be filtered by geographical location, collection date, mutation count, or the presence or absence of a mutation pattern. The geographic or temporal distribution of clusters can be listed.

Results presented here are based on a multiple sequence alignment from GISAID^11,12^ downloaded on February 10, 2021. Additional sequences downloaded from GISAID on February 22, 2021, were aligned with MAFFT v7.464^28^.

### Initial Phylogenetic Analysis

Multiple sequence alignments were performed with MAFFT v7.464^28^. The phylogenetic tree was calculated by IQ-TREE^29^, and the tree diagram was generated using iTOL (Interactive Tree of Life)^30^. The Pango lineage nomenclature system^31^ provides systematic names for SARS-CoV-2 lineages. The Pango lineage designation for B.1.526 was supported by the phylogenetic tree shown in **Supplementary Figure 1**.

### Library preparation and sequencing

RNA was extracted from positive specimens collected at NYC PHL using the EZ1 (Qiagen, CA), NUCLISENS^®^ easyMAG^®^ (bioMérieux Inc., Netherlands), or Kingfisher™ Flex Purification System (Thermo Fisher Scientific, MA). RNA extracts were subjected to annealing reaction with random hexamers and dNTPs (New England Biolabs Inc., NEB, MA), and reverse transcribed with SuperScript IV Reverse Transcriptase at 42ºC for 50 min. The resulting cDNA was amplified using two separate multiplex PCRs with ARTIC V3 primer pools (Integrated DNA Technologies, IA) per sample in the presence of Q5 2X Hot Start Master Mix (NEB) at 98ºC for 30 secs, followed by 35 cycles of 98ºC for 15 secs and 65ºC for 5 min^32,33^. The resulting PCR products per sample were combined and purified using Agencourt Ampure XP magnetic beads (Beckman Coulter, IN), at a ratio of 1:1 sample to bead ratio and quantified using a Qubit 3.0 fluorometer (Thermo Fisher Scientific, MA). The PCR products were normalized to 90 ng as input for the NEBNext Ultra II Library Preparation Kit according to standard protocol (NEB): Briefly, the ARTIC PCR products were subjected to simultaneous end-repair, 5’-phosphorylation, and dA-tailing reaction at 20ºC for 30 min, followed by heat inactivation at 65ºC for 30 min. NEBNext Adaptor was then ligated at 25º for 30 min, and then cleaved by USER Enzyme at 37ºC for 15 min. This product was subjected to bead cleanup at a ratio of 0.6x sample to bed ratio. The eluted product was amplified for 6 cycles using NEBNext Ultra II Q5 Master Mix in the presence of NEBNext Multiplex Oligos for Illumina (NEB). The PCR product was purified with Ampure XP beads at a 0.6x sample to bead ratio. The product was a barcoded library containing Illumina P5 and P7 adapters for sequencing on Illumina instruments. The individual libraries were quantified, normalized and pooled at equimolar concentration and loaded onto the Illumina MiSeq sequencing instrument using V3 600-cycle reagent kits and a V3 flow cell for 250-cycle paired end sequencing (Illumina, CA).

### Genome Assembly

All raw paired end sequence reads are trimmed using Trim Galore version 0.6.4_dev^34^ removing NEB adapters and quality score below 20 from ends of the reads. The trimmed reads were assembled using the Burrows-Wheeler Aligner MEM algorithm (BWA-MEM) version 0.7.12^35^ with SARS-CoV-2 Wuhan-Hu-1 (GenBank accession number MN908947.3) as the reference sequence. Intrahost variant analysis of replicates (iVar)^36^ tool was used to remove primer sequences from the amplicon-based sequencing data. Finally, the mutation calls and consensus genome were built using a combination of samtools mpileup^37^ and iVar consensus, with a minimum quality score of 20, frequency threshold of 0.6, and minimum depth of 15 to optimize high quality variant calls. A sequence mapping quality control tool developed in-house was used to assess depth of coverage across all sequences, percent of ambiguous bases in the consensus genome and percent sequence mapped to the reference genome. Consensus genome with more than 3% ambiguous bases or less than 95% reference mapped were excluded from any further analyses.

### Library preparation and sequencing (PRL)

Positive RNA specimens between cycle threshold of 15-30 were selected from all samples tested at Pandemic Response Labs, NYC and cDNA for each specimen was generated using LunaScript RT SuperMix (NEB, MA) according to manufacturer protocol. To target SARS-CoV-2 specifically, cDNA for each specimen was amplified in two separate pools, 28- and 30-plex respectively, to generate 1200bp of overlapping amplicons^38^ using Q5 2x Hot-Start Master Mix (NEB, MA). The resulting pools are combined in equal volume and enriched for full length 1200 bp product using a SPRI-based magnetic bead cleanup. Enriched amplicons are tagmented (Illumina, CA) and barcoded (IDT, IA) and paired-end sequenced on an Illumina MiSeq or NextSeq 550.

### Genome Assembly (PRL)

For each specimen, sequencing adapters are first trimmed using Trim Galore v0.6.6^34^, then aligned to the SARS-CoV-2 Wuhan-Hu-1 reference genome (NCBI Nucleotide NC_045512.2) using BWA MEM 0.7.17-r1188^35^. Reads that are unmapped or those that have secondary alignments are discarded from the alignment. Consensus and mutations were called using samtools^37^ and Intrahost variant analysis of replicates (iVar)^36^ with a minimum quality score of 20, frequency threshold of 0.6 and a minimum read depth of 10x coverage. A consensus genome with ≥ 90% breath-of-coverage with ≤ 3000 ambiguous bases is considered a successful reconstruction (as per APHL recommendation).

### Genome alignment

Complete genome sequences produced by the NYC PHL and the PRL with reported collection dates on or before 04 March 2021 were analyzed. We restricted our analysis to genomes produced by public health surveillance to NYC to reduce bias due to geography or preferential sequencing of viral variants by academic institutions. Genomes were aligned to the Wuhan-Hu-1 reference genome (GenBank Accession MN908947) using mafft v7.475 (mafft --6merpair --keeplength --addfragments)^28^. Pango lineage designations^31^ for variants were assigned using Pangolin v2.3.2^39^.

### Segmented regression analysis

To estimate the timing and approximate linear slope of increase in B.1.526 and the E484K clade prevalence, we employed a segmented regression analysis (segmented package in R).

### Maximum likelihood phylogenetic inference

Maximum likelihood trees were inferred using IQTree2 for B.1.1.7, B.1.427, B.1.429, and B.1.526 genomes using a GTR+F+Γ4 substitution model^40^. Minimum branch length of 1e-9 was enforced and an expanded NNI search (--allnni) was employed to improve topology search. Preliminary molecular clock analyses were performed in TreeTime v0.8.1 using a fixed substitution rate of 8×10^−4^ substitutions/site/year and a skyline coalescent model^41^. This analysis identified 34 genomes whose root-to-tip genetic distance were flagged as problematic and excluded from subsequent phylodynamic analyses. TreeTime was also used to root and perform ancestral state reconstruction for a tree inferred from the 258 B.1.526 genomes sampled by the NYC PHL used to display the history of spike mutations in B.1.526 (**Figure 1**).

### Bayesian phylodynamic inference

We performed population growth rate inference in coalescence-based framework using an exponential growth model in BEAST 1.10.4^22^. We used a strict molecular clock model with the fixed substitution rate of 8×10^−4^ substitutions/site/year. We applied a GTR+F+Γ_4_ substitution model and specified the following priors for the population growth model: OneOnX distribution prior for the population size parameter and Laplace distribution prior (mean = 0.0, scale = 1.0) for the growth rate prior. Markov chain Monte Carlo analyses were run for 100-300 million generations; the first 10% of samples were discarded as burn-in. Separate inference was performed for B.1.1.7 (n=354), B.1.427 (n=35), B.1.429 (n=69), B.1.526 E484 (n=569), and B.1.526 E484K (n=678). For the B.1.526 phylodynamic inference, we did not include two sequences most closely related to B.1.526 (hCoV-19/USA/NY-NYCPHL-001701/2020 and hCoV-19/USA/NY-NYCPHL-002542/2021).

### Geocoding addresses

To identify areas with the highest density of B.1.526 sequenced genomes in NYC from December 2020 to March 2021, patient addresses were geocoded to be visualized on a map^42^. Geocoding was performed using the NYC DOHMH’s Geoportal application. Once geocoded, a map representing the point locations of individuals with sequenced B.1.526 genomes was created in ArcMap (v. 10.6.1) and exported as a point feature class.

### Point density method

Point density maps of individuals with B.1.526 sequenced genomes were created by using the point density tool in ArcMap. Point density calculates the density-per-unit area from point features (individuals with a SARS-CoV-2 B.1.526 sequenced genome) that fall within a defined neighborhood by totaling the number of points that fall within the neighborhood divided by the neighborhood area. Density calculations result in the observed gradient patterns. The point density map parameters were 4000 ft radius from the center of 250 square foot cells. The symbology class for point density classification was set at equal intervals of 5.

### Human plasma samples

Human plasma samples were among those collected in previously reported studies^8,13,15^. The study visits and blood draws were performed in compliance with all relevant ethical regulations and the protocol for human participants was approved by the Institutional Review Board (IRB) of the Rockefeller University (protocol #DRO-1006).

### Pseudovirus neutralization by human plasma samples

Human plasma samples were assayed for neutralization activity against lentiviruses pseudotyped with SARS-CoV-2 spike containing a 21-amino acid cytoplasmic tail deletion and either D614G or mutations corresponding to lineage B.1.526 (L5F, T95I, D253G, E484K, D614G, and A701V). Pseudotyped lentiviruses were generated and neutralizations assays were conducted as previously described^43,44^. Briefly, lentiviral particles were produced by co-transfecting the gene encoding SARS-CoV-2 spike protein (D614G or B.1.526) and Env-deficient HIV backbone expressing Luciferase-IRES-ZsGreen. Plasma samples were heat inactivated at 56ºC for 1 hour, then 3-fold serial diluted and incubated with SARS-CoV-2 pseudotyped virus for 1 hour at 37ºC. The virus/plasma mixture was added to 293T_ACE2_ target cells, which were seeded the previous day on poly-L-lysine coated plates. After incubating for 48 hours at 37ºC, target cells were lysed with Britelite Plus (Perkin Elmer) and luciferase activity was measured as relative luminesce units (RLUs) and normalized to values derived from cells infected with pseudotyped virus in the absence of plasma. Data were fit to 2-parameter non-linear regression in Antibody database^45^.

## Supporting information

Supp. Table 3 GISAID Acknowledgment part 1

Supp. Table 3 GISAID Acknowledgment part 2

Supp. Table 3 GISAID Acknowledgment part 3

## Data availability

The data analyzed as part of this project were obtained from the GISAID database and through a Data Use Agreement between NYC DOHMH and the University of California San Diego. Sequences analyzed by using the **vdb** tool were downloaded from GISAID. No personally identifying information were included as part of these analyses. SARS-CoV-2 genomes included in these analyses have been deposited in GISAID. See **Supplementary Data 1** for a list of genomes, including which genomes were excluded from the phylogenetic analysis. Data for **Figure 5** are provided in **Supplementary Table 2**.

## Code availability

The source code for the vdb program is available at the Github repository: https://github.com/variant-database/vdb.

## Acknowledgments

We thank the Global Initiative on Sharing Avian Influenza Data (GISAID) and the originating and submitting laboratories for sharing the SARS-CoV-2 genome sequences; see **Supplementary Table 3** for a list of sequence contributors. We thank Andrew Rambaut and Áine O’Toole for lineage designation. This work was supported by the Caltech Merkin Institute for Translational Research (P.J.B.) and the Bill and Melinda Gates Foundation Collaboration for AIDS Vaccine Discovery (CAVD) (INV-002143). J.O.W. acknowledges funding from the National Institutes of Health (AI135992 and AI136056). T.I.V. is funded by a Branco Weiss Fellowship. M.C.N. is an HHMI Investigator.

## Author Contributions

A.P.W., J.O.W., J.L.H., T.I.V., H.H.L., S.H. and J.C.W. analyzed data. J.C.W., M.A.C., E.G. and H.H.L. performed genome sequencing and assembly. J.O.W. curated data. C.G., M. Caskey and M.C.N. provided clinical samples. P.N.P.G. and J.R.K. carried out experiments. A.P.W., C.O.B., Z.Y., S.H., S.S.D., C.E.F. and J.O.W. prepared figures. A.P.W, J.O.W., T.I.V., C.O.B., J.C.W. and S.H. wrote the manuscript with input from all co-authors. A.P.W., P.J.B., J.O.W., J.L.R. and S.H. supervised the study.

## Supplementary Material

### Supplementary Methods

Commands for the program **vdb**, implementing a mutation pattern query language:

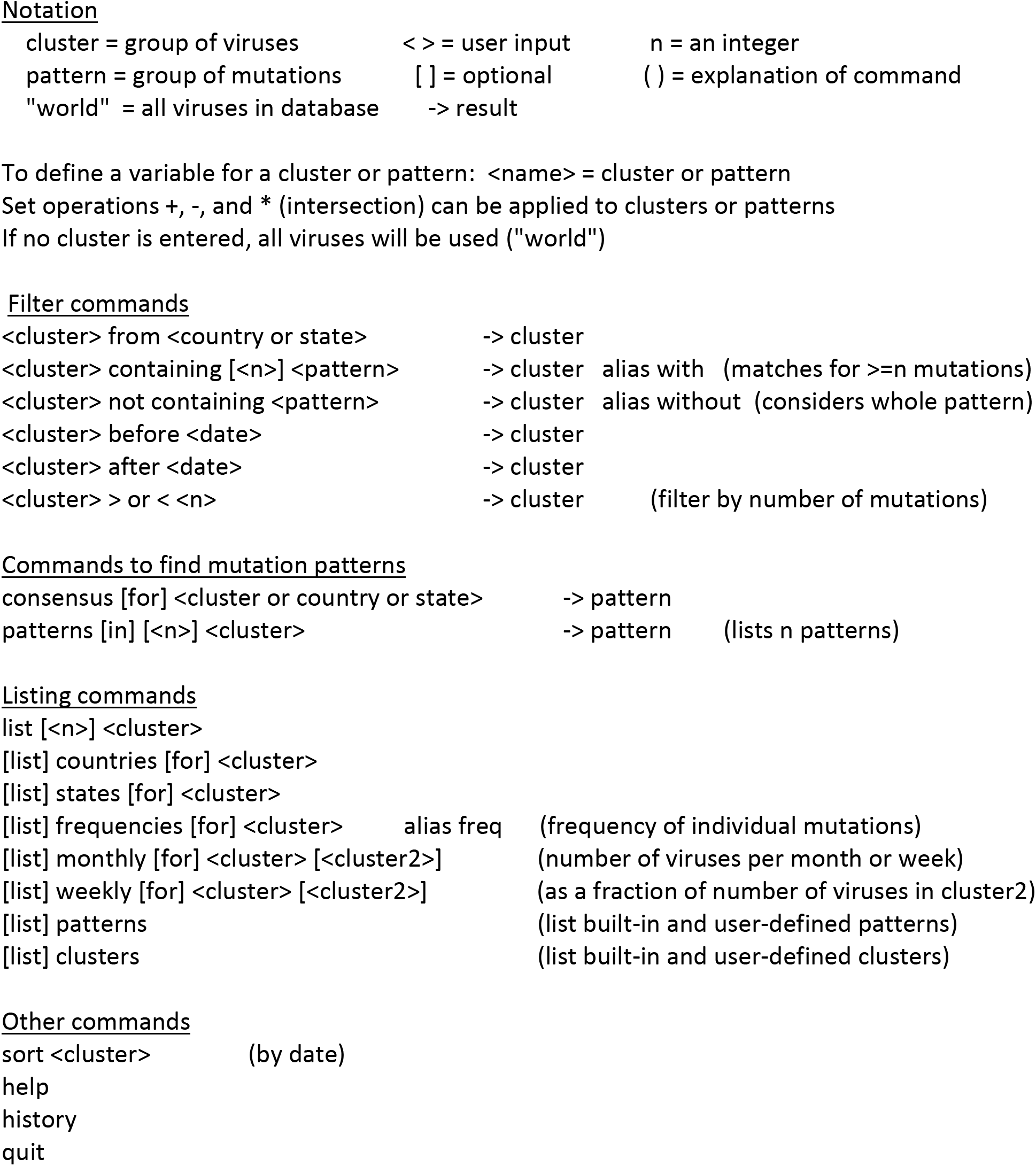

**Supplementary Figure 1.**
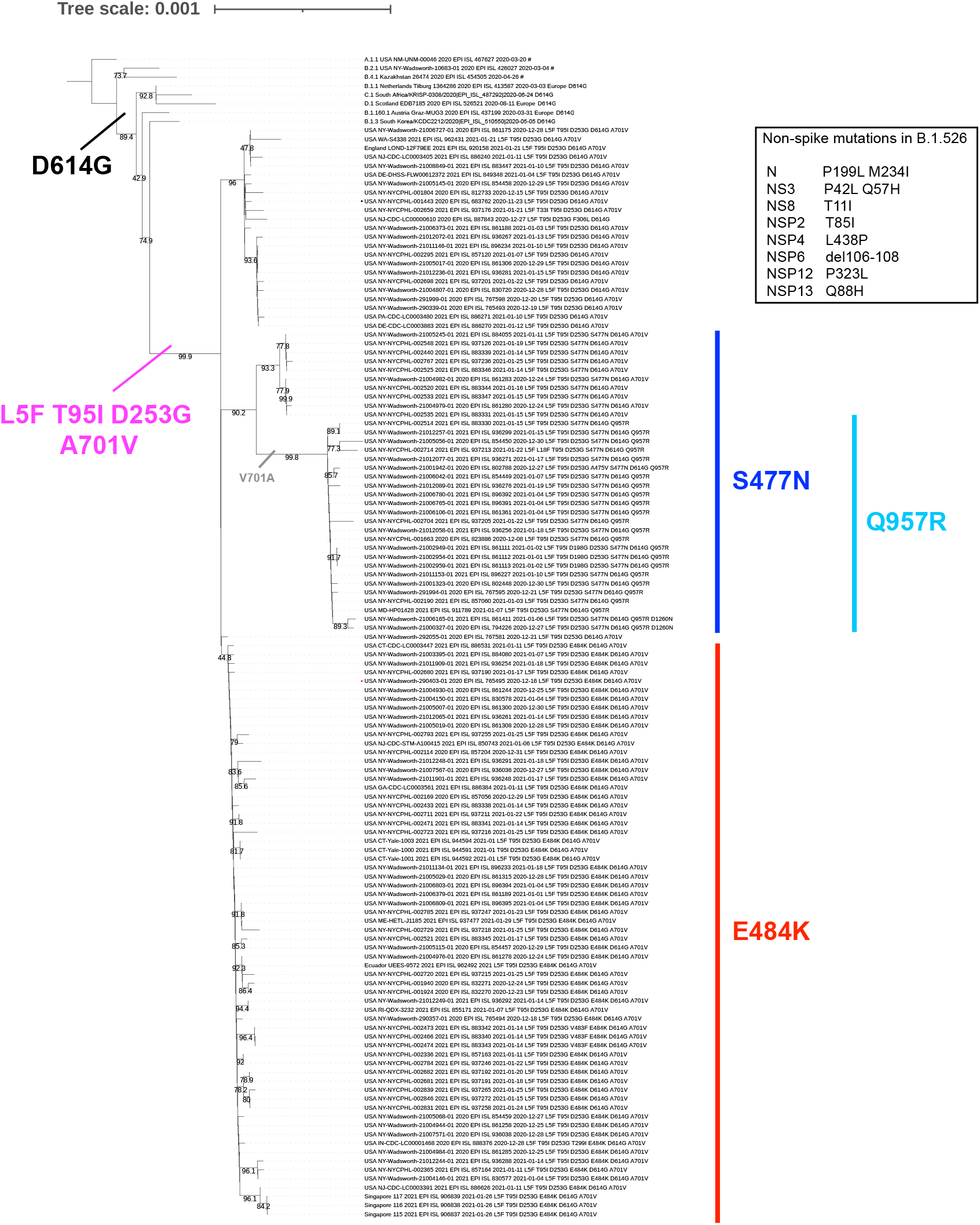
Phylogenetic tree of lineage B.1.526 indicating spike mutations. The inset lists non-spike mutations common in this lineage.

**Supplementary Figure 2.**
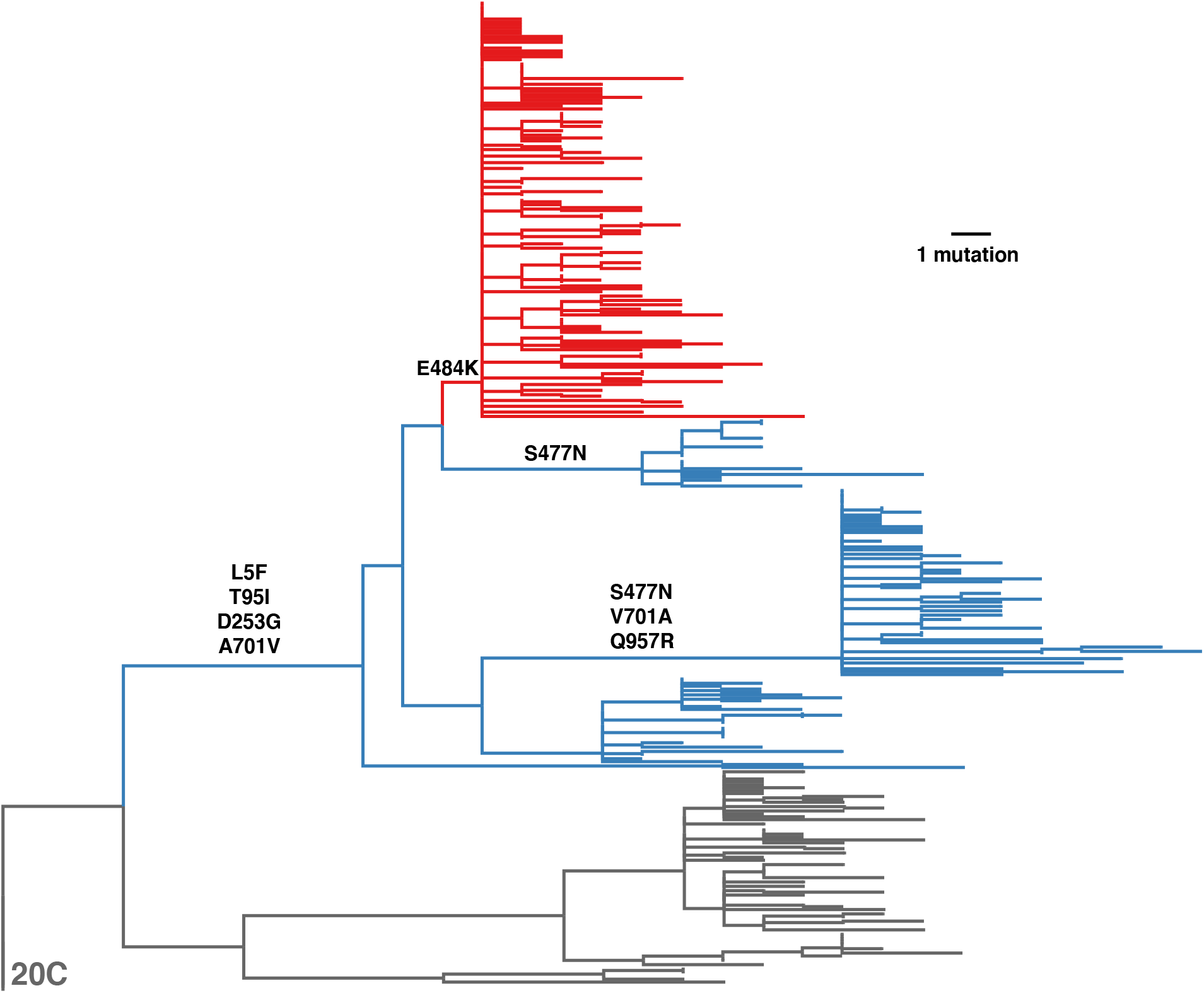
Maximum likelihood phylogenetic tree of the B.1.526 lineage in relation to a sister clade defined by an L452R spike mutation and the 20C ancestral virus (both shown in gray). Tree was rooted using the clade 20C ancestral viruses. Amino acid substitutions in the spike protein occurring on internal branches are labeled leading to and within B.1.526 are labeled. The B.1.526 lineage is colored blue, except for the clade defined by the E484K mutation, which is highlighted in red. The most common pattern of spike mutations in the sister clade is D80G, ΔY144, F157S, L452R, D614G, T859N, and D950H.

**Supplementary Table 1.**
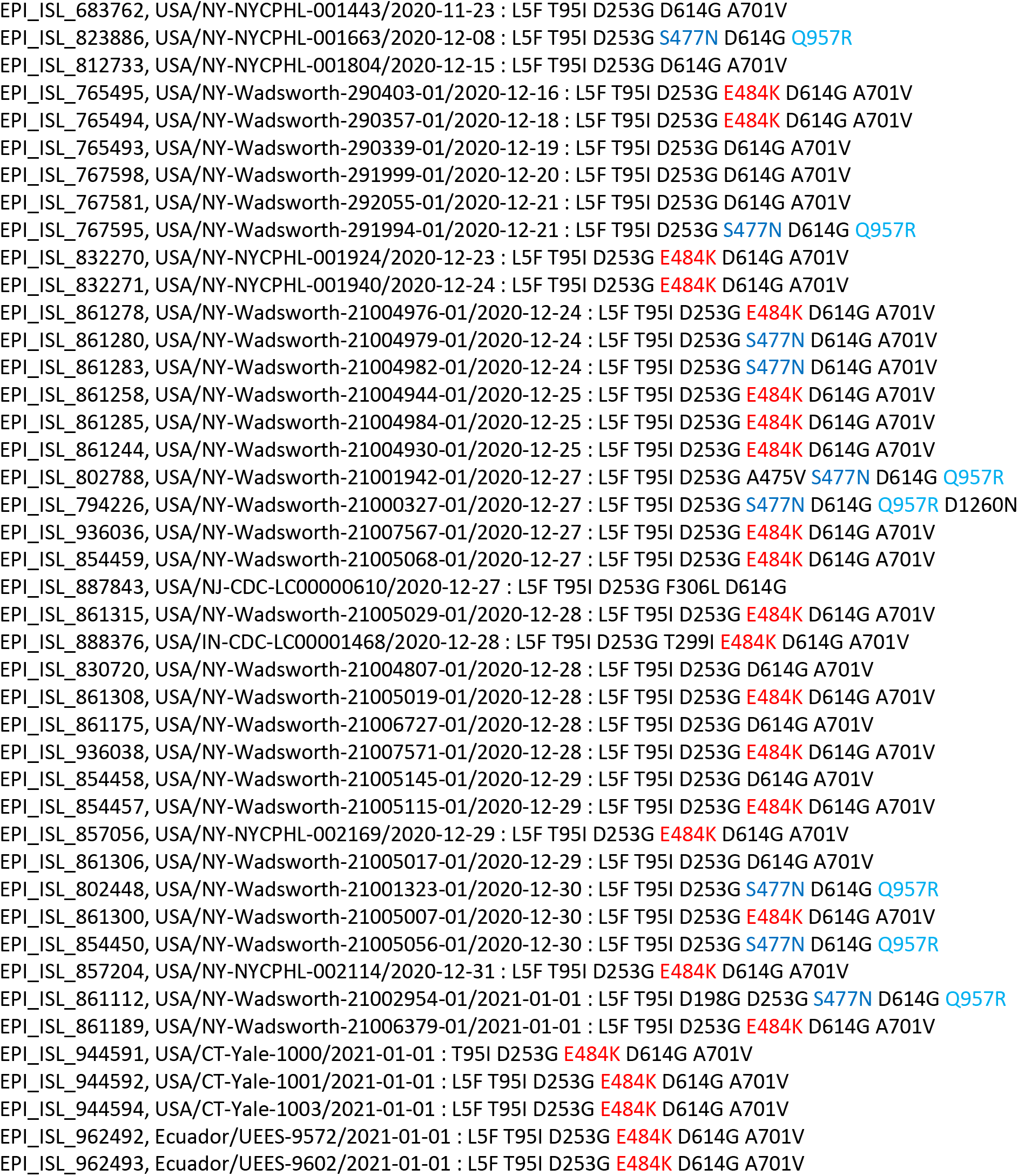
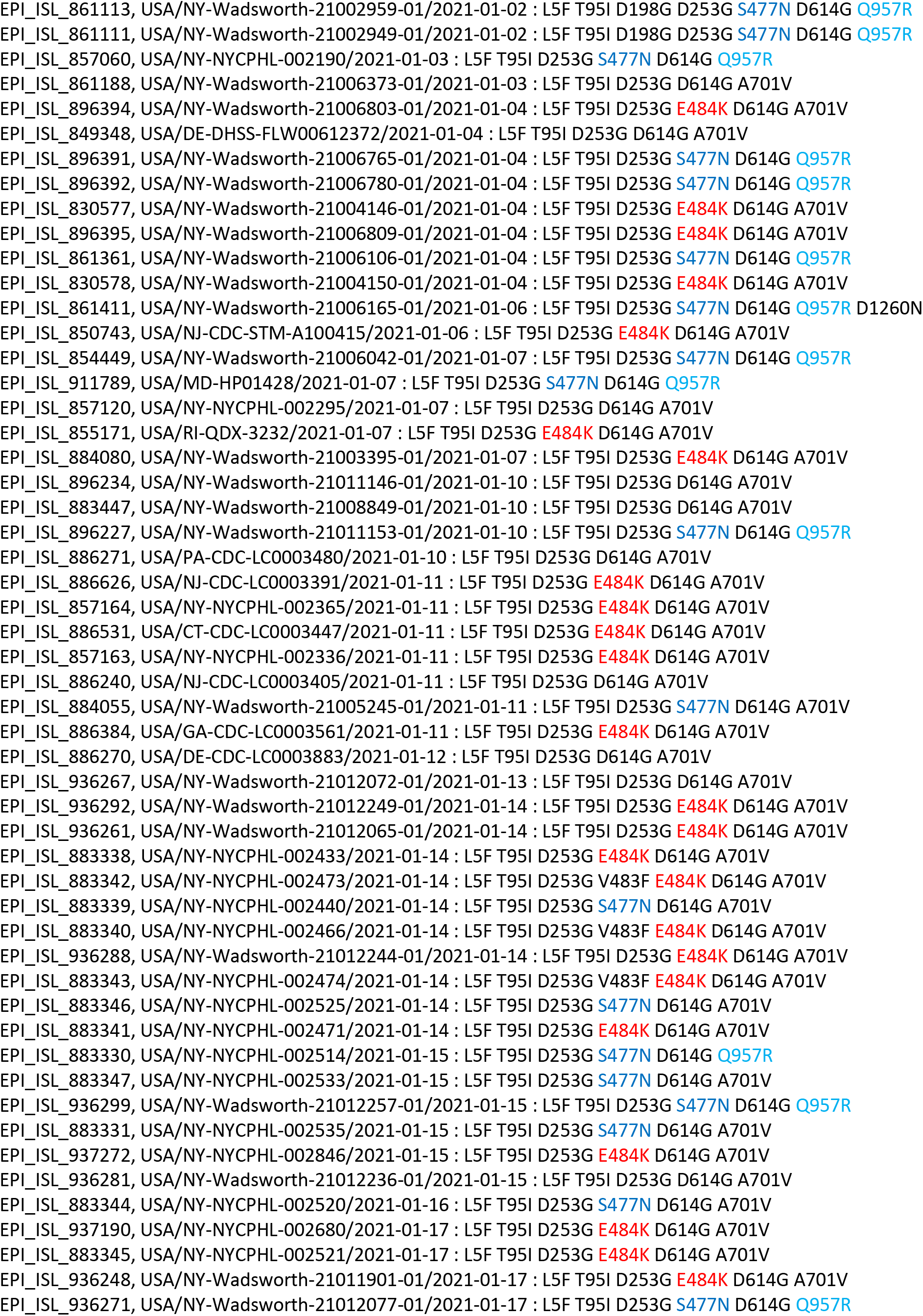
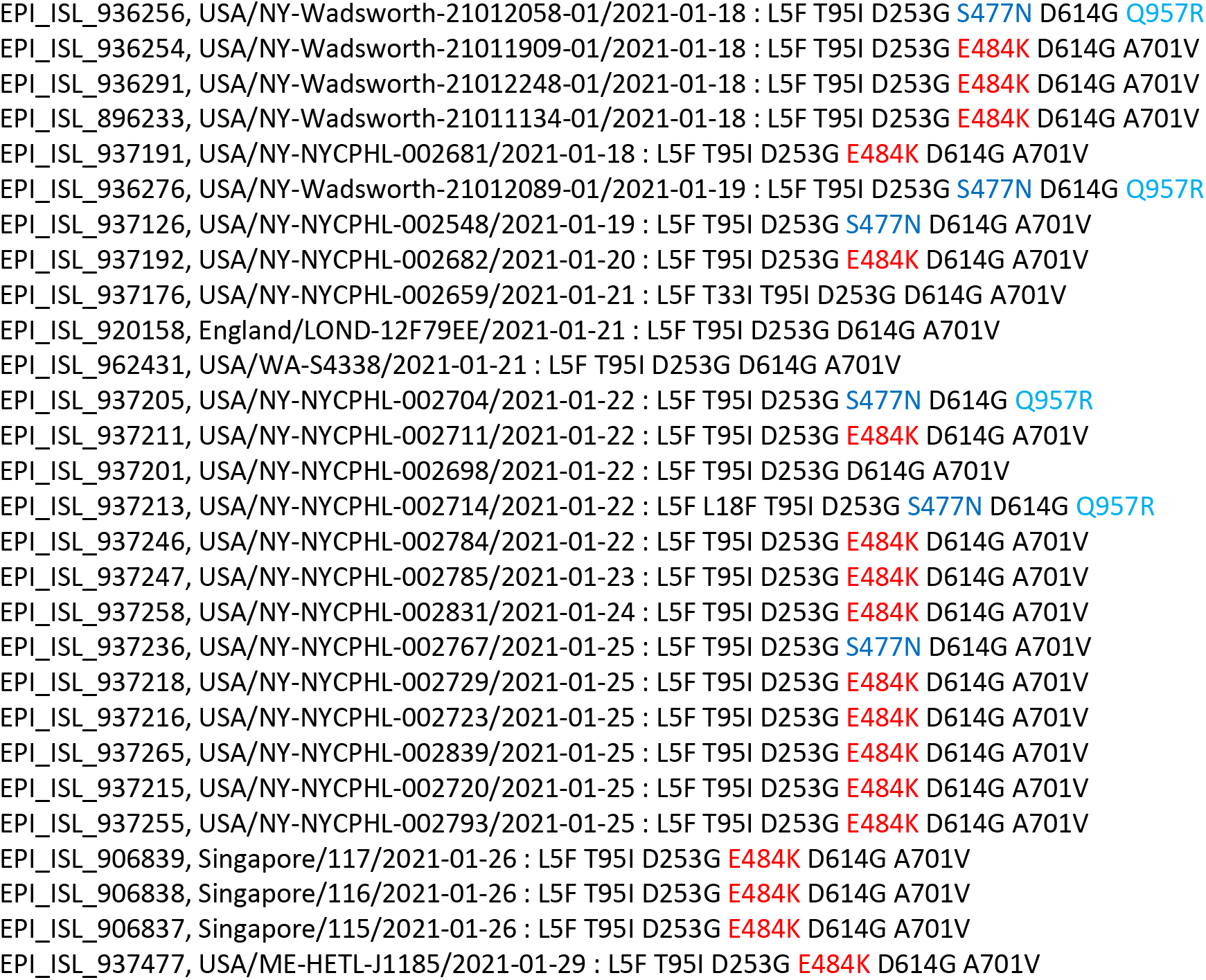
List of 124 viral genomes (with their accession number, location, collection date, and spike mutations) in lineage B.1.526. Mutations E484K, S477N, Q957R are highlighted in red, blue, and cyan, respectively.

