## Supplementary material for "Detection and characterization of the SARS-CoV-2 lineage B.1.526 in New York": Supp. Table 3 GISAID Acknowledgment part 2: gisaid_hcov-19_acknowledgement_table_2021_02_12_16-3.pdf

We gratefully acknowledge the following Authors from the Originating laboratories responsible for obtaining the specimens, as well as the Submitting laboratories where the genome data were generated and shared via GISAID, on which this research is based.

All Submitters of data may be contacted directly via www.gisaid.org

Authors are sorted alphabetically.

| Accession ID | Originating Laboratory | Submitting Laboratory | Authors |
|---|---|---|---|
| EPI_ISL_415710 | WHO National Influenza Centre Russian Federation | WHO National Influenza Centre Russian Federation | Andrey Komissarov, Artem Fadeev, Anna Ivanova, Daria Danilenko |
| EPI_ISL_416426 | Virological Research Group, Szentágothai Research Centre, University of Pécs | Bioinformatics Research Group, Szentágothai Research Centre, University of Pécs | Péter Urbán, Endre Gábor Tóth, Gábor Kemenesi, Róbert Herczeg, Attila Gyenesei, Ferenc Jakab |
| EPI_ISL_416457 | Andersen Lab, The Scripps Research Institute | Andersen Lab, The Scripps Research Institute | Mark Zeller, Catie Anderson, Emily Spender, Sarah Topol, Raphaelle Klitting, Refugio Robles-Sikisaka, Karthik Gangavarapu, Laura Nicholson, Kristian Andersen |
| EPI_ISL_416481 | R. G. Lugar Center for Public Health Research, National Center for Disease Control and Public Health (NCDC) of Georgia. | R. G. Lugar Center for Public Health Research, National Center for Disease Control and Public Health (NCDC) of Georgia. | Gvantsa Chanturia, Marine Murtskhvaladze, Nato Kotaria, Ann Machablishvili, Lela Sabadze, Mari Gavashelidze, Ana Papkiauri, Meri Pantsulaia, Gvantsa Brachveli, Tata Imnadze, Tamar Jashiashvili, Tea Tevdoradze, Ketevan Sidamonidze, Ekaterine Khmaladze, Ekaterine Zhghenti, Roena Sukhiashvili, Mariam Zakalashvili, Lela Urushadze, Magda Dgebuadze, Giorgi Tomashvili, Davit Tsaguria, Ekaterine Zangaladze, Nino Berishvili, Adam Kotorashvili, Maia Alkhazashvili, Irma Burjanadze, Anna Kasradze, Khatuna Zakhashvili, Paata Imnadze, Amiran Gamkrelidze. |
| EPI_ISL_416489, EPI_ISL_416491, EPI_ISL_416492 | University of Wisconsin-Madison AIDS Vaccine Research Laboratories | University of Wisconsin-Madison AIDS Vaccine Research Laboratories | Gage Moreno, Katarina Braun, et al. AIDS Vaccine Research Laboratories |
| EPI_ISL_416514, EPI_ISL_416515, EPI_ISL_416516, EPI_ISL_416517, EPI_ISL_416518 | Victorian Infectious Diseases Reference Laboratory (VIDRL) | Victorian Infectious Diseases Reference Laboratory and Microbiological Diagnostic Unit Public Health Laboratory, Doherty Institute | Caly L., Seemann T., Schultz M., Taiaroa, G., Druce J. |
| EPI_ISL_416538 | Wellington Hospital | Institute of Environmental Science and Research (ESR) | Wellington SCL, Wellington Hospital, Riddiford Street, Newtown, Wellington 6021, New Zealand |
| EPI_ISL_416539 | Wellington Hospital | Institute of Environmental Science and Research (ESR) | Matt Storey, Xiaoyun Ren, Craig Thornley, Maxim Bloomfield, Erasmus Smit, Lauren Jelly, Joep de Ligt |
| EPI_ISL_416683, EPI_ISL_416685, EPI_ISL_416704, EPI_ISL_416711, EPI_ISL_416713, EPI_ISL_416715, EPI_ISL_416717 | UW Virology Lab | UW Virology Lab | Pavitra Roychoudhury, Hong Xie, Keith Jerome, Alexander Greninger |
| EPI_ISL_416830, EPI_ISL_416831, EPI_ISL_416832 | NYU Langone Health | Department of Pathology and Medicine, New York University School of Medicine | John Chen, Dacia Dimartino, Xiaojun Feng, Adriana Heguy, Megan Hogan, Emily Huang, George Jour, Christian Marier, Matt Maurano, Mark Mulligan, Peter Meyn, Marie Samanovic-Golden, Amy Rapkiewicz, Guomiao Shen, Matija Snuderl, Gael Westby, Paul Zappile |
| EPI_ISL_417020, EPI_ISL_417021, EPI_ISL_417022, EPI_ISL_417023, EPI_ISL_417025 | Department of Clinical Microbiology | GIGA Medical Genomics | Durkin Keith, Artesi Maria, Bontems Sébastien, Boreux Raphaël, Meex Cécile, Melin Pierrette, Hayette Marie-Pierre, Bours Vincent. |
| EPI_ISL_417034 | Laboratorio de Ecologia de Doencas Transmissiveis na Amazonia, Instituto Leonidas e Maria Deane - Fiocruz Amazonia | Laboratorio de Ecologia de Doencas Transmissiveis na Amazonia, Instituto Leonidas e Maria Deane - Fiocruz Amazonia | Valdinete Nascimento, André Corado, Fernanda Nascimento, Ágatha Costa, Debora Duarte, Luciana Gonçalves, Michele Jesus, Sérgio Luz, Felipe Naveca |
| EPI_ISL_417374, EPI_ISL_417375, EPI_ISL_417376, EPI_ISL_417377, EPI_ISL_417379, EPI_ISL_417382 | UW Virology Lab | UW Virology Lab | Pavitra Roychoudhury, Hong Xie, Keith Jerome, Alexander Greninger |
| EPI_ISL_417408 | Centre for Infectious Diseases and Microbiology Public Health | NSW Health Pathology - Institute of Clinical Pathology and Medical Research; Westmead Hospital; University of Sydney | Carter I, Rahman H, Holmes EC, O'Sullivan MV, Sintchenko V, Chen SC, Maddocks S, Kok J, Dwyer DE, Rockett R, Eden J-S, Lam C, Gray K, Timms V, Gall M, Arnott A and Sadsad R for the 2019-nCoV Study Group |
| EPI_ISL_417409 | Centre for Infectious Diseases and Microbiology Public Health | NSW Health Pathology - Institute of Clinical Pathology and Medical Research; Westmead Hospital; University of Sydney | Rahman H, Holmes EC, O'Sullivan MV, Sintchenko V, Chen SC, Maddocks S, Kok J, Dwyer DE, Rockett R, Eden J-S, Lam C, Gray K, Timms V, Gall M, Arnott A, Sadsad R and Carter I for the 2019-nCoV Study Group |
| EPI_ISL_417410 | Centre for Infectious Diseases and Microbiology Public Health | NSW Health Pathology - Institute of Clinical Pathology and Medical Research; Westmead Hospital; University of Sydney | Holmes EC, O'Sullivan MV, Sintchenko V, Chen SC, Maddocks S, Kok J, Dwyer DE, Rockett R, Eden J-S, Lam C, Gray K, Timms V, Gall M, Arnott A, Sadsad R, Carter I and Rahman H for the 2019-nCoV Study Group |
| EPI_ISL_417411 | Centre for Infectious Diseases and Microbiology Public Health | NSW Health Pathology - Institute of Clinical Pathology and Medical Research; Westmead Hospital; University of Sydney | O'Sullivan MV, Sintchenko V, Chen SC, Maddocks S, Kok J, Dwyer DE, Rockett R, Eden J-S, Lam C, Gray K, Timms V, Gall M, Arnott A, Sadsad R, Carter I, Rahman H and Holmes EC for the 2019-nCoV Study Group |
| EPI_ISL_417412 | Centre for Infectious Diseases and Microbiology Public Health | NSW Health Pathology - Institute of Clinical Pathology and Medical Research; Westmead Hospital; University of Sydney | Sintchenko V, Chen SC, Maddocks S, Kok J, Dwyer DE, Rockett R, Eden J-S, Lam C, Gray K, Timms V, Gall M, Arnott A, Sadsad R, Carter I, Rahman H, Holmes EC and O'Sullivan MV for the 2019-nCoV Study Group |
| EPI_ISL_417413 | Ministry of Health Turkey | Ministry of Health Turkey | Fatma Bayrakdar,Aye Baak Alta,Yasemin Cogun,Gülay Korukluolu,Selçuk Klç |
| EPI_ISL_417435, EPI_ISL_417436, EPI_ISL_417437, EPI_ISL_417438, EPI_ISL_417439, EPI_ISL_417440, EPI_ISL_417441, EPI_ISL_417442 | Viral Respiratory Lab, National Institute for Biomedical Research (INRB) | Pathogen Sequencing Lab, National Institute for Biomedical Research (INRB) | Placide Mbala-Kingebeni, Edith Nkwembe, Eddy Kinganda-Lusamaki, Amuri Aziza, Catherine Pratt, Matthias Pauthner, Josh Quick, Allison Black, James Hadfield, Trevor Bedford, Ian Goodfellow, Nick Loman, Kristian Andersen, Michael Wiley, Steve Ahuka-Mundeke, Jean-Jacques Muyembe Tamfum |
| EPI_ISL_417467, EPI_ISL_417468 | Center of Medical Microbiology, Virology, and Hospital Hygiene, University of Duesseldorf | Center of Medical Microbiology, Virology, and Hospital Hygiene, University of Duesseldorf | Ortwin Adams, Marcel Andree, Alexander Dilthey, Torsten Feldt, Sandra Hauka, Torsten Houwaart, Björn-Erik Jensen, Detlef Kindgen-Milles, Malte Kohns Vasconcelos, Klaus Pfeffer, Tina Senff, Daniel Strelow, Jörg Timm, Andreas Walker, Tobias Wienemann |
| EPI_ISL_417504, EPI_ISL_417505, EPI_ISL_417506, EPI_ISL_417507, EPI_ISL_417508, EPI_ISL_417510, EPI_ISL_417512, EPI_ISL_417513, EPI_ISL_417515, EPI_ISL_417516 | University of Wisconsin-Madison AIDS Vaccine Research Laboratories | University of Wisconsin-Madison AIDS Vaccine Research Laboratories | Gage Moreno, Katarina Braun, et al. AIDS Vaccine Research Laboratories |
| EPI_ISL_417526, EPI_ISL_417527, EPI_ISL_417528, EPI_ISL_417529, EPI_ISL_417530, EPI_ISL_417531, EPI_ISL_417532, EPI_ISL_417533, EPI_ISL_417534 | Laboratoire Nationale de Santé, Microbiology, Virology | Laboratoire Nationale de Santé, Microbiology, Epidemiology and Microbial Genomics | Anke Wienecke-Baldacchino, Ardashel Latsuzbaia, Jessica Tapp, Catherine Ragimbeau, Guillaume Fournier, Tamir Abdelrahman, Trung Nguyen Nguyen, Joel Mossong |
| EPI_ISL_417536, EPI_ISL_417537, EPI_ISL_417538, EPI_ISL_417539, EPI_ISL_417540, EPI_ISL_417541, EPI_ISL_417542, EPI_ISL_417543, EPI_ISL_417544, EPI_ISL_417545, EPI_ISL_417546, EPI_ISL_417547, EPI_ISL_417548 | | | |
| see above | deCODE genetics | deCODE genetics | Daniel F Gudbjartsson; Agnar Helgason; Hakon Jonsson; Olafur T Magnusson; Pall Melsted; Gudmundur L Norddahl; Jona Saemundsdottir; Asgeir Sigurdsson; Patrick Sulem; Arna B Agustsdottir; Berglind Eiriksdottir; Run Fridriksdottir; Elisabet E Gardarsdottir; Gudmundur Georgsson; Olafia S Gretarsdottir; Kjartan R Gudmundsson; Thora R Gunnarsdottir; Arnaldur Gylfason; Hilma Holm; Brynjar O Jensson; Aslaug Jonasdottir; Kamilla S Josefsdottir; Thordur Kristjansson; Droplaug N Magnusdottir; Louise le Roux; Gudrun Sigmundsdottir; Gardar Sveinbjornsson; Kristin E Sveinsdottir; Maney |

| | | | |
|---|---|---|---|
| | | | Sveinsdottir; Emil A Thorarensen; Bjarni Thorbjornsson; Gisli Masson; Ingileif Jonsdottir; Alma Moller; Thorolfur Gudnason; Karl G Kristinsson; Unnur Thorsteinsdottir; Kari Stefansson |
| EPI_ISL_417550, EPI_ISL_417551 | The National University Hospital of Iceland | deCODE genetics | Daniel F Gudbjartsson; Agnar Helgason; Hakon Jonsson; Olafur T Magnusson; Pall Melsted; Gudmundur L Norddahl; Jona Saemundsdottir; Asgeir Sigurdsson; Patrick Sulem; Arna B Agustsdottir; Berglind Eiriksdottir; Run Fridriksdottir; Elisabet E Gardarsdottir; Gudmundur Georgsson; Olafia S Gretarsdottir; Kjartan R Gudmundsson; Thora R Gunnarsdottir; Arnaldur Gylfason; Hilma Holm; Brynjar O Jensson; Aslaug Jonasdottir; Kamilla S Josefsdottir; Thordur Kristjansson; Droplaug N Magnusdottir; Louise le Roux; Gudrun Sigmundsdottir; Gardar Sveinbjornsson; Kristin E Sveinsdottir; Maney Sveinsdottir; Emil A Thorarensen; Bjarni Thorbjornsson; Gisli Masson; Ingileif Jonsdottir; Alma Moller; Thorolfur Gudnason; Karl G Kristinsson; Unnur Thorsteinsdottir; Kari Stefansson |
| EPI_ISL_417552 | deCODE genetics | deCODE genetics | Daniel F Gudbjartsson; Agnar Helgason; Hakon Jonsson; Olafur T Magnusson; Pall Melsted; Gudmundur L Norddahl; Jona Saemundsdottir; Asgeir Sigurdsson; Patrick Sulem; Arna B Agustsdottir; Berglind Eiriksdottir; Run Fridriksdottir; Elisabet E Gardarsdottir; Gudmundur Georgsson; Olafia S Gretarsdottir; Kjartan R Gudmundsson; Thora R Gunnarsdottir; Arnaldur Gylfason; Hilma Holm; Brynjar O Jensson; Aslaug Jonasdottir; Kamilla S Josefsdottir; Thordur Kristjansson; Droplaug N Magnusdottir; Louise le Roux; Gudrun Sigmundsdottir; Gardar Sveinbjornsson; Kristin E Sveinsdottir; Maney Sveinsdottir; Emil A Thorarensen; Bjarni Thorbjornsson; Gisli Masson; Ingileif Jonsdottir; Alma Moller; Thorolfur Gudnason; Karl G Kristinsson; Unnur Thorsteinsdottir; Kari Stefansson |
| EPI_ISL_417553, EPI_ISL_417554, EPI_ISL_417555, EPI_ISL_417556, EPI_ISL_417557, EPI_ISL_417558, EPI_ISL_417559, EPI_ISL_417560, EPI_ISL_417561, EPI_ISL_417562, EPI_ISL_417563, EPI_ISL_417564, EPI_ISL_417565, EPI_ISL_417566, EPI_ISL_417567, EPI_ISL_417568, EPI_ISL_417569, EPI_ISL_417570, EPI_ISL_417571, EPI_ISL_417572, EPI_ISL_417573, EPI_ISL_417574, EPI_ISL_417575, EPI_ISL_417576, EPI_ISL_417577, EPI_ISL_417578, EPI_ISL_417579, EPI_ISL_417580, EPI_ISL_417581, EPI_ISL_417582, EPI_ISL_417583, EPI_ISL_417584, EPI_ISL_417585, EPI_ISL_417586, EPI_ISL_417587, EPI_ISL_417588, EPI_ISL_417589 | | | |
| see above | The National University Hospital of Iceland | deCODE genetics | Daniel F Gudbjartsson; Agnar Helgason; Hakon Jonsson; Olafur T Magnusson; Pall Melsted; Gudmundur L Norddahl; Jona Saemundsdottir; Asgeir Sigurdsson; Patrick Sulem; Arna B Agustsdottir; Berglind Eiriksdottir; Run Fridriksdottir; Elisabet E Gardarsdottir; Gudmundur Georgsson; Olafia S Gretarsdottir; Kjartan R Gudmundsson; Thora R Gunnarsdottir; Arnaldur Gylfason; Hilma Holm; Brynjar O Jensson; Aslaug Jonasdottir; Kamilla S Josefsdottir; Thordur Kristjansson; Droplaug N Magnusdottir; Louise le Roux; Gudrun Sigmundsdottir; Gardar Sveinbjornsson; Kristin E Sveinsdottir; Maney Sveinsdottir; Emil A Thorarensen; Bjarni Thorbjornsson; Gisli Masson; Ingileif Jonsdottir; Alma Moller; Thorolfur Gudnason; Karl G Kristinsson; Unnur Thorsteinsdottir; Kari Stefansson |
| EPI_ISL_417590 | deCODE genetics | deCODE genetics | Daniel F Gudbjartsson; Agnar Helgason; Hakon Jonsson; Olafur T Magnusson; Pall Melsted; Gudmundur L Norddahl; Jona Saemundsdottir; Asgeir Sigurdsson; Patrick Sulem; Arna B Agustsdottir; Berglind Eiriksdottir; Run Fridriksdottir; Elisabet E Gardarsdottir; Gudmundur Georgsson; Olafia S Gretarsdottir; Kjartan R Gudmundsson; Thora R Gunnarsdottir; Arnaldur Gylfason; Hilma Holm; Brynjar O Jensson; Aslaug Jonasdottir; Kamilla S Josefsdottir; Thordur Kristjansson; Droplaug N Magnusdottir; Louise le Roux; Gudrun Sigmundsdottir; Gardar Sveinbjornsson; Kristin E Sveinsdottir; Maney Sveinsdottir; Emil A Thorarensen; Bjarni Thorbjornsson; Gisli Masson; Ingileif Jonsdottir; Alma Moller; Thorolfur Gudnason; Karl G Kristinsson; Unnur Thorsteinsdottir; Kari Stefansson |
| EPI_ISL_417591, EPI_ISL_417592, EPI_ISL_417593, EPI_ISL_417594, EPI_ISL_417595, EPI_ISL_417596, EPI_ISL_417597, EPI_ISL_417598, EPI_ISL_417599, EPI_ISL_417600, EPI_ISL_417601, EPI_ISL_417602, EPI_ISL_417603, EPI_ISL_417604, EPI_ISL_417605, EPI_ISL_417606, EPI_ISL_417607, EPI_ISL_417608, EPI_ISL_417609, EPI_ISL_417610, EPI_ISL_417611, EPI_ISL_417612, EPI_ISL_417613, EPI_ISL_417614, EPI_ISL_417615, EPI_ISL_417616, EPI_ISL_417617 | | | |
| see above | The National University Hospital of Iceland | deCODE genetics | Daniel F Gudbjartsson; Agnar Helgason; Hakon Jonsson; Olafur T Magnusson; Pall Melsted; Gudmundur L Norddahl; Jona Saemundsdottir; Asgeir Sigurdsson; Patrick Sulem; Arna B Agustsdottir; Berglind Eiriksdottir; Run Fridriksdottir; Elisabet E Gardarsdottir; Gudmundur Georgsson; Olafia S Gretarsdottir; Kjartan R Gudmundsson; Thora R Gunnarsdottir; Arnaldur Gylfason; Hilma Holm; Brynjar O Jensson; Aslaug Jonasdottir; Kamilla S Josefsdottir; Thordur Kristjansson; Droplaug N Magnusdottir; Louise le Roux; Gudrun Sigmundsdottir; Gardar Sveinbjornsson; Kristin E Sveinsdottir; Maney Sveinsdottir; Emil A Thorarensen; Bjarni Thorbjornsson; Gisli Masson; Ingileif Jonsdottir; Alma Moller; Thorolfur Gudnason; Karl G Kristinsson; Unnur Thorsteinsdottir; Kari Stefansson |
| EPI_ISL_417618 | deCODE genetics | deCODE genetics | Daniel F Gudbjartsson; Agnar Helgason; Hakon Jonsson; Olafur T Magnusson; Pall Melsted; Gudmundur L Norddahl; Jona Saemundsdottir; Asgeir Sigurdsson; Patrick Sulem; Arna B Agustsdottir; Berglind Eiriksdottir; Run Fridriksdottir; Elisabet E Gardarsdottir; Gudmundur Georgsson; Olafia S Gretarsdottir; Kjartan R Gudmundsson; Thora R Gunnarsdottir; Arnaldur Gylfason; Hilma Holm; Brynjar O Jensson; Aslaug Jonasdottir; Kamilla S Josefsdottir; Thordur Kristjansson; Droplaug N Magnusdottir; Louise le Roux; Gudrun Sigmundsdottir; Gardar Sveinbjornsson; Kristin E Sveinsdottir; Maney Sveinsdottir; Emil A Thorarensen; Bjarni Thorbjornsson; Gisli Masson; Ingileif Jonsdottir; Alma Moller; Thorolfur Gudnason; Karl G Kristinsson; Unnur Thorsteinsdottir; Kari Stefansson |
| EPI_ISL_417619, EPI_ISL_417620, EPI_ISL_417621, EPI_ISL_417622, EPI_ISL_417623, EPI_ISL_417624, EPI_ISL_417625, EPI_ISL_417626, EPI_ISL_417627, EPI_ISL_417628, EPI_ISL_417629, EPI_ISL_417630, EPI_ISL_417631, EPI_ISL_417632, EPI_ISL_417633, EPI_ISL_417634, EPI_ISL_417635, EPI_ISL_417636, EPI_ISL_417637, EPI_ISL_417638, EPI_ISL_417639, EPI_ISL_417640, EPI_ISL_417641, EPI_ISL_417642, EPI_ISL_417643, EPI_ISL_417644, EPI_ISL_417645, EPI_ISL_417646, EPI_ISL_417647, EPI_ISL_417648, EPI_ISL_417649, EPI_ISL_417650, EPI_ISL_417651, EPI_ISL_417652, EPI_ISL_417653, EPI_ISL_417654 | | | |
| see above | The National University Hospital of Iceland | deCODE genetics | Daniel F Gudbjartsson; Agnar Helgason; Hakon Jonsson; Olafur T Magnusson; Pall Melsted; Gudmundur L Norddahl; Jona Saemundsdottir; Asgeir Sigurdsson; Patrick Sulem; Arna B Agustsdottir; Berglind Eiriksdottir; Run Fridriksdottir; Elisabet E Gardarsdottir; Gudmundur Georgsson; Olafia S Gretarsdottir; Kjartan R Gudmundsson; Thora R Gunnarsdottir; Arnaldur Gylfason; Hilma Holm; Brynjar O Jensson; Aslaug Jonasdottir; Kamilla S Josefsdottir; Thordur Kristjansson; Droplaug N Magnusdottir; Louise le Roux; Gudrun Sigmundsdottir; Gardar Sveinbjornsson; Kristin E Sveinsdottir; Maney Sveinsdottir; Emil A Thorarensen; Bjarni Thorbjornsson; Gisli Masson; Ingileif Jonsdottir; Alma Moller; Thorolfur Gudnason; Karl G Kristinsson; Unnur Thorsteinsdottir; Kari Stefansson |
| EPI_ISL_417666, EPI_ISL_417667, EPI_ISL_417668, EPI_ISL_417669, EPI_ISL_417670, EPI_ISL_417671, EPI_ISL_417672, EPI_ISL_417676 | deCODE genetics | deCODE genetics | Daniel F Gudbjartsson; Agnar Helgason; Hakon Jonsson; Olafur T Magnusson; Pall Melsted; Gudmundur L Norddahl; Jona Saemundsdottir; Asgeir Sigurdsson; Patrick Sulem; Arna B Agustsdottir; Berglind Eiriksdottir; Run Fridriksdottir; Elisabet E Gardarsdottir; Gudmundur Georgsson; Olafia S Gretarsdottir; Kjartan R Gudmundsson; Thora R Gunnarsdottir; Arnaldur Gylfason; Hilma Holm; Brynjar O Jensson; Aslaug Jonasdottir; Kamilla S Josefsdottir; Thordur Kristjansson; Droplaug N Magnusdottir; Louise le Roux; Gudrun Sigmundsdottir; Gardar Sveinbjornsson; Kristin E Sveinsdottir; Maney Sveinsdottir; Emil A Thorarensen; Bjarni Thorbjornsson; Gisli Masson; Ingileif Jonsdottir; Alma Moller; Thorolfur Gudnason; Karl G Kristinsson; Unnur Thorsteinsdottir; Kari Stefansson |
| EPI_ISL_417678, EPI_ISL_417680, EPI_ISL_417685, EPI_ISL_417699, EPI_ISL_417700, EPI_ISL_417703, EPI_ISL_417706, EPI_ISL_417709, EPI_ISL_417712, EPI_ISL_417716, EPI_ISL_417717, EPI_ISL_417724, EPI_ISL_417733, EPI_ISL_417737, EPI_ISL_417740, EPI_ISL_417742, EPI_ISL_417743, EPI_ISL_417746, EPI_ISL_417750, EPI_ISL_417752, EPI_ISL_417753, EPI_ISL_417754, EPI_ISL_417762, EPI_ISL_417763, EPI_ISL_417764, EPI_ISL_417766, EPI_ISL_417774, EPI_ISL_417808, EPI_ISL_417809, EPI_ISL_417813, EPI_ISL_417814, EPI_ISL_417815, EPI_ISL_417816, EPI_ISL_417818, EPI_ISL_417819, EPI_ISL_417820, EPI_ISL_417821, EPI_ISL_417822, EPI_ISL_417823, EPI_ISL_417824, EPI_ISL_417825, EPI_ISL_417826, EPI_ISL_417827, EPI_ISL_417829, EPI_ISL_417830, EPI_ISL_417831, EPI_ISL_417832, EPI_ISL_417833, EPI_ISL_417834, EPI_ISL_417835, EPI_ISL_417836, EPI_ISL_417837, EPI_ISL_417838, EPI_ISL_417839, EPI_ISL_417864 | | | |
| see above | The National University Hospital of Iceland | deCODE genetics | Daniel F Gudbjartsson; Agnar Helgason; Hakon Jonsson; Olafur T Magnusson; Pall Melsted; Gudmundur L Norddahl; Jona Saemundsdottir; Asgeir Sigurdsson; Patrick Sulem; Arna B Agustsdottir; Berglind Eiriksdottir; Run Fridriksdottir; Elisabet E Gardarsdottir; Gudmundur Georgsson; Olafia S Gretarsdottir; Kjartan R Gudmundsson; Thora R Gunnarsdottir; Arnaldur Gylfason; Hilma Holm; Brynjar O Jensson; Aslaug Jonasdottir; Kamilla S Josefsdottir; Thordur Kristjansson; Droplaug N Magnusdottir; Louise le Roux; Gudrun Sigmundsdottir; Gardar Sveinbjornsson; Kristin E Sveinsdottir; Maney Sveinsdottir; Emil A Thorarensen; Bjarni Thorbjornsson; Gisli Masson; Ingileif Jonsdottir; Alma Moller; Thorolfur Gudnason; Karl G Kristinsson; Unnur Thorsteinsdottir; Kari Stefansson |
| EPI_ISL_417918 | Department of Medical Microbiology, University Malaya Medical Centre | Department of Medical Microbiology, Faculty of Medicine, University of Malaya | Yoong Min CHONG, Sasheela PONNAMPALAVANAR, Sharifah Faridah SYED OMAR, Adeeba KAMARULZAMAN,Vijayan MUNUSAMY, Chee Kuan WONG, Cindy Shuan Ju TEH, I-Ching SAM, Yoke Fun Chan, University Malaya Medical Centre COVID Team |
| EPI_ISL_417925 | Laboratório Simili | Bioinformatics Laboratory / LNCC | Filipe Romero, Ana Paula Guimarães, Mariane Talon, Luiz Gonzaga Paula de Almeida, Ronaldo da Silva, Francisco Junior, Diana Mariani, Lídia Boullosa, Alexandra Gerber, Jaqueline Goes de Jesus, Ingra Morales Claro, Ester Cerdeira Sabino, Nuno Rodrigues Faria, Terezinha Marta Pereira, Pinto Castiñeiras, Isabela de Carvalho Leitão, Rafael de Mello Galliez, Cássia Cristina Alves Gonçalves, Érica Ramos dos Santos Nascimento, Richard Araújo Maia, Mauro Teixeira,Cristiano Xavier Lima, Orlando Ferreira Jr., Rodrigo Brindeiro, Luciana Jesus Costa e André Felipe Santos, Laboratorio Hermes Pardini, Laboratorio Simile, Amilcar Tanuri, Renato Santana Aguiar e Ana Tereza Vasconcelos |
| EPI_ISL_417926 | Laboratório Simili | Bioinformatics Laboratory / LNCC | Filipe Romero, Ana Paula Guimarães, Mariane Talon, Luiz Gonzaga Paula de Almeida, Ronaldo da Silva Francisco Junior, Diana Mariani, Lídia Boullosa, Alexandra Gerber, Jaqueline Goes de Jesus, Ingra Morales Claro, Ester Cerdeira Sabino, Nuno Rodrigues Faria, Terezinha Marta Pereira, Pinto Castiñeiras, Isabela de Carvalho Leitão, Rafael de Mello Galliez, Cássia Cristina Alves Gonçalves, Érica Ramos dos Santos Nascimento, Richard Araújo Maia, Mauro Teixeira,Cristiano Xavier Lima, Orlando Ferreira Jr., Rodrigo Brindeiro, Luciana Jesus Costa e André Felipe Santos, Laboratorio Hermes Pardini, Laboratorio Simile, Amilcar Tanuri, Renato Santana Aguiar e Ana Tereza Vasconcelos |

| | | | |
|---|---|---|---|
| EPI_ISL_417931, EPI_ISL_417932, EPI_ISL_417933, EPI_ISL_417935 | UCSF Clinical Microbiology Laboratory | Chan-Zuckerberg Biohub | Shaun Arevalo, Josh Batson, Olga Botvinnik, Gloria Castaneda, Angela Detweiler, David Dynerman, Samantha Hao, Jack Kamm, Amy Kistler, G. Renuka Kumar, Chaz Langelier, Lucy Li, Steve Miller, Lusajo Mwakibete, Norma Neff, Angela Pisco, Maira Phelps, Michelle Tan, Chunyu Zhao |
| EPI_ISL_417936 | Laboratório Hermes Pardini | Bioinformatics Laboratory - LNCC | Filipe Romero, Ana Paula Guimarães, Mariane Talon, Luiz Gonzaga Paula de Almeida, Ronaldo da Silva Francisco Junior, Diana Mariani, Lídia Boullosa, Alexandra Gerber, Jaqueline Goes de Jesus, Ingra Morales Claro, Ester Cerdeira Sabino, Nuno Rodrigues Faria, Terezinha Marta Pereira, Pinto Castiñeiras, Isabela de Carvalho Leitão, Rafael de Mello Galliez, Cássia Cristina Alves Gonçalves, Érica Ramos dos Santos Nascimento, Richard Araújo Maia, Mauro Teixeira,Cristiano Xavier Lima, Orlando Ferreira Jr., Rodrigo Brindeiro, Luciana Jesus Costa e André Felipe Santos, Laboratorio Hermes Pardini, Laboratorio Simile, Amilcar Tanuri, Renato Santana Aguiar e Ana Tereza Vasconcelos |
| EPI_ISL_417937, EPI_ISL_417938, EPI_ISL_417939 | UCSF Clinical Microbiology Laboratory | Chan-Zuckerberg Biohub | Shaun Arevalo, Josh Batson, Olga Botvinnik, Gloria Castaneda, Angela Detweiler, David Dynerman, Samantha Hao, Jack Kamm, Amy Kistler, G. Renuka Kumar, Chaz Langelier, Lucy Li, Steve Miller, Lusajo Mwakibete, Norma Neff, Angela Pisco, Maira Phelps, Michelle Tan, Chunyu Zhao |
| EPI_ISL_417940 | Laboratório Hermes Pardini | Bioinformatics Laboratory - LNCC | Filipe Romero, Ana Paula Guimarães, Mariane Talon, Luiz Gonzaga Paula de Almeida, Ronaldo da Silva Francisco Junior, Diana Mariani, Lídia Boullosa,Alexandra Gerber, Jaqueline Goes de Jesus, Ingra Morales Claro, Ester Cerdeira Sabino, Nuno Rodrigues Faria, Terezinha Marta Pereira, Pinto Castiñeiras, Isabela de Carvalho Leitão, Rafael de Mello Galliez, Cássia Alves Gonçalves, Érica Ramos dos Santos Nascimento, Richard Araújo Maia, Mauro Teixeira,Cristiano Xavier Lima, Orlando Ferreira Jr., Rodrigo Brindeiro, Luciana Jesus Costa e André Felipe Santos, Laboratorio Hermes Pardini, Laboratorio Simile, Amilcar Tanuri, Renato Santana Aguiar e Ana Tereza Vasconcelos |
| EPI_ISL_417941, EPI_ISL_417942 | Viral Respiratory Lab, National Institute for Biomedical Research (INRB) | Pathogen Sequencing Lab, National Institute for Biomedical Research (INRB) | Placide Mbala-Kingebeni, Edith Nkwembe, Eddy Kinganda-Lusamaki, Amuri Aziza, Catherine Pratt, Matthias Pauthner, Josh Quick, Allison Black, James Hadfield, Trevor Bedford, Ian Goodfellow, Nick Loman, Kristian Andersen, Michael Wiley, Steve Ahuka-Mundeke, Jean-Jacques Muyembe Tamfum |
| EPI_ISL_417943 | Laboratório Hermes Pardini | Bioinformatics Laboratory | Filipe Romero, Ana Paula Guimarães, Mariane Talon, Luiz Gonzaga Paula de Almeida, Ronaldo da Silva Francisco Junior, Diana Mariani, Lídia Boullosa,Alexandra Gerber, Jaqueline Goes de Jesus, Ingra Morales Claro, Ester Cerdeira Sabino, Nuno Rodrigues Faria, Terezinha Marta Pereira, Pinto Castiñeiras, Isabela de Carvalho Leitão, Rafael de Mello Galliez, Cássia Alves Gonçalves, Érica Ramos dos Santos Nascimento, Richard Araújo Maia, Mauro Teixeira,Cristiano Xavier Lima, Orlando Ferreira Jr., Rodrigo Brindeiro, Luciana Jesus Costa e André Felipe Santos, Laboratorio Hermes Pardini, Laboratorio Simile, Amilcar Tanuri, Renato Santana Aguiar e Ana Tereza Vasconcelos |
| EPI_ISL_417944 | Viral Respiratory Lab, National Institute for Biomedical Research (INRB) | Pathogen Sequencing Lab, National Institute for Biomedical Research (INRB) | Placide Mbala-Kingebeni, Edith Nkwembe, Eddy Kinganda-Lusamaki, Amuri Aziza, Catherine Pratt, Matthias Pauthner, Josh Quick, Allison Black, James Hadfield, Trevor Bedford, Ian Goodfellow, Nick Loman, Kristian Andersen, Michael Wiley, Steve Ahuka-Mundeke, Jean-Jacques Muyembe Tamfum |
| EPI_ISL_417945 | Laboratório Hermes Pardini | Bioinformatics Laboratory - LNCC | Filipe Romero, Ana Paula Guimarães, Mariane Talon, Luiz Gonzaga Paula de Almeida, Ronaldo da Silva Francisco Junior, Diana Mariani, Lídia Boullosa,Alexandra Gerber, Jaqueline Goes de Jesus, Ingra Morales Claro, Ester Cerdeira Sabino, Nuno Rodrigues Faria, Terezinha Marta Pereira, Pinto Castiñeiras, Isabela de Carvalho Leitão, Rafael de Mello Galliez, Cássia Alves Gonçalves, Érica Ramos dos Santos Nascimento, Richard Araújo Maia, Mauro Teixeira,Cristiano Xavier Lima, Orlando Ferreira Jr., Rodrigo Brindeiro, Luciana Jesus Costa e André Felipe Santos, Laboratorio Hermes Pardini, Laboratorio Simile, Amilcar Tanuri, Renato Santana Aguiar e Ana Tereza Vasconcelos |
| EPI_ISL_417946 | Viral Respiratory Lab, National Institute for Biomedical Research (INRB) | Pathogen Sequencing Lab, National Institute for Biomedical Research (INRB) | Placide Mbala-Kingebeni, Edith Nkwembe, Eddy Kinganda-Lusamaki, Amuri Aziza, Catherine Pratt, Matthias Pauthner, Josh Quick, Allison Black, James Hadfield, Trevor Bedford, Ian Goodfellow, Nick Loman, Kristian Andersen, Michael Wiley, Steve Ahuka-Mundeke, Jean-Jacques Muyembe Tamfum |
| EPI_ISL_417949 | Laboratório Hermes Pardini | Bioinformatics Laboratory - LNCC | Filipe Romero, Ana Paula Guimarães, Mariane Talon, Luiz Gonzaga Paula de Almeida, Ronaldo da Silva Francisco Junior, Diana Mariani, Lídia Boullosa,Alexandra Gerber, Jaqueline Goes de Jesus, Ingra Morales Claro, Ester Cerdeira Sabino, Nuno Rodrigues Faria, Terezinha Marta Pereira, Pinto Castiñeiras, Isabela de Carvalho Leitão, Rafael de Mello Galliez, Cássia Alves Gonçalves, Érica Ramos dos Santos Nascimento, Richard Araújo Maia, Mauro Teixeira,Cristiano Xavier Lima, Orlando Ferreira Jr., Rodrigo Brindeiro, Luciana Jesus Costa e André Felipe Santos, Laboratorio Hermes Pardini, Laboratorio Simile, Amilcar Tanuri, Renato Santana Aguiar e Ana Tereza Vasconcelos |
| EPI_ISL_417951, EPI_ISL_417953 | Universidade Federal do Rio de Janeiro | Bioinformatics Laboratory - LNCC | Filipe Romero, Ana Paula Guimarães, Mariane Talon, Luiz Gonzaga Paula de Almeida, Ronaldo da Silva Francisco Junior, Diana Mariani, Lídia Boullosa,Alexandra Gerber, Jaqueline Goes de Jesus, Ingra Morales Claro, Ester Cerdeira Sabino, Nuno Rodrigues Faria, Terezinha Marta Pereira, Pinto Castiñeiras, Isabela de Carvalho Leitão, Rafael de Mello Galliez, Cássia Alves Gonçalves, Érica Ramos dos Santos Nascimento, Richard Araújo Maia, Mauro Teixeira,Cristiano Xavier Lima, Orlando Ferreira Jr., Rodrigo Brindeiro, Luciana Jesus Costa e André Felipe Santos, Laboratorio Hermes Pardini, Laboratorio Simile, Amilcar Tanuri, Renato Santana Aguiar e Ana Tereza Vasconcelos |
| EPI_ISL_417970 | Utah Public Health Laboratory | Utah Public Health Laboratory | Erin Young, Kelly Oakeson |
| EPI_ISL_418009 | HSE Ilha Terceira - Angra do Heroismo | Instituto Nacional de Saude (INSA) | Guiomar et al |
| EPI_ISL_418017 | CHMT | Instituto Nacional de Saude (INSA) | Guiomar et al |
| EPI_ISL_418018 | H Garcia de Orta | Instituto Nacional de Saude (INSA) | Guiomar et al |
| EPI_ISL_418019, EPI_ISL_418020, EPI_ISL_418021, EPI_ISL_418022 | H Braga | Instituto Nacional de Saude (INSA) | Guiomar et al |
| EPI_ISL_418023 | H Evora | Instituto Nacional de Saude (INSA) | Guiomar et al |
| EPI_ISL_418024 | CHUA - Faro | Instituto Nacional de Saude (INSA) | Guiomar et al |
| EPI_ISL_418025 | H Santarem | Instituto Nacional de Saude (INSA) | Guiomar et al |
| EPI_ISL_418026 | H Dr. Nelio Mendonca - Funchal | Instituto Nacional de Saude (INSA) | Guiomar et al |
| EPI_ISL_418027 | CHTMAD | Instituto Nacional de Saude (INSA) | Guiomar et al |
| EPI_ISL_418029, EPI_ISL_418030, EPI_ISL_418031, EPI_ISL_418032, EPI_ISL_418033, EPI_ISL_418034, EPI_ISL_418037, EPI_ISL_418038, EPI_ISL_418040, EPI_ISL_418046, EPI_ISL_418047, EPI_ISL_418048, EPI_ISL_418050, EPI_ISL_418052, EPI_ISL_418053, EPI_ISL_418054, EPI_ISL_418063, EPI_ISL_418064, EPI_ISL_418067, EPI_ISL_418071, EPI_ISL_418072, EPI_ISL_418073, EPI_ISL_418074, EPI_ISL_418075, EPI_ISL_418076, EPI_ISL_418077, EPI_ISL_418078, EPI_ISL_418079, EPI_ISL_418080, EPI_ISL_418081, EPI_ISL_418082 | | | |
| see above | UW Virology Lab | UW Virology Lab | Pavitra Roychoudhury, Hong Xie, Keith Jerome, Alexander Greninger |
| EPI_ISL_418101, EPI_ISL_418102, EPI_ISL_418103, EPI_ISL_418104, EPI_ISL_418105, EPI_ISL_418126, EPI_ISL_418127, EPI_ISL_418128, EPI_ISL_418129, EPI_ISL_418130, EPI_ISL_418131, EPI_ISL_418132, EPI_ISL_418133, EPI_ISL_418134, EPI_ISL_418135, EPI_ISL_418136, EPI_ISL_418137, EPI_ISL_418138, EPI_ISL_418139, EPI_ISL_418140, EPI_ISL_418148, EPI_ISL_418149, EPI_ISL_418150, EPI_ISL_418151, EPI_ISL_418152, EPI_ISL_418153, EPI_ISL_418154, EPI_ISL_418155, EPI_ISL_418156, EPI_ISL_418157, EPI_ISL_418158, EPI_ISL_418159, EPI_ISL_418160, EPI_ISL_418161, EPI_ISL_418162, EPI_ISL_418163, EPI_ISL_418164, EPI_ISL_418165 | | | |
| see above | Wales Specialist Virology Centre | Public Health Wales Microbiology Cardiff | Catherine Moore, Joanne Watkins, Sally Corden, Sara Rey, Matt Bull, Tom Connor |
| EPI_ISL_418183 | Virological Research Group, Szentágothai Research Centre | Bioinformatics Research Group, Szentágothai Research Centre | Péter Urbán, Endre Gábor Tóth, Gábor Kemenesi, Róbert Herczeg, Attila Gyenesei, Ferenc Jakab |
| EPI_ISL_418184 | Gundersen Molecular Diagnostics Laboratory | Kabara Cancer Research Institute | Craig S. Richmond & Paraic A. Kenny |
| EPI_ISL_418185 | Gundersen Molecular Diagnostic Laboratory | Kabara Cancer Research Institute | Craig S. Richmond & Paraic A. Kenny |
| EPI_ISL_418190, EPI_ISL_418191, EPI_ISL_418192, EPI_ISL_418193, EPI_ISL_418194, EPI_ISL_418195, EPI_ISL_418197, EPI_ISL_418198, EPI_ISL_418199, EPI_ISL_418200, EPI_ISL_418201, EPI_ISL_418202, EPI_ISL_418203, EPI_ISL_418204 | | | |
| see above | NYU Langone Health | Department of Pathology and Medicine, New York University School of Medicine | Margaret Black, John Cadley, Paolo Cotzia, John Chen, Dacia Dimartino, Xiaojun Feng, Adriana Heguy, Megan Hogan, Emily Huang, George Jour, Christian Marier, Matthew T. Maurano, Mark J. Mulligan, Peter Meyn, Jared Pinnell, Amy Rapkiewicz, Marie Samanovic-Golden, Antonio Serrano, Guomiao Shen, Matija Snuderl, Nick Vulpescu, Gael Westby, Paul Zappile |
| EPI_ISL_418231 | Centre Hospitalier Compiègne Laboratoire de Biologie | National Reference Center for Viruses of Respiratory Infections, Institut Pasteur, Paris | Mélanie Albert, Marion Barbet, Sylvie Behillil, Méline Bizard, Angela Brisebarre, Flora Donati, Etienne Simon-Lorière, Vincent Enouf, Maud Vanpeene, Sylvie van der Werf, Raulin Olivia |
| EPI_ISL_418232, EPI_ISL_418233 | Service des Urgences | National Reference Center for Viruses of Respiratory Infections, Institut Pasteur, Paris | Mélanie Albert, Marion Barbet, Sylvie Behillil, Méline Bizard, Angela Brisebarre, Flora Donati, Etienne Simon-Lorière, Vincent Enouf, Maud Vanpeene, Sylvie van der Werf, Boubkeur |
| EPI_ISL_418235 | Cabinet médical | National Reference Center for Viruses of Respiratory Infections, Institut Pasteur, Paris | Mélanie Albert, Marion Barbet, Sylvie Behillil, Méline Bizard, Angela Brisebarre, Flora Donati, Etienne Simon-Lorière, Vincent Enouf, Maud Vanpeene, Sylvie van der Werf |

| | | | |
|---|---|---|---|
| EPI_ISL_418236, EPI_ISL_418237, EPI_ISL_418238, EPI_ISL_418239 | Centre Hospitalier Compiègne Laboratoire de Biologie | National Reference Center for Viruses of Respiratory Infections, Institut Pasteur, Paris | Mélanie Albert, Marion Barbet, Sylvie Behillil, Méline Bizard, Angela Brisebarre, Flora Donati, Etienne Simon-Lorière, Vincent Enouf, Maud Vanpeene, Sylvie van der Werf, Raulin Olivia |
| EPI_ISL_418240 | LABM GH nord Essonne | National Reference Center for Viruses of Respiratory Infections, Institut Pasteur, Paris | Mélanie Albert, Marion Barbet, Sylvie Behillil, Méline Bizard, Angela Brisebarre, Flora Donati, Etienne Simon-Lorière, Vincent Enouf, Maud Vanpeene, Sylvie van der Werf, Christine Lambert |
| EPI_ISL_418254 | NYU Langone Health | Department of Pathology and Medicine, New York University School of Medicine | Margaret Black, John Cadley, Paolo Cotzia, John Chen, Dacia Dimartino, Xiaojun Feng, Adriana Heguy, Megan Hogan, Emily Huang, George Jour, Christian Marier, Matthew T. Maurano, Mark J. Mulligan, Peter Meyn, Jared Pinnell, Amy Rapkiewicz, Marie Samanovic-Golden, Antonio Serrano, Guomiao Shen, Matija Snuderl, Nick Vulpescu, Gael Westby, Paul Zappile |
| EPI_ISL_418257 | Ospedale Civile Giuseppe Mazzini, Teramo | Istituto Zooprofilattico Sperimentale dell'Abruzzo e Molise "G.Caporale" | Lorusso A, Marcacci M, Di Domenico M, Puglia I, Curini V, Ancora M, Di Pasquale A, Rinaldi A, Mangone I, Cammà C, Savini G. |
| EPI_ISL_418260, EPI_ISL_418261 | Ospedale Civile Giuseppe Mazzini | Istituto Zooprofilattico Sperimentale dell'Abruzzo e Molise "G. Caporale" | Lorusso A, Marcacci M, Di Domenico M, Puglia I, Curini V, Ancora M, Di Pasquale A, Rinaldi A, Mangone I, Cammà C, Savini G. |
| EPI_ISL_418263, EPI_ISL_418264, EPI_ISL_418265 | Laboratory of Microbiology, Department of Medicine, National and Kapodistrian University of Athens, Greece | Laboratory of Biology, Department of Medicine, Democritus University of Thrace, Greece | Maria Bampali, Elisavet Gatzidou, Nikolaos Dovrolis, Stavroula Veletza, Nikolaos Spanakis, Ioannis Karakasiliotis |
| EPI_ISL_418286, EPI_ISL_418287, EPI_ISL_418288, EPI_ISL_418289, EPI_ISL_418290, EPI_ISL_418291, EPI_ISL_418292, EPI_ISL_418293, EPI_ISL_418294, EPI_ISL_418295, EPI_ISL_418296, EPI_ISL_418297, EPI_ISL_418298, EPI_ISL_418299, EPI_ISL_418300 | | | |
| see above | Virology Department, Sheffield Teaching Hospitals NHS Foundation Trust | Department of Infection, Immunity and Cardiovascular Disease, The Florey Institute, The Medical School, University of Sheffield | Thushan de Silva, Matthew Parker, Adri Angyal, Rebecca Brown, Rachel Tucker, Paul Parsons, Danielle Groves, Alex Keeley, Dave Partridge, Matthew Wyles, Benjamin Lindsey, Mehmet Yavuz, Mohammad Raza, Cariad Evans |
| EPI_ISL_418412 | Centre Hospitalier des Vals d'Ardeche | CNR Virus des Infections Respiratoires - France SUD | Antonin Bal, Gregory Destras, Gwendolyne Burfin, Solenne Brun, Carine Moustaud, Raphaelle Lamy, Alexandre Gaymard, Maude Bouscambert-Duchamp, Florence Morfin-Sherpa, Martine Valette, Bruno Lina, Laurence Josset |
| EPI_ISL_418413 | Centre Hospitalier de Macon | CNR Virus des Infections Respiratoires - France SUD | Antonin Bal, Gregory Destras, Gwendolyne Burfin, Solenne Brun, Carine Moustaud, Raphaelle Lamy, Alexandre Gaymard, Maude Bouscambert-Duchamp, Florence Morfin-Sherpa, Martine Valette, Bruno Lina, Laurence Josset |
| EPI_ISL_418414, EPI_ISL_418415 | Centre Hospitalier de Valence | CNR Virus des Infections Respiratoires - France SUD | Antonin Bal, Gregory Destras, Gwendolyne Burfin, Solenne Brun, Carine Moustaud, Raphaelle Lamy, Alexandre Gaymard, Maude Bouscambert-Duchamp, Florence Morfin-Sherpa, Martine Valette, Bruno Lina, Laurence Josset |
| EPI_ISL_418416 | GH Les Portes du Sud | CNR Virus des Infections Respiratoires - France SUD | Antonin Bal, Gregory Destras, Gwendolyne Burfin, Solenne Brun, Carine Moustaud, Raphaelle Lamy, Alexandre Gaymard, Maude Bouscambert-Duchamp, Florence Morfin-Sherpa, Martine Valette, Bruno Lina, Laurence Josset |
| EPI_ISL_418417 | Centre Hospitalier de Valence | CNR Virus des Infections Respiratoires - France SUD | Antonin Bal, Gregory Destras, Gwendolyne Burfin, Solenne Brun, Carine Moustaud, Raphaelle Lamy, Alexandre Gaymard, Maude Bouscambert-Duchamp, Florence Morfin-Sherpa, Martine Valette, Bruno Lina, Laurence Josset |
| EPI_ISL_418418, EPI_ISL_418419 | Centre Hospitalier Saint Joseph Saint Luc | CNR Virus des Infections Respiratoires - France SUD | Antonin Bal, Gregory Destras, Gwendolyne Burfin, Solenne Brun, Carine Moustaud, Raphaelle Lamy, Alexandre Gaymard, Maude Bouscambert-Duchamp, Florence Morfin-Sherpa, Martine Valette, Bruno Lina, Laurence Josset |
| EPI_ISL_418420, EPI_ISL_418421, EPI_ISL_418422, EPI_ISL_418423, EPI_ISL_418424, EPI_ISL_418425 | Institut des Agents Infectieux (IAI), Hospices Civils de Lyon | CNR Virus des Infections Respiratoires - France SUD | Antonin Bal, Gregory Destras, Gwendolyne Burfin, Solenne Brun, Carine Moustaud, Raphaelle Lamy, Alexandre Gaymard, Maude Bouscambert-Duchamp, Florence Morfin-Sherpa, Martine Valette, Bruno Lina, Laurence Josset |
| EPI_ISL_418426 | Centre Hospitalier de Bourg en Bresse | CNR Virus des Infections Respiratoires - France SUD | Antonin Bal, Gregory Destras, Gwendolyne Burfin, Solenne Brun, Carine Moustaud, Raphaelle Lamy, Alexandre Gaymard, Maude Bouscambert-Duchamp, Florence Morfin-Sherpa, Martine Valette, Bruno Lina, Laurence Josset |
| EPI_ISL_418427 | Hopital Privé de l'Est Lyonnais | CNR Virus des Infections Respiratoires - France SUD | Antonin Bal, Gregory Destras, Gwendolyne Burfin, Solenne Brun, Carine Moustaud, Raphaelle Lamy, Alexandre Gaymard, Maude Bouscambert-Duchamp, Florence Morfin-Sherpa, Martine Valette, Bruno Lina, Laurence Josset |
| EPI_ISL_418428 | Centre Hospitalier Lucien Hussel | CNR Virus des Infections Respiratoires - France SUD | Antonin Bal, Gregory Destras, Gwendolyne Burfin, Solenne Brun, Carine Moustaud, Raphaelle Lamy, Alexandre Gaymard, Maude Bouscambert-Duchamp, Florence Morfin-Sherpa, Martine Valette, Bruno Lina, Laurence Josset |
| EPI_ISL_418429, EPI_ISL_418430, EPI_ISL_418431, EPI_ISL_418432 | Institut des Agents Infectieux (IAI), Hospices Civils de Lyon | CNR Virus des Infections Respiratoires - France SUD | Antonin Bal, Gregory Destras, Gwendolyne Burfin, Solenne Brun, Carine Moustaud, Raphaelle Lamy, Alexandre Gaymard, Maude Bouscambert-Duchamp, Florence Morfin-Sherpa, Martine Valette, Bruno Lina, Laurence Josset |
| EPI_ISL_418624, EPI_ISL_418626, EPI_ISL_418628, EPI_ISL_418629, EPI_ISL_418630, EPI_ISL_418632, EPI_ISL_418633 | Department of Clinical Microbiology | GIGA Medical Genomics | Keith Durkin, Maria Artesi, Sébastien Bontems, Raphaël Boreux, Cécile Meex, Pierrette Melin, Marie-Pierre Hayette, Vincent Bours. |
| EPI_ISL_418672, EPI_ISL_418673, EPI_ISL_418674, EPI_ISL_418675, EPI_ISL_418677, EPI_ISL_418678, EPI_ISL_418679, EPI_ISL_418680, EPI_ISL_418681, EPI_ISL_418682, EPI_ISL_418683, EPI_ISL_418684, EPI_ISL_418685, EPI_ISL_418686, EPI_ISL_418687, EPI_ISL_418688, EPI_ISL_418689, EPI_ISL_418690, EPI_ISL_418691, EPI_ISL_418692, EPI_ISL_418693, EPI_ISL_418694, EPI_ISL_418695, EPI_ISL_418696, EPI_ISL_418697, EPI_ISL_418698, EPI_ISL_418699, EPI_ISL_418700, EPI_ISL_418701, EPI_ISL_418702, EPI_ISL_418703, EPI_ISL_418704, EPI_ISL_418705, EPI_ISL_418706, EPI_ISL_418707, EPI_ISL_418708, EPI_ISL_418709, EPI_ISL_418710, EPI_ISL_418711, EPI_ISL_418712, EPI_ISL_418713, EPI_ISL_418714, EPI_ISL_418715, EPI_ISL_418716, EPI_ISL_418717, EPI_ISL_418718, EPI_ISL_418719, EPI_ISL_418720, EPI_ISL_418721, EPI_ISL_418722, EPI_ISL_418723, EPI_ISL_418724, EPI_ISL_418725, EPI_ISL_418728, EPI_ISL_418729, EPI_ISL_418730, EPI_ISL_418733, EPI_ISL_418735, EPI_ISL_418736, EPI_ISL_418737, EPI_ISL_418738, EPI_ISL_418739, EPI_ISL_418740, EPI_ISL_418741, EPI_ISL_418743, EPI_ISL_418745, EPI_ISL_418746, EPI_ISL_418748, EPI_ISL_418749, EPI_ISL_418750, EPI_ISL_418751, EPI_ISL_418752, EPI_ISL_418754, EPI_ISL_418755, EPI_ISL_418756, EPI_ISL_418757, EPI_ISL_418758, EPI_ISL_418759, EPI_ISL_418761, EPI_ISL_418763, EPI_ISL_418764, EPI_ISL_418765, EPI_ISL_418766, EPI_ISL_418767, EPI_ISL_418768, EPI_ISL_418769, EPI_ISL_418770 | | | |
| see above | Respiratory Virus Unit, Microbiology Services Colindale, Public Health England | Respiratory Virus Unit, Microbiology Services Colindale, Public Health England | Monica Galiano, Shahjahan Miah, Angie Lackenby, Omolola Akinbami, Tiina Talts, Leena Bhaw, Richard Myers, Steven Platt, Kirstin Edwards, Jonathan Hubb, Joanna Ellis, Maria Zambon |
| EPI_ISL_418802, EPI_ISL_418803, EPI_ISL_418804 | Pathology Queensland | Public Health Virology Laboratory | Bixing Huang, Alyssa Pyke, Amanda De Jong, Andrew Van Den Hurk, Carmel Taylor, David Warrilow, Doris Genge, Elisabeth Gamez, Glen Hewitson, Ian Maxwell Mackay, Inga Sultana, Jamie McMahon, Jean Barcelon, Judy Northill, Mitchell Finger, Natalie Simpson, Neelima Nair, Peter Burtonclay, Peter Moore, Sarah Wheatley, Sean Moody, Sonja Hall-Mendelin, Timothy Gardam, and Frederick Moore |
| EPI_ISL_418860 | Hospital Universitari Vall d'Hebron (HUVH) - Vall d'Hebron Research Institute (VHIR) | Hospital Universitari Vall d'Hebron (HUVH) - Vall d'Hebron Research Institute (VHIR) | Cristina Andrés, Dàmir Garcia-Cehic, Maria Piñana, Mercedes Guerrero-Murillo, Ariadna Rando, Tomàs Pumarola, Maria Gema Codina, Andrés Antón, Josep Quer |
| EPI_ISL_418864 | Virginia DCLS | Virginia DCLS | Virginia DCLS |
| EPI_ISL_418869, EPI_ISL_418870, EPI_ISL_418872, EPI_ISL_418873, EPI_ISL_418874, EPI_ISL_418875, EPI_ISL_418876, EPI_ISL_418884, EPI_ISL_418886, EPI_ISL_418888, EPI_ISL_418889, EPI_ISL_418890, EPI_ISL_418892, EPI_ISL_418897, EPI_ISL_418899, EPI_ISL_418900, EPI_ISL_418903, EPI_ISL_418904, EPI_ISL_418905, EPI_ISL_418906, EPI_ISL_418907, EPI_ISL_418908, EPI_ISL_418909, EPI_ISL_418910, EPI_ISL_418911, EPI_ISL_418912, EPI_ISL_418913, EPI_ISL_418914, EPI_ISL_418921, EPI_ISL_418922, EPI_ISL_418923, EPI_ISL_418927, EPI_ISL_418928, EPI_ISL_418929, EPI_ISL_418930, EPI_ISL_418931, EPI_ISL_418932, EPI_ISL_418933, EPI_ISL_418934, EPI_ISL_418935, EPI_ISL_418936, EPI_ISL_418937, EPI_ISL_418938, EPI_ISL_418939, EPI_ISL_418940, EPI_ISL_418942, EPI_ISL_418943, EPI_ISL_418944, EPI_ISL_418945, EPI_ISL_418947, EPI_ISL_418948, EPI_ISL_418949, EPI_ISL_418950, EPI_ISL_418951, EPI_ISL_418952, EPI_ISL_418953, EPI_ISL_418954, EPI_ISL_418955 | | | |
| see above | UW Virology Lab | UW Virology Lab | Pavitra Roychoudhury, Hong Xie, Keith Jerome, Alexander Greninger |
| EPI_ISL_418956, EPI_ISL_418957, EPI_ISL_418958 | Virginia DCLS | Virginia DCLS | Virginia DCLS |
| EPI_ISL_418959 | Universidade Federal do Rio de Janeiro - UFRJ | Bioinformatics Laboratory - LNCC | Filipe Romero, Ana Paula Guimarães, Mariane Talon, Luiz Gonzaga Paula de Almeida, Ronaldo da Silva Francisco Junior, Diana Mariani, Lídia Boullosa, Alexandra Gerber, Jaqueline Goes de Jesus, Ingra Morales Claro, Ester Cerdeira Sabino, Nuno Rodrigues Faria, Terezinha Marta Pereira, Pinto Castiñeiras, Isabela de Carvalho Leitão, Rafael de Mello Galliez, Cássia Cristina Alves Gonçalves, Érica Ramos dos Santos Nascimento, Richard Araújo Maia, Mauro Teixeira,Cristiano Xavier Lima, Orlando Ferreira Jr., Rodrigo Brindeiro, Luciana Jesus Costa e André Felipe Santos, Laboratorio Hermes Pardini, Laboratorio Simile, Amilcar Tanuri, Renato Santana Aguiar, e Ana Tereza Vasconcelos |
| EPI_ISL_418968, EPI_ISL_418970, EPI_ISL_418971, EPI_ISL_418972, EPI_ISL_418973, EPI_ISL_418974, EPI_ISL_418975, EPI_ISL_418976, EPI_ISL_418977, EPI_ISL_418978, EPI_ISL_418979, EPI_ISL_418980 | | | |
| see above | NYU Langone Health | Department of Pathology and Medicine, New York University School of Medicine | Maria Aguero-Rosenfeld, Margaret Black, John Cadley, Paolo Cotzia, John Chen, Dacia Dimartino, Xiaojun Feng, Adriana Heguy, Megan Hogan, Emily Huang, George Jour, Christian Marier, Matthew T. Maurano, Mark J. Mulligan, Peter Meyn, Jared Pinnell, Sitharam Ramaswami, Amy Rapkiewicz, Marie |

| | | | |
|---|---|---|---|
| | | | Samanovic-Golden, Antonio Serrano, Guomiao Shen, Matija Snuderl, Nick Vulpescu, Gael Westby, Paul Zappile, Yutong Zhang |
| EPI_ISL_419168 | Centre Hospitalier de Valence | CNR Virus des Infections Respiratoires - France SUD | Antonin Bal, Gregory Destras, Gwendolyne Burfin, Solenne Brun, Carine Moustaud, Raphaelle Lamy, Alexandre Gaymard, Maude Bouscambert-Duchamp, Florence Morfin-Sherpa, Martine Valette, Bruno Lina, Laurence Josset |
| EPI_ISL_419256 | Virginia DCLS | Virginia DCLS | Virginia DCLS |
| EPI_ISL_419304, EPI_ISL_419305 | Saitama Prefectural Institute of Public Health | Pathogen Genomics Center, National Institute of Infectious Diseases | Tsuyoshi Sekizuka, Miho Shinohara, Tsuyoshi Kishimoto, Kentaro Itokawa, Rina Tanaka, Masanori Hashino, Hajime Kamiya, Motoi Suzuki, Makoto Kuroda |
| EPI_ISL_419386, EPI_ISL_419387 | Hospital Prof. Doutor Fernando Fonseca, EPE | Instituto Gulbenkian de Ciência | João Costa, Cathy Paulino, Joao Sobral, Susana Ladeiro, Ricardo Leite |
| EPI_ISL_419398, EPI_ISL_419399, EPI_ISL_419400, EPI_ISL_419401, EPI_ISL_419402, EPI_ISL_419403, EPI_ISL_419404, EPI_ISL_419405, EPI_ISL_419406, EPI_ISL_419407, EPI_ISL_419408, EPI_ISL_419409, EPI_ISL_419410, EPI_ISL_419411, EPI_ISL_419412, EPI_ISL_419413, EPI_ISL_419414, EPI_ISL_419415, EPI_ISL_419416, EPI_ISL_419417, EPI_ISL_419418, EPI_ISL_419419, EPI_ISL_419420, EPI_ISL_419421, EPI_ISL_419422, EPI_ISL_419423, EPI_ISL_419424, EPI_ISL_419425, EPI_ISL_419426, EPI_ISL_419427, EPI_ISL_419428, EPI_ISL_419429, EPI_ISL_419430, EPI_ISL_419431, EPI_ISL_419432, EPI_ISL_419433, EPI_ISL_419434, EPI_ISL_419435, EPI_ISL_419436, EPI_ISL_419438, EPI_ISL_419439, EPI_ISL_419440, EPI_ISL_419441, EPI_ISL_419442, EPI_ISL_419443, EPI_ISL_419444, EPI_ISL_419448, EPI_ISL_419449, EPI_ISL_419450, EPI_ISL_419451 | | | |
| see above | Wales Specialist Virology Centre | Public Health Wales Microbiology Cardiff | Catherine Moore, Joanne Watkins, Sally Corden, Sara Rey, Matt Bull, Tom Connor |
| EPI_ISL_419521, EPI_ISL_419522, EPI_ISL_419523, EPI_ISL_419524, EPI_ISL_419525, EPI_ISL_419526, EPI_ISL_419527 | Yale Clinical Virology Laboratory | Grubaugh Lab - Yale School of Public Health | Joseph Fauver, Anderson Brito, Tara Alpert, Chantal Vogels, Ellen Foxman, Albert Ko, Marie Landry, Nathan Grubaugh |
| EPI_ISL_419542, EPI_ISL_419543, EPI_ISL_419544, EPI_ISL_419545, EPI_ISL_419546, EPI_ISL_419547, EPI_ISL_419548, EPI_ISL_419549, EPI_ISL_419550, EPI_ISL_419551, EPI_ISL_419552 | | | |
| see above | Center of Medical Microbiology, Virology, and Hospital Hygiene, University of Duesseldorf | Center of Medical Microbiology, Virology, and Hospital Hygiene, University of Duesseldorf | Ortwin Adams, Marcel Andree, Alexander Dilthey, Torsten Feldt, Sandra Hauka, Torsten Houwaart, Björn-Erik Jensen, Detlef Kindgen-Milles, Malte Kohns Vasconcelos, Klaus Pfeffer, Tina Senff, Daniel Strelow, Jörg Timm, Andreas Walker, Tobias Wienemann |
| EPI_ISL_419564, EPI_ISL_419567, EPI_ISL_419574, EPI_ISL_419576, EPI_ISL_419583, EPI_ISL_419588, EPI_ISL_419589, EPI_ISL_419592, EPI_ISL_419593, EPI_ISL_419594, EPI_ISL_419597, EPI_ISL_419599, EPI_ISL_419601, EPI_ISL_419603, EPI_ISL_419604 | | | |
| see above | Laboratoire National de Santé, Microbiology, Virology | Laboratoire National de Santé, Microbiology, Epidemiology and Microbial Genomics | Anke Wienecke-Baldacchino, Ardashel Latsuzbaia, Jessica Tapp, Catherine Ragimbeau, Guillaume Fournier, Tamir Abdelrahman, Trung Nguyen Nguyen, Joel Mossong |
| EPI_ISL_419663, EPI_ISL_419669, EPI_ISL_419670 | Center for Virology, Medical University of Vienna | Bergthaler laboratory, CeMM Research Center for Molecular Medicine of the Austrian Academy of Sciences | Alexandra Popa, Benedikt Agerer, Henrique Colaco, Lukas Endler, Jakob-Wendelin Genger, Alexander Lercher, Mark Smyth, Thomas Penz, Michael Schuster, Judith Aberle, Stephan Aberle, Elisabeth Puchhammer-Stöckl, Christoph Bock, Andreas Bergthaler |
| EPI_ISL_419696, EPI_ISL_419697, EPI_ISL_419698, EPI_ISL_419699, EPI_ISL_419700, EPI_ISL_419701, EPI_ISL_419702, EPI_ISL_419703, EPI_ISL_419704, EPI_ISL_419705 | NYU Langone Health | Departments of Pathology and Medicine, New York University School of Medicine | Maria Aguero-Rosenfeld, Margaret Black, John Cadley, Paolo Cotzia, John Chen, Dacia Dimartino, Xiaojun Feng, Adriana Heguy, Megan Hogan, Emily Huang, George Jour, Christian Marier, Matthew T. Maurano, Mark J. Mulligan, Peter Meyn, Jared Pinnell, Sitharam Ramaswami, Amy Rapkiewicz, Marie Samanovic-Golden, Antonio Serrano, Guomiao Shen, Matija Snuderl, Nick Vulpescu, Gael Westby, Paul Zappile, Yutong Zhang |
| EPI_ISL_419714, EPI_ISL_419715, EPI_ISL_419716, EPI_ISL_419717, EPI_ISL_419718, EPI_ISL_419719, EPI_ISL_419724 | Microbiological Diagnostic Unit Public Health Laboratory | Microbiological Diagnostic Unit Public Health Laboratory | Seemann T., Schultz M., Sait, M., Sherry, N. |
| EPI_ISL_419795, EPI_ISL_419796, EPI_ISL_419797, EPI_ISL_419798, EPI_ISL_419800, EPI_ISL_419801, EPI_ISL_419804, EPI_ISL_419805, EPI_ISL_419806, EPI_ISL_419807, EPI_ISL_419808, EPI_ISL_419809, EPI_ISL_419810, EPI_ISL_419811, EPI_ISL_419812, EPI_ISL_419813, EPI_ISL_419814, EPI_ISL_419815, EPI_ISL_419817, EPI_ISL_419818, EPI_ISL_419819, EPI_ISL_419820, EPI_ISL_419821, EPI_ISL_419822, EPI_ISL_419836, EPI_ISL_419837, EPI_ISL_419838, EPI_ISL_419839, EPI_ISL_419840, EPI_ISL_419841, EPI_ISL_419842, EPI_ISL_419843, EPI_ISL_419844, EPI_ISL_419845, EPI_ISL_419846, EPI_ISL_419847, EPI_ISL_419848, EPI_ISL_419849, EPI_ISL_419850, EPI_ISL_419851, EPI_ISL_419852, EPI_ISL_419853, EPI_ISL_419854, EPI_ISL_419855, EPI_ISL_419856, EPI_ISL_419857, EPI_ISL_419858, EPI_ISL_419859, EPI_ISL_419860, EPI_ISL_419861, EPI_ISL_419862, EPI_ISL_419863, EPI_ISL_419864, EPI_ISL_419865, EPI_ISL_419866, EPI_ISL_419867, EPI_ISL_419868, EPI_ISL_419869, EPI_ISL_419870, EPI_ISL_419871, EPI_ISL_419872, EPI_ISL_419873, EPI_ISL_419874, EPI_ISL_419875 | | | |
| see above | Victorian Infectious Diseases Reference Laboratory (VIDRL) | Victorian Infectious Diseases Reference Laboratory and Microbiological Diagnostic Unit Public Health Laboratory, Doherty Institute | Caly L., Seemann T., Sait, M., Schultz M., Druce J., Sherry, N. |
| EPI_ISL_420018 | Virginia DCLS | Virginia DCLS | Virginia DCLS |
| EPI_ISL_420038 | Sentinelles network | National Reference Center for Viruses of Respiratory Infections, Institut Pasteur, Paris | Mélanie Albert, Marion Barbet, Sylvie Behillil, Méline Bizard, Angela Brisebarre, Flora Donati, Etienne Simon-Lorière, Vincent Enouf, Maud Vanpeene, Sylvie van der Werf |
| EPI_ISL_420041 | CH Compiègne Laboratoire de Biologie | National Reference Center for Viruses of Respiratory Infections, Institut Pasteur, Paris | Mélanie Albert, Marion Barbet, Sylvie Behillil, Méline Bizard, Angela Brisebarre, Flora Donati, Etienne Simon-Lorière, Vincent Enouf, Maud Vanpeene, Sylvie van der Werf, Raulin Olivia |
| EPI_ISL_420042 | Service de Biologie clinique | National Reference Center for Viruses of Respiratory Infections, Institut Pasteur, Paris | Mélanie Albert, Marion Barbet, Sylvie Behillil, Méline Bizard, Angela Brisebarre, Flora Donati, Etienne Simon-Lorière, Vincent Enouf, Maud Vanpeene, Sylvie van der Werf |
| EPI_ISL_420043 | CMIP | National Reference Center for Viruses of Respiratory Infections, Institut Pasteur, Paris | Mélanie Albert, Marion Barbet, Sylvie Behillil, Méline Bizard, Angela Brisebarre, Flora Donati, Etienne Simon-Lorière, Vincent Enouf, Maud Vanpeene, Sylvie van der Werf |
| EPI_ISL_420044 | CH Jean de Navarre Laboratoire de Biologie | National Reference Center for Viruses of Respiratory Infections, Institut Pasteur, Paris | Mélanie Albert, Marion Barbet, Sylvie Behillil, Méline Bizard, Angela Brisebarre, Flora Donati, Etienne Simon-Lorière, Vincent Enouf, Maud Vanpeene, Sylvie van der Werf |
| EPI_ISL_420045 | Sentinelles network | National Reference Center for Viruses of Respiratory Infections, Institut Pasteur, Paris | Mélanie Albert, Marion Barbet, Sylvie Behillil, Méline Bizard, Angela Brisebarre, Flora Donati, Etienne Simon-Lorière, Vincent Enouf, Maud Vanpeene, Sylvie van der Werf |
| EPI_ISL_420046, EPI_ISL_420047 | Résidence Villa Caroline | National Reference Center for Viruses of Respiratory Infections, Institut Pasteur, Paris | Mélanie Albert, Marion Barbet, Sylvie Behillil, Méline Bizard, Angela Brisebarre, Flora Donati, Etienne Simon-Lorière, Vincent Enouf, Maud Vanpeene, Sylvie van der Werf |
| EPI_ISL_420048 | Service de Biologie Médicale - BP 125 | National Reference Center for Viruses of Respiratory Infections, Institut Pasteur, Paris | Mélanie Albert, Marion Barbet, Sylvie Behillil, Méline Bizard, Angela Brisebarre, Flora Donati, Etienne Simon-Lorière, Vincent Enouf, Maud Vanpeene, Sylvie van der Werf, Christine Lambert |
| EPI_ISL_420055 | Sentinelles network | National Reference Center for Viruses of Respiratory Infections, Institut Pasteur, Paris | Mélanie Albert, Marion Barbet, Sylvie Behillil, Méline Bizard, Angela Brisebarre, Flora Donati, Etienne Simon-Lorière, Vincent Enouf, Maud Vanpeene, Sylvie van der Werf |
| EPI_ISL_420069, EPI_ISL_420070, EPI_ISL_420071 | Institut Pasteur Dakar | Institut Pasteur de Dakar | Ndongo Dia, Moussa Moise Diagne, Mamadou Diop, Ousmane Faye, Amadou Alpha Sall |
| EPI_ISL_420072 | Institut Pasteur Dakar | Institut Pasteur de Dakar | Ndongo Dia, Moussa Moise Diagne, Mamadou Diop, Ousmane Faye , Amadou Alpha Sall |
| EPI_ISL_420080 | WHO National Influenza Centre Russian Federation | WHO National Influenza Centre Russian Federation | Andrey Komissarov, Artem Fadeev, Anna Ivanova, Daria Danilenko |
| EPI_ISL_420081 | WHO National Influenza Centre Russian Federation | WHO National Influenza Centre Russian Federation | Andrey Komissarov, Artem Fadeev, Mariia Sergeeva, Anna Ivanova, Daria Danilenko |
| EPI_ISL_420083 | Centers for Disease Control, R.O.C. (Taiwan) | Centers for Disease Control, R.O.C. (Taiwan) | Ji-Rong Yang, Yu-Chi Lin, Jung-Jung Mu, Ming-Tsan Liu |
| EPI_ISL_420090, EPI_ISL_420091, EPI_ISL_420092, EPI_ISL_420096, EPI_ISL_420097 | Yale Clinical Virology Laboratory | Grubaugh Lab - Yale School of Public Health | Joseph Fauver, Anderson Brito, Tara Alpert, Chantal Vogels, Ellen Foxman, Albert Ko, Marie Landry, Nathan Grubaugh |
| EPI_ISL_420130 | Servicio de Microbiologia. Hospital Clinico Universitario de Valencia | Sequencing and Bioinformatics Service and Molecular Epidemiology Research Group. FISABIO-Public Health | David Navarro, Loreto Ferrús Abad, Maria Alma Bracho, Griselda De Marco, Beatriz Beamud, Lidia Ruiz Roldan, Marta Pla Diaz, Neris Garcia-Gonzalez, Inma Galán Vendrell, Paula Ruiz-Hueso, Mariana Reyes-Prieto, Vicente Soriano Chirona, Sandra Carbo, Ivan Ansari, Llúcia Martínez-Priego, Giuseppe D'Auria, Fernando Gonzalez-Candelas |
| EPI_ISL_420131 | Servicio de Microbiologia. Hospital Clinico Universitario de | Sequencing and Bioinformatics Service and Molecular | Paula Ruiz-Hueso,Loreto Ferrús Abad, Maria Alma Bracho, Griselda De Marco, Beatriz Beamud, Sandra Carbo, Lidia Ruiz Roldan, Marta Pla Diaz, Neris |

| | | | |
|---|---|---|---|
| | Valencia | Epidemiology Research Group. FISABIO-Public Health | Garcia-Gonzalez, Inma Galán Vendrell, Mariana Reyes-Prieto, Vicente Soriano Chirona, Ivan Ansari, David Navarro, Llúcia Martínez-Priego, Giuseppe D'Auria, Fernando Gonzalez-Candelas |
| EPI_ISL_420132 | Servicio de Microbiologia. Hospital Clinico Universitario de Valencia | Sequencing and Bioinformatics Service and Molecular Epidemiology Research Group. FISABIO-Public Health | Giuseppe D'Auria,Sandra Carbo, Loreto Ferrús Abad, Maria Alma Bracho, Griselda De Marco, Beatriz Beamud, Lidia Ruiz Roldan, Marta Pla Diaz, Neris Garcia-Gonzalez, Inma Galán Vendrell, Paula Ruiz-Hueso, Mariana Reyes-Prieto, Vicente Soriano Chirona, Ivan Ansari, David Navarro, Llúcia Martínez-Priego, Fernando Gonzalez-Candelas |
| EPI_ISL_420151 | Nordland Hospital - Bodo, Laboratory Department, Molecular Biology Unit | Norwegian Institute of Public Health, Department of Virology | Kathrine Stene-Johansen, Kamilla Heddeland Instefjord, Hilde Elshaug, Karoline Bragstad, Olav Hungnes |
| EPI_ISL_420152, EPI_ISL_420153 | University Hospital of Northern Norway, Department for Microbiology and Infectious Disease Control | Norwegian Institute of Public Health, Department of Virology | Kathrine Stene-Johansen, Kamilla Heddeland Instefjord, Hilde Elshaug, Karoline Bragstad, Olav Hungnes |
| EPI_ISL_420177, EPI_ISL_420288, EPI_ISL_420289, EPI_ISL_420290, EPI_ISL_420291 | Virology Department, Sheffield Teaching Hospitals NHS Foundation Trust | Department of Infection, Immunity and Cardiovascular Disease, The Florey Institute, The Medical School, University of Sheffield | Thushan de Silva, Matthew Parker, Adri Angyal, Rebecca Brown, Rachel Tucker, Paul Parsons, Luke Green, Danielle Groves, Alex Keeley, Dave Partridge, Matthew Wyles, Benjamin Lindsey, Mehmet Yavuz, Mohammad Raza, Cariad Evans |
| EPI_ISL_420296, EPI_ISL_420297, EPI_ISL_420299, EPI_ISL_420300, EPI_ISL_420301, EPI_ISL_420302 | NYU Langone Health | Departments of Pathology and Medicine, New York University School of Medicine | Maria Aguero-Rosenfeld, Margaret Black, John Cadley, Paolo Cotzia, John Chen, Dacia Dimartino, Xiaojun Feng, Adriana Heguy, Megan Hogan, Emily Huang, George Jour, Christian Marier, Matthew T. Maurano, Mark J. Mulligan, Peter Meyn, Jared Pinnell, Sitharam Ramaswami, Amy Rapkiewicz, Marie Samanovic-Golden, Antonio Serrano, Guomiao Shen, Matija Snuderl, Nick Vulpescu, Gael Westby, Paul Zappile, Yutong Zhang |
| EPI_ISL_420303 | Alaska State Virology Laboratory | Alaska State Virology Laboratory | Chen, J |
| EPI_ISL_420304, EPI_ISL_420305, EPI_ISL_420306 | Alaska State Virology Laboratory | Alaska State Virology Laboratory | Chen, J. |
| EPI_ISL_420310 | University Hospital of Northern Norway, Department for Microbiology and Infectious Disease Control | Norwegian Institute of Public Health, Department of Virology | Kathrine Stene-Johansen, Kamilla Heddeland Instefjord, Hilde Elshaug, Karoline Bragstad, Olav Hungnes |
| EPI_ISL_420311, EPI_ISL_420312 | Akershus University Hospital, Department for Microbiology and Infectious Disease Control | Norwegian Institute of Public Health, Department of Virology | Kathrine Stene-Johansen, Kamilla Heddeland Instefjord, Hilde Elshaug, Karoline Bragstad, Olav Hungnes |
| EPI_ISL_420358, EPI_ISL_420359, EPI_ISL_420360, EPI_ISL_420361, EPI_ISL_420362 | KU Leuven, Clinical and Epidemiological Virology | KU Leuven, Clinical and Epidemiological Virology | Joan Marti-Carreras, Bert Vanmechelen, Tony Wawina, Piet Maes |
| EPI_ISL_420465, EPI_ISL_420466, EPI_ISL_420467, EPI_ISL_420468, EPI_ISL_420469, EPI_ISL_420470, EPI_ISL_420471, EPI_ISL_420472, EPI_ISL_420473, EPI_ISL_420474, EPI_ISL_420476, EPI_ISL_420477, EPI_ISL_420480, EPI_ISL_420481, EPI_ISL_420482, EPI_ISL_420483, EPI_ISL_420485 | | | |
| see above | Respiratory Virus Unit, Microbiology Services Colindale, Public Health England | Respiratory Virus Unit, Microbiology Services Colindale, Public Health England | Monica Galiano, Shahjahan Miah, Angie Lackenby, Omolola Akinbami, Tiina Talts, Leena Bhaw, Richard Myers, Steven Platt, Kirstin Edwards, Jonathan Hubb, Joanna Ellis, Maria Zambon |
| EPI_ISL_420536 | Department of Microbiology, PathWest QEII Medical Centre | Department of Microbiology, PathWest QEII Medical Centre | Chisha Sikazwe, Jurissa Lang, Avram Levy, David Speers and David Smith |
| EPI_ISL_420563 | Ospedale Civile Giuseppe Mazzini | Istituto Zooprofilattico Sperimentale dell'Abruzzo e Molise "G. Caporale" | Lorusso A, Marcacci M, Di Domenico M, Ancora M, Curini V, Mangone I, Rinaldi A, Di Pasquale A, Cammà C, Puglia I, Savini G |
| EPI_ISL_420570, EPI_ISL_420571, EPI_ISL_420572, EPI_ISL_420573, EPI_ISL_420574, EPI_ISL_420575, EPI_ISL_420576, EPI_ISL_420577, EPI_ISL_420578, EPI_ISL_420579, EPI_ISL_420580, EPI_ISL_420581, EPI_ISL_420582, EPI_ISL_420584, EPI_ISL_420585, EPI_ISL_420587, EPI_ISL_420588, EPI_ISL_420589, EPI_ISL_420590, EPI_ISL_420591 | | | |
| see above | NYU Langone Health | Departments of Pathology and Medicine, New York University School of Medicine | Maria Aguero-Rosenfeld, Brendan Belovarac, Margaret Black, Ludovic Boytard, John Cadley, Paolo Cotzia, John Chen, Dacia Dimartino, Xiaojun Feng, Tatyana Gindin, Adriana Heguy, Megan Hogan, Emily Huang, George Jour, Andrew Lytle, Christian Marier, Matthew T. Maurano, Mark J. Mulligan, Peter Meyn, Iman Osman, Jared Pinnell, Sitharam Ramaswami, Amy Rapkiewicz, Marie Samanovic-Golden, Antonio Serrano, Guomiao Shen, Matija Snuderl, Theodore Vougiouklakis, Nick Vulpescu, Gael Westby, Paul Zappile, Yutong Zhang |
| EPI_ISL_420599 | Servicio Virosis Respiratorias-Departamento Virología-INEI | Instituto Nacional Enfermedades Infecciosas C.G.Malbran | Baumeister E., Avaro M., Benedetti E., Russo M., Dattero ME, Pontoriero A., Cisterna D., Molina V., Perandones C., Tuduri E., Lorenzo F., Poklepovich T., Campos J. |
| EPI_ISL_420904, EPI_ISL_420905, EPI_ISL_420906, EPI_ISL_420907 | Max von Pettenkofer Institute, Virology, National Reference Center for Retroviruses, LMU Munich | Laboratory for Functional Genome Analysis, Dept. Genomics, Gene Center of the LMU Munich | Max Muenchhoff, Stefan Krebs, Alexander Graf, Ashok Varadharajan, Oliver Keppler, Helmut Blum |
| EPI_ISL_420917 | Wales Specialist Virology Centre | Public Health Wales Microbiology Cardiff | Catherine Moore, Joanne Watkins, Sally Corden, Malorie Perry, Simon Cottrell Sara Rey, Matt Bull, Tom Connor |
| EPI_ISL_421221 | Hangzhou Center for Diseases Control and Prevention | Hangzhou Center for Diseases Control and Prevention | Jun Li, Haoqiu Wang, Lingfeng Mao, Hua Yu, Xinfen Yu, Zhou Sun, Xin Qian, Shuchang Chen, Junfang Chen, Xuchu Wang |
| EPI_ISL_421275 | Russian State Collection of Viruses | Pathogenic Microorganisms Variability Laboratory | Alexey Shchetinin, Maria Nikiforova, Nadezhda Kuznetsova, Ekaterina Aksenova, Marina Kunda, Natalia Ryzhova, Olga Voronina, Inna Dolzhikova, Daria Grousova, Andrey Botikov, Denis Logunov, Alexander Gintsburg, Vladimir Gushchin |
| EPI_ISL_421300, EPI_ISL_421302, EPI_ISL_421310, EPI_ISL_421311, EPI_ISL_421314 | University of Wisconsin-Madison AIDS Vaccine Research Laboratories | University of Wisconsin-Madison AIDS Vaccine Research Laboratories | Gage Moreno, Katarina Braun, et al. AIDS Vaccine Research Laboratories |
| EPI_ISL_421347 | Wyoming Public Health Laboratory | Center for Global Health, University of New Mexico Health Sciences Center | Daryl Domman, Kurt Schwalm, Rob Christensen, Wanda Manley, Cari Sloma, Noah Hull, Darrell Dinwiddie |
| EPI_ISL_421348, EPI_ISL_421349, EPI_ISL_421350, EPI_ISL_421357, EPI_ISL_421358, EPI_ISL_421359, EPI_ISL_421361, EPI_ISL_421362, EPI_ISL_421363, EPI_ISL_421364, EPI_ISL_421366, EPI_ISL_421367, EPI_ISL_421368, EPI_ISL_421369, EPI_ISL_421370, EPI_ISL_421371, EPI_ISL_421372, EPI_ISL_421373, EPI_ISL_421374, EPI_ISL_421375, EPI_ISL_421376, EPI_ISL_421377, EPI_ISL_421378, EPI_ISL_421379, EPI_ISL_421380, EPI_ISL_421381, EPI_ISL_421383, EPI_ISL_421384, EPI_ISL_421388, EPI_ISL_421389, EPI_ISL_421390, EPI_ISL_421391, EPI_ISL_421392, EPI_ISL_421393, EPI_ISL_421394, EPI_ISL_421395, EPI_ISL_421396, EPI_ISL_421397, EPI_ISL_421398, EPI_ISL_421399, EPI_ISL_421400, EPI_ISL_421401, EPI_ISL_421402, EPI_ISL_421403, EPI_ISL_421404, EPI_ISL_421405, EPI_ISL_421406, EPI_ISL_421407, EPI_ISL_421408, EPI_ISL_421409, EPI_ISL_421410, EPI_ISL_421411, EPI_ISL_421412, EPI_ISL_421413, EPI_ISL_421414, EPI_ISL_421416, EPI_ISL_421417, EPI_ISL_421418, EPI_ISL_421419, EPI_ISL_421420, EPI_ISL_421421, EPI_ISL_421422, EPI_ISL_421423, EPI_ISL_421424, EPI_ISL_421425, EPI_ISL_421426, EPI_ISL_421427, EPI_ISL_421428, EPI_ISL_421429, EPI_ISL_421430, EPI_ISL_421431, EPI_ISL_421432, EPI_ISL_421433, EPI_ISL_421434, EPI_ISL_421435 | | | |
| see above | MSHS Clinical Microbiology Laboratories | MSHS Pathogen Surveillance Program | Ana S. Gonzalez-Reiche, Mitchell Sullivan, Ajay Obla, Gopi Patel, Emilia Sordillo, Melissa Gitman, Alberto Paniz-mondolfi, Matthew Hernandez, Shelcie Fabre, Jose Polanco, Zenab Khan, Bremy Albuquerque, Jayeeta Dutta, Juan Soto, Shwetha Sridhar Hara, Ying-Chih Wang, Melissa Smith, Robert Sebra, Lisa Miorin, Wen-chun Liu, Randy Albrecht, Judith Aberg, Florian Krammer, Adolfo Garcia-Sarstre, Viviana Simon, Harm van Bakel |
| EPI_ISL_421446, EPI_ISL_421447, EPI_ISL_421448 | H Guimaraes | Instituto Nacional de Saude (INSA) | Guiomar et al |
| EPI_ISL_421449 | H Dr. Nelio Mendonca - Funchal | Instituto Nacional de Saude (INSA) | Guiomar et al |
| EPI_ISL_421450, EPI_ISL_421451, EPI_ISL_421452 | Instituto Nacional de Saude (INSA) | Instituto Nacional de Saude (INSA) | Guiomar et al |
| EPI_ISL_421453 | CHTMAD | Instituto Nacional de Saude (INSA) | Guiomar et al |
| EPI_ISL_421454 | H Beatriz Angelo | Instituto Nacional de Saude (INSA) | Guiomar et al |
| EPI_ISL_421455 | CH Barreiro Montijo | Instituto Nacional de Saude (INSA) | Guiomar et al |
| EPI_ISL_421457 | H Dr Nelio Mendonca - Funchal | Instituto Nacional de Saude (INSA) | Guiomar et al |
| EPI_ISL_421464 | CHTMAD | Instituto Nacional de Saude (INSA) | Guiomar et al |
| EPI_ISL_421482, EPI_ISL_421483 | CH VN Gaia - Espinho | Instituto Nacional de Saude (INSA) | Guiomar et al |
| EPI_ISL_421505 | Service de Biologie Médicale - BP 125 | National Reference Center for Viruses of Respiratory | Mélanie Albert, Marion Barbet, Sylvie Behillil, Méline Bizard, Angela Brisebarre, Flora Donati, Etienne Simon-Lorière, Vincent Enouf, Maud Vanpeene, |

| | | | |
|---|---|---|---|
| | | Infections, Institut Pasteur, Paris | Sylvie van der Werf, Christine Lambert |
| EPI_ISL_421516 | Servicio de Microbiologia. Hospital Clinico Universitario de Valencia | Sequencing and Bioinformatics Service and Molecular Epidemiology Research Group. FISABIO-Public Health | Loreto Ferrús Abad, Maria Alma Bracho, Griselda De Marco, Beatriz Beamud, Lidia Ruiz Roldan, Marta Pla Diaz, Neris Garcia-Gonzalez, Inma Galán Vendrell, Paula Ruiz-Hueso, Mariana Reyes-Prieto, Vicente Soriano Chirona, David Navarro, Llúcia Martínez-Priego, Giuseppe D'Auria, Fernando Gonzalez-Candelas |
| EPI_ISL_421517 | Servicio de Microbiologia. Hospital Clinico Universitario de Valencia | Sequencing and Bioinformatics Service and Molecular Epidemiology Research Group. FISABIO-Public Health | Sandra Carbo, Loreto Ferrús Abad, Maria Alma Bracho, Griselda De Marco, Beatriz Beamud, Lidia Ruiz Roldan, Marta Pla Diaz, Neris Garcia-Gonzalez, Inma Galán Vendrell, Paula Ruiz-Hueso, Mariana Reyes-Prieto, Vicente Soriano Chirona, Ivan Ansari, David Navarro, Llúcia Martínez-Priego, Giuseppe D'Auria, Fernando Gonzalez-Candelas |
| EPI_ISL_421518 | Servicio de Microbiologia. Hospital Clinico Universitario de Valencia | Sequencing and Bioinformatics Service and Molecular Epidemiology Research Group. FISABIO-Public Health | Inma Galán Vendrell, Paula Ruiz-Hueso, Sandra Carbo, Loreto Ferrús Abad, Maria Alma Bracho, Griselda De Marco, Beatriz Beamud, Lidia Ruiz Roldan, Marta Pla Diaz, Neris Garcia-Gonzalez, Mariana Reyes-Prieto, Vicente Soriano Chirona, Ivan Ansari, David Navarro, Llúcia Martínez-Priego, Giuseppe D'Auria, Fernando Gonzalez-Candelas |
| EPI_ISL_421531 | State Key Laboratory of Agricultural Microbiology | State Key Laboratory of Agricultural Microbiology | Meilin Jin |
| EPI_ISL_421543, EPI_ISL_421544, EPI_ISL_421545, EPI_ISL_421546, EPI_ISL_421547, EPI_ISL_421550, EPI_ISL_421551 | Wyoming Public Health Laboratory | Center for Global Health, University of New Mexico Health Sciences Center | Daryl Domman, Kurt Schwalm, Rob Christensen, Wanda Manley, Cari Sloma, Noah Hull, Darrell Dinwiddie |
| EPI_ISL_421560, EPI_ISL_421561 | Utah Public Health Laboratory | Utah Public Health Laboratory | Erin Young, Kelly Oakeson |
| EPI_ISL_421577, EPI_ISL_421578, EPI_ISL_421579, EPI_ISL_421580, EPI_ISL_421581, EPI_ISL_421582, EPI_ISL_421583, EPI_ISL_421584, EPI_ISL_421585, EPI_ISL_421586, EPI_ISL_421587, EPI_ISL_421588, EPI_ISL_421589, EPI_ISL_421590, EPI_ISL_421591, EPI_ISL_421592 | | | |
| see above | NYU Langone Health | Departments of Pathology and Medicine, New York University School of Medicine | Maria Aguero-Rosenfeld, Brendan Belovarac, Margaret Black, Ludovic Boytard, John Cadley, Paolo Cotzia, John Chen, Dacia Dimartino, Xiaojun Feng, Tatyana Gindin, Adriana Heguy, Megan Hogan, Emily Huang, George Jour, Andrew Lytle, Christian Marier, Matthew T. Maurano, Mark J. Mulligan, Peter Meyn, Iman Osman, Jared Pinnell, Sitharam Ramaswami, Amy Rapkiewicz, Marie Samanovic-Golden, Antonio Serrano, Guomiao Shen, Matija Snuderl, Theodore Vougiouklakis, Nick Vulpescu, Gael Westby, Paul Zappile, Yutong Zhang |
| EPI_ISL_421627, EPI_ISL_421630, EPI_ISL_421631, EPI_ISL_421633 | MSHS Clinical Microbiology Laboratories | MSHS Pathogen Surveillance Program | Ana S. Gonzalez-Reiche, Mitchell Sullivan, Ajay Obla, Gopi Patel, Emilia Sordillo, Melissa Gitman, Alberto Paniz-mondolfi, Matthew Hernandez, Shelcie Fabre, Jose Polanco, Zenab Khan, Bremy Albuquerque, Jayeeta Dutta, Juan Soto, Shwetha Sridhar Hara, Ying-Chih Wang, Melissa Smith, Robert Sebra, Lisa Miorin, Wen-chun Liu, Randy Albrecht, Judith Aberg, Florian Krammer, Adolfo Garcia-Sarstre, Viviana Simon, Harm van Bakel |
| EPI_ISL_421652 | Dasman Diabetes Institute | Dasman Diabetes Institute | Fahd Al-Mulla, Rasheeba Iqbal, Sumi John, Ebaa Al-Ozairi, Qais Al-Duwairi |
| EPI_ISL_421683, EPI_ISL_421684, EPI_ISL_421685, EPI_ISL_421686, EPI_ISL_421687, EPI_ISL_421688, EPI_ISL_421689, EPI_ISL_421690, EPI_ISL_421691, EPI_ISL_421692 | Minnesota Department of Health, Public Health Laboratory | Minnesota Department of Health, Public Health Laboratory | Matt Plumb, Jacob Garfin, Xiong Wang |
| EPI_ISL_421704, EPI_ISL_421705, EPI_ISL_421706, EPI_ISL_421707, EPI_ISL_421708, EPI_ISL_421709, EPI_ISL_421710, EPI_ISL_421711, EPI_ISL_421712, EPI_ISL_421713, EPI_ISL_421714, EPI_ISL_421715, EPI_ISL_421716, EPI_ISL_421717, EPI_ISL_421718, EPI_ISL_421719, EPI_ISL_421720, EPI_ISL_421721, EPI_ISL_421722, EPI_ISL_421723, EPI_ISL_421724, EPI_ISL_421725, EPI_ISL_421726, EPI_ISL_421727, EPI_ISL_421728, EPI_ISL_421729, EPI_ISL_421730, EPI_ISL_421731, EPI_ISL_421732, EPI_ISL_421733 | | | |
| see above | NYU Langone Health | Departments of Pathology and Medicine, New York University School of Medicine | Maria Aguero-Rosenfeld, Brendan Belovarac, Margaret Black, Ludovic Boytard, John Cadley, Paolo Cotzia, John Chen, Dacia Dimartino, Xiaojun Feng, Tatyana Gindin, Adriana Heguy, Megan Hogan, Emily Huang, George Jour, Andrew Lytle, Christian Marier, Matthew T. Maurano, Mark J. Mulligan, Peter Meyn, Iman Osman, Jared Pinnell, Sitharam Ramaswami, Amy Rapkiewicz, Marie Samanovic-Golden, Antonio Serrano, Guomiao Shen, Matija Snuderl, Theodore Vougiouklakis, Nick Vulpescu, Gael Westby, Paul Zappile, Yutong Zhang |
| EPI_ISL_421734, EPI_ISL_421735, EPI_ISL_421736, EPI_ISL_421737, EPI_ISL_421738, EPI_ISL_421739, EPI_ISL_421740, EPI_ISL_421741, EPI_ISL_421742, EPI_ISL_421743, EPI_ISL_421744, EPI_ISL_421745, EPI_ISL_421746, EPI_ISL_421747, EPI_ISL_421748, EPI_ISL_421749, EPI_ISL_421750, EPI_ISL_421751, EPI_ISL_421752, EPI_ISL_421753, EPI_ISL_421754, EPI_ISL_421755, EPI_ISL_421756, EPI_ISL_421757, EPI_ISL_421758, EPI_ISL_421759, EPI_ISL_421760, EPI_ISL_421761, EPI_ISL_421762, EPI_ISL_421763 | | | |
| see above | Laboratoire National de Sante, Microbiology, Virology | Laboratoire National de Sante, Microbiology, Epidemiology and Microbial Genomics | Anke Wienecke-Baldacchino, Ardashel Latsuzbaia, Jessica Tapp, Catherine Ragimbeau, Guillaume Fournier, Tamir Abdelrahman, Trung Nguyen Nguyen, Joel Mossong |
| EPI_ISL_422413, EPI_ISL_422414, EPI_ISL_422415, EPI_ISL_422416, EPI_ISL_422417 | Department of Laboratory Medicine, National Taiwan University Hospital | Microbial Genomics Core Lab, National Taiwan University Centers of Genomic and Precision Medicine | Shiou-Hwei Yeh, You-Yu Lin, Ya-Yun Lai, Chiao-Ling Li, Shan-Chwen Chang, Pei-Jer Chen, Sui-Yuan Chang |
| EPI_ISL_422424 | Jaber Al Ahmad Al Sabah Hospital | Dasman diabetes Institute | Fahd Al-Mulla, Rasheeba Iqbal, Sumi John, Ebaa Al-Ozairi, Qais Al-Duwairi |
| EPI_ISL_422426, EPI_ISL_422427 | JABER AL AHMAD AL SABAH HOSPITAL - KUWAIT CITY | Dasman Diabetes Institute | Fahd Al-Mulla, Rasheeba Iqbal, Sumi John, Ebaa Al-Ozairi, Qais Al-Duwairi |
| EPI_ISL_422645, EPI_ISL_422649, EPI_ISL_422650, EPI_ISL_422651, EPI_ISL_422652, EPI_ISL_422653, EPI_ISL_422654, EPI_ISL_422655, EPI_ISL_422656, EPI_ISL_422657, EPI_ISL_422658, EPI_ISL_422659, EPI_ISL_422664, EPI_ISL_422665, EPI_ISL_422666, EPI_ISL_422667, EPI_ISL_422668, EPI_ISL_422669, EPI_ISL_422670, EPI_ISL_422671, EPI_ISL_422672, EPI_ISL_422673, EPI_ISL_422674, EPI_ISL_422703, EPI_ISL_422704, EPI_ISL_422705, EPI_ISL_422814, EPI_ISL_422815, EPI_ISL_422816, EPI_ISL_422817, EPI_ISL_422846, EPI_ISL_422847, EPI_ISL_422848, EPI_ISL_422849, EPI_ISL_422850, EPI_ISL_422851, EPI_ISL_422852, EPI_ISL_422853, EPI_ISL_422854, EPI_ISL_422855, EPI_ISL_422856, EPI_ISL_422857, EPI_ISL_422858, EPI_ISL_422888, EPI_ISL_422889, EPI_ISL_422890, EPI_ISL_422901, EPI_ISL_422906, EPI_ISL_422907, EPI_ISL_422908, EPI_ISL_422909, EPI_ISL_422910, EPI_ISL_422911, EPI_ISL_422912, EPI_ISL_422913 | | | |
| see above | Dutch COVID-19 response team | Erasmus Medical Center | Bas Oude Munnink, David Nieuwenhuijse, Reina Sikkema, Claudia Schapendonk, Irina Chestakova, Anne van der Linden, Theo Bestebroer, Stefan van Nieuwkoop, Mark Pronk, Pascal Lexmond, Corien Swaan, Manon Haverkate, Madelief Mollers, Mart Stein, Sandra Kengne Kamga Mobou, Jeroen van Kampen, Jolanda Voermans, Aura Timen, Corine GeurtsvanKessel, Annemiek van der Eijk, Richard Molenkamp, Marion Koopmans, on behalf of the Dutch national COVID-19 response team. |
| EPI_ISL_422962, EPI_ISL_422963, EPI_ISL_422964, EPI_ISL_422965, EPI_ISL_422966, EPI_ISL_422967, EPI_ISL_422969, EPI_ISL_422970, EPI_ISL_422971, EPI_ISL_422972, EPI_ISL_422973, EPI_ISL_422974, EPI_ISL_422975, EPI_ISL_422976, EPI_ISL_422977, EPI_ISL_422978, EPI_ISL_422979, EPI_ISL_422980, EPI_ISL_422981, EPI_ISL_422982, EPI_ISL_422983, EPI_ISL_422984, EPI_ISL_422985, EPI_ISL_422986, EPI_ISL_422987, EPI_ISL_422988, EPI_ISL_422989, EPI_ISL_422990, EPI_ISL_422991, EPI_ISL_422992, EPI_ISL_422993, EPI_ISL_422994, EPI_ISL_422995, EPI_ISL_422996, EPI_ISL_422997, EPI_ISL_422998, EPI_ISL_422999, EPI_ISL_423000, EPI_ISL_423008, EPI_ISL_423009, EPI_ISL_423010, EPI_ISL_423024, EPI_ISL_423025, EPI_ISL_423026, EPI_ISL_423027, EPI_ISL_423028, EPI_ISL_423029, EPI_ISL_423030, EPI_ISL_423031, EPI_ISL_423032, EPI_ISL_423033 | | | |
| see above | UW Virology Lab | UW Virology Lab | Pavitra Roychoudhury, Hong Xie, Keith Jerome, Alexander Greninger |
| EPI_ISL_423043 | Ramathibodi Hospital | COVID-19 Network Investigations (CONI) Alliance | Elizabeth Batty, Wasun Chantratita, Thanat Chookajorn, Stefan Fernandez, Angkana Huang, Anthony R. Jones, Khajohn Joonsalak, Chonticha Klungtong, Theerarat Kochakarn, Namfon Kotanan, Krittikorn Kumpornsin, Wudtichai Manasatienkij, Bhakbhoom Panthan, Ekawat Pasomsub, Insee Sensorn, Arporn Wangwiwatsin |
| EPI_ISL_423053, EPI_ISL_423054, EPI_ISL_423637, EPI_ISL_423854, EPI_ISL_423855, EPI_ISL_423856, EPI_ISL_423857, EPI_ISL_423858, EPI_ISL_423859, EPI_ISL_423860, EPI_ISL_423861, EPI_ISL_423862, EPI_ISL_423863, EPI_ISL_423864, EPI_ISL_423867, EPI_ISL_423868, EPI_ISL_423869, EPI_ISL_423870, EPI_ISL_423871, EPI_ISL_423872, EPI_ISL_423873, EPI_ISL_423874, EPI_ISL_423875, EPI_ISL_423876, EPI_ISL_423877, EPI_ISL_423878, EPI_ISL_423881, EPI_ISL_423883, EPI_ISL_423884, EPI_ISL_423887, EPI_ISL_423888, EPI_ISL_423890, EPI_ISL_423891, EPI_ISL_423892, EPI_ISL_423893, EPI_ISL_423894, EPI_ISL_423895, EPI_ISL_423896, EPI_ISL_423897, EPI_ISL_423898, EPI_ISL_423900, EPI_ISL_423901, EPI_ISL_423902, EPI_ISL_423903, EPI_ISL_423904, EPI_ISL_423905, EPI_ISL_423907, EPI_ISL_423909, EPI_ISL_423911, EPI_ISL_423912, EPI_ISL_423913, EPI_ISL_423914, EPI_ISL_423915, EPI_ISL_423916, EPI_ISL_423917, EPI_ISL_423924, EPI_ISL_423925, EPI_ISL_423926, EPI_ISL_423927, EPI_ISL_423935, EPI_ISL_423936, EPI_ISL_423940, EPI_ISL_423941, EPI_ISL_423948, EPI_ISL_423953, EPI_ISL_423955, EPI_ISL_423956, EPI_ISL_423964 | | | |
| see above | Respiratory Virus Unit, Microbiology Services Colindale, Public Health England | Respiratory Virus Unit, Microbiology Services Colindale, Public Health England | Monica Galiano, Shahjahan Miah, Angie Lackenby, Omolola Akinbami, Tiina Talts, Leena Bhaw, Richard Myers, Steven Platt, Kirstin Edwards, Jonathan Hubb, Joanna Ellis, Maria Zambon |
| EPI_ISL_424175, EPI_ISL_424176, EPI_ISL_424177, EPI_ISL_424178, EPI_ISL_424179, EPI_ISL_424180, EPI_ISL_424181, EPI_ISL_424182, EPI_ISL_424183, EPI_ISL_424184, EPI_ISL_424185, EPI_ISL_424255 | | | |
| see above | UW Virology Lab | UW Virology Lab | Pavitra Roychoudhury, Hong Xie, Keith Jerome, Alexander Greninger |
| EPI_ISL_424364, EPI_ISL_424365 | National Influenza Center, Indian Council of Medical Research - National Institute of Virology | Indian Council of Medical Research-National Institute of Virology, Microbial Containment Complex | Pragya D. Yadav, Varsha Potdar, Savita Patil, Dimpal A. Nyayanit, Triparna Majumdar, Manohar. L. Chaudhary, Gururaj Deshpande, Padinjaremattathil Thankappan Ullas, Anita Shete-Aich, Hitesh Dighe, Sreelekshmy Mohandas, Gajanan Sapkal, Atanu Basu, Amita Jain, Bharti Malhotra, Deepika Chaudhary, Sarah Cherian, Priya Abraham |
| EPI_ISL_424366 | Vaccine Research, Development and Application Center, Erciyes University | Gen Era Diagnostics Inc. | Shaikh Terkis Islam Pavel, Hazel Yetiskin, Gunsu Aydin, Can Holyavkin, Muhammet Ali Uygut, Zehra B Dursun, lhami Celik, Alper Iseri, Aykut Ozdarendeli |

| | | | |
|---|---|---|---|
| EPI_ISL_424377 | deCODE genetics | deCODE genetics | Daniel F Gudbjartsson; Agnar Helgason; Hakon Jonsson; Olafur T Magnusson; Pall Melsted; Gudmundur L Norddahl; Jona Saemundsdottir; Asgeir Sigurdsson; Patrick Sulem; Arna B Agustsdottir; Berglind Eiriksdottir; Run Fridriksdottir; Elisabet E Gardarsdottir; Gudmundur Georgsson; Olafia S Gretarsdottir; Kjartan R Gudmundsson; Thora R Gunnarsdottir; Arnaldur Gylfason; Hilma Holm; Brynjar O Jensson; Aslaug Jonasdottir; Kamilla S Josefsdottir; Thordur Kristjansson; Droplaug N Magnusdottir; Louise le Roux; Gudrun Sigmundsdottir; Gardar Sveinbjornsson; Kristin E Sveinsdottir; Maney Sveinsdottir; Emil A Thorarensen; Bjarni Thorbjornsson; Gisli Masson; Ingileif Jonsdottir; Alma Moller; Thorolfur Gudnason; Karl G Kristinsson; Unnur Thorsteinsdottir; Kari Stefansson |
| EPI_ISL_424403 | The National University Hospital of Iceland | deCODE genetics | Daniel F Gudbjartsson; Agnar Helgason; Hakon Jonsson; Olafur T Magnusson; Pall Melsted; Gudmundur L Norddahl; Jona Saemundsdottir; Asgeir Sigurdsson; Patrick Sulem; Arna B Agustsdottir; Berglind Eiriksdottir; Run Fridriksdottir; Elisabet E Gardarsdottir; Gudmundur Georgsson; Olafia S Gretarsdottir; Kjartan R Gudmundsson; Thora R Gunnarsdottir; Arnaldur Gylfason; Hilma Holm; Brynjar O Jensson; Aslaug Jonasdottir; Kamilla S Josefsdottir; Thordur Kristjansson; Droplaug N Magnusdottir; Louise le Roux; Gudrun Sigmundsdottir; Gardar Sveinbjornsson; Kristin E Sveinsdottir; Maney Sveinsdottir; Emil A Thorarensen; Bjarni Thorbjornsson; Gisli Masson; Ingileif Jonsdottir; Alma Moller; Thorolfur Gudnason; Karl G Kristinsson; Unnur Thorsteinsdottir; Kari Stefansson |
| EPI_ISL_424626, EPI_ISL_424627 | Instituto Nacional de Enfermedades Respiratorias | Instituto Nacional de Enfermedades Respiratorias | Joel Armando Vázquez Pérez, Celia Boukadida, Santiago Avila Ríos, Mario Mújica Sánchez, José Arturo Martínez Orozco, Eduardo Becerril Vargas, Jorge Salas Hernández, Irma López Martínez, Lucia Hernández Rivas, Gisela Barrera Badillo, Edgar Mendieta Condado, Fabiola Garcés Ayala, Adnan Araiza Rodríguez, José Ernesto Ramírez González, Victor Hugo Borja Aburto, Concepción Grajales Muñiz, Cesar Raúl González Bonilla, Carolina González Torres, Francisco Javier Gaytán Cervantes, José Esteban Muñoz Medina, Guillermo M. Ruiz-Palacios, Pilar Ramos Cervantes, Violeta Ibarra Gonzalez, Fernando Ledesma Barrientos, Luis Alberto García Andrade, Alfredo Ponce de León Garduño, Blanca Taboada, Alejandro Sánchez, Pavel Isa, Ricardo Grande, Gloria Vázquez, Francisco Pulido, Carlos F. Arias. |
| EPI_ISL_424628 | Department of Clinical Microbiology | GIGA Medical Genomics | Keith Durkin, Maria Artesi, Sébastien Bontems, Raphaël Boreux, Cécile Meex, Pierrette Melin, Marie-Pierre Hayette, Vincent Bours. |
| EPI_ISL_424668, EPI_ISL_424669, EPI_ISL_424671 | Arizona State University Health Services | Arizona State University | Rabia Maqsood, LaRinda A. Holland, Emily A. Kaelin, Bereket Estifanos, Nicholas J. Mellor, Jason Steel, Lily I. Wu, Arvind Varsani, Rolf U. Halden, Brenda G. Hogue, Matthew Scotch, Efrem S. Lim |
| EPI_ISL_424703 | Klinisk mikrobiologi, Region Västerbotten | Unit for Biological Agents, Department for CBRN Defence and Security, Swedish Defence Research Agency | FOI Bioinformatics team |
| EPI_ISL_424969, EPI_ISL_424970, EPI_ISL_424971, EPI_ISL_424972, EPI_ISL_424973, EPI_ISL_424974, EPI_ISL_424975, EPI_ISL_424978 | Laboratory Medicine | Department of Laboratory Medicine, Lin-Kou Chang Gung Memorial Hospital, Taoyuan, Taiwan | Kuo-Chien Tsao, Yu-Nong Gong, Shu-Li Yang, Yi-Chun Liu, Chung-Guei Huang, Mei-Jen Hsiao, Po-Wei Huang, Cheng-Ta Yang, Cheng-Hsun Chiu, Peng-Nien Huang, Kuo-Ming Lee, Guang-Wu Chen, Shin-Ru Shih |
| EPI_ISL_425120, EPI_ISL_425121, EPI_ISL_425122, EPI_ISL_425123, EPI_ISL_425124, EPI_ISL_425125, EPI_ISL_425126 | Center of Medical Microbiology, Virology, and Hospital Hygiene, University of Duesseldorf | Center of Medical Microbiology, Virology, and Hospital Hygiene, University of Duesseldorf | Ortwin Adams, Marcel Andree, Alexander Dilthey, Torsten Feldt, Sandra Hauka, Torsten Houwaart, Björn-Erik Jensen, Detlef Kindgen-Milles, Malte Kohns Vasconcelos, Klaus Pfeffer, Tina Senff, Daniel Strelow, Jörg Timm, Andreas Walker, Tobias Wienemann |
| EPI_ISL_425187 | Servicio de Microbiología. Consorcio Hospital General Universitario de Valencia | Sequencing and Bioinformatics Service and Molecular Epidemiology Research Group. FISABIO-Public Health | Loreto Ferrús Abad, Paula Ruiz-Hueso, Mariana Reyes-Prieto, Vicente Soriano Chirona, Ivan Ansari, David Navarro, Maria Alma Bracho, Griselda De Marco, Beatriz Beamud, Lidia Ruiz Roldan, Marta Pla Diaz, Neris Garcia-Gonzalez, Inma Galán Vendrell, Sandra Carbo, Llúcia Martínez-Priego, Giuseppe D'Auria, Fernando Gonzalez-Candelas |
| EPI_ISL_425188 | Servicio de Microbiología. Consorcio Hospital General Universitario de Valencia | Sequencing and Bioinformatics Service and Molecular Epidemiology Research Group. FISABIO-Public Health | Paula Ruiz-Hueso, Mariana Reyes-Prieto, Vicente Soriano Chirona, Ivan Ansari, David Navarro, Maria Alma Bracho, Griselda De Marco, Beatriz Beamud, Lidia Ruiz Roldan, Marta Pla Diaz, Neris Garcia-Gonzalez, Inma Galán Vendrell, Sandra Carbo, Loreto Ferrús Abad, Llúcia Martínez-Priego, Giuseppe D'Auria, Fernando Gonzalez-Candelas |
| EPI_ISL_425189 | Servicio de Microbiología. Consorcio Hospital General Universitario de Valencia | Sequencing and Bioinformatics Service and Molecular Epidemiology Research Group. FISABIO-Public Health | Mariana Reyes-Prieto, Vicente Soriano Chirona, Ivan Ansari, David Navarro, Maria Alma Bracho, Griselda De Marco, Beatriz Beamud, Lidia Ruiz Roldan, Marta Pla Diaz, Neris Garcia-Gonzalez, Inma Galán Vendrell, Sandra Carbo, Loreto Ferrús Abad, Paula Ruiz-Hueso, Llúcia Martínez-Priego, Giuseppe D'Auria, Fernando Gonzalez-Candelas |
| EPI_ISL_425190 | Servicio de Microbiología. Consorcio Hospital General Universitario de Valencia | Sequencing and Bioinformatics Service and Molecular Epidemiology Research Group. FISABIO-Public Health | Vicente Soriano Chirona, Ivan Ansari, David Navarro, Maria Alma Bracho, Griselda De Marco, Beatriz Beamud, Lidia Ruiz Roldan, Marta Pla Diaz, Neris Garcia-Gonzalez, Inma Galán Vendrell, Sandra Carbo, Loreto Ferrús Abad, Paula Ruiz-Hueso, Mariana Reyes-Prieto, Llúcia Martínez-Priego, Giuseppe D'Auria, Fernando Gonzalez-Candelas |
| EPI_ISL_425191 | Servicio de Microbiología. Consorcio Hospital General Universitario de Valencia | Sequencing and Bioinformatics Service and Molecular Epidemiology Research Group. FISABIO-Public Health | Ivan Ansari, David Navarro, Maria Alma Bracho, Griselda De Marco, Beatriz Beamud, Lidia Ruiz Roldan, Marta Pla Diaz, Neris Garcia-Gonzalez, Inma Galán Vendrell, Sandra Carbo, Loreto Ferrús Abad, Paula Ruiz-Hueso, Mariana Reyes-Prieto, Vicente Soriano Chirona, Llúcia Martínez-Priego, Giuseppe D'Auria, Fernando Gonzalez-Candelas |
| EPI_ISL_425192 | Servicio de Microbiología. Consorcio Hospital General Universitario de Valencia | Sequencing and Bioinformatics Service and Molecular Epidemiology Research Group. FISABIO-Public Health | David Navarro, Maria Alma Bracho, Griselda De Marco, Beatriz Beamud, Lidia Ruiz Roldan, Marta Pla Diaz, Neris Garcia-Gonzalez, Inma Galán Vendrell, Sandra Carbo, Loreto Ferrús Abad, Paula Ruiz-Hueso, Mariana Reyes-Prieto, Vicente Soriano Chirona, Ivan Ansari, Llúcia Martínez-Priego, Giuseppe D'Auria, Fernando Gonzalez-Candelas |
| EPI_ISL_425193 | Servicio de Microbiología. Consorcio Hospital General Universitario de Valencia | Sequencing and Bioinformatics Service and Molecular Epidemiology Research Group. FISABIO-Public Health | Maria Alma Bracho, Griselda De Marco, Beatriz Beamud, Lidia Ruiz Roldan, Marta Pla Diaz, Neris Garcia-Gonzalez, Inma Galán Vendrell, Sandra Carbo, Loreto Ferrús Abad, Paula Ruiz-Hueso, Mariana Reyes-Prieto, Vicente Soriano Chirona, Ivan Ansari, David Navarro, Llúcia Martínez-Priego, Giuseppe D'Auria, Fernando Gonzalez-Candelas |
| EPI_ISL_425194 | Servicio de Microbiología. Consorcio Hospital General Universitario de Valencia | Sequencing and Bioinformatics Service and Molecular Epidemiology Research Group. FISABIO-Public Health | Griselda De Marco, Beatriz Beamud, Lidia Ruiz Roldan, Marta Pla Diaz, Neris Garcia-Gonzalez, Inma Galán Vendrell, Sandra Carbo, Loreto Ferrús Abad, Paula Ruiz-Hueso, Mariana Reyes-Prieto, Vicente Soriano Chirona, Ivan Ansari, David Navarro, Maria Alma Bracho, Llúcia Martínez-Priego, Giuseppe D'Auria, Fernando Gonzalez-Candelas |
| EPI_ISL_425195 | Servicio de Microbiología. Consorcio Hospital General Universitario de Valencia | Sequencing and Bioinformatics Service and Molecular Epidemiology Research Group. FISABIO-Public Health | Beatriz Beamud, Lidia Ruiz Roldan, Marta Pla Diaz, Neris Garcia-Gonzalez, Inma Galán Vendrell, Sandra Carbo, Loreto Ferrús Abad, Paula Ruiz-Hueso, Mariana Reyes-Prieto, Vicente Soriano Chirona, Ivan Ansari, David Navarro, Maria Alma Bracho, Griselda De Marco, Llúcia Martínez-Priego, Giuseppe D'Auria, Fernando Gonzalez-Candelas |
| EPI_ISL_425196 | Servicio de Microbiología. Consorcio Hospital General Universitario de Valencia | Sequencing and Bioinformatics Service and Molecular Epidemiology Research Group. FISABIO-Public Health | Lidia Ruiz Roldan, Marta Pla Diaz, Neris Garcia-Gonzalez, Inma Galán Vendrell, Sandra Carbo, Loreto Ferrús Abad, Paula Ruiz-Hueso, Mariana Reyes-Prieto, Vicente Soriano Chirona, Ivan Ansari, David Navarro, Maria Alma Bracho, Griselda De Marco, Beatriz Beamud, Llúcia Martínez-Priego, Giuseppe D'Auria, Fernando Gonzalez-Candelas |
| EPI_ISL_425197 | Servicio de Microbiología. Consorcio Hospital General Universitario de Valencia | Sequencing and Bioinformatics Service and Molecular Epidemiology Research Group. FISABIO-Public Health | Marta Pla Diaz, Neris Garcia-Gonzalez, Inma Galán Vendrell, Sandra Carbo, Loreto Ferrús Abad, Paula Ruiz-Hueso, Mariana Reyes-Prieto, Vicente Soriano Chirona, Ivan Ansari, David Navarro, Maria Alma Bracho, Griselda De Marco, Beatriz Beamud, Lidia Ruiz Roldan, Llúcia Martínez-Priego, Giuseppe D'Auria, Fernando Gonzalez-Candelas |
| EPI_ISL_425198 | Servicio de Microbiología. Consorcio Hospital General Universitario de Valencia | Sequencing and Bioinformatics Service and Molecular Epidemiology Research Group. FISABIO-Public Health | Neris Garcia-Gonzalez, Inma Galán Vendrell, Sandra Carbo, Loreto Ferrús Abad, Paula Ruiz-Hueso, Mariana Reyes-Prieto, Vicente Soriano Chirona, Ivan Ansari, David Navarro, Maria Alma Bracho, Griselda De Marco, Beatriz Beamud, Lidia Ruiz Roldan, Marta Pla Diaz, Llúcia Martínez-Priego, Giuseppe D'Auria, Fernando Gonzalez-Candelas |
| EPI_ISL_425200 | Servicio de Microbiologia. Hospital Clinico Universitario de Valencia | Sequencing and Bioinformatics Service and Molecular Epidemiology Research Group. FISABIO-Public Health | Paula Ruiz-Hueso, Mariana Reyes-Prieto, Vicente Soriano Chirona, Maria Alma Bracho, Griselda De Marco, Beatriz Beamud, Lidia Ruiz Roldan, Marta Pla Diaz,Neris Garcia-Gonzalez, Loreto Ferrús Abad, Maria Dolores Ocete, Llúcia Martínez-Priego, Inma Galán Vendrell, Concepcion Gimeno, Giuseppe D'Auria, Fernando Gonzalez-Candelas |
| EPI_ISL_425215 | Servicio de Microbiología. Consorcio Hospital General Universitario de Valencia | Sequencing and Bioinformatics Service and Molecular Epidemiology Research Group. FISABIO-Public Health | Beatriz Beamud, Lidia Ruiz Roldan, Marta Pla Diaz, Neris Garcia-Gonzalez, Inma Galán Vendrell, Sandra Carbo, Loreto Ferrús Abad, Paula Ruiz-Hueso, Mariana Reyes-Prieto, Vicente Soriano Chirona, Ivan Ansari, David Navarro, Maria Alma Bracho, Griselda De Marco, Llúcia Martínez-Priego, Giuseppe D'Auria, Fernando Gonzalez-Candelas |
| EPI_ISL_425216 | Servicio de Microbiología. Consorcio Hospital General Universitario de Valencia | Sequencing and Bioinformatics Service and Molecular Epidemiology Research Group. FISABIO-Public Health | Lidia Ruiz Roldan, Marta Pla Diaz, Neris Garcia-Gonzalez, Inma Galán Vendrell, Sandra Carbo, Loreto Ferrús Abad, Paula Ruiz-Hueso, Mariana Reyes-Prieto, Vicente Soriano Chirona, Ivan Ansari, David Navarro, Maria Alma Bracho, Griselda De Marco, Beatriz Beamud, Llúcia Martínez-Priego, Giuseppe D'Auria, Fernando Gonzalez-Candelas |

| | | | |
|---|---|---|---|
| EPI_ISL_425217 | Servicio de Microbiología. Consorcio Hospital General Universitario de Valencia | Sequencing and Bioinformatics Service and Molecular Epidemiology Research Group. FISABIO-Public Health | Marta Pla Diaz, Neris Garcia-Gonzalez, Inma Galán Vendrell, Sandra Carbo, Loreto Ferrús Abad, Paula Ruiz-Hueso, Mariana Reyes-Prieto, Vicente Soriano Chirona, Ivan Ansari, David Navarro, Maria Alma Bracho, Griselda De Marco, Beatriz Beamud, Lidia Ruiz Roldan, Llúcia Martínez-Priego, Giuseppe D'Auria, Fernando Gonzalez-Candelas |
| EPI_ISL_425218, EPI_ISL_425219 | Servicio de Microbiología. Consorcio Hospital General Universitario de Valencia | Sequencing and Bioinformatics Service and Molecular Epidemiology Research Group. FISABIO-Public Health | Neris Garcia-Gonzalez, Inma Galán Vendrell, Sandra Carbo, Loreto Ferrús Abad, Paula Ruiz-Hueso, Mariana Reyes-Prieto, Vicente Soriano Chirona, Ivan Ansari, David Navarro, Maria Alma Bracho, Griselda De Marco, Beatriz Beamud, Lidia Ruiz Roldan, Marta Pla Diaz, Llúcia Martínez-Priego, Giuseppe D'Auria, Fernando Gonzalez-Candelas |
| EPI_ISL_425220 | Servicio de Microbiología. Consorcio Hospital General Universitario de Valencia | Sequencing and Bioinformatics Service and Molecular Epidemiology Research Group. FISABIO-Public Health | Inma Galán Vendrell, Sandra Carbo, Loreto Ferrús Abad, Paula Ruiz-Hueso, Mariana Reyes-Prieto, Vicente Soriano Chirona, Ivan Ansari, David Navarro, Maria Alma Bracho, Griselda De Marco, Beatriz Beamud, Lidia Ruiz Roldan, Marta Pla Diaz, Neris Garcia-Gonzalez, Llúcia Martínez-Priego, Giuseppe D'Auria, Fernando Gonzalez-Candelas |
| EPI_ISL_425221 | Servicio de Microbiología. Consorcio Hospital General Universitario de Valencia | Sequencing and Bioinformatics Service and Molecular Epidemiology Research Group. FISABIO-Public Health | Sandra Carbo, Loreto Ferrús Abad, Paula Ruiz-Hueso, Mariana Reyes-Prieto, Vicente Soriano Chirona, Ivan Ansari, David Navarro, Maria Alma Bracho, Griselda De Marco, Beatriz Beamud, Lidia Ruiz Roldan, Marta Pla Diaz, Neris Garcia-Gonzalez, Inma Galán Vendrell, Llúcia Martínez-Priego, Giuseppe D'Auria, Fernando Gonzalez-Candelas |
| EPI_ISL_425222 | Servicio de Microbiología. Consorcio Hospital General Universitario de Valencia | Sequencing and Bioinformatics Service and Molecular Epidemiology Research Group. FISABIO-Public Health | Loreto Ferrús Abad, Paula Ruiz-Hueso, Mariana Reyes-Prieto, Vicente Soriano Chirona, Ivan Ansari, David Navarro, Maria Alma Bracho, Griselda De Marco, Beatriz Beamud, Lidia Ruiz Roldan, Marta Pla Diaz, Neris Garcia-Gonzalez, Inma Galán Vendrell, Sandra Carbo, Llúcia Martínez-Priego, Giuseppe D'Auria, Fernando Gonzalez-Candelas |
| EPI_ISL_425223 | Servicio de Microbiología. Consorcio Hospital General Universitario de Valencia | Sequencing and Bioinformatics Service and Molecular Epidemiology Research Group. FISABIO-Public Health | Paula Ruiz-Hueso, Mariana Reyes-Prieto, Vicente Soriano Chirona, Ivan Ansari, David Navarro, Maria Alma Bracho, Griselda De Marco, Beatriz Beamud, Lidia Ruiz Roldan, Marta Pla Diaz, Neris Garcia-Gonzalez, Inma Galán Vendrell, Sandra Carbo, Loreto Ferrús Abad, Llúcia Martínez-Priego, Giuseppe D'Auria, Fernando Gonzalez-Candelas |
| EPI_ISL_425284, EPI_ISL_425341, EPI_ISL_425342, EPI_ISL_425343, EPI_ISL_425344, EPI_ISL_425345, EPI_ISL_425347, EPI_ISL_425348, EPI_ISL_425349, EPI_ISL_425350, EPI_ISL_425351, EPI_ISL_425352, EPI_ISL_425353, EPI_ISL_425354, EPI_ISL_425355, EPI_ISL_425356, EPI_ISL_425357, EPI_ISL_425358, EPI_ISL_425359, EPI_ISL_425362, EPI_ISL_425363, EPI_ISL_425364, EPI_ISL_425365, EPI_ISL_425366, EPI_ISL_425367, EPI_ISL_425368, EPI_ISL_425375, EPI_ISL_425376, EPI_ISL_425377, EPI_ISL_425378, EPI_ISL_425379, EPI_ISL_425380, EPI_ISL_425381, EPI_ISL_425382, EPI_ISL_425383, EPI_ISL_425384, EPI_ISL_425385, EPI_ISL_425389, EPI_ISL_425393, EPI_ISL_425395, EPI_ISL_425396, EPI_ISL_425397, EPI_ISL_425403, EPI_ISL_425406, EPI_ISL_425408, EPI_ISL_425409, EPI_ISL_425411, EPI_ISL_425412, EPI_ISL_425413, EPI_ISL_425416, EPI_ISL_425417, EPI_ISL_425418, EPI_ISL_425420 | | | |
| see above | Department of Pathology, University of Cambridge | COVID-19 Genomics UK (COG-UK) Consortium | Luke W Meredith, M. Estee Torok , Myra Hosmillo, William L. Hamilton, Martin D. Curran, Theresa Feltwell, Anna Yakovleva, Charlotte J. Houldcroft, Aminu S. Jahun, Sarah L. Caddy, Ian Goodfellow |
| EPI_ISL_425478, EPI_ISL_425479, EPI_ISL_425480, EPI_ISL_425481, EPI_ISL_425482, EPI_ISL_425483, EPI_ISL_425484, EPI_ISL_425485, EPI_ISL_425486, EPI_ISL_425487, EPI_ISL_425488, EPI_ISL_425489, EPI_ISL_425490, EPI_ISL_425491, EPI_ISL_425492, EPI_ISL_425493, EPI_ISL_425494, EPI_ISL_425495, EPI_ISL_425501, EPI_ISL_425502, EPI_ISL_425503, EPI_ISL_425504, EPI_ISL_425505, EPI_ISL_425644, EPI_ISL_425645 | | | |
| see above | Queens Medical Centre, Clinical Microbiology Department / DeepSeq Nottingham | COVID-19 Genomics UK (COG-UK) Consortium | Gemma Clark, Wendy Smith, Manjinder Khakh, Hannah Howson-Wells, Jonathan Ball, Patrick McClure, Joseph Chappell, Theocharis Tsoleridis, Nadine Holmes, Matthew Carlisle, Christopher Moore, Fei Sang, Johnny Debebe, Victoria Wright, Matthew Loose |
| EPI_ISL_425657, EPI_ISL_425658, EPI_ISL_425665, EPI_ISL_425667, EPI_ISL_425669, EPI_ISL_425673, EPI_ISL_425675, EPI_ISL_425676, EPI_ISL_425677, EPI_ISL_425678, EPI_ISL_425679, EPI_ISL_425680, EPI_ISL_425681, EPI_ISL_425682, EPI_ISL_425684, EPI_ISL_425685, EPI_ISL_425686, EPI_ISL_425687, EPI_ISL_425688, EPI_ISL_425691, EPI_ISL_425693, EPI_ISL_425694, EPI_ISL_425695, EPI_ISL_425696, EPI_ISL_425697, EPI_ISL_425698, EPI_ISL_425700, EPI_ISL_425701, EPI_ISL_425702, EPI_ISL_425703, EPI_ISL_425704, EPI_ISL_425705, EPI_ISL_425706, EPI_ISL_425707, EPI_ISL_425708, EPI_ISL_425710, EPI_ISL_425711, EPI_ISL_425712, EPI_ISL_425713, EPI_ISL_425714, EPI_ISL_425715, EPI_ISL_425716, EPI_ISL_425717, EPI_ISL_425718, EPI_ISL_425719, EPI_ISL_425721, EPI_ISL_425722, EPI_ISL_425723, EPI_ISL_425725, EPI_ISL_425726, EPI_ISL_425727, EPI_ISL_425729, EPI_ISL_425731, EPI_ISL_425732, EPI_ISL_425734, EPI_ISL_425736, EPI_ISL_425737, EPI_ISL_425740, EPI_ISL_425741, EPI_ISL_425743, EPI_ISL_425747 | | | |
| see above | West of Scotland Specialist Virology Centre, NHSGGC / MRC-University of Glasgow Centre for Virus Research | COVID-19 Genomics UK (COG-UK) Consortium | Ana da Silva Filipe, Kathy Smollett, Stephen Carmichael, Natasha Johnson, Daniel Mair, Lily Tong, Jenna Nichols; Sarah McDonald; Richard Orton, Joseph Hughes, Sreenu Vattipally, David L Robertson; Kathy Li, Natasha Jesudason, Rajiv Shah, James Shepherd, Antonia Ho, Emma Thomson; Alasdair MacLean, Rory Gunson. |
| EPI_ISL_425831, EPI_ISL_425850, EPI_ISL_425851, EPI_ISL_425852, EPI_ISL_425853, EPI_ISL_425854, EPI_ISL_425855, EPI_ISL_425856, EPI_ISL_425857, EPI_ISL_425858, EPI_ISL_425860, EPI_ISL_425861, EPI_ISL_425862, EPI_ISL_425865, EPI_ISL_425906 | | | |
| see above | Virology Department, Royal Infirmary of Edinburgh, NHS Lothian / School of Biological Sciences, University of Edinburgh / Institute of Genetics and Molecular Medicine, University of Edinburgh | COVID-19 Genomics UK (COG-UK) Consortium | McHugh M, Dewar R, Rooke S, Gallagher M, Balcaza C, O'Toole A, Hill V, McCrone JT, Colquhoun R, Yu X, Jackson B, Scher E, Rambaut A, Williams TC, Templeton K |
| EPI_ISL_426297, EPI_ISL_426306, EPI_ISL_426316, EPI_ISL_426317, EPI_ISL_426319, EPI_ISL_426323 | Wadsworth Center, New York State Department.of Health | Wadsworth Center, New York State Department.of Health | Kirsten St. George, Daryl M. Lamson, Sara Griesemer, Jonathan Plitnick, Navjot Singh, Matthew D. Shudt, Erica Lasek-Nesselquist |
| EPI_ISL_426357, EPI_ISL_426359, EPI_ISL_426360 | Laboratory of Molecular Genetics, 2nd Faculty of Medicine, Charles University in Prague, Prague, Czech Republic | Laboratory of Molecular Genetics, 2nd Faculty of Medicine, Charles University in Prague, Prague, Czech Republic | Lenka Kramna, Katerina Polackova, Ondrej Cinek |
| EPI_ISL_426405 | Queen Mary Hospital | Hong Kong Department of Health | Mak Gannon C.K., Cheng Peter K.C., Lam Edman T.K., Chan Rickjason C.W., Tsang Dominic N.C. |
| EPI_ISL_426406 | Queen Elizabeth Hospital | Hong Kong Department of Health | Mak Gannon C.K., Cheng Peter K.C., Lam Edman T.K., Chan Rickjason C.W., Tsang Dominic N.C. |
| EPI_ISL_426407 | Caritas Medical Centre | Hong Kong Department of Health | Mak Gannon C.K., Cheng Peter K.C., Lam Edman T.K., Chan Rickjason C.W., Tsang Dominic N.C. |
| EPI_ISL_426408 | Ruttonjee Hospital | Hong Kong Department of Health | Mak Gannon C.K., Cheng Peter K.C., Lam Edman T.K., Chan Rickjason C.W., Tsang Dominic N.C. |
| EPI_ISL_426476, EPI_ISL_426477, EPI_ISL_426478 | Microbial Genomics Laboratory, Institut Pasteur Montevideo | Microbial Genomics Laboratory, Institut Pasteur Montevideo, Uruguay | Cecilia Salazar, Florencia Díaz-Viraqué, Marianoel Pereira, Pilar Moreno, Gonzalo Moratorio, Gregorio Iraola |
| EPI_ISL_426479, EPI_ISL_426480 | Microbial Genomics Laboratory, Institut Pasteur Montevideo | Microbial Genomics Laboratory, Institut Pasteur Montevideo | Cecilia Salazar, Florencia Díaz-Viraqué, Marianoel Pereira, Pilar Moreno, Gonzalo Moratorio, Gregorio Iraola |
| EPI_ISL_426481, EPI_ISL_426482 | Microbial Genomics Laboratory, Institut Pasteur Montevideo, Uruguay | Microbial Genomics Laboratory, Institut Pasteur Montevideo | Cecilia Salazar, Florencia Díaz-Viraqué, Marianoel Pereira, Pilar Moreno, Gonzalo Moratorio, Gregorio Iraola |
| EPI_ISL_426499, EPI_ISL_426500, EPI_ISL_426501, EPI_ISL_426502, EPI_ISL_426503, EPI_ISL_426504, EPI_ISL_426505, EPI_ISL_426506, EPI_ISL_426507, EPI_ISL_426509, EPI_ISL_426510 | | | |
| see above | TGen North | TGen North | Jolene Bowers, Megan Folkerts, Darrin Lemmer, Dave Engelthaler |
| EPI_ISL_426527, EPI_ISL_426529, EPI_ISL_426530 | AZ SPHL, Arizona Department of Health Services | TGen North | Jolene Bowers, Megan Folkerts, Darrin Lemmer, Dave Engelthaler |
| EPI_ISL_426535 | TGen North | TGen North | Jolene Bowers, Megan Folkerts, Darrin Lemmer, Dave Engelthaler |
| EPI_ISL_426537 | AZ SPHL, Arizona Department of Health Services | TGen North | Jolene Bowers, Megan Folkerts, Darrin Lemmer, Dave Engelthaler |
| EPI_ISL_426583 | Microbial Genomics Laboratory, Institut Pasteur Monteivdeo | Microbial Genomics Laboratory, Institut Pasteur Montevideo, Uruguay | Cecilia Salazar, Florencia Díaz-Viraqué, Marianoel Pereira, Pilar Moreno, Gonzalo Moratorio, Gregorio Iraola |
| EPI_ISL_426630 | TSGH-CP molecular lab | TSGH-CP molecular lab | Cherng-Lih Perng, Ming-Jr Jian, Chih-Kai Chang, Jung-Chung Lin, Kuo-Ming Yeh, Chien-Wen Chen, Sheng-Kang Chiu, Hsing-Yi Chung, Shih-Hung Tsai, Kuo-Sheng Hung, Feng-Yee Chang, Hung-Sheng Shang |
| EPI_ISL_426631, EPI_ISL_426632 | TSGH-CP molecular lab | TSGH-CP molecular lab | Cherng-Lih Perng, Ming-Jr Jian, Chih-Kai Chang, Jung-Chung Lin, Kuo-Ming Yeh, Chien-Wen Chen, Sheng-Kang Chiu, Hsing-Yi Chung, Shih-Hung Tsai, Kuo-Sheng Hung, Tien-Yao Chang, Feng-Yee Chang, Hung-Sheng Shang |
| EPI_ISL_426637, EPI_ISL_426638, EPI_ISL_426639, EPI_ISL_426640, EPI_ISL_426641, EPI_ISL_426642, EPI_ISL_427042 | Victorian Infectious Diseases Reference Laboratory (VIDRL) | Microbiological Diagnostic Unit Public Health Laboratory and Victorian Infectious Diseases Reference Laboratory, Doherty Institute | Caly L., Seemann T., Sait, M., Schultz M., Druce J., Sherry, N. |
| EPI_ISL_427043 | Laboratory of Microbiology, Medical School, National and Kapodistrian University of Athens | Laboratory of Biology, Department of Medicine, Democritus University of Thrace | Bampali,M., Dovrolis,N., Gatzidou,E., Froukala,E., Stavropoulou,A., Veletza,S., Tsakris,A., Spanakis,N. and Karakasiliotis,I. |

| | | | |
|---|---|---|---|
| EPI_ISL_427047, EPI_ISL_427050, EPI_ISL_427079 | Victorian Infectious Diseases Reference Laboratory (VIDRL) | Microbiological Diagnostic Unit Public Health Laboratory and Victorian Infectious Diseases Reference Laboratory, Doherty Institute | Caly L., Seemann T., Sait, M., Schultz M., Druce J., Sherry, N. |
| EPI_ISL_427281 | Minnesota Department of Health, Public Health Laboratory | Minnesota Department of Health, Public Health Laboratory | Matt Plumb, Jacob Garfin and Xiong Wang |
| EPI_ISL_427292 | LACEN-AL - Laboratorio Central de Alagoas | Instituto Oswaldo Cruz FIOCRUZ - Laboratory of Respiratory Viruses and Measles (LVRS) | Paola Resende, Fernando Motta, Luciana Appolinario, Sunando Roy, Aline Mattos, Milene Miranda, Cristiana Garcia, Braulia Caetano, Maria Ogrzewalska, Priscila Born, Jonathan Lopes, Marilda Siqueira |
| EPI_ISL_427393 | TSGH-CP molecular lab | TSGH-CP molecular lab | Cherng-Lih Perng, Ming-Jr Jian, Chih-Kai Chang, Jung-Chung Lin, Kuo-Ming Yeh, Chien-Wen Chen, Sheng-Kang Chiu, Hsing-Yi Chung, Shih-Hung Tsai, Kuo-Sheng Hung, Tien-Yao Chang, Feng-Yee Chang, Hung-Sheng Shang |
| EPI_ISL_427515, EPI_ISL_427516, EPI_ISL_427517, EPI_ISL_427518, EPI_ISL_427519, EPI_ISL_427520, EPI_ISL_427521 | NYU Langone Health | Departments of Pathology and Medicine, New York University School of Medicine | Maria Aguero-Rosenfeld, Brendan Belovarac, Margaret Black, Ludovic Boytard, John Cadley, Paolo Cotzia, John Chen, Dacia Dimartino, Xiaojun Feng, Tatyana Gindin, Emily Guzman, Adriana Heguy, Megan Hogan, Emily Huang, George Jour, Andrew Lytle, Christian Marier, Matthew T. Maurano, Mark J. Mulligan, Peter Meyn, Iman Osman, Jared Pinnell, Vanessa Raabe, Sitharam Ramaswami, Amy Rapkiewicz, Marie Samanovic-Golden, Antonio Serrano, Guomiao Shen, Matija Snuderl, Theodore Vougiouklakis, Nick Vulpescu, Gael Westby, Paul Zappile, Yutong Zhang |
| EPI_ISL_427553, EPI_ISL_427554, EPI_ISL_427555, EPI_ISL_427556, EPI_ISL_427557, EPI_ISL_427558, EPI_ISL_427559, EPI_ISL_427560, EPI_ISL_427561, EPI_ISL_427562, EPI_ISL_427563, EPI_ISL_427564, EPI_ISL_427565, EPI_ISL_427566, EPI_ISL_427567, EPI_ISL_427568, EPI_ISL_427569, EPI_ISL_427570, EPI_ISL_427571, EPI_ISL_427572, EPI_ISL_427573, EPI_ISL_427574, EPI_ISL_427575, EPI_ISL_427576, EPI_ISL_427577, EPI_ISL_427578, EPI_ISL_427579, EPI_ISL_427580, EPI_ISL_427581, EPI_ISL_427582, EPI_ISL_427583, EPI_ISL_427584, EPI_ISL_427585, EPI_ISL_427586, EPI_ISL_427587, EPI_ISL_427588, EPI_ISL_427589, EPI_ISL_427590, EPI_ISL_427591, EPI_ISL_427592, EPI_ISL_427593, EPI_ISL_427594, EPI_ISL_427595, EPI_ISL_427596, EPI_ISL_427597, EPI_ISL_427598, EPI_ISL_427599, EPI_ISL_427600, EPI_ISL_427601, EPI_ISL_427602, EPI_ISL_427603, EPI_ISL_427604, EPI_ISL_427605, EPI_ISL_427606, EPI_ISL_427607, EPI_ISL_427608, EPI_ISL_427609, EPI_ISL_427610, EPI_ISL_427611, EPI_ISL_427612, EPI_ISL_427613, EPI_ISL_427614, EPI_ISL_427615, EPI_ISL_427616, EPI_ISL_427617, EPI_ISL_427618 | | | |
| see above | NewYork-Presbyterian & Mason Lab | Mason Lab | Daniel J. Butler, Christopher Mozsary, Cem Meydan, David Danko, Jonathan Foox, Joel Rosiene, Alon Shaiber, Matthew MacKay, Ebrahim Afshinnekoo, Fritz J. Sedlazeck, Nikolay A. Ivanov, Maria Sierra, Craig D. Westover, Krista Ryon, Benjamin Young, Chandrima Bhattacharya, Phyllis Ruggiero, Justyna Gawrys, Iman Hajirasouliha, Dmitry Meleshko, Mirella Salvatore, Dong Xu, Jenny Xiang, John Sipley, Lin Cong, Arryn Craney, Priya Velu, Lars F. Westblade, Massimo Loda, Shawn Levy, Melissa Cushing, Marcin Imielinski, Hanna Rennert, Christopher E. Mason |
| EPI_ISL_427661 | Centre for Infectious Diseases and Microbiology Public Health | NSW Health Pathology - Institute of Clinical Pathology and Medical Research; Westmead Hospital; University of Sydney | Sim E, Bachmann N, Rockett R, Lam C, Gray K, Timms V, Gall M, Arnott A, Sadsad R, Draper J, Carter I, Holmes EC, O'Sullivan MV, Byun R, Sintchenko V, Chen SC, Eden JS, Maddocks S, Kok J, Propenko M, Sorrell T, Chang S, Basile K, Dwyer DE for the 2019-nCoV Study Group |
| EPI_ISL_427662 | Centre for Infectious Diseases and Microbiology Public Health | NSW Health Pathology - Institute of Clinical Pathology and Medical Research; Westmead Hospital; University of Sydney | Lam C, Gray K, Timms, V, Gall M, Arnott A, Sadsad R, Draper J, Sim E, Bachmann N, Rockett R, Carter I, Holmes EC, O'Sullivan MV, Byun R, Sintchenko V, Chen SC, Eden JS, Maddocks S, Kok J, Propenko M, Sorrell T, Chang S, Basile K, Dwyer DE for the 2019-nCoV Study Group |
| EPI_ISL_427663 | Centre for Infectious Diseases and Microbiology Public Health | NSW Health Pathology - Institute of Clinical Pathology and Medical Research; Westmead Hospital; University of Sydney | Timms V, Gall M, Arnott A, Sadsad R, Draper J, Sim E, Bachmann N, Rockett R, Lam C, Gray K, Carter I, Holmes EC, O'Sullivan MV, Byun R, Sintchenko V, Chen SC, Eden JS, Maddocks S, Kok J, Propenko M, Sorrell T, Chang S, Basile K, Dwyer DE for the 2019-nCoV Study Group |
| EPI_ISL_427666, EPI_ISL_427675 | Centre for Infectious Diseases and Microbiology Public Health | NSW Health Pathology - Institute of Clinical Pathology and Medical Research; Westmead Hospital; University of Sydney | Bachmann N, Rockett R, Lam C, Gray K, Timms V, Gall M, Arnott A, Sadsad R, Draper J, Sim E, Carter I, Holmes EC, O'Sullivan MV, Byun R, Sintchenko V, Chen SC, Eden JS, Maddocks S, Kok J, Propenko M, Sorrell T, Chang S, Basile K, Dwyer DE for the 2019-nCoV Study Group |
| EPI_ISL_427677 | Centre for Infectious Diseases and Microbiology Public Health | NSW Health Pathology - Institute of Clinical Pathology and Medical Research; Westmead Hospital; University of Sydney | Lam C, Gray K, Timms, V, Gall M, Arnott A, Sadsad R, Draper J, Sim E, Bachmann N, Rockett R, Carter I, Holmes EC, O'Sullivan MV, Byun R, Sintchenko V, Chen SC, Eden JS, Maddocks S, Kok J, Propenko M, Sorrell T, Chang S, Basile K, Dwyer DE for the 2019-nCoV Study Group |
| EPI_ISL_427687 | Centre for Infectious Diseases and Microbiology Public Health | NSW Health Pathology - Institute of Clinical Pathology and Medical Research; Westmead Hospital; University of Sydney | Gray K, Timms V, Gall M, Arnott A, Sadsad R, Draper J, Sim E, Bachmann N, Rockett R, Lam C, Carter I, Holmes EC, O'Sullivan MV, Byun R, Sintchenko V, Chen SC, Eden JS, Maddocks S, Kok J, Propenko M, Sorrell T, Chang S, Basile K, Dwyer DE for the 2019-nCoV Study Group |
| EPI_ISL_427702 | Centre for Infectious Diseases and Microbiology Public Health | NSW Health Pathology - Institute of Clinical Pathology and Medical Research; Westmead Hospital; University of Sydney | Arnott A, Sadsad R, Draper J, Sim E, Bachmann N, Rockett R, Lam C, Gray K, Timms V, Gall M, Carter I, Holmes EC, O'Sullivan MV, Byun R, Sintchenko V, Chen SC, Eden JS, Maddocks S, Kok J, Propenko M, Sorrell T, Chang S, Basile K, Dwyer DE for the 2019-nCoV Study Group |
| EPI_ISL_427703 | Centre for Infectious Diseases and Microbiology Public Health | NSW Health Pathology - Institute of Clinical Pathology and Medical Research; Westmead Hospital; University of Sydney | Gray K, Timms V, Gall M, Arnott A, Sadsad R, Draper J, Sim E, Bachmann N, Rockett R, Lam C, Carter I, Holmes EC, O'Sullivan MV, Byun R, Sintchenko V, Chen SC, Eden JS, Maddocks S, Kok J, Propenko M, Sorrell T, Chang S, Basile K, Dwyer DE for the 2019-nCoV Study Group |
| EPI_ISL_427716 | Centre for Infectious Diseases and Microbiology Public Health | NSW Health Pathology - Institute of Clinical Pathology and Medical Research; Westmead Hospital; University of Sydney | Lam C, Gray K, Timms, V, Gall M, Arnott A, Sadsad R, Draper J, Sim E, Bachmann N, Rockett R, Carter I, Holmes EC, O'Sullivan MV, Byun R, Sintchenko V, Chen SC, Eden JS, Maddocks S, Kok J, Propenko M, Sorrell T, Chang S, Basile K, Dwyer DE for the 2019-nCoV Study Group |
| EPI_ISL_427717 | ACT Pathology, The Canberra Hospital | NSW Health Pathology - Institute of Clinical Pathology and Medical Research; Westmead Hospital; University of Sydney | Sim E, Bachmann N, Rockett R, Lam C, Gray K, Timms V, Gall M, Arnott A, Sadsad R, Draper J, Carter I, Holmes EC, O'Sullivan MV, Byun R, Sintchenko V, Chen SC, Eden JS, Maddocks S, Kok J, Propenko M, Sorrell T, Chang S, Basile K, Dwyer DE for the 2019-nCoV Study Group |
| EPI_ISL_427719 | ACT Pathology, The Canberra Hospital | NSW Health Pathology - Institute of Clinical Pathology and Medical Research; Westmead Hospital; University of Sydney | Timms V, Gall M, Arnott A, Sadsad R, Draper J, Sim E, Bachmann N, Rockett R, Lam C, Gray K, Carter I, Holmes EC, O'Sullivan MV, Byun R, Sintchenko V, Chen SC, Eden JS, Maddocks S, Kok J, Propenko M, Sorrell T, Chang S, Basile K, Dwyer DE for the 2019-nCoV Study Group |
| EPI_ISL_427721 | ACT Pathology, The Canberra Hospital | NSW Health Pathology - Institute of Clinical Pathology and Medical Research; Westmead Hospital; University of Sydney | Sim E, Bachmann N, Rockett R, Lam C, Gray K, Timms V, Gall M, Arnott A, Sadsad R, Draper J, Carter I, Holmes EC, O'Sullivan MV, Byun R, Sintchenko V, Chen SC, Eden JS, Maddocks S, Kok J, Propenko M, Sorrell T, Chang S, Basile K, Dwyer DE for the 2019-nCoV Study Group |
| EPI_ISL_427726 | Centre for Infectious Diseases and Microbiology Public Health | NSW Health Pathology - Institute of Clinical Pathology and Medical Research; Westmead Hospital; University of Sydney | Rockett R, Lam C, Gray K, Timms V, Gall M, Arnott A, Sadsad R, Draper J, Sim E, Bachmann N, Carter I, Holmes EC, O'Sullivan MV, Byun R, Sintchenko V, Chen SC, Eden JS, Maddocks S, Kok J, Propenko M, Sorrell T, Chang S, Basile K, Dwyer DE for the 2019-nCoV Study Group |
| EPI_ISL_427727 | Centre for Infectious Diseases and Microbiology Public Health | NSW Health Pathology - Institute of Clinical Pathology and Medical Research; Westmead Hospital; University of Sydney | Lam C, Gray K, Timms, V, Gall M, Arnott A, Sadsad R, Draper J, Sim E, Bachmann N, Rockett R, Carter I, Holmes EC, O'Sullivan MV, Byun R, Sintchenko V, Chen SC, Eden JS, Maddocks S, Kok J, Propenko M, Sorrell T, Chang S, Basile K, Dwyer DE for the 2019-nCoV Study Group |
| EPI_ISL_427728 | Centre for Infectious Diseases and Microbiology Public Health | NSW Health Pathology - Institute of Clinical Pathology and Medical Research; Westmead Hospital; University of Sydney | Rockett R, Lam C, Gray K, Timms V, Gall M, Arnott A, Sadsad R, Draper J, Sim E, Bachmann N, Carter I, Holmes EC, O'Sullivan MV, Byun R, Sintchenko V, Chen SC, Eden JS, Maddocks S, Kok J, Propenko M, Sorrell T, Chang S, Basile K, Dwyer DE for the 2019-nCoV Study Group |
| EPI_ISL_427729 | Centre for Infectious Diseases and Microbiology Public Health | NSW Health Pathology - Institute of Clinical Pathology and Medical Research; Westmead Hospital; University of Sydney | Gall M, Arnott A, Sadsad R, Draper J, Sim E, Bachmann N, Rockett R, Lam C, Gray K, Timms V, Carter I, Holmes EC, O'Sullivan MV, Byun R, Sintchenko V, Chen SC, Eden JS, Maddocks S, Kok J, Propenko M, Sorrell T, Chang S, Basile K, Dwyer DE for the 2019-nCoV Study Group |
| EPI_ISL_427730, EPI_ISL_427731, EPI_ISL_427732 | Centre for Infectious Diseases and Microbiology Public Health | NSW Health Pathology - Institute of Clinical Pathology and Medical Research; Westmead Hospital; University of Sydney | Gray K, Timms V, Gall M, Arnott A, Sadsad R, Draper J, Sim E, Bachmann N, Rockett R, Lam C, Carter I, Holmes EC, O'Sullivan MV, Byun R, Sintchenko V, Chen SC, Eden JS, Maddocks S, Kok J, Propenko M, Sorrell T, Chang S, Basile K, Dwyer DE for the 2019-nCoV Study Group |
| EPI_ISL_427733 | Centre for Infectious Diseases and Microbiology Public Health | NSW Health Pathology - Institute of Clinical Pathology and Medical Research; Westmead Hospital; University of Sydney | Gall M, Arnott A, Sadsad R, Draper J, Sim E, Bachmann N, Rockett R, Lam C, Gray K, Timms V, Carter I, Holmes EC, O'Sullivan MV, Byun R, Sintchenko V, Chen SC, Eden JS, Maddocks S, Kok J, Propenko M, Sorrell T, Chang S, Basile K, Dwyer DE for the 2019-nCoV Study Group |
| EPI_ISL_427734 | Centre for Infectious Diseases and Microbiology Public Health | NSW Health Pathology - Institute of Clinical Pathology and Medical Research; Westmead Hospital; University of Sydney | Timms V, Gall M, Arnott A, Sadsad R, Draper J, Sim E, Bachmann N, Rockett R, Lam C, Gray K, Carter I, Holmes EC, O'Sullivan MV, Byun R, Sintchenko V, Chen SC, Eden JS, Maddocks S, Kok J, Propenko M, Sorrell T, Chang S, Basile K, Dwyer DE for the 2019-nCoV Study Group |
| EPI_ISL_427735 | Centre for Infectious Diseases and Microbiology Public Health | NSW Health Pathology - Institute of Clinical Pathology and Medical Research; Westmead Hospital; University of Sydney | Arnott A, Sadsad R, Draper J, Sim E, Bachmann N, Rockett R, Lam C, Gray K, Timms V, Gall M, Carter I, Holmes EC, O'Sullivan MV, Byun R, Sintchenko V, Chen SC, Eden JS, Maddocks S, Kok J, Propenko M, Sorrell T, Chang S, Basile K, Dwyer DE for the 2019-nCoV Study Group |
| EPI_ISL_428148 | Klinisk mikrobiologi, Region Västerbotten | Unit for Biological Agents, Department for CBRN Defence and Security, Swedish Defence Research Agency | FOI bioinformatics team |
| EPI_ISL_428229, EPI_ISL_428230 | TSGH-CP molecular lab | TSGH-CP molecular lab | Cherng-Lih Perng, Ming-Jr Jian, Chih-Kai Chang, Jung-Chung Lin, Kuo-Ming Yeh, Chien-Wen Chen, Sheng-Kang Chiu, Hsing-Yi Chung, Shih-Hung Tsai, Kuo-Sheng Hung, Tien-Yao Chang, Feng-Yee Chang, Hung-Sheng Shang |
| EPI_ISL_428232, EPI_ISL_428233 | Hematology Laboratory, Section of Molecular Diagnostics, University Clinical Centre, Medical University of Gdansk | Department of Virology, Faculty of Medicine, University of Helsinki, Helsinki, Finland | Marlena Robakowska, Aneta Szulc, Maciej Grzybek, Olli Vapalahti, Teemu Smura |
| EPI_ISL_428253, EPI_ISL_428254, EPI_ISL_428255, EPI_ISL_428256, EPI_ISL_428259, EPI_ISL_428260, EPI_ISL_428261, EPI_ISL_428262, EPI_ISL_428263, EPI_ISL_428345 | University of Wisconsin-Madison AIDS Vaccine Research Laboratories | University of Wisconsin-Madison AIDS Vaccine Research Laboratories | Gage Moreno, Katarina Braun, et al. AIDS Vaccine Research Laboratories |
| EPI_ISL_428370, EPI_ISL_428371, | Yale Clinical Virology Laboratory | Grubaugh Lab - Yale School of Public Health | Joseph Fauver, Anderson Brito, Tara Alpert, Chantal Vogels, Ellen Foxman, Albert Ko, Marie Landry, Nathan Grubaugh |

| | | | |
|---|---|---|---|
| EPI_ISL_428372, EPI_ISL_428381 | | | |
| EPI_ISL_428490, EPI_ISL_428491 | Centers for Disease Control, R.O.C. (Taiwan) | Centers for Disease Control, R.O.C. (Taiwan) | Ji-Rong Yang, Yu-Chi Lin, Jung-Jung Mu, Ming-Tsan Liu |
| EPI_ISL_428670 | Centre for Dengue Research | Centre for Dengue Research | Chandima Jeewandara, Dinuka Ariyaratne, Laksiri Gomes, Deshni Jayathilaka, Ananda Wijewickrama, Eranga Narangoda, Damayanthi Idampitiya, Neelika Malaige |
| EPI_ISL_428712, EPI_ISL_428713, EPI_ISL_428714, EPI_ISL_428715, EPI_ISL_428716 | Ministry of Health Turkey | Ministry of Health Turkey | Fatma Bayrakdar,Aye Baak Alta,Yasemin Cogun,Gülay Korukluolu,Selçuk Klç |
| EPI_ISL_428824, EPI_ISL_428825, EPI_ISL_428826, EPI_ISL_428827 | National Public Health Laboratory, National Centre for Infectious Diseases | National Public Health Laboratory, National Centre for Infectious Diseases | Mak TM, Octavia S, Chavatte JM, Cui L, Lin RTP |
| EPI_ISL_428855 | MRCG at LSHTM Geomics lab | MRCG at LSHTM Genomics lab | Sesay et al |
| EPI_ISL_428875, EPI_ISL_428881 | State Research Center of Virology and Biotechnology VECTOR, Department of Collection of Microorganisms | State Research Center of Virology and Biotechnology VECTOR, Department of Collection of Microorganisms | Sergey A. Bodnev, Oleg V. Pyankov, Tatyana V. Tregubchak, Alexander N. Shvalov, Elena V. Gavrilova, Rinat A. Maksyutov |
| EPI_ISL_428882 | State Research Center of Virology and Biotechnology VECTOR, Department of Collection of Microorganisms | State Research Center of Virology and Biotechnology VECTOR, Department of Collection of Microorganisms | Oleg V. Pyankov, Sergey A. Bodnev, Tatyana V. Tregubchak, Alexander N. Shvalov, Elena V. Gavrilova, Rinat A. Maksyutov |
| EPI_ISL_429007 | UCSF Clinical Microbiology Laboratory | Chan-Zuckerberg Biohub | CZB Cliahub Consortium |
| EPI_ISL_429116 | Klinisk mikrobiologi och vardhygien Halmstad | The Public Health Agency of Sweden | Arne Kotz, Olov Svartstrom, Maria Lind Karlberg, Anna-Malin Linde, Oskar Karlsson Lindsjo, Anna Risberg, Shaman Muradrasoli, Karin Tegmark-Wisell |
| EPI_ISL_429162, EPI_ISL_429163 | The Public Health Agency of Sweden | The Public Health Agency of Sweden | Olov Svartstrom, Maria Lind Karlberg, Anna-Malin Linde, Oskar Karlsson Lindsjo, Anna Risberg, Shaman Muradrasoli, Karin Tegmark-Wisell |
| EPI_ISL_429168, EPI_ISL_429169, EPI_ISL_429170, EPI_ISL_429171 | Ramathibodi Hospital | COVID-19 Network Investigations (CONI) Alliance | Elizabeth Batty, Wasun Chantratita, Thanat Chookajorn, Stefan Fernandez, Angkana Huang, Poramate Jiaranai, Anthony R. Jones, Khajohn Joonsalak, Chonticha Klungtong, Theerarat Kochakarn, Namfon Kotanan, Krittikorn Kumpornsin, Wudtichai Manasatienkij, Bhakbhoom Panthan, Ekawat Pasomsub, Kingkan Rakmanee, Insee Sensorn, Janjira Thaipadungpanit, Arporn Wangwiwatsin,Treewat Watthanachockchai |
| EPI_ISL_429196, EPI_ISL_429208, EPI_ISL_429209, EPI_ISL_429210, EPI_ISL_429213 | University Hospitals of Geneva Laboratory of Virology | University Hospitals of Geneva Laboratory of Virology | Laubscher F. |
| EPI_ISL_429226, EPI_ISL_429227 | Presidio Ospedaliero Santo Spirito | Istituto Zooprofilattico Sperimentale dell'Abruzzo e Molise "G. Caporale" | Lorusso A, Marcacci M, Di Domenico M, Ancora M, Curini V, Mangone I, Rinaldi A, Di Pasquale A, Camma C, Puglia I, Savini G |
| EPI_ISL_429298, EPI_ISL_429301, EPI_ISL_429302, EPI_ISL_429303, EPI_ISL_429304, EPI_ISL_429305, EPI_ISL_429306, EPI_ISL_429307, EPI_ISL_429308, EPI_ISL_429309, EPI_ISL_429310, EPI_ISL_429311, EPI_ISL_429312, EPI_ISL_429313, EPI_ISL_429314, EPI_ISL_429315, EPI_ISL_429316, EPI_ISL_429317, EPI_ISL_429318, EPI_ISL_429320, EPI_ISL_429321, EPI_ISL_429322, EPI_ISL_429323, EPI_ISL_429324, EPI_ISL_429325, EPI_ISL_429326, EPI_ISL_429327, EPI_ISL_429328, EPI_ISL_429329, EPI_ISL_429330, EPI_ISL_429331, EPI_ISL_429332 | | | |
| see above | Department of Clinical Microbiology, Copenhagen University Hospital, Hvidovre, Kettegaard Alle 30, 2650 Hvidovre. | Albertsen lab, Department of Chemistry and Bioscience, Aalborg University, Denmark | Rasmus Kirkegaard |
| EPI_ISL_429582 | Department of Virus and Microbiological Special Diagnostics, Statens Serum Institut, Copenhagen, Denmark, Artillerivej 5, 2300 Copenahgen S | Albertsen lab, Department of Chemistry and Bioscience, Aalborg University, Denmark | Rasmus Kirkegaard |
| EPI_ISL_429673, EPI_ISL_429675, EPI_ISL_429676, EPI_ISL_429678, EPI_ISL_429679, EPI_ISL_429680, EPI_ISL_429681, EPI_ISL_429682, EPI_ISL_429683, EPI_ISL_429684, EPI_ISL_429685, EPI_ISL_429686, EPI_ISL_429687, EPI_ISL_429688, EPI_ISL_429689, EPI_ISL_429690, EPI_ISL_429691, EPI_ISL_429692, EPI_ISL_429694, EPI_ISL_429696, EPI_ISL_429697 | | | |
| see above | Central Public Health Laboratory/Octávio Magalhães Institute (IOM) from the Ezequiel Dias Foundation (FUNED) | Instituto Octávio Magalhães / Fundação Ezequiel Dias (IOM/Funed) | Talita Adelino, Joilson Xavier, Marta Giovanetti, Vagner Fonseca, Marcos Vinícius Silva, Luiz Carlos Junior Alcantara, Marluce Aparecida Assunção Oliveira |
| EPI_ISL_429707, EPI_ISL_429738, EPI_ISL_429748, EPI_ISL_429750 | Laboratoire National de Sante, Microbiology, Virology | Laboratoire National de Sante, Microbiology, Epidemiology and Microbial Genomics | Anke Wienecke-Baldacchino, Ardashel Latsuzbaia, Jessica Tapp, Catherine Ragimbeau, Guillaume Fournier, Tamir Abdelrahman, Trung Nguyen Nguyen, Joel Mossong |
| EPI_ISL_429813, EPI_ISL_429814 | Queen Elizabeth II Health Science Centre | National Microbiology Laboratory | Anna Majer, Shari Tyson, Grace Seo, Kristyn Burak, Philip Mabon, Elsie Grudeski, Rhiannon Huzarewich, Russell Mandes, Jennifer Tanner, Natalie Knox, Morag Graham, Gary Van Domselaar, Todd Hatchette, Jason LeBlanc, Nathalie Bastien, Yan Li, Timothy Booth, Matthew Gilmour |
| EPI_ISL_429815 | Public Health Laboratory | National Microbiology Laboratory | Anna Majer, Shari Tyson, Grace Seo, Kristyn Burak, Philip Mabon, Elsie Grudeski, Rhiannon Huzarewich, Russell Mandes, Jennifer Tanner, Natalie Knox, Morag Graham, Gary Van Domselaar, Robert Needle, Yang Yu, Adel Malek, Laura Gilbert, George Zahariadis, Nathalie Bastien, Yan Li, Timothy Booth, Matthew Gilmour |
| EPI_ISL_429818, EPI_ISL_429819 | Cadham Provincial Laboratory | National Microbiology Laboratory | Anna Majer, Shari Tyson, Grace Seo, Kristyn Burak, Philip Mabon, Elsie Grudeski, Rhiannon Huzarewich, Russell Mandes, Jennifer Tanner, Natalie Knox, Morag Graham, Gary Van Domselaar, Paul Van Caeseele, Jared Bullard, David Alexander, Kerry Dust, Nathalie Bastien, Yan Li, Timothy Booth, Matthew Gilmour |
| EPI_ISL_429865, EPI_ISL_429866, EPI_ISL_429867, EPI_ISL_429868, EPI_ISL_429869 | Ministry of Health Turkey | Ministry of Health Turkey | Fatma Bayrakdar,Aye Baak Alta,Yasemin Cogun,Gülay Korukluolu,Selçuk Klç |
| EPI_ISL_429993, EPI_ISL_429994, EPI_ISL_429997, EPI_ISL_429999, EPI_ISL_430001, EPI_ISL_430002, EPI_ISL_430003, EPI_ISL_430004, EPI_ISL_430014 | Biolab Diagnostic Laboratories | Andersen lab at Scripps Research | Issa Abu-Dayyeh, Ahmad Tibi, Lama Hussein, Lina Mohammad, Zein Naber, Amid Abdelnour with SEARCH Alliance San Diego |
| EPI_ISL_430019, EPI_ISL_430020, EPI_ISL_430021, EPI_ISL_430022, EPI_ISL_430023, EPI_ISL_430024, EPI_ISL_430025, EPI_ISL_430026, EPI_ISL_430027, EPI_ISL_430028, EPI_ISL_430029, EPI_ISL_430030, EPI_ISL_430031, EPI_ISL_430032, EPI_ISL_430033, EPI_ISL_430034, EPI_ISL_430035, EPI_ISL_430038 | | | |
| see above | Utah Public Health Laboratory | Utah Public Health Laboratory | Erin Young, Kelly Oakeson |
| EPI_ISL_430116, EPI_ISL_430132, EPI_ISL_430140 | Seattle Flu Study | Seattle Flu Study | Chu et al |
| EPI_ISL_430180, EPI_ISL_430181, EPI_ISL_430182, EPI_ISL_430188, EPI_ISL_430189, EPI_ISL_430190, EPI_ISL_430191, EPI_ISL_430192, EPI_ISL_430193, EPI_ISL_430194, EPI_ISL_430196, EPI_ISL_430197, EPI_ISL_430198, EPI_ISL_430199, EPI_ISL_430203, EPI_ISL_430204, EPI_ISL_430205, EPI_ISL_430206, EPI_ISL_430207, EPI_ISL_430208 | | | |
| see above | Washington State Department of Health | Seattle Flu Study | Chu et al |
| EPI_ISL_430819 | Center of Scientific Excellence for Influenza Viruses,National Research Centre (NRC), Egypt. | Center of Scientific Excellence for Influenza Viruses,National Research Centre (NRC), Egypt. | Mohamed Ahmed Ali, Ahmed Kandeil, Ahmed Mostafa, Rabeh El-Shesheny, Mahmoud Shehata, Wael Roshdy, Shymaa Showky Ahmed , Amal Naguib, Nancy M. El Guindy, Mokhtar Gomaa, Ahmed El-Taweel, Ahmed E Kayed, Yassmin Moatasim, Omnia Kutkat, Sara Mahmoud, Mina Kamel, Abo Shama, M Noura, Mohamed El Sayes |
| EPI_ISL_430820 | Center of Scientific Excellence for Influenza Viruses, National Research Centre (NRC), Egypt. | Center of Scientific Excellence for Influenza Viruses, National Research Centre (NRC), Egypt. | Mohamed Ahmed Ali, Ahmed Kandeil, Ahmed Mostafa, Rabeh El-Shesheny, Mahmoud Shehata, Wael Roshdy, Shymaa Showky Ahmed , Amal Naguib, Mokhtar Gomaa, Ahmed El-Taweel, Ahmed E Kayed, Yassmin Moatasim, Omnia Kutkat, Sara Mahmoud, Mina Kamel, Abo Shama, M Noura, Mohamed El Sayes, Nancy M. El Guindy |
| EPI_ISL_430841 | Praram 9 Hospital | National Institute of Health. Department of medical Sciences, Ministry of Public Health, Thailand | Pilailuk,Okada; Siripaporn,Phuygun; Thanutsapa,Thanadachakul; Sittiporn,Parnmen;Warawan,Wongboot; Sunthareeya,Waicharoen; Malinee,Chittaganpitch |
| EPI_ISL_430842 | Central chest Institute of Thailand | National Institute of Health. Department of medical Sciences, Ministry of Public Health, Thailand | Pilailuk,Okada; Siripaporn,Phuygun; Thanutsapa,Thanadachakul; Sittiporn,Parnmen;Warawan,Wongboot; Sunthareeya,Waicharoen; Malinee,Chittaganpitch |

| | | | |
|---|---|---|---|
| EPI_ISL_430864 | The Public Health Agency of Sweden | The Public Health Agency of Sweden | Oskar Karlsson Lindsjo, Maria Lind Karlberg, Anna-Malin Linde, Olov Svartstrom, Anna Risberg, Shaman Muradrasoli, Karin Tegmark-Wisell |
| EPI_ISL_431013 | Alaska State Virology Laboratory | Alaska State Virology Laboratory | Jack Chen |
| EPI_ISL_431014, EPI_ISL_431016 | Alaska State Virology Laboratory | Alaska State Virology Laboratory | Jack Chen, Ph.D. |
| EPI_ISL_431103 | Department of Microbiology, Gandhi Medical College and Hospital, Secendrabad, Hyderabad, India | Department of Microbiology, Gandhi Medical College and Hospital, Secendrabad, Hyderabad, India | Nagamani K, Muttineni Radhakrishna, Thrilok Chander B, Raja Rao M, Kalyani Putty, Ravikumar P, Sunitha P, Pankaj Singh D, Anand Kumar K, Amit A. Upadhyay, Steven E. Bosinger, Rama Amara |
| EPI_ISL_431180, EPI_ISL_431240, EPI_ISL_431292, EPI_ISL_431780, EPI_ISL_431784 | Fujian Center for Disease Control and Prevention | Fujian Center for Disease Control and Prevention | Lin Qi, Huang Zhimiao, Zhang Yanhua, Weng Yuwei |
| EPI_ISL_432640, EPI_ISL_432653, EPI_ISL_432710, EPI_ISL_432794, EPI_ISL_432810 | Virology Department, Sheffield Teaching Hospitals NHS Foundation Trust / Virology Department, Sheffield Teaching Hospitals NHS Foundation Trust | COVID-19 Genomics UK (COG-UK) Consortium | Thushan de Silva, Matthew Parker,Adri Angyal, Rebecca Brown, Luke Green, Rachel Tucker, Paul Parsons, Danielle Groves, Alex Keeley, Dave Partridge, Matthew Wyles, Benjamin Lindsey, Mehmet Yavuz, Mohammad Raza, Cariad Evans |
| EPI_ISL_433274 | West of Scotland Specialist Virology Centre, NHSGGC / MRC-University of Glasgow Centre for Virus Research | COVID-19 Genomics UK (COG-UK) Consortium | Ana da Silva Filipe, Natasha Johnson, Kathy Smollett, Daniel Mair, Stephen Carmichael, Lily Tong, Jenna Nichols, Elihu Aranday-Cortes, Kirstyn Brunker, Yasmin Parr, Kyriaki Nomikou; Sarah McDonald, Marc Niebel, Patawee Asamaphan; Richard Orton, Joseph Hughes, Sreenu Vattipally, David L Robertson; Alasdair MacLean, Rory Gunson; Kathy Li, Natasha Jesudason, Rajiv Shah, James Shepherd, Antonia Ho, Emma Thomson |
| EPI_ISL_434464, EPI_ISL_434477, EPI_ISL_434482, EPI_ISL_434483 | Laboratory of Microbiology, Medical School, National and Kapodistrian University of Athens | Laboratory of Biology, Department of Medicine, Democritus University of Thrace | Kassela K., Bampali,M., Dovrolis,N., Gatzidou,E., Froukala,E., Stavropoulou,A., Veletza,S., Tsakris,A., Spanakis,N. and Karakasiliotis,I. |
| EPI_ISL_434516 | Biolab Diagnostic Laboratories | Andersen lab at Scripps Research | Issa Abu-Dayyeh, Ahmad Tibi, Lama Hussein, Lina Mohammad, Zein Naber, Amid Abdelnour with SEARCH Alliance San Diego |
| EPI_ISL_434533, EPI_ISL_434535 | Area de Salud Alajuela Sur | Inciensa, Instituto Costarricense de Investigación y Enseñanza en Nutrición y Salud | Francisco Duarte, Hebleen Porras, Claudio Soto-Garita, Estela Cordero, Adriana Godinez & Melany Calderon |
| EPI_ISL_434536 | Hospital San Vicente de Paul | Inciensa, Instituto Costarricense de Investigación y Enseñanza en Nutrición y Salud | Francisco Duarte, Hebleen Porras, Claudio Soto-Garita, Estela Cordero, Adriana Godinez & Melany Calderon |
| EPI_ISL_434540 | EBAIS Concepción Norte | Inciensa, Instituto Costarricense de Investigación y Enseñanza en Nutrición y Salud | Francisco Duarte, Hebleen Porras, Claudio Soto-Garita, Estela Cordero, Adriana Godinez & Melany Calderon |
| EPI_ISL_434647, EPI_ISL_434648 | Victoria Vard och Halsa | The Public Health Agency of Sweden | Sarah Henriksson, Oskar Karlsson Lindsjo, Maria Lind Karlberg, Anna-Malin Linde, Olov Svartstrom, Anna Risberg, Theresa Enkirch, Mia Brytting, Karin Tegmark-Wisell |
| EPI_ISL_434649 | Svardsjo VC | The Public Health Agency of Sweden | Tommy Janers, Oskar Karlsson Lindsjo, Maria Lind Karlberg, Anna-Malin Linde, Olov Svartstrom, Anna Risberg, Theresa Enkirch, Mia Brytting, Karin Tegmark-Wisell |
| EPI_ISL_434689, EPI_ISL_434690 | Johns Hopkins Hospital Department of Pathology | Johns Hopkins Hospital Department of Pathology | Peter M. Thielen, Thomas Mehoke, Shirlee Wohl, Srividya Ramakrishnan, Melanie Kirsche, Amanda Ernlund, Oluwaseun Falade-Nwulia, Timothy Gilpatrick, Paul Morris, Norah Sadowski, Nídiá Trovao, Victoria Gniazdowski, Michael Schatz, Stuart C. Ray, Winston Timp, Heba Mostafa |
| EPI_ISL_434704, EPI_ISL_434705, EPI_ISL_434706 | Praram 9 Hospital | National Institute of Health. Department of medical Sciences, Ministry of Public Health, Thailand | Pilailuk,Okada; Siripaporn,Phuygun; Thanutsapa,Thanadachakul; Sittiporn,Parnmen;Warawan,Wongboot; Sunthareeya,Waicharoen; Malinee,Chittaganpitch |
| EPI_ISL_434707 | Thammasat University Hospital | National Institute of Health. Department of medical Sciences, Ministry of Public Health, Thailand | Pilailuk,Okada; Siripaporn,Phuygun; Thanutsapa,Thanadachakul; Sittiporn,Parnmen;Warawan,Wongboot; Sunthareeya,Waicharoen; Malinee,Chittaganpitch |
| EPI_ISL_434708 | unknown | National Institute of Health. Department of medical Sciences, Ministry of Public Health, Thailand | Pilailuk,Okada; Siripaporn,Phuygun; Thanutsapa,Thanadachakul; Sittiporn,Parnmen;Warawan,Wongboot; Sunthareeya,Waicharoen; Malinee,Chittaganpitch |
| EPI_ISL_434712, EPI_ISL_434713, EPI_ISL_434714, EPI_ISL_434715, EPI_ISL_434716, EPI_ISL_434717, EPI_ISL_434718, EPI_ISL_434719, EPI_ISL_434720, EPI_ISL_434721, EPI_ISL_434723, EPI_ISL_434725, EPI_ISL_434728, EPI_ISL_434729, EPI_ISL_434731, EPI_ISL_434732, EPI_ISL_434754, EPI_ISL_434756, EPI_ISL_434757, EPI_ISL_434758, EPI_ISL_434759, EPI_ISL_434760, EPI_ISL_434761, EPI_ISL_434762, EPI_ISL_434763, EPI_ISL_434764, EPI_ISL_434765, EPI_ISL_434766, EPI_ISL_434767, EPI_ISL_434768, EPI_ISL_434769, EPI_ISL_434770, EPI_ISL_434772, EPI_ISL_434773, EPI_ISL_434774, EPI_ISL_434775, EPI_ISL_434776, EPI_ISL_434777, EPI_ISL_434778, EPI_ISL_434779, EPI_ISL_434780, EPI_ISL_434781, EPI_ISL_434782, EPI_ISL_434783, EPI_ISL_434786 | | | |
| see above | Houston Methodist Hospital | Houston Methodist Hospital | S. Wesley Long, Randall J. Olsen, Paul A. Christensen, David W. Bernard, James J. Davis, Maulik Shukla, Marcus Nguyen, Matthew Ojeda Saavedra, Concepcion C. Cantu, Prasanti Yerramilli, Layne Pruitt, Sishir Subedi, Heather Hendrickson, Ghazaleh Eskandari, Muthiah Kumaraswami, Jason S. McLellan, Hakon Jonsson, Kari Stefansson, and James M. Musser |
| EPI_ISL_435061, EPI_ISL_435062, EPI_ISL_435064, EPI_ISL_435065, EPI_ISL_435066, EPI_ISL_435067, EPI_ISL_435068, EPI_ISL_435069, EPI_ISL_435070, EPI_ISL_435101, EPI_ISL_435102, EPI_ISL_435103, EPI_ISL_435104, EPI_ISL_435105, EPI_ISL_435106, EPI_ISL_435108, EPI_ISL_435109, EPI_ISL_435110 | | | |
| see above | National Centre for Disease control (NCDC), CSIR-Institute of Genomics and Integrative Biology (CSIR-IGIB) | NCDC/CSIR-IGIB | Pramod Kumar, Rajesh Pandey, Pooja Sharma, Mahesh Dhar, Vivekanand A, Bharathram Uppili, Himanshu Vashisht, Saruchi Wadhwa, Nishu Tyagi, Uma Sharma, Priyanka Singh, Hemlata Lall, Meena Datta, Poonam Gupta, Nidhi Saini, Aarti Tewari, Bibhash Nandi, Dhirendra Kumar, Satyabrata Bag, Varun Jaiswal, Hema Gogia, Preeti Madan, Simrita Singh, Prateek Singh, Debasis Dash, Mitali Mukerji, Manju Bala, Sandhya Kabra, Sujeet Singh, Mohammed Faruq, Anurag Agrawal, Partha Rakshit |
| EPI_ISL_435124, EPI_ISL_435125, EPI_ISL_435127, EPI_ISL_435128, EPI_ISL_435132, EPI_ISL_435143 | Mohammed Bin Rashid University of Medicine and Health Sciences | Al Jalila Genomics Center | Ahmad Abou Tayoun, Tom Loney, Hamda Khansaheb, Sathishkumar Ramaswamy, Divinlal Harilal, Zulfa Omar Deesi, Rupa Murthy Varghese, Hanan Al Suwaidi, Abdulmajeed Alkhaja, Mohammed Uddin, Rifat Hamoudi, Rabih Halwani, Abiola Catherine Senok, Qutayba Hamid, Norbert Nowotny, Alawi Alsheikh-Ali |
| EPI_ISL_435281 | Medistra Hospital Jakarta | Eijkman Institute for Molecular Biology, Ministry of Research and Technology/National Agency for Research and Innovation | Edison Johar, Frilasita A Yudhaputri, Hidayat Trimarsanto, David H Muljono, Safarina G Malik, Khin Saw Myint, Amin Soebandrio |
| EPI_ISL_435315, EPI_ISL_435316, EPI_ISL_435317 | National Hospital of Tropical Diseases | Oxford University Clinical Research Unit, Hanoi, Vietnam | Nguyen Thi Tam, Van Dinh Trang, Nguyen Thu Trang, Nguyen Thi Ngoc Diep, Le Nguyen Minh Hoa, Pham Ngoc Thach, H. Rogier van Doorn, on behalf of the OUCRU COVID-19 research group |
| EPI_ISL_435658, EPI_ISL_435659, EPI_ISL_435660, EPI_ISL_435661, EPI_ISL_435662, EPI_ISL_435663, EPI_ISL_435664, EPI_ISL_435665 | Santa Clara County Public Health Department | Chiu Laboratory, University of California, San Francisco | Xianding Deng, Scot Federman, Wei Gu, Elsa Villarino, Brandon Bonin, Debra A. Wadford, and Charles Y. Chiu |
| EPI_ISL_435697, EPI_ISL_435698, EPI_ISL_435699, EPI_ISL_435700 | National Public Health Laboratory, National Centre for Infectious Diseases | National Public Health Laboratory, National Centre for Infectious Diseases | Mak Tze Minn, Octavia Sophie, Chavatte Jean-Marc, Cui Lin, Lin Raymond Tzer Pin |
| EPI_ISL_436057, EPI_ISL_436058, EPI_ISL_436059, EPI_ISL_436064, EPI_ISL_436065, EPI_ISL_436071, EPI_ISL_436073, EPI_ISL_436080, EPI_ISL_436081 | NYC Department of Health and Mental Hygiene | Pathogen Discovery, Respiratory Viruses Branch, Division of Viral Diseases, Centers for Disease Control and Prevention | Ying Tao, Krista Queen, Christy Harrison, Jennifer Rakeman, Clinton R. Paden, Jing Zhang, Anna Uehara, Yan Li, Haibin Wang, Jasmine Padilla, Justin Lee, Bettina Bankamp, Zachary Weiner, Suxiang Tong |
| EPI_ISL_436196 | Servicio de Microbiología. Consorcio Hospital General Universitario de Valencia | Sequencing and Bioinformatics Service and Molecular Epidemiology Research Group. FISABIO-Public Health | Griselda De Marco, Beatriz Beamud, Lidia Ruiz Roldan, Marta Pla Diaz,Neris Garcia-Gonzalez, Loreto Ferrús Abad, Maria Dolores Ocete, Inma Galán Vendrell, Paula Ruiz-Hueso, Mariana Reyes-Prieto, Vicente Soriano Chirona, Maria Alma Bracho, Llúcia Martínez-Priego, Concepcion Gimeno, Giuseppe D'Auria, Fernando Gonzalez-Candelas |
| EPI_ISL_436197 | Servicio de Microbiología. Consorcio Hospital General Universitario de Valencia | Sequencing and Bioinformatics Service and Molecular Epidemiology Research Group. FISABIO-Public Health | Beatriz Beamud, Lidia Ruiz Roldan, Marta Pla Diaz,Neris Garcia-Gonzalez, Loreto Ferrús Abad, Maria Dolores Ocete, Inma Galán Vendrell, Paula Ruiz-Hueso, Mariana Reyes-Prieto, Vicente Soriano Chirona, Maria Alma Bracho, Griselda De Marco, Llúcia Martínez-Priego, Concepcion Gimeno, Giuseppe D'Auria, Fernando Gonzalez-Candelas |
| EPI_ISL_436198 | Servicio de Microbiología. Consorcio Hospital General Universitario de Valencia | Sequencing and Bioinformatics Service and Molecular Epidemiology Research Group. FISABIO-Public Health | Lidia Ruiz Roldan, Marta Pla Diaz,Neris Garcia-Gonzalez, Loreto Ferrús Abad, Maria Dolores Ocete, Inma Galán Vendrell, Paula Ruiz-Hueso, Mariana Reyes-Prieto, Vicente Soriano Chirona, Maria Alma Bracho, Griselda De Marco, Beatriz Beamud, Llúcia Martínez-Priego, Concepcion Gimeno, Giuseppe D'Auria, Fernando Gonzalez-Candelas |

| | | | |
|---|---|---|---|
| EPI_ISL_436199 | Servicio de Microbiología. Consorcio Hospital General Universitario de Valencia | Sequencing and Bioinformatics Service and Molecular Epidemiology Research Group. FISABIO-Public Health | Marta Pla Diaz,Neris Garcia-Gonzalez, Loreto Ferrús Abad, Maria Dolores Ocete, Inma Galán Vendrell, Paula Ruiz-Hueso, Mariana Reyes-Prieto, Vicente Soriano Chirona, Maria Alma Bracho, Griselda De Marco, Beatriz Beamud, Lidia Ruiz Roldan, Llúcia Martínez-Priego, Concepcion Gimeno, Giuseppe D'Auria, Fernando Gonzalez-Candelas |
| EPI_ISL_436200 | Servicio de Microbiología. Consorcio Hospital General Universitario de Valencia | Sequencing and Bioinformatics Service and Molecular Epidemiology Research Group. FISABIO-Public Health | Neris Garcia-Gonzalez, Loreto Ferrús Abad, Maria Dolores Ocete, Inma Galán Vendrell, Paula Ruiz-Hueso, Mariana Reyes-Prieto, Vicente Soriano Chirona, Maria Alma Bracho, Griselda De Marco, Beatriz Beamud, Lidia Ruiz Roldan, Marta Pla Diaz, Llúcia Martínez-Priego, Concepcion Gimeno, Giuseppe D'Auria, Fernando Gonzalez-Candelas |
| EPI_ISL_436201 | Servicio de Microbiología. Consorcio Hospital General Universitario de Valencia | Sequencing and Bioinformatics Service and Molecular Epidemiology Research Group. FISABIO-Public Health | Loreto Ferrús Abad, Maria Dolores Ocete, Inma Galán Vendrell, Paula Ruiz-Hueso, Mariana Reyes-Prieto, Vicente Soriano Chirona, Maria Alma Bracho, Griselda De Marco, Beatriz Beamud, Lidia Ruiz Roldan, Marta Pla Diaz,Neris Garcia-Gonzalez, Llúcia Martínez-Priego, Concepcion Gimeno, Giuseppe D'Auria, Fernando Gonzalez-Candelas |
| EPI_ISL_436202 | Servicio de Microbiología. Consorcio Hospital General Universitario de Valencia | Sequencing and Bioinformatics Service and Molecular Epidemiology Research Group. FISABIO-Public Health | Loreto Ferrús Abad, Maria Dolores Ocete,Inma Galán Vendrell, Paula Ruiz-Hueso, Mariana Reyes-Prieto, Vicente Soriano Chirona, Maria Alma Bracho, Griselda De Marco, Beatriz Beamud, Lidia Ruiz Roldan, Marta Pla Diaz,Neris Garcia-Gonzalez, Llúcia Martínez-Priego, Concepcion Gimeno, Giuseppe D'Auria, Fernando Gonzalez-Candelas |
| EPI_ISL_436203 | Servicio de Microbiología. Consorcio Hospital General Universitario de Valencia | Sequencing and Bioinformatics Service and Molecular Epidemiology Research Group. FISABIO-Public Health | Maria Dolores Ocete, Inma Galán Vendrell, Paula Ruiz-Hueso, Mariana Reyes-Prieto, Vicente Soriano Chirona, Maria Alma Bracho, Griselda De Marco, Beatriz Beamud, Lidia Ruiz Roldan, Marta Pla Diaz,Neris Garcia-Gonzalez, Loreto Ferrús Abad, Llúcia Martínez-Priego, Concepcion Gimeno, Giuseppe D'Auria, Fernando Gonzalez-Candelas |
| EPI_ISL_436224 | Servicio de Microbiología. Consorcio Hospital General Universitario de Valencia | Sequencing and Bioinformatics Service and Molecular Epidemiology Research Group. FISABIO-Public Health | Neris Garcia-Gonzalez, Loreto Ferrús Abad, Maria Dolores Ocete, Inma Galán Vendrell, Paula Ruiz-Hueso, Mariana Reyes-Prieto, Vicente Soriano Chirona, Maria Alma Bracho, Griselda De Marco, Beatriz Beamud, Lidia Ruiz Roldan, Marta Pla Diaz, Llúcia Martínez-Priego, Concepcion Gimeno, Giuseppe D'Auria, Fernando Gonzalez-Candelas |
| EPI_ISL_436225 | Servicio de Microbiología. Consorcio Hospital General Universitario de Valencia | Sequencing and Bioinformatics Service and Molecular Epidemiology Research Group. FISABIO-Public Health | Loreto Ferrús Abad, Maria Dolores Ocete, Inma Galán Vendrell, Paula Ruiz-Hueso, Mariana Reyes-Prieto, Vicente Soriano Chirona, Maria Alma Bracho, Griselda De Marco, Beatriz Beamud, Lidia Ruiz Roldan, Marta Pla Diaz,Neris Garcia-Gonzalez, Llúcia Martínez-Priego, Concepcion Gimeno, Giuseppe D'Auria, Fernando Gonzalez-Candelas |
| EPI_ISL_436226 | Servicio de Microbiología. Consorcio Hospital General Universitario de Valencia | Sequencing and Bioinformatics Service and Molecular Epidemiology Research Group. FISABIO-Public Health | Maria Dolores Ocete, Inma Galán Vendrell, Paula Ruiz-Hueso, Mariana Reyes-Prieto, Vicente Soriano Chirona, Maria Alma Bracho, Griselda De Marco, Beatriz Beamud, Lidia Ruiz Roldan, Marta Pla Diaz,Neris Garcia-Gonzalez, Loreto Ferrús Abad, Llúcia Martínez-Priego, Concepcion Gimeno, Giuseppe D'Auria, Fernando Gonzalez-Candelas |
| EPI_ISL_436227 | Servicio de Microbiología. Consorcio Hospital General Universitario de Valencia | Sequencing and Bioinformatics Service and Molecular Epidemiology Research Group. FISABIO-Public Health | Griselda De Marco, Beatriz Beamud, Lidia Ruiz Roldan, Marta Pla Diaz,Neris Garcia-Gonzalez, Loreto Ferrús Abad, Maria Dolores Ocete, Inma Galán Vendrell, Paula Ruiz-Hueso, Mariana Reyes-Prieto, Vicente Soriano Chirona, Maria Alma Bracho, Llúcia Martínez-Priego, Concepcion Gimeno, Giuseppe D'Auria, Fernando Gonzalez-Candelas |
| EPI_ISL_436228 | Servicio de Microbiología. Consorcio Hospital General Universitario de Valencia | Sequencing and Bioinformatics Service and Molecular Epidemiology Research Group. FISABIO-Public Health | Beatriz Beamud, Lidia Ruiz Roldan, Marta Pla Diaz,Neris Garcia-Gonzalez, Loreto Ferrús Abad, Maria Dolores Ocete, Inma Galán Vendrell, Paula Ruiz-Hueso, Mariana Reyes-Prieto, Vicente Soriano Chirona, Maria Alma Bracho, Griselda De Marco, Llúcia Martínez-Priego, Concepcion Gimeno, Giuseppe D'Auria, Fernando Gonzalez-Candelas |
| EPI_ISL_436229 | Servicio de Microbiología. Consorcio Hospital General Universitario de Valencia | Sequencing and Bioinformatics Service and Molecular Epidemiology Research Group. FISABIO-Public Health | Lidia Ruiz Roldan, Marta Pla Diaz,Neris Garcia-Gonzalez, Loreto Ferrús Abad, Maria Dolores Ocete, Inma Galán Vendrell, Paula Ruiz-Hueso, Mariana Reyes-Prieto, Vicente Soriano Chirona, Maria Alma Bracho, Griselda De Marco, Beatriz Beamud, Llúcia Martínez-Priego, Concepcion Gimeno, Giuseppe D'Auria, Fernando Gonzalez-Candelas |
| EPI_ISL_436230 | Servicio de Microbiología. Consorcio Hospital General Universitario de Valencia | Sequencing and Bioinformatics Service and Molecular Epidemiology Research Group. FISABIO-Public Health | Marta Pla Diaz,Neris Garcia-Gonzalez, Loreto Ferrús Abad, Maria Dolores Ocete, Inma Galán Vendrell, Paula Ruiz-Hueso, Mariana Reyes-Prieto, Vicente Soriano Chirona, Maria Alma Bracho, Griselda De Marco, Beatriz Beamud, Lidia Ruiz Roldan, Llúcia Martínez-Priego, Concepcion Gimeno, Giuseppe D'Auria, Fernando Gonzalez-Candelas |
| EPI_ISL_436231 | Servicio de Microbiología. Consorcio Hospital General Universitario de Valencia | Sequencing and Bioinformatics Service and Molecular Epidemiology Research Group. FISABIO-Public Health | Neris Garcia-Gonzalez, Loreto Ferrús Abad, Maria Dolores Ocete, Inma Galán Vendrell, Paula Ruiz-Hueso, Mariana Reyes-Prieto, Vicente Soriano Chirona, Maria Alma Bracho, Griselda De Marco, Beatriz Beamud, Lidia Ruiz Roldan, Marta Pla Diaz, Llúcia Martínez-Priego, Concepcion Gimeno, Giuseppe D'Auria, Fernando Gonzalez-Candelas |
| EPI_ISL_436232 | Servicio de Microbiología. Consorcio Hospital General Universitario de Valencia | Sequencing and Bioinformatics Service and Molecular Epidemiology Research Group. FISABIO-Public Health | Loreto Ferrús Abad, Maria Dolores Ocete, Inma Galán Vendrell, Paula Ruiz-Hueso, Mariana Reyes-Prieto, Vicente Soriano Chirona, Maria Alma Bracho, Griselda De Marco, Beatriz Beamud, Lidia Ruiz Roldan, Marta Pla Diaz,Neris Garcia-Gonzalez, Llúcia Martínez-Priego, Concepcion Gimeno, Giuseppe D'Auria, Fernando Gonzalez-Candelas |
| EPI_ISL_436233 | Servicio de Microbiología. Consorcio Hospital General Universitario de Valencia | Sequencing and Bioinformatics Service and Molecular Epidemiology Research Group. FISABIO-Public Health | Loreto Ferrús Abad, Maria Dolores Ocete,Inma Galán Vendrell, Paula Ruiz-Hueso, Mariana Reyes-Prieto, Vicente Soriano Chirona, Maria Alma Bracho, Griselda De Marco, Beatriz Beamud, Lidia Ruiz Roldan, Marta Pla Diaz,Neris Garcia-Gonzalez, Llúcia Martínez-Priego, Concepcion Gimeno, Giuseppe D'Auria, Fernando Gonzalez-Candelas |
| EPI_ISL_436234 | Servicio de Microbiología. Consorcio Hospital General Universitario de Valencia | Sequencing and Bioinformatics Service and Molecular Epidemiology Research Group. FISABIO-Public Health | Maria Dolores Ocete, Inma Galán Vendrell, Paula Ruiz-Hueso, Mariana Reyes-Prieto, Vicente Soriano Chirona, Maria Alma Bracho, Griselda De Marco, Beatriz Beamud, Lidia Ruiz Roldan, Marta Pla Diaz,Neris Garcia-Gonzalez, Loreto Ferrús Abad, Llúcia Martínez-Priego, Concepcion Gimeno, Giuseppe D'Auria, Fernando Gonzalez-Candelas |
| EPI_ISL_436235, EPI_ISL_436236, EPI_ISL_436237, EPI_ISL_436238, EPI_ISL_436239, EPI_ISL_436240, EPI_ISL_436241, EPI_ISL_436242, EPI_ISL_436243 | Servicio de Microbiologia. Hospital Universitario Doctor Peset | Sequencing and Bioinformatics Service and Molecular Epidemiology Research Group. FISABIO-Public Health | Juan Alberola Enguídanos, Juan Jose Camarena Miñana, Rosa González Pellicer, Neris Garcia-Gonzalez, Inma Galán Vendrell, Sandra Carbo, Loreto Ferrús Abad, Paula Ruiz-Hueso, Mariana Reyes-Prieto, Vicente Soriano Chirona, Ivan Ansari, Maria Alma Bracho, Griselda De Marco, Beatriz Beamud, Lidia Ruiz Roldan, Marta Pla Diaz, Llúcia Martínez-Priego, Giuseppe D'Auria, Jose Miguel Nogueira Coito, Fernando Gonzalez-Candelas |
| EPI_ISL_436277 | Servicio de Microbiología. Consorcio Hospital General Universitario de Valencia | Sequencing and Bioinformatics Service and Molecular Epidemiology Research Group. FISABIO-Public Health | Neris Garcia-Gonzalez, Loreto Ferrús Abad, Maria Dolores Ocete, Inma Galán Vendrell, Paula Ruiz-Hueso, Mariana Reyes-Prieto, Vicente Soriano Chirona, Maria Alma Bracho, Griselda De Marco, Beatriz Beamud, Lidia Ruiz Roldan, Marta Pla Diaz, Llúcia Martínez-Priego, Concepcion Gimeno, Giuseppe D'Auria, Fernando Gonzalez-Candelas |
| EPI_ISL_436279 | Servicio de Microbiologia. Hospital Clinico Universitario de Valencia | Sequencing and Bioinformatics Service and Molecular Epidemiology Research Group. FISABIO-Public Health | Griselda De Marco, Beatriz Beamud, Lidia Ruiz Roldan, Marta Pla Diaz, Neris Garcia-Gonzalez, Inma Galán Vendrell, Sandra Carbo, Loreto Ferrús Abad, Paula Ruiz-Hueso, Mariana Reyes-Prieto, Vicente Soriano Chirona, Ivan Ansari, David Navarro, Maria Alma Bracho, Llúcia Martínez-Priego, Giuseppe D'Auria, Fernando Gonzalez-Candelas |
| EPI_ISL_436280 | Servicio de Microbiologia. Hospital Clinico Universitario de Valencia | Sequencing and Bioinformatics Service and Molecular Epidemiology Research Group. FISABIO-Public Health | Beatriz Beamud, Lidia Ruiz Roldan, Marta Pla Diaz, Neris Garcia-Gonzalez, Inma Galán Vendrell, Sandra Carbo, Loreto Ferrús Abad, Paula Ruiz-Hueso, Mariana Reyes-Prieto, Vicente Soriano Chirona, Ivan Ansari, David Navarro, Maria Alma Bracho, Griselda De Marco, Llúcia Martínez-Priego, Giuseppe D'Auria, Fernando Gonzalez-Candelas |
| EPI_ISL_436283 | Servicio de Microbiologia. Hospital Clinico Universitario de Valencia | Sequencing and Bioinformatics Service and Molecular Epidemiology Research Group. FISABIO-Public Health | Neris Garcia-Gonzalez, Inma Galán Vendrell, Sandra Carbo, Loreto Ferrús Abad, Paula Ruiz-Hueso, Mariana Reyes-Prieto, Vicente Soriano Chirona, Ivan Ansari, David Navarro, Maria Alma Bracho, Griselda De Marco, Beatriz Beamud, Lidia Ruiz Roldan, Marta Pla Diaz, Llúcia Martínez-Priego, Giuseppe D'Auria, Fernando Gonzalez-Candelas |
| EPI_ISL_436291 | Servicio de Microbiologia. Hospital Clinico Universitario de Valencia | Sequencing and Bioinformatics Service and Molecular Epidemiology Research Group. FISABIO-Public Health | David Navarro, Maria Alma Bracho, Griselda De Marco, Beatriz Beamud, Lidia Ruiz Roldan, Marta Pla Diaz, Neris Garcia-Gonzalez, Inma Galán Vendrell, Sandra Carbo, Loreto Ferrús Abad, Paula Ruiz-Hueso, Mariana Reyes-Prieto, Vicente Soriano Chirona, Ivan Ansari, Llúcia Martínez-Priego, Giuseppe D'Auria, Fernando Gonzalez-Candelas |
| EPI_ISL_436301 | Servicio de Microbiologia. Hospital Clinico Universitario de Valencia | Sequencing and Bioinformatics Service and Molecular Epidemiology Research Group. FISABIO-Public Health | Lidia Ruiz Roldan, Marta Pla Diaz, Neris Garcia-Gonzalez, Inma Galán Vendrell, Sandra Carbo, Loreto Ferrús Abad, Paula Ruiz-Hueso, Mariana Reyes-Prieto, Vicente Soriano Chirona, Ivan Ansari, David Navarro, Maria Alma Bracho, Griselda De Marco, Beatriz Beamud, Llúcia Martínez-Priego, Giuseppe D'Auria, Fernando Gonzalez-Candelas |
| EPI_ISL_436362 | Servicio de Microbiologia. Hospital Clinico Universitario de Valencia | Sequencing and Bioinformatics Service and Molecular Epidemiology Research Group. FISABIO-Public Health | David Navarro, Maria Alma Bracho, Griselda De Marco, Beatriz Beamud, Lidia Ruiz Roldan, Marta Pla Diaz, Neris Garcia-Gonzalez, Inma Galán Vendrell, Sandra Carbo, Loreto Ferrús Abad, Paula Ruiz-Hueso, Mariana Reyes-Prieto, Vicente Soriano Chirona, Ivan Ansari, David Navarro, Llúcia Martínez-Priego, Giuseppe D'Auria, Fernando Gonzalez-Candelas |
| EPI_ISL_436363 | Servicio de Microbiologia. Hospital Clinico Universitario de | Sequencing and Bioinformatics Service and Molecular | Griselda De Marco, Beatriz Beamud, Lidia Ruiz Roldan, Marta Pla Diaz, Neris Garcia-Gonzalez, Inma Galán Vendrell, Sandra Carbo, Loreto Ferrús Abad, |

| | | | |
|---|---|---|---|
| | Valencia | Epidemiology Research Group. FISABIO-Public Health | Paula Ruiz-Hueso, Mariana Reyes-Prieto, Vicente Soriano Chirona, Ivan Ansari, David Navarro, Maria Alma Bracho, Llúcia Martínez-Priego, Giuseppe D'Auria, Fernando Gonzalez-Candelas |
| EPI_ISL_436372, EPI_ISL_436373, EPI_ISL_436374, EPI_ISL_436375, EPI_ISL_436376, EPI_ISL_436377, EPI_ISL_436378, EPI_ISL_436379, EPI_ISL_436380, EPI_ISL_436381 | Servicio de Microbiologia. Hospital Universitario Doctor Peset | Sequencing and Bioinformatics Service and Molecular Epidemiology Research Group. FISABIO-Public Health | Juan Alberola Enguídanos, Juan Jose Camarena Miñana, Rosa González Pellicer, Neris Garcia-Gonzalez, Inma Galán Vendrell, Sandra Carbo, Loreto Ferrús Abad, Paula Ruiz-Hueso, Mariana Reyes-Prieto, Vicente Soriano Chirona, Ivan Ansari, Maria Alma Bracho, Griselda De Marco, Beatriz Beamud, Lidia Ruiz Roldan, Marta Pla Diaz, Llúcia Martínez-Priego, Giuseppe D'Auria, Jose Miguel Nogueira Coito, Fernando Gonzalez-Candelas |
| EPI_ISL_436382 | Servicio de Microbiología. Consorcio Hospital General Universitario de Valencia | Sequencing and Bioinformatics Service and Molecular Epidemiology Research Group. FISABIO-Public Health | Loreto Ferrús Abad, Maria Dolores Ocete,Inma Galán Vendrell, Paula Ruiz-Hueso, Mariana Reyes-Prieto, Vicente Soriano Chirona, Maria Alma Bracho, Griselda De Marco, Beatriz Beamud, Lidia Ruiz Roldan, Marta Pla Diaz,Neris Garcia-Gonzalez, Llúcia Martínez-Priego, Concepcion Gimeno, Giuseppe D'Auria, Fernando Gonzalez-Candelas |
| EPI_ISL_436383 | Servicio de Microbiología. Consorcio Hospital General Universitario de Valencia | Sequencing and Bioinformatics Service and Molecular Epidemiology Research Group. FISABIO-Public Health | Maria Dolores Ocete, Inma Galán Vendrell, Paula Ruiz-Hueso, Mariana Reyes-Prieto, Vicente Soriano Chirona, Maria Alma Bracho, Griselda De Marco, Beatriz Beamud, Lidia Ruiz Roldan, Marta Pla Diaz,Neris Garcia-Gonzalez, Loreto Ferrús Abad, Llúcia Martínez-Priego, Concepcion Gimeno, Giuseppe D'Auria, Fernando Gonzalez-Candelas |
| EPI_ISL_436384 | Servicio de Microbiología. Consorcio Hospital General Universitario de Valencia | Sequencing and Bioinformatics Service and Molecular Epidemiology Research Group. FISABIO-Public Health | Griselda De Marco, Beatriz Beamud, Lidia Ruiz Roldan, Marta Pla Diaz,Neris Garcia-Gonzalez, Loreto Ferrús Abad, Maria Dolores Ocete, Inma Galán Vendrell, Paula Ruiz-Hueso, Mariana Reyes-Prieto, Vicente Soriano Chirona, Maria Alma Bracho, Llúcia Martínez-Priego, Concepcion Gimeno, Giuseppe D'Auria, Fernando Gonzalez-Candelas |
| EPI_ISL_436385 | Servicio de Microbiología. Consorcio Hospital General Universitario de Valencia | Sequencing and Bioinformatics Service and Molecular Epidemiology Research Group. FISABIO-Public Health | Beatriz Beamud, Lidia Ruiz Roldan, Marta Pla Diaz,Neris Garcia-Gonzalez, Loreto Ferrús Abad, Maria Dolores Ocete, Inma Galán Vendrell, Paula Ruiz-Hueso, Mariana Reyes-Prieto, Vicente Soriano Chirona, Maria Alma Bracho, Griselda De Marco, Llúcia Martínez-Priego, Concepcion Gimeno, Giuseppe D'Auria, Fernando Gonzalez-Candelas |
| EPI_ISL_436386 | Servicio de Microbiología. Consorcio Hospital General Universitario de Valencia | Sequencing and Bioinformatics Service and Molecular Epidemiology Research Group. FISABIO-Public Health | Lidia Ruiz Roldan, Marta Pla Diaz,Neris Garcia-Gonzalez, Loreto Ferrús Abad, Maria Dolores Ocete, Inma Galán Vendrell, Paula Ruiz-Hueso, Mariana Reyes-Prieto, Vicente Soriano Chirona, Maria Alma Bracho, Griselda De Marco, Beatriz Beamud, Llúcia Martínez-Priego, Concepcion Gimeno, Giuseppe D'Auria, Fernando Gonzalez-Candelas |
| EPI_ISL_436387 | Servicio de Microbiología. Consorcio Hospital General Universitario de Valencia | Sequencing and Bioinformatics Service and Molecular Epidemiology Research Group. FISABIO-Public Health | Marta Pla Diaz,Neris Garcia-Gonzalez, Loreto Ferrús Abad, Maria Dolores Ocete, Inma Galán Vendrell, Paula Ruiz-Hueso, Mariana Reyes-Prieto, Vicente Soriano Chirona, Maria Alma Bracho, Griselda De Marco, Beatriz Beamud, Lidia Ruiz Roldan, Llúcia Martínez-Priego, Concepcion Gimeno, Giuseppe D'Auria, Fernando Gonzalez-Candelas |
| EPI_ISL_436388 | Servicio de Microbiología. Consorcio Hospital General Universitario de Valencia | Sequencing and Bioinformatics Service and Molecular Epidemiology Research Group. FISABIO-Public Health | Neris Garcia-Gonzalez, Loreto Ferrús Abad, Maria Dolores Ocete, Inma Galán Vendrell, Paula Ruiz-Hueso, Mariana Reyes-Prieto, Vicente Soriano Chirona, Maria Alma Bracho, Griselda De Marco, Beatriz Beamud, Lidia Ruiz Roldan, Marta Pla Diaz, Llúcia Martínez-Priego, Concepcion Gimeno, Giuseppe D'Auria, Fernando Gonzalez-Candelas |
| EPI_ISL_436389 | Servicio de Microbiología. Consorcio Hospital General Universitario de Valencia | Sequencing and Bioinformatics Service and Molecular Epidemiology Research Group. FISABIO-Public Health | Loreto Ferrús Abad, Maria Dolores Ocete, Inma Galán Vendrell, Paula Ruiz-Hueso, Mariana Reyes-Prieto, Vicente Soriano Chirona, Maria Alma Bracho, Griselda De Marco, Beatriz Beamud, Lidia Ruiz Roldan, Marta Pla Diaz,Neris Garcia-Gonzalez, Llúcia Martínez-Priego, Concepcion Gimeno, Giuseppe D'Auria, Fernando Gonzalez-Candelas |
| EPI_ISL_436390 | Servicio de Microbiología. Consorcio Hospital General Universitario de Valencia | Sequencing and Bioinformatics Service and Molecular Epidemiology Research Group. FISABIO-Public Health | Maria Dolores Ocete, Inma Galán Vendrell, Paula Ruiz-Hueso, Mariana Reyes-Prieto, Vicente Soriano Chirona, Maria Alma Bracho, Griselda De Marco, Beatriz Beamud, Lidia Ruiz Roldan, Marta Pla Diaz,Neris Garcia-Gonzalez, Loreto Ferrús Abad, Llúcia Martínez-Priego, Concepcion Gimeno, Giuseppe D'Auria, Fernando Gonzalez-Candelas |
| EPI_ISL_436391 | Servicio de Microbiología. Consorcio Hospital General Universitario de Valencia | Sequencing and Bioinformatics Service and Molecular Epidemiology Research Group. FISABIO-Public Health | Griselda De Marco, Beatriz Beamud, Lidia Ruiz Roldan, Marta Pla Diaz,Neris Garcia-Gonzalez, Loreto Ferrús Abad, Maria Dolores Ocete, Inma Galán Vendrell, Paula Ruiz-Hueso, Mariana Reyes-Prieto, Vicente Soriano Chirona, Maria Alma Bracho, Llúcia Martínez-Priego, Concepcion Gimeno, Giuseppe D'Auria, Fernando Gonzalez-Candelas |
| EPI_ISL_436392 | Servicio de Microbiología. Consorcio Hospital General Universitario de Valencia | Sequencing and Bioinformatics Service and Molecular Epidemiology Research Group. FISABIO-Public Health | Beatriz Beamud, Lidia Ruiz Roldan, Marta Pla Diaz,Neris Garcia-Gonzalez, Loreto Ferrús Abad, Maria Dolores Ocete, Inma Galán Vendrell, Paula Ruiz-Hueso, Mariana Reyes-Prieto, Vicente Soriano Chirona, Maria Alma Bracho, Griselda De Marco, Llúcia Martínez-Priego, Concepcion Gimeno, Giuseppe D'Auria, Fernando Gonzalez-Candelas |
| EPI_ISL_436393 | Servicio de Microbiología. Consorcio Hospital General Universitario de Valencia | Sequencing and Bioinformatics Service and Molecular Epidemiology Research Group. FISABIO-Public Health | Lidia Ruiz Roldan, Marta Pla Diaz,Neris Garcia-Gonzalez, Loreto Ferrús Abad, Maria Dolores Ocete, Inma Galán Vendrell, Paula Ruiz-Hueso, Mariana Reyes-Prieto, Vicente Soriano Chirona, Maria Alma Bracho, Griselda De Marco, Beatriz Beamud, Llúcia Martínez-Priego, Concepcion Gimeno, Giuseppe D'Auria, Fernando Gonzalez-Candelas |
| EPI_ISL_436394 | Servicio de Microbiología. Consorcio Hospital General Universitario de Valencia | Sequencing and Bioinformatics Service and Molecular Epidemiology Research Group. FISABIO-Public Health | Marta Pla Diaz,Neris Garcia-Gonzalez, Loreto Ferrús Abad, Maria Dolores Ocete, Inma Galán Vendrell, Paula Ruiz-Hueso, Mariana Reyes-Prieto, Vicente Soriano Chirona, Maria Alma Bracho, Griselda De Marco, Beatriz Beamud, Lidia Ruiz Roldan, Llúcia Martínez-Priego, Concepcion Gimeno, Giuseppe D'Auria, Fernando Gonzalez-Candelas |
| EPI_ISL_436395 | Servicio de Microbiología. Consorcio Hospital General Universitario de Valencia | Sequencing and Bioinformatics Service and Molecular Epidemiology Research Group. FISABIO-Public Health | Neris Garcia-Gonzalez, Loreto Ferrús Abad, Maria Dolores Ocete, Inma Galán Vendrell, Paula Ruiz-Hueso, Mariana Reyes-Prieto, Vicente Soriano Chirona, Maria Alma Bracho, Griselda De Marco, Beatriz Beamud, Lidia Ruiz Roldan, Marta Pla Diaz, Llúcia Martínez-Priego, Concepcion Gimeno, Giuseppe D'Auria, Fernando Gonzalez-Candelas |
| EPI_ISL_436733, EPI_ISL_436734, EPI_ISL_436735, EPI_ISL_436736, EPI_ISL_436737, EPI_ISL_436738, EPI_ISL_436739, EPI_ISL_436740, EPI_ISL_436741, EPI_ISL_436742, EPI_ISL_436743 | | | |
| see above | NYU Langone Health | Departments of Pathology and Medicine, New York University School of Medicine | Maria Aguero-Rosenfeld, Brendan Belovarac, Margaret Black, Ludovic Boytard, John Cadley, Paolo Cotzia, John Chen, Dacia Dimartino, Xiaojun Feng, Tatyana Gindin, Emily Guzman, Adriana Heguy, Megan Hogan, Emily Huang, George Jour, Alireza Khodadadi-Jamayran, Lawrence H. Lin, Raven Luther, Andrew Lytle, Christian Marier, Matthew T. Maurano, Mark J. Mulligan, Peter Meyn, Raquel Ordonez Ciriza, Iman Osman, Jared Pinnell, Vanessa Raabe, Sitharam Ramaswami, Amy Rapkiewicz, Andre M. Ribeiro-dos-Santos, Marie Samanovic-Golden, Antonio Serrano, Guomiao Shen, Matija Snuderl, Theodore Vougiouklakis, Nick Vulpescu, Gael Westby, Paul Zappile, Yutong Zhang |
| EPI_ISL_436817, EPI_ISL_436818, EPI_ISL_436819, EPI_ISL_436820, EPI_ISL_436821, EPI_ISL_436822, EPI_ISL_436823, EPI_ISL_436824, EPI_ISL_436889 | Michigan Department of Health and Human Services, Bureau of Laboratories | Michigan Department of Health and Human Services, Bureau of Laboratories | Blankenship HM, Riner D, Soehnlen MK |
| EPI_ISL_437034 | Department of Virus and Microbiological Special Diagnostics, Statens Serum Institut, Copenhagen, Denmark, Artillerivej 5, 2300 Copenahgen S | Albertsen lab, Department of Chemistry and Bioscience, Aalborg University, Denmark | Rasmus Kirkegaard |
| EPI_ISL_437108, EPI_ISL_437109, EPI_ISL_437110, EPI_ISL_437111, EPI_ISL_437113, EPI_ISL_437114, EPI_ISL_437115, EPI_ISL_437116, EPI_ISL_437117, EPI_ISL_437118, EPI_ISL_437119, EPI_ISL_437120, EPI_ISL_437122, EPI_ISL_437123, EPI_ISL_437131, EPI_ISL_437138 | | | |
| see above | Michigan Department of Health and Human Services, Bureau of Laboratories | Michigan Department of Health and Human Services, Bureau of Laboratories | Blankenship HM, Riner D, Soehnlen MK |
| EPI_ISL_437205, EPI_ISL_437206, EPI_ISL_437234, EPI_ISL_437252, EPI_ISL_437253, EPI_ISL_437254 | Max von Pettenkofer Institute, Virology, National Reference Center for Retroviruses, LMU München | Laboratory for Functional Genome Analysis, Dept. Genomics, Gene Center of the LMU Munich | Max Muenchhoff, Stefan Krebs, Alexander Graf, Oliver Keppler, Helmut Blum |
| EPI_ISL_437332 | Ministry of Health Turkey | Ministry of Health Turkey | Fatma Bayrakdar,Aye Baak Alta,Yasemin Cogun,Süleyman Yalçn, Gülay Korukluolu,Selçuk Klç |
| EPI_ISL_437481 | Pathogen Genomics Lab King Abdullah University of Science and Technology(KAUST) | Pathogen Genomics Lab King Abdullah University of Science and Technology(KAUST) | Sara Mfarrej,Raeece Naeem,Sharif Hala,Amit Subudhi,Fathia Rached,Arnab Pain |
| EPI_ISL_437519 | The National Institute of Public Health Center for Epidemiology and Microbiology | The National Institute of Public Health Center for Epidemiology and Microbiology | Alexander Nagy, Helena Jirincova, Ludmila Novakova, Dusan Trnka, Jaromira Vecerova |

| | | | |
|---|---|---|---|
| EPI_ISL_437561, EPI_ISL_437584 | Scripps Medical Laboratory | Andersen lab at Scripps Research | SEARCH Alliance San Diego with Michael Quigley, Ellen Stefanski, Ian Mchardy |
| EPI_ISL_437605, EPI_ISL_437606, EPI_ISL_437607 | unknown | Faculty of Medicine | Rodpan,A., Joyjinda,Y., Wacharapluesadee,S., Buathong,R., Ghai,S., Petcharat,S., Bunprakob,S., Sirichan,N., Prasithsirikul,W., Mungaomklang,A., Plipat,T. and Hemachudha,T. |
| EPI_ISL_437625 | Laboratory of Genomics & Bioinformatics, Institute of Immunology and Experimental Therapy, Polish Academy of Sciences Oddzial Mikrobiologii Wojewodzkiej Stacji Sanitarno-Epidemiologicznej. | Laboratory of Genomics & Bioinformatics, Institute of Immunology and Experimental Therapy, Polish Academy of Sciences | Dorota Kujawa, Aleksandra Herud, Dariusz Martynowski, Krzysztof Jakub Pawlik, Joanna Sikorska, Paulina Zebrowska, Grazyna Zalewska, Oskar Karpinski and Lukasz Laczmanski |
| EPI_ISL_437635, EPI_ISL_437680 | Department of Virus and Microbiological Special Diagnostics, Statens Serum Institut, Copenhagen, Denmark, Artillerivej 5, 2300 Copenahgen S | Albertsen lab, Department of Chemistry and Bioscience, Aalborg University, Denmark | Rasmus Kirkegaard |
| EPI_ISL_437883, EPI_ISL_437884, EPI_ISL_437885, EPI_ISL_437886, EPI_ISL_437887, EPI_ISL_437889, EPI_ISL_437890, EPI_ISL_437891 | Laboratory of Microbiology, Medical School, National and Kapodistrian University of Athens | Laboratory of Biology, Department of Medicine, Democritus University of Thrace | Kassela K., Dovrolis,N., Bampali,M., Gatzidou,E., Froukala,E., Stavropoulou,A., Veletza,S., Tsakris,A., Spanakis,N. and Karakasiliotis,I. |
| EPI_ISL_437943, EPI_ISL_437944, EPI_ISL_437945, EPI_ISL_437966, EPI_ISL_437970, EPI_ISL_437971, EPI_ISL_437972, EPI_ISL_437973 | Universitaetsklinik für Innere Medizin II Innsbruck | Bergthaler laboratory, CeMM Research Center for Molecular Medicine of the Austrian Academy of Sciences | Alexandra Popa, Benedikt Agerer, Henrique Colaco, Lukas Endler, Jakob-Wendelin Genger, Alexander Lercher, Mark Smyth, Thomas Penz, Michael Schuster, Jan Laine, Martin Senekowitsch, Judith Aberle, Stephan Aberle, Elisabeth Puchhammer-Stoeckl, Manfred Nairz, Guenter Weiss, Wegene Borena, Dorothee von Laer, Christoph Bock, Andreas Bergthaler |
| EPI_ISL_437977 | Institut für Virologie am Department für Hygiene, Mikrobiologie und Public Health | Bergthaler laboratory, CeMM Research Center for Molecular Medicine of the Austrian Academy of Sciences | Alexandra Popa, Benedikt Agerer, Henrique Colaco, Lukas Endler, Jakob-Wendelin Genger, Alexander Lercher, Mark Smyth, Thomas Penz, Michael Schuster, Jan Laine, Martin Senekowitsch, Judith Aberle, Stephan Aberle, Elisabeth Puchhammer-Stoeckl, Manfred Nairz, Guenter Weiss, Wegene Borena, Dorothee von Laer, Christoph Bock, Andreas Bergthaler |
| EPI_ISL_438040, EPI_ISL_438041, EPI_ISL_438042, EPI_ISL_438043, EPI_ISL_438044, EPI_ISL_438045, EPI_ISL_438046, EPI_ISL_438047, EPI_ISL_438048, EPI_ISL_438049, EPI_ISL_438050, EPI_ISL_438051, EPI_ISL_438052, EPI_ISL_438053, EPI_ISL_438054, EPI_ISL_438055, EPI_ISL_438056, EPI_ISL_438057, EPI_ISL_438058, EPI_ISL_438059, EPI_ISL_438060, EPI_ISL_438061, EPI_ISL_438062, EPI_ISL_438063, EPI_ISL_438064, EPI_ISL_438065 | | | |
| see above | Center for Virology, Medical University of Vienna | Bergthaler laboratory, CeMM Research Center for Molecular Medicine of the Austrian Academy of Sciences | Alexandra Popa, Benedikt Agerer, Henrique Colaco, Lukas Endler, Jakob-Wendelin Genger, Alexander Lercher, Mark Smyth, Thomas Penz, Michael Schuster, Jan Laine, Martin Senekowitsch, Judith Aberle, Stephan Aberle, Elisabeth Puchhammer-Stoeckl, Manfred Nairz, Guenter Weiss, Wegene Borena, Dorothee von Laer, Christoph Bock, Andreas Bergthaler |
| EPI_ISL_438222, EPI_ISL_438226, EPI_ISL_438229, EPI_ISL_438231 | Johns Hopkins Hospital Department of Pathology | Johns Hopkins Hospital Department of Pathology | Peter M. Thielen, Thomas Mehoke, Shirlee Wohl, Srividya Ramakrishnan, Melanie Kirsche,_Amanda Ernlund,_Oluwaseun Falade-Nwulia, Timothy Gilpatrick, Paul Morris, Norah Sadowski, N_di_ Trovao, Victoria Gniazdowski, Michael Schatz, Stuart C. Ray, Winston Timp, Heba Mostafa |
| EPI_ISL_438236, EPI_ISL_438246, EPI_ISL_438247 | Johns Hopkins Hospital Department of Pathology | Johns Hopkins Hospital Department of Pathology | Peter M. Thielen, Thomas Mehoke, Shirlee Wohl, Srividya Ramakrishnan, Melanie Kirsche, Amanda Ernlund, Oluwaseun Falade-Nwulia, Timothy Gilpatrick, Paul Morris, Norah Sadowski, Nidia Trovao, Victoria Gniazdowski, Michael Schatz, Stuart C. Ray, Winston Timp, Heba Mostafa |
| EPI_ISL_438883, EPI_ISL_438885, EPI_ISL_438906 | West of Scotland Specialist Virology Centre, NHSGGC / MRC-University of Glasgow Centre for Virus Research | COVID-19 Genomics UK (COG-UK) Consortium | Ana da Silva Filipe, Natasha Johnson, Kathy Smollett, Daniel Mair, Stephen Carmichael, Lily Tong, Jenna Nichols, Elihu Aranday-Cortes, Kirstyn Brunker, Yasmin Parr, Kyriaki Nomikou; Sarah McDonald, Marc Niebel, Patawee Asamaphan; Richard Orton, Joseph Hughes, Sreenu Vattipally, David L Robertson; Alasdair MacLean, Rory Gunson; Kathy Li, Natasha Jesudason, Rajiv Shah, James Shepherd, Antonia Ho, Emma Thomson |
| EPI_ISL_440006, EPI_ISL_440037, EPI_ISL_440098, EPI_ISL_440105, EPI_ISL_440119, EPI_ISL_440131, EPI_ISL_440134, EPI_ISL_440136, EPI_ISL_440139, EPI_ISL_440140, EPI_ISL_440148, EPI_ISL_440149, EPI_ISL_440151, EPI_ISL_440165, EPI_ISL_440168, EPI_ISL_440174, EPI_ISL_440175, EPI_ISL_440194, EPI_ISL_440196, EPI_ISL_440198, EPI_ISL_440200, EPI_ISL_440201, EPI_ISL_440204, EPI_ISL_440206, EPI_ISL_440215, EPI_ISL_440218, EPI_ISL_440220, EPI_ISL_440232, EPI_ISL_440233, EPI_ISL_440241, EPI_ISL_440242, EPI_ISL_440243, EPI_ISL_440248, EPI_ISL_440251, EPI_ISL_440253, EPI_ISL_440255, EPI_ISL_440266, EPI_ISL_440273, EPI_ISL_440276, EPI_ISL_440277 | | | |
| see above | PHE South West Regional Laboratory, National Infection Service | Wellcome Sanger Institute for the COVID-19 Genomics UK (COG-UK) consortium | Stephanie Hutchings, Hannah Pymont, Dr Peter Muir, Barry Vipond, Rich Hopes, Alex Alderton, Roberto Amato, Sonia Goncalves, Ewan Harrison, David K. Jackson, Ian Johnston, Dominic Kwiatkowski, Cordelia Langford, John Sillitoe on behalf of the Wellcome Sanger Institute COVID-19 Surveillance Team (http://www.sanger.ac.uk/covid-team) |
| EPI_ISL_440288 | Department of Pathology, University of Cambridge | Wellcome Sanger Institute for the COVID-19 Genomics UK (COG-UK) consortium | Luke W Meredith, M. Estée Török , Myra Hosmillo, William L. Hamilton, Martin D. Curran, Theresa Feltwell, Grant Hall, Anna Yakovleva, Fahad A Khokhar, Charlotte J. Houldcroft, Laura G Caller, Aminu S. Jahun, Sarah L. Caddy, Ian Goodfellow, Alex Alderton, Roberto Amato, Sonia Goncalves, Ewan Harrison, David K. Jackson, Ian Johnston, Dominic Kwiatkowski, Cordelia Langford, John Sillitoe on behalf of the Wellcome Sanger Institute COVID-19 Surveillance Team (http://www.sanger.ac.uk/covid-team) |
| EPI_ISL_440291, EPI_ISL_440299 | PHE South West Regional Laboratory, National Infection Service | Wellcome Sanger Institute for the COVID-19 Genomics UK (COG-UK) consortium | Stephanie Hutchings, Hannah Pymont, Dr Peter Muir, Barry Vipond, Rich Hopes, Alex Alderton, Roberto Amato, Sonia Goncalves, Ewan Harrison, David K. Jackson, Ian Johnston, Dominic Kwiatkowski, Cordelia Langford, John Sillitoe on behalf of the Wellcome Sanger Institute COVID-19 Surveillance Team (http://www.sanger.ac.uk/covid-team) |
| EPI_ISL_440303, EPI_ISL_440308, EPI_ISL_440315, EPI_ISL_440317, EPI_ISL_440320, EPI_ISL_440326, EPI_ISL_440328, EPI_ISL_440329, EPI_ISL_440330, EPI_ISL_440334, EPI_ISL_440337, EPI_ISL_440340, EPI_ISL_440341, EPI_ISL_440349, EPI_ISL_440353, EPI_ISL_440356, EPI_ISL_440358, EPI_ISL_440368, EPI_ISL_440370, EPI_ISL_440376, EPI_ISL_440377, EPI_ISL_440379, EPI_ISL_440382, EPI_ISL_440384, EPI_ISL_440385, EPI_ISL_440386, EPI_ISL_440388, EPI_ISL_440390, EPI_ISL_440391, EPI_ISL_440392, EPI_ISL_440393, EPI_ISL_440394, EPI_ISL_440395, EPI_ISL_440398, EPI_ISL_440400, EPI_ISL_440401, EPI_ISL_440402, EPI_ISL_440405, EPI_ISL_440406, EPI_ISL_440410, EPI_ISL_440412, EPI_ISL_440414, EPI_ISL_440416, EPI_ISL_440417, EPI_ISL_440418, EPI_ISL_440420, EPI_ISL_440422, EPI_ISL_440424, EPI_ISL_440425, EPI_ISL_440428, EPI_ISL_440429, EPI_ISL_440430, EPI_ISL_440431, EPI_ISL_440432, EPI_ISL_440433, EPI_ISL_440434, EPI_ISL_440436, EPI_ISL_440437, EPI_ISL_440438, EPI_ISL_440441, EPI_ISL_440454 | | | |
| see above | Department of Pathology, University of Cambridge | Wellcome Sanger Institute for the COVID-19 Genomics UK (COG-UK) consortium | Luke W Meredith, M. Estée Török , Myra Hosmillo, William L. Hamilton, Martin D. Curran, Theresa Feltwell, Grant Hall, Anna Yakovleva, Fahad A Khokhar, Charlotte J. Houldcroft, Laura G Caller, Aminu S. Jahun, Sarah L. Caddy, Ian Goodfellow, Alex Alderton, Roberto Amato, Sonia Goncalves, Ewan Harrison, David K. Jackson, Ian Johnston, Dominic Kwiatkowski, Cordelia Langford, John Sillitoe on behalf of the Wellcome Sanger Institute COVID-19 Surveillance Team (http://www.sanger.ac.uk/covid-team) |
| EPI_ISL_441409, EPI_ISL_441413, EPI_ISL_441414, EPI_ISL_441419, EPI_ISL_441420, EPI_ISL_441421, EPI_ISL_441422, EPI_ISL_441423, EPI_ISL_441425, EPI_ISL_441426, EPI_ISL_441428, EPI_ISL_441429, EPI_ISL_441430, EPI_ISL_441431, EPI_ISL_441432, EPI_ISL_441433, EPI_ISL_441434, EPI_ISL_441435, EPI_ISL_441436 | | | |
| see above | Regional Virus Laboratory, Belfast Health and Social Care Trust | COVID-19 Genomics UK (COG-UK) Consortium | Conall McCaughey, James McKenna, Tanya Curran, Susan Feeney, Alison Watt, Ciara Cox, Mairead Connor, Zoltan Molnar, David Simpson, Derek Fairley |
| EPI_ISL_443188, EPI_ISL_443189, EPI_ISL_443193, EPI_ISL_443194, EPI_ISL_443197, EPI_ISL_443199, EPI_ISL_443200, EPI_ISL_443202, EPI_ISL_443203, EPI_ISL_443204, EPI_ISL_443205, EPI_ISL_443206, EPI_ISL_443207, EPI_ISL_443208, EPI_ISL_443209, EPI_ISL_443210, EPI_ISL_443211, EPI_ISL_443214, EPI_ISL_443215, EPI_ISL_443226, EPI_ISL_443227, EPI_ISL_443228, EPI_ISL_443229, EPI_ISL_443230, EPI_ISL_443231, EPI_ISL_443232, EPI_ISL_443233, EPI_ISL_443234, EPI_ISL_443235, EPI_ISL_443236, EPI_ISL_443237, EPI_ISL_443238, EPI_ISL_443239, EPI_ISL_443240, EPI_ISL_443241, EPI_ISL_443242, EPI_ISL_443243 | | | |
| see above | National Public Health Laboratory, National Centre for Infectious Diseases | National Public Health Laboratory, National Centre for Infectious Diseases | Mak Tze Minn, Octavia Sophie, Chavatte Jean-Marc, Cui Lin, Lin Raymond Tzer Pin |
| EPI_ISL_443266, EPI_ISL_443267, EPI_ISL_443268 | CHU - Hôpital Cavale Blanche - Labo. de Virologie | National Reference Center for Viruses of Respiratory Infections, Institut Pasteur, Paris | Mélanie Albert, Marion Barbet, Sylvie Behillil, Méline Bizard, Angela Brisebarre, Flora Donati, Etienne Simon-Lorière, Vincent Enouf, Maud Vanpeene, Sylvie van der Werf, Léa Pilorge |
| EPI_ISL_443300 | Cabinet Médical | National Reference Center for Viruses of Respiratory Infections, Institut Pasteur, Paris | Mélanie Albert, Marion Barbet, Sylvie Behillil, Méline Bizard, Angela Brisebarre, Flora Donati, Etienne Simon-Lorière, Vincent Enouf, Maud Vanpeene, Sylvie van der Werf |
| EPI_ISL_444022 | Baylor College of Medicine | Baylor College of Medicine: HGSC | Vasanthi Avadhanula, Erin Nicholson, David Henke, Pedro Piedra, Harsha Doddapaneni, Donna Muzny, Qingchang Meng, Hsu Chao, Zeineen Momin, Hua Shen, George Weissenberger, Kavya Kottapalli, Yimiti Meiheerguli, Sejal Salvi, Ginger Metcalf, Vipin Menon, Sara J.J. Cregeen, Matthew C. Ross, Tulin Ayvaz, Richard Sucgang, Kristi L. Hoffman, Matthew Wong, Joseph F. Petrosino |
| EPI_ISL_444275 | Laboratory Medicine | Department of Laboratory Medicine, Lin-Kou Chang Gung Memorial Hospital, Taoyuan, Taiwan | Kuo-Chien Tsao, Yu-Nong Gong, Shu-Li Yang, Yi-Chun Liu, Chung-Guei Huang, Mei-Jen Hsiao, Po-Wei Huang, Cheng-Ta Yang, Cheng-Hsun Chiu, Peng-Nien Huang, Kuo-Ming Lee, Guang-Wu Chen, Shin-Ru Shih |
| EPI_ISL_444520, EPI_ISL_444521, EPI_ISL_444522, EPI_ISL_444523, EPI_ISL_444524, EPI_ISL_444525, EPI_ISL_444526, EPI_ISL_444527, EPI_ISL_444528, EPI_ISL_444529, EPI_ISL_444530, EPI_ISL_444531, EPI_ISL_444532, EPI_ISL_444535, EPI_ISL_444536, EPI_ISL_444537, EPI_ISL_444538, EPI_ISL_444539, EPI_ISL_444540, EPI_ISL_444541, EPI_ISL_444542, EPI_ISL_444549, EPI_ISL_444550, EPI_ISL_444552, EPI_ISL_444553, EPI_ISL_444554, EPI_ISL_444555, EPI_ISL_444556, EPI_ISL_444557, EPI_ISL_444558, EPI_ISL_444559, EPI_ISL_444560, EPI_ISL_444561, EPI_ISL_444562, EPI_ISL_444563, EPI_ISL_444564, | | | |

EPI_ISL_444565, EPI_ISL_444566, EPI_ISL_444567, EPI_ISL_444568, EPI_ISL_444569, EPI_ISL_444570, EPI_ISL_444584, EPI_ISL_444585

| | | | |
|---|---|---|---|
| see above | Northwestern Memorial Hospital | Ozer Lab | Ramon Lorenzo-Redondo, Hannah H. Nam, Scott C. Roberts, Lacy M. Simons, Chad J. Achenbach, Lawrence J. Jennings, Chao Qi, Alan R. Hauser, Michael G. Ison, Judd F. Hultquist, Egon A. Ozer |

EPI_ISL_444773, EPI_ISL_444774, EPI_ISL_444775, EPI_ISL_444776, EPI_ISL_444777, EPI_ISL_444778, EPI_ISL_444779, EPI_ISL_444780, EPI_ISL_444781, EPI_ISL_444782, EPI_ISL_444783, EPI_ISL_444784, EPI_ISL_444785, EPI_ISL_444786, EPI_ISL_444788, EPI_ISL_444789, EPI_ISL_444790, EPI_ISL_444791, EPI_ISL_444792

| | | | |
|---|---|---|---|
| see above | NYU Langone Health | Departments of Pathology and Medicine, New York University School of Medicine | Maria Aguero-Rosenfeld, Brendan Belovarac, Margaret Black, Ludovic Boytard, John Cadley, Paolo Cotzia, John Chen, Dacia Dimartino, Xiaojun Feng, Tatyana Gindin, Emily Guzman, Adriana Heguy, Megan Hogan, Emily Huang, George Jour, Alireza Khodadadi-Jamayran, Lawrence H. Lin, Raven Luther, Andrew Lytle, Christian Marier, Matthew T. Maurano, Mark J. Mulligan, Peter Meyn, Raquel Ordonez Ciriza, Iman Osman, Jared Pinnell, Vanessa Raabe, Sitharam Ramaswami, Amy Rapkiewicz, Andre M. Ribeiro-dos-Santos, Marie Samanovic-Golden, Antonio Serrano, Guomiao Shen, Matija Snuderl, Theodore Vougiouklakis, Nick Vulpescu, Gael Westby, Paul Zappile, Yutong Zhang |
| EPI_ISL_444897, EPI_ISL_444898, EPI_ISL_444899 | Department of Virus and Microbiological Special Diagnostics, Statens Serum Institut, Copenhagen, Denmark, Artillerivej 5, 2300 Copenahgen S | Albertsen lab, Department of Chemistry and Bioscience, Aalborg University, Denmark | Rasmus Kirkegaard |
| EPI_ISL_444977 | Hospital Universitari Vall d'Hebron - Vall d'Hebron Institut de Recerca | Hospital Universitari Vall d'Hebron | Cristina Andrés, Maria Piñana, Damir Garcia-Cehic, Mercedes Guerrero-Murillo, Ariadna Rando, Juliana Esperalba, Maria Gema Codina, Tomàs Pumarola, Josep Quer, Andrés Antón |
| EPI_ISL_445094, EPI_ISL_445100, EPI_ISL_445101 | UC San Diego Center for Advanced Laboratory Medicine | Andersen lab at Scripps Research | SEARCH Alliance San Diego with David Pride, Ji H Shin |
| EPI_ISL_445165 | Scripps Medical Laboratory | Andersen lab at Scripps Research | SEARCH Alliance San Diego with Michael Quigley, Ellen Stefanski, Ian Mchardy |
| EPI_ISL_445222 | Saroledens Familjelakare | The Public Health Agency of Sweden | Katarina Jarbur, Oskar Karlsson Lindsjo, Maria Lind Karlberg, Anna-Malin Linde, Olov Svartstrom, Anna Risberg, Theresa Enkirch, Mia Brytting, Karin Tegmark-Wisell |
| EPI_ISL_445269 | HOSPITAL REG.LAUTARO NAVARRO AVARIA | Instituto de Salud Publica de Chile | Andrés E Castillo, Bárbara Parra,Paz Tapia, Jaime Lagos, Loredana Arata, Alejandra Acevedo, Winston Andrade, Gabriel Leal, Carolina Tambley, Patricia Bustos, Rodrigo Fasce, Jorge Fernandez |
| EPI_ISL_445272 | CLINICA CIUDAD DEL MAR | Instituto de Salud Publica de Chile | Andrés E Castillo, Bárbara Parra,Paz Tapia, Jaime Lagos, Loredana Arata, Alejandra Acevedo, Winston Andrade, Gabriel Leal, Carolina Tambley, Patricia Bustos, Rodrigo Fasce, Jorge Fernandez |
| EPI_ISL_445273, EPI_ISL_445274 | LABORATORIO TORRE MEDICA LTDA. | Instituto de Salud Publica de Chile | Andrés E Castillo, Bárbara Parra,Paz Tapia, Jaime Lagos, Loredana Arata, Alejandra Acevedo, Winston Andrade, Gabriel Leal, Carolina Tambley, Patricia Bustos, Rodrigo Fasce, Jorge Fernandez |
| EPI_ISL_445275, EPI_ISL_445276 | HOSPITAL CLINICO FUSAT | Instituto de Salud Publica de Chile | Andrés E Castillo, Bárbara Parra,Paz Tapia, Jaime Lagos, Loredana Arata, Alejandra Acevedo, Winston Andrade, Gabriel Leal, Carolina Tambley, Patricia Bustos, Rodrigo Fasce, Jorge Fernandez |
| EPI_ISL_445277 | FUNDACION DE SALUD EL TENIENTE | Instituto de Salud Publica de Chile | Andrés E Castillo, Bárbara Parra,Paz Tapia, Jaime Lagos, Loredana Arata, Alejandra Acevedo, Winston Andrade, Gabriel Leal, Carolina Tambley, Patricia Bustos, Rodrigo Fasce, Jorge Fernandez |
| EPI_ISL_445278 | LABORATORIO TORRE MEDICA LTDA. | Instituto de Salud Publica de Chile | Andrés E Castillo, Bárbara Parra,Paz Tapia, Jaime Lagos, Loredana Arata, Alejandra Acevedo, Winston Andrade, Gabriel Leal, Carolina Tambley, Patricia Bustos, Rodrigo Fasce, Jorge Fernandez |
| EPI_ISL_445279 | LABORATORIO INMUNOLAB SPA | Instituto de Salud Publica de Chile | Andrés E Castillo, Bárbara Parra,Paz Tapia, Jaime Lagos, Loredana Arata, Alejandra Acevedo, Winston Andrade, Gabriel Leal, Carolina Tambley, Patricia Bustos, Rodrigo Fasce, Jorge Fernandez |
| EPI_ISL_445280 | HOSPITAL REG.LAUTARO NAVARRO AVARIA | Instituto de Salud Publica de Chile | Andrés E Castillo, Bárbara Parra,Paz Tapia, Jaime Lagos, Loredana Arata, Alejandra Acevedo, Winston Andrade, Gabriel Leal, Carolina Tambley, Patricia Bustos, Rodrigo Fasce, Jorge Fernandez |
| EPI_ISL_445281 | HOSPITAL CLINICO DEL SUR | Instituto de Salud Publica de Chile | Andrés E Castillo, Bárbara Parra,Paz Tapia, Jaime Lagos, Loredana Arata, Alejandra Acevedo, Winston Andrade, Gabriel Leal, Carolina Tambley, Patricia Bustos, Rodrigo Fasce, Jorge Fernandez |
| EPI_ISL_445304 | HOSPITAL SAN JUAN DE DIOS | Instituto de Salud Publica de Chile | Andrés E Castillo, Bárbara Parra,Paz Tapia, Jaime Lagos, Loredana Arata, Alejandra Acevedo, Winston Andrade, Gabriel Leal, Carolina Tambley, Patricia Bustos, Rodrigo Fasce, Jorge Fernandez |
| EPI_ISL_445305 | HOSPITAL DE CARABINEROS | Instituto de Salud Publica de Chile | Andrés E Castillo, Bárbara Parra,Paz Tapia, Jaime Lagos, Loredana Arata, Alejandra Acevedo, Winston Andrade, Gabriel Leal, Carolina Tambley, Patricia Bustos, Rodrigo Fasce, Jorge Fernandez |
| EPI_ISL_445306 | CLINICA UC SAN CARLOS DE APOQUINDO | Instituto de Salud Publica de Chile | Andrés E Castillo, Bárbara Parra,Paz Tapia, Jaime Lagos, Loredana Arata, Alejandra Acevedo, Winston Andrade, Gabriel Leal, Carolina Tambley, Patricia Bustos, Rodrigo Fasce, Jorge Fernandez |
| EPI_ISL_445307 | HOSPITAL SAN JOSE DE MAIPO | Instituto de Salud Publica de Chile | Andrés E Castillo, Bárbara Parra,Paz Tapia, Jaime Lagos, Loredana Arata, Alejandra Acevedo, Winston Andrade, Gabriel Leal, Carolina Tambley, Patricia Bustos, Rodrigo Fasce, Jorge Fernandez |
| EPI_ISL_445308 | HOSPITAL EL CARMEN DR.LUIS VALENTIN F. | Instituto de Salud Publica de Chile | Andrés E Castillo, Bárbara Parra,Paz Tapia, Jaime Lagos, Loredana Arata, Alejandra Acevedo, Winston Andrade, Gabriel Leal, Carolina Tambley, Patricia Bustos, Rodrigo Fasce, Jorge Fernandez |
| EPI_ISL_445309 | MUTUAL DE SEGURIDAD C.CH.C. | Instituto de Salud Publica de Chile | Andrés E Castillo, Bárbara Parra,Paz Tapia, Jaime Lagos, Loredana Arata, Alejandra Acevedo, Winston Andrade, Gabriel Leal, Carolina Tambley, Patricia Bustos, Rodrigo Fasce, Jorge Fernandez |
| EPI_ISL_445310 | HOSPITAL DR.SOTERO DEL RIO | Instituto de Salud Publica de Chile | Andrés E Castillo, Bárbara Parra,Paz Tapia, Jaime Lagos, Loredana Arata, Alejandra Acevedo, Winston Andrade, Gabriel Leal, Carolina Tambley, Patricia Bustos, Rodrigo Fasce, Jorge Fernandez |
| EPI_ISL_445311 | MEGASALUD S.A. | Instituto de Salud Publica de Chile | Andrés E Castillo, Bárbara Parra,Paz Tapia, Jaime Lagos, Loredana Arata, Alejandra Acevedo, Winston Andrade, Gabriel Leal, Carolina Tambley, Patricia Bustos, Rodrigo Fasce, Jorge Fernandez |
| EPI_ISL_445312 | CLINICA UC SAN CARLOS DE APOQUINDO | Instituto de Salud Publica de Chile | Andrés E Castillo, Bárbara Parra,Paz Tapia, Jaime Lagos, Loredana Arata, Alejandra Acevedo, Winston Andrade, Gabriel Leal, Carolina Tambley, Patricia Bustos, Rodrigo Fasce, Jorge Fernandez |
| EPI_ISL_445313, EPI_ISL_445314 | HOSPITAL EL CARMEN DR.LUIS VALENTIN F. | Instituto de Salud Publica de Chile | Andrés E Castillo, Bárbara Parra,Paz Tapia, Jaime Lagos, Loredana Arata, Alejandra Acevedo, Winston Andrade, Gabriel Leal, Carolina Tambley, Patricia Bustos, Rodrigo Fasce, Jorge Fernandez |
| EPI_ISL_445316 | CESFAM BALMACEDA DE RENCA | Instituto de Salud Publica de Chile | Andrés E Castillo, Bárbara Parra,Paz Tapia, Jaime Lagos, Loredana Arata, Alejandra Acevedo, Winston Andrade, Gabriel Leal, Carolina Tambley, Patricia Bustos, Rodrigo Fasce, Jorge Fernandez |
| EPI_ISL_445317 | UNIV.DE CHILE HOSP.CLINICO | Instituto de Salud Publica de Chile | Andrés E Castillo, Bárbara Parra,Paz Tapia, Jaime Lagos, Loredana Arata, Alejandra Acevedo, Winston Andrade, Gabriel Leal, Carolina Tambley, Patricia Bustos, Rodrigo Fasce, Jorge Fernandez |
| EPI_ISL_445319 | HOSPITAL FELIX BULNES | Instituto de Salud Publica de Chile | Andrés E Castillo, Bárbara Parra,Paz Tapia, Jaime Lagos, Loredana Arata, Alejandra Acevedo, Winston Andrade, Gabriel Leal, Carolina Tambley, Patricia Bustos, Rodrigo Fasce, Jorge Fernandez |
| EPI_ISL_445320 | C.C.SALUD FAMILIAR PADRE FELIX DONOSO G. | Instituto de Salud Publica de Chile | Andrés E Castillo, Bárbara Parra,Paz Tapia, Jaime Lagos, Loredana Arata, Alejandra Acevedo, Winston Andrade, Gabriel Leal, Carolina Tambley, Patricia Bustos, Rodrigo Fasce, Jorge Fernandez |
| EPI_ISL_445321 | HOSPITAL FELIX BULNES | Instituto de Salud Publica de Chile | Andrés E Castillo, Bárbara Parra,Paz Tapia, Jaime Lagos, Loredana Arata, Alejandra Acevedo, Winston Andrade, Gabriel Leal, Carolina Tambley, Patricia Bustos, Rodrigo Fasce, Jorge Fernandez |
| EPI_ISL_445322 | UNIV.DE CHILE HOSP.CLINICO | Instituto de Salud Publica de Chile | Andrés E Castillo, Bárbara Parra,Paz Tapia, Jaime Lagos, Loredana Arata, Alejandra Acevedo, Winston Andrade, Gabriel Leal, Carolina Tambley, Patricia Bustos, Rodrigo Fasce, Jorge Fernandez |
| EPI_ISL_445323 | HOSP.ENFERMEDADES INFECCIOSAS | Instituto de Salud Publica de Chile | Andrés E Castillo, Bárbara Parra,Paz Tapia, Jaime Lagos, Loredana Arata, Alejandra Acevedo, Winston Andrade, Gabriel Leal, Carolina Tambley, Patricia |

| | | | |
|---|---|---|---|
| | | | Bustos, Rodrigo Fasce, Jorge Fernandez |
| EPI_ISL_445359 | HOSP. SANTIAGO ORIENTE DR. LUIS TISNE B. | Instituto de Salud Publica de Chile | Andrés E Castillo, Bárbara Parra,Paz Tapia, Jaime Lagos, Loredana Arata, Alejandra Acevedo, Winston Andrade, Gabriel Leal, Carolina Tambley, Patricia Bustos, Rodrigo Fasce, Jorge Fernandez |
| EPI_ISL_445360 | HOSPITAL DEL PROFESOR | Instituto de Salud Publica de Chile | Andrés E Castillo, Bárbara Parra,Paz Tapia, Jaime Lagos, Loredana Arata, Alejandra Acevedo, Winston Andrade, Gabriel Leal, Carolina Tambley, Patricia Bustos, Rodrigo Fasce, Jorge Fernandez |
| EPI_ISL_445361 | CLINICA UC SAN CARLOS DE APOQUINDO | Instituto de Salud Publica de Chile | Andrés E Castillo, Bárbara Parra,Paz Tapia, Jaime Lagos, Loredana Arata, Alejandra Acevedo, Winston Andrade, Gabriel Leal, Carolina Tambley, Patricia Bustos, Rodrigo Fasce, Jorge Fernandez |
| EPI_ISL_445362 | BUPA SERVICIOS CLINICOS S.A | Instituto de Salud Publica de Chile | Andrés E Castillo, Bárbara Parra,Paz Tapia, Jaime Lagos, Loredana Arata, Alejandra Acevedo, Winston Andrade, Gabriel Leal, Carolina Tambley, Patricia Bustos, Rodrigo Fasce, Jorge Fernandez |
| EPI_ISL_445364 | HOSPITAL EL CARMEN DR.LUIS VALENTIN F. | Instituto de Salud Publica de Chile | Andrés E Castillo, Bárbara Parra,Paz Tapia, Jaime Lagos, Loredana Arata, Alejandra Acevedo, Winston Andrade, Gabriel Leal, Carolina Tambley, Patricia Bustos, Rodrigo Fasce, Jorge Fernandez |
| EPI_ISL_447059, EPI_ISL_447060, EPI_ISL_447092, EPI_ISL_447101, EPI_ISL_447114 | Michigan Department of Health and Human Services, Bureau of Laboratories | Michigan Department of Health and Human Services, Bureau of Laboratories | Blankenship HM, Riner D, Soehnlen MK |
| EPI_ISL_447136 | Department of Clinical Microbiology | GIGA Medical Genomics | Keith Durkin, Maria Artesi, Sébastien Bontems, Raphaël Boreux, Cécile Meex, Pierrette Melin, Marie-Pierre Hayette, Vincent Bours. |
| EPI_ISL_447169, EPI_ISL_447175, EPI_ISL_447177, EPI_ISL_447180, EPI_ISL_447183, EPI_ISL_447186, EPI_ISL_447191, EPI_ISL_447192, EPI_ISL_447194, EPI_ISL_447196, EPI_ISL_447200, EPI_ISL_447206, EPI_ISL_447214, EPI_ISL_447218, EPI_ISL_447228 | | | |
| see above | Michigan Department of Health and Human Services, Bureau of Laboratories | Michigan Department of Health and Human Services, Bureau of Laboratories | Blankenship HM, Riner D, Soehnlen MK |
| EPI_ISL_447250 | Central Virology Laboratory | Central Virology Laboratory | Neta Zuckerman, Efrat Bucris, Oran Erster, Danit Sofer, Orna Mor, Ella Mendelson, Michal Mandelboim |
| EPI_ISL_447254, EPI_ISL_447255 | TSGH-CP molecular lab | TSGH-CP molecular lab | Cherng-Lih Perng, Ming-Jr JIAN, Chih-Kai Chang, Jung-Chung Lin, Kuo-Ming Yeh, Chien-Wen Chen, Sheng-Kang Chiu, Hsing-Yi Chung, Shih-Hung Tsai, Kuo-Sheng Hung, Tien-Yao Chang, Feng-Yee Chang, Hung-Sheng Shang |
| EPI_ISL_447324, EPI_ISL_447325, EPI_ISL_447326, EPI_ISL_447327, EPI_ISL_447328, EPI_ISL_447329 | Clinical Virology Laboratory, Soroka Medical Center and the Faculty of Health Sciences, Ben-Gurion University of the Negev | Stern Lab | Stern Lab |
| EPI_ISL_447407, EPI_ISL_447408, EPI_ISL_447409, EPI_ISL_447410, EPI_ISL_447411, EPI_ISL_447412, EPI_ISL_447413, EPI_ISL_447414, EPI_ISL_447415, EPI_ISL_447416 | Clinical Virology Unit, Hadassah Hebrew University Medical Center | Stern Lab | Stern Lab |
| EPI_ISL_447532 | Hospital Universitari Vall d'Hebron - Vall d'Hebron Institut de Recerca | Hospital Universitari Vall d'Hebron | Cristina Andrés, Maria Piñana, Damir Garcia-Cehic, Mercedes Guerrero-Murillo, Ariadna Rando, Juliana Esperalba, Maria Gema Codina, Tomàs Pumarola, Josep Quer, Andrés Antón |
| EPI_ISL_447614 | Department of Laboratory Medicine, National Taiwan University Hospital | Microbial Genomics Core Lab, National Taiwan University Centers of Genomic and Precision Medicine | Shiou-Hwei Yeh, You-Yu Lin, Ya-Yun Lai, Chiao-Ling Li, Shan-Chwen Chang, Pei-Jer Chen, Sui-Yuan Chang |
| EPI_ISL_447640, EPI_ISL_447641, EPI_ISL_447836 | unknown | Department of Medicine | Kassela,K., Dovrolis,N., Bampali,M., Gatzidou,E., Froukala,E., Stavropoulou,A., Veletza,S., Tsakris,A., Spanakis,N. and Karakasiliotis,I. |
| EPI_ISL_448222 | Pasteur Insitute Ho Chi Minh City | National Key Laboratory of Gene Technology, Institute of Biotechnology, Vietnam Academy of Science and Technology | Le Tung Lam, Nguyen Trung Hieu, Nguyen Hong Trang, Ho Thi Thuong, Nguyen Thi Ngoc Thao, Huynh Thi Kim Loan, Luu Thuy Tien, Tran Huyen Linh, Pham Duy Quang, Luong Chan Quang, Cao Minh Thang, Nguyen Vu Thuong, Hoang Ha, Chu Hoang Ha, Phan Trong Lan, Truong Nam Hai |
| EPI_ISL_448815, EPI_ISL_448816, EPI_ISL_448818, EPI_ISL_448824, EPI_ISL_448825 | Oxford Viromics, NDM, University of Oxford; Oxford University Hospitals; Basingstoke and North Hampshire Hospital | COVID-19 Genomics UK (COG-UK) Consortium | Tanya Golubchik, David Bonsall, George Macintyre, Amy Trebes, Mariateresa de Cesare, Catrin Moore, Alex Mobbs, Anita Justice, Robert Shaw, Monique Andersson, Emma Wise, Nathan Moore, Jessica Lynch, Nick Cortes, Stephen Kidd, David Buck, John Todd, Christophe Fraser |
| EPI_ISL_450011, EPI_ISL_450012, EPI_ISL_450013, EPI_ISL_450014, EPI_ISL_450015, EPI_ISL_450016, EPI_ISL_450017, EPI_ISL_450018, EPI_ISL_450019, EPI_ISL_450020, EPI_ISL_450021, EPI_ISL_450022, EPI_ISL_450023, EPI_ISL_450024, EPI_ISL_450025, EPI_ISL_450043, EPI_ISL_450044, EPI_ISL_450045, EPI_ISL_450046, EPI_ISL_450047, EPI_ISL_450048, EPI_ISL_450049, EPI_ISL_450050, EPI_ISL_450051, EPI_ISL_450052, EPI_ISL_450053, EPI_ISL_450054 | | | |
| see above | MSHS Clinical Microbiology Laboratories | MSHS Pathogen Surveillance Program | Ana S. Gonzalez-Reiche, Mitchell Sullivan, Ajay Obla, Gopi Patel, Emilia Sordillo, Melissa Gitman, Alberto Paniz-mondolfi, Matthew Hernandez, Shelcie Fabre, Jose Polanco, Zenab Khan, Bremy Alburquerque, Jayeeta Dutta, Juan Soto, Shwetha Sridhar Hara, Ying-Chih Wang, Melissa Smith, Robert Sebra, Lisa Miorin, Wen-chun Liu, Randy Albrecht, Judith Aberg, Florian Krammer, Adolfo Garcia-Sastre, Viviana Simon, Harm van Bakel |
| EPI_ISL_450193 | SA Pathology | SA Pathology | Lex Leong |
| EPI_ISL_450214 | unknown | Microbiological Diagnostic Unit Public Health Laboratory (MDU-PHL) and Victorian Infectious Disease Reference Laboratory (VIDRL) | Seemann,T., Lane,C.R., Sherry,N.L., Duchene,S., Goncalves da Silva,A., Caly,L., Sait,M., Ballard,S.A., Horan,K., Schultz,M.B., Hoang,T., Easton,M., Dougal,S., Stinear,T.P., Druce,J., Catton,M., Sutton,B., van Diemen,A., Alpren,C., Williamson,D.A., Howden,B.P. |
| EPI_ISL_450232, EPI_ISL_450238 | UCSF Clinical Microbiology Laboratory | Chiu Laboratory, University of California, San Francisco | Xianding Deng, Scot Federman, Wei Gu, and Charles Y. Chiu |
| EPI_ISL_450296 | National Institute for Communicable Diseases of the National Health Laboratory Service | National Institute for Communicable Diseases of the National Health Laboratory Service | Allam M, Ismail A, Khumalo Z, Kwenda S, van Heusden P, Mtshali P, Mnyameni F, Mohale T, Subramoney K, Bhiman JN |
| EPI_ISL_450325 | NIV Pune | CSIR-Centre for Cellular and Molecular Biology | Dr V A Potdar, Dr ML Choudhary,Dr Priya Abraham,V. Vipat, S. Jadhav, U. Saha, H. Kengle, A. Awhale, A. Jagtap, A. Gondhalikar, V Malik, N Srivastava, S. Digraskar, P. Malsane, S. Hundekar, K. Patel, Yogesh Balakartik, M. Kakade, S. Jadhav, R. Gunjikar, V. Awtade, S. Bhorekar, P Shinde, S. Salve, B. Minhas S. Bharadwaj, H Kaushal Y. Gurav, S. Tomar,Payel Mukherjee, Sofia Banu, Priya Singh, Dhiviya Vedagiri, Divya Gupta, Vishal Sah, Santosh Kumar Kuncha, Krishnan Harinivas Harshan, Archana Bharadwaj Siva, Karthik Bharadwaj Tallapaka, Shagufta Khan, Lamuk Zaveri, Namami Gaur, Sakshi Shambhavi, Tulasi Nagabandi, Purushotham Vodnala,G. Aditya Kumar, Koushick Sivakumar, Pooja Ramesh Gupta, Rajan Kumar Jha, Shraddha Vijay Lahoti, Deepak Kumar, Devi Prasad Vijayashankara, Disha Nanda, Divya Das, Jotin Gogoi, Manish |
| EPI_ISL_450403 | School of Public Health, The University of Hong Kong | School of Public Health, The University of Hong Kong | Sit,T.H.S., Brackman,C.J., Sims,L.D., Tsang,D.N.C., Chu,D.K.W., Perera,R.A.P.M., Poon,L.L.M. and Peiris,M. |
| EPI_ISL_450404 | unknown | School of Public Health, The University of Hong Kong | Sit,T.H.S., Brackman,C.J., Sims,L.D., Tsang,D.N.C., Chu,D.K.W., Perera,R.A.P.M., Poon,L.L.M. and Peiris,M. |
| EPI_ISL_450468, EPI_ISL_450469, EPI_ISL_450474, EPI_ISL_450477, EPI_ISL_450478, EPI_ISL_450479, EPI_ISL_450480, EPI_ISL_450481 | Stanford clinical virology lab | Chan-Zuckerberg Biohub | Benjamin Pinksy, Katharine Walter, Victoria N. Parikh, John Gorzynski, Hannah N. DeJong, Matthew T. Wheeler, Jason Andrews, Manuel Rivas, Carlos Bustamante, Euan Ashley, with CZB Cliahub Consortium |
| EPI_ISL_450506 | Clinical Laboratory, Hospital Israelita Albert Einstein | Clinical Laboratory, Hospital Israelita Albert Einstein | Malta,F., Amgarten,D., de Oliveira,D.B.L., Araujo,D.B., Machado,R.R.G., Santana,R.A.F., Mangueira,C.L.P., Durigon,E.L. and Pinho,J.R.R. |
| EPI_ISL_450509, EPI_ISL_450516 | Rafik Hariri University Hospital | Rafik Hariri University Hospital | Rita Feghali |
| EPI_ISL_450539 | Utah Public Health Laboratory | Utah Public Health Laboratory | Erin Young, Kelly Oakeson |
| EPI_ISL_450578, EPI_ISL_450580, EPI_ISL_450581, EPI_ISL_450582, EPI_ISL_450583, EPI_ISL_450587 | Michigan Department of Health and Human Services, Bureau of Laboratories | Michigan Department of Health and Human Services, Bureau of Laboratories | Blankenship HM; Riner D; Soehnlen MK |
| EPI_ISL_450602, EPI_ISL_450604, | Michigan Department of Health and Human Services, Bureau | Michigan Department of Health and Human Services, Bureau | Blankenship HM, Riner D, Soehnlen MK |

| | | | |
|---|---|---|---|
| EPI_ISL_450606, EPI_ISL_450607, EPI_ISL_450608, EPI_ISL_450610 | of Laboratories | of Laboratories | |
| EPI_ISL_450700 | University of Wisconsin-Madison AIDS Vaccine Research Laboratories | University of Wisconsin-Madison AIDS Vaccine Research Laboratories | Gage Moreno, Katarina Braun, et al. AIDS Vaccine Research Laboratories |
| EPI_ISL_450739 | OUCRU/HTD | OUCRU/HTD | Nguyen Van Vinh Chau, Nguyen Thi Thu Hong, Nguyen Thi Han Ny, Le Nguyen Truc Nhu, Nghiem My Ngoc, Vo Thanh Lam, Nguyen Thanh Dung, Lam Minh Yen, Ngo Ngoc Quang Minh, Le Manh Hung, Nguyen Tri Dung, Dinh Nguyen Huy Man, Lam Anh Nguyet, Tran Chanh Xuan, Tran Tinh Hien, Nguyen Thanh Phong, Tran Nguyen Hoang Tu, Tran Tan Thanh, Nguyen Thanh Truong, Nguyen Tan Binh, Tang Chi Thuong, Guy Thwaites, and Le Van Tan, for OUCRU COVID-19 research group* |
| EPI_ISL_450796, EPI_ISL_450797 | Jamaica Ministry of Health and Wellness | Pathogen Discovery, Respiratory Viruses Branch, Division of Viral Diseases, Centers for Disease Control and Prevention | Yan Li, Anna Montmayeur, Ying Tao, Krista Queen, Jing Zhang, Anna Uehara, Clinton R. Paden, Rachel Marine, Haibin Wang, Zachary Weiner, Bettina Bankamp, Suxiang Tong |
| EPI_ISL_450798 | Jamaica Ministry of Health and Wellness | Pathogen Discovery, Respiratory Viruses Branch, Division of Viral Diseases, Centers for Disease Control and Prevention | Krista Queen, Yan Li, Anna Montmayeur, Ying Tao, Jing Zhang, Anna Uehara, Clinton R. Paden, Rachel Marine, Haibin Wang, Jasmine Padilla, Justin Lee, Zachary Weiner, Bettina Bankamp, Suxiang Tong |
| EPI_ISL_450799 | Jamaica Ministry of Health and Wellness | Pathogen Discovery, Respiratory Viruses Branch, Division of Viral Diseases, Centers for Disease Control and Prevention | Yan Li, Anna Montmayeur, Ying Tao, Krista Queen, Jing Zhang, Anna Uehara, Clinton R. Paden, Rachel Marine, Haibin Wang, Zachary Weiner, Bettina Bankamp, Suxiang Tong |
| EPI_ISL_450803 | PA Department of Health, Bureau of Laboratories | Pathogen Discovery, Respiratory Viruses Branch, Division of Viral Diseases, Centers for Disease Control and Prevention | Yan Li, Anna Montmayeur, Ying Tao, Krista Queen, Jing Zhang, Anna Uehara, Clinton R. Paden, Rachel Marine, Haibin Wang, Zachary Weiner, Bettina Bankamp, Suxiang Tong |
| EPI_ISL_450873 | Evandro Chagas Institute | Evandro Chagas Institute | Santos, M.C.; Silva, A.M.; Junior, W.D.C.; Barbagelata, L.S.; Ferreira, J.A.; Sousa, E.M.A.; da Silva, P.S.; Martins, L.C.; Sousa Junior, E.C.; Viana, G.M.R |
| EPI_ISL_450878, EPI_ISL_450879, EPI_ISL_450880, EPI_ISL_450881, EPI_ISL_450886, EPI_ISL_450887, EPI_ISL_450888, EPI_ISL_450889, EPI_ISL_450894, EPI_ISL_450895, EPI_ISL_450907, EPI_ISL_450918, EPI_ISL_450919, EPI_ISL_450920, EPI_ISL_450921, EPI_ISL_450926, EPI_ISL_450927, EPI_ISL_450928, EPI_ISL_450929, EPI_ISL_450934, EPI_ISL_450935, EPI_ISL_450947, EPI_ISL_450958, EPI_ISL_450959, EPI_ISL_450960, EPI_ISL_450961, EPI_ISL_450966, EPI_ISL_450967, EPI_ISL_450968, EPI_ISL_450969, EPI_ISL_450974, EPI_ISL_450975, EPI_ISL_450987, EPI_ISL_450998, EPI_ISL_450999, EPI_ISL_451000, EPI_ISL_451001, EPI_ISL_451006, EPI_ISL_451007, EPI_ISL_451008, EPI_ISL_451009, EPI_ISL_451014, EPI_ISL_451015, EPI_ISL_451027, EPI_ISL_451038, EPI_ISL_451039, EPI_ISL_451040, EPI_ISL_451041, EPI_ISL_451046, EPI_ISL_451047, EPI_ISL_451048, EPI_ISL_451049, EPI_ISL_451054, EPI_ISL_451055, EPI_ISL_451067 | | | |
| see above | Center of Excellence in Clinical Virology | Center of Excellence in Clinical Virology | Puenpa,J., Chansaenroj,J., Nilyanimit,P., Auphimai,C., Yorsaeng,R., Suwannakarn,K., Poovorawan,Y. |
| EPI_ISL_451077, EPI_ISL_451130, EPI_ISL_451131, EPI_ISL_451132, EPI_ISL_451147 | SA Pathology | SA Pathology | Lex Leong, Chuan Kok Lim, Mark Turra, Ivan Bastian, Geoff Higgins |
| EPI_ISL_451694, EPI_ISL_451695, EPI_ISL_451696, EPI_ISL_451697, EPI_ISL_451698, EPI_ISL_451699, EPI_ISL_451700, EPI_ISL_451701, EPI_ISL_451702, EPI_ISL_451703, EPI_ISL_451704, EPI_ISL_451705, EPI_ISL_451706, EPI_ISL_451707, EPI_ISL_451708, EPI_ISL_451709, EPI_ISL_451710, EPI_ISL_451711, EPI_ISL_451712, EPI_ISL_451713, EPI_ISL_451714, EPI_ISL_451715, EPI_ISL_451716, EPI_ISL_451717, EPI_ISL_451718, EPI_ISL_451719, EPI_ISL_451720, EPI_ISL_451721, EPI_ISL_451722, EPI_ISL_451723, EPI_ISL_451724, EPI_ISL_451725, EPI_ISL_451726, EPI_ISL_451727, EPI_ISL_451728, EPI_ISL_451729, EPI_ISL_451730, EPI_ISL_451731, EPI_ISL_451732, EPI_ISL_451733, EPI_ISL_451734, EPI_ISL_451735, EPI_ISL_451736, EPI_ISL_451737, EPI_ISL_451738, EPI_ISL_451739 | | | |
| see above | Viollier AG | Department of Biosystems Science and Engineering, ETH Zürich | Christian Beisel, Sarah Nadeau, Ivan Topolsky, Pedro Ferreira, Philipp Jablonski, Susana Posada-Céspedes, Tobias Schär, Ina Nissen, Natascha Santacroce, Elodie Burcklen, Christiane Beckmann, Maurice Redondo, Olivier Kobel, Christoph Noppen, Sophie Seidel, Noemie Santamaria de Souza, Niko Beerenwinkel, Tanja Stadler |
| EPI_ISL_451935 | CUB Hopital Erasme Laboratoire d'Anatomie Pathologique | CUB Hopital Erasme Laboratoire d'Anatomie Pathologique | Prof. Isabelle Salmon, Dr. Nicky D'Haene |
| EPI_ISL_451937 | Max von Pettenkofer Institute, Virology, National Reference Center for Retroviruses, LMU München | Laboratory for Functional Genome Analysis, Dept. Genomics, Gene Center of the LMU Munich | Max Muenchhoff, Stefan Krebs, Alexander Graf, Oliver Keppler, Helmut Blum |
| EPI_ISL_451958 | Jamil-ur-Rahman Center for Genome Research, Dr. Panjwani Center for Molecular Medicine and Drug Research, International Center for Chemical and Biological Sciences, University of Karachi | Jamil-ur-Rahman Center for Genome Research, Dr. Panjwani Center for Molecular Medicine and Drug Research, International Center for Chemical and Biological Sciences, University of Karachi | Shakeel,M., Raza,S.A., Khan,S., Khan,B.A., Zahid,M., Qureshi,M.A.and Khan,I.A |
| EPI_ISL_451968 | Federal Budget Institution of Science, State Research Center for Applied Microbiology & Biotechnology | Federal Budget Institution of Science, State Research Center for Applied Microbiology & Biotechnology | Dyatlov I, Shemyakin I, Khramov M, Bogun A, Kislichkina A, Frolov V, Shishkina L, Sizova A, Chekan L, Blagodatskikh S, Podkopaev Y, Kosilova I, Koroleva-Ushakova A, Tyurin E, Galkina E, Slukina N, Shaikhutdinova R, Kalmantayev T, Kalmantayeva O, Fursova N, Silkina M, Gorbatov A, Titareva G, Firstova V, Makarova M, Gapelchenkova T, Solovieva A, Slukin P, Dentovskaya S, Detushev K, Vagayskaya A, Kartsev N, Detusheva E, Zeninskaya N, Ivanov S, Kartseva A, Platonov M, Hlyntseva A, Khomyakov A, Chernysh S, Krasilnikova E, Ryabko A, Solomentsev V, Teymurazov M, Bakhteeva I, Borzilov A, Skryabin Y, Kanashenko M, Abaimova A, Kolchanova A, Novikova T, Goncharova J, Timofeev V, Kuzina E, Fursov M, Zhumakaev R, Marin M, Denisenko E, Trunyakova A, Kuzin V |
| EPI_ISL_452034 | Department of Clinical Microbiology, Copenhagen University Hospital, Hvidovre, Kettegaard Alle 30, 2650 Hvidovre. | Albertsen lab, Department of Chemistry and Bioscience, Aalborg University, Denmark | Rasmus Kirkegaard |
| EPI_ISL_452119 | VI-US Virgin Islands Department of Health | Pathogen Discovery, Respiratory Viruses Branch, Division of Viral Diseases, Centers for Disease Control and Prevention | Yan Li, Anna Montmayeur, Ying Tao, Krista Queen, Jing Zhang, Anna Uehara, Clinton R. Paden, Rachel Marine, Mary S. Keckler, Alison S. Laufer Halpin, Haibin Wang, Christopher A. Elkins, Zachary Weiner, Suxiang Tong |
| EPI_ISL_452202 | NIV Influenza | NIV Influenza | Potdar V |
| EPI_ISL_452229 | Narhalsan Backa vardcentral | The Public Health Agency of Sweden | Mats Olsson, Anna-Malin Linde, Maria Lind Karlberg, Oskar Karlsson Lindsjo, Olov Svartstrom, Anna Risberg, Theresa Enkirch, Mia Brytting, Karin Tegmark-Wisell |
| EPI_ISL_452285, EPI_ISL_452286, EPI_ISL_452288, EPI_ISL_452289, EPI_ISL_452290, EPI_ISL_452291, EPI_ISL_452292, EPI_ISL_452294, EPI_ISL_452295, EPI_ISL_452296, EPI_ISL_452297, EPI_ISL_452298, EPI_ISL_452299, EPI_ISL_452300, EPI_ISL_452301, EPI_ISL_452303, EPI_ISL_452315, EPI_ISL_452317, EPI_ISL_452318, EPI_ISL_452319, EPI_ISL_452321, EPI_ISL_452322, EPI_ISL_452323, EPI_ISL_452324 | | | |
| see above | Michigan Department of Health and Human Services, Bureau of Laboratories | Michigan Department of Health and Human Services, Bureau of Laboratories | Blankenship HM, Riner D, Soehnlen MK |
| EPI_ISL_452328, EPI_ISL_452335, EPI_ISL_452343, EPI_ISL_452347, EPI_ISL_452348 | Laboratory of Infectious Diseases Center of Beijing Ditan Hospital | Laboratory of Infectious Diseases Center of Beijing Ditan Hospital | Siyuan Yang, Chengjie Jie, Fengting Yu, Yunxia Tang, Liting Yan, Linghang Wang |
| EPI_ISL_452374, EPI_ISL_452375, EPI_ISL_452376, EPI_ISL_452377, EPI_ISL_452379, EPI_ISL_452380, EPI_ISL_452381, EPI_ISL_452382, EPI_ISL_452383, EPI_ISL_452384, EPI_ISL_452385, EPI_ISL_452386, EPI_ISL_452387, EPI_ISL_452388, EPI_ISL_452389, EPI_ISL_452390, EPI_ISL_452391, EPI_ISL_452392, EPI_ISL_452393, EPI_ISL_452394, EPI_ISL_452397, EPI_ISL_452398, EPI_ISL_452399, EPI_ISL_452400, EPI_ISL_452401, EPI_ISL_452402, EPI_ISL_452403, EPI_ISL_452404, EPI_ISL_452405, EPI_ISL_452406, EPI_ISL_452407, EPI_ISL_452408 | | | |
| see above | Servicio de Microbiología. HRU de Málaga. Servicio Andaluz de Salud | SeqCOVID-SPAIN consortium/IBV(CSIC) | Inmaculada de Toro Peinado, Maria Concepción Mediavilla Gradolph, Begoña Palop Borrás and SeqCOVID-SPAIN consortium |
| EPI_ISL_452453 | Hospital Universitario Puerta del Mar de Cádiz - INIBICA | SeqCOVID-SPAIN consortium/IBV(CSIC) | Salud Rodríguez-Pallares, Fátima-Galán-Sánchez, Manuel Rodrí-guez-Iglesias and SeqCOVID-SPAIN consortium |
| EPI_ISL_452500, EPI_ISL_452501, EPI_ISL_452502, EPI_ISL_452503, EPI_ISL_452504, EPI_ISL_452505, EPI_ISL_452506, EPI_ISL_452507, EPI_ISL_452508, EPI_ISL_452509, EPI_ISL_452510, EPI_ISL_452511, EPI_ISL_452512, EPI_ISL_452513, EPI_ISL_452514, EPI_ISL_452515, EPI_ISL_452516, EPI_ISL_452517, EPI_ISL_452518, EPI_ISL_452519 | | | |
| see above | Clínica Universidad de Navarra. Servicio de Enfermedades Infecciosas y Microbiología clínica | SeqCOVID-SPAIN consortium/IBV(CSIC) | Mirian Fernández-Alonso, Jose Luis del Pozo and SeqCOVID-SPAIN consortium |
| EPI_ISL_453825, EPI_ISL_454002, EPI_ISL_454018, EPI_ISL_454024, EPI_ISL_454029, EPI_ISL_454030, EPI_ISL_454033, EPI_ISL_454034, EPI_ISL_454035, EPI_ISL_454036, EPI_ISL_454037, EPI_ISL_454038, EPI_ISL_454039, EPI_ISL_454040, EPI_ISL_454041, EPI_ISL_454042, EPI_ISL_454043, EPI_ISL_454044, EPI_ISL_454045, EPI_ISL_454046, EPI_ISL_454047, EPI_ISL_454048, EPI_ISL_454049, EPI_ISL_454050, EPI_ISL_454051, EPI_ISL_454052, EPI_ISL_454053, EPI_ISL_454054, EPI_ISL_454055, EPI_ISL_454056, EPI_ISL_454057, EPI_ISL_454059, EPI_ISL_454060, EPI_ISL_454061, EPI_ISL_454062, EPI_ISL_454063, EPI_ISL_454064, EPI_ISL_454065, EPI_ISL_454067, EPI_ISL_454068, EPI_ISL_454069, EPI_ISL_454070, EPI_ISL_454071, EPI_ISL_454127, EPI_ISL_454128, EPI_ISL_454225, EPI_ISL_454226, EPI_ISL_454227, EPI_ISL_454228, EPI_ISL_454229, EPI_ISL_454334, EPI_ISL_454335, EPI_ISL_454336, EPI_ISL_454337, EPI_ISL_454338, EPI_ISL_454339, EPI_ISL_454340, EPI_ISL_454341, EPI_ISL_454342 | | | |
| see above | unknown | Instituto Nacional de Saude (INSA) | Borges et al |

| | | | |
|---|---|---|---|
| EPI_ISL_454434, EPI_ISL_454435 | Maryland Department of Health Laboratories Administration | Maryland Department of Health Laboratories Administration | MDH Laboratories Administration |
| EPI_ISL_454528 | NIV Influenza | NIV Influenza | Potdar V |
| EPI_ISL_454581, EPI_ISL_454583 | University Hospital for Infectious Diseases "Dr. Fran Mihaljevi", Research Unit | University of Zagreb, Centre for research and knowledge transfer in biotechnology | Ivan-Christian Kurolt, Jelena Ivancic Jelecki, Anamarija Slovic |
| EPI_ISL_454635 | County Of San Luis Obispo Public Health Laboratory | Chan-Zuckerberg Biohub | CZB Cliahub Consortium |
| EPI_ISL_454762, EPI_ISL_454763, EPI_ISL_454764, EPI_ISL_454765, EPI_ISL_454788 | Dutch COVID-19 response team | National Institute for Public Health and the Environment (RIVM) | Adam Meijer, Harry Vennema, Jeroen Cremer, Sharon van den Brink, Pieter Overduin, Florian Zwagemaker, Dennis Schmitz, Chantal Reusken, on behalf of the national COVID-19 response team |
| EPI_ISL_454945, EPI_ISL_454946, EPI_ISL_454953, EPI_ISL_454954, EPI_ISL_454960, EPI_ISL_454969, EPI_ISL_454970, EPI_ISL_454971, EPI_ISL_454972, EPI_ISL_454977, EPI_ISL_454979, EPI_ISL_454984 | | | |
| see above | Wuhan Chain Medical Labs (CMLabs) | State Key Laboratory of Biotherapy of Sichuan University | Baowen Du, Minjin Wang, Chao Tang, Chuan Chen, Yongzhao Zhou, Mingxia Yu, Hancheng Wei, Weimin Li, Jing-wen Lin, Jia Geng, Binwu Ying, Lu Chen |
| EPI_ISL_455207, EPI_ISL_455208, EPI_ISL_455209, EPI_ISL_455283 | Dutch COVID-19 response team | Erasmus Medical Center | Bas Oude Munnink, David Nieuwenhuijse, Reina Sikkema, Claudia Schapendonk, Irina Chestakova, Anne van der Linden, Theo Bestebroer, Stefan van Nieuwkoop, Mark Pronk, Pascal Lexmond, Corien Swaan, Manon Haverkate, Madelief Mollers, Mart Stein, Sandra Kengne Kamga Mobou, Jeroen van Kampen, Jolanda Voermans, Aura Timen, Corine GeurtsvanKessel, Annemiek van der Eijk, Richard Molenkamp, Marion Koopmans, on behalf of the Dutch national COVID-19 response team. |
| EPI_ISL_455381, EPI_ISL_455384, EPI_ISL_455387, EPI_ISL_455388, EPI_ISL_455392, EPI_ISL_455393, EPI_ISL_455394, EPI_ISL_455395, EPI_ISL_455397, EPI_ISL_455398 | Wuhan Chain Medical Labs (CMLabs) | State Key Laboratory of Biotherapy of Sichuan University | Baowen Du, Minjin Wang, Chao Tang, Chuan Chen, Yongzhao Zhou, Mingxia Yu, Hancheng Wei, Weimin Li, Jing-wen Lin, Jia Geng, Binwu Ying, Lu Chen |
| EPI_ISL_455442, EPI_ISL_455449, EPI_ISL_455451 | 1. ViroGenetics - BSL3 Laboratory of Virology, Maopolska Centre of Biotechnology, Jagiellonian University; 2. II Department of Internal Medicine, Faculty of Medicine, Jagiellonian University Medical College; 3. Narodowy Instytut Zdrowia Publicznego - Pastwowy Zakad Higieny (NIZP-PZH) | 1. ViroGenetics - BSL3 Laboratory of Virology, Maopolska Centre of Biotechnology, Jagiellonian University; 2. II Department of Internal Medicine, Faculty of Medicine, Jagiellonian University Medical College; 3. Narodowy Instytut Zdrowia Publicznego - Pastwowy Zakad Higieny (NIZP-PZH). | Katarzyna Pancer, Marek Sanak, Aleksandra A. Zasada, Magdalena Rzeczkowska, Tomasz Wokowicz, Katarzyna Zacharczuk, Agnieszka Koakowska-Kulesza, Katarzyna Owczarek, Aleksandra Milewska, Natalia Wolaniuk, Ewelina Hallman-Szeliska, Pawe P abaj, Wojciech Branicki, Krzysztof Pyr |
| EPI_ISL_455454 | Instituto de Diagnostico y Referencia Epidemiologicos (INDRE) | Instituto de Diagnostico y Referencia Epidemiologicos (INDRE) | Mendieta-Condado Edgar, Araiza-Rodriguez Adnan, Garces-Ayala Fabiola, , Rodriguez-Maldonado Abril, Wong-Arambula Claudia, Barrera-Badillo Gisela, Taboada Ramírez Blanca, Ramirez-Gonzalez Ernesto, Hernandez-Rivas Lucia, Lopez-Martinez Irma. |
| EPI_ISL_455583 | Central Chest Institute of Thailand | National Institute of Health. Department of medical Sciences, Ministry of Public Health, Thailand | Pilailuk,Okada; Siripaporn,Phuygun; Thanutsapa,Thanadachakul; Sittiporn,Parnmen;Warawan,Wongboot; Sunthareeya,Waicharoen; Malinee,Chittaganpitch |
| EPI_ISL_455584 | National Institute of Health. Department of medical Sciences, Ministry of Public Health, Thailand | National Institute of Health. Department of medical Sciences, Ministry of Public Health, Thailand | Pilailuk,Okada; Siripaporn,Phuygun; Thanutsapa,Thanadachakul; Sittiporn,Parnmen;Warawan,Wongboot; Sunthareeya,Waicharoen; Malinee,Chittaganpitch |
| EPI_ISL_455586 | Siriraj hospital | National Institute of Health. Department of medical Sciences, Ministry of Public Health, Thailand | Pilailuk,Okada; Siripaporn,Phuygun; Thanutsapa,Thanadachakul; Sittiporn,Parnmen;Warawan,Wongboot; Sunthareeya,Waicharoen; Malinee,Chittaganpitch |
| EPI_ISL_455594 | Central Chest Institute of Thailand | National Institute of Health. Department of medical Sciences, Ministry of Public Health, Thailand | Pilailuk,Okada; Siripaporn,Phuygun; Thanutsapa,Thanadachakul; Sittiporn,Parnmen;Warawan,Wongboot; Sunthareeya,Waicharoen; Malinee,Chittaganpitch |
| EPI_ISL_455694, EPI_ISL_455695, EPI_ISL_455696 | National Hospital of Tropical Diseases | Oxford University Clinical Research Unit, Hanoi, Vietnam | Nguyen Thi Tam, Van Dinh Trang, Nguyen Thu Trang, Nguyen Thi Ngoc Diep, Le Nguyen Minh Hoa, Pham Ngoc Thach, H. Rogier van Doorn, on behalf of the OUCRU COVID-19 research group |
| EPI_ISL_455716, EPI_ISL_455717 | National Hospital of Tropical Diseases | Oxford University Clinical Research Unit, Hanoi, Vietnam | Nguyen Thi Tam, Van Dinh Trang, Nguyen Thi Hong Thuong, Vu Thi Ngoc Bich, Nguyen Thu Trang, Nguyen Thi Ngoc Diep, Le Nguyen Minh Hoa, Pham Ngoc Thach, H. Rogier van Doorn, on behalf of the OUCRU COVID-19 research group |
| EPI_ISL_455937, EPI_ISL_455938, EPI_ISL_455939, EPI_ISL_455940, EPI_ISL_455941, EPI_ISL_455942, EPI_ISL_455943, EPI_ISL_455944, EPI_ISL_455945, EPI_ISL_455946, EPI_ISL_455947, EPI_ISL_455948 | | | |
| see above | Ramathibodi Hospital | COVID-19 Network Investigations (CONI) Alliance | Elizabeth Batty, Wasun Chantratita, Thanat Chookajorn, Stefan Fernandez, Angkana Huang, Anthony R. Jones, Khajohn Joonsalak, Chonticha Klungtong, Theerarat Kochakarn, Namfon Kotanan, Krittikorn Kumpornsin, Wudtichai Manasatienkij, Bhakbhoom Panthan, Ekawat Pasomsub, Kingkan Rakmanee, Insee Sensorn, Janjira Thaipadungpanit, Arporn Wangwiwatsin,Treewat Watthanachockchai |
| EPI_ISL_456118, EPI_ISL_456119, EPI_ISL_456122, EPI_ISL_456124, EPI_ISL_456126, EPI_ISL_456127, EPI_ISL_456128, EPI_ISL_456132, EPI_ISL_456138, EPI_ISL_456139, EPI_ISL_456141, EPI_ISL_456142, EPI_ISL_456148, EPI_ISL_456149, EPI_ISL_456150, EPI_ISL_456155 | | | |
| see above | Instituto Nacional de Salud - Unidad de Secuenciación y Análisis Genómico | Instituto Nacional de Salud, Universidad Cooperativa de Colombia, Instituto Alexander von Humboldt, Imperial College-London, London School of Hygiene & Tropical Medicine | Katherine Laiton-Donato, Diego A. Álvarez-Díaz, Carlos Franco-Muñoz, Jose A. Usme-Ciro, Gloria Puerto, Nicolas D. Franco-Sierra, Mailyn A.Gonzalez, Zulma M. Cucunubá, Christian Julian VillabonaArenas, Liz Villabona-Arenas, Sussy Echeverria, Astrid C. Flórez, Sergio Gomez-Rangel, Luz Dary Rodriguez, Juliana Barbosa, Erika Ospitia, Diana Marcela Walteros-Acero, Martha Lucia Ospina Martinez, Marcela Mercado-Reyes. |
| EPI_ISL_456159 | Southern Community Labs Dunedin | Institute of Environmental Science and Research (ESR) | Matt Storey, Xiaoyun Ren, Anja Werno, Antje van der Linden, Arlo Upton, Chris Mansell, David Hammer, Dragana Drinkovic, Erasmus Smit, Gary McAuliffe, Hana Sofia Andersson, James Ussher, Jill Sherwood, Josh Freeman, Julia Howard, Juliet Elvy, Mary DeAlmeida, Matt Blakiston, Matthew Rogers, Max Bloomfield, Michael Addidle, Michelle Balm, Sally Roberts, Sarah Jefferies, Sharmini Muttaiyah, Susan Morpeth, Susan Taylor, Timothy Blackmore, Vani Sathyendran, Veronica Playle, Virginia Hope, Erasmus Smit, Lauren Jelly, Joep de Ligt |
| EPI_ISL_456160 | Waikato Hospital | Institute of Environmental Science and Research (ESR) | Matt Storey, Xiaoyun Ren, Anja Werno, Antje van der Linden, Arlo Upton, Chris Mansell, David Hammer, Dragana Drinkovic, Erasmus Smit, Gary McAuliffe, Hana Sofia Andersson, James Ussher, Jill Sherwood, Josh Freeman, Julia Howard, Juliet Elvy, Mary DeAlmeida, Matt Blakiston, Matthew Rogers, Max Bloomfield, Michael Addidle, Michelle Balm, Sally Roberts, Sarah Jefferies, Sharmini Muttaiyah, Susan Morpeth, Susan Taylor, Timothy Blackmore, Vani Sathyendran, Veronica Playle, Virginia Hope, Erasmus Smit, Lauren Jelly, Joep de Ligt |
| EPI_ISL_456161 | Wellington SCL | Institute of Environmental Science and Research (ESR) | Matt Storey, Xiaoyun Ren, Anja Werno, Antje van der Linden, Arlo Upton, Chris Mansell, David Hammer, Dragana Drinkovic, Erasmus Smit, Gary McAuliffe, Hana Sofia Andersson, James Ussher, Jill Sherwood, Josh Freeman, Julia Howard, Juliet Elvy, Mary DeAlmeida, Matt Blakiston, Matthew Rogers, Max Bloomfield, Michael Addidle, Michelle Balm, Sally Roberts, Sarah Jefferies, Sharmini Muttaiyah, Susan Morpeth, Susan Taylor, Timothy Blackmore, Vani Sathyendran, Veronica Playle, Virginia Hope, Erasmus Smit, Lauren Jelly, Joep de Ligt |
| EPI_ISL_456162 | Southern Community Labs Dunedin | Institute of Environmental Science and Research (ESR) | Matt Storey, Xiaoyun Ren, Anja Werno, Antje van der Linden, Arlo Upton, Chris Mansell, David Hammer, Dragana Drinkovic, Erasmus Smit, Gary McAuliffe, Hana Sofia Andersson, James Ussher, Jill Sherwood, Josh Freeman, Julia Howard, Juliet Elvy, Mary DeAlmeida, Matt Blakiston, Matthew Rogers, Max Bloomfield, Michael Addidle, Michelle Balm, Sally Roberts, Sarah Jefferies, Sharmini Muttaiyah, Susan Morpeth, Susan Taylor, Timothy Blackmore, Vani Sathyendran, Veronica Playle, Virginia Hope, Erasmus Smit, Lauren Jelly, Joep de Ligt |
| EPI_ISL_456164, EPI_ISL_456165, EPI_ISL_456166 | Wellington SCL | Institute of Environmental Science and Research (ESR) | Matt Storey, Xiaoyun Ren, Anja Werno, Antje van der Linden, Arlo Upton, Chris Mansell, David Hammer, Dragana Drinkovic, Erasmus Smit, Gary McAuliffe, Hana Sofia Andersson, James Ussher, Jill Sherwood, Josh Freeman, Julia Howard, Juliet Elvy, Mary DeAlmeida, Matt Blakiston, Matthew Rogers, Max Bloomfield, Michael Addidle, Michelle Balm, Sally Roberts, Sarah Jefferies, Sharmini Muttaiyah, Susan Morpeth, Susan Taylor, Timothy Blackmore, Vani Sathyendran, Veronica Playle, Virginia Hope, Erasmus Smit, Lauren Jelly, Joep de Ligt |
| EPI_ISL_456167 | Southern Community Labs Dunedin | Institute of Environmental Science and Research (ESR) | Matt Storey, Xiaoyun Ren, Anja Werno, Antje van der Linden, Arlo Upton, Chris Mansell, David Hammer, Dragana Drinkovic, Erasmus Smit, Gary McAuliffe, Hana Sofia Andersson, James Ussher, Jill Sherwood, Josh Freeman, Julia Howard, Juliet Elvy, Mary DeAlmeida, Matt Blakiston, Matthew Rogers, Max Bloomfield, Michael Addidle, Michelle Balm, Sally Roberts, Sarah Jefferies, Sharmini Muttaiyah, Susan Morpeth, Susan Taylor, Timothy Blackmore, Vani Sathyendran, Veronica Playle, Virginia Hope, Erasmus Smit, Lauren Jelly, Joep de Ligt |
| EPI_ISL_456168 | Waikato Hospital | Institute of Environmental Science and Research (ESR) | Matt Storey, Xiaoyun Ren, Anja Werno, Antje van der Linden, Arlo Upton, Chris Mansell, David Hammer, Dragana Drinkovic, Erasmus Smit, Gary McAuliffe, Hana Sofia Andersson, James Ussher, Jill Sherwood, Josh Freeman, Julia Howard, Juliet Elvy, Mary DeAlmeida, Matt Blakiston, Matthew Rogers, Max Bloomfield, Michael Addidle, Michelle Balm, Sally Roberts, Sarah Jefferies, Sharmini Muttaiyah, Susan Morpeth, Susan Taylor, Timothy |

| | | | |
|---|---|---|---|
| | | | Blackmore, Vani Sathyendran, Veronica Playle, Virginia Hope, Erasmus Smit, Lauren Jelly, Joep de Ligt |
| EPI_ISL_456172, EPI_ISL_456173, EPI_ISL_456174, EPI_ISL_456175, EPI_ISL_456176, EPI_ISL_456177, EPI_ISL_456178, EPI_ISL_456179, EPI_ISL_456180, EPI_ISL_456209 | LabPLUS | Institute of Environmental Science and Research (ESR) | Matt Storey, Xiaoyun Ren, Anja Werno, Antje van der Linden, Arlo Upton, Chris Mansell, David Hammer, Dragana Drinkovic, Erasmus Smit, Gary McAuliffe, Hana Sofia Andersson, James Ussher, Jill Sherwood, Josh Freeman, Julia Howard, Juliet Elvy, Mary DeAlmeida, Matt Blakiston, Matthew Rogers, Max Bloomfield, Michael Addidle, Michelle Balm, Sally Roberts, Sarah Jefferies, Sharmini Muttaiyah, Susan Morpeth, Susan Taylor, Timothy Blackmore, Vani Sathyendran, Veronica Playle, Virginia Hope, Erasmus Smit, Lauren Jelly, Joep de Ligt |
| EPI_ISL_457124, EPI_ISL_457140 | University of Exeter | COVID-19 Genomics UK (COG-UK) Consortium | Ben Temperton,Aaron Jeffries,Michelle Michelsen,Joanna Warwick-Dugdale,Audrey Farbos,Robyn Manley,Stephen Michell,Jane Masoli |
| EPI_ISL_457724 | Department of Infectious Diseases, Istituto Superiore di Sanità, Roma , Italy | Army Medical and Veterinary Research Center | Paola Stefanelli, Alessandra Lo Presti, Stefano Fiore, Antonella Marchi, Eleonora Benedetti, Concetta Fabiani Silvia Fillo, Giovanni Faggioni, Riccardo De Sanctis, Antonella Fortunato, Anna Anselmo, Francesco Giordani, Vanessa Vera Fain, Nino D'Amore, Florigio Lista |
| EPI_ISL_457752, EPI_ISL_457753, EPI_ISL_457754, EPI_ISL_457755, EPI_ISL_457763, EPI_ISL_457770, EPI_ISL_457774, EPI_ISL_457785, EPI_ISL_457786, EPI_ISL_457789, EPI_ISL_457791, EPI_ISL_457792, EPI_ISL_457803, EPI_ISL_457806, EPI_ISL_457807, EPI_ISL_457812, EPI_ISL_457814, EPI_ISL_457819, EPI_ISL_457820 | | | |
| see above | Johns Hopkins Hospital Department of Pathology | Johns Hopkins Hospital Department of Pathology | Peter M. Thielen, Thomas Mehoke, Shirlee Wohl, Srividya Ramakrishnan, Melanie Kirsche, Amanda Ernlund, Craig Howser, Kristina Zudock, Oluwaseun Falade-Nwulia, Norah Sadowski, Paul Morris, Mark Hopkins, Yunfan Fan, Nidia Trovao, Victoria Gniazdowski, Michael C. Schatz, Stuart C. Ray, Winston Timp, Heba H. Mostafa |
| EPI_ISL_457845 | KEMRI-CGMR-C | KEMRI-Wellcome Trust Research Programme/KEMRI-CGMR-C Kilifi | Githinji G. et al 2020 |
| EPI_ISL_457937, EPI_ISL_457938, EPI_ISL_457939 | Oman-NIC | Oman-NIC | Samira Al-Maruqi, Fahad Zadjali, Amina Al Jardani, Khulood Al-Mammary, Hanan Al-kindi, Fatma BaAlawi, Hamida AL Barwani, Zeyana AL-Dahmani, Intisar Al-Shukri, Aisha Al-Busaidi, Aisha Al-Amri, Ahlam Al-Amri, Mohammed Al-Tobi, Samiha Al Kharusi, Abdulla Balkhair |
| EPI_ISL_457940, EPI_ISL_457941, EPI_ISL_457942, EPI_ISL_457943, EPI_ISL_457944, EPI_ISL_457945, EPI_ISL_457946, EPI_ISL_457947, EPI_ISL_457948, EPI_ISL_457949 | Laboratorio de Biología Molecular Asociación Española Primera en Salud | Departments of Pathology and Medicine, New York University School of Medicine | Maria Victoria Elizondo, Maria Noel Zubillaga, Gonzalo Manrique, Paul Zappile, Gael Westby, Matthew T Maurano, Christian Marier, Adriana Heguy |
| EPI_ISL_457975, EPI_ISL_457976, EPI_ISL_457977, EPI_ISL_457978, EPI_ISL_457979, EPI_ISL_457980, EPI_ISL_457985 | Oman-NIC | Oman-NIC | Samira Al-Maruqi, Fahad Zadjali, Amina Al Jardani, Khulood Al-Mammary, Hanan Al-kindi, Fatma BaAlawi, Hamida AL Barwani, Zeyana AL-Dahmani, Intisar Al-Shukri, Aisha Al-Busaidi, Aisha Al-Amri, Ahlam Al-Amri, Mohammed Al-Tobi, Samiha Al Kharusi, Abdulla Balkhair |
| EPI_ISL_458000 | Centre For Biotechnology Research and Development | Centre For Biotechnology Research and Development | Matoke-Muhia,D.K., Symeker,S.L., Muuo,S.N., Ochwoto,M., Zablon,J.O., Kimotho,J., Waruhiu,C.N. and Michuki,G.N. |
| EPI_ISL_458017, EPI_ISL_458018, EPI_ISL_458019, EPI_ISL_458020, EPI_ISL_458021 | NYU Langone Health | Departments of Pathology and Medicine, New York University School of Medicine | Maria Aguero-Rosenfeld, Brendan Belovarac, Margaret Black, Ludovic Boytard, John Cadley, Paolo Cotzia, John Chen, Dacia Dimartino, Xiaojun Feng, Tatyana Gindin, Emily Guzman, Adriana Heguy, Megan Hogan, Emily Huang, George Jour, Alireza Khodadadi-Jamayran, Lawrence H. Lin, Raven Luther, Andrew Lytle, Christian Marier, Matthew T. Maurano, Mark J. Mulligan, Peter Meyn, Raquel Ordonez Ciriza, Iman Osman, Jared Pinnell, Vanessa Raabe, Sitharam Ramaswami, Amy Rapkiewicz, Andre M. Ribeiro-dos-Santos, Marie Samanovic-Golden, Antonio Serrano, Guomiao Shen, Matija Snuderl, Theodore Vougiouklakis, Nick Vulpescu, Gael Westby, Paul Zappile, Yutong Zhang |
| EPI_ISL_458139 | Evandro Chagas Institute | Evandro Chagas Institute | Santos, M.C.; Silva, A.M.; Junior, W.D.C.; Barbagelata, L.S.; Ferreira, J.A.; Sousa, E.M.A.; da Silva, P.S.; Resque, H.R; Martins, L.C.; Sousa Junior, E.C.;Viana, G.M.R |
| EPI_ISL_458230 | KU Leuven, Rega Institute, Clinical and Epidemiological Virology | KU Leuven, Rega Institute, Clinical and Epidemiological Virology | Tony Wawina-Bokalanga, Bert Vanmechelen, Joan Marti-Carerras, Piet Maes |
| EPI_ISL_459856, EPI_ISL_459857, EPI_ISL_459858, EPI_ISL_459862 | Center for Genome Regulation (CRG) | Center for Mathematical Modeling and Center for Genome Regulation. Santiago, Chile | Gaete A, Travisany D, Palma R, Urra C, Varas M, Allende ML, Maass A, González M. |
| EPI_ISL_459866 | Kingston Health Sciences Center | Queen's Genomics Lab at Ongwanada (Q-GLO) | Sjaarda CP, Rustom N, Huang D, Perez-Patrigeon S, Hudson ML, Wong H,Guan H, Ayub M, Soares CN, Colautti R, Evans GA, Sheth P |
| EPI_ISL_459953 | Institute for Medical Research, Infectious Disease Research Centre, National Institutes of Health, Ministry of Health Malaysia | Institute for Medical Research Infectious Disease Research Centre, National Institutes of Health, Ministry of Health Malaysia | Suppiah J, Mohd-Zawawi Z, Kamel KA, Ellan K, Kalyanasundram J, Mohd-Zain R, Thayan R |
| EPI_ISL_459954, EPI_ISL_459955 | Institute for Medical Research, Infectious Disease Research Centre, National Institutes of Health, Ministry of Health Malaysia | Institute for Medical Research, Infectious Disease Research Centre, National Institutes of Health, Ministry of Health Malaysia | Suppiah J, Mohd-Zawawi Z, Kamel KA, Ellan K, Kalyanasundram J, Mohd-Zain R, Thayan R |
| EPI_ISL_459966, EPI_ISL_459967, EPI_ISL_459968, EPI_ISL_459970, EPI_ISL_459971, EPI_ISL_459976 | Institut Pasteur du Maroc | Institut Pasteur du Maroc | Marion Barbet, Sylvie Behillil, Méline Bizard, Angela Brisebarre, Camille Capel, Etienne Simon-Lorière, Vincent Enouf, Maud Vanpeene, Sylvie van der Werf, Latifa Anga, Abdellah Faouzi, Anass Abbad, Mjid Eloualid, Jalal Nourlil, Anderrahmane Maaroufi |
| EPI_ISL_460045, EPI_ISL_460046, EPI_ISL_460047, EPI_ISL_460048, EPI_ISL_460049, EPI_ISL_460050, EPI_ISL_460051, EPI_ISL_460062, EPI_ISL_460063, EPI_ISL_460064, EPI_ISL_460065, EPI_ISL_460066, EPI_ISL_460068, EPI_ISL_460069, EPI_ISL_460070, EPI_ISL_460071, EPI_ISL_460072, EPI_ISL_460073, EPI_ISL_460075, EPI_ISL_460076, EPI_ISL_460077, EPI_ISL_460078 | | | |
| see above | Minnesota Department of Health, Public Health Laboratory | Minnesota Department of Health, Public Health Laboratory | Matt Plumb, Jacob Garfin, and Xiong Wang |
| EPI_ISL_460178, EPI_ISL_460211, EPI_ISL_460224, EPI_ISL_460262, EPI_ISL_460344, EPI_ISL_460381, EPI_ISL_460413, EPI_ISL_460458 | Massachusetts General Hospital | Infectious Disease Program, Broad Institute of Harvard and MIT | Lemieux,J.E., Siddle,K.J., Shaw,B., Adams,G., Pierce,V., Turbett,S., Anahtar,M., Branda,J., Slater,D., Harris,J., Lin,A.E., Gladden-Young,A., Lagerborg,K., Rudy,M., DeRuff,K., Carter,A., Normandin,E., Bauer,M., Reilly,S., Tomkins-Tinch,C., Loreth,C., Chaluvadi,S., Neumann,A., Cusick,C., Chapman,S.B., Gnirke,A., Flowers,K., Cerrato,F., Birren,B.W., Gallagher,G., Smole,S., Park,D.J., MacInnis,B.L., Ryan,E., LaRocque,R., Rosenberg,E., Sabeti,P.C. |
| EPI_ISL_460655, EPI_ISL_460666, EPI_ISL_460667, EPI_ISL_460698, EPI_ISL_460714, EPI_ISL_460720, EPI_ISL_460778, EPI_ISL_460779, EPI_ISL_460815, EPI_ISL_460816, EPI_ISL_460817, EPI_ISL_460818, EPI_ISL_461116, EPI_ISL_461259, EPI_ISL_461287 | | | |
| see above | Dutch COVID-19 response team | Erasmus Medical Center | Bas Oude Munnink, David Nieuwenhuijse, Reina Sikkema, Claudia Schapendonk, Irina Chestakova, Anne van der Linden, Theo Bestebroer, Stefan van Nieuwkoop, Mark Pronk, Pascal Lexmond, Corien Swaan, Manon Haverkate, Madelief Mollers, Mart Stein, Sandra Kengne Kamga Mobou, Jeroen van Kampen, Jolanda Voermans, Aura Timen, Corine GeurtsvanKessel, Annemiek van der Eijk, Richard Molenkamp, Marion Koopmans, on behalf of the Dutch national COVID-19 response team. |
| EPI_ISL_462178, EPI_ISL_462179, EPI_ISL_462180, EPI_ISL_462181 | KU Leuven, Rega Institute, Clinical and Epidemiological Virology | KU Leuven, Rega Institute, Clinical and Epidemiological Virology | Tony Wawina-Bokalanga, Bert Vanmechelen, Joan Marti-Carerras, Piet Maes |
| EPI_ISL_462284, EPI_ISL_462292, EPI_ISL_462387, EPI_ISL_462401, EPI_ISL_462405, EPI_ISL_462409, EPI_ISL_462416, EPI_ISL_462429, EPI_ISL_462432 | National Public Health Laboratory, National Centre for Infectious Diseases | National Public Health Laboratory, National Centre for Infectious Diseases | Mak TM, Octavia S, Chavatte JM, Cui L, Lin RTP |
| EPI_ISL_462450 | Clinical Center, University of Sarajevo | Charite Universitatsmedizin Berlin, Institute of Virology | Victor M Corman, Jorn Beheim-Schwarzbach, Barbara Muehlemann, Talitha Veith, Julia Schneider, Terry Jones, Amela Dedeic-Ljubovic, Irma Salimovic-Besic, Suzana Arapcic, Almedina Hadzihasanovic-Moro, Selma Mutevelic, Christian Drosten |
| EPI_ISL_463001 | unknown | Clinical virology | Fares,W., Triki,H. |
| EPI_ISL_463311, EPI_ISL_463312 | Queen Elizabeth Hospital | Hong Kong Department of Health | Mak Gannon C.K., Cheng Peter K.C., Lam Edman T.K., Chan Rickjason C.W., Tsang Dominic N.C. |
| EPI_ISL_463313 | Central Health Medical Practice | Hong Kong Department of Health | Mak Gannon C.K., Cheng Peter K.C., Lam Edman T.K., Chan Rickjason C.W., Tsang Dominic N.C. |

| | | | |
|---|---|---|---|
| EPI_ISL_463315 | Princess Margaret Hospital | Hong Kong Department of Health | Mak Gannon C.K., Cheng Peter K.C., Lam Edman T.K., Chan Rickjason C.W., Tsang Dominic N.C. |
| EPI_ISL_463975, EPI_ISL_463977, EPI_ISL_463984, EPI_ISL_463987, EPI_ISL_463989, EPI_ISL_463990 | Toronto Invasive Bacterial Diseases Network | McMaster University | Allison McGeer, Patryk Aftanas, Angel Li, Kuganya Nirmalarajah, Samira Mubareka, Andrew G. McArthur |
| EPI_ISL_464009, EPI_ISL_464051 | Unity Health Toronto | Ontario Institute for Cancer Research | Ramzi Fattouh,Larissa M. Matukas,Mark Downing,Annette Gower,Karel Boissinot,Samira Mubareka,TIBDN,Ilinca Lungu,Bernard Lam,Jeremy Johns,Paul Krzyzanowski,Richard de Borja,Philip Zuzarte,Jared Simpson |
| EPI_ISL_464092 | Laboratory Medicine | Department of Laboratory Medicine, Lin-Kou Chang Gung Memorial Hospital, Taoyuan, Taiwan | Kuo-Chien Tsao, Yu-Nong Gong, Shu-Li Yang, Yi-Chun Liu, Chung-Guei Huang, Mei-Jen Hsiao, Po-Wei Huang, Cheng-Ta Yang, Cheng-Hsun Chiu, Peng-Nien Huang, Kuo-Ming Lee, Guang-Wu Chen, Shin-Ru Shih |
| EPI_ISL_464112, EPI_ISL_464116 | National Health Laboratory Service (NHLS), Tygerberg | Division of Medical Virology, Stellenbosch University and National Health Laboratory Service (NHLS) | Susan Engelbrecht, Kayla Delaney, Bronwyn Kleinhans, Houriiyah Tegally, Eduan Wilkindon, Gert van Zyl, Wolfgang Preiser, Tulio de Oliveira |
| EPI_ISL_464877, EPI_ISL_465096, EPI_ISL_465099, EPI_ISL_465102, EPI_ISL_465114, EPI_ISL_465115, EPI_ISL_465116, EPI_ISL_465117, EPI_ISL_465118, EPI_ISL_465121, EPI_ISL_465133, EPI_ISL_465135, EPI_ISL_465136, EPI_ISL_465137, EPI_ISL_465138, EPI_ISL_465139, EPI_ISL_465142, EPI_ISL_465145, EPI_ISL_465146, EPI_ISL_465149, EPI_ISL_465154, EPI_ISL_465155, EPI_ISL_465156, EPI_ISL_465157, EPI_ISL_465158, EPI_ISL_465159, EPI_ISL_465160, EPI_ISL_465161, EPI_ISL_465721, EPI_ISL_465722, EPI_ISL_465723, EPI_ISL_465724, EPI_ISL_465725, EPI_ISL_465726, EPI_ISL_465727, EPI_ISL_465728, EPI_ISL_465729, EPI_ISL_465730, EPI_ISL_465731, EPI_ISL_465732, EPI_ISL_465733, EPI_ISL_465734, EPI_ISL_465735, EPI_ISL_465736, EPI_ISL_465737, EPI_ISL_465738, EPI_ISL_465739, EPI_ISL_465740, EPI_ISL_465741, EPI_ISL_465742, EPI_ISL_465743, EPI_ISL_465744, EPI_ISL_465745, EPI_ISL_465746, EPI_ISL_465747, EPI_ISL_465748, EPI_ISL_465749, EPI_ISL_465750, EPI_ISL_465751, EPI_ISL_465752, EPI_ISL_465753, EPI_ISL_465754, EPI_ISL_465755, EPI_ISL_465756, EPI_ISL_465757, EPI_ISL_465758, EPI_ISL_465759, EPI_ISL_465760, EPI_ISL_465761, EPI_ISL_465762, EPI_ISL_465763, EPI_ISL_465764, EPI_ISL_465765, EPI_ISL_465766, EPI_ISL_465767, EPI_ISL_465768, EPI_ISL_465770, EPI_ISL_465771, EPI_ISL_465772, EPI_ISL_465773, EPI_ISL_465774, EPI_ISL_465775, EPI_ISL_465776, EPI_ISL_465777, EPI_ISL_465778, EPI_ISL_465779, EPI_ISL_465780, EPI_ISL_465781, EPI_ISL_465782, EPI_ISL_465783, EPI_ISL_465784, EPI_ISL_465785, EPI_ISL_465786, EPI_ISL_465787, EPI_ISL_465788, EPI_ISL_465789, EPI_ISL_465790, EPI_ISL_465792, EPI_ISL_465793, EPI_ISL_465795, EPI_ISL_465797, EPI_ISL_465798 | | | |
| see above | Respiratory Virus Unit, Microbiology Services Colindale, Public Health England | Respiratory Virus Unit, Microbiology Services Colindale, Public Health England | PHE Covid Sequencing Team |
| EPI_ISL_467055 | Servicio de Microbiología, Hospital Universitario Son Espases | SeqCOVID-SPAIN consortium/IBV(CSIC) | Carla López-Causapé, Jordi Reina y Antonio Oliver and SeqCOVID-SPAIN consortium |
| EPI_ISL_467062 | Hospital Universitario Virgen de las Nieves de Granada-SAS | SeqCOVID-SPAIN consortium/IBV(CSIC) | Mercedes Pérez Ruiz, Sara Sanbonmatsu Gámez, Irene Pedrosa Corral, José M. Navarro-Marí and SeqCOVID-SPAIN consortium |
| EPI_ISL_467064, EPI_ISL_467066, EPI_ISL_467067, EPI_ISL_467068, EPI_ISL_467069, EPI_ISL_467070, EPI_ISL_467071, EPI_ISL_467072, EPI_ISL_467073, EPI_ISL_467074, EPI_ISL_467075, EPI_ISL_467076, EPI_ISL_467077, EPI_ISL_467078, EPI_ISL_467079, EPI_ISL_467080, EPI_ISL_467081, EPI_ISL_467083, EPI_ISL_467084, EPI_ISL_467085 | | | |
| see above | Hospital Universitario Puerta del Mar de Cádiz - INIBICA | SeqCOVID-SPAIN consortium/IBV(CSIC) | Salud Rodríguez-Pallares, Fátima-Galán-Sánchez, Manuel Rodríguez-Iglesias and SeqCOVID-SPAIN consortium |
| EPI_ISL_467086, EPI_ISL_467087, EPI_ISL_467088, EPI_ISL_467089 | Hospital Universitario de Gran Canaria Dr. Negrín | SeqCOVID-SPAIN consortium/IBV(CSIC) | M. Carmen Pérez González, Francisco J. Chamizo López, Ana Bordes Benítez and SeqCOVID-SPAIN consortium |
| EPI_ISL_467092, EPI_ISL_467094, EPI_ISL_467095, EPI_ISL_467098, EPI_ISL_467099, EPI_ISL_467101, EPI_ISL_467102, EPI_ISL_467106, EPI_ISL_467108, EPI_ISL_467109, EPI_ISL_467110, EPI_ISL_467111, EPI_ISL_467115, EPI_ISL_467117, EPI_ISL_467118, EPI_ISL_467119, EPI_ISL_467120, EPI_ISL_467122, EPI_ISL_467123, EPI_ISL_467125, EPI_ISL_467127, EPI_ISL_467128, EPI_ISL_467130, EPI_ISL_467131, EPI_ISL_467133, EPI_ISL_467137, EPI_ISL_467138, EPI_ISL_467140, EPI_ISL_467143, EPI_ISL_467144, EPI_ISL_467145, EPI_ISL_467148, EPI_ISL_467149, EPI_ISL_467151, EPI_ISL_467152, EPI_ISL_467154, EPI_ISL_467155, EPI_ISL_467163, EPI_ISL_467166, EPI_ISL_467167, EPI_ISL_467168, EPI_ISL_467169, EPI_ISL_467170, EPI_ISL_467173, EPI_ISL_467174, EPI_ISL_467176, EPI_ISL_467178, EPI_ISL_467179, EPI_ISL_467181, EPI_ISL_467182, EPI_ISL_467183 | | | |
| see above | Hospital Universitario Araba. Vitoria-Gasteiz | SeqCOVID-SPAIN consortium/IBV(CSIC) | Silvia Hernáez Crespo, Carmen Gómez González, Amaia Aguirre Quiñonero, Marina Fernández Torres, Mª Rosario Almela Ferrer, Mª Concepción Lecaroz Agara, Andrés Canut Blasco. and SeqCOVID-SPAIN consortium |
| EPI_ISL_467372 | Arizona State University Health Services | Arizona State University | Peter T. Skidmore, Rabia Maqsood, LaRinda A. Holland, Emily A. Kaelin, Lily I. Wu, Arvind Varsani, Rolf U. Halden, Brenda G. Hogue, Matthew Scotch, Efrem S. Lim |
| EPI_ISL_467374 | Dinkes Samarinda | Eijkman Institute for Molecular Biology, Ministry of Research and Technology/National Agency for Research and Innovation | Edison Johar, Frilasita A Yudhaputri, Hidayat Trimarsanto, David H Muljono, Safarina G Malik, Khin Saw Myint, Amin Soebandrio |
| EPI_ISL_467411, EPI_ISL_467412, EPI_ISL_467413, EPI_ISL_467414, EPI_ISL_467415, EPI_ISL_467416, EPI_ISL_467417, EPI_ISL_467418, EPI_ISL_467419, EPI_ISL_467420, EPI_ISL_467421, EPI_ISL_467422 | | | |
| see above | NYU Langone Health | Departments of Pathology and Medicine, New York University School of Medicine | Maria Aguero-Rosenfeld, Brendan Belovarac, Margaret Black, Ludovic Boytard, John Cadley, Paolo Cotzia, John Chen, Dacia Dimartino, Xiaojun Feng, Tatyana Gindin, Emily Guzman, Adriana Heguy, Megan Hogan, Emily Huang, George Jour, Alireza Khodadadi-Jamayran, Lawrence H. Lin, Raven Luther, Andrew Lytle, Christian Marier, Matthew T. Maurano, Mark J. Mulligan, Peter Meyn, Raquel Ordonez Ciriza, Iman Osman, Jared Pinnell, Vanessa Raabe, Sitharam Ramaswami, Amy Rapkiewicz, Andre M. Ribeiro-dos-Santos, Marie Samanovic-Golden, Antonio Serrano, Guomiao Shen, Matija Snuderl, Theodore Vougiouklakis, Nick Vulpescu, Gael Westby, Paul Zappile, Yutong Zhang |
| EPI_ISL_467526, EPI_ISL_467527, EPI_ISL_467532, EPI_ISL_467594, EPI_ISL_467595, EPI_ISL_467601, EPI_ISL_467602, EPI_ISL_467603, EPI_ISL_467604, EPI_ISL_467605, EPI_ISL_467606, EPI_ISL_467607, EPI_ISL_467608, EPI_ISL_467609, EPI_ISL_467610, EPI_ISL_467611, EPI_ISL_467612, EPI_ISL_467613, EPI_ISL_467614, EPI_ISL_467615, EPI_ISL_467616, EPI_ISL_467619, EPI_ISL_467620, EPI_ISL_467621, EPI_ISL_467622, EPI_ISL_467623, EPI_ISL_467624 | | | |
| see above | New Mexico Department of Health Scientific Laboratory Division | Center for Global Health, University of New Mexico Health Sciences Center | Daryl Domman, Kurt Schwalm, Twila Kunde, Joseph Hicks, Michael Edwards, Darrell Dinwiddie |
| EPI_ISL_467811, EPI_ISL_467812, EPI_ISL_467813, EPI_ISL_467814, EPI_ISL_467815, EPI_ISL_467816, EPI_ISL_467817, EPI_ISL_467818, EPI_ISL_467819, EPI_ISL_467820, EPI_ISL_467821, EPI_ISL_467822, EPI_ISL_467823, EPI_ISL_467824, EPI_ISL_467825, EPI_ISL_467826, EPI_ISL_467827, EPI_ISL_467828, EPI_ISL_467829, EPI_ISL_467830, EPI_ISL_467831, EPI_ISL_467833, EPI_ISL_467834, EPI_ISL_467835, EPI_ISL_467836, EPI_ISL_467837, EPI_ISL_467838, EPI_ISL_467840, EPI_ISL_467841, EPI_ISL_467842, EPI_ISL_467843, EPI_ISL_467858, EPI_ISL_467859, EPI_ISL_467860, EPI_ISL_467861, EPI_ISL_467863, EPI_ISL_467865, EPI_ISL_467869, EPI_ISL_467870, EPI_ISL_467871, EPI_ISL_467872, EPI_ISL_467875, EPI_ISL_467877, EPI_ISL_467879, EPI_ISL_467884, EPI_ISL_467885, EPI_ISL_467886, EPI_ISL_467887, EPI_ISL_467888, EPI_ISL_467889, EPI_ISL_467890, EPI_ISL_467891, EPI_ISL_467892, EPI_ISL_467893, EPI_ISL_467894, EPI_ISL_467895, EPI_ISL_467896, EPI_ISL_467897, EPI_ISL_467898, EPI_ISL_467899, EPI_ISL_467900, EPI_ISL_467901, EPI_ISL_467903, EPI_ISL_467906, EPI_ISL_467907 | | | |
| see above | Quest Diagnostics | Quest Diagnostics | Anderson,B.P., Rosenthal,S.H., Gerasimova,A., Kagan,R.M. and Owen, R. |
| EPI_ISL_467952, EPI_ISL_467955, EPI_ISL_467969, EPI_ISL_467978 | San Diego County Public Health Laboratory | Andersen lab at Scripps Research | SEARCH Alliance San Diego with Tracy Basler, Jovan Shephard, Brett Austin |
| EPI_ISL_467995, EPI_ISL_468000, EPI_ISL_468016 | SA Pathology | SA Pathology | Lex Leong, Chuan Kok Lim, Mark Turra, Ivan Bastian, Geoff Higgins |
| EPI_ISL_468063, EPI_ISL_468064, EPI_ISL_468065 | unknown | Computer Science and Engineering | Rouchka,E.C., Chariker,J.H., Chung,D., Ramirez,J., Palmer,K.E., Lasnik,A.B., Carrico,R., Arnold,F.W., Adcock,R.S., Zhang,M., Alejandro,B., Wolf,L.A., Hwang,J.Y., Park,J.W., Waigel,S., Zacharias,W. |
| EPI_ISL_468389, EPI_ISL_468390, EPI_ISL_468392, EPI_ISL_468393 | County of San Luis Obispo Public Health Laboratory | Chan-Zuckerberg Biohub | CZB Cliahub Consortium |
| EPI_ISL_468495 | Ventura County Public Health Lab | Chan-Zuckerberg Biohub | CZB Cliahub Consortium |
| EPI_ISL_468509, EPI_ISL_468510 | San Joaquin County Public Health Lab | Chan-Zuckerberg Biohub | CZB Cliahub Consortium |
| EPI_ISL_468737, EPI_ISL_468743, EPI_ISL_468744, EPI_ISL_468745, EPI_ISL_468746 | Lab voor klinische biologie | Onderzoeksgroep Virologie | Nick Vereecke, Laurens Lambrechts, Marthe Pauwels, Bruno Verhasselt, Linos Vandekerckhove, Hans Nauwynck, Sebastiaan Theuns |
| EPI_ISL_468760 | Center for Genome Regulation (CRG) | Center for Mathematical Modeling and Center for Genome Regulation. Santiago, Chile | Gaete A, Travisany D, Palma R, Urra C, Varas M, Allende ML, Maass A, González M. |
| EPI_ISL_468765, EPI_ISL_468769, EPI_ISL_468778, EPI_ISL_468797, EPI_ISL_468801, EPI_ISL_468803, EPI_ISL_468818, EPI_ISL_468830, EPI_ISL_468834, EPI_ISL_468836, EPI_ISL_468840, EPI_ISL_468856 | | | |
| see above | Servicio de Microbiología, Hospital Miguel Servet, Zaragoza | SeqCOVID-SPAIN consortium/IBV(CSIC) | Antonio Rezusta López, Alexander Tristancho Baró, Ana Milagro, Yolanda Gracia Grataloup, Nieves Martínez Cameo and SeqCOVID-SPAIN consortium |
| EPI_ISL_468914 | Istituto Zooprofilattico Sperimentale Puglia e Basilicata; Dipartimento di Bioscienze, Biotecnologie e Biofarmaceutica dell'Università degli Studi di Bari "A.Moro"; Istituto di Biomembrane. Bioenergetica e Biotecnologie Molecolari del Consiglio Nazionale delle Ricerche di Bari | Beaconlab (Bioinformatics, Evolution and Comparative Genomics lab), Dept of Biosciences, University on Milan | Parisi A.,Pesole G., Manzari C., Chiara M. |

| | | | |
|---|---|---|---|
| EPI_ISL_468961, EPI_ISL_468970, EPI_ISL_468982, EPI_ISL_468985, EPI_ISL_468989, EPI_ISL_468990, EPI_ISL_469005 | Servicio de Microbiología, Hospital Universitario Son Espases | SeqCOVID-SPAIN consortium/IBV(CSIC) | Carla López-Causapé, Jordi Reina, Antonio Oliver and SeqCOVID-SPAIN consortium |
| EPI_ISL_469016 | Istituto Zooprofilattico Sperimentale Puglia e Basilicata; Dipartimento di Bioscienze, Biotecnologie e Biofarmaceutica dell'Università degli Studi di Bari "A.Moro"; Istituto di Biomembrane. Bioenergetica e Biotecnologie Molecolari del Consiglio Nazionale delle Ricerche di Bari | Beaconlab (Bioinformatics, Evolution and Comparative Genomics lab), Dept of Biosciences, University on Milan | Parisi A.,Pesole G., Manzari C., Chiara M. |
| EPI_ISL_469241, EPI_ISL_469242, EPI_ISL_469243, EPI_ISL_469244, EPI_ISL_469245, EPI_ISL_469246, EPI_ISL_469247, EPI_ISL_469248, EPI_ISL_469249, EPI_ISL_469250, EPI_ISL_469251, EPI_ISL_469252 | | | |
| see above | Special Infectious Agents Unit | Special Infectious Agents Unit | Azhar,E.I., Hassan,A.M., Tolah,A.M., Uthman,N.A., Al-Sobahy,T.L., Farraj,S.A., El-Kafrawy,S.A. |
| EPI_ISL_469283, EPI_ISL_469284 | Service de Virologie Hôpital Saint-Louis | Laboratory Cell Biology of Viral Infection-INSERM unit 944 | Laurent Meertens, Lucie Bonnet-Madin, Séverine Mercier-Delarue, Maud SALMONA, Constance Delaugerre, Ali Amara |
| EPI_ISL_470540 | Wisconsin State Laboratory of Hygiene Communicable Disease Division | Wisconsin State Laboratory of Hygiene Communicable Disease Division | Kelsey R Florek |
| EPI_ISL_470575, EPI_ISL_470580, EPI_ISL_470598, EPI_ISL_470652, EPI_ISL_470653, EPI_ISL_470654 | Hermes Pardini | Bioinformatics Laboratory / LNCC | Alexandra Gerber, Ana Paula Guimarães, Luiz Gonzaga Paula de Almeida, Ronaldo da Silva Francisco Junior, Mariane Talon, Filipe Romero, Átila Duque Rossi, Terezinha Marta Pereira, working group UFRJ, Jaqueline Goes de Jesus, Ingra Morales Claro, Ester Cerdeira Sabino, Nuno Rodrigues Faria, CADDE-group, Laboratorio Hermes Pardini, Laboratorio Simile, working group UFMG, Amilcar Tanuri, Carolina Voloch, Renato Santana Aguiar e Ana Tereza Vasconcelos |
| EPI_ISL_471205, EPI_ISL_471207, EPI_ISL_471235, EPI_ISL_471249, EPI_ISL_471250, EPI_ISL_471251, EPI_ISL_471252 | Wisconsin State Laboratory of Hygiene Communicable Disease Division | Wisconsin State Laboratory of Hygiene Communicable Disease Division | Kelsey R. Florek, Abigail C. Shockey |
| EPI_ISL_471472 | Hospital Universitari Germans Trias i Pujol(HUGTiP)/Fundació Lluita contra la SIDA (FLSida)/IRTA-CReSA | IrsiCaixa AIDS Research Lab | Marc Noguera-Julian, Pilar Armengol, Jordi Rodón, Julia Vergara, Lidia Ruiz, Nuria Izquierdo, Jorge Carrillo, Roger Paredes, Albert Bensaid, Julia Blanco, Joaquim Segalés, Bonaventura Clotet |
| EPI_ISL_471553, EPI_ISL_471555 | The National Institute of Public Health | State Veterinary Institute Prague and The National Institute of Public Health | Nagy,A;Jirincova,H;Novakova,L;Trnka,D;Vecerova,J |
| EPI_ISL_471677 | Michigan Department of Health and Human Services, Bureau of Laboratories | Michigan Department of Health and Human Services, Bureau of Laboratories | Blankenship HM, Riner D, Soehnlen MK |
| EPI_ISL_471970 | University of Exeter | COVID-19 Genomics UK (COG-UK) Consortium | Ben Temperton,Aaron Jeffries,Michelle Michelsen,Joanna Warwick-Dugdale,Audrey Farbos,Robyn Manley,Stephen Michell,Jane Masoli |
| EPI_ISL_472165, EPI_ISL_472166 | Northumbria University / South Tees Hospitals NHS Foundation Trust / North Cumbria Integrated Care NHS Foundation Trust / North Tees and Hartlepool NHS Foundation Trust / Newcastle Hospitals NHS Foundation Trust | COVID-19 Genomics UK (COG-UK) Consortium | Darren L Smith,Andrew Nelson,Matthew Bashton,Greg R Young,Joshua Loh,John Allan,Mohammad A Tariq,Giles S Holt,Gary Black,Wen C Yew,Lynn Dover,Paul Baker,Steve Liggett,Sarah Essex,Jane Greenaway,Debra Padgett,Clive Graham,Garren Scott,Edward Barton,Emma Swindells,Brendan Payne,Jennifer Collins,Yusri Taha,Gary Eltringham |
| EPI_ISL_474224, EPI_ISL_474225, EPI_ISL_474226, EPI_ISL_474227, EPI_ISL_474228, EPI_ISL_474229, EPI_ISL_474230 | Wales Specialist Virology Centre Sequencing lab: Pathogen Genomics Unit | COVID-19 Genomics UK (COG-UK) Consortium | Catherine Moore, Johnathan Evans, Laura Gifford, Malorie Perry, Simon Cottrell, Angela Marchbank, Alec Birchley, Alexander Adams, Amy Gaskin, Bree Gatica-Wilcox, Jason Coombes, Joel Southgate, Lauren Gilbert, Lee Graham, Nicole Pacchiarini, Sara Kumziene-Summerhayes, Sarah Taylor, Sophie Jones, Sara Rey, Matthew Bull, Joanne Watkins, Sally Corden, Tom Connor |
| EPI_ISL_474833, EPI_ISL_474834, EPI_ISL_474844, EPI_ISL_474852, EPI_ISL_474860, EPI_ISL_474861, EPI_ISL_474862, EPI_ISL_474863, EPI_ISL_474864, EPI_ISL_474865, EPI_ISL_474866, EPI_ISL_474867, EPI_ISL_474868, EPI_ISL_474869, EPI_ISL_474897, EPI_ISL_474909 | | | |
| see above | Hospital Universitario Virgen de las Nieves de Granada-SAS | SeqCOVID-SPAIN consortium/IBV(CSIC) | Mercedes Pérez Ruiz, Sara Sanbonmatsu Gámez, Irene Pedrosa Corral, José M. Navarro-Marí and SeqCOVID-SPAIN consortium |
| EPI_ISL_474912, EPI_ISL_474913, EPI_ISL_474914, EPI_ISL_474915, EPI_ISL_474916, EPI_ISL_474917, EPI_ISL_474918 | Hospital Universitario de Gran Canaria Dr. Negrín | SeqCOVID-SPAIN consortium/IBV(CSIC) | M. Carmen Pérez González, Francisco J. Chamizo López, Ana Bordes Benítez and SeqCOVID-SPAIN consortium |
| EPI_ISL_474920, EPI_ISL_474934, EPI_ISL_474936, EPI_ISL_474937, EPI_ISL_474938, EPI_ISL_474942, EPI_ISL_474948 | Hospital Universitario Virgen de las Nieves de Granada-SAS | SeqCOVID-SPAIN consortium/IBV(CSIC) | Mercedes Pérez Ruiz, Sara Sanbonmatsu Gámez, Irene Pedrosa Corral, José M. Navarro-Marí and SeqCOVID-SPAIN consortium |
| EPI_ISL_474960, EPI_ISL_474965, EPI_ISL_474969, EPI_ISL_474973, EPI_ISL_474974, EPI_ISL_474975, EPI_ISL_474979, EPI_ISL_474980, EPI_ISL_474981, EPI_ISL_474986, EPI_ISL_474987, EPI_ISL_474988, EPI_ISL_474992, EPI_ISL_474993, EPI_ISL_474994, EPI_ISL_474999, EPI_ISL_475000, EPI_ISL_475001, EPI_ISL_475002, EPI_ISL_475003, EPI_ISL_475010, EPI_ISL_475013, EPI_ISL_475014, EPI_ISL_475015, EPI_ISL_475018, EPI_ISL_475019, EPI_ISL_475021, EPI_ISL_475022 | | | |
| see above | Israel Central Virology laboratory | Israel Central Virology laboratory | Neta Zuckerman, Efrat Dahan Bucris, Oran Erster, Ella Mendelson, Michal Mandelboim |
| EPI_ISL_475060, EPI_ISL_475061, EPI_ISL_475062, EPI_ISL_475063, EPI_ISL_475064, EPI_ISL_475065, EPI_ISL_475066, EPI_ISL_475067, EPI_ISL_475068, EPI_ISL_475069, EPI_ISL_475070 | | | |
| see above | Lab voor klinische biologie | Onderzoeksgroep Virologie | Laurens Lambrechts, Nick Vereecke, Marthe Pauwels, Bruno Verhasselt, Linos Vandekerckhove, Hans Nauwynck, Sebastiaan Theuns |
| EPI_ISL_475071, EPI_ISL_475072 | Lab voor klinische biologie | Onderzoeksgroep Virologie | Nick Vereecke, Laurens Lambrechts, Marthe Pauwels, Bruno Verhasselt, Linos Vandekerckhove, Hans Nauwynck, Sebastiaan Theuns |
| EPI_ISL_475092 | Skovde/Unilabs | The Public Health Agency of Sweden | Oskar Karlsson Lindsjo, Maria Lind Karlberg, Mattias Haukland, Reza Advani, Olov Svartstrom, Anna-Malin Linde, Sandra Broddesson, Petra Edquist, Shamam Muradrasoli, Anna Risberg, Karin Tegmark-Wisell |
| EPI_ISL_475093, EPI_ISL_475094 | Halmstad klinisk mikrobiologi | The Public Health Agency of Sweden | Oskar Karlsson Lindsjo, Maria Lind Karlberg, Mattias Haukland, Reza Advani, Olov Svartstrom, Anna-Malin Linde, Sandra Broddesson, Petra Edquist, Shamam Muradrasoli, Anna Risberg, Karin Tegmark-Wisell |
| EPI_ISL_475095 | Karolinska Universitetslaboratoriet | The Public Health Agency of Sweden | Oskar Karlsson Lindsjo, Maria Lind Karlberg, Mattias Haukland, Reza Advani, Olov Svartstrom, Anna-Malin Linde, Sandra Broddesson, Petra Edquist, Shamam Muradrasoli, Anna Risberg, Karin Tegmark-Wisell |
| EPI_ISL_475137, EPI_ISL_475138 | Kalmar klinisk mikrobiologi | The Public Health Agency of Sweden | Oskar Karlsson Lindsjo, Maria Lind Karlberg, Mattias Haukland, Reza Advani, Olov Svartstrom, Anna-Malin Linde, Sandra Broddesson, Petra Edquist, Shamam Muradrasoli, Anna Risberg, Karin Tegmark-Wisell |
| EPI_ISL_475139 | Orebro klinisk mikrobiologi | The Public Health Agency of Sweden | Oskar Karlsson Lindsjo, Maria Lind Karlberg, Mattias Haukland, Reza Advani, Olov Svartstrom, Anna-Malin Linde, Sandra Broddesson, Petra Edquist, Shamam Muradrasoli, Anna Risberg, Karin Tegmark-Wisell |
| EPI_ISL_475562, EPI_ISL_475563 | Din Klinik | The Public Health Agency of Sweden | Oskar Karlsson Lindsjo, Maria Lind Karlberg, Mattias Haukland, Reza Advani, Olov Svartstrom, Anna-Malin Linde, Sandra Broddesson, Mia Brytting, Anna Risberg, Karin Tegmark-Wisell |
| EPI_ISL_475572 | Imperial College London | Imperial College London | Jie Zhou, Wendy Barclay |
| EPI_ISL_475768 | Universitaetsklinik für Innere Medizin II Innsbruck | Bergthaler laboratory, CeMM Research Center for Molecular Medicine of the Austrian Academy of Sciences | Alexandra Popa, Benedikt Agerer, Henrique Colaco, Lukas Endler, Jakob-Wendelin Genger, Alexander Lercher, Mark Smyth, Thomas Penz, Michael Schuster, Jan Laine, Martin Senekowitsch, Judith Aberle, Stephan Aberle, Peter Hufnagl, Daniela Schmid, Franz Allerberger, Elisabeth Puchhammer-Stoeckl, Manfred Nairz, Guenter Weiss, Gregor Hörmann, Kinga Rigler-Hohenwarter, Rainer Gattringer, Wegene Borena, Dorothee von Laer, Christoph Bock, Andreas Bergthaler |
| EPI_ISL_475771 | Center for Virology, Medical University of Vienna | Bergthaler laboratory, CeMM Research Center for Molecular Medicine of the Austrian Academy of Sciences | Alexandra Popa, Benedikt Agerer, Henrique Colaco, Lukas Endler, Jakob-Wendelin Genger, Alexander Lercher, Mark Smyth, Thomas Penz, Michael Schuster, Jan Laine, Martin Senekowitsch, Judith Aberle, Stephan Aberle, Peter Hufnagl, Daniela Schmid, Franz Allerberger, Elisabeth |

| | | | |
|---|---|---|---|
| | | | Puchhammer-Stoeckl, Manfred Nairz, Guenter Weiss, Gregor Hörmann, Kinga Rigler-Hohenwarter, Rainer Gattringer, Wegene Borena, Dorothee von Laer, Christoph Bock, Andreas Bergthaler |
| EPI_ISL_475813, EPI_ISL_475814 | Institut für Virologie am Department für Hygiene, Mikrobiologie und Public Health | Bergthaler laboratory, CeMM Research Center for Molecular Medicine of the Austrian Academy of Sciences | Alexandra Popa, Benedikt Agerer, Henrique Colaco, Lukas Endler, Jakob-Wendelin Genger, Alexander Lercher, Mark Smyth, Thomas Penz, Michael Schuster, Jan Laine, Martin Senekowitsch, Judith Aberle, Stephan Aberle, Peter Hufnagl, Daniela Schmid, Franz Allerberger, Elisabeth Puchhammer-Stoeckl, Manfred Nairz, Guenter Weiss, Gregor Hörmann, Kinga Rigler-Hohenwarter, Rainer Gattringer, Wegene Borena, Dorothee von Laer, Christoph Bock, Andreas Bergthaler |
| EPI_ISL_475863, EPI_ISL_475867, EPI_ISL_475874, EPI_ISL_475880 | Austrian Agency for Health and Food Safety (AGES) | Bergthaler laboratory, CeMM Research Center for Molecular Medicine of the Austrian Academy of Sciences | Alexandra Popa, Benedikt Agerer, Henrique Colaco, Lukas Endler, Jakob-Wendelin Genger, Alexander Lercher, Mark Smyth, Thomas Penz, Michael Schuster, Jan Laine, Martin Senekowitsch, Judith Aberle, Stephan Aberle, Peter Hufnagl, Daniela Schmid, Franz Allerberger, Elisabeth Puchhammer-Stoeckl, Manfred Nairz, Guenter Weiss, Gregor Hörmann, Kinga Rigler-Hohenwarter, Rainer Gattringer, Wegene Borena, Dorothee von Laer, Christoph Bock, Andreas Bergthaler |
| EPI_ISL_475905, EPI_ISL_475906, EPI_ISL_475907, EPI_ISL_475908, EPI_ISL_475909 | Zentralinstitut für medizinische und chemische Labordiagnostik, Universitätskliniken Insbruck | Bergthaler laboratory, CeMM Research Center for Molecular Medicine of the Austrian Academy of Sciences | Alexandra Popa, Benedikt Agerer, Henrique Colaco, Lukas Endler, Jakob-Wendelin Genger, Alexander Lercher, Mark Smyth, Thomas Penz, Michael Schuster, Jan Laine, Martin Senekowitsch, Judith Aberle, Stephan Aberle, Peter Hufnagl, Daniela Schmid, Franz Allerberger, Elisabeth Puchhammer-Stoeckl, Manfred Nairz, Guenter Weiss, Gregor Hörmann, Kinga Rigler-Hohenwarter, Rainer Gattringer, Wegene Borena, Dorothee von Laer, Christoph Bock, Andreas Bergthaler |
| EPI_ISL_475924 | Institut für Virologie am Department für Hygiene, Mikrobiologie und Public Health | Bergthaler laboratory, CeMM Research Center for Molecular Medicine of the Austrian Academy of Sciences | Alexandra Popa, Benedikt Agerer, Henrique Colaco, Lukas Endler, Jakob-Wendelin Genger, Alexander Lercher, Mark Smyth, Thomas Penz, Michael Schuster, Jan Laine, Martin Senekowitsch, Judith Aberle, Stephan Aberle, Peter Hufnagl, Daniela Schmid, Franz Allerberger, Elisabeth Puchhammer-Stoeckl, Manfred Nairz, Guenter Weiss, Gregor Hörmann, Kinga Rigler-Hohenwarter, Rainer Gattringer, Wegene Borena, Dorothee von Laer, Christoph Bock, Andreas Bergthaler |
| EPI_ISL_475948, EPI_ISL_475949, EPI_ISL_475950, EPI_ISL_475961, EPI_ISL_475962, EPI_ISL_475963, EPI_ISL_475964 | National Public Health Laboratory, National Centre for Infectious Diseases | National Public Health Laboratory, National Centre for Infectious Diseases | Mak TM, Octavia S, Chavatte JM, Cui L, Lin RTP |
| EPI_ISL_476175, EPI_ISL_476178, EPI_ISL_476179, EPI_ISL_476180, EPI_ISL_476181, EPI_ISL_476182, EPI_ISL_476183, EPI_ISL_476185, EPI_ISL_476186, EPI_ISL_476187, EPI_ISL_476189, EPI_ISL_476190, EPI_ISL_476191, EPI_ISL_476192, EPI_ISL_476193, EPI_ISL_476194, EPI_ISL_476195, EPI_ISL_476196, EPI_ISL_476197, EPI_ISL_476202, EPI_ISL_476211, EPI_ISL_476212, EPI_ISL_476213, EPI_ISL_476214, EPI_ISL_476224, EPI_ISL_476225, EPI_ISL_476226, EPI_ISL_476227, EPI_ISL_476228, EPI_ISL_476229 | | | |
| see above | DB Diagnósticos do Brasil | Instituto de Medicina Tropical da Univesidade de São Paulo | Samples: Nelson Gaburo Jr; Sequencing: Ingra Morales Claro, Jaqueline Goes de Jesus, Erika Regina Manuli, Flavia Cristina da Silva Sales, Thais de Moura Coletti, Camila Alves Maia da Silva, Mariana Severo Ramundo, Giula Magalhaes Ferreira, Darlan da Silva Candido, Julien Theze, Nuno Faria, Ester Sabino |
| EPI_ISL_476428, EPI_ISL_476429, EPI_ISL_476433, EPI_ISL_476458, EPI_ISL_476471, EPI_ISL_476488 | Hospital da Clínicas da Faculdade de Medicina da Universidade de São Paulo | Instituto de Medicina Tropical da Univesidade de São Paulo | Samples: Ingra Morales Claro, Erika Regina Manuli, Cecilia Salete Alencar, Carolina S. Lazar, Silvia F. Costa; Sequencing: Ingra Morales Claro, Jaqueline Goes de Jesus, Erika Regina Manuli, Flavia Cristina da Silva Sales, Thais de Moura Coletti, Camila Alves Maia da Silva, Mariana Severo Ramundo, Giula Magalhaes Ferreira, Darlan da Silva Candido, Julien Theze, Nuno Faria, Ester Sabino |
| EPI_ISL_476777, EPI_ISL_476779, EPI_ISL_476780, EPI_ISL_476793 | Stanford clinical virology lab | Chan-Zuckerberg Biohub | Benjamin Pinksy, Katharine Walter, Victoria N. Parikh, John Gorzynski, Hannah N. DeJong, Matthew T. Wheeler, Jason Andrews, Manuel Rivas, Carlos Bustamante, Euan Ashley, with CZB Cliahub Consortium |
| EPI_ISL_476801, EPI_ISL_476803 | Hong Kong Department of Health | School of Public Health, The University of Hong Kong | Dominic N.C. Tsang, Daniel K.W. Chu, Leo L.M. Poon, Malik Peiris |
| EPI_ISL_476806 | Department of Laboratory Medicine Tan Tock Seng Hospital | Department of Laboratory Medicine Tan Tock Seng Hospital | Chen YYC, Zair X, Li C, Tang WY, Maurer-Stroh S, Barkham TMS, Nagarajan N, Sessions OM |
| EPI_ISL_476816, EPI_ISL_476817 | Department of Laboratory Medicine, Tan Tock Seng Hospital | Department of Laboratory Medicine, Tan Tock Seng Hospital | Chen YYC, Zair X, Li C, Tang WY, Maurer-Stroh S, Barkham TMS, Nagarajan N, Sessions OM |
| EPI_ISL_476822, EPI_ISL_476823, EPI_ISL_476824, EPI_ISL_476833, EPI_ISL_476834 | Laboratoire des Fièvres Hémorragiques Virales du Benin | Charité-Universitätsmedizin Berlin | Yadouleton, Anges; Sander Anna-Lena; Moreira-Soto Andres; Drexler, Jan Felix |
| EPI_ISL_476902, EPI_ISL_476905 | UW Virology Lab | UW Virology Lab | Pavitra Roychoudhury, Hong Xie, Lasata Shrestha, Amin Addetia, Truong Nguyen, Victoria M Rachleff, Meei-Li Huang, Keith R Jerome, Alexander Greninger |
| EPI_ISL_477171 | Department of Laboratory, Medicine Tan Tock Seng Hospital | Department of Laboratory, Medicine Tan Tock Seng Hospital | Chen YYC, Zair X, Li C, Tang WY, Maurer-Stroh S, Barkham TMS, Nagarajan N, Sessions OM |
| EPI_ISL_477172 | Department of Laboratory Medicine Tan Tock Seng Hospital | Department of Laboratory Medicine Tan Tock Seng Hospital | Chen YYC, Zair X, Li C, Tang WY, Maurer-Stroh S, Barkham TMS, Nagarajan N, Sessions OM |
| EPI_ISL_477193, EPI_ISL_477194 | Istituto Zooprofilattico Sperimentale Puglia e Basilicata; | Beaconlab (Bioinformatics, Evolution and Comparative Genomics lab), Dept of Biosciences, University on Mila | Parisi A.,Pesole G., Manzari C., Chiara M. |
| EPI_ISL_477291 | Mayo Clinic & Mayo Clinic Laboratories | Minnesota Department of Health, Public Health Laboratory | Matt Plumb, Jacob Garfin, Kelly Pung, and Xiong Wang |
| EPI_ISL_477685, EPI_ISL_477686, EPI_ISL_477687, EPI_ISL_477690 | UW Virology Lab | UW Virology Lab | Pavitra Roychoudhury, Hong Xie, Lasata Shrestha, Amin Addetia, Truong Nguyen, Victoria M Rachleff, Meei-Li Huang, Keith R Jerome, Alexander Greninger |
| EPI_ISL_477824 | West of Scotland Specialist Virology Centre, NHSGGC / MRC-University of Glasgow Centre for Virus Research | COVID-19 Genomics UK (COG-UK) Consortium | Ana da Silva Filipe, Natasha Johnson, Kathy Smollett, Daniel Mair, Stephen Carmichael, Lily Tong, Jenna Nichols, Elihu Aranday-Cortes, Kirstyn Brunker, Yasmin Parr, Alice Broos, Kyriaki Nomikou; Sarah McDonald, Marc Niebel, Patawee Asamaphan; Richard Orton, Joseph Hughes, Sreenu Vattipally, David L Robertson; Alasdair MacLean, Rory Gunson; Kathy Li, Natasha Jesudason, Rajiv Shah, James Shepherd, Antonia Ho, Emma Thomson |
| EPI_ISL_478536, EPI_ISL_478537, EPI_ISL_478538, EPI_ISL_478539, EPI_ISL_478540, EPI_ISL_478541 | Northumbria University / South Tees Hospitals NHS Foundation Trust / North Cumbria Integrated Care NHS Foundation Trust / North Tees and Hartlepool NHS Foundation Trust / Newcastle Hospitals NHS Foundation Trust | COVID-19 Genomics UK (COG-UK) Consortium | Darren L Smith,Andrew Nelson,Matthew Bashton,Greg R Young,Joshua Loh,John Allan,Mohammad A Tariq,Giles S Holt,Gary Black,Wen C Yew,Lynn Dover,Paul Baker,Steve Liggett,Sarah Essex,Jane Greenaway,Debra Padgett,Clive Graham,Garren Scott,Edward Barton,Emma Swindells,Brendan Payne,Jennifer Collins,Yusri Taha,Gary Eltringham |
| EPI_ISL_478696, EPI_ISL_478697, EPI_ISL_478698 | South Eastern Area Laboratory Services (SEALS) | NSW Health Pathology - Institute of Clinical Pathology and Medical Research; Westmead Hospital; University of Sydney | CIDM-PH et al. |
| EPI_ISL_479214, EPI_ISL_479218 | Virology Department, Sheffield Teaching Hospitals NHS Foundation Trust/Department of Infection, Immunity and Cardiovascular Disease, The Medical School, University of Sheffield | COVID-19 Genomics UK (COG-UK) Consortium | Thushan de Silva, Matthew Parker, Nikki Smith, Adri Angyal, Rebecca Brown, Luke Green, Rachel Tucker, Paul Parsons, Danielle Groves, Katie Johnson, Laura Carrilero, Alex Keeley, Dave Partridge, Matthew Wyles, Benjamin Lindsey, Mehmet Yavuz, Mohammad Raza, Cariad Evans |
| EPI_ISL_479664, EPI_ISL_479665, EPI_ISL_479666 | Center for Genomics and System Biology, New York University | Center for Genomics and System Biology, New York University | Roder,A., Banakis,S., Johnson,K., Khalfan,M., Borenstein,E.S., Samanovic,M., Cornelius,A., Herati,R., Ulrich,R., Fleming,A., Kottkamp,A., Raabe,V., Mulligan,M.J., Gresham,D., Ghedin,E. |
| EPI_ISL_479861, EPI_ISL_479868 | Department of Infectious Diseases, Kobe Institute of Health | Pathogen Genomics Center, National Institute of Infectious Diseases | Tsuyoshi Sekizuka, Ryohei Nomoto, Kentaro Itokawa, Rina Tanaka, Masanori Hashino, Hajime Kamiya, Motoi Suzuki, Makoto Kuroda |
| EPI_ISL_479881, EPI_ISL_479882, EPI_ISL_479883, EPI_ISL_479884 | Sapporo City Institute of Public Health | Pathogen Genomics Center, National Institute of Infectious Diseases | Tsuyoshi Sekizuka, Asami Ohnishi, Kentaro Itokawa, Rina Tanaka, Masanori Hashino, Hajime Kamiya, Motoi Suzuki, Makoto Kuroda |
| EPI_ISL_479891, EPI_ISL_479892, EPI_ISL_479893, EPI_ISL_479894, EPI_ISL_479895 | Tokyo Metropolitan Institute of Public Health | Pathogen Genomics Center, National Institute of Infectious Diseases | Tsuyoshi Sekizuka, Kenji Sadamasu, Takashi Chiba, Mami Nagashima, Kentaro Itokawa, Rina Tanaka, Masanori Hashino, Hajime Kamiya, Motoi Suzuki, Makoto Kuroda |
| EPI_ISL_479898, EPI_ISL_479899, EPI_ISL_479900 | Gunma Prefectural Institute of Public Health and Environmental Sciences | Pathogen Genomics Center, National Institute of Infectious Diseases | Tsuyoshi Sekizuka, Hiroyuki Tsukagoshi, Kentaro Itokawa, Rina Tanaka, Masanori Hashino, Hajime Kamiya, Motoi Suzuki, Makoto Kuroda |
| EPI_ISL_479908 | Himeji City Institute of Environment and Health | Pathogen Genomics Center, National Institute of Infectious Diseases | Tsuyoshi Sekizuka, Kentaro Itokawa, Rina Tanaka, Masanori Hashino, Hajime Kamiya, Motoi Suzuki, Makoto Kuroda |

| | | | |
|---|---|---|---|
| EPI_ISL_479924 | Niigata City Public Health Research Institute | Pathogen Genomics Center, National Institute of Infectious Diseases | Tsuyoshi Sekizuka, Yurie Takahashi, Kentaro Itokawa, Rina Tanaka, Masanori Hashino, Hajime Kamiya, Motoi Suzuki, Makoto Kuroda |
| EPI_ISL_479933, EPI_ISL_479934, EPI_ISL_479935 | Saitama Prefectural Institute of Public Health | Pathogen Genomics Center, National Institute of Infectious Diseases | Tsuyoshi Sekizuka, Hayato Ehara, Kentaro Itokawa, Rina Tanaka, Masanori Hashino, Hajime Kamiya, Motoi Suzuki, Makoto Kuroda |
| EPI_ISL_479937, EPI_ISL_479938, EPI_ISL_479939, EPI_ISL_479940 | Ibaraki Prefectural Institute of Public Health | Pathogen Genomics Center, National Institute of Infectious Diseases | Tsuyoshi Sekizuka, Keiko Goto, Kentaro Itokawa, Rina Tanaka, Masanori Hashino, Hajime Kamiya, Motoi Suzuki, Makoto Kuroda |
| EPI_ISL_479967 | Fukui Prefectural Institute of Public Health and Environmental Science | Pathogen Genomics Center, National Institute of Infectious Diseases | Tsuyoshi Sekizuka, Miho Toho, Kentaro Itokawa, Rina Tanaka, Masanori Hashino, Hajime Kamiya, Motoi Suzuki, Makoto Kuroda |
| EPI_ISL_480227 | Tokyo Metropolitan Institute of Public Health | Pathogen Genomics Center, National Institute of Infectious Diseases | Tsuyoshi Sekizuka, Kenji Sadamasu, Takashi Chiba, Mami Nagashima, Kentaro Itokawa, Rina Tanaka, Masanori Hashino, Hajime Kamiya, Motoi Suzuki, Makoto Kuroda |
| EPI_ISL_480312, EPI_ISL_480313, EPI_ISL_480314 | Hospital Mexico | Charité Virology-University of Costa Rica | Andres Moreira-Soto, Eugenia Corrales-Aguilar, Ignacio Postigo-Hidalgo, Teresita Somogyi, Jan Felix Drexler |
| EPI_ISL_480322, EPI_ISL_480323 | Hospital Nacional de Niños | Charité Virology-University of Costa Rica | Andres Moreira-Soto, Eugenia Corrales-Aguilar, Ignacio Postigo-Hidalgo, Cristian Pérez Corrales, Andrei Montero Bonilla, Jan Felix Drexler |
| EPI_ISL_480377, EPI_ISL_480379, EPI_ISL_480380, EPI_ISL_480381, EPI_ISL_480382, EPI_ISL_480383, EPI_ISL_480384, EPI_ISL_480385, EPI_ISL_480387 | University of Wisconsin-Madison AIDS Vaccine Research Laboratories | University of Wisconsin-Madison AIDS Vaccine Research Laboratories | Gage Moreno, Katarina Braun, et al. AIDS Vaccine Research Laboratories |
| EPI_ISL_480438 | Laboratorio de Biología Molecular Asociación Española Primera en Salud | Departments of Pathology and Medicine, New York University School of Medicine | Maria Victoria Elizondo, Maria Noel Zubillaga, Gonzalo Manrique, Paul Zappile, Gael Westby, Matthew T Maurano, Christian Marier, Adriana Heguy |
| EPI_ISL_480953, EPI_ISL_480955, EPI_ISL_480956, EPI_ISL_480957, EPI_ISL_480958, EPI_ISL_480965, EPI_ISL_480966, EPI_ISL_480967, EPI_ISL_480973, EPI_ISL_480976, EPI_ISL_480977, EPI_ISL_480978, EPI_ISL_480983, EPI_ISL_480984, EPI_ISL_480987, EPI_ISL_480988, EPI_ISL_480991, EPI_ISL_480993, EPI_ISL_480994, EPI_ISL_480996, EPI_ISL_480997, EPI_ISL_480998, EPI_ISL_481002, EPI_ISL_481004, EPI_ISL_481006, EPI_ISL_481008, EPI_ISL_481011, EPI_ISL_481012, EPI_ISL_481014, EPI_ISL_481019, EPI_ISL_481021, EPI_ISL_481022, EPI_ISL_481023, EPI_ISL_481028, EPI_ISL_481032, EPI_ISL_481037, EPI_ISL_481039, EPI_ISL_481040 | | | |
| see above | Servicio de Microbiología. Hospital Universitario Donostia. OSI Donostialdea. Área de Enfermedades Infecciosas, Grupo de Infección Respiratoria y Resistencia Antimicrobiana. Instituto de Investigación Sanitaria Biodonostia | SeqCOVID-SPAIN consortium/IBV(CSIC) | Gustavo Cilla, Milagrosa Montes, Luis Piñeiro, Jose Maria Marimón and SeqCOVID-SPAIN consortium |
| EPI_ISL_481043, EPI_ISL_481054 | Hospital General Universitario Gregorio Marañón | SeqCOVID-SPAIN consortium/IBV(CSIC) | Laura Pérez-Lago, Marta Herranz, Jon Sicilia, Julia Suárez, Pilar Catalán, Patricia Muñoz, Darío García de Viedma and SeqCOVID-SPAIN consortium |
| EPI_ISL_481569, EPI_ISL_481570 | Department of Virology and Immunology, University of Helsinki and Helsinki University Hospital, Huslab Finland | Department of Virology, Faculty of Medicine, University of Helsinki, Helsinki, Finland | Teemu Smura, Hannimari Kallio-Kokko, Jenni Virtanen, Maija Suvanto, Sari Hannula, Harri Kangas, Pekka Ellonen, Olli Vapalahti |
| EPI_ISL_482323, EPI_ISL_482463, EPI_ISL_482464 | Providence St. Joseph Health Molecular Genomics Laboratory | Providence St. Joseph Health Molecular Genomics Laboratory | Alexa K Dowdell, Brian D Piening, Fred L Robinson, Carlo B Bifulco, Mary Campbell |
| EPI_ISL_482470 | Queen Elizabeth II Health Science Centre | National Microbiology Laboratory | Anna Majer, Shari Tyson, Grace Seo, Kristyn Burak, Philip Mabon, Elsie Grudeski, Rhiannon Huzarewich, Russell Mandes, Jennifer Tanner, Natalie Knox, Morag Graham, Gary Van Domselaar, Todd Hatchette, Jason LeBlanc, Nathalie Bastien, Yan Li, Timothy Booth |
| EPI_ISL_482471, EPI_ISL_482472, EPI_ISL_482473 | Dr. Georges-L.-Dumont University Hospital Centre | National Microbiology Laboratory | Anna Majer, Shari Tyson, Grace Seo, Kristyn Burak, Philip Mabon, Elsie Grudeski, Rhiannon Huzarewich, Russell Mandes, Jennifer Tanner, Natalie Knox, Morag Graham, Gary Van Domselaar, Richard Garceau, Guillaume Desnoyers, Nathalie Bastien, Yan Li, Timothy Booth |
| EPI_ISL_482475 | Cadham Provincial Laboratory | National Microbiology Laboratory | Anna Majer, Shari Tyson, Grace Seo, Kristyn Burak, Philip Mabon, Elsie Grudeski, Rhiannon Huzarewich, Russell Mandes, Jennifer Tanner, Natalie Knox, Morag Graham, Gary Van Domselaar, Paul Van Caeseele, Jared Bullard, David Alexander, Kerry Dust, Nathalie Bastien, Yan Li, Timothy Booth, |
| EPI_ISL_482476, EPI_ISL_482477 | Queen Elizabeth II Health Science Centre | National Microbiology Laboratory | Anna Majer, Shari Tyson, Grace Seo, Kristyn Burak, Philip Mabon, Elsie Grudeski, Rhiannon Huzarewich, Russell Mandes, Jennifer Tanner, Natalie Knox, Morag Graham, Gary Van Domselaar, Todd Hatchette, Jason LeBlanc, Nathalie Bastien, Yan Li, Timothy Booth |
| EPI_ISL_482478, EPI_ISL_482481, EPI_ISL_482482 | Cadham Provincial Laboratory | National Microbiology Laboratory | Anna Majer, Shari Tyson, Grace Seo, Kristyn Burak, Philip Mabon, Elsie Grudeski, Rhiannon Huzarewich, Russell Mandes, Jennifer Tanner, Natalie Knox, Morag Graham, Gary Van Domselaar, Paul Van Caeseele, Jared Bullard, David Alexander, Kerry Dust, Nathalie Bastien, Yan Li, Timothy Booth, |
| EPI_ISL_482579, EPI_ISL_482580 | Hangzhou Center for Diseases Control and Prevention | Hangzhou Center for Diseases Control and Prevention | Jun Li, Haoqiu Wang, Lingfeng Mao, Hua Yu, Xinfen Yu, Zhou Sun, Xin Qian, Shuchang Chen, Junfang Chen, Xuchu Wang |
| EPI_ISL_482677 | Singapore General Hospital | Department of Microbiology | Nurdyana Abdul Rahman, Kun Lee Lim, Chenhao Li, Kian Sing Chan, Lynette Oon, Kern Rei Chng, Niranjan Nagarajan, Karrie Ko |
| EPI_ISL_482999, EPI_ISL_483000, EPI_ISL_483001, EPI_ISL_483003 | Minnesota Department of Health, Public Health Laboratory | Minnesota Department of Health, Public Health Laboratory | Matt Plumb, Jacob Garfin, and Xiong Wang |
| EPI_ISL_483059 | Hospital Universitari Germans Trias i Pujol | IrsiCaixa AIDS Research Lab | J. Segalés, M. Puig, J. Rodon, C. Avila-Nieto, J. Carrillo, G. Cantero, M.T. Terrón, S. Cruz, M. Parera ,M. Noguera-Julián, N. Izquierdo-Useros, V. Guallar, E. Vidal, A. Valencia, I. Blanco, J. Blanco, B. Clotet, J. Vergara-Alert |
| EPI_ISL_483077 | SA Pathology | SA Pathology | Lex Leong, Chuan Kok Lim, Mark Turra, Ivan Bastian, Geoff Higgins |
| EPI_ISL_483544, EPI_ISL_483546, EPI_ISL_483549, EPI_ISL_483550 | Kingdom of Bahrein Ministry of Health | Erasmus Medical Center | Bas Oude Munnink, David Nieuwenhuijse, Reina Sikkema, Fatema, Ebrahim Shehad, Amjad Ghanem Mohamed, Hashmeya Al Wasti, Claudia Schapendonk, Irina Chestakova, Anne van der Linden, Theo Bestebroer, Stefan van Nieuwkoop, Mark Pronk, Pascal Lexmond, Richard Molenkamp, Marion Koopmans, on behalf of the Dutch national COVID-19 response team. |
| EPI_ISL_483637 | National Laboratory of Virology, Szentágothai Research Centre | National Laboratory of Virology, Szentágothai Research Centre | Endre Gábor Tóth, Balázs Somogyi, Ferenc Jakab, Gábor Kemenesi |
| EPI_ISL_483704, EPI_ISL_483708, EPI_ISL_483715, EPI_ISL_483717 | Israel Central Virology laboratory | Israel Central Virology laboratory | Neta Zuckerman, Efrat Dahan Bucris, Oran Erster, Ella Mendelson, Michal Mandelboim |
| EPI_ISL_483922, EPI_ISL_483923, EPI_ISL_483924, EPI_ISL_483925, EPI_ISL_483926, EPI_ISL_483927, EPI_ISL_483928, EPI_ISL_483929, EPI_ISL_483930, EPI_ISL_483931, EPI_ISL_483932, EPI_ISL_483933, EPI_ISL_483934, EPI_ISL_483935, EPI_ISL_483936, EPI_ISL_483937, EPI_ISL_483938, EPI_ISL_483939, EPI_ISL_483940, EPI_ISL_483941, EPI_ISL_483942, EPI_ISL_483943, EPI_ISL_483944, EPI_ISL_483945, EPI_ISL_483946, EPI_ISL_483947, EPI_ISL_483948, EPI_ISL_483949, EPI_ISL_483950, EPI_ISL_483951, EPI_ISL_483952, EPI_ISL_483953, EPI_ISL_483954, EPI_ISL_483955, EPI_ISL_483957, EPI_ISL_483958, EPI_ISL_483969, EPI_ISL_484111, EPI_ISL_484197 | | | |
| see above | Centre for Clinical Infection and Diagnostics Research and Genomics Innovation Unit, Guy's and St. Thomas' NHS Trust | COVID-19 Genomics UK (COG-UK) Consortium | Chloe Fisher, Luke Snell, Penny Cliff, Rahul Batra, Jonathan Edgeworth, Ali Raza Awan |
| EPI_ISL_484339 | Queens Medical Centre, Clinical Microbiology Department / DeepSeq Nottingham | COVID-19 Genomics UK (COG-UK) Consortium | Gemma Clark, Wendy Smith, Manjinder Khakh, Vicki M Fleming, Michelle M Lister, Hannah Howson-Wells, Jonathan Ball, Patrick McClure, Joseph Chappell, Theocharis Tsoleridis, Nadine Holmes, Matthew Carlisle, Christopher Moore, Fei Sang, Johnny Debebe, Victoria Wright, Matthew Loose |
| EPI_ISL_484701, EPI_ISL_484705 | Department of Clinical Microbiology | GIGA Medical Genomics | Keith Durkin, Maria Artesi, Sébastien Bontems, Raphaël Boreux, Cécile Meex, Axelle Chaslain, Céline Fombellida-Lopez, Pierrette Melin, Marie-Pierre Hayette, Vincent Bours. |
| EPI_ISL_485848, EPI_ISL_485856, EPI_ISL_485857, EPI_ISL_485858, EPI_ISL_485859, EPI_ISL_485860, EPI_ISL_485861, EPI_ISL_485862, EPI_ISL_485863, EPI_ISL_485864, EPI_ISL_485865, EPI_ISL_485866, EPI_ISL_485867, EPI_ISL_485868, EPI_ISL_485869, EPI_ISL_485870 | | | |
| see above | Virginia DCLS | Virginia DCLS | Virginia DCLS |
| EPI_ISL_485971, EPI_ISL_486113 | UW Virology Lab | UW Virology Lab | Pavitra Roychoudhury, Hong Xie, Lasata Shrestha, Amin Addetia, Truong Nguyen, Victoria M Rachleff, Meei-Li Huang, Keith R Jerome, Alexander Greninger |
| EPI_ISL_486422, EPI_ISL_486423, EPI_ISL_486431, EPI_ISL_486432, | Latvijas Infektoloijas centrs | Latvian Biomedical Research and Study Centre | Ivars Silamielis, Kaspars Megnis, Monta Ustinova, ikita Zrelovs, Vita Rovte, Jeena Storoženko, Tatjana Kolupajeva, Oksana Savicka, Uga Dumpis, Jnis Kloviš |

| | | | |
|---|---|---|---|
| EPI_ISL_486433, EPI_ISL_486434 | | | |
| EPI_ISL_486650 | Microbiology, Virology and Biemergency Laboratory-ASST FBF Sacco | Microbiology, Virology and Biemergency Laboratory-ASST FBF Sacco | Romeri F, Comandatore F, Mancon A, Micheli V, Rimoldi SG |
| EPI_ISL_488335 | PHE South West Regional Laboratory, National Infection Service | Wellcome Sanger Institute for the COVID-19 Genomics UK (COG-UK) consortium | Stephanie Hutchings, Hannah Pymont, Dr Peter Muir, Barry Vipond, Rich Hopes; and Alex Alderton, Roberto Amato, Sonia Goncalves, Ewan Harrison, David K. Jackson, Ian Johnston, Dominic Kwiatkowski, Cordelia Langford, John Sillitoe on behalf of the Wellcome Sanger Institute COVID-19 Surveillance Team (http://www.sanger.ac.uk/covid-team) |
| EPI_ISL_488485, EPI_ISL_488508, EPI_ISL_488541, EPI_ISL_488588, EPI_ISL_488628, EPI_ISL_488707, EPI_ISL_488744, EPI_ISL_488749, EPI_ISL_488750, EPI_ISL_488757, EPI_ISL_488762, EPI_ISL_488766, EPI_ISL_488767, EPI_ISL_488772, EPI_ISL_488774, EPI_ISL_488775, EPI_ISL_488779, EPI_ISL_488785, EPI_ISL_488786, EPI_ISL_488792, EPI_ISL_488794, EPI_ISL_488797, EPI_ISL_488799, EPI_ISL_488801, EPI_ISL_488805, EPI_ISL_488809, EPI_ISL_488818, EPI_ISL_488823, EPI_ISL_488828 | | | |
| see above | NU-OMICS DNA Sequencing research facility, Northumbria University | Wellcome Sanger Institute for the COVID-19 Genomics UK (COG-UK) consortium | Chris Duncan, Sheia Waugh, Shirelle Burton-Fanning, Gary Eltringham, Jennifer Collins, Brendan Payne, Yusri Taha, Emma Swindells, Jane Greenaway, Edward Barton, Garren Scott, Debra Padgett, Clive Graham, Sarah Essex, Steve Ligget, Paul Baker, Lynn Dover, Wen Yew, Gary Black, John Allan, Joshua Loh, Greg Young, Matthew Bashton, Andrew Nelson, Darren Smith and Alex Alderton, Roberto Amato, Sonia Goncalves, Ewan Harrison, David K. Jackson, Ian Johnston, Dominic Kwiatkowski, Cordelia Langford, John Sillitoe on behalf of the Wellcome Sanger Institute COVID-19 Surveillance Team (http://www.sanger.ac.uk/covid-team) |
| EPI_ISL_489626, EPI_ISL_489630, EPI_ISL_489662, EPI_ISL_489688, EPI_ISL_489692, EPI_ISL_489700 | NHSGGC West of Scotland Specialist Virology Centre / MRC-University of Glasgow Centre for Virus Research | Wellcome Sanger Institute for the COVID-19 Genomics UK (COG-UK) consortium | Ana da Silva Filipe, Natasha Johnson, Kathy Smollett, Daniel Mair, Stephen Carmichael, Lily Tong, Jenna Nichols, Elihu Aranday-Cortes, Kirstyn Brunker, Yasmin Parr, Kyriaki Nomikou; Sarah McDonald, Marc Niebel, Patawee Asamaphan; Richard Orton, Joseph Hughes, Sreenu Vattipally, David L Robertson; Alasdair MacLean, Rory Gunson; Kathy Li, Natasha Jesudason, Rajiv Shah, James Shepherd, Antonia Ho, Alice Broos, Emma Thomson and Alex Alderton, Roberto Amato, Sonia Goncalves, Ewan Harrison, David K. Jackson, Ian Johnston, Dominic Kwiatkowski, Cordelia Langford, John Sillitoe on behalf of the Wellcome Sanger Institute COVID-19 Surveillance Team (http://www.sanger.ac.uk/covid-team) |
| EPI_ISL_489800, EPI_ISL_489801, EPI_ISL_489802, EPI_ISL_489803, EPI_ISL_489805, EPI_ISL_489808 | Florida Bureau of Public Health Laboratories | Florida Bureau of Public Health Laboratories | Sarah Schmedes, Jason Blanton |
| EPI_ISL_490018 | South Eastern Area Laboratory Services (SEALS) | NSW Health Pathology - Institute of Clinical Pathology and Medical Research; Westmead Hospital; University of Sydney | CIDM-PH et al. |
| EPI_ISL_490112 | The National Institute of Public Health | The National Institute of Public Health and State Veterinary Institute Prague | Nagy,A;Jirincova,H;Novakova,L;Trnka,D;Vecerova,J |
| EPI_ISL_490988, EPI_ISL_490997, EPI_ISL_490998, EPI_ISL_491000 | Mayo Clinic Laboratories | UW Virology Lab | Pavitra Roychoudhury, Hong Xie, Lasata Shrestha, Amin Addetia, Truong Nguyen, Victoria M Rachleff, Meei-Li Huang, Keith R Jerome, Alexander Greninger |
| EPI_ISL_491093 | The National Institute of Public Health | The National Institute of Public Health and State Veterinary Institute Prague | Nagy,A; Jirincova,H; Novakova,L; Trnka,D; Vecerova,J |
| EPI_ISL_491100 | SC Department of Health and Environmental Control | SC Department of Health and Environmental Control | Flores,H. |
| EPI_ISL_491117 | The National Institute of Public Health | The National Institute of Public Health and State Veterinary Institute Prague | Nagy,A;Jirincova,H;Novakova,L;Trnka,D;Vecerova,J |
| EPI_ISL_491444 | Area de Salud Escazu (Coopesana) | Inciensa, Instituto Costarricense de Investigación y Enseñanza en Nutrición y Salud | Francisco Duarte, Hebleen Brenes, Claudio Soto-Garita, Estela Cordero, Adriana Godinez & Melany Calderon |
| EPI_ISL_491445 | Area de Salud Mata Redonda | Inciensa, Instituto Costarricense de Investigación y Enseñanza en Nutrición y Salud | Francisco Duarte, Hebleen Brenes, Claudio Soto-Garita, Estela Cordero, Adriana Godinez & Melany Calderon |
| EPI_ISL_491447 | Hospital Fernando Escalante Pradilla | Inciensa, Instituto Costarricense de Investigación y Enseñanza en Nutrición y Salud | Francisco Duarte, Hebleen Brenes, Claudio Soto-Garita, Estela Cordero, Adriana Godinez & Melany Calderon |
| EPI_ISL_491448 | Hospital San Rafael de Alajuela | Inciensa, Instituto Costarricense de Investigación y Enseñanza en Nutrición y Salud | Francisco Duarte, Hebleen Brenes, Claudio Soto-Garita, Estela Cordero, Adriana Godinez & Melany Calderon |
| EPI_ISL_491449 | Area de Salud Alajuela Sur | Inciensa, Instituto Costarricense de Investigación y Enseñanza en Nutrición y Salud | Francisco Duarte, Hebleen Brenes, Claudio Soto-Garita, Estela Cordero, Adriana Godinez & Melany Calderon |
| EPI_ISL_491450, EPI_ISL_491454 | Hospital San Juan de Dios | Inciensa, Instituto Costarricense de Investigación y Enseñanza en Nutrición y Salud | Francisco Duarte, Hebleen Brenes, Claudio Soto-Garita, Estela Cordero, Adriana Godinez & Melany Calderon |
| EPI_ISL_491476 | BSL3 Lab, Pendik Veterinary Control Enstitue | Genomic Laboratory (GLAB), Istanbul Technical University | Mustafa HASOKSUZ, Fahriye SARAC, Osman ERGANIS, Serdar UZAR, Hakan ENUL, Cumhur ADIAY, Ahmet SAIT, Orbay SAYI, Kadir YESILBAG, Oguz KARABEY |
| EPI_ISL_492114, EPI_ISL_492115, EPI_ISL_492116, EPI_ISL_492121, EPI_ISL_492122, EPI_ISL_492172, EPI_ISL_492175, EPI_ISL_492176 | SA Pathology | SA Pathology | Lex Leong, Chuan Kok Lim, Mark Turra, Ivan Bastian, Geoff Higgins |
| EPI_ISL_492536, EPI_ISL_492544, EPI_ISL_492606, EPI_ISL_492659 | PHE South West Regional Laboratory, National Infection Service | Wellcome Sanger Institute for the COVID-19 Genomics UK (COG-UK) consortium | Stephanie Hutchings, Hannah Pymont, Dr Peter Muir, Barry Vipond, Rich Hopes; and Alex Alderton, Roberto Amato, Sonia Goncalves, Ewan Harrison, David K. Jackson, Ian Johnston, Dominic Kwiatkowski, Cordelia Langford, John Sillitoe on behalf of the Wellcome Sanger Institute COVID-19 Surveillance Team (http://www.sanger.ac.uk/covid-team) |
| EPI_ISL_492862 | Royal Free Hospital / Health Services Laboratories | Wellcome Sanger Institute for the COVID-19 Genomics UK (COG-UK) Consortium | Tanzina Haque, Tabitha Mahungu, Dianne Irish, Cate Goodlad, Jenny Cross, Judith Heaney and Alex Alderton, Roberto Amato, Sonia Goncalves, Ewan Harrison, David K. Jackson, Ian Johnston, Dominic Kwiatkowski, Cordelia Langford, John Sillitoe on behalf of the Wellcome Sanger Institute COVID-19 Surveillance Team |
| EPI_ISL_492978 | Department of Laboratory Medicine Tan Tock Seng Hospital | Department of Laboratory Medicine Tan Tock Seng Hospital | Chen YYC, Zair X, Li C, Tang WY, Maurer-Stroh S, Barkham TMS, Nagarajan N, Sessions OM |
| EPI_ISL_492980 | IRCCS Sacro Cuore Don Calabria Hospital, Department of Infectious, Tropical Diseases & Microbiology | University of Verona, Department of Biotechnology | Antonio Mori, Michela Deiana, Elena Pomari, Chiara Piubelli; Giulia Lopatriello, Luca Marcolungo, Cristina Beltrami, Chiara Degli Esposti, Emanuela Cosentino, Massimo Delledonne |
| EPI_ISL_493198, EPI_ISL_493199, EPI_ISL_493200 | Virology Lab,Department of Pathology, National Cheng Kung University Hospital | Virology Lab,Department of Pathology, National Cheng Kung University Hospital | Huey-Pin Tsai, et al |
| EPI_ISL_493328 | INMI Lazzaro Spallanzani IRCCS | INMI Lazzaro Spallanzani IRCCS | Martina Rueca, Cesare E.M. Gruber, Barbara Bartolini, Francesco Messina, Maria R. Capobianchi, Antonino Di Caro |
| EPI_ISL_493741 | West of Scotland Specialist Virology Centre, NHSGGC / MRC-University of Glasgow Centre for Virus Research | COVID-19 Genomics UK (COG-UK) Consortium | Ana da Silva Filipe, Natasha Johnson, Kathy Smollett, Daniel Mair, Stephen Carmichael, Lily Tong, Jenna Nichols, Elihu Aranday-Cortes, Kirstyn Brunker, Yasmin Parr, Alice Broos, Kyriaki Nomikou; Sarah McDonald, Marc Niebel, Patawee Asamaphan; Richard Orton, Joseph Hughes, Sreenu Vattipally, David L Robertson; Alasdair MacLean, Rory Gunson; Kathy Li, Natasha Jesudason, Rajiv Shah, James Shepherd, Antonia Ho, Emma Thomson |
| EPI_ISL_494575, EPI_ISL_494592 | San Diego County Public Health Laboratory | Andersen lab at Scripps Research | SEARCH Alliance San Diego with Tracy Basler, Jovan Shephard, Brett Austin |
| EPI_ISL_495389 | Florida Bureau of Public Health Laboratories | Florida Bureau of Public Health Laboratories | Sarah Schmedes, Jason Blanton |
| EPI_ISL_495610 | Minnesota Department of Health, Public Health Laboratory | Minnesota Department of Health, Public Health Laboratory | Matt Plumb, Jacob Garfin, and Xiong Wang |
| EPI_ISL_496482 | Dept. Infectious, Tropical Diseases & Microbiology, IRCCS Sacro Cuore Don Calabria Hospital | 1) Dept. Infectious, Tropical Diseases & Microbiology, IRCCS Sacro Cuore Don Calabria Hospital; 2) Centro Piattaforme Tecnologiche, University of Verona; 3) Dept. Neurosciences, Biomedicine and Movement Sciences, University of Verona. | 1) Antonio Mori, Michela Deiana, Elena Pomari, Chiara Piubelli; 2) Monica Castellucci and Francesca Griggio; 3) Giovanni Malerba |

| | | | |
|---|---|---|---|
| EPI_ISL_496644, EPI_ISL_496645, EPI_ISL_496657, EPI_ISL_496659, EPI_ISL_496668, EPI_ISL_496669, EPI_ISL_496674, EPI_ISL_496678, EPI_ISL_496680, EPI_ISL_496684, EPI_ISL_496689, EPI_ISL_496690, EPI_ISL_496694, EPI_ISL_496696, EPI_ISL_496704, EPI_ISL_496714, EPI_ISL_496718, EPI_ISL_496739, EPI_ISL_496747, EPI_ISL_496748 | | | |
| see above | Gorgas Memorial Laboratory of Health Studies | Gorgas Memorial Laboratory of Health Studies | Danilo Franco, Claudia Gonzalez Sandra Lopez-Verges, Alexander A Martinez |
| EPI_ISL_496919 | Minnesota Department of Health, Public Health Laboratory | Minnesota Department of Health, Public Health Laboratory | Matt Plumb, Jacob Garfin, and Xiong Wang |
| EPI_ISL_497736 | Instituto Nacional de Salud, Bogotá, Colombia | Instituto Nacional de Salud, Bogotá, Colombia | Katherine Laiton-Donato, Diego A. Álvarez-Díaz, Carlos Franco-Muñoz, Jonathan Reales, Diego Andrés Prada, Jose A. Usme-Ciro, Nicolas D. Franco-Sierra, Zulma M. Cucunubá, Christian Julian VillabonaArenas, Liz Villabona-Arenas, Sussy Echeverria, Astrid C. Flórez, Carolina Ferro, Diana Marcela Walteros-Acero, Franklin Prieto, Carlos Andrés Durán, Martha Lucia Ospina Martinez, Marcela Mercado-Reyes |
| EPI_ISL_497773, EPI_ISL_497796, EPI_ISL_497799, EPI_ISL_497802, EPI_ISL_497805, EPI_ISL_497810, EPI_ISL_497820, EPI_ISL_497840, EPI_ISL_497845, EPI_ISL_497850, EPI_ISL_497864, EPI_ISL_497870 | | | |
| see above | Department of Microbiology, The University of Hong Kong | Department of Microbiology, The University of Hong Kong | Kelvin K.W. To, Kwok-Yung Yuen |
| EPI_ISL_498018, EPI_ISL_498019, EPI_ISL_498020, EPI_ISL_498021, EPI_ISL_498022, EPI_ISL_498023, EPI_ISL_498024, EPI_ISL_498025, EPI_ISL_498026, EPI_ISL_498027, EPI_ISL_498028, EPI_ISL_498029, EPI_ISL_498030, EPI_ISL_498031, EPI_ISL_498032, EPI_ISL_498033 | | | |
| see above | Division of Viral Diseases, Center for Laboratory Control of Infectious Diseases, Korea Centers for Diseases Control and Prevention | Division of Viral Diseases, Center for Laboratory Control of Infectious Diseases, Korea Centers for Diseases Control and Prevention | Jeong-Min Kim, Yoon-Seok Chung, Namjoo Lee, Sang Hee Woo, Hye-Jun Jo, Heui Man Kim, Jun-Sub Kim, Dong Hyun Song, Daesang Lee, Seong Tae Jeong, Myung Guk Han |
| EPI_ISL_498036 | Division of Viral Diseases, Center for Laboratory Control of Infectious Diseases, Korea Centers for Diseases Control and Prevention | Division of Viral Diseases, Center for Laboratory Control of Infectious Diseases, Korea Centers for Diseases Control and Prevention | Jeong-Min Kim, Yoon-Seok Chung, Namjoo Lee, Sang Hee Woo, Hye-Jun Jo, Heui Man Kim, Jun-Sub Kim, Myung Guk Han |
| EPI_ISL_498037 | Division of Viral Diseases, Center for Laboratory Control of Infectious Diseases, Korea Centers for Diseases Control and Prevention | Division of Viral Diseases, Center for Laboratory Control of Infectious Diseases, Korea Centers for Diseases Control and Prevention | Jeong-Min Kim, Yoon-Seok Chung, Namjoo Lee, Sang Hee Woo, Hye-Jun Jo, Heui Man Kim, Jun-Sub Kim, Dong Hyun Song, Daesang Lee, Seong Tae Jeong, Myung Guk Han |
| EPI_ISL_498176, EPI_ISL_498181, EPI_ISL_498186 | OUCRU | OUCRU | Nguyen Van Vinh Chau, Nguyen Thi Thu Hong, Nguyen Thi Han Ny, Le Nguyen Truc Nhu, Nghiem My Ngoc, Vo Thanh Lam, Nguyen Thanh Dung, Lam Minh Yen, Ngo Ngoc Quang Minh, Le Manh Hung, Nguyen Tri Dung, Dinh Nguyen Huy Man, Lam Anh Nguyet, Tran Chanh Xuan, Tran Tinh Hien, Nguyen Thanh Phong, Tran Nguyen Hoang Tu, Tran Tan Thanh, Nguyen Thanh Truong, Nguyen Tan Binh, Tang Chi Thuong, Guy Thwaites, and Le Van Tan, for OUCRU COVID-19 research group* |
| EPI_ISL_498266 | Ramathibodi Hospital | COVID-19 Network Investigations (CONI) Alliance | Elizabeth Batty, Wasun Chantratita, Thanat Chookajorn, Stefan Fernandez, Angkana Huang, Anthony R. Jones, Khajohn Joonsalak, Chonticha Klungtong, Theerarat Kochakarn, Namfon Kotanan, Krittikorn Kumpornsin, Wudtichai Manasatienkij, Bhakbhoom Panthan, Ekawat Pasomsub, Kingkan Rakmanee, Insee Sensorn, Janjira Thaipadungpanit, Arporn Wangwiwatsin,Treewat Watthanachockchai |
| EPI_ISL_498481, EPI_ISL_498492, EPI_ISL_498514, EPI_ISL_498529, EPI_ISL_498538, EPI_ISL_498542 | ACT Pathology | Schwessinger Lab | Ashley Jones, Benjamin Schwessinger, Robert Lanfear, Robyn N Hall, Megan McDonald, Ming-Dao Chia, Kevin Murray, Craig Kennedy, Karina Kennedy |
| EPI_ISL_498552 | Lebanese American University | Lebanese American University | Abi Habib,W., Abdallah,J., El Shesheny,R., Mokhbat,J., Webby,R.J., Goldstein,J. and Kayali,G. |
| EPI_ISL_499301, EPI_ISL_499302 | Centre for Enzyme Innovation, University of Portsmouth / Translational Research Laboratory, Portsmouth Hospitals NHS Trust | COVID-19 Genomics UK (COG-UK) Consortium | Angela Beckett,Yann Bourgeois,Garry Scarlett,Sharon Glaysher,Scott Elliott,Kelly Bicknell,Robert Impey,Allyson Lloyd,Sarah Wyllie,Ethan Butcher,Anoop Chauhan,Samuel Robson |
| EPI_ISL_499460, EPI_ISL_499462, EPI_ISL_499463, EPI_ISL_499464, EPI_ISL_499467, EPI_ISL_499469, EPI_ISL_499473, EPI_ISL_499475, EPI_ISL_499476, EPI_ISL_499477, EPI_ISL_499481, EPI_ISL_499483, EPI_ISL_499486, EPI_ISL_499487, EPI_ISL_499488, EPI_ISL_499489, EPI_ISL_499496, EPI_ISL_499498, EPI_ISL_499499, EPI_ISL_499500, EPI_ISL_499501, EPI_ISL_499509, EPI_ISL_499511, EPI_ISL_499513, EPI_ISL_499514, EPI_ISL_499516, EPI_ISL_499518, EPI_ISL_499521, EPI_ISL_499522, EPI_ISL_499526, EPI_ISL_499527, EPI_ISL_499528, EPI_ISL_499529, EPI_ISL_499531, EPI_ISL_499532, EPI_ISL_499533, EPI_ISL_499540, EPI_ISL_499542, EPI_ISL_499544, EPI_ISL_499566, EPI_ISL_499567, EPI_ISL_499569, EPI_ISL_499570, EPI_ISL_499571, EPI_ISL_499573, EPI_ISL_499574, EPI_ISL_499575, EPI_ISL_499576, EPI_ISL_499577, EPI_ISL_499580, EPI_ISL_499581, EPI_ISL_499582, EPI_ISL_499583, EPI_ISL_499584, EPI_ISL_499590, EPI_ISL_499594, EPI_ISL_499600, EPI_ISL_499605, EPI_ISL_499607, EPI_ISL_499608, EPI_ISL_499611, EPI_ISL_499613, EPI_ISL_499616, EPI_ISL_499617, EPI_ISL_499621, EPI_ISL_499624, EPI_ISL_499627, EPI_ISL_499628, EPI_ISL_499632, EPI_ISL_499633, EPI_ISL_499634, EPI_ISL_499635, EPI_ISL_499636, EPI_ISL_499637, EPI_ISL_499639, EPI_ISL_499644, EPI_ISL_499646, EPI_ISL_499647, EPI_ISL_499648, EPI_ISL_499649, EPI_ISL_499650, EPI_ISL_499651, EPI_ISL_499652, EPI_ISL_499653, EPI_ISL_499654, EPI_ISL_499655, EPI_ISL_499656, EPI_ISL_499657, EPI_ISL_499658, EPI_ISL_499660, EPI_ISL_499661, EPI_ISL_499662, EPI_ISL_499700, EPI_ISL_499702, EPI_ISL_499703, EPI_ISL_499705, EPI_ISL_499707, EPI_ISL_499708, EPI_ISL_499709, EPI_ISL_499710, EPI_ISL_499711, EPI_ISL_499712, EPI_ISL_499713, EPI_ISL_499714, EPI_ISL_499715, EPI_ISL_499716, EPI_ISL_499717, EPI_ISL_499718, EPI_ISL_499719, EPI_ISL_499720, EPI_ISL_499721, EPI_ISL_499722, EPI_ISL_499723, EPI_ISL_499728, EPI_ISL_499729, EPI_ISL_499734, EPI_ISL_499735, EPI_ISL_499736, EPI_ISL_499737, EPI_ISL_499740, EPI_ISL_499746, EPI_ISL_499747, EPI_ISL_499751, EPI_ISL_499752, EPI_ISL_499753, EPI_ISL_499754, EPI_ISL_499757, EPI_ISL_499758, EPI_ISL_499760, EPI_ISL_499761, EPI_ISL_499762, EPI_ISL_499763, EPI_ISL_499764, EPI_ISL_499765, EPI_ISL_499767, EPI_ISL_499768, EPI_ISL_499769, EPI_ISL_499771, EPI_ISL_499772, EPI_ISL_499773, EPI_ISL_499774, EPI_ISL_499775 | | | |
| see above | Liverpool Clinical Laboratories | COVID-19 Genomics UK (COG-UK) Consortium | Sam Haldenby, Anita Lucaci, Steve Paterson, Julian Hiscox, Alistair Darby, M Almsaud, A Alrezaihi, Muhannad Alruwaili, Stuart D Armstrong, Jones Benjamin, Eleanor G Bentley, Anu Chawla, Jordan J Clark, Angela Cowell, Richard Eccles, Isabel García-Dorival, Matthew Gemmell, Alessandro Gerada, PKF Gilmore, Richard Gregory, Ximeng Han, Catherine Hartley, Margaret Hughes, Miren Iturriza-Gomara, James Johnson, L Luu, Jenifer Manson, Charlotte Nelson, Elaine O'Toole, Cassie Olateju, Rebekah Penrice-Randal , Lucille Rainbow, N.P Randle, Trevor Ian Robinson, Parul Sharma, Ghada T Shawli, James P Stewart, Neil Swainston, Ecaterina Vamos, Joanne Watts, Mark Whitehead |
| EPI_ISL_500219 | Hospital Clínico Universitario de Santiago de Compostela | SeqCOVID-SPAIN consortium/IBV(CSIC) | José Javier Costa Alcalde, Antonio Aguilera Guirao, Mª Luisa Pérez del Molino Bernal, Amparo Coira Nieto, Gema Barbeito Castiñeiras, Rocio Trastoy Pena and SeqCOVID-SPAIN consortium |
| EPI_ISL_500503, EPI_ISL_500504, EPI_ISL_500508, EPI_ISL_500509, EPI_ISL_500511, EPI_ISL_500513, EPI_ISL_500514, EPI_ISL_500516, EPI_ISL_500519, EPI_ISL_500520, EPI_ISL_500521, EPI_ISL_500526, EPI_ISL_500528, EPI_ISL_500537, EPI_ISL_500538 | | | |
| see above | Mayo Clinic Laboratories | University of Washington Virology Lab | Pavitra Roychoudhury, Hong Xie, Lasata Shrestha, Amin Addetia, Truong Nguyen, Victoria M Rachleff, Meei-Li Huang, Keith R Jerome, Alexander Greninger |
| EPI_ISL_500596, EPI_ISL_500629, EPI_ISL_500630, EPI_ISL_500631, EPI_ISL_500643, EPI_ISL_500644, EPI_ISL_500646, EPI_ISL_500676 | Area of Virology, Serology and Virology Division (SAViD), New South Wales Health Pathology Randwick | Area of Virology, Serology and Virology Division (SAViD), New South Wales Health Pathology Randwick | Rawlinson, W. |
| EPI_ISL_500716 | BSL3 Lab Pendik Veterinary Control Institute | Department of Medicinal Genetics, Bursa Uluda University, Faculty of medicine By Sehime Gülsün Temel, Adem Alemdar, Kadir Yeilba | Mustafa HASOKSUZ, Fahriye SARAC, Osman ERGANIS, Serdar UZAR, Hakan ENUL, Cumhur ADIAY, Ahmet SAIT, Orbay SAYI, Kadir YESILBAG, Oguz KARABEY |
| EPI_ISL_500831 | Virginia DCLS | Virginia DCLS | Virginia DCLS |
| EPI_ISL_501073, EPI_ISL_501078 | Mayo Clinic Laboratories | University of Washington Virology Lab | Pavitra Roychoudhury, Hong Xie, Lasata Shrestha, Amin Addetia, Truong Nguyen, Victoria M Rachleff, Meei-Li Huang, Keith R Jerome, Alexander Greninger |
| EPI_ISL_501221, EPI_ISL_501223, EPI_ISL_501224 | Department of Medical Microbiology, University Malaya Medical Centre | Department of Medical Microbiology, Faculty of Medicine, University of Malaya | Yoong Min CHONG, Jennifer Chong, I-Ching SAM, Yoke Fun CHAN, University Malaya Medical Centre COVID Team |
| EPI_ISL_507016, EPI_ISL_507017, EPI_ISL_507018, EPI_ISL_507019 | unknown | Infectious Diseases Research, King Abdullah International Medical Research Center (KAIMRC) | Alghoribi,M.F. |
| EPI_ISL_507052, EPI_ISL_507053, EPI_ISL_507054, EPI_ISL_507055, EPI_ISL_507056, EPI_ISL_507085 | University College London Hospital | COVID-19 Genomics UK (COG-UK) Consortium | Judith Heaney, Matthew Byott, Catherine Houlihan, Dan Frampton, Stuart Kirk, Moira Spyer and Eleni Nastouli |
| EPI_ISL_507291 | WHO National Influenza Centre Russian Federation | WHO National Influenza Centre Russian Federation | Andrey Komissarov, Artem Fadeev, Mariia Sergeeva, Anna Ivanova, Daria Danilenko |
| EPI_ISL_507707, EPI_ISL_507721, EPI_ISL_507743 | Michigan Department of Health and Human Services, Bureau of Laboratories | Michigan Department of Health and Human Services, Bureau of Laboratories | Blankenship HM, Riner D, Soehnlen MK |
| EPI_ISL_508124, EPI_ISL_508130, EPI_ISL_508143, EPI_ISL_508144 | SA Pathology | SA Pathology | Lex Leong, Chuan Kok Lim, Mark Turra, Ivan Bastian, Geoff Higgins |

| | | | |
|---|---|---|---|
| EPI_ISL_508615, EPI_ISL_508616, EPI_ISL_508617, EPI_ISL_508618, EPI_ISL_508619, EPI_ISL_508620, EPI_ISL_508621, EPI_ISL_508622, EPI_ISL_508623, EPI_ISL_508624, EPI_ISL_508634, EPI_ISL_508636, EPI_ISL_508637, EPI_ISL_508638, EPI_ISL_508639, EPI_ISL_508641, EPI_ISL_508642, EPI_ISL_508643, EPI_ISL_508644, EPI_ISL_508645, EPI_ISL_508675, EPI_ISL_508678, EPI_ISL_508681, EPI_ISL_508682, EPI_ISL_508684 | | | |
| see above | Departamento de Microbiología, CDB, Hospital Clínic, Barcelona | SeqCOVID-SPAIN consortium/IBV(CSIC) | Andrea Vergara, Mikel Martínez, Elisa Rubio, Jéssica Navero, Aida Peiró and SeqCOVID-SPAIN consortium |
| EPI_ISL_508770, EPI_ISL_508771, EPI_ISL_508773, EPI_ISL_508805, EPI_ISL_508807 | Florida Bureau of Public Health Laboratories | Florida Bureau of Public Health Laboratories | Sarah Schmedes, Jason Blanton |
| EPI_ISL_508864 | Division of Infectious Diseases and Hospital Epidemiology, University Hospital Zürich | Institute of Medical Virology, University of Zurich | Verena Kufner, Maryam Zaheri, Dana Weissberg, Jürg Böni, Silvana K. Rampini, Peter W. Schreiber, Irene A. Abela, Hugo Sax, Aline Wolfensberger, Michael Huber |
| EPI_ISL_508865, EPI_ISL_508867 | Division of Infectious Diseases and Hospital Epidemiology, University Hospital Zürich | Institute of Medical Virology, University of Zurich | Maryam Zaheri, Verena Kufner, Dana Weissberg, Jürg Böni, Silvana K. Rampini, Peter W. Schreiber, Irene A. Abela, Hugo Sax, Aline Wolfensberger, Michael Huber |
| EPI_ISL_508868 | Division of Infectious Diseases and Hospital Epidemiology, University Hospital Zürich | Institute of Medical Virology, University of Zurich | Verena Kufner, Maryam Zaheri, Dana Weissberg, Jürg Böni, Silvana K. Rampini, Peter W. Schreiber, Irene A. Abela, Hugo Sax, Aline Wolfensberger, Michael Huber |
| EPI_ISL_508869 | Division of Infectious Diseases and Hospital Epidemiology, University Hospital Zürich | Institute of Medical Virology, University of Zurich | Maryam Zaheri, Verena Kufner, Dana Weissberg, Jürg Böni, Silvana K. Rampini, Peter W. Schreiber, Irene A. Abela, Hugo Sax, Aline Wolfensberger, Michael Huber |
| EPI_ISL_508871, EPI_ISL_508877 | Institut des Agents Infectieux (IAI), Hospices Civils de Lyon | CNR Virus des Infections Respiratoires - France SUD | Antonin Bal, Gregory Destras, Gwendolyne Burfin, Solenne Brun, Carine Moustaud, Raphaelle Lamy, Alexandre Gaymard, Maude Bouscambert-Duchamp, Florence Morfin-Sherpa, Martine Valette, Bruno Lina, Laurence Josset |
| EPI_ISL_508878 | GH Les Portes du Sud | CNR Virus des Infections Respiratoires - France SUD | Antonin Bal, Gregory Destras, Gwendolyne Burfin, Solenne Brun, Carine Moustaud, Raphaelle Lamy, Alexandre Gaymard, Maude Bouscambert-Duchamp, Florence Morfin-Sherpa, Martine Valette, Bruno Lina, Laurence Josset |
| EPI_ISL_508879, EPI_ISL_508880 | Centre Hospitalier Saint Joseph Saint Luc | CNR Virus des Infections Respiratoires - France SUD | Antonin Bal, Gregory Destras, Gwendolyne Burfin, Solenne Brun, Carine Moustaud, Raphaelle Lamy, Alexandre Gaymard, Maude Bouscambert-Duchamp, Florence Morfin-Sherpa, Martine Valette, Bruno Lina, Laurence Josset |
| EPI_ISL_508881 | Centre Hospitalier de Valence | CNR Virus des Infections Respiratoires - France SUD | Antonin Bal, Gregory Destras, Gwendolyne Burfin, Solenne Brun, Carine Moustaud, Raphaelle Lamy, Alexandre Gaymard, Maude Bouscambert-Duchamp, Florence Morfin-Sherpa, Martine Valette, Bruno Lina, Laurence Josset |
| EPI_ISL_508882, EPI_ISL_508883, EPI_ISL_508909 | Institut des Agents Infectieux (IAI), Hospices Civils de Lyon | CNR Virus des Infections Respiratoires - France SUD | Antonin Bal, Gregory Destras, Gwendolyne Burfin, Solenne Brun, Carine Moustaud, Raphaelle Lamy, Alexandre Gaymard, Maude Bouscambert-Duchamp, Florence Morfin-Sherpa, Martine Valette, Bruno Lina, Laurence Josset |
| EPI_ISL_508998 | Centre Hospitalier Pierre Oudot | CNR Virus des Infections Respiratoires - France SUD | Antonin Bal, Gregory Destras, Gwendolyne Burfin, Solenne Brun, Carine Moustaud, Raphaelle Lamy, Alexandre Gaymard, Maude Bouscambert-Duchamp, Florence Morfin-Sherpa, Martine Valette, Bruno Lina, Laurence Josset |
| EPI_ISL_508999, EPI_ISL_509000, EPI_ISL_509001 | CNR Virus des Infections Respiratoires - France SUD | CNR Virus des Infections Respiratoires - France SUD | Antonin Bal, Gregory Destras, Gwendolyne Burfin, Solenne Brun, Carine Moustaud, Raphaelle Lamy, Alexandre Gaymard, Maude Bouscambert-Duchamp, Florence Morfin-Sherpa, Martine Valette, Bruno Lina, Laurence Josset |
| EPI_ISL_509003, EPI_ISL_509004 | GH Les Portes du Sud | CNR Virus des Infections Respiratoires - France SUD | Antonin Bal, Gregory Destras, Gwendolyne Burfin, Solenne Brun, Carine Moustaud, Raphaelle Lamy, Alexandre Gaymard, Maude Bouscambert-Duchamp, Florence Morfin-Sherpa, Martine Valette, Bruno Lina, Laurence Josset |
| EPI_ISL_509501, EPI_ISL_509502, EPI_ISL_509504 | Area of Virology, Serology and Virology Division (SAViD), New South Wales Health Pathology Randwick | Area of Virology, Serology and Virology Division (SAViD), New South Wales Health Pathology Randwick | Rawlinson, W. |
| EPI_ISL_509603, EPI_ISL_509604, EPI_ISL_509605, EPI_ISL_509606, EPI_ISL_509608, EPI_ISL_509609, EPI_ISL_509611 | Servicio de Microbiología. HRU de Málaga. Servicio Andaluz de Salud | SeqCOVID-SPAIN consortium/IBV(CSIC) | Inmaculada de Toro Peinado. MªConcepción Mediavilla Gradolph. Begoña Palop Borrás and SeqCOVID-SPAIN consortium |
| EPI_ISL_509616, EPI_ISL_509617, EPI_ISL_509618 | Hospital Universitario Araba. Vitoria-Gasteiz | SeqCOVID-SPAIN consortium/IBV(CSIC) | Silvia Hernáez Crespo, Carmen Gómez González, Amaia Aguirre Quiñonero, Marina Fernández Torres, Mª Rosario Almela Ferrer, Mª Concepción Lecaroz Agara, Andrés Canut Blasco and SeqCOVID-SPAIN consortium |
| EPI_ISL_509619, EPI_ISL_509620, EPI_ISL_509621, EPI_ISL_509622, EPI_ISL_509623, EPI_ISL_509624, EPI_ISL_509625, EPI_ISL_509626, EPI_ISL_509627, EPI_ISL_509628, EPI_ISL_509629, EPI_ISL_509630, EPI_ISL_509631, EPI_ISL_509632 | | | |
| see above | Servicio de Microbiología. HRU de Málaga. Servicio Andaluz de Salud | SeqCOVID-SPAIN consortium/IBV(CSIC) | Inmaculada de Toro Peinado. MªConcepción Mediavilla Gradolph. Begoña Palop Borrás and SeqCOVID-SPAIN consortium |
| EPI_ISL_509696, EPI_ISL_509697, EPI_ISL_509698, EPI_ISL_509699, EPI_ISL_509700 | Guatemala Ministry of Public Health | Pathogen Discovery, Respiratory Viruses Branch, Division of Viral Diseases, Centers for Disease Control and Prevention | Ying Tao, Jing Zhang, Krista Queen, Anna Uehara, Yan Li, Clinton Paden, Haibin Wang, Suxiang Tong |
| EPI_ISL_509710 | Guatemala Ministry of Public Health | Pathogen Discovery, Respiratory Viruses Branch, Division of Viral Diseases, Centers for Disease Control and Prevention | Jing Zhang, Ying Tao, Krista Queen, Anna Uehara, Yan Li, Clinton Paden, Haibin Wang, Suxiang Tong |
| EPI_ISL_509723, EPI_ISL_509733, EPI_ISL_509736 | Florida Bureau of Public Health Laboratories | Florida Bureau of Public Health Laboratories | Sarah Schmedes, Jason Blanton |
| EPI_ISL_510149, EPI_ISL_510171, EPI_ISL_510172, EPI_ISL_510173, EPI_ISL_510174, EPI_ISL_510175, EPI_ISL_510176, EPI_ISL_510177, EPI_ISL_510178, EPI_ISL_510179, EPI_ISL_510180, EPI_ISL_510181, EPI_ISL_510182, EPI_ISL_510183, EPI_ISL_510184, EPI_ISL_510185, EPI_ISL_510189, EPI_ISL_510190, EPI_ISL_510191, EPI_ISL_510192, EPI_ISL_510193, EPI_ISL_510194, EPI_ISL_510195, EPI_ISL_510196, EPI_ISL_510197, EPI_ISL_510198, EPI_ISL_510199, EPI_ISL_510200, EPI_ISL_510201, EPI_ISL_510202, EPI_ISL_510203, EPI_ISL_510204, EPI_ISL_510205, EPI_ISL_510206, EPI_ISL_510207, EPI_ISL_510208, EPI_ISL_510209, EPI_ISL_510210, EPI_ISL_510211, EPI_ISL_510212, EPI_ISL_510213, EPI_ISL_510214, EPI_ISL_510215, EPI_ISL_510216, EPI_ISL_510217, EPI_ISL_510218, EPI_ISL_510219 | | | |
| see above | Hospital General Universitario Gregorio Marañón | SeqCOVID-SPAIN consortium/IBV(CSIC) | Laura Pérez-Lago, Marta Herranz, Jon Sicilia, Julia Suárez, Pilar Catalán, Patricia Muñoz, Darío García de Viedma and SeqCOVID-SPAIN consortium |
| EPI_ISL_510247, EPI_ISL_510248, EPI_ISL_510249, EPI_ISL_510250, EPI_ISL_510251, EPI_ISL_510252, EPI_ISL_510253, EPI_ISL_510254, EPI_ISL_510255, EPI_ISL_510256, EPI_ISL_510257, EPI_ISL_510258, EPI_ISL_510259, EPI_ISL_510260, EPI_ISL_510261, EPI_ISL_510262, EPI_ISL_510263, EPI_ISL_510264 | | | |
| see above | Hospital de la Santa Creu i Sant Pau. Servicio de Microbiología | SeqCOVID-SPAIN consortium/IBV(CSIC) | Ferran Navarro, Núria Rabella, Elisenda Miró and SeqCOVID-SPAIN consortium |
| EPI_ISL_510277, EPI_ISL_510278, EPI_ISL_510279, EPI_ISL_510280, EPI_ISL_510281, EPI_ISL_510282, EPI_ISL_510283, EPI_ISL_510284, EPI_ISL_510285, EPI_ISL_510286, EPI_ISL_510287, EPI_ISL_510288, EPI_ISL_510289, EPI_ISL_510290, EPI_ISL_510291, EPI_ISL_510292, EPI_ISL_510293, EPI_ISL_510294, EPI_ISL_510295, EPI_ISL_510296, EPI_ISL_510297, EPI_ISL_510298, EPI_ISL_510299, EPI_ISL_510300, EPI_ISL_510301, EPI_ISL_510302, EPI_ISL_510303, EPI_ISL_510304 | | | |
| see above | Hospital Clínico Universitario de Santiago de Compostela | SeqCOVID-SPAIN consortium/IBV(CSIC) | José Javier Costa Alcalde, Antonio Aguilera Guirao, Mª Luisa Pérez del Molino Bernal, Amparo Coira Nieto, Gema Barbeito Castiñeiras, Rocio Trastoy Pena and SeqCOVID-SPAIN consortium |
| EPI_ISL_510444 | Hospital Universitario Virgen de las Nieves de Granada-SAS | SeqCOVID-SPAIN consortium/IBV(CSIC) | Mercedes Pérez Ruiz, Sara Sanbonmatsu Gámez, Irene Pedrosa Corral, José M. Navarro-Marí and SeqCOVID-SPAIN consortium |
| EPI_ISL_510529 | School of Veterinary Medicine, Disease Control | School of Veterinary Medicine, Disease Control | Simulundu,E., Kapata,N., Mupeta,F., Kapata,P.C., Saasa,N., Changula,K., Muleya,W., Chitanga,S., Chambaro,H., Mubemba,B., Masahiro,K., Chanda,D., Mulenga,L., Fwoloshi,S., Shibemba,A.L., Kapaya,F., Zulu,P., Musonda,K., Monze,M., Sinyange,N., Liwewe,M.M., Kapin'a,M., Chipimo,P.J., Ngosa,W., Morales,A.N., Kayeyi,N., Malama,K., Tembo,J., Bates,M., Sawa,H., Takada,A., Nalubamba,K.S., Mukonka,V., Chilufya,C. and Zumla,A. |
| EPI_ISL_510959, EPI_ISL_510960, EPI_ISL_510961, EPI_ISL_510962, EPI_ISL_510988, EPI_ISL_511037, EPI_ISL_511040, EPI_ISL_511041, EPI_ISL_511042, EPI_ISL_511043, EPI_ISL_511044, EPI_ISL_511045, EPI_ISL_511046, EPI_ISL_511047, EPI_ISL_511048, EPI_ISL_511049, EPI_ISL_511050, EPI_ISL_511051, EPI_ISL_511052, EPI_ISL_511189 | | | |
| see above | Instituto Nacional de Saude (INSA) | Instituto Nacional de Saude (INSA) | Borges et al |
| EPI_ISL_511210, EPI_ISL_511211, EPI_ISL_511212, EPI_ISL_511213, EPI_ISL_511214, EPI_ISL_511215, EPI_ISL_511216, EPI_ISL_511217, EPI_ISL_511218, EPI_ISL_511219, EPI_ISL_511220, EPI_ISL_511221, EPI_ISL_511257, EPI_ISL_511258, EPI_ISL_511259, EPI_ISL_511260, EPI_ISL_511261, EPI_ISL_511262, EPI_ISL_511263, EPI_ISL_511264, EPI_ISL_511265, EPI_ISL_511266, EPI_ISL_511267, EPI_ISL_511268, EPI_ISL_511269, EPI_ISL_511270, EPI_ISL_511271, EPI_ISL_511272, EPI_ISL_511273, EPI_ISL_511274, EPI_ISL_511275, EPI_ISL_511276, EPI_ISL_511277, EPI_ISL_511278, EPI_ISL_511279, EPI_ISL_511280, EPI_ISL_511281, EPI_ISL_511282, EPI_ISL_511283, EPI_ISL_511284, EPI_ISL_511285, EPI_ISL_511286, EPI_ISL_511287, EPI_ISL_511288, EPI_ISL_511289, EPI_ISL_511290, EPI_ISL_511293, EPI_ISL_511294, EPI_ISL_511295 | | | |
| see above | Instituto Nacional de Saude (INSA) and Instituto Gulbenkian | Instituto Nacional de Saude (INSA) and Instituto Gulbenkian | Borges et al |

| | | | |
|---|---|---|---|
| | de Ciencia (IGC) | de Ciencia (IGC) | |
| EPI_ISL_511372, EPI_ISL_511374, EPI_ISL_511385, EPI_ISL_511389, EPI_ISL_511412, EPI_ISL_511417, EPI_ISL_511426, EPI_ISL_511427, EPI_ISL_511428, EPI_ISL_511429, EPI_ISL_511434, EPI_ISL_511441, EPI_ISL_511444, EPI_ISL_511447, EPI_ISL_511450, EPI_ISL_511452, EPI_ISL_511455, EPI_ISL_511459, EPI_ISL_511460, EPI_ISL_511462, EPI_ISL_511463, EPI_ISL_511465, EPI_ISL_511467, EPI_ISL_511471, EPI_ISL_511472, EPI_ISL_511473, EPI_ISL_511474, EPI_ISL_511477, EPI_ISL_511479, EPI_ISL_511642, EPI_ISL_511643, EPI_ISL_511645, EPI_ISL_511646 | | | |
| see above | Instituto Nacional de Saude (INSA) | Instituto Nacional de Saude (INSA) | Borges et al |
| EPI_ISL_511893, EPI_ISL_511894, EPI_ISL_511895, EPI_ISL_511896 | National Hospital of Tropical Diseases | Oxford University Clinical Research Unit, Hanoi, Vietnam | Nguyen Thi Tam, Van Dinh Trang, Nguyen Thi Hong Thuong, Vu Thi Ngoc Bich, Nguyen Thu Trang, Nguyen Thi Ngoc Diep, Le Nguyen Minh Hoa, Pham Ngoc Thach, H. Rogier van Doorn, on behalf of the OUCRU COVID-19 research group |
| EPI_ISL_512384, EPI_ISL_512385, EPI_ISL_512386, EPI_ISL_512387, EPI_ISL_512388 | Centre for Enzyme Innovation, University of Portsmouth / Translational Research Laboratory, Portsmouth Hospitals NHS Trust | COVID-19 Genomics UK (COG-UK) Consortium | Angela Beckett,Yann Bourgeois,Garry Scarlett,Sharon Glaysher,Scott Elliott,Kelly Bicknell,Robert Impey,Allyson Lloyd,Sarah Wyllie,Ethan Butcher,Anoop Chauhan,Samuel Robson |
| EPI_ISL_512654 | Hospital Dr. Rafael A. Calderon Guardia [San Jose/San Jose] | Inciensa, Instituto Costarricense de Investigación y Enseñanza en Nutrición y Salud | Francisco Duarte, Hebleen Porras, Claudio Soto-Garita, Estela Cordero, Adriana Godinez & Melany Calderon |
| EPI_ISL_512759, EPI_ISL_512760, EPI_ISL_512761 | PathWest Laboratory Medicine WA | PathWest Laboratory Medicine WA Microbial Surveillance Unit | PathWest Laboratory Medicine WA Microbial Surveillance Unit |
| EPI_ISL_512848, EPI_ISL_512851, EPI_ISL_512852, EPI_ISL_512853, EPI_ISL_512854, EPI_ISL_512855, EPI_ISL_512856, EPI_ISL_512857, EPI_ISL_512858, EPI_ISL_512859, EPI_ISL_512860, EPI_ISL_512861, EPI_ISL_512862, EPI_ISL_512863, EPI_ISL_512864 | | | |
| see above | Ramathibodi Hospital | COVID-19 Network Investigations (CONI) Alliance | Elizabeth Batty, Wasun Chantratita, Thanat Chookajorn, Stefan Fernandez, Angkana Huang, Anthony R. Jones, Khajohn Joonsalak, Chonticha Klungtong, Theerarat Kochakarn, Namfon Kotanan, Krittikorn Kumpornsin, Wudtichai Manasatienkij, Bhakbhoom Panthan, Ekawat Pasomsub, Kingkan Rakmanee, Insee Sensorn, Janjira Thaipadungpanit, Arporn Wangwiwatsin,Treewat Watthanachockchai |
| EPI_ISL_513298 | Department of Infection Prevention and Infectious Diseases, University Hospital Regensburg | University Hospital Regensburg | Fritsch,J., Holzmann,T., Schneider-Brachert,W. |
| EPI_ISL_513310 | Public Health, United States Air Force School of Aerospace Medicine | Public Health, United States Air Force School of Aerospace Medicine | Fries,A.C., Purves,S.M., Meyer,J.R., Javorina,A.K., Connors,B.C., Macias,E.A., Lambert,A.W., Chapleau,R.R. and Starr,C.R. |
| EPI_ISL_513428, EPI_ISL_513431, EPI_ISL_513433, EPI_ISL_513436, EPI_ISL_513437, EPI_ISL_513438, EPI_ISL_513439, EPI_ISL_513440, EPI_ISL_513441, EPI_ISL_513442, EPI_ISL_513443, EPI_ISL_513444, EPI_ISL_513445, EPI_ISL_513446, EPI_ISL_513447, EPI_ISL_513448, EPI_ISL_513449, EPI_ISL_513450, EPI_ISL_513451, EPI_ISL_513452, EPI_ISL_513453, EPI_ISL_513454, EPI_ISL_513455, EPI_ISL_513456, EPI_ISL_513457, EPI_ISL_513458, EPI_ISL_513459, EPI_ISL_513463, EPI_ISL_513466, EPI_ISL_513467, EPI_ISL_513468, EPI_ISL_513469, EPI_ISL_513470, EPI_ISL_513474, EPI_ISL_513475, EPI_ISL_513487, EPI_ISL_513488, EPI_ISL_513492 | | | |
| see above | Maine HETL | Tewhey Lab, The Jackson Laboratory | Matluk,N., Dewey,H., Barter,M., Lynch,R., Munger,H. and Tewhey,R. |
| EPI_ISL_513511, EPI_ISL_513512 | The National Institute of Public Health | The National Institute of Public Health and State Veterinary Institute Prague | Nagy,A;Jirincova,H;Novakova,L;Trnka,D;Vecerova,J |
| EPI_ISL_513773 | County of Santa Clara Public Health Department | Chan-Zuckerberg Biohub | CZB Cliahub Consortium |
| EPI_ISL_514140 | Florida Bureau of Public Health Laboratories | Florida Bureau of Public Health Laboratories | Sarah Schmedes, Jason Blanton |
| EPI_ISL_514308, EPI_ISL_514309, EPI_ISL_514310, EPI_ISL_514311 | Israel Central Virology laboratory | Israel Central Virology laboratory | Neta Zuckerman, Efrat Dahan Bucris, Oran Erster, Ella Mendelson, Michal Mandelboim |
| EPI_ISL_514456, EPI_ISL_514457, EPI_ISL_514459, EPI_ISL_514460, EPI_ISL_514462, EPI_ISL_514463 | Centre for Enzyme Innovation, University of Portsmouth / Translational Research Laboratory, Portsmouth Hospitals NHS Trust | COVID-19 Genomics UK (COG-UK) Consortium | Angela Beckett,Yann Bourgeois,Garry Scarlett,Sharon Glaysher,Scott Elliott,Kelly Bicknell,Robert Impey,Allyson Lloyd,Sarah Wyllie,Ethan Butcher,Anoop Chauhan,Samuel Robson |
| EPI_ISL_514637 | M Health Fairview St. Joseph's Hospital | Minnesota Department of Health, Public Health Laboratory | Matt Plumb, Jacob Garfin, and Xiong Wang |
| EPI_ISL_514638 | Mayo Clinic & Mayo Clinic Laboratories | Minnesota Department of Health, Public Health Laboratory | Matt Plumb, Jacob Garfin, and Xiong Wang |
| EPI_ISL_515300, EPI_ISL_515301, EPI_ISL_515302, EPI_ISL_515303, EPI_ISL_515304, EPI_ISL_515305, EPI_ISL_515306, EPI_ISL_515307, EPI_ISL_515308, EPI_ISL_515309, EPI_ISL_515310, EPI_ISL_515311, EPI_ISL_515312 | | | |
| see above | Nevada State Public Health Laboratory | Nevada State Public Health Laboratory | Richard Tillett, Joel R. Sevinsky, Paul Hartley, Heather Kerwin, David Jackson, Subhash C. Verma, Cyprian Rossetto, Andrew Gorzalski, Chris Laverdure, Natalie Crawford, Stephanie Van Hooser, and Mark Pandori |
| EPI_ISL_515527 | Hospital Santa Clara | Instituto Adolfo Lutz, Interdiciplinary Procedures Center, Strategic Laboratory | Claudio Tavares Sacchi, Claudia Regina Gonçalves, Erica Valessa Ramos Gomes |
| EPI_ISL_515894, EPI_ISL_515895 | California Department of Public Health | California Department of Public Health | CDPH IDLB COVIDNet |
| EPI_ISL_516082 | Biomedical Sciences and Public Health, Polytechnic University of Marche | Biomedical Sciences and Public Health, Polytechnic University of Marche | Bagnarelli,P., Caucci,S., Di Sante,L., Menzo,S., Alessandrini,F., Onofri,V., Turchi,C., Melchionda,F., Tagliabracci,A. |
| EPI_ISL_516269, EPI_ISL_516281, EPI_ISL_516282 | Michigan Department of Health and Human Services, Bureau of Laboratories | Michigan Department of Health and Human Services, Bureau of Laboratories | Blankenship HM, Riner D, Soehnlen MK |
| EPI_ISL_516833, EPI_ISL_516835, EPI_ISL_516837, EPI_ISL_516838, EPI_ISL_516840, EPI_ISL_516841, EPI_ISL_516845, EPI_ISL_516850, EPI_ISL_516851, EPI_ISL_516852, EPI_ISL_516855, EPI_ISL_516856, EPI_ISL_516857, EPI_ISL_516859, EPI_ISL_516862, EPI_ISL_516863, EPI_ISL_516868 | | | |
| see above | North West London Pathology, Imperial College Healthcare NHS Trust | Wellcome Sanger Institute for the COVID-19 Genomics UK (COG-UK) consortium | Ling Li, Paul Randell, David Muir, Frankie Bolt, Alison Holmes, James Price, Aileen Rowan, Graham Taylor, Anjna Badhan, Carolina Herrera and Alex Alderton, Roberto Amato, Sonia Goncalves, Ewan Harrison, David K. Jackson, Ian Johnston, Dominic Kwiatkowski, Cordelia Langford, John Sillitoe on behalf of the Wellcome Sanger Institute COVID-19 Surveillance Team (http://www.sanger.ac.uk/covid-team) |
| EPI_ISL_516922, EPI_ISL_516923 | Department for Molecular Diagnostics, Centre for Medical Microbiology, Institute of Public Health of Montenegro | Charite Universitatsmedizin Berlin, Institut fur Virologie | Victor M Corman, Terry Jones, Jörn Beheim-Schwarzbach, Barbara Muehlemann, Talitha Veith, Julia Schneider, Marija Govedarica and Danijela Vujoševi, Christian Drosten |
| EPI_ISL_516948 | King Georges Medical University | CSIR-National Botanical Research Institute | Priti Prasad, Shantanu Prakash, Kishan Sahu, Babita Singh, Suruchi Shukla, Hricha Mishra, Danish Nasar Khan , Om Prakash, MLB Bhatt, SK Barik, Mehar H.Asif,Samir V. Sawant,Amita Jain, Sumit Kr. Bag |
| EPI_ISL_516987 | Laboratorio de Referencia Nacional de Virus Respiratorio. Centro Nacional de Salud Publica. Instituto Nacional de Salud Peru. | Laboratorio de Referencia Nacional de Biotecnología y Biología Molecular. Centro Nacional de Salud Publica. Instituto Nacional de Salud Peru. | Carlos Padilla Rojas, Karolyn Vega Chozo, Priscila Lope Pari, Omar Caceres Rey, Marco Galarza Perez, Maribel Huaringa Nuñez, Johanna Balbuena Torres, Henri Bailon Calderon, Nancy Rojas Serrano. |
| EPI_ISL_518868, EPI_ISL_518874, EPI_ISL_518875, EPI_ISL_518877, EPI_ISL_518879, EPI_ISL_518880, EPI_ISL_518881, EPI_ISL_518892, EPI_ISL_521153 | Mayo Clinic & Mayo Clinic Laboratories | Minnesota Department of Health, Public Health Laboratory | Matt Plumb, Jacob Garfin, and Xiong Wang |
| EPI_ISL_521885 | Victorian Infectious Diseases Reference Laboratory (VIDRL) | VIDRL and MDU-PHL | Caly L., Seemann T., Sait, M., Schultz M., Druce J., Sherry, N. |
| EPI_ISL_522566, EPI_ISL_522737, EPI_ISL_522742, EPI_ISL_522743 | Royal Hobart Hospital Microbiology Department | MDU-PHL | Cooley L., van Haeften R., Seemann T., Sait M., Schultz, M.B., Sherry N. |
| EPI_ISL_523332, EPI_ISL_523470, EPI_ISL_523525 | Dutch COVID-19 response team | Erasmus Medical Center | Bas Oude Munnink, David Nieuwenhuijse, Reina Sikkema, Claudia Schapendonk, Irina Chestakova, Anne van der Linden, Theo Bestebroer, Stefan van Nieuwkoop, Mark Pronk, Pascal Lexmond, Corien Swaan, Manon Haverkate, Madelief Mollers, Mart Stein, Sandra Kengne Kamga Mobou, Jeroen van Kampen, Jolanda Voermans, Aura Timen, Corine GeurtsvanKessel, Annemiek van der Eijk, Richard Molenkamp, Marion Koopmans, on behalf of the Dutch national COVID-19 response team. |
| EPI_ISL_523933, EPI_ISL_523939 | Center of Medical Microbiology, Virology, and Hospital | Center of Medical Microbiology, Virology, and Hospital | Maximilian Damagnez, Alexander Dilthey, Torsten Houwaart, Malte Kohns Vasconcelos, Marek Korencak, Jessica Nicolai, Klaus Pfeffer, Hendrik Streeck, |

| | | | |
|---|---|---|---|
| | Hygiene, University of Duesseldorf | Hygiene, University of Duesseldorf | Daniel Strelow, Jörg Timm, Andreas Walker, Tobias Wienemann |
| EPI_ISL_523954, EPI_ISL_523979 | Laboratorio de Referencia Nacional de Virus Respiratorio. Centro Nacional de Salud Publica. Instituto Nacional de Salud Peru. | Laboratorio de Referencia Nacional de Biotecnología y Biología Molecular. Centro Nacional de Salud Publica. Instituto Nacional de Salud Peru. | Carlos Padilla Rojas, Karolyn Vega Chozo, Priscila Lope Pari, Omar Caceres Rey, Marco Galarza Perez, Maribel Huaringa Nuñez, Johanna Balbuena Torres, Henri Bailon Calderon, Nancy Rojas Serrano. |
| EPI_ISL_524060, EPI_ISL_524061, EPI_ISL_524062, EPI_ISL_524063, EPI_ISL_524064, EPI_ISL_524065, EPI_ISL_524066 | Texas Department of State Health Services | Texas Department of State Health Services | Rashmi Tuladhar, Bonnie Oh, Cara Akrout, Jenny Zhang, Maliha Rahman, Anita Pokharel, Myong Koag, Chun Wang, Rachel Lee, Grace Kubin |
| EPI_ISL_524462 | Hospital Metropolitano | Instituto Adolfo Lutz, Interdiciplinary Procedures Center, Strategic Laboratory | Claudio Tavares Sacchi, Claudia Regina Gonçalves, Erica Valessa Ramos Gomes |
| EPI_ISL_524471 | Laboratorio de Referencia Nacional de Virus Respiratorio. Centro Nacional de Salud Publica. Instituto Nacional de Salud Peru. | Laboratorio de Referencia Nacional de Biotecnología y Biología Molecular. Centro Nacional de Salud Publica. Instituto Nacional de Salud Peru. | Carlos Padilla Rojas, Karolyn Vega Chozo, Priscila Lope Pari, Omar Caceres Rey, Marco Galarza Perez, Maribel Huaringa Nuñez, Johanna Balbuena Torres, Henri Bailon Calderon, Nancy Rojas Serrano. |
| EPI_ISL_524627, EPI_ISL_524638, EPI_ISL_524645, EPI_ISL_524651, EPI_ISL_524652, EPI_ISL_524653, EPI_ISL_524654, EPI_ISL_524658, EPI_ISL_524660, EPI_ISL_524661, EPI_ISL_524662, EPI_ISL_524663, EPI_ISL_524668, EPI_ISL_524669, EPI_ISL_524670, EPI_ISL_524671, EPI_ISL_524673, EPI_ISL_524676 | | | |
| see above | North West London Pathology, Imperial College Healthcare NHS Trust | Wellcome Sanger Institute for the COVID-19 Genomics UK (COG-UK) consortium | Ling Li, Paul Randell, David Muir, Frankie Bolt, Alison Holmes, James Price, Aileen Rowan, Graham Taylor, Anjna Badhan, Carolina Herrera and Alex Alderton, Roberto Amato, Sonia Goncalves, Ewan Harrison, David K. Jackson, Ian Johnston, Dominic Kwiatkowski, Cordelia Langford, John Sillitoe on behalf of the Wellcome Sanger Institute COVID-19 Surveillance Team (http://www.sanger.ac.uk/covid-team) |
| EPI_ISL_525537, EPI_ISL_525538 | CNR Virus des Infections Respiratoires - France SUD | CNR Virus des Infections Respiratoires - France SUD | Antonin Bal, Gregory Destras, Gwendolyne Burfin, Solenne Brun, Alexandre Gaymard, Maude Bouscambert-Duchamp, Florence Morfin-Sherpa, Martine Valette, Bruno Lina, Laurence Josset |
| EPI_ISL_525539 | Centre Hospitalier de Bourg en Bresse | CNR Virus des Infections Respiratoires - France SUD | Antonin Bal, Gregory Destras, Gwendolyne Burfin, Solenne Brun, Alexandre Gaymard, Maude Bouscambert-Duchamp, Florence Morfin-Sherpa, Martine Valette, Bruno Lina, Laurence Josset |
| EPI_ISL_525540, EPI_ISL_525541, EPI_ISL_525542 | CNR Virus des Infections Respiratoires - France SUD | CNR Virus des Infections Respiratoires - France SUD | Antonin Bal, Gregory Destras, Gwendolyne Burfin, Solenne Brun, Alexandre Gaymard, Maude Bouscambert-Duchamp, Florence Morfin-Sherpa, Martine Valette, Bruno Lina, Laurence Josset |
| EPI_ISL_525553, EPI_ISL_525556, EPI_ISL_525557, EPI_ISL_525570 | Istituto Zooprofilattico Sperimentale Puglia e Basilicata; Dipartimento di Bioscienze, Biotecnologie e Biofarmaceutica dell'Università degli Studi di Bari "A.Moro"; Istituto di Biomembrane. Bioenergetica e Biotecnologie Molecolari del Consiglio Nazionale delle Ricerche di Bari | Beaconlab (Bioinformatics, Evolution and Comparative Genomics lab), Dept of Biosciences, University on Milan | Parisi A.,Pesole G., Manzari C., Chiara M |
| EPI_ISL_525614, EPI_ISL_525615, EPI_ISL_525616, EPI_ISL_525617, EPI_ISL_525618, EPI_ISL_525619, EPI_ISL_525620, EPI_ISL_525621, EPI_ISL_525622 | Wadsworth Center, New York State Department of Health | Wadsworth Center, New York State Department of Health | Kirsten St. George, Daryl M. Lamson, Sara Griesemer, Jonathan Plitnick, Navjot Singh, Matthew D. Shudt, Erica Lasek-Nesselquist |
| EPI_ISL_525764, EPI_ISL_525765, EPI_ISL_525766 | Texas Department of State Health Services | Texas Department of State Health Services | Jenny Zhang, Rashmi Tuladhar, Bonnie Oh, Maliha Rahman, Anita Pokharel, Myong Koag, Chun Wang, Rachel Lee, Grace Kubin |
| EPI_ISL_526558, EPI_ISL_526581 | Florida Bureau of Public Health Laboratories | Florida Bureau of Public Health Laboratories | Sarah Schmedes, Jason Blanton |
| EPI_ISL_526797, EPI_ISL_526798, EPI_ISL_526799, EPI_ISL_526801, EPI_ISL_526802, EPI_ISL_526803, EPI_ISL_526804, EPI_ISL_526805, EPI_ISL_526806, EPI_ISL_526807 | Virginia DCLS | Virginia DCLS | Virginia DCLS |
| EPI_ISL_526942, EPI_ISL_526943, EPI_ISL_526944, EPI_ISL_526945, EPI_ISL_526946, EPI_ISL_526947, EPI_ISL_526948 | Faroese National Reference Laboratory for Fish and Animal Diseases | Faroese National Reference Laboratory for Fish and Animal Diseases | Maria Marjunardóttir Dahl, Petra Elisabeth Petersen, Debes Hammershaimb Christiansen |
| EPI_ISL_527008 | Area of Virology, Serology and Virology Division (SAViD), New South Wales Health Pathology Randwick | Area of Virology, Serology and Virology Division (SAViD), New South Wales Health Pathology Randwick | Rawlinson, W. |
| EPI_ISL_527759 | Clinica Biblica | Inciensa, Instituto Costarricense de Investigación y Enseñanza en Nutrición y Salud | Francisco Duarte, Hebleen Porras, Claudio Soto-Garita, Estela Cordero, Adriana Godinez & Melany Calderon |
| EPI_ISL_528095, EPI_ISL_528096, EPI_ISL_528097, EPI_ISL_528098, EPI_ISL_528099, EPI_ISL_528100, EPI_ISL_528101, EPI_ISL_528102, EPI_ISL_528103, EPI_ISL_528104, EPI_ISL_528105, EPI_ISL_528106, EPI_ISL_528109, EPI_ISL_528110, EPI_ISL_528111, EPI_ISL_528112, EPI_ISL_528114, EPI_ISL_528115, EPI_ISL_528116, EPI_ISL_528117, EPI_ISL_528118, EPI_ISL_528119, EPI_ISL_528120, EPI_ISL_528121, EPI_ISL_528122, EPI_ISL_528123, EPI_ISL_528124, EPI_ISL_528125, EPI_ISL_528126, EPI_ISL_528127, EPI_ISL_528128, EPI_ISL_528129, EPI_ISL_528130, EPI_ISL_528131, EPI_ISL_528132, EPI_ISL_528133, EPI_ISL_528134, EPI_ISL_528135, EPI_ISL_528136, EPI_ISL_528137, EPI_ISL_528138, EPI_ISL_528139, EPI_ISL_528140, EPI_ISL_528141, EPI_ISL_528142, EPI_ISL_528143, EPI_ISL_528144, EPI_ISL_528145, EPI_ISL_528146, EPI_ISL_528147, EPI_ISL_528148, EPI_ISL_528149, EPI_ISL_528150, EPI_ISL_528151, EPI_ISL_528152, EPI_ISL_528153, EPI_ISL_528154, EPI_ISL_528155, EPI_ISL_528156, EPI_ISL_528157, EPI_ISL_528158, EPI_ISL_528159, EPI_ISL_528160, EPI_ISL_528161, EPI_ISL_528162, EPI_ISL_528163, EPI_ISL_528164, EPI_ISL_528165, EPI_ISL_528166, EPI_ISL_528167, EPI_ISL_528168, EPI_ISL_528169, EPI_ISL_528170, EPI_ISL_528171, EPI_ISL_528172, EPI_ISL_528173, EPI_ISL_528174, EPI_ISL_528175, EPI_ISL_528176, EPI_ISL_528177, EPI_ISL_528178, EPI_ISL_528179, EPI_ISL_528180, EPI_ISL_528181, EPI_ISL_528182, EPI_ISL_528183, EPI_ISL_528184, EPI_ISL_528185, EPI_ISL_528186, EPI_ISL_528187, EPI_ISL_528188, EPI_ISL_528189, EPI_ISL_528190, EPI_ISL_528191, EPI_ISL_528192, EPI_ISL_528193, EPI_ISL_528194, EPI_ISL_528195, EPI_ISL_528196, EPI_ISL_528197, EPI_ISL_528198, EPI_ISL_528199, EPI_ISL_528200, EPI_ISL_528201, EPI_ISL_528202, EPI_ISL_528203, EPI_ISL_528204, EPI_ISL_528205, EPI_ISL_528206, EPI_ISL_528207, EPI_ISL_528208, EPI_ISL_528209, EPI_ISL_528210, EPI_ISL_528211, EPI_ISL_528212, EPI_ISL_528213, EPI_ISL_528214, EPI_ISL_528215, EPI_ISL_528216, EPI_ISL_528217, EPI_ISL_528218, EPI_ISL_528219, EPI_ISL_528377, EPI_ISL_528378 | | | |
| see above | University Hospital Basel, Clinical Virology | University Hospital Basel, Clinical Bacteriology | Madlen Stange, Alfredo Mari, Tim Roloff, Helena MB Seth-Smith, Michael Schweitzer, Myrta Brunner, Karoline Leuzinger, Kirstine K. Soegaard, Alexander Gensch, Sarah Tschudin-Sutter, Simon Fuchs, Julia Bielicki, Hans Pargger, Martin Siegemund, Christian Nickel, Roland Bingisser, Michael Osthoff, Stefano Bassetti, Rita Schneider-Sliwa, Manuel Battegay, Hans Hirsch, Adrian Egli |
| EPI_ISL_528637, EPI_ISL_528638 | LVM/UFRJ | Bioinformatics Laboratory / LNCC | Gustavo D. P. Silva; M. Romário M. de Souza; Bruno B. Bezerra; Lucio A. Caldas; Fabio Limonte; Elena Cobos; Sharton V. A. Coelho; Luiz Almeida; Luiza Higga; Isadora A. Correa; Diana Marianni; Luciana B. Arruda; Marcelo Bozza; Orlando Ferreira; Wanderley de Souza; Ana Teresa R. Vasconcelos; Terezinha M. Castineiras; Amilcar Tanuri; Luciana J. Costa |
| EPI_ISL_528919 | Ospedale Civile S. Liberatore-Atri | Istituto Zooprofilattico Sperimentale dell'Abruzzo e Molise "G.Caporale" | Lorusso A, Marcacci M, Di Domenico M, Curini V, Ancora M, Cammà C, Rinaldi A, Mangone I, Di Pasquale A, Puglia I, Savini G. |
| EPI_ISL_528920, EPI_ISL_528921 | Presidio Ospedaliero "Santo Spirito"-Pescara | Istituto Zooprofilattico Sperimentale dell'Abruzzo e Molise "G.Caporale" | Lorusso A, Marcacci M, Di Domenico M, Curini V, Ancora M, Cammà C, Rinaldi A, Mangone I, Di Pasquale A, Puglia I, Savini G. |
| EPI_ISL_528922 | Ospedale "Giuseppe Mazzini"-Teramo | Istituto Zooprofilattico Sperimentale dell'Abruzzo e Molise "G.Caporale" | Lorusso A, Marcacci M, Di Domenico M, Curini V, Ancora M, Cammà C, Rinaldi A, Mangone I, Di Pasquale A, Puglia I, Savini G. |
| EPI_ISL_528923 | Ospedale Civile S. Liberatore-Atri | Istituto Zooprofilattico Sperimentale dell'Abruzzo e Molise "G.Caporale" | Lorusso A, Marcacci M, Di Domenico M, Curini V, Ancora M, Cammà C, Rinaldi A, Mangone I, Di Pasquale A, Puglia I, Savini G. |
| EPI_ISL_528924 | Ospedale "Giuseppe Mazzini"-Teramo | Istituto Zooprofilattico Sperimentale dell'Abruzzo e Molise "G.Caporale" | Lorusso A, Marcacci M, Di Domenico M, Curini V, Ancora M, Cammà C, Rinaldi A, Mangone I, Di Pasquale A, Puglia I, Savini G. |
| EPI_ISL_528925 | Ospedale Regionale San Salvatore-L'Aquila | Istituto Zooprofilattico Sperimentale dell'Abruzzo e Molise "G.Caporale" | Lorusso A, Marcacci M, Di Domenico M, Curini V, Ancora M, Cammà C, Rinaldi A, Mangone I, Di Pasquale A, Puglia I, Savini G. |

| | | | |
|---|---|---|---|
| EPI_ISL_529148 | Democritus University of Thrace, Department of Medicine | Democritus University of Thrace, Department of Medicine | Kassela,K., Dovrolis,N., Bampali,M., Gatzidou,E., Froukala,E., Stavropoulou,A., Veletza,S., Tsakris,A., Spanakis,N., Karakasiliotis,I. |
| EPI_ISL_529172 | South Carolina Department of Health and Environmental Control | South Carolina Department of Health and Environmental Control | Haley V. Flores |
| EPI_ISL_529202 | Utah Public Health Laboratory | Utah Public Health Laboratory | Erin Young, Kelly Oakeson |
| EPI_ISL_529962 | Universitas Airlangga Hospital | Institute of Tropical Disease, Universitas Airlangga | Jezzy R Dewantari, Rima R Prasetya, Krisnoadi Rahardjo, Aldise M Nastri, Nasronudin, Gatot Soegiarto, Laksmi Wulandari, Retno A Setyoningrum, Resti Yudhawati, Yohko K Shimizu, Mitsuhiro Nishimura, Yasuko Mori, Soetjipto, Kazufumi Shimizu, Maria I Lusida |
| EPI_ISL_530109, EPI_ISL_530110, EPI_ISL_530112, EPI_ISL_530113 | Hospital Universitario Ramón y Cajal | Hospital Universitario La Paz | Raúl Recio, Sara González, Elias Dahdouh, Fernando Lázaro, Esther Viedma, Natalia Stella, Julio García, Juan Carlos Galán, Rafael Cantón, Mª Dolores Folgueira, Rafael Delgado, Jesús Mingorance |
| EPI_ISL_530347, EPI_ISL_530350 | The National Institute of Public Health | State Veterinary Institute Prague | Nagy,A;Jirincova,H;Novakova,L;Trnka,D;Vecerova,J |
| EPI_ISL_534722 | Respiratory Virus Unit, Microbiology Services Colindale, Public Health England | Respiratory Virus Unit, Microbiology Services Colindale, Public Health England | PHE Covid Sequencing Team |
| EPI_ISL_534891 | Oxford Viromics, NDM, University of Oxford; Oxford University Hospitals; Basingstoke and North Hampshire Hospital | COVID-19 Genomics UK (COG-UK) Consortium | Tanya Golubchik, David Bonsall, George Macintyre, Amy Trebes, Mariateresa de Cesare, Catrin Moore, Alex Mobbs, Anita Justice, Robert Shaw, Monique Andersson, Timothy Peto, Emma Wise, Nathan Moore, Jessica Lynch, Nick Cortes, Matilde Mori, Stephen Kidd, David Buck, John Todd, Christophe Fraser |
| EPI_ISL_535793 | Hôpital de Saint-Eustache | Laboratoire de santé publique du Québec | Sandrine Moreira, Ioannis Ragoussis, Guillaume Bourque, Jesse Shapiro, Mark Lathrop and Michel Roger |
| EPI_ISL_535795 | Hôpital Notre-Dame | Laboratoire de santé publique du Québec | Sandrine Moreira, Ioannis Ragoussis, Guillaume Bourque, Jesse Shapiro, Mark Lathrop and Michel Roger |
| EPI_ISL_535796 | Hôpital Charles-LeMoyne | Laboratoire de santé publique du Québec | Sandrine Moreira, Ioannis Ragoussis, Guillaume Bourque, Jesse Shapiro, Mark Lathrop and Michel Roger |
| EPI_ISL_535797 | Hôpital Honoré-Mercier | Laboratoire de santé publique du Québec | Sandrine Moreira, Ioannis Ragoussis, Guillaume Bourque, Jesse Shapiro, Mark Lathrop and Michel Roger |
| EPI_ISL_535798 | Hôpital régional de Saint-Jérôme | Laboratoire de santé publique du Québec | Sandrine Moreira, Ioannis Ragoussis, Guillaume Bourque, Jesse Shapiro, Mark Lathrop and Michel Roger |
| EPI_ISL_535799, EPI_ISL_535800 | CSSS Haut-Richelieu/Rouville (Hôpital) | Laboratoire de santé publique du Québec | Sandrine Moreira, Ioannis Ragoussis, Guillaume Bourque, Jesse Shapiro, Mark Lathrop and Michel Roger |
| EPI_ISL_535801 | CHU Sainte-Justine | Laboratoire de santé publique du Québec | Sandrine Moreira, Ioannis Ragoussis, Guillaume Bourque, Jesse Shapiro, Mark Lathrop and Michel Roger |
| EPI_ISL_535802, EPI_ISL_535806 | CHUM - Microbiologie - Hôpital Saint-Luc | Laboratoire de santé publique du Québec | Sandrine Moreira, Ioannis Ragoussis, Guillaume Bourque, Jesse Shapiro, Mark Lathrop and Michel Roger |
| EPI_ISL_535807 | Centre hospitalier régional du Grand Portage | Laboratoire de santé publique du Québec | Sandrine Moreira, Ioannis Ragoussis, Guillaume Bourque, Jesse Shapiro, Mark Lathrop and Michel Roger |
| EPI_ISL_535809 | Hôpital de Hull | Laboratoire de santé publique du Québec | Sandrine Moreira, Ioannis Ragoussis, Guillaume Bourque, Jesse Shapiro, Mark Lathrop and Michel Roger |
| EPI_ISL_535810 | Hôpital Sainte-Croix | Laboratoire de santé publique du Québec | Sandrine Moreira, Ioannis Ragoussis, Guillaume Bourque, Jesse Shapiro, Mark Lathrop and Michel Roger |
| EPI_ISL_535811 | CUSM-Site Glen-LAB Microbiologie | Laboratoire de santé publique du Québec | Sandrine Moreira, Ioannis Ragoussis, Guillaume Bourque, Jesse Shapiro, Mark Lathrop and Michel Roger |
| EPI_ISL_535812 | Hôpital du Sacré-Coeur de Montréal | Laboratoire de santé publique du Québec | Sandrine Moreira, Ioannis Ragoussis, Guillaume Bourque, Jesse Shapiro, Mark Lathrop and Michel Roger |
| EPI_ISL_535813 | CSSS Haut-Richelieu/Rouville (Hôpital) | Laboratoire de santé publique du Québec | Sandrine Moreira, Ioannis Ragoussis, Guillaume Bourque, Jesse Shapiro, Mark Lathrop and Michel Roger |
| EPI_ISL_535814 | CHUM - Microbiologie - Hôpital Saint-Luc | Laboratoire de santé publique du Québec | Sandrine Moreira, Ioannis Ragoussis, Guillaume Bourque, Jesse Shapiro, Mark Lathrop and Michel Roger |
| EPI_ISL_535815 | CSSS Haut-Richelieu/Rouville (Hôpital) | Laboratoire de santé publique du Québec | Sandrine Moreira, Ioannis Ragoussis, Guillaume Bourque, Jesse Shapiro, Mark Lathrop and Michel Roger |
| EPI_ISL_535817 | Hôpital Maisonneuve-Rosemont | Laboratoire de santé publique du Québec | Sandrine Moreira, Ioannis Ragoussis, Guillaume Bourque, Jesse Shapiro, Mark Lathrop and Michel Roger |
| EPI_ISL_535818 | Hôpital Pierre-Boucher | Laboratoire de santé publique du Québec | Sandrine Moreira, Ioannis Ragoussis, Guillaume Bourque, Jesse Shapiro, Mark Lathrop and Michel Roger |
| EPI_ISL_535819 | Hôpital Charles-LeMoyne | Laboratoire de santé publique du Québec | Sandrine Moreira, Ioannis Ragoussis, Guillaume Bourque, Jesse Shapiro, Mark Lathrop and Michel Roger |
| EPI_ISL_535820, EPI_ISL_535821 | Hôtel-Dieu de Lévis | Laboratoire de santé publique du Québec | Sandrine Moreira, Ioannis Ragoussis, Guillaume Bourque, Jesse Shapiro, Mark Lathrop and Michel Roger |
| EPI_ISL_535822, EPI_ISL_535824, EPI_ISL_535825 | CHUL-LABO MULTI / MICRO | Laboratoire de santé publique du Québec | Sandrine Moreira, Ioannis Ragoussis, Guillaume Bourque, Jesse Shapiro, Mark Lathrop and Michel Roger |
| EPI_ISL_535826, EPI_ISL_535827 | CHUM - Microbiologie - Hôpital Saint-Luc | Laboratoire de santé publique du Québec | Sandrine Moreira, Ioannis Ragoussis, Guillaume Bourque, Jesse Shapiro, Mark Lathrop and Michel Roger |
| EPI_ISL_535828 | CHUL-LABO MULTI / MICRO | Laboratoire de santé publique du Québec | Sandrine Moreira, Ioannis Ragoussis, Guillaume Bourque, Jesse Shapiro, Mark Lathrop and Michel Roger |
| EPI_ISL_535829 | Hôpital de Chicoutimi | Laboratoire de santé publique du Québec | Sandrine Moreira, Ioannis Ragoussis, Guillaume Bourque, Jesse Shapiro, Mark Lathrop and Michel Roger |
| EPI_ISL_535830 | CHUM - Microbiologie - Hôpital Saint-Luc | Laboratoire de santé publique du Québec | Sandrine Moreira, Ioannis Ragoussis, Guillaume Bourque, Jesse Shapiro, Mark Lathrop and Michel Roger |
| EPI_ISL_535831 | Hôpital Maisonneuve-Rosemont | Laboratoire de santé publique du Québec | Sandrine Moreira, Ioannis Ragoussis, Guillaume Bourque, Jesse Shapiro, Mark Lathrop and Michel Roger |
| EPI_ISL_535832 | CHUM - Microbiologie - Hôpital Saint-Luc | Laboratoire de santé publique du Québec | Sandrine Moreira, Ioannis Ragoussis, Guillaume Bourque, Jesse Shapiro, Mark Lathrop and Michel Roger |
| EPI_ISL_535833, EPI_ISL_535834, EPI_ISL_535835, EPI_ISL_535836 | Hôtel-Dieu de Lévis | Laboratoire de santé publique du Québec | Sandrine Moreira, Ioannis Ragoussis, Guillaume Bourque, Jesse Shapiro, Mark Lathrop and Michel Roger |
| EPI_ISL_535837, EPI_ISL_535838 | Hôpital régional de Saint-Jérôme | Laboratoire de santé publique du Québec | Sandrine Moreira, Ioannis Ragoussis, Guillaume Bourque, Jesse Shapiro, Mark Lathrop and Michel Roger |
| EPI_ISL_535839 | Hôpital Cité de la Santé | Laboratoire de santé publique du Québec | Sandrine Moreira, Ioannis Ragoussis, Guillaume Bourque, Jesse Shapiro, Mark Lathrop and Michel Roger |
| EPI_ISL_535840 | Hôpital Pierre-Boucher | Laboratoire de santé publique du Québec | Sandrine Moreira, Ioannis Ragoussis, Guillaume Bourque, Jesse Shapiro, Mark Lathrop and Michel Roger |
| EPI_ISL_535841 | CSSS Haut-Richelieu/Rouville (Hôpital) | Laboratoire de santé publique du Québec | Sandrine Moreira, Ioannis Ragoussis, Guillaume Bourque, Jesse Shapiro, Mark Lathrop and Michel Roger |
| EPI_ISL_535842, EPI_ISL_535843, EPI_ISL_535844 | Centre Hospitalier Régional de Lanaudière | Laboratoire de santé publique du Québec | Sandrine Moreira, Ioannis Ragoussis, Guillaume Bourque, Jesse Shapiro, Mark Lathrop and Michel Roger |
| EPI_ISL_535845 | Centre de SSS D'Arthabaska-et-de-l'érable - Hôtel-Dieu | Laboratoire de santé publique du Québec | Sandrine Moreira, Ioannis Ragoussis, Guillaume Bourque, Jesse Shapiro, Mark Lathrop and Michel Roger |
| EPI_ISL_535846 | CSSS Haut-Richelieu/Rouville (Hôpital) | Laboratoire de santé publique du Québec | Sandrine Moreira, Ioannis Ragoussis, Guillaume Bourque, Jesse Shapiro, Mark Lathrop and Michel Roger |
| EPI_ISL_535847 | Hôpital du Suroît | Laboratoire de santé publique du Québec | Sandrine Moreira, Ioannis Ragoussis, Guillaume Bourque, Jesse Shapiro, Mark Lathrop and Michel Roger |
| EPI_ISL_535848 | Hôpital Charles-LeMoyne | Laboratoire de santé publique du Québec | Sandrine Moreira, Ioannis Ragoussis, Guillaume Bourque, Jesse Shapiro, Mark Lathrop and Michel Roger |
| EPI_ISL_535849 | Centre de SSS de Trois-Rivières | Laboratoire de santé publique du Québec | Sandrine Moreira, Ioannis Ragoussis, Guillaume Bourque, Jesse Shapiro, Mark Lathrop and Michel Roger |
| EPI_ISL_535850 | Hôpital Pierre-Le Gardeur | Laboratoire de santé publique du Québec | Sandrine Moreira, Ioannis Ragoussis, Guillaume Bourque, Jesse Shapiro, Mark Lathrop and Michel Roger |
| EPI_ISL_535851 | Hôpital Charles-LeMoyne | Laboratoire de santé publique du Québec | Sandrine Moreira, Ioannis Ragoussis, Guillaume Bourque, Jesse Shapiro, Mark Lathrop and Michel Roger |
| EPI_ISL_535852 | Hôpital général du Lakeshore | Laboratoire de santé publique du Québec | Sandrine Moreira, Ioannis Ragoussis, Guillaume Bourque, Jesse Shapiro, Mark Lathrop and Michel Roger |
| EPI_ISL_535853 | Hôpital Charles-LeMoyne | Laboratoire de santé publique du Québec | Sandrine Moreira, Ioannis Ragoussis, Guillaume Bourque, Jesse Shapiro, Mark Lathrop and Michel Roger |
| EPI_ISL_535854, EPI_ISL_535855 | Centre de SSS de la Haute-Yamaska | Laboratoire de santé publique du Québec | Sandrine Moreira, Ioannis Ragoussis, Guillaume Bourque, Jesse Shapiro, Mark Lathrop and Michel Roger |
| EPI_ISL_535856 | Hôpital de Verdun | Laboratoire de santé publique du Québec | Sandrine Moreira, Ioannis Ragoussis, Guillaume Bourque, Jesse Shapiro, Mark Lathrop and Michel Roger |
| EPI_ISL_535857 | Hôpital Pierre-Boucher | Laboratoire de santé publique du Québec | Sandrine Moreira, Ioannis Ragoussis, Guillaume Bourque, Jesse Shapiro, Mark Lathrop and Michel Roger |

| | | | |
|---|---|---|---|
| EPI_ISL_535858 | Hôpital de Gatineau | Laboratoire de santé publique du Québec | Sandrine Moreira, Ioannis Ragoussis, Guillaume Bourque, Jesse Shapiro, Mark Lathrop and Michel Roger |
| EPI_ISL_535859, EPI_ISL_535860 | Hôpital Sainte-Croix | Laboratoire de santé publique du Québec | Sandrine Moreira, Ioannis Ragoussis, Guillaume Bourque, Jesse Shapiro, Mark Lathrop and Michel Roger |
| EPI_ISL_535861, EPI_ISL_535862 | Hôpital Pierre-Le Gardeur | Laboratoire de santé publique du Québec | Sandrine Moreira, Ioannis Ragoussis, Guillaume Bourque, Jesse Shapiro, Mark Lathrop and Michel Roger |
| EPI_ISL_535870 | Centre Hospitalier Régional de Lanaudière | Laboratoire de santé publique du Québec | Sandrine Moreira, Ioannis Ragoussis, Guillaume Bourque, Jesse Shapiro, Mark Lathrop and Michel Roger |
| EPI_ISL_535872 | Centre hospitalier régional du Grand Portage | Laboratoire de santé publique du Québec | Sandrine Moreira, Ioannis Ragoussis, Guillaume Bourque, Jesse Shapiro, Mark Lathrop and Michel Roger |
| EPI_ISL_536527, EPI_ISL_536531, EPI_ISL_536533, EPI_ISL_536534, EPI_ISL_536535, EPI_ISL_536536, EPI_ISL_536537, EPI_ISL_536538, EPI_ISL_536539, EPI_ISL_536540, EPI_ISL_536541, EPI_ISL_536543, EPI_ISL_536544, EPI_ISL_536546, EPI_ISL_536551, EPI_ISL_536554, EPI_ISL_536560, EPI_ISL_536565 | | | |
| see above | Instituto Nacional de Salud | Laboratorio de Infecciones Respiratorias Agudas | Eduardo Juscamayta Lopez, David Tarazona, Faviola Valdivia Guerrero, Nancy Rojas Serrano, Dennis Carhuaricra, Lenin Maturrano Hernandez, Ronnie Gavilan Chavez |
| EPI_ISL_537507, EPI_ISL_537568, EPI_ISL_537569, EPI_ISL_537571, EPI_ISL_537572, EPI_ISL_537581, EPI_ISL_537582, EPI_ISL_537586, EPI_ISL_537587, EPI_ISL_537595, EPI_ISL_537596, EPI_ISL_537598, EPI_ISL_537599, EPI_ISL_537603 | | | |
| see above | UCLA Pathology Clinical Microbiology Lab | Kruglyak Lab | Guo et al. |
| EPI_ISL_537608 | Servicio de Microbiología. Consorcio Hospital General Universitario de Valencia | SeqCOVID-SPAIN consortium/IBV(CSIC) | Paula Ruiz-Hueso, Mariana Reyes-Prieto, Vicente Soriano Chirona, Ivan Ansari, David Navarro, Maria Alma Bracho, Griselda De Marco, Beatriz Beamud, Lidia Ruiz Roldan, Marta Pla Diaz, Neris Garcia-Gonzalez, Inma Galán Vendrell, Sandra Carbo, Loreto Ferrús Abad, Llúcia Martínez-Priego, Giuseppe D'Auria, Fernando Gonzalez-Candelas and SeqCOVID-SPAIN consortium |
| EPI_ISL_537788, EPI_ISL_537789 | Hospital Universitario de Gran Canaria Dr. Negrín | SeqCOVID-SPAIN consortium/IBV(CSIC) | M. Carmen Pérez González, Francisco J. Chamizo López, Ana Bordes Benítez and SeqCOVID-SPAIN consortium |
| EPI_ISL_537874, EPI_ISL_537875, EPI_ISL_537876, EPI_ISL_537877, EPI_ISL_537878, EPI_ISL_537880, EPI_ISL_537881, EPI_ISL_537882, EPI_ISL_537883, EPI_ISL_537884, EPI_ISL_537885, EPI_ISL_537886, EPI_ISL_537887, EPI_ISL_537888, EPI_ISL_537889, EPI_ISL_537890, EPI_ISL_537891, EPI_ISL_537892, EPI_ISL_537893, EPI_ISL_537894, EPI_ISL_537895, EPI_ISL_537896, EPI_ISL_537897, EPI_ISL_537898, EPI_ISL_537899, EPI_ISL_537900, EPI_ISL_537901, EPI_ISL_537902, EPI_ISL_537903, EPI_ISL_537905, EPI_ISL_537906, EPI_ISL_537907, EPI_ISL_537908, EPI_ISL_537909, EPI_ISL_537910, EPI_ISL_537911, EPI_ISL_537912, EPI_ISL_537913, EPI_ISL_537914, EPI_ISL_537915, EPI_ISL_537916, EPI_ISL_537917, EPI_ISL_537920, EPI_ISL_537928, EPI_ISL_537933, EPI_ISL_537937, EPI_ISL_537938, EPI_ISL_537939, EPI_ISL_537940, EPI_ISL_537941, EPI_ISL_537942, EPI_ISL_537943, EPI_ISL_537944, EPI_ISL_537945 | | | |
| see above | Hospital Universitario Araba. Vitoria-Gasteiz | SeqCOVID-SPAIN consortium/IBV(CSIC) | Silvia Hernáez Crespo, Carmen Gómez González, Amaia Aguirre Quiñonero, Marina Fernández Torres, Ma Rosario Almela Ferrer, Ma Concepción Lecaroz Agara, Andrés Canut Blasco and SeqCOVID-SPAIN consortium |
| EPI_ISL_537985 | Servicio de Microbiología. Hospital General Universitario de Castellón | SeqCOVID-SPAIN consortium/IBV(CSIC) | Rosario Moreno, María Dolores Tirado and SeqCOVID-SPAIN consortium |
| EPI_ISL_538032, EPI_ISL_538033, EPI_ISL_538034, EPI_ISL_538035, EPI_ISL_538036, EPI_ISL_538037, EPI_ISL_538038, EPI_ISL_538039, EPI_ISL_538040, EPI_ISL_538041, EPI_ISL_538042, EPI_ISL_538043, EPI_ISL_538044, EPI_ISL_538045, EPI_ISL_538046, EPI_ISL_538047, EPI_ISL_538048, EPI_ISL_538049, EPI_ISL_538050, EPI_ISL_538051, EPI_ISL_538052, EPI_ISL_538053, EPI_ISL_538054 | | | |
| see above | Hospital Universitari i Politècnic La Fe de València | SeqCOVID-SPAIN consortium/IBV(CSIC) | María Dolores Gómez Ruiz, Eva González Barbera, Ana Gil Brusola, Salvador Giner Almaraz, José Luis López Hontangas and SeqCOVID-SPAIN consortium |
| EPI_ISL_538118 | Hospital Universitario Araba. Vitoria-Gasteiz | SeqCOVID-SPAIN consortium/IBV(CSIC) | Silvia Hernáez Crespo, Carmen Gómez González, Amaia Aguirre Quiñonero, Marina Fernández Torres, Mª Rosario Almela Ferrer, Mª Concepción Lecaroz Agara, Andrés Canut Blasco and SeqCOVID-SPAIN consortium |
| EPI_ISL_538437, EPI_ISL_538438, EPI_ISL_538439, EPI_ISL_538440, EPI_ISL_538447, EPI_ISL_538448 | Department of Laboratory Medicine, Tan Tock Seng Hospital | Department of Laboratory Medicine, Tan Tock Seng Hospital | Chen YYC, Zair X, Lim JX, Li C, Tang WY, Maurer-Stroh S, Barkham TMS, Nagarajan N, Sessions OM |
| EPI_ISL_538540, EPI_ISL_538548 | Department of Microbiology, The University of Hong Kong | Department of Microbiology, The University of Hong Kong | Kelvin K.W. To, Kwok-Yung Yuen |
| EPI_ISL_538617, EPI_ISL_538618, EPI_ISL_538619, EPI_ISL_538620, EPI_ISL_538622, EPI_ISL_538623, EPI_ISL_538624, EPI_ISL_538625, EPI_ISL_538626, EPI_ISL_538627, EPI_ISL_538628, EPI_ISL_538629, EPI_ISL_538630, EPI_ISL_538631, EPI_ISL_538632, EPI_ISL_538633, EPI_ISL_538634, EPI_ISL_538635, EPI_ISL_538636, EPI_ISL_538637 | | | |
| see above | Servicio de Microbiología. Hospital General Universitario de Castellón | SeqCOVID-SPAIN consortium/IBV(CSIC) | Rosario Moreno, María Dolores Tirado and SeqCOVID-SPAIN consortium |
| EPI_ISL_538673, EPI_ISL_538674, EPI_ISL_538675, EPI_ISL_538676, EPI_ISL_538677, EPI_ISL_538678, EPI_ISL_538679, EPI_ISL_538680, EPI_ISL_538681, EPI_ISL_538686 | Hospital Universitario Virgen de las Nieves de Granada-SAS | SeqCOVID-SPAIN consortium/IBV(CSIC) | Mercedes Pérez Ruiz, Sara Sanbonmatsu Gámez, Irene Pedrosa Corral, José M. Navarro-Marí and SeqCOVID-SPAIN consortium |
| EPI_ISL_538881, EPI_ISL_539049, EPI_ISL_539174 | Leeds Teaching Hospitals NHS Trust and Public Health England, National Infection Service (Leeds laboratory) | Wellcome Sanger Institute for the COVID-19 Genomics UK (COG-UK) consortium | Louissa Macfarlane-Smith, Holli Carden, Katherine L. Harper, Antony Hale and Alex Alderton, Roberto Amato, Sonia Goncalves, Ewan Harrison, David K. Jackson, Ian Johnston, Dominic Kwiatkowski, Cordelia Langford, John Sillitoe on behalf of the Wellcome Sanger Institute COVID-19 Surveillance Team |
| EPI_ISL_539230, EPI_ISL_539243 | Servicio de Microbiologia. Hospital Clinico Universitario de Valencia | SeqCOVID-SPAIN consortium/IBV(CSIC) | Maria Dolores Ocete, Inma Galán Vendrell, Paula Ruiz-Hueso, Mariana Reyes-Prieto, Vicente Soriano Chirona, Maria Alma Bracho, Griselda De Marco, Beatriz Beamud, Lidia Ruiz Roldan, Marta Pla Diaz,Neris Garcia-Gonzalez, Loreto Ferrús Abad, Llúcia Martínez-Priego, Concepcion Gimeno, Giuseppe D'Auria, Fernando Gonzalez-Candelas and SeqCOVID-SPAIN consortium |
| EPI_ISL_539245 | Hospital Universitari i Politècnic La Fe de València | SeqCOVID-SPAIN consortium/IBV(CSIC) | María Dolores Gómez Ruiz, Eva González Barbera, Ana Gil Brusola, Salvador Giner Almaraz, José Luis López Hontangas and SeqCOVID-SPAIN consortium |
| EPI_ISL_539291, EPI_ISL_539292, EPI_ISL_539293, EPI_ISL_539294, EPI_ISL_539295, EPI_ISL_539296, EPI_ISL_539297, EPI_ISL_539298 | Servicio de Microbiología. Consorcio Hospital General Universitario de Valencia | SeqCOVID-SPAIN consortium/IBV(CSIC) | Paula Ruiz-Hueso, Mariana Reyes-Prieto, Vicente Soriano Chirona, Ivan Ansari, David Navarro, Maria Alma Bracho, Griselda De Marco, Beatriz Beamud, Lidia Ruiz Roldan, Marta Pla Diaz, Neris Garcia-Gonzalez, Inma Galán Vendrell, Sandra Carbo, Loreto Ferrús Abad, Llúcia Martínez-Priego, Giuseppe D'Auria, Fernando Gonzalez-Candelas and SeqCOVID-SPAIN consortium |
| EPI_ISL_539521 | Hospital Universitario de Ceuta | Instituto de Salud Carlos III | Iglesias-Caballero, M. Molinero Calamita, M. González-Esguevillas, M. Camarero, S. Pozo, F. Casas, I. Jiménez, P. Jiménez, M. Zaballos, A. Monzón, S. Varona, S. Juliá, M. Cuesta, I, J. López |
| EPI_ISL_539522 | Hospital Universitario de Ceuta | Instituto de Salud Carlos III | Iglesias-Caballero, M. Molinero Calamita, M. González-Esguevillas, M. Camarero, S. Pozo, F. Casas, I. Jiménez, P. Jiménez, M. Zaballos, A. Monzón, S. Varona, S. Juliá, M. Cuesta, I, G. Sánchez |
| EPI_ISL_539524 | Gerencia de Asistencia Sanitaria de Soria | Instituto de Salud Carlos III | Iglesias-Caballero, M. Molinero Calamita, M. González-Esguevillas, M. Camarero, S. Pozo, F. Casas, I. Jiménez, P. Jiménez, M. Zaballos, A. Monzón, S. Varona, S. Juliá, M. Cuesta, I, C. Aldea |
| EPI_ISL_539781 | National Institute of Public Health (Czech Republic) | State Veterinary Institute Prague | Nagy, A; Jirincova, H; Novakova, L; Trnka, D; Vecerova, J. |
| EPI_ISL_541072, EPI_ISL_541073 | Hospital de la Santa Creu i Sant Pau. Servicio de Microbiología | SeqCOVID-SPAIN consortium/Institute of Biomedicine of Valencia, IBV-CSIC | Ferran Navarro, Núria Rabella, Elisenda Miró and SeqCOVID-SPAIN consortium |
| EPI_ISL_541242 | Florida Bureau of Public Health Laboratories, Florida Department of Health | Florida Bureau of Public Health Laboratories, Florida Department of Health | Schmedes,S., Blanton,J. |
| EPI_ISL_541340, EPI_ISL_541341, EPI_ISL_541342 | LACEN/PR | Laboratory of Respiratory Viruses and Measles, Oswaldo Cruz Institute, FIOCRUZ | Paola Resende, Luciana Appolinario, Fernando Motta, Anna Carolina Paixão, Ana Carolina Mendonça, Jonathan Lopes, Irina Riediger, Maria do Carmo Debur, Marilda Siqueira |
| EPI_ISL_541942 | Health and Environmental Research Institute of Gwangju | Health & Environment Institute of Gwangju | Min Ji Kim, Ji-eun Lee |
| EPI_ISL_541958, EPI_ISL_541960, EPI_ISL_541963, EPI_ISL_541966 | Servicio de Microbiología, Hospital Universitario Son Espases | SeqCOVID-SPAIN consortium/IBV(CSIC) | Carla López-Causapé, Jordi Reina, Antonio Oliver and SeqCOVID-SPAIN consortium |
| EPI_ISL_541970 | Influenza Centre, University of Bergen | Norwegian Institute of Public Health, Department of Virology | Fan Zhou, Rebecca J Cox, Karl A Brokstad, Bjørn Blomberg, Kathrine Stene-Johansen, Kamilla Heddeland Instefjord, Hilde Elshaug, Rasmus Riis Kopperud, Hilde Synnøve Vollan, Karoline Bragstad, Olav Hungnes |
| EPI_ISL_542114, EPI_ISL_542117, EPI_ISL_542122, EPI_ISL_542123, EPI_ISL_542129, EPI_ISL_542130, EPI_ISL_542134, EPI_ISL_542135, EPI_ISL_542147, EPI_ISL_542148, EPI_ISL_542150, EPI_ISL_542159, EPI_ISL_542161, EPI_ISL_542169, EPI_ISL_542171, EPI_ISL_542178, EPI_ISL_542179, EPI_ISL_542204, | | | |

| | | | |
|---|---|---|---|
| EPI_ISL_542206 | | | |
| see above | ASST GOM Niguarda | Dep. Of Oncology and Hemato-Oncology University of Milan | Claudia Alteri, Valeria Cento, Antonio Piralla, Valentino Costabile, Monica Tallarita, Luna Colagrossi, Silvia Renica, Federica Giardina, Federica Novazzi, Stefano Gaiarsa, Elisa Matarazzo, Maria Antonello, Chiara Vismara, Roberto Fumagalli, Oscar Massimiliano Epis, Massimo Puoti, Carlo Federico Perno, Fausto Baldanti |
| EPI_ISL_542325, EPI_ISL_542327, EPI_ISL_542333, EPI_ISL_542334, EPI_ISL_542335, EPI_ISL_542336, EPI_ISL_542337, EPI_ISL_542339, EPI_ISL_542340, EPI_ISL_542341, EPI_ISL_542343, EPI_ISL_542346, EPI_ISL_542348, EPI_ISL_542366, EPI_ISL_542378, EPI_ISL_542385, EPI_ISL_542393, EPI_ISL_542394 | | | |
| see above | San Matteo Hospital Pavia | Dep. Of Oncology and Hemato-Oncology University of Milan | Claudia Alteri, Valeria Cento, Antonio Piralla, Valentino Costabile, Monica Tallarita, Luna Colagrossi, Silvia Renica, Federica Giardina, Federica Novazzi, Stefano Gaiarsa, Elisa Matarazzo, Maria Antonello, Chiara Vismara, Roberto Fumagalli, Oscar Massimiliano Epis, Massimo Puoti, Carlo Federico Perno, Fausto Baldanti |
| EPI_ISL_545707, EPI_ISL_545714 | Houston Methodist Hospital | Houston Methodist Hospital | S. Wesley Long, Randall J. Olsen, Paul A. Christensen, David W. Bernard, James J. Davis, Maulik Shukla, Marcus Nguyen, Matthew Ojeda Saavedra, Concepcion C. Cantu, Prasanti Yerramilli, Layne Pruitt, Sishir Subedi, Hung-Che Kuo, Heather Hendrickson, Ghazaleh Eskandari, Hoang A. T. Nguyen, J. Hunter Long, Muthiah Kumaraswami, Jule Goike, Daniel Boutz, Jimmy Gollihar, Jason S. McLellan, Chia-Wei Chou, Kamyab Javanmardi, Ilya J. Finkelstein, and James M. Musser |
| EPI_ISL_547446, EPI_ISL_547447, EPI_ISL_547463, EPI_ISL_547464, EPI_ISL_547482, EPI_ISL_547492, EPI_ISL_547503, EPI_ISL_547504, EPI_ISL_547529 | Dutch COVID-19 response team | National Institute for Public Health and the Environment (RIVM) | Adam Meijer, Harry Vennema, Jeroen Cremer, Sharon van den Brink, Bas van der Veer, AnneMarie van den Brandt, Florian Zwagemaker, Dennis Schmitz, Chantal Reusken, on behalf of the national COVID-19 response team |
| EPI_ISL_548349, EPI_ISL_548356 | Ventura County Public Health Lab | Chan-Zuckerberg Biohub | CZB Cliahub Consortium |
| EPI_ISL_548961 | Max von Pettenkofer Institute, Virology, National Reference Center for Retroviruses, LMU München | Laboratory for Functional Genome Analysis, Dept. Genomics, Gene Center of the LMU Munich | Max Muenchhoff, Stefan Krebs, Alexander Graf, Oliver Keppler, Helmut Blum |
| EPI_ISL_548984, EPI_ISL_548989, EPI_ISL_548998, EPI_ISL_549005 | National Public Health Laboratory, National Centre for Infectious Diseases | National Public Health Laboratory, National Centre for Infectious Diseases | Mak TM, Octavia S, Zhou Z, Cui L, Lin RTP |
| EPI_ISL_549026 | Klinisk mikrobiologi, Region Västerbotten | Unit for Biological Agents, Department for CBRN Defence and Security, Swedish Defence Research Agency | FOI Bioinformatics team |
| EPI_ISL_549359 | Quadram Institute Bioscience | COVID-19 Genomics UK (COG-UK) Consortium | Dave J. Baker, Gemma L. Kay, Alp Aydin, Thanh Le-Viet, Steven Rudder, Ana P. Tedim, Anastasia Kolyva, Maria Diaz, Leonardo de Oliveira Martins, Nabil-Fareed Alikhan, Lizzie Meadows, Rachael Stanley, Ngozi Elumogo, Muhammed Yasir, Nicholas M. Thomson, Alexander J Trotter, Rachel Gilroy, Samuel Bloomfield, Claire Stuart, Andrew Bell, Reenesh Prakash, Samir Dervisevic, Alison E. Mather, John Wain, Mark Webber, Andrew J. Page, Justin O'Grady |
| EPI_ISL_560623 | Hospital | National Reference Center for Viruses of Respiratory Infections, Institut Pasteur, Paris | Sylvie Behillil, Fabiana Gambaro, Etienne Simon-Lorière, Vincent Enouf, Maud Vanpeene, Sylvie van der Werf |
| EPI_ISL_560651, EPI_ISL_560652, EPI_ISL_560653, EPI_ISL_560654, EPI_ISL_560655, EPI_ISL_560656, EPI_ISL_560657, EPI_ISL_560658, EPI_ISL_560659, EPI_ISL_560660, EPI_ISL_560662, EPI_ISL_560664 | | | |
| see above | Centre for Clinical Infection and Diagnostics Research and Genomics Innovation Unit, Guy's and St. Thomas' NHS Trust | Centre for Clinical Infection and Diagnostics Research and Genomics Innovation Unit, Guy's and St. Thomas' NHS Trust | Chloe Fisher, Luke Snell, Rahul Batra, Jonathan Edgeworth, Ali Raza Awan |
| EPI_ISL_561356, EPI_ISL_561357, EPI_ISL_561358, EPI_ISL_561359, EPI_ISL_561360 | Delaware Public Health Lab | Delaware Public Health Lab | Gregory Hovan |
| EPI_ISL_561363 | Centre for Clinical Infection and Diagnostics Research and Genomics Innovation Unit, Guy's and St. Thomas' NHS Trust | Centre for Clinical Infection and Diagnostics Research and Genomics Innovation Unit, Guy's and St. Thomas' NHS Trust | Chloe Fisher, Luke Snell, Rahul Batra, Jonathan Edgeworth, Ali Raza Awan |
| EPI_ISL_565863, EPI_ISL_565864, EPI_ISL_565865, EPI_ISL_565867, EPI_ISL_565868, EPI_ISL_565869 | Michigan Department of Health and Human Services, Bureau of Laboratories | Michigan Department of Health and Human Services, Bureau of Laboratories | Blankenship HM, Riner D, Soehnlen MK |
| EPI_ISL_565912, EPI_ISL_565913, EPI_ISL_565914, EPI_ISL_565917, EPI_ISL_565918, EPI_ISL_565919 | Servicio de Microbiología. HRU de Málaga. Servicio Andaluz de Salud | SeqCOVID-SPAIN consortium/IBV(CSIC) | Inmaculada de Toro Peinado, Ma Concepción Mediavilla Gradolph, Begoña Palop Borrás and SeqCOVID-SPAIN consortium |
| EPI_ISL_565961, EPI_ISL_566024, EPI_ISL_566028 | Michigan Department of Health and Human Services, Bureau of Laboratories | Michigan Department of Health and Human Services, Bureau of Laboratories | Blankenship HM, Riner D, Soehnlen MK |
| EPI_ISL_568580 | Ramathibodi Hospital | COVID-19 Network Investigations (CONI) Alliance | Elizabeth Batty, Wasun Chantratita, Thanat Chookajorn, Stefan Fernandez, Angkana Huang, Anthony R. Jones, Khajohn Joonsalak, Chonticha Klungtong, Theerarat Kochakarn, Namfon Kotanan, Krittikorn Kumpornsin, Wudtichai Manasatienkij, Bhakbhoom Panthan, Ekawat Pasomsub, Kingkan Rakmanee, Insee Sensorn, Janjira Thaipadungpanit, Arporn Wangwiwatsin,Treewat Watthanachockchai |
| EPI_ISL_568914, EPI_ISL_568919, EPI_ISL_568920, EPI_ISL_568921, EPI_ISL_568922, EPI_ISL_568956, EPI_ISL_568989, EPI_ISL_569254, EPI_ISL_569255, EPI_ISL_569256, EPI_ISL_569257, EPI_ISL_569258, EPI_ISL_569259, EPI_ISL_569260, EPI_ISL_569261, EPI_ISL_569262, EPI_ISL_569263, EPI_ISL_569264, EPI_ISL_569265, EPI_ISL_569266, EPI_ISL_569267, EPI_ISL_569268, EPI_ISL_569269, EPI_ISL_569270, EPI_ISL_569271, EPI_ISL_569331, EPI_ISL_569332, EPI_ISL_569338 | | | |
| see above | MEPHI, Aix Marseille University | MEPHI, Aix Marseille University | Anthony LEVASSEUR |
| EPI_ISL_570201, EPI_ISL_570536, EPI_ISL_570537, EPI_ISL_570547, EPI_ISL_570806, EPI_ISL_570816, EPI_ISL_570895, EPI_ISL_570901, EPI_ISL_570912 | UW Virology Lab | UW Virology Lab | Pavitra Roychoudhury, Hong Xie, Lasata Shrestha, Amin Addetia, Victoria M Rachleff, Meei-Li Huang, Keith R Jerome, Alexander Greninger |
| EPI_ISL_571027, EPI_ISL_571028, EPI_ISL_571032, EPI_ISL_571033, EPI_ISL_571036, EPI_ISL_571037, EPI_ISL_571078, EPI_ISL_571149, EPI_ISL_571150, EPI_ISL_571152, EPI_ISL_571156, EPI_ISL_571157, EPI_ISL_571178, EPI_ISL_571179, EPI_ISL_571180, EPI_ISL_571181, EPI_ISL_571182, EPI_ISL_571183, EPI_ISL_571184, EPI_ISL_571185, EPI_ISL_571186, EPI_ISL_571187, EPI_ISL_571188, EPI_ISL_571248, EPI_ISL_571249, EPI_ISL_571250, EPI_ISL_571251, EPI_ISL_571253, EPI_ISL_571255, EPI_ISL_571256, EPI_ISL_571259, EPI_ISL_571261, EPI_ISL_571262, EPI_ISL_571263, EPI_ISL_571264, EPI_ISL_571265, EPI_ISL_571266, EPI_ISL_571267, EPI_ISL_571268, EPI_ISL_571270, EPI_ISL_571286, EPI_ISL_571287, EPI_ISL_571288, EPI_ISL_571289, EPI_ISL_571290, EPI_ISL_571291, EPI_ISL_571292, EPI_ISL_571293, EPI_ISL_571294, EPI_ISL_571295, EPI_ISL_571296, EPI_ISL_571297, EPI_ISL_571299, EPI_ISL_571300, EPI_ISL_571301, EPI_ISL_571302, EPI_ISL_571303, EPI_ISL_571304, EPI_ISL_571305, EPI_ISL_571306, EPI_ISL_571307, EPI_ISL_571309, EPI_ISL_571310, EPI_ISL_571311, EPI_ISL_571312, EPI_ISL_571313, EPI_ISL_571314, EPI_ISL_571315, EPI_ISL_571316, EPI_ISL_571317, EPI_ISL_571318, EPI_ISL_571319, EPI_ISL_571321, EPI_ISL_571322, EPI_ISL_571324, EPI_ISL_571325, EPI_ISL_571327, EPI_ISL_571328, EPI_ISL_571329, EPI_ISL_571330, EPI_ISL_571332, EPI_ISL_571333, EPI_ISL_571334, EPI_ISL_571340, EPI_ISL_571341, EPI_ISL_571342, EPI_ISL_571343, EPI_ISL_571344, EPI_ISL_571345, EPI_ISL_571346, EPI_ISL_571347, EPI_ISL_571348, EPI_ISL_571349, EPI_ISL_571350, EPI_ISL_571351, EPI_ISL_571353, EPI_ISL_571354, EPI_ISL_571356, EPI_ISL_571357, EPI_ISL_571358, EPI_ISL_571359, EPI_ISL_571360, EPI_ISL_571361, EPI_ISL_571362, EPI_ISL_571363, EPI_ISL_571364, EPI_ISL_571365, EPI_ISL_571366, EPI_ISL_571367, EPI_ISL_571368, EPI_ISL_571369, EPI_ISL_571370, EPI_ISL_571371, EPI_ISL_571376, EPI_ISL_571377, EPI_ISL_571381, EPI_ISL_571382, EPI_ISL_571383, EPI_ISL_571384, EPI_ISL_571385, EPI_ISL_571386, EPI_ISL_571387, EPI_ISL_571388, EPI_ISL_571389, EPI_ISL_571390, EPI_ISL_571391, EPI_ISL_571392, EPI_ISL_571393, EPI_ISL_571394, EPI_ISL_571395, EPI_ISL_571396, EPI_ISL_571398, EPI_ISL_571399, EPI_ISL_571400, EPI_ISL_571401, EPI_ISL_571402, EPI_ISL_571403, EPI_ISL_571404, EPI_ISL_571405, EPI_ISL_571406, EPI_ISL_571407, EPI_ISL_571408, EPI_ISL_571409, EPI_ISL_571424, EPI_ISL_571425, EPI_ISL_571426, EPI_ISL_571427, EPI_ISL_571428, EPI_ISL_571429, EPI_ISL_571430, EPI_ISL_571431, EPI_ISL_571432, EPI_ISL_571434, EPI_ISL_571435, EPI_ISL_571436, EPI_ISL_571437, EPI_ISL_571438, EPI_ISL_571440, EPI_ISL_571473, EPI_ISL_571474, EPI_ISL_571475, EPI_ISL_571476, EPI_ISL_571477, EPI_ISL_571478, EPI_ISL_571479, EPI_ISL_571480, EPI_ISL_571481, EPI_ISL_571482, EPI_ISL_571483, EPI_ISL_571509, EPI_ISL_571510, EPI_ISL_571511, EPI_ISL_571512, EPI_ISL_571513, EPI_ISL_571514, EPI_ISL_571515, EPI_ISL_571516, EPI_ISL_571517, EPI_ISL_571518, EPI_ISL_571528, EPI_ISL_571529, EPI_ISL_571530, EPI_ISL_571531, EPI_ISL_571532, EPI_ISL_571545, EPI_ISL_571546, EPI_ISL_571548, EPI_ISL_571551, EPI_ISL_571552, EPI_ISL_571553, EPI_ISL_571554, EPI_ISL_571555, EPI_ISL_571556, EPI_ISL_571563, EPI_ISL_571565, EPI_ISL_571567, EPI_ISL_571568, EPI_ISL_571573, EPI_ISL_571592, EPI_ISL_571602, EPI_ISL_571603, EPI_ISL_571604, EPI_ISL_571606, EPI_ISL_571607, EPI_ISL_571608, EPI_ISL_571609, EPI_ISL_571610, EPI_ISL_571611, EPI_ISL_571614, EPI_ISL_571641, EPI_ISL_571642, EPI_ISL_571643, EPI_ISL_571644, EPI_ISL_571645, EPI_ISL_571646, EPI_ISL_571647, EPI_ISL_571648, EPI_ISL_571649, EPI_ISL_571650, EPI_ISL_571651, EPI_ISL_571652, EPI_ISL_571653, EPI_ISL_571654, EPI_ISL_571655, EPI_ISL_571656, EPI_ISL_571657, EPI_ISL_571658, EPI_ISL_571659, EPI_ISL_571660, EPI_ISL_571661, EPI_ISL_571662, EPI_ISL_571663, EPI_ISL_571664, EPI_ISL_571665, EPI_ISL_571666, EPI_ISL_571667, EPI_ISL_571668, EPI_ISL_571669, EPI_ISL_571670, EPI_ISL_571671, EPI_ISL_571672, EPI_ISL_571673, EPI_ISL_571674, EPI_ISL_571675, EPI_ISL_571676, EPI_ISL_571677, EPI_ISL_571678, EPI_ISL_571679, EPI_ISL_571680, EPI_ISL_571681, EPI_ISL_571682, EPI_ISL_571683, EPI_ISL_571684, EPI_ISL_571687, EPI_ISL_571688, EPI_ISL_571689, EPI_ISL_571690, EPI_ISL_571692, EPI_ISL_571693, EPI_ISL_571694, EPI_ISL_571695, EPI_ISL_571696, EPI_ISL_571697, EPI_ISL_571698, EPI_ISL_571699, EPI_ISL_571700, EPI_ISL_571701, EPI_ISL_571702, EPI_ISL_571703, EPI_ISL_571704, EPI_ISL_571705, EPI_ISL_571706, EPI_ISL_571707, EPI_ISL_571708, EPI_ISL_571709, EPI_ISL_571710, EPI_ISL_571711, EPI_ISL_571712, EPI_ISL_571713, EPI_ISL_571714, EPI_ISL_571715, EPI_ISL_571716, EPI_ISL_571717, EPI_ISL_571718, EPI_ISL_571719, EPI_ISL_571720, EPI_ISL_571721, EPI_ISL_571722, EPI_ISL_571723, EPI_ISL_571724, EPI_ISL_571725, EPI_ISL_571726, EPI_ISL_571728, EPI_ISL_571731, EPI_ISL_571732, EPI_ISL_571733, EPI_ISL_571734, EPI_ISL_571735, EPI_ISL_571738, EPI_ISL_571739, EPI_ISL_571740, EPI_ISL_571743, EPI_ISL_571746, EPI_ISL_571747, EPI_ISL_571748, EPI_ISL_571749, EPI_ISL_571750, EPI_ISL_571751, EPI_ISL_571753, EPI_ISL_571754, EPI_ISL_571755, EPI_ISL_571756, EPI_ISL_571757, EPI_ISL_571758, EPI_ISL_571759, EPI_ISL_571762, EPI_ISL_571764, EPI_ISL_571765, EPI_ISL_571768, EPI_ISL_571769, EPI_ISL_571773, EPI_ISL_571774, EPI_ISL_571777, EPI_ISL_571779, EPI_ISL_571780, EPI_ISL_571781, EPI_ISL_571782, EPI_ISL_571783, EPI_ISL_571784, EPI_ISL_571789, EPI_ISL_571797, EPI_ISL_571798, EPI_ISL_571799, EPI_ISL_571800, EPI_ISL_571801, EPI_ISL_571802, EPI_ISL_571821, EPI_ISL_571823, EPI_ISL_571843, EPI_ISL_571844, EPI_ISL_571901, EPI_ISL_571911, | | | |

| | | | |
|---|---|---|---|
| EPI_ISL_571912, EPI_ISL_571913, EPI_ISL_571916, EPI_ISL_571917, EPI_ISL_571918, EPI_ISL_571919, EPI_ISL_571920, EPI_ISL_571922, EPI_ISL_571923, EPI_ISL_571924, EPI_ISL_571927, EPI_ISL_571928, EPI_ISL_571929, EPI_ISL_571930, EPI_ISL_571931, EPI_ISL_571932, EPI_ISL_571933, EPI_ISL_571934, EPI_ISL_571935, EPI_ISL_571936, EPI_ISL_571937, EPI_ISL_571938, EPI_ISL_571939, EPI_ISL_571940, EPI_ISL_571941, EPI_ISL_571942, EPI_ISL_571943, EPI_ISL_571944, EPI_ISL_571945, EPI_ISL_571946, EPI_ISL_571947, EPI_ISL_571948, EPI_ISL_571949, EPI_ISL_571950, EPI_ISL_571951, EPI_ISL_571952, EPI_ISL_571953, EPI_ISL_571954, EPI_ISL_571955, EPI_ISL_571956, EPI_ISL_571957, EPI_ISL_571958, EPI_ISL_571959, EPI_ISL_571960, EPI_ISL_571961, EPI_ISL_571962, EPI_ISL_571963, EPI_ISL_571964, EPI_ISL_571965, EPI_ISL_571966, EPI_ISL_571968, EPI_ISL_571970, EPI_ISL_571971, EPI_ISL_571972, EPI_ISL_571973, EPI_ISL_571974, EPI_ISL_571975, EPI_ISL_571976, EPI_ISL_571977, EPI_ISL_571978, EPI_ISL_571979, EPI_ISL_571980, EPI_ISL_571981, EPI_ISL_571982, EPI_ISL_571983, EPI_ISL_571990, EPI_ISL_571994, EPI_ISL_571995, EPI_ISL_571996, EPI_ISL_571997, EPI_ISL_571998, EPI_ISL_571999, EPI_ISL_572000, EPI_ISL_572001, EPI_ISL_572002, EPI_ISL_572003, EPI_ISL_572004, EPI_ISL_572005, EPI_ISL_572006, EPI_ISL_572007, EPI_ISL_572008, EPI_ISL_572009, EPI_ISL_572010, EPI_ISL_572011, EPI_ISL_572012, EPI_ISL_572013, EPI_ISL_572014, EPI_ISL_572015, EPI_ISL_572016, EPI_ISL_572017, EPI_ISL_572018, EPI_ISL_572019, EPI_ISL_572020, EPI_ISL_572021, EPI_ISL_572022, EPI_ISL_572023, EPI_ISL_572024, EPI_ISL_572025, EPI_ISL_572026, EPI_ISL_572027, EPI_ISL_572028, EPI_ISL_572029, EPI_ISL_572030, EPI_ISL_572031, EPI_ISL_572032, EPI_ISL_572033, EPI_ISL_572034, EPI_ISL_572035, EPI_ISL_572036, EPI_ISL_572037, EPI_ISL_572038, EPI_ISL_572039, EPI_ISL_572040, EPI_ISL_572042, EPI_ISL_572044, EPI_ISL_572045, EPI_ISL_572046, EPI_ISL_572047, EPI_ISL_572049, EPI_ISL_572050, EPI_ISL_572051, EPI_ISL_572052, EPI_ISL_572053, EPI_ISL_572054, EPI_ISL_572055, EPI_ISL_572056, EPI_ISL_572057, EPI_ISL_572058, EPI_ISL_572059, EPI_ISL_572060, EPI_ISL_572061, EPI_ISL_572062, EPI_ISL_572063, EPI_ISL_572064, EPI_ISL_572065, EPI_ISL_572066, EPI_ISL_572067, EPI_ISL_572068, EPI_ISL_572069, EPI_ISL_572070, EPI_ISL_572071, EPI_ISL_572072, EPI_ISL_572073, EPI_ISL_572074, EPI_ISL_572075, EPI_ISL_572076, EPI_ISL_572077, EPI_ISL_572078, EPI_ISL_572079, EPI_ISL_572080, EPI_ISL_572081, EPI_ISL_572082, EPI_ISL_572083, EPI_ISL_572084, EPI_ISL_572085, EPI_ISL_572086, EPI_ISL_572087, EPI_ISL_572088, EPI_ISL_572089, EPI_ISL_572090, EPI_ISL_572091, EPI_ISL_572092, EPI_ISL_572093, EPI_ISL_572094, EPI_ISL_572095, EPI_ISL_572096, EPI_ISL_572097, EPI_ISL_572098, EPI_ISL_572099, EPI_ISL_572100, EPI_ISL_572101, EPI_ISL_572102, EPI_ISL_572103, EPI_ISL_572104, EPI_ISL_572107, EPI_ISL_572109, EPI_ISL_572111, EPI_ISL_572112, EPI_ISL_572113, EPI_ISL_572114, EPI_ISL_572115, EPI_ISL_572116, EPI_ISL_572117, EPI_ISL_572118, EPI_ISL_572119, EPI_ISL_572120, EPI_ISL_572121, EPI_ISL_572122, EPI_ISL_572123, EPI_ISL_572127, EPI_ISL_572128, EPI_ISL_572129, EPI_ISL_572130, EPI_ISL_572131, EPI_ISL_572132, EPI_ISL_572133, EPI_ISL_572134, EPI_ISL_572135, EPI_ISL_572137, EPI_ISL_572142, EPI_ISL_572147, EPI_ISL_572151, EPI_ISL_572153, EPI_ISL_572156, EPI_ISL_572157, EPI_ISL_572159, EPI_ISL_572160, EPI_ISL_572161, EPI_ISL_572162, EPI_ISL_572163, EPI_ISL_572164, EPI_ISL_572165, EPI_ISL_572166, EPI_ISL_572167, EPI_ISL_572170, EPI_ISL_572171, EPI_ISL_572173, EPI_ISL_572174, EPI_ISL_572175, EPI_ISL_572176, EPI_ISL_572178, EPI_ISL_572180, EPI_ISL_572181, EPI_ISL_572182, EPI_ISL_572183 | | | |
| see above | Quest Diagnostics | Quest Diagnostics | Rosenthal,S.H., Gerasimova,A., Kagan,R.M., Anderson, B., Grover, D., Livingston, K.E., Hua, M., Liu Y., Shalhout, D.F., Owen, R., Lacbawan, F. |
| EPI_ISL_574292 | Instituto Nacional de Investigación en Salud Pública | Instituto Nacional de Investigación en Salud Pública | Leandro Patino Patino, Domenica de Mora Coloma, Denisse Portugal, Alfredo Bruno Caicedo, Andrés Carrasco, Orson Mestanza, Manuel Gonzalez, Alberto Orlando Narvaez. |
| EPI_ISL_574581 | Delaware Public Health Lab | Delaware Public Health Lab | Gregory Hovan |
| EPI_ISL_574800, EPI_ISL_574801, EPI_ISL_574802, EPI_ISL_574803, EPI_ISL_574804, EPI_ISL_574805, EPI_ISL_574806, EPI_ISL_574807, EPI_ISL_574808, EPI_ISL_574809, EPI_ISL_574810, EPI_ISL_574811, EPI_ISL_574812 | | | |
| see above | Institute for Infectious Diseases, University of Bern | Institute for Infectious Diseases, University of Bern | Michel C Koch, Christian Baumann, Miguel A Terrazos Miani, Cora Sägesser, Stephen L Leib, Peter Keller, Franziska Suter-Riniker, Alban Ramette |
| EPI_ISL_574876, EPI_ISL_574877, EPI_ISL_574878, EPI_ISL_574883, EPI_ISL_574884, EPI_ISL_574885, EPI_ISL_574886, EPI_ISL_574887, EPI_ISL_574888, EPI_ISL_574905, EPI_ISL_574906, EPI_ISL_574907, EPI_ISL_574908, EPI_ISL_574909, EPI_ISL_574910, EPI_ISL_574913, EPI_ISL_574914, EPI_ISL_574915, EPI_ISL_574916, EPI_ISL_574917, EPI_ISL_574918, EPI_ISL_574919, EPI_ISL_574920, EPI_ISL_574931, EPI_ISL_574932, EPI_ISL_574933, EPI_ISL_574934, EPI_ISL_574936, EPI_ISL_574937, EPI_ISL_574938, EPI_ISL_574944, EPI_ISL_574945, EPI_ISL_574946, EPI_ISL_574947, EPI_ISL_574948, EPI_ISL_574949, EPI_ISL_574950, EPI_ISL_574951, EPI_ISL_574954, EPI_ISL_574955, EPI_ISL_574956, EPI_ISL_574957, EPI_ISL_574958, EPI_ISL_574959, EPI_ISL_574960, EPI_ISL_574961, EPI_ISL_574962, EPI_ISL_574963, EPI_ISL_574964, EPI_ISL_574965, EPI_ISL_574966, EPI_ISL_574967, EPI_ISL_574968, EPI_ISL_574969, EPI_ISL_574970, EPI_ISL_574971, EPI_ISL_574972, EPI_ISL_574977, EPI_ISL_574978, EPI_ISL_574979, EPI_ISL_574980, EPI_ISL_574981, EPI_ISL_574982, EPI_ISL_574983, EPI_ISL_574984, EPI_ISL_574985, EPI_ISL_574986, EPI_ISL_574987, EPI_ISL_574988, EPI_ISL_574989, EPI_ISL_574990, EPI_ISL_574991, EPI_ISL_574992, EPI_ISL_574993, EPI_ISL_574994, EPI_ISL_574995, EPI_ISL_574996, EPI_ISL_574997, EPI_ISL_574998, EPI_ISL_574999, EPI_ISL_575000, EPI_ISL_575003, EPI_ISL_575004, EPI_ISL_575005, EPI_ISL_575006, EPI_ISL_575007, EPI_ISL_575008, EPI_ISL_575009, EPI_ISL_575010, EPI_ISL_575011, EPI_ISL_575013, EPI_ISL_575014, EPI_ISL_575015, EPI_ISL_575016, EPI_ISL_575017, EPI_ISL_575018 | | | |
| see above | Viollier AG | Department of Biosystems Science and Engineering, ETH Zürich | Christian Beisel, Sarah Nadeau, Ivan Topolsky, Pedro Ferreira, Philipp Jablonski, Susana Posada-Céspedes, Tobias Schär, Ina Nissen, Natascha Santacroce, Elodie Burcklen, Christiane Beckmann, Maurice Redondo, Olivier Kobel, Christoph Noppen, Sophie Seidel, Noemie Santamaria de Souza, Niko Beerenwinkel, Tanja Stadler |
| EPI_ISL_576171 | PA Department of Health, Bureau of Laboratories | Pathogen Discovery, Respiratory Viruses Branch, Division of Viral Diseases, Centers for Disease Control and Prevention | Yan Li, Anna Montmayeur, Brian Lynch, Jing Zhang, Krista Queen, Ying Tao, Anna Uehara, Rachel Marine, Clinton R. Paden, Peter Cook, Haibin Wang, Suxiang Tong |
| EPI_ISL_578420, EPI_ISL_578422, EPI_ISL_578423, EPI_ISL_578424, EPI_ISL_578425, EPI_ISL_578470, EPI_ISL_578471 | Wisconsin State Laboratory of Hygiene Communicable Disease Division | Wisconsin State Laboratory of Hygiene Communicable Disease Division | Kelsey R. Florek, Abigail C. Shockey |
| EPI_ISL_579060, EPI_ISL_579061 | Canterbury Health Laboratories | Institute of Environmental Science and Research (ESR) | Xiaoyun Ren, Matt Storey, Nikki Freed, Muhammad Faisal, Jing Wang, Hermes Perez, Anja Werno, Antje van der Linden, Arlo Upton, Chris Mansell, David Hammer, Dragana Drinkovic, Gary McAuliffe, Hana Sofia Andersson, James Ussher, Jill Sherwood, Josh Freeman, Julia Howard, Juliet Elvy, Mary DeAlmeida, Matt Blakiston, Matthew Rogers, Max Bloomfield, Michael Addidle, Michelle Balm, Sally Roberts, Sarah Jefferies, Sharmini Muttaiyah, Susan Morpeth, Susan Taylor, Timothy Blackmore, Vani Sathyendran, Veronica Playle, Virginia Hope, Erasmus Smit, Lauren Jelly, Olin Silander, Joep de Ligt |
| EPI_ISL_579118 | Waikato Hospital | Institute of Environmental Science and Research (ESR) | Xiaoyun Ren, Matt Storey, Nikki Freed, Muhammad Faisal, Jing Wang, Hermes Perez, Anja Werno, Antje van der Linden, Arlo Upton, Chris Mansell, David Hammer, Dragana Drinkovic, Gary McAuliffe, Hana Sofia Andersson, James Ussher, Jill Sherwood, Josh Freeman, Julia Howard, Juliet Elvy, Mary DeAlmeida, Matt Blakiston, Matthew Rogers, Max Bloomfield, Michael Addidle, Michelle Balm, Sally Roberts, Sarah Jefferies, Sharmini Muttaiyah, Susan Morpeth, Susan Taylor, Timothy Blackmore, Vani Sathyendran, Veronica Playle, Virginia Hope, Erasmus Smit, Lauren Jelly, Olin Silander, Joep de Ligt |
| EPI_ISL_579119, EPI_ISL_579120 | LabPLUS | Institute of Environmental Science and Research (ESR) | Xiaoyun Ren, Matt Storey, Nikki Freed, Muhammad Faisal, Jing Wang, Hermes Perez, Anja Werno, Antje van der Linden, Arlo Upton, Chris Mansell, David Hammer, Dragana Drinkovic, Gary McAuliffe, Hana Sofia Andersson, James Ussher, Jill Sherwood, Josh Freeman, Julia Howard, Juliet Elvy, Mary DeAlmeida, Matt Blakiston, Matthew Rogers, Max Bloomfield, Michael Addidle, Michelle Balm, Sally Roberts, Sarah Jefferies, Sharmini Muttaiyah, Susan Morpeth, Susan Taylor, Timothy Blackmore, Vani Sathyendran, Veronica Playle, Virginia Hope, Erasmus Smit, Lauren Jelly, Olin Silander, Joep de Ligt |
| EPI_ISL_579517, EPI_ISL_579537 | QEII Health Sciences Centre | National Microbiology Laboratory (NML) | Anna Majer, Shari Tyson, Grace Seo, Philip Mabon, Darian Hole, Elsie Grudeski, Rhiannon Huzarewich, Russell Mandes, Anneliese Landgraff, Jennifer Tanner, Natalie Knox, Morag Graham, Gary Van Domselaar, Todd Hatchette, Jason LeBlanc, Nathalie Bastien, Yan Li, Timothy Booth, CanCOGeN's metadata curation team, Public Health Agency of Canada's CanCOGeN team |
| EPI_ISL_581452 | Medizinische Klinik Innere Medizin I, Universitätsklinikum Tübingen | NGS Competence Center Tübingen, Institut für Medizinische Mikrobiologie und Hygiene, Universitätsklinikum Tübingen | Angel Angelov |
| EPI_ISL_581719 | University Hospital Basel, Clinical Virology | University Hospital Basel, Clinical Bacteriology | Madlen Stange, Alfredo Mari, Tim Roloff, Helena MB Seth-Smith, Michael Schweitzer, Myrta Brunner, Karoline Leuzinger, Kirstine K. Soegaard, Alexander Gensch, Sarah Tschudin-Sutter, Simon Fuchs, Julia Bielicki, Hans Pargger, Martin Siegemund, Christian Nickel, Roland Bingisser, Michael Osthoff, Stefano Bassetti, Rita Schneider-Sliwa, Manuel Battegay, Hans Hirsch, Adrian Egli |
| EPI_ISL_582127 | Antwerp University Hospital | Institute of Tropical Medicine | Philippe Selhorst, Colin Anthony |
| EPI_ISL_582303, EPI_ISL_582304, EPI_ISL_582305 | Cadham Provincial Laboratory | National Microbiology Laboratory (NML) | Anna Majer, Shari Tyson, Grace Seo, Philip Mabon, Elsie Grudeski, Rhiannon Huzarewich, Russell Mandes, Anneliese Landgraff, Jennifer Tanner, Natalie Knox, Morag Graham, Gary Van Domselaar, Paul Van Caeseele, Jared Bullard, David Alexander, Kerry Dust, Nathalie Bastien, Yan Li, Timothy Booth, Darian Hole, Madison Chapel, CanCOGeN's metadata curation team, Public Health Agency of Canada CanCOGeN team |
| EPI_ISL_582703, EPI_ISL_582707 | Servicio de Microbiología, Hospital Universitario Cruces, Bilbao | SeqCOVID-SPAIN consortium/IBV(CSIC) | Maitane Aranzamendi Zaldumbide and SeqCOVID-SPAIN consortium |
| EPI_ISL_583461 | Memorial Sloan Kettering Cancer Center | van Bakel Laboratory, Genetics and Genomics Sciences, Icahn School of Medicine at Mount Sinai | Teresa Aydillo, Ana S. Gonzalez-Reiche, Sadaf Aslam, Adriana van de Guchte, Zenab Khan, Ajay Obla, Jayeeta Dutta, Harm van Bakel, Judith Aberg, Adolfo García-Sastre, Gunjan Shah, Tobias Hohl, Genovefa Papanicolaou, Miguel-Angel Perales, Kent Sepkowitz, Ngolela Esther Babady, and Mini Kamboj |
| EPI_ISL_583463 | Garcia-Sastre Laboratory, Department of Microbiology, Icahn School of Medicine at Mount Sinai | van Bakel Laboratory, Genetics and Genomics Sciences, Icahn School of Medicine at Mount Sinai | Teresa Aydillo, Ana S. Gonzalez-Reiche, Sadaf Aslam, Adriana van de Guchte, Zenab Khan, Ajay Obla, Jayeeta Dutta, Harm van Bakel, Judith Aberg, Adolfo García-Sastre, Gunjan Shah, Tobias Hohl, Genovefa Papanicolaou, Miguel-Angel Perales, Kent Sepkowitz, Ngolela Esther Babady, and Mini Kamboj |
| EPI_ISL_583469 | Memorial Sloan Kettering Cancer Center | van Bakel Laboratory, Genetics and Genomics Sciences, Icahn School of Medicine at Mount Sinai | Teresa Aydillo, Ana S. Gonzalez-Reiche, Sadaf Aslam, Adriana van de Guchte, Zenab Khan, Ajay Obla, Jayeeta Dutta, Harm van Bakel, Judith Aberg, Adolfo García-Sastre, Gunjan Shah, Tobias Hohl, Genovefa Papanicolaou, Miguel-Angel Perales, Kent Sepkowitz, Ngolela Esther Babady, and Mini Kamboj |
| EPI_ISL_583558 | Universitaetsklinik für Innere Medizin II Innsbruck | Bergthaler laboratory, CeMM Research Center for Molecular Medicine of the Austrian Academy of Sciences | Alexandra Popa, Benedikt Agerer, Henrique Colaco, Lukas Endler, Jakob-Wendelin Genger, Alexander Lercher, Mark Smyth, Thomas Penz, Michael Schuster, Jan Laine, Martin Senekowitsch, Judith Aberle, Stephan Aberle, Peter Hufnagl, Daniela Schmid, Franz Allerberger, Elisabeth Puchhammer-Stoeckl, Manfred Nairz, Guenter Weiss, Gregor Hörmann, Kinga Rigler-Hohenwarter, Rainer Gattringer, Wegene Borena, Dorothee von Laer, Gernot Walder, Peter Obrist, Christian Paar, Sabine Sussitz-Rack, Gunther Vogl, Adi Steinrigl, Christoph Bock, Andreas Bergthaler |
| EPI_ISL_583566 | Center for Virology, Medical University of Vienna | Bergthaler laboratory, CeMM Research Center for Molecular Medicine of the Austrian Academy of Sciences | Alexandra Popa, Benedikt Agerer, Henrique Colaco, Lukas Endler, Jakob-Wendelin Genger, Alexander Lercher, Mark Smyth, Thomas Penz, Michael Schuster, Jan Laine, Martin Senekowitsch, Judith Aberle, Stephan Aberle, Peter Hufnagl, Daniela Schmid, Franz Allerberger, Elisabeth Puchhammer-Stoeckl, Manfred Nairz, Guenter Weiss, Gregor Hörmann, Kinga Rigler-Hohenwarter, Rainer Gattringer, Wegene Borena, Dorothee von Laer, |

| | | | |
|---|---|---|---|
| | | | Gernot Walder, Peter Obrist, Christian Paar, Sabine Sussitz-Rack, Gunther Vogl, Adi Steinrigl, Christoph Bock, Andreas Bergthaler |
| EPI_ISL_583577 | Austrian Agency for Health and Food Safety (AGES) | Bergthaler laboratory, CeMM Research Center for Molecular Medicine of the Austrian Academy of Sciences | Alexandra Popa, Benedikt Agerer, Henrique Colaco, Lukas Endler, Jakob-Wendelin Genger, Alexander Lercher, Mark Smyth, Thomas Penz, Michael Schuster, Jan Laine, Martin Senekowitsch, Judith Aberle, Stephan Aberle, Peter Hufnagl, Daniela Schmid, Franz Allerberger, Elisabeth Puchhammer-Stoeckl, Manfred Nairz, Guenter Weiss, Gregor Hörmann, Kinga Rigler-Hohenwarter, Rainer Gattringer, Wegene Borena, Dorothee von Laer, Gernot Walder, Peter Obrist, Christian Paar, Sabine Sussitz-Rack, Gunther Vogl, Adi Steinrigl, Christoph Bock, Andreas Bergthaler |
| EPI_ISL_583583, EPI_ISL_583585, EPI_ISL_583596, EPI_ISL_583598 | Institute for Medical and Chemical Laboratory Diagnostics, Kepler Universitätsklinikum | Bergthaler laboratory, CeMM Research Center for Molecular Medicine of the Austrian Academy of Sciences | Alexandra Popa, Benedikt Agerer, Henrique Colaco, Lukas Endler, Jakob-Wendelin Genger, Alexander Lercher, Mark Smyth, Thomas Penz, Michael Schuster, Jan Laine, Martin Senekowitsch, Judith Aberle, Stephan Aberle, Peter Hufnagl, Daniela Schmid, Franz Allerberger, Elisabeth Puchhammer-Stoeckl, Manfred Nairz, Guenter Weiss, Gregor Hörmann, Kinga Rigler-Hohenwarter, Rainer Gattringer, Wegene Borena, Dorothee von Laer, Gernot Walder, Peter Obrist, Christian Paar, Sabine Sussitz-Rack, Gunther Vogl, Adi Steinrigl, Christoph Bock, Andreas Bergthaler |
| EPI_ISL_583605, EPI_ISL_583609, EPI_ISL_583610, EPI_ISL_583611, EPI_ISL_583612 | Institut für Virologie am Department für Hygiene, Mikrobiologie und Public Health | Bergthaler laboratory, CeMM Research Center for Molecular Medicine of the Austrian Academy of Sciences | Alexandra Popa, Benedikt Agerer, Henrique Colaco, Lukas Endler, Jakob-Wendelin Genger, Alexander Lercher, Mark Smyth, Thomas Penz, Michael Schuster, Jan Laine, Martin Senekowitsch, Judith Aberle, Stephan Aberle, Peter Hufnagl, Daniela Schmid, Franz Allerberger, Elisabeth Puchhammer-Stoeckl, Manfred Nairz, Guenter Weiss, Gregor Hörmann, Kinga Rigler-Hohenwarter, Rainer Gattringer, Wegene Borena, Dorothee von Laer, Gernot Walder, Peter Obrist, Christian Paar, Sabine Sussitz-Rack, Gunther Vogl, Adi Steinrigl, Christoph Bock, Andreas Bergthaler |
| EPI_ISL_583744, EPI_ISL_583745, EPI_ISL_583746, EPI_ISL_583747, EPI_ISL_583766, EPI_ISL_583767, EPI_ISL_583768, EPI_ISL_583769, EPI_ISL_583770, EPI_ISL_583771, EPI_ISL_583772, EPI_ISL_583773, EPI_ISL_583774, EPI_ISL_583775, EPI_ISL_583776, EPI_ISL_583777, EPI_ISL_583778, EPI_ISL_583779 | | | |
| see above | Dr. Gernot Walder GmbH | Bergthaler laboratory, CeMM Research Center for Molecular Medicine of the Austrian Academy of Sciences | Alexandra Popa, Benedikt Agerer, Henrique Colaco, Lukas Endler, Jakob-Wendelin Genger, Alexander Lercher, Mark Smyth, Thomas Penz, Michael Schuster, Jan Laine, Martin Senekowitsch, Judith Aberle, Stephan Aberle, Peter Hufnagl, Daniela Schmid, Franz Allerberger, Elisabeth Puchhammer-Stoeckl, Manfred Nairz, Guenter Weiss, Gregor Hörmann, Kinga Rigler-Hohenwarter, Rainer Gattringer, Wegene Borena, Dorothee von Laer, Gernot Walder, Peter Obrist, Christian Paar, Sabine Sussitz-Rack, Gunther Vogl, Adi Steinrigl, Christoph Bock, Andreas Bergthaler |
| EPI_ISL_583879, EPI_ISL_583880, EPI_ISL_583881 | Center for Virology, Medical University of Vienna | Bergthaler laboratory, CeMM Research Center for Molecular Medicine of the Austrian Academy of Sciences | Alexandra Popa, Benedikt Agerer, Henrique Colaco, Lukas Endler, Jakob-Wendelin Genger, Alexander Lercher, Mark Smyth, Thomas Penz, Michael Schuster, Jan Laine, Martin Senekowitsch, Judith Aberle, Stephan Aberle, Peter Hufnagl, Daniela Schmid, Franz Allerberger, Elisabeth Puchhammer-Stoeckl, Manfred Nairz, Guenter Weiss, Gregor Hörmann, Kinga Rigler-Hohenwarter, Rainer Gattringer, Wegene Borena, Dorothee von Laer, Gernot Walder, Peter Obrist, Christian Paar, Sabine Sussitz-Rack, Gunther Vogl, Adi Steinrigl, Christoph Bock, Andreas Bergthaler |
| EPI_ISL_586297, EPI_ISL_586298, EPI_ISL_586299, EPI_ISL_586356, EPI_ISL_586361, EPI_ISL_586369, EPI_ISL_586370, EPI_ISL_586373, EPI_ISL_586381, EPI_ISL_586393, EPI_ISL_586402, EPI_ISL_586406, EPI_ISL_586407, EPI_ISL_586409, EPI_ISL_586410, EPI_ISL_586411, EPI_ISL_586412, EPI_ISL_586413, EPI_ISL_586416, EPI_ISL_586418, EPI_ISL_586419, EPI_ISL_586422 | | | |
| see above | Toronto Invasive Bacterial Diseases Network | McMaster University | Allison McGeer, Patryk Aftanas, Hooman Derakhshani, Angel Li, Kuganya Nirmalarajah, Emily Panousis, Ahmed Draia, Jalees Nasir, Michael Surette, Samira Mubareka, Andrew G. McArthur |
| EPI_ISL_590917 | Unilabs Laboratory Medicine | Norwegian Institute of Public Health, Department of Virology | Kathrine Stene-Johansen, Kamilla Heddeland Instefjord, Hilde Elshaug, Rasmus Riis Kopperud, Hilde Vollan, Karoline Bragstad, Olav Hungnes |
| EPI_ISL_591124, EPI_ISL_591127, EPI_ISL_591128, EPI_ISL_591130, EPI_ISL_591138 | Toronto Invasive Bacterial Diseases Network | McMaster University | Allison McGeer, Patryk Aftanas, Hooman Derakhshani, Angel Li, Kuganya Nirmalarajah, Emily Panousis, Ahmed Draia, Jalees Nasir, Michael Surette, Samira Mubareka, Andrew G. McArthur |
| EPI_ISL_593756, EPI_ISL_593757, EPI_ISL_593758, EPI_ISL_593759 | Sydney South West Pathology Service (SSWPS) - Liverpool Hospital - NSW Health Pathology | NSW Health Pathology - Institute of Clinical Pathology and Medical Research; Westmead Hospital; University of Sydney | CIDM-PH et al. |
| EPI_ISL_594190, EPI_ISL_594191, EPI_ISL_594192 | Michigan Department of Health and Human Services, Bureau of Laboratories | Michigan Department of Health and Human Services, Bureau of Laboratories | Blankenship HM, Riner D, Soehnlen MK |
| EPI_ISL_596505, EPI_ISL_596506 | Palestinian Ministry of Health | Molecular Genetics Lab | Nouar Qutob, Zaidoun Salah, Damien Richard, Hisham Darwish, Husam Sallam, Issa Shtayeh, Osama Najjar, Mahmoud Ruzayqat, Dana Najjar, Francois Balloux, Lucy van Dorp |
| EPI_ISL_596824, EPI_ISL_596827, EPI_ISL_596829, EPI_ISL_596846, EPI_ISL_596847, EPI_ISL_596854, EPI_ISL_596865, EPI_ISL_596874, EPI_ISL_596877, EPI_ISL_596880 | PathWest Laboratory Medicine WA | PathWest Laboratory Medicine WA Microbial Surveillance Unit | PathWest Laboratory Medicine WA Microbial Surveillance Unit |
| EPI_ISL_600436, EPI_ISL_600440 | Institute of Epidemiology Disease Control And Research | Institute for Developing Science and Health Initiatives | Lauren Cowley, Mokibul Hassan Afrad, Sadia Isfat Ara Rahman, Md. Mahfuz-Al-mamun, Firadausi Qadri, Tahmina Shirin |
| EPI_ISL_602486, EPI_ISL_602487, EPI_ISL_602488, EPI_ISL_602489 | Institute for Virology, University Hospital Essen | Center of Medical Microbiology, Virology, and Hospital Hygiene, University of Duesseldorf | Olympia E. Anastasiou, Ulf Dittmer, Maximilian Damagnez, Alexander Dilthey, Torsten Houwaart, Lisanna Hülse, Malte Kohns Vasconcelos, Nadine Lübke, Jessica Nicolai, Klaus Pfeffer, Daniel Strelow, Jörg Timm, Andreas Walker, Tobias Wienemann |
| EPI_ISL_602516 | Center of Medical Microbiology, Virology, and Hospital Hygiene, University of Duesseldorf | Center of Medical Microbiology, Virology, and Hospital Hygiene, University of Duesseldorf | Maximilian Damagnez, Alexander Dilthey, Torsten Houwaart, Lisanna Hülse, Malte Kohns Vasconcelos, Marek Korencak, Nadine Lübke, Jessica Nicolai, Klaus Pfeffer, Hendrik Streeck, Daniel Strelow, Jörg Timm, Andreas Walker, Tobias Wienemann |
| EPI_ISL_602532 | Institute for Virology, University Hospital Essen | Center of Medical Microbiology, Virology, and Hospital Hygiene, University of Duesseldorf | Olympia E. Anastasiou, Ulf Dittmer, Maximilian Damagnez, Alexander Dilthey, Torsten Houwaart, Lisanna Hülse, Malte Kohns Vasconcelos, Nadine Lübke, Jessica Nicolai, Klaus Pfeffer, Daniel Strelow, Jörg Timm, Andreas Walker, Tobias Wienemann |
| EPI_ISL_602534 | Center of Medical Microbiology, Virology, and Hospital Hygiene, University of Duesseldorf | Center of Medical Microbiology, Virology, and Hospital Hygiene, University of Duesseldorf | Maximilian Damagnez, Alexander Dilthey, Torsten Houwaart, Malte Kohns Vasconcelos, Marek Korencak, Nadine Lübke, Jessica Nicolai, Klaus Pfeffer, Hendrik Streeck, Daniel Strelow, Jörg Timm, Andreas Walker, Tobias Wienemann |
| EPI_ISL_603759, EPI_ISL_603760, EPI_ISL_603761, EPI_ISL_603762, EPI_ISL_603763, EPI_ISL_603764, EPI_ISL_603765, EPI_ISL_603766, EPI_ISL_603767, EPI_ISL_603768, EPI_ISL_603769, EPI_ISL_603770, EPI_ISL_603771, EPI_ISL_603772, EPI_ISL_603773, EPI_ISL_603774, EPI_ISL_603775, EPI_ISL_603776, EPI_ISL_603777, EPI_ISL_603778, EPI_ISL_603779, EPI_ISL_603780, EPI_ISL_603781, EPI_ISL_603782, EPI_ISL_603783, EPI_ISL_603784, EPI_ISL_603785, EPI_ISL_603786, EPI_ISL_603787, EPI_ISL_603788, EPI_ISL_603789, EPI_ISL_603790, EPI_ISL_603791, EPI_ISL_603792, EPI_ISL_603793, EPI_ISL_603794, EPI_ISL_603795, EPI_ISL_603796, EPI_ISL_603797, EPI_ISL_603798, EPI_ISL_603799, EPI_ISL_603800, EPI_ISL_603801, EPI_ISL_603802, EPI_ISL_603803, EPI_ISL_603804, EPI_ISL_603805, EPI_ISL_603806, EPI_ISL_603807, EPI_ISL_603808, EPI_ISL_603809, EPI_ISL_603810, EPI_ISL_603811, EPI_ISL_603812, EPI_ISL_603813, EPI_ISL_603814, EPI_ISL_603815, EPI_ISL_603816, EPI_ISL_603817, EPI_ISL_603818, EPI_ISL_603819, EPI_ISL_603820, EPI_ISL_603821, EPI_ISL_603822, EPI_ISL_603823, EPI_ISL_603826, EPI_ISL_603827, EPI_ISL_603828, EPI_ISL_603829, EPI_ISL_603830, EPI_ISL_603831, EPI_ISL_603832, EPI_ISL_603833, EPI_ISL_603834, EPI_ISL_603835, EPI_ISL_603836, EPI_ISL_603837, EPI_ISL_603838, EPI_ISL_603839, EPI_ISL_603840, EPI_ISL_603841, EPI_ISL_603842, EPI_ISL_603843, EPI_ISL_603844, EPI_ISL_603845, EPI_ISL_603847, EPI_ISL_603849, EPI_ISL_603850, EPI_ISL_603851, EPI_ISL_603852, EPI_ISL_603853, EPI_ISL_603854, EPI_ISL_603855, EPI_ISL_603856, EPI_ISL_603857, EPI_ISL_603859, EPI_ISL_603862, EPI_ISL_603866, EPI_ISL_603872, EPI_ISL_603873, EPI_ISL_603874, EPI_ISL_603877, EPI_ISL_603878, EPI_ISL_603879, EPI_ISL_603880, EPI_ISL_603881, EPI_ISL_603882, EPI_ISL_603883, EPI_ISL_603884, EPI_ISL_603886, EPI_ISL_603888, EPI_ISL_603889, EPI_ISL_603890, EPI_ISL_603891, EPI_ISL_603893, EPI_ISL_603894, EPI_ISL_603895, EPI_ISL_603898, EPI_ISL_603899, EPI_ISL_603901, EPI_ISL_603903, EPI_ISL_603906, EPI_ISL_603908, EPI_ISL_603909, EPI_ISL_603910, EPI_ISL_603911, EPI_ISL_603914, EPI_ISL_603919, EPI_ISL_603920, EPI_ISL_603924, EPI_ISL_603927, EPI_ISL_603928, EPI_ISL_603934, EPI_ISL_603936, EPI_ISL_603937, EPI_ISL_603938, EPI_ISL_603939, EPI_ISL_603940, EPI_ISL_603941, EPI_ISL_603942, EPI_ISL_603943, EPI_ISL_603944, EPI_ISL_603945, EPI_ISL_603946, EPI_ISL_603947, EPI_ISL_603948, EPI_ISL_603949, EPI_ISL_603950, EPI_ISL_603951, EPI_ISL_603952, EPI_ISL_603953, EPI_ISL_603954, EPI_ISL_603955, EPI_ISL_603956, EPI_ISL_603957, EPI_ISL_603958, EPI_ISL_603959, EPI_ISL_603960, EPI_ISL_603961, EPI_ISL_603962, EPI_ISL_603963, EPI_ISL_603964, EPI_ISL_603965, EPI_ISL_603966, EPI_ISL_603967, EPI_ISL_603968, EPI_ISL_603969, EPI_ISL_603970, EPI_ISL_603971, EPI_ISL_603972, EPI_ISL_603973, EPI_ISL_603974, EPI_ISL_603975, EPI_ISL_603976, EPI_ISL_603977, EPI_ISL_603978, EPI_ISL_603979, EPI_ISL_603980, EPI_ISL_603981, EPI_ISL_603982, EPI_ISL_603983, EPI_ISL_603984, EPI_ISL_603985, EPI_ISL_603986, EPI_ISL_603987, EPI_ISL_603988, EPI_ISL_603989, EPI_ISL_603990, EPI_ISL_603991, EPI_ISL_603992, EPI_ISL_603993, EPI_ISL_603994, EPI_ISL_603995, EPI_ISL_603996, EPI_ISL_603997, EPI_ISL_603998, EPI_ISL_603999, EPI_ISL_604000, EPI_ISL_604001, EPI_ISL_604002, EPI_ISL_604003, EPI_ISL_604004, EPI_ISL_604005, EPI_ISL_604006, EPI_ISL_604007, EPI_ISL_604008, EPI_ISL_604009, EPI_ISL_604010, EPI_ISL_604011, EPI_ISL_604012, EPI_ISL_604013, EPI_ISL_604014, EPI_ISL_604015, EPI_ISL_604016, EPI_ISL_604017, EPI_ISL_604018, EPI_ISL_604020, EPI_ISL_604021, EPI_ISL_604022, EPI_ISL_604023, EPI_ISL_604024, EPI_ISL_604025, EPI_ISL_604026, EPI_ISL_604027, EPI_ISL_604028, EPI_ISL_604029, EPI_ISL_604030, EPI_ISL_604031, EPI_ISL_604032, EPI_ISL_604033, EPI_ISL_604034, EPI_ISL_604035, EPI_ISL_604036, EPI_ISL_604037, EPI_ISL_604038, EPI_ISL_604039, EPI_ISL_604040, EPI_ISL_604041, EPI_ISL_604042, EPI_ISL_604043, EPI_ISL_604044, EPI_ISL_604045, EPI_ISL_604046, EPI_ISL_604047, EPI_ISL_604048, EPI_ISL_604049, EPI_ISL_604050, EPI_ISL_604051, EPI_ISL_604052, EPI_ISL_604053, EPI_ISL_604054, EPI_ISL_604055, EPI_ISL_604056, EPI_ISL_604057, EPI_ISL_604058, EPI_ISL_604059, EPI_ISL_604060, EPI_ISL_604061, EPI_ISL_604063, EPI_ISL_604064, EPI_ISL_604065, EPI_ISL_604066, EPI_ISL_604067, EPI_ISL_604068, EPI_ISL_604069, EPI_ISL_604070, EPI_ISL_604071, EPI_ISL_604072, EPI_ISL_604073, EPI_ISL_604074, EPI_ISL_604075, EPI_ISL_604076, EPI_ISL_604077, EPI_ISL_604078, EPI_ISL_604079, EPI_ISL_604080, EPI_ISL_604081, EPI_ISL_604082, EPI_ISL_604083, EPI_ISL_604084, EPI_ISL_604085, EPI_ISL_604086, EPI_ISL_604087, EPI_ISL_604088, EPI_ISL_604089, EPI_ISL_604091, EPI_ISL_604092, EPI_ISL_604094, EPI_ISL_604095, EPI_ISL_604096, EPI_ISL_604097, EPI_ISL_604098, EPI_ISL_604099, EPI_ISL_604100, EPI_ISL_604102, EPI_ISL_604103, EPI_ISL_604104, EPI_ISL_604105, EPI_ISL_604106, EPI_ISL_604107, EPI_ISL_604108, EPI_ISL_604109, EPI_ISL_604110, EPI_ISL_604111, EPI_ISL_604112, EPI_ISL_604113, EPI_ISL_604114, EPI_ISL_604115, EPI_ISL_604116, EPI_ISL_604118, EPI_ISL_604120, EPI_ISL_604121, EPI_ISL_604124, EPI_ISL_604125, EPI_ISL_604126, EPI_ISL_604127, EPI_ISL_604128, EPI_ISL_604130, EPI_ISL_604133, EPI_ISL_604135, EPI_ISL_604137, EPI_ISL_604138, EPI_ISL_604140, EPI_ISL_604141, EPI_ISL_604142, EPI_ISL_604143, EPI_ISL_604146, EPI_ISL_604147, EPI_ISL_604148, EPI_ISL_604155, EPI_ISL_604158, EPI_ISL_604159, EPI_ISL_604160, EPI_ISL_604162, EPI_ISL_604164, EPI_ISL_604165, EPI_ISL_604166, EPI_ISL_604167, EPI_ISL_604168, EPI_ISL_604169, EPI_ISL_604171, EPI_ISL_604172, EPI_ISL_604175, EPI_ISL_604176, EPI_ISL_604177, EPI_ISL_604179, EPI_ISL_604180, EPI_ISL_604182, EPI_ISL_604183, EPI_ISL_604185, EPI_ISL_604186, EPI_ISL_604188, EPI_ISL_604192, EPI_ISL_604194, EPI_ISL_604195, EPI_ISL_604196, EPI_ISL_604197, EPI_ISL_604198, EPI_ISL_604200, EPI_ISL_604201, EPI_ISL_604202, EPI_ISL_604203, EPI_ISL_604204, EPI_ISL_604205, EPI_ISL_604206, EPI_ISL_604207, EPI_ISL_604209, EPI_ISL_604211, EPI_ISL_604212, EPI_ISL_604213, EPI_ISL_604214, EPI_ISL_604215, EPI_ISL_604216, EPI_ISL_604217, EPI_ISL_604218, EPI_ISL_604220, EPI_ISL_604221, EPI_ISL_604224, EPI_ISL_604226, EPI_ISL_604227, EPI_ISL_604230, EPI_ISL_604232, EPI_ISL_604234, EPI_ISL_604235, EPI_ISL_604236, EPI_ISL_604237, EPI_ISL_604238, EPI_ISL_604239, EPI_ISL_604240, EPI_ISL_604241, EPI_ISL_604243, EPI_ISL_604244, EPI_ISL_604245, EPI_ISL_604246, EPI_ISL_604247, EPI_ISL_604248, EPI_ISL_604249, EPI_ISL_604250, EPI_ISL_604251, EPI_ISL_604252, EPI_ISL_604253, EPI_ISL_604254, EPI_ISL_604255, EPI_ISL_604256, EPI_ISL_604257, EPI_ISL_604259, EPI_ISL_604260, EPI_ISL_604261, EPI_ISL_604262, EPI_ISL_604263, EPI_ISL_604265, EPI_ISL_604266, EPI_ISL_604267, EPI_ISL_604268, EPI_ISL_604269, EPI_ISL_604270, EPI_ISL_604271, EPI_ISL_604272, EPI_ISL_604273, EPI_ISL_604274, EPI_ISL_604275, EPI_ISL_604276, EPI_ISL_604277, EPI_ISL_604278, EPI_ISL_604293, EPI_ISL_604294, EPI_ISL_604295, EPI_ISL_604296, EPI_ISL_604297, EPI_ISL_604298, EPI_ISL_604299, EPI_ISL_604300, EPI_ISL_604301, EPI_ISL_604302, EPI_ISL_604303, EPI_ISL_604304, EPI_ISL_604306, EPI_ISL_604307, EPI_ISL_604309, EPI_ISL_604312, EPI_ISL_604313, EPI_ISL_604314, EPI_ISL_604315, EPI_ISL_604316, EPI_ISL_604317, EPI_ISL_604318, EPI_ISL_604319, EPI_ISL_604322, EPI_ISL_604323, EPI_ISL_604324, EPI_ISL_604325, EPI_ISL_604326, EPI_ISL_604327, EPI_ISL_604328, EPI_ISL_604329, EPI_ISL_604330, EPI_ISL_604331, EPI_ISL_604332, EPI_ISL_604333, EPI_ISL_604334, EPI_ISL_604335, EPI_ISL_604336, EPI_ISL_604337, EPI_ISL_604338, EPI_ISL_604339, EPI_ISL_604340, EPI_ISL_604341, EPI_ISL_604342, EPI_ISL_604343, EPI_ISL_604344, EPI_ISL_604345, EPI_ISL_604346, EPI_ISL_604347, EPI_ISL_604349, EPI_ISL_604350, EPI_ISL_604351, EPI_ISL_604353, EPI_ISL_604354, EPI_ISL_604355, EPI_ISL_604357, EPI_ISL_604358, EPI_ISL_604359, EPI_ISL_604360, EPI_ISL_604361, EPI_ISL_604362, EPI_ISL_604363, | | | |

| | | | |
|---|---|---|---|
| EPI_ISL_604364, EPI_ISL_604365, EPI_ISL_604366, EPI_ISL_604367, EPI_ISL_604368, EPI_ISL_604369, EPI_ISL_604370, EPI_ISL_604371, EPI_ISL_604372, EPI_ISL_604373, EPI_ISL_604374, EPI_ISL_604375, EPI_ISL_604376, EPI_ISL_604377, EPI_ISL_604378, EPI_ISL_604379, EPI_ISL_604381, EPI_ISL_604382, EPI_ISL_604384, EPI_ISL_604385, EPI_ISL_604386, EPI_ISL_604387, EPI_ISL_604388, EPI_ISL_604389, EPI_ISL_604392, EPI_ISL_604400, EPI_ISL_604403, EPI_ISL_604406, EPI_ISL_604407, EPI_ISL_604408, EPI_ISL_604410, EPI_ISL_604412, EPI_ISL_604413, EPI_ISL_604414, EPI_ISL_604415, EPI_ISL_604418, EPI_ISL_604420, EPI_ISL_604421, EPI_ISL_604423, EPI_ISL_604427, EPI_ISL_604428, EPI_ISL_604429, EPI_ISL_604431, EPI_ISL_604432, EPI_ISL_604435, EPI_ISL_604437, EPI_ISL_604438, EPI_ISL_604441, EPI_ISL_604443, EPI_ISL_604444, EPI_ISL_604446, EPI_ISL_604455, EPI_ISL_604457, EPI_ISL_604458, EPI_ISL_604459, EPI_ISL_604460, EPI_ISL_604461, EPI_ISL_604462, EPI_ISL_604463, EPI_ISL_604464, EPI_ISL_604465, EPI_ISL_604466, EPI_ISL_604467, EPI_ISL_604468, EPI_ISL_604469, EPI_ISL_604470, EPI_ISL_604471, EPI_ISL_604472, EPI_ISL_604473, EPI_ISL_604474, EPI_ISL_604475, EPI_ISL_604476, EPI_ISL_604477, EPI_ISL_604478, EPI_ISL_604479, EPI_ISL_604480, EPI_ISL_604481, EPI_ISL_604482, EPI_ISL_604483, EPI_ISL_604488, EPI_ISL_604496, EPI_ISL_604497, EPI_ISL_604501, EPI_ISL_604503, EPI_ISL_604504, EPI_ISL_604505, EPI_ISL_604506, EPI_ISL_604507, EPI_ISL_604509, EPI_ISL_604510, EPI_ISL_604511, EPI_ISL_604514, EPI_ISL_604515, EPI_ISL_604516, EPI_ISL_604517, EPI_ISL_604518, EPI_ISL_604519, EPI_ISL_604520, EPI_ISL_604521, EPI_ISL_604522, EPI_ISL_604523, EPI_ISL_604528, EPI_ISL_604529, EPI_ISL_604530, EPI_ISL_604534, EPI_ISL_604539, EPI_ISL_604541, EPI_ISL_604542, EPI_ISL_604544, EPI_ISL_604547, EPI_ISL_604549, EPI_ISL_604553, EPI_ISL_604554, EPI_ISL_604561, EPI_ISL_604565, EPI_ISL_604566, EPI_ISL_604567, EPI_ISL_604568, EPI_ISL_604569, EPI_ISL_604570, EPI_ISL_604571, EPI_ISL_604573, EPI_ISL_604574, EPI_ISL_604575, EPI_ISL_604576, EPI_ISL_604577, EPI_ISL_604621, EPI_ISL_604622, EPI_ISL_604626, EPI_ISL_604627, EPI_ISL_604628, EPI_ISL_604629, EPI_ISL_604630, EPI_ISL_604631, EPI_ISL_604632, EPI_ISL_604633, EPI_ISL_604634, EPI_ISL_604635, EPI_ISL_604636, EPI_ISL_604637, EPI_ISL_604638, EPI_ISL_604639, EPI_ISL_604640, EPI_ISL_604642, EPI_ISL_604643, EPI_ISL_604644, EPI_ISL_604645, EPI_ISL_604647, EPI_ISL_604648, EPI_ISL_604649, EPI_ISL_604650, EPI_ISL_604651, EPI_ISL_604652, EPI_ISL_604653, EPI_ISL_604654, EPI_ISL_604655, EPI_ISL_604656, EPI_ISL_604657, EPI_ISL_604658, EPI_ISL_604659, EPI_ISL_604660, EPI_ISL_604662, EPI_ISL_604663, EPI_ISL_604664, EPI_ISL_604665, EPI_ISL_604666, EPI_ISL_604667, EPI_ISL_604668, EPI_ISL_604669, EPI_ISL_604670, EPI_ISL_604671, EPI_ISL_604672, EPI_ISL_604673, EPI_ISL_604674, EPI_ISL_604675, EPI_ISL_604676, EPI_ISL_604677, EPI_ISL_604678, EPI_ISL_604679, EPI_ISL_604680, EPI_ISL_604681, EPI_ISL_604682, EPI_ISL_604683, EPI_ISL_604684, EPI_ISL_604685, EPI_ISL_604686, EPI_ISL_604687, EPI_ISL_604688, EPI_ISL_604689, EPI_ISL_604690, EPI_ISL_604691, EPI_ISL_604692, EPI_ISL_604693, EPI_ISL_604694, EPI_ISL_604695, EPI_ISL_604696, EPI_ISL_604697, EPI_ISL_604698, EPI_ISL_604699, EPI_ISL_604700, EPI_ISL_604701, EPI_ISL_604702, EPI_ISL_604703, EPI_ISL_604704, EPI_ISL_604705, EPI_ISL_604706, EPI_ISL_604707, EPI_ISL_604709, EPI_ISL_604710, EPI_ISL_604711, EPI_ISL_604712, EPI_ISL_604713, EPI_ISL_604714, EPI_ISL_604715, EPI_ISL_604716, EPI_ISL_604717, EPI_ISL_604718, EPI_ISL_604719, EPI_ISL_604720, EPI_ISL_604721, EPI_ISL_604722, EPI_ISL_604723, EPI_ISL_604724, EPI_ISL_604727, EPI_ISL_604729, EPI_ISL_604731, EPI_ISL_604732, EPI_ISL_604733, EPI_ISL_604734, EPI_ISL_604735, EPI_ISL_604736, EPI_ISL_604737, EPI_ISL_604738, EPI_ISL_604739, EPI_ISL_604740, EPI_ISL_604741, EPI_ISL_604742, EPI_ISL_604743, EPI_ISL_604744, EPI_ISL_604745, EPI_ISL_604746, EPI_ISL_604747, EPI_ISL_604748, EPI_ISL_604749, EPI_ISL_604750, EPI_ISL_604752, EPI_ISL_604753, EPI_ISL_604754, EPI_ISL_604755, EPI_ISL_604756, EPI_ISL_604757, EPI_ISL_604759, EPI_ISL_604760, EPI_ISL_604761, EPI_ISL_604763, EPI_ISL_604764, EPI_ISL_604765, EPI_ISL_604766, EPI_ISL_604767, EPI_ISL_604768, EPI_ISL_604769, EPI_ISL_604770, EPI_ISL_604772, EPI_ISL_604773, EPI_ISL_604774, EPI_ISL_604775, EPI_ISL_604776, EPI_ISL_604786, EPI_ISL_604792, EPI_ISL_604794, EPI_ISL_604796, EPI_ISL_604797, EPI_ISL_604798, EPI_ISL_604799, EPI_ISL_604802, EPI_ISL_604803, EPI_ISL_604804, EPI_ISL_604806, EPI_ISL_604807, EPI_ISL_604808, EPI_ISL_604810, EPI_ISL_604812, EPI_ISL_604813, EPI_ISL_604815, EPI_ISL_604816, EPI_ISL_604817, EPI_ISL_604818, EPI_ISL_604820, EPI_ISL_604821, EPI_ISL_604822, EPI_ISL_604824, EPI_ISL_604825, EPI_ISL_604826, EPI_ISL_604827, EPI_ISL_604828, EPI_ISL_604830, EPI_ISL_604833, EPI_ISL_604834, EPI_ISL_604840, EPI_ISL_604868, EPI_ISL_604869, EPI_ISL_604870, EPI_ISL_604871, EPI_ISL_604872, EPI_ISL_604873, EPI_ISL_604874, EPI_ISL_604875, EPI_ISL_604876, EPI_ISL_604877, EPI_ISL_604878, EPI_ISL_604879, EPI_ISL_604880, EPI_ISL_604881, EPI_ISL_604882, EPI_ISL_604883, EPI_ISL_604884, EPI_ISL_604885, EPI_ISL_604886, EPI_ISL_604887, EPI_ISL_604888, EPI_ISL_604889, EPI_ISL_604890, EPI_ISL_604891, EPI_ISL_604892, EPI_ISL_604893, EPI_ISL_604894, EPI_ISL_604895, EPI_ISL_604896, EPI_ISL_604897, EPI_ISL_604898, EPI_ISL_604899, EPI_ISL_604900, EPI_ISL_604901, EPI_ISL_604902, EPI_ISL_604903, EPI_ISL_604904, EPI_ISL_604905, EPI_ISL_604906, EPI_ISL_604907, EPI_ISL_604908, EPI_ISL_604909, EPI_ISL_604910, EPI_ISL_604911, EPI_ISL_604912, EPI_ISL_604913, EPI_ISL_604914, EPI_ISL_604915, EPI_ISL_604916, EPI_ISL_604917, EPI_ISL_604918, EPI_ISL_604921, EPI_ISL_604922, EPI_ISL_604923, EPI_ISL_604925, EPI_ISL_604926, EPI_ISL_604927, EPI_ISL_604928, EPI_ISL_604929, EPI_ISL_604930, EPI_ISL_604934, EPI_ISL_604935, EPI_ISL_604936, EPI_ISL_604938, EPI_ISL_604939, EPI_ISL_604940, EPI_ISL_604941, EPI_ISL_604942, EPI_ISL_604943, EPI_ISL_604945, EPI_ISL_604947, EPI_ISL_604948, EPI_ISL_604950, EPI_ISL_604951, EPI_ISL_604952, EPI_ISL_604953, EPI_ISL_604954, EPI_ISL_604955, EPI_ISL_604956, EPI_ISL_604957, EPI_ISL_604958, EPI_ISL_604959, EPI_ISL_604960, EPI_ISL_604992, EPI_ISL_604993, EPI_ISL_604994, EPI_ISL_604995, EPI_ISL_604996, EPI_ISL_604997, EPI_ISL_604998, EPI_ISL_604999, EPI_ISL_605000, EPI_ISL_605001, EPI_ISL_605002, EPI_ISL_605003, EPI_ISL_605004, EPI_ISL_605005, EPI_ISL_605006, EPI_ISL_605007, EPI_ISL_605008, EPI_ISL_605009, EPI_ISL_605010, EPI_ISL_605011, EPI_ISL_605012, EPI_ISL_605013, EPI_ISL_605014, EPI_ISL_605015, EPI_ISL_605016, EPI_ISL_605017, EPI_ISL_605018, EPI_ISL_605019, EPI_ISL_605020, EPI_ISL_605021, EPI_ISL_605022, EPI_ISL_605023, EPI_ISL_605024, EPI_ISL_605025, EPI_ISL_605037, EPI_ISL_605038, EPI_ISL_605039, EPI_ISL_605040, EPI_ISL_605042, EPI_ISL_605043, EPI_ISL_605046, EPI_ISL_605047, EPI_ISL_605048, EPI_ISL_605050, EPI_ISL_605053, EPI_ISL_605054, EPI_ISL_605055, EPI_ISL_605056, EPI_ISL_605057 | | | |
| see above | Quest Diagnostics | Quest Diagnostics | Rosenthal,S.H., Gerasimova,A., Kagan,R.M., Anderson, B., Grover, D., Livingston, K.E., Hua, M., Liu Y., Shalhout, D.F., Owen, R., Lacbawan, F. |
| EPI_ISL_605059, EPI_ISL_605064, EPI_ISL_605068, EPI_ISL_605073, EPI_ISL_605079 | National Virus Reference Laboratory | Irish Coronavirus Sequencing Consortium - Helixworks | Sachin Chalapati, Conor Crosbie, Nimesh Pinnamaneni |
| EPI_ISL_605926 | Emergence des Maladies Virales, Centre Interdisciplinaire de Recherches Medicales de Franceville | Emergence des Maladies Virales, Centre Interdisciplinaire de Recherches Medicales de Franceville | Lekana-Douki,S.E., N'dilimabaka,N., Mbongo kama,E., Kandet Yattara,M., Mintsa Ndong,A., Ngonga Dikongo,A.M., Andeko,J.C., Zong Minko,O., Lekana-Douki,J.-B. |
| EPI_ISL_610164, EPI_ISL_610165, EPI_ISL_610166 | Department of Health Technology and Informatics, The Hong Kong Polytechnic University | Department of Health Technology and Informatics, The Hong Kong Polytechnic University | Siu,G.K.-H., Lee,L.-K., Leung,K.S.-S., Leung,J.S.-L., Ng,T.T.-L., Chan,C.T.-M., Tam,K.K.-G., Lao,H.-Y., Wu,A.K.-L., Yau,M.C.-Y., Lai,Y.W.-M., Fung,K.S.-C., Chau,S.K.-Y., Wong,B.K.-C., To,W.-K., Luk,K., Ho,A.Y.-M., Que,T.-L., Yip,K.-T., Yam,W.C., Shum,D.H.-K., Yip,S.P. |
| EPI_ISL_613419, EPI_ISL_613422, EPI_ISL_613423, EPI_ISL_613425, EPI_ISL_613426, EPI_ISL_613427, EPI_ISL_613428, EPI_ISL_613430, EPI_ISL_613431 | Institut Pasteur de la Guadeloupe | Institut Pasteur de la Guadeloupe | Marion Barbet, Sylvie Behillil, Méline Bizard, Angela Brisebarre, Camille Capel, Etienne Simon-Lorière, Vincent Enouf, Maud Vanpeene, Sylvie van der Werf, Stéphanie Guyomard, Sébastien Breurec, Antoine Talarmin |
| EPI_ISL_614403, EPI_ISL_614415, EPI_ISL_614416, EPI_ISL_614420, EPI_ISL_614421, EPI_ISL_614422, EPI_ISL_614439, EPI_ISL_614473, EPI_ISL_614600, EPI_ISL_614601, EPI_ISL_614602, EPI_ISL_614603, EPI_ISL_614604, EPI_ISL_614605, EPI_ISL_614606, EPI_ISL_614607, EPI_ISL_614608, EPI_ISL_614609, EPI_ISL_614610, EPI_ISL_614611, EPI_ISL_614612, EPI_ISL_614613, EPI_ISL_614614, EPI_ISL_614615, EPI_ISL_614616, EPI_ISL_614617, EPI_ISL_614618, EPI_ISL_614619, EPI_ISL_614620, EPI_ISL_614621, EPI_ISL_614622, EPI_ISL_614623, EPI_ISL_614624, EPI_ISL_614625, EPI_ISL_614626, EPI_ISL_614627, EPI_ISL_614628, EPI_ISL_614630, EPI_ISL_614633, EPI_ISL_614634, EPI_ISL_614635, EPI_ISL_614637, EPI_ISL_614638, EPI_ISL_614639, EPI_ISL_614640, EPI_ISL_614641, EPI_ISL_614642, EPI_ISL_614643, EPI_ISL_614644, EPI_ISL_614645, EPI_ISL_614646, EPI_ISL_614647, EPI_ISL_614648, EPI_ISL_614649, EPI_ISL_614650, EPI_ISL_614651, EPI_ISL_614652, EPI_ISL_614653, EPI_ISL_614654, EPI_ISL_614655, EPI_ISL_614658, EPI_ISL_614659, EPI_ISL_614660, EPI_ISL_614661, EPI_ISL_614662, EPI_ISL_614664, EPI_ISL_614665, EPI_ISL_614666, EPI_ISL_614667, EPI_ISL_614668, EPI_ISL_614675, EPI_ISL_614677, EPI_ISL_614678, EPI_ISL_614679, EPI_ISL_614681, EPI_ISL_614682, EPI_ISL_614683, EPI_ISL_614684, EPI_ISL_614685, EPI_ISL_614686, EPI_ISL_614687, EPI_ISL_614689, EPI_ISL_614690, EPI_ISL_614691, EPI_ISL_614692, EPI_ISL_614755, EPI_ISL_614757, EPI_ISL_615467, EPI_ISL_619455, EPI_ISL_619456, EPI_ISL_619492, EPI_ISL_619534, EPI_ISL_619556, EPI_ISL_619657, EPI_ISL_620145, EPI_ISL_620146, EPI_ISL_620147, EPI_ISL_620286, EPI_ISL_620302, EPI_ISL_622500, EPI_ISL_622511, EPI_ISL_622524, EPI_ISL_622679, EPI_ISL_622724 | | | |
| see above | Department of Virus and Microbiological Special Diagnostics, Statens Serum Institut, Denmark | Albertsen lab, Department of Chemistry and Bioscience, Aalborg University, Denmark | Danish Covid-19 Genome Consortia |
| EPI_ISL_623104 | Simile Medicina Diagnóstica | Bioinformatics Laboratory / LNCC | Carolina M Voloch, Ronaldo S Francisco Jr, Luiz G P de Almeida, Otavio J. Brustolini, Cynthia C Cardoso, Alexandra L Gerber, Ana Paula de C Guimarães, Diana Mariani, Covid19-UFRJ Workgroup, Luís Cristóvão Pôrto, Renato S Aguiar, Terezinha M P P Castiñeiras, Orlando C. Ferreira, Amilcar Tanuri, Ana Tereza R de Vasconcelos |
| EPI_ISL_626225, EPI_ISL_626226 | Institute for Virology, University Hospital Essen | Center of Medical Microbiology, Virology, and Hospital Hygiene, University of Duesseldorf | Olympia E. Anastasiou, Ulf Dittmer, Maximilian Damagnez, Alexander Dilthey, Torsten Houwaart, Lisanna Hülse, Malte Kohns Vasconcelos, Nadine Lübke, Jessica Nicolai, Klaus Pfeffer, Daniel Strelow, Jörg Timm, Andreas Walker, Tobias Wienemann |
| EPI_ISL_626357, EPI_ISL_626358, EPI_ISL_626359, EPI_ISL_626360, EPI_ISL_626361, EPI_ISL_626362 | Northwestern Memorial Hospital | Ozer Lab | Ramon Lorenzo-Redondo, Hannah H. Nam, Scott C. Roberts, Lacy M. Simons, Chad J. Achenbach, Lawrence J. Jennings, Chao Qi, Alan R. Hauser, Michael G. Ison, Judd F. Hultquist, Egon A. Ozer |
| EPI_ISL_626566 | National Centre for Communication Disease (NCCD) National Influenza Center | National Centre for Communication Disease (NCCD) National Influenza Center | Naranzul Ts,Darmaa B,Bayasgalan N,Ankhbayar S,Tsogtbaatar B,Erden-Ochir Ts,Nymadawa P,Chang-Seon Song |
| EPI_ISL_629024 | Centro de Biotecnología Vegetal, Universidad Andrés Bello, Center for Genome Regulation | Center for Mathematical Modeling and Center for Genome Regulation. Santiago, Chile | Bastias M, Sanhueza D, Travisany D, Allende ML, Maass A, González M, Bustos F, Arriagada G, Montecino, M, Orellana A, Castro E, Meneses C. |
| EPI_ISL_629088 | Laboratoire du Centre Hospitalier Annecy Genevois | CNR Virus des Infections Respiratoires - France SUD | Antonin Bal, Gregory Destras, Gwendolyne Burfin, Hadrien Règue, Quentin Semanas, Martine Valette, Bruno Lina, Hélène Petitprez, Bruno Chanzy, Laurence Josset |
| EPI_ISL_631550, EPI_ISL_631551 | Jamaica Hospital Medical Center | New York City Public Health Laboratory | Jade Wang, et al. |
| EPI_ISL_631552, EPI_ISL_631553 | Brookdale University Hospital Medical Center | New York City Public Health Laboratory | Jade Wang, et al. |
| EPI_ISL_631554 | Jamaica Hospital Medical Center | New York City Public Health Laboratory | Jade Wang, et al. |
| EPI_ISL_631555 | Richmond University Medical Center | New York City Public Health Laboratory | Jade Wang, et al. |
| EPI_ISL_631556 | NYC HH Jacobi Medical Center | New York City Public Health Laboratory | Jade Wang, et al. |
| EPI_ISL_631573 | Brookdale University Hospital Medical Center | New York City Public Health Laboratory | Jade Wang, et al. |
| EPI_ISL_631574, EPI_ISL_631575 | Jamaica Hospital Medical Center | New York City Public Health Laboratory | Jade Wang, et al. |
| EPI_ISL_631576 | Richmond University Medical Center | New York City Public Health Laboratory | Jade Wang, et al. |
| EPI_ISL_631654, EPI_ISL_631655, EPI_ISL_631656, EPI_ISL_631657, EPI_ISL_631658, EPI_ISL_631659, EPI_ISL_631660, EPI_ISL_631661, EPI_ISL_631662, EPI_ISL_631663, EPI_ISL_631664, EPI_ISL_631665, EPI_ISL_631666, EPI_ISL_631667, EPI_ISL_631668, EPI_ISL_631669, EPI_ISL_631670, EPI_ISL_631671, EPI_ISL_631672 | | | |
| see above | Texas Department of State Health Services | Texas Department of State Health Services | Rashmi Tuladhar, Bonnie Oh, Jenny Zhang, Maliha Rahman, Anita Pokharel, Myong Koag, Chung Wang, Rachel Lee, Grace Kubin, Mayela Pedrueza |
| EPI_ISL_631752 | NYC HH Lincoln Medical And Mental Health Center | New York City Public Health Laboratory | Jade Wang, et al. |
| EPI_ISL_631764 | Mount Sinai Hospital | New York City Public Health Laboratory | Jade Wang, et al. |
| EPI_ISL_631765 | Bellevue Hospital Center | New York City Public Health Laboratory | Jade Wang, et al. |
| EPI_ISL_631785, EPI_ISL_631786 | Mount Sinai Hospital | New York City Public Health Laboratory | Jade Wang, et al. |

| | | | |
|---|---|---|---|
| EPI_ISL_631787 | Flushing Hospital Medical Center | New York City Public Health Laboratory | Jade Wang, et al. |
| EPI_ISL_631788, EPI_ISL_631789, EPI_ISL_631790, EPI_ISL_631791 | Jamaica Hospital Medical Center | New York City Public Health Laboratory | Jade Wang, et al. |
| EPI_ISL_631792 | Richmond University Medical Center | New York City Public Health Laboratory | Jade Wang, et al. |
| EPI_ISL_631793, EPI_ISL_631794 | SUNY Downstate Medical Center | New York City Public Health Laboratory | Jade Wang, et al. |
| EPI_ISL_631795 | Jamaica Hospital Medical Center | New York City Public Health Laboratory | Jade Wang, et al. |
| EPI_ISL_631796 | Maimonides Medical Center | New York City Public Health Laboratory | Jade Wang, et al. |
| EPI_ISL_631797 | OCME Office Of Chief Medical Examiner | New York City Public Health Laboratory | Jade Wang, et al. |
| EPI_ISL_631798 | Jamaica Hospital Medical Center | New York City Public Health Laboratory | Jade Wang, et al. |
| EPI_ISL_631799 | NYC HH Lincoln Medical And Mental Health Center | New York City Public Health Laboratory | Jade Wang, et al. |
| EPI_ISL_631800 | Brookdale University Hospital Medical Center | New York City Public Health Laboratory | Jade Wang, et al. |
| EPI_ISL_631801 | Richmond University Medical Center | New York City Public Health Laboratory | Jade Wang, et al. |
| EPI_ISL_631832 | Metro Urgicare | New York City Public Health Laboratory | Jade Wang, et al. |
| EPI_ISL_631833 | Jamaica Hospital Medical Center | New York City Public Health Laboratory | Jade Wang, et al. |
| EPI_ISL_631834 | Mount Sinai Hospital | New York City Public Health Laboratory | Jade Wang, et al. |
| EPI_ISL_631835, EPI_ISL_631836 | Jamaica Hospital Medical Center | New York City Public Health Laboratory | Jade Wang, et al. |
| EPI_ISL_631837 | Richmond University Medical Center | New York City Public Health Laboratory | Jade Wang, et al. |
| EPI_ISL_631838 | Bellevue Hospital Center | New York City Public Health Laboratory | Jade Wang, et al. |
| EPI_ISL_631839 | Mount Sinai Hospital | New York City Public Health Laboratory | Jade Wang, et al. |
| EPI_ISL_631868, EPI_ISL_631869 | Richmond University Medical Center | New York City Public Health Laboratory | Jade Wang, et al. |
| EPI_ISL_631915 | Maimonides Medical Center | New York City Public Health Laboratory | Jade Wang, et al. |
| EPI_ISL_631978 | Brookdale University Hospital Medical Center | New York City Public Health Laboratory | Jade Wang, et al. |
| EPI_ISL_631979 | Jamaica Hospital Medical Center | New York City Public Health Laboratory | Jade Wang, et al. |
| EPI_ISL_631980 | St Barnabas Hospital | New York City Public Health Laboratory | Jade Wang, et al. |
| EPI_ISL_631981 | Brookdale University Hospital Medical Center | New York City Public Health Laboratory | Jade Wang, et al. |
| EPI_ISL_631994 | Jamaica Hospital Medical Center | New York City Public Health Laboratory | Jade Wang, et al. |
| EPI_ISL_631995 | Wyckoff Heights Medical Center | New York City Public Health Laboratory | Jade Wang, et al. |
| EPI_ISL_631996, EPI_ISL_631997 | Jamaica Hospital Medical Center | New York City Public Health Laboratory | Jade Wang, et al. |
| EPI_ISL_631998, EPI_ISL_632085 | Richmond University Medical Center | New York City Public Health Laboratory | Jade Wang, et al. |
| EPI_ISL_632086 | Jamaica Hospital Medical Center | New York City Public Health Laboratory | Jade Wang, et al. |
| EPI_ISL_632087 | Flushing Hospital Medical Center | New York City Public Health Laboratory | Jade Wang, et al. |
| EPI_ISL_632088 | Jamaica Hospital Medical Center | New York City Public Health Laboratory | Jade Wang, et al. |
| EPI_ISL_632089 | Brookdale University Hospital Medical Center | New York City Public Health Laboratory | Jade Wang, et al. |
| EPI_ISL_632090, EPI_ISL_632108, EPI_ISL_632109, EPI_ISL_632110 | Jamaica Hospital Medical Center | New York City Public Health Laboratory | Jade Wang, et al. |
| EPI_ISL_632112, EPI_ISL_632113, EPI_ISL_632115 | Richmond University Medical Center | New York City Public Health Laboratory | Jade Wang, et al. |
| EPI_ISL_632116 | NYC HH Lincoln Medical And Mental Health Center | New York City Public Health Laboratory | Jade Wang, et al. |
| EPI_ISL_632119 | Flushing Hospital Medical Center | New York City Public Health Laboratory | Jade Wang, et al. |
| EPI_ISL_632121 | Bellevue Hospital Center | New York City Public Health Laboratory | Jade Wang, et al. |
| EPI_ISL_632123 | Brookdale University Hospital Medical Center | New York City Public Health Laboratory | Jade Wang, et al. |
| EPI_ISL_632129 | Wyckoff Heights Medical Center | New York City Public Health Laboratory | Jade Wang, et al. |
| EPI_ISL_632133, EPI_ISL_632140, EPI_ISL_632144 | Flushing Hospital Medical Center | New York City Public Health Laboratory | Jade Wang, et al. |
| EPI_ISL_632154, EPI_ISL_632163 | Richmond University Medical Center | New York City Public Health Laboratory | Jade Wang, et al. |
| EPI_ISL_632172 | Jamaica Hospital Medical Center | New York City Public Health Laboratory | Jade Wang, et al. |
| EPI_ISL_632174 | Flushing Hospital Medical Center | New York City Public Health Laboratory | Jade Wang, et al. |
| EPI_ISL_632847, EPI_ISL_632848, EPI_ISL_632850, EPI_ISL_632852, EPI_ISL_632853, EPI_ISL_632864, EPI_ISL_632865, EPI_ISL_632871, EPI_ISL_632873, EPI_ISL_632876, EPI_ISL_632881, EPI_ISL_632882 | | | |
| see above | Idaho Bureau of Laboratories | Center for Global Health, University of New Mexico Health Sciences Center | Daryl Domman, Kurt Schwalm, Matthew Burns, Robert Voermans, Christopher Ball, Darrell Dinwiddie |
| EPI_ISL_632926, EPI_ISL_632927, EPI_ISL_632928 | Cadham Provincial laboratory | Cadham Provincial laboratory | Anna Majer, Shari Tyson, Grace Seo, Philip Mabon, Elsie Grudeski, Rhiannon Huzarewich, Russell Mandes, Anneliese Landgraff, Jennifer Tanner, Natalie Knox, Morag Graham, Gary Van Domselaar, Paul Van Caeseele, Jared Bullard, David Alexander, Kerry Dust, Nathalie Bastien, Yan Li, Timothy Booth, Darian Hole, Madison Chapel, CanCOGeN's metadata curation team, Public Health Agency of Canada CanCOGeN team |
| EPI_ISL_632966, EPI_ISL_632967 | Flushing Hospital Medical Center | New York City Public Health Laboratory | Jade Wang, et al. |
| EPI_ISL_635205, EPI_ISL_635220, EPI_ISL_635244, EPI_ISL_635245, EPI_ISL_635246, EPI_ISL_635247, EPI_ISL_635248, EPI_ISL_635249 | Institute of Microbiology and Immunology, Faculty of Medicine, University of Ljubljana | Institute of Microbiology and Immunology, Faculty of Medicine, University of Ljubljana | Samo Zakotnik, Tomaž Mark Zorec, Miša Korva, Mario Poljak, Tatjana Avši - Županc |
| EPI_ISL_635250, EPI_ISL_635251 | Institute of Microbiology and Immunology, Faculty of Medicine, University of Ljubljana | Institute of Microbiology and Immunology, Faculty of Medicine, University of Ljubljana | Tomaž Mark Zorec, Samo Zakotnik, Miša Korva, Tatjana Avši - Županc, Mario Poljak |
| EPI_ISL_636531 | Dutch COVID-19 response team | National Institute for Public Health and the Environment (RIVM) | Adam Meijer, Harry Vennema, Jeroen Cremer, Sharon van den Brink, Bas van der Veer, AnneMarie van den Brandt, Florian Zwagemaker, Dennis Schmitz, Chantal Reusken, on behalf of the national COVID-19 response team |
| EPI_ISL_639794, EPI_ISL_639795, | unknown | Public Health Virology Laboratory, Forensic and Scientific | Son Nguyen et al. |

| | | | |
|---|---|---|---|
| EPI_ISL_639796 | | Services (PHV-FSS) | |
| EPI_ISL_640038 | Knysna Hospital wc KNY | NHLS/UCT | Arash Iranzadeh, Deelan Doolabh, Lynn Tyers, Bruna Galvao, Innocent Mudau, Marvin Hsiao, Kruger Marais, Diana Hardie, Stephen Korsman, Carolyn Williamson |
| EPI_ISL_640060 | Groote Schuur Hospital wc GSH | NHLS/UCT | Arash Iranzadeh, Deelan Doolabh, Lynn Tyers, Bruna Galvao, Innocent Mudau, Marvin Hsiao, Kruger Marais, Diana Hardie, Stephen Korsman, Carolyn Williamson |
| EPI_ISL_641519, EPI_ISL_641543 | CHU de Saint-Étienne Hôpital Nord | CNR Virus des Infections Respiratoires - France SUD | Antonin Bal, Gregory Destras, Gwendolyne Burfin, Hadrien Règue, Quentin Semanas, Martine Valette, Bruno Lina, Issam Bechri, Manon Vogrig, Marine Delorme, Bruno Pozzetto, Thomas Bourlet, Sylvie Gonzalo, Sylvie Pillet, Laurence Josset |
| EPI_ISL_644403, EPI_ISL_644405, EPI_ISL_644409, EPI_ISL_644410 | MEPHI, Aix Marseille University | MEPHI, Aix Marseille University | Anthony LEVASSEUR |
| EPI_ISL_644687 | CHU Montpellier | CNR Virus des Infections Respiratoires - France SUD | Antonin Bal, Gregory Destras, Gwendolyne Burfin, Hadrien Règue, Quentin Semanas, Martine Valette, Bruno Lina, Michel Segondy, Vincent Foulongne, Laurence Josset |
| EPI_ISL_644702 | CHU Nîmes | CNR Virus des Infections Respiratoires - France SUD | Antonin Bal, Gregory Destras, Gwendolyne Burfin, Hadrien Règue, Quentin Semanas, Martine Valette, Bruno Lina, Jean-Philippe Lavigne, Stephan Robin, Maxence Lotellier, Marie-Josée Carles, Laurence Josset |
| EPI_ISL_644710, EPI_ISL_644712 | Unité des Virus Émergents | CNR Virus des Infections Respiratoires - France SUD | Antonin Bal, Gregory Destras, Gwendolyne Burfin, Hadrien Règue, Quentin Semanas, Martine Valette, Bruno Lina, Laetitia Ninove, Léa Luciani, Antoine Nougairède, Laurence Josset |
| EPI_ISL_645174 | CHU de Limoges | CNR Virus des Infections Respiratoires - France SUD | Antonin Bal, Gregory Destras, Gwendolyne Burfin, Hadrien Règue, Quentin Semanas, Martine Valette, Bruno Lina, Sylvie Rogez, Laurence Josset |
| EPI_ISL_645190, EPI_ISL_645191 | CHU Clermont-Ferrand | CNR Virus des Infections Respiratoires - France SUD | Antonin Bal, Gregory Destras, Gwendolyne Burfin, Hadrien Règue, Quentin Semanas, Martine Valette, Bruno Lina, Christine Archimbaud, Amélie Brebion, Hélène Chabrolles, Martine Chambon, Audrey Mirand, Christel Regagnon, Maxime Bisseux, Patricia Combes, Cécile Henquell, Laurence Josset |
| EPI_ISL_645197, EPI_ISL_645198 | CHU Nîmes | CNR Virus des Infections Respiratoires - France SUD | Antonin Bal, Gregory Destras, Gwendolyne Burfin, Hadrien Règue, Quentin Semanas, Martine Valette, Bruno Lina, Jean-Philippe Lavigne, Stephan Robin, Maxence Lotellier, Marie-Josée Carles, Laurence Josset |
| EPI_ISL_648024, EPI_ISL_648025, EPI_ISL_648026 | MS Public Health Laboratory | Pathogen Discovery, Respiratory Viruses Branch, Division of Viral Diseases, Centers for Disease Control and Prevention | Yan Li, Jing Zhang, Ying Tao, Brian Lynch, Krista Queen, Anna Montmayeur, Anna Uehara, Clinton R. Paden, Rachel Marine, Haibin Wang, Suxiang Tong |
| EPI_ISL_648075, EPI_ISL_648076, EPI_ISL_648077, EPI_ISL_648078, EPI_ISL_648079, EPI_ISL_648080 | Department of Laboratory Medicine, Tan Tock Seng Hospital | Department of Laboratory Medicine, Tan Tock Seng Hospital | Chen YYC, Zair X, Lim JX, Li C, Tang WY, Maurer-Stroh S, Barkham TMS, Nagarajan N, Sessions OM |
| EPI_ISL_648380, EPI_ISL_648595 | Laboratorio de Infectología Servicio de Infectología Hospital Universitario Dr. José Eleuterio González - Universidad Autónoma de Nuevo León | Laboratorio de Infectología Molecular Departamento de Bioquímica y Medicina Molecular Facultad de Medicina - Universidad Autónoma de Nuevo León | Kame A. Galán-Huerta, María F. Herrera-Saldivar, Natalia Martínez-Acuña, Sonia A. Lozano-Sepúlveda, Daniel Arellanos-Soto, Ana M. Rivas-Estilla, Paola Bocanegra-Ibarias, Samantha M. Flores-Treviño, Elvira Garza-González, Eduardo Perez-Alba, Laura Nuzzolo-Shihadeh, Adrian Camacho-Ortiz, Roberto Montes-de-Oca, Consuelo Treviño-Garza, Manuel E. de-la-O-Cavazos |
| EPI_ISL_653179, EPI_ISL_653277, EPI_ISL_653278 | Florida Bureau of Public Health Laboratories | Florida Bureau of Public Health Laboratories | Sarah Schmedes, Jason Blanton |
| EPI_ISL_654016 | Laboratory of Microbiology, National Reference Lab, Charles Nicolle Hospital; 2-University of Tunis ElManar, Faculty of Medicine of Tunis, LR99ES09, Tunis, Tunisia | 1-Clinical and Experimental Pharmacology Lab, LR16SP02, National Center of Pharmacovigilance, University of Tunis El Manar, Tunis, Tunisia. 2-Neurodegenerative diseases and psychiatric troubles, LR18SP03, Razi Hospital, University of Tunis El Manar, Tunis, Tunisia. 3- Ministry of Health, National Observatory of New and Emerging Diseases, 1006, Tunis, Tunisia | Ilhem Boutiba-Ben Boubaker, Sameh Trabelsi, Nissaf Ben Alaya, Maher Kharrat, Alia Ben Kahla, Jalila Ben Khelil, Salma Abid, Sana Ferjani, Mouna Ben Sassi, Mouna Safer, Imen Mkada, Imen Kacem, Gaies Emna, Soumaya Rammeh, Riadh Daghfous, Riadh Gouider. |
| EPI_ISL_654187 | Hospital General Universitario Gregorio Marañón | SeqCOVID-SPAIN consortium/IBV(CSIC) | Darío García de Viedma, Laura Pérez-Lago, Marta Herranz, Jon Sicilia, Julia Suárez, Pilar Catalán, Patricia Muñoz and SeqCOVID-SPAIN consortium |
| EPI_ISL_654673 | Servicio de Microbiología, Hospital Universitario Central de Asturias | SeqCOVID-SPAIN consortium/IBV(CSIC) | Cristián Castelló Abietar, Jose A. Boga, Susana Rojo-Alba, Marta Elena Álvarez-Argüelles, Santiago Melón and SeqCOVID-SPAIN consortium |
| EPI_ISL_654754 | Texas Department of State Health Services | Texas Department of State Health Services | Rashmi Tuladhar, Bonnie Oh, Jenny Zhang, Maliha Rahman, Anita Pokharel, Myong Koag, Chung Wang, Rachel Lee, Grace Kubin, Mayela Pedrueza, James Daniel Bonser |
| EPI_ISL_654867 | Pasteur Institute in Ho Chi Minh city | Department of Microbiology and Immunology - Pasteur Institute in Ho Chi Minh city | Hoàng Minh, Cao Minh Thng, Phm Th Thu Hng, V Phm Hng Nhung, ào Huy Mnh, Hunh Th Kim Loan, Nguyn Hoàng Quân, Hunh Phng Tho, Hoàng Nh ào, Nguyn Trung Hiu, Nguyn Hoàng Anh, Nguyn Thu Ngc, Phm Th Nhung, La Hoàng Huy, ng Thanh Giang, Nguyn Th Thanh Thng, Trn Th Hng Kim, Nguyn Thanh Long, Phm Duy Quang, Lng Chn Quang, Hoàng Quc Cng, Phan Trng Lân, Nguyn V Thng |
| EPI_ISL_654868 | Pasteur Institute in Ho Chi Minh city | Department of Microbiology and Immunology - Pasteur Institute in Ho Chi Minh city | ào Huy Mnh, Phm Th Thu Hng, V Phm Hng Nhung, Cao Minh Thng, Hunh Th Kim Loan, Nguyn Hoàng Quân, Hunh Phng Tho, Hoàng Nh ào, Nguyn Trung Hiu, Nguyn Hoàng Anh, Nguyn Thu Ngc, Phm Th Nhung, Nguyn Th Ngc Tho, Hà Bích Ngc, Nguyn Th Thanh Thng, Hoàng Minh, Trn Th Hng Kim, Nguyn Thanh Long, Phm Duy Quang, Lng Chn Quang, Hoàng Quc Cng, Nguyn V Thng, Phan Trng Lân |
| EPI_ISL_654869 | Pasteur Institute in Ho Chi Minh city | Department of Microbiology and Immunology - Pasteur Institute in Ho Chi Minh city | Phm Th Nhung, ào Huy Mnh, Phm Th Thu Hng, V Phm Hng Nhung, Cao Minh Thng, Hunh Th Kim Loan, Nguyn Hoàng Quân, Hunh Phng Tho, Hoàng Nh ào, Nguyn Trung Hiu, Nguyn Hoàng Anh, Nguyn Thu Ngc, Nguyn Th Ngc Tho, ng Thanh Giang, Võ Th Trang ài, Hoàng Minh, Trn Th Hng Kim, Nguyn Thanh Long, Phm Duy Quang, Lng Chn Quang, Hoàng Quc Cng, Phan Trng Lân, Nguyn V Thng |
| EPI_ISL_654870 | Pasteur Institute in Ho Chi Minh city | Department of Microbiology and Immunology - Pasteur Institute in Ho Chi Minh city | Trn Th Hng Kim, Phm Th Thu Hng, ào Huy Mnh, V Phm Hng Nhung, Cao Minh Thng, Hunh Th Kim Loan, Nguyn Hoàng Quân, Hunh Phng Tho, Hoàng Nh ào, Nguyn Trung Hiu, Nguyn Hoàng Anh, Nguyn Thu Ngc, Phm Th Nhung, Nguyn Th Ngc Tho, ng Thanh Giang, Nguyn Th Thanh Thng, Hoàng Kim Loan, Nguyn Thanh Long, Phm Duy Quang, Lng Chn Quang, Hoàng Quc Cng, Nguyn V Thng, Phan Trng Lân |
| EPI_ISL_654878 | Pasteur Institute in Ho Chi Minh city | Department of Microbiology and Immunology - Pasteur Institute in Ho Chi Minh city | Phm Duy Quang, ào Huy Mnh, Phm Th Thu Hng, V Phm Hng Nhung, Cao Minh Thng, Hunh Th Kim Loan, Nguyn Hoàng Quân, Hunh Phng Tho, Hoàng Nh ào, Nguyn Trung Hiu, Nguyn Hoàng Anh, Nguyn Thu Ngc, Phm Th Nhung, Nguyn Th Ngc Tho, Viên ng Khánh Linh, Nguyn Th Thanh Thng, Hoàng Minh, Trn Th Hng Kim, Nguyn Thanh Long, Lng Chn Quang, Hoàng Quc Cng, Phan Trng Lân, Nguyn V Thng |
| EPI_ISL_654885 | Pasteur Institute in Ho Chi Minh city | Department of Microbiology and Immunology - Pasteur Institute in Ho Chi Minh city | Nguyn V Thng, ào Huy Mnh, Phm Th Thu Hng, V Phm Hng Nhung, Cao Minh Thng, Hunh Th Kim Loan, Nguyn Hoàng Quân, Hunh Phng Tho, Hoàng Nh ào, Nguyn Trung Hiu, Nguyn Hoàng Anh, Nguyn Thu Ngc, Phm Th Nhung, Nguyn Th Ngc Tho, ng Thanh Giang, Hoàng Thùy Linh, Hoàng Minh, Trn Th Hng Kim, Nguyn Thanh Long, Phm Duy Quang, Lng Chn Quang, Hoàng Quc Cng, Phan Trng Lân |
| EPI_ISL_654888 | Pasteur Institute in Ho Chi Minh city | Department of Microbiology and Immunology - Pasteur Institute in Ho Chi Minh city | Nguyn Hoàng Anh, Hoàng Quc Cng, ào Huy Mnh, Phm Th Thu Hng, V Phm Hng Nhung, Cao Minh Thng, Hunh Th Kim Loan, Nguyn Hoàng Quân, Hunh Phng Tho, Hoàng Nh ào, Nguyn Trung Hiu, Nguyn Thu Ngc, Phm Th Nhung, Nguyn Th Ngc Tho, ng Thanh Giang, Nguyn Th Thanh Thng, Hoàng Minh, Khu Vn Ngha, Nguyn Thanh Long, Phm Duy Quang, Lng Chn Quang, Phan Trng Lân, Nguyn V Thng |
| EPI_ISL_657460, EPI_ISL_657463 | Servicio de Microbiología. Hospital General Universitario de Castellón | SeqCOVID-SPAIN consortium/IBV(CSIC) | Rosario Moreno Muñoz, María Dolores Tirado Balaguer and SeqCOVID-SPAIN consortium |
| EPI_ISL_660338, EPI_ISL_660339, EPI_ISL_660340, EPI_ISL_660341 | CHU de Saint-Étienne Hôpital Nord | CNR Virus des Infections Respiratoires - France SUD | Antonin Bal, Gregory Destras, Gwendolyne Burfin, Hadrien Règue, Quentin Semanas, Martine Valette, Bruno Lina, Issam Bechri, Manon Vogrig, Marine Delorme, Bruno Pozzetto, Thomas Bourlet, Sylvie Gonzalo, Sylvie Pillet, Laurence Josset |
| EPI_ISL_660356, EPI_ISL_660357, EPI_ISL_660358 | CHU Clermont-Ferrand | CNR Virus des Infections Respiratoires - France SUD | Antonin Bal, Gregory Destras, Gwendolyne Burfin, Hadrien Règue, Quentin Semanas, Martine Valette, Bruno Lina, Christine Archimbaud, Amélie Brebion, Hélène Chabrolles, Martine Chambon, Audrey Mirand, Christel Regagnon, Maxime Bisseux, Patricia Combes, Cécile Henquell, Laurence Josset |
| EPI_ISL_660385 | Klinisk mikrobiologi Linkoping | The Public Health Agency of Sweden | Anna-Malin Linde, Maria Lind Karlberg, Mattias Haukland, Reza Advani, Olov Svartstrom, Oskar Karlsson Lindsjo, Sandra Broddesson, Petra Edquist, Mia Brytting, Anna Risberg, Karin Tegmark-Wisell |
| EPI_ISL_660675, EPI_ISL_660676, EPI_ISL_660677, EPI_ISL_660678, | CHU Montpellier | CNR Virus des Infections Respiratoires - France SUD | Antonin Bal, Gregory Destras, Gwendolyne Burfin, Hadrien Règue, Quentin Semanas, Martine Valette, Bruno Lina, Michel Segondy, Vincent Foulongne, Laurence Josset |

| | | | |
|---|---|---|---|
| EPI_ISL_660679, EPI_ISL_660680 | | | |
| EPI_ISL_660713 | CHU Nîmes | CNR Virus des Infections Respiratoires - France SUD | Antonin Bal, Gregory Destras, Gwendolyne Burfin, Hadrien Règue, Quentin Semanas, Martine Valette, Bruno Lina, Jean-Philippe Lavigne, Stephan Robin, Maxence Lotellier, Marie-Josée Carles, Laurence Josset |
| EPI_ISL_660734, EPI_ISL_660736 | Unité des Virus Émergents | CNR Virus des Infections Respiratoires - France SUD | Antonin Bal, Gregory Destras, Gwendolyne Burfin, Hadrien Règue, Quentin Semanas, Martine Valette, Bruno Lina, Laetitia Ninove, Léa Luciani, Antoine Nougairède, Laurence Josset |
| EPI_ISL_661253, EPI_ISL_661254 | Quest Diagnostics | Quest Diagnostics | Rosenthal,S.H., Gerasimova,A., Kagan,R.M., Anderson, B., Owen, R., Lacbawan, F. |
| EPI_ISL_663206 | CHU de Limoges | CNR Virus des Infections Respiratoires - France SUD | Antonin Bal, Gregory Destras, Gwendolyne Burfin, Hadrien Règue, Quentin Semanas, Martine Valette, Bruno Lina, Sylvie Rogez, Laurence Josset |
| EPI_ISL_663251, EPI_ISL_663252, EPI_ISL_663253 | CHU Nantes | CNR Virus des Infections Respiratoires - France SUD | Antonin Bal, Louise Castain, Gregory Destras, Gwendolyne Burfin, Hadrien Règue, Quentin Semanas, Martine Valette, Bruno Lina, Virginie Ferré, Celine Bressollette, Laurence Josset |
| EPI_ISL_664707 | University College London Hospital | COVID-19 Genomics UK (COG-UK) Consortium | Judith Heaney, Matthew Byott, Catherine Houlihan, Dan Frampton, Stuart Kirk, Moira Spyer and Eleni Nastouli |
| EPI_ISL_666654 | Laboratoire du Centre Hospitalier Annecy Genevois | CNR Virus des Infections Respiratoires - France SUD | Antonin Bal, Gregory Destras, Gwendolyne Burfin, Hadrien Règue, Quentin Semanas, Martine Valette, Bruno Lina, Hélène Petitprez, Bruno Chanzy, Laurence Josset |
| EPI_ISL_666683, EPI_ISL_666684, EPI_ISL_666685 | Centre hospitalier Métropole Savoie | CNR Virus des Infections Respiratoires - France SUD | Antonin Bal, Carine Dumollard, Gregory Destras, Gwendolyne Burfin, Hadrien Règue, Quentin Semanas, Martine Valette, Bruno Lina, Jérôme Grosjean, Laurence Josset |
| EPI_ISL_666964 | Quest Diagnostics | Quest Diagnostics | Rosenthal,S.H., Gerasimova,A., Kagan,R.M., Anderson, B., Owen, R., Lacbawan, F. |
| EPI_ISL_671374 | National Virus Reference Laboratory | Irish Coronavirus Sequencing Consortium - Teagasc Moorepark | Calum Walsh, Fiona Crispie, John Kenny, Paul Cotter |
| EPI_ISL_671765 | Santa Clara County Public Health Department | Chiu Laboratory, University of California, San Francisco | Xianding Deng, Scot Federman, Wei Gu, Elsa Villarino, Brandon Bonin, Debra A. Wadford, and Charles Y. Chiu |
| EPI_ISL_672158, EPI_ISL_672159 | The Ashley Laboratory, Stanford University | Chan-Zuckerberg Biohub | CZB Cliahub Consortium |
| EPI_ISL_676703, EPI_ISL_676704, EPI_ISL_676737, EPI_ISL_676738, EPI_ISL_676739, EPI_ISL_676740, EPI_ISL_676742, EPI_ISL_676743, EPI_ISL_676744, EPI_ISL_676745, EPI_ISL_676746, EPI_ISL_676747 | | | |
| see above | Wadsworth Center, New York State Department.of Health | Wadsworth Center, New York State Department.of Health | Kirsten St. George, Daryl M. Lamson, Alexis Russel, Jonathan Plitnick, Navjot Singh, John Kelly, Sara Griesemer, Erasmus Schneider, Erica Lasek-Nesselquist |
| EPI_ISL_677260 | Colorado Department of Public Health and Environment | Colorado Department of Puplic Health and Environment | Laura Bankers, Molly Hetherington-Rauth, Shannon Ely, Shannon R. Matzinger, Sarah Elizabeth Totten, Emily A. Travanty |
| EPI_ISL_677664 | Wadsworth Center, New York State Department.of Health | Wadsworth Center, New York State Department.of Health | Kirsten St. George, Daryl M. Lamson, Alexis Russel, Jonathan Plitnick, Navjot Singh, John Kelly, Sara Griesemer, Erasmus Schneider, Erica Lasek-Nesselquist |
| EPI_ISL_678275, EPI_ISL_678279, EPI_ISL_678287 | Mikrobiologie, RARI | Mikrobiologie, RARI | Krasnov,Y.M., Naryshkina,E.A., Guseva,N.P., Sosedova,E.A., Fedorov,A.V., Badanin,D.V., Sharapova,N.A., Portenko,S.A., Shcherbakova,S.A., Kutyrev,V.V. |
| EPI_ISL_678300 | Area of Virology, Serology and Virology Division (SAViD), New South Wales Health Pathology Randwick | Virology Research Laboratory; Area of Virology, Serology and Virology Division (SAViD), New South Wales Health Pathology Randwick | Foster, C.; Au, J.; Ruiz Silva, M.; Deveson, I.; Bull, R.; Van Hal, S.; Rawlinson, W. |
| EPI_ISL_678494, EPI_ISL_678495 | CNR Virus des Infections Respiratoires - France SUD | CNR Virus des Infections Respiratoires - France SUD | Antonin Bal, Gregory Destras, Gwendolyne Burfin, Solenne Brun, Martine Valette, Bruno Lina, Laurence Josset |
| EPI_ISL_681999 | UPMC Clinical Microbiology Laboratory | Microbial Genomic Epidemiology Laboratory, University of Pittsburgh | Mustapha M. Mustapha, Jane W. Marsh, Dan Snyder, Marissa P. Griffith, Stephanie L. Mitchell, Vatsala R. Srinivasa, Kady D. Waggle, Chinelo Ezeonwuku, Vaughn S. Cooper, Lee H. Harrison |
| EPI_ISL_683329 | 1-Laboratory of Microbiology, National Reference Lab, Charles Nicolle Hospital; 2-University of Tunis ElManar, Faculty of Medicine of Tunis, LR99ES09, Tunis, Tunisia | 1-Clinical and Experimental Pharmacology Lab, LR16SP02, National Center of Pharmacovigilance, University of Tunis El Manar, Tunis, Tunisia. 2-Neurodegenerative diseases and psychiatric troubles, LR18SP03, Razi Hospital, University of Tunis El Manar, Tunis, Tunisia. 3- Ministry of Health, National Observatory of New and Emerging Diseases, 1006, Tunis, Tunisia | Ilhem Boutiba-Ben Boubaker, Sameh Trabelsi, Nissaf Ben Alaya, Maher Kharrat, Alia Ben Kahla, Jalila Ben Khelil, Salma Abid, Sana Ferjani, Asma Ferjani, Mouna Ben Sassi, Mouna Safer, Guedi Berrabeh,Salwa Mrabet, Hanen ElJebari, Gaies Emna, Riadh Daghfous, Riadh Gouider. |
| EPI_ISL_683665, EPI_ISL_683666, EPI_ISL_683674, EPI_ISL_683675 | IdISSC/Hospital Clínico San Carlos de Madrid | SeqCOVID-SPAIN consortium/IBV(CSIC) | Alberto Delgado-Iribarren, Esther Culebras Lopez, Vicente Estrada Pérez, Jorge Matias-Guiu, Luis Ortega Medina, Silvia Sánchez Ramón, Ulises Gómez-Pinedo and SeqCOVID-SPAIN consortium |
| EPI_ISL_693359, EPI_ISL_693360, EPI_ISL_693361, EPI_ISL_693362, EPI_ISL_693363, EPI_ISL_693364, EPI_ISL_693365, EPI_ISL_693366, EPI_ISL_693367, EPI_ISL_693368, EPI_ISL_693369, EPI_ISL_693370, EPI_ISL_693371, EPI_ISL_693372, EPI_ISL_693373, EPI_ISL_693374, EPI_ISL_693375, EPI_ISL_693376, EPI_ISL_693377, EPI_ISL_693378, EPI_ISL_693379, EPI_ISL_693380, EPI_ISL_693381, EPI_ISL_693382, EPI_ISL_693383, EPI_ISL_693384, EPI_ISL_693385 | | | |
| see above | Ramathibodi Hospital | COVID-19 Network Investigations (CONI) Alliance | Elizabeth Batty, Wasun Chantratita, Thanat Chookajorn, Stefan Fernandez, Angkana Huang, Anthony R. Jones, Khajohn Joonsalak, Chonticha Klungtong, Theerarat Kochakarn, Namfon Kotanan, Krittikorn Kumpornsin, Wudtichai Manasatienkij, Bhakbhoom Panthan, Ekawat Pasomsub, Kingkan Rakmanee, Insee Sensorn, Janjira Thaipadungpanit, Arporn Wangwiwatsin,Treewat Watthanachockchai |
| EPI_ISL_694048 | TGen North | TGen North | Jolene Bowers, Megan Folkerts, Chris French, Hayley Yaglom, Ashlyn Pfeiffer, Darrin Lemmer, Dave Engelthaler, The Arizona COVID Genomics Union (ACGU) |
| EPI_ISL_694049 | AZ SPHL, Arizona Department of Health Services | TGen North | Jolene Bowers, Megan Folkerts, Chris French, Hayley Yaglom, Ashlyn Pfeiffer, Darrin Lemmer, Dave Engelthaler, The Arizona COVID Genomics Union (ACGU) |
| EPI_ISL_694052 | TGen North | TGen North | Jolene Bowers, Megan Folkerts, Chris French, Hayley Yaglom, Ashlyn Pfeiffer, Darrin Lemmer, Dave Engelthaler, The Arizona COVID Genomics Union (ACGU) |
| EPI_ISL_699464 | Regional Medical Sciences Center 11/1 phuket | National Institute of Health, Department of Medical Sciences, Ministry of Public Health, Thailand | Pilailuk Okada; Siripaporn Phuygun; Thanutsapa Thanadachakul; Sittiporn Parnmen; Warawan Wongboot; Sunthareeya Waicharoen; Malinee Chittaganpitch |
| EPI_ISL_699657 | 1-Laboratory of Microbiology, National Reference Lab, Charles Nicolle Hospital; 2-University of Tunis ElManar, Faculty of Medicine of Tunis, LR99ES09, Tunis, Tunisia | 1-Clinical and Experimental Pharmacology Lab, LR16SP02, National Center of Pharmacovigilance, University of Tunis El Manar, Tunis, Tunisia. 2-Neurodegenerative diseases and psychiatric troubles, LR18SP03, Razi Hospital, University of Tunis El Manar, Tunis, Tunisia. 3- Ministry of Health, National Observatory of New and Emerging Diseases, 1006, Tunis, Tunisia | Ilhem Boutiba-Ben Boubaker, Sameh Trabelsi, Nissaf Ben Alaya, Maher Kharrat, Alia Ben Kahla, Jalila Ben Khelil, Salma Abid, Sana Ferjani, Asma Ferjani, Mouna Ben Sassi, Mouna Safer, Guedi Berrabeh,Salwa Mrabet, Hanen ElJebari, Gaies Emna, Riadh Daghfous, Riadh Gouider. |
| EPI_ISL_700326 | CHU Bordeaux | CNR Virus des Infections Respiratoires - France SUD | Antonin Bal, Gregory Destras, Gwendolyne Burfin, Hadrien Règue, Quentin Semanas, Martine Valette, Bruno Lina, Pantxika Bellecave, Camille Ciccone, Isabelle Garrigue, Marie-Edith Lafon, Pascale Trimoulet, Laurence Josset |
| EPI_ISL_700371 | CHU Nantes | CNR Virus des Infections Respiratoires - France SUD | Antonin Bal, Louise Castain, Gregory Destras, Gwendolyne Burfin, Hadrien Règue, Quentin Semanas, Martine Valette, Bruno Lina, Virginie Ferré, Celine Bressollette, Laurence Josset |
| EPI_ISL_707782 | CHU Poitiers | CNR Virus des Infections Respiratoires - France SUD | Antonin Bal, Gregory Destras, Gwendolyne Burfin, Hadrien Règue, Quentin Semanas, Martine Valette, Bruno Lina, Agnès Beby-Defaux, Magali Garcia, Clément Jousselin, Nicolas Lévêque, Laurence Josset |
| EPI_ISL_707949, EPI_ISL_707950, EPI_ISL_707951, EPI_ISL_707952, EPI_ISL_707953, EPI_ISL_707954 | Virology, Universitätsklinikum des Saarlandes | Epigenetics, Saarland University | Kathrin Kattler, Markus Vogelgesang, Stefan Lohse, Sascha Tierling, Sigrun Smola, Jörn Walter |

| | | | |
|---|---|---|---|
| EPI_ISL_723079, EPI_ISL_723085, EPI_ISL_723086 | Institute of Medical Genetics and Applied Genomics | Institute of Medical Genetics and Applied Genomics | Caspar Gross, Tina Ganzenmüller, Siri Göpel, Michaela Pogoda, Daniela Bezdan, Michael Sonnabend, Angel Angelov, Nicolas Casadei, Stephan Ossowski, Thomas Iftner, Michael Bitzer |
| EPI_ISL_729340, EPI_ISL_729466 | A. Krumbholz, Labor Dr. Krause und Kollegen MVZ GmbH, Kiel | Charité Universitätsmedizin Berlin, Institut für Virologie | Victor M Corman, Barbara Mühlemann, Jörn Beheim-Schwarzbach, Talitha Veith, Julia Schneider, Terry Jones, Christian Drosten |
| EPI_ISL_729482, EPI_ISL_729483 | Charité Universitätsmedizin Berlin, Institut für Virologie/Labor Berlin | Charité Universitätsmedizin Berlin, Institut für Virologie | Victor M Corman, Barbara Mühlemann, Jörn Beheim-Schwarzbach, Talitha Veith, Julia Schneider, Terry Jones, Christian Drosten |
| EPI_ISL_729507, EPI_ISL_729508, EPI_ISL_729509, EPI_ISL_729510, EPI_ISL_729511, EPI_ISL_729512, EPI_ISL_729513, EPI_ISL_729515, EPI_ISL_729516, EPI_ISL_729517, EPI_ISL_729518, EPI_ISL_729519, EPI_ISL_729520, EPI_ISL_729521, EPI_ISL_729549 | | | |
| see above | A. Krumbholz, Labor Dr. Krause und Kollegen MVZ GmbH, Kiel | Charité Universitätsmedizin Berlin, Institut für Virologie | Victor M Corman, Barbara Mühlemann, Jörn Beheim-Schwarzbach, Talitha Veith, Julia Schneider, Terry Jones, Christian Drosten |
| EPI_ISL_729805, EPI_ISL_729806, EPI_ISL_729807 | Laboratorio Central de Saude Publica do Estado do Rio Grande do Sul (LACEN-RS) | Laboratory of Respiratory Viruses and Measles, Oswaldo Cruz Institute, FIOCRUZ | Paola Resende, Luciana Appolinario, Fernando Motta, Anna Carolina Paixão, Ana Carolina Mendonça, Tatiana Schaffer Gregianini, Marilda Tereza Mar da Rosa, Marilda Siqueira |
| EPI_ISL_732545, EPI_ISL_732546 | Bundeswehr Institute of Microbiology | Bundeswehr Institute of Microbiology | Markus Antwerpen, Alexandra Rehn, Mathias Walter, Malena Bestehorn-Willmann, Sabine Zange, Enrico Georgi, Roman Wölfel |
| EPI_ISL_734505, EPI_ISL_734506 | UZ Leuven, National Reference Laboratory for Coronaviruses, Laboratory Medicine, Leuven, Belgium | KU Leuven, Rega Institute, Clinical and Epidemiological Virology | Tony Wawina-Bokalanga, Joan Marti-Carerras, Bert Vanmechelen, Piet Maes |
| EPI_ISL_735449, EPI_ISL_736775 | UW Virology Lab | UW Virology Lab | Pavitra Roychoudhury, Hong Xie, Lasata Shrestha, Meei-Li Huang, Keith R Jerome, Alexander Greninger |
| EPI_ISL_738085 | University Hospitals of Geneva, Laboratory of Virology | University Hospitals of Geneva, Laboratory of Virology | Cordey Samuel and Laubscher Florian |
| EPI_ISL_740250, EPI_ISL_740440, EPI_ISL_744478 | Laboratoire national de santé, Microbiology, Virology | Laboratoire national de santé, Microbiology, Microbial Genomics Platform | Anke Wienecke-Baldacchino, Catherine Ragimbeau, Tamir Abdelrahman, Jessica Tapp, Fatu Djabi |
| EPI_ISL_751380, EPI_ISL_751381 | IRCCS Sacro Cuore Don Calabria Hospital, Department of Infectious, Tropical Diseases & Microbiology | University of Verona, Department of Biotechnology | Antonio Mori, Michela Deiana, Elena Pomari, Chiara Piubelli; Giulia Lopatriello, Luca Marcolungo, Cristina Beltrami, Chiara Degli Esposti, Emanuela Cosentino, Massimo Delledonne |
| EPI_ISL_752608, EPI_ISL_752614, EPI_ISL_752615 | State Laboratories Division, Hawaii State Department of Health | State Laboratories Division, Hawaii State Department of Health | Pamela O'Brien, Sabrina Diemert, Drew Kuwazaki, Razvan Sultana, Edward Desmond |
| EPI_ISL_753763, EPI_ISL_753799, EPI_ISL_753921, EPI_ISL_753922, EPI_ISL_754056, EPI_ISL_754058 | Charité Universitätsmedizin Berlin, Institut für Virologie/Labor Berlin | Charité Universitätsmedizin Berlin, Institut für Virologie | Victor M Corman, Jörn Beheim-Schwarzbach, Barbara Mühlemann, Julia Schneider, Talitha Veith, Terry Jones, Christian Drosten |
| EPI_ISL_754749, EPI_ISL_754751, EPI_ISL_754752, EPI_ISL_754753, EPI_ISL_754754, EPI_ISL_754756, EPI_ISL_754757, EPI_ISL_754758, EPI_ISL_754759, EPI_ISL_754760, EPI_ISL_754761, EPI_ISL_754762, EPI_ISL_754763, EPI_ISL_754764, EPI_ISL_754765 | | | |
| see above | Emory Molecular Diagnostics Laboratory, Emory Healthcare | Piantadosi Lab, Emory Department of Pathology | Ahmed Babiker, Anne Piantadosi |
| EPI_ISL_755794, EPI_ISL_755881 | Toronto Invasive Bacterial Diseases Network | McMaster University | Allison McGeer, Patryk Aftanas, Hooman Derakhshani, Angel Li, Kuganya Nirmalarajah, Emily Panousis, Ahmed Draia, Jalees Nasir, Michael Surette, Samira Mubareka, Andrew G. McArthur |
| EPI_ISL_766032 | RSUP Dr. Sardjito | Eijkman Institute for Molecular Biology, Ministry of Research and Technology/National Agency for Research and Innovation | Frilasita A Yudhaputri, Edison Johar, Hidayat Trimarsanto, David H Muljono, Safarina G Malik, Khin Saw Myint, Amin Soebandrio |
| EPI_ISL_775610, EPI_ISL_775615, EPI_ISL_775616, EPI_ISL_775618, EPI_ISL_775620, EPI_ISL_775622, EPI_ISL_775625, EPI_ISL_775629, EPI_ISL_775630, EPI_ISL_775634, EPI_ISL_775635, EPI_ISL_775637, EPI_ISL_775638, EPI_ISL_775639, EPI_ISL_775642, EPI_ISL_775645, EPI_ISL_775646, EPI_ISL_775647, EPI_ISL_775649, EPI_ISL_775651, EPI_ISL_775656, EPI_ISL_775659, EPI_ISL_775664, EPI_ISL_775665, EPI_ISL_775671, EPI_ISL_775672, EPI_ISL_775673, EPI_ISL_775674, EPI_ISL_775676, EPI_ISL_775677, EPI_ISL_775679, EPI_ISL_775680, EPI_ISL_775681, EPI_ISL_775682, EPI_ISL_775688, EPI_ISL_775694, EPI_ISL_775697, EPI_ISL_775699, EPI_ISL_775707, EPI_ISL_775709, EPI_ISL_775714, EPI_ISL_775715, EPI_ISL_775716, EPI_ISL_775718, EPI_ISL_775719, EPI_ISL_775720, EPI_ISL_775725, EPI_ISL_775727, EPI_ISL_775730, EPI_ISL_775736, EPI_ISL_775740, EPI_ISL_775742, EPI_ISL_775743, EPI_ISL_775744, EPI_ISL_775747, EPI_ISL_775748, EPI_ISL_775759, EPI_ISL_775763, EPI_ISL_775764, EPI_ISL_775767, EPI_ISL_775770, EPI_ISL_775775, EPI_ISL_775776, EPI_ISL_775777, EPI_ISL_775780, EPI_ISL_775781, EPI_ISL_775784, EPI_ISL_775792, EPI_ISL_775794, EPI_ISL_775796, EPI_ISL_775799, EPI_ISL_775801, EPI_ISL_775803, EPI_ISL_775804, EPI_ISL_775807, EPI_ISL_775808, EPI_ISL_775809, EPI_ISL_775811, EPI_ISL_775812, EPI_ISL_775814, EPI_ISL_775815, EPI_ISL_775818, EPI_ISL_775819, EPI_ISL_775822, EPI_ISL_775823, EPI_ISL_775824, EPI_ISL_775833, EPI_ISL_775835, EPI_ISL_775838, EPI_ISL_775839, EPI_ISL_775842, EPI_ISL_775846, EPI_ISL_775847, EPI_ISL_775854, EPI_ISL_775858, EPI_ISL_775859, EPI_ISL_775860, EPI_ISL_775861, EPI_ISL_775862, EPI_ISL_775867, EPI_ISL_775871, EPI_ISL_775872, EPI_ISL_775874, EPI_ISL_775875, EPI_ISL_775876, EPI_ISL_775878, EPI_ISL_775879, EPI_ISL_775889, EPI_ISL_775891, EPI_ISL_775895, EPI_ISL_775901, EPI_ISL_775902, EPI_ISL_775905, EPI_ISL_775906, EPI_ISL_775925, EPI_ISL_775926, EPI_ISL_775927, EPI_ISL_775928, EPI_ISL_775929, EPI_ISL_775930, EPI_ISL_775937, EPI_ISL_775938, EPI_ISL_776013, EPI_ISL_776014, EPI_ISL_776015, EPI_ISL_776016, EPI_ISL_776017, EPI_ISL_776018, EPI_ISL_776019, EPI_ISL_776020, EPI_ISL_776021, EPI_ISL_776022, EPI_ISL_776023, EPI_ISL_776025, EPI_ISL_776026, EPI_ISL_776027, EPI_ISL_776028, EPI_ISL_776029, EPI_ISL_776030, EPI_ISL_776031, EPI_ISL_776032, EPI_ISL_776033, EPI_ISL_776034, EPI_ISL_776035, EPI_ISL_776036, EPI_ISL_776037, EPI_ISL_776038, EPI_ISL_776039, EPI_ISL_776040, EPI_ISL_776041, EPI_ISL_776042, EPI_ISL_776043, EPI_ISL_776044, EPI_ISL_776045, EPI_ISL_776046, EPI_ISL_776047, EPI_ISL_776048, EPI_ISL_776049, EPI_ISL_776050, EPI_ISL_776051, EPI_ISL_776052, EPI_ISL_776053, EPI_ISL_776054, EPI_ISL_776055, EPI_ISL_776056, EPI_ISL_776057, EPI_ISL_776058, EPI_ISL_776059, EPI_ISL_776060, EPI_ISL_776061, EPI_ISL_776062, EPI_ISL_776063, EPI_ISL_776064, EPI_ISL_776065, EPI_ISL_776066, EPI_ISL_776067, EPI_ISL_776068, EPI_ISL_776069, EPI_ISL_776070, EPI_ISL_776071, EPI_ISL_776072, EPI_ISL_776073, EPI_ISL_776074, EPI_ISL_776075, EPI_ISL_776076, EPI_ISL_776077, EPI_ISL_776078, EPI_ISL_776079, EPI_ISL_776080, EPI_ISL_776081, EPI_ISL_776082, EPI_ISL_776083, EPI_ISL_776084, EPI_ISL_776085, EPI_ISL_776086, EPI_ISL_776087, EPI_ISL_776088, EPI_ISL_776089, EPI_ISL_776090, EPI_ISL_776091, EPI_ISL_776092, EPI_ISL_776093, EPI_ISL_776094, EPI_ISL_776095, EPI_ISL_776096, EPI_ISL_776097, EPI_ISL_776098, EPI_ISL_776099, EPI_ISL_776100, EPI_ISL_776101, EPI_ISL_776102, EPI_ISL_776103, EPI_ISL_776104, EPI_ISL_776105, EPI_ISL_776106, EPI_ISL_776107, EPI_ISL_776108, EPI_ISL_776109, EPI_ISL_776110, EPI_ISL_776111, EPI_ISL_776115, EPI_ISL_776116, EPI_ISL_776336, EPI_ISL_776418, EPI_ISL_776443, EPI_ISL_776549, EPI_ISL_776550, EPI_ISL_776551, EPI_ISL_776555, EPI_ISL_776559, EPI_ISL_776561 | | | |
| see above | University Medical Center Hamburg Eppendorf | Heinrich Pette Institute, Leibniz Institute for Experimental Virology | Alexis Robitaille, Thomas Günther, Johannes Knobloch, Martin Aepfelbacher, Nicole Fischer, Adam Grundhoff |
| EPI_ISL_779169 | Laboratorio de Infectología, Servicio de Infectología, Hospital Universitario Dr. José Eleuterio González - Universidad Autónoma de Nuevo León | Laboratorio de Infectología Molecular, Departamento de Bioquímica y Medicina Molecular,Facultad de Medicina - Universidad Autónoma de Nuevo León | Kame A. Galán-Huerta, María F. Herrera-Saldivar, Natalia Martínez-Acuña, Sonia A. Lozano-Sepúlveda, Daniel Arellanos-Soto, Ana M. Rivas-Estilla, Paola Bocanegra-Ibarias, Samantha M. Flores-Treviño, Elvira Garza-González, Eduardo Perez-Alba, Laura Nuzzolo-Shihadeh, Adrian Camacho-Ortiz |
| EPI_ISL_779739, EPI_ISL_779741, EPI_ISL_779742, EPI_ISL_779744, EPI_ISL_779746, EPI_ISL_779758, EPI_ISL_779759, EPI_ISL_779760, EPI_ISL_779761, EPI_ISL_779764 | IdISSC/Hospital Clínico San Carlos de Madrid | SeqCOVID-SPAIN consortium/IBV(CSIC) | Alberto Delgado-Iribarren, Esther Culebras López, Vicente Estrada Pérez, Jorge Matias-Guiu, Luis Ortega Medina, Silvia Sánchez Ramón, Ulises Gómez-Pinedo and SeqCOVID-SPAIN consortium |
| EPI_ISL_780376, EPI_ISL_780378, EPI_ISL_780381, EPI_ISL_780411 | Laboratory of Infectious Diseases, Department of Biomedical and Clinical Sciences L. Sacco, University of Milan | Laboratory of Infectious Diseases, Department of Biomedical and Clinical Sciences L. Sacco, University of Milan | Alessia Lai, Annalisa Bergna, Carla Della Ventura, Claudia Balotta, Massimo Galli, Gianguglielmo Zehender on behalf of SARS-CoV-2 ITALIAN RESEARCH ENTERPRISE-(SCIRE) Collaborative Group |
| EPI_ISL_792458 | Laboratorio del Hospital Regional Ushuaia Gdor. Ernesto Campos | Hospital Regional Ushuaia - Centro Austral De Investigaciones Científicas - Universidad Nacional De Tierra Del Fuego on behalf of 'Proyecto Argentino Interinstitucional de genomica de SARS-CoV-2' (PAIS Consortium) | Ceballos, SG; Nardi, CF; Gramundi, ID; Gallego, F; De Roccis, CA; Castro, G; Cáceres, SB; Yulan, CB; Boutureira, MF. |
| EPI_ISL_794606 | Central Public Health Laboratories, Egyptian Ministry of Health and Population | Central Public Health Laboratories, Egyptian Ministry of Health and Population | El-Shesheny,R., Roshdy,W.H., Kayed,A.E., Mostafa,A., Khalifa,M.K., Shehata,M., Shawky,S., Saleh,M., Gomaa,M., El Taweel,A., Mahmoud,S.H., Moatasim,Y., Kutkat,O., Kamel,M.N., Mahrous,N., El Sayes,M.A., El Guindy,N.M., Naguib,A., Kandeil,A., Kayali,G., Ali,M.A. |
| EPI_ISL_794678, EPI_ISL_794679, EPI_ISL_794680, EPI_ISL_794692, EPI_ISL_794694, EPI_ISL_794702, EPI_ISL_794704, EPI_ISL_794712, EPI_ISL_794714, EPI_ISL_794720 | PathWest Laboratory Medicine WA | PathWest Laboratory Medicine WA Microbial Surveillance Unit | PathWest Laboratory Medicine WA Microbial Surveillance Unit |
| EPI_ISL_796067, EPI_ISL_796101, EPI_ISL_796102, EPI_ISL_796135, EPI_ISL_796136 | Servicio de Microbiología. Hospital General Universitario de Castellón | SeqCOVID-SPAIN consortium/IBV(CSIC) | Rosario Moreno Muñoz, María Dolores Tirado Balaguer and SeqCOVID-SPAIN consortium |
| EPI_ISL_801549 | Laboratory of Molecular Virology, Pontificia Universidad Católica de Chile | MSHS Pathogen Surveillance Program | Leonardo I. Almonacid, Ana S. Gonzalez-Reiche, Matthew M. Hernandez, Jorge Levican, Ana Maria Contreras, Carlos Palma, Tamara García-Salum, Zenab Khan, Adriana van De Guchte, Ajay Obla, Jayeeta Dutta, Bremy Alburquerque, Eileen Serrano, Constanza Maldonado, M. Belen Leyton, Erick Salinas, Hala Alshammary, Juan Soto, Shwetha Hara Sridhar, Ying-Chih Wang, Melissa Smith, Robert Sebra, Marcela Ferres, Adolfo Garcia-Sastre, Edward C. Holmes, Viviana Simon, Harm van Bakel, Rafael A. Medina. |

| | | | |
|---|---|---|---|
| EPI_ISL_801587, EPI_ISL_801588, EPI_ISL_801589, EPI_ISL_801590 | Laboratory of Molecular Virology, Pontificia Universidad Católica de Chile | MSHS Pathogen Surveillance Program | Leonardo I. Almonacid, Ana S. Gonzalez-Reiche, Matthew M. Hernandez, Jorge Levican, Tamara García-Salum, Zenab Khan, Adriana van De Guchte, Ajay Obla, Jayeeta Dutta, Bremy Alburquerque, Eileen Serrano, Erick Salinas, Hala Alshammary, Juan Soto, Shwetha Hara Sridhar, Ying-Chih Wang, Melissa Smith, Robert Sebra, Adolfo Garcia-Sastre, Edward C. Holmes, Viviana Simon, Harm van Bakel, Rafael A. Medina |
| EPI_ISL_801600, EPI_ISL_801601, EPI_ISL_801602, EPI_ISL_801603, EPI_ISL_801605, EPI_ISL_801611, EPI_ISL_801612, EPI_ISL_801613 | Laboratory of Molecular Virology, Pontificia Universidad Católica de Chile | MSHS Pathogen Surveillance Program | Leonardo I. Almonacid, Ana S. Gonzalez-Reiche, Matthew M. Hernandez, Jorge Levican, Ana Maria Contreras, Carlos Palma, Tamara García-Salum, Zenab Khan, Adriana van De Guchte, Ajay Obla, Jayeeta Dutta, Bremy Alburquerque, Eileen Serrano, Constanza Maldonado, M. Belen Leyton, Erick Salinas, Hala Alshammary, Juan Soto, Shwetha Hara Sridhar, Ying-Chih Wang, Melissa Smith, Robert Sebra, Marcela Ferres, Adolfo Garcia-Sastre, Edward C. Holmes, Viviana Simon, Harm van Bakel, Rafael A. Medina. |
| EPI_ISL_803306, EPI_ISL_803638 | Wisconsin State Laboratory of Hygiene Communicable Disease Division | Wisconsin State Laboratory of Hygiene Communicable Disease Division | Kelsey R. Florek, Abigail C. Shockey |
| EPI_ISL_803886 | Department of Medical Biotechnologies, University of Siena | Laboratory of Infectious Diseases, Department of Biomedical and Clinical Sciences L. Sacco, University of Milan | Ilaria Vicenti, Filippo Dragoni, Maurizio Zazzi, Maria Grazia Cusi, Alessia Lai, Annalisa Bergna, Carla Della Ventura, Claudia Balotta, Massimo Galli, Gianguglielmo Zehender on behalf of SARS-CoV-2 ITALIAN RESEARCH ENTERPRISE-(SCIRE) Collaborative Group |
| EPI_ISL_804406, EPI_ISL_804459 | Dutch COVID-19 response team | National Institute for Public Health and the Environment (RIVM) | Adam Meijer, Harry Vennema, Dirk Eggink, Matthijs Welkers, Jeroen Cremer, Sharon van den Brink, Bas van der Veer, AnneMarie van den Brandt, Florian Zwagemaker, Dennis Schmitz, Chantal Reusken, on behalf of the national COVID-19 response team |
| EPI_ISL_804851, EPI_ISL_804862, EPI_ISL_804885, EPI_ISL_804888, EPI_ISL_804891, EPI_ISL_804893, EPI_ISL_804898, EPI_ISL_804899 | DC Public Health Lab/ Dept. of Forensic Sciences | DC Public Health Lab/ Dept. of Forensic Sciences | Scott Nguyen, Elizabeth Zelaya, Connie Maza, Monica Mann, Brittany Hamilton, David Payne, Jocelyn Hauser |
| EPI_ISL_805020, EPI_ISL_805581, EPI_ISL_805679, EPI_ISL_805695, EPI_ISL_805801, EPI_ISL_805859, EPI_ISL_805897, EPI_ISL_805922, EPI_ISL_805953, EPI_ISL_806032, EPI_ISL_806197, EPI_ISL_806201, EPI_ISL_806215, EPI_ISL_806230, EPI_ISL_806245, EPI_ISL_806263, EPI_ISL_806284, EPI_ISL_806340, EPI_ISL_806383, EPI_ISL_806389, EPI_ISL_806409, EPI_ISL_806462 | | | |
| see above | Alberta Precision Labs (APL) | Alberta Precision Labs (APL) | Gordon P, Lam LG, Pabbaraju K, Wong A, Ma R, Li V, Melin A, Tipples G, Berenger B, Zelyas N, Kellner J, Bernier F, Chui L, Croxen M |
| EPI_ISL_812237, EPI_ISL_812238, EPI_ISL_812239, EPI_ISL_812240, EPI_ISL_812241, EPI_ISL_812242, EPI_ISL_812243, EPI_ISL_812244 | North Dakota Department of Health, Public Health Laboratory | North Dakota Department of Health, Public Health Laboratory | Lisa Wingerter |
| EPI_ISL_812321 | United States Air Force School of Aerospace Medicine | United States Air Force School of Aerospace Medicine | Anthony Fries, Jennifer Meyer, Amanda Javorina, Sarah Purves, William Gruner, Clarise Starr, Elizabeth Macias |
| EPI_ISL_812957, EPI_ISL_812958, EPI_ISL_812959, EPI_ISL_812960, EPI_ISL_812961 | Division of Pathogen Resource Management, Korea National Institute of Health, Korea Disease Control and Prevention Agency | Division of Pathogen Resource Management, Korea National Institute of Health, Korea Disease Control and Prevention Agency | Kim,S.T., Kim,S.Y., Choi,Y.S., Kim,E.-J., Kim,J.-M., Yun,M.-r., Choi,C. |
| EPI_ISL_812966 | University Clinical Research Center, University of Sciences | University Clinical Research Center, University of Sciences | Diarra,B., Kone,A., Guindo,I., Bane,S., Diakite,M., Dao,S., Iknane,A.A., Doumbia,S. |
| EPI_ISL_812968 | Department of Molecular Medicine, University of Padova | Department of Molecular Medicine, University of Padova | Lavezzo,E., Franchin,E., Barzon,L., Del Vecchio,C., Rossi,L.,Manganelli,R., Loregian,A., Abate,D., Sciro,M., De Canale,E.,Vanuzzo,M.C., Besutti,V., Saluzzo,F., Onelia,F., Pacenti,M.,Manuto,L., Parisi,S.G., Masi,G., Trevisan,M., Toppo,S. and Crisanti,A. |
| EPI_ISL_819301, EPI_ISL_819302 | Hospital Universitari Vall d'Hebron - Vall d'Hebron Institut de Rercerca | Hospital Universitari Vall d'Hebron | Cristina Andrés, Maria Piñana, Josep F Abril, Damir Garcia-Cehic, Ariadna Rando, Juliana Esperalba, Maria Gema Codina, Carla Castillo, Maria Carmen Martin, Tomàs Pumarola, Josep Quer, Andrés Antón |
| EPI_ISL_831975 | Uppsala klinisk mikrobiologi | The Public Health Agency of Sweden | Department of Microbiology, The Public Health Agency of Sweden |
| EPI_ISL_837608 | Instituto Nacional de Enfermedades Respiratorias (INER) | Instituto Nacional de Enfermedades Respiratorias (INER) | Celia Boukadida, Margarita Matías-Florentino, Alma Rincón-Rubio, Hector Esteban Paz-Juárez, Olivia Briceño, Edgar Sevilla-Reyes, Fidencio Mejía-Nepomuceno, Mario Mújica-Sánchez, Eduardo Becerril-Vargas, José Arturo Martínez-Orozco, Alejandra Hernández-Terán, Jorge Salas-Hernández, Santiago Ávila-Ríos, Joel Armando Vázquez-Pérez |
| EPI_ISL_848085, EPI_ISL_848086, EPI_ISL_848096, EPI_ISL_848100, EPI_ISL_848106, EPI_ISL_848108, EPI_ISL_848109, EPI_ISL_848110, EPI_ISL_848111, EPI_ISL_848112, EPI_ISL_848113, EPI_ISL_848114, EPI_ISL_848115, EPI_ISL_848116, EPI_ISL_848117, EPI_ISL_848118, EPI_ISL_848119, EPI_ISL_848120, EPI_ISL_848121, EPI_ISL_848122, EPI_ISL_848123, EPI_ISL_848124, EPI_ISL_848125, EPI_ISL_848126, EPI_ISL_848146, EPI_ISL_848148, EPI_ISL_848149, EPI_ISL_848151, EPI_ISL_848152, EPI_ISL_848153, EPI_ISL_848154, EPI_ISL_848155, EPI_ISL_848156, EPI_ISL_848157, EPI_ISL_848158, EPI_ISL_848159, EPI_ISL_848160, EPI_ISL_848161, EPI_ISL_848162, EPI_ISL_848163 | | | |
| see above | National Virus Reference Laboratory | Irish Coronavirus Sequencing Consortium - National University of Ireland Galway | Kate Reddington, Grainne Mc Andrew, Simone Coughlan |
| EPI_ISL_848205 | Illinois Department of Public Health | Gagnon Lab, Southern Illinois University | Keith Gagnon |
| EPI_ISL_848599, EPI_ISL_848600, EPI_ISL_848601, EPI_ISL_848602 | Evandro Chagas Institute | Evandro Chagas Institute | Santos, M.C.; Silva, A.M.; Junior, W.D.C.; Barbagelata, L.S.; Ferreira, J.A.; Sousa, E.M.A.; da Silva, P.S.; Pinheiro, K.C.; L.C.; Sousa Junior, E.C. |
| EPI_ISL_848634, EPI_ISL_848635, EPI_ISL_848636 | North Dakota Department of Health, Public Health Laboratory | North Dakota Department of Health, Public Health Laboratory | Lisa Wingerter |
| EPI_ISL_849696, EPI_ISL_849698, EPI_ISL_849699, EPI_ISL_849700, EPI_ISL_849701, EPI_ISL_849702, EPI_ISL_849707, EPI_ISL_849708, EPI_ISL_849709, EPI_ISL_849724 | unknown | PHV-FSS | Son Nguyen et al. |
| EPI_ISL_852724, EPI_ISL_852725, EPI_ISL_852726, EPI_ISL_852727, EPI_ISL_852728, EPI_ISL_852729, EPI_ISL_852730, EPI_ISL_852731, EPI_ISL_852732, EPI_ISL_852733, EPI_ISL_852775, EPI_ISL_852776, EPI_ISL_852777, EPI_ISL_852778, EPI_ISL_852779, EPI_ISL_852780, EPI_ISL_852781, EPI_ISL_852782, EPI_ISL_852783, EPI_ISL_852784 | | | |
| see above | Institute of Virology, Medical Center, University of Freiburg, Freiburg, Germany | Institute of Virology, Clinial Virus Genomics, Medical Center, University of Freiburg, Freiburg, Germany | Jonas Fuchs, Lisa Kern, Sandra Reuter, Hajo Grundmann, Marcus Panning |
| EPI_ISL_853759, EPI_ISL_853882 | Center for Virology, Medical University of Vienna | Bergthaler laboratory, CeMM Research Center for Molecular Medicine of the Austrian Academy of Sciences | Lukas Endler, Alexandra Popa, Benedikt Agerer, Jakob-Wendelin Genger, Alexander Lercher, Anna Schedl, Thomas Penz, Michael Schuster, Jan Laine, Martin Senekowitsch, Christoph Bock, Andreas Bergthaler |
| EPI_ISL_854461 | Alberta Precision Labs (APL) | Alberta Precision Labs (APL) | Gordon P, Lam LG, Pabbaraju K, Wong A, Ma R, Li V, Melin A, Tipples G, Berenger B, Zelyas N, Kellner J, Bernier F, Chui L, Croxen M |
| EPI_ISL_855186, EPI_ISL_855187, EPI_ISL_855188, EPI_ISL_855189, EPI_ISL_855190, EPI_ISL_855191, EPI_ISL_855192, EPI_ISL_855193, EPI_ISL_855194, EPI_ISL_855195, EPI_ISL_855196, EPI_ISL_855197, EPI_ISL_855198, EPI_ISL_855199, EPI_ISL_855200, EPI_ISL_855201, EPI_ISL_855202, EPI_ISL_855203, EPI_ISL_855204, EPI_ISL_855205, EPI_ISL_855206, EPI_ISL_855207, EPI_ISL_855208, EPI_ISL_855209, EPI_ISL_855210, EPI_ISL_855211, EPI_ISL_855212, EPI_ISL_855213, EPI_ISL_855214, EPI_ISL_855215, EPI_ISL_855216, EPI_ISL_855217, EPI_ISL_855218, EPI_ISL_855219, EPI_ISL_855220, EPI_ISL_855221, EPI_ISL_855222, EPI_ISL_855223, EPI_ISL_855224, EPI_ISL_855225, EPI_ISL_855226, EPI_ISL_855227, EPI_ISL_855228, EPI_ISL_855229, EPI_ISL_855230, EPI_ISL_855231, EPI_ISL_855232, EPI_ISL_855233, EPI_ISL_855234, EPI_ISL_855235, EPI_ISL_855236, EPI_ISL_855237, EPI_ISL_855238, EPI_ISL_855239, EPI_ISL_855240, EPI_ISL_855241, EPI_ISL_855242, EPI_ISL_855243, EPI_ISL_855244, EPI_ISL_855245, EPI_ISL_855246, EPI_ISL_855247, EPI_ISL_855248, EPI_ISL_855249, EPI_ISL_855250, EPI_ISL_855251, EPI_ISL_855252, EPI_ISL_855253, EPI_ISL_855254, EPI_ISL_855255, EPI_ISL_855256, EPI_ISL_855257, EPI_ISL_855258, EPI_ISL_855259, EPI_ISL_855260, EPI_ISL_855261, EPI_ISL_855262, EPI_ISL_855263, EPI_ISL_855264, EPI_ISL_855265, EPI_ISL_855266, EPI_ISL_855267, EPI_ISL_855268, EPI_ISL_855269, EPI_ISL_855270, EPI_ISL_855271, EPI_ISL_855272, EPI_ISL_855273, EPI_ISL_855274, EPI_ISL_855275, EPI_ISL_855276, EPI_ISL_855277, EPI_ISL_855278, EPI_ISL_855279, EPI_ISL_855280, EPI_ISL_855281, EPI_ISL_855282, EPI_ISL_855283, EPI_ISL_855284, EPI_ISL_855285, EPI_ISL_855286, EPI_ISL_855287, EPI_ISL_855288, EPI_ISL_855289, EPI_ISL_855290, EPI_ISL_855291, EPI_ISL_855292, EPI_ISL_855293, EPI_ISL_855294, EPI_ISL_855295, EPI_ISL_855296, EPI_ISL_855297, EPI_ISL_855298, EPI_ISL_855299, EPI_ISL_855300, EPI_ISL_855301, EPI_ISL_855302, EPI_ISL_855303, EPI_ISL_855304, EPI_ISL_855305, EPI_ISL_855306, EPI_ISL_855307, EPI_ISL_855308, EPI_ISL_855309, EPI_ISL_855310, EPI_ISL_855311, EPI_ISL_855312, EPI_ISL_855313, EPI_ISL_855314, EPI_ISL_855315, EPI_ISL_855316, EPI_ISL_855317, EPI_ISL_855318, EPI_ISL_855319, EPI_ISL_855320, EPI_ISL_855321, EPI_ISL_855322, EPI_ISL_855323, EPI_ISL_855324, EPI_ISL_855325, EPI_ISL_855326, EPI_ISL_855327, EPI_ISL_855328, EPI_ISL_855329, EPI_ISL_855330, EPI_ISL_855331, EPI_ISL_855332, EPI_ISL_855333, EPI_ISL_855334, EPI_ISL_855335, EPI_ISL_855336, EPI_ISL_855337, EPI_ISL_855338, EPI_ISL_855339 | | | |
| see above | Institute for Computational Biomedicine, Weill Cornell Medicine | Institute for Computational Biomedicine, Weill Cornell Medicine | Butler,D.J., Mozsary,C., Meydan,C., Danko,D., Foox,J., Rosiene,J., Shaiber,A., Afshinnekoo,E., MacKay,M., Sedlazeck,F.J., Ivanov,N.A., Sierra,M., Pohle,D., Zietz,M., Gisladottir,U., Ramlall,V., Westover,C.D., Ryon,K., Young,B., Bhattacharya,C., Ruggiero,P., Langhorst,B.W., Tanner,N., Gawrys,J., Meleshko,D., Xu,D., Steel,P.A.D., Shemesh,A.J., Xiang,J., Thierry-Mieg,J., Thierry-Mieg,D., Schwartz,R.E., Iftner,A., Bezdan,D., Sipley,J., Cong,L., Craney,A., Velu,P., Melnick,A.M., Hajirasouliha,I., Horner,S.M., Iftner,T., Salvatore,M., Loda,M., Westblade,L.F., Cushing,M., Levy,S., Wu,S., Tatonetti,N., Imielinski,M., Rennert,H., Mason,C.E. |

| | | | |
|---|---|---|---|
| EPI_ISL_856695 | University of Zambia, School of Veterinary Medicine, Disease Control | University of Zambia, School of Veterinary Medicine, Disease Control | Simulundu,E., Kapata,N., Mupeta,F., Kapata,P.C., Saasa,N., Changula,K., Muleya,W., Chitanga,S., Chambaro,H., Mubemba,B., Masahiro,K., Chanda,D., Mulenga,L., Fwoloshi,S., Shibemba,A.L., Kapaya,F., Zulu,P., Musonda,K., Monze,M., Sinyange,N., Liwewe,M.M., Kapin'a,M., Chipimo,P.J., Ngosa,W., Morales,A.N., Kayeyi,N., Malama,K., Tembo,J., Bates,M., Sawa,H., Takada,A., Nalubamba,K.S., Mukonka,V., Chilufya,C. and Zumla,A. |
| EPI_ISL_856719 | Infectious Diseases Unit, Department of Internal Medicine, Azienda Ospedaliera-Universitaria di Padova | Laboratory of Infectious Diseases, Department of Biomedical and Clinical Sciences L. Sacco, University of Milan | Anna Maria Cattelan, Lolita Sasset, Davide Leoni, Alessia Lai, Annalisa Bergna, Carla Della Ventura, Claudia Balotta, Massimo Galli, Gianguglielmo Zehender on behalf of SARS-CoV-2 ITALIAN RESEARCH ENTERPRISE-(SCIRE) Collaborative Group |
| EPI_ISL_856897 | Department of Infectious Diseases, Istituto Superiore di Sanità, Roma, Italy; PO Cardarelli, Laboratorio Analisi Microbiologia e Virologia, Campobasso, Italy | Virology Laboratory, Scientific Department, Army Medical Center | Paola Stefanelli, Angela Di Martino, Alessandra Lo Presti, Stefano Fiore, Massimiliano Scutellà, Silvia Fillo, Giovanni Faggioni, Riccardo De Sanctis, Antonella Fortunato, Anna Anselmo, Francesco Giordani, Vanessa Vera Fain, Nino D'Amore, Florigio Lista |
| EPI_ISL_856907 | Department of Infectious Diseases, Istituto Superiore di Sanità, Roma, Italy; AUSL Valle d'Aosta, Aosta, Italy | Virology Laboratory, Scientific Department, Army Medical Center | Paola Stefanelli, Angela Di Martino, Alessandra Lo Presti, Stefano Fiore, Massimo Di Benedetto, Silvia Fillo, Giovanni Faggioni, Riccardo De Sanctis, Antonella Fortunato, Anna Anselmo, Francesco Giordani, Vanessa Vera Fain, Nino D'Amore, Florigio Lista |
| EPI_ISL_857039 | The Ashley Laboratory, Stanford University | Chan-Zuckerberg Biohub | CZB Cliahub Consortium |
| EPI_ISL_860711, EPI_ISL_860714 | Ohio Department of Health Laboratory | Ohio Department of Health Laboratory | Holmes, Jennifer; Eric Brandt, Keoni Omura, Glen McGillivary, Caitlin McDonnell, Kirtana Ramadugu, Erica Leasure, Kelsey Florek, Heather Blankenship, Quanta Brown, and Tammy Bannerman |
| EPI_ISL_861532 | Instituto Nacional de Saude (INSA) | Instituto Nacional de Saude (INSA) | Borges et al |
| EPI_ISL_861893 | LATE - Laboratório de Técnicas Especiais - Hospital Israelita Albert Einstein | LATE - Laboratório de Técnicas Especiais - Hospital Israelita Albert Einstein | Deyvid Amgarten, Fernanda de Mello Malta, Raquel Riyuzo, Ana Paula Moreira Salles, Pedro Henrique Sebe Rodrigues, João Renato Rebello Pinho |
| EPI_ISL_862658 | Hospital Universitario de Ceuta | Instituto de Salud Carlos III | Iglesias-Caballero, M. Camarero, S. Molinero Calamita, M. González-Esguevillas, M. Pozo, F. Casas, I. Jiménez, P. Jiménez, M. Zaballos, A. Monzón, S. Varona, S. Juliá, M. Cuesta, I. Hijano, S. |
| EPI_ISL_875676, EPI_ISL_875678, EPI_ISL_875679, EPI_ISL_875680, EPI_ISL_875681 | Kansas Health and Environmental Lab | Kansas Health and Environmental Lab | Mike Grose, Carissa Robertson, Ben Olsen, and Phil Adam |
| EPI_ISL_876975, EPI_ISL_876976, EPI_ISL_876978, EPI_ISL_876983, EPI_ISL_876984, EPI_ISL_876985, EPI_ISL_876986, EPI_ISL_876987, EPI_ISL_876988, EPI_ISL_876989, EPI_ISL_876990, EPI_ISL_876991, EPI_ISL_876992, EPI_ISL_876993, EPI_ISL_876994, EPI_ISL_876995, EPI_ISL_876996, EPI_ISL_877036 | | | |
| see above | 1.AO Universitaria 'S. Giovanni di Dio e Ruggi D'Aragona, Scuola Medica Salernitana' Hospital / 2.UOC di Virologia e Microbiologia, Università della Campania 'L. Vanvitelli' / 3.AO Universitaria 'Federico II' Napoli Hospital / 4.AORN 'San Giuseppe Moscati' Avellino Hospital / 5.AO 'San Pio - presidio G. Rummo' Benevento Hospital / 6.AO 'Sant'Anna e San Sebastiano' Caserta Hospital / 7.PO 'Maria Santissima Addolorata' Eboli Hospital / 8.Biogem Istituto di Ricerche Genetiche | 1. Genome Research Center for Health (CRGS) / 2. Laboratory of Molecular Medicine and Genomics(LMMGe) / 3. Center for Research in Pure and Applied Mathematics (CRMPA) | Giorgio Giurato (Corresponding Author), Francesca Rizzo (Corresponding Author), Alessandro Weisz (Corresponding Author), Gianluigi Franci, Giovanni Nassa, Pasquale Pagliano, Roberta Tarallo, Elena Alexandrova, Ylenia D'Agostino, Carlo Ferravante, Jessica Lamberti, Viola Melone, Domenico Memoli, Valeria Mirici Cappa, Domenico Palumbo, Giovanni Pecoraro, Assunta Sellitto, Oriana Strianese, Ilaria Terenzi, Giuseppe Fenza, Aniello Gentile, Antonello Saccomanno, Sonia Amabile, Teresa Rocco, Annamaria Salvati, Emilia Vaccaro, Massimiliano Galdiero, Michele Cennamo, Giuseppe Portella, Maria Grazia Foti, Mariarosaria Ingino, Maria Landi, Maurizio Fumi, Vincenzo Rocco, Rita Greco, Vittoria Letizia, Arnolfo Petruzziello, Maddalena Schioppa, Gregorio Goffredi, Francesca Marciano, Michele Caraglia, Alessia Cossu, Marianna Scrima |
| EPI_ISL_884531, EPI_ISL_884532 | Molecular Microbiology & Immunology, University of Missouri | Molecular Microbiology & Immunology, University of Missouri | Tang,C.Y., Li,T., Hang,J., Lidl,G.M., Wan,X.-F. |
| EPI_ISL_887173, EPI_ISL_887176, EPI_ISL_887193, EPI_ISL_887198, EPI_ISL_887212, EPI_ISL_887216, EPI_ISL_887217, EPI_ISL_887219, EPI_ISL_887242, EPI_ISL_887252, EPI_ISL_887269, EPI_ISL_887297, EPI_ISL_887319, EPI_ISL_887324 | | | |
| see above | Protzer Lab | Protzer Lab, Gagneur Lab, Robert Koch Institut | Ulrike Protzer, Dieter Hoffmann, Eva Schulte, Andrea Theumer, Oliver Drechsel, Max von Kleist,Aleksandar Radonic,Stephan Fuchs, Alexander Karollus, Julien Gagneur |
| EPI_ISL_888681 | 1.AO Universitaria 'S. Giovanni di Dio e Ruggi D'Aragona, Scuola Medica Salernitana' Hospital / 2.UOC di Virologia e Microbiologia, Università della Campania 'L. Vanvitelli' / 3.AO Universitaria 'Federico II' Napoli Hospital / 4.AORN 'San Giuseppe Moscati' Avellino Hospital / 5.AO 'San Pio - presidio G. Rummo' Benevento Hospital / 6.AO 'Sant'Anna e San Sebastiano' Caserta Hospital / 7.PO 'Maria Santissima Addolorata' Eboli Hospital / 8.Biogem Istituto di Ricerche Genetiche | 1. Genome Research Center for Health (CRGS) / 2. Laboratory of Molecular Medicine and Genomics(LMMGe) / 3. Center for Research in Pure and Applied Mathematics (CRMPA) | Giorgio Giurato, Francesca Rizzo, Alessandro Weisz, Gianluigi Franci, Giovanni Nassa, Pasquale Pagliano, Roberta Tarallo, Elena Alexandrova, Ylenia D'Agostino, Carlo Ferravante, Jessica Lamberti, Viola Melone, Domenico Memoli, Valeria Mirici Cappa, Domenico Palumbo, Giovanni Pecoraro, Assunta Sellitto, Oriana Strianese, Ilaria Terenzi, Giuseppe Fenza, Aniello Gentile, Antonello Saccomanno, Sonia Amabile, Teresa Rocco, Annamaria Salvati, Emilia Vaccaro, Massimiliano Galdiero, Michele Cennamo, Giuseppe Portella, Maria Grazia Foti, Mariarosaria Ingino, Maria Landi, Maurizio Fumi, Vincenzo Rocco, Rita Greco, Vittoria Letizia, Arnolfo Petruzziello, Maddalena Schioppa, Gregorio Goffredi, Francesca Marciano, Michele Caraglia, Alessia Cossu, Marianna Scrima |
| EPI_ISL_888982 | RS Pondok Indah Hospital - Pondok Indah | Eijkman Institute for Molecular Biology, Ministry of Research and Technology/National Agency for Research and Innovation | Edison Johar, Frilasita A Yudhaputri, Hidayat Trimarsanto, Iskandar Adnan, Lydia V. Panggalo, Sukma Oktavianthi, Willy Agustine, Safarina G Malik, Khin Saw Myint, Amin Soebandrio |
| EPI_ISL_889309, EPI_ISL_889310, EPI_ISL_889311, EPI_ISL_889312 | The University Hospital Brno | Institute of Applied Biotechnologies a.s. | Petr Klempt, Ondej Brzo, Martin Kašný, Kateina Kvapilová, Martina Lengerová, Petr Kvapil |
| EPI_ISL_890376, EPI_ISL_890426, EPI_ISL_890430, EPI_ISL_890431, EPI_ISL_890432, EPI_ISL_890433, EPI_ISL_890434, EPI_ISL_890435, EPI_ISL_890436 | Laboratoire de santé publique du Québec | Laboratoire de santé publique du Québec | Sandrine Moreira, Ioannis Ragoussis, Guillaume Bourque, Jesse Shapiro, Mark Lathrop and Michel Roger on behalf of the CoVSeQ research group (http://covseq.ca/researchgroup) |
| EPI_ISL_894218 | 1.AO Universitaria 'S. Giovanni di Dio e Ruggi D'Aragona, Scuola Medica Salernitana' Hospital / 2.UOC di Virologia e Microbiologia, Università della Campania 'L. Vanvitelli' / 3.AO Universitaria 'Federico II' Napoli Hospital / 4.AORN 'San Giuseppe Moscati' Avellino Hospital / 5.AO 'San Pio - presidio G. Rummo' Benevento Hospital / 6.AO 'Sant'Anna e San Sebastiano' Caserta Hospital / 7.PO 'Maria Santissima Addolorata' Eboli Hospital / 8.Biogem Istituto di Ricerche Genetiche | 1. Genome Research Center for Health (CRGS) / 2. Laboratory of Molecular Medicine and Genomics(LMMGe) / 3. Center for Research in Pure and Applied Mathematics (CRMPA) | Giorgio Giurato, Francesca Rizzo, Alessandro Weisz, Gianluigi Franci, Giovanni Nassa, Pasquale Pagliano, Roberta Tarallo, Elena Alexandrova, Ylenia D'Agostino, Carlo Ferravante, Jessica Lamberti, Viola Melone, Domenico Memoli, Valeria Mirici Cappa, Domenico Palumbo, Giovanni Pecoraro, Assunta Sellitto, Oriana Strianese, Ilaria Terenzi, Giuseppe Fenza, Aniello Gentile, Antonello Saccomanno, Sonia Amabile, Teresa Rocco, Annamaria Salvati, Emilia Vaccaro, Massimiliano Galdiero, Michele Cennamo, Giuseppe Portella, Maria Grazia Foti, Mariarosaria Ingino, Maria Landi, Maurizio Fumi, Vincenzo Rocco, Rita Greco, Vittoria Letizia, Arnolfo Petruzziello, Maddalena Schioppa, Gregorio Goffredi, Francesca Marciano, Michele Caraglia, Alessia Cossu, Marianna Scrima |
| EPI_ISL_895735, EPI_ISL_895776, EPI_ISL_895779, EPI_ISL_895835, EPI_ISL_895837, EPI_ISL_895842, EPI_ISL_895847 | Molecular biology division, Institute of Clinical Biochemistry and Diagnostics, Charles University, Faculty of Medicine in Hradec Králové and University Hospital Hradec Králové | Molecular biology division, Institute of Clinical Biochemistry and Diagnostics, Charles University, Faculty of Medicine in Hradec Králové and University Hospital Hradec Králové | Helena Kovaíková, Petr Brož, Ivana Baranová, Kateina Hrochová, Tereza Baková, Jitka Novotná, Kateina Pehlíková, Vladimír Palika. Cooperation project with BioVendor-R&D and bioinformatics company BIOXSYS s r.o. |
| EPI_ISL_900076, EPI_ISL_900111, EPI_ISL_900115, EPI_ISL_900144, EPI_ISL_900204, EPI_ISL_900226, EPI_ISL_900314, EPI_ISL_900359, EPI_ISL_900392, EPI_ISL_900420, EPI_ISL_900446 | | | |
| see above | MEPHI, Aix Marseille University | MEPHI, Aix Marseille University | Anthony LEVASSEUR |
| EPI_ISL_904628 | Servicio de Microbiología, Laboratori Clínic Metropolitana Nord. Hospital Universitari Germans Trias i Pujol. Institut d'Investigació en Ciències de la Salut Germans Trias i Pujol (IGTP) | IrsiCaixa - Can Ruti CovidSeq | Marc Noguera-Julian, Mariona Parera, Maria Casadellà, Pilar Armengol, Francesc Catala-Moll, Roger Paredes, Bonaventura Clotet Elisa Martró, Verónica Saludes, Anna Not, Ana Pérez, Montserrat Giménez, Ignacio Blanco, Cristina Casañ, Antoni E. Bordoy, Adrián Antuori |
| EPI_ISL_906760 | Dr. Leonard A. Miller Centre for Health Services | National Microbiology Laboratory (NML) | Anna Majer, Shari Tyson, Grace Seo, Philip Mabon, Elsie Grudeski, Rhiannon Huzarewich, Russell Mandes, Anneliese Landgraff, Jennifer Tanner, Natalie Knox, Morag Graham, Gary Van Domselaar, Robert Needle, Yang Yu, Adel Malek, Laura Gilbert, George Zahariadis, Nathalie Bastien, Yan Li, Timothy Booth, Darian Hole, Madison Chapel, CanCOGeN's metadata curation team, Public Health Agency of Canada CanCOGeN team |

| | | | |
|---|---|---|---|
| EPI_ISL_913323 | Klinisk mikrobiologi, Viruslab | The Public Health Agency of Sweden | Anna-Malin Linde, Maria Lind Karlberg, Carlo Berg, Oskar Karlsson Lindsjo, Sofia Stamouli, Reza Advani, Mattias Haukland, Petra Holmstrom, Noura Walai, Petra Edquist, Mia Brytting, Anna Risberg, Karin Tegmark-Wisell |
| EPI_ISL_918457, EPI_ISL_918458, EPI_ISL_918459, EPI_ISL_918460, EPI_ISL_918461, EPI_ISL_918462, EPI_ISL_918463, EPI_ISL_918470, EPI_ISL_918476 | 1.AO Universitaria 'S. Giovanni di Dio e Ruggi D'Aragona, Scuola Medica Salernitana' Hospital / 2.UOC di Virologia e Microbiologia, Università della Campania 'L. Vanvitelli' / 3.AO Universitaria 'Federico II' Napoli Hospital / 4.AORN 'San Giuseppe Moscati' Avellino Hospital / 5.AO 'San Pio - presidio G. Rummo' Benevento Hospital / 6.AO 'Sant'Anna e San Sebastiano' Caserta Hospital / 7.PO 'Maria Santissima Addolorata' Eboli Hospital / 8.Biogem Istituto di Ricerche Genetiche | 1. Genome Research Center for Health (CRGS) / 2. Laboratory of Molecular Medicine and Genomics(LMMGe) / 3. Center for Research in Pure and Applied Mathematics (CRMPA) | Giorgio Giurato (Corresponding Author), Francesca Rizzo (Corresponding Author), Alessandro Weisz (Corresponding Author), Gianluigi Franci, Giovanni Nassa, Pasquale Pagliano, Roberta Tarallo, Elena Alexandrova, Ylenia D'Agostino, Carlo Ferravante, Jessica Lamberti, Viola Melone, Domenico Memoli, Valeria Mirici Cappa, Domenico Palumbo, Giovanni Pecoraro, Assunta Sellitto, Oriana Strianese, Ilaria Terenzi, Giuseppe Fenza, Aniello Gentile, Antonello Saccomanno, Sonia Amabile, Teresa Rocco, Annamaria Salvati, Emilia Vaccaro, Massimiliano Galdiero, Michele Cennamo, Giuseppe Portella, Maria Grazia Foti, Mariarosaria Ingino, Maria Landi, Maurizio Fumi, Vincenzo Rocco, Rita Greco, Vittoria Letizia, Arnolfo Petruzziello, Maddalena Schioppa, Gregorio Goffredi, Francesca Marciano, Michele Caraglia, Alessia Cossu, Marianna Scrima |
| EPI_ISL_933604 | Toronto Invasive Bacterial Diseases Network | McMaster University | Allison McGeer, Patryk Aftanas, Hooman Derakhshani, Angel Li, Kuganya Nirmalarajah, Emily Panousis, Ahmed Draia, Jalees Nasir, Michael Surette, Samira Mubareka, Andrew G. McArthur |
| EPI_ISL_934322 | Klinisk mikrobiologi, Viruslab | The Public Health Agency of Sweden | Anna-Malin Linde, Maria Lind Karlberg, Carlo Berg, Oskar Karlsson Lindsjo, Sofia Stamouli, Reza Advani, Mattias Haukland, Petra Holmstrom, Noura Walai, Petra Edquist, Mia Brytting, Anna Risberg, Karin Tegmark-Wisell |
| EPI_ISL_937539, EPI_ISL_937540, EPI_ISL_937541, EPI_ISL_937542, EPI_ISL_937543 | Thai Red Cross Emerging Infectious Diseases Health Science Centre, Chulalongkorn Hospital, Faculty of Medicine, Chulalongkorn University | Thai Red Cross Emerging Infectious Diseases Center and Faculty of Medicine, Chulalongkorn University | Opass Putcharoen, Watsamon Jantarabenjakul, Pattama Torvorapanit, Gompol Suwanpimolkul, Leilani Paitoonpong, Sopon Iamsirithaworn, Sininat Petcharat, Yutthana Joyjinda, Weenassarin Ampoot, Apaporn Rodpan, Thiravat Hemachudha, Supaporn Wacharapluesadee |
| EPI_ISL_940430, EPI_ISL_940431, EPI_ISL_940432, EPI_ISL_940433, EPI_ISL_940434, EPI_ISL_940435, EPI_ISL_940436, EPI_ISL_940437, EPI_ISL_940438, EPI_ISL_940439, EPI_ISL_940440, EPI_ISL_940441, EPI_ISL_940442, EPI_ISL_940443, EPI_ISL_940444, EPI_ISL_940445, EPI_ISL_940446, EPI_ISL_940447, EPI_ISL_940448, EPI_ISL_940449, EPI_ISL_940450, EPI_ISL_940454 | | | |
| see above | Hôpital Bichat Claude Bernard, Laboratoire de Virologie | IAME UMR1137 Inserm, Université de Paris, Hôpital Bichat | Antoine Bridier-Nahmias, Amélie Recoing, Quentin Le Hingrat, Lena Daniel, Siham Hamri, Gilles Collin, Alexandre Storto, Mélanie Bertine, Charlotte Charpentier, Nadhira Houhou-Fidouh, Diane Descamps, Benoit Visseaux |
| EPI_ISL_949249 | Cliniques universitaires Saint-Luc | UCLouvain/IREC/MBLG | Jean Ruelle, Lysa Pinsmaye, Benoit Kabamba Mukadi |
| EPI_ISL_956390, EPI_ISL_956391, EPI_ISL_956392, EPI_ISL_956393, EPI_ISL_956394 | Isolation - Virology Unit, Institut Pasteur du Cambodge; Sequencing - US National Institute of Allergy and Infectious Diseases Cambodia, US Naval Medical Research Unit -2, Cambodia National Institute for Public Health | Virology Unit, Institut Pasteur du Cambodge | Vireak Heang, Jennifer Bohl, Sophana Chea, Sreyngim Lay, Ly Sovann, Kraing Sidonn, Yi Sengdoeurn, Chin Savuth, Chau Darapheak, Veasna Duong, Jose A Garcia-Rivera, Jessica Manning, Erik A Karlsson |
| EPI_ISL_961010 | Viral Respiratory Lab, National Institute for Biomedical Research (INRB) | Pathogen Sequencing Lab, National Institute for Biomedical Research (INRB) | Placide Mbala-Kingebeni, Edith Nkwembe, Eddy Kinganda-Lusamaki, Amuri Aziza, Francisca Muyembe Mawete, Emmanuel Lokilo Lofiko, Jean-Claude Makangara Cigolo, Catherine Pratt, Matthias Pauthner, Josh Quick, Allison Black, James Hadfield, Trevor Bedford, Ian Goodfellow, Andrew Rambaut, Nick Loman, Kristian Andersen, Michael Wiley, Steve Ahuka-Mundeke, Jean-Jacques Muyembe Tamfum |
| EPI_ISL_976894, EPI_ISL_976895 | Thai Red Cross Emerging Infectious Diseases Health Science Centre, Chulalongkorn Hospital, Faculty of Medicine, Chulalongkorn University | Thai Red Cross Emerging Infectious Diseases Center and Faculty of Medicine, Chulalongkorn University | Opass Putcharoen, Watsamon Jantarabenjakul, Pattama Torvorapanit, Gompol Suwanpimolkul, Leilani Paitoonpong, Sopon Iamsirithaworn, Sininat Petcharat, Yutthana Joyjinda, Weenassarin Ampoot, Apaporn Rodpan, Thiravat Hemachudha, Supaporn Wacharapluesadee |
| EPI_ISL_978503, EPI_ISL_978511, EPI_ISL_978519, EPI_ISL_978520, EPI_ISL_978524, EPI_ISL_978533 | Central Public Health Laboratory - LACEN -Bahia, Salvador, Brazil | Central Public Health Laboratory - LACEN -Bahia, Salvador, Brazil | Stephane Tosta, Luciana Oliveira, Vanessa Nardy,Patrícia Cajado,Marcela Gómez, Breno Dominguez, Jaqueline Gomes, Vagner Fonseca,Marta Giovanetti,Luiz Alcantara, Felicidade Pereira, Arabela Leal |
| EPI_ISL_979273, EPI_ISL_979278, EPI_ISL_979280, EPI_ISL_979283, EPI_ISL_979284, EPI_ISL_979287 | Cadham Provincial laboratory | National Microbiology Laboratory (NML) | Anna Majer, Shari Tyson, Grace Seo, Philip Mabon, Elsie Grudeski, Rhiannon Huzarewich, Russell Mandes, Anneliese Landgraff, Jennifer Tanner, Natalie Knox, Morag Graham, Gary Van Domselaar, Paul Van Caeseele, Jared Bullard, David Alexander, Kerry Dust, Nathalie Bastien, Yan Li, Timothy Booth, Darian Hole, Madison Chapel, Kirsten Biggar, CanCOGeN's metadata curation team, Public Health Agency of Canada CanCOGeN team |
