## Supplementary material for "Detection and characterization of the SARS-CoV-2 lineage B.1.526 in New York": Supp. Table 3 GISAID Acknowledgment part 2: gisaid_hcov-19_acknowledgement_table_2021_02_12_16-6.pdf

We gratefully acknowledge the following Authors from the Originating laboratories responsible for obtaining the specimens, as well as the Submitting laboratories where the genome data were generated and shared via GISAID, on which this research is based.

All Submitters of data may be contacted directly via www.gisaid.org

Authors are sorted alphabetically.

| Accession ID | Originating Laboratory | Submitting Laboratory | Authors |
| --- | --- | --- | --- |
| EPI_ISL_419651 | Gundersen Molecular Diagnostics Laboratory | Kabara Cancer Research Institute | Craig S. Richmond & Paraic A. Kenny |
| EPI_ISL_420024, EPI_ISL_420025, EPI_ISL_420026, EPI_ISL_420027 | Virginia DCLS | Virginia DCLS | Virginia DCLS |
| EPI_ISL_420140 | Department for Virology, Molecular Biology and Genome Research, R. G. Lugar Center for Public Health Research, National Center for Disease Control and Public Health (NCDC) of Georgia. | Department for Virology, Molecular Biology and Genome Research, R. G. Lugar Center for Public Health Research, National Center for Disease Control and Public Health (NCDC) of Georgia. | Nato Kotaria, Marine Murtskhvaladze, Ann Machablishvili, Lela Sabadze, Mari Gavashelidze, Ana Papkiauri, Meri Pantsulaia, Gvantsa Brachveli, Tata Imnadze, Tamar Jashiashvili, Tea Tevdoradze, Ketevan Sidamonidze, Ekaterine Khmaladze, Ekaterine Zhghenti, Roena Sukhiashvili, Mariam Zakalashvili, Lela Urushadze, Magda Dgebuadze, Giorgi Tomashvili, Davit Tsaguria, Ekaterine Zangaladze, Nino Berishvili, Gvantsa Chanturia, Adam Kotorashvili, Maia Alkhazashvili, Irma Burjanadze, Anna Kasradze, Khatuna Zakhashvili, Paata Imnadze, Amiran Gamkrelidze |
| EPI_ISL_420142 | Department for Virology, Molecular Biology and Genome Research, R. G. Lugar Center for Public Health Research, National Center for Disease Control and Public Health (NCDC) of Georgia. | Department for Virology, Molecular Biology and Genome Research, R. G. Lugar Center for Public Health Research, National Center for Disease Control and Public Health (NCDC) of Georgia. | Marine Murtskhvaladze, Ann Machablishvili, Lela Sabadze, Mari Gavashelidze, Ana Papkiauri, Meri Pantsulaia, Gvantsa Brachveli, Tata Imnadze, Tamar Jashiashvili, Tea Tevdoradze, Ketevan Sidamonidze, Ekaterine Khmaladze, Ekaterine Zhghenti, Roena Sukhiashvili, Mariam Zakalashvili, Lela Urushadze, Magda Dgebuadze, Giorgi Tomashvili, Davit Tsaguria, Ekaterine Zangaladze, Nino Berishvili, Gvantsa Chanturia, Adam Kotorashvili, Maia Alkhazashvili, Irma Burjanadze, Anna Kasradze, Khatuna Zakhashvili, Paata Imnadze, Amiran Gamkrelidze. |
| EPI_ISL_420154, EPI_ISL_420155, EPI_ISL_420156, EPI_ISL_420157, EPI_ISL_420158, EPI_ISL_420159, EPI_ISL_420160, EPI_ISL_420178, EPI_ISL_420179, EPI_ISL_420180, EPI_ISL_420181, EPI_ISL_420182, EPI_ISL_420183, EPI_ISL_420184, EPI_ISL_420185, EPI_ISL_420186, EPI_ISL_420187, EPI_ISL_420188, EPI_ISL_420189, EPI_ISL_420190, EPI_ISL_420191, EPI_ISL_420192, EPI_ISL_420193, EPI_ISL_420194, EPI_ISL_420195, EPI_ISL_420196, EPI_ISL_420197, EPI_ISL_420198, EPI_ISL_420199, EPI_ISL_420200, EPI_ISL_420201, EPI_ISL_420202, EPI_ISL_420203, EPI_ISL_420204, EPI_ISL_420205, EPI_ISL_420206, EPI_ISL_420207, EPI_ISL_420208, EPI_ISL_420209, EPI_ISL_420210, EPI_ISL_420211, EPI_ISL_420212, EPI_ISL_420213, EPI_ISL_420214, EPI_ISL_420215, EPI_ISL_420216, EPI_ISL_420217, EPI_ISL_420218, EPI_ISL_420219, EPI_ISL_420220, EPI_ISL_420221, EPI_ISL_420222 | | | |
| see above | Virology Department, Sheffield Teaching Hospitals NHS Foundation Trust | Department of Infection, Immunity and Cardiovascular Disease, The Florey Institute, The Medical School, University of Sheffield | Thushan de Silva, Matthew Parker, Adri Angyal, Rebecca Brown, Rachel Tucker, Paul Parsons, Luke Green, Danielle Groves, Alex Keeley, Dave Partridge, Matthew Wyles, Benjamin Lindsey, Mehmet Yavuz, Mohammad Raza, Cariad Evans |
| EPI_ISL_420526, EPI_ISL_420527, EPI_ISL_420528, EPI_ISL_420529, EPI_ISL_420530, EPI_ISL_420658, EPI_ISL_420660, EPI_ISL_420661, EPI_ISL_420662, EPI_ISL_420668, EPI_ISL_420669, EPI_ISL_420670, EPI_ISL_420671, EPI_ISL_420672, EPI_ISL_420673, EPI_ISL_420674, EPI_ISL_420675, EPI_ISL_420676, EPI_ISL_420677, EPI_ISL_420678, EPI_ISL_420679, EPI_ISL_420680, EPI_ISL_420681, EPI_ISL_420682, EPI_ISL_420683, EPI_ISL_420684, EPI_ISL_420685, EPI_ISL_420686, EPI_ISL_420687, EPI_ISL_420688, EPI_ISL_420689, EPI_ISL_420690, EPI_ISL_420691, EPI_ISL_420692, EPI_ISL_420693, EPI_ISL_420694, EPI_ISL_420695, EPI_ISL_420696, EPI_ISL_420697, EPI_ISL_420698, EPI_ISL_420699, EPI_ISL_420700, EPI_ISL_420701, EPI_ISL_420702, EPI_ISL_420703, EPI_ISL_420704, EPI_ISL_420705, EPI_ISL_420706, EPI_ISL_420707, EPI_ISL_420708, EPI_ISL_420709, EPI_ISL_420710, EPI_ISL_420711, EPI_ISL_420712, EPI_ISL_420713, EPI_ISL_420714, EPI_ISL_420715, EPI_ISL_420718, EPI_ISL_420719, EPI_ISL_420720, EPI_ISL_420721, EPI_ISL_420722, EPI_ISL_420723, EPI_ISL_420724, EPI_ISL_420725, EPI_ISL_420726, EPI_ISL_420727, EPI_ISL_420728, EPI_ISL_420729, EPI_ISL_420730, EPI_ISL_420731, EPI_ISL_420732, EPI_ISL_420733, EPI_ISL_420734, EPI_ISL_420735, EPI_ISL_420736, EPI_ISL_420737, EPI_ISL_420738, EPI_ISL_420739, EPI_ISL_420740, EPI_ISL_420741, EPI_ISL_420742, EPI_ISL_420743, EPI_ISL_420744, EPI_ISL_420745, EPI_ISL_420747, EPI_ISL_420748, EPI_ISL_420751, EPI_ISL_420754, EPI_ISL_420755, EPI_ISL_420756, EPI_ISL_420757, EPI_ISL_420758, EPI_ISL_420760, EPI_ISL_420765, EPI_ISL_420766, EPI_ISL_420767, EPI_ISL_420768, EPI_ISL_420770, EPI_ISL_420771, EPI_ISL_420772, EPI_ISL_420773, EPI_ISL_420774, EPI_ISL_420775, EPI_ISL_420776, EPI_ISL_420777 | | | |
| see above | Respiratory Virus Unit, Microbiology Services Colindale, Public Health England | Respiratory Virus Unit, Microbiology Services Colindale, Public Health England | Monica Galiano, Shahjahan Miah, Angie Lackenby, Omolola Akinbami, Tiina Talts, Leena Bhaw, Richard Myers, Steven Platt, Kirstin Edwards, Jonathan Hubb, Joanna Ellis, Maria Zambon |
| EPI_ISL_420810, EPI_ISL_420811, EPI_ISL_420812, EPI_ISL_420813, EPI_ISL_420814, EPI_ISL_420823 | Utah Public Health Laboratory | Utah Public Health Laboratory | Erin Young, Kelly Oakeson |
| EPI_ISL_420849 | Viral Respiratory Lab, National Institute for Biomedical Research (INRB) | Pathogen Sequencing Lab, National Institute for Biomedical Research (INRB) | Placide Mbala-Kingebeni, Edith Nkwembe, Eddy Kinganda-Lusamaki, Amuri Aziza, Catherine Pratt, Matthias Pauthner, Josh Quick, Allison Black, James Hadfield, Trevor Bedford, Ian Goodfellow, Nick Loman, Kristian Andersen, Michael Wiley, Steve Ahuka-Mundeke, Jean-Jacques Muyembe Tamfum |
| EPI_ISL_420877 | Geelong Centre for Emerging Infectious Diseases | Geelong Centre for Emerging Infectious Diseases | Chamings,A., Raj Bhatta T., Alexandersen S. |
| EPI_ISL_421174, EPI_ISL_421175, EPI_ISL_421176, EPI_ISL_421177, EPI_ISL_421178, EPI_ISL_421179, EPI_ISL_421180 | Hospital Universitario 12 de Octubre | Hospital Universitario 12 de Octubre | Esther Viedma, Sara González, Elias Dahdouh, Raúl Recio, Fernando Lázaro, Julio García, Mª Dolores Folgueira, Jesús Mingorance, Rafael Delgado |
| EPI_ISL_421182, EPI_ISL_421183, EPI_ISL_421185, EPI_ISL_421186, EPI_ISL_421187, EPI_ISL_421188, EPI_ISL_421189, EPI_ISL_421191, EPI_ISL_421192, EPI_ISL_421194, EPI_ISL_421195, EPI_ISL_421196, EPI_ISL_421198, EPI_ISL_421202, EPI_ISL_421203, EPI_ISL_421208, EPI_ISL_421214 | | | |
| see above | Department of Clinical Microbiology | GIGA Medical Genomics | Keith Durkin, Maria Artesi, Sébastien Bontems, Raphaël Boreux, Cécile Meex, Pierrette Melin, Marie-Pierre Hayette, Vincent Bours. |
| EPI_ISL_421563 | Utah Public Health Laboratory | Utah Public Health Laboratory | Erin Young, Kelly Oakeson |
| EPI_ISL_421796, EPI_ISL_421798, EPI_ISL_421802, EPI_ISL_421820, EPI_ISL_421837, EPI_ISL_421838, EPI_ISL_421839, EPI_ISL_421848, EPI_ISL_421849, EPI_ISL_421850, EPI_ISL_421851, EPI_ISL_421852, EPI_ISL_421853, EPI_ISL_421854, EPI_ISL_421855, EPI_ISL_421860, EPI_ISL_421861, EPI_ISL_421864, EPI_ISL_421865, EPI_ISL_421866, EPI_ISL_421867, EPI_ISL_421868, EPI_ISL_421869, EPI_ISL_421870, EPI_ISL_421871, EPI_ISL_421872, EPI_ISL_421873, EPI_ISL_421874, EPI_ISL_421877, EPI_ISL_421878, EPI_ISL_421879, EPI_ISL_421882, EPI_ISL_421883, EPI_ISL_421884, EPI_ISL_421885, EPI_ISL_421886, EPI_ISL_421887, EPI_ISL_421888, EPI_ISL_421889, EPI_ISL_421890, EPI_ISL_421891, EPI_ISL_421892, EPI_ISL_421893, EPI_ISL_421894, EPI_ISL_421895, EPI_ISL_421902, EPI_ISL_421906, EPI_ISL_421908, EPI_ISL_421913, EPI_ISL_421914, EPI_ISL_421915, EPI_ISL_421916, EPI_ISL_421917, EPI_ISL_421918, EPI_ISL_421919, EPI_ISL_421920, EPI_ISL_421921, EPI_ISL_421922, EPI_ISL_421923, EPI_ISL_421924, EPI_ISL_421925, EPI_ISL_421926, EPI_ISL_421927, EPI_ISL_421928, EPI_ISL_421929, EPI_ISL_421930, EPI_ISL_421937, EPI_ISL_421938, EPI_ISL_421940, EPI_ISL_421954, EPI_ISL_421955, EPI_ISL_422002 | | | |
| see above | Respiratory Virus Unit, Microbiology Services Colindale, Public Health England | Respiratory Virus Unit, Microbiology Services Colindale, Public Health England | Monica Galiano, Shahjahan Miah, Angie Lackenby, Omolola Akinbami, Tiina Talts, Leena Bhaw, Richard Myers, Steven Platt, Kirstin Edwards, Jonathan Hubb, Joanna Ellis, Maria Zambon |
| EPI_ISL_422016, EPI_ISL_422017, EPI_ISL_422018, EPI_ISL_422019, EPI_ISL_422020, EPI_ISL_422021, EPI_ISL_422022, EPI_ISL_422023, EPI_ISL_422024, EPI_ISL_422025, EPI_ISL_422026, EPI_ISL_422027, EPI_ISL_422028, EPI_ISL_422030, EPI_ISL_422031, EPI_ISL_422032, EPI_ISL_422033, EPI_ISL_422034, EPI_ISL_422035, EPI_ISL_422036, EPI_ISL_422037, EPI_ISL_422040, EPI_ISL_422042, EPI_ISL_422043, EPI_ISL_422044, EPI_ISL_422046, EPI_ISL_422047, EPI_ISL_422048, EPI_ISL_422050, EPI_ISL_422051, EPI_ISL_422052, EPI_ISL_422054, EPI_ISL_422056, EPI_ISL_422057, EPI_ISL_422058, EPI_ISL_422059, EPI_ISL_422061, EPI_ISL_422062, EPI_ISL_422068, EPI_ISL_422069, EPI_ISL_422070, EPI_ISL_422072, EPI_ISL_422074, EPI_ISL_422075, EPI_ISL_422076, EPI_ISL_422077, EPI_ISL_422079, EPI_ISL_422080, EPI_ISL_422081, EPI_ISL_422082, EPI_ISL_422083, EPI_ISL_422084, EPI_ISL_422086, EPI_ISL_422087, EPI_ISL_422088, EPI_ISL_422089, EPI_ISL_422090, EPI_ISL_422091, EPI_ISL_422092, EPI_ISL_422093, EPI_ISL_422095, EPI_ISL_422096, EPI_ISL_422097, EPI_ISL_422098, EPI_ISL_422099, EPI_ISL_422102, EPI_ISL_422103, EPI_ISL_422109, EPI_ISL_422112, EPI_ISL_422113, EPI_ISL_422115, EPI_ISL_422119, EPI_ISL_422120, EPI_ISL_422121, EPI_ISL_422122, EPI_ISL_422123, EPI_ISL_422124, EPI_ISL_422125, EPI_ISL_422126, EPI_ISL_422127, EPI_ISL_422128, EPI_ISL_422129, EPI_ISL_422130, EPI_ISL_422131, EPI_ISL_422132, EPI_ISL_422133, EPI_ISL_422137, EPI_ISL_422140, EPI_ISL_422141, EPI_ISL_422143, EPI_ISL_422144, EPI_ISL_422145, EPI_ISL_422146, EPI_ISL_422147, EPI_ISL_422148, EPI_ISL_422151, EPI_ISL_422152, EPI_ISL_422155, EPI_ISL_422156, EPI_ISL_422157, EPI_ISL_422158, EPI_ISL_422159, EPI_ISL_422160, EPI_ISL_422161, EPI_ISL_422162, EPI_ISL_422163, EPI_ISL_422165, EPI_ISL_422166, EPI_ISL_422167, EPI_ISL_422170, EPI_ISL_422171, EPI_ISL_422173, EPI_ISL_422174, EPI_ISL_422176, EPI_ISL_422177, EPI_ISL_422178, EPI_ISL_422179, EPI_ISL_422184, EPI_ISL_422186, EPI_ISL_422187, EPI_ISL_422189, EPI_ISL_422190, EPI_ISL_422191, EPI_ISL_422192, EPI_ISL_422193, EPI_ISL_422196, EPI_ISL_422197, EPI_ISL_422198, EPI_ISL_422200, EPI_ISL_422201, EPI_ISL_422203, EPI_ISL_422204, EPI_ISL_422206, EPI_ISL_422207, EPI_ISL_422209, EPI_ISL_422211, EPI_ISL_422212, EPI_ISL_422213, EPI_ISL_422214, EPI_ISL_422215, EPI_ISL_422216, EPI_ISL_422217, EPI_ISL_422222, EPI_ISL_422224, EPI_ISL_422225, EPI_ISL_422230, EPI_ISL_422231, EPI_ISL_422232, EPI_ISL_422233, EPI_ISL_422235, EPI_ISL_422236, EPI_ISL_422237, EPI_ISL_422239, EPI_ISL_422242, EPI_ISL_422244, EPI_ISL_422245, EPI_ISL_422246, EPI_ISL_422250, EPI_ISL_422251, EPI_ISL_422253, EPI_ISL_422254, EPI_ISL_422256, EPI_ISL_422260, EPI_ISL_422261, EPI_ISL_422265, EPI_ISL_422267, EPI_ISL_422268, EPI_ISL_422270, EPI_ISL_422273, EPI_ISL_422275, EPI_ISL_422278, EPI_ISL_422279, EPI_ISL_422281, EPI_ISL_422283, EPI_ISL_422288, EPI_ISL_422292, EPI_ISL_422293, EPI_ISL_422294, EPI_ISL_422297, EPI_ISL_422300, EPI_ISL_422302, EPI_ISL_422305, EPI_ISL_422306, EPI_ISL_422308, EPI_ISL_422311, EPI_ISL_422312, EPI_ISL_422315, EPI_ISL_422319, EPI_ISL_422325, EPI_ISL_422327, EPI_ISL_422334, EPI_ISL_422336, EPI_ISL_422340, EPI_ISL_422352, EPI_ISL_422356, EPI_ISL_422357, EPI_ISL_422360, EPI_ISL_422365, EPI_ISL_422366, EPI_ISL_422371, EPI_ISL_422372, EPI_ISL_422373, EPI_ISL_422374, EPI_ISL_422375, EPI_ISL_422376, EPI_ISL_422377 | | | |
| see above | Wales Specialist Virology Centre | Public Health Wales Microbiology Cardiff | Catherine Moore, Johnathan Evans, Malorie Perry, Simon Cottrell, Alec Birchley, Alexander Adams, Amy Gaskin, Bree Gatica-Wilcox, Jason Coombes, Lauren Gilbert, Lee Graham, Nicole Pacchiarini, Sara Kumziene-Summerhayes, Sarah Taylor, Sophie Jones, Sara Rey, Matthew Bull, Joanne Watkins, Sally Corden, Tom Connor |
| EPI_ISL_422394, EPI_ISL_422397, EPI_ISL_422398, EPI_ISL_422399, EPI_ISL_422400, EPI_ISL_422402, EPI_ISL_422403, EPI_ISL_422406 | NMIMR, Department of Virology | WACCBIP, University of Ghana | Joyce M. Ngoi, Bright Adu, Collins M. Morang'a, Selassie Kumordjie, Miriam Eshun, Linda Boatemaa, Vanessa Magnussen, Erasmus Kotey, Fred Tei-Maya, Dominic S. Y. Amuzu, Peter Quashie, Augustina Arjarquah, Ivy Asante, Evelyn Bonney, George B. Kyei, Kofi Bonney, Abraham Kwabena Anang, Gordon A. Awandare, William Ampofo |
| EPI_ISL_422436 | Respiratory Virus Unit, Microbiology Services Colindale, Public Health England | Respiratory Virus Unit, Microbiology Services Colindale, Public Health England | Monica Galiano, Shahjahan Miah, Angie Lackenby, Omolola Akinbami, Tiina Talts, Leena Bhaw, Richard Myers, Steven Platt, Kirstin Edwards, Jonathan Hubb, Joanna Ellis, Maria Zambon |

| | | | |
|---|---|---|---|
| EPI_ISL_422465 | Gundersen Molecular Diagnostics Laboratory | Kabara Cancer Research Institute | Craig S. Richmond, Paraic A. Kenny |
| EPI_ISL_422563 | Institute of Microbiology Universidad San Francisco de Quito | Institute of Microbiology Universidad San Francisco de Quito | Belen Prado-Vivar, Sully Marquez, Juan Jose Guadalupe, Bernardo Gutierrez, Francisco Mora, Juan Gaviria, Alejandra Ramones, Franklin Espinoza, Edison Ligña, Jorge Reyes, Patricio Rojas-Silva, Veronica Barragan, Gabriel Trueba, Michelle Grunauer, Paul Cardenas |
| EPI_ISL_422564 | Institute of Microbiology Universidad San Francisco de Quito | Institute of Microbiology Universidad San Francisco de Quito | Juan Jose Guadalupe, Belen Prado-Vivar, Sully Marquez, Bernardo Gutierrez, Francisco Mora, Juan Gaviria, Alejandra Ramones, Franklin Espinoza, Edison Ligña, Jorge Reyes, Patricio Rojas-Silva, Veronica Barragan, Gabriel Trueba, Michelle Grunauer, Paul Cardenas |
| EPI_ISL_422565 | Institute of Microbiology Universidad San Francisco de Quito | Institute of Microbiology Universidad San Francisco de Quito | Sully Marquez, Belen Prado-Vivar, Juan Jose Guadalupe, Bernardo Gutierrez, Francisco Mora, Juan Gaviria, Alejandra Ramones, Franklin Espinoza, Edison Ligña, Jorge Reyes, Patricio Rojas-Silva, Veronica Barragan, Gabriel Trueba, Michelle Grunauer, Paul Cardenas |
| EPI_ISL_422566, EPI_ISL_422568, EPI_ISL_422569, EPI_ISL_422570, EPI_ISL_422571, EPI_ISL_422572, EPI_ISL_422573, EPI_ISL_422574, EPI_ISL_422575, EPI_ISL_422576, EPI_ISL_422577, EPI_ISL_422578, EPI_ISL_422579, EPI_ISL_422580, EPI_ISL_422581, EPI_ISL_422582, EPI_ISL_422583, EPI_ISL_422584, EPI_ISL_422585, EPI_ISL_422586, EPI_ISL_422632, EPI_ISL_422756, EPI_ISL_422758, EPI_ISL_422760, EPI_ISL_422761, EPI_ISL_422762, EPI_ISL_422763, EPI_ISL_422764, EPI_ISL_422765, EPI_ISL_422766, EPI_ISL_422767, EPI_ISL_422768, EPI_ISL_422769, EPI_ISL_422771, EPI_ISL_422772, EPI_ISL_422773, EPI_ISL_422774, EPI_ISL_422775, EPI_ISL_422776, EPI_ISL_422777, EPI_ISL_422778, EPI_ISL_422779, EPI_ISL_422780, EPI_ISL_422781, EPI_ISL_422782, EPI_ISL_422783, EPI_ISL_422784, EPI_ISL_422785, EPI_ISL_422786, EPI_ISL_422787, EPI_ISL_422788, EPI_ISL_422789, EPI_ISL_422790, EPI_ISL_422791, EPI_ISL_422792, EPI_ISL_422793, EPI_ISL_422794, EPI_ISL_422795, EPI_ISL_422796, EPI_ISL_422896, EPI_ISL_422902, EPI_ISL_422903, EPI_ISL_422948, EPI_ISL_422949, EPI_ISL_422950, EPI_ISL_422951, EPI_ISL_422952, EPI_ISL_422953, EPI_ISL_422954, EPI_ISL_422955, EPI_ISL_422956, EPI_ISL_422957, EPI_ISL_422958, EPI_ISL_422959, EPI_ISL_422960 | | | |
| see above | Dutch COVID-19 response team | Erasmus Medical Center | Bas Oude Munnink, David Nieuwenhuijse, Reina Sikkema, Claudia Schapendonk, Irina Chestakova, Anne van der Linden, Theo Bestebroer, Stefan van Nieuwkoop, Mark Pronk, Pascal Lexmond, Corien Swaan, Manon Haverkate, Madelief Mollers, Mart Stein, Sandra Kengne Kamga Mobou, Jeroen van Kampen, Jolanda Voermans, Aura Timen, Corine GeurtsvanKessel, Annemiek van der Eijk, Richard Molenkamp, Marion Koopmans, on behalf of the Dutch national COVID-19 response team. |
| EPI_ISL_423187, EPI_ISL_423210, EPI_ISL_423269, EPI_ISL_423271, EPI_ISL_423289, EPI_ISL_423326, EPI_ISL_423327, EPI_ISL_423328, EPI_ISL_423329, EPI_ISL_423331, EPI_ISL_423332, EPI_ISL_423333, EPI_ISL_423334, EPI_ISL_423335, EPI_ISL_423336, EPI_ISL_423337, EPI_ISL_423338, EPI_ISL_423339, EPI_ISL_423340, EPI_ISL_423341, EPI_ISL_423342, EPI_ISL_423343, EPI_ISL_423344, EPI_ISL_423345, EPI_ISL_423346, EPI_ISL_423347, EPI_ISL_423348, EPI_ISL_423349, EPI_ISL_423350, EPI_ISL_423351, EPI_ISL_423352, EPI_ISL_423353, EPI_ISL_423354, EPI_ISL_423355, EPI_ISL_423356, EPI_ISL_423357, EPI_ISL_423358, EPI_ISL_423359, EPI_ISL_423360, EPI_ISL_423361, EPI_ISL_423362, EPI_ISL_423363, EPI_ISL_423364, EPI_ISL_423365, EPI_ISL_423366, EPI_ISL_423367, EPI_ISL_423368, EPI_ISL_423369, EPI_ISL_423370, EPI_ISL_423371, EPI_ISL_423372, EPI_ISL_423373, EPI_ISL_423374, EPI_ISL_423375, EPI_ISL_423376, EPI_ISL_423377, EPI_ISL_423378, EPI_ISL_423379, EPI_ISL_423380, EPI_ISL_423381, EPI_ISL_423382, EPI_ISL_423383, EPI_ISL_423384, EPI_ISL_423385, EPI_ISL_423386, EPI_ISL_423387, EPI_ISL_423388, EPI_ISL_423389, EPI_ISL_423390, EPI_ISL_423391, EPI_ISL_423392, EPI_ISL_423393, EPI_ISL_423394, EPI_ISL_423395, EPI_ISL_423396, EPI_ISL_423397, EPI_ISL_423398, EPI_ISL_423399, EPI_ISL_423400, EPI_ISL_423401, EPI_ISL_423402, EPI_ISL_423403, EPI_ISL_423404, EPI_ISL_423405, EPI_ISL_423406, EPI_ISL_423407, EPI_ISL_423408, EPI_ISL_423409, EPI_ISL_423410, EPI_ISL_423411, EPI_ISL_423412, EPI_ISL_423413, EPI_ISL_423414, EPI_ISL_423415, EPI_ISL_423416, EPI_ISL_423417, EPI_ISL_423418, EPI_ISL_423419, EPI_ISL_423420, EPI_ISL_423421, EPI_ISL_423422, EPI_ISL_423423, EPI_ISL_423424, EPI_ISL_423425, EPI_ISL_423426, EPI_ISL_423428, EPI_ISL_423431, EPI_ISL_423432, EPI_ISL_423433, EPI_ISL_423434, EPI_ISL_423435, EPI_ISL_423436, EPI_ISL_423437, EPI_ISL_423438, EPI_ISL_423439, EPI_ISL_423440, EPI_ISL_423441, EPI_ISL_423442, EPI_ISL_423443, EPI_ISL_423444, EPI_ISL_423445, EPI_ISL_423446, EPI_ISL_423447, EPI_ISL_423449, EPI_ISL_423450, EPI_ISL_423451, EPI_ISL_423452, EPI_ISL_423453, EPI_ISL_423455, EPI_ISL_423456, EPI_ISL_423457, EPI_ISL_423458, EPI_ISL_423459, EPI_ISL_423460, EPI_ISL_423461, EPI_ISL_423462, EPI_ISL_423463, EPI_ISL_423464, EPI_ISL_423465, EPI_ISL_423466, EPI_ISL_423467, EPI_ISL_423468, EPI_ISL_423470, EPI_ISL_423471, EPI_ISL_423475, EPI_ISL_423476, EPI_ISL_423477, EPI_ISL_423478, EPI_ISL_423495, EPI_ISL_423496, EPI_ISL_423505, EPI_ISL_423506, EPI_ISL_423507, EPI_ISL_423513, EPI_ISL_423514, EPI_ISL_423515, EPI_ISL_423516, EPI_ISL_423517, EPI_ISL_423518, EPI_ISL_423520, EPI_ISL_423521, EPI_ISL_423522, EPI_ISL_423523, EPI_ISL_423524, EPI_ISL_423526, EPI_ISL_423527, EPI_ISL_423529, EPI_ISL_423542, EPI_ISL_423543, EPI_ISL_423544, EPI_ISL_423554, EPI_ISL_423556, EPI_ISL_423589, EPI_ISL_423841 | | | |
| see above | Respiratory Virus Unit, Microbiology Services Colindale, Public Health England | Respiratory Virus Unit, Microbiology Services Colindale, Public Health England | Monica Galiano, Shahjahan Miah, Angie Lackenby, Omolola Akinbami, Tiina Talts, Leena Bhaw, Richard Myers, Steven Platt, Kirstin Edwards, Jonathan Hubb, Joanna Ellis, Maria Zambon |
| EPI_ISL_424347 | Alaska State Virology Laboratory | Alaska State Virology Laboratory | Chen, J. |
| EPI_ISL_424351 | Environmental and Global Health | Environmental and Global Health | Elbadry,M.A., Subramaniam,K., Waltzek,T.B., Gibson,J.C., Stephenson,C.J., Morris,J.G. Jr. and Lednicky,J.A. |
| EPI_ISL_424552 | The National University Hospital of Iceland | deCODE genetics | Daniel F Gudbjartsson; Agnar Helgason; Hakon Jonsson; Olafur T Magnusson; Pall Melsted; Gudmundur L Norddahl; Jona Saemundsdottir; Asgeir Sigurdsson; Patrick Sulem; Arna B Agustsdottir; Berglind Eiriksdottir; Run Fridriksdottir; Elisabet E Gardarsdottir; Gudmundur Georgsson; Olafia S Gretarsdottir; Kjartan R Gudmundsson; Thora R Gunnarsdottir; Arnaldur Gylfason; Hilma Holm; Brynjar O Jensson; Aslaug Jonasdottir; Kamilla S Josefsdottir; Thordur Kristjansson; Droplaug N Magnusdottir; Louise le Roux; Gudrun Sigmundsdottir; Gardar Sveinbjornsson; Kristin E Sveinsdottir; Maney Sveinsdottir; Emil A Thorarensen; Bjarni Thorbjornsson; Gisli Masson; Ingileif Jonsdottir; Alma Moller; Thorolfur Gudnason; Karl G Kristinsson; Unnur Thorsteinsdottir; Kari Stefansson |
| EPI_ISL_424553 | deCODE genetics | deCODE genetics | Daniel F Gudbjartsson; Agnar Helgason; Hakon Jonsson; Olafur T Magnusson; Pall Melsted; Gudmundur L Norddahl; Jona Saemundsdottir; Asgeir Sigurdsson; Patrick Sulem; Arna B Agustsdottir; Berglind Eiriksdottir; Run Fridriksdottir; Elisabet E Gardarsdottir; Gudmundur Georgsson; Olafia S Gretarsdottir; Kjartan R Gudmundsson; Thora R Gunnarsdottir; Arnaldur Gylfason; Hilma Holm; Brynjar O Jensson; Aslaug Jonasdottir; Kamilla S Josefsdottir; Thordur Kristjansson; Droplaug N Magnusdottir; Louise le Roux; Gudrun Sigmundsdottir; Gardar Sveinbjornsson; Kristin E Sveinsdottir; Maney Sveinsdottir; Emil A Thorarensen; Bjarni Thorbjornsson; Gisli Masson; Ingileif Jonsdottir; Alma Moller; Thorolfur Gudnason; Karl G Kristinsson; Unnur Thorsteinsdottir; Kari Stefansson |
| EPI_ISL_424554, EPI_ISL_424555, EPI_ISL_424556, EPI_ISL_424557, EPI_ISL_424558, EPI_ISL_424559 | The National University Hospital of Iceland | deCODE genetics | Daniel F Gudbjartsson; Agnar Helgason; Hakon Jonsson; Olafur T Magnusson; Pall Melsted; Gudmundur L Norddahl; Jona Saemundsdottir; Asgeir Sigurdsson; Patrick Sulem; Arna B Agustsdottir; Berglind Eiriksdottir; Run Fridriksdottir; Elisabet E Gardarsdottir; Gudmundur Georgsson; Olafia S Gretarsdottir; Kjartan R Gudmundsson; Thora R Gunnarsdottir; Arnaldur Gylfason; Hilma Holm; Brynjar O Jensson; Aslaug Jonasdottir; Kamilla S Josefsdottir; Thordur Kristjansson; Droplaug N Magnusdottir; Louise le Roux; Gudrun Sigmundsdottir; Gardar Sveinbjornsson; Kristin E Sveinsdottir; Maney Sveinsdottir; Emil A Thorarensen; Bjarni Thorbjornsson; Gisli Masson; Ingileif Jonsdottir; Alma Moller; Thorolfur Gudnason; Karl G Kristinsson; Unnur Thorsteinsdottir; Kari Stefansson |
| EPI_ISL_424560, EPI_ISL_424561 | deCODE genetics | deCODE genetics | Daniel F Gudbjartsson; Agnar Helgason; Hakon Jonsson; Olafur T Magnusson; Pall Melsted; Gudmundur L Norddahl; Jona Saemundsdottir; Asgeir Sigurdsson; Patrick Sulem; Arna B Agustsdottir; Berglind Eiriksdottir; Run Fridriksdottir; Elisabet E Gardarsdottir; Gudmundur Georgsson; Olafia S Gretarsdottir; Kjartan R Gudmundsson; Thora R Gunnarsdottir; Arnaldur Gylfason; Hilma Holm; Brynjar O Jensson; Aslaug Jonasdottir; Kamilla S Josefsdottir; Thordur Kristjansson; Droplaug N Magnusdottir; Louise le Roux; Gudrun Sigmundsdottir; Gardar Sveinbjornsson; Kristin E Sveinsdottir; Maney Sveinsdottir; Emil A Thorarensen; Bjarni Thorbjornsson; Gisli Masson; Ingileif Jonsdottir; Alma Moller; Thorolfur Gudnason; Karl G Kristinsson; Unnur Thorsteinsdottir; Kari Stefansson |
| EPI_ISL_424563, EPI_ISL_424564 | The National University Hospital of Iceland | deCODE genetics | Daniel F Gudbjartsson; Agnar Helgason; Hakon Jonsson; Olafur T Magnusson; Pall Melsted; Gudmundur L Norddahl; Jona Saemundsdottir; Asgeir Sigurdsson; Patrick Sulem; Arna B Agustsdottir; Berglind Eiriksdottir; Run Fridriksdottir; Elisabet E Gardarsdottir; Gudmundur Georgsson; Olafia S Gretarsdottir; Kjartan R Gudmundsson; Thora R Gunnarsdottir; Arnaldur Gylfason; Hilma Holm; Brynjar O Jensson; Aslaug Jonasdottir; Kamilla S Josefsdottir; Thordur Kristjansson; Droplaug N Magnusdottir; Louise le Roux; Gudrun Sigmundsdottir; Gardar Sveinbjornsson; Kristin E Sveinsdottir; Maney Sveinsdottir; Emil A Thorarensen; Bjarni Thorbjornsson; Gisli Masson; Ingileif Jonsdottir; Alma Moller; Thorolfur Gudnason; Karl G Kristinsson; Unnur Thorsteinsdottir; Kari Stefansson |
| EPI_ISL_424567, EPI_ISL_424568, EPI_ISL_424569, EPI_ISL_424570 | deCODE genetics | deCODE genetics | Daniel F Gudbjartsson; Agnar Helgason; Hakon Jonsson; Olafur T Magnusson; Pall Melsted; Gudmundur L Norddahl; Jona Saemundsdottir; Asgeir Sigurdsson; Patrick Sulem; Arna B Agustsdottir; Berglind Eiriksdottir; Run Fridriksdottir; Elisabet E Gardarsdottir; Gudmundur Georgsson; Olafia S Gretarsdottir; Kjartan R Gudmundsson; Thora R Gunnarsdottir; Arnaldur Gylfason; Hilma Holm; Brynjar O Jensson; Aslaug Jonasdottir; Kamilla S Josefsdottir; Thordur Kristjansson; Droplaug N Magnusdottir; Louise le Roux; Gudrun Sigmundsdottir; Gardar Sveinbjornsson; Kristin E Sveinsdottir; Maney Sveinsdottir; Emil A Thorarensen; Bjarni Thorbjornsson; Gisli Masson; Ingileif Jonsdottir; Alma Moller; Thorolfur Gudnason; Karl G Kristinsson; Unnur Thorsteinsdottir; Kari Stefansson |
| EPI_ISL_424571, EPI_ISL_424573, EPI_ISL_424574, EPI_ISL_424575, EPI_ISL_424576, EPI_ISL_424577, EPI_ISL_424578, EPI_ISL_424580, EPI_ISL_424581, EPI_ISL_424582, EPI_ISL_424583, EPI_ISL_424584, EPI_ISL_424585, EPI_ISL_424586, EPI_ISL_424587, EPI_ISL_424588, EPI_ISL_424589, EPI_ISL_424590, EPI_ISL_424591, EPI_ISL_424592, EPI_ISL_424593, EPI_ISL_424595, EPI_ISL_424596, EPI_ISL_424597, EPI_ISL_424598, EPI_ISL_424599, EPI_ISL_424600, EPI_ISL_424601, EPI_ISL_424602, EPI_ISL_424603, EPI_ISL_424604, EPI_ISL_424605, EPI_ISL_424606, EPI_ISL_424607 | | | |
| see above | The National University Hospital of Iceland | deCODE genetics | Daniel F Gudbjartsson; Agnar Helgason; Hakon Jonsson; Olafur T Magnusson; Pall Melsted; Gudmundur L Norddahl; Jona Saemundsdottir; Asgeir Sigurdsson; Patrick Sulem; Arna B Agustsdottir; Berglind Eiriksdottir; Run Fridriksdottir; Elisabet E Gardarsdottir; Gudmundur Georgsson; Olafia S Gretarsdottir; Kjartan R Gudmundsson; Thora R Gunnarsdottir; Arnaldur Gylfason; Hilma Holm; Brynjar O Jensson; Aslaug Jonasdottir; Kamilla S Josefsdottir; Thordur Kristjansson; Droplaug N Magnusdottir; Louise le Roux; Gudrun Sigmundsdottir; Gardar Sveinbjornsson; Kristin E Sveinsdottir; Maney Sveinsdottir; Emil A Thorarensen; Bjarni Thorbjornsson; Gisli Masson; Ingileif Jonsdottir; Alma Moller; Thorolfur Gudnason; Karl G Kristinsson; Unnur Thorsteinsdottir; Kari Stefansson |
| EPI_ISL_424608, EPI_ISL_424609 | deCODE genetics | deCODE genetics | Daniel F Gudbjartsson; Agnar Helgason; Hakon Jonsson; Olafur T Magnusson; Pall Melsted; Gudmundur L Norddahl; Jona Saemundsdottir; Asgeir Sigurdsson; Patrick Sulem; Arna B Agustsdottir; Berglind Eiriksdottir; Run Fridriksdottir; Elisabet E Gardarsdottir; Gudmundur Georgsson; Olafia S Gretarsdottir; Kjartan R Gudmundsson; Thora R Gunnarsdottir; Arnaldur Gylfason; Hilma Holm; Brynjar O Jensson; Aslaug Jonasdottir; Kamilla S Josefsdottir; Thordur Kristjansson; Droplaug N Magnusdottir; Louise le Roux; Gudrun Sigmundsdottir; Gardar Sveinbjornsson; Kristin E Sveinsdottir; Maney |

| | | | |
|---|---|---|---|
| | | | Sveinsdottir; Emil A Thorarensen; Bjarni Thorbjornsson; Gisli Masson; Ingileif Jonsdottir; Alma Moller; Thorolfur Gudnason; Karl G Kristinsson; Unnur Thorsteinsdottir; Kari Stefansson |
| EPI_ISL_424611, EPI_ISL_424612, EPI_ISL_424613, EPI_ISL_424614, EPI_ISL_424615, EPI_ISL_424616, EPI_ISL_424617, EPI_ISL_424618, EPI_ISL_424619, EPI_ISL_424620, EPI_ISL_424621, EPI_ISL_424622, EPI_ISL_424623, EPI_ISL_424624 | | | |
| see above | The National University Hospital of Iceland | deCODE genetics | Daniel F Gudbjartsson; Agnar Helgason; Hakon Jonsson; Olafur T Magnusson; Pall Melsted; Gudmundur L Norddahl; Jona Saemundsdottir; Asgeir Sigurdsson; Patrick Sulem; Arna B Agustsdottir; Berglind Eiriksdottir; Run Fridriksdottir; Elisabet E Gardarsdottir; Gudmundur Georgsson; Olafia S Gretarsdottir; Kjartan R Gudmundsson; Thora R Gunnarsdottir; Arnaldur Gylfason; Hilma Holm; Brynjar O Jensson; Aslaug Jonasdottir; Kamilla S Josefsdottir; Thordur Kristjansson; Droplaug N Magnusdottir; Louise le Roux; Gudrun Sigmundsdottir; Gardar Sveinbjornsson; Kristin E Sveinsdottir; Maney Sveinsdottir; Emil A Thorarensen; Bjarni Thorbjornsson; Gisli Masson; Ingileif Jonsdottir; Alma Moller; Thorolfur Gudnason; Karl G Kristinsson; Unnur Thorsteinsdottir; Kari Stefansson |
| EPI_ISL_425149, EPI_ISL_425151, EPI_ISL_425152, EPI_ISL_425154, EPI_ISL_425159, EPI_ISL_425162, EPI_ISL_425163, EPI_ISL_425165, EPI_ISL_425166, EPI_ISL_425169, EPI_ISL_425171, EPI_ISL_425173, EPI_ISL_425174 | | | |
| see above | University of Wisconsin-Madison AIDS Vaccine Research Laboratories | University of Wisconsin-Madison AIDS Vaccine Research Laboratories | Gage Moreno, Katarina Braun, et al. AIDS Vaccine Research Laboratories |
| EPI_ISL_425231, EPI_ISL_425232, EPI_ISL_425233, EPI_ISL_425234, EPI_ISL_425235, EPI_ISL_425236, EPI_ISL_425237, EPI_ISL_425238, EPI_ISL_425247, EPI_ISL_425248, EPI_ISL_425250, EPI_ISL_425253, EPI_ISL_425254, EPI_ISL_425257, EPI_ISL_425260, EPI_ISL_425273, EPI_ISL_425277, EPI_ISL_425279, EPI_ISL_425281, EPI_ISL_425300, EPI_ISL_425301, EPI_ISL_425302, EPI_ISL_425303, EPI_ISL_425304, EPI_ISL_425305, EPI_ISL_425306, EPI_ISL_425307, EPI_ISL_425308, EPI_ISL_425309, EPI_ISL_425310, EPI_ISL_425311, EPI_ISL_425312, EPI_ISL_425313, EPI_ISL_425319, EPI_ISL_425320, EPI_ISL_425321, EPI_ISL_425322, EPI_ISL_425323, EPI_ISL_425324, EPI_ISL_425325, EPI_ISL_425326, EPI_ISL_425327, EPI_ISL_425328, EPI_ISL_425329, EPI_ISL_425330, EPI_ISL_425331, EPI_ISL_425332, EPI_ISL_425333, EPI_ISL_425334, EPI_ISL_425335, EPI_ISL_425336, EPI_ISL_425337, EPI_ISL_425338, EPI_ISL_425339, EPI_ISL_425423, EPI_ISL_425425, EPI_ISL_425429, EPI_ISL_425430, EPI_ISL_425431, EPI_ISL_425433, EPI_ISL_425461 | | | |
| see above | Department of Pathology, University of Cambridge | COVID-19 Genomics UK (COG-UK) Consortium | Luke W Meredith, M. Estee Torok , Myra Hosmillo, William L. Hamilton, Martin D. Curran, Theresa Feltwell, Anna Yakovleva, Charlotte J. Houldcroft, Aminu S. Jahun, Sarah L. Caddy, Ian Goodfellow |
| EPI_ISL_425610, EPI_ISL_425611, EPI_ISL_425612, EPI_ISL_425613, EPI_ISL_425614, EPI_ISL_425615, EPI_ISL_425616, EPI_ISL_425617, EPI_ISL_425618, EPI_ISL_425619, EPI_ISL_425620, EPI_ISL_425621, EPI_ISL_425622, EPI_ISL_425623, EPI_ISL_425624, EPI_ISL_425625, EPI_ISL_425626, EPI_ISL_425627, EPI_ISL_425628, EPI_ISL_425629, EPI_ISL_425630, EPI_ISL_425631, EPI_ISL_425632, EPI_ISL_425633, EPI_ISL_425634, EPI_ISL_425635, EPI_ISL_425636, EPI_ISL_425637, EPI_ISL_425638, EPI_ISL_425639, EPI_ISL_425640, EPI_ISL_425641, EPI_ISL_425642, EPI_ISL_425643 | | | |
| see above | Queens Medical Centre, Clinical Microbiology Department / DeepSeq Nottingham | COVID-19 Genomics UK (COG-UK) Consortium | Gemma Clark, Wendy Smith, Manjinder Khakh, Hannah Howson-Wells, Jonathan Ball, Patrick McClure, Joseph Chappell, Theocharis Tsoleridis, Nadine Holmes, Matthew Carlisle, Christopher Moore, Fei Sang, Johnny Debebe, Victoria Wright, Matthew Loose |
| EPI_ISL_425951, EPI_ISL_425952, EPI_ISL_425954, EPI_ISL_425955, EPI_ISL_425957, EPI_ISL_425958, EPI_ISL_425962, EPI_ISL_425963, EPI_ISL_425964, EPI_ISL_425965, EPI_ISL_425966, EPI_ISL_425967, EPI_ISL_425968, EPI_ISL_425969, EPI_ISL_425970, EPI_ISL_425971, EPI_ISL_425972, EPI_ISL_425973, EPI_ISL_425974, EPI_ISL_425975, EPI_ISL_425976, EPI_ISL_425977, EPI_ISL_425978, EPI_ISL_425979, EPI_ISL_425980, EPI_ISL_425981, EPI_ISL_425982, EPI_ISL_425983, EPI_ISL_425984, EPI_ISL_425985, EPI_ISL_425986, EPI_ISL_425987, EPI_ISL_425988, EPI_ISL_425989, EPI_ISL_425990, EPI_ISL_425991, EPI_ISL_425992, EPI_ISL_425993, EPI_ISL_425994, EPI_ISL_425995, EPI_ISL_425996, EPI_ISL_425997, EPI_ISL_425998, EPI_ISL_425999, EPI_ISL_426000, EPI_ISL_426001, EPI_ISL_426002, EPI_ISL_426003, EPI_ISL_426004, EPI_ISL_426005, EPI_ISL_426006 | | | |
| see above | Virology Department, Royal Infirmary of Edinburgh, NHS Lothian / School of Biological Sciences, University of Edinburgh / Institute of Genetics and Molecular Medicine, University of Edinburgh | COVID-19 Genomics UK (COG-UK) Consortium | McHugh M, Dewar R, Rooke S, Gallagher M, Balcaza C, O'Toole A, Hill V, McCrone JT, Colquhoun R, Yu X, Jackson B, Scher E, Rambaut A, Williams TC, Templeton K |
| EPI_ISL_426052, EPI_ISL_426053, EPI_ISL_426054, EPI_ISL_426055, EPI_ISL_426056, EPI_ISL_426064, EPI_ISL_426065, EPI_ISL_426066, EPI_ISL_426067, EPI_ISL_426068, EPI_ISL_426069, EPI_ISL_426070, EPI_ISL_426071, EPI_ISL_426072, EPI_ISL_426073, EPI_ISL_426074, EPI_ISL_426075, EPI_ISL_426076, EPI_ISL_426135 | | | |
| see above | UW Virology Lab | UW Virology Lab | Pavitra Roychoudhury, Hong Xie, Keith Jerome, Alexander Greninger |
| EPI_ISL_426285, EPI_ISL_426286, EPI_ISL_426287, EPI_ISL_426288, EPI_ISL_426289 | E. Gulbja Laboratorija | Latvian Biomedical Research and Study Centre | Ivars Silamielis, Kaspars Megnis, Monta Ustinova, ikita Zrelovs, Vita Rovte, Mikus Gavars, Dmitrijs Perminovs, Uga Dumpis, Jnis Kloviš |
| EPI_ISL_426298, EPI_ISL_426299, EPI_ISL_426301, EPI_ISL_426303, EPI_ISL_426305, EPI_ISL_426309, EPI_ISL_426312, EPI_ISL_426321 | Wadsworth Center, New York State Department.of Health | Wadsworth Center, New York State Department.of Health | Kirsten St. George, Daryl M. Lamson, Sara Griesemer, Jonathan Plitnick, Navjot Singh, Matthew D. Shudt, Erica Lasek-Nesselquist |
| EPI_ISL_426379 | The National Laboratory of Health, Environment and Food, Maribor, Slovenia | The National Laboratory of Health, Environment and Food, Maribor, Slovenia | Mahnic A., Hedzet S., Janezic S., Duh D., Zavrsnik J., Blazun Vosner H., Rupnik M. |
| EPI_ISL_426455, EPI_ISL_426456, EPI_ISL_426457, EPI_ISL_426458, EPI_ISL_426459, EPI_ISL_426460, EPI_ISL_426461, EPI_ISL_426462, EPI_ISL_426463, EPI_ISL_426469 | Virginia DCLS | Virginia DCLS | Virginia DCLS |
| EPI_ISL_426534 | TGen North | TGen North | Jolene Bowers, Megan Folkerts, Darrin Lemmer, Dave Engelthaler |
| EPI_ISL_426551, EPI_ISL_426552, EPI_ISL_426553, EPI_ISL_426560 | AZ SPHL, Arizona Department of Health Services | TGen North | Jolene Bowers, Megan Folkerts, Darrin Lemmer, Dave Engelthaler |
| EPI_ISL_426738, EPI_ISL_426739, EPI_ISL_426740, EPI_ISL_426741, EPI_ISL_426742, EPI_ISL_426743, EPI_ISL_426754, EPI_ISL_426755, EPI_ISL_426758, EPI_ISL_426759, EPI_ISL_426760, EPI_ISL_426761, EPI_ISL_426860, EPI_ISL_426861, EPI_ISL_426862, EPI_ISL_426863, EPI_ISL_426865, EPI_ISL_426870, EPI_ISL_426875, EPI_ISL_426876, EPI_ISL_426877 | | | |
| see above | Victorian Infectious Diseases Reference Laboratory (VIDRL) | Microbiological Diagnostic Unit Public Health Laboratory and Victorian Infectious Diseases Reference Laboratory, Doherty Institute | Caly L., Seemann T., Sait, M., Schultz M., Druce J., Sherry, N. |
| EPI_ISL_426883, EPI_ISL_426884, EPI_ISL_426885, EPI_ISL_426886, EPI_ISL_426892, EPI_ISL_426893, EPI_ISL_426894, EPI_ISL_426895, EPI_ISL_426896, EPI_ISL_426897 | Motol University Hospital | Institute of Applied Biotechnologies a.s. | Petr Brož, Jan Geryk, Petr Klempt, Martin Kašný, Adam Novotný, Kateina Kvapilová, Pavel Devínek, Petr Kvapil, Milan Macek |
| EPI_ISL_426906, EPI_ISL_426907, EPI_ISL_426908, EPI_ISL_426909, EPI_ISL_426910, EPI_ISL_426911, EPI_ISL_426912, EPI_ISL_426913, EPI_ISL_426914, EPI_ISL_426915, EPI_ISL_426916, EPI_ISL_426917, EPI_ISL_426918, EPI_ISL_426919, EPI_ISL_426920, EPI_ISL_426921, EPI_ISL_426922 | | | |
| see above | Microbiological Diagnostic Unit Public Health Laboratory | Microbiological Diagnostic Unit Public Health Laboratory | Seemann T., Schultz M., Sait, M., Sherry, N. |
| EPI_ISL_426923, EPI_ISL_426924, EPI_ISL_426925, EPI_ISL_426926, EPI_ISL_426927, EPI_ISL_426928, EPI_ISL_426929, EPI_ISL_426930, EPI_ISL_426931, EPI_ISL_426932, EPI_ISL_426933, EPI_ISL_426934, EPI_ISL_426935, EPI_ISL_426936, EPI_ISL_426937, EPI_ISL_426938, EPI_ISL_426939, EPI_ISL_426940, EPI_ISL_426941, EPI_ISL_426943, EPI_ISL_426944, EPI_ISL_426945, EPI_ISL_426946, EPI_ISL_426947, EPI_ISL_426948, EPI_ISL_426949, EPI_ISL_426950, EPI_ISL_426951, EPI_ISL_426952, EPI_ISL_426953, EPI_ISL_426954, EPI_ISL_426955, EPI_ISL_426956, EPI_ISL_426957, EPI_ISL_426958, EPI_ISL_426959, EPI_ISL_426960, EPI_ISL_426961, EPI_ISL_426962, EPI_ISL_426963, EPI_ISL_426964, EPI_ISL_426965, EPI_ISL_426966, EPI_ISL_426967, EPI_ISL_426968, EPI_ISL_426970, EPI_ISL_426971, EPI_ISL_426972, EPI_ISL_426973, EPI_ISL_426974, EPI_ISL_426975, EPI_ISL_426976, EPI_ISL_426977, EPI_ISL_426978, EPI_ISL_426979, EPI_ISL_426980, EPI_ISL_426981, EPI_ISL_426982, EPI_ISL_426983, EPI_ISL_426984, EPI_ISL_426985, EPI_ISL_426986, EPI_ISL_426987, EPI_ISL_426988, EPI_ISL_426989, EPI_ISL_426990, EPI_ISL_426992, EPI_ISL_426993, EPI_ISL_426994, EPI_ISL_426995, EPI_ISL_426996, EPI_ISL_426997, EPI_ISL_426998, EPI_ISL_426999, EPI_ISL_427000, EPI_ISL_427001, EPI_ISL_427002, EPI_ISL_427003, EPI_ISL_427004, EPI_ISL_427005, EPI_ISL_427006, EPI_ISL_427007, EPI_ISL_427008, EPI_ISL_427009, EPI_ISL_427010, EPI_ISL_427011, EPI_ISL_427012, EPI_ISL_427013, EPI_ISL_427014, EPI_ISL_427015, EPI_ISL_427016, EPI_ISL_427017, EPI_ISL_427018, EPI_ISL_427019, EPI_ISL_427020, EPI_ISL_427021, EPI_ISL_427022, EPI_ISL_427023, EPI_ISL_427024, EPI_ISL_427026, EPI_ISL_427028, EPI_ISL_427029, EPI_ISL_427030, EPI_ISL_427031, EPI_ISL_427032, EPI_ISL_427033, EPI_ISL_427035, EPI_ISL_427036, EPI_ISL_427037, EPI_ISL_427038, EPI_ISL_427039, EPI_ISL_427040, EPI_ISL_427041, EPI_ISL_427051, EPI_ISL_427052 | | | |
| see above | Victorian Infectious Diseases Reference Laboratory (VIDRL) | Microbiological Diagnostic Unit Public Health Laboratory and Victorian Infectious Diseases Reference Laboratory, Doherty Institute | Caly L., Seemann T., Sait, M., Schultz M., Druce J., Sherry, N. |
| EPI_ISL_427054, EPI_ISL_427055, EPI_ISL_427056, EPI_ISL_427057, EPI_ISL_427058, EPI_ISL_427059, | Microbiological Diagnostic Unit Public Health Laboratory | Microbiological Diagnostic Unit Public Health Laboratory | Seemann T., Schultz M., Sait, M., Sherry, N. |

| | | | |
|---|---|---|---|
| EPI_ISL_427060, EPI_ISL_427061, EPI_ISL_427062 | | | |
| EPI_ISL_427082, EPI_ISL_427083, EPI_ISL_427152, EPI_ISL_427153, EPI_ISL_427154, EPI_ISL_427155, EPI_ISL_427156, EPI_ISL_427157, EPI_ISL_427158, EPI_ISL_427159 | Victorian Infectious Diseases Reference Laboratory (VIDRL) | Microbiological Diagnostic Unit Public Health Laboratory and Victorian Infectious Diseases Reference Laboratory, Doherty Institute | Caly L., Seemann T., Sait, M., Schultz M., Druce J., Sherry, N. |
| EPI_ISL_427162, EPI_ISL_427165, EPI_ISL_427168, EPI_ISL_427169, EPI_ISL_427170, EPI_ISL_427171, EPI_ISL_427172, EPI_ISL_427178, EPI_ISL_427191, EPI_ISL_427201, EPI_ISL_427202, EPI_ISL_427203, EPI_ISL_427204, EPI_ISL_427205, EPI_ISL_427206, EPI_ISL_427207, EPI_ISL_427208, EPI_ISL_427209, EPI_ISL_427210, EPI_ISL_427211, EPI_ISL_427212, EPI_ISL_427213, EPI_ISL_427214, EPI_ISL_427215, EPI_ISL_427216, EPI_ISL_427217, EPI_ISL_427218, EPI_ISL_427219, EPI_ISL_427220, EPI_ISL_427221, EPI_ISL_427224, EPI_ISL_427225, EPI_ISL_427226, EPI_ISL_427227, EPI_ISL_427228, EPI_ISL_427229, EPI_ISL_427230, EPI_ISL_427231, EPI_ISL_427232, EPI_ISL_427233, EPI_ISL_427234, EPI_ISL_427235, EPI_ISL_427236, EPI_ISL_427237, EPI_ISL_427262, EPI_ISL_427264, EPI_ISL_427266, EPI_ISL_427267, EPI_ISL_427268 | | | |
| see above | UW Virology Lab | UW Virology Lab | Pavitra Roychoudhury, Hong Xie, Keith Jerome, Alexander Greninger |
| EPI_ISL_427310, EPI_ISL_427311, EPI_ISL_427312 | WHO National Influenza Centre Russian Federation | WHO National Influenza Centre Russian Federation | Andrey Komissarov, Artem Fadeev, Mariia Sergeeva, Anna Ivanova, Daria Danilenko |
| EPI_ISL_427341, EPI_ISL_427344, EPI_ISL_427345, EPI_ISL_427346 | Department of Clinical Microbiology | GIGA Medical Genomics | Keith Durkin, Maria Artesi, Sébastien Bontems, Raphaël Boreux, Cécile Meex, Pierrette Melin, Marie-Pierre Hayette, Vincent Bours. |
| EPI_ISL_427629 | NYU Langone Health | Departments of Pathology and Medicine, New York University School of Medicine | Maria Aguero-Rosenfeld, Brendan Belovarac, Margaret Black, Ludovic Boytard, John Cadley, Paolo Cotzia, John Chen, Dacia Dimartino, Xiaojun Feng, Tatyana Gindin, Emily Guzman, Adriana Heguy, Megan Hogan, Emily Huang, George Jour, Andrew Lytle, Christian Marier, Matthew T. Maurano, Mark J. Mulligan, Peter Meyn, Iman Osman, Jared Pinnell, Vanessa Raabe, Sitharam Ramaswami, Amy Rapkiewicz, Marie Samanovic-Golden, Antonio Serrano, Guomiao Shen, Matija Snuderl, Theodore Vougiouklakis, Nick Vulpescu, Gael Westby, Paul Zappile, Yutong Zhang |
| EPI_ISL_428236 | Hematology Laboratory, Section of Molecular Diagnostics, University Clinical Centre, Medical University of Gdansk | Department of Virology, Faculty of Medicine, University of Helsinki, Helsinki, Finland | Marlena Robakowska, Aneta Szulc, Maciej Grzybek, Olli Vapalahti, Teemu Smura |
| EPI_ISL_428257, EPI_ISL_428258, EPI_ISL_428328, EPI_ISL_428329, EPI_ISL_428331, EPI_ISL_428332, EPI_ISL_428333, EPI_ISL_428334, EPI_ISL_428335, EPI_ISL_428336, EPI_ISL_428337, EPI_ISL_428338, EPI_ISL_428339, EPI_ISL_428340, EPI_ISL_428341, EPI_ISL_428342, EPI_ISL_428343, EPI_ISL_428344 | | | |
| see above | University of Wisconsin-Madison AIDS Vaccine Research Laboratories | University of Wisconsin-Madison AIDS Vaccine Research Laboratories | Gage Moreno, Katarina Braun, et al. AIDS Vaccine Research Laboratories |
| EPI_ISL_428366 | CH Jeanne de Navarre Laboratoire de Biologie | National Reference Center for Viruses of Respiratory Infections, Institut Pasteur, Paris | Mélanie Albert, Marion Barbet, Sylvie Behillil, Méline Bizard, Angela Brisebarre, Flora Donati, Etienne Simon-Lorière, Vincent Enouf, Maud Vanpeene, Sylvie van der Werf |
| EPI_ISL_428374, EPI_ISL_428375, EPI_ISL_428376, EPI_ISL_428377, EPI_ISL_428378, EPI_ISL_428379, EPI_ISL_428388 | Yale COVID-19 Biorepository | Grubaugh Lab - Yale School of Public Health | Joseph Fauver, Tara Alpert, Anderson Brito, Anne Wyllie, Chantal Vogels, Mary Petrone, Chaney Kalinich, Isabel Ott, Arnau Casanovas, Catherine Muenker, Adam Moore, Alice Lu, Maria Tokuyama, Patrick Wong, Peiwen Lu, Saad Omer, Richard Martinello, Allison Nelson, Shelli Farhadian, Akiko Iwasaki, Charlese Dela Cruz, Albert Ko, Nathan Grubaugh |
| EPI_ISL_428400 | Yale COVID-19 Biorepository | Grubaugh Lab - Yale School of Public Health | Joseph Fauver, Tara Alpert, Anderson Brito, Anne Wyllie, Chantal Vogels, Mary Petrone, Cole Jensen, Chaney Kalinich, Isabel Ott, Arnau Casanovas, Catherine Muenker, Adam Moore, Alice Lu, Maria Tokuyama, Patrick Wong, Peiwen Lu, Saad Omer, Richard Martinello, Allison Nelson, Shelli Farhadian, Akiko Iwasaki, Charlese Dela Cruz, Albert Ko, Nathan Grubaugh |
| EPI_ISL_428684, EPI_ISL_428685 | Hospital Universitario 12 de Octubre | Hospital Universitario 12 de Octubre | Sara González, Raúl Recio,Elias Dahdouh, Fernando Lázaro, Esther Viedma, Natalia Stella, Julio García, Juan Carlos Galán, Rafael Cantón, Mª Dolores Folgueira, Rafael Delgado, Jesús Mingorance |
| EPI_ISL_428700, EPI_ISL_428701 | Hospital Universitario 12 de Octubre | Hospital Universitario 12 de Octubre | Esther Viedma, Sara González, Raúl Recio, Elias Dahdouh, Fernando Lázaro, Julio García, Mª Dolores Folgueira, Jesús Mingorance, Rafael Delgado |
| EPI_ISL_428757 | NYU Langone Health | Departments of Pathology and Medicine, New York University School of Medicine | Maria Aguero-Rosenfeld, Brendan Belovarac, Margaret Black, Ludovic Boytard, John Cadley, Paolo Cotzia, John Chen, Dacia Dimartino, Xiaojun Feng, Tatyana Gindin, Emily Guzman, Adriana Heguy, Megan Hogan, Emily Huang, George Jour, Andrew Lytle, Christian Marier, Matthew T. Maurano, Mark J. Mulligan, Peter Meyn, Iman Osman, Jared Pinnell, Vanessa Raabe, Sitharam Ramaswami, Amy Rapkiewicz, Marie Samanovic-Golden, Antonio Serrano, Guomiao Shen, Matija Snuderl, Theodore Vougiouklakis, Nick Vulpescu, Gael Westby, Paul Zappile, Yutong Zhang |
| EPI_ISL_428853 | Laboratory of Molecular Virology International Center for Genetic Engineering and Biotechnology (ICGEB) | ARGO Open Lab Platform for Genome Sequencing | Licastro D, Rajasekharan S, Dal Monego S, Segat L, D'Agaro P, Marcello A |
| EPI_ISL_428854 | Laboratory of Molecular Virology International Center for Genetic Engineering and Biotechnology (ICGEB) | ARGO Open Lab Platform for Genome sequencing | Licastro D, Rajasekharan S, Dal Monego S, Segat L, D'Agaro P, Marcello A |
| EPI_ISL_428894 | State Research Center of Virology and Biotechnology VECTOR, Department of Collection of Microorganisms | State Research Center of Virology and Biotechnology VECTOR, Department of Collection of Microorganisms | Oleg V. Pyankov, Sergey A. Bodnev, Tatyana V. Tregubchak, Alexander N. Shvalov, Elena V. Gavrilova, Rinat A. Maksyutov |
| EPI_ISL_428896, EPI_ISL_428899 | State Research Center of Virology and Biotechnology VECTOR, Department of Collection of Microorganisms | State Research Center of Virology and Biotechnology VECTOR, Department of Collection of Microorganisms | Sergey A. Bodnev, Oleg V. Pyankov, Tatyana V. Tregubchak, Alexander N. Shvalov, Elena V. Gavrilova, Rinat A. Maksyutov |
| EPI_ISL_428906, EPI_ISL_428912, EPI_ISL_428913, EPI_ISL_428914 | State Research Center of Virology and Biotechnology VECTOR, Department of Collection of Microorganisms | State Research Center of Virology and Biotechnology VECTOR, Department of Collection of Microorganisms | Oleg V. Pyankov, Sergey A. Bodnev, Tatyana V. Tregubchak, Alexander N. Shvalov, Elena V. Gavrilova, Rinat A. Maksyutov |
| EPI_ISL_428918, EPI_ISL_428919, EPI_ISL_428920, EPI_ISL_428922, EPI_ISL_428923, EPI_ISL_428924 | State Research Center of Virology and Biotechnology VECTOR, Department of Collection of Microorganisms | State Research Center of Virology and Biotechnology VECTOR, Department of Collection of Microorganisms | Sergey A. Bodnev, Oleg V. Pyankov, Tatyana V. Tregubchak, Alexander N. Shvalov, Elena V. Gavrilova, Rinat A. Maksyutov |
| EPI_ISL_428925, EPI_ISL_428926, EPI_ISL_428927, EPI_ISL_428928, EPI_ISL_428929, EPI_ISL_428930, EPI_ISL_428931, EPI_ISL_428932 | ViroGenetics - BSL3 Laboratory of Virology; Human Genome Variation Research Group & Genomics Centre MCB; Bioinformatics Research Group; Wojewódzka Stacja Sanitarno-Epidemiologiczna w Krakowie | ViroGenetics - BSL3 Laboratory of Virology; Human Genome Variation Research Group & Genomics Centre MCB; Bioinformatics Research Group | Wojciech Branicki, Ewelina Popiech, Micha Kowalski, Agnieszka Starowicz, Adrianna Klajmon, Aleksandra Pisarek, Danuta Piniewska-Róg, Kamila Marszaek, Tomasz Gromowski, Katarzyna Kopera, Katarzyna Dudek, Inga Drebot, Katarzyna Gua, Magda Pachota, Aleksandra Synowiec, Marek Sanak, Jarosaw Foremny, Pawe P abaj, Krzysztof Pyr |
| EPI_ISL_428990, EPI_ISL_428991, EPI_ISL_428996, EPI_ISL_428998, EPI_ISL_428999, EPI_ISL_429000, EPI_ISL_429002, EPI_ISL_429004, EPI_ISL_429005, EPI_ISL_429006, EPI_ISL_429008, EPI_ISL_429010, EPI_ISL_429013, EPI_ISL_429015, EPI_ISL_429019, EPI_ISL_429029, EPI_ISL_429030, EPI_ISL_429032, EPI_ISL_429033, EPI_ISL_429036, EPI_ISL_429041, EPI_ISL_429049, EPI_ISL_429050, EPI_ISL_429056, EPI_ISL_429057, EPI_ISL_429058, EPI_ISL_429059, EPI_ISL_429065, EPI_ISL_429066, EPI_ISL_429071 | | | |
| see above | UCSF Clinical Microbiology Laboratory | Chan-Zuckerberg Biohub | CZB Cliahub Consortium |
| EPI_ISL_429129, EPI_ISL_429131, EPI_ISL_429132 | Laboratoriemedicin | The Public Health Agency of Sweden | Olov Svartstrom, Maria Lind Karlberg, Anna-Malin Linde, Oskar Karlsson Lindsjo, Anna Risberg, Shaman Muradrasoli, Karin Tegmark-Wisell |
| EPI_ISL_429175, EPI_ISL_429176, EPI_ISL_429177, EPI_ISL_429178, EPI_ISL_429179, EPI_ISL_429180, EPI_ISL_429181, EPI_ISL_429182, EPI_ISL_429183, EPI_ISL_429184 | Ramathibodi Hospital | COVID-19 Network Investigations (CONI) Alliance | Elizabeth Batty, Wasun Chantratita, Thanat Chookajorn, Stefan Fernandez, Angkana Huang, Poramate Jiaranai, Anthony R. Jones, Khajohn Joonsalak, Chonticha Klungtong, Theerarat Kochakarn, Namfon Kotanan, Krittikorn Kumpornsin, Wudtichai Manasatienkij, Bhakbhoom Panthan, Ekawat Pasomsub, Kingkan Rakmanee, Insee Sensorn, Janjira Thaipadungpanit, Arporn Wangwiwatsin,Treewat Watthanachockchai |
| EPI_ISL_429201 | University Hospitals of Geneva Laboratory of Virology | University Hospitals of Geneva Laboratory of Virology | Laubscher F. |
| EPI_ISL_429236 | Ospedale Civile S. Liberatore di Atri | Istituto Zooprofilattico Sperimentale dell'Abruzzo e Molise "G. Caporale" | Lorusso A, Marcacci M, Di Domenico M, Ancora M, Curini V, Mangone I, Rinaldi A, Di Pasquale A, Camma C, Puglia I, Savini G |
| EPI_ISL_429589, EPI_ISL_429590 | Department of Virus and Microbiological Special Diagnostics, Statens Serum Institut, Copenhagen, Denmark, Artillerivej 5, 2300 Copenahgen S | Albertsen lab, Department of Chemistry and Bioscience, Aalborg University, Denmark | Rasmus Kirkegaard |

| | | | |
|---|---|---|---|
| EPI_ISL_429597, EPI_ISL_429599, EPI_ISL_429601, EPI_ISL_429602, EPI_ISL_429603, EPI_ISL_429604, EPI_ISL_429610, EPI_ISL_429615, EPI_ISL_429616, EPI_ISL_429617, EPI_ISL_429619, EPI_ISL_429620, EPI_ISL_429621, EPI_ISL_429622, EPI_ISL_429623, EPI_ISL_429624, EPI_ISL_429625, EPI_ISL_429626, EPI_ISL_429627 | | | |
| see above | UW Virology Lab | UW Virology Lab | Pavitra Roychoudhury, Hong Xie, Keith Jerome, Alexander Greninger |
| EPI_ISL_429708, EPI_ISL_429710, EPI_ISL_429716, EPI_ISL_429719, EPI_ISL_429725, EPI_ISL_429736, EPI_ISL_429741, EPI_ISL_429743, EPI_ISL_429745, EPI_ISL_429747, EPI_ISL_429751, EPI_ISL_429755, EPI_ISL_429756, EPI_ISL_429761, EPI_ISL_429763, EPI_ISL_429769, EPI_ISL_429770, EPI_ISL_429772, EPI_ISL_429782, EPI_ISL_429783, EPI_ISL_429784, EPI_ISL_429788, EPI_ISL_429790, EPI_ISL_429791, EPI_ISL_429793, EPI_ISL_429797, EPI_ISL_429799 | | | |
| see above | Laboratoire National de Sante, Microbiology, Virology | Laboratoire National de Sante, Microbiology, Epidemiology and Microbial Genomics | Anke Wienecke-Baldacchino, Ardashel Latsuzbaia, Jessica Tapp, Catherine Ragimbeau, Guillaume Fournier, Tamir Abdelrahman, Trung Nguyen Nguyen, Joel Mossong |
| EPI_ISL_429882, EPI_ISL_429884 | Centers for Disease Control, R.O.C. (Taiwan) | Centers for Disease Control, R.O.C. (Taiwan) | Ji-Rong Yang, Yu-Chi Lin, Jung-Jung Mu, Ming-Tsan Liu |
| EPI_ISL_429970, EPI_ISL_429971, EPI_ISL_429972, EPI_ISL_429973, EPI_ISL_429974, EPI_ISL_429975, EPI_ISL_429977, EPI_ISL_429978, EPI_ISL_429983 | Virginia DCLS | Virginia DCLS | Virginia DCLS |
| EPI_ISL_429995, EPI_ISL_429998, EPI_ISL_430000, EPI_ISL_430009, EPI_ISL_430012 | Biolab Diagnostic Laboratories | Andersen lab at Scripps Research | Issa Abu-Dayyeh, Ahmad Tibi, Lama Hussein, Lina Mohammad, Zein Naber, Amid Abdelnour with SEARCH Alliance San Diego |
| EPI_ISL_430039, EPI_ISL_430041, EPI_ISL_430042, EPI_ISL_430043, EPI_ISL_430044 | Utah Public Health Laboratory | Utah Public Health Laboratory | Erin Young, Kelly Oakeson |
| EPI_ISL_430115, EPI_ISL_430139 | Seattle Flu Study | Seattle Flu Study | Chu et al |
| EPI_ISL_430326, EPI_ISL_430327, EPI_ISL_430328, EPI_ISL_430329, EPI_ISL_430330, EPI_ISL_430331, EPI_ISL_430332, EPI_ISL_430333, EPI_ISL_430334, EPI_ISL_430335, EPI_ISL_430336, EPI_ISL_430337, EPI_ISL_430338, EPI_ISL_430339, EPI_ISL_430340, EPI_ISL_430341, EPI_ISL_430342, EPI_ISL_430343, EPI_ISL_430344, EPI_ISL_430345, EPI_ISL_430346, EPI_ISL_430347, EPI_ISL_430348, EPI_ISL_430349 | | | |
| see above | NYU Langone Health | Departments of Pathology and Medicine, New York University School of Medicine | Maria Aguero-Rosenfeld, Brendan Belovarac, Margaret Black, Ludovic Boytard, John Cadley, Paolo Cotzia, John Chen, Dacia Dimartino, Xiaojun Feng, Tatyana Gindin, Emily Guzman, Adriana Heguy, Megan Hogan, Emily Huang, George Jour, Lawrence H. Lin, Raven Luther, Andrew Lytle, Christian Marier, Matthew T. Maurano, Mark J. Mulligan, Peter Meyn, Raquel Ordonez Ciriza, Iman Osman, Jared Pinnell, Vanessa Raabe, Sitharam Ramaswami, Amy Rapkiewicz, Andre M. Ribeiro-dos-Santos, Marie Samanovic-Golden, Antonio Serrano, Guomiao Shen, Matija Snuderl, Theodore Vougiouklakis, Nick Vulpescu, Gael Westby, Paul Zappile, Yutong Zhang |
| EPI_ISL_430465 | ICMR-National Institute of Cholera and Enteric Diseases | National Institute of Biomedical Genomics | Arindam Maitra, Mamta Chawla Sarkar, Sreedhar Chinnaswamy, Hasina Banu, Ananya Chatterjee, Shanta Dutta, Saumitra Das |
| EPI_ISL_430528, EPI_ISL_430529, EPI_ISL_430542, EPI_ISL_430543, EPI_ISL_430545, EPI_ISL_430552, EPI_ISL_430554, EPI_ISL_430557, EPI_ISL_430558 | Victorian Infectious Diseases Reference Laboratory (VIDRL) | Microbiological Diagnostic Unit Public Health Laboratory and Victorian Infectious Diseases Reference Laboratory, The Peter Doherty Institute for Infection and Immunity | Caly L., Seemann T., Sait, M., Schultz M., Druce J., Sherry, N. |
| EPI_ISL_430648, EPI_ISL_430649, EPI_ISL_430650, EPI_ISL_430651, EPI_ISL_430652, EPI_ISL_430653, EPI_ISL_430654, EPI_ISL_430655, EPI_ISL_430656, EPI_ISL_430657, EPI_ISL_430658, EPI_ISL_430659, EPI_ISL_430660, EPI_ISL_430661, EPI_ISL_430662, EPI_ISL_430663 | | | |
| see above | Microbiological Diagnostic Unit Public Health Laboratory | Microbiological Diagnostic Unit Public Health Laboratory | Seemann T., Schultz M., Sait, M., Sherry, N. |
| EPI_ISL_430791 | UCSF Clinical Microbiology Laboratory | Chan-Zuckerberg Biohub | CZB Cliahub Consortium |
| EPI_ISL_430796, EPI_ISL_430797, EPI_ISL_430798 | Departamento de Biología y genética molecular, IACA Laboratorios. | Área de Secuenciación del Laboratorio de Virología del Hospital de Niños Dr. Ricardo Gutierrez on behalf of 'Proyecto Argentino Interinstitucional de genomica de SARS-CoV-2' (PAIS Consortium) | Nabaes Jodar, MS; Goya, S; Natale, MI; Lusso, S; Tittarelli, E; Suárez, A; Masciovecchio MV; Streitenberger ER; Mistchenko, AS; Valinotto, LE; Viegas, M. |
| EPI_ISL_430799, EPI_ISL_430800 | Laboratorio de Virología del Hospital de Niños Dr. Ricardo Gutierrez | Área de Secuenciación del Laboratorio de Virología del Hospital de Niños Dr. Ricardo Gutierrez on behalf of 'Proyecto Argentino Interinstitucional de genomica de SARS-CoV-2' (PAIS Consortium) | Nabaes Jodar, MS; Goya, S; Natale, MI; Lusso, S; Gravis, E; Mistchenko, AS; Valinotto, LE; Viegas, M. |
| EPI_ISL_430896, EPI_ISL_430940, EPI_ISL_430943, EPI_ISL_430944, EPI_ISL_430945, EPI_ISL_430946, EPI_ISL_430947, EPI_ISL_430948, EPI_ISL_430949, EPI_ISL_430950, EPI_ISL_430952 | | | |
| see above | UW Virology Lab | UW Virology Lab | Pavitra Roychoudhury, Hong Xie, Keith Jerome, Alexander Greninger |
| EPI_ISL_431019 | Alaska State Virology Laboratory | Alaska State Virology Laboratory | Jack Chen, Ph.D. |
| EPI_ISL_431909, EPI_ISL_431913, EPI_ISL_431918, EPI_ISL_431921, EPI_ISL_431925, EPI_ISL_431927, EPI_ISL_431933, EPI_ISL_431942, EPI_ISL_431943, EPI_ISL_431946, EPI_ISL_431960, EPI_ISL_431963, EPI_ISL_431966, EPI_ISL_431967, EPI_ISL_431968, EPI_ISL_431973, EPI_ISL_431977, EPI_ISL_431980, EPI_ISL_431995, EPI_ISL_432000, EPI_ISL_432015, EPI_ISL_432018, EPI_ISL_432023, EPI_ISL_432029, EPI_ISL_432031, EPI_ISL_432034, EPI_ISL_432047, EPI_ISL_432050, EPI_ISL_432059, EPI_ISL_432062, EPI_ISL_432067, EPI_ISL_432068, EPI_ISL_432073, EPI_ISL_432074, EPI_ISL_432079, EPI_ISL_432084, EPI_ISL_432089, EPI_ISL_432095, EPI_ISL_432097, EPI_ISL_432100, EPI_ISL_432103, EPI_ISL_432113, EPI_ISL_432116, EPI_ISL_432117, EPI_ISL_432118, EPI_ISL_432121, EPI_ISL_432125, EPI_ISL_432148, EPI_ISL_432153, EPI_ISL_432155, EPI_ISL_432160, EPI_ISL_432180, EPI_ISL_432198, EPI_ISL_432211, EPI_ISL_432222, EPI_ISL_432227, EPI_ISL_432235, EPI_ISL_432250, EPI_ISL_432255, EPI_ISL_432257, EPI_ISL_432262, EPI_ISL_432266, EPI_ISL_432290, EPI_ISL_432293, EPI_ISL_432301, EPI_ISL_432317, EPI_ISL_432319, EPI_ISL_432321, EPI_ISL_432331, EPI_ISL_432337, EPI_ISL_432338, EPI_ISL_432339, EPI_ISL_432341, EPI_ISL_432342, EPI_ISL_432347, EPI_ISL_432352, EPI_ISL_432357, EPI_ISL_432359, EPI_ISL_432360, EPI_ISL_432361, EPI_ISL_432367, EPI_ISL_432369, EPI_ISL_432379, EPI_ISL_432384, EPI_ISL_432387, EPI_ISL_432390, EPI_ISL_432395, EPI_ISL_432396, EPI_ISL_432397, EPI_ISL_432401, EPI_ISL_432402, EPI_ISL_432403, EPI_ISL_432410, EPI_ISL_432411, EPI_ISL_432420, EPI_ISL_432421, EPI_ISL_432422, EPI_ISL_432423, EPI_ISL_432428, EPI_ISL_432432, EPI_ISL_432442 | | | |
| see above | Wales Specialist Virology Centre | Public Health Wales Microbiology Cardiff | Catherine Moore, Johnathan Evans, Malorie Perry, Simon Cottrell, Alec Birchley, Alexander Adams, Amy Gaskin, Bree Gatica-Wilcox, Jason Coombes, Lauren Gilbert, Lee Graham, Nicole Pacchiarini, Sara Kumziene-Summerhayes, Sarah Taylor, Sophie Jones, Sara Rey, Matthew Bull, Joanne Watkins, Sally Corden, Tom Connor |
| EPI_ISL_432458, EPI_ISL_432476, EPI_ISL_432480, EPI_ISL_432481, EPI_ISL_432486, EPI_ISL_432487, EPI_ISL_432490, EPI_ISL_432492, EPI_ISL_432495, EPI_ISL_432498, EPI_ISL_432500, EPI_ISL_432501, EPI_ISL_432510, EPI_ISL_432514, EPI_ISL_432516, EPI_ISL_432517, EPI_ISL_432520, EPI_ISL_432523, EPI_ISL_432530, EPI_ISL_432534, EPI_ISL_432535, EPI_ISL_432539, EPI_ISL_432540, EPI_ISL_432543, EPI_ISL_432548, EPI_ISL_432549, EPI_ISL_432551, EPI_ISL_432553, EPI_ISL_432554, EPI_ISL_432557, EPI_ISL_432562, EPI_ISL_432563, EPI_ISL_432587, EPI_ISL_432588, EPI_ISL_432593, EPI_ISL_432596, EPI_ISL_432600, EPI_ISL_432601, EPI_ISL_432602, EPI_ISL_432604, EPI_ISL_432609, EPI_ISL_432612, EPI_ISL_432619, EPI_ISL_432620, EPI_ISL_432628, EPI_ISL_432631, EPI_ISL_432632, EPI_ISL_432636, EPI_ISL_432638, EPI_ISL_432644, EPI_ISL_432651, EPI_ISL_432652, EPI_ISL_432656, EPI_ISL_432717, EPI_ISL_432720, EPI_ISL_432724, EPI_ISL_432726, EPI_ISL_432730, EPI_ISL_432734, EPI_ISL_432746, EPI_ISL_432753, EPI_ISL_432754, EPI_ISL_432758, EPI_ISL_432759, EPI_ISL_432767, EPI_ISL_432773, EPI_ISL_432777, EPI_ISL_432786, EPI_ISL_432788, EPI_ISL_432796, EPI_ISL_432799, EPI_ISL_432804, EPI_ISL_432813, EPI_ISL_432815, EPI_ISL_432828, EPI_ISL_432831, EPI_ISL_432837, EPI_ISL_432843, EPI_ISL_432844, EPI_ISL_432852, EPI_ISL_432867 | | | |
| see above | Virology Department, Sheffield Teaching Hospitals NHS Foundation Trust / Virology Department, Sheffield Teaching Hospitals NHS Foundation Trust | COVID-19 Genomics UK (COG-UK) Consortium | Thushan de Silva, Matthew Parker,Adri Angyal, Rebecca Brown, Luke Green, Rachel Tucker, Paul Parsons, Danielle Groves, Alex Keeley, Dave Partridge, Matthew Wyles, Benjamin Lindsey, Mehmet Yavuz, Mohammad Raza, Cariad Evans |
| EPI_ISL_432902, EPI_ISL_432903, EPI_ISL_432904, EPI_ISL_432905, EPI_ISL_432906, EPI_ISL_432907, EPI_ISL_432908, EPI_ISL_432909, EPI_ISL_432910, EPI_ISL_432911, EPI_ISL_432912, EPI_ISL_432913, EPI_ISL_432914, EPI_ISL_432915, EPI_ISL_432916, EPI_ISL_432917, EPI_ISL_432918, EPI_ISL_432919, EPI_ISL_432920, EPI_ISL_432921, EPI_ISL_432922, EPI_ISL_432923, EPI_ISL_432924, EPI_ISL_432925, EPI_ISL_432926, EPI_ISL_432927, EPI_ISL_432928, EPI_ISL_432929, EPI_ISL_432930, EPI_ISL_432931, EPI_ISL_432932, EPI_ISL_432933, EPI_ISL_432934, EPI_ISL_432935, EPI_ISL_432936, EPI_ISL_432937, EPI_ISL_432938, EPI_ISL_432939, EPI_ISL_432940, EPI_ISL_432941, EPI_ISL_432942, EPI_ISL_432943, EPI_ISL_432944, EPI_ISL_432945, EPI_ISL_432946, EPI_ISL_432947, EPI_ISL_432948, EPI_ISL_432951, EPI_ISL_432952, EPI_ISL_432953, EPI_ISL_432954, EPI_ISL_432955, EPI_ISL_432956, EPI_ISL_432957, EPI_ISL_432958, EPI_ISL_432959, EPI_ISL_432960, EPI_ISL_432961, EPI_ISL_432964 | | | |
| see above | Queens Medical Centre, Clinical Microbiology Department / DeepSeq Nottingham | COVID-19 Genomics UK (COG-UK) Consortium | Gemma Clark, Wendy Smith, Manjinder Khakh, Hannah Howson-Wells, Jonathan Ball, Patrick McClure, Joseph Chappell, Theocharis Tsoleridis, Nadine Holmes, Matthew Carlisle, Christopher Moore, Fei Sang, Johnny Debebe, Victoria Wright, Matthew Loose |
| EPI_ISL_433198, EPI_ISL_433199, EPI_ISL_433219 | Virology Department, Royal Infirmary of Edinburgh, NHS Lothian / School of Biological Sciences, University of Edinburgh / Institute of Genetics and Molecular Medicine, University of Edinburgh | COVID-19 Genomics UK (COG-UK) Consortium | McHugh M, Dewar R, Rooke S, Gallagher M, Balcaza C, O'Toole A, Hill V, McCrone JT, Colquhoun R, Yu X, Jackson B, Rambaut A, Williams TC, Templeton K |

| | | | |
|---|---|---|---|
| EPI_ISL_433338, EPI_ISL_433342, EPI_ISL_433343, EPI_ISL_433344, EPI_ISL_433345, EPI_ISL_433346, EPI_ISL_433347, EPI_ISL_433348, EPI_ISL_433351, EPI_ISL_433352, EPI_ISL_433354, EPI_ISL_433355, EPI_ISL_433356, EPI_ISL_433357, EPI_ISL_433358, EPI_ISL_433359, EPI_ISL_433360, EPI_ISL_433361, EPI_ISL_433362, EPI_ISL_433363, EPI_ISL_433364, EPI_ISL_433365, EPI_ISL_433366, EPI_ISL_433367, EPI_ISL_433368, EPI_ISL_433369, EPI_ISL_433370, EPI_ISL_433371, EPI_ISL_433372, EPI_ISL_433373, EPI_ISL_433498, EPI_ISL_433499, EPI_ISL_433500, EPI_ISL_433501, EPI_ISL_433502, EPI_ISL_433503, EPI_ISL_433504, EPI_ISL_433505, EPI_ISL_433506, EPI_ISL_433507, EPI_ISL_433508, EPI_ISL_433509, EPI_ISL_433510, EPI_ISL_433511, EPI_ISL_433512, EPI_ISL_433513, EPI_ISL_433514, EPI_ISL_433515, EPI_ISL_433516, EPI_ISL_433517, EPI_ISL_433518, EPI_ISL_433519, EPI_ISL_433520, EPI_ISL_433521, EPI_ISL_433522, EPI_ISL_433523, EPI_ISL_433531, EPI_ISL_433532, EPI_ISL_433534, EPI_ISL_433537, EPI_ISL_433538 | | | |
| see above | West of Scotland Specialist Virology Centre, NHSGGC / MRC-University of Glasgow Centre for Virus Research | COVID-19 Genomics UK (COG-UK) Consortium | Ana da Silva Filipe, Natasha Johnson, Kathy Smollett, Daniel Mair, Stephen Carmichael, Lily Tong, Jenna Nichols, Elihu Aranday-Cortes, Kirstyn Brunker, Yasmin Parr, Kyriaki Nomikou; Sarah McDonald, Marc Niebel, Patawee Asamaphan; Richard Orton, Joseph Hughes, Sreenu Vattipally, David L Robertson; Alasdair MacLean, Rory Gunson; Kathy Li, Natasha Jesudason, Rajiv Shah, James Shepherd, Antonia Ho, Emma Thomson |
| EPI_ISL_433744, EPI_ISL_433746, EPI_ISL_433816, EPI_ISL_433818, EPI_ISL_433819, EPI_ISL_433821, EPI_ISL_433823, EPI_ISL_433825 | Department of Pathology, University of Cambridge | COVID-19 Genomics UK (COG-UK) Consortium | Luke W Meredith, M. Estee Torok , Myra Hosmillo, William L. Hamilton, Martin D. Curran, Theresa Feltwell, Grant Hall, Anna Yakovleva, Fahad A Khokhar, Charlotte J. Houldcroft, Laura G Caller, Aminu S. Jahun, Sarah L. Caddy, Ian Goodfellow |
| EPI_ISL_434081, EPI_ISL_434130, EPI_ISL_434131, EPI_ISL_434133, EPI_ISL_434136, EPI_ISL_434138, EPI_ISL_434139, EPI_ISL_434140, EPI_ISL_434141, EPI_ISL_434142, EPI_ISL_434143, EPI_ISL_434144, EPI_ISL_434145, EPI_ISL_434147, EPI_ISL_434148, EPI_ISL_434149, EPI_ISL_434150, EPI_ISL_434152, EPI_ISL_434153, EPI_ISL_434154, EPI_ISL_434155, EPI_ISL_434158, EPI_ISL_434159, EPI_ISL_434160, EPI_ISL_434161, EPI_ISL_434162, EPI_ISL_434167, EPI_ISL_434168, EPI_ISL_434169, EPI_ISL_434170, EPI_ISL_434171, EPI_ISL_434172, EPI_ISL_434175, EPI_ISL_434176, EPI_ISL_434177, EPI_ISL_434179, EPI_ISL_434180, EPI_ISL_434181, EPI_ISL_434183, EPI_ISL_434184, EPI_ISL_434185, EPI_ISL_434186, EPI_ISL_434187, EPI_ISL_434188, EPI_ISL_434191, EPI_ISL_434192, EPI_ISL_434193, EPI_ISL_434194, EPI_ISL_434213, EPI_ISL_434265, EPI_ISL_434268, EPI_ISL_434269, EPI_ISL_434274, EPI_ISL_434279, EPI_ISL_434309, EPI_ISL_434315, EPI_ISL_434316, EPI_ISL_434319, EPI_ISL_434321, EPI_ISL_434331, EPI_ISL_434336, EPI_ISL_434337, EPI_ISL_434338, EPI_ISL_434341 | | | |
| see above | Washington State Department of Health | Seattle Flu Study | Chu et al |
| EPI_ISL_434351, EPI_ISL_434354, EPI_ISL_434355 | Lab voor klinische biologie | Onderzoeksgroep Virologie | Laurens Lambrechts, Nick Vereecke, Marthe Pauwels, Jozefien De Clercq, Bruno Verhasselt, Linos Vandekerckhove, Hans Nauwynck, Sebastiaan Theuns |
| EPI_ISL_434356, EPI_ISL_434357, EPI_ISL_434359, EPI_ISL_434360, EPI_ISL_434361, EPI_ISL_434362, EPI_ISL_434363, EPI_ISL_434364, EPI_ISL_434365 | Lab voor klinische biologie | Onderzoeksgroep Virologie | Nick Vereecke, Laurens Lambrechts, Marthe Pauwels, Jozefien De Clercq, Bruno Verhasselt, Linos Vandekerckhove, Hans Nauwynck, Sebastiaan Theuns |
| EPI_ISL_434366 | Hospital AZ Rivierenland | Institute of Tropical Medicine | Philippe Selhorst, Colin Anthony |
| EPI_ISL_434373, EPI_ISL_434375 | Hospital AZ Rivierenland | Institute of Tropical Medicine | Philippe Selhorst, Colin Anthony, |
| EPI_ISL_434457, EPI_ISL_434459, EPI_ISL_434461 | Laboratory of Microbiology, Medical School, National and Kapodistrian University of Athens | Laboratory of Biology, Department of Medicine, Democritus University of Thrace | Kassela K., Bampali,M., Dovrolis,N., Gatzidou,E., Froukala,E., Stavropoulou,A., Veletza,S., Tsakris,A., Spanakis,N. and Karakasiliotis,I. |
| EPI_ISL_434557, EPI_ISL_434558 | National Institutes of Health, University of the Philippines Manila | Philippine Genome Center | Carlo M. Lapid, Francis A. Tablizo, Benedict A. Maralit, Jan Michael C. Yap, Raul V. Destura, Marissa M. Alejandria, El King D. Morado, Joshua Gregor A. Dizon, Jo-Hannah S. Llames, Shiela Mae M. Araiza, Kris P. Punayan, Kristianne Arielle D. Gabriel, Shebna Rose D. Fabilloren, Shana F. Genavia, Jarvin E. Nipales, Alessandra C. Sanchez, Haifa L.Gaza, Joy Ann Petronio-Santos, Julius Aaron Mejia, Maribell Dollete, Sonia Salamat, Christina Tan, Bernard Demot, John Mark Velasco, Eva Maria Cutiongco-de la Paz, and Cynthia P. Saloma |
| EPI_ISL_434637 | Lednicky Laboratory, Emerging Pathogens Institute, University of Florida. | Lednicky Laboratory, Emerging Pathogens Institute, University of Florida. | Elbadry,M.A.; Subramaniam,K.; Waltzek,T.B.; Gibson,J.C.; Stephenson,C.J.; Morris,J.G. Jr. and Lednicky,J.A. |
| EPI_ISL_434642 | Laboratoriemedicin | The Public Health Agency of Sweden | Oskar Karlsson Lindsjo, Maria Lind Karlberg, Anna-Malin Linde, Olov Svartstrom, Anna Risberg, Shaman Muradrasoli, Karin Tegmark-Wisell |
| EPI_ISL_434656 | Trollbackens VC | The Public Health Agency of Sweden | Amelie Holmqvist, Oskar Karlsson Lindsjo, Maria Lind Karlberg, Anna-Malin Linde, Olov Svartstrom, Anna Risberg, Theresa Enkirch, Mia Brytting, Karin Tegmark-Wisell |
| EPI_ISL_434902, EPI_ISL_434903, EPI_ISL_434904, EPI_ISL_434905, EPI_ISL_434906, EPI_ISL_434907, EPI_ISL_434908, EPI_ISL_434909, EPI_ISL_434910, EPI_ISL_434911, EPI_ISL_434912, EPI_ISL_434913, EPI_ISL_434914, EPI_ISL_434915, EPI_ISL_434916, EPI_ISL_434917, EPI_ISL_434918, EPI_ISL_434919, EPI_ISL_434920, EPI_ISL_434921, EPI_ISL_434922, EPI_ISL_434923, EPI_ISL_434924, EPI_ISL_434925, EPI_ISL_434926, EPI_ISL_434927, EPI_ISL_434928, EPI_ISL_434929, EPI_ISL_434930, EPI_ISL_434931, EPI_ISL_434932, EPI_ISL_434933, EPI_ISL_434934, EPI_ISL_434935, EPI_ISL_434936, EPI_ISL_434937, EPI_ISL_434938, EPI_ISL_434939, EPI_ISL_434940, EPI_ISL_434945, EPI_ISL_434946, EPI_ISL_434947, EPI_ISL_434948, EPI_ISL_434949, EPI_ISL_434950, EPI_ISL_434951, EPI_ISL_434952, EPI_ISL_434953, EPI_ISL_434954, EPI_ISL_434955, EPI_ISL_434956, EPI_ISL_434957, EPI_ISL_434959, EPI_ISL_434961, EPI_ISL_434963, EPI_ISL_434965, EPI_ISL_434967, EPI_ISL_434968, EPI_ISL_434969, EPI_ISL_434973, EPI_ISL_434974, EPI_ISL_434977, EPI_ISL_434978, EPI_ISL_434989, EPI_ISL_434993, EPI_ISL_434996 | | | |
| see above | Houston Methodist Hospital | Houston Methodist Hospital | S. Wesley Long, Randall J. Olsen, Paul A. Christensen, David W. Bernard, James J. Davis, Maulik Shukla, Marcus Nguyen, Matthew Ojeda Saavedra, Concepcion C. Cantu, Prasanti Yerramilli, Layne Pruitt, Sishir Subedi, Heather Hendrickson, Ghazaleh Eskandari, Muthiah Kumaraswami, Jason S. McLellan, Hakon Jonsson, Kari Stefansson, and James M. Musser |
| EPI_ISL_435074, EPI_ISL_435075, EPI_ISL_435076, EPI_ISL_435077, EPI_ISL_435078, EPI_ISL_435079, EPI_ISL_435080, EPI_ISL_435081, EPI_ISL_435082, EPI_ISL_435083, EPI_ISL_435084, EPI_ISL_435085, EPI_ISL_435086, EPI_ISL_435087, EPI_ISL_435088, EPI_ISL_435089, EPI_ISL_435090, EPI_ISL_435091, EPI_ISL_435092, EPI_ISL_435093, EPI_ISL_435094, EPI_ISL_435095, EPI_ISL_435096, EPI_ISL_435097, EPI_ISL_435098, EPI_ISL_435099, EPI_ISL_435100, EPI_ISL_435107, EPI_ISL_435111, EPI_ISL_435112 | | | |
| see above | National Centre for Disease control (NCDC), CSIR-Institute of Genomics and Integrative Biology (CSIR-IGIB) | NCDC/CSIR-IGIB | Pramod Kumar, Rajesh Pandey, Pooja Sharma, Mahesh Dhar, Vivekanand A, Bharathram Uppili, Himanshu Vashisht, Saruchi Wadhwa, Nishu Tyagi, Uma Sharma, Priyanka Singh, Hemlata Lall, Meena Datta, Poonam Gupta, Nidhi Saini, Aarti Tewari, Bibhash Nandi, Dhirendra Kumar, Satyabrata Bag, Varun Jaiswal, Hema Gogia, Preeti Madan, Simrita Singh, Prateek Singh, Debasis Dash, Mitali Mukerji, Manju Bala, Sandhya Kabra, Sujeet Singh, Mohammed Faruq, Anurag Agrawal, Partha Rakshit |
| EPI_ISL_435283 | RS Pondok Indah Hospital - Pondok Indah | Eijkman Institute for Molecular Biology, Ministry of Research and Technology/National Agency for Research and Innovation | Edison Johar, Frilasita A Yudhaputri, Hidayat Trimarsanto, David H Muljono, Safarina G Malik, Khin Saw Myint, Amin Soebandrio |
| EPI_ISL_435353, EPI_ISL_435354, EPI_ISL_435355, EPI_ISL_435356, EPI_ISL_435357 | Utah Public Health Laboratory | Utah Public Health Laboratory | Erin Young, Kelly Oakeson |
| EPI_ISL_435411, EPI_ISL_435412, EPI_ISL_435413, EPI_ISL_435414, EPI_ISL_435415, EPI_ISL_435416, EPI_ISL_435417, EPI_ISL_435422, EPI_ISL_435423, EPI_ISL_435424, EPI_ISL_435429, EPI_ISL_435430 | | | |
| see above | Virological Research Group, Szentágothai Research Centre | Bioinformatics Research Group, Szentágothai Research Centre | Péter Urbán, Endre Gábor Tóth, Gábor Kemenesi, Róbert Herczeg, Attila Gyenesei, Ferenc Jakab |
| EPI_ISL_435446, EPI_ISL_435448, EPI_ISL_435449, EPI_ISL_435450, EPI_ISL_435451, EPI_ISL_435452, EPI_ISL_435453, EPI_ISL_435463, EPI_ISL_435464, EPI_ISL_435466 | Robert Garry lab | Andersen lab at Scripps Research | Allison Smither, Gilberto Sabino-Santos, Patricia Snarski, Lilia Melnik, Antoinette Bell, Kaylynn Genemaras, Arnaud Drouin, Dahlene Fusco, Robert Garry with SEARCH Alliance San Diego |
| EPI_ISL_435475, EPI_ISL_435476, EPI_ISL_435477, EPI_ISL_435478, EPI_ISL_435479, EPI_ISL_435480, EPI_ISL_435481, EPI_ISL_435482, EPI_ISL_435483, EPI_ISL_435484, EPI_ISL_435485, EPI_ISL_435486 | | | |
| see above | NYU Langone Health | Departments of Pathology and Medicine, New York University School of Medicine | Maria Aguero-Rosenfeld, Brendan Belovarac, Margaret Black, Ludovic Boytard, John Cadley, Paolo Cotzia, John Chen, Dacia Dimartino, Xiaojun Feng, Tatyana Gindin, Emily Guzman, Adriana Heguy, Megan Hogan, Emily Huang, George Jour, Lawrence H. Lin, Raven Luther, Andrew Lytle, Christian Marier, Matthew T. Maurano, Mark J. Mulligan, Peter Meyn, Raquel Ordonez Ciriza, Iman Osman, Jared Pinnell, Vanessa Raabe, Sitharam Ramaswami, Amy Rapkiewicz, Andre M. Ribeiro-dos-Santos, Marie Samanovic-Golden, Antonio Serrano, Guomiao Shen, Matija Snuderl, Theodore Vougiouklakis, Nick Vulpescu, Gael Westby, Paul Zappile, Yutong Zhang |
| EPI_ISL_436042, EPI_ISL_436043 | DC Public Health Lab Dept of Forensic Science | Pathogen Discovery, Respiratory Viruses Branch, Division of Viral Diseases, Centers for Disease Control and Prevention | Ying Tao, Jing Zhang, Krista Queen, Yan Li, Anna Uehara, Clinton R. Paden, Haibin Wang, Zachary Weiner, Bettina Bankamp, Suxiang Tong |
| EPI_ISL_436105 | TSGH-CP molecular lab | TSGH-CP molecular lab | Cherng-Lih Perng, Ming-Jr Jian, Chih-Kai Chang, Jung-Chung Lin, Kuo-Ming Yeh, Chien-Wen Chen, Sheng-Kang Chiu, Hsing-Yi Chung, Shih-Hung Tsai, Kuo-Sheng Hung, Tien-Yao Chang, Feng-Yee Chang, Hung-Sheng Shang |
| EPI_ISL_436106 | TSGH-CP molecular lab | TSGH-CP molecular lab | Cherng-Lih Perng, Ming-Jr JIAN, Chih-Kai Chang, Jung-Chung Lin, Kuo-Ming Yeh, Chien-Wen Chen, Sheng-Kang Chiu, Hsing-Yi Chung, Shih-Hung Tsai, Kuo-Sheng Hung, Tien-Yao Chang, Feng-Yee Chang, Hung-Sheng Shang |

| | | | |
|---|---|---|---|
| EPI_ISL_436303 | Servicio de Microbiologia. Hospital Clinico Universitario de Valencia | Sequencing and Bioinformatics Service and Molecular Epidemiology Research Group. FISABIO-Public Health | Neris Garcia-Gonzalez, Inma Galán Vendrell, Sandra Carbo, Loreto Ferrús Abad, Paula Ruiz-Hueso, Mariana Reyes-Prieto, Vicente Soriano Chirona, Ivan Ansari, David Navarro, Maria Alma Bracho, Griselda De Marco, Beatriz Beamud, Lidia Ruiz Roldan, Marta Pla Diaz, Llúcia Martínez-Priego, Giuseppe D'Auria, Fernando Gonzalez-Candelas |
| EPI_ISL_436306 | Servicio de Microbiologia. Hospital Clinico Universitario de Valencia | Sequencing and Bioinformatics Service and Molecular Epidemiology Research Group. FISABIO-Public Health | Loreto Ferrús Abad, Paula Ruiz-Hueso, Mariana Reyes-Prieto, Vicente Soriano Chirona, Ivan Ansari, David Navarro, Maria Alma Bracho, Griselda De Marco, Beatriz Beamud, Lidia Ruiz Roldan, Marta Pla Diaz, Neris Garcia-Gonzalez, Inma Galán Vendrell, Sandra Carbo, Llúcia Martínez-Priego, Giuseppe D'Auria, Fernando Gonzalez-Candelas |
| EPI_ISL_436307 | Servicio de Microbiologia. Hospital Clinico Universitario de Valencia | Sequencing and Bioinformatics Service and Molecular Epidemiology Research Group. FISABIO-Public Health | Paula Ruiz-Hueso, Mariana Reyes-Prieto, Vicente Soriano Chirona, Ivan Ansari, David Navarro, Maria Alma Bracho, Griselda De Marco, Beatriz Beamud, Lidia Ruiz Roldan, Marta Pla Diaz, Neris Garcia-Gonzalez, Inma Galán Vendrell, Sandra Carbo, Loreto Ferrús Abad, Llúcia Martínez-Priego, Giuseppe D'Auria, Fernando Gonzalez-Candelas |
| EPI_ISL_436308 | Servicio de Microbiologia. Hospital Clinico Universitario de Valencia | Sequencing and Bioinformatics Service and Molecular Epidemiology Research Group. FISABIO-Public Health | Mariana Reyes-Prieto, Vicente Soriano Chirona, Ivan Ansari, David Navarro, Maria Alma Bracho, Griselda De Marco, Beatriz Beamud, Lidia Ruiz Roldan, Marta Pla Diaz, Neris Garcia-Gonzalez, Inma Galán Vendrell, Sandra Carbo, Loreto Ferrús Abad, Paula Ruiz-Hueso, Llúcia Martínez-Priego, Giuseppe D'Auria, Fernando Gonzalez-Candelas |
| EPI_ISL_436309 | Servicio de Microbiologia. Hospital Clinico Universitario de Valencia | Sequencing and Bioinformatics Service and Molecular Epidemiology Research Group. FISABIO-Public Health | Vicente Soriano Chirona, Ivan Ansari, David Navarro, Maria Alma Bracho, Griselda De Marco, Beatriz Beamud, Lidia Ruiz Roldan, Marta Pla Diaz, Neris Garcia-Gonzalez, Inma Galán Vendrell, Sandra Carbo, Loreto Ferrús Abad, Paula Ruiz-Hueso, Mariana Reyes-Prieto, Llúcia Martínez-Priego, Giuseppe D'Auria, Fernando Gonzalez-Candelas |
| EPI_ISL_436310 | Servicio de Microbiologia. Hospital Clinico Universitario de Valencia | Sequencing and Bioinformatics Service and Molecular Epidemiology Research Group. FISABIO-Public Health | Ivan Ansari, David Navarro, Maria Alma Bracho, Griselda De Marco, Beatriz Beamud, Lidia Ruiz Roldan, Marta Pla Diaz, Neris Garcia-Gonzalez, Inma Galán Vendrell, Sandra Carbo, Loreto Ferrús Abad, Paula Ruiz-Hueso, Mariana Reyes-Prieto, Vicente Soriano Chirona, Llúcia Martínez-Priego, Giuseppe D'Auria, Fernando Gonzalez-Candelas |
| EPI_ISL_436311 | Servicio de Microbiologia. Hospital Clinico Universitario de Valencia | Sequencing and Bioinformatics Service and Molecular Epidemiology Research Group. FISABIO-Public Health | David Navarro, Maria Alma Bracho, Griselda De Marco, Beatriz Beamud, Lidia Ruiz Roldan, Marta Pla Diaz, Neris Garcia-Gonzalez, Inma Galán Vendrell, Sandra Carbo, Loreto Ferrús Abad, Paula Ruiz-Hueso, Mariana Reyes-Prieto, Vicente Soriano Chirona, Ivan Ansari, Llúcia Martínez-Priego, Giuseppe D'Auria, Fernando Gonzalez-Candelas |
| EPI_ISL_436312 | Servicio de Microbiologia. Hospital Clinico Universitario de Valencia | Sequencing and Bioinformatics Service and Molecular Epidemiology Research Group. FISABIO-Public Health | Maria Alma Bracho, Griselda De Marco, Beatriz Beamud, Lidia Ruiz Roldan, Marta Pla Diaz, Neris Garcia-Gonzalez, Inma Galán Vendrell, Sandra Carbo, Loreto Ferrús Abad, Paula Ruiz-Hueso, Mariana Reyes-Prieto, Vicente Soriano Chirona, Ivan Ansari, David Navarro, Llúcia Martínez-Priego, Giuseppe D'Auria, Fernando Gonzalez-Candelas |
| EPI_ISL_436315 | Servicio de Microbiologia. Hospital Clinico Universitario de Valencia | Sequencing and Bioinformatics Service and Molecular Epidemiology Research Group. FISABIO-Public Health | Lidia Ruiz Roldan, Marta Pla Diaz, Neris Garcia-Gonzalez, Inma Galán Vendrell, Sandra Carbo, Loreto Ferrús Abad, Paula Ruiz-Hueso, Mariana Reyes-Prieto, Vicente Soriano Chirona, Ivan Ansari, David Navarro, Maria Alma Bracho, Griselda De Marco, Beatriz Beamud, Llúcia Martínez-Priego, Giuseppe D'Auria, Fernando Gonzalez-Candelas |
| EPI_ISL_436319, EPI_ISL_436339 | Servicio de Microbiologia. Hospital Clinico Universitario de Valencia | Sequencing and Bioinformatics Service and Molecular Epidemiology Research Group. FISABIO-Public Health | Griselda De Marco, Beatriz Beamud, Lidia Ruiz Roldan, Marta Pla Diaz, Neris Garcia-Gonzalez, Inma Galán Vendrell, Sandra Carbo, Loreto Ferrús Abad, Paula Ruiz-Hueso, Mariana Reyes-Prieto, Vicente Soriano Chirona, Ivan Ansari, David Navarro, Maria Alma Bracho, Llúcia Martínez-Priego, Giuseppe D'Auria, Fernando Gonzalez-Candelas |
| EPI_ISL_436340 | Servicio de Microbiologia. Hospital Clinico Universitario de Valencia | Sequencing and Bioinformatics Service and Molecular Epidemiology Research Group. FISABIO-Public Health | Beatriz Beamud, Lidia Ruiz Roldan, Marta Pla Diaz, Neris Garcia-Gonzalez, Inma Galán Vendrell, Sandra Carbo, Loreto Ferrús Abad, Paula Ruiz-Hueso, Mariana Reyes-Prieto, Vicente Soriano Chirona, Ivan Ansari, David Navarro, Maria Alma Bracho, Griselda De Marco, Llúcia Martínez-Priego, Giuseppe D'Auria, Fernando Gonzalez-Candelas |
| EPI_ISL_436341 | Servicio de Microbiologia. Hospital Clinico Universitario de Valencia | Sequencing and Bioinformatics Service and Molecular Epidemiology Research Group. FISABIO-Public Health | Lidia Ruiz Roldan, Marta Pla Diaz, Neris Garcia-Gonzalez, Inma Galán Vendrell, Sandra Carbo, Loreto Ferrús Abad, Paula Ruiz-Hueso, Mariana Reyes-Prieto, Vicente Soriano Chirona, Ivan Ansari, David Navarro, Maria Alma Bracho, Griselda De Marco, Beatriz Beamud, Llúcia Martínez-Priego, Giuseppe D'Auria, Fernando Gonzalez-Candelas |
| EPI_ISL_436346 | Servicio de Microbiologia. Hospital Clinico Universitario de Valencia | Sequencing and Bioinformatics Service and Molecular Epidemiology Research Group. FISABIO-Public Health | Loreto Ferrús Abad, Paula Ruiz-Hueso, Mariana Reyes-Prieto, Vicente Soriano Chirona, Ivan Ansari, David Navarro, Maria Alma Bracho, Griselda De Marco, Beatriz Beamud, Lidia Ruiz Roldan, Marta Pla Diaz, Neris Garcia-Gonzalez, Inma Galán Vendrell, Sandra Carbo, Llúcia Martínez-Priego, Giuseppe D'Auria, Fernando Gonzalez-Candelas |
| EPI_ISL_436347 | Servicio de Microbiologia. Hospital Clinico Universitario de Valencia | Sequencing and Bioinformatics Service and Molecular Epidemiology Research Group. FISABIO-Public Health | Paula Ruiz-Hueso, Mariana Reyes-Prieto, Vicente Soriano Chirona, Ivan Ansari, David Navarro, Maria Alma Bracho, Griselda De Marco, Beatriz Beamud, Lidia Ruiz Roldan, Marta Pla Diaz, Neris Garcia-Gonzalez, Inma Galán Vendrell, Sandra Carbo, Loreto Ferrús Abad, Llúcia Martínez-Priego, Giuseppe D'Auria, Fernando Gonzalez-Candelas |
| EPI_ISL_436359 | Servicio de Microbiologia. Hospital Clinico Universitario de Valencia | Sequencing and Bioinformatics Service and Molecular Epidemiology Research Group. FISABIO-Public Health | Marta Pla Diaz, Neris Garcia-Gonzalez, Inma Galán Vendrell, Sandra Carbo, Loreto Ferrús Abad, Paula Ruiz-Hueso, Mariana Reyes-Prieto, Vicente Soriano Chirona, Ivan Ansari, David Navarro, Maria Alma Bracho, Griselda De Marco, Beatriz Beamud, Lidia Ruiz Roldan, Llúcia Martínez-Priego, Giuseppe D'Auria, Fernando Gonzalez-Candelas |
| EPI_ISL_436360 | Servicio de Microbiologia. Hospital Clinico Universitario de Valencia | Sequencing and Bioinformatics Service and Molecular Epidemiology Research Group. FISABIO-Public Health | Neris Garcia-Gonzalez, Inma Galán Vendrell, Sandra Carbo, Loreto Ferrús Abad, Paula Ruiz-Hueso, Mariana Reyes-Prieto, Vicente Soriano Chirona, Ivan Ansari, David Navarro, Maria Alma Bracho, Griselda De Marco, Beatriz Beamud, Lidia Ruiz Roldan, Marta Pla Diaz, Llúcia Martínez-Priego, Giuseppe D'Auria, Fernando Gonzalez-Candelas |
| EPI_ISL_436413 | National Centre for Disease control (NCDC) | NCDC/CSIR-IGIB | Pramod Kumar#, Rajesh Pandey#, Pooja Sharma, Mahesh S Dhar, Vivekanand A, Bharathram Uppili, Himanshu Vashisht, Saruchi Wadhwa, Nishu Tyagi, Uma Sharma, Priyanka Singh, Hemlata Lall, Meena Datta, Poonam Gupta, Nidhi Saini, Aarti Tewari, Bibhash Nandi, Dhirendra Kumar, Satyabrata Bag, Varun Jaiswal, Hema Gogia, Preeti Madan, Simrita Singh, Prateek Singh, Debasis Dash, Mitali Mukerji, Manju Bala, Sandhya Kabra, Sujeet Singh, Mohammed Faruq, Anurag Agrawal*, Partha Rakshit* |
| EPI_ISL_436469, EPI_ISL_436475, EPI_ISL_436483, EPI_ISL_436484, EPI_ISL_436485, EPI_ISL_436487, EPI_ISL_436491 | UPMC Clinical Laboratory | Microbial Genome Sequencing Center, Microbial Genomic Epidemiological Laboratory | Dan Snyder, Stephanie L Mitchell, Mustapha M Mustapha, Marissa P Griffith, Vatsala R Srinivasa, Kady D Waggle, Chinelo Ezeonwuku, Jane W. Marsh, Lee H. Harrison, Vaughn S. Cooper |
| EPI_ISL_436566 | University of Wisconsin-Madison AIDS Vaccine Research Laboratories | University of Wisconsin-Madison AIDS Vaccine Research Laboratories | Gage Moreno, Katarina Braun, et al. AIDS Vaccine Research Laboratories |
| EPI_ISL_436672 | County of Santa Clara Public Health Department | Chan-Zuckerberg Biohub | CZB Cliahub Consortium |
| EPI_ISL_436686, EPI_ISL_436687 | KRISP, KZN Research Innovation and Sequencing Platform | KRISP, KZN Research Innovation and Sequencing Platform | Giandhari J, Pillay S, Lessells R, Chimukangara B, Deforche K, Tegally H, Wilkinson E, de Oliveira T |
| EPI_ISL_436870, EPI_ISL_436871, EPI_ISL_436872, EPI_ISL_436873, EPI_ISL_436874, EPI_ISL_436875, EPI_ISL_436876, EPI_ISL_436877, EPI_ISL_436878, EPI_ISL_436879, EPI_ISL_436880, EPI_ISL_436881, EPI_ISL_436882, EPI_ISL_436883, EPI_ISL_436890 | | | |
| see above | Michigan Department of Health and Human Services, Bureau of Laboratories | Michigan Department of Health and Human Services, Bureau of Laboratories | Blankenship HM, Riner D, Soehnlen MK |
| EPI_ISL_436968, EPI_ISL_436969, EPI_ISL_436971, EPI_ISL_436972, EPI_ISL_436973, EPI_ISL_436974, EPI_ISL_436975, EPI_ISL_436976, EPI_ISL_436977, EPI_ISL_436979, EPI_ISL_436980, EPI_ISL_436981, EPI_ISL_436982, EPI_ISL_436983, EPI_ISL_436984, EPI_ISL_436985, EPI_ISL_436986, EPI_ISL_436987, EPI_ISL_436988, EPI_ISL_436989, EPI_ISL_437019, EPI_ISL_437020, EPI_ISL_437021, EPI_ISL_437022, EPI_ISL_437023, EPI_ISL_437024, EPI_ISL_437025, EPI_ISL_437026, EPI_ISL_437027, EPI_ISL_437028, EPI_ISL_437029, EPI_ISL_437031, EPI_ISL_437032, EPI_ISL_437035, EPI_ISL_437036, EPI_ISL_437037, EPI_ISL_437038, EPI_ISL_437039, EPI_ISL_437040, EPI_ISL_437041, EPI_ISL_437042 | | | |
| see above | Department of Virus and Microbiological Special Diagnostics, Statens Serum Institut, Copenhagen, Denmark, Artillerivej 5, 2300 Copenahgen S | Albertsen lab, Department of Chemistry and Bioscience, Aalborg University, Denmark | Rasmus Kirkegaard |
| EPI_ISL_437171, EPI_ISL_437172, EPI_ISL_437173, EPI_ISL_437175, EPI_ISL_437176, EPI_ISL_437177, EPI_ISL_437179, EPI_ISL_437180, EPI_ISL_437181, EPI_ISL_437182, EPI_ISL_437183, EPI_ISL_437184, EPI_ISL_437185, EPI_ISL_437186 | | | |
| see above | Michigan Department of Health and Human Services, Bureau of Laboratories | Michigan Department of Health and Human Services, Bureau of Laboratories | Blankenship HM, Riner D, Soehnlen MK |
| EPI_ISL_437191 | RS Pondok Indah Hospital - Pondok Indah | Eijkman Institute for Molecular Biology, Ministry of Research and Technology/National Agency for Research and Innovation | Edison Johar, Frilasita A Yudhaputri, Hidayat Trimarsanto, David H Muljono, Safarina G Malik, Khin Saw Myint, Amin Soebandrio |

| | | | |
|---|---|---|---|
| EPI_ISL_437198 | Diagnostic- and Research Institute of Pathology, Medical University of Graz | Diagnostic- and Research Institute of Pathology, Medical University of Graz | Karl Kashofer, Peter Regitnig, Martin Zacharias, Gregor Gorkiewicz |
| EPI_ISL_437224, EPI_ISL_437267, EPI_ISL_437268, EPI_ISL_437269, EPI_ISL_437270, EPI_ISL_437271, EPI_ISL_437272, EPI_ISL_437273, EPI_ISL_437274, EPI_ISL_437275, EPI_ISL_437276 | | | |
| see above | Max von Pettenkofer Institute, Virology, National Reference Center for Retroviruses, LMU München | Laboratory for Functional Genome Analysis, Dept. Genomics, Gene Center of the LMU Munich | Max Muenchhoff, Stefan Krebs, Alexander Graf, Oliver Keppler, Helmut Blum |
| EPI_ISL_437305, EPI_ISL_437306, EPI_ISL_437310, EPI_ISL_437311, EPI_ISL_437313, EPI_ISL_437317 | Ministry of Health Turkey | Ministry of Health Turkey | Fatma Bayrakdar,Tülin Demir,Süleyman Yalçn, Selçuk Klç |
| EPI_ISL_437361, EPI_ISL_437362 | Minnesota Department of Health, Public Health Laboratory | Minnesota Department of Health, Public Health Laboratory | Matt Plumb, Jacob Garfin, and Xiong Wang |
| EPI_ISL_437398 | Virginia DCLS | Virginia DCLS | Virginia DCLS |
| EPI_ISL_437434 | Bozeman Water Reclamation Facility | Wiedenheft lab, Montana State University | Artem Nemudryi, Anna Nemudraia, Kevin Surya, Tanner Wiegand, Murat Buyukyoruk, Royce Wilkinson, Blake Wiedenheft |
| EPI_ISL_437469, EPI_ISL_437470, EPI_ISL_437471, EPI_ISL_437472, EPI_ISL_437473, EPI_ISL_437474 | Pathogen Genomics Lab King Abdullah University of Science and Technology(KAUST) | Pathogen Genomics Lab King Abdullah University of Science and Technology(KAUST) | Sharif Hala,Raeece Naeem,Sara Mfarrej,Arnab Pain |
| EPI_ISL_437484 | Pathogen Genomics Lab King Abdullah University of Science and Technology(KAUST) | Pathogen Genomics Lab King Abdullah University of Science and Technology(KAUST) | Sara Mfarrej,Raeece Naeem,Sharif Hala,Amit Subudhi,Fathia Rached,Arnab Pain |
| EPI_ISL_437515 | Alaska State Virology Laboratory | Alaska State Virology Laboratory | Jack Chen, Ph.D. |
| EPI_ISL_437537 | ICMR-National Institute of Cholera and Enteric Diseases | National Institute of Biomedical Genomics | Arindam Maitra, Mamta Chawla Sarkar, Sreedhar Chinnaswamy, Hasina Banu, Ananya Chatterjee, Shanta Dutta, Saumitra Das |
| EPI_ISL_437545, EPI_ISL_437546 | Robert Garry lab | Andersen lab at Scripps Research | Allison Smither, Gilberto Sabino-Santos, Patricia Snarski, Lilia Melnik, Antoinette Bell, Kaylynn Genemaras, Arnaud Drouin, Dahlene Fusco, Robert Garry with SEARCH Alliance San Diego |
| EPI_ISL_437557, EPI_ISL_437558, EPI_ISL_437563, EPI_ISL_437565, EPI_ISL_437587, EPI_ISL_437590, EPI_ISL_437592 | Scripps Medical Laboratory | Andersen lab at Scripps Research | SEARCH Alliance San Diego with Michael Quigley, Ellen Stefanski, Ian Mchardy |
| EPI_ISL_437640, EPI_ISL_437665, EPI_ISL_437666, EPI_ISL_437667, EPI_ISL_437668, EPI_ISL_437669, EPI_ISL_437670, EPI_ISL_437671, EPI_ISL_437672, EPI_ISL_437673, EPI_ISL_437674, EPI_ISL_437675, EPI_ISL_437676, EPI_ISL_437677, EPI_ISL_437678, EPI_ISL_437679, EPI_ISL_437681 | | | |
| see above | Department of Virus and Microbiological Special Diagnostics, Statens Serum Institut, Copenhagen, Denmark, Artillerivej 5, 2300 Copenahgen S | Albertsen lab, Department of Chemistry and Bioscience, Aalborg University, Denmark | Rasmus Kirkegaard |
| EPI_ISL_437733, EPI_ISL_437736, EPI_ISL_437737, EPI_ISL_437738, EPI_ISL_437739, EPI_ISL_437740, EPI_ISL_437741, EPI_ISL_437742, EPI_ISL_437743, EPI_ISL_437744, EPI_ISL_437745, EPI_ISL_437746, EPI_ISL_437747, EPI_ISL_437748, EPI_ISL_437749, EPI_ISL_437750, EPI_ISL_437751, EPI_ISL_437752, EPI_ISL_437753, EPI_ISL_437754, EPI_ISL_437755, EPI_ISL_437756, EPI_ISL_437757, EPI_ISL_437758, EPI_ISL_437759, EPI_ISL_437760, EPI_ISL_437761, EPI_ISL_437762 | | | |
| see above | Pathogen Genomics Lab King Abdullah University of Science and Technology(KAUST) | Pathogen Genomics Lab King Abdullah University of Science and Technology(KAUST) | Sharif Hala,Fadwa Alofi,Afrah Alsomali, Asim Khogeer, Sara Mfarrej, Khaled Algithami,Raeece Naeem, Amit Kumar Subudhi,Fathia Ben-Rached, Rahul Salunke, Anwar Hashem, Naif Almontashiri, Arnab Pain |
| EPI_ISL_437803, EPI_ISL_437804, EPI_ISL_437805, EPI_ISL_437806, EPI_ISL_437807, EPI_ISL_437808, EPI_ISL_437809, EPI_ISL_437810, EPI_ISL_437811, EPI_ISL_437812, EPI_ISL_437825, EPI_ISL_437826, EPI_ISL_437828 | | | |
| see above | UW Virology Lab | UW Virology Lab | Pavitra Roychoudhury, Hong Xie, Keith Jerome, Alexander Greninger |
| EPI_ISL_437893, EPI_ISL_437898, EPI_ISL_437909 | Laboratory of Microbiology, Medical School, National and Kapodistrian University of Athens | Laboratory of Biology, Department of Medicine, Democritus University of Thrace | Kassela K., Dovrolis,N., Bampali,M., Gatzidou,E., Froukala,E., Stavropoulou,A., Veletza,S., Tsakris,A., Spanakis,N. and Karakasiliotis,I. |
| EPI_ISL_437935, EPI_ISL_437936, EPI_ISL_437937 | Universitaetsklinik für Innere Medizin II Innsbruck | Bergthaler laboratory, CeMM Research Center for Molecular Medicine of the Austrian Academy of Sciences | Alexandra Popa, Benedikt Agerer, Henrique Colaco, Lukas Endler, Jakob-Wendelin Genger, Alexander Lercher, Mark Smyth, Thomas Penz, Michael Schuster, Jan Laine, Martin Senekowitsch, Judith Aberle, Stephan Aberle, Elisabeth Puchhammer-Stoeckl, Manfred Nairz, Guenter Weiss, Wegene Borena, Dorothee von Laer, Christoph Bock, Andreas Bergthaler |
| EPI_ISL_438114, EPI_ISL_438115, EPI_ISL_438116, EPI_ISL_438117, EPI_ISL_438118, EPI_ISL_438119, EPI_ISL_438125 | Center for Virology, Medical University of Vienna | Bergthaler laboratory, CeMM Research Center for Molecular Medicine of the Austrian Academy of Sciences | Alexandra Popa, Benedikt Agerer, Henrique Colaco, Lukas Endler, Jakob-Wendelin Genger, Alexander Lercher, Mark Smyth, Thomas Penz, Michael Schuster, Jan Laine, Martin Senekowitsch, Judith Aberle, Stephan Aberle, Elisabeth Puchhammer-Stoeckl, Manfred Nairz, Guenter Weiss, Wegene Borena, Dorothee von Laer, Christoph Bock, Andreas Bergthaler |
| EPI_ISL_438241, EPI_ISL_438243 | Johns Hopkins Hospital Department of Pathology | Johns Hopkins Hospital Department of Pathology | Peter M. Thielen, Thomas Mehoke, Shirlee Wohl, Srividya Ramakrishnan, Melanie Kirsche, Amanda Ernlund, Oluwaseun Falade-Nwulia, Timothy Gilpatrick, Paul Morris, Norah Sadowski, Nidia Trovao, Victoria Gniazdowski, Michael Schatz, Stuart C. Ray, Winston Timp, Heba Mostafa |
| EPI_ISL_438255, EPI_ISL_438256, EPI_ISL_438257, EPI_ISL_438258, EPI_ISL_438259, EPI_ISL_438260, EPI_ISL_438261, EPI_ISL_438262, EPI_ISL_438263, EPI_ISL_438266, EPI_ISL_438267, EPI_ISL_438268, EPI_ISL_438269, EPI_ISL_438270, EPI_ISL_438271, EPI_ISL_438272, EPI_ISL_438273, EPI_ISL_438274, EPI_ISL_438275, EPI_ISL_438276, EPI_ISL_438277, EPI_ISL_438278, EPI_ISL_438280, EPI_ISL_438281, EPI_ISL_438283, EPI_ISL_438284, EPI_ISL_438285, EPI_ISL_438286, EPI_ISL_438287, EPI_ISL_438288, EPI_ISL_438289, EPI_ISL_438290, EPI_ISL_438292, EPI_ISL_438293, EPI_ISL_438294, EPI_ISL_438295, EPI_ISL_438296, EPI_ISL_438297, EPI_ISL_438298, EPI_ISL_438299, EPI_ISL_438300, EPI_ISL_438301, EPI_ISL_438302, EPI_ISL_438303, EPI_ISL_438304, EPI_ISL_438305, EPI_ISL_438306, EPI_ISL_438307, EPI_ISL_438308, EPI_ISL_438309, EPI_ISL_438310, EPI_ISL_438311, EPI_ISL_438312, EPI_ISL_438313, EPI_ISL_438314, EPI_ISL_438315, EPI_ISL_438317, EPI_ISL_438318, EPI_ISL_438319, EPI_ISL_438320, EPI_ISL_438321, EPI_ISL_438322, EPI_ISL_438323, EPI_ISL_438324, EPI_ISL_438325, EPI_ISL_438326, EPI_ISL_438327, EPI_ISL_438328, EPI_ISL_438329, EPI_ISL_438330, EPI_ISL_438331, EPI_ISL_438332, EPI_ISL_438333, EPI_ISL_438334, EPI_ISL_438335, EPI_ISL_438336, EPI_ISL_438337, EPI_ISL_438338, EPI_ISL_438339, EPI_ISL_438341, EPI_ISL_438342, EPI_ISL_438343, EPI_ISL_438344, EPI_ISL_438345, EPI_ISL_438346, EPI_ISL_438347, EPI_ISL_438348, EPI_ISL_438349, EPI_ISL_438350, EPI_ISL_438351, EPI_ISL_438352, EPI_ISL_438353, EPI_ISL_438354, EPI_ISL_438355, EPI_ISL_438356, EPI_ISL_438357, EPI_ISL_438358, EPI_ISL_438359, EPI_ISL_438360, EPI_ISL_438361, EPI_ISL_438362, EPI_ISL_438363, EPI_ISL_438364, EPI_ISL_438365, EPI_ISL_438366, EPI_ISL_438367, EPI_ISL_438368, EPI_ISL_438369, EPI_ISL_438370, EPI_ISL_438371, EPI_ISL_438372, EPI_ISL_438373, EPI_ISL_438374, EPI_ISL_438375, EPI_ISL_438376, EPI_ISL_438377, EPI_ISL_438378, EPI_ISL_438379, EPI_ISL_438380, EPI_ISL_438383, EPI_ISL_438384, EPI_ISL_438385, EPI_ISL_438386, EPI_ISL_438387, EPI_ISL_438388, EPI_ISL_438389, EPI_ISL_438390, EPI_ISL_438392, EPI_ISL_438393, EPI_ISL_438394, EPI_ISL_438395, EPI_ISL_438396, EPI_ISL_438397, EPI_ISL_438399, EPI_ISL_438400, EPI_ISL_438401, EPI_ISL_438402, EPI_ISL_438403, EPI_ISL_438404, EPI_ISL_438405, EPI_ISL_438406, EPI_ISL_438407, EPI_ISL_438408, EPI_ISL_438409, EPI_ISL_438410, EPI_ISL_438411, EPI_ISL_438412, EPI_ISL_438413, EPI_ISL_438414, EPI_ISL_438415, EPI_ISL_438416, EPI_ISL_438417, EPI_ISL_438418, EPI_ISL_438420, EPI_ISL_438421, EPI_ISL_438422, EPI_ISL_438423, EPI_ISL_438424, EPI_ISL_438425, EPI_ISL_438426, EPI_ISL_438427, EPI_ISL_438428, EPI_ISL_438429, EPI_ISL_438430, EPI_ISL_438432, EPI_ISL_438433, EPI_ISL_438434, EPI_ISL_438435, EPI_ISL_438436, EPI_ISL_438437, EPI_ISL_438438, EPI_ISL_438439, EPI_ISL_438440, EPI_ISL_438441, EPI_ISL_438442, EPI_ISL_438443, EPI_ISL_438444, EPI_ISL_438445, EPI_ISL_438446, EPI_ISL_438447, EPI_ISL_438448, EPI_ISL_438449, EPI_ISL_438451, EPI_ISL_438452, EPI_ISL_438453, EPI_ISL_438454, EPI_ISL_438455, EPI_ISL_438456, EPI_ISL_438457, EPI_ISL_438458, EPI_ISL_438460, EPI_ISL_438461, EPI_ISL_438462, EPI_ISL_438463, EPI_ISL_438464, EPI_ISL_438465, EPI_ISL_438466, EPI_ISL_438467, EPI_ISL_438468, EPI_ISL_438469, EPI_ISL_438471, EPI_ISL_438472, EPI_ISL_438474, EPI_ISL_438475, EPI_ISL_438477, EPI_ISL_438478, EPI_ISL_438479, EPI_ISL_438480, EPI_ISL_438481, EPI_ISL_438482, EPI_ISL_438484, EPI_ISL_438485, EPI_ISL_438486, EPI_ISL_438487, EPI_ISL_438488, EPI_ISL_438490, EPI_ISL_438491, EPI_ISL_438492, EPI_ISL_438493, EPI_ISL_438494, EPI_ISL_438495, EPI_ISL_438496, EPI_ISL_438497, EPI_ISL_438498, EPI_ISL_438499, EPI_ISL_438500, EPI_ISL_438501, EPI_ISL_438502, EPI_ISL_438503, EPI_ISL_438504, EPI_ISL_438505, EPI_ISL_438506, EPI_ISL_438508, EPI_ISL_438512, EPI_ISL_438513, EPI_ISL_438516, EPI_ISL_438518, EPI_ISL_438521, EPI_ISL_438522, EPI_ISL_438526, EPI_ISL_438529, EPI_ISL_438538, EPI_ISL_438541, EPI_ISL_438543 | | | |
| see above | Department of Pathology, University of Cambridge | Wellcome Sanger Institute for the COVID-19 Genomics UK (COG-UK) consortium | Luke W Meredith, M. Estée Török , Myra Hosmillo, William L. Hamilton, Martin D. Curran, Theresa Feltwell, Grant Hall, Anna Yakovleva, Fahad A Khokhar, Charlotte J. Houldcroft, Laura G Caller, Aminu S. Jahun, Sarah L. Caddy, Ian Goodfellow, Alex Alderton, Roberto Amato, Sonia Goncalves, Ewan Harrison, David K. Jackson, Ian Johnston, Dominic Kwiatkowski, Cordelia Langford, John Sillitoe on behalf of the Wellcome Sanger Institute COVID-19 Surveillance Team (http://www.sanger.ac.uk/covid-team) |
| EPI_ISL_438749, EPI_ISL_438750, EPI_ISL_438751, EPI_ISL_438752, EPI_ISL_438753, EPI_ISL_438754, EPI_ISL_438755, EPI_ISL_438756, EPI_ISL_438757, EPI_ISL_438758, EPI_ISL_438759, EPI_ISL_438760, EPI_ISL_438761, EPI_ISL_438762, EPI_ISL_438763, EPI_ISL_438764, EPI_ISL_438765, EPI_ISL_438766, EPI_ISL_438767, EPI_ISL_438768, EPI_ISL_438769, EPI_ISL_438770, EPI_ISL_438771, EPI_ISL_438772, EPI_ISL_438773, EPI_ISL_438775, EPI_ISL_438776, EPI_ISL_438782, EPI_ISL_438786 | | | |
| see above | West of Scotland Specialist Virology Centre, NHSGGC / MRC-University of Glasgow Centre for Virus Research | COVID-19 Genomics UK (COG-UK) Consortium | Ana da Silva Filipe, Natasha Johnson, Kathy Smollett, Daniel Mair, Stephen Carmichael, Lily Tong, Jenna Nichols, Elihu Aranday-Cortes, Kirstyn Brunker, Yasmin Parr, Kyriaki Nomikou; Sarah McDonald, Marc Niebel, Patawee Asamaphan; Richard Orton, Joseph Hughes, Sreenu Vattipally, David L Robertson; Alasdair MacLean, Rory Gunson; Kathy Li, Natasha Jesudason, Rajiv Shah, James Shepherd, Antonia Ho, Emma Thomson |
| EPI_ISL_438972 | Keio University School of Medicine | Keio University School of Medicine | Kenjiro Kosaki |
| EPI_ISL_439110, EPI_ISL_439111, EPI_ISL_439113, EPI_ISL_439114, EPI_ISL_439115, EPI_ISL_439116, EPI_ISL_439117, EPI_ISL_439118, EPI_ISL_439119, EPI_ISL_439120, EPI_ISL_439122, EPI_ISL_439123, EPI_ISL_439124, EPI_ISL_439125, EPI_ISL_439126, EPI_ISL_439127, EPI_ISL_439128, EPI_ISL_439129, EPI_ISL_439130, EPI_ISL_439131, EPI_ISL_439132, EPI_ISL_439133, EPI_ISL_439134, EPI_ISL_439135, EPI_ISL_439136, EPI_ISL_439137, EPI_ISL_439138, EPI_ISL_439139, EPI_ISL_439140, EPI_ISL_439141, EPI_ISL_439142, EPI_ISL_439143 | | | |
| see above | West of Scotland Specialist Virology Centre, NHSGGC / MRC-University of Glasgow Centre for Virus Research | COVID-19 Genomics UK (COG-UK) Consortium | Ana da Silva Filipe, Natasha Johnson, Kathy Smollett, Daniel Mair, Stephen Carmichael, Lily Tong, Jenna Nichols, Elihu Aranday-Cortes, Kirstyn Brunker, Yasmin Parr, Kyriaki Nomikou; Sarah McDonald, Marc Niebel, Patawee Asamaphan; Richard Orton, Joseph Hughes, Sreenu Vattipally, David L |

| | | | |
|---|---|---|---|
| | | | Robertson; Alasdair MacLean, Rory Gunson; Kathy Li, Natasha Jesudason, Rajiv Shah, James Shepherd, Antonia Ho, Emma Thomson |
| EPI_ISL_439379, EPI_ISL_439384, EPI_ISL_439385, EPI_ISL_439394, EPI_ISL_439396, EPI_ISL_439398, EPI_ISL_439407, EPI_ISL_439408, EPI_ISL_439410, EPI_ISL_439411, EPI_ISL_439419, EPI_ISL_439421, EPI_ISL_439422, EPI_ISL_439427, EPI_ISL_439432, EPI_ISL_439442, EPI_ISL_439448, EPI_ISL_439450, EPI_ISL_439457, EPI_ISL_439458, EPI_ISL_439461, EPI_ISL_439465, EPI_ISL_439467, EPI_ISL_439469, EPI_ISL_439471, EPI_ISL_439475, EPI_ISL_439476, EPI_ISL_439482, EPI_ISL_439486, EPI_ISL_439491, EPI_ISL_439493, EPI_ISL_439499, EPI_ISL_439500, EPI_ISL_439505, EPI_ISL_439506, EPI_ISL_439507, EPI_ISL_439508, EPI_ISL_439515, EPI_ISL_439518, EPI_ISL_439520, EPI_ISL_439522, EPI_ISL_439523, EPI_ISL_439527, EPI_ISL_439530, EPI_ISL_439532, EPI_ISL_439534, EPI_ISL_439539, EPI_ISL_439543, EPI_ISL_439544, EPI_ISL_439545, EPI_ISL_439546, EPI_ISL_439548, EPI_ISL_439549, EPI_ISL_439550, EPI_ISL_439551, EPI_ISL_439554, EPI_ISL_439555, EPI_ISL_439557, EPI_ISL_439559, EPI_ISL_439560, EPI_ISL_439564, EPI_ISL_439567, EPI_ISL_439575, EPI_ISL_439576, EPI_ISL_439577, EPI_ISL_439581, EPI_ISL_439582, EPI_ISL_439585, EPI_ISL_439586, EPI_ISL_439587, EPI_ISL_439589, EPI_ISL_439590, EPI_ISL_439592, EPI_ISL_439593, EPI_ISL_439598, EPI_ISL_439602, EPI_ISL_439603, EPI_ISL_439605, EPI_ISL_439608, EPI_ISL_439610, EPI_ISL_439611, EPI_ISL_439618, EPI_ISL_439622, EPI_ISL_439623, EPI_ISL_439624, EPI_ISL_439625, EPI_ISL_439626, EPI_ISL_439627, EPI_ISL_439629, EPI_ISL_439631, EPI_ISL_439633, EPI_ISL_439640, EPI_ISL_439646, EPI_ISL_439649, EPI_ISL_439652, EPI_ISL_439653, EPI_ISL_439654, EPI_ISL_439655, EPI_ISL_439656, EPI_ISL_439657, EPI_ISL_439660, EPI_ISL_439661, EPI_ISL_439662, EPI_ISL_439664, EPI_ISL_439864, EPI_ISL_439867, EPI_ISL_439869, EPI_ISL_439873, EPI_ISL_439876, EPI_ISL_439885, EPI_ISL_439887, EPI_ISL_439891, EPI_ISL_439892, EPI_ISL_439894, EPI_ISL_439896, EPI_ISL_439899, EPI_ISL_439902, EPI_ISL_439907, EPI_ISL_439911, EPI_ISL_439916, EPI_ISL_439917, EPI_ISL_439918, EPI_ISL_439919, EPI_ISL_439923, EPI_ISL_439930, EPI_ISL_439932, EPI_ISL_439933, EPI_ISL_439934, EPI_ISL_439935, EPI_ISL_439936, EPI_ISL_439942, EPI_ISL_439944, EPI_ISL_439947, EPI_ISL_439951, EPI_ISL_439952 | | | |
| see above | Department of Pathology, University of Cambridge | Wellcome Sanger Institute for the COVID-19 Genomics UK (COG-UK) consortium | Luke W Meredith, M. Estée Török , Myra Hosmillo, William L. Hamilton, Martin D. Curran, Theresa Feltwell, Grant Hall, Anna Yakovleva, Fahad A Khokhar, Charlotte J. Houldcroft, Laura G Caller, Aminu S. Jahun, Sarah L. Caddy, Ian Goodfellow, Alex Alderton, Roberto Amato, Sonia Goncalves, Ewan Harrison, David K. Jackson, Ian Johnston, Dominic Kwiatkowski, Cordelia Langford, John Sillitoe on behalf of the Wellcome Sanger Institute COVID-19 Surveillance Team (http://www.sanger.ac.uk/covid-team) |
| EPI_ISL_440058, EPI_ISL_440130, EPI_ISL_440141, EPI_ISL_440145, EPI_ISL_440153, EPI_ISL_440192, EPI_ISL_440238 | PHE South West Regional Laboratory, National Infection Service | Wellcome Sanger Institute for the COVID-19 Genomics UK (COG-UK) consortium | Stephanie Hutchings, Hannah Pymont, Dr Peter Muir, Barry Vipond, Rich Hopes, Alex Alderton, Roberto Amato, Sonia Goncalves, Ewan Harrison, David K. Jackson, Ian Johnston, Dominic Kwiatkowski, Cordelia Langford, John Sillitoe on behalf of the Wellcome Sanger Institute COVID-19 Surveillance Team (http://www.sanger.ac.uk/covid-team) |
| EPI_ISL_440951, EPI_ISL_440952, EPI_ISL_440953, EPI_ISL_440954, EPI_ISL_440955, EPI_ISL_440956, EPI_ISL_440957, EPI_ISL_440958 | University College London, Great Ormond Street Hospital for Children NHS Foundation Trust, Imperial College Healthcare NHS Trust | COVID-19 Genomics UK (COG-UK) Consortium | Sergi Castellano, Rachel Williams, Mark Kristiansen, Paola Resende Silva, Sunando Roy, Tony Brooks, Helena Tutill, Paola Niola, Patricia Dyal, Charlotte Williams, Leysa Forrest, Yasmin Panchbhaya, Jacqueline Findlay, Sam Weeks, Julianne Brown, Kathryn Harris, Paul Randell, James Price, Alison Holmes, Judith Breuer |
| EPI_ISL_441662, EPI_ISL_441663, EPI_ISL_441675, EPI_ISL_441685, EPI_ISL_441690, EPI_ISL_441691, EPI_ISL_441696, EPI_ISL_441697, EPI_ISL_441712, EPI_ISL_441714, EPI_ISL_441716, EPI_ISL_441730, EPI_ISL_441737 | | | |
| see above | Regional Virus Laboratory, Belfast Health and Social Care Trust | Wellcome Sanger Institute for the COVID-19 Genomics UK (COG-UK) consortium | Conall McCaughey, James McKenna, Tanya Curran, Susan Feeney, Alison Watt, Ciara Cox, Mairead Connor, Zoltan Molnar, David Simpson, Derek Fairley, Alex Alderton, Roberto Amato, Sonia Goncalves, Ewan Harrison, David K. Jackson, Ian Johnston, Dominic Kwiatkowski, Cordelia Langford, John Sillitoe on behalf of the Wellcome Sanger Institute COVID-19 Surveillance Team (http://www.sanger.ac.uk/covid-team) |
| EPI_ISL_441910, EPI_ISL_441950, EPI_ISL_441962, EPI_ISL_441977, EPI_ISL_442013, EPI_ISL_442031, EPI_ISL_442034, EPI_ISL_442395, EPI_ISL_442414 | Virology Department, Sheffield Teaching Hospitals NHS Foundation Trust/Department of Infection, Immunity and Cardiovascular Disease, The Medical School, University of Sheffield | COVID-19 Genomics UK (COG-UK) Consortium | Thushan de Silva, Matthew Parker, Nikki Smith, Adri Angyal, Rebecca Brown, Luke Green, Rachel Tucker, Paul Parsons, Danielle Groves, Katie Johnson, Laura Carrilero, Alex Keeley, Dave Partridge, Matthew Wyles, Benjamin Lindsey, Mehmet Yavuz, Mohammad Raza, Cariad Evans |
| EPI_ISL_443185 | UW Virology Lab | UW Virology Lab | Pavitra Roychoudhury, Hong Xie, Keith Jerome, Alexander Greninger |
| EPI_ISL_443257 | M Health Fairview | University of Minnesota Genomics Center | Daryl M. Gohl, John Garbe, Patrick Grady, Jerry Daniel, Ray Watson, Benjamin Auch, Andrew Nelson, Sophia Yohe, and Kenneth B. Beckman |
| EPI_ISL_443275, EPI_ISL_443276, EPI_ISL_443277, EPI_ISL_443278 | CHU - Hôpital Cavale Blanche - Labo. de Virologie | National Reference Center for Viruses of Respiratory Infections, Institut Pasteur, Paris | Mélanie Albert, Marion Barbet, Sylvie Behillil, Méline Bizard, Angela Brisebarre, Flora Donati, Etienne Simon-Lorière, Vincent Enouf, Maud Vanpeene, Sylvie van der Werf, Léa Pilorge |
| EPI_ISL_443289 | CHRU Pontchaillou - Laboratoire de Virologie | National Reference Center for Viruses of Respiratory Infections, Institut Pasteur, Paris | Mélanie Albert, Marion Barbet, Sylvie Behillil, Méline Bizard, Angela Brisebarre, Flora Donati, Etienne Simon-Lorière, Vincent Enouf, Maud Vanpeene, Sylvie van der Werf, Gisèle Lagathu |
| EPI_ISL_443686, EPI_ISL_443687, EPI_ISL_443689, EPI_ISL_443690, EPI_ISL_443691, EPI_ISL_443692, EPI_ISL_443694, EPI_ISL_443697, EPI_ISL_443698, EPI_ISL_443699, EPI_ISL_443700, EPI_ISL_443701, EPI_ISL_443702, EPI_ISL_443703, EPI_ISL_443705, EPI_ISL_443706, EPI_ISL_443707, EPI_ISL_443708, EPI_ISL_443710, EPI_ISL_443711, EPI_ISL_443712, EPI_ISL_443713, EPI_ISL_443715, EPI_ISL_443716, EPI_ISL_443717, EPI_ISL_443718, EPI_ISL_443722, EPI_ISL_443723, EPI_ISL_443726, EPI_ISL_443728, EPI_ISL_443730, EPI_ISL_443731, EPI_ISL_443732, EPI_ISL_443734, EPI_ISL_443739, EPI_ISL_443740, EPI_ISL_443746, EPI_ISL_443748, EPI_ISL_443749, EPI_ISL_443750, EPI_ISL_443751, EPI_ISL_443752, EPI_ISL_443753, EPI_ISL_443756, EPI_ISL_443758, EPI_ISL_443759, EPI_ISL_443760, EPI_ISL_443763, EPI_ISL_443765, EPI_ISL_443766, EPI_ISL_443767, EPI_ISL_443770, EPI_ISL_443774, EPI_ISL_443775, EPI_ISL_443777, EPI_ISL_443778, EPI_ISL_443779, EPI_ISL_443780, EPI_ISL_443781, EPI_ISL_443782, EPI_ISL_443783, EPI_ISL_443784, EPI_ISL_443786, EPI_ISL_443789, EPI_ISL_443790, EPI_ISL_443792, EPI_ISL_443794, EPI_ISL_443795, EPI_ISL_443796, EPI_ISL_443798, EPI_ISL_443800, EPI_ISL_443802, EPI_ISL_443803, EPI_ISL_443804, EPI_ISL_443806, EPI_ISL_443807, EPI_ISL_443808, EPI_ISL_443818, EPI_ISL_443819, EPI_ISL_443820, EPI_ISL_443821, EPI_ISL_443822, EPI_ISL_443824, EPI_ISL_443826, EPI_ISL_443828, EPI_ISL_443830, EPI_ISL_443831, EPI_ISL_443834, EPI_ISL_443837, EPI_ISL_443838, EPI_ISL_443840, EPI_ISL_443842, EPI_ISL_443845, EPI_ISL_443847, EPI_ISL_443849, EPI_ISL_443850, EPI_ISL_443851, EPI_ISL_443853, EPI_ISL_443856, EPI_ISL_443858, EPI_ISL_443860, EPI_ISL_443863, EPI_ISL_443865, EPI_ISL_443866, EPI_ISL_443867, EPI_ISL_443868, EPI_ISL_443869, EPI_ISL_443873, EPI_ISL_443875, EPI_ISL_443876, EPI_ISL_443877, EPI_ISL_443879, EPI_ISL_443880, EPI_ISL_443881, EPI_ISL_443885, EPI_ISL_443886, EPI_ISL_443888, EPI_ISL_443889, EPI_ISL_443893, EPI_ISL_443896, EPI_ISL_443898, EPI_ISL_443899, EPI_ISL_443902, EPI_ISL_443903, EPI_ISL_443906, EPI_ISL_443908, EPI_ISL_443909, EPI_ISL_443913, EPI_ISL_443915, EPI_ISL_443916, EPI_ISL_443917, EPI_ISL_443920, EPI_ISL_443921, EPI_ISL_443922, EPI_ISL_443924, EPI_ISL_443927, EPI_ISL_443928, EPI_ISL_443929, EPI_ISL_443930, EPI_ISL_443931, EPI_ISL_443933, EPI_ISL_443934, EPI_ISL_443935, EPI_ISL_443937, EPI_ISL_443938, EPI_ISL_443943, EPI_ISL_443947, EPI_ISL_443948, EPI_ISL_443949, EPI_ISL_443951, EPI_ISL_443953, EPI_ISL_443955, EPI_ISL_443956, EPI_ISL_443957, EPI_ISL_443959, EPI_ISL_443961, EPI_ISL_443962, EPI_ISL_443963, EPI_ISL_443965, EPI_ISL_443967, EPI_ISL_443968, EPI_ISL_443969, EPI_ISL_443970, EPI_ISL_443971, EPI_ISL_443972, EPI_ISL_443973, EPI_ISL_443978, EPI_ISL_443980, EPI_ISL_443981, EPI_ISL_443983, EPI_ISL_443984, EPI_ISL_443987, EPI_ISL_443989, EPI_ISL_443992, EPI_ISL_443993, EPI_ISL_443994, EPI_ISL_443996, EPI_ISL_444000, EPI_ISL_444003, EPI_ISL_444006, EPI_ISL_444011, EPI_ISL_444013, EPI_ISL_444014, EPI_ISL_444017, EPI_ISL_444018, EPI_ISL_444020 | | | |
| see above | PHE South West Regional Laboratory, National Infection Service | Wellcome Sanger Institute for the COVID-19 Genomics UK (COG-UK) consortium | Stephanie Hutchings, Hannah Pymont, Dr Peter Muir, Barry Vipond, Rich Hopes; and Alex Alderton, Roberto Amato, Sonia Goncalves, Ewan Harrison, David K. Jackson, Ian Johnston, Dominic Kwiatkowski, Cordelia Langford, John Sillitoe on behalf of the Wellcome Sanger Institute COVID-19 Surveillance Team (http://www.sanger.ac.uk/covid-team) |
| EPI_ISL_444616, EPI_ISL_444617, EPI_ISL_444618, EPI_ISL_444619, EPI_ISL_444620, EPI_ISL_444621, EPI_ISL_444622, EPI_ISL_444623, EPI_ISL_444624, EPI_ISL_444625, EPI_ISL_444626, EPI_ISL_444627, EPI_ISL_444628, EPI_ISL_444629, EPI_ISL_444630, EPI_ISL_444631, EPI_ISL_444632, EPI_ISL_444633, EPI_ISL_444634, EPI_ISL_444635, EPI_ISL_444636, EPI_ISL_444637, EPI_ISL_444638, EPI_ISL_444639, EPI_ISL_444640, EPI_ISL_444641, EPI_ISL_444642, EPI_ISL_444643, EPI_ISL_444644, EPI_ISL_444645, EPI_ISL_444646, EPI_ISL_444647, EPI_ISL_444648, EPI_ISL_444649, EPI_ISL_444650, EPI_ISL_444651, EPI_ISL_444652, EPI_ISL_444653, EPI_ISL_444654, EPI_ISL_444655, EPI_ISL_444656, EPI_ISL_444657, EPI_ISL_444658, EPI_ISL_444659, EPI_ISL_444660, EPI_ISL_444661, EPI_ISL_444664, EPI_ISL_444665, EPI_ISL_444666, EPI_ISL_444667, EPI_ISL_444668, EPI_ISL_444670, EPI_ISL_444671, EPI_ISL_444672, EPI_ISL_444674, EPI_ISL_444675, EPI_ISL_444678, EPI_ISL_444679, EPI_ISL_444680, EPI_ISL_444681, EPI_ISL_444682, EPI_ISL_444683, EPI_ISL_444684, EPI_ISL_444685, EPI_ISL_444686, EPI_ISL_444687, EPI_ISL_444688, EPI_ISL_444690, EPI_ISL_444691, EPI_ISL_444692, EPI_ISL_444693, EPI_ISL_444694, EPI_ISL_444695, EPI_ISL_444696, EPI_ISL_444697, EPI_ISL_444698, EPI_ISL_444699, EPI_ISL_444700, EPI_ISL_444701, EPI_ISL_444702, EPI_ISL_444703, EPI_ISL_444704, EPI_ISL_444705, EPI_ISL_444765 | | | |
| see above | NYU Langone Health | Departments of Pathology and Medicine, New York University School of Medicine | Maria Aguero-Rosenfeld, Brendan Belovarac, Margaret Black, Ludovic Boytard, John Cadley, Paolo Cotzia, John Chen, Dacia Dimartino, Xiaojun Feng, Tatyana Gindin, Emily Guzman, Adriana Heguy, Megan Hogan, Emily Huang, George Jour, Alireza Khodadadi-Jamayran, Lawrence H. Lin, Raven Luther, Andrew Lytle, Christian Marier, Matthew T. Maurano, Mark J. Mulligan, Peter Meyn, Raquel Ordonez Ciriza, Iman Osman, Jared Pinnell, Vanessa Raabe, Sitharam Ramaswami, Amy Rapkiewicz, Andre M. Ribeiro-dos-Santos, Marie Samanovic-Golden, Antonio Serrano, Guomiao Shen, Matija Snuderl, Theodore Vougiouklakis, Nick Vulpescu, Gael Westby, Paul Zappile, Yutong Zhang |
| EPI_ISL_444859, EPI_ISL_444860, EPI_ISL_444861, EPI_ISL_444862, EPI_ISL_444863, EPI_ISL_444864, EPI_ISL_444865, EPI_ISL_444866, EPI_ISL_444867, EPI_ISL_444868, EPI_ISL_444869, EPI_ISL_444870, EPI_ISL_444871, EPI_ISL_444872, EPI_ISL_444873, EPI_ISL_444874, EPI_ISL_444875, EPI_ISL_444876, EPI_ISL_444877, EPI_ISL_444878, EPI_ISL_444879, EPI_ISL_444880, EPI_ISL_444881, EPI_ISL_444882, EPI_ISL_444883, EPI_ISL_444884, EPI_ISL_444885, EPI_ISL_444886, EPI_ISL_444887, EPI_ISL_444903, EPI_ISL_444904, EPI_ISL_444905, EPI_ISL_444906, EPI_ISL_444907, EPI_ISL_444908, EPI_ISL_444909, EPI_ISL_444910, EPI_ISL_444911, EPI_ISL_444912, EPI_ISL_444913, EPI_ISL_444914, EPI_ISL_444915, EPI_ISL_444916, EPI_ISL_444917, EPI_ISL_444918, EPI_ISL_444919, EPI_ISL_444920, EPI_ISL_444921, EPI_ISL_444922, EPI_ISL_444923, EPI_ISL_444924, EPI_ISL_444925, EPI_ISL_444926, EPI_ISL_444927, EPI_ISL_444928, EPI_ISL_444929, EPI_ISL_444930, EPI_ISL_444931, EPI_ISL_444932, EPI_ISL_444933, EPI_ISL_444934, EPI_ISL_444935, EPI_ISL_444936, EPI_ISL_444937, EPI_ISL_444938, EPI_ISL_444939, EPI_ISL_444940, EPI_ISL_444941, EPI_ISL_444942, EPI_ISL_444943, EPI_ISL_444944, EPI_ISL_444945, EPI_ISL_444946, EPI_ISL_444947, EPI_ISL_444948, EPI_ISL_444949, EPI_ISL_444950, EPI_ISL_444951, EPI_ISL_444952, EPI_ISL_444953, EPI_ISL_444955, EPI_ISL_444957, EPI_ISL_444961, EPI_ISL_444962, EPI_ISL_444963, EPI_ISL_444965, EPI_ISL_444966, EPI_ISL_444967 | | | |
| see above | Department of Virus and Microbiological Special Diagnostics, Statens Serum Institut, Copenhagen, Denmark, Artillerivej 5, 2300 Copenahgen S | Albertsen lab, Department of Chemistry and Bioscience, Aalborg University, Denmark | Rasmus Kirkegaard |
| EPI_ISL_444988, EPI_ISL_444989, EPI_ISL_444990 | Hospital Universitari Vall d'Hebron - Vall d'Hebron Institut de Recerca | Hospital Universitari Vall d'Hebron | Cristina Andrés, Maria Piñana, Damir Garcia-Cehic, Mercedes Guerrero-Murillo, Ariadna Rando, Juliana Esperalba, Maria Gema Codina, Tomàs Pumarola, Josep Quer, Andrés Antón |
| EPI_ISL_444995, EPI_ISL_444996 | Naval Health Research Center | Naval Medical Research Center Biological Defense Research Directorate | Logan Voegtly, Regina Cer, Dessiree Pena-Gomez, Adrian Paskey,Kyle Long, Roger Pan, Melinda Balansay-Ames, Chris Myers, Ewell Hollis, Nathaniel Christy, Kimberly Bishop-Lilly |
| EPI_ISL_445113, EPI_ISL_445114, EPI_ISL_445115 | UC San Diego Center for Advanced Laboratory Medicine | Andersen lab at Scripps Research | SEARCH Alliance San Diego with David Pride, Ji H Shin |
| EPI_ISL_445167 | Scripps Medical Laboratory | Andersen lab at Scripps Research | SEARCH Alliance San Diego with Michael Quigley, Ellen Stefanski, Ian Mchardy |
| EPI_ISL_445169, EPI_ISL_445170, | UCSF Clinical Microbiology Laboratory | Chan-Zuckerberg Biohub | CZB Cliahub Consortium |

| | | | |
|---|---|---|---|
| EPI_ISL_445171, EPI_ISL_445180 | | | |
| EPI_ISL_445225 | Surbrunns VC | The Public Health Agency of Sweden | Erik Embring, Oskar Karlsson Lindsjo, Maria Lind Karlberg, Anna-Malin Linde, Olov Svartstrom, Anna Risberg, Theresa Enkirch, Mia Brytting, Karin Tegmark-Wisell |
| EPI_ISL_445334 | CLINICA UNIVERSITARIA DE PUERTO MONTT S.A. | Instituto de Salud Publica de Chile | Andrés E Castillo, Bárbara Parra,Paz Tapia, Jaime Lagos, Loredana Arata, Alejandra Acevedo, Winston Andrade, Gabriel Leal, Carolina Tambley, Patricia Bustos, Rodrigo Fasce, Jorge Fernandez |
| EPI_ISL_445335 | HOSPITAL DE CALBUCO | Instituto de Salud Publica de Chile | Andrés E Castillo, Bárbara Parra,Paz Tapia, Jaime Lagos, Loredana Arata, Alejandra Acevedo, Winston Andrade, Gabriel Leal, Carolina Tambley, Patricia Bustos, Rodrigo Fasce, Jorge Fernandez |
| EPI_ISL_445337 | HOSPITAL HANGA ROA | Instituto de Salud Publica de Chile | Andrés E Castillo, Bárbara Parra,Paz Tapia, Jaime Lagos, Loredana Arata, Alejandra Acevedo, Winston Andrade, Gabriel Leal, Carolina Tambley, Patricia Bustos, Rodrigo Fasce, Jorge Fernandez |
| EPI_ISL_445345 | HOSPITAL DR.HERNAN HENRIQUEZ ARAVENA | Instituto de Salud Publica de Chile | Andrés E Castillo, Bárbara Parra,Paz Tapia, Jaime Lagos, Loredana Arata, Alejandra Acevedo, Winston Andrade, Gabriel Leal, Carolina Tambley, Patricia Bustos, Rodrigo Fasce, Jorge Fernandez |
| EPI_ISL_445371 | HOSPITAL DR.SOTERO DEL RIO | Instituto de Salud Publica de Chile | Andrés E Castillo, Bárbara Parra,Paz Tapia, Jaime Lagos, Loredana Arata, Alejandra Acevedo, Winston Andrade, Gabriel Leal, Carolina Tambley, Patricia Bustos, Rodrigo Fasce, Jorge Fernandez |
| EPI_ISL_445372 | HOSPITAL FF.AA. "CIRUJANO C. GUZMAN | Instituto de Salud Publica de Chile | Andrés E Castillo, Bárbara Parra,Paz Tapia, Jaime Lagos, Loredana Arata, Alejandra Acevedo, Winston Andrade, Gabriel Leal, Carolina Tambley, Patricia Bustos, Rodrigo Fasce, Jorge Fernandez |
| EPI_ISL_445380 | Ramathibodi Hospital | COVID-19 Network Investigations (CONI) Alliance | Elizabeth Batty, Wasun Chantratita, Thanat Chookajorn, Stefan Fernandez, Angkana Huang, Anthony R. Jones, Khajohn Joonsalak, Chonticha Klungtong, Theerarat Kochakarn, Namfon Kotanan, Krittikorn Kumpornsin, Wudtichai Manasatienkij, Bhakbhoom Panthan, Ekawat Pasomsub, Kingkan Rakmanee, Insee Sensorn, Janjira Thaipadungpanit, Arporn Wangwiwatsin, Treewat Watthanachockchai |
| EPI_ISL_445786, EPI_ISL_445787, EPI_ISL_445788, EPI_ISL_445789, EPI_ISL_445790, EPI_ISL_445791, EPI_ISL_445792, EPI_ISL_445793, EPI_ISL_445794, EPI_ISL_445795, EPI_ISL_445797, EPI_ISL_445798, EPI_ISL_445799, EPI_ISL_445800, EPI_ISL_445801, EPI_ISL_445802, EPI_ISL_445803, EPI_ISL_445804, EPI_ISL_445805, EPI_ISL_445806, EPI_ISL_445807, EPI_ISL_445808, EPI_ISL_445809, EPI_ISL_445810, EPI_ISL_445811, EPI_ISL_445812, EPI_ISL_445813, EPI_ISL_445814, EPI_ISL_445815, EPI_ISL_445816, EPI_ISL_445817, EPI_ISL_445818, EPI_ISL_445819, EPI_ISL_445820, EPI_ISL_445821, EPI_ISL_445822 | | | |
| see above | Wales Specialist Virology Centre | Public Health Wales Microbiology Cardiff | Catherine Moore, Johnathan Evans, Laura Gifford, Malorie Perry, Simon Cottrell, Alec Birchley, Alexander Adams, Amy Gaskin, Bree Gatica-Wilcox, Jason Coombes, Lauren Gilbert, Lee Graham, Nicole Pacchiarini, Sara Kumziene-Summerhayes, Sarah Taylor, Sophie Jones, Sara Rey, Matthew Bull, Joanne Watkins, Sally Corden, Tom Connor |
| EPI_ISL_446996, EPI_ISL_446997, EPI_ISL_446998, EPI_ISL_446999, EPI_ISL_447000, EPI_ISL_447001, EPI_ISL_447002, EPI_ISL_447003, EPI_ISL_447004, EPI_ISL_447005, EPI_ISL_447006, EPI_ISL_447007, EPI_ISL_447008, EPI_ISL_447009, EPI_ISL_447010, EPI_ISL_447011, EPI_ISL_447012 | | | |
| see above | Ramathibodi Hospital | COVID-19 Network Investigations (CONI) Alliance | Elizabeth Batty, Wasun Chantratita, Thanat Chookajorn, Stefan Fernandez, Angkana Huang, Anthony R. Jones, Khajohn Joonsalak, Chonticha Klungtong, Theerarat Kochakarn, Namfon Kotanan, Krittikorn Kumpornsin, Wudtichai Manasatienkij, Bhakbhoom Panthan, Ekawat Pasomsub, Kingkan Rakmanee, Insee Sensorn, Janjira Thaipadungpanit, Arporn Wangwiwatsin,Treewat Watthanachockchai |
| EPI_ISL_447061, EPI_ISL_447063, EPI_ISL_447066, EPI_ISL_447071, EPI_ISL_447074, EPI_ISL_447076, EPI_ISL_447079, EPI_ISL_447081, EPI_ISL_447083, EPI_ISL_447084, EPI_ISL_447085, EPI_ISL_447086, EPI_ISL_447087, EPI_ISL_447095, EPI_ISL_447118, EPI_ISL_447172, EPI_ISL_447203, EPI_ISL_447204 | | | |
| see above | Michigan Department of Health and Human Services, Bureau of Laboratories | Michigan Department of Health and Human Services, Bureau of Laboratories | Blankenship HM, Riner D, Soehnlen MK |
| EPI_ISL_447258, EPI_ISL_447259, EPI_ISL_447267 | Microbiology laboratory, Assuta Ashdod University-Affiliated Hospital | Stern Lab | Stern Lab |
| EPI_ISL_447281, EPI_ISL_447282, EPI_ISL_447283, EPI_ISL_447284, EPI_ISL_447285 | Microbiology Division, Barzilai University Medical Center | Stern Lab | Stern Lab |
| EPI_ISL_447318 | Clinical Virology Laboratory, Soroka Medical Center and the Faculty of Health Sciences, Ben-Gurion University of the Negev | Stern Lab | Stern Lab |
| EPI_ISL_447385, EPI_ISL_447386, EPI_ISL_447387, EPI_ISL_447388, EPI_ISL_447389, EPI_ISL_447390, EPI_ISL_447391, EPI_ISL_447392, EPI_ISL_447393, EPI_ISL_447399 | Clinical Microbiology Laboratory, The Baruch Padeh Medical Center, Poriya | Stern Lab | Stern Lab |
| EPI_ISL_447441, EPI_ISL_447442, EPI_ISL_447443, EPI_ISL_447444, EPI_ISL_447445, EPI_ISL_447446, EPI_ISL_447447, EPI_ISL_447448, EPI_ISL_447449, EPI_ISL_447450 | Clinical Microbiology Laboratory, Sheba Medical Center | Stern Lab | Stern Lab |
| EPI_ISL_447475 | Servicio de Microbiologia. Hospital Clinico Universitario de Valencia | Sequencing and Bioinformatics Service and Molecular Epidemiology Research Group. FISABIO-Public Health | Neris Garcia-Gonzalez, Inma Galán Vendrell, Sandra Carbo, Loreto Ferrús Abad, Paula Ruiz-Hueso, Mariana Reyes-Prieto, Vicente Soriano Chirona, Ivan Ansari, Llúcia Martínez-Priego, Giuseppe 'Auria, David Navarro, Eliseo Albert, Maria Alma Bracho, Lidia Ruiz Roldan, Fernando Gonzalez-Candelas |
| EPI_ISL_447476 | Servicio de Microbiologia. Hospital Clinico Universitario de Valencia | Sequencing and Bioinformatics Service and Molecular Epidemiology Research Group. FISABIO-Public Health | Inma Galán Vendrell, Sandra Carbo, Loreto Ferrús Abad, Paula Ruiz-Hueso, Mariana Reyes-Prieto, Vicente Soriano Chirona, Ivan Ansari, Llúcia Martínez-Priego, Giuseppe 'Auria, David Navarro, Eliseo Albert, Maria Alma Bracho, Lidia Ruiz Roldan, Neris Garcia-Gonzalez, Fernando Gonzalez-Candelas |
| EPI_ISL_447477 | Servicio de Microbiologia. Hospital Clinico Universitario de Valencia | Sequencing and Bioinformatics Service and Molecular Epidemiology Research Group. FISABIO-Public Health | Sandra Carbo, Loreto Ferrús Abad, Paula Ruiz-Hueso, Mariana Reyes-Prieto, Vicente Soriano Chirona, Ivan Ansari, Llúcia Martínez-Priego, Giuseppe 'Auria, David Navarro, Eliseo Albert, Maria Alma Bracho, Lidia Ruiz Roldan, Neris Garcia-Gonzalez, Inma Galán Vendrell, Fernando Gonzalez-Candelas |
| EPI_ISL_447478 | Servicio de Microbiologia. Hospital Clinico Universitario de Valencia | Sequencing and Bioinformatics Service and Molecular Epidemiology Research Group. FISABIO-Public Health | Loreto Ferrús Abad, Paula Ruiz-Hueso, Mariana Reyes-Prieto, Vicente Soriano Chirona, Ivan Ansari, Llúcia Martínez-Priego, Giuseppe 'Auria, David Navarro, Eliseo Albert, Maria Alma Bracho, Lidia Ruiz Roldan, Neris Garcia-Gonzalez, Inma Galán Vendrell, Sandra Carbo, Fernando Gonzalez-Candelas |
| EPI_ISL_447479 | Servicio de Microbiologia. Hospital Clinico Universitario de Valencia | Sequencing and Bioinformatics Service and Molecular Epidemiology Research Group. FISABIO-Public Health | Paula Ruiz-Hueso, Mariana Reyes-Prieto, Vicente Soriano Chirona, Ivan Ansari, Llúcia Martínez-Priego, Giuseppe 'Auria, David Navarro, Eliseo Albert, Maria Alma Bracho, Lidia Ruiz Roldan, Neris Garcia-Gonzalez, Inma Galán Vendrell, Sandra Carbo, Loreto Ferrús Abad, Fernando Gonzalez-Candelas |
| EPI_ISL_447480 | Servicio de Microbiologia. Hospital Clinico Universitario de Valencia | Sequencing and Bioinformatics Service and Molecular Epidemiology Research Group. FISABIO-Public Health | Mariana Reyes-Prieto, Vicente Soriano Chirona, Ivan Ansari, Llúcia Martínez-Priego, Giuseppe 'Auria, David Navarro, Eliseo Albert, Maria Alma Bracho, Lidia Ruiz Roldan, Neris Garcia-Gonzalez, Inma Galán Vendrell, Sandra Carbo, Loreto Ferrús Abad, Paula Ruiz-Hueso, Fernando Gonzalez-Candelas |
| EPI_ISL_447481 | Servicio de Microbiologia. Hospital Clinico Universitario de Valencia | Sequencing and Bioinformatics Service and Molecular Epidemiology Research Group. FISABIO-Public Health | Vicente Soriano Chirona, Ivan Ansari, Llúcia Martínez-Priego, Giuseppe 'Auria, David Navarro, Eliseo Albert, Maria Alma Bracho, Lidia Ruiz Roldan, Neris Garcia-Gonzalez, Inma Galán Vendrell, Sandra Carbo, Loreto Ferrús Abad, Paula Ruiz-Hueso, Mariana Reyes-Prieto, Fernando Gonzalez-Candelas |
| EPI_ISL_447482 | Servicio de Microbiologia. Hospital Clinico Universitario de Valencia | Sequencing and Bioinformatics Service and Molecular Epidemiology Research Group. FISABIO-Public Health | Giuseppe 'Auria, David Navarro, Eliseo Albert, Maria Alma Bracho, Lidia Ruiz Roldan, Neris Garcia-Gonzalez, Inma Galán Vendrell, Sandra Carbo, Loreto Ferrús Abad, Paula Ruiz-Hueso, Mariana Reyes-Prieto, Vicente Soriano Chirona, Ivan Ansari, Llúcia Martínez-Priego, Fernando Gonzalez-Candelas |
| EPI_ISL_447483 | Servicio de Microbiologia. Hospital Clinico Universitario de Valencia | Sequencing and Bioinformatics Service and Molecular Epidemiology Research Group. FISABIO-Public Health | Llúcia Martínez-Priego, Giuseppe 'Auria, David Navarro, Eliseo Albert, Maria Alma Bracho, Lidia Ruiz Roldan, Neris Garcia-Gonzalez, Inma Galán Vendrell, Sandra Carbo, Loreto Ferrús Abad, Paula Ruiz-Hueso, Mariana Reyes-Prieto, Vicente Soriano Chirona, Ivan Ansari, Fernando Gonzalez-Candelas |
| EPI_ISL_447484 | Servicio de Microbiologia. Hospital Clinico Universitario de Valencia | Sequencing and Bioinformatics Service and Molecular Epidemiology Research Group. FISABIO-Public Health | David Navarro, Eliseo Albert, Maria Alma Bracho, Griselda De Marco, Lidia Ruiz Roldan, Neris Garcia-Gonzalez, Inma Galán Vendrell, Sandra Carbo, Loreto Ferrús Abad, Paula Ruiz-Hueso, Mariana Reyes-Prieto, Vicente Soriano Chirona, Ivan Ansari, Llúcia Martínez-Priego, Giuseppe 'Auria, Fernando Gonzalez-Candelas |
| EPI_ISL_447485 | Servicio de Microbiologia. Hospital Clinico Universitario de Valencia | Sequencing and Bioinformatics Service and Molecular Epidemiology Research Group. FISABIO-Public Health | Eliseo Albert, Maria Alma Bracho, Griselda De Marco, Lidia Ruiz Roldan, Neris Garcia-Gonzalez, Inma Galán Vendrell, Sandra Carbo, Loreto Ferrús Abad, Paula Ruiz-Hueso, Mariana Reyes-Prieto, Vicente Soriano Chirona, Ivan Ansari, Llúcia Martínez-Priego, Giuseppe 'Auria, David Navarro, Fernando |

| | | | |
|---|---|---|---|
| | | | Gonzalez-Candelas |
| EPI_ISL_447486 | Servicio de Microbiologia. Hospital Clinico Universitario de Valencia | Sequencing and Bioinformatics Service and Molecular Epidemiology Research Group. FISABIO-Public Health | Maria Alma Bracho, Griselda De Marco, Lidia Ruiz Roldan, Neris Garcia-Gonzalez, Inma Galán Vendrell, Sandra Carbo, Loreto Ferrús Abad, Paula Ruiz-Hueso, Mariana Reyes-Prieto, Vicente Soriano Chirona, Ivan Ansari, Llúcia Martínez-Priego, Giuseppe 'Auria, David Navarro, Eliseo Albert, Fernando Gonzalez-Candelas |
| EPI_ISL_447487 | Servicio de Microbiologia. Hospital Clinico Universitario de Valencia | Sequencing and Bioinformatics Service and Molecular Epidemiology Research Group. FISABIO-Public Health | Griselda De Marco, Lidia Ruiz Roldan, Neris Garcia-Gonzalez, Inma Galán Vendrell, Sandra Carbo, Loreto Ferrús Abad, Paula Ruiz-Hueso, Mariana Reyes-Prieto, Vicente Soriano Chirona, Ivan Ansari, Llúcia Martínez-Priego, Giuseppe 'Auria, David Navarro, Eliseo Albert, Maria Alma Bracho, Fernando Gonzalez-Candelas |
| EPI_ISL_447495 | Servicio de Microbiologia. Hospital Clinico Universitario de Valencia | Sequencing and Bioinformatics Service and Molecular Epidemiology Research Group. FISABIO-Public Health | Vicente Soriano Chirona, Ivan Ansari, Llúcia Martínez-Priego, Giuseppe 'Auria, David Navarro, Eliseo Albert, Maria Alma Bracho, Lidia Ruiz Roldan, Neris Garcia-Gonzalez, Inma Galán Vendrell, Sandra Carbo, Loreto Ferrús Abad, Paula Ruiz-Hueso, Mariana Reyes-Prieto, Fernando Gonzalez-Candelas |
| EPI_ISL_447496 | Servicio de Microbiologia. Hospital Clinico Universitario de Valencia | Sequencing and Bioinformatics Service and Molecular Epidemiology Research Group. FISABIO-Public Health | Giuseppe 'Auria, David Navarro, Eliseo Albert, Maria Alma Bracho, Lidia Ruiz Roldan, Neris Garcia-Gonzalez, Inma Galán Vendrell, Sandra Carbo, Loreto Ferrús Abad, Paula Ruiz-Hueso, Mariana Reyes-Prieto, Vicente Soriano Chirona, Ivan Ansari, Llúcia Martínez-Priego, Fernando Gonzalez-Candelas |
| EPI_ISL_447497 | Servicio de Microbiologia. Hospital Clinico Universitario de Valencia | Sequencing and Bioinformatics Service and Molecular Epidemiology Research Group. FISABIO-Public Health | Llúcia Martínez-Priego, Giuseppe 'Auria, David Navarro, Eliseo Albert, Maria Alma Bracho, Lidia Ruiz Roldan, Neris Garcia-Gonzalez, Inma Galán Vendrell, Sandra Carbo, Loreto Ferrús Abad, Paula Ruiz-Hueso, Mariana Reyes-Prieto, Vicente Soriano Chirona, Ivan Ansari, Fernando Gonzalez-Candelas |
| EPI_ISL_447498 | Servicio de Microbiologia. Hospital Clinico Universitario de Valencia | Sequencing and Bioinformatics Service and Molecular Epidemiology Research Group. FISABIO-Public Health | David Navarro, Eliseo Albert, Maria Alma Bracho, Griselda De Marco, Lidia Ruiz Roldan, Neris Garcia-Gonzalez, Inma Galán Vendrell, Sandra Carbo, Loreto Ferrús Abad, Paula Ruiz-Hueso, Mariana Reyes-Prieto, Vicente Soriano Chirona, Ivan Ansari, Llúcia Martínez-Priego, Giuseppe 'Auria, Fernando Gonzalez-Candelas |
| EPI_ISL_447499 | Servicio de Microbiologia. Hospital Clinico Universitario de Valencia | Sequencing and Bioinformatics Service and Molecular Epidemiology Research Group. FISABIO-Public Health | Eliseo Albert, Maria Alma Bracho, Griselda De Marco, Lidia Ruiz Roldan, Neris Garcia-Gonzalez, Inma Galán Vendrell, Sandra Carbo, Loreto Ferrús Abad, Paula Ruiz-Hueso, Mariana Reyes-Prieto, Vicente Soriano Chirona, Ivan Ansari, Llúcia Martínez-Priego, Giuseppe 'Auria, David Navarro, Fernando Gonzalez-Candelas |
| EPI_ISL_447500 | Servicio de Microbiologia. Hospital Clinico Universitario de Valencia | Sequencing and Bioinformatics Service and Molecular Epidemiology Research Group. FISABIO-Public Health | Maria Alma Bracho, Griselda De Marco, Lidia Ruiz Roldan, Neris Garcia-Gonzalez, Inma Galán Vendrell, Sandra Carbo, Loreto Ferrús Abad, Paula Ruiz-Hueso, Mariana Reyes-Prieto, Vicente Soriano Chirona, Ivan Ansari, Llúcia Martínez-Priego, Giuseppe 'Auria, David Navarro, Eliseo Albert, Fernando Gonzalez-Candelas |
| EPI_ISL_447501 | Servicio de Microbiologia. Hospital Clinico Universitario de Valencia | Sequencing and Bioinformatics Service and Molecular Epidemiology Research Group. FISABIO-Public Health | Griselda De Marco, Lidia Ruiz Roldan, Neris Garcia-Gonzalez, Inma Galán Vendrell, Sandra Carbo, Loreto Ferrús Abad, Paula Ruiz-Hueso, Mariana Reyes-Prieto, Vicente Soriano Chirona, Ivan Ansari, Llúcia Martínez-Priego, Giuseppe 'Auria, David Navarro, Eliseo Albert, Maria Alma Bracho, Fernando Gonzalez-Candelas |
| EPI_ISL_447502 | Servicio de Microbiologia. Hospital Clinico Universitario de Valencia | Sequencing and Bioinformatics Service and Molecular Epidemiology Research Group. FISABIO-Public Health | Lidia Ruiz Roldan, Neris Garcia-Gonzalez, Inma Galán Vendrell, Sandra Carbo, Loreto Ferrús Abad, Paula Ruiz-Hueso, Mariana Reyes-Prieto, Vicente Soriano Chirona, Ivan Ansari, Llúcia Martínez-Priego, Giuseppe 'Auria, David Navarro, Eliseo Albert, Maria Alma Bracho, Fernando Gonzalez-Candelas |
| EPI_ISL_447503 | Servicio de Microbiologia. Hospital Clinico Universitario de Valencia | Sequencing and Bioinformatics Service and Molecular Epidemiology Research Group. FISABIO-Public Health | Neris Garcia-Gonzalez, Inma Galán Vendrell, Sandra Carbo, Loreto Ferrús Abad, Paula Ruiz-Hueso, Mariana Reyes-Prieto, Vicente Soriano Chirona, Ivan Ansari, Llúcia Martínez-Priego, Giuseppe 'Auria, David Navarro, Eliseo Albert, Maria Alma Bracho, Lidia Ruiz Roldan, Fernando Gonzalez-Candelas |
| EPI_ISL_447504 | Servicio de Microbiologia. Hospital Clinico Universitario de Valencia | Sequencing and Bioinformatics Service and Molecular Epidemiology Research Group. FISABIO-Public Health | Inma Galán Vendrell, Sandra Carbo, Loreto Ferrús Abad, Paula Ruiz-Hueso, Mariana Reyes-Prieto, Vicente Soriano Chirona, Ivan Ansari, Llúcia Martínez-Priego, Giuseppe 'Auria, David Navarro, Eliseo Albert, Maria Alma Bracho, Lidia Ruiz Roldan, Neris Garcia-Gonzalez, Fernando Gonzalez-Candelas |
| EPI_ISL_447505 | Servicio de Microbiologia. Hospital Clinico Universitario de Valencia | Sequencing and Bioinformatics Service and Molecular Epidemiology Research Group. FISABIO-Public Health | Sandra Carbo, Loreto Ferrús Abad, Paula Ruiz-Hueso, Mariana Reyes-Prieto, Vicente Soriano Chirona, Ivan Ansari, Llúcia Martínez-Priego, Giuseppe 'Auria, David Navarro, Eliseo Albert, Maria Alma Bracho, Lidia Ruiz Roldan, Neris Garcia-Gonzalez, Inma Galán Vendrell, Fernando Gonzalez-Candelas |
| EPI_ISL_447506 | Servicio de Microbiologia. Hospital Clinico Universitario de Valencia | Sequencing and Bioinformatics Service and Molecular Epidemiology Research Group. FISABIO-Public Health | Loreto Ferrús Abad, Paula Ruiz-Hueso, Mariana Reyes-Prieto, Vicente Soriano Chirona, Ivan Ansari, Llúcia Martínez-Priego, Giuseppe 'Auria, David Navarro, Eliseo Albert, Maria Alma Bracho, Lidia Ruiz Roldan, Neris Garcia-Gonzalez, Inma Galán Vendrell, Sandra Carbo, Fernando Gonzalez-Candelas |
| EPI_ISL_447507 | Servicio de Microbiologia. Hospital Clinico Universitario de Valencia | Sequencing and Bioinformatics Service and Molecular Epidemiology Research Group. FISABIO-Public Health | Paula Ruiz-Hueso, Mariana Reyes-Prieto, Vicente Soriano Chirona, Ivan Ansari, Llúcia Martínez-Priego, Giuseppe 'Auria, David Navarro, Eliseo Albert, Maria Alma Bracho, Lidia Ruiz Roldan, Neris Garcia-Gonzalez, Inma Galán Vendrell, Sandra Carbo, Loreto Ferrús Abad, Fernando Gonzalez-Candelas |
| EPI_ISL_447508 | Servicio de Microbiologia. Hospital Clinico Universitario de Valencia | Sequencing and Bioinformatics Service and Molecular Epidemiology Research Group. FISABIO-Public Health | Mariana Reyes-Prieto, Vicente Soriano Chirona, Ivan Ansari, Llúcia Martínez-Priego, Giuseppe 'Auria, David Navarro, Eliseo Albert, Maria Alma Bracho, Lidia Ruiz Roldan, Neris Garcia-Gonzalez, Inma Galán Vendrell, Sandra Carbo, Loreto Ferrús Abad, Paula Ruiz-Hueso, Fernando Gonzalez-Candelas |
| EPI_ISL_447509 | Servicio de Microbiologia. Hospital Clinico Universitario de Valencia | Sequencing and Bioinformatics Service and Molecular Epidemiology Research Group. FISABIO-Public Health | Vicente Soriano Chirona, Ivan Ansari, Llúcia Martínez-Priego, Giuseppe 'Auria, David Navarro, Eliseo Albert, Maria Alma Bracho, Lidia Ruiz Roldan, Neris Garcia-Gonzalez, Inma Galán Vendrell, Sandra Carbo, Loreto Ferrús Abad, Paula Ruiz-Hueso, Mariana Reyes-Prieto, Fernando Gonzalez-Candelas |
| EPI_ISL_447510 | Servicio de Microbiologia. Hospital Clinico Universitario de Valencia | Sequencing and Bioinformatics Service and Molecular Epidemiology Research Group. FISABIO-Public Health | Giuseppe 'Auria, David Navarro, Eliseo Albert, Maria Alma Bracho, Lidia Ruiz Roldan, Neris Garcia-Gonzalez, Inma Galán Vendrell, Sandra Carbo, Loreto Ferrús Abad, Paula Ruiz-Hueso, Mariana Reyes-Prieto, Vicente Soriano Chirona, Ivan Ansari, Llúcia Martínez-Priego, Fernando Gonzalez-Candelas |
| EPI_ISL_447511 | Servicio de Microbiologia. Hospital Clinico Universitario de Valencia | Sequencing and Bioinformatics Service and Molecular Epidemiology Research Group. FISABIO-Public Health | Llúcia Martínez-Priego, Giuseppe 'Auria, David Navarro, Eliseo Albert, Maria Alma Bracho, Lidia Ruiz Roldan, Neris Garcia-Gonzalez, Inma Galán Vendrell, Sandra Carbo, Loreto Ferrús Abad, Paula Ruiz-Hueso, Mariana Reyes-Prieto, Vicente Soriano Chirona, Ivan Ansari, Fernando Gonzalez-Candelas |
| EPI_ISL_447512 | Servicio de Microbiologia. Hospital Clinico Universitario de Valencia | Sequencing and Bioinformatics Service and Molecular Epidemiology Research Group. FISABIO-Public Health | David Navarro, Eliseo Albert, Maria Alma Bracho, Griselda De Marco, Lidia Ruiz Roldan, Neris Garcia-Gonzalez, Inma Galán Vendrell, Sandra Carbo, Loreto Ferrús Abad, Paula Ruiz-Hueso, Mariana Reyes-Prieto, Vicente Soriano Chirona, Ivan Ansari, Llúcia Martínez-Priego, Giuseppe 'Auria, Fernando Gonzalez-Candelas |
| EPI_ISL_447513 | Servicio de Microbiologia. Hospital Clinico Universitario de Valencia | Sequencing and Bioinformatics Service and Molecular Epidemiology Research Group. FISABIO-Public Health | Eliseo Albert, Maria Alma Bracho, Griselda De Marco, Lidia Ruiz Roldan, Neris Garcia-Gonzalez, Inma Galán Vendrell, Sandra Carbo, Loreto Ferrús Abad, Paula Ruiz-Hueso, Mariana Reyes-Prieto, Vicente Soriano Chirona, Ivan Ansari, Llúcia Martínez-Priego, Giuseppe 'Auria, David Navarro, Fernando Gonzalez-Candelas |
| EPI_ISL_447514 | Servicio de Microbiologia. Hospital Clinico Universitario de Valencia | Sequencing and Bioinformatics Service and Molecular Epidemiology Research Group. FISABIO-Public Health | Maria Alma Bracho, Griselda De Marco, Lidia Ruiz Roldan, Neris Garcia-Gonzalez, Inma Galán Vendrell, Sandra Carbo, Loreto Ferrús Abad, Paula Ruiz-Hueso, Mariana Reyes-Prieto, Vicente Soriano Chirona, Ivan Ansari, Llúcia Martínez-Priego, Giuseppe 'Auria, David Navarro, Eliseo Albert, Fernando Gonzalez-Candelas |
| EPI_ISL_447515 | Servicio de Microbiologia. Hospital Clinico Universitario de Valencia | Sequencing and Bioinformatics Service and Molecular Epidemiology Research Group. FISABIO-Public Health | Griselda De Marco, Lidia Ruiz Roldan, Neris Garcia-Gonzalez, Inma Galán Vendrell, Sandra Carbo, Loreto Ferrús Abad, Paula Ruiz-Hueso, Mariana Reyes-Prieto, Vicente Soriano Chirona, Ivan Ansari, Llúcia Martínez-Priego, Giuseppe 'Auria, David Navarro, Eliseo Albert, Maria Alma Bracho, Fernando Gonzalez-Candelas |
| EPI_ISL_447556 | CSIR-Centre for Cellular and Molecular Biology | CSIR-Centre for Cellular and Molecular Biology | Sofia Banu, Payel Mukherjee, Priya Singh, Dhiviya Vedagiri, Divya Gupta, Vishal Sah, Santosh Kumar Kuncha, Krishnan Harinivas Harshan, Archana Bharadwaj Siva, Karthik Bharadwaj Tallapaka, Shagufta Khan, Lamuk Zaveri, Namami Gaur, Sakshi Shambhavi, Tulasi Nagabandi, Purushotham Vodnala, Rakesh K Mishra, Divya Tej Sowpati |
| EPI_ISL_447557, EPI_ISL_447558 | CSIR-Centre for Cellular and Molecular Biology | CSIR-Centre for Cellular and Molecular Biology | Payel Mukherjee, Sofia Banu, Priya Singh, Dhiviya Vedagiri, Divya Gupta, Vishal Sah, Santosh Kumar Kuncha, Krishnan Harinivas Harshan, Archana Bharadwaj Siva, Karthik Bharadwaj Tallapaka, Shagufta Khan, Lamuk Zaveri, Namami Gaur, Sakshi Shambhavi, Tulasi Nagabandi, Purushotham Vodnala, Rakesh K Mishra, Divya Tej Sowpati |
| EPI_ISL_447588 | Lednicky Lab | Lednicky lab | Elbadry,M.A., Subramaniam,K., Waltzek,T.B., Gibson,J.C., Stephenson,C.J., Alam,M.M., Morris,J.G. Jr. and Lednicky,J.A. |
| EPI_ISL_447589 | University of Florida, Lednicky Lab | University of Florida, Lednicky Lab | Elbadry,M.A., Subramaniam,K., Waltzek,T.B., Gibson,J.C., Stephenson,C.J., Alam,M.M., Morris,J.G. Jr. and Lednicky,J.A. |
| EPI_ISL_447592 | TSGH-CP molecular lab | TSGH-CP molecular lab | Cherng-Lih Perng, Ming-Jr JIAN, Chih-Kai Chang, Jung-Chung Lin, Kuo-Ming Yeh, Chien-Wen Chen, Sheng-Kang Chiu, Hsing-Yi Chung, Shih-Hung Tsai, Kuo-Sheng Hung, Tien-Yao Chang, Feng-Yee Chang, Hung-Sheng Shang |
| EPI_ISL_447620 | Department of Laboratory Medicine, National Taiwan University Hospital | Microbial Genomics Core Lab, National Taiwan University Centers of Genomic and Precision Medicine | Shiou-Hwei Yeh, You-Yu Lin, Ya-Yun Lai, Chiao-Ling Li, Shan-Chwen Chang, Pei-Jer Chen, Sui-Yuan Chang |
| EPI_ISL_447739, EPI_ISL_447740, EPI_ISL_447741, EPI_ISL_447742, EPI_ISL_447743, EPI_ISL_447744, EPI_ISL_447745, EPI_ISL_447746, EPI_ISL_447747, EPI_ISL_447748, EPI_ISL_447749, EPI_ISL_447750, EPI_ISL_447751, EPI_ISL_447752, EPI_ISL_447753 | | | |

| | | | |
|---|---|---|---|
| see above | Grupo de Investigaciones Microbiológicas-UR (GIMUR), Departamento de Biología, Facultad de Ciencias Naturales, Universidad del Rosario, Bogotá, Colombia | Grupo de Investigaciones Microbiológicas-UR (GIMUR), Departamento de Biología, Facultad de Ciencias Naturales, Universidad del Rosario, Bogotá, Colombia Instituto Nacional de Salud, Bogotá, Colombia Icahn School of Medicine at Mount Sinai, New York, USA | Juan David Ramírez, Carolina Florez, Marina Muñoz, Carolina Hernandez, Adriana Castillo, Sergio Castañeda, Nathalia Ballesteros, David Martínez, Laura Vega, Jesús E. Jaimes, Sergio Gomez, Angelica Rico, Lisseth Pardo, Esther C. Barros, Martha L. Ospina, Aníbal A. Teherán, Ana S. Gonzalez-Reiche, Matthew M. Hernandez, Emilia Mia Sordillo, Viviana Simon, Harm van Bakel, Alberto Paniz-Mondolfi |
| EPI_ISL_447887 | University of California, Davis | Chan-Zuckerberg Biohub | CZB Cliahub Consortium |
| EPI_ISL_448864, EPI_ISL_448866, EPI_ISL_448906 | Virology Laboratory, Castle Hill Hospital, Hull University Teaching Hospitals NHS Trust/Department of Infection, Immunity and Cardiovascular Disease, The Medical School, University of Sheffield | COVID-19 Genomics UK (COG-UK) Consortium | Thushan de Silva, Matthew Parker, Nikki Smith, Adri Angyal, Rebecca Brown, Luke Green, Rachel Tucker, Paul Parsons, Danielle Groves, Katie Johnson, Laura Carrilero, Alex Keeley, Dave Partridge, Matthew Wyles, Benjamin Lindsey, Mehmet Yavuz, Mohammad Raza, Cariad Evans |
| EPI_ISL_448948, EPI_ISL_448951, EPI_ISL_448953, EPI_ISL_448955, EPI_ISL_448957, EPI_ISL_448958, EPI_ISL_448959, EPI_ISL_448960, EPI_ISL_448962, EPI_ISL_448964, EPI_ISL_448965, EPI_ISL_448967, EPI_ISL_448969, EPI_ISL_448971, EPI_ISL_448973, EPI_ISL_448975, EPI_ISL_448977 | | | |
| see above | Regional Virus Laboratory, Belfast Health and Social Care Trust | COVID-19 Genomics UK (COG-UK) Consortium | Conall McCaughey, James McKenna, Tanya Curran, Susan Feeney, Alison Watt, Ciara Cox, Mairead Connor, Zoltan Molnar, David Simpson, Derek Fairley |
| EPI_ISL_449333, EPI_ISL_449334, EPI_ISL_449335, EPI_ISL_449336, EPI_ISL_449354, EPI_ISL_449355, EPI_ISL_449356, EPI_ISL_449357, EPI_ISL_449358, EPI_ISL_449359, EPI_ISL_449360, EPI_ISL_449361, EPI_ISL_449362, EPI_ISL_449363, EPI_ISL_449364, EPI_ISL_449365, EPI_ISL_449366, EPI_ISL_449367, EPI_ISL_449368, EPI_ISL_449369, EPI_ISL_449370, EPI_ISL_449371, EPI_ISL_449372, EPI_ISL_449373, EPI_ISL_449374, EPI_ISL_449375, EPI_ISL_449376, EPI_ISL_449377, EPI_ISL_449378, EPI_ISL_449379, EPI_ISL_449390, EPI_ISL_449391, EPI_ISL_449392, EPI_ISL_449393, EPI_ISL_449394, EPI_ISL_449395, EPI_ISL_449396, EPI_ISL_449397, EPI_ISL_449398, EPI_ISL_449399, EPI_ISL_449400, EPI_ISL_449401, EPI_ISL_449402, EPI_ISL_449403, EPI_ISL_449404, EPI_ISL_449405, EPI_ISL_449406, EPI_ISL_449407, EPI_ISL_449420, EPI_ISL_449421, EPI_ISL_449422, EPI_ISL_449423, EPI_ISL_449427, EPI_ISL_449428, EPI_ISL_449429, EPI_ISL_449430, EPI_ISL_449431, EPI_ISL_449432, EPI_ISL_449433, EPI_ISL_449438, EPI_ISL_449439, EPI_ISL_449440, EPI_ISL_449491, EPI_ISL_449516, EPI_ISL_449524, EPI_ISL_449525, EPI_ISL_449526, EPI_ISL_449527, EPI_ISL_449528, EPI_ISL_449529, EPI_ISL_449533, EPI_ISL_449535, EPI_ISL_449537, EPI_ISL_449538, EPI_ISL_449539, EPI_ISL_449541 | | | |
| see above | Liverpool Clinical Laboratories | COVID-19 Genomics UK (COG-UK) Consortium | Sam Haldenby, Anita Lucaci, Steve Paterson, Julian Hiscox, Alistair Darby, M Almsaud, A Alrezaihi, Muhannad Alruwaili, Stuart D Armstrong, Jones Benjamin , Eleanor G Bentley, Anu Chawla, Jordan J Clark, Angela Cowell, Richard Eccles, Isabel García-Dorival, Matthew Gemmell, Alessandro Gerada, PKF Gilmore, Richard Gregory, Ximeng Han, Catherine Hartley, Margaret Hughes, Miren Iturriza-Gomara, James Johnson, L Luu, Jenifer Manson , Charlotte Nelson, Elaine O'Toole, Cassie Olateju, Rebekah Penrice-Randal , Lucille Rainbow, N.P Randle, Trevor Ian Robinson, Parul Sharma, Ghada T Shawli, James P Stewart , Neil Swainston, Ecaterina Vamos, Joanne Watts, Mark Whitehead |
| EPI_ISL_449789 | Dept. of Medical Microbiology, Stavanger University Hospital, Helse Stavanger HF | Norwegian Institute of Public Health, Department of Virology | Kathrine Stene-Johansen, Kamilla Heddeland Instefjord, Hilde Elshaug, Rasmus Riis Kopperud, Karoline Bragstad, Olav Hungnes |
| EPI_ISL_449805, EPI_ISL_449808, EPI_ISL_449809, EPI_ISL_449810 | Utah Public Health Laboratory | Utah Public Health Laboratory | Erin Young, Kelly Oakeson |
| EPI_ISL_450300 | National Institute for Communicable Diseases of the National Health Laboratory Service | National Institute for Communicable Diseases of the National Health Laboratory Service | Allam M, Ismail A, Khumalo Z, Kwenda S, van Heusden P, Mtshali P, Mnyameni F, Mohale T, Subramoney K, Bhiman JN |
| EPI_ISL_450302, EPI_ISL_450303 | Centre hospitalier Anna-Laberge | Laboratoire de santé publique du Québec | Sandrine Moreira, Ioannis Ragoussis, Guillaume Bourque, Jesse Shapiro, Mark Lathrop and Michel Roger on behalf of the CoVSeQ research group (http://covseq.ca/researchgroup) |
| EPI_ISL_450304, EPI_ISL_450305 | Hôpital Charles-LeMoyne | Laboratoire de santé publique du Québec | Sandrine Moreira, Ioannis Ragoussis, Guillaume Bourque, Jesse Shapiro, Mark Lathrop and Michel Roger on behalf of the CoVSeQ research group (http://covseq.ca/researchgroup) |
| EPI_ISL_450306 | CSSS Haut-Richelieu/Rouville (Hôpital) | Laboratoire de santé publique du Québec | Sandrine Moreira, Ioannis Ragoussis, Guillaume Bourque, Jesse Shapiro, Mark Lathrop and Michel Roger on behalf of the CoVSeQ research group (http://covseq.ca/researchgroup) |
| EPI_ISL_450307 | Hôpital du Suroît | Laboratoire de santé publique du Québec | Sandrine Moreira, Ioannis Ragoussis, Guillaume Bourque, Jesse Shapiro, Mark Lathrop and Michel Roger on behalf of the CoVSeQ research group (http://covseq.ca/researchgroup) |
| EPI_ISL_450308 | Hôpital Charles-LeMoyne | Laboratoire de santé publique du Québec | Sandrine Moreira, Ioannis Ragoussis, Guillaume Bourque, Jesse Shapiro, Mark Lathrop and Michel Roger on behalf of the CoVSeQ research group (http://covseq.ca/researchgroup) |
| EPI_ISL_450309 | Hôpital Pierre-Boucher | Laboratoire de santé publique du Québec | Sandrine Moreira, Ioannis Ragoussis, Guillaume Bourque, Jesse Shapiro, Mark Lathrop and Michel Roger on behalf of the CoVSeQ research group (http://covseq.ca/researchgroup) |
| EPI_ISL_450310 | Hôpital Charles-LeMoyne | Laboratoire de santé publique du Québec | Sandrine Moreira, Ioannis Ragoussis, Guillaume Bourque, Jesse Shapiro, Mark Lathrop and Michel Roger on behalf of the CoVSeQ research group (http://covseq.ca/researchgroup) |
| EPI_ISL_450311, EPI_ISL_450312, EPI_ISL_450313 | Hôpital du Suroît | Laboratoire de santé publique du Québec | Sandrine Moreira, Ioannis Ragoussis, Guillaume Bourque, Jesse Shapiro, Mark Lathrop and Michel Roger on behalf of the CoVSeQ research group (http://covseq.ca/researchgroup) |
| EPI_ISL_450314, EPI_ISL_450315 | Hôpital Pierre-Boucher | Laboratoire de santé publique du Québec | Sandrine Moreira, Ioannis Ragoussis, Guillaume Bourque, Jesse Shapiro, Mark Lathrop and Michel Roger on behalf of the CoVSeQ research group (http://covseq.ca/researchgroup) |
| EPI_ISL_450316 | Hôpital Charles-LeMoyne | Laboratoire de santé publique du Québec | Sandrine Moreira, Ioannis Ragoussis, Guillaume Bourque, Jesse Shapiro, Mark Lathrop and Michel Roger on behalf of the CoVSeQ research group (http://covseq.ca/researchgroup) |
| EPI_ISL_450317, EPI_ISL_450318 | Hôpital Pierre-Boucher | Laboratoire de santé publique du Québec | Sandrine Moreira, Ioannis Ragoussis, Guillaume Bourque, Jesse Shapiro, Mark Lathrop and Michel Roger on behalf of the CoVSeQ research group (http://covseq.ca/researchgroup) |
| EPI_ISL_450323 | NIV Pune | CSIR-Centre for Cellular and Molecular Biology | Dr V A Potdar, Dr ML Choudhary,Dr Priya Abraham,V. Vipat, S. Jadhav, U. Saha, H. Kengle, A. Awhale, A. Jagtap, A. Gondhalikar, V Malik, N Srivastava, S. Digraskar, P. Malsane, S. Hundekar, K. Patel, Yogesh Balakartik, M. Kakade, S. Jadhav, R. Gunjikar, V. Awtade, S. Bhorekar, P Shinde, S. Salve, B. Minhas S. Bharadwaj, H Kaushal Y. Gurav, S. Tomar,Payel Mukherjee, Sofia Banu, Priya Singh, Dhiviya Vedagiri, Divya Gupta, Vishal Sah, Santosh Kumar Kuncha, Krishnan Harinivas Harshan, Archana Bharadwaj Siva, Karthik Bharadwaj Tallapaka, Shagufta Khan, Lamuk Zaveri, Namami Gaur, Sakshi Shambhavi, Tulasi Nagabandi, Purushotham Vodnala,G. Aditya Kumar, Koushick Sivakumar, Pooja Ramesh Gupta, Rajan Kumar Jha, Shraddha Vijay Lahoti, Deepak Kumar, Devi Prasad Vijayashankara, Disha Nanda, Divya Das, Jotin Gogoi, Manish |
| EPI_ISL_450349, EPI_ISL_450350 | St.Olavs hospital/NTNU | Institute of Genomics Core Facility, University of Tartu | Aleksandr Ianevski, Tuuli Reisberg, Janne-Fossum Malmring, Svein Arne Nordbø, Denis Kainov |
| EPI_ISL_450393, EPI_ISL_450394, EPI_ISL_450395, EPI_ISL_450396, EPI_ISL_450397, EPI_ISL_450398, EPI_ISL_450399 | NYU Langone Health | Departments of Pathology and Medicine, New York University School of Medicine | Maria Aguero-Rosenfeld, Brendan Belovarac, Margaret Black, Ludovic Boytard, John Cadley, Paolo Cotzia, John Chen, Dacia Dimartino, Xiaojun Feng, Tatyana Gindin, Emily Guzman, Adriana Heguy, Megan Hogan, Emily Huang, George Jour, Alireza Khodadadi-Jamayran, Lawrence H. Lin, Raven Luther, Andrew Lytle, Christian Marier, Matthew T. Maurano, Mark J. Mulligan, Peter Meyn, Raquel Ordonez Ciriza, Iman Osman, Jared Pinnell, Vanessa Raabe, Sitharam Ramaswami, Amy Rapkiewicz, Andre M. Ribeiro-dos-Santos, Marie Samanovic-Golden, Antonio Serrano, Guomiao Shen, Matija Snuderl, Theodore Vougiouklakis, Nick Vulpescu, Gael Westby, Paul Zappile, Yutong Zhang |
| EPI_ISL_450414 | Microbiology, Regional Medical Research Centre (ICMR) | Microbiology, Regional Medical Research Centre (ICMR) | Borkakoty,B., Bali,N.K., Barua,P., Hazarika,R., Sharma,M.D. and Phukon,P. |
| EPI_ISL_450445, EPI_ISL_450446, EPI_ISL_450447, EPI_ISL_450448, EPI_ISL_450456 | Stanford clinical virology lab | Chan-Zuckerberg Biohub | Benjamin Pinksy, Katharine Walter, Victoria N. Parikh, John Gorzynski, Hannah N. DeJong, Matthew T. Wheeler, Jason Andrews, Manuel Rivas, Carlos Bustamante, Euan Ashley, with CZB Cliahub Consortium |
| EPI_ISL_450490, EPI_ISL_450492 | unknown | National Influenza and other Respiratory Viruses Centre-Tunisia | El Moussi,A., Abid,S., Ben Nasr,M., Landolsi,I., Charaa,L., Ferjeni,A., Arab Ennigrou,D., Boutiba,I. |
| EPI_ISL_450525, EPI_ISL_450526, EPI_ISL_450527, EPI_ISL_450528 | Hematology Laboratory, Section of Molecular Diagnostics, University Clinical Centre, Medical University of Gdansk | Department of Virology, Faculty of Medicine, University of Helsinki, Helsinki, Finland | Maciej Grzybek, Marlena Robakowska, Aneta Szulc, Olli Vapalahti, Teemu Smura |
| EPI_ISL_450543 | Utah Public Health Laboratory | Utah Public Health Laboratory | Erin Young, Kelly Oakeson |
| EPI_ISL_450599 | Michigan Department of Health and Human Services, Bureau | Michigan Department of Health and Human Services, Bureau | Blankenship HM; Riner D; Soehnlen MK |

| | | | |
|---|---|---|---|
| | of Laboratories | of Laboratories | |
| EPI_ISL_450620, EPI_ISL_450622, EPI_ISL_450623, EPI_ISL_450624, EPI_ISL_450625, EPI_ISL_450626, EPI_ISL_450627, EPI_ISL_450628, EPI_ISL_450629, EPI_ISL_450630, EPI_ISL_450631, EPI_ISL_450632, EPI_ISL_450633, EPI_ISL_450634, EPI_ISL_450635, EPI_ISL_450636, EPI_ISL_450637 | | | |
| see above | Michigan Department of Health and Human Services, Bureau of Laboratories | Michigan Department of Health and Human Services, Bureau of Laboratories | Blankenship HM, Riner D, Soehnlen MK |
| EPI_ISL_450727, EPI_ISL_450728, EPI_ISL_450729, EPI_ISL_450730 | Hospital AZ Rivierenland | Institute of Tropical Medicine | Philippe Selhorst, Colin Anthony |
| EPI_ISL_450738, EPI_ISL_450740 | OUCRU/HTD | OUCRU/HTD | Nguyen Van Vinh Chau, Nguyen Thi Thu Hong, Nguyen Thi Han Ny, Le Nguyen Truc Nhu, Nghiem My Ngoc, Vo Thanh Lam, Nguyen Thanh Dung, Lam Minh Yen, Ngo Ngoc Quang Minh, Le Manh Hung, Nguyen Tri Dung, Dinh Nguyen Huy Man, Lam Anh Nguyet, Tran Chanh Xuan, Tran Tinh Hien, Nguyen Thanh Phong, Tran Nguyen Hoang Tu, Tran Tan Thanh, Nguyen Thanh Truong, Nguyen Tan Binh, Tang Chi Thuong, Guy Thwaites, and Le Van Tan, for OUCRU COVID-19 research group* |
| EPI_ISL_450748, EPI_ISL_450749, EPI_ISL_450750, EPI_ISL_450751, EPI_ISL_450752, EPI_ISL_450753, EPI_ISL_450754, EPI_ISL_450755, EPI_ISL_450759, EPI_ISL_450760, EPI_ISL_450761, EPI_ISL_450762, EPI_ISL_450764, EPI_ISL_450766, EPI_ISL_450767, EPI_ISL_450768, EPI_ISL_450771, EPI_ISL_450772, EPI_ISL_450773, EPI_ISL_450774, EPI_ISL_450775, EPI_ISL_450776, EPI_ISL_450777, EPI_ISL_450778, EPI_ISL_450779, EPI_ISL_450780 | | | |
| see above | Minnesota Department of Health, Public Health Laboratory | Minnesota Department of Health, Public Health Laboratory | Matt Plumb, Jacob Garfin, and Xiong Wang |
| EPI_ISL_451095, EPI_ISL_451096, EPI_ISL_451097, EPI_ISL_451098, EPI_ISL_451099, EPI_ISL_451100, EPI_ISL_451101, EPI_ISL_451102, EPI_ISL_451103, EPI_ISL_451104, EPI_ISL_451105, EPI_ISL_451106, EPI_ISL_451107, EPI_ISL_451108, EPI_ISL_451142 | | | |
| see above | SA Pathology | SA Pathology | Lex Leong, Chuan Kok Lim, Mark Turra, Ivan Bastian, Geoff Higgins |
| EPI_ISL_451185, EPI_ISL_451186, EPI_ISL_451187, EPI_ISL_451188, EPI_ISL_451189, EPI_ISL_451190, EPI_ISL_451191, EPI_ISL_451192, EPI_ISL_451193 | Uganda Virus Research Institute | MRC/UVRI & LSHTM Uganda Research Unit | Dan Lule Bugembe, John Kayiwa, My V.T Phan, Phionah Tushabe, Stephen Balinandi, Beatrice Dhaala, Deogratius Ssemwanga, Jonas Lexow, Henry Mwebesa, Jane Aceng, Henry Kyobe, Julius Lutwama, Pontiano Kaleebu, Matthew Cotten |
| EPI_ISL_451310, EPI_ISL_451311 | Hellenic Pasteur Institute, National Influenza Reference laboratory of Southern Greece & Unit of Bioinformatics and Applied Genomics | Hellenic Pasteur Institute, National Influenza Reference laboratory of Southern Greece & Unit of Bioinformatics and Applied Genomics | Vasiliki Pogka, Timokratis Karamitros, Athanasios Kossyvakis, Antonios Kalliaropoulos, Horefti Elina, Evangelidou Maria, Androniki Voulgari-Kokota, Aspasia Kontou, Andreas Mentis |
| EPI_ISL_451490, EPI_ISL_451491, EPI_ISL_451492, EPI_ISL_451493, EPI_ISL_451494, EPI_ISL_451495, EPI_ISL_451497, EPI_ISL_451498, EPI_ISL_451499, EPI_ISL_451500, EPI_ISL_451503, EPI_ISL_451504, EPI_ISL_451509, EPI_ISL_451510, EPI_ISL_451511, EPI_ISL_451512, EPI_ISL_451513, EPI_ISL_451514, EPI_ISL_451515, EPI_ISL_451516, EPI_ISL_451523, EPI_ISL_451524, EPI_ISL_451526, EPI_ISL_451527, EPI_ISL_451528 | | | |
| see above | Pathology West - NSW Health Pathology | NSW Health Pathology - Institute of Clinical Pathology and Medical Research; Westmead Hospital; University of Sydney | CIDM-PH et al. |
| EPI_ISL_451530, EPI_ISL_451531, EPI_ISL_451533, EPI_ISL_451536, EPI_ISL_451542, EPI_ISL_451543 | Pathology Sydney South West - NSW Health Pathology | NSW Health Pathology - Institute of Clinical Pathology and Medical Research; Westmead Hospital; University of Sydney | CIDM-PH et al. |
| EPI_ISL_451552 | Pathology North - NSW Health Pathology | NSW Health Pathology - Institute of Clinical Pathology and Medical Research; Westmead Hospital; University of Sydney | CIDM-PH et al. |
| EPI_ISL_451556 | Pathology West - NSW Health Pathology | NSW Health Pathology - Institute of Clinical Pathology and Medical Research; Westmead Hospital; University of Sydney | CIDM-PH et al. |
| EPI_ISL_451567 | Medlab Pathology | NSW Health Pathology - Institute of Clinical Pathology and Medical Research; Westmead Hospital; University of Sydney | CIDM-PH et al. |
| EPI_ISL_451569, EPI_ISL_451575, EPI_ISL_451576 | Pathology West - NSW Health Pathology | NSW Health Pathology - Institute of Clinical Pathology and Medical Research; Westmead Hospital; University of Sydney | CIDM-PH et al. |
| EPI_ISL_451578 | Pathology Sydney South West - NSW Health Pathology | NSW Health Pathology - Institute of Clinical Pathology and Medical Research; Westmead Hospital; University of Sydney | CIDM-PH et al. |
| EPI_ISL_451579, EPI_ISL_451584, EPI_ISL_451585 | Pathology West - NSW Health Pathology | NSW Health Pathology - Institute of Clinical Pathology and Medical Research; Westmead Hospital; University of Sydney | CIDM-PH et al. |
| EPI_ISL_451590, EPI_ISL_451598 | ACT pathology | NSW Health Pathology - Institute of Clinical Pathology and Medical Research; Westmead Hospital; University of Sydney | CIDM-PH et al. |
| EPI_ISL_451604 | Pathology North - NSW Health Pathology | NSW Health Pathology - Institute of Clinical Pathology and Medical Research; Westmead Hospital; University of Sydney | CIDM-PH et al. |
| EPI_ISL_451607 | Pathology West - NSW Health Pathology | NSW Health Pathology - Institute of Clinical Pathology and Medical Research; Westmead Hospital; University of Sydney | CIDM-PH et al. |
| EPI_ISL_451608 | Pathology Sydney South West - NSW Health Pathology | NSW Health Pathology - Institute of Clinical Pathology and Medical Research; Westmead Hospital; University of Sydney | CIDM-PH et al. |
| EPI_ISL_451632 | South Eastern Area Laboratory Services | NSW Health Pathology - Institute of Clinical Pathology and Medical Research; Westmead Hospital; University of Sydney | CIDM-PH et al. |
| EPI_ISL_451795, EPI_ISL_451796, EPI_ISL_451797, EPI_ISL_451798, EPI_ISL_451799, EPI_ISL_451800, EPI_ISL_451801, EPI_ISL_451802, EPI_ISL_451803, EPI_ISL_451804, EPI_ISL_451845 | | | |
| see above | Viollier AG | Department of Biosystems Science and Engineering, ETH Zürich | Christian Beisel, Sarah Nadeau, Ivan Topolsky, Pedro Ferreira, Philipp Jablonski, Susana Posada-Céspedes, Tobias Schär, Ina Nissen, Natascha Santacroce, Elodie Burcklen, Christiane Beckmann, Maurice Redondo, Olivier Kobel, Christoph Noppen, Sophie Seidel, Noemie Santamaria de Souza, Niko Beerenwinkel, Tanja Stadler |
| EPI_ISL_451942, EPI_ISL_451943 | Max von Pettenkofer Institute, Virology, National Reference Center for Retroviruses, LMU München | Laboratory for Functional Genome Analysis, Dept. Genomics, Gene Center of the LMU Munich | Max Muenchhoff, Stefan Krebs, Alexander Graf, Oliver Keppler, Helmut Blum |
| EPI_ISL_451982, EPI_ISL_451984, EPI_ISL_451985 | 1. ViroGenetics - BSL3 Laboratory of Virology, Maopolska Centre of Biotechnology, Jagiellonian University; 2. II Department of Internal Medicine, Faculty of Medicine, Jagiellonian University Medical College; 3. DIAGNOSTYKA Ltd. | 1. ViroGenetics - BSL3 Laboratory of Virology, Maopolska Centre of Biotechnology, Jagiellonian University; 2. II Department of Internal Medicine, Faculty of Medicine, Jagiellonian University Medical College. | Marek Sanak, Marcin Surmiak, Monika Gsecka-Czapla, Wojciech Branicki, Pawe P abaj, Marta Rogalska-Kupiec, Jakub Swadba, Krzysztof Pyr |
| EPI_ISL_452044, EPI_ISL_452079 | Department of Clinical Microbiology, Copenhagen University Hospital, Hvidovre, Kettegaard Alle 30, 2650 Hvidovre. | Albertsen lab, Department of Chemistry and Bioscience, Aalborg University, Denmark | Rasmus Kirkegaard |
| EPI_ISL_452101 | Department of Virus and Microbiological Special Diagnostics, Statens Serum Institut, Copenhagen, Denmark, Artillerivej 5, 2300 Copenahgen S | Albertsen lab, Department of Chemistry and Bioscience, Aalborg University, Denmark | Rasmus Kirkegaard |
| EPI_ISL_452142 | CUB Hopital Erasme Laboratoire d'Anatomie Pathologique | CUB Hopital Erasme Laboratoire d'Anatomie Pathologique | Isabelle Salmon, Nikcy D'Haene |
| EPI_ISL_452186 | ULSS9 Distretto di Bussolengo | Istituto Zooprofilattico Sperimentale delle Venezie | Adelaide Milani, Alessia Schivo, Annalisa Salviato, Erika Giorgia Quaranta, Ambra Pastori, Bianca Zecchin, Alice Fusaro, Isabella Monne, Calogero Terregino, Antonia Ricci |

| | | | |
|---|---|---|---|
| EPI_ISL_452206 | NIV Influenza | NIV Influenza | Potdar V |
| EPI_ISL_452306, EPI_ISL_452307, EPI_ISL_452310 | Michigan Department of Health and Human Services, Bureau of Laboratories | Michigan Department of Health and Human Services, Bureau of Laboratories | Blankenship HM, Riner D, Soehnlen MK |
| EPI_ISL_452372, EPI_ISL_452373 | Servicio de Microbiología. HRU de Málaga. Servicio Andaluz de Salud | SeqCOVID-SPAIN consortium/IBV(CSIC) | Inmaculada de Toro Peinado, Maria Concepción Mediavilla Gradolph, Begoña Palop Borrás and SeqCOVID-SPAIN consortium |
| EPI_ISL_452578, EPI_ISL_452579, EPI_ISL_452580, EPI_ISL_452581, EPI_ISL_452582, EPI_ISL_452583, EPI_ISL_452584, EPI_ISL_452585, EPI_ISL_452586, EPI_ISL_452587, EPI_ISL_452588, EPI_ISL_452589, EPI_ISL_452590, EPI_ISL_452591, EPI_ISL_452592, EPI_ISL_452594, EPI_ISL_452595, EPI_ISL_452596, EPI_ISL_452599, EPI_ISL_452600, EPI_ISL_452601, EPI_ISL_452608 | | | |
| see above | Servicio de Microbiología y Parasitología clínica. UCEIMP. Hospital Universitario Virgen del Rocío/IBIS/CSIC/US. | SeqCOVID-SPAIN consortium/IBV(CSIC) | Guillermo Martí-n Gutiérrez, Ángel Rodrí-guez Villodres, Lidia Gálvez Benitez, Verónica González Galán, Javier Aznar Martí-n and SeqCOVID-SPAIN consortium |
| EPI_ISL_452617, EPI_ISL_452618, EPI_ISL_452619, EPI_ISL_452620, EPI_ISL_452621, EPI_ISL_452622, EPI_ISL_452623, EPI_ISL_452624, EPI_ISL_452625, EPI_ISL_452626, EPI_ISL_452627, EPI_ISL_452628, EPI_ISL_452629, EPI_ISL_452630, EPI_ISL_452631, EPI_ISL_452632, EPI_ISL_452633, EPI_ISL_452634, EPI_ISL_452635, EPI_ISL_452636, EPI_ISL_452637, EPI_ISL_452638, EPI_ISL_452639, EPI_ISL_452640, EPI_ISL_452641, EPI_ISL_452642, EPI_ISL_452643, EPI_ISL_452644, EPI_ISL_452645, EPI_ISL_452646, EPI_ISL_452647, EPI_ISL_452648, EPI_ISL_452649, EPI_ISL_452650, EPI_ISL_452651, EPI_ISL_452652, EPI_ISL_452653, EPI_ISL_452654, EPI_ISL_452655 | | | |
| see above | Servicio de Microbiología. Hospital Universitario Donostia. OSI Donostialdea. Área de Enfermedades Infecciosas, Grupo de Infección Respiratoria y Resistencia Antimicrobiana. Instituto de Investigación Sanitaria Biodonostia. | SeqCOVID-SPAIN consortium/IBV(CSIC) | Gustavo Cilla, Milagrosa Montes, Luis Piñeiro, Jose Maria Marimón and SeqCOVID-SPAIN consortium |
| EPI_ISL_453184, EPI_ISL_453185, EPI_ISL_453189 | Virology Department, Royal Infirmary of Edinburgh, NHS Lothian / School of Biological Sciences, University of Edinburgh / Institute of Genetics and Molecular Medicine, University of Edinburgh | COVID-19 Genomics UK (COG-UK) Consortium | McHugh M, Dewar R, Rooke S, Gallagher M, Balcaza C, O'Toole Á, Scher E, Hill V, McCrone JT, Colquhoun R, Yu X, Jackson B, Rambaut A, Williams TC, Templeton K |
| EPI_ISL_453497, EPI_ISL_453498, EPI_ISL_453499, EPI_ISL_453500, EPI_ISL_453501, EPI_ISL_453502, EPI_ISL_453503 | Northumbria University / South Tees Hospitals NHS Foundation Trust / North Cumbria Integrated Care NHS Foundation Trust / North Tees and Hartlepool NHS Foundation Trust / Newcastle Hospitals NHS Foundation Trust | COVID-19 Genomics UK (COG-UK) Consortium | Darren L Smith,Andrew Nelson,Matthew Bashton,Greg R Young,Joshua Loh,John Allan,Mohammad A Tariq,Giles S Holt,Gary Black,Wen C Yew,Lynn Dover ,Paul Baker,Steve Liggett,Sarah Essex,Jane Greenaway ,Debra Padgett,Clive Graham,Garren Scott,Edward Barton ,Emma Swindells ,Brendan Payne,Jennifer Collins,Yusri Taha,Gary Eltringham |
| EPI_ISL_453854, EPI_ISL_453855, EPI_ISL_453856, EPI_ISL_453857, EPI_ISL_453858, EPI_ISL_453859, EPI_ISL_453860, EPI_ISL_453861, EPI_ISL_453862, EPI_ISL_453863, EPI_ISL_453864, EPI_ISL_453868, EPI_ISL_453869, EPI_ISL_453870, EPI_ISL_453871, EPI_ISL_453872, EPI_ISL_453873, EPI_ISL_453874, EPI_ISL_453875, EPI_ISL_453876, EPI_ISL_453877, EPI_ISL_453878, EPI_ISL_453879, EPI_ISL_453880, EPI_ISL_453881, EPI_ISL_453882, EPI_ISL_453883, EPI_ISL_453884, EPI_ISL_453885, EPI_ISL_453886, EPI_ISL_453887, EPI_ISL_453888, EPI_ISL_453889, EPI_ISL_453890, EPI_ISL_453891, EPI_ISL_453892, EPI_ISL_453893, EPI_ISL_453894, EPI_ISL_453895, EPI_ISL_453896, EPI_ISL_453897, EPI_ISL_453898, EPI_ISL_453899, EPI_ISL_453900, EPI_ISL_453901, EPI_ISL_453902, EPI_ISL_453903, EPI_ISL_453904, EPI_ISL_453905, EPI_ISL_453906, EPI_ISL_453907, EPI_ISL_453908, EPI_ISL_453909, EPI_ISL_453910, EPI_ISL_453911, EPI_ISL_453912, EPI_ISL_453913, EPI_ISL_453914, EPI_ISL_453915, EPI_ISL_453916, EPI_ISL_453917, EPI_ISL_453918, EPI_ISL_453919, EPI_ISL_453920, EPI_ISL_453921, EPI_ISL_453922, EPI_ISL_453923, EPI_ISL_453924, EPI_ISL_453925, EPI_ISL_453926, EPI_ISL_453927, EPI_ISL_453928, EPI_ISL_453929, EPI_ISL_453931, EPI_ISL_453932, EPI_ISL_453933, EPI_ISL_453934, EPI_ISL_453935, EPI_ISL_453936, EPI_ISL_453937, EPI_ISL_454145, EPI_ISL_454146, EPI_ISL_454147, EPI_ISL_454148, EPI_ISL_454149, EPI_ISL_454150, EPI_ISL_454151, EPI_ISL_454152, EPI_ISL_454170, EPI_ISL_454171, EPI_ISL_454172, EPI_ISL_454173, EPI_ISL_454204, EPI_ISL_454205, EPI_ISL_454206, EPI_ISL_454207, EPI_ISL_454209, EPI_ISL_454210, EPI_ISL_454297, EPI_ISL_454306, EPI_ISL_454307, EPI_ISL_454308, EPI_ISL_454309, EPI_ISL_454310, EPI_ISL_454311, EPI_ISL_454312, EPI_ISL_454313, EPI_ISL_454314, EPI_ISL_454315, EPI_ISL_454319, EPI_ISL_454320, EPI_ISL_454321, EPI_ISL_454322, EPI_ISL_454323, EPI_ISL_454327, EPI_ISL_454348, EPI_ISL_454349 | | | |
| see above | unknown | Instituto Nacional de Saude (INSA) | Borges et al |
| EPI_ISL_454353 | Microbial Genome Sequencing Center; Microbial Genomic Epidemiology Laboratory | Microbial Genomic Epidemiology Laboratory, University of Pittsburgh | Mustapha M. Mustapha, Jane W. Marsh, Dan Snyder, Marissa P. Griffith, Stephanie L. Mitchell, Vatsala R. Srinivasa, Kady D. Waggle, Chinelo Ezeonwuku, Vaughn S. Cooper, Lee H. Harrison |
| EPI_ISL_454354 | UPMC Clinical Microbiology Laboratory | Microbial Genome Sequencing Center; Microbial Genomic Epidemiology Laboratory, University of Pittsburgh | Mustapha M. Mustapha, Jane W. Marsh, Dan Snyder, Marissa P. Griffith, Stephanie L. Mitchell, Vatsala R. Srinivasa, Kady D. Waggle, Chinelo Ezeonwuku, Vaughn S. Cooper, Lee H. Harrison |
| EPI_ISL_454357, EPI_ISL_454358, EPI_ISL_454359, EPI_ISL_454360, EPI_ISL_454362, EPI_ISL_454363, EPI_ISL_454364, EPI_ISL_454365, EPI_ISL_454366, EPI_ISL_454367 | UPMC Clinical Microbiology Laboratory | Microbial Genome Sequencing Center, Microbial Genomic Epidemiological Laboratory | Mustapha M. Mustapha, Jane W. Marsh, Dan Snyder, Marissa P. Griffith, Stephanie L. Mitchell, Vatsala R. Srinivasa, Kady D. Waggle, Chinelo Ezeonwuku, Vaughn S. Cooper, Lee H. Harrison |
| EPI_ISL_454541, EPI_ISL_454542 | NIV Influenza | NIV Influenza | Potdar V |
| EPI_ISL_454606 | Institute for Public Health | Laboratory for advanced genomics | Filip Roki, Lovro Trgovec-Greif, Neven Sui, Tomislav Rukavina, Igor Jurak, Oliver Vugrek |
| EPI_ISL_454636, EPI_ISL_454637 | Humboldt County Public Health Laboratory | Chan-Zuckerberg Biohub | CZB Cliahub Consortium |
| EPI_ISL_454646 | City of El Paso Department of Public Health Laboratory | Pathogen Discovery, Respiratory Viruses Branch, Division of Viral Diseases, Centers for Disease Control and Prevention | Ying Tao, Clinton R. Paden, Jing Zhang, Anna Uehara, Krista Queen, Yan Li, Haibin Wang, Zachary Weiner, Michael Bowen, Suxiang Tong |
| EPI_ISL_454774, EPI_ISL_454792 | Dutch COVID-19 response team | National Institute for Public Health and the Environment (RIVM) | Adam Meijer, Harry Vennema, Jeroen Cremer, Sharon van den Brink, Pieter Overduin, Florian Zwagemaker, Dennis Schmitz, Chantal Reusken, on behalf of the national COVID-19 response team |
| EPI_ISL_454956 | Wuhan Chain Medical Labs (CMLabs) | State Key Laboratory of Biotherapy of Sichuan University | Baowen Du, Minjin Wang, Chao Tang, Chuan Chen, Yongzhao Zhou, Mingxia Yu, Hancheng Wei, Weimin Li, Jing-wen Lin, Jia Geng, Binwu Ying, Lu Chen |
| EPI_ISL_455044, EPI_ISL_455045, EPI_ISL_455046 | Pathology West - NSW Health Pathology | NSW Health Pathology - Institute of Clinical Pathology and Medical Research; Westmead Hospital; University of Sydney | CIDM-PH et al. |
| EPI_ISL_455051, EPI_ISL_455053 | Douglas Hanly Moir Pathology | NSW Health Pathology - Institute of Clinical Pathology and Medical Research; Westmead Hospital; University of Sydney | CIDM-PH et al. |
| EPI_ISL_455077 | South Eastern Area Laboratory Services | NSW Health Pathology - Institute of Clinical Pathology and Medical Research; Westmead Hospital; University of Sydney | CIDM-PH et al. |
| EPI_ISL_455094 | Pathology Sydney South West - NSW Health Pathology | NSW Health Pathology - Institute of Clinical Pathology and Medical Research; Westmead Hospital; University of Sydney | CIDM-PH et al. |
| EPI_ISL_455096 | South Eastern Area Laboratory Services | NSW Health Pathology - Institute of Clinical Pathology and Medical Research; Westmead Hospital; University of Sydney | CIDM-PH et al. |
| EPI_ISL_455165, EPI_ISL_455205, EPI_ISL_455206, EPI_ISL_455220, EPI_ISL_455221, EPI_ISL_455253, EPI_ISL_455254, EPI_ISL_455255, EPI_ISL_455282 | Dutch COVID-19 response team | Erasmus Medical Center | Bas Oude Munnink, David Nieuwenhuijse, Reina Sikkema, Claudia Schapendonk, Irina Chestakova, Anne van der Linden, Theo Bestebroer, Stefan van Nieuwkoop, Mark Pronk, Pascal Lexmond, Corien Swaan, Manon Haverkate, Madelief Mollers, Mart Stein, Sandra Kengne Kamga Mobou, Jeroen van Kampen, Jolanda Voermans, Aura Timen, Corine GeurtsvanKessel, Annemiek van der Eijk, Richard Molenkamp, Marion Koopmans, on behalf of the Dutch national COVID-19 response team. |
| EPI_ISL_455361 | Emory Molecular Diagnostics Laboratory, Emory Healthcare | Piantadosi Lab, Emory Department of Pathology | Ahmed Babiker, Anne Piantadosi |
| EPI_ISL_455389 | Wuhan Chain Medical Labs (CMLabs) | State Key Laboratory of Biotherapy of Sichuan University | Baowen Du, Minjin Wang, Chao Tang, Chuan Chen, Yongzhao Zhou, Mingxia Yu, Hancheng Wei, Weimin Li, Jing-wen Lin, Jia Geng, Binwu Ying, Lu Chen |
| EPI_ISL_455412, EPI_ISL_455413, EPI_ISL_455414 | Nigeria Centre for Disease Control (NCDC) | African Centre of Excellence for Genomics of Infectious Diseases (ACEGID), Redeemer's University, Ede, Osun State, Nigeria | Oluniyi P.E., Ajogbasile F.V., Kayode A., Oguzie J., Olawoye I., Uwanibe J., Olumade T., Folarin O.A., Ihekweazu C., Happi C.T. |
| EPI_ISL_455415 | Nigeria Centre for Disease Control (NCDC) | African Centre of Excellence for Genomics of Infectious Diseases (ACEGID), Redeemer's University, Ede, Osun State, | Oluniyi P.E., Ajogbasile F.V., Kayode A., Oguzie J., Olawoye I., Uwanibe J., Olumade T., Folarin O.A., Ihekweazu C., Happi C.T. |

| | | | |
|---|---|---|---|
| | | Nigeria | |
| EPI_ISL_455418, EPI_ISL_455419 | Nigeria Centre for Disease Control (NCDC) | African Centre of Excellence for Genomics of Infectious Diseases (ACEGID), Redeemer's University, Ede, Osun State, Nigeria | Oluniyi P.E., Ajogbasile F.V., Kayode A., Oguzie J., Olawoye I., Uwanibe J., Olumade T., Folarin O.A., Ihekweazu C., Happi C.T. |
| EPI_ISL_455422 | Nigeria Centre for Disease Control | African Centre of Excellence for Genomics of Infectious Diseases (ACEGID), Redeemer's University, Ede, Osun State, Nigeria | Oluniyi P.E., Ajogbasile F.V., Kayode A., Oguzie J., Olawoye I., Uwanibe J., Olumade T., Folarin O.A., Ihekweazu C., Happi C.T. |
| EPI_ISL_455423, EPI_ISL_455424, EPI_ISL_455425 | Nigeria Centre for Disease Control (NCDC) | African Centre of Excellence for Genomics of Infectious Diseases (ACEGID), Redeemer's University, Ede, Osun State, Nigeria | Oluniyi P.E., Ajogbasile F.V., Kayode A., Oguzie J., Olawoye I., Uwanibe J., Olumade T., Folarin O.A., Ihekweazu C., Happi C.T. |
| EPI_ISL_455426 | Nigeria Centre for Disease Control | African Centre of Excellence for Genomics of Infectious Diseases (ACEGID), Redeemer's University, Ede, Osun State, Nigeria | Oluniyi P.E., Ajogbasile F.V., Kayode A., Oguzie J., Olawoye I., Uwanibe J., Olumade T., Folarin O.A., Ihekweazu C., Happi C.T. |
| EPI_ISL_455427, EPI_ISL_455429 | Nigeria Centre for Disease Control (NCDC) | African Centre of Excellence for Genomics of Infectious Diseases (ACEGID), Redeemer's University, Ede, Osun State, Nigeria | Oluniyi P.E., Ajogbasile F.V., Kayode A., Oguzie J., Olawoye I., Uwanibe J., Olumade T., Folarin O.A., Ihekweazu C., Happi C.T. |
| EPI_ISL_455440, EPI_ISL_455452 | 1. ViroGenetics - BSL3 Laboratory of Virology, Maopolska Centre of Biotechnology, Jagiellonian University; 2. II Department of Internal Medicine, Faculty of Medicine, Jagiellonian University Medical College; 3. Narodowy Instytut Zdrowia Publicznego - Pastwowy Zakad Higieny (NIZP-PZH) | 1. ViroGenetics - BSL3 Laboratory of Virology, Maopolska Centre of Biotechnology, Jagiellonian University; 2. II Department of Internal Medicine, Faculty of Medicine, Jagiellonian University Medical College; 3. Narodowy Instytut Zdrowia Publicznego - Pastwowy Zakad Higieny (NIZP-PZH). | Katarzyna Pancer, Marek Sanak, Aleksandra A. Zasada, Magdalena Rzeczkowska, Tomasz Wokowicz, Katarzyna Zacharczuk, Agnieszka Koakowska-Kulesza, Katarzyna Owczarek, Aleksandra Milewska, Natalia Wolaniuk, Ewelina Hallman-Szeliska, Pawe P abaj, Wojciech Branicki, Krzysztof Pyr |
| EPI_ISL_455566, EPI_ISL_455567 | Institute for Public Health | Laboratory for advanced genomics | Filip Roki, Lovro Trgovec-Greif, Neven Sui, Tomislav Rukavina, Igor Jurak, Oliver Vugrek |
| EPI_ISL_455625, EPI_ISL_455626 | unknown | Instituto Nacional de Saude (INSA) | Borges et al |
| EPI_ISL_455640, EPI_ISL_455643 | ICMR-National Institute of Cholera and Enteric Diseases | National Institute of Biomedical Genomics | Arindam Maitra, Mamta Chawla Sarkar, Sreedhar Chinnaswamy, Hasina Banu, Ananya Chatterjee, Shanta Dutta, Saumitra Das |
| EPI_ISL_455702, EPI_ISL_455703, EPI_ISL_455706 | National Hospital of Tropical Diseases | Oxford University Clinical Research Unit, Hanoi, Vietnam | Nguyen Thi Tam, Van Dinh Trang, Nguyen Thu Trang, Nguyen Thi Ngoc Diep, Le Nguyen Minh Hoa, Pham Ngoc Thach, H. Rogier van Doorn, on behalf of the OUCRU COVID-19 research group |
| EPI_ISL_455722 | Servicio de Microbiologia. Hospital Clinico Universitario de Valencia | Sequencing and Bioinformatics Service and Molecular Epidemiology Research Group. FISABIO-Public Health, and SeqCOVID-Spain Consortium | Giuseppe 'Auria, David Navarro, Eliseo Albert, Maria Alma Bracho, Lidia Ruiz Roldan, Neris Garcia-Gonzalez, Inma Galán Vendrell, Sandra Carbo, Loreto Ferrús Abad, Paula Ruiz-Hueso, Mariana Reyes-Prieto, Vicente Soriano Chirona, Ivan Ansari, Llúcia Martínez-Priego, Fernando Gonzalez-Candelas |
| EPI_ISL_455740 | Servicio de Microbiologia. Hospital Clinico Universitario de Valencia | Sequencing and Bioinformatics Service and Molecular Epidemiology Research Group. FISABIO-Public Health, and SeqCOVID-Spain Consortium | Paula Ruiz-Hueso, Mariana Reyes-Prieto, Vicente Soriano Chirona, Ivan Ansari, Llúcia Martínez-Priego, Giuseppe 'Auria, David Navarro, Eliseo Albert, Maria Alma Bracho, Lidia Ruiz Roldan, Neris Garcia-Gonzalez, Inma Galán Vendrell, Sandra Carbo, Loreto Ferrús Abad, Fernando Gonzalez-Candelas |
| EPI_ISL_455741 | Servicio de Microbiologia. Hospital Clinico Universitario de Valencia | Sequencing and Bioinformatics Service and Molecular Epidemiology Research Group. FISABIO-Public Health, and SeqCOVID-Spain Consortium | David Navarro, Eliseo Albert, Maria Alma Bracho, Griselda De Marco, Lidia Ruiz Roldan, Neris Garcia-Gonzalez, Inma Galán Vendrell, Sandra Carbo, Loreto Ferrús Abad, Paula Ruiz-Hueso, Mariana Reyes-Prieto, Vicente Soriano Chirona, Ivan Ansari, Llúcia Martínez-Priego, Giuseppe 'Auria, Fernando Gonzalez-Candelas |
| EPI_ISL_455743 | Servicio de Microbiologia. Hospital Clinico Universitario de Valencia | Sequencing and Bioinformatics Service and Molecular Epidemiology Research Group. FISABIO-Public Health, and SeqCOVID-Spain Consortium | Eliseo Albert, Maria Alma Bracho, Griselda De Marco, Lidia Ruiz Roldan, Neris Garcia-Gonzalez, Inma Galán Vendrell, Sandra Carbo, Loreto Ferrús Abad, Paula Ruiz-Hueso, Mariana Reyes-Prieto, Vicente Soriano Chirona, Ivan Ansari, Llúcia Martínez-Priego, Giuseppe 'Auria, David Navarro, Fernando Gonzalez-Candelas |
| EPI_ISL_455980, EPI_ISL_455981, EPI_ISL_455982, EPI_ISL_455983, EPI_ISL_455984, EPI_ISL_455985 | LSUHS Emerging Viral Threat Laboratory | Microbial Genome Sequencing Center | Jeremy P. Kamil, John A. Vanchiere, Rona S. Scott, Camille F. Abshire, Abida Siddiqa, Byeong-Jae Lee, Chan-ki Min, Md Maksudul Alam, Monica Gestal-Cartele, Edna Ondari, Adam Greer, Malgorzata Bienkowska-Haba, Katarzyna Zwolinska, Michelle M. Arnold, Jason M. Bodily, Andrew D. Yurochko, Paul M. Weinberger, Christopher G. Kevil, Martin J. Sapp, Daniel J. Snyder, Vaughn S. Cooper |
| EPI_ISL_455992, EPI_ISL_455993 | LSUHS Emerging Viral Threat Laboratory | Microbial Genome Sequencing Center | John A. Vanchiere, Jeremy P. Kamil, Rona S. Scott, Camille F. Abshire, Abida Siddiqa, Byeong-Jae Lee, Chan-ki Min, Md Maksudul Alam, Monica Gestal-Cartele, Edna Ondari, Adam Greer, Malgorzata Bienkowska-Haba, Katarzyna Zwolinska, Michelle M. Arnold, Jason M. Bodily, Andrew D. Yurochko, Paul M. Weinberger, Christopher G. Kevil, Martin J. Sapp, Daniel J. Snyder, Vaughn S. Cooper |
| EPI_ISL_456003, EPI_ISL_456004, EPI_ISL_456006, EPI_ISL_456008, EPI_ISL_456010, EPI_ISL_456011, EPI_ISL_456013, EPI_ISL_456014, EPI_ISL_456016, EPI_ISL_456018, EPI_ISL_456021, EPI_ISL_456025, EPI_ISL_456026, EPI_ISL_456028, EPI_ISL_456030, EPI_ISL_456032, EPI_ISL_456034, EPI_ISL_456035, EPI_ISL_456036, EPI_ISL_456037, EPI_ISL_456038, EPI_ISL_456040, EPI_ISL_456041, EPI_ISL_456043, EPI_ISL_456044, EPI_ISL_456045, EPI_ISL_456047, EPI_ISL_456052, EPI_ISL_456055, EPI_ISL_456058, EPI_ISL_456061, EPI_ISL_456065, EPI_ISL_456070 | | | |
| see above | NYU Langone Health | Departments of Pathology and Medicine, New York University School of Medicine | Maria Aguero-Rosenfeld, Brendan Belovarac, Margaret Black, Ludovic Boytard, John Cadley, Paolo Cotzia, John Chen, Dacia Dimartino, Xiaojun Feng, Tatyana Gindin, Emily Guzman, Adriana Heguy, Megan Hogan, Emily Huang, George Jour, Alireza Khodadadi-Jamayran, Lawrence H. Lin, Raven Luther, Andrew Lytle, Christian Marier, Matthew T. Maurano, Mark J. Mulligan, Peter Meyn, Raquel Ordonez Ciriza, Iman Osman, Jared Pinnell, Vanessa Raabe, Sitharam Ramaswami, Amy Rapkiewicz, Andre M. Ribeiro-dos-Santos, Marie Samanovic-Golden, Antonio Serrano, Guomiao Shen, Matija Snuderl, Theodore Vougiouklakis, Nick Vulpescu, Gael Westby, Paul Zappile, Yutong Zhang |
| EPI_ISL_456071 | Laboratory of Respiratory Viruses and Measles, Oswaldo Cruz Institute, FIOCRUZ | Laboratory of Respiratory Viruses and Measles, Oswaldo Cruz Institute, FIOCRUZ | Paola Resende, Luciana Appolinario, Fernando Motta, Aline Mattos, Milene Miranda, Cristiana Garcia, Braulia Caetano, Maria Ogrzewalska, Jonathan Lopes, Marilda Siqueira |
| EPI_ISL_456234 | Southern Community Labs Dunedin | Institute of Environmental Science and Research (ESR) | Matt Storey, Xiaoyun Ren, Anja Werno, Antje van der Linden, Arlo Upton, Chris Mansell, David Hammer, Dragana Drinkovic, Erasmus Smit, Gary McAuliffe, Hana Sofia Andersson, James Ussher, Jill Sherwood, Josh Freeman, Julia Howard, Juliet Elvy, Mary DeAlmeida, Matt Blakiston, Matthew Rogers, Max Bloomfield, Michael Addidle, Michelle Balm, Sally Roberts, Sarah Jefferies, Sharmini Muttaiyah, Susan Morpeth, Susan Taylor, Timothy Blackmore, Vani Sathyendran, Veronica Playle, Virginia Hope, Erasmus Smit, Lauren Jelly, Joep de Ligt |
| EPI_ISL_456241, EPI_ISL_456259, EPI_ISL_456260 | Waikato Hospital | Institute of Environmental Science and Research (ESR) | Matt Storey, Xiaoyun Ren, Anja Werno, Antje van der Linden, Arlo Upton, Chris Mansell, David Hammer, Dragana Drinkovic, Erasmus Smit, Gary McAuliffe, Hana Sofia Andersson, James Ussher, Jill Sherwood, Josh Freeman, Julia Howard, Juliet Elvy, Mary DeAlmeida, Matt Blakiston, Matthew Rogers, Max Bloomfield, Michael Addidle, Michelle Balm, Sally Roberts, Sarah Jefferies, Sharmini Muttaiyah, Susan Morpeth, Susan Taylor, Timothy Blackmore, Vani Sathyendran, Veronica Playle, Virginia Hope, Erasmus Smit, Lauren Jelly, Joep de Ligt |
| EPI_ISL_456261, EPI_ISL_456262, EPI_ISL_456263, EPI_ISL_456264, EPI_ISL_456265, EPI_ISL_456266, EPI_ISL_456267, EPI_ISL_456268, EPI_ISL_456269, EPI_ISL_456270, EPI_ISL_456271, EPI_ISL_456272, EPI_ISL_456273, EPI_ISL_456274, EPI_ISL_456275, EPI_ISL_456276, EPI_ISL_456277, EPI_ISL_456278, EPI_ISL_456279, EPI_ISL_456280, EPI_ISL_456281, EPI_ISL_456282, EPI_ISL_456283, EPI_ISL_456284, EPI_ISL_456285, EPI_ISL_456286, EPI_ISL_456288, EPI_ISL_456289, EPI_ISL_456290, EPI_ISL_456291, EPI_ISL_456293 | | | |
| see above | Southern Community Labs Dunedin | Institute of Environmental Science and Research (ESR) | Matt Storey, Xiaoyun Ren, Anja Werno, Antje van der Linden, Arlo Upton, Chris Mansell, David Hammer, Dragana Drinkovic, Erasmus Smit, Gary McAuliffe, Hana Sofia Andersson, James Ussher, Jill Sherwood, Josh Freeman, Julia Howard, Juliet Elvy, Mary DeAlmeida, Matt Blakiston, Matthew Rogers, Max Bloomfield, Michael Addidle, Michelle Balm, Sally Roberts, Sarah Jefferies, Sharmini Muttaiyah, Susan Morpeth, Susan Taylor, Timothy Blackmore, Vani Sathyendran, Veronica Playle, Virginia Hope, Erasmus Smit, Lauren Jelly, Joep de Ligt |
| EPI_ISL_456319 | Wellington SCL | Institute of Environmental Science and Research (ESR) | Matt Storey, Xiaoyun Ren, Anja Werno, Antje van der Linden, Arlo Upton, Chris Mansell, David Hammer, Dragana Drinkovic, Erasmus Smit, Gary McAuliffe, Hana Sofia Andersson, James Ussher, Jill Sherwood, Josh Freeman, Julia Howard, Juliet Elvy, Mary DeAlmeida, Matt Blakiston, Matthew Rogers, Max Bloomfield, Michael Addidle, Michelle Balm, Sally Roberts, Sarah Jefferies, Sharmini Muttaiyah, Susan Morpeth, Susan Taylor, Timothy Blackmore, Vani Sathyendran, Veronica Playle, Virginia Hope, Erasmus Smit, Lauren Jelly, Joep de Ligt |
| EPI_ISL_456409 | unknown | Research Center Of Tropical and Infectious Of Medical Sciences | Mollaei,H.R., Aghaei-Afshar,A., Kalantar-Neyestanaki,D. |
| EPI_ISL_456600 | National Health Laboratory, Timor-Leste | Microbiological Diagnostic Unit Public Health Laboratory, The | Soares da Silva, E., Dolores de Jesus da Costa, M., Salles de Sousa, A., Jayanti Pereira Tilman, A., Antonia da Costa, E., Barreto, I., Marr, I., Wapling, J., |

| | | | |
|---|---|---|---|
| | | Peter Doherty Institute for Infection and Immunity | Francis, J., Ximenes, J., Canisia, D., Freeman, K., Dakh, F., Douglas, N., Baird, R., Caly, L., Seemann, T., Sait, M., Schultz, M., Sherry, N. |
| EPI_ISL_456756, EPI_ISL_456889 | West of Scotland Specialist Virology Centre, NHSGGC / MRC-University of Glasgow Centre for Virus Research | COVID-19 Genomics UK (COG-UK) Consortium | Ana da Silva Filipe, Natasha Johnson, Kathy Smollett, Daniel Mair, Stephen Carmichael, Lily Tong, Jenna Nichols, Elihu Aranday-Cortes, Kirstyn Brunker, Yasmin Parr, Kyriaki Nomikou; Sarah McDonald, Marc Niebel, Patawee Asamaphan; Richard Orton, Joseph Hughes, Sreenu Vattipally, David L Robertson; Alasdair MacLean, Rory Gunson; Kathy Li, Natasha Jesudason, Rajiv Shah, James Shepherd, Antonia Ho, Emma Thomson |
| EPI_ISL_456896, EPI_ISL_456897, EPI_ISL_456898, EPI_ISL_456901, EPI_ISL_456902, EPI_ISL_456903, EPI_ISL_456905, EPI_ISL_456906, EPI_ISL_456912, EPI_ISL_456956, EPI_ISL_456966, EPI_ISL_456967, EPI_ISL_456968, EPI_ISL_456969, EPI_ISL_456970, EPI_ISL_456971, EPI_ISL_456972, EPI_ISL_456973, EPI_ISL_457009, EPI_ISL_457010, EPI_ISL_457011 | | | |
| see above | Virology Department, Royal Infirmary of Edinburgh, NHS Lothian / School of Biological Sciences, University of Edinburgh / Institute of Genetics and Molecular Medicine, University of Edinburgh | COVID-19 Genomics UK (COG-UK) Consortium | McHugh M, Dewar R, Rooke S, Gallagher M, Balcaza C, O'Toole Á, Scher E, Hill V, McCrone JT, Colquhoun R, Yu X, Jackson B, Rambaut A, Williams TC, Templeton K |
| EPI_ISL_457082, EPI_ISL_457146, EPI_ISL_457151, EPI_ISL_457156, EPI_ISL_457162, EPI_ISL_457163, EPI_ISL_457167, EPI_ISL_457168, EPI_ISL_457171, EPI_ISL_457176, EPI_ISL_457183, EPI_ISL_457185 | | | |
| see above | University of Exeter | COVID-19 Genomics UK (COG-UK) Consortium | Ben Temperton,Aaron Jeffries,Michelle Michelsen,Joanna Warwick-Dugdale,Audrey Farbos,Robyn Manley,Stephen Michell,Jane Masoli |
| EPI_ISL_457590, EPI_ISL_457632, EPI_ISL_457646, EPI_ISL_457649, EPI_ISL_457656, EPI_ISL_457675 | Virology Department, Sheffield Teaching Hospitals NHS Foundation Trust/Department of Infection, Immunity and Cardiovascular Disease, The Medical School, University of Sheffield | COVID-19 Genomics UK (COG-UK) Consortium | Thushan de Silva, Matthew Parker, Nikki Smith, Adri Angyal, Rebecca Brown, Luke Green, Rachel Tucker, Paul Parsons, Danielle Groves, Katie Johnson, Laura Carrilero, Alex Keeley, Dave Partridge, Matthew Wyles, Benjamin Lindsey, Mehmet Yavuz, Mohammad Raza, Cariad Evans |
| EPI_ISL_457760, EPI_ISL_457764, EPI_ISL_457767, EPI_ISL_457773, EPI_ISL_457776, EPI_ISL_457777, EPI_ISL_457780, EPI_ISL_457781, EPI_ISL_457783, EPI_ISL_457787, EPI_ISL_457810, EPI_ISL_457811, EPI_ISL_457818, EPI_ISL_457821 | | | |
| see above | Johns Hopkins Hospital Department of Pathology | Johns Hopkins Hospital Department of Pathology | Peter M. Thielen, Thomas Mehoke, Shirlee Wohl, Srividya Ramakrishnan, Melanie Kirsche, Amanda Ernlund, Craig Howser, Kristina Zudock, Oluwaseun Falade-Nwulia, Norah Sadowski, Paul Morris, Mark Hopkins, Yunfan Fan, Nidia Trovao, Victoria Gniazdowski, Michael C. Schatz, Stuart C. Ray, Winston Timp, Heba H. Mostafa |
| EPI_ISL_457842, EPI_ISL_457843, EPI_ISL_457844 | National Public Health Laboratory | KEMRI-Wellcome Trust Research Programme/KEMRI-CGMR-C Kilifi | Githinji G. et al 2020 |
| EPI_ISL_457934, EPI_ISL_457935, EPI_ISL_457936 | KEMRI-Centre for Virus Research | KEMRI-Wellcome Trust Research Programme/KEMRI-CGMR-C Kilifi | Githinji G. et al 2020 |
| EPI_ISL_457960, EPI_ISL_457961, EPI_ISL_457962, EPI_ISL_457963 | Laboratorio de Biología Molecular Asociación Española Primera en Salud | Departments of Pathology and Medicine, New York University School of Medicine | Maria Victoria Elizondo, Maria Noel Zubillaga, Gonzalo Manrique, Paul Zappile, Gael Westby, Matthew T Maurano, Christian Marier, Adriana Heguy |
| EPI_ISL_457996, EPI_ISL_457997, EPI_ISL_457998 | Oman-NIC | Oman-NIC | Samira Al-Maruqi, Fahad Zadjali, Amina Al Jardani, Khulood Al-Mammary, Hanan Al-kindi, Fatma BaAlawi, Hamida AL Barwani, Zeyana AL-Dahmani, Intisar Al-Shukri, Aisha Al-Busaidi, Aisha Al-Amri, Ahlam Al-Amri, Mohammed Al-Tobi, Samiha Al Kharusi, Abdulla Balkhair |
| EPI_ISL_458023, EPI_ISL_458024 | Hospital for Tropical Diseases | COVID-19 Network Investigations (CONI) Alliance | Elizabeth Batty, Nantarat Chantawat, Wasun Chantratita, Thanat Chookajorn, Stefan Fernandez, Angkana Huang, Weena Janwitthayanan, Akanitt Jittmittraphap, Anthony R. Jones, Khajohn Joonsalak, Chonticha Klungtong, Theerarat Kochakarn, Namfon Kotanan, Krittikorn Kumpornsin, Pornsawan Leaungwutiwong, Wudtichai Manasatienkij, Bhakbhoom Panthan, Ekawat Pasomsub, Kingkan Rakmanee, Insee Sensorn, Janjira Thaipadungpanit, Arporn Wangwiwatsin,Treewat Watthanachockchai |
| EPI_ISL_458079 | Mitra Keluarga Hospital Kenjeran | Institute of Tropical Disease, Universitas Airlangga | Aldise M Nastri, Jezzy R Dewantari, Rima R Prasetya, Krisnoadi Rahardjo, Anastasia W Jefuna, Gatot Soegiarto, Laksmi Wulandari, Retno A Setyoningrum, Resti Yudhawati, Yohko K Shimizu, Mitsuhiro Nishimura, Yasuko Mori, Soetjipto, Kazufumi Shimizu, Maria I Lusida |
| EPI_ISL_458081 | RSUD Bangil Pasuruan | Institute of Tropical Disease, Universitas Airlangga | Jezzy R Dewantari, Rima R Prasetya, Krisnoadi Rahardjo, Aldise M Nastri, Arma Roosalina, Gatot Soegiarto, Laksmi Wulandari, Retno A Setyoningrum, Resti Yudhawati, Yohko K Shimizu, Mitsuhiro Nishimura, Yasuko Mori, Soetjipto, Kazufumi Shimizu, Maria I Lusida |
| EPI_ISL_458156, EPI_ISL_458157, EPI_ISL_458158, EPI_ISL_458159, EPI_ISL_458160, EPI_ISL_458161, EPI_ISL_458162, EPI_ISL_458163, EPI_ISL_458164, EPI_ISL_458165, EPI_ISL_458166, EPI_ISL_458167, EPI_ISL_458168, EPI_ISL_458169, EPI_ISL_458170, EPI_ISL_458171, EPI_ISL_458172, EPI_ISL_458173, EPI_ISL_458174, EPI_ISL_458175, EPI_ISL_458176, EPI_ISL_458177, EPI_ISL_458178, EPI_ISL_458179, EPI_ISL_458180, EPI_ISL_458181, EPI_ISL_458182, EPI_ISL_458183, EPI_ISL_458184, EPI_ISL_458185, EPI_ISL_458186, EPI_ISL_458187, EPI_ISL_458188, EPI_ISL_458189, EPI_ISL_458190, EPI_ISL_458191, EPI_ISL_458192, EPI_ISL_458193, EPI_ISL_458194, EPI_ISL_458195, EPI_ISL_458196, EPI_ISL_458197, EPI_ISL_458198, EPI_ISL_458199, EPI_ISL_458200, EPI_ISL_458201, EPI_ISL_458202, EPI_ISL_458203, EPI_ISL_458204, EPI_ISL_458205, EPI_ISL_458206, EPI_ISL_458207, EPI_ISL_458208, EPI_ISL_458209, EPI_ISL_458210, EPI_ISL_458211, EPI_ISL_458212, EPI_ISL_458213, EPI_ISL_458214, EPI_ISL_458215, EPI_ISL_458216, EPI_ISL_458217, EPI_ISL_458218, EPI_ISL_458219, EPI_ISL_458220, EPI_ISL_458221, EPI_ISL_458222, EPI_ISL_458223, EPI_ISL_458224, EPI_ISL_458225, EPI_ISL_458226, EPI_ISL_458227 | | | |
| see above | KU Leuven, Rega Institute, Clinical and Epidemiological Virology | KU Leuven, Rega Institute, Clinical and Epidemiological Virology | Tony Wawina-Bokalanga, Bert Vanmechelen, Joan Marti-Carerras, Piet Maes |
| EPI_ISL_458466 | PHE South West Regional Laboratory, National Infection Service | Wellcome Sanger Institute for the COVID-19 Genomics UK (COG-UK) consortium | Stephanie Hutchings, Hannah Pymont, Dr Peter Muir, Barry Vipond, Rich Hopes; and Alex Alderton, Roberto Amato, Sonia Goncalves, Ewan Harrison, David K. Jackson, Ian Johnston, Dominic Kwiatkowski, Cordelia Langford, John Sillitoe on behalf of the Wellcome Sanger Institute COVID-19 Surveillance Team (http://www.sanger.ac.uk/covid-team) |
| EPI_ISL_458583, EPI_ISL_458584, EPI_ISL_458586, EPI_ISL_458588, EPI_ISL_458592, EPI_ISL_458593, EPI_ISL_458594, EPI_ISL_458595, EPI_ISL_458596, EPI_ISL_458597, EPI_ISL_458600, EPI_ISL_458601, EPI_ISL_458606, EPI_ISL_458609, EPI_ISL_458611, EPI_ISL_458612, EPI_ISL_458614, EPI_ISL_458615, EPI_ISL_458617, EPI_ISL_458618, EPI_ISL_458626, EPI_ISL_458632, EPI_ISL_458633, EPI_ISL_458634, EPI_ISL_458640, EPI_ISL_458644, EPI_ISL_458645, EPI_ISL_458646, EPI_ISL_458652, EPI_ISL_458657, EPI_ISL_458661, EPI_ISL_458662, EPI_ISL_458663, EPI_ISL_458667, EPI_ISL_458670, EPI_ISL_458674, EPI_ISL_458677, EPI_ISL_458678, EPI_ISL_458680, EPI_ISL_458682, EPI_ISL_458685, EPI_ISL_458686, EPI_ISL_458687, EPI_ISL_458688, EPI_ISL_458691, EPI_ISL_458693, EPI_ISL_458694, EPI_ISL_458697, EPI_ISL_458698, EPI_ISL_458699, EPI_ISL_458701, EPI_ISL_458703, EPI_ISL_458707, EPI_ISL_458710, EPI_ISL_458711, EPI_ISL_458713, EPI_ISL_458715, EPI_ISL_458717, EPI_ISL_458718 | | | |
| see above | NU-OMICS DNA Sequencing research facility, Northumbria University | Wellcome Sanger Institute for the COVID-19 Genomics UK (COG-UK) consortium | Chris Duncan, Sheia Waugh, Shirelle Burton-Fanning, Gary Eltringham, Jennifer Collins, Brendan Payne, Yusri Taha, Emma Swindells, Jane Greenaway, Edward Barton, Garren Scott, Debra Padgett, Clive Graham, Sarah Essex, Steve Ligget, Paul Baker, Lynn Dover, Wen Yew, Gary Black, John Allan, Joshua Loh, Greg Young, Matthew Bashton, Andrew Nelson, Darren Smith and Alex Alderton, Roberto Amato, Sonia Goncalves, Ewan Harrison, David K. Jackson, Ian Johnston, Dominic Kwiatkowski, Cordelia Langford, John Sillitoe on behalf of the Wellcome Sanger Institute COVID-19 Surveillance Team (http://www.sanger.ac.uk/covid-team) |
| EPI_ISL_458734, EPI_ISL_458768, EPI_ISL_458817, EPI_ISL_458890 | PHE South West Regional Laboratory, National Infection Service | Wellcome Sanger Institute for the COVID-19 Genomics UK (COG-UK) consortium | Stephanie Hutchings, Hannah Pymont, Dr Peter Muir, Barry Vipond, Rich Hopes; and Alex Alderton, Roberto Amato, Sonia Goncalves, Ewan Harrison, David K. Jackson, Ian Johnston, Dominic Kwiatkowski, Cordelia Langford, John Sillitoe on behalf of the Wellcome Sanger Institute COVID-19 Surveillance Team (http://www.sanger.ac.uk/covid-team) |
| EPI_ISL_459509, EPI_ISL_459510, EPI_ISL_459517, EPI_ISL_459520, EPI_ISL_459530, EPI_ISL_459540, EPI_ISL_459543, EPI_ISL_459551, EPI_ISL_459552, EPI_ISL_459565, EPI_ISL_459576, EPI_ISL_459582, EPI_ISL_459596, EPI_ISL_459616, EPI_ISL_459621, EPI_ISL_459629, EPI_ISL_459641, EPI_ISL_459649, EPI_ISL_459656, EPI_ISL_459666, EPI_ISL_459670, EPI_ISL_459682, EPI_ISL_459692, EPI_ISL_459693, EPI_ISL_459695, EPI_ISL_459698, EPI_ISL_459701 | | | |
| see above | NHSGGC West of Scotland Specialist Virology Centre / MRC-University of Glasgow Centre for Virus Research | Wellcome Sanger Institute for the COVID-19 Genomics UK (COG-UK) consortium | Ana da Silva Filipe, Natasha Johnson, Kathy Smollett, Daniel Mair, Stephen Carmichael, Lily Tong, Jenna Nichols, Elihu Aranday-Cortes, Kirstyn Brunker, Yasmin Parr, Kyriaki Nomikou; Sarah McDonald, Marc Niebel, Patawee Asamaphan; Richard Orton, Joseph Hughes, Sreenu Vattipally, David L Robertson; Alasdair MacLean, Rory Gunson; Kathy Li, Natasha Jesudason, Rajiv Shah, James Shepherd, Antonia Ho, Alice Broos, Emma Thomson and Alex Alderton, Roberto Amato, Sonia Goncalves, Ewan Harrison, David K. Jackson, Ian Johnston, Dominic Kwiatkowski, Cordelia Langford, John Sillitoe on behalf of the Wellcome Sanger Institute COVID-19 Surveillance Team (http://www.sanger.ac.uk/covid-team) |
| EPI_ISL_459859 | Center for Genome Regulation (CRG) | Center for Mathematical Modeling and Center for Genome Regulation. Santiago, Chile | Gaete A, Travisany D, Palma R, Urra C, Varas M, Allende ML, Maass A, González M. |
| EPI_ISL_459874, EPI_ISL_459875, EPI_ISL_459877, EPI_ISL_459878, EPI_ISL_459879, EPI_ISL_459880, EPI_ISL_459881, EPI_ISL_459882, EPI_ISL_459883, EPI_ISL_459884, EPI_ISL_459885, EPI_ISL_459886, EPI_ISL_459887, EPI_ISL_459888, EPI_ISL_459889, EPI_ISL_459890, EPI_ISL_459891, EPI_ISL_459892 | | | |
| see above | Kingston Health Sciences Center | Queen's Genomics Lab at Ongwanada (Q-GLO) | Sjaarda CP, Rustom N, Huang D, Perez-Patrigeon S, Hudson ML, Wong H,Guan H, Ayub M, Soares CN, Colautti R, Evans GA, Sheth P |
| EPI_ISL_459904 | Laboratoire National de Sante, Microbiology, Virology | Laboratoire National de Sante, Microbiology, Epidemiology and Microbial Genomics | Anke Wienecke-Baldacchino, Jessica Tapp, Guillaume Fournier, Tamir Abdelrahman, Trung Nguyen Nguyen, Catherine Ragimbeau |
| EPI_ISL_459956 | Institute for Medical Research, Infectious Disease Research Centre, National Institutes of Health, Ministry of Health Malaysia | Institute for Medical Research, Infectious Disease Research Centre, National Institutes of Health, Ministry of Health Malaysia | Suppiah J, Mohd-Zawawi Z, Kamel KA, Ellan K, Kalyanasundram J, Mohd-Zain R, Thayan R |

| | | | |
|---|---|---|---|
| EPI_ISL_459964 | Centogene AG | Centogene AG | Prof. Dr. Peter Bauer, Dr. Krishna Kumar Kandaswamy |
| EPI_ISL_459999, EPI_ISL_460000, EPI_ISL_460001, EPI_ISL_460002, EPI_ISL_460003, EPI_ISL_460004, EPI_ISL_460005, EPI_ISL_460006, EPI_ISL_460007, EPI_ISL_460009, EPI_ISL_460019, EPI_ISL_460029 | | | |
| see above | Michigan Department of Health and Human Services, Bureau of Laboratories | Michigan Department of Health and Human Services, Bureau of Laboratories | Blankenship HM, Riner D, Soehnlen MK |
| EPI_ISL_460096 | Molecular diagnostic laboratory of Federal Budget Institution of Science "Central Research Institute of Epidemiology" of The Federal Service on Customers' Rights Protection and Human Well-being Surveillance | Group of Genomics and Postgenomic Technologies of Central Research Institute of Epidemilology | Speranskaya AS, Kaptelova VV, Samoilov AE, Korneenko EV, Tivanova EV, Shipulina OY, Akimkin VG |
| EPI_ISL_460108, EPI_ISL_460110, EPI_ISL_460113, EPI_ISL_460115, EPI_ISL_460129, EPI_ISL_460134, EPI_ISL_460137, EPI_ISL_460142, EPI_ISL_460156, EPI_ISL_460161, EPI_ISL_460163, EPI_ISL_460185, EPI_ISL_460192, EPI_ISL_460194, EPI_ISL_460204, EPI_ISL_460205, EPI_ISL_460229, EPI_ISL_460242, EPI_ISL_460247, EPI_ISL_460258, EPI_ISL_460269, EPI_ISL_460270, EPI_ISL_460273, EPI_ISL_460276, EPI_ISL_460279, EPI_ISL_460283, EPI_ISL_460286, EPI_ISL_460288, EPI_ISL_460289, EPI_ISL_460295, EPI_ISL_460306, EPI_ISL_460314, EPI_ISL_460317, EPI_ISL_460322, EPI_ISL_460329, EPI_ISL_460337, EPI_ISL_460340, EPI_ISL_460343, EPI_ISL_460350, EPI_ISL_460362, EPI_ISL_460374, EPI_ISL_460376, EPI_ISL_460382, EPI_ISL_460393, EPI_ISL_460396, EPI_ISL_460400, EPI_ISL_460410, EPI_ISL_460425, EPI_ISL_460426, EPI_ISL_460427, EPI_ISL_460437, EPI_ISL_460445, EPI_ISL_460449, EPI_ISL_460459, EPI_ISL_460468, EPI_ISL_460471 | | | |
| see above | Massachusetts General Hospital | Infectious Disease Program, Broad Institute of Harvard and MIT | Lemieux,J.E., Siddle,K.J., Shaw,B., Adams,G., Pierce,V., Turbett,S., Anahtar,M., Branda,J., Slater,D., Harris,J., Lin,A.E., Gladden-Young,A., Lagerborg,K., Rudy,M., DeRuff,K., Carter,A., Normandin,E., Bauer,M., Reilly,S., Tomkins-Tinch,C., Loreth,C., Chaluvadi,S., Neumann,A., Cusick,C., Chapman,S.B., Gnirke,A., Flowers,K., Cerrato,F., Birren,B.W., Gallagher,G., Smole,S., Park,D.J., MacInnis,B.L., Ryan,E., LaRocque,R., Rosenberg,E., Sabeti,P.C. |
| EPI_ISL_460604, EPI_ISL_460605 | Molecular diagnostic laboratory of Federal Budget Institution of Science "Central Research Institute of Epidemiology" of The Federal Service on Customers' Rights Protection and Human Well-being Surveillance | Group of Genomics and Postgenomic Technologies of Central Research Institute of Epidemilology | Speranskaya AS, Kaptelova VV, Samoilov AE, Korneenko EV, Tivanova EV, Shipulina OY, Akimkin VG |
| EPI_ISL_460621, EPI_ISL_460622, EPI_ISL_460624, EPI_ISL_460628, EPI_ISL_460629, EPI_ISL_460630, EPI_ISL_460631, EPI_ISL_460632 | UW Virology Lab | UW Virology Lab | Pavitra Roychoudhury, Amin Addetia, Hong Xie, Lasata Shrestha, Truong Nguyen, Meei-Li Huang, Keith Jerome, Alexander Greninger |
| EPI_ISL_460664, EPI_ISL_460673, EPI_ISL_460687, EPI_ISL_460691, EPI_ISL_460692, EPI_ISL_460693, EPI_ISL_460694, EPI_ISL_460699, EPI_ISL_460704, EPI_ISL_460717, EPI_ISL_460718, EPI_ISL_460737, EPI_ISL_460738, EPI_ISL_460754, EPI_ISL_460755, EPI_ISL_460792, EPI_ISL_460793, EPI_ISL_460794, EPI_ISL_461011, EPI_ISL_461013, EPI_ISL_461018, EPI_ISL_461039, EPI_ISL_461061, EPI_ISL_461065, EPI_ISL_461066, EPI_ISL_461068, EPI_ISL_461123, EPI_ISL_461127, EPI_ISL_461161, EPI_ISL_461163, EPI_ISL_461251, EPI_ISL_461254, EPI_ISL_461264, EPI_ISL_461268, EPI_ISL_461282, EPI_ISL_461286 | | | |
| see above | Dutch COVID-19 response team | Erasmus Medical Center | Bas Oude Munnink, David Nieuwenhuijse, Reina Sikkema, Claudia Schapendonk, Irina Chestakova, Anne van der Linden, Theo Bestebroer, Stefan van Nieuwkoop, Mark Pronk, Pascal Lexmond, Corien Swaan, Manon Haverkate, Madelief Mollers, Mart Stein, Sandra Kengne Kamga Mobou, Jeroen van Kampen, Jolanda Voermans, Aura Timen, Corine GeurtsvanKessel, Annemiek van der Eijk, Richard Molenkamp, Marion Koopmans, on behalf of the Dutch national COVID-19 response team. |
| EPI_ISL_461403, EPI_ISL_461422, EPI_ISL_461424, EPI_ISL_461425, EPI_ISL_461430, EPI_ISL_461431, EPI_ISL_461432 | UW Virology Lab | UW Virology Lab | Pavitra Roychoudhury, Amin Addetia, Hong Xie, Lasata Shrestha, Truong Nguyen, Meei-Li Huang, Keith Jerome, Alexander Greninger |
| EPI_ISL_461705 | West of Scotland Specialist Virology Centre, NHSGGC / MRC-University of Glasgow Centre for Virus Research | COVID-19 Genomics UK (COG-UK) Consortium | Ana da Silva Filipe, Natasha Johnson, Kathy Smollett, Daniel Mair, Stephen Carmichael, Lily Tong, Jenna Nichols, Elihu Aranday-Cortes, Kirstyn Brunker, Yasmin Parr, Kyriaki Nomikou; Sarah McDonald, Marc Niebel, Patawee Asamaphan; Richard Orton, Joseph Hughes, Sreenu Vattipally, David L Robertson; Alasdair MacLean, Rory Gunson; Kathy Li, Natasha Jesudason, Rajiv Shah, James Shepherd, Antonia Ho, Emma Thomson |
| EPI_ISL_461706, EPI_ISL_461707, EPI_ISL_461708, EPI_ISL_461709, EPI_ISL_461710 | Virology Department, Royal Infirmary of Edinburgh, NHS Lothian / School of Biological Sciences, University of Edinburgh / Institute of Genetics and Molecular Medicine, University of Edinburgh | COVID-19 Genomics UK (COG-UK) Consortium | McHugh M, Dewar R, Rooke S, Gallagher M, Balcaza C, O'Toole Á, Scher E, Hill V, McCrone JT, Colquhoun R, Yu X, Jackson B, Rambaut A, Williams TC, Templeton K |
| EPI_ISL_461782, EPI_ISL_461784, EPI_ISL_461785, EPI_ISL_461786, EPI_ISL_461787, EPI_ISL_461788, EPI_ISL_461789, EPI_ISL_461790 | Regional Virus Laboratory, Belfast Health and Social Care Trust | COVID-19 Genomics UK (COG-UK) Consortium | Conall McCaughey, James McKenna, Tanya Curran, Susan Feeney, Alison Watt, Ciara Cox, Mairead Connor, Zoltan Molnar, David Simpson, Derek Fairley |
| EPI_ISL_462087 | Singapore General Hospital | Department of Microbiology | Nurdyana Abdul Rahman, Kun Lee Lim, Chenhao Li, Kian Sing Chan, Lynette Oon, Kern Rei Chng, Niranjan Nagarajan, Karrie Ko |
| EPI_ISL_462149 | Molecular diagnostic laboratory of Federal Budget Institution of Science "Central Research Institute of Epidemiology" of The Federal Service on Customers' Rights Protection and Human Well-being Surveillance | Group of Genomics and Postgenomic Technologies of Central Research Institute of Epidemilology | Speranskaya AS, Kaptelova VV, Samoilov AE, Korneenko EV, Sizova TV, Tivanova EV, Shipulina OY, Akimkin VG |
| EPI_ISL_462158, EPI_ISL_462159, EPI_ISL_462160, EPI_ISL_462161, EPI_ISL_462163, EPI_ISL_462164, EPI_ISL_462165, EPI_ISL_462166, EPI_ISL_462167, EPI_ISL_462168, EPI_ISL_462171, EPI_ISL_462172, EPI_ISL_462173, EPI_ISL_462174, EPI_ISL_462175, EPI_ISL_462182, EPI_ISL_462183, EPI_ISL_462184, EPI_ISL_462185, EPI_ISL_462186, EPI_ISL_462189, EPI_ISL_462192, EPI_ISL_462195, EPI_ISL_462196, EPI_ISL_462197, EPI_ISL_462198, EPI_ISL_462199, EPI_ISL_462200, EPI_ISL_462201, EPI_ISL_462202, EPI_ISL_462203, EPI_ISL_462205, EPI_ISL_462206, EPI_ISL_462207, EPI_ISL_462208, EPI_ISL_462211, EPI_ISL_462212, EPI_ISL_462217, EPI_ISL_462219, EPI_ISL_462221, EPI_ISL_462223, EPI_ISL_462224, EPI_ISL_462238, EPI_ISL_462239, EPI_ISL_462240, EPI_ISL_462241, EPI_ISL_462242, EPI_ISL_462243, EPI_ISL_462244, EPI_ISL_462246 | | | |
| see above | KU Leuven, Rega Institute, Clinical and Epidemiological Virology | KU Leuven, Rega Institute, Clinical and Epidemiological Virology | Tony Wawina-Bokalanga, Bert Vanmechelen, Joan Marti-Carerras, Piet Maes |
| EPI_ISL_462336, EPI_ISL_462341, EPI_ISL_462342, EPI_ISL_462346, EPI_ISL_462349 | National Public Health Laboratory, National Centre for Infectious Diseases | National Public Health Laboratory, National Centre for Infectious Diseases | Mak TM, Octavia S, Chavatte JM, Cui L, Lin RTP |
| EPI_ISL_462468, EPI_ISL_462469, EPI_ISL_462471, EPI_ISL_462473, EPI_ISL_462474, EPI_ISL_462475, EPI_ISL_462476 | Clinical Center, University of Sarajevo | Charite Universitatsmedizin Berlin, Institute of Virology | Victor M Corman, Jorn Beheim-Schwarzbach, Barbara Muehlemann, Talitha Veith, Julia Schneider, Terry Jones, Amela Dedeic-Ljubovic, Irma Salimovic-Besic, Suzana Arapcic, Almedina Hadzihasanovic-Moro, Selma Mutevelic, Christian Drosten |
| EPI_ISL_462913, EPI_ISL_462954, EPI_ISL_462958, EPI_ISL_462959, EPI_ISL_462961, EPI_ISL_462962, EPI_ISL_462963, EPI_ISL_462964, EPI_ISL_462965, EPI_ISL_462966, EPI_ISL_462967, EPI_ISL_462968, EPI_ISL_462969, EPI_ISL_462970, EPI_ISL_462971, EPI_ISL_462972, EPI_ISL_462973, EPI_ISL_462974, EPI_ISL_462975, EPI_ISL_462976, EPI_ISL_462977, EPI_ISL_462978, EPI_ISL_462979, EPI_ISL_462980, EPI_ISL_462981, EPI_ISL_462982, EPI_ISL_462983, EPI_ISL_462984, EPI_ISL_462985, EPI_ISL_462986, EPI_ISL_462987, EPI_ISL_462988, EPI_ISL_462989 | | | |
| see above | Wyoming Public Health Laboratory | Center for Global Health, University of New Mexico Health Sciences Center | Daryl Domman, Kurt Schwalm, Rob Christensen, Wanda Manley, Cari Sloma, Noah Hull, Darrell Dinwiddie |
| EPI_ISL_463002, EPI_ISL_463003, EPI_ISL_463005 | unknown | Clinical virology | Fares,W., Triki,H. |
| EPI_ISL_463138, EPI_ISL_463139, EPI_ISL_463140, EPI_ISL_463141, EPI_ISL_463142, EPI_ISL_463143, EPI_ISL_463144, EPI_ISL_463145, EPI_ISL_463146, EPI_ISL_463147, EPI_ISL_463148, EPI_ISL_463149, EPI_ISL_463150 | | | |
| see above | Yale Clinical Virology Laboratory | Grubaugh Lab - Yale School of Public Health | Joseph Fauver, Tara Alpert, Anderson Brito, Anne Wyllie, Chantal Vogels, Mary Petrone, Cole Jensen, Chaney Kalinich, Isabel Ott, Arnau Casanovas, Catherine Muenker, Adam Moore, Alice Lu, Maria Tokuyama, Patrick Wong, Peiwen Lu, Saad Omer, Richard Martinello, Allison Nelson, Shelli Farhadian, Akiko Iwasaki, Charlese Dela Cruz, Albert Ko, Nathan Grubaugh |
| EPI_ISL_463740 | Mohammed Bin Rashid University of Medicine and Health Sciences | Al Jalila Genomics Center | Ahmad Abou Tayoun, Tom Loney, Hamda Khansaheb, Sathishkumar Ramaswamy, Divinlal Harilal, Zulfa Omar Deesi, Rupa Murthy Varghese, Hanan Al Suwaidi, Abdulmajeed Alkhaja, Mohammed Uddin, Rifat Hamoudi, Rabih Halwani, Abiola Catherine Senok, Qutayba Hamid, Norbert Nowotny, Alawi Alsheikh-Ali |

| | | | |
|---|---|---|---|
| EPI_ISL_463976, EPI_ISL_463978, EPI_ISL_463979, EPI_ISL_463980, EPI_ISL_463981, EPI_ISL_463982, EPI_ISL_463983, EPI_ISL_463985 | Toronto Invasive Bacterial Diseases Network | McMaster University | Allison McGeer, Patryk Aftanas, Angel Li, Kuganya Nirmalarajah, Samira Mubareka, Andrew G. McArthur |
| EPI_ISL_464002, EPI_ISL_464003, EPI_ISL_464006, EPI_ISL_464017, EPI_ISL_464025, EPI_ISL_464027, EPI_ISL_464039, EPI_ISL_464049, EPI_ISL_464053 | Unity Health Toronto | Ontario Institute for Cancer Research | Ramzi Fattouh,Larissa M. Matukas,Mark Downing,Annette Gower,Karel Boissinot,Samira Mubareka,TIBDN,Ilinca Lungu,Bernard Lam,Jeremy Johns,Paul Krzyzanowski,Richard de Borja,Philip Zuzarte,Jared Simpson |
| EPI_ISL_464113, EPI_ISL_464114, EPI_ISL_464141, EPI_ISL_464142, EPI_ISL_464143 | National Health Laboratory Service (NHLS), Tygerberg | Division of Medical Virology, Stellenbosch University and National Health Laboratory Service (NHLS) | Susan Engelbrecht, Kayla Delaney, Bronwyn Kleinhans, Houriiyah Tegally, Eduan Wilkindon, Gert van Zyl, Wolfgang Preiser, Tulio de Oliveira |
| EPI_ISL_465679, EPI_ISL_465680 | Hôpital du Suroît | Laboratoire de santé publique du Québec | Sandrine Moreira, Ioannis Ragoussis, Guillaume Bourque, Jesse Shapiro, Mark Lathrop and Michel Roger on behalf of the CoVSeQ research group (http://covseq.ca/researchgroup) |
| EPI_ISL_465681, EPI_ISL_465682 | Hôpital Charles-LeMoyne | Laboratoire de santé publique du Québec | Sandrine Moreira, Ioannis Ragoussis, Guillaume Bourque, Jesse Shapiro, Mark Lathrop and Michel Roger on behalf of the CoVSeQ research group (http://covseq.ca/researchgroup) |
| EPI_ISL_465685 | CSSS Haut-Richelieu/Rouville (Hôpital) | Laboratoire de santé publique du Québec | Sandrine Moreira, Ioannis Ragoussis, Guillaume Bourque, Jesse Shapiro, Mark Lathrop and Michel Roger on behalf of the CoVSeQ research group (http://covseq.ca/researchgroup) |
| EPI_ISL_465686 | Hôpital de Hull | Laboratoire de santé publique du Québec | Sandrine Moreira, Ioannis Ragoussis, Guillaume Bourque, Jesse Shapiro, Mark Lathrop and Michel Roger on behalf of the CoVSeQ research group (http://covseq.ca/researchgroup) |
| EPI_ISL_465687 | CSSS Haut-Richelieu/Rouville (Hôpital) | Laboratoire de santé publique du Québec | Sandrine Moreira, Ioannis Ragoussis, Guillaume Bourque, Jesse Shapiro, Mark Lathrop and Michel Roger on behalf of the CoVSeQ research group (http://covseq.ca/researchgroup) |
| EPI_ISL_465688, EPI_ISL_465689, EPI_ISL_465690 | Centre hospitalier Anna-Laberge | Laboratoire de santé publique du Québec | Sandrine Moreira, Ioannis Ragoussis, Guillaume Bourque, Jesse Shapiro, Mark Lathrop and Michel Roger on behalf of the CoVSeQ research group (http://covseq.ca/researchgroup) |
| EPI_ISL_465691 | Hôpital de Gatineau | Laboratoire de santé publique du Québec | Sandrine Moreira, Ioannis Ragoussis, Guillaume Bourque, Jesse Shapiro, Mark Lathrop and Michel Roger on behalf of the CoVSeQ research group (http://covseq.ca/researchgroup) |
| EPI_ISL_465692 | CSSS Haut-Richelieu/Rouville (Hôpital) | Laboratoire de santé publique du Québec | Sandrine Moreira, Ioannis Ragoussis, Guillaume Bourque, Jesse Shapiro, Mark Lathrop and Michel Roger on behalf of the CoVSeQ research group (http://covseq.ca/researchgroup) |
| EPI_ISL_465693 | Hôpital du Suroît | Laboratoire de santé publique du Québec | Sandrine Moreira, Ioannis Ragoussis, Guillaume Bourque, Jesse Shapiro, Mark Lathrop and Michel Roger on behalf of the CoVSeQ research group (http://covseq.ca/researchgroup) |
| EPI_ISL_465694 | Hôpital Charles-LeMoyne | Laboratoire de santé publique du Québec | Sandrine Moreira, Ioannis Ragoussis, Guillaume Bourque, Jesse Shapiro, Mark Lathrop and Michel Roger on behalf of the CoVSeQ research group (http://covseq.ca/researchgroup) |
| EPI_ISL_465695, EPI_ISL_465696 | Hôpital Pierre-Boucher | Laboratoire de santé publique du Québec | Sandrine Moreira, Ioannis Ragoussis, Guillaume Bourque, Jesse Shapiro, Mark Lathrop and Michel Roger on behalf of the CoVSeQ research group (http://covseq.ca/researchgroup) |
| EPI_ISL_465698, EPI_ISL_465699 | Hôpital Charles-LeMoyne | Laboratoire de santé publique du Québec | Sandrine Moreira, Ioannis Ragoussis, Guillaume Bourque, Jesse Shapiro, Mark Lathrop and Michel Roger on behalf of the CoVSeQ research group (http://covseq.ca/researchgroup) |
| EPI_ISL_465701 | Hôpital du Suroît | Laboratoire de santé publique du Québec | Sandrine Moreira, Ioannis Ragoussis, Guillaume Bourque, Jesse Shapiro, Mark Lathrop and Michel Roger on behalf of the CoVSeQ research group (http://covseq.ca/researchgroup) |
| EPI_ISL_465702, EPI_ISL_465703 | Hôpital Pierre-Boucher | Laboratoire de santé publique du Québec | Sandrine Moreira, Ioannis Ragoussis, Guillaume Bourque, Jesse Shapiro, Mark Lathrop and Michel Roger on behalf of the CoVSeQ research group (http://covseq.ca/researchgroup) |
| EPI_ISL_465839, EPI_ISL_465840, EPI_ISL_465844, EPI_ISL_465845, EPI_ISL_465846, EPI_ISL_465847, EPI_ISL_465848, EPI_ISL_465849, EPI_ISL_465851, EPI_ISL_465852, EPI_ISL_465853, EPI_ISL_465854, EPI_ISL_465855, EPI_ISL_465856, EPI_ISL_465857, EPI_ISL_465858, EPI_ISL_465859, EPI_ISL_465860, EPI_ISL_465861, EPI_ISL_465862, EPI_ISL_465863, EPI_ISL_465864, EPI_ISL_465865, EPI_ISL_465866, EPI_ISL_465867, EPI_ISL_465868, EPI_ISL_465870, EPI_ISL_465871, EPI_ISL_465872, EPI_ISL_465873, EPI_ISL_465874, EPI_ISL_465875, EPI_ISL_465876, EPI_ISL_465877, EPI_ISL_465878, EPI_ISL_465879, EPI_ISL_465880, EPI_ISL_465881, EPI_ISL_465882, EPI_ISL_465883, EPI_ISL_465884, EPI_ISL_465885, EPI_ISL_465886, EPI_ISL_465887, EPI_ISL_465888, EPI_ISL_465889, EPI_ISL_465890, EPI_ISL_465891, EPI_ISL_465892, EPI_ISL_465893, EPI_ISL_465895, EPI_ISL_465896, EPI_ISL_465897, EPI_ISL_465898, EPI_ISL_465899, EPI_ISL_465900, EPI_ISL_465901, EPI_ISL_465902, EPI_ISL_465903, EPI_ISL_465904, EPI_ISL_465905, EPI_ISL_465906, EPI_ISL_465908, EPI_ISL_465909, EPI_ISL_465910, EPI_ISL_465911, EPI_ISL_465912, EPI_ISL_465913, EPI_ISL_465914, EPI_ISL_465915, EPI_ISL_465916, EPI_ISL_465917, EPI_ISL_465918, EPI_ISL_465920, EPI_ISL_465921, EPI_ISL_465923, EPI_ISL_465924, EPI_ISL_465925, EPI_ISL_465926, EPI_ISL_465927, EPI_ISL_465928, EPI_ISL_465929, EPI_ISL_465930, EPI_ISL_465931, EPI_ISL_465932, EPI_ISL_465933, EPI_ISL_465934, EPI_ISL_465935, EPI_ISL_465936, EPI_ISL_465937, EPI_ISL_465938, EPI_ISL_465939, EPI_ISL_465940, EPI_ISL_465941, EPI_ISL_465942, EPI_ISL_465943, EPI_ISL_465944, EPI_ISL_465945, EPI_ISL_465946, EPI_ISL_465947, EPI_ISL_465948, EPI_ISL_465949, EPI_ISL_465950, EPI_ISL_465951, EPI_ISL_465952, EPI_ISL_465953, EPI_ISL_465954, EPI_ISL_465955, EPI_ISL_465956, EPI_ISL_465957, EPI_ISL_465958, EPI_ISL_465959, EPI_ISL_465960, EPI_ISL_465961, EPI_ISL_465962, EPI_ISL_465963, EPI_ISL_465964, EPI_ISL_465965, EPI_ISL_465966, EPI_ISL_465967, EPI_ISL_465968, EPI_ISL_465969, EPI_ISL_465970, EPI_ISL_465972, EPI_ISL_465974, EPI_ISL_465976, EPI_ISL_465977, EPI_ISL_465978, EPI_ISL_465979, EPI_ISL_465980, EPI_ISL_465981, EPI_ISL_465984, EPI_ISL_465985, EPI_ISL_465986, EPI_ISL_465987, EPI_ISL_465989, EPI_ISL_465990, EPI_ISL_465991, EPI_ISL_465992, EPI_ISL_465993, EPI_ISL_465994, EPI_ISL_465995, EPI_ISL_466005, EPI_ISL_466006, EPI_ISL_466042, EPI_ISL_466046, EPI_ISL_466061, EPI_ISL_466105, EPI_ISL_466120, EPI_ISL_466124, EPI_ISL_466125, EPI_ISL_466126, EPI_ISL_466138, EPI_ISL_466141, EPI_ISL_466142, EPI_ISL_466144, EPI_ISL_466145 | | | |
| see above | Respiratory Virus Unit, Microbiology Services Colindale, Public Health England | Respiratory Virus Unit, Microbiology Services Colindale, Public Health England | PHE Covid Sequencing Team |
| EPI_ISL_467061 | Hospital Universitario Virgen de las Nieves de Granada-SAS | SeqCOVID-SPAIN consortium/IBV(CSIC) | Mercedes Pérez Ruiz, Sara Sanbonmatsu Gámez, Irene Pedrosa Corral, José M. Navarro-Marí and SeqCOVID-SPAIN consortium |
| EPI_ISL_467432 | AMPATH-DBN | KRISP, KZN Research Innovation and Sequencing Platform | Giandhari J, Pillay S, Lessells R, Chimukangara B, Mdlalose K, York D, Khan S, Tegally H, Wilkinson E, de Oliveira T |
| EPI_ISL_467441, EPI_ISL_467442 | NHLS-IALCH | KRISP, KZN Research Innovation and Sequencing Platform | Giandhari J, Pillay S, Lessells R, Chimukangara B, Mdlalose K, York D, Khan S, Tegally H, Wilkinson E, de Oliveira T |
| EPI_ISL_467446 | Molecular Diagnostics Services (MDS) | KRISP, KZN Research Innovation and Sequencing Platform | Giandhari J, Pillay S, Lessells R, Chimukangara B, Mdlalose K, York D, Khan S, Tegally H, Wilkinson E, de Oliveira T |
| EPI_ISL_467538 | New Mexico Department of Health Scientific Laboratory Division | Center for Global Health, University of New Mexico Health Sciences Center | Daryl Domman, Kurt Schwalm, Twila Kunde, Joseph Hicks, Michael Edwards, Darrell Dinwiddie |
| EPI_ISL_467775 | Molecular diagnostic laboratory of Federal Budget Institution of Science "Central Research Institute of Epidemiology" of The Federal Service on Customers' Rights Protection and Human Well-being Surveillance | Group of Genomics and Postgenomic Technologies of Central Research Institute of Epidemilology | Speranskaya AS, Kaptelova VV, Samoilov AE, Korneenko EV, Sizova TV, Tivanova EV, Shipulina OY, Akimkin VG |
| EPI_ISL_467809 | Cedars-Sinai Medical Center, Department of Pathology & Laboratory Medicine, Molecular Pathology Laboratory | Cedars-Sinai Medical Center, Molecular Pathology Laboratory of Department of Pathology & Laboratory Medicine and Genomic Core | Wenjuan Zhang, John Paul Govindavari, Brian Davis, Stephanie Chen, Jong Taek Kim, Jianbo Song, Jean Lopategui, Jasmine T Plummer, Eric Vail |
| EPI_ISL_467953, EPI_ISL_467966 | San Diego County Public Health Laboratory | Andersen lab at Scripps Research | SEARCH Alliance San Diego with Tracy Basler, Jovan Shephard, Brett Austin |
| EPI_ISL_467967 | Scripps Medical Laboratory | Andersen lab at Scripps Research | SEARCH Alliance San Diego with Tracy Basler, Jovan Shephard, Brett Austin |
| EPI_ISL_467971, EPI_ISL_467975, EPI_ISL_467979, EPI_ISL_467983 | San Diego County Public Health Laboratory | Andersen lab at Scripps Research | SEARCH Alliance San Diego with Tracy Basler, Jovan Shephard, Brett Austin |
| EPI_ISL_467997, EPI_ISL_468004, EPI_ISL_468008, EPI_ISL_468013, EPI_ISL_468031, EPI_ISL_468036 | SA Pathology | SA Pathology | Lex Leong, Chuan Kok Lim, Mark Turra, Ivan Bastian, Geoff Higgins |
| EPI_ISL_468145 | [Romania, Bucharest] National Institute for Infectious Diseases "Prof. Dr. Matei Bal" | [Romania, Bucharest] National Institute for Infectious Diseases "Prof. Dr. Matei Bal" | Leontina Banica, Marius Cotic, Corina Casangiu, Marius Surleac, Simona Paraschiv |

| | | | |
|---|---|---|---|
| EPI_ISL_468408, EPI_ISL_468409, EPI_ISL_468410, EPI_ISL_468411, EPI_ISL_468412, EPI_ISL_468413 | County of San Luis Obispo Public Health Laboratory | Chan-Zuckerberg Biohub | CZB Cliahub Consortium |
| EPI_ISL_468446, EPI_ISL_468447, EPI_ISL_468449, EPI_ISL_468452 | Humboldt County Public Health Laboratory | Chan-Zuckerberg Biohub | CZB Cliahub Consortium |
| EPI_ISL_468496, EPI_ISL_468497 | Ventura County Public Health Lab | Chan-Zuckerberg Biohub | CZB Cliahub Consortium |
| EPI_ISL_468523, EPI_ISL_468524, EPI_ISL_468525, EPI_ISL_468526, EPI_ISL_468527, EPI_ISL_468559 | San Joaquin County Public Health Lab | Chan-Zuckerberg Biohub | CZB Cliahub Consortium |
| EPI_ISL_468615, EPI_ISL_468616, EPI_ISL_468617, EPI_ISL_468618, EPI_ISL_468619 | Contra Costa Public Health Lab | Chan-Zuckerberg Biohub | CZB Cliahub Consortium |
| EPI_ISL_468766, EPI_ISL_468770, EPI_ISL_468773, EPI_ISL_468781, EPI_ISL_468782, EPI_ISL_468783, EPI_ISL_468790, EPI_ISL_468793, EPI_ISL_468794, EPI_ISL_468799, EPI_ISL_468800, EPI_ISL_468802, EPI_ISL_468807, EPI_ISL_468809, EPI_ISL_468813, EPI_ISL_468820, EPI_ISL_468821, EPI_ISL_468822, EPI_ISL_468825, EPI_ISL_468827, EPI_ISL_468828, EPI_ISL_468837, EPI_ISL_468839, EPI_ISL_468842, EPI_ISL_468843, EPI_ISL_468849 | | | |
| see above | Servicio de Microbiología, Hospital Miguel Servet, Zaragoza | SeqCOVID-SPAIN consortium/IBV(CSIC) | Antonio Rezusta López, Alexander Tristancho Baró, Ana Milagro, Yolanda Gracia Grataloup, Nieves Martínez Cameo and SeqCOVID-SPAIN consortium |
| EPI_ISL_468968, EPI_ISL_468969, EPI_ISL_468992, EPI_ISL_468998, EPI_ISL_469003 | Servicio de Microbiología, Hospital Universitario Son Espases | SeqCOVID-SPAIN consortium/IBV(CSIC) | Carla López-Causapé, Jordi Reina, Antonio Oliver and SeqCOVID-SPAIN consortium |
| EPI_ISL_469049, EPI_ISL_469052, EPI_ISL_469053 | LNR National Reference Laboratory, Mohammed VI University of Health Sciences | Medical Biotechnology Laboratory, Rabat Medical and Pharmacy School, Mohammed The Vth University in Rabat | Meriem LAAMARTI, Souad KARTTI, Rokaia LAAMRTI , M.W. CHEMAO-ELFIHRI, Loubna ALLAM, Mouna OUADGHIRI, Imane SMYEJ, Jalila RAHOUI, Houda BENRAHMA, Jalil El Atar, Idrissa Diawara, Rachid EL JAOUDI, Laila SBABOU, Chakib NEJJARI, Saaid AMZAZI, Rachid MENTAG, Lahcen BELYAMANI and Azeddine IBRAHIMI |
| EPI_ISL_469106 | National Public Health Laboratory, National Centre for Infectious Diseases | National Public Health Laboratory, National Centre for Infectious Diseases | Mak TM, Octavia S, Chavatte JM, Cui L, Lin RTP |
| EPI_ISL_469276, EPI_ISL_469280, EPI_ISL_469281 | Mohammed Bin Rashid University of Medicine and Health Sciences | Al Jalila Genomics Center | Ahmad Abou Tayoun, Tom Loney, Hamda Khansaheb, Sathishkumar Ramaswamy, Divinlal Harilal, Zulfa Omar Deesi, Rupa Murthy Varghese, Hanan Al Suwaidi, Abdulmajeed Alkhaja, Mohammed Uddin, Rifat Hamoudi, Rabih Halwani, Abiola Catherine Senok, Qutayba Hamid, Norbert Nowotny, Alawi Alsheikh-Ali |
| EPI_ISL_469631, EPI_ISL_469658, EPI_ISL_469749 | PHE South West Regional Laboratory, National Infection Service | Wellcome Sanger Institute for the COVID-19 Genomics UK (COG-UK) consortium | Stephanie Hutchings, Hannah Pymont, Dr Peter Muir, Barry Vipond, Rich Hopes; and Alex Alderton, Roberto Amato, Sonia Goncalves, Ewan Harrison, David K. Jackson, Ian Johnston, Dominic Kwiatkowski, Cordelia Langford, John Sillitoe on behalf of the Wellcome Sanger Institute COVID-19 Surveillance Team (http://www.sanger.ac.uk/covid-team) |
| EPI_ISL_469930, EPI_ISL_469932, EPI_ISL_469933, EPI_ISL_469934, EPI_ISL_469935, EPI_ISL_469936, EPI_ISL_469937, EPI_ISL_469938, EPI_ISL_469940, EPI_ISL_469941, EPI_ISL_469942, EPI_ISL_469943, EPI_ISL_469945, EPI_ISL_469946, EPI_ISL_469947, EPI_ISL_469948, EPI_ISL_469949, EPI_ISL_469951, EPI_ISL_469952, EPI_ISL_469953, EPI_ISL_469954, EPI_ISL_469956, EPI_ISL_469957, EPI_ISL_469958, EPI_ISL_469959, EPI_ISL_469960, EPI_ISL_469961, EPI_ISL_469963, EPI_ISL_469964, EPI_ISL_469965, EPI_ISL_469966, EPI_ISL_469967, EPI_ISL_469968, EPI_ISL_469969, EPI_ISL_469970, EPI_ISL_469971, EPI_ISL_469972, EPI_ISL_469973, EPI_ISL_469974, EPI_ISL_469975, EPI_ISL_469976, EPI_ISL_469977, EPI_ISL_469979, EPI_ISL_469980, EPI_ISL_469981, EPI_ISL_469982, EPI_ISL_469983, EPI_ISL_469984, EPI_ISL_469985, EPI_ISL_469987, EPI_ISL_469988, EPI_ISL_469989, EPI_ISL_469990, EPI_ISL_469991, EPI_ISL_469992, EPI_ISL_469993, EPI_ISL_469994, EPI_ISL_469995, EPI_ISL_469996, EPI_ISL_470000, EPI_ISL_470001, EPI_ISL_470002, EPI_ISL_470003, EPI_ISL_470004, EPI_ISL_470005, EPI_ISL_470006, EPI_ISL_470007, EPI_ISL_470008, EPI_ISL_470009, EPI_ISL_470011, EPI_ISL_470012 | | | |
| see above | NHSGGC West of Scotland Specialist Virology Centre / MRC-University of Glasgow Centre for Virus Research | Wellcome Sanger Institute for the COVID-19 Genomics UK (COG-UK) consortium | Ana da Silva Filipe, Natasha Johnson, Kathy Smollett, Daniel Mair, Stephen Carmichael, Lily Tong, Jenna Nichols, Elihu Aranday-Cortes, Kirstyn Brunker, Yasmin Parr, Kyriaki Nomikou; Sarah McDonald, Marc Niebel, Patawee Asamaphan; Richard Orton, Joseph Hughes, Sreenu Vattipally, David L Robertson; Alasdair MacLean, Rory Gunson; Kathy Li, Natasha Jesudason, Rajiv Shah, James Shepherd, Antonia Ho, Alice Broos, Emma Thomson and Alex Alderton, Roberto Amato, Sonia Goncalves, Ewan Harrison, David K. Jackson, Ian Johnston, Dominic Kwiatkowski, Cordelia Langford, John Sillitoe on behalf of the Wellcome Sanger Institute COVID-19 Surveillance Team (http://www.sanger.ac.uk/covid-team) |
| EPI_ISL_470539 | Molecular diagnostic laboratory of Federal Budget Institution of Science "Central Research Institute of Epidemiology" of The Federal Service on Customers' Rights Protection and Human Well-being Surveillance | Group of Genomics and Postgenomic Technologies of Central Research Institute of Epidemilology | Speranskaya AS, Kaptelova VV, Samoilov AE, Korneenko EV, Sizova TV, Tivanova EV, Shipulina OY, Akimkin VG |
| EPI_ISL_470619, EPI_ISL_470620 | Laboratorio de Virologia Molecular / UFRJ | Bioinformatics Laboratory / LNCC | Alexandra Gerber, Ana Paula Guimarães, Luiz Gonzaga Paula de Almeida, Ronaldo da Silva Francisco Junior, Mariane Talon, Filipe Romero, Átila Duque Rossi, Terezinha Marta Pereira, working group UFRJ, Jaqueline Goes de Jesus, Ingra Morales Claro, Ester Cerdeira Sabino, Nuno Rodrigues Faria, CADDE-group, Laboratorio Hermes Pardini, Laboratorio Simile, working group UFMG, Amilcar Tanuri, Carolina Voloch, Renato Santana Aguiar e Ana Tereza Vasconcelos |
| EPI_ISL_470841, EPI_ISL_470850, EPI_ISL_470863 | PathWest Laboratory Medicine WA | PathWest Laboratory Medicine WA | Chisha Sikazwe, Jurissa Lang, Avram Levy, David Smith and David Speers |
| EPI_ISL_471158 | MRCG at LSHTM Genomics lab | MRCG at LSHTM Genomics lab | Sesay et al |
| EPI_ISL_471180, EPI_ISL_471181, EPI_ISL_471199, EPI_ISL_471201, EPI_ISL_471203, EPI_ISL_471210, EPI_ISL_471221, EPI_ISL_471233, EPI_ISL_471260, EPI_ISL_471261, EPI_ISL_471262, EPI_ISL_471263, EPI_ISL_471264, EPI_ISL_471265 | | | |
| see above | Wisconsin State Laboratory of Hygiene Communicable Disease Division | Wisconsin State Laboratory of Hygiene Communicable Disease Division | Kelsey R. Florek, Abigail C. Shockey |
| EPI_ISL_471554 | Hospital Bosque da Saúde | Instituto Adolfo Lutz, Interdiciplinary Procedures Center, Strategic Laboratory | Claudio Tavares Sacchi, Claudia Regina Gonçalves, Erica Valessa Ramos Gomes |
| EPI_ISL_471958, EPI_ISL_471961, EPI_ISL_471962, EPI_ISL_471964, EPI_ISL_471965, EPI_ISL_471966, EPI_ISL_471967, EPI_ISL_471968 | University of Exeter | COVID-19 Genomics UK (COG-UK) Consortium | Ben Temperton,Aaron Jeffries,Michelle Michelsen,Joanna Warwick-Dugdale,Audrey Farbos,Robyn Manley,Stephen Michell,Jane Masoli |
| EPI_ISL_472145, EPI_ISL_472147, EPI_ISL_472148, EPI_ISL_472149, EPI_ISL_472150, EPI_ISL_472151, EPI_ISL_472152, EPI_ISL_472153, EPI_ISL_472154 | Regional Virus Laboratory, Belfast Health and Social Care Trust | COVID-19 Genomics UK (COG-UK) Consortium | Conall McCaughey, James McKenna, Tanya Curran, Susan Feeney, Alison Watt, Ciara Cox, Mairead Connor, Zoltan Molnar, David Simpson, Derek Fairley |
| EPI_ISL_472204, EPI_ISL_472205, EPI_ISL_472206, EPI_ISL_472207, EPI_ISL_472208, EPI_ISL_472209, EPI_ISL_472210, EPI_ISL_472211, EPI_ISL_472212, EPI_ISL_472213, EPI_ISL_472214, EPI_ISL_472215, EPI_ISL_472216, EPI_ISL_472217, EPI_ISL_472218, EPI_ISL_472219, EPI_ISL_472220, EPI_ISL_472221, EPI_ISL_472222, EPI_ISL_472223, EPI_ISL_472224, EPI_ISL_472225, EPI_ISL_472226, EPI_ISL_472227, EPI_ISL_472228, EPI_ISL_472229, EPI_ISL_472230, EPI_ISL_472231, EPI_ISL_472232, EPI_ISL_472233, EPI_ISL_472234, EPI_ISL_472235, EPI_ISL_472236, EPI_ISL_472237, EPI_ISL_472238, EPI_ISL_472240, EPI_ISL_472241, EPI_ISL_472242, EPI_ISL_472243, EPI_ISL_472244, EPI_ISL_472245 | | | |
| see above | Northumbria University / South Tees Hospitals NHS Foundation Trust / North Cumbria Integrated Care NHS Foundation Trust / North Tees and Hartlepool NHS Foundation Trust / Newcastle Hospitals NHS Foundation Trust | COVID-19 Genomics UK (COG-UK) Consortium | Darren L Smith,Andrew Nelson,Matthew Bashton,Greg R Young,Joshua Loh,John Allan,Mohammad A Tariq,Giles S Holt,Gary Black,Wen C Yew,Lynn Dover,Paul Baker,Steve Liggett,Sarah Essex,Jane Greenaway,Debra Padgett,Clive Graham,Garren Scott,Edward Barton,Emma Swindells,Brendan Payne,Jennifer Collins,Yusri Taha,Gary Eltringham |
| EPI_ISL_472444, EPI_ISL_472449, EPI_ISL_472464, EPI_ISL_472479, EPI_ISL_472486, EPI_ISL_472487, EPI_ISL_472493, EPI_ISL_472495, EPI_ISL_472509, EPI_ISL_472519, EPI_ISL_472524, EPI_ISL_472528, EPI_ISL_472538, EPI_ISL_472547, EPI_ISL_472559, EPI_ISL_472579, EPI_ISL_472581, EPI_ISL_472595, EPI_ISL_472612, EPI_ISL_472646, EPI_ISL_472648, EPI_ISL_472657, EPI_ISL_472658, EPI_ISL_472674, EPI_ISL_472677, EPI_ISL_472679, EPI_ISL_472693, EPI_ISL_472698, EPI_ISL_472706, EPI_ISL_472726 | | | |
| see above | Wales Specialist Virology Centre Sequencing lab: Pathogen Genomics Unit | COVID-19 Genomics UK (COG-UK) Consortium | Catherine Moore, Johnathan Evans, Laura Gifford, Malorie Perry, Simon Cottrell, Angela Marchbank, Alec Birchley, Alexander Adams, Amy Gaskin, Bree Gatica-Wilcox, Jason Coombes, Joel Southgate, Lauren Gilbert, Lee Graham, Nicole Pacchiarini, Sara Kumziene-Summerhayes, Sarah Taylor, Sophie |

| | | | |
|---|---|---|---|
| | | | Jones, Sara Rey, Matthew Bull, Joanne Watkins, Sally Corden, Tom Connor |
| EPI_ISL_473782 | West of Scotland Specialist Virology Centre, NHSGGC / MRC-University of Glasgow Centre for Virus Research | COVID-19 Genomics UK (COG-UK) Consortium | Ana da Silva Filipe, Natasha Johnson, Kathy Smollett, Daniel Mair, Stephen Carmichael, Lily Tong, Jenna Nichols, Elihu Aranday-Cortes, Kirstyn Brunker, Yasmin Parr, Alice Broos, Kyriaki Nomikou; Sarah McDonald, Marc Niebel, Patawee Asamaphan; Richard Orton, Joseph Hughes, Sreenu Vattipally, David L Robertson; Alasdair MacLean, Rory Gunson; Kathy Li, Natasha Jesudason, Rajiv Shah, James Shepherd, Antonia Ho, Emma Thomson |
| EPI_ISL_474305 | Wales Specialist Virology Centre Sequencing lab: Pathogen Genomics Unit | COVID-19 Genomics UK (COG-UK) Consortium | Catherine Moore, Johnathan Evans, Laura Gifford, Malorie Perry, Simon Cottrell, Angela Marchbank, Alec Birchley, Alexander Adams, Amy Gaskin, Bree Gatica-Wilcox, Jason Coombes, Joel Southgate, Lauren Gilbert, Lee Graham, Nicole Pacchiarini, Sara Kumziene-Summerhayes, Sarah Taylor, Sophie Jones, Sara Rey, Matthew Bull, Joanne Watkins, Sally Corden, Tom Connor |
| EPI_ISL_474855, EPI_ISL_474877, EPI_ISL_474878, EPI_ISL_474879, EPI_ISL_474880, EPI_ISL_474881, EPI_ISL_474882, EPI_ISL_474883, EPI_ISL_474886, EPI_ISL_474928, EPI_ISL_474929, EPI_ISL_474930, EPI_ISL_474957 | | | |
| see above | Hospital Universitario Virgen de las Nieves de Granada-SAS | SeqCOVID-SPAIN consortium/IBV(CSIC) | Mercedes Pérez Ruiz, Sara Sanbonmatsu Gámez, Irene Pedrosa Corral, José M. Navarro-Marí and SeqCOVID-SPAIN consortium |
| EPI_ISL_475108, EPI_ISL_475109 | Skovde/Unilabs | The Public Health Agency of Sweden | Oskar Karlsson Lindsjo, Maria Lind Karlberg, Mattias Haukland, Reza Advani, Olov Svartstrom, Anna-Malin Linde, Sandra Broddesson, Petra Edquist, Shamam Muradrasoli, Anna Risberg, Karin Tegmark-Wisell |
| EPI_ISL_475110 | Gavle klinisk mikrobiologi | The Public Health Agency of Sweden | Oskar Karlsson Lindsjo, Maria Lind Karlberg, Mattias Haukland, Reza Advani, Olov Svartstrom, Anna-Malin Linde, Sandra Broddesson, Petra Edquist, Shamam Muradrasoli, Anna Risberg, Karin Tegmark-Wisell |
| EPI_ISL_475111, EPI_ISL_475112, EPI_ISL_475113 | Skovde/Unilabs | The Public Health Agency of Sweden | Oskar Karlsson Lindsjo, Maria Lind Karlberg, Mattias Haukland, Reza Advani, Olov Svartstrom, Anna-Malin Linde, Sandra Broddesson, Petra Edquist, Shamam Muradrasoli, Anna Risberg, Karin Tegmark-Wisell |
| EPI_ISL_475114 | Halmstad klinisk mikrobiologi | The Public Health Agency of Sweden | Oskar Karlsson Lindsjo, Maria Lind Karlberg, Mattias Haukland, Reza Advani, Olov Svartstrom, Anna-Malin Linde, Sandra Broddesson, Petra Edquist, Shamam Muradrasoli, Anna Risberg, Karin Tegmark-Wisell |
| EPI_ISL_475115 | Gavle klinisk mikrobiologi | The Public Health Agency of Sweden | Oskar Karlsson Lindsjo, Maria Lind Karlberg, Mattias Haukland, Reza Advani, Olov Svartstrom, Anna-Malin Linde, Sandra Broddesson, Petra Edquist, Shamam Muradrasoli, Anna Risberg, Karin Tegmark-Wisell |
| EPI_ISL_475144, EPI_ISL_475145 | Klinisk mikrobiologi Vasternorrland | The Public Health Agency of Sweden | Oskar Karlsson Lindsjo, Maria Lind Karlberg, Mattias Haukland, Reza Advani, Olov Svartstrom, Anna-Malin Linde, Sandra Broddesson, Petra Edquist, Shamam Muradrasoli, Anna Risberg, Karin Tegmark-Wisell |
| EPI_ISL_475281, EPI_ISL_475282, EPI_ISL_475283, EPI_ISL_475284, EPI_ISL_475286, EPI_ISL_475287, EPI_ISL_475288, EPI_ISL_475290, EPI_ISL_475291, EPI_ISL_475292, EPI_ISL_475293, EPI_ISL_475295, EPI_ISL_475296, EPI_ISL_475298, EPI_ISL_475300 | | | |
| see above | Centre for Enzyme Innovation, University of Portsmouth / Translational Research Laboratory, Portsmouth Hospitals NHS Trust | COVID-19 Genomics UK (COG-UK) Consortium | Angela Beckett,Yann Bourgeois,Garry Scarlett,Sharon Glaysher,Scott Elliott,Kelly Bicknell,Robert Impey,Allyson Lloyd,Sarah Wyllie,Ethan Butcher,Anoop Chauhan,Samuel Robson |
| EPI_ISL_475556 | Halmstad klinisk mikrobiologi | The Public Health Agency of Sweden | Oskar Karlsson Lindsjo, Maria Lind Karlberg, Mattias Haukland, Reza Advani, Olov Svartstrom, Anna-Malin Linde, Sandra Broddesson, Shamam Muradrasoli, Anna Risberg, Karin Tegmark-Wisell |
| EPI_ISL_475574, EPI_ISL_475596, EPI_ISL_475607, EPI_ISL_475612, EPI_ISL_475624, EPI_ISL_475626, EPI_ISL_475629, EPI_ISL_475640, EPI_ISL_475651, EPI_ISL_475653, EPI_ISL_475673, EPI_ISL_475677, EPI_ISL_475678, EPI_ISL_475679, EPI_ISL_475680, EPI_ISL_475681, EPI_ISL_475682, EPI_ISL_475683, EPI_ISL_475684, EPI_ISL_475695, EPI_ISL_475706, EPI_ISL_475707, EPI_ISL_475708 | | | |
| see above | Cedars-Sinai Medical Center, Department of Pathology & Laboratory Medicine, Molecular Pathology Laboratory | Cedars-Sinai Medical Center, Molecular Pathology Laboratory of Department of Pathology & Laboratory Medicine and Genomic Core | Wenjuan Zhang, John Paul Govindavari, Brian Davis, Stephanie Chen, Jong Taek Kim, Jianbo Song, Jean Lopategui, Jasmine T Plummer, Eric Vail |
| EPI_ISL_475718, EPI_ISL_475721 | Microbiology, University Hospital Donostia | Microbiology, University Hospital Donostia | Cilla,G., Montes,M., Pineiro,L., Marimon,J.M. |
| EPI_ISL_475765 | Universitaetsklinik für Innere Medizin II Innsbruck | Bergthaler laboratory, CeMM Research Center for Molecular Medicine of the Austrian Academy of Sciences | Alexandra Popa, Benedikt Agerer, Henrique Colaco, Lukas Endler, Jakob-Wendelin Genger, Alexander Lercher, Mark Smyth, Thomas Penz, Michael Schuster, Jan Laine, Martin Senekowitsch, Judith Aberle, Stephan Aberle, Peter Hufnagl, Daniela Schmid, Franz Allerberger, Elisabeth Puchhammer-Stoeckl, Manfred Nairz, Guenter Weiss, Gregor Hörmann, Kinga Rigler-Hohenwarter, Rainer Gattringer, Wegene Borena, Dorothee von Laer, Christoph Bock, Andreas Bergthaler |
| EPI_ISL_475774, EPI_ISL_475775, EPI_ISL_475776, EPI_ISL_475777, EPI_ISL_475778, EPI_ISL_475779, EPI_ISL_475780, EPI_ISL_475781, EPI_ISL_475782, EPI_ISL_475783, EPI_ISL_475784, EPI_ISL_475785, EPI_ISL_475786, EPI_ISL_475787, EPI_ISL_475788, EPI_ISL_475789, EPI_ISL_475790, EPI_ISL_475791 | | | |
| see above | Center for Virology, Medical University of Vienna | Bergthaler laboratory, CeMM Research Center for Molecular Medicine of the Austrian Academy of Sciences | Alexandra Popa, Benedikt Agerer, Henrique Colaco, Lukas Endler, Jakob-Wendelin Genger, Alexander Lercher, Mark Smyth, Thomas Penz, Michael Schuster, Jan Laine, Martin Senekowitsch, Judith Aberle, Stephan Aberle, Peter Hufnagl, Daniela Schmid, Franz Allerberger, Elisabeth Puchhammer-Stoeckl, Manfred Nairz, Guenter Weiss, Gregor Hörmann, Kinga Rigler-Hohenwarter, Rainer Gattringer, Wegene Borena, Dorothee von Laer, Christoph Bock, Andreas Bergthaler |
| EPI_ISL_475819, EPI_ISL_475820, EPI_ISL_475821 | Institut für Virologie am Department für Hygiene, Mikrobiologie und Public Health | Bergthaler laboratory, CeMM Research Center for Molecular Medicine of the Austrian Academy of Sciences | Alexandra Popa, Benedikt Agerer, Henrique Colaco, Lukas Endler, Jakob-Wendelin Genger, Alexander Lercher, Mark Smyth, Thomas Penz, Michael Schuster, Jan Laine, Martin Senekowitsch, Judith Aberle, Stephan Aberle, Peter Hufnagl, Daniela Schmid, Franz Allerberger, Elisabeth Puchhammer-Stoeckl, Manfred Nairz, Guenter Weiss, Gregor Hörmann, Kinga Rigler-Hohenwarter, Rainer Gattringer, Wegene Borena, Dorothee von Laer, Christoph Bock, Andreas Bergthaler |
| EPI_ISL_475850, EPI_ISL_475884 | Austrian Agency for Health and Food Safety (AGES) | Bergthaler laboratory, CeMM Research Center for Molecular Medicine of the Austrian Academy of Sciences | Alexandra Popa, Benedikt Agerer, Henrique Colaco, Lukas Endler, Jakob-Wendelin Genger, Alexander Lercher, Mark Smyth, Thomas Penz, Michael Schuster, Jan Laine, Martin Senekowitsch, Judith Aberle, Stephan Aberle, Peter Hufnagl, Daniela Schmid, Franz Allerberger, Elisabeth Puchhammer-Stoeckl, Manfred Nairz, Guenter Weiss, Gregor Hörmann, Kinga Rigler-Hohenwarter, Rainer Gattringer, Wegene Borena, Dorothee von Laer, Christoph Bock, Andreas Bergthaler |
| EPI_ISL_475887, EPI_ISL_475888, EPI_ISL_475889, EPI_ISL_475890, EPI_ISL_475891 | Zentralinstitut für medizinische und chemische Labordiagnostik, Universitätskliniken Insbruck | Bergthaler laboratory, CeMM Research Center for Molecular Medicine of the Austrian Academy of Sciences | Alexandra Popa, Benedikt Agerer, Henrique Colaco, Lukas Endler, Jakob-Wendelin Genger, Alexander Lercher, Mark Smyth, Thomas Penz, Michael Schuster, Jan Laine, Martin Senekowitsch, Judith Aberle, Stephan Aberle, Peter Hufnagl, Daniela Schmid, Franz Allerberger, Elisabeth Puchhammer-Stoeckl, Manfred Nairz, Guenter Weiss, Gregor Hörmann, Kinga Rigler-Hohenwarter, Rainer Gattringer, Wegene Borena, Dorothee von Laer, Christoph Bock, Andreas Bergthaler |
| EPI_ISL_475916, EPI_ISL_475921, EPI_ISL_475925, EPI_ISL_475926, EPI_ISL_475927 | Institut für Virologie am Department für Hygiene, Mikrobiologie und Public Health | Bergthaler laboratory, CeMM Research Center for Molecular Medicine of the Austrian Academy of Sciences | Alexandra Popa, Benedikt Agerer, Henrique Colaco, Lukas Endler, Jakob-Wendelin Genger, Alexander Lercher, Mark Smyth, Thomas Penz, Michael Schuster, Jan Laine, Martin Senekowitsch, Judith Aberle, Stephan Aberle, Peter Hufnagl, Daniela Schmid, Franz Allerberger, Elisabeth Puchhammer-Stoeckl, Manfred Nairz, Guenter Weiss, Gregor Hörmann, Kinga Rigler-Hohenwarter, Rainer Gattringer, Wegene Borena, Dorothee von Laer, Christoph Bock, Andreas Bergthaler |
| EPI_ISL_476069 | University of Debrecen, Department of Medical Microbiology | National Laboratory of Virology, Szentágothai Research Centre | Endre Gábor Tóth, Balázs Somogyi, Brigitta Zana, Eszter Csoma, Ferenc Jakab, Gábor Kemenesi |
| EPI_ISL_476078 | University of Szeged, Institute of Clinical Microbiology | National Laboratory of Virology, Szentágothai Research Centre | Endre Gábor Tóth, Balázs Somogyi, Brigitta Zana, Terhes Gabriella, Ferenc Jakab, Gábor Kemenesi |
| EPI_ISL_476143, EPI_ISL_476144 | Skovde/Unilabs | The Public Health Agency of Sweden | Oskar Karlsson Lindsjo, Maria Lind Karlberg, Mattias Haukland, Reza Advani, Olov Svartstrom, Anna-Malin Linde, Sandra Broddesson, Petra Edquist, Shamam Muradrasoli, Anna Risberg, Karin Tegmark-Wisell |
| EPI_ISL_476246, EPI_ISL_476247, EPI_ISL_476248, EPI_ISL_476249, EPI_ISL_476250, EPI_ISL_476251, EPI_ISL_476252, EPI_ISL_476253, EPI_ISL_476254, EPI_ISL_476255, EPI_ISL_476256, EPI_ISL_476257, EPI_ISL_476258, EPI_ISL_476259, EPI_ISL_476260, EPI_ISL_476261, EPI_ISL_476262, EPI_ISL_476263, EPI_ISL_476264, EPI_ISL_476265, EPI_ISL_476266, EPI_ISL_476267, EPI_ISL_476268, EPI_ISL_476269, EPI_ISL_476270, EPI_ISL_476271, EPI_ISL_476272, EPI_ISL_476273, EPI_ISL_476274, EPI_ISL_476275, EPI_ISL_476276, EPI_ISL_476277 | | | |
| see above | Hospital da Clínicas da Faculdade de Medicina da Universidade de São Paulo | Instituto de Medicina Tropical da Univesidade de São Paulo | Samples: Ingra Morales Claro, Erika Regina Manuli, Cecilia Salete Alencar, Carolina S. Lazar, Silvia F. Costa; Sequencing: Ingra Morales Claro, Jaqueline Goes de Jesus, Erika Regina Manuli, Flavia Cristina da Silva Sales, Thais de Moura Coletti, Camila Alves Maia da Silva, Mariana Severo Ramundo, Giula Magalhaes Ferreira, Darlan da Silva Candido, Julien Theze, Nuno Faria, Ester Sabino |
| EPI_ISL_476281, EPI_ISL_476282, EPI_ISL_476283, EPI_ISL_476284, EPI_ISL_476285, EPI_ISL_476286, EPI_ISL_476287, EPI_ISL_476288, EPI_ISL_476289, EPI_ISL_476290, EPI_ISL_476291, EPI_ISL_476292, EPI_ISL_476298, EPI_ISL_476299, EPI_ISL_476300, EPI_ISL_476301 | | | |
| see above | DB Diagnósticos do Brasil | Instituto de Medicina Tropical da Univesidade de São Paulo | Samples: Nelson Gaburo Jr; Sequencing: Ingra Morales Claro, Jaqueline Goes de Jesus, Erika Regina Manuli, Flavia Cristina da Silva Sales, Thais de Moura Coletti, Camila Alves Maia da Silva, Mariana Severo Ramundo, Giula Magalhaes Ferreira, Darlan da Silva Candido, Julien Theze, Nuno Faria, Ester Sabino |

| | | | |
|---|---|---|---|
| EPI_ISL_476517, EPI_ISL_476518, EPI_ISL_476519, EPI_ISL_476520, EPI_ISL_476521, EPI_ISL_476522, EPI_ISL_476523, EPI_ISL_476524, EPI_ISL_476525, EPI_ISL_476526, EPI_ISL_476527, EPI_ISL_476528, EPI_ISL_476529, EPI_ISL_476530, EPI_ISL_476531, EPI_ISL_476532, EPI_ISL_476533, EPI_ISL_476534, EPI_ISL_476535, EPI_ISL_476536, EPI_ISL_476537, EPI_ISL_476538 | | | |
| see above | Yale Clinical Virology Laboratory | Grubaugh Lab - Yale School of Public Health | Joseph Fauver, Tara Alpert, Anderson Brito, Anne Wyllie, Chantal Vogels, Mary Petrone, Cole Jensen, Chaney Kalinich, Isabel Ott, Arnau Casanovas, Catherine Muenker, Adam Moore, Alice Lu, Maria Tokuyama, Patrick Wong, Peiwen Lu, Saad Omer, Richard Martinello, Allison Nelson, Shelli Farhadian, Akiko Iwasaki, Charlese Dela Cruz, Albert Ko, Nathan Grubaugh |
| EPI_ISL_476767, EPI_ISL_476786, EPI_ISL_476789, EPI_ISL_476794 | Stanford clinical virology lab | Chan-Zuckerberg Biohub | Benjamin Pinksy, Katharine Walter, Victoria N. Parikh, John Gorzynski, Hannah N. DeJong, Matthew T. Wheeler, Jason Andrews, Manuel Rivas, Carlos Bustamante, Euan Ashley, with CZB Cliahub Consortium |
| EPI_ISL_476827 | Laboratoire des Fièvres Hémorragiques Virales du Benin | Charité-Universitätsmedizin Berlin | Yadouleton, Anges; Sander Anna-Lena; Moreira-Soto Andres; Drexler, Jan Felix |
| EPI_ISL_476838 | National Influenza Centre for Northern Greece | National Influenza Centre for Northern Greece | Maria Christoforidi |
| EPI_ISL_476900, EPI_ISL_476901 | UW Virology Lab | UW Virology Lab | Pavitra Roychoudhury, Hong Xie, Lasata Shrestha, Amin Addetia, Truong Nguyen, Victoria M Rachleff, Meei-Li Huang, Keith R Jerome, Alexander Greninger |
| EPI_ISL_477014 | Institute of Microbiology, Universidad San Francisco de Quito | Institute of Microbiology, Universidad San Francisco de Quito | Belen Prado-Vivar, Sully Marquez, Juan Jose Guadalupe, Monica Becerra-Wong, Carla Torres, Bernardo Gutierrez, Francisco Mora, Juan Gaviria, Alejandra Ramones, Franklin Espinoza, Edison Ligña, Jorge Reyes, Patricio Rojas-Silva, Veronica Barragan, Gabriel Trueba, Michelle Grunauer, Paul Cardenas |
| EPI_ISL_477621, EPI_ISL_477622, EPI_ISL_477623, EPI_ISL_477624, EPI_ISL_477625 | University of Szeged, Institute of Clinical Microbiology | National Laboratory of Virology, Szentágothai Research Centre | Endre Gábor Tóth, Balázs Somogyi, Brigitta Zana, Terhes Gabriella, Ferenc Jakab, Gábor Kemenesi |
| EPI_ISL_477673, EPI_ISL_477674, EPI_ISL_477675, EPI_ISL_477676, EPI_ISL_477677, EPI_ISL_477678, EPI_ISL_477679, EPI_ISL_477682, EPI_ISL_477684, EPI_ISL_477705, EPI_ISL_477707, EPI_ISL_477708, EPI_ISL_477709, EPI_ISL_477710, EPI_ISL_477711, EPI_ISL_477712, EPI_ISL_477713, EPI_ISL_477714, EPI_ISL_477715, EPI_ISL_477716, EPI_ISL_477717, EPI_ISL_477718, EPI_ISL_477719, EPI_ISL_477720, EPI_ISL_477721, EPI_ISL_477722, EPI_ISL_477723, EPI_ISL_477724 | | | |
| see above | UW Virology Lab | UW Virology Lab | Pavitra Roychoudhury, Hong Xie, Lasata Shrestha, Amin Addetia, Truong Nguyen, Victoria M Rachleff, Meei-Li Huang, Keith R Jerome, Alexander Greninger |
| EPI_ISL_477817 | Department of Pathology, University of Cambridge | COVID-19 Genomics UK (COG-UK) Consortium | Luke W Meredith, M. Estée Török, Myra Hosmillo, William L. Hamilton, Martin D. Curran, Theresa Feltwell, Grant Hall, Anna Yakovleva, Fahad A Khokhar, Charlotte J. Houldcroft, Laura G Caller, Aminu S. Jahun, Sarah L. Caddy, Yasmin Chaudhry, Malte Pinckert, Ian Goodfellow |
| EPI_ISL_478411, EPI_ISL_478412, EPI_ISL_478413, EPI_ISL_478414, EPI_ISL_478415, EPI_ISL_478416 | University College London, Great Ormond Street Hospital for Children NHS Foundation Trust, Imperial College Healthcare NHS Trust | COVID-19 Genomics UK (COG-UK) Consortium | Sergi Castellano, Rachel Williams, Mark Kristiansen, Paola Resende Silva, Sunando Roy, Tony Brooks, Helena Tutill, Paola Niola, Patricia Dyal, Charlotte Williams, Leysa Forrest, Yasmin Panchbhaya, Jacqueline Findlay, Samuel Weeks, Julianne Brown, Kathryn Harris, Paul Randell, James Price, Alison Holmes, Judith Breuer |
| EPI_ISL_478494, EPI_ISL_478495, EPI_ISL_478591, EPI_ISL_478592, EPI_ISL_478593, EPI_ISL_478594, EPI_ISL_478595, EPI_ISL_478596, EPI_ISL_478608, EPI_ISL_478609, EPI_ISL_478610, EPI_ISL_478611, EPI_ISL_478612, EPI_ISL_478613, EPI_ISL_478614, EPI_ISL_478615, EPI_ISL_478616, EPI_ISL_478617, EPI_ISL_478618, EPI_ISL_478619, EPI_ISL_478620, EPI_ISL_478621, EPI_ISL_478622, EPI_ISL_478624, EPI_ISL_478625, EPI_ISL_478626, EPI_ISL_478627, EPI_ISL_478628, EPI_ISL_478629, EPI_ISL_478630, EPI_ISL_478631, EPI_ISL_478632, EPI_ISL_478634 | | | |
| see above | Northumbria University / South Tees Hospitals NHS Foundation Trust / North Cumbria Integrated Care NHS Foundation Trust / North Tees and Hartlepool NHS Foundation Trust / Newcastle Hospitals NHS Foundation Trust | COVID-19 Genomics UK (COG-UK) Consortium | Darren L Smith,Andrew Nelson,Matthew Bashton,Greg R Young,Joshua Loh,John Allan,Mohammad A Tariq,Giles S Holt,Gary Black,Wen C Yew,Lynn Dover,Paul Baker,Steve Liggett,Sarah Essex,Jane Greenaway,Debra Padgett,Clive Graham,Garren Scott,Edward Barton,Emma Swindells,Brendan Payne,Jennifer Collins,Yusri Taha,Gary Eltringham |
| EPI_ISL_478669 | unknown | Microbiology, Koc University | Can,F., Ozer,B., Nurtop,E., Dogan,O. |
| EPI_ISL_478674 | Pathology North - Royal North Shore Hospital - NSW Health Pathology | NSW Health Pathology - Institute of Clinical Pathology and Medical Research; Westmead Hospital; University of Sydney | CIDM-PH et al. |
| EPI_ISL_478693 | South Eastern Area Laboratory Services (SEALS) | NSW Health Pathology - Institute of Clinical Pathology and Medical Research; Westmead Hospital; University of Sydney | CIDM-PH et al. |
| EPI_ISL_478706 | Sydney South West Pathology Service (SSWPS) - Liverpool Hospital - NSW Health Pathology | NSW Health Pathology - Institute of Clinical Pathology and Medical Research; Westmead Hospital; University of Sydney | CIDM-PH et al. |
| EPI_ISL_478708 | South Eastern Area Laboratory Services (SEALS) | NSW Health Pathology - Institute of Clinical Pathology and Medical Research; Westmead Hospital; University of Sydney | CIDM-PH et al. |
| EPI_ISL_479616, EPI_ISL_479617 | Laboratory of Molecular Virology of the International Centre for Genetic Engineering and Biotechnology (ICGEB) | ARGO Open Lab Platform for Genome Sequencing | Licastro, D, Rajasekharan S, Dal Monego S, Segat L, D'Agaro P, Salton F, Confalonieri P, Confalonieri M Marcello A |
| EPI_ISL_479624 | Molecular diagnostic laboratory of Federal Budget Institution of Science "Central Research Institute of Epidemiology" of The Federal Service on Customers' Rights Protection and Human Well-being Surveillance | Group of Genomics and Postgenomic Technologies of Central Research Institute of Epidemilology | Speranskaya AS, Kaptelova VV,Valdokhina AV, Bulanenko VP, Samoilov AE, Korneenko EV, Sizova TV, Tivanova EV, Shipulina OY, Akimkin VG |
| EPI_ISL_479626, EPI_ISL_479627, EPI_ISL_479628, EPI_ISL_479629, EPI_ISL_479630, EPI_ISL_479631, EPI_ISL_479632, EPI_ISL_479633, EPI_ISL_479643 | Dr. Georges-L.-Dumont University Hospital Centre | National Microbiology Laboratory | Anna Majer, Shari Tyson, Grace Seo, Kristyn Burak, Philip Mabon, Elsie Grudeski, Rhiannon Huzarewich, Russell Mandes, Jennifer Tanner, Natalie Knox, Morag Graham, Gary Van Domselaar, Richard Garceau, Guillaume Desnoyers, Nathalie Bastien, Yan Li, Timothy Booth |
| EPI_ISL_479756, EPI_ISL_479758 | National Institute of Hygiene and Epidemiology (NIHE) | National Key Laboratory of Gene Technology, Institute of Biotechnology (IBT) | Le Tung Lam, Nguyen Hong Trang, Ho Thi Thuong, Tran Huyen Linh, Ung Thi Hong Trang, Le Thi Thanh, Nguyen Vu Son, Vuong Duc Cuong, Tran Thu Huong, Pham Thi Hien, Nguyen Phuong Anh, Nguyen Le Khanh Hang, Hoang Vu Mai Phuong, Hoang Ha, Taichiro Takemura, Futoshi Hasebe, Chu Hoang Ha, Le Quynh Mai, Dang Duc Anh, Truong Nam Hai |
| EPI_ISL_479972, EPI_ISL_479973, EPI_ISL_479974, EPI_ISL_479975, EPI_ISL_479976, EPI_ISL_479977, EPI_ISL_479978 | Fukui Prefectural Institute of Public Health and Environmental Science | Pathogen Genomics Center, National Institute of Infectious Diseases | Tsuyoshi Sekizuka, Miho Toho, Kentaro Itokawa, Rina Tanaka, Masanori Hashino, Hajime Kamiya, Motoi Suzuki, Makoto Kuroda |
| EPI_ISL_479986, EPI_ISL_479987, EPI_ISL_479988, EPI_ISL_479989 | Department of Infectious Diseases, Kobe Institute of Health | Pathogen Genomics Center, National Institute of Infectious Diseases | Tsuyoshi Sekizuka, Ryohei Nomoto, Kentaro Itokawa, Rina Tanaka, Masanori Hashino, Hajime Kamiya, Motoi Suzuki, Makoto Kuroda |
| EPI_ISL_479992, EPI_ISL_479993, EPI_ISL_479994, EPI_ISL_479995, EPI_ISL_479996 | Kumamoto City Public Health Research Institute | Pathogen Genomics Center, National Institute of Infectious Diseases | Tsuyoshi Sekizuka, Kaori Tashiro, Kentaro Itokawa, Rina Tanaka, Masanori Hashino, Hajime Kamiya, Motoi Suzuki, Makoto Kuroda |
| EPI_ISL_480004, EPI_ISL_480005, EPI_ISL_480006, EPI_ISL_480007, EPI_ISL_480008, EPI_ISL_480011, EPI_ISL_480012, EPI_ISL_480013, EPI_ISL_480014 | Chiba Prefectural Institute of Public Health | Pathogen Genomics Center, National Institute of Infectious Diseases | Tsuyoshi Sekizuka, Masakatsu Taira, Kentaro Itokawa, Rina Tanaka, Masanori Hashino, Hajime Kamiya, Motoi Suzuki, Makoto Kuroda |
| EPI_ISL_480015, EPI_ISL_480016, EPI_ISL_480018, EPI_ISL_480019, EPI_ISL_480020 | Gunma Prefectural Institute of Public Health and Environmental Sciences | Pathogen Genomics Center, National Institute of Infectious Diseases | Tsuyoshi Sekizuka, Hiroyuki Tsukagoshi, Kentaro Itokawa, Rina Tanaka, Masanori Hashino, Hajime Kamiya, Motoi Suzuki, Makoto Kuroda |
| EPI_ISL_480021, EPI_ISL_480022, | Ibaraki Prefectural Institute of Public Health | Pathogen Genomics Center, National Institute of Infectious | Tsuyoshi Sekizuka, Keiko Goto, Kentaro Itokawa, Rina Tanaka, Masanori Hashino, Hajime Kamiya, Motoi Suzuki, Makoto Kuroda |

| | | | |
|---|---|---|---|
| EPI_ISL_480023, EPI_ISL_480025 | | Diseases | |
| EPI_ISL_480041 | Tochigi Prefectural Institute of Public Health and Environmental Science | Pathogen Genomics Center, National Institute of Infectious Diseases | Tsuyoshi Sekizuka, Ako Nakajima, Kentaro Itokawa, Rina Tanaka, Masanori Hashino, Hajime Kamiya, Motoi Suzuki, Makoto Kuroda |
| EPI_ISL_480065, EPI_ISL_480066 | Sakai City Institute of Public Health | Pathogen Genomics Center, National Institute of Infectious Diseases | Tsuyoshi Sekizuka, Tatsuya Miyoshi, Kentaro Itokawa, Rina Tanaka, Masanori Hashino, Hajime Kamiya, Motoi Suzuki, Makoto Kuroda |
| EPI_ISL_480074, EPI_ISL_480075, EPI_ISL_480076, EPI_ISL_480077, EPI_ISL_480078, EPI_ISL_480079, EPI_ISL_480080 | Shizuoka City Institute of Environmental Sciences and Public Health | Pathogen Genomics Center, National Institute of Infectious Diseases | Tsuyoshi Sekizuka, Takaharu Maehata,Sou Okamura,Yuji Kanazawa,Kenji Yagi, Kentaro Itokawa, Rina Tanaka, Masanori Hashino, Hajime Kamiya, Motoi Suzuki, Makoto Kuroda |
| EPI_ISL_480086, EPI_ISL_480087, EPI_ISL_480088, EPI_ISL_480089 | Gifu Prefectural Institute of Public Health and Environmental Sciences | Pathogen Genomics Center, National Institute of Infectious Diseases | Tsuyoshi Sekizuka, Yoshihiko Kameyama, Kentaro Itokawa, Rina Tanaka, Masanori Hashino, Hajime Kamiya, Motoi Suzuki, Makoto Kuroda |
| EPI_ISL_480108 | Koshigaya City Public Health Center | Pathogen Genomics Center, National Institute of Infectious Diseases | Tsuyoshi Sekizuka, Yuka Furui, Aya Tamura, Kyohei Sakata, Takumi Daimon, Yoko Togawa, Yoshiko Hamada, Kentaro Itokawa, Rina Tanaka, Masanori Hashino, Hajime Kamiya, Motoi Suzuki, Makoto Kuroda |
| EPI_ISL_480118, EPI_ISL_480119 | Oita Prefectural Institute of Public Health and Environmental Science | Pathogen Genomics Center, National Institute of Infectious Diseases | Tsuyoshi Sekizuka, Mari Sasaki, Kentaro Itokawa, Rina Tanaka, Masanori Hashino, Hajime Kamiya, Motoi Suzuki, Makoto Kuroda |
| EPI_ISL_480120, EPI_ISL_480121, EPI_ISL_480122, EPI_ISL_480123, EPI_ISL_480124 | Fukui Prefectural Institute of Public Health and Environmental Science | Pathogen Genomics Center, National Institute of Infectious Diseases | Tsuyoshi Sekizuka, Miho Toho, Kentaro Itokawa, Rina Tanaka, Masanori Hashino, Hajime Kamiya, Motoi Suzuki, Makoto Kuroda |
| EPI_ISL_480179 | Hiroshima City Institute of Public Health | Pathogen Genomics Center, National Institute of Infectious Diseases | Tsuyoshi Sekizuka, Kota Noritsune, Kentaro Itokawa, Rina Tanaka, Masanori Hashino, Hajime Kamiya, Motoi Suzuki, Makoto Kuroda |
| EPI_ISL_480180, EPI_ISL_480181 | Ibaraki Prefectural Institute of Public Health | Pathogen Genomics Center, National Institute of Infectious Diseases | Tsuyoshi Sekizuka, Keiko Goto, Kentaro Itokawa, Rina Tanaka, Masanori Hashino, Hajime Kamiya, Motoi Suzuki, Makoto Kuroda |
| EPI_ISL_480196, EPI_ISL_480198 | Toyama Institute of Health | Pathogen Genomics Center, National Institute of Infectious Diseases | Tsuyoshi Sekizuka, Masae Itamochi, Kazunori Oishi, Kentaro Itokawa, Rina Tanaka, Masanori Hashino, Hajime Kamiya, Motoi Suzuki, Makoto Kuroda |
| EPI_ISL_480303, EPI_ISL_480307 | National Reference Laboratory "Influenza and acute respiratory diseases" | NRL-HIV | Ivan Ivanov, Ivailo Alexiev, Ivva Philipova |
| EPI_ISL_480317, EPI_ISL_480318, EPI_ISL_480319 | Hospital Clínica Bíblica | Charité Virology-University of Costa Rica | Andres Moreira-Soto, Eugenia Corrales-Aguilar, Ignacio Postigo-Hidalgo, Karla Sofía Gutiérrez, Jan Felix Drexler |
| EPI_ISL_480390 | University of Wisconsin-Madison AIDS Vaccine Research Laboratories | University of Wisconsin-Madison AIDS Vaccine Research Laboratories | Gage Moreno, Katarina Braun, et al. AIDS Vaccine Research Laboratories |
| EPI_ISL_480556 | Institut Pasteur Dakar | Institut Pasteur de Dakar | Ndongo Dia, Moussa Moise Diagne, Mamadou Diop, Marie Henriette Dior Ndione, Mamadou Malado Jallow, Safietou Sanke, Ousmane Faye, Amadou Alpha Sall. |
| EPI_ISL_480565, EPI_ISL_480566, EPI_ISL_480567, EPI_ISL_480568, EPI_ISL_480569 | Victorian Infectious Diseases Reference Laboratory (VIDRL) | VIDRL and MDU-PHL | Caly L., Seemann T., Sait, M., Schultz M., Druce J., Sherry, N. |
| EPI_ISL_480613, EPI_ISL_480614 | Microbiological Diagnostic Unit - Public Health Laboratory (MDU-PHL) | MDU-PHL | Seemann T., Schultz M., Sait, M., Sherry, N. |
| EPI_ISL_480782 | Institut Pasteur Dakar | Institut Pasteur de Dakar | Ndongo Dia, Moussa Moise Diagne, Mamadou Diop, Marie Henriette Dior Ndione, Mamadou Malado Jallow, Safietou Sanke, Ousmane Faye, Amadou Alpha Sall. |
| EPI_ISL_480803, EPI_ISL_480805 | Florida Bureau of Public Health Laboratories | Florida Bureau of Public Health Laboratories | Sarah Schmedes, Jason Blanton |
| EPI_ISL_480989 | ISGlobal, Institut de Salut Global de Barcelona | SeqCOVID-SPAIN consortium/IBV(CSIC) | Alfredo Mayor, Alberto L Garcia-Basteiro, Carlota Dobaño, Gemma Moncunill, Pau Cisteró and SeqCOVID-SPAIN consortium |
| EPI_ISL_481262, EPI_ISL_481264 | Robert Koch Institute, National Reference center for Influenza, Berlin, Germany | Robert Koch Institute, Bioinformatics MF1, Berlin, Germany | Marianne Wedde, Oliver Drechsel, Andrea Thuermer, Rene Kmiecinski, Ralf Duerrwald, Thorsten Wolff, Stephan Fuchs, Max v. Kleist |
| EPI_ISL_481639, EPI_ISL_481640, EPI_ISL_481641, EPI_ISL_481642, EPI_ISL_481643, EPI_ISL_481644, EPI_ISL_481645, EPI_ISL_481646, EPI_ISL_481647, EPI_ISL_481648, EPI_ISL_481656, EPI_ISL_481657, EPI_ISL_481658, EPI_ISL_481659, EPI_ISL_481660, EPI_ISL_481661, EPI_ISL_481662, EPI_ISL_481663 | | | |
| see above | Department of Virology and Immunology, University of Helsinki and Helsinki University Hospital, Huslab Finland | Department of Virology, Faculty of Medicine, University of Helsinki, Helsinki, Finland | Teemu Smura, Hannimari Kallio-Kokko, Jenni Virtanen, Maija Suvanto, Sari Hannula, Harri Kangas, Pekka Ellonen, Olli Vapalahti |
| EPI_ISL_481747 | Dr. Georges-L.-Dumont University Hospital Centre | National Microbiology Laboratory | Anna Majer, Shari Tyson, Grace Seo, Kristyn Burak, Philip Mabon, Elsie Grudeski, Rhiannon Huzarewich, Russell Mandes, Jennifer Tanner, Natalie Knox, Morag Graham, Gary Van Domselaar, Richard Garceau, Guillaume Desnoyers, Nathalie Bastien, Yan Li, Timothy Booth |
| EPI_ISL_482307, EPI_ISL_482308, EPI_ISL_482309, EPI_ISL_482310, EPI_ISL_482311, EPI_ISL_482312, EPI_ISL_482320, EPI_ISL_482321, EPI_ISL_482322, EPI_ISL_482452, EPI_ISL_482455, EPI_ISL_482456, EPI_ISL_482458, EPI_ISL_482459, EPI_ISL_482460, EPI_ISL_482466 | | | |
| see above | Providence St. Joseph Health Molecular Genomics Laboratory | Providence St. Joseph Health Molecular Genomics Laboratory | Alexa K Dowdell, Brian D Piening, Fred L Robinson, Carlo B Bifulco, Mary Campbell |
| EPI_ISL_483076, EPI_ISL_483079, EPI_ISL_483085, EPI_ISL_483088, EPI_ISL_483092, EPI_ISL_483093, EPI_ISL_483100, EPI_ISL_483101, EPI_ISL_483102, EPI_ISL_483103, EPI_ISL_483104, EPI_ISL_483105, EPI_ISL_483106, EPI_ISL_483107, EPI_ISL_483108, EPI_ISL_483109, EPI_ISL_483110, EPI_ISL_483111, EPI_ISL_483112, EPI_ISL_483136, EPI_ISL_483138 | | | |
| see above | SA Pathology | SA Pathology | Lex Leong, Chuan Kok Lim, Mark Turra, Ivan Bastian, Geoff Higgins |
| EPI_ISL_483201, EPI_ISL_483299, EPI_ISL_483300, EPI_ISL_483301, EPI_ISL_483302, EPI_ISL_483303, EPI_ISL_483304, EPI_ISL_483305, EPI_ISL_483306, EPI_ISL_483307, EPI_ISL_483309, EPI_ISL_483310, EPI_ISL_483311, EPI_ISL_483312, EPI_ISL_483313, EPI_ISL_483314, EPI_ISL_483315, EPI_ISL_483316, EPI_ISL_483318, EPI_ISL_483319, EPI_ISL_483320, EPI_ISL_483321, EPI_ISL_483322, EPI_ISL_483323, EPI_ISL_483371, EPI_ISL_483372, EPI_ISL_483373, EPI_ISL_483374, EPI_ISL_483375, EPI_ISL_483377, EPI_ISL_483378, EPI_ISL_483383, EPI_ISL_483424, EPI_ISL_483425, EPI_ISL_483426, EPI_ISL_483427, EPI_ISL_483428, EPI_ISL_483429, EPI_ISL_483430, EPI_ISL_483431, EPI_ISL_483432, EPI_ISL_483435, EPI_ISL_483436, EPI_ISL_483445, EPI_ISL_483446 | | | |
| see above | UC San Diego Center for Advanced Laboratory Medicine | Andersen lab at Scripps Research | SEARCH Alliance San Diego with David Pride, Ji H Shin |
| EPI_ISL_483551, EPI_ISL_483552, EPI_ISL_483553, EPI_ISL_483554 | Kingdom of Bahrein Ministry of Health | Erasmus Medical Center | Bas Oude Munnink, David Nieuwenhuijse, Reina Sikkema, Fatema, Ebrahim Shehad, Amjad Ghanem Mohamed, Hashmeya Al Wasti, Claudia Schapendonk, Irina Chestakova, Anne van der Linden, Theo Bestebroer, Stefan van Nieuwkoop, Mark Pronk, Pascal Lexmond, Richard Molenkamp, Marion Koopmans, on behalf of the Dutch national COVID-19 response team. |
| EPI_ISL_483610, EPI_ISL_483617 | National Public Health Laboratory, National Centre for Infectious Diseases | National Public Health Laboratory, National Centre for Infectious Diseases | Mak TM, Octavia S, Zhou Z, Chavatte JM, Cui L, Lin RTP |
| EPI_ISL_483701, EPI_ISL_483711 | Israel Central Virology laboratory | Israel Central Virology laboratory | Neta Zuckerman, Efrat Dahan Bucris, Oran Erster, Ella Mendelson, Michal Mandelboim |
| EPI_ISL_483961, EPI_ISL_484059, EPI_ISL_484060, EPI_ISL_484061, EPI_ISL_484062, EPI_ISL_484063, EPI_ISL_484064, EPI_ISL_484065, EPI_ISL_484066, EPI_ISL_484067, EPI_ISL_484068, EPI_ISL_484069, EPI_ISL_484070, EPI_ISL_484071, EPI_ISL_484072, EPI_ISL_484073, EPI_ISL_484074, EPI_ISL_484075, EPI_ISL_484076, EPI_ISL_484077, EPI_ISL_484078, EPI_ISL_484079, EPI_ISL_484080, EPI_ISL_484081, EPI_ISL_484099, EPI_ISL_484100, EPI_ISL_484101, EPI_ISL_484102, EPI_ISL_484103, EPI_ISL_484104, EPI_ISL_484105, EPI_ISL_484106, EPI_ISL_484107, EPI_ISL_484108, EPI_ISL_484109, EPI_ISL_484110, EPI_ISL_484112, EPI_ISL_484113, EPI_ISL_484114, EPI_ISL_484115, EPI_ISL_484116, EPI_ISL_484117, EPI_ISL_484118, EPI_ISL_484119, EPI_ISL_484120, EPI_ISL_484121, EPI_ISL_484122, EPI_ISL_484123, EPI_ISL_484124, EPI_ISL_484125, EPI_ISL_484126, EPI_ISL_484127, EPI_ISL_484128, EPI_ISL_484129, EPI_ISL_484130, EPI_ISL_484131, EPI_ISL_484132, EPI_ISL_484133, EPI_ISL_484134, EPI_ISL_484135, EPI_ISL_484136, EPI_ISL_484137, EPI_ISL_484138, EPI_ISL_484139, EPI_ISL_484140, EPI_ISL_484141, EPI_ISL_484142, EPI_ISL_484143, EPI_ISL_484144, EPI_ISL_484145, EPI_ISL_484146, EPI_ISL_484147, EPI_ISL_484148, EPI_ISL_484149, EPI_ISL_484150, EPI_ISL_484151, EPI_ISL_484152, EPI_ISL_484154, EPI_ISL_484155, EPI_ISL_484157, EPI_ISL_484159, EPI_ISL_484161, EPI_ISL_484163, EPI_ISL_484168, EPI_ISL_484174, EPI_ISL_484200, EPI_ISL_484201 | | | |
| see above | Centre for Clinical Infection and Diagnostics Research and Genomics Innovation Unit, Guy's and St. Thomas' NHS Trust | COVID-19 Genomics UK (COG-UK) Consortium | Chloe Fisher, Luke Snell, Penny Cliff, Rahul Batra, Jonathan Edgeworth, Ali Raza Awan |

| | | | |
|---|---|---|---|
| EPI_ISL_484711, EPI_ISL_484712, EPI_ISL_484715, EPI_ISL_484719, EPI_ISL_484721, EPI_ISL_484722, EPI_ISL_484724, EPI_ISL_484725, EPI_ISL_484727, EPI_ISL_484729, EPI_ISL_484731, EPI_ISL_484736, EPI_ISL_484739, EPI_ISL_484740, EPI_ISL_484741, EPI_ISL_484743, EPI_ISL_484744, EPI_ISL_484746, EPI_ISL_484748, EPI_ISL_484750, EPI_ISL_484751, EPI_ISL_484752, EPI_ISL_484754, EPI_ISL_484755, EPI_ISL_484758, EPI_ISL_484759, EPI_ISL_484764, EPI_ISL_484766, EPI_ISL_484767, EPI_ISL_484768, EPI_ISL_484770, EPI_ISL_484772, EPI_ISL_484773, EPI_ISL_484774, EPI_ISL_484775, EPI_ISL_484776, EPI_ISL_484777, EPI_ISL_484778, EPI_ISL_484779, EPI_ISL_484780, EPI_ISL_484781, EPI_ISL_484782, EPI_ISL_484783, EPI_ISL_484784, EPI_ISL_484785, EPI_ISL_484786, EPI_ISL_484791 | | | |
| see above | University of Michigan Clinical Microbiology Laboratory | Lauring Lab, University of Michigan, Department of Microbiology and Immunology | Valesano et al. |
| EPI_ISL_485393 | University of Ulsan College of Medicine and Asan Medical Center | University of Ulsan College of Medicine and Asan Medical Center | Kuenyoul Park, Jaewoong Lee, Kihyun Lee, Jiwon Jung, Sung-Han Kim, Jina Lee, Mauricio Chalita, Seok-Hwan Yoon, Jongsik Chun, Kyu-Hwa Hur, Heungsup Sung, Mi-Na Kim, and Hae Kyung Lee |
| EPI_ISL_485917, EPI_ISL_485918, EPI_ISL_485919, EPI_ISL_485920, EPI_ISL_485921, EPI_ISL_485922, EPI_ISL_485923, EPI_ISL_485924, EPI_ISL_485925, EPI_ISL_485926, EPI_ISL_485927, EPI_ISL_485928, EPI_ISL_485929, EPI_ISL_485930, EPI_ISL_485931, EPI_ISL_485932, EPI_ISL_485933, EPI_ISL_485934, EPI_ISL_485935, EPI_ISL_485936, EPI_ISL_485938, EPI_ISL_485939, EPI_ISL_485940, EPI_ISL_485942, EPI_ISL_485946, EPI_ISL_485981, EPI_ISL_485982, EPI_ISL_485986, EPI_ISL_485988, EPI_ISL_485989, EPI_ISL_485994, EPI_ISL_485995, EPI_ISL_485996, EPI_ISL_485997, EPI_ISL_485998, EPI_ISL_485999, EPI_ISL_486000, EPI_ISL_486001, EPI_ISL_486002, EPI_ISL_486003, EPI_ISL_486004, EPI_ISL_486005, EPI_ISL_486006, EPI_ISL_486007, EPI_ISL_486008, EPI_ISL_486009, EPI_ISL_486010, EPI_ISL_486011, EPI_ISL_486012, EPI_ISL_486013, EPI_ISL_486017, EPI_ISL_486018, EPI_ISL_486020, EPI_ISL_486021, EPI_ISL_486028, EPI_ISL_486030, EPI_ISL_486033, EPI_ISL_486034, EPI_ISL_486035, EPI_ISL_486037, EPI_ISL_486038, EPI_ISL_486039, EPI_ISL_486040, EPI_ISL_486041, EPI_ISL_486042, EPI_ISL_486043, EPI_ISL_486044, EPI_ISL_486045, EPI_ISL_486046, EPI_ISL_486047, EPI_ISL_486048, EPI_ISL_486049, EPI_ISL_486050, EPI_ISL_486051, EPI_ISL_486052, EPI_ISL_486053, EPI_ISL_486054, EPI_ISL_486055, EPI_ISL_486056, EPI_ISL_486057, EPI_ISL_486059, EPI_ISL_486064, EPI_ISL_486066, EPI_ISL_486067, EPI_ISL_486068, EPI_ISL_486069, EPI_ISL_486070, EPI_ISL_486071, EPI_ISL_486072, EPI_ISL_486073, EPI_ISL_486074, EPI_ISL_486075, EPI_ISL_486076, EPI_ISL_486077, EPI_ISL_486078, EPI_ISL_486079, EPI_ISL_486083, EPI_ISL_486084, EPI_ISL_486085, EPI_ISL_486086, EPI_ISL_486087, EPI_ISL_486088, EPI_ISL_486089, EPI_ISL_486090, EPI_ISL_486095, EPI_ISL_486098, EPI_ISL_486099 | | | |
| see above | UW Virology Lab | UW Virology Lab | Pavitra Roychoudhury, Hong Xie, Lasata Shrestha, Amin Addetia, Truong Nguyen, Victoria M Rachleff, Meei-Li Huang, Keith R Jerome, Alexander Greninger |
| EPI_ISL_486815, EPI_ISL_486816, EPI_ISL_486817, EPI_ISL_486818, EPI_ISL_486819 | Molecular diagnostic laboratory of Federal Budget Institution of Science "Central Research Institute of Epidemiology" of The Federal Service on Customers' Rights Protection and Human Well-being Surveillance | Group of Genomics and Postgenomic Technologies of Central Research Institute of Epidemilology | Speranskaya AS, Kaptelova VV, Valdokhina AV, Bulanenko VP, Samoilov AE, Korneenko EV, Tivanova EV, Shipulina OY, Akimkin VG |
| EPI_ISL_486830 | Providence St. Joseph Health Molecular Genomics Laboratory | Providence St. Joseph Health Molecular Genomics Laboratory | Alexa K Dowdell, Brian D Piening, Fred L Robinson, Carlo B Bifulco, Mary Campbell |
| EPI_ISL_487229 | University of Michigan Clinical Microbiology Laboratory | Lauring Lab, University of Michigan, Department of Microbiology and Immunology | Valesano et al. |
| EPI_ISL_487648, EPI_ISL_487654, EPI_ISL_487678, EPI_ISL_487698, EPI_ISL_487728, EPI_ISL_487764, EPI_ISL_487788, EPI_ISL_487794, EPI_ISL_487802, EPI_ISL_487811, EPI_ISL_487824, EPI_ISL_487846, EPI_ISL_487865, EPI_ISL_487869, EPI_ISL_487876, EPI_ISL_487883, EPI_ISL_487918, EPI_ISL_487925, EPI_ISL_487933, EPI_ISL_487957, EPI_ISL_487993 | | | |
| see above | Virology Department, Royal Infirmary of Edinburgh, NHS Lothian / School of Biological Sciences, University of Edinburgh | Wellcome Sanger Institute for the COVID-19 Genomics UK (COG-UK) consortium | McHugh M, Dewar R, Rooke S, O'Toole Á, Scher E, Hill V, McCrone JT, Colquhoun R, Yu X, Jackson B, Rambaut A, Templeton K and Alex Alderton, Roberto Amato, Sonia Goncalves, Ewan Harrison, David K. Jackson, Ian Johnston, Dominic Kwiatkowski, Cordelia Langford, John Sillitoe on behalf of the Wellcome Sanger Institute COVID-19 Surveillance Team (http://www.sanger.ac.uk/covid-team) |
| EPI_ISL_488195, EPI_ISL_488215, EPI_ISL_488312, EPI_ISL_488362, EPI_ISL_488394, EPI_ISL_488399, EPI_ISL_488404 | PHE South West Regional Laboratory, National Infection Service | Wellcome Sanger Institute for the COVID-19 Genomics UK (COG-UK) consortium | Stephanie Hutchings, Hannah Pymont, Dr Peter Muir, Barry Vipond, Rich Hopes; and Alex Alderton, Roberto Amato, Sonia Goncalves, Ewan Harrison, David K. Jackson, Ian Johnston, Dominic Kwiatkowski, Cordelia Langford, John Sillitoe on behalf of the Wellcome Sanger Institute COVID-19 Surveillance Team (http://www.sanger.ac.uk/covid-team) |
| EPI_ISL_488458, EPI_ISL_488459, EPI_ISL_488463, EPI_ISL_488464, EPI_ISL_488467, EPI_ISL_488469, EPI_ISL_488472, EPI_ISL_488473, EPI_ISL_488474, EPI_ISL_488479, EPI_ISL_488480, EPI_ISL_488482, EPI_ISL_488486, EPI_ISL_488487, EPI_ISL_488491, EPI_ISL_488493, EPI_ISL_488494, EPI_ISL_488495, EPI_ISL_488497, EPI_ISL_488498, EPI_ISL_488503, EPI_ISL_488506, EPI_ISL_488507, EPI_ISL_488513, EPI_ISL_488514, EPI_ISL_488515, EPI_ISL_488517, EPI_ISL_488521, EPI_ISL_488535, EPI_ISL_488544, EPI_ISL_488548, EPI_ISL_488549, EPI_ISL_488550, EPI_ISL_488551, EPI_ISL_488552, EPI_ISL_488555, EPI_ISL_488556, EPI_ISL_488560, EPI_ISL_488563, EPI_ISL_488564, EPI_ISL_488565, EPI_ISL_488571, EPI_ISL_488573, EPI_ISL_488574, EPI_ISL_488576, EPI_ISL_488578, EPI_ISL_488581, EPI_ISL_488582, EPI_ISL_488584, EPI_ISL_488585, EPI_ISL_488591, EPI_ISL_488593, EPI_ISL_488596, EPI_ISL_488599, EPI_ISL_488601, EPI_ISL_488603, EPI_ISL_488605, EPI_ISL_488606, EPI_ISL_488613, EPI_ISL_488614, EPI_ISL_488616, EPI_ISL_488617, EPI_ISL_488619, EPI_ISL_488627, EPI_ISL_488631, EPI_ISL_488633, EPI_ISL_488635, EPI_ISL_488638, EPI_ISL_488639, EPI_ISL_488644, EPI_ISL_488646, EPI_ISL_488647, EPI_ISL_488654, EPI_ISL_488655, EPI_ISL_488659, EPI_ISL_488662, EPI_ISL_488671, EPI_ISL_488679, EPI_ISL_488680, EPI_ISL_488686, EPI_ISL_488687, EPI_ISL_488689, EPI_ISL_488695, EPI_ISL_488700, EPI_ISL_488704, EPI_ISL_488706, EPI_ISL_488713, EPI_ISL_488715, EPI_ISL_488717, EPI_ISL_488718, EPI_ISL_488719, EPI_ISL_488720, EPI_ISL_488724, EPI_ISL_488726, EPI_ISL_488727, EPI_ISL_488728, EPI_ISL_488730, EPI_ISL_488732, EPI_ISL_488733, EPI_ISL_488734, EPI_ISL_488740, EPI_ISL_488746, EPI_ISL_488754 | | | |
| see above | NU-OMICS DNA Sequencing research facility, Northumbria University | Wellcome Sanger Institute for the COVID-19 Genomics UK (COG-UK) consortium | Chris Duncan, Sheia Waugh, Shirelle Burton-Fanning, Gary Eltringham, Jennifer Collins, Brendan Payne, Yusri Taha, Emma Swindells, Jane Greenaway, Edward Barton, Garren Scott, Debra Padgett, Clive Graham, Sarah Essex, Steve Ligget, Paul Baker, Lynn Dover, Wen Yew, Gary Black, John Allan, Joshua Loh, Greg Young, Matthew Bashton, Andrew Nelson, Darren Smith and Alex Alderton, Roberto Amato, Sonia Goncalves, Ewan Harrison, David K. Jackson, Ian Johnston, Dominic Kwiatkowski, Cordelia Langford, John Sillitoe on behalf of the Wellcome Sanger Institute COVID-19 Surveillance Team (http://www.sanger.ac.uk/covid-team) |
| EPI_ISL_488887, EPI_ISL_488892, EPI_ISL_488899, EPI_ISL_488901, EPI_ISL_488905, EPI_ISL_488912, EPI_ISL_488924, EPI_ISL_488925, EPI_ISL_488926, EPI_ISL_488937, EPI_ISL_488943, EPI_ISL_488947, EPI_ISL_488955, EPI_ISL_488958, EPI_ISL_488964, EPI_ISL_488969, EPI_ISL_488977, EPI_ISL_488980, EPI_ISL_488984, EPI_ISL_488985, EPI_ISL_488988, EPI_ISL_488996, EPI_ISL_488997, EPI_ISL_489002, EPI_ISL_489003, EPI_ISL_489015, EPI_ISL_489018, EPI_ISL_489020, EPI_ISL_489026, EPI_ISL_489033, EPI_ISL_489039, EPI_ISL_489047, EPI_ISL_489060, EPI_ISL_489062 | | | |
| see above | Virology Department, Royal Infirmary of Edinburgh, NHS Lothian / School of Biological Sciences, University of Edinburgh | Wellcome Sanger Institute for the COVID-19 Genomics UK (COG-UK) consortium | McHugh M, Dewar R, Rooke S, O'Toole Á, Scher E, Hill V, McCrone JT, Colquhoun R, Yu X, Jackson B, Rambaut A, Templeton K and Alex Alderton, Roberto Amato, Sonia Goncalves, Ewan Harrison, David K. Jackson, Ian Johnston, Dominic Kwiatkowski, Cordelia Langford, John Sillitoe on behalf of the Wellcome Sanger Institute COVID-19 Surveillance Team (http://www.sanger.ac.uk/covid-team) |
| EPI_ISL_489380, EPI_ISL_489393, EPI_ISL_489396, EPI_ISL_489401, EPI_ISL_489407, EPI_ISL_489413, EPI_ISL_489418, EPI_ISL_489429, EPI_ISL_489442, EPI_ISL_489449, EPI_ISL_489455, EPI_ISL_489470, EPI_ISL_489480, EPI_ISL_489489, EPI_ISL_489523, EPI_ISL_489530, EPI_ISL_489536, EPI_ISL_489537, EPI_ISL_489543, EPI_ISL_489547, EPI_ISL_489580 | | | |
| see above | NHSGGC West of Scotland Specialist Virology Centre / MRC-University of Glasgow Centre for Virus Research | Wellcome Sanger Institute for the COVID-19 Genomics UK (COG-UK) consortium | Ana da Silva Filipe, Natasha Johnson, Kathy Smollett, Daniel Mair, Stephen Carmichael, Lily Tong, Jenna Nichols, Elihu Aranday-Cortes, Kirstyn Brunker, Yasmin Parr, Kyriaki Nomikou; Sarah McDonald, Marc Niebel, Patawee Asamaphan; Richard Orton, Joseph Hughes, Sreenu Vattipally, David L Robertson; Alasdair MacLean, Rory Gunson; Kathy Li, Natasha Jesudason, Rajiv Shah, James Shepherd, Antonia Ho, Alice Broos, Emma Thomson and Alex Alderton, Roberto Amato, Sonia Goncalves, Ewan Harrison, David K. Jackson, Ian Johnston, Dominic Kwiatkowski, Cordelia Langford, John Sillitoe on behalf of the Wellcome Sanger Institute COVID-19 Surveillance Team (http://www.sanger.ac.uk/covid-team) |
| EPI_ISL_489581 | NHSGGC West of Scotland Specialist Virology Centre / MRC-University of Glasgow Centre for Virus Research | Wellcome Sanger Institute for the COVID-19 Genomics UK (COG-UK) Consortium | Ana da Silva Filipe, Natasha Johnson, Kathy Smollett, Daniel Mair, Stephen Carmichael, Lily Tong, Jenna Nichols, Elihu Aranday-Cortes, Kirstyn Brunker, Yasmin Parr, Kyriaki Nomikou; Sarah McDonald, Marc Niebel, Patawee Asamaphan; Richard Orton, Joseph Hughes, Sreenu Vattipally, David L Robertson; Alasdair MacLean, Rory Gunson; Kathy Li, Natasha Jesudason, Rajiv Shah, James Shepherd, Antonia Ho, Alice Broos, Emma Thomson and Alex Alderton, Roberto Amato, Sonia Goncalves, Ewan Harrison, David K. Jackson, Ian Johnston, Dominic Kwiatkowski, Cordelia Langford, John Sillitoe on behalf of the Wellcome Sanger Institute COVID-19 Surveillance Team |
| EPI_ISL_489586, EPI_ISL_489589, EPI_ISL_489590, EPI_ISL_489591, EPI_ISL_489592, EPI_ISL_489596, EPI_ISL_489599, EPI_ISL_489600, EPI_ISL_489601, EPI_ISL_489602, EPI_ISL_489604, EPI_ISL_489605, EPI_ISL_489606, EPI_ISL_489609, EPI_ISL_489611, EPI_ISL_489612, EPI_ISL_489615, EPI_ISL_489616, EPI_ISL_489617, EPI_ISL_489619, EPI_ISL_489620, EPI_ISL_489622, EPI_ISL_489627, EPI_ISL_489628, EPI_ISL_489629, EPI_ISL_489640, EPI_ISL_489644, EPI_ISL_489645, EPI_ISL_489652, EPI_ISL_489656, EPI_ISL_489658, EPI_ISL_489660, EPI_ISL_489661, EPI_ISL_489664, EPI_ISL_489665, EPI_ISL_489666, EPI_ISL_489671, EPI_ISL_489674, EPI_ISL_489676, EPI_ISL_489677, EPI_ISL_489678, EPI_ISL_489679, EPI_ISL_489684, EPI_ISL_489685, EPI_ISL_489686, EPI_ISL_489687, EPI_ISL_489689, EPI_ISL_489691, EPI_ISL_489695, EPI_ISL_489696, EPI_ISL_489697, EPI_ISL_489698, EPI_ISL_489699, EPI_ISL_489701, EPI_ISL_489702, EPI_ISL_489703, EPI_ISL_489704, EPI_ISL_489705, EPI_ISL_489706 | | | |
| see above | NHSGGC West of Scotland Specialist Virology Centre / MRC-University of Glasgow Centre for Virus Research | Wellcome Sanger Institute for the COVID-19 Genomics UK (COG-UK) consortium | Ana da Silva Filipe, Natasha Johnson, Kathy Smollett, Daniel Mair, Stephen Carmichael, Lily Tong, Jenna Nichols, Elihu Aranday-Cortes, Kirstyn Brunker, Yasmin Parr, Kyriaki Nomikou; Sarah McDonald, Marc Niebel, Patawee Asamaphan; Richard Orton, Joseph Hughes, Sreenu Vattipally, David L Robertson; Alasdair MacLean, Rory Gunson; Kathy Li, Natasha Jesudason, Rajiv Shah, James Shepherd, Antonia Ho, Alice Broos, Emma Thomson and Alex Alderton, Roberto Amato, Sonia Goncalves, Ewan Harrison, David K. Jackson, Ian Johnston, Dominic Kwiatkowski, Cordelia Langford, John Sillitoe on behalf of the Wellcome Sanger Institute COVID-19 Surveillance Team (http://www.sanger.ac.uk/covid-team) |
| EPI_ISL_489994 | Institute for Medical Research, Infectious Disease Research Centre, National Institutes of Health, Ministry of Health Malaysia | Institute for Medical Research, Infectious Disease Research Centre, National Institutes of Health, Ministry of Health Malaysia | Suppiah J, Mohd-Zawawi Z, Kamel K, Kalyanasundram J, Thayan R |
| EPI_ISL_490035 | South Eastern Area Laboratory Services (SEALS) | NSW Health Pathology - Institute of Clinical Pathology and Medical Research; Westmead Hospital; University of Sydney | CIDM-PH et al. |
| EPI_ISL_490446, EPI_ISL_490447, EPI_ISL_490448, EPI_ISL_490449, EPI_ISL_490450, EPI_ISL_490451, EPI_ISL_490452, EPI_ISL_490453, EPI_ISL_490454, EPI_ISL_490455, EPI_ISL_490456, EPI_ISL_490457, EPI_ISL_490458, EPI_ISL_490459, EPI_ISL_490460, EPI_ISL_490461, EPI_ISL_490462, EPI_ISL_490463, EPI_ISL_490464, EPI_ISL_490465, EPI_ISL_490466, EPI_ISL_490467 | | | |
| see above | Northumbria University / South Tees Hospitals NHS | COVID-19 Genomics UK (COG-UK) Consortium | Darren L Smith,Andrew Nelson,Matthew Bashton,Greg R Young,Joshua Loh,John Allan,Mohammad A Tariq,Giles S Holt,Gary Black,Wen C Yew,Lynn |

| | | | |
|---|---|---|---|
| | Foundation Trust / North Cumbria Integrated Care NHS Foundation Trust / North Tees and Hartlepool NHS Foundation Trust / Newcastle Hospitals NHS Foundation Trust | | Dover,Paul Baker,Steve Liggett,Sarah Essex,Jane Greenaway,Debra Padgett,Clive Graham,Garren Scott,Edward Barton,Emma Swindells,Brendan Payne,Jennifer Collins,Yusri Taha,Gary Eltringham |
| EPI_ISL_490976 | Laboratorio de Referencia Nacional de Virus Respiratorio. Instituto Nacional de Salud Perú | Laboratorio de Referencia Nacional de Biotecnología y Biología Molecular. Instituto Nacional de Salud Perú | Carlos Padilla Rojas, Karolyn Chozo Vega, Priscila Lope Pari, Omar Caceres Rey, Marco Galarza Perez, Maribel Huaringa Nuñez, Johanna Balbuena Torres, Henri Bailon Calderon, Nancy Rojas Serrano. |
| EPI_ISL_490979, EPI_ISL_490980, EPI_ISL_491005, EPI_ISL_491006, EPI_ISL_491007, EPI_ISL_491009, EPI_ISL_491010, EPI_ISL_491011, EPI_ISL_491012, EPI_ISL_491013, EPI_ISL_491014, EPI_ISL_491015, EPI_ISL_491020, EPI_ISL_491021, EPI_ISL_491023, EPI_ISL_491024, EPI_ISL_491025, EPI_ISL_491026, EPI_ISL_491027, EPI_ISL_491028, EPI_ISL_491029 | | | |
| see above | UW Virology Lab | UW Virology Lab | Pavitra Roychoudhury, Hong Xie, Lasata Shrestha, Amin Addetia, Truong Nguyen, Victoria M Rachleff, Meei-Li Huang, Keith R Jerome, Alexander Greninger |
| EPI_ISL_491104 | SC Department of Health and Environmental Control | SC Department of Health and Environmental Control | Flores,H. |
| EPI_ISL_491134 | Oman-National Influenza Center | Biotechnology & OMICs Laboratory | Samiha Al-Kharusi, Sajjad Asaf, Abdul Latif Khan, Samira Al-Mahruqi, Adil Khan, Ahmed Al-Rawahi, Amina Al-Jardani, Hanan Al-Kindi, Intisar Al-Shukri, Ahlam Al-Amri, Aisha Al-Amri, Aisha Al-Busaidi, Adil Al-Wahaibi, Seif Al-Abri, Ahmed Al-Harrasi |
| EPI_ISL_491154 | Oman-National Influenza Center | Biotechnology & OMICs Laboratory | Abdul Latif Khan, Samira Al-Mahruqi, Ahmed Al-Harrasi, Samiha Al-Kharusi, Adil Khan, Ahmed Al-Rawahi, Sajjad Asaf, Amina Al-Jardani, Hanan Al-Kindi, Intisar Al-Shukri, Ahlam Al-Amri, Aisha Al-Amri, Aisha Al-Busaidi, Adil Al-Wahaibi, Seif Al-Abri. |
| EPI_ISL_491545 | Virology Department, Royal Infirmary of Edinburgh, NHS Lothian / School of Biological Sciences, University of Edinburgh | Wellcome Sanger Institute for the COVID-19 Genomics UK (COG-UK) consortium | McHugh M, Dewar R, Rooke S, O'Toole Á, Scher E, Hill V, McCrone JT, Colquhoun R, Yu X, Jackson B, Rambaut A, Templeton K and Alex Alderton, Roberto Amato, Sonia Goncalves, Ewan Harrison, David K. Jackson, Ian Johnston, Dominic Kwiatkowski, Cordelia Langford, John Sillitoe on behalf of the Wellcome Sanger Institute COVID-19 Surveillance Team (http://www.sanger.ac.uk/covid-team) |
| EPI_ISL_491916, EPI_ISL_491917, EPI_ISL_491918, EPI_ISL_491919 | Naval Infectious Diseases Diagnostic Laboratory | Naval Medical Research Center Biological Defense Research Directorate | Logan Voegtly, Regina Cer, Lindsay Glang, Victor Sugiharto, Francisco Malgon Bautista, Hua Wei Chen, Dessiree Pena-Gomez, Megan Schilling, Adrian Paskey, Kyle Long, Mark Simons, Kimberly Bishop-Lilly |
| EPI_ISL_492028 | Child Health Research Foundation | Child Health Research Foundation | Senjuti Saha, Md Saiful Islam Sajib, Roly Malaker, Md Hafizur Rahman, Afroza Akter Tanni, Syed Muktadir Al Sium, Maksuda Islam, Samir K Saha |
| EPI_ISL_492053, EPI_ISL_492054, EPI_ISL_492055, EPI_ISL_492056 | Alaska State Virology Laboratory | Alaska State Virology Laboratory | Chen J et al with Pathogenomics group Dagdag R, Redlinger M, Milton E, George W, Kovalenko A, Drown DM, Bortz E |
| EPI_ISL_492067 | 1. ViroGenetics - BSL3 Laboratory of Virology, Maopolska Centre of Biotechnology, Jagiellonian University; 2. II Department of Internal Medicine, Faculty of Medicine, Jagiellonian University Medical College; 3. Narodowy Instytut Zdrowia Publicznego - Pastwowy Zakad Higieny (NIZP-PZH) | 1. ViroGenetics - BSL3 Laboratory of Virology, Maopolska Centre of Biotechnology, Jagiellonian University; 2. II Department of Internal Medicine, Faculty of Medicine, Jagiellonian University Medical College; 3. Narodowy Instytut Zdrowia Publicznego - Pastwowy Zakad Higieny (NIZP-PZH). | Katarzyna Pancer, Marek Sanak, Aleksandra A. Zasada, Magdalena Rzeczkowska, Tomasz Wokowicz, Katarzyna Zacharczuk, Agnieszka Koakowska-Kulesza, Katarzyna Owczarek, Aleksandra Milewska, Natalia Wolaniuk, Ewelina Hallman-Szeliska, Pawe P abaj, Wojciech Branicki, Krzysztof Pyr |
| EPI_ISL_492129, EPI_ISL_492130, EPI_ISL_492131, EPI_ISL_492149, EPI_ISL_492177 | SA Pathology | SA Pathology | Lex Leong, Chuan Kok Lim, Mark Turra, Ivan Bastian, Geoff Higgins |
| EPI_ISL_492259, EPI_ISL_492298, EPI_ISL_492368, EPI_ISL_492422 | PHE South West Regional Laboratory, National Infection Service | Wellcome Sanger Institute for the COVID-19 Genomics UK (COG-UK) consortium | Stephanie Hutchings, Hannah Pymont, Dr Peter Muir, Barry Vipond, Rich Hopes; and Alex Alderton, Roberto Amato, Sonia Goncalves, Ewan Harrison, David K. Jackson, Ian Johnston, Dominic Kwiatkowski, Cordelia Langford, John Sillitoe on behalf of the Wellcome Sanger Institute COVID-19 Surveillance Team (http://www.sanger.ac.uk/covid-team) |
| EPI_ISL_492863, EPI_ISL_492874, EPI_ISL_492876, EPI_ISL_492890 | Royal Free Hospital / Health Services Laboratories | Wellcome Sanger Institute for the COVID-19 Genomics UK (COG-UK) consortium | Tanzina Haque, Tabitha Mahungu, Dianne Irish, Cate Goodlad, Jenny Cross, Judith Heaney and Alex Alderton, Roberto Amato, Sonia Goncalves, Ewan Harrison, David K. Jackson, Ian Johnston, Dominic Kwiatkowski, Cordelia Langford, John Sillitoe on behalf of the Wellcome Sanger Institute COVID-19 Surveillance Team (http://www.sanger.ac.uk/covid-team) |
| EPI_ISL_493061 | Wyoming Public Health Laboratory | Wyoming Public Health Laboratory | Noah Hull, Rob Christensen, Jim Mildenberger, Joel Sevinsky, Cari Sloma, and Wanda Manley |
| EPI_ISL_493205, EPI_ISL_493206, EPI_ISL_493207 | Virology Lab,Department of Pathology, National Cheng Kung University Hospital | Virology Lab,Department of Pathology, National Cheng Kung University Hospital | Huey-Pin Tsai, et al |
| EPI_ISL_493332 | Istituto Zooprofilattico Sperimentale del Mezzogiorno | INMI Lazzaro Spallanzani IRCCS | Cesare E.M. Gruber, Martina Rueca, Barbara Bartolini, Francesco Messina, Antonino Di Caro, Giovanna Fusco, Maurizio Viscardi, Giorgia Borriello, Maria R. Capobianchi |
| EPI_ISL_493333 | Istituto Zooprofilattico Sperimentale del Mezzogiorno | INMI Lazzaro Spallanzani IRCCS | Barbara Bartolini, Martina Rueca, Cesare E.M. Gruber, Francesco Messina, Antonino Di Caro, Giovanna Fusco, Maurizio Viscardi, Giorgia Borriello, Maria R. Capobianchi |
| EPI_ISL_493334 | Instituto de Diagnostico y Referencia Epidemiologicos (INDRE) | Instituto de Diagnostico y Referencia Epidemiologicos (INDRE) | Gisela Barrera-Badillo , Abril Rodriguez-Maldonado, Claudia Wong-Arambula , Natividad Cruz-Ortiz, Tatiana Nunez-Garcia, Dayanira Arellano-Suarez, Fabiola Garces-Ayala, Edgar Mendieta-Condado, Lucia Hernandez-Rivas, Irma Lopez-Martinez, Ernesto Ramirez-Gonzalez. |
| EPI_ISL_493415, EPI_ISL_493417, EPI_ISL_493419 | National Public Health Laboratory, National Centre for Infectious Diseases | National Public Health Laboratory, National Centre for Infectious Diseases | Mak TM, Octavia S, Zhou Z, Chavatte JM, Cui L, Lin RTP |
| EPI_ISL_494286 | Wales Specialist Virology Centre Sequencing lab: Pathogen Genomics Unit | COVID-19 Genomics UK (COG-UK) Consortium | Catherine Moore, Johnathan Evans, Laura Gifford, Malorie Perry, Simon Cottrell, Angela Marchbank, Alec Birchley, Alexander Adams, Amy Gaskin, Bree Gatica-Wilcox, Jason Coombes, Joel Southgate, Lauren Gilbert, Lee Graham, Nicole Pacchiarini, Sara Kumziene-Summerhayes, Sarah Taylor, Sophie Jones, Sara Rey, Matthew Bull, Joanne Watkins, Sally Corden, Tom Connor |
| EPI_ISL_494596, EPI_ISL_494597, EPI_ISL_494598, EPI_ISL_494599, EPI_ISL_494600, EPI_ISL_494601, EPI_ISL_494602, EPI_ISL_494603, EPI_ISL_494604, EPI_ISL_494605, EPI_ISL_494606, EPI_ISL_494607, EPI_ISL_494608, EPI_ISL_494610, EPI_ISL_494612, EPI_ISL_494614, EPI_ISL_494615 | | | |
| see above | Scripps Medical Laboratory | Andersen lab at Scripps Research | SEARCH Alliance San Diego with Michael Quigley, Ellen Stefanski, Ian Mchardy |
| EPI_ISL_495594, EPI_ISL_495596 | University of Michigan Clinical Microbiology Laboratory | Lauring Lab, University of Michigan, Department of Microbiology and Immunology | Valesano et al. |
| EPI_ISL_495597 | Mayo Clinic & Mayo Clinic Laboratories | Minnesota Department of Health, Public Health Laboratory | Matt Plumb, Jacob Garfin, and Xiong Wang |
| EPI_ISL_495658 | Seattle Flu Study | Seattle Flu Study | Deborah A. Nickerson, Chris D. Frazar, Jover Lee, Benjamin Pelle, Matthew Richardson, Amanda Adler, Elisabeth Brandstetter, Peter D. Han, Kairsten Fay, Misja Ilcisin, Kirsten Lacombe, Thomas R. Sibley, Melissa Truong, Caitlin R. Wolf, Karen Cowgill, Stephanie Schrag, Jeff Duchin, Michael Boeckh, Janet A. Englund, Michael Famulare, Barry R. Lutz, Mark J. Rieder, Lea M. Starita, Matthew Thompson, Helen Y. Chu, Trevor Bedford, Jay Shendure |
| EPI_ISL_496370 | Infectolab | Andersen lab at Scripps Research | SEARCH Alliance San Diego with Samuel Navarro Alvarez, Carlos A. Cota Haros, Octavio Renteria Pacheco |
| EPI_ISL_496745, EPI_ISL_496746, EPI_ISL_496751, EPI_ISL_496752, EPI_ISL_496757, EPI_ISL_496759, EPI_ISL_496760, EPI_ISL_496761, EPI_ISL_496763, EPI_ISL_496765, EPI_ISL_496766, EPI_ISL_496768, EPI_ISL_496769, EPI_ISL_496771, EPI_ISL_496772, EPI_ISL_496773, EPI_ISL_496774, EPI_ISL_496775, EPI_ISL_496776, EPI_ISL_496777, EPI_ISL_496778, EPI_ISL_496779, EPI_ISL_496781, EPI_ISL_496782, EPI_ISL_496783, EPI_ISL_496784, EPI_ISL_496785 | | | |
| see above | Gorgas Memorial Laboratory of Health Studies | Gorgas Memorial Laboratory of Health Studies | Danilo Franco, Claudia Gonzalez Sandra Lopez-Verges, Alexander A Martinez |
| EPI_ISL_497745 | Instituto Nacional de Salud, Bogotá, Colombia | Instituto Nacional de Salud, Bogotá, Colombia | Katherine Laiton-Donato, Diego A. Álvarez-Díaz, Carlos Franco-Muñoz, Jonathan Reales, Diego Andrés Prada, Jose A. Usme-Ciro, Nicolas D. Franco-Sierra, Zulma M. Cucunubá, Christian Julian VillabonaArenas, Liz Villabona-Arenas, Sussy Echeverria, Astrid C. Flórez, Carolina Ferro, Diana Marcela Walteros-Acero, Franklin Prieto, Carlos Andrés Durán, Martha Lucia Ospina Martinez, Marcela Mercado-Reyes |
| EPI_ISL_497806, EPI_ISL_497811, EPI_ISL_497819, EPI_ISL_497821, EPI_ISL_497828 | Department of Microbiology, The University of Hong Kong | Department of Microbiology, The University of Hong Kong | Kelvin K.W. To, Kwok-Yung Yuen |
| EPI_ISL_498156, EPI_ISL_498157, EPI_ISL_498161, EPI_ISL_498162 | Instituto Nacional de Salud, Bogotá, Colombia | Instituto Nacional de Salud, Bogotá, Colombia | Katherine Laiton-Donato, Diego A. Álvarez-Díaz, Carlos Franco-Muñoz, Jonathan Reales, Diego Andrés Prada, Jose A. Usme-Ciro, Nicolas D. Franco-Sierra, Zulma M. Cucunubá, Christian Julian VillabonaArenas, Liz Villabona-Arenas, Sussy Echeverria, Astrid C. Flórez, Carolina Ferro, Diana Marcela Walteros-Acero, Franklin Prieto, Carlos Andrés Durán, Martha Lucia Ospina Martinez, Marcela Mercado-Reyes |

| | | | |
|---|---|---|---|
| EPI_ISL_498261 | Hospital for Tropical Diseases | COVID-19 Network Investigations (CONI) Alliance | Elizabeth Batty, Nantarat Chantawat, Wasun Chantratita, Thanat Chookajorn, Stefan Fernandez, Angkana Huang, Weena Janwitthayanan, Akanitt Jittmittraphap, Anthony R. Jones, Khajohn Joonsalak, Chonticha Klungtong, Theerarat Kochakarn, Namfon Kotanan, Krittikorn Kumpornsin, Pornsawan Leaungwutiwong, Wudtichai Manasatienkij, Bhakbhoom Panthan, Ekawat Pasomsub, Kingkan Rakmanee, Insee Sensorn, Janjira Thaipadungpanit, Arporn Wangwiwatsin,Treewat Watthanachockchai |
| EPI_ISL_498470, EPI_ISL_498476, EPI_ISL_498510, EPI_ISL_498516, EPI_ISL_498521, EPI_ISL_498522, EPI_ISL_498527, EPI_ISL_498532, EPI_ISL_498536, EPI_ISL_498540 | ACT Pathology | Schwessinger Lab | Ashley Jones, Benjamin Schwessinger, Robert Lanfear, Robyn N Hall, Megan McDonald, Ming-Dao Chia, Kevin Murray, Craig Kennedy, Karina Kennedy |
| EPI_ISL_500369, EPI_ISL_500375, EPI_ISL_500380, EPI_ISL_500381, EPI_ISL_500383, EPI_ISL_500386, EPI_ISL_500406, EPI_ISL_500433, EPI_ISL_500440, EPI_ISL_500441, EPI_ISL_500442, EPI_ISL_500443, EPI_ISL_500444, EPI_ISL_500445, EPI_ISL_500446, EPI_ISL_500447, EPI_ISL_500452, EPI_ISL_500453, EPI_ISL_500454, EPI_ISL_500455 | | | |
| see above | Centro de Investigación Biomédica de La Rioja - Hospital San Pedro Logroño | SeqCOVID-SPAIN consortium/IBV(CSIC) | María de Toro, José Manuel Azcona Gutiérrez, María Pilar Bea Escudero, Miriam Blasco Alberdi and SeqCOVID-SPAIN consortium |
| EPI_ISL_500500 | University of Washington Virology Lab | University of Washington Virology Lab | Pavitra Roychoudhury, Hong Xie, Lasata Shrestha, Amin Addetia, Truong Nguyen, Victoria M Rachleff, Meei-Li Huang, Keith R Jerome, Alexander Greninger |
| EPI_ISL_500512, EPI_ISL_500533, EPI_ISL_500534 | Mayo Clinic Laboratories | University of Washington Virology Lab | Pavitra Roychoudhury, Hong Xie, Lasata Shrestha, Amin Addetia, Truong Nguyen, Victoria M Rachleff, Meei-Li Huang, Keith R Jerome, Alexander Greninger |
| EPI_ISL_500705 | Area of Virology, Serology and Virology Division (SAViD), New South Wales Health Pathology Randwick | Area of Virology, Serology and Virology Division (SAViD), New South Wales Health Pathology Randwick | Rawlinson, W. |
| EPI_ISL_501080 | Mayo Clinic Laboratories | University of Washington Virology Lab | Pavitra Roychoudhury, Hong Xie, Lasata Shrestha, Amin Addetia, Truong Nguyen, Victoria M Rachleff, Meei-Li Huang, Keith R Jerome, Alexander Greninger |
| EPI_ISL_501168 | Baylor College of Medicine | Baylor College of Medicine: HGSC | Vasanthi Avadhanula, Erin Nicholson, David Henke, Pedro Piedra, Harsha Doddapaneni, Donna Muzny, Qingchang Meng, Hsu Chao, Zeineen Momin, Hua Shen, George Weissenberger, Kavya Kottapalli, Yimiti Meiheerguli, Sejal Salvi, Ginger Metcalf, Vipin Menon, Sara J.J. Cregeen, Matthew C. Ross, Tulin Ayvaz, Richard Sucgang, Kristi L. Hoffman, Matthew Wong, Joseph F. Petrosino |
| EPI_ISL_501193, EPI_ISL_501194, EPI_ISL_501195, EPI_ISL_501196, EPI_ISL_501197, EPI_ISL_501198, EPI_ISL_501199, EPI_ISL_501201, EPI_ISL_501202, EPI_ISL_501203, EPI_ISL_501204, EPI_ISL_501205 | | | |
| see above | Department of Medical Microbiology, University Malaya Medical Centre | Department of Medical Microbiology, Faculty of Medicine, University of Malaya | Yoong Min CHONG, Jennifer Chong, I-Ching SAM, Yoke Fun CHAN, University Malaya Medical Centre COVID Team |
| EPI_ISL_501241, EPI_ISL_501242 | Hellenic Pasteur Institute, National Influenza Reference laboratory of Southern Greece & Unit of Bioinformatics and Applied Genomics | Hellenic Pasteur Institute, National Influenza Reference laboratory of Southern Greece & Unit of Bioinformatics and Applied Genomics | Vasiliki Pogka, Timokratis Karamitros, Athanasios Kossyvakis, Antonios Kalliaropoulos, Horefti Elina, Evangelidou Maria, Androniki Voulgari-Kokota, Aspasia Kontou, Andreas Mentis |
| EPI_ISL_501585 | PHE South West Regional Laboratory, National Infection Service | Wellcome Sanger Institute for the COVID-19 Genomics UK (COG-UK) consortium | Stephanie Hutchings, Hannah Pymont, Dr Peter Muir, Barry Vipond, Rich Hopes; and Alex Alderton, Roberto Amato, Sonia Goncalves, Ewan Harrison, David K. Jackson, Ian Johnston, Dominic Kwiatkowski, Cordelia Langford, John Sillitoe on behalf of the Wellcome Sanger Institute COVID-19 Surveillance Team (http://www.sanger.ac.uk/covid-team) |
| EPI_ISL_501620, EPI_ISL_501628 | Lab Microbiology, Pathology Department, William Harvey Hospital | Wellcome Sanger Institute for the COVID-19 Genomics UK (COG-UK) consortium | Samuel Moses, Hannah Lowe, Felicity Ryan and Alex Alderton, Roberto Amato, Sonia Goncalves, Ewan Harrison, David K. Jackson, Ian Johnston, Dominic Kwiatkowski, Cordelia Langford, John Sillitoe on behalf of the Wellcome Sanger Institute COVID-19 Surveillance Team (http://www.sanger.ac.uk/covid-team) |
| EPI_ISL_506978, EPI_ISL_506981, EPI_ISL_506982, EPI_ISL_506983, EPI_ISL_506984, EPI_ISL_506985, EPI_ISL_506986 | Division of Viral Diseases, Center for Laboratory Control of Infectious Diseases, Korea Centers for Diseases Control and Prevention | Division of Viral Diseases, Center for Laboratory Control of Infectious Diseases, Korea Centers for Diseases Control and Prevention | Jeong-Min Kim, Yoon-Seok Chung, Namjoo Lee, Sang Hee Woo, Hye-Jun Jo, Heui Man Kim, Jun-Sub Kim, Dong Hyun Song, Daesang Lee, Seong Tae Jeong, Myung Guk Han |
| EPI_ISL_506998 | Department of Medical Microbiology, University Malaya Medical Centre | Department of Medical Microbiology, Faculty of Medicine, University of Malaya | Yoong Min CHONG, Jennifer Chong, I-Ching SAM, Yoke Fun CHAN, University Malaya Medical Centre COVID Team |
| EPI_ISL_507008 | National Veterinary Institute | National Veterinary Institute | Siamak Zohari |
| EPI_ISL_507024, EPI_ISL_507036, EPI_ISL_507037, EPI_ISL_507038 | unknown | Infectious Diseases Research, King Abdullah International Medical Research Center (KAIMRC) | Alghoribi,M.F. |
| EPI_ISL_507043, EPI_ISL_507074, EPI_ISL_507075, EPI_ISL_507076, EPI_ISL_507077, EPI_ISL_507078, EPI_ISL_507093 | University College London Hospital | COVID-19 Genomics UK (COG-UK) Consortium | Judith Heaney, Matthew Byott, Catherine Houlihan, Dan Frampton, Stuart Kirk, Moira Spyer and Eleni Nastouli |
| EPI_ISL_507127, EPI_ISL_507129 | Northumbria University / South Tees Hospitals NHS Foundation Trust / North Cumbria Integrated Care NHS Foundation Trust / North Tees and Hartlepool NHS Foundation Trust / Newcastle Hospitals NHS Foundation Trust | COVID-19 Genomics UK (COG-UK) Consortium | Darren L Smith,Andrew Nelson,Matthew Bashton,Greg R Young,Joshua Loh,John Allan,Mohammad A Tariq,Giles S Holt,Gary Black,Wen C Yew,Lynn Dover,Paul Baker,Steve Liggett,Sarah Essex,Jane Greenaway,Debra Padgett,Clive Graham,Garren Scott,Edward Barton,Emma Swindells,Brendan Payne,Jennifer Collins,Yusri Taha,Gary Eltringham |
| EPI_ISL_507449, EPI_ISL_507451, EPI_ISL_507453, EPI_ISL_507454, EPI_ISL_507456, EPI_ISL_507458, EPI_ISL_507464, EPI_ISL_507465, EPI_ISL_507466, EPI_ISL_507467, EPI_ISL_507468, EPI_ISL_507469, EPI_ISL_507752, EPI_ISL_507754, EPI_ISL_507762, EPI_ISL_507763, EPI_ISL_507790, EPI_ISL_507791, EPI_ISL_507792, EPI_ISL_507794, EPI_ISL_507795, EPI_ISL_507796, EPI_ISL_507798, EPI_ISL_507799, EPI_ISL_507800, EPI_ISL_507801, EPI_ISL_507802, EPI_ISL_507803, EPI_ISL_507805, EPI_ISL_507806, EPI_ISL_507808, EPI_ISL_507810, EPI_ISL_507817, EPI_ISL_507822, EPI_ISL_507824, EPI_ISL_507827, EPI_ISL_507828, EPI_ISL_507830, EPI_ISL_507831, EPI_ISL_507832, EPI_ISL_507833, EPI_ISL_507834, EPI_ISL_507836, EPI_ISL_507844, EPI_ISL_507852, EPI_ISL_507854, EPI_ISL_507855, EPI_ISL_507857, EPI_ISL_507858, EPI_ISL_507861, EPI_ISL_507862, EPI_ISL_507865, EPI_ISL_507869, EPI_ISL_507873, EPI_ISL_507874, EPI_ISL_507878, EPI_ISL_507879, EPI_ISL_507883, EPI_ISL_507888, EPI_ISL_507889, EPI_ISL_507891, EPI_ISL_507894, EPI_ISL_507895, EPI_ISL_507896, EPI_ISL_507898, EPI_ISL_507899, EPI_ISL_507903, EPI_ISL_507906, EPI_ISL_507907, EPI_ISL_507910, EPI_ISL_507911, EPI_ISL_507912, EPI_ISL_507915, EPI_ISL_507927, EPI_ISL_507929, EPI_ISL_507933 | | | |
| see above | Michigan Department of Health and Human Services, Bureau of Laboratories | Michigan Department of Health and Human Services, Bureau of Laboratories | Blankenship HM, Riner D, Soehnlen MK |
| EPI_ISL_507977, EPI_ISL_507978, EPI_ISL_507979, EPI_ISL_507980, EPI_ISL_507981, EPI_ISL_507982, EPI_ISL_507983 | Minnesota Department of Health, Public Health Laboratory | Minnesota Department of Health, Public Health Laboratory | Matt Plumb, Jacob Garfin, and Xiong Wang |
| EPI_ISL_508141 | SA Pathology | SA Pathology | Lex Leong, Chuan Kok Lim, Mark Turra, Ivan Bastian, Geoff Higgins |
| EPI_ISL_508338, EPI_ISL_508349 | Institute of Post Graduate Medical Education & Research | National Institute of Biomedical Genomics | Arindam Maitra, Aritra Biswas, Jayeeta Haldar, Raja Ray, Monimoy Banerjee, Saumitra Das |
| EPI_ISL_508627, EPI_ISL_508631, EPI_ISL_508651, EPI_ISL_508661, EPI_ISL_508676, EPI_ISL_508680 | Departamento de Microbiología, CDB, Hospital Clínic, Barcelona | SeqCOVID-SPAIN consortium/IBV(CSIC) | Andrea Vergara, Mikel Martínez, Elisa Rubio, Jéssica Navero, Aida Peiró and SeqCOVID-SPAIN consortium |
| EPI_ISL_508896, EPI_ISL_508897, EPI_ISL_508898, EPI_ISL_508899, EPI_ISL_508900, EPI_ISL_508901, EPI_ISL_508902, EPI_ISL_508903, EPI_ISL_508904, EPI_ISL_508905, EPI_ISL_508906, EPI_ISL_508907, EPI_ISL_508908 | | | |
| see above | Institut des Agents Infectieux (IAI), Hospices Civils de Lyon | CNR Virus des Infections Respiratoires - France SUD | Antonin Bal, Gregory Destras, Gwendolyne Burfin, Solenne Brun, Carine Moustaud, Raphaelle Lamy, Alexandre Gaymard, Maude Bouscambert-Duchamp, Florence Morfin-Sherpa, Martine Valette, Bruno Lina, Laurence Josset |
| EPI_ISL_508958 | Centre Hospitalier Alpes Leman | CNR Virus des Infections Respiratoires - France SUD | Antonin Bal, Gregory Destras, Gwendolyne Burfin, Solenne Brun, Carine Moustaud, Raphaelle Lamy, Alexandre Gaymard, Maude Bouscambert-Duchamp, |

| | | | |
|---|---|---|---|
| | | | Florence Morfin-Sherpa, Martine Valette, Bruno Lina, Laurence Josset |
| EPI_ISL_508987 | CNR Virus des Infections Respiratoires - France SUD | CNR Virus des Infections Respiratoires - France SUD | Antonin Bal, Gregory Destras, Gwendolyne Burfin, Solenne Brun, Alexandre Gaymard, Maude Bouscambert-Duchamp, Florence Morfin-Sherpa, Martine Valette, Bruno Lina, Laurence Josset |
| EPI_ISL_509010 | Institut des Agents Infectieux (IAI), Hospices Civils de Lyon | CNR Virus des Infections Respiratoires - France SUD | Antonin Bal, Gregory Destras, Gwendolyne Burfin, Solenne Brun, Carine Moustaud, Raphaelle Lamy, Alexandre Gaymard, Maude Bouscambert-Duchamp, Florence Morfin-Sherpa, Martine Valette, Bruno Lina, Laurence Josset |
| EPI_ISL_509012 | Centre Hospitalier Alpes Leman | CNR Virus des Infections Respiratoires - France SUD | Antonin Bal, Gregory Destras, Gwendolyne Burfin, Solenne Brun, Carine Moustaud, Raphaelle Lamy, Alexandre Gaymard, Maude Bouscambert-Duchamp, Florence Morfin-Sherpa, Martine Valette, Bruno Lina, Laurence Josset |
| EPI_ISL_509069, EPI_ISL_509070, EPI_ISL_509071 | OHSU Lab Services Molecular Microbiology Lab | Oregon SARS-CoV-2 Genome Sequencing Center | Brendan L. O'Connell, Ruth V. Nichols, Sally B. Grindstaff, Alec J. Hirsch, Guang Fan, Daniel N. Streblow, William B. Messer, Andrew C. Adey, Benjamin N. Bimber, Brian J. O'Roak |
| EPI_ISL_509713 | Belize Ministry of Health | Pathogen Discovery, Respiratory Viruses Branch, Division of Viral Diseases, Centers for Disease Control and Prevention | Jing Zhang, Ying Tao, Krista Queen, Anna Uehara, Yan Li, Clinton Paden, Haibin Wang, Suxiang Tong |
| EPI_ISL_509752, EPI_ISL_509753, EPI_ISL_509754, EPI_ISL_509755, EPI_ISL_509756, EPI_ISL_509757, EPI_ISL_509758, EPI_ISL_509759, EPI_ISL_509760, EPI_ISL_509761, EPI_ISL_509762, EPI_ISL_509763, EPI_ISL_509764, EPI_ISL_509765, EPI_ISL_509766, EPI_ISL_509767, EPI_ISL_509770, EPI_ISL_509771 | | | |
| see above | Florida Bureau of Public Health Laboratories | Florida Bureau of Public Health Laboratories | Sarah Schmedes, Jason Blanton |
| EPI_ISL_510070 | Instituto de Investigaciones Biomédicas de Barcelona (CSIC), Hospital Clinic i Provincial de Barcelona, Instituto de Biomedicina de Valencia (CSIC), Hospital de Sant Pau | SeqCOVID-SPAIN consortium/IBV(CSIC) | Anna M. Planas, Mª Angeles Marcos, Miguel J. Martínez, Andrea Vergara, Alex Soriano, Jordi Pérez Tur, Israel Fernández Cadenas and SeqCOVID-SPAIN consortium |
| EPI_ISL_510114 | Hospital General Universitario Gregorio Marañón | SeqCOVID-SPAIN consortium/IBV(CSIC) | Laura Pérez-Lago, Marta Herranz, Jon Sicilia, Julia Suárez, Pilar Catalán, Patricia Muñoz, Darío García de Viedma and SeqCOVID-SPAIN consortium |
| EPI_ISL_510461, EPI_ISL_510463 | Instituto de Investigaciones Biomédicas de Barcelona (CSIC), Hospital Clinic i Provincial de Barcelona, Instituto de Biomedicina de Valencia (CSIC), Hospital de Sant Pau | SeqCOVID-SPAIN consortium/IBV(CSIC) | Anna M. Planas, Mª Angeles Marcos, Miguel J. Martínez, Andrea Vergara, Alex Soriano, Jordi Pérez Tur, Israel Fernández Cadenas and SeqCOVID-SPAIN consortium |
| EPI_ISL_510818 | Klinisk mikrobiologi Vasternorrland | The Public Health Agency of Sweden | Oskar Karlsson Lindsjo, Maria Lind Karlberg, Mattias Haukland, Reza Advani, Olov Svartstrom, Anna-Malin Linde, Sandra Broddesson, Petra Edquist, Mia Brytting, Anna Risberg, Karin Tegmark-Wisell |
| EPI_ISL_510897, EPI_ISL_510898, EPI_ISL_510899, EPI_ISL_510900, EPI_ISL_510901, EPI_ISL_510902, EPI_ISL_510903, EPI_ISL_510904, EPI_ISL_510905, EPI_ISL_510906, EPI_ISL_510907, EPI_ISL_510908, EPI_ISL_510916, EPI_ISL_510917, EPI_ISL_510918, EPI_ISL_510919, EPI_ISL_510920, EPI_ISL_510926, EPI_ISL_510940, EPI_ISL_510942, EPI_ISL_510943, EPI_ISL_510944, EPI_ISL_510945, EPI_ISL_510981, EPI_ISL_510982, EPI_ISL_510983, EPI_ISL_510984, EPI_ISL_510985, EPI_ISL_510986, EPI_ISL_510987, EPI_ISL_511026, EPI_ISL_511107, EPI_ISL_511108, EPI_ISL_511109, EPI_ISL_511110, EPI_ISL_511111, EPI_ISL_511112, EPI_ISL_511113, EPI_ISL_511114, EPI_ISL_511115, EPI_ISL_511116, EPI_ISL_511117, EPI_ISL_511118, EPI_ISL_511119, EPI_ISL_511120, EPI_ISL_511121, EPI_ISL_511122, EPI_ISL_511123, EPI_ISL_511124, EPI_ISL_511125, EPI_ISL_511126, EPI_ISL_511127, EPI_ISL_511128, EPI_ISL_511129, EPI_ISL_511130, EPI_ISL_511131, EPI_ISL_511132, EPI_ISL_511133, EPI_ISL_511134, EPI_ISL_511135, EPI_ISL_511136, EPI_ISL_511137, EPI_ISL_511138, EPI_ISL_511155, EPI_ISL_511156, EPI_ISL_511157, EPI_ISL_511158, EPI_ISL_511159, EPI_ISL_511160, EPI_ISL_511161, EPI_ISL_511162, EPI_ISL_511163, EPI_ISL_511164, EPI_ISL_511165, EPI_ISL_511166, EPI_ISL_511167, EPI_ISL_511168, EPI_ISL_511169, EPI_ISL_511170, EPI_ISL_511171, EPI_ISL_511172, EPI_ISL_511173, EPI_ISL_511174, EPI_ISL_511175, EPI_ISL_511176, EPI_ISL_511177, EPI_ISL_511179, EPI_ISL_511180 | | | |
| see above | Instituto Nacional de Saude (INSA) | Instituto Nacional de Saude (INSA) | Borges et al |
| EPI_ISL_511190, EPI_ISL_511191, EPI_ISL_511307 | Instituto Nacional de Saude (INSA) and Instituto Gulbenkian de Ciencia (IGC) | Instituto Nacional de Saude (INSA) and Instituto Gulbenkian de Ciencia (IGC) | Borges et al |
| EPI_ISL_511329, EPI_ISL_511340, EPI_ISL_511341, EPI_ISL_511342, EPI_ISL_511343, EPI_ISL_511344, EPI_ISL_511345, EPI_ISL_511346, EPI_ISL_511347, EPI_ISL_511348, EPI_ISL_511349, EPI_ISL_511350, EPI_ISL_511351, EPI_ISL_511352, EPI_ISL_511353, EPI_ISL_511354, EPI_ISL_511355, EPI_ISL_511356, EPI_ISL_511357, EPI_ISL_511358, EPI_ISL_511359, EPI_ISL_511360, EPI_ISL_511365, EPI_ISL_511370, EPI_ISL_511375, EPI_ISL_511376, EPI_ISL_511382, EPI_ISL_511383, EPI_ISL_511384, EPI_ISL_511387, EPI_ISL_511388, EPI_ISL_511395, EPI_ISL_511399, EPI_ISL_511404, EPI_ISL_511406, EPI_ISL_511407, EPI_ISL_511409, EPI_ISL_511414, EPI_ISL_511415, EPI_ISL_511419, EPI_ISL_511420, EPI_ISL_511421, EPI_ISL_511422, EPI_ISL_511423, EPI_ISL_511424, EPI_ISL_511436, EPI_ISL_511438, EPI_ISL_511439, EPI_ISL_511442, EPI_ISL_511449, EPI_ISL_511453, EPI_ISL_511454, EPI_ISL_511457, EPI_ISL_511461, EPI_ISL_511464, EPI_ISL_511466, EPI_ISL_511469, EPI_ISL_511480 | | | |
| see above | Instituto Nacional de Saude (INSA) | Instituto Nacional de Saude (INSA) | Borges et al |
| EPI_ISL_511485, EPI_ISL_511486 | Instituto Nacional de Saude (INSA) | Instituto Nacional de Saude (INSA) and Instituto Gulbenkian de Ciencia (IGC) | Borges et al |
| EPI_ISL_511516, EPI_ISL_511517, EPI_ISL_511518, EPI_ISL_511519, EPI_ISL_511520, EPI_ISL_511521, EPI_ISL_511522, EPI_ISL_511523, EPI_ISL_511524, EPI_ISL_511525, EPI_ISL_511526, EPI_ISL_511527, EPI_ISL_511528, EPI_ISL_511553, EPI_ISL_511554, EPI_ISL_511555, EPI_ISL_511556, EPI_ISL_511557, EPI_ISL_511558, EPI_ISL_511585, EPI_ISL_511586, EPI_ISL_511587, EPI_ISL_511588, EPI_ISL_511589, EPI_ISL_511590, EPI_ISL_511591, EPI_ISL_511592, EPI_ISL_511593, EPI_ISL_511594, EPI_ISL_511595, EPI_ISL_511596, EPI_ISL_511597, EPI_ISL_511598, EPI_ISL_511599, EPI_ISL_511600, EPI_ISL_511601, EPI_ISL_511602, EPI_ISL_511603, EPI_ISL_511604, EPI_ISL_511605, EPI_ISL_511606, EPI_ISL_511607, EPI_ISL_511608, EPI_ISL_511609, EPI_ISL_511610, EPI_ISL_511611, EPI_ISL_511612, EPI_ISL_511613, EPI_ISL_511614, EPI_ISL_511615, EPI_ISL_511616, EPI_ISL_511617, EPI_ISL_511618, EPI_ISL_511619, EPI_ISL_511620, EPI_ISL_511621, EPI_ISL_511622, EPI_ISL_511623, EPI_ISL_511624, EPI_ISL_511625, EPI_ISL_511626, EPI_ISL_511627, EPI_ISL_511628, EPI_ISL_511629, EPI_ISL_511630, EPI_ISL_511631, EPI_ISL_511632, EPI_ISL_511633, EPI_ISL_511634, EPI_ISL_511635, EPI_ISL_511636, EPI_ISL_511637, EPI_ISL_511638, EPI_ISL_511639, EPI_ISL_511640, EPI_ISL_511641, EPI_ISL_511647, EPI_ISL_511648, EPI_ISL_511649, EPI_ISL_511650, EPI_ISL_511651, EPI_ISL_511652, EPI_ISL_511653, EPI_ISL_511654, EPI_ISL_511655, EPI_ISL_511657, EPI_ISL_511658, EPI_ISL_511659, EPI_ISL_511660, EPI_ISL_511661, EPI_ISL_511662, EPI_ISL_511664, EPI_ISL_511665, EPI_ISL_511666, EPI_ISL_511667, EPI_ISL_511670, EPI_ISL_511674, EPI_ISL_511679, EPI_ISL_511680, EPI_ISL_511681, EPI_ISL_511682, EPI_ISL_511686, EPI_ISL_511734, EPI_ISL_511735, EPI_ISL_511736, EPI_ISL_511748, EPI_ISL_511749, EPI_ISL_511750 | | | |
| see above | Instituto Nacional de Saude (INSA) | Instituto Nacional de Saude (INSA) | Borges et al |
| EPI_ISL_511759, EPI_ISL_511760, EPI_ISL_511761, EPI_ISL_511763, EPI_ISL_511764, EPI_ISL_511765, EPI_ISL_511766, EPI_ISL_511767, EPI_ISL_511768, EPI_ISL_511769, EPI_ISL_511771 | | | |
| see above | Instituto Nacional de Saude (INSA) | Instituto Nacional de Saude (INSA) and Instituto Gulbenkian de Ciencia (IGC) | Borges et al |
| EPI_ISL_512436 | West of Scotland Specialist Virology Centre, NHSGGC / MRC-University of Glasgow Centre for Virus Research | COVID-19 Genomics UK (COG-UK) Consortium | Ana da Silva Filipe, Natasha Johnson, Kathy Smollett, Daniel Mair, Stephen Carmichael, Lily Tong, Jenna Nichols, Elihu Aranday-Cortes, Kirstyn Brunker, Yasmin Parr, Alice Broos, Kyriaki Nomikou; Sarah McDonald, Marc Niebel, Patawee Asamaphan; Richard Orton, Joseph Hughes, Sreenu Vattipally, David L Robertson; Alasdair MacLean, Rory Gunson; Kathy Li, Natasha Jesudason, Rajiv Shah, James Shepherd, Antonia Ho, Emma Thomson |
| EPI_ISL_512491, EPI_ISL_512493, EPI_ISL_512500 | Wales Specialist Virology Centre Sequencing lab: Pathogen Genomics Unit | COVID-19 Genomics UK (COG-UK) Consortium | Catherine Moore, Johnathan Evans, Laura Gifford, Malorie Perry, Simon Cottrell, Angela Marchbank, Alec Birchley, Alexander Adams, Amy Gaskin, Bree Gatica-Wilcox, Jason Coombes, Joel Southgate, Lauren Gilbert, Lee Graham, Nicole Pacchiarini, Sara Kumziene-Summerhayes, Sarah Taylor, Sophie Jones, Sara Rey, Matthew Bull, Joanne Watkins, Sally Corden, Tom Connor |
| EPI_ISL_512738, EPI_ISL_512739, EPI_ISL_512749, EPI_ISL_512750, EPI_ISL_512751, EPI_ISL_512756, EPI_ISL_512764 | PathWest Laboratory Medicine WA | PathWest Laboratory Medicine WA Microbial Surveillance Unit | PathWest Laboratory Medicine WA Microbial Surveillance Unit |
| EPI_ISL_512787, EPI_ISL_512788, EPI_ISL_512790, EPI_ISL_512792, EPI_ISL_512798, EPI_ISL_512803, EPI_ISL_512804, EPI_ISL_512805, EPI_ISL_512808 | Public Health, United States Air Force School of Aerospace Medicine | Public Health, United States Air Force School of Aerospace Medicine | Fries,A.C., Purves,S.M., Meyer,J.R., Javorina,A.K., Connors,B.C., Macias,E.A., Lambert,A.W., Chapleau,R.R., Starr,C.R. |
| EPI_ISL_512811 | Kenema Government Hospital, Ministry of Health and Sanitation | Kenema Government Hospital, Ministry of Health and Sanitation | Goba,A., Momoh,M., Sandi,J., Tomkins-Tinch,C., Siddle,K., Mehta,S., Oluniyi,P., Jalloh,S., Park,D., Andersen,K., Garry,R., Happi,C., Grant,D., Olawoye,I. |
| EPI_ISL_512872 | Ramathibodi Hospital | COVID-19 Network Investigations (CONI) Alliance | Elizabeth Batty, Wasun Chantratita, Thanat Chookajorn, Stefan Fernandez, Angkana Huang, Anthony R. Jones, Khajohn Joonsalak, Chonticha Klungtong, Theerarat Kochakarn, Namfon Kotanan, Krittikorn Kumpornsin, Wudtichai Manasatienkij, Bhakbhoom Panthan, Ekawat Pasomsub, Kingkan Rakmanee, Insee Sensorn, Janjira Thaipadungpanit, Arporn Wangwiwatsin,Treewat Watthanachockchai |
| EPI_ISL_513305, EPI_ISL_513306, EPI_ISL_513307 | Department of Infection Prevention and Infectious Diseases, University Hospital Regensburg | University Hospital Regensburg | Fritsch,J., Holzmann,T., Schneider-Brachert,W. |
| EPI_ISL_513633, EPI_ISL_513636, EPI_ISL_513637 | University of Washington Virology Lab | University of Washington Virology Lab | Pavitra Roychoudhury, Hong Xie, Lasata Shrestha, Amin Addetia, Truong Nguyen, Victoria M Rachleff, Meei-Li Huang, Keith R Jerome, Alexander Greninger |
| EPI_ISL_513876 | San Francisco Public Health Laboratory | Chan-Zuckerberg Biohub | CZB Cliahub Consortium |

| | | | |
|---|---|---|---|
| EPI_ISL_514314 | Israel Central Virology laboratory | Israel Central Virology laboratory | Neta Zuckerman, Efrat Dahan Bucris, Oran Erster, Ella Mendelson, Michal Mandelboim |
| EPI_ISL_514488, EPI_ISL_514489, EPI_ISL_514490, EPI_ISL_514491, EPI_ISL_514492, EPI_ISL_514493, EPI_ISL_514494, EPI_ISL_514495, EPI_ISL_514496, EPI_ISL_514497, EPI_ISL_514498, EPI_ISL_514499, EPI_ISL_514500, EPI_ISL_514501, EPI_ISL_514502 | | | |
| see above | Centre for Enzyme Innovation, University of Portsmouth / Translational Research Laboratory, Portsmouth Hospitals NHS Trust | COVID-19 Genomics UK (COG-UK) Consortium | Angela Beckett,Yann Bourgeois,Garry Scarlett,Sharon Glaysher,Scott Elliott,Kelly Bicknell,Robert Impey,Allyson Lloyd,Sarah Wyllie,Ethan Butcher,Anoop Chauhan,Samuel Robson |
| EPI_ISL_514651 | Allina Health Laboratory | Minnesota Department of Health, Public Health Laboratory | Matt Plumb, Jacob Garfin, and Xiong Wang |
| EPI_ISL_514652 | M Health Fairview St. Joseph's Hospital | Minnesota Department of Health, Public Health Laboratory | Matt Plumb, Jacob Garfin, and Xiong Wang |
| EPI_ISL_514665 | Minnesota Department of Health, Public Health Laboratory | Minnesota Department of Health, Public Health Laboratory | Matt Plumb, Jacob Garfin, and Xiong Wang |
| EPI_ISL_515082, EPI_ISL_515083, EPI_ISL_515084, EPI_ISL_515085 | Department of Biochemistry, Cell and Molecular Biology | WACCBIP, University of Ghana | Ngoi,J.M., Quashie,P., Morang'a,C.M., Amuzu,D.S., Adu,B., Kumordjie,S., Eshun,M., Boatemaa,L., Magnussen,V., Kotey,E., Tei-Maya,F., Arjarquah,A., Mutungi,J.K., Bediako,Y., Asante,I., Bonney,E., Kyei,G.B., Bonney,K., Amenga-Etego,L.N., Anang,A.K., Awandare,G.A., Ampofo,W. |
| EPI_ISL_515182, EPI_ISL_515183, EPI_ISL_515184, EPI_ISL_515247, EPI_ISL_515248, EPI_ISL_515249 | Kumasi Centre for Collaborative Research in Tropical Medicine, Kumasi. | Institute of Virology, Charité - Universitätsmedizin Berlin | Augustina Sylverken, Philip El-Duah, Michael Owusu, Julia Schneider, Richmond Yeboah, Richmond Gorman, Eric Adu, Sherihane Aryeetey, Jesse Addo Asamoah,Jörn Beheim-Schwarzbach, Victor Max Corman, Christian Drosten, Richard Phillips. |
| EPI_ISL_515270, EPI_ISL_515271, EPI_ISL_515277, EPI_ISL_515283, EPI_ISL_515284, EPI_ISL_515285 | University of Washington Virology Lab | University of Washington Virology Lab | Pavitra Roychoudhury, Hong Xie, Lasata Shrestha, Amin Addetia, Truong Nguyen, Victoria M Rachleff, Meei-Li Huang, Keith R Jerome, Alexander Greninger |
| EPI_ISL_515335, EPI_ISL_515336, EPI_ISL_515337, EPI_ISL_515338, EPI_ISL_515339, EPI_ISL_515340, EPI_ISL_515341, EPI_ISL_515342, EPI_ISL_515343, EPI_ISL_515344 | Nevada State Public Health Laboratory | Nevada State Public Health Laboratory | Richard Tillett, Joel R. Sevinsky, Paul Hartley, Heather Kerwin, David Jackson, Subhash C. Verma, Cyprian Rossetto, Andrew Gorzalski, Chris Laverdure, Natalie Crawford, Stephanie Van Hooser, and Mark Pandori |
| EPI_ISL_515520 | Hospital Municipal do Tatuape Carmino Caricchio | Instituto Adolfo Lutz, Interdiciplinary Procedures Center, Strategic Laboratory | Claudio Tavares Sacchi, Claudia Regina Gonçalves, Erica Valessa Ramos Gomes |
| EPI_ISL_515521 | Hospital Municipal Dr Waldemar Tebaldi | Instituto Adolfo Lutz, Interdiciplinary Procedures Center, Strategic Laboratory | Claudio Tavares Sacchi, Claudia Regina Gonçalves, Erica Valessa Ramos Gomes |
| EPI_ISL_515902, EPI_ISL_515903, EPI_ISL_515904, EPI_ISL_515907 | California Department of Public Health | California Department of Public Health | CDPH IDLB COVIDNet |
| EPI_ISL_516194, EPI_ISL_516195, EPI_ISL_516196, EPI_ISL_516329, EPI_ISL_516330, EPI_ISL_516331, EPI_ISL_516332, EPI_ISL_516333, EPI_ISL_516334, EPI_ISL_516335, EPI_ISL_516336, EPI_ISL_516337, EPI_ISL_516338, EPI_ISL_516339, EPI_ISL_516340, EPI_ISL_516341, EPI_ISL_516342, EPI_ISL_516343, EPI_ISL_516344, EPI_ISL_516345, EPI_ISL_516346, EPI_ISL_516347, EPI_ISL_516348, EPI_ISL_516349, EPI_ISL_516350, EPI_ISL_516351, EPI_ISL_516352, EPI_ISL_516353, EPI_ISL_516354, EPI_ISL_516355, EPI_ISL_516356, EPI_ISL_516357, EPI_ISL_516358, EPI_ISL_516359, EPI_ISL_516360, EPI_ISL_516361, EPI_ISL_516362, EPI_ISL_516363, EPI_ISL_516364, EPI_ISL_516376, EPI_ISL_516377, EPI_ISL_516378, EPI_ISL_516379 | | | |
| see above | Michigan Department of Health and Human Services, Bureau of Laboratories | Michigan Department of Health and Human Services, Bureau of Laboratories | Blankenship HM, Riner D, Soehnlen MK |
| EPI_ISL_516969 | King Georges Medical University | CSIR-National Botanical Research Institute | Priti Prasad, Shantanu Prakash, Kishan Sahu, Babita Singh, Suruchi Shukla, Hricha Mishra, Danish Nasar Khan , Om Prakash, MLB Bhatt, SK Barik, Mehar H.Asif,Samir V. Sawant,Amita Jain, Sumit Kr. Bag |
| EPI_ISL_517001, EPI_ISL_517002, EPI_ISL_517003, EPI_ISL_517004, EPI_ISL_517005, EPI_ISL_517006, EPI_ISL_517007, EPI_ISL_517008, EPI_ISL_517009, EPI_ISL_517010, EPI_ISL_517011, EPI_ISL_517012, EPI_ISL_517013, EPI_ISL_517014, EPI_ISL_517015, EPI_ISL_517016, EPI_ISL_517017, EPI_ISL_517018, EPI_ISL_517019, EPI_ISL_517020, EPI_ISL_517021, EPI_ISL_517022, EPI_ISL_517023, EPI_ISL_517024, EPI_ISL_517025, EPI_ISL_517026, EPI_ISL_517027, EPI_ISL_517028, EPI_ISL_517029, EPI_ISL_517030, EPI_ISL_517031, EPI_ISL_517032, EPI_ISL_517033, EPI_ISL_517034, EPI_ISL_517035, EPI_ISL_517036, EPI_ISL_517037, EPI_ISL_517038, EPI_ISL_517039, EPI_ISL_517040, EPI_ISL_517041, EPI_ISL_517042, EPI_ISL_517043, EPI_ISL_517044, EPI_ISL_517045, EPI_ISL_517046, EPI_ISL_517047, EPI_ISL_517048, EPI_ISL_517049, EPI_ISL_517050, EPI_ISL_517051, EPI_ISL_517052, EPI_ISL_517053, EPI_ISL_517054, EPI_ISL_517055, EPI_ISL_517056, EPI_ISL_517057, EPI_ISL_517058, EPI_ISL_517059, EPI_ISL_517060, EPI_ISL_517061, EPI_ISL_517062, EPI_ISL_517063, EPI_ISL_517064, EPI_ISL_517065, EPI_ISL_517066, EPI_ISL_517067, EPI_ISL_517068, EPI_ISL_517069, EPI_ISL_517070, EPI_ISL_517071, EPI_ISL_517072, EPI_ISL_517073, EPI_ISL_517074, EPI_ISL_517075, EPI_ISL_517076, EPI_ISL_517077, EPI_ISL_517078, EPI_ISL_517079, EPI_ISL_517080, EPI_ISL_517081, EPI_ISL_517082, EPI_ISL_517083, EPI_ISL_517084, EPI_ISL_517085, EPI_ISL_517086, EPI_ISL_517087, EPI_ISL_517158, EPI_ISL_517166, EPI_ISL_517174, EPI_ISL_517183, EPI_ISL_517191, EPI_ISL_517199, EPI_ISL_517200, EPI_ISL_517206, EPI_ISL_517213, EPI_ISL_517214, EPI_ISL_517222, EPI_ISL_517223 | | | |
| see above | Liverpool Clinical Laboratories | COVID-19 Genomics UK (COG-UK) Consortium | Sam Haldenby, Anita Lucaci, Steve Paterson, Julian Hiscox, Alistair Darby, M Almsaud, A Alrezaihi, Muhannad Alruwaili, Stuart D Armstrong, Jones Benjamin, Eleanor G Bentley, Anu Chawla, Jordan J Clark, Angela Cowell, Richard Eccles, Isabel García-Dorival, Matthew Gemmell, Alessandro Gerada, PKF Gilmore, Richard Gregory, Ximeng Han, Catherine Hartley, Margaret Hughes, Miren Iturriza-Gomara, James Johnson, L Luu, Jenifer Manson, Charlotte Nelson, Elaine O'Toole, Cassie Olateju, Rebekah Penrice-Randal , Lucille Rainbow, N.P Randle, Trevor Ian Robinson, Parul Sharma, Ghada T Shawli, James P Stewart, Neil Swainston, Ecaterina Vamos, Joanne Watts, Mark Whitehead |
| EPI_ISL_518856, EPI_ISL_518857, EPI_ISL_518859, EPI_ISL_518866, EPI_ISL_518867, EPI_ISL_518870, EPI_ISL_518899 | Mayo Clinic & Mayo Clinic Laboratories | Minnesota Department of Health, Public Health Laboratory | Matt Plumb, Jacob Garfin, and Xiong Wang |
| EPI_ISL_520720, EPI_ISL_520721, EPI_ISL_520722, EPI_ISL_520723, EPI_ISL_520724, EPI_ISL_520725 | Mohammed Bin Rashid University of Medicine and Health Sciences | Al Jalila Genomics Center | Ahmad Abou Tayoun, Tom Loney, Hamda Khansaheb, Sathishkumar Ramaswamy, Divinlal Harilal, Zulfa Omar Deesi, Rupa Murthy Varghese, Hanan Al Suwaidi, Abdulmajeed Alkhaja, Mohammed Uddin, Rifat Hamoudi, Rabih Halwani, Abiola Catherine Senok, Qutayba Hamid, Norbert Nowotny, Alawi Alsheikh-Ali |
| EPI_ISL_521890, EPI_ISL_521891 | Victorian Infectious Diseases Reference Laboratory (VIDRL) | VIDRL and MDU-PHL | Caly L., Seemann T., Sait, M., Schultz M., Druce J., Sherry, N. |
| EPI_ISL_522409 | Department of Infection Prevention and Infectious Diseases, University Hospital Regensburg | Department of Infection Prevention and Infectious Diseases, University Hospital Regensburg | Fritsch,J., Holzmann,T., Schneider-Brachert,W. |
| EPI_ISL_522578, EPI_ISL_522590, EPI_ISL_522611, EPI_ISL_522641, EPI_ISL_522644, EPI_ISL_522652, EPI_ISL_522653, EPI_ISL_522654, EPI_ISL_522655, EPI_ISL_522656, EPI_ISL_522690, EPI_ISL_522721, EPI_ISL_522741, EPI_ISL_522753, EPI_ISL_522754 | | | |
| see above | Royal Hobart Hospital Microbiology Department | MDU-PHL | Cooley L., van Haeften R., Seemann T., Sait M., Schultz, M.B., Sherry N. |
| EPI_ISL_523289, EPI_ISL_523475, EPI_ISL_523504, EPI_ISL_523505, EPI_ISL_523604, EPI_ISL_523605 | Dutch COVID-19 response team | Erasmus Medical Center | Bas Oude Munnink, David Nieuwenhuijse, Reina Sikkema, Claudia Schapendonk, Irina Chestakova, Anne van der Linden, Theo Bestebroer, Stefan van Nieuwkoop, Mark Pronk, Pascal Lexmond, Corien Swaan, Manon Haverkate, Madelief Mollers, Mart Stein, Sandra Kengne Kamga Mobou, Jeroen van Kampen, Jolanda Voermans, Aura Timen, Corine GeurtsvanKessel, Annemiek van der Eijk, Richard Molenkamp, Marion Koopmans, on behalf of the Dutch national COVID-19 response team. |
| EPI_ISL_523929, EPI_ISL_523941, EPI_ISL_523944 | Center of Medical Microbiology, Virology, and Hospital Hygiene, University of Duesseldorf | Center of Medical Microbiology, Virology, and Hospital Hygiene, University of Duesseldorf | Maximilian Damagnez, Alexander Dilthey, Torsten Houwaart, Malte Kohns Vasconcelos, Marek Korencak, Jessica Nicolai, Klaus Pfeffer, Hendrik Streeck, Daniel Strelow, Jörg Timm, Andreas Walker, Tobias Wienemann |
| EPI_ISL_523951 | Respiratory Virus Unit, Microbiology Services Colindale, Public Health England | Respiratory Virus Unit, Microbiology Services Colindale, Public Health England | PHE Covid Sequencing Team |
| EPI_ISL_524433, EPI_ISL_524434 | Environmental and Global Health, University of Florida - Gainesville | University of Florida | Elbadry,M.A., Subramaniam,K., Waltzek,T.B., Gibson,J.C., Stephenson,C.J., Alam,M.M., Morris,J.G. Jr., Lednicky,J.A. |
| EPI_ISL_524494, EPI_ISL_524499, EPI_ISL_524500, EPI_ISL_524507, EPI_ISL_524512, EPI_ISL_524513, EPI_ISL_524518, EPI_ISL_524519, EPI_ISL_524520, EPI_ISL_524527, EPI_ISL_524532, EPI_ISL_524533, EPI_ISL_524534, EPI_ISL_524536, EPI_ISL_524539, EPI_ISL_524540, EPI_ISL_524543, EPI_ISL_524545, EPI_ISL_524547, EPI_ISL_524549, EPI_ISL_524550, EPI_ISL_524552, EPI_ISL_524554, EPI_ISL_524555, EPI_ISL_524557, EPI_ISL_524559, EPI_ISL_524561, EPI_ISL_524563, EPI_ISL_524566, EPI_ISL_524567, EPI_ISL_524569, EPI_ISL_524571 | | | |
| see above | NHSGGC West of Scotland Specialist Virology Centre / MRC-University of Glasgow Centre for Virus Research | Wellcome Sanger Institute for the COVID-19 Genomics UK (COG-UK) consortium | Ana da Silva Filipe, Natasha Johnson, Kathy Smollett, Daniel Mair, Stephen Carmichael, Lily Tong, Jenna Nichols, Elihu Aranday-Cortes, Kirstyn Brunker, Yasmin Parr, Kyriaki Nomikou; Sarah McDonald, Marc Niebel, Patawee Asamaphan; Richard Orton, Joseph Hughes, Sreenu Vattipally, David L |

| | | | |
|---|---|---|---|
| | | | Robertson; Alasdair MacLean, Rory Gunson; Kathy Li, Natasha Jesudason, Rajiv Shah, James Shepherd, Antonia Ho, Alice Broos, Emma Thomson and Alex Alderton, Roberto Amato, Sonia Goncalves, Ewan Harrison, David K. Jackson, Ian Johnston, Dominic Kwiatkowski, Cordelia Langford, John Sillitoe on behalf of the Wellcome Sanger Institute COVID-19 Surveillance Team (http://www.sanger.ac.uk/covid-team) |
| EPI_ISL_524648, EPI_ISL_524659, EPI_ISL_524674, EPI_ISL_524679, EPI_ISL_524683, EPI_ISL_524696, EPI_ISL_524701, EPI_ISL_524702, EPI_ISL_524706, EPI_ISL_524707 | North West London Pathology, Imperial College Healthcare NHS Trust | Wellcome Sanger Institute for the COVID-19 Genomics UK (COG-UK) consortium | Ling Li, Paul Randell, David Muir, Frankie Bolt, Alison Holmes, James Price, Aileen Rowan, Graham Taylor, Anjna Badhan, Carolina Herrera and Alex Alderton, Roberto Amato, Sonia Goncalves, Ewan Harrison, David K. Jackson, Ian Johnston, Dominic Kwiatkowski, Cordelia Langford, John Sillitoe on behalf of the Wellcome Sanger Institute COVID-19 Surveillance Team (http://www.sanger.ac.uk/covid-team) |
| EPI_ISL_525571, EPI_ISL_525573 | Istituto Zooprofilattico Sperimentale Puglia e Basilicata; Dipartimento di Bioscienze, Biotecnologie e Biofarmaceutica dell'Università degli Studi di Bari "A.Moro"; Istituto di Biomembrane. Bioenergetica e Biotecnologie Molecolari del Consiglio Nazionale delle Ricerche di Bari | Beaconlab (Bioinformatics, Evolution and Comparative Genomics lab), Dept of Biosciences, University on Milan | Parisi A.,Pesole G., Manzari C., Chiara M |
| EPI_ISL_525635, EPI_ISL_525636, EPI_ISL_525637, EPI_ISL_525638, EPI_ISL_525639, EPI_ISL_525640, EPI_ISL_525641, EPI_ISL_525642 | Wadsworth Center, New York State Department of Health | Wadsworth Center, New York State Department of Health | Kirsten St. George, Daryl M. Lamson, Sara Griesemer, Jonathan Plitnick, Navjot Singh, Matthew D. Shudt, Erica Lasek-Nesselquist |
| EPI_ISL_525771 | Texas Department of State Health Services | Texas Department of State Health Services | Jenny Zhang, Rashmi Tuladhar, Bonnie Oh, Maliha Rahman, Anita Pokharel, Myong Koag, Chun Wang, Rachel Lee, Grace Kubin |
| EPI_ISL_526490, EPI_ISL_526491, EPI_ISL_526492 | Virology Department, Royal Infirmary of Edinburgh, NHS Lothian / School of Biological Sciences, University of Edinburgh / Institute of Genetics and Molecular Medicine, University of Edinburgh | COVID-19 Genomics UK (COG-UK) Consortium | McHugh M, Dewar R, Rooke S, Gallagher M, Balcaza C, O'Toole Á, Scher E, Hill V, McCrone JT, Colquhoun R, Yu X, Jackson B, Rambaut A, Williams TC, Templeton K |
| EPI_ISL_526808, EPI_ISL_526809, EPI_ISL_526811, EPI_ISL_526812, EPI_ISL_526813, EPI_ISL_526814, EPI_ISL_526815, EPI_ISL_526816, EPI_ISL_526817, EPI_ISL_526818, EPI_ISL_526819, EPI_ISL_526820, EPI_ISL_526821, EPI_ISL_526822, EPI_ISL_526823, EPI_ISL_526824, EPI_ISL_526825, EPI_ISL_526826 | | | |
| see above | Virginia DCLS | Virginia DCLS | Virginia DCLS |
| EPI_ISL_527862 | Hospital Municipal de Urgência | Instituto Adolfo Lutz, Interdiciplinary Procedures Center, Strategic Laboratory | Claudio Tavares Sacchi, Claudia Regina Gonçalves, Erica Valessa Ramos Gomes |
| EPI_ISL_527881 | Nigeria Centre for Disease Control (NCDC) | African Centre of Excellence for Genomics of Infectious Diseases (ACEGID), Redeemer's University, Ede, Osun State, Nigeria | Oluniyi P.E. et al |
| EPI_ISL_529026 | Ospedale "Giuseppe Mazzini"-Teramo | Istituto Zooprofilattico Sperimentale dell'Abruzzo e Molise "G.Caporale" | Lorusso A, Marcacci M, Di Domenico M, Curini V, Ancora M, Cammà C, Rinaldi A, Mangone I, Di Pasquale A, Puglia I, Savini G. |
| EPI_ISL_529029, EPI_ISL_529030 | Kingston Health Sciences Center | Queen's Genomics Lab at Ongwanada (Q-GLO) | Sjaarda CP, Rustom N, Huang D, Perez-Patrigeon S, Hudson ML, Wong H,Guan H, Ayub M, Soares CN, Colautti R, Evans GA, Sheth P |
| EPI_ISL_529135 | Department of Virology III, National Institute of Infectious Diseases | Pathogen Genomics Center, National Institute of Infectious Diseases | Tsuyoshi Sekizuka, Shutoku Matsuyama, Kentaro Itokawa, Rina Tanaka, Masanori Hashino, Makoto Takeda, Takaji Wakita, Makoto Kuroda |
| EPI_ISL_529155, EPI_ISL_529156, EPI_ISL_529157, EPI_ISL_529158, EPI_ISL_529159 | University of Washington, Laboratory Medicine | University of Washington, Laboratory Medicine | Roychoudhury,P., Greninger,A., Jerome,K. |
| EPI_ISL_529163 | Department of Immunology, The Scripps Research Institute | Andersen lab at Scripps Research | Allison Smither, Gilberto Sabino-Santos, Patricia Snarski, Lilia Melnik, Antoinette Bell, Kaylynn Genemaras, Arnaud Drouin, Dahlene Fusco, Robert Garry with SEARCH Alliance San Diego |
| EPI_ISL_529170 | University of Washington, Laboratory Medicine | University of Washington, Laboratory Medicine | Roychoudhury,P., Greninger,A., Jerome,K. |
| EPI_ISL_529175 | South Carolina Department of Health and Environmental Control | South Carolina Department of Health and Environmental Control | Haley V. Flores |
| EPI_ISL_529200 | University of Washington, Laboratory Medicine | University of Washington, Laboratory Medicine | Roychoudhury,P., Greninger,A., Jerome,K. |
| EPI_ISL_529208, EPI_ISL_529209, EPI_ISL_529210, EPI_ISL_529211 | Laboratory Medicine, University of Washington | University of Washington, Laboratory Medicine | Roychoudhury,P., Greninger,A., Jerome,K. |
| EPI_ISL_529223, EPI_ISL_529278 | University of Birmingham | COVID-19 Genomics UK (COG-UK) Consortium | Institute of Microbiology, University of Birmingham: Claire McMurray, Joanne Stockton, Samuel Nicholls, Radoslaw Poplawski, Will Rowe, Josh Quick, Nicholas Loman. University of Birmingham Testing Laboratory: Celina M Whalley, Andrew Bosworth, Charlotte Poxon, Kasun Wanigasooriya, Oliver Pickles, Mike Kidd, Alex Richter, Andrew D Beggs PHE Heartlands Lab: Husam Osman, Andrew Bosworth. Queen Elizabeth Hospital: Anna Casey |
| EPI_ISL_529319 | West of Scotland Specialist Virology Centre, NHSGGC / MRC-University of Glasgow Centre for Virus Research | COVID-19 Genomics UK (COG-UK) Consortium | Ana da Silva Filipe, Natasha Johnson, Kathy Smollett, Daniel Mair, Stephen Carmichael, Lily Tong, Jenna Nichols, Elihu Aranday-Cortes, Kirstyn Brunker, Yasmin Parr, Alice Broos, Kyriaki Nomikou; Sarah McDonald, Marc Niebel, Patawee Asamaphan; Richard Orton, Joseph Hughes, Sreenu Vattipally, David L Robertson; Alasdair MacLean, Rory Gunson; Kathy Li, Natasha Jesudason, Rajiv Shah, James Shepherd, Antonia Ho, Emma Thomson |
| EPI_ISL_529358, EPI_ISL_529395, EPI_ISL_529396, EPI_ISL_529405, EPI_ISL_529441, EPI_ISL_529473, EPI_ISL_529615, EPI_ISL_529617 | University of Birmingham | COVID-19 Genomics UK (COG-UK) Consortium | Institute of Microbiology, University of Birmingham: Claire McMurray, Joanne Stockton, Samuel Nicholls, Radoslaw Poplawski, Will Rowe, Josh Quick, Nicholas Loman. University of Birmingham Testing Laboratory: Celina M Whalley, Andrew Bosworth, Charlotte Poxon, Kasun Wanigasooriya, Oliver Pickles, Mike Kidd, Alex Richter, Andrew D Beggs PHE Heartlands Lab: Husam Osman, Andrew Bosworth. Queen Elizabeth Hospital: Anna Casey |
| EPI_ISL_529964 | RSUD Bangil Pasuruan | Institute of Tropical Disease, Universitas Airlangga | Aldise M Nastri, Jezzy R Dewantari, Rima R Prasetya, Krisnoadi Rahardjo, Arma Roosalina, Gatot Soegiarto, Laksmi Wulandari, Retno A Setyoningrum, Resti Yudhawati, Yohko K Shimizu, Mitsuhiro Nishimura, Yasuko Mori, Soetjipto, Kazufumi Shimizu, Maria I Lusida |
| EPI_ISL_529982, EPI_ISL_529983 | Hospital Universitario 12 de Octubre | Hospital Universitario 12 de Octubre | Raúl Recio, Sara González, Esther Viedma, Elias Dahdouh, Fernando Lázaro, Natalia Stella, Julio García, Juan Carlos Galán, Rafael Cantón, Mª Dolores Folgueira, Rafael Delgado, Jesús Mingorance |
| EPI_ISL_530224 | Minnesota Department of Health, Public Health Laboratory | Minnesota Department of Health, Public Health Laboratory | Matt Plumb, Jacob Garfin, and Xiong Wang |
| EPI_ISL_530273, EPI_ISL_530274 | Queensland Health Forensic and Scientific Services, Public Health Virology | Public Health Virology Laboratory, Forensic and Scientific Services, Queensland Health | Son Nguyen et al |
| EPI_ISL_534815, EPI_ISL_534816, EPI_ISL_534822, EPI_ISL_534832, EPI_ISL_534840, EPI_ISL_534846, EPI_ISL_534851, EPI_ISL_534859, EPI_ISL_534862, EPI_ISL_534865, EPI_ISL_534877, EPI_ISL_534882, EPI_ISL_534887, EPI_ISL_534893, EPI_ISL_534900, EPI_ISL_534905, EPI_ISL_534915, EPI_ISL_534929, EPI_ISL_534931, EPI_ISL_534934, EPI_ISL_534935, EPI_ISL_534938, EPI_ISL_534939, EPI_ISL_534940, EPI_ISL_534944, EPI_ISL_534945, EPI_ISL_534946, EPI_ISL_534957, EPI_ISL_534976 | | | |
| see above | Oxford Viromics, NDM, University of Oxford; Oxford University Hospitals; Basingstoke and North Hampshire Hospital | COVID-19 Genomics UK (COG-UK) Consortium | Tanya Golubchik, David Bonsall, George Macintyre, Amy Trebes, Mariateresa de Cesare, Catrin Moore, Alex Mobbs, Anita Justice, Robert Shaw, Monique Andersson, Timothy Peto, Emma Wise, Nathan Moore, Jessica Lynch, Nick Cortes, Matilde Mori, Stephen Kidd, David Buck, John Todd, Christophe Fraser |
| EPI_ISL_536268, EPI_ISL_536269, EPI_ISL_536270 | Hôpital Charles-LeMoyne | Laboratoire de santé publique du Québec | Sandrine Moreira, Ioannis Ragoussis, Guillaume Bourque, Jesse Shapiro, Mark Lathrop and Michel Roger |
| EPI_ISL_536271 | Hôpital Honoré-Mercier | Laboratoire de santé publique du Québec | Sandrine Moreira, Ioannis Ragoussis, Guillaume Bourque, Jesse Shapiro, Mark Lathrop and Michel Roger |
| EPI_ISL_536272 | CSSS Haut-Richelieu/Rouville (Hôpital) | Laboratoire de santé publique du Québec | Sandrine Moreira, Ioannis Ragoussis, Guillaume Bourque, Jesse Shapiro, Mark Lathrop and Michel Roger |
| EPI_ISL_536273, EPI_ISL_536274 | Hôpital du Suroît | Laboratoire de santé publique du Québec | Sandrine Moreira, Ioannis Ragoussis, Guillaume Bourque, Jesse Shapiro, Mark Lathrop and Michel Roger |
| EPI_ISL_536275 | Hôpital de Hull | Laboratoire de santé publique du Québec | Sandrine Moreira, Ioannis Ragoussis, Guillaume Bourque, Jesse Shapiro, Mark Lathrop and Michel Roger |

| | | | |
|---|---|---|---|
| EPI_ISL_536276, EPI_ISL_536277 | Hôpital Pierre-Boucher | Laboratoire de santé publique du Québec | Sandrine Moreira, Ioannis Ragoussis, Guillaume Bourque, Jesse Shapiro, Mark Lathrop and Michel Roger |
| EPI_ISL_536278 | CSSS Haut-Richelieu/Rouville (Hôpital) | Laboratoire de santé publique du Québec | Sandrine Moreira, Ioannis Ragoussis, Guillaume Bourque, Jesse Shapiro, Mark Lathrop and Michel Roger |
| EPI_ISL_536279, EPI_ISL_536280, EPI_ISL_536281, EPI_ISL_536282 | Hôpital Charles-LeMoyne | Laboratoire de santé publique du Québec | Sandrine Moreira, Ioannis Ragoussis, Guillaume Bourque, Jesse Shapiro, Mark Lathrop and Michel Roger |
| EPI_ISL_536283 | Centre hospitalier Anna-Laberge | Laboratoire de santé publique du Québec | Sandrine Moreira, Ioannis Ragoussis, Guillaume Bourque, Jesse Shapiro, Mark Lathrop and Michel Roger |
| EPI_ISL_536284, EPI_ISL_536285 | Hôpital Charles-LeMoyne | Laboratoire de santé publique du Québec | Sandrine Moreira, Ioannis Ragoussis, Guillaume Bourque, Jesse Shapiro, Mark Lathrop and Michel Roger |
| EPI_ISL_536286 | CSSS Haut-Richelieu/Rouville (Hôpital) | Laboratoire de santé publique du Québec | Sandrine Moreira, Ioannis Ragoussis, Guillaume Bourque, Jesse Shapiro, Mark Lathrop and Michel Roger |
| EPI_ISL_536287 | Hôpital du Suroît | Laboratoire de santé publique du Québec | Sandrine Moreira, Ioannis Ragoussis, Guillaume Bourque, Jesse Shapiro, Mark Lathrop and Michel Roger |
| EPI_ISL_536289 | Hôpital de Maria | Laboratoire de santé publique du Québec | Sandrine Moreira, Ioannis Ragoussis, Guillaume Bourque, Jesse Shapiro, Mark Lathrop and Michel Roger |
| EPI_ISL_536290 | Hôpital Charles-LeMoyne | Laboratoire de santé publique du Québec | Sandrine Moreira, Ioannis Ragoussis, Guillaume Bourque, Jesse Shapiro, Mark Lathrop and Michel Roger |
| EPI_ISL_536291 | Hôpital de Hull | Laboratoire de santé publique du Québec | Sandrine Moreira, Ioannis Ragoussis, Guillaume Bourque, Jesse Shapiro, Mark Lathrop and Michel Roger |
| EPI_ISL_536292, EPI_ISL_536293, EPI_ISL_536294 | Hôpital Charles-LeMoyne | Laboratoire de santé publique du Québec | Sandrine Moreira, Ioannis Ragoussis, Guillaume Bourque, Jesse Shapiro, Mark Lathrop and Michel Roger |
| EPI_ISL_536295 | Hôpital Pierre-Boucher | Laboratoire de santé publique du Québec | Sandrine Moreira, Ioannis Ragoussis, Guillaume Bourque, Jesse Shapiro, Mark Lathrop and Michel Roger |
| EPI_ISL_536296, EPI_ISL_536297, EPI_ISL_536298 | Hôpital Charles-LeMoyne | Laboratoire de santé publique du Québec | Sandrine Moreira, Ioannis Ragoussis, Guillaume Bourque, Jesse Shapiro, Mark Lathrop and Michel Roger |
| EPI_ISL_536299 | CSSS Haut-Richelieu/Rouville (Hôpital) | Laboratoire de santé publique du Québec | Sandrine Moreira, Ioannis Ragoussis, Guillaume Bourque, Jesse Shapiro, Mark Lathrop and Michel Roger |
| EPI_ISL_536300 | Hôpital Pierre-Boucher | Laboratoire de santé publique du Québec | Sandrine Moreira, Ioannis Ragoussis, Guillaume Bourque, Jesse Shapiro, Mark Lathrop and Michel Roger |
| EPI_ISL_536301, EPI_ISL_536302 | CSSS Haut-Richelieu/Rouville (Hôpital) | Laboratoire de santé publique du Québec | Sandrine Moreira, Ioannis Ragoussis, Guillaume Bourque, Jesse Shapiro, Mark Lathrop and Michel Roger |
| EPI_ISL_536303 | Hôpital Pierre-Boucher | Laboratoire de santé publique du Québec | Sandrine Moreira, Ioannis Ragoussis, Guillaume Bourque, Jesse Shapiro, Mark Lathrop and Michel Roger |
| EPI_ISL_536304 | CSSS Haut-Richelieu/Rouville (Hôpital) | Laboratoire de santé publique du Québec | Sandrine Moreira, Ioannis Ragoussis, Guillaume Bourque, Jesse Shapiro, Mark Lathrop and Michel Roger |
| EPI_ISL_536305, EPI_ISL_536306, EPI_ISL_536307, EPI_ISL_536308, EPI_ISL_536309 | Hôpital Pierre-Boucher | Laboratoire de santé publique du Québec | Sandrine Moreira, Ioannis Ragoussis, Guillaume Bourque, Jesse Shapiro, Mark Lathrop and Michel Roger |
| EPI_ISL_536310 | CLSC et Centre d'Hébergement la Petite-Nation | Laboratoire de santé publique du Québec | Sandrine Moreira, Ioannis Ragoussis, Guillaume Bourque, Jesse Shapiro, Mark Lathrop and Michel Roger |
| EPI_ISL_536311 | Hôpital de Gatineau | Laboratoire de santé publique du Québec | Sandrine Moreira, Ioannis Ragoussis, Guillaume Bourque, Jesse Shapiro, Mark Lathrop and Michel Roger |
| EPI_ISL_536312 | Hôpital de Hull | Laboratoire de santé publique du Québec | Sandrine Moreira, Ioannis Ragoussis, Guillaume Bourque, Jesse Shapiro, Mark Lathrop and Michel Roger |
| EPI_ISL_536313, EPI_ISL_536314 | Hôpital Charles-LeMoyne | Laboratoire de santé publique du Québec | Sandrine Moreira, Ioannis Ragoussis, Guillaume Bourque, Jesse Shapiro, Mark Lathrop and Michel Roger |
| EPI_ISL_536315 | Centre hospitalier Anna-Laberge | Laboratoire de santé publique du Québec | Sandrine Moreira, Ioannis Ragoussis, Guillaume Bourque, Jesse Shapiro, Mark Lathrop and Michel Roger |
| EPI_ISL_536316, EPI_ISL_536317 | Hôpital Charles-LeMoyne | Laboratoire de santé publique du Québec | Sandrine Moreira, Ioannis Ragoussis, Guillaume Bourque, Jesse Shapiro, Mark Lathrop and Michel Roger |
| EPI_ISL_536318, EPI_ISL_536319, EPI_ISL_536320 | Hôpital du Suroît | Laboratoire de santé publique du Québec | Sandrine Moreira, Ioannis Ragoussis, Guillaume Bourque, Jesse Shapiro, Mark Lathrop and Michel Roger |
| EPI_ISL_536321 | CSSS Haut-Richelieu/Rouville (Hôpital) | Laboratoire de santé publique du Québec | Sandrine Moreira, Ioannis Ragoussis, Guillaume Bourque, Jesse Shapiro, Mark Lathrop and Michel Roger |
| EPI_ISL_536322, EPI_ISL_536323, EPI_ISL_536324 | Hôpital Pierre-Boucher | Laboratoire de santé publique du Québec | Sandrine Moreira, Ioannis Ragoussis, Guillaume Bourque, Jesse Shapiro, Mark Lathrop and Michel Roger |
| EPI_ISL_536325 | Centre hospitalier Anna-Laberge | Laboratoire de santé publique du Québec | Sandrine Moreira, Ioannis Ragoussis, Guillaume Bourque, Jesse Shapiro, Mark Lathrop and Michel Roger |
| EPI_ISL_536326, EPI_ISL_536327, EPI_ISL_536328 | Hôpital Charles-LeMoyne | Laboratoire de santé publique du Québec | Sandrine Moreira, Ioannis Ragoussis, Guillaume Bourque, Jesse Shapiro, Mark Lathrop and Michel Roger |
| EPI_ISL_536329 | Centre hospitalier Anna-Laberge | Laboratoire de santé publique du Québec | Sandrine Moreira, Ioannis Ragoussis, Guillaume Bourque, Jesse Shapiro, Mark Lathrop and Michel Roger |
| EPI_ISL_536330 | Hôpital Honoré-Mercier | Laboratoire de santé publique du Québec | Sandrine Moreira, Ioannis Ragoussis, Guillaume Bourque, Jesse Shapiro, Mark Lathrop and Michel Roger |
| EPI_ISL_536331 | Hôpital Charles-LeMoyne | Laboratoire de santé publique du Québec | Sandrine Moreira, Ioannis Ragoussis, Guillaume Bourque, Jesse Shapiro, Mark Lathrop and Michel Roger |
| EPI_ISL_536332, EPI_ISL_536333, EPI_ISL_536334, EPI_ISL_536335 | Hôpital Pierre-Boucher | Laboratoire de santé publique du Québec | Sandrine Moreira, Ioannis Ragoussis, Guillaume Bourque, Jesse Shapiro, Mark Lathrop and Michel Roger |
| EPI_ISL_536336 | Hôpital Charles-LeMoyne | Laboratoire de santé publique du Québec | Sandrine Moreira, Ioannis Ragoussis, Guillaume Bourque, Jesse Shapiro, Mark Lathrop and Michel Roger |
| EPI_ISL_536337 | CUSM-Site Glen-LAB Microbiologie | Laboratoire de santé publique du Québec | Sandrine Moreira, Ioannis Ragoussis, Guillaume Bourque, Jesse Shapiro, Mark Lathrop and Michel Roger |
| EPI_ISL_536338, EPI_ISL_536339, EPI_ISL_536340, EPI_ISL_536341, EPI_ISL_536342, EPI_ISL_536343 | Hôpital Charles-LeMoyne | Laboratoire de santé publique du Québec | Sandrine Moreira, Ioannis Ragoussis, Guillaume Bourque, Jesse Shapiro, Mark Lathrop and Michel Roger |
| EPI_ISL_536344, EPI_ISL_536345 | CSSS Haut-Richelieu/Rouville (Hôpital) | Laboratoire de santé publique du Québec | Sandrine Moreira, Ioannis Ragoussis, Guillaume Bourque, Jesse Shapiro, Mark Lathrop and Michel Roger |
| EPI_ISL_536346, EPI_ISL_536347 | Hôpital Pierre-Boucher | Laboratoire de santé publique du Québec | Sandrine Moreira, Ioannis Ragoussis, Guillaume Bourque, Jesse Shapiro, Mark Lathrop and Michel Roger |
| EPI_ISL_536348 | Centre hospitalier Anna-Laberge | Laboratoire de santé publique du Québec | Sandrine Moreira, Ioannis Ragoussis, Guillaume Bourque, Jesse Shapiro, Mark Lathrop and Michel Roger |
| EPI_ISL_536349 | Hôpital du Suroît | Laboratoire de santé publique du Québec | Sandrine Moreira, Ioannis Ragoussis, Guillaume Bourque, Jesse Shapiro, Mark Lathrop and Michel Roger |
| EPI_ISL_536350 | Hôpital Charles-LeMoyne | Laboratoire de santé publique du Québec | Sandrine Moreira, Ioannis Ragoussis, Guillaume Bourque, Jesse Shapiro, Mark Lathrop and Michel Roger |
| EPI_ISL_536351, EPI_ISL_536352 | Hôpital Pierre-Boucher | Laboratoire de santé publique du Québec | Sandrine Moreira, Ioannis Ragoussis, Guillaume Bourque, Jesse Shapiro, Mark Lathrop and Michel Roger |
| EPI_ISL_536353 | Centre de santé Inuulitsivik | Laboratoire de santé publique du Québec | Sandrine Moreira, Ioannis Ragoussis, Guillaume Bourque, Jesse Shapiro, Mark Lathrop and Michel Roger |
| EPI_ISL_536354, EPI_ISL_536355, EPI_ISL_536356, EPI_ISL_536357, EPI_ISL_536358, EPI_ISL_536359 | Hôpital du Suroît | Laboratoire de santé publique du Québec | Sandrine Moreira, Ioannis Ragoussis, Guillaume Bourque, Jesse Shapiro, Mark Lathrop and Michel Roger |
| EPI_ISL_536360 | Hôpital Pierre-Boucher | Laboratoire de santé publique du Québec | Sandrine Moreira, Ioannis Ragoussis, Guillaume Bourque, Jesse Shapiro, Mark Lathrop and Michel Roger |
| EPI_ISL_536361 | CSSS Haut-Richelieu/Rouville (Hôpital) | Laboratoire de santé publique du Québec | Sandrine Moreira, Ioannis Ragoussis, Guillaume Bourque, Jesse Shapiro, Mark Lathrop and Michel Roger |
| EPI_ISL_536362 | Hôpital du Suroît | Laboratoire de santé publique du Québec | Sandrine Moreira, Ioannis Ragoussis, Guillaume Bourque, Jesse Shapiro, Mark Lathrop and Michel Roger |
| EPI_ISL_536363, EPI_ISL_536364 | Hôpital de Hull | Laboratoire de santé publique du Québec | Sandrine Moreira, Ioannis Ragoussis, Guillaume Bourque, Jesse Shapiro, Mark Lathrop and Michel Roger |
| EPI_ISL_536365, EPI_ISL_536366 | CUSM-Site Glen-LAB Microbiologie | Laboratoire de santé publique du Québec | Sandrine Moreira, Ioannis Ragoussis, Guillaume Bourque, Jesse Shapiro, Mark Lathrop and Michel Roger |

| | | | |
|---|---|---|---|
| EPI_ISL_536367 | Hôpital de Hull | Laboratoire de santé publique du Québec | Sandrine Moreira, Ioannis Ragoussis, Guillaume Bourque, Jesse Shapiro, Mark Lathrop and Michel Roger |
| EPI_ISL_536375 | Hôpital Pierre-Boucher | Laboratoire de santé publique du Québec | Sandrine Moreira, Ioannis Ragoussis, Guillaume Bourque, Jesse Shapiro, Mark Lathrop and Michel Roger |
| EPI_ISL_536791 | Waikato Hospital | Institute of Environmental Science and Research (ESR) | Xiaoyun Ren, Matt Storey, Nikki Freed, Muhammad Faisal, Jing Wang, Hermes Perez, Anja Werno, Antje van der Linden, Arlo Upton, Chris Mansell, David Hammer, Dragana Drinkovic, Gary McAuliffe, Hana Sofia Andersson, James Ussher, Jill Sherwood, Josh Freeman, Julia Howard, Juliet Elvy, Mary DeAlmeida, Matt Blakiston, Matthew Rogers, Max Bloomfield, Michael Addidle, Michelle Balm, Sally Roberts, Sarah Jefferies, Sharmini Muttaiyah, Susan Morpeth, Susan Taylor, Timothy Blackmore, Vani Sathyendran, Veronica Playle, Virginia Hope, Erasmus Smit, Lauren Jelly, Olin Silander, Joep de Ligt |
| EPI_ISL_537390, EPI_ISL_537391, EPI_ISL_537392, EPI_ISL_537454, EPI_ISL_537455, EPI_ISL_537456, EPI_ISL_537457, EPI_ISL_537459, EPI_ISL_537460, EPI_ISL_537461, EPI_ISL_537465, EPI_ISL_537466 | | | |
| see above | Centro de Investigación Biomédica de La Rioja - Hospital San Pedro Logroño | SeqCOVID-SPAIN consortium/IBV(CSIC) | María de Toro, José Manuel Azcona Gutiérrez, María Pilar Bea Escudero, Miriam Blasco Alberdi and SeqCOVID-SPAIN consortium |
| EPI_ISL_537493, EPI_ISL_537494, EPI_ISL_537495, EPI_ISL_537512, EPI_ISL_537583, EPI_ISL_537589, EPI_ISL_537590, EPI_ISL_537606, EPI_ISL_537607 | UCLA Pathology Clinical Microbiology Lab | Kruglyak Lab | Guo et al. |
| EPI_ISL_537706, EPI_ISL_537707, EPI_ISL_537708, EPI_ISL_537709, EPI_ISL_537710, EPI_ISL_537711, EPI_ISL_537712, EPI_ISL_537713, EPI_ISL_537714, EPI_ISL_537715, EPI_ISL_537716, EPI_ISL_537717, EPI_ISL_537718 | | | |
| see above | Hospital Universitario de Gran Canaria Dr. Negrín | SeqCOVID-SPAIN consortium/IBV(CSIC) | M. Carmen Pérez González, Francisco J. Chamizo López, Ana Bordes Benítez and SeqCOVID-SPAIN consortium |
| EPI_ISL_537851, EPI_ISL_537852, EPI_ISL_537853, EPI_ISL_537854, EPI_ISL_537855, EPI_ISL_537856 | Centro de Investigación Biomédica de La Rioja - Hospital San Pedro Logroño | SeqCOVID-SPAIN consortium/IBV(CSIC) | María de Toro, José Manuel Azcona Gutiérrez, María Pilar Bea Escudero, Miriam Blasco Alberdi and SeqCOVID-SPAIN consortium |
| EPI_ISL_537953, EPI_ISL_537954, EPI_ISL_537955, EPI_ISL_537956, EPI_ISL_537957, EPI_ISL_537958, EPI_ISL_537959, EPI_ISL_537960, EPI_ISL_537961, EPI_ISL_537962, EPI_ISL_537963, EPI_ISL_537964, EPI_ISL_537965, EPI_ISL_537966, EPI_ISL_537967, EPI_ISL_537968, EPI_ISL_537969, EPI_ISL_537970, EPI_ISL_537971, EPI_ISL_537972, EPI_ISL_537973, EPI_ISL_537974, EPI_ISL_537975, EPI_ISL_537976, EPI_ISL_537977, EPI_ISL_537978, EPI_ISL_537979, EPI_ISL_537980, EPI_ISL_537981, EPI_ISL_537992, EPI_ISL_537993 | | | |
| see above | Servicio de Microbiología. Hospital General Universitario de Castellón | SeqCOVID-SPAIN consortium/IBV(CSIC) | Rosario Moreno, María Dolores Tirado and SeqCOVID-SPAIN consortium |
| EPI_ISL_538062, EPI_ISL_538063, EPI_ISL_538064, EPI_ISL_538065, EPI_ISL_538066, EPI_ISL_538067, EPI_ISL_538068, EPI_ISL_538069 | Servicio de Microbiología. Hospital Universitario Donostia. OSI Donostialdea. Área de Enfermedades Infecciosas, Grupo de Infección Respiratoria y Resistencia Antimicrobiana. Instituto de Investigación Sanitaria Biodonostia | SeqCOVID-SPAIN consortium/IBV(CSIC) | Gustavo Cilla, Milagrosa Montes, Luis Piñeiro, Jose Maria Marimón and SeqCOVID-SPAIN consortium |
| EPI_ISL_538111, EPI_ISL_538112, EPI_ISL_538113, EPI_ISL_538114, EPI_ISL_538115, EPI_ISL_538116, EPI_ISL_538117 | Clínica Universidad de Navarra. Servicio de Enfermedades Infecciosas y Microbiología clínica | SeqCOVID-SPAIN consortium/IBV(CSIC) | Mirian Fernández-Alonso, Jose Luis del Pozo and SeqCOVID-SPAIN consortium |
| EPI_ISL_538172, EPI_ISL_538173, EPI_ISL_538174, EPI_ISL_538175, EPI_ISL_538176, EPI_ISL_538177, EPI_ISL_538178, EPI_ISL_538179, EPI_ISL_538180, EPI_ISL_538181, EPI_ISL_538182, EPI_ISL_538183, EPI_ISL_538184, EPI_ISL_538185, EPI_ISL_538186, EPI_ISL_538187, EPI_ISL_538188 | | | |
| see above | Servicio de Microbiología y Parasitología clínica. UCEIMP. Hospital Universitario Virgen del Rocío/IBIS/CSIC/US | SeqCOVID-SPAIN consortium/IBV(CSIC) | Guillermo Martín Gutiérrez, Ángel Rodríguez Villodres, Lidia Gálvez Benitez, Verónica González Galán, Javier Aznar Martín and SeqCOVID-SPAIN consortium |
| EPI_ISL_538327, EPI_ISL_538328, EPI_ISL_538329, EPI_ISL_538330, EPI_ISL_538331, EPI_ISL_538332, EPI_ISL_538333, EPI_ISL_538334, EPI_ISL_538335 | Kingston Health Sciences Centre / Queen's University | Ontario Institute for Cancer Research | Prameet M. Sheth, Calvin Sjaarda, Robert Colautti, Katya Douchant, Ilinca Lungu, Bernard Lam, Paul Krzyzanowski, Michael Laszloffy, Lawrence E Heisler, Richard de Borja, Jared T. Simpson |
| EPI_ISL_538460, EPI_ISL_538461, EPI_ISL_538462, EPI_ISL_538463, EPI_ISL_538464 | Department of Laboratory Medicine, Tan Tock Seng Hospital | Department of Laboratory Medicine, Tan Tock Seng Hospital | Chen YYC, Zair X, Lim JX, Li C, Tang WY, Maurer-Stroh S, Barkham TMS, Nagarajan N, Sessions OM |
| EPI_ISL_538608, EPI_ISL_538609, EPI_ISL_538610, EPI_ISL_538611, EPI_ISL_538612 | Clínica Universidad de Navarra. Servicio de Enfermedades Infecciosas y Microbiología clínica | SeqCOVID-SPAIN consortium/IBV(CSIC) | Mirian Fernández-Alonso, Jose Luis del Pozo and SeqCOVID-SPAIN consortium |
| EPI_ISL_538614 | Servicio de Microbiología. Hospital General Universitario de Castellón | SeqCOVID-SPAIN consortium/IBV(CSIC) | Rosario Moreno, María Dolores Tirado and SeqCOVID-SPAIN consortium |
| EPI_ISL_538683, EPI_ISL_538687 | Hospital Universitario Virgen de las Nieves de Granada-SAS | SeqCOVID-SPAIN consortium/IBV(CSIC) | Mercedes Pérez Ruiz, Sara Sanbonmatsu Gámez, Irene Pedrosa Corral, José M. Navarro-Marí and SeqCOVID-SPAIN consortium |
| EPI_ISL_538787, EPI_ISL_538789, EPI_ISL_538799, EPI_ISL_538802, EPI_ISL_538819, EPI_ISL_538852, EPI_ISL_538869, EPI_ISL_538900, EPI_ISL_538906, EPI_ISL_538912, EPI_ISL_538915, EPI_ISL_538925, EPI_ISL_538928, EPI_ISL_538932, EPI_ISL_538938, EPI_ISL_538939, EPI_ISL_538940, EPI_ISL_538944, EPI_ISL_538947, EPI_ISL_538948, EPI_ISL_538951, EPI_ISL_538955, EPI_ISL_538961, EPI_ISL_538962, EPI_ISL_538966, EPI_ISL_538968, EPI_ISL_538972, EPI_ISL_538974, EPI_ISL_538976, EPI_ISL_538993, EPI_ISL_538995, EPI_ISL_538996, EPI_ISL_538998, EPI_ISL_539003, EPI_ISL_539011, EPI_ISL_539013, EPI_ISL_539019, EPI_ISL_539031, EPI_ISL_539038, EPI_ISL_539041, EPI_ISL_539053, EPI_ISL_539054, EPI_ISL_539055, EPI_ISL_539057, EPI_ISL_539059, EPI_ISL_539077, EPI_ISL_539081, EPI_ISL_539083, EPI_ISL_539086, EPI_ISL_539100, EPI_ISL_539105, EPI_ISL_539121, EPI_ISL_539129, EPI_ISL_539132, EPI_ISL_539151, EPI_ISL_539158, EPI_ISL_539159, EPI_ISL_539160, EPI_ISL_539161, EPI_ISL_539162, EPI_ISL_539169, EPI_ISL_539172, EPI_ISL_539178, EPI_ISL_539181, EPI_ISL_539186, EPI_ISL_539187, EPI_ISL_539190, EPI_ISL_539191, EPI_ISL_539195, EPI_ISL_539196, EPI_ISL_539200, EPI_ISL_539206, EPI_ISL_539211, EPI_ISL_539217 | | | |
| see above | Leeds Teaching Hospitals NHS Trust and Public Health England, National Infection Service (Leeds laboratory) | Wellcome Sanger Institute for the COVID-19 Genomics UK (COG-UK) consortium | Louissa Macfarlane-Smith, Holli Carden, Katherine L. Harper, Antony Hale and Alex Alderton, Roberto Amato, Sonia Goncalves, Ewan Harrison, David K. Jackson, Ian Johnston, Dominic Kwiatkowski, Cordelia Langford, John Sillitoe on behalf of the Wellcome Sanger Institute COVID-19 Surveillance Team |
| EPI_ISL_539222, EPI_ISL_539223 | Servicio de Microbiologia. Hospital Clinico Universitario de Valencia | SeqCOVID-SPAIN consortium/IBV(CSIC) | Maria Dolores Ocete, Inma Galán Vendrell, Paula Ruiz-Hueso, Mariana Reyes-Prieto, Vicente Soriano Chirona, Maria Alma Bracho, Griselda De Marco, Beatriz Beamud, Lidia Ruiz Roldan, Marta Pla Diaz,Neris Garcia-Gonzalez, Loreto Ferrús Abad, Llúcia Martínez-Priego, Concepcion Gimeno, Giuseppe D'Auria, Fernando Gonzalez-Candelas and SeqCOVID-SPAIN consortium |
| EPI_ISL_539299 | Molecular diagnostic laboratory of Federal Budget Institution of Science "Central Research Institute of Epidemiology" of The Federal Service on Customers' Rights Protection and Human Well-being Surveillance | Group of Genomics and Postgenomic Technologies of Central Research Institute of Epidemilology | Speranskaya AS, Kaptelova VV, Valdokhina AV, Bulanenko VP, Samoilov AE, Korneenko EV, Tivanova EV, Shipulina OY, Akimkin VG |
| EPI_ISL_539310, EPI_ISL_539313, EPI_ISL_539314, EPI_ISL_539325 | KWR Watercycle Research Institute | Erasmus Medical Center | Ray Izquierdo-Lara, Goffe Elsinga, Leo Heijnen, Bas B. Oude Munnink, Claudia M. E. Schapendonk, David Nieuwenhuijse, Matthijs Kon, Lu Lu, Frank M. Aarestrup, Samantha Lycett, Gertjan Medema, Marion P.G. Koopmans, Miranda de Graaf |
| EPI_ISL_539574, EPI_ISL_539575, EPI_ISL_539576 | Centre de Recherches Medicales de Lambarene (CERMEL) | Department of Emerging Infectious Diseases, Institute of Tropical Medicine, Nagasaki University | Haruka Abe, Yuri Ushijima, Rodrigue Bikangui, Akim A. Adegnika, Bertrand Lell, Jiro Yasuda |
| EPI_ISL_539776 | National Institute of Public Health (Czech Republic) | State Veterinary Institute Prague | Nagy, A; Jirincova, H; Novakova, L; Trnka, D; Vecerova, J. |
| EPI_ISL_539820 | Queen Mary Hospital | Hong Kong Department of Health | Alan K.L. Tsang, Peter C.W. Yip, Edman T.K. Lam, Rickjason C.W. Chan, Dominic N.C. Tsang |
| EPI_ISL_541237 | Florida Bureau of Public Health Laboratories, Florida Department of Health | Florida Bureau of Public Health Laboratories, Florida Department of Health | Schmedes,S., Blanton,J. |
| EPI_ISL_541345 | LACEN/PR | Laboratory of Respiratory Viruses and Measles, Oswaldo Cruz Institute, FIOCRUZ | Paola Resende, Luciana Appolinario, Fernando Motta, Anna Carolina Paixão, Ana Carolina Mendonça, Jonathan Lopes, Irina Riediger, Maria do Carmo Debur, Marilda Siqueira |

| | | | |
|---|---|---|---|
| EPI_ISL_541372 | LACEN/SE | Laboratory of Respiratory Viruses and Measles, Oswaldo Cruz Institute, FIOCRUZ | Paola Resende, Luciana Appolinario, Fernando Motta, Anna Carolina Paixão, Ana Carolina Mendonça, Jonathan Lopes, Clioma Santos, Marilda Siqueira |
| EPI_ISL_541769, EPI_ISL_541771, EPI_ISL_541782 | Barts Health NHS Trust | Wellcome Sanger Institute for the COVID-19 Genomics UK (COG-UK) consortium | Teresa Cutino-Moguel, Mark Hopkins, Beatrix Kele, David Harrington and Alex Alderton, Roberto Amato, Sonia Goncalves, Ewan Harrison, David K. Jackson, Ian Johnston, Dominic Kwiatkowski, Cordelia Langford, John Sillitoe on behalf of the Wellcome Sanger Institute COVID-19 Surveillance Team |
| EPI_ISL_542358, EPI_ISL_542380, EPI_ISL_542382, EPI_ISL_542384, EPI_ISL_542386, EPI_ISL_542387, EPI_ISL_542388, EPI_ISL_542395 | San Matteo Hospital Pavia | Dep. Of Oncology and Hemato-Oncology University of Milan | Claudia Alteri, Valeria Cento, Antonio Piralla, Valentino Costabile, Monica Tallarita, Luna Colagrossi, Silvia Renica, Federica Giardina, Federica Novazzi, Stefano Gaiarsa, Elisa Matarazzo, Maria Antonello, Chiara Vismara, Roberto Fumagalli, Oscar Massimiliano Epis, Massimo Puoti, Carlo Federico Perno, Fausto Baldanti |
| EPI_ISL_542578, EPI_ISL_542615, EPI_ISL_542617, EPI_ISL_542795, EPI_ISL_542803, EPI_ISL_542805, EPI_ISL_542821, EPI_ISL_543980 | Houston Methodist Hospital | Houston Methodist Hospital | S. Wesley Long, Randall J. Olsen, Paul A. Christensen, David W. Bernard, James J. Davis, Maulik Shukla, Marcus Nguyen, Matthew Ojeda Saavedra, Concepcion C. Cantu, Prasanti Yerramilli, Layne Pruitt, Sishir Subedi, Hung-Che Kuo, Heather Hendrickson, Ghazaleh Eskandari, Hoang A. T. Nguyen, J. Hunter Long, Muthiah Kumaraswami, Jule Goike, Daniel Boutz, Jimmy Gollihar, Jason S. McLellan, Chia-Wei Chou, Kamyab Javanmardi, Ilya J. Finkelstein, and James M. Musser |
| EPI_ISL_547474, EPI_ISL_547475, EPI_ISL_547484, EPI_ISL_547485, EPI_ISL_547493, EPI_ISL_547517, EPI_ISL_547518, EPI_ISL_547533 | Dutch COVID-19 response team | National Institute for Public Health and the Environment (RIVM) | Adam Meijer, Harry Vennema, Jeroen Cremer, Sharon van den Brink, Bas van der Veer, AnneMarie van den Brandt, Florian Zwagemaker, Dennis Schmitz, Chantal Reusken, on behalf of the national COVID-19 response team |
| EPI_ISL_548127 | Middlemore Hospital | Institute of Environmental Science and Research (ESR) | Xiaoyun Ren, Matt Storey, Nikki Freed, Muhammad Faisal, Jing Wang, Hermes Perez, Anja Werno, Antje van der Linden, Arlo Upton, Chris Mansell, David Hammer, Dragana Drinkovic, Gary McAuliffe, Hana Sofia Andersson, James Ussher, Jill Sherwood, Josh Freeman, Julia Howard, Juliet Elvy, Mary DeAlmeida, Matt Blakiston, Matthew Rogers, Max Bloomfield, Michael Addidle, Michelle Balm, Sally Roberts, Sarah Jefferies, Sharmini Muttaiyah, Susan Morpeth, Susan Taylor, Timothy Blackmore, Vani Sathyendran, Veronica Playle, Virginia Hope, Erasmus Smit, Lauren Jelly, Olin Silander, Joep de Ligt |
| EPI_ISL_548353 | Ventura County Public Health Lab | Chan-Zuckerberg Biohub | CZB Cliahub Consortium |
| EPI_ISL_548949 | Max von Pettenkofer Institute, Virology, National Reference Center for Retroviruses, LMU München | Laboratory for Functional Genome Analysis, Dept. Genomics, Gene Center of the LMU Munich | Max Muenchhoff, Stefan Krebs, Alexander Graf, Oliver Keppler, Helmut Blum |
| EPI_ISL_548993 | National Public Health Laboratory, National Centre for Infectious Diseases | National Public Health Laboratory, National Centre for Infectious Diseases | Mak TM, Octavia S, Zhou Z, Cui L, Lin RTP |
| EPI_ISL_549010, EPI_ISL_549014, EPI_ISL_549015 | KWR Watercycle Research Institute | Erasmus Medical Center | Ray Izquierdo-Lara, Goffe Elsinga, Leo Heijnen, Bas B. Oude Munnink, Claudia M. E. Schapendonk, David Nieuwenhuijse, Matthijs Kon, Lu Lu, Frank M. Aarestrup, Samantha Lycett, Gertjan Medema, Marion P.G. Koopmans, Miranda de Graaf |
| EPI_ISL_560386 | National Health Laboratory | Botswana Institute for Technology Research and innovation | Kefentse Arnold Tumedi, Madisa Mine, Dineo Emang Tshiamo. Gape Nyepetsi, Thongbotho Mphoyakgosi, Maitshwarelo Ignatius Matsheka |
| EPI_ISL_560570, EPI_ISL_560573, EPI_ISL_560578 | hôpital | National Reference Center for Viruses of Respiratory Infections, Institut Pasteur, Paris | Sylvie Behillil, Fabiana Gambaro, Etienne Simon-Lorière, Vincent Enouf, Maud Vanpeene, Sylvie van der Werf |
| EPI_ISL_560627 | Hospital | National Reference Center for Viruses of Respiratory Infections, Institut Pasteur, Paris | Sylvie Behillil, Fabiana Gambaro, Etienne Simon-Lorière, Vincent Enouf, Maud Vanpeene, Sylvie van der Werf |
| EPI_ISL_560718, EPI_ISL_560719, EPI_ISL_560720, EPI_ISL_560721, EPI_ISL_560722, EPI_ISL_560723, EPI_ISL_560724, EPI_ISL_560725, EPI_ISL_560726, EPI_ISL_560727, EPI_ISL_560728, EPI_ISL_560729, EPI_ISL_560730, EPI_ISL_560731, EPI_ISL_560732, EPI_ISL_560733, EPI_ISL_560734, EPI_ISL_560735, EPI_ISL_560736, EPI_ISL_560737, EPI_ISL_560738, EPI_ISL_560740, EPI_ISL_561367, EPI_ISL_561368, EPI_ISL_561369 | | | |
| see above | Centre for Clinical Infection and Diagnostics Research and Genomics Innovation Unit, Guy's and St. Thomas' NHS Trust | Centre for Clinical Infection and Diagnostics Research and Genomics Innovation Unit, Guy's and St. Thomas' NHS Trust | Chloe Fisher, Luke Snell, Rahul Batra, Jonathan Edgeworth, Ali Raza Awan |
| EPI_ISL_561420, EPI_ISL_561532, EPI_ISL_561728, EPI_ISL_561777, EPI_ISL_561825, EPI_ISL_561843, EPI_ISL_562055, EPI_ISL_562105 | Victorian Infectious Diseases Reference Laboratory (VIDRL) | VIDRL and MDU-PHL | Caly, L., Seemann, T., Sait, M., Schultz, M. B., Druce J., Sherry, N. |
| EPI_ISL_562167 | Microbiological Diagnostic Unit - Public Health Laboratory (MDU-PHL) | MDU-PHL | Seemann, T., Schultz M. B., Sait, M., Sherry, N. |
| EPI_ISL_562258, EPI_ISL_562431, EPI_ISL_562535, EPI_ISL_563880, EPI_ISL_564270 | Victorian Infectious Diseases Reference Laboratory (VIDRL) | VIDRL and MDU-PHL | Caly, L., Seemann, T., Sait, M., Schultz, M. B., Druce J., Sherry, N. |
| EPI_ISL_565929, EPI_ISL_565956, EPI_ISL_565959, EPI_ISL_565962, EPI_ISL_565972, EPI_ISL_565990, EPI_ISL_565995, EPI_ISL_565999, EPI_ISL_566001 | Michigan Department of Health and Human Services, Bureau of Laboratories | Michigan Department of Health and Human Services, Bureau of Laboratories | Blankenship HM, Riner D, Soehnlen MK |
| EPI_ISL_568485 | Virology, Iran University of Medical Sciences | Virology, Iran University of Medical Sciences | Keyvani,H., Ranjbar,M.M., Soleimani,S., Keyvani,F. |
| EPI_ISL_568500 | Virology, Iran University of Medical Sciences | Virology, Iran University of Medical Sciences | Keyvani,H., Ranjbar,Mm., Soleimani,S., Keyvani,F. |
| EPI_ISL_568693 | RS Freeport Tembagapura | Eijkman Institute for Molecular Biology, Ministry of Research and Technology/National Agency for Research and Innovation | Frilasita A Yudhaputri, Edison Johar, Hidayat Trimarsanto, Iskandar A Adnan, Willy Agustine, David H Muljono, Safarina G Malik, Herawati Sudoyo, Khin Saw Myint, Amin Soebandrio |
| EPI_ISL_568694 | RSUP Prof. Dr. R. Kandou Manado | Eijkman Institute for Molecular Biology, Ministry of Research and Technology/National Agency for Research and Innovation | Frilasita A Yudhaputri, Edison Johar, Hidayat Trimarsanto, Iskandar A Adnan, Willy Agustine, David H Muljono, Safarina G Malik, Herawati Sudoyo, Khin Saw Myint, Amin Soebandrio |
| EPI_ISL_568990, EPI_ISL_569354, EPI_ISL_569355, EPI_ISL_569356, EPI_ISL_569357, EPI_ISL_569358, EPI_ISL_569359, EPI_ISL_569360, EPI_ISL_569361, EPI_ISL_569362, EPI_ISL_569363, EPI_ISL_569364, EPI_ISL_569365, EPI_ISL_569366, EPI_ISL_569367, EPI_ISL_569368, EPI_ISL_569369, EPI_ISL_569370, EPI_ISL_569371, EPI_ISL_569372, EPI_ISL_569373, EPI_ISL_569374, EPI_ISL_569375, EPI_ISL_569376, EPI_ISL_569377, EPI_ISL_569378, EPI_ISL_569379, EPI_ISL_569380, EPI_ISL_569381, EPI_ISL_569382, EPI_ISL_569383, EPI_ISL_569384, EPI_ISL_569385, EPI_ISL_569386, EPI_ISL_569387, EPI_ISL_569388, EPI_ISL_569389, EPI_ISL_569390, EPI_ISL_569391, EPI_ISL_569392, EPI_ISL_569393, EPI_ISL_569394, EPI_ISL_569395, EPI_ISL_569396, EPI_ISL_569397, EPI_ISL_569398, EPI_ISL_569399, EPI_ISL_569400, EPI_ISL_569401, EPI_ISL_569402, EPI_ISL_569403, EPI_ISL_569404, EPI_ISL_569405, EPI_ISL_569406, EPI_ISL_569407, EPI_ISL_569408, EPI_ISL_569409, EPI_ISL_569410, EPI_ISL_569411, EPI_ISL_569412, EPI_ISL_569413, EPI_ISL_569414, EPI_ISL_569415, EPI_ISL_569416, EPI_ISL_569417, EPI_ISL_569421, EPI_ISL_569422, EPI_ISL_569478, EPI_ISL_569479, EPI_ISL_569480, EPI_ISL_569481, EPI_ISL_569482, EPI_ISL_569483, EPI_ISL_569484, EPI_ISL_569485, EPI_ISL_569486, EPI_ISL_569487, EPI_ISL_569488, EPI_ISL_569489, EPI_ISL_569490 | | | |
| see above | MEPHI, Aix Marseille University | MEPHI, Aix Marseille University | Anthony LEVASSEUR |
| EPI_ISL_570047, EPI_ISL_570048, EPI_ISL_570050, EPI_ISL_570052, EPI_ISL_570053, EPI_ISL_570059, EPI_ISL_570060, EPI_ISL_570152, EPI_ISL_570153, EPI_ISL_570154, EPI_ISL_570155, EPI_ISL_570156, EPI_ISL_570612 | | | |
| see above | UW Virology Lab | UW Virology Lab | Pavitra Roychoudhury, Hong Xie, Lasata Shrestha, Amin Addetia, Victoria M Rachleff, Meei-Li Huang, Keith R Jerome, Alexander Greninger |
| EPI_ISL_571176, EPI_ISL_571397, EPI_ISL_571815, EPI_ISL_571816, EPI_ISL_571817, EPI_ISL_571818, EPI_ISL_571819, EPI_ISL_571820 | Quest Diagnostics | Quest Diagnostics | Rosenthal,S.H., Gerasimova,A., Kagan,R.M., Anderson, B., Grover, D., Livingston, K.E., Hua, M., Liu Y., Shalhout, D.F., Owen, R., Lacbawan, F. |
| EPI_ISL_572324 | IZSM | IZSM | Maurizio Viscardi, Lorena Cardillo, Giovanna Fusco |
| EPI_ISL_574605 | RSUP dr. Sardjito | Eijkman Institute for Molecular Biology, Ministry of Research and Technology/National Agency for Research and Innovation | Frilasita A Yudhaputri, Edison Johar, Hidayat Trimarsanto, Iskandar A Adnan, Willy Agustine, David H Muljono, Safarina G Malik, Herawati Sudoyo, Khin Saw Myint, Amin Soebandrio |
| EPI_ISL_574840, EPI_ISL_574841, EPI_ISL_574842 | Institute for Infectious Diseases, University of Bern | Institute for Infectious Diseases, University of Bern | Michel C Koch, Christian Baumann, Miguel A Terrazos Miani, Cora Sägesser, Stephen L Leib, Peter Keller, Franziska Suter-Riniker, Alban Ramette |

| | | | |
|---|---|---|---|
| EPI_ISL_576150, EPI_ISL_576151, EPI_ISL_576161, EPI_ISL_576168, EPI_ISL_576222 | Delaware Public Health Lab | Delaware Public Health Lab | Gregory Hovan |
| EPI_ISL_577565 | Michigan Department of Health and Human Services, Bureau of Laboratories | Michigan Department of Health and Human Services, Bureau of Laboratories | Blankenship HM, Riner D, Soehnlen MK |
| EPI_ISL_578085, EPI_ISL_578088, EPI_ISL_578093, EPI_ISL_578113, EPI_ISL_578114, EPI_ISL_578115, EPI_ISL_578116, EPI_ISL_578117, EPI_ISL_578118, EPI_ISL_578119, EPI_ISL_578120, EPI_ISL_578121, EPI_ISL_578122, EPI_ISL_578123, EPI_ISL_578124, EPI_ISL_578125, EPI_ISL_578126, EPI_ISL_578127, EPI_ISL_578128, EPI_ISL_578129 | | | |
| see above | University of Michigan Clinical Microbiology Laboratory | Lauring Lab, University of Michigan, Department of Microbiology and Immunology | Valesano |
| EPI_ISL_578378, EPI_ISL_578379, EPI_ISL_578380, EPI_ISL_578381, EPI_ISL_578382, EPI_ISL_578387, EPI_ISL_578431, EPI_ISL_578432, EPI_ISL_578433, EPI_ISL_578434, EPI_ISL_578435, EPI_ISL_578436, EPI_ISL_578437 | | | |
| see above | Wisconsin State Laboratory of Hygiene Communicable Disease Division | Wisconsin State Laboratory of Hygiene Communicable Disease Division | Kelsey R. Florek, Abigail C. Shockey |
| EPI_ISL_579081 | Canterbury Health Laboratories | Institute of Environmental Science and Research (ESR) | Xiaoyun Ren, Matt Storey, Nikki Freed, Muhammad Faisal, Jing Wang, Hermes Perez, Anja Werno, Antje van der Linden, Arlo Upton, Chris Mansell, David Hammer, Dragana Drinkovic, Gary McAuliffe, Hana Sofia Andersson, James Ussher, Jill Sherwood, Josh Freeman, Julia Howard, Juliet Elvy, Mary DeAlmeida, Matt Blakiston, Matthew Rogers, Max Bloomfield, Michael Addidle, Michelle Balm, Sally Roberts, Sarah Jefferies, Sharmini Muttaiyah, Susan Morpeth, Susan Taylor, Timothy Blackmore, Vani Sathyendran, Veronica Playle, Virginia Hope, Erasmus Smit, Lauren Jelly, Olin Silander, Joep de Ligt |
| EPI_ISL_579133, EPI_ISL_579134, EPI_ISL_579135, EPI_ISL_579136, EPI_ISL_579137, EPI_ISL_579138, EPI_ISL_579139, EPI_ISL_579141, EPI_ISL_579142, EPI_ISL_579143, EPI_ISL_579144, EPI_ISL_579145, EPI_ISL_579146, EPI_ISL_579147, EPI_ISL_579150, EPI_ISL_579151, EPI_ISL_579152, EPI_ISL_579154 | | | |
| see above | Southern Community Labs Dunedin | Institute of Environmental Science and Research (ESR) | Xiaoyun Ren, Matt Storey, Nikki Freed, Muhammad Faisal, Jing Wang, Hermes Perez, Anja Werno, Antje van der Linden, Arlo Upton, Chris Mansell, David Hammer, Dragana Drinkovic, Gary McAuliffe, Hana Sofia Andersson, James Ussher, Jill Sherwood, Josh Freeman, Julia Howard, Juliet Elvy, Mary DeAlmeida, Matt Blakiston, Matthew Rogers, Max Bloomfield, Michael Addidle, Michelle Balm, Sally Roberts, Sarah Jefferies, Sharmini Muttaiyah, Susan Morpeth, Susan Taylor, Timothy Blackmore, Vani Sathyendran, Veronica Playle, Virginia Hope, Erasmus Smit, Lauren Jelly, Olin Silander, Joep de Ligt |
| EPI_ISL_579311 | Middlemore Hospital | Institute of Environmental Science and Research (ESR) | Xiaoyun Ren, Matt Storey, Nikki Freed, Muhammad Faisal, Jing Wang, Hermes Perez, Anja Werno, Antje van der Linden, Arlo Upton, Chris Mansell, David Hammer, Dragana Drinkovic, Gary McAuliffe, Hana Sofia Andersson, James Ussher, Jill Sherwood, Josh Freeman, Julia Howard, Juliet Elvy, Mary DeAlmeida, Matt Blakiston, Matthew Rogers, Max Bloomfield, Michael Addidle, Michelle Balm, Sally Roberts, Sarah Jefferies, Sharmini Muttaiyah, Susan Morpeth, Susan Taylor, Timothy Blackmore, Vani Sathyendran, Veronica Playle, Virginia Hope, Erasmus Smit, Lauren Jelly, Olin Silander, Joep de Ligt |
| EPI_ISL_579405 | LabPLUS | Institute of Environmental Science and Research (ESR) | Xiaoyun Ren, Matt Storey, Nikki Freed, Muhammad Faisal, Jing Wang, Hermes Perez, Anja Werno, Antje van der Linden, Arlo Upton, Chris Mansell, David Hammer, Dragana Drinkovic, Gary McAuliffe, Hana Sofia Andersson, James Ussher, Jill Sherwood, Josh Freeman, Julia Howard, Juliet Elvy, Mary DeAlmeida, Matt Blakiston, Matthew Rogers, Max Bloomfield, Michael Addidle, Michelle Balm, Sally Roberts, Sarah Jefferies, Sharmini Muttaiyah, Susan Morpeth, Susan Taylor, Timothy Blackmore, Vani Sathyendran, Veronica Playle, Virginia Hope, Erasmus Smit, Lauren Jelly, Olin Silander, Joep de Ligt |
| EPI_ISL_579474, EPI_ISL_579481, EPI_ISL_579482, EPI_ISL_579483, EPI_ISL_579484, EPI_ISL_579485, EPI_ISL_579486 | Canterbury Health Laboratories | Institute of Environmental Science and Research (ESR) | Xiaoyun Ren, Matt Storey, Nikki Freed, Muhammad Faisal, Jing Wang, Hermes Perez, Anja Werno, Antje van der Linden, Arlo Upton, Chris Mansell, David Hammer, Dragana Drinkovic, Gary McAuliffe, Hana Sofia Andersson, James Ussher, Jill Sherwood, Josh Freeman, Julia Howard, Juliet Elvy, Mary DeAlmeida, Matt Blakiston, Matthew Rogers, Max Bloomfield, Michael Addidle, Michelle Balm, Sally Roberts, Sarah Jefferies, Sharmini Muttaiyah, Susan Morpeth, Susan Taylor, Timothy Blackmore, Vani Sathyendran, Veronica Playle, Virginia Hope, Erasmus Smit, Lauren Jelly, Olin Silander, Joep de Ligt |
| EPI_ISL_579583, EPI_ISL_579584, EPI_ISL_579585, EPI_ISL_579586, EPI_ISL_579587, EPI_ISL_579588, EPI_ISL_579589, EPI_ISL_579590, EPI_ISL_579591, EPI_ISL_579592, EPI_ISL_579593 | | | |
| see above | QEII Health Sciences Centre | National Microbiology Laboratory (NML) | Anna Majer, Shari Tyson, Grace Seo, Philip Mabon, Darian Hole, Elsie Grudeski, Rhiannon Huzarewich, Russell Mandes, Anneliese Landgraff, Jennifer Tanner, Natalie Knox, Morag Graham, Gary Van Domselaar, Todd Hatchette, Jason LeBlanc, Nathalie Bastien, Yan Li, Timothy Booth, CanCOGeN's metadata curation team, Public Health Agency of Canada's CanCOGeN team |
| EPI_ISL_581461, EPI_ISL_581463, EPI_ISL_581464, EPI_ISL_581473, EPI_ISL_581476, EPI_ISL_581481 | Medizinische Klinik Innere Medizin I, Universitätsklinikum Tübingen | NGS Competence Center Tübingen, Institut für Medizinische Mikrobiologie und Hygiene, Universitätsklinikum Tübingen | Angel Angelov |
| EPI_ISL_581736, EPI_ISL_581768, EPI_ISL_581774, EPI_ISL_581775, EPI_ISL_581776, EPI_ISL_581777, EPI_ISL_581778, EPI_ISL_581779, EPI_ISL_581780, EPI_ISL_581781, EPI_ISL_581782, EPI_ISL_581798, EPI_ISL_581799, EPI_ISL_581800, EPI_ISL_581801, EPI_ISL_581802, EPI_ISL_581803, EPI_ISL_581804, EPI_ISL_581805, EPI_ISL_581806, EPI_ISL_581807, EPI_ISL_581808, EPI_ISL_581809, EPI_ISL_581810, EPI_ISL_581811, EPI_ISL_581812 | | | |
| see above | University Hospital Basel, Clinical Virology | University Hospital Basel, Clinical Bacteriology | Madlen Stange, Alfredo Mari, Tim Roloff, Helena MB Seth-Smith, Michael Schweitzer, Myrta Brunner, Karoline Leuzinger, Kirstine K. Soegaard, Alexander Gensch, Sarah Tschudin-Sutter, Simon Fuchs, Julia Bielicki, Hans Pargger, Martin Siegemund, Christian Nickel, Roland Bingisser, Michael Osthoff, Stefano Bassetti, Rita Schneider-Sliwa, Manuel Battegay, Hans Hirsch, Adrian Egli |
| EPI_ISL_582243, EPI_ISL_582245, EPI_ISL_582253, EPI_ISL_582254, EPI_ISL_582255, EPI_ISL_582259, EPI_ISL_582265, EPI_ISL_582267, EPI_ISL_582268, EPI_ISL_582269, EPI_ISL_582272, EPI_ISL_582276, EPI_ISL_582277, EPI_ISL_582279, EPI_ISL_582280, EPI_ISL_582282, EPI_ISL_582286, EPI_ISL_582288, EPI_ISL_582291, EPI_ISL_582292, EPI_ISL_582295, EPI_ISL_582301, EPI_ISL_582306, EPI_ISL_582328, EPI_ISL_582330, EPI_ISL_582331, EPI_ISL_582340, EPI_ISL_582342, EPI_ISL_582343, EPI_ISL_582345, EPI_ISL_582347, EPI_ISL_582350, EPI_ISL_582351, EPI_ISL_582353, EPI_ISL_582379, EPI_ISL_582387, EPI_ISL_582394, EPI_ISL_582400 | | | |
| see above | Cadham Provincial Laboratory | National Microbiology Laboratory (NML) | Anna Majer, Shari Tyson, Grace Seo, Philip Mabon, Elsie Grudeski, Rhiannon Huzarewich, Russell Mandes, Anneliese Landgraff, Jennifer Tanner, Natalie Knox, Morag Graham, Gary Van Domselaar, Paul Van Caeseele, Jared Bullard, David Alexander, Kerry Dust, Nathalie Bastien, Yan Li, Timothy Booth, Darian Hole, Madison Chapel, CanCOGeN's metadata curation team, Public Health Agency of Canada CanCOGeN team |
| EPI_ISL_582520 | unknown | Instituto Nacional de Saude (INSA) | Borges et al |
| EPI_ISL_582741, EPI_ISL_582742, EPI_ISL_582743, EPI_ISL_582744, EPI_ISL_582748, EPI_ISL_582749, EPI_ISL_582750, EPI_ISL_582751, EPI_ISL_582752, EPI_ISL_582753, EPI_ISL_582754, EPI_ISL_582755, EPI_ISL_582757, EPI_ISL_582759, EPI_ISL_582760, EPI_ISL_582761, EPI_ISL_582763, EPI_ISL_582764, EPI_ISL_582765, EPI_ISL_582766 | | | |
| see above | Servicio de Microbiología, Hospital Universitario Cruces, Bilbao | SeqCOVID-SPAIN consortium/IBV(CSIC) | Maitane Aranzamendi Zaldumbide and SeqCOVID-SPAIN consortium |
| EPI_ISL_583428 | Memorial Sloan Kettering Cancer Center | van Bakel Laboratory, Genetics and Genomics Sciences, Icahn School of Medicine at Mount Sinai | Teresa Aydillo, Ana S. Gonzalez-Reiche, Sadaf Aslam, Adriana van de Guchte, Zenab Khan, Ajay Obla, Jayeeta Dutta, Harm van Bakel, Judith Aberg, Adolfo García-Sastre, Gunjan Shah, Tobias Hohl, Genovefa Papanicolaou, Miguel-Angel Perales, Kent Sepkowitz, Ngolela Esther Babady, and Mini Kamboj |
| EPI_ISL_583429 | Garcia-Sastre Laboratory, Department of Microbiology, Icahn School of Medicine at Mount Sinai | van Bakel Laboratory, Genetics and Genomics Sciences, Icahn School of Medicine at Mount Sinai | Teresa Aydillo, Ana S. Gonzalez-Reiche, Sadaf Aslam, Adriana van de Guchte, Zenab Khan, Ajay Obla, Jayeeta Dutta, Harm van Bakel, Judith Aberg, Adolfo García-Sastre, Gunjan Shah, Tobias Hohl, Genovefa Papanicolaou, Miguel-Angel Perales, Kent Sepkowitz, Ngolela Esther Babady, and Mini Kamboj |
| EPI_ISL_583479 | Memorial Sloan Kettering Cancer Center | van Bakel Laboratory, Genetics and Genomics Sciences, Icahn School of Medicine at Mount Sinai | Teresa Aydillo, Ana S. Gonzalez-Reiche, Sadaf Aslam, Adriana van de Guchte, Zenab Khan, Ajay Obla, Jayeeta Dutta, Harm van Bakel, Judith Aberg, Adolfo García-Sastre, Gunjan Shah, Tobias Hohl, Genovefa Papanicolaou, Miguel-Angel Perales, Kent Sepkowitz, Ngolela Esther Babady, and Mini Kamboj |
| EPI_ISL_583557 | Universitaetsklinik für Innere Medizin II Innsbruck | Bergthaler laboratory, CeMM Research Center for Molecular Medicine of the Austrian Academy of Sciences | Alexandra Popa, Benedikt Agerer, Henrique Colaco, Lukas Endler, Jakob-Wendelin Genger, Alexander Lercher, Mark Smyth, Thomas Penz, Michael Schuster, Jan Laine, Martin Senekowitsch, Judith Aberle, Stephan Aberle, Peter Hufnagl, Daniela Schmid, Franz Allerberger, Elisabeth Puchhammer-Stoeckl, Manfred Nairz, Guenter Weiss, Gregor Hörmann, Kinga Rigler-Hohenwarter, Rainer Gattringer, Wegene Borena, Dorothee von Laer, Gernot Walder, Peter Obrist, Christian Paar, Sabine Sussitz-Rack, Gunther Vogl, Adi Steinrigl, Christoph Bock, Andreas Bergthaler |
| EPI_ISL_583569 | Center for Virology, Medical University of Vienna | Bergthaler laboratory, CeMM Research Center for Molecular Medicine of the Austrian Academy of Sciences | Alexandra Popa, Benedikt Agerer, Henrique Colaco, Lukas Endler, Jakob-Wendelin Genger, Alexander Lercher, Mark Smyth, Thomas Penz, Michael Schuster, Jan Laine, Martin Senekowitsch, Judith Aberle, Stephan Aberle, Peter Hufnagl, Daniela Schmid, Franz Allerberger, Elisabeth Puchhammer-Stoeckl, Manfred Nairz, Guenter Weiss, Gregor Hörmann, Kinga Rigler-Hohenwarter, Rainer Gattringer, Wegene Borena, Dorothee von Laer, Gernot Walder, Peter Obrist, Christian Paar, Sabine Sussitz-Rack, Gunther Vogl, Adi Steinrigl, Christoph Bock, Andreas Bergthaler |

| | | | |
|---|---|---|---|
| EPI_ISL_583574 | Austrian Agency for Health and Food Safety (AGES) | Bergthaler laboratory, CeMM Research Center for Molecular Medicine of the Austrian Academy of Sciences | Alexandra Popa, Benedikt Agerer, Henrique Colaco, Lukas Endler, Jakob-Wendelin Genger, Alexander Lercher, Mark Smyth, Thomas Penz, Michael Schuster, Jan Laine, Martin Senekowitsch, Judith Aberle, Stephan Aberle, Peter Hufnagl, Daniela Schmid, Franz Allerberger, Elisabeth Puchhammer-Stoeckl, Manfred Nairz, Guenter Weiss, Gregor Hörmann, Kinga Rigler-Hohenwarter, Rainer Gattringer, Wegene Borena, Dorothee von Laer, Gernot Walder, Peter Obrist, Christian Paar, Sabine Sussitz-Rack, Gunther Vogl, Adi Steinrigl, Christoph Bock, Andreas Bergthaler |
| EPI_ISL_583580, EPI_ISL_583591, EPI_ISL_583594 | Institute for Medical and Chemical Laboratory Diagnostics, Kepler Universitätsklinikum | Bergthaler laboratory, CeMM Research Center for Molecular Medicine of the Austrian Academy of Sciences | Alexandra Popa, Benedikt Agerer, Henrique Colaco, Lukas Endler, Jakob-Wendelin Genger, Alexander Lercher, Mark Smyth, Thomas Penz, Michael Schuster, Jan Laine, Martin Senekowitsch, Judith Aberle, Stephan Aberle, Peter Hufnagl, Daniela Schmid, Franz Allerberger, Elisabeth Puchhammer-Stoeckl, Manfred Nairz, Guenter Weiss, Gregor Hörmann, Kinga Rigler-Hohenwarter, Rainer Gattringer, Wegene Borena, Dorothee von Laer, Gernot Walder, Peter Obrist, Christian Paar, Sabine Sussitz-Rack, Gunther Vogl, Adi Steinrigl, Christoph Bock, Andreas Bergthaler |
| EPI_ISL_583600, EPI_ISL_583604, EPI_ISL_583613, EPI_ISL_583614 | Institut für Virologie am Department für Hygiene, Mikrobiologie und Public Health | Bergthaler laboratory, CeMM Research Center for Molecular Medicine of the Austrian Academy of Sciences | Alexandra Popa, Benedikt Agerer, Henrique Colaco, Lukas Endler, Jakob-Wendelin Genger, Alexander Lercher, Mark Smyth, Thomas Penz, Michael Schuster, Jan Laine, Martin Senekowitsch, Judith Aberle, Stephan Aberle, Peter Hufnagl, Daniela Schmid, Franz Allerberger, Elisabeth Puchhammer-Stoeckl, Manfred Nairz, Guenter Weiss, Gregor Hörmann, Kinga Rigler-Hohenwarter, Rainer Gattringer, Wegene Borena, Dorothee von Laer, Gernot Walder, Peter Obrist, Christian Paar, Sabine Sussitz-Rack, Gunther Vogl, Adi Steinrigl, Christoph Bock, Andreas Bergthaler |
| EPI_ISL_583620, EPI_ISL_583621, EPI_ISL_583623, EPI_ISL_583624, EPI_ISL_583627, EPI_ISL_583628 | Pathologie-Labor Dr. Obrist- Dr. Brunhuber | Bergthaler laboratory, CeMM Research Center for Molecular Medicine of the Austrian Academy of Sciences | Alexandra Popa, Benedikt Agerer, Henrique Colaco, Lukas Endler, Jakob-Wendelin Genger, Alexander Lercher, Mark Smyth, Thomas Penz, Michael Schuster, Jan Laine, Martin Senekowitsch, Judith Aberle, Stephan Aberle, Peter Hufnagl, Daniela Schmid, Franz Allerberger, Elisabeth Puchhammer-Stoeckl, Manfred Nairz, Guenter Weiss, Gregor Hörmann, Kinga Rigler-Hohenwarter, Rainer Gattringer, Wegene Borena, Dorothee von Laer, Gernot Walder, Peter Obrist, Christian Paar, Sabine Sussitz-Rack, Gunther Vogl, Adi Steinrigl, Christoph Bock, Andreas Bergthaler |
| EPI_ISL_583753, EPI_ISL_583754, EPI_ISL_583755, EPI_ISL_583797, EPI_ISL_583798, EPI_ISL_583799, EPI_ISL_583800, EPI_ISL_583801, EPI_ISL_583802, EPI_ISL_583803, EPI_ISL_583804, EPI_ISL_583805, EPI_ISL_583806, EPI_ISL_583807, EPI_ISL_583808, EPI_ISL_583809, EPI_ISL_583810, EPI_ISL_583811, EPI_ISL_583812, EPI_ISL_583813, EPI_ISL_583814, EPI_ISL_583851 | | | |
| see above | Dr. Gernot Walder GmbH | Bergthaler laboratory, CeMM Research Center for Molecular Medicine of the Austrian Academy of Sciences | Alexandra Popa, Benedikt Agerer, Henrique Colaco, Lukas Endler, Jakob-Wendelin Genger, Alexander Lercher, Mark Smyth, Thomas Penz, Michael Schuster, Jan Laine, Martin Senekowitsch, Judith Aberle, Stephan Aberle, Peter Hufnagl, Daniela Schmid, Franz Allerberger, Elisabeth Puchhammer-Stoeckl, Manfred Nairz, Guenter Weiss, Gregor Hörmann, Kinga Rigler-Hohenwarter, Rainer Gattringer, Wegene Borena, Dorothee von Laer, Gernot Walder, Peter Obrist, Christian Paar, Sabine Sussitz-Rack, Gunther Vogl, Adi Steinrigl, Christoph Bock, Andreas Bergthaler |
| EPI_ISL_583967 | UOC Microbiologia e Virologia, Azienda Ospedaliera Universitaria Senese, Siena, Italy | Dipartimento di Biotecnologie Mediche | Maria Grazia Cusi, David Pinzauti, Claudia Gandolfo, Gabriele Anichini, Gianni Pozzi, Francesco Santoro |
| EPI_ISL_583995, EPI_ISL_584014 | Delaware Public Health Lab | Delaware Public Health Lab | Gregory Hovan |
| EPI_ISL_585092, EPI_ISL_585094, EPI_ISL_585095, EPI_ISL_585096, EPI_ISL_585097, EPI_ISL_585098, EPI_ISL_585099, EPI_ISL_585100, EPI_ISL_585101, EPI_ISL_585102, EPI_ISL_585103, EPI_ISL_585104, EPI_ISL_585105, EPI_ISL_585106, EPI_ISL_585107, EPI_ISL_585108, EPI_ISL_585109, EPI_ISL_585110, EPI_ISL_585111, EPI_ISL_585112, EPI_ISL_585113, EPI_ISL_585114, EPI_ISL_585115, EPI_ISL_585116, EPI_ISL_585117, EPI_ISL_585118, EPI_ISL_585119, EPI_ISL_585120, EPI_ISL_585121, EPI_ISL_585122, EPI_ISL_585123, EPI_ISL_585124, EPI_ISL_585125, EPI_ISL_585126, EPI_ISL_585127, EPI_ISL_585128, EPI_ISL_585129, EPI_ISL_585131, EPI_ISL_585133, EPI_ISL_585135, EPI_ISL_585136, EPI_ISL_585137, EPI_ISL_585138, EPI_ISL_585139, EPI_ISL_585140, EPI_ISL_585141, EPI_ISL_585142, EPI_ISL_585143, EPI_ISL_585150, EPI_ISL_585154, EPI_ISL_585155, EPI_ISL_585156, EPI_ISL_585157, EPI_ISL_585158, EPI_ISL_585159, EPI_ISL_585160, EPI_ISL_585161, EPI_ISL_585162, EPI_ISL_585163, EPI_ISL_585164, EPI_ISL_585165, EPI_ISL_585166, EPI_ISL_585167, EPI_ISL_585168, EPI_ISL_585169, EPI_ISL_585170, EPI_ISL_585171, EPI_ISL_585172, EPI_ISL_585173 | | | |
| see above | Regional Virus Laboratory, Belfast Health and Social Care Trust | COVID-19 Genomics UK (COG-UK) Consortium | Conall McCaughey, James McKenna, Tanya Curran, Susan Feeney, Alison Watt, Ciara Cox, Mairead Connor, Zoltan Molnar, David Simpson, Derek Fairley |
| EPI_ISL_585720 | Wales Specialist Virology Centre Sequencing lab: Pathogen Genomics Unit | COVID-19 Genomics UK (COG-UK) Consortium | Catherine Moore, Johnathan Evans, Laura Gifford, Malorie Perry, Simon Cottrell, Angela Marchbank, Alec Birchley, Alexander Adams, Amy Gaskin, Bree Gatica-Wilcox, Jason Coombes, Joel Southgate, Lauren Gilbert, Lee Graham, Nicole Pacchiarini, Sara Kumziene-Summerhayes, Sarah Taylor, Sophie Jones, Sara Rey, Matthew Bull, Joanne Watkins, Sally Corden, Tom Connor |
| EPI_ISL_586300, EPI_ISL_586301, EPI_ISL_586386, EPI_ISL_586387, EPI_ISL_586388, EPI_ISL_586389, EPI_ISL_586391, EPI_ISL_586392, EPI_ISL_586394, EPI_ISL_586395, EPI_ISL_586396, EPI_ISL_586397, EPI_ISL_586398, EPI_ISL_586399, EPI_ISL_586400, EPI_ISL_586401, EPI_ISL_586403, EPI_ISL_586404, EPI_ISL_586405 | | | |
| see above | Toronto Invasive Bacterial Diseases Network | McMaster University | Allison McGeer, Patryk Aftanas, Hooman Derakhshani, Angel Li, Kuganya Nirmalarajah, Emily Panousis, Ahmed Draia, Jalees Nasir, Michael Surette, Samira Mubareka, Andrew G. McArthur |
| EPI_ISL_590828, EPI_ISL_590829, EPI_ISL_590830 | Institute of Medical Virology, University of Zurich | Institute of Medical Virology, University of Zurich | Marie O. Pohl, Idoia Busnadiego, Verena Kufner, Stefan Schmutz, Maryam Zaheri, Irene Abela, Alexandra Trkola, Michael Huber, Silke Stertz, Benjamin G. Hale |
| EPI_ISL_590914, EPI_ISL_590918 | Oslo University Hospital, Department of Medical Microbiology | Norwegian Institute of Public Health, Department of Virology | Kathrine Stene-Johansen, Kamilla Heddeland Instefjord, Hilde Elshaug, Rasmus Riis Kopperud, Hilde Vollan, Karoline Bragstad, Olav Hungnes |
| EPI_ISL_591125, EPI_ISL_591126, EPI_ISL_591129, EPI_ISL_591131, EPI_ISL_591132, EPI_ISL_591133, EPI_ISL_591134, EPI_ISL_591136, EPI_ISL_591137, EPI_ISL_591139, EPI_ISL_591140, EPI_ISL_591208 | | | |
| see above | Toronto Invasive Bacterial Diseases Network | McMaster University | Allison McGeer, Patryk Aftanas, Hooman Derakhshani, Angel Li, Kuganya Nirmalarajah, Emily Panousis, Ahmed Draia, Jalees Nasir, Michael Surette, Samira Mubareka, Andrew G. McArthur |
| EPI_ISL_591340 | Dipartimento di Biotecnologie Mediche, University of Siena | Dipartimento di Biotecnologie Mediche, University of Siena | Cusi,M.G., Pinzauti,D., Gandolfo,C., Anichini,G., Pozzi,G., Santoro,F. |
| EPI_ISL_593543 | Eastern Ontario Regional Laboratory Association | McMaster University | Leanne Mortimer, Hooman Derakhshani, Emily Panousis, Ahmed Draia, Jalees Nasir, Robert Slinger, Andrew G. McArthur |
| EPI_ISL_593604, EPI_ISL_593605, EPI_ISL_593607, EPI_ISL_593608, EPI_ISL_593609, EPI_ISL_593611, EPI_ISL_593612, EPI_ISL_593627, EPI_ISL_593629, EPI_ISL_593631, EPI_ISL_593632 | | | |
| see above | unknown | Public Health Virology Laboratory, Forensic and Scientific Services (PHV-FSS) | Son Nguyen et al. |
| EPI_ISL_593663, EPI_ISL_593664 | Pathology West - NSW Health Pathology | NSW Health Pathology - Institute of Clinical Pathology and Medical Research; Westmead Hospital; University of Sydney | CIDM-PH et al. |
| EPI_ISL_593755 | Sydney South West Pathology Service (SSWPS) - Liverpool Hospital - NSW Health Pathology | NSW Health Pathology - Institute of Clinical Pathology and Medical Research; Westmead Hospital; University of Sydney | CIDM-PH et al. |
| EPI_ISL_593937, EPI_ISL_593939, EPI_ISL_593942, EPI_ISL_593943, EPI_ISL_593944, EPI_ISL_593945, EPI_ISL_593946, EPI_ISL_593947, EPI_ISL_593948, EPI_ISL_593949, EPI_ISL_593950, EPI_ISL_593951, EPI_ISL_593952, EPI_ISL_593953, EPI_ISL_593954, EPI_ISL_593955, EPI_ISL_593956, EPI_ISL_593965 | | | |
| see above | Delaware Public Health Lab | Delaware Public Health Lab | Gregory Hovan |
| EPI_ISL_594186 | Department of Pathology, School of Medicine, Imam Khomeini Hospital, Tehran University of Medical Sciences | Genetics Research Center, University of Social Welfare and Rehabilitation Sciences | Zohreh Fattahi, Marzieh Mohseni, Khadijeh Jalalvand, Azam Ghaziasadi, Seyedeh elham Mortazavi, Ali Jafarpour, Azar Hadadi, Alireza Abdollahi, Ali Jafarpour, Azam Ghaziasad, Seyedeh elham Mortazavi, Saber Soltani, Reza Najafipour, Kimia Kahrizi, Seyed Mohammad Jazayeri, Hossein Najmabadi |
| EPI_ISL_594198, EPI_ISL_594199, EPI_ISL_594200, EPI_ISL_594201, EPI_ISL_594202, EPI_ISL_594203, EPI_ISL_594204 | Michigan Department of Health and Human Services, Bureau of Laboratories | Michigan Department of Health and Human Services, Bureau of Laboratories | Blankenship HM, Riner D, Soehnlen MK |
| EPI_ISL_594458 | California Department of Public Health | Pathogen Discovery, Respiratory Viruses Branch, Division of Viral Diseases, Centers for Disease Control and Prevention | Ying Tao, Yan Li, Clinton Paden, Jing Zhang, Krista Queen, Anna Uehara, Haibin Wang, Julu Bhatnagar, Suxiang Tong |
| EPI_ISL_596512 | Palestinian Ministry of Health | Molecular Genetics Lab | Nouar Qutob, Zaidoun Salah, Damien Richard, Hisham Darwish, Husam Sallam, Issa Shtayeh, Osama Najjar, Mahmoud Ruzayqat, Dana Najjar, Francois Balloux, Lucy van Dorp |
| EPI_ISL_596646, EPI_ISL_596649, EPI_ISL_596653, EPI_ISL_596668, EPI_ISL_596670 | St.Vincent's University Hospital | St.Vincent's University Hospital | Mary Lucey, Guerrino Macori, Niamh Mullane, Una Sutton-Fitzpatrick, Gabriel Gonzalez, Suzie Coughlan, Aisling Purcell, Lynda Fenelon, Séamus Fanning, Kirsten Schaffer |
| EPI_ISL_596679, EPI_ISL_596682, EPI_ISL_596686, EPI_ISL_596687, EPI_ISL_596688, EPI_ISL_596701, EPI_ISL_596705, EPI_ISL_596709, EPI_ISL_596710, EPI_ISL_596711, EPI_ISL_596715, EPI_ISL_596739, EPI_ISL_596746, EPI_ISL_596747, EPI_ISL_596762, EPI_ISL_596768, EPI_ISL_596774, EPI_ISL_596779, EPI_ISL_596781, EPI_ISL_596782, EPI_ISL_596786, EPI_ISL_596791, EPI_ISL_596801, EPI_ISL_596806, EPI_ISL_596815, EPI_ISL_596831, EPI_ISL_596858 | | | |
| see above | PathWest Laboratory Medicine WA | PathWest Laboratory Medicine WA Microbial Surveillance Unit | PathWest Laboratory Medicine WA Microbial Surveillance Unit |
| EPI_ISL_602290, EPI_ISL_602292 | Evangelisches Klinikum Bethel, Institut für | Bielefeld University | David Brandt, Tobias Busche, Markus Haak, Jörn Kalinowski, Levin-Joe Klages, Christiane Scherer, Alexander Sczyrba, Marina Simunovic, Svenja Vinke |

| | | | |
|---|---|---|---|
| | Laboratoriumsmedizin, Mikrobiologie und Hygiene | | |
| EPI_ISL_602462, EPI_ISL_602463, EPI_ISL_602464, EPI_ISL_602465, EPI_ISL_602466, EPI_ISL_602503, EPI_ISL_602504, EPI_ISL_602505, EPI_ISL_602506, EPI_ISL_602507, EPI_ISL_602508, EPI_ISL_602509 | | | |
| see above | Institute for Virology, University Hospital Essen | Center of Medical Microbiology, Virology, and Hospital Hygiene, University of Duesseldorf | Olympia E. Anastasiou, Ulf Dittmer, Maximilian Damagnez, Alexander Dilthey, Torsten Houwaart, Lisanna Hülse, Malte Kohns Vasconcelos, Nadine Lübke, Jessica Nicolai, Klaus Pfeffer, Daniel Strelow, Jörg Timm, Andreas Walker, Tobias Wienemann |
| EPI_ISL_602544, EPI_ISL_602551 | Center of Medical Microbiology, Virology, and Hospital Hygiene, University of Duesseldorf | Center of Medical Microbiology, Virology, and Hospital Hygiene, University of Duesseldorf | Maximilian Damagnez, Alexander Dilthey, Torsten Houwaart, Malte Kohns Vasconcelos, Marek Korencak, Nadine Lübke, Jessica Nicolai, Klaus Pfeffer, Hendrik Streeck, Daniel Strelow, Jörg Timm, Andreas Walker, Tobias Wienemann |
| EPI_ISL_603262 | UW Virology Lab | UW Virology Lab | Pavitra Roychoudhury, Hong Xie, Lasata Shrestha, Meei-Li Huang, Keith R Jerome, Alexander Greninger |
| EPI_ISL_605067, EPI_ISL_605070, EPI_ISL_605075, EPI_ISL_605080 | National Virus Reference Laboratory | Irish Coronavirus Sequencing Consortium - Helixworks | Sachin Chalapati, Conor Crosbie, Nimesh Pinnamaneni |
| EPI_ISL_610168, EPI_ISL_610169, EPI_ISL_610170, EPI_ISL_610174, EPI_ISL_610176, EPI_ISL_610177 | Department of Health Technology and Informatics, The Hong Kong Polytechnic University | Department of Health Technology and Informatics, The Hong Kong Polytechnic University | Siu,G.K.-H., Lee,L.-K., Leung,K.S.-S., Leung,J.S.-L., Ng,T.T.-L., Chan,C.T.-M., Tam,K.K.-G., Lao,H.-Y., Wu,A.K.-L., Yau,M.C.-Y., Lai,Y.W.-M., Fung,K.S.-C., Chau,S.K.-Y., Wong,B.K.-C., To,W.-K., Luk,K., Ho,A.Y.-M., Que,T.-L., Yip,K.-T., Yam,W.C., Shum,D.H.-K., Yip,S.P. |
| EPI_ISL_611514, EPI_ISL_611567, EPI_ISL_611575, EPI_ISL_611654, EPI_ISL_611694, EPI_ISL_611734, EPI_ISL_611739, EPI_ISL_611796, EPI_ISL_611811, EPI_ISL_611821, EPI_ISL_611891, EPI_ISL_611893, EPI_ISL_611895, EPI_ISL_611897, EPI_ISL_611899, EPI_ISL_611915, EPI_ISL_611918, EPI_ISL_611929, EPI_ISL_611974, EPI_ISL_611991, EPI_ISL_612051, EPI_ISL_612070, EPI_ISL_612075, EPI_ISL_612108, EPI_ISL_612190, EPI_ISL_612401, EPI_ISL_612402, EPI_ISL_612403, EPI_ISL_612404, EPI_ISL_612405, EPI_ISL_612406, EPI_ISL_612407, EPI_ISL_612408, EPI_ISL_612409, EPI_ISL_612410, EPI_ISL_612411, EPI_ISL_612412, EPI_ISL_612413, EPI_ISL_612414, EPI_ISL_612415, EPI_ISL_612417, EPI_ISL_612418, EPI_ISL_612420, EPI_ISL_612421, EPI_ISL_612423, EPI_ISL_612424, EPI_ISL_612426, EPI_ISL_612427 | | | |
| see above | Liverpool Clinical Laboratories | COVID-19 Genomics UK (COG-UK) Consortium | Sam Haldenby, Anita Lucaci, Steve Paterson, Julian Hiscox, Alistair Darby, M Almsaud, A Alrezaihi, Muhannad Alruwaili, Stuart D Armstrong, Jones Benjamin, Eleanor G Bentley, Anu Chawla, Jordan J Clark, Angela Cowell, Richard Eccles, Isabel García-Dorival, Matthew Gemmell, Alessandro Gerada, PKF Gilmore, Richard Gregory, Ximeng Han, Catherine Hartley, Margaret Hughes, Miren Iturriza-Gomara, James Johnson, L Luu, Jenifer Manson, Charlotte Nelson, Elaine O'Toole, Cassie Olateju, Rebekah Penrice-Randal , Lucille Rainbow, N.P Randle, Trevor Ian Robinson, Parul Sharma, Ghada T Shawli, James P Stewart, Neil Swainston, Ecaterina Vamos, Joanne Watts, Mark Whitehead |
| EPI_ISL_613443, EPI_ISL_613444 | Institut Pasteur de la Guadeloupe | Institut Pasteur de la Guadeloupe | Marion Barbet, Sylvie Behillil, Méline Bizard, Angela Brisebarre, Camille Capel, Etienne Simon-Lorière, Vincent Enouf, Maud Vanpeene, Sylvie van der Werf, Stéphanie Guyomard, Sébastien Breurec, Antoine Talarmin |
| EPI_ISL_613448, EPI_ISL_613449, EPI_ISL_613451, EPI_ISL_613452 | Institut Pasteur de la Guadeloupe | Institut Pasteur de la Guadeloupe | Marion Barbet, Sylvie Behillil, Méline Bizard, Angela Brisebarre, Camille Capel, Etienne Simon-Lorière, Vincent Enouf, Maud Vanpeene, Sylvie van der Werf, Stéphanie Guyomard, Sébastien Breurec, Radjin Steingrover, Cherina Fleming, Antoine Talarmin |
| EPI_ISL_613462 | Microbiology, Koc University | Microbiology, Koc University | Nurtop,E., Ozer,B., Kuskucu,M.A., Dogan,O., Can,F. |
| EPI_ISL_613964 | Laboratory of Molecular Biology, Blood Center of Ribeirão Preto, Faculty of Medicine of Ribeirão Preto, University of São Paulo | Laboratory of Molecular Biology, Blood Center of Ribeirão Preto, Faculty of Medicine of Ribeirão Preto, University of São Paulo | Svetoslav N Slavov, Marta Giovanetti, Vagner Fonseca, Elaine V Santos, Evandra S Rodrigues, Talita Adelino, Joilson Xavier, Glauco de Carvalho Pereira, Aparecida Y Yamamoto, Diego Villa Clé, Rodrigo T Calado; Dimas T Covas, Luiz CJ Alcantara, Simone Kashima |
| EPI_ISL_614408, EPI_ISL_614414, EPI_ISL_614417, EPI_ISL_614435, EPI_ISL_614443, EPI_ISL_614444, EPI_ISL_614475, EPI_ISL_614476, EPI_ISL_614477, EPI_ISL_614478, EPI_ISL_614479, EPI_ISL_614515, EPI_ISL_614516, EPI_ISL_614517, EPI_ISL_614518, EPI_ISL_614519, EPI_ISL_614520, EPI_ISL_614521, EPI_ISL_614522, EPI_ISL_614523, EPI_ISL_614524, EPI_ISL_614525, EPI_ISL_614526, EPI_ISL_614527, EPI_ISL_614528, EPI_ISL_614529, EPI_ISL_614530, EPI_ISL_614531, EPI_ISL_614532, EPI_ISL_614533, EPI_ISL_614534, EPI_ISL_614535, EPI_ISL_614536, EPI_ISL_614537, EPI_ISL_614538, EPI_ISL_614539, EPI_ISL_614877, EPI_ISL_614881, EPI_ISL_614882, EPI_ISL_614883, EPI_ISL_614884, EPI_ISL_614885, EPI_ISL_614886, EPI_ISL_614887, EPI_ISL_614888, EPI_ISL_615404, EPI_ISL_615405, EPI_ISL_615577, EPI_ISL_615617, EPI_ISL_616416, EPI_ISL_616569, EPI_ISL_617814, EPI_ISL_617815, EPI_ISL_621824, EPI_ISL_622201, EPI_ISL_622204, EPI_ISL_622205, EPI_ISL_622206, EPI_ISL_622207, EPI_ISL_622208, EPI_ISL_622209, EPI_ISL_622210, EPI_ISL_622211, EPI_ISL_622212, EPI_ISL_622213, EPI_ISL_622214, EPI_ISL_622215, EPI_ISL_622216, EPI_ISL_622217, EPI_ISL_622218, EPI_ISL_622219, EPI_ISL_622220, EPI_ISL_622221, EPI_ISL_622222, EPI_ISL_622223, EPI_ISL_622224, EPI_ISL_622225, EPI_ISL_622226, EPI_ISL_622227, EPI_ISL_622228, EPI_ISL_622229, EPI_ISL_622230, EPI_ISL_622231, EPI_ISL_622232, EPI_ISL_622233, EPI_ISL_622234, EPI_ISL_622235, EPI_ISL_622236, EPI_ISL_622237, EPI_ISL_622437, EPI_ISL_622438, EPI_ISL_622439, EPI_ISL_622440, EPI_ISL_622461, EPI_ISL_622479, EPI_ISL_622480, EPI_ISL_622481, EPI_ISL_622482, EPI_ISL_622483, EPI_ISL_622484, EPI_ISL_622485, EPI_ISL_622486, EPI_ISL_622487, EPI_ISL_622488, EPI_ISL_622489, EPI_ISL_622490, EPI_ISL_622491, EPI_ISL_622504, EPI_ISL_622505, EPI_ISL_622506, EPI_ISL_622507, EPI_ISL_622508, EPI_ISL_622510, EPI_ISL_622549, EPI_ISL_622550, EPI_ISL_622551, EPI_ISL_622552, EPI_ISL_622596, EPI_ISL_622597, EPI_ISL_622598 | | | |
| see above | Department of Virus and Microbiological Special Diagnostics, Statens Serum Institut, Denmark | Albertsen lab, Department of Chemistry and Bioscience, Aalborg University, Denmark | Danish Covid-19 Genome Consortia |
| EPI_ISL_622790, EPI_ISL_622794, EPI_ISL_622796, EPI_ISL_622797, EPI_ISL_622798, EPI_ISL_622799 | Canterbury Health Laboratories | Institute of Environmental Science and Research (ESR) | Xiaoyun Ren, Matt Storey, Nikki Freed, Muhammad Faisal, Jing Wang, Hermes Perez, Anja Werno, Antje van der Linden, Arlo Upton, Chris Mansell, David Hammer, Dragana Drinkovic, Gary McAuliffe, Hana Sofia Andersson, James Ussher, Jill Sherwood, Josh Freeman, Julia Howard, Juliet Elvy, Mary DeAlmeida, Matt Blakiston, Matthew Rogers, Max Bloomfield, Michael Addidle, Michelle Balm, Sally Roberts, Sarah Jefferies, Sharmini Muttaiyah, Susan Morpeth, Susan Taylor, Timothy Blackmore, Vani Sathyendran, Veronica Playle, Virginia Hope, Erasmus Smit, Lauren Jelly, Olin Silander, Joep de Ligt |
| EPI_ISL_623138 | Laboratorio de Virologia Molecular / UFRJ | Bioinformatics Laboratory / LNCC | Carolina M Voloch, Ronaldo S Francisco Jr, Luiz G P de Almeida, Otavio J. Brustolini, Cynthia C Cardoso, Alexandra L Gerber, Ana Paula de C Guimarães, Diana Mariani, Covid19-UFRJ Workgroup, Luís Cristóvão Pôrto, Renato S Aguiar, Terezinha M P P Castiñeiras, Orlando C. Ferreira, Amilcar Tanuri, Ana Tereza R de Vasconcelos |
| EPI_ISL_626227 | Institute for Virology, University Hospital Essen | Center of Medical Microbiology, Virology, and Hospital Hygiene, University of Duesseldorf | Olympia E. Anastasiou, Ulf Dittmer, Maximilian Damagnez, Alexander Dilthey, Torsten Houwaart, Lisanna Hülse, Malte Kohns Vasconcelos, Nadine Lübke, Jessica Nicolai, Klaus Pfeffer, Daniel Strelow, Jörg Timm, Andreas Walker, Tobias Wienemann |
| EPI_ISL_626381, EPI_ISL_626382, EPI_ISL_626383, EPI_ISL_626384, EPI_ISL_626464, EPI_ISL_626465, EPI_ISL_626466, EPI_ISL_626467, EPI_ISL_626468 | Northwestern Memorial Hospital | Ozer Lab | Ramon Lorenzo-Redondo, Hannah H. Nam, Scott C. Roberts, Lacy M. Simons, Chad J. Achenbach, Lawrence J. Jennings, Chao Qi, Alan R. Hauser, Michael G. Ison, Judd F. Hultquist, Egon A. Ozer |
| EPI_ISL_629093, EPI_ISL_629094, EPI_ISL_629095 | Laboratoire du Centre Hospitalier Annecy Genevois | CNR Virus des Infections Respiratoires - France SUD | Antonin Bal, Gregory Destras, Gwendolyne Burfin, Hadrien Règue, Quentin Semanas, Martine Valette, Bruno Lina, Hélène Petitprez, Bruno Chanzy, Laurence Josset |
| EPI_ISL_631600 | James J. Peters VA Medical Center | New York City Public Health Laboratory | Jade Wang, et al. |
| EPI_ISL_631601 | Jamaica Hospital Medical Center | New York City Public Health Laboratory | Jade Wang, et al. |
| EPI_ISL_631602 | Flushing Hospital Medical Center | New York City Public Health Laboratory | Jade Wang, et al. |
| EPI_ISL_631603 | NYC HH Lincoln Medical And Mental Health Center | New York City Public Health Laboratory | Jade Wang, et al. |
| EPI_ISL_631604 | Jamaica Hospital Medical Center | New York City Public Health Laboratory | Jade Wang, et al. |
| EPI_ISL_631605 | OCME Office Of Chief Medical Examiner | New York City Public Health Laboratory | Jade Wang, et al. |
| EPI_ISL_631606 | Flushing Hospital Medical Center | New York City Public Health Laboratory | Jade Wang, et al. |
| EPI_ISL_631607 | NYC HH Lincoln Medical And Mental Health Center | New York City Public Health Laboratory | Jade Wang, et al. |
| EPI_ISL_631608, EPI_ISL_631609, EPI_ISL_631610 | Jamaica Hospital Medical Center | New York City Public Health Laboratory | Jade Wang, et al. |
| EPI_ISL_631611 | OCME Office Of Chief Medical Examiner | New York City Public Health Laboratory | Jade Wang, et al. |
| EPI_ISL_631612 | Jamaica Hospital Medical Center | New York City Public Health Laboratory | Jade Wang, et al. |
| EPI_ISL_631678, EPI_ISL_631679, EPI_ISL_631680, EPI_ISL_631681, EPI_ISL_631682, EPI_ISL_631684 | Texas Department of State Health Services | Texas Department of State Health Services | Rashmi Tuladhar, Bonnie Oh, Jenny Zhang, Maliha Rahman, Anita Pokharel, Myong Koag, Chung Wang, Rachel Lee, Grace Kubin, Mayela Pedrueza |
| EPI_ISL_631761 | OCME Office Of Chief Medical Examiner | New York City Public Health Laboratory | Jade Wang, et al. |
| EPI_ISL_631874 | James J. Peters VA Medical Center | New York City Public Health Laboratory | Jade Wang, et al. |
| EPI_ISL_631875 | Richmond University Medical Center | New York City Public Health Laboratory | Jade Wang, et al. |

| | | | |
|---|---|---|---|
| EPI_ISL_631896 | Jamaica Hospital Medical Center | New York City Public Health Laboratory | Jade Wang, et al. |
| EPI_ISL_631897 | Richmond University Medical Center | New York City Public Health Laboratory | Jade Wang, et al. |
| EPI_ISL_631898 | OCME Office Of Chief Medical Examiner | New York City Public Health Laboratory | Jade Wang, et al. |
| EPI_ISL_631899 | Richmond University Medical Center | New York City Public Health Laboratory | Jade Wang, et al. |
| EPI_ISL_631900 | St Barnabas Hospital | New York City Public Health Laboratory | Jade Wang, et al. |
| EPI_ISL_631901, EPI_ISL_631902 | Jamaica Hospital Medical Center | New York City Public Health Laboratory | Jade Wang, et al. |
| EPI_ISL_631903 | Richmond University Medical Center | New York City Public Health Laboratory | Jade Wang, et al. |
| EPI_ISL_631952 | St Barnabas Hospital | New York City Public Health Laboratory | Jade Wang, et al. |
| EPI_ISL_631961 | Flushing Hospital Medical Center | New York City Public Health Laboratory | Jade Wang, et al. |
| EPI_ISL_631962 | Richmond University Medical Center | New York City Public Health Laboratory | Jade Wang, et al. |
| EPI_ISL_631963 | Jamaica Hospital Medical Center | New York City Public Health Laboratory | Jade Wang, et al. |
| EPI_ISL_631964 | Flushing Hospital Medical Center | New York City Public Health Laboratory | Jade Wang, et al. |
| EPI_ISL_631965 | Richmond University Medical Center | New York City Public Health Laboratory | Jade Wang, et al. |
| EPI_ISL_632017 | Jamaica Hospital Medical Center | New York City Public Health Laboratory | Jade Wang, et al. |
| EPI_ISL_632018, EPI_ISL_632019, EPI_ISL_632020 | Flushing Hospital Medical Center | New York City Public Health Laboratory | Jade Wang, et al. |
| EPI_ISL_632021 | Jamaica Hospital Medical Center | New York City Public Health Laboratory | Jade Wang, et al. |
| EPI_ISL_632022 | Flushing Hospital Medical Center | New York City Public Health Laboratory | Jade Wang, et al. |
| EPI_ISL_632023 | Jamaica Hospital Medical Center | New York City Public Health Laboratory | Jade Wang, et al. |
| EPI_ISL_632024 | Flushing Hospital Medical Center | New York City Public Health Laboratory | Jade Wang, et al. |
| EPI_ISL_632025 | Jamaica Hospital Medical Center | New York City Public Health Laboratory | Jade Wang, et al. |
| EPI_ISL_632026 | NYC HH Lincoln Medical And Mental Health Center | New York City Public Health Laboratory | Jade Wang, et al. |
| EPI_ISL_632027 | Flushing Hospital Medical Center | New York City Public Health Laboratory | Jade Wang, et al. |
| EPI_ISL_632028, EPI_ISL_632029 | Jamaica Hospital Medical Center | New York City Public Health Laboratory | Jade Wang, et al. |
| EPI_ISL_632030 | Richmond University Medical Center | New York City Public Health Laboratory | Jade Wang, et al. |
| EPI_ISL_632175 | OCME Office Of Chief Medical Examiner | New York City Public Health Laboratory | Jade Wang, et al. |
| EPI_ISL_632192, EPI_ISL_632193 | NYC HH Lincoln Medical And Mental Health Center | New York City Public Health Laboratory | Jade Wang, et al. |
| EPI_ISL_632196 | Flushing Hospital Medical Center | New York City Public Health Laboratory | Jade Wang, et al. |
| EPI_ISL_632200 | James J. Peters VA Medical Center | New York City Public Health Laboratory | Jade Wang, et al. |
| EPI_ISL_632215, EPI_ISL_632216, EPI_ISL_632217, EPI_ISL_632218 | Richmond University Medical Center | New York City Public Health Laboratory | Jade Wang, et al. |
| EPI_ISL_632219, EPI_ISL_632221 | Jamaica Hospital Medical Center | New York City Public Health Laboratory | Jade Wang, et al. |
| EPI_ISL_632222, EPI_ISL_632223, EPI_ISL_632224, EPI_ISL_632225, EPI_ISL_632226 | Flushing Hospital Medical Center | New York City Public Health Laboratory | Jade Wang, et al. |
| EPI_ISL_632227, EPI_ISL_632228, EPI_ISL_632229 | Jamaica Hospital Medical Center | New York City Public Health Laboratory | Jade Wang, et al. |
| EPI_ISL_632233, EPI_ISL_632234 | Richmond University Medical Center | New York City Public Health Laboratory | Jade Wang, et al. |
| EPI_ISL_632851, EPI_ISL_632854, EPI_ISL_632860, EPI_ISL_632861, EPI_ISL_632869, EPI_ISL_632872, EPI_ISL_632888 | Idaho Bureau of Laboratories | Center for Global Health, University of New Mexico Health Sciences Center | Daryl Domman, Kurt Schwalm, Matthew Burns, Robert Voermans, Christopher Ball, Darrell Dinwiddie |
| EPI_ISL_632912, EPI_ISL_632914 | Cadham Provincial laboratory | Cadham Provincial laboratory | Anna Majer, Shari Tyson, Grace Seo, Philip Mabon, Elsie Grudeski, Rhiannon Huzarewich, Russell Mandes, Anneliese Landgraff, Jennifer Tanner, Natalie Knox, Morag Graham, Gary Van Domselaar, Paul Van Caeseele, Jared Bullard, David Alexander, Kerry Dust, Nathalie Bastien, Yan Li, Timothy Booth, Darian Hole, Madison Chapel, CanCOGeN's metadata curation team, Public Health Agency of Canada CanCOGeN team |
| EPI_ISL_632952 | Flushing Hospital Medical Center | New York City Public Health Laboratory | Jade Wang, et al. |
| EPI_ISL_632953 | Jamaica Hospital Medical Center | New York City Public Health Laboratory | Jade Wang, et al. |
| EPI_ISL_632954, EPI_ISL_632955, EPI_ISL_632956 | Flushing Hospital Medical Center | New York City Public Health Laboratory | Jade Wang, et al. |
| EPI_ISL_632957, EPI_ISL_632958 | Jamaica Hospital Medical Center | New York City Public Health Laboratory | Jade Wang, et al. |
| EPI_ISL_632959 | Richmond University Medical Center | New York City Public Health Laboratory | Jade Wang, et al. |
| EPI_ISL_632960 | Flushing Hospital Medical Center | New York City Public Health Laboratory | Jade Wang, et al. |
| EPI_ISL_632961 | Jamaica Hospital Medical Center | New York City Public Health Laboratory | Jade Wang, et al. |
| EPI_ISL_632962, EPI_ISL_632965 | Richmond University Medical Center | New York City Public Health Laboratory | Jade Wang, et al. |
| EPI_ISL_634833 | Laboratoire de virologie, CHU de Grenoble - CS 10217 - 38043 Grenoble cedex 25 | CNR Virus des Infections Respiratoires - France SUD | Antonin Bal, Gregory Destras, Gwendolyne Burfin, Hadrien Règue, Quentin Semanas, Martine Valette, Bruno Lina, Sylvie Larrat, Laurence Josset |
| EPI_ISL_634834 | Laboratoire de virologie, CHU de Grenoble - CS 10217 - 38043 Grenoble cedex 26 | CNR Virus des Infections Respiratoires - France SUD | Antonin Bal, Gregory Destras, Gwendolyne Burfin, Hadrien Règue, Quentin Semanas, Martine Valette, Bruno Lina, Sylvie Larrat, Laurence Josset |
| EPI_ISL_634835 | Laboratoire de virologie, CHU de Grenoble - CS 10217 - 38043 Grenoble cedex 27 | CNR Virus des Infections Respiratoires - France SUD | Antonin Bal, Gregory Destras, Gwendolyne Burfin, Hadrien Règue, Quentin Semanas, Martine Valette, Bruno Lina, Sylvie Larrat, Laurence Josset |
| EPI_ISL_635203 | Institute of Microbiology and Immunology, Faculty of Medicine, University of Ljubljana | Institute of Microbiology and Immunology, Faculty of Medicine, University of Ljubljana | Samo Zakotnik, Tomaž Mark Zorec, Miša Korva, Mario Poljak, Tatjana Avši - Županc |
| EPI_ISL_635269, EPI_ISL_635270, EPI_ISL_635276, EPI_ISL_635282, | Institute of Microbiology and Immunology, Faculty of Medicine, University of Ljubljana | Institute of Microbiology and Immunology, Faculty of Medicine, University of Ljubljana | Tomaž Mark Zorec, Samo Zakotnik, Miša Korva, Tatjana Avši - Županc, Mario Poljak |

| | | | |
|---|---|---|---|
| EPI_ISL_635287, EPI_ISL_635290, EPI_ISL_635291, EPI_ISL_635293 | | | |
| EPI_ISL_636838 | Laboratório de Imunofarmacologia - Instituto Oswaldo Cruz | Laboratório de Imunofarmacologia - Instituto Oswaldo Cruz | Souza,T.M., Fintelman-Rodrigues,N., De Paula,A.D., Saraiva,F.B., Ferreira,M.A. and Sacramento,C.Q. |
| EPI_ISL_637108 | Laboratorio Biologia Molecolare Sars Cov2 - UOC Laboratorio Analisi - Servizio Medicina di Laboratorio, Ospedale "San Francesco" - ATS-ASSL Nuoro | Laboratorio specialistico UOC Ematologia - Ospedale "San Francesco" - ATS-ASSL Nuoro | Piras Giovanna, Fancello Tatiana, Asproni Rosanna, Fiamma Maura, Monne Maria Itria, Toja Alessandro, Sanna Filomena, Floris Anna Rita, Sulis Vincenzo, Palmas Angelo Domenico, Casu Gavino, Lo Maglio Iana, Mameli Giuseppe |
| EPI_ISL_639740 | Respiratory virus Laboratory, Chinese Academy of Medical Science | Respiratory virus Laboratory, Chinese Academy of Medical Science | Li,J., Zhen,H., Chen,Y. and Liu,L. |
| EPI_ISL_639749, EPI_ISL_639751, EPI_ISL_639752, EPI_ISL_639753, EPI_ISL_639754, EPI_ISL_639756, EPI_ISL_639757, EPI_ISL_639776, EPI_ISL_639778, EPI_ISL_639779, EPI_ISL_639780, EPI_ISL_639781, EPI_ISL_639782, EPI_ISL_639783, EPI_ISL_639786, EPI_ISL_639787, EPI_ISL_639788, EPI_ISL_639789, EPI_ISL_639790, EPI_ISL_639791, EPI_ISL_639792, EPI_ISL_639793, EPI_ISL_639814, EPI_ISL_639815 | | | |
| see above | unknown | Public Health Virology Laboratory, Forensic and Scientific Services (PHV-FSS) | Son Nguyen et al. |
| EPI_ISL_640043, EPI_ISL_640058 | Groote Schuur Hospital wc GSH | NHLS/UCT | Arash Iranzadeh, Deelan Doolabh, Lynn Tyers, Bruna Galvao, Innocent Mudau, Marvin Hsiao, Kruger Marais, Diana Hardie, Stephen Korsman, Carolyn Williamson |
| EPI_ISL_640062, EPI_ISL_640063 | CoVid WC Cape Town Metro | NHLS/UCT | Arash Iranzadeh, Deelan Doolabh, Lynn Tyers, Bruna Galvao, Innocent Mudau, Marvin Hsiao, Kruger Marais, Diana Hardie, Stephen Korsman, Carolyn Williamson |
| EPI_ISL_641520 | CHU de Nice - Hôpital Archet 2 | CNR Virus des Infections Respiratoires - France SUD | Antonin Bal, Géraldine Gonfrier, Gregory Destras, Gwendolyne Burfin, Hadrien Règue, Quentin Semanas, Martine Valette, Bruno Lina, Valérie Giordanengo, Laurence Josset |
| EPI_ISL_641521 | CHU de Nice - Hôpital Archet 8 | CNR Virus des Infections Respiratoires - France SUD | Antonin Bal, Géraldine Gonfrier, Gregory Destras, Gwendolyne Burfin, Hadrien Règue, Quentin Semanas, Martine Valette, Bruno Lina, Valérie Giordanengo, Laurence Josset |
| EPI_ISL_641522 | CHU de Nice - Hôpital Archet 3 | CNR Virus des Infections Respiratoires - France SUD | Antonin Bal, Géraldine Gonfrier, Gregory Destras, Gwendolyne Burfin, Hadrien Règue, Quentin Semanas, Martine Valette, Bruno Lina, Valérie Giordanengo, Laurence Josset |
| EPI_ISL_641523 | CHU de Nice - Hôpital Archet 4 | CNR Virus des Infections Respiratoires - France SUD | Antonin Bal, Géraldine Gonfrier, Gregory Destras, Gwendolyne Burfin, Hadrien Règue, Quentin Semanas, Martine Valette, Bruno Lina, Valérie Giordanengo, Laurence Josset |
| EPI_ISL_641524 | CHU de Nice - Hôpital Archet 5 | CNR Virus des Infections Respiratoires - France SUD | Antonin Bal, Géraldine Gonfrier, Gregory Destras, Gwendolyne Burfin, Hadrien Règue, Quentin Semanas, Martine Valette, Bruno Lina, Valérie Giordanengo, Laurence Josset |
| EPI_ISL_641525 | CHU de Nice - Hôpital Archet 6 | CNR Virus des Infections Respiratoires - France SUD | Antonin Bal, Géraldine Gonfrier, Gregory Destras, Gwendolyne Burfin, Hadrien Règue, Quentin Semanas, Martine Valette, Bruno Lina, Valérie Giordanengo, Laurence Josset |
| EPI_ISL_641526 | CHU de Nice - Hôpital Archet 7 | CNR Virus des Infections Respiratoires - France SUD | Antonin Bal, Géraldine Gonfrier, Gregory Destras, Gwendolyne Burfin, Hadrien Règue, Quentin Semanas, Martine Valette, Bruno Lina, Valérie Giordanengo, Laurence Josset |
| EPI_ISL_641527 | CHU de Nice - Hôpital Archet 9 | CNR Virus des Infections Respiratoires - France SUD | Antonin Bal, Géraldine Gonfrier, Gregory Destras, Gwendolyne Burfin, Hadrien Règue, Quentin Semanas, Martine Valette, Bruno Lina, Valérie Giordanengo, Laurence Josset |
| EPI_ISL_644164, EPI_ISL_644166, EPI_ISL_644167, EPI_ISL_644168, EPI_ISL_644179, EPI_ISL_644180, EPI_ISL_644181, EPI_ISL_644182, EPI_ISL_644183, EPI_ISL_644184 | Texas Department of State Health Services | Texas Department of State Health Services | Rashmi Tuladhar, Bonnie Oh, Jenny Zhang, Maliha Rahman, Anita Pokharel, Myong Koag, Chung Wang, Rachel Lee, Grace Kubin, Mayela Pedrueza |
| EPI_ISL_644245, EPI_ISL_644273, EPI_ISL_644310, EPI_ISL_644319, EPI_ISL_644342 | CEPHR / Vincent's Hospital | Irish Coronavirus Sequencing Consortium - National Virus Reference Laboratory | Michael Carr, Gabriel Gonzalez, Alejandro Abner Garcia Leon, Patrick Mallon |
| EPI_ISL_644673, EPI_ISL_644674 | CHU de Limoges | CNR Virus des Infections Respiratoires - France SUD | Antonin Bal, Gregory Destras, Gwendolyne Burfin, Hadrien Règue, Quentin Semanas, Martine Valette, Bruno Lina, Sylvie Rogez, Laurence Josset |
| EPI_ISL_645013, EPI_ISL_645014, EPI_ISL_645015, EPI_ISL_645016, EPI_ISL_645017, EPI_ISL_645018, EPI_ISL_645019, EPI_ISL_645020, EPI_ISL_645021, EPI_ISL_645022, EPI_ISL_645023, EPI_ISL_645024 | | | |
| see above | Human Genome Variation Research Group, Malopolska Centre of Biotechnology | Human Genome Variation Research Group, Malopolska Centre of Biotechnology | Kowalski,M., Pospiech,E., Klajmon,A., Gromowski,T., Pisarek,A., Marszalek,K., Kopera,K., Foremny,J., Swadzba,J., Sanak,M., Owczarek,K., Dabrowska,A., Szczepanski,A., Botwina,P., Labaj,P.P., Pyrc,K., Branicki,W. |
| EPI_ISL_645170, EPI_ISL_645171 | CHU Bordeaux | CNR Virus des Infections Respiratoires - France SUD | Antonin Bal, Gregory Destras, Gwendolyne Burfin, Hadrien Règue, Quentin Semanas, Martine Valette, Bruno Lina, Pantxika Bellecave, Camille Ciccone, Isabelle Garrigue, Marie-Edith Lafon, Pascale Trimoulet, Laurence Josset |
| EPI_ISL_645173 | CHU de Nice | CNR Virus des Infections Respiratoires - France SUD | Antonin Bal, Géraldine Gonfrier, Gregory Destras, Gwendolyne Burfin, Hadrien Règue, Quentin Semanas, Martine Valette, Bruno Lina, Valérie Giordanengo, Laurence Josset |
| EPI_ISL_645183 | CHU de Limoges | CNR Virus des Infections Respiratoires - France SUD | Antonin Bal, Gregory Destras, Gwendolyne Burfin, Hadrien Règue, Quentin Semanas, Martine Valette, Bruno Lina, Sylvie Rogez, Laurence Josset |
| EPI_ISL_645194 | CHU Clermont-Ferrand | CNR Virus des Infections Respiratoires - France SUD | Antonin Bal, Gregory Destras, Gwendolyne Burfin, Hadrien Règue, Quentin Semanas, Martine Valette, Bruno Lina, Christine Archimbaud, Amélie Brebion, Hélène Chabrolles, Martine Chambon, Audrey Mirand, Christel Regagnon, Maxime Bisseux, Patricia Combes, Cécile Henquell, Laurence Josset |
| EPI_ISL_645199, EPI_ISL_645200 | CHU Nîmes | CNR Virus des Infections Respiratoires - France SUD | Antonin Bal, Gregory Destras, Gwendolyne Burfin, Hadrien Règue, Quentin Semanas, Martine Valette, Bruno Lina, Jean-Philippe Lavigne, Stephan Robin, Maxence Lotellier, Marie-Josée Carles, Laurence Josset |
| EPI_ISL_648084, EPI_ISL_648085 | Department of Laboratory Medicine, Tan Tock Seng Hospital | Department of Laboratory Medicine, Tan Tock Seng Hospital | Chen YYC, Zair X, Lim JX, Li C, Tang WY, Maurer-Stroh S, Barkham TMS, Nagarajan N, Sessions OM |
| EPI_ISL_649962, EPI_ISL_649963, EPI_ISL_649964, EPI_ISL_649965, EPI_ISL_649966, EPI_ISL_649967, EPI_ISL_649968, EPI_ISL_649969, EPI_ISL_649970 | CHU Bordeaux | CNR Virus des Infections Respiratoires - France SUD | Antonin Bal, Gregory Destras, Gwendolyne Burfin, Hadrien Règue, Quentin Semanas, Martine Valette, Bruno Lina, Pantxika Bellecave, Camille Ciccone, Isabelle Garrigue, Marie-Edith Lafon, Pascale Trimoulet, Laurence Josset |
| EPI_ISL_653181, EPI_ISL_653182, EPI_ISL_653184 | Florida Bureau of Public Health Laboratories | Florida Bureau of Public Health Laboratories | Sarah Schmedes, Jason Blanton |
| EPI_ISL_654618 | Servicio de Microbiología, Hospital Universitario Cruces, Bilbao | SeqCOVID-SPAIN consortium/IBV(CSIC) | Maitane Aranzamendi Zaldumbide and SeqCOVID-SPAIN consortium |
| EPI_ISL_654631 | Servicio de Microbiología, Hospital Universitario Central de Asturias | SeqCOVID-SPAIN consortium/IBV(CSIC) | Cristián Castelló Abietar, Jose A. Boga, Susana Rojo-Alba, Marta Elena Álvarez-Argüelles, Santiago Melón and SeqCOVID-SPAIN consortium |
| EPI_ISL_654731, EPI_ISL_654765 | Texas Department of State Health Services | Texas Department of State Health Services | Rashmi Tuladhar, Bonnie Oh, Jenny Zhang, Maliha Rahman, Anita Pokharel, Myong Koag, Chung Wang, Rachel Lee, Grace Kubin, Mayela Pedrueza, James Daniel Bonser |
| EPI_ISL_654887 | Pasteur Institute in Ho Chi Minh city | Department of Microbiology and Immunology - Pasteur Institute in Ho Chi Minh city | Phan Trng Lân, ào Huy Mnh, Phm Th Thu Hng, V Phm Hng Nhung, Cao Minh Thng, Hunh Th Kim Loan, Nguyn Hoàng Quân, Hunh Phng Tho, Hoàng Nh ào, Nguyn Trung Hiu, Nguyn Hoàng Anh, Nguyn Thu Ngc, Phm Th Nhung, Nguyn Th Ngc Tho, ng Thanh Giang, Nguyn Th Thanh Thng, Hoàng Minh, Trn Th Hng Kim, Nguyn Thanh Long, Phm Duy Quang, Lng Chn Quang, Hoàng Quc Cng, Nguyn V Thng |
| EPI_ISL_660360 | CHU Clermont-Ferrand | CNR Virus des Infections Respiratoires - France SUD | Antonin Bal, Gregory Destras, Gwendolyne Burfin, Hadrien Règue, Quentin Semanas, Martine Valette, Bruno Lina, Christine Archimbaud, Amélie Brebion, |

| | | | |
|---|---|---|---|
| | | | Hélène Chabrolles, Martine Chambon, Audrey Mirand, Christel Regagnon, Maxime Bisseux, Patricia Combes, Cécile Henquell, Laurence Josset |
| EPI_ISL_660390, EPI_ISL_660392, EPI_ISL_660393, EPI_ISL_660394 | Klinisk mikrobiologi Linkoping | The Public Health Agency of Sweden | Anna-Malin Linde, Maria Lind Karlberg, Mattias Haukland, Reza Advani, Olov Svartstrom, Oskar Karlsson Lindsjo, Sandra Broddesson, Petra Edquist, Mia Brytting, Anna Risberg, Karin Tegmark-Wisell |
| EPI_ISL_660443 | Hospital de Braga | Instituto Nacional de Saude (INSA) | Borges et al |
| EPI_ISL_660691, EPI_ISL_660692 | CHU Montpellier | CNR Virus des Infections Respiratoires - France SUD | Antonin Bal, Gregory Destras, Gwendolyne Burfin, Hadrien Règue, Quentin Semanas, Martine Valette, Bruno Lina, Michel Segondy, Vincent Foulongne, Laurence Josset |
| EPI_ISL_660715, EPI_ISL_660716, EPI_ISL_660717, EPI_ISL_660718 | CHU Nîmes | CNR Virus des Infections Respiratoires - France SUD | Antonin Bal, Gregory Destras, Gwendolyne Burfin, Hadrien Règue, Quentin Semanas, Martine Valette, Bruno Lina, Jean-Philippe Lavigne, Stephan Robin, Maxence Lotellier, Marie-Josée Carles, Laurence Josset |
| EPI_ISL_663260, EPI_ISL_663261, EPI_ISL_663262, EPI_ISL_663263 | CHU Nantes | CNR Virus des Infections Respiratoires - France SUD | Antonin Bal, Louise Castain, Gregory Destras, Gwendolyne Burfin, Hadrien Règue, Quentin Semanas, Martine Valette, Bruno Lina, Virginie Ferré, Celine Bressollette, Laurence Josset |
| EPI_ISL_664101 | Dept. of Microbiology and Infection Control, Akershus University Hospital HF | Dept. of Microbiology and Infection Control, Akershus University Hospital HF | Hege Vangstein Aamot, Alexander Hesselberg Løvestad, Silje Bakken Jørgensen, Nina Handal, Ole Herman Ambur |
| EPI_ISL_664171, EPI_ISL_664189, EPI_ISL_664497, EPI_ISL_664512, EPI_ISL_664701, EPI_ISL_664702, EPI_ISL_664703, EPI_ISL_664744, EPI_ISL_665144, EPI_ISL_665174, EPI_ISL_665175, EPI_ISL_665176, EPI_ISL_665177, EPI_ISL_665178, EPI_ISL_665179, EPI_ISL_665180, EPI_ISL_665181 | | | |
| see above | University College London Hospital | COVID-19 Genomics UK (COG-UK) Consortium | Judith Heaney, Matthew Byott, Catherine Houlihan, Dan Frampton, Stuart Kirk, Moira Spyer and Eleni Nastouli |
| EPI_ISL_666660, EPI_ISL_666661 | Laboratoire du Centre Hospitalier Annecy Genevois | CNR Virus des Infections Respiratoires - France SUD | Antonin Bal, Gregory Destras, Gwendolyne Burfin, Hadrien Règue, Quentin Semanas, Martine Valette, Bruno Lina, Hélène Petitprez, Bruno Chanzy, Laurence Josset |
| EPI_ISL_671267 | Department of Virus and Microbiological Special Diagnostics, Statens Serum Institut, Copenhagen, Denmark | Albertsen Lab, Department of Chemistry and Bioscience, Aalborg University, Denmark | Danish Covid-19 Genome Consortium |
| EPI_ISL_671377, EPI_ISL_671378 | National Virus Reference Laboratory | Irish Coronavirus Sequencing Consortium - Teagasc Moorepark | Calum Walsh, Fiona Crispie, John Kenny, Paul Cotter |
| EPI_ISL_672532, EPI_ISL_672538, EPI_ISL_672542, EPI_ISL_672545, EPI_ISL_672551, EPI_ISL_672557 | Utah Public Health Laboratory | Utah Public Health Laboratory | Erin Young, Kelly Oakeson |
| EPI_ISL_672575 | Infectious Diseases and Tropical Medicine Research Center, Infectious Diseases and Tropical Medicine Research Center | Infectious Diseases and Tropical Medicine Research Center, Infectious Diseases and Tropical Medicine Research Center | Ahangarzadeh,S., Ataei,B., Shariati,L., Haghjooy Javanmard,S. |
| EPI_ISL_676656, EPI_ISL_676669, EPI_ISL_676687, EPI_ISL_676690, EPI_ISL_676710, EPI_ISL_676768, EPI_ISL_676769, EPI_ISL_676772, EPI_ISL_676774, EPI_ISL_676775, EPI_ISL_676776, EPI_ISL_676777, EPI_ISL_676778, EPI_ISL_676779, EPI_ISL_676780, EPI_ISL_676781, EPI_ISL_676782, EPI_ISL_676783, EPI_ISL_676784, EPI_ISL_676785, EPI_ISL_676786, EPI_ISL_676787, EPI_ISL_676788, EPI_ISL_676789, EPI_ISL_676791, EPI_ISL_676793 | | | |
| see above | Wadsworth Center, New York State Department.of Health | Wadsworth Center, New York State Department.of Health | Kirsten St. George, Daryl M. Lamson, Alexis Russel, Jonathan Plitnick, Navjot Singh, John Kelly, Sara Griesemer, Erasmus Schneider, Erica Lasek-Nesselquist |
| EPI_ISL_677246 | Colorado Department of Public Health and Environment | Colorado Department of Puplic Health and Environment | Laura Bankers, Molly Hetherington-Rauth, Shannon Ely, Shannon R. Matzinger, Sarah Elizabeth Totten, Emily A. Travanty |
| EPI_ISL_677964 | Pathogen Genomics Lab King Abdullah University of Science and Technology(KAUST) | Pathogen Genomics Lab King Abdullah University of Science and Technology(KAUST) | Sharif Hala, Fadwa Alofi, Sara Mfarrej, Amit Kumar Subudhi, Fathia Ben Rached, Raeece Naeem, Asim Khogeer, Afrah Alsomali, Jumana Taha, Abdulaziz Alahmadi, Kahled Algithami, Anwar Hashem, Naif Almontashiri, Arnab Pain |
| EPI_ISL_678295, EPI_ISL_678382 | Area of Virology, Serology and Virology Division (SAViD), New South Wales Health Pathology Randwick | Virology Research Laboratory; Area of Virology, Serology and Virology Division (SAViD), New South Wales Health Pathology Randwick | Foster, C.; Au, J.; Ruiz Silva, M.; Deveson, I.; Bull, R.; Van Hal, S.; Rawlinson, W. |
| EPI_ISL_678498, EPI_ISL_678499, EPI_ISL_678500, EPI_ISL_678501 | CNR Virus des Infections Respiratoires - France SUD | CNR Virus des Infections Respiratoires - France SUD | Antonin Bal, Gregory Destras, Gwendolyne Burfin, Solenne Brun, Martine Valette, Bruno Lina, Laurence Josset |
| EPI_ISL_681673 | Molecular Medicine Laboratory, University of Magallanes | Centro Asistencial Docente y de Investigacion, Universidad de Magallanes | Jorge González, Jacqueline Aldridge, Diego Alvarez, Marco Montes de Oca, Hermy Alvarez, Roberto Uribe-Paredes, Marcelo Navarrete |
| EPI_ISL_681888, EPI_ISL_681908, EPI_ISL_681910, EPI_ISL_681912, EPI_ISL_681913, EPI_ISL_681919, EPI_ISL_681921, EPI_ISL_681923, EPI_ISL_681928 | National Virus Reference Laboratory | Irish Coronavirus Sequencing Consortium - Teagasc Moorepark | Paul Cotter, Fiona Crispie, John Kenny, Matthew McCabe, Calum Walsh |
| EPI_ISL_683333 | Laboratoire du Centre Hospitalier Annecy Genevois | CNR Virus des Infections Respiratoires - France SUD | Antonin Bal, Gregory Destras, Gwendolyne Burfin, Hadrien Règue, Quentin Semanas, Martine Valette, Bruno Lina, Hélène Petitprez, Bruno Chanzy, Laurence Josset |
| EPI_ISL_683657, EPI_ISL_683658, EPI_ISL_683659, EPI_ISL_683660, EPI_ISL_683662, EPI_ISL_683663 | IdISSC/Hospital Clínico San Carlos de Madrid | SeqCOVID-SPAIN consortium/IBV(CSIC) | Alberto Delgado-Iribarren, Esther Culebras Lopez, Vicente Estrada Pérez, Jorge Matias-Guiu, Luis Ortega Medina, Silvia Sánchez Ramón, Ulises Gómez-Pinedo and SeqCOVID-SPAIN consortium |
| EPI_ISL_683681, EPI_ISL_683682, EPI_ISL_683683, EPI_ISL_683684, EPI_ISL_683687 | M Health Fairview | Minnesota Department of Health, Public Health Laboratory | Alexandra Lorentz, Jacob Garfin, Matt Plumb, and Xiong Wang |
| EPI_ISL_693288 | unknown | Public Health Virology Laboratory, Forensic and Scientific Services (PHV-FSS) | Son Nguyen et al. |
| EPI_ISL_696192, EPI_ISL_696249, EPI_ISL_696415, EPI_ISL_696418, EPI_ISL_696420, EPI_ISL_696430, EPI_ISL_696434 | Sonora Quest Laboratories, Laboratory Sciences of Arizona | TGen North | Jolene Bowers, Megan Folkerts, Chris French, Hayley Yaglom, Ashlyn Pfeiffer, Darrin Lemmer, Dave Engelthaler, The Arizona COVID Genomics Union (ACGU) |
| EPI_ISL_699647 | Pathology Sydney South West - NSW Health Pathology | NSW Health Pathology - Institute of Clinical Pathology and Medical Research; Westmead Hospital; University of Sydney | CIDM-PH et al. |
| EPI_ISL_700359 | Laboratoire de virologie, CHU de Grenoble - CS 10217 - 38043 Grenoble cedex 20 | CNR Virus des Infections Respiratoires - France SUD | Antonin Bal, Gregory Destras, Gwendolyne Burfin, Hadrien Règue, Quentin Semanas, Martine Valette, Bruno Lina, Sylvie Larrat, Laurence Josset |
| EPI_ISL_700360 | Laboratoire de virologie, CHU de Grenoble - CS 10217 - 38043 Grenoble cedex 21 | CNR Virus des Infections Respiratoires - France SUD | Antonin Bal, Gregory Destras, Gwendolyne Burfin, Hadrien Règue, Quentin Semanas, Martine Valette, Bruno Lina, Sylvie Larrat, Laurence Josset |
| EPI_ISL_700377, EPI_ISL_700386, EPI_ISL_700387 | Centre hospitalier Métropole Savoie | CNR Virus des Infections Respiratoires - France SUD | Antonin Bal, Carine Dumollard, Gregory Destras, Gwendolyne Burfin, Hadrien Règue, Quentin Semanas, Martine Valette, Bruno Lina, Jérôme Grosjean, Laurence Josset |
| EPI_ISL_707959 | Virology, Universitätsklinikum des Saarlandes | Epigenetics, Saarland University | Kathrin Kattler, Markus Vogelgesang, Stefan Lohse, Sascha Tierling, Sigrun Smola, Jörn Walter |
| EPI_ISL_708504 | M Health Fairview | Minnesota Department of Health, Public Health Laboratory | Alexandra Lorentz, Jacob Garfin, Matt Plumb, and Xiong Wang |
| EPI_ISL_708512, EPI_ISL_708516 | National Virus Reference Laboratory | Irish Coronavirus Sequencing Consortium - Teagasc Oakpark | Michele Della Bartola, Matthew McCabe, Ewen Mullins |
| EPI_ISL_708738, EPI_ISL_708739, EPI_ISL_708740, EPI_ISL_708743, EPI_ISL_708745, EPI_ISL_708746, EPI_ISL_708749, EPI_ISL_708750, EPI_ISL_708751, EPI_ISL_708752, EPI_ISL_708756, EPI_ISL_708757, EPI_ISL_708758, EPI_ISL_708759, EPI_ISL_708760, EPI_ISL_708762, EPI_ISL_708763, EPI_ISL_708767, EPI_ISL_708768, EPI_ISL_708769, EPI_ISL_708770, EPI_ISL_708771, EPI_ISL_708778, EPI_ISL_708780, EPI_ISL_708786, EPI_ISL_708787 | | | |

| | | | |
|---|---|---|---|
| see above | PathWest Laboratory Medicine WA | PathWest Laboratory Medicine WA Microbial Surveillance Unit | PathWest Laboratory Medicine WA Microbial Surveillance Unit |
| EPI_ISL_710489, EPI_ISL_710491 | Virology (microbiology), Hospital Universitario Central de Asturias | Virology (microbiology), Hospital Universitario Central de Asturias | Castello,C., Gomez-deOna,J., Martin-Rodriguez,G., Abreu,F., Costales,I., Boga,J.A., Coto,E., Rojo,S., Alvarez-Argueelles,M.E., Perez-Martinez,Z., Sandoval,M., Melon,S. |
| EPI_ISL_710584 | Skovde/Unilabs | The Public Health Agency of Sweden | Department of Microbiology, The Public Health Agency of Sweden |
| EPI_ISL_723104 | Institute of Medical Genetics and Applied Genomics | Institute of Medical Genetics and Applied Genomics | Caspar Gross, Tina Ganzenmüller, Siri Göpel, Michaela Pogoda, Daniela Bezdan, Michael Sonnabend, Angel Angelov, Nicolas Casadei, Stephan Ossowski, Thomas Iftner, Michael Bitzer |
| EPI_ISL_729533 | Charité Universitätsmedizin Berlin, Institut für Virologie/Labor Berlin | Charité Universitätsmedizin Berlin, Institut für Virologie | Victor M Corman, Barbara Mühlemann, Jörn Beheim-Schwarzbach, Talitha Veith, Julia Schneider, Terry Jones, Christian Drosten |
| EPI_ISL_729815 | Laboratorio Central de Saude Publica do Estado do Rio Grande do Sul (LACEN-RS) | Laboratory of Respiratory Viruses and Measles, Oswaldo Cruz Institute, FIOCRUZ | Paola Resende, Luciana Appolinario, Fernando Motta, Anna Carolina Paixão, Ana Carolina Mendonça, Tatiana Schaffer Gregianini, Marilda Tereza Mar da Rosa, Marilda Siqueira |
| EPI_ISL_730051 | Yale COVID-19 Biorepository | Grubaugh Lab - Yale School of Public Health | Joseph Fauver, Tara Alpert, Anderson Brito, Annie Watkins, Anne Wyllie, Chantal Vogels, Mary Petrone, Chaney Kalinich, Isabel Ott, Arnau Casanovas, Catherine Muenker, Adam Moore, Alice Lu, Maria Tokuyama, Patrick Wong, Peiwen Lu, Saad Omer, Richard Martinello, Allison Nelson, Shelli Farhadian, Akiko Iwasaki, Charlese Dela Cruz, Albert Ko, Nathan Grubaugh |
| EPI_ISL_732550, EPI_ISL_732551 | Bundeswehr Institute of Microbiology | Bundeswehr Institute of Microbiology | Markus Antwerpen, Alexandra Rehn, Mathias Walter, Malena Bestehorn-Willmann, Sabine Zange, Enrico Georgi, Roman Wölfel |
| EPI_ISL_734486, EPI_ISL_734524, EPI_ISL_734525 | UZ Leuven, National Reference Laboratory for Coronaviruses, Laboratory Medicine, Leuven, Belgium | KU Leuven, Rega Institute, Clinical and Epidemiological Virology | Tony Wawina-Bokalanga, Joan Marti-Carerras, Bert Vanmechelen, Piet Maes |
| EPI_ISL_737955, EPI_ISL_737956, EPI_ISL_737959 | Uganda Central Public Health Lab and Uganda Virus Research Institute | MRC/UVRI & LSHTM Uganda Research Unit | Matthew Cotten, Dan Lule Bugembe, My V.T. Phan, Pontiano Kaleebu et al. |
| EPI_ISL_744644 | Laboratoire national de santé, Microbiology, Virology | Laboratoire national de santé, Microbiology, Microbial Genomics Platform | Anke Wienecke-Baldacchino, Catherine Ragimbeau, Tamir Abdelrahman, Jessica Tapp, Fatu Djabi |
| EPI_ISL_751319, EPI_ISL_751320, EPI_ISL_751329, EPI_ISL_751412, EPI_ISL_751413, EPI_ISL_751414, EPI_ISL_751415, EPI_ISL_751416, EPI_ISL_751417, EPI_ISL_751418, EPI_ISL_751419, EPI_ISL_751420, EPI_ISL_751421, EPI_ISL_751422, EPI_ISL_751423, EPI_ISL_751424, EPI_ISL_751425, EPI_ISL_751426 | | | |
| see above | IRCCS Sacro Cuore Don Calabria Hospital, Department of Infectious, Tropical Diseases & Microbiology | University of Verona, Department of Biotechnology | Antonio Mori, Michela Deiana, Elena Pomari, Chiara Piubelli; Giulia Lopatriello, Luca Marcolungo, Cristina Beltrami, Chiara Degli Esposti, Emanuela Cosentino, Massimo Delledonne |
| EPI_ISL_752627, EPI_ISL_752628, EPI_ISL_752660, EPI_ISL_752700, EPI_ISL_752710, EPI_ISL_752711, EPI_ISL_752845 | State Laboratories Division, Hawaii State Department of Health | State Laboratories Division, Hawaii State Department of Health | Pamela O'Brien, Sabrina Diemert, Drew Kuwazaki, Razvan Sultana, Edward Desmond |
| EPI_ISL_753704, EPI_ISL_753844, EPI_ISL_753845, EPI_ISL_753917, EPI_ISL_753926, EPI_ISL_753927, EPI_ISL_753928, EPI_ISL_754024, EPI_ISL_754025, EPI_ISL_754027, EPI_ISL_754028, EPI_ISL_754029, EPI_ISL_754030, EPI_ISL_754031, EPI_ISL_754032, EPI_ISL_754033, EPI_ISL_754034, EPI_ISL_754035, EPI_ISL_754037, EPI_ISL_754038 | | | |
| see above | Charité Universitätsmedizin Berlin, Institut für Virologie/Labor Berlin | Charité Universitätsmedizin Berlin, Institut für Virologie | Victor M Corman, Jörn Beheim-Schwarzbach, Barbara Mühlemann, Julia Schneider, Talitha Veith, Terry Jones, Christian Drosten |
| EPI_ISL_754747, EPI_ISL_754750, EPI_ISL_754767, EPI_ISL_754768 | Emory Molecular Diagnostics Laboratory, Emory Healthcare | Piantadosi Lab, Emory Department of Pathology | Ahmed Babiker, Anne Piantadosi |
| EPI_ISL_755022, EPI_ISL_755036, EPI_ISL_755037 | California Department of Public Health | California Department of Public Health | CDPH IDLB COVIDNet |
| EPI_ISL_755740, EPI_ISL_755797, EPI_ISL_755800, EPI_ISL_755820, EPI_ISL_755826, EPI_ISL_755828, EPI_ISL_755829, EPI_ISL_755830, EPI_ISL_755831, EPI_ISL_755832, EPI_ISL_755833, EPI_ISL_755834, EPI_ISL_755835, EPI_ISL_755836, EPI_ISL_755837, EPI_ISL_755838, EPI_ISL_755839, EPI_ISL_755840, EPI_ISL_755841, EPI_ISL_755842, EPI_ISL_755843, EPI_ISL_755844, EPI_ISL_755845, EPI_ISL_755846, EPI_ISL_755847, EPI_ISL_755848, EPI_ISL_755849, EPI_ISL_755850, EPI_ISL_755851, EPI_ISL_755852, EPI_ISL_755853, EPI_ISL_755854, EPI_ISL_755855, EPI_ISL_755856, EPI_ISL_755857, EPI_ISL_755858, EPI_ISL_755859, EPI_ISL_755860, EPI_ISL_755861, EPI_ISL_755862, EPI_ISL_755863, EPI_ISL_755864, EPI_ISL_755865, EPI_ISL_755866, EPI_ISL_755867, EPI_ISL_755868, EPI_ISL_755869, EPI_ISL_755870, EPI_ISL_755871, EPI_ISL_755872, EPI_ISL_755873, EPI_ISL_755874, EPI_ISL_755875, EPI_ISL_755876, EPI_ISL_755877 | | | |
| see above | Toronto Invasive Bacterial Diseases Network | McMaster University | Allison McGeer, Patryk Aftanas, Hooman Derakhshani, Angel Li, Kuganya Nirmalarajah, Emily Panousis, Ahmed Draia, Jalees Nasir, Michael Surette, Samira Mubareka, Andrew G. McArthur |
| EPI_ISL_756358, EPI_ISL_756359 | The Caribbean Public Health Agency | Carrington Lab, Department of PreClinical Sciences, Faculty of Medical Sciences, The University of the West Indies | Nikita S. D. Sahadeo, Gabriel Escobar, Sarah Hill, Vernie Ramkissoon, Risha Singh, SueMin Nathaniel, Arianne Brown-Jordan, Naresh Nandram, Avery Hinds, Jerome Foster, Stanley Giddings, Karla Georges, Marsha Ivey, Rahul Naidu, , Rajini Haraksingh, Jaya Jayaraman, Chinna Chinnadurai, Adesh Ramsubhag, Nuno Faria, Oliver Pybus, Christopher Oura, Christine V. F. Carrington |
| EPI_ISL_759858 | School of Public Health, The University of Hong Kong | School of Public Health, The University of Hong Kong | Barrs,V.R., Peiris,M., Tam,K.W.S., Law,P.Y.L., Brackman,C.J.,To,E.M.W., Yu,V.Y.T., Chu,D.K.W., Perera,R.A.P.M. and Sit,T.H.C. |
| EPI_ISL_759959 | Premier Hospital Surabaya | Institute of Tropical Disease, Universitas Airlangga | Krisnoadi Rahardjo, Aldise M Nastri, Jezzy R Dewantari, Rima R Prasetya, Hartono Tanto, Gatot Soegiarto, Laksmi Wulandari, Resti Yudhawati, Soetjipto, Yasuko Mori, Maria I Lusida, Kazufumi Shimizu |
| EPI_ISL_759961 | Universitas Airlangga Hospital | Institute of Tropical Disease, Universitas Airlangga | Jezzy R Dewantari, Rima R Prasetya, Krisnoadi Rahardjo, Aldise M Nastri, Nasronudin, Gatot Soegiarto, Laksmi Wulandari, Resti Yudhawati, Soetjipto, Yasuko Mori, Maria I Lusida, Kazufumi Shimizu |
| EPI_ISL_763091 | Jena University Hospital, Institute for Infectious Diseases and Infection Control | Institute of infectious medicine & hospital hygiene, CaSe-Group | Spott, Riccardo; Marquet, Mike; Pletz, Matthias W.; Brandt, Christian |
| EPI_ISL_766033 | RSUD Prof. DR. W. Z. Johannes Kupang | Eijkman Institute for Molecular Biology, Ministry of Research and Technology/National Agency for Research and Innovation | Frilasita A Yudhaputri, Edison Johar, Hidayat Trimarsanto, David H Muljono, Safarina G Malik, Khin Saw Myint, Amin Soebandrio |
| EPI_ISL_767769, EPI_ISL_767798 | National Virus Reference Laboratory | Irish Coronavirus Sequencing Consortium - Teagasc Moorepark | Calum Walsh, Genuity Ireland |
| EPI_ISL_770823 | Eastern Ontario Regional Laboratory Association | McMaster University | Leanne Mortimer, Hooman Derakhshani, Emily Panousis, Ahmed Draia, Jalees Nasir, Robert Slinger, Andrew G. McArthur |
| EPI_ISL_775257 | INT Fondazione Pascale | INT Fondazione Pascale | INT Fondazione Pascale |
| EPI_ISL_775613, EPI_ISL_775690, EPI_ISL_775762, EPI_ISL_775765, EPI_ISL_775774, EPI_ISL_775779, EPI_ISL_775783, EPI_ISL_775790, EPI_ISL_775802, EPI_ISL_775816, EPI_ISL_775837, EPI_ISL_775890, EPI_ISL_775899, EPI_ISL_775900, EPI_ISL_775909, EPI_ISL_776267, EPI_ISL_776268, EPI_ISL_776269, EPI_ISL_776270, EPI_ISL_776271, EPI_ISL_776272, EPI_ISL_776273, EPI_ISL_776274, EPI_ISL_776275, EPI_ISL_776276, EPI_ISL_776277, EPI_ISL_776307, EPI_ISL_776308, EPI_ISL_776309, EPI_ISL_776310, EPI_ISL_776311, EPI_ISL_776312, EPI_ISL_776313, EPI_ISL_776314, EPI_ISL_776315, EPI_ISL_776316, EPI_ISL_776376, EPI_ISL_776377, EPI_ISL_776378, EPI_ISL_776379, EPI_ISL_776380, EPI_ISL_776382, EPI_ISL_776383, EPI_ISL_776384, EPI_ISL_776385, EPI_ISL_776386, EPI_ISL_776387, EPI_ISL_776388, EPI_ISL_776389, EPI_ISL_776390, EPI_ISL_776391, EPI_ISL_776392, EPI_ISL_776393, EPI_ISL_776394, EPI_ISL_776395, EPI_ISL_776396, EPI_ISL_776397, EPI_ISL_776398, EPI_ISL_776399, EPI_ISL_776400, EPI_ISL_776401, EPI_ISL_776402, EPI_ISL_776403, EPI_ISL_776406, EPI_ISL_776407, EPI_ISL_776408, EPI_ISL_776409, EPI_ISL_776410, EPI_ISL_776411, EPI_ISL_776412, EPI_ISL_776444, EPI_ISL_776445, EPI_ISL_776446, EPI_ISL_776448, EPI_ISL_776560 | | | |
| see above | University Medical Center Hamburg Eppendorf | Heinrich Pette Institute, Leibniz Institute for Experimental Virology | Alexis Robitaille, Thomas Günther, Johannes Knobloch, Martin Aepfelbacher, Nicole Fischer, Adam Grundhoff |
| EPI_ISL_778647, EPI_ISL_778650, EPI_ISL_778651 | Istituto Zooprofilattico Sperimentale del Mezzogiorno | TIGEM | Patrizia Annunziata, Andrea Ballabio, Valentina Bouche, Davide Cacchiarelli (CorrespAuthor), Pellegrino Cerino, Chiara Colantuono, Lucio Di Filippo, Antonio Grimaldi, Antonio Limone, Gabriella Loconte, Anna Manfredi, Francesco Panariello, Biancamaria Pierri, Marcello Salvi, Lucia Vassallo |
| EPI_ISL_779715, EPI_ISL_779748, EPI_ISL_779749, EPI_ISL_779756, EPI_ISL_779757, EPI_ISL_779763 | IdISSC/Hospital Clínico San Carlos de Madrid | SeqCOVID-SPAIN consortium/IBV(CSIC) | Alberto Delgado-Iribarren, Esther Culebras López, Vicente Estrada Pérez, Jorge Matias-Guiu, Luis Ortega Medina, Silvia Sánchez Ramón, Ulises Gómez-Pinedo and SeqCOVID-SPAIN consortium |
| EPI_ISL_790628 | Dutch COVID-19 response team | National Institute for Public Health and the Environment (RIVM) | Adam Meijer, Harry Vennema, Jeroen Cremer, Sharon van den Brink, Bas van der Veer, AnneMarie van den Brandt, Florian Zwagemaker, Dennis Schmitz, Chantal Reusken, on behalf of the national COVID-19 response team |

| | | | |
|---|---|---|---|
| EPI_ISL_791519, EPI_ISL_791520, EPI_ISL_791521 | Massachusetts State Public Health Laboratory | Infectious Disease Program, Broad Institute of Harvard and MIT | Lemieux,J.E., Siddle,K.J., Shaw,B., Adams,G., Pierce,V., Turbett,S., Anahtar,M., Branda,J., Slater,D., Harris,J., Lin,A.E., Gladden-Young,A., Lagerborg,K., Rudy,M., DeRuff,K., Carter,A., Normandin,E., Bauer,M., Reilly,S., Tomkins-Tinch,C., Loreth,C., Chaluvadi,S., Neumann,A., Cusick,C., Chapman,S.B., Gnirke,A., Flowers,K., Cerrato,F., Birren,B.W., Gallagher,G., Smole,S., Park,D.J., MacInnis,B.L., Ryan,E., LaRocque,R., Rosenberg,E. and Sabeti,P.C. |
| EPI_ISL_792091 | Toronto Invasive Bacterial Diseases Network | McMaster University | Allison McGeer, Patryk Aftanas, Hooman Derakhshani, Angel Li, Kuganya Nirmalarajah, Emily Panousis, Ahmed Draia, Jalees Nasir, Michael Surette, Samira Mubareka, Andrew G. McArthur |
| EPI_ISL_792095, EPI_ISL_792097, EPI_ISL_792099, EPI_ISL_792100 | UW Virology Lab | UW Virology Lab | Pavitra Roychoudhury, Hong Xie, Lasata Shrestha, Meei-Li Huang, Keith R Jerome, Alexander Greninger |
| EPI_ISL_792102, EPI_ISL_792105 | Instituto Adolfo Lutz - Central | Instituto Adolfo Lutz, Interdiciplinary Procedures Center, Strategic Laboratory | Claudio Tavares Sacchi, Claudia Regina Gonçalves, Erica Valessa Ramos Gomes, Karoline Rodrigues Campos |
| EPI_ISL_792303 | Unidad de Virología, Centro de Educación Médica en Investigaciones Clínicas CEMIC | Área de Secuenciación del Laboratorio de Virología del Hospital de Niños Dr. Ricardo Gutierrez on behalf of 'Proyecto Argentino Interinstitucional de genomica de SARS-CoV-2' (PAIS Consortium) | Nabaes Jodar, MS; Goya, S; Natale, MI; Lusso, S; Echavarría, M; Mistchenko, AS; Valinotto, LE; Viegas, M. |
| EPI_ISL_792358 | Laboratorio de Virologia - HIEAyC San Juan de Dios | Área de Secuenciación del Laboratorio de Virología del Hospital de Niños Dr. Ricardo Gutierrez on behalf of 'Proyecto Argentino Interinstitucional de genomica de SARS-CoV-2' (PAIS Consortium) | Nabaes Jodar, MS; Goya, S; Natale, MI; Lusso, S; Ferioli, M; Colmeiro, M; Gatelli, A; Ercole, R; Valinotto, LE; Viegas, M. |
| EPI_ISL_792453, EPI_ISL_792454, EPI_ISL_792460, EPI_ISL_792462, EPI_ISL_792464, EPI_ISL_792471 | Laboratorio del Hospital Regional Ushuaia Gdor. Ernesto Campos | Hospital Regional Ushuaia - Centro Austral De Investigaciones Científicas - Universidad Nacional De Tierra Del Fuego on behalf of 'Proyecto Argentino Interinstitucional de genomica de SARS-CoV-2' (PAIS Consortium) | Ceballos, SG; Nardi, CF; Gramundi, ID; Gallego, F; De Roccis, CA; Castro, G; Cáceres, SB; Yulan, CB; Boutureira, MF. |
| EPI_ISL_792481, EPI_ISL_792482, EPI_ISL_792484 | Laboratorio Central Mg. Luis Alfredo Pianciola | Hospital Regional Ushuaia - Centro Austral De Investigaciones Científicas - Universidad Nacional De Tierra Del Fuego on behalf of 'Proyecto Argentino Interinstitucional de genomica de SARS-CoV-2' (PAIS Consortium) | Ceballos, SG; Nardi, CF; Gramundi, ID; Gallego, F; Pinto, C; Ziehm, MC; Mazzeo, M; Pianciola, L. |
| EPI_ISL_792532, EPI_ISL_792535 | Laboratorio Central, Ministerio de Salud Córdoba | Instituto de Patologia Vegetal (CIAP-INTA) on behalf of 'Proyecto Argentino Interinstitucional de genomica de SARS-CoV-2' (PAIS Consortium) | Fernández, FD; Debat, HJ., Re, V; Pisano, MB; Castro, G.; Barbas, G. |
| EPI_ISL_801581 | Laboratory of Molecular Virology, Pontificia Universidad Católica de Chile | MSHS Pathogen Surveillance Program | Leonardo I. Almonacid, Ana S. Gonzalez-Reiche, Matthew M. Hernandez, Jorge Levican, Tamara García-Salum, Zenab Khan, Adriana van De Guchte, Ajay Obla, Jayeeta Dutta, Bremy Alburquerque, Eileen Serrano, Erick Salinas, Hala Alshammary, Juan Soto, Shwetha Hara Sridhar, Ying-Chih Wang, Melissa Smith, Robert Sebra, Adolfo Garcia-Sastre, Edward C. Holmes, Viviana Simon, Harm van Bakel, Rafael A. Medina |
| EPI_ISL_801626, EPI_ISL_801627, EPI_ISL_801630, EPI_ISL_801631, EPI_ISL_801632, EPI_ISL_801633, EPI_ISL_801634, EPI_ISL_801657, EPI_ISL_801658, EPI_ISL_801659, EPI_ISL_801660, EPI_ISL_801661, EPI_ISL_801662, EPI_ISL_801663, EPI_ISL_801664, EPI_ISL_801665, EPI_ISL_801666, EPI_ISL_801667, EPI_ISL_801668, EPI_ISL_801669, EPI_ISL_801671, EPI_ISL_801672, EPI_ISL_801673, EPI_ISL_801674 | | | |
| see above | Laboratory of Molecular Virology, Pontificia Universidad Católica de Chile | MSHS Pathogen Surveillance Program | Leonardo I. Almonacid, Ana S. Gonzalez-Reiche, Matthew M. Hernandez, Jorge Levican, Ana Maria Contreras, Carlos Palma, Tamara García-Salum, Zenab Khan, Adriana van De Guchte, Ajay Obla, Jayeeta Dutta, Bremy Alburquerque, Eileen Serrano, Constanza Maldonado, M. Belen Leyton, Erick Salinas, Hala Alshammary, Juan Soto, Shwetha Hara Sridhar, Ying-Chih Wang, Melissa Smith, Robert Sebra, Marcela Ferres, Adolfo Garcia-Sastre, Edward C. Holmes, Viviana Simon, Harm van Bakel, Rafael A. Medina. |
| EPI_ISL_803882, EPI_ISL_803887, EPI_ISL_803888 | Department of Medical Biotechnologies, University of Siena | Laboratory of Infectious Diseases, Department of Biomedical and Clinical Sciences L. Sacco, University of Milan | Ilaria Vicenti, Filippo Dragoni, Maurizio Zazzi, Maria Grazia Cusi, Alessia Lai, Annalisa Bergna, Carla Della Ventura, Claudia Balotta, Massimo Galli, Gianguglielmo Zehender on behalf of SARS-CoV-2 ITALIAN RESEARCH ENTERPRISE-(SCIRE) Collaborative Group |
| EPI_ISL_804383, EPI_ISL_804386, EPI_ISL_804397, EPI_ISL_804410, EPI_ISL_804411, EPI_ISL_804421, EPI_ISL_804422, EPI_ISL_804435, EPI_ISL_804443, EPI_ISL_804448, EPI_ISL_804449, EPI_ISL_804455, EPI_ISL_804460 | | | |
| see above | Dutch COVID-19 response team | National Institute for Public Health and the Environment (RIVM) | Adam Meijer, Harry Vennema, Dirk Eggink, Matthijs Welkers, Jeroen Cremer, Sharon van den Brink, Bas van der Veer, AnneMarie van den Brandt, Florian Zwagemaker, Dennis Schmitz, Chantal Reusken, on behalf of the national COVID-19 response team |
| EPI_ISL_804854, EPI_ISL_804864, EPI_ISL_804865, EPI_ISL_804890, EPI_ISL_804919, EPI_ISL_804920, EPI_ISL_804921, EPI_ISL_804922 | DC Public Health Lab/ Dept. of Forensic Sciences | DC Public Health Lab/ Dept. of Forensic Sciences | Scott Nguyen, Elizabeth Zelaya, Connie Maza, Monica Mann, Brittany Hamilton, David Payne, Jocelyn Hauser |
| EPI_ISL_805669, EPI_ISL_805701, EPI_ISL_805746, EPI_ISL_805755, EPI_ISL_805782, EPI_ISL_805872, EPI_ISL_805873, EPI_ISL_805938, EPI_ISL_805960, EPI_ISL_805964, EPI_ISL_805965, EPI_ISL_805985, EPI_ISL_806034, EPI_ISL_806056, EPI_ISL_806112, EPI_ISL_806157, EPI_ISL_806168, EPI_ISL_806193, EPI_ISL_806195, EPI_ISL_806196, EPI_ISL_806211, EPI_ISL_806233, EPI_ISL_806248, EPI_ISL_806250, EPI_ISL_806288, EPI_ISL_806293, EPI_ISL_806317, EPI_ISL_806329, EPI_ISL_806330, EPI_ISL_806344, EPI_ISL_806376, EPI_ISL_806384, EPI_ISL_806405, EPI_ISL_806416, EPI_ISL_806423, EPI_ISL_806461, EPI_ISL_806463, EPI_ISL_806493 | | | |
| see above | Alberta Precision Labs (APL) | Alberta Precision Labs (APL) | Gordon P, Lam LG, Pabbaraju K, Wong A, Ma R, Li V, Melin A, Tipples G, Berenger B, Zelyas N, Kellner J, Bernier F, Chui L, Croxen M |
| EPI_ISL_806526, EPI_ISL_806529 | Charité Universitätsmedizin Berlin, Institut für Virologie/Labor Berlin | Charité Universitätsmedizin Berlin, Institut für Virologie | Victor M Corman, Jörn Beheim-Schwarzbach, Barbara Mühlemann, Julia Schneider, Talitha Veith, Cornelia Schlee, Tomasz Zemojtel, Terry Jones, Christian Drosten |
| EPI_ISL_812309, EPI_ISL_812318, EPI_ISL_812320 | United States Air Force School of Aerospace Medicine | United States Air Force School of Aerospace Medicine | Anthony Fries, Jennifer Meyer, Amanda Javorina, Sarah Purves, William Gruner, Clarise Starr, Elizabeth Macias |
| EPI_ISL_819312, EPI_ISL_819313, EPI_ISL_819314 | Hospital Universitari Vall d'Hebron - Vall d'Hebron Institut de Rercerca | Hospital Universitari Vall d'Hebron | Cristina Andrés, Maria Piñana, Josep F Abril, Damir Garcia-Cehic, Ariadna Rando, Juliana Esperalba, Maria Gema Codina, Carla Castillo, Maria Carmen Martin, Tomàs Pumarola, Josep Quer, Andrés Antón |
| EPI_ISL_825866 | Laboratoire de santé publique du Québec | Laboratoire de santé publique du Québec | Sandrine Moreira, Ioannis Ragoussis, Guillaume Bourque, Jesse Shapiro, Mark Lathrop and Michel Roger on behalf of the CoVSeQ research group (http://covseq.ca/researchgroup) |
| EPI_ISL_826641, EPI_ISL_826642, EPI_ISL_826643, EPI_ISL_826644, EPI_ISL_826645 | Montefiore Medical Center | Albert Einstein College of Medicine, Dept. of Microbiology & Immunology, Chandran lab | J. Maximilian Fels, Saad Khan, Ryan Forster, Karin A. Skalina, Surksha Sirichand, Amy S. Fox, Aviv Bergman, William B. Mitchell, Lucia R. Wolgast, Wendy Szymczak, Robert H. Bortz III, M. Eugenia Dieterle, Catalina Florez, Denise Haslwanter, Rohit K. Jangra, Ethan Laudermilch, Ariel S. Wirchnianski, Jason Barnhill, David L. Goldman, Hnin Khine, D. Yitzchak Goldstein, Johanna P. Daily, Kartik Chandran, Libusha Kelly |
| EPI_ISL_826807, EPI_ISL_826809, EPI_ISL_826811, EPI_ISL_826812, EPI_ISL_826814, EPI_ISL_826816, EPI_ISL_826818 | INSPI-CRN DE INFLUENZA Y OTROS VIRUS RESPIRATORIOS | Instituto de Salud Publica de Chile | Javier Tognarelli, Barbara Parra, Loredana Arata, Jaime Lagos, Gisselle Barra, Alfredo Bruno, Domenica de Mora, Solon Narvaez, Jimmy Garcez, Michelle Paez, Martiza Olmedo, Manuel Gonzalez, Patricia Bustos, Rodrigo Fasce, Andres Castillo, Jorge Fernandez |
| EPI_ISL_831039 | Hospital Universitario La Paz (Madrid) | SeqCOVID-SPAIN consortium/IBV(CSIC) | María Rodríguez-Tejedor, Elias Dahdouh, Fernando Lázaro-Perona, Jesús Mingorance and SeqCOVID-SPAIN consortium |
| EPI_ISL_833153 | Instituto Adolfo Lutz - Central | Instituto Adolfo Lutz, Interdiciplinary Procedures Center, Strategic Laboratory | Claudio Tavares Sacchi, Claudia Regina Gonçalves, Erica Valessa Ramos Gomes, Karoline Rodrigues Campos |
| EPI_ISL_837618, EPI_ISL_837619, EPI_ISL_837620 | Instituto Nacional de Enfermedades Respiratorias (INER) | Instituto Nacional de Enfermedades Respiratorias (INER) | Celia Boukadida, Margarita Matías-Florentino, Alma Rincón-Rubio, Hector Esteban Paz-Juárez, Olivia Briceño, Edgar Sevilla-Reyes, Fidencio Mejía-Nepomuceno, Mario Mújica-Sánchez, Eduardo Becerril-Vargas, José Arturo Martínez-Orozco, Alejandra Hernández-Terán, Jorge Salas-Hernández, Santiago Ávila-Ríos, Joel Armando Vázquez-Pérez |
| EPI_ISL_842812, EPI_ISL_842818, EPI_ISL_842822, EPI_ISL_842823, EPI_ISL_843079 | Barts Health NHS Trust | COVID-19 Genomics UK (COG-UK) Consortium | CUTINO-MOGUEL, Maria-Teresa; HARRINGTON, David; OWOYEMI, Dola; SHYLINI, Raghavendran; BROAD, Claire; KELE, Beatrix |

| | | | |
|---|---|---|---|
| EPI_ISL_848087, EPI_ISL_848090, EPI_ISL_848091, EPI_ISL_848092, EPI_ISL_848093, EPI_ISL_848094, EPI_ISL_848098, EPI_ISL_848099, EPI_ISL_848104, EPI_ISL_848105, EPI_ISL_848107, EPI_ISL_848132, EPI_ISL_848133, EPI_ISL_848134, EPI_ISL_848135, EPI_ISL_848136, EPI_ISL_848137, EPI_ISL_848147, EPI_ISL_848166 | | | |
| see above | National Virus Reference Laboratory | Irish Coronavirus Sequencing Consortium - National University of Ireland Galway | Kate Reddington, Grainne Mc Andrew, Simone Coughlan |
| EPI_ISL_848246, EPI_ISL_848281, EPI_ISL_848282 | Illinois Department of Public Health | Gagnon Lab, Southern Illinois University | Keith Gagnon |
| EPI_ISL_848596, EPI_ISL_848625 | Evandro Chagas Institute | Evandro Chagas Institute | Santos, M.C.; Silva, A.M.; Junior, W.D.C.; Barbagelata, L.S.; Ferreira, J.A.; Sousa, E.M.A.; da Silva, P.S.; Pinheiro, K.C.; L.C.; Sousa Junior, E.C. |
| EPI_ISL_849678, EPI_ISL_849679, EPI_ISL_849680, EPI_ISL_849681, EPI_ISL_849687, EPI_ISL_849688 | unknown | PHV-FSS | Son Nguyen et al. |
| EPI_ISL_852668, EPI_ISL_852669, EPI_ISL_852670, EPI_ISL_852671, EPI_ISL_852672, EPI_ISL_852673, EPI_ISL_852674, EPI_ISL_852675, EPI_ISL_852676, EPI_ISL_852677, EPI_ISL_852764, EPI_ISL_852765, EPI_ISL_852766, EPI_ISL_852767, EPI_ISL_852768 | | | |
| see above | Institute of Virology, Medical Center, University of Freiburg, Freiburg, Germany | Institute of Virology, Clinial Virus Genomics, Medical Center, University of Freiburg, Freiburg, Germany | Jonas Fuchs, Lisa Kern, Sandra Reuter, Hajo Grundmann, Marcus Panning |
| EPI_ISL_854471, EPI_ISL_854472 | Alberta Precision Labs (APL) | Alberta Precision Labs (APL) | Gordon P, Lam LG, Pabbaraju K, Wong A, Ma R, Li V, Melin A, Tipples G, Berenger B, Zelyas N, Kellner J, Bernier F, Chui L, Croxen M |
| EPI_ISL_855404, EPI_ISL_855405, EPI_ISL_855406 | IdISSC/Hospital Clínico San Carlos de Madrid | SeqCOVID-SPAIN consortium/IBV(CSIC) | Alberto Delgado-Iribarren, Esther Culebras López, Vicente Estrada Pérez, Jorge Matias-Guiu, Luis Ortega Medina, Silvia Sánchez Ramón, Ulises Gómez-Pinedo and SeqCOVID-SPAIN consortium |
| EPI_ISL_856869 | Virology Laboratory, Scientific Department, Army Medical Center | Virology Laboratory, Scientific Department, Army Medical Center | Silvia Fillo, Giovanni Faggioni, Riccardo De Sanctis, Antonella Fortunato, Anella Monte, Anna Anselmo, Vanessa Vera Fain, Francesco Giordani, Nino D'Amore, Filippo Molinari, Giancarlo Petralito, Florigio Lista |
| EPI_ISL_856870, EPI_ISL_856872, EPI_ISL_856885 | Virology Laboratory, Scientific Department, Army Medical Center | Virology Laboratory, Scientific Department, Army Medical Center | Silvia Fillo, Giovanni Faggioni, Riccardo De Sanctis, Antonella Fortunato, Anella Monte, Anna Anselmo, Vanessa Vera Fain, Francesco Giordani, Nino D'Amore, Filippo Molinari, Giancarlo Petralito, Florigio Lista |
| EPI_ISL_861530 | Instituto Nacional de Saude (INSA) | Instituto Nacional de Saude (INSA) | Borges et al |
| EPI_ISL_861633 | Instituto Adolfo Lutz - Central | Instituto Adolfo Lutz, Interdiciplinary Procedures Center, Strategic Laboratory | Claudio Tavares Sacchi, Claudia Regina Gonçalves, Erica Valessa Ramos Gomes, Karoline Rodrigues Campos |
| EPI_ISL_861636 | Hospital Geral de Sao Mateus São Paulo | Instituto Adolfo Lutz, Interdiciplinary Procedures Center, Strategic Laboratory | Claudio Tavares Sacchi, Claudia Regina Gonçalves, Erica Valessa Ramos Gomes, Karoline Rodrigues Campos |
| EPI_ISL_861900 | LATE - Laboratório de Técnicas Especiais - Hospital Israelita Albert Einstein | LATE - Laboratório de Técnicas Especiais - Hospital Israelita Albert Einstein | Deyvid Amgarten, Fernanda de Mello Malta, Raquel Riyuzo, Ana Paula Moreira Salles, Pedro Henrique Sebe Rodrigues, João Renato Rebello Pinho |
| EPI_ISL_862841 | Fujita Health University Hospital | Fujita health University, School of Medicine, Department of Microbiology | Masahiro suzuki, Yohei Doi |
| EPI_ISL_862859 | Nara Medical University Hospital | Fujita health University, School of Medicine, Department of Microbiology | Masahiro suzuki, Yohei Doi |
| EPI_ISL_876972, EPI_ISL_877012, EPI_ISL_877013 | 1.AO Universitaria 'S. Giovanni di Dio e Ruggi D'Aragona, Scuola Medica Salernitana' Hospital / 2.UOC di Virologia e Microbiologia, Università della Campania 'L. Vanvitelli' / 3.AO Universitaria 'Federico II' Napoli Hospital / 4.AORN 'San Giuseppe Moscati' Avellino Hospital / 5.AO 'San Pio - presidio G. Rummo' Benevento Hospital / 6.AO 'Sant'Anna e San Sebastiano' Caserta Hospital / 7.PO 'Maria Santissima Addolorata' Eboli Hospital / 8.Biogem Istituto di Ricerche Genetiche | 1. Genome Research Center for Health (CRGS) / 2. Laboratory of Molecular Medicine and Genomics(LMMGe) / 3. Center for Research in Pure and Applied Mathematics (CRMPA) | Giorgio Giurato (Corresponding Author), Francesca Rizzo (Corresponding Author), Alessandro Weisz (Corresponding Author), Gianluigi Franci, Giovanni Nassa, Pasquale Pagliano, Roberta Tarallo, Elena Alexandrova, Ylenia D'Agostino, Carlo Ferravante, Jessica Lamberti, Viola Melone, Domenico Memoli, Valeria Mirici Cappa, Domenico Palumbo, Giovanni Pecoraro, Assunta Sellitto, Oriana Strianese, Ilaria Terenzi, Giuseppe Fenza, Aniello Gentile, Antonello Saccomanno, Sonia Amabile, Teresa Rocco, Annamaria Salvati, Emilia Vaccaro, Massimiliano Galdiero, Michele Cennamo, Giuseppe Portella, Maria Grazia Foti, Mariarosaria Ingino, Maria Landi, Maurizio Fumi, Vincenzo Rocco, Rita Greco, Vittoria Letizia, Arnolfo Petruzziello, Maddalena Schioppa, Gregorio Goffredi, Francesca Marciano, Michele Caraglia, Alessia Cossu, Marianna Scrima |
| EPI_ISL_877590, EPI_ISL_877591 | Clinical Molecular Microbiology Laboratory, UNC Hospital | Dirk Dittmer | Razia Moorad , Justin T. Landis , Brent A. Eason, Melissa B. Miller, Linda Pluta, Dirk Dittmer, Angelica Juarez, Cecilia Thompson , Cameroon Grant, Evelyn Hoffman, Patricio Cano, Jason Wong, Carolina Caro-Vegas, Blossom Damania. |
| EPI_ISL_883060, EPI_ISL_883061 | Kansas Health and Environmental Lab | Kansas Health and Environmental Lab | Mike Grose, Carissa Robertson, Ben Olsen, and Phil Adam |
| EPI_ISL_883163 | Hannover Medical School, Institute of Virology | Hannover Medical School, Institute of Virology | Lars Steinbrück, Jasper Götting |
| EPI_ISL_884549, EPI_ISL_884550, EPI_ISL_884551, EPI_ISL_884552, EPI_ISL_884553, EPI_ISL_884554, EPI_ISL_884555, EPI_ISL_884571, EPI_ISL_884574, EPI_ISL_884575, EPI_ISL_884576 | | | |
| see above | Molecular Microbiology & Immunology, University of Missouri | Molecular Microbiology & Immunology, University of Missouri | Tang,C.Y., Li,T., Hang,J., Lidl,G.M., Wan,X.-F. |
| EPI_ISL_887182, EPI_ISL_887231, EPI_ISL_887235, EPI_ISL_887270, EPI_ISL_887276, EPI_ISL_887327, EPI_ISL_887330, EPI_ISL_887331, EPI_ISL_887336, EPI_ISL_887339, EPI_ISL_887341, EPI_ISL_887344, EPI_ISL_887346, EPI_ISL_887349, EPI_ISL_887350, EPI_ISL_887352, EPI_ISL_887353, EPI_ISL_887354, EPI_ISL_887357, EPI_ISL_887359, EPI_ISL_887361, EPI_ISL_887362, EPI_ISL_887371, EPI_ISL_887374, EPI_ISL_887378, EPI_ISL_887379, EPI_ISL_887381, EPI_ISL_887382, EPI_ISL_887384, EPI_ISL_887386, EPI_ISL_887388, EPI_ISL_887394, EPI_ISL_887395, EPI_ISL_887397, EPI_ISL_887401, EPI_ISL_887408, EPI_ISL_887413, EPI_ISL_887418 | | | |
| see above | Protzer Lab | Protzer Lab, Gagneur Lab, Robert Koch Institut | Ulrike Protzer, Dieter Hoffmann, Eva Schulte, Andrea Theumer, Oliver Drechsel, Max von Kleist,Aleksandar Radonic,Stephan Fuchs, Alexander Karollus, Julien Gagneur |
| EPI_ISL_889307, EPI_ISL_889308 | Motol University Hospital | Institute of Applied Biotechnologies a.s. | Petr Klempt, Ondej Brzo, Martin Kašný, Kateina Kvapilová, Pavel Devínek, Petr Kvapil |
| EPI_ISL_889324, EPI_ISL_889325, EPI_ISL_889326 | The University Hospital Brno | Institute of Applied Biotechnologies a.s. | Petr Klempt, Ondej Brzo, Martin Kašný, Kateina Kvapilová, Martina Lengerová, Petr Kvapil |
| EPI_ISL_889375, EPI_ISL_889376, EPI_ISL_889377, EPI_ISL_889378, EPI_ISL_889379, EPI_ISL_889380 | Olomouc University Hospital | Institute of Applied Biotechnologies a.s. | Petr Klempt, Ondej Brzo, Martin Kašný, Kateina Kvapilová, Marián Hajdúch, Petr Kvapil |
| EPI_ISL_889802, EPI_ISL_889803, EPI_ISL_889807, EPI_ISL_889809, EPI_ISL_889810, EPI_ISL_889811, EPI_ISL_889812, EPI_ISL_889813, EPI_ISL_889814, EPI_ISL_889818, EPI_ISL_889819, EPI_ISL_889820, EPI_ISL_889821, EPI_ISL_889823, EPI_ISL_889824, EPI_ISL_889825, EPI_ISL_889826, EPI_ISL_889827, EPI_ISL_889828, EPI_ISL_889830, EPI_ISL_889831, EPI_ISL_889832, EPI_ISL_889834, EPI_ISL_889836, EPI_ISL_889838, EPI_ISL_889841, EPI_ISL_889857, EPI_ISL_889858, EPI_ISL_889859, EPI_ISL_889861, EPI_ISL_889867, EPI_ISL_889868, EPI_ISL_889869, EPI_ISL_889870, EPI_ISL_889871, EPI_ISL_889873, EPI_ISL_889874, EPI_ISL_889875, EPI_ISL_889876, EPI_ISL_889877, EPI_ISL_889878, EPI_ISL_889879, EPI_ISL_889880, EPI_ISL_889881, EPI_ISL_889882, EPI_ISL_889883, EPI_ISL_889884, EPI_ISL_889886, EPI_ISL_889887, EPI_ISL_889888, EPI_ISL_889890, EPI_ISL_889891, EPI_ISL_889892, EPI_ISL_889893, EPI_ISL_889894, EPI_ISL_889895, EPI_ISL_889896, EPI_ISL_889897, EPI_ISL_889898, EPI_ISL_889899, EPI_ISL_889900, EPI_ISL_889901, EPI_ISL_889902, EPI_ISL_889903, EPI_ISL_889904, EPI_ISL_889905, EPI_ISL_889908, EPI_ISL_889909, EPI_ISL_889910, EPI_ISL_889911, EPI_ISL_889912, EPI_ISL_889915, EPI_ISL_889916, EPI_ISL_889921, EPI_ISL_889922, EPI_ISL_889923, EPI_ISL_889924, EPI_ISL_889925, EPI_ISL_889926, EPI_ISL_889935, EPI_ISL_889937, EPI_ISL_889938, EPI_ISL_889939, EPI_ISL_889940, EPI_ISL_889941, EPI_ISL_889947, EPI_ISL_889948, EPI_ISL_889949, EPI_ISL_889950, EPI_ISL_889951, EPI_ISL_889952, EPI_ISL_889953, EPI_ISL_889955, EPI_ISL_889957, EPI_ISL_889961, EPI_ISL_889962, EPI_ISL_889963, EPI_ISL_889964, EPI_ISL_889965, EPI_ISL_889966, EPI_ISL_889967, EPI_ISL_889968, EPI_ISL_889969, EPI_ISL_889970, EPI_ISL_889971, EPI_ISL_889972, EPI_ISL_889973, EPI_ISL_889976, EPI_ISL_889977, EPI_ISL_889978, EPI_ISL_889979, EPI_ISL_889983, EPI_ISL_889984, EPI_ISL_889985, EPI_ISL_889986, EPI_ISL_889987, EPI_ISL_889990, EPI_ISL_889991, EPI_ISL_890000, EPI_ISL_890001, EPI_ISL_890002, EPI_ISL_890004, EPI_ISL_890006, EPI_ISL_890008, EPI_ISL_890009, EPI_ISL_890010, EPI_ISL_890011, EPI_ISL_890012, EPI_ISL_890013, EPI_ISL_890014 | | | |
| see above | Laboratoire de santé publique du Québec | Laboratoire de santé publique du Québec | Sandrine Moreira, Ioannis Ragoussis, Guillaume Bourque, Jesse Shapiro, Mark Lathrop and Michel Roger on behalf of the CoVSeQ research group |
| EPI_ISL_890373, EPI_ISL_890499, EPI_ISL_890516, EPI_ISL_890518, EPI_ISL_890519, EPI_ISL_890521, EPI_ISL_890522, EPI_ISL_890523, EPI_ISL_890524, EPI_ISL_890525, EPI_ISL_890526, EPI_ISL_890527, EPI_ISL_890528, EPI_ISL_890529, EPI_ISL_890530, EPI_ISL_890531, EPI_ISL_890532, EPI_ISL_890533, EPI_ISL_890536, EPI_ISL_890537, EPI_ISL_890538, EPI_ISL_890539, EPI_ISL_890544, EPI_ISL_890548, EPI_ISL_890549, EPI_ISL_890550, EPI_ISL_890551, EPI_ISL_890553, EPI_ISL_890557, EPI_ISL_890558, EPI_ISL_890559, EPI_ISL_890654, EPI_ISL_890706, EPI_ISL_890711, EPI_ISL_890712, EPI_ISL_890713, EPI_ISL_890714, EPI_ISL_890715, EPI_ISL_890716, EPI_ISL_890717, EPI_ISL_890718, EPI_ISL_890719, EPI_ISL_890721, EPI_ISL_890724, EPI_ISL_890741, EPI_ISL_890742, EPI_ISL_890750, EPI_ISL_890751, EPI_ISL_890752, EPI_ISL_890753, EPI_ISL_890754, EPI_ISL_890761, EPI_ISL_890762, EPI_ISL_890763, EPI_ISL_890764, EPI_ISL_890765, EPI_ISL_890767, EPI_ISL_890768, EPI_ISL_890769, EPI_ISL_890770, EPI_ISL_890771, EPI_ISL_890772, EPI_ISL_890774, EPI_ISL_890775, EPI_ISL_890776, EPI_ISL_890777, EPI_ISL_890778, EPI_ISL_890779, EPI_ISL_890780, EPI_ISL_890781, EPI_ISL_890782, EPI_ISL_890783, EPI_ISL_890784, EPI_ISL_890785, EPI_ISL_890786, EPI_ISL_890787, EPI_ISL_890791, EPI_ISL_890797, EPI_ISL_890798, EPI_ISL_890799, EPI_ISL_890800, EPI_ISL_890801, EPI_ISL_890802, EPI_ISL_890803, EPI_ISL_890804, EPI_ISL_890805, EPI_ISL_890806, EPI_ISL_890807, EPI_ISL_890808, EPI_ISL_890809, EPI_ISL_890810, EPI_ISL_890811, EPI_ISL_890812, EPI_ISL_890813, EPI_ISL_890814, EPI_ISL_890815, EPI_ISL_890824, EPI_ISL_890825, EPI_ISL_890826, EPI_ISL_890827, EPI_ISL_890828, EPI_ISL_890829, EPI_ISL_890830, EPI_ISL_890831, EPI_ISL_890832, EPI_ISL_890833, EPI_ISL_890834, EPI_ISL_890835, | | | |

| | | | |
|---|---|---|---|
| EPI_ISL_890836, EPI_ISL_890837, EPI_ISL_890838, EPI_ISL_890839, EPI_ISL_890840, EPI_ISL_890841, EPI_ISL_890842, EPI_ISL_890843, EPI_ISL_890844, EPI_ISL_890845, EPI_ISL_890846, EPI_ISL_890847, EPI_ISL_890848, EPI_ISL_890850, EPI_ISL_890853, EPI_ISL_890854, EPI_ISL_890855, EPI_ISL_890856, EPI_ISL_890857, EPI_ISL_890858, EPI_ISL_890859, EPI_ISL_890860, EPI_ISL_890861, EPI_ISL_890862, EPI_ISL_890863, EPI_ISL_890864, EPI_ISL_890865, EPI_ISL_890866, EPI_ISL_890867, EPI_ISL_890868, EPI_ISL_890869, EPI_ISL_890870, EPI_ISL_890871, EPI_ISL_890872, EPI_ISL_890873, EPI_ISL_890874, EPI_ISL_890875, EPI_ISL_890876, EPI_ISL_890877, EPI_ISL_890878, EPI_ISL_890879, EPI_ISL_890881, EPI_ISL_890882, EPI_ISL_890883, EPI_ISL_890884, EPI_ISL_890885, EPI_ISL_890886, EPI_ISL_890887, EPI_ISL_890888 | | | |
| see above | Laboratoire de santé publique du Québec | Laboratoire de santé publique du Québec | Sandrine Moreira, Ioannis Ragoussis, Guillaume Bourque, Jesse Shapiro, Mark Lathrop and Michel Roger on behalf of the CoVSeQ research group (http://covseq.ca/researchgroup) |
| EPI_ISL_895728, EPI_ISL_895736, EPI_ISL_895741, EPI_ISL_895742, EPI_ISL_895747, EPI_ISL_895752, EPI_ISL_895759, EPI_ISL_895761, EPI_ISL_895762, EPI_ISL_895770, EPI_ISL_895771, EPI_ISL_895778, EPI_ISL_895780, EPI_ISL_895783, EPI_ISL_895785, EPI_ISL_895791, EPI_ISL_895794, EPI_ISL_895798, EPI_ISL_895815, EPI_ISL_895817, EPI_ISL_895823, EPI_ISL_895824, EPI_ISL_895828, EPI_ISL_895830, EPI_ISL_895832, EPI_ISL_895833, EPI_ISL_895839 | | | |
| see above | Molecular biology division, Institute of Clinical Biochemistry and Diagnostics, Charles University, Faculty of Medicine in Hradec Králové and University Hospital Hradec Králové | Molecular biology division, Institute of Clinical Biochemistry and Diagnostics, Charles University, Faculty of Medicine in Hradec Králové and University Hospital Hradec Králové | Helena Kovaíková, Petr Brož, Ivana Baranová, Kateina Hrochová, Tereza Baková, Jitka Novotná, Kateina Pehlíková, Vladimír Palika. Cooperation project with BioVendor-R&D and bioinformatics company BIOXSYS s r.o. |
| EPI_ISL_896121, EPI_ISL_900071, EPI_ISL_900103, EPI_ISL_900139, EPI_ISL_900153, EPI_ISL_900465 | MEPHI, Aix Marseille University | MEPHI, Aix Marseille University | Anthony LEVASSEUR |
| EPI_ISL_904018 | DB Diagnosticos do Brasil | Laboratório de Parasitologia Médica - Instituto de Medicina Tropical - Universidade de São Paulo | Brazil-UK Centre for Arbovirus Discovery Diagnosis Genomics and Epidemiology (CADDE) Genomic Network - Instituto de Medicina Tropical |
| EPI_ISL_906766, EPI_ISL_906768 | Dr. Leonard A. Miller Centre for Health Services | National Microbiology Laboratory (NML) | Anna Majer, Shari Tyson, Grace Seo, Philip Mabon, Elsie Grudeski, Rhiannon Huzarewich, Russell Mandes, Anneliese Landgraff, Jennifer Tanner, Natalie Knox, Morag Graham, Gary Van Domselaar, Robert Needle, Yang Yu, Laura Gilbert, George Zahariadis, Nathalie Bastien, Yan Li, Timothy Booth, Darian Hole, Madison Chapel, Kerri Smith, CanCOGeN's metadata curation team, Public Health Agency of Canada CanCOGeN team |
| EPI_ISL_911924, EPI_ISL_911928 | Clinical Diagnostics Laboratory, Diagnostic & Experimental Pathology, Lilly Research Laboratories | Clinical Diagnostics Laboratory, Diagnostic & Experimental Pathology, Lilly Research Laboratories | Tim Holzer, Mayuri Vaidya, Angie Fulford, Sam McNeely, Rachael Redmond, Phil Ebert, John Calley, Leslie ONeill Reising, Pat Finnegan, Erin Wray, John McElwee, Jeff Fill, Joe Oakley, Andrew Schade |
| EPI_ISL_914632 | TGen North | TGen North | "Jolene Bowers, Megan Folkerts, Chris French, Hayley Yaglom, Ashlyn Pfeiffer, Darrin Lemmer, Dave Engelthaler, The Arizona COVID Genomics Union (ACGU)" |
| EPI_ISL_914981, EPI_ISL_914982, EPI_ISL_914983, EPI_ISL_914984, EPI_ISL_914994, EPI_ISL_914995, EPI_ISL_914996 | QEII Health Sciences Centre | National Microbiology Laboratory (NML) | Anna Majer, Shari Tyson, Grace Seo, Philip Mabon, Elsie Grudeski, Rhiannon Huzarewich, Russell Mandes, Anneliese Landgraff, Jennifer Tanner, Natalie Knox, Morag Graham, Gary Van Domselaar, Todd Hatchette, Jason LeBlanc, Janice Pettipas, Dan Gaston, Nathalie Bastien, Yan Li, Timothy Booth, Darian Hole, Madison Chapel, CanCOGeN's metadata curation team, Public Health Agency of Canada CanCOGeN team |
| EPI_ISL_918163 | Thai Red Cross Emerging Infectious Diseases Health Science Centre, Chulalongkorn Hospital, Faculty of Medicine, Chulalongkorn University | Thai Red Cross Emerging Infectious Diseases Center and Faculty of Medicine, Chulalongkorn University | Opass Putcharoen, Watsamon Jantarabenjakul, Pattama Torvorapanit, Gompol Suwanpimolkul, Leilani Paitoonpong, Sopon Iamsirithaworn, Sininat Petcharat, Yutthana Joyjinda, Weenassarin Ampoot, Apaporn Rodpan, Thiravat Hemachudha, Supaporn Wacharapluesadee |
| EPI_ISL_918415, EPI_ISL_918467, EPI_ISL_918468 | 1.AO Universitaria 'S. Giovanni di Dio e Ruggi D'Aragona, Scuola Medica Salernitana' Hospital / 2.UOC di Virologia e Microbiologia, Università della Campania 'L. Vanvitelli' / 3.AO Universitaria 'Federico II' Napoli Hospital / 4.AORN 'San Giuseppe Moscati' Avellino Hospital / 5.AO 'San Pio - presidio G. Rummo' Benevento Hospital / 6.AO 'Sant'Anna e San Sebastiano' Caserta Hospital / 7.PO 'Maria Santissima Addolorata' Eboli Hospital / 8.Biogem Istituto di Ricerche Genetiche | 1. Genome Research Center for Health (CRGS) / 2. Laboratory of Molecular Medicine and Genomics(LMMGe) / 3. Center for Research in Pure and Applied Mathematics (CRMPA) | Giorgio Giurato (Corresponding Author), Francesca Rizzo (Corresponding Author), Alessandro Weisz (Corresponding Author), Gianluigi Franci, Giovanni Nassa, Pasquale Pagliano, Roberta Tarallo, Elena Alexandrova, Ylenia D'Agostino, Carlo Ferravante, Jessica Lamberti, Viola Melone, Domenico Memoli, Valeria Mirici Cappa, Domenico Palumbo, Giovanni Pecoraro, Assunta Sellitto, Oriana Strianese, Ilaria Terenzi, Giuseppe Fenza, Aniello Gentile, Antonello Saccomanno, Sonia Amabile, Teresa Rocco, Annamaria Salvati, Emilia Vaccaro, Massimiliano Galdiero, Michele Cennamo, Giuseppe Portella, Maria Grazia Foti, Mariarosaria Ingino, Maria Landi, Maurizio Fumi, Vincenzo Rocco, Rita Greco, Vittoria Letizia, Arnolfo Petruzziello, Maddalena Schioppa, Gregorio Goffredi, Francesca Marciano, Michele Caraglia, Alessia Cossu, Marianna Scrima |
| EPI_ISL_933614, EPI_ISL_933615, EPI_ISL_933616, EPI_ISL_933617 | Toronto Invasive Bacterial Diseases Network | McMaster University | Allison McGeer, Patryk Aftanas, Hooman Derakhshani, Angel Li, Kuganya Nirmalarajah, Emily Panousis, Ahmed Draia, Jalees Nasir, Michael Surette, Samira Mubareka, Andrew G. McArthur |
| EPI_ISL_934421 | 1.AO Universitaria 'S. Giovanni di Dio e Ruggi D'Aragona, Scuola Medica Salernitana' Hospital / 2.UOC di Virologia e Microbiologia, Università della Campania 'L. Vanvitelli' / 3.AO Universitaria 'Federico II' Napoli Hospital / 4.AORN 'San Giuseppe Moscati' Avellino Hospital / 5.AO 'San Pio - presidio G. Rummo' Benevento Hospital / 6.AO 'Sant'Anna e San Sebastiano' Caserta Hospital / 7.PO 'Maria Santissima Addolorata' Eboli Hospital / 8.Biogem Istituto di Ricerche Genetiche | 1. Genome Research Center for Health (CRGS) / 2. Laboratory of Molecular Medicine and Genomics(LMMGe) / 3. Center for Research in Pure and Applied Mathematics (CRMPA) | Giorgio Giurato (Corresponding Author), Francesca Rizzo (Corresponding Author), Alessandro Weisz (Corresponding Author), Gianluigi Franci, Giovanni Nassa, Pasquale Pagliano, Roberta Tarallo, Elena Alexandrova, Ylenia D'Agostino, Carlo Ferravante, Jessica Lamberti, Viola Melone, Domenico Memoli, Valeria Mirici Cappa, Domenico Palumbo, Giovanni Pecoraro, Assunta Sellitto, Oriana Strianese, Ilaria Terenzi, Giuseppe Fenza, Aniello Gentile, Antonello Saccomanno, Sonia Amabile, Teresa Rocco, Annamaria Salvati, Emilia Vaccaro, Massimiliano Galdiero, Michele Cennamo, Giuseppe Portella, Maria Grazia Foti, Mariarosaria Ingino, Maria Landi, Maurizio Fumi, Vincenzo Rocco, Rita Greco, Vittoria Letizia, Arnolfo Petruzziello, Maddalena Schioppa, Gregorio Goffredi, Francesca Marciano, Michele Caraglia, Alessia Cossu, Marianna Scrima, Edmondo Adorisio, Morena D'Avenia, Michela Iacobellis, Rosanna Piluscio, Giorgio Dirani, Vittorio Sambri, Simona Semprini, Silvia Zanoli, Francesco Curcio, Stefania Marzinotto, Andreina Baj, Fausto Sessa. |
| EPI_ISL_935753 | Cadham Provincial laboratory | National Microbiology Laboratory (NML) | Anna Majer, Shari Tyson, Grace Seo, Philip Mabon, Elsie Grudeski, Rhiannon Huzarewich, Russell Mandes, Anneliese Landgraff, Jennifer Tanner, Natalie Knox, Morag Graham, Gary Van Domselaar, Paul Van Caeseele, Jared Bullard, David Alexander, Kerry Dust, Nathalie Bastien, Yan Li, Timothy Booth, Darian Hole, Madison Chapel, Kirsten Biggar, CanCOGeN's metadata curation team, Public Health Agency of Canada CanCOGeN team |
| EPI_ISL_937531 | Thai Red Cross Emerging Infectious Diseases Health Science Centre, Chulalongkorn Hospital, Faculty of Medicine, Chulalongkorn University | Thai Red Cross Emerging Infectious Diseases Center and Faculty of Medicine, Chulalongkorn University | Opass Putcharoen, Watsamon Jantarabenjakul, Pattama Torvorapanit, Gompol Suwanpimolkul, Leilani Paitoonpong, Sopon Iamsirithaworn, Sininat Petcharat, Yutthana Joyjinda, Weenassarin Ampoot, Apaporn Rodpan, Thiravat Hemachudha, Supaporn Wacharapluesadee |
| EPI_ISL_940209, EPI_ISL_940211, EPI_ISL_940212, EPI_ISL_940213, EPI_ISL_940215, EPI_ISL_940216, EPI_ISL_940217, EPI_ISL_940218, EPI_ISL_940219, EPI_ISL_940220, EPI_ISL_940221, EPI_ISL_940222, EPI_ISL_940223, EPI_ISL_940225, EPI_ISL_940226, EPI_ISL_940227, EPI_ISL_940228, EPI_ISL_940229, EPI_ISL_940230, EPI_ISL_940231, EPI_ISL_940232, EPI_ISL_940233, EPI_ISL_940235, EPI_ISL_940236, EPI_ISL_940237, EPI_ISL_940238, EPI_ISL_940239, EPI_ISL_940240, EPI_ISL_940507, EPI_ISL_940508, EPI_ISL_940509, EPI_ISL_940510, EPI_ISL_940511, EPI_ISL_940512, EPI_ISL_940513, EPI_ISL_940514, EPI_ISL_940515, EPI_ISL_940516, EPI_ISL_940517, EPI_ISL_940518, EPI_ISL_940519, EPI_ISL_940520, EPI_ISL_940523, EPI_ISL_940524, EPI_ISL_940525, EPI_ISL_940526, EPI_ISL_940527, EPI_ISL_940528 | | | |
| see above | Hôpital Bichat Claude Bernard, Laboratoire de Virologie | IAME UMR1137 Inserm, Université de Paris, Hôpital Bichat | Antoine Bridier-Nahmias, Amélie Recoing, Quentin Le Hingrat, Lena Daniel, Siham Hamri, Gilles Collin, Alexandre Storto, Mélanie Bertine, Charlotte Charpentier, Nadhira Houhou-Fidouh, Diane Descamps, Benoit Visseaux |
| EPI_ISL_940753 | University of Bari Biomedical Sciences and Human Oncology | University of Bari Biomedical Sciences and Human Oncology | Chironna M., Sallustio A., Loconsole D., Accogli M. |
| EPI_ISL_940848, EPI_ISL_940860, EPI_ISL_941272, EPI_ISL_941274, EPI_ISL_941298, EPI_ISL_941299 | Ohio Department of Health Laboratory | Ohio Department of Health Laboratory | Holmes, Jennifer; Eric Brandt, Keoni Omura, Glen McGillivary, Caitlin McDonnell, Jade Mowery, Stephanie Mccracken, Tyler Payne, Kirtana Ramadugu, Erica Leasure, Brent Lee, Kelsey Florek, Heather Blankenship, Quanta Brown, and Tammy Bannerman |
| EPI_ISL_943808 | Molecular Pathology, Pathology and Laboratory Medicine Institute, Cleveland Clinic, Ohio, USA | Molecular Pathology, Pathology and Laboratory Medicine Institute, Cleveland Clinic, Ohio, USA | Frank P. Esper, Yu-Wei Cheng, Thamali M. Adhikari, Zheng Jin Tu, Daniel H. Farkas, Gary Procop, Jennifer Ko, Timothy A. Chan, Jing Li, Brian P. Rubin |
| EPI_ISL_944634 | Centro de Investigaciones en Microbiología y Biotecnología-UR (CIMBIUR), Facultad de Ciencias Naturales, Universidad del Rosario, Bogotá, Colombia | Centro de Investigaciones en Microbiología y Biotecnología-UR (CIMBIUR), Facultad de Ciencias Naturales, Universidad del Rosario, Bogotá, Colombia Instituto Nacional de Salud, Bogotá, Colombia Icahn School of Medicine at Mount Sinai, New York, USA | Luz Helena Patiño, Marina Muñoz, Nathalia Ballesteros, Carolina Hernández, Carolina Flórez, Sergio Gomez, Adriana van de Guchte, Zenab Khan, Jayeeta Dutta, Hala Alejel Alshammary, Ana S. Gonzalez-Reiche, Matthew M. Hernandez, Emilia Mia Sordillo, Viviana Simon, Harm van Bakel, Alberto Paniz-Mondolfi, Juan David Ramírez |
| EPI_ISL_949244, EPI_ISL_949245, EPI_ISL_949246, EPI_ISL_949247 | Cliniques universitaires Saint-Luc | UCLouvain/IREC/MBLG | Jean Ruelle, Lysa Pinsmaye, Benoit Kabamba Mukadi |
| EPI_ISL_949248 | Cliniques Saint-Pierre Ottignies | UCLouvain/IREC/MBLG | Jean Ruelle, Lysa Pinsmaye, Benoit Kabamba Mukadi |
| EPI_ISL_949250, EPI_ISL_949251, | Cliniques universitaires Saint-Luc | UCLouvain/IREC/MBLG | Jean Ruelle, Lysa Pinsmaye, Benoit Kabamba Mukadi |

| | | | |
|---|---|---|---|
| EPI_ISL_949252 | | | |
| EPI_ISL_956400, EPI_ISL_956401 | Isolation - Virology Unit, Institut Pasteur du Cambodge; Sequencing - US National Institute of Allergy and Infectious Diseases Cambodia, US Naval Medical Research Unit -2, Cambodia National Institute for Public Health | Virology Unit, Institut Pasteur du Cambodge | Vireak Heang, Jennifer Bohl, Sophana Chea, Sreyngim Lay, Ly Sovann, Kraing Sidonn, Yi Sengdoeurn, Chin Savuth, Chau Darapheak, Veasna Duong, Jose A Garcia-Rivera, Jessica Manning, Erik A Karlsson |
| EPI_ISL_959378, EPI_ISL_959379, EPI_ISL_959380, EPI_ISL_959381, EPI_ISL_959382, EPI_ISL_959383, EPI_ISL_959385, EPI_ISL_959387, EPI_ISL_959414, EPI_ISL_959415, EPI_ISL_959417, EPI_ISL_959472, EPI_ISL_959473, EPI_ISL_959474 | | | |
| see above | Ohio Department of Health Laboratory | Ohio Department of Health Laboratory | Holmes, Jennifer; Eric Brandt, Keoni Omura, Glen McGillivary, Caitlin McDonnell, Jade Mowery, Stephanie Mccracken, Tyler Payne, Kirtana Ramadugu, Erica Leasure, Brent Lee, Kelsey Florek, Heather Blankenship, Quanta Brown, and Tammy Bannerman |
| EPI_ISL_960310 | NLZOH, Laboratory for Virology | NLZOH, Laboratory for Virology | Katarina Prosenc (Laboratory for Virology), Cesare Camma (IZSAM), Erik Alm (ECDC) |
| EPI_ISL_961670, EPI_ISL_961671 | Hôpital Georges L. Dumont | National Microbiology Laboratory (NML) | Anna Majer, Shari Tyson, Grace Seo, Philip Mabon, Elsie Grudeski, Rhiannon Huzarewich, Russell Mandes, Anneliese Landgraff, Jennifer Tanner, Natalie Knox, Morag Graham, Gary Van Domselaar, Richard Garceau, Guillaume Desnoyers, Nathalie Bastien, Yan Li, Timothy Booth, Darian Hole, Madison Chapel, Kirsten Biggar, CanCOGeN's metadata curation team, Public Health Agency of Canada CanCOGeN team |
| EPI_ISL_964878 | Institute of Virology Ulm University Medical Center Prof. Dr. Thomas Stamminger | Münch Lab / Ulm University Medical Center Kirchhoff Lab / Ulm University Medical Center Sparrer Lab / Ulm University Medical Center Blum Lab / LMU Munich | Thomas Stamminger, Helmut Blum, Stefan Krebs, Alexander Graf, Rüdiger Gross, Janis Müller, Carina Conzelmann, Jan Münch, Frank Kirchhoff, Konstantin Sparrer |
| EPI_ISL_965815, EPI_ISL_965816, EPI_ISL_965817, EPI_ISL_965818, EPI_ISL_965819, EPI_ISL_965820, EPI_ISL_965821, EPI_ISL_965822, EPI_ISL_965823, EPI_ISL_965824, EPI_ISL_965825, EPI_ISL_965827, EPI_ISL_966355, EPI_ISL_966356, EPI_ISL_966358, EPI_ISL_966359 | | | |
| see above | Kentucky State Public Health Lab | Kentucky State Public Health Lab | Stephanie Lunn, Karim George, Joshua Tobias, William Grooms, Vaneet Arora, Matthew Johnson, Rachel Zinner, Rhonda Lucas |
| EPI_ISL_976902, EPI_ISL_976903, EPI_ISL_976905 | Thai Red Cross Emerging Infectious Diseases Health Science Centre, Chulalongkorn Hospital, Faculty of Medicine, Chulalongkorn University | Thai Red Cross Emerging Infectious Diseases Center and Faculty of Medicine, Chulalongkorn University | Opass Putcharoen, Watsamon Jantarabenjakul, Pattama Torvorapanit, Gompol Suwanpimolkul, Leilani Paitoonpong, Sopon Iamsirithaworn, Sininat Petcharat, Yutthana Joyjinda, Weenassarin Ampoot, Apaporn Rodpan, Thiravat Hemachudha, Supaporn Wacharapluesadee |
| EPI_ISL_977308, EPI_ISL_977422, EPI_ISL_977459, EPI_ISL_977467 | University of Zambia, School of Veterinary Medicine | UNZAVET and PATH | Mulenga Mwenda-Chimfwembe, Ngonda Saasa, Daniel Bridges |
| EPI_ISL_978502, EPI_ISL_978506, EPI_ISL_978507, EPI_ISL_978508, EPI_ISL_978509, EPI_ISL_978513, EPI_ISL_978516, EPI_ISL_978518, EPI_ISL_978530 | Central Public Health Laboratory - LACEN -Bahia, Salvador, Brazil | Central Public Health Laboratory - LACEN -Bahia, Salvador, Brazil | Stephane Tosta, Luciana Oliveira, Vanessa Nardy,Patrícia Cajado,Marcela Gómez, Breno Dominguez, Jaqueline Gomes, Vagner Fonseca,Marta Giovanetti,Luiz Alcantara, Felicidade Pereira, Arabela Leal |
| EPI_ISL_979281, EPI_ISL_979282, EPI_ISL_979285, EPI_ISL_979286 | Cadham Provincial laboratory | National Microbiology Laboratory (NML) | Anna Majer, Shari Tyson, Grace Seo, Philip Mabon, Elsie Grudeski, Rhiannon Huzarewich, Russell Mandes, Anneliese Landgraff, Jennifer Tanner, Natalie Knox, Morag Graham, Gary Van Domselaar, Paul Van Caeseele, Jared Bullard, David Alexander, Kerry Dust, Nathalie Bastien, Yan Li, Timothy Booth, Darian Hole, Madison Chapel, Kirsten Biggar, CanCOGeN's metadata curation team, Public Health Agency of Canada CanCOGeN team |
