## Supplementary material for "Detection and characterization of the SARS-CoV-2 lineage B.1.526 in New York": Supp. Table 3 GISAID Acknowledgment part 2: gisaid_hcov-19_acknowledgement_table_2021_02_12_16-7.pdf

We gratefully acknowledge the following Authors from the Originating laboratories responsible for obtaining the specimens, as well as the Submitting laboratories where the genome data were generated and shared via GISAID, on which this research is based.

All Submitters of data may be contacted directly via www.gisaid.org

Authors are sorted alphabetically.

| Accession ID | Originating Laboratory | Submitting Laboratory | Authors |
|---|---|---|---|
| EPI_ISL_420144 | Department for Virology, Molecular Biology and Genome Research, R. G. Lugar Center for Public Health Research, National Center for Disease Control and Public Health (NCDC) of Georgia. | Department for Virology, Molecular Biology and Genome Research, R. G. Lugar Center for Public Health Research, National Center for Disease Control and Public Health (NCDC) of Georgia. | Gvantsa Chanturia, Ann Machablishvili, Nato Kotaria, Marine Murtskhvaladze, Lela Sabadze, Mari Gavashelidze, Ana Papkiauri, Meri Pantsulaia, Gvantsa Brachveli, Tata Imnadze, Tamar Jashiashvili, Tea Tevdoradze, Ketevan Sidamonidze, Ekaterine Khmaladze, Ekaterine Zhghenti, Roena Sukhiashvili, Mariam Zakalashvili, Lela Urushadze, Magda Dgebuadze, Giorgi Tomashvili, Davit Tsaguria, Ekaterine Zangaladze, Nino Berishvili, Adam Kotorashvili, Maia Alkhazashvili, Irma Burjanadze, Anna Kasradze, Khatuna Zakhashvili, Paata Imnadze, Amiran Gamkrelidze. |
| EPI_ISL_420769 | Respiratory Virus Unit, Microbiology Services Colindale, Public Health England | Respiratory Virus Unit, Microbiology Services Colindale, Public Health England | Monica Galiano, Shahjahan Miah, Angie Lackenby, Omolola Akinbami, Tiina Talts, Leena Bhaw, Richard Myers, Steven Platt, Kirstin Edwards, Jonathan Hubb, Joanna Ellis, Maria Zambon |
| EPI_ISL_421184, EPI_ISL_421190, EPI_ISL_421193, EPI_ISL_421197, EPI_ISL_421199, EPI_ISL_421200, EPI_ISL_421201, EPI_ISL_421204, EPI_ISL_421205, EPI_ISL_421206, EPI_ISL_421207, EPI_ISL_421209, EPI_ISL_421210, EPI_ISL_421211, EPI_ISL_421212, EPI_ISL_421213 | | | |
| see above | Department of Clinical Microbiology | GIGA Medical Genomics | Keith Durkin, Maria Artesi, Sébastien Bontems, Raphaël Boreux, Cécile Meex, Pierrette Melin, Marie-Pierre Hayette, Vincent Bours. |
| EPI_ISL_421573, EPI_ISL_421574, EPI_ISL_421575, EPI_ISL_421576 | Molecular Diagnostic Services | KRISP, KZN Research Innovation and Sequencing Platform | Giandhari J, Pillay S, Ngcapu S, Samsunder N, Lessells R, Chimukangara B, Deforche K, Tegally H, Wilkinson E, de Oliveira T |
| EPI_ISL_421660 | The Ohio State University | The Ohio State University-James Molecular Lab at Polaris | Huolin Tu, Matthew Avenarius, Preeti Panchioli, Sean Caruthers, Joan-Miquel Balada-Llasat, Jason Garee, Matt Hunt, Xiaokang Pan, Dan Jones |
| EPI_ISL_421675 | The Ohio State University Wexner Medical Center | The Ohio State University James Molecular lab | Huolin Tu, Sean Caruthers, Matthew Avenarius, Joan-Miquel Balada-Llasat, Matthew Hunt, Preeti Panchioli, Xiaokang Pen, Jason Garee, Pam Snyder, Dan Jones |
| EPI_ISL_421931, EPI_ISL_421932, EPI_ISL_421933, EPI_ISL_421934, EPI_ISL_421935, EPI_ISL_421936, EPI_ISL_421939, EPI_ISL_421941, EPI_ISL_421942, EPI_ISL_421943, EPI_ISL_421944, EPI_ISL_421945, EPI_ISL_421946, EPI_ISL_421947, EPI_ISL_421948, EPI_ISL_421949, EPI_ISL_421951, EPI_ISL_421952, EPI_ISL_421953, EPI_ISL_421956, EPI_ISL_421958, EPI_ISL_421959, EPI_ISL_421960, EPI_ISL_421961, EPI_ISL_421962, EPI_ISL_421963, EPI_ISL_421964, EPI_ISL_421965, EPI_ISL_421966, EPI_ISL_421967, EPI_ISL_421968, EPI_ISL_421969, EPI_ISL_421970, EPI_ISL_421971, EPI_ISL_421972, EPI_ISL_421973, EPI_ISL_421975, EPI_ISL_421977, EPI_ISL_421978, EPI_ISL_421979, EPI_ISL_421980, EPI_ISL_421981, EPI_ISL_421982, EPI_ISL_421983, EPI_ISL_421984, EPI_ISL_421985, EPI_ISL_421987, EPI_ISL_421988, EPI_ISL_421990, EPI_ISL_421991, EPI_ISL_421992, EPI_ISL_421993, EPI_ISL_421994, EPI_ISL_421995, EPI_ISL_421996, EPI_ISL_421997, EPI_ISL_421999, EPI_ISL_422003, EPI_ISL_422004, EPI_ISL_422005, EPI_ISL_422007 | | | |
| see above | Respiratory Virus Unit, Microbiology Services Colindale, Public Health England | Respiratory Virus Unit, Microbiology Services Colindale, Public Health England | Monica Galiano, Shahjahan Miah, Angie Lackenby, Omolola Akinbami, Tiina Talts, Leena Bhaw, Richard Myers, Steven Platt, Kirstin Edwards, Jonathan Hubb, Joanna Ellis, Maria Zambon |
| EPI_ISL_422105 | Wales Specialist Virology Centre | Public Health Wales Microbiology Cardiff | Catherine Moore, Johnathan Evans, Malorie Perry, Simon Cottrell, Alec Birchley, Alexander Adams, Amy Gaskin, Bree Gatica-Wilcox, Jason Coombes, Lauren Gilbert, Lee Graham, Nicole Pacchiarini, Sara Kumziene-Summerhayes, Sarah Taylor, Sophie Jones, Sara Rey, Matthew Bull, Joanne Watkins, Sally Corden, Tom Connor |
| EPI_ISL_422401 | NMIMR, Department of Virology | WACCBIP, University of Ghana | Joyce M. Ngoi, Bright Adu, Collins M. Morang'a, Selassie Kumordjie, Miriam Eshun, Linda Boatemaa, Vanessa Magnussen, Erasmus Kotey, Fred Tei-Maya, Dominic S. Y. Amuzu, Peter Quashie, Augustina Arjarquah, Ivy Asante, Evelyn Bonney, George B. Kyei, Kofi Bonney, Abraham Kwabena Anang, Gordon A. Awandare, William Ampofo |
| EPI_ISL_422431, EPI_ISL_422432, EPI_ISL_422433, EPI_ISL_422434, EPI_ISL_422435 | National Public Health Laboratory, National Centre for Infectious Diseases | National Public Health Laboratory, National Centre for Infectious Diseases | Mak TM, Octavia S, Cui L, Lin RTP |
| EPI_ISL_422453 | Gundersen Molecular Diagnostics Laboratory | Kabara Cancer Research Institute | Craig S. Richmond & Paraic A. Kenny |
| EPI_ISL_422567, EPI_ISL_422587, EPI_ISL_422588, EPI_ISL_422589, EPI_ISL_422590, EPI_ISL_422591, EPI_ISL_422592, EPI_ISL_422593, EPI_ISL_422594, EPI_ISL_422595, EPI_ISL_422596, EPI_ISL_422597, EPI_ISL_422598, EPI_ISL_422599, EPI_ISL_422600, EPI_ISL_422601, EPI_ISL_422602, EPI_ISL_422603, EPI_ISL_422604, EPI_ISL_422605, EPI_ISL_422606, EPI_ISL_422607, EPI_ISL_422608, EPI_ISL_422609, EPI_ISL_422610, EPI_ISL_422611, EPI_ISL_422612, EPI_ISL_422613, EPI_ISL_422614, EPI_ISL_422615, EPI_ISL_422628, EPI_ISL_422629, EPI_ISL_422630, EPI_ISL_422631, EPI_ISL_422633, EPI_ISL_422634, EPI_ISL_422635 | | | |
| see above | Dutch COVID-19 response team | Erasmus Medical Center | Bas Oude Munnink, David Nieuwenhuijse, Reina Sikkema, Claudia Schapendonk, Irina Chestakova, Anne van der Linden, Theo Bestebroer, Stefan van Nieuwkoop, Mark Pronk, Pascal Lexmond, Corien Swaan, Manon Haverkate, Madelief Mollers, Mart Stein, Sandra Kengne Kamga Mobou, Jeroen van Kampen, Jolanda Voermans, Aura Timen, Corine GeurtsvanKessel, Annemiek van der Eijk, Richard Molenkamp, Marion Koopmans, on behalf of the Dutch national COVID-19 response team. |
| EPI_ISL_423427, EPI_ISL_423429, EPI_ISL_423430, EPI_ISL_423448, EPI_ISL_423454, EPI_ISL_423469, EPI_ISL_423472, EPI_ISL_423473, EPI_ISL_423474, EPI_ISL_423479, EPI_ISL_423480, EPI_ISL_423481, EPI_ISL_423482, EPI_ISL_423483, EPI_ISL_423484, EPI_ISL_423485, EPI_ISL_423486, EPI_ISL_423487, EPI_ISL_423488, EPI_ISL_423489, EPI_ISL_423490, EPI_ISL_423491, EPI_ISL_423492, EPI_ISL_423493, EPI_ISL_423494, EPI_ISL_423497, EPI_ISL_423498, EPI_ISL_423499, EPI_ISL_423500, EPI_ISL_423501, EPI_ISL_423502, EPI_ISL_423503, EPI_ISL_423504, EPI_ISL_423508, EPI_ISL_423509, EPI_ISL_423510, EPI_ISL_423511, EPI_ISL_423512, EPI_ISL_423534, EPI_ISL_423535, EPI_ISL_423536, EPI_ISL_423537, EPI_ISL_423538, EPI_ISL_423539, EPI_ISL_423540, EPI_ISL_423541, EPI_ISL_423545, EPI_ISL_423546, EPI_ISL_423547, EPI_ISL_423548, EPI_ISL_423549, EPI_ISL_423550, EPI_ISL_423551, EPI_ISL_423552, EPI_ISL_423555, EPI_ISL_423557, EPI_ISL_423558, EPI_ISL_423559, EPI_ISL_423560, EPI_ISL_423561, EPI_ISL_423562, EPI_ISL_423563, EPI_ISL_423564, EPI_ISL_423565, EPI_ISL_423566, EPI_ISL_423567, EPI_ISL_423568, EPI_ISL_423569, EPI_ISL_423570, EPI_ISL_423572, EPI_ISL_423573, EPI_ISL_423574, EPI_ISL_423575, EPI_ISL_423576, EPI_ISL_423577, EPI_ISL_423578, EPI_ISL_423579, EPI_ISL_423582, EPI_ISL_423583, EPI_ISL_423584 | | | |
| see above | Respiratory Virus Unit, Microbiology Services Colindale, Public Health England | Respiratory Virus Unit, Microbiology Services Colindale, Public Health England | Monica Galiano, Shahjahan Miah, Angie Lackenby, Omolola Akinbami, Tiina Talts, Leena Bhaw, Richard Myers, Steven Platt, Kirstin Edwards, Jonathan Hubb, Joanna Ellis, Maria Zambon |
| EPI_ISL_424929, EPI_ISL_424930, EPI_ISL_424931, EPI_ISL_424932, EPI_ISL_424933, EPI_ISL_424934, EPI_ISL_424935, EPI_ISL_424936, EPI_ISL_424937, EPI_ISL_424938, EPI_ISL_424939, EPI_ISL_424940, EPI_ISL_424941, EPI_ISL_424942, EPI_ISL_424943, EPI_ISL_424944, EPI_ISL_424945, EPI_ISL_424946, EPI_ISL_424947, EPI_ISL_424948, EPI_ISL_424949, EPI_ISL_424950, EPI_ISL_424951, EPI_ISL_424952, EPI_ISL_424953, EPI_ISL_424954, EPI_ISL_424955, EPI_ISL_424956, EPI_ISL_424957, EPI_ISL_424958, EPI_ISL_424959, EPI_ISL_424960, EPI_ISL_424961, EPI_ISL_424962, EPI_ISL_424963, EPI_ISL_424964, EPI_ISL_424965, EPI_ISL_424966, EPI_ISL_424967, EPI_ISL_424968 | | | |
| see above | NYU Langone Health | Departments of Pathology and Medicine, New York University School of Medicine | Maria Aguero-Rosenfeld, Brendan Belovarac, Margaret Black, Ludovic Boytard, John Cadley, Paolo Cotzia, John Chen, Dacia Dimartino, Xiaojun Feng, Tatyana Gindin, Adriana Heguy, Megan Hogan, Emily Huang, George Jour, Andrew Lytle, Christian Marier, Matthew T. Maurano, Mark J. Mulligan, Peter Meyn, Iman Osman, Jared Pinnell, Sitharam Ramaswami, Amy Rapkiewicz, Marie Samanovic-Golden, Antonio Serrano, Guomiao Shen, Matija Snuderl, Theodore Vougiouklakis, Nick Vulpescu, Gael Westby, Paul Zappile, Yutong Zhang |
| EPI_ISL_425146, EPI_ISL_425148, EPI_ISL_425153, EPI_ISL_425155, EPI_ISL_425158, EPI_ISL_425164, EPI_ISL_425167, EPI_ISL_425168, EPI_ISL_425170, EPI_ISL_425172 | University of Wisconsin-Madison AIDS Vaccine Research Laboratories | University of Wisconsin-Madison AIDS Vaccine Research Laboratories | Gage Moreno, Katarina Braun, et al. AIDS Vaccine Research Laboratories |
| EPI_ISL_425239, EPI_ISL_425240, EPI_ISL_425241, EPI_ISL_425242, EPI_ISL_425243, EPI_ISL_425244, EPI_ISL_425245, EPI_ISL_425246, EPI_ISL_425249, EPI_ISL_425251, EPI_ISL_425252, EPI_ISL_425255, EPI_ISL_425256, EPI_ISL_425262, EPI_ISL_425263, EPI_ISL_425264, EPI_ISL_425265, EPI_ISL_425266, EPI_ISL_425267, EPI_ISL_425268, EPI_ISL_425269, EPI_ISL_425270, EPI_ISL_425271, EPI_ISL_425272, EPI_ISL_425275, EPI_ISL_425276, EPI_ISL_425278, EPI_ISL_425424, EPI_ISL_425426, EPI_ISL_425427, EPI_ISL_425428, EPI_ISL_425432, EPI_ISL_425434, EPI_ISL_425435, EPI_ISL_425436, EPI_ISL_425437, EPI_ISL_425438, EPI_ISL_425441, EPI_ISL_425442, EPI_ISL_425443, EPI_ISL_425444, EPI_ISL_425445, EPI_ISL_425446, EPI_ISL_425448, EPI_ISL_425449, EPI_ISL_425450, EPI_ISL_425451, EPI_ISL_425452, EPI_ISL_425453, EPI_ISL_425454, EPI_ISL_425455, EPI_ISL_425456, EPI_ISL_425457, EPI_ISL_425458, EPI_ISL_425459, EPI_ISL_425460 | | | |
| see above | Department of Pathology, University of Cambridge | COVID-19 Genomics UK (COG-UK) Consortium | Luke W Meredith, M. Estee Torok , Myra Hosmillo, William L. Hamilton, Martin D. Curran, Theresa Feltwell, Anna Yakovleva, Charlotte J. Houldcroft, Aminu S. Jahun, Sarah L. Caddy, Ian Goodfellow |
| EPI_ISL_426007, EPI_ISL_426008, EPI_ISL_426009, EPI_ISL_426010, EPI_ISL_426011, EPI_ISL_426012, EPI_ISL_426013, EPI_ISL_426014, EPI_ISL_426015, EPI_ISL_426016, EPI_ISL_426017, EPI_ISL_426018, EPI_ISL_426019, EPI_ISL_426020, EPI_ISL_426021, EPI_ISL_426022, EPI_ISL_426023, EPI_ISL_426024 | | | |
| see above | Virology Department, Royal Infirmary of Edinburgh, NHS Lothian / School of Biological Sciences, University of | COVID-19 Genomics UK (COG-UK) Consortium | McHugh M, Dewar R, Rooke S, Gallagher M, Balcaza C, O'Toole A, Hill V, McCrone JT, Colquhoun R, Yu X, Jackson B, Scher E, Rambaut A, Williams TC, Templeton K |

| | | | |
|---|---|---|---|
| | Edinburgh / Institute of Genetics and Molecular Medicine, University of Edinburgh | | |
| EPI_ISL_426134, EPI_ISL_426136, EPI_ISL_426137 | UW Virology Lab | UW Virology Lab | Pavitra Roychoudhury, Hong Xie, Keith Jerome, Alexander Greninger |
| EPI_ISL_426464, EPI_ISL_426465, EPI_ISL_426466, EPI_ISL_426467, EPI_ISL_426468, EPI_ISL_426470 | Virginia DCLS | Virginia DCLS | Virginia DCLS |
| EPI_ISL_426556, EPI_ISL_426557 | TGen North | TGen North | Jolene Bowers, Megan Folkerts, Darrin Lemmer, Dave Engelthaler |
| EPI_ISL_426561, EPI_ISL_426562, EPI_ISL_426563, EPI_ISL_426564, EPI_ISL_426565, EPI_ISL_426566, EPI_ISL_426567 | AZ SPHL, Arizona Department of Health Services | TGen North | Jolene Bowers, Megan Folkerts, Darrin Lemmer, Dave Engelthaler |
| EPI_ISL_426617, EPI_ISL_426618, EPI_ISL_426619, EPI_ISL_426620, EPI_ISL_426621, EPI_ISL_426622, EPI_ISL_426623, EPI_ISL_426624, EPI_ISL_426625, EPI_ISL_426626 | NYU Langone Health | Departments of Pathology and Medicine, New York University School of Medicine | Maria Aguero-Rosenfeld, Brendan Belovarac, Margaret Black, Ludovic Boytard, John Cadley, Paolo Cotzia, John Chen, Dacia Dimartino, Xiaojun Feng, Tatyana Gindin, Emily Guzman, Adriana Heguy, Megan Hogan, Emily Huang, George Jour, Andrew Lytle, Christian Marier, Matthew T. Maurano, Mark J. Mulligan, Peter Meyn, Iman Osman, Jared Pinnell, Vanessa Raabe, Sitharam Ramaswami, Amy Rapkiewicz, Marie Samanovic-Golden, Antonio Serrano, Guomiao Shen, Matija Snuderl, Theodore Vougiouklakis, Nick Vulpescu, Gael Westby, Paul Zappile, Yutong Zhang |
| EPI_ISL_426887, EPI_ISL_426888 | Motol University Hospital | Institute of Applied Biotechnologies a.s. | Petr Brož, Jan Geryk, Petr Klempt, Martin Kašný, Adam Novotný, Kateina Kvapilová, Pavel Devínek, Petr Kvapil, Milan Macek |
| EPI_ISL_427025, EPI_ISL_427027, EPI_ISL_427034, EPI_ISL_427053 | Victorian Infectious Diseases Reference Laboratory (VIDRL) | Microbiological Diagnostic Unit Public Health Laboratory and Victorian Infectious Diseases Reference Laboratory, Doherty Institute | Caly L., Seemann T., Sait, M., Schultz M., Druce J., Sherry, N. |
| EPI_ISL_427063, EPI_ISL_427064, EPI_ISL_427065, EPI_ISL_427066, EPI_ISL_427067, EPI_ISL_427068, EPI_ISL_427069, EPI_ISL_427070, EPI_ISL_427071, EPI_ISL_427072, EPI_ISL_427073, EPI_ISL_427074, EPI_ISL_427075, EPI_ISL_427076, EPI_ISL_427077, EPI_ISL_427078 | | | |
| see above | Microbiological Diagnostic Unit Public Health Laboratory | Microbiological Diagnostic Unit Public Health Laboratory | Seemann T., Schultz M., Sait, M., Sherry, N. |
| EPI_ISL_427084, EPI_ISL_427160 | Victorian Infectious Diseases Reference Laboratory (VIDRL) | Microbiological Diagnostic Unit Public Health Laboratory and Victorian Infectious Diseases Reference Laboratory, Doherty Institute | Caly L., Seemann T., Sait, M., Schultz M., Druce J., Sherry, N. |
| EPI_ISL_427163, EPI_ISL_427164, EPI_ISL_427173, EPI_ISL_427222, EPI_ISL_427223, EPI_ISL_427238, EPI_ISL_427239, EPI_ISL_427240, EPI_ISL_427241, EPI_ISL_427242, EPI_ISL_427243, EPI_ISL_427244, EPI_ISL_427245, EPI_ISL_427246, EPI_ISL_427247, EPI_ISL_427248, EPI_ISL_427249, EPI_ISL_427250, EPI_ISL_427251, EPI_ISL_427252, EPI_ISL_427253, EPI_ISL_427254, EPI_ISL_427255, EPI_ISL_427256, EPI_ISL_427257, EPI_ISL_427258, EPI_ISL_427259, EPI_ISL_427260, EPI_ISL_427261, EPI_ISL_427263, EPI_ISL_427265, EPI_ISL_427269, EPI_ISL_427270 | | | |
| see above | UW Virology Lab | UW Virology Lab | Pavitra Roychoudhury, Hong Xie, Keith Jerome, Alexander Greninger |
| EPI_ISL_427288 | The Ohio State University | The Ohio State University-James Molecular Lab at Polaris | Huolin Tu, Preeti Panchioli, Jason Garee, Matthew Hunt, Joan-Miquel Balada-Llasat, Erica Vincent, Weiqiang Zhao, Dan Jones |
| EPI_ISL_427289 | The Ohio State University | The Ohio State University-James Molecular Lab at Polaris | Huolin Tu, Joan-Miquel Balada-Llasat, Jason Garee, Matthew Hunt, Preeti Pancholi, Erica Vincent, Xiaokang Zhao, Dan Jones |
| EPI_ISL_427290 | The Ohio State University | The Ohio State University-James Molecular Lab at Polaris | Huolin Tu, Jason Garee, Matthew Hunt, Joan-Miquel Balada-Llasat, Preeti Pancholi, Erica Vincent, Rongqin Ren, Dan Jones |
| EPI_ISL_427291 | The Ohio State University | The Ohio State University-James Molecular Lab at Polaris | Huolin Tu, Matthew Hunt, Preeti Pancholi, Jason Garee, Joan-Miquel Balada-Llasat, Erica Vincent, Weiqiang Zhao, Dan Jones |
| EPI_ISL_427340, EPI_ISL_427342, EPI_ISL_427343, EPI_ISL_427347 | Department of Clinical Microbiology | GIGA Medical Genomics | Keith Durkin, Maria Artesi, Sébastien Bontems, Raphaël Boreux, Cécile Meex, Pierrette Melin, Marie-Pierre Hayette, Vincent Bours. |
| EPI_ISL_427469, EPI_ISL_427470, EPI_ISL_427471, EPI_ISL_427472, EPI_ISL_427473, EPI_ISL_427474, EPI_ISL_427475, EPI_ISL_427476, EPI_ISL_427477, EPI_ISL_427478, EPI_ISL_427479, EPI_ISL_427480, EPI_ISL_427481, EPI_ISL_427482, EPI_ISL_427483, EPI_ISL_427484, EPI_ISL_427485, EPI_ISL_427486, EPI_ISL_427487, EPI_ISL_427488, EPI_ISL_427489, EPI_ISL_427490, EPI_ISL_427491, EPI_ISL_427492, EPI_ISL_427493, EPI_ISL_427494, EPI_ISL_427495, EPI_ISL_427496, EPI_ISL_427497, EPI_ISL_427498, EPI_ISL_427499, EPI_ISL_427500, EPI_ISL_427501, EPI_ISL_427524, EPI_ISL_427525, EPI_ISL_427627 | | | |
| see above | NYU Langone Health | Departments of Pathology and Medicine, New York University School of Medicine | Maria Aguero-Rosenfeld, Brendan Belovarac, Margaret Black, Ludovic Boytard, John Cadley, Paolo Cotzia, John Chen, Dacia Dimartino, Xiaojun Feng, Tatyana Gindin, Emily Guzman, Adriana Heguy, Megan Hogan, Emily Huang, George Jour, Andrew Lytle, Christian Marier, Matthew T. Maurano, Mark J. Mulligan, Peter Meyn, Iman Osman, Jared Pinnell, Vanessa Raabe, Sitharam Ramaswami, Amy Rapkiewicz, Marie Samanovic-Golden, Antonio Serrano, Guomiao Shen, Matija Snuderl, Theodore Vougiouklakis, Nick Vulpescu, Gael Westby, Paul Zappile, Yutong Zhang |
| EPI_ISL_428380, EPI_ISL_428387, EPI_ISL_428391 | Yale COVID-19 Biorepository | Grubaugh Lab - Yale School of Public Health | Joseph Fauver, Tara Alpert, Anderson Brito, Anne Wyllie, Chantal Vogels, Mary Petrone, Chaney Kalinich, Isabel Ott, Arnau Casanovas, Catherine Muenker, Adam Moore, Alice Lu, Maria Tokuyama, Patrick Wong, Peiwen Lu, Saad Omer, Richard Martinello, Allison Nelson, Shelli Farhadian, Akiko Iwasaki, Charlese Dela Cruz, Albert Ko, Nathan Grubaugh |
| EPI_ISL_428673 | Centre for Dengue Research | Centre for Dengue Research | Chandima Jeewandara, Dinuka Ariyaratne, Laksiri Gomes, Deshni Jayathilaka, Diyanath Ranasinghe, Ananda Wijewickrama, Eranga Narangoda, Damayanthi Idampitiya, Neelika Malavige |
| EPI_ISL_428728 | University of Wisconsin-Madison AIDS Vaccine Research Laboratories | University of Wisconsin-Madison AIDS Vaccine Research Laboratories | Gage Moreno, Katarina Braun, et al. AIDS Vaccine Research Laboratories |
| EPI_ISL_428878 | State Research Center of Virology and Biotechnology VECTOR, Department of Collection of Microorganisms | State Research Center of Virology and Biotechnology VECTOR, Department of Collection of Microorganisms | Sergey A. Bodnev, Oleg V. Pyankov, Tatyana V. Tregubchak, Alexander N. Shvalov, Elena V. Gavrilova, Rinat A. Maksyutov |
| EPI_ISL_428907, EPI_ISL_428910, EPI_ISL_428911 | State Research Center of Virology and Biotechnology VECTOR, Department of Collection of Microorganisms | State Research Center of Virology and Biotechnology VECTOR, Department of Collection of Microorganisms | Oleg V. Pyankov, Sergey A. Bodnev, Tatyana V. Tregubchak, Alexander N. Shvalov, Elena V. Gavrilova, Rinat A. Maksyutov |
| EPI_ISL_428939, EPI_ISL_428943, EPI_ISL_428944, EPI_ISL_428950, EPI_ISL_428952, EPI_ISL_428955, EPI_ISL_428956, EPI_ISL_428959 | Laboratoire National de Sante, Microbiology, Virology | Laboratoire National de Sante, Microbiology, Epidemiology and Microbial Genomics | Anke Wienecke-Baldacchino, Ardashel Latsuzbaia, Jessica Tapp, Catherine Ragimbeau, Guillaume Fournier, Tamir Abdelrahman, Trung Nguyen Nguyen, Joel Mossong |
| EPI_ISL_429020, EPI_ISL_429023, EPI_ISL_429025, EPI_ISL_429026, EPI_ISL_429055, EPI_ISL_429060, EPI_ISL_429067, EPI_ISL_429072 | UCSF Clinical Microbiology Laboratory | Chan-Zuckerberg Biohub | CZB Cliahub Consortium |
| EPI_ISL_429598, EPI_ISL_429600, EPI_ISL_429605, EPI_ISL_429606, EPI_ISL_429607, EPI_ISL_429608, EPI_ISL_429609, EPI_ISL_429611, EPI_ISL_429648 | UW Virology Lab | UW Virology Lab | Pavitra Roychoudhury, Hong Xie, Keith Jerome, Alexander Greninger |
| EPI_ISL_429717, EPI_ISL_429721, EPI_ISL_429722, EPI_ISL_429731, EPI_ISL_429786, EPI_ISL_429789, EPI_ISL_429794 | Laboratoire National de Sante, Microbiology, Virology | Laboratoire National de Sante, Microbiology, Epidemiology and Microbial Genomics | Anke Wienecke-Baldacchino, Ardashel Latsuzbaia, Jessica Tapp, Catherine Ragimbeau, Guillaume Fournier, Tamir Abdelrahman, Trung Nguyen Nguyen, Joel Mossong |
| EPI_ISL_429989 | Virginia DCLS | Virginia DCLS | Virginia DCLS |
| EPI_ISL_430040, EPI_ISL_430045, EPI_ISL_430050 | Utah Public Health Laboratory | Utah Public Health Laboratory | Erin Young, Kelly Oakeson |

| | | | |
|---|---|---|---|
| EPI_ISL_430121, EPI_ISL_430125, EPI_ISL_430130, EPI_ISL_430147, EPI_ISL_430149, EPI_ISL_430153 | Seattle Flu Study | Seattle Flu Study | Chu et al |
| EPI_ISL_430351, EPI_ISL_430352, EPI_ISL_430353 | NYU Langone Health | Departments of Pathology and Medicine, New York University School of Medicine | Maria Aguero-Rosenfeld, Brendan Belovarac, Margaret Black, Ludovic Boytard, John Cadley, Paolo Cotzia, John Chen, Dacia Dimartino, Xiaojun Feng, Tatyana Gindin, Emily Guzman, Adriana Heguy, Megan Hogan, Emily Huang, George Jour, Lawrence H. Lin, Raven Luther, Andrew Lytle, Christian Marier, Matthew T. Maurano, Mark J. Mulligan, Peter Meyn, Raquel Ordonez Ciriza, Iman Osman, Jared Pinnell, Vanessa Raabe, Sitharam Ramaswami, Amy Rapkiewicz, Andre M. Ribeiro-dos-Santos, Marie Samanovic-Golden, Antonio Serrano, Guomiao Shen, Matija Snuderl, Theodore Vougiouklakis, Nick Vulpescu, Gael Westby, Paul Zappile, Yutong Zhang |
| EPI_ISL_430473, EPI_ISL_430474, EPI_ISL_430475, EPI_ISL_430476, EPI_ISL_430477, EPI_ISL_430478, EPI_ISL_430479, EPI_ISL_430480, EPI_ISL_430481, EPI_ISL_430482, EPI_ISL_430483, EPI_ISL_430484, EPI_ISL_430485, EPI_ISL_430486, EPI_ISL_430487, EPI_ISL_430488, EPI_ISL_430489, EPI_ISL_430490, EPI_ISL_430491, EPI_ISL_430492, EPI_ISL_430493, EPI_ISL_430498, EPI_ISL_430499, EPI_ISL_430500, EPI_ISL_430501, EPI_ISL_430502, EPI_ISL_430503, EPI_ISL_430504, EPI_ISL_430505, EPI_ISL_430506, EPI_ISL_430507, EPI_ISL_430508, EPI_ISL_430509, EPI_ISL_430513, EPI_ISL_430514, EPI_ISL_430515, EPI_ISL_430516, EPI_ISL_430517, EPI_ISL_430518, EPI_ISL_430519, EPI_ISL_430522, EPI_ISL_430523, EPI_ISL_430527, EPI_ISL_430544, EPI_ISL_430547, EPI_ISL_430548, EPI_ISL_430551, EPI_ISL_430553, EPI_ISL_430555, EPI_ISL_430556, EPI_ISL_430559, EPI_ISL_430560 | | | |
| see above | Victorian Infectious Diseases Reference Laboratory (VIDRL) | Microbiological Diagnostic Unit Public Health Laboratory and Victorian Infectious Diseases Reference Laboratory, The Peter Doherty Institute for Infection and Immunity | Caly L., Seemann T., Sait, M., Schultz M., Druce J., Sherry, N. |
| EPI_ISL_430801 | Laboratorio de Virología del Hospital de Niños Dr. Ricardo Gutierrez | Área de Secuenciación del Laboratorio de Virología del Hospital de Niños Dr. Ricardo Gutierrez on behalf of 'Proyecto Argentino Interinstitucional de genomica de SARS-CoV-2' (PAIS Consortium) | Nabaes Jodar, MS; Goya, S; Natale, MI; Lusso, S; Gravis, E; Mistchenko, AS; Valinotto, LE; Viegas, M. |
| EPI_ISL_430802 | Departamento de Biología y genética molecular, IACA Laboratorios. | Área de Secuenciación del Laboratorio de Virología del Hospital de Niños Dr. Ricardo Gutierrez on behalf of 'Proyecto Argentino Interinstitucional de genomica de SARS-CoV-2' (PAIS Consortium) | Nabaes Jodar, MS; Goya, S; Natale, MI; Lusso, S; Tittarelli, E; Suárez, A; Masciovecchio MV; Streitenberger ER; Mistchenko, AS; Valinotto, LE; Viegas, M. |
| EPI_ISL_430803, EPI_ISL_430804 | Laboratorio de Virología del Hospital de Niños Dr. Ricardo Gutierrez | Área de Secuenciación del Laboratorio de Virología del Hospital de Niños Dr. Ricardo Gutierrez on behalf of 'Proyecto Argentino Interinstitucional de genomica de SARS-CoV-2' (PAIS Consortium) | Nabaes Jodar, MS; Goya, S; Natale, MI; Lusso, S; Gravis, E; Mistchenko, AS; Valinotto, LE; Viegas, M. |
| EPI_ISL_430805, EPI_ISL_430806 | Departamento de Biología y genética molecular, IACA Laboratorios. | Área de Secuenciación del Laboratorio de Virología del Hospital de Niños Dr. Ricardo Gutierrez on behalf of 'Proyecto Argentino Interinstitucional de genomica de SARS-CoV-2' (PAIS Consortium) | Nabaes Jodar, MS; Goya, S; Natale, MI; Lusso, S; Tittarelli, E; Suárez, A; Masciovecchio MV; Streitenberger ER; Mistchenko, AS; Valinotto, LE; Viegas, M. |
| EPI_ISL_430846 | General Intensive Care Unit, Raymond Poincaré Hospital (AP-HP), Lab Inflammation & Infection, U1173 University Paris Saclay-UVSQ/INSERM, Garches, France. | Institut Pasteur, Laboratory for Urgent Response to biological Threats | Annane Djillali, Vanhomwegen Jessica, Caro Valérie, Manuguerra Jean-Claude |
| EPI_ISL_430868, EPI_ISL_430869, EPI_ISL_430873, EPI_ISL_430874, EPI_ISL_430876, EPI_ISL_430899, EPI_ISL_430933, EPI_ISL_430934, EPI_ISL_430935, EPI_ISL_430936, EPI_ISL_430937, EPI_ISL_430938, EPI_ISL_430939, EPI_ISL_430941, EPI_ISL_430942, EPI_ISL_430951, EPI_ISL_430953, EPI_ISL_430954, EPI_ISL_430955, EPI_ISL_430956, EPI_ISL_430957, EPI_ISL_430958, EPI_ISL_430960, EPI_ISL_430961, EPI_ISL_430962, EPI_ISL_430963, EPI_ISL_430965, EPI_ISL_430968, EPI_ISL_430970, EPI_ISL_430972, EPI_ISL_430973 | | | |
| see above | UW Virology Lab | UW Virology Lab | Pavitra Roychoudhury, Hong Xie, Keith Jerome, Alexander Greninger |
| EPI_ISL_431902, EPI_ISL_431903, EPI_ISL_431905, EPI_ISL_431908, EPI_ISL_431910, EPI_ISL_431912, EPI_ISL_431916, EPI_ISL_431917, EPI_ISL_431922, EPI_ISL_431923, EPI_ISL_431931, EPI_ISL_431936, EPI_ISL_431937, EPI_ISL_431939, EPI_ISL_431940, EPI_ISL_431941, EPI_ISL_431944, EPI_ISL_431954, EPI_ISL_431956, EPI_ISL_431957, EPI_ISL_431958, EPI_ISL_431959, EPI_ISL_431962, EPI_ISL_431972, EPI_ISL_431975, EPI_ISL_431976, EPI_ISL_431985, EPI_ISL_431989, EPI_ISL_431990, EPI_ISL_431991, EPI_ISL_431992, EPI_ISL_431994, EPI_ISL_432001, EPI_ISL_432003, EPI_ISL_432006, EPI_ISL_432007, EPI_ISL_432008, EPI_ISL_432011, EPI_ISL_432013, EPI_ISL_432014, EPI_ISL_432016, EPI_ISL_432022, EPI_ISL_432024, EPI_ISL_432028, EPI_ISL_432032, EPI_ISL_432033, EPI_ISL_432035, EPI_ISL_432036, EPI_ISL_432039, EPI_ISL_432040, EPI_ISL_432041, EPI_ISL_432045, EPI_ISL_432048, EPI_ISL_432049, EPI_ISL_432052, EPI_ISL_432053, EPI_ISL_432054, EPI_ISL_432057, EPI_ISL_432065, EPI_ISL_432066, EPI_ISL_432070, EPI_ISL_432072, EPI_ISL_432075, EPI_ISL_432078, EPI_ISL_432080, EPI_ISL_432081, EPI_ISL_432082, EPI_ISL_432086, EPI_ISL_432092, EPI_ISL_432093, EPI_ISL_432096, EPI_ISL_432099, EPI_ISL_432106, EPI_ISL_432108, EPI_ISL_432119, EPI_ISL_432131, EPI_ISL_432134, EPI_ISL_432136, EPI_ISL_432142, EPI_ISL_432143, EPI_ISL_432158, EPI_ISL_432173, EPI_ISL_432176, EPI_ISL_432185, EPI_ISL_432192, EPI_ISL_432199, EPI_ISL_432200, EPI_ISL_432206, EPI_ISL_432208, EPI_ISL_432210, EPI_ISL_432212, EPI_ISL_432214, EPI_ISL_432217, EPI_ISL_432218, EPI_ISL_432221, EPI_ISL_432223, EPI_ISL_432224, EPI_ISL_432232, EPI_ISL_432233, EPI_ISL_432236, EPI_ISL_432238, EPI_ISL_432239, EPI_ISL_432240, EPI_ISL_432242, EPI_ISL_432245, EPI_ISL_432254, EPI_ISL_432256, EPI_ISL_432258, EPI_ISL_432260, EPI_ISL_432270, EPI_ISL_432277, EPI_ISL_432278, EPI_ISL_432282, EPI_ISL_432285, EPI_ISL_432287, EPI_ISL_432288, EPI_ISL_432289, EPI_ISL_432298, EPI_ISL_432299, EPI_ISL_432302, EPI_ISL_432303, EPI_ISL_432306, EPI_ISL_432307, EPI_ISL_432310, EPI_ISL_432312, EPI_ISL_432314, EPI_ISL_432316, EPI_ISL_432322, EPI_ISL_432325, EPI_ISL_432327, EPI_ISL_432328, EPI_ISL_432329, EPI_ISL_432335, EPI_ISL_432340, EPI_ISL_432346, EPI_ISL_432349, EPI_ISL_432350, EPI_ISL_432351, EPI_ISL_432355, EPI_ISL_432356, EPI_ISL_432362, EPI_ISL_432365, EPI_ISL_432366, EPI_ISL_432370, EPI_ISL_432371, EPI_ISL_432374, EPI_ISL_432380, EPI_ISL_432381, EPI_ISL_432385, EPI_ISL_432389, EPI_ISL_432394, EPI_ISL_432407, EPI_ISL_432408, EPI_ISL_432412, EPI_ISL_432413, EPI_ISL_432416, EPI_ISL_432417, EPI_ISL_432425, EPI_ISL_432426, EPI_ISL_432427, EPI_ISL_432429, EPI_ISL_432430, EPI_ISL_432431, EPI_ISL_432434, EPI_ISL_432435, EPI_ISL_432437, EPI_ISL_432438, EPI_ISL_432441, EPI_ISL_432445, EPI_ISL_432446, EPI_ISL_432448, EPI_ISL_432450 | | | |
| see above | Wales Specialist Virology Centre | Public Health Wales Microbiology Cardiff | Catherine Moore, Johnathan Evans, Malorie Perry, Simon Cottrell, Alec Birchley, Alexander Adams, Amy Gaskin, Bree Gatica-Wilcox, Jason Coombes, Lauren Gilbert, Lee Graham, Nicole Pacchiarini, Sara Kumziene-Summerhayes, Sarah Taylor, Sophie Jones, Sara Rey, Matthew Bull, Joanne Watkins, Sally Corden, Tom Connor |
| EPI_ISL_432454, EPI_ISL_432466, EPI_ISL_432467, EPI_ISL_432484, EPI_ISL_432494, EPI_ISL_432496, EPI_ISL_432504, EPI_ISL_432505, EPI_ISL_432507, EPI_ISL_432512, EPI_ISL_432518, EPI_ISL_432522, EPI_ISL_432529, EPI_ISL_432531, EPI_ISL_432533, EPI_ISL_432542, EPI_ISL_432547, EPI_ISL_432555, EPI_ISL_432571, EPI_ISL_432573, EPI_ISL_432579, EPI_ISL_432584, EPI_ISL_432595, EPI_ISL_432606, EPI_ISL_432614, EPI_ISL_432617, EPI_ISL_432626, EPI_ISL_432633, EPI_ISL_432654, EPI_ISL_432657, EPI_ISL_432666, EPI_ISL_432714, EPI_ISL_432727, EPI_ISL_432732, EPI_ISL_432733, EPI_ISL_432736, EPI_ISL_432738, EPI_ISL_432744, EPI_ISL_432745, EPI_ISL_432751, EPI_ISL_432755, EPI_ISL_432763, EPI_ISL_432764, EPI_ISL_432765, EPI_ISL_432772, EPI_ISL_432775, EPI_ISL_432787, EPI_ISL_432790, EPI_ISL_432792, EPI_ISL_432802, EPI_ISL_432809, EPI_ISL_432812, EPI_ISL_432816, EPI_ISL_432818, EPI_ISL_432824, EPI_ISL_432825, EPI_ISL_432842, EPI_ISL_432849, EPI_ISL_432851, EPI_ISL_432855, EPI_ISL_432857, EPI_ISL_432859, EPI_ISL_432861 | | | |
| see above | Virology Department, Sheffield Teaching Hospitals NHS Foundation Trust / Virology Department, Sheffield Teaching Hospitals NHS Foundation Trust | COVID-19 Genomics UK (COG-UK) Consortium | Thushan de Silva, Matthew Parker,Adri Angyal, Rebecca Brown, Luke Green, Rachel Tucker, Paul Parsons, Danielle Groves, Alex Keeley, Dave Partridge, Matthew Wyles, Benjamin Lindsey, Mehmet Yavuz, Mohammad Raza, Cariad Evans |
| EPI_ISL_432949, EPI_ISL_432950, EPI_ISL_432962, EPI_ISL_432963, EPI_ISL_432965, EPI_ISL_432966, EPI_ISL_432967, EPI_ISL_432968, EPI_ISL_432969, EPI_ISL_432970, EPI_ISL_432971, EPI_ISL_432972, EPI_ISL_432973, EPI_ISL_432974, EPI_ISL_432975, EPI_ISL_432976, EPI_ISL_432977, EPI_ISL_432978, EPI_ISL_432979, EPI_ISL_432980, EPI_ISL_432981, EPI_ISL_432982, EPI_ISL_432983, EPI_ISL_432984, EPI_ISL_432985, EPI_ISL_432986, EPI_ISL_432987, EPI_ISL_432988, EPI_ISL_432989, EPI_ISL_432990, EPI_ISL_432991, EPI_ISL_432992, EPI_ISL_432994 | | | |
| see above | Queens Medical Centre, Clinical Microbiology Department / DeepSeq Nottingham | COVID-19 Genomics UK (COG-UK) Consortium | Gemma Clark, Wendy Smith, Manjinder Khakh, Hannah Howson-Wells, Jonathan Ball, Patrick McClure, Joseph Chappell, Theocharis Tsoleridis, Nadine Holmes, Matthew Carlisle, Christopher Moore, Fei Sang, Johnny Debebe, Victoria Wright, Matthew Loose |
| EPI_ISL_433200, EPI_ISL_433201, EPI_ISL_433202, EPI_ISL_433203, EPI_ISL_433207, EPI_ISL_433208, EPI_ISL_433209, EPI_ISL_433210, EPI_ISL_433211, EPI_ISL_433212, EPI_ISL_433214, EPI_ISL_433216, EPI_ISL_433217, EPI_ISL_433218, EPI_ISL_433220, EPI_ISL_433224, EPI_ISL_433225, EPI_ISL_433226, EPI_ISL_433227 | | | |
| see above | Virology Department, Royal Infirmary of Edinburgh, NHS Lothian / School of Biological Sciences, University of Edinburgh / Institute of Genetics and Molecular Medicine, University of Edinburgh | COVID-19 Genomics UK (COG-UK) Consortium | McHugh M, Dewar R, Rooke S, Gallagher M, Balcaza C, O'Toole A, Hill V, McCrone JT, Colquhoun R, Yu X, Jackson B, Rambaut A, Williams TC, Templeton K |
| EPI_ISL_433524, EPI_ISL_433525, EPI_ISL_433526, EPI_ISL_433527, EPI_ISL_433528, EPI_ISL_433529, EPI_ISL_433530, EPI_ISL_433533, EPI_ISL_433539, EPI_ISL_433540, EPI_ISL_433541, EPI_ISL_433542, EPI_ISL_433543, EPI_ISL_433544, EPI_ISL_433545, EPI_ISL_433546, EPI_ISL_433547, EPI_ISL_433548, EPI_ISL_433549, EPI_ISL_433550, EPI_ISL_433551, EPI_ISL_433552, EPI_ISL_433553, EPI_ISL_433554, EPI_ISL_433555, EPI_ISL_433556, EPI_ISL_433557, EPI_ISL_433558, EPI_ISL_433559, EPI_ISL_433560, EPI_ISL_433561, EPI_ISL_433562, EPI_ISL_433563, EPI_ISL_433564, EPI_ISL_433565, EPI_ISL_433566, EPI_ISL_433567, EPI_ISL_433568, EPI_ISL_433569, EPI_ISL_433570, EPI_ISL_433571, EPI_ISL_433572, EPI_ISL_433573, EPI_ISL_433574, EPI_ISL_433575, EPI_ISL_433577, EPI_ISL_433578, EPI_ISL_433579, EPI_ISL_433582, EPI_ISL_433584, EPI_ISL_433586, EPI_ISL_433588, EPI_ISL_433589, EPI_ISL_433593, EPI_ISL_433599 | | | |
| see above | West of Scotland Specialist Virology Centre, NHSGGC / MRC-University of Glasgow Centre for Virus Research | COVID-19 Genomics UK (COG-UK) Consortium | Ana da Silva Filipe, Natasha Johnson, Kathy Smollett, Daniel Mair, Stephen Carmichael, Lily Tong, Jenna Nichols, Elihu Aranday-Cortes, Kirstyn Brunker, Yasmin Parr, Kyriaki Nomikou; Sarah McDonald, Marc Niebel, Patawee Asamaphan; Richard Orton, Joseph Hughes, Sreenu Vattipally, David L Robertson; Alasdair MacLean, Rory Gunson; Kathy Li, Natasha Jesudason, Rajiv Shah, James Shepherd, Antonia Ho, Emma Thomson |
| EPI_ISL_433684, EPI_ISL_433688, EPI_ISL_433689, EPI_ISL_433691, EPI_ISL_433692, EPI_ISL_433693, EPI_ISL_433694, EPI_ISL_433696, EPI_ISL_433697, EPI_ISL_433698, EPI_ISL_433699 | | | |
| see above | Department of Pathology, University of Cambridge | COVID-19 Genomics UK (COG-UK) Consortium | Luke W Meredith, M. Estee Torok , Myra Hosmillo, William L. Hamilton, Martin D. Curran, Theresa Feltwell, Grant Hall, Anna Yakovleva, Fahad A Khokhar, Charlotte J. Houldcroft, Laura G Caller, Aminu S. Jahun, Sarah L. Caddy, Ian Goodfellow |

| | | | |
|---|---|---|---|
| EPI_ISL_434065, EPI_ISL_434070, EPI_ISL_434072, EPI_ISL_434076, EPI_ISL_434078, EPI_ISL_434080, EPI_ISL_434082, EPI_ISL_434084, EPI_ISL_434086, EPI_ISL_434089, EPI_ISL_434097, EPI_ISL_434107, EPI_ISL_434108, EPI_ISL_434196, EPI_ISL_434197, EPI_ISL_434198, EPI_ISL_434200, EPI_ISL_434201, EPI_ISL_434204, EPI_ISL_434205, EPI_ISL_434206, EPI_ISL_434207, EPI_ISL_434208, EPI_ISL_434209, EPI_ISL_434210, EPI_ISL_434211, EPI_ISL_434212, EPI_ISL_434214, EPI_ISL_434215, EPI_ISL_434225, EPI_ISL_434226, EPI_ISL_434234, EPI_ISL_434235, EPI_ISL_434238, EPI_ISL_434245, EPI_ISL_434246, EPI_ISL_434247, EPI_ISL_434248, EPI_ISL_434250, EPI_ISL_434253, EPI_ISL_434254, EPI_ISL_434255, EPI_ISL_434256, EPI_ISL_434257, EPI_ISL_434258, EPI_ISL_434259, EPI_ISL_434264, EPI_ISL_434266, EPI_ISL_434272, EPI_ISL_434273, EPI_ISL_434275, EPI_ISL_434276, EPI_ISL_434280, EPI_ISL_434282, EPI_ISL_434284, EPI_ISL_434285, EPI_ISL_434286, EPI_ISL_434287, EPI_ISL_434288, EPI_ISL_434298, EPI_ISL_434305, EPI_ISL_434308, EPI_ISL_434313, EPI_ISL_434314, EPI_ISL_434317, EPI_ISL_434323, EPI_ISL_434328, EPI_ISL_434334, EPI_ISL_434335, EPI_ISL_434342, EPI_ISL_434346 | | | |
| see above | Washington State Department of Health | Seattle Flu Study | Chu et al |
| EPI_ISL_434368, EPI_ISL_434371 | Hospital AZ Rivierenland | Institute of Tropical Medicine | Philippe Selhorst, Colin Anthony |
| EPI_ISL_434520, EPI_ISL_434521, EPI_ISL_434522, EPI_ISL_434529, EPI_ISL_434530, EPI_ISL_434531, EPI_ISL_434532 | Robert Garry lab | Andersen lab at Scripps Research | Allison Smither, Gilberto Sabino-Santos, Patricia Snarski, Lilia Melnik, Antoinette Bell, Kaylynn Genemaras, Arnaud Drouin, Dahlene Fusco, Robert Garry with SEARCH Alliance San Diego |
| EPI_ISL_434552, EPI_ISL_434553 | Puerto Rico Department of Health | Centers for Disease Control and Prevention, Dengue Branch | Gilberto A. Santiago, Glenda Gonzalez, Betzabel Flores, Keyla Charriez, Fabiola Cruz, Chaney Kalinich, Joseph Fauver, Jessica I. Falcon, Nathan Grubaugh, Jorge L. Munoz-Jordan |
| EPI_ISL_434657 | Kristianstadkliniken | The Public Health Agency of Sweden | Mia Settergren Hammer, Oskar Karlsson Lindsjo, Maria Lind Karlberg, Anna-Malin Linde, Olov Svartstrom, Anna Risberg, Theresa Enkirch, Mia Brytting, Karin Tegmark-Wisell |
| EPI_ISL_434941, EPI_ISL_434943, EPI_ISL_434964, EPI_ISL_434966, EPI_ISL_434970, EPI_ISL_434971, EPI_ISL_434972, EPI_ISL_434975, EPI_ISL_434976, EPI_ISL_434979, EPI_ISL_434980, EPI_ISL_434981, EPI_ISL_434982, EPI_ISL_434983, EPI_ISL_434984, EPI_ISL_434985, EPI_ISL_434986, EPI_ISL_434987, EPI_ISL_434988, EPI_ISL_434990, EPI_ISL_434991, EPI_ISL_434992, EPI_ISL_434994, EPI_ISL_434995, EPI_ISL_434997, EPI_ISL_434998, EPI_ISL_434999, EPI_ISL_435000, EPI_ISL_435001, EPI_ISL_435002, EPI_ISL_435003, EPI_ISL_435004, EPI_ISL_435005, EPI_ISL_435006, EPI_ISL_435007, EPI_ISL_435008, EPI_ISL_435009 | | | |
| see above | Houston Methodist Hospital | Houston Methodist Hospital | S. Wesley Long, Randall J. Olsen, Paul A. Christensen, David W. Bernard, James J. Davis, Maulik Shukla, Marcus Nguyen, Matthew Ojeda Saavedra, Concepcion C. Cantu, Prasanti Yerramilli, Layne Pruitt, Sishir Subedi, Heather Hendrickson, Ghazaleh Eskandari, Muthiah Kumaraswami, Jason S. McLellan, Hakon Jonsson, Kari Stefansson, and James M. Musser |
| EPI_ISL_435343, EPI_ISL_435344, EPI_ISL_435345, EPI_ISL_435346, EPI_ISL_435347 | Laboratoire de microbiologie, Hopital de Verdun | Smith Laboratory, Centre de Recherche CHU Sainte-Justine | Martin Smith, Marieke Rozendaal, Ivan Pavlov |
| EPI_ISL_435374 | Utah Public Health Laboratory | Utah Public Health Laboratory | Erin Young, Kelly Oakeson |
| EPI_ISL_435445, EPI_ISL_435447, EPI_ISL_435465, EPI_ISL_435467 | Robert Garry lab | Andersen lab at Scripps Research | Allison Smither, Gilberto Sabino-Santos, Patricia Snarski, Lilia Melnik, Antoinette Bell, Kaylynn Genemaras, Arnaud Drouin, Dahlene Fusco, Robert Garry with SEARCH Alliance San Diego |
| EPI_ISL_435489 | NYU Langone Health | Departments of Pathology and Medicine, New York University School of Medicine | Maria Aguero-Rosenfeld, Brendan Belovarac, Margaret Black, Ludovic Boytard, John Cadley, Paolo Cotzia, John Chen, Dacia Dimartino, Xiaojun Feng, Tatyana Gindin, Emily Guzman, Adriana Heguy, Megan Hogan, Emily Huang, George Jour, Lawrence H. Lin, Raven Luther, Andrew Lytle, Christian Marier, Matthew T. Maurano, Mark J. Mulligan, Peter Meyn, Raquel Ordonez Ciriza, Iman Osman, Jared Pinnell, Vanessa Raabe, Sitharam Ramaswami, Amy Rapkiewicz, Andre M. Ribeiro-dos-Santos, Marie Samanovic-Golden, Antonio Serrano, Guomiao Shen, Matija Snuderl, Theodore Vougiouklakis, Nick Vulpescu, Gael Westby, Paul Zappile, Yutong Zhang |
| EPI_ISL_435679, EPI_ISL_435680 | National Public Health Laboratory, National Centre for Infectious Diseases | National Public Health Laboratory, National Centre for Infectious Diseases | Mak Tze Minn, Octavia Sophie, Chavatte Jean-Marc, Cui Lin, Lin Raymond Tzer Pin |
| EPI_ISL_436046 | US VI Department of Health | Pathogen Discovery, Respiratory Viruses Branch, Division of Viral Diseases, Centers for Disease Control and Prevention | Ying Tao, Jing Zhang, Krista Queen, Yan Li, Anna Uehara, Clinton R. Paden, Haibin Wang, Zachary Weiner, Bettina Bankamp, Suxiang Tong |
| EPI_ISL_436107 | TSGH-CP molecular lab | TSGH-CP molecular lab | Cherng-Lih Perng, Ming-Jr JIAN, Chih-Kai Chang, Jung-Chung Lin, Kuo-Ming Yeh, Chien-Wen Chen, Sheng-Kang Chiu, Hsing-Yi Chung, Shih-Hung Tsai, Kuo-Sheng Hung, Tien-Yao Chang, Feng-Yee Chang, Hung-Sheng Shang |
| EPI_ISL_436414, EPI_ISL_436415, EPI_ISL_436416, EPI_ISL_436417, EPI_ISL_436418, EPI_ISL_436419, EPI_ISL_436420, EPI_ISL_436421, EPI_ISL_436422 | National Centre for Disease control (NCDC) | NCDC/CSIR-IGIB | Pramod Kumar#, Rajesh Pandey#, Pooja Sharma, Mahesh S Dhar, Vivekanand A, Bharathram Uppili, Himanshu Vashisht, Saruchi Wadhwa, Nishu Tyagi, Uma Sharma, Priyanka Singh, Hemlata Lall, Meena Datta, Poonam Gupta, Nidhi Saini, Aarti Tewari, Bibhash Nandi, Dhirendra Kumar, Satyabrata Bag, Varun Jaiswal, Hema Gogia, Preeti Madan, Simrita Singh, Prateek Singh, Debasis Dash, Mitali Mukerji, Manju Bala, Sandhya Kabra, Sujeet Singh, Mohammed Faruq, Anurag Agrawal*, Partha Rakshit* |
| EPI_ISL_436567, EPI_ISL_436568, EPI_ISL_436569, EPI_ISL_436570, EPI_ISL_436633, EPI_ISL_436634 | University of Wisconsin-Madison AIDS Vaccine Research Laboratories | University of Wisconsin-Madison AIDS Vaccine Research Laboratories | Gage Moreno, Katarina Braun, et al. AIDS Vaccine Research Laboratories |
| EPI_ISL_436641, EPI_ISL_436642, EPI_ISL_436643, EPI_ISL_436645, EPI_ISL_436673, EPI_ISL_436674, EPI_ISL_436675, EPI_ISL_436676 | County of Santa Clara Public Health Department | Chan-Zuckerberg Biohub | CZB Cliahub Consortium |
| EPI_ISL_436684, EPI_ISL_436685 | KRISP, KZN Research Innovation and Sequencing Platform | KRISP, KZN Research Innovation and Sequencing Platform | Giandhari J, Pillay S, Lessells R, Chimukangara B, Deforche K, Tegally H, Wilkinson E, de Oliveira T |
| EPI_ISL_437043, EPI_ISL_437044 | County of Santa Clara Public Health | Chan-Zuckerberg Biohub | CZB Cliahub Consortium |
| EPI_ISL_437192 | Mitra Keluarga Kelapa Gading Hospital | Eijkman Institute for Molecular Biology, Ministry of Research and Technology/National Agency for Research and Innovation | Edison Johar, Frilasita A Yudhaputri, Hidayat Trimarsanto, David H Muljono, Safarina G Malik, Khin Saw Myint, Amin Soebandrio |
| EPI_ISL_437199, EPI_ISL_437200 | Diagnostic- and Research Institute of Pathology, Medical University of Graz | Diagnostic- and Research Institute of Pathology, Medical University of Graz | Karl Kashofer, Peter Regitnig, Martin Zacharias, Gregor Gorkiewicz |
| EPI_ISL_437225, EPI_ISL_437232, EPI_ISL_437277, EPI_ISL_437278, EPI_ISL_437279, EPI_ISL_437280, EPI_ISL_437281, EPI_ISL_437282 | Max von Pettenkofer Institute, Virology, National Reference Center for Retroviruses, LMU München | Laboratory for Functional Genome Analysis, Dept. Genomics, Gene Center of the LMU Munich | Max Muenchhoff, Stefan Krebs, Alexander Graf, Oliver Keppler, Helmut Blum |
| EPI_ISL_437363 | Minnesota Department of Health, Public Health Laboratory | Minnesota Department of Health, Public Health Laboratory | Matt Plumb, Jacob Garfin, and Xiong Wang |
| EPI_ISL_437399 | Virginia DCLS | Virginia DCLS | Virginia DCLS |
| EPI_ISL_437475 | Pathogen Genomics Lab King Abdullah University of Science and Technology(KAUST) | Pathogen Genomics Lab King Abdullah University of Science and Technology(KAUST) | Sharif Hala,Raeece Naeem,Sara Mfarrej,Arnab Pain |
| EPI_ISL_437547, EPI_ISL_437548 | Robert Garry lab | Andersen lab at Scripps Research | Allison Smither, Gilberto Sabino-Santos, Patricia Snarski, Lilia Melnik, Antoinette Bell, Kaylynn Genemaras, Arnaud Drouin, Dahlene Fusco, Robert Garry with SEARCH Alliance San Diego |
| EPI_ISL_437627, EPI_ISL_437628, EPI_ISL_437629, EPI_ISL_437630, EPI_ISL_437633, EPI_ISL_437634, EPI_ISL_437636, EPI_ISL_437682, EPI_ISL_437683 | Department of Virus and Microbiological Special Diagnostics, Statens Serum Institut, Copenhagen, Denmark, Artillerivej 5, 2300 Copenahgen S | Albertsen lab, Department of Chemistry and Bioscience, Aalborg University, Denmark | Rasmus Kirkegaard |
| EPI_ISL_437684, EPI_ISL_437685, EPI_ISL_437686, EPI_ISL_437687, EPI_ISL_437688 | UCD National Virus Reference Laboratory | UCD National Virus Reference Laboratory | Michael J. Carr, Gabriel Gonzalez, Brendan Crowley, Cillian F De Gascun |

| | | | |
|---|---|---|---|
| EPI_ISL_437734, EPI_ISL_437735 | Pathogen Genomics Lab King Abdullah University of Science and Technology(KAUST) | Pathogen Genomics Lab King Abdullah University of Science and Technology(KAUST) | Sharif Hala,Fadwa Alofi,Afrah Alsomali, Asim Khogeer, Sara Mfarrej, Khaled Algithami,Raeece Naeem, Amit Kumar Subudhi,Fathia Ben-Rached, Rahul Salunke, Anwar Hashem, Naif Almontashiri, Arnab Pain |
| EPI_ISL_437800 | Virginia DCLS | Virginia DCLS | Virginia DCLS |
| EPI_ISL_437813, EPI_ISL_437814, EPI_ISL_437815, EPI_ISL_437816, EPI_ISL_437817, EPI_ISL_437818, EPI_ISL_437819, EPI_ISL_437820, EPI_ISL_437821, EPI_ISL_437822, EPI_ISL_437823, EPI_ISL_437824, EPI_ISL_437827, EPI_ISL_437829 | | | |
| see above | UW Virology Lab | UW Virology Lab | Pavitra Roychoudhury, Hong Xie, Keith Jerome, Alexander Greninger |
| EPI_ISL_437873 | Alaska State Virology Laboratory | Alaska State Virology Laboratory | Jack Chen, Ph.D. |
| EPI_ISL_437910, EPI_ISL_437911 | Laboratory of Microbiology, Medical School, National and Kapodistrian University of Athens | Laboratory of Biology, Department of Medicine, Democritus University of Thrace | Kassela K., Dovrolis,N., Bampali,M., Gatzidou,E., Froukala,E., Stavropoulou,A., Veletza,S., Tsakris,A., Spanakis,N. and Karakasiliotis,I. |
| EPI_ISL_438121 | Center for Virology, Medical University of Vienna | Bergthaler laboratory, CeMM Research Center for Molecular Medicine of the Austrian Academy of Sciences | Alexandra Popa, Benedikt Agerer, Henrique Colaco, Lukas Endler, Jakob-Wendelin Genger, Alexander Lercher, Mark Smyth, Thomas Penz, Michael Schuster, Jan Laine, Martin Senekowitsch, Judith Aberle, Stephan Aberle, Elisabeth Puchhammer-Stoeckl, Manfred Nairz, Guenter Weiss, Wegene Borena, Dorothee von Laer, Christoph Bock, Andreas Bergthaler |
| EPI_ISL_438140 | Seattle Flu Study | Seattle Flu Study | Chu et al |
| EPI_ISL_438242 | Johns Hopkins Hospital Department of Pathology | Johns Hopkins Hospital Department of Pathology | Peter M. Thielen, Thomas Mehoke, Shirlee Wohl, Srividya Ramakrishnan, Melanie Kirsche, Amanda Ernlund, Oluwaseun Falade-Nwulia, Timothy Gilpatrick, Paul Morris, Norah Sadowski, Nidia Trovao, Victoria Gniazdowski, Michael Schatz, Stuart C. Ray, Winston Timp, Heba Mostafa |
| EPI_ISL_438248, EPI_ISL_438249, EPI_ISL_438250, EPI_ISL_438252, EPI_ISL_438254, EPI_ISL_438264, EPI_ISL_438265, EPI_ISL_438279, EPI_ISL_438282, EPI_ISL_438291, EPI_ISL_438316, EPI_ISL_438340, EPI_ISL_438381, EPI_ISL_438382, EPI_ISL_438391, EPI_ISL_438398, EPI_ISL_438419, EPI_ISL_438431, EPI_ISL_438450, EPI_ISL_438459, EPI_ISL_438470, EPI_ISL_438473, EPI_ISL_438476, EPI_ISL_438483, EPI_ISL_438489, EPI_ISL_438507, EPI_ISL_438509, EPI_ISL_438510, EPI_ISL_438511, EPI_ISL_438514, EPI_ISL_438515, EPI_ISL_438517, EPI_ISL_438519, EPI_ISL_438520, EPI_ISL_438523, EPI_ISL_438524, EPI_ISL_438525, EPI_ISL_438527, EPI_ISL_438528, EPI_ISL_438530, EPI_ISL_438531, EPI_ISL_438532, EPI_ISL_438533, EPI_ISL_438534, EPI_ISL_438535, EPI_ISL_438536, EPI_ISL_438537, EPI_ISL_438539, EPI_ISL_438540, EPI_ISL_438542, EPI_ISL_438544, EPI_ISL_438545 | | | |
| see above | Department of Pathology, University of Cambridge | Wellcome Sanger Institute for the COVID-19 Genomics UK (COG-UK) consortium | Luke W Meredith, M. Estée Török , Myra Hosmillo, William L. Hamilton, Martin D. Curran, Theresa Feltwell, Grant Hall, Anna Yakovleva, Fahad A Khokhar, Charlotte J. Houldcroft, Laura G Caller, Aminu S. Jahun, Sarah L. Caddy, Ian Goodfellow, Alex Alderton, Roberto Amato, Sonia Goncalves, Ewan Harrison, David K. Jackson, Ian Johnston, Dominic Kwiatkowski, Cordelia Langford, John Sillitoe on behalf of the Wellcome Sanger Institute COVID-19 Surveillance Team (http://www.sanger.ac.uk/covid-team) |
| EPI_ISL_438774, EPI_ISL_438777, EPI_ISL_438778, EPI_ISL_438779, EPI_ISL_438780, EPI_ISL_438781, EPI_ISL_438783, EPI_ISL_438784, EPI_ISL_438785, EPI_ISL_438787, EPI_ISL_438788, EPI_ISL_438789, EPI_ISL_438790, EPI_ISL_438791, EPI_ISL_438792, EPI_ISL_438793, EPI_ISL_438794, EPI_ISL_438795, EPI_ISL_438796, EPI_ISL_438797, EPI_ISL_438798, EPI_ISL_438799, EPI_ISL_438800, EPI_ISL_438801, EPI_ISL_438802, EPI_ISL_438803, EPI_ISL_438804, EPI_ISL_438807, EPI_ISL_438808, EPI_ISL_438809, EPI_ISL_438810, EPI_ISL_438812, EPI_ISL_438813, EPI_ISL_438814, EPI_ISL_438815, EPI_ISL_438818, EPI_ISL_438819, EPI_ISL_438821, EPI_ISL_438822, EPI_ISL_438823 | | | |
| see above | West of Scotland Specialist Virology Centre, NHSGGC / MRC-University of Glasgow Centre for Virus Research | COVID-19 Genomics UK (COG-UK) Consortium | Ana da Silva Filipe, Natasha Johnson, Kathy Smollett, Daniel Mair, Stephen Carmichael, Lily Tong, Jenna Nichols, Elihu Aranday-Cortes, Kirstyn Brunker, Yasmin Parr, Kyriaki Nomikou; Sarah McDonald, Marc Niebel, Patawee Asamaphan; Richard Orton, Joseph Hughes, Sreenu Vattipally, David L Robertson; Alasdair MacLean, Rory Gunson; Kathy Li, Natasha Jesudason, Rajiv Shah, James Shepherd, Antonia Ho, Emma Thomson |
| EPI_ISL_439368, EPI_ISL_439369, EPI_ISL_439370, EPI_ISL_439371, EPI_ISL_439372, EPI_ISL_439373, EPI_ISL_439374, EPI_ISL_439375, EPI_ISL_439376, EPI_ISL_439377, EPI_ISL_439378, EPI_ISL_439380, EPI_ISL_439381, EPI_ISL_439382, EPI_ISL_439386, EPI_ISL_439387, EPI_ISL_439388, EPI_ISL_439389, EPI_ISL_439390, EPI_ISL_439391, EPI_ISL_439392, EPI_ISL_439393, EPI_ISL_439395, EPI_ISL_439397, EPI_ISL_439399, EPI_ISL_439400, EPI_ISL_439401, EPI_ISL_439404, EPI_ISL_439405, EPI_ISL_439406, EPI_ISL_439409, EPI_ISL_439412, EPI_ISL_439413, EPI_ISL_439414, EPI_ISL_439415, EPI_ISL_439416, EPI_ISL_439417, EPI_ISL_439418, EPI_ISL_439420, EPI_ISL_439423, EPI_ISL_439424, EPI_ISL_439425, EPI_ISL_439426, EPI_ISL_439428, EPI_ISL_439429, EPI_ISL_439430, EPI_ISL_439431, EPI_ISL_439433, EPI_ISL_439434, EPI_ISL_439435, EPI_ISL_439436, EPI_ISL_439437, EPI_ISL_439438, EPI_ISL_439439, EPI_ISL_439440, EPI_ISL_439441, EPI_ISL_439443, EPI_ISL_439444, EPI_ISL_439445, EPI_ISL_439446, EPI_ISL_439447, EPI_ISL_439449, EPI_ISL_439451, EPI_ISL_439452, EPI_ISL_439453, EPI_ISL_439454, EPI_ISL_439455, EPI_ISL_439456, EPI_ISL_439459, EPI_ISL_439460, EPI_ISL_439462, EPI_ISL_439463, EPI_ISL_439464, EPI_ISL_439466, EPI_ISL_439468, EPI_ISL_439470, EPI_ISL_439472, EPI_ISL_439473, EPI_ISL_439474, EPI_ISL_439477, EPI_ISL_439478, EPI_ISL_439479, EPI_ISL_439480, EPI_ISL_439481, EPI_ISL_439483, EPI_ISL_439484, EPI_ISL_439485, EPI_ISL_439487, EPI_ISL_439488, EPI_ISL_439489, EPI_ISL_439490, EPI_ISL_439492, EPI_ISL_439494, EPI_ISL_439495, EPI_ISL_439496, EPI_ISL_439497, EPI_ISL_439498, EPI_ISL_439501, EPI_ISL_439502, EPI_ISL_439503, EPI_ISL_439504, EPI_ISL_439509, EPI_ISL_439510, EPI_ISL_439511, EPI_ISL_439512, EPI_ISL_439513, EPI_ISL_439514, EPI_ISL_439516, EPI_ISL_439517, EPI_ISL_439519, EPI_ISL_439521, EPI_ISL_439524, EPI_ISL_439525, EPI_ISL_439526, EPI_ISL_439528, EPI_ISL_439529, EPI_ISL_439531, EPI_ISL_439536, EPI_ISL_439563, EPI_ISL_439571, EPI_ISL_439578, EPI_ISL_439599, EPI_ISL_439601, EPI_ISL_439607, EPI_ISL_439612, EPI_ISL_439617, EPI_ISL_439619, EPI_ISL_439620, EPI_ISL_439637, EPI_ISL_439647, EPI_ISL_439650, EPI_ISL_439659, EPI_ISL_439866, EPI_ISL_439870, EPI_ISL_439880, EPI_ISL_439901, EPI_ISL_439903, EPI_ISL_439927, EPI_ISL_439929, EPI_ISL_439946, EPI_ISL_439950 | | | |
| see above | Department of Pathology, University of Cambridge | Wellcome Sanger Institute for the COVID-19 Genomics UK (COG-UK) consortium | Luke W Meredith, M. Estée Török , Myra Hosmillo, William L. Hamilton, Martin D. Curran, Theresa Feltwell, Grant Hall, Anna Yakovleva, Fahad A Khokhar, Charlotte J. Houldcroft, Laura G Caller, Aminu S. Jahun, Sarah L. Caddy, Ian Goodfellow, Alex Alderton, Roberto Amato, Sonia Goncalves, Ewan Harrison, David K. Jackson, Ian Johnston, Dominic Kwiatkowski, Cordelia Langford, John Sillitoe on behalf of the Wellcome Sanger Institute COVID-19 Surveillance Team (http://www.sanger.ac.uk/covid-team) |
| EPI_ISL_439998, EPI_ISL_440029, EPI_ISL_440114, EPI_ISL_440158, EPI_ISL_440294 | PHE South West Regional Laboratory, National Infection Service | Wellcome Sanger Institute for the COVID-19 Genomics UK (COG-UK) consortium | Stephanie Hutchings, Hannah Pymont, Dr Peter Muir, Barry Vipond, Rich Hopes, Alex Alderton, Roberto Amato, Sonia Goncalves, Ewan Harrison, David K. Jackson, Ian Johnston, Dominic Kwiatkowski, Cordelia Langford, John Sillitoe on behalf of the Wellcome Sanger Institute COVID-19 Surveillance Team (http://www.sanger.ac.uk/covid-team) |
| EPI_ISL_440564, EPI_ISL_440568, EPI_ISL_440575, EPI_ISL_440601, EPI_ISL_440602, EPI_ISL_440613, EPI_ISL_440617, EPI_ISL_440620 | Department of Pathology, University of Cambridge | Wellcome Sanger Institute for the COVID-19 Genomics UK (COG-UK) consortium | Luke W Meredith, M. Estée Török , Myra Hosmillo, William L. Hamilton, Martin D. Curran, Theresa Feltwell, Grant Hall, Anna Yakovleva, Fahad A Khokhar, Charlotte J. Houldcroft, Laura G Caller, Aminu S. Jahun, Sarah L. Caddy, Ian Goodfellow, Alex Alderton, Roberto Amato, Sonia Goncalves, Ewan Harrison, David K. Jackson, Ian Johnston, Dominic Kwiatkowski, Cordelia Langford, John Sillitoe on behalf of the Wellcome Sanger Institute COVID-19 Surveillance Team (http://www.sanger.ac.uk/covid-team) |
| EPI_ISL_440662, EPI_ISL_440669, EPI_ISL_440678, EPI_ISL_440694, EPI_ISL_440696, EPI_ISL_440700, EPI_ISL_440703, EPI_ISL_440720, EPI_ISL_440736, EPI_ISL_440765, EPI_ISL_440779, EPI_ISL_440786, EPI_ISL_440809 | | | |
| see above | PHE South West Regional Laboratory, National Infection Service | Wellcome Sanger Institute for the COVID-19 Genomics UK (COG-UK) consortium | Stephanie Hutchings, Hannah Pymont, Dr Peter Muir, Barry Vipond, Rich Hopes, Alex Alderton, Roberto Amato, Sonia Goncalves, Ewan Harrison, David K. Jackson, Ian Johnston, Dominic Kwiatkowski, Cordelia Langford, John Sillitoe on behalf of the Wellcome Sanger Institute COVID-19 Surveillance Team (http://www.sanger.ac.uk/covid-team) |
| EPI_ISL_440810, EPI_ISL_440811, EPI_ISL_440813, EPI_ISL_440815, EPI_ISL_440817, EPI_ISL_440818, EPI_ISL_440819, EPI_ISL_440820, EPI_ISL_440821, EPI_ISL_440822, EPI_ISL_440823, EPI_ISL_440824, EPI_ISL_440825, EPI_ISL_440826, EPI_ISL_440827, EPI_ISL_440828, EPI_ISL_440829, EPI_ISL_440831, EPI_ISL_440832, EPI_ISL_440836, EPI_ISL_440837, EPI_ISL_440838, EPI_ISL_440839, EPI_ISL_440840, EPI_ISL_440841, EPI_ISL_440842, EPI_ISL_440844, EPI_ISL_440845, EPI_ISL_440846, EPI_ISL_440848, EPI_ISL_440849, EPI_ISL_440850, EPI_ISL_440851 | | | |
| see above | Department of Pathology, University of Cambridge | Wellcome Sanger Institute for the COVID-19 Genomics UK (COG-UK) consortium | Luke W Meredith, M. Estée Török , Myra Hosmillo, William L. Hamilton, Martin D. Curran, Theresa Feltwell, Grant Hall, Anna Yakovleva, Fahad A Khokhar, Charlotte J. Houldcroft, Laura G Caller, Aminu S. Jahun, Sarah L. Caddy, Ian Goodfellow, Alex Alderton, Roberto Amato, Sonia Goncalves, Ewan Harrison, David K. Jackson, Ian Johnston, Dominic Kwiatkowski, Cordelia Langford, John Sillitoe on behalf of the Wellcome Sanger Institute COVID-19 Surveillance Team (http://www.sanger.ac.uk/covid-team) |
| EPI_ISL_440971, EPI_ISL_440972, EPI_ISL_440973, EPI_ISL_440974, EPI_ISL_440975, EPI_ISL_440976, EPI_ISL_440977, EPI_ISL_440978, EPI_ISL_440979, EPI_ISL_440980, EPI_ISL_440981, EPI_ISL_440982, EPI_ISL_440983, EPI_ISL_440984, EPI_ISL_440985, EPI_ISL_440986, EPI_ISL_440987, EPI_ISL_440988, EPI_ISL_440989, EPI_ISL_440990 | | | |
| see above | University College London, Great Ormond Street Hospital for Children NHS Foundation Trust, Imperial College Healthcare NHS Trust | COVID-19 Genomics UK (COG-UK) Consortium | Sergi Castellano, Rachel Williams, Mark Kristiansen, Paola Resende Silva, Sunando Roy, Tony Brooks, Helena Tutill, Paola Niola, Patricia Dyal, Charlotte Williams, Leysa Forrest, Yasmin Panchbhaya, Jacqueline Findlay, Sam Weeks, Julianne Brown, Kathryn Harris, Paul Randell, James Price, Alison Holmes, Judith Breuer |
| EPI_ISL_441053, EPI_ISL_441065, EPI_ISL_441068, EPI_ISL_441071, EPI_ISL_441076, EPI_ISL_441077, EPI_ISL_441082, EPI_ISL_441090, EPI_ISL_441093, EPI_ISL_441103, EPI_ISL_441107, EPI_ISL_441108, EPI_ISL_441110, EPI_ISL_441112, EPI_ISL_441118, EPI_ISL_441120, EPI_ISL_441121, EPI_ISL_441123, EPI_ISL_441125, EPI_ISL_441127, EPI_ISL_441137, EPI_ISL_441144, EPI_ISL_441145, EPI_ISL_441146, EPI_ISL_441149, EPI_ISL_441152, EPI_ISL_441153, EPI_ISL_441160, EPI_ISL_441162, EPI_ISL_441163, EPI_ISL_441166, EPI_ISL_441167, EPI_ISL_441173, EPI_ISL_441179, EPI_ISL_441182, EPI_ISL_441185, EPI_ISL_441187, EPI_ISL_441204, EPI_ISL_441211, EPI_ISL_441227, EPI_ISL_441229, EPI_ISL_441235, EPI_ISL_441241, EPI_ISL_441243, EPI_ISL_441246, EPI_ISL_441248, EPI_ISL_441250, EPI_ISL_441254, EPI_ISL_441256, EPI_ISL_441257, EPI_ISL_441264, EPI_ISL_441274, EPI_ISL_441283, EPI_ISL_441290, EPI_ISL_441291, EPI_ISL_441293, EPI_ISL_441296, EPI_ISL_441302, EPI_ISL_441303, EPI_ISL_441304, EPI_ISL_441307, EPI_ISL_441312, EPI_ISL_441321, EPI_ISL_441325, EPI_ISL_441328, EPI_ISL_441331, EPI_ISL_441339, EPI_ISL_441341 | | | |
| see above | Department of Pathology, University of Cambridge | Wellcome Sanger Institute for the COVID-19 Genomics UK (COG-UK) consortium | Luke W Meredith, M. Estée Török , Myra Hosmillo, William L. Hamilton, Martin D. Curran, Theresa Feltwell, Grant Hall, Anna Yakovleva, Fahad A Khokhar, Charlotte J. Houldcroft, Laura G Caller, Aminu S. Jahun, Sarah L. Caddy, Ian Goodfellow, Alex Alderton, Roberto Amato, Sonia Goncalves, Ewan Harrison, David K. Jackson, Ian Johnston, Dominic Kwiatkowski, Cordelia Langford, John Sillitoe on behalf of the Wellcome Sanger Institute COVID-19 Surveillance Team (http://www.sanger.ac.uk/covid-team) |
| EPI_ISL_441659, EPI_ISL_441670, EPI_ISL_441704, EPI_ISL_441717, EPI_ISL_441733, EPI_ISL_441766, EPI_ISL_441775 | Regional Virus Laboratory, Belfast Health and Social Care Trust | Wellcome Sanger Institute for the COVID-19 Genomics UK (COG-UK) consortium | Conall McCaughey, James McKenna, Tanya Curran, Susan Feeney, Alison Watt, Ciara Cox, Mairead Connor, Zoltan Molnar, David Simpson, Derek Fairley, Alex Alderton, Roberto Amato, Sonia Goncalves, Ewan Harrison, David K. Jackson, Ian Johnston, Dominic Kwiatkowski, Cordelia Langford, John Sillitoe on behalf of the Wellcome Sanger Institute COVID-19 Surveillance Team (http://www.sanger.ac.uk/covid-team) |
| EPI_ISL_441783, EPI_ISL_441788, EPI_ISL_441789, EPI_ISL_441790, EPI_ISL_441795, EPI_ISL_441796, EPI_ISL_441797, EPI_ISL_441803, EPI_ISL_441805, EPI_ISL_441809, EPI_ISL_441811, EPI_ISL_441816, EPI_ISL_441818, EPI_ISL_441819, EPI_ISL_441831, EPI_ISL_441835, EPI_ISL_441836, EPI_ISL_441837, | | | |

| | | | |
|---|---|---|---|
| EPI_ISL_441840, EPI_ISL_441841, EPI_ISL_441842, EPI_ISL_441844 | | | |
| see above | Department of Pathology, University of Cambridge | Wellcome Sanger Institute for the COVID-19 Genomics UK (COG-UK) consortium | Luke W Meredith, M. Estée Török , Myra Hosmillo, William L. Hamilton, Martin D. Curran, Theresa Feltwell, Grant Hall, Anna Yakovleva, Fahad A Khokhar, Charlotte J. Houldcroft, Laura G Caller, Aminu S. Jahun, Sarah L. Caddy, Ian Goodfellow, Alex Alderton, Roberto Amato, Sonia Goncalves, Ewan Harrison, David K. Jackson, Ian Johnston, Dominic Kwiatkowski, Cordelia Langford, John Sillitoe on behalf of the Wellcome Sanger Institute COVID-19 Surveillance Team (http://www.sanger.ac.uk/covid-team) |
| EPI_ISL_441914, EPI_ISL_441934, EPI_ISL_441963, EPI_ISL_441973 | Virology Department, Sheffield Teaching Hospitals NHS Foundation Trust/Department of Infection, Immunity and Cardiovascular Disease, The Medical School, University of Sheffield | COVID-19 Genomics UK (COG-UK) Consortium | Thushan de Silva, Matthew Parker, Nikki Smith, Adri Angyal, Rebecca Brown, Luke Green, Rachel Tucker, Paul Parsons, Danielle Groves, Katie Johnson, Laura Carrilero, Alex Keeley, Dave Partridge, Matthew Wyles, Benjamin Lindsey, Mehmet Yavuz, Mohammad Raza, Cariad Evans |
| EPI_ISL_442322 | Department of Pathology, University of Cambridge | Wellcome Sanger Institute for the COVID-19 Genomics UK (COG-UK) consortium | Luke W Meredith, M. Estée Török , Myra Hosmillo, William L. Hamilton, Martin D. Curran, Theresa Feltwell, Grant Hall, Anna Yakovleva, Fahad A Khokhar, Charlotte J. Houldcroft, Laura G Caller, Aminu S. Jahun, Sarah L. Caddy, Ian Goodfellow, Alex Alderton, Roberto Amato, Sonia Goncalves, Ewan Harrison, David K. Jackson, Ian Johnston, Dominic Kwiatkowski, Cordelia Langford, John Sillitoe on behalf of the Wellcome Sanger Institute COVID-19 Surveillance Team (http://www.sanger.ac.uk/covid-team) |
| EPI_ISL_442348, EPI_ISL_442363, EPI_ISL_442374, EPI_ISL_442435, EPI_ISL_442461, EPI_ISL_442484 | Virology Department, Sheffield Teaching Hospitals NHS Foundation Trust/Department of Infection, Immunity and Cardiovascular Disease, The Medical School, University of Sheffield | COVID-19 Genomics UK (COG-UK) Consortium | Thushan de Silva, Matthew Parker, Nikki Smith, Adri Angyal, Rebecca Brown, Luke Green, Rachel Tucker, Paul Parsons, Danielle Groves, Katie Johnson, Laura Carrilero, Alex Keeley, Dave Partridge, Matthew Wyles, Benjamin Lindsey, Mehmet Yavuz, Mohammad Raza, Cariad Evans |
| EPI_ISL_443184, EPI_ISL_443186 | UW Virology Lab | UW Virology Lab | Pavitra Roychoudhury, Hong Xie, Keith Jerome, Alexander Greninger |
| EPI_ISL_443255 | M Health Fairview | University of Minnesota Genomics Center | Daryl M. Gohl, John Garbe, Patrick Grady, Jerry Daniel, Ray Watson, Benjamin Auch, Andrew Nelson, Sophia Yohe, and Kenneth B. Beckman |
| EPI_ISL_443279, EPI_ISL_443280 | CHU - Hôpital Cavale Blanche - Labo. de Virologie | National Reference Center for Viruses of Respiratory Infections, Institut Pasteur, Paris | Mélanie Albert, Marion Barbet, Sylvie Behillil, Méline Bizard, Angela Brisebarre, Flora Donati, Etienne Simon-Lorière, Vincent Enouf, Maud Vanpeene, Sylvie van der Werf, Léa Pilorge |
| EPI_ISL_443290 | CHRU Pontchaillou - Laboratoire de Virologie | National Reference Center for Viruses of Respiratory Infections, Institut Pasteur, Paris | Mélanie Albert, Marion Barbet, Sylvie Behillil, Méline Bizard, Angela Brisebarre, Flora Donati, Etienne Simon-Lorière, Vincent Enouf, Maud Vanpeene, Sylvie van der Werf, Gisèle Lagathu |
| EPI_ISL_443304 | Résidence Esterel | National Reference Center for Viruses of Respiratory Infections, Institut Pasteur, Paris | Mélanie Albert, Marion Barbet, Sylvie Behillil, Méline Bizard, Angela Brisebarre, Flora Donati, Etienne Simon-Lorière, Vincent Enouf, Maud Vanpeene, Sylvie van der Werf |
| EPI_ISL_443762, EPI_ISL_443791, EPI_ISL_443827, EPI_ISL_443829, EPI_ISL_443841, EPI_ISL_443855, EPI_ISL_443894 | PHE South West Regional Laboratory, National Infection Service | Wellcome Sanger Institute for the COVID-19 Genomics UK (COG-UK) consortium | Stephanie Hutchings, Hannah Pymont, Dr Peter Muir, Barry Vipond, Rich Hopes; and Alex Alderton, Roberto Amato, Sonia Goncalves, Ewan Harrison, David K. Jackson, Ian Johnston, Dominic Kwiatkowski, Cordelia Langford, John Sillitoe on behalf of the Wellcome Sanger Institute COVID-19 Surveillance Team (http://www.sanger.ac.uk/covid-team) |
| EPI_ISL_444662, EPI_ISL_444663, EPI_ISL_444669, EPI_ISL_444673, EPI_ISL_444676, EPI_ISL_444677, EPI_ISL_444689, EPI_ISL_444706, EPI_ISL_444708, EPI_ISL_444709, EPI_ISL_444710, EPI_ISL_444728, EPI_ISL_444729, EPI_ISL_444731, EPI_ISL_444733 | | | |
| see above | NYU Langone Health | Departments of Pathology and Medicine, New York University School of Medicine | Maria Aguero-Rosenfeld, Brendan Belovarac, Margaret Black, Ludovic Boytard, John Cadley, Paolo Cotzia, John Chen, Dacia Dimartino, Xiaojun Feng, Tatyana Gindin, Emily Guzman, Adriana Heguy, Megan Hogan, Emily Huang, George Jour, Alireza Khodadadi-Jamayran, Lawrence H. Lin, Raven Luther, Andrew Lytle, Christian Marier, Matthew T. Maurano, Mark J. Mulligan, Peter Meyn, Raquel Ordonez Ciriza, Iman Osman, Jared Pinnell, Vanessa Raabe, Sitharam Ramaswami, Amy Rapkiewicz, Andre M. Ribeiro-dos-Santos, Marie Samanovic-Golden, Antonio Serrano, Guomiao Shen, Matija Snuderl, Theodore Vougiouklakis, Nick Vulpescu, Gael Westby, Paul Zappile, Yutong Zhang |
| EPI_ISL_444847, EPI_ISL_444848, EPI_ISL_444954, EPI_ISL_444956, EPI_ISL_444958, EPI_ISL_444959, EPI_ISL_444960, EPI_ISL_444964, EPI_ISL_444968 | Department of Virus and Microbiological Special Diagnostics, Statens Serum Institut, Copenhagen, Denmark, Artillerivej 5, 2300 Copenahgen S | Albertsen lab, Department of Chemistry and Bioscience, Aalborg University, Denmark | Rasmus Kirkegaard |
| EPI_ISL_445116 | UC San Diego Center for Advanced Laboratory Medicine | Andersen lab at Scripps Research | SEARCH Alliance San Diego with David Pride, Ji H Shin |
| EPI_ISL_445119, EPI_ISL_445121, EPI_ISL_445122, EPI_ISL_445123, EPI_ISL_445125, EPI_ISL_445126, EPI_ISL_445127, EPI_ISL_445128, EPI_ISL_445130, EPI_ISL_445131, EPI_ISL_445133, EPI_ISL_445134, EPI_ISL_445135, EPI_ISL_445136, EPI_ISL_445137, EPI_ISL_445138, EPI_ISL_445139, EPI_ISL_445140, EPI_ISL_445141, EPI_ISL_445142, EPI_ISL_445143, EPI_ISL_445144, EPI_ISL_445145 | | | |
| see above | Robert Garry lab | Andersen lab at Scripps Research | Allison Smither, Gilberto Sabino-Santos, Patricia Snarski, Lilia Melnik, Antoinette Bell, Kaylynn Genemaras, Arnaud Drouin, Dahlene Fusco, Robert Garry with SEARCH Alliance San Diego |
| EPI_ISL_445173, EPI_ISL_445174, EPI_ISL_445177, EPI_ISL_445179 | UCSF Clinical Microbiology Laboratory | Chan-Zuckerberg Biohub | CZB Cliahub Consortium |
| EPI_ISL_445226 | Saroledens Familjelakare | The Public Health Agency of Sweden | Katarina Jarbur, Oskar Karlsson Lindsjo, Maria Lind Karlberg, Anna-Malin Linde, Olov Svartstrom, Anna Risberg, Theresa Enkirch, Mia Brytting, Karin Tegmark-Wisell |
| EPI_ISL_445227 | Uppsala Narakut Aleris | The Public Health Agency of Sweden | Annika Nilsson, Oskar Karlsson Lindsjo, Maria Lind Karlberg, Anna-Malin Linde, Olov Svartstrom, Anna Risberg, Theresa Enkirch, Mia Brytting, Karin Tegmark-Wisell |
| EPI_ISL_445228 | Ulltuna Vardcentral | The Public Health Agency of Sweden | Heidi Lindback, Oskar Karlsson Lindsjo, Maria Lind Karlberg, Anna-Malin Linde, Olov Svartstrom, Anna Risberg, Theresa Enkirch, Mia Brytting, Karin Tegmark-Wisell |
| EPI_ISL_445229 | Narhalsan Backa vardcentral | The Public Health Agency of Sweden | Mats Olsson, Oskar Karlsson Lindsjo, Maria Lind Karlberg, Anna-Malin Linde, Olov Svartstrom, Anna Risberg, Theresa Enkirch, Mia Brytting, Karin Tegmark-Wisell |
| EPI_ISL_445349, EPI_ISL_445350 | HOSPITAL SAN JUAN DE DIOS | Instituto de Salud Publica de Chile | Andrés E Castillo, Bárbara Parra,Paz Tapia, Jaime Lagos, Loredana Arata, Alejandra Acevedo, Winston Andrade, Gabriel Leal, Carolina Tambley, Patricia Bustos, Rodrigo Fasce, Jorge Fernandez |
| EPI_ISL_445352 | HOSPITAL DEL PROFESOR | Instituto de Salud Publica de Chile | Andrés E Castillo, Bárbara Parra,Paz Tapia, Jaime Lagos, Loredana Arata, Alejandra Acevedo, Winston Andrade, Gabriel Leal, Carolina Tambley, Patricia Bustos, Rodrigo Fasce, Jorge Fernandez |
| EPI_ISL_445381, EPI_ISL_445384, EPI_ISL_445385, EPI_ISL_445386, EPI_ISL_445387, EPI_ISL_445388, EPI_ISL_445397, EPI_ISL_445418, EPI_ISL_445419, EPI_ISL_445420, EPI_ISL_445421, EPI_ISL_445464, EPI_ISL_445465, EPI_ISL_445466, EPI_ISL_445518, EPI_ISL_445525, EPI_ISL_445568, EPI_ISL_445748, EPI_ISL_445782, EPI_ISL_445875, EPI_ISL_445876, EPI_ISL_445877, EPI_ISL_445878, EPI_ISL_445879, EPI_ISL_445880, EPI_ISL_445881, EPI_ISL_445882, EPI_ISL_445883, EPI_ISL_445884, EPI_ISL_445885, EPI_ISL_445886, EPI_ISL_445887, EPI_ISL_445888, EPI_ISL_445889, EPI_ISL_445890, EPI_ISL_445891, EPI_ISL_445892, EPI_ISL_445893, EPI_ISL_445894, EPI_ISL_445895, EPI_ISL_445896, EPI_ISL_445897, EPI_ISL_445898, EPI_ISL_445899, EPI_ISL_445900, EPI_ISL_445901, EPI_ISL_445902, EPI_ISL_445903, EPI_ISL_445904, EPI_ISL_445905, EPI_ISL_445906, EPI_ISL_445907, EPI_ISL_445908, EPI_ISL_445909, EPI_ISL_445910, EPI_ISL_445911, EPI_ISL_445912, EPI_ISL_445913, EPI_ISL_445914, EPI_ISL_445915, EPI_ISL_445916, EPI_ISL_445917, EPI_ISL_445918, EPI_ISL_445919, EPI_ISL_445920, EPI_ISL_445921, EPI_ISL_445922, EPI_ISL_445923, EPI_ISL_445924, EPI_ISL_445925, EPI_ISL_445926, EPI_ISL_445927, EPI_ISL_445928, EPI_ISL_445929, EPI_ISL_445930, EPI_ISL_445931, EPI_ISL_445932, EPI_ISL_445933, EPI_ISL_445934, EPI_ISL_445935, EPI_ISL_445936, EPI_ISL_445937, EPI_ISL_445938, EPI_ISL_445939, EPI_ISL_445940, EPI_ISL_445941, EPI_ISL_445942, EPI_ISL_445943, EPI_ISL_445944, EPI_ISL_445945, EPI_ISL_445946, EPI_ISL_445947, EPI_ISL_445948, EPI_ISL_445949, EPI_ISL_445950, EPI_ISL_445951, EPI_ISL_445952, EPI_ISL_445953, EPI_ISL_445954, EPI_ISL_445955, EPI_ISL_446271 | | | |
| see above | Wales Specialist Virology Centre | Public Health Wales Microbiology Cardiff | Catherine Moore, Johnathan Evans, Laura Gifford, Malorie Perry, Simon Cottrell, Alec Birchley, Alexander Adams, Amy Gaskin, Bree Gatica-Wilcox, Jason Coombes, Lauren Gilbert, Lee Graham, Nicole Pacchiarini, Sara Kumziene-Summerhayes, Sarah Taylor, Sophie Jones, Sara Rey, Matthew Bull, Joanne Watkins, Sally Corden, Tom Connor |
| EPI_ISL_447015, EPI_ISL_447016, EPI_ISL_447017, EPI_ISL_447018, EPI_ISL_447019, EPI_ISL_447020, EPI_ISL_447021 | Ramathibodi Hospital | COVID-19 Network Investigations (CONI) Alliance | Elizabeth Batty, Wasun Chantratita, Thanat Chookajorn, Stefan Fernandez, Angkana Huang, Anthony R. Jones, Khajohn Joonsalak, Chonticha Klungtong, Theerarat Kochakarn, Namfon Kotanan, Krittikorn Kumpornsin, Wudtichai Manasatienkij, Bhakbhoom Panthan, Ekawat Pasomsub, Kingkan Rakmanee, Insee Sensorn, Janjira Thaipadungpanit, Arporn Wangwiwatsin,Treewat Watthanachockchai |
| EPI_ISL_447090 | Michigan Department of Health and Human Services, Bureau of Laboratories | Michigan Department of Health and Human Services, Bureau of Laboratories | Blankenship HM, Riner D, Soehnlen MK |

| | | | |
|---|---|---|---|
| EPI_ISL_447265, EPI_ISL_447266 | Microbiology laboratory, Assuta Ashdod University-Affiliated Hospital | Stern Lab | Stern Lab |
| EPI_ISL_447317 | Clinical Virology Laboratory, Soroka Medical Center and the Faculty of Health Sciences, Ben-Gurion University of the Negev | Stern Lab | Stern Lab |
| EPI_ISL_447359, EPI_ISL_447360, EPI_ISL_447361, EPI_ISL_447362, EPI_ISL_447363, EPI_ISL_447364, EPI_ISL_447365, EPI_ISL_447366, EPI_ISL_447367, EPI_ISL_447368, EPI_ISL_447369, EPI_ISL_447370, EPI_ISL_447371, EPI_ISL_447372, EPI_ISL_447373, EPI_ISL_447374, EPI_ISL_447375, EPI_ISL_447376, EPI_ISL_447377, EPI_ISL_447378, EPI_ISL_447379, EPI_ISL_447380, EPI_ISL_447381, EPI_ISL_447382 | | | |
| see above | Clinical Virology Unit, Hadassah Hebrew University Medical Center | Stern Lab | Stern Lab |
| EPI_ISL_447383, EPI_ISL_447384, EPI_ISL_447403, EPI_ISL_447404, EPI_ISL_447405, EPI_ISL_447406 | Clinical Microbiology Laboratory, The Baruch Padeh Medical Center, Poriya | Stern Lab | Stern Lab |
| EPI_ISL_447516 | Servicio de Microbiologia. Hospital Clinico Universitario de Valencia | Sequencing and Bioinformatics Service and Molecular Epidemiology Research Group. FISABIO-Public Health | Lidia Ruiz Roldan, Neris Garcia-Gonzalez, Inma Galán Vendrell, Sandra Carbo, Loreto Ferrús Abad, Paula Ruiz-Hueso, Mariana Reyes-Prieto, Vicente Soriano Chirona, Ivan Ansari, Llúcia Martínez-Priego, Giuseppe 'Auria, David Navarro, Eliseo Albert, Maria Alma Bracho, Fernando Gonzalez-Candelas |
| EPI_ISL_447517 | Servicio de Microbiologia. Hospital Clinico Universitario de Valencia | Sequencing and Bioinformatics Service and Molecular Epidemiology Research Group. FISABIO-Public Health | Neris Garcia-Gonzalez, Inma Galán Vendrell, Sandra Carbo, Loreto Ferrús Abad, Paula Ruiz-Hueso, Mariana Reyes-Prieto, Vicente Soriano Chirona, Ivan Ansari, Llúcia Martínez-Priego, Giuseppe 'Auria, David Navarro, Eliseo Albert, Maria Alma Bracho, Lidia Ruiz Roldan, Fernando Gonzalez-Candelas |
| EPI_ISL_447518 | Servicio de Microbiologia. Hospital Clinico Universitario de Valencia | Sequencing and Bioinformatics Service and Molecular Epidemiology Research Group. FISABIO-Public Health | Inma Galán Vendrell, Sandra Carbo, Loreto Ferrús Abad, Paula Ruiz-Hueso, Mariana Reyes-Prieto, Vicente Soriano Chirona, Ivan Ansari, Llúcia Martínez-Priego, Giuseppe 'Auria, David Navarro, Eliseo Albert, Maria Alma Bracho, Lidia Ruiz Roldan, Neris Garcia-Gonzalez, Fernando Gonzalez-Candelas |
| EPI_ISL_447519 | Servicio de Microbiologia. Hospital Clinico Universitario de Valencia | Sequencing and Bioinformatics Service and Molecular Epidemiology Research Group. FISABIO-Public Health | Sandra Carbo, Loreto Ferrús Abad, Paula Ruiz-Hueso, Mariana Reyes-Prieto, Vicente Soriano Chirona, Ivan Ansari, Llúcia Martínez-Priego, Giuseppe 'Auria, David Navarro, Eliseo Albert, Maria Alma Bracho, Lidia Ruiz Roldan, Neris Garcia-Gonzalez, Inma Galán Vendrell, Fernando Gonzalez-Candelas |
| EPI_ISL_447520 | Servicio de Microbiologia. Hospital Clinico Universitario de Valencia | Sequencing and Bioinformatics Service and Molecular Epidemiology Research Group. FISABIO-Public Health | Loreto Ferrús Abad, Paula Ruiz-Hueso, Mariana Reyes-Prieto, Vicente Soriano Chirona, Ivan Ansari, Llúcia Martínez-Priego, Giuseppe 'Auria, David Navarro, Eliseo Albert, Maria Alma Bracho, Lidia Ruiz Roldan, Neris Garcia-Gonzalez, Inma Galán Vendrell, Sandra Carbo, Fernando Gonzalez-Candelas |
| EPI_ISL_447521 | Servicio de Microbiologia. Hospital Clinico Universitario de Valencia | Sequencing and Bioinformatics Service and Molecular Epidemiology Research Group. FISABIO-Public Health | Paula Ruiz-Hueso, Mariana Reyes-Prieto, Vicente Soriano Chirona, Ivan Ansari, Llúcia Martínez-Priego, Giuseppe 'Auria, David Navarro, Eliseo Albert, Maria Alma Bracho, Lidia Ruiz Roldan, Neris Garcia-Gonzalez, Inma Galán Vendrell, Sandra Carbo, Loreto Ferrús Abad, Fernando Gonzalez-Candelas |
| EPI_ISL_447522 | Servicio de Microbiologia. Hospital Clinico Universitario de Valencia | Sequencing and Bioinformatics Service and Molecular Epidemiology Research Group. FISABIO-Public Health | Vicente Soriano Chirona, Ivan Ansari, Llúcia Martínez-Priego, Giuseppe 'Auria, David Navarro, Eliseo Albert, Maria Alma Bracho, Lidia Ruiz Roldan, Neris Garcia-Gonzalez, Inma Galán Vendrell, Sandra Carbo, Loreto Ferrús Abad, Paula Ruiz-Hueso, Mariana Reyes-Prieto, Fernando Gonzalez-Candelas |
| EPI_ISL_447523 | Servicio de Microbiologia. Hospital Clinico Universitario de Valencia | Sequencing and Bioinformatics Service and Molecular Epidemiology Research Group. FISABIO-Public Health | Giuseppe 'Auria, David Navarro, Eliseo Albert, Maria Alma Bracho, Lidia Ruiz Roldan, Neris Garcia-Gonzalez, Inma Galán Vendrell, Sandra Carbo, Loreto Ferrús Abad, Paula Ruiz-Hueso, Mariana Reyes-Prieto, Vicente Soriano Chirona, Ivan Ansari, Llúcia Martínez-Priego, Fernando Gonzalez-Candelas |
| EPI_ISL_447524 | Servicio de Microbiologia. Hospital Clinico Universitario de Valencia | Sequencing and Bioinformatics Service and Molecular Epidemiology Research Group. FISABIO-Public Health | Llúcia Martínez-Priego, Giuseppe 'Auria, David Navarro, Eliseo Albert, Maria Alma Bracho, Lidia Ruiz Roldan, Neris Garcia-Gonzalez, Inma Galán Vendrell, Sandra Carbo, Loreto Ferrús Abad, Paula Ruiz-Hueso, Mariana Reyes-Prieto, Vicente Soriano Chirona, Ivan Ansari, Fernando Gonzalez-Candelas |
| EPI_ISL_447525 | Servicio de Microbiologia. Hospital Clinico Universitario de Valencia | Sequencing and Bioinformatics Service and Molecular Epidemiology Research Group. FISABIO-Public Health | David Navarro, Eliseo Albert, Maria Alma Bracho, Griselda De Marco, Lidia Ruiz Roldan, Neris Garcia-Gonzalez, Inma Galán Vendrell, Sandra Carbo, Loreto Ferrús Abad, Paula Ruiz-Hueso, Mariana Reyes-Prieto, Vicente Soriano Chirona, Ivan Ansari, Llúcia Martínez-Priego, Giuseppe 'Auria, Fernando Gonzalez-Candelas |
| EPI_ISL_447526 | Servicio de Microbiologia. Hospital Clinico Universitario de Valencia | Sequencing and Bioinformatics Service and Molecular Epidemiology Research Group. FISABIO-Public Health | Eliseo Albert, Maria Alma Bracho, Griselda De Marco, Lidia Ruiz Roldan, Neris Garcia-Gonzalez, Inma Galán Vendrell, Sandra Carbo, Loreto Ferrús Abad, Paula Ruiz-Hueso, Mariana Reyes-Prieto, Vicente Soriano Chirona, Ivan Ansari, Llúcia Martínez-Priego, Giuseppe 'Auria, David Navarro, Fernando Gonzalez-Candelas |
| EPI_ISL_447527 | Servicio de Microbiologia. Hospital Clinico Universitario de Valencia | Sequencing and Bioinformatics Service and Molecular Epidemiology Research Group. FISABIO-Public Health | Maria Alma Bracho, Griselda De Marco, Lidia Ruiz Roldan, Neris Garcia-Gonzalez, Inma Galán Vendrell, Sandra Carbo, Loreto Ferrús Abad, Paula Ruiz-Hueso, Mariana Reyes-Prieto, Vicente Soriano Chirona, Ivan Ansari, Llúcia Martínez-Priego, Giuseppe 'Auria, David Navarro, Eliseo Albert, Fernando Gonzalez-Candelas |
| EPI_ISL_447528 | Servicio de Microbiologia. Hospital Clinico Universitario de Valencia | Sequencing and Bioinformatics Service and Molecular Epidemiology Research Group. FISABIO-Public Health | Griselda De Marco, Lidia Ruiz Roldan, Neris Garcia-Gonzalez, Inma Galán Vendrell, Sandra Carbo, Loreto Ferrús Abad, Paula Ruiz-Hueso, Mariana Reyes-Prieto, Vicente Soriano Chirona, Ivan Ansari, Llúcia Martínez-Priego, Giuseppe 'Auria, David Navarro, Eliseo Albert, Maria Alma Bracho, Fernando Gonzalez-Candelas |
| EPI_ISL_447529 | Servicio de Microbiologia. Hospital Clinico Universitario de Valencia | Sequencing and Bioinformatics Service and Molecular Epidemiology Research Group. FISABIO-Public Health | Lidia Ruiz Roldan, Neris Garcia-Gonzalez, Inma Galán Vendrell, Sandra Carbo, Loreto Ferrús Abad, Paula Ruiz-Hueso, Mariana Reyes-Prieto, Vicente Soriano Chirona, Ivan Ansari, Llúcia Martínez-Priego, Giuseppe 'Auria, David Navarro, Eliseo Albert, Maria Alma Bracho, Fernando Gonzalez-Candelas |
| EPI_ISL_447530 | Servicio de Microbiologia. Hospital Clinico Universitario de Valencia | Sequencing and Bioinformatics Service and Molecular Epidemiology Research Group. FISABIO-Public Health | Neris Garcia-Gonzalez, Inma Galán Vendrell, Sandra Carbo, Loreto Ferrús Abad, Paula Ruiz-Hueso, Mariana Reyes-Prieto, Vicente Soriano Chirona, Ivan Ansari, Llúcia Martínez-Priego, Giuseppe 'Auria, David Navarro, Eliseo Albert, Maria Alma Bracho, Lidia Ruiz Roldan, Fernando Gonzalez-Candelas |
| EPI_ISL_447531 | Servicio de Microbiologia. Hospital Clinico Universitario de Valencia | Sequencing and Bioinformatics Service and Molecular Epidemiology Research Group. FISABIO-Public Health | Inma Galán Vendrell, Sandra Carbo, Loreto Ferrús Abad, Paula Ruiz-Hueso, Mariana Reyes-Prieto, Vicente Soriano Chirona, Ivan Ansari, Llúcia Martínez-Priego, Giuseppe 'Auria, David Navarro, Eliseo Albert, Maria Alma Bracho, Lidia Ruiz Roldan, Neris Garcia-Gonzalez, Fernando Gonzalez-Candelas |
| EPI_ISL_447559 | CSIR-Centre for Cellular and Molecular Biology | CSIR-Centre for Cellular and Molecular Biology | Sofia Banu, Payel Mukherjee, Priya Singh, Dhiviya Vedagiri, Divya Gupta, Vishal Sah, Santosh Kumar Kuncha, Krishnan Harinivas Harshan, Archana Bharadwaj Siva, Karthik Bharadwaj Tallapaka, Shagufta Khan, Lamuk Zaveri, Namami Gaur, Sakshi Shambhavi, Tulasi Nagabandi, Purushotham Vodnala, Rakesh K Mishra, Divya Tej Sowpati |
| EPI_ISL_447560 | CSIR-Centre for Cellular and Molecular Biology | CSIR-Centre for Cellular and Molecular Biology | Payel Mukherjee, Sofia Banu, Priya Singh, Dhiviya Vedagiri, Divya Gupta, Vishal Sah, Santosh Kumar Kuncha, Krishnan Harinivas Harshan, Archana Bharadwaj Siva, Karthik Bharadwaj Tallapaka, Shagufta Khan, Lamuk Zaveri, Namami Gaur, Sakshi Shambhavi, Tulasi Nagabandi, Purushotham Vodnala, Rakesh K Mishra, Divya Tej Sowpati |
| EPI_ISL_447561, EPI_ISL_447562 | CSIR-Centre for Cellular and Molecular Biology | CSIR-Centre for Cellular and Molecular Biology | Shagufta Khan, Lamuk Zaveri, Namami Gaur, Sakshi Shambhavi, Tulasi Nagabandi, Purushotham Vodnala, Payel Mukherjee, Sofia Banu, Priya Singh, Dhiviya Vedagiri, Divya Gupta, Vishal Sah, Santosh Kumar Kuncha, Krishnan Harinivas Harshan, Archana Bharadwaj Siva, Karthik Bharadwaj Tallapaka, Rakesh K Mishra, Divya Tej Sowpati |
| EPI_ISL_447563 | CSIR-Centre for Cellular and Molecular Biology | CSIR-Centre for Cellular and Molecular Biology | Sakshi Shambhavi, Lamuk Zaveri, Shagufta Khan, Namami Gaur, Tulasi Nagabandi, Purushotham Vodnala, Payel Mukherjee, Sofia Banu, Priya Singh, Dhiviya Vedagiri, Divya Gupta, Vishal Sah, Santosh Kumar Kuncha, Krishnan Harinivas Harshan, Archana Bharadwaj Siva, Karthik Bharadwaj Tallapaka, Rakesh K Mishra, Divya Tej Sowpati |
| EPI_ISL_447564, EPI_ISL_447565, EPI_ISL_447566 | CSIR-Centre for Cellular and Molecular Biology | CSIR-Centre for Cellular and Molecular Biology | Sofia Banu, Payel Mukherjee, Priya Singh, Dhiviya Vedagiri, Divya Gupta, Vishal Sah, Santosh Kumar Kuncha, Krishnan Harinivas Harshan, Archana Bharadwaj Siva, Karthik Bharadwaj Tallapaka, Shagufta Khan, Lamuk Zaveri, Namami Gaur, Sakshi Shambhavi, Tulasi Nagabandi, Purushotham Vodnala, Rakesh K Mishra, Divya Tej Sowpati |
| EPI_ISL_447567 | CSIR-Centre for Cellular and Molecular Biology | CSIR-Centre for Cellular and Molecular Biology | Payel Mukherjee, Sofia Banu, Priya Singh, Dhiviya Vedagiri, Divya Gupta, Vishal Sah, Santosh Kumar Kuncha, Krishnan Harinivas Harshan, Archana Bharadwaj Siva, Karthik Bharadwaj Tallapaka, Shagufta Khan, Lamuk Zaveri, Namami Gaur, Sakshi Shambhavi, Tulasi Nagabandi, Purushotham Vodnala, Rakesh K Mishra, Divya Tej Sowpati |
| EPI_ISL_447568, EPI_ISL_447569 | CSIR-Centre for Cellular and Molecular Biology | CSIR-Centre for Cellular and Molecular Biology | Shagufta Khan, Lamuk Zaveri, Namami Gaur, Sakshi Shambhavi, Tulasi Nagabandi, Purushotham Vodnala, Payel Mukherjee, Sofia Banu, Priya Singh, Dhiviya Vedagiri, Divya Gupta, Vishal Sah, Santosh Kumar Kuncha, Krishnan Harinivas Harshan, Archana Bharadwaj Siva, Karthik Bharadwaj Tallapaka, Rakesh K Mishra, Divya Tej Sowpati |
| EPI_ISL_447570, EPI_ISL_447571, EPI_ISL_447572 | CSIR-Centre for Cellular and Molecular Biology | CSIR-Centre for Cellular and Molecular Biology | Lamuk Zaveri, Shagufta Khan, Namami Gaur, Sakshi Shambhavi, Tulasi Nagabandi, Purushotham Vodnala, Payel Mukherjee, Sofia Banu, Priya Singh, Dhiviya Vedagiri, Divya Gupta, Vishal Sah, Santosh Kumar Kuncha, Krishnan Harinivas Harshan, Archana Bharadwaj Siva, Karthik Bharadwaj Tallapaka, |

| | | | |
|---|---|---|---|
| | | | Rakesh K Mishra, Divya Tej Sowpati |
| EPI_ISL_447573 | CSIR-Centre for Cellular and Molecular Biology | CSIR-Centre for Cellular and Molecular Biology | Sakshi Shambhavi, Lamuk Zaveri, Shagufta Khan, Namami Gaur, Tulasi Nagabandi, Purushotham Vodnala, Payel Mukherjee, Sofia Banu, Priya Singh, Dhiviya Vedagiri, Divya Gupta, Vishal Sah, Santosh Kumar Kuncha, Krishnan Harinivas Harshan, Archana Bharadwaj Siva, Karthik Bharadwaj Tallapaka, Rakesh K Mishra, Divya Tej Sowpati |
| EPI_ISL_447574 | CSIR-Centre for Cellular and Molecular Biology | CSIR-Centre for Cellular and Molecular Biology | Namami Gaur, Sakshi Shambhavi, Lamuk Zaveri, Shagufta Khan, Tulasi Nagabandi, Purushotham Vodnala, Payel Mukherjee, Sofia Banu, Priya Singh, Dhiviya Vedagiri, Divya Gupta, Vishal Sah, Santosh Kumar Kuncha, Krishnan Harinivas Harshan, Archana Bharadwaj Siva, Karthik Bharadwaj Tallapaka, Rakesh K Mishra, Divya Tej Sowpati |
| EPI_ISL_447593 | TSGH-CP molecular lab | TSGH-CP molecular lab | Cherng-Lih Perng, Ming-Jr JIAN, Chih-Kai Chang, Jung-Chung Lin, Kuo-Ming Yeh, Chien-Wen Chen, Sheng-Kang Chiu, Hsing-Yi Chung, Shih-Hung Tsai, Kuo-Sheng Hung, Tien-Yao Chang, Feng-Yee Chang, Hung-Sheng Shang |
| EPI_ISL_447754 | Grupo de Investigaciones Microbiológicas-UR (GIMUR), Departamento de Biología, Facultad de Ciencias Naturales, Universidad del Rosario, Bogotá, Colombia | Grupo de Investigaciones Microbiológicas-UR (GIMUR), Departamento de Biología, Facultad de Ciencias Naturales, Universidad del Rosario, Bogotá, Colombia Instituto Nacional de Salud, Bogotá, Colombia Icahn School of Medicine at Mount Sinai, New York, USA | Juan David Ramírez, Carolina Florez, Marina Muñoz, Carolina Hernandez, Adriana Castillo, Sergio Castañeda, Nathalia Ballesteros, David Martínez, Laura Vega, Jesús E. Jaimes, Sergio Gomez, Angelica Rico, Lisseth Pardo, Esther C. Barros, Martha L. Ospina, Aníbal A. Teherán, Ana S. Gonzalez-Reiche, Matthew M. Hernandez, Emilia Mia Sordillo, Viviana Simon, Harm van Bakel, Alberto Paniz-Mondolfi |
| EPI_ISL_447755, EPI_ISL_447756, EPI_ISL_447757, EPI_ISL_447758, EPI_ISL_447759, EPI_ISL_447760, EPI_ISL_447761, EPI_ISL_447762, EPI_ISL_447763, EPI_ISL_447764, EPI_ISL_447765, EPI_ISL_447766, EPI_ISL_447767, EPI_ISL_447768, EPI_ISL_447769, EPI_ISL_447771, EPI_ISL_447772, EPI_ISL_447774, EPI_ISL_447775, EPI_ISL_447776, EPI_ISL_447777, EPI_ISL_447778, EPI_ISL_447779, EPI_ISL_447780, EPI_ISL_447781 | | | |
| see above | Instituto Nacional de Salud, Bogotá, Colombia | Grupo de Investigaciones Microbiológicas-UR (GIMUR), Departamento de Biología, Facultad de Ciencias Naturales, Universidad del Rosario, Bogotá, Colombia Instituto Nacional de Salud, Bogotá, Colombia Icahn School of Medicine at Mount Sinai, New York, USA | Juan David Ramírez, Carolina Florez, Marina Muñoz, Carolina Hernandez, Adriana Castillo, Sergio Castañeda, Nathalia Ballesteros, David Martínez, Laura Vega, Jesús E. Jaimes, Sergio Gomez, Angelica Rico, Lisseth Pardo, Esther C. Barros, Martha L. Ospina, Aníbal A. Teherán, Ana S. Gonzalez-Reiche, Matthew M. Hernandez, Emilia Mia Sordillo, Viviana Simon, Harm van Bakel, Alberto Paniz-Mondolfi |
| EPI_ISL_447835 | unknown | Department of Medicine | Kassela,K., Dovrolis,N., Bampali,M., Gatzidou,E., Froukala,E., Stavropoulou,A., Veletza,S., Tsakris,A., Spanakis,N. and Karakasiliotis,I. |
| EPI_ISL_447847 | CSIR-Centre for Cellular and Molecular Biology | CSIR-Centre for Cellular and Molecular Biology | Payel Mukherjee, Sofia Banu, Priya Singh, Dhiviya Vedagiri, Divya Gupta, Vishal Sah, Santosh Kumar Kuncha, Krishnan Harinivas Harshan, Archana Bharadwaj Siva, Karthik Bharadwaj Tallapaka, Shagufta Khan, Lamuk Zaveri, Namami Gaur, Sakshi Shambhavi, Tulasi Nagabandi, Purushotham Vodnala, Rakesh K Mishra, Divya Tej Sowpati |
| EPI_ISL_447848 | CSIR-Centre for Cellular and Molecular Biology | CSIR-Centre for Cellular and Molecular Biology | Sofia Banu, Payel Mukherjee, Priya Singh, Dhiviya Vedagiri, Divya Gupta, Vishal Sah, Santosh Kumar Kuncha, Krishnan Harinivas Harshan, Archana Bharadwaj Siva, Karthik Bharadwaj Tallapaka, Shagufta Khan, Lamuk Zaveri, Namami Gaur, Sakshi Shambhavi, Tulasi Nagabandi, Purushotham Vodnala, Rakesh K Mishra, Divya Tej Sowpati |
| EPI_ISL_447849, EPI_ISL_447850 | CSIR-Centre for Cellular and Molecular Biology | CSIR-Centre for Cellular and Molecular Biology | Shagufta Khan, Lamuk Zaveri, Namami Gaur, Sakshi Shambhavi, Tulasi Nagabandi, Purushotham Vodnala, Payel Mukherjee, Sofia Banu, Priya Singh, Dhiviya Vedagiri, Divya Gupta, Vishal Sah, Santosh Kumar Kuncha, Krishnan Harinivas Harshan, Archana Bharadwaj Siva, Karthik Bharadwaj Tallapaka, Rakesh K Mishra, Divya Tej Sowpati |
| EPI_ISL_447851, EPI_ISL_447852 | CSIR-Centre for Cellular and Molecular Biology | CSIR-Centre for Cellular and Molecular Biology | Lamuk Zaveri, Shagufta Khan, Namami Gaur, Sakshi Shambhavi, Tulasi Nagabandi, Purushotham Vodnala, Payel Mukherjee, Sofia Banu, Priya Singh, Dhiviya Vedagiri, Divya Gupta, Vishal Sah, Santosh Kumar Kuncha, Krishnan Harinivas Harshan, Archana Bharadwaj Siva, Karthik Bharadwaj Tallapaka, Rakesh K Mishra, Divya Tej Sowpati |
| EPI_ISL_447853 | CSIR-Centre for Cellular and Molecular Biology | CSIR-Centre for Cellular and Molecular Biology | Namami Gaur, Sakshi Shambhavi, Lamuk Zaveri, Shagufta Khan, Tulasi Nagabandi, Purushotham Vodnala, Payel Mukherjee, Sofia Banu, Priya Singh, Dhiviya Vedagiri, Divya Gupta, Vishal Sah, Santosh Kumar Kuncha, Krishnan Harinivas Harshan, Archana Bharadwaj Siva, Karthik Bharadwaj Tallapaka, Rakesh K Mishra, Divya Tej Sowpati |
| EPI_ISL_447862 | CSIR-Centre for Cellular and Molecular Biology | CSIR-Centre for Cellular and Molecular Biology | Payel Mukherjee, Sofia Banu, Priya Singh, Dhiviya Vedagiri, Divya Gupta, Vishal Sah, Santosh Kumar Kuncha, Krishnan Harinivas Harshan, Archana Bharadwaj Siva, Karthik Bharadwaj Tallapaka, Shagufta Khan, Lamuk Zaveri, Namami Gaur, Sakshi Shambhavi, Tulasi Nagabandi, Purushotham Vodnala, Rakesh K Mishra, Divya Tej Sowpati |
| EPI_ISL_447889, EPI_ISL_447891 | University of California, Davis | Chan-Zuckerberg Biohub | CZB Cliahub Consortium |
| EPI_ISL_448116 | West of Scotland Specialist Virology Centre, NHSGGC / MRC-University of Glasgow Centre for Virus Research | COVID-19 Genomics UK (COG-UK) Consortium | Ana da Silva Filipe, Natasha Johnson, Kathy Smollett, Daniel Mair, Stephen Carmichael, Lily Tong, Jenna Nichols, Elihu Aranday-Cortes, Kirstyn Brunker, Yasmin Parr, Kyriaki Nomikou, Sarah McDonald, Marc Niebel, Patawee Asamaphan, Richard Orton, Joseph Hughes, Sreenu Vattipally, David L Robertson, Alasdair MacLean, Rory Gunson, Kathy Li, Natasha Jesudason, Rajiv Shah, James Shepherd, Antonia Ho, Emma Thomson |
| EPI_ISL_448894 | Virology Laboratory, Castle Hill Hospital, Hull University Teaching Hospitals NHS Trust/Department of Infection, Immunity and Cardiovascular Disease, The Medical School, University of Sheffield | COVID-19 Genomics UK (COG-UK) Consortium | Thushan de Silva, Matthew Parker, Nikki Smith, Adri Angyal, Rebecca Brown, Luke Green, Rachel Tucker, Paul Parsons, Danielle Groves, Katie Johnson, Laura Carrilero, Alex Keeley, Dave Partridge, Matthew Wyles, Benjamin Lindsey, Mehmet Yavuz, Mohammad Raza, Cariad Evans |
| EPI_ISL_449381, EPI_ISL_449382, EPI_ISL_449383, EPI_ISL_449384, EPI_ISL_449385, EPI_ISL_449386, EPI_ISL_449387, EPI_ISL_449388, EPI_ISL_449389, EPI_ISL_449448, EPI_ISL_449449, EPI_ISL_449450, EPI_ISL_449451, EPI_ISL_449452, EPI_ISL_449504, EPI_ISL_449505, EPI_ISL_449506, EPI_ISL_449507, EPI_ISL_449508, EPI_ISL_449509, EPI_ISL_449510, EPI_ISL_449511, EPI_ISL_449512, EPI_ISL_449513, EPI_ISL_449514, EPI_ISL_449515, EPI_ISL_449517, EPI_ISL_449518, EPI_ISL_449519, EPI_ISL_449520, EPI_ISL_449521, EPI_ISL_449522, EPI_ISL_449523, EPI_ISL_449530, EPI_ISL_449531, EPI_ISL_449532, EPI_ISL_449534, EPI_ISL_449536, EPI_ISL_449540, EPI_ISL_449542, EPI_ISL_449543, EPI_ISL_449567, EPI_ISL_449568, EPI_ISL_449569, EPI_ISL_449570, EPI_ISL_449571, EPI_ISL_449572, EPI_ISL_449573, EPI_ISL_449574, EPI_ISL_449576, EPI_ISL_449577, EPI_ISL_449578, EPI_ISL_449579 | | | |
| see above | Liverpool Clinical Laboratories | COVID-19 Genomics UK (COG-UK) Consortium | Sam Haldenby, Anita Lucaci, Steve Paterson, Julian Hiscox, Alistair Darby, M Almsaud, A Alrezaihi, Muhannad Alruwaili, Stuart D Armstrong, Jones Benjamin , Eleanor G Bentley, Anu Chawla, Jordan J Clark, Angela Cowell, Richard Eccles, Isabel García-Dorival, Matthew Gemmell, Alessandro Gerada, PKF Gilmore, Richard Gregory, Ximeng Han, Catherine Hartley, Margaret Hughes, Miren Iturriza-Gomara, James Johnson, L Luu, Jenifer Manson , Charlotte Nelson, Elaine O'Toole, Cassie Olateju, Rebekah Penrice-Randal , Lucille Rainbow, N.P Randle, Trevor Ian Robinson, Parul Sharma, Ghada T Shawli, James P Stewart , Neil Swainston, Ecaterina Vamos, Joanne Watts, Mark Whitehead |
| EPI_ISL_449790 | Dept. of Medical Microbiology, Stavanger University Hospital, Helse Stavanger HF | Norwegian Institute of Public Health, Department of Virology | Kathrine Stene-Johansen, Kamilla Heddeland Instefjord, Hilde Elshaug, Rasmus Riis Kopperud, Karoline Bragstad, Olav Hungnes |
| EPI_ISL_449800 | HOSPITAL SAN JUAN DE DIOS | Instituto de Salud Publica de Chile | Andrés E Castillo, Bárbara Parra,Paz Tapia, Jaime Lagos, Loredana Arata, Alejandra Acevedo, Winston Andrade, Gabriel Leal, Carolina Tambley, Patricia Bustos, Rodrigo Fasce, Jorge Fernandez |
| EPI_ISL_449811, EPI_ISL_449812, EPI_ISL_449813 | Utah Public Health Laboratory | Utah Public Health Laboratory | Erin Young, Kelly Oakeson |
| EPI_ISL_450217, EPI_ISL_450218, EPI_ISL_450219, EPI_ISL_450220, EPI_ISL_450221, EPI_ISL_450222, EPI_ISL_450223, EPI_ISL_450224, EPI_ISL_450225, EPI_ISL_450226, EPI_ISL_450227, EPI_ISL_450228, EPI_ISL_450229, EPI_ISL_450230 | | | |
| see above | unknown | Hamadan University of Medical Sciences | Teimoori,A., Azizi Jallilian,F., Ansari,N., Jamehdor,S., Nazari,A., Saadat,N., Mazaheri,Z., Zanjani,M. |
| EPI_ISL_450231 | Robert Garry lab | Andersen lab at Scripps Research | Allison Smither, Gilberto Sabino-Santos, Patricia Snarski, Lilia Melnik, Antoinette Bell, Kaylynn Genemaras, Arnaud Drouin, Dahlene Fusco, Robert Garry with SEARCH Alliance San Diego |
| EPI_ISL_450299, EPI_ISL_450301 | National Institute for Communicable Diseases of the National Health Laboratory Service | National Institute for Communicable Diseases of the National Health Laboratory Service | Allam M, Ismail A, Khumalo Z, Kwenda S, van Heusden P, Mtshali P, Mnyameni F, Mohale T, Subramoney K, Bhiman JN |
| EPI_ISL_450326 | CSIR-Centre for Cellular and Molecular Biology | CSIR-Centre for Cellular and Molecular Biology | Payel Mukherjee, Sofia Banu, Priya Singh, Dhiviya Vedagiri, Divya Gupta, Vishal Sah, Santosh Kumar Kuncha, Krishnan Harinivas Harshan, Archana Bharadwaj Siva, Karthik Bharadwaj Tallapaka, Shagufta Khan, Lamuk Zaveri, Namami Gaur, Sakshi Shambhavi, Tulasi Nagabandi, Purushotham Vodnala,G. Aditya Kumar, Koushick Sivakumar, Pooja Ramesh Gupta, Rajan Kumar Jha, Shraddha Vijay Lahoti, Deepak Kumar, Devi Prasad Vijayashankara, Disha Nanda, Divya Das, Jotin Gogoi, Manish Bhattacharjee, Rakesh K Mishra, Divya Tej Sowpati |
| EPI_ISL_450327 | CSIR-Centre for Cellular and Molecular Biology | CSIR-Centre for Cellular and Molecular Biology | Sofia Banu, Payel Mukherjee, Priya Singh, Dhiviya Vedagiri, Divya Gupta, Vishal Sah, Santosh Kumar Kuncha, Krishnan Harinivas Harshan, Archana Bharadwaj Siva, Karthik Bharadwaj Tallapaka, Shagufta Khan, Lamuk Zaveri, Namami Gaur, Sakshi Shambhavi, Tulasi Nagabandi, Purushotham Vodnala, |

| | | | |
|---|---|---|---|
| | | | Disha Nanda, Divya Das, Jotin Gogoi, Manish Bhattacharjee, Ravi Prasad Mukku, Renu Sudhakar, Somesh Gorde, Gangumala Srinivas Reddy, Sujoy Deb, Swati Bayyana, Zeba Rizvi, Rakesh K Mishra, Divya Tej Sowpati |
| EPI_ISL_450328 | CSIR-Centre for Cellular and Molecular Biology | CSIR-Centre for Cellular and Molecular Biology | Shagufta Khan, Lamuk Zaveri, Namami Gaur, Sakshi Shambhavi, Tulasi Nagabandi, Purushotham Vodnala, Payel Mukherjee, Sofia Banu, Priya Singh, Dhiviya Vedagiri, Divya Gupta, Vishal Sah, Santosh Kumar Kuncha, Krishnan Harinivas Harshan, Archana Bharadwaj Siva, Karthik Bharadwaj Tallapaka, Zeba Rizvi, Zuberwasim Sayyad, Kakade Aishwarya Arun, Amrutha H C, Ananga Ghosh, Kezia J Ann, Radhika Khandelwal, Roshan Maku Venkata, Shemin Mansuri, Sonu Uday, Sudipta Mondal, Rakesh K Mishra, Divya Tej Sowpati |
| EPI_ISL_450329 | CSIR-Centre for Cellular and Molecular Biology | CSIR-Centre for Cellular and Molecular Biology | Namami Gaur, Sakshi Shambhavi, Lamuk Zaveri, Shagufta Khan, Tulasi Nagabandi, Purushotham Vodnala, Payel Mukherjee, Sofia Banu, Priya Singh, Dhiviya Vedagiri, Divya Gupta, Vishal Sah, Santosh Kumar Kuncha, Krishnan Harinivas Harshan, Archana Bharadwaj Siva, Karthik Bharadwaj Tallapaka, Sonu Uday, Sudipta Mondal, Annapoorna P Karthyayani, Debabrata Jana, Debrya Saha, Gokulan C G, Gunjan Purohit, Hanuman Tulashiram Kale, Pankaj Kumar, Prachand Issarapu, Preethi Jampala Rakesh K Mishra, Divya Tej Sowpati |
| EPI_ISL_450331 | CSIR-Centre for Cellular and Molecular Biology | CSIR-Centre for Cellular and Molecular Biology | Tulasi Nagabandi, Namami Gaur, Sakshi Shambhavi, Lamuk Zaveri, Shagufta Khan, Purushotham Vodnala, Payel Mukherjee, Sofia Banu, Priya Singh, Dhiviya Vedagiri, Divya Gupta, Vishal Sah, Santosh Kumar Kuncha, Krishnan Harinivas Harshan, Archana Bharadwaj Siva, Karthik Bharadwaj Tallapaka,G. Aditya Kumar, Koushick Sivakumar, Pooja Ramesh Gupta, Rajan Kumar Jha, Shraddha Vijay Lahoti, Deepak Kumar, Devi Prasad Vijayashankara, Disha Nanda, Divya Das, Jotin Gogoi, Manish Bhattacharjee, Rakesh K Mishra, Divya Tej Sowpati |
| EPI_ISL_450332 | CSIR-Centre for Cellular and Molecular Biology | CSIR-Centre for Cellular and Molecular Biology | Payel Mukherjee, Sofia Banu, Priya Singh, Dhiviya Vedagiri, Divya Gupta, Vishal Sah, Santosh Kumar Kuncha, Krishnan Harinivas Harshan, Archana Bharadwaj Siva, Karthik Bharadwaj Tallapaka, Shagufta Khan, Lamuk Zaveri, Namami Gaur, Sakshi Shambhavi, Tulasi Nagabandi, Purushotham Vodnala,G. Aditya Kumar, Koushick Sivakumar, Pooja Ramesh Gupta, Rajan Kumar Jha, Shraddha Vijay Lahoti, Deepak Kumar, Devi Prasad Vijayashankara, Disha Nanda, Divya Das, Jotin Gogoi, Manish Bhattacharjee, Rakesh K Mishra, Divya Tej Sowpati |
| EPI_ISL_450401 | NYU Langone Health | Departments of Pathology and Medicine, New York University School of Medicine | Maria Aguero-Rosenfeld, Brendan Belovarac, Margaret Black, Ludovic Boytard, John Cadley, Paolo Cotzia, John Chen, Dacia Dimartino, Xiaojun Feng, Tatyana Gindin, Emily Guzman, Adriana Heguy, Megan Hogan, Emily Huang, George Jour, Alireza Khodadadi-Jamayran, Lawrence H. Lin, Raven Luther, Andrew Lytle, Christian Marier, Matthew T. Maurano, Mark J. Mulligan, Peter Meyn, Raquel Ordonez Ciriza, Iman Osman, Jared Pinnell, Vanessa Raabe, Sitharam Ramaswami, Amy Rapkiewicz, Andre M. Ribeiro-dos-Santos, Marie Samanovic-Golden, Antonio Serrano, Guomiao Shen, Matija Snuderl, Theodore Vougiouklakis, Nick Vulpescu, Gael Westby, Paul Zappile, Yutong Zhang |
| EPI_ISL_450406 | Molecular Diagnostics, Antech Diagnostics | Molecular Diagnostics, Antech Diagnostics | Leutenegger,C.M., Lozoya,C.E., Tereski,J.L., Wyler,D. and Moroff,S. |
| EPI_ISL_450443 | Institute for Forensic Medicine, Faculty of Medicine, University of Belgrade | Institute for Forensic Medicine, Faculty of Medicine, University of Belgrade | Vidanovic,D., Skadric,I.R., Dordevic,N., Tolic,A., Tesovic,B., Sekler,M., Dmitric,M., Debeljak,Z., Zarkovic,A., Kolarevic,M., Petrovic,T. and Baskic,D. |
| EPI_ISL_450486, EPI_ISL_450487 | unknown | Data Science | Carroll,T.D., Tran,N.K., Cohen,S.H., Miller,C.J. |
| EPI_ISL_450529, EPI_ISL_450530 | Hematology Laboratory, Section of Molecular Diagnostics, University Clinical Centre, Medical University of Gdansk | Department of Virology, Faculty of Medicine, University of Helsinki, Helsinki, Finland | Maciej Grzybek, Marlena Robakowska, Aneta Szulc, Olli Vapalahti, Teemu Smura |
| EPI_ISL_450708, EPI_ISL_450721 | University of Wisconsin-Madison AIDS Vaccine Research Laboratories | University of Wisconsin-Madison AIDS Vaccine Research Laboratories | Gage Moreno, Katarina Braun, et al. AIDS Vaccine Research Laboratories |
| EPI_ISL_450731, EPI_ISL_450732 | Hospital AZ Rivierenland | Institute of Tropical Medicine | Philippe Selhorst, Colin Anthony |
| EPI_ISL_450756, EPI_ISL_450757, EPI_ISL_450758, EPI_ISL_450769 | Minnesota Department of Health, Public Health Laboratory | Minnesota Department of Health, Public Health Laboratory | Matt Plumb, Jacob Garfin, and Xiong Wang |
| EPI_ISL_450800 | AR Dept. of Health-Public Health Lab | Pathogen Discovery, Respiratory Viruses Branch, Division of Viral Diseases, Centers for Disease Control and Prevention | Yan Li, Anna Montmayeur, Ying Tao, Krista Queen, Jing Zhang, Anna Uehara, Clinton R. Paden, Rachel Marine, Haibin Wang, Zachary Weiner, Bettina Bankamp, Suxiang Tong |
| EPI_ISL_450802 | PA Department of Health, Bureau of Laboratories | Pathogen Discovery, Respiratory Viruses Branch, Division of Viral Diseases, Centers for Disease Control and Prevention | Yan Li, Anna Montmayeur, Ying Tao, Krista Queen, Jing Zhang, Anna Uehara, Clinton R. Paden, Rachel Marine, Haibin Wang, Zachary Weiner, Bettina Bankamp, Suxiang Tong |
| EPI_ISL_450808 | Saroledens Familjelakare | The Public Health Agency of Sweden | Katarina Jarbur, Anna-Malin Linde, Maria Lind Karlberg, Oskar Karlsson Lindsjo, Olov Svartstrom, Anna Risberg, Theresa Enkirch, Mia Brytting, Karin Tegmark-Wisell |
| EPI_ISL_450809 | Kungsholmsdoktorn | The Public Health Agency of Sweden | Linus Hammar, Anna-Malin Linde, Maria Lind Karlberg, Oskar Karlsson Lindsjo, Olov Svartstrom, Anna Risberg, Theresa Enkirch, Mia Brytting, Karin Tegmark-Wisell |
| EPI_ISL_450810 | Saroledens Familjelakare | The Public Health Agency of Sweden | Katarina Jarbur, Anna-Malin Linde, Maria Lind Karlberg, Oskar Karlsson Lindsjo, Olov Svartstrom, Anna Risberg, Theresa Enkirch, Mia Brytting, Karin Tegmark-Wisell |
| EPI_ISL_450901, EPI_ISL_450941, EPI_ISL_450981, EPI_ISL_451021, EPI_ISL_451061 | Center of Excellence in Clinical Virology | Center of Excellence in Clinical Virology | Puenpa,J., Chansaenroj,J., Nilyanimit,P., Auphimai,C., Yorsaeng,R., Suwannakarn,K., Poovorawan,Y. |
| EPI_ISL_451109, EPI_ISL_451110, EPI_ISL_451111, EPI_ISL_451112, EPI_ISL_451113, EPI_ISL_451114, EPI_ISL_451115, EPI_ISL_451116, EPI_ISL_451117, EPI_ISL_451118, EPI_ISL_451119, EPI_ISL_451120, EPI_ISL_451121, EPI_ISL_451122, EPI_ISL_451123, EPI_ISL_451124, EPI_ISL_451125, EPI_ISL_451126, EPI_ISL_451127, EPI_ISL_451143 | | | |
| see above | SA Pathology | SA Pathology | Lex Leong, Chuan Kok Lim, Mark Turra, Ivan Bastian, Geoff Higgins |
| EPI_ISL_451165, EPI_ISL_451166, EPI_ISL_451167 | Lab voor klinische biologie | Onderzoeksgroep Virologie | Laurens Lambrechts, Nick Vereecke, Marthe Pauwels, Jozefien De Clercq, Bruno Verhasselt, Linos Vandekerckhove, Hans Nauwynck, Sebastiaan Theuns |
| EPI_ISL_451496, EPI_ISL_451538, EPI_ISL_451546, EPI_ISL_451547 | Pathology West - NSW Health Pathology | NSW Health Pathology - Institute of Clinical Pathology and Medical Research; Westmead Hospital; University of Sydney | CIDM-PH et al. |
| EPI_ISL_451549 | Pathology Sydney South West - NSW Health Pathology | NSW Health Pathology - Institute of Clinical Pathology and Medical Research; Westmead Hospital; University of Sydney | CIDM-PH et al. |
| EPI_ISL_451553 | Medlab Pathology | NSW Health Pathology - Institute of Clinical Pathology and Medical Research; Westmead Hospital; University of Sydney | CIDM-PH et al. |
| EPI_ISL_451557 | Pathology West - NSW Health Pathology | NSW Health Pathology - Institute of Clinical Pathology and Medical Research; Westmead Hospital; University of Sydney | CIDM-PH et al. |
| EPI_ISL_451564 | Pathology Sydney South West - NSW Health Pathology | NSW Health Pathology - Institute of Clinical Pathology and Medical Research; Westmead Hospital; University of Sydney | CIDM-PH et al. |
| EPI_ISL_451570, EPI_ISL_451577, EPI_ISL_451580 | Pathology West - NSW Health Pathology | NSW Health Pathology - Institute of Clinical Pathology and Medical Research; Westmead Hospital; University of Sydney | CIDM-PH et al. |
| EPI_ISL_451581 | Pathology Sydney South West - NSW Health Pathology | NSW Health Pathology - Institute of Clinical Pathology and Medical Research; Westmead Hospital; University of Sydney | CIDM-PH et al. |
| EPI_ISL_451582 | Pathology West - NSW Health Pathology | NSW Health Pathology - Institute of Clinical Pathology and Medical Research; Westmead Hospital; University of Sydney | CIDM-PH et al. |
| EPI_ISL_451591 | ACT pathology | NSW Health Pathology - Institute of Clinical Pathology and Medical Research; Westmead Hospital; University of Sydney | CIDM-PH et al. |
| EPI_ISL_451599 | Australian Clinical Labs | NSW Health Pathology - Institute of Clinical Pathology and Medical Research; Westmead Hospital; University of Sydney | CIDM-PH et al. |

| | | | |
|---|---|---|---|
| EPI_ISL_451603 | Pathology Sydney South West - NSW Health Pathology | NSW Health Pathology - Institute of Clinical Pathology and Medical Research; Westmead Hospital; University of Sydney | CIDM-PH et al. |
| EPI_ISL_451606 | Pathology West - NSW Health Pathology | NSW Health Pathology - Institute of Clinical Pathology and Medical Research; Westmead Hospital; University of Sydney | CIDM-PH et al. |
| EPI_ISL_451805, EPI_ISL_451806, EPI_ISL_451807, EPI_ISL_451808, EPI_ISL_451809, EPI_ISL_451810, EPI_ISL_451811, EPI_ISL_451812, EPI_ISL_451813, EPI_ISL_451814, EPI_ISL_451815, EPI_ISL_451816, EPI_ISL_451817, EPI_ISL_451818, EPI_ISL_451819, EPI_ISL_451820, EPI_ISL_451821, EPI_ISL_451822, EPI_ISL_451823, EPI_ISL_451824, EPI_ISL_451825, EPI_ISL_451826, EPI_ISL_451827, EPI_ISL_451828, EPI_ISL_451829, EPI_ISL_451830, EPI_ISL_451831, EPI_ISL_451832, EPI_ISL_451833, EPI_ISL_451834, EPI_ISL_451835, EPI_ISL_451836, EPI_ISL_451837, EPI_ISL_451838, EPI_ISL_451839, EPI_ISL_451840, EPI_ISL_451841, EPI_ISL_451842, EPI_ISL_451843, EPI_ISL_451844, EPI_ISL_451846, EPI_ISL_451847, EPI_ISL_451848, EPI_ISL_451849, EPI_ISL_451850, EPI_ISL_451851, EPI_ISL_451852, EPI_ISL_451853, EPI_ISL_451854, EPI_ISL_451855 | | | |
| see above | Viollier AG | Department of Biosystems Science and Engineering, ETH Zürich | Christian Beisel, Sarah Nadeau, Ivan Topolsky, Pedro Ferreira, Philipp Jablonski, Susana Posada-Céspedes, Tobias Schär, Ina Nissen, Natascha Santacroce, Elodie Burcklen, Christiane Beckmann, Maurice Redondo, Olivier Kobel, Christoph Noppen, Sophie Seidel, Noemie Santamaria de Souza, Niko Beerenwinkel, Tanja Stadler |
| EPI_ISL_451944, EPI_ISL_451945 | Max von Pettenkofer Institute, Virology, National Reference Center for Retroviruses, LMU München | Laboratory for Functional Genome Analysis, Dept. Genomics, Gene Center of the LMU Munich | Max Muenchhoff, Stefan Krebs, Alexander Graf, Oliver Keppler, Helmut Blum |
| EPI_ISL_451957 | Molecular Pathology Division, Department of Pathology, Hong Kong Sanatorium & Hospital | Molecular Pathology Division, Department of Pathology, Hong Kong Sanatorium & Hospital | Chun Hang AU, Wai Sing CHAN, Ho Yin LAM, Dona N. HO, Simon Y.M. LAM, Jonpaul S.T. ZEE, Tsun Leung CHAN, Edmond S.K. MA |
| EPI_ISL_451959 | Alaska State Virology Laboratory | Alaska State Virology Laboratory | DeRonde,S., Deuling,H. and Chen,J. |
| EPI_ISL_451960 | VI-US Virgin Islands Department of Health | Centers for Disease Control and Prevention | Queen,K., Li,Y., Montmayeur,A., Tao,Y., Zhang,J., Uehara,A., Paden,C.R., Marine,R., Wang,H., Padilla,J., Lee,J., Weiner,Z., Bankamp,B. and Tong,S. |
| EPI_ISL_451971, EPI_ISL_451973, EPI_ISL_451975, EPI_ISL_451978, EPI_ISL_451979, EPI_ISL_451987 | 1. ViroGenetics - BSL3 Laboratory of Virology, Maopolska Centre of Biotechnology, Jagiellonian University; 2. II Department of Internal Medicine, Faculty of Medicine, Jagiellonian University Medical College; 3. DIAGNOSTYKA Ltd. | 1. ViroGenetics - BSL3 Laboratory of Virology, Maopolska Centre of Biotechnology, Jagiellonian University; 2. II Department of Internal Medicine, Faculty of Medicine, Jagiellonian University Medical College. | Marek Sanak, Marcin Surmiak, Monika Gsecka-Czapla, Wojciech Branicki, Pawe P abaj, Marta Rogalska-Kupiec, Jakub Swadba, Krzysztof Pyr |
| EPI_ISL_452148 | CUB Hopital Erasme Laboratoire d'Anatomie Pathologique | CUB Hopital Erasme Laboratoire d'Anatomie Pathologique | Isabelle Salmon, Nicky D'Haene |
| EPI_ISL_452187, EPI_ISL_452188, EPI_ISL_452189 | ULSS9 Distretto di Bussolengo | Istituto Zooprofilattico Sperimentale delle Venezie | Adelaide Milani, Alessia Schivo, Annalisa Salviato, Erika Giorgia Quaranta, Ambra Pastori, Bianca Zecchin, Alice Fusaro, Isabella Monne, Calogero Terregino, Antonia Ricci |
| EPI_ISL_452234 | Din Klinik | The Public Health Agency of Sweden | Helene Warnborg, Anna-Malin Linde, Maria Lind Karlberg, Oskar Karlsson Lindsjo, Olov Svartstrom, Anna Risberg, Theresa Enkirch, Mia Brytting, Karin Tegmark-Wisell |
| EPI_ISL_452308 | Michigan Department of Health and Human Services, Bureau of Laboratories | Michigan Department of Health and Human Services, Bureau of Laboratories | Blankenship HM, Riner D, Soehnlen MK |
| EPI_ISL_452371 | Servicio de Microbiología. HRU de Málaga. Servicio Andaluz de Salud | SeqCOVID-SPAIN consortium/IBV(CSIC) | Inmaculada de Toro Peinado, Maria Concepción Mediavilla Gradolph, Begoña Palop Borrás and SeqCOVID-SPAIN consortium |
| EPI_ISL_452609, EPI_ISL_452610, EPI_ISL_452611, EPI_ISL_452612, EPI_ISL_452613, EPI_ISL_452614, EPI_ISL_452615, EPI_ISL_452616 | Servicio de Microbiología y Parasitología clínica. UCEIMP. Hospital Universitario Virgen del Rocío/IBIS/CSIC/US. | SeqCOVID-SPAIN consortium/IBV(CSIC) | Guillermo Martí-n Gutiérrez, Ángel Rodrí-guez Villodres, Lidia Gálvez Benitez, Verónica González Galán, Javier Aznar Martí-n and SeqCOVID-SPAIN consortium |
| EPI_ISL_452809 | Virginia DCLS | Virginia DCLS | Virginia DCLS |
| EPI_ISL_453006, EPI_ISL_453007 | West of Scotland Specialist Virology Centre, NHSGGC / MRC-University of Glasgow Centre for Virus Research | COVID-19 Genomics UK (COG-UK) Consortium | Ana da Silva Filipe, Natasha Johnson, Kathy Smollett, Daniel Mair, Stephen Carmichael, Lily Tong, Jenna Nichols, Elihu Aranday-Cortes, Kirstyn Brunker, Yasmin Parr, Kyriaki Nomikou; Sarah McDonald, Marc Niebel, Patawee Asamaphan; Richard Orton, Joseph Hughes, Sreenu Vattipally, David L Robertson; Alasdair MacLean, Rory Gunson; Kathy Li, Natasha Jesudason, Rajiv Shah, James Shepherd, Antonia Ho, Emma Thomson |
| EPI_ISL_453157, EPI_ISL_453164, EPI_ISL_453165, EPI_ISL_453166, EPI_ISL_453186, EPI_ISL_453187, EPI_ISL_453188, EPI_ISL_453190 | Virology Department, Royal Infirmary of Edinburgh, NHS Lothian / School of Biological Sciences, University of Edinburgh / Institute of Genetics and Molecular Medicine, University of Edinburgh | COVID-19 Genomics UK (COG-UK) Consortium | McHugh M, Dewar R, Rooke S, Gallagher M, Balcaza C, O'Toole Á, Scher E, Hill V, McCrone JT, Colquhoun R, Yu X, Jackson B, Rambaut A, Williams TC, Templeton K |
| EPI_ISL_453504, EPI_ISL_453505, EPI_ISL_453506 | Northumbria University / South Tees Hospitals NHS Foundation Trust / North Cumbria Integrated Care NHS Foundation Trust / North Tees and Hartlepool NHS Foundation Trust / Newcastle Hospitals NHS Foundation Trust | COVID-19 Genomics UK (COG-UK) Consortium | Darren L Smith,Andrew Nelson,Matthew Bashton,Greg R Young,Joshua Loh,John Allan,Mohammad A Tariq,Giles S Holt,Gary Black,Wen C Yew,Lynn Dover ,Paul Baker,Steve Liggett,Sarah Essex,Jane Greenaway ,Debra Padgett,Clive Graham,Garren Scott,Edward Barton ,Emma Swindells ,Brendan Payne,Jennifer Collins,Yusri Taha,Gary Eltringham |
| EPI_ISL_453938, EPI_ISL_453939, EPI_ISL_453940, EPI_ISL_453941, EPI_ISL_453942, EPI_ISL_453943, EPI_ISL_453944, EPI_ISL_453945, EPI_ISL_453946, EPI_ISL_453947, EPI_ISL_453948, EPI_ISL_453949, EPI_ISL_453950, EPI_ISL_453951, EPI_ISL_453952, EPI_ISL_453953, EPI_ISL_453954, EPI_ISL_453955, EPI_ISL_453956, EPI_ISL_453957, EPI_ISL_453958, EPI_ISL_453959, EPI_ISL_454153, EPI_ISL_454154, EPI_ISL_454155, EPI_ISL_454156, EPI_ISL_454157, EPI_ISL_454174, EPI_ISL_454208, EPI_ISL_454211, EPI_ISL_454212, EPI_ISL_454350 | | | |
| see above | unknown | Instituto Nacional de Saude (INSA) | Borges et al |
| EPI_ISL_454356, EPI_ISL_454368, EPI_ISL_454370, EPI_ISL_454372, EPI_ISL_454373, EPI_ISL_454374, EPI_ISL_454376, EPI_ISL_454378, EPI_ISL_454380, EPI_ISL_454381, EPI_ISL_454382, EPI_ISL_454383, EPI_ISL_454384 | | | |
| see above | UPMC Clinical Microbiology Laboratory | Microbial Genome Sequencing Center, Microbial Genomic Epidemiological Laboratory | Mustapha M. Mustapha, Jane W. Marsh, Dan Snyder, Marissa P. Griffith, Stephanie L. Mitchell, Vatsala R. Srinivasa, Kady D. Waggle, Chinelo Ezeonwuku, Vaughn S. Cooper, Lee H. Harrison |
| EPI_ISL_454540, EPI_ISL_454543, EPI_ISL_454565 | NIV Influenza | NIV Influenza | Potdar V |
| EPI_ISL_454605 | Institute for Public Health | Laboratory for advanced genomics | Filip Roki, Lovro Trgovec-Greif, Neven Sui, Tomislav Rukavina, Igor Jurak, Oliver Vugrek |
| EPI_ISL_454638, EPI_ISL_454639, EPI_ISL_454640 | Humboldt County Public Health Laboratory | Chan-Zuckerberg Biohub | CZB Cliahub Consortium |
| EPI_ISL_454775, EPI_ISL_454777, EPI_ISL_454793 | Dutch COVID-19 response team | National Institute for Public Health and the Environment (RIVM) | Adam Meijer, Harry Vennema, Jeroen Cremer, Sharon van den Brink, Pieter Overduin, Florian Zwagemaker, Dennis Schmitz, Chantal Reusken, on behalf of the national COVID-19 response team |
| EPI_ISL_454809, EPI_ISL_454810, EPI_ISL_454811, EPI_ISL_454812, EPI_ISL_454813, EPI_ISL_454829 | Dirk Dittmer | Dirk Dittmer | Bailey,A.G., Caro-Vegas,C.P., Dittmer,D., Eason,A.B., Juarez,A., Landis,J.T., McNamara,R.P., Miller,M.B., Moorad,R., Pluta,L.J., Seltzer,T.A., Thompson,C., Vahrson,W., Villamor,F. |
| EPI_ISL_455036 | South Eastern Area Laboratory Services | NSW Health Pathology - Institute of Clinical Pathology and Medical Research; Westmead Hospital; University of Sydney | CIDM-PH et al. |
| EPI_ISL_455052 | Douglas Hanly Moir Pathology | NSW Health Pathology - Institute of Clinical Pathology and Medical Research; Westmead Hospital; University of Sydney | CIDM-PH et al. |
| EPI_ISL_455054 | South Eastern Area Laboratory Services | NSW Health Pathology - Institute of Clinical Pathology and Medical Research; Westmead Hospital; University of Sydney | CIDM-PH et al. |
| EPI_ISL_455065 | Douglas Hanly Moir Pathology | NSW Health Pathology - Institute of Clinical Pathology and Medical Research; Westmead Hospital; University of Sydney | CIDM-PH et al. |
| EPI_ISL_455081, EPI_ISL_455082 | South Eastern Area Laboratory Services | NSW Health Pathology - Institute of Clinical Pathology and Medical Research; Westmead Hospital; University of Sydney | CIDM-PH et al. |

| | | | |
|---|---|---|---|
| EPI_ISL_455100 | Orsa VC | The Public Health Agency of Sweden | Anna-Karin Lundqvist, Anna-Malin Linde, Maria Lind Karlberg, Oskar Karlsson Lindsjo, Olov Svartstrom, Anna Risberg, Theresa Enkirch, Mia Brytting, Karin Tegmark-Wisell |
| EPI_ISL_455112, EPI_ISL_455113, EPI_ISL_455114, EPI_ISL_455115, EPI_ISL_455116, EPI_ISL_455117, EPI_ISL_455118, EPI_ISL_455119, EPI_ISL_455196, EPI_ISL_455204, EPI_ISL_455222, EPI_ISL_455249, EPI_ISL_455258, EPI_ISL_455264, EPI_ISL_455280, EPI_ISL_455285, EPI_ISL_455287 | | | |
| see above | Dutch COVID-19 response team | Erasmus Medical Center | Bas Oude Munnink, David Nieuwenhuijse, Reina Sikkema, Claudia Schapendonk, Irina Chestakova, Anne van der Linden, Theo Bestebroer, Stefan van Nieuwkoop, Mark Pronk, Pascal Lexmond, Corien Swaan, Manon Haverkate, Madelief Mollers, Mart Stein, Sandra Kengne Kamga Mobou, Jeroen van Kampen, Jolanda Voermans, Aura Timen, Corine GeurtsvanKessel, Annemiek van der Eijk, Richard Molenkamp, Marion Koopmans, on behalf of the Dutch national COVID-19 response team. |
| EPI_ISL_455441, EPI_ISL_455443, EPI_ISL_455448 | 1. ViroGenetics - BSL3 Laboratory of Virology, Maopolska Centre of Biotechnology, Jagiellonian University; 2. II Department of Internal Medicine, Faculty of Medicine, Jagiellonian University Medical College; 3. Narodowy Instytut Zdrowia Publicznego - Pastwowy Zakad Higieny (NIZP-PZH) | 1. ViroGenetics - BSL3 Laboratory of Virology, Maopolska Centre of Biotechnology, Jagiellonian University; 2. II Department of Internal Medicine, Faculty of Medicine, Jagiellonian University Medical College; 3. Narodowy Instytut Zdrowia Publicznego - Pastwowy Zakad Higieny (NIZP-PZH). | Katarzyna Pancer, Marek Sanak, Aleksandra A. Zasada, Magdalena Rzeczkowska, Tomasz Wokowicz, Katarzyna Zacharczuk, Agnieszka Koakowska-Kulesza, Katarzyna Owczarek, Aleksandra Milewska, Natalia Wolaniuk, Ewelina Hallman-Szeliska, Pawe P abaj, Wojciech Branicki, Krzysztof Pyr |
| EPI_ISL_455598, EPI_ISL_455599, EPI_ISL_455600, EPI_ISL_455601 | SA Pathology | VPRL | Beard, MR., Van Der Hoek, K., Lim, C.K., Leong, L.E.X., Coldbeck-Shackley, R., Shue, B., Kirby, E., Merrett, J., Llamas, B. |
| EPI_ISL_455636, EPI_ISL_455639 | KRISP, KZN Research Innovation and Sequencing Platform | KRISP, KZN Research Innovation and Sequencing Platform | Giandhari J, Pillay S, Lessells R, Chimukangara B, Deforche K, Tegally H, Wilkinson E, de Oliveira T |
| EPI_ISL_455641 | ICMR-National Institute of Cholera and Enteric Diseases | National Institute of Biomedical Genomics | Arindam Maitra, Mamta Chawla Sarkar, Sreedhar Chinnaswamy, Hasina Banu, Ananya Chatterjee, Shanta Dutta, Saumitra Das |
| EPI_ISL_455704, EPI_ISL_455705 | National Hospital of Tropical Diseases | Oxford University Clinical Research Unit, Hanoi, Vietnam | Nguyen Thi Tam, Van Dinh Trang, Nguyen Thu Trang, Nguyen Thi Ngoc Diep, Le Nguyen Minh Hoa, Pham Ngoc Thach, H. Rogier van Doorn, on behalf of the OUCRU COVID-19 research group |
| EPI_ISL_455721 | Servicio de Microbiologia. Hospital Clinico Universitario de Valencia | Sequencing and Bioinformatics Service and Molecular Epidemiology Research Group. FISABIO-Public Health, and SeqCOVID-Spain Consortium | Sandra Carbo, Loreto Ferrús Abad, Paula Ruiz-Hueso, Mariana Reyes-Prieto, Vicente Soriano Chirona, Ivan Ansari, Llúcia Martínez-Priego, Giuseppe 'Auria, David Navarro, Eliseo Albert, Maria Alma Bracho, Lidia Ruiz Roldan, Neris Garcia-Gonzalez, Inma Galán Vendrell, Fernando Gonzalez-Candelas |
| EPI_ISL_455723 | Servicio de Microbiologia. Hospital Clinico Universitario de Valencia | Sequencing and Bioinformatics Service and Molecular Epidemiology Research Group. FISABIO-Public Health, and SeqCOVID-Spain Consortium | Vicente Soriano Chirona, Ivan Ansari, Llúcia Martínez-Priego, Giuseppe 'Auria, David Navarro, Eliseo Albert, Maria Alma Bracho, Lidia Ruiz Roldan, Neris Garcia-Gonzalez, Inma Galán Vendrell, Sandra Carbo, Loreto Ferrús Abad, Paula Ruiz-Hueso, Mariana Reyes-Prieto, Fernando Gonzalez-Candelas |
| EPI_ISL_455724 | Servicio de Microbiologia. Hospital Clinico Universitario de Valencia | Sequencing and Bioinformatics Service and Molecular Epidemiology Research Group. FISABIO-Public Health, and SeqCOVID-Spain Consortium | Loreto Ferrús Abad, Paula Ruiz-Hueso, Mariana Reyes-Prieto, Vicente Soriano Chirona, Ivan Ansari, Llúcia Martínez-Priego, Giuseppe 'Auria, David Navarro, Eliseo Albert, Maria Alma Bracho, Lidia Ruiz Roldan, Neris Garcia-Gonzalez, Inma Galán Vendrell, Sandra Carbo, Fernando Gonzalez-Candelas |
| EPI_ISL_455725 | Servicio de Microbiologia. Hospital Clinico Universitario de Valencia | Sequencing and Bioinformatics Service and Molecular Epidemiology Research Group. FISABIO-Public Health, and SeqCOVID-Spain Consortium | Eliseo Albert, Maria Alma Bracho, Griselda De Marco, Lidia Ruiz Roldan, Neris Garcia-Gonzalez, Inma Galán Vendrell, Sandra Carbo, Loreto Ferrús Abad, Paula Ruiz-Hueso, Mariana Reyes-Prieto, Vicente Soriano Chirona, Ivan Ansari, Llúcia Martínez-Priego, Giuseppe 'Auria, David Navarro, Fernando Gonzalez-Candelas |
| EPI_ISL_455726 | Servicio de Microbiologia. Hospital Clinico Universitario de Valencia | Sequencing and Bioinformatics Service and Molecular Epidemiology Research Group. FISABIO-Public Health, and SeqCOVID-Spain Consortium | Inma Galán Vendrell, Sandra Carbo, Loreto Ferrús Abad, Paula Ruiz-Hueso, Mariana Reyes-Prieto, Vicente Soriano Chirona, Ivan Ansari, Llúcia Martínez-Priego, Giuseppe 'Auria, David Navarro, Eliseo Albert, Maria Alma Bracho, Lidia Ruiz Roldan, Neris Garcia-Gonzalez, Fernando Gonzalez-Candelas |
| EPI_ISL_455727 | Servicio de Microbiologia. Hospital Clinico Universitario de Valencia | Sequencing and Bioinformatics Service and Molecular Epidemiology Research Group. FISABIO-Public Health, and SeqCOVID-Spain Consortium | David Navarro, Eliseo Albert, Maria Alma Bracho, Griselda De Marco, Lidia Ruiz Roldan, Neris Garcia-Gonzalez, Inma Galán Vendrell, Sandra Carbo, Loreto Ferrús Abad, Paula Ruiz-Hueso, Mariana Reyes-Prieto, Vicente Soriano Chirona, Ivan Ansari, Llúcia Martínez-Priego, Giuseppe 'Auria, Fernando Gonzalez-Candelas |
| EPI_ISL_455728 | Servicio de Microbiologia. Hospital Clinico Universitario de Valencia | Sequencing and Bioinformatics Service and Molecular Epidemiology Research Group. FISABIO-Public Health, and SeqCOVID-Spain Consortium | Maria Alma Bracho, Griselda De Marco, Lidia Ruiz Roldan, Neris Garcia-Gonzalez, Inma Galán Vendrell, Sandra Carbo, Loreto Ferrús Abad, Paula Ruiz-Hueso, Mariana Reyes-Prieto, Vicente Soriano Chirona, Ivan Ansari, Llúcia Martínez-Priego, Giuseppe 'Auria, David Navarro, Eliseo Albert, Fernando Gonzalez-Candelas |
| EPI_ISL_455729 | Servicio de Microbiologia. Hospital Clinico Universitario de Valencia | Sequencing and Bioinformatics Service and Molecular Epidemiology Research Group. FISABIO-Public Health, and SeqCOVID-Spain Consortium | Sandra Carbo, Loreto Ferrús Abad, Paula Ruiz-Hueso, Mariana Reyes-Prieto, Vicente Soriano Chirona, Ivan Ansari, Llúcia Martínez-Priego, Giuseppe 'Auria, David Navarro, Eliseo Albert, Maria Alma Bracho, Lidia Ruiz Roldan, Neris Garcia-Gonzalez, Inma Galán Vendrell, Fernando Gonzalez-Candelas |
| EPI_ISL_455730, EPI_ISL_455731 | Servicio de Microbiologia. Hospital Clinico Universitario de Valencia | Sequencing and Bioinformatics Service and Molecular Epidemiology Research Group. FISABIO-Public Health, and SeqCOVID-Spain Consortium | Mariana Reyes-Prieto, Vicente Soriano Chirona, Ivan Ansari, Llúcia Martínez-Priego, Giuseppe 'Auria, David Navarro, Eliseo Albert, Maria Alma Bracho, Lidia Ruiz Roldan, Neris Garcia-Gonzalez, Inma Galán Vendrell, Sandra Carbo, Loreto Ferrús Abad, Paula Ruiz-Hueso, Fernando Gonzalez-Candelas |
| EPI_ISL_455732 | Servicio de Microbiologia. Hospital Clinico Universitario de Valencia | Sequencing and Bioinformatics Service and Molecular Epidemiology Research Group. FISABIO-Public Health, and SeqCOVID-Spain Consortium | Lidia Ruiz Roldan, Neris Garcia-Gonzalez, Inma Galán Vendrell, Sandra Carbo, Loreto Ferrús Abad, Paula Ruiz-Hueso, Mariana Reyes-Prieto, Vicente Soriano Chirona, Ivan Ansari, Llúcia Martínez-Priego, Giuseppe 'Auria, David Navarro, Eliseo Albert, Maria Alma Bracho, Fernando Gonzalez-Candelas |
| EPI_ISL_455733 | Servicio de Microbiologia. Hospital Clinico Universitario de Valencia | Sequencing and Bioinformatics Service and Molecular Epidemiology Research Group. FISABIO-Public Health, and SeqCOVID-Spain Consortium | Loreto Ferrús Abad, Paula Ruiz-Hueso, Mariana Reyes-Prieto, Vicente Soriano Chirona, Ivan Ansari, Llúcia Martínez-Priego, Giuseppe 'Auria, David Navarro, Eliseo Albert, Maria Alma Bracho, Lidia Ruiz Roldan, Neris Garcia-Gonzalez, Inma Galán Vendrell, Sandra Carbo, Fernando Gonzalez-Candelas |
| EPI_ISL_455734, EPI_ISL_455737 | Servicio de Microbiologia. Hospital Clinico Universitario de Valencia | Sequencing and Bioinformatics Service and Molecular Epidemiology Research Group. FISABIO-Public Health, and SeqCOVID-Spain Consortium | Griselda De Marco, Lidia Ruiz Roldan, Neris Garcia-Gonzalez, Inma Galán Vendrell, Sandra Carbo, Loreto Ferrús Abad, Paula Ruiz-Hueso, Mariana Reyes-Prieto, Vicente Soriano Chirona, Ivan Ansari, Llúcia Martínez-Priego, Giuseppe 'Auria, David Navarro, Eliseo Albert, Maria Alma Bracho, Fernando Gonzalez-Candelas |
| EPI_ISL_455738 | Servicio de Microbiologia. Hospital Clinico Universitario de Valencia | Sequencing and Bioinformatics Service and Molecular Epidemiology Research Group. FISABIO-Public Health, and SeqCOVID-Spain Consortium | Maria Alma Bracho, Griselda De Marco, Lidia Ruiz Roldan, Neris Garcia-Gonzalez, Inma Galán Vendrell, Sandra Carbo, Loreto Ferrús Abad, Paula Ruiz-Hueso, Mariana Reyes-Prieto, Vicente Soriano Chirona, Ivan Ansari, Llúcia Martínez-Priego, Giuseppe 'Auria, David Navarro, Eliseo Albert, Fernando Gonzalez-Candelas |
| EPI_ISL_455739 | Servicio de Microbiologia. Hospital Clinico Universitario de Valencia | Sequencing and Bioinformatics Service and Molecular Epidemiology Research Group. FISABIO-Public Health, and SeqCOVID-Spain Consortium | Lidia Ruiz Roldan, Neris Garcia-Gonzalez, Inma Galán Vendrell, Sandra Carbo, Loreto Ferrús Abad, Paula Ruiz-Hueso, Mariana Reyes-Prieto, Vicente Soriano Chirona, Ivan Ansari, Llúcia Martínez-Priego, Giuseppe 'Auria, David Navarro, Eliseo Albert, Maria Alma Bracho, Fernando Gonzalez-Candelas |
| EPI_ISL_455744 | Servicio de Microbiologia. Hospital Clinico Universitario de Valencia | Sequencing and Bioinformatics Service and Molecular Epidemiology Research Group. FISABIO-Public Health, and SeqCOVID-Spain Consortium | Llúcia Martínez-Priego, Giuseppe 'Auria, David Navarro, Eliseo Albert, Maria Alma Bracho, Lidia Ruiz Roldan, Neris Garcia-Gonzalez, Inma Galán Vendrell, Sandra Carbo, Loreto Ferrús Abad, Paula Ruiz-Hueso, Mariana Reyes-Prieto, Vicente Soriano Chirona, Ivan Ansari, Fernando Gonzalez-Candelas |
| EPI_ISL_455994, EPI_ISL_455995, EPI_ISL_455996, EPI_ISL_455997, EPI_ISL_455998, EPI_ISL_455999 | LSUHS Emerging Viral Threat Laboratory | Microbial Genome Sequencing Center | John A. Vanchiere, Jeremy P. Kamil, Rona S. Scott, Camille F. Abshire, Abida Siddiqa, Byeong-Jae Lee, Chan-ki Min, Md Maksudul Alam, Monica Gestal-Cartele, Edna Ondari, Adam Greer, Malgorzata Bienkowska-Haba, Katarzyna Zwolinska, Michelle M. Arnold, Jason M. Bodily, Andrew D. Yurochko, Paul M. Weinberger, Christopher G. Kevil, Martin J. Sapp, Daniel J. Snyder, Vaughn S. Cooper |
| EPI_ISL_456000, EPI_ISL_456001, EPI_ISL_456002 | LSUHS Emerging Viral Threat Laboratory | Microbial Genome Sequencing Center | Rona S. Scott, Jeremy P. Kamil, John A. Vanchiere, Camille F. Abshire, Abida Siddiqa, Byeong-Jae Lee, Chan-ki Min, Md Maksudul Alam, Monica Gestal-Cartele, Edna Ondari, Adam Greer, Malgorzata Bienkowska-Haba, Katarzyna Zwolinska, Michelle M. Arnold, Jason M. Bodily, Andrew D. Yurochko, Paul M. Weinberger, Christopher G. Kevil, Martin J. Sapp, Daniel J. Snyder, Vaughn S. Cooper |
| EPI_ISL_456072, EPI_ISL_456073, EPI_ISL_456074 | Laboratory of Respiratory Viruses and Measles, Oswaldo Cruz Institute, FIOCRUZ | Laboratory of Respiratory Viruses and Measles, Oswaldo Cruz Institute, FIOCRUZ | Paola Resende, Luciana Appolinario, Fernando Motta, Aline Mattos, Milene Miranda, Cristiana Garcia, Braulia Caetano, Maria Ogrzewalska, Jonathan Lopes, Marilda Siqueira |
| EPI_ISL_456147, EPI_ISL_456152 | Instituto Nacional de Salud - Unidad de Secuenciación y Análisis Genómico | Instituto Nacional de Salud, Universidad Cooperativa de Colombia, Instituto Alexander von Humboldt, Imperial College-London, London School of Hygiene & Tropical | Katherine Laiton-Donato, Diego A. Álvarez-Díaz, Carlos Franco-Muñoz, Jose A. Usme-Ciro, Gloria Puerto, Nicolas D. Franco-Sierra, Mailyn A.Gonzalez, Zulma M. Cucunubá, Christian Julian VillabonaArenas, Liz Villabona-Arenas, Sussy Echeverria, Astrid C. Flórez, Sergio Gomez-Rangel, Luz Dary Rodriguez, Juliana Barbosa, Erika Ospitia, Diana Marcela Walteros-Acero, Martha Lucia Ospina Martinez, Marcela Mercado-Reyes. |

| | | | |
|---|---|---|---|
| | | Medicine | |
| EPI_ISL_456287, EPI_ISL_456292, EPI_ISL_456295, EPI_ISL_456296, EPI_ISL_456297, EPI_ISL_456298, EPI_ISL_456299 | Southern Community Labs Dunedin | Institute of Environmental Science and Research (ESR) | Matt Storey, Xiaoyun Ren, Anja Werno, Antje van der Linden, Arlo Upton, Chris Mansell, David Hammer, Dragana Drinkovic, Erasmus Smit, Gary McAuliffe, Hana Sofia Andersson, James Ussher, Jill Sherwood, Josh Freeman, Julia Howard, Juliet Elvy, Mary DeAlmeida, Matt Blakiston, Matthew Rogers, Max Bloomfield, Michael Addidle, Michelle Balm, Sally Roberts, Sarah Jefferies, Sharmini Muttaiyah, Susan Morpeth, Susan Taylor, Timothy Blackmore, Vani Sathyendran, Veronica Playle, Virginia Hope, Erasmus Smit, Lauren Jelly, Joep de Ligt |
| EPI_ISL_456300 | LabPLUS | Institute of Environmental Science and Research (ESR) | Matt Storey, Xiaoyun Ren, Anja Werno, Antje van der Linden, Arlo Upton, Chris Mansell, David Hammer, Dragana Drinkovic, Erasmus Smit, Gary McAuliffe, Hana Sofia Andersson, James Ussher, Jill Sherwood, Josh Freeman, Julia Howard, Juliet Elvy, Mary DeAlmeida, Matt Blakiston, Matthew Rogers, Max Bloomfield, Michael Addidle, Michelle Balm, Sally Roberts, Sarah Jefferies, Sharmini Muttaiyah, Susan Morpeth, Susan Taylor, Timothy Blackmore, Vani Sathyendran, Veronica Playle, Virginia Hope, Erasmus Smit, Lauren Jelly, Joep de Ligt |
| EPI_ISL_456304, EPI_ISL_456307, EPI_ISL_456308, EPI_ISL_456316 | Southern Community Labs Dunedin | Institute of Environmental Science and Research (ESR) | Matt Storey, Xiaoyun Ren, Anja Werno, Antje van der Linden, Arlo Upton, Chris Mansell, David Hammer, Dragana Drinkovic, Erasmus Smit, Gary McAuliffe, Hana Sofia Andersson, James Ussher, Jill Sherwood, Josh Freeman, Julia Howard, Juliet Elvy, Mary DeAlmeida, Matt Blakiston, Matthew Rogers, Max Bloomfield, Michael Addidle, Michelle Balm, Sally Roberts, Sarah Jefferies, Sharmini Muttaiyah, Susan Morpeth, Susan Taylor, Timothy Blackmore, Vani Sathyendran, Veronica Playle, Virginia Hope, Erasmus Smit, Lauren Jelly, Joep de Ligt |
| EPI_ISL_456322 | Canterbury Health Laboratories | Institute of Environmental Science and Research (ESR) | Matt Storey, Xiaoyun Ren, Anja Werno, Antje van der Linden, Arlo Upton, Chris Mansell, David Hammer, Dragana Drinkovic, Erasmus Smit, Gary McAuliffe, Hana Sofia Andersson, James Ussher, Jill Sherwood, Josh Freeman, Julia Howard, Juliet Elvy, Mary DeAlmeida, Matt Blakiston, Matthew Rogers, Max Bloomfield, Michael Addidle, Michelle Balm, Sally Roberts, Sarah Jefferies, Sharmini Muttaiyah, Susan Morpeth, Susan Taylor, Timothy Blackmore, Vani Sathyendran, Veronica Playle, Virginia Hope, Erasmus Smit, Lauren Jelly, Joep de Ligt |
| EPI_ISL_456324, EPI_ISL_456325, EPI_ISL_456326, EPI_ISL_456327, EPI_ISL_456328, EPI_ISL_456329, EPI_ISL_456330, EPI_ISL_456331, EPI_ISL_456332, EPI_ISL_456333, EPI_ISL_456334, EPI_ISL_456335, EPI_ISL_456336, EPI_ISL_456337, EPI_ISL_456338, EPI_ISL_456339, EPI_ISL_456340, EPI_ISL_456341, EPI_ISL_456342, EPI_ISL_456343 | | | |
| see above | Wellington SCL | Institute of Environmental Science and Research (ESR) | Matt Storey, Xiaoyun Ren, Anja Werno, Antje van der Linden, Arlo Upton, Chris Mansell, David Hammer, Dragana Drinkovic, Erasmus Smit, Gary McAuliffe, Hana Sofia Andersson, James Ussher, Jill Sherwood, Josh Freeman, Julia Howard, Juliet Elvy, Mary DeAlmeida, Matt Blakiston, Matthew Rogers, Max Bloomfield, Michael Addidle, Michelle Balm, Sally Roberts, Sarah Jefferies, Sharmini Muttaiyah, Susan Morpeth, Susan Taylor, Timothy Blackmore, Vani Sathyendran, Veronica Playle, Virginia Hope, Erasmus Smit, Lauren Jelly, Joep de Ligt |
| EPI_ISL_456404 | unknown | Research Center Of Tropical and Infectious Of Medical Sciences | Mollaei,H.R., Aghaei-Afshar,A., Kalantar-Neyestanaki,D., Tabatabaeifar,F., Morones Ramirez,J.R. |
| EPI_ISL_456406 | unknown | Research Center Of Tropical and Infectious Of Medical Sciences | Mollaei,H.R., Aghaei-Afshar,A., Kalantar-Neyestanaki,D., Morones Ramirez,J.R. |
| EPI_ISL_456899, EPI_ISL_456900, EPI_ISL_456904, EPI_ISL_456950, EPI_ISL_456953, EPI_ISL_456954, EPI_ISL_456955, EPI_ISL_457012, EPI_ISL_457013, EPI_ISL_457014, EPI_ISL_457015 | | | |
| see above | Virology Department, Royal Infirmary of Edinburgh, NHS Lothian / School of Biological Sciences, University of Edinburgh / Institute of Genetics and Molecular Medicine, University of Edinburgh | COVID-19 Genomics UK (COG-UK) Consortium | McHugh M, Dewar R, Rooke S, Gallagher M, Balcaza C, O'Toole Á, Scher E, Hill V, McCrone JT, Colquhoun R, Yu X, Jackson B, Rambaut A, Williams TC, Templeton K |
| EPI_ISL_457052, EPI_ISL_457089, EPI_ISL_457101, EPI_ISL_457126, EPI_ISL_457134, EPI_ISL_457139, EPI_ISL_457150, EPI_ISL_457181 | University of Exeter | COVID-19 Genomics UK (COG-UK) Consortium | Ben Temperton,Aaron Jeffries,Michelle Michelsen,Joanna Warwick-Dugdale,Audrey Farbos,Robyn Manley,Stephen Michell,Jane Masoli |
| EPI_ISL_457608, EPI_ISL_457627, EPI_ISL_457630, EPI_ISL_457638, EPI_ISL_457642, EPI_ISL_457659, EPI_ISL_457668, EPI_ISL_457673, EPI_ISL_457680, EPI_ISL_457682 | Virology Department, Sheffield Teaching Hospitals NHS Foundation Trust/Department of Infection, Immunity and Cardiovascular Disease, The Medical School, University of Sheffield | COVID-19 Genomics UK (COG-UK) Consortium | Thushan de Silva, Matthew Parker, Nikki Smith, Adri Angyal, Rebecca Brown, Luke Green, Rachel Tucker, Paul Parsons, Danielle Groves, Katie Johnson, Laura Carrilero, Alex Keeley, Dave Partridge, Matthew Wyles, Benjamin Lindsey, Mehmet Yavuz, Mohammad Raza, Cariad Evans |
| EPI_ISL_457758, EPI_ISL_457759 | Johns Hopkins Hospital Department of Pathology | Johns Hopkins Hospital Department of Pathology | Peter M. Thielen, Thomas Mehoke, Shirlee Wohl, Srividya Ramakrishnan, Melanie Kirsche, Amanda Ernlund, Craig Howser, Kristina Zudock, Oluwaseun Falade-Nwulia, Norah Sadowski, Paul Morris, Mark Hopkins, Yunfan Fan, Nidia Trovao, Victoria Gniazdowski, Michael C. Schatz, Stuart C. Ray, Winston Timp, Heba H. Mostafa |
| EPI_ISL_457855 | KEMRI-CGMR-C | KEMRI-Wellcome Trust Research Programme/KEMRI-CGMR-C Kilifi | Githinji G. et al 2020 |
| EPI_ISL_457964 | Laboratorio de Biología Molecular Asociación Española Primera en Salud | Departments of Pathology and Medicine, New York University School of Medicine | Maria Victoria Elizondo, Maria Noel Zubillaga, Gonzalo Manrique, Paul Zappile, Gael Westby, Matthew T Maurano, Christian Marier, Adriana Heguy |
| EPI_ISL_458001, EPI_ISL_458002, EPI_ISL_458003, EPI_ISL_458004, EPI_ISL_458005, EPI_ISL_458006 | Dirk Dittmer | Dirk Dittmer | Bailey,A.G., Caro-Vegas,C.P., Dittmer,D., Eason,A.B., Juarez,A., Landis,J.T., McNamara,R.P., Miller,M.B., Moorad,R., Pluta,L.J., Seltzer,T.A., Thompson,C., Vahrson,W., Villamor,F. |
| EPI_ISL_458012, EPI_ISL_458013, EPI_ISL_458015 | Department of Food Safety, Nutrition and Veterinary public health, Istituto Superiore di Sanita' | Department of Food Safety, Nutrition and Veterinary public health, Istituto Superiore di Sanita' | La Rosa,G., Iaconelli,M., Mancini,P., Bonanno Ferraro,G., Veneri,C., Bonadonna,L., Lucentini,L., Suffredini,E. |
| EPI_ISL_458022 | NYU Langone Health | Departments of Pathology and Medicine, New York University School of Medicine | Maria Aguero-Rosenfeld, Brendan Belovarac, Margaret Black, Ludovic Boytard, John Cadley, Paolo Cotzia, John Chen, Dacia Dimartino, Xiaojun Feng, Tatyana Gindin, Emily Guzman, Adriana Heguy, Megan Hogan, Emily Huang, George Jour, Alireza Khodadadi-Jamayran, Lawrence H. Lin, Raven Luther, Andrew Lytle, Christian Marier, Matthew T. Maurano, Mark J. Mulligan, Peter Meyn, Raquel Ordonez Ciriza, Iman Osman, Jared Pinnell, Vanessa Raabe, Sitharam Ramaswami, Amy Rapkiewicz, Andre M. Ribeiro-dos-Santos, Marie Samanovic-Golden, Antonio Serrano, Guomiao Shen, Matija Snuderl, Theodore Vougiouklakis, Nick Vulpescu, Gael Westby, Paul Zappile, Yutong Zhang |
| EPI_ISL_458025 | Hospital for Tropical Diseases | COVID-19 Network Investigations (CONI) Alliance | Elizabeth Batty, Nantarat Chantawat, Wasun Chantratita, Thanat Chookajorn, Stefan Fernandez, Angkana Huang, Weena Janwitthayanan, Akanitt Jittmittraphap, Anthony R. Jones, Khajohn Joonsalak, Chonticha Klungtong, Theerarat Kochakarn, Namfon Kotanan, Krittikorn Kumpornsin, Pornsawan Leaungwutiwong, Wudtichai Manasatienkij, Bhakbhoom Panthan, Ekawat Pasomsub, Kingkan Rakmanee, Insee Sensorn, Janjira Thaipadungpanit, Arporn Wangwiwatsin,Treewat Watthanachockchai |
| EPI_ISL_458071 | CSIR-Centre for Cellular and Molecular Biology | CSIR-Centre for Cellular and Molecular Biology | Sakshi Shambhavi, Lamuk Zaveri, Shagufta Khan, Namami Gaur, Tulasi Nagabandi, Purushotham Vodnala, Payel Mukherjee, Sofia Banu, Priya Singh, Dhiviya Vedagiri, Divya Gupta, Vishal Sah, Santosh Kumar Kuncha, Krishnan Harinivas Harshan, Archana Bharadwaj Siva, Karthik Bharadwaj Tallapaka,Nikhil Hajirnis, Pratheusa Maccha, M Soujanya Reddy,G. Aditya Kumar, Koushick Sivakumar, Rakesh K Mishra, Divya Tej Sowpati |
| EPI_ISL_458233, EPI_ISL_458234 | KU Leuven, Rega Institute, Clinical and Epidemiological Virology | KU Leuven, Rega Institute, Clinical and Epidemiological Virology | Tony Wawina-Bokalanga, Bert Vanmechelen, Joan Marti-Carerras, Piet Maes |
| EPI_ISL_458262 | Scripps Medical Laboratory | Andersen lab at Scripps Research | SEARCH Alliance San Diego with Michael Quigley, Ellen Stefanski, Ian Mchardy |
| EPI_ISL_458291, EPI_ISL_458294, EPI_ISL_458295, EPI_ISL_458296, EPI_ISL_458297 | Dirk Dittmer | Dirk Dittmer | Bailey,A.G., Caro-Vegas,C.P., Dittmer,D., Eason,A.B., Juarez,A., Landis,J.T., McNamara,R.P., Miller,M.B., Moorad,R., Pluta,L.J., Seltzer,T.A., Thompson,C., Vahrson,W., Villamor,F. |
| EPI_ISL_458302, EPI_ISL_458340, EPI_ISL_458357, EPI_ISL_458365, EPI_ISL_458372, EPI_ISL_458378, EPI_ISL_458417, EPI_ISL_458425, EPI_ISL_458426, EPI_ISL_458430, EPI_ISL_458433, EPI_ISL_458436, EPI_ISL_458440, EPI_ISL_458448, EPI_ISL_458455, EPI_ISL_458499, EPI_ISL_458503 | | | |
| see above | PHE South West Regional Laboratory, National Infection Service | Wellcome Sanger Institute for the COVID-19 Genomics UK (COG-UK) consortium | Stephanie Hutchings, Hannah Pymont, Dr Peter Muir, Barry Vipond, Rich Hopes; and Alex Alderton, Roberto Amato, Sonia Goncalves, Ewan Harrison, David K. Jackson, Ian Johnston, Dominic Kwiatkowski, Cordelia Langford, John Sillitoe on behalf of the Wellcome Sanger Institute COVID-19 Surveillance Team (http://www.sanger.ac.uk/covid-team) |

| | | | |
|---|---|---|---|
| EPI_ISL_458581, EPI_ISL_458582, EPI_ISL_458585, EPI_ISL_458587, EPI_ISL_458589, EPI_ISL_458590, EPI_ISL_458598, EPI_ISL_458599, EPI_ISL_458605, EPI_ISL_458608, EPI_ISL_458616, EPI_ISL_458620, EPI_ISL_458624, EPI_ISL_458627, EPI_ISL_458630, EPI_ISL_458635, EPI_ISL_458637, EPI_ISL_458643, EPI_ISL_458647, EPI_ISL_458648, EPI_ISL_458649, EPI_ISL_458655, EPI_ISL_458656, EPI_ISL_458658, EPI_ISL_458664, EPI_ISL_458665, EPI_ISL_458671, EPI_ISL_458673, EPI_ISL_458675, EPI_ISL_458676, EPI_ISL_458679, EPI_ISL_458681, EPI_ISL_458683, EPI_ISL_458689, EPI_ISL_458692, EPI_ISL_458695, EPI_ISL_458702, EPI_ISL_458706 | | | |
| see above | NU-OMICS DNA Sequencing research facility, Northumbria University | Wellcome Sanger Institute for the COVID-19 Genomics UK (COG-UK) consortium | Chris Duncan, Sheia Waugh, Shirelle Burton-Fanning, Gary Eltringham, Jennifer Collins, Brendan Payne, Yusri Taha, Emma Swindells, Jane Greenaway, Edward Barton, Garren Scott, Debra Padgett, Clive Graham, Sarah Essex, Steve Ligget, Paul Baker, Lynn Dover, Wen Yew, Gary Black, John Allan, Joshua Loh, Greg Young, Matthew Bashton, Andrew Nelson, Darren Smith and Alex Alderton, Roberto Amato, Sonia Goncalves, Ewan Harrison, David K. Jackson, Ian Johnston, Dominic Kwiatkowski, Cordelia Langford, John Sillitoe on behalf of the Wellcome Sanger Institute COVID-19 Surveillance Team (http://www.sanger.ac.uk/covid-team) |
| EPI_ISL_458712 | NU-OMICS DNA Sequencing research facility, Northumbria University | Wellcome Sanger Institute for the COVID-19 Genomics UK (COG-UK) Consortium | Chris Duncan, Sheia Waugh, Shirelle Burton-Fanning, Gary Eltringham, Jennifer Collins, Brendan Payne, Yusri Taha, Emma Swindells, Jane Greenaway, Edward Barton, Garren Scott, Debra Padgett, Clive Graham, Sarah Essex, Steve Ligget, Paul Baker, Lynn Dover, Wen Yew, Gary Black, John Allan, Joshua Loh, Greg Young, Matthew Bashton, Andrew Nelson, Darren Smith and Alex Alderton, Roberto Amato, Sonia Goncalves, Ewan Harrison, David K. Jackson, Ian Johnston, Dominic Kwiatkowski, Cordelia Langford, John Sillitoe on behalf of the Wellcome Sanger Institute COVID-19 Surveillance Team |
| EPI_ISL_458714, EPI_ISL_458716 | NU-OMICS DNA Sequencing research facility, Northumbria University | Wellcome Sanger Institute for the COVID-19 Genomics UK (COG-UK) consortium | Chris Duncan, Sheia Waugh, Shirelle Burton-Fanning, Gary Eltringham, Jennifer Collins, Brendan Payne, Yusri Taha, Emma Swindells, Jane Greenaway, Edward Barton, Garren Scott, Debra Padgett, Clive Graham, Sarah Essex, Steve Ligget, Paul Baker, Lynn Dover, Wen Yew, Gary Black, John Allan, Joshua Loh, Greg Young, Matthew Bashton, Andrew Nelson, Darren Smith and Alex Alderton, Roberto Amato, Sonia Goncalves, Ewan Harrison, David K. Jackson, Ian Johnston, Dominic Kwiatkowski, Cordelia Langford, John Sillitoe on behalf of the Wellcome Sanger Institute COVID-19 Surveillance Team (http://www.sanger.ac.uk/covid-team) |
| EPI_ISL_458770, EPI_ISL_458778, EPI_ISL_458801, EPI_ISL_458802, EPI_ISL_458821, EPI_ISL_458823, EPI_ISL_458825, EPI_ISL_458828, EPI_ISL_458830, EPI_ISL_458837, EPI_ISL_458841, EPI_ISL_458851, EPI_ISL_458855, EPI_ISL_458862, EPI_ISL_458875, EPI_ISL_458895, EPI_ISL_458899, EPI_ISL_458902, EPI_ISL_458904 | | | |
| see above | PHE South West Regional Laboratory, National Infection Service | Wellcome Sanger Institute for the COVID-19 Genomics UK (COG-UK) consortium | Stephanie Hutchings, Hannah Pymont, Dr Peter Muir, Barry Vipond, Rich Hopes; and Alex Alderton, Roberto Amato, Sonia Goncalves, Ewan Harrison, David K. Jackson, Ian Johnston, Dominic Kwiatkowski, Cordelia Langford, John Sillitoe on behalf of the Wellcome Sanger Institute COVID-19 Surveillance Team (http://www.sanger.ac.uk/covid-team) |
| EPI_ISL_459957 | Institute for Medical Research, Infectious Disease Research Centre, National Institutes of Health, Minis | Institute for Medical Research, Infectious Disease Research Centre, National Institutes of Health, Minis | Suppiah J, Mohd-Zawawi Z, Kamel KA, Ellan K, Kalyanasundram J, Mohd-Zain R, Thayan R |
| EPI_ISL_459962, EPI_ISL_459963 | Centogene AG | Centogene AG | Prof. Dr. Peter Bauer, Dr. Krishna Kumar Kandaswamy |
| EPI_ISL_460010, EPI_ISL_460011, EPI_ISL_460012, EPI_ISL_460013, EPI_ISL_460014, EPI_ISL_460015, EPI_ISL_460016, EPI_ISL_460017, EPI_ISL_460018, EPI_ISL_460020, EPI_ISL_460021, EPI_ISL_460022, EPI_ISL_460023, EPI_ISL_460024, EPI_ISL_460025, EPI_ISL_460026, EPI_ISL_460027, EPI_ISL_460028, EPI_ISL_460030 | | | |
| see above | Michigan Department of Health and Human Services, Bureau of Laboratories | Michigan Department of Health and Human Services, Bureau of Laboratories | Blankenship HM, Riner D, Soehnlen MK |
| EPI_ISL_460098, EPI_ISL_460100, EPI_ISL_460120, EPI_ISL_460131, EPI_ISL_460133, EPI_ISL_460141, EPI_ISL_460144, EPI_ISL_460148, EPI_ISL_460151, EPI_ISL_460155, EPI_ISL_460157, EPI_ISL_460160, EPI_ISL_460164, EPI_ISL_460169, EPI_ISL_460171, EPI_ISL_460172, EPI_ISL_460181, EPI_ISL_460193, EPI_ISL_460195, EPI_ISL_460196, EPI_ISL_460198, EPI_ISL_460210, EPI_ISL_460234, EPI_ISL_460243, EPI_ISL_460249, EPI_ISL_460255, EPI_ISL_460260, EPI_ISL_460265, EPI_ISL_460274, EPI_ISL_460282, EPI_ISL_460287, EPI_ISL_460294, EPI_ISL_460308, EPI_ISL_460342, EPI_ISL_460352, EPI_ISL_460379, EPI_ISL_460408, EPI_ISL_460416, EPI_ISL_460430, EPI_ISL_460432, EPI_ISL_460461, EPI_ISL_460463, EPI_ISL_460464 | | | |
| see above | Massachusetts General Hospital | Infectious Disease Program, Broad Institute of Harvard and MIT | Lemieux,J.E., Siddle,K.J., Shaw,B., Adams,G., Pierce,V., Turbett,S., Anahtar,M., Branda,J., Slater,D., Harris,J., Lin,A.E., Gladden-Young,A., Lagerborg,K., Rudy,M., DeRuff,K., Carter,A., Normandin,E., Bauer,M., Reilly,S., Tomkins-Tinch,C., Loreth,C., Chaluvadi,S., Neumann,A., Cusick,C., Chapman,S.B., Gnirke,A., Flowers,K., Cerrato,F., Birren,B.W., Gallagher,G., Smole,S., Park,D.J., MacInnis,B.L., Ryan,E., LaRocque,R., Rosenberg,E., Sabeti,P.C. |
| EPI_ISL_460623, EPI_ISL_460625, EPI_ISL_460626, EPI_ISL_460627 | UW Virology Lab | UW Virology Lab | Pavitra Roychoudhury, Amin Addetia, Hong Xie, Lasata Shrestha, Truong Nguyen, Meei-Li Huang, Keith Jerome, Alexander Greninger |
| EPI_ISL_460638, EPI_ISL_460675, EPI_ISL_460685, EPI_ISL_460688, EPI_ISL_460719, EPI_ISL_460791, EPI_ISL_460796, EPI_ISL_460797, EPI_ISL_460798, EPI_ISL_461076, EPI_ISL_461077, EPI_ISL_461078, EPI_ISL_461079, EPI_ISL_461080, EPI_ISL_461081, EPI_ISL_461082, EPI_ISL_461128, EPI_ISL_461164, EPI_ISL_461250, EPI_ISL_461261, EPI_ISL_461337 | | | |
| see above | Dutch COVID-19 response team | Erasmus Medical Center | Bas Oude Munnink, David Nieuwenhuijse, Reina Sikkema, Claudia Schapendonk, Irina Chestakova, Anne van der Linden, Theo Bestebroer, Stefan van Nieuwkoop, Mark Pronk, Pascal Lexmond, Corien Swaan, Manon Haverkate, Madelief Mollers, Mart Stein, Sandra Kengne Kamga Mobou, Jeroen van Kampen, Jolanda Voermans, Aura Timen, Corine GeurtsvanKessel, Annemiek van der Eijk, Richard Molenkamp, Marion Koopmans, on behalf of the Dutch national COVID-19 response team. |
| EPI_ISL_461423, EPI_ISL_461443, EPI_ISL_461444, EPI_ISL_461445 | UW Virology Lab | UW Virology Lab | Pavitra Roychoudhury, Amin Addetia, Hong Xie, Lasata Shrestha, Truong Nguyen, Meei-Li Huang, Keith Jerome, Alexander Greninger |
| EPI_ISL_461732, EPI_ISL_461733, EPI_ISL_461734, EPI_ISL_461761 | Virology Department, Royal Infirmary of Edinburgh, NHS Lothian / School of Biological Sciences, University of Edinburgh / Institute of Genetics and Molecular Medicine, University of Edinburgh | COVID-19 Genomics UK (COG-UK) Consortium | McHugh M, Dewar R, Rooke S, Gallagher M, Balcaza C, O'Toole Á, Scher E, Hill V, McCrone JT, Colquhoun R, Yu X, Jackson B, Rambaut A, Williams TC, Templeton K |
| EPI_ISL_461773, EPI_ISL_461774, EPI_ISL_461775, EPI_ISL_461776, EPI_ISL_461777, EPI_ISL_461778, EPI_ISL_461779, EPI_ISL_461780, EPI_ISL_461781, EPI_ISL_461783 | Regional Virus Laboratory, Belfast Health and Social Care Trust | COVID-19 Genomics UK (COG-UK) Consortium | Conall McCaughey, James McKenna, Tanya Curran, Susan Feeney, Alison Watt, Ciara Cox, Mairead Connor, Zoltan Molnar, David Simpson, Derek Fairley |
| EPI_ISL_462150 | Molecular diagnostic laboratory of Federal Budget Institution of Science "Central Research Institute of Epidemiology" of The Federal Service on Customers' Rights Protection and Human Well-being Surveillance | Group of Genomics and Postgenomic Technologies of Central Research Institute of Epidemilology | Speranskaya AS, Kaptelova VV, Samoilov AE, Korneenko EV, Sizova TV, Tivanova EV, Shipulina OY, Akimkin VG |
| EPI_ISL_462213, EPI_ISL_462214, EPI_ISL_462215, EPI_ISL_462216, EPI_ISL_462218, EPI_ISL_462220, EPI_ISL_462222, EPI_ISL_462225, EPI_ISL_462226, EPI_ISL_462227, EPI_ISL_462229, EPI_ISL_462230, EPI_ISL_462231, EPI_ISL_462232, EPI_ISL_462233 | | | |
| see above | KU Leuven, Rega Institute, Clinical and Epidemiological Virology | KU Leuven, Rega Institute, Clinical and Epidemiological Virology | Tony Wawina-Bokalanga, Bert Vanmechelen, Joan Marti-Carerras, Piet Maes |
| EPI_ISL_462335, EPI_ISL_462350, EPI_ISL_462368, EPI_ISL_462418 | National Public Health Laboratory, National Centre for Infectious Diseases | National Public Health Laboratory, National Centre for Infectious Diseases | Mak TM, Octavia S, Chavatte JM, Cui L, Lin RTP |
| EPI_ISL_462434 | unknown | Laboratory Diagnostic | Vidanovic,D., Tesovic,B., Banovic Djeri,B., Knezevic,A., Jankovic,M., Sekler,M., Dmitric,M., Petrovic,T., Volkening,J., Afonso,C.L. |
| EPI_ISL_462439, EPI_ISL_462440, EPI_ISL_462441, EPI_ISL_462442, EPI_ISL_462443, EPI_ISL_462444, EPI_ISL_462445 | unknown | Ryota Kumagai Tokyo Metropolitan Institute of Public Health | Asakura,H., Kumagai,R., Yoshida,I., Nagashima,M., Chiba,T., Sadamasu,K. |
| EPI_ISL_462472 | Clinical Center, University of Sarajevo | Charite Universitatsmedizin Berlin, Institute of Virology | Victor M Corman, Jorn Beheim-Schwarzbach, Barbara Muehlemann, Talitha Veith, Julia Schneider, Terry Jones, Amela Dedeic-Ljubovic, Irma Salimovic-Besic, Suzana Arapcic, Almedina Hadzihasanovic-Moro, Selma Mutevelic, Christian Drosten |
| EPI_ISL_463004, EPI_ISL_463006 | unknown | Clinical virology | Fares,W., Triki,H. |
| EPI_ISL_463151, EPI_ISL_463152, EPI_ISL_463153, EPI_ISL_463154, EPI_ISL_463155, EPI_ISL_463156, EPI_ISL_463157, EPI_ISL_463158, EPI_ISL_463159, EPI_ISL_463160, EPI_ISL_463161, EPI_ISL_463162, EPI_ISL_463163, EPI_ISL_463164, EPI_ISL_463165, EPI_ISL_463166, EPI_ISL_463167, EPI_ISL_463168, EPI_ISL_463169, EPI_ISL_463170, EPI_ISL_463171, EPI_ISL_463172, EPI_ISL_463173, EPI_ISL_463174 | | | |
| see above | Yale Clinical Virology Laboratory | Grubaugh Lab - Yale School of Public Health | Joseph Fauver, Tara Alpert, Anderson Brito, Anne Wyllie, Chantal Vogels, Mary Petrone, Cole Jensen, Chaney Kalinich, Isabel Ott, Arnau Casanovas, |

| | | | |
|---|---|---|---|
| | | | Catherine Muenker, Adam Moore, Alice Lu, Maria Tokuyama, Patrick Wong, Peiwen Lu, Saad Omer, Richard Martinello, Allison Nelson, Shelli Farhadian, Akiko Iwasaki, Charlese Dela Cruz, Albert Ko, Nathan Grubaugh |
| EPI_ISL_463745 | Department of Molecular Virology, Cyprus Institute of Neurology and Genetics | Department of Molecular Virology, Cyprus Institute of Neurology and Genetics | Jan Richter, George Krashias, Christina Tryfonos, Stavros Bashiardes, Dana Koptides, Christina Christodoulou |
| EPI_ISL_463928, EPI_ISL_463933, EPI_ISL_463940, EPI_ISL_463952 | Laboratoire de microbiologie, Hopital de Verdun | Smith Laboratory, Centre de Recherche CHU Sainte-Justine | Martin Smith, Marieke Rozendaal, Ivan Pavlov |
| EPI_ISL_463986 | Toronto Invasive Bacterial Diseases Network | McMaster University | Allison McGeer, Patryk Aftanas, Angel Li, Kuganya Nirmalarajah, Samira Mubareka, Andrew G. McArthur |
| EPI_ISL_464000, EPI_ISL_464005, EPI_ISL_464016, EPI_ISL_464032, EPI_ISL_464035, EPI_ISL_464044, EPI_ISL_464045, EPI_ISL_464048, EPI_ISL_464055, EPI_ISL_464060 | Unity Health Toronto | Ontario Institute for Cancer Research | Ramzi Fattouh,Larissa M. Matukas,Mark Downing,Annette Gower,Karel Boissinot,Samira Mubareka,TIBDN,Ilinca Lungu,Bernard Lam,Jeremy Johns,Paul Krzyzanowski,Richard de Borja,Philip Zuzarte,Jared Simpson |
| EPI_ISL_465894, EPI_ISL_465907, EPI_ISL_465919, EPI_ISL_465922, EPI_ISL_465971, EPI_ISL_465973, EPI_ISL_465975, EPI_ISL_465982, EPI_ISL_465983, EPI_ISL_465998, EPI_ISL_465999, EPI_ISL_466000, EPI_ISL_466001, EPI_ISL_466002, EPI_ISL_466003, EPI_ISL_466004, EPI_ISL_466007, EPI_ISL_466008, EPI_ISL_466009, EPI_ISL_466010, EPI_ISL_466011, EPI_ISL_466012, EPI_ISL_466013, EPI_ISL_466014, EPI_ISL_466015, EPI_ISL_466016, EPI_ISL_466017, EPI_ISL_466018, EPI_ISL_466019, EPI_ISL_466020, EPI_ISL_466021, EPI_ISL_466022, EPI_ISL_466023, EPI_ISL_466024, EPI_ISL_466025, EPI_ISL_466026, EPI_ISL_466027, EPI_ISL_466028, EPI_ISL_466029, EPI_ISL_466030, EPI_ISL_466031, EPI_ISL_466032, EPI_ISL_466033, EPI_ISL_466034, EPI_ISL_466035, EPI_ISL_466038, EPI_ISL_466039, EPI_ISL_466040, EPI_ISL_466041, EPI_ISL_466043, EPI_ISL_466044, EPI_ISL_466045, EPI_ISL_466047, EPI_ISL_466051, EPI_ISL_466052, EPI_ISL_466053, EPI_ISL_466054, EPI_ISL_466055, EPI_ISL_466057, EPI_ISL_466062, EPI_ISL_466064, EPI_ISL_466065, EPI_ISL_466072, EPI_ISL_466106, EPI_ISL_466108, EPI_ISL_466109, EPI_ISL_466110, EPI_ISL_466111, EPI_ISL_466112, EPI_ISL_466113, EPI_ISL_466114, EPI_ISL_466115, EPI_ISL_466116, EPI_ISL_466117, EPI_ISL_466118, EPI_ISL_466119, EPI_ISL_466121, EPI_ISL_466122, EPI_ISL_466123, EPI_ISL_466127, EPI_ISL_466128, EPI_ISL_466129, EPI_ISL_466130, EPI_ISL_466131, EPI_ISL_466132, EPI_ISL_466133, EPI_ISL_466134, EPI_ISL_466135, EPI_ISL_466136, EPI_ISL_466137, EPI_ISL_466139, EPI_ISL_466140, EPI_ISL_466143, EPI_ISL_466146, EPI_ISL_466147, EPI_ISL_466148, EPI_ISL_466149, EPI_ISL_466229, EPI_ISL_466230, EPI_ISL_466239 | | | |
| see above | Respiratory Virus Unit, Microbiology Services Colindale, Public Health England | Respiratory Virus Unit, Microbiology Services Colindale, Public Health England | PHE Covid Sequencing Team |
| EPI_ISL_466696, EPI_ISL_466697, EPI_ISL_466698, EPI_ISL_466699, EPI_ISL_466700, EPI_ISL_466701, EPI_ISL_466702, EPI_ISL_466703, EPI_ISL_466704, EPI_ISL_466705, EPI_ISL_466706, EPI_ISL_466707, EPI_ISL_466708, EPI_ISL_466709, EPI_ISL_466710, EPI_ISL_466711, EPI_ISL_466712, EPI_ISL_466713, EPI_ISL_466714, EPI_ISL_466715, EPI_ISL_466716, EPI_ISL_466717, EPI_ISL_466718, EPI_ISL_466719, EPI_ISL_466720, EPI_ISL_466721, EPI_ISL_466722, EPI_ISL_466723, EPI_ISL_466724, EPI_ISL_466725, EPI_ISL_466726, EPI_ISL_466727, EPI_ISL_466728, EPI_ISL_466729, EPI_ISL_466730, EPI_ISL_466731, EPI_ISL_466732, EPI_ISL_466733, EPI_ISL_466734, EPI_ISL_466735, EPI_ISL_466736, EPI_ISL_466737, EPI_ISL_466738, EPI_ISL_466739, EPI_ISL_466740, EPI_ISL_466741, EPI_ISL_466742, EPI_ISL_466743, EPI_ISL_466744, EPI_ISL_466745, EPI_ISL_466746, EPI_ISL_466747, EPI_ISL_466748, EPI_ISL_466749, EPI_ISL_466750, EPI_ISL_466751, EPI_ISL_466752, EPI_ISL_466753, EPI_ISL_466754, EPI_ISL_466755, EPI_ISL_466756, EPI_ISL_466757, EPI_ISL_466758, EPI_ISL_466759, EPI_ISL_466760, EPI_ISL_466761, EPI_ISL_466762, EPI_ISL_466763, EPI_ISL_466764, EPI_ISL_466765, EPI_ISL_466766, EPI_ISL_466767, EPI_ISL_466768, EPI_ISL_466769, EPI_ISL_466770, EPI_ISL_466771, EPI_ISL_466772, EPI_ISL_466773, EPI_ISL_466774, EPI_ISL_466775, EPI_ISL_466776, EPI_ISL_466777, EPI_ISL_466778, EPI_ISL_466779, EPI_ISL_466780, EPI_ISL_466781, EPI_ISL_466782, EPI_ISL_466783, EPI_ISL_466784, EPI_ISL_466785, EPI_ISL_466786, EPI_ISL_466787, EPI_ISL_466788, EPI_ISL_466789, EPI_ISL_466790, EPI_ISL_466791, EPI_ISL_466792, EPI_ISL_466793, EPI_ISL_466794, EPI_ISL_466795, EPI_ISL_466796, EPI_ISL_466797, EPI_ISL_466798, EPI_ISL_466799, EPI_ISL_466800, EPI_ISL_466801, EPI_ISL_466802, EPI_ISL_466803, EPI_ISL_466804, EPI_ISL_466805, EPI_ISL_466806, EPI_ISL_466807, EPI_ISL_466808, EPI_ISL_466809, EPI_ISL_466810, EPI_ISL_466811, EPI_ISL_466812, EPI_ISL_466813, EPI_ISL_466814, EPI_ISL_466815, EPI_ISL_466816, EPI_ISL_466817, EPI_ISL_466818, EPI_ISL_466819, EPI_ISL_466820, EPI_ISL_466821, EPI_ISL_466822, EPI_ISL_466823, EPI_ISL_466824, EPI_ISL_466825, EPI_ISL_466826, EPI_ISL_466827, EPI_ISL_466828, EPI_ISL_466829, EPI_ISL_466830, EPI_ISL_466831, EPI_ISL_466832, EPI_ISL_466833, EPI_ISL_466834, EPI_ISL_466835, EPI_ISL_466836, EPI_ISL_466837, EPI_ISL_466838 | | | |
| see above | BCCDC Public Health Laboratory | BCCDC Public Health Laboratory | Richard Harrigan, Hope Lapointe, Jinny Choi, Kimia Kamelian, John Tyson,Terry Snutch, Linda Hoang, Inna Sekirov, Paul Levett, Mel Krajden, Natalie Prystajecky |
| EPI_ISL_466873 | Centre for Clinical Infection and Diagnostics Research and Genomics Innovation Unit | Respiratory Virus Unit, Microbiology Services Colindale, Public Health England | PHE Covid Sequencing Team, Chloe Fisher, Luke Snell, Gaia Nebbia, Ali Awan |
| EPI_ISL_467059, EPI_ISL_467063 | Hospital Universitario Virgen de las Nieves de Granada-SAS | SeqCOVID-SPAIN consortium/IBV(CSIC) | Mercedes Pérez Ruiz, Sara Sanbonmatsu Gámez, Irene Pedrosa Corral, José M. Navarro-Marí and SeqCOVID-SPAIN consortium |
| EPI_ISL_467187, EPI_ISL_467188, EPI_ISL_467189, EPI_ISL_467193, EPI_ISL_467196, EPI_ISL_467197, EPI_ISL_467201, EPI_ISL_467204, EPI_ISL_467205, EPI_ISL_467206, EPI_ISL_467208, EPI_ISL_467209, EPI_ISL_467213, EPI_ISL_467214, EPI_ISL_467215, EPI_ISL_467218, EPI_ISL_467221, EPI_ISL_467222, EPI_ISL_467223, EPI_ISL_467225, EPI_ISL_467226, EPI_ISL_467228, EPI_ISL_467232, EPI_ISL_467239, EPI_ISL_467244, EPI_ISL_467245, EPI_ISL_467247, EPI_ISL_467250, EPI_ISL_467253, EPI_ISL_467254, EPI_ISL_467255 | | | |
| see above | Hospital General Universitario Gregorio Marañón | SeqCOVID-SPAIN consortium/IBV(CSIC) | Laura Pérez-Lago, Marta Herranz, Jon Sicilia, Julia Suárez, Pilar Catalán, Patricia Muñoz, Darío García de Viedma and SeqCOVID-SPAIN consortium |
| EPI_ISL_467303, EPI_ISL_467305, EPI_ISL_467306, EPI_ISL_467307, EPI_ISL_467308, EPI_ISL_467309, EPI_ISL_467310, EPI_ISL_467311, EPI_ISL_467312, EPI_ISL_467313, EPI_ISL_467314, EPI_ISL_467315, EPI_ISL_467316, EPI_ISL_467317, EPI_ISL_467318, EPI_ISL_467319, EPI_ISL_467320, EPI_ISL_467321, EPI_ISL_467322, EPI_ISL_467323, EPI_ISL_467324, EPI_ISL_467325, EPI_ISL_467326, EPI_ISL_467327, EPI_ISL_467328, EPI_ISL_467329, EPI_ISL_467330, EPI_ISL_467331, EPI_ISL_467332, EPI_ISL_467333, EPI_ISL_467334, EPI_ISL_467335, EPI_ISL_467336, EPI_ISL_467337, EPI_ISL_467338, EPI_ISL_467339, EPI_ISL_467340, EPI_ISL_467341, EPI_ISL_467342, EPI_ISL_467343, EPI_ISL_467423, EPI_ISL_467424, EPI_ISL_467425, EPI_ISL_467426, EPI_ISL_467427, EPI_ISL_467428, EPI_ISL_467429 | | | |
| see above | BCCDC Public Health Laboratory | BCCDC Public Health Laboratory | Richard Harrigan, Hope Lapointe, Jinny Choi, Kimia Kamelian, John Tyson,Terry Snutch, Linda Hoang, Inna Sekirov, Paul Levett, Mel Krajden, Natalie Prystajecky |
| EPI_ISL_467436, EPI_ISL_467437, EPI_ISL_467440 | NHLS-IALCH | KRISP, KZN Research Innovation and Sequencing Platform | Giandhari J, Pillay S, Lessells R, Chimukangara B, Mdlalose K, York D, Khan S, Tegally H, Wilkinson E, de Oliveira T |
| EPI_ISL_467448 | Molecular Diagnostics Services (MDS) | KRISP, KZN Research Innovation and Sequencing Platform | Giandhari J, Pillay S, Lessells R, Chimukangara B, Mdlalose K, York D, Khan S, Tegally H, Wilkinson E, de Oliveira T |
| EPI_ISL_467454, EPI_ISL_467461, EPI_ISL_467462, EPI_ISL_467463, EPI_ISL_467464, EPI_ISL_467465, EPI_ISL_467470, EPI_ISL_467471, EPI_ISL_467472, EPI_ISL_467473, EPI_ISL_467474 | | | |
| see above | AMPATH-DBN | KRISP, KZN Research Innovation and Sequencing Platform | Giandhari J, Pillay S, Lessells R, Chimukangara B, Mdlalose K, York D, Khan S, Tegally H, Wilkinson E, de Oliveira T |
| EPI_ISL_467539, EPI_ISL_467540, EPI_ISL_467541, EPI_ISL_467542, EPI_ISL_467543, EPI_ISL_467544, EPI_ISL_467545, EPI_ISL_467546, EPI_ISL_467547, EPI_ISL_467629, EPI_ISL_467630, EPI_ISL_467631, EPI_ISL_467632, EPI_ISL_467633, EPI_ISL_467634, EPI_ISL_467635, EPI_ISL_467636, EPI_ISL_467637, EPI_ISL_467638, EPI_ISL_467639, EPI_ISL_467640 | | | |
| see above | New Mexico Department of Health Scientific Laboratory Division | Center for Global Health, University of New Mexico Health Sciences Center | Daryl Domman, Kurt Schwalm, Twila Kunde, Joseph Hicks, Michael Edwards, Darrell Dinwiddie |
| EPI_ISL_467774 | Molecular diagnostic laboratory of Federal Budget Institution of Science "Central Research Institute of Epidemiology" of The Federal Service on Customers' Rights Protection and Human Well-being Surveillance | Group of Genomics and Postgenomic Technologies of Central Research Institute of Epidemilology | Speranskaya AS, Kaptelova VV, Samoilov AE, Korneenko EV, Sizova TV, Tivanova EV, Shipulina OY, Akimkin VG |
| EPI_ISL_467993, EPI_ISL_467999, EPI_ISL_468005, EPI_ISL_468006, EPI_ISL_468010, EPI_ISL_468035, EPI_ISL_468040, EPI_ISL_468041, EPI_ISL_468043 | SA Pathology | SA Pathology | Lex Leong, Chuan Kok Lim, Mark Turra, Ivan Bastian, Geoff Higgins |
| EPI_ISL_468359 | Alameda County Public Health Lab | Chan-Zuckerberg Biohub | CZB Cliahub Consortium |
| EPI_ISL_468453, EPI_ISL_468454, EPI_ISL_468455, EPI_ISL_468456, EPI_ISL_468457, EPI_ISL_468458 | Humboldt County Public Health Laboratory | Chan-Zuckerberg Biohub | CZB Cliahub Consortium |
| EPI_ISL_468498 | Ventura County Public Health Lab | Chan-Zuckerberg Biohub | CZB Cliahub Consortium |
| EPI_ISL_468528, EPI_ISL_468529, EPI_ISL_468531 | San Joaquin County Public Health Lab | Chan-Zuckerberg Biohub | CZB Cliahub Consortium |
| EPI_ISL_468620, EPI_ISL_468621, EPI_ISL_468622 | Contra Costa Public Health Lab | Chan-Zuckerberg Biohub | CZB Cliahub Consortium |
| EPI_ISL_468657, EPI_ISL_468658, EPI_ISL_468659, EPI_ISL_468660, EPI_ISL_468661, EPI_ISL_468662, EPI_ISL_468663, EPI_ISL_468664, EPI_ISL_468665, EPI_ISL_468666, EPI_ISL_468667, EPI_ISL_468668, EPI_ISL_468669, EPI_ISL_468670, EPI_ISL_468671, EPI_ISL_468672, EPI_ISL_468673, EPI_ISL_468674, EPI_ISL_468675, EPI_ISL_468676, EPI_ISL_468677, EPI_ISL_468678, EPI_ISL_468679, EPI_ISL_468680, EPI_ISL_468681, EPI_ISL_468682, EPI_ISL_468683, EPI_ISL_468684, EPI_ISL_468685, EPI_ISL_468686, EPI_ISL_468687, EPI_ISL_468688, EPI_ISL_468689, EPI_ISL_468690, EPI_ISL_468691, EPI_ISL_468692, EPI_ISL_468693, EPI_ISL_468694, EPI_ISL_468695, EPI_ISL_468696, EPI_ISL_468697, EPI_ISL_468698, EPI_ISL_468699, EPI_ISL_468700 | | | |
| see above | BCCDC Public Health Laboratory | BCCDC Public Health Laboratory | Richard Harrigan, Hope Lapointe, Jinny Choi, Kimia Kamelian, John Tyson,Terry Snutch, Linda Hoang, Inna Sekirov, Paul Levett, Mel Krajden, Natalie Prystajecky |

| | | | |
|---|---|---|---|
| EPI_ISL_469054 | LNR National Reference Laboratory, Mohammed VI University of Health Sciences | Medical Biotechnology Laboratory, Rabat Medical and Pharmacy School, Mohammed The Vth University in Rabat | Meriem LAAMARTI, Souad KARTTI, Rokaia LAAMRTI , M.W. CHEMAO-ELFIHRI, Loubna ALLAM, Mouna OUADGHIRI, Imane SMYEJ, Jalila RAHOUI, Houda BENRAHMA, Jalil El Atar, Idrissa Diawara, Rachid EL JAOUDI, Laila SBABOU, Chakib NEJJARI, Saaid AMZAZI, Rachid MENTAG, Lahcen BELYAMANI and Azeddine IBRAHIMI |
| EPI_ISL_469098, EPI_ISL_469103, EPI_ISL_469104, EPI_ISL_469121 | National Public Health Laboratory, National Centre for Infectious Diseases | National Public Health Laboratory, National Centre for Infectious Diseases | Mak TM, Octavia S, Chavatte JM, Cui L, Lin RTP |
| EPI_ISL_469171, EPI_ISL_469172, EPI_ISL_469173, EPI_ISL_469174, EPI_ISL_469175, EPI_ISL_469176, EPI_ISL_469177, EPI_ISL_469178, EPI_ISL_469179, EPI_ISL_469180, EPI_ISL_469181, EPI_ISL_469182, EPI_ISL_469183, EPI_ISL_469184, EPI_ISL_469185, EPI_ISL_469186, EPI_ISL_469187, EPI_ISL_469188 | | | |
| see above | Yale Clinical Virology Laboratory | Grubaugh Lab - Yale School of Public Health | Joseph Fauver, Tara Alpert, Anderson Brito, Anne Wyllie, Chantal Vogels, Mary Petrone, Cole Jensen, Chaney Kalinich, Isabel Ott, Arnau Casanovas, Catherine Muenker, Adam Moore, Alice Lu, Maria Tokuyama, Patrick Wong, Peiwen Lu, Saad Omer, Richard Martinello, Allison Nelson, Shelli Farhadian, Akiko Iwasaki, Charlese Dela Cruz, Albert Ko, Nathan Grubaugh |
| EPI_ISL_469209, EPI_ISL_469210, EPI_ISL_469211, EPI_ISL_469212, EPI_ISL_469213, EPI_ISL_469214, EPI_ISL_469215, EPI_ISL_469216, EPI_ISL_469217, EPI_ISL_469218, EPI_ISL_469219, EPI_ISL_469220, EPI_ISL_469221, EPI_ISL_469222, EPI_ISL_469223 | | | |
| see above | BCCDC Public Health Laboratory | BCCDC Public Health Laboratory | Richard Harrigan, Hope Lapointe, Jinny Choi, Kimia Kamelian, John Tyson,Terry Snutch, Linda Hoang, Inna Sekirov, Paul Levett, Mel Krajden, Natalie Prystajecky |
| EPI_ISL_469279 | Mohammed Bin Rashid University of Medicine and Health Sciences | Al Jalila Genomics Center | Ahmad Abou Tayoun, Tom Loney, Hamda Khansaheb, Sathishkumar Ramaswamy, Divinlal Harilal, Zulfa Omar Deesi, Rupa Murthy Varghese, Hanan Al Suwaidi, Abdulmajeed Alkhaja, Mohammed Uddin, Rifat Hamoudi, Rabih Halwani, Abiola Catherine Senok, Qutayba Hamid, Norbert Nowotny, Alawi Alsheikh-Ali |
| EPI_ISL_469289 | Keio University Hospital | Keio University Hospital | Kenjiro Kosaki |
| EPI_ISL_469541, EPI_ISL_469548, EPI_ISL_469551, EPI_ISL_469553, EPI_ISL_469556, EPI_ISL_469561, EPI_ISL_469565, EPI_ISL_469568, EPI_ISL_469573, EPI_ISL_469577, EPI_ISL_469594, EPI_ISL_469597, EPI_ISL_469601, EPI_ISL_469603, EPI_ISL_469609, EPI_ISL_469617, EPI_ISL_469620, EPI_ISL_469634, EPI_ISL_469642, EPI_ISL_469645, EPI_ISL_469648, EPI_ISL_469650, EPI_ISL_469662, EPI_ISL_469670, EPI_ISL_469681, EPI_ISL_469698, EPI_ISL_469708, EPI_ISL_469751, EPI_ISL_469771, EPI_ISL_469778, EPI_ISL_469784, EPI_ISL_469793, EPI_ISL_469800 | | | |
| see above | PHE South West Regional Laboratory, National Infection Service | Wellcome Sanger Institute for the COVID-19 Genomics UK (COG-UK) consortium | Stephanie Hutchings, Hannah Pymont, Dr Peter Muir, Barry Vipond, Rich Hopes; and Alex Alderton, Roberto Amato, Sonia Goncalves, Ewan Harrison, David K. Jackson, Ian Johnston, Dominic Kwiatkowski, Cordelia Langford, John Sillitoe on behalf of the Wellcome Sanger Institute COVID-19 Surveillance Team (http://www.sanger.ac.uk/covid-team) |
| EPI_ISL_469915 | NU-OMICS DNA Sequencing research facility, Northumbria University | Wellcome Sanger Institute for the COVID-19 Genomics UK (COG-UK) consortium | Chris Duncan, Sheia Waugh, Shirelle Burton-Fanning, Gary Eltringham, Jennifer Collins, Brendan Payne, Yusri Taha, Emma Swindells, Jane Greenaway, Edward Barton, Garren Scott, Debra Padgett, Clive Graham, Sarah Essex, Steve Ligget, Paul Baker, Lynn Dover, Wen Yew, Gary Black, John Allan, Joshua Loh, Greg Young, Matthew Bashton, Andrew Nelson, Darren Smith and Alex Alderton, Roberto Amato, Sonia Goncalves, Ewan Harrison, David K. Jackson, Ian Johnston, Dominic Kwiatkowski, Cordelia Langford, John Sillitoe on behalf of the Wellcome Sanger Institute COVID-19 Surveillance Team (http://www.sanger.ac.uk/covid-team) |
| EPI_ISL_469931, EPI_ISL_469939, EPI_ISL_469944, EPI_ISL_469950, EPI_ISL_469955, EPI_ISL_469962, EPI_ISL_469978, EPI_ISL_469986, EPI_ISL_469997, EPI_ISL_469998, EPI_ISL_469999, EPI_ISL_470010 | | | |
| see above | NHSGGC West of Scotland Specialist Virology Centre / MRC-University of Glasgow Centre for Virus Research | Wellcome Sanger Institute for the COVID-19 Genomics UK (COG-UK) consortium | Ana da Silva Filipe, Natasha Johnson, Kathy Smollett, Daniel Mair, Stephen Carmichael, Lily Tong, Jenna Nichols, Elihu Aranday-Cortes, Kirstyn Brunker, Yasmin Parr, Kyriaki Nomikou; Sarah McDonald, Marc Niebel, Patawee Asamaphan; Richard Orton, Joseph Hughes, Sreenu Vattipally, David L Robertson; Alasdair MacLean, Rory Gunson; Kathy Li, Natasha Jesudason, Rajiv Shah, James Shepherd, Antonia Ho, Alice Broos, Emma Thomson and Alex Alderton, Roberto Amato, Sonia Goncalves, Ewan Harrison, David K. Jackson, Ian Johnston, Dominic Kwiatkowski, Cordelia Langford, John Sillitoe on behalf of the Wellcome Sanger Institute COVID-19 Surveillance Team (http://www.sanger.ac.uk/covid-team) |
| EPI_ISL_470534 | Regional Virus Laboratory, Belfast Health and Social Care Trust | Wellcome Sanger Institute for the COVID-19 Genomics UK (COG-UK) consortium | Conall McCaughey, James McKenna, Tanya Curran, Susan Feeney, Alison Watt, Ciara Cox, Mairead Connor, Zoltan Molnar, David Simpson, Derek Fairley; and Alex Alderton, Roberto Amato, Sonia Goncalves, Ewan Harrison, David K. Jackson, Ian Johnston, Dominic Kwiatkowski, Cordelia Langford, John Sillitoe on behalf of the Wellcome Sanger Institute COVID-19 Surveillance Team (http://www.sanger.ac.uk/covid-team) |
| EPI_ISL_470568, EPI_ISL_470569, EPI_ISL_470600, EPI_ISL_470601, EPI_ISL_470602, EPI_ISL_470603 | Hermes Pardini | Bioinformatics Laboratory / LNCC | Alexandra Gerber, Ana Paula Guimarães, Luiz Gonzaga Paula de Almeida, Ronaldo da Silva Francisco Junior, Mariane Talon, Filipe Romero, Átila Duque Rossi, Terezinha Marta Pereira, working group UFRJ, Jaqueline Goes de Jesus, Ingra Morales Claro, Ester Cerdeira Sabino, Nuno Rodrigues Faria, CADDE-group, Laboratorio Hermes Pardini, Laboratorio Simile, working group UFMG, Amilcar Tanuri, Carolina Voloch, Renato Santana Aguiar e Ana Tereza Vasconcelos |
| EPI_ISL_470621, EPI_ISL_470622 | Laboratorio de Virologia Molecular / UFRJ | Bioinformatics Laboratory / LNCC | Alexandra Gerber, Ana Paula Guimarães, Luiz Gonzaga Paula de Almeida, Ronaldo da Silva Francisco Junior, Mariane Talon, Filipe Romero, Átila Duque Rossi, Terezinha Marta Pereira, working group UFRJ, Jaqueline Goes de Jesus, Ingra Morales Claro, Ester Cerdeira Sabino, Nuno Rodrigues Faria, CADDE-group, Laboratorio Hermes Pardini, Laboratorio Simile, working group UFMG, Amilcar Tanuri, Carolina Voloch, Renato Santana Aguiar e Ana Tereza Vasconcelos |
| EPI_ISL_470830, EPI_ISL_470842, EPI_ISL_470843, EPI_ISL_470866 | PathWest Laboratory Medicine WA | PathWest Laboratory Medicine WA | Chisha Sikazwe, Jurissa Lang, Avram Levy, David Smith and David Speers |
| EPI_ISL_471182, EPI_ISL_471223, EPI_ISL_471266 | Wisconsin State Laboratory of Hygiene Communicable Disease Division | Wisconsin State Laboratory of Hygiene Communicable Disease Division | Kelsey R. Florek, Abigail C. Shockey |
| EPI_ISL_471456 | Centre de Virologie des Maladies Tropicales | Functional Genomic Platform/Service Analyses Biologique/UATRS/ Centre National Pour la Recherche Scientifique Et Technique (CNRST) | Hicham EL ANNAZ, Elmostafa EL FAHIME, Marouane MELLOUL, Youssef AKHOUAD, Mly Abdelaziz ELALAOUI, Ahmed REGGAD, Sanaa ALAOUI-Amine, Rachid ABI, Rida TAGAJDID, Zohour KASMY, Safae ELKOCHRI, Nadia TOUIL, Farida HILALI, Abdelkader LAATIRIS, Abdelillah LARAQUI, Tahar BAJJOU , Yassine SEKHSOKH , Idriss-Amine LAHLOU, Mostafa ELOUENNASS, Khalid ENNIBI |
| EPI_ISL_471552 | Hospital Sancta Maggiore | Instituto Adolfo Lutz, Interdiciplinary Procedures Center, Strategic Laboratory | Claudio Tavares Sacchi, Claudia Regina Gonçalves, Erica Valessa Ramos Gomes |
| EPI_ISL_471957, EPI_ISL_471959, EPI_ISL_471960, EPI_ISL_471963 | University of Exeter | COVID-19 Genomics UK (COG-UK) Consortium | Ben Temperton,Aaron Jeffries,Michelle Michelsen,Joanna Warwick-Dugdale,Audrey Farbos,Robyn Manley,Stephen Michell,Jane Masoli |
| EPI_ISL_472136, EPI_ISL_472137, EPI_ISL_472138, EPI_ISL_472139, EPI_ISL_472140, EPI_ISL_472141, EPI_ISL_472142, EPI_ISL_472143, EPI_ISL_472144, EPI_ISL_472146 | Regional Virus Laboratory, Belfast Health and Social Care Trust | COVID-19 Genomics UK (COG-UK) Consortium | Conall McCaughey, James McKenna, Tanya Curran, Susan Feeney, Alison Watt, Ciara Cox, Mairead Connor, Zoltan Molnar, David Simpson, Derek Fairley |
| EPI_ISL_472246, EPI_ISL_472262 | Northumbria University / South Tees Hospitals NHS Foundation Trust / North Cumbria Integrated Care NHS Foundation Trust / North Tees and Hartlepool NHS Foundation Trust / Newcastle Hospitals NHS Foundation Trust | COVID-19 Genomics UK (COG-UK) Consortium | Darren L Smith,Andrew Nelson,Matthew Bashton,Greg R Young,Joshua Loh,John Allan,Mohammad A Tariq,Giles S Holt,Gary Black,Wen C Yew,Lynn Dover,Paul Baker,Steve Liggett,Sarah Essex,Jane Greenaway,Debra Padgett,Clive Graham,Garren Scott,Edward Barton,Emma Swindells,Brendan Payne,Jennifer Collins,Yusri Taha,Gary Eltringham |
| EPI_ISL_472432, EPI_ISL_472434, EPI_ISL_472435, EPI_ISL_472456, EPI_ISL_472458, EPI_ISL_472478, EPI_ISL_472499, EPI_ISL_472521, EPI_ISL_472526, EPI_ISL_472527, EPI_ISL_472530, EPI_ISL_472533, EPI_ISL_472539, EPI_ISL_472552, EPI_ISL_472565, EPI_ISL_472569, EPI_ISL_472572, EPI_ISL_472578, EPI_ISL_472591, EPI_ISL_472592, EPI_ISL_472593, EPI_ISL_472600, EPI_ISL_472619, EPI_ISL_472632, EPI_ISL_472654, EPI_ISL_472690, EPI_ISL_472704, EPI_ISL_472709, EPI_ISL_472714, EPI_ISL_473244, EPI_ISL_474234, EPI_ISL_474238, EPI_ISL_474257, EPI_ISL_474309, EPI_ISL_474310, EPI_ISL_474311, EPI_ISL_474312, EPI_ISL_474314 | | | |
| see above | Wales Specialist Virology Centre Sequencing lab: Pathogen Genomics Unit | COVID-19 Genomics UK (COG-UK) Consortium | Catherine Moore, Johnathan Evans, Laura Gifford, Malorie Perry, Simon Cottrell, Angela Marchbank, Alec Birchley, Alexander Adams, Amy Gaskin, Bree Gatica-Wilcox, Jason Coombes, Joel Southgate, Lauren Gilbert, Lee Graham, Nicole Pacchiarini, Sara Kumziene-Summerhayes, Sarah Taylor, Sophie Jones, Sara Rey, Matthew Bull, Joanne Watkins, Sally Corden, Tom Connor |
| EPI_ISL_474798, EPI_ISL_474799, EPI_ISL_474806 | Complejo Hospitalario Universitario de Albacete | SeqCOVID-SPAIN consortium/IBV(CSIC) | Encarnacion Simarro Córdoba, Julia Lozano Serra, Lorena Robles Fonseca , Monica Parra Grandes, Caridad Sainz de Baranda Camino and SeqCOVID-SPAIN consortium |
| EPI_ISL_474832, EPI_ISL_474835 | Hospital Universitario Virgen de las Nieves de Granada-SAS | SeqCOVID-SPAIN consortium/IBV(CSIC) | Mercedes Pérez Ruiz, Sara Sanbonmatsu Gámez, Irene Pedrosa Corral, José M. Navarro-Marí and SeqCOVID-SPAIN consortium |
| EPI_ISL_474837 | Complejo Hospitalario Universitario de Albacete | SeqCOVID-SPAIN consortium/IBV(CSIC) | Encarnacion Simarro Córdoba, Julia Lozano Serra, Lorena Robles Fonseca , Monica Parra Grandes, Caridad Sainz de Baranda Camino and SeqCOVID-SPAIN consortium |

| | | | |
|---|---|---|---|
| EPI_ISL_474846, EPI_ISL_474856, EPI_ISL_474884, EPI_ISL_474885, EPI_ISL_474887, EPI_ISL_474888, EPI_ISL_474898, EPI_ISL_474899 | Hospital Universitario Virgen de las Nieves de Granada-SAS | SeqCOVID-SPAIN consortium/IBV(CSIC) | Mercedes Pérez Ruiz, Sara Sanbonmatsu Gámez, Irene Pedrosa Corral, José M. Navarro-Marí and SeqCOVID-SPAIN consortium |
| EPI_ISL_474904 | Complejo Hospitalario Universitario de Albacete | SeqCOVID-SPAIN consortium/IBV(CSIC) | Encarnacion Simarro Córdoba, Julia Lozano Serra, Lorena Robles Fonseca , Monica Parra Grandes, Caridad Sainz de Baranda Camino and SeqCOVID-SPAIN consortium |
| EPI_ISL_474908, EPI_ISL_474931, EPI_ISL_474943, EPI_ISL_474944 | Hospital Universitario Virgen de las Nieves de Granada-SAS | SeqCOVID-SPAIN consortium/IBV(CSIC) | Mercedes Pérez Ruiz, Sara Sanbonmatsu Gámez, Irene Pedrosa Corral, José M. Navarro-Marí and SeqCOVID-SPAIN consortium |
| EPI_ISL_474952, EPI_ISL_474953, EPI_ISL_474955 | Complejo Hospitalario Universitario de Albacete | SeqCOVID-SPAIN consortium/IBV(CSIC) | Encarnacion Simarro Córdoba, Julia Lozano Serra, Lorena Robles Fonseca , Monica Parra Grandes, Caridad Sainz de Baranda Camino and SeqCOVID-SPAIN consortium |
| EPI_ISL_474962, EPI_ISL_474968 | Israel Central Virology laboratory | Israel Central Virology laboratory | Neta Zuckerman, Efrat Dahan Bucris, Oran Erster, Ella Mendelson, Michal Mandelboim |
| EPI_ISL_475116, EPI_ISL_475117 | Halmstad klinisk mikrobiologi | The Public Health Agency of Sweden | Oskar Karlsson Lindsjo, Maria Lind Karlberg, Mattias Haukland, Reza Advani, Olov Svartstrom, Anna-Malin Linde, Sandra Broddesson, Petra Edquist, Shamam Muradrasoli, Anna Risberg, Karin Tegmark-Wisell |
| EPI_ISL_475118 | Uppsala klinisk mikrobiologi | The Public Health Agency of Sweden | Oskar Karlsson Lindsjo, Maria Lind Karlberg, Mattias Haukland, Reza Advani, Olov Svartstrom, Anna-Malin Linde, Sandra Broddesson, Petra Edquist, Shamam Muradrasoli, Anna Risberg, Karin Tegmark-Wisell |
| EPI_ISL_475146, EPI_ISL_475147, EPI_ISL_475148 | Klinisk mikrobiologi Vasternorrland | The Public Health Agency of Sweden | Oskar Karlsson Lindsjo, Maria Lind Karlberg, Mattias Haukland, Reza Advani, Olov Svartstrom, Anna-Malin Linde, Sandra Broddesson, Petra Edquist, Shamam Muradrasoli, Anna Risberg, Karin Tegmark-Wisell |
| EPI_ISL_475279, EPI_ISL_475280, EPI_ISL_475285, EPI_ISL_475289, EPI_ISL_475297, EPI_ISL_475299 | Centre for Enzyme Innovation, University of Portsmouth / Translational Research Laboratory, Portsmouth Hospitals NHS Trust | COVID-19 Genomics UK (COG-UK) Consortium | Angela Beckett,Yann Bourgeois,Garry Scarlett,Sharon Glaysher,Scott Elliott,Kelly Bicknell,Robert Impey,Allyson Lloyd,Sarah Wyllie,Ethan Butcher,Anoop Chauhan,Samuel Robson |
| EPI_ISL_475585, EPI_ISL_475642, EPI_ISL_475646, EPI_ISL_475654, EPI_ISL_475655, EPI_ISL_475656, EPI_ISL_475657, EPI_ISL_475661, EPI_ISL_475666, EPI_ISL_475669, EPI_ISL_475670, EPI_ISL_475671, EPI_ISL_475674, EPI_ISL_475675, EPI_ISL_475676 | | | |
| see above | Cedars-Sinai Medical Center, Department of Pathology & Laboratory Medicine, Molecular Pathology Laboratory | Cedars-Sinai Medical Center, Molecular Pathology Laboratory of Department of Pathology & Laboratory Medicine and Genomic Core | Wenjuan Zhang, John Paul Govindavari, Brian Davis, Stephanie Chen, Jong Taek Kim, Jianbo Song, Jean Lopategui, Jasmine T Plummer, Eric Vail |
| EPI_ISL_475792, EPI_ISL_475793, EPI_ISL_475794, EPI_ISL_475795, EPI_ISL_475796, EPI_ISL_475797, EPI_ISL_475798, EPI_ISL_475799 | Center for Virology, Medical University of Vienna | Bergthaler laboratory, CeMM Research Center for Molecular Medicine of the Austrian Academy of Sciences | Alexandra Popa, Benedikt Agerer, Henrique Colaco, Lukas Endler, Jakob-Wendelin Genger, Alexander Lercher, Mark Smyth, Thomas Penz, Michael Schuster, Jan Laine, Martin Senekowitsch, Judith Aberle, Stephan Aberle, Peter Hufnagl, Daniela Schmid, Franz Allerberger, Elisabeth Puchhammer-Stoeckl, Manfred Nairz, Guenter Weiss, Gregor Hörmann, Kinga Rigler-Hohenwarter, Rainer Gattringer, Wegene Borena, Dorothee von Laer, Christoph Bock, Andreas Bergthaler |
| EPI_ISL_475822, EPI_ISL_475823, EPI_ISL_475824, EPI_ISL_475825, EPI_ISL_475826, EPI_ISL_475827 | Institut für Virologie am Department für Hygiene, Mikrobiologie und Public Health | Bergthaler laboratory, CeMM Research Center for Molecular Medicine of the Austrian Academy of Sciences | Alexandra Popa, Benedikt Agerer, Henrique Colaco, Lukas Endler, Jakob-Wendelin Genger, Alexander Lercher, Mark Smyth, Thomas Penz, Michael Schuster, Jan Laine, Martin Senekowitsch, Judith Aberle, Stephan Aberle, Peter Hufnagl, Daniela Schmid, Franz Allerberger, Elisabeth Puchhammer-Stoeckl, Manfred Nairz, Guenter Weiss, Gregor Hörmann, Kinga Rigler-Hohenwarter, Rainer Gattringer, Wegene Borena, Dorothee von Laer, Christoph Bock, Andreas Bergthaler |
| EPI_ISL_475892, EPI_ISL_475893 | Zentralinstitut für medizinische und chemische Labordiagnostik, Universitätskliniken Insbruck | Bergthaler laboratory, CeMM Research Center for Molecular Medicine of the Austrian Academy of Sciences | Alexandra Popa, Benedikt Agerer, Henrique Colaco, Lukas Endler, Jakob-Wendelin Genger, Alexander Lercher, Mark Smyth, Thomas Penz, Michael Schuster, Jan Laine, Martin Senekowitsch, Judith Aberle, Stephan Aberle, Peter Hufnagl, Daniela Schmid, Franz Allerberger, Elisabeth Puchhammer-Stoeckl, Manfred Nairz, Guenter Weiss, Gregor Hörmann, Kinga Rigler-Hohenwarter, Rainer Gattringer, Wegene Borena, Dorothee von Laer, Christoph Bock, Andreas Bergthaler |
| EPI_ISL_475911, EPI_ISL_475914 | Klinikum Wels-Grieskirchen | Bergthaler laboratory, CeMM Research Center for Molecular Medicine of the Austrian Academy of Sciences | Alexandra Popa, Benedikt Agerer, Henrique Colaco, Lukas Endler, Jakob-Wendelin Genger, Alexander Lercher, Mark Smyth, Thomas Penz, Michael Schuster, Jan Laine, Martin Senekowitsch, Judith Aberle, Stephan Aberle, Peter Hufnagl, Daniela Schmid, Franz Allerberger, Elisabeth Puchhammer-Stoeckl, Manfred Nairz, Guenter Weiss, Gregor Hörmann, Kinga Rigler-Hohenwarter, Rainer Gattringer, Wegene Borena, Dorothee von Laer, Christoph Bock, Andreas Bergthaler |
| EPI_ISL_475923 | Institut für Virologie am Department für Hygiene, Mikrobiologie und Public Health | Bergthaler laboratory, CeMM Research Center for Molecular Medicine of the Austrian Academy of Sciences | Alexandra Popa, Benedikt Agerer, Henrique Colaco, Lukas Endler, Jakob-Wendelin Genger, Alexander Lercher, Mark Smyth, Thomas Penz, Michael Schuster, Jan Laine, Martin Senekowitsch, Judith Aberle, Stephan Aberle, Peter Hufnagl, Daniela Schmid, Franz Allerberger, Elisabeth Puchhammer-Stoeckl, Manfred Nairz, Guenter Weiss, Gregor Hörmann, Kinga Rigler-Hohenwarter, Rainer Gattringer, Wegene Borena, Dorothee von Laer, Christoph Bock, Andreas Bergthaler |
| EPI_ISL_476023 | Defence Research & Development Establishment (DRDE) | Defence Research & Development Establishment (DRDE) | Shashi Sharma, Paban Kumar Dash, Sushil Kumar Sharma, Ambuj Shrivastava, Jyoti S. Kumar |
| EPI_ISL_476161, EPI_ISL_476164 | Laboratório de Patologia Clínica - UNICAMP | Laboratório de Estudos de Vírus Emergentes - UNICAMP | José Luiz Proença-Modena, Magnun Nueldo Nunes dos Santos, Angelica Schreiber, Julia Forato,Camila Simeoni, Marcilio Jorge Fumagalli, Mariene Ribeiro Amorim, Darlan da Silva Candido, Nuno Rodrigues Faria, Julien Theze, Luiz Gonzaga,Jaqueline Goes Jesus e William Marciel de Souza |
| EPI_ISL_476302, EPI_ISL_476303, EPI_ISL_476304, EPI_ISL_476305, EPI_ISL_476306, EPI_ISL_476307, EPI_ISL_476308, EPI_ISL_476309, EPI_ISL_476310, EPI_ISL_476311, EPI_ISL_476312, EPI_ISL_476313, EPI_ISL_476314, EPI_ISL_476315, EPI_ISL_476316, EPI_ISL_476317, EPI_ISL_476318, EPI_ISL_476319, EPI_ISL_476320, EPI_ISL_476323, EPI_ISL_476324, EPI_ISL_476325, EPI_ISL_476326, EPI_ISL_476354 | | | |
| see above | DB Diagnósticos do Brasil | Instituto de Medicina Tropical da Univesidade de São Paulo | Samples: Nelson Gaburo Jr; Sequencing: Ingra Morales Claro, Jaqueline Goes de Jesus, Erika Regina Manuli, Flavia Cristina da Silva Sales, Thais de Moura Coletti, Camila Alves Maia da Silva, Mariana Severo Ramundo, Giula Magalhaes Ferreira, Darlan da Silva Candido, Julien Theze, Nuno Faria, Ester Sabino |
| EPI_ISL_476372, EPI_ISL_476373 | Hospital da Clínicas da Faculdade de Medicina da Universidade de São Paulo | Instituto de Medicina Tropical da Univesidade de São Paulo | Samples: Ingra Morales Claro, Erika Regina Manuli, Cecilia Salete Alencar, Carolina S. Lazar, Silvia F. Costa; Sequencing: Ingra Morales Claro, Jaqueline Goes de Jesus, Erika Regina Manuli, Flavia Cristina da Silva Sales, Thais de Moura Coletti, Camila Alves Maia da Silva, Mariana Severo Ramundo, Giula Magalhaes Ferreira, Darlan da Silva Candido, Julien Theze, Nuno Faria, Ester Sabino |
| EPI_ISL_476539, EPI_ISL_476540, EPI_ISL_476541, EPI_ISL_476542, EPI_ISL_476543, EPI_ISL_476544, EPI_ISL_476545, EPI_ISL_476546, EPI_ISL_476547, EPI_ISL_476548, EPI_ISL_476549, EPI_ISL_476550, EPI_ISL_476551, EPI_ISL_476552, EPI_ISL_476553, EPI_ISL_476554, EPI_ISL_476555, EPI_ISL_476556, EPI_ISL_476557 | | | |
| see above | Yale Clinical Virology Laboratory | Grubaugh Lab - Yale School of Public Health | Joseph Fauver, Tara Alpert, Anderson Brito, Anne Wyllie, Chantal Vogels, Mary Petrone, Cole Jensen, Chaney Kalinich, Isabel Ott, Arnau Casanovas, Catherine Muenker, Adam Moore, Alice Lu, Maria Tokuyama, Patrick Wong, Peiwen Lu, Saad Omer, Richard Martinello, Allison Nelson, Shelli Farhadian, Akiko Iwasaki, Charlese Dela Cruz, Albert Ko, Nathan Grubaugh |
| EPI_ISL_476792 | Stanford clinical virology lab | Chan-Zuckerberg Biohub | Benjamin Pinksy, Katharine Walter, Victoria N. Parikh, John Gorzynski, Hannah N. DeJong, Matthew T. Wheeler, Jason Andrews, Manuel Rivas, Carlos Bustamante, Euan Ashley, with CZB Cliahub Consortium |
| EPI_ISL_476846 | Defence Research & Development Establishment (DRDE) | Defence Research & Development Establishment (DRDE) | Shashi Sharma, Paban Kumar Dash, Sushil Kumar Sharma, Ambuj Shrivastava, Jyoti S. Kumar |
| EPI_ISL_477017, EPI_ISL_477018, EPI_ISL_477019, EPI_ISL_477020, EPI_ISL_477021, EPI_ISL_477022, EPI_ISL_477023, EPI_ISL_477024, EPI_ISL_477025, EPI_ISL_477026, EPI_ISL_477027, EPI_ISL_477028, EPI_ISL_477029, EPI_ISL_477030, EPI_ISL_477031, EPI_ISL_477032, EPI_ISL_477033, EPI_ISL_477034, EPI_ISL_477035, EPI_ISL_477036, EPI_ISL_477037, EPI_ISL_477038, EPI_ISL_477064, EPI_ISL_477076, EPI_ISL_477090, EPI_ISL_477091, EPI_ISL_477092, EPI_ISL_477093, EPI_ISL_477094, EPI_ISL_477095, EPI_ISL_477096, EPI_ISL_477097, EPI_ISL_477098, EPI_ISL_477099, EPI_ISL_477100, EPI_ISL_477101, EPI_ISL_477102, EPI_ISL_477103, EPI_ISL_477104, EPI_ISL_477105, EPI_ISL_477106, EPI_ISL_477107, EPI_ISL_477108, EPI_ISL_477109, EPI_ISL_477110, EPI_ISL_477111, EPI_ISL_477112, EPI_ISL_477113, EPI_ISL_477114, EPI_ISL_477115, EPI_ISL_477116, EPI_ISL_477117, EPI_ISL_477118, EPI_ISL_477119 | | | |
| see above | BCCDC Public Health Laboratory | BCCDC Public Health Laboratory | Richard Harrigan, Hope Lapointe, Jinny Choi, Kimia Kamelian, John Tyson,Terry Snutch, Linda Hoang, Inna Sekirov, Paul Levett, Mel Krajden, Natalie Prystajecky |
| EPI_ISL_477706 | UW Virology Lab | UW Virology Lab | Pavitra Roychoudhury, Hong Xie, Lasata Shrestha, Amin Addetia, Truong Nguyen, Victoria M Rachleff, Meei-Li Huang, Keith R Jerome, Alexander Greninger |
| EPI_ISL_477821 | West of Scotland Specialist Virology Centre, NHSGGC / MRC-University of Glasgow Centre for Virus Research | COVID-19 Genomics UK (COG-UK) Consortium | Ana da Silva Filipe, Natasha Johnson, Kathy Smollett, Daniel Mair, Stephen Carmichael, Lily Tong, Jenna Nichols, Elihu Aranday-Cortes, Kirstyn Brunker, Yasmin Parr, Alice Broos, Kyriaki Nomikou; Sarah McDonald, Marc Niebel, Patawee Asamaphan; Richard Orton, Joseph Hughes, Sreenu Vattipally, David L Robertson; Alasdair MacLean, Rory Gunson; Kathy Li, Natasha Jesudason, Rajiv Shah, James Shepherd, Antonia Ho, Emma Thomson |

| | | | |
|---|---|---|---|
| EPI_ISL_478417, EPI_ISL_478418, EPI_ISL_478419, EPI_ISL_478420, EPI_ISL_478421, EPI_ISL_478422, EPI_ISL_478423, EPI_ISL_478424, EPI_ISL_478425, EPI_ISL_478426, EPI_ISL_478427, EPI_ISL_478428, EPI_ISL_478429, EPI_ISL_478430, EPI_ISL_478431 | | | |
| see above | University College London, Great Ormond Street Hospital for Children NHS Foundation Trust, Imperial College Healthcare NHS Trust | COVID-19 Genomics UK (COG-UK) Consortium | Sergi Castellano, Rachel Williams, Mark Kristiansen, Paola Resende Silva, Sunando Roy, Tony Brooks, Helena Tutill, Paola Niola, Patricia Dyal, Charlotte Williams, Leysa Forrest, Yasmin Panchbhaya, Jacqueline Findlay, Samuel Weeks, Julianne Brown, Kathryn Harris, Paul Randell, James Price, Alison Holmes, Judith Breuer |
| EPI_ISL_478496, EPI_ISL_478497, EPI_ISL_478498, EPI_ISL_478500, EPI_ISL_478501, EPI_ISL_478502, EPI_ISL_478503, EPI_ISL_478504, EPI_ISL_478505, EPI_ISL_478506, EPI_ISL_478507, EPI_ISL_478508, EPI_ISL_478509, EPI_ISL_478510, EPI_ISL_478511, EPI_ISL_478633, EPI_ISL_478635, EPI_ISL_478636, EPI_ISL_478637, EPI_ISL_478638, EPI_ISL_478663, EPI_ISL_478664, EPI_ISL_478665, EPI_ISL_478666, EPI_ISL_478667 | | | |
| see above | Northumbria University / South Tees Hospitals NHS Foundation Trust / North Cumbria Integrated Care NHS Foundation Trust / North Tees and Hartlepool NHS Foundation Trust / Newcastle Hospitals NHS Foundation Trust | COVID-19 Genomics UK (COG-UK) Consortium | Darren L Smith,Andrew Nelson,Matthew Bashton,Greg R Young,Joshua Loh,John Allan,Mohammad A Tariq,Giles S Holt,Gary Black,Wen C Yew,Lynn Dover,Paul Baker,Steve Liggett,Sarah Essex,Jane Greenaway,Debra Padgett,Clive Graham,Garren Scott,Edward Barton,Emma Swindells,Brendan Payne,Jennifer Collins,Yusri Taha,Gary Eltringham |
| EPI_ISL_478673 | Pathology North - Royal North Shore Hospital - NSW Health Pathology | NSW Health Pathology - Institute of Clinical Pathology and Medical Research; Westmead Hospital; University of Sydney | CIDM-PH et al. |
| EPI_ISL_478700, EPI_ISL_478701 | South Eastern Area Laboratory Services (SEALS) | NSW Health Pathology - Institute of Clinical Pathology and Medical Research; Westmead Hospital; University of Sydney | CIDM-PH et al. |
| EPI_ISL_479620, EPI_ISL_479621, EPI_ISL_479622, EPI_ISL_479623 | Molecular diagnostic laboratory of Federal Budget Institution of Science "Central Research Institute of Epidemiology" of The Federal Service on Customers' Rights Protection and Human Well-being Surveillance | Group of Genomics and Postgenomic Technologies of Central Research Institute of Epidemilology | Speranskaya AS, Kaptelova VV,Valdokhina AV, Bulanenko VP, Samoilov AE, Korneenko EV, Sizova TV, Tivanova EV, Shipulina OY, Akimkin VG |
| EPI_ISL_479648, EPI_ISL_479649, EPI_ISL_479650 | Dr. Georges-L.-Dumont University Hospital Centre | National Microbiology Laboratory | Anna Majer, Shari Tyson, Grace Seo, Kristyn Burak, Philip Mabon, Elsie Grudeski, Rhiannon Huzarewich, Russell Mandes, Jennifer Tanner, Natalie Knox, Morag Graham, Gary Van Domselaar, Richard Garceau, Guillaume Desnoyers, Nathalie Bastien, Yan Li, Timothy Booth |
| EPI_ISL_479673 | Center for Genomics and System Biology, New York University | Center for Genomics and System Biology, New York University | Roder,A., Banakis,S., Johnson,K., Khalfan,M., Borenstein,E.S., Samanovic,M., Cornelius,A., Herati,R., Ulrich,R., Fleming,A., Kottkamp,A., Raabe,V., Mulligan,M.J., Gresham,D., Ghedin,E. |
| EPI_ISL_479676, EPI_ISL_479677, EPI_ISL_479678, EPI_ISL_479679, EPI_ISL_479680, EPI_ISL_479681, EPI_ISL_479682, EPI_ISL_479683, EPI_ISL_479684, EPI_ISL_479685 | unknown | Contact:Hiroyuki Asakura Tokyo Metropolitan Institute of Public Health, Department of Microbiology | Asakura,H., Yoshida,I., Kumagai,R., Nagashima,M., Chiba,T., Sadamasu,K. |
| EPI_ISL_480009, EPI_ISL_480010 | Chiba Prefectural Institute of Public Health | Pathogen Genomics Center, National Institute of Infectious Diseases | Tsuyoshi Sekizuka, Masakatsu Taira, Kentaro Itokawa, Rina Tanaka, Masanori Hashino, Hajime Kamiya, Motoi Suzuki, Makoto Kuroda |
| EPI_ISL_480026, EPI_ISL_480027, EPI_ISL_480028, EPI_ISL_480029 | Ibaraki Prefectural Institute of Public Health | Pathogen Genomics Center, National Institute of Infectious Diseases | Tsuyoshi Sekizuka, Keiko Goto, Kentaro Itokawa, Rina Tanaka, Masanori Hashino, Hajime Kamiya, Motoi Suzuki, Makoto Kuroda |
| EPI_ISL_480067, EPI_ISL_480068, EPI_ISL_480069 | Sakai City Institute of Public Health | Pathogen Genomics Center, National Institute of Infectious Diseases | Tsuyoshi Sekizuka, Tatsuya Miyoshi, Kentaro Itokawa, Rina Tanaka, Masanori Hashino, Hajime Kamiya, Motoi Suzuki, Makoto Kuroda |
| EPI_ISL_480081 | Shizuoka City Institute of Environmental Sciences and Public Health | Pathogen Genomics Center, National Institute of Infectious Diseases | Tsuyoshi Sekizuka, Takaharu Maehata,Sou Okamura,Yuji Kanazawa,Kenji Yagi, Kentaro Itokawa, Rina Tanaka, Masanori Hashino, Hajime Kamiya, Motoi Suzuki, Makoto Kuroda |
| EPI_ISL_480090 | Department of Infectious Diseases, Kobe Institute of Health | Pathogen Genomics Center, National Institute of Infectious Diseases | Tsuyoshi Sekizuka, Ryohei Nomoto, Kentaro Itokawa, Rina Tanaka, Masanori Hashino, Hajime Kamiya, Motoi Suzuki, Makoto Kuroda |
| EPI_ISL_480125, EPI_ISL_480127, EPI_ISL_480128, EPI_ISL_480129, EPI_ISL_480130, EPI_ISL_480131, EPI_ISL_480132, EPI_ISL_480133, EPI_ISL_480134 | Fukui Prefectural Institute of Public Health and Environmental Science | Pathogen Genomics Center, National Institute of Infectious Diseases | Tsuyoshi Sekizuka, Miho Toho, Kentaro Itokawa, Rina Tanaka, Masanori Hashino, Hajime Kamiya, Motoi Suzuki, Makoto Kuroda |
| EPI_ISL_480177 | Gunma Prefectural Institute of Public Health and Environmental Sciences | Pathogen Genomics Center, National Institute of Infectious Diseases | Tsuyoshi Sekizuka, Hiroyuki Tsukagoshi, Kentaro Itokawa, Rina Tanaka, Masanori Hashino, Hajime Kamiya, Motoi Suzuki, Makoto Kuroda |
| EPI_ISL_480182 | Ibaraki Prefectural Institute of Public Health | Pathogen Genomics Center, National Institute of Infectious Diseases | Tsuyoshi Sekizuka, Keiko Goto, Kentaro Itokawa, Rina Tanaka, Masanori Hashino, Hajime Kamiya, Motoi Suzuki, Makoto Kuroda |
| EPI_ISL_480190, EPI_ISL_480191, EPI_ISL_480192, EPI_ISL_480193, EPI_ISL_480194, EPI_ISL_480195 | Ota Health Center Welfare Section | Pathogen Genomics Center, National Institute of Infectious Diseases | Tsuyoshi Sekizuka, Chika Takahashi, Kentaro Itokawa, Rina Tanaka, Masanori Hashino, Hajime Kamiya, Motoi Suzuki, Makoto Kuroda |
| EPI_ISL_480199, EPI_ISL_480200, EPI_ISL_480201, EPI_ISL_480202 | Toyama Institute of Health | Pathogen Genomics Center, National Institute of Infectious Diseases | Tsuyoshi Sekizuka, Masae Itamochi, Kazunori Oishi, Kentaro Itokawa, Rina Tanaka, Masanori Hashino, Hajime Kamiya, Motoi Suzuki, Makoto Kuroda |
| EPI_ISL_480221 | Koshigaya City Public Health Center | Pathogen Genomics Center, National Institute of Infectious Diseases | Tsuyoshi Sekizuka, Yuka Furui, Aya Tamura, Kyohei Sakata, Takumi Daimon, Yoko Togawa, Yoshiko Hamada, Kentaro Itokawa, Rina Tanaka, Masanori Hashino, Hajime Kamiya, Motoi Suzuki, Makoto Kuroda |
| EPI_ISL_480320 | Hospital Clínica Bíblica | Charité Virology-University of Costa Rica | Andres Moreira-Soto, Eugenia Corrales-Aguilar, Ignacio Postigo-Hidalgo, Karla Sofía Gutiérrez, Jan Felix Drexler |
| EPI_ISL_480326 | Hospital Nacional de Niños | Charité Virology-University of Costa Rica | Andres Moreira-Soto, Eugenia Corrales-Aguilar, Ignacio Postigo-Hidalgo, Cristian Pérez Corrales, Andrei Montero Bonilla, Jan Felix Drexler |
| EPI_ISL_480391, EPI_ISL_480392, EPI_ISL_480393 | University of Wisconsin-Madison AIDS Vaccine Research Laboratories | University of Wisconsin-Madison AIDS Vaccine Research Laboratories | Gage Moreno, Katarina Braun, et al. AIDS Vaccine Research Laboratories |
| EPI_ISL_480570, EPI_ISL_480572 | Victorian Infectious Diseases Reference Laboratory (VIDRL) | VIDRL and MDU-PHL | Caly L., Seemann T., Sait, M., Schultz M., Druce J., Sherry, N. |
| EPI_ISL_480615 | Microbiological Diagnostic Unit - Public Health Laboratory (MDU-PHL) | MDU-PHL | Seemann T., Schultz M., Sait, M., Sherry, N. |
| EPI_ISL_480783, EPI_ISL_480786, EPI_ISL_480787, EPI_ISL_480788, EPI_ISL_480789 | Institut Pasteur Dakar | Institut Pasteur de Dakar | Ndongo Dia, Moussa Moise Diagne, Mamadou Diop, Marie Henriette Dior Ndione, Mamadou Malado Jallow, Safietou Sanke, Ousmane Faye, Amadou Alpha Sall. |
| EPI_ISL_480801, EPI_ISL_480802, EPI_ISL_480804, EPI_ISL_480806, EPI_ISL_480807, EPI_ISL_480808, EPI_ISL_480809, EPI_ISL_480810, EPI_ISL_480811, EPI_ISL_480812, EPI_ISL_480813, EPI_ISL_480814, EPI_ISL_480816 | | | |
| see above | Florida Bureau of Public Health Laboratories | Florida Bureau of Public Health Laboratories | Sarah Schmedes, Jason Blanton |
| EPI_ISL_480961 | ISGlobal, Institut de Salut Global de Barcelona | SeqCOVID-SPAIN consortium/IBV(CSIC) | Alfredo Mayor, Alberto L Garcia-Basteiro, Carlota Dobaño, Gemma Moncunill, Pau Cisteró and SeqCOVID-SPAIN consortium |
| EPI_ISL_481254 | Department of Emerging Infectious Diseases, Institute of Tropical Medicine, Nagasaki University | Department of Emerging Infectious Diseases, Institute of Tropical Medicine, Nagasaki University | Jiro Yasuda, Rokusuke Yoshikawa, Yuichiro Furusato, Haruka Abe |
| EPI_ISL_481265, EPI_ISL_481266, EPI_ISL_481267, EPI_ISL_481268, EPI_ISL_481269, EPI_ISL_481270 | Maryland Department of Health | Maryland Department of Health | Keller,E. |
| EPI_ISL_481284 | Department of Experimental Modeling and Pathogenesis of | Department of Experimental Modeling and Pathogenesis of | Sobolev,I.A., Shanshin,D.V., Chepurnov,A.A., Kononova,J.V., Bondar,A.A., Alekseev,A.Y. and Shestopalov,A.M. |

| | | | |
|---|---|---|---|
| | Infectious Diseases, Federal Research Center of Fundamental and Translational Medicine | Infectious Diseases, Federal Research Center of Fundamental and Translational Medicine | |
| EPI_ISL_481744, EPI_ISL_481745, EPI_ISL_481746, EPI_ISL_481749 | Dr. Georges-L.-Dumont University Hospital Centre | National Microbiology Laboratory | Anna Majer, Shari Tyson, Grace Seo, Kristyn Burak, Philip Mabon, Elsie Grudeski, Rhiannon Huzarewich, Russell Mandes, Jennifer Tanner, Natalie Knox, Morag Graham, Gary Van Domselaar, Richard Garceau, Guillaume Desnoyers, Nathalie Bastien, Yan Li, Timothy Booth |
| EPI_ISL_482292, EPI_ISL_482293, EPI_ISL_482298, EPI_ISL_482299, EPI_ISL_482300, EPI_ISL_482301 | Providence St. Joseph Health Molecular Genomics Laboratory | Providence St. Joseph Health Molecular Genomics Laboratory | Alexa K Dowdell, Brian D Piening, Fred L Robinson, Carlo B Bifulco, Mary Campbell |
| EPI_ISL_482733, EPI_ISL_482734 | LNR National Reference Laboratory, Mohammed VI University of Health Sciences | Medical Biotechnology Laboratory, Rabat Medical and Pharmacy School, Mohammed The Vth University in Rabat | Meriem LAAMARTI, Souad KARTTI, Rokia LAAMARTI , M.W. CHEMAO-ELFIHRI, Loubna ALLAM, Mouna OUADGHIRI, Imane SMYEJ, Jalila RAHOUI, Houda BENRHAMA, Jalil El ATAR, Idrissa DIAWARA, Rachid EL JAOUDI, Laila SBABOU, Chakib NEJJARI, Saaid AMZAZI, Rachid MENTAG, Lahcen BELYAMANI and Azeddine IBRAHIMI |
| EPI_ISL_483072, EPI_ISL_483086, EPI_ISL_483091 | SA Pathology | SA Pathology | Lex Leong, Chuan Kok Lim, Mark Turra, Ivan Bastian, Geoff Higgins |
| EPI_ISL_483167, EPI_ISL_483170, EPI_ISL_483171, EPI_ISL_483172, EPI_ISL_483177, EPI_ISL_483179, EPI_ISL_483183, EPI_ISL_483188, EPI_ISL_483189, EPI_ISL_483190, EPI_ISL_483193, EPI_ISL_483196, EPI_ISL_483197, EPI_ISL_483317, EPI_ISL_483367, EPI_ISL_483368, EPI_ISL_483370, EPI_ISL_483376, EPI_ISL_483379, EPI_ISL_483386, EPI_ISL_483388, EPI_ISL_483392, EPI_ISL_483434, EPI_ISL_483439, EPI_ISL_483441, EPI_ISL_483442, EPI_ISL_483443, EPI_ISL_483447 | | | |
| see above | UC San Diego Center for Advanced Laboratory Medicine | Andersen lab at Scripps Research | SEARCH Alliance San Diego with David Pride, Ji H Shin |
| EPI_ISL_483621 | National Public Health Laboratory, National Centre for Infectious Diseases | National Public Health Laboratory, National Centre for Infectious Diseases | Mak TM, Octavia S, Zhou Z, Chavatte JM, Cui L, Lin RTP |
| EPI_ISL_484153, EPI_ISL_484156, EPI_ISL_484158, EPI_ISL_484160, EPI_ISL_484162, EPI_ISL_484164, EPI_ISL_484165, EPI_ISL_484166, EPI_ISL_484167, EPI_ISL_484169, EPI_ISL_484170, EPI_ISL_484171, EPI_ISL_484172, EPI_ISL_484173, EPI_ISL_484175, EPI_ISL_484176, EPI_ISL_484177, EPI_ISL_484178, EPI_ISL_484179, EPI_ISL_484180, EPI_ISL_484181, EPI_ISL_484182, EPI_ISL_484202, EPI_ISL_484203, EPI_ISL_484204, EPI_ISL_484205, EPI_ISL_484206, EPI_ISL_484207, EPI_ISL_484208, EPI_ISL_484209, EPI_ISL_484210, EPI_ISL_484211, EPI_ISL_484212, EPI_ISL_484213, EPI_ISL_484214, EPI_ISL_484215, EPI_ISL_484216, EPI_ISL_484217 | | | |
| see above | Centre for Clinical Infection and Diagnostics Research and Genomics Innovation Unit, Guy's and St. Thomas' NHS Trust | COVID-19 Genomics UK (COG-UK) Consortium | Chloe Fisher, Luke Snell, Penny Cliff, Rahul Batra, Jonathan Edgeworth, Ali Raza Awan |
| EPI_ISL_484415, EPI_ISL_484416, EPI_ISL_484417 | Centre for Enzyme Innovation, University of Portsmouth / Translational Research Laboratory, Portsmouth Hospitals NHS Trust | COVID-19 Genomics UK (COG-UK) Consortium | Angela Beckett,Yann Bourgeois,Garry Scarlett,Sharon Glaysher,Scott Elliott,Kelly Bicknell,Robert Impey,Allyson Lloyd,Sarah Wyllie,Ethan Butcher,Anoop Chauhan,Samuel Robson |
| EPI_ISL_484710, EPI_ISL_484713, EPI_ISL_484717, EPI_ISL_484718, EPI_ISL_484723, EPI_ISL_484742, EPI_ISL_484745, EPI_ISL_484760, EPI_ISL_484762, EPI_ISL_484787, EPI_ISL_484788, EPI_ISL_484793, EPI_ISL_484794, EPI_ISL_484801, EPI_ISL_484806 | | | |
| see above | University of Michigan Clinical Microbiology Laboratory | Lauring Lab, University of Michigan, Department of Microbiology and Immunology | Valesano et al. |
| EPI_ISL_485388, EPI_ISL_485394, EPI_ISL_485395, EPI_ISL_485396 | University of Ulsan College of Medicine and Asan Medical Center | University of Ulsan College of Medicine and Asan Medical Center | Kuenyoul Park, Jaewoong Lee, Kihyun Lee, Jiwon Jung, Sung-Han Kim, Jina Lee, Mauricio Chalita, Seok-Hwan Yoon, Jongsik Chun, Kyu-Hwa Hur, Heungsup Sung, Mi-Na Kim, and Hae Kyung Lee |
| EPI_ISL_485871 | Virginia DCLS | Virginia DCLS | Virginia DCLS |
| EPI_ISL_485987 | UW Virology Lab | UW Virology Lab | Pavitra Roychoudhury, Hong Xie, Lasata Shrestha, Amin Addetia, Truong Nguyen, Victoria M Rachleff, Meei-Li Huang, Keith R Jerome, Alexander Greninger |
| EPI_ISL_486442, EPI_ISL_486443, EPI_ISL_486444, EPI_ISL_486445, EPI_ISL_486446, EPI_ISL_486447, EPI_ISL_486448, EPI_ISL_486449, EPI_ISL_486450, EPI_ISL_486451, EPI_ISL_486452, EPI_ISL_486453, EPI_ISL_486454, EPI_ISL_486455, EPI_ISL_486456, EPI_ISL_486457, EPI_ISL_486458, EPI_ISL_486459, EPI_ISL_486460, EPI_ISL_486461, EPI_ISL_486462, EPI_ISL_486463, EPI_ISL_486464, EPI_ISL_486465, EPI_ISL_486466, EPI_ISL_486467, EPI_ISL_486468, EPI_ISL_486469, EPI_ISL_486470, EPI_ISL_486471, EPI_ISL_486472, EPI_ISL_486473, EPI_ISL_486474, EPI_ISL_486475, EPI_ISL_486476, EPI_ISL_486477, EPI_ISL_486478, EPI_ISL_486479, EPI_ISL_486480, EPI_ISL_486481, EPI_ISL_486482, EPI_ISL_486483, EPI_ISL_486484, EPI_ISL_486485, EPI_ISL_486486, EPI_ISL_486487, EPI_ISL_486488, EPI_ISL_486489, EPI_ISL_486490, EPI_ISL_486491, EPI_ISL_486492, EPI_ISL_486493, EPI_ISL_486494, EPI_ISL_486495, EPI_ISL_486496, EPI_ISL_486497, EPI_ISL_486498, EPI_ISL_486499, EPI_ISL_486500 | | | |
| see above | Viollier AG | Department of Biosystems Science and Engineering, ETH Zürich | Christian Beisel, Sarah Nadeau, Ivan Topolsky, Pedro Ferreira, Philipp Jablonski, Susana Posada-Céspedes, Tobias Schär, Ina Nissen, Natascha Santacroce, Elodie Burcklen, Christiane Beckmann, Maurice Redondo, Olivier Kobel, Christoph Noppen, Sophie Seidel, Noemie Santamaria de Souza, Niko Beerenwinkel, Tanja Stadler |
| EPI_ISL_486658 | Microbiology, Virology and Biemergency Laboratory-ASST FBF Sacco | Microbiology, Virology and Biemergency Laboratory-ASST FBF Sacco | Romeri F, Comandatore F, Mancon A, Micheli V, Rimoldi SG |
| EPI_ISL_486820, EPI_ISL_486821, EPI_ISL_486822, EPI_ISL_486823, EPI_ISL_486824, EPI_ISL_486825, EPI_ISL_486826, EPI_ISL_486827, EPI_ISL_486828, EPI_ISL_486829 | Molecular diagnostic laboratory of Federal Budget Institution of Science "Central Research Institute of Epidemiology" of The Federal Service on Customers' Rights Protection and Human Well-being Surveillance | Group of Genomics and Postgenomic Technologies of Central Research Institute of Epidemilology | Speranskaya AS, Kaptelova VV, Valdokhina AV, Bulanenko VP, Samoilov AE, Korneenko EV, Tivanova EV, Shipulina OY, Akimkin VG |
| EPI_ISL_486890, EPI_ISL_486891, EPI_ISL_486892, EPI_ISL_486893, EPI_ISL_486894, EPI_ISL_486895, EPI_ISL_486896 | Tokyo Metoropolitan Institute of Public Health | Tokyo Metoropolitan Institute of Public Health | Asakura,H., Yoshida,I., Kumagai,R., Nagashima,M., Chiba,T., Sadamasu,K. |
| EPI_ISL_486912, EPI_ISL_486913, EPI_ISL_486914 | Maryland Department of Health | Maryland Department of Health | Keller,E. |
| EPI_ISL_487684, EPI_ISL_487703, EPI_ISL_487707, EPI_ISL_487710, EPI_ISL_487721, EPI_ISL_487789, EPI_ISL_487805, EPI_ISL_487809, EPI_ISL_487812, EPI_ISL_487815, EPI_ISL_487817, EPI_ISL_487845, EPI_ISL_487856, EPI_ISL_487934, EPI_ISL_487991 | | | |
| see above | Virology Department, Royal Infirmary of Edinburgh, NHS Lothian / School of Biological Sciences, University of Edinburgh | Wellcome Sanger Institute for the COVID-19 Genomics UK (COG-UK) consortium | McHugh M, Dewar R, Rooke S, O'Toole Á, Scher E, Hill V, McCrone JT, Colquhoun R, Yu X, Jackson B, Rambaut A, Templeton K and Alex Alderton, Roberto Amato, Sonia Goncalves, Ewan Harrison, David K. Jackson, Ian Johnston, Dominic Kwiatkowski, Cordelia Langford, John Sillitoe on behalf of the Wellcome Sanger Institute COVID-19 Surveillance Team (http://www.sanger.ac.uk/covid-team) |
| EPI_ISL_488214, EPI_ISL_488410 | PHE South West Regional Laboratory, National Infection Service | Wellcome Sanger Institute for the COVID-19 Genomics UK (COG-UK) consortium | Stephanie Hutchings, Hannah Pymont, Dr Peter Muir, Barry Vipond, Rich Hopes; and Alex Alderton, Roberto Amato, Sonia Goncalves, Ewan Harrison, David K. Jackson, Ian Johnston, Dominic Kwiatkowski, Cordelia Langford, John Sillitoe on behalf of the Wellcome Sanger Institute COVID-19 Surveillance Team (http://www.sanger.ac.uk/covid-team) |
| EPI_ISL_488465, EPI_ISL_488477, EPI_ISL_488488, EPI_ISL_488496, EPI_ISL_488501, EPI_ISL_488516, EPI_ISL_488522, EPI_ISL_488523, EPI_ISL_488524, EPI_ISL_488526, EPI_ISL_488529, EPI_ISL_488531, EPI_ISL_488533, EPI_ISL_488534, EPI_ISL_488537, EPI_ISL_488539, EPI_ISL_488540, EPI_ISL_488545, EPI_ISL_488546, EPI_ISL_488553, EPI_ISL_488558, EPI_ISL_488559, EPI_ISL_488561, EPI_ISL_488566, EPI_ISL_488568, EPI_ISL_488569, EPI_ISL_488577, EPI_ISL_488583, EPI_ISL_488597, EPI_ISL_488604, EPI_ISL_488607, EPI_ISL_488615, EPI_ISL_488623, EPI_ISL_488625, EPI_ISL_488642, EPI_ISL_488643, EPI_ISL_488645, EPI_ISL_488649, EPI_ISL_488651, EPI_ISL_488652, EPI_ISL_488653, EPI_ISL_488660, EPI_ISL_488663, EPI_ISL_488665, EPI_ISL_488668, EPI_ISL_488672, EPI_ISL_488677, EPI_ISL_488684, EPI_ISL_488691, EPI_ISL_488692, EPI_ISL_488693, EPI_ISL_488696, EPI_ISL_488702, EPI_ISL_488703, EPI_ISL_488705, EPI_ISL_488708, EPI_ISL_488709, EPI_ISL_488711, EPI_ISL_488712, EPI_ISL_488721, EPI_ISL_488735, EPI_ISL_488736, EPI_ISL_488738, EPI_ISL_488741 | | | |
| see above | NU-OMICS DNA Sequencing research facility, Northumbria University | Wellcome Sanger Institute for the COVID-19 Genomics UK (COG-UK) consortium | Chris Duncan, Sheia Waugh, Shirelle Burton-Fanning, Gary Eltringham, Jennifer Collins, Brendan Payne, Yusri Taha, Emma Swindells, Jane Greenaway, Edward Barton, Garren Scott, Debra Padgett, Clive Graham, Sarah Essex, Steve Ligget, Paul Baker, Lynn Dover, Wen Yew, Gary Black, John Allan, Joshua Loh, Greg Young, Matthew Bashton, Andrew Nelson, Darren Smith and Alex Alderton, Roberto Amato, Sonia Goncalves, Ewan Harrison, David K. Jackson, Ian Johnston, Dominic Kwiatkowski, Cordelia Langford, John Sillitoe on behalf of the Wellcome Sanger Institute COVID-19 Surveillance Team (http://www.sanger.ac.uk/covid-team) |
| EPI_ISL_488884, EPI_ISL_488886, EPI_ISL_488895, EPI_ISL_488897, EPI_ISL_488902, EPI_ISL_488909, EPI_ISL_488915, EPI_ISL_488918, EPI_ISL_488930, EPI_ISL_488931, EPI_ISL_488940, EPI_ISL_488945, EPI_ISL_488957, EPI_ISL_488961, EPI_ISL_488965, EPI_ISL_488970, EPI_ISL_488973, EPI_ISL_488995, EPI_ISL_489010, EPI_ISL_489016, EPI_ISL_489025, EPI_ISL_489027, EPI_ISL_489028, EPI_ISL_489030, EPI_ISL_489041 | | | |
| see above | Virology Department, Royal Infirmary of Edinburgh, NHS Lothian / School of Biological Sciences, University of | Wellcome Sanger Institute for the COVID-19 Genomics UK (COG-UK) consortium | McHugh M, Dewar R, Rooke S, O'Toole Á, Scher E, Hill V, McCrone JT, Colquhoun R, Yu X, Jackson B, Rambaut A, Templeton K and Alex Alderton, Roberto Amato, Sonia Goncalves, Ewan Harrison, David K. Jackson, Ian Johnston, Dominic Kwiatkowski, Cordelia Langford, John Sillitoe on behalf of the |

| | | | |
|---|---|---|---|
| | Edinburgh | | Wellcome Sanger Institute COVID-19 Surveillance Team (http://www.sanger.ac.uk/covid-team) |
| EPI_ISL_490031 | South Eastern Area Laboratory Services (SEALS) | NSW Health Pathology - Institute of Clinical Pathology and Medical Research; Westmead Hospital; University of Sydney | CIDM-PH et al. |
| EPI_ISL_490058 | National Public Health Laboratory, National Centre for Infectious Diseases | National Public Health Laboratory, National Centre for Infectious Diseases | Mak TM, Octavia S, Zhou Z, Chavatte JM, Cui L, Lin RTP |
| EPI_ISL_490105 | South Eastern Area Laboratory Services (SEALS) | NSW Health Pathology - Institute of Clinical Pathology and Medical Research; Westmead Hospital; University of Sydney | CIDM-PH et al. |
| EPI_ISL_490210, EPI_ISL_490211, EPI_ISL_490212, EPI_ISL_490213, EPI_ISL_490214, EPI_ISL_490215, EPI_ISL_490216, EPI_ISL_490217, EPI_ISL_490218, EPI_ISL_490219, EPI_ISL_490220, EPI_ISL_490221, EPI_ISL_490222, EPI_ISL_490223 | | | |
| see above | Quest Diagnostics | Q Squared Solutions - QRTP facility | Victor J Weigman |
| EPI_ISL_491118 | The National Institute of Public Health | The National Institute of Public Health and State Veterinary Institute Prague | Nagy,A;Jirincova,H;Novakova,L;Trnka,D;Vecerova,J |
| EPI_ISL_491459, EPI_ISL_491460, EPI_ISL_491461, EPI_ISL_491462, EPI_ISL_491464 | Laboratorio de Referencia Nacional de Virus Respiratorio. Instituto Nacional de Salud Perú | Laboratorio de Referencia Nacional de Biotecnología y Biología Molecular. Instituto Nacional de Salud Perú | Carlos Padilla Rojas, Karolyn Vega Chozo, Priscila Lope Pari, Omar Caceres Rey, Marco Galarza Perez, Maribel Huaringa Nuñez, Johanna Balbuena Torrez, Henri Bailon Calderon, Nancy Rojas Serrano |
| EPI_ISL_491504, EPI_ISL_491558, EPI_ISL_491574, EPI_ISL_491605, EPI_ISL_491668, EPI_ISL_491689 | Virology Department, Royal Infirmary of Edinburgh, NHS Lothian / School of Biological Sciences, University of Edinburgh | Wellcome Sanger Institute for the COVID-19 Genomics UK (COG-UK) consortium | McHugh M, Dewar R, Rooke S, O'Toole Á, Scher E, Hill V, McCrone JT, Colquhoun R, Yu X, Jackson B, Rambaut A, Templeton K and Alex Alderton, Roberto Amato, Sonia Goncalves, Ewan Harrison, David K. Jackson, Ian Johnston, Dominic Kwiatkowski, Cordelia Langford, John Sillitoe on behalf of the Wellcome Sanger Institute COVID-19 Surveillance Team (http://www.sanger.ac.uk/covid-team) |
| EPI_ISL_491920 | Naval Infectious Diseases Diagnostic Laboratory | Naval Medical Research Center Biological Defense Research Directorate | Logan Voegtly, Regina Cer, Lindsay Glang, Victor Sugiharto, Francisco Malgon Bautista, Hua Wei Chen, Dessiree Pena-Gomez, Megan Schilling, Adrian Paskey, Kyle Long, Mark Simons, Kimberly Bishop-Lilly |
| EPI_ISL_492068 | 1. ViroGenetics - BSL3 Laboratory of Virology, Maopolska Centre of Biotechnology, Jagiellonian University; 2. II Department of Internal Medicine, Faculty of Medicine, Jagiellonian University Medical College; 3. Narodowy Instytut Zdrowia Publicznego - Pastwowy Zakad Higieny (NIZP-PZH) | 1. ViroGenetics - BSL3 Laboratory of Virology, Maopolska Centre of Biotechnology, Jagiellonian University; 2. II Department of Internal Medicine, Faculty of Medicine, Jagiellonian University Medical College; 3. Narodowy Instytut Zdrowia Publicznego - Pastwowy Zakad Higieny (NIZP-PZH). | Katarzyna Pancer, Marek Sanak, Aleksandra A. Zasada, Magdalena Rzeczkowska, Tomasz Wokowicz, Katarzyna Zacharczuk, Agnieszka Koakowska-Kulesza, Katarzyna Owczarek, Aleksandra Milewska, Natalia Wolaniuk, Ewelina Hallman-Szeliska, Pawe P abaj, Wojciech Branicki, Krzysztof Pyr |
| EPI_ISL_492152, EPI_ISL_492166 | SA Pathology | SA Pathology | Lex Leong, Chuan Kok Lim, Mark Turra, Ivan Bastian, Geoff Higgins |
| EPI_ISL_492214, EPI_ISL_492216, EPI_ISL_492224, EPI_ISL_492241, EPI_ISL_492263, EPI_ISL_492285, EPI_ISL_492291, EPI_ISL_492309, EPI_ISL_492313, EPI_ISL_492333, EPI_ISL_492341, EPI_ISL_492364, EPI_ISL_492371, EPI_ISL_492376, EPI_ISL_492407, EPI_ISL_492414, EPI_ISL_492415, EPI_ISL_492425, EPI_ISL_492444 | | | |
| see above | PHE South West Regional Laboratory, National Infection Service | Wellcome Sanger Institute for the COVID-19 Genomics UK (COG-UK) consortium | Stephanie Hutchings, Hannah Pymont, Dr Peter Muir, Barry Vipond, Rich Hopes; and Alex Alderton, Roberto Amato, Sonia Goncalves, Ewan Harrison, David K. Jackson, Ian Johnston, Dominic Kwiatkowski, Cordelia Langford, John Sillitoe on behalf of the Wellcome Sanger Institute COVID-19 Surveillance Team (http://www.sanger.ac.uk/covid-team) |
| EPI_ISL_492852 | Royal Free Hospital / Health Services Laboratories | Wellcome Sanger Institute for the COVID-19 Genomics UK (COG-UK) consortium | Tanzina Haque, Tabitha Mahungu, Dianne Irish, Cate Goodlad, Jenny Cross, Judith Heaney and Alex Alderton, Roberto Amato, Sonia Goncalves, Ewan Harrison, David K. Jackson, Ian Johnston, Dominic Kwiatkowski, Cordelia Langford, John Sillitoe on behalf of the Wellcome Sanger Institute COVID-19 Surveillance Team (http://www.sanger.ac.uk/covid-team) |
| EPI_ISL_493062, EPI_ISL_493063 | Wyoming Public Health Laboratory | Wyoming Public Health Laboratory | Noah Hull, Rob Christensen, Jim Mildenberger, Joel Sevinsky, Cari Sloma, and Wanda Manley |
| EPI_ISL_493646, EPI_ISL_493647, EPI_ISL_493653, EPI_ISL_493656, EPI_ISL_493662 | Centre for Enzyme Innovation, University of Portsmouth / Translational Research Laboratory, Portsmouth Hospitals NHS Trust | COVID-19 Genomics UK (COG-UK) Consortium | Angela Beckett,Yann Bourgeois,Garry Scarlett,Sharon Glaysher,Scott Elliott,Kelly Bicknell,Robert Impey,Allyson Lloyd,Sarah Wyllie,Ethan Butcher,Anoop Chauhan,Samuel Robson |
| EPI_ISL_494185, EPI_ISL_494188, EPI_ISL_494198, EPI_ISL_494209, EPI_ISL_494213, EPI_ISL_494218, EPI_ISL_494231, EPI_ISL_494239, EPI_ISL_494253, EPI_ISL_494276, EPI_ISL_494285, EPI_ISL_494294, EPI_ISL_494296, EPI_ISL_494317, EPI_ISL_494344, EPI_ISL_494345, EPI_ISL_494358, EPI_ISL_494364 | | | |
| see above | Wales Specialist Virology Centre Sequencing lab: Pathogen Genomics Unit | COVID-19 Genomics UK (COG-UK) Consortium | Catherine Moore, Johnathan Evans, Laura Gifford, Malorie Perry, Simon Cottrell, Angela Marchbank, Alec Birchley, Alexander Adams, Amy Gaskin, Bree Gatica-Wilcox, Jason Coombes, Joel Southgate, Lauren Gilbert, Lee Graham, Nicole Pacchiarini, Sara Kumziene-Summerhayes, Sarah Taylor, Sophie Jones, Sara Rey, Matthew Bull, Joanne Watkins, Sally Corden, Tom Connor |
| EPI_ISL_494395, EPI_ISL_494400, EPI_ISL_494408, EPI_ISL_494412, EPI_ISL_494420, EPI_ISL_494572, EPI_ISL_494622, EPI_ISL_494623 | San Diego County Public Health Laboratory | Andersen lab at Scripps Research | SEARCH Alliance San Diego with Tracy Basler, Jovan Shephard, Brett Austin |
| EPI_ISL_494667 | Scripps Medical Laboratory | Andersen lab at Scripps Research | SEARCH Alliance San Diego with Michael Quigley, Ellen Stefanski, Ian Mchardy |
| EPI_ISL_494715, EPI_ISL_494717, EPI_ISL_494719, EPI_ISL_494720, EPI_ISL_494725, EPI_ISL_494736, EPI_ISL_494739, EPI_ISL_494742 | San Diego County Public Health Laboratory | Andersen lab at Scripps Research | SEARCH Alliance San Diego with Tracy Basler, Jovan Shephard, Brett Austin |
| EPI_ISL_495374, EPI_ISL_495379 | Florida Bureau of Public Health Laboratories | Florida Bureau of Public Health Laboratories | Sarah Schmedes, Jason Blanton |
| EPI_ISL_495564, EPI_ISL_495570 | University of Michigan Clinical Microbiology Laboratory | Lauring Lab, University of Michigan, Department of Microbiology and Immunology | Valesano et al. |
| EPI_ISL_496786, EPI_ISL_496787, EPI_ISL_496788, EPI_ISL_496789, EPI_ISL_496790, EPI_ISL_496791, EPI_ISL_496792, EPI_ISL_496793, EPI_ISL_496794, EPI_ISL_496795, EPI_ISL_496796, EPI_ISL_496797, EPI_ISL_496798, EPI_ISL_496799, EPI_ISL_496800, EPI_ISL_496801, EPI_ISL_496802, EPI_ISL_496803, EPI_ISL_496804, EPI_ISL_496805, EPI_ISL_496806, EPI_ISL_496807, EPI_ISL_496808, EPI_ISL_496809 | | | |
| see above | Gorgas Memorial Laboratory of Health Studies | Gorgas Memorial Laboratory of Health Studies | Danilo Franco, Claudia Gonzalez Sandra Lopez-Verges, Alexander A Martinez |
| EPI_ISL_497818, EPI_ISL_497858 | Department of Microbiology, The University of Hong Kong | Department of Microbiology, The University of Hong Kong | Kelvin K.W. To, Kwok-Yung Yuen |
| EPI_ISL_497872 | UW Virology Lab | UW Virology Lab | Pavitra Roychoudhury, Amin Addetia, Hong Xie, Lasata Shrestha, Truong Nguyen, Meei-Li Huang, Keith Jerome, Alexander Greninger |
| EPI_ISL_498159, EPI_ISL_498160, EPI_ISL_498163, EPI_ISL_498166 | Instituto Nacional de Salud, Bogotá, Colombia | Instituto Nacional de Salud, Bogotá, Colombia | Katherine Laiton-Donato, Diego A. Álvarez-Díaz, Carlos Franco-Muñoz, Jonathan Reales, Diego Andrés Prada, Jose A. Usme-Ciro, Nicolas D. Franco-Sierra, Zulma M. Cucunubá, Christian Julian VillabonaArenas, Liz Villabona-Arenas, Sussy Echeverria, Astrid C. Flórez, Carolina Ferro, Diana Marcela Walteros-Acero, Franklin Prieto, Carlos Andrés Durán, Martha Lucia Ospina Martinez, Marcela Mercado-Reyes |
| EPI_ISL_498191, EPI_ISL_498192 | OUCRU | OUCRU | Nguyen Van Vinh Chau, Nguyen Thi Thu Hong, Nguyen Thi Han Ny, Le Nguyen Truc Nhu, Nghiem My Ngoc, Vo Thanh Lam, Nguyen Thanh Dung, Lam Minh Yen, Ngo Ngoc Quang Minh, Le Manh Hung, Nguyen Tri Dung, Dinh Nguyen Huy Man, Lam Anh Nguyet, Tran Chanh Xuan, Tran Tinh Hien, Nguyen Thanh Phong, Tran Nguyen Hoang Tu, Tran Tan Thanh, Nguyen Thanh Truong, Nguyen Tan Binh, Tang Chi Thuong, Guy Thwaites, and Le Van Tan, for OUCRU COVID-19 research group* |
| EPI_ISL_498508, EPI_ISL_498524, EPI_ISL_498526, EPI_ISL_498530 | ACT Pathology | Schwessinger Lab | Ashley Jones, Benjamin Schwessinger, Robert Lanfear, Robyn N Hall, Megan McDonald, Ming-Dao Chia, Kevin Murray, Craig Kennedy, Karina Kennedy |
| EPI_ISL_499281 | Centre for Enzyme Innovation, University of Portsmouth / Translational Research Laboratory, Portsmouth Hospitals NHS Trust | COVID-19 Genomics UK (COG-UK) Consortium | Angela Beckett,Yann Bourgeois,Garry Scarlett,Sharon Glaysher,Scott Elliott,Kelly Bicknell,Robert Impey,Allyson Lloyd,Sarah Wyllie,Ethan Butcher,Anoop Chauhan,Samuel Robson |
| EPI_ISL_499333 | Virology Department, Sheffield Teaching Hospitals NHS Foundation Trust/Department of Infection, Immunity and | COVID-19 Genomics UK (COG-UK) Consortium | Thushan de Silva, Matthew Parker, Nikki Smith, Adri Angyal, Rebecca Brown, Luke Green, Rachel Tucker, Paul Parsons, Danielle Groves, Katie Johnson, Laura Carrilero, Alex Keeley, Dave Partridge, Matthew Wyles, Benjamin Lindsey, Mehmet Yavuz, Mohammad Raza, Cariad Evans |

| | | | |
|---|---|---|---|
| | Cardiovascular Disease, The Medical School, University of Sheffield | | |
| EPI_ISL_499358, EPI_ISL_499369, EPI_ISL_499374, EPI_ISL_499403, EPI_ISL_499413, EPI_ISL_499429, EPI_ISL_499433, EPI_ISL_499442, EPI_ISL_499447, EPI_ISL_499451 | Wales Specialist Virology Centre Sequencing lab: Pathogen Genomics Unit | COVID-19 Genomics UK (COG-UK) Consortium | Catherine Moore, Johnathan Evans, Laura Gifford, Malorie Perry, Simon Cottrell, Angela Marchbank, Alec Birchley, Alexander Adams, Amy Gaskin, Bree Gatica-Wilcox, Jason Coombes, Joel Southgate, Lauren Gilbert, Lee Graham, Nicole Pacchiarini, Sara Kumziene-Summerhayes, Sarah Taylor, Sophie Jones, Sara Rey, Matthew Bull, Joanne Watkins, Sally Corden, Tom Connor |
| EPI_ISL_500371, EPI_ISL_500389, EPI_ISL_500426, EPI_ISL_500448, EPI_ISL_500456, EPI_ISL_500457 | Centro de Investigación Biomédica de La Rioja - Hospital San Pedro Logroño | SeqCOVID-SPAIN consortium/IBV(CSIC) | María de Toro, José Manuel Azcona Gutiérrez, María Pilar Bea Escudero, Miriam Blasco Alberdi and SeqCOVID-SPAIN consortium |
| EPI_ISL_500832 | Virginia DCLS | Virginia DCLS | Virginia DCLS |
| EPI_ISL_501191, EPI_ISL_501206, EPI_ISL_501207, EPI_ISL_501208, EPI_ISL_501209, EPI_ISL_501210 | Department of Medical Microbiology, University Malaya Medical Centre | Department of Medical Microbiology, Faculty of Medicine, University of Malaya | Yoong Min CHONG, Jennifer Chong, I-Ching SAM, Yoke Fun CHAN, University Malaya Medical Centre COVID Team |
| EPI_ISL_501246, EPI_ISL_501251 | Hellenic Pasteur Institute, National Influenza Reference laboratory of Southern Greece & Unit of Bioinformatics and Applied Genomics | Hellenic Pasteur Institute, National Influenza Reference laboratory of Southern Greece & Unit of Bioinformatics and Applied Genomics | Vasiliki Pogka, Timokratis Karamitros, Athanasios Kossyvakis, Antonios Kalliaropoulos, Horefti Elina, Evangelidou Maria, Androniki Voulgari-Kokota, Aspasia Kontou, Andreas Mentis |
| EPI_ISL_506987, EPI_ISL_506990 | Division of Viral Diseases, Center for Laboratory Control of Infectious Diseases, Korea Centers for Diseases Control and Prevention | Division of Viral Diseases, Center for Laboratory Control of Infectious Diseases, Korea Centers for Diseases Control and Prevention | Jeong-Min Kim, Yoon-Seok Chung, Namjoo Lee, Sang Hee Woo, Hye-Jun Jo, Heui Man Kim, Jun-Sub Kim, Dong Hyun Song, Daesang Lee, Seong Tae Jeong, Myung Guk Han |
| EPI_ISL_507025, EPI_ISL_507026 | unknown | Infectious Diseases Research, King Abdullah International Medical Research Center (KAIMRC) | Alghoribi,M.F. |
| EPI_ISL_507079, EPI_ISL_507080 | University College London Hospital | COVID-19 Genomics UK (COG-UK) Consortium | Judith Heaney, Matthew Byott, Catherine Houlihan, Dan Frampton, Stuart Kirk, Moira Spyer and Eleni Nastouli |
| EPI_ISL_507135 | Centre for Enzyme Innovation, University of Portsmouth / Translational Research Laboratory, Portsmouth Hospitals NHS Trust | COVID-19 Genomics UK (COG-UK) Consortium | Angela Beckett,Yann Bourgeois,Garry Scarlett,Sharon Glaysher,Scott Elliott,Kelly Bicknell,Robert Impey,Allyson Lloyd,Sarah Wyllie,Ethan Butcher,Anoop Chauhan,Samuel Robson |
| EPI_ISL_507431, EPI_ISL_507447, EPI_ISL_507448, EPI_ISL_507450, EPI_ISL_507452, EPI_ISL_507455, EPI_ISL_507457, EPI_ISL_507460, EPI_ISL_507461, EPI_ISL_507462, EPI_ISL_507463, EPI_ISL_507470, EPI_ISL_507471, EPI_ISL_507560, EPI_ISL_507564, EPI_ISL_507566, EPI_ISL_507567, EPI_ISL_507570, EPI_ISL_507572, EPI_ISL_507573, EPI_ISL_507575, EPI_ISL_507578, EPI_ISL_507580, EPI_ISL_507584, EPI_ISL_507585, EPI_ISL_507586, EPI_ISL_507595, EPI_ISL_507597, EPI_ISL_507600, EPI_ISL_507601, EPI_ISL_507602, EPI_ISL_507604, EPI_ISL_507609, EPI_ISL_507612, EPI_ISL_507613, EPI_ISL_507614, EPI_ISL_507619, EPI_ISL_507804, EPI_ISL_507812, EPI_ISL_507813, EPI_ISL_507826, EPI_ISL_507835, EPI_ISL_507837, EPI_ISL_507840, EPI_ISL_507841, EPI_ISL_507843, EPI_ISL_507847, EPI_ISL_507849, EPI_ISL_507853, EPI_ISL_507863, EPI_ISL_507880, EPI_ISL_507884, EPI_ISL_507904, EPI_ISL_507913, EPI_ISL_507920, EPI_ISL_507921, EPI_ISL_507922, EPI_ISL_507931 | | | |
| see above | Michigan Department of Health and Human Services, Bureau of Laboratories | Michigan Department of Health and Human Services, Bureau of Laboratories | Blankenship HM, Riner D, Soehnlen MK |
| EPI_ISL_508145 | SA Pathology | SA Pathology | Lex Leong, Chuan Kok Lim, Mark Turra, Ivan Bastian, Geoff Higgins |
| EPI_ISL_508368, EPI_ISL_508376 | Institute of Post Graduate Medical Education & Research | National Institute of Biomedical Genomics | Arindam Maitra, Aritra Biswas, Jayeeta Haldar, Raja Ray, Monimoy Banerjee, Saumitra Das |
| EPI_ISL_508745, EPI_ISL_508746, EPI_ISL_508747, EPI_ISL_508748, EPI_ISL_508749 | Florida Bureau of Public Health Laboratories | Florida Bureau of Public Health Laboratories | Sarah Schmedes, Jason Blanton |
| EPI_ISL_508959, EPI_ISL_508960 | Centre Hospitalier Saint Joseph Saint Luc | CNR Virus des Infections Respiratoires - France SUD | Antonin Bal, Gregory Destras, Gwendolyne Burfin, Solenne Brun, Carine Moustaud, Raphaelle Lamy, Alexandre Gaymard, Maude Bouscambert-Duchamp, Florence Morfin-Sherpa, Martine Valette, Bruno Lina, Laurence Josset |
| EPI_ISL_508961, EPI_ISL_508962, EPI_ISL_508963, EPI_ISL_508976 | CNR Virus des Infections Respiratoires - France SUD | CNR Virus des Infections Respiratoires - France SUD | Antonin Bal, Gregory Destras, Gwendolyne Burfin, Solenne Brun, Carine Moustaud, Raphaelle Lamy, Alexandre Gaymard, Maude Bouscambert-Duchamp, Florence Morfin-Sherpa, Martine Valette, Bruno Lina, Laurence Josset |
| EPI_ISL_509011, EPI_ISL_509013, EPI_ISL_509014 | Institut des Agents Infectieux (IAI), Hospices Civils de Lyon | CNR Virus des Infections Respiratoires - France SUD | Antonin Bal, Gregory Destras, Gwendolyne Burfin, Solenne Brun, Carine Moustaud, Raphaelle Lamy, Alexandre Gaymard, Maude Bouscambert-Duchamp, Florence Morfin-Sherpa, Martine Valette, Bruno Lina, Laurence Josset |
| EPI_ISL_509015 | Centre Hospitalier de Villefranche | CNR Virus des Infections Respiratoires - France SUD | Antonin Bal, Gregory Destras, Gwendolyne Burfin, Solenne Brun, Carine Moustaud, Raphaelle Lamy, Alexandre Gaymard, Maude Bouscambert-Duchamp, Florence Morfin-Sherpa, Martine Valette, Bruno Lina, Laurence Josset |
| EPI_ISL_509016 | Centre Hospitalier du Haut-Bugey | CNR Virus des Infections Respiratoires - France SUD | Antonin Bal, Gregory Destras, Gwendolyne Burfin, Solenne Brun, Carine Moustaud, Raphaelle Lamy, Alexandre Gaymard, Maude Bouscambert-Duchamp, Florence Morfin-Sherpa, Martine Valette, Bruno Lina, Laurence Josset |
| EPI_ISL_509587, EPI_ISL_509588, EPI_ISL_509589, EPI_ISL_509590, EPI_ISL_509591, EPI_ISL_509597, EPI_ISL_509598, EPI_ISL_509600 | Utah Public Health Laboratory | Utah Public Health Laboratory | Heidi Butz, Erin Young, Kelly Oakeson |
| EPI_ISL_509769, EPI_ISL_509772, EPI_ISL_509773, EPI_ISL_509774 | Florida Bureau of Public Health Laboratories | Florida Bureau of Public Health Laboratories | Sarah Schmedes, Jason Blanton |
| EPI_ISL_510071 | Instituto de Investigaciones Biomédicas de Barcelona (CSIC), Hospital Clinic i Provincial de Barcelona, Instituto de Biomedicina de Valencia (CSIC), Hospital de Sant Pau | SeqCOVID-SPAIN consortium/IBV(CSIC) | Anna M. Planas, Mª Angeles Marcos, Miguel J. Martínez, Andrea Vergara, Alex Soriano, Jordi Pérez Tur, Israel Fernández Cadenas and SeqCOVID-SPAIN consortium |
| EPI_ISL_510119, EPI_ISL_510134, EPI_ISL_510141, EPI_ISL_510239, EPI_ISL_510240, EPI_ISL_510241, EPI_ISL_510242 | Hospital General Universitario Gregorio Marañón | SeqCOVID-SPAIN consortium/IBV(CSIC) | Laura Pérez-Lago, Marta Herranz, Jon Sicilia, Julia Suárez, Pilar Catalán, Patricia Muñoz, Darío García de Viedma and SeqCOVID-SPAIN consortium |
| EPI_ISL_510464 | Instituto de Investigaciones Biomédicas de Barcelona (CSIC), Hospital Clinic i Provincial de Barcelona, Instituto de Biomedicina de Valencia (CSIC), Hospital de Sant Pau | SeqCOVID-SPAIN consortium/IBV(CSIC) | Anna M. Planas, Mª Angeles Marcos, Miguel J. Martínez, Andrea Vergara, Alex Soriano, Jordi Pérez Tur, Israel Fernández Cadenas and SeqCOVID-SPAIN consortium |
| EPI_ISL_510530 | UMR 190 - Faculte de medecine, UMR 'Emergence des Pathologies Virales' (EPV: Aix-Marseille University - IRD 190 - Inserm 1207 - EHE | UMR 190 - Faculte de medecine, UMR 'Emergence des Pathologies Virales' (EPV: Aix-Marseille University - IRD 190 - Inserm 1207 - EHE | Durand, G.A., Piorkowski,G., Pommier De Santi,V., De Laval,F., Leparc Goffart,I., Gilles,M., Geulen,M., Peduzzi,F., Le Flem,F.-X. |
| EPI_ISL_510537 | Microbiology Division, SC DHEC | Microbiology Division, SC DHEC | Flores,H. |
| EPI_ISL_510909, EPI_ISL_510910, EPI_ISL_510911, EPI_ISL_510912, EPI_ISL_510927, EPI_ISL_510946, EPI_ISL_511028, EPI_ISL_511178 | Instituto Nacional de Saude (INSA) | Instituto Nacional de Saude (INSA) | Borges et al |
| EPI_ISL_511192, EPI_ISL_511193, EPI_ISL_511194, EPI_ISL_511195, EPI_ISL_511196, EPI_ISL_511197, EPI_ISL_511198, EPI_ISL_511199, EPI_ISL_511200, EPI_ISL_511201, EPI_ISL_511202, EPI_ISL_511203, EPI_ISL_511204, EPI_ISL_511205, EPI_ISL_511206, EPI_ISL_511207, EPI_ISL_511208, EPI_ISL_511209 | | | |
| see above | Instituto Nacional de Saude (INSA) and Instituto Gulbenkian | Instituto Nacional de Saude (INSA) and Instituto Gulbenkian | Borges et al |

| | | | |
|---|---|---|---|
| | de Ciencia (IGC) | de Ciencia (IGC) | |
| EPI_ISL_511362, EPI_ISL_511398, EPI_ISL_511431, EPI_ISL_511432, EPI_ISL_511435, EPI_ISL_511445, EPI_ISL_511451, EPI_ISL_511456, EPI_ISL_511470, EPI_ISL_511529, EPI_ISL_511656, EPI_ISL_511663, EPI_ISL_511668, EPI_ISL_511669, EPI_ISL_511675, EPI_ISL_511676, EPI_ISL_511687, EPI_ISL_511688, EPI_ISL_511689, EPI_ISL_511705, EPI_ISL_511707, EPI_ISL_511737, EPI_ISL_511738, EPI_ISL_511739, EPI_ISL_511740, EPI_ISL_511741, EPI_ISL_511742, EPI_ISL_511743, EPI_ISL_511744, EPI_ISL_511745, EPI_ISL_511746, EPI_ISL_511747 | | | |
| see above | Instituto Nacional de Saude (INSA) | Instituto Nacional de Saude (INSA) | Borges et al |
| EPI_ISL_511865, EPI_ISL_511873 | Johns Hopkins Hospital Department of Pathology | Johns Hopkins Hospital Department of Pathology | Peter M. Thielen, Thomas Mehoke, Shirlee Wohl, Srividya Ramakrishnan, Melanie Kirsche, Amanda Ernlund, Craig Howser, Kristina Zudock, Oluwaseun Falade-Nwulia, Norah Sadowski, Paul Morris, Mark Hopkins, Yunfan Fan, Nidia Trovao, Victoria Gniazdowski, Michael C. Schatz, Stuart C. Ray, Winston Timp, Heba H. Mostafa |
| EPI_ISL_511900 | Institute of Post Graduate Medical Education & Research | National Institute of Biomedical Genomics - DBT's PAN-INDIA 1000 SARS--CoV-2 RNA Genome Sequencing Consortium | Arindam Maitra, Aritra Biswas, Jayeeta Haldar, Raja Ray, Monimoy Banerjee, Saumitra Das |
| EPI_ISL_512393, EPI_ISL_512394, EPI_ISL_512395, EPI_ISL_512396, EPI_ISL_512397, EPI_ISL_512398, EPI_ISL_512399, EPI_ISL_512400, EPI_ISL_512405 | Centre for Enzyme Innovation, University of Portsmouth / Translational Research Laboratory, Portsmouth Hospitals NHS Trust | COVID-19 Genomics UK (COG-UK) Consortium | Angela Beckett,Yann Bourgeois,Garry Scarlett,Sharon Glaysher,Scott Elliott,Kelly Bicknell,Robert Impey,Allyson Lloyd,Sarah Wyllie,Ethan Butcher,Anoop Chauhan,Samuel Robson |
| EPI_ISL_512437 | West of Scotland Specialist Virology Centre, NHSGGC / MRC-University of Glasgow Centre for Virus Research | COVID-19 Genomics UK (COG-UK) Consortium | Ana da Silva Filipe, Natasha Johnson, Kathy Smollett, Daniel Mair, Stephen Carmichael, Lily Tong, Jenna Nichols, Elihu Aranday-Cortes, Kirstyn Brunker, Yasmin Parr, Alice Broos, Kyriaki Nomikou; Sarah McDonald, Marc Niebel, Patawee Asamaphan; Richard Orton, Joseph Hughes, Sreenu Vattipally, David L Robertson; Alasdair MacLean, Rory Gunson; Kathy Li, Natasha Jesudason, Rajiv Shah, James Shepherd, Antonia Ho, Emma Thomson |
| EPI_ISL_512482 | Wales Specialist Virology Centre Sequencing lab: Pathogen Genomics Unit | COVID-19 Genomics UK (COG-UK) Consortium | Catherine Moore, Johnathan Evans, Laura Gifford, Malorie Perry, Simon Cottrell, Angela Marchbank, Alec Birchley, Alexander Adams, Amy Gaskin, Bree Gatica-Wilcox, Jason Coombes, Joel Southgate, Lauren Gilbert, Lee Graham, Nicole Pacchiarini, Sara Kumziene-Summerhayes, Sarah Taylor, Sophie Jones, Sara Rey, Matthew Bull, Joanne Watkins, Sally Corden, Tom Connor |
| EPI_ISL_512745, EPI_ISL_512746 | PathWest Laboratory Medicine WA | PathWest Laboratory Medicine WA Microbial Surveillance Unit | PathWest Laboratory Medicine WA Microbial Surveillance Unit |
| EPI_ISL_512793, EPI_ISL_512799, EPI_ISL_512800, EPI_ISL_512809 | Public Health, United States Air Force School of Aerospace Medicine | Public Health, United States Air Force School of Aerospace Medicine | Fries,A.C., Purves,S.M., Meyer,J.R., Javorina,A.K., Connors,B.C., Macias,E.A., Lambert,A.W., Chapleau,R.R., Starr,C.R. |
| EPI_ISL_512812 | Kenema Government Hospital, Ministry of Health and Sanitation | Kenema Government Hospital, Ministry of Health and Sanitation | Goba,A., Momoh,M., Sandi,J., Tomkins-Tinch,C., Siddle,K., Mehta,S., Oluniyi,P., Jalloh,S., Park,D., Andersen,K., Garry,R., Happi,C., Grant,D., Olawoye,I. |
| EPI_ISL_512928, EPI_ISL_512929 | Pathogen Genomics Lab King Abdullah University of Science and Technology(KAUST) | Pathogen Genomics Lab King Abdullah University of Science and Technology(KAUST) | Fadwa Alofi, Sharif Hala, Rahul P Salunke, Sara Mfarrej, Amit Kumar Subudhi, Fathia Ben Rached, Amanda, Luke, Afrah Alsomali, Asim Khogeer, Jumana Taha, Abdulaziz Alahmadi, Kahled Algithami, Raeece Naeem, Anwar Hashem, Naif Almontashiri, Arnab Pain |
| EPI_ISL_512989, EPI_ISL_512993, EPI_ISL_512994, EPI_ISL_513000, EPI_ISL_513001 | Pathogen Genomics Lab King Abdullah University of Science and Technology(KAUST) | Pathogen Genomics Lab King Abdullah University of Science and Technology(KAUST) | Amit Kumar Subudhi, Rahul P Salunke, Sara Mfarrej, Sharif Hala, Fadwa Alofi, Fathia Ben Rached, Afrah Alsomali, Asim Khogeer, Nashwa Al-khotani, Raeece Naeem, Anwar Hashem, Naif Almontashiri, Arnab Pain |
| EPI_ISL_513121, EPI_ISL_513122, EPI_ISL_513123, EPI_ISL_513124, EPI_ISL_513125, EPI_ISL_513126 | Pathogen Genomics Lab King Abdullah University of Science and Technology(KAUST) | Pathogen Genomics Lab King Abdullah University of Science and Technology(KAUST) | Sharif Hala, Fadwa Alofi, Sara Mfarrej, Amit Kumar Subudhi, Rahul P Salunke, Fathia Ben Rached, Amanda Ooi, Luke Esau, Afrah Alsomali, Asim Khogeer, Jumana Taha, Abdulaziz Alahmadi, Kahled Algithami, Raeece Naeem, Anwar Hashem, Naif Almontashiri, Arnab Pain |
| EPI_ISL_513182, EPI_ISL_513183, EPI_ISL_513184, EPI_ISL_513185, EPI_ISL_513186, EPI_ISL_513187, EPI_ISL_513188, EPI_ISL_513189, EPI_ISL_513232, EPI_ISL_513233 | Pathogen Genomics Lab King Abdullah University of Science and Technology(KAUST) | Pathogen Genomics Lab King Abdullah University of Science and Technology(KAUST) | Amit Kumar Subudhi, Rahul P Salunke, Sara Mfarrej, Sharif Hala, Fadwa Alofi, Fathia Ben Rached, Afrah Alsomali, Asim Khogeer, Nashwa Al-khotani, Raeece Naeem, Anwar Hashem, Naif Almontashiri, Arnab Pain |
| EPI_ISL_513632 | University of Washington Virology Lab | University of Washington Virology Lab | Pavitra Roychoudhury, Hong Xie, Lasata Shrestha, Amin Addetia, Truong Nguyen, Victoria M Rachleff, Meei-Li Huang, Keith R Jerome, Alexander Greninger |
| EPI_ISL_514264 | Laboratorio de Referencia Nacional de Virus Respiratorio. Instituto Nacional de Salud. Perú | Laboratorio de Referencia Nacional de Biotecnologia y Biologia Molecular.Instituto Nacional de Salud.Perú | Carlos Padilla Rojas, Karolyn Vega Chozo, Priscila Lope Pari, Omar Caceres Rey, Marco Galarza Perez, Maribel Huaringa Nuñez, Johanna Balbuena Torrez, Henri Bailon Calderon, Nancy Rojas Serrano |
| EPI_ISL_514653, EPI_ISL_514654, EPI_ISL_514655, EPI_ISL_514656 | Mayo Clinic & Mayo Clinic Laboratories | Minnesota Department of Health, Public Health Laboratory | Matt Plumb, Jacob Garfin, and Xiong Wang |
| EPI_ISL_514657 | M Health Fairview St. Joseph's Hospital | Minnesota Department of Health, Public Health Laboratory | Matt Plumb, Jacob Garfin, and Xiong Wang |
| EPI_ISL_515272, EPI_ISL_515273, EPI_ISL_515274, EPI_ISL_515275, EPI_ISL_515276, EPI_ISL_515278 | University of Washington Virology Lab | University of Washington Virology Lab | Pavitra Roychoudhury, Hong Xie, Lasata Shrestha, Amin Addetia, Truong Nguyen, Victoria M Rachleff, Meei-Li Huang, Keith R Jerome, Alexander Greninger |
| EPI_ISL_515345 | Nevada State Public Health Laboratory | Nevada State Public Health Laboratory | Richard Tillett, Joel R. Sevinsky, Paul Hartley, Heather Kerwin, David Jackson, Subhash C. Verma, Cyprian Rossetto, Andrew Gorzalski, Chris Laverdure, Natalie Crawford, Stephanie Van Hooser, and Mark Pandori |
| EPI_ISL_515522 | UPA 24HS de Itatiba | Instituto Adolfo Lutz, Interdiciplinary Procedures Center, Strategic Laboratory | Claudio Tavares Sacchi, Claudia Regina Gonçalves, Erica Valessa Ramos Gomes |
| EPI_ISL_515524 | PS Municipal Dr Lauro Ribas Braga | Instituto Adolfo Lutz, Interdiciplinary Procedures Center, Strategic Laboratory | Claudio Tavares Sacchi, Claudia Regina Gonçalves, Erica Valessa Ramos Gomes |
| EPI_ISL_515543 | Serviço de Vigilância Sanitária e Epidemiológica | Instituto Adolfo Lutz, Interdiciplinary Procedures Center, Strategic Laboratory | Claudio Tavares Sacchi, Claudia Regina Gonçalves, Erica Valessa Ramos Gomes |
| EPI_ISL_516075 | BIMS | Department of Neurovirology, National Institute of Mental Health and Neuroscience (NIMHANS) | Chitra Pattabiraman,Vijayalakshmi Reddy, Harsha PK, Risha Rasheed, Pramada Prasad, Shafeeq S Hameed, Manjunatha Venkataswamy, Anita Desai, Ravi Vasanthapuram |
| EPI_ISL_516197, EPI_ISL_516226, EPI_ISL_516227, EPI_ISL_516228, EPI_ISL_516229, EPI_ISL_516230, EPI_ISL_516231, EPI_ISL_516232, EPI_ISL_516233, EPI_ISL_516234, EPI_ISL_516236, EPI_ISL_516237, EPI_ISL_516239, EPI_ISL_516240, EPI_ISL_516241, EPI_ISL_516365, EPI_ISL_516366, EPI_ISL_516367, EPI_ISL_516368, EPI_ISL_516369, EPI_ISL_516370, EPI_ISL_516371, EPI_ISL_516372, EPI_ISL_516373, EPI_ISL_516374, EPI_ISL_516375, EPI_ISL_516380 | | | |
| see above | Michigan Department of Health and Human Services, Bureau of Laboratories | Michigan Department of Health and Human Services, Bureau of Laboratories | Blankenship HM, Riner D, Soehnlen MK |
| EPI_ISL_516945 | King Georges Medical University | CSIR-National Botanical Research Institute | Priti Prasad, Shantanu Prakash, Kishan Sahu, Babita Singh, Suruchi Shukla, Hricha Mishra, Danish Nasar Khan , Om Prakash, MLB Bhatt, SK Barik, Mehar H.Asif,Samir V. Sawant,Amita Jain, Sumit Kr. Bag |
| EPI_ISL_517713 | Laboratorio de Referencia Nacional de Virus Respiratorio. Centro Nacional de Salud Publica. Instituto Nacional de Salud Peru. | Laboratorio de Referencia Nacional de Biotecnología y Biología Molecular. Centro Nacional de Salud Publica. Instituto Nacional de Salud Peru. | Carlos Padilla Rojas, Karolyn Vega Chozo, Priscila Lope Pari, Omar Caceres Rey, Marco Galarza Perez, Maribel Huaringa Nuñez, Johanna Balbuena Torres, Henri Bailon Calderon, Nancy Rojas Serrano. |
| EPI_ISL_520726, EPI_ISL_520727, EPI_ISL_520728 | Mohammed Bin Rashid University of Medicine and Health Sciences | Al Jalila Genomics Center | Ahmad Abou Tayoun, Tom Loney, Hamda Khansaheb, Sathishkumar Ramaswamy, Divinlal Harilal, Zulfa Omar Deesi, Rupa Murthy Varghese, Hanan Al Suwaidi, Abdulmajeed Alkhaja, Mohammed Uddin, Rifat Hamoudi, Rabih Halwani, Abiola Catherine Senok, Qutayba Hamid, Norbert Nowotny, Alawi Alsheikh-Ali |
| EPI_ISL_521867, EPI_ISL_521868 | Victorian Infectious Diseases Reference Laboratory (VIDRL) | VIDRL and MDU-PHL | Caly L., Seemann T., Sait, M., Schultz M., Druce J., Sherry, N. |
| EPI_ISL_521910 | Microbiological Diagnostic Unit - Public Health Laboratory | MDU-PHL | Seemann T., Schultz M., Sait, M., Sherry, N. |

| | | | |
|---|---|---|---|
| | (MDU-PHL) | | |
| EPI_ISL_522549 | Félix Guyon Hospital | UMR PIMIT Université de La Réunion | David Wilkinson, Camille Lebarbenchon, Patrick Mavingui |
| EPI_ISL_522577, EPI_ISL_522657, EPI_ISL_522761 | Royal Hobart Hospital Microbiology Department | MDU-PHL | Cooley L., van Haeften R., Seemann T., Sait M., Schultz, M.B., Sherry N. |
| EPI_ISL_523476 | Dutch COVID-19 response team | Erasmus Medical Center | Bas Oude Munnink, David Nieuwenhuijse, Reina Sikkema, Claudia Schapendonk, Irina Chestakova, Anne van der Linden, Theo Bestebroer, Stefan van Nieuwkoop, Mark Pronk, Pascal Lexmond, Corien Swaan, Manon Haverkate, Madelief Mollers, Mart Stein, Sandra Kengne Kamga Mobou, Jeroen van Kampen, Jolanda Voermans, Aura Timen, Corine GeurtsvanKessel, Annemiek van der Eijk, Richard Molenkamp, Marion Koopmans, on behalf of the Dutch national COVID-19 response team. |
| EPI_ISL_523936, EPI_ISL_523945, EPI_ISL_523946 | Center of Medical Microbiology, Virology, and Hospital Hygiene, University of Duesseldorf | Center of Medical Microbiology, Virology, and Hospital Hygiene, University of Duesseldorf | Maximilian Damagnez, Alexander Dilthey, Torsten Houwaart, Malte Kohns Vasconcelos, Marek Korencak, Jessica Nicolai, Klaus Pfeffer, Hendrik Streeck, Daniel Strelow, Jörg Timm, Andreas Walker, Tobias Wienemann |
| EPI_ISL_523956 | Hospital Regional de Assis | Instituto Adolfo Lutz, Interdiciplinary Procedures Center, Strategic Laboratory | Claudio Tavares Sacchi, Claudia Regina Gonçalves, Erica Valessa Ramos Gomes |
| EPI_ISL_523969 | Hospital Sao Paulo de Ensino da Unifesp | Instituto Adolfo Lutz, Interdiciplinary Procedures Center, Strategic Laboratory | Claudio Tavares Sacchi, Claudia Regina Gonçalves, Erica Valessa Ramos Gomes |
| EPI_ISL_524450, EPI_ISL_524451, EPI_ISL_524452, EPI_ISL_524453, EPI_ISL_524454 | Microbiology & Immunology, University of North Carolina | Microbiology & Immunology, University of North Carolina | Bailey,A.G., Caro-Vegas,C.P., Dittmer,D., Eason,A.B., Juarez,A., Landis,J.T., McNamara,R.P., Miller,M.B., Moorad,R., Pluta,L.J., Seltzer,T.A., Thompson,C., Vahrson,W., Villamor,F. |
| EPI_ISL_524473 | Laboratorio de Referencia Nacional de Virus Respiratorio. Centro Nacional de Salud Publica. Instituto Nacional de Salud Peru. | Laboratorio de Referencia Nacional de Biotecnología y Biología Molecular. Centro Nacional de Salud Publica. Instituto Nacional de Salud Peru. | Carlos Padilla Rojas, Karolyn Vega Chozo, Priscila Lope Pari, Omar Caceres Rey, Marco Galarza Perez, Maribel Huaringa Nuñez, Johanna Balbuena Torres, Henri Bailon Calderon, Nancy Rojas Serrano. |
| EPI_ISL_525472 | Institute of Clinical Microbiology and Hygiene, University Hospital Regensburg | Institute of Clinical Microbiology and Hygiene, University Hospital Regensburg | Hiergeist, A. |
| EPI_ISL_525479 | Centre for Dengue Research | Centre for Dengue Research | Chandima Jeewandara, Deshni Jayathilaka, Dinuka Ariyaratne, Laksiri Gomes, Diyanath Ranasinghe, Ananda Wijewickrama, Eranga Narangoda, Damayanthi Idampitiya, Gathsaurie Neelika Malavige |
| EPI_ISL_525643, EPI_ISL_525644, EPI_ISL_525645, EPI_ISL_525646, EPI_ISL_525647, EPI_ISL_525656 | Wadsworth Center, New York State Department of Health | Wadsworth Center, New York State Department of Health | Kirsten St. George, Daryl M. Lamson, Sara Griesemer, Jonathan Plitnick, Navjot Singh, Matthew D. Shudt, Erica Lasek-Nesselquist |
| EPI_ISL_525772, EPI_ISL_525773, EPI_ISL_525774, EPI_ISL_525775, EPI_ISL_525776, EPI_ISL_525777, EPI_ISL_525778 | Texas Department of State Health Services | Texas Department of State Health Services | Jenny Zhang, Rashmi Tuladhar, Bonnie Oh, Maliha Rahman, Anita Pokharel, Myong Koag, Chun Wang, Rachel Lee, Grace Kubin |
| EPI_ISL_526451, EPI_ISL_526452, EPI_ISL_526453, EPI_ISL_526454, EPI_ISL_526455, EPI_ISL_526456 | West of Scotland Specialist Virology Centre, NHSGGC / MRC-University of Glasgow Centre for Virus Research | COVID-19 Genomics UK (COG-UK) Consortium | Ana da Silva Filipe, Natasha Johnson, Kathy Smollett, Daniel Mair, Stephen Carmichael, Lily Tong, Jenna Nichols, Elihu Aranday-Cortes, Kirstyn Brunker, Yasmin Parr, Alice Broos, Kyriaki Nomikou; Sarah McDonald, Marc Niebel, Patawee Asamaphan; Richard Orton, Joseph Hughes, Sreenu Vattipally, David L Robertson; Alasdair MacLean, Rory Gunson; Kathy Li, Natasha Jesudason, Rajiv Shah, James Shepherd, Antonia Ho, Emma Thomson |
| EPI_ISL_526470 | Virology Department, Royal Infirmary of Edinburgh, NHS Lothian / School of Biological Sciences, University of Edinburgh / Institute of Genetics and Molecular Medicine, University of Edinburgh | COVID-19 Genomics UK (COG-UK) Consortium | McHugh M, Dewar R, Rooke S, Gallagher M, Balcaza C, O'Toole Á, Scher E, Hill V, McCrone JT, Colquhoun R, Yu X, Jackson B, Rambaut A, Williams TC, Templeton K |
| EPI_ISL_526827, EPI_ISL_526828, EPI_ISL_526829, EPI_ISL_526830, EPI_ISL_526831, EPI_ISL_526832, EPI_ISL_526833, EPI_ISL_526834, EPI_ISL_526835 | Virginia DCLS | Virginia DCLS | Virginia DCLS |
| EPI_ISL_527878 | Nigeria Centre for Disease Control (NCDC) | African Centre of Excellence for Genomics of Infectious Diseases (ACEGID), Redeemer's University, Ede, Osun State, Nigeria | Oluniyi P.E. et al |
| EPI_ISL_528636 | National Genomics Core-Center for DNA Fingerprinting and Diagnostics | National Genomics Core- Center for DNA Fingerprinting and Diagnostics (NGC-CDFD)- DBT's PAN-INDIA-1000 Genome consortium | Heena Shah, G Shashikanth, Bala Pratyusha, Vinay Donipadi, K.Manohar, Muthumohan Rao, Shruti Dasgupta, Kandali Sreethi Sreenivasulu Reddy, Chandra Shekhar Singh, Sunke Vijayakumar, R Lakshmi Vaishna, Jenige Aravindh Kumar, Muthulakshmi, V Naga Sailaja, R Harinarayanan, Rashna Bhandari, Murali Dharan Bashyam, Debashish Mitra, Divya Vashisht, Ashwin Dalal |
| EPI_ISL_528990 | Ospedale Civile Maria SS. dello Splendore | Istitituto Zooprofilattico Sperimentale dell'Abruzzo e Molise "G.Caporale" | Lorusso A, Marcacci M, Di Domenico M, Curini V, Ancora M, Cammà C, Rinaldi A, Mangone I, Di Pasquale A, Puglia I, Savini G. |
| EPI_ISL_529009 | Ospedale Civile S. Liberatore-Atri | Istitituto Zooprofilattico Sperimentale dell'Abruzzo e Molise "G.Caporale" | Lorusso A, Marcacci M, Di Domenico M, Curini V, Ancora M, Cammà C, Rinaldi A, Mangone I, Di Pasquale A, Puglia I, Savini G. |
| EPI_ISL_529162 | Department of Immunology, The Scripps Research Institute | Andersen lab at Scripps Research | Allison Smither, Gilberto Sabino-Santos, Patricia Snarski, Lilia Melnik, Antoinette Bell, Kaylynn Genemaras, Arnaud Drouin, Dahlene Fusco, Robert Garry with SEARCH Alliance San Diego |
| EPI_ISL_529176 | South Carolina Department of Health and Environmental Control | South Carolina Department of Health and Environmental Control | Haley V. Flores |
| EPI_ISL_529972, EPI_ISL_529973 | Hospital Universitario 12 de Octubre | Hospital Universitario 12 de Octubre | Raúl Recio, Sara González, Esther Viedma, Elias Dahdouh, Fernando Lázaro, Natalia Stella, Julio García, Juan Carlos Galán, Rafael Cantón, Mª Dolores Folgueira, Rafael Delgado, Jesús Mingorance |
| EPI_ISL_530005, EPI_ISL_530006 | Hospital Universitario 12 de Octubre | Hospital Universitario 12 de Octubre | Sara González, Esther Viedma, Raúl Recio, Elias Dahdouh, Fernando Lázaro, Natalia Stella, Julio García, Juan Carlos Galán, Rafael Cantón, Mª Dolores Folgueira, Rafael Delgado, Jesús Mingorance |
| EPI_ISL_530220, EPI_ISL_530222, EPI_ISL_530223 | Minnesota Department of Health, Public Health Laboratory | Minnesota Department of Health, Public Health Laboratory | Matt Plumb, Jacob Garfin, and Xiong Wang |
| EPI_ISL_530226, EPI_ISL_530227, EPI_ISL_530228, EPI_ISL_530229, EPI_ISL_530230 | Queensland Health Forensic and Scientific Services, Public Health Virology | Public Health Virology Laboratory, Forensic and Scientific Services, Queensland Health | Son Nguyen et al |
| EPI_ISL_534346, EPI_ISL_534347, EPI_ISL_534348, EPI_ISL_534349 | Molecular diagnostic laboratory of Federal Budget Institution of Science "Central Research Institute of Epidemiology" of The Federal Service on Customers' Rights Protection and Human Well-being Surveillance | Group of Genomics and Postgenomic Technologies of Central Research Institute of Epidemilology | Speranskaya AS, Kaptelova VV, Valdokhina AV, Bulanenko VP, Samoilov AE, Korneenko EV, Tivanova EV, Shipulina OY, Akimkin VG |
| EPI_ISL_534817, EPI_ISL_534821, EPI_ISL_534824, EPI_ISL_534841, EPI_ISL_534852, EPI_ISL_534868, EPI_ISL_534883, EPI_ISL_534884, EPI_ISL_534899, EPI_ISL_534904, EPI_ISL_534918, EPI_ISL_534924, EPI_ISL_534948, EPI_ISL_534954, EPI_ISL_534955, EPI_ISL_534971, EPI_ISL_534980, EPI_ISL_534991, EPI_ISL_534999, EPI_ISL_535001, EPI_ISL_535006, EPI_ISL_535010, EPI_ISL_535016 | | | |
| see above | Oxford Viromics, NDM, University of Oxford; Oxford University | COVID-19 Genomics UK (COG-UK) Consortium | Tanya Golubchik, David Bonsall, George Macintyre, Amy Trebes, Mariateresa de Cesare, Catrin Moore, Alex Mobbs, Anita Justice, Robert Shaw, Monique |

| | | | |
|---|---|---|---|
| | Hospitals; Basingstoke and North Hampshire Hospital | | Andersson, Timothy Peto, Emma Wise, Nathan Moore, Jessica Lynch, Nick Cortes, Matilde Mori, Stephen Kidd, David Buck, John Todd, Christophe Fraser |
| EPI_ISL_536368, EPI_ISL_536369, EPI_ISL_536370 | Centre hospitalier Anna-Laberge | Laboratoire de santé publique du Québec | Sandrine Moreira, Ioannis Ragoussis, Guillaume Bourque, Jesse Shapiro, Mark Lathrop and Michel Roger |
| EPI_ISL_536371, EPI_ISL_536372, EPI_ISL_536373 | Hôpital Charles-LeMoyne | Laboratoire de santé publique du Québec | Sandrine Moreira, Ioannis Ragoussis, Guillaume Bourque, Jesse Shapiro, Mark Lathrop and Michel Roger |
| EPI_ISL_536374 | Hôpital Pierre-Boucher | Laboratoire de santé publique du Québec | Sandrine Moreira, Ioannis Ragoussis, Guillaume Bourque, Jesse Shapiro, Mark Lathrop and Michel Roger |
| EPI_ISL_536376, EPI_ISL_536377, EPI_ISL_536378 | Centre hospitalier Anna-Laberge | Laboratoire de santé publique du Québec | Sandrine Moreira, Ioannis Ragoussis, Guillaume Bourque, Jesse Shapiro, Mark Lathrop and Michel Roger |
| EPI_ISL_536379 | Hôpital Honoré-Mercier | Laboratoire de santé publique du Québec | Sandrine Moreira, Ioannis Ragoussis, Guillaume Bourque, Jesse Shapiro, Mark Lathrop and Michel Roger |
| EPI_ISL_536380 | Hôpital Pierre-Boucher | Laboratoire de santé publique du Québec | Sandrine Moreira, Ioannis Ragoussis, Guillaume Bourque, Jesse Shapiro, Mark Lathrop and Michel Roger |
| EPI_ISL_536381 | Centre Hospitalier Régional de Lanaudière | Laboratoire de santé publique du Québec | Sandrine Moreira, Ioannis Ragoussis, Guillaume Bourque, Jesse Shapiro, Mark Lathrop and Michel Roger |
| EPI_ISL_536382, EPI_ISL_536383 | CSSS Haut-Richelieu/Rouville (Hôpital) | Laboratoire de santé publique du Québec | Sandrine Moreira, Ioannis Ragoussis, Guillaume Bourque, Jesse Shapiro, Mark Lathrop and Michel Roger |
| EPI_ISL_536384 | Hôpital de Gatineau | Laboratoire de santé publique du Québec | Sandrine Moreira, Ioannis Ragoussis, Guillaume Bourque, Jesse Shapiro, Mark Lathrop and Michel Roger |
| EPI_ISL_536385, EPI_ISL_536386, EPI_ISL_536387 | Hôpital de Hull | Laboratoire de santé publique du Québec | Sandrine Moreira, Ioannis Ragoussis, Guillaume Bourque, Jesse Shapiro, Mark Lathrop and Michel Roger |
| EPI_ISL_536388, EPI_ISL_536389 | Hôpital Pierre-Boucher | Laboratoire de santé publique du Québec | Sandrine Moreira, Ioannis Ragoussis, Guillaume Bourque, Jesse Shapiro, Mark Lathrop and Michel Roger |
| EPI_ISL_536390 | Hôpital du Suroît | Laboratoire de santé publique du Québec | Sandrine Moreira, Ioannis Ragoussis, Guillaume Bourque, Jesse Shapiro, Mark Lathrop and Michel Roger |
| EPI_ISL_536391 | Hôpital Pierre-Boucher | Laboratoire de santé publique du Québec | Sandrine Moreira, Ioannis Ragoussis, Guillaume Bourque, Jesse Shapiro, Mark Lathrop and Michel Roger |
| EPI_ISL_536392, EPI_ISL_536393, EPI_ISL_536394 | Hôpital de Hull | Laboratoire de santé publique du Québec | Sandrine Moreira, Ioannis Ragoussis, Guillaume Bourque, Jesse Shapiro, Mark Lathrop and Michel Roger |
| EPI_ISL_536395 | Hôpital Pierre-Boucher | Laboratoire de santé publique du Québec | Sandrine Moreira, Ioannis Ragoussis, Guillaume Bourque, Jesse Shapiro, Mark Lathrop and Michel Roger |
| EPI_ISL_536396 | Hôpital de Hull | Laboratoire de santé publique du Québec | Sandrine Moreira, Ioannis Ragoussis, Guillaume Bourque, Jesse Shapiro, Mark Lathrop and Michel Roger |
| EPI_ISL_536397 | Hôpital Pierre-Boucher | Laboratoire de santé publique du Québec | Sandrine Moreira, Ioannis Ragoussis, Guillaume Bourque, Jesse Shapiro, Mark Lathrop and Michel Roger |
| EPI_ISL_536523, EPI_ISL_536524, EPI_ISL_536525, EPI_ISL_536526, EPI_ISL_536528, EPI_ISL_536529, EPI_ISL_536530, EPI_ISL_536562, EPI_ISL_536563, EPI_ISL_536564 | Instituto Nacional de Salud | Laboratorio de Infecciones Respiratorias Agudas | Eduardo Juscamayta Lopez, David Tarazona, Faviola Valdivia Guerrero, Nancy Rojas Serrano, Dennis Carhuaricra, Lenin Maturrano Hernandez, Ronnie Gavilan Chavez |
| EPI_ISL_537288, EPI_ISL_537289, EPI_ISL_537290, EPI_ISL_537291, EPI_ISL_537292, EPI_ISL_537293, EPI_ISL_537294, EPI_ISL_537295 | Universidad de León | SeqCOVID-SPAIN consortium/IBV(CSIC) | Ana Carvajal, Vicente Martín, Héctor Argüello, Juan M. Fregeneda, Tania Fernández-Villa, Antonio J. Molina and SeqCOVID-SPAIN consortium |
| EPI_ISL_537393, EPI_ISL_537394, EPI_ISL_537401, EPI_ISL_537438 | Centro de Investigación Biomédica de La Rioja - Hospital San Pedro Logroño | SeqCOVID-SPAIN consortium/IBV(CSIC) | María de Toro, José Manuel Azcona Gutiérrez, María Pilar Bea Escudero, Miriam Blasco Alberdi and SeqCOVID-SPAIN consortium |
| EPI_ISL_537496, EPI_ISL_537497, EPI_ISL_537573, EPI_ISL_537574, EPI_ISL_537591, EPI_ISL_537592, EPI_ISL_537593 | UCLA Pathology Clinical Microbiology Lab | Kruglyak Lab | Guo et al. |
| EPI_ISL_537719 | Hospital Universitario de Gran Canaria Dr. Negrín | SeqCOVID-SPAIN consortium/IBV(CSIC) | M. Carmen Pérez González, Francisco J. Chamizo López, Ana Bordes Benítez and SeqCOVID-SPAIN consortium |
| EPI_ISL_537857, EPI_ISL_537858, EPI_ISL_537869, EPI_ISL_537870 | Centro de Investigación Biomédica de La Rioja - Hospital San Pedro Logroño | SeqCOVID-SPAIN consortium/IBV(CSIC) | María de Toro, José Manuel Azcona Gutiérrez, María Pilar Bea Escudero, Miriam Blasco Alberdi and SeqCOVID-SPAIN consortium |
| EPI_ISL_537994, EPI_ISL_537995 | Servicio de Microbiología. Hospital General Universitario de Castellón | SeqCOVID-SPAIN consortium/IBV(CSIC) | Rosario Moreno, María Dolores Tirado and SeqCOVID-SPAIN consortium |
| EPI_ISL_538070, EPI_ISL_538071, EPI_ISL_538072, EPI_ISL_538073, EPI_ISL_538074, EPI_ISL_538075, EPI_ISL_538076, EPI_ISL_538077, EPI_ISL_538078, EPI_ISL_538079, EPI_ISL_538080, EPI_ISL_538081 | | | |
| see above | Servicio de Microbiología. Hospital Universitario Donostia. OSI Donostialdea. Área de Enfermedades Infecciosas, Grupo de Infección Respiratoria y Resistencia Antimicrobiana. Instituto de Investigación Sanitaria Biodonostia | SeqCOVID-SPAIN consortium/IBV(CSIC) | Gustavo Cilla, Milagrosa Montes, Luis Piñeiro, Jose Maria Marimón and SeqCOVID-SPAIN consortium |
| EPI_ISL_538158, EPI_ISL_538159, EPI_ISL_538160, EPI_ISL_538163, EPI_ISL_538166, EPI_ISL_538167, EPI_ISL_538168, EPI_ISL_538169, EPI_ISL_538170, EPI_ISL_538171 | Servicio de Microbiología y Parasitología clínica. UCEIMP. Hospital Universitario Virgen del Rocío/IBIS/CSIC/US | SeqCOVID-SPAIN consortium/IBV(CSIC) | Guillermo Martín Gutiérrez, Ángel Rodríguez Villodres, Lidia Gálvez Benitez, Verónica González Galán, Javier Aznar Martín and SeqCOVID-SPAIN consortium |
| EPI_ISL_538336, EPI_ISL_538337, EPI_ISL_538338, EPI_ISL_538339, EPI_ISL_538340, EPI_ISL_538341, EPI_ISL_538342, EPI_ISL_538343 | Kingston Health Sciences Centre / Queen's University | Ontario Institute for Cancer Research | Prameet M. Sheth, Calvin Sjaarda, Robert Colautti, Katya Douchant, Ilinca Lungu, Bernard Lam, Paul Krzyzanowski, Michael Laszloffy, Lawrence E Heisler, Richard de Borja, Jared T. Simpson |
| EPI_ISL_538465, EPI_ISL_538466, EPI_ISL_538467, EPI_ISL_538468 | Department of Laboratory Medicine, Tan Tock Seng Hospital | Department of Laboratory Medicine, Tan Tock Seng Hospital | Chen YYC, Zair X, Lim JX, Li C, Tang WY, Maurer-Stroh S, Barkham TMS, Nagarajan N, Sessions OM |
| EPI_ISL_538613, EPI_ISL_538615, EPI_ISL_538616 | Servicio de Microbiología. Hospital General Universitario de Castellón | SeqCOVID-SPAIN consortium/IBV(CSIC) | Rosario Moreno, María Dolores Tirado and SeqCOVID-SPAIN consortium |
| EPI_ISL_538688 | Hospital Universitario Virgen de las Nieves de Granada-SAS | SeqCOVID-SPAIN consortium/IBV(CSIC) | Mercedes Pérez Ruiz, Sara Sanbonmatsu Gámez, Irene Pedrosa Corral, José M. Navarro-Marí and SeqCOVID-SPAIN consortium |
| EPI_ISL_538791, EPI_ISL_538816, EPI_ISL_538820, EPI_ISL_538832, EPI_ISL_538844, EPI_ISL_538847, EPI_ISL_538884, EPI_ISL_538901, EPI_ISL_538979, EPI_ISL_538983, EPI_ISL_538986, EPI_ISL_538988, EPI_ISL_538989, EPI_ISL_538990, EPI_ISL_539002, EPI_ISL_539004, EPI_ISL_539009, EPI_ISL_539023, EPI_ISL_539034, EPI_ISL_539047, EPI_ISL_539051, EPI_ISL_539068, EPI_ISL_539069, EPI_ISL_539072, EPI_ISL_539088, EPI_ISL_539090, EPI_ISL_539095, EPI_ISL_539099, EPI_ISL_539111, EPI_ISL_539112, EPI_ISL_539118, EPI_ISL_539119, EPI_ISL_539134, EPI_ISL_539137, EPI_ISL_539139 | | | |
| see above | Leeds Teaching Hospitals NHS Trust and Public Health England, National Infection Service (Leeds laboratory) | Wellcome Sanger Institute for the COVID-19 Genomics UK (COG-UK) consortium | Louissa Macfarlane-Smith, Holli Carden, Katherine L. Harper, Antony Hale and Alex Alderton, Roberto Amato, Sonia Goncalves, Ewan Harrison, David K. Jackson, Ian Johnston, Dominic Kwiatkowski, Cordelia Langford, John Sillitoe on behalf of the Wellcome Sanger Institute COVID-19 Surveillance Team |
| EPI_ISL_539152 | Leeds Teaching Hospitals NHS Trust and Public Health England, National Infection Service (Leeds laboratory) | Wellcome Sanger Institute for the COVID-19 Genomics UK (COG-UK) Consortium | Louissa Macfarlane-Smith, Holli Carden, Katherine L. Harper, Antony Hale and Alex Alderton, Roberto Amato, Sonia Goncalves, Ewan Harrison, David K. Jackson, Ian Johnston, Dominic Kwiatkowski, Cordelia Langford, John Sillitoe on behalf of the Wellcome Sanger Institute COVID-19 Surveillance Team |
| EPI_ISL_539163, EPI_ISL_539171, EPI_ISL_539176, EPI_ISL_539194, | Leeds Teaching Hospitals NHS Trust and Public Health England, National Infection Service (Leeds laboratory) | Wellcome Sanger Institute for the COVID-19 Genomics UK (COG-UK) consortium | Louissa Macfarlane-Smith, Holli Carden, Katherine L. Harper, Antony Hale and Alex Alderton, Roberto Amato, Sonia Goncalves, Ewan Harrison, David K. Jackson, Ian Johnston, Dominic Kwiatkowski, Cordelia Langford, John Sillitoe on behalf of the Wellcome Sanger Institute COVID-19 Surveillance Team |

| | | | |
|---|---|---|---|
| EPI_ISL_539203, EPI_ISL_539219 | | | |
| EPI_ISL_539246, EPI_ISL_539247, EPI_ISL_539248 | Hospital Universitario de La Ribera (Alzira, València) | SeqCOVID-SPAIN consortium/IBV(CSIC) | Olalla Martínez Macias, Julia González and SeqCOVID-SPAIN consortium |
| EPI_ISL_539306, EPI_ISL_539308, EPI_ISL_539309, EPI_ISL_539317, EPI_ISL_539318, EPI_ISL_539319, EPI_ISL_539320 | KWR Watercycle Research Institute | Erasmus Medical Center | Ray Izquierdo-Lara, Goffe Elsinga, Leo Heijnen, Bas B. Oude Munnink, Claudia M. E. Schapendonk, David Nieuwenhuijse, Matthijs Kon, Lu Lu, Frank M. Aarestrup, Samantha Lycett, Gertjan Medema, Marion P.G. Koopmans, Miranda de Graaf |
| EPI_ISL_539494, EPI_ISL_539777 | The National Institute of Public Health | State Veterinary Institute Prague | Nagy,A;Jirincova,H;Novakova,L;Trnka,D;Vecerova,J |
| EPI_ISL_541037 | Hospital Clínico Universitario de Santiago de Compostela | SeqCOVID-SPAIN consortium/Institute of Biomedicine of Valencia, IBV-CSIC | José Javier Costa Alcalde, Antonio Aguilera Guirao, Mª Luisa Pérez del Molino Bernal, Amparo Coira Nieto, Gema Barbeito Castiñeiras, Rocio Trastoy Pena and SeqCOVID-SPAIN consortium |
| EPI_ISL_541761, EPI_ISL_541770 | Barts Health NHS Trust | Wellcome Sanger Institute for the COVID-19 Genomics UK (COG-UK) consortium | Teresa Cutino-Moguel, Mark Hopkins, Beatrix Kele, David Harrington and Alex Alderton, Roberto Amato, Sonia Goncalves, Ewan Harrison, David K. Jackson, Ian Johnston, Dominic Kwiatkowski, Cordelia Langford, John Sillitoe on behalf of the Wellcome Sanger Institute COVID-19 Surveillance Team |
| EPI_ISL_541953 | Servicio de Microbiología, Hospital Universitario Son Espases | SeqCOVID-SPAIN consortium/IBV(CSIC) | Carla López-Causapé, Jordi Reina, Antonio Oliver and SeqCOVID-SPAIN consortium |
| EPI_ISL_542389, EPI_ISL_542390, EPI_ISL_542392 | San Matteo Hospital Pavia | Dep. Of Oncology and Hemato-Oncology University of Milan | Claudia Alteri, Valeria Cento, Antonio Piralla, Valentino Costabile, Monica Tallarita, Luna Colagrossi, Silvia Renica, Federica Giardina, Federica Novazzi, Stefano Gaiarsa, Elisa Matarazzo, Maria Antonello, Chiara Vismara, Roberto Fumagalli, Oscar Massimiliano Epis, Massimo Puoti, Carlo Federico Perno, Fausto Baldanti |
| EPI_ISL_542571, EPI_ISL_542573, EPI_ISL_542581, EPI_ISL_542582, EPI_ISL_542585, EPI_ISL_542587, EPI_ISL_542590, EPI_ISL_542599, EPI_ISL_542610, EPI_ISL_542692, EPI_ISL_542801, EPI_ISL_544456, EPI_ISL_544465, EPI_ISL_544471, EPI_ISL_544479, EPI_ISL_545240, EPI_ISL_545867, EPI_ISL_545882 | | | |
| see above | Houston Methodist Hospital | Houston Methodist Hospital | S. Wesley Long, Randall J. Olsen, Paul A. Christensen, David W. Bernard, James J. Davis, Maulik Shukla, Marcus Nguyen, Matthew Ojeda Saavedra, Concepcion C. Cantu, Prasanti Yerramilli, Layne Pruitt, Sishir Subedi, Hung-Che Kuo, Heather Hendrickson, Ghazaleh Eskandari, Hoang A. T. Nguyen, J. Hunter Long, Muthiah Kumaraswami, Jule Goike, Daniel Boutz, Jimmy Gollihar, Jason S. McLellan, Chia-Wei Chou, Kamyab Javanmardi, Ilya J. Finkelstein, and James M. Musser |
| EPI_ISL_545955 | Laboratorio de Infecciones Respiratorias Agudas. Centro Nacional de Salud Publica,Instituto Nacional de Salud | Laboratorio de Infecciones Respiratorias Agudas. Centro Nacional de Salud Publica,Instituto Nacional de Salud | Juscamayta,E. |
| EPI_ISL_546738 | Houston Methodist Hospital | Houston Methodist Hospital | S. Wesley Long, Randall J. Olsen, Paul A. Christensen, David W. Bernard, James J. Davis, Maulik Shukla, Marcus Nguyen, Matthew Ojeda Saavedra, Concepcion C. Cantu, Prasanti Yerramilli, Layne Pruitt, Sishir Subedi, Hung-Che Kuo, Heather Hendrickson, Ghazaleh Eskandari, Hoang A. T. Nguyen, J. Hunter Long, Muthiah Kumaraswami, Jule Goike, Daniel Boutz, Jimmy Gollihar, Jason S. McLellan, Chia-Wei Chou, Kamyab Javanmardi, Ilya J. Finkelstein, and James M. Musser |
| EPI_ISL_547494, EPI_ISL_547505, EPI_ISL_547534 | Dutch COVID-19 response team | National Institute for Public Health and the Environment (RIVM) | Adam Meijer, Harry Vennema, Jeroen Cremer, Sharon van den Brink, Bas van der Veer, AnneMarie van den Brandt, Florian Zwagemaker, Dennis Schmitz, Chantal Reusken, on behalf of the national COVID-19 response team |
| EPI_ISL_548364 | Ventura County Public Health Lab | Chan-Zuckerberg Biohub | CZB Cliahub Consortium |
| EPI_ISL_548781, EPI_ISL_548931 | Public Health Ontario Laboratory | Public Health Ontario Laboratory | Vanessa G Allen, Philip Banh, Richard de Borja, Yao Chen, Alireza Eshaghi, Nahuel Fittipaldi, Christine Frantz, Jonathan B Gubbay, Jennifer L Guthrie, Lawrence Heisler, Esha Joshi, Michael Laszloffy, Aimin Li, Michael CY Li, Dean Maxwell, Sandeep Nagra, Samir N Patel, Heather Rilkoff, Jared Simpson, Karthikeyan Sivaraman, Yogi Sundaravadanam, Sarah Teatero, Andre Villegas, Sandra Zittermann |
| EPI_ISL_549016, EPI_ISL_549021 | KWR Watercycle Research Institute | Erasmus Medical Center | Ray Izquierdo-Lara, Goffe Elsinga, Leo Heijnen, Bas B. Oude Munnink, Claudia M. E. Schapendonk, David Nieuwenhuijse, Matthijs Kon, Lu Lu, Frank M. Aarestrup, Samantha Lycett, Gertjan Medema, Marion P.G. Koopmans, Miranda de Graaf |
| EPI_ISL_549336, EPI_ISL_549349, EPI_ISL_549353, EPI_ISL_549368, EPI_ISL_549369, EPI_ISL_549370, EPI_ISL_549373, EPI_ISL_549377, EPI_ISL_549378, EPI_ISL_549379, EPI_ISL_549383, EPI_ISL_549477, EPI_ISL_549482, EPI_ISL_549483, EPI_ISL_549484, EPI_ISL_549485, EPI_ISL_549489, EPI_ISL_549490 | | | |
| see above | Quadram Institute Bioscience | COVID-19 Genomics UK (COG-UK) Consortium | Dave J. Baker, Gemma L. Kay, Alp Aydin, Thanh Le-Viet, Steven Rudder, Ana P. Tedim, Anastasia Kolyva, Maria Diaz, Leonardo de Oliveira Martins, Nabil-Fareed Alikhan, Lizzie Meadows, Rachael Stanley, Ngozi Elumogo, Muhammed Yasir, Nicholas M. Thomson, Alexander J Trotter, Rachel Gilroy, Samuel Bloomfield, Claire Stuart, Andrew Bell, Reenesh Prakash, Samir Dervisevic, Alison E. Mather, John Wain, Mark Webber, Andrew J. Page, Justin O'Grady |
| EPI_ISL_560569 | hôpital | National Reference Center for Viruses of Respiratory Infections, Institut Pasteur, Paris | Sylvie Behillil, Fabiana Gambaro, Etienne Simon-Lorière, Vincent Enouf, Maud Vanpeene, Sylvie van der Werf |
| EPI_ISL_560582, EPI_ISL_560583 | Hopital | National Reference Center for Viruses of Respiratory Infections, Institut Pasteur, Paris | Sylvie Behillil, Fabiana Gambaro, Etienne Simon-Lorière, Vincent Enouf, Maud Vanpeene, Sylvie van der Werf |
| EPI_ISL_560596 | hopital | National Reference Center for Viruses of Respiratory Infections, Institut Pasteur, Paris | Sylvie Behillil, Fabiana Gambaro, Etienne Simon-Lorière, Vincent Enouf, Maud Vanpeene, Sylvie van der Werf |
| EPI_ISL_560739 | Centre for Clinical Infection and Diagnostics Research and Genomics Innovation Unit, Guy's and St. Thomas' NHS Trust | Centre for Clinical Infection and Diagnostics Research and Genomics Innovation Unit, Guy's and St. Thomas' NHS Trust | Chloe Fisher, Luke Snell, Rahul Batra, Jonathan Edgeworth, Ali Raza Awan |
| EPI_ISL_560743, EPI_ISL_560744, EPI_ISL_560745 | Minnesota Department of Health, Public Health Laboratory | Minnesota Department of Health, Public Health Laboratory | Matt Plumb, Jacob Garfin, and Xiong Wang |
| EPI_ISL_560841, EPI_ISL_560847, EPI_ISL_560861 | Utah Public Health Laboratory | Utah Public Health Laboratory | Erin Young, Kelly Oakeson |
| EPI_ISL_562588 | Victorian Infectious Diseases Reference Laboratory (VIDRL) | VIDRL and MDU-PHL | Caly, L., Seemann, T., Sait, M., Schultz, M. B., Druce J., Sherry, N. |
| EPI_ISL_565930, EPI_ISL_565931, EPI_ISL_565953, EPI_ISL_565973, EPI_ISL_565974, EPI_ISL_565978, EPI_ISL_565997, EPI_ISL_566002, EPI_ISL_566004, EPI_ISL_566011 | Michigan Department of Health and Human Services, Bureau of Laboratories | Michigan Department of Health and Human Services, Bureau of Laboratories | Blankenship HM, Riner D, Soehnlen MK |
| EPI_ISL_568558 | Department of Infectious Diseases and Immunology, National Hospital Organization Nagoya Medical Center | Clinical Research Center, National Hospital Organization Nagoya Medical Center | Yoshihiro Nakata, Hirotaka Ode, Mai Kubota, Masakazu Matsuda, Kazuhiro Matsuoka, Nakasuji Miho, Mikiko Mori, Mayumi Imahashi, Yoshiyuki Yokomaku, Yasumasa Iwatani |
| EPI_ISL_569418, EPI_ISL_569419, EPI_ISL_569420, EPI_ISL_569423, EPI_ISL_569424, EPI_ISL_569425, EPI_ISL_569426, EPI_ISL_569427, EPI_ISL_569428, EPI_ISL_569429, EPI_ISL_569430, EPI_ISL_569431, EPI_ISL_569432, EPI_ISL_569433, EPI_ISL_569434, EPI_ISL_569435, EPI_ISL_569436, EPI_ISL_569437, EPI_ISL_569438, EPI_ISL_569439, EPI_ISL_569440, EPI_ISL_569441, EPI_ISL_569442, EPI_ISL_569443, EPI_ISL_569444, EPI_ISL_569445, EPI_ISL_569446, EPI_ISL_569447, EPI_ISL_569448, EPI_ISL_569449, EPI_ISL_569451, EPI_ISL_569452, EPI_ISL_569453, EPI_ISL_569454, EPI_ISL_569455, EPI_ISL_569456, EPI_ISL_569457, EPI_ISL_569458, EPI_ISL_569459, EPI_ISL_569460, EPI_ISL_569461, EPI_ISL_569462, EPI_ISL_569463, EPI_ISL_569464 | | | |
| see above | MEPHI, Aix Marseille University | MEPHI, Aix Marseille University | Anthony LEVASSEUR |
| EPI_ISL_569864, EPI_ISL_569868, EPI_ISL_569869, EPI_ISL_569870, EPI_ISL_569871, EPI_ISL_569872, EPI_ISL_569873, EPI_ISL_569874, EPI_ISL_569875, EPI_ISL_569876, EPI_ISL_569877, EPI_ISL_569878, EPI_ISL_569879, EPI_ISL_569880, EPI_ISL_569881, EPI_ISL_569882 | | | |
| see above | Amedeo di savoia | Crosetto lab, Karolinska Institutet, SciLifeLab | Michele Simonetti, Maria Grazia Milia, Luuk Harbers, Ning Zhang, Anna Sapino, Valeria Ghisetti, Nicola Crosetto |
| EPI_ISL_570045, EPI_ISL_570046, EPI_ISL_570051, EPI_ISL_570082, EPI_ISL_570083, EPI_ISL_570084, EPI_ISL_570085, EPI_ISL_570086, EPI_ISL_570087, EPI_ISL_570088, EPI_ISL_570089, EPI_ISL_570092, EPI_ISL_570093, EPI_ISL_570094, EPI_ISL_570095, EPI_ISL_570099, EPI_ISL_570100, EPI_ISL_570104, EPI_ISL_570105, EPI_ISL_570106, EPI_ISL_570107, EPI_ISL_570109, EPI_ISL_570110 | | | |
| see above | UW Virology Lab | UW Virology Lab | Pavitra Roychoudhury, Hong Xie, Lasata Shrestha, Amin Addetia, Victoria M Rachleff, Meei-Li Huang, Keith R Jerome, Alexander Greninger |
| EPI_ISL_574541, EPI_ISL_574542 | National Public Health Laboratory, National Centre for Infectious Diseases | National Public Health Laboratory, National Centre for Infectious Diseases | Tze Minn Mak, Sophie Octavia, Zhenyang Zhou, Lin Cui, Raymond Tzer Pin Lin |

| | | | |
|---|---|---|---|
| EPI_ISL_574591, EPI_ISL_574592 | Hospital Domingos Leonardo Ceravolo Presidente Prudente | Instituto Adolfo Lutz, Interdiciplinary Procedures Center, Strategic Laboratory | Claudio Tavares Sacchi, Claudia Regina Gonçalves, Erica Valessa Ramos Gomes, Karoline Rodrigues Campos |
| EPI_ISL_574843, EPI_ISL_574844, EPI_ISL_574845, EPI_ISL_574846 | Institute for Infectious Diseases, University of Bern | Institute for Infectious Diseases, University of Bern | Michel C Koch, Christian Baumann, Miguel A Terrazos Miani, Cora Sägesser, Stephen L Leib, Peter Keller, Franziska Suter-Riniker, Alban Ramette |
| EPI_ISL_575038, EPI_ISL_575039, EPI_ISL_575040, EPI_ISL_575041, EPI_ISL_575044, EPI_ISL_575045, EPI_ISL_575046, EPI_ISL_575047, EPI_ISL_575057, EPI_ISL_575058, EPI_ISL_575059, EPI_ISL_575105, EPI_ISL_575106, EPI_ISL_575107, EPI_ISL_575108 | | | |
| see above | Utah Public Health Laboratory | Utah Public Health Laboratory | Erin Young, Kelly Oakeson |
| EPI_ISL_577742 | Institute of Virology, Biomedical Research Center of the Slovak Academy of Sciences, Bratislava | Faculty of Natural Sciences, Comenius University, Bratislava | Broa Brejová, Viktória Hodorová, Kristína Boršová, Viktória abanová, Dominika Friová, Sabina Fumaová Havlíková, Juraj Kopáek, Martina Liková, ubomíra Lukáiková, Martina Neboháová, Monika Sláviková, Edita Staroová, Elena Tichá, Tomáš Vina, Jozef Nosek, Boris Klempa |
| EPI_ISL_578084, EPI_ISL_578094, EPI_ISL_578130, EPI_ISL_578131, EPI_ISL_578132, EPI_ISL_578133, EPI_ISL_578134, EPI_ISL_578135 | University of Michigan Clinical Microbiology Laboratory | Lauring Lab, University of Michigan, Department of Microbiology and Immunology | Valesano |
| EPI_ISL_578335, EPI_ISL_578336, EPI_ISL_578337, EPI_ISL_578383, EPI_ISL_578384, EPI_ISL_578385, EPI_ISL_578386, EPI_ISL_578388, EPI_ISL_578389, EPI_ISL_578390, EPI_ISL_578391, EPI_ISL_578392, EPI_ISL_578393, EPI_ISL_578394, EPI_ISL_578395, EPI_ISL_578396, EPI_ISL_578397 | | | |
| see above | Wisconsin State Laboratory of Hygiene Communicable Disease Division | Wisconsin State Laboratory of Hygiene Communicable Disease Division | Kelsey R. Florek, Abigail C. Shockey |
| EPI_ISL_579087, EPI_ISL_579088, EPI_ISL_579089 | Canterbury Health Laboratories | Institute of Environmental Science and Research (ESR) | Xiaoyun Ren, Matt Storey, Nikki Freed, Muhammad Faisal, Jing Wang, Hermes Perez, Anja Werno, Antje van der Linden, Arlo Upton, Chris Mansell, David Hammer, Dragana Drinkovic, Gary McAuliffe, Hana Sofia Andersson, James Ussher, Jill Sherwood, Josh Freeman, Julia Howard, Juliet Elvy, Mary DeAlmeida, Matt Blakiston, Matthew Rogers, Max Bloomfield, Michael Addidle, Michelle Balm, Sally Roberts, Sarah Jefferies, Sharmini Muttaiyah, Susan Morpeth, Susan Taylor, Timothy Blackmore, Vani Sathyendran, Veronica Playle, Virginia Hope, Erasmus Smit, Lauren Jelly, Olin Silander, Joep de Ligt |
| EPI_ISL_579153, EPI_ISL_579155, EPI_ISL_579157, EPI_ISL_579158, EPI_ISL_579163, EPI_ISL_579174 | Southern Community Labs Dunedin | Institute of Environmental Science and Research (ESR) | Xiaoyun Ren, Matt Storey, Nikki Freed, Muhammad Faisal, Jing Wang, Hermes Perez, Anja Werno, Antje van der Linden, Arlo Upton, Chris Mansell, David Hammer, Dragana Drinkovic, Gary McAuliffe, Hana Sofia Andersson, James Ussher, Jill Sherwood, Josh Freeman, Julia Howard, Juliet Elvy, Mary DeAlmeida, Matt Blakiston, Matthew Rogers, Max Bloomfield, Michael Addidle, Michelle Balm, Sally Roberts, Sarah Jefferies, Sharmini Muttaiyah, Susan Morpeth, Susan Taylor, Timothy Blackmore, Vani Sathyendran, Veronica Playle, Virginia Hope, Erasmus Smit, Lauren Jelly, Olin Silander, Joep de Ligt |
| EPI_ISL_579229 | Middlemore Hospital | Institute of Environmental Science and Research (ESR) | Xiaoyun Ren, Matt Storey, Nikki Freed, Muhammad Faisal, Jing Wang, Hermes Perez, Anja Werno, Antje van der Linden, Arlo Upton, Chris Mansell, David Hammer, Dragana Drinkovic, Gary McAuliffe, Hana Sofia Andersson, James Ussher, Jill Sherwood, Josh Freeman, Julia Howard, Juliet Elvy, Mary DeAlmeida, Matt Blakiston, Matthew Rogers, Max Bloomfield, Michael Addidle, Michelle Balm, Sally Roberts, Sarah Jefferies, Sharmini Muttaiyah, Susan Morpeth, Susan Taylor, Timothy Blackmore, Vani Sathyendran, Veronica Playle, Virginia Hope, Erasmus Smit, Lauren Jelly, Olin Silander, Joep de Ligt |
| EPI_ISL_579418 | Wellington SCL (WN) | Institute of Environmental Science and Research (ESR) | Xiaoyun Ren, Matt Storey, Nikki Freed, Muhammad Faisal, Jing Wang, Hermes Perez, Anja Werno, Antje van der Linden, Arlo Upton, Chris Mansell, David Hammer, Dragana Drinkovic, Gary McAuliffe, Hana Sofia Andersson, James Ussher, Jill Sherwood, Josh Freeman, Julia Howard, Juliet Elvy, Mary DeAlmeida, Matt Blakiston, Matthew Rogers, Max Bloomfield, Michael Addidle, Michelle Balm, Sally Roberts, Sarah Jefferies, Sharmini Muttaiyah, Susan Morpeth, Susan Taylor, Timothy Blackmore, Vani Sathyendran, Veronica Playle, Virginia Hope, Erasmus Smit, Lauren Jelly, Olin Silander, Joep de Ligt |
| EPI_ISL_579487, EPI_ISL_579488, EPI_ISL_579489, EPI_ISL_579490, EPI_ISL_579491, EPI_ISL_579492, EPI_ISL_579493, EPI_ISL_579494, EPI_ISL_579495 | Canterbury Health Laboratories | Institute of Environmental Science and Research (ESR) | Xiaoyun Ren, Matt Storey, Nikki Freed, Muhammad Faisal, Jing Wang, Hermes Perez, Anja Werno, Antje van der Linden, Arlo Upton, Chris Mansell, David Hammer, Dragana Drinkovic, Gary McAuliffe, Hana Sofia Andersson, James Ussher, Jill Sherwood, Josh Freeman, Julia Howard, Juliet Elvy, Mary DeAlmeida, Matt Blakiston, Matthew Rogers, Max Bloomfield, Michael Addidle, Michelle Balm, Sally Roberts, Sarah Jefferies, Sharmini Muttaiyah, Susan Morpeth, Susan Taylor, Timothy Blackmore, Vani Sathyendran, Veronica Playle, Virginia Hope, Erasmus Smit, Lauren Jelly, Olin Silander, Joep de Ligt |
| EPI_ISL_579594, EPI_ISL_579595, EPI_ISL_579596, EPI_ISL_579597, EPI_ISL_579598 | QEII Health Sciences Centre | National Microbiology Laboratory (NML) | Anna Majer, Shari Tyson, Grace Seo, Philip Mabon, Darian Hole, Elsie Grudeski, Rhiannon Huzarewich, Russell Mandes, Anneliese Landgraff, Jennifer Tanner, Natalie Knox, Morag Graham, Gary Van Domselaar, Todd Hatchette, Jason LeBlanc, Nathalie Bastien, Yan Li, Timothy Booth, CanCOGeN's metadata curation team, Public Health Agency of Canada's CanCOGeN team |
| EPI_ISL_581462 | Fondation Congolaise pour la recherche medicale (FCRM) | NGS Competence Center Tübingen, Institut für Medizinische Mikrobiologie und Hygiene, Universitätsklinikum Tübingen | Angel Angelov |
| EPI_ISL_581482, EPI_ISL_581483 | Medizinische Klinik Innere Medizin I, Universitätsklinikum Tübingen | NGS Competence Center Tübingen, Institut für Medizinische Mikrobiologie und Hygiene, Universitätsklinikum Tübingen | Angel Angelov |
| EPI_ISL_581486 | Fondation Congolaise pour la recherche medicale (FCRM) | NGS Competence Center Tübingen, Institut für Medizinische Mikrobiologie und Hygiene, Universitätsklinikum Tübingen | Angel Angelov |
| EPI_ISL_581668, EPI_ISL_581669, EPI_ISL_581670, EPI_ISL_581671, EPI_ISL_581672, EPI_ISL_581673, EPI_ISL_581674, EPI_ISL_581813, EPI_ISL_581814, EPI_ISL_581815, EPI_ISL_581816 | | | |
| see above | University Hospital Basel, Clinical Virology | University Hospital Basel, Clinical Bacteriology | Madlen Stange, Alfredo Mari, Tim Roloff, Helena MB Seth-Smith, Michael Schweitzer, Myrta Brunner, Karoline Leuzinger, Kirstine K. Soegaard, Alexander Gensch, Sarah Tschudin-Sutter, Simon Fuchs, Julia Bielicki, Hans Pargger, Martin Siegemund, Christian Nickel, Roland Bingisser, Michael Osthoff, Stefano Bassetti, Rita Schneider-Sliwa, Manuel Battegay, Hans Hirsch, Adrian Egli |
| EPI_ISL_582242, EPI_ISL_582244, EPI_ISL_582256, EPI_ISL_582257, EPI_ISL_582258, EPI_ISL_582273, EPI_ISL_582278, EPI_ISL_582308, EPI_ISL_582354 | Cadham Provincial Laboratory | National Microbiology Laboratory (NML) | Anna Majer, Shari Tyson, Grace Seo, Philip Mabon, Elsie Grudeski, Rhiannon Huzarewich, Russell Mandes, Anneliese Landgraff, Jennifer Tanner, Natalie Knox, Morag Graham, Gary Van Domselaar, Paul Van Caeseele, Jared Bullard, David Alexander, Kerry Dust, Nathalie Bastien, Yan Li, Timothy Booth, Darian Hole, Madison Chapel, CanCOGeN's metadata curation team, Public Health Agency of Canada CanCOGeN team |
| EPI_ISL_582521 | unknown | Instituto Nacional de Saude (INSA) | Borges et al |
| EPI_ISL_583472 | Memorial Sloan Kettering Cancer Center | van Bakel Laboratory, Genetics and Genomics Sciences, Icahn School of Medicine at Mount Sinai | Teresa Aydillo, Ana S. Gonzalez-Reiche, Sadaf Aslam, Adriana van de Guchte, Zenab Khan, Ajay Obla, Jayeeta Dutta, Harm van Bakel, Judith Aberg, Adolfo García-Sastre, Gunjan Shah, Tobias Hohl, Genovefa Papanicolaou, Miguel-Angel Perales, Kent Sepkowitz, Ngolela Esther Babady, and Mini Kamboj |
| EPI_ISL_583570 | Center for Virology, Medical University of Vienna | Bergthaler laboratory, CeMM Research Center for Molecular Medicine of the Austrian Academy of Sciences | Alexandra Popa, Benedikt Agerer, Henrique Colaco, Lukas Endler, Jakob-Wendelin Genger, Alexander Lercher, Mark Smyth, Thomas Penz, Michael Schuster, Jan Laine, Martin Senekowitsch, Judith Aberle, Stephan Aberle, Peter Hufnagl, Daniela Schmid, Franz Allerberger, Elisabeth Puchhammer-Stoeckl, Manfred Nairz, Guenter Weiss, Gregor Hörmann, Kinga Rigler-Hohenwarter, Rainer Gattringer, Wegene Borena, Dorothee von Laer, Gernot Walder, Peter Obrist, Christian Paar, Sabine Sussitz-Rack, Gunther Vogl, Adi Steinrigl, Christoph Bock, Andreas Bergthaler |
| EPI_ISL_583589 | Institute for Medical and Chemical Laboratory Diagnostics, Kepler Universitätsklinikum | Bergthaler laboratory, CeMM Research Center for Molecular Medicine of the Austrian Academy of Sciences | Alexandra Popa, Benedikt Agerer, Henrique Colaco, Lukas Endler, Jakob-Wendelin Genger, Alexander Lercher, Mark Smyth, Thomas Penz, Michael Schuster, Jan Laine, Martin Senekowitsch, Judith Aberle, Stephan Aberle, Peter Hufnagl, Daniela Schmid, Franz Allerberger, Elisabeth Puchhammer-Stoeckl, Manfred Nairz, Guenter Weiss, Gregor Hörmann, Kinga Rigler-Hohenwarter, Rainer Gattringer, Wegene Borena, Dorothee von Laer, Gernot Walder, Peter Obrist, Christian Paar, Sabine Sussitz-Rack, Gunther Vogl, Adi Steinrigl, Christoph Bock, Andreas Bergthaler |
| EPI_ISL_583599 | Zentralinstitut für medizinische und chemische Labordiagnostik, Universitätskliniken Insbruck | Bergthaler laboratory, CeMM Research Center for Molecular Medicine of the Austrian Academy of Sciences | Alexandra Popa, Benedikt Agerer, Henrique Colaco, Lukas Endler, Jakob-Wendelin Genger, Alexander Lercher, Mark Smyth, Thomas Penz, Michael Schuster, Jan Laine, Martin Senekowitsch, Judith Aberle, Stephan Aberle, Peter Hufnagl, Daniela Schmid, Franz Allerberger, Elisabeth Puchhammer-Stoeckl, Manfred Nairz, Guenter Weiss, Gregor Hörmann, Kinga Rigler-Hohenwarter, Rainer Gattringer, Wegene Borena, Dorothee von Laer, Gernot Walder, Peter Obrist, Christian Paar, Sabine Sussitz-Rack, Gunther Vogl, Adi Steinrigl, Christoph Bock, Andreas Bergthaler |
| EPI_ISL_583633, EPI_ISL_583640, EPI_ISL_583648, EPI_ISL_583649, EPI_ISL_583658, EPI_ISL_583678, EPI_ISL_583686 | Austrian Agency for Health and Food Safety (AGES) | Bergthaler laboratory, CeMM Research Center for Molecular Medicine of the Austrian Academy of Sciences | Alexandra Popa, Benedikt Agerer, Henrique Colaco, Lukas Endler, Jakob-Wendelin Genger, Alexander Lercher, Mark Smyth, Thomas Penz, Michael Schuster, Jan Laine, Martin Senekowitsch, Judith Aberle, Stephan Aberle, Peter Hufnagl, Daniela Schmid, Franz Allerberger, Elisabeth Puchhammer-Stoeckl, Manfred Nairz, Guenter Weiss, Gregor Hörmann, Kinga Rigler-Hohenwarter, Rainer Gattringer, Wegene Borena, Dorothee von Laer, Gernot Walder, Peter Obrist, Christian Paar, Sabine Sussitz-Rack, Gunther Vogl, Adi Steinrigl, Christoph Bock, Andreas Bergthaler |

| | | | |
|---|---|---|---|
| EPI_ISL_583756, EPI_ISL_583757, EPI_ISL_583758, EPI_ISL_583759, EPI_ISL_583760, EPI_ISL_583815, EPI_ISL_583816, EPI_ISL_583817, EPI_ISL_583818, EPI_ISL_583819, EPI_ISL_583820, EPI_ISL_583821, EPI_ISL_583822 | | | |
| see above | Dr. Gernot Walder GmbH | Bergthaler laboratory, CeMM Research Center for Molecular Medicine of the Austrian Academy of Sciences | Alexandra Popa, Benedikt Agerer, Henrique Colaco, Lukas Endler, Jakob-Wendelin Genger, Alexander Lercher, Mark Smyth, Thomas Penz, Michael Schuster, Jan Laine, Martin Senekowitsch, Judith Aberle, Stephan Aberle, Peter Hufnagl, Daniela Schmid, Franz Allerberger, Elisabeth Puchhammer-Stoeckl, Manfred Nairz, Guenter Weiss, Gregor Hörmann, Kinga Rigler-Hohenwarter, Rainer Gattringer, Wegene Borena, Dorothee von Laer, Gernot Walder, Peter Obrist, Christian Paar, Sabine Sussitz-Rack, Gunther Vogl, Adi Steinrigl, Christoph Bock, Andreas Bergthaler |
| EPI_ISL_584079 | The National Institute of Public Health | State Veterinary Institute Prague | Nagy,A;Jirincova,H;Novakova,L;Trnka,D;Vecerova,J |
| EPI_ISL_585093, EPI_ISL_585130, EPI_ISL_585132, EPI_ISL_585134, EPI_ISL_585144, EPI_ISL_585145, EPI_ISL_585146, EPI_ISL_585147, EPI_ISL_585148, EPI_ISL_585149, EPI_ISL_585151, EPI_ISL_585152, EPI_ISL_585153, EPI_ISL_585174, EPI_ISL_585175, EPI_ISL_585176, EPI_ISL_585177, EPI_ISL_585178, EPI_ISL_585179, EPI_ISL_585180, EPI_ISL_585181, EPI_ISL_585182, EPI_ISL_585183, EPI_ISL_585184, EPI_ISL_585246, EPI_ISL_585248, EPI_ISL_585250, EPI_ISL_585252, EPI_ISL_585254, EPI_ISL_585256 | | | |
| see above | Regional Virus Laboratory, Belfast Health and Social Care Trust | COVID-19 Genomics UK (COG-UK) Consortium | Conall McCaughey, James McKenna, Tanya Curran, Susan Feeney, Alison Watt, Ciara Cox, Mairead Connor, Zoltan Molnar, David Simpson, Derek Fairley |
| EPI_ISL_586408, EPI_ISL_586429, EPI_ISL_586439, EPI_ISL_586450, EPI_ISL_586451, EPI_ISL_586454, EPI_ISL_586457, EPI_ISL_586462, EPI_ISL_586464, EPI_ISL_586465 | Toronto Invasive Bacterial Diseases Network | McMaster University | Allison McGeer, Patryk Aftanas, Hooman Derakhshani, Angel Li, Kuganya Nirmalarajah, Emily Panousis, Ahmed Draia, Jalees Nasir, Michael Surette, Samira Mubareka, Andrew G. McArthur |
| EPI_ISL_591081, EPI_ISL_591082, EPI_ISL_591093 | Department of Pathology, University of Cambridge | Wellcome Sanger Institute for the COVID-19 Genomics UK (COG-UK) consortium | Luke W Meredith, M. Estée Török , Myra Hosmillo, William L. Hamilton, Martin D. Curran, Theresa Feltwell, Grant Hall, Anna Yakovleva, Fahad A Khokhar, Charlotte J. Houldcroft, Laura G Caller, Aminu S. Jahun, Sarah L. Caddy, Ian Goodfellow; and Alex Alderton, Roberto Amato, Sonia Goncalves, Ewan Harrison, David K. Jackson, Ian Johnston, Dominic Kwiatkowski, Cordelia Langford, John Sillitoe on behalf of the Wellcome Sanger Institute COVID-19 Surveillance Team |
| EPI_ISL_591135, EPI_ISL_591142, EPI_ISL_591144, EPI_ISL_591145, EPI_ISL_591146, EPI_ISL_591147, EPI_ISL_591149, EPI_ISL_591151, EPI_ISL_591153, EPI_ISL_591161, EPI_ISL_591162, EPI_ISL_591163, EPI_ISL_591167, EPI_ISL_591171, EPI_ISL_591172, EPI_ISL_591173, EPI_ISL_591174, EPI_ISL_591213, EPI_ISL_591214, EPI_ISL_591215, EPI_ISL_591217, EPI_ISL_591218, EPI_ISL_591219, EPI_ISL_591221, EPI_ISL_591226, EPI_ISL_591227, EPI_ISL_591228, EPI_ISL_591229, EPI_ISL_591230, EPI_ISL_591231, EPI_ISL_591232, EPI_ISL_591233, EPI_ISL_591234, EPI_ISL_591235, EPI_ISL_591236, EPI_ISL_591239, EPI_ISL_591241, EPI_ISL_591242, EPI_ISL_591243, EPI_ISL_591244, EPI_ISL_591248, EPI_ISL_591250, EPI_ISL_591251, EPI_ISL_591252 | | | |
| see above | Toronto Invasive Bacterial Diseases Network | McMaster University | Allison McGeer, Patryk Aftanas, Hooman Derakhshani, Angel Li, Kuganya Nirmalarajah, Emily Panousis, Ahmed Draia, Jalees Nasir, Michael Surette, Samira Mubareka, Andrew G. McArthur |
| EPI_ISL_591316, EPI_ISL_591317, EPI_ISL_591318, EPI_ISL_591319, EPI_ISL_591324 | Virology, Iran University of Medical Sciences | Virology, Iran University of Medical Sciences | Keyvani,H., Ranjbar,Mm., Keyvani,F., Soleimani,S. |
| EPI_ISL_591351, EPI_ISL_591361, EPI_ISL_591362, EPI_ISL_591363, EPI_ISL_591364, EPI_ISL_591365, EPI_ISL_591366, EPI_ISL_591367, EPI_ISL_591368, EPI_ISL_591369, EPI_ISL_591377, EPI_ISL_591378 | | | |
| see above | Pathogen Genomics Center, National Institute of Infectious Diseases | Pathogen Genomics Center, National Institute of Infectious Diseases | Tsuyoshi Sekizuka, Kentaro Itokawa, Rina Tanaka, Masanori Hashino, Makoto Kuroda |
| EPI_ISL_591479, EPI_ISL_591480, EPI_ISL_591481, EPI_ISL_591482, EPI_ISL_591483 | Pathogen Genomics Center, National Institute of Infectious Diseases | Pathogen Genomics Center, National Institute of Infectious Diseases | Tsuyoshi Sekizuka, Kentaro Itokawa, Rina Tanaka, Masanori Hashino, Hajime Kamiya, Tomoe Shimada, Makoto Kuroda |
| EPI_ISL_591536 | Pathogen Genomics Center, National Institute of Infectious Diseases | Pathogen Genomics Center, National Institute of Infectious Diseases | Tsuyoshi Sekizuka, Kentaro Itokawa, Rina Tanaka, Masanori Hashino, Makoto Kuroda |
| EPI_ISL_593613, EPI_ISL_593633 | unknown | Public Health Virology Laboratory, Forensic and Scientific Services (PHV-FSS) | Son Nguyen et al. |
| EPI_ISL_593957, EPI_ISL_593958, EPI_ISL_593959, EPI_ISL_593960, EPI_ISL_593961, EPI_ISL_593962, EPI_ISL_593963, EPI_ISL_593964, EPI_ISL_593966, EPI_ISL_593967, EPI_ISL_593968, EPI_ISL_593969, EPI_ISL_593970, EPI_ISL_593971, EPI_ISL_593973, EPI_ISL_593974, EPI_ISL_593975 | | | |
| see above | Delaware Public Health Lab | Delaware Public Health Lab | Gregory Hovan |
| EPI_ISL_596378, EPI_ISL_596380, EPI_ISL_596381, EPI_ISL_596383 | Virology, Iran University of Medical Sciences | Virology, Iran University of Medical Sciences | Keyvani,H., Ranjbar,Mm., Keyvani,F., Soleimani,S. |
| EPI_ISL_596513, EPI_ISL_596515, EPI_ISL_596516, EPI_ISL_596517, EPI_ISL_596518 | Palestinian Ministry of Health | Molecular Genetics Lab | Nouar Qutob, Zaidoun Salah, Damien Richard, Hisham Darwish, Husam Sallam, Issa Shtayeh, Osama Najjar, Mahmoud Ruzayqat, Dana Najjar, Francois Balloux, Lucy van Dorp |
| EPI_ISL_596656, EPI_ISL_596657, EPI_ISL_596665, EPI_ISL_596675 | St.Vincent's University Hospital | St.Vincent's University Hospital | Mary Lucey, Guerrino Macori, Niamh Mullane, Una Sutton-Fitzpatrick, Gabriel Gonzalez, Suzie Coughlan, Aisling Purcell, Lynda Fenelon, Séamus Fanning, Kirsten Schaffer |
| EPI_ISL_596677, EPI_ISL_596684, EPI_ISL_596707, EPI_ISL_596754, EPI_ISL_596755, EPI_ISL_596761, EPI_ISL_596776, EPI_ISL_596794, EPI_ISL_596796, EPI_ISL_596797, EPI_ISL_596798, EPI_ISL_596812, EPI_ISL_596816, EPI_ISL_596825, EPI_ISL_596849, EPI_ISL_596852, EPI_ISL_596870 | | | |
| see above | PathWest Laboratory Medicine WA | PathWest Laboratory Medicine WA Microbial Surveillance Unit | PathWest Laboratory Medicine WA Microbial Surveillance Unit |
| EPI_ISL_602282, EPI_ISL_602283, EPI_ISL_602287 | Evangelisches Klinikum Bethel, Institut für Laboratoriumsmedizin, Mikrobiologie und Hygiene | Bielefeld University | David Brandt, Tobias Busche, Markus Haak, Jörn Kalinowski, Levin-Joe Klages, Christiane Scherer, Alexander Sczyrba, Marina Simunovic, Svenja Vinke |
| EPI_ISL_602467 | Institute for Virology, University Hospital Essen | Center of Medical Microbiology, Virology, and Hospital Hygiene, University of Duesseldorf | Olympia E. Anastasiou, Ulf Dittmer, Maximilian Damagnez, Alexander Dilthey, Torsten Houwaart, Lisanna Hülse, Malte Kohns Vasconcelos, Nadine Lübke, Jessica Nicolai, Klaus Pfeffer, Daniel Strelow, Jörg Timm, Andreas Walker, Tobias Wienemann |
| EPI_ISL_605058, EPI_ISL_605061, EPI_ISL_605062, EPI_ISL_605063, EPI_ISL_605065, EPI_ISL_605066, EPI_ISL_605072, EPI_ISL_605078 | National Virus Reference Laboratory | Irish Coronavirus Sequencing Consortium - Helixworks | Sachin Chalapati, Conor Crosbie, Nimesh Pinnamaneni |
| EPI_ISL_605793 | Department of Experimental Modeling and Pathogenesis of Infectious Diseases | WHO National Influenza Centre Russian Federation | Andrey Komissarov, Artem Fadeev, Anna Ivanova, Kseniya Komissarova, Sobolev I.A., Alekseev A.Yu., Chepurnov A.A., Kononova Yu.V., Shestopalov A.M. |
| EPI_ISL_605794 | Department of Experimental Modeling and Pathogenesis of Infectious Diseases | WHO National Influenza Centre Russian Federation | Andrey Komissarov, Artem Fadeev, Anna Ivanova, Kseniya Komissarova, Sobolev I.A., Alekseev A.Yu., Kononova Yu.V., Shestopalov A.M. |
| EPI_ISL_605795, EPI_ISL_605796 | Department of Experimental Modeling and Pathogenesis of Infectious Diseases | WHO National Influenza Centre Russian Federation | Andrey Komissarov, Artem Fadeev, Anna Ivanova, Kseniya Komissarova, Sobolev I.A., Alekseev A.Yu., Chepurnov A.A., Kononova Yu.V., Shestopalov A.M. |
| EPI_ISL_605803, EPI_ISL_605811, EPI_ISL_605812, EPI_ISL_605816 | Clinical Virology Laboratory, Institute of Liver and Biliary Sciences | ILBS - IGIB | Ekta Gupta, Sheetalnath Rooge, Abhishek Padhi, Reshu Agarwal, Jaswinder Singh Maras, Shridhar Sivasubbu, Vinod Scaria, Shvetank Sharma |
| EPI_ISL_605868 | PathWest Laboratory Medicine WA | PathWest Laboratory Medicine WA Microbial Surveillance Unit | PathWest Laboratory Medicine WA Microbial Surveillance Unit |
| EPI_ISL_610237, EPI_ISL_610238 | Molecular diagnostic laboratory of Federal Budget Institution of Science "Central Research Institute of Epidemiology" of The Federal Service on Customers' Rights Protection and Human Well-being Surveillance | Group of Genomics and Postgenomic Technologies of Central Research Institute of Epidemilology | Samoilov AE, Kaptelova VV, Bukharina AY, Speranskaya AS, Tivanova EV, Shipulina OY, Akimkin VG |
| EPI_ISL_611653, EPI_ISL_611672, EPI_ISL_611845, EPI_ISL_611853, EPI_ISL_612017, EPI_ISL_612191, EPI_ISL_612416, EPI_ISL_612419, EPI_ISL_612422, EPI_ISL_612425, EPI_ISL_612428, EPI_ISL_612429, EPI_ISL_612430, EPI_ISL_612431 | | | |
| see above | Liverpool Clinical Laboratories | COVID-19 Genomics UK (COG-UK) Consortium | Sam Haldenby, Anita Lucaci, Steve Paterson, Julian Hiscox, Alistair Darby, M Almsaud, A Alrezaihi, Muhannad Alruwaili, Stuart D Armstrong, Jones Benjamin, Eleanor G Bentley, Anu Chawla, Jordan J Clark, Angela Cowell, Richard Eccles, Isabel García-Dorival, Matthew Gemmell, Alessandro Gerada, |

| | | | |
|---|---|---|---|
| | | | PKF Gilmore, Richard Gregory, Ximeng Han, Catherine Hartley, Margaret Hughes, Miren Iturriza-Gomara, James Johnson, L Luu, Jenifer Manson, Charlotte Nelson, Elaine O'Toole, Cassie Olateju, Rebekah Penrice-Randal , Lucille Rainbow, N.P Randle, Trevor Ian Robinson, Parul Sharma, Ghada T Shawli, James P Stewart, Neil Swainston, Ecaterina Vamos, Joanne Watts, Mark Whitehead |
| EPI_ISL_613445 | Institut Pasteur de la Guadeloupe | Institut Pasteur de la Guadeloupe | Marion Barbet, Sylvie Behillil, Méline Bizard, Angela Brisebarre, Camille Capel, Etienne Simon-Lorière, Vincent Enouf, Maud Vanpeene, Sylvie van der Werf, Stéphanie Guyomard, Sébastien Breurec, Antoine Talarmin |
| EPI_ISL_613563 | Laboratory of Molecular Biology, Blood Center of Ribeirão Preto | Laboratory of Molecular Biology, Blood Center of Ribeirão Preto, Faculty of Medicine of Ribeirão Preto, University of São Paulo | Svetoslav N Slavov, Marta Giovanetti, Vagner Fonseca, Elaine V Santos, Evandra S Rodrigues, Talita Adelino, Joilson Xavier, Glauco de Carvalho Pereira, Aparecida Y Yamamoto, Diego Villa Clé, Rodrigo T Calado; Dimas T Covas, Luiz CJ Alcantara, Simone Kashima |
| EPI_ISL_613711, EPI_ISL_613951, EPI_ISL_613965, EPI_ISL_614011, EPI_ISL_614156 | Laboratory of Molecular Biology, Blood Center of Ribeirão Preto, Faculty of Medicine of Ribeirão Preto, University of São Paulo | Laboratory of Molecular Biology, Blood Center of Ribeirão Preto, Faculty of Medicine of Ribeirão Preto, University of São Paulo | Svetoslav N Slavov, Marta Giovanetti, Vagner Fonseca, Elaine V Santos, Evandra S Rodrigues, Talita Adelino, Joilson Xavier, Glauco de Carvalho Pereira, Aparecida Y Yamamoto, Diego Villa Clé, Rodrigo T Calado; Dimas T Covas, Luiz CJ Alcantara, Simone Kashima |
| EPI_ISL_622788, EPI_ISL_622800, EPI_ISL_622801 | Canterbury Health Laboratories | Institute of Environmental Science and Research (ESR) | Xiaoyun Ren, Matt Storey, Nikki Freed, Muhammad Faisal, Jing Wang, Hermes Perez, Anja Werno, Antje van der Linden, Arlo Upton, Chris Mansell, David Hammer, Dragana Drinkovic, Gary McAuliffe, Hana Sofia Andersson, James Ussher, Jill Sherwood, Josh Freeman, Julia Howard, Juliet Elvy, Mary DeAlmeida, Matt Blakiston, Matthew Rogers, Max Bloomfield, Michael Addidle, Michelle Balm, Sally Roberts, Sarah Jefferies, Sharmini Muttaiyah, Susan Morpeth, Susan Taylor, Timothy Blackmore, Vani Sathyendran, Veronica Playle, Virginia Hope, Erasmus Smit, Lauren Jelly, Olin Silander, Joep de Ligt |
| EPI_ISL_623126 | Laboratorio de Virologia Molecular / UFRJ | Bioinformatics Laboratory / LNCC | Carolina M Voloch, Ronaldo S Francisco Jr, Luiz G P de Almeida, Otavio J. Brustolini, Cynthia C Cardoso, Alexandra L Gerber, Ana Paula de C Guimarães, Diana Mariani, Covid19-UFRJ Workgroup, Luís Cristóvão Pôrto, Renato S Aguiar, Terezinha M P P Castiñeiras, Orlando C. Ferreira, Amilcar Tanuri, Ana Tereza R de Vasconcelos |
| EPI_ISL_623181, EPI_ISL_623182 | Utah Public Health Laboratory | Utah Public Health Laboratory | Erin Young, Kelly Oakeson |
| EPI_ISL_626228, EPI_ISL_626229 | Institute for Virology, University Hospital Essen | Center of Medical Microbiology, Virology, and Hospital Hygiene, University of Duesseldorf | Olympia E. Anastasiou, Ulf Dittmer, Maximilian Damagnez, Alexander Dilthey, Torsten Houwaart, Lisanna Hülse, Malte Kohns Vasconcelos, Nadine Lübke, Jessica Nicolai, Klaus Pfeffer, Daniel Strelow, Jörg Timm, Andreas Walker, Tobias Wienemann |
| EPI_ISL_626513, EPI_ISL_626514, EPI_ISL_626515, EPI_ISL_626516 | Northwestern Memorial Hospital | Ozer Lab | Ramon Lorenzo-Redondo, Hannah H. Nam, Scott C. Roberts, Lacy M. Simons, Chad J. Achenbach, Lawrence J. Jennings, Chao Qi, Alan R. Hauser, Michael G. Ison, Judd F. Hultquist, Egon A. Ozer |
| EPI_ISL_629096 | Laboratoire du Centre Hospitalier Annecy Genevois | CNR Virus des Infections Respiratoires - France SUD | Antonin Bal, Gregory Destras, Gwendolyne Burfin, Hadrien Règue, Quentin Semanas, Martine Valette, Bruno Lina, Hélène Petitprez, Bruno Chanzy, Laurence Josset |
| EPI_ISL_631390 | Wisconsin State Laboratory of Hygiene Communicable Disease Division | Wisconsin State Laboratory of Hygiene Communicable Disease Division | Kelsey R. Florek, Abigail C. Shockey |
| EPI_ISL_631613 | Jamaica Hospital Medical Center | New York City Public Health Laboratory | Jade Wang, et al. |
| EPI_ISL_631614 | Flushing Hospital Medical Center | New York City Public Health Laboratory | Jade Wang, et al. |
| EPI_ISL_631615 | Jamaica Hospital Medical Center | New York City Public Health Laboratory | Jade Wang, et al. |
| EPI_ISL_631616 | Flushing Hospital Medical Center | New York City Public Health Laboratory | Jade Wang, et al. |
| EPI_ISL_631617, EPI_ISL_631762 | OCME Office Of Chief Medical Examiner | New York City Public Health Laboratory | Jade Wang, et al. |
| EPI_ISL_631904 | Jamaica Hospital Medical Center | New York City Public Health Laboratory | Jade Wang, et al. |
| EPI_ISL_631905 | Richmond University Medical Center | New York City Public Health Laboratory | Jade Wang, et al. |
| EPI_ISL_631906 | Flushing Hospital Medical Center | New York City Public Health Laboratory | Jade Wang, et al. |
| EPI_ISL_631917 | St Barnabas Hospital | New York City Public Health Laboratory | Jade Wang, et al. |
| EPI_ISL_631918 | Flushing Hospital Medical Center | New York City Public Health Laboratory | Jade Wang, et al. |
| EPI_ISL_631919, EPI_ISL_631920 | Richmond University Medical Center | New York City Public Health Laboratory | Jade Wang, et al. |
| EPI_ISL_631953 | NYC HH Lincoln Medical And Mental Health Center | New York City Public Health Laboratory | Jade Wang, et al. |
| EPI_ISL_631954, EPI_ISL_631992, EPI_ISL_632031, EPI_ISL_632032, EPI_ISL_632033 | Richmond University Medical Center | New York City Public Health Laboratory | Jade Wang, et al. |
| EPI_ISL_632220 | Flushing Hospital Medical Center | New York City Public Health Laboratory | Jade Wang, et al. |
| EPI_ISL_632866, EPI_ISL_632867, EPI_ISL_632889 | Idaho Bureau of Laboratories | Center for Global Health, University of New Mexico Health Sciences Center | Daryl Domman, Kurt Schwalm, Matthew Burns, Robert Voermans, Christopher Ball, Darrell Dinwiddie |
| EPI_ISL_632908 | Genomic Sciences, Rehman Medical Institute | Genomic Sciences, Rehman Medical Institute | Ali,J., Afridi,U.K., Haider,S.A., Sabiha,B., Jan,H. and Jehanzeb,V. |
| EPI_ISL_634829 | Laboratoire de virologie, CHU de Grenoble - CS 10217 - 38043 Grenoble cedex 21 | CNR Virus des Infections Respiratoires - France SUD | Antonin Bal, Gregory Destras, Gwendolyne Burfin, Hadrien Règue, Quentin Semanas, Martine Valette, Bruno Lina, Sylvie Larrat, Laurence Josset |
| EPI_ISL_634830 | Laboratoire de virologie, CHU de Grenoble - CS 10217 - 38043 Grenoble cedex 22 | CNR Virus des Infections Respiratoires - France SUD | Antonin Bal, Gregory Destras, Gwendolyne Burfin, Hadrien Règue, Quentin Semanas, Martine Valette, Bruno Lina, Sylvie Larrat, Laurence Josset |
| EPI_ISL_634836 | Laboratoire de virologie, CHU de Grenoble - CS 10217 - 38043 Grenoble cedex 28 | CNR Virus des Infections Respiratoires - France SUD | Antonin Bal, Gregory Destras, Gwendolyne Burfin, Hadrien Règue, Quentin Semanas, Martine Valette, Bruno Lina, Sylvie Larrat, Laurence Josset |
| EPI_ISL_635278, EPI_ISL_635280, EPI_ISL_635281, EPI_ISL_635283, EPI_ISL_635284 | Institute of Microbiology and Immunology, Faculty of Medicine, University of Ljubljana | Institute of Microbiology and Immunology, Faculty of Medicine, University of Ljubljana | Tomaž Mark Zorec, Samo Zakotnik, Miša Korva, Tatjana Avši - Županc, Mario Poljak |
| EPI_ISL_636542, EPI_ISL_636565 | Dutch COVID-19 response team | National Institute for Public Health and the Environment (RIVM) | Adam Meijer, Harry Vennema, Jeroen Cremer, Sharon van den Brink, Bas van der Veer, AnneMarie van den Brandt, Florian Zwagemaker, Dennis Schmitz, Chantal Reusken, on behalf of the national COVID-19 response team |
| EPI_ISL_636946, EPI_ISL_636947 | Public Health Ontario Laboratory | Public Health Ontario Laboratory | Vanessa G Allen, Philip Banh, Richard de Borja, Yao Chen, Alireza Eshaghi, Nahuel Fittipaldi, Christine Frantz, Jonathan B Gubbay, Jennifer L Guthrie, Lawrence Heisler, Esha Joshi, Michael Laszloffy, Aimin Li, Michael CY Li, Dean Maxwell, Sandeep Nagra, Samir N Patel, Heather Rilkoff, Jared Simpson, Karthikeyan Sivaraman, Yogi Sundaravadanam, Sarah Teatero, Andre Villegas, Sandra Zittermann |
| EPI_ISL_636969, EPI_ISL_636970 | Etlik Veterinary Control Central Research Institute | Etlik Veterinary Control Central Research Institute | Sabri Hacioglu, Ahu Pakdemirli, Dilek Dulger, Erdem Danyer, Ummu Sena Sari, Cevdet Yarali, Ozcan Yildirim |
| EPI_ISL_636990 | Department of Infectious Diseases and Immunology, National Hospital Organization Nagoya Medical Center | Clinical Research Center, National Hospital Organization Nagoya Medical Center | Yoshihiro Nakata, Hirotaka Ode, Mai Kubota, Masakazu Matsuda, Kazuhiro Matsuoka, Miho Nakasuji, Mikiko Mori, Mayumi Imahashi, Yoshiyuki Yokomaku, Yasumasa Iwatani |
| EPI_ISL_637333 | Wales Specialist Virology Centre Sequencing lab: Pathogen Genomics Unit | COVID-19 Genomics UK (COG-UK) Consortium | Catherine Moore, Johnathan Evans, Laura Gifford, Malorie Perry, Simon Cottrell, Angela Marchbank, Alec Birchley, Alexander Adams, Amy Gaskin, Bree Gatica-Wilcox, Jason Coombes, Joel Southgate, Lauren Gilbert, Lee Graham, Nicole Pacchiarini, Sara Kumziene-Summerhayes, Sarah Taylor, Sophie Jones, Sara Rey, Matthew Bull, Joanne Watkins, Sally Corden, Tom Connor |
| EPI_ISL_639727, EPI_ISL_639728 | Public Health Ontario Laboratory | Public Health Ontario Laboratory | Vanessa G Allen, Philip Banh, Richard de Borja, Yao Chen, Alireza Eshaghi, Nahuel Fittipaldi, Christine Frantz, Jonathan B Gubbay, Jennifer L Guthrie, Lawrence Heisler, Esha Joshi, Michael Laszloffy, Aimin Li, Michael CY Li, Dean Maxwell, Sandeep Nagra, Samir N Patel, Heather Rilkoff, Jared Simpson, Karthikeyan Sivaraman, Yogi Sundaravadanam, Sarah Teatero, Andre Villegas, Sandra Zittermann |

| | | | |
|---|---|---|---|
| EPI_ISL_639755, EPI_ISL_639758, EPI_ISL_639759, EPI_ISL_639760, EPI_ISL_639761, EPI_ISL_639762, EPI_ISL_639816, EPI_ISL_639817 | unknown | Public Health Virology Laboratory, Forensic and Scientific Services (PHV-FSS) | Son Nguyen et al. |
| EPI_ISL_640044 | District 6 CDC wc DSI | NHLS/UCT | Arash Iranzadeh, Deelan Doolabh, Lynn Tyers, Bruna Galvao, Innocent Mudau, Marvin Hsiao, Kruger Marais, Diana Hardie, Stephen Korsman, Carolyn Williamson |
| EPI_ISL_640045 | Groote Schuur Hospital wc GSH | NHLS/UCT | Arash Iranzadeh, Deelan Doolabh, Lynn Tyers, Bruna Galvao, Innocent Mudau, Marvin Hsiao, Kruger Marais, Diana Hardie, Stephen Korsman, Carolyn Williamson |
| EPI_ISL_641309, EPI_ISL_641310 | unknown | Public Health Virology Laboratory, Forensic and Scientific Services (PHV-FSS) | Son Nguyen et al. |
| EPI_ISL_641319 | Environmental and Global Health, University of Florida | Environmental and Global Health, University of Florida | Elbadry,M.A., Subramaniam,K., Waltzek,T.B., Loeb,J.C., Stephenson,C.J., Lauzardo,M., Morris,J.G., Lednicky,J.A. |
| EPI_ISL_644214, EPI_ISL_644225, EPI_ISL_644227, EPI_ISL_644228 | CEPHR / Vincent's Hospital | Irish Coronavirus Sequencing Consortium - National Virus Reference Laboratory | Michael Carr, Gabriel Gonzalez, Alejandro Abner Garcia Leon, Patrick Mallon |
| EPI_ISL_644675 | CHU de Limoges | CNR Virus des Infections Respiratoires - France SUD | Antonin Bal, Gregory Destras, Gwendolyne Burfin, Hadrien Règue, Quentin Semanas, Martine Valette, Bruno Lina, Sylvie Rogez, Laurence Josset |
| EPI_ISL_644753, EPI_ISL_644757, EPI_ISL_644772, EPI_ISL_644773, EPI_ISL_644774, EPI_ISL_644781, EPI_ISL_644783, EPI_ISL_644784, EPI_ISL_644785, EPI_ISL_644786, EPI_ISL_644787, EPI_ISL_644788, EPI_ISL_644789, EPI_ISL_644822 | | | |
| see above | National Microbiology Reference Laboratory | Quadram Institute Bioscience | Thanh Le Viet, Andrew J. Page, Justin O'Grady, Gemma Kay, David Baker, Gaetan Thilliez, Ana-Victoria Gutierrez, Robert Kingsley, Leonardo de Oliveira Martins, Sekesai Zinyowera, Tatenda Takawira, Muchaneta Mugabe, Gibson Mhlanga, Portia Manangazira, Andrew Tarupiwa, Hlanai Gumbo, Agnes Juru, Charles Nyagupe, Alexander Goredema, Isaac Phiri, Barbra Murwira, Beuty Makamure, Tapfumanei Mashe |
| EPI_ISL_645182 | CHU de Limoges | CNR Virus des Infections Respiratoires - France SUD | Antonin Bal, Gregory Destras, Gwendolyne Burfin, Hadrien Règue, Quentin Semanas, Martine Valette, Bruno Lina, Sylvie Rogez, Laurence Josset |
| EPI_ISL_648030, EPI_ISL_648031, EPI_ISL_648032, EPI_ISL_648033, EPI_ISL_648041 | MS Public Health Laboratory | Pathogen Discovery, Respiratory Viruses Branch, Division of Viral Diseases, Centers for Disease Control and Prevention | Yan Li, Jing Zhang, Ying Tao, Brian Lynch, Krista Queen, Anna Montmayeur, Anna Uehara, Clinton R. Paden, Rachel Marine, Haibin Wang, Suxiang Tong |
| EPI_ISL_648219, EPI_ISL_648220, EPI_ISL_648221, EPI_ISL_648222, EPI_ISL_648223, EPI_ISL_648224, EPI_ISL_648225, EPI_ISL_648226, EPI_ISL_648227, EPI_ISL_648228, EPI_ISL_648229, EPI_ISL_648230, EPI_ISL_648231, EPI_ISL_648232, EPI_ISL_648233, EPI_ISL_648234, EPI_ISL_648236, EPI_ISL_648237, EPI_ISL_648238, EPI_ISL_648239, EPI_ISL_648240, EPI_ISL_648241, EPI_ISL_648242, EPI_ISL_648243, EPI_ISL_648244, EPI_ISL_648245, EPI_ISL_648246, EPI_ISL_648247, EPI_ISL_648248, EPI_ISL_648249, EPI_ISL_648250, EPI_ISL_648251, EPI_ISL_648252, EPI_ISL_648253, EPI_ISL_648254, EPI_ISL_648255, EPI_ISL_648256, EPI_ISL_648257, EPI_ISL_648258, EPI_ISL_648259, EPI_ISL_648260, EPI_ISL_648261, EPI_ISL_648262, EPI_ISL_648263 | | | |
| see above | Texas Department of State Health Services | Texas Department of State Health Services | Rashmi Tuladhar, Bonnie Oh, Jenny Zhang, Maliha Rahman, Anita Pokharel, Myong Koag, Chung Wang, Rachel Lee, Grace Kubin, Mayela Pedrueza, James Daniel Bonser |
| EPI_ISL_648555, EPI_ISL_648556 | Utah Public Health Laboratory | Utah Public Health Laboratory | Erin Young, Kelly Oakeson |
| EPI_ISL_649176, EPI_ISL_649177, EPI_ISL_649178, EPI_ISL_649179 | CHU Nantes | CNR Virus des Infections Respiratoires - France SUD | Antonin Bal, Louise Castain, Gregory Destras, Gwendolyne Burfin, Hadrien Règue, Quentin Semanas, Martine Valette, Bruno Lina, Celine Bressollette, Laurence Josset |
| EPI_ISL_653185, EPI_ISL_653186 | Florida Bureau of Public Health Laboratories | Florida Bureau of Public Health Laboratories | Sarah Schmedes, Jason Blanton |
| EPI_ISL_654625, EPI_ISL_654663, EPI_ISL_654664, EPI_ISL_654665, EPI_ISL_654666, EPI_ISL_654667 | Servicio de Microbiología, Hospital Universitario Central de Asturias | SeqCOVID-SPAIN consortium/IBV(CSIC) | Cristián Castelló Abietar, Jose A. Boga, Susana Rojo-Alba, Marta Elena Álvarez-Argüelles, Santiago Melón and SeqCOVID-SPAIN consortium |
| EPI_ISL_654881 | Pasteur Institute in Ho Chi Minh city | Department of Microbiology and Immunology - Pasteur Institute in Ho Chi Minh city | Lng Chn Quang, ào Huy Mnh, Phm Th Thu Hng, V Phm Hng Nhung, Cao Minh Thng, Hunh Th Kim Loan, Nguyn Hoàng Quân, Hunh Phng Tho, Hoàng Nh ào, Nguyn Trung Hiu, Nguyn Hoàng Anh, Nguyn Thu Ngc, Phm Th Nhung, Nguyn Th Ngc Tho, ng Thanh Giang, Nguyn Th Thanh Thng, Hoàng Minh, Nguyn Th Phng Thúy, Nguyn Thanh Long, Phm Duy Quang, Hoàng Quc Cng, Nguyn V Thng, Phan Trng Lân |
| EPI_ISL_654900, EPI_ISL_654901, EPI_ISL_654902, EPI_ISL_654903, EPI_ISL_654904, EPI_ISL_654905, EPI_ISL_654906, EPI_ISL_654907, EPI_ISL_654908, EPI_ISL_654909, EPI_ISL_654910, EPI_ISL_654913 | | | |
| see above | Texas Department of State Health Services | Texas Department of State Health Services | Rashmi Tuladhar, Bonnie Oh, Jenny Zhang, Maliha Rahman, Anita Pokharel, Myong Koag, Chung Wang, Rachel Lee, Grace Kubin, Mayela Pedrueza, James Daniel Bonser |
| EPI_ISL_657450, EPI_ISL_657452, EPI_ISL_657455, EPI_ISL_657458 | Servicio de Microbiología. Hospital General Universitario de Castellón | SeqCOVID-SPAIN consortium/IBV(CSIC) | Rosario Moreno Muñoz, María Dolores Tirado Balaguer and SeqCOVID-SPAIN consortium |
| EPI_ISL_660378, EPI_ISL_660383 | Orebro klinisk mikrobiologi | The Public Health Agency of Sweden | Anna-Malin Linde, Maria Lind Karlberg, Mattias Haukland, Reza Advani, Olov Svartstrom, Oskar Karlsson Lindsjo, Sandra Broddesson, Petra Edquist, Mia Brytting, Anna Risberg, Karin Tegmark-Wisell |
| EPI_ISL_660395, EPI_ISL_660396, EPI_ISL_660397, EPI_ISL_660398, EPI_ISL_660399 | Klinisk mikrobiologi Linkoping | The Public Health Agency of Sweden | Anna-Malin Linde, Maria Lind Karlberg, Mattias Haukland, Reza Advani, Olov Svartstrom, Oskar Karlsson Lindsjo, Sandra Broddesson, Petra Edquist, Mia Brytting, Anna Risberg, Karin Tegmark-Wisell |
| EPI_ISL_660737 | Unité des Virus Émergents | CNR Virus des Infections Respiratoires - France SUD | Antonin Bal, Gregory Destras, Gwendolyne Burfin, Hadrien Règue, Quentin Semanas, Martine Valette, Bruno Lina, Laetitia Ninove, Léa Luciani, Antoine Nougairède, Laurence Josset |
| EPI_ISL_661200 | Scientific Veterinary Institute Novi Sad | Veterinary Specialized Institute "Kraljevo", Serbia | Vidanovic,D., Tesovic,B., Knezevic,A., Jovanovic,T., Jankovic,M., Sekler,M., Banovic Djeri,B., Petrovic,T., Volkening,J., Afonso,C. |
| EPI_ISL_663264, EPI_ISL_663265, EPI_ISL_663266, EPI_ISL_663267, EPI_ISL_663268, EPI_ISL_663269, EPI_ISL_663270 | CHU Nantes | CNR Virus des Infections Respiratoires - France SUD | Antonin Bal, Louise Castain, Gregory Destras, Gwendolyne Burfin, Hadrien Règue, Quentin Semanas, Martine Valette, Bruno Lina, Virginie Ferré, Celine Bressollette, Laurence Josset |
| EPI_ISL_664188, EPI_ISL_664196, EPI_ISL_664236, EPI_ISL_664651, EPI_ISL_664728, EPI_ISL_665207 | University College London Hospital | COVID-19 Genomics UK (COG-UK) Consortium | Judith Heaney, Matthew Byott, Catherine Houlihan, Dan Frampton, Stuart Kirk, Moira Spyer and Eleni Nastouli |
| EPI_ISL_666662, EPI_ISL_666663, EPI_ISL_666664, EPI_ISL_666665 | Laboratoire du Centre Hospitalier Annecy Genevois | CNR Virus des Infections Respiratoires - France SUD | Antonin Bal, Gregory Destras, Gwendolyne Burfin, Hadrien Règue, Quentin Semanas, Martine Valette, Bruno Lina, Hélène Petitprez, Bruno Chanzy, Laurence Josset |
| EPI_ISL_666694 | National Virus Reference Lab (NVRL) | Irish Coronavirus Sequencing Consortium-Teagasc Grange | Matthew McCabe, Calum Walsh, Fiona Crispie, Paul Cotter, Michael Carr, Aljandro Abner Garcia Leon |
| EPI_ISL_671379, EPI_ISL_671380 | National Virus Reference Laboratory | Irish Coronavirus Sequencing Consortium - Teagasc Moorepark | Calum Walsh, Fiona Crispie, John Kenny, Paul Cotter |
| EPI_ISL_672172, EPI_ISL_672184 | The Ashley Laboratory, Stanford University | Chan-Zuckerberg Biohub | CZB Cliahub Consortium |
| EPI_ISL_672540, EPI_ISL_672541, EPI_ISL_672550, EPI_ISL_672552, EPI_ISL_672553, EPI_ISL_672554, EPI_ISL_672555, EPI_ISL_672556, EPI_ISL_672558, EPI_ISL_672559 | Utah Public Health Laboratory | Utah Public Health Laboratory | Erin Young, Kelly Oakeson |
| EPI_ISL_672672, EPI_ISL_672686 | DB Diagnosticos do Brasil | Laboratório de Parasitologia Médica - Instituto de Medicina Tropical - Universidade de São Paulo | Brazil-UK Centre for Arbovirus Discovery Diagnosis Genomics and Epidemiology (CADDE) Genomic Network - Instituto de Medicina Tropical |
| EPI_ISL_676576 | Scientific Veterinary Institute Novi Sad | Veterinary Specialized Institute "Kraljevo", Serbia | Vidanovic,D., Tesovic,B., Knezevic,A., Jovanovic,T., Jankovic,M., Sekler,M., Banovic Djeri,B., Petrovic,T., Volkening,J., Afonso,C. |

| | | | |
|---|---|---|---|
| EPI_ISL_676655, EPI_ISL_676792, EPI_ISL_676794, EPI_ISL_676795, EPI_ISL_676796 | Wadsworth Center, New York State Department.of Health | Wadsworth Center, New York State Department.of Health | Kirsten St. George, Daryl M. Lamson, Alexis Russel, Jonathan Plitnick, Navjot Singh, John Kelly, Sara Griesemer, Erasmus Schneider, Erica Lasek-Nesselquist |
| EPI_ISL_677944 | Pathogen Genomics Lab King Abdullah University of Science and Technology(KAUST) | Pathogen Genomics Lab King Abdullah University of Science and Technology(KAUST) | Sara Mfarrej, Olga Douvropoulou, Raushan Nugmanova, Sharif Hala, Raeece Naeem, Afrah Alsomali, Fadwa Alofi, Asim Khogeer, Afrah Alsomali, Jumana Taha, Abdulaziz Alahmadi, Kahled Algithami, Anwar Hashem, Naif Almontashiri, Arnab Pain |
| EPI_ISL_677946 | Pathogen Genomics Lab King Abdullah University of Science and Technology(KAUST) | Pathogen Genomics Lab King Abdullah University of Science and Technology(KAUST) | Fathia Ben Rached, Sharif Hala, Amit Kumar Subudhi, Sara Mfarrej, Raeece Naeem, Rahul P Salunke, Fadwa Alofi, Afrah Alsomali, Asim Khogeer, Jumana Taha, Abdulaziz Alahmadi, Kahled Algithami, Anwar Hashem, Naif Almontashiri, Arnab Pain |
| EPI_ISL_677947 | Pathogen Genomics Lab King Abdullah University of Science and Technology(KAUST) | Pathogen Genomics Lab King Abdullah University of Science and Technology(KAUST) | Sara Mfarrej, Amanda Ooi, Luke Esau, Sharif Hala, Raeece Naeem, Fadwa Alofi, Afrah Alsomali, Asim Khogeer, Jumana Taha, Abdulaziz Alahmadi, Kahled Algithami, Anwar Hashem, Naif Almontashiri, Arnab Pain |
| EPI_ISL_677955 | Pathogen Genomics Lab King Abdullah University of Science and Technology(KAUST) | Pathogen Genomics Lab King Abdullah University of Science and Technology(KAUST) | Sara Mfarrej, Olga Douvropoulou, Raushan Nugmanova, Sharif Hala, Raeece Naeem, Amanda Ooi, Luke Esau, Fadwa Alofi, Afrah Alsomali, Asim Khogeer, Jumana Taha, Abdulaziz Alahmadi, Kahled Algithami, Anwar Hashem, Naif Almontashiri, Arnab Pain |
| EPI_ISL_677958, EPI_ISL_677960, EPI_ISL_677963 | Pathogen Genomics Lab King Abdullah University of Science and Technology(KAUST) | Pathogen Genomics Lab King Abdullah University of Science and Technology(KAUST) | Fathia Ben Rached, Sharif Hala, Amit Kumar Subudhi, Sara Mfarrej, Raeece Naeem, Rahul P Salunke, Fadwa Alofi, Afrah Alsomali, Asim Khogeer, Jumana Taha, Abdulaziz Alahmadi, Kahled Algithami, Anwar Hashem, Naif Almontashiri, Arnab Pain |
| EPI_ISL_677966 | Pathogen Genomics Lab King Abdullah University of Science and Technology(KAUST) | Pathogen Genomics Lab King Abdullah University of Science and Technology(KAUST) | Sara Mfarrej, Amanda Ooi, Luke Esau, Sharif Hala, Raeece Naeem, Fadwa Alofi, Afrah Alsomali, Asim Khogeer, Jumana Taha, Abdulaziz Alahmadi, Kahled Algithami, Anwar Hashem, Naif Almontashiri, Arnab Pain |
| EPI_ISL_677968, EPI_ISL_677970 | Pathogen Genomics Lab King Abdullah University of Science and Technology(KAUST) | Pathogen Genomics Lab King Abdullah University of Science and Technology(KAUST) | Fathia Ben Rached, Sharif Hala, Amit Kumar Subudhi, Sara Mfarrej, Raeece Naeem, Rahul P Salunke, Fadwa Alofi, Afrah Alsomali, Asim Khogeer, Jumana Taha, Abdulaziz Alahmadi, Kahled Algithami, Anwar Hashem, Naif Almontashiri, Arnab Pain |
| EPI_ISL_677973 | Pathogen Genomics Lab King Abdullah University of Science and Technology(KAUST) | Pathogen Genomics Lab King Abdullah University of Science and Technology(KAUST) | Sara Mfarrej, Amanda Ooi, Luke Esau, Sharif Hala, Raeece Naeem, Fadwa Alofi, Afrah Alsomali, Asim Khogeer, Jumana Taha, Abdulaziz Alahmadi, Kahled Algithami, Anwar Hashem, Naif Almontashiri, Arnab Pain |
| EPI_ISL_678026 | Pathogen Genomics Lab King Abdullah University of Science and Technology(KAUST) | Pathogen Genomics Lab King Abdullah University of Science and Technology(KAUST) | Sharif Hala, Sara Mfarrej, Raeece Naeem, Amit Kumar Subudhi, Rahul P Salunke, Fadwa Alofi, Asim Khogeer, Afrah Alsomali, Jumana Taha, Abdulaziz Alahmadi, Kahled Algithami, Anwar Hashem, Naif Almontashiri, Arnab Pain |
| EPI_ISL_678028 | Pathogen Genomics Lab King Abdullah University of Science and Technology(KAUST) | Pathogen Genomics Lab King Abdullah University of Science and Technology(KAUST) | Fathia Ben Rached, Sharif Hala, Amit Kumar Subudhi, Sara Mfarrej, Raeece Naeem, Rahul P Salunke, Fadwa Alofi, Afrah Alsomali, Asim Khogeer, Jumana Taha, Abdulaziz Alahmadi, Kahled Algithami, Anwar Hashem, Naif Almontashiri, Arnab Pain |
| EPI_ISL_678029 | Pathogen Genomics Lab King Abdullah University of Science and Technology(KAUST) | Pathogen Genomics Lab King Abdullah University of Science and Technology(KAUST) | Sharif Hala, Sara Mfarrej, Raushan Nugmanova, Olga Douvropoulou, Raeece Naeem, Fadwa Alofi, Asim Khogeer, Afrah Alsomali, Jumana Taha, Abdulaziz Alahmadi, Kahled Algithami, Anwar Hashem, Naif Almontashiri, Arnab Pain |
| EPI_ISL_678035, EPI_ISL_678036, EPI_ISL_678038 | Pathogen Genomics Lab King Abdullah University of Science and Technology(KAUST) | Pathogen Genomics Lab King Abdullah University of Science and Technology(KAUST) | Sara Mfarrej, Amanda Ooi, Luke Esau, Sharif Hala, Raeece Naeem, Fadwa Alofi, Afrah Alsomali, Asim Khogeer, Jumana Taha, Abdulaziz Alahmadi, Kahled Algithami, Anwar Hashem, Naif Almontashiri, Arnab Pain |
| EPI_ISL_678234 | Pathogen Genomics Lab King Abdullah University of Science and Technology(KAUST) | Pathogen Genomics Lab King Abdullah University of Science and Technology(KAUST) | Sharif Hala, Sara Mfarrej, Raeece Naeem, Amit Kumar Subudhi, Rahul P Salunke, Fadwa Alofi, Asim Khogeer, Afrah Alsomali, Jumana Taha, Abdulaziz Alahmadi, Kahled Algithami, Anwar Hashem, Naif Almontashiri, Arnab Pain |
| EPI_ISL_678235 | Pathogen Genomics Lab King Abdullah University of Science and Technology(KAUST) | Pathogen Genomics Lab King Abdullah University of Science and Technology(KAUST) | Fathia Ben Rached, Sharif Hala, Amit Kumar Subudhi, Sara Mfarrej, Raeece Naeem, Rahul P Salunke, Fadwa Alofi, Afrah Alsomali, Asim Khogeer, Jumana Taha, Abdulaziz Alahmadi, Kahled Algithami, Anwar Hashem, Naif Almontashiri, Arnab Pain |
| EPI_ISL_678502 | CNR Virus des Infections Respiratoires - France SUD | CNR Virus des Infections Respiratoires - France SUD | Antonin Bal, Gregory Destras, Gwendolyne Burfin, Solenne Brun, Martine Valette, Bruno Lina, Laurence Josset |
| EPI_ISL_680283, EPI_ISL_680284, EPI_ISL_680285 | Regional Virus Laboratory, Belfast Health and Social Care Trust | COVID-19 Genomics UK (COG-UK) Consortium | Conall McCaughey, James McKenna, Tanya Curran, Susan Feeney, Alison Watt, Ciara Cox, Mairead Connor, Zoltan Molnar, David Simpson, Derek Fairley |
| EPI_ISL_681721, EPI_ISL_681728, EPI_ISL_681731, EPI_ISL_681752, EPI_ISL_681754, EPI_ISL_681759, EPI_ISL_681760, EPI_ISL_681764, EPI_ISL_681767 | University Hospital Limerick | Irish Coronavirus Sequencing Consortium - Teagasc Moorepark | Paul Cotter, Fiona Crispie, John Kenny, Calum Walsh |
| EPI_ISL_681920, EPI_ISL_682057 | National Virus Reference Laboratory | Irish Coronavirus Sequencing Consortium - Teagasc Moorepark | Paul Cotter, Fiona Crispie, John Kenny, Matthew McCabe, Calum Walsh |
| EPI_ISL_684166, EPI_ISL_684167, EPI_ISL_684174, EPI_ISL_684175, EPI_ISL_684176, EPI_ISL_684177, EPI_ISL_684178, EPI_ISL_684179, EPI_ISL_684180, EPI_ISL_684181, EPI_ISL_684182, EPI_ISL_684183, EPI_ISL_684184, EPI_ISL_684185, EPI_ISL_684186, EPI_ISL_684187, EPI_ISL_684188, EPI_ISL_684189, EPI_ISL_684190, EPI_ISL_684191, EPI_ISL_684192, EPI_ISL_684193, EPI_ISL_684194, EPI_ISL_684195, EPI_ISL_684196, EPI_ISL_684197, EPI_ISL_684198, EPI_ISL_684199, EPI_ISL_684200, EPI_ISL_684201, EPI_ISL_684202, EPI_ISL_684203, EPI_ISL_684204, EPI_ISL_684205, EPI_ISL_684206, EPI_ISL_684207, EPI_ISL_684208, EPI_ISL_684209, EPI_ISL_684210, EPI_ISL_684211, EPI_ISL_684212, EPI_ISL_684213, EPI_ISL_684214, EPI_ISL_684215, EPI_ISL_684216, EPI_ISL_684217, EPI_ISL_684218 | | | |
| see above | Pathogen Genomics Center, National Institute of Infectious Diseases | Pathogen Genomics Center, National Institute of Infectious Diseases | Tsuyoshi Sekizuka, Kentaro Itokawa, Rina Tanaka, Masanori Hashino, Makoto Kuroda |
| EPI_ISL_684220, EPI_ISL_684221, EPI_ISL_684222, EPI_ISL_684223, EPI_ISL_684224, EPI_ISL_684225, EPI_ISL_684226, EPI_ISL_684227, EPI_ISL_684228, EPI_ISL_684229, EPI_ISL_684230, EPI_ISL_684231, EPI_ISL_684232, EPI_ISL_684233, EPI_ISL_684234, EPI_ISL_684235, EPI_ISL_684236, EPI_ISL_684237, EPI_ISL_684238, EPI_ISL_684239, EPI_ISL_684240, EPI_ISL_684241, EPI_ISL_684242, EPI_ISL_684243, EPI_ISL_684244, EPI_ISL_684245, EPI_ISL_684246, EPI_ISL_684247, EPI_ISL_684248, EPI_ISL_684249, EPI_ISL_684250, EPI_ISL_684251, EPI_ISL_684252, EPI_ISL_684253, EPI_ISL_684254, EPI_ISL_684255, EPI_ISL_684256, EPI_ISL_684257, EPI_ISL_684258, EPI_ISL_684259, EPI_ISL_684260, EPI_ISL_684261, EPI_ISL_684262, EPI_ISL_684263, EPI_ISL_684264, EPI_ISL_684265, EPI_ISL_684266, EPI_ISL_684267, EPI_ISL_684268, EPI_ISL_684269, EPI_ISL_684270, EPI_ISL_684271, EPI_ISL_684272, EPI_ISL_684273, EPI_ISL_684274, EPI_ISL_684275, EPI_ISL_684276, EPI_ISL_684277, EPI_ISL_684278, EPI_ISL_684279, EPI_ISL_684280, EPI_ISL_684281, EPI_ISL_684282, EPI_ISL_684283, EPI_ISL_684284, EPI_ISL_684285, EPI_ISL_684286, EPI_ISL_684287, EPI_ISL_684288 | | | |
| see above | Gunma Prefectural Institute of Public Health and Environmental Sciences | Pathogen Genomics Center, National Institute of Infectious Diseases | Tsuyoshi Sekizuka, Kentaro Itokawa, Rina Tanaka, Masanori Hashino, Makoto Kuroda |
| EPI_ISL_684289, EPI_ISL_684290, EPI_ISL_684291, EPI_ISL_684292, EPI_ISL_684293, EPI_ISL_684294, EPI_ISL_684295, EPI_ISL_684297, EPI_ISL_684298, EPI_ISL_684299, EPI_ISL_684300, EPI_ISL_684301, EPI_ISL_684302, EPI_ISL_684303, EPI_ISL_684305, EPI_ISL_684306, EPI_ISL_684307, EPI_ISL_684308, EPI_ISL_684309, EPI_ISL_684310, EPI_ISL_684311, EPI_ISL_684312, EPI_ISL_684313, EPI_ISL_684314, EPI_ISL_684315, EPI_ISL_684316, EPI_ISL_684317, EPI_ISL_684318, EPI_ISL_684319, EPI_ISL_684320, EPI_ISL_684321, EPI_ISL_684322, EPI_ISL_684323, EPI_ISL_684324 | | | |
| see above | Pathogen Genomics Center, National Institute of Infectious Diseases | Pathogen Genomics Center, National Institute of Infectious Diseases | Tsuyoshi Sekizuka, Kentaro Itokawa, Rina Tanaka, Masanori Hashino, Makoto Kuroda |
| EPI_ISL_684325, EPI_ISL_684326, EPI_ISL_684327, EPI_ISL_684328, EPI_ISL_684329, EPI_ISL_684330, EPI_ISL_684331, EPI_ISL_684332, EPI_ISL_684333, EPI_ISL_684334, EPI_ISL_684335, EPI_ISL_684336, EPI_ISL_684337, EPI_ISL_684338 | | | |
| see above | Gunma Prefectural Institute of Public Health and Environmental Sciences | Pathogen Genomics Center, National Institute of Infectious Diseases | Tsuyoshi Sekizuka, Kentaro Itokawa, Rina Tanaka, Masanori Hashino, Makoto Kuroda |
| EPI_ISL_684339, EPI_ISL_684340, EPI_ISL_684341, EPI_ISL_684342, EPI_ISL_684343, EPI_ISL_684344, EPI_ISL_684345, EPI_ISL_684346, EPI_ISL_684347, EPI_ISL_684348, EPI_ISL_684349, EPI_ISL_684350, EPI_ISL_684351, EPI_ISL_684352, EPI_ISL_684353, EPI_ISL_684354, EPI_ISL_684355, EPI_ISL_684356, EPI_ISL_684357, EPI_ISL_684358, EPI_ISL_684359, EPI_ISL_684360, EPI_ISL_684362, EPI_ISL_684363, EPI_ISL_684364, EPI_ISL_684365, EPI_ISL_684366, EPI_ISL_684367, EPI_ISL_684368, EPI_ISL_684369, EPI_ISL_684385, EPI_ISL_684386, EPI_ISL_684387, EPI_ISL_684388, EPI_ISL_684389, EPI_ISL_684390, EPI_ISL_684391, EPI_ISL_684392, EPI_ISL_684393, EPI_ISL_684394, EPI_ISL_684395, EPI_ISL_684396, EPI_ISL_684397, EPI_ISL_684398, EPI_ISL_684399, EPI_ISL_684400, EPI_ISL_684401, EPI_ISL_684402, EPI_ISL_684403, EPI_ISL_684404, EPI_ISL_684405, EPI_ISL_684406, EPI_ISL_684407 | | | |
| see above | Pathogen Genomics Center, National Institute of Infectious Diseases | Pathogen Genomics Center, National Institute of Infectious Diseases | Tsuyoshi Sekizuka, Kentaro Itokawa, Rina Tanaka, Masanori Hashino, Makoto Kuroda |
| EPI_ISL_684408, EPI_ISL_684409, EPI_ISL_684410, EPI_ISL_684411, EPI_ISL_684412, EPI_ISL_684413, EPI_ISL_684414, EPI_ISL_684415, EPI_ISL_684416, EPI_ISL_684417, EPI_ISL_684418, EPI_ISL_684419, EPI_ISL_684420, EPI_ISL_684423, EPI_ISL_684424, EPI_ISL_684425, EPI_ISL_684426, EPI_ISL_684427, EPI_ISL_684428, EPI_ISL_684430, EPI_ISL_684431, EPI_ISL_684432, EPI_ISL_684433, EPI_ISL_684440, EPI_ISL_684441, EPI_ISL_684442, EPI_ISL_684443, EPI_ISL_684444, EPI_ISL_684445, EPI_ISL_684446, EPI_ISL_684447, EPI_ISL_684448, EPI_ISL_684449, EPI_ISL_684450, EPI_ISL_684451, EPI_ISL_684452, EPI_ISL_684453, EPI_ISL_684454, EPI_ISL_684455, EPI_ISL_684456, EPI_ISL_684457, EPI_ISL_684458, EPI_ISL_684459, EPI_ISL_684460, EPI_ISL_684461, EPI_ISL_684462, EPI_ISL_684463, EPI_ISL_684464, EPI_ISL_684465, EPI_ISL_684466, EPI_ISL_684467, EPI_ISL_684468, EPI_ISL_684469, EPI_ISL_684470 | | | |
| see above | Fukuoka Institute of Health and Environmental Sciences | Pathogen Genomics Center, National Institute of Infectious Diseases | Tsuyoshi Sekizuka, Kentaro Itokawa, Rina Tanaka, Masanori Hashino, Makoto Kuroda |
| EPI_ISL_684484, EPI_ISL_684485, EPI_ISL_684489, EPI_ISL_684490, EPI_ISL_684491, EPI_ISL_684494, EPI_ISL_684495, EPI_ISL_684496, EPI_ISL_684497, EPI_ISL_684498, EPI_ISL_684499, EPI_ISL_684502, EPI_ISL_684503, EPI_ISL_684504, EPI_ISL_684505 | | | |
| see above | Pathogen Genomics Center, National Institute of Infectious Diseases | Pathogen Genomics Center, National Institute of Infectious Diseases | Tsuyoshi Sekizuka, Kentaro Itokawa, Rina Tanaka, Masanori Hashino, Makoto Kuroda |

| | | | |
|---|---|---|---|
| EPI_ISL_684515, EPI_ISL_684516, EPI_ISL_684518 | Tokyo Metropolitan Institute of Public Health | Pathogen Genomics Center, National Institute of Infectious Diseases | Tsuyoshi Sekizuka, Kentaro Itokawa, Rina Tanaka, Masanori Hashino, Makoto Kuroda |
| EPI_ISL_684523, EPI_ISL_684524, EPI_ISL_684525, EPI_ISL_684526, EPI_ISL_684527, EPI_ISL_684528, EPI_ISL_684529, EPI_ISL_684530, EPI_ISL_684531, EPI_ISL_684532, EPI_ISL_684533, EPI_ISL_684534, EPI_ISL_684535, EPI_ISL_684536, EPI_ISL_684537, EPI_ISL_684538, EPI_ISL_684539, EPI_ISL_684540, EPI_ISL_684541, EPI_ISL_684542, EPI_ISL_684543, EPI_ISL_684544, EPI_ISL_684545, EPI_ISL_684546, EPI_ISL_684547, EPI_ISL_684548, EPI_ISL_684549, EPI_ISL_684550, EPI_ISL_684551, EPI_ISL_684552, EPI_ISL_684553, EPI_ISL_684554, EPI_ISL_684555 | | | |
| see above | Pathogen Genomics Center, National Institute of Infectious Diseases | Pathogen Genomics Center, National Institute of Infectious Diseases | Tsuyoshi Sekizuka, Kentaro Itokawa, Rina Tanaka, Masanori Hashino, Makoto Kuroda |
| EPI_ISL_684557 | Tokyo Metropolitan Institute of Public Health | Pathogen Genomics Center, National Institute of Infectious Diseases | Tsuyoshi Sekizuka, Kentaro Itokawa, Rina Tanaka, Masanori Hashino, Makoto Kuroda |
| EPI_ISL_684558, EPI_ISL_684559, EPI_ISL_684560, EPI_ISL_684561, EPI_ISL_684562, EPI_ISL_684563, EPI_ISL_684564, EPI_ISL_684565 | Pathogen Genomics Center, National Institute of Infectious Diseases | Pathogen Genomics Center, National Institute of Infectious Diseases | Tsuyoshi Sekizuka, Kentaro Itokawa, Rina Tanaka, Masanori Hashino, Makoto Kuroda |
| EPI_ISL_684566, EPI_ISL_684567, EPI_ISL_684568, EPI_ISL_684569, EPI_ISL_684570, EPI_ISL_684571, EPI_ISL_684572, EPI_ISL_684573, EPI_ISL_684574, EPI_ISL_684575, EPI_ISL_684576, EPI_ISL_684577 | | | |
| see above | Gunma Prefectural Institute of Public Health and Environmental Sciences | Pathogen Genomics Center, National Institute of Infectious Diseases | Tsuyoshi Sekizuka, Kentaro Itokawa, Rina Tanaka, Masanori Hashino, Makoto Kuroda |
| EPI_ISL_684578, EPI_ISL_684579, EPI_ISL_684580, EPI_ISL_684581, EPI_ISL_684582, EPI_ISL_684583, EPI_ISL_684584, EPI_ISL_684585, EPI_ISL_684586, EPI_ISL_684587, EPI_ISL_684588, EPI_ISL_684589, EPI_ISL_684590, EPI_ISL_684591, EPI_ISL_684592, EPI_ISL_684593, EPI_ISL_684594, EPI_ISL_684595, EPI_ISL_684596, EPI_ISL_684597, EPI_ISL_684598, EPI_ISL_684599, EPI_ISL_684600, EPI_ISL_684601, EPI_ISL_684602, EPI_ISL_684603, EPI_ISL_684604, EPI_ISL_684605, EPI_ISL_684606, EPI_ISL_684607, EPI_ISL_684608, EPI_ISL_684609, EPI_ISL_684610, EPI_ISL_684611, EPI_ISL_684612, EPI_ISL_684613, EPI_ISL_684614 | | | |
| see above | Pathogen Genomics Center, National Institute of Infectious Diseases | Pathogen Genomics Center, National Institute of Infectious Diseases | Tsuyoshi Sekizuka, Kentaro Itokawa, Rina Tanaka, Masanori Hashino, Makoto Kuroda |
| EPI_ISL_684632, EPI_ISL_684633 | Tokyo Metropolitan Institute of Public Health | Pathogen Genomics Center, National Institute of Infectious Diseases | Tsuyoshi Sekizuka, Kentaro Itokawa, Rina Tanaka, Masanori Hashino, Makoto Kuroda |
| EPI_ISL_684634, EPI_ISL_684635, EPI_ISL_684636, EPI_ISL_684637, EPI_ISL_684638, EPI_ISL_684639, EPI_ISL_684640, EPI_ISL_684641, EPI_ISL_684642, EPI_ISL_684643, EPI_ISL_684644, EPI_ISL_684645, EPI_ISL_684646, EPI_ISL_684647, EPI_ISL_684648, EPI_ISL_684649, EPI_ISL_684650, EPI_ISL_684651, EPI_ISL_684652, EPI_ISL_684653, EPI_ISL_684654, EPI_ISL_684655, EPI_ISL_684656, EPI_ISL_684657, EPI_ISL_684658, EPI_ISL_684659, EPI_ISL_684660, EPI_ISL_684661, EPI_ISL_684662, EPI_ISL_684663, EPI_ISL_684664, EPI_ISL_684665, EPI_ISL_684666, EPI_ISL_684667, EPI_ISL_684668, EPI_ISL_684669, EPI_ISL_684670, EPI_ISL_684671, EPI_ISL_684672, EPI_ISL_684673, EPI_ISL_684674, EPI_ISL_684675, EPI_ISL_684676, EPI_ISL_684677, EPI_ISL_684678, EPI_ISL_684679, EPI_ISL_684680, EPI_ISL_684681, EPI_ISL_684682, EPI_ISL_684683, EPI_ISL_684684, EPI_ISL_684685, EPI_ISL_684686, EPI_ISL_684687, EPI_ISL_684688, EPI_ISL_684689, EPI_ISL_684690, EPI_ISL_684691, EPI_ISL_684692 | | | |
| see above | Fukuoka Institute of Health and Environmental Sciences | Pathogen Genomics Center, National Institute of Infectious Diseases | Tsuyoshi Sekizuka, Kentaro Itokawa, Rina Tanaka, Masanori Hashino, Makoto Kuroda |
| EPI_ISL_684695, EPI_ISL_684696, EPI_ISL_684697, EPI_ISL_684698, EPI_ISL_684699, EPI_ISL_684700, EPI_ISL_684701, EPI_ISL_684702, EPI_ISL_684703, EPI_ISL_684704, EPI_ISL_684705, EPI_ISL_684706, EPI_ISL_684707, EPI_ISL_684708, EPI_ISL_684709, EPI_ISL_684710, EPI_ISL_684711, EPI_ISL_684712, EPI_ISL_684713, EPI_ISL_684714, EPI_ISL_684715, EPI_ISL_684719, EPI_ISL_684720, EPI_ISL_684721, EPI_ISL_684722 | | | |
| see above | Pathogen Genomics Center, National Institute of Infectious Diseases | Pathogen Genomics Center, National Institute of Infectious Diseases | Tsuyoshi Sekizuka, Kentaro Itokawa, Rina Tanaka, Masanori Hashino, Makoto Kuroda |
| EPI_ISL_684723, EPI_ISL_684724, EPI_ISL_684725, EPI_ISL_684726, EPI_ISL_684727 | Fukuoka Institute of Health and Environmental Sciences | Pathogen Genomics Center, National Institute of Infectious Diseases | Tsuyoshi Sekizuka, Kentaro Itokawa, Rina Tanaka, Masanori Hashino, Makoto Kuroda |
| EPI_ISL_684728, EPI_ISL_684729, EPI_ISL_684730, EPI_ISL_684731, EPI_ISL_684732, EPI_ISL_684733, EPI_ISL_684734, EPI_ISL_684735 | Tokyo Metropolitan Institute of Public Health | Pathogen Genomics Center, National Institute of Infectious Diseases | Tsuyoshi Sekizuka, Kentaro Itokawa, Rina Tanaka, Masanori Hashino, Makoto Kuroda |
| EPI_ISL_684737, EPI_ISL_684738, EPI_ISL_684739, EPI_ISL_684740, EPI_ISL_684741, EPI_ISL_684742, EPI_ISL_684743, EPI_ISL_684744, EPI_ISL_684745, EPI_ISL_684746, EPI_ISL_684747, EPI_ISL_684748, EPI_ISL_684749, EPI_ISL_684750, EPI_ISL_684751, EPI_ISL_684752, EPI_ISL_684753, EPI_ISL_684754, EPI_ISL_684755, EPI_ISL_684756, EPI_ISL_684757, EPI_ISL_684758, EPI_ISL_684759, EPI_ISL_684760, EPI_ISL_684761, EPI_ISL_684764, EPI_ISL_684765, EPI_ISL_684766, EPI_ISL_684767, EPI_ISL_684768, EPI_ISL_684769, EPI_ISL_684770, EPI_ISL_684771, EPI_ISL_684772, EPI_ISL_684773, EPI_ISL_684774, EPI_ISL_684775, EPI_ISL_684776, EPI_ISL_684777, EPI_ISL_684778, EPI_ISL_684779, EPI_ISL_684780, EPI_ISL_684781, EPI_ISL_684782, EPI_ISL_684783, EPI_ISL_684784, EPI_ISL_684785, EPI_ISL_684786, EPI_ISL_684787, EPI_ISL_684788, EPI_ISL_684789, EPI_ISL_684790, EPI_ISL_684791, EPI_ISL_684792, EPI_ISL_684793, EPI_ISL_684794, EPI_ISL_684795, EPI_ISL_684796, EPI_ISL_684797 | | | |
| see above | Pathogen Genomics Center, National Institute of Infectious Diseases | Pathogen Genomics Center, National Institute of Infectious Diseases | Tsuyoshi Sekizuka, Kentaro Itokawa, Rina Tanaka, Masanori Hashino, Makoto Kuroda |
| EPI_ISL_684800, EPI_ISL_684801, EPI_ISL_684802, EPI_ISL_684803, EPI_ISL_684804, EPI_ISL_684805, EPI_ISL_684806, EPI_ISL_684807, EPI_ISL_684808, EPI_ISL_684809, EPI_ISL_684810, EPI_ISL_684811, EPI_ISL_684812, EPI_ISL_684813, EPI_ISL_684814, EPI_ISL_684815, EPI_ISL_684816, EPI_ISL_684817, EPI_ISL_684818, EPI_ISL_684819, EPI_ISL_684820, EPI_ISL_684821, EPI_ISL_684822, EPI_ISL_684823, EPI_ISL_684824, EPI_ISL_684825, EPI_ISL_684826, EPI_ISL_684827, EPI_ISL_684828, EPI_ISL_684829, EPI_ISL_684830, EPI_ISL_684831, EPI_ISL_684832, EPI_ISL_684833, EPI_ISL_684834, EPI_ISL_684835, EPI_ISL_684836, EPI_ISL_684837, EPI_ISL_684838, EPI_ISL_684839, EPI_ISL_684840, EPI_ISL_684841, EPI_ISL_684842, EPI_ISL_684843, EPI_ISL_684844, EPI_ISL_684845, EPI_ISL_684846, EPI_ISL_684847, EPI_ISL_684848, EPI_ISL_684849, EPI_ISL_684850, EPI_ISL_684851, EPI_ISL_684852, EPI_ISL_684853, EPI_ISL_684854, EPI_ISL_684855, EPI_ISL_684856 | | | |
| see above | Yamagata Prefectural Institute of Public Health | Pathogen Genomics Center, National Institute of Infectious Diseases | Tsuyoshi Sekizuka, Kentaro Itokawa, Rina Tanaka, Masanori Hashino, Makoto Kuroda |
| EPI_ISL_684910, EPI_ISL_684911, EPI_ISL_684912, EPI_ISL_684913, EPI_ISL_684914, EPI_ISL_684915, EPI_ISL_684916, EPI_ISL_684917, EPI_ISL_684918, EPI_ISL_684919, EPI_ISL_684920, EPI_ISL_684921, EPI_ISL_684922, EPI_ISL_684923, EPI_ISL_684924, EPI_ISL_684925, EPI_ISL_684926, EPI_ISL_684927, EPI_ISL_684928, EPI_ISL_684929, EPI_ISL_684930, EPI_ISL_684931, EPI_ISL_684932, EPI_ISL_684933, EPI_ISL_684934, EPI_ISL_684935, EPI_ISL_684936, EPI_ISL_684937, EPI_ISL_684938, EPI_ISL_684939, EPI_ISL_684940, EPI_ISL_684941, EPI_ISL_684942, EPI_ISL_684943, EPI_ISL_684944, EPI_ISL_684945, EPI_ISL_684946, EPI_ISL_684947, EPI_ISL_684948, EPI_ISL_684949, EPI_ISL_684950, EPI_ISL_684951, EPI_ISL_684952, EPI_ISL_684953, EPI_ISL_684954, EPI_ISL_684955, EPI_ISL_684956, EPI_ISL_684957, EPI_ISL_684958, EPI_ISL_684959, EPI_ISL_684960, EPI_ISL_684961, EPI_ISL_684962, EPI_ISL_684963, EPI_ISL_684964, EPI_ISL_684965, EPI_ISL_684966, EPI_ISL_684967, EPI_ISL_684968, EPI_ISL_684969, EPI_ISL_684970, EPI_ISL_684971, EPI_ISL_684972, EPI_ISL_684973, EPI_ISL_684974, EPI_ISL_684975, EPI_ISL_684976, EPI_ISL_684977, EPI_ISL_684978, EPI_ISL_684979, EPI_ISL_684980, EPI_ISL_684981, EPI_ISL_684982, EPI_ISL_684983, EPI_ISL_684984, EPI_ISL_684985, EPI_ISL_684986, EPI_ISL_684987, EPI_ISL_684988, EPI_ISL_684989, EPI_ISL_684990, EPI_ISL_684991, EPI_ISL_684992, EPI_ISL_684993, EPI_ISL_684994, EPI_ISL_684995, EPI_ISL_684996, EPI_ISL_684997, EPI_ISL_684998, EPI_ISL_684999, EPI_ISL_685000, EPI_ISL_685001, EPI_ISL_685002, EPI_ISL_685003, EPI_ISL_685004, EPI_ISL_685005, EPI_ISL_685006, EPI_ISL_685007, EPI_ISL_685008, EPI_ISL_685009, EPI_ISL_685010, EPI_ISL_685011, EPI_ISL_685012, EPI_ISL_685013, EPI_ISL_685014, EPI_ISL_685015, EPI_ISL_685016, EPI_ISL_685017, EPI_ISL_685018, EPI_ISL_685019, EPI_ISL_685020, EPI_ISL_685021, EPI_ISL_685022, EPI_ISL_685023, EPI_ISL_685024, EPI_ISL_685025, EPI_ISL_685026, EPI_ISL_685027, EPI_ISL_685028, EPI_ISL_685029, EPI_ISL_685030, EPI_ISL_685031, EPI_ISL_685032, EPI_ISL_685033, EPI_ISL_685034, EPI_ISL_685035, EPI_ISL_685036, EPI_ISL_685037, EPI_ISL_685038, EPI_ISL_685039, EPI_ISL_685040, EPI_ISL_685041, EPI_ISL_685042, EPI_ISL_685043, EPI_ISL_685044, EPI_ISL_685045, EPI_ISL_685046, EPI_ISL_685047, EPI_ISL_685048, EPI_ISL_685049, EPI_ISL_685050, EPI_ISL_685051, EPI_ISL_685052, EPI_ISL_685053, EPI_ISL_685054, EPI_ISL_685055, EPI_ISL_685056, EPI_ISL_685057, EPI_ISL_685058, EPI_ISL_685059, EPI_ISL_685060, EPI_ISL_685061, EPI_ISL_685062, EPI_ISL_685063, EPI_ISL_685064, EPI_ISL_685065, EPI_ISL_685066, EPI_ISL_685067, EPI_ISL_685068, EPI_ISL_685069, EPI_ISL_685070, EPI_ISL_685071, EPI_ISL_685072, EPI_ISL_685073, EPI_ISL_685074, EPI_ISL_685075, EPI_ISL_685076, EPI_ISL_685077, EPI_ISL_685078, EPI_ISL_685079, EPI_ISL_685080, EPI_ISL_685081, EPI_ISL_685082, EPI_ISL_685083, EPI_ISL_685084, EPI_ISL_685085, EPI_ISL_685086, EPI_ISL_685087, EPI_ISL_685088, EPI_ISL_685089, EPI_ISL_685090, EPI_ISL_685091, EPI_ISL_685092, EPI_ISL_685093, EPI_ISL_685094, EPI_ISL_685095, EPI_ISL_685096, EPI_ISL_685097, EPI_ISL_685098, EPI_ISL_685099, EPI_ISL_685100, EPI_ISL_685101, EPI_ISL_685102, EPI_ISL_685103, EPI_ISL_685104, EPI_ISL_685105, EPI_ISL_685106, EPI_ISL_685107, EPI_ISL_685108, EPI_ISL_685109, EPI_ISL_685110, EPI_ISL_685111, EPI_ISL_685112, EPI_ISL_685113, EPI_ISL_685114, EPI_ISL_685115, EPI_ISL_685116, EPI_ISL_685117, EPI_ISL_685118, EPI_ISL_685119, EPI_ISL_685120, EPI_ISL_685121, EPI_ISL_685122, EPI_ISL_685123, EPI_ISL_685124, EPI_ISL_685125, EPI_ISL_685126, EPI_ISL_685127, EPI_ISL_685128, EPI_ISL_685129, EPI_ISL_685130, EPI_ISL_685131 | | | |
| see above | Saitama Prefectural Institute of Public Health | Pathogen Genomics Center, National Institute of Infectious Diseases | Tsuyoshi Sekizuka, Kentaro Itokawa, Rina Tanaka, Masanori Hashino, Makoto Kuroda |
| EPI_ISL_685136, EPI_ISL_685137, EPI_ISL_685138, EPI_ISL_685139, EPI_ISL_685140, EPI_ISL_685141, EPI_ISL_685142, EPI_ISL_685143, EPI_ISL_685144, EPI_ISL_685145, EPI_ISL_685146, EPI_ISL_685147, EPI_ISL_685148, EPI_ISL_685149, EPI_ISL_685150, EPI_ISL_685151, EPI_ISL_685152, EPI_ISL_685153, EPI_ISL_685154, EPI_ISL_685155, EPI_ISL_685156, EPI_ISL_685157, EPI_ISL_685158, EPI_ISL_685159, EPI_ISL_685160, EPI_ISL_685161, EPI_ISL_685162, EPI_ISL_685163, EPI_ISL_685164, EPI_ISL_685165, EPI_ISL_685166, EPI_ISL_685167, EPI_ISL_685168, EPI_ISL_685169, EPI_ISL_685170, EPI_ISL_685171, EPI_ISL_685172, EPI_ISL_685173, EPI_ISL_685174, EPI_ISL_685175, EPI_ISL_685176, EPI_ISL_685177, EPI_ISL_685178, EPI_ISL_685179, EPI_ISL_685180, EPI_ISL_685181, EPI_ISL_685182, EPI_ISL_685183, EPI_ISL_685184, EPI_ISL_685185, EPI_ISL_685186, EPI_ISL_685187, EPI_ISL_685188, EPI_ISL_685189, EPI_ISL_685190, EPI_ISL_685191, EPI_ISL_685192, EPI_ISL_685193, EPI_ISL_685194, EPI_ISL_685195, EPI_ISL_685196, EPI_ISL_685197, EPI_ISL_685198, EPI_ISL_685199, EPI_ISL_685200, EPI_ISL_685201, EPI_ISL_685202, EPI_ISL_685204, EPI_ISL_685205, EPI_ISL_685206, EPI_ISL_685207, EPI_ISL_685208, EPI_ISL_685209, EPI_ISL_685210, EPI_ISL_685211, EPI_ISL_685212, EPI_ISL_685213, EPI_ISL_685214, EPI_ISL_685215, EPI_ISL_685216, EPI_ISL_685217, EPI_ISL_685218, EPI_ISL_685219 | | | |
| see above | Pathogen Genomics Center, National Institute of Infectious Diseases | Pathogen Genomics Center, National Institute of Infectious Diseases | Tsuyoshi Sekizuka, Kentaro Itokawa, Rina Tanaka, Masanori Hashino, Makoto Kuroda |
| EPI_ISL_685221, EPI_ISL_685222, EPI_ISL_685223, EPI_ISL_685224, EPI_ISL_685225, EPI_ISL_685226, EPI_ISL_685227, EPI_ISL_685228, EPI_ISL_685229, EPI_ISL_685230, EPI_ISL_685231, EPI_ISL_685232, EPI_ISL_685233, EPI_ISL_685234, EPI_ISL_685235, EPI_ISL_685236, EPI_ISL_685237 | | | |

| | | | |
|---|---|---|---|
| see above | Tokyo Metropolitan Institute of Public Health | Pathogen Genomics Center, National Institute of Infectious Diseases | Tsuyoshi Sekizuka, Kentaro Itokawa, Rina Tanaka, Masanori Hashino, Makoto Kuroda |
| EPI_ISL_685244, EPI_ISL_685245, EPI_ISL_685246, EPI_ISL_685247, EPI_ISL_685248, EPI_ISL_685249, EPI_ISL_685250, EPI_ISL_685251, EPI_ISL_685252, EPI_ISL_685253, EPI_ISL_685254, EPI_ISL_685255, EPI_ISL_685256, EPI_ISL_685257, EPI_ISL_685258, EPI_ISL_685259, EPI_ISL_685260, EPI_ISL_685261, EPI_ISL_685262, EPI_ISL_685263, EPI_ISL_685264, EPI_ISL_685265, EPI_ISL_685266, EPI_ISL_685267, EPI_ISL_685268, EPI_ISL_685269, EPI_ISL_685270, EPI_ISL_685271, EPI_ISL_685272, EPI_ISL_685273, EPI_ISL_685274, EPI_ISL_685275, EPI_ISL_685276, EPI_ISL_685277, EPI_ISL_685278 | | | |
| see above | Pathogen Genomics Center, National Institute of Infectious Diseases | Pathogen Genomics Center, National Institute of Infectious Diseases | Tsuyoshi Sekizuka, Kentaro Itokawa, Rina Tanaka, Masanori Hashino, Makoto Kuroda |
| EPI_ISL_685279, EPI_ISL_685280 | Gunma Prefectural Institute of Public Health and Environmental Sciences | Pathogen Genomics Center, National Institute of Infectious Diseases | Tsuyoshi Sekizuka, Kentaro Itokawa, Rina Tanaka, Masanori Hashino, Makoto Kuroda |
| EPI_ISL_685282 | Pathogen Genomics Center, National Institute of Infectious Diseases | Pathogen Genomics Center, National Institute of Infectious Diseases | Tsuyoshi Sekizuka, Kentaro Itokawa, Rina Tanaka, Masanori Hashino, Makoto Kuroda |
| EPI_ISL_685299, EPI_ISL_685300, EPI_ISL_685301, EPI_ISL_685302, EPI_ISL_685303, EPI_ISL_685304, EPI_ISL_685305, EPI_ISL_685306, EPI_ISL_685307, EPI_ISL_685308, EPI_ISL_685309, EPI_ISL_685310 | | | |
| see above | Niigata Prefectural Institute of Public Health and Environmental Sciences | Pathogen Genomics Center, National Institute of Infectious Diseases | Tsuyoshi Sekizuka, Kentaro Itokawa, Rina Tanaka, Masanori Hashino, Makoto Kuroda |
| EPI_ISL_685313, EPI_ISL_685314, EPI_ISL_685315, EPI_ISL_685316, EPI_ISL_685317, EPI_ISL_685318, EPI_ISL_685319, EPI_ISL_685320, EPI_ISL_685321, EPI_ISL_685322, EPI_ISL_685323, EPI_ISL_685324, EPI_ISL_685325, EPI_ISL_685326, EPI_ISL_685327, EPI_ISL_685328, EPI_ISL_685329, EPI_ISL_685330, EPI_ISL_685331, EPI_ISL_685332, EPI_ISL_685333, EPI_ISL_685334, EPI_ISL_685335, EPI_ISL_685336, EPI_ISL_685337, EPI_ISL_685338, EPI_ISL_685339, EPI_ISL_685340, EPI_ISL_685341, EPI_ISL_685342, EPI_ISL_685343, EPI_ISL_685354, EPI_ISL_685355, EPI_ISL_685356, EPI_ISL_685357, EPI_ISL_685358, EPI_ISL_685359, EPI_ISL_685360, EPI_ISL_685361, EPI_ISL_685362, EPI_ISL_685363, EPI_ISL_685364, EPI_ISL_685365, EPI_ISL_685366, EPI_ISL_685367, EPI_ISL_685368, EPI_ISL_685369, EPI_ISL_685370, EPI_ISL_685371, EPI_ISL_685372, EPI_ISL_685373, EPI_ISL_685374, EPI_ISL_685375, EPI_ISL_685376, EPI_ISL_685377, EPI_ISL_685378, EPI_ISL_685379, EPI_ISL_685380, EPI_ISL_685381, EPI_ISL_685382, EPI_ISL_685383, EPI_ISL_685384, EPI_ISL_685385, EPI_ISL_685386, EPI_ISL_685387, EPI_ISL_685388, EPI_ISL_685389 | | | |
| see above | Pathogen Genomics Center, National Institute of Infectious Diseases | Pathogen Genomics Center, National Institute of Infectious Diseases | Tsuyoshi Sekizuka, Kentaro Itokawa, Rina Tanaka, Masanori Hashino, Makoto Kuroda |
| EPI_ISL_685407, EPI_ISL_685408, EPI_ISL_685409, EPI_ISL_685410, EPI_ISL_685411, EPI_ISL_685412, EPI_ISL_685413, EPI_ISL_685414, EPI_ISL_685415, EPI_ISL_685416, EPI_ISL_685417, EPI_ISL_685418, EPI_ISL_685419, EPI_ISL_685420, EPI_ISL_685421, EPI_ISL_685422, EPI_ISL_685423, EPI_ISL_685424, EPI_ISL_685425, EPI_ISL_685426, EPI_ISL_685427, EPI_ISL_685428, EPI_ISL_685429, EPI_ISL_685430, EPI_ISL_685431, EPI_ISL_685432, EPI_ISL_685433, EPI_ISL_685434, EPI_ISL_685435, EPI_ISL_685436, EPI_ISL_685437, EPI_ISL_685438, EPI_ISL_685439, EPI_ISL_685440, EPI_ISL_685441, EPI_ISL_685442, EPI_ISL_685443, EPI_ISL_685444, EPI_ISL_685445, EPI_ISL_685446, EPI_ISL_685447, EPI_ISL_685448, EPI_ISL_685449, EPI_ISL_685450, EPI_ISL_685451, EPI_ISL_685452, EPI_ISL_685453 | | | |
| see above | Tokyo Metropolitan Institute of Public Health | Pathogen Genomics Center, National Institute of Infectious Diseases | Tsuyoshi Sekizuka, Kentaro Itokawa, Rina Tanaka, Masanori Hashino, Makoto Kuroda |
| EPI_ISL_685498, EPI_ISL_685499, EPI_ISL_685500, EPI_ISL_685501, EPI_ISL_685503, EPI_ISL_685504 | Yamagata Prefectural Institute of Public Health | Pathogen Genomics Center, National Institute of Infectious Diseases | Tsuyoshi Sekizuka, Kentaro Itokawa, Rina Tanaka, Masanori Hashino, Makoto Kuroda |
| EPI_ISL_685506, EPI_ISL_685507, EPI_ISL_685508, EPI_ISL_685509, EPI_ISL_685510, EPI_ISL_685511, EPI_ISL_685512, EPI_ISL_685513, EPI_ISL_685514, EPI_ISL_685515, EPI_ISL_685516, EPI_ISL_685517, EPI_ISL_685518, EPI_ISL_685519, EPI_ISL_685520, EPI_ISL_685528, EPI_ISL_685529, EPI_ISL_685530, EPI_ISL_685531, EPI_ISL_685532, EPI_ISL_685533, EPI_ISL_685534, EPI_ISL_685535, EPI_ISL_685536, EPI_ISL_685537, EPI_ISL_685538, EPI_ISL_685539, EPI_ISL_685540, EPI_ISL_685541, EPI_ISL_685542, EPI_ISL_685543, EPI_ISL_685544, EPI_ISL_685545, EPI_ISL_685546, EPI_ISL_685547, EPI_ISL_685548, EPI_ISL_685549, EPI_ISL_685550, EPI_ISL_685551, EPI_ISL_685552, EPI_ISL_685553, EPI_ISL_685554, EPI_ISL_685555, EPI_ISL_685556, EPI_ISL_685557, EPI_ISL_685558, EPI_ISL_685559, EPI_ISL_685560, EPI_ISL_685561, EPI_ISL_685562, EPI_ISL_685563, EPI_ISL_685564, EPI_ISL_685565, EPI_ISL_685566, EPI_ISL_685567, EPI_ISL_685568, EPI_ISL_685569, EPI_ISL_685570, EPI_ISL_685571, EPI_ISL_685572, EPI_ISL_685573, EPI_ISL_685587, EPI_ISL_685588, EPI_ISL_685589, EPI_ISL_685590, EPI_ISL_685591, EPI_ISL_685592, EPI_ISL_685593, EPI_ISL_685594, EPI_ISL_685595, EPI_ISL_685596, EPI_ISL_685597, EPI_ISL_685598, EPI_ISL_685599, EPI_ISL_685600, EPI_ISL_685601, EPI_ISL_685602, EPI_ISL_685603 | | | |
| see above | Pathogen Genomics Center, National Institute of Infectious Diseases | Pathogen Genomics Center, National Institute of Infectious Diseases | Tsuyoshi Sekizuka, Kentaro Itokawa, Rina Tanaka, Masanori Hashino, Makoto Kuroda |
| EPI_ISL_685607, EPI_ISL_685608, EPI_ISL_685609, EPI_ISL_685610, EPI_ISL_685611, EPI_ISL_685612, EPI_ISL_685613, EPI_ISL_685614, EPI_ISL_685615, EPI_ISL_685616 | Tokyo Metropolitan Institute of Public Health | Pathogen Genomics Center, National Institute of Infectious Diseases | Tsuyoshi Sekizuka, Kentaro Itokawa, Rina Tanaka, Masanori Hashino, Makoto Kuroda |
| EPI_ISL_685623, EPI_ISL_685624, EPI_ISL_685625, EPI_ISL_685626, EPI_ISL_685627, EPI_ISL_685628, EPI_ISL_685629, EPI_ISL_685630, EPI_ISL_685631, EPI_ISL_685632, EPI_ISL_685633, EPI_ISL_685634, EPI_ISL_685635, EPI_ISL_685636, EPI_ISL_685637, EPI_ISL_685649, EPI_ISL_685650, EPI_ISL_685651, EPI_ISL_685652, EPI_ISL_685653, EPI_ISL_685654, EPI_ISL_685655, EPI_ISL_685656, EPI_ISL_685657, EPI_ISL_685658, EPI_ISL_685659, EPI_ISL_685660, EPI_ISL_685661, EPI_ISL_685662, EPI_ISL_685663, EPI_ISL_685664, EPI_ISL_685665 | | | |
| see above | Pathogen Genomics Center, National Institute of Infectious Diseases | Pathogen Genomics Center, National Institute of Infectious Diseases | Tsuyoshi Sekizuka, Kentaro Itokawa, Rina Tanaka, Masanori Hashino, Makoto Kuroda |
| EPI_ISL_685666, EPI_ISL_685667, EPI_ISL_685668, EPI_ISL_685669, EPI_ISL_685670, EPI_ISL_685671, EPI_ISL_685672, EPI_ISL_685673, EPI_ISL_685674, EPI_ISL_685675, EPI_ISL_685676, EPI_ISL_685677, EPI_ISL_685678, EPI_ISL_685679, EPI_ISL_685680, EPI_ISL_685681, EPI_ISL_685682, EPI_ISL_685683, EPI_ISL_685684, EPI_ISL_685685, EPI_ISL_685686, EPI_ISL_685687, EPI_ISL_685688, EPI_ISL_685689, EPI_ISL_685690, EPI_ISL_685691, EPI_ISL_685692, EPI_ISL_685693, EPI_ISL_685694 | | | |
| see above | Tokyo Metropolitan Institute of Public Health | Pathogen Genomics Center, National Institute of Infectious Diseases | Tsuyoshi Sekizuka, Kentaro Itokawa, Rina Tanaka, Masanori Hashino, Makoto Kuroda |
| EPI_ISL_685698, EPI_ISL_685699, EPI_ISL_685701, EPI_ISL_685702, EPI_ISL_685703, EPI_ISL_685704, EPI_ISL_685705, EPI_ISL_685706, EPI_ISL_685707, EPI_ISL_685708, EPI_ISL_685709, EPI_ISL_685710, EPI_ISL_685711, EPI_ISL_685734, EPI_ISL_685735, EPI_ISL_685736, EPI_ISL_685737, EPI_ISL_685738, EPI_ISL_685739, EPI_ISL_685740, EPI_ISL_685741, EPI_ISL_685742, EPI_ISL_685743, EPI_ISL_685746, EPI_ISL_685747, EPI_ISL_685748, EPI_ISL_685749, EPI_ISL_685750, EPI_ISL_685751, EPI_ISL_685752, EPI_ISL_685753, EPI_ISL_685754, EPI_ISL_685755, EPI_ISL_685756, EPI_ISL_685757, EPI_ISL_685758, EPI_ISL_685759, EPI_ISL_685760, EPI_ISL_685761, EPI_ISL_685762, EPI_ISL_685763, EPI_ISL_685764, EPI_ISL_685765, EPI_ISL_685766, EPI_ISL_685767, EPI_ISL_685768, EPI_ISL_685769, EPI_ISL_685770, EPI_ISL_685771, EPI_ISL_685772, EPI_ISL_685773, EPI_ISL_685774, EPI_ISL_685786, EPI_ISL_685787, EPI_ISL_685788, EPI_ISL_685789, EPI_ISL_685790, EPI_ISL_685791, EPI_ISL_685792, EPI_ISL_685793, EPI_ISL_685794, EPI_ISL_685795, EPI_ISL_685796, EPI_ISL_685797, EPI_ISL_685798, EPI_ISL_685799, EPI_ISL_685800, EPI_ISL_685801, EPI_ISL_685802, EPI_ISL_685803, EPI_ISL_685804, EPI_ISL_685805, EPI_ISL_685806, EPI_ISL_685807, EPI_ISL_685808, EPI_ISL_685809, EPI_ISL_685810, EPI_ISL_685811, EPI_ISL_685812, EPI_ISL_685813, EPI_ISL_685814, EPI_ISL_685815, EPI_ISL_685816, EPI_ISL_685817, EPI_ISL_685818, EPI_ISL_685819, EPI_ISL_685820, EPI_ISL_685821, EPI_ISL_685822, EPI_ISL_685823, EPI_ISL_685824, EPI_ISL_685825, EPI_ISL_685826, EPI_ISL_685827, EPI_ISL_685828, EPI_ISL_685829, EPI_ISL_685830, EPI_ISL_685831, EPI_ISL_685832, EPI_ISL_685833, EPI_ISL_685834, EPI_ISL_685835, EPI_ISL_685836, EPI_ISL_685837, EPI_ISL_685838, EPI_ISL_685839, EPI_ISL_685840, EPI_ISL_685841, EPI_ISL_685842, EPI_ISL_685843, EPI_ISL_685844, EPI_ISL_685845, EPI_ISL_685846, EPI_ISL_685847, EPI_ISL_685848, EPI_ISL_685849, EPI_ISL_685850, EPI_ISL_685851, EPI_ISL_685852, EPI_ISL_685853, EPI_ISL_685854, EPI_ISL_685855, EPI_ISL_685856, EPI_ISL_685857, EPI_ISL_685858, EPI_ISL_685859, EPI_ISL_685860, EPI_ISL_685861, EPI_ISL_685862, EPI_ISL_685863, EPI_ISL_685864, EPI_ISL_685865, EPI_ISL_685866, EPI_ISL_685867, EPI_ISL_685868, EPI_ISL_685869, EPI_ISL_685870, EPI_ISL_685871, EPI_ISL_685872, EPI_ISL_685873, EPI_ISL_685874, EPI_ISL_685875, EPI_ISL_685876, EPI_ISL_685877, EPI_ISL_685878, EPI_ISL_685879, EPI_ISL_685880, EPI_ISL_685881, EPI_ISL_685882, EPI_ISL_685883, EPI_ISL_685884, EPI_ISL_685885, EPI_ISL_685886, EPI_ISL_685887, EPI_ISL_685888, EPI_ISL_685889, EPI_ISL_685890, EPI_ISL_685891, EPI_ISL_685892, EPI_ISL_685893, EPI_ISL_685894, EPI_ISL_685895, EPI_ISL_685896, EPI_ISL_685897, EPI_ISL_685898, EPI_ISL_685899, EPI_ISL_685900, EPI_ISL_685901, EPI_ISL_685902, EPI_ISL_685903, EPI_ISL_685906, EPI_ISL_685907, EPI_ISL_685908 | | | |
| see above | Pathogen Genomics Center, National Institute of Infectious Diseases | Pathogen Genomics Center, National Institute of Infectious Diseases | Tsuyoshi Sekizuka, Kentaro Itokawa, Rina Tanaka, Masanori Hashino, Makoto Kuroda |
| EPI_ISL_685918, EPI_ISL_685919, EPI_ISL_685920, EPI_ISL_685921 | Fukuoka Institute of Health and Environmental Sciences | Pathogen Genomics Center, National Institute of Infectious Diseases | Tsuyoshi Sekizuka, Kentaro Itokawa, Rina Tanaka, Masanori Hashino, Makoto Kuroda |
| EPI_ISL_685925, EPI_ISL_685926, EPI_ISL_685927, EPI_ISL_685929, EPI_ISL_685931, EPI_ISL_685932, EPI_ISL_685933, EPI_ISL_685934, EPI_ISL_685935, EPI_ISL_685936, EPI_ISL_685947, EPI_ISL_685953, EPI_ISL_685954, EPI_ISL_685955, EPI_ISL_685957, EPI_ISL_685958, EPI_ISL_685959, EPI_ISL_685960, EPI_ISL_685961 | | | |
| see above | Pathogen Genomics Center, National Institute of Infectious Diseases | Pathogen Genomics Center, National Institute of Infectious Diseases | Tsuyoshi Sekizuka, Kentaro Itokawa, Rina Tanaka, Masanori Hashino, Makoto Kuroda |
| EPI_ISL_685987 | Tokyo Metropolitan Institute of Public Health | Pathogen Genomics Center, National Institute of Infectious Diseases | Tsuyoshi Sekizuka, Kentaro Itokawa, Rina Tanaka, Masanori Hashino, Makoto Kuroda |
| EPI_ISL_685992, EPI_ISL_685993, EPI_ISL_685994, EPI_ISL_685995, EPI_ISL_685996, EPI_ISL_685997, EPI_ISL_685998, EPI_ISL_685999, EPI_ISL_686000, EPI_ISL_686001, EPI_ISL_686002, EPI_ISL_686003, EPI_ISL_686004, EPI_ISL_686005, EPI_ISL_686006, EPI_ISL_686007, EPI_ISL_686008, EPI_ISL_686009, EPI_ISL_686010, EPI_ISL_686011, EPI_ISL_686012, EPI_ISL_686013, EPI_ISL_686014, EPI_ISL_686015, EPI_ISL_686016, EPI_ISL_686017, EPI_ISL_686018, EPI_ISL_686019, EPI_ISL_686020, EPI_ISL_686025, EPI_ISL_686026, EPI_ISL_686027, EPI_ISL_686028, EPI_ISL_686029, EPI_ISL_686030, EPI_ISL_686031, EPI_ISL_686032, EPI_ISL_686033, EPI_ISL_686034, EPI_ISL_686035, EPI_ISL_686036, EPI_ISL_686037, EPI_ISL_686044, EPI_ISL_686045, EPI_ISL_686046, EPI_ISL_686047, EPI_ISL_686048, EPI_ISL_686049, EPI_ISL_686050, EPI_ISL_686051, EPI_ISL_686052, EPI_ISL_686053, EPI_ISL_686054, EPI_ISL_686055, EPI_ISL_686056, EPI_ISL_686057, EPI_ISL_686058, EPI_ISL_686059, EPI_ISL_686060, EPI_ISL_686061, EPI_ISL_686062, EPI_ISL_686063, EPI_ISL_686064, EPI_ISL_686065, EPI_ISL_686066, EPI_ISL_686067, EPI_ISL_686068, EPI_ISL_686069, EPI_ISL_686070, EPI_ISL_686071, EPI_ISL_686072, EPI_ISL_686073 | | | |
| see above | Pathogen Genomics Center, National Institute of Infectious Diseases | Pathogen Genomics Center, National Institute of Infectious Diseases | Tsuyoshi Sekizuka, Kentaro Itokawa, Rina Tanaka, Masanori Hashino, Makoto Kuroda |
| EPI_ISL_686083, EPI_ISL_686084, EPI_ISL_686085, EPI_ISL_686086, EPI_ISL_686087, EPI_ISL_686088, EPI_ISL_686089, EPI_ISL_686090, EPI_ISL_686091, EPI_ISL_686092, EPI_ISL_686093, EPI_ISL_686094, EPI_ISL_686095, EPI_ISL_686096, EPI_ISL_686097, EPI_ISL_686098, EPI_ISL_686099, EPI_ISL_686100, | | | |

| | | | |
|---|---|---|---|
| EPI_ISL_686101, EPI_ISL_686102, EPI_ISL_686103, EPI_ISL_686104, EPI_ISL_686105, EPI_ISL_686106, EPI_ISL_686107 | | | |
| see above | Saitama Prefectural Institute of Public Health | Pathogen Genomics Center, National Institute of Infectious Diseases | Tsuyoshi Sekizuka, Kentaro Itokawa, Rina Tanaka, Masanori Hashino, Makoto Kuroda |
| EPI_ISL_686849, EPI_ISL_686850, EPI_ISL_686851, EPI_ISL_686852, EPI_ISL_686853, EPI_ISL_686854, EPI_ISL_686855, EPI_ISL_686856, EPI_ISL_686857, EPI_ISL_686858, EPI_ISL_686859, EPI_ISL_686860, EPI_ISL_686861, EPI_ISL_686862, EPI_ISL_686863, EPI_ISL_686864, EPI_ISL_686865, EPI_ISL_686866, EPI_ISL_686867, EPI_ISL_686868, EPI_ISL_686869, EPI_ISL_686870, EPI_ISL_686871, EPI_ISL_686872, EPI_ISL_686873, EPI_ISL_687618, EPI_ISL_687921, EPI_ISL_687922, EPI_ISL_687924, EPI_ISL_687925, EPI_ISL_687926, EPI_ISL_687927, EPI_ISL_687928, EPI_ISL_687929, EPI_ISL_687930, EPI_ISL_688084, EPI_ISL_688085, EPI_ISL_688762, EPI_ISL_688763, EPI_ISL_688764, EPI_ISL_688765, EPI_ISL_688766, EPI_ISL_688767, EPI_ISL_688768, EPI_ISL_688769, EPI_ISL_688770, EPI_ISL_688771, EPI_ISL_688772, EPI_ISL_688773, EPI_ISL_688774, EPI_ISL_688775, EPI_ISL_688776, EPI_ISL_688777, EPI_ISL_688778, EPI_ISL_688779, EPI_ISL_688780, EPI_ISL_688781, EPI_ISL_688782, EPI_ISL_688783, EPI_ISL_688784, EPI_ISL_688785, EPI_ISL_688786, EPI_ISL_688787, EPI_ISL_688788, EPI_ISL_688789, EPI_ISL_688790, EPI_ISL_688791, EPI_ISL_688792, EPI_ISL_688793, EPI_ISL_688794, EPI_ISL_688795, EPI_ISL_688796, EPI_ISL_688797, EPI_ISL_688798, EPI_ISL_688799, EPI_ISL_688800, EPI_ISL_688801, EPI_ISL_688802, EPI_ISL_688803, EPI_ISL_688804, EPI_ISL_688805, EPI_ISL_688806, EPI_ISL_688807, EPI_ISL_688808, EPI_ISL_688809, EPI_ISL_688810, EPI_ISL_688811, EPI_ISL_690614, EPI_ISL_690615, EPI_ISL_690617, EPI_ISL_690620, EPI_ISL_690621, EPI_ISL_690622, EPI_ISL_690623, EPI_ISL_690625 | | | |
| see above | Pathogen Genomics Center, National Institute of Infectious Diseases | Pathogen Genomics Center, National Institute of Infectious Diseases | Tsuyoshi Sekizuka, Kentaro Itokawa, Rina Tanaka, Masanori Hashino, Makoto Kuroda |
| EPI_ISL_690626, EPI_ISL_690627, EPI_ISL_690628 | Sakai City Institute of Public Health | Pathogen Genomics Center, National Institute of Infectious Diseases | Tsuyoshi Sekizuka, Kentaro Itokawa, Rina Tanaka, Masanori Hashino, Makoto Kuroda |
| EPI_ISL_690629 | Pathogen Genomics Center, National Institute of Infectious Diseases | Pathogen Genomics Center, National Institute of Infectious Diseases | Tsuyoshi Sekizuka, Kentaro Itokawa, Rina Tanaka, Masanori Hashino, Makoto Kuroda |
| EPI_ISL_690630, EPI_ISL_690631 | Saitama Prefectural Institute of Public Health | Pathogen Genomics Center, National Institute of Infectious Diseases | Tsuyoshi Sekizuka, Kentaro Itokawa, Rina Tanaka, Masanori Hashino, Makoto Kuroda |
| EPI_ISL_690636, EPI_ISL_690637 | Pathogen Genomics Center, National Institute of Infectious Diseases | Pathogen Genomics Center, National Institute of Infectious Diseases | Tsuyoshi Sekizuka, Kentaro Itokawa, Rina Tanaka, Masanori Hashino, Makoto Kuroda |
| EPI_ISL_690641, EPI_ISL_690642, EPI_ISL_690643, EPI_ISL_690644, EPI_ISL_690645, EPI_ISL_690646, EPI_ISL_690647, EPI_ISL_690648, EPI_ISL_690649, EPI_ISL_690650, EPI_ISL_690651, EPI_ISL_690652, EPI_ISL_690653, EPI_ISL_690654, EPI_ISL_690655, EPI_ISL_690657 | | | |
| see above | Saitama Prefectural Institute of Public Health | Pathogen Genomics Center, National Institute of Infectious Diseases | Tsuyoshi Sekizuka, Kentaro Itokawa, Rina Tanaka, Masanori Hashino, Makoto Kuroda |
| EPI_ISL_690684, EPI_ISL_690685, EPI_ISL_690686, EPI_ISL_690711, EPI_ISL_690712, EPI_ISL_690713, EPI_ISL_690714, EPI_ISL_690715, EPI_ISL_690716, EPI_ISL_690717, EPI_ISL_690718, EPI_ISL_690719, EPI_ISL_690720, EPI_ISL_690721, EPI_ISL_690722, EPI_ISL_690723, EPI_ISL_690724, EPI_ISL_690725, EPI_ISL_690726, EPI_ISL_690727, EPI_ISL_690728, EPI_ISL_690729, EPI_ISL_690730, EPI_ISL_690731, EPI_ISL_690732, EPI_ISL_690733, EPI_ISL_690734, EPI_ISL_690735, EPI_ISL_690736, EPI_ISL_690737, EPI_ISL_690738, EPI_ISL_690739, EPI_ISL_690740, EPI_ISL_690752, EPI_ISL_690753, EPI_ISL_690754, EPI_ISL_690755, EPI_ISL_690756, EPI_ISL_690757, EPI_ISL_690758, EPI_ISL_690759, EPI_ISL_690760, EPI_ISL_690761, EPI_ISL_690762, EPI_ISL_690763, EPI_ISL_690764, EPI_ISL_690765, EPI_ISL_690766, EPI_ISL_690767, EPI_ISL_690768, EPI_ISL_690769, EPI_ISL_690770, EPI_ISL_690771, EPI_ISL_690772, EPI_ISL_690773, EPI_ISL_690774, EPI_ISL_690775, EPI_ISL_690776, EPI_ISL_690777, EPI_ISL_690779, EPI_ISL_690780, EPI_ISL_690781, EPI_ISL_690782, EPI_ISL_690783, EPI_ISL_690784, EPI_ISL_690785, EPI_ISL_690786, EPI_ISL_690787, EPI_ISL_690788, EPI_ISL_690789, EPI_ISL_690790, EPI_ISL_690791 | | | |
| see above | Pathogen Genomics Center, National Institute of Infectious Diseases | Pathogen Genomics Center, National Institute of Infectious Diseases | Tsuyoshi Sekizuka, Kentaro Itokawa, Rina Tanaka, Masanori Hashino, Makoto Kuroda |
| EPI_ISL_690792, EPI_ISL_690793 | Tokyo Metropolitan Institute of Public Health | Pathogen Genomics Center, National Institute of Infectious Diseases | Tsuyoshi Sekizuka, Kentaro Itokawa, Rina Tanaka, Masanori Hashino, Makoto Kuroda |
| EPI_ISL_690827, EPI_ISL_690828, EPI_ISL_690829, EPI_ISL_690830, EPI_ISL_690831, EPI_ISL_690832, EPI_ISL_690833, EPI_ISL_690834 | Kanagawa Prefectural Institute of Public Health | Pathogen Genomics Center, National Institute of Infectious Diseases | Tsuyoshi Sekizuka, Kentaro Itokawa, Rina Tanaka, Masanori Hashino, Makoto Kuroda |
| EPI_ISL_690837, EPI_ISL_690838, EPI_ISL_690839, EPI_ISL_690840, EPI_ISL_690841 | Sakai City Institute of Public Health | Pathogen Genomics Center, National Institute of Infectious Diseases | Tsuyoshi Sekizuka, Kentaro Itokawa, Rina Tanaka, Masanori Hashino, Makoto Kuroda |
| EPI_ISL_690874, EPI_ISL_690875, EPI_ISL_690876, EPI_ISL_690877, EPI_ISL_690878, EPI_ISL_690879, EPI_ISL_690880, EPI_ISL_690881, EPI_ISL_690882, EPI_ISL_690883, EPI_ISL_690884, EPI_ISL_690885, EPI_ISL_690886, EPI_ISL_690887, EPI_ISL_690888, EPI_ISL_690889, EPI_ISL_690890, EPI_ISL_690891, EPI_ISL_690892, EPI_ISL_690893, EPI_ISL_690894, EPI_ISL_690895, EPI_ISL_690896, EPI_ISL_690897, EPI_ISL_690898, EPI_ISL_690899, EPI_ISL_690900, EPI_ISL_690901, EPI_ISL_690909, EPI_ISL_690910, EPI_ISL_690911, EPI_ISL_690912, EPI_ISL_690913, EPI_ISL_690914, EPI_ISL_690915, EPI_ISL_690916, EPI_ISL_690917, EPI_ISL_690918, EPI_ISL_690919, EPI_ISL_690920, EPI_ISL_690921, EPI_ISL_690922, EPI_ISL_690923, EPI_ISL_690924, EPI_ISL_690925, EPI_ISL_690926, EPI_ISL_690927, EPI_ISL_690928, EPI_ISL_690929, EPI_ISL_690930, EPI_ISL_690931, EPI_ISL_690932, EPI_ISL_690933, EPI_ISL_690934, EPI_ISL_690935, EPI_ISL_690936, EPI_ISL_690937, EPI_ISL_690938, EPI_ISL_690939, EPI_ISL_690940, EPI_ISL_690941, EPI_ISL_690942, EPI_ISL_690943, EPI_ISL_690944, EPI_ISL_690945, EPI_ISL_690946, EPI_ISL_690947, EPI_ISL_690948, EPI_ISL_690949, EPI_ISL_690950, EPI_ISL_690951, EPI_ISL_690952, EPI_ISL_690953, EPI_ISL_690954, EPI_ISL_690955, EPI_ISL_690956, EPI_ISL_690957, EPI_ISL_690958, EPI_ISL_690959, EPI_ISL_690960, EPI_ISL_690961, EPI_ISL_690962, EPI_ISL_690963, EPI_ISL_690964, EPI_ISL_690965, EPI_ISL_690966, EPI_ISL_690967, EPI_ISL_690968, EPI_ISL_690969, EPI_ISL_690970, EPI_ISL_690971, EPI_ISL_690972, EPI_ISL_690978, EPI_ISL_690979, EPI_ISL_690980 | | | |
| see above | Pathogen Genomics Center, National Institute of Infectious Diseases | Pathogen Genomics Center, National Institute of Infectious Diseases | Tsuyoshi Sekizuka, Kentaro Itokawa, Rina Tanaka, Masanori Hashino, Makoto Kuroda |
| EPI_ISL_690982, EPI_ISL_690983, EPI_ISL_690984, EPI_ISL_690985, EPI_ISL_690986 | Fukuoka Institute of Health and Environmental Sciences | Pathogen Genomics Center, National Institute of Infectious Diseases | Tsuyoshi Sekizuka, Kentaro Itokawa, Rina Tanaka, Masanori Hashino, Makoto Kuroda |
| EPI_ISL_690991, EPI_ISL_690992, EPI_ISL_690993, EPI_ISL_690994, EPI_ISL_690995, EPI_ISL_690996, EPI_ISL_690997, EPI_ISL_690998, EPI_ISL_690999, EPI_ISL_691000, EPI_ISL_691001, EPI_ISL_691002, EPI_ISL_691003, EPI_ISL_691004, EPI_ISL_691005, EPI_ISL_691006, EPI_ISL_691007, EPI_ISL_691008, EPI_ISL_691009, EPI_ISL_691010, EPI_ISL_691011, EPI_ISL_691033, EPI_ISL_691034, EPI_ISL_691035, EPI_ISL_691036, EPI_ISL_691037, EPI_ISL_691038, EPI_ISL_691039, EPI_ISL_691040, EPI_ISL_691041, EPI_ISL_691042, EPI_ISL_691043, EPI_ISL_691044, EPI_ISL_691045, EPI_ISL_691046, EPI_ISL_691047, EPI_ISL_691048, EPI_ISL_691049, EPI_ISL_691050, EPI_ISL_691051, EPI_ISL_691052, EPI_ISL_691053, EPI_ISL_691054, EPI_ISL_691055, EPI_ISL_691056, EPI_ISL_691057, EPI_ISL_691058, EPI_ISL_691059, EPI_ISL_691060, EPI_ISL_691061, EPI_ISL_691062, EPI_ISL_691063, EPI_ISL_691064, EPI_ISL_691065, EPI_ISL_691066, EPI_ISL_691067, EPI_ISL_691068, EPI_ISL_691069, EPI_ISL_691070, EPI_ISL_691071, EPI_ISL_691072, EPI_ISL_691073, EPI_ISL_691074, EPI_ISL_691075, EPI_ISL_691076, EPI_ISL_691077, EPI_ISL_691078, EPI_ISL_691079, EPI_ISL_691080, EPI_ISL_691081, EPI_ISL_691082, EPI_ISL_691083, EPI_ISL_691084, EPI_ISL_691085, EPI_ISL_691086, EPI_ISL_691087, EPI_ISL_691088, EPI_ISL_691089, EPI_ISL_691090, EPI_ISL_691091, EPI_ISL_691092, EPI_ISL_691093, EPI_ISL_691094, EPI_ISL_691095, EPI_ISL_691096, EPI_ISL_691097, EPI_ISL_691098, EPI_ISL_691099, EPI_ISL_691100, EPI_ISL_691101, EPI_ISL_691102, EPI_ISL_691103, EPI_ISL_691104, EPI_ISL_691109, EPI_ISL_691110, EPI_ISL_691111, EPI_ISL_691112, EPI_ISL_691113, EPI_ISL_691114, EPI_ISL_691115, EPI_ISL_691116, EPI_ISL_691117, EPI_ISL_691118, EPI_ISL_691119, EPI_ISL_691120, EPI_ISL_691121, EPI_ISL_691122, EPI_ISL_691123, EPI_ISL_691124, EPI_ISL_691125, EPI_ISL_691126, EPI_ISL_691127, EPI_ISL_691128, EPI_ISL_691129, EPI_ISL_691130, EPI_ISL_691131, EPI_ISL_691132, EPI_ISL_691133, EPI_ISL_691134, EPI_ISL_691135, EPI_ISL_691136, EPI_ISL_691137, EPI_ISL_691138, EPI_ISL_691139, EPI_ISL_691140, EPI_ISL_691141, EPI_ISL_691142, EPI_ISL_691143, EPI_ISL_691144, EPI_ISL_691145, EPI_ISL_691146, EPI_ISL_691147, EPI_ISL_691148, EPI_ISL_691149, EPI_ISL_691150, EPI_ISL_691151, EPI_ISL_691152, EPI_ISL_691153, EPI_ISL_691154, EPI_ISL_691155, EPI_ISL_691156, EPI_ISL_691157, EPI_ISL_691158, EPI_ISL_691159, EPI_ISL_691160, EPI_ISL_691161, EPI_ISL_691162, EPI_ISL_691163, EPI_ISL_691164, EPI_ISL_691165, EPI_ISL_691166, EPI_ISL_691167, EPI_ISL_691168, EPI_ISL_691169, EPI_ISL_691170, EPI_ISL_691171, EPI_ISL_691172, EPI_ISL_691173, EPI_ISL_691174, EPI_ISL_691175, EPI_ISL_691176, EPI_ISL_691177, EPI_ISL_691178, EPI_ISL_691179, EPI_ISL_691180, EPI_ISL_691181, EPI_ISL_691182, EPI_ISL_691183, EPI_ISL_691184, EPI_ISL_691185, EPI_ISL_691186, EPI_ISL_691187, EPI_ISL_691188, EPI_ISL_691189, EPI_ISL_691190, EPI_ISL_691191, EPI_ISL_691192, EPI_ISL_691193, EPI_ISL_691194, EPI_ISL_691195, EPI_ISL_691196, EPI_ISL_691197, EPI_ISL_691198, EPI_ISL_691199, EPI_ISL_691200, EPI_ISL_691201, EPI_ISL_691202, EPI_ISL_691203, EPI_ISL_691204, EPI_ISL_691205, EPI_ISL_691206, EPI_ISL_691207, EPI_ISL_691208, EPI_ISL_691209, EPI_ISL_691210, EPI_ISL_691211, EPI_ISL_691212, EPI_ISL_691213, EPI_ISL_691214, EPI_ISL_691215, EPI_ISL_691216, EPI_ISL_691217, EPI_ISL_691218, EPI_ISL_691219, EPI_ISL_691220, EPI_ISL_691221, EPI_ISL_691222, EPI_ISL_691223, EPI_ISL_691224, EPI_ISL_691225, EPI_ISL_691228, EPI_ISL_691229, EPI_ISL_691230, EPI_ISL_691231, EPI_ISL_691243, EPI_ISL_691244, EPI_ISL_691245, EPI_ISL_691246, EPI_ISL_691247, EPI_ISL_691249, EPI_ISL_691250, EPI_ISL_691251, EPI_ISL_691257, EPI_ISL_691258, EPI_ISL_691300, EPI_ISL_691301, EPI_ISL_691302, EPI_ISL_691303, EPI_ISL_691304, EPI_ISL_691305, EPI_ISL_691306, EPI_ISL_691307, EPI_ISL_691308, EPI_ISL_691309, EPI_ISL_691310, EPI_ISL_691311, EPI_ISL_691312, EPI_ISL_691313, EPI_ISL_691314, EPI_ISL_691315, EPI_ISL_691316, EPI_ISL_691317, EPI_ISL_691318, EPI_ISL_691319, EPI_ISL_691320, EPI_ISL_691321, EPI_ISL_691322, EPI_ISL_691323, EPI_ISL_691324, EPI_ISL_691325, EPI_ISL_691326, EPI_ISL_691327, EPI_ISL_691328, EPI_ISL_691329, EPI_ISL_691330, EPI_ISL_691331, EPI_ISL_691332, EPI_ISL_691333, EPI_ISL_691334, EPI_ISL_691335, EPI_ISL_691336, EPI_ISL_691337, EPI_ISL_691338, EPI_ISL_691339, EPI_ISL_691340, EPI_ISL_691341, EPI_ISL_691342, EPI_ISL_691343, EPI_ISL_691344, EPI_ISL_691345, EPI_ISL_691346, EPI_ISL_691347, EPI_ISL_691348, EPI_ISL_691349, EPI_ISL_691350, EPI_ISL_691351, EPI_ISL_691352, EPI_ISL_691353, EPI_ISL_691354, EPI_ISL_691355, EPI_ISL_691356, EPI_ISL_691357, EPI_ISL_691358, EPI_ISL_691359, EPI_ISL_691360, EPI_ISL_691361, EPI_ISL_691362, EPI_ISL_691363, EPI_ISL_691384, EPI_ISL_691385, EPI_ISL_691396, EPI_ISL_691397, EPI_ISL_691398, EPI_ISL_691399, EPI_ISL_691400, EPI_ISL_691401, EPI_ISL_691402, EPI_ISL_691403, EPI_ISL_691404, EPI_ISL_691405, EPI_ISL_691406, EPI_ISL_691407, EPI_ISL_691408, EPI_ISL_691409, EPI_ISL_691410, EPI_ISL_691411, EPI_ISL_691412, EPI_ISL_691413, EPI_ISL_691414, EPI_ISL_691415, EPI_ISL_691416, EPI_ISL_691417, EPI_ISL_691418, EPI_ISL_691419, EPI_ISL_691420, EPI_ISL_691421, EPI_ISL_691422, EPI_ISL_691423, EPI_ISL_691424, EPI_ISL_691425, EPI_ISL_691426, EPI_ISL_691427, EPI_ISL_691428, EPI_ISL_691429, EPI_ISL_691430, EPI_ISL_691431, EPI_ISL_691432, EPI_ISL_691433, EPI_ISL_691434, EPI_ISL_691435, EPI_ISL_691436, EPI_ISL_691437, EPI_ISL_691438, EPI_ISL_691439, EPI_ISL_691440, EPI_ISL_691441, EPI_ISL_691442, EPI_ISL_691443, EPI_ISL_691444, EPI_ISL_691445, EPI_ISL_691446, EPI_ISL_691447, EPI_ISL_691448, EPI_ISL_691449, EPI_ISL_691450, EPI_ISL_691451, EPI_ISL_691452, EPI_ISL_691453, EPI_ISL_691454, EPI_ISL_691455, EPI_ISL_691456, EPI_ISL_691457, EPI_ISL_691458, EPI_ISL_691459, EPI_ISL_691460, EPI_ISL_691461, EPI_ISL_691462, EPI_ISL_691463, EPI_ISL_691464, EPI_ISL_691465, EPI_ISL_691466, EPI_ISL_691467, EPI_ISL_691468, EPI_ISL_691469, EPI_ISL_691470, EPI_ISL_691471, EPI_ISL_691472, EPI_ISL_691473, EPI_ISL_691474, EPI_ISL_691475, EPI_ISL_691476, EPI_ISL_691477, EPI_ISL_691478, EPI_ISL_691479, EPI_ISL_691480, EPI_ISL_691481, EPI_ISL_691482, EPI_ISL_691483, EPI_ISL_691484, EPI_ISL_691485, EPI_ISL_691486, EPI_ISL_691487, EPI_ISL_691488, EPI_ISL_691489, EPI_ISL_691490, EPI_ISL_691491, EPI_ISL_691492, EPI_ISL_691493, EPI_ISL_691494, EPI_ISL_691495, EPI_ISL_691496, EPI_ISL_691497, EPI_ISL_691498, EPI_ISL_691499, EPI_ISL_691507, EPI_ISL_691508 | | | |
| see above | Pathogen Genomics Center, National Institute of Infectious Diseases | Pathogen Genomics Center, National Institute of Infectious Diseases | Tsuyoshi Sekizuka, Kentaro Itokawa, Rina Tanaka, Masanori Hashino, Makoto Kuroda |
| EPI_ISL_691509, EPI_ISL_691510, EPI_ISL_691511, EPI_ISL_691512, EPI_ISL_691513, EPI_ISL_691514, EPI_ISL_691515, EPI_ISL_691516, EPI_ISL_691517, EPI_ISL_691518, EPI_ISL_691519, EPI_ISL_691520, EPI_ISL_691521, EPI_ISL_691522, EPI_ISL_691523, EPI_ISL_691524, EPI_ISL_691525, EPI_ISL_691526, EPI_ISL_691527, EPI_ISL_691528, EPI_ISL_691529, EPI_ISL_691530, EPI_ISL_691531, EPI_ISL_691532, EPI_ISL_691533, EPI_ISL_691534, EPI_ISL_691535, EPI_ISL_691536, EPI_ISL_691537, EPI_ISL_691538, EPI_ISL_691539, EPI_ISL_691540, EPI_ISL_691541, EPI_ISL_691542 | | | |
| see above | Sakai City Institute of Public Health | Pathogen Genomics Center, National Institute of Infectious Diseases | Tsuyoshi Sekizuka, Kentaro Itokawa, Rina Tanaka, Masanori Hashino, Makoto Kuroda |

| | | | |
|---|---|---|---|
| EPI_ISL_691757, EPI_ISL_691758, EPI_ISL_691767, EPI_ISL_691872, EPI_ISL_691873, EPI_ISL_691874, EPI_ISL_691875, EPI_ISL_691876, EPI_ISL_691877, EPI_ISL_691878, EPI_ISL_691879, EPI_ISL_691880, EPI_ISL_691881, EPI_ISL_691882, EPI_ISL_691883, EPI_ISL_691884, EPI_ISL_691885, EPI_ISL_691886, EPI_ISL_691887, EPI_ISL_691888, EPI_ISL_691889, EPI_ISL_691890, EPI_ISL_691891, EPI_ISL_691892, EPI_ISL_691893, EPI_ISL_691894, EPI_ISL_691895, EPI_ISL_691896, EPI_ISL_691897, EPI_ISL_691898, EPI_ISL_691899, EPI_ISL_691900, EPI_ISL_691901, EPI_ISL_691902, EPI_ISL_691903, EPI_ISL_691904, EPI_ISL_691905, EPI_ISL_691906, EPI_ISL_691907, EPI_ISL_691908, EPI_ISL_691909, EPI_ISL_691910, EPI_ISL_691911, EPI_ISL_691912, EPI_ISL_691913, EPI_ISL_691914, EPI_ISL_691915, EPI_ISL_691916, EPI_ISL_691917, EPI_ISL_691918, EPI_ISL_691919, EPI_ISL_691920, EPI_ISL_691921, EPI_ISL_691922, EPI_ISL_691923, EPI_ISL_691924, EPI_ISL_691925, EPI_ISL_691926, EPI_ISL_691927, EPI_ISL_691928, EPI_ISL_691929, EPI_ISL_691930, EPI_ISL_691931, EPI_ISL_691932, EPI_ISL_691933, EPI_ISL_691934, EPI_ISL_691935, EPI_ISL_691936, EPI_ISL_691937, EPI_ISL_691938, EPI_ISL_691939, EPI_ISL_691940, EPI_ISL_691941, EPI_ISL_691942, EPI_ISL_691943, EPI_ISL_691944, EPI_ISL_691945, EPI_ISL_691946, EPI_ISL_691947, EPI_ISL_691948, EPI_ISL_691949, EPI_ISL_691950, EPI_ISL_691951, EPI_ISL_691952, EPI_ISL_691953, EPI_ISL_691954, EPI_ISL_691955, EPI_ISL_691956, EPI_ISL_691957, EPI_ISL_691958, EPI_ISL_691959, EPI_ISL_691960, EPI_ISL_691961, EPI_ISL_691962, EPI_ISL_691963, EPI_ISL_691964, EPI_ISL_691965, EPI_ISL_691966, EPI_ISL_691967, EPI_ISL_691968, EPI_ISL_691969, EPI_ISL_691970, EPI_ISL_691971, EPI_ISL_691972, EPI_ISL_691973, EPI_ISL_691974, EPI_ISL_691975, EPI_ISL_691976, EPI_ISL_691977, EPI_ISL_691978, EPI_ISL_691979, EPI_ISL_691980, EPI_ISL_691981, EPI_ISL_691982, EPI_ISL_691983, EPI_ISL_691984, EPI_ISL_691985, EPI_ISL_691986, EPI_ISL_691987, EPI_ISL_691988, EPI_ISL_691989, EPI_ISL_691990, EPI_ISL_691991, EPI_ISL_691992, EPI_ISL_691993, EPI_ISL_691994, EPI_ISL_691995, EPI_ISL_691996, EPI_ISL_691997, EPI_ISL_691998, EPI_ISL_691999, EPI_ISL_692000, EPI_ISL_692001, EPI_ISL_692002, EPI_ISL_692003, EPI_ISL_692004, EPI_ISL_692005, EPI_ISL_692006, EPI_ISL_692007, EPI_ISL_692008, EPI_ISL_692009, EPI_ISL_692010, EPI_ISL_692011, EPI_ISL_692012, EPI_ISL_692013, EPI_ISL_692014, EPI_ISL_692015, EPI_ISL_692016, EPI_ISL_692017, EPI_ISL_692018, EPI_ISL_692019, EPI_ISL_692020, EPI_ISL_692021, EPI_ISL_692022, EPI_ISL_692023, EPI_ISL_692024, EPI_ISL_692025, EPI_ISL_692026, EPI_ISL_692027, EPI_ISL_692028, EPI_ISL_692029, EPI_ISL_692030, EPI_ISL_692031, EPI_ISL_692032, EPI_ISL_692033, EPI_ISL_692034, EPI_ISL_692035, EPI_ISL_692036, EPI_ISL_692037, EPI_ISL_692038, EPI_ISL_692039, EPI_ISL_692040, EPI_ISL_692041, EPI_ISL_692042, EPI_ISL_692043, EPI_ISL_692197 | | | |
| see above | Pathogen Genomics Center, National Institute of Infectious Diseases | Pathogen Genomics Center, National Institute of Infectious Diseases | Tsuyoshi Sekizuka, Kentaro Itokawa, Rina Tanaka, Masanori Hashino, Makoto Kuroda |
| EPI_ISL_693201 | Hospital Sao Paulo de Ensino da Unifesp | Instituto Adolfo Lutz, Interdiciplinary Procedures Center, Strategic Laboratory | Claudio Tavares Sacchi, Claudia Regina Gonçalves, Erica Valessa Ramos Gomes, Karoline Rodrigues Campos |
| EPI_ISL_693202 | Pronto Socorro Municipal Prof. Joao Catarin Mezomo | Instituto Adolfo Lutz, Interdiciplinary Procedures Center, Strategic Laboratory | Claudio Tavares Sacchi, Claudia Regina Gonçalves, Erica Valessa Ramos Gomes, Karoline Rodrigues Campos |
| EPI_ISL_693287 | unknown | Public Health Virology Laboratory, Forensic and Scientific Services (PHV-FSS) | Son Nguyen et al. |
| EPI_ISL_694055 | AZ SPHL, Arizona Department of Health Services | TGen North | Jolene Bowers, Megan Folkerts, Chris French, Hayley Yaglom, Ashlyn Pfeiffer, Darrin Lemmer, Dave Engelthaler, The Arizona COVID Genomics Union (ACGU) |
| EPI_ISL_696015, EPI_ISL_696017, EPI_ISL_696019, EPI_ISL_696022, EPI_ISL_696023, EPI_ISL_696350, EPI_ISL_696379 | Sonora Quest Laboratories, Laboratory Sciences of Arizona | TGen North | Jolene Bowers, Megan Folkerts, Chris French, Hayley Yaglom, Ashlyn Pfeiffer, Darrin Lemmer, Dave Engelthaler, The Arizona COVID Genomics Union (ACGU) |
| EPI_ISL_699641 | Sydney South West Pathology Service (SSWPS) - Liverpool Hospital - NSW Health Pathology | NSW Health Pathology - Institute of Clinical Pathology and Medical Research; Westmead Hospital; University of Sydney | CIDM-PH et al. |
| EPI_ISL_700361 | Laboratoire de virologie, CHU de Grenoble - CS 10217 - 38043 Grenoble cedex 22 | CNR Virus des Infections Respiratoires - France SUD | Antonin Bal, Gregory Destras, Gwendolyne Burfin, Hadrien Règue, Quentin Semanas, Martine Valette, Bruno Lina, Sylvie Larrat, Laurence Josset |
| EPI_ISL_707792 | 1-Laboratory of Microbiology, National Reference Lab, Charles Nicolle Hospital; 2-University of Tunis ElManar, Faculty of Medicine of Tunis, LR99ES09, Tunis, Tunisia | 1-Clinical and Experimental Pharmacology Lab, LR16SP02, National Center of Pharmacovigilance, University of Tunis El Manar, Tunis, Tunisia. 2-Neurodegenerative diseases and psychiatric troubles, LR18SP03, Razi Hospital, University of Tunis El Manar, Tunis, Tunisia. 3- Ministry of Health, National Observatory of New and Emerging Diseases, 1006, Tunis, Tunisia | Ilhem Boutiba-Ben Boubaker, Sameh Trabelsi, Nissaf Ben Alaya, Maher Kharrat, Alia Ben Kahla, Jalila Ben Khelil, Salma Abid, Sana Ferjani, Mouna Ben Sassi, Mouna Safer, Awatef El MOussi, Habiba Ben Romdhane, Souissi Amira, Ines Mdini, Hanen El Jebari, Asma Ferjani, Gaies Emna, Riadh Daghfous, Riadh Gouider. |
| EPI_ISL_707960, EPI_ISL_707961, EPI_ISL_707962, EPI_ISL_708016, EPI_ISL_708017 | Virology, Universitätsklinikum des Saarlandes | Epigenetics, Saarland University | Kathrin Kattler, Markus Vogelgesang, Stefan Lohse, Sascha Tierling, Sigrun Smola, Jörn Walter |
| EPI_ISL_708741, EPI_ISL_708744, EPI_ISL_708765, EPI_ISL_708766, EPI_ISL_708772, EPI_ISL_708775, EPI_ISL_708782, EPI_ISL_708784 | PathWest Laboratory Medicine WA | PathWest Laboratory Medicine WA Microbial Surveillance Unit | PathWest Laboratory Medicine WA Microbial Surveillance Unit |
| EPI_ISL_718140, EPI_ISL_718142, EPI_ISL_718150, EPI_ISL_718152, EPI_ISL_718157, EPI_ISL_718162, EPI_ISL_718163, EPI_ISL_718164 | Ministry of Health Hospitals | Institute of Health and Community Medicine | David Perera, Ooi Mong How, Chua Hock Hin, Tonnii Sia Loong Loong, Wong Jyn Shan, Wong Kiing Aik, Chan Chia Jui |
| EPI_ISL_722208 | Triemli Hospital Zurich | Institute of Medical Virology, University of Zurich | Stefan Schmutz, Verena Kufner, Maryam Zaheri, Gabriela Ziltener, Gerhard Eich, Daniel Röösli, Jürg Böni, Michael Huber, Alexandra Trkola |
| EPI_ISL_723105, EPI_ISL_723106, EPI_ISL_723107 | Institute of Medical Genetics and Applied Genomics | Institute of Medical Genetics and Applied Genomics | Caspar Gross, Tina Ganzenmüller, Siri Göpel, Michaela Pogoda, Daniela Bezdan, Michael Sonnabend, Angel Angelov, Nicolas Casadei, Stephan Ossowski, Thomas Iftner, Michael Bitzer |
| EPI_ISL_728221, EPI_ISL_728222, EPI_ISL_728223, EPI_ISL_728224, EPI_ISL_728235, EPI_ISL_728236, EPI_ISL_728237 | LNR National Reference Laboratory, Mohammed VI University of Health Sciences | Medical Biotechnology Laboratory, Rabat Medical and Pharmacy School, Mohammed The Vth University in Rabat | Souad KARTTI, Housna ARROUCHI, Loubna ALLAM, Mouneem Essabbar, Mouna OUADGHIRI, Tarek Aanniz, Nabila Soara, Adib Ghassan, Saaid AMZAZI, Lahcen BELYAMANI and Azeddine IBRAHIMI |
| EPI_ISL_728358 | Servicio de Microbiología. Hospital Clínico Universitario de Valencia | SeqCOVID-SPAIN consortium/IBV(CSIC) | David Navarro Ortega, Eliseo Albert Vicent, Ignacio Torres and SeqCOVID-SPAIN consortium |
| EPI_ISL_729405, EPI_ISL_729534, EPI_ISL_729548, EPI_ISL_729558 | Charité Universitätsmedizin Berlin, Institut für Virologie/Labor Berlin | Charité Universitätsmedizin Berlin, Institut für Virologie | Victor M Corman, Barbara Mühlemann, Jörn Beheim-Schwarzbach, Talitha Veith, Julia Schneider, Terry Jones, Christian Drosten |
| EPI_ISL_730153 | Scripps Medical Laboratory | Andersen lab at Scripps Research | SEARCH Alliance San Diego with Michael Quigley, Ellen Stefanski, Ian Mchardy |
| EPI_ISL_732552 | Bundeswehr Institute of Microbiology | Bundeswehr Institute of Microbiology | Markus Antwerpen, Alexandra Rehn, Mathias Walter, Malena Bestehorn-Willmann, Sabine Zange, Enrico Georgi, Roman Wölfel |
| EPI_ISL_734526, EPI_ISL_734527, EPI_ISL_734528, EPI_ISL_734529, EPI_ISL_734530 | UZ Leuven, National Reference Laboratory for Coronaviruses, Laboratory Medicine, Leuven, Belgium | KU Leuven, Rega Institute, Clinical and Epidemiological Virology | Tony Wawina-Bokalanga, Joan Marti-Carerras, Bert Vanmechelen, Piet Maes |
| EPI_ISL_738286, EPI_ISL_738287, EPI_ISL_738288, EPI_ISL_738300, EPI_ISL_738301, EPI_ISL_738302, EPI_ISL_738303, EPI_ISL_738304, EPI_ISL_738305, EPI_ISL_738306 | Connecticut Veterans' Affairs Hospital | Grubaugh Lab - Yale School of Public Health | Joseph Fauver, Tara Alpert, Chantal Vogels, Mary Petrone, Isabel Ott, Ellen Foxman, Shaili Gupta, Danielle Plank, Nathan Grubaugh |
| EPI_ISL_751212 | Pathogen Genomics Lab King Abdullah University of Science and Technology(KAUST) | Pathogen Genomics Lab King Abdullah University of Science and Technology(KAUST) | Sharif Hala, Sara Mfarrej, Raushan Nugmanova, Olga Douvropoulou, Rahul P Salunke, Raeece Naeem, Fadwa Alofi, Asim Khogeer, Afrah Alsomali, Jumana Taha, Abdulaziz Alahmadi, Kahled Algithami, Anwar Hashem, Naif Almontashiri, Arnab Pain |
| EPI_ISL_751321, EPI_ISL_751328, EPI_ISL_751427, EPI_ISL_751428 | IRCCS Sacro Cuore Don Calabria Hospital, Department of Infectious, Tropical Diseases & Microbiology | University of Verona, Department of Biotechnology | Antonio Mori, Michela Deiana, Elena Pomari, Chiara Piubelli; Giulia Lopatriello, Luca Marcolungo, Cristina Beltrami, Chiara Degli Esposti, Emanuela Cosentino, Massimo Delledonne |
| EPI_ISL_753839, EPI_ISL_753840, EPI_ISL_753846, EPI_ISL_753847, EPI_ISL_753848, EPI_ISL_753850, EPI_ISL_753851, EPI_ISL_753929, EPI_ISL_753936, EPI_ISL_753937, EPI_ISL_753945 | | | |
| see above | Charité Universitätsmedizin Berlin, Institut für Virologie/Labor Berlin | Charité Universitätsmedizin Berlin, Institut für Virologie | Victor M Corman, Jörn Beheim-Schwarzbach, Barbara Mühlemann, Julia Schneider, Talitha Veith, Terry Jones, Christian Drosten |

| | | | |
|---|---|---|---|
| EPI_ISL_755878, EPI_ISL_755879, EPI_ISL_755880 | Toronto Invasive Bacterial Diseases Network | McMaster University | Allison McGeer, Patryk Aftanas, Hooman Derakhshani, Angel Li, Kuganya Nirmalarajah, Emily Panousis, Ahmed Draia, Jalees Nasir, Michael Surette, Samira Mubareka, Andrew G. McArthur |
| EPI_ISL_759967 | RSUD Dr. Iskak Tulungagung | Institute of Tropical Disease, Universitas Airlangga | Rima R Prasetya, Krisnoadi Rahardjo, Aldise M Nastri, Jezzy R Dewantari, Supriyanto Dharmoredjo, Gatot Soegiarto, Laksmi Wulandari, Resti Yudhawati, Soetjipto, Yasuko Mori, Maria I Lusida, Kazufumi Shimizu |
| EPI_ISL_763065 | 1-Laboratory of Microbiology, National Reference Lab, Charles Nicolle Hospital; 2-University of Tunis ElManar, Faculty of Medicine of Tunis, LR99ES09, Tunis, Tunisia | 1-Clinical and Experimental Pharmacology Lab, LR16SP02, National Center of Pharmacovigilance, University of Tunis El Manar, Tunis, Tunisia. 2-Neurodegenerative diseases and psychiatric troubles, LR18SP03, Razi Hospital, University of Tunis El Manar, Tunis, Tunisia. 3- Ministry of Health, National Observatory of New and Emerging Diseases, 1006, Tunis, Tunisia | Ilhem Boutiba-Ben Boubaker, Sameh Trabelsi, Nissaf Ben Alaya, Maher Kharrat, Alia Ben Kahla, Jalila Ben Khelil, Salma Abid, Sana Ferjani, Mouna Ben Sassi, Mouna Safer, Awatef El Moussi, Habiba Ben Romdhane, Souissi Amira, Ines Mdini, Hanen El Jebari, Asma Ferjani, Gaies Emna, Riadh Daghfous, Riadh Gouider. |
| EPI_ISL_763092 | Jena University Hospital, Institute for Infectious Diseases and Infection Control | Institute of infectious medicine & hospital hygiene, CaSe-Group | Spott, Riccardo; Marquet, Mike; Pletz, Matthias W.; Brandt, Christian |
| EPI_ISL_763351 | University Hospital Limerick | Irish Coronavirus Sequencing Consortium - Teagasc Moorepark | Paul Cotter, Fiona Crispie, John Kenny, Calum Walsh |
| EPI_ISL_770829 | Toronto Invasive Bacterial Diseases Network | McMaster University | Allison McGeer, Patryk Aftanas, Hooman Derakhshani, Angel Li, Kuganya Nirmalarajah, Emily Panousis, Ahmed Draia, Jalees Nasir, Michael Surette, Samira Mubareka, Andrew G. McArthur |
| EPI_ISL_775243, EPI_ISL_775244 | INT Fondazione Pascale | INT Fondazione Pascale | INT Fondazione Pascale |
| EPI_ISL_776753, EPI_ISL_776754 | Instituto Adolfo Lutz - Central | Instituto Adolfo Lutz, Interdiciplinary Procedures Center, Strategic Laboratory | Claudio Tavares Sacchi, Claudia Regina Gonçalves, Erica Valessa Ramos Gomes, Karoline Rodrigues Campos |
| EPI_ISL_778686, EPI_ISL_778687, EPI_ISL_778688, EPI_ISL_778689, EPI_ISL_778690 | Istituto Zooprofilattico Sperimentale del Mezzogiorno | TIGEM | Patrizia Annunziata, Andrea Ballabio, Valentina Bouche, Davide Cacchiarelli (CorrespAuthor), Pellegrino Cerino, Chiara Colantuono, Lucio Di Filippo, Antonio Grimaldi, Antonio Limone, Gabriella Loconte, Anna Manfredi, Francesco Panariello, Biancamaria Pierri, Marcello Salvi, Lucia Vassallo |
| EPI_ISL_791522, EPI_ISL_791523, EPI_ISL_791524, EPI_ISL_791525, EPI_ISL_791526 | Massachusetts State Public Health Laboratory | Infectious Disease Program, Broad Institute of Harvard and MIT | Lemieux,J.E., Siddle,K.J., Shaw,B., Adams,G., Pierce,V., Turbett,S., Anahtar,M., Branda,J., Slater,D., Harris,J., Lin,A.E., Gladden-Young,A., Lagerborg,K., Rudy,M., DeRuff,K., Carter,A., Normandin,E., Bauer,M., Reilly,S., Tomkins-Tinch,C., Loreth,C., Chaluvadi,S., Neumann,A., Cusick,C., Chapman,S.B., Gnirke,A., Flowers,K., Cerrato,F., Birren,B.W., Gallagher,G., Smole,S., Park,D.J., MacInnis,B.L., Ryan,E., LaRocque,R., Rosenberg,E. and Sabeti,P.C. |
| EPI_ISL_792092, EPI_ISL_792094 | UW Virology Lab | UW Virology Lab | Pavitra Roychoudhury, Hong Xie, Lasata Shrestha, Meei-Li Huang, Keith R Jerome, Alexander Greninger |
| EPI_ISL_792103 | Instituto Adolfo Lutz - Regional de Santo Andre | Instituto Adolfo Lutz, Interdiciplinary Procedures Center, Strategic Laboratory | Claudio Tavares Sacchi, Claudia Regina Gonçalves, Erica Valessa Ramos Gomes, Karoline Rodrigues Campos |
| EPI_ISL_792107 | Instituto Adolfo Lutz - Central | Instituto Adolfo Lutz, Interdiciplinary Procedures Center, Strategic Laboratory | Claudio Tavares Sacchi, Claudia Regina Gonçalves, Erica Valessa Ramos Gomes, Karoline Rodrigues Campos |
| EPI_ISL_792364 | Laboratorio de Virologia - HIEAyC San Juan de Dios | Área de Secuenciación del Laboratorio de Virología del Hospital de Niños Dr. Ricardo Gutierrez on behalf of 'Proyecto Argentino Interinstitucional de genomica de SARS-CoV-2' (PAIS Consortium) | Nabaes Jodar, MS; Goya, S; Natale, MI; Lusso, S; Ferioli, M; Colmeiro, M; Gatelli, A; Ercole, R; Valinotto, LE; Viegas, M. |
| EPI_ISL_792424 | Laboratorio de Inmunología del Hospital Perrando e Instituto de Medicina Regional de la UNNE | Instituto de Biotecnología, IABIMO (CONICET), Instituto de Virología, IVIT(CONICET), Instituto de Patobiología, IPVET(CONICET), CICVyA, INTA on behalf of 'Proyecto Argentino Interinstitucional de genomica de SARS-CoV-2' (PAIS Consortium) | König, GA; Peralta, AV; Distéfano, AJ; Zavallo, D; Muñoz Hidalgo, MG; Vera, PA; Fass, M; Farber, MD; Cacciabué, MPD; Pedroarias, VC; Lozano Calderón, LC; Bengoa Luoni, S; Asurmendi, S; Foussal, MD; Deluca, G; Ayala, NA; Gómez, MV; Giusiano, G; Lucero, H; Marin, M; Lescano, L; Cayré, A; Paniego, NB; Rivarola, M; Puebla, AF; Viegas, M. |
| EPI_ISL_792455, EPI_ISL_792468, EPI_ISL_792469, EPI_ISL_792473 | Laboratorio del Hospital Regional Ushuaia Gdor. Ernesto Campos | Hospital Regional Ushuaia - Centro Austral De Investigaciones Científicas - Universidad Nacional De Tierra Del Fuego on behalf of 'Proyecto Argentino Interinstitucional de genomica de SARS-CoV-2' (PAIS Consortium) | Ceballos, SG; Nardi, CF; Gramundi, ID; Gallego, F; De Roccis, CA; Castro, G; Cáceres, SB; Yulan, CB; Boutureira, MF. |
| EPI_ISL_792485, EPI_ISL_792486 | Laboratorio Central Mg. Luis Alfredo Pianciola | Hospital Regional Ushuaia - Centro Austral De Investigaciones Científicas - Universidad Nacional De Tierra Del Fuego on behalf of 'Proyecto Argentino Interinstitucional de genomica de SARS-CoV-2' (PAIS Consortium) | Ceballos, SG; Nardi, CF; Gramundi, ID; Gallego, F; Pinto, C; Ziehm, MC; Mazzeo, M; Pianciola, L. |
| EPI_ISL_792528, EPI_ISL_792533, EPI_ISL_792537 | Laboratorio Central, Ministerio de Salud Córdoba | Instituto de Patologia Vegetal (CIAP-INTA) on behalf of 'Proyecto Argentino Interinstitucional de genomica de SARS-CoV-2' (PAIS Consortium) | Fernández, FD; Debat, HJ., Re, V; Pisano, MB; Castro, G.; Barbas, G. |
| EPI_ISL_793013 | Laboratorio de Virología del Hospital de Niños Dr. Ricardo Gutierrez | Área de Secuenciación del Laboratorio de Virología del Hospital de Niños Dr. Ricardo Gutierrez on behalf of 'Proyecto Argentino Interinstitucional de genomica de SARS-CoV-2' (PAIS Consortium) | Nabaes Jodar, MS; Goya, S; Natale, MI; Lusso, S; Gravis, E; Mistchenko, AS; Valinotto, LE; Viegas, M. |
| EPI_ISL_794745 | Istituto Zooprofilattico Sperimentale della Puglia e della Basilicata | Istituto Zooprofilattico Sperimentale della Puglia e della Basilicata | Parisi A., Bianco A., Capozzi L., Del Sambro L., Manzulli V, Rondinone V., Pace L., Cipolletta D., Galante D. |
| EPI_ISL_796069, EPI_ISL_796100, EPI_ISL_796110, EPI_ISL_796113 | Servicio de Microbiología. Hospital General Universitario de Castellón | SeqCOVID-SPAIN consortium/IBV(CSIC) | Rosario Moreno Muñoz, María Dolores Tirado Balaguer and SeqCOVID-SPAIN consortium |
| EPI_ISL_801582 | Laboratory of Molecular Virology, Pontificia Universidad Católica de Chile | MSHS Pathogen Surveillance Program | Leonardo I. Almonacid, Ana S. Gonzalez-Reiche, Matthew M. Hernandez, Jorge Levican, Tamara García-Salum, Zenab Khan, Adriana van De Guchte, Ajay Obla, Jayeeta Dutta, Bremy Alburquerque, Eileen Serrano, Erick Salinas, Hala Alshammary, Juan Soto, Shwetha Hara Sridhar, Ying-Chih Wang, Melissa Smith, Robert Sebra, Adolfo Garcia-Sastre, Edward C. Holmes, Viviana Simon, Harm van Bakel, Rafael A. Medina |
| EPI_ISL_801670, EPI_ISL_801675, EPI_ISL_801676, EPI_ISL_801677, EPI_ISL_801678, EPI_ISL_801679, EPI_ISL_801680, EPI_ISL_801681, EPI_ISL_801682, EPI_ISL_801683 | Laboratory of Molecular Virology, Pontificia Universidad Católica de Chile | MSHS Pathogen Surveillance Program | Leonardo I. Almonacid, Ana S. Gonzalez-Reiche, Matthew M. Hernandez, Jorge Levican, Ana Maria Contreras, Carlos Palma, Tamara García-Salum, Zenab Khan, Adriana van de Guchte, Ajay Obla, Jayeeta Dutta, Bremy Alburquerque, Eileen Serrano, Constanza Maldonado, M. Belen Leyton, Erick Salinas, Hala Alshammary, Juan Soto, Shwetha Hara Sridhar, Ying-Chih Wang, Melissa Smith, Robert Sebra, Marcela Ferres, Adolfo Garcia-Sastre, Edward C. Holmes, Viviana Simon, Harm van Bakel, Rafael A. Medina. |
| EPI_ISL_801877, EPI_ISL_801906, EPI_ISL_801935, EPI_ISL_801981, EPI_ISL_801982, EPI_ISL_801983, EPI_ISL_801984, EPI_ISL_801985, EPI_ISL_801986 | MSHS Clinical Microbiology Laboratories | MSHS Pathogen Surveillance Program | Ana S. Gonzalez-Reiche, Hala Alshammary, Mitchell J. Sullivan, Brianne Ciferri, Ajay Obla, Angela Amoako, Mahmoud Awawda, Elena Hirsch, Ashley S. Salimbangon, Levy Sominsky, Katherine Beach, Kayla Russo, Charles Gleason, Shelcie Fabre, Giulio Kleiner, Zenab Khan, Bremy Alburquerque, Adriana van de Guchte, Komal Srivastava, Matthew M. Hernandez, Jayeeta Dutta, Denise Jurczyszak, Emily Ferreri, Rachel Chernet, Nancy Francoeur, Betsaida Salom Melo, Irina Oussenko, Gintaras Deikus, Juan Soto, Shwetha Hara Sridhar, Ying-Chih Wang, Kathryn Twyman, Andrew Kasarskis, Deena R. Altman, Robert Sebra, Adolfo García-Sastre, Marta Luksza, Gopi Patel, Sarah Schaefer, Melissa Gitman, Michael D. Nowak, Alberto Paniz-Mondolfi, Emilia Mia Sordillo, Viviana Simon, Harm van Bakel |
| EPI_ISL_804380, EPI_ISL_804396, EPI_ISL_804399, EPI_ISL_804409, | Dutch COVID-19 response team | National Institute for Public Health and the Environment (RIVM) | Adam Meijer, Harry Vennema, Dirk Eggink, Matthijs Welkers, Jeroen Cremer, Sharon van den Brink, Bas van der Veer, AnneMarie van den Brandt, Florian Zwagemaker, Dennis Schmitz, Chantal Reusken, on behalf of the national COVID-19 response team |

| | | | |
|---|---|---|---|
| EPI_ISL_804424, EPI_ISL_804432, EPI_ISL_804456, EPI_ISL_804458, EPI_ISL_804462 | | | |
| EPI_ISL_806092, EPI_ISL_806094, EPI_ISL_806172, EPI_ISL_806224, EPI_ISL_806290, EPI_ISL_806321, EPI_ISL_806372, EPI_ISL_806404, EPI_ISL_806417 | Alberta Precision Labs (APL) | Alberta Precision Labs (APL) | Gordon P, Lam LG, Pabbaraju K, Wong A, Ma R, Li V, Melin A, Tipples G, Berenger B, Zelyas N, Kellner J, Bernier F, Chui L, Croxen M |
| EPI_ISL_806532, EPI_ISL_806538 | Charité Universitätsmedizin Berlin, Institut für Virologie/Labor Berlin | Charité Universitätsmedizin Berlin, Institut für Virologie | Victor M Corman, Jörn Beheim-Schwarzbach, Barbara Mühlemann, Julia Schneider, Talitha Veith, Cornelia Schlee, Tomasz Zemojtel, Terry Jones, Christian Drosten |
| EPI_ISL_812314, EPI_ISL_812323, EPI_ISL_812333, EPI_ISL_812344, EPI_ISL_812345 | United States Air Force School of Aerospace Medicine | United States Air Force School of Aerospace Medicine | Anthony Fries, Jennifer Meyer, Amanda Javorina, Sarah Purves, William Gruner, Clarise Starr, Elizabeth Macias |
| EPI_ISL_812967 | University Clinical Research Center, University of Sciences | University Clinical Research Center, University of Sciences | Diarra,B., Kone,A., Guindo,I., Bane,S., Diakite,M., Dao,S., Iknane,A.A., Doumbia,S. |
| EPI_ISL_819293, EPI_ISL_819315, EPI_ISL_819316, EPI_ISL_819317 | Hospital Universitari Vall d'Hebron - Vall d'Hebron Institut de Rercerca | Hospital Universitari Vall d'Hebron | Cristina Andrés, Maria Piñana, Josep F Abril, Damir Garcia-Cehic, Ariadna Rando, Juliana Esperalba, Maria Gema Codina, Carla Castillo, Maria Carmen Martin, Tomàs Pumarola, Josep Quer, Andrés Antón |
| EPI_ISL_831334, EPI_ISL_831335, EPI_ISL_831336, EPI_ISL_831337, EPI_ISL_831338, EPI_ISL_831339, EPI_ISL_831340, EPI_ISL_831341, EPI_ISL_831342, EPI_ISL_831343, EPI_ISL_831344, EPI_ISL_831345, EPI_ISL_831346, EPI_ISL_831347, EPI_ISL_831348, EPI_ISL_831349, EPI_ISL_831350, EPI_ISL_831351, EPI_ISL_831352, EPI_ISL_831353, EPI_ISL_831354, EPI_ISL_831355, EPI_ISL_831356, EPI_ISL_831357, EPI_ISL_831358, EPI_ISL_831359, EPI_ISL_831360, EPI_ISL_832082, EPI_ISL_832083, EPI_ISL_832084, EPI_ISL_832085, EPI_ISL_832086, EPI_ISL_832087, EPI_ISL_832088, EPI_ISL_832089, EPI_ISL_832091, EPI_ISL_832092, EPI_ISL_832093, EPI_ISL_832094 | | | |
| see above | Clinical Molecular Microbiology Laboratory, UNC Hospitals | Jeremy Wang | Jeremy Wang, Alexander Rubinsteyn, Cecilia Thompson, Clark Cunningham, Kaylee Gentry, Jonathan Juliano, Benjamin Vincent, Melissa Miller, Robert Hagan, Corbin Jones |
| EPI_ISL_833576 | Veterinary Specialized Instute "Nis" | Veterinary Specialized Institute "Nis", Serbia | Vidanovic,D., Tesovic,B., Manic,M., Petrovic,M.,Knezevic,A., Jovanovic,T., Jankovic,M., Sekler,M., Banovic Djeri,B., Petrovic,T., Volkening,J., Afonso,C. |
| EPI_ISL_837576 | Centro Nacional de Enfermedades Tropicales (CENETROP) | Laboratory of Respiratory Viruses and Measles, Oswaldo Cruz Institute, FIOCRUZ | Paola Resende, Roxana Loayza, Cinthia Avila, Luciana Appolinario, Fernando Motta, Anna Carolina Paixao, Ana Carolina Mendonca, Marilda Siqueira |
| EPI_ISL_837621, EPI_ISL_837622, EPI_ISL_837623 | Instituto Nacional de Enfermedades Respiratorias (INER) | Instituto Nacional de Enfermedades Respiratorias (INER) | Celia Boukadida, Margarita Matías-Florentino, Alma Rincón-Rubio, Hector Esteban Paz-Juárez, Olivia Briceño, Edgar Sevilla-Reyes, Fidencio Mejía-Nepomuceno, Mario Mújica-Sánchez, Eduardo Becerril-Vargas, José Arturo Martínez-Orozco, Alejandra Hernández-Terán, Jorge Salas-Hernández, Santiago Ávila-Ríos, Joel Armando Vázquez-Pérez |
| EPI_ISL_842830, EPI_ISL_842831 | Barts Health NHS Trust | COVID-19 Genomics UK (COG-UK) Consortium | CUTINO-MOGUEL, Maria-Teresa; HARRINGTON, David; OWOYEMI, Dola; SHYLINI, Raghavendran; BROAD, Claire; KELE, Beatrix |
| EPI_ISL_848624 | Evandro Chagas Institute | Evandro Chagas Institute | Santos, M.C.; Silva, A.M.; Junior, W.D.C.; Barbagelata, L.S.; Ferreira, J.A.; Sousa, E.M.A.; da Silva, P.S.; Pinheiro, K.C.; L.C.; Sousa Junior, E.C. |
| EPI_ISL_852763 | Institute of Virology, Medical Center, University of Freiburg, Freiburg, Germany | Institute of Virology, Clinial Virus Genomics, Medical Center, University of Freiburg, Freiburg, Germany | Jonas Fuchs, Lisa Kern, Sandra Reuter, Hajo Grundmann, Marcus Panning |
| EPI_ISL_856720 | Infectious Diseases Unit, Department of Internal Medicine, Azienda Ospedaliera-Universitaria di Padova | Laboratory of Infectious Diseases, Department of Biomedical and Clinical Sciences L. Sacco, University of Milan | Anna Maria Cattelan, Lolita Sasset, Davide Leoni, Alessia Lai, Annalisa Bergna, Carla Della Ventura, Claudia Balotta, Massimo Galli, Gianguglielmo Zehnder on behalf of SARS-CoV-2 ITALIAN RESEARCH ENTERPRISE-(SCIRE) Collaborative Group |
| EPI_ISL_861915 | LATE - Laboratório de Técnicas Especiais - Hospital Israelita Albert Einstein | LATE - Laboratório de Técnicas Especiais - Hospital Israelita Albert Einstein | Deyvid Amgarten, Fernanda de Mello Malta, Raquel Riyuzo, Ana Paula Moreira Salles, Pedro Henrique Sebe Rodrigues, João Renato Rebello Pinho |
| EPI_ISL_862849 | Ehime Prefectural Central Hospital | Fujita health University, School of Medicine, Department of Microbiology | Masahiro suzuki, Yohei Doi |
| EPI_ISL_862853 | Tosei General Hospital | Fujita health University, School of Medicine, Department of Microbiology | Masahiro suzuki, Yohei Doi |
| EPI_ISL_862857 | Fujita Health University Bantane Hospital | Fujita health University, School of Medicine, Department of Microbiology | Masahiro suzuki, Yohei Doi |
| EPI_ISL_862860 | Nara Medical University Hospital | Fujita health University, School of Medicine, Department of Microbiology | Masahiro suzuki, Yohei Doi |
| EPI_ISL_872089 | Instituto de Diagnostico y Referencia Epidemiologicos (INDRE) | Instituto de Diagnostico y Referencia Epidemiologicos (INDRE) | Abril Rodriguez-Maldonado, Gisela Barrera-Badillo ,Claudia Wong-Arambula , Natividad Cruz-Ortiz, Tatiana Nunez-Garcia, Dayanira Arellano-Suarez, Fabiola Garces-Ayala, Adnan-Araiza Rodríguez, Edgar Mendieta-Condado, Lucia Hernandez-Rivas, Irma Lopez-Martinez, Ernesto Ramirez-Gonzalez. |
| EPI_ISL_877014, EPI_ISL_877015 | 1.AO Universitaria 'S. Giovanni di Dio e Ruggi D'Aragona, Scuola Medica Salernitana' Hospital / 2.UOC di Virologia e Microbiologia, Università della Campania 'L. Vanvitelli' / 3.AO Universitaria 'Federico II' Napoli Hospital / 4.AORN 'San Giuseppe Moscati' Avellino Hospital / 5.AO 'San Pio - presidio G. Rummo' Benevento Hospital / 6.AO 'Sant'Anna e San Sebastiano' Caserta Hospital / 7.PO 'Maria Santissima Addolorata' Eboli Hospital / 8.Biogem Istituto di Ricerche Genetiche | 1. Genome Research Center for Health (CRGS) / 2. Laboratory of Molecular Medicine and Genomics(LMMGe) / 3. Center for Research in Pure and Applied Mathematics (CRMPA) | Giorgio Giurato (Corresponding Author), Francesca Rizzo (Corresponding Author), Alessandro Weisz (Corresponding Author), Gianluigi Franci, Giovanni Nassa, Pasquale Pagliano, Roberta Tarallo, Elena Alexandrova, Ylenia D'Agostino, Carlo Ferravante, Jessica Lamberti, Viola Melone, Domenico Memoli, Valeria Mirici Cappa, Domenico Palumbo, Giovanni Pecoraro, Assunta Sellitto, Oriana Strianese, Ilaria Terenzi, Giuseppe Fenza, Aniello Gentile, Antonello Saccomanno, Sonia Amabile, Teresa Rocco, Annamaria Salvati, Emilia Vaccaro, Massimiliano Galdiero, Michele Cennamo, Giuseppe Portella, Maria Grazia Foti, Mariarosaria Ingino, Maria Landi, Maurizio Fumi, Vincenzo Rocco, Rita Greco, Vittoria Letizia, Arnolfo Petruzziello, Maddalena Schioppa, Gregorio Goffredi, Francesca Marciano, Michele Caraglia, Alessia Cossu, Marianna Scrima |
| EPI_ISL_877592, EPI_ISL_877761 | Clinical Molecular Microbiology Laboratory, UNC Hospital | Dirk Dittmer | Razia Moorad , Justin T. Landis , Brent A. Eason, Melissa B. Miller, Linda Pluta, Dirk Dittmer, Angelica Juarez, Cecilia Thompson , Cameroon Grant, Evelyn Hoffman, Patricio Cano, Jason Wong, Carolina Caro-Vegas, Blossom Damania. |
| EPI_ISL_878539, EPI_ISL_878541 | Robert Garry lab | Andersen lab at Scripps Research | Allison Smither, Gilberto Sabino-Santos, Patricia Snarski, Lilia Melnik, Antoinette Bell, Kaylynn Genemaras, Arnaud Drouin, Dahlene Fusco, Robert Garry with SEARCH Alliance San Diego |
| EPI_ISL_883038, EPI_ISL_883039, EPI_ISL_883040, EPI_ISL_883041, EPI_ISL_883042, EPI_ISL_883043, EPI_ISL_883044, EPI_ISL_883045, EPI_ISL_883046, EPI_ISL_883047, EPI_ISL_883048, EPI_ISL_883049, EPI_ISL_883062, EPI_ISL_883063, EPI_ISL_883064, EPI_ISL_883065 | | | |
| see above | Kansas Health and Environmental Lab | Kansas Health and Environmental Lab | Mike Grose, Carissa Robertson, Ben Olsen, and Phil Adam |
| EPI_ISL_884556, EPI_ISL_884557 | Molecular Microbiology & Immunology, University of Missouri | Molecular Microbiology & Immunology, University of Missouri | Tang,C.Y., Li,T., Hang,J., Lidl,G.M., Wan,X.-F. |
| EPI_ISL_887230, EPI_ISL_887246, EPI_ISL_887261, EPI_ISL_887275, EPI_ISL_887289, EPI_ISL_887338, EPI_ISL_887347, EPI_ISL_887364, EPI_ISL_887389, EPI_ISL_887393, EPI_ISL_887396, EPI_ISL_887399, EPI_ISL_887411, EPI_ISL_887414 | | | |
| see above | Protzer Lab | Protzer Lab, Gagneur Lab, Robert Koch Institut | Ulrike Protzer, Dieter Hoffmann, Eva Schulte, Andrea Theumer, Oliver Drechsel, Max von Kleist,Aleksandar Radonic,Stephan Fuchs, Alexander Karollus, Julien Gagneur |
| EPI_ISL_887506 | 1.AO Universitaria 'S. Giovanni di Dio e Ruggi D'Aragona, Scuola Medica Salernitana' Hospital / 2.UOC di Virologia e Microbiologia, Università della Campania 'L. Vanvitelli' / 3.AO Universitaria 'Federico II' Napoli Hospital / 4.AORN 'San Giuseppe Moscati' Avellino Hospital / 5.AO 'San Pio - presidio G. Rummo' Benevento Hospital / 6.AO 'Sant'Anna e San Sebastiano' Caserta Hospital / 7.PO 'Maria Santissima Addolorata' Eboli Hospital / 8.Biogem Istituto di Ricerche Genetiche | 1. Genome Research Center for Health (CRGS) / 2. Laboratory of Molecular Medicine and Genomics(LMMGe) / 3. Center for Research in Pure and Applied Mathematics (CRMPA) | Giorgio Giurato, Francesca Rizzo, Alessandro Weisz, Gianluigi Franci, Giovanni Nassa, Pasquale Pagliano, Roberta Tarallo, Elena Alexandrova, Ylenia D'Agostino, Carlo Ferravante, Jessica Lamberti, Viola Melone, Domenico Memoli, Valeria Mirici Cappa, Domenico Palumbo, Giovanni Pecoraro, Assunta Sellitto, Oriana Strianese, Ilaria Terenzi, Giuseppe Fenza, Aniello Gentile, Antonello Saccomanno, Sonia Amabile, Teresa Rocco, Annamaria Salvati, Emilia Vaccaro, Massimiliano Galdiero, Michele Cennamo, Giuseppe Portella, Maria Grazia Foti, Mariarosaria Ingino, Maria Landi, Maurizio Fumi, Vincenzo Rocco, Rita Greco, Vittoria Letizia, Arnolfo Petruzziello, Maddalena Schioppa, Gregorio Goffredi, Francesca Marciano, Michele Caraglia, Alessia Cossu, Marianna Scrima |

| | | | |
|---|---|---|---|
| EPI_ISL_889008 | RS Sentra Medika Cibinong | Eijkman Institute for Molecular Biology, Ministry of Research and Technology/National Agency for Research and Innovation | Edison Johar, Frilasita A Yudhaputri, Hidayat Trimarsanto, Iskandar Adnan, Lydia V. Panggalo, Sukma Oktavianthi, Willy Agustine, Safarina G Malik, Khin Saw Myint, Amin Soebandrio |
| EPI_ISL_889381, EPI_ISL_889382, EPI_ISL_889383 | Olomouc University Hospital | Institute of Applied Biotechnologies a.s. | Petr Klempt, Ondej Brzo, Martin Kašný, Kateina Kvapilová, Marián Hajdúch, Petr Kvapil |
| EPI_ISL_889808, EPI_ISL_889815, EPI_ISL_889822, EPI_ISL_889829, EPI_ISL_889833, EPI_ISL_889835, EPI_ISL_889837, EPI_ISL_889839, EPI_ISL_889840, EPI_ISL_889842, EPI_ISL_889843, EPI_ISL_889845, EPI_ISL_889848, EPI_ISL_889849, EPI_ISL_889850, EPI_ISL_889851, EPI_ISL_889852, EPI_ISL_889853, EPI_ISL_889854, EPI_ISL_889855, EPI_ISL_889856, EPI_ISL_889860, EPI_ISL_889862, EPI_ISL_889863, EPI_ISL_889864, EPI_ISL_889865, EPI_ISL_889866, EPI_ISL_889872, EPI_ISL_889889, EPI_ISL_889906, EPI_ISL_889907, EPI_ISL_889913, EPI_ISL_889914, EPI_ISL_889918, EPI_ISL_889919, EPI_ISL_889920, EPI_ISL_889927, EPI_ISL_889928, EPI_ISL_889929, EPI_ISL_889930, EPI_ISL_889931, EPI_ISL_889932, EPI_ISL_889933, EPI_ISL_889934, EPI_ISL_889936, EPI_ISL_889942, EPI_ISL_889943, EPI_ISL_889944, EPI_ISL_889945, EPI_ISL_889946, EPI_ISL_889954, EPI_ISL_889956, EPI_ISL_889959, EPI_ISL_889960, EPI_ISL_889974, EPI_ISL_889975, EPI_ISL_889980, EPI_ISL_889981, EPI_ISL_889982, EPI_ISL_889988, EPI_ISL_889989, EPI_ISL_889992, EPI_ISL_889993, EPI_ISL_889994, EPI_ISL_889995, EPI_ISL_889996, EPI_ISL_889997, EPI_ISL_889998, EPI_ISL_889999, EPI_ISL_890003, EPI_ISL_890005, EPI_ISL_890007, EPI_ISL_890015, EPI_ISL_890016, EPI_ISL_890017, EPI_ISL_890018, EPI_ISL_890019, EPI_ISL_890020, EPI_ISL_890021, EPI_ISL_890022, EPI_ISL_890023, EPI_ISL_890024, EPI_ISL_890025, EPI_ISL_890026, EPI_ISL_890027, EPI_ISL_890030, EPI_ISL_890031, EPI_ISL_890032 | | | |
| see above | Laboratoire de santé publique du Québec | Laboratoire de santé publique du Québec | Sandrine Moreira, Ioannis Ragoussis, Guillaume Bourque, Jesse Shapiro, Mark Lathrop and Michel Roger on behalf of the CoVSeQ research group |
| EPI_ISL_890851 | Laboratoire de santé publique du Québec | Laboratoire de santé publique du Québec | Sandrine Moreira, Ioannis Ragoussis, Guillaume Bourque, Jesse Shapiro, Mark Lathrop and Michel Roger on behalf of the CoVSeQ research group (http://covseq.ca/researchgroup) |
| EPI_ISL_892515, EPI_ISL_892516, EPI_ISL_892517, EPI_ISL_892518, EPI_ISL_892519, EPI_ISL_892520, EPI_ISL_892521, EPI_ISL_892522, EPI_ISL_892523, EPI_ISL_892524, EPI_ISL_892525, EPI_ISL_892526, EPI_ISL_892527, EPI_ISL_892528, EPI_ISL_892529, EPI_ISL_892530, EPI_ISL_892531, EPI_ISL_892532, EPI_ISL_892533 | | | |
| see above | Osaka Institute of Public Health, Morinomiya Center | Pathogen Genomics Center, National Institute of Infectious Diseases | Tsuyoshi Sekizuka, Kentaro Itokawa, Rina Tanaka, Masanori Hashino, Makoto Kuroda |
| EPI_ISL_895749, EPI_ISL_895774, EPI_ISL_895782, EPI_ISL_895804, EPI_ISL_895812, EPI_ISL_895820, EPI_ISL_895821 | Molecular biology division, Institute of Clinical Biochemistry and Diagnostics, Charles University, Faculty of Medicine in Hradec Králové and University Hospital Hradec Králové | Molecular biology division, Institute of Clinical Biochemistry and Diagnostics, Charles University, Faculty of Medicine in Hradec Králové and University Hospital Hradec Králové | Helena Kovaíková, Petr Brož, Ivana Baranová, Kateina Hrochová, Tereza Baková, Jitka Novotná, Kateina Pehlíková, Vladimír Palika. Cooperation project with BioVendor-R&D and bioinformatics company BIOXSYS s r.o. |
| EPI_ISL_896572, EPI_ISL_896573, EPI_ISL_896574, EPI_ISL_896575 | Department of Medical Research | DMR | Myat Htut Nyunt |
| EPI_ISL_897529, EPI_ISL_897530, EPI_ISL_897531, EPI_ISL_897532, EPI_ISL_897533, EPI_ISL_897534, EPI_ISL_897535, EPI_ISL_897536, EPI_ISL_898569, EPI_ISL_898570, EPI_ISL_898571, EPI_ISL_898572, EPI_ISL_898573, EPI_ISL_898574, EPI_ISL_898575, EPI_ISL_898576, EPI_ISL_898577, EPI_ISL_898578, EPI_ISL_898579, EPI_ISL_898580, EPI_ISL_898581, EPI_ISL_898582, EPI_ISL_898583, EPI_ISL_898584 | | | |
| see above | Pathogen Genomics Center, National Institute of Infectious Diseases | Pathogen Genomics Center, National Institute of Infectious Diseases | Tsuyoshi Sekizuka, Kentaro Itokawa, Rina Tanaka, Masanori Hashino, Makoto Kuroda |
| EPI_ISL_900117, EPI_ISL_900142, EPI_ISL_900197, EPI_ISL_900240, EPI_ISL_900264, EPI_ISL_900286, EPI_ISL_900292, EPI_ISL_900327, EPI_ISL_900329 | MEPHI, Aix Marseille University | MEPHI, Aix Marseille University | Anthony LEVASSEUR |
| EPI_ISL_901227, EPI_ISL_901228, EPI_ISL_901229, EPI_ISL_901230, EPI_ISL_901231, EPI_ISL_901232, EPI_ISL_901233, EPI_ISL_901234, EPI_ISL_901235, EPI_ISL_901236, EPI_ISL_901237, EPI_ISL_901238, EPI_ISL_901239, EPI_ISL_901240, EPI_ISL_901241, EPI_ISL_901242, EPI_ISL_901243, EPI_ISL_901244, EPI_ISL_901245, EPI_ISL_901246, EPI_ISL_901247, EPI_ISL_901248, EPI_ISL_901249, EPI_ISL_901250, EPI_ISL_901328, EPI_ISL_901329, EPI_ISL_901330, EPI_ISL_901331, EPI_ISL_901332, EPI_ISL_901333, EPI_ISL_901334, EPI_ISL_901335, EPI_ISL_901336, EPI_ISL_901337, EPI_ISL_901338, EPI_ISL_901339, EPI_ISL_901340, EPI_ISL_901341, EPI_ISL_901342, EPI_ISL_901343, EPI_ISL_901344, EPI_ISL_901345, EPI_ISL_901346, EPI_ISL_901347, EPI_ISL_901348, EPI_ISL_901349, EPI_ISL_901350, EPI_ISL_901351, EPI_ISL_901352, EPI_ISL_901353, EPI_ISL_901354, EPI_ISL_901355, EPI_ISL_901356, EPI_ISL_901364, EPI_ISL_901365, EPI_ISL_901366, EPI_ISL_901367, EPI_ISL_902308, EPI_ISL_902473, EPI_ISL_902475, EPI_ISL_902476 | | | |
| see above | Pathogen Genomics Center, National Institute of Infectious Diseases | Pathogen Genomics Center, National Institute of Infectious Diseases | Tsuyoshi Sekizuka, Kentaro Itokawa, Rina Tanaka, Masanori Hashino, Makoto Kuroda |
| EPI_ISL_904019 | DB Diagnosticos do Brasil | Laboratório de Parasitologia Médica - Instituto de Medicina Tropical - Universidade de São Paulo | Brazil-UK Centre for Arbovirus Discovery Diagnosis Genomics and Epidemiology (CADDE) Genomic Network - Instituto de Medicina Tropical |
| EPI_ISL_906476 | Pathogen Genomics Center, National Institute of Infectious Diseases | Pathogen Genomics Center, National Institute of Infectious Diseases | Tsuyoshi Sekizuka, Kentaro Itokawa, Rina Tanaka, Masanori Hashino, Makoto Kuroda |
| EPI_ISL_906769, EPI_ISL_906770, EPI_ISL_906771 | Dr. Leonard A. Miller Centre for Health Services | National Microbiology Laboratory (NML) | Anna Majer, Shari Tyson, Grace Seo, Philip Mabon, Elsie Grudeski, Rhiannon Huzarewich, Russell Mandes, Anneliese Landgraff, Jennifer Tanner, Natalie Knox, Morag Graham, Gary Van Domselaar, Robert Needle, Yang Yu, Laura Gilbert, George Zahariadis, Nathalie Bastien, Yan Li, Timothy Booth, Darian Hole, Madison Chapel, Kerri Smith, CanCOGeN's metadata curation team, Public Health Agency of Canada CanCOGeN team |
| EPI_ISL_914997 | QEII Health Sciences Centre | National Microbiology Laboratory (NML) | Anna Majer, Shari Tyson, Grace Seo, Philip Mabon, Elsie Grudeski, Rhiannon Huzarewich, Russell Mandes, Anneliese Landgraff, Jennifer Tanner, Natalie Knox, Morag Graham, Gary Van Domselaar, Todd Hatchette, Jason LeBlanc, Janice Pettipas, Dan Gaston, Nathalie Bastien, Yan Li, Timothy Booth, Darian Hole, Madison Chapel, CanCOGeN's metadata curation team, Public Health Agency of Canada CanCOGeN team |
| EPI_ISL_925537, EPI_ISL_925543, EPI_ISL_925544, EPI_ISL_925545, EPI_ISL_925546, EPI_ISL_925547, EPI_ISL_925548, EPI_ISL_925549, EPI_ISL_925550, EPI_ISL_925551, EPI_ISL_925552, EPI_ISL_925553, EPI_ISL_925554, EPI_ISL_925555, EPI_ISL_925556, EPI_ISL_925557, EPI_ISL_925558, EPI_ISL_925559, EPI_ISL_925560, EPI_ISL_925561, EPI_ISL_925562, EPI_ISL_925563, EPI_ISL_925564, EPI_ISL_925565, EPI_ISL_925566, EPI_ISL_925567, EPI_ISL_925568, EPI_ISL_925569, EPI_ISL_925570, EPI_ISL_925571, EPI_ISL_925572, EPI_ISL_925573, EPI_ISL_925574, EPI_ISL_925575, EPI_ISL_925576, EPI_ISL_925577, EPI_ISL_925578, EPI_ISL_925579, EPI_ISL_925580, EPI_ISL_925581, EPI_ISL_925582, EPI_ISL_925583, EPI_ISL_925678, EPI_ISL_925679, EPI_ISL_925680, EPI_ISL_925681, EPI_ISL_925682, EPI_ISL_925683, EPI_ISL_925684, EPI_ISL_925685, EPI_ISL_925686, EPI_ISL_925687, EPI_ISL_925688, EPI_ISL_925689, EPI_ISL_925690, EPI_ISL_925691, EPI_ISL_925692, EPI_ISL_925693, EPI_ISL_925694, EPI_ISL_925695, EPI_ISL_925696, EPI_ISL_925697, EPI_ISL_925726, EPI_ISL_925727, EPI_ISL_925731, EPI_ISL_925732, EPI_ISL_925733, EPI_ISL_925734, EPI_ISL_925735, EPI_ISL_925736, EPI_ISL_925737, EPI_ISL_925738, EPI_ISL_925739, EPI_ISL_925740, EPI_ISL_925741, EPI_ISL_925742, EPI_ISL_925743, EPI_ISL_925744, EPI_ISL_925745, EPI_ISL_925746, EPI_ISL_925747, EPI_ISL_925751, EPI_ISL_925752, EPI_ISL_925753, EPI_ISL_925754, EPI_ISL_925755, EPI_ISL_925756, EPI_ISL_925757, EPI_ISL_925758, EPI_ISL_925759, EPI_ISL_925760, EPI_ISL_925761 | | | |
| see above | Public Health Ontario Laboratory | Public Health Ontario Laboratory | Vanessa G Allen, Philip Banh, Yao Chen, Richard de Borja, Alireza Eshaghi, Nahuel Fittipaldi, Christine Frantz, Jonathan B Gubbay, Jennifer L Guthrie, Lawrence Heisler, Esha Joshi, Michael Laszloffy, Aimin Li, Michael CY Li, Dean Maxwell, Sandeep Nagra, Samir N Patel, Jared Simpson, Karthikeyan Sivaraman, Ashleigh Sullivan, Yogi Sundaravadanam, Sarah Teatero, Matthew Watson, Andre Villegas, Sandra Zittermann |
| EPI_ISL_933618, EPI_ISL_933619 | Toronto Invasive Bacterial Diseases Network | McMaster University | Allison McGeer, Patryk Aftanas, Hooman Derakhshani, Angel Li, Kuganya Nirmalarajah, Emily Panousis, Ahmed Draia, Jalees Nasir, Michael Surette, Samira Mubareka, Andrew G. McArthur |
| EPI_ISL_934422, EPI_ISL_934423 | 1.AO Universitaria 'S. Giovanni di Dio e Ruggi D'Aragona, Scuola Medica Salernitana' Hospital / 2.UOC di Virologia e Microbiologia, Università della Campania 'L. Vanvitelli' / 3.AO Universitaria 'Federico II' Napoli Hospital / 4.AORN 'San Giuseppe Moscati' Avellino Hospital / 5.AO 'San Pio - presidio G. Rummo' Benevento Hospital / 6.AO 'Sant'Anna e San Sebastiano' Caserta Hospital / 7.PO 'Maria Santissima Addolorata' Eboli Hospital / 8.Biogem Istituto di Ricerche Genetiche | 1. Genome Research Center for Health (CRGS) / 2. Laboratory of Molecular Medicine and Genomics(LMMGe) / 3. Center for Research in Pure and Applied Mathematics (CRMPA) | Giorgio Giurato (Corresponding Author), Francesca Rizzo (Corresponding Author), Alessandro Weisz (Corresponding Author), Gianluigi Franci, Giovanni Nassa, Pasquale Pagliano, Roberta Tarallo, Elena Alexandrova, Ylenia D'Agostino, Carlo Ferravante, Jessica Lamberti, Viola Melone, Domenico Memoli, Valeria Mirici Cappa, Domenico Palumbo, Giovanni Pecoraro, Assunta Sellitto, Oriana Strianese, Ilaria Terenzi, Giuseppe Fenza, Aniello Gentile, Antonello Saccomanno, Sonia Amabile, Teresa Rocco, Annamaria Salvati, Emilia Vaccaro, Massimiliano Galdiero, Michele Cennamo, Giuseppe Portella, Maria Grazia Foti, Mariarosaria Ingino, Maria Landi, Maurizio Fumi, Vincenzo Rocco, Rita Greco, Vittoria Letizia, Arnolfo Petruzziello, Maddalena Schioppa, Gregorio Goffredi, Francesca Marciano, Michele Caraglia, Alessia Cossu, Marianna Scrima, Edmondo Adorisio, Morena D'Avenia, Michela Iacobellis, Rosanna Piluscio, Giorgio Dirani, Vittorio Sambri, Simona Semprini, Silvia Zanoli, Francesco Curcio, Stefania Marzinotto, Andreina Baj, Fausto Sessa. |
| EPI_ISL_939660, EPI_ISL_939661 | Division of Infectious Diseases, University Hospital Zürich | Institute of Medical Virology, University of Zurich | Stefan Schmutz, Maryam Zaheri, Verena Kufner, Annette Audigé, Maria Grünberg, Kevin Steiner, Jon Huder, Cyril Shah, Riccarda Capaul, Guido Bloemberg, Jürg Böni, Michael Huber, Alexandra Trkola |
| EPI_ISL_940163, EPI_ISL_940164, EPI_ISL_940166, EPI_ISL_940167, EPI_ISL_940168, EPI_ISL_940169, EPI_ISL_940170, EPI_ISL_940183, EPI_ISL_940184, EPI_ISL_940242, EPI_ISL_940247, EPI_ISL_940248, EPI_ISL_940249, EPI_ISL_940250, EPI_ISL_940251, EPI_ISL_940252, EPI_ISL_940253, EPI_ISL_940254, EPI_ISL_940255, EPI_ISL_940256, EPI_ISL_940257, EPI_ISL_940352, EPI_ISL_940353, EPI_ISL_940357, EPI_ISL_940358, EPI_ISL_940359, EPI_ISL_940361, EPI_ISL_940362, EPI_ISL_940363, EPI_ISL_940364, EPI_ISL_940370 | | | |
| see above | Hôpital Bichat Claude Bernard, Laboratoire de Virologie | IAME UMR1137 Inserm, Université de Paris, Hôpital Bichat | Antoine Bridier-Nahmias, Amélie Recoing, Quentin Le Hingrat, Lena Daniel, Siham Hamri, Gilles Collin, Alexandre Storto, Mélanie Bertine, Charlotte Charpentier, Nadhira Houhou-Fidouh, Diane Descamps, Benoit Visseaux |
| EPI_ISL_949240 | Departamento de Microbiología, CDB, Hospital Clínic, Barcelona | SeqCOVID-SPAIN consortium/IBV(CSIC) | Andrea Vergara, Mikel Martínez, Elisa Rubio, Jéssica Navero, Aida Peiró and SeqCOVID-SPAIN consortium |

| | | | |
|---|---|---|---|
| EPI_ISL_949345 | University of Birmingham | COVID-19 Genomics UK (COG-UK) Consortium | Institute of Microbiology, University of Birmingham: Claire McMurray, Joanne Stockton, Samuel Nicholls, Radoslaw Poplawski, Will Rowe, Josh Quick, Nicholas Loman. University of Birmingham Testing Laboratory: Celina M Whalley, Andrew Bosworth, Charlotte Poxon, Kasun Wanigasooriya, Oliver Pickles, Mike Kidd, Alex Richter, Andrew D Beggs PHE Heartlands Lab: Husam Osman, Andrew Bosworth. Queen Elizabeth Hospital: Anna Casey |
| EPI_ISL_954727, EPI_ISL_954728, EPI_ISL_954729, EPI_ISL_954730, EPI_ISL_954731, EPI_ISL_954732, EPI_ISL_954733, EPI_ISL_954734, EPI_ISL_954735, EPI_ISL_954736, EPI_ISL_954737, EPI_ISL_954738, EPI_ISL_954739, EPI_ISL_954740, EPI_ISL_954741, EPI_ISL_954742, EPI_ISL_954743, EPI_ISL_954744 | | | |
| see above | Laboratoire de santé publique du Québec | Laboratoire de santé publique du Québec | Sandrine Moreira, Ioannis Ragoussis, Guillaume Bourque, Jesse Shapiro, Mark Lathrop and Michel Roger on behalf of the CoVSeQ research group (http://covseq.ca/researchgroup) |
| EPI_ISL_955257 | HGZ 32 VILLA COAPA | BIOBANCO / COCTI | Borja-Aburto VH, Grajales-Muñiz C, Santacruz Tinoco CE, Rojas-Mendoza T, Ochoa Carrera LA, Sánchez A, Grande R, Isa P, Taboada B, López S, Arias C, Muñoz-Medina JE |
| EPI_ISL_955982, EPI_ISL_955983, EPI_ISL_955984, EPI_ISL_955985, EPI_ISL_955987, EPI_ISL_955989, EPI_ISL_955991, EPI_ISL_955992, EPI_ISL_955993, EPI_ISL_955994, EPI_ISL_955995, EPI_ISL_955996, EPI_ISL_955997, EPI_ISL_955998, EPI_ISL_955999, EPI_ISL_956000, EPI_ISL_956005, EPI_ISL_956007, EPI_ISL_956008, EPI_ISL_956010, EPI_ISL_956011, EPI_ISL_956012, EPI_ISL_956013, EPI_ISL_956015, EPI_ISL_956016, EPI_ISL_956018, EPI_ISL_956019, EPI_ISL_956022, EPI_ISL_956024, EPI_ISL_956029, EPI_ISL_956033, EPI_ISL_956034, EPI_ISL_956036, EPI_ISL_956039, EPI_ISL_956041, EPI_ISL_956046, EPI_ISL_956047, EPI_ISL_956048, EPI_ISL_956049, EPI_ISL_956051, EPI_ISL_956052, EPI_ISL_956053, EPI_ISL_956056, EPI_ISL_956061, EPI_ISL_956062, EPI_ISL_956070, EPI_ISL_956074, EPI_ISL_956075, EPI_ISL_956076, EPI_ISL_956080, EPI_ISL_956081, EPI_ISL_956082, EPI_ISL_956083, EPI_ISL_956084, EPI_ISL_956086, EPI_ISL_956089, EPI_ISL_956090, EPI_ISL_956092, EPI_ISL_956093, EPI_ISL_956097, EPI_ISL_956098, EPI_ISL_956100, EPI_ISL_956102, EPI_ISL_956104, EPI_ISL_956105, EPI_ISL_956107, EPI_ISL_956109, EPI_ISL_956113, EPI_ISL_956115, EPI_ISL_956120, EPI_ISL_956121, EPI_ISL_956123, EPI_ISL_956124, EPI_ISL_956125, EPI_ISL_956126, EPI_ISL_956130, EPI_ISL_956131, EPI_ISL_956132, EPI_ISL_956133, EPI_ISL_956137, EPI_ISL_956141, EPI_ISL_956144, EPI_ISL_956145, EPI_ISL_956149, EPI_ISL_956152, EPI_ISL_956153, EPI_ISL_956154, EPI_ISL_956155, EPI_ISL_956157, EPI_ISL_956158, EPI_ISL_956159, EPI_ISL_956160, EPI_ISL_956161, EPI_ISL_956163, EPI_ISL_956165, EPI_ISL_956166, EPI_ISL_956167, EPI_ISL_956170, EPI_ISL_956172, EPI_ISL_956174, EPI_ISL_956176, EPI_ISL_956177, EPI_ISL_956178, EPI_ISL_956179, EPI_ISL_956180, EPI_ISL_956181, EPI_ISL_956182, EPI_ISL_956183, EPI_ISL_956184, EPI_ISL_956185, EPI_ISL_956186, EPI_ISL_956187, EPI_ISL_956188, EPI_ISL_956189, EPI_ISL_956190, EPI_ISL_956192, EPI_ISL_956193, EPI_ISL_956194, EPI_ISL_956195, EPI_ISL_956196, EPI_ISL_956199, EPI_ISL_956200, EPI_ISL_956203, EPI_ISL_956205, EPI_ISL_956207, EPI_ISL_956209, EPI_ISL_956210, EPI_ISL_956211, EPI_ISL_956212, EPI_ISL_956213, EPI_ISL_956214, EPI_ISL_956216, EPI_ISL_956217, EPI_ISL_956218, EPI_ISL_956219, EPI_ISL_956220, EPI_ISL_956221, EPI_ISL_956222, EPI_ISL_956224, EPI_ISL_956225, EPI_ISL_956227, EPI_ISL_956228, EPI_ISL_956229, EPI_ISL_956230, EPI_ISL_956231, EPI_ISL_956232, EPI_ISL_956234, EPI_ISL_956236, EPI_ISL_956237, EPI_ISL_956238, EPI_ISL_956239, EPI_ISL_956240, EPI_ISL_956242, EPI_ISL_956243, EPI_ISL_956244, EPI_ISL_956246, EPI_ISL_956247, EPI_ISL_956248, EPI_ISL_956250, EPI_ISL_956253, EPI_ISL_956254, EPI_ISL_956255, EPI_ISL_956256, EPI_ISL_956257, EPI_ISL_956258, EPI_ISL_956259, EPI_ISL_956260, EPI_ISL_956262, EPI_ISL_956268 | | | |
| see above | Pathology and Laboratory Medicine Institute, Cleveland Clinic, Ohio, USA | Pathology and Laboratory Medicine Institute, Cleveland Clinic, Ohio, USA | Frank P. Esper, Yu-Wei Cheng, Thamali M. Adhikari, Zheng Jin Tu, Daniel H. Farkas, Gary Procop, Jennifer Ko, Timothy A. Chan, Jing Li, Brian P. Rubin |
| EPI_ISL_960309 | NLZOH, Laboratory for Virology | NLZOH, Laboratory for Virology | Katarina Prosenc (Laboratory for Virology), Cesare Camma (IZSAM), Erik Alm (ECDC) |
| EPI_ISL_961290, EPI_ISL_961292, EPI_ISL_961293, EPI_ISL_961294, EPI_ISL_961295, EPI_ISL_961296, EPI_ISL_961297, EPI_ISL_961298, EPI_ISL_961299, EPI_ISL_961300, EPI_ISL_961301, EPI_ISL_961302, EPI_ISL_961303, EPI_ISL_961304, EPI_ISL_961305, EPI_ISL_961306, EPI_ISL_961307, EPI_ISL_961308, EPI_ISL_961309, EPI_ISL_961310, EPI_ISL_961311, EPI_ISL_961312, EPI_ISL_961313, EPI_ISL_961314, EPI_ISL_961315 | | | |
| see above | Biotia | Biotia | Dorottya Nagy-Szakal, Mara Couto-Rodriguez, Heather Wells, Joseph Barrows, Marilyne Debieu, Kristin Butcher, Siyuan Chen, Agnes Berki, Courteny Hager, Robert Boorstein, Mariah Taylor, Colleen Jonsson, Christopher Mason, Niamh O'Hara |
| EPI_ISL_961669 | Hôpital Georges L. Dumont | National Microbiology Laboratory (NML) | Anna Majer, Shari Tyson, Grace Seo, Philip Mabon, Elsie Grudeski, Rhiannon Huzarewich, Russell Mandes, Anneliese Landgraff, Jennifer Tanner, Natalie Knox, Morag Graham, Gary Van Domselaar, Richard Garceau, Guillaume Desnoyers, Nathalie Bastien, Yan Li, Timothy Booth, Darian Hole, Madison Chapel, Kirsten Biggar, CanCOGeN's metadata curation team, Public Health Agency of Canada CanCOGeN team |
| EPI_ISL_964879 | Institute of Virology Ulm University Medical Center Prof. Dr. Thomas Stamminger | Münch Lab / Ulm University Medical Center Kirchhoff Lab / Ulm University Medical Center Sparrer Lab / Ulm University Medical Center Blum Lab / LMU Munich | Thomas Stamminger, Helmut Blum, Stefan Krebs, Alexander Graf, Rüdiger Gross, Janis Müller, Carina Conzelmann, Jan Münch, Frank Kirchhoff, Konstantin Sparrer |
| EPI_ISL_965583 | Dutch COVID-19 response team | Medical Microbiology, Maastricht University Medical Centre | Jozef Dingemans*, Brian van der Veer*, Erik Beuken, Carmen Reumkens, Lieke van Alphen, Christian Hoebe, Paul Savelkoul |
| EPI_ISL_965983, EPI_ISL_965999, EPI_ISL_966001, EPI_ISL_966010, EPI_ISL_966015, EPI_ISL_966018, EPI_ISL_966024, EPI_ISL_966026, EPI_ISL_966031, EPI_ISL_966176, EPI_ISL_966177, EPI_ISL_966178, EPI_ISL_966181, EPI_ISL_966183, EPI_ISL_966185, EPI_ISL_966186, EPI_ISL_966187, EPI_ISL_966188, EPI_ISL_966189, EPI_ISL_966190, EPI_ISL_966191, EPI_ISL_966192, EPI_ISL_966193, EPI_ISL_966194, EPI_ISL_966195, EPI_ISL_966196, EPI_ISL_966197, EPI_ISL_966198, EPI_ISL_966199, EPI_ISL_966200, EPI_ISL_966201, EPI_ISL_966202, EPI_ISL_966203, EPI_ISL_966205, EPI_ISL_966206, EPI_ISL_966207, EPI_ISL_966208, EPI_ISL_966209, EPI_ISL_966210, EPI_ISL_966211, EPI_ISL_966212, EPI_ISL_966213, EPI_ISL_966214, EPI_ISL_966215, EPI_ISL_966216, EPI_ISL_966217, EPI_ISL_966218, EPI_ISL_966219, EPI_ISL_966220, EPI_ISL_966221, EPI_ISL_966222, EPI_ISL_966223, EPI_ISL_966224, EPI_ISL_966225, EPI_ISL_966226, EPI_ISL_966236, EPI_ISL_966237, EPI_ISL_966239, EPI_ISL_966240, EPI_ISL_966242, EPI_ISL_966243, EPI_ISL_966244, EPI_ISL_966247, EPI_ISL_966248, EPI_ISL_966249, EPI_ISL_966250, EPI_ISL_966251, EPI_ISL_966252, EPI_ISL_966254, EPI_ISL_966255, EPI_ISL_966256, EPI_ISL_966257, EPI_ISL_966258, EPI_ISL_966259 | | | |
| see above | Public Health Ontario Laboratory | Public Health Ontario Laboratory | Vanessa G Allen, Philip Banh, Yao Chen, Richard de Borja, Alireza Eshaghi, Nahuel Fittipaldi, Christine Frantz, Jonathan B Gubbay, Jennifer L Guthrie, Lawrence Heisler, Esha Joshi, Michael Laszloffy, Aimin Li, Michael CY Li, Dean Maxwell, Sandeep Nagra, Samir N Patel, Jared Simpson, Karthikeyan Sivaraman, Ashleigh Sullivan, Yogi Sundaravadanam, Sarah Teatero, Andre Villegas, Matthew Watson, Sandra Zittermann |
| EPI_ISL_968174 | Clinical Molecular Microbiology Laboratory, UNC Hospital | Dirk Dittmer | Justin T. Landis , Razia Moorad , Brent A. Eason, Melissa B. Miller, Linda Pluta, Dirk Dittmer, Angelica Juarez, Cecilia Thompson, Shawn Hawken, Cameroon Grant, Evelyn Hoffman, Patricio Cano, Jason Wong, Carolina Caro-Vegas, Ryan McNamara, Blossom Damania. |
| EPI_ISL_978494 | Central Public Health Laboratory - LACEN -Bahia, Salvador, Brazil | Central Public Health Laboratory - LACEN -Bahia, Salvador, Brazil | Stephane Tosta, Luciana Oliveira, Vanessa Nardy,Patrícia Cajado,Marcela Gómez, Breno Dominguez, Jaqueline Gomes, Vagner Fonseca,Marta Giovanetti,Luiz Alcantara, Felicidade Pereira, Arabela Leal |
