## Supplementary material for "Detection and characterization of the SARS-CoV-2 lineage B.1.526 in New York": Supp. Table 3 GISAID Acknowledgment part 2: gisaid_hcov-19_acknowledgement_table_2021_02_12_16-8.pdf

We gratefully acknowledge the following Authors from the Originating laboratories responsible for obtaining the specimens, as well as the Submitting laboratories where the genome data were generated and shared via GISAID, on which this research is based.

All Submitters of data may be contacted directly via www.gisaid.org

Authors are sorted alphabetically.

| Accession ID | Originating Laboratory | Submitting Laboratory | Authors |
|---|---|---|---|
| EPI_ISL_420293 | Wildlife Conservation Society, Bronx Zoo | Diagnostic Virology Laboratory, United States Department of Agriculture, National Veterinary Services Laboratories | Patrick K. Mitchell, Renee R. Anderson, Brittany Chilson, Roopa Venugopalan, D. G. Diel, Laura B. Goodman, L. Wang, F. Yuan, Y. Fang, Mary Lea Killian, Kerrie Franzen, Nichole Hines Bergeson, Ivan Kuzmin, Melinda Jenkins-Moore, Tod P. Stuber |
| EPI_ISL_421974, EPI_ISL_421976, EPI_ISL_421986, EPI_ISL_421989, EPI_ISL_421998, EPI_ISL_422000, EPI_ISL_422001, EPI_ISL_422006, EPI_ISL_422008, EPI_ISL_422009, EPI_ISL_422010, EPI_ISL_422011, EPI_ISL_422012 | | | |
| see above | Respiratory Virus Unit, Microbiology Services Colindale, Public Health England | Respiratory Virus Unit, Microbiology Services Colindale, Public Health England | Monica Galiano, Shahjahan Miah, Angie Lackenby, Omolola Akinbami, Tiina Talts, Leena Bhaw, Richard Myers, Steven Platt, Kirstin Edwards, Jonathan Hubb, Joanna Ellis, Maria Zambon |
| EPI_ISL_422459 | Gundersen Molecular Diagnostics Laboratory | Kabara Cancer Research Institute | Craig S. Richmond & Paraic A. Kenny |
| EPI_ISL_422461, EPI_ISL_422462, EPI_ISL_422463 | Gundersen Molecular Diagnostics Laboratory | Kabara Cancer Research Institute | Craig S. Richmond, Paraic A. Kenny |
| EPI_ISL_422616, EPI_ISL_422617, EPI_ISL_422618, EPI_ISL_422619, EPI_ISL_422620, EPI_ISL_422621, EPI_ISL_422622, EPI_ISL_422623, EPI_ISL_422624, EPI_ISL_422625, EPI_ISL_422626, EPI_ISL_422627 | | | |
| see above | Dutch COVID-19 response team | Erasmus Medical Center | Bas Oude Munnink, David Nieuwenhuijse, Reina Sikkema, Claudia Schapendonk, Irina Chestakova, Anne van der Linden, Theo Bestebroer, Stefan van Nieuwkoop, Mark Pronk, Pascal Lexmond, Corien Swaan, Manon Haverkate, Madelief Mollers, Mart Stein, Sandra Kengne Kamga Mobou, Jeroen van Kampen, Jolanda Voermans, Aura Timen, Corine GeurtsvanKessel, Annemiek van der Eijk, Richard Molenkamp, Marion Koopmans, on behalf of the Dutch national COVID-19 response team. |
| EPI_ISL_423571, EPI_ISL_423580, EPI_ISL_423581, EPI_ISL_423585, EPI_ISL_423586, EPI_ISL_423587, EPI_ISL_423588, EPI_ISL_423590, EPI_ISL_423591, EPI_ISL_423592, EPI_ISL_423593 | | | |
| see above | Respiratory Virus Unit, Microbiology Services Colindale, Public Health England | Respiratory Virus Unit, Microbiology Services Colindale, Public Health England | Monica Galiano, Shahjahan Miah, Angie Lackenby, Omolola Akinbami, Tiina Talts, Leena Bhaw, Richard Myers, Steven Platt, Kirstin Edwards, Jonathan Hubb, Joanna Ellis, Maria Zambon |
| EPI_ISL_424629, EPI_ISL_424630, EPI_ISL_424631, EPI_ISL_424632, EPI_ISL_424633, EPI_ISL_424634, EPI_ISL_424635, EPI_ISL_424636, EPI_ISL_424637, EPI_ISL_424638, EPI_ISL_424639, EPI_ISL_424640, EPI_ISL_424641, EPI_ISL_424642, EPI_ISL_424643, EPI_ISL_424644, EPI_ISL_424645, EPI_ISL_424646, EPI_ISL_424647, EPI_ISL_424648, EPI_ISL_424649, EPI_ISL_424650, EPI_ISL_424651, EPI_ISL_424652, EPI_ISL_424653, EPI_ISL_424654, EPI_ISL_424655, EPI_ISL_424656, EPI_ISL_424657, EPI_ISL_424658, EPI_ISL_424659, EPI_ISL_424660, EPI_ISL_424661, EPI_ISL_424662, EPI_ISL_424663, EPI_ISL_424664, EPI_ISL_424665 | | | |
| see above | Department of Clinical Microbiology | GIGA Medical Genomics | Keith Durkin, Maria Artesi, Sébastien Bontems, Raphaël Boreux, Cécile Meex, Pierrette Melin, Marie-Pierre Hayette, Vincent Bours. |
| EPI_ISL_425142, EPI_ISL_425143, EPI_ISL_425144, EPI_ISL_425145, EPI_ISL_425147, EPI_ISL_425150, EPI_ISL_425156, EPI_ISL_425157, EPI_ISL_425160, EPI_ISL_425161 | University of Wisconsin-Madison AIDS Vaccine Research Laboratories | University of Wisconsin-Madison AIDS Vaccine Research Laboratories | Gage Moreno, Katarina Braun, et al. AIDS Vaccine Research Laboratories |
| EPI_ISL_425447 | Department of Pathology, University of Cambridge | COVID-19 Genomics UK (COG-UK) Consortium | Luke W Meredith, M. Estee Torok , Myra Hosmillo, William L. Hamilton, Martin D. Curran, Theresa Feltwell, Anna Yakovleva, Charlotte J. Houldcroft, Aminu S. Jahun, Sarah L. Caddy, Ian Goodfellow |
| EPI_ISL_426159, EPI_ISL_426160, EPI_ISL_426161 | Gundersen Molecular Diagnostics Laboratory | Kabara Cancer Research Institute | Craig S. Richmond, Paraic A. Kenny |
| EPI_ISL_426412 | Pok Oi Hospital | Hong Kong Department of Health | Mak Gannon C.K., Cheng Peter K.C., Lam Edman T.K., Chan Rickjason C.W., Tsang Dominic N.C. |
| EPI_ISL_426414 | Sir M P Shah Government Medical College | Gujarat Biotechnology Research Centre | Ramesh Pandit, Tejas Shah, Ankit Hinsu, Pritesh Sabara, Apurvasinh Puvar, Janvi Raval, Monika Gandhi, Pinal Trivedi, Maharshi Pandya, Amit Kanani, Akanksha Verma, Nitin Savaliya, Raghawendra Kumar, Dinesh Kumar, Zubair Saiyed, Dipa Kinariwala, Disha Patel, Binita Aring, Geeta Vaghela, Sonia Barve, Bhavesh Modi, Kairavi Joshi, Nidhi Sood, Pranay Shah, R D Dixit, Snehal Bagatharia, Madhvi Joshi, Chaitanya Joshi |
| EPI_ISL_426415 | Sir M P Shah Government Medical College, Jamnagar | Gujarat Biotechnology Research Centre, Gandhinagar | Ramesh Pandit, Tejas Shah, Ankit Hinsu, Pritesh Sabara, Apurvasinh Puvar, Janvi Raval, Monika Gandhi, Pinal Trivedi, Maharshi Pandya, Amit Kanani, Akanksha Verma, Nitin Savaliya, Raghvendra Kumar, Dinesh Kumar, Zuber Saiyed, Dipa Kinariwala, Disha Patel, Binita Aring, Geeta Vaghela, Sonia Barve, Bhavesh Modi, Kairavi Joshi, Nidhi Sood, Pranay Shah, R D Dixit, Snehal Bagatharia, Madhvi Joshi, Chaitanya Joshi |
| EPI_ISL_426471, EPI_ISL_426472, EPI_ISL_426473, EPI_ISL_426474, EPI_ISL_426475 | Virginia DCLS | Virginia DCLS | Virginia DCLS |
| EPI_ISL_426568, EPI_ISL_426569 | AZ SPHL, Arizona Department of Health Services | TGen North | Jolene Bowers, Megan Folkerts, Darrin Lemmer, Dave Engelthaler |
| EPI_ISL_426627, EPI_ISL_426628 | Ochsner Health | BioInfoExperts, LLC | Amy Feehan, David Nolan, Rebecca Rose, Susanna Lamers, Sissy Cross, Julia-Garcia-Diaz, Tong Yang, Luke Caruso, David Moraga Amador, Wayra Navia, Lydia Von Borstel, Xiao Hui Zhou |
| EPI_ISL_426889 | Motol University Hospital | Institute of Applied Biotechnologies a.s. | Petr Brož, Jan Geryk, Petr Klempt, Martin Kašný, Adam Novotný, Kateina Kvapilová, Pavel Devínek, Petr Kvapil, Milan Macek |
| EPI_ISL_427085, EPI_ISL_427086, EPI_ISL_427087, EPI_ISL_427088, EPI_ISL_427089, EPI_ISL_427090, EPI_ISL_427091, EPI_ISL_427092, EPI_ISL_427093, EPI_ISL_427094, EPI_ISL_427095, EPI_ISL_427096, EPI_ISL_427097, EPI_ISL_427098, EPI_ISL_427099, EPI_ISL_427100, EPI_ISL_427101, EPI_ISL_427102, EPI_ISL_427103, EPI_ISL_427104, EPI_ISL_427105, EPI_ISL_427106, EPI_ISL_427107, EPI_ISL_427108, EPI_ISL_427109, EPI_ISL_427110, EPI_ISL_427111, EPI_ISL_427112, EPI_ISL_427113, EPI_ISL_427114, EPI_ISL_427115, EPI_ISL_427116, EPI_ISL_427117, EPI_ISL_427118, EPI_ISL_427119, EPI_ISL_427120, EPI_ISL_427121, EPI_ISL_427122, EPI_ISL_427123, EPI_ISL_427124, EPI_ISL_427125, EPI_ISL_427126, EPI_ISL_427127, EPI_ISL_427128, EPI_ISL_427129, EPI_ISL_427130, EPI_ISL_427131, EPI_ISL_427132 | | | |
| see above | Victorian Infectious Diseases Reference Laboratory (VIDRL) | Microbiological Diagnostic Unit Public Health Laboratory and Victorian Infectious Diseases Reference Laboratory, Doherty Institute | Caly L., Seemann T., Sait, M., Schultz M., Druce J., Sherry, N. |
| EPI_ISL_427133 | Microbiological Diagnostic Unit Public Health Laboratory | Microbiological Diagnostic Unit Public Health Laboratory | Seemann T., Schultz M., Sait, M., Sherry, N. |
| EPI_ISL_427134, EPI_ISL_427135, EPI_ISL_427136, EPI_ISL_427137, EPI_ISL_427138, EPI_ISL_427139, EPI_ISL_427140, EPI_ISL_427141, EPI_ISL_427142, EPI_ISL_427143, EPI_ISL_427144, EPI_ISL_427145, EPI_ISL_427146, EPI_ISL_427147 | | | |
| see above | Victorian Infectious Diseases Reference Laboratory (VIDRL) | Microbiological Diagnostic Unit Public Health Laboratory and Victorian Infectious Diseases Reference Laboratory, Doherty Institute | Caly L., Seemann T., Sait, M., Schultz M., Druce J., Sherry, N. |
| EPI_ISL_427313, EPI_ISL_427314, EPI_ISL_427316, EPI_ISL_427317, EPI_ISL_427318, EPI_ISL_427319, EPI_ISL_427320, EPI_ISL_427321, EPI_ISL_427322, EPI_ISL_427323 | WHO National Influenza Centre Russian Federation | WHO National Influenza Centre Russian Federation | Andrey Komissarov, Artem Fadeev, Mariia Sergeeva, Anna Ivanova, Daria Danilenko |
| EPI_ISL_427348, EPI_ISL_427349, EPI_ISL_427350, EPI_ISL_427352, EPI_ISL_427353, EPI_ISL_427354, EPI_ISL_427355, EPI_ISL_427356, EPI_ISL_427358, EPI_ISL_427359, EPI_ISL_427362, EPI_ISL_427389 | | | |
| see above | Department of Clinical Microbiology | GIGA Medical Genomics | Keith Durkin, Maria Artesi, Sébastien Bontems, Raphaël Boreux, Cécile Meex, Pierrette Melin, Marie-Pierre Hayette, Vincent Bours. |

| | | | |
|---|---|---|---|
| EPI_ISL_427436, EPI_ISL_427451, EPI_ISL_427453, EPI_ISL_427459 | University of Wisconsin-Madison AIDS Vaccine Research Laboratories | University of Wisconsin-Madison AIDS Vaccine Research Laboratories | Gage Moreno, Katarina Braun, et al. AIDS Vaccine Research Laboratories |
| EPI_ISL_427502, EPI_ISL_427503, EPI_ISL_427504, EPI_ISL_427505, EPI_ISL_427506, EPI_ISL_427507, EPI_ISL_427508, EPI_ISL_427509, EPI_ISL_427510, EPI_ISL_427511, EPI_ISL_427512, EPI_ISL_427513, EPI_ISL_427514 | | | |
| see above | NYU Langone Health | Departments of Pathology and Medicine, New York University School of Medicine | Maria Aguero-Rosenfeld, Brendan Belovarac, Margaret Black, Ludovic Boytard, John Cadley, Paolo Cotzia, John Chen, Dacia Dimartino, Xiaojun Feng, Tatyana Gindin, Emily Guzman, Adriana Heguy, Megan Hogan, Emily Huang, George Jour, Andrew Lytle, Christian Marier, Matthew T. Maurano, Mark J. Mulligan, Peter Meyn, Iman Osman, Jared Pinnell, Vanessa Raabe, Sitharam Ramaswami, Amy Rapkiewicz, Marie Samanovic-Golden, Antonio Serrano, Guomiao Shen, Matija Snuderl, Theodore Vougiouklakis, Nick Vulpescu, Gael Westby, Paul Zappile, Yutong Zhang |
| EPI_ISL_427619, EPI_ISL_427620 | Alaska State Virology Laboratory | Alaska State Virology Laboratory | Chen, J. |
| EPI_ISL_427628, EPI_ISL_427630, EPI_ISL_427631, EPI_ISL_427632, EPI_ISL_427633, EPI_ISL_427634, EPI_ISL_427635, EPI_ISL_427636, EPI_ISL_427637, EPI_ISL_427638, EPI_ISL_427639, EPI_ISL_427640, EPI_ISL_427641, EPI_ISL_427642 | | | |
| see above | NYU Langone Health | Departments of Pathology and Medicine, New York University School of Medicine | Maria Aguero-Rosenfeld, Brendan Belovarac, Margaret Black, Ludovic Boytard, John Cadley, Paolo Cotzia, John Chen, Dacia Dimartino, Xiaojun Feng, Tatyana Gindin, Emily Guzman, Adriana Heguy, Megan Hogan, Emily Huang, George Jour, Andrew Lytle, Christian Marier, Matthew T. Maurano, Mark J. Mulligan, Peter Meyn, Iman Osman, Jared Pinnell, Vanessa Raabe, Sitharam Ramaswami, Amy Rapkiewicz, Marie Samanovic-Golden, Antonio Serrano, Guomiao Shen, Matija Snuderl, Theodore Vougiouklakis, Nick Vulpescu, Gael Westby, Paul Zappile, Yutong Zhang |
| EPI_ISL_428209 | Laboratory of Molecular Biology, Diagnostyka sp. z o.o. | Laboratory of Recombinant Vaccines | Lukasz Rabalski, Anna Piotrowska-Mietelska, Boguslaw Szewczyk, Krystyna Bienkowska-Szewczyk |
| EPI_ISL_428384, EPI_ISL_428385, EPI_ISL_428386, EPI_ISL_428389, EPI_ISL_428392, EPI_ISL_428393 | Yale COVID-19 Biorepository | Grubaugh Lab - Yale School of Public Health | Joseph Fauver, Tara Alpert, Anderson Brito, Anne Wyllie, Chantal Vogels, Mary Petrone, Chaney Kalinich, Isabel Ott, Arnau Casanovas, Catherine Muenker, Adam Moore, Alice Lu, Maria Tokuyama, Patrick Wong, Peiwen Lu, Saad Omer, Richard Martinello, Allison Nelson, Shelli Farhadian, Akiko Iwasaki, Charlese Dela Cruz, Albert Ko, Nathan Grubaugh |
| EPI_ISL_428399, EPI_ISL_428401, EPI_ISL_428402 | Yale COVID-19 Biorepository | Grubaugh Lab - Yale School of Public Health | Joseph Fauver, Tara Alpert, Anderson Brito, Anne Wyllie, Chantal Vogels, Mary Petrone, Cole Jensen, Chaney Kalinich, Isabel Ott, Arnau Casanovas, Catherine Muenker, Adam Moore, Alice Lu, Maria Tokuyama, Patrick Wong, Peiwen Lu, Saad Omer, Richard Martinello, Allison Nelson, Shelli Farhadian, Akiko Iwasaki, Charlese Dela Cruz, Albert Ko, Nathan Grubaugh |
| EPI_ISL_428479, EPI_ISL_428480, EPI_ISL_428481 | District Surveillence Unit | Department of Neurovirology, National Institute of Mental Health and Neuroscience (NIMHANS) | Chitra Pattabiraman, Vijayalakshmi Reddy, Harsha PK, Risha Rasheed, Shafeeq S Hameed, Manjunatha Venkataswamy, Anita Desai, Ravi Vasanthapuram |
| EPI_ISL_428729, EPI_ISL_428730, EPI_ISL_428731, EPI_ISL_428732 | University of Wisconsin-Madison AIDS Vaccine Research Laboratories | University of Wisconsin-Madison AIDS Vaccine Research Laboratories | Gage Moreno, Katarina Braun, et al. AIDS Vaccine Research Laboratories |
| EPI_ISL_428747, EPI_ISL_428749 | Yale COVID-19 Biorepository | Grubaugh Lab - Yale School of Public Health | Joseph Fauver, Tara Alpert, Anderson Brito, Anne Wyllie, Chantal Vogels, Mary Petrone, Cole Jensen, Chaney Kalinich, Isabel Ott, Arnau Casanovas, Catherine Muenker, Adam Moore, Alice Lu, Maria Tokuyama, Patrick Wong, Peiwen Lu, Saad Omer, Richard Martinello, Allison Nelson, Shelli Farhadian, Akiko Iwasaki, Charlese Dela Cruz, Albert Ko, Nathan Grubaugh |
| EPI_ISL_428758, EPI_ISL_428759, EPI_ISL_428760, EPI_ISL_428761, EPI_ISL_428762, EPI_ISL_428763, EPI_ISL_428764, EPI_ISL_428765, EPI_ISL_428766, EPI_ISL_428767, EPI_ISL_428768, EPI_ISL_428769, EPI_ISL_428770, EPI_ISL_428771, EPI_ISL_428772, EPI_ISL_428773, EPI_ISL_428774, EPI_ISL_428775, EPI_ISL_428776, EPI_ISL_428777, EPI_ISL_428778, EPI_ISL_428779, EPI_ISL_428780, EPI_ISL_428781, EPI_ISL_428782, EPI_ISL_428783, EPI_ISL_428784, EPI_ISL_428785, EPI_ISL_428786, EPI_ISL_428787, EPI_ISL_428788, EPI_ISL_428789, EPI_ISL_428790, EPI_ISL_428791, EPI_ISL_428792, EPI_ISL_428793, EPI_ISL_428794, EPI_ISL_428795, EPI_ISL_428796, EPI_ISL_428797, EPI_ISL_428798, EPI_ISL_428799, EPI_ISL_428800, EPI_ISL_428801, EPI_ISL_428802, EPI_ISL_428803, EPI_ISL_428804, EPI_ISL_428805 | | | |
| see above | NYU Langone Health | Departments of Pathology and Medicine, New York University School of Medicine | Maria Aguero-Rosenfeld, Brendan Belovarac, Margaret Black, Ludovic Boytard, John Cadley, Paolo Cotzia, John Chen, Dacia Dimartino, Xiaojun Feng, Tatyana Gindin, Emily Guzman, Adriana Heguy, Megan Hogan, Emily Huang, George Jour, Andrew Lytle, Christian Marier, Matthew T. Maurano, Mark J. Mulligan, Peter Meyn, Iman Osman, Jared Pinnell, Vanessa Raabe, Sitharam Ramaswami, Amy Rapkiewicz, Marie Samanovic-Golden, Antonio Serrano, Guomiao Shen, Matija Snuderl, Theodore Vougiouklakis, Nick Vulpescu, Gael Westby, Paul Zappile, Yutong Zhang |
| EPI_ISL_428851, EPI_ISL_428852 | FSBSI "Chumakov Federal Scientific Center for Research and Development of Immune-and-Biological Products of Russian Academy of Sciences" | FSBSI "Chumakov Federal Scientific Center for Research and Development of Immune-and-Biological Products of Russian Academy of Sciences" & NRC "Kurchatov institute" | Liubov Kozlovskaya, Anastasia Piniaeva, Georgy Ignatyev, Anna Shishova, Aydar Ishmukhametov, Mikhail Rychev, Egor Prokhorchuk, Denis Protsenko, Anastasia Berestovskaya |
| EPI_ISL_428940, EPI_ISL_428941, EPI_ISL_428942, EPI_ISL_428945, EPI_ISL_428946, EPI_ISL_428947, EPI_ISL_428948, EPI_ISL_428949, EPI_ISL_428951, EPI_ISL_428953, EPI_ISL_428954, EPI_ISL_428960, EPI_ISL_428961, EPI_ISL_428962 | | | |
| see above | Laboratoire National de Sante, Microbiology, Virology | Laboratoire National de Sante, Microbiology, Epidemiology and Microbial Genomics | Anke Wienecke-Baldacchino, Ardashel Latsuzbaia, Jessica Tapp, Catherine Ragimbeau, Guillaume Fournier, Tamir Abdelrahman, Trung Nguyen Nguyen, Joel Mossong |
| EPI_ISL_429018, EPI_ISL_429028, EPI_ISL_429051 | UCSF Clinical Microbiology Laboratory | Chan-Zuckerberg Biohub | CZB Cliahub Consortium |
| EPI_ISL_429128, EPI_ISL_429135 | Laboratoriemedicin | The Public Health Agency of Sweden | Olov Svartstrom, Maria Lind Karlberg, Anna-Malin Linde, Oskar Karlsson Lindsjo, Anna Risberg, Shaman Muradrasoli, Karin Tegmark-Wisell |
| EPI_ISL_429204, EPI_ISL_429207, EPI_ISL_429217, EPI_ISL_429218, EPI_ISL_429219, EPI_ISL_429220, EPI_ISL_429221 | University Hospitals of Geneva Laboratory of Virology | University Hospitals of Geneva Laboratory of Virology | Laubscher F. |
| EPI_ISL_429254, EPI_ISL_429255, EPI_ISL_429258, EPI_ISL_429259 | Viral Respiratory Lab, National Institute for Biomedical Research (INRB) | Pathogen Sequencing Lab, National Institute for Biomedical Research (INRB) | Placide Mbala-Kingebeni, Edith Nkwembe, Eddy Kinganda-Lusamaki, Amuri Aziza, Catherine Pratt, Matthias Pauthner, Josh Quick, Allison Black, James Hadfield, Trevor Bedford, Ian Goodfellow, Nick Loman, Kristian Andersen, Michael Wiley, Steve Ahuka-Mundeke, Jean-Jacques Muyembe Tamfum |
| EPI_ISL_429628, EPI_ISL_429629, EPI_ISL_429630, EPI_ISL_429631, EPI_ISL_429632, EPI_ISL_429633, EPI_ISL_429641, EPI_ISL_429642, EPI_ISL_429645, EPI_ISL_429646, EPI_ISL_429649, EPI_ISL_429650, EPI_ISL_429651, EPI_ISL_429652, EPI_ISL_429653, EPI_ISL_429654, EPI_ISL_429655, EPI_ISL_429656 | | | |
| see above | UW Virology Lab | UW Virology Lab | Pavitra Roychoudhury, Hong Xie, Keith Jerome, Alexander Greninger |
| EPI_ISL_429848 | Gundersen Molecular Diagnostics Laboratory | Kabara Cancer Research Institute | Craig S. Richmond, Paraic A. Kenny |
| EPI_ISL_429976, EPI_ISL_429980, EPI_ISL_429981, EPI_ISL_429982, EPI_ISL_429984, EPI_ISL_429985, EPI_ISL_429986, EPI_ISL_429987, EPI_ISL_429988 | Virginia DCLS | Virginia DCLS | Virginia DCLS |
| EPI_ISL_430008, EPI_ISL_430013 | Biolab Diagnostic Laboratories | Andersen lab at Scripps Research | Issa Abu-Dayyeh, Ahmad Tibi, Lama Hussein, Lina Mohammad, Zein Naber, Amid Abdelnour with SEARCH Alliance San Diego |
| EPI_ISL_430046, EPI_ISL_430047, EPI_ISL_430048, EPI_ISL_430049, EPI_ISL_430051, EPI_ISL_430052, EPI_ISL_430053, EPI_ISL_430054, EPI_ISL_430055, EPI_ISL_430056, EPI_ISL_430057, EPI_ISL_430058, EPI_ISL_430059, EPI_ISL_430060, EPI_ISL_430061, EPI_ISL_430062 | | | |
| see above | Utah Public Health Laboratory | Utah Public Health Laboratory | Erin Young, Kelly Oakeson |
| EPI_ISL_430319, EPI_ISL_430320, EPI_ISL_430321, EPI_ISL_430322, EPI_ISL_430350, EPI_ISL_430354, EPI_ISL_430355, EPI_ISL_430356, EPI_ISL_430357 | NYU Langone Health | Departments of Pathology and Medicine, New York University School of Medicine | Maria Aguero-Rosenfeld, Brendan Belovarac, Margaret Black, Ludovic Boytard, John Cadley, Paolo Cotzia, John Chen, Dacia Dimartino, Xiaojun Feng, Tatyana Gindin, Emily Guzman, Adriana Heguy, Megan Hogan, Emily Huang, George Jour, Lawrence H. Lin, Raven Luther, Andrew Lytle, Christian Marier, Matthew T. Maurano, Mark J. Mulligan, Peter Meyn, Raquel Ordonez Ciriza, Iman Osman, Jared Pinnell, Vanessa Raabe, Sitharam Ramaswami, Amy Rapkiewicz, Andre M. Ribeiro-dos-Santos, Marie Samanovic-Golden, Antonio Serrano, Guomiao Shen, Matija Snuderl, Theodore Vougiouklakis, Nick Vulpescu, Gael Westby, Paul Zappile, Yutong Zhang |
| EPI_ISL_430467 | ICMR-National Institute of Cholera and Enteric Diseases | National Institute of Biomedical Genomics | Arindam Maitra, Mamta Chawla Sarkar, Sreedhar Chinnaswamy, Hasina Banu, Ananya Chatterjee, Shanta Dutta, Saumitra Das |
| EPI_ISL_430510, EPI_ISL_430511, EPI_ISL_430512, EPI_ISL_430520, EPI_ISL_430524, EPI_ISL_430525, EPI_ISL_430526, EPI_ISL_430530, EPI_ISL_430531, EPI_ISL_430532, EPI_ISL_430533, EPI_ISL_430534, EPI_ISL_430535, EPI_ISL_430536, EPI_ISL_430537, EPI_ISL_430538, EPI_ISL_430539, EPI_ISL_430540, EPI_ISL_430541, EPI_ISL_430546, EPI_ISL_430549, EPI_ISL_430550, EPI_ISL_430561, EPI_ISL_430562, EPI_ISL_430563, EPI_ISL_430564, EPI_ISL_430565, EPI_ISL_430566, EPI_ISL_430574, EPI_ISL_430578, EPI_ISL_430579, EPI_ISL_430580, EPI_ISL_430581, EPI_ISL_430587, EPI_ISL_430597, EPI_ISL_430599 | | | |
| see above | Victorian Infectious Diseases Reference Laboratory (VIDRL) | Microbiological Diagnostic Unit Public Health Laboratory and Victorian Infectious Diseases Reference Laboratory, The Peter Doherty Institute for Infection and Immunity | Caly L., Seemann T., Sait, M., Schultz M., Druce J., Sherry, N. |
| EPI_ISL_430664, EPI_ISL_430665, EPI_ISL_430666, EPI_ISL_430667, EPI_ISL_430668, EPI_ISL_430669, EPI_ISL_430670, EPI_ISL_430671, EPI_ISL_430672, EPI_ISL_430673, EPI_ISL_430685 | | | |

| | | | |
|---|---|---|---|
| see above | Microbiological Diagnostic Unit Public Health Laboratory | Microbiological Diagnostic Unit Public Health Laboratory | Seemann T., Schultz M., Sait, M., Sherry, N. |
| EPI_ISL_430807 | Laboratorio de Virología del Hospital de Niños Dr. Ricardo Gutierrez | Área de Secuenciación del Laboratorio de Virología del Hospital de Niños Dr. Ricardo Gutierrez on behalf of 'Proyecto Argentino Interinstitucional de genomica de SARS-CoV-2' (PAIS Consortium) | Nabaes Jodar, MS; Goya, S; Natale, MI; Lusso, S; Gravis, E; Mistchenko, AS; Valinotto, LE; Viegas, M. |
| EPI_ISL_430808 | Departamento de Biología y genética molecular, IACA Laboratorios. | Área de Secuenciación del Laboratorio de Virología del Hospital de Niños Dr. Ricardo Gutierrez on behalf of 'Proyecto Argentino Interinstitucional de genomica de SARS-CoV-2' (PAIS Consortium) | Nabaes Jodar, MS; Goya, S; Natale, MI; Lusso, S; Tittarelli, E; Suárez, A; Masciovecchio MV; Streitenberger ER; Mistchenko, AS; Valinotto, LE; Viegas, M. |
| EPI_ISL_430809 | Laboratorio de Virología del Hospital de Niños Dr. Ricardo Gutierrez | Área de Secuenciación del Laboratorio de Virología del Hospital de Niños Dr. Ricardo Gutierrez on behalf of 'Proyecto Argentino Interinstitucional de genomica de SARS-CoV-2' (PAIS Consortium) | Nabaes Jodar, MS; Goya, S; Natale, MI; Lusso, S; Gravis, E; Mistchenko, AS; Valinotto, LE; Viegas, M. |
| EPI_ISL_430858, EPI_ISL_430859 | Laboratoriemedicin | The Public Health Agency of Sweden | Oskar Karlsson Lindsjo, Maria Lind Karlberg, Anna-Malin Linde, Olov Svartstrom, Anna Risberg, Shaman Muradrasoli, Karin Tegmark-Wisell |
| EPI_ISL_430871, EPI_ISL_430872, EPI_ISL_430875, EPI_ISL_430897, EPI_ISL_430898, EPI_ISL_430900, EPI_ISL_430901, EPI_ISL_430902, EPI_ISL_430903, EPI_ISL_430904, EPI_ISL_430905, EPI_ISL_430906, EPI_ISL_430913, EPI_ISL_430914, EPI_ISL_430915, EPI_ISL_430917, EPI_ISL_430918, EPI_ISL_430919, EPI_ISL_430921, EPI_ISL_430922, EPI_ISL_430923, EPI_ISL_430925, EPI_ISL_430926, EPI_ISL_430927, EPI_ISL_430959, EPI_ISL_430964, EPI_ISL_430966, EPI_ISL_430967, EPI_ISL_430969, EPI_ISL_430971, EPI_ISL_430974, EPI_ISL_430975, EPI_ISL_430976, EPI_ISL_430977, EPI_ISL_430978, EPI_ISL_430979, EPI_ISL_430980 | | | |
| see above | UW Virology Lab | UW Virology Lab | Pavitra Roychoudhury, Hong Xie, Keith Jerome, Alexander Greninger |
| EPI_ISL_431081 | Yale COVID-19 Biorepository | Grubaugh Lab - Yale School of Public Health | Joseph Fauver, Tara Alpert, Anderson Brito, Anne Wyllie, Chantal Vogels, Mary Petrone, Cole Jensen, Chaney Kalinich, Isabel Ott, Arnau Casanovas, Catherine Muenker, Adam Moore, Alice Lu, Maria Tokuyama, Patrick Wong, Peiwen Lu, Saad Omer, Richard Martinello, Allison Nelson, Shelli Farhadian, Akiko Iwasaki, Charlese Dela Cruz, Albert Ko, Nathan Grubaugh |
| EPI_ISL_431901, EPI_ISL_431911, EPI_ISL_431914, EPI_ISL_431920, EPI_ISL_431929, EPI_ISL_431930, EPI_ISL_431932, EPI_ISL_431934, EPI_ISL_431935, EPI_ISL_431951, EPI_ISL_431952, EPI_ISL_431953, EPI_ISL_431964, EPI_ISL_431965, EPI_ISL_431974, EPI_ISL_431978, EPI_ISL_431982, EPI_ISL_431988, EPI_ISL_431998, EPI_ISL_431999, EPI_ISL_432002, EPI_ISL_432012, EPI_ISL_432017, EPI_ISL_432021, EPI_ISL_432037, EPI_ISL_432046, EPI_ISL_432051, EPI_ISL_432056, EPI_ISL_432090, EPI_ISL_432094, EPI_ISL_432128, EPI_ISL_432129, EPI_ISL_432133, EPI_ISL_432139, EPI_ISL_432152, EPI_ISL_432154, EPI_ISL_432156, EPI_ISL_432157, EPI_ISL_432159, EPI_ISL_432163, EPI_ISL_432168, EPI_ISL_432171, EPI_ISL_432172, EPI_ISL_432178, EPI_ISL_432181, EPI_ISL_432184, EPI_ISL_432186, EPI_ISL_432187, EPI_ISL_432189, EPI_ISL_432193, EPI_ISL_432194, EPI_ISL_432195, EPI_ISL_432196, EPI_ISL_432197, EPI_ISL_432201, EPI_ISL_432202, EPI_ISL_432203, EPI_ISL_432204, EPI_ISL_432205, EPI_ISL_432209, EPI_ISL_432215, EPI_ISL_432216, EPI_ISL_432219, EPI_ISL_432226, EPI_ISL_432229, EPI_ISL_432230, EPI_ISL_432234, EPI_ISL_432237, EPI_ISL_432243, EPI_ISL_432244, EPI_ISL_432247, EPI_ISL_432248, EPI_ISL_432249, EPI_ISL_432251, EPI_ISL_432252, EPI_ISL_432253, EPI_ISL_432259, EPI_ISL_432267, EPI_ISL_432274, EPI_ISL_432279, EPI_ISL_432292, EPI_ISL_432294, EPI_ISL_432297, EPI_ISL_432313, EPI_ISL_432320, EPI_ISL_432330, EPI_ISL_432333, EPI_ISL_432334, EPI_ISL_432343, EPI_ISL_432344, EPI_ISL_432345, EPI_ISL_432353, EPI_ISL_432363, EPI_ISL_432368, EPI_ISL_432372, EPI_ISL_432373, EPI_ISL_432383, EPI_ISL_432388, EPI_ISL_432391, EPI_ISL_432393, EPI_ISL_432398, EPI_ISL_432399, EPI_ISL_432409, EPI_ISL_432414, EPI_ISL_432433, EPI_ISL_432436, EPI_ISL_432440, EPI_ISL_432443, EPI_ISL_432447 | | | |
| see above | Wales Specialist Virology Centre | Public Health Wales Microbiology Cardiff | Catherine Moore, Johnathan Evans, Malorie Perry, Simon Cottrell, Alec Birchley, Alexander Adams, Amy Gaskin, Bree Gatica-Wilcox, Jason Coombes, Lauren Gilbert, Lee Graham, Nicole Pacchiarini, Sara Kumziene-Summerhayes, Sarah Taylor, Sophie Jones, Sara Rey, Matthew Bull, Joanne Watkins, Sally Corden, Tom Connor |
| EPI_ISL_432452, EPI_ISL_432455, EPI_ISL_432456, EPI_ISL_432457, EPI_ISL_432459, EPI_ISL_432461, EPI_ISL_432462, EPI_ISL_432463, EPI_ISL_432468, EPI_ISL_432469, EPI_ISL_432470, EPI_ISL_432473, EPI_ISL_432474, EPI_ISL_432477, EPI_ISL_432479, EPI_ISL_432482, EPI_ISL_432483, EPI_ISL_432488, EPI_ISL_432489, EPI_ISL_432491, EPI_ISL_432493, EPI_ISL_432497, EPI_ISL_432499, EPI_ISL_432503, EPI_ISL_432511, EPI_ISL_432519, EPI_ISL_432524, EPI_ISL_432525, EPI_ISL_432526, EPI_ISL_432527, EPI_ISL_432528, EPI_ISL_432532, EPI_ISL_432536, EPI_ISL_432538, EPI_ISL_432541, EPI_ISL_432545, EPI_ISL_432552, EPI_ISL_432556, EPI_ISL_432558, EPI_ISL_432559, EPI_ISL_432565, EPI_ISL_432566, EPI_ISL_432567, EPI_ISL_432569, EPI_ISL_432570, EPI_ISL_432572, EPI_ISL_432575, EPI_ISL_432576, EPI_ISL_432577, EPI_ISL_432578, EPI_ISL_432580, EPI_ISL_432582, EPI_ISL_432583, EPI_ISL_432586, EPI_ISL_432589, EPI_ISL_432598, EPI_ISL_432599, EPI_ISL_432607, EPI_ISL_432608, EPI_ISL_432610, EPI_ISL_432613, EPI_ISL_432615, EPI_ISL_432616, EPI_ISL_432618, EPI_ISL_432622, EPI_ISL_432625, EPI_ISL_432630, EPI_ISL_432634, EPI_ISL_432635, EPI_ISL_432637, EPI_ISL_432641, EPI_ISL_432642, EPI_ISL_432643, EPI_ISL_432645, EPI_ISL_432646, EPI_ISL_432648, EPI_ISL_432649, EPI_ISL_432655, EPI_ISL_432659, EPI_ISL_432660, EPI_ISL_432661, EPI_ISL_432662, EPI_ISL_432665 | | | |
| see above | Virology Department, Sheffield Teaching Hospitals NHS Foundation Trust / Virology Department, Sheffield Teaching Hospitals NHS Foundation Trust | COVID-19 Genomics UK (COG-UK) Consortium | Thushan de Silva, Matthew Parker,Adri Angyal, Rebecca Brown, Luke Green, Rachel Tucker, Paul Parsons, Danielle Groves, Alex Keeley, Dave Partridge, Matthew Wyles, Benjamin Lindsey, Mehmet Yavuz, Mohammad Raza, Cariad Evans |
| EPI_ISL_432668, EPI_ISL_432669, EPI_ISL_432670, EPI_ISL_432671, EPI_ISL_432672, EPI_ISL_432673, EPI_ISL_432674, EPI_ISL_432675, EPI_ISL_432676, EPI_ISL_432677, EPI_ISL_432678, EPI_ISL_432679, EPI_ISL_432680, EPI_ISL_432700, EPI_ISL_432707 | | | |
| see above | Queens Medical Centre, Clinical Microbiology Department / DeepSeq Nottingham | COVID-19 Genomics UK (COG-UK) Consortium | Gemma Clark, Wendy Smith, Manjinder Khakh, Hannah Howson-Wells, Jonathan Ball, Patrick McClure, Joseph Chappell, Theocharis Tsoleridis, Nadine Holmes, Matthew Carlisle, Christopher Moore, Fei Sang, Johnny Debebe, Victoria Wright, Matthew Loose |
| EPI_ISL_432715, EPI_ISL_432721, EPI_ISL_432731, EPI_ISL_432737, EPI_ISL_432741, EPI_ISL_432742, EPI_ISL_432743, EPI_ISL_432748, EPI_ISL_432749, EPI_ISL_432750, EPI_ISL_432756, EPI_ISL_432757, EPI_ISL_432766, EPI_ISL_432774, EPI_ISL_432776, EPI_ISL_432778, EPI_ISL_432779, EPI_ISL_432781, EPI_ISL_432782, EPI_ISL_432783, EPI_ISL_432785, EPI_ISL_432789, EPI_ISL_432793, EPI_ISL_432798, EPI_ISL_432800, EPI_ISL_432801, EPI_ISL_432803, EPI_ISL_432805, EPI_ISL_432808, EPI_ISL_432814, EPI_ISL_432822, EPI_ISL_432826, EPI_ISL_432827, EPI_ISL_432830, EPI_ISL_432833, EPI_ISL_432834, EPI_ISL_432835, EPI_ISL_432836, EPI_ISL_432839, EPI_ISL_432840, EPI_ISL_432845, EPI_ISL_432847, EPI_ISL_432848, EPI_ISL_432850, EPI_ISL_432854, EPI_ISL_432858 | | | |
| see above | Virology Department, Sheffield Teaching Hospitals NHS Foundation Trust / Virology Department, Sheffield Teaching Hospitals NHS Foundation Trust | COVID-19 Genomics UK (COG-UK) Consortium | Thushan de Silva, Matthew Parker,Adri Angyal, Rebecca Brown, Luke Green, Rachel Tucker, Paul Parsons, Danielle Groves, Alex Keeley, Dave Partridge, Matthew Wyles, Benjamin Lindsey, Mehmet Yavuz, Mohammad Raza, Cariad Evans |
| EPI_ISL_432868, EPI_ISL_432869, EPI_ISL_432870, EPI_ISL_432871, EPI_ISL_432872, EPI_ISL_432873, EPI_ISL_432874, EPI_ISL_432875, EPI_ISL_432876, EPI_ISL_432877, EPI_ISL_432878, EPI_ISL_432879, EPI_ISL_432880, EPI_ISL_432881, EPI_ISL_432882, EPI_ISL_432883, EPI_ISL_432884, EPI_ISL_432885, EPI_ISL_432886, EPI_ISL_432887, EPI_ISL_432888, EPI_ISL_432889, EPI_ISL_432890, EPI_ISL_432891, EPI_ISL_432892, EPI_ISL_432893, EPI_ISL_432894, EPI_ISL_432895, EPI_ISL_432896, EPI_ISL_432897, EPI_ISL_432898, EPI_ISL_432899 | | | |
| see above | Virology Department, Royal Infirmary of Edinburgh, NHS Lothian / School of Biological Sciences, University of Edinburgh / Institute of Genetics and Molecular Medicine, University of Edinburgh | COVID-19 Genomics UK (COG-UK) Consortium | McHugh M, Dewar R, Rooke S, Gallagher M, Balcaza C, O'Toole A, Hill V, McCrone JT, Colquhoun R, Yu X, Jackson B, Rambaut A, Williams TC, Templeton K |
| EPI_ISL_432993, EPI_ISL_432995, EPI_ISL_432996, EPI_ISL_432997, EPI_ISL_432998, EPI_ISL_432999, EPI_ISL_433000, EPI_ISL_433001, EPI_ISL_433002, EPI_ISL_433003, EPI_ISL_433004, EPI_ISL_433005, EPI_ISL_433006, EPI_ISL_433007, EPI_ISL_433008, EPI_ISL_433009, EPI_ISL_433010, EPI_ISL_433011, EPI_ISL_433012, EPI_ISL_433013, EPI_ISL_433014, EPI_ISL_433015, EPI_ISL_433016, EPI_ISL_433017, EPI_ISL_433018, EPI_ISL_433019, EPI_ISL_433020, EPI_ISL_433021, EPI_ISL_433022, EPI_ISL_433023, EPI_ISL_433024, EPI_ISL_433025, EPI_ISL_433026, EPI_ISL_433027, EPI_ISL_433028, EPI_ISL_433029, EPI_ISL_433030, EPI_ISL_433031, EPI_ISL_433032, EPI_ISL_433033, EPI_ISL_433034, EPI_ISL_433035, EPI_ISL_433036, EPI_ISL_433037, EPI_ISL_433038, EPI_ISL_433039, EPI_ISL_433040, EPI_ISL_433041, EPI_ISL_433042, EPI_ISL_433043, EPI_ISL_433044, EPI_ISL_433045, EPI_ISL_433046, EPI_ISL_433047, EPI_ISL_433048, EPI_ISL_433049, EPI_ISL_433050, EPI_ISL_433051, EPI_ISL_433052, EPI_ISL_433053, EPI_ISL_433054, EPI_ISL_433055, EPI_ISL_433056, EPI_ISL_433057, EPI_ISL_433058, EPI_ISL_433059, EPI_ISL_433060, EPI_ISL_433061, EPI_ISL_433062, EPI_ISL_433063, EPI_ISL_433064, EPI_ISL_433065, EPI_ISL_433066, EPI_ISL_433067 | | | |
| see above | Queens Medical Centre, Clinical Microbiology Department / DeepSeq Nottingham | COVID-19 Genomics UK (COG-UK) Consortium | Gemma Clark, Wendy Smith, Manjinder Khakh, Hannah Howson-Wells, Jonathan Ball, Patrick McClure, Joseph Chappell, Theocharis Tsoleridis, Nadine Holmes, Matthew Carlisle, Christopher Moore, Fei Sang, Johnny Debebe, Victoria Wright, Matthew Loose |
| EPI_ISL_433204, EPI_ISL_433205, EPI_ISL_433206, EPI_ISL_433213, EPI_ISL_433215, EPI_ISL_433221, EPI_ISL_433222, EPI_ISL_433223, EPI_ISL_433228, EPI_ISL_433229, EPI_ISL_433230, EPI_ISL_433231, EPI_ISL_433232, EPI_ISL_433233, EPI_ISL_433234, EPI_ISL_433235, EPI_ISL_433236, EPI_ISL_433237, EPI_ISL_433238, EPI_ISL_433239, EPI_ISL_433240, EPI_ISL_433241, EPI_ISL_433242, EPI_ISL_433243, EPI_ISL_433244, EPI_ISL_433245, EPI_ISL_433246, EPI_ISL_433247, EPI_ISL_433248, EPI_ISL_433249, EPI_ISL_433250, EPI_ISL_433251, EPI_ISL_433252, EPI_ISL_433253, EPI_ISL_433254, EPI_ISL_433255, EPI_ISL_433256, EPI_ISL_433257, EPI_ISL_433258, EPI_ISL_433259, EPI_ISL_433260, EPI_ISL_433261, EPI_ISL_433262, EPI_ISL_433263, EPI_ISL_433264, EPI_ISL_433265, EPI_ISL_433266, EPI_ISL_433267, EPI_ISL_433374, EPI_ISL_433375, EPI_ISL_433376, EPI_ISL_433377, EPI_ISL_433378, EPI_ISL_433379, EPI_ISL_433380, EPI_ISL_433381, EPI_ISL_433382, EPI_ISL_433383, EPI_ISL_433399, EPI_ISL_433400, EPI_ISL_433401, EPI_ISL_433402, EPI_ISL_433403, EPI_ISL_433405, EPI_ISL_433406 | | | |
| see above | Virology Department, Royal Infirmary of Edinburgh, NHS Lothian / School of Biological Sciences, University of Edinburgh / Institute of Genetics and Molecular Medicine, University of Edinburgh | COVID-19 Genomics UK (COG-UK) Consortium | McHugh M, Dewar R, Rooke S, Gallagher M, Balcaza C, O'Toole A, Hill V, McCrone JT, Colquhoun R, Yu X, Jackson B, Rambaut A, Williams TC, Templeton K |
| EPI_ISL_433576, EPI_ISL_433580, EPI_ISL_433581, EPI_ISL_433583, EPI_ISL_433585, EPI_ISL_433587, EPI_ISL_433590, EPI_ISL_433591, EPI_ISL_433592, EPI_ISL_433594, EPI_ISL_433595, EPI_ISL_433596, EPI_ISL_433597, EPI_ISL_433598, EPI_ISL_433600, EPI_ISL_433601, EPI_ISL_433602, EPI_ISL_433603, EPI_ISL_433604, EPI_ISL_433605, EPI_ISL_433606, EPI_ISL_433607, EPI_ISL_433608, EPI_ISL_433609, EPI_ISL_433610, EPI_ISL_433611, EPI_ISL_433612, EPI_ISL_433613, EPI_ISL_433614, EPI_ISL_433615, EPI_ISL_433616, EPI_ISL_433617, EPI_ISL_433618, EPI_ISL_433619, EPI_ISL_433620, EPI_ISL_433621, EPI_ISL_433622, EPI_ISL_433623, EPI_ISL_433624, EPI_ISL_433625, EPI_ISL_433626, EPI_ISL_433627, EPI_ISL_433628, EPI_ISL_433629, EPI_ISL_433630, EPI_ISL_433631, EPI_ISL_433632, EPI_ISL_433633, EPI_ISL_433634, EPI_ISL_433635, EPI_ISL_433636, EPI_ISL_433637, EPI_ISL_433638, EPI_ISL_433639, EPI_ISL_433640, EPI_ISL_433641, EPI_ISL_433642, EPI_ISL_433643, EPI_ISL_433644, EPI_ISL_433645, EPI_ISL_433646, EPI_ISL_433647, EPI_ISL_433648, EPI_ISL_433649, EPI_ISL_433650, EPI_ISL_433651, EPI_ISL_433652, EPI_ISL_433653, EPI_ISL_433654, EPI_ISL_433655, EPI_ISL_433656, EPI_ISL_433657, EPI_ISL_433658 | | | |
| see above | West of Scotland Specialist Virology Centre, NHSGGC / | COVID-19 Genomics UK (COG-UK) Consortium | Ana da Silva Filipe, Natasha Johnson, Kathy Smollett, Daniel Mair, Stephen Carmichael, Lily Tong, Jenna Nichols, Elihu Aranday-Cortes, Kirstyn Brunker, |

| | | | |
|---|---|---|---|
| | MRC-University of Glasgow Centre for Virus Research | | Yasmin Parr, Kyriaki Nomikou; Sarah McDonald, Marc Niebel, Patawee Asamaphan; Richard Orton, Joseph Hughes, Sreenu Vattipally, David L Robertson; Alasdair MacLean, Rory Gunson; Kathy Li, Natasha Jesudason, Rajiv Shah, James Shepherd, Antonia Ho, Emma Thomson |
| EPI_ISL_433666, EPI_ISL_433667, EPI_ISL_433668, EPI_ISL_433669, EPI_ISL_433670, EPI_ISL_433671, EPI_ISL_433672, EPI_ISL_433673, EPI_ISL_433674, EPI_ISL_433675, EPI_ISL_433676, EPI_ISL_433677, EPI_ISL_433678, EPI_ISL_433679, EPI_ISL_433680, EPI_ISL_433681, EPI_ISL_433682, EPI_ISL_433683, EPI_ISL_433685, EPI_ISL_433686, EPI_ISL_433690, EPI_ISL_433695, EPI_ISL_433701, EPI_ISL_433702, EPI_ISL_433703, EPI_ISL_433704, EPI_ISL_433705, EPI_ISL_433706, EPI_ISL_433707, EPI_ISL_433709, EPI_ISL_433710, EPI_ISL_433711, EPI_ISL_433712, EPI_ISL_433713, EPI_ISL_433714, EPI_ISL_433715, EPI_ISL_433717, EPI_ISL_433718, EPI_ISL_433719, EPI_ISL_433720, EPI_ISL_433721, EPI_ISL_433722, EPI_ISL_433723, EPI_ISL_433724, EPI_ISL_433725, EPI_ISL_433726, EPI_ISL_433727, EPI_ISL_433728, EPI_ISL_433729, EPI_ISL_433730, EPI_ISL_433731, EPI_ISL_433732, EPI_ISL_433733, EPI_ISL_433734, EPI_ISL_433735, EPI_ISL_433736, EPI_ISL_433737, EPI_ISL_433738, EPI_ISL_433742, EPI_ISL_433745, EPI_ISL_433748, EPI_ISL_433749, EPI_ISL_433750, EPI_ISL_433751, EPI_ISL_433752, EPI_ISL_433753, EPI_ISL_433754, EPI_ISL_433755, EPI_ISL_433756, EPI_ISL_433757, EPI_ISL_433758, EPI_ISL_433759, EPI_ISL_433760, EPI_ISL_433761, EPI_ISL_433762, EPI_ISL_433763, EPI_ISL_433764, EPI_ISL_433765, EPI_ISL_433767, EPI_ISL_433768, EPI_ISL_433769, EPI_ISL_433770, EPI_ISL_433771, EPI_ISL_433773, EPI_ISL_433780, EPI_ISL_433781, EPI_ISL_433782, EPI_ISL_433783, EPI_ISL_433785, EPI_ISL_433787, EPI_ISL_433788, EPI_ISL_433793, EPI_ISL_433794, EPI_ISL_433800, EPI_ISL_433801, EPI_ISL_433802, EPI_ISL_433804, EPI_ISL_433806, EPI_ISL_433808, EPI_ISL_433810, EPI_ISL_433811, EPI_ISL_433842, EPI_ISL_433848, EPI_ISL_433850, EPI_ISL_433862 | | | |
| see above | Department of Pathology, University of Cambridge | COVID-19 Genomics UK (COG-UK) Consortium | Luke W Meredith, M. Estee Torok , Myra Hosmillo, William L. Hamilton, Martin D. Curran, Theresa Feltwell, Grant Hall, Anna Yakovleva, Fahad A Khokhar, Charlotte J. Houldcroft, Laura G Caller, Aminu S. Jahun, Sarah L. Caddy, Ian Goodfellow |
| EPI_ISL_434063, EPI_ISL_434064, EPI_ISL_434066, EPI_ISL_434068, EPI_ISL_434069, EPI_ISL_434071, EPI_ISL_434073, EPI_ISL_434074, EPI_ISL_434075, EPI_ISL_434077, EPI_ISL_434079, EPI_ISL_434083, EPI_ISL_434085, EPI_ISL_434087, EPI_ISL_434088, EPI_ISL_434090, EPI_ISL_434091, EPI_ISL_434092, EPI_ISL_434093, EPI_ISL_434094, EPI_ISL_434095, EPI_ISL_434096, EPI_ISL_434098, EPI_ISL_434099, EPI_ISL_434100, EPI_ISL_434101, EPI_ISL_434102, EPI_ISL_434103, EPI_ISL_434104, EPI_ISL_434105, EPI_ISL_434106, EPI_ISL_434109, EPI_ISL_434110, EPI_ISL_434199, EPI_ISL_434216, EPI_ISL_434217, EPI_ISL_434218, EPI_ISL_434219, EPI_ISL_434220, EPI_ISL_434221, EPI_ISL_434222, EPI_ISL_434223, EPI_ISL_434224, EPI_ISL_434227, EPI_ISL_434228, EPI_ISL_434229, EPI_ISL_434230, EPI_ISL_434231, EPI_ISL_434232, EPI_ISL_434233, EPI_ISL_434236, EPI_ISL_434237, EPI_ISL_434239, EPI_ISL_434240, EPI_ISL_434241, EPI_ISL_434242, EPI_ISL_434243, EPI_ISL_434244, EPI_ISL_434249, EPI_ISL_434251 | | | |
| see above | Washington State Department of Health | Seattle Flu Study | Chu et al |
| EPI_ISL_434367, EPI_ISL_434369, EPI_ISL_434370 | Hospital AZ Rivierenland | Institute of Tropical Medicine | Philippe Selhorst, Colin Anthony |
| EPI_ISL_434379, EPI_ISL_434380 | Hospital AZ Rivierenland | Institute of Tropical Medicine | Philippe Selhorst, Colin Anthony, |
| EPI_ISL_434466 | Laboratory of Microbiology, Medical School, National and Kapodistrian University of Athens | Laboratory of Biology, Department of Medicine, Democritus University of Thrace | Kassela K., Bampali,M., Dovrolis,N., Gatzidou,E., Froukala,E., Stavropoulou,A., Veletza,S., Tsakris,A., Spanakis,N. and Karakasiliotis,I. |
| EPI_ISL_434614, EPI_ISL_434615 | University of Wisconsin-Madison AIDS Vaccine Research Laboratories | University of Wisconsin-Madison AIDS Vaccine Research Laboratories | Gage Moreno, Katarina Braun, et al. AIDS Vaccine Research Laboratories |
| EPI_ISL_434658 | Lundens VC | The Public Health Agency of Sweden | Marita Dagner, Oskar Karlsson Lindsjo, Maria Lind Karlberg, Anna-Malin Linde, Olov Svartstrom, Anna Risberg, Theresa Enkirch, Mia Brytting, Karin Tegmark-Wisell |
| EPI_ISL_434659 | Knivsta VC | The Public Health Agency of Sweden | Johanna Carlson, Oskar Karlsson Lindsjo, Maria Lind Karlberg, Anna-Malin Linde, Olov Svartstrom, Anna Risberg, Theresa Enkirch, Mia Brytting, Karin Tegmark-Wisell |
| EPI_ISL_434660 | Narhalsan Backa vardcentral | The Public Health Agency of Sweden | Mats Olsson, Oskar Karlsson Lindsjo, Maria Lind Karlberg, Anna-Malin Linde, Olov Svartstrom, Anna Risberg, Theresa Enkirch, Mia Brytting, Karin Tegmark-Wisell |
| EPI_ISL_434663 | Surbrunns VC | The Public Health Agency of Sweden | Erik Embring, Oskar Karlsson Lindsjo, Maria Lind Karlberg, Anna-Malin Linde, Olov Svartstrom, Anna Risberg, Theresa Enkirch, Mia Brytting, Karin Tegmark-Wisell |
| EPI_ISL_434664 | Omtanken Grimmered | The Public Health Agency of Sweden | Bernd Sengpiel, Oskar Karlsson Lindsjo, Maria Lind Karlberg, Anna-Malin Linde, Olov Svartstrom, Anna Risberg, Theresa Enkirch, Mia Brytting, Karin Tegmark-Wisell |
| EPI_ISL_434665 | Uppsala Narakut Aleris | The Public Health Agency of Sweden | Annika Nilsson, Oskar Karlsson Lindsjo, Maria Lind Karlberg, Anna-Malin Linde, Olov Svartstrom, Anna Risberg, Theresa Enkirch, Mia Brytting, Karin Tegmark-Wisell |
| EPI_ISL_434942, EPI_ISL_434944, EPI_ISL_435010, EPI_ISL_435011, EPI_ISL_435012, EPI_ISL_435013, EPI_ISL_435014, EPI_ISL_435015, EPI_ISL_435016, EPI_ISL_435017, EPI_ISL_435018, EPI_ISL_435019, EPI_ISL_435020, EPI_ISL_435021, EPI_ISL_435022, EPI_ISL_435023, EPI_ISL_435024, EPI_ISL_435025, EPI_ISL_435026, EPI_ISL_435027, EPI_ISL_435028, EPI_ISL_435029, EPI_ISL_435030, EPI_ISL_435031 | | | |
| see above | Houston Methodist Hospital | Houston Methodist Hospital | S. Wesley Long, Randall J. Olsen, Paul A. Christensen, David W. Bernard, James J. Davis, Maulik Shukla, Marcus Nguyen, Matthew Ojeda Saavedra, Concepcion C. Cantu, Prasanti Yerramilli, Layne Pruitt, Sishir Subedi, Heather Hendrickson, Ghazaleh Eskandari, Muthiah Kumaraswami, Jason S. McLellan, Hakon Jonsson, Kari Stefansson, and James M. Musser |
| EPI_ISL_435358, EPI_ISL_435359, EPI_ISL_435360, EPI_ISL_435361, EPI_ISL_435362, EPI_ISL_435363, EPI_ISL_435364, EPI_ISL_435365, EPI_ISL_435366, EPI_ISL_435367, EPI_ISL_435368, EPI_ISL_435369, EPI_ISL_435370, EPI_ISL_435371, EPI_ISL_435372, EPI_ISL_435373, EPI_ISL_435375, EPI_ISL_435376, EPI_ISL_435377, EPI_ISL_435378, EPI_ISL_435379, EPI_ISL_435380, EPI_ISL_435381, EPI_ISL_435382, EPI_ISL_435383, EPI_ISL_435384, EPI_ISL_435385, EPI_ISL_435386, EPI_ISL_435387, EPI_ISL_435388, EPI_ISL_435389, EPI_ISL_435390, EPI_ISL_435391 | | | |
| see above | Utah Public Health Laboratory | Utah Public Health Laboratory | Erin Young, Kelly Oakeson |
| EPI_ISL_435418 | Virological Research Group, Szentágothai Research Centre | Bioinformatics Research Group, Szentágothai Research Centre | Péter Urbán, Endre Gábor Tóth, Gábor Kemenesi, Róbert Herczeg, Attila Gyenesei, Ferenc Jakab |
| EPI_ISL_435454, EPI_ISL_435455, EPI_ISL_435456, EPI_ISL_435457, EPI_ISL_435458, EPI_ISL_435459, EPI_ISL_435460, EPI_ISL_435461, EPI_ISL_435462, EPI_ISL_435468, EPI_ISL_435469, EPI_ISL_435470, EPI_ISL_435471, EPI_ISL_435472 | | | |
| see above | Robert Garry lab | Andersen lab at Scripps Research | Allison Smither, Gilberto Sabino-Santos, Patricia Snarski, Lilia Melnik, Antoinette Bell, Kaylynn Genemaras, Arnaud Drouin, Dahlene Fusco, Robert Garry with SEARCH Alliance San Diego |
| EPI_ISL_435702 | Yale COVID-19 Biorepository | Grubaugh Lab - Yale School of Public Health | Joseph Fauver, Tara Alpert, Anderson Brito, Anne Wyllie, Chantal Vogels, Mary Petrone, Cole Jensen, Chaney Kalinich, Isabel Ott, Arnau Casanovas, Catherine Muenker, Adam Moore, Alice Lu, Maria Tokuyama, Patrick Wong, Peiwen Lu, Saad Omer, Richard Martinello, Allison Nelson, Shelli Farhadian, Akiko Iwasaki, Charlese Dela Cruz, Albert Ko, Nathan Grubaugh |
| EPI_ISL_436099, EPI_ISL_436108 | TSGH-CP molecular lab | TSGH-CP molecular lab | Cherng-Lih Perng, Ming-Jr JIAN, Chih-Kai Chang, Jung-Chung Lin, Kuo-Ming Yeh, Chien-Wen Chen, Sheng-Kang Chiu, Hsing-Yi Chung, Shih-Hung Tsai, Kuo-Sheng Hung, Tien-Yao Chang, Feng-Yee Chang, Hung-Sheng Shang |
| EPI_ISL_436423, EPI_ISL_436424, EPI_ISL_436425, EPI_ISL_436426, EPI_ISL_436427, EPI_ISL_436428, EPI_ISL_436429, EPI_ISL_436430, EPI_ISL_436431, EPI_ISL_436432, EPI_ISL_436433, EPI_ISL_436434, EPI_ISL_436436 | | | |
| see above | National Centre for Disease control (NCDC) | NCDC/CSIR-IGIB | Pramod Kumar#, Rajesh Pandey#, Pooja Sharma, Mahesh S Dhar, Vivekanand A, Bharathram Uppili, Himanshu Vashisht, Saruchi Wadhwa, Nishu Tyagi, Uma Sharma, Priyanka Singh, Hemlata Lall, Meena Datta, Poonam Gupta, Nidhi Saini, Aarti Tewari, Bibhash Nandi, Dhirendra Kumar, Satyabrata Bag, Varun Jaiswal, Hema Gogia, Preeti Madan, Simrita Singh, Prateek Singh, Debasis Dash, Mitali Mukerji, Manju Bala, Sandhya Kabra, Sujeet Singh, Mohammed Faruq, Anurag Agrawal*, Partha Rakshit* |
| EPI_ISL_436505, EPI_ISL_436506, EPI_ISL_436507, EPI_ISL_436508, EPI_ISL_436509, EPI_ISL_436510, EPI_ISL_436511, EPI_ISL_436512, EPI_ISL_436513, EPI_ISL_436514, EPI_ISL_436515, EPI_ISL_436516 | | | |
| see above | Florida Bureau of Public Health Laboratories | Florida Bureau of Public Health Laboratories | Sarah Schmedes, Jason Blanton |
| EPI_ISL_436571, EPI_ISL_436572, EPI_ISL_436573, EPI_ISL_436574, EPI_ISL_436575, EPI_ISL_436576, EPI_ISL_436577, EPI_ISL_436579, EPI_ISL_436580, EPI_ISL_436581, EPI_ISL_436635, EPI_ISL_436636, EPI_ISL_436637, EPI_ISL_436638 | | | |
| see above | University of Wisconsin-Madison AIDS Vaccine Research Laboratories | University of Wisconsin-Madison AIDS Vaccine Research Laboratories | Gage Moreno, Katarina Braun, et al. AIDS Vaccine Research Laboratories |
| EPI_ISL_436644, EPI_ISL_436646, EPI_ISL_436647, EPI_ISL_436648, EPI_ISL_436649, EPI_ISL_436650, EPI_ISL_436651, EPI_ISL_436652, EPI_ISL_436677, EPI_ISL_436678, EPI_ISL_436679, EPI_ISL_436680, EPI_ISL_436681 | | | |
| see above | County of Santa Clara Public Health Department | Chan-Zuckerberg Biohub | CZB Cliahub Consortium |
| EPI_ISL_436688 | Victorian Infectious Diseases Reference Laboratory (VIDRL) | Microbiological Diagnostic Unit Public Health Laboratory and Victorian Infectious Diseases Reference Laboratory, The Peter Doherty Institute for Infection & Immunity | Caly L., Seemann T., Sait, M., Schultz M., Druce J., Sherry, N. |
| EPI_ISL_436904 | Utah Public Health Laboratory | Utah Public Health Laboratory | Erin Young, Kelly Oakeson |
| EPI_ISL_436939, EPI_ISL_436943, EPI_ISL_436944, EPI_ISL_436945, EPI_ISL_436947, EPI_ISL_436948, EPI_ISL_436949, EPI_ISL_436950, EPI_ISL_436951, EPI_ISL_436952, EPI_ISL_436953, EPI_ISL_436956, EPI_ISL_436957, EPI_ISL_436958, EPI_ISL_436959, EPI_ISL_436960, EPI_ISL_436961 | | | |
| see above | Ochsner Health | Bioinfoexperts, LLC | Amy Feehan, David J. Nolan, Rebecca Rose, Sissy Cross, David Moraga Amador, Tong Yang, Luke Caruso, Wayra Navia, Lydia Von Borstel, Xiao Hui |

| | | | |
|---|---|---|---|
| | | | Zhou, Julia-Garcia-Diaz, Susanna L. Lamers |
| EPI_ISL_436962 | Department of Virus and Microbiological Special Diagnostics, Statens Serum Institut, Copenhagen, Denmark, Artillerivej 5, 2300 Copenahgen S | Albertsen lab, Department of Chemistry and Bioscience, Aalborg University, Denmark | Rasmus Kirkegaard |
| EPI_ISL_437045, EPI_ISL_437046, EPI_ISL_437047, EPI_ISL_437048, EPI_ISL_437049, EPI_ISL_437050, EPI_ISL_437051, EPI_ISL_437052, EPI_ISL_437053, EPI_ISL_437054, EPI_ISL_437055, EPI_ISL_437058, EPI_ISL_437080, EPI_ISL_437086, EPI_ISL_437087, EPI_ISL_437088 | | | |
| see above | County of Santa Clara Public Health | Chan-Zuckerberg Biohub | CZB Cliahub Consortium |
| EPI_ISL_437201, EPI_ISL_437202, EPI_ISL_437203 | Diagnostic- and Research Institute of Pathology, Medical University of Graz | Diagnostic- and Research Institute of Pathology, Medical University of Graz | Karl Kashofer, Peter Regitnig, Martin Zacharias, Gregor Gorkiewicz |
| EPI_ISL_437213, EPI_ISL_437219, EPI_ISL_437220, EPI_ISL_437221, EPI_ISL_437239, EPI_ISL_437240, EPI_ISL_437283, EPI_ISL_437284, EPI_ISL_437285, EPI_ISL_437286, EPI_ISL_437287, EPI_ISL_437288, EPI_ISL_437289, EPI_ISL_437290, EPI_ISL_437291 | | | |
| see above | Max von Pettenkofer Institute, Virology, National Reference Center for Retroviruses, LMU München | Laboratory for Functional Genome Analysis, Dept. Genomics, Gene Center of the LMU Munich | Max Muenchhoff, Stefan Krebs, Alexander Graf, Oliver Keppler, Helmut Blum |
| EPI_ISL_437364, EPI_ISL_437365, EPI_ISL_437366 | Minnesota Department of Health, Public Health Laboratory | Minnesota Department of Health, Public Health Laboratory | Matt Plumb, Jacob Garfin, and Xiong Wang |
| EPI_ISL_437393, EPI_ISL_437394, EPI_ISL_437395, EPI_ISL_437396, EPI_ISL_437397, EPI_ISL_437426, EPI_ISL_437427, EPI_ISL_437428, EPI_ISL_437429, EPI_ISL_437431 | Virginia DCLS | Virginia DCLS | Virginia DCLS |
| EPI_ISL_437435, EPI_ISL_437436 | Veterinary Specialized Institue Kraljevo | Veterinary Specialized Institue Kraljevo | Dejan Vidanovic. Bojana Tesovic, Milanko Sekler, Marko Dmitric, Kazimir Matovic, Zoran Debeljak, Nikola Vaskovic, Tamas Petrovic, Jeremy Volkening, Claudio L Afonso |
| EPI_ISL_437456, EPI_ISL_437457 | Clinical Diagnostics Laboratory, Diagnostic & Experimental Pathology, Lilly Research Laboratories | Clinical Diagnostics Laboratory, Diagnostic & Experimental Pathology, Lilly Research Laboratories | Tim Holzer, Mayuri Vaidya, Angie Fulford, Sam McNeely, Rachael Redmond, Phil Ebert, John Calley, Leslie ONeill Reising, Pat Finnegan, Erin Wray, John McElwee, Jeff Fill, Joe Oakley, Andrew Schade |
| EPI_ISL_437476, EPI_ISL_437477, EPI_ISL_437478, EPI_ISL_437479 | Pathogen Genomics Lab King Abdullah University of Science and Technology(KAUST) | Pathogen Genomics Lab King Abdullah University of Science and Technology(KAUST) | Sharif Hala,Raeece Naeem,Sara Mfarrej,Arnab Pain |
| EPI_ISL_437538, EPI_ISL_437539 | ICMR-National Institute of Cholera and Enteric Diseases | National Institute of Biomedical Genomics | Arindam Maitra, Mamta Chawla Sarkar, Sreedhar Chinnaswamy, Hasina Banu, Ananya Chatterjee, Shanta Dutta, Saumitra Das |
| EPI_ISL_437541, EPI_ISL_437542, EPI_ISL_437544 | Robert Garry lab | Andersen lab at Scripps Research | Allison Smither, Gilberto Sabino-Santos, Patricia Snarski, Lilia Melnik, Antoinette Bell, Kaylynn Genemaras, Arnaud Drouin, Dahlene Fusco, Robert Garry with SEARCH Alliance San Diego |
| EPI_ISL_437575, EPI_ISL_437576, EPI_ISL_437579 | Scripps Medical Laboratory | Andersen lab at Scripps Research | SEARCH Alliance San Diego with Michael Quigley, Ellen Stefanski, Ian Mchardy |
| EPI_ISL_437601 | Keio University School of Medicine | Keio University School of Medicine | Kenjiro Kosaki, Yuka Iwasaki, Toshiki Takenouchi, Haruhiko Siomi, |
| EPI_ISL_437631, EPI_ISL_437632, EPI_ISL_437637, EPI_ISL_437639, EPI_ISL_437641, EPI_ISL_437643, EPI_ISL_437644, EPI_ISL_437645, EPI_ISL_437646, EPI_ISL_437651, EPI_ISL_437654, EPI_ISL_437655, EPI_ISL_437656 | | | |
| see above | Department of Virus and Microbiological Special Diagnostics, Statens Serum Institut, Copenhagen, Denmark, Artillerivej 5, 2300 Copenahgen S | Albertsen lab, Department of Chemistry and Bioscience, Aalborg University, Denmark | Rasmus Kirkegaard |
| EPI_ISL_437696, EPI_ISL_437697, EPI_ISL_437698, EPI_ISL_437699, EPI_ISL_437700, EPI_ISL_437701, EPI_ISL_437702, EPI_ISL_437703, EPI_ISL_437704, EPI_ISL_437705, EPI_ISL_437706, EPI_ISL_437707, EPI_ISL_437708, EPI_ISL_437709, EPI_ISL_437710, EPI_ISL_437711, EPI_ISL_437712, EPI_ISL_437713, EPI_ISL_437714, EPI_ISL_437715, EPI_ISL_437716, EPI_ISL_437717, EPI_ISL_437718, EPI_ISL_437719, EPI_ISL_437720, EPI_ISL_437721, EPI_ISL_437722, EPI_ISL_437723, EPI_ISL_437724, EPI_ISL_437725, EPI_ISL_437726, EPI_ISL_437727, EPI_ISL_437728, EPI_ISL_437729, EPI_ISL_437730, EPI_ISL_437731, EPI_ISL_437732 | | | |
| see above | Pathogen Genomics Lab King Abdullah University of Science and Technology(KAUST) | Pathogen Genomics Lab King Abdullah University of Science and Technology(KAUST) | Sharif Hala,Fadwa Alofi,Afrah Alsomali, Asim Khogeer, Sara Mfarrej, Khaled Algithami,Raeece Naeem, Amit Kumar Subudhi,Fathia Ben-Rached, Rahul Salunke, Anwar Hashem, Naif Almontashiri, Arnab Pain |
| EPI_ISL_437830, EPI_ISL_437831, EPI_ISL_437832, EPI_ISL_437833, EPI_ISL_437834, EPI_ISL_437835, EPI_ISL_437836, EPI_ISL_437837, EPI_ISL_437838, EPI_ISL_437839, EPI_ISL_437840, EPI_ISL_437841, EPI_ISL_437842, EPI_ISL_437843, EPI_ISL_437844, EPI_ISL_437845, EPI_ISL_437846, EPI_ISL_437847, EPI_ISL_437848, EPI_ISL_437849, EPI_ISL_437850, EPI_ISL_437851, EPI_ISL_437852, EPI_ISL_437853, EPI_ISL_437854, EPI_ISL_437855, EPI_ISL_437856, EPI_ISL_437857, EPI_ISL_437858, EPI_ISL_437859, EPI_ISL_437860, EPI_ISL_437861, EPI_ISL_437862, EPI_ISL_437863, EPI_ISL_437864, EPI_ISL_437865, EPI_ISL_437866, EPI_ISL_437867, EPI_ISL_437868, EPI_ISL_437869, EPI_ISL_437870, EPI_ISL_437871, EPI_ISL_437872 | | | |
| see above | UW Virology Lab | UW Virology Lab | Pavitra Roychoudhury, Hong Xie, Keith Jerome, Alexander Greninger |
| EPI_ISL_437938, EPI_ISL_437939 | Universitaetsklinik für Innere Medizin II Innsbruck | Bergthaler laboratory, CeMM Research Center for Molecular Medicine of the Austrian Academy of Sciences | Alexandra Popa, Benedikt Agerer, Henrique Colaco, Lukas Endler, Jakob-Wendelin Genger, Alexander Lercher, Mark Smyth, Thomas Penz, Michael Schuster, Jan Laine, Martin Senekowitsch, Judith Aberle, Stephan Aberle, Elisabeth Puchhammer-Stoeckl, Manfred Nairz, Guenter Weiss, Wegene Borena, Dorothee von Laer, Christoph Bock, Andreas Bergthaler |
| EPI_ISL_438122, EPI_ISL_438123, EPI_ISL_438126, EPI_ISL_438127 | Center for Virology, Medical University of Vienna | Bergthaler laboratory, CeMM Research Center for Molecular Medicine of the Austrian Academy of Sciences | Alexandra Popa, Benedikt Agerer, Henrique Colaco, Lukas Endler, Jakob-Wendelin Genger, Alexander Lercher, Mark Smyth, Thomas Penz, Michael Schuster, Jan Laine, Martin Senekowitsch, Judith Aberle, Stephan Aberle, Elisabeth Puchhammer-Stoeckl, Manfred Nairz, Guenter Weiss, Wegene Borena, Dorothee von Laer, Christoph Bock, Andreas Bergthaler |
| EPI_ISL_438141, EPI_ISL_438142, EPI_ISL_438143, EPI_ISL_438144, EPI_ISL_438145, EPI_ISL_438146 | Seattle Flu Study | Seattle Flu Study | Chu et al |
| EPI_ISL_438251, EPI_ISL_438253 | Department of Pathology, University of Cambridge | Wellcome Sanger Institute for the COVID-19 Genomics UK (COG-UK) consortium | Luke W Meredith, M. Estée Török , Myra Hosmillo, William L. Hamilton, Martin D. Curran, Theresa Feltwell, Grant Hall, Anna Yakovleva, Fahad A Khokhar, Charlotte J. Houldcroft, Laura G Caller, Aminu S. Jahun, Sarah L. Caddy, Ian Goodfellow, Alex Alderton, Roberto Amato, Sonia Goncalves, Ewan Harrison, David K. Jackson, Ian Johnston, Dominic Kwiatkowski, Cordelia Langford, John Sillitoe on behalf of the Wellcome Sanger Institute COVID-19 Surveillance Team (http://www.sanger.ac.uk/covid-team) |
| EPI_ISL_438805, EPI_ISL_438806, EPI_ISL_438811, EPI_ISL_438816, EPI_ISL_438817, EPI_ISL_438820, EPI_ISL_438824, EPI_ISL_438825, EPI_ISL_438826, EPI_ISL_438827, EPI_ISL_438828, EPI_ISL_438829, EPI_ISL_438830, EPI_ISL_438831, EPI_ISL_438832, EPI_ISL_438833, EPI_ISL_438834, EPI_ISL_438835, EPI_ISL_438836, EPI_ISL_438837, EPI_ISL_438838, EPI_ISL_438839, EPI_ISL_438840, EPI_ISL_438841, EPI_ISL_438842, EPI_ISL_438843, EPI_ISL_438844, EPI_ISL_438845, EPI_ISL_438846, EPI_ISL_438847, EPI_ISL_438848, EPI_ISL_438850 | | | |
| see above | West of Scotland Specialist Virology Centre, NHSGGC / MRC-University of Glasgow Centre for Virus Research | COVID-19 Genomics UK (COG-UK) Consortium | Ana da Silva Filipe, Natasha Johnson, Kathy Smollett, Daniel Mair, Stephen Carmichael, Lily Tong, Jenna Nichols, Elihu Aranday-Cortes, Kirstyn Brunker, Yasmin Parr, Kyriaki Nomikou; Sarah McDonald, Marc Niebel, Patawee Asamaphan; Richard Orton, Joseph Hughes, Sreenu Vattipally, David L Robertson; Alasdair MacLean, Rory Gunson; Kathy Li, Natasha Jesudason, Rajiv Shah, James Shepherd, Antonia Ho, Emma Thomson |
| EPI_ISL_438947, EPI_ISL_438948, EPI_ISL_438949, EPI_ISL_438951, EPI_ISL_438952, EPI_ISL_438953, EPI_ISL_438954 | Keio University School of Medicine | Keio University School of Medicine | Kenjiro Kosaki |
| EPI_ISL_439145 | Virology Department, Royal Infirmary of Edinburgh, NHS Lothian / School of Biological Sciences, University of Edinburgh / Institute of Genetics and Molecular Medicine, University of Edinburgh | COVID-19 Genomics UK (COG-UK) Consortium | McHugh M, Dewar R, Rooke S, Gallagher M, Balcaza C, O'ÄôToole Å, Scher E, Hill V, McCrone JT, Colquhoun R, Yu X, Jackson B, Rambaut A, Williams TC, Templeton K |
| EPI_ISL_439383, EPI_ISL_439402, EPI_ISL_439403 | Department of Pathology, University of Cambridge | Wellcome Sanger Institute for the COVID-19 Genomics UK (COG-UK) consortium | Luke W Meredith, M. Estée Török , Myra Hosmillo, William L. Hamilton, Martin D. Curran, Theresa Feltwell, Grant Hall, Anna Yakovleva, Fahad A Khokhar, Charlotte J. Houldcroft, Laura G Caller, Aminu S. Jahun, Sarah L. Caddy, Ian Goodfellow, Alex Alderton, Roberto Amato, Sonia Goncalves, Ewan Harrison, David K. Jackson, Ian Johnston, Dominic Kwiatkowski, Cordelia Langford, John Sillitoe on behalf of the Wellcome Sanger Institute COVID-19 Surveillance Team (http://www.sanger.ac.uk/covid-team) |

| | | | |
|---|---|---|---|
| EPI_ISL_439671 | Virology Department, Royal Infirmary of Edinburgh, NHS Lothian / School of Biological Sciences, University of Edinburgh / Institute of Genetics and Molecular Medicine, University of Edinburgh | COVID-19 Genomics UK (COG-UK) Consortium | McHugh M, Dewar R, Rooke S, Gallagher M, Balcaza C, O'ÄôToole Å, Scher E, Hill V, McCrone JT, Colquhoun R, Yu X, Jackson B, Rambaut A, Williams TC, Templeton K |
| EPI_ISL_439672, EPI_ISL_439680, EPI_ISL_439692, EPI_ISL_439701, EPI_ISL_439706, EPI_ISL_439708, EPI_ISL_439737 | Liverpool Clinical Laboratories | COVID-19 Genomics UK (COG-UK) Consortium | Sam Haldenby, Anita Lucaci, Steve Paterson, Julian Hiscox, Alistair Darby, M Almsaud, A Alrezaihi, Muhannad Alruwaili, Stuart D Armstrong, Jones Benjamin , Eleanor G Bentley, Anu Chawla, Jordan J Clark, Angela Cowell, Richard Eccles, Isabel Garca-Dorival, Matthew Gemmell, Alessandro Gerada, PKF Gilmore, Richard Gregory, Ximeng Han, Catherine Hartley, Margaret Hughes, Miren Iturriza-Gomara, James Johnson, L Luu, Jenifer Manson , Charlotte Nelson, Elaine O'ÄôToole, Cassie Olateju, Rebekah Penrice-Randal¬†, Lucille Rainbow, N.P Randle, Trevor Ian Robinson, Parul Sharma, Ghada T Shawli, James P Stewart , Neil Swainston, Ecaterina Vamos, Joanne Watts, Mark Whitehead |
| EPI_ISL_440555, EPI_ISL_440556, EPI_ISL_440557, EPI_ISL_440558, EPI_ISL_440559, EPI_ISL_440560, EPI_ISL_440561, EPI_ISL_440562, EPI_ISL_440563, EPI_ISL_440565, EPI_ISL_440566, EPI_ISL_440567, EPI_ISL_440569, EPI_ISL_440570, EPI_ISL_440571, EPI_ISL_440572, EPI_ISL_440573, EPI_ISL_440574, EPI_ISL_440576, EPI_ISL_440577, EPI_ISL_440578, EPI_ISL_440579, EPI_ISL_440580, EPI_ISL_440581, EPI_ISL_440582, EPI_ISL_440583, EPI_ISL_440584, EPI_ISL_440585, EPI_ISL_440586, EPI_ISL_440587, EPI_ISL_440588, EPI_ISL_440589, EPI_ISL_440590, EPI_ISL_440591, EPI_ISL_440592, EPI_ISL_440593, EPI_ISL_440594, EPI_ISL_440595, EPI_ISL_440596, EPI_ISL_440597, EPI_ISL_440598, EPI_ISL_440599, EPI_ISL_440600, EPI_ISL_440603, EPI_ISL_440604, EPI_ISL_440605, EPI_ISL_440606, EPI_ISL_440607, EPI_ISL_440608, EPI_ISL_440609, EPI_ISL_440610, EPI_ISL_440611, EPI_ISL_440612, EPI_ISL_440614, EPI_ISL_440615, EPI_ISL_440616, EPI_ISL_440618, EPI_ISL_440619, EPI_ISL_440621, EPI_ISL_440622 | | | |
| see above | Department of Pathology, University of Cambridge | Wellcome Sanger Institute for the COVID-19 Genomics UK (COG-UK) consortium | Luke W Meredith, M. Estée Török , Myra Hosmillo, William L. Hamilton, Martin D. Curran, Theresa Feltwell, Grant Hall, Anna Yakovleva, Fahad A Khokhar, Charlotte J. Houldcroft, Laura G Caller, Aminu S. Jahun, Sarah L. Caddy, Ian Goodfellow, Alex Alderton, Roberto Amato, Sonia Goncalves, Ewan Harrison, David K. Jackson, Ian Johnston, Dominic Kwiatkowski, Cordelia Langford, John Sillitoe on behalf of the Wellcome Sanger Institute COVID-19 Surveillance Team (http://www.sanger.ac.uk/covid-team) |
| EPI_ISL_440623, EPI_ISL_440624, EPI_ISL_440625, EPI_ISL_440626, EPI_ISL_440627, EPI_ISL_440628, EPI_ISL_440629, EPI_ISL_440630, EPI_ISL_440631, EPI_ISL_440632, EPI_ISL_440633, EPI_ISL_440634, EPI_ISL_440635, EPI_ISL_440636, EPI_ISL_440637, EPI_ISL_440638, EPI_ISL_440639, EPI_ISL_440640, EPI_ISL_440641, EPI_ISL_440642, EPI_ISL_440643, EPI_ISL_440644, EPI_ISL_440645, EPI_ISL_440646, EPI_ISL_440647, EPI_ISL_440648, EPI_ISL_440649, EPI_ISL_440650, EPI_ISL_440651, EPI_ISL_440652, EPI_ISL_440653, EPI_ISL_440654, EPI_ISL_440655, EPI_ISL_440656, EPI_ISL_440657, EPI_ISL_440658, EPI_ISL_440659, EPI_ISL_440660, EPI_ISL_440661, EPI_ISL_440663, EPI_ISL_440664, EPI_ISL_440665, EPI_ISL_440666, EPI_ISL_440667, EPI_ISL_440668, EPI_ISL_440670, EPI_ISL_440671, EPI_ISL_440672, EPI_ISL_440673, EPI_ISL_440674, EPI_ISL_440675, EPI_ISL_440676, EPI_ISL_440677, EPI_ISL_440679, EPI_ISL_440680, EPI_ISL_440681, EPI_ISL_440682, EPI_ISL_440683, EPI_ISL_440684, EPI_ISL_440685, EPI_ISL_440686, EPI_ISL_440687, EPI_ISL_440688, EPI_ISL_440689, EPI_ISL_440690, EPI_ISL_440691, EPI_ISL_440692, EPI_ISL_440693, EPI_ISL_440695, EPI_ISL_440697, EPI_ISL_440698, EPI_ISL_440699, EPI_ISL_440701, EPI_ISL_440702, EPI_ISL_440704, EPI_ISL_440705, EPI_ISL_440706, EPI_ISL_440707, EPI_ISL_440708, EPI_ISL_440709, EPI_ISL_440710, EPI_ISL_440711, EPI_ISL_440712, EPI_ISL_440713, EPI_ISL_440714, EPI_ISL_440715, EPI_ISL_440716, EPI_ISL_440717, EPI_ISL_440718, EPI_ISL_440719, EPI_ISL_440721, EPI_ISL_440722, EPI_ISL_440723, EPI_ISL_440724, EPI_ISL_440725, EPI_ISL_440726, EPI_ISL_440727, EPI_ISL_440728, EPI_ISL_440729, EPI_ISL_440730, EPI_ISL_440731, EPI_ISL_440732, EPI_ISL_440733, EPI_ISL_440734, EPI_ISL_440735, EPI_ISL_440737, EPI_ISL_440738, EPI_ISL_440739, EPI_ISL_440740, EPI_ISL_440741, EPI_ISL_440742, EPI_ISL_440743, EPI_ISL_440744, EPI_ISL_440745, EPI_ISL_440746, EPI_ISL_440747, EPI_ISL_440748, EPI_ISL_440749, EPI_ISL_440750, EPI_ISL_440751, EPI_ISL_440752, EPI_ISL_440753, EPI_ISL_440754, EPI_ISL_440755, EPI_ISL_440756, EPI_ISL_440757, EPI_ISL_440758, EPI_ISL_440759, EPI_ISL_440760, EPI_ISL_440761, EPI_ISL_440762, EPI_ISL_440763, EPI_ISL_440764, EPI_ISL_440766, EPI_ISL_440767, EPI_ISL_440768, EPI_ISL_440769, EPI_ISL_440770, EPI_ISL_440771, EPI_ISL_440772, EPI_ISL_440773, EPI_ISL_440774, EPI_ISL_440775, EPI_ISL_440776, EPI_ISL_440777, EPI_ISL_440778, EPI_ISL_440780, EPI_ISL_440781, EPI_ISL_440782, EPI_ISL_440783, EPI_ISL_440784, EPI_ISL_440785, EPI_ISL_440787, EPI_ISL_440788, EPI_ISL_440789, EPI_ISL_440790, EPI_ISL_440791, EPI_ISL_440792, EPI_ISL_440793, EPI_ISL_440794, EPI_ISL_440795, EPI_ISL_440796, EPI_ISL_440797, EPI_ISL_440798, EPI_ISL_440799, EPI_ISL_440800, EPI_ISL_440801, EPI_ISL_440802, EPI_ISL_440803, EPI_ISL_440804, EPI_ISL_440805, EPI_ISL_440806, EPI_ISL_440807, EPI_ISL_440808 | | | |
| see above | PHE South West Regional Laboratory, National Infection Service | Wellcome Sanger Institute for the COVID-19 Genomics UK (COG-UK) consortium | Stephanie Hutchings, Hannah Pymont, Dr Peter Muir, Barry Vipond, Rich Hopes, Alex Alderton, Roberto Amato, Sonia Goncalves, Ewan Harrison, David K. Jackson, Ian Johnston, Dominic Kwiatkowski, Cordelia Langford, John Sillitoe on behalf of the Wellcome Sanger Institute COVID-19 Surveillance Team (http://www.sanger.ac.uk/covid-team) |
| EPI_ISL_440812, EPI_ISL_440814, EPI_ISL_440816, EPI_ISL_440830, EPI_ISL_440833, EPI_ISL_440834, EPI_ISL_440835, EPI_ISL_440843, EPI_ISL_440847 | Department of Pathology, University of Cambridge | Wellcome Sanger Institute for the COVID-19 Genomics UK (COG-UK) consortium | Luke W Meredith, M. Estée Török , Myra Hosmillo, William L. Hamilton, Martin D. Curran, Theresa Feltwell, Grant Hall, Anna Yakovleva, Fahad A Khokhar, Charlotte J. Houldcroft, Laura G Caller, Aminu S. Jahun, Sarah L. Caddy, Ian Goodfellow, Alex Alderton, Roberto Amato, Sonia Goncalves, Ewan Harrison, David K. Jackson, Ian Johnston, Dominic Kwiatkowski, Cordelia Langford, John Sillitoe on behalf of the Wellcome Sanger Institute COVID-19 Surveillance Team (http://www.sanger.ac.uk/covid-team) |
| EPI_ISL_440959, EPI_ISL_440960, EPI_ISL_440961, EPI_ISL_440962, EPI_ISL_440963, EPI_ISL_440964, EPI_ISL_440965, EPI_ISL_440966, EPI_ISL_440967, EPI_ISL_440968, EPI_ISL_440969, EPI_ISL_440970, EPI_ISL_440991, EPI_ISL_440992, EPI_ISL_440993, EPI_ISL_440994, EPI_ISL_440995, EPI_ISL_440996, EPI_ISL_440997, EPI_ISL_440998, EPI_ISL_440999, EPI_ISL_441000, EPI_ISL_441001, EPI_ISL_441002, EPI_ISL_441003, EPI_ISL_441004, EPI_ISL_441005, EPI_ISL_441006, EPI_ISL_441007, EPI_ISL_441008, EPI_ISL_441009, EPI_ISL_441010, EPI_ISL_441011, EPI_ISL_441012, EPI_ISL_441013, EPI_ISL_441014, EPI_ISL_441015, EPI_ISL_441016, EPI_ISL_441017, EPI_ISL_441018, EPI_ISL_441019, EPI_ISL_441020, EPI_ISL_441021, EPI_ISL_441022, EPI_ISL_441023, EPI_ISL_441024, EPI_ISL_441025, EPI_ISL_441026, EPI_ISL_441027, EPI_ISL_441028, EPI_ISL_441029, EPI_ISL_441030, EPI_ISL_441031, EPI_ISL_441032, EPI_ISL_441033, EPI_ISL_441034, EPI_ISL_441035, EPI_ISL_441036, EPI_ISL_441037, EPI_ISL_441038, EPI_ISL_441039, EPI_ISL_441040, EPI_ISL_441041, EPI_ISL_441042, EPI_ISL_441043, EPI_ISL_441044, EPI_ISL_441045, EPI_ISL_441046, EPI_ISL_441047, EPI_ISL_441048, EPI_ISL_441049, EPI_ISL_441050, EPI_ISL_441051 | | | |
| see above | University College London, Great Ormond Street Hospital for Children NHS Foundation Trust, Imperial College Healthcare NHS Trust | COVID-19 Genomics UK (COG-UK) Consortium | Sergi Castellano, Rachel Williams, Mark Kristiansen, Paola Resende Silva, Sunando Roy, Tony Brooks, Helena Tutill, Paola Niola, Patricia Dyal, Charlotte Williams, Leysa Forrest, Yasmin Panchbhaya, Jacqueline Findlay, Sam Weeks, Julianne Brown, Kathryn Harris, Paul Randell, James Price, Alison Holmes, Judith Breuer |
| EPI_ISL_441052, EPI_ISL_441054, EPI_ISL_441055, EPI_ISL_441056, EPI_ISL_441057, EPI_ISL_441058, EPI_ISL_441059, EPI_ISL_441060, EPI_ISL_441061, EPI_ISL_441062, EPI_ISL_441063, EPI_ISL_441064, EPI_ISL_441066, EPI_ISL_441067, EPI_ISL_441069, EPI_ISL_441070, EPI_ISL_441072, EPI_ISL_441073, EPI_ISL_441074, EPI_ISL_441075, EPI_ISL_441078, EPI_ISL_441079, EPI_ISL_441080, EPI_ISL_441081, EPI_ISL_441083, EPI_ISL_441084, EPI_ISL_441085, EPI_ISL_441086, EPI_ISL_441087, EPI_ISL_441088, EPI_ISL_441089, EPI_ISL_441091, EPI_ISL_441092, EPI_ISL_441094, EPI_ISL_441095, EPI_ISL_441096, EPI_ISL_441097, EPI_ISL_441098, EPI_ISL_441099, EPI_ISL_441100, EPI_ISL_441101, EPI_ISL_441102, EPI_ISL_441104, EPI_ISL_441105, EPI_ISL_441106, EPI_ISL_441109, EPI_ISL_441111, EPI_ISL_441113, EPI_ISL_441114, EPI_ISL_441115, EPI_ISL_441116, EPI_ISL_441117, EPI_ISL_441119, EPI_ISL_441122, EPI_ISL_441124, EPI_ISL_441126, EPI_ISL_441128, EPI_ISL_441129, EPI_ISL_441130, EPI_ISL_441131, EPI_ISL_441132, EPI_ISL_441133, EPI_ISL_441134, EPI_ISL_441135, EPI_ISL_441136, EPI_ISL_441138, EPI_ISL_441139, EPI_ISL_441140, EPI_ISL_441141, EPI_ISL_441142, EPI_ISL_441143, EPI_ISL_441147, EPI_ISL_441148, EPI_ISL_441150, EPI_ISL_441151, EPI_ISL_441154, EPI_ISL_441155, EPI_ISL_441156, EPI_ISL_441157, EPI_ISL_441158, EPI_ISL_441159, EPI_ISL_441161, EPI_ISL_441164, EPI_ISL_441165, EPI_ISL_441168, EPI_ISL_441169, EPI_ISL_441170, EPI_ISL_441171, EPI_ISL_441172, EPI_ISL_441174, EPI_ISL_441175, EPI_ISL_441176, EPI_ISL_441177, EPI_ISL_441178, EPI_ISL_441180, EPI_ISL_441181, EPI_ISL_441183, EPI_ISL_441184, EPI_ISL_441186, EPI_ISL_441188, EPI_ISL_441189, EPI_ISL_441190, EPI_ISL_441191, EPI_ISL_441192, EPI_ISL_441193, EPI_ISL_441194, EPI_ISL_441195, EPI_ISL_441196, EPI_ISL_441197, EPI_ISL_441198, EPI_ISL_441199, EPI_ISL_441200, EPI_ISL_441201, EPI_ISL_441202, EPI_ISL_441203, EPI_ISL_441205, EPI_ISL_441206, EPI_ISL_441207, EPI_ISL_441208, EPI_ISL_441209, EPI_ISL_441210, EPI_ISL_441212, EPI_ISL_441213, EPI_ISL_441214, EPI_ISL_441215, EPI_ISL_441216, EPI_ISL_441217, EPI_ISL_441218, EPI_ISL_441219, EPI_ISL_441220, EPI_ISL_441221, EPI_ISL_441222, EPI_ISL_441223, EPI_ISL_441224, EPI_ISL_441225, EPI_ISL_441226, EPI_ISL_441228, EPI_ISL_441230, EPI_ISL_441231, EPI_ISL_441232, EPI_ISL_441233, EPI_ISL_441234, EPI_ISL_441236, EPI_ISL_441237, EPI_ISL_441238, EPI_ISL_441239, EPI_ISL_441240, EPI_ISL_441242, EPI_ISL_441244, EPI_ISL_441245, EPI_ISL_441247, EPI_ISL_441249, EPI_ISL_441251, EPI_ISL_441252, EPI_ISL_441253, EPI_ISL_441255, EPI_ISL_441258, EPI_ISL_441259, EPI_ISL_441260, EPI_ISL_441261, EPI_ISL_441262, EPI_ISL_441263, EPI_ISL_441265, EPI_ISL_441266, EPI_ISL_441267, EPI_ISL_441268, EPI_ISL_441269, EPI_ISL_441270, EPI_ISL_441271, EPI_ISL_441272, EPI_ISL_441273, EPI_ISL_441275, EPI_ISL_441276, EPI_ISL_441277, EPI_ISL_441278, EPI_ISL_441279, EPI_ISL_441280, EPI_ISL_441281, EPI_ISL_441282, EPI_ISL_441284, EPI_ISL_441285, EPI_ISL_441286, EPI_ISL_441287, EPI_ISL_441288, EPI_ISL_441289, EPI_ISL_441292, EPI_ISL_441294, EPI_ISL_441295, EPI_ISL_441297, EPI_ISL_441298, EPI_ISL_441299, EPI_ISL_441300, EPI_ISL_441301, EPI_ISL_441305, EPI_ISL_441306, EPI_ISL_441308, EPI_ISL_441309, EPI_ISL_441310, EPI_ISL_441311, EPI_ISL_441313, EPI_ISL_441314, EPI_ISL_441315, EPI_ISL_441316, EPI_ISL_441317, EPI_ISL_441318, EPI_ISL_441319, EPI_ISL_441320, EPI_ISL_441322, EPI_ISL_441323, EPI_ISL_441324, EPI_ISL_441326, EPI_ISL_441327, EPI_ISL_441329, EPI_ISL_441330, EPI_ISL_441332, EPI_ISL_441333, EPI_ISL_441334, EPI_ISL_441335, EPI_ISL_441336, EPI_ISL_441337, EPI_ISL_441338, EPI_ISL_441340, EPI_ISL_441342, EPI_ISL_441343, EPI_ISL_441344, EPI_ISL_441345, EPI_ISL_441346, EPI_ISL_441347, EPI_ISL_441348, EPI_ISL_441349 | | | |
| see above | Department of Pathology, University of Cambridge | Wellcome Sanger Institute for the COVID-19 Genomics UK (COG-UK) consortium | Luke W Meredith, M. Estée Török , Myra Hosmillo, William L. Hamilton, Martin D. Curran, Theresa Feltwell, Grant Hall, Anna Yakovleva, Fahad A Khokhar, Charlotte J. Houldcroft, Laura G Caller, Aminu S. Jahun, Sarah L. Caddy, Ian Goodfellow, Alex Alderton, Roberto Amato, Sonia Goncalves, Ewan Harrison, David K. Jackson, Ian Johnston, Dominic Kwiatkowski, Cordelia Langford, John Sillitoe on behalf of the Wellcome Sanger Institute COVID-19 Surveillance Team (http://www.sanger.ac.uk/covid-team) |
| EPI_ISL_441460 | Queens Medical Centre, Clinical Microbiology Department / DeepSeq Nottingham | COVID-19 Genomics UK (COG-UK) Consortium | Gemma Clark, Wendy Smith, Manjinder Khakh, Hannah Howson-Wells, Jonathan Ball, Patrick McClure, Joseph Chappell, Theocharis Tsoleridis, Nadine Holmes, Matthew Carlisle, Christopher Moore, Fei Sang, Johnny Debebe, Victoria Wright, Matthew Loose |
| EPI_ISL_441547, EPI_ISL_441548, EPI_ISL_441549, EPI_ISL_441550, EPI_ISL_441551, EPI_ISL_441552, EPI_ISL_441553, EPI_ISL_441554, EPI_ISL_441555, EPI_ISL_441556, EPI_ISL_441557, EPI_ISL_441558, EPI_ISL_441559, EPI_ISL_441560, EPI_ISL_441561, EPI_ISL_441562, EPI_ISL_441563, EPI_ISL_441564, EPI_ISL_441565, EPI_ISL_441567, EPI_ISL_441568, EPI_ISL_441569, EPI_ISL_441570, EPI_ISL_441571, EPI_ISL_441572, EPI_ISL_441573, EPI_ISL_441574, EPI_ISL_441575, EPI_ISL_441577, EPI_ISL_441578, EPI_ISL_441579, EPI_ISL_441580, EPI_ISL_441581, EPI_ISL_441583, EPI_ISL_441584, EPI_ISL_441585, EPI_ISL_441586, EPI_ISL_441587, EPI_ISL_441588, EPI_ISL_441589, EPI_ISL_441590, EPI_ISL_441591, EPI_ISL_441593, EPI_ISL_441594, EPI_ISL_441595, EPI_ISL_441596, EPI_ISL_441597, EPI_ISL_441598, EPI_ISL_441599, EPI_ISL_441600, EPI_ISL_441601, EPI_ISL_441602, EPI_ISL_441603, EPI_ISL_441604, EPI_ISL_441605, EPI_ISL_441606, EPI_ISL_441607, EPI_ISL_441608, EPI_ISL_441609, EPI_ISL_441611, EPI_ISL_441612, EPI_ISL_441613, EPI_ISL_441614, EPI_ISL_441615, EPI_ISL_441616, EPI_ISL_441618, EPI_ISL_441619, EPI_ISL_441620, EPI_ISL_441621, EPI_ISL_441622, EPI_ISL_441623, EPI_ISL_441624, EPI_ISL_441625, EPI_ISL_441626, EPI_ISL_441627, EPI_ISL_441628, EPI_ISL_441629, EPI_ISL_441630, EPI_ISL_441631, EPI_ISL_441632, EPI_ISL_441634, EPI_ISL_441635, EPI_ISL_441636, EPI_ISL_441638, EPI_ISL_441639, EPI_ISL_441640, EPI_ISL_441641, EPI_ISL_441642, EPI_ISL_441643, EPI_ISL_441644, EPI_ISL_441645, EPI_ISL_441646, EPI_ISL_441647, EPI_ISL_441648, EPI_ISL_441649, EPI_ISL_441650, EPI_ISL_441651, EPI_ISL_441652, EPI_ISL_441653, EPI_ISL_441654, EPI_ISL_441655, EPI_ISL_441656, EPI_ISL_441657, EPI_ISL_441658, EPI_ISL_441661, EPI_ISL_441665, EPI_ISL_441666, EPI_ISL_441667, EPI_ISL_441669, EPI_ISL_441671, EPI_ISL_441672, EPI_ISL_441673, EPI_ISL_441674, EPI_ISL_441677, EPI_ISL_441678, EPI_ISL_441679, EPI_ISL_441680, EPI_ISL_441681, EPI_ISL_441682, EPI_ISL_441683, EPI_ISL_441684, EPI_ISL_441686, EPI_ISL_441688, EPI_ISL_441689 | | | |
| see above | Department of Pathology, University of Cambridge | Wellcome Sanger Institute for the COVID-19 Genomics UK (COG-UK) consortium | Luke W Meredith, M. Estée Török , Myra Hosmillo, William L. Hamilton, Martin D. Curran, Theresa Feltwell, Grant Hall, Anna Yakovleva, Fahad A Khokhar, Charlotte J. Houldcroft, Laura G Caller, Aminu S. Jahun, Sarah L. Caddy, Ian Goodfellow, Alex Alderton, Roberto Amato, Sonia Goncalves, Ewan Harrison, David K. Jackson, Ian Johnston, Dominic Kwiatkowski, Cordelia Langford, John Sillitoe on behalf of the Wellcome Sanger Institute COVID-19 Surveillance Team (http://www.sanger.ac.uk/covid-team) |
| EPI_ISL_441693, EPI_ISL_441694 | Regional Virus Laboratory, Belfast Health and Social Care Trust | Wellcome Sanger Institute for the COVID-19 Genomics UK (COG-UK) consortium | Conall McCaughey, James McKenna, Tanya Curran, Susan Feeney, Alison Watt, Ciara Cox, Mairead Connor, Zoltan Molnar, David Simpson, Derek Fairley, Alex Alderton, Roberto Amato, Sonia Goncalves, Ewan Harrison, David K. Jackson, Ian Johnston, Dominic Kwiatkowski, Cordelia Langford, John Sillitoe on behalf of the Wellcome Sanger Institute COVID-19 Surveillance Team (http://www.sanger.ac.uk/covid-team) |
| EPI_ISL_441695 | Department of Pathology, University of Cambridge | Wellcome Sanger Institute for the COVID-19 Genomics UK (COG-UK) consortium | Luke W Meredith, M. Estée Török , Myra Hosmillo, William L. Hamilton, Martin D. Curran, Theresa Feltwell, Grant Hall, Anna Yakovleva, Fahad A Khokhar, Charlotte J. Houldcroft, Laura G Caller, Aminu S. Jahun, Sarah L. Caddy, Ian Goodfellow, Alex Alderton, Roberto Amato, Sonia Goncalves, Ewan Harrison, David K. Jackson, Ian Johnston, Dominic Kwiatkowski, Cordelia Langford, John Sillitoe on behalf of the Wellcome Sanger Institute COVID-19 Surveillance |

| | | | |
|---|---|---|---|
| | | | Team (http://www.sanger.ac.uk/covid-team) |
| EPI_ISL_441698, EPI_ISL_441699 | Regional Virus Laboratory, Belfast Health and Social Care Trust | Wellcome Sanger Institute for the COVID-19 Genomics UK (COG-UK) consortium | Conall McCaughey, James McKenna, Tanya Curran, Susan Feeney, Alison Watt, Ciara Cox, Mairead Connor, Zoltan Molnar, David Simpson, Derek Fairley, Alex Alderton, Roberto Amato, Sonia Goncalves, Ewan Harrison, David K. Jackson, Ian Johnston, Dominic Kwiatkowski, Cordelia Langford, John Sillitoe on behalf of the Wellcome Sanger Institute COVID-19 Surveillance Team (http://www.sanger.ac.uk/covid-team) |
| EPI_ISL_441700, EPI_ISL_441701, EPI_ISL_441702, EPI_ISL_441703, EPI_ISL_441706, EPI_ISL_441707 | Department of Pathology, University of Cambridge | Wellcome Sanger Institute for the COVID-19 Genomics UK (COG-UK) consortium | Luke W Meredith, M. Estée Török , Myra Hosmillo, William L. Hamilton, Martin D. Curran, Theresa Feltwell, Grant Hall, Anna Yakovleva, Fahad A Khokhar, Charlotte J. Houldcroft, Laura G Caller, Aminu S. Jahun, Sarah L. Caddy, Ian Goodfellow, Alex Alderton, Roberto Amato, Sonia Goncalves, Ewan Harrison, David K. Jackson, Ian Johnston, Dominic Kwiatkowski, Cordelia Langford, John Sillitoe on behalf of the Wellcome Sanger Institute COVID-19 Surveillance Team (http://www.sanger.ac.uk/covid-team) |
| EPI_ISL_441708 | Regional Virus Laboratory, Belfast Health and Social Care Trust | Wellcome Sanger Institute for the COVID-19 Genomics UK (COG-UK) consortium | Conall McCaughey, James McKenna, Tanya Curran, Susan Feeney, Alison Watt, Ciara Cox, Mairead Connor, Zoltan Molnar, David Simpson, Derek Fairley, Alex Alderton, Roberto Amato, Sonia Goncalves, Ewan Harrison, David K. Jackson, Ian Johnston, Dominic Kwiatkowski, Cordelia Langford, John Sillitoe on behalf of the Wellcome Sanger Institute COVID-19 Surveillance Team (http://www.sanger.ac.uk/covid-team) |
| EPI_ISL_441709, EPI_ISL_441710, EPI_ISL_441711, EPI_ISL_441713, EPI_ISL_441715, EPI_ISL_441718, EPI_ISL_441720, EPI_ISL_441721, EPI_ISL_441722, EPI_ISL_441723, EPI_ISL_441724, EPI_ISL_441725, EPI_ISL_441726, EPI_ISL_441727, EPI_ISL_441729, EPI_ISL_441731, EPI_ISL_441732, EPI_ISL_441735, EPI_ISL_441736, EPI_ISL_441738, EPI_ISL_441739, EPI_ISL_441740, EPI_ISL_441743, EPI_ISL_441744, EPI_ISL_441745, EPI_ISL_441746 | | | |
| see above | Department of Pathology, University of Cambridge | Wellcome Sanger Institute for the COVID-19 Genomics UK (COG-UK) consortium | Luke W Meredith, M. Estée Török , Myra Hosmillo, William L. Hamilton, Martin D. Curran, Theresa Feltwell, Grant Hall, Anna Yakovleva, Fahad A Khokhar, Charlotte J. Houldcroft, Laura G Caller, Aminu S. Jahun, Sarah L. Caddy, Ian Goodfellow, Alex Alderton, Roberto Amato, Sonia Goncalves, Ewan Harrison, David K. Jackson, Ian Johnston, Dominic Kwiatkowski, Cordelia Langford, John Sillitoe on behalf of the Wellcome Sanger Institute COVID-19 Surveillance Team (http://www.sanger.ac.uk/covid-team) |
| EPI_ISL_441747 | Regional Virus Laboratory, Belfast Health and Social Care Trust | Wellcome Sanger Institute for the COVID-19 Genomics UK (COG-UK) consortium | Conall McCaughey, James McKenna, Tanya Curran, Susan Feeney, Alison Watt, Ciara Cox, Mairead Connor, Zoltan Molnar, David Simpson, Derek Fairley, Alex Alderton, Roberto Amato, Sonia Goncalves, Ewan Harrison, David K. Jackson, Ian Johnston, Dominic Kwiatkowski, Cordelia Langford, John Sillitoe on behalf of the Wellcome Sanger Institute COVID-19 Surveillance Team (http://www.sanger.ac.uk/covid-team) |
| EPI_ISL_441748, EPI_ISL_441749 | Department of Pathology, University of Cambridge | Wellcome Sanger Institute for the COVID-19 Genomics UK (COG-UK) consortium | Luke W Meredith, M. Estée Török , Myra Hosmillo, William L. Hamilton, Martin D. Curran, Theresa Feltwell, Grant Hall, Anna Yakovleva, Fahad A Khokhar, Charlotte J. Houldcroft, Laura G Caller, Aminu S. Jahun, Sarah L. Caddy, Ian Goodfellow, Alex Alderton, Roberto Amato, Sonia Goncalves, Ewan Harrison, David K. Jackson, Ian Johnston, Dominic Kwiatkowski, Cordelia Langford, John Sillitoe on behalf of the Wellcome Sanger Institute COVID-19 Surveillance Team (http://www.sanger.ac.uk/covid-team) |
| EPI_ISL_441751 | Regional Virus Laboratory, Belfast Health and Social Care Trust | Wellcome Sanger Institute for the COVID-19 Genomics UK (COG-UK) consortium | Conall McCaughey, James McKenna, Tanya Curran, Susan Feeney, Alison Watt, Ciara Cox, Mairead Connor, Zoltan Molnar, David Simpson, Derek Fairley, Alex Alderton, Roberto Amato, Sonia Goncalves, Ewan Harrison, David K. Jackson, Ian Johnston, Dominic Kwiatkowski, Cordelia Langford, John Sillitoe on behalf of the Wellcome Sanger Institute COVID-19 Surveillance Team (http://www.sanger.ac.uk/covid-team) |
| EPI_ISL_441752, EPI_ISL_441754, EPI_ISL_441755, EPI_ISL_441756, EPI_ISL_441757, EPI_ISL_441758, EPI_ISL_441759, EPI_ISL_441760 | Department of Pathology, University of Cambridge | Wellcome Sanger Institute for the COVID-19 Genomics UK (COG-UK) consortium | Luke W Meredith, M. Estée Török , Myra Hosmillo, William L. Hamilton, Martin D. Curran, Theresa Feltwell, Grant Hall, Anna Yakovleva, Fahad A Khokhar, Charlotte J. Houldcroft, Laura G Caller, Aminu S. Jahun, Sarah L. Caddy, Ian Goodfellow, Alex Alderton, Roberto Amato, Sonia Goncalves, Ewan Harrison, David K. Jackson, Ian Johnston, Dominic Kwiatkowski, Cordelia Langford, John Sillitoe on behalf of the Wellcome Sanger Institute COVID-19 Surveillance Team (http://www.sanger.ac.uk/covid-team) |
| EPI_ISL_441761 | Regional Virus Laboratory, Belfast Health and Social Care Trust | Wellcome Sanger Institute for the COVID-19 Genomics UK (COG-UK) consortium | Conall McCaughey, James McKenna, Tanya Curran, Susan Feeney, Alison Watt, Ciara Cox, Mairead Connor, Zoltan Molnar, David Simpson, Derek Fairley, Alex Alderton, Roberto Amato, Sonia Goncalves, Ewan Harrison, David K. Jackson, Ian Johnston, Dominic Kwiatkowski, Cordelia Langford, John Sillitoe on behalf of the Wellcome Sanger Institute COVID-19 Surveillance Team (http://www.sanger.ac.uk/covid-team) |
| EPI_ISL_441763, EPI_ISL_441764, EPI_ISL_441768, EPI_ISL_441772, EPI_ISL_441773, EPI_ISL_441774, EPI_ISL_441777, EPI_ISL_441778, EPI_ISL_441779, EPI_ISL_441780, EPI_ISL_441784, EPI_ISL_441785, EPI_ISL_441786, EPI_ISL_441787, EPI_ISL_441791, EPI_ISL_441792, EPI_ISL_441793, EPI_ISL_441794, EPI_ISL_441798, EPI_ISL_441799, EPI_ISL_441800, EPI_ISL_441801, EPI_ISL_441802, EPI_ISL_441804, EPI_ISL_441806, EPI_ISL_441807, EPI_ISL_441808, EPI_ISL_441810, EPI_ISL_441812, EPI_ISL_441813, EPI_ISL_441814, EPI_ISL_441815, EPI_ISL_441817, EPI_ISL_441820, EPI_ISL_441821, EPI_ISL_441822, EPI_ISL_441823, EPI_ISL_441824, EPI_ISL_441825, EPI_ISL_441826, EPI_ISL_441827, EPI_ISL_441828, EPI_ISL_441829, EPI_ISL_441830, EPI_ISL_441832, EPI_ISL_441833, EPI_ISL_441834, EPI_ISL_441838, EPI_ISL_441839, EPI_ISL_441843, EPI_ISL_441845, EPI_ISL_441846 | | | |
| see above | Department of Pathology, University of Cambridge | Wellcome Sanger Institute for the COVID-19 Genomics UK (COG-UK) consortium | Luke W Meredith, M. Estée Török , Myra Hosmillo, William L. Hamilton, Martin D. Curran, Theresa Feltwell, Grant Hall, Anna Yakovleva, Fahad A Khokhar, Charlotte J. Houldcroft, Laura G Caller, Aminu S. Jahun, Sarah L. Caddy, Ian Goodfellow, Alex Alderton, Roberto Amato, Sonia Goncalves, Ewan Harrison, David K. Jackson, Ian Johnston, Dominic Kwiatkowski, Cordelia Langford, John Sillitoe on behalf of the Wellcome Sanger Institute COVID-19 Surveillance Team (http://www.sanger.ac.uk/covid-team) |
| EPI_ISL_441906, EPI_ISL_441919, EPI_ISL_441961, EPI_ISL_441965, EPI_ISL_441978, EPI_ISL_441995, EPI_ISL_441997, EPI_ISL_442015, EPI_ISL_442022, EPI_ISL_442037 | Virology Department, Sheffield Teaching Hospitals NHS Foundation Trust/Department of Infection, Immunity and Cardiovascular Disease, The Medical School, University of Sheffield | COVID-19 Genomics UK (COG-UK) Consortium | Thushan de Silva, Matthew Parker, Nikki Smith, Adri Angyal, Rebecca Brown, Luke Green, Rachel Tucker, Paul Parsons, Danielle Groves, Katie Johnson, Laura Carrilero, Alex Keeley, Dave Partridge, Matthew Wyles, Benjamin Lindsey, Mehmet Yavuz, Mohammad Raza, Cariad Evans |
| EPI_ISL_442045, EPI_ISL_442046, EPI_ISL_442047, EPI_ISL_442048, EPI_ISL_442049, EPI_ISL_442050, EPI_ISL_442051, EPI_ISL_442052, EPI_ISL_442053, EPI_ISL_442054, EPI_ISL_442055, EPI_ISL_442057, EPI_ISL_442058, EPI_ISL_442059, EPI_ISL_442060, EPI_ISL_442061, EPI_ISL_442062, EPI_ISL_442063, EPI_ISL_442064, EPI_ISL_442065, EPI_ISL_442066, EPI_ISL_442067, EPI_ISL_442068, EPI_ISL_442069, EPI_ISL_442070, EPI_ISL_442071, EPI_ISL_442072, EPI_ISL_442073, EPI_ISL_442074, EPI_ISL_442075, EPI_ISL_442076, EPI_ISL_442077, EPI_ISL_442078, EPI_ISL_442079, EPI_ISL_442080, EPI_ISL_442082, EPI_ISL_442083, EPI_ISL_442085, EPI_ISL_442086, EPI_ISL_442087, EPI_ISL_442088, EPI_ISL_442089, EPI_ISL_442090, EPI_ISL_442091, EPI_ISL_442092, EPI_ISL_442093, EPI_ISL_442094, EPI_ISL_442095, EPI_ISL_442096, EPI_ISL_442097, EPI_ISL_442098, EPI_ISL_442099, EPI_ISL_442100, EPI_ISL_442101, EPI_ISL_442102, EPI_ISL_442103, EPI_ISL_442104, EPI_ISL_442105, EPI_ISL_442107, EPI_ISL_442108, EPI_ISL_442109, EPI_ISL_442110, EPI_ISL_442111, EPI_ISL_442112, EPI_ISL_442113, EPI_ISL_442114, EPI_ISL_442115, EPI_ISL_442116, EPI_ISL_442117, EPI_ISL_442118, EPI_ISL_442119, EPI_ISL_442120, EPI_ISL_442121, EPI_ISL_442122, EPI_ISL_442123, EPI_ISL_442124, EPI_ISL_442125, EPI_ISL_442126, EPI_ISL_442127, EPI_ISL_442128, EPI_ISL_442130, EPI_ISL_442131, EPI_ISL_442132, EPI_ISL_442133, EPI_ISL_442134, EPI_ISL_442135, EPI_ISL_442136, EPI_ISL_442137, EPI_ISL_442138, EPI_ISL_442139, EPI_ISL_442140, EPI_ISL_442141, EPI_ISL_442142, EPI_ISL_442143, EPI_ISL_442144, EPI_ISL_442145, EPI_ISL_442146, EPI_ISL_442147, EPI_ISL_442150, EPI_ISL_442151, EPI_ISL_442153, EPI_ISL_442154, EPI_ISL_442155, EPI_ISL_442156, EPI_ISL_442157, EPI_ISL_442158, EPI_ISL_442159, EPI_ISL_442160, EPI_ISL_442162, EPI_ISL_442163, EPI_ISL_442164, EPI_ISL_442165, EPI_ISL_442166, EPI_ISL_442167, EPI_ISL_442168, EPI_ISL_442170, EPI_ISL_442171, EPI_ISL_442173, EPI_ISL_442174, EPI_ISL_442176, EPI_ISL_442177, EPI_ISL_442178, EPI_ISL_442180, EPI_ISL_442181, EPI_ISL_442182, EPI_ISL_442183, EPI_ISL_442184, EPI_ISL_442185, EPI_ISL_442186, EPI_ISL_442187, EPI_ISL_442188, EPI_ISL_442189, EPI_ISL_442190, EPI_ISL_442191, EPI_ISL_442192, EPI_ISL_442193, EPI_ISL_442194, EPI_ISL_442195, EPI_ISL_442196, EPI_ISL_442197, EPI_ISL_442198, EPI_ISL_442199, EPI_ISL_442202, EPI_ISL_442206, EPI_ISL_442207, EPI_ISL_442208, EPI_ISL_442209, EPI_ISL_442210, EPI_ISL_442211, EPI_ISL_442212, EPI_ISL_442213, EPI_ISL_442214, EPI_ISL_442215, EPI_ISL_442217, EPI_ISL_442218, EPI_ISL_442220, EPI_ISL_442222, EPI_ISL_442223, EPI_ISL_442224, EPI_ISL_442226, EPI_ISL_442227, EPI_ISL_442229, EPI_ISL_442230, EPI_ISL_442231, EPI_ISL_442232, EPI_ISL_442233, EPI_ISL_442234, EPI_ISL_442235, EPI_ISL_442237, EPI_ISL_442238, EPI_ISL_442239, EPI_ISL_442240, EPI_ISL_442242, EPI_ISL_442243, EPI_ISL_442244, EPI_ISL_442245, EPI_ISL_442246, EPI_ISL_442247, EPI_ISL_442248, EPI_ISL_442249, EPI_ISL_442250, EPI_ISL_442251, EPI_ISL_442252, EPI_ISL_442253, EPI_ISL_442254, EPI_ISL_442255, EPI_ISL_442256, EPI_ISL_442257, EPI_ISL_442258, EPI_ISL_442259, EPI_ISL_442260, EPI_ISL_442262, EPI_ISL_442263, EPI_ISL_442264, EPI_ISL_442265, EPI_ISL_442266, EPI_ISL_442267, EPI_ISL_442268, EPI_ISL_442269, EPI_ISL_442270, EPI_ISL_442271, EPI_ISL_442272, EPI_ISL_442274, EPI_ISL_442275, EPI_ISL_442276, EPI_ISL_442277, EPI_ISL_442278, EPI_ISL_442279, EPI_ISL_442280, EPI_ISL_442281, EPI_ISL_442282, EPI_ISL_442283, EPI_ISL_442284, EPI_ISL_442285, EPI_ISL_442286, EPI_ISL_442287, EPI_ISL_442288, EPI_ISL_442289, EPI_ISL_442290, EPI_ISL_442291, EPI_ISL_442292, EPI_ISL_442293, EPI_ISL_442295, EPI_ISL_442296, EPI_ISL_442297, EPI_ISL_442298, EPI_ISL_442299, EPI_ISL_442300, EPI_ISL_442301, EPI_ISL_442302, EPI_ISL_442303, EPI_ISL_442304, EPI_ISL_442305, EPI_ISL_442306, EPI_ISL_442307, EPI_ISL_442308, EPI_ISL_442309, EPI_ISL_442310, EPI_ISL_442312, EPI_ISL_442313, EPI_ISL_442314, EPI_ISL_442315, EPI_ISL_442316, EPI_ISL_442317, EPI_ISL_442318, EPI_ISL_442319, EPI_ISL_442320, EPI_ISL_442321, EPI_ISL_442323, EPI_ISL_442324, EPI_ISL_442325, EPI_ISL_442326, EPI_ISL_442328, EPI_ISL_442329, EPI_ISL_442330, EPI_ISL_442331, EPI_ISL_442333, EPI_ISL_442334, EPI_ISL_442335, EPI_ISL_442336, EPI_ISL_442337, EPI_ISL_442338, EPI_ISL_442340, EPI_ISL_442341, EPI_ISL_442342, EPI_ISL_442343 | | | |
| see above | Department of Pathology, University of Cambridge | Wellcome Sanger Institute for the COVID-19 Genomics UK (COG-UK) consortium | Luke W Meredith, M. Estée Török , Myra Hosmillo, William L. Hamilton, Martin D. Curran, Theresa Feltwell, Grant Hall, Anna Yakovleva, Fahad A Khokhar, Charlotte J. Houldcroft, Laura G Caller, Aminu S. Jahun, Sarah L. Caddy, Ian Goodfellow, Alex Alderton, Roberto Amato, Sonia Goncalves, Ewan Harrison, David K. Jackson, Ian Johnston, Dominic Kwiatkowski, Cordelia Langford, John Sillitoe on behalf of the Wellcome Sanger Institute COVID-19 Surveillance Team (http://www.sanger.ac.uk/covid-team) |
| EPI_ISL_442356, EPI_ISL_442359, EPI_ISL_442371, EPI_ISL_442388, EPI_ISL_442410, EPI_ISL_442412, EPI_ISL_442415, EPI_ISL_442433, EPI_ISL_442434, EPI_ISL_442442, EPI_ISL_442443, EPI_ISL_442452, EPI_ISL_442463, EPI_ISL_442478, EPI_ISL_442483, EPI_ISL_442490, EPI_ISL_442491, EPI_ISL_442507, EPI_ISL_442512, EPI_ISL_442515 | | | |
| see above | Virology Department, Sheffield Teaching Hospitals NHS Foundation Trust/Department of Infection, Immunity and Cardiovascular Disease, The Medical School, University of Sheffield | COVID-19 Genomics UK (COG-UK) Consortium | Thushan de Silva, Matthew Parker, Nikki Smith, Adri Angyal, Rebecca Brown, Luke Green, Rachel Tucker, Paul Parsons, Danielle Groves, Katie Johnson, Laura Carrilero, Alex Keeley, Dave Partridge, Matthew Wyles, Benjamin Lindsey, Mehmet Yavuz, Mohammad Raza, Cariad Evans |
| EPI_ISL_442530, EPI_ISL_442600, EPI_ISL_442607 | Department of Pathology, University of Cambridge | Wellcome Sanger Institute for the COVID-19 Genomics UK (COG-UK) consortium | Luke W Meredith, M. Estée Török , Myra Hosmillo, William L. Hamilton, Martin D. Curran, Theresa Feltwell, Grant Hall, Anna Yakovleva, Fahad A Khokhar, Charlotte J. Houldcroft, Laura G Caller, Aminu S. Jahun, Sarah L. Caddy, Ian Goodfellow, Alex Alderton, Roberto Amato, Sonia Goncalves, Ewan Harrison, David K. Jackson, Ian Johnston, Dominic Kwiatkowski, Cordelia Langford, John Sillitoe on behalf of the Wellcome Sanger Institute COVID-19 Surveillance Team (http://www.sanger.ac.uk/covid-team) |
| EPI_ISL_442624, EPI_ISL_442625, EPI_ISL_442626, EPI_ISL_442627, EPI_ISL_442628, EPI_ISL_442629, EPI_ISL_442630, EPI_ISL_442631, EPI_ISL_442632, EPI_ISL_442634, EPI_ISL_442635, EPI_ISL_442636, EPI_ISL_442638, EPI_ISL_442641, EPI_ISL_442642, EPI_ISL_442643, EPI_ISL_442645, EPI_ISL_442646, EPI_ISL_442647, EPI_ISL_442650, EPI_ISL_442651, EPI_ISL_442653, EPI_ISL_442655, EPI_ISL_442657, EPI_ISL_442658, EPI_ISL_442659, EPI_ISL_442660, EPI_ISL_442662, EPI_ISL_442663, EPI_ISL_442664, EPI_ISL_442665, EPI_ISL_442666, EPI_ISL_442668, EPI_ISL_442671, EPI_ISL_442674, EPI_ISL_442675, EPI_ISL_442676, EPI_ISL_442677, EPI_ISL_442678, EPI_ISL_442679, EPI_ISL_442680, EPI_ISL_442681, EPI_ISL_442682, EPI_ISL_442683, EPI_ISL_442686, EPI_ISL_442687, EPI_ISL_442688, EPI_ISL_442689, EPI_ISL_442690, EPI_ISL_442691, EPI_ISL_442692, EPI_ISL_442694, EPI_ISL_442696, EPI_ISL_442698, | | | |

| | | | |
|---|---|---|---|
| EPI_ISL_442700, EPI_ISL_442701, EPI_ISL_442703, EPI_ISL_442704, EPI_ISL_442705, EPI_ISL_442707, EPI_ISL_442708, EPI_ISL_442709, EPI_ISL_442710, EPI_ISL_442711, EPI_ISL_442713, EPI_ISL_442715, EPI_ISL_442717, EPI_ISL_442718, EPI_ISL_442719, EPI_ISL_442720, EPI_ISL_442721, EPI_ISL_442723, EPI_ISL_442724, EPI_ISL_442725, EPI_ISL_442726, EPI_ISL_442727, EPI_ISL_442728, EPI_ISL_442730, EPI_ISL_442731, EPI_ISL_442732, EPI_ISL_442733, EPI_ISL_442734, EPI_ISL_442735, EPI_ISL_442736, EPI_ISL_442737, EPI_ISL_442738, EPI_ISL_442739, EPI_ISL_442740, EPI_ISL_442741, EPI_ISL_442742, EPI_ISL_442743, EPI_ISL_442744, EPI_ISL_442746, EPI_ISL_442748, EPI_ISL_442749, EPI_ISL_442750, EPI_ISL_442751, EPI_ISL_442753, EPI_ISL_442754, EPI_ISL_442755, EPI_ISL_442756, EPI_ISL_442757, EPI_ISL_442758, EPI_ISL_442759, EPI_ISL_442760, EPI_ISL_442761, EPI_ISL_442762, EPI_ISL_442763, EPI_ISL_442765, EPI_ISL_442766, EPI_ISL_442767, EPI_ISL_442770, EPI_ISL_442773, EPI_ISL_442775, EPI_ISL_442776, EPI_ISL_442778, EPI_ISL_442781, EPI_ISL_442783, EPI_ISL_442784, EPI_ISL_442785, EPI_ISL_442786, EPI_ISL_442787, EPI_ISL_442788, EPI_ISL_442791, EPI_ISL_442792, EPI_ISL_442794, EPI_ISL_442795, EPI_ISL_442798, EPI_ISL_442799, EPI_ISL_442801, EPI_ISL_442803, EPI_ISL_442804, EPI_ISL_442805, EPI_ISL_442806, EPI_ISL_442807, EPI_ISL_442808, EPI_ISL_442809, EPI_ISL_442810 | | | |
| see above | PHE South West Regional Laboratory, National Infection Service | Wellcome Sanger Institute for the COVID-19 Genomics UK (COG-UK) consortium | Stephanie Hutchings, Hannah Pymont, Dr Peter Muir, Barry Vipond, Rich Hopes, Alex Alderton, Roberto Amato, Sonia Goncalves, Ewan Harrison, David K. Jackson, Ian Johnston, Dominic Kwiatkowski, Cordelia Langford, John Sillitoe on behalf of the Wellcome Sanger Institute COVID-19 Surveillance Team (http://www.sanger.ac.uk/covid-team) |
| EPI_ISL_442811, EPI_ISL_442813, EPI_ISL_442814, EPI_ISL_442815, EPI_ISL_442816, EPI_ISL_442817, EPI_ISL_442818, EPI_ISL_442819, EPI_ISL_442820, EPI_ISL_442824, EPI_ISL_442826, EPI_ISL_442834, EPI_ISL_442860, EPI_ISL_442881, EPI_ISL_442893, EPI_ISL_442920, EPI_ISL_442954, EPI_ISL_442972, EPI_ISL_442994, EPI_ISL_443025, EPI_ISL_443032, EPI_ISL_443061, EPI_ISL_443063, EPI_ISL_443067 | | | |
| see above | Department of Pathology, University of Cambridge | Wellcome Sanger Institute for the COVID-19 Genomics UK (COG-UK) consortium | Luke W Meredith, M. Estée Török , Myra Hosmillo, William L. Hamilton, Martin D. Curran, Theresa Feltwell, Grant Hall, Anna Yakovleva, Fahad A Khokhar, Charlotte J. Houldcroft, Laura G Caller, Aminu S. Jahun, Sarah L. Caddy, Ian Goodfellow, Alex Alderton, Roberto Amato, Sonia Goncalves, Ewan Harrison, David K. Jackson, Ian Johnston, Dominic Kwiatkowski, Cordelia Langford, John Sillitoe on behalf of the Wellcome Sanger Institute COVID-19 Surveillance Team (http://www.sanger.ac.uk/covid-team) |
| EPI_ISL_443183, EPI_ISL_443253, EPI_ISL_443254, EPI_ISL_443256 | M Health Fairview | University of Minnesota Genomics Center | Daryl M. Gohl, John Garbe, Patrick Grady, Jerry Daniel, Ray Watson, Benjamin Auch, Andrew Nelson, Sophia Yohe, and Kenneth B. Beckman |
| EPI_ISL_443258, EPI_ISL_443259 | Résidence Ornano | National Reference Center for Viruses of Respiratory Infections, Institut Pasteur, Paris | Mélanie Albert, Marion Barbet, Sylvie Behillil, Méline Bizard, Angela Brisebarre, Flora Donati, Etienne Simon-Lorière, Vincent Enouf, Maud Vanpeene, Sylvie van der Werf |
| EPI_ISL_443260 | LABM GH nord Essonne de Longjumeau - BP 125 | National Reference Center for Viruses of Respiratory Infections, Institut Pasteur, Paris | Mélanie Albert, Marion Barbet, Sylvie Behillil, Méline Bizard, Angela Brisebarre, Flora Donati, Etienne Simon-Lorière, Vincent Enouf, Maud Vanpeene, Sylvie van der Werf |
| EPI_ISL_443261, EPI_ISL_443262, EPI_ISL_443263, EPI_ISL_443264 | CHU de Dijon - Laboratoire de Virologie | National Reference Center for Viruses of Respiratory Infections, Institut Pasteur, Paris | Mélanie Albert, Marion Barbet, Sylvie Behillil, Méline Bizard, Angela Brisebarre, Flora Donati, Etienne Simon-Lorière, Vincent Enouf, Maud Vanpeene, Sylvie van der Werf, Jean-Baptiste Bour |
| EPI_ISL_443265, EPI_ISL_443281 | CHU - Hôpital Cavale Blanche - Labo. de Virologie | National Reference Center for Viruses of Respiratory Infections, Institut Pasteur, Paris | Mélanie Albert, Marion Barbet, Sylvie Behillil, Méline Bizard, Angela Brisebarre, Flora Donati, Etienne Simon-Lorière, Vincent Enouf, Maud Vanpeene, Sylvie van der Werf, Léa Pilorge |
| EPI_ISL_443284, EPI_ISL_443285, EPI_ISL_443286, EPI_ISL_443287, EPI_ISL_443288 | Laboratoire de Microbiologie - Bât A - CH René Dubois | National Reference Center for Viruses of Respiratory Infections, Institut Pasteur, Paris | Mélanie Albert, Marion Barbet, Sylvie Behillil, Méline Bizard, Angela Brisebarre, Flora Donati, Etienne Simon-Lorière, Vincent Enouf, Maud Vanpeene, Sylvie van der Werf, Pascale Martres |
| EPI_ISL_443291, EPI_ISL_443292, EPI_ISL_443293, EPI_ISL_443294 | CHRU Pontchaillou - Laboratoire de Virologie | National Reference Center for Viruses of Respiratory Infections, Institut Pasteur, Paris | Mélanie Albert, Marion Barbet, Sylvie Behillil, Méline Bizard, Angela Brisebarre, Flora Donati, Etienne Simon-Lorière, Vincent Enouf, Maud Vanpeene, Sylvie van der Werf, Gisèle Lagathu |
| EPI_ISL_443306 | Cabinet Médical | National Reference Center for Viruses of Respiratory Infections, Institut Pasteur, Paris | Mélanie Albert, Marion Barbet, Sylvie Behillil, Méline Bizard, Angela Brisebarre, Flora Donati, Etienne Simon-Lorière, Vincent Enouf, Maud Vanpeene, Sylvie van der Werf |
| EPI_ISL_443315 | Château de la Source | National Reference Center for Viruses of Respiratory Infections, Institut Pasteur, Paris | Mélanie Albert, Marion Barbet, Sylvie Behillil, Méline Bizard, Angela Brisebarre, Flora Donati, Etienne Simon-Lorière, Vincent Enouf, Maud Vanpeene, Sylvie van der Werf |
| EPI_ISL_443627 | Department of Pathology, University of Cambridge | Wellcome Sanger Institute for the COVID-19 Genomics UK (COG-UK) consortium | Luke W Meredith, M. Estée Török , Myra Hosmillo, William L. Hamilton, Martin D. Curran, Theresa Feltwell, Grant Hall, Anna Yakovleva, Fahad A Khokhar, Charlotte J. Houldcroft, Laura G Caller, Aminu S. Jahun, Sarah L. Caddy, Ian Goodfellow; and Alex Alderton, Roberto Amato, Sonia Goncalves, Ewan Harrison, David K. Jackson, Ian Johnston, Dominic Kwiatkowski, Cordelia Langford, John Sillitoe on behalf of the Wellcome Sanger Institute COVID-19 Surveillance Team (http://www.sanger.ac.uk/covid-team) |
| EPI_ISL_444027 | Pamela Youde Nethersole Eastern Hospital | Hong Kong Department of Health | Mak Gannon C.K., Cheng Peter K.C., Lam Edman T.K., Chan Rickjason C.W., Tsang Dominic N.C. |
| EPI_ISL_444028 | Queen Elizabeth Hospital | Hong Kong Department of Health | Mak Gannon C.K., Cheng Peter K.C., Lam Edman T.K., Chan Rickjason C.W., Tsang Dominic N.C. |
| EPI_ISL_444029 | Prince of Wales Hospital | Hong Kong Department of Health | Mak Gannon C.K., Cheng Peter K.C., Lam Edman T.K., Chan Rickjason C.W., Tsang Dominic N.C. |
| EPI_ISL_444030 | United Christian Hospital | Hong Kong Department of Health | Mak Gannon C.K., Cheng Peter K.C., Lam Edman T.K., Chan Rickjason C.W., Tsang Dominic N.C. |
| EPI_ISL_444031 | Queen Mary Hospital | Hong Kong Department of Health | Mak Gannon C.K., Cheng Peter K.C., Lam Edman T.K., Chan Rickjason C.W., Tsang Dominic N.C. |
| EPI_ISL_444032 | North Lantau Hospital | Hong Kong Department of Health | Mak Gannon C.K., Cheng Peter K.C., Lam Edman T.K., Chan Rickjason C.W., Tsang Dominic N.C. |
| EPI_ISL_444033 | Queen Mary Hospital | Hong Kong Department of Health | Mak Gannon C.K., Cheng Peter K.C., Lam Edman T.K., Chan Rickjason C.W., Tsang Dominic N.C. |
| EPI_ISL_444034 | Prince of Wales Hospital | Hong Kong Department of Health | Mak Gannon C.K., Cheng Peter K.C., Lam Edman T.K., Chan Rickjason C.W., Tsang Dominic N.C. |
| EPI_ISL_444035 | Princess Margaret Hospital | Hong Kong Department of Health | Mak Gannon C.K., Cheng Peter K.C., Lam Edman T.K., Chan Rickjason C.W., Tsang Dominic N.C. |
| EPI_ISL_444036 | North Lantau Hospital | Hong Kong Department of Health | Mak Gannon C.K., Cheng Peter K.C., Lam Edman T.K., Chan Rickjason C.W., Tsang Dominic N.C. |
| EPI_ISL_444037 | Hong Kong Adventist Hospital | Hong Kong Department of Health | Mak Gannon C.K., Cheng Peter K.C., Lam Edman T.K., Chan Rickjason C.W., Tsang Dominic N.C. |
| EPI_ISL_444038 | Princess Margaret Hospital | Hong Kong Department of Health | Mak Gannon C.K., Cheng Peter K.C., Lam Edman T.K., Chan Rickjason C.W., Tsang Dominic N.C. |
| EPI_ISL_444047 | United Christian Hospital | Hong Kong Department of Health | Mak Gannon C.K., Cheng Peter K.C., Lam Edman T.K., Chan Rickjason C.W., Tsang Dominic N.C. |
| EPI_ISL_444048 | Queen Elizabeth Hospital | Hong Kong Department of Health | Mak Gannon C.K., Cheng Peter K.C., Lam Edman T.K., Chan Rickjason C.W., Tsang Dominic N.C. |
| EPI_ISL_444049 | North Lantau Hospital | Hong Kong Department of Health | Mak Gannon C.K., Cheng Peter K.C., Lam Edman T.K., Chan Rickjason C.W., Tsang Dominic N.C. |
| EPI_ISL_444079, EPI_ISL_444080, EPI_ISL_444081, EPI_ISL_444082, EPI_ISL_444083, EPI_ISL_444084, EPI_ISL_444085, EPI_ISL_444086, EPI_ISL_444087, EPI_ISL_444088, EPI_ISL_444089, EPI_ISL_444090, EPI_ISL_444091, EPI_ISL_444092, EPI_ISL_444093, EPI_ISL_444094, EPI_ISL_444095, EPI_ISL_444096, EPI_ISL_444097, EPI_ISL_444098, EPI_ISL_444099, EPI_ISL_444100, EPI_ISL_444101, EPI_ISL_444102, EPI_ISL_444103, EPI_ISL_444104, EPI_ISL_444105, EPI_ISL_444106, EPI_ISL_444107, EPI_ISL_444108, EPI_ISL_444109, EPI_ISL_444110, EPI_ISL_444111, EPI_ISL_444112, EPI_ISL_444113, EPI_ISL_444114, EPI_ISL_444115, EPI_ISL_444116, EPI_ISL_444117, EPI_ISL_444118, EPI_ISL_444119, EPI_ISL_444120 | | | |
| see above | University College London, Great Ormond Street Hospital for Children NHS Foundation Trust, Imperial College Healthcare NHS Trust | COVID-19 Genomics UK (COG-UK) Consortium | Sergi Castellano, Rachel Williams, Mark Kristiansen, Paola Resende Silva, Sunando Roy, Tony Brooks, Helena Tutill, Paola Niola, Patricia Dyal, Charlotte Williams, Leysa Forrest, Yasmin Panchbhaya, Jacqueline Findlay, Sam Weeks, Julianne Brown, Kathryn Harris, Paul Randell, James Price, Alison Holmes, Judith Breuer |
| EPI_ISL_444495, EPI_ISL_444498, EPI_ISL_444499, EPI_ISL_444501, EPI_ISL_444502, EPI_ISL_444503, EPI_ISL_444504, EPI_ISL_444505, EPI_ISL_444506, EPI_ISL_444507, EPI_ISL_444508, EPI_ISL_444509, EPI_ISL_444510, EPI_ISL_444512, EPI_ISL_444513, EPI_ISL_444515, EPI_ISL_444516 | | | |
| see above | Laboratoire de microbiologie, Hopital de Verdun | Smith Laboratory, Centre de Recherche CHU Sainte-Justine | Martin Smith, Marieke Rozendaal, Ivan Pavlov |
| EPI_ISL_444613, EPI_ISL_444614, EPI_ISL_444615, EPI_ISL_444707, EPI_ISL_444711, EPI_ISL_444712, EPI_ISL_444713, EPI_ISL_444714, EPI_ISL_444715, EPI_ISL_444716, EPI_ISL_444717, EPI_ISL_444718, EPI_ISL_444719, EPI_ISL_444720, EPI_ISL_444721, EPI_ISL_444722, EPI_ISL_444723, EPI_ISL_444724, EPI_ISL_444725, EPI_ISL_444726, EPI_ISL_444727, EPI_ISL_444730, EPI_ISL_444732, EPI_ISL_444734, EPI_ISL_444735, EPI_ISL_444736, EPI_ISL_444737, EPI_ISL_444738, EPI_ISL_444739, EPI_ISL_444740, EPI_ISL_444741, EPI_ISL_444742, EPI_ISL_444743, EPI_ISL_444744, EPI_ISL_444745, EPI_ISL_444746, EPI_ISL_444747, EPI_ISL_444748, EPI_ISL_444749, EPI_ISL_444750, EPI_ISL_444751, EPI_ISL_444752, EPI_ISL_444753, EPI_ISL_444754, EPI_ISL_444755, EPI_ISL_444756, EPI_ISL_444757, EPI_ISL_444758, EPI_ISL_444759, EPI_ISL_444760, EPI_ISL_444761, EPI_ISL_444762, EPI_ISL_444763, EPI_ISL_444764, EPI_ISL_444766, EPI_ISL_444767, EPI_ISL_444768, EPI_ISL_444769 | | | |
| see above | NYU Langone Health | Departments of Pathology and Medicine, New York University School of Medicine | Maria Aguero-Rosenfeld, Brendan Belovarac, Margaret Black, Ludovic Boytard, John Cadley, Paolo Cotzia, John Chen, Dacia Dimartino, Xiaojun Feng, Tatyana Gindin, Emily Guzman, Adriana Heguy, Megan Hogan, Emily Huang, George Jour, Alireza Khodadadi-Jamayran, Lawrence H. Lin, Raven Luther, Andrew Lytle, Christian Marier, Matthew T. Maurano, Mark J. Mulligan, Peter Meyn, Raquel Ordonez Ciriza, Iman Osman, Jared Pinnell, Vanessa Raabe, Sitharam Ramaswami, Amy Rapkiewicz, Andre M. Ribeiro-dos-Santos, Marie Samanovic-Golden, Antonio Serrano, Guomiao Shen, Matija Snuderl, Theodore Vougiouklakis, Nick Vulpescu, Gael Westby, Paul Zappile, Yutong Zhang |
| EPI_ISL_444843, EPI_ISL_444844, EPI_ISL_444845, EPI_ISL_444846, EPI_ISL_444849, EPI_ISL_444850, EPI_ISL_444851, EPI_ISL_444852, EPI_ISL_444853, EPI_ISL_444854, EPI_ISL_444855, EPI_ISL_444856, EPI_ISL_444857, EPI_ISL_444858 | | | |

| | | | |
|---|---|---|---|
| see above | Department of Virus and Microbiological Special Diagnostics, Statens Serum Institut, Copenhagen, Denmark, Artillerivej 5, 2300 Copenahgen S | Albertsen lab, Department of Chemistry and Bioscience, Aalborg University, Denmark | Rasmus Kirkegaard |
| EPI_ISL_444971 | Hospital Universitari Vall d'Hebron - Vall d'Hebron Institut de Recerca | Hospital Universitari Vall d'Hebron | Cristina Andrés, Maria Piñana, Damir Garcia-Cehic, Mercedes Guerrero-Murillo, Ariadna Rando, Juliana Esperalba, Maria Gema Codina, Tomàs Pumarola, Josep Quer, Andrés Antón |
| EPI_ISL_444972 | Hospital Universitari Vall d'Hebron - Vall d'Hebron Institut de Recerca | Hospital Universitari Vall d'Hebron | Cristina Andrés, Maria Piñana, DAmir Garcia-Cehic, Mercedes Guerrero-Murillo, Ariadna Rando, Juliana Esperalba, Maria Gema Codina, Tomàs Pumarola, Josep Quer, Andrés Antón |
| EPI_ISL_444973 | Hospital Universitari Vall d`Hebron - Vall d'Hebron Institut de Recerca | Hospital Universitari Vall d'Hebron | Cristina Andrés, Maria Piñana, DAmir Garcia-Cehic, Mercedes Guerrero-Murillo, Ariadna Rando, Juliana Esperalba, Maria Gema Codina, Tomàs Pumarola, Josep Quer, Andrés Antón |
| EPI_ISL_444974, EPI_ISL_444976 | Hospital Universitari Vall d'Hebron - Vall d'Hebron Institut de Recerca | Hospital Universitari Vall d'Hebron | Cristina Andrés, Maria Piñana, Damir Garcia-Cehic, Mercedes Guerrero-Murillo, Ariadna Rando, Juliana Esperalba, Maria Gema Codina, Tomàs Pumarola, Josep Quer, Andrés Antón |
| EPI_ISL_445085 | Virology Unit, Agrobiodiversity and Biotechnology Project, CIAT - International Center for Tropical Agriculture | Virology Unit, Agrobiodiversity and Biotechnology Project, CIAT - International Center for Tropical Agriculture | Lopez,D., Parra,B. and Cuellar,W.J. |
| EPI_ISL_445087 | Laboratory Diagnostic, Veterinary Specialized Institute Kraljevo | Laboratory Diagnostic, Veterinary Specialized Institute Kraljevo | Vidanovic,D., Tesovic,B., Sekler,M., Dmitric,M., Debeljak,Z., Matovic,K., Vaskovic,N., Petrovic,T., Volkening,J. and Alfonso,C. |
| EPI_ISL_445120, EPI_ISL_445124, EPI_ISL_445129, EPI_ISL_445132, EPI_ISL_445146, EPI_ISL_445162 | Robert Garry lab | Andersen lab at Scripps Research | Allison Smither, Gilberto Sabino-Santos, Patricia Snarski, Lilia Melnik, Antoinette Bell, Kaylynn Genemaras, Arnaud Drouin, Dahlene Fusco, Robert Garry with SEARCH Alliance San Diego |
| EPI_ISL_445175, EPI_ISL_445176, EPI_ISL_445178 | UCSF Clinical Microbiology Laboratory | Chan-Zuckerberg Biohub | CZB Cliahub Consortium |
| EPI_ISL_445219 | Universidad del Valle, Laboratorio de Microbiologia, VIREM | Universidad del Valle, Universidad Nacional de Colombia-Sede Palmira, International Center for Tropical Agriculture | Beatriz Parra, Diana López-Alvarez, Wilmer J. Cuellar |
| EPI_ISL_445230 | Uppsala Narakut Aleris | The Public Health Agency of Sweden | Annika Nilsson, Oskar Karlsson Lindsjo, Maria Lind Karlberg, Anna-Malin Linde, Olov Svartstrom, Anna Risberg, Theresa Enkirch, Mia Brytting, Karin Tegmark-Wisell |
| EPI_ISL_445351 | HOSPITAL SAN JUAN DE DIOS | Instituto de Salud Publica de Chile | Andrés E Castillo, Bárbara Parra,Paz Tapia, Jaime Lagos, Loredana Arata, Alejandra Acevedo, Winston Andrade, Gabriel Leal, Carolina Tambley, Patricia Bustos, Rodrigo Fasce, Jorge Fernandez |
| EPI_ISL_445353 | HOSPITAL PADRE HURTADO | Instituto de Salud Publica de Chile | Andrés E Castillo, Bárbara Parra,Paz Tapia, Jaime Lagos, Loredana Arata, Alejandra Acevedo, Winston Andrade, Gabriel Leal, Carolina Tambley, Patricia Bustos, Rodrigo Fasce, Jorge Fernandez |
| EPI_ISL_445354, EPI_ISL_445369, EPI_ISL_445370 | HOSPITAL DE CARABINEROS | Instituto de Salud Publica de Chile | Andrés E Castillo, Bárbara Parra,Paz Tapia, Jaime Lagos, Loredana Arata, Alejandra Acevedo, Winston Andrade, Gabriel Leal, Carolina Tambley, Patricia Bustos, Rodrigo Fasce, Jorge Fernandez |
| EPI_ISL_445373, EPI_ISL_445374, EPI_ISL_445375, EPI_ISL_445376, EPI_ISL_445377 | HOSPITAL SAN JUAN DE DIOS | Instituto de Salud Publica de Chile | Andrés E Castillo, Bárbara Parra,Paz Tapia, Jaime Lagos, Loredana Arata, Alejandra Acevedo, Winston Andrade, Gabriel Leal, Carolina Tambley, Patricia Bustos, Rodrigo Fasce, Jorge Fernandez |
| EPI_ISL_445378 | HOSPITAL DE BULNES | Instituto de Salud Publica de Chile | Andrés E Castillo, Bárbara Parra,Paz Tapia, Jaime Lagos, Loredana Arata, Alejandra Acevedo, Winston Andrade, Gabriel Leal, Carolina Tambley, Patricia Bustos, Rodrigo Fasce, Jorge Fernandez |
| EPI_ISL_445379 | IMALAB- HOSPITAL FACH | Instituto de Salud Publica de Chile | Andrés E Castillo, Bárbara Parra,Paz Tapia, Jaime Lagos, Loredana Arata, Alejandra Acevedo, Winston Andrade, Gabriel Leal, Carolina Tambley, Patricia Bustos, Rodrigo Fasce, Jorge Fernandez |
| EPI_ISL_445382, EPI_ISL_445383, EPI_ISL_445389, EPI_ISL_445390, EPI_ISL_445391, EPI_ISL_445392, EPI_ISL_445393, EPI_ISL_445394, EPI_ISL_445395, EPI_ISL_445396, EPI_ISL_445398, EPI_ISL_445399, EPI_ISL_445400, EPI_ISL_445401, EPI_ISL_445402, EPI_ISL_445403, EPI_ISL_445404, EPI_ISL_445405, EPI_ISL_445406, EPI_ISL_445407, EPI_ISL_445408, EPI_ISL_445409, EPI_ISL_445410, EPI_ISL_445411, EPI_ISL_445412, EPI_ISL_445413, EPI_ISL_445414, EPI_ISL_445415, EPI_ISL_445416, EPI_ISL_445417, EPI_ISL_445422, EPI_ISL_445423, EPI_ISL_445424, EPI_ISL_445425, EPI_ISL_445426, EPI_ISL_445427, EPI_ISL_445428, EPI_ISL_445429, EPI_ISL_445430, EPI_ISL_445431, EPI_ISL_445432, EPI_ISL_445433, EPI_ISL_445434, EPI_ISL_445435, EPI_ISL_445436, EPI_ISL_445437, EPI_ISL_445438, EPI_ISL_445439, EPI_ISL_445440, EPI_ISL_445441, EPI_ISL_445442, EPI_ISL_445443, EPI_ISL_445444, EPI_ISL_445445, EPI_ISL_445446, EPI_ISL_445447, EPI_ISL_445448, EPI_ISL_445449, EPI_ISL_445450, EPI_ISL_445451, EPI_ISL_445452, EPI_ISL_445453, EPI_ISL_445454, EPI_ISL_445455, EPI_ISL_445456, EPI_ISL_445457, EPI_ISL_445458, EPI_ISL_445459, EPI_ISL_445460, EPI_ISL_445461, EPI_ISL_445462, EPI_ISL_445463, EPI_ISL_445467, EPI_ISL_445468, EPI_ISL_445469, EPI_ISL_445470, EPI_ISL_445471, EPI_ISL_445472, EPI_ISL_445473, EPI_ISL_445474, EPI_ISL_445475, EPI_ISL_445476, EPI_ISL_445477, EPI_ISL_445478, EPI_ISL_445479, EPI_ISL_445480, EPI_ISL_445481, EPI_ISL_445482, EPI_ISL_445483, EPI_ISL_445484, EPI_ISL_445485, EPI_ISL_445486, EPI_ISL_445487, EPI_ISL_445488, EPI_ISL_445489, EPI_ISL_445490, EPI_ISL_445491, EPI_ISL_445492, EPI_ISL_445493, EPI_ISL_445494, EPI_ISL_445495, EPI_ISL_445496, EPI_ISL_445497, EPI_ISL_445498, EPI_ISL_445499, EPI_ISL_445500, EPI_ISL_445501, EPI_ISL_445502, EPI_ISL_445503, EPI_ISL_445504, EPI_ISL_445505, EPI_ISL_445506, EPI_ISL_445507, EPI_ISL_445508, EPI_ISL_445509, EPI_ISL_445510, EPI_ISL_445511, EPI_ISL_445512, EPI_ISL_445513, EPI_ISL_445514, EPI_ISL_445515, EPI_ISL_445516, EPI_ISL_445517, EPI_ISL_445519, EPI_ISL_445520, EPI_ISL_445521, EPI_ISL_445522, EPI_ISL_445523, EPI_ISL_445524, EPI_ISL_445526, EPI_ISL_445527, EPI_ISL_445528, EPI_ISL_445529, EPI_ISL_445530, EPI_ISL_445531, EPI_ISL_445532, EPI_ISL_445533, EPI_ISL_445534, EPI_ISL_445535, EPI_ISL_445536, EPI_ISL_445537, EPI_ISL_445538, EPI_ISL_445539, EPI_ISL_445540, EPI_ISL_445541, EPI_ISL_445542, EPI_ISL_445543, EPI_ISL_445544, EPI_ISL_445545, EPI_ISL_445546, EPI_ISL_445547, EPI_ISL_445548, EPI_ISL_445549, EPI_ISL_445550, EPI_ISL_445551, EPI_ISL_445553, EPI_ISL_445554, EPI_ISL_445555, EPI_ISL_445556, EPI_ISL_445557, EPI_ISL_445558, EPI_ISL_445559, EPI_ISL_445560, EPI_ISL_445561, EPI_ISL_445562, EPI_ISL_445563, EPI_ISL_445564, EPI_ISL_445565, EPI_ISL_445566, EPI_ISL_445567, EPI_ISL_445569, EPI_ISL_445570, EPI_ISL_445571, EPI_ISL_445572, EPI_ISL_445573, EPI_ISL_445574, EPI_ISL_445575, EPI_ISL_445576, EPI_ISL_445577, EPI_ISL_445578, EPI_ISL_445579, EPI_ISL_445580, EPI_ISL_445581, EPI_ISL_445587, EPI_ISL_445588, EPI_ISL_445589, EPI_ISL_445590, EPI_ISL_445591, EPI_ISL_445592, EPI_ISL_445593, EPI_ISL_445594, EPI_ISL_445595, EPI_ISL_445596, EPI_ISL_445597, EPI_ISL_445598, EPI_ISL_445599, EPI_ISL_445600, EPI_ISL_445603, EPI_ISL_445606, EPI_ISL_445607, EPI_ISL_445608, EPI_ISL_445609, EPI_ISL_445610, EPI_ISL_445611, EPI_ISL_445612, EPI_ISL_445613, EPI_ISL_445614, EPI_ISL_445615, EPI_ISL_445642, EPI_ISL_445643, EPI_ISL_445644, EPI_ISL_445645, EPI_ISL_445646, EPI_ISL_445648, EPI_ISL_445650, EPI_ISL_445651, EPI_ISL_445652, EPI_ISL_445653, EPI_ISL_445660, EPI_ISL_445661, EPI_ISL_445662, EPI_ISL_445663, EPI_ISL_445664, EPI_ISL_445665, EPI_ISL_445666, EPI_ISL_445667, EPI_ISL_445668, EPI_ISL_445669, EPI_ISL_445674, EPI_ISL_445675, EPI_ISL_445676, EPI_ISL_445677, EPI_ISL_445678, EPI_ISL_445679, EPI_ISL_445680, EPI_ISL_445681, EPI_ISL_445682, EPI_ISL_445683, EPI_ISL_445684, EPI_ISL_445685, EPI_ISL_445686, EPI_ISL_445687, EPI_ISL_445688, EPI_ISL_445689, EPI_ISL_445690, EPI_ISL_445691, EPI_ISL_445692, EPI_ISL_445693, EPI_ISL_445694, EPI_ISL_445695, EPI_ISL_445696, EPI_ISL_445697, EPI_ISL_445698, EPI_ISL_445956, EPI_ISL_445957, EPI_ISL_445958, EPI_ISL_445959, EPI_ISL_445960, EPI_ISL_445961, EPI_ISL_445962, EPI_ISL_445963, EPI_ISL_445964, EPI_ISL_445965, EPI_ISL_445966, EPI_ISL_445967, EPI_ISL_445968, EPI_ISL_445969, EPI_ISL_445970, EPI_ISL_445971, EPI_ISL_445972, EPI_ISL_445973, EPI_ISL_445974, EPI_ISL_445975, EPI_ISL_445976, EPI_ISL_445977, EPI_ISL_445978, EPI_ISL_445979, EPI_ISL_445980, EPI_ISL_445981, EPI_ISL_445982, EPI_ISL_445983, EPI_ISL_445984, EPI_ISL_445985, EPI_ISL_445986, EPI_ISL_445987, EPI_ISL_445988, EPI_ISL_445989, EPI_ISL_445990, EPI_ISL_445991, EPI_ISL_445992, EPI_ISL_445993, EPI_ISL_445994, EPI_ISL_445995, EPI_ISL_445996, EPI_ISL_445997, EPI_ISL_445998, EPI_ISL_445999, EPI_ISL_446000, EPI_ISL_446001, EPI_ISL_446002, EPI_ISL_446003, EPI_ISL_446004, EPI_ISL_446005, EPI_ISL_446006, EPI_ISL_446007, EPI_ISL_446008, EPI_ISL_446009, EPI_ISL_446010, EPI_ISL_446011, EPI_ISL_446012, EPI_ISL_446013, EPI_ISL_446014, EPI_ISL_446015, EPI_ISL_446016, EPI_ISL_446017, EPI_ISL_446018, EPI_ISL_446019, EPI_ISL_446020, EPI_ISL_446021, EPI_ISL_446022, EPI_ISL_446032, EPI_ISL_446033, EPI_ISL_446119, EPI_ISL_446120, EPI_ISL_446122, EPI_ISL_446126, EPI_ISL_446129, EPI_ISL_446132, EPI_ISL_446133, EPI_ISL_446134, EPI_ISL_446139, EPI_ISL_446141, EPI_ISL_446142, EPI_ISL_446143, EPI_ISL_446146, EPI_ISL_446147, EPI_ISL_446149, EPI_ISL_446161, EPI_ISL_446164, EPI_ISL_446169, EPI_ISL_446170, EPI_ISL_446175, EPI_ISL_446176, EPI_ISL_446177, EPI_ISL_446178, EPI_ISL_446179, EPI_ISL_446180, EPI_ISL_446181, EPI_ISL_446182, EPI_ISL_446183, EPI_ISL_446184, EPI_ISL_446185, EPI_ISL_446186, EPI_ISL_446187, EPI_ISL_446188, EPI_ISL_446189, EPI_ISL_446190, EPI_ISL_446191, EPI_ISL_446192, EPI_ISL_446193, EPI_ISL_446194, EPI_ISL_446195, EPI_ISL_446196, EPI_ISL_446197, EPI_ISL_446198, EPI_ISL_446199, EPI_ISL_446200, EPI_ISL_446201, EPI_ISL_446202, EPI_ISL_446203, EPI_ISL_446204, EPI_ISL_446205, EPI_ISL_446206, EPI_ISL_446207, EPI_ISL_446208, EPI_ISL_446209, EPI_ISL_446210, EPI_ISL_446211, EPI_ISL_446212, EPI_ISL_446213, EPI_ISL_446214, EPI_ISL_446215, EPI_ISL_446216, EPI_ISL_446217, EPI_ISL_446218, EPI_ISL_446219, EPI_ISL_446220, EPI_ISL_446221, EPI_ISL_446222, EPI_ISL_446223, EPI_ISL_446224, EPI_ISL_446225, EPI_ISL_446226, EPI_ISL_446228, EPI_ISL_446229, EPI_ISL_446230, EPI_ISL_446231, EPI_ISL_446232, EPI_ISL_446233, EPI_ISL_446234, EPI_ISL_446235, EPI_ISL_446236, EPI_ISL_446237, EPI_ISL_446238, EPI_ISL_446239, EPI_ISL_446240, EPI_ISL_446241, EPI_ISL_446242, EPI_ISL_446243, EPI_ISL_446244, EPI_ISL_446266, EPI_ISL_446267, EPI_ISL_446268, EPI_ISL_446269, EPI_ISL_446270, EPI_ISL_446560, EPI_ISL_446577, EPI_ISL_446582, EPI_ISL_446583, EPI_ISL_446674, EPI_ISL_446678, EPI_ISL_446679 | | | |
| see above | Wales Specialist Virology Centre | Public Health Wales Microbiology Cardiff | Catherine Moore, Johnathan Evans, Laura Gifford, Malorie Perry, Simon Cottrell, Alec Birchley, Alexander Adams, Amy Gaskin, Bree Gatica-Wilcox, Jason Coombes, Lauren Gilbert, Lee Graham, Nicole Pacchiarini, Sara Kumziene-Summerhayes, Sarah Taylor, Sophie Jones, Sara Rey, Matthew Bull, Joanne Watkins, Sally Corden, Tom Connor |
| EPI_ISL_447022, EPI_ISL_447023, EPI_ISL_447024, EPI_ISL_447025, EPI_ISL_447026, EPI_ISL_447027 | Ramathibodi Hospital | COVID-19 Network Investigations (CONI) Alliance | Elizabeth Batty, Wasun Chantratita, Thanat Chookajorn, Stefan Fernandez, Angkana Huang, Anthony R. Jones, Khajohn Joonsalak, Chonticha Klungtong, Theerarat Kochakarn, Namfon Kotanan, Krittikorn Kumpornsin, Wudtichai Manasatienkij, Bhakbhoom Panthan, Ekawat Pasomsub, Kingkan Rakmanee, Insee Sensorn, Janjira Thaipadungpanit, Arporn Wangwiwatsin,Treewat Watthanachockchai |
| EPI_ISL_447058, EPI_ISL_447064, EPI_ISL_447068, EPI_ISL_447082, EPI_ISL_447088, EPI_ISL_447098, EPI_ISL_447100, EPI_ISL_447102, EPI_ISL_447104, EPI_ISL_447105, EPI_ISL_447108, EPI_ISL_447109, EPI_ISL_447110, EPI_ISL_447112, EPI_ISL_447113, EPI_ISL_447115 | | | |
| see above | Michigan Department of Health and Human Services, Bureau of Laboratories | Michigan Department of Health and Human Services, Bureau of Laboratories | Blankenship HM, Riner D, Soehnlen MK |
| EPI_ISL_447264 | Microbiology laboratory, Assuta Ashdod University-Affiliated Hospital | Stern Lab | Stern Lab |
| EPI_ISL_447311, EPI_ISL_447312, EPI_ISL_447313, EPI_ISL_447314, EPI_ISL_447315, EPI_ISL_447316 | Clinical Virology Laboratory, Soroka Medical Center and the Faculty of Health Sciences, Ben-Gurion University of the Negev | Stern Lab | Stern Lab |

| | | | |
|---|---|---|---|
| EPI_ISL_447353, EPI_ISL_447354, EPI_ISL_447355, EPI_ISL_447356, EPI_ISL_447357, EPI_ISL_447358 | Clinical Virology Unit, Hadassah Hebrew University Medical Center | Stern Lab | Stern Lab |
| EPI_ISL_447400, EPI_ISL_447401, EPI_ISL_447402 | Clinical Microbiology Laboratory, The Baruch Padeh Medical Center, Poriya | Stern Lab | Stern Lab |
| EPI_ISL_447419, EPI_ISL_447420, EPI_ISL_447421, EPI_ISL_447422, EPI_ISL_447423, EPI_ISL_447424, EPI_ISL_447425, EPI_ISL_447426, EPI_ISL_447427, EPI_ISL_447428, EPI_ISL_447429, EPI_ISL_447430, EPI_ISL_447431, EPI_ISL_447432, EPI_ISL_447433, EPI_ISL_447434, EPI_ISL_447435, EPI_ISL_447436, EPI_ISL_447437, EPI_ISL_447438, EPI_ISL_447439, EPI_ISL_447440 | | | |
| see above | Clinical Microbiology Laboratory, Sheba Medical Center | Stern Lab | Stern Lab |
| EPI_ISL_447575 | CSIR-Centre for Cellular and Molecular Biology | CSIR-Centre for Cellular and Molecular Biology | Sofia Banu, Payel Mukherjee, Priya Singh, Dhiviya Vedagiri, Divya Gupta, Vishal Sah, Santosh Kumar Kuncha, Krishnan Harinivas Harshan, Archana Bharadwaj Siva, Karthik Bharadwaj Tallapaka, Shagufta Khan, Lamuk Zaveri, Namami Gaur, Sakshi Shambhavi, Tulasi Nagabandi, Purushotham Vodnala, Rakesh K Mishra, Divya Tej Sowpati |
| EPI_ISL_447576, EPI_ISL_447577, EPI_ISL_447578 | CSIR-Centre for Cellular and Molecular Biology | CSIR-Centre for Cellular and Molecular Biology | Namami Gaur, Sakshi Shambhavi, Lamuk Zaveri, Shagufta Khan, Tulasi Nagabandi, Purushotham Vodnala, Payel Mukherjee, Sofia Banu, Priya Singh, Dhiviya Vedagiri, Divya Gupta, Vishal Sah, Santosh Kumar Kuncha, Krishnan Harinivas Harshan, Archana Bharadwaj Siva, Karthik Bharadwaj Tallapaka, Rakesh K Mishra, Divya Tej Sowpati |
| EPI_ISL_447596 | Viral Respiratory Lab, National Institute for Biomedical Research (INRB) | Pathogen Sequencing Lab, National Institute for Biomedical Research (INRB) | Placide Mbala-Kingebeni, Edith Nkwembe, Eddy Kinganda-Lusamaki, Amuri Aziza, Francisca Muyembe Mawete, Catherine Pratt, Matthias Pauthner, Josh Quick, Allison Black, James Hadfield, Trevor Bedford, Ian Goodfellow, Andrew Rambaut, Nick Loman, Kristian Andersen, Michael Wiley, Steve Ahuka-Mundeke, Jean-Jacques Muyembe Tamfum |
| EPI_ISL_447782, EPI_ISL_447783, EPI_ISL_447784, EPI_ISL_447785, EPI_ISL_447786, EPI_ISL_447787, EPI_ISL_447789, EPI_ISL_447790, EPI_ISL_447791, EPI_ISL_447792, EPI_ISL_447793, EPI_ISL_447794, EPI_ISL_447795, EPI_ISL_447796, EPI_ISL_447797, EPI_ISL_447798, EPI_ISL_447799, EPI_ISL_447800, EPI_ISL_447801, EPI_ISL_447802, EPI_ISL_447803, EPI_ISL_447804, EPI_ISL_447805, EPI_ISL_447806, EPI_ISL_447807, EPI_ISL_447808, EPI_ISL_447809, EPI_ISL_447810, EPI_ISL_447811, EPI_ISL_447812, EPI_ISL_447813, EPI_ISL_447814, EPI_ISL_447815 | | | |
| see above | Instituto Nacional de Salud, Bogotá, Colombia | Grupo de Investigaciones Microbiológicas-UR (GIMUR), Departamento de Biología, Facultad de Ciencias Naturales, Universidad del Rosario, Bogotá, Colombia Instituto Nacional de Salud, Bogotá, Colombia Icahn School of Medicine at Mount Sinai, New York, USA | Juan David Ramírez, Carolina Florez, Marina Muñoz, Carolina Hernandez, Adriana Castillo, Sergio Castañeda, Nathalia Ballesteros, David Martínez, Laura Vega, Jesús E. Jaimes, Sergio Gomez, Angelica Rico, Lisseth Pardo, Esther C. Barros, Martha L. Ospina, Aníbal A. Teherán, Ana S. Gonzalez-Reiche, Matthew M. Hernandez, Emilia Mia Sordillo, Viviana Simon, Harm van Bakel, Alberto Paniz-Mondolfi |
| EPI_ISL_447854 | CSIR-Centre for Cellular and Molecular Biology | CSIR-Centre for Cellular and Molecular Biology | Payel Mukherjee, Sofia Banu, Priya Singh, Dhiviya Vedagiri, Divya Gupta, Vishal Sah, Santosh Kumar Kuncha, Krishnan Harinivas Harshan, Archana Bharadwaj Siva, Karthik Bharadwaj Tallapaka, Shagufta Khan, Lamuk Zaveri, Namami Gaur, Sakshi Shambhavi, Tulasi Nagabandi, Purushotham Vodnala, Rakesh K Mishra, Divya Tej Sowpati |
| EPI_ISL_447855 | CSIR-Centre for Cellular and Molecular Biology | CSIR-Centre for Cellular and Molecular Biology | Lamuk Zaveri, Shagufta Khan, Namami Gaur, Sakshi Shambhavi, Tulasi Nagabandi, Purushotham Vodnala, Payel Mukherjee, Sofia Banu, Priya Singh, Dhiviya Vedagiri, Divya Gupta, Vishal Sah, Santosh Kumar Kuncha, Krishnan Harinivas Harshan, Archana Bharadwaj Siva, Karthik Bharadwaj Tallapaka, Rakesh K Mishra, Divya Tej Sowpati |
| EPI_ISL_447856, EPI_ISL_447857, EPI_ISL_447858 | CSIR-Centre for Cellular and Molecular Biology | CSIR-Centre for Cellular and Molecular Biology | Sakshi Shambhavi, Lamuk Zaveri, Shagufta Khan, Namami Gaur, Tulasi Nagabandi, Purushotham Vodnala, Payel Mukherjee, Sofia Banu, Priya Singh, Dhiviya Vedagiri, Divya Gupta, Vishal Sah, Santosh Kumar Kuncha, Krishnan Harinivas Harshan, Archana Bharadwaj Siva, Karthik Bharadwaj Tallapaka, Rakesh K Mishra, Divya Tej Sowpati |
| EPI_ISL_447890, EPI_ISL_447892, EPI_ISL_447893, EPI_ISL_447894 | University of California, Davis | Chan-Zuckerberg Biohub | CZB Cliahub Consortium |
| EPI_ISL_448117, EPI_ISL_448118, EPI_ISL_448119, EPI_ISL_448120, EPI_ISL_448121, EPI_ISL_448122, EPI_ISL_448123, EPI_ISL_448124, EPI_ISL_448125, EPI_ISL_448126, EPI_ISL_448127, EPI_ISL_448128 | | | |
| see above | West of Scotland Specialist Virology Centre, NHSGGC / MRC-University of Glasgow Centre for Virus Research | COVID-19 Genomics UK (COG-UK) Consortium | Ana da Silva Filipe, Natasha Johnson, Kathy Smollett, Daniel Mair, Stephen Carmichael, Lily Tong, Jenna Nichols, Elihu Aranday-Cortes, Kirstyn Brunker, Yasmin Parr, Kyriaki Nomikou, Sarah McDonald, Marc Niebel, Patawee Asamaphan, Richard Orton, Joseph Hughes, Sreenu Vattipally, David L Robertson, Alasdair MacLean, Rory Gunson, Kathy Li, Natasha Jesudason, Rajiv Shah, James Shepherd, Antonia Ho, Emma Thomson |
| EPI_ISL_448450, EPI_ISL_448453, EPI_ISL_448455, EPI_ISL_448458, EPI_ISL_448460, EPI_ISL_448462, EPI_ISL_448464, EPI_ISL_448466, EPI_ISL_448468, EPI_ISL_448470, EPI_ISL_448472, EPI_ISL_448474, EPI_ISL_448476, EPI_ISL_448477, EPI_ISL_448478, EPI_ISL_448480, EPI_ISL_448483, EPI_ISL_448485, EPI_ISL_448486, EPI_ISL_448487, EPI_ISL_448489, EPI_ISL_448490, EPI_ISL_448491, EPI_ISL_448492, EPI_ISL_448493, EPI_ISL_448494, EPI_ISL_448496, EPI_ISL_448497, EPI_ISL_448500, EPI_ISL_448503, EPI_ISL_448504, EPI_ISL_448505, EPI_ISL_448506, EPI_ISL_448507, EPI_ISL_448508, EPI_ISL_448509, EPI_ISL_448510, EPI_ISL_448511, EPI_ISL_448512, EPI_ISL_448513, EPI_ISL_448514, EPI_ISL_448515, EPI_ISL_448516, EPI_ISL_448517, EPI_ISL_448518, EPI_ISL_448519, EPI_ISL_448520, EPI_ISL_448521, EPI_ISL_448522, EPI_ISL_448523, EPI_ISL_448524, EPI_ISL_448525, EPI_ISL_448526, EPI_ISL_448527, EPI_ISL_448528, EPI_ISL_448529, EPI_ISL_448530, EPI_ISL_448531, EPI_ISL_448532, EPI_ISL_448533, EPI_ISL_448534, EPI_ISL_448535, EPI_ISL_448536, EPI_ISL_448537, EPI_ISL_448538, EPI_ISL_448539, EPI_ISL_448540, EPI_ISL_448541, EPI_ISL_448542, EPI_ISL_448543, EPI_ISL_448544, EPI_ISL_448545, EPI_ISL_448546, EPI_ISL_448547, EPI_ISL_448551 | | | |
| see above | Oxford Viromics, NDM, University of Oxford; Oxford University Hospitals; Basingstoke and North Hampshire Hospital | COVID-19 Genomics UK (COG-UK) Consortium | Tanya Golubchik, David Bonsall, George Macintyre, Amy Trebes, Mariateresa de Cesare, Catrin Moore, Alex Mobbs, Anita Justice, Robert Shaw, Monique Andersson, Emma Wise, Nathan Moore, Jessica Lynch, Nick Cortes, Stephen Kidd, David Buck, John Todd, Christophe Fraser |
| EPI_ISL_448850 | Virology Laboratory, Castle Hill Hospital, Hull University Teaching Hospitals NHS Trust/Department of Infection, Immunity and Cardiovascular Disease, The Medical School, University of Sheffield | COVID-19 Genomics UK (COG-UK) Consortium | Thushan de Silva, Matthew Parker, Nikki Smith, Adri Angyal, Rebecca Brown, Luke Green, Rachel Tucker, Paul Parsons, Danielle Groves, Katie Johnson, Laura Carrilero, Alex Keeley, Dave Partridge, Matthew Wyles, Benjamin Lindsey, Mehmet Yavuz, Mohammad Raza, Cariad Evans |
| EPI_ISL_449380, EPI_ISL_449441, EPI_ISL_449442, EPI_ISL_449443, EPI_ISL_449444, EPI_ISL_449445, EPI_ISL_449446, EPI_ISL_449453, EPI_ISL_449454, EPI_ISL_449455, EPI_ISL_449456, EPI_ISL_449457, EPI_ISL_449458, EPI_ISL_449459, EPI_ISL_449460, EPI_ISL_449461, EPI_ISL_449462, EPI_ISL_449463, EPI_ISL_449464, EPI_ISL_449465, EPI_ISL_449575, EPI_ISL_449597, EPI_ISL_449598, EPI_ISL_449599, EPI_ISL_449600, EPI_ISL_449601, EPI_ISL_449602, EPI_ISL_449603 | | | |
| see above | Liverpool Clinical Laboratories | COVID-19 Genomics UK (COG-UK) Consortium | Sam Haldenby, Anita Lucaci, Steve Paterson, Julian Hiscox, Alistair Darby, M Almsaud, A Alrezaihi, Muhannad Alruwaili, Stuart D Armstrong, Jones Benjamin , Eleanor G Bentley, Anu Chawla, Jordan J Clark, Angela Cowell, Richard Eccles, Isabel García-Dorival, Matthew Gemmell, Alessandro Gerada, PKF Gilmore, Richard Gregory, Ximeng Han, Catherine Hartley, Margaret Hughes, Miren Iturriza-Gomara, James Johnson, L Luu, Jenifer Manson , Charlotte Nelson, Elaine O'Toole, Cassie Olateju, Rebekah Penrice-Randal , Lucille Rainbow, N.P Randle, Trevor Ian Robinson, Parul Sharma, Ghada T Shawli, James P Stewart , Neil Swainston, Ecaterina Vamos, Joanne Watts, Mark Whitehead |
| EPI_ISL_449626, EPI_ISL_449627, EPI_ISL_449628, EPI_ISL_449629, EPI_ISL_449630, EPI_ISL_449631, EPI_ISL_449632, EPI_ISL_449633, EPI_ISL_449634, EPI_ISL_449635, EPI_ISL_449636, EPI_ISL_449637, EPI_ISL_449638, EPI_ISL_449639, EPI_ISL_449640, EPI_ISL_449641, EPI_ISL_449642, EPI_ISL_449643, EPI_ISL_449644, EPI_ISL_449645, EPI_ISL_449646, EPI_ISL_449647, EPI_ISL_449648, EPI_ISL_449649, EPI_ISL_449650, EPI_ISL_449651, EPI_ISL_449652, EPI_ISL_449653, EPI_ISL_449654, EPI_ISL_449655 | | | |
| see above | University College London, Great Ormond Street Hospital for Children NHS Foundation Trust, Imperial College Healthcare NHS Trust | COVID-19 Genomics UK (COG-UK) Consortium | Sergi Castellano, Rachel Williams, Mark Kristiansen, Paola Resende Silva, Sunando Roy, Tony Brooks, Helena Tutill, Paola Niola, Patricia Dyal, Charlotte Williams, Leysa Forrest, Yasmin Panchbhaya, Jacqueline Findlay, Sam Weeks, Julianne Brown, Kathryn Harris, Paul Randell, James Price, Alison Holmes, Judith Breuer |
| EPI_ISL_449791, EPI_ISL_449792, EPI_ISL_449793, EPI_ISL_449794 | Dept. of Medical Microbiology, Stavanger University Hospital, Helse Stavanger HF | Norwegian Institute of Public Health, Department of Virology | Kathrine Stene-Johansen, Kamilla Heddeland Instefjord, Hilde Elshaug, Rasmus Riis Kopperud, Karoline Bragstad, Olav Hungnes |
| EPI_ISL_449833, EPI_ISL_449834, EPI_ISL_449835, EPI_ISL_449836, EPI_ISL_449837, EPI_ISL_449838 | Utah Public Health Laboratory | Utah Public Health Laboratory | Erin Young, Kelly Oakeson |
| EPI_ISL_450173, EPI_ISL_450174, EPI_ISL_450175, EPI_ISL_450176, EPI_ISL_450178, EPI_ISL_450179, EPI_ISL_450181, EPI_ISL_450182, EPI_ISL_450183, EPI_ISL_450184, EPI_ISL_450185 | | | |
| see above | Robert Garry lab | Andersen lab at Scripps Research | Allison Smither, Gilberto Sabino-Santos, Patricia Snarski, Lilia Melnik, Antoinette Bell, Kaylynn Genemaras, Arnaud Drouin, Dahlene Fusco, Robert Garry with SEARCH Alliance San Diego |
| EPI_ISL_450188, EPI_ISL_450189 | Biolab Diagnostic Laboratories | Andersen lab at Scripps Research | Issa Abu-Dayyeh, Ahmad Tibi, Lama Hussein, Lina Mohammad, Zein Naber, Amid Abdelnour with SEARCH Alliance San Diego |
| EPI_ISL_450215, EPI_ISL_450216 | unknown | Microbiological Diagnostic Unit Public Health Laboratory (MDU-PHL) and Victorian Infectious Disease Reference Laboratory (VIDRL) | Seemann,T., Lane,C.R., Sherry,N.L., Duchene,S., Goncalves da Silva,A., Caly,L., Sait,M., Ballard,S.A., Horan,K., Schultz,M.B., Hoang,T., Easton,M., Dougal,S., Stinear,T.P., Druce,J., Catton,M., Sutton,B., van Diemen,A., Alpren,C., Williamson,D.A., Howden,B.P. |

| | | | |
|---|---|---|---|
| EPI_ISL_450321 | NIV Pune | CSIR-Centre for Cellular and Molecular Biology | Dr V A Potdar, Dr ML Choudhary,Dr Priya Abraham,V. Vipat, S. Jadhav, U. Saha, H. Kengle, A. Awhale, A. Jagtap, A. Gondhalikar, V Malik, N Srivastava, S. Digraskar, P. Malsane, S. Hundekar, K. Patel, Yogesh Balakartik, M. Kakade, S. Jadhav, R. Gunjikar, V. Awtade, S. Bhorekar, P Shinde, S. Salve, B. Minhas S. Bharadwaj, H Kaushal Y. Gurav, S. Tomar,Payel Mukherjee, Sofia Banu, Priya Singh, Dhiviya Vedagiri, Divya Gupta, Vishal Sah, Santosh Kumar Kuncha, Krishnan Harinivas Harshan, Archana Bharadwaj Siva, Karthik Bharadwaj Tallapaka, Shagufta Khan, Lamuk Zaveri, Namami Gaur, Sakshi Shambhavi, Tulasi Nagabandi, Purushotham Vodnala,G. Aditya Kumar, Koushick Sivakumar, Pooja Ramesh Gupta, Rajan Kumar Jha, Shraddha Vijay Lahoti, Deepak Kumar, Devi Prasad Vijayashankara, Disha Nanda, Divya Das, Jotin Gogoi, Manish |
| EPI_ISL_450322 | NIV Pune | CSIR-Centre for Cellular and Molecular Biology | Dr V A Potdar, Dr ML Choudhary,Dr Priya Abraham,V. Vipat, S. Jadhav, U. Saha, H. Kengle, A. Awhale, A. Jagtap, A. Gondhalikar, V Malik, N Srivastava, S. Digraskar, P. Malsane, S. Hundekar, K. Patel, Yogesh Balakartik, M. Kakade, S. Jadhav, R. Gunjikar, V. Awtade, S. Bhorekar, P Shinde, S. Salve, B. Minhas S. Bharadwaj, H Kaushal Y. Gurav, S. Tomar,Sofia Banu, Payel Mukherjee, Priya Singh, Dhiviya Vedagiri, Divya Gupta, Vishal Sah, Santosh Kumar Kuncha, Krishnan Harinivas Harshan, Archana Bharadwaj Siva, Karthik Bharadwaj Tallapaka, Shagufta Khan, Lamuk Zaveri, Namami Gaur, Sakshi Shambhavi, Tulasi Nagabandi, Purushotham Vodnala, Disha Nanda, Divya Das, Jotin Gogoi, Manish Bhattacharjee, Ravi Prasad Mukku, Renu Sudhakar, Somesh Gorde, Gangumala Srinivas Reddy, Sujoy Deb, Swati Bayyana, Zeba Rizvi, Rakesh K Mishra |
| EPI_ISL_450330 | CSIR-Centre for Cellular and Molecular Biology | CSIR-Centre for Cellular and Molecular Biology | Sakshi Shambhavi, Lamuk Zaveri, Shagufta Khan, Namami Gaur, Tulasi Nagabandi, Purushotham Vodnala, Payel Mukherjee, Sofia Banu, Priya Singh, Dhiviya Vedagiri, Divya Gupta, Vishal Sah, Santosh Kumar Kuncha, Krishnan Harinivas Harshan, Archana Bharadwaj Siva, Karthik Bharadwaj Tallapaka,Preethi Jampala, Sharada Ravi Iyer, Sulagana Mukherjee, Swetha Sundar, Peddapuvala Sai Uday Kiran, Umesh Kumar, Unis Ahmad Bhat, Ajay Sarawagi, Priyanka Pant, Rajkanwar Nathawat, Nikhil Hajirnis, Pratheusa Maccha, M Soujanya Reddy Rakesh K Mishra, Divya Tej Sowpati |
| EPI_ISL_450400, EPI_ISL_450402 | NYU Langone Health | Departments of Pathology and Medicine, New York University School of Medicine | Maria Aguero-Rosenfeld, Brendan Belovarac, Margaret Black, Ludovic Boytard, John Cadley, Paolo Cotzia, John Chen, Dacia Dimartino, Xiaojun Feng, Tatyana Gindin, Emily Guzman, Adriana Heguy, Megan Hogan, Emily Huang, George Jour, Alireza Khodadadi-Jamayran, Lawrence H. Lin, Raven Luther, Andrew Lytle, Christian Marier, Matthew T. Maurano, Mark J. Mulligan, Peter Meyn, Raquel Ordonez Ciriza, Iman Osman, Jared Pinnell, Vanessa Raabe, Sitharam Ramaswami, Amy Rapkiewicz, Andre M. Ribeiro-dos-Santos, Marie Samanovic-Golden, Antonio Serrano, Guomiao Shen, Matija Snuderl, Theodore Vougiouklakis, Nick Vulpescu, Gael Westby, Paul Zappile, Yutong Zhang |
| EPI_ISL_450407 | Molecular Diagnostics, Antech Diagnostics | Molecular Diagnostics, Antech Diagnostics | Leutenegger,C.M., Lozoya,C.E., Tereski,J.L. and Moroff,S. |
| EPI_ISL_450415 | Laboratory Diagnostic, Veterinary Specialized Institute Kraljevo | Laboratory Diagnostic, Veterinary Specialized Institute Kraljevo | Vidanovic,D., Skadric,I., Tesovic,B., Tolic,A., Sekler,M., Petrovic,T., Matovic,K., Dmitric,M., Debeljak,Z. and Vaskovic,N. |
| EPI_ISL_450482 | National Influenza and other Respiratory Viruses Centre-Tunisia, Virology Unit, Microbiology Laboratory, Charles Nicolle Hospital | National Influenza and other Respiratory Viruses Centre-Tunisia, Virology Unit, Microbiology Laboratory, Charles Nicolle Hospital | El Moussi,A., Abid,S., Ben Nasr,M., Landolsi,I., Charaa,L., Enigrou,D. and Boutiba,I. |
| EPI_ISL_450483 | Molecular Genetic, Immuno Gene Center | Molecular Genetic, Immuno Gene Center | Dlovan,M.F., Haval,F.M., Hazha,H.J. and Ariamand,A. |
| EPI_ISL_450491, EPI_ISL_450493, EPI_ISL_450494 | unknown | National Influenza and other Respiratory Viruses Centre-Tunisia | El Moussi,A., Abid,S., Ben Nasr,M., Landolsi,I., Charaa,L., Ferjeni,A., Arab Ennigrou,D., Boutiba,I. |
| EPI_ISL_450544, EPI_ISL_450545, EPI_ISL_450546, EPI_ISL_450547, EPI_ISL_450548, EPI_ISL_450549 | Utah Public Health Laboratory | Utah Public Health Laboratory | Erin Young, Kelly Oakeson |
| EPI_ISL_450649 | Laboratoire de microbiologie, Hopital de Verdun | Smith Laboratory, Centre de Recherche CHU Sainte-Justine | Martin Smith, Marieke Rozendaal, Ivan Pavlov |
| EPI_ISL_450733, EPI_ISL_450734 | Hospital AZ Rivierenland | Institute of Tropical Medicine | Philippe Selhorst, Colin Anthony |
| EPI_ISL_450801 | Georgia Department of Health | Pathogen Discovery, Respiratory Viruses Branch, Division of Viral Diseases, Centers for Disease Control and Prevention | Yan Li, Anna Montmayeur, Ying Tao, Krista Queen, Jing Zhang, Anna Uehara, Clinton R. Paden, Rachel Marine, Haibin Wang, Zachary Weiner, Bettina Bankamp, Suxiang Tong |
| EPI_ISL_450804 | VI-US Virgin Islands Department of Health | Pathogen Discovery, Respiratory Viruses Branch, Division of Viral Diseases, Centers for Disease Control and Prevention | Yan Li, Anna Montmayeur, Ying Tao, Krista Queen, Jing Zhang, Anna Uehara, Clinton R. Paden, Rachel Marine, Haibin Wang, Zachary Weiner, Bettina Bankamp, Suxiang Tong |
| EPI_ISL_450805 | VI-US Virgin Islands Department of Health | Pathogen Discovery, Respiratory Viruses Branch, Division of Viral Diseases, Centers for Disease Control and Prevention | Krista Queen, Yan Li, Anna Montmayeur, Ying Tao, Jing Zhang, Anna Uehara, Clinton R. Paden, Rachel Marine, Haibin Wang, Jasmine Padilla, Justin Lee, Zachary Weiner, Bettina Bankamp, Suxiang Tong |
| EPI_ISL_450811 | Knivsta VC | The Public Health Agency of Sweden | Johanna Carlson, Anna-Malin Linde, Maria Lind Karlberg, Oskar Karlsson Lindsjo, Olov Svartstrom, Anna Risberg, Theresa Enkirch, Mia Brytting, Karin Tegmark-Wisell |
| EPI_ISL_450812 | Uppsala Narakut Aleris | The Public Health Agency of Sweden | Annika Nilsson, Anna-Malin Linde, Maria Lind Karlberg, Oskar Karlsson Lindsjo, Olov Svartstrom, Anna Risberg, Theresa Enkirch, Mia Brytting, Karin Tegmark-Wisell |
| EPI_ISL_450836, EPI_ISL_450837 | Laboratoriemedicin | The Public Health Agency of Sweden | Anna-Malin Linde, Maria Lind Karlberg, Oskar Karlsson Lindsjo, Olov Svartstrom, Anna Risberg, Theresa Enkirch, Mia Brytting, Karin Tegmark-Wisell |
| EPI_ISL_450844, EPI_ISL_450845, EPI_ISL_450846 | Florida Bureau of Public Health Laboratories | Florida Bureau of Public Health Laboratories | Sarah Schmedes, Jason Blanton |
| EPI_ISL_451128 | SA Pathology | SA Pathology | Lex Leong, Chuan Kok Lim, Mark Turra, Ivan Bastian, Geoff Higgins |
| EPI_ISL_451168, EPI_ISL_451169, EPI_ISL_451170, EPI_ISL_451171, EPI_ISL_451173 | Lab voor klinische biologie | Onderzoeksgroep Virologie | Laurens Lambrechts, Nick Vereecke, Marthe Pauwels, Jozefien De Clercq, Bruno Verhasselt, Linos Vandekerckhove, Hans Nauwynck, Sebastiaan Theuns |
| EPI_ISL_451174 | Lab voor klinische biologie | Onderzoeksgroep Virologie | Nick Vereecke, Laurens Lambrechts, Marthe Pauwels, Jozefien De Clercq, Bruno Verhasselt, Linos Vandekerckhove, Hans Nauwynck, Sebastiaan Theuns |
| EPI_ISL_451532 | Pathology West - NSW Health Pathology | NSW Health Pathology - Institute of Clinical Pathology and Medical Research; Westmead Hospital; University of Sydney | CIDM-PH et al. |
| EPI_ISL_451534 | Medlab Pathology | NSW Health Pathology - Institute of Clinical Pathology and Medical Research; Westmead Hospital; University of Sydney | CIDM-PH et al. |
| EPI_ISL_451535, EPI_ISL_451537, EPI_ISL_451539, EPI_ISL_451541 | Pathology West - NSW Health Pathology | NSW Health Pathology - Institute of Clinical Pathology and Medical Research; Westmead Hospital; University of Sydney | CIDM-PH et al. |
| EPI_ISL_451548 | Childrens Hospital Westmead | NSW Health Pathology - Institute of Clinical Pathology and Medical Research; Westmead Hospital; University of Sydney | CIDM-PH et al. |
| EPI_ISL_451550, EPI_ISL_451551 | Pathology West - NSW Health Pathology | NSW Health Pathology - Institute of Clinical Pathology and Medical Research; Westmead Hospital; University of Sydney | CIDM-PH et al. |
| EPI_ISL_451554, EPI_ISL_451555 | Medlab Pathology | NSW Health Pathology - Institute of Clinical Pathology and Medical Research; Westmead Hospital; University of Sydney | CIDM-PH et al. |
| EPI_ISL_451559, EPI_ISL_451560, EPI_ISL_451561, EPI_ISL_451562, EPI_ISL_451565 | Pathology Sydney South West - NSW Health Pathology | NSW Health Pathology - Institute of Clinical Pathology and Medical Research; Westmead Hospital; University of Sydney | CIDM-PH et al. |
| EPI_ISL_451568, EPI_ISL_451571, EPI_ISL_451573 | Pathology West - NSW Health Pathology | NSW Health Pathology - Institute of Clinical Pathology and Medical Research; Westmead Hospital; University of Sydney | CIDM-PH et al. |
| EPI_ISL_451583 | Medlab Pathology | NSW Health Pathology - Institute of Clinical Pathology and Medical Research; Westmead Hospital; University of Sydney | CIDM-PH et al. |

| | | | |
|---|---|---|---|
| EPI_ISL_451589, EPI_ISL_451600 | Pathology North Hunter- NSW Health Pathology | NSW Health Pathology - Institute of Clinical Pathology and Medical Research; Westmead Hospital; University of Sydney | CIDM-PH et al. |
| EPI_ISL_451605 | Childrens Hospital Westmead | NSW Health Pathology - Institute of Clinical Pathology and Medical Research; Westmead Hospital; University of Sydney | CIDM-PH et al. |
| EPI_ISL_451610 | Medlab Pathology | NSW Health Pathology - Institute of Clinical Pathology and Medical Research; Westmead Hospital; University of Sydney | CIDM-PH et al. |
| EPI_ISL_451641 | Laverty Pathology | NSW Health Pathology - Institute of Clinical Pathology and Medical Research; Westmead Hospital; University of Sydney | CIDM-PH et al. |
| EPI_ISL_451856, EPI_ISL_451884, EPI_ISL_451885, EPI_ISL_451886, EPI_ISL_451887, EPI_ISL_451888, EPI_ISL_451889, EPI_ISL_451891 | Viollier AG | Department of Biosystems Science and Engineering, ETH Zürich | Christian Beisel, Sarah Nadeau, Ivan Topolsky, Pedro Ferreira, Philipp Jablonski, Susana Posada-Céspedes, Tobias Schär, Ina Nissen, Natascha Santacroce, Elodie Burcklen, Christiane Beckmann, Maurice Redondo, Olivier Kobel, Christoph Noppen, Sophie Seidel, Noemie Santamaria de Souza, Niko Beerenwinkel, Tanja Stadler |
| EPI_ISL_451946 | Max von Pettenkofer Institute, Virology, National Reference Center for Retroviruses, LMU München | Laboratory for Functional Genome Analysis, Dept. Genomics, Gene Center of the LMU Munich | Max Muenchhoff, Stefan Krebs, Alexander Graf, Oliver Keppler, Helmut Blum |
| EPI_ISL_451972, EPI_ISL_451974, EPI_ISL_451977, EPI_ISL_451980, EPI_ISL_451983 | 1. ViroGenetics - BSL3 Laboratory of Virology, Maopolska Centre of Biotechnology, Jagiellonian University; 2. II Department of Internal Medicine, Faculty of Medicine, Jagiellonian University Medical College; 3. DIAGNOSTYKA Ltd. | 1. ViroGenetics - BSL3 Laboratory of Virology, Maopolska Centre of Biotechnology, Jagiellonian University; 2. II Department of Internal Medicine, Faculty of Medicine, Jagiellonian University Medical College. | Marek Sanak, Marcin Surmiak, Monika Gsecka-Czapla, Wojciech Branicki, Pawe P abaj, Marta Rogalska-Kupiec, Jakub Swadba, Krzysztof Pyr |
| EPI_ISL_452104 | Max von Pettenkofer Institute, Virology, National Reference Center for Retroviruses, LMU München | Laboratory for Functional Genome Analysis, Dept. Genomics, Gene Center of the LMU Munich | Max Muenchhoff, Stefan Krebs, Alexander Graf, Oliver Keppler, Helmut Blum |
| EPI_ISL_452140 | CUB Hopital Erasme Laboratoire d'Anatomie Pathologique | CUB Hopital Erasme Laboratoire d'Anatomie Pathologique | Isabelle Salmon, Nicky D'Haene |
| EPI_ISL_452190, EPI_ISL_452191 | ULSS9 Distretto di San Bonifacio | Istituto Zooprofilattico Sperimentale delle Venezie | Adelaide Milani, Alessia Schivo, Annalisa Salviato, Erika Giorgia Quaranta, Ambra Pastori, Bianca Zecchin, Alice Fusaro, Isabella Monne, Calogero Terregino, Antonia Ricci |
| EPI_ISL_452207, EPI_ISL_452208, EPI_ISL_452209, EPI_ISL_452210 | NIV Influenza | NIV Influenza | Potdar V |
| EPI_ISL_452367, EPI_ISL_452370 | Servicio de Microbiología. HRU de Málaga. Servicio Andaluz de Salud | SeqCOVID-SPAIN consortium/IBV(CSIC) | Inmaculada de Toro Peinado, Maria Concepción Mediavilla Gradolph, Begoña Palop Borrás and SeqCOVID-SPAIN consortium |
| EPI_ISL_452797, EPI_ISL_452798, EPI_ISL_452799, EPI_ISL_452800, EPI_ISL_452801, EPI_ISL_452802, EPI_ISL_452803, EPI_ISL_452804, EPI_ISL_452805, EPI_ISL_452806, EPI_ISL_452807, EPI_ISL_452808, EPI_ISL_452810 | | | |
| see above | Virginia DCLS | Virginia DCLS | Virginia DCLS |
| EPI_ISL_453008, EPI_ISL_453009, EPI_ISL_453010, EPI_ISL_453011, EPI_ISL_453012 | West of Scotland Specialist Virology Centre, NHSGGC / MRC-University of Glasgow Centre for Virus Research | COVID-19 Genomics UK (COG-UK) Consortium | Ana da Silva Filipe, Natasha Johnson, Kathy Smollett, Daniel Mair, Stephen Carmichael, Lily Tong, Jenna Nichols, Elihu Aranday-Cortes, Kirstyn Brunker, Yasmin Parr, Kyriaki Nomikou; Sarah McDonald, Marc Niebel, Patawee Asamaphan; Richard Orton, Joseph Hughes, Sreenu Vattipally, David L Robertson; Alasdair MacLean, Rory Gunson; Kathy Li, Natasha Jesudason, Rajiv Shah, James Shepherd, Antonia Ho, Emma Thomson |
| EPI_ISL_453100, EPI_ISL_453156, EPI_ISL_453158, EPI_ISL_453159, EPI_ISL_453160, EPI_ISL_453161, EPI_ISL_453162, EPI_ISL_453163 | Virology Department, Royal Infirmary of Edinburgh, NHS Lothian / School of Biological Sciences, University of Edinburgh / Institute of Genetics and Molecular Medicine, University of Edinburgh | COVID-19 Genomics UK (COG-UK) Consortium | McHugh M, Dewar R, Rooke S, Gallagher M, Balcaza C, O'Toole Á, Scher E, Hill V, McCrone JT, Colquhoun R, Yu X, Jackson B, Rambaut A, Williams TC, Templeton K |
| EPI_ISL_453240, EPI_ISL_453241, EPI_ISL_453242, EPI_ISL_453243, EPI_ISL_453244, EPI_ISL_453245, EPI_ISL_453246, EPI_ISL_453247, EPI_ISL_453248, EPI_ISL_453254, EPI_ISL_453255, EPI_ISL_453256, EPI_ISL_453257, EPI_ISL_453259, EPI_ISL_453260, EPI_ISL_453261, EPI_ISL_453262, EPI_ISL_453263, EPI_ISL_453264, EPI_ISL_453265, EPI_ISL_453266, EPI_ISL_453267, EPI_ISL_453268, EPI_ISL_453269, EPI_ISL_453270, EPI_ISL_453271, EPI_ISL_453272, EPI_ISL_453273, EPI_ISL_453274, EPI_ISL_453275, EPI_ISL_453276, EPI_ISL_453277, EPI_ISL_453278, EPI_ISL_453279, EPI_ISL_453280, EPI_ISL_453281, EPI_ISL_453282, EPI_ISL_453283, EPI_ISL_453284, EPI_ISL_453285, EPI_ISL_453286, EPI_ISL_453287, EPI_ISL_453288, EPI_ISL_453289, EPI_ISL_453290, EPI_ISL_453291, EPI_ISL_453292, EPI_ISL_453293, EPI_ISL_453294, EPI_ISL_453295, EPI_ISL_453296, EPI_ISL_453297, EPI_ISL_453298, EPI_ISL_453299, EPI_ISL_453300, EPI_ISL_453301, EPI_ISL_453302, EPI_ISL_453303, EPI_ISL_453304, EPI_ISL_453305, EPI_ISL_453306, EPI_ISL_453307, EPI_ISL_453308, EPI_ISL_453309, EPI_ISL_453310, EPI_ISL_453311, EPI_ISL_453312, EPI_ISL_453313, EPI_ISL_453314, EPI_ISL_453328, EPI_ISL_453329, EPI_ISL_453331, EPI_ISL_453360, EPI_ISL_453361, EPI_ISL_453362, EPI_ISL_453363, EPI_ISL_453364, EPI_ISL_453365, EPI_ISL_453366, EPI_ISL_453367, EPI_ISL_453368, EPI_ISL_453369, EPI_ISL_453370, EPI_ISL_453371, EPI_ISL_453372, EPI_ISL_453373, EPI_ISL_453374, EPI_ISL_453375, EPI_ISL_453376, EPI_ISL_453377, EPI_ISL_453378, EPI_ISL_453379, EPI_ISL_453380, EPI_ISL_453381, EPI_ISL_453382, EPI_ISL_453383, EPI_ISL_453384, EPI_ISL_453385, EPI_ISL_453386, EPI_ISL_453387, EPI_ISL_453388, EPI_ISL_453389, EPI_ISL_453390, EPI_ISL_453391, EPI_ISL_453392, EPI_ISL_453394, EPI_ISL_453395, EPI_ISL_453396, EPI_ISL_453397, EPI_ISL_453398, EPI_ISL_453399, EPI_ISL_453400, EPI_ISL_453401, EPI_ISL_453426, EPI_ISL_453427, EPI_ISL_453428, EPI_ISL_453429, EPI_ISL_453430 | | | |
| see above | Liverpool Clinical Laboratories | COVID-19 Genomics UK (COG-UK) Consortium | Sam Haldenby, Anita Lucaci, Steve Paterson, Julian Hiscox, Alistair Darby, M Almsaud, A Alrezaihi, Muhannad Alruwaili, Stuart D Armstrong, Jones Benjamin , Eleanor G Bentley, Anu Chawla, Jordan J Clark, Angela Cowell, Richard Eccles, Isabel García-Dorival, Matthew Gemmell, Alessandro Gerada, PKF Gilmore, Richard Gregory, Ximeng Han, Catherine Hartley, Margaret Hughes, Miren Iturriza-Gomara, James Johnson, L Luu, Jenifer Manson , Charlotte Nelson, Elaine O'Toole, Cassie Olateju, Rebekah Penrice-Randal , Lucille Rainbow, N.P Randle, Trevor Ian Robinson, Parul Sharma, Ghada T Shawli, James P Stewart , Neil Swainston, Ecaterina Vamos, Joanne Watts, Mark Whitehead |
| EPI_ISL_453507, EPI_ISL_453508, EPI_ISL_453509, EPI_ISL_453510, EPI_ISL_453511, EPI_ISL_453512, EPI_ISL_453513, EPI_ISL_453514, EPI_ISL_453515, EPI_ISL_453516, EPI_ISL_453517, EPI_ISL_453518 | | | |
| see above | Northumbria University / South Tees Hospitals NHS Foundation Trust / North Cumbria Integrated Care NHS Foundation Trust / North Tees and Hartlepool NHS Foundation Trust / Newcastle Hospitals NHS Foundation Trust | COVID-19 Genomics UK (COG-UK) Consortium | Darren L Smith,Andrew Nelson,Matthew Bashton,Greg R Young,Joshua Loh,John Allan,Mohammad A Tariq,Giles S Holt,Gary Black,Wen C Yew,Lynn Dover ,Paul Baker,Steve Liggett,Sarah Essex,Jane Greenaway ,Debra Padgett,Clive Graham,Garren Scott,Edward Barton ,Emma Swindells ,Brendan Payne,Jennifer Collins,Yusri Taha,Gary Eltringham |
| EPI_ISL_453960, EPI_ISL_453961, EPI_ISL_453962, EPI_ISL_453963, EPI_ISL_453964, EPI_ISL_453965, EPI_ISL_453966, EPI_ISL_453967, EPI_ISL_453968, EPI_ISL_453969, EPI_ISL_453970, EPI_ISL_453971, EPI_ISL_453972, EPI_ISL_453973, EPI_ISL_453974, EPI_ISL_453975, EPI_ISL_453976, EPI_ISL_453977, EPI_ISL_453978, EPI_ISL_453979, EPI_ISL_453980, EPI_ISL_453981, EPI_ISL_453982, EPI_ISL_453983, EPI_ISL_453984, EPI_ISL_453985, EPI_ISL_453986, EPI_ISL_453987, EPI_ISL_453988, EPI_ISL_453989, EPI_ISL_453990, EPI_ISL_453991, EPI_ISL_453992, EPI_ISL_453993, EPI_ISL_453994, EPI_ISL_453995, EPI_ISL_453996, EPI_ISL_454175, EPI_ISL_454176, EPI_ISL_454177, EPI_ISL_454250, EPI_ISL_454251, EPI_ISL_454252, EPI_ISL_454253, EPI_ISL_454254, EPI_ISL_454271, EPI_ISL_454317, EPI_ISL_454324, EPI_ISL_454351, EPI_ISL_454352 | | | |
| see above | unknown | Instituto Nacional de Saude (INSA) | Borges et al |
| EPI_ISL_454371, EPI_ISL_454375, EPI_ISL_454379, EPI_ISL_454387, EPI_ISL_454401, EPI_ISL_454402, EPI_ISL_454410 | UPMC Clinical Microbiology Laboratory | Microbial Genome Sequencing Center, Microbial Genomic Epidemiological Laboratory | Mustapha M. Mustapha, Jane W. Marsh, Dan Snyder, Marissa P. Griffith, Stephanie L. Mitchell, Vatsala R. Srinivasa, Kady D. Waggle, Chinelo Ezeonwuku, Vaughn S. Cooper, Lee H. Harrison |
| EPI_ISL_454544, EPI_ISL_454545, EPI_ISL_454546, EPI_ISL_454547, EPI_ISL_454548, EPI_ISL_454549, EPI_ISL_454550, EPI_ISL_454552, EPI_ISL_454553, EPI_ISL_454554, EPI_ISL_454560, EPI_ISL_454564 | | | |
| see above | NIV Influenza | NIV Influenza | Potdar V |
| EPI_ISL_454641 | Humboldt County Public Health Laboratory | Chan-Zuckerberg Biohub | CZB Cliahub Consortium |
| EPI_ISL_454732 | Russian State Collection of Viruses | Pathogenic Microorganisms Variability Laboratory | Denis Protsenko, Alexey Shchetinin, Maria Nikiforova, Elena Shidlovskaya, Nadezhda Kuznetsova, Vladimir Gushchin, Inna Dolzhikova, Daria Grousova, Andrey Botikov, Denis Logunov, Alexander Gintsburg, Alexey Mazus |
| EPI_ISL_454776, EPI_ISL_454778, EPI_ISL_454779, EPI_ISL_454780, EPI_ISL_454781 | Dutch COVID-19 response team | National Institute for Public Health and the Environment (RIVM) | Adam Meijer, Harry Vennema, Jeroen Cremer, Sharon van den Brink, Pieter Overduin, Florian Zwagemaker, Dennis Schmitz, Chantal Reusken, on behalf of the national COVID-19 response team |
| EPI_ISL_455047 | Pathology West - NSW Health Pathology | NSW Health Pathology - Institute of Clinical Pathology and Medical Research; Westmead Hospital; University of Sydney | CIDM-PH et al. |
| EPI_ISL_455083, EPI_ISL_455084, | South Eastern Area Laboratory Services | NSW Health Pathology - Institute of Clinical Pathology and | CIDM-PH et al. |

| | | | |
|---|---|---|---|
| EPI_ISL_455086 | | Medical Research; Westmead Hospital; University of Sydney | |
| EPI_ISL_455120, EPI_ISL_455121, EPI_ISL_455124, EPI_ISL_455125, EPI_ISL_455126, EPI_ISL_455127, EPI_ISL_455128, EPI_ISL_455129, EPI_ISL_455130, EPI_ISL_455131, EPI_ISL_455132, EPI_ISL_455133, EPI_ISL_455134, EPI_ISL_455135, EPI_ISL_455136, EPI_ISL_455137, EPI_ISL_455138, EPI_ISL_455139, EPI_ISL_455140, EPI_ISL_455141, EPI_ISL_455142, EPI_ISL_455143, EPI_ISL_455144, EPI_ISL_455145, EPI_ISL_455146, EPI_ISL_455147, EPI_ISL_455148, EPI_ISL_455149, EPI_ISL_455150, EPI_ISL_455151, EPI_ISL_455152, EPI_ISL_455153, EPI_ISL_455154, EPI_ISL_455155, EPI_ISL_455156, EPI_ISL_455157, EPI_ISL_455195, EPI_ISL_455197, EPI_ISL_455198, EPI_ISL_455199, EPI_ISL_455200, EPI_ISL_455203, EPI_ISL_455226, EPI_ISL_455227, EPI_ISL_455228, EPI_ISL_455229, EPI_ISL_455230, EPI_ISL_455231, EPI_ISL_455232, EPI_ISL_455233, EPI_ISL_455234, EPI_ISL_455235, EPI_ISL_455250, EPI_ISL_455251, EPI_ISL_455252, EPI_ISL_455261, EPI_ISL_455265, EPI_ISL_455266, EPI_ISL_455290, EPI_ISL_455291, EPI_ISL_455294, EPI_ISL_455295, EPI_ISL_455296 | | | |
| see above | Dutch COVID-19 response team | Erasmus Medical Center | Bas Oude Munnink, David Nieuwenhuijse, Reina Sikkema, Claudia Schapendonk, Irina Chestakova, Anne van der Linden, Theo Bestebroer, Stefan van Nieuwkoop, Mark Pronk, Pascal Lexmond, Corien Swaan, Manon Haverkate, Madelief Mollers, Mart Stein, Sandra Kengne Kamga Mobou, Jeroen van Kampen, Jolanda Voermans, Aura Timen, Corine GeurtsvanKessel, Annemiek van der Eijk, Richard Molenkamp, Marion Koopmans, on behalf of the Dutch national COVID-19 response team. |
| EPI_ISL_455308, EPI_ISL_455309 | REGIONAL VRDL,ICMR-RMRC BBSR | Immunogenomics group, Institute of Life Sciences, Bhubaneswar | Sunil Raghav, Jyotirmayee Turuk, Arup Ghosh, Atimukta Jha, Viplov K. Biswas, Swati Madhulika, Manasi Priyadarshini, Shuchi Smita, Jaya Singh Khastri, Rupesh Dash, Soma Chattopadhyay, Ghulam Hussain Syed, Shanti Senapati, Tushar K. Beuria, Debdutta Bhattacharya, Rajeeb Swain, Punit Prasad, COVID-19 team of ILS & RMRC, Orissa COVID-19 study group, DBT's PAN-INDIA 1000 SARS-CoV2 RNA genome sequencing consortium, Sanghamitra Pati, Ajay Parida |
| EPI_ISL_455313 | Microbiology Unit, Department of Pathology & Laboratory Medicine, IIUM Medical Centre | SEA Microbiome Unit, Faculty of Industrial Sciences & Technology, Universiti Malaysia Pahang | Hajar Fauzan Ahmad, Norhidayah Kamarudin, Ahmad Hafiz Zulkifly, IIUM Medical Centre Covid19 Taskforce, UMP Covid19 Team |
| EPI_ISL_455358 | Emory Molecular Diagnostics Laboratory, Emory Healthcare | Piantadosi Lab, Emory Department of Pathology | Ahmed Babiker, Anne Piantadosi |
| EPI_ISL_455430, EPI_ISL_455431 | Nigeria Centre for Disease Control (NCDC) | African Centre of Excellence for Genomics of Infectious Diseases (ACEGID), Redeemer's University, Ede, Osun State, Nigeria | Oluniyi P.E., Ajogbasile F.V., Kayode A., Oguzie J., Olawoye I., Uwanibe J., Olumade T., Folarin O.A., Ihekweazu C., Happi C.T. |
| EPI_ISL_455597 | SA Pathology | VPRL | Beard, MR., Van Der Hoek, K., Lim, C.K., Leong, L.E.X., Coldbeck-Shackley, R., Shue, B., Kirby, E., Merrett, J., Llamas, B. |
| EPI_ISL_455609, EPI_ISL_455611, EPI_ISL_455615, EPI_ISL_455617, EPI_ISL_455619, EPI_ISL_455620, EPI_ISL_455621, EPI_ISL_455623 | Ochsner Health | Bioinfoexperts, LLC | Susanna L. Lamers, David J. Nolan, Rebecca Rose, Sissy Cross, David Moraga Amador, Tong Yang, Luke Caruso, Wayra Navia, Lydia Von Borstel, Xiao Hui Zhou, Amy Feehan, Julia-Garcia-Diaz |
| EPI_ISL_455627 | unknown | Instituto Nacional de Saude (INSA) | Borges et al |
| EPI_ISL_455642, EPI_ISL_455644, EPI_ISL_455645, EPI_ISL_455646 | ICMR-National Institute of Cholera and Enteric Diseases | National Institute of Biomedical Genomics | Arindam Maitra, Mamta Chawla Sarkar, Sreedhar Chinnaswamy, Hasina Banu, Ananya Chatterjee, Shanta Dutta, Saumitra Das |
| EPI_ISL_455707 | National Hospital of Tropical Diseases | Oxford University Clinical Research Unit, Hanoi, Vietnam | Nguyen Thi Tam, Van Dinh Trang, Nguyen Thu Trang, Nguyen Thi Ngoc Diep, Le Nguyen Minh Hoa, Pham Ngoc Thach, H. Rogier van Doorn, on behalf of the OUCRU COVID-19 research group |
| EPI_ISL_455772, EPI_ISL_455773, EPI_ISL_455774, EPI_ISL_455775, EPI_ISL_455776, EPI_ISL_455777, EPI_ISL_455778, EPI_ISL_455779, EPI_ISL_455782, EPI_ISL_455783, EPI_ISL_455784 | | | |
| see above | REGIONAL VRDL,ICMR-RMRC BBSR | Immunogenomics lab, Institute of Life Sciences, Bhubaneswar | Sunil Raghav, Jyotirmayee Turuk, Arup Ghosh, Atimukta Jha, Viplov K. Biswas, Swati Madhulika, Manasi Priyadarshini, Shuchi Smita, Jaya Singh Khastri, Rupesh Dash, Soma Chattopadhyay, Ghulam Hussain Syed, Shanti Senapati, Tushar K. Beuria, Debdutta Bhattacharya, Rajeeb Swain, Punit Prasad, COVID-19 team of ILS & RMRC, Orissa COVID-19 study group, DBT's PAN-INDIA 1000 SARS-CoV2 RNA genome sequencing consortium, Sanghamitra Pati, Ajay Parida |
| EPI_ISL_456075, EPI_ISL_456079, EPI_ISL_456080, EPI_ISL_456081 | Laboratory of Respiratory Viruses and Measles, Oswaldo Cruz Institute, FIOCRUZ | Laboratory of Respiratory Viruses and Measles, Oswaldo Cruz Institute, FIOCRUZ | Paola Resende, Luciana Appolinario, Fernando Motta, Aline Mattos, Milene Miranda, Cristiana Garcia, Braulia Caetano, Maria Ogrzewalska, Jonathan Lopes, Marilda Siqueira |
| EPI_ISL_456082, EPI_ISL_456083, EPI_ISL_456088 | LACEN RJ - Laboratório Central de Saúde Pública Noel Nutels | Laboratory of Respiratory Viruses and Measles, Oswaldo Cruz Institute, FIOCRUZ | Paola Resende, Luciana Appolinario, Fernando Motta, Aline Mattos, Milene Miranda, Cristiana Garcia, Braulia Caetano, Maria Ogrzewalska, Jonathan Lopes, Marilda Siqueira |
| EPI_ISL_456156 | Instituto Nacional de Salud - Unidad de Secuenciación y Análisis Genómico | Instituto Nacional de Salud, Universidad Cooperativa de Colombia, Instituto Alexander von Humboldt, Imperial College-London, London School of Hygiene & Tropical Medicine | Katherine Laiton-Donato, Diego A. Álvarez-Díaz, Carlos Franco-Muñoz, Jose A. Usme-Ciro, Gloria Puerto, Nicolas D. Franco-Sierra, Mailyn A.Gonzalez, Zulma M. Cucunubá, Christian Julian VillabonaArenas, Liz Villabona-Arenas, Sussy Echeverria, Astrid C. Flórez, Sergio Gomez-Rangel, Luz Dary Rodriguez, Juliana Barbosa, Erika Ospitia, Diana Marcela Walteros-Acero, Martha Lucia Ospina Martinez, Marcela Mercado-Reyes. |
| EPI_ISL_456294 | North Shore Hospital | Institute of Environmental Science and Research (ESR) | Matt Storey, Xiaoyun Ren, Anja Werno, Antje van der Linden, Arlo Upton, Chris Mansell, David Hammer, Dragana Drinkovic, Erasmus Smit, Gary McAuliffe, Hana Sofia Andersson, James Ussher, Jill Sherwood, Josh Freeman, Julia Howard, Juliet Elvy, Mary DeAlmeida, Matt Blakiston, Matthew Rogers, Max Bloomfield, Michael Addidle, Michelle Balm, Sally Roberts, Sarah Jefferies, Sharmini Muttaiyah, Susan Morpeth, Susan Taylor, Timothy Blackmore, Vani Sathyendran, Veronica Playle, Virginia Hope, Erasmus Smit, Lauren Jelly, Joep de Ligt |
| EPI_ISL_456301 | PathLab Bay of Plenty | Institute of Environmental Science and Research (ESR) | Matt Storey, Xiaoyun Ren, Anja Werno, Antje van der Linden, Arlo Upton, Chris Mansell, David Hammer, Dragana Drinkovic, Erasmus Smit, Gary McAuliffe, Hana Sofia Andersson, James Ussher, Jill Sherwood, Josh Freeman, Julia Howard, Juliet Elvy, Mary DeAlmeida, Matt Blakiston, Matthew Rogers, Max Bloomfield, Michael Addidle, Michelle Balm, Sally Roberts, Sarah Jefferies, Sharmini Muttaiyah, Susan Morpeth, Susan Taylor, Timothy Blackmore, Vani Sathyendran, Veronica Playle, Virginia Hope, Erasmus Smit, Lauren Jelly, Joep de Ligt |
| EPI_ISL_456302, EPI_ISL_456303, EPI_ISL_456305, EPI_ISL_456306, EPI_ISL_456309, EPI_ISL_456310, EPI_ISL_456311, EPI_ISL_456312, EPI_ISL_456313, EPI_ISL_456314, EPI_ISL_456315, EPI_ISL_456317, EPI_ISL_456318 | | | |
| see above | Southern Community Labs Dunedin | Institute of Environmental Science and Research (ESR) | Matt Storey, Xiaoyun Ren, Anja Werno, Antje van der Linden, Arlo Upton, Chris Mansell, David Hammer, Dragana Drinkovic, Erasmus Smit, Gary McAuliffe, Hana Sofia Andersson, James Ussher, Jill Sherwood, Josh Freeman, Julia Howard, Juliet Elvy, Mary DeAlmeida, Matt Blakiston, Matthew Rogers, Max Bloomfield, Michael Addidle, Michelle Balm, Sally Roberts, Sarah Jefferies, Sharmini Muttaiyah, Susan Morpeth, Susan Taylor, Timothy Blackmore, Vani Sathyendran, Veronica Playle, Virginia Hope, Erasmus Smit, Lauren Jelly, Joep de Ligt |
| EPI_ISL_456348, EPI_ISL_456349, EPI_ISL_456350, EPI_ISL_456351, EPI_ISL_456352, EPI_ISL_456353, EPI_ISL_456354, EPI_ISL_456355, EPI_ISL_456356, EPI_ISL_456357, EPI_ISL_456358, EPI_ISL_456373 | | | |
| see above | Canterbury Health Laboratories | Institute of Environmental Science and Research (ESR) | Matt Storey, Xiaoyun Ren, Anja Werno, Antje van der Linden, Arlo Upton, Chris Mansell, David Hammer, Dragana Drinkovic, Erasmus Smit, Gary McAuliffe, Hana Sofia Andersson, James Ussher, Jill Sherwood, Josh Freeman, Julia Howard, Juliet Elvy, Mary DeAlmeida, Matt Blakiston, Matthew Rogers, Max Bloomfield, Michael Addidle, Michelle Balm, Sally Roberts, Sarah Jefferies, Sharmini Muttaiyah, Susan Morpeth, Susan Taylor, Timothy Blackmore, Vani Sathyendran, Veronica Playle, Virginia Hope, Erasmus Smit, Lauren Jelly, Joep de Ligt |
| EPI_ISL_456376 | Middlemore Hospital | Institute of Environmental Science and Research (ESR) | Matt Storey, Xiaoyun Ren, Anja Werno, Antje van der Linden, Arlo Upton, Chris Mansell, David Hammer, Dragana Drinkovic, Erasmus Smit, Gary McAuliffe, Hana Sofia Andersson, James Ussher, Jill Sherwood, Josh Freeman, Julia Howard, Juliet Elvy, Mary DeAlmeida, Matt Blakiston, Matthew Rogers, Max Bloomfield, Michael Addidle, Michelle Balm, Sally Roberts, Sarah Jefferies, Sharmini Muttaiyah, Susan Morpeth, Susan Taylor, Timothy Blackmore, Vani Sathyendran, Veronica Playle, Virginia Hope, Erasmus Smit, Lauren Jelly, Joep de Ligt |
| EPI_ISL_456890, EPI_ISL_456907, EPI_ISL_456908, EPI_ISL_456909, EPI_ISL_456910, EPI_ISL_456911, EPI_ISL_456913, EPI_ISL_456951, EPI_ISL_456952, EPI_ISL_457016, EPI_ISL_457017, EPI_ISL_457018, EPI_ISL_457019, EPI_ISL_457020, EPI_ISL_457021, EPI_ISL_457022, EPI_ISL_457023 | | | |
| see above | Virology Department, Royal Infirmary of Edinburgh, NHS Lothian / School of Biological Sciences, University of Edinburgh / Institute of Genetics and Molecular Medicine, University of Edinburgh | COVID-19 Genomics UK (COG-UK) Consortium | McHugh M, Dewar R, Rooke S, Gallagher M, Balcaza C, O'Toole Á, Scher E, Hill V, McCrone JT, Colquhoun R, Yu X, Jackson B, Rambaut A, Williams TC, Templeton K |
| EPI_ISL_457031, EPI_ISL_457032, EPI_ISL_457033, EPI_ISL_457034, EPI_ISL_457035, EPI_ISL_457036, EPI_ISL_457037, EPI_ISL_457038, EPI_ISL_457039, EPI_ISL_457040, EPI_ISL_457041, EPI_ISL_457042, EPI_ISL_457043, EPI_ISL_457044, EPI_ISL_457045, EPI_ISL_457046, EPI_ISL_457047, EPI_ISL_457048, EPI_ISL_457054, EPI_ISL_457058, EPI_ISL_457063, EPI_ISL_457078, EPI_ISL_457080, EPI_ISL_457084, EPI_ISL_457096, EPI_ISL_457131, EPI_ISL_457145, EPI_ISL_457155, EPI_ISL_457160, EPI_ISL_457161, EPI_ISL_457165, EPI_ISL_457166, EPI_ISL_457180, EPI_ISL_457184 | | | |
| see above | University of Exeter | COVID-19 Genomics UK (COG-UK) Consortium | Ben Temperton,Aaron Jeffries,Michelle Michelsen,Joanna Warwick-Dugdale,Audrey Farbos,Robyn Manley,Stephen Michell,Jane Masoli |
| EPI_ISL_457587, EPI_ISL_457595, | Virology Department, Sheffield Teaching Hospitals NHS | COVID-19 Genomics UK (COG-UK) Consortium | Thushan de Silva, Matthew Parker, Nikki Smith, Adri Angyal, Rebecca Brown, Luke Green, Rachel Tucker, Paul Parsons, Danielle Groves, Katie Johnson, |

| | | | |
|---|---|---|---|
| EPI_ISL_457601, EPI_ISL_457643, EPI_ISL_457653, EPI_ISL_457681 | Foundation Trust/Department of Infection, Immunity and Cardiovascular Disease, The Medical School, University of Sheffield | | Laura Carrilero, Alex Keeley, Dave Partridge, Matthew Wyles, Benjamin Lindsey, Mehmet Yavuz, Mohammad Raza, Cariad Evans |
| EPI_ISL_457868, EPI_ISL_457871, EPI_ISL_457873, EPI_ISL_457874, EPI_ISL_457875, EPI_ISL_457876, EPI_ISL_457886, EPI_ISL_457890, EPI_ISL_457895 | KEMRI-CGMR-C | KEMRI-Wellcome Trust Research Programme/KEMRI-CGMR-C Kilifi | Githinji G. et al 2020 |
| EPI_ISL_457965, EPI_ISL_457966 | Laboratorio de Biología Molecular Asociación Española Primera en Salud | Departments of Pathology and Medicine, New York University School of Medicine | Maria Victoria Elizondo, Maria Noel Zubillaga, Gonzalo Manrique, Paul Zappile, Gael Westby, Matthew T Maurano, Christian Marier, Adriana Heguy |
| EPI_ISL_457981 | Oman-NIC | Department of Microbiology and Immunology-SQUH | Fahad Zadjali, Samira Al-Maruqi, Amina Al Jardani, Khulood Al-Mammary, Hanan Al-kindi, Fatma BaAlawi, Hamida AL Barwani, Zeyana AL-Dahmani, Intisar Al-Shukri, Aisha Al-Busaidi, Aisha Al-Amri, Ahlam Al-Amri, Mohammed Al-Tobi, Samiha Al Kharusi, Abdulla Balkhair |
| EPI_ISL_458026, EPI_ISL_458027, EPI_ISL_458028 | Hospital for Tropical Diseases | COVID-19 Network Investigations (CONI) Alliance | Elizabeth Batty, Nantarat Chantawat, Wasun Chantratita, Thanat Chookajorn, Stefan Fernandez, Angkana Huang, Weena Janwitthayanan, Akanitt Jittmittraphap, Anthony R. Jones, Khajohn Joonsalak, Chonticha Klungtong, Theerarat Kochakarn, Namfon Kotanan, Krittikorn Kumpornsin, Pornsawan Leaungwutiwong, Wudtichai Manasatienkij, Bhakbhoom Panthan, Ekawat Pasomsub, Kingkan Rakmanee, Insee Sensorn, Janjira Thaipadungpanit, Arporn Wangwiwatsin,Treewat Watthanachockchai |
| EPI_ISL_458069 | Osmania Medical College | CSIR-Centre for Cellular and Molecular Biology | Shashikala Reddy, Mahboob Khan,Payel Mukherjee, Sofia Banu, Priya Singh, Dhiviya Vedagiri, Divya Gupta, Vishal Sah, Santosh Kumar Kuncha, Krishnan Harinivas Harshan, Archana Bharadwaj Siva, Karthik Bharadwaj Tallapaka, Shagufta Khan, Lamuk Zaveri, Namami Gaur, Sakshi Shambhavi, Tulasi Nagabandi, Purushotham Vodnala, Rakesh K Mishra, Divya Tej Sowpati |
| EPI_ISL_458080 | CSIR-Centre for Cellular and Molecular Biology | CSIR-Centre for Cellular and Molecular Biology | Sakshi Shambhavi, Lamuk Zaveri, Shagufta Khan, Namami Gaur, Tulasi Nagabandi, Purushotham Vodnala, Payel Mukherjee, Sofia Banu, Priya Singh, Dhiviya Vedagiri, Divya Gupta, Vishal Sah, Santosh Kumar Kuncha, Krishnan Harinivas Harshan, Archana Bharadwaj Siva, Karthik Bharadwaj Tallapaka, G. Aditya Kumar, Koushick Sivakumar, Pooja Ramesh Gupta, Rajan Kumar Jha, Shraddha Vijay Lahoti, Rakesh K Mishra, Divya Tej Sowpati |
| EPI_ISL_458084 | Laboratorio Biologia Molecolare Sars Cov2 - UOC Laboratorio Analisi - Servizio Medicina di Laboratorio, Ospedale "San Francesco" - ATS-ASSL Nuoro | Laboratorio specialistico UOC Ematologia - Ospedale "San Francesco" - ATS-ASSL Nuoro | Piras Giovanna, Fancello Tatiana, Asproni Rosanna, Fiamma Maura, Monne Maria Itria, Toja Alessandro, Sanna Filomena, Floris Anna Rita, Sulis Vincenzo, Palmas Angelo Domenico, Casu Gavino, Lo Maglio Iana, Mameli Giuseppe. |
| EPI_ISL_458123, EPI_ISL_458127 | Oman National Influenza Centre | Department of Microbiology and Immunology-SQUH | Fahad Zadjali, Samira Al-Maruqi, Amina Al Jardani, Khulood Al-Mammary, Hanan Al-kindi, Fatma BaAlawi, Hamida AL Barwani, Zeyana AL-Dahmani, Intisar Al-Shukri, Aisha Al-Busaidi, Aisha Al-Amri, Ahlam Al-Amri, Mohammed Al-Tobi, Samiha Al Kharusi, Abdulla Balkhair |
| EPI_ISL_458138, EPI_ISL_458142, EPI_ISL_458149 | Evandro Chagas Institute | Evandro Chagas Institute | Santos, M.C.; Silva, A.M.; Junior, W.D.C.; Barbagelata, L.S.; Ferreira, J.A.; Sousa, E.M.A.; da Silva, P.S.; Resque, H.R; Martins, L.C.; Sousa Junior, E.C.;Viana, G.M.R |
| EPI_ISL_458240, EPI_ISL_458245, EPI_ISL_458250, EPI_ISL_458252, EPI_ISL_458261, EPI_ISL_458270, EPI_ISL_458276, EPI_ISL_458281 | Scripps Medical Laboratory | Andersen lab at Scripps Research | SEARCH Alliance San Diego with Michael Quigley, Ellen Stefanski, Ian Mchardy |
| EPI_ISL_458298 | CSIR-Centre for Cellular and Molecular Biology | CSIR-Centre for Cellular and Molecular Biology | Sakshi Shambhavi, Lamuk Zaveri, Shagufta Khan, Namami Gaur, Tulasi Nagabandi, Purushotham Vodnala, Payel Mukherjee, Sofia Banu, Priya Singh, Dhiviya Vedagiri, Divya Gupta, Vishal Sah, Santosh Kumar Kuncha, Krishnan Harinivas Harshan, Archana Bharadwaj Siva, Karthik Bharadwaj Tallapaka, Deepak Kumar, Devi Prasad Vijayashankar, Disha Nanda, Divya Das, Jotin Gogoi, Manish Bhattacharjee, Rakesh K Mishra, Divya Tej Sowpati |
| EPI_ISL_458578, EPI_ISL_458610, EPI_ISL_458672 | NU-OMICS DNA Sequencing research facility, Northumbria University | Wellcome Sanger Institute for the COVID-19 Genomics UK (COG-UK) consortium | Chris Duncan, Sheia Waugh, Shirelle Burton-Fanning, Gary Eltringham, Jennifer Collins, Brendan Payne, Yusri Taha, Emma Swindells, Jane Greenaway, Edward Barton, Garren Scott, Debra Padgett, Clive Graham, Sarah Essex, Steve Ligget, Paul Baker, Lynn Dover, Wen Yew, Gary Black, John Allan, Joshua Loh, Greg Young, Matthew Bashton, Andrew Nelson, Darren Smith and Alex Alderton, Roberto Amato, Sonia Goncalves, Ewan Harrison, David K. Jackson, Ian Johnston, Dominic Kwiatkowski, Cordelia Langford, John Sillitoe on behalf of the Wellcome Sanger Institute COVID-19 Surveillance Team (http://www.sanger.ac.uk/covid-team) |
| EPI_ISL_458965 | PHE South West Regional Laboratory, National Infection Service | Wellcome Sanger Institute for the COVID-19 Genomics UK (COG-UK) consortium | Stephanie Hutchings, Hannah Pymont, Dr Peter Muir, Barry Vipond, Rich Hopes; and Alex Alderton, Roberto Amato, Sonia Goncalves, Ewan Harrison, David K. Jackson, Ian Johnston, Dominic Kwiatkowski, Cordelia Langford, John Sillitoe on behalf of the Wellcome Sanger Institute COVID-19 Surveillance Team (http://www.sanger.ac.uk/covid-team) |
| EPI_ISL_459095, EPI_ISL_459096, EPI_ISL_459099, EPI_ISL_459100, EPI_ISL_459101, EPI_ISL_459102, EPI_ISL_459103, EPI_ISL_459104 | NHSGGC West of Scotland Specialist Virology Centre / MRC-University of Glasgow Centre for Virus Research | Wellcome Sanger Institute for the COVID-19 Genomics UK (COG-UK) consortium | Ana da Silva Filipe, Natasha Johnson, Kathy Smollett, Daniel Mair, Stephen Carmichael, Lily Tong, Jenna Nichols, Elihu Aranday-Cortes, Kirstyn Brunker, Yasmin Parr, Kyriaki Nomikou; Sarah McDonald, Marc Niebel, Patawee Asamaphan; Richard Orton, Joseph Hughes, Sreenu Vattipally, David L Robertson; Alasdair MacLean, Rory Gunson; Kathy Li, Natasha Jesudason, Rajiv Shah, James Shepherd, Antonia Ho, Alice Broos, Emma Thomson and Alex Alderton, Roberto Amato, Sonia Goncalves, Ewan Harrison, David K. Jackson, Ian Johnston, Dominic Kwiatkowski, Cordelia Langford, John Sillitoe on behalf of the Wellcome Sanger Institute COVID-19 Surveillance Team (http://www.sanger.ac.uk/covid-team) |
| EPI_ISL_459108 | NHSGGC West of Scotland Specialist Virology Centre / MRC-University of Glasgow Centre for Virus Research | Wellcome Sanger Institute for the COVID-19 Genomics UK (COG-UK) Consortium | Ana da Silva Filipe, Natasha Johnson, Kathy Smollett, Daniel Mair, Stephen Carmichael, Lily Tong, Jenna Nichols, Elihu Aranday-Cortes, Kirstyn Brunker, Yasmin Parr, Kyriaki Nomikou; Sarah McDonald, Marc Niebel, Patawee Asamaphan; Richard Orton, Joseph Hughes, Sreenu Vattipally, David L Robertson; Alasdair MacLean, Rory Gunson; Kathy Li, Natasha Jesudason, Rajiv Shah, James Shepherd, Antonia Ho, Alice Broos, Emma Thomson and Alex Alderton, Roberto Amato, Sonia Goncalves, Ewan Harrison, David K. Jackson, Ian Johnston, Dominic Kwiatkowski, Cordelia Langford, John Sillitoe on behalf of the Wellcome Sanger Institute COVID-19 Surveillance Team |
| EPI_ISL_459109, EPI_ISL_459115, EPI_ISL_459116, EPI_ISL_459117, EPI_ISL_459118, EPI_ISL_459120, EPI_ISL_459123, EPI_ISL_459124, EPI_ISL_459125, EPI_ISL_459126, EPI_ISL_459131, EPI_ISL_459134, EPI_ISL_459135, EPI_ISL_459137, EPI_ISL_459139, EPI_ISL_459141, EPI_ISL_459145, EPI_ISL_459150, EPI_ISL_459151, EPI_ISL_459153, EPI_ISL_459154, EPI_ISL_459156, EPI_ISL_459157, EPI_ISL_459158, EPI_ISL_459160, EPI_ISL_459162, EPI_ISL_459163, EPI_ISL_459164 | | | |
| see above | NHSGGC West of Scotland Specialist Virology Centre / MRC-University of Glasgow Centre for Virus Research | Wellcome Sanger Institute for the COVID-19 Genomics UK (COG-UK) consortium | Ana da Silva Filipe, Natasha Johnson, Kathy Smollett, Daniel Mair, Stephen Carmichael, Lily Tong, Jenna Nichols, Elihu Aranday-Cortes, Kirstyn Brunker, Yasmin Parr, Kyriaki Nomikou; Sarah McDonald, Marc Niebel, Patawee Asamaphan; Richard Orton, Joseph Hughes, Sreenu Vattipally, David L Robertson; Alasdair MacLean, Rory Gunson; Kathy Li, Natasha Jesudason, Rajiv Shah, James Shepherd, Antonia Ho, Alice Broos, Emma Thomson and Alex Alderton, Roberto Amato, Sonia Goncalves, Ewan Harrison, David K. Jackson, Ian Johnston, Dominic Kwiatkowski, Cordelia Langford, John Sillitoe on behalf of the Wellcome Sanger Institute COVID-19 Surveillance Team (http://www.sanger.ac.uk/covid-team) |
| EPI_ISL_459325, EPI_ISL_459326, EPI_ISL_459329, EPI_ISL_459335, EPI_ISL_459336, EPI_ISL_459360, EPI_ISL_459375, EPI_ISL_459391, EPI_ISL_459393 | Regional Virus Laboratory, Belfast Health and Social Care Trust | Wellcome Sanger Institute for the COVID-19 Genomics UK (COG-UK) consortium | Conall McCaughey, James McKenna, Tanya Curran, Susan Feeney, Alison Watt, Ciara Cox, Mairead Connor, Zoltan Molnar, David Simpson, Derek Fairley; and Alex Alderton, Roberto Amato, Sonia Goncalves, Ewan Harrison, David K. Jackson, Ian Johnston, Dominic Kwiatkowski, Cordelia Langford, John Sillitoe on behalf of the Wellcome Sanger Institute COVID-19 Surveillance Team (http://www.sanger.ac.uk/covid-team) |
| EPI_ISL_459984 | Institut Pasteur du Maroc | Institut Pasteur du Maroc | Marion Barbet, Sylvie Behillil, Méline Bizard, Angela Brisebarre, Camille Capel, Etienne Simon-Lorière, Vincent Enouf, Maud Vanpeene, Sylvie van der Werf, Latifa Anga, Abdellah Faouzi, Anass Abbad, Mjid Eloualid, Jalal Nourlil, Anderrahmane Maaroufi |
| EPI_ISL_460099, EPI_ISL_460121, EPI_ISL_460127, EPI_ISL_460132, EPI_ISL_460135, EPI_ISL_460139, EPI_ISL_460147, EPI_ISL_460152, EPI_ISL_460153, EPI_ISL_460159, EPI_ISL_460162, EPI_ISL_460165, EPI_ISL_460168, EPI_ISL_460173, EPI_ISL_460176, EPI_ISL_460177, EPI_ISL_460187, EPI_ISL_460188, EPI_ISL_460189, EPI_ISL_460190, EPI_ISL_460191, EPI_ISL_460197, EPI_ISL_460201, EPI_ISL_460208, EPI_ISL_460213, EPI_ISL_460215, EPI_ISL_460217, EPI_ISL_460220, EPI_ISL_460223, EPI_ISL_460225, EPI_ISL_460226, EPI_ISL_460227, EPI_ISL_460230, EPI_ISL_460233, EPI_ISL_460237, EPI_ISL_460244, EPI_ISL_460248, EPI_ISL_460254, EPI_ISL_460256, EPI_ISL_460257, EPI_ISL_460259, EPI_ISL_460268, EPI_ISL_460272, EPI_ISL_460284, EPI_ISL_460293, EPI_ISL_460296, EPI_ISL_460298, EPI_ISL_460299, EPI_ISL_460305, EPI_ISL_460307, EPI_ISL_460309, EPI_ISL_460318, EPI_ISL_460325, EPI_ISL_460327, EPI_ISL_460328, EPI_ISL_460330, EPI_ISL_460332, EPI_ISL_460336, EPI_ISL_460341, EPI_ISL_460345, EPI_ISL_460346, EPI_ISL_460351, EPI_ISL_460356, EPI_ISL_460357, EPI_ISL_460358, EPI_ISL_460360, EPI_ISL_460361, EPI_ISL_460365, EPI_ISL_460370, EPI_ISL_460373, EPI_ISL_460377, EPI_ISL_460378, EPI_ISL_460383, EPI_ISL_460385, EPI_ISL_460386, EPI_ISL_460389, EPI_ISL_460390, EPI_ISL_460392, EPI_ISL_460402, EPI_ISL_460403, EPI_ISL_460404, EPI_ISL_460405, EPI_ISL_460407, EPI_ISL_460412, EPI_ISL_460414, EPI_ISL_460418, EPI_ISL_460419, EPI_ISL_460420, EPI_ISL_460422, EPI_ISL_460429, EPI_ISL_460431, EPI_ISL_460435, EPI_ISL_460439, EPI_ISL_460441, EPI_ISL_460443, EPI_ISL_460450, EPI_ISL_460454, EPI_ISL_460455 | | | |
| see above | Massachusetts General Hospital | Infectious Disease Program, Broad Institute of Harvard and MIT | Lemieux,J.E., Siddle,K.J., Shaw,B., Adams,G., Pierce,V., Turbett,S., Anahtar,M., Branda,J., Slater,D., Harris,J., Lin,A.E., Gladden-Young,A., Lagerborg,K., Rudy,M., DeRuff,K., Carter,A., Normandin,E., Bauer,M., Reilly,S., Tomkins-Tinch,C., Loreth,C., Chaluvadi,S., Neumann,A., Cusick,C., Chapman,S.B., Gnirke,A., Flowers,K., Cerrato,F., Birren,B.W., Gallagher,G., Smole,S., Park,D.J., MacInnis,B.L., Ryan,E., LaRocque,R., Rosenberg,E., Sabeti,P.C. |

| | | | |
|---|---|---|---|
| EPI_ISL_460593 | Michigan Department of Health and Human Services, Bureau of Laboratories | Michigan Department of Health and Human Services, Bureau of Laboratories | Blankenship HM, Riner D, Soehnlen MK |
| EPI_ISL_460633, EPI_ISL_460634 | UW Virology Lab | UW Virology Lab | Pavitra Roychoudhury, Amin Addetia, Hong Xie, Lasata Shrestha, Truong Nguyen, Meei-Li Huang, Keith Jerome, Alexander Greninger |
| EPI_ISL_460636, EPI_ISL_460637, EPI_ISL_460643, EPI_ISL_460645, EPI_ISL_460651, EPI_ISL_460652, EPI_ISL_460660, EPI_ISL_460663, EPI_ISL_460670, EPI_ISL_460676, EPI_ISL_460679, EPI_ISL_460683, EPI_ISL_460705, EPI_ISL_460753, EPI_ISL_460795, EPI_ISL_460824, EPI_ISL_460825, EPI_ISL_460829, EPI_ISL_460830, EPI_ISL_460930, EPI_ISL_460941, EPI_ISL_461032, EPI_ISL_461035, EPI_ISL_461036, EPI_ISL_461038, EPI_ISL_461067, EPI_ISL_461071, EPI_ISL_461120, EPI_ISL_461129, EPI_ISL_461130, EPI_ISL_461131, EPI_ISL_461132, EPI_ISL_461181, EPI_ISL_461252, EPI_ISL_461253, EPI_ISL_461256, EPI_ISL_461265, EPI_ISL_461280, EPI_ISL_461288, EPI_ISL_461339 | | | |
| see above | Dutch COVID-19 response team | Erasmus Medical Center | Bas Oude Munnink, David Nieuwenhuijse, Reina Sikkema, Claudia Schapendonk, Irina Chestakova, Anne van der Linden, Theo Bestebroer, Stefan van Nieuwkoop, Mark Pronk, Pascal Lexmond, Corien Swaan, Manon Haverkate, Madelief Mollers, Mart Stein, Sandra Kengne Kamga Mobou, Jeroen van Kampen, Jolanda Voermans, Aura Timen, Corine GeurtsvanKessel, Annemiek van der Eijk, Richard Molenkamp, Marion Koopmans, on behalf of the Dutch national COVID-19 response team. |
| EPI_ISL_461399, EPI_ISL_461400, EPI_ISL_461401, EPI_ISL_461402, EPI_ISL_461408, EPI_ISL_461409, EPI_ISL_461410, EPI_ISL_461411, EPI_ISL_461412, EPI_ISL_461413, EPI_ISL_461414, EPI_ISL_461415, EPI_ISL_461416, EPI_ISL_461417, EPI_ISL_461418, EPI_ISL_461419, EPI_ISL_461420, EPI_ISL_461421, EPI_ISL_461426, EPI_ISL_461427, EPI_ISL_461428, EPI_ISL_461429, EPI_ISL_461435, EPI_ISL_461439, EPI_ISL_461440, EPI_ISL_461441, EPI_ISL_461442 | | | |
| see above | UW Virology Lab | UW Virology Lab | Pavitra Roychoudhury, Amin Addetia, Hong Xie, Lasata Shrestha, Truong Nguyen, Meei-Li Huang, Keith Jerome, Alexander Greninger |
| EPI_ISL_461735, EPI_ISL_461736, EPI_ISL_461737, EPI_ISL_461738, EPI_ISL_461762 | Virology Department, Royal Infirmary of Edinburgh, NHS Lothian / School of Biological Sciences, University of Edinburgh / Institute of Genetics and Molecular Medicine, University of Edinburgh | COVID-19 Genomics UK (COG-UK) Consortium | McHugh M, Dewar R, Rooke S, Gallagher M, Balcaza C, O'Toole Á, Scher E, Hill V, McCrone JT, Colquhoun R, Yu X, Jackson B, Rambaut A, Williams TC, Templeton K |
| EPI_ISL_462228, EPI_ISL_462235, EPI_ISL_462236, EPI_ISL_462237, EPI_ISL_462252, EPI_ISL_462254 | KU Leuven, Rega Institute, Clinical and Epidemiological Virology | KU Leuven, Rega Institute, Clinical and Epidemiological Virology | Tony Wawina-Bokalanga, Bert Vanmechelen, Joan Marti-Carerras, Piet Maes |
| EPI_ISL_462348, EPI_ISL_462355, EPI_ISL_462357, EPI_ISL_462360, EPI_ISL_462378, EPI_ISL_462407, EPI_ISL_462419, EPI_ISL_462421, EPI_ISL_462428 | National Public Health Laboratory, National Centre for Infectious Diseases | National Public Health Laboratory, National Centre for Infectious Diseases | Mak TM, Octavia S, Chavatte JM, Cui L, Lin RTP |
| EPI_ISL_463277, EPI_ISL_463278, EPI_ISL_463279, EPI_ISL_463280, EPI_ISL_463281, EPI_ISL_463282, EPI_ISL_463283, EPI_ISL_463284, EPI_ISL_463285, EPI_ISL_463286, EPI_ISL_463287, EPI_ISL_463288, EPI_ISL_463289, EPI_ISL_463291, EPI_ISL_463294, EPI_ISL_463295, EPI_ISL_463296, EPI_ISL_463297, EPI_ISL_463298, EPI_ISL_463299, EPI_ISL_463300 | | | |
| see above | Ochsner Health | Bioinfoexperts, LLC | Susanna L. Lamers, David J. Nolan, Rebecca Rose, Sissy Cross, David Moraga Amador, Tong Yang, Luke Caruso, Wayra Navia, Lydia Von Borstel, Xiao Hui Zhou, Amy Feehan, Julia-Garcia-Diaz |
| EPI_ISL_463903, EPI_ISL_463904, EPI_ISL_463906, EPI_ISL_463908, EPI_ISL_463910, EPI_ISL_463911, EPI_ISL_463912, EPI_ISL_463924, EPI_ISL_463927, EPI_ISL_463929, EPI_ISL_463930, EPI_ISL_463931, EPI_ISL_463932, EPI_ISL_463934, EPI_ISL_463935, EPI_ISL_463936, EPI_ISL_463937, EPI_ISL_463938, EPI_ISL_463939, EPI_ISL_463941, EPI_ISL_463942, EPI_ISL_463943, EPI_ISL_463944, EPI_ISL_463946, EPI_ISL_463947, EPI_ISL_463948, EPI_ISL_463949, EPI_ISL_463950, EPI_ISL_463951 | | | |
| see above | Laboratoire de microbiologie, Hopital de Verdun | Smith Laboratory, Centre de Recherche CHU Sainte-Justine | Martin Smith, Marieke Rozendaal, Ivan Pavlov |
| EPI_ISL_464001, EPI_ISL_464010, EPI_ISL_464012, EPI_ISL_464013, EPI_ISL_464014, EPI_ISL_464018, EPI_ISL_464020, EPI_ISL_464024, EPI_ISL_464028, EPI_ISL_464037, EPI_ISL_464046, EPI_ISL_464047, EPI_ISL_464056, EPI_ISL_464063 | | | |
| see above | Unity Health Toronto | Ontario Institute for Cancer Research | Ramzi Fattouh,Larissa M. Matukas,Mark Downing,Annette Gower,Karel Boissinot,Samira Mubareka,TIBDN,Ilinca Lungu,Bernard Lam,Jeremy Johns,Paul Krzyzanowski,Richard de Borja,Philip Zuzarte,Jared Simpson |
| EPI_ISL_464067, EPI_ISL_464069, EPI_ISL_464070, EPI_ISL_464071, EPI_ISL_464072, EPI_ISL_464073, EPI_ISL_464079, EPI_ISL_464083, EPI_ISL_464084 | KU Leuven, Rega Institute, Clinical and Epidemiological Virology | KU Leuven, Rega Institute, Clinical and Epidemiological Virology | Tony Wawina-Bokalanga, Bert Vanmechelen, Joan Marti-Carerras, Piet Maes |
| EPI_ISL_464093 | Laboratory Medicine | Department of Laboratory Medicine, Lin-Kou Chang Gung Memorial Hospital, Taoyuan, Taiwan | Kuo-Chien Tsao, Yu-Nong Gong, Shu-Li Yang, Yi-Chun Liu, Chung-Guei Huang, Mei-Jen Hsiao, Po-Wei Huang, Cheng-Ta Yang, Cheng-Hsun Chiu, Peng-Nien Huang, Kuo-Ming Lee, Guang-Wu Chen, Shin-Ru Shih |
| EPI_ISL_464144, EPI_ISL_464145, EPI_ISL_464146, EPI_ISL_464147 | National Health Laboratory Service (NHLS), Tygerberg | Division of Medical Virology, Stellenbosch University and National Health Laboratory Service (NHLS) | Susan Engelbrecht, Kayla Delaney, Bronwyn Kleinhans, Houriiyah Tegally, Eduan Wilkindon, Gert van Zyl, Wolfgang Preiser, Tulio de Oliveira |
| EPI_ISL_465281, EPI_ISL_465499, EPI_ISL_466036, EPI_ISL_466037, EPI_ISL_466048, EPI_ISL_466049, EPI_ISL_466050, EPI_ISL_466056, EPI_ISL_466058, EPI_ISL_466059, EPI_ISL_466060, EPI_ISL_466063, EPI_ISL_466066, EPI_ISL_466067, EPI_ISL_466068, EPI_ISL_466069, EPI_ISL_466070, EPI_ISL_466071, EPI_ISL_466073, EPI_ISL_466074, EPI_ISL_466075, EPI_ISL_466076, EPI_ISL_466077, EPI_ISL_466078, EPI_ISL_466079, EPI_ISL_466080, EPI_ISL_466081, EPI_ISL_466082, EPI_ISL_466083, EPI_ISL_466084, EPI_ISL_466085, EPI_ISL_466086, EPI_ISL_466087, EPI_ISL_466088, EPI_ISL_466089, EPI_ISL_466090, EPI_ISL_466091, EPI_ISL_466092, EPI_ISL_466093, EPI_ISL_466094, EPI_ISL_466095, EPI_ISL_466096, EPI_ISL_466097, EPI_ISL_466098, EPI_ISL_466099, EPI_ISL_466100, EPI_ISL_466101, EPI_ISL_466102, EPI_ISL_466103, EPI_ISL_466104, EPI_ISL_466107, EPI_ISL_466150, EPI_ISL_466151, EPI_ISL_466152, EPI_ISL_466153, EPI_ISL_466154, EPI_ISL_466155, EPI_ISL_466156, EPI_ISL_466157, EPI_ISL_466158, EPI_ISL_466159, EPI_ISL_466160, EPI_ISL_466161, EPI_ISL_466162, EPI_ISL_466163, EPI_ISL_466164, EPI_ISL_466165, EPI_ISL_466166, EPI_ISL_466167, EPI_ISL_466168, EPI_ISL_466169, EPI_ISL_466170, EPI_ISL_466171, EPI_ISL_466172, EPI_ISL_466173, EPI_ISL_466174, EPI_ISL_466175, EPI_ISL_466176, EPI_ISL_466177, EPI_ISL_466178, EPI_ISL_466179, EPI_ISL_466180, EPI_ISL_466181, EPI_ISL_466182, EPI_ISL_466183, EPI_ISL_466184, EPI_ISL_466185, EPI_ISL_466186, EPI_ISL_466187, EPI_ISL_466188, EPI_ISL_466189, EPI_ISL_466190, EPI_ISL_466191, EPI_ISL_466192, EPI_ISL_466193, EPI_ISL_466194, EPI_ISL_466195, EPI_ISL_466196, EPI_ISL_466197, EPI_ISL_466198, EPI_ISL_466199, EPI_ISL_466200, EPI_ISL_466201, EPI_ISL_466202, EPI_ISL_466203, EPI_ISL_466204, EPI_ISL_466205, EPI_ISL_466206, EPI_ISL_466207, EPI_ISL_466208, EPI_ISL_466209, EPI_ISL_466210, EPI_ISL_466211, EPI_ISL_466212, EPI_ISL_466213, EPI_ISL_466214, EPI_ISL_466215, EPI_ISL_466216, EPI_ISL_466217, EPI_ISL_466218, EPI_ISL_466219, EPI_ISL_466220, EPI_ISL_466221, EPI_ISL_466222, EPI_ISL_466223, EPI_ISL_466224, EPI_ISL_466225, EPI_ISL_466226, EPI_ISL_466227, EPI_ISL_466228, EPI_ISL_466231, EPI_ISL_466232, EPI_ISL_466233, EPI_ISL_466234, EPI_ISL_466235, EPI_ISL_466236, EPI_ISL_466237, EPI_ISL_466238, EPI_ISL_466240, EPI_ISL_466241, EPI_ISL_466242, EPI_ISL_466243, EPI_ISL_466244, EPI_ISL_466245, EPI_ISL_466246, EPI_ISL_466247, EPI_ISL_466249, EPI_ISL_466250, EPI_ISL_466251, EPI_ISL_466252, EPI_ISL_466253, EPI_ISL_466254, EPI_ISL_466255, EPI_ISL_466263, EPI_ISL_466264, EPI_ISL_466265, EPI_ISL_466269, EPI_ISL_466271, EPI_ISL_466274, EPI_ISL_466275, EPI_ISL_466276, EPI_ISL_466321, EPI_ISL_466323, EPI_ISL_466325, EPI_ISL_466330, EPI_ISL_466331, EPI_ISL_466334, EPI_ISL_466336, EPI_ISL_466340, EPI_ISL_466414, EPI_ISL_466418, EPI_ISL_466559 | | | |
| see above | Respiratory Virus Unit, Microbiology Services Colindale, Public Health England | Respiratory Virus Unit, Microbiology Services Colindale, Public Health England | PHE Covid Sequencing Team |
| EPI_ISL_466874 | Max von Pettenkofer Institute, Virology, National Reference Center for Retroviruses, LMU München | Laboratory for Functional Genome Analysis, Dept. Genomics, Gene Center of the LMU Munich | Max Muenchhoff, Stefan Krebs, Alexander Graf, Oliver Keppler, Helmut Blum |
| EPI_ISL_467184, EPI_ISL_467185, EPI_ISL_467186, EPI_ISL_467191, EPI_ISL_467192, EPI_ISL_467194, EPI_ISL_467198, EPI_ISL_467199, EPI_ISL_467200, EPI_ISL_467202, EPI_ISL_467203, EPI_ISL_467207, EPI_ISL_467210, EPI_ISL_467211, EPI_ISL_467212, EPI_ISL_467216, EPI_ISL_467217, EPI_ISL_467219, EPI_ISL_467220, EPI_ISL_467224, EPI_ISL_467227, EPI_ISL_467229, EPI_ISL_467230, EPI_ISL_467231, EPI_ISL_467233, EPI_ISL_467235, EPI_ISL_467236, EPI_ISL_467237, EPI_ISL_467238, EPI_ISL_467240, EPI_ISL_467241, EPI_ISL_467242, EPI_ISL_467243, EPI_ISL_467248, EPI_ISL_467249, EPI_ISL_467251, EPI_ISL_467252, EPI_ISL_467256, EPI_ISL_467257, EPI_ISL_467258, EPI_ISL_467259, EPI_ISL_467260, EPI_ISL_467261 | | | |
| see above | Hospital General Universitario Gregorio Marañón | SeqCOVID-SPAIN consortium/IBV(CSIC) | Laura Pérez-Lago, Marta Herranz, Jon Sicilia, Julia Suárez, Pilar Catalán, Patricia Muñoz, Darío García de Viedma and SeqCOVID-SPAIN consortium |
| EPI_ISL_467344, EPI_ISL_467345, EPI_ISL_467346 | Laboratory of Respiratory Viruses and Measles, Oswaldo Cruz Institute, FIOCRUZ | Laboratory of Respiratory Viruses and Measles, Oswaldo Cruz Institute, FIOCRUZ | Paola Resende, Luciana Appolinario, Fernando Motta, Ana Carolina Paixão, Ana Carolina Mendonça, Aline Mattos, Milene Miranda, Cristiana Garcia, Braulia Caetano, Maria Ogrzewalska, Jonathan Lopes, Marilda Siqueira |
| EPI_ISL_467381, EPI_ISL_467383 | NYU Langone Health | Departments of Pathology and Medicine, New York University School of Medicine | Maria Aguero-Rosenfeld, Brendan Belovarac, Margaret Black, Ludovic Boytard, John Cadley, Paolo Cotzia, John Chen, Dacia Dimartino, Xiaojun Feng, Tatyana Gindin, Emily Guzman, Adriana Heguy, Megan Hogan, Emily Huang, George Jour, Alireza Khodadadi-Jamayran, Lawrence H. Lin, Raven Luther, Andrew Lytle, Christian Marier, Matthew T. Maurano, Mark J. Mulligan, Peter Meyn, Raquel Ordonez Ciriza, Iman Osman, Jared Pinnell, Vanessa Raabe, Sitharam Ramaswami, Amy Rapkiewicz, Andre M. Ribeiro-dos-Santos, Marie Samanovic-Golden, Antonio Serrano, Guomiao Shen, Matija Snuderl, Theodore Vougiouklakis, Nick Vulpescu, Gael Westby, Paul Zappile, Yutong Zhang |
| EPI_ISL_467433, EPI_ISL_467434, EPI_ISL_467435 | AMPATH-DBN | KRISP, KZN Research Innovation and Sequencing Platform | Giandhari J, Pillay S, Lessells R, Chimukangara B, Mdlalose K, York D, Khan S, Tegally H, Wilkinson E, de Oliveira T |
| EPI_ISL_467438, EPI_ISL_467439 | NHLS-IALCH | KRISP, KZN Research Innovation and Sequencing Platform | Giandhari J, Pillay S, Lessells R, Chimukangara B, Mdlalose K, York D, Khan S, Tegally H, Wilkinson E, de Oliveira T |
| EPI_ISL_467445, EPI_ISL_467447 | Molecular Diagnostics Services (MDS) | KRISP, KZN Research Innovation and Sequencing Platform | Giandhari J, Pillay S, Lessells R, Chimukangara B, Mdlalose K, York D, Khan S, Tegally H, Wilkinson E, de Oliveira T |

| | | | |
|---|---|---|---|
| EPI_ISL_467449, EPI_ISL_467450, EPI_ISL_467451, EPI_ISL_467452, EPI_ISL_467453, EPI_ISL_467455, EPI_ISL_467456, EPI_ISL_467458, EPI_ISL_467459, EPI_ISL_467460, EPI_ISL_467466, EPI_ISL_467467, EPI_ISL_467468, EPI_ISL_467469 | | | |
| see above | AMPATH-DBN | KRISP, KZN Research Innovation and Sequencing Platform | Giandhari J, Pillay S, Lessells R, Chimukangara B, Mdlalose K, York D, Khan S, Tegally H, Wilkinson E, de Oliveira T |
| EPI_ISL_467548, EPI_ISL_467549, EPI_ISL_467550, EPI_ISL_467551, EPI_ISL_467552, EPI_ISL_467553, EPI_ISL_467554, EPI_ISL_467555, EPI_ISL_467556, EPI_ISL_467557, EPI_ISL_467558, EPI_ISL_467559, EPI_ISL_467560, EPI_ISL_467561, EPI_ISL_467562, EPI_ISL_467563, EPI_ISL_467564, EPI_ISL_467565, EPI_ISL_467566, EPI_ISL_467567, EPI_ISL_467568, EPI_ISL_467569, EPI_ISL_467570, EPI_ISL_467571, EPI_ISL_467572, EPI_ISL_467573, EPI_ISL_467574, EPI_ISL_467575, EPI_ISL_467576, EPI_ISL_467577, EPI_ISL_467578, EPI_ISL_467579, EPI_ISL_467580, EPI_ISL_467581, EPI_ISL_467582, EPI_ISL_467583, EPI_ISL_467584, EPI_ISL_467585, EPI_ISL_467586, EPI_ISL_467641, EPI_ISL_467642, EPI_ISL_467643, EPI_ISL_467644, EPI_ISL_467645, EPI_ISL_467646, EPI_ISL_467647, EPI_ISL_467648, EPI_ISL_467649, EPI_ISL_467650, EPI_ISL_467651, EPI_ISL_467652, EPI_ISL_467653, EPI_ISL_467654, EPI_ISL_467655, EPI_ISL_467656, EPI_ISL_467657, EPI_ISL_467658, EPI_ISL_467659, EPI_ISL_467660, EPI_ISL_467661, EPI_ISL_467662, EPI_ISL_467663, EPI_ISL_467664 | | | |
| see above | New Mexico Department of Health Scientific Laboratory Division | Center for Global Health, University of New Mexico Health Sciences Center | Daryl Domman, Kurt Schwalm, Twila Kunde, Joseph Hicks, Michael Edwards, Darrell Dinwiddie |
| EPI_ISL_467980 | San Diego County Public Health Laboratory | Andersen lab at Scripps Research | SEARCH Alliance San Diego with Tracy Basler, Jovan Shephard, Brett Austin |
| EPI_ISL_467987 | SA Pathology | SA Pathology | Lex Leong, Chuan Kok Lim, Mark Turra, Ivan Bastian, Geoff Higgins |
| EPI_ISL_467990 | SA Pathology | SA Pathology | Leong, LEX, Lim, CK, Turra, M, Bastian, I, Higgins, G |
| EPI_ISL_467994, EPI_ISL_467996, EPI_ISL_467998, EPI_ISL_468003, EPI_ISL_468009, EPI_ISL_468011, EPI_ISL_468012, EPI_ISL_468017, EPI_ISL_468019, EPI_ISL_468024 | SA Pathology | SA Pathology | Lex Leong, Chuan Kok Lim, Mark Turra, Ivan Bastian, Geoff Higgins |
| EPI_ISL_468358 | Alameda County Public Health Lab | Chan-Zuckerberg Biohub | CZB Cliahub Consortium |
| EPI_ISL_468414 | County of San Luis Obispo Public Health Laboratory | Chan-Zuckerberg Biohub | CZB Cliahub Consortium |
| EPI_ISL_468459 | Humboldt County Public Health Laboratory | Chan-Zuckerberg Biohub | CZB Cliahub Consortium |
| EPI_ISL_468499, EPI_ISL_468500, EPI_ISL_468501, EPI_ISL_468502 | Ventura County Public Health Lab | Chan-Zuckerberg Biohub | CZB Cliahub Consortium |
| EPI_ISL_468530, EPI_ISL_468532, EPI_ISL_468533, EPI_ISL_468534, EPI_ISL_468535, EPI_ISL_468536, EPI_ISL_468537 | San Joaquin County Public Health Lab | Chan-Zuckerberg Biohub | CZB Cliahub Consortium |
| EPI_ISL_468623, EPI_ISL_468624, EPI_ISL_468625, EPI_ISL_468626, EPI_ISL_468627, EPI_ISL_468628, EPI_ISL_468629, EPI_ISL_468630, EPI_ISL_468631, EPI_ISL_468632, EPI_ISL_468633, EPI_ISL_468634, EPI_ISL_468635, EPI_ISL_468636, EPI_ISL_468637, EPI_ISL_468638, EPI_ISL_468639, EPI_ISL_468640, EPI_ISL_468641, EPI_ISL_468642, EPI_ISL_468643, EPI_ISL_468644, EPI_ISL_468645, EPI_ISL_468646, EPI_ISL_468647, EPI_ISL_468648, EPI_ISL_468649, EPI_ISL_468650 | | | |
| see above | Contra Costa Public Health Lab | Chan-Zuckerberg Biohub | CZB Cliahub Consortium |
| EPI_ISL_468701, EPI_ISL_468702, EPI_ISL_468703, EPI_ISL_468712, EPI_ISL_468713, EPI_ISL_468714, EPI_ISL_468715, EPI_ISL_468717 | Ochsner Health | Bioinfoexperts, LLC | Rebecca Rose, Amy Feehan, David J. Nolan, Sissy Cross, David Moraga Amador, Tong Yang, Luke Caruso, Wayra Navia, Lydia Von Borstel, Xiao Hui Zhou, Julia-Garcia-Diaz, Susanna L. Lamers |
| EPI_ISL_468761 | Centro de Investigación Biomédica de La Rioja - Hospital San Pedro Logroño | SeqCOVID-SPAIN consortium/IBV(CSIC) | María de Toro, José Manuel Azcona Gutiérrez, María Pilar Bea Escudero, Miriam Blasco Alberdi and SeqCOVID-SPAIN consortium |
| EPI_ISL_468986, EPI_ISL_468997 | Servicio de Microbiología, Hospital Universitario Son Espases | SeqCOVID-SPAIN consortium/IBV(CSIC) | Carla López-Causapé, Jordi Reina, Antonio Oliver and SeqCOVID-SPAIN consortium |
| EPI_ISL_469051 | LNR National Reference Laboratory, Mohammed VI University of Health Sciences | Medical Biotechnology Laboratory, Rabat Medical and Pharmacy School, Mohammed The Vth University in Rabat | Meriem LAAMARTI, Souad KARTTI, Rokaia LAAMRTI , M.W. CHEMAO-ELFIHRI, Loubna ALLAM, Mouna OUADGHIRI, Imane SMYEJ, Jalila RAHOUI, Houda BENRAHMA, Jalil El Atar, Idrissa Diawara, Rachid EL JAOUDI, Laila SBABOU, Chakib NEJJARI, Saaid AMZAZI, Rachid MENTAG, Lahcen BELYAMANI and Azeddine IBRAHIMI |
| EPI_ISL_469105, EPI_ISL_469113, EPI_ISL_469114, EPI_ISL_469120, EPI_ISL_469122, EPI_ISL_469123, EPI_ISL_469125, EPI_ISL_469129 | National Public Health Laboratory, National Centre for Infectious Diseases | National Public Health Laboratory, National Centre for Infectious Diseases | Mak TM, Octavia S, Chavatte JM, Cui L, Lin RTP |
| EPI_ISL_469189, EPI_ISL_469190, EPI_ISL_469191, EPI_ISL_469192, EPI_ISL_469193, EPI_ISL_469194, EPI_ISL_469195, EPI_ISL_469196, EPI_ISL_469197, EPI_ISL_469198, EPI_ISL_469199, EPI_ISL_469200, EPI_ISL_469201, EPI_ISL_469202, EPI_ISL_469203, EPI_ISL_469204, EPI_ISL_469205, EPI_ISL_469206, EPI_ISL_469207, EPI_ISL_469208 | | | |
| see above | Yale Clinical Virology Laboratory | Grubaugh Lab - Yale School of Public Health | Joseph Fauver, Tara Alpert, Anderson Brito, Anne Wyllie, Chantal Vogels, Mary Petrone, Cole Jensen, Chaney Kalinich, Isabel Ott, Arnau Casanovas, Catherine Muenker, Adam Moore, Alice Lu, Maria Tokuyama, Patrick Wong, Peiwen Lu, Saad Omer, Richard Martinello, Allison Nelson, Shelli Farhadian, Akiko Iwasaki, Charlese Dela Cruz, Albert Ko, Nathan Grubaugh |
| EPI_ISL_469274 | National Public Health Laboratory, National Centre for Infectious Diseases | National Public Health Laboratory, National Centre for Infectious Diseases | Mak TM, Octavia S, Chavatte JM, Cui L, Lin RTP |
| EPI_ISL_469293 | Keio University Hospital | Keio University Hospital | Kenjiro Kosaki |
| EPI_ISL_469528, EPI_ISL_469605, EPI_ISL_469608, EPI_ISL_469622, EPI_ISL_469627, EPI_ISL_469636, EPI_ISL_469669, EPI_ISL_469675, EPI_ISL_469693, EPI_ISL_469704, EPI_ISL_469752, EPI_ISL_469797 | | | |
| see above | PHE South West Regional Laboratory, National Infection Service | Wellcome Sanger Institute for the COVID-19 Genomics UK (COG-UK) consortium | Stephanie Hutchings, Hannah Pymont, Dr Peter Muir, Barry Vipond, Rich Hopes; and Alex Alderton, Roberto Amato, Sonia Goncalves, Ewan Harrison, David K. Jackson, Ian Johnston, Dominic Kwiatkowski, Cordelia Langford, John Sillitoe on behalf of the Wellcome Sanger Institute COVID-19 Surveillance Team (http://www.sanger.ac.uk/covid-team) |
| EPI_ISL_470019, EPI_ISL_470035, EPI_ISL_470037, EPI_ISL_470069, EPI_ISL_470080, EPI_ISL_470088 | Regional Virus Laboratory, Belfast Health and Social Care Trust | Wellcome Sanger Institute for the COVID-19 Genomics UK (COG-UK) consortium | Conall McCaughey, James McKenna, Tanya Curran, Susan Feeney, Alison Watt, Ciara Cox, Mairead Connor, Zoltan Molnar, David Simpson, Derek Fairley; and Alex Alderton, Roberto Amato, Sonia Goncalves, Ewan Harrison, David K. Jackson, Ian Johnston, Dominic Kwiatkowski, Cordelia Langford, John Sillitoe on behalf of the Wellcome Sanger Institute COVID-19 Surveillance Team (http://www.sanger.ac.uk/covid-team) |
| EPI_ISL_470362, EPI_ISL_470368, EPI_ISL_470372, EPI_ISL_470374, EPI_ISL_470380, EPI_ISL_470383, EPI_ISL_470385, EPI_ISL_470391, EPI_ISL_470403, EPI_ISL_470415, EPI_ISL_470419, EPI_ISL_470435, EPI_ISL_470441, EPI_ISL_470452, EPI_ISL_470459, EPI_ISL_470460, EPI_ISL_470471, EPI_ISL_470481, EPI_ISL_470484, EPI_ISL_470485, EPI_ISL_470493, EPI_ISL_470497, EPI_ISL_470509, EPI_ISL_470513, EPI_ISL_470515, EPI_ISL_470520 | | | |
| see above | Department of Pathology, University of Cambridge | Wellcome Sanger Institute for the COVID-19 Genomics UK (COG-UK) consortium | Luke W Meredith, M. Estée Török , Myra Hosmillo, William L. Hamilton, Martin D. Curran, Theresa Feltwell, Grant Hall, Anna Yakovleva, Fahad A Khokhar, Charlotte J. Houldcroft, Laura G Caller, Aminu S. Jahun, Sarah L. Caddy, Ian Goodfellow; and Alex Alderton, Roberto Amato, Sonia Goncalves, Ewan Harrison, David K. Jackson, Ian Johnston, Dominic Kwiatkowski, Cordelia Langford, John Sillitoe on behalf of the Wellcome Sanger Institute COVID-19 Surveillance Team (http://www.sanger.ac.uk/covid-team) |
| EPI_ISL_470530 | PHE South West Regional Laboratory, National Infection Service | Wellcome Sanger Institute for the COVID-19 Genomics UK (COG-UK) consortium | Stephanie Hutchings, Hannah Pymont, Dr Peter Muir, Barry Vipond, Rich Hopes; and Alex Alderton, Roberto Amato, Sonia Goncalves, Ewan Harrison, David K. Jackson, Ian Johnston, Dominic Kwiatkowski, Cordelia Langford, John Sillitoe on behalf of the Wellcome Sanger Institute COVID-19 Surveillance Team (http://www.sanger.ac.uk/covid-team) |
| EPI_ISL_470531, EPI_ISL_470532, EPI_ISL_470533, EPI_ISL_470535, EPI_ISL_470536, EPI_ISL_470537, EPI_ISL_470538 | Department of Pathology, University of Cambridge | Wellcome Sanger Institute for the COVID-19 Genomics UK (COG-UK) consortium | Luke W Meredith, M. Estée Török , Myra Hosmillo, William L. Hamilton, Martin D. Curran, Theresa Feltwell, Grant Hall, Anna Yakovleva, Fahad A Khokhar, Charlotte J. Houldcroft, Laura G Caller, Aminu S. Jahun, Sarah L. Caddy, Ian Goodfellow; and Alex Alderton, Roberto Amato, Sonia Goncalves, Ewan Harrison, David K. Jackson, Ian Johnston, Dominic Kwiatkowski, Cordelia Langford, John Sillitoe on behalf of the Wellcome Sanger Institute COVID-19 Surveillance Team (http://www.sanger.ac.uk/covid-team) |
| EPI_ISL_470570, EPI_ISL_470571, EPI_ISL_470572, EPI_ISL_470573, | Hermes Pardini | Bioinformatics Laboratory / LNCC | Alexandra Gerber, Ana Paula Guimarães, Luiz Gonzaga Paula de Almeida, Ronaldo da Silva Francisco Junior, Mariane Talon, Filipe Romero, Átila Duque Rossi, Terezinha Marta Pereira, working group UFRJ, Jaqueline Goes de Jesus, Ingra Morales Claro, Ester Cerdeira Sabino, Nuno Rodrigues Faria, |

| | | | |
|---|---|---|---|
| EPI_ISL_470576, EPI_ISL_470577, EPI_ISL_470585, EPI_ISL_470586, EPI_ISL_470587, EPI_ISL_470588 | | | CADDE-group, Laboratorio Hermes Pardini, Laboratorio Simile, working group UFMG, Amilcar Tanuri, Carolina Voloch, Renato Santana Aguiar e Ana Tereza Vasconcelos |
| EPI_ISL_470595, EPI_ISL_470597 | Simile | Bioinformatics Laboratory / LNCC | Alexandra Gerber, Ana Paula Guimarães, Luiz Gonzaga Paula de Almeida, Ronaldo da Silva Francisco Junior, Mariane Talon, Filipe Romero, Átila Duque Rossi, Terezinha Marta Pereira, working group UFRJ, Jaqueline Goes de Jesus, Ingra Morales Claro, Ester Cerdeira Sabino, Nuno Rodrigues Faria, CADDE-group, Laboratorio Hermes Pardini, Laboratorio Simile, working group UFMG, Amilcar Tanuri, Carolina Voloch, Renato Santana Aguiar e Ana Tereza Vasconcelos |
| EPI_ISL_470604, EPI_ISL_470605, EPI_ISL_470606, EPI_ISL_470607, EPI_ISL_470608, EPI_ISL_470609, EPI_ISL_470610, EPI_ISL_470611, EPI_ISL_470612, EPI_ISL_470614 | Hermes Pardini | Bioinformatics Laboratory / LNCC | Alexandra Gerber, Ana Paula Guimarães, Luiz Gonzaga Paula de Almeida, Ronaldo da Silva Francisco Junior, Mariane Talon, Filipe Romero, Átila Duque Rossi, Terezinha Marta Pereira, working group UFRJ, Jaqueline Goes de Jesus, Ingra Morales Claro, Ester Cerdeira Sabino, Nuno Rodrigues Faria, CADDE-group, Laboratorio Hermes Pardini, Laboratorio Simile, working group UFMG, Amilcar Tanuri, Carolina Voloch, Renato Santana Aguiar e Ana Tereza Vasconcelos |
| EPI_ISL_470623, EPI_ISL_470624, EPI_ISL_470625 | Laboratorio de Virologia Molecular / UFRJ | Bioinformatics Laboratory / LNCC | Alexandra Gerber, Ana Paula Guimarães, Luiz Gonzaga Paula de Almeida, Ronaldo da Silva Francisco Junior, Mariane Talon, Filipe Romero, Átila Duque Rossi, Terezinha Marta Pereira, working group UFRJ, Jaqueline Goes de Jesus, Ingra Morales Claro, Ester Cerdeira Sabino, Nuno Rodrigues Faria, CADDE-group, Laboratorio Hermes Pardini, Laboratorio Simile, working group UFMG, Amilcar Tanuri, Carolina Voloch, Renato Santana Aguiar e Ana Tereza Vasconcelos |
| EPI_ISL_470655 | Hermes Pardini | Bioinformatics Laboratory / LNCC | Alexandra Gerber, Ana Paula Guimarães, Luiz Gonzaga Paula de Almeida, Ronaldo da Silva Francisco Junior, Mariane Talon, Filipe Romero, Átila Duque Rossi, Terezinha Marta Pereira, working group UFRJ, Jaqueline Goes de Jesus, Ingra Morales Claro, Ester Cerdeira Sabino, Nuno Rodrigues Faria, CADDE-group, Laboratorio Hermes Pardini, Laboratorio Simile, working group UFMG, Amilcar Tanuri, Carolina Voloch, Renato Santana Aguiar e Ana Tereza Vasconcelos |
| EPI_ISL_470712, EPI_ISL_470713, EPI_ISL_470714, EPI_ISL_470716, EPI_ISL_470717, EPI_ISL_470718 | Utah Public Health Laboratory | Utah Public Health Laboratory | Erin Young, Kelly Oakeson |
| EPI_ISL_470724, EPI_ISL_470725, EPI_ISL_470727 | Utah Public Health Laboratory | Utah Public Health Laboratory | Heidi Butz, Erin Young, Kelly Oakeson |
| EPI_ISL_470831, EPI_ISL_470832, EPI_ISL_470833, EPI_ISL_470859, EPI_ISL_470874 | PathWest Laboratory Medicine WA | PathWest Laboratory Medicine WA | Chisha Sikazwe, Jurissa Lang, Avram Levy, David Smith and David Speers |
| EPI_ISL_470901 | Influenza etiology and epidemiology laboratory | Pathogenic Microorganisms Variability Laboratory | Alexey Shchetinin, Maria Nikiforova, Elena Shidlovskaya, Nadezhda Kuznetsova, Vladimir Gushchin, Inna Dolzhikova, Daria Grousova, Andrey Botikov, Denis Logunov, Kirill Krasnoslobotsev, Svetlana Trushakova, Elena Burtseva, Ludmila Kolobukhina, Svetlana Smetanina, Alexander Gintsburg |
| EPI_ISL_471183, EPI_ISL_471206, EPI_ISL_471208, EPI_ISL_471214, EPI_ISL_471217, EPI_ISL_471224, EPI_ISL_471227, EPI_ISL_471237, EPI_ISL_471238, EPI_ISL_471240, EPI_ISL_471241, EPI_ISL_471243, EPI_ISL_471244, EPI_ISL_471248 | | | |
| see above | Wisconsin State Laboratory of Hygiene Communicable Disease Division | Wisconsin State Laboratory of Hygiene Communicable Disease Division | Kelsey R. Florek, Abigail C. Shockey |
| EPI_ISL_471541 | Hospital Geral Santa Marcelina | Instituto Adolfo Lutz, Interdiciplinary Procedures Center, Strategic Laboratory | Claudio Tavares Sacchi, Claudia Regina Gonçalves, Erica Valessa Ramos Gomes |
| EPI_ISL_471542 | Secretaria de Saude de Mogi das Cruzes | Instituto Adolfo Lutz, Interdiciplinary Procedures Center, Strategic Laboratory | Claudio Tavares Sacchi, Claudia Regina Gonçalves, Erica Valessa Ramos Gomes |
| EPI_ISL_471543 | Centro de Saude I Tacito Leite de Carvalho e Silva | Instituto Adolfo Lutz, Interdiciplinary Procedures Center, Strategic Laboratory | Claudio Tavares Sacchi, Claudia Regina Gonçalves, Erica Valessa Ramos Gomes |
| EPI_ISL_471916, EPI_ISL_471929, EPI_ISL_471956 | University of Exeter | COVID-19 Genomics UK (COG-UK) Consortium | Ben Temperton,Aaron Jeffries,Michelle Michelsen,Joanna Warwick-Dugdale,Audrey Farbos,Robyn Manley,Stephen Michell,Jane Masoli |
| EPI_ISL_472247, EPI_ISL_472248, EPI_ISL_472249, EPI_ISL_472250, EPI_ISL_472251, EPI_ISL_472252, EPI_ISL_472253, EPI_ISL_472254, EPI_ISL_472255, EPI_ISL_472256, EPI_ISL_472257, EPI_ISL_472258, EPI_ISL_472263, EPI_ISL_472264 | | | |
| see above | Northumbria University / South Tees Hospitals NHS Foundation Trust / North Cumbria Integrated Care NHS Foundation Trust / North Tees and Hartlepool NHS Foundation Trust / Newcastle Hospitals NHS Foundation Trust | COVID-19 Genomics UK (COG-UK) Consortium | Darren L Smith,Andrew Nelson,Matthew Bashton,Greg R Young,Joshua Loh,John Allan,Mohammad A Tariq,Giles S Holt,Gary Black,Wen C Yew,Lynn Dover,Paul Baker,Steve Liggett,Sarah Essex,Jane Greenaway,Debra Padgett,Clive Graham,Garren Scott,Edward Barton,Emma Swindells,Brendan Payne,Jennifer Collins,Yusri Taha,Gary Eltringham |
| EPI_ISL_472440, EPI_ISL_472450, EPI_ISL_472468, EPI_ISL_472534, EPI_ISL_472549, EPI_ISL_472568, EPI_ISL_472577, EPI_ISL_472639, EPI_ISL_472667, EPI_ISL_472700, EPI_ISL_472701, EPI_ISL_472728, EPI_ISL_472736, EPI_ISL_472741, EPI_ISL_472756, EPI_ISL_472788, EPI_ISL_472802 | | | |
| see above | Wales Specialist Virology Centre Sequencing lab: Pathogen Genomics Unit | COVID-19 Genomics UK (COG-UK) Consortium | Catherine Moore, Johnathan Evans, Laura Gifford, Malorie Perry, Simon Cottrell, Angela Marchbank, Alec Birchley, Alexander Adams, Amy Gaskin, Bree Gatica-Wilcox, Jason Coombes, Joel Southgate, Lauren Gilbert, Lee Graham, Nicole Pacchiarini, Sara Kumziene-Summerhayes, Sarah Taylor, Sophie Jones, Sara Rey, Matthew Bull, Joanne Watkins, Sally Corden, Tom Connor |
| EPI_ISL_473778, EPI_ISL_473779 | West of Scotland Specialist Virology Centre, NHSGGC / MRC-University of Glasgow Centre for Virus Research | COVID-19 Genomics UK (COG-UK) Consortium | Ana da Silva Filipe, Natasha Johnson, Kathy Smollett, Daniel Mair, Stephen Carmichael, Lily Tong, Jenna Nichols, Elihu Aranday-Cortes, Kirstyn Brunker, Yasmin Parr, Alice Broos, Kyriaki Nomikou; Sarah McDonald, Marc Niebel, Patawee Asamaphan; Richard Orton, Joseph Hughes, Sreenu Vattipally, David L Robertson; Alasdair MacLean, Rory Gunson; Kathy Li, Natasha Jesudason, Rajiv Shah, James Shepherd, Antonia Ho, Emma Thomson |
| EPI_ISL_473783 | Virology Department, Royal Infirmary of Edinburgh, NHS Lothian / School of Biological Sciences, University of Edinburgh / Institute of Genetics and Molecular Medicine, University of Edinburgh | COVID-19 Genomics UK (COG-UK) Consortium | McHugh M, Dewar R, Rooke S, Gallagher M, Balcaza C, O'Toole Á, Scher E, Hill V, McCrone JT, Colquhoun R, Yu X, Jackson B, Rambaut A, Williams TC, Templeton K |
| EPI_ISL_474232, EPI_ISL_474233, EPI_ISL_474235, EPI_ISL_474236, EPI_ISL_474237, EPI_ISL_474239, EPI_ISL_474240, EPI_ISL_474241, EPI_ISL_474242, EPI_ISL_474243, EPI_ISL_474244, EPI_ISL_474245, EPI_ISL_474246, EPI_ISL_474247, EPI_ISL_474248, EPI_ISL_474249, EPI_ISL_474250, EPI_ISL_474251, EPI_ISL_474252, EPI_ISL_474253, EPI_ISL_474254, EPI_ISL_474255, EPI_ISL_474256, EPI_ISL_474258, EPI_ISL_474259, EPI_ISL_474260, EPI_ISL_474261, EPI_ISL_474262, EPI_ISL_474263, EPI_ISL_474264, EPI_ISL_474265, EPI_ISL_474266, EPI_ISL_474267, EPI_ISL_474268, EPI_ISL_474269, EPI_ISL_474270, EPI_ISL_474271, EPI_ISL_474272, EPI_ISL_474273, EPI_ISL_474274, EPI_ISL_474275, EPI_ISL_474276, EPI_ISL_474277, EPI_ISL_474278, EPI_ISL_474279, EPI_ISL_474280, EPI_ISL_474281, EPI_ISL_474282, EPI_ISL_474283, EPI_ISL_474284, EPI_ISL_474285, EPI_ISL_474286, EPI_ISL_474287, EPI_ISL_474288, EPI_ISL_474289, EPI_ISL_474290, EPI_ISL_474291, EPI_ISL_474292, EPI_ISL_474293, EPI_ISL_474294, EPI_ISL_474295, EPI_ISL_474296, EPI_ISL_474297, EPI_ISL_474298, EPI_ISL_474299, EPI_ISL_474300, EPI_ISL_474301, EPI_ISL_474302, EPI_ISL_474303, EPI_ISL_474304, EPI_ISL_474306, EPI_ISL_474307, EPI_ISL_474308, EPI_ISL_474315, EPI_ISL_474316, EPI_ISL_474317, EPI_ISL_474327 | | | |
| see above | Wales Specialist Virology Centre Sequencing lab: Pathogen Genomics Unit | COVID-19 Genomics UK (COG-UK) Consortium | Catherine Moore, Johnathan Evans, Laura Gifford, Malorie Perry, Simon Cottrell, Angela Marchbank, Alec Birchley, Alexander Adams, Amy Gaskin, Bree Gatica-Wilcox, Jason Coombes, Joel Southgate, Lauren Gilbert, Lee Graham, Nicole Pacchiarini, Sara Kumziene-Summerhayes, Sarah Taylor, Sophie Jones, Sara Rey, Matthew Bull, Joanne Watkins, Sally Corden, Tom Connor |
| EPI_ISL_474800, EPI_ISL_474801, EPI_ISL_474802, EPI_ISL_474803, EPI_ISL_474804, EPI_ISL_474805, EPI_ISL_474807, EPI_ISL_474808, EPI_ISL_474809, EPI_ISL_474810, EPI_ISL_474811, EPI_ISL_474812, EPI_ISL_474813, EPI_ISL_474815, EPI_ISL_474816, EPI_ISL_474817, EPI_ISL_474818, EPI_ISL_474822, EPI_ISL_474823 | | | |
| see above | Complejo Hospitalario Universitario de Albacete | SeqCOVID-SPAIN consortium/IBV(CSIC) | Encarnacion Simarro Córdoba, Julia Lozano Serra, Lorena Robles Fonseca , Monica Parra Grandes, Caridad Sainz de Baranda Camino and SeqCOVID-SPAIN consortium |
| EPI_ISL_474836 | Hospital Universitario Virgen de las Nieves de Granada-SAS | SeqCOVID-SPAIN consortium/IBV(CSIC) | Mercedes Pérez Ruiz, Sara Sanbonmatsu Gámez, Irene Pedrosa Corral, José M. Navarro-Marí and SeqCOVID-SPAIN consortium |
| EPI_ISL_474838, EPI_ISL_474839, EPI_ISL_474847 | Complejo Hospitalario Universitario de Albacete | SeqCOVID-SPAIN consortium/IBV(CSIC) | Encarnacion Simarro Córdoba, Julia Lozano Serra, Lorena Robles Fonseca , Monica Parra Grandes, Caridad Sainz de Baranda Camino and SeqCOVID-SPAIN consortium |
| EPI_ISL_474850 | Hospital Universitario Virgen de las Nieves de Granada-SAS | SeqCOVID-SPAIN consortium/IBV(CSIC) | Mercedes Pérez Ruiz, Sara Sanbonmatsu Gámez, Irene Pedrosa Corral, José M. Navarro-Marí and SeqCOVID-SPAIN consortium |

| | | | |
|---|---|---|---|
| EPI_ISL_474853 | Complejo Hospitalario Universitario de Albacete | SeqCOVID-SPAIN consortium/IBV(CSIC) | Encarnacion Simarro Córdoba, Julia Lozano Serra, Lorena Robles Fonseca , Monica Parra Grandes, Caridad Sainz de Baranda Camino and SeqCOVID-SPAIN consortium |
| EPI_ISL_474889, EPI_ISL_474890, EPI_ISL_474891, EPI_ISL_474892, EPI_ISL_474893, EPI_ISL_474894, EPI_ISL_474895, EPI_ISL_474896 | Hospital Universitario Virgen de las Nieves de Granada-SAS | SeqCOVID-SPAIN consortium/IBV(CSIC) | Mercedes Pérez Ruiz, Sara Sanbonmatsu Gámez, Irene Pedrosa Corral, José M. Navarro-Marí and SeqCOVID-SPAIN consortium |
| EPI_ISL_474901, EPI_ISL_474902, EPI_ISL_474903, EPI_ISL_474919, EPI_ISL_474921 | Complejo Hospitalario Universitario de Albacete | SeqCOVID-SPAIN consortium/IBV(CSIC) | Encarnacion Simarro Córdoba, Julia Lozano Serra, Lorena Robles Fonseca , Monica Parra Grandes, Caridad Sainz de Baranda Camino and SeqCOVID-SPAIN consortium |
| EPI_ISL_474932 | Hospital Universitario Virgen de las Nieves de Granada-SAS | SeqCOVID-SPAIN consortium/IBV(CSIC) | Mercedes Pérez Ruiz, Sara Sanbonmatsu Gámez, Irene Pedrosa Corral, José M. Navarro-Marí and SeqCOVID-SPAIN consortium |
| EPI_ISL_474933 | Complejo Hospitalario Universitario de Albacete | SeqCOVID-SPAIN consortium/IBV(CSIC) | Encarnacion Simarro Córdoba, Julia Lozano Serra, Lorena Robles Fonseca , Monica Parra Grandes, Caridad Sainz de Baranda Camino and SeqCOVID-SPAIN consortium |
| EPI_ISL_474935 | Hospital Universitario Virgen de las Nieves de Granada-SAS | SeqCOVID-SPAIN consortium/IBV(CSIC) | Mercedes Pérez Ruiz, Sara Sanbonmatsu Gámez, Irene Pedrosa Corral, José M. Navarro-Marí and SeqCOVID-SPAIN consortium |
| EPI_ISL_474940, EPI_ISL_474945, EPI_ISL_474946, EPI_ISL_474947 | Complejo Hospitalario Universitario de Albacete | SeqCOVID-SPAIN consortium/IBV(CSIC) | Encarnacion Simarro Córdoba, Julia Lozano Serra, Lorena Robles Fonseca , Monica Parra Grandes, Caridad Sainz de Baranda Camino and SeqCOVID-SPAIN consortium |
| EPI_ISL_474949 | Hospital Universitario Virgen de las Nieves de Granada-SAS | SeqCOVID-SPAIN consortium/IBV(CSIC) | Mercedes Pérez Ruiz, Sara Sanbonmatsu Gámez, Irene Pedrosa Corral, José M. Navarro-Marí and SeqCOVID-SPAIN consortium |
| EPI_ISL_474951, EPI_ISL_474954 | Complejo Hospitalario Universitario de Albacete | SeqCOVID-SPAIN consortium/IBV(CSIC) | Encarnacion Simarro Córdoba, Julia Lozano Serra, Lorena Robles Fonseca , Monica Parra Grandes, Caridad Sainz de Baranda Camino and SeqCOVID-SPAIN consortium |
| EPI_ISL_475119 | Uppsala klinisk mikrobiologi | The Public Health Agency of Sweden | Oskar Karlsson Lindsjo, Maria Lind Karlberg, Mattias Haukland, Reza Advani, Olov Svartstrom, Anna-Malin Linde, Sandra Broddesson, Petra Edquist, Shamam Muradrasoli, Anna Risberg, Karin Tegmark-Wisell |
| EPI_ISL_475120, EPI_ISL_475121 | Halmstad klinisk mikrobiologi | The Public Health Agency of Sweden | Oskar Karlsson Lindsjo, Maria Lind Karlberg, Mattias Haukland, Reza Advani, Olov Svartstrom, Anna-Malin Linde, Sandra Broddesson, Petra Edquist, Shamam Muradrasoli, Anna Risberg, Karin Tegmark-Wisell |
| EPI_ISL_475149, EPI_ISL_475150 | Karolinska Universitetslaboratoriet | The Public Health Agency of Sweden | Oskar Karlsson Lindsjo, Maria Lind Karlberg, Mattias Haukland, Reza Advani, Olov Svartstrom, Anna-Malin Linde, Sandra Broddesson, Petra Edquist, Shamam Muradrasoli, Anna Risberg, Karin Tegmark-Wisell |
| EPI_ISL_475151 | Uppsala klinisk mikrobiologi | The Public Health Agency of Sweden | Oskar Karlsson Lindsjo, Maria Lind Karlberg, Mattias Haukland, Reza Advani, Olov Svartstrom, Anna-Malin Linde, Sandra Broddesson, Petra Edquist, Shamam Muradrasoli, Anna Risberg, Karin Tegmark-Wisell |
| EPI_ISL_475152 | Folkhalsomyndigheten | The Public Health Agency of Sweden | Oskar Karlsson Lindsjo, Maria Lind Karlberg, Mattias Haukland, Reza Advani, Olov Svartstrom, Anna-Malin Linde, Sandra Broddesson, Petra Edquist, Shamam Muradrasoli, Anna Risberg, Karin Tegmark-Wisell |
| EPI_ISL_475153, EPI_ISL_475154, EPI_ISL_475155 | Klinisk mikrobiologi Vasternorrland | The Public Health Agency of Sweden | Oskar Karlsson Lindsjo, Maria Lind Karlberg, Mattias Haukland, Reza Advani, Olov Svartstrom, Anna-Malin Linde, Sandra Broddesson, Petra Edquist, Shamam Muradrasoli, Anna Risberg, Karin Tegmark-Wisell |
| EPI_ISL_475537 | Narhalsan Oden VC | The Public Health Agency of Sweden | Oskar Karlsson Lindsjo, Maria Lind Karlberg, Mattias Haukland, Reza Advani, Olov Svartstrom, Anna-Malin Linde, Sandra Broddesson, Mia Brytting, Anna Risberg, Karin Tegmark-Wisell |
| EPI_ISL_475538 | Omtanken Grimmered | The Public Health Agency of Sweden | Oskar Karlsson Lindsjo, Maria Lind Karlberg, Mattias Haukland, Reza Advani, Olov Svartstrom, Anna-Malin Linde, Sandra Broddesson, Mia Brytting, Anna Risberg, Karin Tegmark-Wisell |
| EPI_ISL_475565 | Narhalsan Backa vardcentral | The Public Health Agency of Sweden | Oskar Karlsson Lindsjo, Maria Lind Karlberg, Mattias Haukland, Reza Advani, Olov Svartstrom, Anna-Malin Linde, Sandra Broddesson, Mia Brytting, Anna Risberg, Karin Tegmark-Wisell |
| EPI_ISL_475575, EPI_ISL_475576, EPI_ISL_475577, EPI_ISL_475579, EPI_ISL_475580, EPI_ISL_475581, EPI_ISL_475582, EPI_ISL_475583, EPI_ISL_475587, EPI_ISL_475588, EPI_ISL_475590, EPI_ISL_475593, EPI_ISL_475594, EPI_ISL_475595, EPI_ISL_475597, EPI_ISL_475598, EPI_ISL_475599, EPI_ISL_475601, EPI_ISL_475602, EPI_ISL_475604, EPI_ISL_475606, EPI_ISL_475614, EPI_ISL_475616, EPI_ISL_475621, EPI_ISL_475622, EPI_ISL_475644, EPI_ISL_475645, EPI_ISL_475647, EPI_ISL_475648, EPI_ISL_475649, EPI_ISL_475650, EPI_ISL_475652, EPI_ISL_475663, EPI_ISL_475664, EPI_ISL_475665, EPI_ISL_475686, EPI_ISL_475690, EPI_ISL_475691, EPI_ISL_475697, EPI_ISL_475701, EPI_ISL_475702, EPI_ISL_475709, EPI_ISL_475712, EPI_ISL_475713, EPI_ISL_475714 | | | |
| see above | Cedars-Sinai Medical Center, Department of Pathology & Laboratory Medicine, Molecular Pathology Laboratory | Cedars-Sinai Medical Center, Molecular Pathology Laboratory of Department of Pathology & Laboratory Medicine and Genomic Core | Wenjuan Zhang, John Paul Govindavari, Brian Davis, Stephanie Chen, Jong Taek Kim, Jianbo Song, Jean Lopategui, Jasmine T Plummer, Eric Vail |
| EPI_ISL_475766, EPI_ISL_475767 | Universitaetsklinik für Innere Medizin II Innsbruck | Bergthaler laboratory, CeMM Research Center for Molecular Medicine of the Austrian Academy of Sciences | Alexandra Popa, Benedikt Agerer, Henrique Colaco, Lukas Endler, Jakob-Wendelin Genger, Alexander Lercher, Mark Smyth, Thomas Penz, Michael Schuster, Jan Laine, Martin Senekowitsch, Judith Aberle, Stephan Aberle, Peter Hufnagl, Daniela Schmid, Franz Allerberger, Elisabeth Puchhammer-Stoeckl, Manfred Nairz, Guenter Weiss, Gregor Hörmann, Kinga Rigler-Hohenwarter, Rainer Gattringer, Wegene Borena, Dorothee von Laer, Christoph Bock, Andreas Bergthaler |
| EPI_ISL_475800, EPI_ISL_475801, EPI_ISL_475802, EPI_ISL_475803, EPI_ISL_475804, EPI_ISL_475805, EPI_ISL_475806, EPI_ISL_475807, EPI_ISL_475808, EPI_ISL_475809, EPI_ISL_475810, EPI_ISL_475811 | | | |
| see above | Center for Virology, Medical University of Vienna | Bergthaler laboratory, CeMM Research Center for Molecular Medicine of the Austrian Academy of Sciences | Alexandra Popa, Benedikt Agerer, Henrique Colaco, Lukas Endler, Jakob-Wendelin Genger, Alexander Lercher, Mark Smyth, Thomas Penz, Michael Schuster, Jan Laine, Martin Senekowitsch, Judith Aberle, Stephan Aberle, Peter Hufnagl, Daniela Schmid, Franz Allerberger, Elisabeth Puchhammer-Stoeckl, Manfred Nairz, Guenter Weiss, Gregor Hörmann, Kinga Rigler-Hohenwarter, Rainer Gattringer, Wegene Borena, Dorothee von Laer, Christoph Bock, Andreas Bergthaler |
| EPI_ISL_475828 | Institut für Virologie am Department für Hygiene, Mikrobiologie und Public Health | Bergthaler laboratory, CeMM Research Center for Molecular Medicine of the Austrian Academy of Sciences | Alexandra Popa, Benedikt Agerer, Henrique Colaco, Lukas Endler, Jakob-Wendelin Genger, Alexander Lercher, Mark Smyth, Thomas Penz, Michael Schuster, Jan Laine, Martin Senekowitsch, Judith Aberle, Stephan Aberle, Peter Hufnagl, Daniela Schmid, Franz Allerberger, Elisabeth Puchhammer-Stoeckl, Manfred Nairz, Guenter Weiss, Gregor Hörmann, Kinga Rigler-Hohenwarter, Rainer Gattringer, Wegene Borena, Dorothee von Laer, Christoph Bock, Andreas Bergthaler |
| EPI_ISL_475870 | Austrian Agency for Health and Food Safety (AGES) | Bergthaler laboratory, CeMM Research Center for Molecular Medicine of the Austrian Academy of Sciences | Alexandra Popa, Benedikt Agerer, Henrique Colaco, Lukas Endler, Jakob-Wendelin Genger, Alexander Lercher, Mark Smyth, Thomas Penz, Michael Schuster, Jan Laine, Martin Senekowitsch, Judith Aberle, Stephan Aberle, Peter Hufnagl, Daniela Schmid, Franz Allerberger, Elisabeth Puchhammer-Stoeckl, Manfred Nairz, Guenter Weiss, Gregor Hörmann, Kinga Rigler-Hohenwarter, Rainer Gattringer, Wegene Borena, Dorothee von Laer, Christoph Bock, Andreas Bergthaler |
| EPI_ISL_475894, EPI_ISL_475895, EPI_ISL_475896, EPI_ISL_475897, EPI_ISL_475898 | Zentralinstitut für medizinische und chemische Labordiagnostik, Universitätskliniken Insbruck | Bergthaler laboratory, CeMM Research Center for Molecular Medicine of the Austrian Academy of Sciences | Alexandra Popa, Benedikt Agerer, Henrique Colaco, Lukas Endler, Jakob-Wendelin Genger, Alexander Lercher, Mark Smyth, Thomas Penz, Michael Schuster, Jan Laine, Martin Senekowitsch, Judith Aberle, Stephan Aberle, Peter Hufnagl, Daniela Schmid, Franz Allerberger, Elisabeth Puchhammer-Stoeckl, Manfred Nairz, Guenter Weiss, Gregor Hörmann, Kinga Rigler-Hohenwarter, Rainer Gattringer, Wegene Borena, Dorothee von Laer, Christoph Bock, Andreas Bergthaler |
| EPI_ISL_475929, EPI_ISL_475932, EPI_ISL_475933, EPI_ISL_475934, EPI_ISL_475935 | Universitaetsklinik für Innere Medizin II Innsbruck | Bergthaler laboratory, CeMM Research Center for Molecular Medicine of the Austrian Academy of Sciences | Alexandra Popa, Benedikt Agerer, Henrique Colaco, Lukas Endler, Jakob-Wendelin Genger, Alexander Lercher, Mark Smyth, Thomas Penz, Michael Schuster, Jan Laine, Martin Senekowitsch, Judith Aberle, Stephan Aberle, Peter Hufnagl, Daniela Schmid, Franz Allerberger, Elisabeth Puchhammer-Stoeckl, Manfred Nairz, Guenter Weiss, Gregor Hörmann, Kinga Rigler-Hohenwarter, Rainer Gattringer, Wegene Borena, Dorothee von Laer, Christoph Bock, Andreas Bergthaler |
| EPI_ISL_476070 | University of Debrecen, Department of Medical Microbiology | National Laboratory of Virology, Szentágothai Research Centre | Endre Gábor Tóth, Balázs Somogyi, Brigitta Zana, Eszter Csoma, Ferenc Jakab, Gábor Kemenesi |
| EPI_ISL_476152, EPI_ISL_476168, EPI_ISL_476169, EPI_ISL_476170 | Laboratório de Patologia Clínica - UNICAMP | Laboratório de Estudos de Vírus Emergentes - UNICAMP | José Luiz Proença-Modena, Magnun Nueldo Nunes dos Santos, Angelica Schreiber, Julia Forato,Camila Simeoni, Marcilio Jorge Fumagalli, Mariene Ribeiro Amorim, Darlan da Silva Candido, Nuno Rodrigues Faria, Julien Theze, Luiz Gonzaga,Jaqueline Goes Jesus e William Marciel de Souza |
| EPI_ISL_476293, EPI_ISL_476294, EPI_ISL_476295, EPI_ISL_476296, EPI_ISL_476321, EPI_ISL_476322, EPI_ISL_476327, EPI_ISL_476328, EPI_ISL_476329, EPI_ISL_476330, EPI_ISL_476331, EPI_ISL_476332, EPI_ISL_476333, EPI_ISL_476334, EPI_ISL_476335, EPI_ISL_476336, EPI_ISL_476350, EPI_ISL_476351, EPI_ISL_476352, EPI_ISL_476353, EPI_ISL_476355, EPI_ISL_476356, EPI_ISL_476357, EPI_ISL_476358, EPI_ISL_476359, EPI_ISL_476360, EPI_ISL_476361, EPI_ISL_476362, EPI_ISL_476363, EPI_ISL_476364, EPI_ISL_476365, EPI_ISL_476366, EPI_ISL_476367, EPI_ISL_476368, EPI_ISL_476369, EPI_ISL_476370, | | | |

| | | | |
|---|---|---|---|
| EPI_ISL_476371 | | | |
| see above | DB Diagnósticos do Brasil | Instituto de Medicina Tropical da Univesidade de São Paulo | Samples: Nelson Gaburo Jr; Sequencing: Ingra Morales Claro, Jaqueline Goes de Jesus, Erika Regina Manuli, Flavia Cristina da Silva Sales, Thais de Moura Coletti, Camila Alves Maia da Silva, Mariana Severo Ramundo, Giula Magalhaes Ferreira, Darlan da Silva Candido, Julien Theze, Nuno Faria, Ester Sabino |
| EPI_ISL_476374, EPI_ISL_476375, EPI_ISL_476376, EPI_ISL_476377, EPI_ISL_476378, EPI_ISL_476379, EPI_ISL_476380, EPI_ISL_476381, EPI_ISL_476382, EPI_ISL_476383, EPI_ISL_476384, EPI_ISL_476385, EPI_ISL_476386 | | | |
| see above | Hospital da Clínicas da Faculdade de Medicina da Universidade de São Paulo | Instituto de Medicina Tropical da Univesidade de São Paulo | Samples: Ingra Morales Claro, Erika Regina Manuli, Cecilia Salete Alencar, Carolina S. Lazar, Silvia F. Costa; Sequencing: Ingra Morales Claro, Jaqueline Goes de Jesus, Erika Regina Manuli, Flavia Cristina da Silva Sales, Thais de Moura Coletti, Camila Alves Maia da Silva, Mariana Severo Ramundo, Giula Magalhaes Ferreira, Darlan da Silva Candido, Julien Theze, Nuno Faria, Ester Sabino |
| EPI_ISL_476391, EPI_ISL_476402 | Laboratório de Patologia Clínica - UNICAMP | Laboratório de Estudos de Vírus Emergentes - UNICAMP | José Luiz Proença-Modena, Magnun Nueldo Nunes dos Santos, Angelica Schreiber, Julia Forato,Camila Simeoni, Marcilio Jorge Fumagalli, Mariene Ribeiro Amorim, Darlan da Silva Candido, Nuno Rodrigues Faria, Julien Theze, Luiz Gonzaga,Jaqueline Goes Jesus e William Marciel de Souza |
| EPI_ISL_476498, EPI_ISL_476499, EPI_ISL_476500, EPI_ISL_476501, EPI_ISL_476511, EPI_ISL_476512, EPI_ISL_476513 | Laboratoire de microbiologie, Hopital de Verdun | Smith Laboratory, Centre de Recherche CHU Sainte-Justine | Martin Smith, Marieke Rozendaal, Ivan Pavlov |
| EPI_ISL_476702 | Incubadora Venezolana de Ciencia, Venezuela | Incubadora Venezolana de Ciencia, Venezuela / Instituto Nacional de Salud, Bogotá, Colombia / Grupo de Investigaciones Microbiológicas-UR (GIMUR), Departamento de Biología, Facultad de Ciencias Naturales, Universidad del Rosario, Bogotá, Colombia / Icahn School of Medicine at Mount Sinai, New York, USA | Alberto Paniz-Mondolfi, Marina Muñoz, Luis Perez-Garcia, Lourdes Delgado, Carolina Florez, Sergio Gomez, Angelica Rico, Lisseth Pardo, Esther C. Barros, Carolina Hernández, Jesús E. Jaimes, Aníbal A. Teherán, Ana S. Gonzalez-Reiche, Matthew M. Hernandez, Emilia Mia Sordillo, Viviana Simon, Harm van Bakel, Juan David Ramírez |
| EPI_ISL_476790, EPI_ISL_476791 | Stanford clinical virology lab | Chan-Zuckerberg Biohub | Benjamin Pinksy, Katharine Walter, Victoria N. Parikh, John Gorzynski, Hannah N. DeJong, Matthew T. Wheeler, Jason Andrews, Manuel Rivas, Carlos Bustamante, Euan Ashley, with CZB Cliahub Consortium |
| EPI_ISL_476825, EPI_ISL_476828 | Laboratoire des Fièvres Hémorragiques Virales du Benin | Charité-Universitätsmedizin Berlin | Yadouleton, Anges; Sander Anna-Lena; Moreira-Soto Andres; Drexler, Jan Felix |
| EPI_ISL_476835 | National Influenza Centre for Northern Greece | National Influenza Centre for Northern Greece | Maria Christoforidi |
| EPI_ISL_477196, EPI_ISL_477197 | Istituto Zooprofilattico Sperimentale Puglia e Basilicata; | Beaconlab (Bioinformatics, Evolution and Comparative Genomics lab), Dept of Biosciences, University on Milan | Parisi A.,Pesole G., Manzari C., Chiara M. |
| EPI_ISL_477816, EPI_ISL_477818 | Department of Pathology, University of Cambridge | COVID-19 Genomics UK (COG-UK) Consortium | Luke W Meredith, M. Estée Török, Myra Hosmillo, William L. Hamilton, Martin D. Curran, Theresa Feltwell, Grant Hall, Anna Yakovleva, Fahad A Khokhar, Charlotte J. Houldcroft, Laura G Caller, Aminu S. Jahun, Sarah L. Caddy, Yasmin Chaudhry, Malte Pinckert, Ian Goodfellow |
| EPI_ISL_477822, EPI_ISL_477823 | West of Scotland Specialist Virology Centre, NHSGGC / MRC-University of Glasgow Centre for Virus Research | COVID-19 Genomics UK (COG-UK) Consortium | Ana da Silva Filipe, Natasha Johnson, Kathy Smollett, Daniel Mair, Stephen Carmichael, Lily Tong, Jenna Nichols, Elihu Aranday-Cortes, Kirstyn Brunker, Yasmin Parr, Alice Broos, Kyriaki Nomikou; Sarah McDonald, Marc Niebel, Patawee Asamaphan; Richard Orton, Joseph Hughes, Sreenu Vattipally, David L Robertson; Alasdair MacLean, Rory Gunson; Kathy Li, Natasha Jesudason, Rajiv Shah, James Shepherd, Antonia Ho, Emma Thomson |
| EPI_ISL_478274, EPI_ISL_478279, EPI_ISL_478287 | University of Exeter | COVID-19 Genomics UK (COG-UK) Consortium | Ben Temperton,Aaron Jeffries,Michelle Michelsen,Joanna Warwick-Dugdale,Audrey Farbos,Robyn Manley,Stephen Michell,Jane Masoli |
| EPI_ISL_478372 | Liverpool Clinical Laboratories | COVID-19 Genomics UK (COG-UK) Consortium | Sam Haldenby, Anita Lucaci, Steve Paterson, Julian Hiscox, Alistair Darby, M Almsaud, A Alrezaihi, Muhannad Alruwaili, Stuart D Armstrong, Jones Benjamin, Eleanor G Bentley, Anu Chawla, Jordan J Clark, Angela Cowell, Richard Eccles, Isabel García-Dorival, Matthew Gemmell, Alessandro Gerada, PKF Gilmore, Richard Gregory, Ximeng Han, Catherine Hartley, Margaret Hughes, Miren Iturriza-Gomara, James Johnson, L Luu, Jenifer Manson, Charlotte Nelson, Elaine O'Toole, Cassie Olateju, Rebekah Penrice-Randal , Lucille Rainbow, N.P Randle, Trevor Ian Robinson, Parul Sharma, Ghada T Shawli, James P Stewart, Neil Swainston, Ecaterina Vamos, Joanne Watts, Mark Whitehead |
| EPI_ISL_478404, EPI_ISL_478432, EPI_ISL_478433, EPI_ISL_478434, EPI_ISL_478435, EPI_ISL_478436, EPI_ISL_478437, EPI_ISL_478438, EPI_ISL_478439, EPI_ISL_478440, EPI_ISL_478441, EPI_ISL_478442, EPI_ISL_478443, EPI_ISL_478444, EPI_ISL_478445, EPI_ISL_478446, EPI_ISL_478447, EPI_ISL_478448, EPI_ISL_478449, EPI_ISL_478450, EPI_ISL_478451, EPI_ISL_478452, EPI_ISL_478453, EPI_ISL_478454, EPI_ISL_478455, EPI_ISL_478456, EPI_ISL_478457, EPI_ISL_478458 | | | |
| see above | University College London, Great Ormond Street Hospital for Children NHS Foundation Trust, Imperial College Healthcare NHS Trust | COVID-19 Genomics UK (COG-UK) Consortium | Sergi Castellano, Rachel Williams, Mark Kristiansen, Paola Resende Silva, Sunando Roy, Tony Brooks, Helena Tutill, Paola Niola, Patricia Dyal, Charlotte Williams, Leysa Forrest, Yasmin Panchbhaya, Jacqueline Findlay, Samuel Weeks, Julianne Brown, Kathryn Harris, Paul Randell, James Price, Alison Holmes, Judith Breuer |
| EPI_ISL_478513, EPI_ISL_478516, EPI_ISL_478517, EPI_ISL_478518, EPI_ISL_478520, EPI_ISL_478521, EPI_ISL_478522, EPI_ISL_478523, EPI_ISL_478524, EPI_ISL_478525, EPI_ISL_478526, EPI_ISL_478527, EPI_ISL_478528, EPI_ISL_478529, EPI_ISL_478530, EPI_ISL_478639, EPI_ISL_478640, EPI_ISL_478641, EPI_ISL_478642, EPI_ISL_478643, EPI_ISL_478644, EPI_ISL_478645, EPI_ISL_478646, EPI_ISL_478647, EPI_ISL_478648, EPI_ISL_478649, EPI_ISL_478650, EPI_ISL_478651, EPI_ISL_478652, EPI_ISL_478653, EPI_ISL_478654, EPI_ISL_478655, EPI_ISL_478656, EPI_ISL_478657, EPI_ISL_478658, EPI_ISL_478659, EPI_ISL_478660, EPI_ISL_478661 | | | |
| see above | Northumbria University / South Tees Hospitals NHS Foundation Trust / North Cumbria Integrated Care NHS Foundation Trust / North Tees and Hartlepool NHS Foundation Trust / Newcastle Hospitals NHS Foundation Trust | COVID-19 Genomics UK (COG-UK) Consortium | Darren L Smith,Andrew Nelson,Matthew Bashton,Greg R Young,Joshua Loh,John Allan,Mohammad A Tariq,Giles S Holt,Gary Black,Wen C Yew,Lynn Dover,Paul Baker,Steve Liggett,Sarah Essex,Jane Greenaway,Debra Padgett,Clive Graham,Garren Scott,Edward Barton,Emma Swindells,Brendan Payne,Jennifer Collins,Yusri Taha,Gary Eltringham |
| EPI_ISL_478723, EPI_ISL_478724 | Queens Medical Centre, Clinical Microbiology Department / DeepSeq Nottingham | COVID-19 Genomics UK (COG-UK) Consortium | Gemma Clark, Wendy Smith, Manjinder Khakh, Vicki M Fleming, Michelle M Lister, Hannah Howson-Wells, Jonathan Ball, Patrick McClure, Joseph Chappell, Theocharis Tsoleridis, Nadine Holmes, Matthew Carlisle, Christopher Moore, Fei Sang, Johnny Debebe, Victoria Wright, Matthew Loose |
| EPI_ISL_478807, EPI_ISL_478971, EPI_ISL_479137, EPI_ISL_479141, EPI_ISL_479142, EPI_ISL_479148, EPI_ISL_479150, EPI_ISL_479156, EPI_ISL_479162, EPI_ISL_479169 | Oxford Viromics, NDM, University of Oxford; Oxford University Hospitals; Basingstoke and North Hampshire Hospital | COVID-19 Genomics UK (COG-UK) Consortium | Tanya Golubchik, David Bonsall, George Macintyre, Amy Trebes, Mariateresa de Cesare, Catrin Moore, Alex Mobbs, Anita Justice, Robert Shaw, Monique Andersson, Timothy Peto, Emma Wise, Nathan Moore, Jessica Lynch, Nick Cortes, Matilde Mori, Stephen Kidd, David Buck, John Todd, Christophe Fraser |
| EPI_ISL_479572 | NIV Influenza | NIV Influenza | Potdar V |
| EPI_ISL_479579, EPI_ISL_479580, EPI_ISL_479581 | National Public Health Laboratory, National Centre for Infectious Diseases | National Public Health Laboratory, National Centre for Infectious Diseases | Mak TM, Octavia S, Zhou Z, Chavatte JM, Cui L, Lin RTP |
| EPI_ISL_479759, EPI_ISL_479760 | University of Miami Immunology and Histocompatibility Laboratory | University of Miami Immunology and Histocompatibility Laboratory | Emilio Margolles-Clark, PhD and Phillip Ruiz, MD, PhD |
| EPI_ISL_480070, EPI_ISL_480071, EPI_ISL_480072 | Sakai City Institute of Public Health | Pathogen Genomics Center, National Institute of Infectious Diseases | Tsuyoshi Sekizuka, Tatsuya Miyoshi, Kentaro Itokawa, Rina Tanaka, Masanori Hashino, Hajime Kamiya, Motoi Suzuki, Makoto Kuroda |
| EPI_ISL_480082 | Shizuoka City Institute of Environmental Sciences and Public Health | Pathogen Genomics Center, National Institute of Infectious Diseases | Tsuyoshi Sekizuka, Takaharu Maehata,Sou Okamura,Yuji Kanazawa,Kenji Yagi, Kentaro Itokawa, Rina Tanaka, Masanori Hashino, Hajime Kamiya, Motoi Suzuki, Makoto Kuroda |
| EPI_ISL_480091, EPI_ISL_480092, EPI_ISL_480093, EPI_ISL_480094, EPI_ISL_480095, EPI_ISL_480096, EPI_ISL_480097, EPI_ISL_480098, EPI_ISL_480099, EPI_ISL_480100, EPI_ISL_480101, EPI_ISL_480102 | | | |
| see above | Department of Infectious Diseases, Kobe Institute of Health | Pathogen Genomics Center, National Institute of Infectious Diseases | Tsuyoshi Sekizuka, Ryohei Nomoto, Kentaro Itokawa, Rina Tanaka, Masanori Hashino, Hajime Kamiya, Motoi Suzuki, Makoto Kuroda |
| EPI_ISL_480135, EPI_ISL_480136, EPI_ISL_480137, EPI_ISL_480138, EPI_ISL_480139, EPI_ISL_480140, EPI_ISL_480141, EPI_ISL_480142, EPI_ISL_480143, EPI_ISL_480144, EPI_ISL_480145, EPI_ISL_480146, EPI_ISL_480147, EPI_ISL_480148, EPI_ISL_480149, EPI_ISL_480150, EPI_ISL_480151, EPI_ISL_480152, EPI_ISL_480154, EPI_ISL_480155, EPI_ISL_480156, EPI_ISL_480158, EPI_ISL_480159, EPI_ISL_480160, EPI_ISL_480161, EPI_ISL_480162, EPI_ISL_480163 | | | |
| see above | Fukui Prefectural Institute of Public Health and Environmental Science | Pathogen Genomics Center, National Institute of Infectious Diseases | Tsuyoshi Sekizuka, Miho Toho, Kentaro Itokawa, Rina Tanaka, Masanori Hashino, Hajime Kamiya, Motoi Suzuki, Makoto Kuroda |

| | | | |
|---|---|---|---|
| EPI_ISL_480169, EPI_ISL_480170, EPI_ISL_480171, EPI_ISL_480172, EPI_ISL_480173, EPI_ISL_480174, EPI_ISL_480175, EPI_ISL_480176 | Gunma Prefectural Institute of Public Health and Environmental Sciences | Pathogen Genomics Center, National Institute of Infectious Diseases | Tsuyoshi Sekizuka, Hiroyuki Tsukagoshi, Kentaro Itokawa, Rina Tanaka, Masanori Hashino, Hajime Kamiya, Motoi Suzuki, Makoto Kuroda |
| EPI_ISL_480183, EPI_ISL_480184, EPI_ISL_480185, EPI_ISL_480186, EPI_ISL_480187, EPI_ISL_480188, EPI_ISL_480189 | Ibaraki Prefectural Institute of Public Health | Pathogen Genomics Center, National Institute of Infectious Diseases | Tsuyoshi Sekizuka, Keiko Goto, Kentaro Itokawa, Rina Tanaka, Masanori Hashino, Hajime Kamiya, Motoi Suzuki, Makoto Kuroda |
| EPI_ISL_480197, EPI_ISL_480203 | Toyama Institute of Health | Pathogen Genomics Center, National Institute of Infectious Diseases | Tsuyoshi Sekizuka, Masae Itamochi, Kazunori Oishi, Kentaro Itokawa, Rina Tanaka, Masanori Hashino, Hajime Kamiya, Motoi Suzuki, Makoto Kuroda |
| EPI_ISL_480217 | Department of Infectious Diseases, Kobe Institute of Health | Pathogen Genomics Center, National Institute of Infectious Diseases | Tsuyoshi Sekizuka, Ryohei Nomoto, Kentaro Itokawa, Rina Tanaka, Masanori Hashino, Hajime Kamiya, Motoi Suzuki, Makoto Kuroda |
| EPI_ISL_480222, EPI_ISL_480223 | Koshigaya City Public Health Center | Pathogen Genomics Center, National Institute of Infectious Diseases | Tsuyoshi Sekizuka, Yuka Furui, Aya Tamura, Kyohei Sakata, Takumi Daimon, Yoko Togawa, Yoshiko Hamada, Kentaro Itokawa, Rina Tanaka, Masanori Hashino, Hajime Kamiya, Motoi Suzuki, Makoto Kuroda |
| EPI_ISL_480327 | Hospital Nacional de Niños | Charité Virology-University of Costa Rica | Andres Moreira-Soto, Eugenia Corrales-Aguilar, Ignacio Postigo-Hidalgo, Cristian Pérez Corrales, Andrei Montero Bonilla, Jan Felix Drexler |
| EPI_ISL_480328 | Laboratorio LABIN | Charité Virology-University of Costa Rica | Andres Moreira-Soto, Eugenia Corrales-Aguilar, Ignacio Postigo-Hidalgo, Ignacio Soto Pacheco, Jan Felix Drexler |
| EPI_ISL_480394, EPI_ISL_480396 | University of Wisconsin-Madison AIDS Vaccine Research Laboratories | University of Wisconsin-Madison AIDS Vaccine Research Laboratories | Gage Moreno, Katarina Braun, et al. AIDS Vaccine Research Laboratories |
| EPI_ISL_480571, EPI_ISL_480574, EPI_ISL_480576, EPI_ISL_480577, EPI_ISL_480579 | Victorian Infectious Diseases Reference Laboratory (VIDRL) | VIDRL and MDU-PHL | Caly L., Seemann T., Sait, M., Schultz M., Druce J., Sherry, N. |
| EPI_ISL_480616, EPI_ISL_480617 | Microbiological Diagnostic Unit - Public Health Laboratory (MDU-PHL) | MDU-PHL | Seemann T., Schultz M., Sait, M., Sherry, N. |
| EPI_ISL_480784, EPI_ISL_480785 | NYC Department of Health and Mental Hygiene | Pathogen Discovery, Respiratory Viruses Branch, Division of Viral Diseases, Centers for Disease Control and Prevention | Krista Queen, Christine Mahl, Jennifer Rakeman, Anna Uehara, Ying Tao, Jing Zhang, Yan Li, Clinton R. Paden, Haibin Wang, Jasmine Padilla, Justin Lee, Sally Slavinski, Suxiang Tong |
| EPI_ISL_480815, EPI_ISL_480817, EPI_ISL_480818, EPI_ISL_480819, EPI_ISL_480820, EPI_ISL_480821, EPI_ISL_480822, EPI_ISL_480823, EPI_ISL_480824, EPI_ISL_480825, EPI_ISL_480826, EPI_ISL_480827, EPI_ISL_480828, EPI_ISL_480829, EPI_ISL_480830, EPI_ISL_480831, EPI_ISL_480832, EPI_ISL_480833, EPI_ISL_480834, EPI_ISL_480835, EPI_ISL_480836, EPI_ISL_480837, EPI_ISL_480838, EPI_ISL_480839, EPI_ISL_480840, EPI_ISL_480841, EPI_ISL_480842, EPI_ISL_480843, EPI_ISL_480844, EPI_ISL_480845, EPI_ISL_480846, EPI_ISL_480847, EPI_ISL_480848, EPI_ISL_480849, EPI_ISL_480851, EPI_ISL_480852 | | | |
| see above | Florida Bureau of Public Health Laboratories | Florida Bureau of Public Health Laboratories | Sarah Schmedes, Jason Blanton |
| EPI_ISL_480981, EPI_ISL_480995, EPI_ISL_481025, EPI_ISL_481035 | ISGlobal, Institut de Salut Global de Barcelona | SeqCOVID-SPAIN consortium/IBV(CSIC) | Alfredo Mayor, Alberto L Garcia-Basteiro, Carlota Dobaño, Gemma Moncunill, Pau Cisteró and SeqCOVID-SPAIN consortium |
| EPI_ISL_481080, EPI_ISL_481088, EPI_ISL_481102 | Hospital General Universitario Gregorio Marañón | SeqCOVID-SPAIN consortium/IBV(CSIC) | Laura Pérez-Lago, Marta Herranz, Jon Sicilia, Julia Suárez, Pilar Catalán, Patricia Muñoz, Darío García de Viedma and SeqCOVID-SPAIN consortium |
| EPI_ISL_481217 | Oslo University Hospital, Department of Medical Microbiology | Norwegian Institute of Public Health, Department of Virology | Kathrine Stene-Johansen, Kamilla Heddeland Instefjord, Hilde Elshaug, Rasmus Riis Kopperud, Karoline Bragstad, Olav Hungnes |
| EPI_ISL_481220 | Institut Pasteur Dakar | Institut Pasteur de Dakar | Ndongo Dia, Moussa Moise Diagne, Mamadou Diop, Marie Henriette Dior Ndione, Mamadou Malado Jallow, Safietou Sanke, Ousmane Faye, Amadou Alpha Sall. |
| EPI_ISL_481252 | Department of Emerging Infectious Diseases, Institute of Tropical Medicine, Nagasaki University | Department of Emerging Infectious Diseases, Institute of Tropical Medicine, Nagasaki University | Jiro Yasuda, Rokusuke Yoshikawa, Yuichiro Furusato, Haruka Abe |
| EPI_ISL_481271, EPI_ISL_481272, EPI_ISL_481273, EPI_ISL_481274, EPI_ISL_481275 | Maryland Department of Health | Maryland Department of Health | Keller,E. |
| EPI_ISL_481649, EPI_ISL_481650, EPI_ISL_481651, EPI_ISL_481652, EPI_ISL_481653, EPI_ISL_481654, EPI_ISL_481655, EPI_ISL_481664, EPI_ISL_481665, EPI_ISL_481666, EPI_ISL_481667, EPI_ISL_481668, EPI_ISL_481669, EPI_ISL_481670, EPI_ISL_481671, EPI_ISL_481672, EPI_ISL_481673, EPI_ISL_481682, EPI_ISL_481683, EPI_ISL_481684, EPI_ISL_481685, EPI_ISL_481686, EPI_ISL_481687, EPI_ISL_481688, EPI_ISL_481689, EPI_ISL_481690, EPI_ISL_481691, EPI_ISL_481692, EPI_ISL_481693, EPI_ISL_481707, EPI_ISL_481708, EPI_ISL_481709, EPI_ISL_481710, EPI_ISL_481711, EPI_ISL_481712, EPI_ISL_481713 | | | |
| see above | Department of Virology and Immunology, University of Helsinki and Helsinki University Hospital, Huslab Finland | Department of Virology, Faculty of Medicine, University of Helsinki, Helsinki, Finland | Teemu Smura, Hannimari Kallio-Kokko, Jenni Virtanen, Maija Suvanto, Sari Hannula, Harri Kangas, Pekka Ellonen, Olli Vapalahti |
| EPI_ISL_481748, EPI_ISL_481750, EPI_ISL_481751, EPI_ISL_481752, EPI_ISL_481753 | Dr. Georges-L.-Dumont University Hospital Centre | National Microbiology Laboratory | Anna Majer, Shari Tyson, Grace Seo, Kristyn Burak, Philip Mabon, Elsie Grudeski, Rhiannon Huzarewich, Russell Mandes, Jennifer Tanner, Natalie Knox, Morag Graham, Gary Van Domselaar, Richard Garceau, Guillaume Desnoyers, Nathalie Bastien, Yan Li, Timothy Booth |
| EPI_ISL_481989, EPI_ISL_482006 | PHE South West Regional Laboratory, National Infection Service | Wellcome Sanger Institute for the COVID-19 Genomics UK (COG-UK) consortium | Stephanie Hutchings, Hannah Pymont, Dr Peter Muir, Barry Vipond, Rich Hopes; and Alex Alderton, Roberto Amato, Sonia Goncalves, Ewan Harrison, David K. Jackson, Ian Johnston, Dominic Kwiatkowski, Cordelia Langford, John Sillitoe on behalf of the Wellcome Sanger Institute COVID-19 Surveillance Team (http://www.sanger.ac.uk/covid-team) |
| EPI_ISL_482294, EPI_ISL_482295, EPI_ISL_482302, EPI_ISL_482303, EPI_ISL_482304, EPI_ISL_482305, EPI_ISL_482448, EPI_ISL_482450, EPI_ISL_482453, EPI_ISL_482454 | Providence St. Joseph Health Molecular Genomics Laboratory | Providence St. Joseph Health Molecular Genomics Laboratory | Alexa K Dowdell, Brian D Piening, Fred L Robinson, Carlo B Bifulco, Mary Campbell |
| EPI_ISL_482673, EPI_ISL_482674, EPI_ISL_482675, EPI_ISL_482676, EPI_ISL_482681 | Singapore General Hospital | Department of Microbiology | Nurdyana Abdul Rahman, Kun Lee Lim, Chenhao Li, Kian Sing Chan, Lynette Oon, Kern Rei Chng, Niranjan Nagarajan, Karrie Ko |
| EPI_ISL_482732 | LNR National Reference Laboratory, Mohammed VI University of Health Sciences | Medical Biotechnology Laboratory, Rabat Medical and Pharmacy School, Mohammed The Vth University in Rabat | Meriem LAAMARTI, Souad KARTTI, Rokia LAAMARTI , M.W. CHEMAO-ELFIHRI, Loubna ALLAM, Mouna OUADGHIRI, Imane SMYEJ, Jalila RAHOUI, Houda BENRAHMA, Jalil El ATAR, Idrissa DIAWARA, Rachid EL JAOUDI, Laila SBABOU, Chakib NEJJARI, Saaid AMZAZI, Rachid MENTAG, Lahcen BELYAMANI and Azeddine IBRAHIMI |
| EPI_ISL_482744 | Laboratory Diagnostic, Veterinary Specialized Institute Kraljevo | Laboratory Diagnostic, Veterinary Specialized Institute Kraljevo | Vidanovic,D., Tesovic,B., Banovic Djeri,B., Knezevic,A., Vidanovic,D., Tesovic,B., Banovic Djeri,B., Knezevic,A., Afonso,C. |
| EPI_ISL_482777 | Queen Elizabeth Hospital | Hong Kong Department of Health | Mak Gannon C.K., Cheng Peter K.C., Lam Edman T.K., Chan Rickjason C.W., Tsang Dominic N.C. |
| EPI_ISL_483081 | SA Pathology | SA Pathology | Lex Leong, Chuan Kok Lim, Mark Turra, Ivan Bastian, Geoff Higgins |
| EPI_ISL_483165, EPI_ISL_483166, EPI_ISL_483169, EPI_ISL_483176, EPI_ISL_483178, EPI_ISL_483180, EPI_ISL_483184, EPI_ISL_483187, EPI_ISL_483191, EPI_ISL_483192, EPI_ISL_483198, EPI_ISL_483203, EPI_ISL_483258, EPI_ISL_483259, EPI_ISL_483261, EPI_ISL_483343, EPI_ISL_483354, EPI_ISL_483360, EPI_ISL_483361, EPI_ISL_483362, EPI_ISL_483363, EPI_ISL_483364, EPI_ISL_483365, EPI_ISL_483366, EPI_ISL_483369, EPI_ISL_483380, EPI_ISL_483381, EPI_ISL_483382, EPI_ISL_483384, EPI_ISL_483385, EPI_ISL_483387, EPI_ISL_483390, EPI_ISL_483437, EPI_ISL_483438, EPI_ISL_483440, EPI_ISL_483444, EPI_ISL_483448 | | | |
| see above | UC San Diego Center for Advanced Laboratory Medicine | Andersen lab at Scripps Research | SEARCH Alliance San Diego with David Pride, Ji H Shin |
| EPI_ISL_483556, EPI_ISL_483562 | Kingdom of Bahrein Ministry of Health | Erasmus Medical Center | Bas Oude Munnink, David Nieuwenhuijse, Reina Sikkema, Fatema, Ebrahim Shehad, Amjad Ghanem Mohamed, Hashmeya Al Wasti, Claudia Schapendonk, Irina Chestakova, Anne van der Linden, Theo Bestebroer, Stefan van Nieuwkoop, Mark Pronk, Pascal Lexmond, Richard Molenkamp, Marion Koopmans, on behalf of the Dutch national COVID-19 response team. |

| | | | |
|---|---|---|---|
| EPI_ISL_483585, EPI_ISL_483587, EPI_ISL_483589, EPI_ISL_483602, EPI_ISL_483605, EPI_ISL_483612 | National Public Health Laboratory, National Centre for Infectious Diseases | National Public Health Laboratory, National Centre for Infectious Diseases | Mak TM, Octavia S, Zhou Z, Chavatte JM, Cui L, Lin RTP |
| EPI_ISL_483640 | Kingdom of Bahrein Ministry of Health | Erasmus Medical Center | Bas Oude Munnink, David Nieuwenhuijse, Reina Sikkema, Fatema, Ebrahim Shehad, Amjad Ghanem Mohamed, Hashmeya Al Wasti, Claudia Schapendonk, Irina Chestakova, Anne van der Linden, Theo Bestebroer, Stefan van Nieuwkoop, Mark Pronk, Pascal Lexmond, Richard Molenkamp, Marion Koopmans, on behalf of the Dutch national COVID-19 response team. |
| EPI_ISL_483714, EPI_ISL_483716, EPI_ISL_483719, EPI_ISL_483725 | Israel Central Virology laboratory | Israel Central Virology laboratory | Neta Zuckerman, Efrat Dahan Bucris, Oran Erster, Ella Mendelson, Michal Mandelboim |
| EPI_ISL_484769, EPI_ISL_484792, EPI_ISL_484796, EPI_ISL_484798, EPI_ISL_484799, EPI_ISL_484800, EPI_ISL_484802, EPI_ISL_484803, EPI_ISL_484805 | University of Michigan Clinical Microbiology Laboratory | Lauring Lab, University of Michigan, Department of Microbiology and Immunology | Valesano et al. |
| EPI_ISL_485251, EPI_ISL_485257, EPI_ISL_485258, EPI_ISL_485259, EPI_ISL_485273, EPI_ISL_485279, EPI_ISL_485283, EPI_ISL_485284, EPI_ISL_485298, EPI_ISL_485330, EPI_ISL_485355, EPI_ISL_485356, EPI_ISL_485357, EPI_ISL_485358, EPI_ISL_485359, EPI_ISL_485360, EPI_ISL_485361, EPI_ISL_485362, EPI_ISL_485363, EPI_ISL_485364, EPI_ISL_485365, EPI_ISL_485366, EPI_ISL_485367, EPI_ISL_485368, EPI_ISL_485369, EPI_ISL_485370, EPI_ISL_485371, EPI_ISL_485372, EPI_ISL_485373, EPI_ISL_485374, EPI_ISL_485375, EPI_ISL_485376, EPI_ISL_485377, EPI_ISL_485378, EPI_ISL_485379, EPI_ISL_485380, EPI_ISL_485381, EPI_ISL_485382, EPI_ISL_485383, EPI_ISL_485384, EPI_ISL_485385, EPI_ISL_485386, EPI_ISL_485387 | | | |
| see above | River Road Testing Lab | Ginkgo Bioworks Clinical Laboratory | Rebecca C. Christofferson, Stephania A. Cormier, Luan V. Dinh, E. Handly Mayton, Hollis R. O'Neil, Thaya Stoufflet, Malaika Mckenzie-Bennett, James McGann, Jim Griffin, Keith Robison, Alex Plocik, Becky Schilling, Rebecca Littlefield, Michelle Spencer, Birgitte Simen |
| EPI_ISL_485389, EPI_ISL_485390 | University of Ulsan College of Medicine and Asan Medical Center | University of Ulsan College of Medicine and Asan Medical Center | Kuenyoul Park, Jaewoong Lee, Kihyun Lee, Jiwon Jung, Sung-Han Kim, Jina Lee, Mauricio Chalita, Seok-Hwan Yoon, Jongsik Chun, Kyu-Hwa Hur, Heungsup Sung, Mi-Na Kim, and Hae Kyung Lee |
| EPI_ISL_485578 | Instituto de diagnóstico y Referencia Epidemiologicos (INDRE) | Instituto de diagnóstico y Referencia Epidemiologicos (INDRE) | Barrera-Badillo,G., Ramirez-Gonzalez,E. |
| EPI_ISL_485872 | New Mexico Department of Health Scientific Laboratory Division | Center for Global Health, University of New Mexico Health Sciences Center | Daryl Domman, Kurt Schwalm, Twila Kunde, Joseph Hicks, Michael Edwards, Darrell Dinwiddie |
| EPI_ISL_486097, EPI_ISL_486102 | UW Virology Lab | UW Virology Lab | Pavitra Roychoudhury, Hong Xie, Lasata Shrestha, Amin Addetia, Truong Nguyen, Victoria M Rachleff, Meei-Li Huang, Keith R Jerome, Alexander Greninger |
| EPI_ISL_486288 | San Joaquin County Public Health Lab | Chan-Zuckerberg Biohub | CZB Cliahub Consortium |
| EPI_ISL_486652 | Microbiology, Virology and Biemergency Laboratory-ASST FBF Sacco | Microbiology, Virology and Biemergency Laboratory-ASST FBF Sacco | Micheli V, Comandatore F, Romeri F, Mancon A, Rimoldi SG |
| EPI_ISL_486654 | Microbiology, Virology and Biemergency Laboratory-ASST FBF Sacco | Microbiology, Virology and Biemergency Laboratory-ASST FBF Sacco | Romeri F, Comandatore F, Mancon A, Micheli V, Rimoldi SG |
| EPI_ISL_486655 | Microbiology, Virology and Biemergency Laboratory-ASST FBF Sacco | Microbiology, Virology and Biemergency Laboratory-ASST FBF Sacco | Mancon A, Comandatore F, Romeri F, Micheli V, Rimoldi SG |
| EPI_ISL_486865 | Institut Pasteur Dakar | Institut Pasteur de Dakar | Ndongo Dia, Moussa Moise Diagne, Mamadou Diop, Marie Henriette Dior Ndione, Mamadou Malado Jallow, Safietou Sanke, Ousmane Faye, Amadou Alpha Sall. |
| EPI_ISL_487276 | Department of Food Safety, Nutrition and Veterinary public health, Istituto Superiore di Sanita' | Department of Biomedical, Surgical and Dental Sciences and Department of Biomedical Sciences for Health | Delbue,S., Ferrante,P., Basilico,N., Parapini,S., Binda,S., D'Alessandro,S., Galli,C., Signorini,L., Primache,V., Anselmi,G., Pariani,E. |
| EPI_ISL_487370, EPI_ISL_487377, EPI_ISL_487379, EPI_ISL_487381 | Hellenic Pasteur Institute, National Influenza Reference laboratory of Southern Greece & Unit of Bioinformatics and Applied Genomics | Hellenic Pasteur Institute, National Influenza Reference laboratory of Southern Greece & Unit of Bioinformatics and Applied Genomics | Vasiliki Pogka, Timokratis Karamitros, Athanasios Kossyvakis, Antonios Kalliaropoulos, Horefti Elina, Evangelidou Maria, Androniki Voulgari-Kokota, Aspasia Kontou, Andreas Mentis |
| EPI_ISL_487446, EPI_ISL_487447, EPI_ISL_487448, EPI_ISL_487449 | CICM-Mali | Bundeswehr Institut of Microbiology | Kouriba, Dürr, Sangaré, Rehn, Traoré, Bestehorn-Willmann, Walter, Quedraogo, Zimmermann, Maiga, Heitzer, Sogodogo, Antwerpen, Wölfel |
| EPI_ISL_487689, EPI_ISL_487699, EPI_ISL_487709, EPI_ISL_487716, EPI_ISL_487717, EPI_ISL_487724, EPI_ISL_487733, EPI_ISL_487740, EPI_ISL_487762, EPI_ISL_487791, EPI_ISL_487804, EPI_ISL_487808, EPI_ISL_487813, EPI_ISL_487816, EPI_ISL_487838, EPI_ISL_487850, EPI_ISL_487851, EPI_ISL_487858, EPI_ISL_487893, EPI_ISL_487897, EPI_ISL_487914, EPI_ISL_487922, EPI_ISL_487962, EPI_ISL_487966, EPI_ISL_487968, EPI_ISL_487970, EPI_ISL_487971, EPI_ISL_487975, EPI_ISL_487982, EPI_ISL_487983, EPI_ISL_487987 | | | |
| see above | Virology Department, Royal Infirmary of Edinburgh, NHS Lothian / School of Biological Sciences, University of Edinburgh | Wellcome Sanger Institute for the COVID-19 Genomics UK (COG-UK) consortium | McHugh M, Dewar R, Rooke S, O'Toole Á, Scher E, Hill V, McCrone JT, Colquhoun R, Yu X, Jackson B, Rambaut A, Templeton K and Alex Alderton, Roberto Amato, Sonia Goncalves, Ewan Harrison, David K. Jackson, Ian Johnston, Dominic Kwiatkowski, Cordelia Langford, John Sillitoe on behalf of the Wellcome Sanger Institute COVID-19 Surveillance Team (http://www.sanger.ac.uk/covid-team) |
| EPI_ISL_488187, EPI_ISL_488192, EPI_ISL_488193, EPI_ISL_488196, EPI_ISL_488211, EPI_ISL_488242, EPI_ISL_488251, EPI_ISL_488258, EPI_ISL_488261, EPI_ISL_488267, EPI_ISL_488268, EPI_ISL_488279, EPI_ISL_488280, EPI_ISL_488288, EPI_ISL_488293, EPI_ISL_488315, EPI_ISL_488325, EPI_ISL_488329, EPI_ISL_488341 | | | |
| see above | PHE South West Regional Laboratory, National Infection Service | Wellcome Sanger Institute for the COVID-19 Genomics UK (COG-UK) consortium | Stephanie Hutchings, Hannah Pymont, Dr Peter Muir, Barry Vipond, Rich Hopes; and Alex Alderton, Roberto Amato, Sonia Goncalves, Ewan Harrison, David K. Jackson, Ian Johnston, Dominic Kwiatkowski, Cordelia Langford, John Sillitoe on behalf of the Wellcome Sanger Institute COVID-19 Surveillance Team (http://www.sanger.ac.uk/covid-team) |
| EPI_ISL_488342 | PHE South West Regional Laboratory, National Infection Service | Wellcome Sanger Institute for the COVID-19 Genomics UK (COG-UK) Consortium | Stephanie Hutchings, Hannah Pymont, Dr Peter Muir, Barry Vipond, Rich Hopes; and Alex Alderton, Roberto Amato, Sonia Goncalves, Ewan Harrison, David K. Jackson, Ian Johnston, Dominic Kwiatkowski, Cordelia Langford, John Sillitoe on behalf of the Wellcome Sanger Institute COVID-19 Surveillance Team |
| EPI_ISL_488347, EPI_ISL_488382, EPI_ISL_488389, EPI_ISL_488390, EPI_ISL_488392, EPI_ISL_488419, EPI_ISL_488423, EPI_ISL_488432, EPI_ISL_488439 | PHE South West Regional Laboratory, National Infection Service | Wellcome Sanger Institute for the COVID-19 Genomics UK (COG-UK) consortium | Stephanie Hutchings, Hannah Pymont, Dr Peter Muir, Barry Vipond, Rich Hopes; and Alex Alderton, Roberto Amato, Sonia Goncalves, Ewan Harrison, David K. Jackson, Ian Johnston, Dominic Kwiatkowski, Cordelia Langford, John Sillitoe on behalf of the Wellcome Sanger Institute COVID-19 Surveillance Team (http://www.sanger.ac.uk/covid-team) |
| EPI_ISL_488456, EPI_ISL_488457, EPI_ISL_488460, EPI_ISL_488461, EPI_ISL_488462, EPI_ISL_488468, EPI_ISL_488476, EPI_ISL_488484, EPI_ISL_488489, EPI_ISL_488492, EPI_ISL_488499, EPI_ISL_488500, EPI_ISL_488502, EPI_ISL_488504, EPI_ISL_488505, EPI_ISL_488530, EPI_ISL_488532, EPI_ISL_488547, EPI_ISL_488567, EPI_ISL_488570, EPI_ISL_488586, EPI_ISL_488587, EPI_ISL_488589, EPI_ISL_488590, EPI_ISL_488592, EPI_ISL_488595, EPI_ISL_488598, EPI_ISL_488610, EPI_ISL_488612, EPI_ISL_488618, EPI_ISL_488621, EPI_ISL_488624, EPI_ISL_488629, EPI_ISL_488630, EPI_ISL_488634, EPI_ISL_488637, EPI_ISL_488640, EPI_ISL_488641, EPI_ISL_488648, EPI_ISL_488650, EPI_ISL_488656, EPI_ISL_488661, EPI_ISL_488678, EPI_ISL_488685, EPI_ISL_488688, EPI_ISL_488690, EPI_ISL_488694, EPI_ISL_488697, EPI_ISL_488698, EPI_ISL_488699, EPI_ISL_488710, EPI_ISL_488722, EPI_ISL_488742, EPI_ISL_488745, EPI_ISL_488751, EPI_ISL_488761, EPI_ISL_488784, EPI_ISL_488808, EPI_ISL_488827, EPI_ISL_488835, EPI_ISL_488836 | | | |
| see above | NU-OMICS DNA Sequencing research facility, Northumbria University | Wellcome Sanger Institute for the COVID-19 Genomics UK (COG-UK) consortium | Chris Duncan, Sheia Waugh, Shirelle Burton-Fanning, Gary Eltringham, Jennifer Collins, Brendan Payne, Yusri Taha, Emma Swindells, Jane Greenaway, Edward Barton, Garren Scott, Debra Padgett, Clive Graham, Sarah Essex, Steve Ligget, Paul Baker, Lynn Dover, Wen Yew, Gary Black, John Allan, Joshua Loh, Greg Young, Matthew Bashton, Andrew Nelson, Darren Smith and Alex Alderton, Roberto Amato, Sonia Goncalves, Ewan Harrison, David K. Jackson, Ian Johnston, Dominic Kwiatkowski, Cordelia Langford, John Sillitoe on behalf of the Wellcome Sanger Institute COVID-19 Surveillance Team (http://www.sanger.ac.uk/covid-team) |
| EPI_ISL_488879, EPI_ISL_488880, EPI_ISL_488883, EPI_ISL_488888, EPI_ISL_488889, EPI_ISL_488890, EPI_ISL_488891, EPI_ISL_488893, EPI_ISL_488894, EPI_ISL_488903, EPI_ISL_488910, EPI_ISL_488911, EPI_ISL_488913, EPI_ISL_488914, EPI_ISL_488916, EPI_ISL_488917, EPI_ISL_488919, EPI_ISL_488920, EPI_ISL_488922, EPI_ISL_488923, EPI_ISL_488927, EPI_ISL_488929, EPI_ISL_488932, EPI_ISL_488933, EPI_ISL_488934, EPI_ISL_488935, EPI_ISL_488938, EPI_ISL_488939, EPI_ISL_488941, EPI_ISL_488942, EPI_ISL_488948, EPI_ISL_488949, EPI_ISL_488951, EPI_ISL_488954, EPI_ISL_488956, EPI_ISL_488959, EPI_ISL_488962, EPI_ISL_488963, EPI_ISL_488967, EPI_ISL_488972, EPI_ISL_488974, EPI_ISL_488976, EPI_ISL_488979, EPI_ISL_488981, EPI_ISL_488982, EPI_ISL_488986, EPI_ISL_488989, EPI_ISL_488990, EPI_ISL_488991, EPI_ISL_488992, EPI_ISL_488993, EPI_ISL_488994, EPI_ISL_488999, EPI_ISL_489000, EPI_ISL_489001, EPI_ISL_489004, EPI_ISL_489005, EPI_ISL_489007, EPI_ISL_489008, EPI_ISL_489009, EPI_ISL_489013, EPI_ISL_489014, EPI_ISL_489019, EPI_ISL_489021, EPI_ISL_489022, EPI_ISL_489023, EPI_ISL_489029, EPI_ISL_489031, EPI_ISL_489032, EPI_ISL_489034, EPI_ISL_489035, EPI_ISL_489037, EPI_ISL_489038, EPI_ISL_489040, EPI_ISL_489042, EPI_ISL_489045, EPI_ISL_489046, EPI_ISL_489048, EPI_ISL_489050, EPI_ISL_489052, EPI_ISL_489053, EPI_ISL_489055, EPI_ISL_489056, EPI_ISL_489057, EPI_ISL_489058, EPI_ISL_489061, EPI_ISL_489063 | | | |
| see above | Virology Department, Royal Infirmary of Edinburgh, NHS | Wellcome Sanger Institute for the COVID-19 Genomics UK | McHugh M, Dewar R, Rooke S, O'Toole Á, Scher E, Hill V, McCrone JT, Colquhoun R, Yu X, Jackson B, Rambaut A, Templeton K and Alex Alderton, |

| | | | |
|---|---|---|---|
| | Lothian / School of Biological Sciences, University of Edinburgh | (COG-UK) consortium | Roberto Amato, Sonia Goncalves, Ewan Harrison, David K. Jackson, Ian Johnston, Dominic Kwiatkowski, Cordelia Langford, John Sillitoe on behalf of the Wellcome Sanger Institute COVID-19 Surveillance Team (http://www.sanger.ac.uk/covid-team) |
| EPI_ISL_490033 | South Eastern Area Laboratory Services (SEALS) | NSW Health Pathology - Institute of Clinical Pathology and Medical Research; Westmead Hospital; University of Sydney | CIDM-PH et al. |
| EPI_ISL_490775, EPI_ISL_490842 | Wales Specialist Virology Centre Sequencing lab: Pathogen Genomics Unit | COVID-19 Genomics UK (COG-UK) Consortium | Catherine Moore, Johnathan Evans, Laura Gifford, Malorie Perry, Simon Cottrell, Angela Marchbank, Alec Birchley, Alexander Adams, Amy Gaskin, Bree Gatica-Wilcox, Jason Coombes, Joel Southgate, Lauren Gilbert, Lee Graham, Nicole Pacchiarini, Sara Kumziene-Summerhayes, Sarah Taylor, Sophie Jones, Sara Rey, Matthew Bull, Joanne Watkins, Sally Corden, Tom Connor |
| EPI_ISL_491085 | Suceava County Emergency Hospital | "Stefan cel Mare" University Metagenomics Lab | Lobiuc Andrei, Antoniadis Panagiotis et al. |
| EPI_ISL_491121, EPI_ISL_491128 | Oman-National Influenza Center | Biotechnology & OMICs Laboratory | Samira Al-Mahruqi, Abdul Latif Khan, Samiha Al-Kharusi, Adil Khan , Ahmed Al-Rawahi, Sajjad Asaf, Amina Al-Jardani, Hanan Al-Kindi, Intisar Al-Shukri, Ahlam Al-Amri, Aisha Al-Amri, Aisha Al-Busaidi, Adil Al-Wahaibi, Seif Al-Abri, Ahmed Al-Harrasi |
| EPI_ISL_491135 | Oman-National Influenza Center | Biotechnology & OMICs Laboratory | Samiha Al-Kharusi, Sajjad Asaf, Abdul Latif Khan, Samira Al-Mahruqi, Adil Khan, Ahmed Al-Rawahi, Amina Al-Jardani, Hanan Al-Kindi, Intisar Al-Shukri, Ahlam Al-Amri, Aisha Al-Amri, Aisha Al-Busaidi, Adil Al-Wahaibi, Seif Al-Abri, Ahmed Al-Harrasi |
| EPI_ISL_491161 | Oman-National Influenza Center | Biotechnology & OMICs Laboratory | Sajjad Asaf, Samiha Al-Kharusi, Ahmed Al-Harrasi, Samira Al-Mahruqi, Adil Khan, Ahmed Al-Rawahi, Abdul Latif Khan, Amina Al-Jardani, Hanan Al-Kindi, Intisar Al-Shukri, Ahlam Al-Amri, Aisha Al-Amri, Aisha Al-Busaidi, Adil Al-Wahaibi, Seif Al-Abri. |
| EPI_ISL_491481 | Functional Genomics Core University of South Carolina / Prisma Health-Midlands | Functional Genomics Core, Center For Targeted Therapeutics, | Hao Ji, Diego Altomare, B.Celia Cui, Mengqian Chen, Alyssa Clay-Glimour, Michael Wyatt, Phillip Buckhaults, Helmut Albrecht, Michael Shtutman |
| EPI_ISL_491501, EPI_ISL_491537, EPI_ISL_491541, EPI_ISL_491546, EPI_ISL_491552, EPI_ISL_491568, EPI_ISL_491569, EPI_ISL_491576, EPI_ISL_491578, EPI_ISL_491601, EPI_ISL_491602, EPI_ISL_491604, EPI_ISL_491637, EPI_ISL_491640 | | | |
| see above | Virology Department, Royal Infirmary of Edinburgh, NHS Lothian / School of Biological Sciences, University of Edinburgh | Wellcome Sanger Institute for the COVID-19 Genomics UK (COG-UK) consortium | McHugh M, Dewar R, Rooke S, O'Toole Á, Scher E, Hill V, McCrone JT, Colquhoun R, Yu X, Jackson B, Rambaut A, Templeton K and Alex Alderton, Roberto Amato, Sonia Goncalves, Ewan Harrison, David K. Jackson, Ian Johnston, Dominic Kwiatkowski, Cordelia Langford, John Sillitoe on behalf of the Wellcome Sanger Institute COVID-19 Surveillance Team (http://www.sanger.ac.uk/covid-team) |
| EPI_ISL_491907, EPI_ISL_491921, EPI_ISL_491922, EPI_ISL_491923, EPI_ISL_491924, EPI_ISL_491925 | Naval Infectious Diseases Diagnostic Laboratory | Naval Medical Research Center Biological Defense Research Directorate | Logan Voegtly, Regina Cer, Lindsay Glang, Victor Sugiharto, Francisco Malgon Bautista, Hua Wei Chen, Dessiree Pena-Gomez, Megan Schilling, Adrian Paskey, Kyle Long, Mark Simons, Kimberly Bishop-Lilly |
| EPI_ISL_492057, EPI_ISL_492058, EPI_ISL_492059, EPI_ISL_492060 | Alaska State Virology Laboratory | Alaska State Virology Laboratory | Chen J et al with Pathogenomics group Dagdag R, Redlinger M, Milton E, George W, Kovalenko A, Drown DM, Bortz E |
| EPI_ISL_492066, EPI_ISL_492069 | 1. ViroGenetics - BSL3 Laboratory of Virology, Maopolska Centre of Biotechnology, Jagiellonian University; 2. II Department of Internal Medicine, Faculty of Medicine, Jagiellonian University Medical College; 3. Narodowy Instytut Zdrowia Publicznego - Pastwowy Zakad Higieny (NIZP-PZH) | 1. ViroGenetics - BSL3 Laboratory of Virology, Maopolska Centre of Biotechnology, Jagiellonian University; 2. II Department of Internal Medicine, Faculty of Medicine, Jagiellonian University Medical College; 3. Narodowy Instytut Zdrowia Publicznego - Pastwowy Zakad Higieny (NIZP-PZH). | Katarzyna Pancer, Marek Sanak, Aleksandra A. Zasada, Magdalena Rzeczkowska, Tomasz Wokowicz, Katarzyna Zacharczuk, Agnieszka Koakowska-Kulesza, Katarzyna Owczarek, Aleksandra Milewska, Natalia Wolaniuk, Ewelina Hallman-Szeliska, Pawe P abaj, Wojciech Branicki, Krzysztof Pyr |
| EPI_ISL_492187, EPI_ISL_492203, EPI_ISL_492213, EPI_ISL_492215, EPI_ISL_492223, EPI_ISL_492231, EPI_ISL_492242, EPI_ISL_492251, EPI_ISL_492257, EPI_ISL_492261, EPI_ISL_492272, EPI_ISL_492275, EPI_ISL_492280, EPI_ISL_492286, EPI_ISL_492295, EPI_ISL_492302, EPI_ISL_492312, EPI_ISL_492323, EPI_ISL_492326, EPI_ISL_492334, EPI_ISL_492343, EPI_ISL_492345, EPI_ISL_492347, EPI_ISL_492356, EPI_ISL_492369, EPI_ISL_492374, EPI_ISL_492382, EPI_ISL_492387, EPI_ISL_492402, EPI_ISL_492403, EPI_ISL_492418, EPI_ISL_492419, EPI_ISL_492435, EPI_ISL_492436, EPI_ISL_492572, EPI_ISL_492580, EPI_ISL_492588 | | | |
| see above | PHE South West Regional Laboratory, National Infection Service | Wellcome Sanger Institute for the COVID-19 Genomics UK (COG-UK) consortium | Stephanie Hutchings, Hannah Pymont, Dr Peter Muir, Barry Vipond, Rich Hopes; and Alex Alderton, Roberto Amato, Sonia Goncalves, Ewan Harrison, David K. Jackson, Ian Johnston, Dominic Kwiatkowski, Cordelia Langford, John Sillitoe on behalf of the Wellcome Sanger Institute COVID-19 Surveillance Team (http://www.sanger.ac.uk/covid-team) |
| EPI_ISL_492596 | PHE South West Regional Laboratory, National Infection Service | Wellcome Sanger Institute for the COVID-19 Genomics UK (COG-UK) Consortium | Stephanie Hutchings, Hannah Pymont, Dr Peter Muir, Barry Vipond, Rich Hopes; and Alex Alderton, Roberto Amato, Sonia Goncalves, Ewan Harrison, David K. Jackson, Ian Johnston, Dominic Kwiatkowski, Cordelia Langford, John Sillitoe on behalf of the Wellcome Sanger Institute COVID-19 Surveillance Team |
| EPI_ISL_492624, EPI_ISL_492633, EPI_ISL_492635, EPI_ISL_492636, EPI_ISL_492638, EPI_ISL_492738, EPI_ISL_492741, EPI_ISL_492770, EPI_ISL_492773, EPI_ISL_492809 | PHE South West Regional Laboratory, National Infection Service | Wellcome Sanger Institute for the COVID-19 Genomics UK (COG-UK) consortium | Stephanie Hutchings, Hannah Pymont, Dr Peter Muir, Barry Vipond, Rich Hopes; and Alex Alderton, Roberto Amato, Sonia Goncalves, Ewan Harrison, David K. Jackson, Ian Johnston, Dominic Kwiatkowski, Cordelia Langford, John Sillitoe on behalf of the Wellcome Sanger Institute COVID-19 Surveillance Team (http://www.sanger.ac.uk/covid-team) |
| EPI_ISL_492869, EPI_ISL_492872 | Royal Free Hospital / Health Services Laboratories | Wellcome Sanger Institute for the COVID-19 Genomics UK (COG-UK) consortium | Tanzina Haque, Tabitha Mahungu, Dianne Irish, Cate Goodlad, Jenny Cross, Judith Heaney and Alex Alderton, Roberto Amato, Sonia Goncalves, Ewan Harrison, David K. Jackson, Ian Johnston, Dominic Kwiatkowski, Cordelia Langford, John Sillitoe on behalf of the Wellcome Sanger Institute COVID-19 Surveillance Team (http://www.sanger.ac.uk/covid-team) |
| EPI_ISL_493001, EPI_ISL_493064, EPI_ISL_493125, EPI_ISL_493126, EPI_ISL_493127, EPI_ISL_493128 | Wyoming Public Health Laboratory | Wyoming Public Health Laboratory | Noah Hull, Rob Christensen, Jim Mildenberger, Joel Sevinsky, Cari Sloma, and Wanda Manley |
| EPI_ISL_493129 | Functional Genomics Core University of South Carolina / Prisma Health-Midlands | Functional Genomics Core, University of South Carolina, | Hao Ji, Diego Altomare, B.Celia Cui, Mengqian Chen, Alyssa Clay-Glimour, Michael Wyatt, Phillip Buckhaults, Helmut Albrecht, Michael Shtutman |
| EPI_ISL_493335 | Instituto de Diagnostico y Referencia Epidemiologicos (INDRE) | Instituto de Diagnostico y Referencia Epidemiologicos (INDRE) | Gisela Barrera-Badillo , Abril Rodriguez-Maldonado, Claudia Wong-Arambula , Natividad Cruz-Ortiz, Tatiana Nunez-Garcia, Dayanira Arellano-Suarez, Fabiola Garces-Ayala, Edgar Mendieta-Condado, Lucia Hernandez-Rivas, Irma Lopez-Martinez, Ernesto Ramirez-Gonzalez. |
| EPI_ISL_493351 | Oslo University Hospital, Department of Medical Microbiology | Norwegian Institute of Public Health, Department of Virology | Kathrine Stene-Johansen, Kamilla Heddeland Instefjord, Hilde Elshaug, Rasmus Riis Kopperud, Karoline Bragstad, Olav Hungnes |
| EPI_ISL_493610, EPI_ISL_493611 | Lincolnshire Hospitals and DeepSeq Nottingham | COVID-19 Genomics UK (COG-UK) Consortium | Nichola Duckworth, Tim Sloan, Sarah Walsh, Jonathan Ball, Patrick McClure, Joeseph Chappell, Nadine Holmes, Matthew Carlisle, Christopher Moore, Fei Sang, Johnny Debebe, Victoria Wright, Matthew Loose |
| EPI_ISL_493638, EPI_ISL_493639, EPI_ISL_493640, EPI_ISL_493641, EPI_ISL_493642, EPI_ISL_493643, EPI_ISL_493644, EPI_ISL_493645, EPI_ISL_493648, EPI_ISL_493649, EPI_ISL_493650, EPI_ISL_493651, EPI_ISL_493652, EPI_ISL_493654, EPI_ISL_493655, EPI_ISL_493657, EPI_ISL_493658, EPI_ISL_493659, EPI_ISL_493660, EPI_ISL_493661, EPI_ISL_493663, EPI_ISL_493664 | | | |
| see above | Centre for Enzyme Innovation, University of Portsmouth / Translational Research Laboratory, Portsmouth Hospitals NHS Trust | COVID-19 Genomics UK (COG-UK) Consortium | Angela Beckett,Yann Bourgeois,Garry Scarlett,Sharon Glaysher,Scott Elliott,Kelly Bicknell,Robert Impey,Allyson Lloyd,Sarah Wyllie,Ethan Butcher,Anoop Chauhan,Samuel Robson |
| EPI_ISL_493742, EPI_ISL_493743, EPI_ISL_493745 | West of Scotland Specialist Virology Centre, NHSGGC / MRC-University of Glasgow Centre for Virus Research | COVID-19 Genomics UK (COG-UK) Consortium | Ana da Silva Filipe, Natasha Johnson, Kathy Smollett, Daniel Mair, Stephen Carmichael, Lily Tong, Jenna Nichols, Elihu Aranday-Cortes, Kirstyn Brunker, Yasmin Parr, Alice Broos, Kyriaki Nomikou; Sarah McDonald, Marc Niebel, Patawee Asamaphan; Richard Orton, Joseph Hughes, Sreenu Vattipally, David L Robertson; Alasdair MacLean, Rory Gunson; Kathy Li, Natasha Jesudason, Rajiv Shah, James Shepherd, Antonia Ho, Emma Thomson |
| EPI_ISL_494186, EPI_ISL_494192, EPI_ISL_494230, EPI_ISL_494238, EPI_ISL_494271, EPI_ISL_494371 | Wales Specialist Virology Centre Sequencing lab: Pathogen Genomics Unit | COVID-19 Genomics UK (COG-UK) Consortium | Catherine Moore, Johnathan Evans, Laura Gifford, Malorie Perry, Simon Cottrell, Angela Marchbank, Alec Birchley, Alexander Adams, Amy Gaskin, Bree Gatica-Wilcox, Jason Coombes, Joel Southgate, Lauren Gilbert, Lee Graham, Nicole Pacchiarini, Sara Kumziene-Summerhayes, Sarah Taylor, Sophie Jones, Sara Rey, Matthew Bull, Joanne Watkins, Sally Corden, Tom Connor |
| EPI_ISL_494372, EPI_ISL_494373, EPI_ISL_494374, EPI_ISL_494377, EPI_ISL_494378, EPI_ISL_494379, EPI_ISL_494381, EPI_ISL_494385, EPI_ISL_494392, EPI_ISL_494394, EPI_ISL_494398, EPI_ISL_494402, EPI_ISL_494404, EPI_ISL_494405, EPI_ISL_494406, EPI_ISL_494407, EPI_ISL_494409, EPI_ISL_494410, EPI_ISL_494411, EPI_ISL_494416, EPI_ISL_494417, EPI_ISL_494418, EPI_ISL_494419, EPI_ISL_494421, EPI_ISL_494427, EPI_ISL_494627, EPI_ISL_494629, EPI_ISL_494630, EPI_ISL_494631, EPI_ISL_494632 | | | |
| see above | San Diego County Public Health Laboratory | Andersen lab at Scripps Research | SEARCH Alliance San Diego with Tracy Basler, Jovan Shephard, Brett Austin |
| EPI_ISL_494665, EPI_ISL_494668 | Scripps Medical Laboratory | Andersen lab at Scripps Research | SEARCH Alliance San Diego with Michael Quigley, Ellen Stefanski, Ian Mchardy |
| EPI_ISL_494722, EPI_ISL_494724, | San Diego County Public Health Laboratory | Andersen lab at Scripps Research | SEARCH Alliance San Diego with Tracy Basler, Jovan Shephard, Brett Austin |

| | | | |
|---|---|---|---|
| EPI_ISL_494726, EPI_ISL_494730, EPI_ISL_494731, EPI_ISL_494737, EPI_ISL_494738, EPI_ISL_494741 | | | |
| EPI_ISL_495375, EPI_ISL_495376, EPI_ISL_495377, EPI_ISL_495378, EPI_ISL_495380, EPI_ISL_495381, EPI_ISL_495382 | Florida Bureau of Public Health Laboratories | Florida Bureau of Public Health Laboratories | Sarah Schmedes, Jason Blanton |
| EPI_ISL_495593 | University of Michigan Clinical Microbiology Laboratory | Lauring Lab, University of Michigan, Department of Microbiology and Immunology | Valesano et al. |
| EPI_ISL_495598, EPI_ISL_495601, EPI_ISL_495603, EPI_ISL_495606, EPI_ISL_495607, EPI_ISL_495608 | Mayo Clinic & Mayo Clinic Laboratories | Minnesota Department of Health, Public Health Laboratory | Matt Plumb, Jacob Garfin, and Xiong Wang |
| EPI_ISL_496810, EPI_ISL_496811, EPI_ISL_496812, EPI_ISL_496813, EPI_ISL_496814, EPI_ISL_496815, EPI_ISL_496816, EPI_ISL_496817, EPI_ISL_496818, EPI_ISL_496819, EPI_ISL_496820, EPI_ISL_496821, EPI_ISL_496822, EPI_ISL_496823, EPI_ISL_496824, EPI_ISL_496825, EPI_ISL_496826, EPI_ISL_496827, EPI_ISL_496828, EPI_ISL_496829, EPI_ISL_496830, EPI_ISL_496831, EPI_ISL_496832, EPI_ISL_496833, EPI_ISL_496834, EPI_ISL_496835, EPI_ISL_496836, EPI_ISL_496837, EPI_ISL_496838, EPI_ISL_496839, EPI_ISL_496840, EPI_ISL_496841, EPI_ISL_496842, EPI_ISL_496843, EPI_ISL_496844, EPI_ISL_496845, EPI_ISL_496846, EPI_ISL_496847, EPI_ISL_496848, EPI_ISL_496849, EPI_ISL_496850, EPI_ISL_496851, EPI_ISL_496852, EPI_ISL_496853, EPI_ISL_496855, EPI_ISL_496856 | | | |
| see above | Gorgas Memorial Laboratory of Health Studies | Gorgas Memorial Laboratory of Health Studies | Danilo Franco, Claudia Gonzalez Sandra Lopez-Verges, Alexander A Martinez |
| EPI_ISL_498167 | Instituto Nacional de Salud, Bogotá, Colombia | Instituto Nacional de Salud, Bogotá, Colombia | Katherine Laiton-Donato, Diego A. Álvarez-Díaz, Carlos Franco-Muñoz, Jonathan Reales, Diego Andrés Prada, Jose A. Usme-Ciro, Nicolas D. Franco-Sierra, Zulma M. Cucunubá, Christian Julian VillabonaArenas, Liz Villabona-Arenas, Sussy Echeverria, Astrid C. Flórez, Carolina Ferro, Diana Marcela Walteros-Acero, Franklin Prieto, Carlos Andrés Durán, Martha Lucia Ospina Martinez, Marcela Mercado-Reyes |
| EPI_ISL_498520, EPI_ISL_498528, EPI_ISL_498533, EPI_ISL_498535 | ACT Pathology | Schwessinger Lab | Ashley Jones, Benjamin Schwessinger, Robert Lanfear, Robyn N Hall, Megan McDonald, Ming-Dao Chia, Kevin Murray, Craig Kennedy, Karina Kennedy |
| EPI_ISL_499270, EPI_ISL_499271, EPI_ISL_499272, EPI_ISL_499273, EPI_ISL_499274, EPI_ISL_499275, EPI_ISL_499276, EPI_ISL_499277, EPI_ISL_499278, EPI_ISL_499279, EPI_ISL_499280, EPI_ISL_499282, EPI_ISL_499283, EPI_ISL_499284, EPI_ISL_499285, EPI_ISL_499286, EPI_ISL_499287, EPI_ISL_499288, EPI_ISL_499289, EPI_ISL_499290, EPI_ISL_499291, EPI_ISL_499292, EPI_ISL_499293, EPI_ISL_499294, EPI_ISL_499295, EPI_ISL_499296, EPI_ISL_499297, EPI_ISL_499298, EPI_ISL_499299, EPI_ISL_499300 | | | |
| see above | Centre for Enzyme Innovation, University of Portsmouth / Translational Research Laboratory, Portsmouth Hospitals NHS Trust | COVID-19 Genomics UK (COG-UK) Consortium | Angela Beckett,Yann Bourgeois,Garry Scarlett,Sharon Glaysher,Scott Elliott,Kelly Bicknell,Robert Impey,Allyson Lloyd,Sarah Wyllie,Ethan Butcher,Anoop Chauhan,Samuel Robson |
| EPI_ISL_499331, EPI_ISL_499332, EPI_ISL_499335, EPI_ISL_499336, EPI_ISL_499337, EPI_ISL_499338, EPI_ISL_499340, EPI_ISL_499341, EPI_ISL_499342, EPI_ISL_499343, EPI_ISL_499344, EPI_ISL_499345, EPI_ISL_499346, EPI_ISL_499347, EPI_ISL_499349, EPI_ISL_499350, EPI_ISL_499351 | | | |
| see above | Virology Department, Sheffield Teaching Hospitals NHS Foundation Trust/Department of Infection, Immunity and Cardiovascular Disease, The Medical School, University of Sheffield | COVID-19 Genomics UK (COG-UK) Consortium | Thushan de Silva, Matthew Parker, Nikki Smith, Adri Angyal, Rebecca Brown, Luke Green, Rachel Tucker, Paul Parsons, Danielle Groves, Katie Johnson, Laura Carrilero, Alex Keeley, Dave Partridge, Matthew Wyles, Benjamin Lindsey, Mehmet Yavuz, Mohammad Raza, Cariad Evans |
| EPI_ISL_499366, EPI_ISL_499370, EPI_ISL_499384, EPI_ISL_499409, EPI_ISL_499428, EPI_ISL_499440, EPI_ISL_499448 | Wales Specialist Virology Centre Sequencing lab: Pathogen Genomics Unit | COVID-19 Genomics UK (COG-UK) Consortium | Catherine Moore, Johnathan Evans, Laura Gifford, Malorie Perry, Simon Cottrell, Angela Marchbank, Alec Birchley, Alexander Adams, Amy Gaskin, Bree Gatica-Wilcox, Jason Coombes, Joel Southgate, Lauren Gilbert, Lee Graham, Nicole Pacchiarini, Sara Kumziene-Summerhayes, Sarah Taylor, Sophie Jones, Sara Rey, Matthew Bull, Joanne Watkins, Sally Corden, Tom Connor |
| EPI_ISL_500292, EPI_ISL_500293, EPI_ISL_500294, EPI_ISL_500295, EPI_ISL_500296, EPI_ISL_500297, EPI_ISL_500298, EPI_ISL_500299, EPI_ISL_500300, EPI_ISL_500301, EPI_ISL_500302, EPI_ISL_500303, EPI_ISL_500304, EPI_ISL_500305, EPI_ISL_500306, EPI_ISL_500307, EPI_ISL_500308, EPI_ISL_500309, EPI_ISL_500310, EPI_ISL_500311, EPI_ISL_500312, EPI_ISL_500313, EPI_ISL_500314, EPI_ISL_500315, EPI_ISL_500316, EPI_ISL_500317, EPI_ISL_500318, EPI_ISL_500319, EPI_ISL_500320, EPI_ISL_500321, EPI_ISL_500322, EPI_ISL_500323 | | | |
| see above | Servicio de Microbiología. Hospital Universitario Donostia. OSI Donostialdea. Área de Enfermedades Infecciosas, Grupo de Infección Respiratoria y Resistencia Antimicrobiana. Instituto de Investigación Sanitaria Biodonostia | SeqCOVID-SPAIN consortium/IBV(CSIC) | Gustavo Cilla, Milagrosa Montes, Luis Piñeiro, Jose Maria Marimón and SeqCOVID-SPAIN consortium |
| EPI_ISL_500374, EPI_ISL_500376 | Centro de Investigación Biomédica de La Rioja - Hospital San Pedro Logroño | SeqCOVID-SPAIN consortium/IBV(CSIC) | María de Toro, José Manuel Azcona Gutiérrez, María Pilar Bea Escudero, Miriam Blasco Alberdi and SeqCOVID-SPAIN consortium |
| EPI_ISL_500471 | LACEN/PE | WallauLab, Aggeu Magalhaes Institute | Marcelo Henrique Santos Paiva, Duschinka Ribeiro Duarte Guedes, Cássia Docena, Matheus Filgueira Bezerra, Filipe Zimmer Dezordi, Laís Ceschini Machado, Larissa Krokovsky, Elisama Helvecio, Alexandre Freitas da Silva, Luydson Richardson Silva Vasconcelos, Antonio Mauro Rezende, Severino Jefferson Ribeiro da Silva, Kamila Gaudêncio da Silva Sales, Bruna Santos Lima Figueiredo de Sá, Derciliano Lopes da Cruz, Claudio Eduardo Cavalcanti, Armando de Menezes Neto, Caroline Targino Alves da Silva, Renata Pessôa Germano Mendes, Maria Almerice Lopes da Silva, Tiago Gräf, Paola Cristina Resende, Gonzalo Bello, Michelle da Silva Barros, Wheverton Ricardo Correia do Nascimento, Rodrigo Moraes Loyo Arcoverde, Luciane Caroline Albuquerque Bezerra, Sinval Pinto Brandão Filho, Constância Flávia Junqueira Ayres, Gabriel Luz Wallau |
| EPI_ISL_500517, EPI_ISL_500518 | Mayo Clinic Laboratories | University of Washington Virology Lab | Pavitra Roychoudhury, Hong Xie, Lasata Shrestha, Amin Addetia, Truong Nguyen, Victoria M Rachleff, Meei-Li Huang, Keith R Jerome, Alexander Greninger |
| EPI_ISL_500624, EPI_ISL_500625 | Area of Virology, Serology and Virology Division (SAViD), New South Wales Health Pathology Randwick | Area of Virology, Serology and Virology Division (SAViD), New South Wales Health Pathology Randwick | Rawlinson, W. |
| EPI_ISL_501169 | Baylor College of Medicine | Baylor College of Medicine: HGSC | Vasanthi Avadhanula, Erin Nicholson, David Henke, Pedro Piedra, Harsha Doddapaneni, Donna Muzny, Qingchang Meng, Hsu Chao, Zeineen Momin, Hua Shen, George Weissenberger, Kavya Kottapalli, Yimiti Meiheerguli, Sejal Salvi, Ginger Metcalf, Vipin Menon, Sara J.J. Cregeen, Matthew C. Ross, Tulin Ayvaz, Richard Sucgang, Kristi L. Hoffman, Matthew Wong, Joseph F. Petrosino |
| EPI_ISL_501211, EPI_ISL_501212, EPI_ISL_501213, EPI_ISL_501214, EPI_ISL_501215, EPI_ISL_501216, EPI_ISL_501217 | Department of Medical Microbiology, University Malaya Medical Centre | Department of Medical Microbiology, Faculty of Medicine, University of Malaya | Yoong Min CHONG, Jennifer Chong, I-Ching SAM, Yoke Fun CHAN, University Malaya Medical Centre COVID Team |
| EPI_ISL_506988, EPI_ISL_506989, EPI_ISL_506991, EPI_ISL_506992, EPI_ISL_506993 | Division of Viral Diseases, Center for Laboratory Control of Infectious Diseases, Korea Centers for Diseases Control and Prevention | Division of Viral Diseases, Center for Laboratory Control of Infectious Diseases, Korea Centers for Diseases Control and Prevention | Jeong-Min Kim, Yoon-Seok Chung, Namjoo Lee, Sang Hee Woo, Hye-Jun Jo, Heui Man Kim, Jun-Sub Kim, Dong Hyun Song, Daesang Lee, Seong Tae Jeong, Myung Guk Han |
| EPI_ISL_507027, EPI_ISL_507028, EPI_ISL_507029, EPI_ISL_507030, EPI_ISL_507031 | unknown | Infectious Diseases Research, King Abdullah International Medical Research Center (KAIMRC) | Alghoribi,M.F. |
| EPI_ISL_507081 | University College London Hospital | COVID-19 Genomics UK (COG-UK) Consortium | Judith Heaney, Matthew Byott, Catherine Houlihan, Dan Frampton, Stuart Kirk, Moira Spyer and Eleni Nastouli |
| EPI_ISL_507131, EPI_ISL_507132, EPI_ISL_507133, EPI_ISL_507134 | Centre for Enzyme Innovation, University of Portsmouth / Translational Research Laboratory, Portsmouth Hospitals NHS Trust | COVID-19 Genomics UK (COG-UK) Consortium | Angela Beckett,Yann Bourgeois,Garry Scarlett,Sharon Glaysher,Scott Elliott,Kelly Bicknell,Robert Impey,Allyson Lloyd,Sarah Wyllie,Ethan Butcher,Anoop Chauhan,Samuel Robson |
| EPI_ISL_507161, EPI_ISL_507163, EPI_ISL_507165, EPI_ISL_507166, EPI_ISL_507168, EPI_ISL_507169, EPI_ISL_507172, EPI_ISL_507177, EPI_ISL_507179 | Virology Department, Sheffield Teaching Hospitals NHS Foundation Trust/Department of Infection, Immunity and Cardiovascular Disease, The Medical School, University of Sheffield | COVID-19 Genomics UK (COG-UK) Consortium | Thushan de Silva, Matthew Parker, Nikki Smith, Adri Angyal, Rebecca Brown, Luke Green, Rachel Tucker, Paul Parsons, Danielle Groves, Katie Johnson, Laura Carrilero, Alex Keeley, Dave Partridge, Matthew Wyles, Benjamin Lindsey, Mehmet Yavuz, Mohammad Raza, Cariad Evans |

| | | | |
|---|---|---|---|
| EPI_ISL_507415, EPI_ISL_507416, EPI_ISL_507417, EPI_ISL_507419, EPI_ISL_507421, EPI_ISL_507422, EPI_ISL_507423, EPI_ISL_507424, EPI_ISL_507425, EPI_ISL_507426, EPI_ISL_507428, EPI_ISL_507429, EPI_ISL_507430, EPI_ISL_507433, EPI_ISL_507436, EPI_ISL_507437, EPI_ISL_507439, EPI_ISL_507440, EPI_ISL_507443, EPI_ISL_507444, EPI_ISL_507446, EPI_ISL_507472, EPI_ISL_507476, EPI_ISL_507477, EPI_ISL_507478, EPI_ISL_507479, EPI_ISL_507487, EPI_ISL_507491, EPI_ISL_507492, EPI_ISL_507494, EPI_ISL_507497, EPI_ISL_507498, EPI_ISL_507499, EPI_ISL_507500, EPI_ISL_507501, EPI_ISL_507502, EPI_ISL_507503, EPI_ISL_507504, EPI_ISL_507505, EPI_ISL_507506, EPI_ISL_507507, EPI_ISL_507508, EPI_ISL_507509, EPI_ISL_507510, EPI_ISL_507511, EPI_ISL_507512, EPI_ISL_507513, EPI_ISL_507514, EPI_ISL_507515, EPI_ISL_507516, EPI_ISL_507517, EPI_ISL_507518, EPI_ISL_507519, EPI_ISL_507520, EPI_ISL_507521, EPI_ISL_507522, EPI_ISL_507523, EPI_ISL_507524, EPI_ISL_507525, EPI_ISL_507526, EPI_ISL_507527, EPI_ISL_507528, EPI_ISL_507532, EPI_ISL_507550, EPI_ISL_507553, EPI_ISL_507554, EPI_ISL_507557, EPI_ISL_507559, EPI_ISL_507561, EPI_ISL_507562, EPI_ISL_507563, EPI_ISL_507565, EPI_ISL_507568, EPI_ISL_507569, EPI_ISL_507571, EPI_ISL_507574, EPI_ISL_507576, EPI_ISL_507577, EPI_ISL_507579, EPI_ISL_507581, EPI_ISL_507582, EPI_ISL_507583, EPI_ISL_507587, EPI_ISL_507588, EPI_ISL_507589, EPI_ISL_507590, EPI_ISL_507591, EPI_ISL_507592, EPI_ISL_507593, EPI_ISL_507594, EPI_ISL_507596, EPI_ISL_507598, EPI_ISL_507599, EPI_ISL_507603, EPI_ISL_507605, EPI_ISL_507606, EPI_ISL_507607, EPI_ISL_507608, EPI_ISL_507610, EPI_ISL_507611, EPI_ISL_507615, EPI_ISL_507616, EPI_ISL_507617, EPI_ISL_507618, EPI_ISL_507620, EPI_ISL_507621, EPI_ISL_507622, EPI_ISL_507623, EPI_ISL_507661, EPI_ISL_507753, EPI_ISL_507809, EPI_ISL_507846, EPI_ISL_507871, EPI_ISL_507885, EPI_ISL_507928 | | | |
| see above | Michigan Department of Health and Human Services, Bureau of Laboratories | Michigan Department of Health and Human Services, Bureau of Laboratories | Blankenship HM, Riner D, Soehnlen MK |
| EPI_ISL_508005, EPI_ISL_508077 | New Mexico Department of Health Scientific Laboratory Division | Center for Global Health, University of New Mexico Health Sciences Center | Daryl Domman, Kurt Schwalm, Twila Kunde, Joseph Hicks, Michael Edwards, Darrell Dinwiddie |
| EPI_ISL_508128, EPI_ISL_508132 | SA Pathology | SA Pathology | Lex Leong, Chuan Kok Lim, Mark Turra, Ivan Bastian, Geoff Higgins |
| EPI_ISL_508385, EPI_ISL_508394 | Institute of Post Graduate Medical Education & Research | National Institute of Biomedical Genomics | Arindam Maitra, Aritra Biswas, Jayeeta Haldar, Raja Ray, Monimoy Banerjee, Saumitra Das |
| EPI_ISL_508808, EPI_ISL_508809, EPI_ISL_508811, EPI_ISL_508812, EPI_ISL_508813, EPI_ISL_508814, EPI_ISL_508815, EPI_ISL_508816, EPI_ISL_508817, EPI_ISL_508818, EPI_ISL_508819, EPI_ISL_508820, EPI_ISL_508821, EPI_ISL_508822, EPI_ISL_508823, EPI_ISL_508824, EPI_ISL_508825, EPI_ISL_508826, EPI_ISL_508827, EPI_ISL_508828, EPI_ISL_508829, EPI_ISL_508830, EPI_ISL_508831, EPI_ISL_508832, EPI_ISL_508833, EPI_ISL_508834, EPI_ISL_508835, EPI_ISL_508836, EPI_ISL_508837, EPI_ISL_508838, EPI_ISL_508839, EPI_ISL_508840, EPI_ISL_508841, EPI_ISL_508842, EPI_ISL_508843, EPI_ISL_508844, EPI_ISL_508845, EPI_ISL_508846, EPI_ISL_508847, EPI_ISL_508848, EPI_ISL_508849, EPI_ISL_508850, EPI_ISL_508851, EPI_ISL_508852, EPI_ISL_508853, EPI_ISL_508854, EPI_ISL_508855, EPI_ISL_508856, EPI_ISL_508857 | | | |
| see above | Florida Bureau of Public Health Laboratories | Florida Bureau of Public Health Laboratories | Sarah Schmedes, Jason Blanton |
| EPI_ISL_508912, EPI_ISL_508913, EPI_ISL_508914, EPI_ISL_508915, EPI_ISL_508964, EPI_ISL_508965 | CNR Virus des Infections Respiratoires - France SUD | CNR Virus des Infections Respiratoires - France SUD | Antonin Bal, Gregory Destras, Gwendolyne Burfin, Solenne Brun, Carine Moustaud, Raphaelle Lamy, Alexandre Gaymard, Maude Bouscambert-Duchamp, Florence Morfin-Sherpa, Martine Valette, Bruno Lina, Laurence Josset |
| EPI_ISL_508966 | Centre Hospitalier Pierre Oudot | CNR Virus des Infections Respiratoires - France SUD | Antonin Bal, Gregory Destras, Gwendolyne Burfin, Solenne Brun, Carine Moustaud, Raphaelle Lamy, Alexandre Gaymard, Maude Bouscambert-Duchamp, Florence Morfin-Sherpa, Martine Valette, Bruno Lina, Laurence Josset |
| EPI_ISL_508967 | CNR Virus des Infections Respiratoires - France SUD | CNR Virus des Infections Respiratoires - France SUD | Antonin Bal, Gregory Destras, Gwendolyne Burfin, Solenne Brun, Carine Moustaud, Raphaelle Lamy, Alexandre Gaymard, Maude Bouscambert-Duchamp, Florence Morfin-Sherpa, Martine Valette, Bruno Lina, Laurence Josset |
| EPI_ISL_508968 | Centre Hospitalier de Bourg en Bresse | CNR Virus des Infections Respiratoires - France SUD | Antonin Bal, Gregory Destras, Gwendolyne Burfin, Solenne Brun, Carine Moustaud, Raphaelle Lamy, Alexandre Gaymard, Maude Bouscambert-Duchamp, Florence Morfin-Sherpa, Martine Valette, Bruno Lina, Laurence Josset |
| EPI_ISL_508969, EPI_ISL_508970, EPI_ISL_508971, EPI_ISL_508972, EPI_ISL_508973, EPI_ISL_508974 | CNR Virus des Infections Respiratoires - France SUD | CNR Virus des Infections Respiratoires - France SUD | Antonin Bal, Gregory Destras, Gwendolyne Burfin, Solenne Brun, Carine Moustaud, Raphaelle Lamy, Alexandre Gaymard, Maude Bouscambert-Duchamp, Florence Morfin-Sherpa, Martine Valette, Bruno Lina, Laurence Josset |
| EPI_ISL_508975 | Centre Hospitalier Pierre Oudot | CNR Virus des Infections Respiratoires - France SUD | Antonin Bal, Gregory Destras, Gwendolyne Burfin, Solenne Brun, Carine Moustaud, Raphaelle Lamy, Alexandre Gaymard, Maude Bouscambert-Duchamp, Florence Morfin-Sherpa, Martine Valette, Bruno Lina, Laurence Josset |
| EPI_ISL_508977 | CNR Virus des Infections Respiratoires - France SUD | CNR Virus des Infections Respiratoires - France SUD | Antonin Bal, Gregory Destras, Gwendolyne Burfin, Solenne Brun, Carine Moustaud, Raphaelle Lamy, Alexandre Gaymard, Maude Bouscambert-Duchamp, Florence Morfin-Sherpa, Martine Valette, Bruno Lina, Laurence Josset |
| EPI_ISL_508978 | Centre Hospitalier de Villefranche | CNR Virus des Infections Respiratoires - France SUD | Antonin Bal, Gregory Destras, Gwendolyne Burfin, Solenne Brun, Carine Moustaud, Raphaelle Lamy, Alexandre Gaymard, Maude Bouscambert-Duchamp, Florence Morfin-Sherpa, Martine Valette, Bruno Lina, Laurence Josset |
| EPI_ISL_508979, EPI_ISL_508980, EPI_ISL_508981, EPI_ISL_508982, EPI_ISL_508983, EPI_ISL_508984, EPI_ISL_508985 | CNR Virus des Infections Respiratoires - France SUD | CNR Virus des Infections Respiratoires - France SUD | Antonin Bal, Gregory Destras, Gwendolyne Burfin, Solenne Brun, Carine Moustaud, Raphaelle Lamy, Alexandre Gaymard, Maude Bouscambert-Duchamp, Florence Morfin-Sherpa, Martine Valette, Bruno Lina, Laurence Josset |
| EPI_ISL_508988 | CNR Virus des Infections Respiratoires - France SUD | CNR Virus des Infections Respiratoires - France SUD | Antonin Bal, Gregory Destras, Gwendolyne Burfin, Solenne Brun, Alexandre Gaymard, Maude Bouscambert-Duchamp, Florence Morfin-Sherpa, Martine Valette, Bruno Lina, Laurence Josset |
| EPI_ISL_508990 | CNR Virus des Infections Respiratoires - France SUD | CNR Virus des Infections Respiratoires - France SUD | Antonin Bal, Gregory Destras, Gwendolyne Burfin, Solenne Brun, Carine Moustaud, Raphaelle Lamy, Alexandre Gaymard, Maude Bouscambert-Duchamp, Florence Morfin-Sherpa, Martine Valette, Bruno Lina, Laurence Josset |
| EPI_ISL_509072, EPI_ISL_509073, EPI_ISL_509074, EPI_ISL_509075, EPI_ISL_509076, EPI_ISL_509077, EPI_ISL_509078, EPI_ISL_509079, EPI_ISL_509080, EPI_ISL_509081 | OHSU Lab Services Molecular Microbiology Lab | Oregon SARS-CoV-2 Genome Sequencing Center | Brendan L. O'Connell, Ruth V. Nichols, Sally B. Grindstaff, Alec J. Hirsch, Guang Fan, Daniel N. Streblow, William B. Messer, Andrew C. Adey, Benjamin N. Bimber, Brian J. O'Roak |
| EPI_ISL_509413, EPI_ISL_509415 | Acibadem Labcell Cellular Therapy Laboratory | Acibadem Mehmet Ali Aydinlar University School of Medicine, Medical Genetics Department | Ozden Hatirnaz Ng, Sezer Akyoney, Ilayda Sahin, Gunseli Bayram Akcapinar, Ozkan Ozdemir, Derya Dilek Kancagi, Gozde Sir Karakus, Bulut Yurtsever, Cihan Tastan, Ercument Ovali, Ugur Ozbek |
| EPI_ISL_509444, EPI_ISL_509445, EPI_ISL_509446, EPI_ISL_509447, EPI_ISL_509448, EPI_ISL_509449 | Maryland Department of Health | Maryland Department of Health | Keller,E. |
| EPI_ISL_509519 | Area of Virology, Serology and Virology Division (SAViD), New South Wales Health Pathology Randwick | Area of Virology, Serology and Virology Division (SAViD), New South Wales Health Pathology Randwick | Rawlinson, W. |
| EPI_ISL_509579, EPI_ISL_509580, EPI_ISL_509581, EPI_ISL_509582, EPI_ISL_509583, EPI_ISL_509584, EPI_ISL_509585, EPI_ISL_509586, EPI_ISL_509592, EPI_ISL_509593, EPI_ISL_509594, EPI_ISL_509595, EPI_ISL_509596, EPI_ISL_509599 | | | |
| see above | Utah Public Health Laboratory | Utah Public Health Laboratory | Heidi Butz, Erin Young, Kelly Oakeson |
| EPI_ISL_509715 | Wisconsin Department of Health Services | Pathogen Discovery, Respiratory Viruses Branch, Division of Viral Diseases, Centers for Disease Control and Prevention | Jing Zhang, Ying Tao, Krista Queen, Anna Uehara, Yan Li, Clinton Paden, Haibin Wang, Suxiang Tong |
| EPI_ISL_510108, EPI_ISL_510111, EPI_ISL_510128, EPI_ISL_510131, EPI_ISL_510135, EPI_ISL_510144, EPI_ISL_510145, EPI_ISL_510151, EPI_ISL_510243 | Hospital General Universitario Gregorio Marañón | SeqCOVID-SPAIN consortium/IBV(CSIC) | Laura Pérez-Lago, Marta Herranz, Jon Sicilia, Julia Suárez, Pilar Catalán, Patricia Muñoz, Darío García de Viedma and SeqCOVID-SPAIN consortium |
| EPI_ISL_510462, EPI_ISL_510465 | Instituto de Investigaciones Biomédicas de Barcelona (CSIC), Hospital Clinic i Provincial de Barcelona, Instituto de Biomedicina de Valencia (CSIC), Hospital de Sant Pau | SeqCOVID-SPAIN consortium/IBV(CSIC) | Anna M. Planas, Mª Angeles Marcos, Miguel J. Martínez, Andrea Vergara, Alex Soriano, Jordi Pérez Tur, Israel Fernández Cadenas and SeqCOVID-SPAIN consortium |
| EPI_ISL_510467, EPI_ISL_510468, EPI_ISL_510469, EPI_ISL_510470, EPI_ISL_510471, EPI_ISL_510472, EPI_ISL_510473, EPI_ISL_510474, EPI_ISL_510475, EPI_ISL_510476, EPI_ISL_510477, EPI_ISL_510478, EPI_ISL_510479, EPI_ISL_510480, EPI_ISL_510481, EPI_ISL_510482, EPI_ISL_510483, EPI_ISL_510484, EPI_ISL_510485, EPI_ISL_510486, EPI_ISL_510487, EPI_ISL_510488, EPI_ISL_510489, EPI_ISL_510490, EPI_ISL_510491, EPI_ISL_510492, EPI_ISL_510493, EPI_ISL_510494, EPI_ISL_510495, EPI_ISL_510496, EPI_ISL_510497, EPI_ISL_510498, EPI_ISL_510499, EPI_ISL_510500, EPI_ISL_510501, EPI_ISL_510502, EPI_ISL_510503, EPI_ISL_510504, EPI_ISL_510505, EPI_ISL_510506, EPI_ISL_510507, EPI_ISL_510508, EPI_ISL_510509 | | | |
| see above | Servicio de Microbiología. Hospital Universitario Donostia. OSI Donostialdea. Área de Enfermedades Infecciosas, Grupo de Infección Respiratoria y Resistencia Antimicrobiana. Instituto | SeqCOVID-SPAIN consortium/IBV(CSIC) | Gustavo Cilla, Milagrosa Montes, Luis Piñeiro, Jose Maria Marimón and SeqCOVID-SPAIN consortium |

| | | | |
|---|---|---|---|
| | de Investigación Sanitaria Biodonostia | | |
| EPI_ISL_510921, EPI_ISL_510922, EPI_ISL_510930, EPI_ISL_511032, EPI_ISL_511033, EPI_ISL_511181 | Instituto Nacional de Saude (INSA) | Instituto Nacional de Saude (INSA) | Borges et al |
| EPI_ISL_511185 | Instituto Nacional de Saude (INSA) and Instituto Gulbenkian de Ciencia (IGC) | Instituto Nacional de Saude (INSA) and Instituto Gulbenkian de Ciencia (IGC) | Borges et al |
| EPI_ISL_511361 | Instituto Nacional de Saude (INSA) | Instituto Nacional de Saude (INSA) | Borges et al |
| EPI_ISL_511487, EPI_ISL_511488, EPI_ISL_511489 | Instituto Nacional de Saude (INSA) | Instituto Nacional de Saude (INSA) and Instituto Gulbenkian de Ciencia (IGC) | Borges et al |
| EPI_ISL_511690, EPI_ISL_511706, EPI_ISL_511708, EPI_ISL_511709, EPI_ISL_511710, EPI_ISL_511711, EPI_ISL_511712, EPI_ISL_511713, EPI_ISL_511714, EPI_ISL_511715 | Instituto Nacional de Saude (INSA) | Instituto Nacional de Saude (INSA) | Borges et al |
| EPI_ISL_511868, EPI_ISL_511869, EPI_ISL_511871, EPI_ISL_511872, EPI_ISL_511874 | Johns Hopkins Hospital Department of Pathology | Johns Hopkins Hospital Department of Pathology | Peter M. Thielen, Thomas Mehoke, Shirlee Wohl, Srividya Ramakrishnan, Melanie Kirsche, Amanda Ernlund, Craig Howser, Kristina Zudock, Oluwaseun Falade-Nwulia, Norah Sadowski, Paul Morris, Mark Hopkins, Yunfan Fan, Nidia Trovao, Victoria Gniazdowski, Michael C. Schatz, Stuart C. Ray, Winston Timp, Heba H. Mostafa |
| EPI_ISL_511901 | Institute of Post Graduate Medical Education & Research | National Institute of Biomedical Genomics - DBT's PAN-INDIA 1000 SARS--CoV-2 RNA Genome Sequencing Consortium | Arindam Maitra, Aritra Biswas, Jayeeta Haldar, Raja Ray, Monimoy Banerjee, Saumitra Das |
| EPI_ISL_512419, EPI_ISL_512420, EPI_ISL_512421 | Centre for Enzyme Innovation, University of Portsmouth / Translational Research Laboratory, Portsmouth Hospitals NHS Trust | COVID-19 Genomics UK (COG-UK) Consortium | Angela Beckett,Yann Bourgeois,Garry Scarlett,Sharon Glaysher,Scott Elliott,Kelly Bicknell,Robert Impey,Allyson Lloyd,Sarah Wyllie,Ethan Butcher,Anoop Chauhan,Samuel Robson |
| EPI_ISL_512438, EPI_ISL_512439, EPI_ISL_512440, EPI_ISL_512441, EPI_ISL_512442, EPI_ISL_512443, EPI_ISL_512444, EPI_ISL_512445, EPI_ISL_512446, EPI_ISL_512447, EPI_ISL_512448, EPI_ISL_512449 | | | |
| see above | West of Scotland Specialist Virology Centre, NHSGGC / MRC-University of Glasgow Centre for Virus Research | COVID-19 Genomics UK (COG-UK) Consortium | Ana da Silva Filipe, Natasha Johnson, Kathy Smollett, Daniel Mair, Stephen Carmichael, Lily Tong, Jenna Nichols, Elihu Aranday-Cortes, Kirstyn Brunker, Yasmin Parr, Alice Broos, Kyriaki Nomikou; Sarah McDonald, Marc Niebel, Patawee Asamaphan; Richard Orton, Joseph Hughes, Sreenu Vattipally, David L Robertson; Alasdair MacLean, Rory Gunson; Kathy Li, Natasha Jesudason, Rajiv Shah, James Shepherd, Antonia Ho, Emma Thomson |
| EPI_ISL_512596 | Florida Bureau of Public Health Laboratories | Florida Bureau of Public Health Laboratories | Sarah Schmedes, Jason Blanton |
| EPI_ISL_512712, EPI_ISL_512736, EPI_ISL_512740, EPI_ISL_512747, EPI_ISL_512752, EPI_ISL_512753, EPI_ISL_512754 | PathWest Laboratory Medicine WA | PathWest Laboratory Medicine WA Microbial Surveillance Unit | PathWest Laboratory Medicine WA Microbial Surveillance Unit |
| EPI_ISL_512782, EPI_ISL_512783, EPI_ISL_512784 | Utah Public Health Laboratory, Utah Public Health Laboratory Infectious Disease submission group | Utah Public Health Laboratory, Utah Public Health Laboratory Infectious Disease submission group | Butz,H.A., Young,E.L., Oakeson,K. |
| EPI_ISL_512814, EPI_ISL_512815 | Kenema Government Hospital, Ministry of Health and Sanitation | Kenema Government Hospital, Ministry of Health and Sanitation | Goba,A., Momoh,M., Sandi,J., Tomkins-Tinch,C., Siddle,K., Mehta,S., Oluniyi,P., Jalloh,S., Park,D., Andersen,K., Garry,R., Happi,C., Grant,D., Olawoye,I. |
| EPI_ISL_512823, EPI_ISL_512824, EPI_ISL_512825 | National Public Health Laboratory, National Centre for Infectious Diseases | National Public Health Laboratory, National Centre for Infectious Diseases | Mak TM, Octavia S, Zhou Z, Chavatte JM, Cui L, Lin RTP |
| EPI_ISL_512930, EPI_ISL_512932 | Pathogen Genomics Lab King Abdullah University of Science and Technology(KAUST) | Pathogen Genomics Lab King Abdullah University of Science and Technology(KAUST) | Fadwa Alofi, Sharif Hala, Rahul P Salunke, Sara Mfarrej, Amit Kumar Subudhi, Fathia Ben Rached, Amanda, Luke, Afrah Alsomali, Asim Khogeer, Jumana Taha, Abdulaziz Alahmadi, Kahled Algithami, Raeece Naeem, Anwar Hashem, Naif Almontashiri, Arnab Pain |
| EPI_ISL_512995, EPI_ISL_512996, EPI_ISL_512997, EPI_ISL_512998, EPI_ISL_512999 | Pathogen Genomics Lab King Abdullah University of Science and Technology(KAUST) | Pathogen Genomics Lab King Abdullah University of Science and Technology(KAUST) | Amit Kumar Subudhi, Rahul P Salunke, Sara Mfarrej, Sharif Hala, Fadwa Alofi, Fathia Ben Rached, Afrah Alsomali, Asim Khogeer, Nashwa Al-khotani, Raeece Naeem, Anwar Hashem, Naif Almontashiri, Arnab Pain |
| EPI_ISL_513041, EPI_ISL_513042, EPI_ISL_513043, EPI_ISL_513044, EPI_ISL_513045, EPI_ISL_513046, EPI_ISL_513047, EPI_ISL_513048, EPI_ISL_513049, EPI_ISL_513050, EPI_ISL_513051, EPI_ISL_513052, EPI_ISL_513053, EPI_ISL_513054, EPI_ISL_513055, EPI_ISL_513056, EPI_ISL_513057, EPI_ISL_513058, EPI_ISL_513059, EPI_ISL_513060, EPI_ISL_513061, EPI_ISL_513062 | | | |
| see above | Pathogen Genomics Lab King Abdullah University of Science and Technology(KAUST) | Pathogen Genomics Lab King Abdullah University of Science and Technology(KAUST) | Rahul P Salunke, Sharif Hala, Raeece Naeem, Sara Mfarrej, Amit Kumar Subudhi, Amanda Ooi, Luke Esau, Fadwa Alofi, Fathia Ben Rached, Afrah Alsomali, Asim Khogeer, Ahmad Bakur Mahmoud, Anwar Hashem, Naif Almontashiri, Arnab Pain |
| EPI_ISL_513165, EPI_ISL_513166, EPI_ISL_513167 | Pathogen Genomics Lab King Abdullah University of Science and Technology(KAUST) | Pathogen Genomics Lab King Abdullah University of Science and Technology(KAUST) | Fadwa Alofi, Sharif Hala, Rahul P Salunke, Sara Mfarrej, Amit Kumar Subudhi, Fathia Ben Rached, Amanda, Luke, Afrah Alsomali, Asim Khogeer, Jumana Taha, Abdulaziz Alahmadi, Kahled Algithami, Raeece Naeem, Anwar Hashem, Naif Almontashiri, Arnab Pain |
| EPI_ISL_513315 | South Eastern Area Laboratory Services (SEALS) | NSW Health Pathology - Institute of Clinical Pathology and Medical Research; Westmead Hospital; University of Sydney | CIDM-PH et al. |
| EPI_ISL_513634, EPI_ISL_513635 | University of Washington Virology Lab | University of Washington Virology Lab | Pavitra Roychoudhury, Hong Xie, Lasata Shrestha, Amin Addetia, Truong Nguyen, Victoria M Rachleff, Meei-Li Huang, Keith R Jerome, Alexander Greninger |
| EPI_ISL_514317 | Laboratorio de Referencia Nacional de Virus Respiratorio. Instituto Nacional de Salud Perú | Laboratorio de Referencia Nacional de Biotecnología y Biología Molecular. Instituto Nacional de Salud Perú | Carlos Padilla Rojas, Karolyn Vega Chozo, Priscila Lope Pari, Omar Caceres Rey, Marco Galarza Perez, Maribel Huaringa Nuñez, Johanna Balbuena Torres, Henri Bailon Calderon, Nancy Rojas Serrano. |
| EPI_ISL_514464, EPI_ISL_514468, EPI_ISL_514471, EPI_ISL_514475, EPI_ISL_514480 | Centre for Enzyme Innovation, University of Portsmouth / Translational Research Laboratory, Portsmouth Hospitals NHS Trust | COVID-19 Genomics UK (COG-UK) Consortium | Angela Beckett,Yann Bourgeois,Garry Scarlett,Sharon Glaysher,Scott Elliott,Kelly Bicknell,Robert Impey,Allyson Lloyd,Sarah Wyllie,Ethan Butcher,Anoop Chauhan,Samuel Robson |
| EPI_ISL_515197, EPI_ISL_515198, EPI_ISL_515199, EPI_ISL_515200, EPI_ISL_515201, EPI_ISL_515202, EPI_ISL_515225, EPI_ISL_515226, EPI_ISL_515227, EPI_ISL_515228, EPI_ISL_515229, EPI_ISL_515230, EPI_ISL_515231 | | | |
| see above | Laboratoire de microbiologie, Hopital de Verdun | Smith Laboratory, Centre de Recherche CHU Sainte-Justine | Martin Smith, Marieke Rozendaal, Ivan Pavlov |
| EPI_ISL_515346, EPI_ISL_515347 | Nevada State Public Health Laboratory | Nevada State Public Health Laboratory | Richard Tillett, Joel R. Sevinsky, Paul Hartley, Heather Kerwin, David Jackson, Subhash C. Verma, Cyprian Rossetto, Andrew Gorzalski, Chris Laverdure, Natalie Crawford, Stephanie Van Hooser, and Mark Pandori |
| EPI_ISL_515523 | PS Municipal Dr Lauro Ribas Braga | Instituto Adolfo Lutz, Interdiciplinary Procedures Center, Strategic Laboratory | Claudio Tavares Sacchi, Claudia Regina Gonçalves, Erica Valessa Ramos Gomes |
| EPI_ISL_515932 | DH | Department of Neurovirology, National Institute of Mental Health and Neuroscience (NIMHANS) | Chitra Pattabiraman,Vijayalakshmi Reddy, Harsha PK, Risha Rasheed, Pramada Prasad, Shafeeq S Hameed, Manjunatha Venkataswamy, Anita Desai, Ravi Vasanthapuram |
| EPI_ISL_516235, EPI_ISL_516238, EPI_ISL_516242, EPI_ISL_516243, EPI_ISL_516244, EPI_ISL_516245, EPI_ISL_516246, EPI_ISL_516247, EPI_ISL_516248, EPI_ISL_516249, EPI_ISL_516250, EPI_ISL_516251, EPI_ISL_516252, EPI_ISL_516253, EPI_ISL_516254, EPI_ISL_516255, EPI_ISL_516256, EPI_ISL_516257, EPI_ISL_516258, EPI_ISL_516259, EPI_ISL_516260, EPI_ISL_516261, EPI_ISL_516264, EPI_ISL_516265, EPI_ISL_516266 | | | |
| see above | Michigan Department of Health and Human Services, Bureau of Laboratories | Michigan Department of Health and Human Services, Bureau of Laboratories | Blankenship HM, Riner D, Soehnlen MK |
| EPI_ISL_516832, EPI_ISL_516834, EPI_ISL_516843, EPI_ISL_516846, EPI_ISL_516858, EPI_ISL_516860, EPI_ISL_516861, EPI_ISL_516866, EPI_ISL_516869, EPI_ISL_516871, EPI_ISL_516873, EPI_ISL_516874, EPI_ISL_516875, EPI_ISL_516876, EPI_ISL_516877, EPI_ISL_516878, EPI_ISL_516879, EPI_ISL_516880, EPI_ISL_516881, EPI_ISL_516882, EPI_ISL_516883 | | | |

| | | | |
|---|---|---|---|
| see above | North West London Pathology, Imperial College Healthcare NHS Trust | Wellcome Sanger Institute for the COVID-19 Genomics UK (COG-UK) consortium | Ling Li, Paul Randell, David Muir, Frankie Bolt, Alison Holmes, James Price, Aileen Rowan, Graham Taylor, Anjna Badhan, Carolina Herrera and Alex Alderton, Roberto Amato, Sonia Goncalves, Ewan Harrison, David K. Jackson, Ian Johnston, Dominic Kwiatkowski, Cordelia Langford, John Sillitoe on behalf of the Wellcome Sanger Institute COVID-19 Surveillance Team (http://www.sanger.ac.uk/covid-team) |
| EPI_ISL_516940, EPI_ISL_516941, EPI_ISL_516942, EPI_ISL_516943, EPI_ISL_516944 | King Georges Medical University | CSIR-National Botanical Research Institute | Priti Prasad, Shantanu Prakash, Kishan Sahu, Babita Singh, Suruchi Shukla, Hricha Mishra, Danish Nasar Khan , Om Prakash, MLB Bhatt, SK Barik, Mehar H.Asif,Samir V. Sawant,Amita Jain, Sumit Kr. Bag |
| EPI_ISL_517510, EPI_ISL_517511, EPI_ISL_517512, EPI_ISL_517513, EPI_ISL_517514, EPI_ISL_517515, EPI_ISL_517516, EPI_ISL_517517, EPI_ISL_517518, EPI_ISL_517519 | Centre for Enzyme Innovation, University of Portsmouth / Translational Research Laboratory, Portsmouth Hospitals NHS Trust | COVID-19 Genomics UK (COG-UK) Consortium | Angela Beckett,Yann Bourgeois,Garry Scarlett,Sharon Glaysher,Scott Elliott,Kelly Bicknell,Robert Impey,Allyson Lloyd,Sarah Wyllie,Ethan Butcher,Anoop Chauhan,Samuel Robson |
| EPI_ISL_517687 | Laboratorio de Referencia Nacional de Virus Respiratorio. Centro Nacional de Salud Publica. Instituto Nacional de Salud Peru. | Laboratorio de Referencia Nacional de Biotecnología y Biología Molecular. Centro Nacional de Salud Publica. Instituto Nacional de Salud Peru. | Carlos Padilla Rojas, Karolyn Vega Chozo, Priscila Lope Pari, Omar Caceres Rey, Marco Galarza Perez, Maribel Huaringa Nuñez, Johanna Balbuena Torres, Henri Bailon Calderon, Nancy Rojas Serrano. |
| EPI_ISL_517771, EPI_ISL_517772, EPI_ISL_517773, EPI_ISL_517774, EPI_ISL_517783, EPI_ISL_517784, EPI_ISL_517785 | Florida Bureau of Public Health Laboratories | Florida Bureau of Public Health Laboratories | Sarah Schmedes, Jason Blanton |
| EPI_ISL_520729, EPI_ISL_520730, EPI_ISL_520731 | Mohammed Bin Rashid University of Medicine and Health Sciences | Al Jalila Genomics Center | Ahmad Abou Tayoun, Tom Loney, Hamda Khansaheb, Sathishkumar Ramaswamy, Divinlal Harilal, Zulfa Omar Deesi, Rupa Murthy Varghese, Hanan Al Suwaidi, Abdulmajeed Alkhaja, Mohammed Uddin, Rifat Hamoudi, Rabih Halwani, Abiola Catherine Senok, Qutayba Hamid, Norbert Nowotny, Alawi Alsheikh-Ali |
| EPI_ISL_521450 | Microbiological Diagnostic Unit - Public Health Laboratory (MDU-PHL) | MDU-PHL | Seemann T., Schultz M., Sait, M., Sherry, N. |
| EPI_ISL_521864 | Victorian Infectious Diseases Reference Laboratory (VIDRL) | VIDRL and MDU-PHL | Caly L., Seemann T., Sait, M., Schultz M., Druce J., Sherry, N. |
| EPI_ISL_522571, EPI_ISL_522593, EPI_ISL_522594, EPI_ISL_522604, EPI_ISL_522626, EPI_ISL_522628, EPI_ISL_522635, EPI_ISL_522659, EPI_ISL_522691, EPI_ISL_522723, EPI_ISL_522732, EPI_ISL_522747, EPI_ISL_522759, EPI_ISL_522766, EPI_ISL_522767, EPI_ISL_522768 | | | |
| see above | Royal Hobart Hospital Microbiology Department | MDU-PHL | Cooley L., van Haeften R., Seemann T., Sait M., Schultz, M.B., Sherry N. |
| EPI_ISL_522856 | ULSS9 Distretto di San Bonifacio | Istituto Zooprofilattico Sperimentale delle Venezie | Adelaide Milani, Alessia Schivo, Annalisa Salviato, Erika Giorgia Quaranta, Ambra Pastori, Bianca Zecchin, Alice Fusaro, Isabella Monne, Calogero Terregino, Antonia Ricci |
| EPI_ISL_523477, EPI_ISL_523478, EPI_ISL_523507 | Dutch COVID-19 response team | Erasmus Medical Center | Bas Oude Munnink, David Nieuwenhuijse, Reina Sikkema, Claudia Schapendonk, Irina Chestakova, Anne van der Linden, Theo Bestebroer, Stefan van Nieuwkoop, Mark Pronk, Pascal Lexmond, Corien Swaan, Manon Haverkate, Madelief Mollers, Mart Stein, Sandra Kengne Kamga Mobou, Jeroen van Kampen, Jolanda Voermans, Aura Timen, Corine GeurtsvanKessel, Annemiek van der Eijk, Richard Molenkamp, Marion Koopmans, on behalf of the Dutch national COVID-19 response team. |
| EPI_ISL_523947 | Center of Medical Microbiology, Virology, and Hospital Hygiene, University of Duesseldorf | Center of Medical Microbiology, Virology, and Hospital Hygiene, University of Duesseldorf | Maximilian Damagnez, Alexander Dilthey, Torsten Houwaart, Malte Kohns Vasconcelos, Marek Korencak, Jessica Nicolai, Klaus Pfeffer, Hendrik Streeck, Daniel Strelow, Jörg Timm, Andreas Walker, Tobias Wienemann |
| EPI_ISL_523955 | Hospital Municipal do Tatuape Carmino Caricchio | Instituto Adolfo Lutz, Interdiciplinary Procedures Center, Strategic Laboratory | Claudio Tavares Sacchi, Claudia Regina Gonçalves, Erica Valessa Ramos Gomes |
| EPI_ISL_523957 | Hospital Itamaraty | Instituto Adolfo Lutz, Interdiciplinary Procedures Center, Strategic Laboratory | Claudio Tavares Sacchi, Claudia Regina Gonçalves, Erica Valessa Ramos Gomes |
| EPI_ISL_523970 | Conjunto Hospitalar do Mandaqui | Instituto Adolfo Lutz, Interdiciplinary Procedures Center, Strategic Laboratory | Claudio Tavares Sacchi, Claudia Regina Gonçalves, Erica Valessa Ramos Gomes |
| EPI_ISL_523973 | PS Municipal Dona Maria Antonieta Ferreira de Barros | Instituto Adolfo Lutz, Interdiciplinary Procedures Center, Strategic Laboratory | Claudio Tavares Sacchi, Claudia Regina Gonçalves, Erica Valessa Ramos Gomes |
| EPI_ISL_523974 | Hospital Municipal do Tatuape Carmino Caricchio | Instituto Adolfo Lutz, Interdiciplinary Procedures Center, Strategic Laboratory | Claudio Tavares Sacchi, Claudia Regina Gonçalves, Erica Valessa Ramos Gomes |
| EPI_ISL_523975, EPI_ISL_523980 | UPA Tito Lopes | Instituto Adolfo Lutz, Interdiciplinary Procedures Center, Strategic Laboratory | Claudio Tavares Sacchi, Claudia Regina Gonçalves, Erica Valessa Ramos Gomes |
| EPI_ISL_524432 | Dirk Dittmer | Dirk Dittmer | Bailey,A.G., Caro-Vegas,C.P., Dittmer,D., Eason,A.B., Juarez,A., Landis,J.T., McNamara,R.P., Miller,M.B., Moorad,R., Pluta,L.J.,Seltzer,T.A., Thompson,C., Vahrson,W. and Villamor,F. |
| EPI_ISL_524496, EPI_ISL_524504, EPI_ISL_524510, EPI_ISL_524511, EPI_ISL_524514, EPI_ISL_524516, EPI_ISL_524517 | NHSGGC West of Scotland Specialist Virology Centre / MRC-University of Glasgow Centre for Virus Research | Wellcome Sanger Institute for the COVID-19 Genomics UK (COG-UK) consortium | Ana da Silva Filipe, Natasha Johnson, Kathy Smollett, Daniel Mair, Stephen Carmichael, Lily Tong, Jenna Nichols, Elihu Aranday-Cortes, Kirstyn Brunker, Yasmin Parr, Kyriaki Nomikou; Sarah McDonald, Marc Niebel, Patawee Asamaphan; Richard Orton, Joseph Hughes, Sreenu Vattipally, David L Robertson; Alasdair MacLean, Rory Gunson; Kathy Li, Natasha Jesudason, Rajiv Shah, James Shepherd, Antonia Ho, Alice Broos, Emma Thomson and Alex Alderton, Roberto Amato, Sonia Goncalves, Ewan Harrison, David K. Jackson, Ian Johnston, Dominic Kwiatkowski, Cordelia Langford, John Sillitoe on behalf of the Wellcome Sanger Institute COVID-19 Surveillance Team (http://www.sanger.ac.uk/covid-team) |
| EPI_ISL_524521 | Department of Pathology, University of Cambridge | Wellcome Sanger Institute for the COVID-19 Genomics UK (COG-UK) consortium | Luke W Meredith, M. Estée Török , Myra Hosmillo, William L. Hamilton, Martin D. Curran, Theresa Feltwell, Grant Hall, Anna Yakovleva, Fahad A Khokhar, Charlotte J. Houldcroft, Laura G Caller, Aminu S. Jahun, Sarah L. Caddy, Ian Goodfellow; and Alex Alderton, Roberto Amato, Sonia Goncalves, Ewan Harrison, David K. Jackson, Ian Johnston, Dominic Kwiatkowski, Cordelia Langford, John Sillitoe on behalf of the Wellcome Sanger Institute COVID-19 Surveillance Team (http://www.sanger.ac.uk/covid-team) |
| EPI_ISL_524523, EPI_ISL_524524, EPI_ISL_524526, EPI_ISL_524535, EPI_ISL_524542, EPI_ISL_524556, EPI_ISL_524562, EPI_ISL_524565, EPI_ISL_524572 | NHSGGC West of Scotland Specialist Virology Centre / MRC-University of Glasgow Centre for Virus Research | Wellcome Sanger Institute for the COVID-19 Genomics UK (COG-UK) consortium | Ana da Silva Filipe, Natasha Johnson, Kathy Smollett, Daniel Mair, Stephen Carmichael, Lily Tong, Jenna Nichols, Elihu Aranday-Cortes, Kirstyn Brunker, Yasmin Parr, Kyriaki Nomikou; Sarah McDonald, Marc Niebel, Patawee Asamaphan; Richard Orton, Joseph Hughes, Sreenu Vattipally, David L Robertson; Alasdair MacLean, Rory Gunson; Kathy Li, Natasha Jesudason, Rajiv Shah, James Shepherd, Antonia Ho, Alice Broos, Emma Thomson and Alex Alderton, Roberto Amato, Sonia Goncalves, Ewan Harrison, David K. Jackson, Ian Johnston, Dominic Kwiatkowski, Cordelia Langford, John Sillitoe on behalf of the Wellcome Sanger Institute COVID-19 Surveillance Team (http://www.sanger.ac.uk/covid-team) |
| EPI_ISL_524619, EPI_ISL_524620, EPI_ISL_524621, EPI_ISL_524623, EPI_ISL_524624, EPI_ISL_524625, EPI_ISL_524626, EPI_ISL_524628, EPI_ISL_524629, EPI_ISL_524630, EPI_ISL_524631, EPI_ISL_524632, EPI_ISL_524633, EPI_ISL_524634, EPI_ISL_524635, EPI_ISL_524636, EPI_ISL_524637, EPI_ISL_524640, EPI_ISL_524641, EPI_ISL_524642, EPI_ISL_524643, EPI_ISL_524644, EPI_ISL_524646, EPI_ISL_524647, EPI_ISL_524650, EPI_ISL_524655, EPI_ISL_524666, EPI_ISL_524677, EPI_ISL_524678, EPI_ISL_524680, EPI_ISL_524681, EPI_ISL_524682, EPI_ISL_524684, EPI_ISL_524685, EPI_ISL_524686, EPI_ISL_524687, EPI_ISL_524688, EPI_ISL_524689, EPI_ISL_524690, EPI_ISL_524691, EPI_ISL_524692, EPI_ISL_524693, EPI_ISL_524694, EPI_ISL_524695, EPI_ISL_524697, EPI_ISL_524698, EPI_ISL_524699, EPI_ISL_524700, EPI_ISL_524704, EPI_ISL_524705, EPI_ISL_524708, EPI_ISL_524709, EPI_ISL_524710, EPI_ISL_524711, EPI_ISL_524712 | | | |
| see above | North West London Pathology, Imperial College Healthcare NHS Trust | Wellcome Sanger Institute for the COVID-19 Genomics UK (COG-UK) consortium | Ling Li, Paul Randell, David Muir, Frankie Bolt, Alison Holmes, James Price, Aileen Rowan, Graham Taylor, Anjna Badhan, Carolina Herrera and Alex Alderton, Roberto Amato, Sonia Goncalves, Ewan Harrison, David K. Jackson, Ian Johnston, Dominic Kwiatkowski, Cordelia Langford, John Sillitoe on behalf of the Wellcome Sanger Institute COVID-19 Surveillance Team (http://www.sanger.ac.uk/covid-team) |
| EPI_ISL_524792 | Evandro Chagas Institute | Evandro Chagas Institute | Santos, M.C.; Silva, A.M.; Junior, W.D.C.; Barbagelata, L.S.; Ferreira, J.A.; Sousa, E.M.A.; da Silva, P.S.; Resque, H.R; Martins, L.C.; Sousa Junior, E.C.;Viana, G.M.R |
| EPI_ISL_524873 | MD PHL | MD PHL | Maryland Department of Health Laboratories Administration |
| EPI_ISL_525648, EPI_ISL_525649, EPI_ISL_525650, EPI_ISL_525651, | Wadsworth Center, New York State Department of Health | Wadsworth Center, New York State Department of Health | Kirsten St. George, Daryl M. Lamson, Sara Griesemer, Jonathan Plitnick, Navjot Singh, Matthew D. Shudt, Erica Lasek-Nesselquist |

| | | | |
|---|---|---|---|
| EPI_ISL_525652, EPI_ISL_525653, EPI_ISL_525654, EPI_ISL_525655, EPI_ISL_525657, EPI_ISL_525659 | | | |
| EPI_ISL_525779, EPI_ISL_525780, EPI_ISL_525781 | Texas Department of State Health Services | Texas Department of State Health Services | Jenny Zhang, Rashmi Tuladhar, Bonnie Oh, Maliha Rahman, Anita Pokharel, Myong Koag, Chun Wang, Rachel Lee, Grace Kubin |
| EPI_ISL_526128, EPI_ISL_526130 | Pathology North - Royal North Shore Hospital - NSW Health Pathology | NSW Health Pathology - Institute of Clinical Pathology and Medical Research; Westmead Hospital; University of Sydney | CIDM-PH et al. |
| EPI_ISL_526146 | South Eastern Area Laboratory Services (SEALS) | NSW Health Pathology - Institute of Clinical Pathology and Medical Research; Westmead Hospital; University of Sydney | CIDM-PH et al. |
| EPI_ISL_526457 | West of Scotland Specialist Virology Centre, NHSGGC / MRC-University of Glasgow Centre for Virus Research | COVID-19 Genomics UK (COG-UK) Consortium | Ana da Silva Filipe, Natasha Johnson, Kathy Smollett, Daniel Mair, Stephen Carmichael, Lily Tong, Jenna Nichols, Elihu Aranday-Cortes, Kirstyn Brunker, Yasmin Parr, Alice Broos, Kyriaki Nomikou; Sarah McDonald, Marc Niebel, Patawee Asamaphan; Richard Orton, Joseph Hughes, Sreenu Vattipally, David L Robertson; Alasdair MacLean, Rory Gunson; Kathy Li, Natasha Jesudason, Rajiv Shah, James Shepherd, Antonia Ho, Emma Thomson |
| EPI_ISL_526471, EPI_ISL_526493, EPI_ISL_526494 | Virology Department, Royal Infirmary of Edinburgh, NHS Lothian / School of Biological Sciences, University of Edinburgh / Institute of Genetics and Molecular Medicine, University of Edinburgh | COVID-19 Genomics UK (COG-UK) Consortium | McHugh M, Dewar R, Rooke S, Gallagher M, Balcaza C, O'Toole Á, Scher E, Hill V, McCrone JT, Colquhoun R, Yu X, Jackson B, Rambaut A, Williams TC, Templeton K |
| EPI_ISL_527015, EPI_ISL_527039, EPI_ISL_527040, EPI_ISL_527041, EPI_ISL_527042, EPI_ISL_527043, EPI_ISL_527044, EPI_ISL_527045, EPI_ISL_527046, EPI_ISL_527047, EPI_ISL_527048, EPI_ISL_527049, EPI_ISL_527050, EPI_ISL_527051, EPI_ISL_527052, EPI_ISL_527053, EPI_ISL_527054, EPI_ISL_527055, EPI_ISL_527056, EPI_ISL_527062 | | | |
| see above | Area of Virology, Serology and Virology Division (SAViD), New South Wales Health Pathology Randwick | Area of Virology, Serology and Virology Division (SAViD), New South Wales Health Pathology Randwick | Rawlinson, W. |
| EPI_ISL_527381 | University of Miami Immunology and Histocompatibility Laboratory | University of Miami Immunology and Histocompatibility Laboratory | Emilio Margolles-Clark, PhD and Phillip Ruiz, MD, PhD |
| EPI_ISL_527408, EPI_ISL_527410, EPI_ISL_527416, EPI_ISL_527431, EPI_ISL_527433, EPI_ISL_527436, EPI_ISL_527441, EPI_ISL_527442, EPI_ISL_527446, EPI_ISL_527449, EPI_ISL_527450, EPI_ISL_527451, EPI_ISL_527453, EPI_ISL_527454, EPI_ISL_527456, EPI_ISL_527458, EPI_ISL_527459, EPI_ISL_527465, EPI_ISL_527468, EPI_ISL_527472, EPI_ISL_527474, EPI_ISL_527477, EPI_ISL_527478 | | | |
| see above | Colorado State University - Ebel Lab | Colorado State University - Ebel Lab | Greg Ebel et al. |
| EPI_ISL_527710 | MN PHL Division, Minnesota Department of Health | Pathogen Discovery, Respiratory Viruses Branch, Division of Viral Diseases, Centers for Disease Control and Prevention | Yan Li, Anna Montmayeur, Jing Zhang, Krista Queen, Ying Tao, Anna Uehara, Rachel Marine, Clinton R. Paden, Haibin Wang, Suxiang Tong |
| EPI_ISL_528538 | Alsafar | Alsafar | Andreas Henschel, Gihan Elsir Ahmed Daw Elbait, Samuel Feng, Rifat, Ernesto Damiani, Guan Tay, Habiba Alsafar |
| EPI_ISL_528739, EPI_ISL_528740, EPI_ISL_528741 | Malaysia Genome Institute | Malaysia Genome Institute | Mohd Noor Mat Isa, Irni Suhayu Sapian, Yusuf Muhammad Noor, Nurhezreen Md Iqbal, Mohd Faizal Abu Bakar, Enizza Kasim, Shamsidar Sopie, Siti Noraini Othman, Azrin Ahmad, Nor Azfa Johari, Shahrul Hisham Zainal Ariffin |
| EPI_ISL_528742 | Malaysia Genome Institute | Malaysia Genome Institute | Mohd Noor Mat Isa, Irni Suhayu Sapian, Gan Han Ming, Yusuf Muhammad Noor, Tan Ju Lin, Nurhezreen Md Iqbal, Mohd Faizal Abu Bakar, Enizza Kasim, Shamsidar Sopie, Siti Noraini Othman, Azrin Ahmad, Nor Azfa Johari, Shahrul Hisham Zainal Ariffin |
| EPI_ISL_528991, EPI_ISL_529016 | Ospedale SS Annunziata-Sulmona | Istituto Zooprofilattico Sperimentale dell'Abruzzo e Molise "G.Caporale" | Lorusso A, Marcacci M, Di Domenico M, Curini V, Ancora M, Cammà C, Rinaldi A, Mangone I, Di Pasquale A, Puglia I, Savini G. |
| EPI_ISL_529165, EPI_ISL_529166 | Department of Immunology, The Scripps Research Institute | Andersen lab at Scripps Research | Allison Smither, Gilberto Sabino-Santos, Patricia Snarski, Lilia Melnik, Antoinette Bell, Kaylynn Genemaras, Arnaud Drouin, Dahlene Fusco, Robert Garry with SEARCH Alliance San Diego |
| EPI_ISL_529205 | Utah Public Health Laboratory | Utah Public Health Laboratory | Erin Young, Kelly Oakeson |
| EPI_ISL_529341, EPI_ISL_529342, EPI_ISL_529668 | Centre for Enzyme Innovation, University of Portsmouth / Translational Research Laboratory, Portsmouth Hospitals NHS Trust | COVID-19 Genomics UK (COG-UK) Consortium | Angela Beckett,Yann Bourgeois,Garry Scarlett,Sharon Glaysher,Scott Elliott,Kelly Bicknell,Robert Impey,Allyson Lloyd,Sarah Wyllie,Ethan Butcher,Anoop Chauhan,Samuel Robson |
| EPI_ISL_529691, EPI_ISL_529692 | Virology Department, Royal Infirmary of Edinburgh, NHS Lothian / School of Biological Sciences, University of Edinburgh / Institute of Genetics and Molecular Medicine, University of Edinburgh | COVID-19 Genomics UK (COG-UK) Consortium | McHugh M, Dewar R, Rooke S, Gallagher M, Balcaza C, O'Toole Á, Scher E, Hill V, McCrone JT, Colquhoun R, Yu X, Jackson B, Rambaut A, Williams TC, Templeton K |
| EPI_ISL_529821 | Michigan Department of Health and Human Services, Bureau of Laboratories | Michigan Department of Health and Human Services, Bureau of Laboratories | Blankenship HM, Riner D, Soehnlen MK |
| EPI_ISL_529974, EPI_ISL_529975, EPI_ISL_529976, EPI_ISL_529977, EPI_ISL_529978 | Hospital Universitario 12 de Octubre | Hospital Universitario 12 de Octubre | Raúl Recio, Sara González, Esther Viedma, Elias Dahdouh, Fernando Lázaro, Natalia Stella, Julio García, Juan Carlos Galán, Rafael Cantón, Mª Dolores Folgueira, Rafael Delgado, Jesús Mingorance |
| EPI_ISL_529989, EPI_ISL_529990, EPI_ISL_529991 | Hospital Universitario 12 de Octubre | Hospital Universitario 12 de Octubre | Sara González, Esther Viedma, Raúl Recio, Elias Dahdouh, Fernando Lázaro, Natalia Stella, Julio García, Juan Carlos Galán, Rafael Cantón, Mª Dolores Folgueira, Rafael Delgado, Jesús Mingorance |
| EPI_ISL_530219, EPI_ISL_530221 | Minnesota Department of Health, Public Health Laboratory | Minnesota Department of Health, Public Health Laboratory | Matt Plumb, Jacob Garfin, and Xiong Wang |
| EPI_ISL_530231, EPI_ISL_530232, EPI_ISL_530237, EPI_ISL_530275, EPI_ISL_530276 | Queensland Health Forensic and Scientific Services, Public Health Virology | Public Health Virology Laboratory, Forensic and Scientific Services, Queensland Health | Son Nguyen et al |
| EPI_ISL_534313 | Hospital da Sta Casa de Sto Amaro | Instituto Adolfo Lutz, Interdiciplinary Procedures Center, Strategic Laboratory | Claudio Tavares Sacchi, Claudia Regina Gonçalves, Erica Valessa Ramos Gomes |
| EPI_ISL_534773, EPI_ISL_534782, EPI_ISL_534788, EPI_ISL_534798, EPI_ISL_534811, EPI_ISL_534831, EPI_ISL_534837, EPI_ISL_534863, EPI_ISL_534874, EPI_ISL_534875, EPI_ISL_534880, EPI_ISL_534894, EPI_ISL_534901, EPI_ISL_534914, EPI_ISL_534923, EPI_ISL_534926, EPI_ISL_534952, EPI_ISL_534966, EPI_ISL_534978 | | | |
| see above | Oxford Viromics, NDM, University of Oxford; Oxford University Hospitals; Basingstoke and North Hampshire Hospital | COVID-19 Genomics UK (COG-UK) Consortium | Tanya Golubchik, David Bonsall, George Macintyre, Amy Trebes, Mariateresa de Cesare, Catrin Moore, Alex Mobbs, Anita Justice, Robert Shaw, Monique Andersson, Timothy Peto, Emma Wise, Nathan Moore, Jessica Lynch, Nick Cortes, Matilde Mori, Stephen Kidd, David Buck, John Todd, Christophe Fraser |
| EPI_ISL_536412, EPI_ISL_536414 | National Public Health Laboratory, National Centre for Infectious Diseases | National Public Health Laboratory, National Centre for Infectious Diseases | Mak TM, Octavia S, Zhou Z, Cui L, Lin RTP |
| EPI_ISL_536792 | Canterbury Health Laboratories | Institute of Environmental Science and Research (ESR) | Xiaoyun Ren, Matt Storey, Nikki Freed, Muhammad Faisal, Jing Wang, Hermes Perez, Anja Werno, Antje van der Linden, Arlo Upton, Chris Mansell, David Hammer, Dragana Drinkovic, Gary McAuliffe, Hana Sofia Andersson, James Ussher, Jill Sherwood, Josh Freeman, Julia Howard, Juliet Elvy, Mary DeAlmeida, Matt Blakiston, Matthew Rogers, Max Bloomfield, Michael Addidle, Michelle Balm, Sally Roberts, Sarah Jefferies, Sharmini Muttaiyah, Susan Morpeth, Susan Taylor, Timothy Blackmore, Vani Sathyendran, Veronica Playle, Virginia Hope, Erasmus Smit, Lauren Jelly, Olin Silander, Joep de Ligt |
| EPI_ISL_537296, EPI_ISL_537297, EPI_ISL_537298, EPI_ISL_537299, EPI_ISL_537300, EPI_ISL_537301, EPI_ISL_537302, EPI_ISL_537303, EPI_ISL_537304, EPI_ISL_537305, EPI_ISL_537306, EPI_ISL_537307, EPI_ISL_537308, EPI_ISL_537309, EPI_ISL_537310, EPI_ISL_537311, EPI_ISL_537312, EPI_ISL_537313, EPI_ISL_537314, EPI_ISL_537315, EPI_ISL_537316, EPI_ISL_537317, EPI_ISL_537318, EPI_ISL_537319, EPI_ISL_537320, EPI_ISL_537321, EPI_ISL_537322, EPI_ISL_537323, EPI_ISL_537324, EPI_ISL_537325 | | | |
| see above | Universidad de León | SeqCOVID-SPAIN consortium/IBV(CSIC) | Ana Carvajal, Vicente Martín, Héctor Argüello, Juan M. Fregeneda, Tania Fernández-Villa, Antonio J. Molina and SeqCOVID-SPAIN consortium |
| EPI_ISL_537384, EPI_ISL_537385, EPI_ISL_537441, EPI_ISL_537442, | Centro de Investigación Biomédica de La Rioja - Hospital San Pedro Logroño | SeqCOVID-SPAIN consortium/IBV(CSIC) | María de Toro, José Manuel Azcona Gutiérrez, María Pilar Bea Escudero, Miriam Blasco Alberdi and SeqCOVID-SPAIN consortium |

| | | | |
|---|---|---|---|
| EPI_ISL_537443 | | | |
| EPI_ISL_537469, EPI_ISL_537498, EPI_ISL_537575 | UCLA Pathology Clinical Microbiology Lab | Kruglyak Lab | Guo et al. |
| EPI_ISL_537720, EPI_ISL_537721, EPI_ISL_537722, EPI_ISL_537723, EPI_ISL_537724, EPI_ISL_537725, EPI_ISL_537726 | Hospital Universitario de Gran Canaria Dr. Negrín | SeqCOVID-SPAIN consortium/IBV(CSIC) | M. Carmen Pérez González, Francisco J. Chamizo López, Ana Bordes Benítez and SeqCOVID-SPAIN consortium |
| EPI_ISL_537859, EPI_ISL_537871 | Centro de Investigación Biomédica de La Rioja - Hospital San Pedro Logroño | SeqCOVID-SPAIN consortium/IBV(CSIC) | María de Toro, José Manuel Azcona Gutiérrez, María Pilar Bea Escudero, Miriam Blasco Alberdi and SeqCOVID-SPAIN consortium |
| EPI_ISL_538119, EPI_ISL_538124, EPI_ISL_538125, EPI_ISL_538138, EPI_ISL_538143, EPI_ISL_538144, EPI_ISL_538148, EPI_ISL_538149, EPI_ISL_538150, EPI_ISL_538151, EPI_ISL_538152, EPI_ISL_538153, EPI_ISL_538154, EPI_ISL_538155, EPI_ISL_538156, EPI_ISL_538157, EPI_ISL_538161, EPI_ISL_538162, EPI_ISL_538164, EPI_ISL_538165 | | | |
| see above | Servicio de Microbiología y Parasitología clínica. UCEIMP. Hospital Universitario Virgen del Rocío/IBIS/CSIC/US | SeqCOVID-SPAIN consortium/IBV(CSIC) | Guillermo Martín Gutiérrez, Ángel Rodríguez Villodres, Lidia Gálvez Benitez, Verónica González Galán, Javier Aznar Martín and SeqCOVID-SPAIN consortium |
| EPI_ISL_538250 | TriCore Reference Laboratories | Center for Global Health, University of New Mexico Health Sciences Center | Daryl Domman, Kurt Schwalm, Twila Kunde, Joseph Hicks, Michael Edwards, Darrell Dinwiddie |
| EPI_ISL_538344, EPI_ISL_538345, EPI_ISL_538346, EPI_ISL_538347, EPI_ISL_538348, EPI_ISL_538349, EPI_ISL_538350, EPI_ISL_538351, EPI_ISL_538352, EPI_ISL_538353, EPI_ISL_538354, EPI_ISL_538355, EPI_ISL_538356, EPI_ISL_538357, EPI_ISL_538358, EPI_ISL_538359, EPI_ISL_538360, EPI_ISL_538361, EPI_ISL_538362, EPI_ISL_538363, EPI_ISL_538364, EPI_ISL_538365, EPI_ISL_538366, EPI_ISL_538367, EPI_ISL_538368, EPI_ISL_538369, EPI_ISL_538370 | | | |
| see above | Kingston Health Sciences Centre / Queen's University | Ontario Institute for Cancer Research | Prameet M. Sheth, Calvin Sjaarda, Robert Colautti, Katya Douchant, Ilinca Lungu, Bernard Lam, Paul Krzyzanowski, Michael Laszloffy, Lawrence E Heisler, Richard de Borja, Jared T. Simpson |
| EPI_ISL_538469, EPI_ISL_538470, EPI_ISL_538471, EPI_ISL_538472, EPI_ISL_538473, EPI_ISL_538474, EPI_ISL_538475, EPI_ISL_538476, EPI_ISL_538477, EPI_ISL_538478, EPI_ISL_538479, EPI_ISL_538480, EPI_ISL_538481, EPI_ISL_538482, EPI_ISL_538483, EPI_ISL_538484 | | | |
| see above | Department of Laboratory Medicine, Tan Tock Seng Hospital | Department of Laboratory Medicine, Tan Tock Seng Hospital | Chen YYC, Zair X, Lim JX, Li C, Tang WY, Maurer-Stroh S, Barkham TMS, Nagarajan N, Sessions OM |
| EPI_ISL_538577, EPI_ISL_538578, EPI_ISL_538579, EPI_ISL_538580, EPI_ISL_538581, EPI_ISL_538583, EPI_ISL_538588, EPI_ISL_538591, EPI_ISL_538596, EPI_ISL_538597, EPI_ISL_538598, EPI_ISL_538599, EPI_ISL_538600, EPI_ISL_538603, EPI_ISL_538604, EPI_ISL_538605, EPI_ISL_538606, EPI_ISL_538607 | | | |
| see above | Servicio de Microbiología y Parasitología clínica. UCEIMP. Hospital Universitario Virgen del Rocío/IBIS/CSIC/US | SeqCOVID-SPAIN consortium/IBV(CSIC) | Guillermo Martín Gutiérrez, Ángel Rodríguez Villodres, Lidia Gálvez Benitez, Verónica González Galán, Javier Aznar Martín and SeqCOVID-SPAIN consortium |
| EPI_ISL_538689 | Hospital Universitario Virgen de las Nieves de Granada-SAS | SeqCOVID-SPAIN consortium/IBV(CSIC) | Mercedes Pérez Ruiz, Sara Sanbonmatsu Gámez, Irene Pedrosa Corral, José M. Navarro-Marí and SeqCOVID-SPAIN consortium |
| EPI_ISL_538786, EPI_ISL_538792, EPI_ISL_538795, EPI_ISL_538804, EPI_ISL_538805, EPI_ISL_538807, EPI_ISL_538814, EPI_ISL_538815, EPI_ISL_538825, EPI_ISL_538827, EPI_ISL_538830, EPI_ISL_538840 | | | |
| see above | Leeds Teaching Hospitals NHS Trust and Public Health England, National Infection Service (Leeds laboratory) | Wellcome Sanger Institute for the COVID-19 Genomics UK (COG-UK) consortium | Louissa Macfarlane-Smith, Holli Carden, Katherine L. Harper, Antony Hale and Alex Alderton, Roberto Amato, Sonia Goncalves, Ewan Harrison, David K. Jackson, Ian Johnston, Dominic Kwiatkowski, Cordelia Langford, John Sillitoe on behalf of the Wellcome Sanger Institute COVID-19 Surveillance Team |
| EPI_ISL_538851 | Leeds Teaching Hospitals NHS Trust and Public Health England, National Infection Service (Leeds laboratory) | Wellcome Sanger Institute for the COVID-19 Genomics UK (COG-UK) Consortium | Louissa Macfarlane-Smith, Holli Carden, Katherine L. Harper, Antony Hale and Alex Alderton, Roberto Amato, Sonia Goncalves, Ewan Harrison, David K. Jackson, Ian Johnston, Dominic Kwiatkowski, Cordelia Langford, John Sillitoe on behalf of the Wellcome Sanger Institute COVID-19 Surveillance Team |
| EPI_ISL_538857 | Leeds Teaching Hospitals NHS Trust and Public Health England, National Infection Service (Leeds laboratory) | Wellcome Sanger Institute for the COVID-19 Genomics UK (COG-UK) consortium | Louissa Macfarlane-Smith, Holli Carden, Katherine L. Harper, Antony Hale and Alex Alderton, Roberto Amato, Sonia Goncalves, Ewan Harrison, David K. Jackson, Ian Johnston, Dominic Kwiatkowski, Cordelia Langford, John Sillitoe on behalf of the Wellcome Sanger Institute COVID-19 Surveillance Team |
| EPI_ISL_538863 | Leeds Teaching Hospitals NHS Trust and Public Health England, National Infection Service (Leeds laboratory) | Wellcome Sanger Institute for the COVID-19 Genomics UK (COG-UK) Consortium | Louissa Macfarlane-Smith, Holli Carden, Katherine L. Harper, Antony Hale and Alex Alderton, Roberto Amato, Sonia Goncalves, Ewan Harrison, David K. Jackson, Ian Johnston, Dominic Kwiatkowski, Cordelia Langford, John Sillitoe on behalf of the Wellcome Sanger Institute COVID-19 Surveillance Team |
| EPI_ISL_538873, EPI_ISL_538880, EPI_ISL_538882, EPI_ISL_538886, EPI_ISL_538893, EPI_ISL_538894, EPI_ISL_538895, EPI_ISL_538897, EPI_ISL_538903, EPI_ISL_538904, EPI_ISL_538905, EPI_ISL_538926, EPI_ISL_538952, EPI_ISL_538958, EPI_ISL_538980 | | | |
| see above | Leeds Teaching Hospitals NHS Trust and Public Health England, National Infection Service (Leeds laboratory) | Wellcome Sanger Institute for the COVID-19 Genomics UK (COG-UK) consortium | Louissa Macfarlane-Smith, Holli Carden, Katherine L. Harper, Antony Hale and Alex Alderton, Roberto Amato, Sonia Goncalves, Ewan Harrison, David K. Jackson, Ian Johnston, Dominic Kwiatkowski, Cordelia Langford, John Sillitoe on behalf of the Wellcome Sanger Institute COVID-19 Surveillance Team |
| EPI_ISL_538987, EPI_ISL_538994 | Leeds Teaching Hospitals NHS Trust and Public Health England, National Infection Service (Leeds laboratory) | Wellcome Sanger Institute for the COVID-19 Genomics UK (COG-UK) Consortium | Louissa Macfarlane-Smith, Holli Carden, Katherine L. Harper, Antony Hale and Alex Alderton, Roberto Amato, Sonia Goncalves, Ewan Harrison, David K. Jackson, Ian Johnston, Dominic Kwiatkowski, Cordelia Langford, John Sillitoe on behalf of the Wellcome Sanger Institute COVID-19 Surveillance Team |
| EPI_ISL_538997, EPI_ISL_539000, EPI_ISL_539005, EPI_ISL_539007, EPI_ISL_539010, EPI_ISL_539012, EPI_ISL_539016, EPI_ISL_539017, EPI_ISL_539018, EPI_ISL_539020, EPI_ISL_539021, EPI_ISL_539022, EPI_ISL_539024, EPI_ISL_539025, EPI_ISL_539026, EPI_ISL_539027 | | | |
| see above | Leeds Teaching Hospitals NHS Trust and Public Health England, National Infection Service (Leeds laboratory) | Wellcome Sanger Institute for the COVID-19 Genomics UK (COG-UK) consortium | Louissa Macfarlane-Smith, Holli Carden, Katherine L. Harper, Antony Hale and Alex Alderton, Roberto Amato, Sonia Goncalves, Ewan Harrison, David K. Jackson, Ian Johnston, Dominic Kwiatkowski, Cordelia Langford, John Sillitoe on behalf of the Wellcome Sanger Institute COVID-19 Surveillance Team |
| EPI_ISL_539030 | Leeds Teaching Hospitals NHS Trust and Public Health England, National Infection Service (Leeds laboratory) | Wellcome Sanger Institute for the COVID-19 Genomics UK (COG-UK) Consortium | Louissa Macfarlane-Smith, Holli Carden, Katherine L. Harper, Antony Hale and Alex Alderton, Roberto Amato, Sonia Goncalves, Ewan Harrison, David K. Jackson, Ian Johnston, Dominic Kwiatkowski, Cordelia Langford, John Sillitoe on behalf of the Wellcome Sanger Institute COVID-19 Surveillance Team |
| EPI_ISL_539036, EPI_ISL_539037, EPI_ISL_539040, EPI_ISL_539042, EPI_ISL_539043, EPI_ISL_539044, EPI_ISL_539045, EPI_ISL_539046, EPI_ISL_539050, EPI_ISL_539056, EPI_ISL_539062, EPI_ISL_539063, EPI_ISL_539065, EPI_ISL_539066, EPI_ISL_539067 | | | |
| see above | Leeds Teaching Hospitals NHS Trust and Public Health England, National Infection Service (Leeds laboratory) | Wellcome Sanger Institute for the COVID-19 Genomics UK (COG-UK) consortium | Louissa Macfarlane-Smith, Holli Carden, Katherine L. Harper, Antony Hale and Alex Alderton, Roberto Amato, Sonia Goncalves, Ewan Harrison, David K. Jackson, Ian Johnston, Dominic Kwiatkowski, Cordelia Langford, John Sillitoe on behalf of the Wellcome Sanger Institute COVID-19 Surveillance Team |
| EPI_ISL_539070 | Leeds Teaching Hospitals NHS Trust and Public Health England, National Infection Service (Leeds laboratory) | Wellcome Sanger Institute for the COVID-19 Genomics UK (COG-UK) Consortium | Louissa Macfarlane-Smith, Holli Carden, Katherine L. Harper, Antony Hale and Alex Alderton, Roberto Amato, Sonia Goncalves, Ewan Harrison, David K. Jackson, Ian Johnston, Dominic Kwiatkowski, Cordelia Langford, John Sillitoe on behalf of the Wellcome Sanger Institute COVID-19 Surveillance Team |
| EPI_ISL_539071, EPI_ISL_539073, EPI_ISL_539074, EPI_ISL_539075, EPI_ISL_539076, EPI_ISL_539082, EPI_ISL_539084, EPI_ISL_539092, EPI_ISL_539093, EPI_ISL_539094, EPI_ISL_539096, EPI_ISL_539098, EPI_ISL_539101, EPI_ISL_539102, EPI_ISL_539103, EPI_ISL_539104, EPI_ISL_539106, EPI_ISL_539107, EPI_ISL_539108, EPI_ISL_539110, EPI_ISL_539113, EPI_ISL_539115 | | | |
| see above | Leeds Teaching Hospitals NHS Trust and Public Health England, National Infection Service (Leeds laboratory) | Wellcome Sanger Institute for the COVID-19 Genomics UK (COG-UK) consortium | Louissa Macfarlane-Smith, Holli Carden, Katherine L. Harper, Antony Hale and Alex Alderton, Roberto Amato, Sonia Goncalves, Ewan Harrison, David K. Jackson, Ian Johnston, Dominic Kwiatkowski, Cordelia Langford, John Sillitoe on behalf of the Wellcome Sanger Institute COVID-19 Surveillance Team |
| EPI_ISL_539116, EPI_ISL_539117, EPI_ISL_539120 | Leeds Teaching Hospitals NHS Trust and Public Health England, National Infection Service (Leeds laboratory) | Wellcome Sanger Institute for the COVID-19 Genomics UK (COG-UK) Consortium | Louissa Macfarlane-Smith, Holli Carden, Katherine L. Harper, Antony Hale and Alex Alderton, Roberto Amato, Sonia Goncalves, Ewan Harrison, David K. Jackson, Ian Johnston, Dominic Kwiatkowski, Cordelia Langford, John Sillitoe on behalf of the Wellcome Sanger Institute COVID-19 Surveillance Team |
| EPI_ISL_539123, EPI_ISL_539124, EPI_ISL_539128, EPI_ISL_539133, EPI_ISL_539135, EPI_ISL_539136, EPI_ISL_539138, EPI_ISL_539140, EPI_ISL_539141, EPI_ISL_539142 | Leeds Teaching Hospitals NHS Trust and Public Health England, National Infection Service (Leeds laboratory) | Wellcome Sanger Institute for the COVID-19 Genomics UK (COG-UK) consortium | Louissa Macfarlane-Smith, Holli Carden, Katherine L. Harper, Antony Hale and Alex Alderton, Roberto Amato, Sonia Goncalves, Ewan Harrison, David K. Jackson, Ian Johnston, Dominic Kwiatkowski, Cordelia Langford, John Sillitoe on behalf of the Wellcome Sanger Institute COVID-19 Surveillance Team |
| EPI_ISL_539143 | Leeds Teaching Hospitals NHS Trust and Public Health England, National Infection Service (Leeds laboratory) | Wellcome Sanger Institute for the COVID-19 Genomics UK (COG-UK) Consortium | Louissa Macfarlane-Smith, Holli Carden, Katherine L. Harper, Antony Hale and Alex Alderton, Roberto Amato, Sonia Goncalves, Ewan Harrison, David K. Jackson, Ian Johnston, Dominic Kwiatkowski, Cordelia Langford, John Sillitoe on behalf of the Wellcome Sanger Institute COVID-19 Surveillance Team |
| EPI_ISL_539145, EPI_ISL_539146 | Leeds Teaching Hospitals NHS Trust and Public Health England, National Infection Service (Leeds laboratory) | Wellcome Sanger Institute for the COVID-19 Genomics UK (COG-UK) consortium | Louissa Macfarlane-Smith, Holli Carden, Katherine L. Harper, Antony Hale and Alex Alderton, Roberto Amato, Sonia Goncalves, Ewan Harrison, David K. Jackson, Ian Johnston, Dominic Kwiatkowski, Cordelia Langford, John Sillitoe on behalf of the Wellcome Sanger Institute COVID-19 Surveillance Team |
| EPI_ISL_539150 | Leeds Teaching Hospitals NHS Trust and Public Health England, National Infection Service (Leeds laboratory) | Wellcome Sanger Institute for the COVID-19 Genomics UK (COG-UK) Consortium | Louissa Macfarlane-Smith, Holli Carden, Katherine L. Harper, Antony Hale and Alex Alderton, Roberto Amato, Sonia Goncalves, Ewan Harrison, David K. Jackson, Ian Johnston, Dominic Kwiatkowski, Cordelia Langford, John Sillitoe on behalf of the Wellcome Sanger Institute COVID-19 Surveillance Team |
| EPI_ISL_539155, EPI_ISL_539167, EPI_ISL_539179, EPI_ISL_539182, EPI_ISL_539184 | Leeds Teaching Hospitals NHS Trust and Public Health England, National Infection Service (Leeds laboratory) | Wellcome Sanger Institute for the COVID-19 Genomics UK (COG-UK) consortium | Louissa Macfarlane-Smith, Holli Carden, Katherine L. Harper, Antony Hale and Alex Alderton, Roberto Amato, Sonia Goncalves, Ewan Harrison, David K. Jackson, Ian Johnston, Dominic Kwiatkowski, Cordelia Langford, John Sillitoe on behalf of the Wellcome Sanger Institute COVID-19 Surveillance Team |
| EPI_ISL_539185 | Leeds Teaching Hospitals NHS Trust and Public Health | Wellcome Sanger Institute for the COVID-19 Genomics UK | Louissa Macfarlane-Smith, Holli Carden, Katherine L. Harper, Antony Hale and Alex Alderton, Roberto Amato, Sonia Goncalves, Ewan Harrison, David K. |

| | | | |
|---|---|---|---|
| | England, National Infection Service (Leeds laboratory) | (COG-UK) Consortium | Jackson, Ian Johnston, Dominic Kwiatkowski, Cordelia Langford, John Sillitoe on behalf of the Wellcome Sanger Institute COVID-19 Surveillance Team |
| EPI_ISL_539192 | Leeds Teaching Hospitals NHS Trust and Public Health England, National Infection Service (Leeds laboratory) | Wellcome Sanger Institute for the COVID-19 Genomics UK (COG-UK) consortium | Louissa Macfarlane-Smith, Holli Carden, Katherine L. Harper, Antony Hale and Alex Alderton, Roberto Amato, Sonia Goncalves, Ewan Harrison, David K. Jackson, Ian Johnston, Dominic Kwiatkowski, Cordelia Langford, John Sillitoe on behalf of the Wellcome Sanger Institute COVID-19 Surveillance Team |
| EPI_ISL_539197 | Leeds Teaching Hospitals NHS Trust and Public Health England, National Infection Service (Leeds laboratory) | Wellcome Sanger Institute for the COVID-19 Genomics UK (COG-UK) Consortium | Louissa Macfarlane-Smith, Holli Carden, Katherine L. Harper, Antony Hale and Alex Alderton, Roberto Amato, Sonia Goncalves, Ewan Harrison, David K. Jackson, Ian Johnston, Dominic Kwiatkowski, Cordelia Langford, John Sillitoe on behalf of the Wellcome Sanger Institute COVID-19 Surveillance Team |
| EPI_ISL_539198, EPI_ISL_539202, EPI_ISL_539207, EPI_ISL_539209, EPI_ISL_539212, EPI_ISL_539218, EPI_ISL_539220 | Leeds Teaching Hospitals NHS Trust and Public Health England, National Infection Service (Leeds laboratory) | Wellcome Sanger Institute for the COVID-19 Genomics UK (COG-UK) consortium | Louissa Macfarlane-Smith, Holli Carden, Katherine L. Harper, Antony Hale and Alex Alderton, Roberto Amato, Sonia Goncalves, Ewan Harrison, David K. Jackson, Ian Johnston, Dominic Kwiatkowski, Cordelia Langford, John Sillitoe on behalf of the Wellcome Sanger Institute COVID-19 Surveillance Team |
| EPI_ISL_539249, EPI_ISL_539250, EPI_ISL_539251, EPI_ISL_539252, EPI_ISL_539253, EPI_ISL_539254, EPI_ISL_539255, EPI_ISL_539256, EPI_ISL_539257, EPI_ISL_539258, EPI_ISL_539259, EPI_ISL_539260, EPI_ISL_539261, EPI_ISL_539262, EPI_ISL_539263, EPI_ISL_539264, EPI_ISL_539265, EPI_ISL_539266, EPI_ISL_539267, EPI_ISL_539268, EPI_ISL_539269, EPI_ISL_539270, EPI_ISL_539271 | | | |
| see above | Hospital Universitario de La Ribera (Alzira, València) | SeqCOVID-SPAIN consortium/IBV(CSIC) | Olalla Martínez Macias, Julia González and SeqCOVID-SPAIN consortium |
| EPI_ISL_539300, EPI_ISL_539301, EPI_ISL_539302, EPI_ISL_539303, EPI_ISL_539304, EPI_ISL_539311 | KWR Watercycle Research Institute | Erasmus Medical Center | Ray Izquierdo-Lara, Goffe Elsinga, Leo Heijnen, Bas B. Oude Munnink, Claudia M. E. Schapendonk, David Nieuwenhuijse, Matthijs Kon, Lu Lu, Frank M. Aarestrup, Samantha Lycett, Gertjan Medema, Marion P.G. Koopmans, Miranda de Graaf |
| EPI_ISL_539851 | Pok Oi Hospital | Hong Kong Department of Health | Alan K.L. Tsang, Peter C.W. Yip, Edman T.K. Lam, Rickjason C.W. Chan, Dominic N.C. Tsang |
| EPI_ISL_540217 | Lighthouse Lab in Glasgow | Wellcome Sanger Institute for the COVID-19 Genomics UK (COG-UK) consortium | Harper VanSteenhouse, Yumi Kasai, David Gray, Carol Clugston, Anna Dominiczak and Alex Alderton, Roberto Amato, Sonia Goncalves, Ewan Harrison, David K. Jackson, Ian Johnston, Dominic Kwiatkowski, Cordelia Langford, John Sillitoe on behalf of the Wellcome Sanger Institute COVID-19 Surveillance Team |
| EPI_ISL_540431 | IN State Department of Health Laboratory Services | Pathogen Discovery, Respiratory Viruses Branch, Division of Viral Diseases, Centers for Disease Control and Prevention | Yan Li, Jing Zhang, Anna Montmayeur, Krista Queen,Ying Tao, Anna Uehara, Clinton R. Paden, Rachel Marine, Haibin Wang, Suxiang Tong |
| EPI_ISL_541042, EPI_ISL_541043, EPI_ISL_541044, EPI_ISL_541045, EPI_ISL_541046, EPI_ISL_541047, EPI_ISL_541048, EPI_ISL_541049, EPI_ISL_541050, EPI_ISL_541051, EPI_ISL_541052, EPI_ISL_541053 | | | |
| see above | Hospital Clínico Universitario de Santiago de Compostela | SeqCOVID-SPAIN consortium/Institute of Biomedicine of Valencia, IBV-CSIC | José Javier Costa Alcalde, Antonio Aguilera Guirao, Mª Luisa Pérez del Molino Bernal, Amparo Coira Nieto, Gema Barbeito Castiñeiras, Rocio Trastoy Pena and SeqCOVID-SPAIN consortium |
| EPI_ISL_541238, EPI_ISL_541239 | Florida Bureau of Public Health Laboratories, Florida Department of Health | Florida Bureau of Public Health Laboratories, Florida Department of Health | Schmedes,S., Blanton,J. |
| EPI_ISL_541347, EPI_ISL_541348, EPI_ISL_541349, EPI_ISL_541350, EPI_ISL_541351 | Laboratory of Respiratory Viruses and Measles, Oswaldo Cruz Institute, FIOCRUZ | Laboratory of Respiratory Viruses and Measles, Oswaldo Cruz Institute, FIOCRUZ | Paola Resende, Luciana Appolinario, Fernando Motta, Anna Carolina Paixão, Ana Carolina Mendonça, Jonathan Lopes, Marilda Siqueira |
| EPI_ISL_541758, EPI_ISL_541772 | Barts Health NHS Trust | Wellcome Sanger Institute for the COVID-19 Genomics UK (COG-UK) consortium | Teresa Cutino-Moguel, Mark Hopkins, Beatrix Kele, David Harrington and Alex Alderton, Roberto Amato, Sonia Goncalves, Ewan Harrison, David K. Jackson, Ian Johnston, Dominic Kwiatkowski, Cordelia Langford, John Sillitoe on behalf of the Wellcome Sanger Institute COVID-19 Surveillance Team |
| EPI_ISL_541848, EPI_ISL_541849, EPI_ISL_541850, EPI_ISL_541851, EPI_ISL_541852, EPI_ISL_541853, EPI_ISL_541854, EPI_ISL_541855, EPI_ISL_541868, EPI_ISL_541870 | Lithuanian University of Health Sciences Hospital, Department of Laboratory Medicine | Lithuanian University of Health Sciences, Laboratory of Molecular Cardiology | Lukas Zemaitis, Arnoldas Pautienius, Kamile Tamusauskaite, Dovydas Gecys, Laura Pareckaite, Vaiva Lesauskaite, Astra Vitkauskiene |
| EPI_ISL_542490, EPI_ISL_542495, EPI_ISL_542498, EPI_ISL_542500, EPI_ISL_542502, EPI_ISL_542503, EPI_ISL_542506, EPI_ISL_542507, EPI_ISL_542509, EPI_ISL_542510, EPI_ISL_542511, EPI_ISL_542515, EPI_ISL_542521, EPI_ISL_542572, EPI_ISL_542594, EPI_ISL_542600, EPI_ISL_542605, EPI_ISL_542606, EPI_ISL_542611, EPI_ISL_542616, EPI_ISL_542618, EPI_ISL_542681, EPI_ISL_542777, EPI_ISL_542778, EPI_ISL_542796, EPI_ISL_542799, EPI_ISL_542800, EPI_ISL_542806, EPI_ISL_542807 | | | |
| see above | Houston Methodist Hospital | Houston Methodist Hospital | S. Wesley Long, Randall J. Olsen, Paul A. Christensen, David W. Bernard, James J. Davis, Maulik Shukla, Marcus Nguyen, Matthew Ojeda Saavedra, Concepcion C. Cantu, Prasanti Yerramilli, Layne Pruitt, Sishir Subedi, Hung-Che Kuo, Heather Hendrickson, Ghazaleh Eskandari, Hoang A. T. Nguyen, J. Hunter Long, Muthiah Kumaraswami, Jule Goike, Daniel Boutz, Jimmy Gollihar, Jason S. McLellan, Chia-Wei Chou, Kamyab Javanmardi, Ilya J. Finkelstein, and James M. Musser |
| EPI_ISL_542938, EPI_ISL_542948, EPI_ISL_542971 | TriCore Reference Laboratories | Center for Global Health, University of New Mexico Health Sciences Center | Daryl Domman, Kurt Schwalm, Twila Kunde, Joseph Hicks, Michael Edwards, Darrell Dinwiddie |
| EPI_ISL_543614, EPI_ISL_543695, EPI_ISL_543703, EPI_ISL_543724, EPI_ISL_543745, EPI_ISL_543753, EPI_ISL_543767, EPI_ISL_543776, EPI_ISL_543786, EPI_ISL_544243, EPI_ISL_544251, EPI_ISL_544259, EPI_ISL_544268, EPI_ISL_544278, EPI_ISL_544285, EPI_ISL_544296, EPI_ISL_544311, EPI_ISL_544332, EPI_ISL_544360, EPI_ISL_544368, EPI_ISL_544381, EPI_ISL_544392, EPI_ISL_544398, EPI_ISL_544406, EPI_ISL_544407, EPI_ISL_544408, EPI_ISL_544409, EPI_ISL_544415, EPI_ISL_544424, EPI_ISL_544433, EPI_ISL_544485, EPI_ISL_544517, EPI_ISL_544532, EPI_ISL_544538, EPI_ISL_544545, EPI_ISL_544579, EPI_ISL_544585, EPI_ISL_544588, EPI_ISL_544595, EPI_ISL_544604, EPI_ISL_544608, EPI_ISL_544617, EPI_ISL_544632, EPI_ISL_544656, EPI_ISL_544662, EPI_ISL_544678, EPI_ISL_544687, EPI_ISL_544696, EPI_ISL_544705, EPI_ISL_544713, EPI_ISL_544812, EPI_ISL_544885, EPI_ISL_544925, EPI_ISL_544934, EPI_ISL_544950, EPI_ISL_545659, EPI_ISL_545691, EPI_ISL_545724, EPI_ISL_545826, EPI_ISL_545855, EPI_ISL_545858, EPI_ISL_545866, EPI_ISL_545868, EPI_ISL_545869, EPI_ISL_545870, EPI_ISL_545871, EPI_ISL_545872, EPI_ISL_545873, EPI_ISL_545874, EPI_ISL_545876, EPI_ISL_545877, EPI_ISL_545878, EPI_ISL_545879, EPI_ISL_545880, EPI_ISL_545883, EPI_ISL_545884, EPI_ISL_545885, EPI_ISL_545886, EPI_ISL_545889, EPI_ISL_545891, EPI_ISL_545911, EPI_ISL_545912, EPI_ISL_545952, EPI_ISL_545953, EPI_ISL_546656, EPI_ISL_546719, EPI_ISL_546797, EPI_ISL_546810, EPI_ISL_546824, EPI_ISL_546834, EPI_ISL_546843, EPI_ISL_546868, EPI_ISL_546913, EPI_ISL_546952, EPI_ISL_546959, EPI_ISL_547001, EPI_ISL_547069, EPI_ISL_547109 | | | |
| see above | Houston Methodist Hospital | Houston Methodist Hospital | S. Wesley Long, Randall J. Olsen, Paul A. Christensen, David W. Bernard, James J. Davis, Maulik Shukla, Marcus Nguyen, Matthew Ojeda Saavedra, Concepcion C. Cantu, Prasanti Yerramilli, Layne Pruitt, Sishir Subedi, Hung-Che Kuo, Heather Hendrickson, Ghazaleh Eskandari, Hoang A. T. Nguyen, J. Hunter Long, Muthiah Kumaraswami, Jule Goike, Daniel Boutz, Jimmy Gollihar, Jason S. McLellan, Chia-Wei Chou, Kamyab Javanmardi, Ilya J. Finkelstein, and James M. Musser |
| EPI_ISL_547495, EPI_ISL_547536, EPI_ISL_547537 | Dutch COVID-19 response team | National Institute for Public Health and the Environment (RIVM) | Adam Meijer, Harry Vennema, Jeroen Cremer, Sharon van den Brink, Bas van der Veer, AnneMarie van den Brandt, Florian Zwagemaker, Dennis Schmitz, Chantal Reusken, on behalf of the national COVID-19 response team |
| EPI_ISL_548123, EPI_ISL_548124, EPI_ISL_548125 | Middlemore Hospital | Institute of Environmental Science and Research (ESR) | Xiaoyun Ren, Matt Storey, Nikki Freed, Muhammad Faisal, Jing Wang, Hermes Perez, Anja Werno, Antje van der Linden, Arlo Upton, Chris Mansell, David Hammer, Dragana Drinkovic, Gary McAuliffe, Hana Sofia Andersson, James Ussher, Jill Sherwood, Josh Freeman, Julia Howard, Juliet Elvy, Mary DeAlmeida, Matt Blakiston, Matthew Rogers, Max Bloomfield, Michael Addidle, Michelle Balm, Sally Roberts, Sarah Jefferies, Sharmini Muttaiyah, Susan Morpeth, Susan Taylor, Timothy Blackmore, Vani Sathyendran, Veronica Playle, Virginia Hope, Erasmus Smit, Lauren Jelly, Olin Silander, Joep de Ligt |
| EPI_ISL_549006, EPI_ISL_549007, EPI_ISL_549008, EPI_ISL_549009, EPI_ISL_549022 | KWR Watercycle Research Institute | Erasmus Medical Center | Ray Izquierdo-Lara, Goffe Elsinga, Leo Heijnen, Bas B. Oude Munnink, Claudia M. E. Schapendonk, David Nieuwenhuijse, Matthijs Kon, Lu Lu, Frank M. Aarestrup, Samantha Lycett, Gertjan Medema, Marion P.G. Koopmans, Miranda de Graaf |
| EPI_ISL_549384, EPI_ISL_549487 | Quadram Institute Bioscience | COVID-19 Genomics UK (COG-UK) Consortium | Dave J. Baker, Gemma L. Kay, Alp Aydin, Thanh Le-Viet, Steven Rudder, Ana P. Tedim, Anastasia Kolyva, Maria Diaz, Leonardo de Oliveira Martins, Nabil-Fareed Alikhan, Lizzie Meadows, Rachael Stanley, Ngozi Elumogo, Muhammed Yasir, Nicholas M. Thomson, Alexander J Trotter, Rachel Gilroy, Samuel Bloomfield, Claire Stuart, Andrew Bell, Reenesh Prakash, Samir Dervisevic, Alison E. Mather, John Wain, Mark Webber, Andrew J. Page, Justin O'Grady |
| EPI_ISL_559971 | Virology Department, Royal Infirmary of Edinburgh, NHS Lothian / School of Biological Sciences, University of Edinburgh / Institute of Genetics and Molecular Medicine, University of Edinburgh | COVID-19 Genomics UK (COG-UK) Consortium | McHugh M, Dewar R, Rooke S, Gallagher M, Balcaza C, O'Toole Á, Scher E, Hill V, McCrone JT, Colquhoun R, Yu X, Jackson B, Rambaut A, Williams TC, Templeton K |
| EPI_ISL_560387 | National Health Laboratory | Botswana Institute for Technology Research and Innovation | Kefentse Arnold Tumedi, Madisa Mine, Dineo Emang Tshiamo. Gape Nyepetsi, Thongbotho Mphoyakgosi, Maitshwarelo Ignatius Matsheka |
| EPI_ISL_560571, EPI_ISL_560576 | hôpital | National Reference Center for Viruses of Respiratory Infections, Institut Pasteur, Paris | Sylvie Behillil, Fabiana Gambaro, Etienne Simon-Lorière, Vincent Enouf, Maud Vanpeene, Sylvie van der Werf |

| | | | |
|---|---|---|---|
| EPI_ISL_560595, EPI_ISL_560597 | hopital | National Reference Center for Viruses of Respiratory Infections, Institut Pasteur, Paris | Sylvie Behillil, Fabiana Gambaro, Etienne Simon-Lorière, Vincent Enouf, Maud Vanpeene, Sylvie van der Werf |
| EPI_ISL_560628 | Hospital | National Reference Center for Viruses of Respiratory Infections, Institut Pasteur, Paris | Sylvie Behillil, Fabiana Gambaro, Etienne Simon-Lorière, Vincent Enouf, Maud Vanpeene, Sylvie van der Werf |
| EPI_ISL_560746, EPI_ISL_560747, EPI_ISL_560748, EPI_ISL_560749, EPI_ISL_560750, EPI_ISL_560751, EPI_ISL_560752, EPI_ISL_560753, EPI_ISL_560754, EPI_ISL_560755, EPI_ISL_560756, EPI_ISL_560757, EPI_ISL_560758, EPI_ISL_560759, EPI_ISL_560760, EPI_ISL_560761, EPI_ISL_560762, EPI_ISL_560767 | | | |
| see above | Minnesota Department of Health, Public Health Laboratory | Minnesota Department of Health, Public Health Laboratory | Matt Plumb, Jacob Garfin, and Xiong Wang |
| EPI_ISL_560842, EPI_ISL_560843, EPI_ISL_560844, EPI_ISL_560845, EPI_ISL_560846, EPI_ISL_560848, EPI_ISL_560920, EPI_ISL_560921, EPI_ISL_560922, EPI_ISL_560923, EPI_ISL_560924, EPI_ISL_560925 | | | |
| see above | Utah Public Health Laboratory | Utah Public Health Laboratory | Erin Young, Kelly Oakeson |
| EPI_ISL_561371 | Hospital Universitario de Gran Canaria Dr. Negrín | SeqCOVID-SPAIN consortium/IBV(CSIC) | M. Carmen Pérez González, Francisco J. Chamizo López, Ana Bordes Benítez and SeqCOVID-SPAIN consortium |
| EPI_ISL_561788, EPI_ISL_561844, EPI_ISL_561957, EPI_ISL_562482 | Victorian Infectious Diseases Reference Laboratory (VIDRL) | VIDRL and MDU-PHL | Caly, L., Seemann, T., Sait, M., Schultz, M. B., Druce J., Sherry, N. |
| EPI_ISL_562590 | Microbiological Diagnostic Unit - Public Health Laboratory (MDU-PHL) | MDU-PHL | Seemann, T., Schultz M. B., Sait, M., Sherry, N. |
| EPI_ISL_563551 | Victorian Infectious Diseases Reference Laboratory (VIDRL) | VIDRL and MDU-PHL | Caly, L., Seemann, T., Sait, M., Schultz, M. B., Druce J., Sherry, N. |
| EPI_ISL_563594 | Microbiological Diagnostic Unit - Public Health Laboratory (MDU-PHL) | MDU-PHL | Seemann, T., Schultz M. B., Sait, M., Sherry, N. |
| EPI_ISL_565928, EPI_ISL_565937, EPI_ISL_565951, EPI_ISL_565952, EPI_ISL_565954, EPI_ISL_565955, EPI_ISL_565957, EPI_ISL_565963, EPI_ISL_565966, EPI_ISL_565967, EPI_ISL_565975, EPI_ISL_565976, EPI_ISL_565980, EPI_ISL_565982, EPI_ISL_565987, EPI_ISL_565989, EPI_ISL_566019 | | | |
| see above | Michigan Department of Health and Human Services, Bureau of Laboratories | Michigan Department of Health and Human Services, Bureau of Laboratories | Blankenship HM, Riner D, Soehnlen MK |
| EPI_ISL_566036, EPI_ISL_566037, EPI_ISL_566038, EPI_ISL_566039, EPI_ISL_566040, EPI_ISL_566041, EPI_ISL_566042, EPI_ISL_566043 | Wildlife Conservation Society, Bronx Zoo | Diagnostic Virology Laboratory, United States Department of Agriculture, National Veterinary Services Laboratories, Population Medicine and Diagnostic Sciences, Cornell University | McAloose,D., Laverack,M., Wang,L., Killian,M.L., Caserta,L.C., Yuan,F., Mitchell,P.K., Queen,K., Mauldin,M.R., Cronk,B.D., Bartlett,S.L., Sykes,J.M., Zec,S., Stokol,T., Ingerman,K., Delaney,M.A., Frederikson,R., Ivancic,M., Jenkins-Moore,M., Mozingo,K., Farnzen,K., Kuzmin,I., Hines Bergeson,N., Goodman,L., Wang,H., Tong,S., Fang,Y., Olmstead,C., McCann,C., Thomas,P., Goodrich,E., Elvinger,F., Smith,D.C., Slavinski,S., Calle,P.P., Terio,K., Torchetti,M.K., Diel,D.G., Cardia Caserta,L. |
| EPI_ISL_566044 | Wildlife Conservation Society, Bronx Zoo | Diagnostic Virology Laboratory, United States Department of Agriculture, National Veterinary Services Laboratories, Population Medicine and Diagnostic Sciences, Cornell University | McAloose,D., Laverack,M., Wang,L., Killian,M.L., Caserta,L.C., Yuan,F., Mitchell,P.K., Queen,K., Mauldin,M.R., Cronk,B.D., Bartlett,S.L., Sykes,J.M., Zec,S., Stokol,T., Ingerman,K., Delaney,M.A., Frederikson,R., Ivancic,M., Jenkins-Moore,M., Mozingo,K., Farnzen,K., Kuzmin,I., Hines Bergeson,N., Goodman,L., Wang,H., Tong,S., Fang,Y., Olmstead,C., McCann,C., Thomas,P., Goodrich,E., Elvinger,F., Smith,D.C., Slavinski,S., Calle,P.P., Terio,K., Torchetti,M.K., Diel,D.G., Cardia Caserta,L., Mauldin,M. |
| EPI_ISL_568695, EPI_ISL_568696, EPI_ISL_568697 | KEMRI-Wellcome Trust Research Programme/KEMRI-CGMR-C Kilifi | KEMRI-Wellcome Trust Research Programme/KEMRI-CGMR-C Kilifi | Githinji et al 2020 |
| EPI_ISL_568873 | Malaysia Genome Institute | Malaysia Genome Institute | Mohd Noor Mat Isa, Irni Suhayu Sapian, Yusuf Muhammad Noor, Nurhezreen Md Iqbal, Mohd Faizal Abu Bakar, Enizza Kasim, Shamsidar Sopie, Siti Noraini Othman, Azrin Ahmad, Nor Azfa Johari, Shahrul Hisham Zainal Ariffin |
| EPI_ISL_568909, EPI_ISL_568991, EPI_ISL_568992, EPI_ISL_569491, EPI_ISL_569492, EPI_ISL_569493, EPI_ISL_569494, EPI_ISL_569495, EPI_ISL_569496, EPI_ISL_569497, EPI_ISL_569498, EPI_ISL_569499, EPI_ISL_569500, EPI_ISL_569501, EPI_ISL_569502, EPI_ISL_569503, EPI_ISL_569504, EPI_ISL_569505, EPI_ISL_569506, EPI_ISL_569507, EPI_ISL_569508, EPI_ISL_569509, EPI_ISL_569510, EPI_ISL_569511, EPI_ISL_569512, EPI_ISL_569513, EPI_ISL_569514, EPI_ISL_569515, EPI_ISL_569516, EPI_ISL_569517, EPI_ISL_569518, EPI_ISL_569519, EPI_ISL_569520, EPI_ISL_569521, EPI_ISL_569522, EPI_ISL_569523, EPI_ISL_569524, EPI_ISL_569525, EPI_ISL_569526, EPI_ISL_569527, EPI_ISL_569528, EPI_ISL_569529, EPI_ISL_569530, EPI_ISL_569531, EPI_ISL_569532, EPI_ISL_569533, EPI_ISL_569534, EPI_ISL_569535, EPI_ISL_569536, EPI_ISL_569537, EPI_ISL_569539, EPI_ISL_569540, EPI_ISL_569541, EPI_ISL_569542, EPI_ISL_569543, EPI_ISL_569544, EPI_ISL_569545, EPI_ISL_569546, EPI_ISL_569547, EPI_ISL_569548, EPI_ISL_569549, EPI_ISL_569550, EPI_ISL_569551, EPI_ISL_569552, EPI_ISL_569553, EPI_ISL_569554, EPI_ISL_569555, EPI_ISL_569556, EPI_ISL_569557, EPI_ISL_569558, EPI_ISL_569559, EPI_ISL_569560, EPI_ISL_569561, EPI_ISL_569562, EPI_ISL_569563, EPI_ISL_569564, EPI_ISL_569565, EPI_ISL_569566, EPI_ISL_569567, EPI_ISL_569568, EPI_ISL_569569, EPI_ISL_569570, EPI_ISL_569584 | | | |
| see above | MEPHI, Aix Marseille University | MEPHI, Aix Marseille University | Anthony LEVASSEUR |
| EPI_ISL_569609 | Bennet County Hospital | South Dakota Public Health Laboratory | Matt Plumb, Jacob Garfin, Xiong Wang, and Chris Carlson |
| EPI_ISL_570054, EPI_ISL_570055, EPI_ISL_570056, EPI_ISL_570057, EPI_ISL_570061, EPI_ISL_570063, EPI_ISL_570066, EPI_ISL_570067, EPI_ISL_570073, EPI_ISL_570076, EPI_ISL_570079, EPI_ISL_570090, EPI_ISL_570091, EPI_ISL_570096, EPI_ISL_570097, EPI_ISL_570098, EPI_ISL_570101, EPI_ISL_570102, EPI_ISL_570103, EPI_ISL_570108, EPI_ISL_570111, EPI_ISL_570112, EPI_ISL_570113, EPI_ISL_570114, EPI_ISL_570115, EPI_ISL_570116, EPI_ISL_570117, EPI_ISL_570118, EPI_ISL_570119, EPI_ISL_570120, EPI_ISL_570121, EPI_ISL_570122, EPI_ISL_570123, EPI_ISL_570124, EPI_ISL_570125, EPI_ISL_570126, EPI_ISL_570127, EPI_ISL_570128, EPI_ISL_570129, EPI_ISL_570130, EPI_ISL_570131, EPI_ISL_570132, EPI_ISL_570157, EPI_ISL_570158, EPI_ISL_570159, EPI_ISL_570160, EPI_ISL_570161, EPI_ISL_570162, EPI_ISL_570163, EPI_ISL_570165, EPI_ISL_570166, EPI_ISL_570178, EPI_ISL_570179, EPI_ISL_570180, EPI_ISL_570181, EPI_ISL_570182, EPI_ISL_570183, EPI_ISL_570184, EPI_ISL_570185, EPI_ISL_570186, EPI_ISL_570187, EPI_ISL_570188 | | | |
| see above | UW Virology Lab | UW Virology Lab | Pavitra Roychoudhury, Hong Xie, Lasata Shrestha, Amin Addetia, Victoria M Rachleff, Meei-Li Huang, Keith R Jerome, Alexander Greninger |
| EPI_ISL_572270 | Virginia DCLS | Virginia DCLS | Virginia DCLS |
| EPI_ISL_572329 | Public Health Authority of the Slovak Republic, Bratislava | Faculty of Natural Sciences, Comenius University in Bratislava | Dominika Friová, Viktória Hodorová, Kristína Boršová, Broa Brejová, Viktória abanová, Sabina Fumaová Havlíková, Juraj Kopáek, Martina Liková, ubomíra Lukáiková, Martina Neboháová, Monika Sláviková, Edita Staroová, Elena Tichá, Tomáš Vina, Boris Klempa, Jozef Nosek |
| EPI_ISL_574485 | Programme in Emerging Infectious Diseases, Duke-NUS Medical School | National Public Health Laboratory, National Centre for Infectious Diseases | Tze Minn Mak, Sophie Octavia, Zhenyang Zhou, Danielle E Anderson, Adrian Eng Zheng Kang, Lin Cui, Raymond Tzer Pin Lin |
| EPI_ISL_574606 | RS Siloam Purwakarta | Eijkman Institute for Molecular Biology, Ministry of Research and Technology/National Agency for Research and Innovation | Frilasita A Yudhaputri, Edison Johar, Hidayat Trimarsanto, Iskandar A Adnan, Willy Agustine, David H Muljono, Safarina G Malik, Herawati Sudoyo, Khin Saw Myint, Amin Soebandrio |
| EPI_ISL_574847, EPI_ISL_574848, EPI_ISL_574849, EPI_ISL_574850, EPI_ISL_574851, EPI_ISL_574852 | Institute for Infectious Diseases, University of Bern | Institute for Infectious Diseases, University of Bern | Michel C Koch, Christian Baumann, Miguel A Terrazos Miani, Cora Sägesser, Stephen L Leib, Peter Keller, Franziska Suter-Riniker, Alban Ramette |
| EPI_ISL_575042, EPI_ISL_575043, EPI_ISL_575048, EPI_ISL_575049, EPI_ISL_575050, EPI_ISL_575051, EPI_ISL_575052, EPI_ISL_575053, EPI_ISL_575054, EPI_ISL_575055, EPI_ISL_575056, EPI_ISL_575060, EPI_ISL_575098, EPI_ISL_575099, EPI_ISL_575100, EPI_ISL_575101, EPI_ISL_575102, EPI_ISL_575103, EPI_ISL_575104, EPI_ISL_575109, EPI_ISL_575110, EPI_ISL_575111, EPI_ISL_575112, EPI_ISL_575113, EPI_ISL_575114, EPI_ISL_575116 | | | |
| see above | Utah Public Health Laboratory | Utah Public Health Laboratory | Erin Young, Kelly Oakeson |
| EPI_ISL_576174 | VA-Division of Consolidated Laboratory Services | Pathogen Discovery, Respiratory Viruses Branch, Division of Viral Diseases, Centers for Disease Control and Prevention | Ying Tao, Jing Zhang, Brian Lynch, Yan Li, Krista Queen, Anna Uehara, Clinton R. Paden, Peter Cook, Haibin Wang, Suxiang Tong |
| EPI_ISL_576188 | WI State Laboratory of Hygiene | Pathogen Discovery, Respiratory Viruses Branch, Division of Viral Diseases, Centers for Disease Control and Prevention | Ying Tao, Jing Zhang, Brian Lynch, Yan Li, Krista Queen, Anna Uehara, Clinton R. Paden, Peter Cook, Haibin Wang, Suxiang Tong |
| EPI_ISL_577629 | The National Institute of Public Health | State Veterinary Institute Prague | Nagy,A;Jirincova,H;Novakova,L;Trnka,D;Vecerova,J |
| EPI_ISL_577735 | Institute of Virology, Biomedical Research Center of the Slovak Academy of Sciences, Bratislava | Faculty of Natural Sciences, Comenius University, Bratislava | Kristína Boršová, Viktória Hodorová, Broa Brejová, Viktória abanová, Dominika Friová, Sabina Fumaová Havlíková, Juraj Kopáek, Martina Liková, ubomíra Lukáiková, Martina Neboháová, Monika Sláviková, Edita Staroová, Elena Tichá, Tomáš Vina, Boris Klempa, Jozef Nosek |
| EPI_ISL_577736 | Institute of Virology, Biomedical Research Center of the Slovak Academy of Sciences, Bratislava | Faculty of Natural Sciences, Comenius University, Bratislava | Viktória Hodorová, Kristína Boršová, Broa Brejová, Viktória abanová, Dominika Friová, Sabina Fumaová Havlíková, Juraj Kopáek, Martina Liková, ubomíra Lukáiková, Martina Neboháová, Monika Sláviková, Edita Staroová, Elena Tichá, Tomáš Vina, Jozef Nosek, Boris Klempa |
| EPI_ISL_578083, EPI_ISL_578086, EPI_ISL_578087, EPI_ISL_578089, EPI_ISL_578091, EPI_ISL_578092, EPI_ISL_578136, EPI_ISL_578137, EPI_ISL_578138, EPI_ISL_578139, EPI_ISL_578140, EPI_ISL_578141, EPI_ISL_578142, EPI_ISL_578143 | | | |
| see above | University of Michigan Clinical Microbiology Laboratory | Lauring Lab, University of Michigan, Department of Microbiology and Immunology | Valesano |
| EPI_ISL_578338, EPI_ISL_578339, EPI_ISL_578340, EPI_ISL_578341, EPI_ISL_578342, EPI_ISL_578343, EPI_ISL_578344, EPI_ISL_578398, EPI_ISL_578399, EPI_ISL_578400, EPI_ISL_578401, EPI_ISL_578402, EPI_ISL_578403, EPI_ISL_578404, EPI_ISL_578405, EPI_ISL_578406, EPI_ISL_578407, EPI_ISL_578408, EPI_ISL_578409, EPI_ISL_578469 | | | |

| | | | |
|---|---|---|---|
| see above | Wisconsin State Laboratory of Hygiene Communicable Disease Division | Wisconsin State Laboratory of Hygiene Communicable Disease Division | Kelsey R. Florek, Abigail C. Shockey |
| EPI_ISL_578697 | Clinical Virology Laboratory, Institute of Liver and Biliary Sciences | ILBS - IGIB | Ekta Gupta, Sheetalnath Rooge, Abhishek Padhi, Reshu Agarwal, Jaswinder Singh Maras, Shridhar Sivasubbu, Vinod Scaria, Shvetank Sharma |
| EPI_ISL_579090, EPI_ISL_579091 | Canterbury Health Laboratories | Institute of Environmental Science and Research (ESR) | Xiaoyun Ren, Matt Storey, Nikki Freed, Muhammad Faisal, Jing Wang, Hermes Perez, Anja Werno, Antje van der Linden, Arlo Upton, Chris Mansell, David Hammer, Dragana Drinkovic, Gary McAuliffe, Hana Sofia Andersson, James Ussher, Jill Sherwood, Josh Freeman, Julia Howard, Juliet Elvy, Mary DeAlmeida, Matt Blakiston, Matthew Rogers, Max Bloomfield, Michael Addidle, Michelle Balm, Sally Roberts, Sarah Jefferies, Sharmini Muttaiyah, Susan Morpeth, Susan Taylor, Timothy Blackmore, Vani Sathyendran, Veronica Playle, Virginia Hope, Erasmus Smit, Lauren Jelly, Olin Silander, Joep de Ligt |
| EPI_ISL_579159, EPI_ISL_579160, EPI_ISL_579161, EPI_ISL_579162, EPI_ISL_579164, EPI_ISL_579165, EPI_ISL_579166, EPI_ISL_579167, EPI_ISL_579168, EPI_ISL_579169, EPI_ISL_579170, EPI_ISL_579171, EPI_ISL_579172, EPI_ISL_579173, EPI_ISL_579175, EPI_ISL_579176, EPI_ISL_579177, EPI_ISL_579178, EPI_ISL_579179, EPI_ISL_579180, EPI_ISL_579181, EPI_ISL_579182, EPI_ISL_579183, EPI_ISL_579184 | | | |
| see above | Southern Community Labs Dunedin | Institute of Environmental Science and Research (ESR) | Xiaoyun Ren, Matt Storey, Nikki Freed, Muhammad Faisal, Jing Wang, Hermes Perez, Anja Werno, Antje van der Linden, Arlo Upton, Chris Mansell, David Hammer, Dragana Drinkovic, Gary McAuliffe, Hana Sofia Andersson, James Ussher, Jill Sherwood, Josh Freeman, Julia Howard, Juliet Elvy, Mary DeAlmeida, Matt Blakiston, Matthew Rogers, Max Bloomfield, Michael Addidle, Michelle Balm, Sally Roberts, Sarah Jefferies, Sharmini Muttaiyah, Susan Morpeth, Susan Taylor, Timothy Blackmore, Vani Sathyendran, Veronica Playle, Virginia Hope, Erasmus Smit, Lauren Jelly, Olin Silander, Joep de Ligt |
| EPI_ISL_579220, EPI_ISL_579221, EPI_ISL_579222, EPI_ISL_579223 | Canterbury Health Laboratories | Institute of Environmental Science and Research (ESR) | Xiaoyun Ren, Matt Storey, Nikki Freed, Muhammad Faisal, Jing Wang, Hermes Perez, Anja Werno, Antje van der Linden, Arlo Upton, Chris Mansell, David Hammer, Dragana Drinkovic, Gary McAuliffe, Hana Sofia Andersson, James Ussher, Jill Sherwood, Josh Freeman, Julia Howard, Juliet Elvy, Mary DeAlmeida, Matt Blakiston, Matthew Rogers, Max Bloomfield, Michael Addidle, Michelle Balm, Sally Roberts, Sarah Jefferies, Sharmini Muttaiyah, Susan Morpeth, Susan Taylor, Timothy Blackmore, Vani Sathyendran, Veronica Playle, Virginia Hope, Erasmus Smit, Lauren Jelly, Olin Silander, Joep de Ligt |
| EPI_ISL_579228, EPI_ISL_579232, EPI_ISL_579312 | Middlemore Hospital | Institute of Environmental Science and Research (ESR) | Xiaoyun Ren, Matt Storey, Nikki Freed, Muhammad Faisal, Jing Wang, Hermes Perez, Anja Werno, Antje van der Linden, Arlo Upton, Chris Mansell, David Hammer, Dragana Drinkovic, Gary McAuliffe, Hana Sofia Andersson, James Ussher, Jill Sherwood, Josh Freeman, Julia Howard, Juliet Elvy, Mary DeAlmeida, Matt Blakiston, Matthew Rogers, Max Bloomfield, Michael Addidle, Michelle Balm, Sally Roberts, Sarah Jefferies, Sharmini Muttaiyah, Susan Morpeth, Susan Taylor, Timothy Blackmore, Vani Sathyendran, Veronica Playle, Virginia Hope, Erasmus Smit, Lauren Jelly, Olin Silander, Joep de Ligt |
| EPI_ISL_579497, EPI_ISL_579498, EPI_ISL_579499, EPI_ISL_579500, EPI_ISL_579501 | Canterbury Health Laboratories | Institute of Environmental Science and Research (ESR) | Xiaoyun Ren, Matt Storey, Nikki Freed, Muhammad Faisal, Jing Wang, Hermes Perez, Anja Werno, Antje van der Linden, Arlo Upton, Chris Mansell, David Hammer, Dragana Drinkovic, Gary McAuliffe, Hana Sofia Andersson, James Ussher, Jill Sherwood, Josh Freeman, Julia Howard, Juliet Elvy, Mary DeAlmeida, Matt Blakiston, Matthew Rogers, Max Bloomfield, Michael Addidle, Michelle Balm, Sally Roberts, Sarah Jefferies, Sharmini Muttaiyah, Susan Morpeth, Susan Taylor, Timothy Blackmore, Vani Sathyendran, Veronica Playle, Virginia Hope, Erasmus Smit, Lauren Jelly, Olin Silander, Joep de Ligt |
| EPI_ISL_579599, EPI_ISL_579600, EPI_ISL_579601, EPI_ISL_579602, EPI_ISL_579603, EPI_ISL_579604, EPI_ISL_579605, EPI_ISL_579606, EPI_ISL_579607, EPI_ISL_579608, EPI_ISL_579609, EPI_ISL_579610, EPI_ISL_579611, EPI_ISL_579612, EPI_ISL_579613, EPI_ISL_579614, EPI_ISL_579615, EPI_ISL_579616, EPI_ISL_579617, EPI_ISL_579618, EPI_ISL_579619, EPI_ISL_579620, EPI_ISL_579621, EPI_ISL_579622, EPI_ISL_579623, EPI_ISL_579624, EPI_ISL_579625, EPI_ISL_579626, EPI_ISL_579627, EPI_ISL_579628, EPI_ISL_579629, EPI_ISL_579630, EPI_ISL_579631, EPI_ISL_579632, EPI_ISL_579633, EPI_ISL_579634, EPI_ISL_579635, EPI_ISL_579636, EPI_ISL_579637, EPI_ISL_579638, EPI_ISL_579639, EPI_ISL_579640, EPI_ISL_579641, EPI_ISL_579642, EPI_ISL_579643, EPI_ISL_579644, EPI_ISL_579645, EPI_ISL_579646, EPI_ISL_579647, EPI_ISL_579648, EPI_ISL_579649, EPI_ISL_579650 | | | |
| see above | QEII Health Sciences Centre | National Microbiology Laboratory (NML) | Anna Majer, Shari Tyson, Grace Seo, Philip Mabon, Darian Hole, Elsie Grudeski, Rhiannon Huzarewich, Russell Mandes, Anneliese Landgraff, Jennifer Tanner, Natalie Knox, Morag Graham, Gary Van Domselaar, Todd Hatchette, Jason LeBlanc, Nathalie Bastien, Yan Li, Timothy Booth, CanCOGeN's metadata curation team, Public Health Agency of Canada's CanCOGeN team |
| EPI_ISL_581457, EPI_ISL_581470, EPI_ISL_581471, EPI_ISL_581477, EPI_ISL_581484 | Medizinische Klinik Innere Medizin I, Universitätsklinikum Tübingen | NGS Competence Center Tübingen, Institut für Medizinische Mikrobiologie und Hygiene, Universitätsklinikum Tübingen | Angel Angelov |
| EPI_ISL_581675, EPI_ISL_581676, EPI_ISL_581677, EPI_ISL_581678, EPI_ISL_581679, EPI_ISL_581680, EPI_ISL_581681, EPI_ISL_581682, EPI_ISL_581683, EPI_ISL_581684, EPI_ISL_581685, EPI_ISL_581686, EPI_ISL_581817, EPI_ISL_581818, EPI_ISL_581819, EPI_ISL_581820, EPI_ISL_581821, EPI_ISL_581822, EPI_ISL_581823, EPI_ISL_581824, EPI_ISL_581825, EPI_ISL_581826, EPI_ISL_581827, EPI_ISL_581828, EPI_ISL_581829, EPI_ISL_581830, EPI_ISL_581831, EPI_ISL_581832, EPI_ISL_581833, EPI_ISL_581834, EPI_ISL_581835, EPI_ISL_581836, EPI_ISL_581837, EPI_ISL_581838, EPI_ISL_581839, EPI_ISL_581840, EPI_ISL_581841, EPI_ISL_581842, EPI_ISL_581843, EPI_ISL_581844, EPI_ISL_581845, EPI_ISL_581846, EPI_ISL_581847, EPI_ISL_581848, EPI_ISL_581849, EPI_ISL_581850, EPI_ISL_581851, EPI_ISL_581852, EPI_ISL_581853, EPI_ISL_581854, EPI_ISL_581855, EPI_ISL_581856, EPI_ISL_581857, EPI_ISL_581858, EPI_ISL_581859, EPI_ISL_581860, EPI_ISL_581861, EPI_ISL_581863, EPI_ISL_581864, EPI_ISL_581865, EPI_ISL_581866, EPI_ISL_581867, EPI_ISL_581868, EPI_ISL_581869, EPI_ISL_581870, EPI_ISL_581871, EPI_ISL_581872, EPI_ISL_581873 | | | |
| see above | University Hospital Basel, Clinical Virology | University Hospital Basel, Clinical Bacteriology | Madlen Stange, Alfredo Mari, Tim Roloff, Helena MB Seth-Smith, Michael Schweitzer, Myrta Brunner, Karoline Leuzinger, Kirstine K. Soegaard, Alexander Gensch, Sarah Tschudin-Sutter, Simon Fuchs, Julia Bielicki, Hans Pargger, Martin Siegemund, Christian Nickel, Roland Bingisser, Michael Osthoff, Stefano Bassetti, Rita Schneider-Sliwa, Manuel Battegay, Hans Hirsch, Adrian Egli |
| EPI_ISL_582033 | Department of Pathology, School of Medicine, Imam Khomeini Hospital, Tehran University of Medical Sciences | Genetics Research Center. University Of Social Welfare And Rehabilitation Sciences | Zohreh Fattahi, Marzieh Mohseni, Khadijeh Jalalvand, Azam Ghaziasadi, Seyedeh elham Mortazavi, Ali Jafarpour, Azar Hadadi, Alireza Abdollahi, Ali Jafarpour, Azam Ghaziasad, Seyedeh elham Mortazavi, Saber Soltani, Reza Najafipour , Kimia Kahrizi, Seyed Mohammad Jazayeri, Hossein Najmabadi |
| EPI_ISL_582109 | Hospital Universitario Marqués de Valdecilla - IDIVAL (Santander, Cantabria) | SeqCOVID-SPAIN consortium/IBV(CSIC) | María Eliecer Cano García, Mónica Gozalo Margüello, Jose Manuel Méndez Legaza, Daniel Pablo Marcos, Jesús Rodríguez Rodríguez, María Siller Ruiz and SeqCOVID-SPAIN consortium |
| EPI_ISL_582251, EPI_ISL_582252, EPI_ISL_582294, EPI_ISL_582296, EPI_ISL_582300, EPI_ISL_582302, EPI_ISL_582311, EPI_ISL_582312 | Cadham Provincial Laboratory | National Microbiology Laboratory (NML) | Anna Majer, Shari Tyson, Grace Seo, Philip Mabon, Elsie Grudeski, Rhiannon Huzarewich, Russell Mandes, Anneliese Landgraff, Jennifer Tanner, Natalie Knox, Morag Graham, Gary Van Domselaar, Paul Van Caeseele, Jared Bullard, David Alexander, Kerry Dust, Nathalie Bastien, Yan Li, Timothy Booth, Darian Hole, Madison Chapel, CanCOGeN's metadata curation team, Public Health Agency of Canada CanCOGeN team |
| EPI_ISL_582518 | unknown | Instituto Nacional de Saude (INSA) | Borges et al |
| EPI_ISL_582524, EPI_ISL_582525, EPI_ISL_582526 | Veterinary Specialized Institute "Kraljevo", Serbia | Veterinary Specialized Institute "Kraljevo", Serbia | Vidanovic,D., Tesovic,B., Knezevic,A., Jovanovic,T., Jankovic,M., Sekler,M., Banovic Djeri,B., Petrovic,T., Volkening,J., Afonso,C. |
| EPI_ISL_582627, EPI_ISL_582628 | Sheikh Khalifa Medical City | Molecular/Surveillance lab Sheikh Khalifa Medical City | Amirtharaj Francis, Sajeed Abdul, Hala Imambaccus, Sahar Almarzooqi, Hiba Saud, Stefan Weber |
| EPI_ISL_583432 | Memorial Sloan Kettering Cancer Center | van Bakel Laboratory, Genetics and Genomics Sciences, Icahn School of Medicine at Mount Sinai | Teresa Aydillo, Ana S. Gonzalez-Reiche, Sadaf Aslam, Adriana van de Guchte, Zenab Khan, Ajay Obla, Jayeeta Dutta, Harm van Bakel, Judith Aberg, Adolfo García-Sastre, Gunjan Shah, Tobias Hohl, Genovefa Papanicolaou, Miguel-Angel Perales, Kent Sepkowitz, Ngolela Esther Babady, and Mini Kamboj |
| EPI_ISL_583435 | Garcia-Sastre Laboratory, Department of Microbiology, Icahn School of Medicine at Mount Sinai | van Bakel Laboratory, Genetics and Genomics Sciences, Icahn School of Medicine at Mount Sinai | Teresa Aydillo, Ana S. Gonzalez-Reiche, Sadaf Aslam, Adriana van de Guchte, Zenab Khan, Ajay Obla, Jayeeta Dutta, Harm van Bakel, Judith Aberg, Adolfo García-Sastre, Gunjan Shah, Tobias Hohl, Genovefa Papanicolaou, Miguel-Angel Perales, Kent Sepkowitz, Ngolela Esther Babady, and Mini Kamboj |
| EPI_ISL_583475 | Memorial Sloan Kettering Cancer Center | van Bakel Laboratory, Genetics and Genomics Sciences, Icahn School of Medicine at Mount Sinai | Teresa Aydillo, Ana S. Gonzalez-Reiche, Sadaf Aslam, Adriana van de Guchte, Zenab Khan, Ajay Obla, Jayeeta Dutta, Harm van Bakel, Judith Aberg, Adolfo García-Sastre, Gunjan Shah, Tobias Hohl, Genovefa Papanicolaou, Miguel-Angel Perales, Kent Sepkowitz, Ngolela Esther Babady, and Mini Kamboj |
| EPI_ISL_583571 | Center for Virology, Medical University of Vienna | Bergthaler laboratory, CeMM Research Center for Molecular Medicine of the Austrian Academy of Sciences | Alexandra Popa, Benedikt Agerer, Henrique Colaco, Lukas Endler, Jakob-Wendelin Genger, Alexander Lercher, Mark Smyth, Thomas Penz, Michael Schuster, Jan Laine, Martin Senekowitsch, Judith Aberle, Stephan Aberle, Peter Hufnagl, Daniela Schmid, Franz Allerberger, Elisabeth Puchhammer-Stoeckl, Manfred Nairz, Guenter Weiss, Gregor Hörmann, Kinga Rigler-Hohenwarter, Rainer Gattringer, Wegene Borena, Dorothee von Laer, Gernot Walder, Peter Obrist, Christian Paar, Sabine Sussitz-Rack, Gunther Vogl, Adi Steinrigl, Christoph Bock, Andreas Bergthaler |
| EPI_ISL_583607, EPI_ISL_583608 | Institut für Virologie am Department für Hygiene, Mikrobiologie und Public Health | Bergthaler laboratory, CeMM Research Center for Molecular Medicine of the Austrian Academy of Sciences | Alexandra Popa, Benedikt Agerer, Henrique Colaco, Lukas Endler, Jakob-Wendelin Genger, Alexander Lercher, Mark Smyth, Thomas Penz, Michael Schuster, Jan Laine, Martin Senekowitsch, Judith Aberle, Stephan Aberle, Peter Hufnagl, Daniela Schmid, Franz Allerberger, Elisabeth Puchhammer-Stoeckl, Manfred Nairz, Guenter Weiss, Gregor Hörmann, Kinga Rigler-Hohenwarter, Rainer Gattringer, Wegene Borena, Dorothee von Laer, Gernot Walder, Peter Obrist, Christian Paar, Sabine Sussitz-Rack, Gunther Vogl, Adi Steinrigl, Christoph Bock, Andreas Bergthaler |
| EPI_ISL_583712 | Center for Virology, Medical University of Vienna | Bergthaler laboratory, CeMM Research Center for Molecular Medicine of the Austrian Academy of Sciences | Alexandra Popa, Benedikt Agerer, Henrique Colaco, Lukas Endler, Jakob-Wendelin Genger, Alexander Lercher, Mark Smyth, Thomas Penz, Michael Schuster, Jan Laine, Martin Senekowitsch, Judith Aberle, Stephan Aberle, Peter Hufnagl, Daniela Schmid, Franz Allerberger, Elisabeth Puchhammer-Stoeckl, Manfred Nairz, Guenter Weiss, Gregor Hörmann, Kinga Rigler-Hohenwarter, Rainer Gattringer, Wegene Borena, Dorothee von Laer, Gernot Walder, Peter Obrist, Christian Paar, Sabine Sussitz-Rack, Gunther Vogl, Adi Steinrigl, Christoph Bock, Andreas Bergthaler |

| | | | |
|---|---|---|---|
| EPI_ISL_583727, EPI_ISL_583728, EPI_ISL_583761, EPI_ISL_583762, EPI_ISL_583763, EPI_ISL_583823, EPI_ISL_583824, EPI_ISL_583825, EPI_ISL_583826, EPI_ISL_583827 | Dr. Gernot Walder GmbH | Bergthaler laboratory, CeMM Research Center for Molecular Medicine of the Austrian Academy of Sciences | Alexandra Popa, Benedikt Agerer, Henrique Colaco, Lukas Endler, Jakob-Wendelin Genger, Alexander Lercher, Mark Smyth, Thomas Penz, Michael Schuster, Jan Laine, Martin Senekowitsch, Judith Aberle, Stephan Aberle, Peter Hufnagl, Daniela Schmid, Franz Allerberger, Elisabeth Puchhammer-Stoeckl, Manfred Nairz, Guenter Weiss, Gregor Hörmann, Kinga Rigler-Hohenwarter, Rainer Gattringer, Wegene Borena, Dorothee von Laer, Gernot Walder, Peter Obrist, Christian Paar, Sabine Sussitz-Rack, Gunther Vogl, Adi Steinrigl, Christoph Bock, Andreas Bergthaler |
| EPI_ISL_583853, EPI_ISL_583854, EPI_ISL_583862 | Institute for Laboratory Diagnostics and Microbiology, Klinikum Klagenfurt am Worthersee | Bergthaler laboratory, CeMM Research Center for Molecular Medicine of the Austrian Academy of Sciences | Alexandra Popa, Benedikt Agerer, Henrique Colaco, Lukas Endler, Jakob-Wendelin Genger, Alexander Lercher, Mark Smyth, Thomas Penz, Michael Schuster, Jan Laine, Martin Senekowitsch, Judith Aberle, Stephan Aberle, Peter Hufnagl, Daniela Schmid, Franz Allerberger, Elisabeth Puchhammer-Stoeckl, Manfred Nairz, Guenter Weiss, Gregor Hörmann, Kinga Rigler-Hohenwarter, Rainer Gattringer, Wegene Borena, Dorothee von Laer, Gernot Walder, Peter Obrist, Christian Paar, Sabine Sussitz-Rack, Gunther Vogl, Adi Steinrigl, Christoph Bock, Andreas Bergthaler |
| EPI_ISL_584051 | Laboratory of Molecular Virology, Department of Biomedical, Surgical and Dental Sciences University of Milano | Laboratory of Molecular Virology, Department of Biomedical, Surgical and Dental Sciences University of Milano | Delbue,S., Ferrante,P., Basilico,N., Parapini,S., Binda,S., D'Alessandro,S., Galli,C., Signorini,L., Primache,V., Anselmi,G. and Pariani,E |
| EPI_ISL_584052 | Laboratory of Molecular Virology, Department of Biomedical, Surgical and Dental Sciences University of Milano | Laboratory of Molecular Virology, Department of Biomedical, Surgical and Dental Sciences University of Milano | Delbue,S., D'Alessandro,S., Modenese,A., Signorini,L., Parapini,S.,Dolci,M., Binda,S., Primache,V., Taramelli,D., Incorvaia,B. and Ferrante,P. |
| EPI_ISL_586302, EPI_ISL_586414, EPI_ISL_586415, EPI_ISL_586420 | Toronto Invasive Bacterial Diseases Network | McMaster University | Allison McGeer, Patryk Aftanas, Hooman Derakhshani, Angel Li, Kuganya Nirmalarajah, Emily Panousis, Ahmed Draia, Jalees Nasir, Michael Surette, Samira Mubareka, Andrew G. McArthur |
| EPI_ISL_593544, EPI_ISL_593545, EPI_ISL_593546 | Eastern Ontario Regional Laboratory Association | McMaster University | Leanne Mortimer, Hooman Derakhshani, Emily Panousis, Ahmed Draia, Jalees Nasir, Robert Slinger, Andrew G. McArthur |
| EPI_ISL_593614, EPI_ISL_593615, EPI_ISL_593634 | unknown | Public Health Virology Laboratory, Forensic and Scientific Services (PHV-FSS) | Son Nguyen et al. |
| EPI_ISL_593972, EPI_ISL_593976, EPI_ISL_593977, EPI_ISL_593978, EPI_ISL_593979 | Delaware Public Health Lab | Delaware Public Health Lab | Gregory Hovan |
| EPI_ISL_594086 | Utah Public Health Laboratory | Utah Public Health Laboratory | Erin Young, Kelly Oakeson |
| EPI_ISL_594205 | Michigan Department of Health and Human Services, Bureau of Laboratories | Michigan Department of Health and Human Services, Bureau of Laboratories | Blankenship HM, Riner D, Soehnlen MK |
| EPI_ISL_594394, EPI_ISL_594395, EPI_ISL_594396, EPI_ISL_594397, EPI_ISL_594398, EPI_ISL_594399, EPI_ISL_594400, EPI_ISL_594401, EPI_ISL_594402, EPI_ISL_594403, EPI_ISL_594404, EPI_ISL_594405, EPI_ISL_594406, EPI_ISL_594407, EPI_ISL_594408, EPI_ISL_594417 | | | |
| see above | Utah Public Health Laboratory | Utah Public Health Laboratory | Erin Young, Kelly Oakeson |
| EPI_ISL_596641, EPI_ISL_596643, EPI_ISL_596644, EPI_ISL_596645, EPI_ISL_596650, EPI_ISL_596654, EPI_ISL_596658, EPI_ISL_596659, EPI_ISL_596660, EPI_ISL_596661, EPI_ISL_596664, EPI_ISL_596669, EPI_ISL_596673, EPI_ISL_596674 | | | |
| see above | St.Vincent's University Hospital | St.Vincent's University Hospital | Mary Lucey, Guerrino Macori, Niamh Mullane, Una Sutton-Fitzpatrick, Gabriel Gonzalez, Suzie Coughlan, Aisling Purcell, Lynda Fenelon, Séamus Fanning, Kirsten Schaffer |
| EPI_ISL_596683, EPI_ISL_596692, EPI_ISL_596696, EPI_ISL_596725, EPI_ISL_596751, EPI_ISL_596792, EPI_ISL_596805, EPI_ISL_596808, EPI_ISL_596810, EPI_ISL_596814, EPI_ISL_596826, EPI_ISL_596832, EPI_ISL_596833, EPI_ISL_596837, EPI_ISL_596843, EPI_ISL_596844, EPI_ISL_596851, EPI_ISL_596866, EPI_ISL_596869 | | | |
| see above | PathWest Laboratory Medicine WA | PathWest Laboratory Medicine WA Microbial Surveillance Unit | PathWest Laboratory Medicine WA Microbial Surveillance Unit |
| EPI_ISL_600439, EPI_ISL_600441 | Institute of Epidemiology Disease Control And Research | Institute for Developing Science and Health Initiatives | Lauren Cowley, Mokibul Hassan Afrad, Sadia Isfat Ara Rahman, Md. Mahfuz-Al-mamun, Firadausi Qadri, Tahmina Shirin |
| EPI_ISL_602285, EPI_ISL_602288, EPI_ISL_602289, EPI_ISL_602291, EPI_ISL_602293, EPI_ISL_602300, EPI_ISL_602301 | Evangelisches Klinikum Bethel, Institut für Laboratoriumsmedizin, Mikrobiologie und Hygiene | Bielefeld University | David Brandt, Tobias Busche, Markus Haak, Jörn Kalinowski, Levin-Joe Klages, Christiane Scherer, Alexander Sczyrba, Marina Simunovic, Svenja Vinke |
| EPI_ISL_602468, EPI_ISL_602469, EPI_ISL_602470, EPI_ISL_602471, EPI_ISL_602472, EPI_ISL_602473, EPI_ISL_602474, EPI_ISL_602475, EPI_ISL_602476 | Institute for Virology, University Hospital Essen | Center of Medical Microbiology, Virology, and Hospital Hygiene, University of Duesseldorf | Olympia E. Anastasiou, Ulf Dittmer, Maximilian Damagnez, Alexander Dilthey, Torsten Houwaart, Lisanna Hülse, Malte Kohns Vasconcelos, Nadine Lübke, Jessica Nicolai, Klaus Pfeffer, Daniel Strelow, Jörg Timm, Andreas Walker, Tobias Wienemann |
| EPI_ISL_603082, EPI_ISL_603083, EPI_ISL_603084, EPI_ISL_603085, EPI_ISL_603086, EPI_ISL_603087, EPI_ISL_603088, EPI_ISL_603089, EPI_ISL_603090, EPI_ISL_603091, EPI_ISL_603093, EPI_ISL_603118 | | | |
| see above | Lithuanian University of Health Sciences Hospital, Department of Laboratory Medicine | Lithuanian University of Health Sciences, Molecular cardiology lab. | Lukas Zemaitis, Ingrida Olendrait, Arnoldas Pautienius, Kamile Tamusauskaite, Dovydas Gecys, Laura Pareckaite, Vaiva Lesauskaite, Astra Vitkauskiene |
| EPI_ISL_605781 | CEIRS Data Processing and Coordinating Center, St. Jude Center of Excellence for Influenza Research and Surveillance (CEIRS) | CEIRS Data Processing and Coordinating Center, St. Jude Center of Excellence for Influenza Research and Surveillance (CEIRS) | Roshdy,W.H., Kayed,A.E., Naguib,A., Kamel,M.N., El-Taweel,A., El-Shesheny,R., Kandeil,A., Mostafa,A., Shehata,M., Gomaa,M., Mahmoud,S.H., Moatasim,Y., Kutkat,O., Mahrous,N., El-Sayes,M., Showky,S., El-Guindy,N.M., Webby,R., Kayali,G., Ali,M.A. |
| EPI_ISL_605810, EPI_ISL_605814, EPI_ISL_605815 | Clinical Virology Laboratory, Institute of Liver and Biliary Sciences | ILBS - IGIB | Ekta Gupta, Sheetalnath Rooge, Abhishek Padhi, Reshu Agarwal, Jaswinder Singh Maras, Shridhar Sivasubbu, Vinod Scaria, Shvetank Sharma |
| EPI_ISL_605827, EPI_ISL_605849, EPI_ISL_605852, EPI_ISL_605858, EPI_ISL_605870 | PathWest Laboratory Medicine WA | PathWest Laboratory Medicine WA Microbial Surveillance Unit | PathWest Laboratory Medicine WA Microbial Surveillance Unit |
| EPI_ISL_605928 | Emergence des Maladies Virales, Centre Interdisciplinaire de Recherches Medicales de Franceville | Emergence des Maladies Virales, Centre Interdisciplinaire de Recherches Medicales de Franceville | Lekana-Douki,S.E., N'dilimabaka,N., Mbongo kama,E., Kandet Yattara,M., Mintsa Ndong,A., Ngonga Dikongo,A.M., Andeko,J.C., Zong Minko,O., Lekana-Douki,J.-B. |
| EPI_ISL_611569, EPI_ISL_611621 | Liverpool Clinical Laboratories | COVID-19 Genomics UK (COG-UK) Consortium | Sam Haldenby, Anita Lucaci, Steve Paterson, Julian Hiscox, Alistair Darby, M Almsaud, A Alrezaihi, Muhannad Alruwaili, Stuart D Armstrong, Jones Benjamin, Eleanor G Bentley, Anu Chawla, Jordan J Clark, Angela Cowell, Richard Eccles, Isabel García-Dorival, Matthew Gemmell, Alessandro Gerada, PKF Gilmore, Richard Gregory, Ximeng Han, Catherine Hartley, Margaret Hughes, Miren Iturriza-Gomara, James Johnson, L Luu, Jenifer Manson, Charlotte Nelson, Elaine O'Toole, Cassie Olateju, Rebekah Penrice-Randal , Lucille Rainbow, N.P Randle, Trevor Ian Robinson, Parul Sharma, Ghada T Shawli, James P Stewart, Neil Swainston, Ecaterina Vamos, Joanne Watts, Mark Whitehead |
| EPI_ISL_611925 | Centre for Enzyme Innovation, University of Portsmouth / Translational Research Laboratory, Portsmouth Hospitals NHS Trust | COVID-19 Genomics UK (COG-UK) Consortium | Angela Beckett,Yann Bourgeois,Garry Scarlett,Sharon Glaysher,Scott Elliott,Kelly Bicknell,Robert Impey,Allyson Lloyd,Sarah Wyllie,Ethan Butcher,Anoop Chauhan,Samuel Robson |
| EPI_ISL_612432 | Liverpool Clinical Laboratories | COVID-19 Genomics UK (COG-UK) Consortium | Sam Haldenby, Anita Lucaci, Steve Paterson, Julian Hiscox, Alistair Darby, M Almsaud, A Alrezaihi, Muhannad Alruwaili, Stuart D Armstrong, Jones Benjamin, Eleanor G Bentley, Anu Chawla, Jordan J Clark, Angela Cowell, Richard Eccles, Isabel García-Dorival, Matthew Gemmell, Alessandro Gerada, PKF Gilmore, Richard Gregory, Ximeng Han, Catherine Hartley, Margaret Hughes, Miren Iturriza-Gomara, James Johnson, L Luu, Jenifer Manson, Charlotte Nelson, Elaine O'Toole, Cassie Olateju, Rebekah Penrice-Randal , Lucille Rainbow, N.P Randle, Trevor Ian Robinson, Parul Sharma, Ghada T Shawli, James P Stewart, Neil Swainston, Ecaterina Vamos, Joanne Watts, Mark Whitehead |
| EPI_ISL_613453, EPI_ISL_613454, EPI_ISL_613455 | Institut Pasteur de la Guadeloupe | Institut Pasteur de la Guadeloupe | Marion Barbet, Sylvie Behillil, Méline Bizard, Angela Brisebarre, Camille Capel, Etienne Simon-Lorière, Vincent Enouf, Maud Vanpeene, Sylvie van der Werf, Stéphanie Guyomard, Sébastien Breurec, Radjin Steingrover, Cherina Fleming, Antoine Talarmin |
| EPI_ISL_613560 | Laboratorio Biologia Molecolare Sars Cov2 - UOC Laboratorio Analisi - Servizio Medicina di Laboratorio, Ospedale "San | Laboratorio specialistico UOC Ematologia - Ospedale "San Francesco" - ATS-ASSL Nuoro Nuoro | Piras Giovanna, Fancello Tatiana, Asproni Rosanna, Fiamma Maura, Monne Maria Itria, Toja Alessandro, Sanna Filomena, Floris Anna Rita, Sulis Vincenzo, Palmas Angelo Domenico, Casu Gavino, Lo Maglio Iana, Mameli Giuseppe |

| | | | |
|---|---|---|---|
| | Francesco" - ATS-ASSL Nuoro Via Mannironi 1, 08100 Nuoro | | |
| EPI_ISL_613564, EPI_ISL_613708 | Laboratory of Molecular Biology, Blood Center of Ribeirão Preto | Laboratory of Molecular Biology, Blood Center of Ribeirão Preto, Faculty of Medicine of Ribeirão Preto, University of São Paulo | Svetoslav N Slavov, Marta Giovanetti, Vagner Fonseca, Elaine V Santos, Evandra S Rodrigues, Talita Adelino, Joilson Xavier, Glauco de Carvalho Pereira, Aparecida Y Yamamoto, Diego Villa Clé, Rodrigo T Calado; Dimas T Covas, Luiz CJ Alcantara, Simone Kashima |
| EPI_ISL_613717, EPI_ISL_613719, EPI_ISL_613776, EPI_ISL_613779, EPI_ISL_613784, EPI_ISL_613815, EPI_ISL_613818, EPI_ISL_613819, EPI_ISL_613821, EPI_ISL_613916, EPI_ISL_613932, EPI_ISL_613934 | | | |
| see above | Florida Bureau of Public Health Laboratories | Florida Bureau of Public Health Laboratories | Sarah Schmedes, Jason Blanton |
| EPI_ISL_613952, EPI_ISL_613963, EPI_ISL_614155 | Laboratory of Molecular Biology, Blood Center of Ribeirão Preto, Faculty of Medicine of Ribeirão Preto, University of São Paulo | Laboratory of Molecular Biology, Blood Center of Ribeirão Preto, Faculty of Medicine of Ribeirão Preto, University of São Paulo | Svetoslav N Slavov, Marta Giovanetti, Vagner Fonseca, Elaine V Santos, Evandra S Rodrigues, Talita Adelino, Joilson Xavier, Glauco de Carvalho Pereira, Aparecida Y Yamamoto, Diego Villa Clé, Rodrigo T Calado; Dimas T Covas, Luiz CJ Alcantara, Simone Kashima |
| EPI_ISL_614398 | Laboratorio Biologia Molecolare Sars Cov2 - UOC Laboratorio Analisi - Servizio Medicina di Laboratorio, Ospedale "San Francesco" - ATS-ASSL Nuoro | Laboratorio specialistico UOC Ematologia - Ospedale "San Francesco" - ATS-ASSL Nuoro | Piras Giovanna, Fancello Tatiana, Asproni Rosanna, Fiamma Maura, Monne Maria Itria, Toja Alessandro, Sanna Filomena, Floris Anna Rita, Sulis Vincenzo, Palmas Angelo Domenico, Casu Gavino, Lo Maglio Iana, Mameli Giuseppe |
| EPI_ISL_614480, EPI_ISL_614481, EPI_ISL_614482, EPI_ISL_614483, EPI_ISL_614543, EPI_ISL_614693, EPI_ISL_614694, EPI_ISL_616371, EPI_ISL_618643, EPI_ISL_618644, EPI_ISL_618645, EPI_ISL_618646, EPI_ISL_618805, EPI_ISL_618962, EPI_ISL_618965, EPI_ISL_618969, EPI_ISL_618972, EPI_ISL_618978, EPI_ISL_618979, EPI_ISL_618980, EPI_ISL_619063, EPI_ISL_619133, EPI_ISL_619141, EPI_ISL_619145, EPI_ISL_619151, EPI_ISL_619163, EPI_ISL_619170, EPI_ISL_619176, EPI_ISL_619185, EPI_ISL_619211, EPI_ISL_619226, EPI_ISL_619233, EPI_ISL_619256, EPI_ISL_619260, EPI_ISL_619261, EPI_ISL_619262, EPI_ISL_619263, EPI_ISL_619264, EPI_ISL_619265, EPI_ISL_619266, EPI_ISL_619267, EPI_ISL_619268, EPI_ISL_619269, EPI_ISL_619270, EPI_ISL_619271, EPI_ISL_619292, EPI_ISL_619318, EPI_ISL_619885, EPI_ISL_619895, EPI_ISL_619905, EPI_ISL_622454, EPI_ISL_622455, EPI_ISL_622456, EPI_ISL_622457, EPI_ISL_622458, EPI_ISL_622580 | | | |
| see above | Department of Virus and Microbiological Special Diagnostics, Statens Serum Institut, Denmark | Albertsen lab, Department of Chemistry and Bioscience, Aalborg University, Denmark | Danish Covid-19 Genome Consortia |
| EPI_ISL_623170, EPI_ISL_623171, EPI_ISL_623172, EPI_ISL_623173, EPI_ISL_623174, EPI_ISL_623175, EPI_ISL_623176, EPI_ISL_623177, EPI_ISL_623178, EPI_ISL_623180, EPI_ISL_623183, EPI_ISL_623184, EPI_ISL_623185, EPI_ISL_623187, EPI_ISL_623194, EPI_ISL_623197, EPI_ISL_623202 | | | |
| see above | Utah Public Health Laboratory | Utah Public Health Laboratory | Erin Young, Kelly Oakeson |
| EPI_ISL_626230 | Institute for Virology, University Hospital Essen | Center of Medical Microbiology, Virology, and Hospital Hygiene, University of Duesseldorf | Olympia E. Anastasiou, Ulf Dittmer, Maximilian Damagnez, Alexander Dilthey, Torsten Houwaart, Lisanna Hülse, Malte Kohns Vasconcelos, Nadine Lübke, Jessica Nicolai, Klaus Pfeffer, Daniel Strelow, Jörg Timm, Andreas Walker, Tobias Wienemann |
| EPI_ISL_626570, EPI_ISL_626593 | The National Institute of Public Health | State Veterinary Institute Prague | Nagy,A;Jirincova,H;Novakova,L;Trnka,D;Vecerova,J |
| EPI_ISL_629097 | Laboratoire du Centre Hospitalier Annecy Genevois | CNR Virus des Infections Respiratoires - France SUD | Antonin Bal, Gregory Destras, Gwendolyne Burfin, Hadrien Règue, Quentin Semanas, Martine Valette, Bruno Lina, Hélène Petitprez, Bruno Chanzy, Laurence Josset |
| EPI_ISL_631391, EPI_ISL_631392 | Wisconsin State Laboratory of Hygiene Communicable Disease Division | Wisconsin State Laboratory of Hygiene Communicable Disease Division | Kelsey R. Florek, Abigail C. Shockey |
| EPI_ISL_631618, EPI_ISL_631619 | OCME Office Of Chief Medical Examiner | New York City Public Health Laboratory | Jade Wang, et al. |
| EPI_ISL_631620 | Flushing Hospital Medical Center | New York City Public Health Laboratory | Jade Wang, et al. |
| EPI_ISL_631621 | Jamaica Hospital Medical Center | New York City Public Health Laboratory | Jade Wang, et al. |
| EPI_ISL_631622, EPI_ISL_631623, EPI_ISL_631624, EPI_ISL_631625, EPI_ISL_631626, EPI_ISL_631627, EPI_ISL_631628, EPI_ISL_631629, EPI_ISL_631630, EPI_ISL_631631, EPI_ISL_631632, EPI_ISL_631633 | | | |
| see above | Richmond University Medical Center | New York City Public Health Laboratory | Jade Wang, et al. |
| EPI_ISL_631634 | Jamaica Hospital Medical Center | New York City Public Health Laboratory | Jade Wang, et al. |
| EPI_ISL_631635 | OCME Office Of Chief Medical Examiner | New York City Public Health Laboratory | Jade Wang, et al. |
| EPI_ISL_631636, EPI_ISL_631637 | Jamaica Hospital Medical Center | New York City Public Health Laboratory | Jade Wang, et al. |
| EPI_ISL_631638 | Flushing Hospital Medical Center | New York City Public Health Laboratory | Jade Wang, et al. |
| EPI_ISL_631639, EPI_ISL_631685 | OCME Office Of Chief Medical Examiner | New York City Public Health Laboratory | Jade Wang, et al. |
| EPI_ISL_631686 | NYC HH Lincoln Medical And Mental Health Center | New York City Public Health Laboratory | Jade Wang, et al. |
| EPI_ISL_631687 | OCME Office Of Chief Medical Examiner | New York City Public Health Laboratory | Jade Wang, et al. |
| EPI_ISL_631688 | Wildlife Conservation Society - Bronx Zoo | New York City Public Health Laboratory | Jade Wang, et al. |
| EPI_ISL_631689, EPI_ISL_631690 | Lincoln Medical Center | New York City Public Health Laboratory | Jade Wang, et al. |
| EPI_ISL_631756 | Flushing Hospital Medical Center | New York City Public Health Laboratory | Jade Wang, et al. |
| EPI_ISL_631763 | OCME Office Of Chief Medical Examiner | New York City Public Health Laboratory | Jade Wang, et al. |
| EPI_ISL_631823, EPI_ISL_631825 | Jamaica Hospital Medical Center | New York City Public Health Laboratory | Jade Wang, et al. |
| EPI_ISL_631826 | Flushing Hospital Medical Center | New York City Public Health Laboratory | Jade Wang, et al. |
| EPI_ISL_631876, EPI_ISL_631877, EPI_ISL_631882, EPI_ISL_631921, EPI_ISL_631922 | Jamaica Hospital Medical Center | New York City Public Health Laboratory | Jade Wang, et al. |
| EPI_ISL_631923 | Flushing Hospital Medical Center | New York City Public Health Laboratory | Jade Wang, et al. |
| EPI_ISL_631924 | St Barnabas Hospital | New York City Public Health Laboratory | Jade Wang, et al. |
| EPI_ISL_631926, EPI_ISL_631927, EPI_ISL_631928 | Jamaica Hospital Medical Center | New York City Public Health Laboratory | Jade Wang, et al. |
| EPI_ISL_631929 | Flushing Hospital Medical Center | New York City Public Health Laboratory | Jade Wang, et al. |
| EPI_ISL_631930 | Jamaica Hospital Medical Center | New York City Public Health Laboratory | Jade Wang, et al. |
| EPI_ISL_631931, EPI_ISL_631955 | Flushing Hospital Medical Center | New York City Public Health Laboratory | Jade Wang, et al. |
| EPI_ISL_631958, EPI_ISL_631987, EPI_ISL_631988 | St Barnabas Hospital | New York City Public Health Laboratory | Jade Wang, et al. |
| EPI_ISL_631989 | Flushing Hospital Medical Center | New York City Public Health Laboratory | Jade Wang, et al. |
| EPI_ISL_631990 | St Barnabas Hospital | New York City Public Health Laboratory | Jade Wang, et al. |
| EPI_ISL_631991, EPI_ISL_632034 | Flushing Hospital Medical Center | New York City Public Health Laboratory | Jade Wang, et al. |
| EPI_ISL_632035 | Jamaica Hospital Medical Center | New York City Public Health Laboratory | Jade Wang, et al. |
| EPI_ISL_632036 | Richmond University Medical Center | New York City Public Health Laboratory | Jade Wang, et al. |
| EPI_ISL_632037 | Jamaica Hospital Medical Center | New York City Public Health Laboratory | Jade Wang, et al. |
| EPI_ISL_632183 | OCME Office Of Chief Medical Examiner | New York City Public Health Laboratory | Jade Wang, et al. |

| | | | |
|---|---|---|---|
| EPI_ISL_632195 | Jamaica Hospital Medical Center | New York City Public Health Laboratory | Jade Wang, et al. |
| EPI_ISL_632862 | Idaho Bureau of Laboratories | Center for Global Health, University of New Mexico Health Sciences Center | Daryl Domman, Kurt Schwalm, Matthew Burns, Robert Voermans, Christopher Ball, Darrell Dinwiddie |
| EPI_ISL_632913, EPI_ISL_632917 | Cadham Provincial laboratory | Cadham Provincial laboratory | Anna Majer, Shari Tyson, Grace Seo, Philip Mabon, Elsie Grudeski, Rhiannon Huzarewich, Russell Mandes, Anneliese Landgraff, Jennifer Tanner, Natalie Knox, Morag Graham, Gary Van Domselaar, Paul Van Caeseele, Jared Bullard, David Alexander, Kerry Dust, Nathalie Bastien, Yan Li, Timothy Booth, Darian Hole, Madison Chapel, CanCOGeN's metadata curation team, Public Health Agency of Canada CanCOGeN team |
| EPI_ISL_634831 | Laboratoire de virologie, CHU de Grenoble - CS 10217 - 38043 Grenoble cedex 23 | CNR Virus des Infections Respiratoires - France SUD | Antonin Bal, Gregory Destras, Gwendolyne Burfin, Hadrien Règue, Quentin Semanas, Martine Valette, Bruno Lina, Sylvie Larrat, Laurence Josset |
| EPI_ISL_634832 | Laboratoire de virologie, CHU de Grenoble - CS 10217 - 38043 Grenoble cedex 24 | CNR Virus des Infections Respiratoires - France SUD | Antonin Bal, Gregory Destras, Gwendolyne Burfin, Hadrien Règue, Quentin Semanas, Martine Valette, Bruno Lina, Sylvie Larrat, Laurence Josset |
| EPI_ISL_635285, EPI_ISL_635286, EPI_ISL_635289, EPI_ISL_635292, EPI_ISL_636461 | Institute of Microbiology and Immunology, Faculty of Medicine, University of Ljubljana | Institute of Microbiology and Immunology, Faculty of Medicine, University of Ljubljana | Tomaž Mark Zorec, Samo Zakotnik, Miša Korva, Tatjana Avši - Županc, Mario Poljak |
| EPI_ISL_636473 | ULSS6 Piove di Sacco | Istituto Zooprofilattico Sperimentale delle Venezie | Adelaide Milani, Alessia Schivo, Annalisa Salviato, Erika Giorgia Quaranta, Ambra Pastori, Bianca Zecchin, Alice Fusaro, Isabella Monne, Calogero Terregino, Antonia Ricci |
| EPI_ISL_636564 | Dutch COVID-19 response team | National Institute for Public Health and the Environment (RIVM) | Adam Meijer, Harry Vennema, Jeroen Cremer, Sharon van den Brink, Bas van der Veer, AnneMarie van den Brandt, Florian Zwagemaker, Dennis Schmitz, Chantal Reusken, on behalf of the national COVID-19 response team |
| EPI_ISL_636991 | Department of Infectious Diseases and Immunology, National Hospital Organization Nagoya Medical Center | Clinical Research Center, National Hospital Organization Nagoya Medical Center | Yoshihiro Nakata, Hirotaka Ode, Mai Kubota, Masakazu Matsuda, Kazuhiro Matsuoka, Miho Nakasuji, Mikiko Mori, Mayumi Imahashi, Yoshiyuki Yokomaku, Yasumasa Iwatani |
| EPI_ISL_640059 | Du Noon CDC wc DNC | NHLS/UCT | Arash Iranzadeh, Deelan Doolabh, Lynn Tyers, Bruna Galvao, Innocent Mudau, Marvin Hsiao, Kruger Marais, Diana Hardie, Stephen Korsman, Carolyn Williamson |
| EPI_ISL_641528 | CHU de Nice - Hôpital Archet 10 | CNR Virus des Infections Respiratoires - France SUD | Antonin Bal, Géraldine Gonfrier, Gregory Destras, Gwendolyne Burfin, Hadrien Règue, Quentin Semanas, Martine Valette, Bruno Lina, Valérie Giordanengo, Laurence Josset |
| EPI_ISL_641529 | CHU de Nice - Hôpital Archet 11 | CNR Virus des Infections Respiratoires - France SUD | Antonin Bal, Géraldine Gonfrier, Gregory Destras, Gwendolyne Burfin, Hadrien Règue, Quentin Semanas, Martine Valette, Bruno Lina, Valérie Giordanengo, Laurence Josset |
| EPI_ISL_644217, EPI_ISL_644223, EPI_ISL_644278, EPI_ISL_644282, EPI_ISL_644283, EPI_ISL_644285, EPI_ISL_644320, EPI_ISL_644326, EPI_ISL_644340 | CEPHR / Vincent's Hospital | Irish Coronavirus Sequencing Consortium - National Virus Reference Laboratory | Michael Carr, Gabriel Gonzalez, Alejandro Abner Garcia Leon, Patrick Mallon |
| EPI_ISL_644391 | MEPHI, Aix Marseille University | MEPHI, Aix Marseille University | Anthony LEVASSEUR |
| EPI_ISL_644582, EPI_ISL_644583 | Veterinary Specialized Institute "Kraljevo", Serbia | Veterinary Specialized Institute "Kraljevo", Serbia | Vidanovic,D., Tesovic,B., Knezevic,A., Jovanovic,T., Jankovic,M., Sekler,M., Banovic Djeri,B., Petrovic,T., Volkening,J., Afonso,C. |
| EPI_ISL_644676, EPI_ISL_644677 | CHU de Limoges | CNR Virus des Infections Respiratoires - France SUD | Antonin Bal, Gregory Destras, Gwendolyne Burfin, Hadrien Règue, Quentin Semanas, Martine Valette, Bruno Lina, Sylvie Rogez, Laurence Josset |
| EPI_ISL_644713, EPI_ISL_644714 | Unité des Virus Émergents | CNR Virus des Infections Respiratoires - France SUD | Antonin Bal, Gregory Destras, Gwendolyne Burfin, Hadrien Règue, Quentin Semanas, Martine Valette, Bruno Lina, Laetitia Ninove, Léa Luciani, Antoine Nougairède, Laurence Josset |
| EPI_ISL_645195, EPI_ISL_645196 | CHU Clermont-Ferrand | CNR Virus des Infections Respiratoires - France SUD | Antonin Bal, Gregory Destras, Gwendolyne Burfin, Hadrien Règue, Quentin Semanas, Martine Valette, Bruno Lina, Christine Archimbaud, Amélie Brebion, Hélène Chabrolles, Martine Chambon, Audrey Mirand, Christel Regagnon, Maxime Bisseux, Patricia Combes, Cécile Henquell, Laurence Josset |
| EPI_ISL_645201, EPI_ISL_645202, EPI_ISL_645203, EPI_ISL_645204, EPI_ISL_645205, EPI_ISL_645206 | CHU Nîmes | CNR Virus des Infections Respiratoires - France SUD | Antonin Bal, Gregory Destras, Gwendolyne Burfin, Hadrien Règue, Quentin Semanas, Martine Valette, Bruno Lina, Jean-Philippe Lavigne, Stephan Robin, Maxence Lotellier, Marie-Josée Carles, Laurence Josset |
| EPI_ISL_648035 | WI State Laboratory of Hygiene | Pathogen Discovery, Respiratory Viruses Branch, Division of Viral Diseases, Centers for Disease Control and Prevention | Yan Li, Jing Zhang, Ying Tao, Brian Lynch, Krista Queen, Anna Montmayeur, Anna Uehara, Clinton R. Paden, Rachel Marine, Haibin Wang, Suxiang Tong |
| EPI_ISL_648086, EPI_ISL_648087, EPI_ISL_648088, EPI_ISL_648089, EPI_ISL_648090, EPI_ISL_648091, EPI_ISL_648092 | Department of Laboratory Medicine, Tan Tock Seng Hospital | Department of Laboratory Medicine, Tan Tock Seng Hospital | Chen YYC, Zair X, Lim JX, Li C, Tang WY, Maurer-Stroh S, Barkham TMS, Nagarajan N, Sessions OM |
| EPI_ISL_648557, EPI_ISL_648564, EPI_ISL_648565, EPI_ISL_648566, EPI_ISL_648567, EPI_ISL_648568, EPI_ISL_648570, EPI_ISL_648571, EPI_ISL_648572, EPI_ISL_648573, EPI_ISL_648574, EPI_ISL_648576, EPI_ISL_648577 | | | |
| see above | Utah Public Health Laboratory | Utah Public Health Laboratory | Erin Young, Kelly Oakeson |
| EPI_ISL_649180, EPI_ISL_649181, EPI_ISL_649182 | CHU Nantes | CNR Virus des Infections Respiratoires - France SUD | Antonin Bal, Louise Castain, Gregory Destras, Gwendolyne Burfin, Hadrien Règue, Quentin Semanas, Martine Valette, Bruno Lina, Celine Bressollette, Laurence Josset |
| EPI_ISL_654676, EPI_ISL_654677, EPI_ISL_654684 | Servicio de Microbiología, Hospital Universitario Central de Asturias | SeqCOVID-SPAIN consortium/IBV(CSIC) | Cristián Castelló Abietar, Jose A. Boga, Susana Rojo-Alba, Marta Elena Álvarez-Argüelles, Santiago Melón and SeqCOVID-SPAIN consortium |
| EPI_ISL_654911, EPI_ISL_654912, EPI_ISL_654914, EPI_ISL_654915, EPI_ISL_654916, EPI_ISL_654917, EPI_ISL_654918, EPI_ISL_654919, EPI_ISL_654920, EPI_ISL_654921, EPI_ISL_654922, EPI_ISL_654924, EPI_ISL_654925 | | | |
| see above | Texas Department of State Health Services | Texas Department of State Health Services | Rashmi Tuladhar, Bonnie Oh, Jenny Zhang, Maliha Rahman, Anita Pokharel, Myong Koag, Chung Wang, Rachel Lee, Grace Kubin, Mayela Pedrueza, James Daniel Bonser |
| EPI_ISL_660361 | CHU Clermont-Ferrand | CNR Virus des Infections Respiratoires - France SUD | Antonin Bal, Gregory Destras, Gwendolyne Burfin, Hadrien Règue, Quentin Semanas, Martine Valette, Bruno Lina, Christine Archimbaud, Amélie Brebion, Hélène Chabrolles, Martine Chambon, Audrey Mirand, Christel Regagnon, Maxime Bisseux, Patricia Combes, Cécile Henquell, Laurence Josset |
| EPI_ISL_660380, EPI_ISL_660381, EPI_ISL_660382 | Orebro klinisk mikrobiologi | The Public Health Agency of Sweden | Anna-Malin Linde, Maria Lind Karlberg, Mattias Haukland, Reza Advani, Olov Svartstrom, Oskar Karlsson Lindsjo, Sandra Broddesson, Petra Edquist, Mia Brytting, Anna Risberg, Karin Tegmark-Wisell |
| EPI_ISL_660400, EPI_ISL_660401, EPI_ISL_660402, EPI_ISL_660403, EPI_ISL_660404, EPI_ISL_660405, EPI_ISL_660406 | Klinisk mikrobiologi Linkoping | The Public Health Agency of Sweden | Anna-Malin Linde, Maria Lind Karlberg, Mattias Haukland, Reza Advani, Olov Svartstrom, Oskar Karlsson Lindsjo, Sandra Broddesson, Petra Edquist, Mia Brytting, Anna Risberg, Karin Tegmark-Wisell |
| EPI_ISL_660693, EPI_ISL_660694, EPI_ISL_660695 | CHU Montpellier | CNR Virus des Infections Respiratoires - France SUD | Antonin Bal, Gregory Destras, Gwendolyne Burfin, Hadrien Règue, Quentin Semanas, Martine Valette, Bruno Lina, Michel Segondy, Vincent Foulongne, Laurence Josset |
| EPI_ISL_660719, EPI_ISL_660720, EPI_ISL_660721, EPI_ISL_660722, EPI_ISL_660723, EPI_ISL_660724, EPI_ISL_660725 | CHU Nîmes | CNR Virus des Infections Respiratoires - France SUD | Antonin Bal, Gregory Destras, Gwendolyne Burfin, Hadrien Règue, Quentin Semanas, Martine Valette, Bruno Lina, Jean-Philippe Lavigne, Stephan Robin, Maxence Lotellier, Marie-Josée Carles, Laurence Josset |
| EPI_ISL_661279 | Klinisk mikrobiologi Linkoping | The Public Health Agency of Sweden | Department of Microbiology, The Public Health Agency of Sweden |
| EPI_ISL_663271, EPI_ISL_663272, EPI_ISL_663273, EPI_ISL_663274, EPI_ISL_663275, EPI_ISL_663276, EPI_ISL_663277, EPI_ISL_663278, EPI_ISL_663279, EPI_ISL_663280, EPI_ISL_663281 | | | |

| | | | |
|---|---|---|---|
| see above | CHU Nantes | CNR Virus des Infections Respiratoires - France SUD | Antonin Bal, Louise Castain, Gregory Destras, Gwendolyne Burfin, Hadrien Règue, Quentin Semanas, Martine Valette, Bruno Lina, Virginie Ferré, Celine Bressollette, Laurence Josset |
| EPI_ISL_664103 | Dept. of Microbiology and Infection Control, Akershus University Hospital HF | Dept. of Microbiology and Infection Control, Akershus University Hospital HF | Hege Vangstein Aamot, Alexander Hesselberg Løvestad, Silje Bakken Jørgensen, Nina Handal, Ole Herman Ambur |
| EPI_ISL_664447, EPI_ISL_664493, EPI_ISL_664499, EPI_ISL_664694, EPI_ISL_664712, EPI_ISL_665145 | University College London Hospital | COVID-19 Genomics UK (COG-UK) Consortium | Judith Heaney, Matthew Byott, Catherine Houlihan, Dan Frampton, Stuart Kirk, Moira Spyer and Eleni Nastouli |
| EPI_ISL_666604, EPI_ISL_666606, EPI_ISL_666607 | Dept. of Microbiology and Infection Control, Akershus University Hospital HF | Dept. of Microbiology and Infection Control, Akershus University Hospital HF | Hege Vangstein Aamot, Alexander Hesselberg Løvestad, Silje Bakken Jørgensen, Nina Handal, Ole Herman Ambur |
| EPI_ISL_666666, EPI_ISL_666667, EPI_ISL_666668 | Laboratoire du Centre Hospitalier Annecy Genevois | CNR Virus des Infections Respiratoires - France SUD | Antonin Bal, Gregory Destras, Gwendolyne Burfin, Hadrien Règue, Quentin Semanas, Martine Valette, Bruno Lina, Hélène Petitprez, Bruno Chanzy, Laurence Josset |
| EPI_ISL_666688, EPI_ISL_666689, EPI_ISL_666690, EPI_ISL_666691, EPI_ISL_666692, EPI_ISL_666693, EPI_ISL_666695, EPI_ISL_666696, EPI_ISL_666697, EPI_ISL_666698, EPI_ISL_666699, EPI_ISL_666700, EPI_ISL_666701, EPI_ISL_666702, EPI_ISL_666703, EPI_ISL_666704, EPI_ISL_666705, EPI_ISL_666706, EPI_ISL_666707, EPI_ISL_666708, EPI_ISL_666709, EPI_ISL_666710 | | | |
| see above | National Virus Reference Lab (NVRL) | Irish Coronavirus Sequencing Consortium-Teagasc Grange | Matthew McCabe, Calum Walsh, Fiona Crispie, Paul Cotter, Michael Carr, Aljandro Abner Garcia Leon |
| EPI_ISL_667080 | OHSU Lab Services Molecular Microbiology Lab | Oregon SARS-CoV-2 Genome Sequencing Center | Brendan L. O'Connell, Ruth V. Nichols, Sally Grindstaff, Alec J. Hirsch, Donna Hansel, Guang Fan, Daniel N. Streblow, William B. Messer, Andrew C. Adey, Benjamin N. Bimber, Brian J. O'Roak |
| EPI_ISL_672114, EPI_ISL_672115, EPI_ISL_672123, EPI_ISL_672124, EPI_ISL_672125 | The Ashley Laboratory, Stanford University | Chan-Zuckerberg Biohub | CZB Cliahub Consortium |
| EPI_ISL_672531, EPI_ISL_672533, EPI_ISL_672534, EPI_ISL_672535, EPI_ISL_672536, EPI_ISL_672537, EPI_ISL_672539, EPI_ISL_672543, EPI_ISL_672546, EPI_ISL_672547, EPI_ISL_672548, EPI_ISL_672549, EPI_ISL_672560, EPI_ISL_672562, EPI_ISL_672563 | | | |
| see above | Utah Public Health Laboratory | Utah Public Health Laboratory | Erin Young, Kelly Oakeson |
| EPI_ISL_672665, EPI_ISL_672666, EPI_ISL_672667, EPI_ISL_672668, EPI_ISL_672683, EPI_ISL_672685 | DB Diagnosticos do Brasil | Laboratório de Parasitologia Médica - Instituto de Medicina Tropical - Universidade de São Paulo | Brazil-UK Centre for Arbovirus Discovery Diagnosis Genomics and Epidemiology (CADDE) Genomic Network - Instituto de Medicina Tropical |
| EPI_ISL_676574, EPI_ISL_676577, EPI_ISL_676580 | Scientific Veterinary Institute Novi Sad | Veterinary Specialized Institute "Kraljevo", Serbia | Vidanovic,D., Tesovic,B., Knezevic,A., Jovanovic,T., Jankovic,M., Sekler,M., Banovic Djeri,B., Petrovic,T., Volkening,J., Afonso,C. |
| EPI_ISL_676681, EPI_ISL_676790, EPI_ISL_676797, EPI_ISL_676798, EPI_ISL_676799, EPI_ISL_676800, EPI_ISL_676801, EPI_ISL_676802, EPI_ISL_676803, EPI_ISL_676804, EPI_ISL_676805, EPI_ISL_676806, EPI_ISL_676807, EPI_ISL_676808, EPI_ISL_676818, EPI_ISL_676819, EPI_ISL_676962 | | | |
| see above | Wadsworth Center, New York State Department.of Health | Wadsworth Center, New York State Department.of Health | Kirsten St. George, Daryl M. Lamson, Alexis Russel, Jonathan Plitnick, Navjot Singh, John Kelly, Sara Griesemer, Erasmus Schneider, Erica Lasek-Nesselquist |
| EPI_ISL_677943 | Pathogen Genomics Lab King Abdullah University of Science and Technology(KAUST) | Pathogen Genomics Lab King Abdullah University of Science and Technology(KAUST) | Sara Mfarrej, Luke Esau, Amanda Ooi, Sharif Hala, Raeece Naeem, Afrah Alsomali, Fadwa Alofi, Asim Khogeer, Jumana Taha, Abdulaziz Alahmadi, Kahled Algithami, Anwar Hashem, Naif Almontashiri, Arnab Pain |
| EPI_ISL_678027 | Pathogen Genomics Lab King Abdullah University of Science and Technology(KAUST) | Pathogen Genomics Lab King Abdullah University of Science and Technology(KAUST) | Fathia Ben Rached, Sharif Hala, Amit Kumar Subudhi, Sara Mfarrej, Raeece Naeem, Rahul P Salunke, Fadwa Alofi, Afrah Alsomali, Asim Khogeer, Jumana Taha, Abdulaziz Alahmadi, Kahled Algithami, Anwar Hashem, Naif Almontashiri, Arnab Pain |
| EPI_ISL_678039 | Pathogen Genomics Lab King Abdullah University of Science and Technology(KAUST) | Pathogen Genomics Lab King Abdullah University of Science and Technology(KAUST) | Sharif Hala, Sara Mfarrej, Amit Kumar Subudhi, Raeece Naeem, Rahul P Salunke, Asim Khogeer, Fadwa Alofi, Afrah Alsomali, Jumana Taha, Abdulaziz Alahmadi, Kahled Algithami, Anwar Hashem, Naif Almontashiri, Arnab Pain |
| EPI_ISL_678040 | Pathogen Genomics Lab King Abdullah University of Science and Technology(KAUST) | Pathogen Genomics Lab King Abdullah University of Science and Technology(KAUST) | Amit Kumar Subudhi, Rahul P Salunke, Raeece Naeem, Sharif Hala, Sara Mfarrej, Asim Khogeer, Fadwa Alofi, Afrah Alsomali, Jumana Taha, Abdulaziz Alahmadi, Kahled Algithami, Anwar Hashem, Naif Almontashiri, Arnab Pain |
| EPI_ISL_678046 | Pathogen Genomics Lab King Abdullah University of Science and Technology(KAUST) | Pathogen Genomics Lab King Abdullah University of Science and Technology(KAUST) | Amanda Ooi, Luke Esau, Sara Mfarrej, Raeece Naeem, Sharif Hala, Asim Khogeer, Fadwa Alofi, Afrah Alsomali, Jumana Taha, Abdulaziz Alahmadi, Kahled Algithami, Anwar Hashem, Naif Almontashiri, Arnab Pain |
| EPI_ISL_678047 | Pathogen Genomics Lab King Abdullah University of Science and Technology(KAUST) | Pathogen Genomics Lab King Abdullah University of Science and Technology(KAUST) | Sharif Hala, Sara Mfarrej, Raeece Naeem, Amit Kumar Subudhi, Rahul P Salunke, Asim Khogeer, Fadwa Alofi, Afrah Alsomali, Jumana Taha, Abdulaziz Alahmadi, Kahled Algithami, Anwar Hashem, Naif Almontashiri, Arnab Pain |
| EPI_ISL_678048 | Pathogen Genomics Lab King Abdullah University of Science and Technology(KAUST) | Pathogen Genomics Lab King Abdullah University of Science and Technology(KAUST) | Sharif Hala, Sara Mfarrej, Amit Kumar Subudhi, Raeece Naeem, Rahul P Salunke, Asim Khogeer, Fadwa Alofi, Afrah Alsomali, Jumana Taha, Abdulaziz Alahmadi, Kahled Algithami, Anwar Hashem, Naif Almontashiri, Arnab Pain |
| EPI_ISL_678055 | Pathogen Genomics Lab King Abdullah University of Science and Technology(KAUST) | Pathogen Genomics Lab King Abdullah University of Science and Technology(KAUST) | Sharif Hala, Sara Mfarrej, Raeece Naeem, Amit Kumar Subudhi, Rahul P Salunke, Asim Khogeer, Fadwa Alofi, Afrah Alsomali, Jumana Taha, Abdulaziz Alahmadi, Kahled Algithami, Anwar Hashem, Naif Almontashiri, Arnab Pain |
| EPI_ISL_678056 | Pathogen Genomics Lab King Abdullah University of Science and Technology(KAUST) | Pathogen Genomics Lab King Abdullah University of Science and Technology(KAUST) | Fathia Ben Rached, Sharif Hala, Amit Kumar Subudhi, Sara Mfarrej, Raeece Naeem, Rahul P Salunke, Asim Khogeer, Fadwa Alofi, Afrah Alsomali, Jumana Taha, Abdulaziz Alahmadi, Kahled Algithami, Anwar Hashem, Naif Almontashiri, Arnab Pain |
| EPI_ISL_678058 | Pathogen Genomics Lab King Abdullah University of Science and Technology(KAUST) | Pathogen Genomics Lab King Abdullah University of Science and Technology(KAUST) | Sharif Hala, Sara Mfarrej, Amit Kumar Subudhi, Raeece Naeem, Rahul P Salunke, Asim Khogeer, Fadwa Alofi, Afrah Alsomali, Jumana Taha, Abdulaziz Alahmadi, Kahled Algithami, Anwar Hashem, Naif Almontashiri, Arnab Pain |
| EPI_ISL_678059 | Pathogen Genomics Lab King Abdullah University of Science and Technology(KAUST) | Pathogen Genomics Lab King Abdullah University of Science and Technology(KAUST) | Amit Kumar Subudhi, Rahul P Salunke, Raeece Naeem, Sharif Hala, Sara Mfarrej, Asim Khogeer, Fadwa Alofi, Afrah Alsomali, Jumana Taha, Abdulaziz Alahmadi, Kahled Algithami, Anwar Hashem, Naif Almontashiri, Arnab Pain |
| EPI_ISL_678060 | Pathogen Genomics Lab King Abdullah University of Science and Technology(KAUST) | Pathogen Genomics Lab King Abdullah University of Science and Technology(KAUST) | Fathia Ben Rached, Sharif Hala, Amit Kumar Subudhi, Sara Mfarrej, Raeece Naeem, Rahul P Salunke, Asim Khogeer, Fadwa Alofi, Afrah Alsomali, Jumana Taha, Abdulaziz Alahmadi, Kahled Algithami, Anwar Hashem, Naif Almontashiri, Arnab Pain |
| EPI_ISL_678065 | Pathogen Genomics Lab King Abdullah University of Science and Technology(KAUST) | Pathogen Genomics Lab King Abdullah University of Science and Technology(KAUST) | Amit Kumar Subudhi, Rahul P Salunke, Raeece Naeem, Sharif Hala, Sara Mfarrej, Asim Khogeer, Fadwa Alofi, Afrah Alsomali, Jumana Taha, Abdulaziz Alahmadi, Kahled Algithami, Anwar Hashem, Naif Almontashiri, Arnab Pain |
| EPI_ISL_678066 | Pathogen Genomics Lab King Abdullah University of Science and Technology(KAUST) | Pathogen Genomics Lab King Abdullah University of Science and Technology(KAUST) | Fathia Ben Rached, Sharif Hala, Amit Kumar Subudhi, Sara Mfarrej, Raeece Naeem, Rahul P Salunke, Asim Khogeer, Fadwa Alofi, Afrah Alsomali, Jumana Taha, Abdulaziz Alahmadi, Kahled Algithami, Anwar Hashem, Naif Almontashiri, Arnab Pain |
| EPI_ISL_678068 | Pathogen Genomics Lab King Abdullah University of Science and Technology(KAUST) | Pathogen Genomics Lab King Abdullah University of Science and Technology(KAUST) | Sharif Hala, Sara Mfarrej, Raeece Naeem, Amit Kumar Subudhi, Rahul P Salunke, Asim Khogeer, Fadwa Alofi, Afrah Alsomali, Jumana Taha, Abdulaziz Alahmadi, Kahled Algithami, Anwar Hashem, Naif Almontashiri, Arnab Pain |
| EPI_ISL_678069 | Pathogen Genomics Lab King Abdullah University of Science and Technology(KAUST) | Pathogen Genomics Lab King Abdullah University of Science and Technology(KAUST) | Fathia Ben Rached, Sharif Hala, Amit Kumar Subudhi, Sara Mfarrej, Raeece Naeem, Rahul P Salunke, Asim Khogeer, Fadwa Alofi, Afrah Alsomali, Jumana Taha, Abdulaziz Alahmadi, Kahled Algithami, Anwar Hashem, Naif Almontashiri, Arnab Pain |
| EPI_ISL_678071 | Pathogen Genomics Lab King Abdullah University of Science and Technology(KAUST) | Pathogen Genomics Lab King Abdullah University of Science and Technology(KAUST) | Sharif Hala, Sara Mfarrej, Raeece Naeem, Amit Kumar Subudhi, Rahul P Salunke, Asim Khogeer, Fadwa Alofi, Afrah Alsomali, Jumana Taha, Abdulaziz Alahmadi, Kahled Algithami, Anwar Hashem, Naif Almontashiri, Arnab Pain |
| EPI_ISL_678073, EPI_ISL_678074 | Pathogen Genomics Lab King Abdullah University of Science and Technology(KAUST) | Pathogen Genomics Lab King Abdullah University of Science and Technology(KAUST) | Amit Kumar Subudhi, Rahul P Salunke, Raeece Naeem, Sharif Hala, Sara Mfarrej, Asim Khogeer, Fadwa Alofi, Afrah Alsomali, Jumana Taha, Abdulaziz Alahmadi, Kahled Algithami, Anwar Hashem, Naif Almontashiri, Arnab Pain |
| EPI_ISL_678075 | Pathogen Genomics Lab King Abdullah University of Science and Technology(KAUST) | Pathogen Genomics Lab King Abdullah University of Science and Technology(KAUST) | Fathia Ben Rached, Sharif Hala, Amit Kumar Subudhi, Sara Mfarrej, Raeece Naeem, Rahul P Salunke, Asim Khogeer, Fadwa Alofi, Afrah Alsomali, Jumana Taha, Abdulaziz Alahmadi, Kahled Algithami, Anwar Hashem, Naif Almontashiri, Arnab Pain |
| EPI_ISL_678077 | Pathogen Genomics Lab King Abdullah University of Science and Technology(KAUST) | Pathogen Genomics Lab King Abdullah University of Science and Technology(KAUST) | Sharif Hala, Sara Mfarrej, Raeece Naeem, Amit Kumar Subudhi, Rahul P Salunke, Asim Khogeer, Fadwa Alofi, Afrah Alsomali, Jumana Taha, Abdulaziz Alahmadi, Kahled Algithami, Anwar Hashem, Naif Almontashiri, Arnab Pain |
| EPI_ISL_678100, EPI_ISL_678101, | Pathogen Genomics Lab King Abdullah University of Science | Pathogen Genomics Lab King Abdullah University of Science | Amanda Ooi, Luke Esau, Sara Mfarrej, Raeece Naeem, Sharif Hala, Asim Khogeer, Fadwa Alofi, Afrah Alsomali, Jumana Taha, Abdulaziz Alahmadi, |

| | | | |
|---|---|---|---|
| EPI_ISL_678102, EPI_ISL_678103 | and Technology(KAUST) | and Technology(KAUST) | Kahled Algithami, Anwar Hashem, Naif Almontashiri, Arnab Pain |
| EPI_ISL_678116, EPI_ISL_678117 | Pathogen Genomics Lab King Abdullah University of Science and Technology(KAUST) | Pathogen Genomics Lab King Abdullah University of Science and Technology(KAUST) | Amit Kumar Subudhi, Raeece Naeem, Sharif Hala, Sara Mfarrej, Rahul P Salunke, Asim Khogeer, Fadwa Alofi, Afrah Alsomali, Jumana Taha, Abdulaziz Alahmadi, Kahled Algithami, Anwar Hashem, Naif Almontashiri, Arnab Pain |
| EPI_ISL_678119, EPI_ISL_678120 | Pathogen Genomics Lab King Abdullah University of Science and Technology(KAUST) | Pathogen Genomics Lab King Abdullah University of Science and Technology(KAUST) | Luke Esau, Amanda Ooi, Sharif Hala, Raeece Naeem, Sara Mfarrej, Asim Khogeer, Fadwa Alofi, Afrah Alsomali, Jumana Taha, Abdulaziz Alahmadi, Kahled Algithami, Anwar Hashem, Naif Almontashiri, Arnab Pain |
| EPI_ISL_678127 | Pathogen Genomics Lab King Abdullah University of Science and Technology(KAUST) | Pathogen Genomics Lab King Abdullah University of Science and Technology(KAUST) | Amit Kumar Subudhi, Raeece Naeem, Sharif Hala, Sara Mfarrej, Rahul P Salunke, Asim Khogeer, Fadwa Alofi, Afrah Alsomali, Jumana Taha, Abdulaziz Alahmadi, Kahled Algithami, Anwar Hashem, Naif Almontashiri, Arnab Pain |
| EPI_ISL_678135, EPI_ISL_678136 | Pathogen Genomics Lab King Abdullah University of Science and Technology(KAUST) | Pathogen Genomics Lab King Abdullah University of Science and Technology(KAUST) | Sharif Hala, Sara Mfarrej, Amit Kumar Subudhi, Raeece Naeem, Rahul P Salunke, Asim Khogeer, Fadwa Alofi, Afrah Alsomali, Jumana Taha, Abdulaziz Alahmadi, Kahled Algithami, Anwar Hashem, Naif Almontashiri, Arnab Pain |
| EPI_ISL_678137, EPI_ISL_678138 | Pathogen Genomics Lab King Abdullah University of Science and Technology(KAUST) | Pathogen Genomics Lab King Abdullah University of Science and Technology(KAUST) | Sharif Hala, Sara Mfarrej, Raeece Naeem, Amit Kumar Subudhi, Rahul P Salunke, Asim Khogeer, Fadwa Alofi, Afrah Alsomali, Jumana Taha, Abdulaziz Alahmadi, Kahled Algithami, Anwar Hashem, Naif Almontashiri, Arnab Pain |
| EPI_ISL_678139, EPI_ISL_678140 | Pathogen Genomics Lab King Abdullah University of Science and Technology(KAUST) | Pathogen Genomics Lab King Abdullah University of Science and Technology(KAUST) | Fathia Ben Rached, Sharif Hala, Amit Kumar Subudhi, Sara Mfarrej, Raeece Naeem, Rahul P Salunke, Asim Khogeer, Fadwa Alofi, Afrah Alsomali, Jumana Taha, Abdulaziz Alahmadi, Kahled Algithami, Anwar Hashem, Naif Almontashiri, Arnab Pain |
| EPI_ISL_678230, EPI_ISL_678231, EPI_ISL_678232 | Pathogen Genomics Lab King Abdullah University of Science and Technology(KAUST) | Pathogen Genomics Lab King Abdullah University of Science and Technology(KAUST) | Amit Kumar Subudhi, Rahul P Salunke, Raeece Naeem, Sharif Hala, Sara Mfarrej, Asim Khogeer, Fadwa Alofi, Afrah Alsomali, Jumana Taha, Abdulaziz Alahmadi, Kahled Algithami, Anwar Hashem, Naif Almontashiri, Arnab Pain |
| EPI_ISL_678233 | Pathogen Genomics Lab King Abdullah University of Science and Technology(KAUST) | Pathogen Genomics Lab King Abdullah University of Science and Technology(KAUST) | Sharif Hala, Sara Mfarrej, Raeece Naeem, Amit Kumar Subudhi, Rahul P Salunke, Asim Khogeer, Fadwa Alofi, Afrah Alsomali, Jumana Taha, Abdulaziz Alahmadi, Kahled Algithami, Anwar Hashem, Naif Almontashiri, Arnab Pain |
| EPI_ISL_678314, EPI_ISL_678315, EPI_ISL_678322, EPI_ISL_678323, EPI_ISL_678341 | Area of Virology, Serology and Virology Division (SAViD), New South Wales Health Pathology Randwick | Virology Research Laboratory; Area of Virology, Serology and Virology Division (SAViD), New South Wales Health Pathology Randwick | Foster, C.; Au, J.; Ruiz Silva, M.; Deveson, I.; Bull, R.; Van Hal, S.; Rawlinson, W. |
| EPI_ISL_678503, EPI_ISL_678504, EPI_ISL_678505 | CNR Virus des Infections Respiratoires - France SUD | CNR Virus des Infections Respiratoires - France SUD | Antonin Bal, Gregory Destras, Gwendolyne Burfin, Solenne Brun, Martine Valette, Bruno Lina, Laurence Josset |
| EPI_ISL_678509, EPI_ISL_678511, EPI_ISL_678514, EPI_ISL_678515, EPI_ISL_678516, EPI_ISL_678519, EPI_ISL_678520, EPI_ISL_678521, EPI_ISL_678522, EPI_ISL_678523, EPI_ISL_678524, EPI_ISL_678529, EPI_ISL_678530, EPI_ISL_678531, EPI_ISL_678532, EPI_ISL_678533 | | | |
| see above | Utah Public Health Laboratory | Utah Public Health Laboratory | Erin Young, Kelly Oakeson |
| EPI_ISL_680286, EPI_ISL_680287 | Regional Virus Laboratory, Belfast Health and Social Care Trust | COVID-19 Genomics UK (COG-UK) Consortium | Conall McCaughey, James McKenna, Tanya Curran, Susan Feeney, Alison Watt, Ciara Cox, Mairead Connor, Zoltan Molnar, David Simpson, Derek Fairley |
| EPI_ISL_681674 | Molecular Medicine Laboratory, University of Magallanes | Centro Asistencial Docente y de Investigacion, Universidad de Magallanes | Jorge González, Jacqueline Aldridge, Diego Alvarez, Marco Montes de Oca, Hermy Alvarez, Roberto Uribe-Paredes, Marcelo Navarrete |
| EPI_ISL_681719 | University Hospital Limerick | Irish Coronavirus Sequencing Consortium - Teagasc Moorepark | Paul Cotter, Fiona Crispie, John Kenny, Calum Walsh |
| EPI_ISL_681722, EPI_ISL_681724 | University Hospital Limerick | Irish Coronavirus Sequencing Consortium - Teagasc Moorepark | Paul Cotter, Fiona Crispie, Amy Fitzpatrick, John Kenny, Elaine Lawton, Carolyn Meaney, Patrick Stapleton, Calum Walsh |
| EPI_ISL_681736, EPI_ISL_681737 | University Hospital Limerick | Irish Coronavirus Sequencing Consortium - Teagasc Moorepark | Paul Cotter, Fiona Crispie, John Kenny, Calum Walsh |
| EPI_ISL_681738, EPI_ISL_681739 | University Hospital Limerick | Irish Coronavirus Sequencing Consortium - Teagasc Moorepark | Paul Cotter, Fiona Crispie, Amy Fitzpatrick, John Kenny, Carolyn Meaney, Patrick Stapleton, Calum Walsh |
| EPI_ISL_681757, EPI_ISL_681768 | University Hospital Limerick | Irish Coronavirus Sequencing Consortium - Teagasc Moorepark | Paul Cotter, Fiona Crispie, John Kenny, Calum Walsh |
| EPI_ISL_681769 | University Hospital Limerick | Irish Coronavirus Sequencing Consortium - Teagasc Moorepark | Paul Cotter, Fiona Crispie, Amy Fitzpatrick, John Kenny, Carolyn Meaney, Patrick Stapleton, Calum Walsh |
| EPI_ISL_681885, EPI_ISL_681887, EPI_ISL_681893, EPI_ISL_681906, EPI_ISL_681909, EPI_ISL_681911, EPI_ISL_681914, EPI_ISL_681915, EPI_ISL_681916, EPI_ISL_681917, EPI_ISL_681924, EPI_ISL_681926 | | | |
| see above | National Virus Reference Laboratory | Irish Coronavirus Sequencing Consortium - Teagasc Moorepark | Paul Cotter, Fiona Crispie, John Kenny, Matthew McCabe, Calum Walsh |
| EPI_ISL_681952, EPI_ISL_681955 | UPMC Clinical Microbiology Laboratory | Microbial Genomic Epidemiology Laboratory, University of Pittsburgh | Mustapha M. Mustapha, Jane W. Marsh, Dan Snyder, Marissa P. Griffith, Stephanie L. Mitchell, Vatsala R. Srinivasa, Kady D. Waggle, Chinelo Ezeonwuku, Vaughn S. Cooper, Lee H. Harrison |
| EPI_ISL_683595 | Hospital Clínico Universitario Lozano Blesa de Zaragoza (España) | SeqCOVID-SPAIN consortium/IBV(CSIC) | Rafael Benito, Sonia Algarate, Jessica Bueno and SeqCOVID-SPAIN consortium |
| EPI_ISL_692966 | Massachusetts State Public Health Laboratory | Massachusetts State Public Health Laboratory | Andrew Lang, Timelia Fink, Glen Gallagher, Sandra Smole |
| EPI_ISL_694053, EPI_ISL_694054, EPI_ISL_694056, EPI_ISL_694057, EPI_ISL_694058, EPI_ISL_694059, EPI_ISL_694060, EPI_ISL_694061, EPI_ISL_694062, EPI_ISL_694063, EPI_ISL_694064, EPI_ISL_694065, EPI_ISL_694071, EPI_ISL_694072 | | | |
| see above | AZ SPHL, Arizona Department of Health Services | TGen North | Jolene Bowers, Megan Folkerts, Chris French, Hayley Yaglom, Ashlyn Pfeiffer, Darrin Lemmer, Dave Engelthaler, The Arizona COVID Genomics Union (ACGU) |
| EPI_ISL_695828, EPI_ISL_696021, EPI_ISL_696024, EPI_ISL_696026, EPI_ISL_696359, EPI_ISL_696360, EPI_ISL_696362, EPI_ISL_696364, EPI_ISL_696367, EPI_ISL_696373 | Sonora Quest Laboratories, Laboratory Sciences of Arizona | TGen North | Jolene Bowers, Megan Folkerts, Chris French, Hayley Yaglom, Ashlyn Pfeiffer, Darrin Lemmer, Dave Engelthaler, The Arizona COVID Genomics Union (ACGU) |
| EPI_ISL_699645, EPI_ISL_699648 | Pathology West - NSW Health Pathology | NSW Health Pathology - Institute of Clinical Pathology and Medical Research; Westmead Hospital; University of Sydney | CIDM-PH et al. |
| EPI_ISL_700355 | Laboratoire de virologie, CHU de Grenoble - CS 10217 - 38043 Grenoble cedex 16 | CNR Virus des Infections Respiratoires - France SUD | Antonin Bal, Gregory Destras, Gwendolyne Burfin, Hadrien Règue, Quentin Semanas, Martine Valette, Bruno Lina, Sylvie Larrat, Laurence Josset |
| EPI_ISL_700356 | Laboratoire de virologie, CHU de Grenoble - CS 10217 - 38043 Grenoble cedex 17 | CNR Virus des Infections Respiratoires - France SUD | Antonin Bal, Gregory Destras, Gwendolyne Burfin, Hadrien Règue, Quentin Semanas, Martine Valette, Bruno Lina, Sylvie Larrat, Laurence Josset |
| EPI_ISL_700357 | Laboratoire de virologie, CHU de Grenoble - CS 10217 - 38043 Grenoble cedex 18 | CNR Virus des Infections Respiratoires - France SUD | Antonin Bal, Gregory Destras, Gwendolyne Burfin, Hadrien Règue, Quentin Semanas, Martine Valette, Bruno Lina, Sylvie Larrat, Laurence Josset |
| EPI_ISL_700358 | Laboratoire de virologie, CHU de Grenoble - CS 10217 - 38043 Grenoble cedex 19 | CNR Virus des Infections Respiratoires - France SUD | Antonin Bal, Gregory Destras, Gwendolyne Burfin, Hadrien Règue, Quentin Semanas, Martine Valette, Bruno Lina, Sylvie Larrat, Laurence Josset |
| EPI_ISL_700373, EPI_ISL_700375, EPI_ISL_700376, EPI_ISL_700378 | Centre hospitalier Métropole Savoie | CNR Virus des Infections Respiratoires - France SUD | Antonin Bal, Carine Dumollard, Gregory Destras, Gwendolyne Burfin, Hadrien Règue, Quentin Semanas, Martine Valette, Bruno Lina, Jérôme Grosjean, Laurence Josset |
| EPI_ISL_702679, EPI_ISL_703297, EPI_ISL_705113, EPI_ISL_705416, | Virology Department, Royal Infirmary of Edinburgh, NHS Lothian / School of Biological Sciences, University of | COVID-19 Genomics UK (COG-UK) Consortium | McHugh M, Dewar R, Rooke S, Gallagher M, Balcaza C, O'Toole Á, Scher E, Hill V, McCrone JT, Colquhoun R, Yu X, Jackson B, Rambaut A, Williams TC, Templeton K |

| | | | |
|---|---|---|---|
| EPI_ISL_705741, EPI_ISL_705796, EPI_ISL_705797, EPI_ISL_705802 | Edinburgh / Institute of Genetics and Molecular Medicine, University of Edinburgh | | |
| EPI_ISL_707963, EPI_ISL_707964, EPI_ISL_707965, EPI_ISL_707966, EPI_ISL_707967, EPI_ISL_707968, EPI_ISL_708018 | Virology, Universitätsklinikum des Saarlandes | Epigenetics, Saarland University | Kathrin Kattler, Markus Vogelgesang, Stefan Lohse, Sascha Tierling, Sigrun Smola, Jörn Walter |
| EPI_ISL_708508, EPI_ISL_708509, EPI_ISL_708510, EPI_ISL_708511, EPI_ISL_708513, EPI_ISL_708514, EPI_ISL_708515, EPI_ISL_708517, EPI_ISL_708518, EPI_ISL_708519, EPI_ISL_708520, EPI_ISL_708521, EPI_ISL_708522, EPI_ISL_708523, EPI_ISL_708524, EPI_ISL_708525, EPI_ISL_708526, EPI_ISL_708527 | | | |
| see above | National Virus Reference Laboratory | Irish Coronavirus Sequencing Consortium - Teagasc Oakpark | Michele Della Bartola, Matthew McCabe, Ewen Mullins |
| EPI_ISL_708755, EPI_ISL_708761, EPI_ISL_708773, EPI_ISL_708777, EPI_ISL_708785 | PathWest Laboratory Medicine WA | PathWest Laboratory Medicine WA Microbial Surveillance Unit | PathWest Laboratory Medicine WA Microbial Surveillance Unit |
| EPI_ISL_710119 | National Virus Reference Laboratory | Irish Coronavirus Sequencing Consortium - Teagasc Oakpark | Michele Della Bartola, Matthew McCabe, Ewen Mullins |
| EPI_ISL_710580 | Uppsala klinisk mikrobiologi | The Public Health Agency of Sweden | Department of Microbiology, The Public Health Agency of Sweden |
| EPI_ISL_714937 | Department of Virus and Microbiological Special Diagnostics, Statens Serum Institut, Copenhagen, Denmark | Albertsen Lab, Department of Chemistry and Bioscience, Aalborg University, Denmark | Danish Covid-19 Genome Consortium |
| EPI_ISL_718138, EPI_ISL_718148 | Ministry of Health Hospitals | Institute of Health and Community Medicine | David Perera, Ooi Hong How, Chua Hock Hin, Tonnii Sia Loong Loong, Wong Jyn Shan, Wong Kiing Aik, Chan Chia Jui |
| EPI_ISL_721987, EPI_ISL_722004, EPI_ISL_722012, EPI_ISL_722023, EPI_ISL_722029, EPI_ISL_722057, EPI_ISL_722063, EPI_ISL_722065, EPI_ISL_722078, EPI_ISL_722079, EPI_ISL_722080, EPI_ISL_722082, EPI_ISL_722084, EPI_ISL_722099, EPI_ISL_722108, EPI_ISL_722119, EPI_ISL_722122 | | | |
| see above | Hospital das Clínicas Universidade de São Paulo Medical School | Laboratório de Parasitologia Médica - Instituto de Medicina Tropical - Universidade de São Paulo | Brazil-UK Centre for Arbovirus Discovery Diagnosis Genomics and Epidemiology (CADDE) Genomic Network - Instituto de Medicina Tropical |
| EPI_ISL_723108, EPI_ISL_723109, EPI_ISL_723110, EPI_ISL_723111, EPI_ISL_723112 | Institute of Medical Genetics and Applied Genomics | Institute of Medical Genetics and Applied Genomics | Caspar Gross, Tina Ganzenmüller, Siri Göpel, Michaela Pogoda, Daniela Bezdan, Michael Sonnabend, Angel Angelov, Nicolas Casadei, Stephan Ossowski, Thomas Iftner, Michael Bitzer |
| EPI_ISL_728288 | Jena University Hospital, Institute for Infectious Diseases and Infection Control | Institute for infectious medicine & hospital hygiene, CaSe-Group | Spott, Riccardo; Marquet, Mike; Pletz, Matthias W.; Brandt, Christian |
| EPI_ISL_728292, EPI_ISL_728293 | Jena University Hospital, Institute for Infectious Diseases and Infection Control | Institute of infectious medicine & hospital hygiene, CaSe-Group | Spott, Riccardo; Marquet, Mike; Pletz, Matthias W.; Brandt, Christian |
| EPI_ISL_729816, EPI_ISL_729817, EPI_ISL_729818 | Laboratorio Central de Saude Publica do Estado do Rio Grande do Sul (LACEN-RS) | Laboratory of Respiratory Viruses and Measles, Oswaldo Cruz Institute, FIOCRUZ | Paola Resende, Luciana Appolinario, Fernando Motta, Anna Carolina Paixão, Ana Carolina Mendonça, Tatiana Schaffer Gregianini, Marilda Tereza Mar da Rosa, Marilda Siqueira |
| EPI_ISL_729946 | Nigeria Centre for Disease Control (NCDC) | African Centre of Excellence for Genomics of Infectious Diseases (ACEGID), Redeemer's University, Ede, Osun State, Nigeria | Oluniyi P.E. et al |
| EPI_ISL_730138, EPI_ISL_730156, EPI_ISL_730157, EPI_ISL_730158, EPI_ISL_730160 | Scripps Medical Laboratory | Andersen lab at Scripps Research | SEARCH Alliance San Diego with Michael Quigley, Ellen Stefanski, Ian Mchardy |
| EPI_ISL_732441 | National Virus Reference Laboratory | National Virus Reference Laboratory | Michael Carr, Gabriel Gonzalez, Jonathan Dean, Daniel Hare, Cillian F De Gascun |
| EPI_ISL_732758 | Centro de Investigación Biomédica de La Rioja - Hospital San Pedro Logroño | SeqCOVID-SPAIN consortium/IBV(CSIC) | María de Toro, José Manuel Azcona Gutiérrez, María Pilar Bea Escudero, Miriam Blasco Alberdi and SeqCOVID-SPAIN consortium |
| EPI_ISL_734531, EPI_ISL_734532, EPI_ISL_734533, EPI_ISL_734534, EPI_ISL_734535, EPI_ISL_734536, EPI_ISL_734537, EPI_ISL_734538, EPI_ISL_734539 | UZ Leuven, National Reference Laboratory for Coronaviruses, Laboratory Medicine, Leuven, Belgium | KU Leuven, Rega Institute, Clinical and Epidemiological Virology | Tony Wawina-Bokalanga, Joan Marti-Carreras, Bert Vanmechelen, Piet Maes |
| EPI_ISL_735399 | Hospital Municiplal Dr Ignacio de gouvea | Instituto Adolfo Lutz, Interdiciplinary Procedures Center, Strategic Laboratory | Claudio Tavares Sacchi, Claudia Regina Gonçalves, Erica Valessa Ramos Gomes, Karoline Rodrigues Campos |
| EPI_ISL_737935, EPI_ISL_737951 | Uganda Central Public Health Lab and Uganda Virus Research Institute | MRC/UVRI & LSHTM Uganda Research Unit | Matthew Cotten, Dan Lule Bugembe, My V.T. Phan, Pontiano Kaleebu et al. |
| EPI_ISL_738284, EPI_ISL_738285 | Connecticut Veterans' Affairs Hospital | Grubaugh Lab - Yale School of Public Health | Joseph Fauver, Tara Alpert, Chantal Vogels, Mary Petrone, Isabel Ott, Ellen Foxman, Shaili Gupta, Danielle Plank, Nathan Grubaugh |
| EPI_ISL_738290 | Yale Pathology Lab | Grubaugh Lab - Yale School of Public Health | Joseph Fauver, Tara Alpert, Chantal Vogels, Mary Petrone, Isabel Ott, Ellen Foxman, Shaili Gupta, Danielle Plank, Nathan Grubaugh |
| EPI_ISL_738299 | Connecticut Veterans' Affairs Hospital | Grubaugh Lab - Yale School of Public Health | Joseph Fauver, Tara Alpert, Chantal Vogels, Mary Petrone, Isabel Ott, Ellen Foxman, Shaili Gupta, Danielle Plank, Nathan Grubaugh |
| EPI_ISL_751204 | Pathogen Genomics Lab King Abdullah University of Science and Technology(KAUST) | Pathogen Genomics Lab King Abdullah University of Science and Technology(KAUST) | Sharif Hala, Sara Mfarrej, Raushan Nugmanova, Olga Douvropoulou, Rahul P Salunke, Raeece Naeem, Fadwa Alofi, Asim Khogeer, Afrah Alsomali, Jumana Taha, Abdulaziz Alahmadi, Kahled Algithami, Anwar Hashem, Naif Almontashiri, Arnab Pain |
| EPI_ISL_751206 | Pathogen Genomics Lab King Abdullah University of Science and Technology(KAUST) | Pathogen Genomics Lab King Abdullah University of Science and Technology(KAUST) | Sara Mfarrej, Luke Esau, Amanda Ooi, Sharif Hala, Raeece Naeem, Asim Khogeer, Fadwa Alofi, Afrah Alsomali, Jumana Taha, Abdulaziz Alahmadi, Kahled Algithami, Anwar Hashem, Naif Almontashiri, Arnab Pain |
| EPI_ISL_751213 | Pathogen Genomics Lab King Abdullah University of Science and Technology(KAUST) | Pathogen Genomics Lab King Abdullah University of Science and Technology(KAUST) | Sara Mfarrej, Amanda Ooi, Luke Esau, Sharif Hala, Raeece Naeem, Asim Khogeer, Fadwa Alofi, Afrah Alsomali, Jumana Taha, Abdulaziz Alahmadi, Kahled Algithami, Anwar Hashem, Naif Almontashiri, Arnab Pain |
| EPI_ISL_751429, EPI_ISL_751430, EPI_ISL_751431, EPI_ISL_751432, EPI_ISL_751433, EPI_ISL_751434, EPI_ISL_751435, EPI_ISL_751436, EPI_ISL_751437, EPI_ISL_751438, EPI_ISL_751439, EPI_ISL_751440, EPI_ISL_751441 | | | |
| see above | IRCCS Sacro Cuore Don Calabria Hospital, Department of Infectious, Tropical Diseases & Microbiology | University of Verona, Department of Biotechnology | Antonio Mori, Michela Deiana, Elena Pomari, Chiara Piubelli; Giulia Lopatriello, Luca Marcolungo, Cristina Beltrami, Chiara Degli Esposti, Emanuela Cosentino, Massimo Delledonne |
| EPI_ISL_752629, EPI_ISL_752630 | State Laboratories Division, Hawaii State Department of Health | State Laboratories Division, Hawaii State Department of Health | Pamela O'Brien, Sabrina Diemert, Drew Kuwazaki, Razvan Sultana, Edward Desmond |
| EPI_ISL_753729, EPI_ISL_753754, EPI_ISL_753764, EPI_ISL_753774, EPI_ISL_753811, EPI_ISL_753849, EPI_ISL_753930, EPI_ISL_753931, EPI_ISL_753932, EPI_ISL_753933, EPI_ISL_753934, EPI_ISL_753935, EPI_ISL_753938, EPI_ISL_753940, EPI_ISL_754048, EPI_ISL_754049, EPI_ISL_754051 | | | |
| see above | Charité Universitätsmedizin Berlin, Institut für Virologie/Labor Berlin | Charité Universitätsmedizin Berlin, Institut für Virologie | Victor M Corman, Jörn Beheim-Schwarzbach, Barbara Mühlemann, Julia Schneider, Talitha Veith, Terry Jones, Christian Drosten |
| EPI_ISL_754769 | Emory Molecular Diagnostics Laboratory, Emory Healthcare | Piantadosi Lab, Emory Department of Pathology | Ahmed Babiker, Anne Piantadosi |
| EPI_ISL_759862, EPI_ISL_759863, EPI_ISL_759864 | Molecular, ProgenaBiome | Molecular, ProgenaBiome | Papoutsis,A.J. |
| EPI_ISL_760151 | Division of Emerging Infectious Diseases, Bureau of Infectious Diseases Diagnosis Control, Korea Disease Control and Prevention Agency | Division of Emerging Infectious Diseases, Bureau of Infectious Diseases Diagnosis Control, Korea Disease Control and Prevention Agency | Ae Kyung Park, Il-Hwan Kim, Heui Man Kim, Jeong-Min Kim, Namjoo Lee, Chaeyoung Lee, Sang Hee Woo, Eun-Jin Kim |
| EPI_ISL_763067 | 1-Laboratory of Microbiology, National Reference Lab, Charles | 1-Clinical and Experimental Pharmacology Lab, LR16SP02, | Ilhem Boutiba-Ben Boubaker, Sameh Trabelsi, Nissaf Ben Alaya, Maher Kharrat, Alia Ben Kahla, Jalila Ben Khelil, Salma Abid, Sana Ferjani, Mouna Ben |

| | | | |
|---|---|---|---|
| | Nicolle Hospital; 2-University of Tunis ElManar, Faculty of Medicine of Tunis, LR99ES09, Tunis, Tunisia | National Center of Pharmacovigilance, University of Tunis El Manar, Tunis, Tunisia. 2-Neurodegenerative diseases and psychiatric troubles, LR18SP03, Razi Hospital, University of Tunis El Manar, Tunis, Tunisia. 3- Ministry of Health, National Observatory of New and Emerging Diseases, 1006, Tunis, Tunisia | Sassi, Mouna Safer, Awatef El MOussi, Habiba Ben Romdhane, Souissi Amira, Ines Mdini, Hanen El Jebari, Asma Ferjani, Gaies Emna, Riadh Daghfous, Riadh Gouider. |
| EPI_ISL_763093 | Jena University Hospital, Institute for Infectious Diseases and Infection Control | Institute of infectious medicine & hospital hygiene, CaSe-Group | Spott, Riccardo; Marquet, Mike; Pletz, Matthias W.; Brandt, Christian |
| EPI_ISL_763350 | National Virus Reference Laboratory | Irish Coronavirus Sequencing Consortium - Teagasc Moorepark | Paul Cotter, Fiona Crispie, John Kenny, Matthew McCabe, Calum Walsh |
| EPI_ISL_763363 | AIID | Irish Coronavirus Sequencing Consortium - National Virus Reference Laboratory | Michael Carr, Gabriel Gonzalez, Alejandro Abner Garcia Leon, Patrick Mallon |
| EPI_ISL_766259 | Respiratory Virus Unit, National Infection Service, Public Health England | COVID-19 Genomics UK (COG-UK) Consortium | PHE Covid Sequencing Team |
| EPI_ISL_767404, EPI_ISL_767405, EPI_ISL_767407, EPI_ISL_767408 | Ochsner Health | BioInfo Experts, LLC | Rebecca Rose, Amy Feehan, David J. Nolan, Sissy Cross, David Moraga Amador, Tong Yang, Luke Caruso, Wayra Navia, Lydia Von Borstel, Xiao Hui Zhou, Julia-Garcia-Diaz, Susanna L. Lamers |
| EPI_ISL_767764, EPI_ISL_767765, EPI_ISL_767766, EPI_ISL_767767, EPI_ISL_767820, EPI_ISL_767838 | National Virus Reference Laboratory | Irish Coronavirus Sequencing Consortium - Teagasc Moorepark | Calum Walsh, Genuity Ireland |
| EPI_ISL_770822, EPI_ISL_770824, EPI_ISL_770825 | Eastern Ontario Regional Laboratory Association | McMaster University | Leanne Mortimer, Hooman Derakhshani, Emily Panousis, Ahmed Draia, Jalees Nasir, Robert Slinger, Andrew G. McArthur |
| EPI_ISL_775239, EPI_ISL_775240, EPI_ISL_775259 | INT Fondazione Pascale | INT Fondazione Pascale | INT Fondazione Pascale |
| EPI_ISL_778648, EPI_ISL_778649, EPI_ISL_778696, EPI_ISL_778697, EPI_ISL_778698, EPI_ISL_778699, EPI_ISL_778700, EPI_ISL_778701, EPI_ISL_778702, EPI_ISL_778703 | Istituto Zooprofilattico Sperimentale del Mezzogiorno | TIGEM | Patrizia Annunziata, Andrea Ballabio, Valentina Bouche, Davide Cacchiarelli (CorrespAuthor), Pellegrino Cerino, Chiara Colantuono, Lucio Di Filippo, Antonio Grimaldi, Antonio Limone, Gabriella Loconte, Anna Manfredi, Francesco Panariello, Biancamaria Pierri, Marcello Salvi, Lucia Vassallo |
| EPI_ISL_788966 | Ospedale "Di Venere" | Beaconlab (Bioinformatics, Evolution and Comparative Genomics lab), Dept of Biosciences, University on Milan | Iacobellis M, d'Avenia M, Piluscio R, Parisi A, Chiara M, Manzari C, Pesole G |
| EPI_ISL_791527, EPI_ISL_791528, EPI_ISL_791529, EPI_ISL_791530, EPI_ISL_791531, EPI_ISL_791532, EPI_ISL_791533, EPI_ISL_791534, EPI_ISL_791535, EPI_ISL_791536, EPI_ISL_791537, EPI_ISL_791538, EPI_ISL_791539, EPI_ISL_791540, EPI_ISL_791541, EPI_ISL_791542, EPI_ISL_791543, EPI_ISL_791544, EPI_ISL_791545, EPI_ISL_791546, EPI_ISL_791547, EPI_ISL_791548, EPI_ISL_791549, EPI_ISL_791550, EPI_ISL_791551, EPI_ISL_791552, EPI_ISL_791553, EPI_ISL_791554, EPI_ISL_791555, EPI_ISL_791556, EPI_ISL_791557, EPI_ISL_791558, EPI_ISL_791559, EPI_ISL_791560, EPI_ISL_791561, EPI_ISL_791562, EPI_ISL_791563, EPI_ISL_791564, EPI_ISL_791565, EPI_ISL_791566, EPI_ISL_791567, EPI_ISL_791568, EPI_ISL_791569, EPI_ISL_791570, EPI_ISL_791571, EPI_ISL_791572, EPI_ISL_791573, EPI_ISL_791574, EPI_ISL_791575, EPI_ISL_791576, EPI_ISL_791577, EPI_ISL_791578, EPI_ISL_791579, EPI_ISL_791580, EPI_ISL_791581, EPI_ISL_791582, EPI_ISL_791583, EPI_ISL_791584, EPI_ISL_791585, EPI_ISL_791586, EPI_ISL_791587, EPI_ISL_791588, EPI_ISL_791589, EPI_ISL_791590, EPI_ISL_791591, EPI_ISL_791592, EPI_ISL_791593, EPI_ISL_791594, EPI_ISL_791595, EPI_ISL_791596, EPI_ISL_791597, EPI_ISL_791598, EPI_ISL_791599, EPI_ISL_791600, EPI_ISL_791601, EPI_ISL_791602, EPI_ISL_791603, EPI_ISL_791604, EPI_ISL_791605, EPI_ISL_791606, EPI_ISL_791607, EPI_ISL_791608, EPI_ISL_791609, EPI_ISL_791610, EPI_ISL_791611, EPI_ISL_791612, EPI_ISL_791613, EPI_ISL_791614, EPI_ISL_791615, EPI_ISL_791616, EPI_ISL_791617, EPI_ISL_791618, EPI_ISL_791619, EPI_ISL_791620, EPI_ISL_791621, EPI_ISL_791622, EPI_ISL_791623, EPI_ISL_791624, EPI_ISL_791625, EPI_ISL_791626, EPI_ISL_791627, EPI_ISL_791628, EPI_ISL_791629, EPI_ISL_791630, EPI_ISL_791631, EPI_ISL_791632, EPI_ISL_791633, EPI_ISL_791634, EPI_ISL_791635, EPI_ISL_791636, EPI_ISL_791637, EPI_ISL_791638, EPI_ISL_791639, EPI_ISL_791640, EPI_ISL_791641, EPI_ISL_791642, EPI_ISL_791643, EPI_ISL_791644, EPI_ISL_791645, EPI_ISL_791646, EPI_ISL_791647, EPI_ISL_791648, EPI_ISL_791649, EPI_ISL_791650 | | | |
| see above | Massachusetts State Public Health Laboratory | Infectious Disease Program, Broad Institute of Harvard and MIT | Lemieux,J.E., Siddle,K.J., Shaw,B., Adams,G., Pierce,V., Turbett,S., Anahtar,M., Branda,J., Slater,D., Harris,J., Lin,A.E., Gladden-Young,A., Lagerborg,K., Rudy,M., DeRuff,K., Carter,A., Normandin,E., Bauer,M., Reilly,S., Tomkins-Tinch,C., Loreth,C., Chaluvadi,S., Neumann,A., Cusick,C., Chapman,S.B., Gnirke,A., Flowers,K., Cerrato,F., Birren,B.W., Gallagher,G., Smole,S., Park,D.J., MacInnis,B.L., Ryan,E., LaRocque,R., Rosenberg,E. and Sabeti,P.C. |
| EPI_ISL_791815 | Massachusetts General Hospital | Infectious Disease Program, Broad Institute of Harvard and MIT | Lemieux,J.E., Siddle,K.J., Shaw,B., Adams,G., Pierce,V., Turbett,S., Anahtar,M., Branda,J., Slater,D., Harris,J., Lin,A.E., Gladden-Young,A., Lagerborg,K., Rudy,M., DeRuff,K., Carter,A., Normandin,E., Bauer,M., Reilly,S., Tomkins-Tinch,C., Loreth,C., Chaluvadi,S., Neumann,A., Cusick,C., Chapman,S.B., Gnirke,A., Flowers,K., Cerrato,F., Birren,B.W., Gallagher,G., Smole,S., Park,D.J., MacInnis,B.L., Ryan,E., LaRocque,R., Rosenberg,E. and Sabeti,P.C. |
| EPI_ISL_792098 | UW Virology Lab | UW Virology Lab | Pavitra Roychoudhury, Hong Xie, Lasata Shrestha, Meei-Li Huang, Keith R Jerome, Alexander Greninger |
| EPI_ISL_792483, EPI_ISL_792487, EPI_ISL_792488, EPI_ISL_792489, EPI_ISL_792490, EPI_ISL_792492 | Laboratorio Central Mg. Luis Alfredo Pianciola | Hospital Regional Ushuaia - Centro Austral De Investigaciones Científicas - Universidad Nacional De Tierra Del Fuego on behalf of 'Proyecto Argentino Interinstitucional de genomica de SARS-CoV-2' (PAIS Consortium) | Ceballos, SG; Nardi, CF; Gramundi, ID; Gallego, F; Pinto, C; Ziehm, MC; Mazzeo, M; Pianciola, L. |
| EPI_ISL_792500 | Laboratorio de Inmunología del Hospital Perrando e Instituto de Medicina Regional de la UNNE | Grupo de Genómica y Bioinformática del Instituto de Investigación de la Cadena Láctea CONICET-INTA on behalf of 'Proyecto Argentino Interinstitucional de genomica de SARS-CoV-2' (PAIS Consortium) | Eberhardt, MF; Irazoqui, JM; Foussal, MD; Deluca, G; Ayala, NA; Gómez, MV; Giusiano, G; Lucero, H; Marin, M; Lescano, L; Cayré, A; Amadio, AF |
| EPI_ISL_792529, EPI_ISL_792534, EPI_ISL_792536, EPI_ISL_792538 | Laboratorio Central, Ministerio de Salud Córdoba | Instituto de Patologia Vegetal (CIAP-INTA) on behalf of 'Proyecto Argentino Interinstitucional de genomica de SARS-CoV-2' (PAIS Consortium) | Fernández, FD; Debat, HJ., Re, V; Pisano, MB; Castro, G.; Barbas, G. |
| EPI_ISL_792572 | LACEN-PB | Laboratory of Respiratory Viruses and Measles, Oswaldo Cruz Institute, FIOCRUZ | Paola Resende, Luciana Appolinario, Fernando Motta, Anna Carolina Paixao, Ana Carolina Mendonca, João Felipe Bezerra, Romero Henrique Teixeira de Vasconcelos, Dalane Loudal Florentino Teixeira, Thiago Franco de Oliveira Carneiro, Marilda Siqueira |
| EPI_ISL_794600 | Central Laboratories, Egyptian Ministry of Health and Population | Central Laboratories, Egyptian Ministry of Health and Population | Kayed,A.E., Roshdy,W.H., El-Shesheny,R., Mostafa,A., Khalifa,M.K., Shehata,M., Shawky,S., Saleh,M., Gomaa,M., El Taweel,A., Mahmoud,S.H., Moatasim,Y., Kutkat,O., Kamel,M.N., Mahrous,N., El Sayes,M.A., El Guindy,N.M., Naguib,A., Kandeil,A., Kayali,G., Ali,M.A. |
| EPI_ISL_796064, EPI_ISL_796066, EPI_ISL_796071, EPI_ISL_796072, EPI_ISL_796111, EPI_ISL_796112, EPI_ISL_796114, EPI_ISL_796115, EPI_ISL_796116, EPI_ISL_796117, EPI_ISL_796118, EPI_ISL_796119, EPI_ISL_796120, EPI_ISL_796121, EPI_ISL_796122, EPI_ISL_796123, EPI_ISL_796124, EPI_ISL_796125, EPI_ISL_796126 | | | |
| see above | Servicio de Microbiología. Hospital General Universitario de Castellón | SeqCOVID-SPAIN consortium/IBV(CSIC) | Rosario Moreno Muñoz, María Dolores Tirado Balaguer and SeqCOVID-SPAIN consortium |
| EPI_ISL_801542, EPI_ISL_801553, EPI_ISL_801684, EPI_ISL_801685, EPI_ISL_801686, EPI_ISL_801687, EPI_ISL_801688, EPI_ISL_801689, EPI_ISL_801690, EPI_ISL_801691, EPI_ISL_801692, EPI_ISL_801693, EPI_ISL_801694, EPI_ISL_801695, EPI_ISL_801696, EPI_ISL_801697, EPI_ISL_801705, EPI_ISL_801736 | | | |
| see above | Laboratory of Molecular Virology, Pontificia Universidad Católica de Chile | MSHS Pathogen Surveillance Program | Leonardo I. Almonacid, Ana S. Gonzalez-Reiche, Matthew M. Hernandez, Jorge Levican, Ana Maria Contreras, Carlos Palma, Tamara García-Salum, Zenab Khan, Adriana van De Guchte, Ajay Obla, Jayeeta Dutta, Bremy Alburquerque, Eileen Serrano, Constanza Maldonado, M. Belen Leyton, Erick Salinas, Hala Alshammary, Juan Soto, Shwetha Hara Sridhar, Ying-Chih Wang, Melissa Smith, Robert Sebra, Marcela Ferres, Adolfo Garcia-Sastre, Edward C. Holmes, Viviana Simon, Harm van Bakel, Rafael A. Medina. |
| EPI_ISL_801875, EPI_ISL_801879, EPI_ISL_801907, EPI_ISL_801932, EPI_ISL_801948, EPI_ISL_801957, EPI_ISL_801987, EPI_ISL_801988, EPI_ISL_801989, EPI_ISL_801990, EPI_ISL_801991, EPI_ISL_801992, EPI_ISL_801993, EPI_ISL_801994, EPI_ISL_801995, EPI_ISL_801996, EPI_ISL_801997, EPI_ISL_801998, EPI_ISL_801999, EPI_ISL_802000, EPI_ISL_802001 | | | |
| see above | MSHS Clinical Microbiology Laboratories | MSHS Pathogen Surveillance Program | Ana S. Gonzalez-Reiche, Hala Alshammary, Mitchell J. Sullivan, Brianne Ciferri, Ajay Obla, Angela Amoako, Mahmoud Awawda, Elena Hirsch, Ashley S. Salimbangon, Levy Sominsky, Katherine Beach, Kayla Russo, Charles Gleason, Shelcie Fabre, Giulio Kleiner, Zenab Khan, Bremy Alburquerque, Adriana van de Guchte, Komal Srivastava, Matthew M. Hernandez, Jayeeta Dutta, Denise Jurczyszak, Emily Ferreri, Rachel Chernet, Nancy Francoeur, Betsaida |

| | | | |
|---|---|---|---|
| | | | Salom Melo, Irina Oussenko, Gintaras Deikus, Juan Soto, Shwetha Hara Sridhar, Ying-Chih Wang, Kathryn Twyman, Andrew Kasarskis, Deena R. Altman, Robert Sebra, Adolfo García-Sastre, Marta Luksza, Gopi Patel, Sarah Schaefer, Melissa Gitman, Michael D. Nowak, Alberto Paniz-Mondolfi, Emilia Mia Sordillo, Viviana Simon, Harm van Bakel |
| EPI_ISL_804389, EPI_ISL_804395, EPI_ISL_804401, EPI_ISL_804402, EPI_ISL_804412, EPI_ISL_804419, EPI_ISL_804428, EPI_ISL_804429, EPI_ISL_804450, EPI_ISL_804451, EPI_ISL_804452 | | | |
| see above | Dutch COVID-19 response team | National Institute for Public Health and the Environment (RIVM) | Adam Meijer, Harry Vennema, Dirk Eggink, Matthijs Welkers, Jeroen Cremer, Sharon van den Brink, Bas van der Veer, AnneMarie van den Brandt, Florian Zwagemaker, Dennis Schmitz, Chantal Reusken, on behalf of the national COVID-19 response team |
| EPI_ISL_805274, EPI_ISL_805481, EPI_ISL_805496, EPI_ISL_805653, EPI_ISL_805723, EPI_ISL_805764, EPI_ISL_805874, EPI_ISL_805980, EPI_ISL_806021, EPI_ISL_806033, EPI_ISL_806040, EPI_ISL_806063, EPI_ISL_806066, EPI_ISL_806081, EPI_ISL_806126, EPI_ISL_806150, EPI_ISL_806180, EPI_ISL_806181, EPI_ISL_806194, EPI_ISL_806203, EPI_ISL_806210, EPI_ISL_806223, EPI_ISL_806254, EPI_ISL_806294, EPI_ISL_806343, EPI_ISL_806354, EPI_ISL_806360, EPI_ISL_806429, EPI_ISL_806442, EPI_ISL_806455, EPI_ISL_806464, EPI_ISL_806491 | | | |
| see above | Alberta Precision Labs (APL) | Alberta Precision Labs (APL) | Gordon P, Lam LG, Pabbaraju K, Wong A, Ma R, Li V, Melin A, Tipples G, Berenger B, Zelyas N, Kellner J, Bernier F, Chui L, Croxen M |
| EPI_ISL_806527, EPI_ISL_806530, EPI_ISL_806531, EPI_ISL_806533, EPI_ISL_806539, EPI_ISL_806540, EPI_ISL_806541 | Charité Universitätsmedizin Berlin, Institut für Virologie/Labor Berlin | Charité Universitätsmedizin Berlin, Institut für Virologie | Victor M Corman, Jörn Beheim-Schwarzbach, Barbara Mühlemann, Julia Schneider, Talitha Veith, Cornelia Schlee, Tomasz Zemojtel, Terry Jones, Christian Drosten |
| EPI_ISL_812319, EPI_ISL_812324, EPI_ISL_812331, EPI_ISL_812334, EPI_ISL_812339 | United States Air Force School of Aerospace Medicine | United States Air Force School of Aerospace Medicine | Anthony Fries, Jennifer Meyer, Amanda Javorina, Sarah Purves, William Gruner, Clarise Starr, Elizabeth Macias |
| EPI_ISL_814076 | Ministry of Health Turkey | Ministry of Health Turkey | Fatma Bayrakdar, Yasemin Cogun, Süleyman Yalcin, Aye Baak Alta, Gülay Korukluolu |
| EPI_ISL_819318, EPI_ISL_819319 | Hospital Universitari Vall d'Hebron - Vall d'Hebron Institut de Rercerca | Hospital Universitari Vall d'Hebron | Cristina Andrés, Maria Piñana, Josep F Abril, Damir Garcia-Cehic, Ariadna Rando, Juliana Esperalba, Maria Gema Codina, Carla Castillo, Maria Carmen Martin, Tomàs Pumarola, Josep Quer, Andrés Antón |
| EPI_ISL_825033 | Center for Genome Sciences, US Army Medical Research Institute of Infectious Disease (USAMRIID) | Center for Genome Sciences, US Army Medical Research Institute of Infectious Disease (USAMRIID) | Kugelman,J. and Beitzel,B. |
| EPI_ISL_826581, EPI_ISL_826603, EPI_ISL_826604, EPI_ISL_826605, EPI_ISL_826606, EPI_ISL_826607, EPI_ISL_826608, EPI_ISL_826609, EPI_ISL_826621, EPI_ISL_826622, EPI_ISL_826623, EPI_ISL_826624, EPI_ISL_826625, EPI_ISL_826626, EPI_ISL_826627, EPI_ISL_826628, EPI_ISL_826637, EPI_ISL_826639, EPI_ISL_826666 | | | |
| see above | Montefiore Medical Center | Albert Einstein College of Medicine, Dept. of Microbiology & Immunology, Chandran lab | J. Maximilian Fels, Saad Khan, Ryan Forster, Karin A. Skalina, Surksha Sirichand, Amy S. Fox, Aviv Bergman, William B. Mitchell, Lucia R. Wolgast, Wendy Szymczak, Robert H. Bortz III, M. Eugenia Dieterle, Catalina Florez, Denise Haslwanter, Rohit K. Jangra, Ethan Laudermilch, Ariel S. Wirchnianski, Jason Barnhill, David L. Goldman, Hnin Khine, D. Yitzchak Goldstein, Johanna P. Daily, Kartik Chandran, Libusha Kelly |
| EPI_ISL_831040, EPI_ISL_831041 | Hospital Universitario La Paz (Madrid) | SeqCOVID-SPAIN consortium/IBV(CSIC) | María Rodríguez-Tejedor, Elias Dahdouh, Fernando Lázaro-Perona, Jesús Mingorance and SeqCOVID-SPAIN consortium |
| EPI_ISL_831902 | New Mexico Department of Health Scientific Laboratory | New Mexico Department of Health Scientific Laboratory | Ellie Johnson, Anastacia Griego-Fisher, D'eldra Malone |
| EPI_ISL_833513 | Veterinary Specialized Instute "Nis" | Veterinary Specialized Institute "Kraljevo", Serbia | Vidanovic,D., Tesovic,B., Manic,M., Petrovic,M.,Knezevic,A., Jovanovic,T., Jankovic,M., Sekler,M., Banovic Djeri,B., Petrovic,T., Volkening,J., Afonso,C. |
| EPI_ISL_833537, EPI_ISL_833538, EPI_ISL_833539, EPI_ISL_833540, EPI_ISL_833541 | Istituto Zooprofilattico Sperimentale del Mezzogiorno | TIGEM | Patrizia Annunziata, Andrea Ballabio, Valentina Bouche, Davide Cacchiarelli (CorrespAuthor), Pellegrino Cerino, Chiara Colantuono, Lucio Di Filippo, Antonio Grimaldi, Antonio Limone, Gabriella Loconte, Anna Manfredi, Francesco Panariello, Biancamaria Pierri, Marcello Salvi, Lucia Vassallo |
| EPI_ISL_833577 | Veterinary Specialized Instute "Nis" | Veterinary Specialized Institute "Nis", Serbia | Vidanovic,D., Tesovic,B., Manic,M., Petrovic,M.,Knezevic,A., Jovanovic,T., Jankovic,M., Sekler,M., Banovic Djeri,B., Petrovic,T., Volkening,J., Afonso,C. |
| EPI_ISL_837624, EPI_ISL_837625, EPI_ISL_837626, EPI_ISL_837627, EPI_ISL_837628, EPI_ISL_837629, EPI_ISL_837630, EPI_ISL_837631, EPI_ISL_837632, EPI_ISL_837633 | Instituto Nacional de Enfermedades Respiratorias (INER) | Instituto Nacional de Enfermedades Respiratorias (INER) | Celia Boukadida, Margarita Matías-Florentino, Alma Rincón-Rubio, Hector Esteban Paz-Juárez, Olivia Briceño, Edgar Sevilla-Reyes, Fidencio Mejía-Nepomuceno, Mario Mújica-Sánchez, Eduardo Becerril-Vargas, José Arturo Martínez-Orozco, Alejandra Hernández-Terán, Jorge Salas-Hernández, Santiago Ávila-Ríos, Joel Armando Vázquez-Pérez |
| EPI_ISL_842815, EPI_ISL_842816, EPI_ISL_842820, EPI_ISL_842828, EPI_ISL_842832 | Barts Health NHS Trust | COVID-19 Genomics UK (COG-UK) Consortium | CUTINO-MOGUEL, Maria-Teresa; HARRINGTON, David; OWOYEMI, Dola; SHYLINI, Raghavendran; BROAD, Claire; KELE, Beatrix |
| EPI_ISL_848088, EPI_ISL_848101, EPI_ISL_848138, EPI_ISL_848139, EPI_ISL_848140, EPI_ISL_848141, EPI_ISL_848142, EPI_ISL_848143, EPI_ISL_848144, EPI_ISL_848145, EPI_ISL_848150, EPI_ISL_848167, EPI_ISL_848168, EPI_ISL_848169, EPI_ISL_848170, EPI_ISL_848171 | | | |
| see above | National Virus Reference Laboratory | Irish Coronavirus Sequencing Consortium - National University of Ireland Galway | Kate Reddington, Grainne Mc Andrew, Simone Coughlan |
| EPI_ISL_848283, EPI_ISL_848457, EPI_ISL_848467 | Illinois Department of Public Health | Gagnon Lab, Southern Illinois University | Keith Gagnon |
| EPI_ISL_848595 | Evandro Chagas Institute | Evandro Chagas Institute | Santos, M.C.; Silva, A.M.; Junior, W.D.C.; Barbagelata, L.S.; Ferreira, J.A.; Sousa, E.M.A.; da Silva, P.S.; Pinheiro, K.C.; L.C.; Sousa Junior, E.C. |
| EPI_ISL_849689 | unknown | PHV-FSS | Son Nguyen et al. |
| EPI_ISL_852667, EPI_ISL_852762 | Institute of Virology, Medical Center, University of Freiburg, Freiburg, Germany | Institute of Virology, Clinial Virus Genomics, Medical Center, University of Freiburg, Freiburg, Germany | Jonas Fuchs, Lisa Kern, Sandra Reuter, Hajo Grundmann, Marcus Panning |
| EPI_ISL_854473 | Alberta Precision Labs (APL) | Alberta Precision Labs (APL) | Gordon P, Lam LG, Pabbaraju K, Wong A, Ma R, Li V, Melin A, Tipples G, Berenger B, Zelyas N, Kellner J, Bernier F, Chui L, Croxen M |
| EPI_ISL_855549 | Barts Health NHS Trust | Barts Health NHS Trust | CUTINO-MOGUEL, Maria-Teresa; HARRINGTON, David; OWOYEMI, Dola; KULASEGARAN-SHYLINI, Raghavendran; BROAD, Claire; KELE, Beatrix |
| EPI_ISL_860631, EPI_ISL_860632 | Ampath-Netcare | KRISP, KZn Research Innovation and Sequencing Platform | Giandhari J, Pillay S, Lessells R, Mdlalose K, York D, Khan S, Tegally H, Wilkinson E, de Oliveira T |
| EPI_ISL_861905 | LATE - Laboratório de Técnicas Especiais - Hospital Israelita Albert Einstein | LATE - Laboratório de Técnicas Especiais - Hospital Israelita Albert Einstein | Deyvid Amgarten, Fernanda de Mello Malta, Raquel Riyuzo, Ana Paula Moreira Salles, Pedro Henrique Sebe Rodrigues, João Renato Rebello Pinho |
| EPI_ISL_862842 | Fujita Health University Hospital | Fujita health University, School of Medicine, Department of Microbiology | Masahiro suzuki, Yohei Doi |
| EPI_ISL_862850 | Ehime Prefectural Central Hospital | Fujita health University, School of Medicine, Department of Microbiology | Masahiro suzuki, Yohei Doi |
| EPI_ISL_862852 | Tosei General Hospital | Fujita health University, School of Medicine, Department of Microbiology | Masahiro suzuki, Yohei Doi |
| EPI_ISL_872034, EPI_ISL_872035, EPI_ISL_872036 | Ospedale "Di Venere" | Beaconlab (Bioinformatics, Evolution and Comparative Genomics lab), Dept of Biosciences, University on Milan | Iacobellis M, d'Avenia M, Piluscio R, Parisi A, Chiara M, Manzari C, Pesole G |
| EPI_ISL_872090 | Instituto de Diagnostico y Referencia Epidemiologicos (INDRE) | Instituto de Diagnostico y Referencia Epidemiologicos (INDRE) | Claudia Wong-Arambula, Gisela Barrera-Badillo , Abril Rodriguez-Maldonado, Natividad Cruz-Ortiz, Tatiana Nunez-Garcia, Dayanira Arellano-Suarez, Fabiola Garces-Ayala, Adnan-Araiza Rodríguez, Edgar Mendieta-Condado, Lucia Hernandez-Rivas, Irma Lopez-Martinez, Ernesto Ramirez-Gonzalez. |
| EPI_ISL_872091 | Instituto de Diagnostico y Referencia Epidemiologicos (INDRE) | Instituto de Diagnostico y Referencia Epidemiologicos (INDRE) | Adnan-Araiza Rodríguez,Claudia Wong-Arambula, Gisela Barrera-Badillo , Abril Rodriguez-Maldonado, Natividad Cruz-Ortiz, Tatiana Nunez-Garcia, Dayanira Arellano-Suarez, Fabiola Garces-Ayala, Edgar Mendieta-Condado, Lucia Hernandez-Rivas, Irma Lopez-Martinez, Ernesto Ramirez-Gonzalez. |
| EPI_ISL_872093 | Instituto de Diagnostico y Referencia Epidemiologicos (INDRE) | Instituto de Diagnostico y Referencia Epidemiologicos (INDRE) | Edgar Mendieta-Condado, Ernesto Ramirez-Gonzalez, Adnan-Araiza Rodríguez,Claudia Wong-Arambula, Gisela Barrera-Badillo , Abril Rodriguez-Maldonado, Natividad Cruz-Ortiz, Tatiana Nunez-Garcia, Dayanira Arellano-Suarez, Fabiola Garces-Ayala, Lucia Hernandez-Rivas, Irma Lopez-Martinez. |

| | | | |
|---|---|---|---|
| EPI_ISL_876032 | Montefiore Medical Center | Albert Einstein College of Medicine, Dept. of Microbiology & Immunology, Chandran lab | J. Maximilian Fels, Saad Khan, Ryan Forster, Karin A. Skalina, Surksha Sirichand, Amy S. Fox, Aviv Bergman, William B. Mitchell, Lucia R. Wolgast, Wendy Szymczak, Robert H. Bortz III, M. Eugenia Dieterle, Catalina Florez, Denise Haslwanter, Rohit K. Jangra, Ethan Laudermilch, Ariel S. Wirchnianski, Jason Barnhill, David L. Goldman, Hnin Khine, D. Yitzchak Goldstein, Johanna P. Daily, Kartik Chandran, Libusha Kelly |
| EPI_ISL_876973, EPI_ISL_876974, EPI_ISL_877016, EPI_ISL_877017, EPI_ISL_877018, EPI_ISL_877019, EPI_ISL_877020, EPI_ISL_877021, EPI_ISL_877022, EPI_ISL_877023 | 1.AO Universitaria 'S. Giovanni di Dio e Ruggi D'Aragona, Scuola Medica Salernitana' Hospital / 2.UOC di Virologia e Microbiologia, Università della Campania 'L. Vanvitelli' / 3.AO Universitaria 'Federico II' Napoli Hospital / 4.AORN 'San Giuseppe Moscati' Avellino Hospital / 5.AO 'San Pio - presidio G. Rummo' Benevento Hospital / 6.AO 'Sant'Anna e San Sebastiano' Caserta Hospital / 7.PO 'Maria Santissima Addolorata' Eboli Hospital / 8.Biogem Istituto di Ricerche Genetiche | 1. Genome Research Center for Health (CRGS) / 2. Laboratory of Molecular Medicine and Genomics(LMMGe) / 3. Center for Research in Pure and Applied Mathematics (CRMPA) | Giorgio Giurato (Corresponding Author), Francesca Rizzo (Corresponding Author), Alessandro Weisz (Corresponding Author), Gianluigi Franci, Giovanni Nassa, Pasquale Pagliano, Roberta Tarallo, Elena Alexandrova, Ylenia D'Agostino, Carlo Ferravante, Jessica Lamberti, Viola Melone, Domenico Memoli, Valeria Mirici Cappa, Domenico Palumbo, Giovanni Pecoraro, Assunta Sellitto, Oriana Strianese, Ilaria Terenzi, Giuseppe Fenza, Aniello Gentile, Antonello Saccomanno, Sonia Amabile, Teresa Rocco, Annamaria Salvati, Emilia Vaccaro, Massimiliano Galdiero, Michele Cennamo, Giuseppe Portella, Maria Grazia Foti, Mariarosaria Ingino, Maria Landi, Maurizio Fumi, Vincenzo Rocco, Rita Greco, Vittoria Letizia, Arnolfo Petruzziello, Maddalena Schioppa, Gregorio Goffredi, Francesca Marciano, Michele Caraglia, Alessia Cossu, Marianna Scrima |
| EPI_ISL_877593, EPI_ISL_877594 | Clinical Molecular Microbiology Laboratory, UNC Hospital | Dirk Dittmer | Razia Moorad , Justin T. Landis , Brent A. Eason, Melissa B. Miller, Linda Pluta, Dirk Dittmer, Angelica Juarez, Cecilia Thompson , Cameroon Grant, Evelyn Hoffman, Patricio Cano, Jason Wong, Carolina Caro-Vegas, Blossom Damania. |
| EPI_ISL_878529, EPI_ISL_878531, EPI_ISL_878534, EPI_ISL_878537 | Robert Garry lab | Andersen lab at Scripps Research | Allison Smither, Gilberto Sabino-Santos, Patricia Snarski, Lilia Melnik, Antoinette Bell, Kaylynn Genemaras, Arnaud Drouin, Dahlene Fusco, Robert Garry with SEARCH Alliance San Diego |
| EPI_ISL_883037, EPI_ISL_883050, EPI_ISL_883051, EPI_ISL_883052, EPI_ISL_883053, EPI_ISL_883054, EPI_ISL_883055 | Kansas Health and Environmental Lab | Kansas Health and Environmental Lab | Mike Grose, Carissa Robertson, Ben Olsen, and Phil Adam |
| EPI_ISL_883164, EPI_ISL_883165 | Hannover Medical School, Institute of Virology | Hannover Medical School, Institute of Virology | Lars Steinbrück, Jasper Götting |
| EPI_ISL_886157, EPI_ISL_887107 | Institute of Medical Microbiology and Virology, University Hospital Carl Gustav Carus, TU Dresden | DRESDEN-concept Genome Center, CMCB, TU Dresden | Beil J., Brown T., Büttner L., Gscheidel N., Hochauf-Stange K., Klemroth S., Lindemann D., Mehnert G., Petzold A., Reinhardt S., Rost F., Sameith K., Winkler S. |
| EPI_ISL_887179, EPI_ISL_887202, EPI_ISL_887273, EPI_ISL_887279, EPI_ISL_887303, EPI_ISL_887305, EPI_ISL_887307, EPI_ISL_887308, EPI_ISL_887329, EPI_ISL_887333, EPI_ISL_887335, EPI_ISL_887337, EPI_ISL_887342, EPI_ISL_887348, EPI_ISL_887351, EPI_ISL_887366, EPI_ISL_887372, EPI_ISL_887376, EPI_ISL_887377, EPI_ISL_887385, EPI_ISL_887387, EPI_ISL_887391, EPI_ISL_887400, EPI_ISL_887402, EPI_ISL_887405, EPI_ISL_887407, EPI_ISL_887409, EPI_ISL_887415, EPI_ISL_887417 | | | |
| see above | Protzer Lab | Protzer Lab, Gagneur Lab, Robert Koch Institut | Ulrike Protzer, Dieter Hoffmann, Eva Schulte, Andrea Theumer, Oliver Drechsel, Max von Kleist,Aleksandar Radonic,Stephan Fuchs, Alexander Karollus, Julien Gagneur |
| EPI_ISL_888678 | Laboratory for Respiratory Viruses, Cantacuzino National Military-Medical Institute for Research and Development | Cantacuzino Institute Virology | Luiza Ustea, Nicoleta Paraschiv, Mihaela Lazar |
| EPI_ISL_889327 | The University Hospital Brno | Institute of Applied Biotechnologies a.s. | Petr Klempt, Ondej Brzo, Martin Kašný, Kateina Kvapilová, Martina Lengerová, Petr Kvapil |
| EPI_ISL_889384, EPI_ISL_889385, EPI_ISL_889386, EPI_ISL_889387, EPI_ISL_889388 | Olomouc University Hospital | Institute of Applied Biotechnologies a.s. | Petr Klempt, Ondej Brzo, Martin Kašný, Kateina Kvapilová, Marián Hajdúch, Petr Kvapil |
| EPI_ISL_889844, EPI_ISL_889846, EPI_ISL_889847, EPI_ISL_889917, EPI_ISL_889958, EPI_ISL_890028, EPI_ISL_890029 | Laboratoire de santé publique du Québec | Laboratoire de santé publique du Québec | Sandrine Moreira, Ioannis Ragoussis, Guillaume Bourque, Jesse Shapiro, Mark Lathrop and Michel Roger on behalf of the CoVSeQ research group |
| EPI_ISL_895746, EPI_ISL_895760, EPI_ISL_895767, EPI_ISL_895773, EPI_ISL_895797, EPI_ISL_895801, EPI_ISL_895802, EPI_ISL_895808, EPI_ISL_895846 | Molecular biology division, Institute of Clinical Biochemistry and Diagnostics, Charles University, Faculty of Medicine in Hradec Králové and University Hospital Hradec Králové | Molecular biology division, Institute of Clinical Biochemistry and Diagnostics, Charles University, Faculty of Medicine in Hradec Králové and University Hospital Hradec Králové | Helena Kovaíková, Petr Brož, Ivana Baranová, Kateina Hrochová, Tereza Baková, Jitka Novotná, Kateina Pehlíková, Vladimír Palika. Cooperation project with BioVendor-R&D and bioinformatics company BIOXSYS s r.o. |
| EPI_ISL_896123, EPI_ISL_896172, EPI_ISL_900088, EPI_ISL_900094, EPI_ISL_900126, EPI_ISL_900201, EPI_ISL_900357, EPI_ISL_900360, EPI_ISL_900397, EPI_ISL_900417, EPI_ISL_900474, EPI_ISL_900488 | | | |
| see above | MEPHI, Aix Marseille University | MEPHI, Aix Marseille University | Anthony LEVASSEUR |
| EPI_ISL_904012, EPI_ISL_904013, EPI_ISL_904014 | Veterinary Specialized Institute Nis | Veterinary Specialized Institute "Kraljevo", Serbia | Vidanovic,D., Tesovic,B., Knezevic,A., Jovanovic,T., Jankovic,M., Sekler,M., Banovic Djeri,B., Petrovic,M.M., Manic, M., Volkening,J., Afonso,C. |
| EPI_ISL_904020, EPI_ISL_904021, EPI_ISL_904022 | DB Diagnosticos do Brasil | Laboratório de Parasitologia Médica - Instituto de Medicina Tropical - Universidade de São Paulo | Brazil-UK Centre for Arbovirus Discovery Diagnosis Genomics and Epidemiology (CADDE) Genomic Network - Instituto de Medicina Tropical |
| EPI_ISL_906772, EPI_ISL_906773, EPI_ISL_906774 | Dr. Leonard A. Miller Centre for Health Services | National Microbiology Laboratory (NML) | Anna Majer, Shari Tyson, Grace Seo, Philip Mabon, Elsie Grudeski, Rhiannon Huzarewich, Russell Mandes, Anneliese Landgraff, Jennifer Tanner, Natalie Knox, Morag Graham, Gary Van Domselaar, Robert Needle, Yang Yu, Laura Gilbert, George Zahariadis, Nathalie Bastien, Yan Li, Timothy Booth, Darian Hole, Madison Chapel, Kerri Smith, CanCOGeN's metadata curation team, Public Health Agency of Canada CanCOGeN team |
| EPI_ISL_914985, EPI_ISL_914986, EPI_ISL_914987, EPI_ISL_914988, EPI_ISL_914989, EPI_ISL_914998, EPI_ISL_914999, EPI_ISL_915000, EPI_ISL_915001, EPI_ISL_915002, EPI_ISL_915003, EPI_ISL_915004, EPI_ISL_915005, EPI_ISL_915006 | | | |
| see above | QEII Health Sciences Centre | National Microbiology Laboratory (NML) | Anna Majer, Shari Tyson, Grace Seo, Philip Mabon, Elsie Grudeski, Rhiannon Huzarewich, Russell Mandes, Anneliese Landgraff, Jennifer Tanner, Natalie Knox, Morag Graham, Gary Van Domselaar, Todd Hatchette, Jason LeBlanc, Janice Pettipas, Dan Gaston, Nathalie Bastien, Yan Li, Timothy Booth, Darian Hole, Madison Chapel, CanCOGeN's metadata curation team, Public Health Agency of Canada CanCOGeN team |
| EPI_ISL_918128, EPI_ISL_918417, EPI_ISL_918422, EPI_ISL_918469 | 1.AO Universitaria 'S. Giovanni di Dio e Ruggi D'Aragona, Scuola Medica Salernitana' Hospital / 2.UOC di Virologia e Microbiologia, Università della Campania 'L. Vanvitelli' / 3.AO Universitaria 'Federico II' Napoli Hospital / 4.AORN 'San Giuseppe Moscati' Avellino Hospital / 5.AO 'San Pio - presidio G. Rummo' Benevento Hospital / 6.AO 'Sant'Anna e San Sebastiano' Caserta Hospital / 7.PO 'Maria Santissima Addolorata' Eboli Hospital / 8.Biogem Istituto di Ricerche Genetiche | 1. Genome Research Center for Health (CRGS) / 2. Laboratory of Molecular Medicine and Genomics(LMMGe) / 3. Center for Research in Pure and Applied Mathematics (CRMPA) | Giorgio Giurato (Corresponding Author), Francesca Rizzo (Corresponding Author), Alessandro Weisz (Corresponding Author), Gianluigi Franci, Giovanni Nassa, Pasquale Pagliano, Roberta Tarallo, Elena Alexandrova, Ylenia D'Agostino, Carlo Ferravante, Jessica Lamberti, Viola Melone, Domenico Memoli, Valeria Mirici Cappa, Domenico Palumbo, Giovanni Pecoraro, Assunta Sellitto, Oriana Strianese, Ilaria Terenzi, Giuseppe Fenza, Aniello Gentile, Antonello Saccomanno, Sonia Amabile, Teresa Rocco, Annamaria Salvati, Emilia Vaccaro, Massimiliano Galdiero, Michele Cennamo, Giuseppe Portella, Maria Grazia Foti, Mariarosaria Ingino, Maria Landi, Maurizio Fumi, Vincenzo Rocco, Rita Greco, Vittoria Letizia, Arnolfo Petruzziello, Maddalena Schioppa, Gregorio Goffredi, Francesca Marciano, Michele Caraglia, Alessia Cossu, Marianna Scrima |
| EPI_ISL_933620, EPI_ISL_933621, EPI_ISL_933622, EPI_ISL_933623 | Toronto Invasive Bacterial Diseases Network | McMaster University | Allison McGeer, Patryk Aftanas, Hooman Derakhshani, Angel Li, Kuganya Nirmalarajah, Emily Panousis, Ahmed Draia, Jalees Nasir, Michael Surette, Samira Mubareka, Andrew G. McArthur |
| EPI_ISL_937532, EPI_ISL_937535 | Thai Red Cross Emerging Infectious Diseases Health Science Centre, Chulalongkorn Hospital, Faculty of Medicine, Chulalongkorn University | Thai Red Cross Emerging Infectious Diseases Center and Faculty of Medicine, Chulalongkorn University | Opass Putcharoen, Watsamon Jantarabenjakul, Pattama Torvorapanit, Gompol Suwanpimolkul, Leilani Paitoonpong, Sopon Iamsirithaworn, Sininat Petcharat, Yutthana Joyjinda, Weenassarin Ampoot, Apaporn Rodpan, Thiravat Hemachudha, Supaporn Wacharapluesadee |
| EPI_ISL_939662, EPI_ISL_939663, EPI_ISL_939664, EPI_ISL_939665, EPI_ISL_939666, EPI_ISL_939667 | Division of Infectious Diseases, University Hospital Zürich | Institute of Medical Virology, University of Zurich | Stefan Schmutz, Maryam Zaheri, Verena Kufner, Annette Audigé, Maria Grünberg, Kevin Steiner, Jon Huder, Cyril Shah, Riccarda Capaul, Guido Bloemberg, Jürg Böni, Michael Huber, Alexandra Trkola |
| EPI_ISL_940180, EPI_ISL_940181, EPI_ISL_940182, EPI_ISL_940185, EPI_ISL_940194, EPI_ISL_940214, EPI_ISL_940224, EPI_ISL_940246, EPI_ISL_940260, EPI_ISL_940261, EPI_ISL_940264, EPI_ISL_940265, EPI_ISL_940280, EPI_ISL_940281, EPI_ISL_940282, EPI_ISL_940287, EPI_ISL_940288, EPI_ISL_940289, EPI_ISL_940290, EPI_ISL_940292, EPI_ISL_940293, EPI_ISL_940294, EPI_ISL_940295, EPI_ISL_940296, EPI_ISL_940297, EPI_ISL_940298, EPI_ISL_940299, EPI_ISL_940300, EPI_ISL_940301, EPI_ISL_940302, EPI_ISL_940303, EPI_ISL_940304, EPI_ISL_940305, EPI_ISL_940306, EPI_ISL_940307, EPI_ISL_940310, EPI_ISL_940311, EPI_ISL_940312, EPI_ISL_940313, EPI_ISL_940314, EPI_ISL_940315, EPI_ISL_940316, EPI_ISL_940318, EPI_ISL_940319, EPI_ISL_940371, EPI_ISL_940472, EPI_ISL_940473, EPI_ISL_940474, EPI_ISL_940475, EPI_ISL_940476, EPI_ISL_940478, EPI_ISL_940479, EPI_ISL_940480, EPI_ISL_940482, | | | |

| | | | |
|---|---|---|---|
| EPI_ISL_940484, EPI_ISL_940485, EPI_ISL_940492, EPI_ISL_940495 | | | |
| see above | Hôpital Bichat Claude Bernard, Laboratoire de Virologie | IAME UMR1137 Inserm, Université de Paris, Hôpital Bichat | Antoine Bridier-Nahmias, Amélie Recoing, Quentin Le Hingrat, Lena Daniel, Siham Hamri, Gilles Collin, Alexandre Storto, Mélanie Bertine, Charlotte Charpentier, Nadhira Houhou-Fidouh, Diane Descamps, Benoit Visseaux |
| EPI_ISL_941134 | Centro de Investigaciones en Microbiología y Biotecnología-UR (CIMBIUR), Facultad de Ciencias Naturales, Universidad del Rosario, Bogotá, Colombia | Centro de Investigaciones en Microbiología y Biotecnología-UR (CIMBIUR), Facultad de Ciencias Naturales, Universidad del Rosario, Bogotá, Colombia Icahn School of Medicine at Mount Sinai, New York, USA | Nathalia Ballesteros, Marina Muñoz, Luz Helena Patiño, Carolina Hernández, Felipe González-Casabianca, Iván Carroll, Mauricio Santos-Vega, Jaime Cascante, Andrés Angel, Alejandro Feged-Rivadeneira, Mónica Palma-Cuero, Carolina Flórez, Sergio Gomez, Adriana van de Guchte, Zenab Khan, Jayeeta Dutta, Hala Alejel Alshammary, Ana S. Gonzalez-Reiche, Matthew M. Hernandez, Emilia Mia Sordillo, Viviana Simon, Harm van Bakel, Alberto Paniz-Mondolfi, Juan David Ramírez |
| EPI_ISL_943620 | Utah Public Health Laboratory | Utah Public Health Laboratory | Erin L. Young, Kelly F. Oakeson, Tara Gallagher |
| EPI_ISL_954568, EPI_ISL_954569, EPI_ISL_954570, EPI_ISL_954571, EPI_ISL_954572, EPI_ISL_954573, EPI_ISL_954574, EPI_ISL_954575, EPI_ISL_954576, EPI_ISL_954577, EPI_ISL_954578, EPI_ISL_954579, EPI_ISL_954580, EPI_ISL_954581, EPI_ISL_954582, EPI_ISL_954583, EPI_ISL_954584, EPI_ISL_954585, EPI_ISL_954586, EPI_ISL_954587, EPI_ISL_954588, EPI_ISL_954589, EPI_ISL_954590, EPI_ISL_954591, EPI_ISL_954592, EPI_ISL_954593, EPI_ISL_954594, EPI_ISL_954595, EPI_ISL_954596, EPI_ISL_954597, EPI_ISL_954598, EPI_ISL_954599, EPI_ISL_954600, EPI_ISL_954601, EPI_ISL_954602, EPI_ISL_954603, EPI_ISL_954604, EPI_ISL_954605, EPI_ISL_954606, EPI_ISL_954607, EPI_ISL_954608, EPI_ISL_954609, EPI_ISL_954610, EPI_ISL_954612, EPI_ISL_954613, EPI_ISL_954614, EPI_ISL_954615, EPI_ISL_954616, EPI_ISL_954617, EPI_ISL_954618, EPI_ISL_954619, EPI_ISL_954620, EPI_ISL_954621, EPI_ISL_954622, EPI_ISL_954623, EPI_ISL_954624, EPI_ISL_954625, EPI_ISL_954626, EPI_ISL_954627, EPI_ISL_954628, EPI_ISL_954629, EPI_ISL_954630, EPI_ISL_954631, EPI_ISL_954632, EPI_ISL_954633, EPI_ISL_954634, EPI_ISL_954635, EPI_ISL_954636, EPI_ISL_954637, EPI_ISL_954638, EPI_ISL_954639, EPI_ISL_954640, EPI_ISL_954641, EPI_ISL_954642, EPI_ISL_954643, EPI_ISL_954644, EPI_ISL_954645, EPI_ISL_954646, EPI_ISL_954647, EPI_ISL_954648, EPI_ISL_954649, EPI_ISL_954650, EPI_ISL_954651, EPI_ISL_954652, EPI_ISL_954653, EPI_ISL_954654, EPI_ISL_954655, EPI_ISL_954656, EPI_ISL_954657, EPI_ISL_954658, EPI_ISL_954659, EPI_ISL_954660, EPI_ISL_954661, EPI_ISL_954662, EPI_ISL_954663, EPI_ISL_954664, EPI_ISL_954665, EPI_ISL_954666, EPI_ISL_954667, EPI_ISL_954668, EPI_ISL_954669, EPI_ISL_954670, EPI_ISL_954671, EPI_ISL_954672, EPI_ISL_954673, EPI_ISL_954674, EPI_ISL_954675, EPI_ISL_954676, EPI_ISL_954677, EPI_ISL_954678, EPI_ISL_954679, EPI_ISL_954680, EPI_ISL_954681, EPI_ISL_954682, EPI_ISL_954683, EPI_ISL_954684, EPI_ISL_954685, EPI_ISL_954686, EPI_ISL_954687, EPI_ISL_954688, EPI_ISL_954689, EPI_ISL_954690, EPI_ISL_954691, EPI_ISL_954692, EPI_ISL_954693, EPI_ISL_954694, EPI_ISL_954695, EPI_ISL_954696, EPI_ISL_954697, EPI_ISL_954698, EPI_ISL_954699, EPI_ISL_954700, EPI_ISL_954701, EPI_ISL_954702, EPI_ISL_954703, EPI_ISL_954704, EPI_ISL_954705, EPI_ISL_954706, EPI_ISL_954707, EPI_ISL_954708, EPI_ISL_954709, EPI_ISL_954710, EPI_ISL_954711, EPI_ISL_954712, EPI_ISL_954713, EPI_ISL_954714, EPI_ISL_954715, EPI_ISL_954716, EPI_ISL_954717, EPI_ISL_954718, EPI_ISL_954719, EPI_ISL_954720, EPI_ISL_954721, EPI_ISL_954722, EPI_ISL_954723, EPI_ISL_954724, EPI_ISL_954725, EPI_ISL_954726 | | | |
| see above | Laboratoire de santé publique du Québec | Laboratoire de santé publique du Québec | Sandrine Moreira, Ioannis Ragoussis, Guillaume Bourque, Jesse Shapiro, Mark Lathrop and Michel Roger on behalf of the CoVSeQ research group (http://covseq.ca/researchgroup) |
| EPI_ISL_955255 | HGSMF 26 CABO SAN LUCAS | BIOBANCO / COCTI | Borja-Aburto VH, Grajales-Muñiz C, Santacruz Tinoco CE, Rojas-Mendoza T, Ochoa Carrera LA, Sánchez A, Grande R, Isa P, Taboada B, López S, Arias C, Muñoz-Medina JE |
| EPI_ISL_955256 | HGZ 1 CARLOS MC GREGOR | BIOBANCO / COCTI | Borja-Aburto VH, Grajales-Muñiz C, Santacruz Tinoco CE, Rojas-Mendoza T, Ochoa Carrera LA, Sánchez A, Grande R, Isa P, Taboada B, López S, Arias C, Muñoz-Medina JE |
| EPI_ISL_959275 | National Influenza Center, Virology Department | National Influenza Center | V Salimi,NZ Shafiei Jandaghi, J Yavarian, A Nejati, K Sadeghi, N Ghavvami,F Ajaminejad and T Mokhtari Azad |
| EPI_ISL_959276 | National Influenza Center, Virology Department | National Influenza Center | NZ Shafiei Jandaghi, V Salimi, A Nejati, K Sadeghi, J Yavarian, N Ghavvami,F Ajaminejad and T Mokhtari Azad |
| EPI_ISL_959280 | National Influenza Center, Virology Department | National Influenza Center | J Yavarian, NZ Shafiei Jandaghi, V Salimi, A Nejati, K Sadeghi, F Ajaminejad,N Ghavvami and T Mokhtari Azad |
| EPI_ISL_959283 | National Influenza Center, Virology Department | National Influenza Center | NZ Shafiei Jandaghi, V Salimi, A Nejati, K Sadeghi, J Yavarian, F Ajaminejad,N Ghavvami and T Mokhtari Azad |
| EPI_ISL_961668 | Hôpital Georges L. Dumont | National Microbiology Laboratory (NML) | Anna Majer, Shari Tyson, Grace Seo, Philip Mabon, Elsie Grudeski, Rhiannon Huzarewich, Russell Mandes, Anneliese Landgraff, Jennifer Tanner, Natalie Knox, Morag Graham, Gary Van Domselaar, Richard Garceau, Guillaume Desnoyers, Nathalie Bastien, Yan Li, Timothy Booth, Darian Hole, Madison Chapel, Kirsten Biggar, CanCOGeN's metadata curation team, Public Health Agency of Canada CanCOGeN team |
| EPI_ISL_961759 | Viral Respiratory Lab, National Institute for Biomedical Research (INRB) | Pathogen Sequencing Lab, National Institute for Biomedical Research (INRB) | Placide Mbala-Kingebeni, Edith Nkwembe, Eddy Kinganda-Lusamaki, Amuri Aziza, Francisca Muyembe Mawete, Emmanuel Lokilo Lofiko, Jean-Claude Makangara Cigolo, Catherine Pratt, Matthias Pauthner, Josh Quick, Allison Black, James Hadfield, Trevor Bedford, Ian Goodfellow, Andrew Rambaut, Nick Loman, Kristian Andersen, Michael Wiley, Steve Ahuka-Mundeke, Jean-Jacques Muyembe Tamfum |
| EPI_ISL_965595, EPI_ISL_965596, EPI_ISL_965655, EPI_ISL_965693 | Dutch COVID-19 response team | Medical Microbiology, Maastricht University Medical Centre | Jozef Dingemans*, Brian van der Veer*, Erik Beuken, Carmen Reumkens, Lieke van Alphen, Christian Hoebe, Paul Savelkoul |
| EPI_ISL_968175 | Clinical Molecular Microbiology Laboratory, UNC Hospital | Dirk Dittmer | Justin T. Landis , Razia Moorad , Brent A. Eason, Melissa B. Miller, Linda Pluta, Dirk Dittmer, Angelica Juarez, Cecilia Thompson, Shawn Hawken, Cameroon Grant, Evelyn Hoffman, Patricio Cano, Jason Wong, Carolina Caro-Vegas, Ryan McNamara, Blossom Damania. |
