## Supplementary material for "Detection and characterization of the SARS-CoV-2 lineage B.1.526 in New York": Supp. Table 3 GISAID Acknowledgment part 2: gisaid_hcov-19_acknowledgement_table_2021_02_12_16-9.pdf

We gratefully acknowledge the following Authors from the Originating laboratories responsible for obtaining the specimens, as well as the Submitting laboratories where the genome data were generated and shared via GISAID, on which this research is based.

All Submitters of data may be contacted directly via www.gisaid.org

Authors are sorted alphabetically.

| Accession ID | Originating Laboratory | Submitting Laboratory | Authors |
|---|---|---|---|
| EPI_ISL_427148, EPI_ISL_427149 | Microbiological Diagnostic Unit Public Health Laboratory | Microbiological Diagnostic Unit Public Health Laboratory | Seemann T., Schultz M., Sait, M., Sherry, N. |
| EPI_ISL_427324, EPI_ISL_427325, EPI_ISL_427326, EPI_ISL_427327, EPI_ISL_427328, EPI_ISL_427329, EPI_ISL_427330, EPI_ISL_427331, EPI_ISL_427332, EPI_ISL_427333, EPI_ISL_427334, EPI_ISL_427335, EPI_ISL_427336 | | | |
| see above | WHO National Influenza Centre Russian Federation | WHO National Influenza Centre Russian Federation | Andrey Komissarov, Artem Fadeev, Mariia Sergeeva, Anna Ivanova, Daria Danilenko |
| EPI_ISL_427351, EPI_ISL_427357, EPI_ISL_427360, EPI_ISL_427361, EPI_ISL_427363, EPI_ISL_427364, EPI_ISL_427365, EPI_ISL_427366, EPI_ISL_427367, EPI_ISL_427368, EPI_ISL_427369, EPI_ISL_427370, EPI_ISL_427371, EPI_ISL_427372, EPI_ISL_427373, EPI_ISL_427374, EPI_ISL_427375, EPI_ISL_427376, EPI_ISL_427377, EPI_ISL_427378, EPI_ISL_427379, EPI_ISL_427380, EPI_ISL_427381, EPI_ISL_427382, EPI_ISL_427383, EPI_ISL_427384, EPI_ISL_427385, EPI_ISL_427386, EPI_ISL_427387, EPI_ISL_427388, EPI_ISL_427390 | | | |
| see above | Department of Clinical Microbiology | GIGA Medical Genomics | Keith Durkin, Maria Artesi, Sébastien Bontems, Raphaël Boreux, Cécile Meex, Pierrette Melin, Marie-Pierre Hayette, Vincent Bours. |
| EPI_ISL_427428, EPI_ISL_427429, EPI_ISL_427430, EPI_ISL_427431, EPI_ISL_427432, EPI_ISL_427433, EPI_ISL_427434, EPI_ISL_427435, EPI_ISL_427438, EPI_ISL_427439, EPI_ISL_427440, EPI_ISL_427442, EPI_ISL_427443, EPI_ISL_427444, EPI_ISL_427445, EPI_ISL_427446, EPI_ISL_427447, EPI_ISL_427448, EPI_ISL_427449, EPI_ISL_427452, EPI_ISL_427454, EPI_ISL_427455, EPI_ISL_427456, EPI_ISL_427458, EPI_ISL_427460, EPI_ISL_427461, EPI_ISL_427462 | | | |
| see above | University of Wisconsin-Madison AIDS Vaccine Research Laboratories | University of Wisconsin-Madison AIDS Vaccine Research Laboratories | Gage Moreno, Katarina Braun, et al. AIDS Vaccine Research Laboratories |
| EPI_ISL_427815 | WHO National Influenza Centre Russian Federation | WHO National Influenza Centre Russian Federation | Andrey Komissarov, Artem Fadeev, Anna Ivanova, Daria Danilenko |
| EPI_ISL_428394, EPI_ISL_428395, EPI_ISL_428396, EPI_ISL_428397, EPI_ISL_428398 | Yale COVID-19 Biorepository | Grubaugh Lab - Yale School of Public Health | Joseph Fauver, Tara Alpert, Anderson Brito, Anne Wyllie, Chantal Vogels, Mary Petrone, Chaney Kalinich, Isabel Ott, Arnau Casanovas, Catherine Muenker, Adam Moore, Alice Lu, Maria Tokuyama, Patrick Wong, Peiwen Lu, Saad Omer, Richard Martinello, Allison Nelson, Shelli Farhadian, Akiko Iwasaki, Charlese Dela Cruz, Albert Ko, Nathan Grubaugh |
| EPI_ISL_428403, EPI_ISL_428404, EPI_ISL_428405 | Yale COVID-19 Biorepository | Grubaugh Lab - Yale School of Public Health | Joseph Fauver, Tara Alpert, Anderson Brito, Anne Wyllie, Chantal Vogels, Mary Petrone, Cole Jensen, Chaney Kalinich, Isabel Ott, Arnau Casanovas, Catherine Muenker, Adam Moore, Alice Lu, Maria Tokuyama, Patrick Wong, Peiwen Lu, Saad Omer, Richard Martinello, Allison Nelson, Shelli Farhadian, Akiko Iwasaki, Charlese Dela Cruz, Albert Ko, Nathan Grubaugh |
| EPI_ISL_428482, EPI_ISL_428483, EPI_ISL_428484, EPI_ISL_428485 | District Surveillence Unit | Department of Neurovirology, National Institute of Mental Health and Neuroscience (NIMHANS) | Chitra Pattabiraman, Vijayalakshmi Reddy, Harsha PK, Risha Rasheed, Shafeeq S Hameed, Manjunatha Venkataswamy, Anita Desai, Ravi Vasanthapuram |
| EPI_ISL_428746, EPI_ISL_428748, EPI_ISL_428750, EPI_ISL_428751, EPI_ISL_428752 | Yale COVID-19 Biorepository | Grubaugh Lab - Yale School of Public Health | Joseph Fauver, Tara Alpert, Anderson Brito, Anne Wyllie, Chantal Vogels, Mary Petrone, Cole Jensen, Chaney Kalinich, Isabel Ott, Arnau Casanovas, Catherine Muenker, Adam Moore, Alice Lu, Maria Tokuyama, Patrick Wong, Peiwen Lu, Saad Omer, Richard Martinello, Allison Nelson, Shelli Farhadian, Akiko Iwasaki, Charlese Dela Cruz, Albert Ko, Nathan Grubaugh |
| EPI_ISL_428832, EPI_ISL_428833, EPI_ISL_428834, EPI_ISL_428835, EPI_ISL_428836, EPI_ISL_428837, EPI_ISL_428838, EPI_ISL_428839, EPI_ISL_428840, EPI_ISL_428841, EPI_ISL_428842, EPI_ISL_428843, EPI_ISL_428844, EPI_ISL_428845 | | | |
| see above | National Public Health Laboratory, National Centre for Infectious Diseases | National Public Health Laboratory, National Centre for Infectious Diseases | Mak TM, Octavia S, Chavatte JM, Cui L, Lin RTP |
| EPI_ISL_428957, EPI_ISL_428958 | Laboratoire National de Sante, Microbiology, Virology | Laboratoire National de Sante, Microbiology, Epidemiology and Microbial Genomics | Anke Wienecke-Baldacchino, Ardashel Latsuzbaia, Jessica Tapp, Catherine Ragimbeau, Guillaume Fournier, Tamir Abdelrahman, Trung Nguyen Nguyen, Joel Mossong |
| EPI_ISL_428992, EPI_ISL_428993, EPI_ISL_428995, EPI_ISL_429003, EPI_ISL_429009, EPI_ISL_429011, EPI_ISL_429014, EPI_ISL_429027, EPI_ISL_429031, EPI_ISL_429043, EPI_ISL_429044, EPI_ISL_429046, EPI_ISL_429047, EPI_ISL_429053, EPI_ISL_429054, EPI_ISL_429061, EPI_ISL_429062 | | | |
| see above | UCSF Clinical Microbiology Laboratory | Chan-Zuckerberg Biohub | CZB Cliahub Consortium |
| EPI_ISL_429206, EPI_ISL_429222, EPI_ISL_429223 | University Hospitals of Geneva Laboratory of Virology | University Hospitals of Geneva Laboratory of Virology | Laubscher F. |
| EPI_ISL_429634, EPI_ISL_429635, EPI_ISL_429636, EPI_ISL_429637, EPI_ISL_429638, EPI_ISL_429639, EPI_ISL_429640, EPI_ISL_429643, EPI_ISL_429644, EPI_ISL_429647 | UW Virology Lab | UW Virology Lab | Pavitra Roychoudhury, Hong Xie, Keith Jerome, Alexander Greninger |
| EPI_ISL_429706, EPI_ISL_429713, EPI_ISL_429714, EPI_ISL_429730, EPI_ISL_429739, EPI_ISL_429742, EPI_ISL_429764, EPI_ISL_429773, EPI_ISL_429785, EPI_ISL_429796 | Laboratoire National de Sante, Microbiology, Virology | Laboratoire National de Sante, Microbiology, Epidemiology and Microbial Genomics | Anke Wienecke-Baldacchino, Ardashel Latsuzbaia, Jessica Tapp, Catherine Ragimbeau, Guillaume Fournier, Tamir Abdelrahman, Trung Nguyen Nguyen, Joel Mossong |
| EPI_ISL_429843 | Gundersen Molecular Diagnostics Laboratory | Kabara Cancer Research Institute | Craig S. Richmond, Paraic A. Kenny |
| EPI_ISL_429844 | Gundersen Molecular Diagnostics Laboratory | Kabara Cancer Research Institute | Craig S. Richmond; Paraic A. Kenny |
| EPI_ISL_430010 | OSU Wexner Medical Center | James Molecular Lab - OSUWMC | Huolin Tu, Preeti Pancholi, Matt Avenarius, Erica Vincent, Matt Hunt, Dan Jones |
| EPI_ISL_430065, EPI_ISL_430066 | Geelong Centre for Emerging Infectious Diseases | Geelong Centre for Emerging Infectious Diseases | Chamings A., Bhatta T.R., Alexandersen S. |
| EPI_ISL_430067, EPI_ISL_430068, EPI_ISL_430069, EPI_ISL_430070, EPI_ISL_430071, EPI_ISL_430072, EPI_ISL_430073, EPI_ISL_430074, EPI_ISL_430075, EPI_ISL_430076, EPI_ISL_430077, EPI_ISL_430078, EPI_ISL_430079, EPI_ISL_430080, EPI_ISL_430081, EPI_ISL_430082, EPI_ISL_430083, EPI_ISL_430084, EPI_ISL_430085, EPI_ISL_430086, EPI_ISL_430087, EPI_ISL_430088, EPI_ISL_430089, EPI_ISL_430090, EPI_ISL_430091 | | | |
| see above | WHO National Influenza Centre Russian Federation | WHO National Influenza Centre Russian Federation | Andrey Komissarov, Artem Fadeev, Mariia Sergeeva, Anna Ivanova, Daria Danilenko |
| EPI_ISL_430323, EPI_ISL_430324, EPI_ISL_430325, EPI_ISL_430359, EPI_ISL_430360, EPI_ISL_430361, EPI_ISL_430362, EPI_ISL_430363, EPI_ISL_430364, EPI_ISL_430365, EPI_ISL_430366, EPI_ISL_430367, EPI_ISL_430368, EPI_ISL_430369, EPI_ISL_430370, EPI_ISL_430371, EPI_ISL_430372, EPI_ISL_430373, EPI_ISL_430374, EPI_ISL_430375, EPI_ISL_430376, EPI_ISL_430377, EPI_ISL_430378, EPI_ISL_430379, EPI_ISL_430380, EPI_ISL_430381, EPI_ISL_430382, EPI_ISL_430383, EPI_ISL_430384, EPI_ISL_430385, EPI_ISL_430386, EPI_ISL_430387, EPI_ISL_430388, EPI_ISL_430389, EPI_ISL_430390, EPI_ISL_430391, EPI_ISL_430392, EPI_ISL_430393, EPI_ISL_430394, EPI_ISL_430395, EPI_ISL_430396, EPI_ISL_430397, EPI_ISL_430398, EPI_ISL_430399, EPI_ISL_430400, EPI_ISL_430401, EPI_ISL_430402, EPI_ISL_430403, EPI_ISL_430404, EPI_ISL_430405, EPI_ISL_430406, EPI_ISL_430407, EPI_ISL_430408, EPI_ISL_430409, EPI_ISL_430410, EPI_ISL_430411, EPI_ISL_430412, EPI_ISL_430413, EPI_ISL_430414, EPI_ISL_430415, EPI_ISL_430423, EPI_ISL_430434 | | | |
| see above | NYU Langone Health | Departments of Pathology and Medicine, New York University School of Medicine | Maria Aguero-Rosenfeld, Brendan Belovarac, Margaret Black, Ludovic Boytard, John Cadley, Paolo Cotzia, John Chen, Dacia Dimartino, Xiaojun Feng, Tatyana Gindin, Emily Guzman, Adriana Heguy, Megan Hogan, Emily Huang, George Jour, Lawrence H. Lin, Raven Luther, Andrew Lytle, Christian Marier, Matthew T. Maurano, Mark J. Mulligan, Peter Meyn, Raquel Ordonez Ciriza, Iman Osman, Jared Pinnell, Vanessa Raabe, Sitharam Ramaswami, Amy Rapkiewicz, Andre M. Ribeiro-dos-Santos, Marie Samanovic-Golden, Antonio Serrano, Guomiao Shen, Matija Snuderl, Theodore Vougiouklakis, Nick Vulpescu, Gael Westby, Paul Zappile, Yutong Zhang |
| EPI_ISL_430567, EPI_ISL_430568, EPI_ISL_430569, EPI_ISL_430570, EPI_ISL_430571, EPI_ISL_430572, EPI_ISL_430573, EPI_ISL_430575, EPI_ISL_430576, EPI_ISL_430577, EPI_ISL_430582, EPI_ISL_430583, EPI_ISL_430584, EPI_ISL_430585, EPI_ISL_430586, EPI_ISL_430588, EPI_ISL_430589, EPI_ISL_430590, EPI_ISL_430591, EPI_ISL_430592, EPI_ISL_430593, EPI_ISL_430594, EPI_ISL_430595, EPI_ISL_430596, EPI_ISL_430598, EPI_ISL_430600, EPI_ISL_430601, EPI_ISL_430602, EPI_ISL_430603, EPI_ISL_430604, EPI_ISL_430605, EPI_ISL_430606, EPI_ISL_430607, EPI_ISL_430608, EPI_ISL_430609, EPI_ISL_430610, EPI_ISL_430611, EPI_ISL_430612, EPI_ISL_430613, EPI_ISL_430614, EPI_ISL_430615, EPI_ISL_430616, EPI_ISL_430617, EPI_ISL_430618, EPI_ISL_430619, EPI_ISL_430620, EPI_ISL_430621, EPI_ISL_430622, EPI_ISL_430623, EPI_ISL_430624, EPI_ISL_430625, EPI_ISL_430626, EPI_ISL_430627, EPI_ISL_430628, EPI_ISL_430629, EPI_ISL_430630, EPI_ISL_430637, EPI_ISL_430638 | | | |

| | | | |
|---|---|---|---|
| see above | Victorian Infectious Diseases Reference Laboratory (VIDRL) | Microbiological Diagnostic Unit Public Health Laboratory and Victorian Infectious Diseases Reference Laboratory, The Peter Doherty Institute for Infection and Immunity | Caly L., Seemann T., Sait, M., Schultz M., Druce J., Sherry, N. |
| EPI_ISL_430674, EPI_ISL_430675, EPI_ISL_430676, EPI_ISL_430677, EPI_ISL_430678, EPI_ISL_430679, EPI_ISL_430680, EPI_ISL_430681 | Microbiological Diagnostic Unit Public Health Laboratory | Microbiological Diagnostic Unit Public Health Laboratory | Seemann T., Schultz M., Sait, M., Sherry, N. |
| EPI_ISL_430688, EPI_ISL_430689, EPI_ISL_430690, EPI_ISL_430691, EPI_ISL_430693, EPI_ISL_430694, EPI_ISL_430695, EPI_ISL_430696, EPI_ISL_430700, EPI_ISL_430701, EPI_ISL_430702, EPI_ISL_430703, EPI_ISL_430704, EPI_ISL_430705, EPI_ISL_430709, EPI_ISL_430710, EPI_ISL_430711, EPI_ISL_430712, EPI_ISL_430713, EPI_ISL_430714 | | | |
| see above | Victorian Infectious Diseases Reference Laboratory (VIDRL) | Microbiological Diagnostic Unit Public Health Laboratory and Victorian Infectious Diseases Reference Laboratory, The Peter Doherty Institute for Infection and Immunity | Caly L., Seemann T., Sait, M., Schultz M., Druce J., Sherry, N. |
| EPI_ISL_430716, EPI_ISL_430717 | Microbiological Diagnostic Unit Public Health Laboratory | Microbiological Diagnostic Unit Public Health Laboratory | Seemann T., Schultz M., Sait, M., Sherry, N. |
| EPI_ISL_430810, EPI_ISL_430811, EPI_ISL_430812, EPI_ISL_430813 | Laboratorio de Virología del Hospital de Niños Dr. Ricardo Gutierrez | Área de Secuenciación del Laboratorio de Virología del Hospital de Niños Dr. Ricardo Gutierrez on behalf of 'Proyecto Argentino Interinstitucional de genomica de SARS-CoV-2' (PAIS Consortium) | Nabaes Jodar, MS; Goya, S; Natale, MI; Lusso, S; Gravis, E; Mistchenko, AS; Valinotto, LE; Viegas, M. |
| EPI_ISL_430907, EPI_ISL_430908, EPI_ISL_430909, EPI_ISL_430910, EPI_ISL_430911, EPI_ISL_430912, EPI_ISL_430916, EPI_ISL_430920, EPI_ISL_430924, EPI_ISL_430928, EPI_ISL_430929, EPI_ISL_430930, EPI_ISL_430931, EPI_ISL_430932 | | | |
| see above | UW Virology Lab | UW Virology Lab | Pavitra Roychoudhury, Hong Xie, Keith Jerome, Alexander Greninger |
| EPI_ISL_431011, EPI_ISL_431012 | Viral Respiratory Lab, National Institute for Biomedical Research (INRB) | Pathogen Sequencing Lab, National Institute for Biomedical Research (INRB) | Placide Mbala-Kingebeni, Edith Nkwembe, Eddy Kinganda-Lusamaki, Amuri Aziza, Francisca Muyembe Mawete, Catherine Pratt, Matthias Pauthner, Josh Quick, Allison Black, James Hadfield, Trevor Bedford, Ian Goodfellow, Andrew Rambaut, Nick Loman, Kristian Andersen, Michael Wiley, Steve Ahuka-Mundeke, Jean-Jacques Muyembe Tamfum |
| EPI_ISL_431082, EPI_ISL_431083, EPI_ISL_431084, EPI_ISL_431085, EPI_ISL_431086, EPI_ISL_431088, EPI_ISL_431095 | Yale COVID-19 Biorepository | Grubaugh Lab - Yale School of Public Health | Joseph Fauver, Tara Alpert, Anderson Brito, Anne Wyllie, Chantal Vogels, Mary Petrone, Cole Jensen, Chaney Kalinich, Isabel Ott, Arnau Casanovas, Catherine Muenker, Adam Moore, Alice Lu, Maria Tokuyama, Patrick Wong, Peiwen Lu, Saad Omer, Richard Martinello, Allison Nelson, Shelli Farhadian, Akiko Iwasaki, Charlese Dela Cruz, Albert Ko, Nathan Grubaugh |
| EPI_ISL_431904, EPI_ISL_431907, EPI_ISL_431928, EPI_ISL_431945, EPI_ISL_431948, EPI_ISL_431981, EPI_ISL_431984, EPI_ISL_431986, EPI_ISL_431997, EPI_ISL_432005, EPI_ISL_432009, EPI_ISL_432019, EPI_ISL_432026, EPI_ISL_432027, EPI_ISL_432038, EPI_ISL_432042, EPI_ISL_432043, EPI_ISL_432044, EPI_ISL_432055, EPI_ISL_432058, EPI_ISL_432071, EPI_ISL_432077, EPI_ISL_432105, EPI_ISL_432115, EPI_ISL_432127, EPI_ISL_432162, EPI_ISL_432174, EPI_ISL_432191, EPI_ISL_432213, EPI_ISL_432225, EPI_ISL_432228, EPI_ISL_432241, EPI_ISL_432246, EPI_ISL_432261, EPI_ISL_432263, EPI_ISL_432264, EPI_ISL_432265, EPI_ISL_432268, EPI_ISL_432269, EPI_ISL_432271, EPI_ISL_432272, EPI_ISL_432273, EPI_ISL_432275, EPI_ISL_432276, EPI_ISL_432280, EPI_ISL_432281, EPI_ISL_432283, EPI_ISL_432284, EPI_ISL_432286, EPI_ISL_432291, EPI_ISL_432295, EPI_ISL_432296, EPI_ISL_432300, EPI_ISL_432304, EPI_ISL_432305, EPI_ISL_432309, EPI_ISL_432315, EPI_ISL_432318, EPI_ISL_432324, EPI_ISL_432332, EPI_ISL_432354, EPI_ISL_432358, EPI_ISL_432364, EPI_ISL_432378, EPI_ISL_432382, EPI_ISL_432392, EPI_ISL_432404, EPI_ISL_432418, EPI_ISL_432439, EPI_ISL_432444, EPI_ISL_432449 | | | |
| see above | Wales Specialist Virology Centre | Public Health Wales Microbiology Cardiff | Catherine Moore, Johnathan Evans, Malorie Perry, Simon Cottrell, Alec Birchley, Alexander Adams, Amy Gaskin, Bree Gatica-Wilcox, Jason Coombes, Lauren Gilbert, Lee Graham, Nicole Pacchiarini, Sara Kumziene-Summerhayes, Sarah Taylor, Sophie Jones, Sara Rey, Matthew Bull, Joanne Watkins, Sally Corden, Tom Connor |
| EPI_ISL_432478, EPI_ISL_432544, EPI_ISL_432581, EPI_ISL_432591, EPI_ISL_432594, EPI_ISL_432623, EPI_ISL_432647 | Virology Department, Sheffield Teaching Hospitals NHS Foundation Trust / Virology Department, Sheffield Teaching Hospitals NHS Foundation Trust | COVID-19 Genomics UK (COG-UK) Consortium | Thushan de Silva, Matthew Parker,Adri Angyal, Rebecca Brown, Luke Green, Rachel Tucker, Paul Parsons, Danielle Groves, Alex Keeley, Dave Partridge, Matthew Wyles, Benjamin Lindsey, Mehmet Yavuz, Mohammad Raza, Cariad Evans |
| EPI_ISL_432681, EPI_ISL_432682, EPI_ISL_432683, EPI_ISL_432684, EPI_ISL_432685, EPI_ISL_432686, EPI_ISL_432687, EPI_ISL_432688, EPI_ISL_432689, EPI_ISL_432690, EPI_ISL_432691, EPI_ISL_432692, EPI_ISL_432693, EPI_ISL_432694, EPI_ISL_432695, EPI_ISL_432696, EPI_ISL_432697, EPI_ISL_432698, EPI_ISL_432699, EPI_ISL_432701, EPI_ISL_432702, EPI_ISL_432703, EPI_ISL_432704, EPI_ISL_432705, EPI_ISL_432706, EPI_ISL_432708, EPI_ISL_432709 | | | |
| see above | Queens Medical Centre, Clinical Microbiology Department / DeepSeq Nottingham | COVID-19 Genomics UK (COG-UK) Consortium | Gemma Clark, Wendy Smith, Manjinder Khakh, Hannah Howson-Wells, Jonathan Ball, Patrick McClure, Joseph Chappell, Theocharis Tsoleridis, Nadine Holmes, Matthew Carlisle, Christopher Moore, Fei Sang, Johnny Debebe, Victoria Wright, Matthew Loose |
| EPI_ISL_432722, EPI_ISL_432723, EPI_ISL_432841, EPI_ISL_432853, EPI_ISL_432865 | Virology Department, Sheffield Teaching Hospitals NHS Foundation Trust / Virology Department, Sheffield Teaching Hospitals NHS Foundation Trust | COVID-19 Genomics UK (COG-UK) Consortium | Thushan de Silva, Matthew Parker,Adri Angyal, Rebecca Brown, Luke Green, Rachel Tucker, Paul Parsons, Danielle Groves, Alex Keeley, Dave Partridge, Matthew Wyles, Benjamin Lindsey, Mehmet Yavuz, Mohammad Raza, Cariad Evans |
| EPI_ISL_433068, EPI_ISL_433072, EPI_ISL_433075, EPI_ISL_433077, EPI_ISL_433078, EPI_ISL_433079, EPI_ISL_433081, EPI_ISL_433082, EPI_ISL_433084 | Virology Department, Royal Infirmary of Edinburgh, NHS Lothian / School of Biological Sciences, University of Edinburgh / Institute of Genetics and Molecular Medicine, University of Edinburgh | COVID-19 Genomics UK (COG-UK) Consortium | McHugh M, Dewar R, Rooke S, Gallagher M, Balcaza C, O'Toole A, Hill V, McCrone JT, Colquhoun R, Yu X, Jackson B, Rambaut A, Williams TC, Templeton K |
| EPI_ISL_433268, EPI_ISL_433269, EPI_ISL_433270, EPI_ISL_433271, EPI_ISL_433272 | West of Scotland Specialist Virology Centre, NHSGGC / MRC-University of Glasgow Centre for Virus Research | COVID-19 Genomics UK (COG-UK) Consortium | Ana da Silva Filipe, Natasha Johnson, Kathy Smollett, Daniel Mair, Stephen Carmichael, Lily Tong, Jenna Nichols, Elihu Aranday-Cortes, Kirstyn Brunker, Yasmin Parr, Kyriaki Nomikou; Sarah McDonald, Marc Niebel, Patawee Asamaphan; Richard Orton, Joseph Hughes, Sreenu Vattipally, David L Robertson; Alasdair MacLean, Rory Gunson; Kathy Li, Natasha Jesudason, Rajiv Shah, James Shepherd, Antonia Ho, Emma Thomson |
| EPI_ISL_433384, EPI_ISL_433385, EPI_ISL_433386, EPI_ISL_433387, EPI_ISL_433388, EPI_ISL_433389, EPI_ISL_433390, EPI_ISL_433391, EPI_ISL_433392, EPI_ISL_433393, EPI_ISL_433394, EPI_ISL_433395, EPI_ISL_433396, EPI_ISL_433397, EPI_ISL_433398, EPI_ISL_433404, EPI_ISL_433408, EPI_ISL_433409, EPI_ISL_433410, EPI_ISL_433411, EPI_ISL_433412, EPI_ISL_433413, EPI_ISL_433414, EPI_ISL_433415, EPI_ISL_433416, EPI_ISL_433417, EPI_ISL_433418, EPI_ISL_433419, EPI_ISL_433420, EPI_ISL_433421, EPI_ISL_433422, EPI_ISL_433423, EPI_ISL_433424, EPI_ISL_433425, EPI_ISL_433426, EPI_ISL_433427, EPI_ISL_433428, EPI_ISL_433429, EPI_ISL_433430, EPI_ISL_433431, EPI_ISL_433432, EPI_ISL_433434, EPI_ISL_433435, EPI_ISL_433436, EPI_ISL_433437, EPI_ISL_433438, EPI_ISL_433439, EPI_ISL_433440, EPI_ISL_433441, EPI_ISL_433442, EPI_ISL_433443, EPI_ISL_433444, EPI_ISL_433445, EPI_ISL_433446, EPI_ISL_433447, EPI_ISL_433448, EPI_ISL_433451, EPI_ISL_433455, EPI_ISL_433456, EPI_ISL_433457, EPI_ISL_433458, EPI_ISL_433459, EPI_ISL_433460, EPI_ISL_433461, EPI_ISL_433463, EPI_ISL_433465 | | | |
| see above | Virology Department, Royal Infirmary of Edinburgh, NHS Lothian / School of Biological Sciences, University of Edinburgh / Institute of Genetics and Molecular Medicine, University of Edinburgh | COVID-19 Genomics UK (COG-UK) Consortium | McHugh M, Dewar R, Rooke S, Gallagher M, Balcaza C, O'Toole A, Hill V, McCrone JT, Colquhoun R, Yu X, Jackson B, Rambaut A, Williams TC, Templeton K |
| EPI_ISL_433659, EPI_ISL_433660, EPI_ISL_433661, EPI_ISL_433662, EPI_ISL_433663, EPI_ISL_433664, EPI_ISL_433665 | West of Scotland Specialist Virology Centre, NHSGGC / MRC-University of Glasgow Centre for Virus Research | COVID-19 Genomics UK (COG-UK) Consortium | Ana da Silva Filipe, Natasha Johnson, Kathy Smollett, Daniel Mair, Stephen Carmichael, Lily Tong, Jenna Nichols, Elihu Aranday-Cortes, Kirstyn Brunker, Yasmin Parr, Kyriaki Nomikou; Sarah McDonald, Marc Niebel, Patawee Asamaphan; Richard Orton, Joseph Hughes, Sreenu Vattipally, David L Robertson; Alasdair MacLean, Rory Gunson; Kathy Li, Natasha Jesudason, Rajiv Shah, James Shepherd, Antonia Ho, Emma Thomson |
| EPI_ISL_433772, EPI_ISL_433774, EPI_ISL_433775, EPI_ISL_433776, EPI_ISL_433777, EPI_ISL_433778, EPI_ISL_433779, EPI_ISL_433784, EPI_ISL_433786, EPI_ISL_433789, EPI_ISL_433790, EPI_ISL_433791, EPI_ISL_433792, EPI_ISL_433795, EPI_ISL_433796, EPI_ISL_433797, EPI_ISL_433798, EPI_ISL_433799, EPI_ISL_433803, EPI_ISL_433805, EPI_ISL_433809, EPI_ISL_433812, EPI_ISL_433826, EPI_ISL_433827, EPI_ISL_433828, EPI_ISL_433829, EPI_ISL_433830, EPI_ISL_433831, EPI_ISL_433832, EPI_ISL_433833, EPI_ISL_433834, EPI_ISL_433835, EPI_ISL_433836, EPI_ISL_433837, EPI_ISL_433839, EPI_ISL_433840, EPI_ISL_433841, EPI_ISL_433843, EPI_ISL_433844, EPI_ISL_433845, EPI_ISL_433846, EPI_ISL_433847, EPI_ISL_433849, EPI_ISL_433851, EPI_ISL_433852, EPI_ISL_433853, EPI_ISL_433854, EPI_ISL_433855, EPI_ISL_433856, EPI_ISL_433857, EPI_ISL_433858, EPI_ISL_433859, EPI_ISL_433860, EPI_ISL_433861, EPI_ISL_433863, EPI_ISL_433864, EPI_ISL_433865, EPI_ISL_433866, EPI_ISL_433867, EPI_ISL_433868, EPI_ISL_433869, EPI_ISL_433870, EPI_ISL_433871, EPI_ISL_433872, EPI_ISL_433873, EPI_ISL_433874, EPI_ISL_433875, EPI_ISL_433876, EPI_ISL_433877, EPI_ISL_433878, EPI_ISL_433879, EPI_ISL_433880, EPI_ISL_433881, EPI_ISL_433882, EPI_ISL_433883, EPI_ISL_433884, EPI_ISL_433885, EPI_ISL_433886, EPI_ISL_433887, EPI_ISL_433888, EPI_ISL_433889, EPI_ISL_433890, EPI_ISL_433891, EPI_ISL_433892, EPI_ISL_433893, EPI_ISL_433894, EPI_ISL_433895, EPI_ISL_433896, EPI_ISL_433897, EPI_ISL_433898, EPI_ISL_433899, EPI_ISL_433900, EPI_ISL_433901, EPI_ISL_433902, EPI_ISL_433903, EPI_ISL_433904, EPI_ISL_433905, EPI_ISL_433906, EPI_ISL_433907, EPI_ISL_433908, EPI_ISL_433909, EPI_ISL_433910, EPI_ISL_433911, EPI_ISL_433912, EPI_ISL_433913, EPI_ISL_433914, EPI_ISL_433915, EPI_ISL_433916, EPI_ISL_433917, EPI_ISL_433918, EPI_ISL_433919, EPI_ISL_433920, EPI_ISL_433921, EPI_ISL_433922, EPI_ISL_433923, EPI_ISL_433924, EPI_ISL_433925, EPI_ISL_433926, EPI_ISL_433927, EPI_ISL_433928, EPI_ISL_433929, EPI_ISL_433930, EPI_ISL_433931, EPI_ISL_433932, EPI_ISL_433933, EPI_ISL_433934, EPI_ISL_433935, EPI_ISL_433936, EPI_ISL_433937, EPI_ISL_433938, EPI_ISL_433939, EPI_ISL_433940, EPI_ISL_433941, EPI_ISL_433942, EPI_ISL_433943, EPI_ISL_433944, EPI_ISL_433945, EPI_ISL_433946, EPI_ISL_433947, EPI_ISL_433948, EPI_ISL_433949, EPI_ISL_433950, EPI_ISL_433951, EPI_ISL_433952, EPI_ISL_433954, EPI_ISL_433957, EPI_ISL_433959, EPI_ISL_433960, EPI_ISL_433961, EPI_ISL_433962, EPI_ISL_433963, EPI_ISL_433964, EPI_ISL_433965, EPI_ISL_433966, EPI_ISL_433967, EPI_ISL_433968, EPI_ISL_433969, EPI_ISL_433979, EPI_ISL_433987, EPI_ISL_433993, EPI_ISL_433994, EPI_ISL_434002, EPI_ISL_434014 | | | |
| see above | Department of Pathology, University of Cambridge | COVID-19 Genomics UK (COG-UK) Consortium | Luke W Meredith, M. Estee Torok , Myra Hosmillo, William L. Hamilton, Martin D. Curran, Theresa Feltwell, Grant Hall, Anna Yakovleva, Fahad A Khokhar, Charlotte J. Houldcroft, Laura G Caller, Aminu S. Jahun, Sarah L. Caddy, Ian Goodfellow |

| | | | |
|---|---|---|---|
| EPI_ISL_434374, EPI_ISL_434377, EPI_ISL_434382, EPI_ISL_434383 | Hospital AZ Rivierenland | Institute of Tropical Medicine | Philippe Selhorst, Colin Anthony, |
| EPI_ISL_434385, EPI_ISL_434386 | Hospital AZ Rivierenland | Institute of Tropical Medicine | Philippe Selhorst, Colin Anthony |
| EPI_ISL_434490, EPI_ISL_434492, EPI_ISL_434493, EPI_ISL_434494, EPI_ISL_434495, EPI_ISL_434496, EPI_ISL_434497, EPI_ISL_434503, EPI_ISL_434505, EPI_ISL_434507, EPI_ISL_434508, EPI_ISL_434509 | | | |
| see above | Laboratoire National de Sante, Microbiology, Virology | Laboratoire National de Sante, Microbiology, Epidemiology and Microbial Genomics | Anke Wienecke-Baldacchino, Ardashel Latsuzbaia, Jessica Tapp, Catherine Ragimbeau, Guillaume Fournier, Tamir Abdelrahman, Trung Nguyen Nguyen, Joel Mossong |
| EPI_ISL_434590, EPI_ISL_434591, EPI_ISL_434592, EPI_ISL_434593, EPI_ISL_434594, EPI_ISL_434595, EPI_ISL_434596, EPI_ISL_434601, EPI_ISL_434602, EPI_ISL_434603, EPI_ISL_434604, EPI_ISL_434605, EPI_ISL_434606 | | | |
| see above | Virginia DCLS | Virginia DCLS | Virginia DCLS |
| EPI_ISL_434608, EPI_ISL_434609, EPI_ISL_434610, EPI_ISL_434611, EPI_ISL_434612, EPI_ISL_434613 | University of Wisconsin-Madison AIDS Vaccine Research Laboratories | University of Wisconsin-Madison AIDS Vaccine Research Laboratories | Gage Moreno, Katarina Braun, et al. AIDS Vaccine Research Laboratories |
| EPI_ISL_434628, EPI_ISL_434629, EPI_ISL_434630, EPI_ISL_434631, EPI_ISL_434632, EPI_ISL_434633, EPI_ISL_434634, EPI_ISL_434635 | CHU Purpan - Laboratoire de Virologie - Institut Fédératif de Biologie | Laboratoire de virologie - École Nationale Vétérinaire de Toulouse | Guillaume Croville, Jean-Luc Guérin, Jacques Izopet |
| EPI_ISL_434661, EPI_ISL_434662 | Omtanken Grimmered | The Public Health Agency of Sweden | Bernd Sengpiel, Oskar Karlsson Lindsjo, Maria Lind Karlberg, Anna-Malin Linde, Olov Svartstrom, Anna Risberg, Theresa Enkirch, Mia Brytting, Karin Tegmark-Wisell |
| EPI_ISL_434666 | Hornefors Halsocentral | The Public Health Agency of Sweden | Camilla Eiback, Oskar Karlsson Lindsjo, Maria Lind Karlberg, Anna-Malin Linde, Olov Svartstrom, Anna Risberg, Theresa Enkirch, Mia Brytting, Karin Tegmark-Wisell |
| EPI_ISL_434678, EPI_ISL_434679, EPI_ISL_434680, EPI_ISL_434681 | Viral Respiratory Lab, National Institute for Biomedical Research (INRB) | Pathogen Sequencing Lab, National Institute for Biomedical Research (INRB) | Placide Mbala-Kingebeni; Edith Nkwembe; Eddy Kinganda-Lusamaki; Amuri Aziza; Francisca Muyembe Mawete; Catherine Pratt; Matthias Pauthner; Josh Quick; Allison Black; James Hadfield; Trevor Bedford; Ian Goodfellow; Andrew Rambaut; Nick Loman; Kristian Andersen; Michael Wiley; Steve Ahuka-Mundeke; Jean-Jacques Muyembe Tamfum |
| EPI_ISL_434710, EPI_ISL_434711, EPI_ISL_435032, EPI_ISL_435033 | Viral Respiratory Lab, National Institute for Biomedical Research (INRB) | Pathogen Sequencing Lab, National Institute for Biomedical Research (INRB) | Placide Mbala-Kingebeni, Edith Nkwembe, Eddy Kinganda-Lusamaki, Adrienne Amuri Aziza, Francisca Muyembe Mawete, Catherine Pratt, Matthias Pauthner, Josh Quick, Allison Black, James Hadfield, Trevor Bedford, Ian Goodfellow, Andrew Rambaut, Nick Loman, Kristian Andersen, Michael Wiley, Steve Ahuka-Mundeke, Jean-Jacques Muyembe Tamfum |
| EPI_ISL_435034 | LSUHS Emerging Viral Threat Laboratory | Microbial Genome Sequencing Center | Jeremy P. Kamil, John A. Vanchiere, Rona S. Scott, Camille F. Abshire, Abida Siddiqa, Byeong-Jae Lee, Chan-ki Min, Md Maksudul Alam, Monica Gestal-Cartele, Edna Ondari, Adam Greer, Malgorzata Bienkowska-Haba, Katarzyna Zwolinska, Michelle M. Arnold, Jason M. Bodily, Andrew D. Yurochko, Paul M. Weinberger, Christopher G. Kevil, Martin J. Sapp, Daniel J. Snyder, Vaughn S. Cooper |
| EPI_ISL_435035, EPI_ISL_435036 | LSUHS Emerging Viral Threat Laboratory | Microbial Genome Sequencing Center | Jeremy P. Kamil, John A. Vanchiere, Rona S. Scott, Camille F. Abshire, Abida Siddiqa, Byeong-Jae Lee, Chan-ki Min, Md Maksudul Alam, Monica Gestal-Cartele, Edna Ondari, Adam Greer, Malgorzata Bienkowska-Haba, Katarzyna Zwolinska, Jason M. Bodily, Andrew D. Yurochko, Paul M. Weinberger, Christopher G. Kevil, Martin J. Sapp, Daniel J. Snyder, Vaughn S. Cooper |
| EPI_ISL_435053 | B.J. Medical College and Civil hospital | Gujarat Biotechnology Research Centre | Janvi Raval, Monika Gandhi, Pinal Trivedi, Maharshi Pandya, Amit Kanani, Akanksha Verma, Nitin Savaliya, Raghawendra Kumar, Dinesh Kumar, Zuber Saiyed, Dipa Kinariwala, Disha Patel, Binita Aring, Geeta Vaghela, Sonia Barve, Bhavesh Modi, Kairavi Joshi, Gaurishankar Shrimali, Nidhi Sood, Pranay Shah, R D Dixit, Snehal Bagatharia, Kamlesh J Upadhyay, Ramesh Pandit, Tejas Shah, Ankit Hinsu, Pritesh Sabara, Apurvasinh Puvar, Nidhi Patel, Chaitanya Joshi, Madhvi Joshi |
| EPI_ISL_435057 | T.C. Salk Bakanl Adyaman l Salk Müdürlüü Adyaman Eitim Ve Aratrma Hastanesi | VETAL Animal Health Products Company, BSL3+ Production Laboratuary, Turkey | Fatma Nilay Tutak, Haluk Uluca, Fethiye Sevimli, O. Ugur Sezerman |
| EPI_ISL_435113, EPI_ISL_435114, EPI_ISL_435116, EPI_ISL_435117, EPI_ISL_435118 | Viral Respiratory Lab, National Institute for Biomedical Research (INRB) | Pathogen Sequencing Lab, National Institute for Biomedical Research (INRB) | Placide Mbala-Kingebeni, Edith Nkwembe, Eddy Kinganda-Lusamaki, Adrienne Amuri Aziza, Francisca Muyembe Mawete, Catherine Pratt, Matthias Pauthner, Josh Quick, Allison Black, James Hadfield, Trevor Bedford, Ian Goodfellow, Andrew Rambaut, Nick Loman, Kristian Andersen, Michael Wiley, Steve Ahuka-Mundeke, Jean-Jacques Muyembe Tamfum |
| EPI_ISL_435146, EPI_ISL_435147 | Villa Serena del Dr. Leonardo Petruzzi | Istituto Zooprofilattico Sperimentale dell'Abruzzo e Molise "G.Caporale" | Lorusso A, Marcacci M, Di Domenico M, Ancora M, Curini V, Mangone I, Rinaldi A, Di Pasquale A, Cammà C, Puglia I, Savini G |
| EPI_ISL_435148 | Ospedale SS Annunziata | Istituto Zooprofilattico Sperimentale dell'Abruzzo e Molise "G.Caporale" | Lorusso A, Marcacci M, Di Domenico M, Ancora M, Curini V, Mangone I, Rinaldi A, Di Pasquale A, Cammà C, Puglia I, Savini G |
| EPI_ISL_435149 | SERVIZIO DI IGIENE E SANITÀ PUBBLICA ASL Teramo | Istituto Zooprofilattico Sperimentale dell'Abruzzo e Molise "G.Caporale" | Lorusso A, Marcacci M, Di Domenico M, Ancora M, Curini V, Mangone I, Rinaldi A, Di Pasquale A, Cammà C, Puglia I, Savini G |
| EPI_ISL_435150, EPI_ISL_435151 | Ospedale SS Annunziata | Istituto Zooprofilattico Sperimentale dell'Abruzzo e Molise "G.Caporale" | Lorusso A, Marcacci M, Di Domenico M, Ancora M, Curini V, Mangone I, Rinaldi A, Di Pasquale A, Cammà C, Puglia I, Savini G |
| EPI_ISL_435152 | Servizio di Igiene, Epidemiologia e Sanità Pubblica (SIESP) Avezzano | Istituto Zooprofilattico Sperimentale dell'Abruzzo e Molise "G.Caporale" | Lorusso A, Marcacci M, Di Domenico M, Ancora M, Curini V, Mangone I, Rinaldi A, Di Pasquale A, Cammà C, Puglia I, Savini G |
| EPI_ISL_435153, EPI_ISL_435154, EPI_ISL_435155 | SERVIZIO DI IGIENE E SANITÀ PUBBLICA ASL Teramo | Istituto Zooprofilattico Sperimentale dell'Abruzzo e Molise "G.Caporale" | Lorusso A, Marcacci M, Di Domenico M, Ancora M, Curini V, Mangone I, Rinaldi A, Di Pasquale A, Cammà C, Puglia I, Savini G |
| EPI_ISL_435156, EPI_ISL_435157, EPI_ISL_435158, EPI_ISL_435159, EPI_ISL_435160, EPI_ISL_435161, EPI_ISL_435162, EPI_ISL_435163, EPI_ISL_435164 | Viral Respiratory Lab, National Institute for Biomedical Research (INRB) | Pathogen Sequencing Lab, National Institute for Biomedical Research (INRB) | Placide Mbala-Kingebeni, Edith Nkwembe, Eddy Kinganda-Lusamaki, Amuri Aziza, Francisca Muyembe Mawete, Catherine Pratt, Matthias Pauthner, Josh Quick, Allison Black, James Hadfield, Trevor Bedford, Ian Goodfellow, Andrew Rambaut, Nick Loman, Kristian Andersen, Michael Wiley, Steve Ahuka-Mundeke, Jean-Jacques Muyembe Tamfum |
| EPI_ISL_435392, EPI_ISL_435393 | Utah Public Health Laboratory | Utah Public Health Laboratory | Erin Young, Kelly Oakeson |
| EPI_ISL_435441, EPI_ISL_435442, EPI_ISL_435443, EPI_ISL_435444 | Alaska State Virology Laboratory | Alaska State Virology Laboratory | Jack Chen, Ph.D. |
| EPI_ISL_435498 | NYU Langone Health | Departments of Pathology and Medicine, New York University School of Medicine | Maria Aguero-Rosenfeld, Brendan Belovarac, Margaret Black, Ludovic Boytard, John Cadley, Paolo Cotzia, John Chen, Dacia Dimartino, Xiaojun Feng, Tatyana Gindin, Emily Guzman, Adriana Heguy, Megan Hogan, Emily Huang, George Jour, Lawrence H. Lin, Raven Luther, Andrew Lytle, Christian Marier, Matthew T. Maurano, Mark J. Mulligan, Peter Meyn, Raquel Ordonez Ciriza, Iman Osman, Jared Pinnell, Vanessa Raabe, Sitharam Ramaswami, Amy Rapkiewicz, Andre M. Ribeiro-dos-Santos, Marie Samanovic-Golden, Antonio Serrano, Guomiao Shen, Matija Snuderl, Theodore Vougiouklakis, Nick Vulpescu, Gael Westby, Paul Zappile, Yutong Zhang |
| EPI_ISL_435681, EPI_ISL_435682, EPI_ISL_435683, EPI_ISL_435684, EPI_ISL_435685 | National Public Health Laboratory, National Centre for Infectious Diseases | National Public Health Laboratory, National Centre for Infectious Diseases | Mak Tze Minn, Octavia Sophie, Chavatte Jean-Marc, Cui Lin, Lin Raymond Tzer Pin |
| EPI_ISL_435703, EPI_ISL_435704, EPI_ISL_435705, EPI_ISL_435706, EPI_ISL_435707, EPI_ISL_435708 | Yale COVID-19 Biorepository | Grubaugh Lab - Yale School of Public Health | Joseph Fauver, Tara Alpert, Anderson Brito, Anne Wyllie, Chantal Vogels, Mary Petrone, Cole Jensen, Chaney Kalinich, Isabel Ott, Arnau Casanovas, Catherine Muenker, Adam Moore, Alice Lu, Maria Tokuyama, Patrick Wong, Peiwen Lu, Saad Omer, Richard Martinello, Allison Nelson, Shelli Farhadian, Akiko Iwasaki, Charlese Dela Cruz, Albert Ko, Nathan Grubaugh |
| EPI_ISL_436111, EPI_ISL_436112, EPI_ISL_436113, EPI_ISL_436114, EPI_ISL_436115, EPI_ISL_436116, EPI_ISL_436117, EPI_ISL_436118, EPI_ISL_436119, EPI_ISL_436120, EPI_ISL_436121, EPI_ISL_436122, EPI_ISL_436123, EPI_ISL_436124, EPI_ISL_436125, EPI_ISL_436126 | | | |
| see above | Victorian Infectious Diseases Reference Laboratory (VIDRL) | Microbiological Diagnostic Unit Public Health Laboratory and | Caly L., Seemann T., Sait, M., Schultz M., Druce J., Sherry, N. |

| | | | |
|---|---|---|---|
| | Victorian Infectious Diseases Reference Laboratory, Doherty Institute | | |
| EPI_ISL_436435, EPI_ISL_436437, EPI_ISL_436438, EPI_ISL_436439, EPI_ISL_436440, EPI_ISL_436441, EPI_ISL_436442, EPI_ISL_436443, EPI_ISL_436444, EPI_ISL_436445, EPI_ISL_436446, EPI_ISL_436447, EPI_ISL_436448, EPI_ISL_436449 | | | |
| see above | National Centre for Disease control (NCDC) | NCDC/CSIR-IGIB | Pramod Kumar#, Rajesh Pandey#, Pooja Sharma, Mahesh S Dhar, Vivekanand A, Bharathram Uppili, Himanshu Vashisht, Saruchi Wadhwa, Nishu Tyagi, Uma Sharma, Priyanka Singh, Hemlata Lall, Meena Datta, Poonam Gupta, Nidhi Saini, Aarti Tewari, Bibhash Nandi, Dhirendra Kumar, Satyabrata Bag, Varun Jaiswal, Hema Gogia, Preeti Madan, Simrita Singh, Prateek Singh, Debasis Dash, Mitali Mukerji, Manju Bala, Sandhya Kabra, Sujeet Singh, Mohammed Faruq, Anurag Agrawal*, Partha Rakshit* |
| EPI_ISL_436464 | Alaska State Virology Laboratory | Alaska State Virology Laboratory | Jack Chen |
| EPI_ISL_436517, EPI_ISL_436518, EPI_ISL_436519, EPI_ISL_436520, EPI_ISL_436521, EPI_ISL_436522 | Florida Bureau of Public Health Laboratories | Florida Bureau of Public Health Laboratories | Sarah Schmedes, Jason Blanton |
| EPI_ISL_436578, EPI_ISL_436582, EPI_ISL_436583, EPI_ISL_436584, EPI_ISL_436585, EPI_ISL_436586, EPI_ISL_436587, EPI_ISL_436588, EPI_ISL_436589, EPI_ISL_436590, EPI_ISL_436639 | | | |
| see above | University of Wisconsin-Madison AIDS Vaccine Research Laboratories | University of Wisconsin-Madison AIDS Vaccine Research Laboratories | Gage Moreno, Katarina Braun, et al. AIDS Vaccine Research Laboratories |
| EPI_ISL_436653, EPI_ISL_436654, EPI_ISL_436655, EPI_ISL_436656, EPI_ISL_436657, EPI_ISL_436658, EPI_ISL_436659, EPI_ISL_436660, EPI_ISL_436661, EPI_ISL_436662, EPI_ISL_436663, EPI_ISL_436664, EPI_ISL_436665, EPI_ISL_436666, EPI_ISL_436667, EPI_ISL_436682 | | | |
| see above | County of Santa Clara Public Health Department | Chan-Zuckerberg Biohub | CZB Cliahub Consortium |
| EPI_ISL_436901, EPI_ISL_436902, EPI_ISL_436903, EPI_ISL_436905, EPI_ISL_436906, EPI_ISL_436907, EPI_ISL_436908, EPI_ISL_436909, EPI_ISL_436910, EPI_ISL_436911, EPI_ISL_436912, EPI_ISL_436913, EPI_ISL_436914, EPI_ISL_436915, EPI_ISL_436916, EPI_ISL_436917 | | | |
| see above | Utah Public Health Laboratory | Utah Public Health Laboratory | Erin Young, Kelly Oakeson |
| EPI_ISL_436940, EPI_ISL_436941, EPI_ISL_436942, EPI_ISL_436946, EPI_ISL_436954, EPI_ISL_436955 | Ochsner Health | Bioinfoexperts, LLC | Amy Feehan, David J. Nolan, Rebecca Rose, Sissy Cross, David Moraga Amador, Tong Yang, Luke Caruso, Wayra Navia, Lydia Von Borstel, Xiao Hui Zhou, Julia-Garcia-Diaz, Susanna L. Lamers |
| EPI_ISL_437056, EPI_ISL_437057, EPI_ISL_437059, EPI_ISL_437060, EPI_ISL_437061, EPI_ISL_437062, EPI_ISL_437063, EPI_ISL_437064, EPI_ISL_437076, EPI_ISL_437077, EPI_ISL_437078, EPI_ISL_437083, EPI_ISL_437084, EPI_ISL_437085 | | | |
| see above | County of Santa Clara Public Health | Chan-Zuckerberg Biohub | CZB Cliahub Consortium |
| EPI_ISL_437207, EPI_ISL_437208, EPI_ISL_437214, EPI_ISL_437215, EPI_ISL_437216, EPI_ISL_437217, EPI_ISL_437222, EPI_ISL_437226, EPI_ISL_437241, EPI_ISL_437242, EPI_ISL_437243, EPI_ISL_437292, EPI_ISL_437293, EPI_ISL_437294, EPI_ISL_437295 | | | |
| see above | Max von Pettenkofer Institute, Virology, National Reference Center for Retroviruses, LMU München | Laboratory for Functional Genome Analysis, Dept. Genomics, Gene Center of the LMU Munich | Max Muenchhoff, Stefan Krebs, Alexander Graf, Oliver Keppler, Helmut Blum |
| EPI_ISL_437298, EPI_ISL_437299 | Diagnostic- and Research Institute of Pathology, Medical University of Graz | Diagnostic- and Research Institute of Pathology, Medical University of Graz | Karl Kashofer, Peter Regitnig, Martin Zacharias, Gregor Gorkiewicz |
| EPI_ISL_437367, EPI_ISL_437368, EPI_ISL_437369, EPI_ISL_437370, EPI_ISL_437371, EPI_ISL_437372, EPI_ISL_437373, EPI_ISL_437374, EPI_ISL_437375, EPI_ISL_437376, EPI_ISL_437377 | | | |
| see above | Minnesota Department of Health, Public Health Laboratory | Minnesota Department of Health, Public Health Laboratory | Matt Plumb, Jacob Garfin, and Xiong Wang |
| EPI_ISL_437400, EPI_ISL_437401, EPI_ISL_437402, EPI_ISL_437411, EPI_ISL_437412, EPI_ISL_437413, EPI_ISL_437414, EPI_ISL_437415, EPI_ISL_437416, EPI_ISL_437421, EPI_ISL_437422, EPI_ISL_437423, EPI_ISL_437432 | | | |
| see above | Virginia DCLS | Virginia DCLS | Virginia DCLS |
| EPI_ISL_437437 | Alaska State Virology Laboratory | Alaska State Virology Laboratory | Jack Chen, Ph.D. |
| EPI_ISL_437458 | Clinical Diagnostics Laboratory, Diagnostic & Experimental Pathology, Lilly Research Laboratories | Clinical Diagnostics Laboratory, Diagnostic & Experimental Pathology, Lilly Research Laboratories | Tim Holzer, Mayuri Vaidya, Angie Fulford, Sam McNeely, Rachael Redmond, Phil Ebert, John Calley, Leslie ONeill Reising, Pat Finnegan, Erin Wray, John McElwee, Jeff Fill, Joe Oakley, Andrew Schade |
| EPI_ISL_437517 | Alaska State Virology Laboratory | Alaska State Virology Laboratory | Jack Chen, Ph.D. |
| EPI_ISL_437520, EPI_ISL_437521, EPI_ISL_437522, EPI_ISL_437523, EPI_ISL_437529, EPI_ISL_437535 | OHSU Lab Services Molecular Microbiology Lab | Oregon SARS-CoV-2 Genome Sequencing Center | Brendan L. O'Connell, Ruth V. Nichols, Alec J. Hirsch, Guang Fan, Daniel N. Streblow, William B. Messer, Andrew C. Adey, Benjamin N. Bimber, Brian J. O'Roak |
| EPI_ISL_437540, EPI_ISL_437543 | Robert Garry lab | Andersen lab at Scripps Research | Allison Smither, Gilberto Sabino-Santos, Patricia Snarski, Lilia Melnik, Antoinette Bell, Kaylynn Genemaras, Arnaud Drouin, Dahlene Fusco, Robert Garry with SEARCH Alliance San Diego |
| EPI_ISL_437574 | Scripps Medical Laboratory | Andersen lab at Scripps Research | SEARCH Alliance San Diego with Michael Quigley, Ellen Stefanski, Ian Mchardy |
| EPI_ISL_437763, EPI_ISL_437764, EPI_ISL_437767, EPI_ISL_437768, EPI_ISL_437786, EPI_ISL_437787, EPI_ISL_437788, EPI_ISL_437789, EPI_ISL_437796, EPI_ISL_437797, EPI_ISL_437798, EPI_ISL_437799 | | | |
| see above | Virginia DCLS | Virginia DCLS | Virginia DCLS |
| EPI_ISL_438128 | Center for Virology, Medical University of Vienna | Bergthaler laboratory, CeMM Research Center for Molecular Medicine of the Austrian Academy of Sciences | Alexandra Popa, Benedikt Agerer, Henrique Colaco, Lukas Endler, Jakob-Wendelin Genger, Alexander Lercher, Mark Smyth, Thomas Penz, Michael Schuster, Jan Laine, Martin Senekowitsch, Judith Aberle, Stephan Aberle, Elisabeth Puchhammer-Stoeckl, Manfred Nairz, Guenter Weiss, Wegene Borena, Dorothee von Laer, Christoph Bock, Andreas Bergthaler |
| EPI_ISL_438147, EPI_ISL_438148 | Seattle Flu Study | Seattle Flu Study | Chu et al |
| EPI_ISL_438849, EPI_ISL_438851, EPI_ISL_438852, EPI_ISL_438853, EPI_ISL_438854, EPI_ISL_438855, EPI_ISL_438856, EPI_ISL_438857, EPI_ISL_438858, EPI_ISL_438859, EPI_ISL_438860, EPI_ISL_438861, EPI_ISL_438862, EPI_ISL_438863, EPI_ISL_438864, EPI_ISL_438865, EPI_ISL_438866, EPI_ISL_438867, EPI_ISL_438868, EPI_ISL_438869, EPI_ISL_438870, EPI_ISL_438871, EPI_ISL_438872, EPI_ISL_438874, EPI_ISL_438875, EPI_ISL_438876, EPI_ISL_438877, EPI_ISL_438878, EPI_ISL_438879, EPI_ISL_438880, EPI_ISL_438881, EPI_ISL_438882, EPI_ISL_438884, EPI_ISL_438886 | | | |
| see above | West of Scotland Specialist Virology Centre, NHSGGC / MRC-University of Glasgow Centre for Virus Research | COVID-19 Genomics UK (COG-UK) Consortium | Ana da Silva Filipe, Natasha Johnson, Kathy Smollett, Daniel Mair, Stephen Carmichael, Lily Tong, Jenna Nichols, Elihu Aranday-Cortes, Kirstyn Brunker, Yasmin Parr, Kyriaki Nomikou; Sarah McDonald, Marc Niebel, Patawee Asamaphan; Richard Orton, Joseph Hughes, Sreenu Vattipally, David L Robertson; Alasdair MacLean, Rory Gunson; Kathy Li, Natasha Jesudason, Rajiv Shah, James Shepherd, Antonia Ho, Emma Thomson |
| EPI_ISL_438955, EPI_ISL_438956, EPI_ISL_438958, EPI_ISL_438959, EPI_ISL_438961 | Keio University School of Medicine | Keio University School of Medicine | Kenjiro Kosaki |
| EPI_ISL_439144, EPI_ISL_439146, EPI_ISL_439147, EPI_ISL_439265, EPI_ISL_439266, EPI_ISL_439267, EPI_ISL_439270 | Virology Department, Royal Infirmary of Edinburgh, NHS Lothian / School of Biological Sciences, University of Edinburgh / Institute of Genetics and Molecular Medicine, University of Edinburgh | COVID-19 Genomics UK (COG-UK) Consortium | McHugh M, Dewar R, Rooke S, Gallagher M, Balcaza C, O'ÄôToole Å, Scher E, Hill V, McCrone JT, Colquhoun R, Yu X, Jackson B, Rambaut A, Williams TC, Templeton K |
| EPI_ISL_439673, EPI_ISL_439674, EPI_ISL_439675, EPI_ISL_439676, EPI_ISL_439677, EPI_ISL_439678, EPI_ISL_439679, EPI_ISL_439681, EPI_ISL_439682, EPI_ISL_439683, EPI_ISL_439684, EPI_ISL_439685, EPI_ISL_439686, EPI_ISL_439687, EPI_ISL_439688, EPI_ISL_439689, EPI_ISL_439690, EPI_ISL_439691, EPI_ISL_439693, EPI_ISL_439694, EPI_ISL_439695, EPI_ISL_439696, EPI_ISL_439697, EPI_ISL_439698, EPI_ISL_439699, EPI_ISL_439700, EPI_ISL_439702, EPI_ISL_439703, EPI_ISL_439704, EPI_ISL_439705, EPI_ISL_439707, EPI_ISL_439709, EPI_ISL_439710, EPI_ISL_439711, EPI_ISL_439712, EPI_ISL_439713, EPI_ISL_439714, EPI_ISL_439715, EPI_ISL_439716, EPI_ISL_439717, EPI_ISL_439718, EPI_ISL_439719, EPI_ISL_439720, EPI_ISL_439721, EPI_ISL_439722, EPI_ISL_439723, EPI_ISL_439724, EPI_ISL_439725, EPI_ISL_439726, EPI_ISL_439727, EPI_ISL_439728, EPI_ISL_439729, EPI_ISL_439730, EPI_ISL_439731, EPI_ISL_439732, EPI_ISL_439733, EPI_ISL_439734, EPI_ISL_439735, EPI_ISL_439736, EPI_ISL_439738, EPI_ISL_439739, EPI_ISL_439740, EPI_ISL_439741, EPI_ISL_439742, EPI_ISL_439743, EPI_ISL_439744, EPI_ISL_439745, EPI_ISL_439746, EPI_ISL_439747, EPI_ISL_439748, EPI_ISL_439749, EPI_ISL_439750, EPI_ISL_439751, EPI_ISL_439752, EPI_ISL_439753, EPI_ISL_439754, EPI_ISL_439755, EPI_ISL_439756, EPI_ISL_439757, EPI_ISL_439758, EPI_ISL_439759, EPI_ISL_439760, EPI_ISL_439761, EPI_ISL_439762, EPI_ISL_439763, EPI_ISL_439764, EPI_ISL_439765, EPI_ISL_439766, EPI_ISL_439767, EPI_ISL_439768, EPI_ISL_439769, EPI_ISL_439770, EPI_ISL_439771, EPI_ISL_439772, EPI_ISL_439773, EPI_ISL_439774, EPI_ISL_439775, EPI_ISL_439776, EPI_ISL_439777, EPI_ISL_439778, EPI_ISL_439779, EPI_ISL_439780, EPI_ISL_439781, EPI_ISL_439782, EPI_ISL_439783, EPI_ISL_439784, EPI_ISL_439785, EPI_ISL_439787, EPI_ISL_439788, EPI_ISL_439789, EPI_ISL_439792, EPI_ISL_439793, EPI_ISL_439796, EPI_ISL_439797, EPI_ISL_439798, EPI_ISL_439799, EPI_ISL_439800, EPI_ISL_439802, EPI_ISL_439805, EPI_ISL_439807, EPI_ISL_439809, EPI_ISL_439812 | | | |
| see above | Liverpool Clinical Laboratories | COVID-19 Genomics UK (COG-UK) Consortium | Sam Haldenby, Anita Lucaci, Steve Paterson, Julian Hiscox, Alistair Darby, M Almsaud, A Alrezaihi, Muhannad Alruwaili, Stuart D Armstrong, Jones Benjamin , Eleanor G Bentley, Anu Chawla, Jordan J Clark, Angela Cowell, Richard Eccles, Isabel Garca-Dorival, Matthew Gemmell, Alessandro Gerada, PKF Gilmore, Richard Gregory, Ximeng Han, Catherine Hartley, Margaret Hughes, Miren Iturriza-Gomara, James Johnson, L Luu, Jenifer Manson , |

| | | | |
|---|---|---|---|
| | | | Charlotte Nelson, Elaine O'ÄôToole, Cassie Olateju, Rebekah Penrice-Randal¬†, Lucille Rainbow, N.P Randle, Trevor Ian Robinson, Parul Sharma, Ghada T Shawli, James P Stewart , Neil Swainston, Ecaterina Vamos, Joanne Watts, Mark Whitehead |
| EPI_ISL_441437, EPI_ISL_441438, EPI_ISL_441439, EPI_ISL_441440, EPI_ISL_441441, EPI_ISL_441442, EPI_ISL_441443, EPI_ISL_441444, EPI_ISL_441445, EPI_ISL_441446, EPI_ISL_441447, EPI_ISL_441448, EPI_ISL_441449, EPI_ISL_441450, EPI_ISL_441451, EPI_ISL_441452, EPI_ISL_441453, EPI_ISL_441454, EPI_ISL_441455, EPI_ISL_441456, EPI_ISL_441457, EPI_ISL_441458, EPI_ISL_441459, EPI_ISL_441466, EPI_ISL_441467, EPI_ISL_441468, EPI_ISL_441469, EPI_ISL_441470, EPI_ISL_441471, EPI_ISL_441472, EPI_ISL_441473, EPI_ISL_441474, EPI_ISL_441475, EPI_ISL_441476, EPI_ISL_441477, EPI_ISL_441478, EPI_ISL_441479, EPI_ISL_441480, EPI_ISL_441481, EPI_ISL_441482, EPI_ISL_441483, EPI_ISL_441484, EPI_ISL_441485, EPI_ISL_441486, EPI_ISL_441487, EPI_ISL_441488, EPI_ISL_441489, EPI_ISL_441490, EPI_ISL_441491, EPI_ISL_441492, EPI_ISL_441493, EPI_ISL_441494, EPI_ISL_441495, EPI_ISL_441496, EPI_ISL_441497, EPI_ISL_441498, EPI_ISL_441499, EPI_ISL_441500, EPI_ISL_441501, EPI_ISL_441502, EPI_ISL_441503 | | | |
| see above | Queens Medical Centre, Clinical Microbiology Department / DeepSeq Nottingham | COVID-19 Genomics UK (COG-UK) Consortium | Gemma Clark, Wendy Smith, Manjinder Khakh, Hannah Howson-Wells, Jonathan Ball, Patrick McClure, Joseph Chappell, Theocharis Tsoleridis, Nadine Holmes, Matthew Carlisle, Christopher Moore, Fei Sang, Johnny Debebe, Victoria Wright, Matthew Loose |
| EPI_ISL_441566, EPI_ISL_441576, EPI_ISL_441582, EPI_ISL_441592, EPI_ISL_441610, EPI_ISL_441617, EPI_ISL_441633, EPI_ISL_441637, EPI_ISL_441660, EPI_ISL_441664, EPI_ISL_441668, EPI_ISL_441687, EPI_ISL_441692, EPI_ISL_441705, EPI_ISL_441728, EPI_ISL_441734, EPI_ISL_441741, EPI_ISL_441742, EPI_ISL_441750, EPI_ISL_441762, EPI_ISL_441767, EPI_ISL_441769, EPI_ISL_441782 | | | |
| see above | Department of Pathology, University of Cambridge | Wellcome Sanger Institute for the COVID-19 Genomics UK (COG-UK) consortium | Luke W Meredith, M. Estée Török , Myra Hosmillo, William L. Hamilton, Martin D. Curran, Theresa Feltwell, Grant Hall, Anna Yakovleva, Fahad A Khokhar, Charlotte J. Houldcroft, Laura G Caller, Aminu S. Jahun, Sarah L. Caddy, Ian Goodfellow, Alex Alderton, Roberto Amato, Sonia Goncalves, Ewan Harrison, David K. Jackson, Ian Johnston, Dominic Kwiatkowski, Cordelia Langford, John Sillitoe on behalf of the Wellcome Sanger Institute COVID-19 Surveillance Team (http://www.sanger.ac.uk/covid-team) |
| EPI_ISL_441848 | Queens Medical Centre, Clinical Microbiology Department / DeepSeq Nottingham | COVID-19 Genomics UK (COG-UK) Consortium | Gemma Clark, Wendy Smith, Manjinder Khakh, Hannah Howson-Wells, Jonathan Ball, Patrick McClure, Joseph Chappell, Theocharis Tsoleridis, Nadine Holmes, Matthew Carlisle, Christopher Moore, Fei Sang, Johnny Debebe, Victoria Wright, Matthew Loose |
| EPI_ISL_441898, EPI_ISL_441899, EPI_ISL_441900, EPI_ISL_441902, EPI_ISL_441903, EPI_ISL_441904, EPI_ISL_441905, EPI_ISL_441907, EPI_ISL_441908, EPI_ISL_441909, EPI_ISL_441913, EPI_ISL_441915, EPI_ISL_441916, EPI_ISL_441918, EPI_ISL_441924, EPI_ISL_441925, EPI_ISL_441927, EPI_ISL_441928, EPI_ISL_441932, EPI_ISL_441933, EPI_ISL_441935, EPI_ISL_441936, EPI_ISL_441939, EPI_ISL_441941, EPI_ISL_441942, EPI_ISL_441943, EPI_ISL_441946, EPI_ISL_441947, EPI_ISL_441951, EPI_ISL_441952, EPI_ISL_441954, EPI_ISL_441955, EPI_ISL_441956, EPI_ISL_441957, EPI_ISL_441960, EPI_ISL_441966, EPI_ISL_441967, EPI_ISL_441970, EPI_ISL_441971, EPI_ISL_441974, EPI_ISL_441979, EPI_ISL_441980, EPI_ISL_441982, EPI_ISL_441983, EPI_ISL_441987, EPI_ISL_441988, EPI_ISL_441991, EPI_ISL_441993, EPI_ISL_441996, EPI_ISL_441998, EPI_ISL_441999, EPI_ISL_442000, EPI_ISL_442005, EPI_ISL_442006, EPI_ISL_442007, EPI_ISL_442008, EPI_ISL_442009, EPI_ISL_442011, EPI_ISL_442014, EPI_ISL_442017, EPI_ISL_442018, EPI_ISL_442021, EPI_ISL_442023, EPI_ISL_442024, EPI_ISL_442025, EPI_ISL_442027, EPI_ISL_442028, EPI_ISL_442030, EPI_ISL_442033, EPI_ISL_442035, EPI_ISL_442043 | | | |
| see above | Virology Department, Sheffield Teaching Hospitals NHS Foundation Trust/Department of Infection, Immunity and Cardiovascular Disease, The Medical School, University of Sheffield | COVID-19 Genomics UK (COG-UK) Consortium | Thushan de Silva, Matthew Parker, Nikki Smith, Adri Angyal, Rebecca Brown, Luke Green, Rachel Tucker, Paul Parsons, Danielle Groves, Katie Johnson, Laura Carrilero, Alex Keeley, Dave Partridge, Matthew Wyles, Benjamin Lindsey, Mehmet Yavuz, Mohammad Raza, Cariad Evans |
| EPI_ISL_442056, EPI_ISL_442081, EPI_ISL_442084, EPI_ISL_442106, EPI_ISL_442129, EPI_ISL_442148, EPI_ISL_442149, EPI_ISL_442152, EPI_ISL_442161, EPI_ISL_442169, EPI_ISL_442172, EPI_ISL_442175, EPI_ISL_442179, EPI_ISL_442200, EPI_ISL_442201, EPI_ISL_442203, EPI_ISL_442204, EPI_ISL_442205, EPI_ISL_442216, EPI_ISL_442219, EPI_ISL_442221, EPI_ISL_442225, EPI_ISL_442228, EPI_ISL_442236, EPI_ISL_442241, EPI_ISL_442261, EPI_ISL_442273, EPI_ISL_442294, EPI_ISL_442311, EPI_ISL_442327, EPI_ISL_442332, EPI_ISL_442339 | | | |
| see above | Department of Pathology, University of Cambridge | Wellcome Sanger Institute for the COVID-19 Genomics UK (COG-UK) consortium | Luke W Meredith, M. Estée Török , Myra Hosmillo, William L. Hamilton, Martin D. Curran, Theresa Feltwell, Grant Hall, Anna Yakovleva, Fahad A Khokhar, Charlotte J. Houldcroft, Laura G Caller, Aminu S. Jahun, Sarah L. Caddy, Ian Goodfellow, Alex Alderton, Roberto Amato, Sonia Goncalves, Ewan Harrison, David K. Jackson, Ian Johnston, Dominic Kwiatkowski, Cordelia Langford, John Sillitoe on behalf of the Wellcome Sanger Institute COVID-19 Surveillance Team (http://www.sanger.ac.uk/covid-team) |
| EPI_ISL_442345, EPI_ISL_442346, EPI_ISL_442350, EPI_ISL_442351, EPI_ISL_442353, EPI_ISL_442354, EPI_ISL_442355, EPI_ISL_442357, EPI_ISL_442358, EPI_ISL_442360, EPI_ISL_442364, EPI_ISL_442366, EPI_ISL_442368, EPI_ISL_442370, EPI_ISL_442372, EPI_ISL_442373, EPI_ISL_442375, EPI_ISL_442376, EPI_ISL_442377, EPI_ISL_442378, EPI_ISL_442379, EPI_ISL_442380, EPI_ISL_442381, EPI_ISL_442384, EPI_ISL_442387, EPI_ISL_442389, EPI_ISL_442390, EPI_ISL_442391, EPI_ISL_442392, EPI_ISL_442393, EPI_ISL_442394, EPI_ISL_442396, EPI_ISL_442400, EPI_ISL_442402, EPI_ISL_442403, EPI_ISL_442404, EPI_ISL_442405, EPI_ISL_442407, EPI_ISL_442409, EPI_ISL_442413, EPI_ISL_442416, EPI_ISL_442418, EPI_ISL_442421, EPI_ISL_442423, EPI_ISL_442425, EPI_ISL_442426, EPI_ISL_442430, EPI_ISL_442432, EPI_ISL_442436, EPI_ISL_442437, EPI_ISL_442438, EPI_ISL_442439, EPI_ISL_442440, EPI_ISL_442441, EPI_ISL_442447, EPI_ISL_442448, EPI_ISL_442450, EPI_ISL_442451, EPI_ISL_442453, EPI_ISL_442454, EPI_ISL_442455, EPI_ISL_442456, EPI_ISL_442462, EPI_ISL_442464, EPI_ISL_442467, EPI_ISL_442468, EPI_ISL_442471, EPI_ISL_442472, EPI_ISL_442475, EPI_ISL_442476, EPI_ISL_442477, EPI_ISL_442479, EPI_ISL_442481, EPI_ISL_442489, EPI_ISL_442492, EPI_ISL_442493, EPI_ISL_442495, EPI_ISL_442502, EPI_ISL_442503, EPI_ISL_442504, EPI_ISL_442505, EPI_ISL_442508, EPI_ISL_442511, EPI_ISL_442513, EPI_ISL_442516, EPI_ISL_442517 | | | |
| see above | Virology Department, Sheffield Teaching Hospitals NHS Foundation Trust/Department of Infection, Immunity and Cardiovascular Disease, The Medical School, University of Sheffield | COVID-19 Genomics UK (COG-UK) Consortium | Thushan de Silva, Matthew Parker, Nikki Smith, Adri Angyal, Rebecca Brown, Luke Green, Rachel Tucker, Paul Parsons, Danielle Groves, Katie Johnson, Laura Carrilero, Alex Keeley, Dave Partridge, Matthew Wyles, Benjamin Lindsey, Mehmet Yavuz, Mohammad Raza, Cariad Evans |
| EPI_ISL_442524, EPI_ISL_442525, EPI_ISL_442527, EPI_ISL_442528, EPI_ISL_442529, EPI_ISL_442531, EPI_ISL_442532, EPI_ISL_442533, EPI_ISL_442534, EPI_ISL_442535, EPI_ISL_442536, EPI_ISL_442537, EPI_ISL_442538, EPI_ISL_442540, EPI_ISL_442541, EPI_ISL_442542, EPI_ISL_442543, EPI_ISL_442544, EPI_ISL_442545, EPI_ISL_442546, EPI_ISL_442547, EPI_ISL_442548, EPI_ISL_442549, EPI_ISL_442550, EPI_ISL_442551, EPI_ISL_442552, EPI_ISL_442553, EPI_ISL_442554, EPI_ISL_442555, EPI_ISL_442556, EPI_ISL_442557, EPI_ISL_442558, EPI_ISL_442559, EPI_ISL_442560, EPI_ISL_442561, EPI_ISL_442562, EPI_ISL_442563, EPI_ISL_442564, EPI_ISL_442565, EPI_ISL_442566, EPI_ISL_442568, EPI_ISL_442569, EPI_ISL_442570, EPI_ISL_442571, EPI_ISL_442572, EPI_ISL_442573, EPI_ISL_442574, EPI_ISL_442575, EPI_ISL_442576, EPI_ISL_442577, EPI_ISL_442578, EPI_ISL_442579, EPI_ISL_442580, EPI_ISL_442581, EPI_ISL_442582, EPI_ISL_442583, EPI_ISL_442584, EPI_ISL_442585, EPI_ISL_442586, EPI_ISL_442587, EPI_ISL_442588, EPI_ISL_442589, EPI_ISL_442591, EPI_ISL_442592, EPI_ISL_442593, EPI_ISL_442594, EPI_ISL_442595, EPI_ISL_442596, EPI_ISL_442597, EPI_ISL_442598, EPI_ISL_442599, EPI_ISL_442601, EPI_ISL_442604, EPI_ISL_442605, EPI_ISL_442606, EPI_ISL_442608, EPI_ISL_442609, EPI_ISL_442610, EPI_ISL_442611, EPI_ISL_442612, EPI_ISL_442613, EPI_ISL_442614, EPI_ISL_442615, EPI_ISL_442616, EPI_ISL_442617, EPI_ISL_442618, EPI_ISL_442620, EPI_ISL_442621, EPI_ISL_442622, EPI_ISL_442623 | | | |
| see above | Department of Pathology, University of Cambridge | Wellcome Sanger Institute for the COVID-19 Genomics UK (COG-UK) consortium | Luke W Meredith, M. Estée Török , Myra Hosmillo, William L. Hamilton, Martin D. Curran, Theresa Feltwell, Grant Hall, Anna Yakovleva, Fahad A Khokhar, Charlotte J. Houldcroft, Laura G Caller, Aminu S. Jahun, Sarah L. Caddy, Ian Goodfellow, Alex Alderton, Roberto Amato, Sonia Goncalves, Ewan Harrison, David K. Jackson, Ian Johnston, Dominic Kwiatkowski, Cordelia Langford, John Sillitoe on behalf of the Wellcome Sanger Institute COVID-19 Surveillance Team (http://www.sanger.ac.uk/covid-team) |
| EPI_ISL_442633, EPI_ISL_442637, EPI_ISL_442639, EPI_ISL_442640, EPI_ISL_442644, EPI_ISL_442648, EPI_ISL_442649, EPI_ISL_442652, EPI_ISL_442654, EPI_ISL_442656, EPI_ISL_442661, EPI_ISL_442667, EPI_ISL_442669, EPI_ISL_442670, EPI_ISL_442672, EPI_ISL_442673, EPI_ISL_442684, EPI_ISL_442685, EPI_ISL_442693, EPI_ISL_442695, EPI_ISL_442697, EPI_ISL_442699, EPI_ISL_442702, EPI_ISL_442706, EPI_ISL_442712, EPI_ISL_442714, EPI_ISL_442716, EPI_ISL_442722, EPI_ISL_442729, EPI_ISL_442745, EPI_ISL_442747, EPI_ISL_442752, EPI_ISL_442764, EPI_ISL_442768, EPI_ISL_442769, EPI_ISL_442771, EPI_ISL_442772, EPI_ISL_442774, EPI_ISL_442777, EPI_ISL_442779, EPI_ISL_442780, EPI_ISL_442782, EPI_ISL_442789, EPI_ISL_442790, EPI_ISL_442793, EPI_ISL_442796, EPI_ISL_442797, EPI_ISL_442800, EPI_ISL_442802 | | | |
| see above | PHE South West Regional Laboratory, National Infection Service | Wellcome Sanger Institute for the COVID-19 Genomics UK (COG-UK) consortium | Stephanie Hutchings, Hannah Pymont, Dr Peter Muir, Barry Vipond, Rich Hopes, Alex Alderton, Roberto Amato, Sonia Goncalves, Ewan Harrison, David K. Jackson, Ian Johnston, Dominic Kwiatkowski, Cordelia Langford, John Sillitoe on behalf of the Wellcome Sanger Institute COVID-19 Surveillance Team (http://www.sanger.ac.uk/covid-team) |
| EPI_ISL_442812, EPI_ISL_442821, EPI_ISL_442822, EPI_ISL_442823, EPI_ISL_442825, EPI_ISL_442827, EPI_ISL_442828, EPI_ISL_442829, EPI_ISL_442830, EPI_ISL_442831, EPI_ISL_442832, EPI_ISL_442833, EPI_ISL_442835, EPI_ISL_442836, EPI_ISL_442837, EPI_ISL_442838, EPI_ISL_442839, EPI_ISL_442840, EPI_ISL_442841, EPI_ISL_442842, EPI_ISL_442843, EPI_ISL_442844, EPI_ISL_442845, EPI_ISL_442846, EPI_ISL_442847, EPI_ISL_442848, EPI_ISL_442849, EPI_ISL_442850, EPI_ISL_442851, EPI_ISL_442852, EPI_ISL_442853, EPI_ISL_442854, EPI_ISL_442855, EPI_ISL_442856, EPI_ISL_442857, EPI_ISL_442858, EPI_ISL_442859, EPI_ISL_442861, EPI_ISL_442862, EPI_ISL_442863, EPI_ISL_442864, EPI_ISL_442865, EPI_ISL_442866, EPI_ISL_442867, EPI_ISL_442868, EPI_ISL_442869, EPI_ISL_442870, EPI_ISL_442871, EPI_ISL_442872, EPI_ISL_442873, EPI_ISL_442874, EPI_ISL_442875, EPI_ISL_442876, EPI_ISL_442877, EPI_ISL_442878, EPI_ISL_442879, EPI_ISL_442880, EPI_ISL_442882, EPI_ISL_442883, EPI_ISL_442884, EPI_ISL_442885, EPI_ISL_442886, EPI_ISL_442887, EPI_ISL_442888, EPI_ISL_442889, EPI_ISL_442890, EPI_ISL_442891, EPI_ISL_442892, EPI_ISL_442894, EPI_ISL_442895, EPI_ISL_442896, EPI_ISL_442897, EPI_ISL_442898, EPI_ISL_442899, EPI_ISL_442900, EPI_ISL_442901, EPI_ISL_442902, EPI_ISL_442903, EPI_ISL_442904, EPI_ISL_442905, EPI_ISL_442906, EPI_ISL_442907, EPI_ISL_442908, EPI_ISL_442909, EPI_ISL_442910, EPI_ISL_442911, EPI_ISL_442912, EPI_ISL_442913, EPI_ISL_442914, EPI_ISL_442915, EPI_ISL_442916, EPI_ISL_442917, EPI_ISL_442918, EPI_ISL_442919, EPI_ISL_442921, EPI_ISL_442922, EPI_ISL_442923, EPI_ISL_442924, EPI_ISL_442925, EPI_ISL_442926, EPI_ISL_442927, EPI_ISL_442928, EPI_ISL_442929, EPI_ISL_442930, EPI_ISL_442931, EPI_ISL_442932, EPI_ISL_442933, EPI_ISL_442934, EPI_ISL_442935, EPI_ISL_442936, EPI_ISL_442937, EPI_ISL_442938, EPI_ISL_442939, EPI_ISL_442940, EPI_ISL_442941, EPI_ISL_442942, EPI_ISL_442943, EPI_ISL_442944, EPI_ISL_442945, EPI_ISL_442946, EPI_ISL_442947, EPI_ISL_442948, EPI_ISL_442949, EPI_ISL_442950, EPI_ISL_442951, EPI_ISL_442952, EPI_ISL_442953, EPI_ISL_442955, EPI_ISL_442956, EPI_ISL_442957, EPI_ISL_442958, EPI_ISL_442959, EPI_ISL_442960, EPI_ISL_442961, EPI_ISL_442962, EPI_ISL_442963, EPI_ISL_442964, EPI_ISL_442965, EPI_ISL_442966, EPI_ISL_442967, EPI_ISL_442968, EPI_ISL_442969, EPI_ISL_442970, EPI_ISL_442971, EPI_ISL_442973, EPI_ISL_442974, EPI_ISL_442975, EPI_ISL_442976, EPI_ISL_442977, EPI_ISL_442978, EPI_ISL_442979, EPI_ISL_442980, EPI_ISL_442981, EPI_ISL_442982, EPI_ISL_442983, EPI_ISL_442984, EPI_ISL_442985, EPI_ISL_442986, EPI_ISL_442987, EPI_ISL_442988, EPI_ISL_442989, EPI_ISL_442990, EPI_ISL_442991, EPI_ISL_442992, EPI_ISL_442993, EPI_ISL_442995, EPI_ISL_442996, EPI_ISL_442997, EPI_ISL_442998, EPI_ISL_442999, EPI_ISL_443000, EPI_ISL_443001, EPI_ISL_443002, EPI_ISL_443003, EPI_ISL_443004, EPI_ISL_443005, EPI_ISL_443006, EPI_ISL_443007, EPI_ISL_443008, EPI_ISL_443009, EPI_ISL_443010, EPI_ISL_443011, EPI_ISL_443012, EPI_ISL_443013, EPI_ISL_443014, EPI_ISL_443015, EPI_ISL_443016, EPI_ISL_443017, EPI_ISL_443018, EPI_ISL_443019, EPI_ISL_443020, EPI_ISL_443021, EPI_ISL_443022, EPI_ISL_443023, EPI_ISL_443024, EPI_ISL_443026, EPI_ISL_443027, EPI_ISL_443028, EPI_ISL_443029, EPI_ISL_443030, EPI_ISL_443031, EPI_ISL_443033, EPI_ISL_443034, EPI_ISL_443035, EPI_ISL_443036, EPI_ISL_443037, EPI_ISL_443038, EPI_ISL_443039, EPI_ISL_443040, EPI_ISL_443041, EPI_ISL_443042, EPI_ISL_443043, EPI_ISL_443044, EPI_ISL_443045, EPI_ISL_443046, EPI_ISL_443047, EPI_ISL_443048, EPI_ISL_443049, EPI_ISL_443050, EPI_ISL_443051, EPI_ISL_443052, EPI_ISL_443053, EPI_ISL_443054, EPI_ISL_443056, EPI_ISL_443057, EPI_ISL_443058, EPI_ISL_443059, EPI_ISL_443060, EPI_ISL_443062, EPI_ISL_443064, EPI_ISL_443065, EPI_ISL_443066, EPI_ISL_443068, EPI_ISL_443069, EPI_ISL_443070, EPI_ISL_443071, EPI_ISL_443072, EPI_ISL_443073, EPI_ISL_443074, EPI_ISL_443076, EPI_ISL_443077, EPI_ISL_443078, EPI_ISL_443079, EPI_ISL_443080, EPI_ISL_443081, EPI_ISL_443082, EPI_ISL_443083, EPI_ISL_443084, EPI_ISL_443085, EPI_ISL_443086, EPI_ISL_443087, EPI_ISL_443088, EPI_ISL_443089, EPI_ISL_443090, EPI_ISL_443091, EPI_ISL_443092, EPI_ISL_443093, EPI_ISL_443094, EPI_ISL_443095, EPI_ISL_443096, EPI_ISL_443097, EPI_ISL_443098, EPI_ISL_443099, EPI_ISL_443100, EPI_ISL_443102, EPI_ISL_443103, EPI_ISL_443104, EPI_ISL_443105, EPI_ISL_443107, EPI_ISL_443108, EPI_ISL_443109, EPI_ISL_443110, EPI_ISL_443111, EPI_ISL_443112, EPI_ISL_443113, EPI_ISL_443114, EPI_ISL_443115, EPI_ISL_443116, EPI_ISL_443118 | | | |
| see above | Department of Pathology, University of Cambridge | Wellcome Sanger Institute for the COVID-19 Genomics UK (COG-UK) consortium | Luke W Meredith, M. Estée Török , Myra Hosmillo, William L. Hamilton, Martin D. Curran, Theresa Feltwell, Grant Hall, Anna Yakovleva, Fahad A Khokhar, Charlotte J. Houldcroft, Laura G Caller, Aminu S. Jahun, Sarah L. Caddy, Ian Goodfellow, Alex Alderton, Roberto Amato, Sonia Goncalves, Ewan Harrison, David K. Jackson, Ian Johnston, Dominic Kwiatkowski, Cordelia Langford, John Sillitoe on behalf of the Wellcome Sanger Institute COVID-19 Surveillance Team (http://www.sanger.ac.uk/covid-team) |
| EPI_ISL_443190, EPI_ISL_443212, EPI_ISL_443213, EPI_ISL_443244, EPI_ISL_443245, EPI_ISL_443246 | National Public Health Laboratory, National Centre for Infectious Diseases | National Public Health Laboratory, National Centre for Infectious Diseases | Mak Tze Minn, Octavia Sophie, Chavatte Jean-Marc, Cui Lin, Lin Raymond Tzer Pin |
| EPI_ISL_443303 | Résidence Les Marines | National Reference Center for Viruses of Respiratory | Mélanie Albert, Marion Barbet, Sylvie Behillil, Méline Bizard, Angela Brisebarre, Flora Donati, Etienne Simon-Lorière, Vincent Enouf, Maud Vanpeene, |

| | | | |
|---|---|---|---|
| | | Infections, Institut Pasteur, Paris | Sylvie van der Werf |
| EPI_ISL_443307 | La Villa Papyri | National Reference Center for Viruses of Respiratory Infections, Institut Pasteur, Paris | Mélanie Albert, Marion Barbet, Sylvie Behillil, Méline Bizard, Angela Brisebarre, Flora Donati, Etienne Simon-Lorière, Vincent Enouf, Maud Vanpeene, Sylvie van der Werf |
| EPI_ISL_443320, EPI_ISL_443322, EPI_ISL_443333, EPI_ISL_443353, EPI_ISL_443356, EPI_ISL_443366, EPI_ISL_443367, EPI_ISL_443373, EPI_ISL_443381, EPI_ISL_443400, EPI_ISL_443425, EPI_ISL_443427, EPI_ISL_443428, EPI_ISL_443430, EPI_ISL_443437, EPI_ISL_443440, EPI_ISL_443461, EPI_ISL_443465, EPI_ISL_443467, EPI_ISL_443487, EPI_ISL_443504, EPI_ISL_443521, EPI_ISL_443540, EPI_ISL_443545, EPI_ISL_443552, EPI_ISL_443563, EPI_ISL_443587, EPI_ISL_443592, EPI_ISL_443626, EPI_ISL_443628, EPI_ISL_443670, EPI_ISL_443673 | | | |
| see above | Department of Pathology, University of Cambridge | Wellcome Sanger Institute for the COVID-19 Genomics UK (COG-UK) consortium | Luke W Meredith, M. Estée Török , Myra Hosmillo, William L. Hamilton, Martin D. Curran, Theresa Feltwell, Grant Hall, Anna Yakovleva, Fahad A Khokhar, Charlotte J. Houldcroft, Laura G Caller, Aminu S. Jahun, Sarah L. Caddy, Ian Goodfellow; and Alex Alderton, Roberto Amato, Sonia Goncalves, Ewan Harrison, David K. Jackson, Ian Johnston, Dominic Kwiatkowski, Cordelia Langford, John Sillitoe on behalf of the Wellcome Sanger Institute COVID-19 Surveillance Team (http://www.sanger.ac.uk/covid-team) |
| EPI_ISL_444039 | United Christian Hospital | Hong Kong Department of Health | Mak Gannon C.K., Cheng Peter K.C., Lam Edman T.K., Chan Rickjason C.W., Tsang Dominic N.C. |
| EPI_ISL_444040 | Queen Elizabeth Hospital | Hong Kong Department of Health | Mak Gannon C.K., Cheng Peter K.C., Lam Edman T.K., Chan Rickjason C.W., Tsang Dominic N.C. |
| EPI_ISL_444041 | Tuen Mun Hospital | Hong Kong Department of Health | Mak Gannon C.K., Cheng Peter K.C., Lam Edman T.K., Chan Rickjason C.W., Tsang Dominic N.C. |
| EPI_ISL_444042, EPI_ISL_444043 | Pamela Youde Nethersole Eastern Hospital | Hong Kong Department of Health | Mak Gannon C.K., Cheng Peter K.C., Lam Edman T.K., Chan Rickjason C.W., Tsang Dominic N.C. |
| EPI_ISL_444044 | Yan Chai Hospital | Hong Kong Department of Health | Mak Gannon C.K., Cheng Peter K.C., Lam Edman T.K., Chan Rickjason C.W., Tsang Dominic N.C. |
| EPI_ISL_444045, EPI_ISL_444046 | Queen Elizabeth Hospital | Hong Kong Department of Health | Mak Gannon C.K., Cheng Peter K.C., Lam Edman T.K., Chan Rickjason C.W., Tsang Dominic N.C. |
| EPI_ISL_444050 | Shek Wu Hui Jockey Club General Out-patient Clinic | Hong Kong Department of Health | Mak Gannon C.K., Cheng Peter K.C., Lam Edman T.K., Chan Rickjason C.W., Tsang Dominic N.C. |
| EPI_ISL_444121, EPI_ISL_444122, EPI_ISL_444123, EPI_ISL_444124, EPI_ISL_444125, EPI_ISL_444126, EPI_ISL_444127, EPI_ISL_444128, EPI_ISL_444129, EPI_ISL_444130, EPI_ISL_444131, EPI_ISL_444132, EPI_ISL_444133, EPI_ISL_444134, EPI_ISL_444135, EPI_ISL_444136, EPI_ISL_444137, EPI_ISL_444138, EPI_ISL_444139, EPI_ISL_444140, EPI_ISL_444141, EPI_ISL_444142, EPI_ISL_444143, EPI_ISL_444144, EPI_ISL_444145, EPI_ISL_444146, EPI_ISL_444147, EPI_ISL_444148, EPI_ISL_444149, EPI_ISL_444150, EPI_ISL_444151, EPI_ISL_444152, EPI_ISL_444153, EPI_ISL_444154, EPI_ISL_444157, EPI_ISL_444158, EPI_ISL_444159, EPI_ISL_444160, EPI_ISL_444161, EPI_ISL_444162, EPI_ISL_444163, EPI_ISL_444164, EPI_ISL_444165, EPI_ISL_444166, EPI_ISL_444167, EPI_ISL_444168, EPI_ISL_444169, EPI_ISL_444170, EPI_ISL_444171, EPI_ISL_444172, EPI_ISL_444173, EPI_ISL_444174, EPI_ISL_444175, EPI_ISL_444176, EPI_ISL_444177, EPI_ISL_444178, EPI_ISL_444179, EPI_ISL_444180, EPI_ISL_444181, EPI_ISL_444182 | | | |
| see above | University College London, Great Ormond Street Hospital for Children NHS Foundation Trust, Imperial College Healthcare NHS Trust | COVID-19 Genomics UK (COG-UK) Consortium | Sergi Castellano, Rachel Williams, Mark Kristiansen, Paola Resende Silva, Sunando Roy, Tony Brooks, Helena Tutill, Paola Niola, Patricia Dyal, Charlotte Williams, Leysa Forrest, Yasmin Panchbhaya, Jacqueline Findlay, Sam Weeks, Julianne Brown, Kathryn Harris, Paul Randell, James Price, Alison Holmes, Judith Breuer |
| EPI_ISL_444394, EPI_ISL_444395, EPI_ISL_444396 | Department of Pathology, University of Cambridge | COVID-19 Genomics UK (COG-UK) Consortium | Luke W Meredith, M. Estée Török , Myra Hosmillo, William L. Hamilton, Martin D. Curran, Theresa Feltwell, Grant Hall, Anna Yakovleva, Fahad A Khokhar, Charlotte J. Houldcroft, Laura G Caller, Aminu S. Jahun, Sarah L. Caddy, Ian Goodfellow |
| EPI_ISL_444494, EPI_ISL_444496, EPI_ISL_444497, EPI_ISL_444500, EPI_ISL_444511, EPI_ISL_444514 | Laboratoire de microbiologie, Hopital de Verdun | Smith Laboratory, Centre de Recherche CHU Sainte-Justine | Martin Smith, Marieke Rozendaal, Ivan Pavlov |
| EPI_ISL_444770, EPI_ISL_444771, EPI_ISL_444772 | NYU Langone Health | Departments of Pathology and Medicine, New York University School of Medicine | Maria Aguero-Rosenfeld, Brendan Belovarac, Margaret Black, Ludovic Boytard, John Cadley, Paolo Cotzia, John Chen, Dacia Dimartino, Xiaojun Feng, Tatyana Gindin, Emily Guzman, Adriana Heguy, Megan Hogan, Emily Huang, George Jour, Alireza Khodadadi-Jamayran, Lawrence H. Lin, Raven Luther, Andrew Lytle, Christian Marier, Matthew T. Maurano, Mark J. Mulligan, Peter Meyn, Raquel Ordonez Ciriza, Iman Osman, Jared Pinnell, Vanessa Raabe, Sitharam Ramaswami, Amy Rapkiewicz, Andre M. Ribeiro-dos-Santos, Marie Samanovic-Golden, Antonio Serrano, Guomiao Shen, Matija Snuderl, Theodore Vougiouklakis, Nick Vulpescu, Gael Westby, Paul Zappile, Yutong Zhang |
| EPI_ISL_444975 | Hospital Universitari Vall d'Hebron - Vall d'Hebron Institut de Recerca | Hospital Universitari Vall d'Hebron | Cristina Andrés, Maria Piñana, Damir Garcia-Cehic, Mercedes Guerrero-Murillo, Ariadna Rando, Juliana Esperalba, Maria Gema Codina, Tomàs Pumarola, Josep Quer, Andrés Antón |
| EPI_ISL_445147, EPI_ISL_445148, EPI_ISL_445149, EPI_ISL_445150, EPI_ISL_445151, EPI_ISL_445152, EPI_ISL_445153, EPI_ISL_445154, EPI_ISL_445155, EPI_ISL_445156, EPI_ISL_445157, EPI_ISL_445158, EPI_ISL_445159, EPI_ISL_445160, EPI_ISL_445161, EPI_ISL_445163 | | | |
| see above | Robert Garry lab | Andersen lab at Scripps Research | Allison Smither, Gilberto Sabino-Santos, Patricia Snarski, Lilia Melnik, Antoinette Bell, Kaylynn Genemaras, Arnaud Drouin, Dahlene Fusco, Robert Garry with SEARCH Alliance San Diego |
| EPI_ISL_445181, EPI_ISL_445182 | UCSF Clinical Microbiology Laboratory | Chan-Zuckerberg Biohub | CZB Cliahub Consortium |
| EPI_ISL_445552, EPI_ISL_445582, EPI_ISL_445583, EPI_ISL_445584, EPI_ISL_445585, EPI_ISL_445586, EPI_ISL_445601, EPI_ISL_445602, EPI_ISL_445604, EPI_ISL_445605, EPI_ISL_445616, EPI_ISL_445617, EPI_ISL_445618, EPI_ISL_445619, EPI_ISL_445620, EPI_ISL_445621, EPI_ISL_445622, EPI_ISL_445623, EPI_ISL_445624, EPI_ISL_445625, EPI_ISL_445626, EPI_ISL_445627, EPI_ISL_445628, EPI_ISL_445629, EPI_ISL_445630, EPI_ISL_445631, EPI_ISL_445632, EPI_ISL_445633, EPI_ISL_445634, EPI_ISL_445635, EPI_ISL_445636, EPI_ISL_445637, EPI_ISL_445638, EPI_ISL_445639, EPI_ISL_445640, EPI_ISL_445641, EPI_ISL_445647, EPI_ISL_445649, EPI_ISL_445654, EPI_ISL_445655, EPI_ISL_445656, EPI_ISL_445657, EPI_ISL_445658, EPI_ISL_445659, EPI_ISL_445670, EPI_ISL_445671, EPI_ISL_445672, EPI_ISL_445673, EPI_ISL_445699, EPI_ISL_445700, EPI_ISL_445701, EPI_ISL_445702, EPI_ISL_445703, EPI_ISL_445704, EPI_ISL_445705, EPI_ISL_445706, EPI_ISL_445707, EPI_ISL_445708, EPI_ISL_445709, EPI_ISL_445710, EPI_ISL_445711, EPI_ISL_445712, EPI_ISL_445713, EPI_ISL_445714, EPI_ISL_445715, EPI_ISL_445716, EPI_ISL_445717, EPI_ISL_445718, EPI_ISL_445719, EPI_ISL_445720, EPI_ISL_445721, EPI_ISL_445722, EPI_ISL_445723, EPI_ISL_445724, EPI_ISL_445725, EPI_ISL_445726, EPI_ISL_445727, EPI_ISL_445728, EPI_ISL_445729, EPI_ISL_445730, EPI_ISL_445731, EPI_ISL_445732, EPI_ISL_445733, EPI_ISL_445737, EPI_ISL_445738, EPI_ISL_445740, EPI_ISL_445741, EPI_ISL_445744, EPI_ISL_445745, EPI_ISL_445746, EPI_ISL_445747, EPI_ISL_445750, EPI_ISL_445751, EPI_ISL_445752, EPI_ISL_445756, EPI_ISL_445757, EPI_ISL_445758, EPI_ISL_445761, EPI_ISL_445762, EPI_ISL_445763, EPI_ISL_445764, EPI_ISL_445765, EPI_ISL_445766, EPI_ISL_445767, EPI_ISL_445768, EPI_ISL_445769, EPI_ISL_445770, EPI_ISL_445771, EPI_ISL_445772, EPI_ISL_445773, EPI_ISL_445774, EPI_ISL_445775, EPI_ISL_445776, EPI_ISL_445777, EPI_ISL_445778, EPI_ISL_445779, EPI_ISL_445780, EPI_ISL_445781, EPI_ISL_445783, EPI_ISL_445784, EPI_ISL_445785, EPI_ISL_445823, EPI_ISL_445824, EPI_ISL_445825, EPI_ISL_445826, EPI_ISL_445827, EPI_ISL_445828, EPI_ISL_445829, EPI_ISL_445830, EPI_ISL_445831, EPI_ISL_445832, EPI_ISL_445833, EPI_ISL_445834, EPI_ISL_445835, EPI_ISL_445836, EPI_ISL_445837, EPI_ISL_445838, EPI_ISL_445839, EPI_ISL_445840, EPI_ISL_445841, EPI_ISL_445842, EPI_ISL_445843, EPI_ISL_445844, EPI_ISL_445845, EPI_ISL_445846, EPI_ISL_445847, EPI_ISL_445848, EPI_ISL_445849, EPI_ISL_445850, EPI_ISL_445851, EPI_ISL_445852, EPI_ISL_445853, EPI_ISL_445854, EPI_ISL_445855, EPI_ISL_445856, EPI_ISL_445857, EPI_ISL_445858, EPI_ISL_445859, EPI_ISL_445860, EPI_ISL_445861, EPI_ISL_445862, EPI_ISL_445863, EPI_ISL_445864, EPI_ISL_445865, EPI_ISL_445866, EPI_ISL_445867, EPI_ISL_445868, EPI_ISL_445869, EPI_ISL_445870, EPI_ISL_445871, EPI_ISL_445872, EPI_ISL_445873, EPI_ISL_445874, EPI_ISL_446034, EPI_ISL_446035, EPI_ISL_446036, EPI_ISL_446037, EPI_ISL_446038, EPI_ISL_446039, EPI_ISL_446040, EPI_ISL_446041, EPI_ISL_446042, EPI_ISL_446043, EPI_ISL_446044, EPI_ISL_446045, EPI_ISL_446046, EPI_ISL_446047, EPI_ISL_446048, EPI_ISL_446049, EPI_ISL_446050, EPI_ISL_446051, EPI_ISL_446052, EPI_ISL_446053, EPI_ISL_446054, EPI_ISL_446055, EPI_ISL_446056, EPI_ISL_446057, EPI_ISL_446058, EPI_ISL_446059, EPI_ISL_446060, EPI_ISL_446061, EPI_ISL_446062, EPI_ISL_446063, EPI_ISL_446064, EPI_ISL_446065, EPI_ISL_446066, EPI_ISL_446067, EPI_ISL_446068, EPI_ISL_446069, EPI_ISL_446070, EPI_ISL_446071, EPI_ISL_446072, EPI_ISL_446073, EPI_ISL_446074, EPI_ISL_446075, EPI_ISL_446076, EPI_ISL_446077, EPI_ISL_446078, EPI_ISL_446079, EPI_ISL_446080, EPI_ISL_446092, EPI_ISL_446093, EPI_ISL_446094, EPI_ISL_446095, EPI_ISL_446096, EPI_ISL_446097, EPI_ISL_446098, EPI_ISL_446099, EPI_ISL_446100, EPI_ISL_446101, EPI_ISL_446102, EPI_ISL_446103, EPI_ISL_446104, EPI_ISL_446105, EPI_ISL_446106, EPI_ISL_446107, EPI_ISL_446108, EPI_ISL_446109, EPI_ISL_446110, EPI_ISL_446111, EPI_ISL_446112, EPI_ISL_446113, EPI_ISL_446114, EPI_ISL_446115, EPI_ISL_446116, EPI_ISL_446117, EPI_ISL_446118, EPI_ISL_446121, EPI_ISL_446123, EPI_ISL_446124, EPI_ISL_446125, EPI_ISL_446127, EPI_ISL_446128, EPI_ISL_446130, EPI_ISL_446131, EPI_ISL_446135, EPI_ISL_446136, EPI_ISL_446137, EPI_ISL_446138, EPI_ISL_446140, EPI_ISL_446144, EPI_ISL_446145, EPI_ISL_446148, EPI_ISL_446150, EPI_ISL_446151, EPI_ISL_446152, EPI_ISL_446153, EPI_ISL_446154, EPI_ISL_446155, EPI_ISL_446156, EPI_ISL_446157, EPI_ISL_446158, EPI_ISL_446159, EPI_ISL_446160, EPI_ISL_446162, EPI_ISL_446163, EPI_ISL_446165, EPI_ISL_446166, EPI_ISL_446167, EPI_ISL_446168, EPI_ISL_446171, EPI_ISL_446172, EPI_ISL_446173, EPI_ISL_446174, EPI_ISL_446227, EPI_ISL_446245, EPI_ISL_446246, EPI_ISL_446247, EPI_ISL_446248, EPI_ISL_446249, EPI_ISL_446250, EPI_ISL_446251, EPI_ISL_446252, EPI_ISL_446253, EPI_ISL_446254, EPI_ISL_446272, EPI_ISL_446273, EPI_ISL_446274, EPI_ISL_446275, EPI_ISL_446276, EPI_ISL_446277, EPI_ISL_446278, EPI_ISL_446279, EPI_ISL_446280, EPI_ISL_446281, EPI_ISL_446282, EPI_ISL_446283, EPI_ISL_446284, EPI_ISL_446285, EPI_ISL_446286, EPI_ISL_446287, EPI_ISL_446288, EPI_ISL_446289, EPI_ISL_446290, EPI_ISL_446291, EPI_ISL_446292, EPI_ISL_446293, EPI_ISL_446294, EPI_ISL_446295, EPI_ISL_446296, EPI_ISL_446297, EPI_ISL_446298, EPI_ISL_446299, EPI_ISL_446300, EPI_ISL_446301, EPI_ISL_446302, EPI_ISL_446303, EPI_ISL_446304, EPI_ISL_446305, EPI_ISL_446306, EPI_ISL_446307, EPI_ISL_446308, EPI_ISL_446309, EPI_ISL_446310, EPI_ISL_446311, EPI_ISL_446312, EPI_ISL_446313, EPI_ISL_446314, EPI_ISL_446315, EPI_ISL_446316, EPI_ISL_446317, EPI_ISL_446318, EPI_ISL_446319, EPI_ISL_446320, EPI_ISL_446321, EPI_ISL_446322, EPI_ISL_446323, EPI_ISL_446324, EPI_ISL_446325, EPI_ISL_446326, EPI_ISL_446327, EPI_ISL_446328, EPI_ISL_446329, EPI_ISL_446330, EPI_ISL_446331, EPI_ISL_446332, EPI_ISL_446333, EPI_ISL_446334, EPI_ISL_446335, EPI_ISL_446336, EPI_ISL_446337, EPI_ISL_446338, EPI_ISL_446339, EPI_ISL_446340, EPI_ISL_446341, EPI_ISL_446342, EPI_ISL_446343, EPI_ISL_446344, EPI_ISL_446345, EPI_ISL_446346, EPI_ISL_446347, EPI_ISL_446348, EPI_ISL_446349, EPI_ISL_446350, EPI_ISL_446351, EPI_ISL_446352, EPI_ISL_446353, EPI_ISL_446354, EPI_ISL_446355, EPI_ISL_446356, EPI_ISL_446357, EPI_ISL_446358, EPI_ISL_446359, EPI_ISL_446360, EPI_ISL_446361, EPI_ISL_446362, EPI_ISL_446363, EPI_ISL_446364, EPI_ISL_446365, EPI_ISL_446366, EPI_ISL_446367, EPI_ISL_446368, EPI_ISL_446369, EPI_ISL_446370, EPI_ISL_446371, EPI_ISL_446372, EPI_ISL_446373, EPI_ISL_446374, EPI_ISL_446375, EPI_ISL_446376, EPI_ISL_446377, EPI_ISL_446378, EPI_ISL_446379, EPI_ISL_446380, EPI_ISL_446381, EPI_ISL_446382, EPI_ISL_446383, EPI_ISL_446384, EPI_ISL_446385, EPI_ISL_446386, EPI_ISL_446387, EPI_ISL_446388, EPI_ISL_446389, EPI_ISL_446390, EPI_ISL_446391, EPI_ISL_446392, EPI_ISL_446393, EPI_ISL_446394, EPI_ISL_446395, EPI_ISL_446396, EPI_ISL_446397, EPI_ISL_446398, EPI_ISL_446399, EPI_ISL_446400, EPI_ISL_446401, EPI_ISL_446402, EPI_ISL_446403, EPI_ISL_446404, EPI_ISL_446405, EPI_ISL_446406, EPI_ISL_446407, EPI_ISL_446408, EPI_ISL_446409, EPI_ISL_446410, EPI_ISL_446411, EPI_ISL_446412, EPI_ISL_446413, EPI_ISL_446414, EPI_ISL_446415, EPI_ISL_446416, EPI_ISL_446417, EPI_ISL_446418, EPI_ISL_446419, EPI_ISL_446420, EPI_ISL_446421, EPI_ISL_446422, EPI_ISL_446423, EPI_ISL_446424, EPI_ISL_446425, EPI_ISL_446426, EPI_ISL_446427, EPI_ISL_446428, EPI_ISL_446429, EPI_ISL_446430, EPI_ISL_446431, EPI_ISL_446432, EPI_ISL_446433, EPI_ISL_446434, EPI_ISL_446435, EPI_ISL_446436, EPI_ISL_446437, EPI_ISL_446438, EPI_ISL_446439, EPI_ISL_446440, EPI_ISL_446442, EPI_ISL_446443, EPI_ISL_446445, EPI_ISL_446446, EPI_ISL_446447, EPI_ISL_446448, EPI_ISL_446449, EPI_ISL_446450, EPI_ISL_446451, EPI_ISL_446452, EPI_ISL_446453, EPI_ISL_446454, EPI_ISL_446455, EPI_ISL_446456, EPI_ISL_446457, EPI_ISL_446458, EPI_ISL_446459, EPI_ISL_446460, EPI_ISL_446461, EPI_ISL_446462, EPI_ISL_446463, EPI_ISL_446464, EPI_ISL_446466, EPI_ISL_446468, EPI_ISL_446469, EPI_ISL_446470, EPI_ISL_446472, EPI_ISL_446474, EPI_ISL_446475, EPI_ISL_446476, EPI_ISL_446477, EPI_ISL_446479, EPI_ISL_446484, EPI_ISL_446485, EPI_ISL_446486, EPI_ISL_446492, EPI_ISL_446493, EPI_ISL_446494, EPI_ISL_446495, EPI_ISL_446496, EPI_ISL_446497, EPI_ISL_446498, EPI_ISL_446499, EPI_ISL_446500, EPI_ISL_446501, EPI_ISL_446502, EPI_ISL_446503, EPI_ISL_446504, EPI_ISL_446505, EPI_ISL_446506, EPI_ISL_446507, EPI_ISL_446508, EPI_ISL_446509, EPI_ISL_446510, EPI_ISL_446511, EPI_ISL_446512, EPI_ISL_446513, EPI_ISL_446514, EPI_ISL_446515, EPI_ISL_446516, EPI_ISL_446517, EPI_ISL_446518, EPI_ISL_446519, EPI_ISL_446520, EPI_ISL_446521, EPI_ISL_446522, EPI_ISL_446523, EPI_ISL_446524, EPI_ISL_446525, EPI_ISL_446526, EPI_ISL_446527, EPI_ISL_446528, EPI_ISL_446529, EPI_ISL_446530, EPI_ISL_446531, EPI_ISL_446532, EPI_ISL_446533, EPI_ISL_446534, EPI_ISL_446535, EPI_ISL_446536, EPI_ISL_446537, EPI_ISL_446538, EPI_ISL_446539, EPI_ISL_446540, EPI_ISL_446541, EPI_ISL_446542, EPI_ISL_446543, EPI_ISL_446544, EPI_ISL_446545, EPI_ISL_446556, EPI_ISL_446557, EPI_ISL_446558, EPI_ISL_446559, EPI_ISL_446561, EPI_ISL_446562, EPI_ISL_446563, EPI_ISL_446564, EPI_ISL_446565, EPI_ISL_446566, EPI_ISL_446567, EPI_ISL_446568, EPI_ISL_446569, EPI_ISL_446570, EPI_ISL_446571, EPI_ISL_446572, EPI_ISL_446573, EPI_ISL_446574, EPI_ISL_446575, EPI_ISL_446576, EPI_ISL_446578, EPI_ISL_446579, EPI_ISL_446580, EPI_ISL_446581, EPI_ISL_446584, EPI_ISL_446585, EPI_ISL_446586, EPI_ISL_446587, EPI_ISL_446608, EPI_ISL_446619, EPI_ISL_446620, EPI_ISL_446621, EPI_ISL_446622, EPI_ISL_446623, EPI_ISL_446624, EPI_ISL_446625, EPI_ISL_446626, EPI_ISL_446627, EPI_ISL_446628, EPI_ISL_446629, EPI_ISL_446630, EPI_ISL_446631, EPI_ISL_446632, EPI_ISL_446633, EPI_ISL_446634, EPI_ISL_446635, EPI_ISL_446636, EPI_ISL_446637, EPI_ISL_446638, EPI_ISL_446639, EPI_ISL_446640, EPI_ISL_446641, EPI_ISL_446642, EPI_ISL_446643, EPI_ISL_446644, EPI_ISL_446645, EPI_ISL_446646, EPI_ISL_446647, EPI_ISL_446648, EPI_ISL_446649, EPI_ISL_446650, EPI_ISL_446651, EPI_ISL_446672, EPI_ISL_446673, EPI_ISL_446675, EPI_ISL_446676, EPI_ISL_446677, EPI_ISL_446680, EPI_ISL_446681, EPI_ISL_446682, EPI_ISL_446683, EPI_ISL_446684, EPI_ISL_446705, EPI_ISL_446906 | | | |
| see above | Wales Specialist Virology Centre | Public Health Wales Microbiology Cardiff | Catherine Moore, Johnathan Evans, Laura Gifford, Malorie Perry, Simon Cottrell, Alec Birchley, Alexander Adams, Amy Gaskin, Bree Gatica-Wilcox, Jason Coombes, Lauren Gilbert, Lee Graham, Nicole Pacchiarini, Sara Kumziene-Summerhayes, Sarah Taylor, Sophie Jones, Sara Rey, Matthew Bull, Joanne Watkins, Sally Corden, Tom Connor |

| | | | |
|---|---|---|---|
| EPI_ISL_447028, EPI_ISL_447029 | Ramathibodi Hospital | COVID-19 Network Investigations (CONI) Alliance | Elizabeth Batty, Wasun Chantratita, Thanat Chookajorn, Stefan Fernandez, Angkana Huang, Anthony R. Jones, Khajohn Joonsalak, Chonticha Klungtong, Theerarat Kochakarn, Namfon Kotanan, Krittikorn Kumpornsin, Wudtichai Manasatienkij, Bhakbhoom Panthan, Ekawat Pasomsub, Kingkan Rakmanee, Insee Sensorn, Janjira Thaipadungpanit, Arporn Wangwiwatsin,Treewat Watthanachockchai |
| EPI_ISL_447139 | Department of Clinical Microbiology | GIGA Medical Genomics | Keith Durkin, Maria Artesi, Sébastien Bontems, Raphaël Boreux, Cécile Meex, Pierrette Melin, Marie-Pierre Hayette, Vincent Bours. |
| EPI_ISL_447224 | Michigan Department of Health and Human Services, Bureau of Laboratories | Michigan Department of Health and Human Services, Bureau of Laboratories | Blankenship HM, Riner D, Soehnlen MK |
| EPI_ISL_447579 | CSIR-Centre for Cellular and Molecular Biology | CSIR-Centre for Cellular and Molecular Biology | Tulasi Nagabandi, Namami Gaur, Sakshi Shambhavi, Lamuk Zaveri, Shagufta Khan, Purushotham Vodnala, Payel Mukherjee, Sofia Banu, Priya Singh, Dhiviya Vedagiri, Divya Gupta, Vishal Sah, Santosh Kumar Kuncha, Krishnan Harinivas Harshan, Archana Bharadwaj Siva, Karthik Bharadwaj Tallapaka, Rakesh K Mishra, Divya Tej Sowpati |
| EPI_ISL_447816, EPI_ISL_447817 | Instituto Nacional de Salud, Bogotá, Colombia | Grupo de Investigaciones Microbiológicas-UR (GIMUR), Departamento de Biología, Facultad de Ciencias Naturales, Universidad del Rosario, Bogotá, Colombia Instituto Nacional de Salud, Bogotá, Colombia Icahn School of Medicine at Mount Sinai, New York, USA | Juan David Ramírez, Carolina Florez, Marina Muñoz, Carolina Hernandez, Adriana Castillo, Sergio Castañeda, Nathalia Ballesteros, David Martínez, Laura Vega, Jesús E. Jaimes, Sergio Gomez, Angelica Rico, Lisseth Pardo, Esther C. Barros, Martha L. Ospina, Aníbal A. Teherán, Ana S. Gonzalez-Reiche, Matthew M. Hernandez, Emilia Mia Sordillo, Viviana Simon, Harm van Bakel, Alberto Paniz-Mondolfi |
| EPI_ISL_447895, EPI_ISL_447896 | University of California, Davis | Chan-Zuckerberg Biohub | CZB Cliahub Consortium |
| EPI_ISL_447903 | University of Florida | University of Florida | Elbadry,M.A., Subramaniam,K., Waltzek,T.B., Lauzardo,M., Gibson,J.C., Stephenson,C.J., Alam,M.M., Morris,J.G. Jr. and Lednicky,J.A. |
| EPI_ISL_448129, EPI_ISL_448130, EPI_ISL_448131, EPI_ISL_448132, EPI_ISL_448133, EPI_ISL_448134, EPI_ISL_448135, EPI_ISL_448136, EPI_ISL_448137, EPI_ISL_448138, EPI_ISL_448139, EPI_ISL_448140, EPI_ISL_448141, EPI_ISL_448142, EPI_ISL_448143, EPI_ISL_448144, EPI_ISL_448145, EPI_ISL_448146, EPI_ISL_448147, EPI_ISL_448148, EPI_ISL_448149, EPI_ISL_448150, EPI_ISL_448151, EPI_ISL_448152, EPI_ISL_448153, EPI_ISL_448154, EPI_ISL_448155, EPI_ISL_448156, EPI_ISL_448157, EPI_ISL_448158, EPI_ISL_448159, EPI_ISL_448160, EPI_ISL_448161, EPI_ISL_448162, EPI_ISL_448163, EPI_ISL_448164, EPI_ISL_448165, EPI_ISL_448166, EPI_ISL_448167, EPI_ISL_448168, EPI_ISL_448169, EPI_ISL_448170, EPI_ISL_448171, EPI_ISL_448172, EPI_ISL_448173, EPI_ISL_448176, EPI_ISL_448177 | | | |
| see above | West of Scotland Specialist Virology Centre, NHSGGC / MRC-University of Glasgow Centre for Virus Research | COVID-19 Genomics UK (COG-UK) Consortium | Ana da Silva Filipe, Natasha Johnson, Kathy Smollett, Daniel Mair, Stephen Carmichael, Lily Tong, Jenna Nichols, Elihu Aranday-Cortes, Kirstyn Brunker, Yasmin Parr, Kyriaki Nomikou, Sarah McDonald, Marc Niebel, Patawee Asamaphan, Richard Orton, Joseph Hughes, Sreenu Vattipally, David L Robertson, Alasdair MacLean, Rory Gunson, Kathy Li, Natasha Jesudason, Rajiv Shah, James Shepherd, Antonia Ho, Emma Thomson |
| EPI_ISL_448228, EPI_ISL_448231, EPI_ISL_448232, EPI_ISL_448233, EPI_ISL_448234, EPI_ISL_448235, EPI_ISL_448236, EPI_ISL_448237, EPI_ISL_448238, EPI_ISL_448240, EPI_ISL_448242, EPI_ISL_448243, EPI_ISL_448244, EPI_ISL_448246, EPI_ISL_448247, EPI_ISL_448248, EPI_ISL_448249, EPI_ISL_448250, EPI_ISL_448251, EPI_ISL_448252, EPI_ISL_448253, EPI_ISL_448254, EPI_ISL_448255, EPI_ISL_448256, EPI_ISL_448258, EPI_ISL_448259, EPI_ISL_448260, EPI_ISL_448261, EPI_ISL_448262, EPI_ISL_448263, EPI_ISL_448264, EPI_ISL_448265, EPI_ISL_448266, EPI_ISL_448267, EPI_ISL_448270, EPI_ISL_448284, EPI_ISL_448285, EPI_ISL_448294, EPI_ISL_448307 | | | |
| see above | Quadram Institute Bioscience | COVID-19 Genomics UK (COG-UK) Consortium | Dave J. Baker, Gemma L. Kay, Alp Aydin, Thanh Le-Viet, Steven Rudder, Ana P. Tedim, Anastasia Kolyva, Maria Diaz, Leonardo de Oliveira Martins, Nabil-Fareed Alikhan, Lizzie Meadows, Rachael Stanley, Ngozi Elumogo, Muhammed Yasir, Nicholas M. Thomson, Alexander J Trotter, Rachel Gilroy, Samuel Bloomfield, Claire Stuart, Andrew Bell, Reenesh Prakash, Samir Dervisevic, Alison E. Mather, John Wain, Mark Webber, Andrew J. Page, Justin O'Grady |
| EPI_ISL_448415, EPI_ISL_448416 | Queens Medical Centre, Clinical Microbiology Department / DeepSeq Nottingham | COVID-19 Genomics UK (COG-UK) Consortium | Gemma Clark, Wendy Smith, Manjinder Khakh, Hannah Howson-Wells, Jonathan Ball, Patrick McClure, Joseph Chappell, Theocharis Tsoleridis, Nadine Holmes, Matthew Carlisle, Christopher Moore, Fei Sang, Johnny Debebe, Victoria Wright, Matthew Loose |
| EPI_ISL_448449, EPI_ISL_448451, EPI_ISL_448452, EPI_ISL_448454, EPI_ISL_448456, EPI_ISL_448457, EPI_ISL_448459, EPI_ISL_448461, EPI_ISL_448463, EPI_ISL_448465, EPI_ISL_448467, EPI_ISL_448469, EPI_ISL_448471, EPI_ISL_448473, EPI_ISL_448475, EPI_ISL_448479, EPI_ISL_448481, EPI_ISL_448482, EPI_ISL_448484, EPI_ISL_448488, EPI_ISL_448495, EPI_ISL_448498, EPI_ISL_448499, EPI_ISL_448501, EPI_ISL_448502, EPI_ISL_448549, EPI_ISL_448553, EPI_ISL_448555, EPI_ISL_448557, EPI_ISL_448559, EPI_ISL_448561, EPI_ISL_448564, EPI_ISL_448565, EPI_ISL_448774, EPI_ISL_448775, EPI_ISL_448776, EPI_ISL_448777, EPI_ISL_448778, EPI_ISL_448779, EPI_ISL_448780, EPI_ISL_448781, EPI_ISL_448782, EPI_ISL_448783, EPI_ISL_448784, EPI_ISL_448785, EPI_ISL_448786, EPI_ISL_448787, EPI_ISL_448788, EPI_ISL_448789, EPI_ISL_448790, EPI_ISL_448791, EPI_ISL_448792, EPI_ISL_448793, EPI_ISL_448794, EPI_ISL_448795, EPI_ISL_448796, EPI_ISL_448797, EPI_ISL_448798, EPI_ISL_448799, EPI_ISL_448800, EPI_ISL_448801, EPI_ISL_448802, EPI_ISL_448803, EPI_ISL_448804, EPI_ISL_448805, EPI_ISL_448806, EPI_ISL_448807, EPI_ISL_448808, EPI_ISL_448809, EPI_ISL_448810, EPI_ISL_448811, EPI_ISL_448812 | | | |
| see above | Oxford Viromics, NDM, University of Oxford; Oxford University Hospitals; Basingstoke and North Hampshire Hospital | COVID-19 Genomics UK (COG-UK) Consortium | Tanya Golubchik, David Bonsall, George Macintyre, Amy Trebes, Mariateresa de Cesare, Catrin Moore, Alex Mobbs, Anita Justice, Robert Shaw, Monique Andersson, Emma Wise, Nathan Moore, Jessica Lynch, Nick Cortes, Stephen Kidd, David Buck, John Todd, Christophe Fraser |
| EPI_ISL_448844 | Virology Laboratory, Castle Hill Hospital, Hull University Teaching Hospitals NHS Trust/Department of Infection, Immunity and Cardiovascular Disease, The Medical School, University of Sheffield | COVID-19 Genomics UK (COG-UK) Consortium | Thushan de Silva, Matthew Parker, Nikki Smith, Adri Angyal, Rebecca Brown, Luke Green, Rachel Tucker, Paul Parsons, Danielle Groves, Katie Johnson, Laura Carrilero, Alex Keeley, Dave Partridge, Matthew Wyles, Benjamin Lindsey, Mehmet Yavuz, Mohammad Raza, Cariad Evans |
| EPI_ISL_448846, EPI_ISL_448865 | Virology Department, Sheffield Teaching Hospitals NHS Foundation Trust/Department of Infection, Immunity and Cardiovascular Disease, The Medical School, University of Sheffield | COVID-19 Genomics UK (COG-UK) Consortium | Thushan de Silva, Matthew Parker, Nikki Smith, Adri Angyal, Rebecca Brown, Luke Green, Rachel Tucker, Paul Parsons, Danielle Groves, Katie Johnson, Laura Carrilero, Alex Keeley, Dave Partridge, Matthew Wyles, Benjamin Lindsey, Mehmet Yavuz, Mohammad Raza, Cariad Evans |
| EPI_ISL_448981, EPI_ISL_448982, EPI_ISL_448987, EPI_ISL_448993, EPI_ISL_448996, EPI_ISL_449013, EPI_ISL_449022, EPI_ISL_449024, EPI_ISL_449099, EPI_ISL_449100, EPI_ISL_449102, EPI_ISL_449104, EPI_ISL_449105, EPI_ISL_449106, EPI_ISL_449107, EPI_ISL_449109, EPI_ISL_449110, EPI_ISL_449111, EPI_ISL_449112, EPI_ISL_449113, EPI_ISL_449114, EPI_ISL_449115, EPI_ISL_449116, EPI_ISL_449117, EPI_ISL_449118, EPI_ISL_449119, EPI_ISL_449120, EPI_ISL_449121, EPI_ISL_449122, EPI_ISL_449123, EPI_ISL_449124, EPI_ISL_449125, EPI_ISL_449126, EPI_ISL_449127, EPI_ISL_449128, EPI_ISL_449129, EPI_ISL_449130, EPI_ISL_449131, EPI_ISL_449132, EPI_ISL_449133, EPI_ISL_449134, EPI_ISL_449136, EPI_ISL_449138, EPI_ISL_449139, EPI_ISL_449140, EPI_ISL_449141, EPI_ISL_449142, EPI_ISL_449143, EPI_ISL_449144, EPI_ISL_449145, EPI_ISL_449146, EPI_ISL_449147 | | | |
| see above | Quadram Institute Bioscience | COVID-19 Genomics UK (COG-UK) Consortium | Dave J. Baker, Gemma L. Kay, Alp Aydin, Thanh Le-Viet, Steven Rudder, Ana P. Tedim, Anastasia Kolyva, Maria Diaz, Leonardo de Oliveira Martins, Nabil-Fareed Alikhan, Lizzie Meadows, Rachael Stanley, Ngozi Elumogo, Muhammed Yasir, Nicholas M. Thomson, Alexander J Trotter, Rachel Gilroy, Samuel Bloomfield, Claire Stuart, Andrew Bell, Reenesh Prakash, Samir Dervisevic, Alison E. Mather, John Wain, Mark Webber, Andrew J. Page, Justin O'Grady |
| EPI_ISL_449656, EPI_ISL_449657, EPI_ISL_449658, EPI_ISL_449659, EPI_ISL_449660 | University College London, Great Ormond Street Hospital for Children NHS Foundation Trust, Imperial College Healthcare NHS Trust | COVID-19 Genomics UK (COG-UK) Consortium | Sergi Castellano, Rachel Williams, Mark Kristiansen, Paola Resende Silva, Sunando Roy, Tony Brooks, Helena Tutill, Paola Niola, Patricia Dyal, Charlotte Williams, Leysa Forrest, Yasmin Panchbhaya, Jacqueline Findlay, Sam Weeks, Julianne Brown, Kathryn Harris, Paul Randell, James Price, Alison Holmes, Judith Breuer |
| EPI_ISL_449786 | Ostfold Hospital Trust - Kalnes, Centre for Laboratory Medicine, Section for gene technology and infection serology | Norwegian Institute of Public Health, Department of Virology | Kathrine Stene-Johansen, Kamilla Heddeland Instefjord, Hilde Elshaug, Rasmus Riis Kopperud, Karoline Bragstad, Olav Hungnes |
| EPI_ISL_449787, EPI_ISL_449788 | Furst Medical Laboratory | Norwegian Institute of Public Health, Department of Virology | Kathrine Stene-Johansen, Kamilla Heddeland Instefjord, Hilde Elshaug, Rasmus Riis Kopperud, Karoline Bragstad, Olav Hungnes |
| EPI_ISL_449815, EPI_ISL_449816, EPI_ISL_449817, EPI_ISL_449818, EPI_ISL_449819, EPI_ISL_449820, EPI_ISL_449821, EPI_ISL_449822, EPI_ISL_449823, EPI_ISL_449824, EPI_ISL_449825, EPI_ISL_449826, EPI_ISL_449827, EPI_ISL_449828, EPI_ISL_449829, EPI_ISL_449830, EPI_ISL_449831, EPI_ISL_449832 | | | |
| see above | Utah Public Health Laboratory | Utah Public Health Laboratory | Erin Young, Kelly Oakeson |
| EPI_ISL_449839, EPI_ISL_449840, EPI_ISL_449841, EPI_ISL_449842, EPI_ISL_449843, EPI_ISL_449844, EPI_ISL_449845, EPI_ISL_449846, EPI_ISL_449847, EPI_ISL_449848, EPI_ISL_449849, EPI_ISL_449850, EPI_ISL_449851, EPI_ISL_449852, EPI_ISL_449853, EPI_ISL_449854, EPI_ISL_449855, EPI_ISL_449856, EPI_ISL_449857, EPI_ISL_449858, EPI_ISL_449859, EPI_ISL_449860, EPI_ISL_449861, EPI_ISL_449862, EPI_ISL_449863, EPI_ISL_449864, EPI_ISL_449865, EPI_ISL_449866, EPI_ISL_449867, EPI_ISL_449868, EPI_ISL_449869, EPI_ISL_449870, EPI_ISL_449871, EPI_ISL_449872, EPI_ISL_449873, EPI_ISL_449874, EPI_ISL_449875, EPI_ISL_449876, EPI_ISL_449877, EPI_ISL_449878, EPI_ISL_449879, EPI_ISL_449880, EPI_ISL_449881, EPI_ISL_449882, EPI_ISL_449883, EPI_ISL_449884, EPI_ISL_449885, EPI_ISL_449886, EPI_ISL_449887, EPI_ISL_449888, EPI_ISL_449889, EPI_ISL_449890, EPI_ISL_449891, EPI_ISL_449892, EPI_ISL_449893, EPI_ISL_449894, EPI_ISL_449895, EPI_ISL_449896, EPI_ISL_449897, EPI_ISL_449898, EPI_ISL_449899, EPI_ISL_449900, EPI_ISL_449901, EPI_ISL_449914, EPI_ISL_449919, EPI_ISL_449921 | | | |
| see above | Washington State Department of Health | Seattle Flu Study | Chu et al |
| EPI_ISL_450177, EPI_ISL_450180 | Robert Garry lab | Andersen lab at Scripps Research | Allison Smither, Gilberto Sabino-Santos, Patricia Snarski, Lilia Melnik, Antoinette Bell, Kaylynn Genemaras, Arnaud Drouin, Dahlene Fusco, Robert Garry with SEARCH Alliance San Diego |
| EPI_ISL_450186, EPI_ISL_450187 | Biolab Diagnostic Laboratories | Andersen lab at Scripps Research | Issa Abu-Dayyeh, Ahmad Tibi, Lama Hussein, Lina Mohammad, Zein Naber, Amid Abdelnour with SEARCH Alliance San Diego |
| EPI_ISL_450294 | Institute of Human Genetics, Polish Academy of Sciences Sanitary and Epidemiological Station in Pozna | Institute of Human Genetics, Polish Academy of Sciences | Szymon Hryhorowicz, Adam Ustaszewski, Emilia Lis, Marta Kaczmarek-Ry, Micha Witt, Andrzej Pawski |
| EPI_ISL_450295 | Institute of Human Genetics, Polish Academy of Sciences; Sanitary and Epidemiological Station in Pozna | Institute of Human Genetics, Polish Academy of Sciences | Szymon Hryhorowicz, Adam Ustaszewski, Emilia Lis, Marta Kaczmarek-Ry, Micha Witt, Andrzej Pawski |
| EPI_ISL_450338 | Institute of Human Genetics, Polish Academy of Sciences; | Institute of Human Genetics, Polish Academy of Sciences | Szymon Hryhorowicz, Adam Ustaszewski, Emilia Lis, Marta Kaczmarek-Ry, Micha Witt, Andrzej Pawski |

| | | | |
|---|---|---|---|
| | Sanitary and Epidemiological Station in Pozna | | |
| EPI_ISL_450507 | Nigerian Institute of Medical Research | Nigerian Institute of Medical Research | Shaibu,J.O., Onwuamah,C.K., James,A.B., Okwuraiwe,A.P., Amoo,O.S., Salu,O.B., Ige,F.A., Okoli,L.C., Ahmed,R.A., Sokei,J., Oyefolu,A.O., Omilabu,S.A., Salako,B.L. and Audu,R.A. |
| EPI_ISL_450550, EPI_ISL_450551, EPI_ISL_450552, EPI_ISL_450553, EPI_ISL_450554, EPI_ISL_450555, EPI_ISL_450556, EPI_ISL_450557, EPI_ISL_450559, EPI_ISL_450560, EPI_ISL_450561 | | | |
| see above | Utah Public Health Laboratory | Utah Public Health Laboratory | Erin Young, Kelly Oakeson |
| EPI_ISL_450638, EPI_ISL_450639, EPI_ISL_450640, EPI_ISL_450641, EPI_ISL_450642, EPI_ISL_450643, EPI_ISL_450644, EPI_ISL_450645, EPI_ISL_450646, EPI_ISL_450647, EPI_ISL_450650, EPI_ISL_450651, EPI_ISL_450652 | | | |
| see above | Laboratoire de microbiologie, Hopital de Verdun | Smith Laboratory, Centre de Recherche CHU Sainte-Justine | Martin Smith, Marieke Rozendaal, Ivan Pavlov |
| EPI_ISL_450735, EPI_ISL_450736, EPI_ISL_450737 | Hospital AZ Rivierenland | Institute of Tropical Medicine | Philippe Selhorst, Colin Anthony |
| EPI_ISL_450782 | Government Medical College-Bhavnagar | Gujarat Biotechnology Research Centre | Saklain Malek, Shirish Patel, Kairavi Desai, Tejas Shah, Ankit Hinsu, Pritesh Sabara, Apurvasinh Puvar, Janvi Raval, Zarna Patel, Monika Gandhi, Pinal Trivedi, Maharshi Pandya, Amit Kanani, Nidhi Patel, Nitin Savaliya, Raghawendra Kumar, Dinesh Kumar, Zuber Saiyed, Komal Patel, Labdhi Pandya, Snehal Bagatharia, Ramesh Pandit, Bhavesh Modi, Gaurishankar Shrimali, R D Dixit, A M Kadri, Priti Pandita, Chaitanya Joshi, Madhvi Joshi |
| EPI_ISL_450783 | Government Medical College-Bhavnagar | Gujarat Biotechnology Research Centre | Shirish Patel, Kairavi Desai, Saklain Malek, Ankit Hinsu, Pritesh Sabara, Apurvasinh Puvar, Janvi Raval, Zarna Patel, Monika Gandhi, Pinal Trivedi, Maharshi Pandya, Amit Kanani, Nidhi Patel, Nitin Savaliya, Raghawendra Kumar, Dinesh Kumar, Zuber Saiyed, Komal Patel, Labdhi Pandya, Snehal Bagatharia, Ramesh Pandit, Tejas Shah, Bhavesh Modi, Gaurishankar Shrimali, R D Dixit, A M Kadri, Neha Rajpara, Chaitanya Joshi, Madhvi Joshi |
| EPI_ISL_450806 | VI-US Virgin Islands Department of Health | Pathogen Discovery, Respiratory Viruses Branch, Division of Viral Diseases, Centers for Disease Control and Prevention | Yan Li, Anna Montmayeur, Ying Tao, Krista Queen, Jing Zhang, Anna Uehara, Clinton R. Paden, Rachel Marine, Haibin Wang, Zachary Weiner, Bettina Bankamp, Suxiang Tong |
| EPI_ISL_450838 | Utah Public Health Laboratory | Utah Public Health Laboratory | Erin Young, Kelly Oakeson |
| EPI_ISL_450847, EPI_ISL_450848, EPI_ISL_450849, EPI_ISL_450850, EPI_ISL_450851, EPI_ISL_450852, EPI_ISL_450853, EPI_ISL_450854, EPI_ISL_450855, EPI_ISL_450856, EPI_ISL_450857, EPI_ISL_450858, EPI_ISL_450859, EPI_ISL_450860, EPI_ISL_450861, EPI_ISL_450862, EPI_ISL_450863, EPI_ISL_450864, EPI_ISL_450865, EPI_ISL_450866, EPI_ISL_450867, EPI_ISL_450868, EPI_ISL_450869, EPI_ISL_450870, EPI_ISL_450871, EPI_ISL_450872 | | | |
| see above | Florida Bureau of Public Health Laboratories | Florida Bureau of Public Health Laboratories | Sarah Schmedes, Jason Blanton |
| EPI_ISL_450908, EPI_ISL_450909, EPI_ISL_450910, EPI_ISL_450911, EPI_ISL_450912, EPI_ISL_450948, EPI_ISL_450949, EPI_ISL_450950, EPI_ISL_450951, EPI_ISL_450952, EPI_ISL_450988, EPI_ISL_450989, EPI_ISL_450990, EPI_ISL_450991, EPI_ISL_450992, EPI_ISL_451028, EPI_ISL_451029, EPI_ISL_451030, EPI_ISL_451031, EPI_ISL_451032, EPI_ISL_451068, EPI_ISL_451069, EPI_ISL_451070, EPI_ISL_451071, EPI_ISL_451072 | | | |
| see above | Center of Excellence in Clinical Virology | Center of Excellence in Clinical Virology | Puenpa,J., Chansaenroj,J., Nilyanimit,P., Auphimai,C., Yorsaeng,R., Suwannakarn,K., Poovorawan,Y. |
| EPI_ISL_451129, EPI_ISL_451144, EPI_ISL_451145 | SA Pathology | SA Pathology | Lex Leong, Chuan Kok Lim, Mark Turra, Ivan Bastian, Geoff Higgins |
| EPI_ISL_451172 | Lab voor klinische biologie | Onderzoeksgroep Virologie | Laurens Lambrechts, Nick Vereecke, Marthe Pauwels, Jozefien De Clercq, Bruno Verhasselt, Linos Vandekerckhove, Hans Nauwynck, Sebastiaan Theuns |
| EPI_ISL_451175, EPI_ISL_451176, EPI_ISL_451177, EPI_ISL_451178, EPI_ISL_451179 | Lab voor klinische biologie | Onderzoeksgroep Virologie | Nick Vereecke, Laurens Lambrechts, Marthe Pauwels, Jozefien De Clercq, Bruno Verhasselt, Linos Vandekerckhove, Hans Nauwynck, Sebastiaan Theuns |
| EPI_ISL_451558 | Medlab Pathology | NSW Health Pathology - Institute of Clinical Pathology and Medical Research; Westmead Hospital; University of Sydney | CIDM-PH et al. |
| EPI_ISL_451566 | Pathology West - NSW Health Pathology | NSW Health Pathology - Institute of Clinical Pathology and Medical Research; Westmead Hospital; University of Sydney | CIDM-PH et al. |
| EPI_ISL_451587 | Pathology North Hunter- NSW Health Pathology | NSW Health Pathology - Institute of Clinical Pathology and Medical Research; Westmead Hospital; University of Sydney | CIDM-PH et al. |
| EPI_ISL_451594 | Childrens Hospital Westmead | NSW Health Pathology - Institute of Clinical Pathology and Medical Research; Westmead Hospital; University of Sydney | CIDM-PH et al. |
| EPI_ISL_451597 | Medlab Pathology | NSW Health Pathology - Institute of Clinical Pathology and Medical Research; Westmead Hospital; University of Sydney | CIDM-PH et al. |
| EPI_ISL_451613 | Pathology West - NSW Health Pathology | NSW Health Pathology - Institute of Clinical Pathology and Medical Research; Westmead Hospital; University of Sydney | CIDM-PH et al. |
| EPI_ISL_451643 | Pathology Sydney South West - NSW Health Pathology | NSW Health Pathology - Institute of Clinical Pathology and Medical Research; Westmead Hospital; University of Sydney | CIDM-PH et al. |
| EPI_ISL_451857, EPI_ISL_451858, EPI_ISL_451859, EPI_ISL_451860, EPI_ISL_451861, EPI_ISL_451862, EPI_ISL_451863, EPI_ISL_451864, EPI_ISL_451865, EPI_ISL_451866, EPI_ISL_451867, EPI_ISL_451868, EPI_ISL_451869, EPI_ISL_451870, EPI_ISL_451871, EPI_ISL_451872, EPI_ISL_451873, EPI_ISL_451874, EPI_ISL_451875, EPI_ISL_451876, EPI_ISL_451877, EPI_ISL_451878, EPI_ISL_451879, EPI_ISL_451880, EPI_ISL_451881, EPI_ISL_451882, EPI_ISL_451883, EPI_ISL_451890, EPI_ISL_451892, EPI_ISL_451893, EPI_ISL_451894, EPI_ISL_451895, EPI_ISL_451896, EPI_ISL_451897, EPI_ISL_451898, EPI_ISL_451899, EPI_ISL_451900, EPI_ISL_451901, EPI_ISL_451902, EPI_ISL_451903, EPI_ISL_451904, EPI_ISL_451905, EPI_ISL_451906, EPI_ISL_451907, EPI_ISL_451908, EPI_ISL_451909, EPI_ISL_451910, EPI_ISL_451911, EPI_ISL_451912, EPI_ISL_451913, EPI_ISL_451914, EPI_ISL_451915, EPI_ISL_451916 | | | |
| see above | Viollier AG | Department of Biosystems Science and Engineering, ETH Zürich | Christian Beisel, Sarah Nadeau, Ivan Topolsky, Pedro Ferreira, Philipp Jablonski, Susana Posada-Céspedes, Tobias Schär, Ina Nissen, Natascha Santacroce, Elodie Burcklen, Christiane Beckmann, Maurice Redondo, Olivier Kobel, Christoph Noppen, Sophie Seidel, Noemie Santamaria de Souza, Niko Beerenwinkel, Tanja Stadler |
| EPI_ISL_451947 | Max von Pettenkofer Institute, Virology, National Reference Center for Retroviruses, LMU München | Laboratory for Functional Genome Analysis, Dept. Genomics, Gene Center of the LMU Munich | Max Muenchhoff, Stefan Krebs, Alexander Graf, Oliver Keppler, Helmut Blum |
| EPI_ISL_451976, EPI_ISL_451981, EPI_ISL_451986 | 1. ViroGenetics - BSL3 Laboratory of Virology, Maopolska Centre of Biotechnology, Jagiellonian University; 2. II Department of Internal Medicine, Faculty of Medicine, Jagiellonian University Medical College; 3. DIAGNOSTYKA Ltd. | 1. ViroGenetics - BSL3 Laboratory of Virology, Maopolska Centre of Biotechnology, Jagiellonian University; 2. II Department of Internal Medicine, Faculty of Medicine, Jagiellonian University Medical College. | Marek Sanak, Marcin Surmiak, Monika Gsecka-Czapla, Wojciech Branicki, Pawe P abaj, Marta Rogalska-Kupiec, Jakub Swadba, Krzysztof Pyr |
| EPI_ISL_452038 | Department of Clinical Microbiology, Copenhagen University Hospital, Hvidovre, Kettegaard Alle 30, 2650 Hvidovre. | Albertsen lab, Department of Chemistry and Bioscience, Aalborg University, Denmark | Rasmus Kirkegaard |
| EPI_ISL_452149 | CUB Hopital Erasme Laboratoire d'Anatomie Pathologique | CUB Hopital Erasme Laboratoire d'Anatomie Pathologique | Isabelle Salmon, Nicky D'Haene |
| EPI_ISL_452150 | CUB Hopital Erasme Laboratoire d'Anatomie Pathologique | CUB Hopital Erasme Laboratoire d'Anatomie Pathologique | Prof. Isabelle Salmon, Dr Nicky D'Haene |
| EPI_ISL_452153, EPI_ISL_452154, EPI_ISL_452155, EPI_ISL_452156, EPI_ISL_452157, EPI_ISL_452158, EPI_ISL_452159 | Utah Public Health Laboratory | Utah Public Health Laboratory | Erin Young, Kelly Oakeson |
| EPI_ISL_452214 | NIV Influenza | NIV Influenza | Potdar V |
| EPI_ISL_452365, EPI_ISL_452366, EPI_ISL_452369 | Servicio de Microbiología. HRU de Málaga. Servicio Andaluz de Salud | SeqCOVID-SPAIN consortium/IBV(CSIC) | Inmaculada de Toro Peinado, Maria Concepción Mediavilla Gradolph, Begoña Palop Borrás and SeqCOVID-SPAIN consortium |
| EPI_ISL_452796, EPI_ISL_452811, EPI_ISL_452812, EPI_ISL_452814, | Virginia DCLS | Virginia DCLS | Virginia DCLS |

| | | | |
|---|---|---|---|
| EPI_ISL_452815 | | | |
| EPI_ISL_453013, EPI_ISL_453014, EPI_ISL_453015, EPI_ISL_453016, EPI_ISL_453017, EPI_ISL_453018, EPI_ISL_453019, EPI_ISL_453020, EPI_ISL_453021, EPI_ISL_453022, EPI_ISL_453023, EPI_ISL_453024, EPI_ISL_453025, EPI_ISL_453026, EPI_ISL_453027, EPI_ISL_453028, EPI_ISL_453029, EPI_ISL_453030, EPI_ISL_453031, EPI_ISL_453032, EPI_ISL_453033, EPI_ISL_453034, EPI_ISL_453035, EPI_ISL_453036, EPI_ISL_453037, EPI_ISL_453038, EPI_ISL_453039, EPI_ISL_453040, EPI_ISL_453042, EPI_ISL_453043, EPI_ISL_453045 | | | |
| see above | West of Scotland Specialist Virology Centre, NHSGGC / MRC-University of Glasgow Centre for Virus Research | COVID-19 Genomics UK (COG-UK) Consortium | Ana da Silva Filipe, Natasha Johnson, Kathy Smollett, Daniel Mair, Stephen Carmichael, Lily Tong, Jenna Nichols, Elihu Aranday-Cortes, Kirstyn Brunker, Yasmin Parr, Kyriaki Nomikou; Sarah McDonald, Marc Niebel, Patawee Asamaphan; Richard Orton, Joseph Hughes, Sreenu Vattipally, David L Robertson; Alasdair MacLean, Rory Gunson; Kathy Li, Natasha Jesudason, Rajiv Shah, James Shepherd, Antonia Ho, Emma Thomson |
| EPI_ISL_453215, EPI_ISL_453216, EPI_ISL_453217, EPI_ISL_453218, EPI_ISL_453219, EPI_ISL_453220, EPI_ISL_453221, EPI_ISL_453222, EPI_ISL_453223, EPI_ISL_453224, EPI_ISL_453225, EPI_ISL_453226, EPI_ISL_453227, EPI_ISL_453228, EPI_ISL_453229, EPI_ISL_453230, EPI_ISL_453249, EPI_ISL_453250, EPI_ISL_453251, EPI_ISL_453252, EPI_ISL_453253, EPI_ISL_453258, EPI_ISL_453315, EPI_ISL_453316, EPI_ISL_453317, EPI_ISL_453318, EPI_ISL_453319, EPI_ISL_453320, EPI_ISL_453321, EPI_ISL_453322, EPI_ISL_453323, EPI_ISL_453324, EPI_ISL_453325, EPI_ISL_453326, EPI_ISL_453327, EPI_ISL_453330, EPI_ISL_453332, EPI_ISL_453333, EPI_ISL_453335, EPI_ISL_453336, EPI_ISL_453337, EPI_ISL_453338, EPI_ISL_453339, EPI_ISL_453340, EPI_ISL_453341, EPI_ISL_453342, EPI_ISL_453343, EPI_ISL_453344, EPI_ISL_453345, EPI_ISL_453346, EPI_ISL_453347, EPI_ISL_453348, EPI_ISL_453349, EPI_ISL_453350, EPI_ISL_453351, EPI_ISL_453352, EPI_ISL_453353, EPI_ISL_453354, EPI_ISL_453355, EPI_ISL_453356, EPI_ISL_453357, EPI_ISL_453358, EPI_ISL_453359, EPI_ISL_453402, EPI_ISL_453403, EPI_ISL_453404, EPI_ISL_453405, EPI_ISL_453406, EPI_ISL_453407, EPI_ISL_453408, EPI_ISL_453409, EPI_ISL_453410, EPI_ISL_453411, EPI_ISL_453412, EPI_ISL_453413, EPI_ISL_453414, EPI_ISL_453415, EPI_ISL_453416 | | | |
| see above | Liverpool Clinical Laboratories | COVID-19 Genomics UK (COG-UK) Consortium | Sam Haldenby, Anita Lucaci, Steve Paterson, Julian Hiscox, Alistair Darby, M Almsaud, A Alrezaihi, Muhannad Alruwaili, Stuart D Armstrong, Jones Benjamin , Eleanor G Bentley, Anu Chawla, Jordan J Clark, Angela Cowell, Richard Eccles, Isabel García-Dorival, Matthew Gemmell, Alessandro Gerada, PKF Gilmore, Richard Gregory, Ximeng Han, Catherine Hartley, Margaret Hughes, Miren Iturriza-Gomara, James Johnson, L Luu, Jenifer Manson , Charlotte Nelson, Elaine O'Toole, Cassie Olateju, Rebekah Penrice-Randal , Lucille Rainbow, N.P Randle, Trevor Ian Robinson, Parul Sharma, Ghada T Shawli, James P Stewart , Neil Swainston, Ecaterina Vamos, Joanne Watts, Mark Whitehead |
| EPI_ISL_453519, EPI_ISL_453520, EPI_ISL_453521, EPI_ISL_453522, EPI_ISL_453523, EPI_ISL_453524, EPI_ISL_453525, EPI_ISL_453526, EPI_ISL_453527, EPI_ISL_453528, EPI_ISL_453529, EPI_ISL_453530, EPI_ISL_453531, EPI_ISL_453532 | | | |
| see above | Northumbria University / South Tees Hospitals NHS Foundation Trust / North Cumbria Integrated Care NHS Foundation Trust / North Tees and Hartlepool NHS Foundation Trust / Newcastle Hospitals NHS Foundation Trust | COVID-19 Genomics UK (COG-UK) Consortium | Darren L Smith,Andrew Nelson,Matthew Bashton,Greg R Young,Joshua Loh,John Allan,Mohammad A Tariq,Giles S Holt,Gary Black,Wen C Yew,Lynn Dover ,Paul Baker,Steve Liggett,Sarah Essex,Jane Greenaway ,Debra Padgett,Clive Graham,Garren Scott,Edward Barton ,Emma Swindells ,Brendan Payne,Jennifer Collins,Yusri Taha,Gary Eltringham |
| EPI_ISL_453637 | Queens Medical Centre, Clinical Microbiology Department / DeepSeq Nottingham | COVID-19 Genomics UK (COG-UK) Consortium | Gemma Clark, Wendy Smith, Manjinder Khakh, Hannah Howson-Wells, Jonathan Ball, Patrick McClure, Joseph Chappell, Theocharis Tsoleridis, Nadine Holmes, Matthew Carlisle, Christopher Moore, Fei Sang, Johnny Debebe, Victoria Wright, Matthew Loose |
| EPI_ISL_454255, EPI_ISL_454256, EPI_ISL_454257, EPI_ISL_454258, EPI_ISL_454259, EPI_ISL_454260, EPI_ISL_454261, EPI_ISL_454262, EPI_ISL_454263, EPI_ISL_454264, EPI_ISL_454265, EPI_ISL_454266, EPI_ISL_454267, EPI_ISL_454268, EPI_ISL_454269, EPI_ISL_454274, EPI_ISL_454275, EPI_ISL_454276, EPI_ISL_454277, EPI_ISL_454278, EPI_ISL_454289, EPI_ISL_454296, EPI_ISL_454298, EPI_ISL_454325 | | | |
| see above | unknown | Instituto Nacional de Saude (INSA) | Borges et al |
| EPI_ISL_454399, EPI_ISL_454403, EPI_ISL_454404, EPI_ISL_454406, EPI_ISL_454407, EPI_ISL_454408, EPI_ISL_454409, EPI_ISL_454411 | UPMC Clinical Microbiology Laboratory | Microbial Genome Sequencing Center, Microbial Genomic Epidemiological Laboratory | Mustapha M. Mustapha, Jane W. Marsh, Dan Snyder, Marissa P. Griffith, Stephanie L. Mitchell, Vatsala R. Srinivasa, Kady D. Waggle, Chinelo Ezeonwuku, Vaughn S. Cooper, Lee H. Harrison |
| EPI_ISL_454539, EPI_ISL_454551, EPI_ISL_454555, EPI_ISL_454556, EPI_ISL_454561, EPI_ISL_454562, EPI_ISL_454563 | NIV Influenza | NIV Influenza | Potdar V |
| EPI_ISL_454574 | nstitute for Public Health | Laboratory for advanced genomics | Filip Roki, Lovro Trgovec-Greif, Neven Sui, Tomislav Rukavina, Igor Jurak, Oliver Vugrek |
| EPI_ISL_454642 | CT-Dr. Katherine A. Kelley State Public Health Lab | Pathogen Discovery, Respiratory Viruses Branch, Division of Viral Diseases, Centers for Disease Control and Prevention | Jing Zhang, Ying Tao, Clinton R. Paden, Anna Uehara, Krista Queen, Yan Li, Haibin Wang, Zachary Weiner, Bettina Bankamp, Suxiang Tong |
| EPI_ISL_454653, EPI_ISL_454654, EPI_ISL_454655, EPI_ISL_454658, EPI_ISL_454667, EPI_ISL_454668, EPI_ISL_454670, EPI_ISL_454671, EPI_ISL_454673, EPI_ISL_454674, EPI_ISL_454679, EPI_ISL_454684, EPI_ISL_454686 | | | |
| see above | County of Santa Clara Public Health Department | Chan-Zuckerberg Biohub | CZB Cliahub Consortium |
| EPI_ISL_454782, EPI_ISL_454794 | Dutch COVID-19 response team | National Institute for Public Health and the Environment (RIVM) | Adam Meijer, Harry Vennema, Jeroen Cremer, Sharon van den Brink, Pieter Overduin, Florian Zwagemaker, Dennis Schmitz, Chantal Reusken, on behalf of the national COVID-19 response team |
| EPI_ISL_454858, EPI_ISL_454859, EPI_ISL_454860, EPI_ISL_454861, EPI_ISL_454862 | Translational Health Science and Technology Institute -ESIC medical college and hospital, Faridabad | THSTI Bioassay laboratory | Saurabh Kumar, Jigme Wangchuk, Anil Kumar Pandey, Asim Das, Guruprasad R. Medigeshi |
| EPI_ISL_455034 | Pathology West - NSW Health Pathology | NSW Health Pathology - Institute of Clinical Pathology and Medical Research; Westmead Hospital; University of Sydney | CIDM-PH et al. |
| EPI_ISL_455037 | Pathology Sydney South West - NSW Health Pathology | NSW Health Pathology - Institute of Clinical Pathology and Medical Research; Westmead Hospital; University of Sydney | CIDM-PH et al. |
| EPI_ISL_455085 | South Eastern Area Laboratory Services | NSW Health Pathology - Institute of Clinical Pathology and Medical Research; Westmead Hospital; University of Sydney | CIDM-PH et al. |
| EPI_ISL_455158, EPI_ISL_455159, EPI_ISL_455160, EPI_ISL_455161, EPI_ISL_455162, EPI_ISL_455163, EPI_ISL_455166, EPI_ISL_455167, EPI_ISL_455168, EPI_ISL_455169, EPI_ISL_455170, EPI_ISL_455171, EPI_ISL_455172, EPI_ISL_455173, EPI_ISL_455174, EPI_ISL_455175, EPI_ISL_455176, EPI_ISL_455177, EPI_ISL_455178, EPI_ISL_455179, EPI_ISL_455180, EPI_ISL_455194, EPI_ISL_455236, EPI_ISL_455237, EPI_ISL_455238, EPI_ISL_455239, EPI_ISL_455240, EPI_ISL_455241, EPI_ISL_455242, EPI_ISL_455243, EPI_ISL_455244, EPI_ISL_455245, EPI_ISL_455246, EPI_ISL_455247, EPI_ISL_455248, EPI_ISL_455263, EPI_ISL_455267, EPI_ISL_455268, EPI_ISL_455281, EPI_ISL_455284, EPI_ISL_455289, EPI_ISL_455297, EPI_ISL_455298, EPI_ISL_455299, EPI_ISL_455300, EPI_ISL_455301, EPI_ISL_455302, EPI_ISL_455303, EPI_ISL_455304, EPI_ISL_455305, EPI_ISL_455306, EPI_ISL_455307 | | | |
| see above | Dutch COVID-19 response team | Erasmus Medical Center | Bas Oude Munnink, David Nieuwenhuijse, Reina Sikkema, Claudia Schapendonk, Irina Chestakova, Anne van der Linden, Theo Bestebroer, Stefan van Nieuwkoop, Mark Pronk, Pascal Lexmond, Corien Swaan, Manon Haverkate, Madelief Mollers, Mart Stein, Sandra Kengne Kamga Mobou, Jeroen van Kampen, Jolanda Voermans, Aura Timen, Corine GeurtsvanKessel, Annemiek van der Eijk, Richard Molenkamp, Marion Koopmans, on behalf of the Dutch national COVID-19 response team. |
| EPI_ISL_455310 | REGIONAL VRDL,ICMR-RMRC BBSR | Immunogenomics group, Institute of Life Sciences, Bhubaneswar | Sunil Raghav, Jyotirmayee Turuk, Arup Ghosh, Atimukta Jha, Viplov K. Biswas, Swati Madhulika, Manasi Priyadarshini, Shuchi Smita, Jaya Singh Khastri, Rupesh Dash, Soma Chattopadhyay, Ghulam Hussain Syed, Shanti Senapati, Tushar K. Beuria, Debdutta Bhattacharya, Rajeeb Swain, Punit Prasad, COVID-19 team of ILS & RMRC, Orissa COVID-19 study group, DBT's PAN-INDIA 1000 SARS-CoV2 RNA genome sequencing consortium, Sanghamitra Pati, Ajay Parida |
| EPI_ISL_455312 | Microbiology Unit, Department of Pathology & Laboratory Medicine, IIUM Medical Centre | SEA Microbiome Unit, Faculty of Industrial Sciences & Technology, Universiti Malaysia Pahang | Norhidayah Binti Kamarudin, Ahmad Hafiz Bin Zulkifly, Hajar Fauzan Ahmad, Muhammad Adam Lee Abdullah, Mohd Fazli Farida Asras, Ahmad Mahfuz Gazali, Mohd Nazli Bin Kamarulzaman, IIUM Medical Centre Covid19 Taskforce, UMP Covid19 Team |
| EPI_ISL_455362 | Nigeria Centre for Disease Control (NCDC) | African Centre of Excellence for Genomics of Infectious Diseases (ACEGID), Redeemer's University, Ede, Osun State, Nigeria | Oluniyi P.E., Ajogbasile F.V., Kayode A., Olawoye I., Uwanibe J., Oguzie J., Olumade T., Folarin O.A., Ihekweazu C., Happi C.T. |
| EPI_ISL_455478 | REGIONAL VRDL,ICMR-RMRC BBSR | Immunogenomics group, Institute of Life Sciences, Bhubaneswar | Sunil Raghav, Jyotirmayee Turuk, Arup Ghosh, Atimukta Jha, Viplov K. Biswas, Swati Madhulika, Manasi Priyadarshini, Shuchi Smita, Jaya Singh Khastri, Rupesh Dash, Soma Chattopadhyay, Ghulam Hussain Syed, Shanti Senapati, Tushar K. Beuria, Debdutta Bhattacharya, Rajeeb Swain, Punit Prasad, COVID-19 team of ILS & RMRC, Orissa COVID-19 study group, DBT's PAN-INDIA 1000 SARS-CoV2 RNA genome sequencing consortium, Sanghamitra Pati, Ajay Parida |
| EPI_ISL_455480 | Veterinary Specialized Institute Kraljevo | Veterinary Specialized Institute Kraljevo | Vidanovic,D., Tesovic,B., Banovic Djeri,B., Sekler,M., Dmitric,M., Debeljak,Z., Matovic,K., Vaskovic,N., Petrovic,T., Volkening,J. and Alfonso,C.L. |
| EPI_ISL_455610, EPI_ISL_455612, EPI_ISL_455613, EPI_ISL_455614, | Ochsner Health | Bioinfoexperts, LLC | Susanna L. Lamers, David J. Nolan, Rebecca Rose, Sissy Cross, David Moraga Amador, Tong Yang, Luke Caruso, Wayra Navia, Lydia Von Borstel, Xiao Hui Zhou, Amy Feehan, Julia-Garcia-Diaz |

| | | | |
|---|---|---|---|
| EPI_ISL_455616, EPI_ISL_455618, EPI_ISL_455622 | | | |
| EPI_ISL_455708, EPI_ISL_455709, EPI_ISL_455710, EPI_ISL_455711, EPI_ISL_455712 | National Hospital of Tropical Diseases | Oxford University Clinical Research Unit, Hanoi, Vietnam | Nguyen Thi Tam, Van Dinh Trang, Nguyen Thu Trang, Nguyen Thi Ngoc Diep, Le Nguyen Minh Hoa, Pham Ngoc Thach, H. Rogier van Doorn, on behalf of the OUCRU COVID-19 research group |
| EPI_ISL_455718 | National Hospital of Tropical Diseases | Oxford University Clinical Research Unit, Hanoi, Vietnam | Nguyen Thi Tam, Van Dinh Trang, Nguyen Thi Hong Thuong, Vu Thi Ngoc Bich, Nguyen Thu Trang, Nguyen Thi Ngoc Diep, Le Nguyen Minh Hoa, Pham Ngoc Thach, H. Rogier van Doorn, on behalf of the OUCRU COVID-19 research group |
| EPI_ISL_455719 | T.C. Salk Bakanl Adyaman l Salk Müdürlüü Adyaman Eitim Ve Aratrma Hastanesi | VETAL Animal Health Products Company, BSL3+ Production Laboratuary /Turkey | Mehmet Turgut, Muhittin Önderci, Fatma Nilay Tutak, Abidin Ercan Yonucu, Murat Dönen, Haluk Uluca, Fethiye Sevimli, O. Ugur Sezerman |
| EPI_ISL_455780, EPI_ISL_455781 | REGIONAL VRDL,ICMR-RMRC BBSR | Immunogenomics lab, Institute of Life Sciences, Bhubaneswar | Sunil Raghav, Jyotirmayee Turuk, Arup Ghosh, Atimukta Jha, Viplov K. Biswas, Swati Madhulika, Manasi Priyadarshini, Shuchi Smita, Jaya Singh Khastri, Rupesh Dash, Soma Chattopadhyay, Ghulam Hussain Syed, Shanti Senapati, Tushar K. Beuria, Debdutta Bhattacharya, Rajeeb Swain, Punit Prasad, COVID-19 team of ILS & RMRC, Orissa COVID-19 study group, DBT's PAN-INDIA 1000 SARS-CoV2 RNA genome sequencing consortium, Sanghamitra Pati, Ajay Parida |
| EPI_ISL_456084, EPI_ISL_456085, EPI_ISL_456086, EPI_ISL_456087, EPI_ISL_456089, EPI_ISL_456090, EPI_ISL_456091, EPI_ISL_456092, EPI_ISL_456093 | Laboratory of Respiratory Viruses and Measles, Oswaldo Cruz Institute, FIOCRUZ | Laboratory of Respiratory Viruses and Measles, Oswaldo Cruz Institute, FIOCRUZ | Paola Resende, Luciana Appolinario, Fernando Motta, Aline Mattos, Milene Miranda, Cristiana Garcia, Braulia Caetano, Maria Ogrzewalska, Jonathan Lopes, Marilda Siqueira |
| EPI_ISL_456323 | North Shore Hospital | Institute of Environmental Science and Research (ESR) | Matt Storey, Xiaoyun Ren, Anja Werno, Antje van der Linden, Arlo Upton, Chris Mansell, David Hammer, Dragana Drinkovic, Erasmus Smit, Gary McAuliffe, Hana Sofia Andersson, James Ussher, Jill Sherwood, Josh Freeman, Julia Howard, Juliet Elvy, Mary DeAlmeida, Matt Blakiston, Matthew Rogers, Max Bloomfield, Michael Addidle, Michelle Balm, Sally Roberts, Sarah Jefferies, Sharmini Muttaiyah, Susan Morpeth, Susan Taylor, Timothy Blackmore, Vani Sathyendran, Veronica Playle, Virginia Hope, Erasmus Smit, Lauren Jelly, Joep de Ligt |
| EPI_ISL_456344, EPI_ISL_456345, EPI_ISL_456346, EPI_ISL_456347 | Southern Community Labs Dunedin | Institute of Environmental Science and Research (ESR) | Matt Storey, Xiaoyun Ren, Anja Werno, Antje van der Linden, Arlo Upton, Chris Mansell, David Hammer, Dragana Drinkovic, Erasmus Smit, Gary McAuliffe, Hana Sofia Andersson, James Ussher, Jill Sherwood, Josh Freeman, Julia Howard, Juliet Elvy, Mary DeAlmeida, Matt Blakiston, Matthew Rogers, Max Bloomfield, Michael Addidle, Michelle Balm, Sally Roberts, Sarah Jefferies, Sharmini Muttaiyah, Susan Morpeth, Susan Taylor, Timothy Blackmore, Vani Sathyendran, Veronica Playle, Virginia Hope, Erasmus Smit, Lauren Jelly, Joep de Ligt |
| EPI_ISL_456359, EPI_ISL_456360, EPI_ISL_456361, EPI_ISL_456362, EPI_ISL_456363, EPI_ISL_456364, EPI_ISL_456365, EPI_ISL_456366, EPI_ISL_456367, EPI_ISL_456368, EPI_ISL_456369, EPI_ISL_456370, EPI_ISL_456371, EPI_ISL_456374, EPI_ISL_456375 | | | |
| see above | Canterbury Health Laboratories | Institute of Environmental Science and Research (ESR) | Matt Storey, Xiaoyun Ren, Anja Werno, Antje van der Linden, Arlo Upton, Chris Mansell, David Hammer, Dragana Drinkovic, Erasmus Smit, Gary McAuliffe, Hana Sofia Andersson, James Ussher, Jill Sherwood, Josh Freeman, Julia Howard, Juliet Elvy, Mary DeAlmeida, Matt Blakiston, Matthew Rogers, Max Bloomfield, Michael Addidle, Michelle Balm, Sally Roberts, Sarah Jefferies, Sharmini Muttaiyah, Susan Morpeth, Susan Taylor, Timothy Blackmore, Vani Sathyendran, Veronica Playle, Virginia Hope, Erasmus Smit, Lauren Jelly, Joep de Ligt |
| EPI_ISL_456421, EPI_ISL_456427 | Victorian Infectious Diseases Reference Laboratory (VIDRL) | Microbiological Diagnostic Unit Public Health Laboratory and Victorian Infectious Diseases Reference Laboratory, Doherty Institute | Caly L., Seemann T., Sait, M., Schultz M., Druce J., Sherry, N. |
| EPI_ISL_456597, EPI_ISL_456598, EPI_ISL_456601, EPI_ISL_456602, EPI_ISL_456603, EPI_ISL_456604, EPI_ISL_456605, EPI_ISL_456606, EPI_ISL_456608, EPI_ISL_456609, EPI_ISL_456610 | | | |
| see above | National Health Laboratory, Timor-Leste | Microbiological Diagnostic Unit Public Health Laboratory, The Peter Doherty Institute for Infection and Immunity | Soares da Silva, E., Dolores de Jesus da Costa, M., Salles de Sousa, A., Jayanti Pereira Tilman, A., Antonia da Costa, E., Barreto, I., Marr, I., Wapling, J., Francis, J., Ximenes, J., Canisia, D., Freeman, K., Dakh, F., Douglas, N., Baird, R., Caly, L., Seemann, T., Sait, M., Schultz, M., Sherry, N. |
| EPI_ISL_456757, EPI_ISL_456758, EPI_ISL_456759, EPI_ISL_456760, EPI_ISL_456761, EPI_ISL_456762 | West of Scotland Specialist Virology Centre, NHSGGC / MRC-University of Glasgow Centre for Virus Research | COVID-19 Genomics UK (COG-UK) Consortium | Ana da Silva Filipe, Natasha Johnson, Kathy Smollett, Daniel Mair, Stephen Carmichael, Lily Tong, Jenna Nichols, Elihu Aranday-Cortes, Kirstyn Brunker, Yasmin Parr, Kyriaki Nomikou; Sarah McDonald, Marc Niebel, Patawee Asamaphan; Richard Orton, Joseph Hughes, Sreenu Vattipally, David L Robertson; Alasdair MacLean, Rory Gunson; Kathy Li, Natasha Jesudason, Rajiv Shah, James Shepherd, Antonia Ho, Emma Thomson |
| EPI_ISL_456891, EPI_ISL_456892, EPI_ISL_456893, EPI_ISL_456914, EPI_ISL_456915 | Virology Department, Royal Infirmary of Edinburgh, NHS Lothian / School of Biological Sciences, University of Edinburgh / Institute of Genetics and Molecular Medicine, University of Edinburgh | COVID-19 Genomics UK (COG-UK) Consortium | McHugh M, Dewar R, Rooke S, Gallagher M, Balcaza C, O'Toole Á, Scher E, Hill V, McCrone JT, Colquhoun R, Yu X, Jackson B, Rambaut A, Williams TC, Templeton K |
| EPI_ISL_457028, EPI_ISL_457029, EPI_ISL_457049, EPI_ISL_457050, EPI_ISL_457051, EPI_ISL_457059, EPI_ISL_457060, EPI_ISL_457065, EPI_ISL_457066, EPI_ISL_457067, EPI_ISL_457068, EPI_ISL_457069, EPI_ISL_457071, EPI_ISL_457072, EPI_ISL_457074, EPI_ISL_457076, EPI_ISL_457077, EPI_ISL_457079, EPI_ISL_457085, EPI_ISL_457086, EPI_ISL_457087, EPI_ISL_457088, EPI_ISL_457091, EPI_ISL_457092, EPI_ISL_457093, EPI_ISL_457097, EPI_ISL_457098, EPI_ISL_457103, EPI_ISL_457105, EPI_ISL_457122, EPI_ISL_457123, EPI_ISL_457129, EPI_ISL_457130, EPI_ISL_457133, EPI_ISL_457135, EPI_ISL_457136, EPI_ISL_457138, EPI_ISL_457141, EPI_ISL_457142, EPI_ISL_457143, EPI_ISL_457149, EPI_ISL_457153, EPI_ISL_457158, EPI_ISL_457159, EPI_ISL_457164, EPI_ISL_457169, EPI_ISL_457173, EPI_ISL_457174, EPI_ISL_457175, EPI_ISL_457179, EPI_ISL_457215, EPI_ISL_457216, EPI_ISL_457217, EPI_ISL_457222, EPI_ISL_457223, EPI_ISL_457224, EPI_ISL_457225 | | | |
| see above | University of Exeter | COVID-19 Genomics UK (COG-UK) Consortium | Ben Temperton,Aaron Jeffries,Michelle Michelsen,Joanna Warwick-Dugdale,Audrey Farbos,Robyn Manley,Stephen Michell,Jane Masoli |
| EPI_ISL_457304, EPI_ISL_457305, EPI_ISL_457306, EPI_ISL_457307 | Northumbria University / South Tees Hospitals NHS Foundation Trust / North Cumbria Integrated Care NHS Foundation Trust / North Tees and Hartlepool NHS Foundation Trust / Newcastle Hospitals NHS Foundation Trust | COVID-19 Genomics UK (COG-UK) Consortium | Darren L Smith,Andrew Nelson,Matthew Bashton,Greg R Young,Joshua Loh,John Allan,Mohammad A Tariq,Giles S Holt,Gary Black,Wen C Yew,Lynn Dover,Paul Baker,Steve Liggett,Sarah Essex,Jane Greenaway,Debra Padgett,Clive Graham,Garren Scott,Edward Barton,Emma Swindells,Brendan Payne,Jennifer Collins,Yusri Taha,Gary Eltringham |
| EPI_ISL_457610, EPI_ISL_457614, EPI_ISL_457619, EPI_ISL_457625, EPI_ISL_457650, EPI_ISL_457671 | Virology Department, Sheffield Teaching Hospitals NHS Foundation Trust/Department of Infection, Immunity and Cardiovascular Disease, The Medical School, University of Sheffield | COVID-19 Genomics UK (COG-UK) Consortium | Thushan de Silva, Matthew Parker, Nikki Smith, Adri Angyal, Rebecca Brown, Luke Green, Rachel Tucker, Paul Parsons, Danielle Groves, Katie Johnson, Laura Carrilero, Alex Keeley, Dave Partridge, Matthew Wyles, Benjamin Lindsey, Mehmet Yavuz, Mohammad Raza, Cariad Evans |
| EPI_ISL_457702 | Oman-NIC | Microbiology laboratory- Sultan Qaboos University Hospital | Fahad Zadjali, Samira Al-Maruqi, Amina Al Jardani, Khulood Al-Mammary, Hanan Al-kindi, Fatma BaAlawi, Hamida AL Barwani, Zeyana AL-Dahmani, Intisar Al-Shukri, Aisha Al-Busaidi, Aisha Al-Amri, Ahlam Al-Amri, Mohammed Al-Tobi, Samiha Al Kharusi, Abdulla Balkhair |
| EPI_ISL_457905, EPI_ISL_457908, EPI_ISL_457909, EPI_ISL_457910, EPI_ISL_457911, EPI_ISL_457912, EPI_ISL_457913, EPI_ISL_457914 | KEMRI-CGMR-C | KEMRI-Wellcome Trust Research Programme/KEMRI-CGMR-C Kilifi | Githinji G. et al 2020 |
| EPI_ISL_457967, EPI_ISL_457968 | Laboratorio de Biología Molecular Asociación Española Primera en Salud | Departments of Pathology and Medicine, New York University School of Medicine | Maria Victoria Elizondo, Maria Noel Zubillaga, Gonzalo Manrique, Paul Zappile, Gael Westby, Matthew T Maurano, Christian Marier, Adriana Heguy |
| EPI_ISL_458070 | CSIR-Centre for Cellular and Molecular Biology | CSIR-Centre for Cellular and Molecular Biology | Sakshi Shambhavi, Lamuk Zaveri, Shagufta Khan, Namami Gaur, Tulasi Nagabandi, Purushotham Vodnala, Payel Mukherjee, Sofia Banu, Priya Singh, Dhiviya Vedagiri, Divya Gupta, Vishal Sah, Santosh Kumar Kuncha, Krishnan Harinivas Harshan, Archana Bharadwaj Siva, Karthik Bharadwaj Tallapaka,Nikhil Hajirnis, Pratheusa Maccha, M Soujanya Reddy,G. Aditya Kumar, Koushick Sivakumar,Disha Nanda, Divya Das, Jotin Gogoi, Manish Bhattacharjee, Ravi Prasad Mukku, Rakesh K Mishra, Divya Tej Sowpati |
| EPI_ISL_458072 | CSIR-Centre for Cellular and Molecular Biology | CSIR-Centre for Cellular and Molecular Biology | Dhiviya Vedagiri, Divya Gupta, Vishal Sah, Payel Mukherjee, Sofia Banu, Priya Singh, Santosh Kumar Kuncha, Archana Bharadwaj Siva, Karthik Bharadwaj Tallapaka, Shagufta Khan, Lamuk Zaveri, Namami Gaur, Sakshi Shambhavi, Tulasi Nagabandi, Purushotham Vodnala, Rakesh K Mishra, Divya Tej Sowpati, Krishnan Harinivas Harshan |
| EPI_ISL_458082 | Universitas Airlangga Hospital | Institute of Tropical Disease, Universitas Airlangga | Kazufumi Shimizu, Krisnoadi Rahardjo, Aldise M Nastri, Jezzy R Dewantari, Rima R Prasetya, Nasronudin, Gatot Soegiarto, Laksmi Wulandari, Retno A Setyoningrum, Resti Yudhawati, Yohko K Shimizu, Mitsuhiro Nishimura, Yasuko Mori, Soetjipto, Maria I Lusida |
| EPI_ISL_458083 | Adi Husada Undaan Hospital | Institute of Tropical Disease, Universitas Airlangga | Rima R Prasetya, Krisnoadi Rahardjo, Aldise M Nastri, Jezzy R Dewantari, Irawati Marga, Gatot Soegiarto, Laksmi Wulandari, Retno A Setyoningrum, |

| | | | |
|---|---|---|---|
| | | | Resti Yudhawati, Yohko K Shimizu, Mitsuhiro Nishimura, Yasuko Mori, Soetjipto, Kazufumi Shimizu, Maria I Lusida |
| EPI_ISL_458085 | Laboratorio Biologia Molecolare Sars Cov2 - UOC Laboratorio Analisi - Servizio Medicina di Laboratorio , Ospedale "San Francesco" - ATS- ASSL Nuoro | Laboratorio specialistico UOC Ematologia - Ospedale "San Francesco" - ATS-ASSL Nuoro | Piras Giovanna, Fancello Tatiana, Asproni Rosanna, Fiamma Maura, Monne Maria Itria, Toja Alessandro, Sanna Filomena, Floris Anna Rita, Sulis Vincenzo, Palmas Angelo Domenico, Casu Gavino, Lo Maglio Iana, Mameli Giuseppe. |
| EPI_ISL_458119, EPI_ISL_458122, EPI_ISL_458125 | Oman National Influenza Centre | Department of Microbiology and Immunology-SQUH | Fahad Zadjali, Samira Al-Maruqi, Amina Al Jardani, Khulood Al-Mammary, Hanan Al-kindi, Fatma BaAlawi, Hamida AL Barwani, Zeyana AL-Dahmani, Intisar Al-Shukri, Aisha Al-Busaidi, Aisha Al-Amri, Ahlam Al-Amri, Mohammed Al-Tobi, Samiha Al Kharusi, Abdulla Balkhair |
| EPI_ISL_458131 | Hospital Universitari Vall d'Hebron - Vall d'Hebron Institut de Recerca | Hospital Universitari Vall d'Hebron | Cristina Andrés, Maria Piñana, Damir Garcia-Cehic, Mercedes Guerrero-Murillo, Ariadna Rando, Josep Gregori, Juliana Esperalba, Maria Gema Codina, Maria Carmen Martin, Tomàs Pumarola, Josep Quer, Andrés Antón |
| EPI_ISL_458140 | Evandro Chagas Institute | Evandro Chagas Institute | Santos, M.C.; Silva, A.M.; Junior, W.D.C.; Barbagelata, L.S.; Ferreira, J.A.; Sousa, E.M.A.; da Silva, P.S.; Resque, H.R; Martins, L.C.; Sousa Junior, E.C.;Viana, G.M.R |
| EPI_ISL_458239, EPI_ISL_458241, EPI_ISL_458243, EPI_ISL_458256, EPI_ISL_458257, EPI_ISL_458260, EPI_ISL_458265, EPI_ISL_458267, EPI_ISL_458274, EPI_ISL_458275, EPI_ISL_458279 | | | |
| see above | Scripps Medical Laboratory | Andersen lab at Scripps Research | SEARCH Alliance San Diego with Michael Quigley, Ellen Stefanski, Ian Mchardy |
| EPI_ISL_458407, EPI_ISL_458412, EPI_ISL_458415, EPI_ISL_458435, EPI_ISL_458754, EPI_ISL_458912, EPI_ISL_458913, EPI_ISL_458914, EPI_ISL_458915, EPI_ISL_458916, EPI_ISL_458917, EPI_ISL_458918, EPI_ISL_458919, EPI_ISL_458920, EPI_ISL_458921, EPI_ISL_458922, EPI_ISL_458923, EPI_ISL_458924, EPI_ISL_458925, EPI_ISL_458926, EPI_ISL_458927, EPI_ISL_458928, EPI_ISL_458929, EPI_ISL_458930, EPI_ISL_458931, EPI_ISL_458932, EPI_ISL_458933, EPI_ISL_458934, EPI_ISL_458935, EPI_ISL_458936, EPI_ISL_458937, EPI_ISL_458938, EPI_ISL_458939, EPI_ISL_458940, EPI_ISL_458941, EPI_ISL_458942, EPI_ISL_458943, EPI_ISL_458944, EPI_ISL_458945, EPI_ISL_458946, EPI_ISL_458947, EPI_ISL_458948, EPI_ISL_458949, EPI_ISL_458950, EPI_ISL_458951, EPI_ISL_458952, EPI_ISL_458953, EPI_ISL_458954, EPI_ISL_458955, EPI_ISL_458956, EPI_ISL_458957, EPI_ISL_458958, EPI_ISL_458959, EPI_ISL_458960, EPI_ISL_458961, EPI_ISL_458962, EPI_ISL_458963, EPI_ISL_458964, EPI_ISL_458966, EPI_ISL_458967, EPI_ISL_458968, EPI_ISL_458969, EPI_ISL_458970, EPI_ISL_458971, EPI_ISL_458972, EPI_ISL_458973, EPI_ISL_458974, EPI_ISL_458975, EPI_ISL_458976, EPI_ISL_458977, EPI_ISL_458978, EPI_ISL_458979, EPI_ISL_458980, EPI_ISL_458981, EPI_ISL_458982, EPI_ISL_458983, EPI_ISL_458984, EPI_ISL_458985, EPI_ISL_458986, EPI_ISL_458987, EPI_ISL_458988, EPI_ISL_458989, EPI_ISL_458990, EPI_ISL_458991, EPI_ISL_458992, EPI_ISL_458993, EPI_ISL_458994, EPI_ISL_458995, EPI_ISL_458996, EPI_ISL_458997, EPI_ISL_458998, EPI_ISL_458999, EPI_ISL_459000, EPI_ISL_459001, EPI_ISL_459002, EPI_ISL_459003, EPI_ISL_459004, EPI_ISL_459005, EPI_ISL_459006, EPI_ISL_459007, EPI_ISL_459008, EPI_ISL_459009, EPI_ISL_459010, EPI_ISL_459011, EPI_ISL_459012, EPI_ISL_459013, EPI_ISL_459014, EPI_ISL_459015, EPI_ISL_459016, EPI_ISL_459017, EPI_ISL_459018, EPI_ISL_459019, EPI_ISL_459020, EPI_ISL_459021, EPI_ISL_459022, EPI_ISL_459023, EPI_ISL_459024, EPI_ISL_459025, EPI_ISL_459026, EPI_ISL_459027, EPI_ISL_459028, EPI_ISL_459029, EPI_ISL_459030, EPI_ISL_459031, EPI_ISL_459032, EPI_ISL_459033, EPI_ISL_459034, EPI_ISL_459035, EPI_ISL_459036, EPI_ISL_459037, EPI_ISL_459038 | | | |
| see above | PHE South West Regional Laboratory, National Infection Service | Wellcome Sanger Institute for the COVID-19 Genomics UK (COG-UK) consortium | Stephanie Hutchings, Hannah Pymont, Dr Peter Muir, Barry Vipond, Rich Hopes; and Alex Alderton, Roberto Amato, Sonia Goncalves, Ewan Harrison, David K. Jackson, Ian Johnston, Dominic Kwiatkowski, Cordelia Langford, John Sillitoe on behalf of the Wellcome Sanger Institute COVID-19 Surveillance Team (http://www.sanger.ac.uk/covid-team) |
| EPI_ISL_459039 | PHE South West Regional Laboratory, National Infection Service | Wellcome Sanger Institute for the COVID-19 Genomics UK (COG-UK) Consortium | Stephanie Hutchings, Hannah Pymont, Dr Peter Muir, Barry Vipond, Rich Hopes; and Alex Alderton, Roberto Amato, Sonia Goncalves, Ewan Harrison, David K. Jackson, Ian Johnston, Dominic Kwiatkowski, Cordelia Langford, John Sillitoe on behalf of the Wellcome Sanger Institute COVID-19 Surveillance Team |
| EPI_ISL_459040, EPI_ISL_459041, EPI_ISL_459042, EPI_ISL_459043, EPI_ISL_459044, EPI_ISL_459045, EPI_ISL_459046, EPI_ISL_459047, EPI_ISL_459048, EPI_ISL_459049, EPI_ISL_459050, EPI_ISL_459051, EPI_ISL_459052, EPI_ISL_459053, EPI_ISL_459054, EPI_ISL_459055, EPI_ISL_459056, EPI_ISL_459057, EPI_ISL_459058, EPI_ISL_459059, EPI_ISL_459060, EPI_ISL_459061, EPI_ISL_459062, EPI_ISL_459063, EPI_ISL_459064, EPI_ISL_459065, EPI_ISL_459066, EPI_ISL_459067, EPI_ISL_459068, EPI_ISL_459069, EPI_ISL_459070, EPI_ISL_459071, EPI_ISL_459072, EPI_ISL_459073, EPI_ISL_459074, EPI_ISL_459075, EPI_ISL_459076, EPI_ISL_459077, EPI_ISL_459078, EPI_ISL_459079, EPI_ISL_459080, EPI_ISL_459081, EPI_ISL_459082, EPI_ISL_459083, EPI_ISL_459084, EPI_ISL_459085, EPI_ISL_459086, EPI_ISL_459087, EPI_ISL_459088, EPI_ISL_459089, EPI_ISL_459090, EPI_ISL_459091, EPI_ISL_459092, EPI_ISL_459093, EPI_ISL_459094 | | | |
| see above | PHE South West Regional Laboratory, National Infection Service | Wellcome Sanger Institute for the COVID-19 Genomics UK (COG-UK) consortium | Stephanie Hutchings, Hannah Pymont, Dr Peter Muir, Barry Vipond, Rich Hopes; and Alex Alderton, Roberto Amato, Sonia Goncalves, Ewan Harrison, David K. Jackson, Ian Johnston, Dominic Kwiatkowski, Cordelia Langford, John Sillitoe on behalf of the Wellcome Sanger Institute COVID-19 Surveillance Team (http://www.sanger.ac.uk/covid-team) |
| EPI_ISL_459097, EPI_ISL_459098, EPI_ISL_459105, EPI_ISL_459106, EPI_ISL_459107, EPI_ISL_459110, EPI_ISL_459111, EPI_ISL_459112, EPI_ISL_459113, EPI_ISL_459114, EPI_ISL_459119, EPI_ISL_459121, EPI_ISL_459122, EPI_ISL_459127, EPI_ISL_459128, EPI_ISL_459129, EPI_ISL_459130, EPI_ISL_459132, EPI_ISL_459133, EPI_ISL_459136, EPI_ISL_459138, EPI_ISL_459140, EPI_ISL_459142, EPI_ISL_459143, EPI_ISL_459144, EPI_ISL_459146, EPI_ISL_459147, EPI_ISL_459148, EPI_ISL_459149, EPI_ISL_459152, EPI_ISL_459155, EPI_ISL_459159, EPI_ISL_459161, EPI_ISL_459165 | | | |
| see above | NHSGGC West of Scotland Specialist Virology Centre / MRC-University of Glasgow Centre for Virus Research | Wellcome Sanger Institute for the COVID-19 Genomics UK (COG-UK) consortium | Ana da Silva Filipe, Natasha Johnson, Kathy Smollett, Daniel Mair, Stephen Carmichael, Lily Tong, Jenna Nichols, Elihu Aranday-Cortes, Kirstyn Brunker, Yasmin Parr, Kyriaki Nomikou; Sarah McDonald, Marc Niebel, Patawee Asamaphan; Richard Orton, Joseph Hughes, Sreenu Vattipally, David L Robertson; Alasdair MacLean, Rory Gunson; Kathy Li, Natasha Jesudason, Rajiv Shah, James Shepherd, Antonia Ho, Alice Broos, Emma Thomson and Alex Alderton, Roberto Amato, Sonia Goncalves, Ewan Harrison, David K. Jackson, Ian Johnston, Dominic Kwiatkowski, Cordelia Langford, John Sillitoe on behalf of the Wellcome Sanger Institute COVID-19 Surveillance Team (http://www.sanger.ac.uk/covid-team) |
| EPI_ISL_459327, EPI_ISL_459328, EPI_ISL_459331, EPI_ISL_459333, EPI_ISL_459339, EPI_ISL_459343, EPI_ISL_459345, EPI_ISL_459348, EPI_ISL_459353, EPI_ISL_459355 | Regional Virus Laboratory, Belfast Health and Social Care Trust | Wellcome Sanger Institute for the COVID-19 Genomics UK (COG-UK) consortium | Conall McCaughey, James McKenna, Tanya Curran, Susan Feeney, Alison Watt, Ciara Cox, Mairead Connor, Zoltan Molnar, David Simpson, Derek Fairley; and Alex Alderton, Roberto Amato, Sonia Goncalves, Ewan Harrison, David K. Jackson, Ian Johnston, Dominic Kwiatkowski, Cordelia Langford, John Sillitoe on behalf of the Wellcome Sanger Institute COVID-19 Surveillance Team (http://www.sanger.ac.uk/covid-team) |
| EPI_ISL_459358 | Regional Virus Laboratory, Belfast Health and Social Care Trust | Wellcome Sanger Institute for the COVID-19 Genomics UK (COG-UK) Consortium | Conall McCaughey, James McKenna, Tanya Curran, Susan Feeney, Alison Watt, Ciara Cox, Mairead Connor, Zoltan Molnar, David Simpson, Derek Fairley; and Alex Alderton, Roberto Amato, Sonia Goncalves, Ewan Harrison, David K. Jackson, Ian Johnston, Dominic Kwiatkowski, Cordelia Langford, John Sillitoe on behalf of the Wellcome Sanger Institute COVID-19 Surveillance Team |
| EPI_ISL_459359, EPI_ISL_459363, EPI_ISL_459364, EPI_ISL_459370 | Regional Virus Laboratory, Belfast Health and Social Care Trust | Wellcome Sanger Institute for the COVID-19 Genomics UK (COG-UK) consortium | Conall McCaughey, James McKenna, Tanya Curran, Susan Feeney, Alison Watt, Ciara Cox, Mairead Connor, Zoltan Molnar, David Simpson, Derek Fairley; and Alex Alderton, Roberto Amato, Sonia Goncalves, Ewan Harrison, David K. Jackson, Ian Johnston, Dominic Kwiatkowski, Cordelia Langford, John Sillitoe on behalf of the Wellcome Sanger Institute COVID-19 Surveillance Team (http://www.sanger.ac.uk/covid-team) |
| EPI_ISL_459372 | Regional Virus Laboratory, Belfast Health and Social Care Trust | Wellcome Sanger Institute for the COVID-19 Genomics UK (COG-UK) Consortium | Conall McCaughey, James McKenna, Tanya Curran, Susan Feeney, Alison Watt, Ciara Cox, Mairead Connor, Zoltan Molnar, David Simpson, Derek Fairley; and Alex Alderton, Roberto Amato, Sonia Goncalves, Ewan Harrison, David K. Jackson, Ian Johnston, Dominic Kwiatkowski, Cordelia Langford, John Sillitoe on behalf of the Wellcome Sanger Institute COVID-19 Surveillance Team |
| EPI_ISL_459377, EPI_ISL_459380, EPI_ISL_459381, EPI_ISL_459383, EPI_ISL_459385, EPI_ISL_459386, EPI_ISL_459388, EPI_ISL_459389, EPI_ISL_459390, EPI_ISL_459398, EPI_ISL_459402, EPI_ISL_459403, EPI_ISL_459405 | | | |
| see above | Regional Virus Laboratory, Belfast Health and Social Care Trust | Wellcome Sanger Institute for the COVID-19 Genomics UK (COG-UK) consortium | Conall McCaughey, James McKenna, Tanya Curran, Susan Feeney, Alison Watt, Ciara Cox, Mairead Connor, Zoltan Molnar, David Simpson, Derek Fairley; and Alex Alderton, Roberto Amato, Sonia Goncalves, Ewan Harrison, David K. Jackson, Ian Johnston, Dominic Kwiatkowski, Cordelia Langford, John Sillitoe on behalf of the Wellcome Sanger Institute COVID-19 Surveillance Team (http://www.sanger.ac.uk/covid-team) |
| EPI_ISL_459725, EPI_ISL_459726, EPI_ISL_459727 | PHE South West Regional Laboratory, National Infection Service | Wellcome Sanger Institute for the COVID-19 Genomics UK (COG-UK) consortium | Stephanie Hutchings, Hannah Pymont, Dr Peter Muir, Barry Vipond, Rich Hopes; and Alex Alderton, Roberto Amato, Sonia Goncalves, Ewan Harrison, David K. Jackson, Ian Johnston, Dominic Kwiatkowski, Cordelia Langford, John Sillitoe on behalf of the Wellcome Sanger Institute COVID-19 Surveillance Team (http://www.sanger.ac.uk/covid-team) |
| EPI_ISL_460106, EPI_ISL_460112, EPI_ISL_460123, EPI_ISL_460130, EPI_ISL_460246, EPI_ISL_460277, EPI_ISL_460278, EPI_ISL_460315, EPI_ISL_460323 | Massachusetts General Hospital | Infectious Disease Program, Broad Institute of Harvard and MIT | Lemieux,J.E., Siddle,K.J., Shaw,B., Adams,G., Pierce,V., Turbett,S., Anahtar,M., Branda,J., Slater,D., Harris,J., Lin,A.E., Gladden-Young,A., Lagerborg,K., Rudy,M., DeRuff,K., Carter,A., Normandin,E., Bauer,M., Reilly,S., Tomkins-Tinch,C., Loreth,C., Chaluvadi,S., Neumann,A., Cusick,C., Chapman,S.B., Gnirke,A., Flowers,K., Cerrato,F., Birren,B.W., Gallagher,G., Smole,S., Park,D.J., MacInnis,B.L., Ryan,E., LaRocque,R., Rosenberg,E., Sabeti,P.C. |
| EPI_ISL_460562, EPI_ISL_460564 | Michigan Department of Health and Human Services, Bureau of Laboratories | Michigan Department of Health and Human Services, Bureau of Laboratories | Blankenship HM, Riner D, Soehnlen MK |
| EPI_ISL_460635 | UHCW Pathology / University of Warwick | University of Warwick, for the COVID-19 Genomics (COG) UK Consortium | Richard Stark, Chrystala Constantinidou, Meera Unnikrishnan, Laura Baxter, Jeff Cheng, Grace Taylor-Joyce, Hannah Elizabeth Bridgewater, Lucy Frost, Sarojini Pandey, Paul Brown, Tauqeer Alam, Sascha Ott, Dimitris Grammatopoulos |
| EPI_ISL_460641, EPI_ISL_460642, EPI_ISL_460644, EPI_ISL_460646, EPI_ISL_460647, EPI_ISL_460648, EPI_ISL_460649, EPI_ISL_460650, EPI_ISL_460653, EPI_ISL_460654, EPI_ISL_460669, EPI_ISL_460702, EPI_ISL_460723, EPI_ISL_460724, EPI_ISL_460725, EPI_ISL_460726, EPI_ISL_460727, EPI_ISL_460728, EPI_ISL_460729, EPI_ISL_460730, EPI_ISL_460731, EPI_ISL_460732, EPI_ISL_460733, EPI_ISL_460734, EPI_ISL_460789, EPI_ISL_460799, EPI_ISL_460800, EPI_ISL_460801, EPI_ISL_460802, EPI_ISL_460826, EPI_ISL_460827, EPI_ISL_460828, EPI_ISL_460836, EPI_ISL_460837, EPI_ISL_460929, EPI_ISL_460931, EPI_ISL_460932, EPI_ISL_460933, EPI_ISL_460934, EPI_ISL_460937, EPI_ISL_461033, EPI_ISL_461034, EPI_ISL_461037, EPI_ISL_461069, EPI_ISL_461072, EPI_ISL_461119, EPI_ISL_461121, EPI_ISL_461122, EPI_ISL_461165, EPI_ISL_461166, EPI_ISL_461167, EPI_ISL_461234, EPI_ISL_461235, EPI_ISL_461236, EPI_ISL_461242, EPI_ISL_461249, EPI_ISL_461279 | | | |
| see above | Dutch COVID-19 response team | Erasmus Medical Center | Bas Oude Munnink, David Nieuwenhuijse, Reina Sikkema, Claudia Schapendonk, Irina Chestakova, Anne van der Linden, Theo Bestebroer, Stefan van |

| | | | |
|---|---|---|---|
| | | | Nieuwkoop, Mark Pronk, Pascal Lexmond, Corien Swaan, Manon Haverkate, Madelief Mollers, Mart Stein, Sandra Kengne Kamga Mobou, Jeroen van Kampen, Jolanda Voermans, Aura Timen, Corine GeurtsvanKessel, Annemiek van der Eijk, Richard Molenkamp, Marion Koopmans, on behalf of the Dutch national COVID-19 response team. |
| EPI_ISL_461404, EPI_ISL_461405, EPI_ISL_461433, EPI_ISL_461434, EPI_ISL_461437, EPI_ISL_461438, EPI_ISL_461446, EPI_ISL_461447, EPI_ISL_461448, EPI_ISL_461449 | UW Virology Lab | UW Virology Lab | Pavitra Roychoudhury, Amin Addetia, Hong Xie, Lasata Shrestha, Truong Nguyen, Meei-Li Huang, Keith Jerome, Alexander Greninger |
| EPI_ISL_462247, EPI_ISL_462248, EPI_ISL_462249, EPI_ISL_462250, EPI_ISL_462251, EPI_ISL_462253, EPI_ISL_462257, EPI_ISL_462258, EPI_ISL_462259, EPI_ISL_462260, EPI_ISL_462261, EPI_ISL_462262, EPI_ISL_462266, EPI_ISL_462267, EPI_ISL_462268, EPI_ISL_462269 | | | |
| see above | KU Leuven, Rega Institute, Clinical and Epidemiological Virology | KU Leuven, Rega Institute, Clinical and Epidemiological Virology | Tony Wawina-Bokalanga, Bert Vanmechelen, Joan Marti-Carerras, Piet Maes |
| EPI_ISL_462334, EPI_ISL_462337, EPI_ISL_462340, EPI_ISL_462352, EPI_ISL_462353, EPI_ISL_462354, EPI_ISL_462356, EPI_ISL_462363, EPI_ISL_462365, EPI_ISL_462374, EPI_ISL_462375, EPI_ISL_462377, EPI_ISL_462379, EPI_ISL_462381, EPI_ISL_462383, EPI_ISL_462384, EPI_ISL_462395, EPI_ISL_462398, EPI_ISL_462400, EPI_ISL_462402, EPI_ISL_462413, EPI_ISL_462414 | | | |
| see above | National Public Health Laboratory, National Centre for Infectious Diseases | National Public Health Laboratory, National Centre for Infectious Diseases | Mak TM, Octavia S, Chavatte JM, Cui L, Lin RTP |
| EPI_ISL_462461 | Clinical Center, University of Sarajevo | Charite Universitatsmedizin Berlin, Institute of Virology | Victor M Corman, Jorn Beheim-Schwarzbach, Barbara Muehlemann, Talitha Veith, Julia Schneider, Terry Jones, Amela Dedeic-Ljubovic, Irma Salimovic-Besic, Suzana Arapcic, Almedina Hadzihasanovic-Moro, Selma Mutevelic, Christian Drosten |
| EPI_ISL_463007 | Department of Laboratory Medicine, National Taiwan University Hospital | Microbial Genomics Core Lab, National Taiwan University Centers of Genomic and Precision Medicine | Shiou-Hwei Yeh, You-Yu Lin, Ya-Yun Lai, Chiao-Ling Li, Shan-Chwen Chang, Pei-Jer Chen, Sui-Yuan Chang |
| EPI_ISL_463290, EPI_ISL_463292, EPI_ISL_463293 | Ochsner Health | Bioinfoexperts, LLC | Susanna L. Lamers, David J. Nolan, Rebecca Rose, Sissy Cross, David Moraga Amador, Tong Yang, Luke Caruso, Wayra Navia, Lydia Von Borstel, Xiao Hui Zhou, Amy Feehan, Julia-Garcia-Diaz |
| EPI_ISL_463301 | Tuen Mun Hospital | Hong Kong Department of Health | Mak Gannon C.K., Cheng Peter K.C., Lam Edman T.K., Chan Rickjason C.W., Tsang Dominic N.C. |
| EPI_ISL_463320 | Tseung Kwan O Hospital | Hong Kong Department of Health | Mak Gannon C.K., Cheng Peter K.C., Lam Edman T.K., Chan Rickjason C.W., Tsang Dominic N.C. |
| EPI_ISL_463321 | Queen Mary Hospital | Hong Kong Department of Health | Mak Gannon C.K., Cheng Peter K.C., Lam Edman T.K., Chan Rickjason C.W., Tsang Dominic N.C. |
| EPI_ISL_463322 | North Lantau Hospital | Hong Kong Department of Health | Mak Gannon C.K., Cheng Peter K.C., Lam Edman T.K., Chan Rickjason C.W., Tsang Dominic N.C. |
| EPI_ISL_463323 | Yuen Long JC GOPC | Hong Kong Department of Health | Mak Gannon C.K., Cheng Peter K.C., Lam Edman T.K., Chan Rickjason C.W., Tsang Dominic N.C. |
| EPI_ISL_463746 | Department of Molecular Virology, Cyprus Institute of Neurology and Genetics | Department of Molecular Virology, Cyprus Institute of Neurology and Genetics | Jan Richter, George Krashias, Christina Tryfonos, Stavros Bashiardes, Dana Koptides, Christina Christodoulou |
| EPI_ISL_463905, EPI_ISL_463907, EPI_ISL_463909, EPI_ISL_463913, EPI_ISL_463915, EPI_ISL_463916, EPI_ISL_463917, EPI_ISL_463918, EPI_ISL_463919, EPI_ISL_463920, EPI_ISL_463921, EPI_ISL_463922, EPI_ISL_463923, EPI_ISL_463925, EPI_ISL_463945, EPI_ISL_463954, EPI_ISL_463955, EPI_ISL_463956, EPI_ISL_463957, EPI_ISL_463958, EPI_ISL_463959, EPI_ISL_463960, EPI_ISL_463962, EPI_ISL_463967, EPI_ISL_463968 | | | |
| see above | Laboratoire de microbiologie, Hopital de Verdun | Smith Laboratory, Centre de Recherche CHU Sainte-Justine | Martin Smith, Marieke Rozendaal, Ivan Pavlov |
| EPI_ISL_463992 | Toronto Invasive Bacterial Diseases Network | McMaster University | Allison McGeer, Patryk Aftanas, Angel Li, Kuganya Nirmalarajah, Samira Mubareka, Andrew G. McArthur |
| EPI_ISL_463997, EPI_ISL_463999, EPI_ISL_464004, EPI_ISL_464007, EPI_ISL_464011, EPI_ISL_464022, EPI_ISL_464026, EPI_ISL_464033, EPI_ISL_464036, EPI_ISL_464040, EPI_ISL_464042, EPI_ISL_464050, EPI_ISL_464059 | | | |
| see above | Unity Health Toronto | Ontario Institute for Cancer Research | Ramzi Fattouh,Larissa M. Matukas,Mark Downing,Annette Gower,Karel Boissinot,Samira Mubareka,TIBDN,Ilinca Lungu,Bernard Lam,Jeremy Johns,Paul Krzyzanowski,Richard de Borja,Philip Zuzarte,Jared Simpson |
| EPI_ISL_464065, EPI_ISL_464066, EPI_ISL_464068, EPI_ISL_464074, EPI_ISL_464075, EPI_ISL_464076, EPI_ISL_464077, EPI_ISL_464078, EPI_ISL_464080, EPI_ISL_464081, EPI_ISL_464082, EPI_ISL_464085, EPI_ISL_464086, EPI_ISL_464087, EPI_ISL_464088, EPI_ISL_464089, EPI_ISL_464090 | | | |
| see above | KU Leuven, Rega Institute, Clinical and Epidemiological Virology | KU Leuven, Rega Institute, Clinical and Epidemiological Virology | Tony Wawina-Bokalanga, Bert Vanmechelen, Joan Marti-Carerras, Piet Maes |
| EPI_ISL_464094 | Laboratory Medicine | Department of Laboratory Medicine, Lin-Kou Chang Gung Memorial Hospital, Taoyuan, Taiwan | Kuo-Chien Tsao, Yu-Nong Gong, Shu-Li Yang, Yi-Chun Liu, Chung-Guei Huang, Mei-Jen Hsiao, Po-Wei Huang, Cheng-Ta Yang, Cheng-Hsun Chiu, Peng-Nien Huang, Kuo-Ming Lee, Guang-Wu Chen, Shin-Ru Shih |
| EPI_ISL_464118, EPI_ISL_464119, EPI_ISL_464120, EPI_ISL_464121, EPI_ISL_464122, EPI_ISL_464123, EPI_ISL_464124, EPI_ISL_464125, EPI_ISL_464126, EPI_ISL_464127, EPI_ISL_464128, EPI_ISL_464129, EPI_ISL_464130, EPI_ISL_464131, EPI_ISL_464132, EPI_ISL_464133 | | | |
| see above | National Health Laboratory Service (NHLS), Tygerberg | Division of Medical Virology, Stellenbosch University and National Health Laboratory Service (NHLS) | Susan Engelbrecht, Kayla Delaney, Bronwyn Kleinhans, Houriiyah Tegally, Eduan Wilkindon, Gert van Zyl, Wolfgang Preiser, Tulio de Oliveira |
| EPI_ISL_464134 | National Health Laboratory Service (NHLS), Tygerberg | Stellenbosch University and NHLS | Susan Engelbrecht, Kayla Delaney, Bronwyn Kleinhans, Houriiyah Tegally, Eduan Wilkindon, Gert van Zyl, Wolfgang Preiser, Tulio de Oliveira |
| EPI_ISL_464135, EPI_ISL_464136, EPI_ISL_464137, EPI_ISL_464138, EPI_ISL_464139, EPI_ISL_464148, EPI_ISL_464149, EPI_ISL_464150, EPI_ISL_464151, EPI_ISL_464152, EPI_ISL_464153, EPI_ISL_464154, EPI_ISL_464155, EPI_ISL_464156, EPI_ISL_464157, EPI_ISL_464158 | | | |
| see above | National Health Laboratory Service (NHLS), Tygerberg | Division of Medical Virology, Stellenbosch University and National Health Laboratory Service (NHLS) | Susan Engelbrecht, Kayla Delaney, Bronwyn Kleinhans, Houriiyah Tegally, Eduan Wilkindon, Gert van Zyl, Wolfgang Preiser, Tulio de Oliveira |
| EPI_ISL_465224, EPI_ISL_465275, EPI_ISL_465276, EPI_ISL_465277, EPI_ISL_465278, EPI_ISL_465287, EPI_ISL_465288, EPI_ISL_465289, EPI_ISL_465290, EPI_ISL_465291, EPI_ISL_465294, EPI_ISL_465348, EPI_ISL_465349, EPI_ISL_466248, EPI_ISL_466256, EPI_ISL_466257, EPI_ISL_466258, EPI_ISL_466259, EPI_ISL_466260, EPI_ISL_466261, EPI_ISL_466262, EPI_ISL_466266, EPI_ISL_466267, EPI_ISL_466268, EPI_ISL_466270, EPI_ISL_466272, EPI_ISL_466273, EPI_ISL_466277, EPI_ISL_466278, EPI_ISL_466279, EPI_ISL_466280, EPI_ISL_466281, EPI_ISL_466282, EPI_ISL_466283, EPI_ISL_466284, EPI_ISL_466285, EPI_ISL_466286, EPI_ISL_466287, EPI_ISL_466288, EPI_ISL_466289, EPI_ISL_466290, EPI_ISL_466291, EPI_ISL_466292, EPI_ISL_466293, EPI_ISL_466294, EPI_ISL_466295, EPI_ISL_466296, EPI_ISL_466297, EPI_ISL_466298, EPI_ISL_466299, EPI_ISL_466300, EPI_ISL_466301, EPI_ISL_466302, EPI_ISL_466303, EPI_ISL_466304, EPI_ISL_466305, EPI_ISL_466306, EPI_ISL_466307, EPI_ISL_466308, EPI_ISL_466309, EPI_ISL_466310, EPI_ISL_466311, EPI_ISL_466312, EPI_ISL_466313, EPI_ISL_466314, EPI_ISL_466315, EPI_ISL_466316, EPI_ISL_466317, EPI_ISL_466318, EPI_ISL_466319, EPI_ISL_466320, EPI_ISL_466322, EPI_ISL_466324, EPI_ISL_466326, EPI_ISL_466327, EPI_ISL_466328, EPI_ISL_466329, EPI_ISL_466332, EPI_ISL_466333, EPI_ISL_466335, EPI_ISL_466337, EPI_ISL_466338, EPI_ISL_466339, EPI_ISL_466341, EPI_ISL_466342, EPI_ISL_466343, EPI_ISL_466344, EPI_ISL_466345, EPI_ISL_466346, EPI_ISL_466347, EPI_ISL_466348, EPI_ISL_466349, EPI_ISL_466350, EPI_ISL_466351, EPI_ISL_466352, EPI_ISL_466353, EPI_ISL_466354, EPI_ISL_466355, EPI_ISL_466356, EPI_ISL_466357, EPI_ISL_466358, EPI_ISL_466359, EPI_ISL_466360, EPI_ISL_466361, EPI_ISL_466362, EPI_ISL_466363, EPI_ISL_466364, EPI_ISL_466365, EPI_ISL_466366, EPI_ISL_466367, EPI_ISL_466368, EPI_ISL_466369, EPI_ISL_466370, EPI_ISL_466371, EPI_ISL_466372, EPI_ISL_466373, EPI_ISL_466374, EPI_ISL_466375, EPI_ISL_466393, EPI_ISL_466394, EPI_ISL_466400, EPI_ISL_466401, EPI_ISL_466407, EPI_ISL_466408, EPI_ISL_466409, EPI_ISL_466410, EPI_ISL_466411, EPI_ISL_466412, EPI_ISL_466413, EPI_ISL_466415, EPI_ISL_466416, EPI_ISL_466417, EPI_ISL_466419, EPI_ISL_466421, EPI_ISL_466422, EPI_ISL_466423, EPI_ISL_466424, EPI_ISL_466428, EPI_ISL_466429, EPI_ISL_466430, EPI_ISL_466431, EPI_ISL_466459, EPI_ISL_466467, EPI_ISL_466468, EPI_ISL_466469, EPI_ISL_466470, EPI_ISL_466471, EPI_ISL_466472, EPI_ISL_466473, EPI_ISL_466474, EPI_ISL_466475, EPI_ISL_466476, EPI_ISL_466477, EPI_ISL_466539, EPI_ISL_466564, EPI_ISL_466613 | | | |
| see above | Respiratory Virus Unit, Microbiology Services Colindale, Public Health England | Respiratory Virus Unit, Microbiology Services Colindale, Public Health England | PHE Covid Sequencing Team |
| EPI_ISL_467263, EPI_ISL_467264, EPI_ISL_467270, EPI_ISL_467271, EPI_ISL_467273, EPI_ISL_467275, EPI_ISL_467278, EPI_ISL_467279, EPI_ISL_467286, EPI_ISL_467287, EPI_ISL_467293, EPI_ISL_467294, EPI_ISL_467295, EPI_ISL_467297 | | | |
| see above | Hospital Clínico Universitario de Santiago de Compostela | SeqCOVID-SPAIN consortium/IBV(CSIC) | José Javier Costa Alcalde, Antonio Aguilera Guirao, Mª Luisa Pérez del Molino Bernal, Amparo Coira Nieto, Gema Barbeito Castiñeiras, Rocio Trastoy Pena and SeqCOVID-SPAIN consortium |
| EPI_ISL_467300 | General Hospital "Abdulah Nakas" | Alea Genetic Center | Rijad Konjhodzic; Lana Salihefendic; Teufik Goletic; Sead Jazic; Dino Pecar; Nihad Fejzic; Damir Marjanovic; Enis Kandic |
| EPI_ISL_467385 | NYU Langone Health | Departments of Pathology and Medicine, New York University School of Medicine | Maria Aguero-Rosenfeld, Brendan Belovarac, Margaret Black, Ludovic Boytard, John Cadley, Paolo Cotzia, John Chen, Dacia Dimartino, Xiaojun Feng, Tatyana Gindin, Emily Guzman, Adriana Heguy, Megan Hogan, Emily Huang, George Jour, Alireza Khodadadi-Jamayran, Lawrence H. Lin, Raven Luther, Andrew Lytle, Christian Marier, Matthew T. Maurano, Mark J. Mulligan, Peter Meyn, Raquel Ordonez Ciriza, Iman Osman, Jared Pinnell, Vanessa Raabe, Sitharam Ramaswami, Amy Rapkiewicz, Andre M. Ribeiro-dos-Santos, Marie Samanovic-Golden, Antonio Serrano, Guomiao Shen, Matija Snuderl, Theodore Vougiouklakis, Nick Vulpescu, Gael Westby, Paul Zappile, Yutong Zhang |
| EPI_ISL_467457 | AMPATH-DBN | KRISP, KZN Research Innovation and Sequencing Platform | Giandhari J, Pillay S, Lessells R, Chimukangara B, Mdlalose K, York D, Khan S, Tegally H, Wilkinson E, de Oliveira T |
| EPI_ISL_467986, EPI_ISL_468007, EPI_ISL_468025, EPI_ISL_468027 | SA Pathology | SA Pathology | Lex Leong, Chuan Kok Lim, Mark Turra, Ivan Bastian, Geoff Higgins |

| | | | |
|---|---|---|---|
| EPI_ISL_468077 | Child Health Research Foundation | Child Health Research Foundation | Senjuti Saha, Roly Malaker, Md Saiful Islam Sajib, Hafizur Rahman, Afroza Akter Tanni, Syed Muktadir Al Sium, Maksuda Islam, Samir K Saha |
| EPI_ISL_468146, EPI_ISL_468147, EPI_ISL_468148, EPI_ISL_468149, EPI_ISL_468150 | [Romania, Bucharest] National Institute for Infectious Diseases "Prof. Dr. Matei Bal" | [Romania, Bucharest] National Institute for Infectious Diseases "Prof. Dr. Matei Bal" | Leontina Banica, Marius Cotic, Corina Casangiu, Marius Surleac, Simona Paraschiv |
| EPI_ISL_468305, EPI_ISL_468307 | Centro de Vigilancia a Saude de Diadema | Instituto Adolfo Lutz, Interdiciplinary Procedures Center, Strategic Laboratory | Claudio Tavares Sacchi, Claudia Regina Gonçalves, Erica Valessa Ramos Gomes |
| EPI_ISL_468315 | Hospital Municipal do Tatuape Carmino Caricchio | Instituto Adolfo Lutz, Interdiciplinary Procedures Center, Strategic Laboratory | Claudio Tavares Sacchi, Claudia Regina Gonçalves, Erica Valessa Ramos Gomes |
| EPI_ISL_468360, EPI_ISL_468361, EPI_ISL_468362, EPI_ISL_468363, EPI_ISL_468364, EPI_ISL_468365, EPI_ISL_468366, EPI_ISL_468367, EPI_ISL_468368, EPI_ISL_468369, EPI_ISL_468370, EPI_ISL_468371, EPI_ISL_468372 | | | |
| see above | Alameda County Public Health Lab | Chan-Zuckerberg Biohub | CZB Cliahub Consortium |
| EPI_ISL_468415, EPI_ISL_468416, EPI_ISL_468417, EPI_ISL_468418 | County of San Luis Obispo Public Health Laboratory | Chan-Zuckerberg Biohub | CZB Cliahub Consortium |
| EPI_ISL_468503, EPI_ISL_468504, EPI_ISL_468505 | Ventura County Public Health Lab | Chan-Zuckerberg Biohub | CZB Cliahub Consortium |
| EPI_ISL_468538, EPI_ISL_468539, EPI_ISL_468540, EPI_ISL_468541, EPI_ISL_468542 | San Joaquin County Public Health Lab | Chan-Zuckerberg Biohub | CZB Cliahub Consortium |
| EPI_ISL_468591 | Institute for Public Health | Laboratory for advanced genomics | Filip Roki, Lovro Trgovec-Greif, Neven Sui, Tomislav Rukavina, Igor Jurak, Oliver Vugrek |
| EPI_ISL_468651, EPI_ISL_468652, EPI_ISL_468653 | Contra Costa Public Health Lab | Chan-Zuckerberg Biohub | CZB Cliahub Consortium |
| EPI_ISL_468656 | Institute for Public Health | Laboratory for advanced genomics | Filip Roki, Lovro Trgovec-Greif, Neven Sui, Tomislav Rukavina, Igor Jurak, Oliver Vugrek |
| EPI_ISL_468704, EPI_ISL_468705, EPI_ISL_468706, EPI_ISL_468707, EPI_ISL_468708, EPI_ISL_468709, EPI_ISL_468710, EPI_ISL_468711, EPI_ISL_468716 | Ochsner Health | Bioinfoexperts, LLC | Rebecca Rose, Amy Feehan, David J. Nolan, Sissy Cross, David Moraga Amador, Tong Yang, Luke Caruso, Wayra Navia, Lydia Von Borstel, Xiao Hui Zhou, Julia-Garcia-Diaz, Susanna L. Lamers |
| EPI_ISL_468722 | University of Florida | University of Florida | Elbadry,M.A., Subramaniam,K., Waltzek,T.B., Lauzardo,M., Morris,J.G., Lednicky,J.A. |
| EPI_ISL_468860, EPI_ISL_468861, EPI_ISL_468862, EPI_ISL_468863, EPI_ISL_468864, EPI_ISL_468865, EPI_ISL_468866, EPI_ISL_468867, EPI_ISL_468868, EPI_ISL_468869, EPI_ISL_468870, EPI_ISL_468871, EPI_ISL_468872, EPI_ISL_468873, EPI_ISL_468874, EPI_ISL_468875, EPI_ISL_468876, EPI_ISL_468877, EPI_ISL_468878, EPI_ISL_468879, EPI_ISL_468880, EPI_ISL_468881, EPI_ISL_468882, EPI_ISL_468883, EPI_ISL_468884, EPI_ISL_468885, EPI_ISL_468886, EPI_ISL_468887, EPI_ISL_468888, EPI_ISL_468889, EPI_ISL_468890, EPI_ISL_468891, EPI_ISL_468892, EPI_ISL_468893, EPI_ISL_468894, EPI_ISL_468895, EPI_ISL_468896, EPI_ISL_468897, EPI_ISL_468898, EPI_ISL_468899, EPI_ISL_468900, EPI_ISL_468901, EPI_ISL_468902, EPI_ISL_468903, EPI_ISL_468904, EPI_ISL_468905, EPI_ISL_468906, EPI_ISL_468907, EPI_ISL_468908, EPI_ISL_468909, EPI_ISL_468910, EPI_ISL_468911, EPI_ISL_468912, EPI_ISL_468913, EPI_ISL_468915, EPI_ISL_468916, EPI_ISL_468917, EPI_ISL_468918, EPI_ISL_468919, EPI_ISL_468920, EPI_ISL_468921, EPI_ISL_468922, EPI_ISL_468923, EPI_ISL_468924, EPI_ISL_468925, EPI_ISL_468926, EPI_ISL_468927, EPI_ISL_468928, EPI_ISL_468929, EPI_ISL_468930, EPI_ISL_468931, EPI_ISL_468932, EPI_ISL_468933, EPI_ISL_468934, EPI_ISL_468935, EPI_ISL_468936, EPI_ISL_468937, EPI_ISL_468938, EPI_ISL_468939, EPI_ISL_468940, EPI_ISL_468941, EPI_ISL_468942, EPI_ISL_468943, EPI_ISL_468944, EPI_ISL_468945, EPI_ISL_468946, EPI_ISL_468947, EPI_ISL_468948, EPI_ISL_468949, EPI_ISL_468950, EPI_ISL_468951 | | | |
| see above | Servicio de Microbiología. Hospital Universitario Donostia. OSI Donostialdea. Área de Enfermedades Infecciosas, Grupo de Infección Respiratoria y Resistencia Antimicrobiana. Instituto de Investigación Sanitaria Biodonostia. | SeqCOVID-SPAIN consortium/IBV(CSIC) | Gustavo Cilla, Milagrosa Montes, Luis Piñeiro, Jose Maria Marimón and SeqCOVID-SPAIN consortium |
| EPI_ISL_468964, EPI_ISL_468980, EPI_ISL_469006 | Servicio de Microbiología, Hospital Universitario Son Espases | SeqCOVID-SPAIN consortium/IBV(CSIC) | Carla López-Causapé, Jordi Reina, Antonio Oliver and SeqCOVID-SPAIN consortium |
| EPI_ISL_469097, EPI_ISL_469099, EPI_ISL_469101, EPI_ISL_469109, EPI_ISL_469110, EPI_ISL_469111, EPI_ISL_469112, EPI_ISL_469118, EPI_ISL_469126, EPI_ISL_469133 | National Public Health Laboratory, National Centre for Infectious Diseases | National Public Health Laboratory, National Centre for Infectious Diseases | Mak TM, Octavia S, Chavatte JM, Cui L, Lin RTP |
| EPI_ISL_469818, EPI_ISL_469825, EPI_ISL_469917 | PHE South West Regional Laboratory, National Infection Service | Wellcome Sanger Institute for the COVID-19 Genomics UK (COG-UK) consortium | Stephanie Hutchings, Hannah Pymont, Dr Peter Muir, Barry Vipond, Rich Hopes; and Alex Alderton, Roberto Amato, Sonia Goncalves, Ewan Harrison, David K. Jackson, Ian Johnston, Dominic Kwiatkowski, Cordelia Langford, John Sillitoe on behalf of the Wellcome Sanger Institute COVID-19 Surveillance Team (http://www.sanger.ac.uk/covid-team) |
| EPI_ISL_469920, EPI_ISL_469923, EPI_ISL_469924, EPI_ISL_469925, EPI_ISL_469928, EPI_ISL_469929 | Virology Department, Sheffield Teaching Hospitals NHS Foundation Trust / Department of Infection, Immunity and Cardiovascular Disease, The Medical School, University of Sheffield | Wellcome Sanger Institute for the COVID-19 Genomics UK (COG-UK) consortium | Thushan de Silva, Matthew Parker,Adri Angyal, Rebecca Brown, Luke Green, Rachel Tucker, Paul Parsons, Danielle Groves, Alex Keeley, Dave Partridge, Matthew Wyles, Benjamin Lindsey, Mehmet Yavuz, Mohammad Raza, Cariad Evans and Alex Alderton, Roberto Amato, Sonia Goncalves, Ewan Harrison, David K. Jackson, Ian Johnston, Dominic Kwiatkowski, Cordelia Langford, John Sillitoe on behalf of the Wellcome Sanger Institute COVID-19 Surveillance Team (http://www.sanger.ac.uk/covid-team) |
| EPI_ISL_470014, EPI_ISL_470016, EPI_ISL_470021, EPI_ISL_470025, EPI_ISL_470026, EPI_ISL_470028, EPI_ISL_470031, EPI_ISL_470032, EPI_ISL_470034, EPI_ISL_470036, EPI_ISL_470038, EPI_ISL_470041, EPI_ISL_470042, EPI_ISL_470044, EPI_ISL_470045, EPI_ISL_470046, EPI_ISL_470047, EPI_ISL_470049, EPI_ISL_470050, EPI_ISL_470051, EPI_ISL_470052, EPI_ISL_470055, EPI_ISL_470056, EPI_ISL_470058, EPI_ISL_470061, EPI_ISL_470063, EPI_ISL_470065, EPI_ISL_470066, EPI_ISL_470068, EPI_ISL_470071, EPI_ISL_470072, EPI_ISL_470073, EPI_ISL_470075, EPI_ISL_470076, EPI_ISL_470079, EPI_ISL_470081, EPI_ISL_470084, EPI_ISL_470085, EPI_ISL_470086, EPI_ISL_470089 | | | |
| see above | Regional Virus Laboratory, Belfast Health and Social Care Trust | Wellcome Sanger Institute for the COVID-19 Genomics UK (COG-UK) consortium | Conall McCaughey, James McKenna, Tanya Curran, Susan Feeney, Alison Watt, Ciara Cox, Mairead Connor, Zoltan Molnar, David Simpson, Derek Fairley; and Alex Alderton, Roberto Amato, Sonia Goncalves, Ewan Harrison, David K. Jackson, Ian Johnston, Dominic Kwiatkowski, Cordelia Langford, John Sillitoe on behalf of the Wellcome Sanger Institute COVID-19 Surveillance Team (http://www.sanger.ac.uk/covid-team) |
| EPI_ISL_470358, EPI_ISL_470359, EPI_ISL_470360, EPI_ISL_470361, EPI_ISL_470363, EPI_ISL_470364, EPI_ISL_470365, EPI_ISL_470366, EPI_ISL_470367, EPI_ISL_470369, EPI_ISL_470370, EPI_ISL_470371, EPI_ISL_470373, EPI_ISL_470375, EPI_ISL_470376, EPI_ISL_470377, EPI_ISL_470378, EPI_ISL_470379, EPI_ISL_470381, EPI_ISL_470382, EPI_ISL_470384, EPI_ISL_470386, EPI_ISL_470387, EPI_ISL_470388, EPI_ISL_470389, EPI_ISL_470390, EPI_ISL_470392, EPI_ISL_470393, EPI_ISL_470394, EPI_ISL_470395, EPI_ISL_470396, EPI_ISL_470397, EPI_ISL_470398, EPI_ISL_470399, EPI_ISL_470400, EPI_ISL_470401, EPI_ISL_470402, EPI_ISL_470404, EPI_ISL_470405, EPI_ISL_470406, EPI_ISL_470407, EPI_ISL_470408, EPI_ISL_470409, EPI_ISL_470410, EPI_ISL_470411, EPI_ISL_470412, EPI_ISL_470413, EPI_ISL_470414, EPI_ISL_470416, EPI_ISL_470417, EPI_ISL_470418, EPI_ISL_470420, EPI_ISL_470421, EPI_ISL_470422, EPI_ISL_470423, EPI_ISL_470424, EPI_ISL_470425, EPI_ISL_470426, EPI_ISL_470427, EPI_ISL_470428, EPI_ISL_470429, EPI_ISL_470430, EPI_ISL_470431, EPI_ISL_470432, EPI_ISL_470433, EPI_ISL_470434, EPI_ISL_470436, EPI_ISL_470437, EPI_ISL_470438, EPI_ISL_470439, EPI_ISL_470440, EPI_ISL_470442, EPI_ISL_470443, EPI_ISL_470444, EPI_ISL_470445, EPI_ISL_470446, EPI_ISL_470447, EPI_ISL_470448, EPI_ISL_470449, EPI_ISL_470450, EPI_ISL_470451, EPI_ISL_470453, EPI_ISL_470454, EPI_ISL_470455, EPI_ISL_470456, EPI_ISL_470457, EPI_ISL_470458, EPI_ISL_470461, EPI_ISL_470462, EPI_ISL_470463, EPI_ISL_470464, EPI_ISL_470465, EPI_ISL_470466, EPI_ISL_470467, EPI_ISL_470468, EPI_ISL_470469, EPI_ISL_470470, EPI_ISL_470472, EPI_ISL_470473, EPI_ISL_470474, EPI_ISL_470475, EPI_ISL_470476, EPI_ISL_470477, EPI_ISL_470478, EPI_ISL_470479, EPI_ISL_470480, EPI_ISL_470482, EPI_ISL_470483, EPI_ISL_470486, EPI_ISL_470487, EPI_ISL_470488, EPI_ISL_470489, EPI_ISL_470490, EPI_ISL_470491, EPI_ISL_470492, EPI_ISL_470494, EPI_ISL_470495, EPI_ISL_470496, EPI_ISL_470498, EPI_ISL_470499, EPI_ISL_470500, EPI_ISL_470501, EPI_ISL_470502, EPI_ISL_470503, EPI_ISL_470504, EPI_ISL_470505, EPI_ISL_470506, EPI_ISL_470507, EPI_ISL_470508, EPI_ISL_470510, EPI_ISL_470511, EPI_ISL_470512, EPI_ISL_470514, EPI_ISL_470516, EPI_ISL_470517, EPI_ISL_470518, EPI_ISL_470519, EPI_ISL_470521, EPI_ISL_470522, EPI_ISL_470523, EPI_ISL_470524, EPI_ISL_470525, EPI_ISL_470526, EPI_ISL_470527, EPI_ISL_470528 | | | |
| see above | Department of Pathology, University of Cambridge | Wellcome Sanger Institute for the COVID-19 Genomics UK (COG-UK) consortium | Luke W Meredith, M. Estée Török , Myra Hosmillo, William L. Hamilton, Martin D. Curran, Theresa Feltwell, Grant Hall, Anna Yakovleva, Fahad A Khokhar, Charlotte J. Houldcroft, Laura G Caller, Aminu S. Jahun, Sarah L. Caddy, Ian Goodfellow; and Alex Alderton, Roberto Amato, Sonia Goncalves, Ewan Harrison, David K. Jackson, Ian Johnston, Dominic Kwiatkowski, Cordelia Langford, John Sillitoe on behalf of the Wellcome Sanger Institute COVID-19 Surveillance Team (http://www.sanger.ac.uk/covid-team) |
| EPI_ISL_470529 | PHE South West Regional Laboratory, National Infection Service | Wellcome Sanger Institute for the COVID-19 Genomics UK (COG-UK) consortium | Stephanie Hutchings, Hannah Pymont, Dr Peter Muir, Barry Vipond, Rich Hopes; and Alex Alderton, Roberto Amato, Sonia Goncalves, Ewan Harrison, David K. Jackson, Ian Johnston, Dominic Kwiatkowski, Cordelia Langford, John Sillitoe on behalf of the Wellcome Sanger Institute COVID-19 Surveillance Team (http://www.sanger.ac.uk/covid-team) |
| EPI_ISL_470590, EPI_ISL_470593, EPI_ISL_470594 | Simile | Bioinformatics Laboratory / LNCC | Alexandra Gerber, Ana Paula Guimarães, Luiz Gonzaga Paula de Almeida, Ronaldo da Silva Francisco Junior, Mariane Talon, Filipe Romero, Átila Duque Rossi, Terezinha Marta Pereira, working group UFRJ, Jaqueline Goes de Jesus, Ingra Morales Claro, Ester Cerdeira Sabino, Nuno Rodrigues Faria, |

| | | | |
|---|---|---|---|
| | | | CADDE-group, Laboratorio Hermes Pardini, Laboratorio Simile, working group UFMG, Amilcar Tanuri, Carolina Voloch, Renato Santana Aguiar e Ana Tereza Vasconcelos |
| EPI_ISL_470626, EPI_ISL_470627, EPI_ISL_470628, EPI_ISL_470629 | Laboratorio de Virologia Molecular / UFRJ | Bioinformatics Laboratory / LNCC | Alexandra Gerber, Ana Paula Guimarães, Luiz Gonzaga Paula de Almeida, Ronaldo da Silva Francisco Junior, Mariane Talon, Filipe Romero, Átila Duque Rossi, Terezinha Marta Pereira, working group UFRJ, Jaqueline Goes de Jesus, Ingra Morales Claro, Ester Cerdeira Sabino, Nuno Rodrigues Faria, CADDE-group, Laboratorio Hermes Pardini, Laboratorio Simile, working group UFMG, Amilcar Tanuri, Carolina Voloch, Renato Santana Aguiar e Ana Tereza Vasconcelos |
| EPI_ISL_470683, EPI_ISL_470715 | Utah Public Health Laboratory | Utah Public Health Laboratory | Erin Young, Kelly Oakeson |
| EPI_ISL_470720, EPI_ISL_470721, EPI_ISL_470722, EPI_ISL_470723, EPI_ISL_470726, EPI_ISL_470728, EPI_ISL_470745, EPI_ISL_470746 | Utah Public Health Laboratory | Utah Public Health Laboratory | Heidi Butz, Erin Young, Kelly Oakeson |
| EPI_ISL_470834, EPI_ISL_470835, EPI_ISL_470836, EPI_ISL_470860, EPI_ISL_470861, EPI_ISL_470862 | PathWest Laboratory Medicine WA | PathWest Laboratory Medicine WA | Chisha Sikazwe, Jurissa Lang, Avram Levy, David Smith and David Speers |
| EPI_ISL_470898 | Pathogenic Microorganisms Variability Laboratory | Pathogenic Microorganisms Variability Laboratory | Alexey Shchetinin, Maria Nikiforova, Elena Shidlovskaya, Nadezhda Kuznetsova, Andrey Botikov, Alexander Gintsburg, Vladimir Gushchin |
| EPI_ISL_470902 | Influenza etiology and epidemiology laboratory | Pathogenic Microorganisms Variability Laboratory | Alexey Shchetinin, Maria Nikiforova, Elena Shidlovskaya, Nadezhda Kuznetsova, Vladimir Gushchin, Inna Dolzhikova, Daria Grousova, Andrey Botikov, Denis Logunov, Kirill Krasnoslobotsev, Svetlana Trushakova, Elena Burtseva, Ludmila Kolobukhina, Svetlana Smetanina, Alexander Gintsburg |
| EPI_ISL_470903, EPI_ISL_470904 | Influenza etiology and epidemiology laboratory | Pathogenic Microorganisms Variability Laboratory | Alexey Shchetinin, Maria Nikiforova, Elena Shidlovskaya, Nadezhda Kuznetsova, Vladimir Gushchin, Inna Dolzhikova, Daria Grousova, Andrey Botikov, Denis Logunov, Anna Ignatjeva, Evgeniya Mukasheva, Elena Burtseva, Ludmila Kolobukhina, Svetlana Smetanina, Alexander Gintsburg |
| EPI_ISL_471184, EPI_ISL_471185, EPI_ISL_471186, EPI_ISL_471187, EPI_ISL_471188, EPI_ISL_471245, EPI_ISL_471246 | Wisconsin State Laboratory of Hygiene Communicable Disease Division | Wisconsin State Laboratory of Hygiene Communicable Disease Division | Kelsey R. Florek, Abigail C. Shockey |
| EPI_ISL_471418 | Laboratory for Respiratory Viruses, National Influenza Centre, Cantacuzino National Military-Medical Institute for Research and Development | Cantacuzino Institute | Luiza Ustea, Nicoleta Paraschiv, Tim Durfee, Mihaela Lazar |
| EPI_ISL_471548 | Hospital do Servidor Público Estadual Francisco Morato de Oliveira | Instituto Adolfo Lutz, Interdiciplinary Procedures Center, Strategic Laboratory | Claudio Tavares Sacchi, Claudia Regina Gonçalves, Erica Valessa Ramos Gomes |
| EPI_ISL_471551 | Hospital Sao Paulo de Ensino da Unifesp | Instituto Adolfo Lutz, Interdiciplinary Procedures Center, Strategic Laboratory | Claudio Tavares Sacchi, Claudia Regina Gonçalves, Erica Valessa Ramos Gomes |
| EPI_ISL_471585 | CSIR-Centre for Cellular and Molecular Biology | CSIR-Centre for Cellular and Molecular Biology | Dhiviya Vedagiri, Divya Gupta, Vishal Sah, Payel Mukherjee, Sofia Banu, Priya Singh, Santosh Kumar Kuncha, Archana Bharadwaj Siva, Karthik Bharadwaj Tallapaka, Shagufta Khan, Lamuk Zaveri, Namami Gaur, Sakshi Shambhavi, Tulasi Nagabandi, Purushotham Vodnala, Rakesh K Mishra, Divya Tej Sowpati, Krishnan Harinivas Harshan |
| EPI_ISL_471923, EPI_ISL_471924, EPI_ISL_471954 | University of Exeter | COVID-19 Genomics UK (COG-UK) Consortium | Ben Temperton,Aaron Jeffries,Michelle Michelsen,Joanna Warwick-Dugdale,Audrey Farbos,Robyn Manley,Stephen Michell,Jane Masoli |
| EPI_ISL_472442, EPI_ISL_472546, EPI_ISL_472573, EPI_ISL_472613, EPI_ISL_472642, EPI_ISL_472649, EPI_ISL_472652, EPI_ISL_472681, EPI_ISL_472715, EPI_ISL_472940, EPI_ISL_472961, EPI_ISL_473004, EPI_ISL_473047, EPI_ISL_473048, EPI_ISL_473052, EPI_ISL_473069 | | | |
| see above | Wales Specialist Virology Centre Sequencing lab: Pathogen Genomics Unit | COVID-19 Genomics UK (COG-UK) Consortium | Catherine Moore, Johnathan Evans, Laura Gifford, Malorie Perry, Simon Cottrell, Angela Marchbank, Alec Birchley, Alexander Adams, Amy Gaskin, Bree Gatica-Wilcox, Jason Coombes, Joel Southgate, Lauren Gilbert, Lee Graham, Nicole Pacchiarini, Sara Kumziene-Summerhayes, Sarah Taylor, Sophie Jones, Sara Rey, Matthew Bull, Joanne Watkins, Sally Corden, Tom Connor |
| EPI_ISL_473509, EPI_ISL_473511, EPI_ISL_473512, EPI_ISL_473780, EPI_ISL_473781 | West of Scotland Specialist Virology Centre, NHSGGC / MRC-University of Glasgow Centre for Virus Research | COVID-19 Genomics UK (COG-UK) Consortium | Ana da Silva Filipe, Natasha Johnson, Kathy Smollett, Daniel Mair, Stephen Carmichael, Lily Tong, Jenna Nichols, Elihu Aranday-Cortes, Kirstyn Brunker, Yasmin Parr, Alice Broos, Kyriaki Nomikou; Sarah McDonald, Marc Niebel, Patawee Asamaphan; Richard Orton, Joseph Hughes, Sreenu Vattipally, David L Robertson; Alasdair MacLean, Rory Gunson; Kathy Li, Natasha Jesudason, Rajiv Shah, James Shepherd, Antonia Ho, Emma Thomson |
| EPI_ISL_473817, EPI_ISL_473818, EPI_ISL_473819, EPI_ISL_473820, EPI_ISL_473821, EPI_ISL_473822, EPI_ISL_473823, EPI_ISL_473826, EPI_ISL_473827, EPI_ISL_473828, EPI_ISL_473829, EPI_ISL_473830, EPI_ISL_473831, EPI_ISL_473838, EPI_ISL_473839, EPI_ISL_473840, EPI_ISL_473841, EPI_ISL_473842, EPI_ISL_473843, EPI_ISL_473844, EPI_ISL_473845, EPI_ISL_473846, EPI_ISL_473847, EPI_ISL_473848, EPI_ISL_473849, EPI_ISL_473850, EPI_ISL_473851 | | | |
| see above | Virology Department, Royal Infirmary of Edinburgh, NHS Lothian / School of Biological Sciences, University of Edinburgh / Institute of Genetics and Molecular Medicine, University of Edinburgh | COVID-19 Genomics UK (COG-UK) Consortium | McHugh M, Dewar R, Rooke S, Gallagher M, Balcaza C, O'Toole Á, Scher E, Hill V, McCrone JT, Colquhoun R, Yu X, Jackson B, Rambaut A, Williams TC, Templeton K |
| EPI_ISL_474318, EPI_ISL_474319, EPI_ISL_474321, EPI_ISL_474322, EPI_ISL_474323, EPI_ISL_474324, EPI_ISL_474325, EPI_ISL_474326, EPI_ISL_474328, EPI_ISL_474329 | Wales Specialist Virology Centre Sequencing lab: Pathogen Genomics Unit | COVID-19 Genomics UK (COG-UK) Consortium | Catherine Moore, Johnathan Evans, Laura Gifford, Malorie Perry, Simon Cottrell, Angela Marchbank, Alec Birchley, Alexander Adams, Amy Gaskin, Bree Gatica-Wilcox, Jason Coombes, Joel Southgate, Lauren Gilbert, Lee Graham, Nicole Pacchiarini, Sara Kumziene-Summerhayes, Sarah Taylor, Sophie Jones, Sara Rey, Matthew Bull, Joanne Watkins, Sally Corden, Tom Connor |
| EPI_ISL_474958 | Israeli Central Virology laboratory | Israel Central Virology laboratory | Neta Zuckerman, Efrat Dahan Bucris, Oran Erster, Ella Mendelson, Michal Mandelboim |
| EPI_ISL_474959, EPI_ISL_474961, EPI_ISL_474963, EPI_ISL_474964, EPI_ISL_474976, EPI_ISL_474977, EPI_ISL_474978, EPI_ISL_474989, EPI_ISL_474996, EPI_ISL_474997, EPI_ISL_475004, EPI_ISL_475005, EPI_ISL_475011, EPI_ISL_475012, EPI_ISL_475023, EPI_ISL_475024, EPI_ISL_475025 | | | |
| see above | Israel Central Virology laboratory | Israel Central Virology laboratory | Neta Zuckerman, Efrat Dahan Bucris, Oran Erster, Ella Mendelson, Michal Mandelboim |
| EPI_ISL_475122 | Umea klinisk mikrobiologi | The Public Health Agency of Sweden | Oskar Karlsson Lindsjo, Maria Lind Karlberg, Mattias Haukland, Reza Advani, Olov Svartstrom, Anna-Malin Linde, Sandra Broddesson, Petra Edquist, Shamam Muradrasoli, Anna Risberg, Karin Tegmark-Wisell |
| EPI_ISL_475123 | Halmstad klinisk mikrobiologi | The Public Health Agency of Sweden | Oskar Karlsson Lindsjo, Maria Lind Karlberg, Mattias Haukland, Reza Advani, Olov Svartstrom, Anna-Malin Linde, Sandra Broddesson, Petra Edquist, Shamam Muradrasoli, Anna Risberg, Karin Tegmark-Wisell |
| EPI_ISL_475124 | Ostersund klinisk mikrobiologi | The Public Health Agency of Sweden | Oskar Karlsson Lindsjo, Maria Lind Karlberg, Mattias Haukland, Reza Advani, Olov Svartstrom, Anna-Malin Linde, Sandra Broddesson, Petra Edquist, Shamam Muradrasoli, Anna Risberg, Karin Tegmark-Wisell |
| EPI_ISL_475156 | Halmstad klinisk mikrobiologi | The Public Health Agency of Sweden | Oskar Karlsson Lindsjo, Maria Lind Karlberg, Mattias Haukland, Reza Advani, Olov Svartstrom, Anna-Malin Linde, Sandra Broddesson, Petra Edquist, Shamam Muradrasoli, Anna Risberg, Karin Tegmark-Wisell |
| EPI_ISL_475539 | Narhalsan Sjobo vardcentral | The Public Health Agency of Sweden | Oskar Karlsson Lindsjo, Maria Lind Karlberg, Mattias Haukland, Reza Advani, Olov Svartstrom, Anna-Malin Linde, Sandra Broddesson, Mia Brytting, Anna Risberg, Karin Tegmark-Wisell |
| EPI_ISL_475557 | Halmstad klinisk mikrobiologi | The Public Health Agency of Sweden | Oskar Karlsson Lindsjo, Maria Lind Karlberg, Mattias Haukland, Reza Advani, Olov Svartstrom, Anna-Malin Linde, Sandra Broddesson, Shaman Muradrasoli, Anna Risberg, Karin Tegmark-Wisell |
| EPI_ISL_475591, EPI_ISL_475592, EPI_ISL_475603, EPI_ISL_475605, EPI_ISL_475615, EPI_ISL_475617, EPI_ISL_475619, EPI_ISL_475620, EPI_ISL_475623, EPI_ISL_475658, EPI_ISL_475659, EPI_ISL_475660, EPI_ISL_475667, EPI_ISL_475668, EPI_ISL_475698, EPI_ISL_475710, EPI_ISL_475711, EPI_ISL_475715 | | | |
| see above | Cedars-Sinai Medical Center, Department of Pathology & Laboratory Medicine, Molecular Pathology Laboratory | Cedars-Sinai Medical Center, Molecular Pathology Laboratory of Department of Pathology & Laboratory Medicine and Genomic Core | Wenjuan Zhang, John Paul Govindavari, Brian Davis, Stephanie Chen, Jong Taek Kim, Jianbo Song, Jean Lopategui, Jasmine T Plummer, Eric Vail |

| | | | |
|---|---|---|---|
| EPI_ISL_475812 | Center for Virology, Medical University of Vienna | Bergthaler laboratory, CeMM Research Center for Molecular Medicine of the Austrian Academy of Sciences | Alexandra Popa, Benedikt Agerer, Henrique Colaco, Lukas Endler, Jakob-Wendelin Genger, Alexander Lercher, Mark Smyth, Thomas Penz, Michael Schuster, Jan Laine, Martin Senekowitsch, Judith Aberle, Stephan Aberle, Peter Hufnagl, Daniela Schmid, Franz Allerberger, Elisabeth Puchhammer-Stoeckl, Manfred Nairz, Guenter Weiss, Gregor Hörmann, Kinga Rigler-Hohenwarter, Rainer Gattringer, Wegene Borena, Dorothee von Laer, Christoph Bock, Andreas Bergthaler |
| EPI_ISL_475899, EPI_ISL_475900, EPI_ISL_475901, EPI_ISL_475902 | Zentralinstitut für medizinische und chemische Labordiagnostik, Universitätskliniken Insbruck | Bergthaler laboratory, CeMM Research Center for Molecular Medicine of the Austrian Academy of Sciences | Alexandra Popa, Benedikt Agerer, Henrique Colaco, Lukas Endler, Jakob-Wendelin Genger, Alexander Lercher, Mark Smyth, Thomas Penz, Michael Schuster, Jan Laine, Martin Senekowitsch, Judith Aberle, Stephan Aberle, Peter Hufnagl, Daniela Schmid, Franz Allerberger, Elisabeth Puchhammer-Stoeckl, Manfred Nairz, Guenter Weiss, Gregor Hörmann, Kinga Rigler-Hohenwarter, Rainer Gattringer, Wegene Borena, Dorothee von Laer, Christoph Bock, Andreas Bergthaler |
| EPI_ISL_475930, EPI_ISL_475931 | Universitaetsklinik für Innere Medizin II Innsbruck | Bergthaler laboratory, CeMM Research Center for Molecular Medicine of the Austrian Academy of Sciences | Alexandra Popa, Benedikt Agerer, Henrique Colaco, Lukas Endler, Jakob-Wendelin Genger, Alexander Lercher, Mark Smyth, Thomas Penz, Michael Schuster, Jan Laine, Martin Senekowitsch, Judith Aberle, Stephan Aberle, Peter Hufnagl, Daniela Schmid, Franz Allerberger, Elisabeth Puchhammer-Stoeckl, Manfred Nairz, Guenter Weiss, Gregor Hörmann, Kinga Rigler-Hohenwarter, Rainer Gattringer, Wegene Borena, Dorothee von Laer, Christoph Bock, Andreas Bergthaler |
| EPI_ISL_475951, EPI_ISL_475952, EPI_ISL_475953, EPI_ISL_475967, EPI_ISL_475971, EPI_ISL_475972 | National Public Health Laboratory, National Centre for Infectious Diseases | National Public Health Laboratory, National Centre for Infectious Diseases | Mak TM, Octavia S, Chavatte JM, Cui L, Lin RTP |
| EPI_ISL_476071, EPI_ISL_476072, EPI_ISL_476073, EPI_ISL_476074, EPI_ISL_476075, EPI_ISL_476076 | University of Debrecen, Department of Medical Microbiology | National Laboratory of Virology, Szentágothai Research Centre | Endre Gábor Tóth, Balázs Somogyi, Brigitta Zana, Eszter Csoma, Ferenc Jakab, Gábor Kemenesi |
| EPI_ISL_476145, EPI_ISL_476146, EPI_ISL_476147 | Ostersund klinisk mikrobiologi | The Public Health Agency of Sweden | Oskar Karlsson Lindsjo, Maria Lind Karlberg, Mattias Haukland, Reza Advani, Olov Svartstrom, Anna-Malin Linde, Sandra Broddesson, Petra Edquist, Shamam Muradrasoli, Anna Risberg, Karin Tegmark-Wisell |
| EPI_ISL_476153, EPI_ISL_476154, EPI_ISL_476155, EPI_ISL_476156, EPI_ISL_476157, EPI_ISL_476158, EPI_ISL_476159, EPI_ISL_476160, EPI_ISL_476162, EPI_ISL_476387, EPI_ISL_476388, EPI_ISL_476389 | | | |
| see above | Laboratório de Patologia Clínica - UNICAMP | Laboratório de Estudos de Vírus Emergentes - UNICAMP | José Luiz Proença-Modena, Magnun Nueldo Nunes dos Santos, Angelica Schreiber, Julia Forato,Camila Simeoni, Marcilio Jorge Fumagalli, Mariene Ribeiro Amorim, Darlan da Silva Candido, Nuno Rodrigues Faria, Julien Theze, Luiz Gonzaga,Jaqueline Goes Jesus e William Marciel de Souza |
| EPI_ISL_476502, EPI_ISL_476504, EPI_ISL_476505, EPI_ISL_476506, EPI_ISL_476507, EPI_ISL_476508, EPI_ISL_476510 | Laboratoire de microbiologie, Hopital de Verdun | Smith Laboratory, Centre de Recherche CHU Sainte-Justine | Martin Smith, Marieke Rozendaal, Ivan Pavlov |
| EPI_ISL_476703, EPI_ISL_476704 | Incubadora Venezolana de Ciencia, Venezuela | Incubadora Venezolana de Ciencia, Venezuela / Instituto Nacional de Salud, Bogotá, Colombia / Grupo de Investigaciones Microbiológicas-UR (GIMUR), Departamento de Biología, Facultad de Ciencias Naturales, Universidad del Rosario, Bogotá, Colombia / Icahn School of Medicine at Mount Sinai, New York, USA | Alberto Paniz-Mondolfi, Marina Muñoz, Luis Perez-Garcia, Lourdes Delgado, Carolina Florez, Sergio Gomez, Angelica Rico, Lisseth Pardo, Esther C. Barros, Carolina Hernández, Jesús E. Jaimes, Aníbal A. Teherán, Ana S. Gonzalez-Reiche, Matthew M. Hernandez, Emilia Mia Sordillo, Viviana Simon, Harm van Bakel, Juan David Ramírez |
| EPI_ISL_476839 | National Influenza Centre for Northern Greece | National Influenza Centre for Northern Greece | Maria Christoforidi |
| EPI_ISL_476840 | Defence Research & Development Establishment (DRDE) | Defence Research & Development Establishment (DRDE) | Shashi Sharma, Paban Kumar Dash, Sushil Kumar Sharma, Ambuj Shrivastava, Jyoti S. Kumar |
| EPI_ISL_476847 | National Influenza Centre for Northern Greece | National Influenza Centre for Northern Greece | Maria Christoforidi |
| EPI_ISL_476848 | Defence Research & Development Establishment (DRDE) | Defence Research & Development Establishment (DRDE) | Shashi Sharma, Paban Kumar Dash, Sushil Kumar Sharma, Ambuj Shrivastava, Jyoti S. Kumar |
| EPI_ISL_476851 | National Influenza Centre for Northern Greece | National Influenza Centre for Northern Greece | Maria Christoforidi |
| EPI_ISL_476941, EPI_ISL_476942, EPI_ISL_476943, EPI_ISL_476944, EPI_ISL_476945, EPI_ISL_476946, EPI_ISL_476947, EPI_ISL_476948, EPI_ISL_476949, EPI_ISL_476950, EPI_ISL_476951, EPI_ISL_476952, EPI_ISL_476953, EPI_ISL_476954, EPI_ISL_476955, EPI_ISL_476956, EPI_ISL_476957, EPI_ISL_476958, EPI_ISL_476959, EPI_ISL_476960, EPI_ISL_476961, EPI_ISL_476962, EPI_ISL_476963, EPI_ISL_476964, EPI_ISL_476965, EPI_ISL_476966, EPI_ISL_476967, EPI_ISL_476968, EPI_ISL_476969, EPI_ISL_476970, EPI_ISL_476971, EPI_ISL_476972, EPI_ISL_476973, EPI_ISL_476974, EPI_ISL_476975, EPI_ISL_476976, EPI_ISL_476977, EPI_ISL_476978, EPI_ISL_476979, EPI_ISL_476980, EPI_ISL_476981, EPI_ISL_476982 | | | |
| see above | KU Leuven, Rega Institute, Clinical and Epidemiological Virology | KU Leuven, Rega Institute, Clinical and Epidemiological Virology | Tony Wawina-Bokalanga, Joan Marti-Carerras, Bert Vanmechelen, Piet Maes |
| EPI_ISL_477010, EPI_ISL_477011, EPI_ISL_477012 | University of Debrecen, Department of Medical Microbiology | National Laboratory of Virology, Szentágothai Research Centre | Endre Gábor Tóth, Balázs Somogyi, Brigitta Zana, Eszter Csoma, Ferenc Jakab, Gábor Kemenesi |
| EPI_ISL_477195 | Istituto Zooprofilattico Sperimentale Puglia e Basilicata; | Beaconlab (Bioinformatics, Evolution and Comparative Genomics lab), Dept of Biosciences, University on Milan | Parisi A.,Pesole G., Manzari C., Chiara M. |
| EPI_ISL_477825, EPI_ISL_477826, EPI_ISL_477827, EPI_ISL_477828, EPI_ISL_477829, EPI_ISL_477830, EPI_ISL_477831, EPI_ISL_477832, EPI_ISL_477833, EPI_ISL_477834, EPI_ISL_477835, EPI_ISL_477836, EPI_ISL_477837, EPI_ISL_477838, EPI_ISL_477839, EPI_ISL_477840, EPI_ISL_477841, EPI_ISL_477842, EPI_ISL_477843, EPI_ISL_477844, EPI_ISL_477845, EPI_ISL_477846, EPI_ISL_477847, EPI_ISL_477848, EPI_ISL_477849, EPI_ISL_477850, EPI_ISL_477851, EPI_ISL_477852, EPI_ISL_477853, EPI_ISL_477854, EPI_ISL_477855, EPI_ISL_477856, EPI_ISL_477857, EPI_ISL_477858, EPI_ISL_477859, EPI_ISL_477860, EPI_ISL_477861, EPI_ISL_477862, EPI_ISL_477863, EPI_ISL_477864, EPI_ISL_477865, EPI_ISL_477866, EPI_ISL_477867, EPI_ISL_477868, EPI_ISL_477869, EPI_ISL_477870, EPI_ISL_477871, EPI_ISL_477872, EPI_ISL_477873, EPI_ISL_477874, EPI_ISL_477875, EPI_ISL_477876, EPI_ISL_477877, EPI_ISL_477878, EPI_ISL_477879, EPI_ISL_477880, EPI_ISL_477881, EPI_ISL_477882, EPI_ISL_477883, EPI_ISL_477884, EPI_ISL_477885, EPI_ISL_477886, EPI_ISL_477887, EPI_ISL_477888, EPI_ISL_477889, EPI_ISL_477892, EPI_ISL_477893, EPI_ISL_477894, EPI_ISL_477895, EPI_ISL_477896, EPI_ISL_477900, EPI_ISL_477907, EPI_ISL_477913 | | | |
| see above | West of Scotland Specialist Virology Centre, NHSGGC / MRC-University of Glasgow Centre for Virus Research | COVID-19 Genomics UK (COG-UK) Consortium | Ana da Silva Filipe, Natasha Johnson, Kathy Smollett, Daniel Mair, Stephen Carmichael, Lily Tong, Jenna Nichols, Elihu Aranday-Cortes, Kirstyn Brunker, Yasmin Parr, Alice Broos, Kyriaki Nomikou; Sarah McDonald, Marc Niebel, Patawee Asamaphan; Richard Orton, Joseph Hughes, Sreenu Vattipally, David L Robertson; Alasdair MacLean, Rory Gunson; Kathy Li, Natasha Jesudason, Rajiv Shah, James Shepherd, Antonia Ho, Emma Thomson |
| EPI_ISL_478275, EPI_ISL_478283, EPI_ISL_478284, EPI_ISL_478285, EPI_ISL_478286 | University of Exeter | COVID-19 Genomics UK (COG-UK) Consortium | Ben Temperton,Aaron Jeffries,Michelle Michelsen,Joanna Warwick-Dugdale,Audrey Farbos,Robyn Manley,Stephen Michell,Jane Masoli |
| EPI_ISL_478373 | Liverpool Clinical Laboratories | COVID-19 Genomics UK (COG-UK) Consortium | Sam Haldenby, Anita Lucaci, Steve Paterson, Julian Hiscox, Alistair Darby, M Almsaud, A Alrezaihi, Muhannad Alruwaili, Stuart D Armstrong, Jones Benjamin, Eleanor G Bentley, Anu Chawla, Jordan J Clark, Angela Cowell, Richard Eccles, Isabel García-Dorival, Matthew Gemmell, Alessandro Gerada, PKF Gilmore, Richard Gregory, Ximeng Han, Catherine Hartley, Margaret Hughes, Miren Iturriza-Gomara, James Johnson, L Luu, Jenifer Manson, Charlotte Nelson, Elaine O'Toole, Cassie Olateju, Rebekah Penrice-Randal , Lucille Rainbow, N.P Randle, Trevor Ian Robinson, Parul Sharma, Ghada T Shawli, James P Stewart, Neil Swainston, Ecaterina Vamos, Joanne Watts, Mark Whitehead |
| EPI_ISL_478459, EPI_ISL_478460, EPI_ISL_478461, EPI_ISL_478462, EPI_ISL_478463, EPI_ISL_478464, EPI_ISL_478465, EPI_ISL_478466, EPI_ISL_478467, EPI_ISL_478468, EPI_ISL_478469, EPI_ISL_478470 | | | |
| see above | University College London, Great Ormond Street Hospital for Children NHS Foundation Trust, Imperial College Healthcare NHS Trust | COVID-19 Genomics UK (COG-UK) Consortium | Sergi Castellano, Rachel Williams, Mark Kristiansen, Paola Resende Silva, Sunando Roy, Tony Brooks, Helena Tutill, Paola Niola, Patricia Dyal, Charlotte Williams, Leysa Forrest, Yasmin Panchbhaya, Jacqueline Findlay, Samuel Weeks, Julianne Brown, Kathryn Harris, Paul Randell, James Price, Alison Holmes, Judith Breuer |
| EPI_ISL_478670 | unknown | Microbiology, Koc University | Can,F., Ozer,B., Nurtop,E., Dogan,O. |
| EPI_ISL_478707, EPI_ISL_478714, EPI_ISL_478715, EPI_ISL_478716 | Sydney South West Pathology Service (SSWPS) - Liverpool Hospital - NSW Health Pathology | NSW Health Pathology - Institute of Clinical Pathology and Medical Research; Westmead Hospital; University of Sydney | CIDM-PH et al. |
| EPI_ISL_478729, EPI_ISL_478735, EPI_ISL_478739, EPI_ISL_478742, EPI_ISL_478743, EPI_ISL_478749, EPI_ISL_478752, EPI_ISL_478753, EPI_ISL_478756, EPI_ISL_478758, EPI_ISL_478763, EPI_ISL_478764, EPI_ISL_478767, EPI_ISL_478770, EPI_ISL_478772, EPI_ISL_478780, EPI_ISL_478782, EPI_ISL_478784, EPI_ISL_478791, EPI_ISL_478794, EPI_ISL_478798, EPI_ISL_478803, EPI_ISL_478811, EPI_ISL_478815, EPI_ISL_478822, EPI_ISL_478829, EPI_ISL_478837, EPI_ISL_478840, EPI_ISL_478842, EPI_ISL_478856, EPI_ISL_478870, EPI_ISL_478871, EPI_ISL_478878, EPI_ISL_478904, EPI_ISL_478914, EPI_ISL_478915, EPI_ISL_478927, EPI_ISL_478928, EPI_ISL_478964, EPI_ISL_478968, EPI_ISL_478969, EPI_ISL_478974, EPI_ISL_478975, EPI_ISL_478977, EPI_ISL_478979, EPI_ISL_478980, EPI_ISL_478984, EPI_ISL_478986, EPI_ISL_479134, EPI_ISL_479139, EPI_ISL_479152, EPI_ISL_479158, EPI_ISL_479173 | | | |

| | | | |
|---|---|---|---|
| see above | Oxford Viromics, NDM, University of Oxford; Oxford University Hospitals; Basingstoke and North Hampshire Hospital | COVID-19 Genomics UK (COG-UK) Consortium | Tanya Golubchik, David Bonsall, George Macintyre, Amy Trebes, Mariateresa de Cesare, Catrin Moore, Alex Mobbs, Anita Justice, Robert Shaw, Monique Andersson, Timothy Peto, Emma Wise, Nathan Moore, Jessica Lynch, Nick Cortes, Matilde Mori, Stephen Kidd, David Buck, John Todd, Christophe Fraser |
| EPI_ISL_479573 | NIV Influenza | NIV Influenza | Potdar V |
| EPI_ISL_479618, EPI_ISL_479619 | Laboratory of Molecular Virology of the International Centre for Genetic Engineering and Biotechnology (ICGEB) | ARGO Open Lab Platform for Genome Sequencing | Licastro, D, Rajasekharan S, Dal Monego S, Segat L, D'Agaro P, Salton F, Confalonieri P, Confalonieri M, Marcello A |
| EPI_ISL_479674, EPI_ISL_479675 | Center for Genomics and System Biology, New York University | Center for Genomics and System Biology, New York University | Roder,A., Banakis,S., Johnson,K., Khalfan,M., Borenstein,E.S., Samanovic,M., Cornelius,A., Herati,R., Ulrich,R., Fleming,A., Kottkamp,A., Raabe,V., Mulligan,M.J., Gresham,D., Ghedin,E. |
| EPI_ISL_479761, EPI_ISL_479762, EPI_ISL_479766, EPI_ISL_479767 | University of Miami Immunology and Histocompatibility Laboratory | University of Miami Immunology and Histocompatibility Laboratory | Emilio Margolles-Clark, PhD and Phillip Ruiz, MD, PhD |
| EPI_ISL_480126, EPI_ISL_480153, EPI_ISL_480157, EPI_ISL_480164, EPI_ISL_480165, EPI_ISL_480166, EPI_ISL_480167, EPI_ISL_480168 | Fukui Prefectural Institute of Public Health and Environmental Science | Pathogen Genomics Center, National Institute of Infectious Diseases | Tsuyoshi Sekizuka, Miho Toho, Kentaro Itokawa, Rina Tanaka, Masanori Hashino, Hajime Kamiya, Motoi Suzuki, Makoto Kuroda |
| EPI_ISL_480204 | Akita City Public Health Center | Pathogen Genomics Center, National Institute of Infectious Diseases | Tsuyoshi Sekizuka, Koichi Ito, Kentaro Itokawa, Rina Tanaka, Masanori Hashino, Hajime Kamiya, Motoi Suzuki, Makoto Kuroda |
| EPI_ISL_480205, EPI_ISL_480206, EPI_ISL_480207 | Department of Infectious Diseases, Kobe Institute of Health | Pathogen Genomics Center, National Institute of Infectious Diseases | Tsuyoshi Sekizuka, Ryohei Nomoto, Kentaro Itokawa, Rina Tanaka, Masanori Hashino, Hajime Kamiya, Motoi Suzuki, Makoto Kuroda |
| EPI_ISL_480208 | Pathogen Genomics Center, National Institute of Infectious Diseases | Pathogen Genomics Center, National Institute of Infectious Diseases | Tsuyoshi Sekizuka, Ryohei Nomoto, Kentaro Itokawa, Rina Tanaka, Masanori Hashino, Hajime Kamiya, Motoi Suzuki, Makoto Kuroda |
| EPI_ISL_480209, EPI_ISL_480210, EPI_ISL_480211, EPI_ISL_480212, EPI_ISL_480213, EPI_ISL_480214, EPI_ISL_480215, EPI_ISL_480216, EPI_ISL_480218, EPI_ISL_480219, EPI_ISL_480220 | | | |
| see above | Department of Infectious Diseases, Kobe Institute of Health | Pathogen Genomics Center, National Institute of Infectious Diseases | Tsuyoshi Sekizuka, Ryohei Nomoto, Kentaro Itokawa, Rina Tanaka, Masanori Hashino, Hajime Kamiya, Motoi Suzuki, Makoto Kuroda |
| EPI_ISL_480321 | Laboratorio Clínico San José | Charité Virology-University of Costa Rica | Andres Moreira-Soto, Eugenia Corrales-Aguilar, Ignacio Postigo-Hidalgo, Hugo Núñez Navas, Jan Felix Drexler |
| EPI_ISL_480329 | Quadram Institute Bioscience | COVID-19 Genomics UK (COG-UK) Consortium | Dave J. Baker, Gemma L. Kay, Alp Aydin, Thanh Le-Viet, Steven Rudder, Ana P. Tedim, Anastasia Kolyva, Maria Diaz, Leonardo de Oliveira Martins, Nabil-Fareed Alikhan, Lizzie Meadows, Rachael Stanley, Ngozi Elumogo, Muhammed Yasir, Nicholas M. Thomson, Alexander J Trotter, Rachel Gilroy, Samuel Bloomfield, Claire Stuart, Andrew Bell, Reenesh Prakash, Samir Dervisevic, Alison E. Mather, John Wain, Mark Webber, Andrew J. Page, Justin O'Grady |
| EPI_ISL_480332, EPI_ISL_480333, EPI_ISL_480334, EPI_ISL_480335, EPI_ISL_480344 | Microbial Genomics Laboratory, Institut Pasteur de Montevideo | Microbial Genomics Laboratory, Institut Pasteur de Montevideo | Cecilia Salazar, Marianoel Pereira, Ignacio Ferrés, Gonzalo Moratorio, Pilar Moreno, Gregorio Iraola |
| EPI_ISL_480395, EPI_ISL_480397, EPI_ISL_480398 | University of Wisconsin-Madison AIDS Vaccine Research Laboratories | University of Wisconsin-Madison AIDS Vaccine Research Laboratories | Gage Moreno, Katarina Braun, et al. AIDS Vaccine Research Laboratories |
| EPI_ISL_480429 | Laboratorio de Biología Molecular Asociación Española Primera en Salud | Departments of Pathology and Medicine, New York University School of Medicine | Maria Victoria Elizondo, Maria Noel Zubillaga, Gonzalo Manrique, Paul Zappile, Gael Westby, Matthew T Maurano, Christian Marier, Adriana Heguy |
| EPI_ISL_480581, EPI_ISL_480582, EPI_ISL_480583, EPI_ISL_480584, EPI_ISL_480585, EPI_ISL_480586, EPI_ISL_480588, EPI_ISL_480589 | Victorian Infectious Diseases Reference Laboratory (VIDRL) | VIDRL and MDU-PHL | Caly L., Seemann T., Sait, M., Schultz M., Druce J., Sherry, N. |
| EPI_ISL_480601, EPI_ISL_480602 | National Health Laboratory, Timor-Leste | MDU-PHL | Soares da Silva, E., Dolores de Jesus da Costa, M., Salles de Sousa, A., Jayanti Pereira Tilman, A., Antonia da Costa, E., Barreto, I., Marr, I., Wapling, J., Francis, J., Ximenes, J., Canisia, D., Freeman, K., Dakh, F., Douglas, N., Baird, R., Caly, L., Seemann, T., Sait, M., Schultz, M., Sherry, N. |
| EPI_ISL_480850, EPI_ISL_480853, EPI_ISL_480854, EPI_ISL_480855, EPI_ISL_480856 | Florida Bureau of Public Health Laboratories | Florida Bureau of Public Health Laboratories | Sarah Schmedes, Jason Blanton |
| EPI_ISL_480975, EPI_ISL_481003, EPI_ISL_481017, EPI_ISL_481029, EPI_ISL_481034 | ISGlobal, Institut de Salut Global de Barcelona | SeqCOVID-SPAIN consortium/IBV(CSIC) | Alfredo Mayor, Alberto L Garcia-Basteiro, Carlota Dobaño, Gemma Moncunill, Pau Cisteró and SeqCOVID-SPAIN consortium |
| EPI_ISL_481048, EPI_ISL_481062, EPI_ISL_481064, EPI_ISL_481065, EPI_ISL_481072, EPI_ISL_481074, EPI_ISL_481078, EPI_ISL_481081, EPI_ISL_481090, EPI_ISL_481092, EPI_ISL_481104, EPI_ISL_481106, EPI_ISL_481109 | | | |
| see above | Hospital General Universitario Gregorio Marañón | SeqCOVID-SPAIN consortium/IBV(CSIC) | Laura Pérez-Lago, Marta Herranz, Jon Sicilia, Julia Suárez, Pilar Catalán, Patricia Muñoz, Darío García de Viedma and SeqCOVID-SPAIN consortium |
| EPI_ISL_481234, EPI_ISL_481235 | Institut Pasteur Dakar | Institut Pasteur de Dakar | Ndongo Dia, Moussa Moise Diagne, Mamadou Diop, Marie Henriette Dior Ndione, Mamadou Malado Jallow, Safietou Sanke, Ousmane Faye, Amadou Alpha Sall. |
| EPI_ISL_481276, EPI_ISL_481277, EPI_ISL_481278, EPI_ISL_481279, EPI_ISL_481280, EPI_ISL_481281, EPI_ISL_481282 | Maryland Department of Health | Maryland Department of Health | Keller,E. |
| EPI_ISL_481514, EPI_ISL_481515, EPI_ISL_481516, EPI_ISL_481517, EPI_ISL_481518, EPI_ISL_481519, EPI_ISL_481520, EPI_ISL_481521, EPI_ISL_481522, EPI_ISL_481523, EPI_ISL_481524, EPI_ISL_481525, EPI_ISL_481526, EPI_ISL_481527, EPI_ISL_481528, EPI_ISL_481529, EPI_ISL_481530, EPI_ISL_481531, EPI_ISL_481532, EPI_ISL_481533, EPI_ISL_481534, EPI_ISL_481535, EPI_ISL_481538, EPI_ISL_481539, EPI_ISL_481540, EPI_ISL_481541, EPI_ISL_481542, EPI_ISL_481543, EPI_ISL_481544, EPI_ISL_481545, EPI_ISL_481546, EPI_ISL_481547, EPI_ISL_481548, EPI_ISL_481730, EPI_ISL_481731, EPI_ISL_481732, EPI_ISL_481733, EPI_ISL_481734, EPI_ISL_481735, EPI_ISL_481736, EPI_ISL_481737, EPI_ISL_481738, EPI_ISL_481739, EPI_ISL_481740 | | | |
| see above | Department of Virology and Immunology, University of Helsinki and Helsinki University Hospital, Huslab Finland | Department of Virology, Faculty of Medicine, University of Helsinki, Helsinki, Finland | Teemu Smura, Hannimari Kallio-Kokko, Jenni Virtanen, Maija Suvanto, Sari Hannula, Harri Kangas, Pekka Ellonen, Olli Vapalahti |
| EPI_ISL_481754, EPI_ISL_481755, EPI_ISL_481756, EPI_ISL_481757 | Dr. Georges-L.-Dumont University Hospital Centre | National Microbiology Laboratory | Anna Majer, Shari Tyson, Grace Seo, Kristyn Burak, Philip Mabon, Elsie Grudeski, Rhiannon Huzarewich, Russell Mandes, Jennifer Tanner, Natalie Knox, Morag Graham, Gary Van Domselaar, Richard Garceau, Guillaume Desnoyers, Nathalie Bastien, Yan Li, Timothy Booth |
| EPI_ISL_481976, EPI_ISL_481981, EPI_ISL_482013, EPI_ISL_482020 | PHE South West Regional Laboratory, National Infection Service | Wellcome Sanger Institute for the COVID-19 Genomics UK (COG-UK) consortium | Stephanie Hutchings, Hannah Pymont, Dr Peter Muir, Barry Vipond, Rich Hopes; and Alex Alderton, Roberto Amato, Sonia Goncalves, Ewan Harrison, David K. Jackson, Ian Johnston, Dominic Kwiatkowski, Cordelia Langford, John Sillitoe on behalf of the Wellcome Sanger Institute COVID-19 Surveillance Team (http://www.sanger.ac.uk/covid-team) |
| EPI_ISL_482032, EPI_ISL_482041, EPI_ISL_482042, EPI_ISL_482046, EPI_ISL_482049, EPI_ISL_482052, EPI_ISL_482053, EPI_ISL_482056, EPI_ISL_482134, EPI_ISL_482136, EPI_ISL_482137, EPI_ISL_482139 | | | |
| see above | Regional Virus Laboratory, Belfast Health and Social Care Trust | Wellcome Sanger Institute for the COVID-19 Genomics UK (COG-UK) consortium | Conall McCaughey, James McKenna, Tanya Curran, Susan Feeney, Alison Watt, Ciara Cox, Mairead Connor, Zoltan Molnar, David Simpson, Derek Fairley; and Alex Alderton, Roberto Amato, Sonia Goncalves, Ewan Harrison, David K. Jackson, Ian Johnston, Dominic Kwiatkowski, Cordelia Langford, John Sillitoe on behalf of the Wellcome Sanger Institute COVID-19 Surveillance Team (http://www.sanger.ac.uk/covid-team) |
| EPI_ISL_482296, EPI_ISL_482297, EPI_ISL_482334, EPI_ISL_482335, EPI_ISL_482336, EPI_ISL_482337, EPI_ISL_482338, EPI_ISL_482343, EPI_ISL_482344, EPI_ISL_482345, EPI_ISL_482461 | | | |
| see above | Providence St. Joseph Health Molecular Genomics Laboratory | Providence St. Joseph Health Molecular Genomics Laboratory | Alexa K Dowdell, Brian D Piening, Fred L Robinson, Carlo B Bifulco, Mary Campbell |
| EPI_ISL_482738 | LNR National Reference Laboratory, Mohammed VI University of Health Sciences | Medical Biotechnology Laboratory, Rabat Medical and Pharmacy School, Mohammed The Vth University in Rabat | Meriem LAAMARTI, Souad KARTTI, Rokia LAAMARTI , M.W. CHEMAO-ELFIHRI, Loubna ALLAM, Mouna OUADGHIRI, Imane SMYEJ, Jalila RAHOUI, Houda BENRHAMA, Jalil El ATAR, Idrissa DIAWARA, Rachid EL JAOUDI, Laila SBABOU, Chakib NEJJARI, Saaid AMZAZI, Rachid MENTAG, Lahcen |

| | | | |
|---|---|---|---|
| | | | BELYAMANI and Azeddine IBRAHIMI |
| EPI_ISL_483075, EPI_ISL_483095, EPI_ISL_483096 | SA Pathology | SA Pathology | Lex Leong, Chuan Kok Lim, Mark Turra, Ivan Bastian, Geoff Higgins |
| EPI_ISL_483159, EPI_ISL_483160, EPI_ISL_483161 | San Diego County Public Health Laboratory | Andersen lab at Scripps Research | SEARCH Alliance San Diego with Tracy Basler, Jovan Shephard, Brett Austin |
| EPI_ISL_483168, EPI_ISL_483173, EPI_ISL_483174, EPI_ISL_483175, EPI_ISL_483181, EPI_ISL_483195, EPI_ISL_483199, EPI_ISL_483200, EPI_ISL_483202, EPI_ISL_483260, EPI_ISL_483339, EPI_ISL_483340, EPI_ISL_483341, EPI_ISL_483342, EPI_ISL_483346, EPI_ISL_483347, EPI_ISL_483348, EPI_ISL_483350, EPI_ISL_483351, EPI_ISL_483352, EPI_ISL_483355, EPI_ISL_483356, EPI_ISL_483357, EPI_ISL_483358, EPI_ISL_483359, EPI_ISL_483389, EPI_ISL_483433, EPI_ISL_483459, EPI_ISL_483471 | | | |
| see above | UC San Diego Center for Advanced Laboratory Medicine | Andersen lab at Scripps Research | SEARCH Alliance San Diego with David Pride, Ji H Shin |
| EPI_ISL_483555, EPI_ISL_483557, EPI_ISL_483558, EPI_ISL_483559, EPI_ISL_483560, EPI_ISL_483561, EPI_ISL_483563, EPI_ISL_483564, EPI_ISL_483565 | Kingdom of Bahrein Ministry of Health | Erasmus Medical Center | Bas Oude Munnink, David Nieuwenhuijse, Reina Sikkema, Fatema, Ebrahim Shehad, Amjad Ghanem Mohamed, Hashmeya Al Wasti, Claudia Schapendonk, Irina Chestakova, Anne van der Linden, Theo Bestebroer, Stefan van Nieuwkoop, Mark Pronk, Pascal Lexmond, Richard Molenkamp, Marion Koopmans, on behalf of the Dutch national COVID-19 response team. |
| EPI_ISL_483576, EPI_ISL_483579, EPI_ISL_483581, EPI_ISL_483583, EPI_ISL_483600 | National Public Health Laboratory, National Centre for Infectious Diseases | National Public Health Laboratory, National Centre for Infectious Diseases | Mak TM, Octavia S, Zhou Z, Chavatte JM, Cui L, Lin RTP |
| EPI_ISL_483638, EPI_ISL_483639 | Kingdom of Bahrein Ministry of Health | Erasmus Medical Center | Bas Oude Munnink, David Nieuwenhuijse, Reina Sikkema, Fatema, Ebrahim Shehad, Amjad Ghanem Mohamed, Hashmeya Al Wasti, Claudia Schapendonk, Irina Chestakova, Anne van der Linden, Theo Bestebroer, Stefan van Nieuwkoop, Mark Pronk, Pascal Lexmond, Richard Molenkamp, Marion Koopmans, on behalf of the Dutch national COVID-19 response team. |
| EPI_ISL_483668, EPI_ISL_483669, EPI_ISL_483671, EPI_ISL_483674, EPI_ISL_483678, EPI_ISL_483680, EPI_ISL_483683 | University Hospital Zurich | Department of Biosystems Science and Engineering, ETH Zürich | Christian Beisel, Sarah Nadeau, Ivan Topolsky, Pedro Ferreira, Philipp Jablonski, Susana Posada-Céspedes, Tobias Schär, Ina Nissen, Natascha Santacroce, Elodie Burcklen, Julia Martinez-Gomez, Phil Cheng, Mitch Levesque, Philipp Bosshard, Niko Beerenwinkel, Tanja Stadler |
| EPI_ISL_483712, EPI_ISL_483713 | Israel Central Virology laboratory | Israel Central Virology laboratory | Neta Zuckerman, Efrat Dahan Bucris, Oran Erster, Ella Mendelson, Michal Mandelboim |
| EPI_ISL_484375 | Queens Medical Centre, Clinical Microbiology Department / DeepSeq Nottingham | COVID-19 Genomics UK (COG-UK) Consortium | Gemma Clark, Wendy Smith, Manjinder Khakh, Vicki M Fleming, Michelle M Lister, Hannah Howson-Wells, Jonathan Ball, Patrick McClure, Joseph Chappell, Theocharis Tsoleridis, Nadine Holmes, Matthew Carlisle, Christopher Moore, Fei Sang, Johnny Debebe, Victoria Wright, Matthew Loose |
| EPI_ISL_484419, EPI_ISL_484420, EPI_ISL_484421, EPI_ISL_484422, EPI_ISL_484424, EPI_ISL_484425, EPI_ISL_484426, EPI_ISL_484427, EPI_ISL_484428, EPI_ISL_484429, EPI_ISL_484430, EPI_ISL_484431, EPI_ISL_484432 | | | |
| see above | Centre for Enzyme Innovation, University of Portsmouth / Translational Research Laboratory, Portsmouth Hospitals NHS Trust | COVID-19 Genomics UK (COG-UK) Consortium | Angela Beckett,Yann Bourgeois,Garry Scarlett,Sharon Glaysher,Scott Elliott,Kelly Bicknell,Robert Impey,Allyson Lloyd,Sarah Wyllie,Ethan Butcher,Anoop Chauhan,Samuel Robson |
| EPI_ISL_484519, EPI_ISL_484520, EPI_ISL_484521, EPI_ISL_484522, EPI_ISL_484523, EPI_ISL_484524, EPI_ISL_484525, EPI_ISL_484526, EPI_ISL_484527, EPI_ISL_484528, EPI_ISL_484529, EPI_ISL_484530, EPI_ISL_484531, EPI_ISL_484532, EPI_ISL_484533, EPI_ISL_484534, EPI_ISL_484535, EPI_ISL_484536, EPI_ISL_484537, EPI_ISL_484538, EPI_ISL_484539, EPI_ISL_484540, EPI_ISL_484541, EPI_ISL_484542, EPI_ISL_484543, EPI_ISL_484544, EPI_ISL_484545, EPI_ISL_484546, EPI_ISL_484547, EPI_ISL_484548, EPI_ISL_484549, EPI_ISL_484550, EPI_ISL_484551, EPI_ISL_484552, EPI_ISL_484553, EPI_ISL_484554, EPI_ISL_484555, EPI_ISL_484556, EPI_ISL_484557, EPI_ISL_484558, EPI_ISL_484559, EPI_ISL_484560, EPI_ISL_484561, EPI_ISL_484562, EPI_ISL_484563, EPI_ISL_484564, EPI_ISL_484565, EPI_ISL_484566, EPI_ISL_484567, EPI_ISL_484568, EPI_ISL_484569, EPI_ISL_484570, EPI_ISL_484571, EPI_ISL_484572, EPI_ISL_484573, EPI_ISL_484574 | | | |
| see above | West of Scotland Specialist Virology Centre, NHSGGC / MRC-University of Glasgow Centre for Virus Research | COVID-19 Genomics UK (COG-UK) Consortium | Ana da Silva Filipe, Natasha Johnson, Kathy Smollett, Daniel Mair, Stephen Carmichael, Lily Tong, Jenna Nichols, Elihu Aranday-Cortes, Kirstyn Brunker, Yasmin Parr, Alice Broos, Kyriaki Nomikou; Sarah McDonald, Marc Niebel, Patawee Asamaphan; Richard Orton, Joseph Hughes, Sreenu Vattipally, David L Robertson; Alasdair MacLean, Rory Gunson; Kathy Li, Natasha Jesudason, Rajiv Shah, James Shepherd, Antonia Ho, Emma Thomson |
| EPI_ISL_484789, EPI_ISL_484790, EPI_ISL_484795, EPI_ISL_484797, EPI_ISL_484804 | University of Michigan Clinical Microbiology Laboratory | Lauring Lab, University of Michigan, Department of Microbiology and Immunology | Valesano et al. |
| EPI_ISL_485003, EPI_ISL_485004, EPI_ISL_485005, EPI_ISL_485006, EPI_ISL_485007, EPI_ISL_485008, EPI_ISL_485009, EPI_ISL_485010, EPI_ISL_485011, EPI_ISL_485012, EPI_ISL_485013, EPI_ISL_485014, EPI_ISL_485015, EPI_ISL_485016, EPI_ISL_485017, EPI_ISL_485018, EPI_ISL_485249, EPI_ISL_485250, EPI_ISL_485252, EPI_ISL_485253, EPI_ISL_485254, EPI_ISL_485255, EPI_ISL_485256, EPI_ISL_485263, EPI_ISL_485264, EPI_ISL_485265, EPI_ISL_485266, EPI_ISL_485267, EPI_ISL_485268, EPI_ISL_485269, EPI_ISL_485270, EPI_ISL_485271, EPI_ISL_485272, EPI_ISL_485274, EPI_ISL_485275, EPI_ISL_485276, EPI_ISL_485277, EPI_ISL_485278, EPI_ISL_485280, EPI_ISL_485281, EPI_ISL_485282, EPI_ISL_485285, EPI_ISL_485286, EPI_ISL_485287, EPI_ISL_485288, EPI_ISL_485289, EPI_ISL_485290, EPI_ISL_485291, EPI_ISL_485292, EPI_ISL_485293, EPI_ISL_485294, EPI_ISL_485295, EPI_ISL_485296, EPI_ISL_485297, EPI_ISL_485299, EPI_ISL_485300, EPI_ISL_485301, EPI_ISL_485302, EPI_ISL_485303, EPI_ISL_485304, EPI_ISL_485305, EPI_ISL_485306, EPI_ISL_485307, EPI_ISL_485308, EPI_ISL_485309, EPI_ISL_485310, EPI_ISL_485311, EPI_ISL_485312, EPI_ISL_485313, EPI_ISL_485314, EPI_ISL_485315, EPI_ISL_485316, EPI_ISL_485317, EPI_ISL_485318, EPI_ISL_485319, EPI_ISL_485320, EPI_ISL_485321, EPI_ISL_485322, EPI_ISL_485323, EPI_ISL_485324, EPI_ISL_485325, EPI_ISL_485326, EPI_ISL_485327, EPI_ISL_485328, EPI_ISL_485329, EPI_ISL_485331, EPI_ISL_485332, EPI_ISL_485333, EPI_ISL_485334, EPI_ISL_485335, EPI_ISL_485336, EPI_ISL_485337, EPI_ISL_485338, EPI_ISL_485339, EPI_ISL_485340, EPI_ISL_485341, EPI_ISL_485342, EPI_ISL_485343, EPI_ISL_485344, EPI_ISL_485345, EPI_ISL_485346, EPI_ISL_485347, EPI_ISL_485348, EPI_ISL_485349, EPI_ISL_485350, EPI_ISL_485351, EPI_ISL_485352, EPI_ISL_485353, EPI_ISL_485354 | | | |
| see above | River Road Testing Lab | Ginkgo Bioworks Clinical Laboratory | Rebecca C. Christofferson, Stephania A. Cormier, Luan V. Dinh, E. Handly Mayton, Hollis R. O'Neil, Thaya Stoufflet, Malaika Mckenzie-Bennett, James McGann, Jim Griffin, Keith Robison, Alex Plocik, Becky Schilling, Rebecca Littlefield, Michelle Spencer, Birgitte Simen |
| EPI_ISL_485582, EPI_ISL_485602 | Instituto de diagnóstico y Referencia Epidemiologicos (INDRE) | Instituto de diagnóstico y Referencia Epidemiologicos (INDRE) | Barrera-Badillo,G., Ramirez-Gonzalez,E. |
| EPI_ISL_485897, EPI_ISL_485898, EPI_ISL_485901, EPI_ISL_485902, EPI_ISL_485903, EPI_ISL_485904, EPI_ISL_485905, EPI_ISL_485906 | River Road Testing Lab | Ginkgo Bioworks Clinical Laboratory | Rebecca C. Christofferson, Stephania A. Cormier, Luan V. Dinh, E. Handly Mayton, Hollis R. O'Neil, Thaya Stoufflet, Malaika Mckenzie-Bennett, James McGann, Jim Griffin, Keith Robison, Alex Plocik, Becky Schilling, Rebecca Littlefield, Michelle Spencer, Birgitte Simen |
| EPI_ISL_486289, EPI_ISL_486290, EPI_ISL_486291 | San Joaquin County Public Health Lab | Chan-Zuckerberg Biohub | CZB Cliahub Consortium |
| EPI_ISL_486872 | Institut Pasteur Dakar | Institut Pasteur de Dakar | Ndongo Dia, Moussa Moise Diagne, Mamadou Diop, Marie Henriette Dior Ndione, Mamadou Malado Jallow, Safietou Sanke, Ousmane Faye, Amadou Alpha Sall. |
| EPI_ISL_486915, EPI_ISL_486916, EPI_ISL_486917 | Maryland Department of Health | Maryland Department of Health | Keller,E. |
| EPI_ISL_487230 | University of Michigan Clinical Microbiology Laboratory | Lauring Lab, University of Michigan, Department of Microbiology and Immunology | Valesano et al. |
| EPI_ISL_487450, EPI_ISL_487451, EPI_ISL_487452, EPI_ISL_487453, EPI_ISL_487454, EPI_ISL_487455, EPI_ISL_487456, EPI_ISL_487457, EPI_ISL_487458, EPI_ISL_487459, EPI_ISL_487460, EPI_ISL_487461 | | | |
| see above | CICM-Mali | Bundeswehr Institut of Microbiology | Kouriba, Dürr, Sangaré, Rehn, Traoré, Bestehorn-Willmann, Walter, Quedraogo, Zimmermann, Maiga, Heitzer, Sogodogo, Antwerpen, Wölfel |
| EPI_ISL_487635, EPI_ISL_487644 | University College London, Great Ormond Street Hospital for Children NHS Foundation Trust, Imperial College Healthcare NHS Trust | Wellcome Sanger Institute for the COVID-19 Genomics UK (COG-UK) consortium | Sergi Castellano, Rachel Williams, Mark Kristiansen, Paola Resende Silva, Sunando Roy, Tony Brooks, Helena Tutill, Paola Niola, Patricia Dyal, Charlotte Williams, Leysa Forrest, Yasmin Panchbhaya, Jacqueline Findlay, Sam Weeks, Julianne Brown, Kathryn Harris, Paul Randell, James Price, Alison Holmes, Judith Breuer and Alex Alderton, Roberto Amato, Sonia Goncalves, Ewan Harrison, David K. Jackson, Ian Johnston, Dominic Kwiatkowski, Cordelia Langford, John Sillitoe on behalf of the Wellcome Sanger Institute COVID-19 Surveillance Team (http://www.sanger.ac.uk/covid-team) |
| EPI_ISL_487653, EPI_ISL_487655, EPI_ISL_487661, EPI_ISL_487663, EPI_ISL_487670, EPI_ISL_487671, EPI_ISL_487686, EPI_ISL_487688, EPI_ISL_487697, EPI_ISL_487700, EPI_ISL_487702, EPI_ISL_487706, EPI_ISL_487708, EPI_ISL_487712, EPI_ISL_487714, EPI_ISL_487718, EPI_ISL_487719, EPI_ISL_487720, EPI_ISL_487725, EPI_ISL_487726, EPI_ISL_487727, EPI_ISL_487730, EPI_ISL_487741, EPI_ISL_487742, EPI_ISL_487743, EPI_ISL_487744, EPI_ISL_487745, EPI_ISL_487746, EPI_ISL_487757, EPI_ISL_487759, EPI_ISL_487770, EPI_ISL_487771, EPI_ISL_487782, EPI_ISL_487790, EPI_ISL_487806, EPI_ISL_487807, EPI_ISL_487819, EPI_ISL_487821, EPI_ISL_487823, EPI_ISL_487825, EPI_ISL_487826, EPI_ISL_487829, EPI_ISL_487831, EPI_ISL_487832, EPI_ISL_487834, EPI_ISL_487852, EPI_ISL_487867, EPI_ISL_487871, EPI_ISL_487874, EPI_ISL_487875, EPI_ISL_487882, EPI_ISL_487886, EPI_ISL_487888, EPI_ISL_487889, EPI_ISL_487892, EPI_ISL_487898, EPI_ISL_487907, EPI_ISL_487908, EPI_ISL_487911, EPI_ISL_487912, EPI_ISL_487915, EPI_ISL_487916, EPI_ISL_487923, EPI_ISL_487924, EPI_ISL_487927, EPI_ISL_487928, EPI_ISL_487930, EPI_ISL_487935, EPI_ISL_487936, EPI_ISL_487949, EPI_ISL_487952, EPI_ISL_487958, EPI_ISL_487959, EPI_ISL_487960, EPI_ISL_487963, EPI_ISL_487967, EPI_ISL_487973, EPI_ISL_487985, EPI_ISL_487988, EPI_ISL_487995, EPI_ISL_487996, EPI_ISL_487999 | | | |

| | | | |
|---|---|---|---|
| see above | Virology Department, Royal Infirmary of Edinburgh, NHS Lothian / School of Biological Sciences, University of Edinburgh | Wellcome Sanger Institute for the COVID-19 Genomics UK (COG-UK) consortium | McHugh M, Dewar R, Rooke S, O'Toole Á, Scher E, Hill V, McCrone JT, Colquhoun R, Yu X, Jackson B, Rambaut A, Templeton K and Alex Alderton, Roberto Amato, Sonia Goncalves, Ewan Harrison, David K. Jackson, Ian Johnston, Dominic Kwiatkowski, Cordelia Langford, John Sillitoe on behalf of the Wellcome Sanger Institute COVID-19 Surveillance Team (http://www.sanger.ac.uk/covid-team) |
| EPI_ISL_488038, EPI_ISL_488043, EPI_ISL_488055 | NU-OMICS DNA Sequencing research facility, Northumbria University | Wellcome Sanger Institute for the COVID-19 Genomics UK (COG-UK) consortium | Chris Duncan, Sheia Waugh, Shirelle Burton-Fanning, Gary Eltringham, Jennifer Collins, Brendan Payne, Yusri Taha, Emma Swindells, Jane Greenaway, Edward Barton, Garren Scott, Debra Padgett, Clive Graham, Sarah Essex, Steve Ligget, Paul Baker, Lynn Dover, Wen Yew, Gary Black, John Allan, Joshua Loh, Greg Young, Matthew Bashton, Andrew Nelson, Darren Smith and Alex Alderton, Roberto Amato, Sonia Goncalves, Ewan Harrison, David K. Jackson, Ian Johnston, Dominic Kwiatkowski, Cordelia Langford, John Sillitoe on behalf of the Wellcome Sanger Institute COVID-19 Surveillance Team (http://www.sanger.ac.uk/covid-team) |
| EPI_ISL_488232, EPI_ISL_488233, EPI_ISL_488240, EPI_ISL_488303, EPI_ISL_488305, EPI_ISL_488321, EPI_ISL_488331, EPI_ISL_488351, EPI_ISL_488358, EPI_ISL_488363 | PHE South West Regional Laboratory, National Infection Service | Wellcome Sanger Institute for the COVID-19 Genomics UK (COG-UK) consortium | Stephanie Hutchings, Hannah Pymont, Dr Peter Muir, Barry Vipond, Rich Hopes; and Alex Alderton, Roberto Amato, Sonia Goncalves, Ewan Harrison, David K. Jackson, Ian Johnston, Dominic Kwiatkowski, Cordelia Langford, John Sillitoe on behalf of the Wellcome Sanger Institute COVID-19 Surveillance Team (http://www.sanger.ac.uk/covid-team) |
| EPI_ISL_488411 | PHE South West Regional Laboratory, National Infection Service | Wellcome Sanger Institute for the COVID-19 Genomics UK (COG-UK) Consortium | Stephanie Hutchings, Hannah Pymont, Dr Peter Muir, Barry Vipond, Rich Hopes; and Alex Alderton, Roberto Amato, Sonia Goncalves, Ewan Harrison, David K. Jackson, Ian Johnston, Dominic Kwiatkowski, Cordelia Langford, John Sillitoe on behalf of the Wellcome Sanger Institute COVID-19 Surveillance Team |
| EPI_ISL_488417, EPI_ISL_488425, EPI_ISL_488427, EPI_ISL_488428, EPI_ISL_488435 | PHE South West Regional Laboratory, National Infection Service | Wellcome Sanger Institute for the COVID-19 Genomics UK (COG-UK) consortium | Stephanie Hutchings, Hannah Pymont, Dr Peter Muir, Barry Vipond, Rich Hopes; and Alex Alderton, Roberto Amato, Sonia Goncalves, Ewan Harrison, David K. Jackson, Ian Johnston, Dominic Kwiatkowski, Cordelia Langford, John Sillitoe on behalf of the Wellcome Sanger Institute COVID-19 Surveillance Team (http://www.sanger.ac.uk/covid-team) |
| EPI_ISL_488881, EPI_ISL_488882, EPI_ISL_488885, EPI_ISL_488896, EPI_ISL_488898, EPI_ISL_488904, EPI_ISL_488907, EPI_ISL_488908, EPI_ISL_488921, EPI_ISL_488928, EPI_ISL_488936, EPI_ISL_488944, EPI_ISL_488946, EPI_ISL_488950, EPI_ISL_488952, EPI_ISL_488953, EPI_ISL_488960, EPI_ISL_488966, EPI_ISL_488968, EPI_ISL_488971, EPI_ISL_488975, EPI_ISL_488978, EPI_ISL_488983, EPI_ISL_488987, EPI_ISL_488998, EPI_ISL_489011, EPI_ISL_489012, EPI_ISL_489017, EPI_ISL_489024, EPI_ISL_489036 | | | |
| see above | Virology Department, Royal Infirmary of Edinburgh, NHS Lothian / School of Biological Sciences, University of Edinburgh | Wellcome Sanger Institute for the COVID-19 Genomics UK (COG-UK) consortium | McHugh M, Dewar R, Rooke S, O'Toole Á, Scher E, Hill V, McCrone JT, Colquhoun R, Yu X, Jackson B, Rambaut A, Templeton K and Alex Alderton, Roberto Amato, Sonia Goncalves, Ewan Harrison, David K. Jackson, Ian Johnston, Dominic Kwiatkowski, Cordelia Langford, John Sillitoe on behalf of the Wellcome Sanger Institute COVID-19 Surveillance Team (http://www.sanger.ac.uk/covid-team) |
| EPI_ISL_489043 | Virology Department, Royal Infirmary of Edinburgh, NHS Lothian / School of Biological Sciences, University of Edinburgh | Wellcome Sanger Institute for the COVID-19 Genomics UK (COG-UK) Consortium | McHugh M, Dewar R, Rooke S, O'Toole Á, Scher E, Hill V, McCrone JT, Colquhoun R, Yu X, Jackson B, Rambaut A, Templeton K and Alex Alderton, Roberto Amato, Sonia Goncalves, Ewan Harrison, David K. Jackson, Ian Johnston, Dominic Kwiatkowski, Cordelia Langford, John Sillitoe on behalf of the Wellcome Sanger Institute COVID-19 Surveillance Team |
| EPI_ISL_489044, EPI_ISL_489049, EPI_ISL_489051, EPI_ISL_489059 | Virology Department, Royal Infirmary of Edinburgh, NHS Lothian / School of Biological Sciences, University of Edinburgh | Wellcome Sanger Institute for the COVID-19 Genomics UK (COG-UK) consortium | McHugh M, Dewar R, Rooke S, O'Toole Á, Scher E, Hill V, McCrone JT, Colquhoun R, Yu X, Jackson B, Rambaut A, Templeton K and Alex Alderton, Roberto Amato, Sonia Goncalves, Ewan Harrison, David K. Jackson, Ian Johnston, Dominic Kwiatkowski, Cordelia Langford, John Sillitoe on behalf of the Wellcome Sanger Institute COVID-19 Surveillance Team (http://www.sanger.ac.uk/covid-team) |
| EPI_ISL_489731, EPI_ISL_489733, EPI_ISL_489752, EPI_ISL_489753, EPI_ISL_489754, EPI_ISL_489755, EPI_ISL_489763, EPI_ISL_489764, EPI_ISL_489775 | Florida Bureau of Public Health Laboratories | Florida Bureau of Public Health Laboratories | Sarah Schmedes, Jason Blanton |
| EPI_ISL_490014, EPI_ISL_490015, EPI_ISL_490016 | Institute for Medical Research, Infectious Disease Research Centre, National Institutes of Health, Ministry of Health Malaysia | Institute for Medical Research, Infectious Disease Research Centre, National Institutes of Health, Ministry of Health Malaysia | Suppiah J, Mohd-Zawawi Z, Kamel K, Kalyanasundram J, Thayan R |
| EPI_ISL_490020 | Sydney South West Pathology Service (SSWPS) - Liverpool Hospital - NSW Health Pathology | NSW Health Pathology - Institute of Clinical Pathology and Medical Research; Westmead Hospital; University of Sydney | CIDM-PH et al. |
| EPI_ISL_490047, EPI_ISL_490048 | Institute for Medical Research, Infectious Disease Research Centre, National Institutes of Health, Ministry of Health Malaysia | Institute for Medical Research, Infectious Disease Research Centre, National Institutes of Health, Ministry of Health Malaysia | Suppiah J, Mohd-Zawawi Z, Kamel K, Kalyanasundram J, Thayan R |
| EPI_ISL_490071, EPI_ISL_490074 | National Public Health Laboratory, National Centre for Infectious Diseases | National Public Health Laboratory, National Centre for Infectious Diseases | Mak TM, Octavia S, Zhou Z, Chavatte JM, Cui L, Lin RTP |
| EPI_ISL_490646, EPI_ISL_490647, EPI_ISL_490648, EPI_ISL_490649, EPI_ISL_490650, EPI_ISL_490651, EPI_ISL_490652, EPI_ISL_490653, EPI_ISL_490654, EPI_ISL_490655, EPI_ISL_490656, EPI_ISL_490657, EPI_ISL_490658, EPI_ISL_490659, EPI_ISL_490660, EPI_ISL_490661, EPI_ISL_490662, EPI_ISL_490663, EPI_ISL_490664, EPI_ISL_490665, EPI_ISL_490666, EPI_ISL_490667, EPI_ISL_490668, EPI_ISL_490669, EPI_ISL_490670, EPI_ISL_490671, EPI_ISL_490672, EPI_ISL_490673, EPI_ISL_490674, EPI_ISL_490675, EPI_ISL_490676, EPI_ISL_490677, EPI_ISL_490678, EPI_ISL_490679, EPI_ISL_490680, EPI_ISL_490681, EPI_ISL_490682 | | | |
| see above | West of Scotland Specialist Virology Centre, NHSGGC / MRC-University of Glasgow Centre for Virus Research | COVID-19 Genomics UK (COG-UK) Consortium | Ana da Silva Filipe, Natasha Johnson, Kathy Smollett, Daniel Mair, Stephen Carmichael, Lily Tong, Jenna Nichols, Elihu Aranday-Cortes, Kirstyn Brunker, Yasmin Parr, Alice Broos, Kyriaki Nomikou; Sarah McDonald, Marc Niebel, Patawee Asamaphan; Richard Orton, Joseph Hughes, Sreenu Vattipally, David L Robertson; Alasdair MacLean, Rory Gunson; Kathy Li, Natasha Jesudason, Rajiv Shah, James Shepherd, Antonia Ho, Emma Thomson |
| EPI_ISL_490761, EPI_ISL_490847 | Wales Specialist Virology Centre Sequencing lab: Pathogen Genomics Unit | COVID-19 Genomics UK (COG-UK) Consortium | Catherine Moore, Johnathan Evans, Laura Gifford, Malorie Perry, Simon Cottrell, Angela Marchbank, Alec Birchley, Alexander Adams, Amy Gaskin, Bree Gatica-Wilcox, Jason Coombes, Joel Southgate, Lauren Gilbert, Lee Graham, Nicole Pacchiarini, Sara Kumziene-Summerhayes, Sarah Taylor, Sophie Jones, Sara Rey, Matthew Bull, Joanne Watkins, Sally Corden, Tom Connor |
| EPI_ISL_491050, EPI_ISL_491052, EPI_ISL_491054, EPI_ISL_491055, EPI_ISL_491056, EPI_ISL_491057 | Suceava County Emergency Hospital | "Stefan cel Mare" University Metagenomics Lab | Lobiuc Andrei et al. |
| EPI_ISL_491119 | The National Institute of Public Health | The National Institute of Public Health and State Veterinary Institute Prague | Nagy,A;Jirincova,H;Novakova,L;Trnka,D;Vecerova,J |
| EPI_ISL_491147, EPI_ISL_491149, EPI_ISL_491150, EPI_ISL_491152, EPI_ISL_491155, EPI_ISL_491156, EPI_ISL_491157 | Oman-National Influenza Center | Biotechnology & OMICs Laboratory | Abdul Latif Khan, Samira Al-Mahruqi, Ahmed Al-Harrasi, Samiha Al-Kharusi, Adil Khan, Ahmed Al-Rawahi, Sajjad Asaf, Amina Al-Jardani, Hanan Al-Kindi, Intisar Al-Shukri, Ahlam Al-Amri, Aisha Al-Amri, Aisha Al-Busaidi, Adil Al-Wahaibi, Seif Al-Abri. |
| EPI_ISL_491159, EPI_ISL_491160 | Oman-National Influenza Center | Biotechnology & OMICs Laboratory | Sajjad Asaf, Samiha Al-Kharusi, Ahmed Al-Harrasi, Samira Al-Mahruqi, Adil Khan, Ahmed Al-Rawahi, Abdul Latif Khan, Amina Al-Jardani, Hanan Al-Kindi, Intisar Al-Shukri, Ahlam Al-Amri, Aisha Al-Amri, Aisha Al-Busaidi, Adil Al-Wahaibi, Seif Al-Abri. |
| EPI_ISL_491495, EPI_ISL_491500, EPI_ISL_491508, EPI_ISL_491512, EPI_ISL_491522, EPI_ISL_491526, EPI_ISL_491528, EPI_ISL_491529, EPI_ISL_491530, EPI_ISL_491539, EPI_ISL_491540, EPI_ISL_491544, EPI_ISL_491554, EPI_ISL_491559, EPI_ISL_491561, EPI_ISL_491562, EPI_ISL_491564, EPI_ISL_491582, EPI_ISL_491584, EPI_ISL_491585, EPI_ISL_491589, EPI_ISL_491590, EPI_ISL_491592, EPI_ISL_491596, EPI_ISL_491600, EPI_ISL_491606, EPI_ISL_491607, EPI_ISL_491615, EPI_ISL_491624, EPI_ISL_491629, EPI_ISL_491633, EPI_ISL_491634, EPI_ISL_491636, EPI_ISL_491643, EPI_ISL_491644, EPI_ISL_491648, EPI_ISL_491650, EPI_ISL_491653, EPI_ISL_491662, EPI_ISL_491666, EPI_ISL_491667, EPI_ISL_491672, EPI_ISL_491675, EPI_ISL_491694, EPI_ISL_491697, EPI_ISL_491698, EPI_ISL_491699 | | | |
| see above | Virology Department, Royal Infirmary of Edinburgh, NHS Lothian / School of Biological Sciences, University of Edinburgh | Wellcome Sanger Institute for the COVID-19 Genomics UK (COG-UK) consortium | McHugh M, Dewar R, Rooke S, O'Toole Á, Scher E, Hill V, McCrone JT, Colquhoun R, Yu X, Jackson B, Rambaut A, Templeton K and Alex Alderton, Roberto Amato, Sonia Goncalves, Ewan Harrison, David K. Jackson, Ian Johnston, Dominic Kwiatkowski, Cordelia Langford, John Sillitoe on behalf of the Wellcome Sanger Institute COVID-19 Surveillance Team (http://www.sanger.ac.uk/covid-team) |
| EPI_ISL_491908, EPI_ISL_491926, EPI_ISL_491927, EPI_ISL_491928, EPI_ISL_491929, EPI_ISL_491930, EPI_ISL_491931 | Naval Infectious Diseases Diagnostic Laboratory | Naval Medical Research Center Biological Defense Research Directorate | Logan Voegtly, Regina Cer, Lindsay Glang, Victor Sugiharto, Francisco Malgon Bautista, Hua Wei Chen, Dessiree Pena-Gomez, Megan Schilling, Adrian Paskey, Kyle Long, Mark Simons, Kimberly Bishop-Lilly |

| | | | |
|---|---|---|---|
| EPI_ISL_491933, EPI_ISL_491934 | Centro de Investigaciones, Universidad de Especialidades Espíritu Santo | Institute of Microbiology, Universidad San Francisco de Quito | Derly Andrade, Juan Carlos Fernandez, Belén Prado-Vivar, Sully Márquez, Juan José Guadalupe, Monica Becerra-Wong, Bernardo Gutiérrez, Gabriel Morey, Ruben Armas, Jose Pedro Barberan, Fernando Espinoza, Edith Lopez, Verónica Barragán, Patricio Rojas-Silva, Gabriel Trueba, Michelle Grunauer, Paúl Cárdenas |
| EPI_ISL_491942 | Naval Infectious Diseases Diagnostic Laboratory | Naval Medical Research Center Biological Defense Research Directorate | Logan Voegtly, Regina Cer, Lindsay Glang, Victor Sugiharto, Francisco Malgon Bautista, Hua Wei Chen, Dessiree Pena-Gomez, Megan Schilling, Adrian Paskey, Kyle Long, Mark Simons, Kimberly Bishop-Lilly |
| EPI_ISL_492061 | Alaska State Virology Laboratory | Alaska State Virology Laboratory | Chen J et al with Pathogenomics group Dagdag R, Redlinger M, Milton E, George W, Kovalenko A, Drown DM, Bortz E |
| EPI_ISL_492228, EPI_ISL_492237, EPI_ISL_492281, EPI_ISL_492282, EPI_ISL_492287, EPI_ISL_492290, EPI_ISL_492296, EPI_ISL_492301, EPI_ISL_492304, EPI_ISL_492311, EPI_ISL_492317, EPI_ISL_492321, EPI_ISL_492324, EPI_ISL_492344, EPI_ISL_492351, EPI_ISL_492354, EPI_ISL_492357, EPI_ISL_492413, EPI_ISL_492420, EPI_ISL_492426, EPI_ISL_492431, EPI_ISL_492506, EPI_ISL_492510, EPI_ISL_492511, EPI_ISL_492513, EPI_ISL_492516, EPI_ISL_492517, EPI_ISL_492518, EPI_ISL_492519, EPI_ISL_492520, EPI_ISL_492521, EPI_ISL_492522, EPI_ISL_492524, EPI_ISL_492525, EPI_ISL_492526, EPI_ISL_492527, EPI_ISL_492528, EPI_ISL_492529, EPI_ISL_492530, EPI_ISL_492531, EPI_ISL_492532, EPI_ISL_492533, EPI_ISL_492534, EPI_ISL_492535, EPI_ISL_492537, EPI_ISL_492538, EPI_ISL_492540, EPI_ISL_492541, EPI_ISL_492542, EPI_ISL_492543, EPI_ISL_492545, EPI_ISL_492546, EPI_ISL_492548, EPI_ISL_492549, EPI_ISL_492550, EPI_ISL_492551, EPI_ISL_492552, EPI_ISL_492553, EPI_ISL_492554, EPI_ISL_492555, EPI_ISL_492556, EPI_ISL_492558, EPI_ISL_492559, EPI_ISL_492560, EPI_ISL_492562, EPI_ISL_492564, EPI_ISL_492565, EPI_ISL_492566, EPI_ISL_492568, EPI_ISL_492571, EPI_ISL_492573, EPI_ISL_492574, EPI_ISL_492576, EPI_ISL_492578, EPI_ISL_492579, EPI_ISL_492581, EPI_ISL_492582, EPI_ISL_492583, EPI_ISL_492584, EPI_ISL_492586, EPI_ISL_492587, EPI_ISL_492589, EPI_ISL_492593 | | | |
| see above | PHE South West Regional Laboratory, National Infection Service | Wellcome Sanger Institute for the COVID-19 Genomics UK (COG-UK) consortium | Stephanie Hutchings, Hannah Pymont, Dr Peter Muir, Barry Vipond, Rich Hopes; and Alex Alderton, Roberto Amato, Sonia Goncalves, Ewan Harrison, David K. Jackson, Ian Johnston, Dominic Kwiatkowski, Cordelia Langford, John Sillitoe on behalf of the Wellcome Sanger Institute COVID-19 Surveillance Team (http://www.sanger.ac.uk/covid-team) |
| EPI_ISL_492594 | PHE South West Regional Laboratory, National Infection Service | Wellcome Sanger Institute for the COVID-19 Genomics UK (COG-UK) Consortium | Stephanie Hutchings, Hannah Pymont, Dr Peter Muir, Barry Vipond, Rich Hopes; and Alex Alderton, Roberto Amato, Sonia Goncalves, Ewan Harrison, David K. Jackson, Ian Johnston, Dominic Kwiatkowski, Cordelia Langford, John Sillitoe on behalf of the Wellcome Sanger Institute COVID-19 Surveillance Team |
| EPI_ISL_492595, EPI_ISL_492597, EPI_ISL_492598, EPI_ISL_492599, EPI_ISL_492600, EPI_ISL_492602, EPI_ISL_492603, EPI_ISL_492605, EPI_ISL_492608, EPI_ISL_492609, EPI_ISL_492611, EPI_ISL_492612, EPI_ISL_492613, EPI_ISL_492614, EPI_ISL_492616, EPI_ISL_492617, EPI_ISL_492618, EPI_ISL_492621, EPI_ISL_492622, EPI_ISL_492625, EPI_ISL_492626, EPI_ISL_492628, EPI_ISL_492632, EPI_ISL_492637, EPI_ISL_492639, EPI_ISL_492640, EPI_ISL_492641, EPI_ISL_492643, EPI_ISL_492644, EPI_ISL_492645, EPI_ISL_492646, EPI_ISL_492647, EPI_ISL_492648, EPI_ISL_492649, EPI_ISL_492651, EPI_ISL_492652, EPI_ISL_492653, EPI_ISL_492654, EPI_ISL_492656, EPI_ISL_492657, EPI_ISL_492658, EPI_ISL_492661, EPI_ISL_492663, EPI_ISL_492666, EPI_ISL_492667, EPI_ISL_492668, EPI_ISL_492669, EPI_ISL_492670, EPI_ISL_492671, EPI_ISL_492673, EPI_ISL_492674, EPI_ISL_492677, EPI_ISL_492678, EPI_ISL_492679, EPI_ISL_492680, EPI_ISL_492681, EPI_ISL_492684, EPI_ISL_492685, EPI_ISL_492686, EPI_ISL_492687, EPI_ISL_492688, EPI_ISL_492689, EPI_ISL_492690, EPI_ISL_492691, EPI_ISL_492692, EPI_ISL_492693, EPI_ISL_492695, EPI_ISL_492696, EPI_ISL_492697, EPI_ISL_492698, EPI_ISL_492699, EPI_ISL_492700, EPI_ISL_492701, EPI_ISL_492702, EPI_ISL_492703, EPI_ISL_492704, EPI_ISL_492705, EPI_ISL_492706, EPI_ISL_492707, EPI_ISL_492708, EPI_ISL_492709, EPI_ISL_492711, EPI_ISL_492712, EPI_ISL_492713, EPI_ISL_492714, EPI_ISL_492715, EPI_ISL_492716, EPI_ISL_492717, EPI_ISL_492718, EPI_ISL_492720, EPI_ISL_492721, EPI_ISL_492722, EPI_ISL_492723, EPI_ISL_492724, EPI_ISL_492725, EPI_ISL_492728, EPI_ISL_492729, EPI_ISL_492731, EPI_ISL_492732, EPI_ISL_492733, EPI_ISL_492734, EPI_ISL_492737, EPI_ISL_492739, EPI_ISL_492740, EPI_ISL_492742, EPI_ISL_492743, EPI_ISL_492744, EPI_ISL_492745, EPI_ISL_492764, EPI_ISL_492765, EPI_ISL_492766, EPI_ISL_492767, EPI_ISL_492768, EPI_ISL_492769, EPI_ISL_492771, EPI_ISL_492772, EPI_ISL_492774, EPI_ISL_492775, EPI_ISL_492776, EPI_ISL_492777, EPI_ISL_492778, EPI_ISL_492779, EPI_ISL_492780, EPI_ISL_492781, EPI_ISL_492782, EPI_ISL_492783, EPI_ISL_492784, EPI_ISL_492785, EPI_ISL_492786, EPI_ISL_492787, EPI_ISL_492788, EPI_ISL_492789, EPI_ISL_492790, EPI_ISL_492791, EPI_ISL_492792, EPI_ISL_492793, EPI_ISL_492794, EPI_ISL_492795, EPI_ISL_492796, EPI_ISL_492797, EPI_ISL_492798, EPI_ISL_492799, EPI_ISL_492800, EPI_ISL_492801, EPI_ISL_492802, EPI_ISL_492803, EPI_ISL_492804, EPI_ISL_492805, EPI_ISL_492806, EPI_ISL_492807, EPI_ISL_492808, EPI_ISL_492810, EPI_ISL_492811, EPI_ISL_492812, EPI_ISL_492813, EPI_ISL_492814, EPI_ISL_492815, EPI_ISL_492816, EPI_ISL_492817, EPI_ISL_492818, EPI_ISL_492819, EPI_ISL_492820, EPI_ISL_492821, EPI_ISL_492822, EPI_ISL_492823, EPI_ISL_492824, EPI_ISL_492825, EPI_ISL_492826, EPI_ISL_492827, EPI_ISL_492828, EPI_ISL_492829, EPI_ISL_492830, EPI_ISL_492831, EPI_ISL_492832, EPI_ISL_492833, EPI_ISL_492834, EPI_ISL_492835 | | | |
| see above | PHE South West Regional Laboratory, National Infection Service | Wellcome Sanger Institute for the COVID-19 Genomics UK (COG-UK) consortium | Stephanie Hutchings, Hannah Pymont, Dr Peter Muir, Barry Vipond, Rich Hopes; and Alex Alderton, Roberto Amato, Sonia Goncalves, Ewan Harrison, David K. Jackson, Ian Johnston, Dominic Kwiatkowski, Cordelia Langford, John Sillitoe on behalf of the Wellcome Sanger Institute COVID-19 Surveillance Team (http://www.sanger.ac.uk/covid-team) |
| EPI_ISL_492837, EPI_ISL_492840, EPI_ISL_492845, EPI_ISL_492847, EPI_ISL_492851, EPI_ISL_492865, EPI_ISL_492868, EPI_ISL_492880, EPI_ISL_492885, EPI_ISL_492887, EPI_ISL_492906, EPI_ISL_492914 | | | |
| see above | Royal Free Hospital / Health Services Laboratories | Wellcome Sanger Institute for the COVID-19 Genomics UK (COG-UK) consortium | Tanzina Haque, Tabitha Mahungu, Dianne Irish, Cate Goodlad, Jenny Cross, Judith Heaney and Alex Alderton, Roberto Amato, Sonia Goncalves, Ewan Harrison, David K. Jackson, Ian Johnston, Dominic Kwiatkowski, Cordelia Langford, John Sillitoe on behalf of the Wellcome Sanger Institute COVID-19 Surveillance Team (http://www.sanger.ac.uk/covid-team) |
| EPI_ISL_492981, EPI_ISL_492982 | IRCCS Sacro Cuore Don Calabria Hospital, Department of Infectious, Tropical Diseases & Microbiology | University of Verona, Department of Biotechnology | Antonio Mori, Michela Deiana, Elena Pomari, Chiara Piubelli; Giulia Lopatriello, Luca Marcolungo, Cristina Beltrami, Chiara Degli Esposti, Emanuela Cosentino, Massimo Delledonne |
| EPI_ISL_493612, EPI_ISL_493613 | Lincolnshire Hospitals and DeepSeq Nottingham | COVID-19 Genomics UK (COG-UK) Consortium | Nichola Duckworth, Tim Sloan, Sarah Walsh, Jonathan Ball, Patrick McClure, Joeseph Chappell, Nadine Holmes, Matthew Carlisle, Christopher Moore, Fei Sang, Johnny Debebe, Victoria Wright, Matthew Loose |
| EPI_ISL_493665 | Centre for Enzyme Innovation, University of Portsmouth / Translational Research Laboratory, Portsmouth Hospitals NHS Trust | COVID-19 Genomics UK (COG-UK) Consortium | Angela Beckett,Yann Bourgeois,Garry Scarlett,Sharon Glaysher,Scott Elliott,Kelly Bicknell,Robert Impey,Allyson Lloyd,Sarah Wyllie,Ethan Butcher,Anoop Chauhan,Samuel Robson |
| EPI_ISL_493744, EPI_ISL_493746, EPI_ISL_493747, EPI_ISL_493748, EPI_ISL_493749, EPI_ISL_493750, EPI_ISL_493751, EPI_ISL_493752, EPI_ISL_493753, EPI_ISL_493754, EPI_ISL_493755, EPI_ISL_493756, EPI_ISL_493757, EPI_ISL_493758, EPI_ISL_493759, EPI_ISL_493760, EPI_ISL_493761, EPI_ISL_493762, EPI_ISL_493763, EPI_ISL_493764, EPI_ISL_493765, EPI_ISL_493766, EPI_ISL_493767, EPI_ISL_493768, EPI_ISL_493769, EPI_ISL_493770, EPI_ISL_493771, EPI_ISL_493772, EPI_ISL_493773, EPI_ISL_493774, EPI_ISL_493775, EPI_ISL_493776, EPI_ISL_493777, EPI_ISL_493778, EPI_ISL_493779, EPI_ISL_493780, EPI_ISL_493781, EPI_ISL_493782, EPI_ISL_493783, EPI_ISL_493784, EPI_ISL_493785, EPI_ISL_493786, EPI_ISL_493790 | | | |
| see above | West of Scotland Specialist Virology Centre, NHSGGC / MRC-University of Glasgow Centre for Virus Research | COVID-19 Genomics UK (COG-UK) Consortium | Ana da Silva Filipe, Natasha Johnson, Kathy Smollett, Daniel Mair, Stephen Carmichael, Lily Tong, Jenna Nichols, Elihu Aranday-Cortes, Kirstyn Brunker, Yasmin Parr, Alice Broos, Kyriaki Nomikou; Sarah McDonald, Marc Niebel, Patawee Asamaphan; Richard Orton, Joseph Hughes, Sreenu Vattipally, David L Robertson; Alasdair MacLean, Rory Gunson; Kathy Li, Natasha Jesudason, Rajiv Shah, James Shepherd, Antonia Ho, Emma Thomson |
| EPI_ISL_494380, EPI_ISL_494390, EPI_ISL_494391, EPI_ISL_494413, EPI_ISL_494438, EPI_ISL_494439, EPI_ISL_494440, EPI_ISL_494447, EPI_ISL_494449 | San Diego County Public Health Laboratory | Andersen lab at Scripps Research | SEARCH Alliance San Diego with Tracy Basler, Jovan Shephard, Brett Austin |
| EPI_ISL_494521, EPI_ISL_494538, EPI_ISL_494539, EPI_ISL_494540, EPI_ISL_494541 | Quest Diagnostics | Quest Diagnostics | Anderson,B.P., Rosenthal,S.H., Gerasimova,A., Kagan,R.M. and Owen, R. |
| EPI_ISL_495298, EPI_ISL_495299, EPI_ISL_495300, EPI_ISL_495301, EPI_ISL_495302, EPI_ISL_495303, EPI_ISL_495304, EPI_ISL_495305, EPI_ISL_495306, EPI_ISL_495307, EPI_ISL_495308, EPI_ISL_495309, EPI_ISL_495310, EPI_ISL_495311, EPI_ISL_495312, EPI_ISL_495313, EPI_ISL_495314, EPI_ISL_495315, EPI_ISL_495316, EPI_ISL_495317, EPI_ISL_495318, EPI_ISL_495319, EPI_ISL_495320, EPI_ISL_495321, EPI_ISL_495322, EPI_ISL_495323, EPI_ISL_495324, EPI_ISL_495325, EPI_ISL_495326, EPI_ISL_495327, EPI_ISL_495328, EPI_ISL_495329 | | | |
| see above | Virginia DCLS | Virginia DCLS | Virginia DCLS |
| EPI_ISL_495571, EPI_ISL_495573, EPI_ISL_495575 | University of Michigan Clinical Microbiology Laboratory | Lauring Lab, University of Michigan, Department of Microbiology and Immunology | Valesano et al. |
| EPI_ISL_495599, EPI_ISL_495602, EPI_ISL_495604 | Mayo Clinic & Mayo Clinic Laboratories | Minnesota Department of Health, Public Health Laboratory | Matt Plumb, Jacob Garfin, and Xiong Wang |
| EPI_ISL_496553, EPI_ISL_496554 | B.J. Govt. Medical College | National Centre For Cell Science | Dhiraj Paul, Kunal Jani, Radha Chauhan, Janesh Kumar, Vasudevan Seshadri, Girdhari Lal, Rajesh Karyakarte, Suvarna Joshi, Murlidhar Tambe, Sourav Sen, Santosh Karade, Kavita Bala Anand, Shelinder Pal Singh Shergill, Rajiv Mohan Gupta, Manoj Kumar Bhat, Arvind Sahu, Maharashtra COVID-19 Study Group, DBT's PAN-INDIA 1000 SARS-CoV2 RNA genome sequencing consortium, Yogesh S Shouche |
| EPI_ISL_496857, EPI_ISL_496858, EPI_ISL_496859, EPI_ISL_496865, EPI_ISL_496866, EPI_ISL_496867, EPI_ISL_496868, EPI_ISL_496869, EPI_ISL_496870, EPI_ISL_496871, EPI_ISL_496872, EPI_ISL_496873, EPI_ISL_496874, EPI_ISL_496875, EPI_ISL_496876, EPI_ISL_496877, EPI_ISL_496878, EPI_ISL_496879, EPI_ISL_496880, EPI_ISL_496881, EPI_ISL_496882, EPI_ISL_496883, EPI_ISL_496884, EPI_ISL_496885, EPI_ISL_496886, EPI_ISL_496887, EPI_ISL_496888, EPI_ISL_496889, EPI_ISL_496890, EPI_ISL_496891, EPI_ISL_496892, EPI_ISL_496893, EPI_ISL_496894, EPI_ISL_496895, EPI_ISL_496896, EPI_ISL_496897, EPI_ISL_496898, EPI_ISL_496899, EPI_ISL_496900 | | | |
| see above | Gorgas Memorial Laboratory of Health Studies | Gorgas Memorial Laboratory of Health Studies | Danilo Franco, Claudia Gonzalez Sandra Lopez-Verges, Alexander A Martinez |
| EPI_ISL_496918 | Mayo Clinic & Mayo Clinic Laboratories | Minnesota Department of Health, Public Health Laboratory | Matt Plumb, Jacob Garfin, and Xiong Wang |
| EPI_ISL_497823 | Department of Microbiology, The University of Hong Kong | Department of Microbiology, The University of Hong Kong | Kelvin K.W. To, Kwok-Yung Yuen |
| EPI_ISL_498164, EPI_ISL_498165 | Instituto Nacional de Salud, Bogotá, Colombia | Instituto Nacional de Salud, Bogotá, Colombia | Katherine Laiton-Donato, Diego A. Álvarez-Díaz, Carlos Franco-Muñoz, Jonathan Reales, Diego Andrés Prada, Jose A. Usme-Ciro, Nicolas D. Franco-Sierra, Zulma M. Cucunubá, Christian Julian VillabonaArenas, Liz Villabona-Arenas, Sussy Echeverria, Astrid C. Flórez, Carolina Ferro, Diana Marcela Walteros-Acero, Franklin Prieto, Carlos Andrés Durán, Martha Lucia Ospina Martinez, Marcela Mercado-Reyes |

| | | | |
|---|---|---|---|
| EPI_ISL_498518, EPI_ISL_498523, EPI_ISL_498539 | ACT Pathology | Schwessinger Lab | Ashley Jones, Benjamin Schwessinger, Robert Lanfear, Robyn N Hall, Megan McDonald, Ming-Dao Chia, Kevin Murray, Craig Kennedy, Karina Kennedy |
| EPI_ISL_498635, EPI_ISL_498639 | Utah Public Health Laboratory | Utah Public Health Laboratory | Heidi Butz, Erin Young, Kelly Oakeson |
| EPI_ISL_498697, EPI_ISL_498698, EPI_ISL_498699, EPI_ISL_498736, EPI_ISL_498738, EPI_ISL_498739 | Quest Diagnostics | Quest Diagnostics | Rosenthal,S.H., Gerasimova,A., Kagan,R.M. and Owen, R. |
| EPI_ISL_499330, EPI_ISL_499334, EPI_ISL_499339, EPI_ISL_499348 | Virology Department, Sheffield Teaching Hospitals NHS Foundation Trust/Department of Infection, Immunity and Cardiovascular Disease, The Medical School, University of Sheffield | COVID-19 Genomics UK (COG-UK) Consortium | Thushan de Silva, Matthew Parker, Nikki Smith, Adri Angyal, Rebecca Brown, Luke Green, Rachel Tucker, Paul Parsons, Danielle Groves, Katie Johnson, Laura Carrilero, Alex Keeley, Dave Partridge, Matthew Wyles, Benjamin Lindsey, Mehmet Yavuz, Mohammad Raza, Cariad Evans |
| EPI_ISL_500163, EPI_ISL_500164, EPI_ISL_500168, EPI_ISL_500169, EPI_ISL_500170, EPI_ISL_500172, EPI_ISL_500174, EPI_ISL_500175, EPI_ISL_500176, EPI_ISL_500177, EPI_ISL_500178, EPI_ISL_500179, EPI_ISL_500180, EPI_ISL_500181, EPI_ISL_500182, EPI_ISL_500183, EPI_ISL_500184, EPI_ISL_500185, EPI_ISL_500186, EPI_ISL_500202, EPI_ISL_500203, EPI_ISL_500211, EPI_ISL_500212, EPI_ISL_500213, EPI_ISL_500214, EPI_ISL_500215, EPI_ISL_500216, EPI_ISL_500224, EPI_ISL_500225 | | | |
| see above | Hospital Universitario Virgen de las Nieves de Granada-SAS | SeqCOVID-SPAIN consortium/IBV(CSIC) | Mercedes Pérez Ruiz, Sara Sanbonmatsu Gámez, Irene Pedrosa Corral, José M. Navarro-Marí and SeqCOVID-SPAIN consortium |
| EPI_ISL_500324, EPI_ISL_500325, EPI_ISL_500326, EPI_ISL_500327, EPI_ISL_500328, EPI_ISL_500329, EPI_ISL_500330, EPI_ISL_500331, EPI_ISL_500332, EPI_ISL_500333, EPI_ISL_500334, EPI_ISL_500335 | | | |
| see above | Servicio de Microbiología. Hospital Universitario Donostia. OSI Donostialdea. Área de Enfermedades Infecciosas, Grupo de Infección Respiratoria y Resistencia Antimicrobiana. Instituto de Investigación Sanitaria Biodonostia | SeqCOVID-SPAIN consortium/IBV(CSIC) | Gustavo Cilla, Milagrosa Montes, Luis Piñeiro, Jose Maria Marimón and SeqCOVID-SPAIN consortium |
| EPI_ISL_500459 | Quest Diagnostics | Quest Diagnostics | Rosenthal,S.H., Gerasimova,A., Kagan,R.M., Owen, R. AND Lacbawan, F. |
| EPI_ISL_500460, EPI_ISL_500461, EPI_ISL_500462, EPI_ISL_500463, EPI_ISL_500464, EPI_ISL_500465, EPI_ISL_500466, EPI_ISL_500467, EPI_ISL_500468, EPI_ISL_500469, EPI_ISL_500470, EPI_ISL_500472, EPI_ISL_500473, EPI_ISL_500474, EPI_ISL_500475, EPI_ISL_500476, EPI_ISL_500477, EPI_ISL_500481, EPI_ISL_500482, EPI_ISL_500483, EPI_ISL_500484, EPI_ISL_500485, EPI_ISL_500486 | | | |
| see above | LACEN/PE | WallauLab, Aggeu Magalhaes Institute | Marcelo Henrique Santos Paiva, Duschinka Ribeiro Duarte Guedes, Cássia Docena, Matheus Filgueira Bezerra, Filipe Zimmer Dezordi, Laís Ceschini Machado, Larissa Krokovsky, Elisama Helvecio, Alexandre Freitas da Silva, Luydson Richardson Silva Vasconcelos, Antonio Mauro Rezende, Severino Jefferson Ribeiro da Silva, Kamila Gaudêncio da Silva Sales, Bruna Santos Lima Figueiredo de Sá, Derciliano Lopes da Cruz, Claudio Eduardo Cavalcanti, Armando de Menezes Neto, Caroline Targino Alves da Silva, Renata Pessôa Germano Mendes, Maria Almerice Lopes da Silva, Tiago Gräf, Paola Cristina Resende, Gonzalo Bello, Michelle da Silva Barros, Wheverton Ricardo Correia do Nascimento, Rodrigo Moraes Loyo Arcoverde, Luciane Caroline Albuquerque Bezerra, Sinval Pinto Brandão Filho, Constância Flávia Junqueira Ayres, Gabriel Luz Wallau |
| EPI_ISL_500523, EPI_ISL_500524, EPI_ISL_500532 | Mayo Clinic Laboratories | University of Washington Virology Lab | Pavitra Roychoudhury, Hong Xie, Lasata Shrestha, Amin Addetia, Truong Nguyen, Victoria M Rachleff, Meei-Li Huang, Keith R Jerome, Alexander Greninger |
| EPI_ISL_500702 | Area of Virology, Serology and Virology Division (SAViD), New South Wales Health Pathology Randwick | Area of Virology, Serology and Virology Division (SAViD), New South Wales Health Pathology Randwick | Rawlinson, W. |
| EPI_ISL_500865, EPI_ISL_500866, EPI_ISL_500867, EPI_ISL_500868, EPI_ISL_500869, EPI_ISL_500870, EPI_ISL_500871, EPI_ISL_500872 | LACEN/PE | WallauLab, Aggeu Magalhaes Institute | Marcelo Henrique Santos Paiva, Duschinka Ribeiro Duarte Guedes, Cássia Docena, Matheus Filgueira Bezerra, Filipe Zimmer Dezordi, Laís Ceschini Machado, Larissa Krokovsky, Elisama Helvecio, Alexandre Freitas da Silva, Luydson Richardson Silva Vasconcelos, Antonio Mauro Rezende, Severino Jefferson Ribeiro da Silva, Kamila Gaudêncio da Silva Sales, Bruna Santos Lima Figueiredo de Sá, Derciliano Lopes da Cruz, Claudio Eduardo Cavalcanti, Armando de Menezes Neto, Caroline Targino Alves da Silva, Renata Pessôa Germano Mendes, Maria Almerice Lopes da Silva, Tiago Gräf, Paola Cristina Resende, Gonzalo Bello, Michelle da Silva Barros, Wheverton Ricardo Correia do Nascimento, Rodrigo Moraes Loyo Arcoverde, Luciane Caroline Albuquerque Bezerra, Sinval Pinto Brandão Filho, Constância Flávia Junqueira Ayres, Gabriel Luz Wallau |
| EPI_ISL_501072 | Mayo Clinic Laboratories | University of Washington Virology Lab | Pavitra Roychoudhury, Hong Xie, Lasata Shrestha, Amin Addetia, Truong Nguyen, Victoria M Rachleff, Meei-Li Huang, Keith R Jerome, Alexander Greninger |
| EPI_ISL_501170, EPI_ISL_501171 | Baylor College of Medicine | Baylor College of Medicine: HGSC | Vasanthi Avadhanula, Erin Nicholson, David Henke, Pedro Piedra, Harsha Doddapaneni, Donna Muzny, Qingchang Meng, Hsu Chao, Zeineen Momin, Hua Shen, George Weissenberger, Kavya Kottapalli, Yimiti Meiheerguli, Sejal Salvi, Ginger Metcalf, Vipin Menon, Sara J.J. Cregeen, Matthew C. Ross, Tulin Ayvaz, Richard Sucgang, Kristi L. Hoffman, Matthew Wong, Joseph F. Petrosino |
| EPI_ISL_501218, EPI_ISL_501219, EPI_ISL_501222 | Department of Medical Microbiology, University Malaya Medical Centre | Department of Medical Microbiology, Faculty of Medicine, University of Malaya | Yoong Min CHONG, Jennifer Chong, I-Ching SAM, Yoke Fun CHAN, University Malaya Medical Centre COVID Team |
| EPI_ISL_501621 | Lab Microbiology, Pathology Department, William Harvey Hospital | Wellcome Sanger Institute for the COVID-19 Genomics UK (COG-UK) consortium | Samuel Moses, Hannah Lowe, Felicity Ryan and Alex Alderton, Roberto Amato, Sonia Goncalves, Ewan Harrison, David K. Jackson, Ian Johnston, Dominic Kwiatkowski, Cordelia Langford, John Sillitoe on behalf of the Wellcome Sanger Institute COVID-19 Surveillance Team (http://www.sanger.ac.uk/covid-team) |
| EPI_ISL_505003 | Biology Dpt | Microbiology and Infections Diseases | Emmanuelle Billon-Denis, Audrey Ferrier-Rembert, Annabelle Garnier, Laurence Cheutin, Clarisse Vigne, Emilie Tessier, Jessica Denis, Olivier Gorgé, Flora Nolent, Isabelle Drouet, Noémie Verguet, Olivier Ferraris, Jean-Nicolas Tournier |
| EPI_ISL_506041 | Biology Dpt, HIA Percy | Microbiology and Infectious Diseases Dpt | Emmanuelle Billon-Denis, Audrey Ferrier-Rembert, Annabelle Garnier, Laurence Cheutin, Clarisse Vigne, Emilie Tessier, Jessica Denis, Olivier Gorgé, Flora Nolent, Isabelle Drouet, Noémie Verguet, Olivier Ferraris, Jean-Nicolas Tournier |
| EPI_ISL_506994, EPI_ISL_506995 | Division of Viral Diseases, Center for Laboratory Control of Infectious Diseases, Korea Centers for Diseases Control and Prevention | Division of Viral Diseases, Center for Laboratory Control of Infectious Diseases, Korea Centers for Diseases Control and Prevention | Jeong-Min Kim, Yoon-Seok Chung, Namjoo Lee, Sang Hee Woo, Hye-Jun Jo, Heui Man Kim, Jun-Sub Kim, Myung Guk Han |
| EPI_ISL_506996, EPI_ISL_506997, EPI_ISL_507000 | Department of Medical Microbiology, University Malaya Medical Centre | Department of Medical Microbiology, Faculty of Medicine, University of Malaya | Yoong Min CHONG, Jennifer Chong, I-Ching SAM, Yoke Fun CHAN, University Malaya Medical Centre COVID Team |
| EPI_ISL_507044, EPI_ISL_507082, EPI_ISL_507083 | University College London Hospital | COVID-19 Genomics UK (COG-UK) Consortium | Judith Heaney, Matthew Byott, Catherine Houlihan, Dan Frampton, Stuart Kirk, Moira Spyer and Eleni Nastouli |
| EPI_ISL_507156 | Centre for Enzyme Innovation, University of Portsmouth / Translational Research Laboratory, Portsmouth Hospitals NHS Trust | COVID-19 Genomics UK (COG-UK) Consortium | Angela Beckett,Yann Bourgeois,Garry Scarlett,Sharon Glaysher,Scott Elliott,Kelly Bicknell,Robert Impey,Allyson Lloyd,Sarah Wyllie,Ethan Butcher,Anoop Chauhan,Samuel Robson |
| EPI_ISL_507178 | Virology Department, Sheffield Teaching Hospitals NHS Foundation Trust/Department of Infection, Immunity and Cardiovascular Disease, The Medical School, University of Sheffield | COVID-19 Genomics UK (COG-UK) Consortium | Thushan de Silva, Matthew Parker, Nikki Smith, Adri Angyal, Rebecca Brown, Luke Green, Rachel Tucker, Paul Parsons, Danielle Groves, Katie Johnson, Laura Carrilero, Alex Keeley, Dave Partridge, Matthew Wyles, Benjamin Lindsey, Mehmet Yavuz, Mohammad Raza, Cariad Evans |
| EPI_ISL_507180 | Virology Department, Royal Infirmary of Edinburgh, NHS Lothian / School of Biological Sciences, University of Edinburgh / Institute of Genetics and Molecular Medicine, University of Edinburgh | COVID-19 Genomics UK (COG-UK) Consortium | McHugh M, Dewar R, Rooke S, Gallagher M, Balcaza C, O'Toole Á, Scher E, Hill V, McCrone JT, Colquhoun R, Yu X, Jackson B, Rambaut A, Williams TC, Templeton K |
| EPI_ISL_507473, EPI_ISL_507474, EPI_ISL_507475, EPI_ISL_507480, EPI_ISL_507481, EPI_ISL_507482, EPI_ISL_507483, EPI_ISL_507484, EPI_ISL_507485, EPI_ISL_507486, EPI_ISL_507488, EPI_ISL_507489, EPI_ISL_507490, EPI_ISL_507493, EPI_ISL_507495, EPI_ISL_507496, EPI_ISL_507529, EPI_ISL_507530, EPI_ISL_507531, EPI_ISL_507533, EPI_ISL_507534, EPI_ISL_507535, EPI_ISL_507536, EPI_ISL_507537, EPI_ISL_507538, EPI_ISL_507539, EPI_ISL_507540, EPI_ISL_507541, EPI_ISL_507542, EPI_ISL_507543, EPI_ISL_507544, EPI_ISL_507545, EPI_ISL_507546, EPI_ISL_507547, EPI_ISL_507548, EPI_ISL_507549, EPI_ISL_507551, EPI_ISL_507552, EPI_ISL_507555, EPI_ISL_507556, EPI_ISL_507558, EPI_ISL_507624, EPI_ISL_507626, EPI_ISL_507657, EPI_ISL_507674, EPI_ISL_507703, EPI_ISL_507711, EPI_ISL_507714, EPI_ISL_507715 | | | |
| see above | Michigan Department of Health and Human Services, Bureau of Laboratories | Michigan Department of Health and Human Services, Bureau of Laboratories | Blankenship HM, Riner D, Soehnlen MK |

| | | | |
|---|---|---|---|
| EPI_ISL_508035, EPI_ISL_508075 | New Mexico Department of Health Scientific Laboratory Division | Center for Global Health, University of New Mexico Health Sciences Center | Daryl Domman, Kurt Schwalm, Twila Kunde, Joseph Hicks, Michael Edwards, Darrell Dinwiddie |
| EPI_ISL_508663 | Departamento de Microbiología, CDB, Hospital Clínic, Barcelona | SeqCOVID-SPAIN consortium/IBV(CSIC) | Andrea Vergara, Mikel Martínez, Elisa Rubio, Jéssica Navero, Aida Peiró and SeqCOVID-SPAIN consortium |
| EPI_ISL_508858, EPI_ISL_508859, EPI_ISL_508860, EPI_ISL_508861 | Florida Bureau of Public Health Laboratories | Florida Bureau of Public Health Laboratories | Sarah Schmedes, Jason Blanton |
| EPI_ISL_508916, EPI_ISL_508917, EPI_ISL_508918, EPI_ISL_508919, EPI_ISL_508920, EPI_ISL_508921, EPI_ISL_508922, EPI_ISL_508923, EPI_ISL_508924, EPI_ISL_508925, EPI_ISL_508926, EPI_ISL_508927, EPI_ISL_508928, EPI_ISL_508929, EPI_ISL_508930 | | | |
| see above | CNR Virus des Infections Respiratoires - France SUD | CNR Virus des Infections Respiratoires - France SUD | Antonin Bal, Gregory Destras, Gwendolyne Burfin, Solenne Brun, Carine Moustaud, Raphaelle Lamy, Alexandre Gaymard, Maude Bouscambert-Duchamp, Florence Morfin-Sherpa, Martine Valette, Bruno Lina, Laurence Josset |
| EPI_ISL_508931 | L'Hôpital Nord-Ouest Tarare-Grandris | CNR Virus des Infections Respiratoires - France SUD | Antonin Bal, Gregory Destras, Gwendolyne Burfin, Solenne Brun, Carine Moustaud, Raphaelle Lamy, Alexandre Gaymard, Maude Bouscambert-Duchamp, Florence Morfin-Sherpa, Martine Valette, Bruno Lina, Laurence Josset |
| EPI_ISL_508932 | Centre Hospitalier de Villefranche | CNR Virus des Infections Respiratoires - France SUD | Antonin Bal, Gregory Destras, Gwendolyne Burfin, Solenne Brun, Carine Moustaud, Raphaelle Lamy, Alexandre Gaymard, Maude Bouscambert-Duchamp, Florence Morfin-Sherpa, Martine Valette, Bruno Lina, Laurence Josset |
| EPI_ISL_508989 | CNR Virus des Infections Respiratoires - France SUD | CNR Virus des Infections Respiratoires - France SUD | Antonin Bal, Gregory Destras, Gwendolyne Burfin, Solenne Brun, Alexandre Gaymard, Maude Bouscambert-Duchamp, Florence Morfin-Sherpa, Martine Valette, Bruno Lina, Laurence Josset |
| EPI_ISL_508991, EPI_ISL_508992, EPI_ISL_508993, EPI_ISL_508994, EPI_ISL_508995, EPI_ISL_508996, EPI_ISL_508997 | CNR Virus des Infections Respiratoires - France SUD | CNR Virus des Infections Respiratoires - France SUD | Antonin Bal, Gregory Destras, Gwendolyne Burfin, Solenne Brun, Carine Moustaud, Raphaelle Lamy, Alexandre Gaymard, Maude Bouscambert-Duchamp, Florence Morfin-Sherpa, Martine Valette, Bruno Lina, Laurence Josset |
| EPI_ISL_509082, EPI_ISL_509083, EPI_ISL_509084, EPI_ISL_509086, EPI_ISL_509087 | OHSU Lab Services Molecular Microbiology Lab | Oregon SARS-CoV-2 Genome Sequencing Center | Brendan L. O'Connell, Ruth V. Nichols, Sally B. Grindstaff, Alec J. Hirsch, Guang Fan, Daniel N. Streblow, William B. Messer, Andrew C. Adey, Benjamin N. Bimber, Brian J. O'Roak |
| EPI_ISL_509414 | Acibadem Labcell Cellular Therapy Laboratory | Acibadem Mehmet Ali Aydinlar University School of Medicine, Medical Genetics Department | Ozden Hatirnaz Ng, Sezer Akyoney, Ilayda Sahin, Gunseli Bayram Akcapinar, Ozkan Ozdemir, Derya Dilek Kancagi, Gozde Sir Karakus, Bulut Yurtsever, Cihan Tastan, Ercument Ovali, Ugur Ozbek |
| EPI_ISL_509450, EPI_ISL_509451, EPI_ISL_509452 | Maryland Department of Health | Maryland Department of Health | Keller,E. |
| EPI_ISL_509567, EPI_ISL_509568, EPI_ISL_509569, EPI_ISL_509570, EPI_ISL_509571, EPI_ISL_509572, EPI_ISL_509573, EPI_ISL_509574 | Utah Public Health Laboratory | Utah Public Health Laboratory | Heidi Butz, Erin Young, Kelly Oakeson |
| EPI_ISL_509714 | Belize Ministry of Health | Pathogen Discovery, Respiratory Viruses Branch, Division of Viral Diseases, Centers for Disease Control and Prevention | Jing Zhang, Ying Tao, Krista Queen, Anna Uehara, Yan Li, Clinton Paden, Haibin Wang, Suxiang Tong |
| EPI_ISL_510068, EPI_ISL_510072 | Instituto de Investigaciones Biomédicas de Barcelona (CSIC), Hospital Clinic i Provincial de Barcelona, Instituto de Biomedicina de Valencia (CSIC), Hospital de Sant Pau | SeqCOVID-SPAIN consortium/IBV(CSIC) | Anna M. Planas, Mª Angeles Marcos, Miguel J. Martínez, Andrea Vergara, Alex Soriano, Jordi Pérez Tur, Israel Fernández Cadenas and SeqCOVID-SPAIN consortium |
| EPI_ISL_510109, EPI_ISL_510112, EPI_ISL_510120, EPI_ISL_510138, EPI_ISL_510142, EPI_ISL_510244, EPI_ISL_510245 | Hospital General Universitario Gregorio Marañón | SeqCOVID-SPAIN consortium/IBV(CSIC) | Laura Pérez-Lago, Marta Herranz, Jon Sicilia, Julia Suárez, Pilar Catalán, Patricia Muñoz, Darío García de Viedma and SeqCOVID-SPAIN consortium |
| EPI_ISL_510391, EPI_ISL_510392, EPI_ISL_510393, EPI_ISL_510394, EPI_ISL_510395, EPI_ISL_510396, EPI_ISL_510397, EPI_ISL_510398, EPI_ISL_510399, EPI_ISL_510400, EPI_ISL_510401, EPI_ISL_510402, EPI_ISL_510403, EPI_ISL_510404, EPI_ISL_510405, EPI_ISL_510406, EPI_ISL_510407, EPI_ISL_510408, EPI_ISL_510409, EPI_ISL_510410, EPI_ISL_510411, EPI_ISL_510412, EPI_ISL_510413, EPI_ISL_510414, EPI_ISL_510415, EPI_ISL_510416, EPI_ISL_510417, EPI_ISL_510418, EPI_ISL_510419, EPI_ISL_510420, EPI_ISL_510421, EPI_ISL_510422 | | | |
| see above | Servicio de Microbiología. Hospital Universitario Donostia. OSI Donostialdea. Área de Enfermedades Infecciosas, Grupo de Infección Respiratoria y Resistencia Antimicrobiana. Instituto de Investigación Sanitaria Biodonostia | SeqCOVID-SPAIN consortium/IBV(CSIC) | Gustavo Cilla, Milagrosa Montes, Luis Piñeiro, Jose Maria Marimón and SeqCOVID-SPAIN consortium |
| EPI_ISL_510451 | Hospital Universitario Virgen de las Nieves de Granada-SAS | SeqCOVID-SPAIN consortium/IBV(CSIC) | Mercedes Pérez Ruiz, Sara Sanbonmatsu Gámez, Irene Pedrosa Corral, José M. Navarro-Marí and SeqCOVID-SPAIN consortium |
| EPI_ISL_510466 | Instituto de Investigaciones Biomédicas de Barcelona (CSIC), Hospital Clinic i Provincial de Barcelona, Instituto de Biomedicina de Valencia (CSIC), Hospital de Sant Pau | SeqCOVID-SPAIN consortium/IBV(CSIC) | Anna M. Planas, Mª Angeles Marcos, Miguel J. Martínez, Andrea Vergara, Alex Soriano, Jordi Pérez Tur, Israel Fernández Cadenas and SeqCOVID-SPAIN consortium |
| EPI_ISL_511029, EPI_ISL_511481 | Instituto Nacional de Saude (INSA) | Instituto Nacional de Saude (INSA) | Borges et al |
| EPI_ISL_511490, EPI_ISL_511491, EPI_ISL_511492, EPI_ISL_511493, EPI_ISL_511494, EPI_ISL_511495, EPI_ISL_511496, EPI_ISL_511497, EPI_ISL_511498, EPI_ISL_511499, EPI_ISL_511500 | | | |
| see above | Instituto Nacional de Saude (INSA) | Instituto Nacional de Saude (INSA) and Instituto Gulbenkian de Ciencia (IGC) | Borges et al |
| EPI_ISL_511577 | Hospital de Braga | Instituto Nacional de Saude (INSA) | Borges et al |
| EPI_ISL_511644 | Life and Health Sciences Research Institute, University of Minho, Braga | Instituto Nacional de Saude (INSA) | Borges et al |
| EPI_ISL_511677, EPI_ISL_511678, EPI_ISL_511691, EPI_ISL_511692, EPI_ISL_511693, EPI_ISL_511694, EPI_ISL_511695, EPI_ISL_511696 | Instituto Nacional de Saude (INSA) | Instituto Nacional de Saude (INSA) | Borges et al |
| EPI_ISL_511864, EPI_ISL_511875 | Johns Hopkins Hospital Department of Pathology | Johns Hopkins Hospital Department of Pathology | Peter M. Thielen, Thomas Mehoke, Shirlee Wohl, Srividya Ramakrishnan, Melanie Kirsche, Amanda Ernlund, Craig Howser, Kristina Zudock, Oluwaseun Falade-Nwulia, Norah Sadowski, Paul Morris, Mark Hopkins, Yunfan Fan, Nidia Trovao, Victoria Gniazdowski, Michael C. Schatz, Stuart C. Ray, Winston Timp, Heba H. Mostafa |
| EPI_ISL_511903 | Institute of Post Graduate Medical Education & Research | National Institute of Biomedical Genomics - DBT's PAN-INDIA 1000 SARS--CoV-2 RNA Genome Sequencing Consortium | Arindam Maitra, Aritra Biswas, Jayeeta Haldar, Raja Ray, Monimoy Banerjee, Saumitra Das |
| EPI_ISL_512422, EPI_ISL_512423, EPI_ISL_512424 | Centre for Enzyme Innovation, University of Portsmouth / Translational Research Laboratory, Portsmouth Hospitals NHS Trust | COVID-19 Genomics UK (COG-UK) Consortium | Angela Beckett,Yann Bourgeois,Garry Scarlett,Sharon Glaysher,Scott Elliott,Kelly Bicknell,Robert Impey,Allyson Lloyd,Sarah Wyllie,Ethan Butcher,Anoop Chauhan,Samuel Robson |
| EPI_ISL_512450, EPI_ISL_512451, EPI_ISL_512452, EPI_ISL_512453, EPI_ISL_512454, EPI_ISL_512455 | West of Scotland Specialist Virology Centre, NHSGGC / MRC-University of Glasgow Centre for Virus Research | COVID-19 Genomics UK (COG-UK) Consortium | Ana da Silva Filipe, Natasha Johnson, Kathy Smollett, Daniel Mair, Stephen Carmichael, Lily Tong, Jenna Nichols, Elihu Aranday-Cortes, Kirstyn Brunker, Yasmin Parr, Alice Broos, Kyriaki Nomikou; Sarah McDonald, Marc Niebel, Patawee Asamaphan; Richard Orton, Joseph Hughes, Sreenu Vattipally, David L Robertson; Alasdair MacLean, Rory Gunson; Kathy Li, Natasha Jesudason, Rajiv Shah, James Shepherd, Antonia Ho, Emma Thomson |
| EPI_ISL_512502 | Wales Specialist Virology Centre Sequencing lab: Pathogen | COVID-19 Genomics UK (COG-UK) Consortium | Catherine Moore, Johnathan Evans, Laura Gifford, Malorie Perry, Simon Cottrell, Angela Marchbank, Alec Birchley, Alexander Adams, Amy Gaskin, Bree |

| | | | |
|---|---|---|---|
| | Genomics Unit | | Gatica-Wilcox, Jason Coombes, Joel Southgate, Lauren Gilbert, Lee Graham, Nicole Pacchiarini, Sara Kumziene-Summerhayes, Sarah Taylor, Sophie Jones, Sara Rey, Matthew Bull, Joanne Watkins, Sally Corden, Tom Connor |
| EPI_ISL_512580, EPI_ISL_512581, EPI_ISL_512582, EPI_ISL_512583, EPI_ISL_512584, EPI_ISL_512585, EPI_ISL_512586, EPI_ISL_512587, EPI_ISL_512588, EPI_ISL_512589, EPI_ISL_512590, EPI_ISL_512591, EPI_ISL_512592, EPI_ISL_512593, EPI_ISL_512594, EPI_ISL_512595 | | | |
| see above | Florida Bureau of Public Health Laboratories | Florida Bureau of Public Health Laboratories | Sarah Schmedes, Jason Blanton |
| EPI_ISL_512713, EPI_ISL_512714, EPI_ISL_512748, EPI_ISL_512757 | PathWest Laboratory Medicine WA | PathWest Laboratory Medicine WA Microbial Surveillance Unit | PathWest Laboratory Medicine WA Microbial Surveillance Unit |
| EPI_ISL_512777, EPI_ISL_512778 | Utah Public Health Laboratory, Utah Public Health Laboratory Infectious Disease submission group | Utah Public Health Laboratory, Utah Public Health Laboratory Infectious Disease submission group | Butz,H.A., Young,E.L., Oakeson,K. |
| EPI_ISL_512816, EPI_ISL_512817, EPI_ISL_512818, EPI_ISL_512819 | Kenema Government Hospital, Ministry of Health and Sanitation | Kenema Government Hospital, Ministry of Health and Sanitation | Goba,A., Momoh,M., Sandi,J., Tomkins-Tinch,C., Siddle,K., Mehta,S., Oluniyi,P., Jalloh,S., Park,D., Andersen,K., Garry,R., Happi,C., Grant,D., Olawoye,I. |
| EPI_ISL_512822, EPI_ISL_512826, EPI_ISL_512830 | National Public Health Laboratory, National Centre for Infectious Diseases | National Public Health Laboratory, National Centre for Infectious Diseases | Mak TM, Octavia S, Zhou Z, Chavatte JM, Cui L, Lin RTP |
| EPI_ISL_512915 | Pathogen Genomics Lab King Abdullah University of Science and Technology(KAUST) | Pathogen Genomics Lab King Abdullah University of Science and Technology(KAUST) | Sharif Hala, Fadwa Alofi, Sara Mfarrej, Amit Kumar Subudhi, Rahul P Salunke, Fathia Ben Rached, Amanda Ooi, Luke Esau, Afrah Alsomali, Asim Khogeer, Jumana Taha, Abdulaziz Alahmadi, Kahled Algithami, Raeece Naeem, Anwar Hashem, Naif Almontashiri, Arnab Pain |
| EPI_ISL_512931, EPI_ISL_512933, EPI_ISL_512934, EPI_ISL_512935, EPI_ISL_512936, EPI_ISL_512937, EPI_ISL_512938, EPI_ISL_512939, EPI_ISL_512940, EPI_ISL_513162, EPI_ISL_513163, EPI_ISL_513164 | | | |
| see above | Pathogen Genomics Lab King Abdullah University of Science and Technology(KAUST) | Pathogen Genomics Lab King Abdullah University of Science and Technology(KAUST) | Fadwa Alofi, Sharif Hala, Rahul P Salunke, Sara Mfarrej, Amit Kumar Subudhi, Fathia Ben Rached, Amanda, Luke, Afrah Alsomali, Asim Khogeer, Jumana Taha, Abdulaziz Alahmadi, Kahled Algithami, Raeece Naeem, Anwar Hashem, Naif Almontashiri, Arnab Pain |
| EPI_ISL_513316 | South Eastern Area Laboratory Services (SEALS) | NSW Health Pathology - Institute of Clinical Pathology and Medical Research; Westmead Hospital; University of Sydney | CIDM-PH et al. |
| EPI_ISL_513513, EPI_ISL_513514 | Programa de Oncovirologia, Instituto Nacional de Câncer | Programa de Oncovirologia, Instituto Nacional de Câncer | Juliana D. Siqueira, Livia R. Goes, Brunna M. Alves, Claudia Cicala,James Arthos, João P.B. Viola, Andreia C. de Melo, Marcelo A. Soares |
| EPI_ISL_513638 | University of Washington Virology Lab | University of Washington Virology Lab | Pavitra Roychoudhury, Hong Xie, Lasata Shrestha, Amin Addetia, Truong Nguyen, Victoria M Rachleff, Meei-Li Huang, Keith R Jerome, Alexander Greninger |
| EPI_ISL_514103 | Viral Respiratory Lab, National Institute for Biomedical Research (INRB) | Pathogen Sequencing Lab, National Institute for Biomedical Research (INRB) | Placide Mbala-Kingebeni, Edith Nkwembe, Eddy Kinganda-Lusamaki, Amuri Aziza, Francisca Muyembe Mawete, Emmanuel Lokilo Lofiko, Catherine Pratt, Matthias Pauthner, Josh Quick, Allison Black, James Hadfield, Trevor Bedford, Ian Goodfellow, Andrew Rambaut, Nick Loman, Kristian Andersen, Michael Wiley, Steve Ahuka-Mundeke, Jean-Jacques Muyembe Tamfum |
| EPI_ISL_514226 | Laboratorio de Referencia Nacional de Virus Respiratorio. Instituto Nacional de Salud Perú | Laboratorio de Referencia Nacional de Biotecnología y Biología Molecular. Instituto Nacional de Salud Perú | Carlos Padilla Rojas, Karolyn Vega Chozo, Priscila Lope Pari, Omar Caceres Rey, Marco Galarza Perez, Maribel Huaringa Nuñez, Johanna Balbuena Torrez, Henri Bailon Calderon, Nancy Rojas Serrano |
| EPI_ISL_514227 | Laboratorio de Referencia Nacional de Virus Respiratorio. Instituto Nacional de Salud. Peru | Laboratorio de Referencia Nacional de Biotecnologia y Biologia Molecular.Instituto Nacional de Salud.Peru | Carlos Padilla Rojas, Karolyn Vega Chozo, Priscila Lope Pari, Omar Caceres Rey, Marco Galarza Perez, Maribel Huaringa Nuñez, Johanna Balbuena Torrez, Henri Bailon Calderon, Nancy Rojas Serrano |
| EPI_ISL_514265 | Laboratorio de Referencia Nacional de Virus Respiratorio. Instituto Nacional de Salud. Perú | Laboratorio de Referencia Nacional de Biotecnologia y Biologia Molecular.Instituto Nacional de Salud.Perú | Carlos Padilla Rojas, Karolyn Vega Chozo, Priscila Lope Pari, Omar Caceres Rey, Marco Galarza Perez, Maribel Huaringa Nuñez, Johanna Balbuena Torrez, Henri Bailon Calderon, Nancy Rojas Serrano |
| EPI_ISL_514315, EPI_ISL_514316 | Israel Central Virology laboratory | Israel Central Virology laboratory | Neta Zuckerman, Efrat Dahan Bucris, Oran Erster, Ella Mendelson, Michal Mandelboim |
| EPI_ISL_514469, EPI_ISL_514470, EPI_ISL_514472, EPI_ISL_514473, EPI_ISL_514474, EPI_ISL_514476, EPI_ISL_514477, EPI_ISL_514478, EPI_ISL_514479, EPI_ISL_514482 | Centre for Enzyme Innovation, University of Portsmouth / Translational Research Laboratory, Portsmouth Hospitals NHS Trust | COVID-19 Genomics UK (COG-UK) Consortium | Angela Beckett,Yann Bourgeois,Garry Scarlett,Sharon Glaysher,Scott Elliott,Kelly Bicknell,Robert Impey,Allyson Lloyd,Sarah Wyllie,Ethan Butcher,Anoop Chauhan,Samuel Robson |
| EPI_ISL_515203, EPI_ISL_515206, EPI_ISL_515208, EPI_ISL_515211, EPI_ISL_515214, EPI_ISL_515215, EPI_ISL_515216, EPI_ISL_515217, EPI_ISL_515218, EPI_ISL_515220, EPI_ISL_515222, EPI_ISL_515234, EPI_ISL_515235, EPI_ISL_515236, EPI_ISL_515243, EPI_ISL_515245, EPI_ISL_515246 | | | |
| see above | Laboratoire de microbiologie, Hopital de Verdun | Smith Laboratory, Centre de Recherche CHU Sainte-Justine | Martin Smith, Marieke Rozendaal, Ivan Pavlov |
| EPI_ISL_515348, EPI_ISL_515349, EPI_ISL_515350, EPI_ISL_515351, EPI_ISL_515352, EPI_ISL_515353, EPI_ISL_515354 | Nevada State Public Health Laboratory | Nevada State Public Health Laboratory | Richard Tillett, Joel R. Sevinsky, Paul Hartley, Heather Kerwin, David Jackson, Subhash C. Verma, Cyprian Rossetto, Andrew Gorzalski, Chris Laverdure, Natalie Crawford, Stephanie Van Hooser, and Mark Pandori |
| EPI_ISL_515908 | California Department of Public Health | California Department of Public Health | CDPH IDLB COVIDNet |
| EPI_ISL_516262, EPI_ISL_516263 | Michigan Department of Health and Human Services, Bureau of Laboratories | Michigan Department of Health and Human Services, Bureau of Laboratories | Blankenship HM, Riner D, Soehnlen MK |
| EPI_ISL_516628, EPI_ISL_516646, EPI_ISL_516647, EPI_ISL_516653, EPI_ISL_516654, EPI_ISL_516685, EPI_ISL_516686, EPI_ISL_516721, EPI_ISL_516722 | Laboratorio de Referencia Nacional de Virus Respiratorio. Centro Nacional de Salud Publica. Instituto Nacional de Salud Peru. | Laboratorio de Referencia Nacional de Biotecnología y Biología Molecular. Centro Nacional de Salud Publica. Instituto Nacional de Salud Peru. | Carlos Padilla Rojas, Karolyn Vega Chozo, Priscila Lope Pari, Omar Caceres Rey, Marco Galarza Perez, Maribel Huaringa Nuñez, Johanna Balbuena Torres, Henri Bailon Calderon, Nancy Rojas Serrano. |
| EPI_ISL_516807 | National Public Health Laboratory, National Centre for Infectious Diseases | National Public Health Laboratory, National Centre for Infectious Diseases | Mak TM, Octavia S, Zhou Z, Cui L, Lin RTP |
| EPI_ISL_516844 | North West London Pathology, Imperial College Healthcare NHS Trust | Wellcome Sanger Institute for the COVID-19 Genomics UK (COG-UK) consortium | Ling Li, Paul Randell, David Muir, Frankie Bolt, Alison Holmes, James Price, Aileen Rowan, Graham Taylor, Anjna Badhan, Carolina Herrera and Alex Alderton, Roberto Amato, Sonia Goncalves, Ewan Harrison, David K. Jackson, Ian Johnston, Dominic Kwiatkowski, Cordelia Langford, John Sillitoe on behalf of the Wellcome Sanger Institute COVID-19 Surveillance Team (http://www.sanger.ac.uk/covid-team) |
| EPI_ISL_517520, EPI_ISL_517521, EPI_ISL_517522, EPI_ISL_517523, EPI_ISL_517524, EPI_ISL_517525, EPI_ISL_517526, EPI_ISL_517527 | Centre for Enzyme Innovation, University of Portsmouth / Translational Research Laboratory, Portsmouth Hospitals NHS Trust | COVID-19 Genomics UK (COG-UK) Consortium | Angela Beckett,Yann Bourgeois,Garry Scarlett,Sharon Glaysher,Scott Elliott,Kelly Bicknell,Robert Impey,Allyson Lloyd,Sarah Wyllie,Ethan Butcher,Anoop Chauhan,Samuel Robson |
| EPI_ISL_517531, EPI_ISL_517686 | Laboratorio de Referencia Nacional de Virus Respiratorio. Centro Nacional de Salud Publica. Instituto Nacional de Salud Peru. | Laboratorio de Referencia Nacional de Biotecnología y Biología Molecular. Centro Nacional de Salud Publica. Instituto Nacional de Salud Peru. | Carlos Padilla Rojas, Karolyn Vega Chozo, Priscila Lope Pari, Omar Caceres Rey, Marco Galarza Perez, Maribel Huaringa Nuñez, Johanna Balbuena Torres, Henri Bailon Calderon, Nancy Rojas Serrano. |
| EPI_ISL_517786 | Florida Bureau of Public Health Laboratories | Florida Bureau of Public Health Laboratories | Sarah Schmedes, Jason Blanton |
| EPI_ISL_517959 | Laboratorio de Referencia Nacional de Virus Respiratorio. Centro Nacional de Salud Publica. Instituto Nacional de Salud Peru | Laboratorio de Referencia Nacional de Biotecnología y Biología Molecular. Centro Nacional de Salud Publica. Instituto Nacional de Salud Peru. | Carlos Padilla Rojas, Karolyn Vega Chozo, Priscila Lope Pari, Omar Caceres Rey, Marco Galarza Perez, Maribel Huaringa Nuñez, Johanna Balbuena Torres, Henri Bailon Calderon, Nancy Rojas Serrano. |
| EPI_ISL_520456 | Microbiological Diagnostic Unit - Public Health Laboratory (MDU-PHL) | MDU-PHL | Seemann T., Schultz M., Sait, M., Sherry, N. |
| EPI_ISL_520680, EPI_ISL_520718, EPI_ISL_520732, EPI_ISL_520733 | Mohammed Bin Rashid University of Medicine and Health Sciences | Al Jalila Genomics Center | Ahmad Abou Tayoun, Tom Loney, Hamda Khansaheb, Sathishkumar Ramaswamy, Divinlal Harilal, Zulfa Omar Deesi, Rupa Murthy Varghese, Hanan Al Suwaidi, Abdulmajeed Alkhaja, Mohammed Uddin, Rifat Hamoudi, Rabih Halwani, Abiola Catherine Senok, Qutayba Hamid, Norbert Nowotny, Alawi Alsheikh-Ali |

| | | | |
|---|---|---|---|
| EPI_ISL_521865, EPI_ISL_521866 | Victorian Infectious Diseases Reference Laboratory (VIDRL) | VIDRL and MDU-PHL | Caly L., Seemann T., Sait, M., Schultz M., Druce J., Sherry, N. |
| EPI_ISL_522556, EPI_ISL_522557, EPI_ISL_522558, EPI_ISL_522559, EPI_ISL_522560, EPI_ISL_522561, EPI_ISL_522576, EPI_ISL_522581, EPI_ISL_522608, EPI_ISL_522610, EPI_ISL_522616, EPI_ISL_522618, EPI_ISL_522640, EPI_ISL_522649, EPI_ISL_522660, EPI_ISL_522661, EPI_ISL_522662, EPI_ISL_522663, EPI_ISL_522664, EPI_ISL_522665, EPI_ISL_522677, EPI_ISL_522687, EPI_ISL_522689, EPI_ISL_522692, EPI_ISL_522693, EPI_ISL_522694, EPI_ISL_522695, EPI_ISL_522696, EPI_ISL_522697, EPI_ISL_522706, EPI_ISL_522708, EPI_ISL_522709, EPI_ISL_522710, EPI_ISL_522711, EPI_ISL_522712, EPI_ISL_522716, EPI_ISL_522717, EPI_ISL_522720, EPI_ISL_522722, EPI_ISL_522725, EPI_ISL_522726, EPI_ISL_522733, EPI_ISL_522734, EPI_ISL_522740, EPI_ISL_522748, EPI_ISL_522750, EPI_ISL_522751, EPI_ISL_522752, EPI_ISL_522760, EPI_ISL_522765 | | | |
| see above | Royal Hobart Hospital Microbiology Department | MDU-PHL | Cooley L., van Haeften R., Seemann T., Sait M., Schultz, M.B., Sherry N. |
| EPI_ISL_522858 | ULSS9 Distretto di San Bonifacio | Istituto Zooprofilattico Sperimentale delle Venezie | Adelaide Milani, Alessia Schivo, Annalisa Salviato, Erika Giorgia Quaranta, Ambra Pastori, Bianca Zecchin, Alice Fusaro, Isabella Monne, Calogero Terregino, Antonia Ricci |
| EPI_ISL_522860, EPI_ISL_522861, EPI_ISL_522862, EPI_ISL_522863, EPI_ISL_522864, EPI_ISL_522865, EPI_ISL_522866, EPI_ISL_522867 | ULSS9 Distretto di Bussolengo | Istituto Zooprofilattico Sperimentale delle Venezie | Adelaide Milani, Alessia Schivo, Annalisa Salviato, Erika Giorgia Quaranta, Ambra Pastori, Bianca Zecchin, Alice Fusaro, Isabella Monne, Calogero Terregino, Antonia Ricci |
| EPI_ISL_523972 | Hospital do Servidor Público Estadual Francisco Morato de Oliveira | Instituto Adolfo Lutz, Interdiciplinary Procedures Center, Strategic Laboratory | Claudio Tavares Sacchi, Claudia Regina Gonçalves, Erica Valessa Ramos Gomes |
| EPI_ISL_523976 | Hospital Municipal do Tatuape Carmino Caricchio | Instituto Adolfo Lutz, Interdiciplinary Procedures Center, Strategic Laboratory | Claudio Tavares Sacchi, Claudia Regina Gonçalves, Erica Valessa Ramos Gomes |
| EPI_ISL_523977 | Hosp. Municipal Prof. Dr. Alípio Corrêa Netto | Instituto Adolfo Lutz, Interdiciplinary Procedures Center, Strategic Laboratory | Claudio Tavares Sacchi, Claudia Regina Gonçalves, Erica Valessa Ramos Gomes |
| EPI_ISL_523978 | Hospital do Servidor Público Estadual Francisco Morato de Oliveira | Instituto Adolfo Lutz, Interdiciplinary Procedures Center, Strategic Laboratory | Claudio Tavares Sacchi, Claudia Regina Gonçalves, Erica Valessa Ramos Gomes |
| EPI_ISL_523981 | Hospital Sao Paulo de Ensino da Unifesp | Instituto Adolfo Lutz, Interdiciplinary Procedures Center, Strategic Laboratory | Claudio Tavares Sacchi, Claudia Regina Gonçalves, Erica Valessa Ramos Gomes |
| EPI_ISL_524067, EPI_ISL_524068 | Texas Department of State Health Services | Texas Department of State Health Services | Rashmi Tuladhar, Bonnie Oh, Cara Akrout, Jenny Zhang, Maliha Rahman, Anita Pokharel, Myong Koag, Chun Wang, Rachel Lee, Grace Kubin |
| EPI_ISL_524428, EPI_ISL_524429, EPI_ISL_524431 | Dirk Dittmer | Dirk Dittmer | Bailey,A.G., Caro-Vegas,C.P., Dittmer,D., Eason,A.B., Juarez,A., Landis,J.T., McNamara,R.P., Miller,M.B., Moorad,R., Pluta,L.J.,Seltzer,T.A., Thompson,C., Vahrson,W. and Villamor,F. |
| EPI_ISL_524493, EPI_ISL_524495, EPI_ISL_524497, EPI_ISL_524498, EPI_ISL_524501, EPI_ISL_524502, EPI_ISL_524503, EPI_ISL_524505, EPI_ISL_524506, EPI_ISL_524508, EPI_ISL_524509, EPI_ISL_524515, EPI_ISL_524522, EPI_ISL_524525, EPI_ISL_524528, EPI_ISL_524529, EPI_ISL_524530, EPI_ISL_524531, EPI_ISL_524537, EPI_ISL_524538, EPI_ISL_524541, EPI_ISL_524544, EPI_ISL_524546, EPI_ISL_524548, EPI_ISL_524551, EPI_ISL_524553, EPI_ISL_524558, EPI_ISL_524560, EPI_ISL_524564, EPI_ISL_524568, EPI_ISL_524570 | | | |
| see above | Department of Pathology, University of Cambridge | Wellcome Sanger Institute for the COVID-19 Genomics UK (COG-UK) consortium | Luke W Meredith, M. Estée Török , Myra Hosmillo, William L. Hamilton, Martin D. Curran, Theresa Feltwell, Grant Hall, Anna Yakovleva, Fahad A Khokhar, Charlotte J. Houldcroft, Laura G Caller, Aminu S. Jahun, Sarah L. Caddy, Ian Goodfellow; and Alex Alderton, Roberto Amato, Sonia Goncalves, Ewan Harrison, David K. Jackson, Ian Johnston, Dominic Kwiatkowski, Cordelia Langford, John Sillitoe on behalf of the Wellcome Sanger Institute COVID-19 Surveillance Team (http://www.sanger.ac.uk/covid-team) |
| EPI_ISL_524639, EPI_ISL_524649, EPI_ISL_524657, EPI_ISL_524664, EPI_ISL_524667, EPI_ISL_524672, EPI_ISL_524703 | North West London Pathology, Imperial College Healthcare NHS Trust | Wellcome Sanger Institute for the COVID-19 Genomics UK (COG-UK) consortium | Ling Li, Paul Randell, David Muir, Frankie Bolt, Alison Holmes, James Price, Aileen Rowan, Graham Taylor, Anjna Badhan, Carolina Herrera and Alex Alderton, Roberto Amato, Sonia Goncalves, Ewan Harrison, David K. Jackson, Ian Johnston, Dominic Kwiatkowski, Cordelia Langford, John Sillitoe on behalf of the Wellcome Sanger Institute COVID-19 Surveillance Team (http://www.sanger.ac.uk/covid-team) |
| EPI_ISL_524795 | Evandro Chagas Institute | Evandro Chagas Institute | Santos, M.C.; Silva, A.M.; Junior, W.D.C.; Barbagelata, L.S.; Ferreira, J.A.; Sousa, E.M.A.; da Silva, P.S.; Resque, H.R; Martins, L.C.; Sousa Junior, E.C.;Viana, G.M.R |
| EPI_ISL_524874, EPI_ISL_524875, EPI_ISL_524876 | MD PHL | MD PHL | Maryland Department of Health Laboratories Administration |
| EPI_ISL_525423 | Oman-National Influenza Center | Biotechnology & OMICs Laboratory | Sajjad Asaf, Samiha Al-Kharusi, Ahmed Al-Harrasi, Samira Al-Mahruqi, Adil Khan, Ahmed Al-Rawahi, Abdul Latif Khan, Amina Al-Jardani, Hanan Al-Kindi, Intisar Al-Shukri, Ahlam Al-Amri, Aisha Al-Amri, Aisha Al-Busaidi, Adil Al-Wahaibi, Seif Al-Abri. |
| EPI_ISL_525473 | Institute of Clinical Microbiology and Hygiene, University Hospital Regensburg | Institute of Clinical Microbiology and Hygiene, University Hospital Regensburg | Hiergeist, A. |
| EPI_ISL_525555, EPI_ISL_525558, EPI_ISL_525559, EPI_ISL_525560, EPI_ISL_525561, EPI_ISL_525562, EPI_ISL_525563, EPI_ISL_525564, EPI_ISL_525565, EPI_ISL_525566, EPI_ISL_525567, EPI_ISL_525568, EPI_ISL_525569, EPI_ISL_525574 | | | |
| see above | Istituto Zooprofilattico Sperimentale Puglia e Basilicata; Dipartimento di Bioscienze, Biotecnologie e Biofarmaceutica dell'Università degli Studi di Bari "A.Moro"; Istituto di Biomembrane. Bioenergetica e Biotecnologie Molecolari del Consiglio Nazionale delle Ricerche di Bari | Beaconlab (Bioinformatics, Evolution and Comparative Genomics lab), Dept of Biosciences, University on Milan | Parisi A.,Pesole G., Manzari C., Chiara M |
| EPI_ISL_525578, EPI_ISL_525658, EPI_ISL_525660, EPI_ISL_525661, EPI_ISL_525662, EPI_ISL_525663, EPI_ISL_525664, EPI_ISL_525665, EPI_ISL_525666, EPI_ISL_525667, EPI_ISL_525668, EPI_ISL_525669, EPI_ISL_525670, EPI_ISL_525671, EPI_ISL_525672, EPI_ISL_525673, EPI_ISL_525677, EPI_ISL_525678, EPI_ISL_525680 | | | |
| see above | Wadsworth Center, New York State Department of Health | Wadsworth Center, New York State Department of Health | Kirsten St. George, Daryl M. Lamson, Sara Griesemer, Jonathan Plitnick, Navjot Singh, Matthew D. Shudt, Erica Lasek-Nesselquist |
| EPI_ISL_525782, EPI_ISL_525783, EPI_ISL_525784, EPI_ISL_525785 | Texas Department of State Health Services | Texas Department of State Health Services | Jenny Zhang, Rashmi Tuladhar, Bonnie Oh, Maliha Rahman, Anita Pokharel, Myong Koag, Chun Wang, Rachel Lee, Grace Kubin |
| EPI_ISL_526147, EPI_ISL_526148 | South Eastern Area Laboratory Services (SEALS) | NSW Health Pathology - Institute of Clinical Pathology and Medical Research; Westmead Hospital; University of Sydney | CIDM-PH et al. |
| EPI_ISL_526215 | Hungarian Defence Forces Military Medical Centre | National Laboratory of Virology, Szentágothai Research Centre | Endre Gábor Tóth, Balázs Somogyi, Bálint Eszenyi, Ferenc Jakab, Gábor Kemenesi |
| EPI_ISL_526459, EPI_ISL_526460, EPI_ISL_526461, EPI_ISL_526462 | Virology Department, Royal Infirmary of Edinburgh, NHS Lothian / School of Biological Sciences, University of Edinburgh / Institute of Genetics and Molecular Medicine, University of Edinburgh | COVID-19 Genomics UK (COG-UK) Consortium | McHugh M, Dewar R, Rooke S, Gallagher M, Balcaza C, O'Toole Á, Scher E, Hill V, McCrone JT, Colquhoun R, Yu X, Jackson B, Rambaut A, Williams TC, Templeton K |
| EPI_ISL_526836, EPI_ISL_526837, EPI_ISL_526838, EPI_ISL_526839, EPI_ISL_526840 | Virginia DCLS | Virginia DCLS | Virginia DCLS |
| EPI_ISL_526959 | Instituto Nacional de Salud, Bogotá, Colombia | Instituto Nacional de Salud, Bogotá, Colombia | Katherine Laiton-Donato, Diego A. Álvarez-Díaz, Carlos Franco-Muñoz, Mauricio Pacheco-Montealegre, Jonathan Reales, Diego Andrés Prada, Jose A. Usme-Ciro, Zulma M. Cucunubá, Christian Julian VillabonaArenas, Liz Villabona-Arenas, Sussy Echeverria, Astrid C. Flórez, Carolina Ferro, Diana Marcela Walteros-Acero, Franklin Prieto, Carlos Andrés Durán, Martha Lucia Ospina Martinez, Marcela Mercado-Reyes |
| EPI_ISL_527017, EPI_ISL_527037, EPI_ISL_527038, EPI_ISL_527057, EPI_ISL_527058, EPI_ISL_527061 | Area of Virology, Serology and Virology Division (SAViD), New South Wales Health Pathology Randwick | Area of Virology, Serology and Virology Division (SAViD), New South Wales Health Pathology Randwick | Rawlinson, W. |
| EPI_ISL_527370, EPI_ISL_527371, EPI_ISL_527372 | National Public Health Laboratory, National Centre for Infectious Diseases | National Public Health Laboratory, National Centre for Infectious Diseases | Mak TM, Octavia S, Zhou Z, Cui L, Lin RTP |

| | | | |
|---|---|---|---|
| EPI_ISL_527380 | Istituto Zooprofilattico Sperimentale Puglia e Basilicata; Dipartimento di Bioscienze, Biotecnologie e Biofarmaceutica dell'Università degli Studi di Bari "A.Moro"; Istituto di Biomembrane. Bioenergetica e Biotecnologie Molecolari del Consiglio Nazionale delle Ricerche di Bari | Beaconlab (Bioinformatics, Evolution and Comparative Genomics lab), Dept of Biosciences, University on Milan | Parisi A.,Pesole G., Manzari C., Chiara M |
| EPI_ISL_527402, EPI_ISL_527404, EPI_ISL_527405, EPI_ISL_527409, EPI_ISL_527411, EPI_ISL_527414, EPI_ISL_527415, EPI_ISL_527417, EPI_ISL_527421, EPI_ISL_527428, EPI_ISL_527434, EPI_ISL_527437, EPI_ISL_527439, EPI_ISL_527443, EPI_ISL_527445, EPI_ISL_527447, EPI_ISL_527452, EPI_ISL_527455, EPI_ISL_527457, EPI_ISL_527460, EPI_ISL_527466, EPI_ISL_527469, EPI_ISL_527473, EPI_ISL_527475, EPI_ISL_527483 | | | |
| see above | Colorado State University - Ebel Lab | Colorado State University - Ebel Lab | Greg Ebel et al. |
| EPI_ISL_527711, EPI_ISL_527712, EPI_ISL_527713, EPI_ISL_527715 | MN PHL Division, Minnesota Department of Health | Pathogen Discovery, Respiratory Viruses Branch, Division of Viral Diseases, Centers for Disease Control and Prevention | Yan Li, Anna Montmayeur, Jing Zhang, Krista Queen, Ying Tao, Anna Uehara, Rachel Marine, Clinton R. Paden, Haibin Wang, Suxiang Tong |
| EPI_ISL_527737 | WI State Laboratory of Hygiene | Pathogen Discovery, Respiratory Viruses Branch, Division of Viral Diseases, Centers for Disease Control and Prevention | Ying Tao, Jing Zhang, Yan Li, Krista Queen, Anna Uehara, Clinton Paden, Haibin Wang, Suxiang Tong |
| EPI_ISL_527819 | Centro de Investigaciones, Universidad de Especialidades Espíritu Santo | Institute of Microbiology, Universidad San Francisco de Quito | Derly Andrade, Juan Carlos Fernandez, Belén Prado-Vivar, Sully Márquez, Juan José Guadalupe, Monica Becerra-Wong, Bernardo Gutiérrez, Gabriel Morey, Ruben Armas, Jose Pedro Barberan, Fernando Espinoza, Edith Lopez, Verónica Barragán, Patricio Rojas-Silva, Gabriel Trueba, Michelle Grunauer, Paúl Cárdenas |
| EPI_ISL_527863 | Hospital Municipal do Tatuape Carmino Caricchio | Instituto Adolfo Lutz, Interdiciplinary Procedures Center, Strategic Laboratory | Claudio Tavares Sacchi, Claudia Regina Gonçalves, Erica Valessa Ramos Gomes |
| EPI_ISL_528605, EPI_ISL_528606 | National Genomics Core-Center for DNA Fingerprinting and Diagnostics | National Genomics Core- Center for DNA Fingerprinting and Diagnostics (NGC-CDFD)- DBT's PAN-INDIA-1000 Genome consortium | Ashwin Dalal, Bala Pratyusha, Heena Shah, G Shashikanth, Vinay Donipadi, K.Manohar, Madhumohan Rao,Neeraj Kumar, Niteen Pathak, Pradipta Hore, Rahul Baroi, Sayantan Goswami, Shaffiqu T S, Shalini Aricthota, Sobhan Babu, R Harinarayanan, Rashna Bhandari, Murali Dharan Bashyam, Debashish Mitra, Divya Vashisht |
| EPI_ISL_528738 | Malaysia Genome Institute | Malaysia Genome Institute | Mohd Noor Mat Isa, Irni Suhayu Sapian, Yusuf Muhammad Noor, Nurhezreen Md Iqbal, Mohd Faizal Abu Bakar, Enizza Kasim, Shamsidar Sopie, Siti Noraini Othman, Azrin Ahmad, Nor Azfa Johari, Shahrul Hisham Zainal Ariffin |
| EPI_ISL_529017 | RSA San Raffaele Sulmona | Istitituto Zooprofilattico Sperimentale dell'Abruzzo e Molise "G.Caporale" | Lorusso A, Marcacci M, Di Domenico M, Curini V, Ancora M, Cammà C, Rinaldi A, Mangone I, Di Pasquale A, Puglia I, Savini G. |
| EPI_ISL_529036 | Wadsworth Center, New York State Department of Health | Wadsworth Center, New York State Department of Health | Kirsten St. George, Daryl M. Lamson, Sara Griesemer, Jonathan Plitnick, Navjot Singh, Matthew D. Shudt, Erica Lasek-Nesselquist |
| EPI_ISL_529164 | Department of Immunology, The Scripps Research Institute | Andersen lab at Scripps Research | Allison Smither, Gilberto Sabino-Santos, Patricia Snarski, Lilia Melnik, Antoinette Bell, Kaylynn Genemaras, Arnaud Drouin, Dahlene Fusco, Robert Garry with SEARCH Alliance San Diego |
| EPI_ISL_529303, EPI_ISL_529343, EPI_ISL_529491, EPI_ISL_529669, EPI_ISL_529670, EPI_ISL_529671 | Centre for Enzyme Innovation, University of Portsmouth / Translational Research Laboratory, Portsmouth Hospitals NHS Trust | COVID-19 Genomics UK (COG-UK) Consortium | Angela Beckett,Yann Bourgeois,Garry Scarlett,Sharon Glaysher,Scott Elliott,Kelly Bicknell,Robert Impey,Allyson Lloyd,Sarah Wyllie,Ethan Butcher,Anoop Chauhan,Samuel Robson |
| EPI_ISL_529979, EPI_ISL_529980, EPI_ISL_529981, EPI_ISL_529984, EPI_ISL_529985 | Hospital Universitario 12 de Octubre | Hospital Universitario 12 de Octubre | Raúl Recio, Sara González, Esther Viedma, Elias Dahdouh, Fernando Lázaro, Natalia Stella, Julio García, Juan Carlos Galán, Rafael Cantón, Mª Dolores Folgueira, Rafael Delgado, Jesús Mingorance |
| EPI_ISL_529993, EPI_ISL_529994, EPI_ISL_529995, EPI_ISL_529998, EPI_ISL_529999, EPI_ISL_530000, EPI_ISL_530001, EPI_ISL_530002 | Hospital Universitario 12 de Octubre | Hospital Universitario 12 de Octubre | Sara González, Esther Viedma, Raúl Recio, Elias Dahdouh, Fernando Lázaro, Natalia Stella, Julio García, Juan Carlos Galán, Rafael Cantón, Mª Dolores Folgueira, Rafael Delgado, Jesús Mingorance |
| EPI_ISL_530044, EPI_ISL_530045, EPI_ISL_530046, EPI_ISL_530047, EPI_ISL_530048, EPI_ISL_530049, EPI_ISL_530050, EPI_ISL_530051 | Hospital Universitario La Paz | Hospital Universitario La Paz | Elias Dahdouh, Sara González, Raúl Recio, Fernando Lázaro, Esther Viedma, Natalia Stella, Julio García, Juan Carlos Galán, Rafael Cantón, Mª Dolores Folgueira, Rafael Delgado, Jesús Mingorance |
| EPI_ISL_530216, EPI_ISL_530217, EPI_ISL_530218 | Minnesota Department of Health, Public Health Laboratory | Minnesota Department of Health, Public Health Laboratory | Matt Plumb, Jacob Garfin, and Xiong Wang |
| EPI_ISL_530233, EPI_ISL_530234, EPI_ISL_530235, EPI_ISL_530236 | Queensland Health Forensic and Scientific Services, Public Health Virology | Public Health Virology Laboratory, Forensic and Scientific Services, Queensland Health | Son Nguyen et al |
| EPI_ISL_534769, EPI_ISL_534775, EPI_ISL_534780, EPI_ISL_534783, EPI_ISL_534784, EPI_ISL_534787, EPI_ISL_534789, EPI_ISL_534791, EPI_ISL_534794, EPI_ISL_534795, EPI_ISL_534796, EPI_ISL_534799, EPI_ISL_534800, EPI_ISL_534804, EPI_ISL_534812, EPI_ISL_534819, EPI_ISL_534838, EPI_ISL_534864, EPI_ISL_534869, EPI_ISL_534873, EPI_ISL_534876, EPI_ISL_534888, EPI_ISL_534892, EPI_ISL_534917, EPI_ISL_534919, EPI_ISL_534936, EPI_ISL_534956, EPI_ISL_534979, EPI_ISL_534987, EPI_ISL_534989, EPI_ISL_535002, EPI_ISL_535018 | | | |
| see above | Oxford Viromics, NDM, University of Oxford; Oxford University Hospitals; Basingstoke and North Hampshire Hospital | COVID-19 Genomics UK (COG-UK) Consortium | Tanya Golubchik, David Bonsall, George Macintyre, Amy Trebes, Mariateresa de Cesare, Catrin Moore, Alex Mobbs, Anita Justice, Robert Shaw, Monique Andersson, Timothy Peto, Emma Wise, Nathan Moore, Jessica Lynch, Nick Cortes, Matilde Mori, Stephen Kidd, David Buck, John Todd, Christophe Fraser |
| EPI_ISL_535364 | Oklahoma Animal Disease Diagnostic Laboratory | Oklahoma Animal Disease Diagnostic Laboratory | Sai Narayanan, John C Ritchey, Girish Patil, Teluguakula Narasaraju, Sunil More, Jerry Malayer, Jeremiah Saliki, Anil Kaul, Akhilesh Ramachandran |
| EPI_ISL_536448, EPI_ISL_536450 | National Public Health Laboratory, National Centre for Infectious Diseases | National Public Health Laboratory, National Centre for Infectious Diseases | Mak TM, Octavia S, Zhou Z, Cui L, Lin RTP |
| EPI_ISL_537326 | Universidad de León | SeqCOVID-SPAIN consortium/IBV(CSIC) | Ana Carvajal, Vicente Martín, Héctor Argüello, Juan M. Fregeneda, Tania Fernández-Villa, Antonio J. Molina and SeqCOVID-SPAIN consortium |
| EPI_ISL_537467, EPI_ISL_537471, EPI_ISL_537472, EPI_ISL_537473, EPI_ISL_537475, EPI_ISL_537476, EPI_ISL_537477, EPI_ISL_537479, EPI_ISL_537480, EPI_ISL_537481, EPI_ISL_537482, EPI_ISL_537483, EPI_ISL_537485, EPI_ISL_537486, EPI_ISL_537487, EPI_ISL_537488, EPI_ISL_537489, EPI_ISL_537490, EPI_ISL_537500, EPI_ISL_537501, EPI_ISL_537502, EPI_ISL_537503, EPI_ISL_537504, EPI_ISL_537509, EPI_ISL_537513 | | | |
| see above | UCLA Pathology Clinical Microbiology Lab | Kruglyak Lab | Guo et al. |
| EPI_ISL_537727 | Hospital Universitario de Gran Canaria Dr. Negrín | SeqCOVID-SPAIN consortium/IBV(CSIC) | M. Carmen Pérez González, Francisco J. Chamizo López, Ana Bordes Benítez and SeqCOVID-SPAIN consortium |
| EPI_ISL_538120, EPI_ISL_538121, EPI_ISL_538122, EPI_ISL_538123, EPI_ISL_538126, EPI_ISL_538127, EPI_ISL_538128, EPI_ISL_538132, EPI_ISL_538134, EPI_ISL_538135, EPI_ISL_538136, EPI_ISL_538137, EPI_ISL_538139, EPI_ISL_538140, EPI_ISL_538141, EPI_ISL_538142, EPI_ISL_538145, EPI_ISL_538146, EPI_ISL_538147 | | | |
| see above | Servicio de Microbiología y Parasitología clínica. UCEIMP. Hospital Universitario Virgen del Rocío/IBIS/CSIC/US | SeqCOVID-SPAIN consortium/IBV(CSIC) | Guillermo Martín Gutiérrez, Ángel Rodríguez Villodres, Lidia Gálvez Benitez, Verónica González Galán, Javier Aznar Martín and SeqCOVID-SPAIN consortium |
| EPI_ISL_538371, EPI_ISL_538372, EPI_ISL_538373, EPI_ISL_538374, EPI_ISL_538375, EPI_ISL_538376, EPI_ISL_538377, EPI_ISL_538378, EPI_ISL_538379, EPI_ISL_538380, EPI_ISL_538381, EPI_ISL_538382, EPI_ISL_538383, EPI_ISL_538384, EPI_ISL_538385, EPI_ISL_538386 | | | |
| see above | Kingston Health Sciences Centre / Queen's University | Ontario Institute for Cancer Research | Prameet M. Sheth, Calvin Sjaarda, Robert Colautti, Katya Douchant, Ilinca Lungu, Bernard Lam, Paul Krzyzanowski, Michael Laszloffy, Lawrence E Heisler, Richard de Borja, Jared T. Simpson |
| EPI_ISL_538445, EPI_ISL_538485, EPI_ISL_538486, EPI_ISL_538487, EPI_ISL_538488, EPI_ISL_538489, EPI_ISL_538490, EPI_ISL_538491, EPI_ISL_538492, EPI_ISL_538493, EPI_ISL_538494, EPI_ISL_538495, EPI_ISL_538496, EPI_ISL_538497 | | | |
| see above | Department of Laboratory Medicine, Tan Tock Seng Hospital | Department of Laboratory Medicine, Tan Tock Seng Hospital | Chen YYC, Zair X, Lim JX, Li C, Tang WY, Maurer-Stroh S, Barkham TMS, Nagarajan N, Sessions OM |
| EPI_ISL_538568, EPI_ISL_538569, EPI_ISL_538570, EPI_ISL_538571, EPI_ISL_538572, EPI_ISL_538573, EPI_ISL_538574, EPI_ISL_538575, EPI_ISL_538576, EPI_ISL_538582, EPI_ISL_538584, EPI_ISL_538585, EPI_ISL_538586, EPI_ISL_538587, EPI_ISL_538589, EPI_ISL_538590, EPI_ISL_538592, EPI_ISL_538593, EPI_ISL_538594, EPI_ISL_538595, EPI_ISL_538602 | | | |
| see above | Servicio de Microbiología y Parasitología clínica. UCEIMP. Hospital Universitario Virgen del Rocío/IBIS/CSIC/US | SeqCOVID-SPAIN consortium/IBV(CSIC) | Guillermo Martín Gutiérrez, Ángel Rodríguez Villodres, Lidia Gálvez Benitez, Verónica González Galán, Javier Aznar Martín and SeqCOVID-SPAIN consortium |
| EPI_ISL_538748, EPI_ISL_538749, EPI_ISL_538750, EPI_ISL_538752, EPI_ISL_538753, EPI_ISL_538755, EPI_ISL_538756, EPI_ISL_538758, EPI_ISL_538759, EPI_ISL_538760, EPI_ISL_538761, EPI_ISL_538763, EPI_ISL_538764, EPI_ISL_538767, EPI_ISL_538768, EPI_ISL_538770, EPI_ISL_538771, EPI_ISL_538772, | | | |

| | | | |
|---|---|---|---|
| EPI_ISL_538774, EPI_ISL_538775, EPI_ISL_538776, EPI_ISL_538777, EPI_ISL_538778, EPI_ISL_538780, EPI_ISL_538781, EPI_ISL_538782, EPI_ISL_538783, EPI_ISL_538784, EPI_ISL_538785, EPI_ISL_538788, EPI_ISL_538790, EPI_ISL_538793, EPI_ISL_538796, EPI_ISL_538797, EPI_ISL_538800, EPI_ISL_538801, EPI_ISL_538803, EPI_ISL_538806, EPI_ISL_538809, EPI_ISL_538810 | | | |
| see above | Leeds Teaching Hospitals NHS Trust and Public Health England, National Infection Service (Leeds laboratory) | Wellcome Sanger Institute for the COVID-19 Genomics UK (COG-UK) consortium | Louissa Macfarlane-Smith, Holli Carden, Katherine L. Harper, Antony Hale and Alex Alderton, Roberto Amato, Sonia Goncalves, Ewan Harrison, David K. Jackson, Ian Johnston, Dominic Kwiatkowski, Cordelia Langford, John Sillitoe on behalf of the Wellcome Sanger Institute COVID-19 Surveillance Team |
| EPI_ISL_538811 | Leeds Teaching Hospitals NHS Trust and Public Health England, National Infection Service (Leeds laboratory) | Wellcome Sanger Institute for the COVID-19 Genomics UK (COG-UK) Consortium | Louissa Macfarlane-Smith, Holli Carden, Katherine L. Harper, Antony Hale and Alex Alderton, Roberto Amato, Sonia Goncalves, Ewan Harrison, David K. Jackson, Ian Johnston, Dominic Kwiatkowski, Cordelia Langford, John Sillitoe on behalf of the Wellcome Sanger Institute COVID-19 Surveillance Team |
| EPI_ISL_538813, EPI_ISL_538817, EPI_ISL_538824, EPI_ISL_538826, EPI_ISL_538828, EPI_ISL_538829, EPI_ISL_538833, EPI_ISL_538834, EPI_ISL_538836, EPI_ISL_538838, EPI_ISL_538839 | | | |
| see above | Leeds Teaching Hospitals NHS Trust and Public Health England, National Infection Service (Leeds laboratory) | Wellcome Sanger Institute for the COVID-19 Genomics UK (COG-UK) consortium | Louissa Macfarlane-Smith, Holli Carden, Katherine L. Harper, Antony Hale and Alex Alderton, Roberto Amato, Sonia Goncalves, Ewan Harrison, David K. Jackson, Ian Johnston, Dominic Kwiatkowski, Cordelia Langford, John Sillitoe on behalf of the Wellcome Sanger Institute COVID-19 Surveillance Team |
| EPI_ISL_538841 | Leeds Teaching Hospitals NHS Trust and Public Health England, National Infection Service (Leeds laboratory) | Wellcome Sanger Institute for the COVID-19 Genomics UK (COG-UK) Consortium | Louissa Macfarlane-Smith, Holli Carden, Katherine L. Harper, Antony Hale and Alex Alderton, Roberto Amato, Sonia Goncalves, Ewan Harrison, David K. Jackson, Ian Johnston, Dominic Kwiatkowski, Cordelia Langford, John Sillitoe on behalf of the Wellcome Sanger Institute COVID-19 Surveillance Team |
| EPI_ISL_538843, EPI_ISL_538845, EPI_ISL_538846, EPI_ISL_538850, EPI_ISL_538853, EPI_ISL_538854, EPI_ISL_538855, EPI_ISL_538856, EPI_ISL_538858, EPI_ISL_538859, EPI_ISL_538860, EPI_ISL_538861, EPI_ISL_538862, EPI_ISL_538864, EPI_ISL_538865 | | | |
| see above | Leeds Teaching Hospitals NHS Trust and Public Health England, National Infection Service (Leeds laboratory) | Wellcome Sanger Institute for the COVID-19 Genomics UK (COG-UK) consortium | Louissa Macfarlane-Smith, Holli Carden, Katherine L. Harper, Antony Hale and Alex Alderton, Roberto Amato, Sonia Goncalves, Ewan Harrison, David K. Jackson, Ian Johnston, Dominic Kwiatkowski, Cordelia Langford, John Sillitoe on behalf of the Wellcome Sanger Institute COVID-19 Surveillance Team |
| EPI_ISL_538866 | Leeds Teaching Hospitals NHS Trust and Public Health England, National Infection Service (Leeds laboratory) | Wellcome Sanger Institute for the COVID-19 Genomics UK (COG-UK) Consortium | Louissa Macfarlane-Smith, Holli Carden, Katherine L. Harper, Antony Hale and Alex Alderton, Roberto Amato, Sonia Goncalves, Ewan Harrison, David K. Jackson, Ian Johnston, Dominic Kwiatkowski, Cordelia Langford, John Sillitoe on behalf of the Wellcome Sanger Institute COVID-19 Surveillance Team |
| EPI_ISL_538867, EPI_ISL_538870, EPI_ISL_538872, EPI_ISL_538874, EPI_ISL_538875, EPI_ISL_538876, EPI_ISL_538877, EPI_ISL_538878, EPI_ISL_538883, EPI_ISL_538885, EPI_ISL_538887, EPI_ISL_538888, EPI_ISL_538889, EPI_ISL_538890, EPI_ISL_538892, EPI_ISL_538896, EPI_ISL_538898 | | | |
| see above | Leeds Teaching Hospitals NHS Trust and Public Health England, National Infection Service (Leeds laboratory) | Wellcome Sanger Institute for the COVID-19 Genomics UK (COG-UK) consortium | Louissa Macfarlane-Smith, Holli Carden, Katherine L. Harper, Antony Hale and Alex Alderton, Roberto Amato, Sonia Goncalves, Ewan Harrison, David K. Jackson, Ian Johnston, Dominic Kwiatkowski, Cordelia Langford, John Sillitoe on behalf of the Wellcome Sanger Institute COVID-19 Surveillance Team |
| EPI_ISL_538899 | Leeds Teaching Hospitals NHS Trust and Public Health England, National Infection Service (Leeds laboratory) | Wellcome Sanger Institute for the COVID-19 Genomics UK (COG-UK) Consortium | Louissa Macfarlane-Smith, Holli Carden, Katherine L. Harper, Antony Hale and Alex Alderton, Roberto Amato, Sonia Goncalves, Ewan Harrison, David K. Jackson, Ian Johnston, Dominic Kwiatkowski, Cordelia Langford, John Sillitoe on behalf of the Wellcome Sanger Institute COVID-19 Surveillance Team |
| EPI_ISL_538902, EPI_ISL_538908, EPI_ISL_538909, EPI_ISL_538910, EPI_ISL_538911, EPI_ISL_538914, EPI_ISL_538918, EPI_ISL_538919, EPI_ISL_538921, EPI_ISL_538922, EPI_ISL_538923, EPI_ISL_538924, EPI_ISL_538929, EPI_ISL_538931, EPI_ISL_538937, EPI_ISL_538942, EPI_ISL_538943 | | | |
| see above | Leeds Teaching Hospitals NHS Trust and Public Health England, National Infection Service (Leeds laboratory) | Wellcome Sanger Institute for the COVID-19 Genomics UK (COG-UK) consortium | Louissa Macfarlane-Smith, Holli Carden, Katherine L. Harper, Antony Hale and Alex Alderton, Roberto Amato, Sonia Goncalves, Ewan Harrison, David K. Jackson, Ian Johnston, Dominic Kwiatkowski, Cordelia Langford, John Sillitoe on behalf of the Wellcome Sanger Institute COVID-19 Surveillance Team |
| EPI_ISL_538945 | Leeds Teaching Hospitals NHS Trust and Public Health England, National Infection Service (Leeds laboratory) | Wellcome Sanger Institute for the COVID-19 Genomics UK (COG-UK) Consortium | Louissa Macfarlane-Smith, Holli Carden, Katherine L. Harper, Antony Hale and Alex Alderton, Roberto Amato, Sonia Goncalves, Ewan Harrison, David K. Jackson, Ian Johnston, Dominic Kwiatkowski, Cordelia Langford, John Sillitoe on behalf of the Wellcome Sanger Institute COVID-19 Surveillance Team |
| EPI_ISL_538949, EPI_ISL_538959, EPI_ISL_538964, EPI_ISL_538967, EPI_ISL_538969, EPI_ISL_538970, EPI_ISL_538973, EPI_ISL_538977, EPI_ISL_538978, EPI_ISL_538981, EPI_ISL_538982, EPI_ISL_538984, EPI_ISL_538985, EPI_ISL_539052, EPI_ISL_539060, EPI_ISL_539061, EPI_ISL_539079, EPI_ISL_539091, EPI_ISL_539125, EPI_ISL_539126 | | | |
| see above | Leeds Teaching Hospitals NHS Trust and Public Health England, National Infection Service (Leeds laboratory) | Wellcome Sanger Institute for the COVID-19 Genomics UK (COG-UK) consortium | Louissa Macfarlane-Smith, Holli Carden, Katherine L. Harper, Antony Hale and Alex Alderton, Roberto Amato, Sonia Goncalves, Ewan Harrison, David K. Jackson, Ian Johnston, Dominic Kwiatkowski, Cordelia Langford, John Sillitoe on behalf of the Wellcome Sanger Institute COVID-19 Surveillance Team |
| EPI_ISL_539157 | Leeds Teaching Hospitals NHS Trust and Public Health England, National Infection Service (Leeds laboratory) | Wellcome Sanger Institute for the COVID-19 Genomics UK (COG-UK) Consortium | Louissa Macfarlane-Smith, Holli Carden, Katherine L. Harper, Antony Hale and Alex Alderton, Roberto Amato, Sonia Goncalves, Ewan Harrison, David K. Jackson, Ian Johnston, Dominic Kwiatkowski, Cordelia Langford, John Sillitoe on behalf of the Wellcome Sanger Institute COVID-19 Surveillance Team |
| EPI_ISL_539165, EPI_ISL_539168, EPI_ISL_539175, EPI_ISL_539177, EPI_ISL_539180, EPI_ISL_539183, EPI_ISL_539188, EPI_ISL_539193, EPI_ISL_539199, EPI_ISL_539204, EPI_ISL_539205, EPI_ISL_539210 | | | |
| see above | Leeds Teaching Hospitals NHS Trust and Public Health England, National Infection Service (Leeds laboratory) | Wellcome Sanger Institute for the COVID-19 Genomics UK (COG-UK) consortium | Louissa Macfarlane-Smith, Holli Carden, Katherine L. Harper, Antony Hale and Alex Alderton, Roberto Amato, Sonia Goncalves, Ewan Harrison, David K. Jackson, Ian Johnston, Dominic Kwiatkowski, Cordelia Langford, John Sillitoe on behalf of the Wellcome Sanger Institute COVID-19 Surveillance Team |
| EPI_ISL_539213 | Leeds Teaching Hospitals NHS Trust and Public Health England, National Infection Service (Leeds laboratory) | Wellcome Sanger Institute for the COVID-19 Genomics UK (COG-UK) Consortium | Louissa Macfarlane-Smith, Holli Carden, Katherine L. Harper, Antony Hale and Alex Alderton, Roberto Amato, Sonia Goncalves, Ewan Harrison, David K. Jackson, Ian Johnston, Dominic Kwiatkowski, Cordelia Langford, John Sillitoe on behalf of the Wellcome Sanger Institute COVID-19 Surveillance Team |
| EPI_ISL_539214, EPI_ISL_539216 | Leeds Teaching Hospitals NHS Trust and Public Health England, National Infection Service (Leeds laboratory) | Wellcome Sanger Institute for the COVID-19 Genomics UK (COG-UK) consortium | Louissa Macfarlane-Smith, Holli Carden, Katherine L. Harper, Antony Hale and Alex Alderton, Roberto Amato, Sonia Goncalves, Ewan Harrison, David K. Jackson, Ian Johnston, Dominic Kwiatkowski, Cordelia Langford, John Sillitoe on behalf of the Wellcome Sanger Institute COVID-19 Surveillance Team |
| EPI_ISL_539272, EPI_ISL_539273, EPI_ISL_539274, EPI_ISL_539275, EPI_ISL_539276, EPI_ISL_539277, EPI_ISL_539278, EPI_ISL_539279, EPI_ISL_539280, EPI_ISL_539281, EPI_ISL_539282, EPI_ISL_539283 | | | |
| see above | Hospital Universitario de La Ribera (Alzira, València) | SeqCOVID-SPAIN consortium/IBV(CSIC) | Olalla Martínez Macias, Julia González and SeqCOVID-SPAIN consortium |
| EPI_ISL_539312, EPI_ISL_539323 | KWR Watercycle Research Institute | Erasmus Medical Center | Ray Izquierdo-Lara, Goffe Elsinga, Leo Heijnen, Bas B. Oude Munnink, Claudia M. E. Schapendonk, David Nieuwenhuijse, Matthijs Kon, Lu Lu, Frank M. Aarestrup, Samantha Lycett, Gertjan Medema, Marion P.G. Koopmans, Miranda de Graaf |
| EPI_ISL_539778, EPI_ISL_539779 | National Institute of Public Health (Czech Republic) | State Veterinary Institute Prague | Nagy, A; Jirincova, H; Novakova, L; Trnka, D; Vecerova, J. |
| EPI_ISL_539794 | Wyoming Public Health Laboratory | Wyoming Public Health Laboratory | Noah Hull, Rob Christensen, Jim Mildenberger, Joel Sevinsky, Cari Sloma, and Wanda Manley |
| EPI_ISL_541240 | Florida Bureau of Public Health Laboratories, Florida Department of Health | Florida Bureau of Public Health Laboratories, Florida Department of Health | Schmedes,S., Blanton,J. |
| EPI_ISL_541354 | Laboratory of Respiratory Viruses and Measles, Oswaldo Cruz Institute, FIOCRUZ | Laboratory of Respiratory Viruses and Measles, Oswaldo Cruz Institute, FIOCRUZ | Paola Resende, Luciana Appolinario, Fernando Motta, Anna Carolina Paixão, Ana Carolina Mendonça, Jonathan Lopes, Marilda Siqueira |
| EPI_ISL_541752, EPI_ISL_541755, EPI_ISL_541762, EPI_ISL_541766, EPI_ISL_541775, EPI_ISL_541779, EPI_ISL_541783 | Barts Health NHS Trust | Wellcome Sanger Institute for the COVID-19 Genomics UK (COG-UK) consortium | Teresa Cutino-Moguel, Mark Hopkins, Beatrix Kele, David Harrington and Alex Alderton, Roberto Amato, Sonia Goncalves, Ewan Harrison, David K. Jackson, Ian Johnston, Dominic Kwiatkowski, Cordelia Langford, John Sillitoe on behalf of the Wellcome Sanger Institute COVID-19 Surveillance Team |
| EPI_ISL_541856, EPI_ISL_541857, EPI_ISL_541858, EPI_ISL_541859, EPI_ISL_541860, EPI_ISL_541861, EPI_ISL_541862, EPI_ISL_541863, EPI_ISL_541864, EPI_ISL_541865, EPI_ISL_541866, EPI_ISL_541867, EPI_ISL_541869 | | | |
| see above | Lithuanian University of Health Sciences Hospital, Department of Laboratory Medicine | Lithuanian University of Health Sciences, Laboratory of Molecular Cardiology | Lukas Zemaitis, Arnoldas Pautienius, Kamile Tamusauskaite, Dovydas Gecys, Laura Pareckaite, Vaiva Lesauskaite, Astra Vitkauskiene |
| EPI_ISL_542487, EPI_ISL_542488, EPI_ISL_542489, EPI_ISL_542492, EPI_ISL_542494, EPI_ISL_542497, EPI_ISL_542499, EPI_ISL_542501, EPI_ISL_542504, EPI_ISL_542505, EPI_ISL_542508, EPI_ISL_542512, EPI_ISL_542513, EPI_ISL_542514, EPI_ISL_542517, EPI_ISL_542518, EPI_ISL_542519, EPI_ISL_542520, EPI_ISL_542522, EPI_ISL_542523, EPI_ISL_542612, EPI_ISL_542621, EPI_ISL_542650, EPI_ISL_542660, EPI_ISL_542662, EPI_ISL_542664, EPI_ISL_542668, EPI_ISL_542675, EPI_ISL_542680, EPI_ISL_542686, EPI_ISL_542757, EPI_ISL_542775, EPI_ISL_542776, EPI_ISL_542793 | | | |
| see above | Houston Methodist Hospital | Houston Methodist Hospital | S. Wesley Long, Randall J. Olsen, Paul A. Christensen, David W. Bernard, James J. Davis, Maulik Shukla, Marcus Nguyen, Matthew Ojeda Saavedra, Concepcion C. Cantu, Prasanti Yerramilli, Layne Pruitt, Sishir Subedi, Hung-Che Kuo, Heather Hendrickson, Ghazaleh Eskandari, Hoang A. T. Nguyen, J. Hunter Long, Muthiah Kumaraswami, Jule Goike, Daniel Boutz, Jimmy Gollihar, Jason S. McLellan, Chia-Wei Chou, Kamyab Javanmardi, Ilya J. Finkelstein, and James M. Musser |
| EPI_ISL_542979 | TriCore Reference Laboratories | Center for Global Health, University of New Mexico Health Sciences Center | Daryl Domman, Kurt Schwalm, Twila Kunde, Joseph Hicks, Michael Edwards, Darrell Dinwiddie |
| EPI_ISL_544339, EPI_ISL_544345, EPI_ISL_544353, EPI_ISL_544410, EPI_ISL_544434, EPI_ISL_544435, EPI_ISL_544796, EPI_ISL_544805, EPI_ISL_544823, EPI_ISL_544833, EPI_ISL_544868, EPI_ISL_544916, EPI_ISL_545341, EPI_ISL_545590, EPI_ISL_545601, EPI_ISL_545610, EPI_ISL_545627, EPI_ISL_545637, EPI_ISL_545640, EPI_ISL_545645, EPI_ISL_545648, EPI_ISL_545653, EPI_ISL_545668, EPI_ISL_545732, EPI_ISL_545780, EPI_ISL_545788, EPI_ISL_545801, EPI_ISL_545810, EPI_ISL_545817, EPI_ISL_545835, EPI_ISL_545844, EPI_ISL_545846, EPI_ISL_545847, EPI_ISL_545848, EPI_ISL_545849, EPI_ISL_545850, EPI_ISL_545851, EPI_ISL_545852, EPI_ISL_545853, EPI_ISL_545854, EPI_ISL_545856, EPI_ISL_545857, EPI_ISL_545859, EPI_ISL_545860, EPI_ISL_545861, EPI_ISL_545862, EPI_ISL_545863, EPI_ISL_545864, EPI_ISL_545865, EPI_ISL_545875, EPI_ISL_545887, EPI_ISL_545888, EPI_ISL_545890, EPI_ISL_545892, EPI_ISL_545907, EPI_ISL_545908, EPI_ISL_545909, EPI_ISL_545933, EPI_ISL_545934, EPI_ISL_545935, EPI_ISL_545936, EPI_ISL_545937, EPI_ISL_545938, EPI_ISL_545939, EPI_ISL_545940, EPI_ISL_545941, EPI_ISL_545942, EPI_ISL_545943, EPI_ISL_545944, EPI_ISL_545945, EPI_ISL_545946, EPI_ISL_545947, | | | |

| | | | |
|---|---|---|---|
| EPI_ISL_545948, EPI_ISL_545949, EPI_ISL_545950, EPI_ISL_545951, EPI_ISL_545990, EPI_ISL_546019, EPI_ISL_546198, EPI_ISL_546214, EPI_ISL_546555, EPI_ISL_546565, EPI_ISL_546575, EPI_ISL_546593, EPI_ISL_546678, EPI_ISL_546705, EPI_ISL_546728, EPI_ISL_546747, EPI_ISL_546753, EPI_ISL_546775, EPI_ISL_546786, EPI_ISL_546940, EPI_ISL_546965, EPI_ISL_547094, EPI_ISL_547132, EPI_ISL_547140, EPI_ISL_547156, EPI_ISL_547165, EPI_ISL_547172, EPI_ISL_547179, EPI_ISL_547201, EPI_ISL_547217, EPI_ISL_547228, EPI_ISL_547245, EPI_ISL_547255, EPI_ISL_547272, EPI_ISL_547288, EPI_ISL_547300, EPI_ISL_547340, EPI_ISL_547416 | | | |
| see above | Houston Methodist Hospital | Houston Methodist Hospital | S. Wesley Long, Randall J. Olsen, Paul A. Christensen, David W. Bernard, James J. Davis, Maulik Shukla, Marcus Nguyen, Matthew Ojeda Saavedra, Concepcion C. Cantu, Prasanti Yerramilli, Layne Pruitt, Sishir Subedi, Hung-Che Kuo, Heather Hendrickson, Ghazaleh Eskandari, Hoang A. T. Nguyen, J. Hunter Long, Muthiah Kumaraswami, Jule Goike, Daniel Boutz, Jimmy Gollihar, Jason S. McLellan, Chia-Wei Chou, Kamyab Javanmardi, Ilya J. Finkelstein, and James M. Musser |
| EPI_ISL_547519, EPI_ISL_547558 | Dutch COVID-19 response team | National Institute for Public Health and the Environment (RIVM) | Adam Meijer, Harry Vennema, Jeroen Cremer, Sharon van den Brink, Bas van der Veer, AnneMarie van den Brandt, Florian Zwagemaker, Dennis Schmitz, Chantal Reusken, on behalf of the national COVID-19 response team |
| EPI_ISL_548243 | Faith Laboratory, Immunology Institute, Icahn School of Medicine at Mount Sinai | van Bakel Laboratory, Genetics and Genomics Sciences, Icahn School of Medicine at Mount Sinai | Graham J. Britton, Alice Chen-Liaw, Francesca Cossarini, Alexandra Livanos, Matthew P. Spindler, Tamar Plitt, Joseph Eggers, Ilaria Mogno, Ana S. Gonzalez-Reiche, Sophia Sui, Michael Tankelevich, Lauren Tal Grinspan, Rebekah E. Dixon, Divya Jha, Gustavo Martinez-Delgado, Fatima Amanat, Daisy Hoagland, Benjamin R. tenOever, Marla C. Dubinsky, Miriam Merad, Harm Van Bakel, Florian Krammer, Gerold Bongers, Saurabh Mehandru and Jeremiah J. Faith |
| EPI_ISL_548248 | Ostersund klinisk mikrobiologi | The Public Health Agency of Sweden | Anna-Malin Linde, Maria Lind Karlberg, Mattias Haukland, Reza Advani, Olov Svartstrom, Oskar Karlsson Lindsjo, Sandra Broddesson, Petra Edquist, Mia Brytting, Anna Risberg, Karin Tegmark-Wisell |
| EPI_ISL_548352 | Ventura County Public Health Lab | Chan-Zuckerberg Biohub | CZB Cliahub Consortium |
| EPI_ISL_548952 | Max von Pettenkofer Institute, Virology, National Reference Center for Retroviruses, LMU München | Laboratory for Functional Genome Analysis, Dept. Genomics, Gene Center of the LMU Munich | Max Muenchhoff, Stefan Krebs, Alexander Graf, Oliver Keppler, Helmut Blum |
| EPI_ISL_549017, EPI_ISL_549018, EPI_ISL_549023 | KWR Watercycle Research Institute | Erasmus Medical Center | Ray Izquierdo-Lara, Goffe Elsinga, Leo Heijnen, Bas B. Oude Munnink, Claudia M. E. Schapendonk, David Nieuwenhuijse, Matthijs Kon, Lu Lu, Frank M. Aarestrup, Samantha Lycett, Gertjan Medema, Marion P.G. Koopmans, Miranda de Graaf |
| EPI_ISL_549182, EPI_ISL_549183, EPI_ISL_549228, EPI_ISL_549229, EPI_ISL_549230, EPI_ISL_549231, EPI_ISL_549232, EPI_ISL_549233, EPI_ISL_549234, EPI_ISL_549235, EPI_ISL_549236, EPI_ISL_549237, EPI_ISL_549238, EPI_ISL_549239, EPI_ISL_549240, EPI_ISL_549241, EPI_ISL_549242, EPI_ISL_549243 | | | |
| see above | Florida Bureau of Public Health Laboratories | Florida Bureau of Public Health Laboratories | Sarah Schmedes, Jason Blanton |
| EPI_ISL_549478, EPI_ISL_549486 | Quadram Institute Bioscience | COVID-19 Genomics UK (COG-UK) Consortium | Dave J. Baker, Gemma L. Kay, Alp Aydin, Thanh Le-Viet, Steven Rudder, Ana P. Tedim, Anastasia Kolyva, Maria Diaz, Leonardo de Oliveira Martins, Nabil-Fareed Alikhan, Lizzie Meadows, Rachael Stanley, Ngozi Elumogo, Muhammed Yasir, Nicholas M. Thomson, Alexander J Trotter, Rachel Gilroy, Samuel Bloomfield, Claire Stuart, Andrew Bell, Reenesh Prakash, Samir Dervisevic, Alison E. Mather, John Wain, Mark Webber, Andrew J. Page, Justin O'Grady |
| EPI_ISL_559972 | Virology Department, Royal Infirmary of Edinburgh, NHS Lothian / School of Biological Sciences, University of Edinburgh / Institute of Genetics and Molecular Medicine, University of Edinburgh | COVID-19 Genomics UK (COG-UK) Consortium | McHugh M, Dewar R, Rooke S, Gallagher M, Balcaza C, O'Toole Á, Scher E, Hill V, McCrone JT, Colquhoun R, Yu X, Jackson B, Rambaut A, Williams TC, Templeton K |
| EPI_ISL_560385 | National Health Laboratory | Botswana Institute for Technology Research and Innovation | Kefentse Arnold Tumedi, Madisa Mine, Dineo Emang Tshiamo, Gape Nyepetsi, Thongbotho Mphoyakgosi, Maitshwarelo Ignatius Matsheka |
| EPI_ISL_560390 | National Health Laboratory | Botswana Institute for Technology Research and Innovation | Kefentse Arnold Tumedi, Madisa Mine, Dineo Emang Tshiamo. Gape Nyepetsi, Thongbotho Mphoyakgosi, Maitshwarelo Ignatius Matsheka |
| EPI_ISL_560568, EPI_ISL_560575, EPI_ISL_560579 | hôpital | National Reference Center for Viruses of Respiratory Infections, Institut Pasteur, Paris | Sylvie Behillil, Fabiana Gambaro, Etienne Simon-Lorière, Vincent Enouf, Maud Vanpeene, Sylvie van der Werf |
| EPI_ISL_560629 | Hospital | National Reference Center for Viruses of Respiratory Infections, Institut Pasteur, Paris | Sylvie Behillil, Fabiana Gambaro, Etienne Simon-Lorière, Vincent Enouf, Maud Vanpeene, Sylvie van der Werf |
| EPI_ISL_560763, EPI_ISL_560764, EPI_ISL_560765, EPI_ISL_560766, EPI_ISL_560768, EPI_ISL_560769, EPI_ISL_560770, EPI_ISL_560771, EPI_ISL_560772, EPI_ISL_560773, EPI_ISL_560774, EPI_ISL_560775, EPI_ISL_560776, EPI_ISL_560777, EPI_ISL_560778, EPI_ISL_560779, EPI_ISL_560780, EPI_ISL_560781, EPI_ISL_560782, EPI_ISL_560783, EPI_ISL_560784, EPI_ISL_560785, EPI_ISL_560786, EPI_ISL_560787, EPI_ISL_560788, EPI_ISL_560789, EPI_ISL_560790, EPI_ISL_560791 | | | |
| see above | Minnesota Department of Health, Public Health Laboratory | Minnesota Department of Health, Public Health Laboratory | Matt Plumb, Jacob Garfin, and Xiong Wang |
| EPI_ISL_560849, EPI_ISL_560850, EPI_ISL_560851, EPI_ISL_560852, EPI_ISL_560853, EPI_ISL_560854, EPI_ISL_560855, EPI_ISL_560856, EPI_ISL_560857, EPI_ISL_560858, EPI_ISL_560859, EPI_ISL_560860 | | | |
| see above | Utah Public Health Laboratory | Utah Public Health Laboratory | Erin Young, Kelly Oakeson |
| EPI_ISL_561517, EPI_ISL_562136, EPI_ISL_562348, EPI_ISL_562518, EPI_ISL_562540, EPI_ISL_562585, EPI_ISL_563609 | Victorian Infectious Diseases Reference Laboratory (VIDRL) | VIDRL and MDU-PHL | Caly, L., Seemann, T., Sait, M., Schultz, M. B., Druce J., Sherry, N. |
| EPI_ISL_565833, EPI_ISL_565834, EPI_ISL_565839, EPI_ISL_565842, EPI_ISL_565906, EPI_ISL_565907, EPI_ISL_565932, EPI_ISL_565934, EPI_ISL_565939, EPI_ISL_565948, EPI_ISL_565960, EPI_ISL_565968, EPI_ISL_565969, EPI_ISL_565979, EPI_ISL_565983, EPI_ISL_565985, EPI_ISL_565986, EPI_ISL_565988, EPI_ISL_566003, EPI_ISL_566015 | | | |
| see above | Michigan Department of Health and Human Services, Bureau of Laboratories | Michigan Department of Health and Human Services, Bureau of Laboratories | Blankenship HM, Riner D, Soehnlen MK |
| EPI_ISL_568559, EPI_ISL_568560 | Department of Infectious Diseases and Immunology, National Hospital Organization Nagoya Medical Center | Clinical Research Center, National Hospital Organization Nagoya Medical Center | Yoshihiro Nakata, Hirotaka Ode, Mai Kubota, Masakazu Matsuda, Kazuhiro Matsuoka, Nakasuji Miho, Mikiko Mori, Mayumi Imahashi, Yoshiyuki Yokomaku, Yasumasa Iwatani |
| EPI_ISL_568687 | RSUP Fatmawati | Eijkman Institute for Molecular Biology, Ministry of Research and Technology/National Agency for Research and Innovation | Frilasita A Yudhaputri, Edison Johar, Hidayat Trimarsanto, Iskandar A Adnan, Willy Agustine, David H Muljono, Safarina G Malik, Herawati Sudoyo, Khin Saw Myint, Amin Soebandrio |
| EPI_ISL_568698, EPI_ISL_568699, EPI_ISL_568700, EPI_ISL_568702 | KEMRI-Wellcome Trust Research Programme/KEMRI-CGMR-C Kilifi | KEMRI-Wellcome Trust Research Programme/KEMRI-CGMR-C Kilifi | Githinji et al 2020 |
| EPI_ISL_568874 | Malaysia Genome Institute | Malaysia Genome Institute | Mohd Noor Mat Isa, Irni Suhayu Sapian, Yusuf Muhammad Noor, Nurhezreen Md Iqbal, Mohd Faizal Abu Bakar, Enizza Kasim, Shamsidar Sopie, Siti Noraini Othman, Azrin Ahmad, Nor Azfa Johari, Shahrul Hisham Zainal Ariffin |
| EPI_ISL_568877, EPI_ISL_568878, EPI_ISL_568879, EPI_ISL_568880, EPI_ISL_568881, EPI_ISL_568882, EPI_ISL_568883, EPI_ISL_568884, EPI_ISL_568885, EPI_ISL_568886, EPI_ISL_568887, EPI_ISL_568888, EPI_ISL_568889, EPI_ISL_568890, EPI_ISL_568891, EPI_ISL_568892, EPI_ISL_568893, EPI_ISL_568894, EPI_ISL_568895, EPI_ISL_568896, EPI_ISL_568897, EPI_ISL_569571, EPI_ISL_569572, EPI_ISL_569573, EPI_ISL_569574, EPI_ISL_569575, EPI_ISL_569576, EPI_ISL_569577, EPI_ISL_569578, EPI_ISL_569579, EPI_ISL_569580, EPI_ISL_569581, EPI_ISL_569582, EPI_ISL_569583, EPI_ISL_569585, EPI_ISL_569586, EPI_ISL_569587, EPI_ISL_569588, EPI_ISL_569589, EPI_ISL_569590, EPI_ISL_569591, EPI_ISL_569592, EPI_ISL_569593, EPI_ISL_569594, EPI_ISL_569595, EPI_ISL_569596, EPI_ISL_569597, EPI_ISL_569598, EPI_ISL_569599, EPI_ISL_569600, EPI_ISL_569602, EPI_ISL_569603, EPI_ISL_569604, EPI_ISL_569605, EPI_ISL_569606, EPI_ISL_569607, EPI_ISL_569608 | | | |
| see above | MEPHI, Aix Marseille University | MEPHI, Aix Marseille University | Anthony LEVASSEUR |
| EPI_ISL_569826 | Omsk Research Institute of Natural Focal Infections | WHO National Influenza Centre Russian Federation | Artem Fadeev, Ekaterina Gradoboeva, Ekaterina Savkina, Daria Nashatyreva, Elena Poleshchuk, Aleksei Vasilenko, Valery Yakimenko, Andrey Komissarov |
| EPI_ISL_569865, EPI_ISL_569883, EPI_ISL_569884, EPI_ISL_569885, EPI_ISL_569886 | Amedeo di savoia | Crosetto lab, Karolinska Institutet, SciLifeLab | Michele Simonetti, Maria Grazia Milia, Luuk Harbers, Ning Zhang, Anna Sapino, Valeria Ghisetti, Nicola Crosetto |
| EPI_ISL_570070, EPI_ISL_570071, EPI_ISL_570072, EPI_ISL_570074, EPI_ISL_570075, EPI_ISL_570077, EPI_ISL_570078, EPI_ISL_570133, EPI_ISL_570134, EPI_ISL_570135, EPI_ISL_570136, EPI_ISL_570137, EPI_ISL_570138, EPI_ISL_570139, EPI_ISL_570140, EPI_ISL_570141, EPI_ISL_570142, EPI_ISL_570143, EPI_ISL_570144, EPI_ISL_570145, EPI_ISL_570146, EPI_ISL_570147, EPI_ISL_570148, EPI_ISL_570149, EPI_ISL_570150, EPI_ISL_570151, EPI_ISL_570164, EPI_ISL_570167, EPI_ISL_570168, EPI_ISL_570169, EPI_ISL_570170, EPI_ISL_570171, EPI_ISL_570172, EPI_ISL_570173, EPI_ISL_570175, EPI_ISL_570177 | | | |
| see above | UW Virology Lab | UW Virology Lab | Pavitra Roychoudhury, Hong Xie, Lasata Shrestha, Amin Addetia, Victoria M Rachleff, Meei-Li Huang, Keith R Jerome, Alexander Greninger |
| EPI_ISL_572334 | LACEN/PE | WallauLab, Aggeu Magalhaes Institute | Marcelo Henrique Santos Paiva, Duschinka Ribeiro Duarte Guedes, Cássia Docena, Matheus Filgueira Bezerra, Filipe Zimmer Dezordi, Laís Ceschini |

| | | | |
|---|---|---|---|
| | | | Machado, Larissa Krokovsky, Elisama Helvecio, Alexandre Freitas da Silva, Luydson Richardson Silva Vasconcelos, Antonio Mauro Rezende, Severino Jefferson Ribeiro da Silva, Kamila Gaudêncio da Silva Sales, Bruna Santos Lima Figueiredo de Sá, Derciliano Lopes da Cruz, Claudio Eduardo Cavalcanti, Armando de Menezes Neto, Caroline Targino Alves da Silva, Renata Pessôa Germano Mendes, Maria Almerice Lopes da Silva, Tiago Gräf, Paola Cristina Resende, Gonzalo Bello0, Michelle da Silva Barros, Wheverton Ricardo Correia do Nascimento,, Rodrigo Moraes Loyo Arcoverde, Luciane Caroline Albuquerque Bezerra, Sinval Pinto Brandão Filho, Constância Flávia Junqueira Ayres, Gabriel Luz Wallau |
| EPI_ISL_574607 | RS Husada | Eijkman Institute for Molecular Biology, Ministry of Research and Technology/National Agency for Research and Innovation | Frilasita A Yudhaputri, Edison Johar, Hidayat Trimarsanto, Iskandar A Adnan, Willy Agustine, David H Muljono, Safarina G Malik, Herawati Sudoyo, Khin Saw Myint, Amin Soebandrio |
| EPI_ISL_574608 | RS Mitra Keluarga Kelapa Gading | Eijkman Institute for Molecular Biology, Ministry of Research and Technology/National Agency for Research and Innovation | Frilasita A Yudhaputri, Edison Johar, Hidayat Trimarsanto, Iskandar A Adnan, Willy Agustine, David H Muljono, Safarina G Malik, Herawati Sudoyo, Khin Saw Myint, Amin Soebandrio |
| EPI_ISL_574853, EPI_ISL_574854, EPI_ISL_574855, EPI_ISL_574856, EPI_ISL_574857, EPI_ISL_574858 | Institute for Infectious Diseases, University of Bern | Institute for Infectious Diseases, University of Bern | Michel C Koch, Christian Baumann, Miguel A Terrazos Miani, Cora Sägesser, Stephen L Leib, Peter Keller, Franziska Suter-Riniker, Alban Ramette |
| EPI_ISL_575061, EPI_ISL_575062, EPI_ISL_575063, EPI_ISL_575064, EPI_ISL_575065, EPI_ISL_575066, EPI_ISL_575067, EPI_ISL_575068, EPI_ISL_575069, EPI_ISL_575070, EPI_ISL_575071, EPI_ISL_575072 | | | |
| see above | Utah Public Health Laboratory | Utah Public Health Laboratory | Erin Young, Kelly Oakeson |
| EPI_ISL_576237, EPI_ISL_576239, EPI_ISL_576243, EPI_ISL_576252, EPI_ISL_576253, EPI_ISL_576254, EPI_ISL_576255 | Minnesota Department of Health, Public Health Laboratory | Minnesota Department of Health, Public Health Laboratory | Matt Plumb, Jacob Garfin, Alexandra Lorentz, and Xiong Wang |
| EPI_ISL_576489, EPI_ISL_576490, EPI_ISL_576491 | UW Virology Lab | UW Virology Lab | Pavitra Roychoudhury, Hong Xie, Lasata Shrestha, Amin Addetia, Victoria M Rachleff, Meei-Li Huang, Keith R Jerome, Alexander Greninger |
| EPI_ISL_577554 | Michigan Department of Health and Human Services, Bureau of Laboratories | Michigan Department of Health and Human Services, Bureau of Laboratories | Blankenship HM, Riner D, Soehnlen MK |
| EPI_ISL_578144, EPI_ISL_578145, EPI_ISL_578146, EPI_ISL_578147, EPI_ISL_578148, EPI_ISL_578149, EPI_ISL_578150, EPI_ISL_578151, EPI_ISL_578152, EPI_ISL_578153, EPI_ISL_578154, EPI_ISL_578155 | | | |
| see above | University of Michigan Clinical Microbiology Laboratory | Lauring Lab, University of Michigan, Department of Microbiology and Immunology | Valesano |
| EPI_ISL_578345, EPI_ISL_578346, EPI_ISL_578347, EPI_ISL_578348, EPI_ISL_578349, EPI_ISL_578410, EPI_ISL_578411, EPI_ISL_578412, EPI_ISL_578413, EPI_ISL_578414, EPI_ISL_578415, EPI_ISL_578416, EPI_ISL_578417, EPI_ISL_578418, EPI_ISL_578438, EPI_ISL_578439, EPI_ISL_578440, EPI_ISL_578441, EPI_ISL_578442, EPI_ISL_578443, EPI_ISL_578444, EPI_ISL_578451 | | | |
| see above | Wisconsin State Laboratory of Hygiene Communicable Disease Division | Wisconsin State Laboratory of Hygiene Communicable Disease Division | Kelsey R. Florek, Abigail C. Shockey |
| EPI_ISL_579185, EPI_ISL_579186, EPI_ISL_579187, EPI_ISL_579188, EPI_ISL_579189, EPI_ISL_579219 | Southern Community Labs Dunedin | Institute of Environmental Science and Research (ESR) | Xiaoyun Ren, Matt Storey, Nikki Freed, Muhammad Faisal, Jing Wang, Hermes Perez, Anja Werno, Antje van der Linden, Arlo Upton, Chris Mansell, David Hammer, Dragana Drinkovic, Gary McAuliffe, Hana Sofia Andersson, James Ussher, Jill Sherwood, Josh Freeman, Julia Howard, Juliet Elvy, Mary DeAlmeida, Matt Blakiston, Matthew Rogers, Max Bloomfield, Michael Addidle, Michelle Balm, Sally Roberts, Sarah Jefferies, Sharmini Muttaiyah, Susan Morpeth, Susan Taylor, Timothy Blackmore, Vani Sathyendran, Veronica Playle, Virginia Hope, Erasmus Smit, Lauren Jelly, Olin Silander, Joep de Ligt |
| EPI_ISL_579224, EPI_ISL_579225, EPI_ISL_579226, EPI_ISL_579227 | Canterbury Health Laboratories | Institute of Environmental Science and Research (ESR) | Xiaoyun Ren, Matt Storey, Nikki Freed, Muhammad Faisal, Jing Wang, Hermes Perez, Anja Werno, Antje van der Linden, Arlo Upton, Chris Mansell, David Hammer, Dragana Drinkovic, Gary McAuliffe, Hana Sofia Andersson, James Ussher, Jill Sherwood, Josh Freeman, Julia Howard, Juliet Elvy, Mary DeAlmeida, Matt Blakiston, Matthew Rogers, Max Bloomfield, Michael Addidle, Michelle Balm, Sally Roberts, Sarah Jefferies, Sharmini Muttaiyah, Susan Morpeth, Susan Taylor, Timothy Blackmore, Vani Sathyendran, Veronica Playle, Virginia Hope, Erasmus Smit, Lauren Jelly, Olin Silander, Joep de Ligt |
| EPI_ISL_579230, EPI_ISL_579231 | Middlemore Hospital | Institute of Environmental Science and Research (ESR) | Xiaoyun Ren, Matt Storey, Nikki Freed, Muhammad Faisal, Jing Wang, Hermes Perez, Anja Werno, Antje van der Linden, Arlo Upton, Chris Mansell, David Hammer, Dragana Drinkovic, Gary McAuliffe, Hana Sofia Andersson, James Ussher, Jill Sherwood, Josh Freeman, Julia Howard, Juliet Elvy, Mary DeAlmeida, Matt Blakiston, Matthew Rogers, Max Bloomfield, Michael Addidle, Michelle Balm, Sally Roberts, Sarah Jefferies, Sharmini Muttaiyah, Susan Morpeth, Susan Taylor, Timothy Blackmore, Vani Sathyendran, Veronica Playle, Virginia Hope, Erasmus Smit, Lauren Jelly, Olin Silander, Joep de Ligt |
| EPI_ISL_579651, EPI_ISL_579652, EPI_ISL_579653, EPI_ISL_579654, EPI_ISL_579657, EPI_ISL_579658, EPI_ISL_579659, EPI_ISL_579660, EPI_ISL_579661, EPI_ISL_579664, EPI_ISL_579666, EPI_ISL_579667, EPI_ISL_579668, EPI_ISL_579669, EPI_ISL_579670, EPI_ISL_579671, EPI_ISL_579672, EPI_ISL_579673, EPI_ISL_579674, EPI_ISL_579675, EPI_ISL_579676, EPI_ISL_579677, EPI_ISL_579678, EPI_ISL_579679, EPI_ISL_579680, EPI_ISL_579681, EPI_ISL_579682, EPI_ISL_579683, EPI_ISL_579684, EPI_ISL_579685, EPI_ISL_579686, EPI_ISL_579687, EPI_ISL_579688, EPI_ISL_579689, EPI_ISL_579690, EPI_ISL_579691, EPI_ISL_579692, EPI_ISL_579693, EPI_ISL_579694, EPI_ISL_579695, EPI_ISL_579696, EPI_ISL_579697, EPI_ISL_579698, EPI_ISL_579699, EPI_ISL_579700, EPI_ISL_579701, EPI_ISL_579702, EPI_ISL_579703, EPI_ISL_579704, EPI_ISL_579705, EPI_ISL_579706, EPI_ISL_579707, EPI_ISL_579708, EPI_ISL_579709, EPI_ISL_579710, EPI_ISL_579711, EPI_ISL_579712, EPI_ISL_579713, EPI_ISL_579714, EPI_ISL_579715, EPI_ISL_579716, EPI_ISL_579717, EPI_ISL_579718, EPI_ISL_579719, EPI_ISL_579720 | | | |
| see above | QEII Health Sciences Centre | National Microbiology Laboratory (NML) | Anna Majer, Shari Tyson, Grace Seo, Philip Mabon, Darian Hole, Elsie Grudeski, Rhiannon Huzarewich, Russell Mandes, Anneliese Landgraff, Jennifer Tanner, Natalie Knox, Morag Graham, Gary Van Domselaar, Todd Hatchette, Jason LeBlanc, Nathalie Bastien, Yan Li, Timothy Booth, CanCOGeN's metadata curation team, Public Health Agency of Canada's CanCOGeN team |
| EPI_ISL_581454, EPI_ISL_581474 | Medizinische Klinik Innere Medizin I, Universitätsklinikum Tübingen | NGS Competence Center Tübingen, Institut für Medizinische Mikrobiologie und Hygiene, Universitätsklinikum Tübingen | Angel Angelov |
| EPI_ISL_581687, EPI_ISL_581688, EPI_ISL_581689, EPI_ISL_581690, EPI_ISL_581691, EPI_ISL_581692, EPI_ISL_581693, EPI_ISL_581694, EPI_ISL_581695, EPI_ISL_581696, EPI_ISL_581697, EPI_ISL_581698 | | | |
| see above | University Hospital Basel, Clinical Virology | University Hospital Basel, Clinical Bacteriology | Madlen Stange, Alfredo Mari, Tim Roloff, Helena MB Seth-Smith, Michael Schweitzer, Myrta Brunner, Karoline Leuzinger, Kirstine K. Soegaard, Alexander Gensch, Sarah Tschudin-Sutter, Simon Fuchs, Julia Bielicki, Hans Pargger, Martin Siegemund, Christian Nickel, Roland Bingisser, Michael Osthoff, Stefano Bassetti, Rita Schneider-Sliwa, Manuel Battegay, Hans Hirsch, Adrian Egli |
| EPI_ISL_582004 | Infectious Diseases and Tropical Medicine Research Center, Infectious Diseases and Tropical Medicine Research Center | Infectious Diseases and Tropical Medicine Research Center, Infectious Diseases and Tropical Medicine Research Center | Haghjoo javanmard,S., Shariati,l., Ahangarzadeh,S., Ataei,B., Ranjbar,M.M., Shoaei,P. |
| EPI_ISL_582247, EPI_ISL_582248, EPI_ISL_582249, EPI_ISL_582271, EPI_ISL_582506 | Cadham Provincial Laboratory | National Microbiology Laboratory (NML) | Anna Majer, Shari Tyson, Grace Seo, Philip Mabon, Elsie Grudeski, Rhiannon Huzarewich, Russell Mandes, Anneliese Landgraff, Jennifer Tanner, Natalie Knox, Morag Graham, Gary Van Domselaar, Paul Van Caeseele, Jared Bullard, David Alexander, Kerry Dust, Nathalie Bastien, Yan Li, Timothy Booth, Darian Hole, Madison Chapel, CanCOGeN's metadata curation team, Public Health Agency of Canada CanCOGeN team |
| EPI_ISL_582514 | unknown | Instituto Nacional de Saude (INSA) and Instituto Gulbenkian de Ciencia (IGC) | Borges et al |
| EPI_ISL_582527 | Veterinary Specialized Institute "Kraljevo", Serbia | Veterinary Specialized Institute "Kraljevo", Serbia | Vidanovic,D., Tesovic,B., Knezevic,A., Jovanovic,T., Jankovic,M., Sekler,M., Banovic Djeri,B., Petrovic,T., Volkening,J., Afonso,C. |
| EPI_ISL_583444 | Memorial Sloan Kettering Cancer Center | van Bakel Laboratory, Genetics and Genomics Sciences, Icahn School of Medicine at Mount Sinai | Teresa Aydillo, Ana S. Gonzalez-Reiche, Sadaf Aslam, Adriana van de Guchte, Zenab Khan, Ajay Obla, Jayeeta Dutta, Harm van Bakel, Judith Aberg, Adolfo García-Sastre, Gunjan Shah, Tobias Hohl, Genovefa Papanicolaou, Miguel-Angel Perales, Kent Sepkowitz, Ngolela Esther Babady, and Mini Kamboj |
| EPI_ISL_583590 | Institute for Medical and Chemical Laboratory Diagnostics, Kepler Universitätsklinikum | Bergthaler laboratory, CeMM Research Center for Molecular Medicine of the Austrian Academy of Sciences | Alexandra Popa, Benedikt Agerer, Henrique Colaco, Lukas Endler, Jakob-Wendelin Genger, Alexander Lercher, Mark Smyth, Thomas Penz, Michael Schuster, Jan Laine, Martin Senekowitsch, Judith Aberle, Stephan Aberle, Peter Hufnagl, Daniela Schmid, Franz Allerberger, Elisabeth Puchhammer-Stoeckl, Manfred Nairz, Guenter Weiss, Gregor Hörmann, Kinga Rigler-Hohenwarter, Rainer Gattringer, Wegene Borena, Dorothee von Laer, Gernot Walder, Peter Obrist, Christian Paar, Sabine Sussitz-Rack, Gunther Vogl, Adi Steinrigl, Christoph Bock, Andreas Bergthaler |
| EPI_ISL_583617, EPI_ISL_583618 | Universitaetsklinik für Innere Medizin II Innsbruck | Bergthaler laboratory, CeMM Research Center for Molecular Medicine of the Austrian Academy of Sciences | Alexandra Popa, Benedikt Agerer, Henrique Colaco, Lukas Endler, Jakob-Wendelin Genger, Alexander Lercher, Mark Smyth, Thomas Penz, Michael Schuster, Jan Laine, Martin Senekowitsch, Judith Aberle, Stephan Aberle, Peter Hufnagl, Daniela Schmid, Franz Allerberger, Elisabeth Puchhammer-Stoeckl, Manfred Nairz, Guenter Weiss, Gregor Hörmann, Kinga Rigler-Hohenwarter, Rainer Gattringer, Wegene Borena, Dorothee von Laer, Gernot Walder, Peter Obrist, Christian Paar, Sabine Sussitz-Rack, Gunther Vogl, Adi Steinrigl, Christoph Bock, Andreas Bergthaler |

| | | | |
|---|---|---|---|
| EPI_ISL_583713, EPI_ISL_583714, EPI_ISL_583722 | Center for Virology, Medical University of Vienna | Bergthaler laboratory, CeMM Research Center for Molecular Medicine of the Austrian Academy of Sciences | Alexandra Popa, Benedikt Agerer, Henrique Colaco, Lukas Endler, Jakob-Wendelin Genger, Alexander Lercher, Mark Smyth, Thomas Penz, Michael Schuster, Jan Laine, Martin Senekowitsch, Judith Aberle, Stephan Aberle, Peter Hufnagl, Daniela Schmid, Franz Allerberger, Elisabeth Puchhammer-Stoeckl, Manfred Nairz, Guenter Weiss, Gregor Hörmann, Kinga Rigler-Hohenwarter, Rainer Gattringer, Wegene Borena, Dorothee von Laer, Gernot Walder, Peter Obrist, Christian Paar, Sabine Sussitz-Rack, Gunther Vogl, Adi Steinrigl, Christoph Bock, Andreas Bergthaler |
| EPI_ISL_583764, EPI_ISL_583828, EPI_ISL_583829, EPI_ISL_583830, EPI_ISL_583831, EPI_ISL_583832, EPI_ISL_583833, EPI_ISL_583834, EPI_ISL_583835, EPI_ISL_583836, EPI_ISL_583837, EPI_ISL_583838, EPI_ISL_583839 | | | |
| see above | Dr. Gernot Walder GmbH | Bergthaler laboratory, CeMM Research Center for Molecular Medicine of the Austrian Academy of Sciences | Alexandra Popa, Benedikt Agerer, Henrique Colaco, Lukas Endler, Jakob-Wendelin Genger, Alexander Lercher, Mark Smyth, Thomas Penz, Michael Schuster, Jan Laine, Martin Senekowitsch, Judith Aberle, Stephan Aberle, Peter Hufnagl, Daniela Schmid, Franz Allerberger, Elisabeth Puchhammer-Stoeckl, Manfred Nairz, Guenter Weiss, Gregor Hörmann, Kinga Rigler-Hohenwarter, Rainer Gattringer, Wegene Borena, Dorothee von Laer, Gernot Walder, Peter Obrist, Christian Paar, Sabine Sussitz-Rack, Gunther Vogl, Adi Steinrigl, Christoph Bock, Andreas Bergthaler |
| EPI_ISL_583857, EPI_ISL_583858, EPI_ISL_583859, EPI_ISL_583860, EPI_ISL_583861 | Institute for Laboratory Diagnostics and Microbiology, Klinikum Klagenfurt am Worthersee | Bergthaler laboratory, CeMM Research Center for Molecular Medicine of the Austrian Academy of Sciences | Alexandra Popa, Benedikt Agerer, Henrique Colaco, Lukas Endler, Jakob-Wendelin Genger, Alexander Lercher, Mark Smyth, Thomas Penz, Michael Schuster, Jan Laine, Martin Senekowitsch, Judith Aberle, Stephan Aberle, Peter Hufnagl, Daniela Schmid, Franz Allerberger, Elisabeth Puchhammer-Stoeckl, Manfred Nairz, Guenter Weiss, Gregor Hörmann, Kinga Rigler-Hohenwarter, Rainer Gattringer, Wegene Borena, Dorothee von Laer, Gernot Walder, Peter Obrist, Christian Paar, Sabine Sussitz-Rack, Gunther Vogl, Adi Steinrigl, Christoph Bock, Andreas Bergthaler |
| EPI_ISL_583920 | Servicio de Microbiología. Hospital General Universitario de Castellón | SeqCOVID-SPAIN consortium/IBV(CSIC) | Rosario Moreno, María Dolores Tirado and SeqCOVID-SPAIN consortium |
| EPI_ISL_584440, EPI_ISL_584441, EPI_ISL_584442, EPI_ISL_584577, EPI_ISL_584586, EPI_ISL_584587 | UHCW / University of Warwick | COVID-19 Genomics UK (COG-UK) Consortium | Richard Stark, Chrystala Constantinidou, Meera Unnikrishnan, Laura Baxter, Jeff Cheng, Grace Taylor-Joyce, Hannah Elizabeth Bridgewater, Lucy Frost, Sarojini Pandey, Paul Brown, Tauqeer Alam, Sascha Ott, Dimitris Grammatopoulos |
| EPI_ISL_586271, EPI_ISL_586303, EPI_ISL_586306, EPI_ISL_586307, EPI_ISL_586330, EPI_ISL_586344, EPI_ISL_586345, EPI_ISL_586349, EPI_ISL_586350, EPI_ISL_586431, EPI_ISL_586432, EPI_ISL_586434, EPI_ISL_586435, EPI_ISL_586437, EPI_ISL_586438, EPI_ISL_586442, EPI_ISL_586443, EPI_ISL_586444, EPI_ISL_586445, EPI_ISL_586446, EPI_ISL_586452, EPI_ISL_586453, EPI_ISL_586455, EPI_ISL_586459, EPI_ISL_586460, EPI_ISL_586461, EPI_ISL_586463, EPI_ISL_586467, EPI_ISL_586469, EPI_ISL_586470, EPI_ISL_586471, EPI_ISL_586473, EPI_ISL_586474, EPI_ISL_586475, EPI_ISL_586476, EPI_ISL_586477, EPI_ISL_586478, EPI_ISL_586479 | | | |
| see above | Toronto Invasive Bacterial Diseases Network | McMaster University | Allison McGeer, Patryk Aftanas, Hooman Derakhshani, Angel Li, Kuganya Nirmalarajah, Emily Panousis, Ahmed Draia, Jalees Nasir, Michael Surette, Samira Mubareka, Andrew G. McArthur |
| EPI_ISL_590753, EPI_ISL_590754, EPI_ISL_590755, EPI_ISL_590756, EPI_ISL_590757, EPI_ISL_590758, EPI_ISL_590759, EPI_ISL_590760, EPI_ISL_590761, EPI_ISL_590762, EPI_ISL_590763, EPI_ISL_590764, EPI_ISL_590765, EPI_ISL_590766, EPI_ISL_590767, EPI_ISL_590768, EPI_ISL_590769, EPI_ISL_590770, EPI_ISL_590771, EPI_ISL_590772, EPI_ISL_590773, EPI_ISL_590774, EPI_ISL_590775, EPI_ISL_590776 | | | |
| see above | Minnesota Department of Health, Public Health Laboratory | Minnesota Department of Health, Public Health Laboratory | Matt Plumb, Jacob Garfin, Alexandra Lorentz, and Xiong Wang |
| EPI_ISL_590831 | Institute of Medical Virology, University of Zurich | Institute of Medical Virology, University of Zurich | Marie O. Pohl, Idoia Busnadiego, Verena Kufner, Stefan Schmutz, Maryam Zaheri, Irene Abela, Alexandra Trkola, Michael Huber, Silke Stertz, Benjamin G. Hale |
| EPI_ISL_591060, EPI_ISL_591061 | Infectious Diseases and Tropical Medicine Research Center, Infectious Diseases and Tropical Medicine Research Center | Infectious Diseases and Tropical Medicine Research Center, Infectious Diseases and Tropical Medicine Research Center | Haghjooy Javanmard,S., Ahangarzadeh,S., Shariati,L., Ataei,B., Shoaei,P., Ranjbar,M.M. |
| EPI_ISL_591080, EPI_ISL_591092 | Department of Pathology, University of Cambridge | Wellcome Sanger Institute for the COVID-19 Genomics UK (COG-UK) consortium | Luke W Meredith, M. Estée Török , Myra Hosmillo, William L. Hamilton, Martin D. Curran, Theresa Feltwell, Grant Hall, Anna Yakovleva, Fahad A Khokhar, Charlotte J. Houldcroft, Laura G Caller, Aminu S. Jahun, Sarah L. Caddy, Ian Goodfellow; and Alex Alderton, Roberto Amato, Sonia Goncalves, Ewan Harrison, David K. Jackson, Ian Johnston, Dominic Kwiatkowski, Cordelia Langford, John Sillitoe on behalf of the Wellcome Sanger Institute COVID-19 Surveillance Team |
| EPI_ISL_591123, EPI_ISL_591141, EPI_ISL_591148, EPI_ISL_591150, EPI_ISL_591152, EPI_ISL_591154, EPI_ISL_591155, EPI_ISL_591156, EPI_ISL_591157, EPI_ISL_591158, EPI_ISL_591159, EPI_ISL_591160, EPI_ISL_591164, EPI_ISL_591165, EPI_ISL_591166, EPI_ISL_591169, EPI_ISL_591170, EPI_ISL_591175, EPI_ISL_591176, EPI_ISL_591177, EPI_ISL_591178, EPI_ISL_591209, EPI_ISL_591210, EPI_ISL_591211, EPI_ISL_591212, EPI_ISL_591216, EPI_ISL_591220, EPI_ISL_591222, EPI_ISL_591223, EPI_ISL_591224, EPI_ISL_591225, EPI_ISL_591237, EPI_ISL_591238, EPI_ISL_591240, EPI_ISL_591245, EPI_ISL_591246, EPI_ISL_591247, EPI_ISL_591249, EPI_ISL_591253, EPI_ISL_591254, EPI_ISL_591255, EPI_ISL_591256, EPI_ISL_591257 | | | |
| see above | Toronto Invasive Bacterial Diseases Network | McMaster University | Allison McGeer, Patryk Aftanas, Hooman Derakhshani, Angel Li, Kuganya Nirmalarajah, Emily Panousis, Ahmed Draia, Jalees Nasir, Michael Surette, Samira Mubareka, Andrew G. McArthur |
| EPI_ISL_591519 | Programa de Virología, Facultad de Medicina UC | Center for Mathematical Modeling and Center for Genome Regulation. Santiago, Chile | Gaggero A, Valiente F, Gaete A, Travisany D, Palma R, Urra C, Varas M, Allende ML, Maass A, González M, Ferres M. |
| EPI_ISL_593481, EPI_ISL_593547 | Eastern Ontario Regional Laboratory Association | McMaster University | Leanne Mortimer, Hooman Derakhshani, Emily Panousis, Ahmed Draia, Jalees Nasir, Robert Slinger, Andrew G. McArthur |
| EPI_ISL_593771 | South Eastern Area Laboratory Services (SEALS) | NSW Health Pathology - Institute of Clinical Pathology and Medical Research; Westmead Hospital; University of Sydney | CIDM-PH et al. |
| EPI_ISL_593772, EPI_ISL_593773, EPI_ISL_593774, EPI_ISL_593775, EPI_ISL_593776 | HOSPITAL REGIONAL LAMBAYEQUE | GENOMA MAYOR | Franklin R. Aguilar-Gamboa, Luis M. López-Serquén, Heber Silva-Díaz, Percy O. Tullume-Vergara, Luis Salcedo-Mejía, Juan J. Bonifacio-Briceño, Ramsés Salas-Asencios, Marco E. Mechán-Llontop and Juan P. Cárdenas. |
| EPI_ISL_594087, EPI_ISL_594088, EPI_ISL_594089, EPI_ISL_594090, EPI_ISL_594091, EPI_ISL_594092, EPI_ISL_594093, EPI_ISL_594094, EPI_ISL_594095, EPI_ISL_594096, EPI_ISL_594097, EPI_ISL_594098, EPI_ISL_594099, EPI_ISL_594100, EPI_ISL_594101 | | | |
| see above | Utah Public Health Laboratory | Utah Public Health Laboratory | Erin Young, Kelly Oakeson |
| EPI_ISL_594187 | Department of Pathology, School of Medicine, Imam Khomeini Hospital, Tehran University of Medical Sciences | Genetics Research Center, University of Social Welfare and Rehabilitation Sciences | Zohreh Fattahi, Marzieh Mohseni, Khadijeh Jalalvand, Azam Ghaziasadi, Seyedeh elham Mortazavi, Ali Jafarpour, Azar Hadadi, Alireza Abdollahi, Ali Jafarpour, Azam Ghaziasad, Seyedeh elham Mortazavi, Saber Soltani, Reza Najafipour, Kimia Kahrizi, Seyed Mohammad Jazayeri, Hossein Najmabadi |
| EPI_ISL_594409, EPI_ISL_594410, EPI_ISL_594411, EPI_ISL_594412, EPI_ISL_594413, EPI_ISL_594414, EPI_ISL_594415, EPI_ISL_594416 | Utah Public Health Laboratory | Utah Public Health Laboratory | Erin Young, Kelly Oakeson |
| EPI_ISL_594461 | FL Bureau of Public Health Laboratories | Pathogen Discovery, Respiratory Viruses Branch, Division of Viral Diseases, Centers for Disease Control and Prevention | Ying Tao, Yan Li, Clinton Paden, Jing Zhang, Krista Queen, Anna Uehara, Haibin Wang, Julu Bhatnagar, Suxiang Tong |
| EPI_ISL_596447, EPI_ISL_596448 | New Mexico Department of Health Scientific Laboratory Division | Los Alamos National Laboratory Bioscience Division | Chien-Chi Lo, Migun Shakya, Cheryl Gleasner, Kim McMurry, Alina Deshpande, Twila Kunde, Joseph Hicks, Michael Edwards, Patrick Chain |
| EPI_ISL_596465, EPI_ISL_596472 | National Public Health Laboratory, National Centre for Infectious Diseases | National Public Health Laboratory, National Centre for Infectious Diseases | Tze Minn Mak, Sophie Octavia, Zhenyang Zhou, Lin Cui, Raymond Tzer Pin Lin |
| EPI_ISL_596651, EPI_ISL_596652, EPI_ISL_596655, EPI_ISL_596663, EPI_ISL_596666, EPI_ISL_596671, EPI_ISL_596672 | St.Vincent's University Hospital | St.Vincent's University Hospital | Mary Lucey, Guerrino Macori, Niamh Mullane, Una Sutton-Fitzpatrick, Gabriel Gonzalez, Suzie Coughlan, Aisling Purcell, Lynda Fenelon, Séamus Fanning, Kirsten Schaffer |
| EPI_ISL_596678, EPI_ISL_596680, EPI_ISL_596690, EPI_ISL_596713, EPI_ISL_596729, EPI_ISL_596731, EPI_ISL_596733, EPI_ISL_596738, EPI_ISL_596750, EPI_ISL_596773, EPI_ISL_596800, EPI_ISL_596809, EPI_ISL_596811, EPI_ISL_596813, EPI_ISL_596828, EPI_ISL_596838, EPI_ISL_596840, EPI_ISL_596841, EPI_ISL_596857, EPI_ISL_596861, EPI_ISL_596862, EPI_ISL_596867, EPI_ISL_596872 | | | |
| see above | PathWest Laboratory Medicine WA | PathWest Laboratory Medicine WA Microbial Surveillance Unit | PathWest Laboratory Medicine WA Microbial Surveillance Unit |
| EPI_ISL_600428, EPI_ISL_600429, EPI_ISL_600432, EPI_ISL_600433, EPI_ISL_600434, EPI_ISL_600435, EPI_ISL_600567 | Institute of Epidemiology Disease Control And Research | Institute for Developing Science and Health Initiatives | Lauren Cowley, Mokibul Hassan Afrad, Sadia Isfat Ara Rahman, Md. Mahfuz-Al-mamun, Firadausi Qadri, Tahmina Shirin |
| EPI_ISL_602208, EPI_ISL_602209, | New Mexico Department of Health Scientific Laboratory | Los Alamos National Laboratory Bioscience Division | Chien-Chi Lo, Migun Shakya, Cheryl Gleasner, Kim McMurry, Alina Deshpande, Twila Kunde, Joseph Hicks, Michael Edwards, Patrick Chain |

| | | | |
|---|---|---|---|
| EPI_ISL_602278, EPI_ISL_602279, EPI_ISL_602280, EPI_ISL_602281 | Division | | |
| EPI_ISL_602286, EPI_ISL_602297, EPI_ISL_602299, EPI_ISL_602302 | Evangelisches Klinikum Bethel, Institut für Laboratoriumsmedizin, Mikrobiologie und Hygiene | Bielefeld University | David Brandt, Tobias Busche, Markus Haak, Jörn Kalinowski, Levin-Joe Klages, Christiane Scherer, Alexander Sczyrba, Marina Simunovic, Svenja Vinke |
| EPI_ISL_602477, EPI_ISL_602478, EPI_ISL_602479, EPI_ISL_602480, EPI_ISL_602481, EPI_ISL_602482, EPI_ISL_602483, EPI_ISL_602484 | Institute for Virology, University Hospital Essen | Center of Medical Microbiology, Virology, and Hospital Hygiene, University of Duesseldorf | Olympia E. Anastasiou, Ulf Dittmer, Maximilian Damagnez, Alexander Dilthey, Torsten Houwaart, Lisanna Hülse, Malte Kohns Vasconcelos, Nadine Lübke, Jessica Nicolai, Klaus Pfeffer, Daniel Strelow, Jörg Timm, Andreas Walker, Tobias Wienemann |
| EPI_ISL_602514, EPI_ISL_602515 | Center of Medical Microbiology, Virology, and Hospital Hygiene, University of Duesseldorf | Center of Medical Microbiology, Virology, and Hospital Hygiene, University of Duesseldorf | Maximilian Damagnez, Alexander Dilthey, Torsten Houwaart, Lisanna Hülse, Malte Kohns Vasconcelos, Marek Korencak, Nadine Lübke, Jessica Nicolai, Klaus Pfeffer, Hendrik Streeck, Daniel Strelow, Jörg Timm, Andreas Walker, Tobias Wienemann |
| EPI_ISL_602536, EPI_ISL_602539, EPI_ISL_602545, EPI_ISL_602547 | Center of Medical Microbiology, Virology, and Hospital Hygiene, University of Duesseldorf | Center of Medical Microbiology, Virology, and Hospital Hygiene, University of Duesseldorf | Maximilian Damagnez, Alexander Dilthey, Torsten Houwaart, Malte Kohns Vasconcelos, Marek Korencak, Nadine Lübke, Jessica Nicolai, Klaus Pfeffer, Hendrik Streeck, Daniel Strelow, Jörg Timm, Andreas Walker, Tobias Wienemann |
| EPI_ISL_603092, EPI_ISL_603094, EPI_ISL_603095, EPI_ISL_603096, EPI_ISL_603119, EPI_ISL_603120, EPI_ISL_603121 | Lithuanian University of Health Sciences Hospital, Department of Laboratory Medicine | Lithuanian University of Health Sciences, Molecular cardiology lab. | Lukas Zemaitis, Ingrida Olendrait, Arnoldas Pautienius, Kamile Tamusauskaite, Dovydas Gecys, Laura Pareckaite, Vaiva Lesauskaite, Astra Vitkauskiene |
| EPI_ISL_603256, EPI_ISL_603259 | UW Virology Lab | UW Virology Lab | Pavitra Roychoudhury, Hong Xie, Lasata Shrestha, Meei-Li Huang, Keith R Jerome, Alexander Greninger |
| EPI_ISL_605800, EPI_ISL_605813 | Clinical Virology Laboratory, Institute of Liver and Biliary Sciences | ILBS - IGIB | Ekta Gupta, Sheetalnath Rooge, Abhishek Padhi, Reshu Agarwal, Jaswinder Singh Maras, Shridhar Sivasubbu, Vinod Scaria, Shvetank Sharma |
| EPI_ISL_613456 | Institut Pasteur de la Guadeloupe | Institut Pasteur de la Guadeloupe | Marion Barbet, Sylvie Behillil, Méline Bizard, Angela Brisebarre, Camille Capel, Etienne Simon-Lorière, Vincent Enouf, Maud Vanpeene, Sylvie van der Werf, Stéphanie Guyomard, Sébastien Breurec, Radjin Steingrover, Cherina Fleming, Antoine Talarmin |
| EPI_ISL_613710 | Laboratorio Biologia Molecolare Sars Cov2 - UOC Laboratorio Analisi - Servizio Medicina di Laboratorio, Ospedale "San Francesco" - ATS-ASSL Nuoro | Laboratorio specialistico UOC Ematologia - Ospedale "San Francesco" - ATS-ASSL Nuoro | Piras Giovanna, Fancello Tatiana, Asproni Rosanna, Fiamma Maura, Monne Maria Itria, Toja Alessandro, Sanna Filomena, Floris Anna Rita, Sulis Vincenzo, Palmas Angelo Domenico, Casu Gavino, Lo Maglio Iana, Mameli Giuseppe. |
| EPI_ISL_613714, EPI_ISL_613731, EPI_ISL_613734, EPI_ISL_613736, EPI_ISL_613738, EPI_ISL_613744, EPI_ISL_613746, EPI_ISL_613757, EPI_ISL_613758, EPI_ISL_613765, EPI_ISL_613767, EPI_ISL_613775, EPI_ISL_613781, EPI_ISL_613782, EPI_ISL_613788, EPI_ISL_613798, EPI_ISL_613800, EPI_ISL_613802, EPI_ISL_613803, EPI_ISL_613804, EPI_ISL_613806, EPI_ISL_613810, EPI_ISL_613811, EPI_ISL_613814, EPI_ISL_613817, EPI_ISL_613823, EPI_ISL_613824, EPI_ISL_613825, EPI_ISL_613904, EPI_ISL_613905, EPI_ISL_613906, EPI_ISL_613907, EPI_ISL_613908, EPI_ISL_613909, EPI_ISL_613910, EPI_ISL_613911, EPI_ISL_613912, EPI_ISL_613913, EPI_ISL_613914, EPI_ISL_613915, EPI_ISL_613917, EPI_ISL_613918, EPI_ISL_613919, EPI_ISL_613920, EPI_ISL_613921, EPI_ISL_613922, EPI_ISL_613923, EPI_ISL_613924, EPI_ISL_613925, EPI_ISL_613926, EPI_ISL_613927, EPI_ISL_613928, EPI_ISL_613929, EPI_ISL_613930, EPI_ISL_613931, EPI_ISL_613933, EPI_ISL_613935, EPI_ISL_613936, EPI_ISL_613937, EPI_ISL_613938, EPI_ISL_613939, EPI_ISL_613940, EPI_ISL_613941, EPI_ISL_613942, EPI_ISL_613943, EPI_ISL_613944, EPI_ISL_613945, EPI_ISL_613946, EPI_ISL_613947, EPI_ISL_613948 | | | |
| see above | Florida Bureau of Public Health Laboratories | Florida Bureau of Public Health Laboratories | Sarah Schmedes, Jason Blanton |
| EPI_ISL_613953, EPI_ISL_613955 | Laboratorio Biologia Molecolare Sars Cov2 - UOC Laboratorio Analisi - Servizio Medicina di Laboratorio, Ospedale "San Francesco" - ATS-ASSL Nuoro | Laboratorio specialistico UOC Ematologia - Ospedale "San Francesco" - ATS-ASSL Nuoro | Piras Giovanna, Fancello Tatiana, Asproni Rosanna, Fiamma Maura, Monne Maria Itria, Toja Alessandro, Sanna Filomena, Floris Anna Rita, Sulis Vincenzo, Palmas Angelo Domenico, Casu Gavino, Lo Maglio Iana, Mameli Giuseppe |
| EPI_ISL_614120, EPI_ISL_614121, EPI_ISL_614122 | Virginia DCLS | Virginia DCLS | Virginia DCLS |
| EPI_ISL_614247 | Florida Bureau of Public Health Laboratories | Florida Bureau of Public Health Laboratories | Sarah Schmedes, Jason Blanton |
| EPI_ISL_623116, EPI_ISL_623128, EPI_ISL_623132, EPI_ISL_623168 | Laboratorio de Virologia Molecular / UFRJ | Bioinformatics Laboratory / LNCC | Carolina M Voloch, Ronaldo S Francisco Jr, Luiz G P de Almeida, Otavio J. Brustolini, Cynthia C Cardoso, Alexandra L Gerber, Ana Paula de C Guimarães, Diana Mariani, Covid19-UFRJ Workgroup, Luís Cristóvão Pôrto, Renato S Aguiar, Terezinha M P P Castiñeiras, Orlando C. Ferreira, Amilcar Tanuri, Ana Tereza R de Vasconcelos |
| EPI_ISL_623186, EPI_ISL_623188, EPI_ISL_623189, EPI_ISL_623190, EPI_ISL_623191, EPI_ISL_623192, EPI_ISL_623193, EPI_ISL_623195, EPI_ISL_623196, EPI_ISL_623198, EPI_ISL_623199, EPI_ISL_623200, EPI_ISL_623201, EPI_ISL_623203, EPI_ISL_623204, EPI_ISL_623205 | | | |
| see above | Utah Public Health Laboratory | Utah Public Health Laboratory | Erin Young, Kelly Oakeson |
| EPI_ISL_625594, EPI_ISL_625595, EPI_ISL_625596, EPI_ISL_625597, EPI_ISL_625598, EPI_ISL_625599 | Santa Clara County Public Health Laboratory | Chan-Zuckerberg Biohub | CZB Cliahub Consortium |
| EPI_ISL_626403, EPI_ISL_626404, EPI_ISL_626405, EPI_ISL_626406, EPI_ISL_626407, EPI_ISL_626408 | Northwestern Memorial Hospital | Ozer Lab | Ramon Lorenzo-Redondo, Hannah H. Nam, Scott C. Roberts, Lacy M. Simons, Chad J. Achenbach, Lawrence J. Jennings, Chao Qi, Alan R. Hauser, Michael G. Ison, Judd F. Hultquist, Egon A. Ozer |
| EPI_ISL_626690, EPI_ISL_627150, EPI_ISL_628312, EPI_ISL_628313, EPI_ISL_628314, EPI_ISL_628315, EPI_ISL_628316, EPI_ISL_628317, EPI_ISL_628318, EPI_ISL_628319 | Centre for Enzyme Innovation, University of Portsmouth / Translational Research Laboratory, Portsmouth Hospitals NHS Trust | COVID-19 Genomics UK (COG-UK) Consortium | Angela Beckett,Yann Bourgeois,Garry Scarlett,Sharon Glaysher,Scott Elliott,Kelly Bicknell,Robert Impey,Allyson Lloyd,Sarah Wyllie,Ethan Butcher,Anoop Chauhan,Samuel Robson |
| EPI_ISL_628962, EPI_ISL_628963 | Utah Public Health Laboratory | Utah Public Health Laboratory | Erin Young, Kelly Oakeson |
| EPI_ISL_629098, EPI_ISL_629099, EPI_ISL_629100, EPI_ISL_629101 | Laboratoire du Centre Hospitalier Annecy Genevois | CNR Virus des Infections Respiratoires - France SUD | Antonin Bal, Gregory Destras, Gwendolyne Burfin, Hadrien Règue, Quentin Semanas, Martine Valette, Bruno Lina, Hélène Petitprez, Bruno Chanzy, Laurence Josset |
| EPI_ISL_631691 | NYC HH Lincoln Medical And Mental Health Center | New York City Public Health Laboratory | Jade Wang, et al. |
| EPI_ISL_631692 | Flushing Hospital Medical Center | New York City Public Health Laboratory | Jade Wang, et al. |
| EPI_ISL_631693 | NYC HH Lincoln Medical And Mental Health Center | New York City Public Health Laboratory | Jade Wang, et al. |
| EPI_ISL_631694, EPI_ISL_631695 | OCME Office Of Chief Medical Examiner | New York City Public Health Laboratory | Jade Wang, et al. |
| EPI_ISL_631696 | Flushing Hospital Medical Center | New York City Public Health Laboratory | Jade Wang, et al. |
| EPI_ISL_631697, EPI_ISL_631698 | Jamaica Hospital Medical Center | New York City Public Health Laboratory | Jade Wang, et al. |
| EPI_ISL_631699, EPI_ISL_631700, EPI_ISL_631701 | Flushing Hospital Medical Center | New York City Public Health Laboratory | Jade Wang, et al. |
| EPI_ISL_631702, EPI_ISL_631703, EPI_ISL_631704, EPI_ISL_631705, EPI_ISL_631706 | Jamaica Hospital Medical Center | New York City Public Health Laboratory | Jade Wang, et al. |
| EPI_ISL_631707, EPI_ISL_631708 | Flushing Hospital Medical Center | New York City Public Health Laboratory | Jade Wang, et al. |
| EPI_ISL_631709 | Jamaica Hospital Medical Center | New York City Public Health Laboratory | Jade Wang, et al. |
| EPI_ISL_631710 | Flushing Hospital Medical Center | New York City Public Health Laboratory | Jade Wang, et al. |
| EPI_ISL_631711, EPI_ISL_631722, EPI_ISL_631767, EPI_ISL_631768 | Jamaica Hospital Medical Center | New York City Public Health Laboratory | Jade Wang, et al. |

| | | | |
|---|---|---|---|
| EPI_ISL_631769, EPI_ISL_631770, EPI_ISL_631771 | Flushing Hospital Medical Center | New York City Public Health Laboratory | Jade Wang, et al. |
| EPI_ISL_631847 | Jamaica Hospital Medical Center | New York City Public Health Laboratory | Jade Wang, et al. |
| EPI_ISL_631848 | Flushing Hospital Medical Center | New York City Public Health Laboratory | Jade Wang, et al. |
| EPI_ISL_631849, EPI_ISL_631850 | Jamaica Hospital Medical Center | New York City Public Health Laboratory | Jade Wang, et al. |
| EPI_ISL_631851 | Flushing Hospital Medical Center | New York City Public Health Laboratory | Jade Wang, et al. |
| EPI_ISL_631852, EPI_ISL_631853, EPI_ISL_631854 | Jamaica Hospital Medical Center | New York City Public Health Laboratory | Jade Wang, et al. |
| EPI_ISL_631855 | Flushing Hospital Medical Center | New York City Public Health Laboratory | Jade Wang, et al. |
| EPI_ISL_631878, EPI_ISL_631879 | Jamaica Hospital Medical Center | New York City Public Health Laboratory | Jade Wang, et al. |
| EPI_ISL_631893, EPI_ISL_631916 | Flushing Hospital Medical Center | New York City Public Health Laboratory | Jade Wang, et al. |
| EPI_ISL_631950, EPI_ISL_631951 | Jamaica Hospital Medical Center | New York City Public Health Laboratory | Jade Wang, et al. |
| EPI_ISL_631959, EPI_ISL_631960 | Flushing Hospital Medical Center | New York City Public Health Laboratory | Jade Wang, et al. |
| EPI_ISL_631985 | St Barnabas Hospital | New York City Public Health Laboratory | Jade Wang, et al. |
| EPI_ISL_631986, EPI_ISL_632197 | Flushing Hospital Medical Center | New York City Public Health Laboratory | Jade Wang, et al. |
| EPI_ISL_632198 | Jamaica Hospital Medical Center | New York City Public Health Laboratory | Jade Wang, et al. |
| EPI_ISL_634824 | Laboratoire de virologie, CHU de Grenoble - CS 10217 - 38043 Grenoble cedex 16 | CNR Virus des Infections Respiratoires - France SUD | Antonin Bal, Gregory Destras, Gwendolyne Burfin, Hadrien Règue, Quentin Semanas, Martine Valette, Bruno Lina, Sylvie Larrat, Laurence Josset |
| EPI_ISL_634825 | Laboratoire de virologie, CHU de Grenoble - CS 10217 - 38043 Grenoble cedex 17 | CNR Virus des Infections Respiratoires - France SUD | Antonin Bal, Gregory Destras, Gwendolyne Burfin, Hadrien Règue, Quentin Semanas, Martine Valette, Bruno Lina, Sylvie Larrat, Laurence Josset |
| EPI_ISL_634826 | Laboratoire de virologie, CHU de Grenoble - CS 10217 - 38043 Grenoble cedex 18 | CNR Virus des Infections Respiratoires - France SUD | Antonin Bal, Gregory Destras, Gwendolyne Burfin, Hadrien Règue, Quentin Semanas, Martine Valette, Bruno Lina, Sylvie Larrat, Laurence Josset |
| EPI_ISL_634827 | Laboratoire de virologie, CHU de Grenoble - CS 10217 - 38043 Grenoble cedex 19 | CNR Virus des Infections Respiratoires - France SUD | Antonin Bal, Gregory Destras, Gwendolyne Burfin, Hadrien Règue, Quentin Semanas, Martine Valette, Bruno Lina, Sylvie Larrat, Laurence Josset |
| EPI_ISL_634828 | Laboratoire de virologie, CHU de Grenoble - CS 10217 - 38043 Grenoble cedex 20 | CNR Virus des Infections Respiratoires - France SUD | Antonin Bal, Gregory Destras, Gwendolyne Burfin, Hadrien Règue, Quentin Semanas, Martine Valette, Bruno Lina, Sylvie Larrat, Laurence Josset |
| EPI_ISL_635294, EPI_ISL_635295, EPI_ISL_635296, EPI_ISL_635297, EPI_ISL_635298 | Institute of Microbiology and Immunology, Faculty of Medicine, University of Ljubljana | Institute of Microbiology and Immunology, Faculty of Medicine, University of Ljubljana | Tomaž Mark Zorec, Samo Zakotnik, Miša Korva, Tatjana Avši - Županc, Mario Poljak |
| EPI_ISL_636503 | Dutch COVID-19 response team | National Institute for Public Health and the Environment (RIVM) | Adam Meijer, Harry Vennema, Jeroen Cremer, Sharon van den Brink, Bas van der Veer, AnneMarie van den Brandt, Florian Zwagemaker, Dennis Schmitz, Chantal Reusken, on behalf of the national COVID-19 response team |
| EPI_ISL_636992, EPI_ISL_636993 | Department of Infectious Diseases and Immunology, National Hospital Organization Nagoya Medical Center | Clinical Research Center, National Hospital Organization Nagoya Medical Center | Yoshihiro Nakata, Hirotaka Ode, Mai Kubota, Masakazu Matsuda, Kazuhiro Matsuoka, Miho Nakasuji, Mikiko Mori, Mayumi Imahashi, Yoshiyuki Yokomaku, Yasumasa Iwatani |
| EPI_ISL_640220, EPI_ISL_640224, EPI_ISL_640225, EPI_ISL_640227, EPI_ISL_640234, EPI_ISL_640236, EPI_ISL_640242, EPI_ISL_640243, EPI_ISL_640249, EPI_ISL_640251, EPI_ISL_640254, EPI_ISL_640258, EPI_ISL_640260, EPI_ISL_640262, EPI_ISL_640263, EPI_ISL_640266, EPI_ISL_640271, EPI_ISL_640272 | | | |
| see above | MVZ Laborärzte Singen | MVZ Laborärzte Singen | Jonas Schmidt, Frithjof Blessing, Sandro Berghaus, Folker Wenzel |
| EPI_ISL_641530 | CHU de Nice - Hôpital Archet 12 | CNR Virus des Infections Respiratoires - France SUD | Antonin Bal, Géraldine Gonfrier, Gregory Destras, Gwendolyne Burfin, Hadrien Règue, Quentin Semanas, Martine Valette, Bruno Lina, Valérie Giordanengo, Laurence Josset |
| EPI_ISL_641535, EPI_ISL_641536, EPI_ISL_641544 | CHU de Saint-Étienne Hôpital Nord | CNR Virus des Infections Respiratoires - France SUD | Antonin Bal, Gregory Destras, Gwendolyne Burfin, Hadrien Règue, Quentin Semanas, Martine Valette, Bruno Lina, Issam Bechri, Manon Vogrig, Marine Delorme, Bruno Pozzetto, Thomas Bourlet, Sylvie Gonzalo, Sylvie Pillet, Laurence Josset |
| EPI_ISL_641546 | CHU Clermont-Ferrand | CNR Virus des Infections Respiratoires - France SUD | Antonin Bal, Gregory Destras, Gwendolyne Burfin, Hadrien Règue, Quentin Semanas, Martine Valette, Bruno Lina, Christine Archimbaud, Amélie Brebion, Hélène Chabrolles, Martine Chambon, Audrey Mirand, Christel Regagnon, Maxime Bisseux, Patricia Combes, Cécile Henquell, Laurence Josset |
| EPI_ISL_644169 | Texas Department of State Health Services | Texas Department of State Health Services | Rashmi Tuladhar, Bonnie Oh, Jenny Zhang, Maliha Rahman, Anita Pokharel, Myong Koag, Chung Wang, Rachel Lee, Grace Kubin, Mayela Pedrueza |
| EPI_ISL_644234, EPI_ISL_644238, EPI_ISL_644239, EPI_ISL_644244, EPI_ISL_644257, EPI_ISL_644259, EPI_ISL_644260, EPI_ISL_644263, EPI_ISL_644274, EPI_ISL_644276, EPI_ISL_644287, EPI_ISL_644292, EPI_ISL_644300, EPI_ISL_644303, EPI_ISL_644306, EPI_ISL_644309, EPI_ISL_644316, EPI_ISL_644332, EPI_ISL_644335, EPI_ISL_644339 | | | |
| see above | CEPHR / Vincent's Hospital | Irish Coronavirus Sequencing Consortium - National Virus Reference Laboratory | Michael Carr, Gabriel Gonzalez, Alejandro Abner Garcia Leon, Patrick Mallon |
| EPI_ISL_644678, EPI_ISL_644679 | CHU de Limoges | CNR Virus des Infections Respiratoires - France SUD | Antonin Bal, Gregory Destras, Gwendolyne Burfin, Hadrien Règue, Quentin Semanas, Martine Valette, Bruno Lina, Sylvie Rogez, Laurence Josset |
| EPI_ISL_645004, EPI_ISL_645005, EPI_ISL_645006, EPI_ISL_645007, EPI_ISL_645008, EPI_ISL_645009, EPI_ISL_645011, EPI_ISL_645012, EPI_ISL_645036, EPI_ISL_645037, EPI_ISL_645042, EPI_ISL_645077 | | | |
| see above | Human Genome Variation Research Group, Malopolska Centre of Biotechnology | Human Genome Variation Research Group, Malopolska Centre of Biotechnology | Kowalski,M., Pospiech,E., Klajmon,A., Gromowski,T., Pisarek,A., Marszalek,K., Kopera,K., Foremny,J., Swadzba,J., Sanak,M., Owczarek,K., Dabrowska,A., Szczepanski,A., Botwina,P., Labaj,P.P., Pyrc,K., Branicki,W. |
| EPI_ISL_648036 | WI State Laboratory of Hygiene | Pathogen Discovery, Respiratory Viruses Branch, Division of Viral Diseases, Centers for Disease Control and Prevention | Yan Li, Jing Zhang, Ying Tao, Brian Lynch, Krista Queen, Anna Montmayeur, Anna Uehara, Clinton R. Paden, Rachel Marine, Haibin Wang, Suxiang Tong |
| EPI_ISL_648093, EPI_ISL_648094, EPI_ISL_648095, EPI_ISL_648096, EPI_ISL_648097, EPI_ISL_648098, EPI_ISL_648099, EPI_ISL_648100, EPI_ISL_648101, EPI_ISL_648102, EPI_ISL_648103, EPI_ISL_648104, EPI_ISL_648105, EPI_ISL_648106, EPI_ISL_648107, EPI_ISL_648108, EPI_ISL_648109, EPI_ISL_648110, EPI_ISL_648111 | | | |
| see above | Department of Laboratory Medicine, Tan Tock Seng Hospital | Department of Laboratory Medicine, Tan Tock Seng Hospital | Chen YYC, Zair X, Lim JX, Li C, Tang WY, Maurer-Stroh S, Barkham TMS, Nagarajan N, Sessions OM |
| EPI_ISL_648314 | Laboratorio de Investigaciones de Baney | University Hospital Basel, Clinical Bacteriology | Carlos Cortes, Claudia Daubenberger, Adrian Egli, Guillermo Garcia, Salome Hosch, Bonifacio Manguire Nlavo, Alfredo Mari, Maximilian Mpina, Elizabeth Nyakarungu, Diosdado Odjama Nseng Ada, Mitoha Ondo O Ayekaba, Tim Roloff, Tobias Schindler, Helena Seth-Smith, Madlen Stange, Philip Wonder Phiri |
| EPI_ISL_648558, EPI_ISL_648559, EPI_ISL_648560, EPI_ISL_648561, EPI_ISL_648562, EPI_ISL_648563, EPI_ISL_648569, EPI_ISL_648575 | Utah Public Health Laboratory | Utah Public Health Laboratory | Erin Young, Kelly Oakeson |
| EPI_ISL_649183 | CHU Nantes | CNR Virus des Infections Respiratoires - France SUD | Antonin Bal, Louise Castain, Gregory Destras, Gwendolyne Burfin, Hadrien Règue, Quentin Semanas, Martine Valette, Bruno Lina, Celine Bressollette, Laurence Josset |
| EPI_ISL_649189 | Istituto Zooprofilattico Sperimentale della Puglia e della Basilicata | Istituto Zooprofilattico Sperimentale della Puglia e della Basilicata | Parisi A., Bianco A., Capozzi L., Del Sambro L., Manzulli V, Rondinone V., Pace L., Galante D., Cipolletta D. |
| EPI_ISL_649947, EPI_ISL_649948 | CHU de Saint-Étienne Hôpital Nord | CNR Virus des Infections Respiratoires - France SUD | Antonin Bal, Gregory Destras, Gwendolyne Burfin, Hadrien Règue, Quentin Semanas, Martine Valette, Bruno Lina, Issam Bechri, Manon Vogrig, Marine Delorme, Bruno Pozzetto, Thomas Bourlet, Sylvie Gonzalo, Sylvie Pillet, Laurence Josset |

| | | | |
|---|---|---|---|
| EPI_ISL_653098, EPI_ISL_653099, EPI_ISL_653100, EPI_ISL_653101, EPI_ISL_653102, EPI_ISL_653103, EPI_ISL_653104, EPI_ISL_653105, EPI_ISL_653106, EPI_ISL_653107, EPI_ISL_653108, EPI_ISL_653109, EPI_ISL_653113, EPI_ISL_653117, EPI_ISL_653143, EPI_ISL_653279, EPI_ISL_653280, EPI_ISL_653281, EPI_ISL_653282, EPI_ISL_653283 | | | |
| see above | Florida Bureau of Public Health Laboratories | Florida Bureau of Public Health Laboratories | Sarah Schmedes, Jason Blanton |
| EPI_ISL_653813 | Istituto Zooprofilattico Sperimentale della Puglia e della Basilicata | Istituto Zooprofilattico Sperimentale della Puglia e della Basilicata | Parisi A., Bianco A., Capozzi L., Del Sambro L., Manzulli V, Rondinone V., Pace L., Cipolletta D., Galante D. |
| EPI_ISL_653814, EPI_ISL_653815, EPI_ISL_653816, EPI_ISL_653817, EPI_ISL_653818 | I.R.C.C.S. "S. De Bellis" - Ente Ospedaliero | Istituto Zooprofilattico Sperimentale della Puglia e della Basilicata | Parisi A., Bianco A., Capozzi L., Del Sambro L., Lippolis A., Notarnicola M., Cipolletta D., Galante D. |
| EPI_ISL_654535 | Servicio de Microbiología, Laboratori Clínic Metropolitana Nord. Hospital Universitari Germans Trias i Pujol. Institut d'Investigació en Ciències de la Salut Germans Trias i Pujol (IGTP) | SeqCOVID-SPAIN consortium/IBV(CSIC) | Elisa Martró, Antoni E. Bordoy, Anna Not, Adrián Antuori, Anabel Fernández, Nona Romaní and SeqCOVID-SPAIN consortium |
| EPI_ISL_654622, EPI_ISL_654668, EPI_ISL_654669, EPI_ISL_654670, EPI_ISL_654678, EPI_ISL_654679 | Servicio de Microbiología, Hospital Universitario Central de Asturias | SeqCOVID-SPAIN consortium/IBV(CSIC) | Cristián Castelló Abietar, Jose A. Boga, Susana Rojo-Alba, Marta Elena Álvarez-Argüelles, Santiago Melón and SeqCOVID-SPAIN consortium |
| EPI_ISL_654923, EPI_ISL_654926, EPI_ISL_654927, EPI_ISL_654928, EPI_ISL_654929 | Texas Department of State Health Services | Texas Department of State Health Services | Rashmi Tuladhar, Bonnie Oh, Jenny Zhang, Maliha Rahman, Anita Pokharel, Myong Koag, Chung Wang, Rachel Lee, Grace Kubin, Mayela Pedrueza, James Daniel Bonser |
| EPI_ISL_658904 | Instituto de Diagnostico y Referencia Epidemiologicos (INDRE) | Instituto de Diagnostico y Referencia Epidemiologicos (INDRE) | Ernesto Ramirez-Gonzalez, Abril Rodriguez-Maldonado, Claudia Wong-Arambula , Natividad Cruz-Ortiz, Tatiana Nunez-Garcia, Dayanira Arellano-Suarez, Fabiola Garces-Ayala, Lucia Hernandez-Rivas, Irma Lopez-Martinez, Gisela Barrera-Badillo, Marisol Galindo-Galindo, Linda Andarade-Sanchez, Octavio Ruiz-Muñiz, Pamela Ramirez-Medina, Roman Canul-Aguilar, Gaudalupe Herrera-Ramirez, Jazmin Galicia-Hernandez. |
| EPI_ISL_660333, EPI_ISL_660342 | CHU de Saint-Étienne Hôpital Nord | CNR Virus des Infections Respiratoires - France SUD | Antonin Bal, Gregory Destras, Gwendolyne Burfin, Hadrien Règue, Quentin Semanas, Martine Valette, Bruno Lina, Issam Bechri, Manon Vogrig, Marine Delorme, Bruno Pozzetto, Thomas Bourlet, Sylvie Gonzalo, Sylvie Pillet, Laurence Josset |
| EPI_ISL_660362 | CHU Clermont-Ferrand | CNR Virus des Infections Respiratoires - France SUD | Antonin Bal, Gregory Destras, Gwendolyne Burfin, Hadrien Règue, Quentin Semanas, Martine Valette, Bruno Lina, Christine Archimbaud, Amélie Brebion, Hélène Chabrolles, Martine Chambon, Audrey Mirand, Christel Regagnon, Maxime Bisseux, Patricia Combes, Cécile Henquell, Laurence Josset |
| EPI_ISL_660374, EPI_ISL_660375, EPI_ISL_660376, EPI_ISL_660377 | CHU Toulouse | CNR Virus des Infections Respiratoires - France SUD | Antonin Bal, Gregory Destras, Gwendolyne Burfin, Hadrien Règue, Quentin Semanas, Martine Valette, Bruno Lina, Jean Michel Mansuy, Laurence Josset |
| EPI_ISL_660407, EPI_ISL_660408, EPI_ISL_660409, EPI_ISL_660410, EPI_ISL_660411, EPI_ISL_660412, EPI_ISL_660413, EPI_ISL_660414 | Klinisk mikrobiologi Linkoping | The Public Health Agency of Sweden | Anna-Malin Linde, Maria Lind Karlberg, Mattias Haukland, Reza Advani, Olov Svartstrom, Oskar Karlsson Lindsjo, Sandra Broddesson, Petra Edquist, Mia Brytting, Anna Risberg, Karin Tegmark-Wisell |
| EPI_ISL_660540 | MVZ Laborärzte Singen | MVZ Laborärzte Singen | Jonas Schmidt, Frithjof Blessing, Sandro Berghaus, Folker Wenzel |
| EPI_ISL_663224, EPI_ISL_663225, EPI_ISL_663226 | CHU de Limoges | CNR Virus des Infections Respiratoires - France SUD | Antonin Bal, Gregory Destras, Gwendolyne Burfin, Hadrien Règue, Quentin Semanas, Martine Valette, Bruno Lina, Sylvie Rogez, Laurence Josset |
| EPI_ISL_663282, EPI_ISL_663283 | CHU Nantes | CNR Virus des Infections Respiratoires - France SUD | Antonin Bal, Louise Castain, Gregory Destras, Gwendolyne Burfin, Hadrien Règue, Quentin Semanas, Martine Valette, Bruno Lina, Virginie Ferré, Celine Bressollette, Laurence Josset |
| EPI_ISL_664427, EPI_ISL_664438, EPI_ISL_664490, EPI_ISL_664525, EPI_ISL_664669, EPI_ISL_664673, EPI_ISL_664723, EPI_ISL_665190, EPI_ISL_665191, EPI_ISL_665192 | University College London Hospital | COVID-19 Genomics UK (COG-UK) Consortium | Judith Heaney, Matthew Byott, Catherine Houlihan, Dan Frampton, Stuart Kirk, Moira Spyer and Eleni Nastouli |
| EPI_ISL_665260 | Dept. of Microbiology and Infection Control, Akershus University Hospital HF | Dept. of Microbiology and Infection Control, Akershus University Hospital HF | Hege Vangstein Aamot, Alexander Hesselberg Løvestad, Silje Bakken Jørgensen, Nina Handal, Ole Herman Ambur |
| EPI_ISL_666669, EPI_ISL_666670, EPI_ISL_666671, EPI_ISL_666672 | Laboratoire du Centre Hospitalier Annecy Genevois | CNR Virus des Infections Respiratoires - France SUD | Antonin Bal, Gregory Destras, Gwendolyne Burfin, Hadrien Règue, Quentin Semanas, Martine Valette, Bruno Lina, Hélène Petitprez, Bruno Chanzy, Laurence Josset |
| EPI_ISL_672093, EPI_ISL_672094, EPI_ISL_672116, EPI_ISL_672117, EPI_ISL_672118, EPI_ISL_672119, EPI_ISL_672120, EPI_ISL_672121, EPI_ISL_672122, EPI_ISL_672126, EPI_ISL_672189, EPI_ISL_672235 | | | |
| see above | The Ashley Laboratory, Stanford University | Chan-Zuckerberg Biohub | CZB Cliahub Consortium |
| EPI_ISL_672275, EPI_ISL_672276 | Santa Clara County Public Health Laboratory | Chan-Zuckerberg Biohub | CZB Cliahub Consortium |
| EPI_ISL_672544, EPI_ISL_672561 | Utah Public Health Laboratory | Utah Public Health Laboratory | Erin Young, Kelly Oakeson |
| EPI_ISL_672576 | Infectious Diseases and Tropical Medicine Research Center, Infectious Diseases and Tropical Medicine Research Center | Infectious Diseases and Tropical Medicine Research Center, Infectious Diseases and Tropical Medicine Research Center | Ahangarzadeh,S., Ataei,B., Shariati,L., Haghjooy Javanmard,S. |
| EPI_ISL_672585 | Infectious Diseases and Tropical Medicine Research Center, Infectious Diseases and Tropical Medicine Research Center | Infectious Diseases and Tropical Medicine Research Center, Infectious Diseases and Tropical Medicine Research Center | Haghjooy Javanmard,S., Shoaei,P., Ataei,B., Shariati,L., Ahangarzadeh,S. |
| EPI_ISL_672664, EPI_ISL_672670, EPI_ISL_672671, EPI_ISL_672673, EPI_ISL_672680, EPI_ISL_672681, EPI_ISL_672682, EPI_ISL_672684 | DB Diagnosticos do Brasil | Laboratório de Parasitologia Médica - Instituto de Medicina Tropical - Universidade de São Paulo | Brazil-UK Centre for Arbovirus Discovery Diagnosis Genomics and Epidemiology (CADDE) Genomic Network - Instituto de Medicina Tropical |
| EPI_ISL_672687 | Hospital das Clínicas da Faculdade de Medicina da Universidade de São Paulo (HC-FMUSP) | Laboratório de Parasitologia Médica - Instituto de Medicina Tropical - Universidade de São Paulo | Brazil-UK Centre for Arbovirus Discovery Diagnosis Genomics and Epidemiology (CADDE) Genomic Network - Instituto de Medicina Tropical |
| EPI_ISL_672701, EPI_ISL_672702, EPI_ISL_672703, EPI_ISL_672704, EPI_ISL_672705, EPI_ISL_672706, EPI_ISL_672707, EPI_ISL_672708, EPI_ISL_672709, EPI_ISL_672710, EPI_ISL_672711, EPI_ISL_672712 | | | |
| see above | Institute of Tropical Medicine at the University of São Paulo (IMT-USP) | Laboratório de Parasitologia Médica - Instituto de Medicina Tropical - Universidade de São Paulo | Brazil-UK Centre for Arbovirus Discovery Diagnosis Genomics and Epidemiology (CADDE) Genomic Network - Instituto de Medicina Tropical |
| EPI_ISL_676533 | Karolinska Universitetslaboratoriet | The Public Health Agency of Sweden | Department of Microbiology, The Public Health Agency of Sweden |
| EPI_ISL_676542 | CNR Virus des Infections Respiratoires - France SUD | CNR Virus des Infections Respiratoires - France SUD | Antonin Bal, Gregory Destras, Gwendolyne Burfin, Solenne Brun, Alexandre Gaymard, Maude Bouscambert-Duchamp, Florence Morfin-Sherpa, Martine Valette, Bruno Lina, Laurence Josset |
| EPI_ISL_676581, EPI_ISL_676602, EPI_ISL_676603 | Scientific Veterinary Institute Novi Sad | Veterinary Specialized Institute "Kraljevo", Serbia | Vidanovic,D., Tesovic,B., Knezevic,A., Jovanovic,T., Jankovic,M., Sekler,M., Banovic Djeri,B., Petrovic,T., Volkening,J., Afonso,C. |
| EPI_ISL_676650 | Wadsworth Center, New York State Department of Health | Wadsworth Center, New York State Department of Health | Kirsten St. George, Daryl M. Lamson, Sara Griesemer, Jonathan Plitnick, Navjot Singh, Matthew D. Shudt, Erica Lasek-Nesselquist |
| EPI_ISL_676686, EPI_ISL_676697, EPI_ISL_676711, EPI_ISL_676723, EPI_ISL_676733, EPI_ISL_676809, EPI_ISL_676810, EPI_ISL_676811, EPI_ISL_676812, EPI_ISL_676813, EPI_ISL_676814, EPI_ISL_676815, EPI_ISL_676816, EPI_ISL_676817, EPI_ISL_676820, EPI_ISL_676821, EPI_ISL_676822, EPI_ISL_676823, EPI_ISL_676824, EPI_ISL_676825, EPI_ISL_676826, EPI_ISL_676827, EPI_ISL_676828, EPI_ISL_676829, EPI_ISL_676830, EPI_ISL_676831, EPI_ISL_676832, EPI_ISL_676833, EPI_ISL_676834, EPI_ISL_676835, EPI_ISL_676836, EPI_ISL_676837, EPI_ISL_676838, EPI_ISL_676839, EPI_ISL_676844, EPI_ISL_676960, EPI_ISL_676961, EPI_ISL_676963, EPI_ISL_676964, EPI_ISL_676966, EPI_ISL_676968, EPI_ISL_676971, EPI_ISL_676972, EPI_ISL_676973, EPI_ISL_676974, EPI_ISL_676975, EPI_ISL_676976, EPI_ISL_676977 | | | |

| | | | |
|---|---|---|---|
| see above | Wadsworth Center, New York State Department.of Health | Wadsworth Center, New York State Department.of Health | Kirsten St. George, Daryl M. Lamson, Alexis Russel, Jonathan Plitnick, Navjot Singh, John Kelly, Sara Griesemer, Erasmus Schneider, Erica Lasek-Nesselquist |
| EPI_ISL_677681 | Vanda Pharmaceuticals Clinical Site | Vanda Pharmaceuticals | Vanda Pharmaceuticals |
| EPI_ISL_678142 | Pathogen Genomics Lab King Abdullah University of Science and Technology(KAUST) | Pathogen Genomics Lab King Abdullah University of Science and Technology(KAUST) | Sara Mfarrej, Luke Esau, Amanda Ooi, Sharif Hala, Raeece Naeem, Asim Khogeer, Fadwa Alofi, Afrah Alsomali, Jumana Taha, Abdulaziz Alahmadi, Kahled Algithami, Anwar Hashem, Naif Almontashiri, Arnab Pain |
| EPI_ISL_678276, EPI_ISL_678288 | Mikrobiologie, RARI | Mikrobiologie, RARI | Krasnov,Y.M., Naryshkina,E.A., Guseva,N.P., Sosedova,E.A., Fedorov,A.V., Badanin,D.V., Sharapova,N.A., Portenko,S.A., Shcherbakova,S.A., Kutyrev,V.V. |
| EPI_ISL_678316, EPI_ISL_678324, EPI_ISL_678334, EPI_ISL_678354, EPI_ISL_678355, EPI_ISL_678356, EPI_ISL_678357 | Area of Virology, Serology and Virology Division (SAViD), New South Wales Health Pathology Randwick | Virology Research Laboratory; Area of Virology, Serology and Virology Division (SAViD), New South Wales Health Pathology Randwick | Foster, C.; Au, J.; Ruiz Silva, M.; Deveson, I.; Bull, R.; Van Hal, S.; Rawlinson, W. |
| EPI_ISL_678398 | Suceava County Emergency Hospital | "Stefan cel Mare" University Metagenomics Lab | Lobiuc Andrei, Puscaselu Roxana |
| EPI_ISL_678506 | CNR Virus des Infections Respiratoires - France SUD | CNR Virus des Infections Respiratoires - France SUD | Antonin Bal, Gregory Destras, Gwendolyne Burfin, Solenne Brun, Martine Valette, Bruno Lina, Laurence Josset |
| EPI_ISL_678510, EPI_ISL_678512, EPI_ISL_678513, EPI_ISL_678517, EPI_ISL_678518, EPI_ISL_678525, EPI_ISL_678526, EPI_ISL_678527, EPI_ISL_678528 | Utah Public Health Laboratory | Utah Public Health Laboratory | Erin Young, Kelly Oakeson |
| EPI_ISL_681675, EPI_ISL_681676 | Molecular Medicine Laboratory, University of Magallanes | Centro Asistencial Docente y de Investigacion, Universidad de Magallanes | Jorge González, Jacqueline Aldridge, Diego Alvarez, Marco Montes de Oca, Hermy Alvarez, Roberto Uribe-Paredes, Marcelo Navarrete |
| EPI_ISL_681706 | University Hospital Limerick | Irish Coronavirus Sequencing Consortium - Teagasc Moorepark | Paul Cotter, Fiona Crispie, Amy Fitzpatrick, John Kenny, Elaine Lawton, Carolyn Meaney, Patrick Stapleton, Calum Walsh |
| EPI_ISL_681707 | University Hospital Limerick | Irish Coronavirus Sequencing Consortium - Teagasc Moorepark | Paul Cotter, Fiona Crispie, Amy Fitzpatrick, John Kenny, Carolyn Meaney, Patrick Stapleton, Calum Walsh |
| EPI_ISL_681708, EPI_ISL_681709, EPI_ISL_681710 | University Hospital Limerick | Irish Coronavirus Sequencing Consortium - Teagasc Moorepark | Paul Cotter, Fiona Crispie, Amy Fitzpatrick, John Kenny, Elaine Lawton, Carolyn Meaney, Patrick Stapleton, Calum Walsh |
| EPI_ISL_681711 | University Hospital Limerick | Irish Coronavirus Sequencing Consortium - Teagasc Moorepark | Paul Cotter, Fiona Crispie, John Kenny, Calum Walsh |
| EPI_ISL_681720, EPI_ISL_681723, EPI_ISL_681725 | University Hospital Limerick | Irish Coronavirus Sequencing Consortium - Teagasc Moorepark | Paul Cotter, Fiona Crispie, Amy Fitzpatrick, John Kenny, Elaine Lawton, Carolyn Meaney, Patrick Stapleton, Calum Walsh |
| EPI_ISL_681726, EPI_ISL_681727 | University Hospital Limerick | Irish Coronavirus Sequencing Consortium - Teagasc Moorepark | Paul Cotter, Fiona Crispie, Amy Fitzpatrick, John Kenny, Carolyn Meaney, Patrick Stapleton, Calum Walsh |
| EPI_ISL_681729, EPI_ISL_681730, EPI_ISL_681732, EPI_ISL_681733, EPI_ISL_681734, EPI_ISL_681735 | University Hospital Limerick | Irish Coronavirus Sequencing Consortium - Teagasc Moorepark | Paul Cotter, Fiona Crispie, Amy Fitzpatrick, John Kenny, Elaine Lawton, Carolyn Meaney, Patrick Stapleton, Calum Walsh |
| EPI_ISL_681744, EPI_ISL_681749, EPI_ISL_681750, EPI_ISL_681751 | University Hospital Limerick | Irish Coronavirus Sequencing Consortium - Teagasc Moorepark | Paul Cotter, Fiona Crispie, Amy Fitzpatrick, John Kenny, Carolyn Meaney, Patrick Stapleton, Calum Walsh |
| EPI_ISL_681765, EPI_ISL_681766 | University Hospital Limerick | Irish Coronavirus Sequencing Consortium - Teagasc Moorepark | Paul Cotter, Fiona Crispie, Amy Fitzpatrick, John Kenny, Elaine Lawton, Carolyn Meaney, Patrick Stapleton, Calum Walsh |
| EPI_ISL_681771, EPI_ISL_681772, EPI_ISL_681773 | University Hospital Limerick | Irish Coronavirus Sequencing Consortium - Teagasc Moorepark | Paul Cotter, Fiona Crispie, Amy Fitzpatrick, John Kenny, Carolyn Meaney, Patrick Stapleton, Calum Walsh |
| EPI_ISL_681886, EPI_ISL_681892, EPI_ISL_681907, EPI_ISL_681918, EPI_ISL_681927 | National Virus Reference Laboratory | Irish Coronavirus Sequencing Consortium - Teagasc Moorepark | Paul Cotter, Fiona Crispie, John Kenny, Matthew McCabe, Calum Walsh |
| EPI_ISL_681947, EPI_ISL_681948, EPI_ISL_681950, EPI_ISL_681951, EPI_ISL_681953, EPI_ISL_681954, EPI_ISL_681956, EPI_ISL_681957, EPI_ISL_681960, EPI_ISL_681965, EPI_ISL_681966, EPI_ISL_681968 | | | |
| see above | UPMC Clinical Microbiology Laboratory | Microbial Genomic Epidemiology Laboratory, University of Pittsburgh | Mustapha M. Mustapha, Jane W. Marsh, Dan Snyder, Marissa P. Griffith, Stephanie L. Mitchell, Vatsala R. Srinivasa, Kady D. Waggle, Chinelo Ezeonwuku, Vaughn S. Cooper, Lee H. Harrison |
| EPI_ISL_683671 | IdISSC/Hospital Clínico San Carlos de Madrid | SeqCOVID-SPAIN consortium/IBV(CSIC) | Alberto Delgado-Iribarren, Esther Culebras Lopez, Vicente Estrada Pérez, Jorge Matias-Guiu, Luis Ortega Medina, Silvia Sánchez Ramón, Ulises Gómez-Pinedo and SeqCOVID-SPAIN consortium |
| EPI_ISL_692963, EPI_ISL_692964, EPI_ISL_692965 | Massachusetts State Public Health Laboratory | Massachusetts State Public Health Laboratory | Andrew Lang, Timelia Fink, Glen Gallagher, Sandra Smole |
| EPI_ISL_693543, EPI_ISL_693544 | Instituto Nacional de Saude (INSA) | Instituto Nacional de Saude (INSA) | Borges et al |
| EPI_ISL_694066, EPI_ISL_694067, EPI_ISL_694068, EPI_ISL_694069, EPI_ISL_694070, EPI_ISL_694073, EPI_ISL_694074, EPI_ISL_694075, EPI_ISL_694076, EPI_ISL_694077, EPI_ISL_694078, EPI_ISL_694079, EPI_ISL_694080, EPI_ISL_694081, EPI_ISL_694082, EPI_ISL_694083, EPI_ISL_694084, EPI_ISL_694085, EPI_ISL_694086, EPI_ISL_694087, EPI_ISL_694088, EPI_ISL_694089, EPI_ISL_694090, EPI_ISL_694091, EPI_ISL_694092, EPI_ISL_694093, EPI_ISL_694094, EPI_ISL_694095, EPI_ISL_694096, EPI_ISL_694097, EPI_ISL_694098, EPI_ISL_694099, EPI_ISL_694100, EPI_ISL_694101, EPI_ISL_694102, EPI_ISL_694103, EPI_ISL_694104 | | | |
| see above | AZ SPHL, Arizona Department of Health Services | TGen North | Jolene Bowers, Megan Folkerts, Chris French, Hayley Yaglom, Ashlyn Pfeiffer, Darrin Lemmer, Dave Engelthaler, The Arizona COVID Genomics Union (ACGU) |
| EPI_ISL_695815, EPI_ISL_695818, EPI_ISL_695820, EPI_ISL_695823, EPI_ISL_695841, EPI_ISL_695893, EPI_ISL_695894, EPI_ISL_695896, EPI_ISL_695898, EPI_ISL_695899, EPI_ISL_695910, EPI_ISL_695911, EPI_ISL_695913, EPI_ISL_695915, EPI_ISL_695917, EPI_ISL_695918, EPI_ISL_695919, EPI_ISL_695921, EPI_ISL_695923, EPI_ISL_695926, EPI_ISL_695927, EPI_ISL_695930, EPI_ISL_695936, EPI_ISL_695937, EPI_ISL_695939, EPI_ISL_695940, EPI_ISL_695941, EPI_ISL_695944, EPI_ISL_695948, EPI_ISL_695949, EPI_ISL_695951, EPI_ISL_695955, EPI_ISL_695956, EPI_ISL_695962, EPI_ISL_695965, EPI_ISL_695967, EPI_ISL_695969, EPI_ISL_695971, EPI_ISL_695972, EPI_ISL_695974, EPI_ISL_695979, EPI_ISL_695983, EPI_ISL_695985, EPI_ISL_695986, EPI_ISL_695988, EPI_ISL_695991, EPI_ISL_695995, EPI_ISL_695997, EPI_ISL_695999, EPI_ISL_696000, EPI_ISL_696001, EPI_ISL_696007, EPI_ISL_696008, EPI_ISL_696014 | | | |
| see above | Sonora Quest Laboratories, Laboratory Sciences of Arizona | TGen North | Jolene Bowers, Megan Folkerts, Chris French, Hayley Yaglom, Ashlyn Pfeiffer, Darrin Lemmer, Dave Engelthaler, The Arizona COVID Genomics Union (ACGU) |
| EPI_ISL_699651 | Sydney South West Pathology Service (SSWPS) - Liverpool Hospital - NSW Health Pathology | NSW Health Pathology - Institute of Clinical Pathology and Medical Research; Westmead Hospital; University of Sydney | CIDM-PH et al. |
| EPI_ISL_699652 | Pathology West - NSW Health Pathology | NSW Health Pathology - Institute of Clinical Pathology and Medical Research; Westmead Hospital; University of Sydney | CIDM-PH et al. |
| EPI_ISL_700352 | Laboratoire de virologie, CHU de Grenoble - CS 10217 - 38043 Grenoble cedex 13 | CNR Virus des Infections Respiratoires - France SUD | Antonin Bal, Gregory Destras, Gwendolyne Burfin, Hadrien Règue, Quentin Semanas, Martine Valette, Bruno Lina, Sylvie Larrat, Laurence Josset |
| EPI_ISL_700353 | Laboratoire de virologie, CHU de Grenoble - CS 10217 - 38043 Grenoble cedex 14 | CNR Virus des Infections Respiratoires - France SUD | Antonin Bal, Gregory Destras, Gwendolyne Burfin, Hadrien Règue, Quentin Semanas, Martine Valette, Bruno Lina, Sylvie Larrat, Laurence Josset |
| EPI_ISL_700354 | Laboratoire de virologie, CHU de Grenoble - CS 10217 - | CNR Virus des Infections Respiratoires - France SUD | Antonin Bal, Gregory Destras, Gwendolyne Burfin, Hadrien Règue, Quentin Semanas, Martine Valette, Bruno Lina, Sylvie Larrat, Laurence Josset |

| | | | |
|---|---|---|---|
| | 38043 Grenoble cedex 15 | | |
| EPI_ISL_700372, EPI_ISL_700374, EPI_ISL_700380, EPI_ISL_700381, EPI_ISL_700396 | Centre hospitalier Métropole Savoie | CNR Virus des Infections Respiratoires - France SUD | Antonin Bal, Carine Dumollard, Gregory Destras, Gwendolyne Burfin, Hadrien Règue, Quentin Semanas, Martine Valette, Bruno Lina, Jérôme Grosjean, Laurence Josset |
| EPI_ISL_702620, EPI_ISL_705795 | Virology Department, Royal Infirmary of Edinburgh, NHS Lothian / School of Biological Sciences, University of Edinburgh / Institute of Genetics and Molecular Medicine, University of Edinburgh | COVID-19 Genomics UK (COG-UK) Consortium | McHugh M, Dewar R, Rooke S, Gallagher M, Balcaza C, O'Toole Á, Scher E, Hill V, McCrone JT, Colquhoun R, Yu X, Jackson B, Rambaut A, Williams TC, Templeton K |
| EPI_ISL_707793 | 1-Laboratory of Microbiology, National Reference Lab, Charles Nicolle Hospital; 2-University of Tunis ElManar, Faculty of Medicine of Tunis, LR99ES09, Tunis, Tunisia | 1-Clinical and Experimental Pharmacology Lab, LR16SP02, National Center of Pharmacovigilance, University of Tunis El Manar, Tunis, Tunisia. 2-Neurodegenerative diseases and psychiatric troubles, LR18SP03, Razi Hospital, University of Tunis El Manar, Tunis, Tunisia. 3- Ministry of Health, National Observatory of New and Emerging Diseases, 1006, Tunis, Tunisia | Ilhem Boutiba-Ben Boubaker, Sameh Trabelsi, Nissaf Ben Alaya, Maher Kharrat, Alia Ben Kahla, Jalila Ben Khelil, Salma Abid, Sana Ferjani, Mouna Ben Sassi, Mouna Safer, Awatef El MOussi, Habiba Ben Romdhane, Souissi Amira, Ines Mdini, Hanen El Jebari, Asma Ferjani, Gaies Emna, Riadh Daghfous, Riadh Gouider. |
| EPI_ISL_708381 | University Hospitals of Geneva, Laboratory of Virology | University Hospitals of Geneva, Laboratory of Virology | Cordey Samuel, Laubscher Florian |
| EPI_ISL_708742 | PathWest Laboratory Medicine WA | PathWest Laboratory Medicine WA Microbial Surveillance Unit | PathWest Laboratory Medicine WA Microbial Surveillance Unit |
| EPI_ISL_717746 | Kingston Health Sciences Centre and Queen's University | Ontario Institute for Cancer Research | Prameet M. Sheth,Calvin Sjaarda,Robert Colautti,Katya Douchant,Ilinca Lungu,Bernard Lam,Paul Krzyzanowski,Michael Laszloffy,Lawrence E. Heisler,Richard de Borja,Jared T. Simpson |
| EPI_ISL_717837, EPI_ISL_717838, EPI_ISL_717839 | Laboratorio de Virologia Molecular / UFRJ | Bioinformatics Laboratory / LNCC | Carolina M Voloch, Ronaldo da Silva F Jr, Luiz G P de Almeida, Cynthia C Cardoso, Otavio Bustrolini, Alexandra L Gerber, Ana Paula de C Guimarães, Diana Mariani, Andréa Cony Cavalcanti, Claudia dos Santos Rodrigues, Terezinha M P P Castiñeira, Amilcar Tanuri, Ana Tereza R de Vasconcelos |
| EPI_ISL_718105 | Kingston Health Sciences Centre and Queen's University | Ontario Institute for Cancer Research | Prameet M. Sheth,Calvin Sjaarda,Robert Colautti,Katya Douchant,Ilinca Lungu,Bernard Lam,Paul Krzyzanowski,Michael Laszloffy,Lawrence E. Heisler,Richard de Borja,Jared T. Simpson |
| EPI_ISL_718132, EPI_ISL_718136, EPI_ISL_718137, EPI_ISL_718147, EPI_ISL_718149, EPI_ISL_718161 | Ministry of Health Hospitals | Institute of Health and Community Medicine | David Perera, Ooi Mong How, Chua Hock Hin, Tonnii Sia Loong Loong, Wong Jyn Shan, Wong Kiing Aik, Chan Chia Jui |
| EPI_ISL_722000, EPI_ISL_722002, EPI_ISL_722005, EPI_ISL_722013, EPI_ISL_722018, EPI_ISL_722030, EPI_ISL_722031, EPI_ISL_722042, EPI_ISL_722043, EPI_ISL_722055, EPI_ISL_722059, EPI_ISL_722067, EPI_ISL_722068, EPI_ISL_722075, EPI_ISL_722090, EPI_ISL_722118 | | | |
| see above | Hospital das Clínicas Universidade de São Paulo Medical School | Laboratório de Parasitologia Médica - Instituto de Medicina Tropical - Universidade de São Paulo | Brazil-UK Centre for Arbovirus Discovery Diagnosis Genomics and Epidemiology (CADDE) Genomic Network - Instituto de Medicina Tropical |
| EPI_ISL_722134, EPI_ISL_722136, EPI_ISL_722137, EPI_ISL_722139, EPI_ISL_722140, EPI_ISL_722142, EPI_ISL_722143, EPI_ISL_722146, EPI_ISL_722147, EPI_ISL_722148, EPI_ISL_722155, EPI_ISL_722156, EPI_ISL_722159, EPI_ISL_722161, EPI_ISL_722164, EPI_ISL_722169 | | | |
| see above | DB Diagnosticos do Brasil | Laboratório de Parasitologia Médica - Instituto de Medicina Tropical - Universidade de São Paulo | Brazil-UK Centre for Arbovirus Discovery Diagnosis Genomics and Epidemiology (CADDE) Genomic Network - Instituto de Medicina Tropical |
| EPI_ISL_723080, EPI_ISL_723113, EPI_ISL_723114, EPI_ISL_723115 | Institute of Medical Genetics and Applied Genomics | Institute of Medical Genetics and Applied Genomics | Caspar Gross, Tina Ganzenmüller, Siri Göpel, Michaela Pogoda, Daniela Bezdan, Michael Sonnabend, Angel Angelov, Nicolas Casadei, Stephan Ossowski, Thomas Iftner, Michael Bitzer |
| EPI_ISL_729530 | Charité Universitätsmedizin Berlin, Institut für Virologie/Labor Berlin | Charité Universitätsmedizin Berlin, Institut für Virologie | Victor M Corman, Barbara Mühlemann, Jörn Beheim-Schwarzbach, Talitha Veith, Julia Schneider, Terry Jones, Christian Drosten |
| EPI_ISL_729550, EPI_ISL_729551, EPI_ISL_729552 | A. Krumbholz, Labor Dr. Krause und Kollegen MVZ GmbH, Kiel | Charité Universitätsmedizin Berlin, Institut für Virologie | Victor M Corman, Barbara Mühlemann, Jörn Beheim-Schwarzbach, Talitha Veith, Julia Schneider, Terry Jones, Christian Drosten |
| EPI_ISL_729941, EPI_ISL_730031 | Nigeria Centre for Disease Control (NCDC) | African Centre of Excellence for Genomics of Infectious Diseases (ACEGID), Redeemer's University, Ede, Osun State, Nigeria | Oluniyi P.E. et al |
| EPI_ISL_732553 | Bundeswehr Institute of Microbiology | Bundeswehr Institute of Microbiology | Markus Antwerpen, Alexandra Rehn, Mathias Walter, Malena Bestehorn-Willmann, Sabine Zange, Enrico Georgi, Roman Wölfel |
| EPI_ISL_734540, EPI_ISL_734541, EPI_ISL_734542, EPI_ISL_734543, EPI_ISL_734544, EPI_ISL_734545, EPI_ISL_734546, EPI_ISL_734547, EPI_ISL_734548, EPI_ISL_734549, EPI_ISL_734550, EPI_ISL_734551, EPI_ISL_734552, EPI_ISL_734553, EPI_ISL_734554, EPI_ISL_734555, EPI_ISL_734556, EPI_ISL_734557, EPI_ISL_734558, EPI_ISL_734559, EPI_ISL_734560, EPI_ISL_734561, EPI_ISL_734562, EPI_ISL_734563, EPI_ISL_734564, EPI_ISL_734565, EPI_ISL_734566, EPI_ISL_734567 | | | |
| see above | UZ Leuven, National Reference Laboratory for Coronaviruses, Laboratory Medicine, Leuven, Belgium | KU Leuven, Rega Institute, Clinical and Epidemiological Virology | Tony Wawina-Bokalanga, Joan Marti-Carerras, Bert Vanmechelen, Piet Maes |
| EPI_ISL_738086 | National Public Health Laboratory | Department of Medical Microbiology, Hospital Pengajar Universiti Putra Malaysia | Hui-Yee Chee, Syafinaz Amin-Nordin, Narcisse Joseph, Muhammad MI, Nor Zahrin Hasran, Yien-Ping Wong, Jia-Yong Lam |
| EPI_ISL_738208, EPI_ISL_738209, EPI_ISL_738210 | UZ Leuven, National Reference Laboratory for Coronaviruses, Laboratory Medicine, Leuven, Belgium | KU Leuven, Rega Institute, Clinical and Epidemiological Virology | Tony Wawina-Bokalanga, Joan Marti-Carerras, Bert Vanmechelen, Piet Maes |
| EPI_ISL_738249, EPI_ISL_738255, EPI_ISL_738256, EPI_ISL_738257, EPI_ISL_738258, EPI_ISL_738259, EPI_ISL_738260, EPI_ISL_738261, EPI_ISL_738262, EPI_ISL_738263, EPI_ISL_738264, EPI_ISL_738265, EPI_ISL_738266, EPI_ISL_738267, EPI_ISL_738268, EPI_ISL_738269, EPI_ISL_738270, EPI_ISL_738283 | | | |
| see above | Connecticut Veterans' Affairs Hospital | Grubaugh Lab - Yale School of Public Health | Joseph Fauver, Tara Alpert, Chantal Vogels, Mary Petrone, Isabel Ott, Ellen Foxman, Shaili Gupta, Danielle Plank, Nathan Grubaugh |
| EPI_ISL_751224 | Pathogen Genomics Lab King Abdullah University of Science and Technology(KAUST) | Pathogen Genomics Lab King Abdullah University of Science and Technology(KAUST) | Sharif Hala, Sara Mfarrej, Raushan Nugmanova, Olga Douvropoulou, Rahul P Salunke, Raeece Naeem, Afrah Alsomali, Fadwa Alofi, Asim Khogeer,Jumana Taha, Abdulaziz Alahmadi, Kahled Algithami, Anwar Hashem, Naif Almontashiri, Arnab Pain |
| EPI_ISL_751327, EPI_ISL_751442, EPI_ISL_751443, EPI_ISL_751444, EPI_ISL_751445 | IRCCS Sacro Cuore Don Calabria Hospital, Department of Infectious, Tropical Diseases & Microbiology | University of Verona, Department of Biotechnology | Antonio Mori, Michela Deiana, Elena Pomari, Chiara Piubelli; Giulia Lopatriello, Luca Marcolungo, Cristina Beltrami, Chiara Degli Esposti, Emanuela Cosentino, Massimo Delledonne |
| EPI_ISL_752661 | State Laboratories Division, Hawaii State Department of Health | State Laboratories Division, Hawaii State Department of Health | Pamela O'Brien, Sabrina Diemert, Drew Kuwazaki, Razvan Sultana, Edward Desmond |
| EPI_ISL_753718, EPI_ISL_753721, EPI_ISL_753732, EPI_ISL_753761, EPI_ISL_753852, EPI_ISL_753939, EPI_ISL_753941, EPI_ISL_753942, EPI_ISL_753943, EPI_ISL_753944, EPI_ISL_754040, EPI_ISL_754050 | | | |
| see above | Charité Universitätsmedizin Berlin, Institut für Virologie/Labor Berlin | Charité Universitätsmedizin Berlin, Institut für Virologie | Victor M Corman, Jörn Beheim-Schwarzbach, Barbara Mühlemann, Julia Schneider, Talitha Veith, Terry Jones, Christian Drosten |
| EPI_ISL_754748 | Emory Molecular Diagnostics Laboratory, Emory Healthcare | Piantadosi Lab, Emory Department of Pathology | Ahmed Babiker, Anne Piantadosi |
| EPI_ISL_755741 | Toronto Invasive Bacterial Diseases Network | McMaster University | Allison McGeer, Patryk Aftanas, Hooman Derakhshani, Angel Li, Kuganya Nirmalarajah, Emily Panousis, Ahmed Draia, Jalees Nasir, Michael Surette, Samira Mubareka, Andrew G. McArthur |
| EPI_ISL_759965 | RSUD Dr. Soetomo | Institute of Tropical Disease, Universitas Airlangga | Rima R Prasetya, Krisnoadi Rahardjo, Aldise M Nastri, Jezzy R Dewantari, Joni Wahyuhadi, Gatot Soegiarto, Laksmi Wulandari, Resti Yudhawati, Yasuko Mori, Soetjipto, Kazufumi Shimizu ,Maria I Lusida |
| EPI_ISL_766034 | RS Mitra Keluarga Bintaro | Eijkman Institute for Molecular Biology, Ministry of Research and Technology/National Agency for Research and Innovation | Frilasita A Yudhaputri, Edison Johar, Hidayat Trimarsanto, David H Muljono, Safarina G Malik, Khin Saw Myint, Amin Soebandrio |
| EPI_ISL_766035 | RS Bhayangkara Brimob Kelapa Dua | Eijkman Institute for Molecular Biology, Ministry of Research and Technology/National Agency for Research and Innovation | Frilasita A Yudhaputri, Edison Johar, Hidayat Trimarsanto, David H Muljono, Safarina G Malik, Khin Saw Myint, Amin Soebandrio |

| | | | |
|---|---|---|---|
| EPI_ISL_766036 | RS Mitra Keluarga Bintaro | Eijkman Institute for Molecular Biology, Ministry of Research and Technology/National Agency for Research and Innovation | Frilasita A Yudhaputri, Edison Johar, Hidayat Trimarsanto, David H Muljono, Safarina G Malik, Khin Saw Myint, Amin Soebandrio |
| EPI_ISL_767721, EPI_ISL_767725, EPI_ISL_767726, EPI_ISL_767727, EPI_ISL_767736, EPI_ISL_767737, EPI_ISL_767738, EPI_ISL_767739, EPI_ISL_767740, EPI_ISL_767741, EPI_ISL_767742, EPI_ISL_767743, EPI_ISL_767744, EPI_ISL_767745, EPI_ISL_767746, EPI_ISL_767747, EPI_ISL_767748, EPI_ISL_767749, EPI_ISL_767751, EPI_ISL_767755, EPI_ISL_767756, EPI_ISL_767757, EPI_ISL_767758, EPI_ISL_767759, EPI_ISL_767763, EPI_ISL_767777, EPI_ISL_767778, EPI_ISL_767782, EPI_ISL_767783, EPI_ISL_767784, EPI_ISL_767785, EPI_ISL_767787, EPI_ISL_767788, EPI_ISL_767789, EPI_ISL_767790, EPI_ISL_767796, EPI_ISL_767797, EPI_ISL_767803, EPI_ISL_767804, EPI_ISL_767805, EPI_ISL_767815, EPI_ISL_767817, EPI_ISL_767818, EPI_ISL_767819, EPI_ISL_767821, EPI_ISL_767825, EPI_ISL_767827, EPI_ISL_767828, EPI_ISL_767829, EPI_ISL_767830, EPI_ISL_767831, EPI_ISL_767834, EPI_ISL_767835, EPI_ISL_768744, EPI_ISL_768749, EPI_ISL_768750 | | | |
| see above | National Virus Reference Laboratory | Irish Coronavirus Sequencing Consortium - Teagasc Moorepark | Calum Walsh, Genuity Ireland |
| EPI_ISL_768754, EPI_ISL_768758, EPI_ISL_768761 | AIID | Irish Coronavirus Sequencing Consortium - National Virus Reference Laboratory | Michael Carr, Gabriel Gonzalez, Alejandro Abner Garcia Leon, Patrick Mallon |
| EPI_ISL_776733 | UW Virology Lab | UW Virology Lab | Pavitra Roychoudhury, Hong Xie, Lasata Shrestha, Meei-Li Huang, Keith R Jerome, Alexander Greninger |
| EPI_ISL_778646, EPI_ISL_778652, EPI_ISL_778653, EPI_ISL_778654, EPI_ISL_778655, EPI_ISL_778656, EPI_ISL_778657, EPI_ISL_778658, EPI_ISL_778722, EPI_ISL_778723, EPI_ISL_778724, EPI_ISL_778725, EPI_ISL_778726, EPI_ISL_778727, EPI_ISL_778728 | | | |
| see above | Istituto Zooprofilattico Sperimentale del Mezzogiorno | TIGEM | Patrizia Annunziata, Andrea Ballabio, Valentina Bouche, Davide Cacchiarelli (CorrespAuthor), Pellegrino Cerino, Chiara Colantuono, Lucio Di Filippo, Antonio Grimaldi, Antonio Limone, Gabriella Loconte, Anna Manfredi, Francesco Panariello, Biancamaria Pierri, Marcello Salvi, Lucia Vassallo |
| EPI_ISL_791651, EPI_ISL_791652, EPI_ISL_791653, EPI_ISL_791654, EPI_ISL_791655, EPI_ISL_791656, EPI_ISL_791657, EPI_ISL_791658, EPI_ISL_791659, EPI_ISL_791660, EPI_ISL_791661, EPI_ISL_791662, EPI_ISL_791663 | | | |
| see above | Massachusetts State Public Health Laboratory | Infectious Disease Program, Broad Institute of Harvard and MIT | Lemieux,J.E., Siddle,K.J., Shaw,B., Adams,G., Pierce,V., Turbett,S., Anahtar,M., Branda,J., Slater,D., Harris,J., Lin,A.E., Gladden-Young,A., Lagerborg,K., Rudy,M., DeRuff,K., Carter,A., Normandin,E., Bauer,M., Reilly,S., Tomkins-Tinch,C., Loreth,C., Chaluvadi,S., Neumann,A., Cusick,C., Chapman,S.B., Gnirke,A., Flowers,K., Cerrato,F., Birren,B.W., Gallagher,G., Smole,S., Park,D.J., MacInnis,B.L., Ryan,E., LaRocque,R., Rosenberg,E. and Sabeti,P.C. |
| EPI_ISL_791729, EPI_ISL_791730, EPI_ISL_791731, EPI_ISL_791732, EPI_ISL_791733, EPI_ISL_791734, EPI_ISL_791735, EPI_ISL_791736, EPI_ISL_791737, EPI_ISL_791738, EPI_ISL_791739, EPI_ISL_791740, EPI_ISL_791741, EPI_ISL_791742, EPI_ISL_791743, EPI_ISL_791744, EPI_ISL_791745, EPI_ISL_791746, EPI_ISL_791747, EPI_ISL_791748, EPI_ISL_791749, EPI_ISL_791750, EPI_ISL_791751, EPI_ISL_791752, EPI_ISL_791753, EPI_ISL_791754, EPI_ISL_791755, EPI_ISL_791756, EPI_ISL_791757, EPI_ISL_791758, EPI_ISL_791759, EPI_ISL_791760, EPI_ISL_791761, EPI_ISL_791762, EPI_ISL_791763, EPI_ISL_791816, EPI_ISL_791818 | | | |
| see above | Massachusetts General Hospital | Infectious Disease Program, Broad Institute of Harvard and MIT | Lemieux,J.E., Siddle,K.J., Shaw,B., Adams,G., Pierce,V., Turbett,S., Anahtar,M., Branda,J., Slater,D., Harris,J., Lin,A.E., Gladden-Young,A., Lagerborg,K., Rudy,M., DeRuff,K., Carter,A., Normandin,E., Bauer,M., Reilly,S., Tomkins-Tinch,C., Loreth,C., Chaluvadi,S., Neumann,A., Cusick,C., Chapman,S.B., Gnirke,A., Flowers,K., Cerrato,F., Birren,B.W., Gallagher,G., Smole,S., Park,D.J., MacInnis,B.L., Ryan,E., LaRocque,R., Rosenberg,E. and Sabeti,P.C. |
| EPI_ISL_792365, EPI_ISL_792366 | Laboratorio de Virologia - HIEAyC San Juan de Dios | Área de Secuenciación del Laboratorio de Virología del Hospital de Niños Dr. Ricardo Gutierrez on behalf of 'Proyecto Argentino Interinstitucional de genomica de SARS-CoV-2' (PAIS Consortium) | Nabaes Jodar, MS; Goya, S; Natale, MI; Lusso, S; Ferioli, M; Colmeiro, M; Gatelli, A; Ercole, R; Valinotto, LE; Viegas, M. |
| EPI_ISL_792443, EPI_ISL_792456 | Laboratorio del Hospital Regional Ushuaia Gdor. Ernesto Campos | Hospital Regional Ushuaia - Centro Austral De Investigaciones Científicas - Universidad Nacional De Tierra Del Fuego on behalf of 'Proyecto Argentino Interinstitucional de genomica de SARS-CoV-2' (PAIS Consortium) | Ceballos, SG; Nardi, CF; Gramundi, ID; Gallego, F; De Roccis, CA; Castro, G; Cáceres, SB; Yulan, CB; Boutureira, MF. |
| EPI_ISL_792491, EPI_ISL_792493, EPI_ISL_792494 | Laboratorio Central Mg. Luis Alfredo Pianciola | Hospital Regional Ushuaia - Centro Austral De Investigaciones Científicas - Universidad Nacional De Tierra Del Fuego on behalf of 'Proyecto Argentino Interinstitucional de genomica de SARS-CoV-2' (PAIS Consortium) | Ceballos, SG; Nardi, CF; Gramundi, ID; Gallego, F; Pinto, C; Ziehm, MC; Mazzeo, M; Pianciola, L. |
| EPI_ISL_792501, EPI_ISL_792502, EPI_ISL_792503 | Laboratorio de Inmunología del Hospital Perrando e Instituto de Medicina Regional de la UNNE | Grupo de Genómica y Bioinformática del Instituto de Investigación de la Cadena Láctea CONICET-INTA on behalf of 'Proyecto Argentino Interinstitucional de genomica de SARS-CoV-2' (PAIS Consortium) | Eberhardt, MF; Irazoqui, JM; Foussal, MD; Deluca, G; Ayala, NA; Gómez, MV; Giusiano, G; Lucero, H; Marin, M; Lescano, L; Cayré, A; Amadio, AF |
| EPI_ISL_792530, EPI_ISL_792541, EPI_ISL_792542, EPI_ISL_792543 | Laboratorio Central, Ministerio de Salud Córdoba | Instituto de Patologia Vegetal (CIAP-INTA) on behalf of 'Proyecto Argentino Interinstitucional de genomica de SARS-CoV-2' (PAIS Consortium) | Fernández, FD; Debat, HJ., Re, V; Pisano, MB; Castro, G.; Barbas, G. |
| EPI_ISL_792591, EPI_ISL_792592 | LACEN-PB | Laboratory of Respiratory Viruses and Measles, Oswaldo Cruz Institute, FIOCRUZ | Paola Resende, Luciana Appolinario, Fernando Motta, Anna Carolina Paixao, Ana Carolina Mendonca, João Felipe Bezerra, Romero Henrique Teixeira de Vasconcelos, Dalane Loudal Florentino Teixeira, Thiago Franco de Oliveira Carneiro, Marilda Siqueira |
| EPI_ISL_794599 | Central Laboratories, Egyptian Ministry of Health and Population | Central Laboratories, Egyptian Ministry of Health and Population | Roshdy,W.H., Kayed,A.E., Mostafa,A., El-Shesheny,R., Khalifa,M.K., Shehata,M., Shawky,S., Saleh,M., Gomaa,M., El Taweel,A., Mahmoud,S.H., Moatasim,Y., Kutkat,O., Kamel,M.N., Mahrous,N., El Sayes,M.A., El Guindy,N.M., Naguib,A., Kandeil,A., Kayali,G., Ali,M.A. |
| EPI_ISL_794608 | Central Public Health Laboratories, Egyptian Ministry of Health and Population | Central Public Health Laboratories, Egyptian Ministry of Health and Population | El-Shesheny,R., Roshdy,W.H., Kayed,A.E., Mostafa,A., Khalifa,M.K., Shehata,M., Shawky,S., Saleh,M., Gomaa,M., El Taweel,A., Mahmoud,S.H., Moatasim,Y., Kutkat,O., Kamel,M.N., Mahrous,N., El Sayes,M.A., El Guindy,N.M., Naguib,A., Kandeil,A., Kayali,G., Ali,M.A. |
| EPI_ISL_796073, EPI_ISL_796074, EPI_ISL_796075, EPI_ISL_796076, EPI_ISL_796077, EPI_ISL_796078, EPI_ISL_796079, EPI_ISL_796080, EPI_ISL_796081, EPI_ISL_796082, EPI_ISL_796083, EPI_ISL_796084, EPI_ISL_796085, EPI_ISL_796086, EPI_ISL_796087, EPI_ISL_796088, EPI_ISL_796089 | | | |
| see above | Servicio de Microbiología. Hospital General Universitario de Castellón | SeqCOVID-SPAIN consortium/IBV(CSIC) | Rosario Moreno Muñoz, María Dolores Tirado Balaguer and SeqCOVID-SPAIN consortium |
| EPI_ISL_801550, EPI_ISL_801698, EPI_ISL_801699, EPI_ISL_801700, EPI_ISL_801701, EPI_ISL_801702, EPI_ISL_801703, EPI_ISL_801704 | Laboratory of Molecular Virology, Pontificia Universidad Católica de Chile | MSHS Pathogen Surveillance Program | Leonardo I. Almonacid, Ana S. Gonzalez-Reiche, Matthew M. Hernandez, Jorge Levican, Ana Maria Contreras, Carlos Palma, Tamara García-Salum, Zenab Khan, Adriana van De Guchte, Ajay Obla, Jayeeta Dutta, Bremy Alburquerque, Eileen Serrano, Constanza Maldonado, M. Belen Leyton, Erick Salinas, Hala Alshammary, Juan Soto, Shwetha Hara Sridhar, Ying-Chih Wang, Melissa Smith, Robert Sebra, Marcela Ferres, Adolfo García-Sastre, Edward C. Holmes, Viviana Simon, Harm van Bakel, Rafael A. Medina. |
| EPI_ISL_801863, EPI_ISL_801898, EPI_ISL_801908, EPI_ISL_801913, EPI_ISL_801949, EPI_ISL_802002, EPI_ISL_802003, EPI_ISL_802004, EPI_ISL_802005, EPI_ISL_802006, EPI_ISL_802007, EPI_ISL_802008, EPI_ISL_802009, EPI_ISL_802010, EPI_ISL_802011, EPI_ISL_802012, EPI_ISL_802013, EPI_ISL_802014, EPI_ISL_802015 | | | |
| see above | MSHS Clinical Microbiology Laboratories | MSHS Pathogen Surveillance Program | Ana S. Gonzalez-Reiche, Hala Alshammary, Mitchell J. Sullivan, Brianne Ciferri, Ajay Obla, Angela Amoako, Mahmoud Awawda, Elena Hirsch, Ashley S. Salimbangon, Levy Sominsky, Katherine Beach, Kayla Russo, Charles Gleason, Shelcie Fabre, Giulio Kleiner, Zenab Khan, Bremy Alburquerque, Adriana van de Guchte, Komal Srivastava, Matthew M. Hernandez, Jayeeta Dutta, Denise Jurczyszak, Emily Ferreri, Rachel Chernet, Nancy Francoeur, Betsaida Salom Melo, Irina Oussenko, Gintaras Deikus, Juan Soto, Shwetha Hara Sridhar, Ying-Chih Wang, Kathryn Twyman, Andrew Kasarskis, Deena R. Altman, Robert Sebra, Adolfo García-Sastre, Marta Luksza, Gopi Patel, Sarah Schaefer, Melissa Gitman, Michael D. Nowak, Alberto Paniz-Mondolfi, Emilia Mia Sordillo, Viviana Simon, Harm van Bakel |
| EPI_ISL_803817 | Laboratory of Microbiology, National Reference Lab, Charles Nicolle Hospital; 2-University of Tunis ElManar, Faculty of Medicine of Tunis, LR99ES09, Tunis, Tunisia | Clinical and Experimental Pharmacology Lab, LR16SP02, National Center of Pharmacovigilance, University of Tunis El Manar, Tunis, Tunisia. 2-Neurodegenerative diseases and psychiatric troubles, LR18SP03, Razi Hospital, University of Tunis El Manar, Tunis, Tunisia. 3- Ministry of Health, National Observatory of New and Emerging Diseases, 1006, Tunis, Tunisia | Ilhem Boutiba-Ben Boubaker, Sameh Trabelsi, Nissaf Ben Alaya, Maher Kharrat, Alia BenKahla, Jalila Ben Khelil, Salma Abid, Sana Ferjani, Mouna Ben Sassi, Mouna Safer, Zaineb Hamzaoui, Habiba Ben Romdhane, Souissi Amira, Sarra Chamman, Hanen El Jebari, Ahmed Fakhfakh, Gaies Emna, Riadh Daghfous, Riadh Gouider. |
| EPI_ISL_804381, EPI_ISL_804384, EPI_ISL_804390, EPI_ISL_804392, EPI_ISL_804398, EPI_ISL_804400, EPI_ISL_804403, EPI_ISL_804405, EPI_ISL_804415, EPI_ISL_804418, EPI_ISL_804420, EPI_ISL_804426, EPI_ISL_804431, EPI_ISL_804438, EPI_ISL_804444, EPI_ISL_804445, EPI_ISL_804447, EPI_ISL_804457 | | | |
| see above | Dutch COVID-19 response team | National Institute for Public Health and the Environment (RIVM) | Adam Meijer, Harry Vennema, Dirk Eggink, Matthijs Welkers, Jeroen Cremer, Sharon van den Brink, Bas van der Veer, AnneMarie van den Brandt, Florian Zwagemaker, Dennis Schmitz, Chantal Reusken, on behalf of the national COVID-19 response team |

| | | | |
|---|---|---|---|
| EPI_ISL_805306, EPI_ISL_805616, EPI_ISL_805730, EPI_ISL_805787, EPI_ISL_805821, EPI_ISL_805823, EPI_ISL_805866, EPI_ISL_805875, EPI_ISL_805885, EPI_ISL_805941, EPI_ISL_805951, EPI_ISL_805957, EPI_ISL_805977, EPI_ISL_805979, EPI_ISL_806002, EPI_ISL_806003, EPI_ISL_806064, EPI_ISL_806067, EPI_ISL_806068, EPI_ISL_806072, EPI_ISL_806090, EPI_ISL_806095, EPI_ISL_806130, EPI_ISL_806148, EPI_ISL_806151, EPI_ISL_806153, EPI_ISL_806159, EPI_ISL_806165, EPI_ISL_806175, EPI_ISL_806179, EPI_ISL_806189, EPI_ISL_806191, EPI_ISL_806205, EPI_ISL_806226, EPI_ISL_806235, EPI_ISL_806237, EPI_ISL_806259, EPI_ISL_806264, EPI_ISL_806287, EPI_ISL_806291, EPI_ISL_806303, EPI_ISL_806304, EPI_ISL_806309, EPI_ISL_806312, EPI_ISL_806313, EPI_ISL_806314, EPI_ISL_806318, EPI_ISL_806319, EPI_ISL_806323, EPI_ISL_806325, EPI_ISL_806333, EPI_ISL_806334, EPI_ISL_806342, EPI_ISL_806347, EPI_ISL_806350, EPI_ISL_806351, EPI_ISL_806353, EPI_ISL_806355, EPI_ISL_806365, EPI_ISL_806366, EPI_ISL_806373, EPI_ISL_806379, EPI_ISL_806380, EPI_ISL_806381, EPI_ISL_806398, EPI_ISL_806399, EPI_ISL_806410, EPI_ISL_806420, EPI_ISL_806432, EPI_ISL_806435, EPI_ISL_806439, EPI_ISL_806444, EPI_ISL_806448, EPI_ISL_806450, EPI_ISL_806465, EPI_ISL_806469, EPI_ISL_806470, EPI_ISL_806472, EPI_ISL_806501, EPI_ISL_806507, EPI_ISL_806510, EPI_ISL_806515, EPI_ISL_806522 | | | |
| see above | Alberta Precision Labs (APL) | Alberta Precision Labs (APL) | Gordon P, Lam LG, Pabbaraju K, Wong A, Ma R, Li V, Melin A, Tipples G, Berenger B, Zelyas N, Kellner J, Bernier F, Chui L, Croxen M |
| EPI_ISL_806809 | Seattle Flu Study | Seattle Flu Study | Deborah A. Nickerson, Chris D. Frazar, Jover Lee, Benjamin Pelle, Matthew Richardson, Amanda Adler, Elisabeth Brandstetter, Peter D. Han, Kairsten Fay, Misja Ilcisin, Kirsten Lacombe, Thomas R. Sibley, Melissa Truong, Caitlin R. Wolf, Michael Boeckh, Janet A. Englund, Michael Famulare, Barry R. Lutz, Mark J. Rieder, Lea M. Starita, Matthew Thompson, Jay Shendure, Trevor Bedford, Helen Y. Chu |
| EPI_ISL_807152 | Makmal Kesihatan Awam Kebangsaan, Kementerian Kesihatan Malaysia | Integrative Pharmacogenomics Institute (iPROMISE) | Mohd Zaki Salleh, Teh Lay Kek, Ariza Adnan, Sazzli Shahlan Kassim, Siti Hamimah Sheikh Abdul Kadir, Fadzilah Mohd Nor, Mohd Nur Fakhruzzaman Noorizhab, Siti Farah Alwani Mohd Nawi, Wang Seok Mui, Hani Mat Hussin, Nor Zahrin Hasran, Lim Wai Feng |
| EPI_ISL_812332 | United States Air Force School of Aerospace Medicine | United States Air Force School of Aerospace Medicine | Anthony Fries, Jennifer Meyer, Amanda Javorina, Sarah Purves, William Gruner, Clarise Starr, Elizabeth Macias |
| EPI_ISL_814075 | Ministry of Health Turkey | Ministry of Health Turkey | Fatma Bayrakdar, Yasemin Cogun, Süleyman Yalcin, Aye Baak Alta, Gülay Korukluolu |
| EPI_ISL_816736, EPI_ISL_816737, EPI_ISL_816738, EPI_ISL_816739, EPI_ISL_816740, EPI_ISL_816741, EPI_ISL_816742, EPI_ISL_816743, EPI_ISL_816744, EPI_ISL_816745, EPI_ISL_816746, EPI_ISL_816747, EPI_ISL_816748, EPI_ISL_816749, EPI_ISL_816750, EPI_ISL_816751, EPI_ISL_816752, EPI_ISL_816753, EPI_ISL_816754, EPI_ISL_816755, EPI_ISL_816756, EPI_ISL_816802, EPI_ISL_816803, EPI_ISL_816804, EPI_ISL_816805, EPI_ISL_816806, EPI_ISL_816807, EPI_ISL_816808, EPI_ISL_816809, EPI_ISL_816810, EPI_ISL_816811, EPI_ISL_816812, EPI_ISL_816813, EPI_ISL_816814, EPI_ISL_816815, EPI_ISL_816816, EPI_ISL_816817, EPI_ISL_816818, EPI_ISL_816821, EPI_ISL_816822, EPI_ISL_816823, EPI_ISL_816824 | | | |
| see above | Bioinformatics and Biostatistics Lab, Advanced Sequencing Facility | COVID-19 Genomics UK (COG-UK) Consortium | Aengus Stewart,Jerome Nicod,Chelsea Sawyer,Laura Cubitt,Harshil Patel,Margaret Crawford |
| EPI_ISL_819320, EPI_ISL_819321, EPI_ISL_819322, EPI_ISL_819323, EPI_ISL_819324 | Hospital Universitari Vall d'Hebron - Vall d'Hebron Institut de Rercerca | Hospital Universitari Vall d'Hebron | Cristina Andrés, Maria Piñana, Josep F Abril, Damir Garcia-Cehic, Ariadna Rando, Juliana Esperalba, Maria Gema Codina, Carla Castillo, Maria Carmen Martin, Tomàs Pumarola, Josep Quer, Andrés Antón |
| EPI_ISL_825015 | UW Virology Lab | UW Virology Lab | Pavitra Roychoudhury, Hong Xie, Lasata Shrestha, Meei-Li Huang, Keith R Jerome, Alexander Greninger |
| EPI_ISL_825835, EPI_ISL_825836 | Laboratoire de santé publique du Québec | Laboratoire de santé publique du Québec | Sandrine Moreira, Ioannis Ragoussis, Guillaume Bourque, Jesse Shapiro, Mark Lathrop and Michel Roger on behalf of the CoVSeQ research group (http://covseq.ca/researchgroup) |
| EPI_ISL_826635, EPI_ISL_826636, EPI_ISL_826638, EPI_ISL_826640 | Montefiore Medical Center | Albert Einstein College of Medicine, Dept. of Microbiology & Immunology, Chandran lab | J. Maximilian Fels, Saad Khan, Ryan Forster, Karin A. Skalina, Surksha Sirichand, Amy S. Fox, Aviv Bergman, William B. Mitchell, Lucia R. Wolgast, Wendy Szymczak, Robert H. Bortz III, M. Eugenia Dieterle, Catalina Florez, Denise Haslwanter, Rohit K. Jangra, Ethan Laudermilch, Ariel S. Wirchnianski, Jason Barnhill, David L. Goldman, Hnin Khine, D. Yitzchak Goldstein, Johanna P. Daily, Kartik Chandran, Libusha Kelly |
| EPI_ISL_831036, EPI_ISL_831042, EPI_ISL_831049, EPI_ISL_831050, EPI_ISL_831051, EPI_ISL_831052, EPI_ISL_831053 | Hospital Universitario La Paz (Madrid) | SeqCOVID-SPAIN consortium/IBV(CSIC) | María Rodríguez-Tejedor, Elias Dahdouh, Fernando Lázaro-Perona, Jesús Mingorance and SeqCOVID-SPAIN consortium |
| EPI_ISL_831912 | New Mexico Department of Health Scientific Laboratory | New Mexico Department of Health Scientific Laboratory | Ellie Johnson, Anastacia Griego-Fisher, D'eldra Malone |
| EPI_ISL_833574 | Veterinary Specialized Instute "Nis" | Veterinary Specialized Institute "Sabac", Serbia | Vidanovic,D., Tesovic,B., Mrkovacki, S., Vujinovic,S., .,Knezevic,A., Jovanovic,T., Jankovic,M., Sekler,M., Banovic Djeri,B., Petrovic,T., Volkening,J., Afonso,C. |
| EPI_ISL_837606, EPI_ISL_837634, EPI_ISL_837635, EPI_ISL_837636, EPI_ISL_837637, EPI_ISL_837638, EPI_ISL_837639, EPI_ISL_837640, EPI_ISL_837641, EPI_ISL_837642, EPI_ISL_837643, EPI_ISL_837644, EPI_ISL_837645, EPI_ISL_837646 | | | |
| see above | Instituto Nacional de Enfermedades Respiratorias (INER) | Instituto Nacional de Enfermedades Respiratorias (INER) | Celia Boukadida, Margarita Matías-Florentino, Alma Rincón-Rubio, Hector Esteban Paz-Juárez, Olivia Briceño, Edgar Sevilla-Reyes, Fidencio Mejía-Nepomuceno, Mario Mújica-Sánchez, Eduardo Becerril-Vargas, José Arturo Martínez-Orozco, Alejandra Hernández-Terán, Jorge Salas-Hernández, Santiago Ávila-Ríos, Joel Armando Vázquez-Pérez |
| EPI_ISL_842821 | Barts Health NHS Trust | COVID-19 Genomics UK (COG-UK) Consortium | CUTINO-MOGUEL, Maria-Teresa; HARRINGTON, David; OWOYEMI, Dola; SHYLINI, Raghavendran; BROAD, Claire; KELE, Beatrix |
| EPI_ISL_845756 | Emory Molecular Diagnostics Laboratory, Emory Healthcare | Piantadosi Lab, Emory Department of Pathology | Ahmed Babiker, Anne Piantadosi |
| EPI_ISL_851048 | Gandhi Medical College and Hospital | CSIR-Centre for Cellular and Molecular Biology | Nagamani Kammili, Winnie Thomas, Madhavi Latha Manolla, Shailaja VV, Sudhamadhuri Devara, Vanisree Rajoli, Archana GJ, Sushma Rajyalakshmi Gudiseva, Sunitha Pakalapati, Manisha Rani, Amrithesh Kumar, Raja Rao Mesepogu, Vinay Shekar Reddy, Thrilok Chander Bingi, Sofia Banu, Divya Tej Sowpati |
| EPI_ISL_851049 | Gandhi Medical College and Hospital | CSIR-Centre for Cellular and Molecular Biology | Nagamani Kammili, Madhavi Latha Manolla, Winnie Thomas, Shailaja VV, Sudhamadhuri Devara, Vanisree Rajoli, Archana GJ,Sushma Rajyalakshmi Gudiseva, Sunitha Pakalapati, Manisha Rani, Amrithesh Kumar, Raja Rao Mesepogu, Vinay Shekar Reddy, Thrilok Chander Bingi, Sofia Banu, Divya Tej Sowpati |
| EPI_ISL_852662, EPI_ISL_852663, EPI_ISL_852664, EPI_ISL_852665, EPI_ISL_852666, EPI_ISL_852761 | Institute of Virology, Medical Center, University of Freiburg, Freiburg, Germany | Institute of Virology, Clinial Virus Genomics, Medical Center, University of Freiburg, Freiburg, Germany | Jonas Fuchs, Lisa Kern, Sandra Reuter, Hajo Grundmann, Marcus Panning |
| EPI_ISL_853277, EPI_ISL_853278, EPI_ISL_853284, EPI_ISL_853285 | UPMC Clinical Microbiology Laboratory | Microbial Genome Sequencing Center; Microbial Genomic Epidemiology Laboratory | Mustapha M. Mustapha, Jane W. Marsh, Dan Snyder, Marissa P. Griffith, Stephanie L. Mitchell, Vatsala R. Srinivasa, Kady D. Waggle, Chinelo Ezeonwuku, Vaughn S. Cooper, Lee H. Harrison |
| EPI_ISL_853416, EPI_ISL_853551, EPI_ISL_853552, EPI_ISL_853553, EPI_ISL_853554, EPI_ISL_853555, EPI_ISL_853556, EPI_ISL_853557, EPI_ISL_853558, EPI_ISL_853559 | Santa Clara County Public Health Laboratory | Chan-Zuckerberg Biohub | CZB Cliahub Consortium |
| EPI_ISL_853834 | Center for Virology, Medical University of Vienna | Bergthaler laboratory, CeMM Research Center for Molecular Medicine of the Austrian Academy of Sciences | Lukas Endler, Alexandra Popa, Benedikt Agerer, Jakob-Wendelin Genger, Alexander Lercher, Anna Schedl, Thomas Penz, Michael Schuster, Jan Laine, Martin Senekowitsch, Christoph Bock, Andreas Bergthaler |
| EPI_ISL_854474, EPI_ISL_854475, EPI_ISL_854476 | Alberta Precision Labs (APL) | Alberta Precision Labs (APL) | Gordon P, Lam LG, Pabbaraju K, Wong A, Ma R, Li V, Melin A, Tipples G, Berenger B, Zelyas N, Kellner J, Bernier F, Chui L, Croxen M |
| EPI_ISL_855346, EPI_ISL_855347 | Department of Epidemiology, Infectious Disease Control and Prevention, Hiroshima University, Japan | Department of Epidemiology, Infectious Disease Control and Prevention, Hiroshima University, Japan | Junko Tanaka, Kazuaki Takahashi, Masao Kuwabara, Eisaku Kishita, Shintaro Nagashima, Ko Ko |
| EPI_ISL_861419 | Instituto de Diagnostico y Referencia Epidemiologicos (INDRE) | Instituto de Diagnostico y Referencia Epidemiologicos (INDRE) | Fabiola Garces-Ayala, Gisela Barrera-Badillo , Abril Rodriguez-Maldonado, Claudia Wong-Arambula , Natividad Cruz-Ortiz, Tatiana Nunez-Garcia, Dayanira Arellano-Suarez, Adnan-Araiza Rodríguez, Edgar Mendieta-Condado, Lucia Hernandez-Rivas, Irma Lopez-Martinez, Ernesto Ramirez-Gonzalez. |
| EPI_ISL_861634 | Instituto Adolfo Lutz - Central | Instituto Adolfo Lutz, Interdiciplinary Procedures Center, Strategic Laboratory | Claudio Tavares Sacchi, Claudia Regina Gonçalves, Erica Valessa Ramos Gomes, Karoline Rodrigues Campos |
| EPI_ISL_862638, EPI_ISL_862639 | Hospital Nuestra Señora de Sonsoles | Instituto de Salud Carlos III | Iglesias-Caballero, M. Molinero Calamita, M. González-Esguevillas, M. Camarero, S. Pozo, F. Casas, I. Jiménez, P. Jiménez, M. Zaballos, A. Monzón, S. Varona, S. Juliá, M. Cuesta, I, A. San Pedro |
| EPI_ISL_862843, EPI_ISL_862844, EPI_ISL_862845, EPI_ISL_862846, EPI_ISL_862847 | Nara Medical University Hospital | Fujita health University, School of Medicine, Department of Microbiology | Masahiro suzuki, Yohei Doi |
| EPI_ISL_862848 | Japanese Red Cross Narita Hospital | Fujita health University, School of Medicine, Department of | Masahiro suzuki, Yohei Doi |

| | | | |
|---|---|---|---|
| | | Microbiology | |
| EPI_ISL_862851 | Ehime Prefectural Central Hospital | Fujita health University, School of Medicine, Department of Microbiology | Masahiro suzuki, Yohei Doi |
| EPI_ISL_862854 | Tosei General Hospital | Fujita health University, School of Medicine, Department of Microbiology | Masahiro suzuki, Yohei Doi |
| EPI_ISL_862855 | National Hospital Organization Kanazawa Medical Center | Fujita health University, School of Medicine, Department of Microbiology | Masahiro suzuki, Yohei Doi |
| EPI_ISL_862856 | Japanese Red Cross Narita Hospital | Fujita health University, School of Medicine, Department of Microbiology | Masahiro suzuki, Yohei Doi |
| EPI_ISL_862861, EPI_ISL_862862 | Sagamihara Kyodo Hospital | Fujita health University, School of Medicine, Department of Microbiology | Masahiro suzuki, Yohei Doi |
| EPI_ISL_868356 | Centre for Enzyme Innovation, University of Portsmouth / Translational Research Laboratory, Portsmouth Hospitals NHS Trust | COVID-19 Genomics UK (COG-UK) Consortium | Angela Beckett,Yann Bourgeois,Garry Scarlett,Sharon Glaysher,Scott Elliott,Kelly Bicknell,Robert Impey,Allyson Lloyd,Sarah Wyllie,Ethan Butcher,Anoop Chauhan,Samuel Robson |
| EPI_ISL_872033 | Ospedale "Di Venere" | Beaconlab (Bioinformatics, Evolution and Comparative Genomics lab), Dept of Biosciences, University on Milan | Iacobellis M, d'Avenia M, Piluscio R, Parisi A, Chiara M, Manzari C, Pesole G |
| EPI_ISL_872088 | Instituto de Diagnostico y Referencia Epidemiologicos (INDRE) | Instituto de Diagnostico y Referencia Epidemiologicos (INDRE) | Fabiola Garces-Ayala, Gisela Barrera-Badillo , Abril Rodriguez-Maldonado, Claudia Wong-Arambula , Natividad Cruz-Ortiz, Tatiana Nunez-Garcia, Dayanira Arellano-Suarez, Adnan-Araiza Rodríguez, Edgar Mendieta-Condado, Lucia Hernandez-Rivas, Irma Lopez-Martinez, Ernesto Ramirez-Gonzalez. |
| EPI_ISL_876033, EPI_ISL_876037 | Montefiore Medical Center | Albert Einstein College of Medicine, Dept. of Microbiology & Immunology, Chandran lab | J. Maximilian Fels, Saad Khan, Ryan Forster, Karin A. Skalina, Surksha Sirichand, Amy S. Fox, Aviv Bergman, William B. Mitchell, Lucia R. Wolgast, Wendy Szymczak, Robert H. Bortz III, M. Eugenia Dieterle, Catalina Florez, Denise Haslwanter, Rohit K. Jangra, Ethan Laudermilch, Ariel S. Wirchnianski, Jason Barnhill, David L. Goldman, Hnin Khine, D. Yitzchak Goldstein, Johanna P. Daily, Kartik Chandran, Libusha Kelly |
| EPI_ISL_876739, EPI_ISL_876813 | Istituto Zooprofilattico Sperimentale della Puglia e della Basilicata | Istituto Zooprofilattico Sperimentale della Puglia e della Basilicata | Parisi A., Bianco A., Capozzi L., Del Sambro L., Manzulli V, Rondinone V., Pace L., Cipolletta D., Galante D. |
| EPI_ISL_877595, EPI_ISL_877596 | Clinical Molecular Microbiology Laboratory, UNC Hospital | Dirk Dittmer | Razia Moorad , Justin T. Landis , Brent A. Eason, Melissa B. Miller, Linda Pluta, Dirk Dittmer, Angelica Juarez, Cecilia Thompson , Cameroon Grant, Evelyn Hoffman, Patricio Cano, Jason Wong, Carolina Caro-Vegas, Blossom Damania. |
| EPI_ISL_883166, EPI_ISL_883167, EPI_ISL_883168 | Hannover Medical School, Institute of Virology | Hannover Medical School, Institute of Virology | Lars Steinbrück, Jasper Götting |
| EPI_ISL_884291 | Clinical Molecular Microbiology Laboratory, UNC Hospital | Dirk Dittmer | Razia Moorad , Justin T. Landis , Brent A. Eason, Melissa B. Miller, Linda Pluta, Dirk Dittmer, Angelica Juarez, Cecilia Thompson , Cameroon Grant, Evelyn Hoffman, Patricio Cano, Jason Wong, Carolina Caro-Vegas, Blossom Damania. |
| EPI_ISL_884558, EPI_ISL_884559, EPI_ISL_884560, EPI_ISL_884561, EPI_ISL_884562, EPI_ISL_884563, EPI_ISL_884577, EPI_ISL_884578 | Molecular Microbiology & Immunology, University of Missouri | Molecular Microbiology & Immunology, University of Missouri | Tang,C.Y., Li,T., Hang,J., Lidl,G.M., Wan,X.-F. |
| EPI_ISL_887106, EPI_ISL_887108 | Institute of Medical Microbiology and Virology, University Hospital Carl Gustav Carus, TU Dresden | DRESDEN-concept Genome Center, CMCB, TU Dresden | Beil J., Brown T., Büttner L., Gscheidel N., Hochauf-Stange K., Klemroth S., Lindemann D., Mehnert G., Petzold A., Reinhardt S., Rost F., Sameith K., Winkler S. |
| EPI_ISL_887178, EPI_ISL_887185, EPI_ISL_887190, EPI_ISL_887195, EPI_ISL_887204, EPI_ISL_887221, EPI_ISL_887222, EPI_ISL_887228, EPI_ISL_887232, EPI_ISL_887250, EPI_ISL_887271, EPI_ISL_887281, EPI_ISL_887285, EPI_ISL_887299, EPI_ISL_887326, EPI_ISL_887334, EPI_ISL_887340, EPI_ISL_887343, EPI_ISL_887356, EPI_ISL_887358, EPI_ISL_887360, EPI_ISL_887363, EPI_ISL_887367, EPI_ISL_887368, EPI_ISL_887375, EPI_ISL_887390, EPI_ISL_887392, EPI_ISL_887398, EPI_ISL_887403, EPI_ISL_887404, EPI_ISL_887412, EPI_ISL_887416 | | | |
| see above | Protzer Lab | Protzer Lab, Gagneur Lab, Robert Koch Institut | Ulrike Protzer, Dieter Hoffmann, Eva Schulte, Andrea Theumer, Oliver Drechsel, Max von Kleist,Aleksandar Radonic,Stephan Fuchs, Alexander Karollus, Julien Gagneur |
| EPI_ISL_888679 | 1.AO Universitaria 'S. Giovanni di Dio e Ruggi D'Aragona, Scuola Medica Salernitana' Hospital / 2.UOC di Virologia e Microbiologia, Università della Campania 'L. Vanvitelli' / 3.AO Universitaria 'Federico II' Napoli Hospital / 4.AORN 'San Giuseppe Moscati' Avellino Hospital / 5.AO 'San Pio - presidio G. Rummo' Benevento Hospital / 6.AO 'Sant'Anna e San Sebastiano' Caserta Hospital / 7.PO 'Maria Santissima Addolorata' Eboli Hospital / 8.Biogem Istituto di Ricerche Genetiche | 1. Genome Research Center for Health (CRGS) / 2. Laboratory of Molecular Medicine and Genomics(LMMGe) / 3. Center for Research in Pure and Applied Mathematics (CRMPA) | Giorgio Giurato, Francesca Rizzo, Alessandro Weisz, Gianluigi Franci, Giovanni Nassa, Pasquale Pagliano, Roberta Tarallo, Elena Alexandrova, Ylenia D'Agostino, Carlo Ferravante, Jessica Lamberti, Viola Melone, Domenico Memoli, Valeria Mirici Cappa, Domenico Palumbo, Giovanni Pecoraro, Assunta Sellitto, Oriana Strianese, Ilaria Terenzi, Giuseppe Fenza, Aniello Gentile, Antonello Saccomanno, Sonia Amabile, Teresa Rocco, Annamaria Salvati, Emilia Vaccaro, Massimiliano Galdiero, Michele Cennamo, Giuseppe Portella, Maria Grazia Foti, Mariarosaria Ingino, Maria Landi, Maurizio Fumi, Vincenzo Rocco, Rita Greco, Vittoria Letizia, Arnolfo Petruzziello, Maddalena Schioppa, Gregorio Goffredi, Francesca Marciano, Michele Caraglia, Alessia Cossu, Marianna Scrima |
| EPI_ISL_889389, EPI_ISL_889390, EPI_ISL_889391, EPI_ISL_889392, EPI_ISL_889393 | Olomouc University Hospital | Institute of Applied Biotechnologies a.s. | Petr Klempt, Ondej Brzo, Martin Kašný, Kateina Kvapilová, Marián Hajdúch, Petr Kvapil |
| EPI_ISL_895737, EPI_ISL_895740, EPI_ISL_895764, EPI_ISL_895768, EPI_ISL_895796, EPI_ISL_895803, EPI_ISL_895809, EPI_ISL_895827, EPI_ISL_895836, EPI_ISL_895841, EPI_ISL_895844, EPI_ISL_895845 | | | |
| see above | Molecular biology division, Institute of Clinical Biochemistry and Diagnostics, Charles University, Faculty of Medicine in Hradec Králové and University Hospital Hradec Králové | Molecular biology division, Institute of Clinical Biochemistry and Diagnostics, Charles University, Faculty of Medicine in Hradec Králové and University Hospital Hradec Králové | Helena Kovaíková, Petr Brož, Ivana Baranová, Kateina Hrochová, Tereza Baková, Jitka Novotná, Kateina Pehlíková, Vladimír Palika. Cooperation project with BioVendor-R&D and bioinformatics company BIOXSYS s r.o. |
| EPI_ISL_900101, EPI_ISL_900145, EPI_ISL_900245, EPI_ISL_900304, EPI_ISL_900369, EPI_ISL_900382, EPI_ISL_900479, EPI_ISL_900485 | MEPHI, Aix Marseille University | MEPHI, Aix Marseille University | Anthony LEVASSEUR |
| EPI_ISL_904023, EPI_ISL_904024, EPI_ISL_904032, EPI_ISL_904033 | DB Diagnosticos do Brasil | Laboratório de Parasitologia Médica - Instituto de Medicina Tropical - Universidade de São Paulo | Brazil-UK Centre for Arbovirus Discovery Diagnosis Genomics and Epidemiology (CADDE) Genomic Network - Instituto de Medicina Tropical |
| EPI_ISL_904635 | Servicio de Microbiología, Laboratori Clínic Metropolitana Nord. Hospital Universitari Germans Trias i Pujol. Institut d'Investigació en Ciències de la Salut Germans Trias i Pujol (IGTP) | IrsiCaixa - Can Ruti CovidSeq | Marc Noguera-Julian, Mariona Parera, Maria Casadellà, Pilar Armengol, Francesc Catala-Moll, Roger Paredes, Bonaventura Clotet Elisa Martró, Verónica Saludes, Anna Not, Ana Pérez, Montserrat Giménez, Ignacio Blanco, Cristina Casañ, Antoni E. Bordoy, Adrián Antuori |
| EPI_ISL_906775 | Dr. Leonard A. Miller Centre for Health Services | National Microbiology Laboratory (NML) | Anna Majer, Shari Tyson, Grace Seo, Philip Mabon, Elsie Grudeski, Rhiannon Huzarewich, Russell Mandes, Anneliese Landgraff, Jennifer Tanner, Natalie Knox, Morag Graham, Gary Van Domselaar, Robert Needle, Yang Yu, Laura Gilbert, George Zahariadis, Nathalie Bastien, Yan Li, Timothy Booth, Darian Hole, Madison Chapel, Kerri Smith, CanCOGeN's metadata curation team, Public Health Agency of Canada CanCOGeN team |
| EPI_ISL_913324 | Klinisk mikrobiologi, Viruslab | The Public Health Agency of Sweden | Anna-Malin Linde, Maria Lind Karlberg, Carlo Berg, Oskar Karlsson Lindsjo, Sofia Stamouli, Reza Advani, Mattias Haukland, Petra Holmstrom, Noura Walai, Petra Edquist, Mia Brytting, Anna Risberg, Karin Tegmark-Wisell |
| EPI_ISL_913472 | Klinisk mikrobiologi | The Public Health Agency of Sweden | Anna-Malin Linde, Maria Lind Karlberg, Carlo Berg, Oskar Karlsson Lindsjo, Sofia Stamouli, Reza Advani, Mattias Haukland, Petra Holmstrom, Noura Walai, Petra Edquist, Mia Brytting, Anna Risberg, Karin Tegmark-Wisell |
| EPI_ISL_914991, EPI_ISL_914992, | QEII Health Sciences Centre | National Microbiology Laboratory (NML) | Anna Majer, Shari Tyson, Grace Seo, Philip Mabon, Elsie Grudeski, Rhiannon Huzarewich, Russell Mandes, Anneliese Landgraff, Jennifer Tanner, Natalie |

| | | | |
|---|---|---|---|
| EPI_ISL_915007, EPI_ISL_915008 | | | Knox, Morag Graham, Gary Van Domselaar, Todd Hatchette, Jason LeBlanc, Janice Pettipas, Dan Gaston, Nathalie Bastien, Yan Li, Timothy Booth, Darian Hole, Madison Chapel, CanCOGeN's metadata curation team, Public Health Agency of Canada CanCOGeN team |
| EPI_ISL_915386 | Keio University School of Medicine | Keio University School of Medicine | Kenjiro Kosaki, Yuka Iwasaki, Hirotsugu Ishizu, Haruhiko Siomi, Kodai Abe |
| EPI_ISL_931486 | Maryland Public Health Laboratory (MD PHL) | Maryland Public Health Laboratory (MD PHL) | Maryland Department of Health Laboratories Administration |
| EPI_ISL_933624, EPI_ISL_933625, EPI_ISL_933626, EPI_ISL_933627, EPI_ISL_933628, EPI_ISL_933629 | Toronto Invasive Bacterial Diseases Network | McMaster University | Allison McGeer, Patryk Aftanas, Hooman Derakhshani, Angel Li, Kuganya Nirmalarajah, Emily Panousis, Ahmed Draia, Jalees Nasir, Michael Surette, Samira Mubareka, Andrew G. McArthur |
| EPI_ISL_937536 | Thai Red Cross Emerging Infectious Diseases Health Science Centre, Chulalongkorn Hospital, Faculty of Medicine, Chulalongkorn University | Thai Red Cross Emerging Infectious Diseases Center and Faculty of Medicine, Chulalongkorn University | Opass Putcharoen, Watsamon Jantarabenjakul, Pattama Torvorapanit, Gompol Suwanpimolkul, Leilani Paitoonpong, Sopon Iamsirithaworn, Sininat Petcharat, Yutthana Joyjinda, Weenassarin Ampoot, Apaporn Rodpan, Thiravat Hemachudha, Supaporn Wacharapluesadee |
| EPI_ISL_939668, EPI_ISL_939669 | Division of Infectious Diseases, University Hospital Zürich | Institute of Medical Virology, University of Zurich | Stefan Schmutz, Maryam Zaheri, Verena Kufner, Annette Audigé, Maria Grünberg, Kevin Steiner, Jon Huder, Cyril Shah, Riccarda Capaul, Guido Bloemberg, Jürg Böni, Michael Huber, Alexandra Trkola |
| EPI_ISL_940322, EPI_ISL_940323, EPI_ISL_940324, EPI_ISL_940325, EPI_ISL_940326, EPI_ISL_940327, EPI_ISL_940328, EPI_ISL_940329, EPI_ISL_940330, EPI_ISL_940332, EPI_ISL_940333, EPI_ISL_940336, EPI_ISL_940337, EPI_ISL_940338, EPI_ISL_940343, EPI_ISL_940344, EPI_ISL_940349, EPI_ISL_940354, EPI_ISL_940355, EPI_ISL_940356, EPI_ISL_940365, EPI_ISL_940367, EPI_ISL_940368, EPI_ISL_940381, EPI_ISL_940383, EPI_ISL_940548 | | | |
| see above | Hôpital Bichat Claude Bernard, Laboratoire de Virologie | IAME UMR1137 Inserm, Université de Paris, Hôpital Bichat | Antoine Bridier-Nahmias, Amélie Recoing, Quentin Le Hingrat, Lena Daniel, Siham Hamri, Gilles Collin, Alexandre Storto, Mélanie Bertine, Charlotte Charpentier, Nadhira Houhou-Fidouh, Diane Descamps, Benoit Visseaux |
| EPI_ISL_943617, EPI_ISL_943618, EPI_ISL_943619 | Utah Public Health Laboratory | Utah Public Health Laboratory | Erin L. Young, Kelly F. Oakeson, Tara Gallagher |
| EPI_ISL_944638 | Centro de Investigaciones en Microbiología y Biotecnología-UR (CIMBIUR), Facultad de Ciencias Naturales, Universidad del Rosario, Bogotá, Colombia | Centro de Investigaciones en Microbiología y Biotecnología-UR (CIMBIUR), Facultad de Ciencias Naturales, Universidad del Rosario, Bogotá, Colombia Instituto Nacional de Salud, Bogotá, Colombia Icahn School of Medicine at Mount Sinai, New York, USA | Luz Helena Patiño, Marina Muñoz, Nathalia Ballesteros, Carolina Hernández, Carolina Flórez, Sergio Gomez, Adriana van de Guchte, Zenab Khan, Jayeeta Dutta, Hala Alejel Alshammary, Ana S. Gonzalez-Reiche, Matthew M. Hernandez, Emilia Mia Sordillo, Viviana Simon, Harm van Bakel, Alberto Paniz-Mondolfi, Juan David Ramírez |
| EPI_ISL_953290, EPI_ISL_953291, EPI_ISL_953292, EPI_ISL_953293, EPI_ISL_953294, EPI_ISL_953296, EPI_ISL_953297, EPI_ISL_953298, EPI_ISL_953299, EPI_ISL_953301, EPI_ISL_953302, EPI_ISL_953309, EPI_ISL_953310, EPI_ISL_953311, EPI_ISL_953312, EPI_ISL_953313, EPI_ISL_953314, EPI_ISL_953315, EPI_ISL_953316, EPI_ISL_953319, EPI_ISL_953321, EPI_ISL_953322, EPI_ISL_953324, EPI_ISL_953325, EPI_ISL_953326, EPI_ISL_953327, EPI_ISL_953330, EPI_ISL_953331, EPI_ISL_953333, EPI_ISL_953335, EPI_ISL_953336, EPI_ISL_953337, EPI_ISL_953338, EPI_ISL_953341, EPI_ISL_953344, EPI_ISL_953350, EPI_ISL_953351, EPI_ISL_953352, EPI_ISL_953353, EPI_ISL_953354, EPI_ISL_953355, EPI_ISL_953356, EPI_ISL_953357, EPI_ISL_953360, EPI_ISL_953365, EPI_ISL_953366, EPI_ISL_953371, EPI_ISL_953375 | | | |
| see above | Bioinformatics and Biostatistics Lab, Advanced Sequencing Facility | COVID-19 Genomics UK (COG-UK) Consortium | Aengus Stewart,Jerome Nicod,Chelsea Sawyer,Laura Cubitt,Harshil Patel,Margaret Crawford |
| EPI_ISL_953418 | Laboratorio de Investigaciones de Baney | "Swiss Tropical and Public Health Institute" | "Carlos Cortes, Claudia Daubenberger, Guillermo Garcia, Salome Hosch, Bonifacio Manguire Nlavo, Maximilian Mpina, Elizabeth Nyakarungu, Diosdado Odjama Nseng Ada, Mitoha Ondo O Ayekaba, Tobias Schindler, Philip Wonder Phiri" |
| EPI_ISL_954216 | 1.AO Universitaria 'S. Giovanni di Dio e Ruggi D'Aragona, Scuola Medica Salernitana' Hospital / 2.UOC di Virologia e Microbiologia, Università della Campania 'L. Vanvitelli' / 3.AO Universitaria 'Federico II' Napoli Hospital / 4.AORN 'San Giuseppe Moscati' Avellino Hospital / 5.AO 'San Pio - presidio G. Rummo' Benevento Hospital / 6.AO 'Sant'Anna e San Sebastiano' Caserta Hospital / 7.PO 'Maria Santissima Addolorata' Eboli Hospital / 8.Biogem Istituto di Ricerche Genetiche | 1. Genome Research Center for Health (CRGS) / 2. Laboratory of Molecular Medicine and Genomics(LMMGe) / 3. Center for Research in Pure and Applied Mathematics (CRMPA) | Giorgio Giurato, Francesca Rizzo, Alessandro Weisz, Gianluigi Franci, Giovanni Nassa, Pasquale Pagliano, Roberta Tarallo, Elena Alexandrova, Ylenia D'Agostino, Carlo Ferravante, Jessica Lamberti, Viola Melone, Domenico Memoli, Valeria Mirici Cappa, Domenico Palumbo, Giovanni Pecoraro, Assunta Sellitto, Oriana Strianese, Ilaria Terenzi, Giuseppe Fenza, Aniello Gentile, Antonello Saccomanno, Sonia Amabile, Teresa Rocco, Annamaria Salvati, Emilia Vaccaro, Massimiliano Galdiero, Michele Cennamo, Giuseppe Portella, Maria Grazia Foti, Mariarosaria Ingino, Maria Landi, Maurizio Fumi, Vincenzo Rocco, Rita Greco, Vittoria Letizia, Arnolfo Petruzziello, Maddalena Schioppa, Gregorio Goffredi, Francesca Marciano, Michele Caraglia, Alessia Cossu, Marianna Scrima, Edmondo Adorisio, Morena D'Avenia, Michela Iacobellis, Rosanna Piluscio, Giorgio Dirani, Vittorio Sambri, Simona Semprini, Silvia Zanoli, Francesco Curcio, Stefania Marzinotto, Andreina Baj, Fausto Sessa. |
| EPI_ISL_954365, EPI_ISL_954366, EPI_ISL_954367, EPI_ISL_954368, EPI_ISL_954369, EPI_ISL_954370, EPI_ISL_954371, EPI_ISL_954372, EPI_ISL_954373, EPI_ISL_954374, EPI_ISL_954375, EPI_ISL_954376, EPI_ISL_954377, EPI_ISL_954378, EPI_ISL_954379, EPI_ISL_954380, EPI_ISL_954381, EPI_ISL_954382, EPI_ISL_954383, EPI_ISL_954384, EPI_ISL_954385, EPI_ISL_954386, EPI_ISL_954387, EPI_ISL_954388, EPI_ISL_954389, EPI_ISL_954390, EPI_ISL_954391, EPI_ISL_954392, EPI_ISL_954393, EPI_ISL_954394, EPI_ISL_954395, EPI_ISL_954396, EPI_ISL_954397, EPI_ISL_954398, EPI_ISL_954399, EPI_ISL_954400, EPI_ISL_954401, EPI_ISL_954402, EPI_ISL_954403, EPI_ISL_954404, EPI_ISL_954405, EPI_ISL_954406, EPI_ISL_954407, EPI_ISL_954408, EPI_ISL_954409, EPI_ISL_954410, EPI_ISL_954411, EPI_ISL_954412, EPI_ISL_954413, EPI_ISL_954414, EPI_ISL_954415, EPI_ISL_954416, EPI_ISL_954417, EPI_ISL_954418, EPI_ISL_954419, EPI_ISL_954420, EPI_ISL_954421, EPI_ISL_954422, EPI_ISL_954423, EPI_ISL_954424, EPI_ISL_954425, EPI_ISL_954426, EPI_ISL_954427, EPI_ISL_954428, EPI_ISL_954429, EPI_ISL_954430, EPI_ISL_954431, EPI_ISL_954432, EPI_ISL_954433, EPI_ISL_954434, EPI_ISL_954435, EPI_ISL_954436, EPI_ISL_954437, EPI_ISL_954438, EPI_ISL_954439, EPI_ISL_954440, EPI_ISL_954441, EPI_ISL_954442, EPI_ISL_954443, EPI_ISL_954444, EPI_ISL_954445, EPI_ISL_954446, EPI_ISL_954447, EPI_ISL_954448, EPI_ISL_954449, EPI_ISL_954450, EPI_ISL_954451, EPI_ISL_954452, EPI_ISL_954453, EPI_ISL_954454, EPI_ISL_954455, EPI_ISL_954456, EPI_ISL_954457, EPI_ISL_954458, EPI_ISL_954459, EPI_ISL_954460, EPI_ISL_954461, EPI_ISL_954462, EPI_ISL_954463, EPI_ISL_954464, EPI_ISL_954465, EPI_ISL_954466, EPI_ISL_954467, EPI_ISL_954468, EPI_ISL_954469, EPI_ISL_954470, EPI_ISL_954471, EPI_ISL_954472, EPI_ISL_954473, EPI_ISL_954474, EPI_ISL_954475, EPI_ISL_954476, EPI_ISL_954477, EPI_ISL_954478, EPI_ISL_954479, EPI_ISL_954480, EPI_ISL_954481, EPI_ISL_954482, EPI_ISL_954483, EPI_ISL_954484, EPI_ISL_954485, EPI_ISL_954486, EPI_ISL_954487, EPI_ISL_954488, EPI_ISL_954489, EPI_ISL_954490, EPI_ISL_954491, EPI_ISL_954492, EPI_ISL_954493, EPI_ISL_954494, EPI_ISL_954495, EPI_ISL_954496, EPI_ISL_954497, EPI_ISL_954498, EPI_ISL_954499, EPI_ISL_954500, EPI_ISL_954501, EPI_ISL_954502, EPI_ISL_954503, EPI_ISL_954504, EPI_ISL_954505, EPI_ISL_954506, EPI_ISL_954507, EPI_ISL_954508, EPI_ISL_954509, EPI_ISL_954510, EPI_ISL_954511, EPI_ISL_954512, EPI_ISL_954513, EPI_ISL_954514, EPI_ISL_954515, EPI_ISL_954516, EPI_ISL_954517, EPI_ISL_954518, EPI_ISL_954519, EPI_ISL_954520, EPI_ISL_954521, EPI_ISL_954522, EPI_ISL_954523, EPI_ISL_954524, EPI_ISL_954525, EPI_ISL_954526, EPI_ISL_954527, EPI_ISL_954528, EPI_ISL_954529, EPI_ISL_954530, EPI_ISL_954531, EPI_ISL_954532, EPI_ISL_954533, EPI_ISL_954534, EPI_ISL_954535, EPI_ISL_954536, EPI_ISL_954537, EPI_ISL_954538, EPI_ISL_954539, EPI_ISL_954540, EPI_ISL_954541, EPI_ISL_954542, EPI_ISL_954543, EPI_ISL_954544, EPI_ISL_954545, EPI_ISL_954546, EPI_ISL_954547, EPI_ISL_954548, EPI_ISL_954549, EPI_ISL_954550, EPI_ISL_954551, EPI_ISL_954552, EPI_ISL_954553, EPI_ISL_954554, EPI_ISL_954555, EPI_ISL_954556, EPI_ISL_954557, EPI_ISL_954558, EPI_ISL_954559, EPI_ISL_954560, EPI_ISL_954561, EPI_ISL_954562, EPI_ISL_954563, EPI_ISL_954564, EPI_ISL_954565, EPI_ISL_954566, EPI_ISL_954567 | | | |
| see above | Laboratoire de santé publique du Québec | Laboratoire de santé publique du Québec | Sandrine Moreira, Ioannis Ragoussis, Guillaume Bourque, Jesse Shapiro, Mark Lathrop and Michel Roger on behalf of the CoVSeQ research group (http://covseq.ca/researchgroup) |
| EPI_ISL_955233 | HGZMF 1 LA PAZ | BIOBANCO / COCTI | Borja-Aburto VH, Grajales-Muñiz C, Santacruz Tinoco CE, Rojas-Mendoza T, Ochoa Carrera LA, Sánchez A, Grande R, Isa P, Taboada B, López S, Arias C, Muñoz-Medina JE |
| EPI_ISL_955240, EPI_ISL_955241 | HGZ 30 MEXICALI | BIOBANCO / COCTI | Borja-Aburto VH, Grajales-Muñiz C, Santacruz Tinoco CE, Rojas-Mendoza T, Ochoa Carrera LA, Sánchez A, Grande R, Isa P, Taboada B, López S, Arias C, Muñoz-Medina JE |
| EPI_ISL_955242, EPI_ISL_955243 | HGSMF 26 CABO SAN LUCAS | BIOBANCO / COCTI | Borja-Aburto VH, Grajales-Muñiz C, Santacruz Tinoco CE, Rojas-Mendoza T, Ochoa Carrera LA, Sánchez A, Grande R, Isa P, Taboada B, López S, Arias C, Muñoz-Medina JE |
| EPI_ISL_955244 | UMF 26 ACAPULCO | BIOBANCO / COCTI | Borja-Aburto VH, Grajales-Muñiz C, Santacruz Tinoco CE, Rojas-Mendoza T, Ochoa Carrera LA, Sánchez A, Grande R, Isa P, Taboada B, López S, Arias C, Muñoz-Medina JE |
| EPI_ISL_955245 | HGR 1 VICENTE GUERRERO | BIOBANCO / COCTI | Borja-Aburto VH, Grajales-Muñiz C, Santacruz Tinoco CE, Rojas-Mendoza T, Ochoa Carrera LA, Sánchez A, Grande R, Isa P, Taboada B, López S, Arias C, Muñoz-Medina JE |
| EPI_ISL_955246 | HGOMF 60 TLANEPANTLA | BIOBANCO / COCTI | Borja-Aburto VH, Grajales-Muñiz C, Santacruz Tinoco CE, Rojas-Mendoza T, Ochoa Carrera LA, Sánchez A, Grande R, Isa P, Taboada B, López S, Arias C, Muñoz-Medina JE |
| EPI_ISL_955247 | HGR 200 TECAMAC | BIOBANCO / COCTI | Borja-Aburto VH, Grajales-Muñiz C, Santacruz Tinoco CE, Rojas-Mendoza T, Ochoa Carrera LA, Sánchez A, Grande R, Isa P, Taboada B, López S, Arias C, Muñoz-Medina JE |
| EPI_ISL_955248 | BCO SANGRE CM LA RAZA | BIOBANCO / COCTI | Borja-Aburto VH, Grajales-Muñiz C, Santacruz Tinoco CE, Rojas-Mendoza T, Ochoa Carrera LA, Sánchez A, Grande R, Isa P, Taboada B, López S, Arias C, Muñoz-Medina JE |
| EPI_ISL_955249 | HGZ 2A PASO-TRONCOSO | BIOBANCO / COCTI | Borja-Aburto VH, Grajales-Muñiz C, Santacruz Tinoco CE, Rojas-Mendoza T, Ochoa Carrera LA, Sánchez A, Grande R, Isa P, Taboada B, López S, Arias C, Muñoz-Medina JE |
| EPI_ISL_955250, EPI_ISL_955251, EPI_ISL_955252, EPI_ISL_955253 | HGSMF 26 CABO SAN LUCAS | BIOBANCO / COCTI | Borja-Aburto VH, Grajales-Muñiz C, Santacruz Tinoco CE, Rojas-Mendoza T, Ochoa Carrera LA, Sánchez A, Grande R, Isa P, Taboada B, López S, Arias C, Muñoz-Medina JE |
| EPI_ISL_955254 | UMF 28 DEL VALLE | BIOBANCO / COCTI | Borja-Aburto VH, Grajales-Muñiz C, Santacruz Tinoco CE, Rojas-Mendoza T, Ochoa Carrera LA, Sánchez A, Grande R, Isa P, Taboada B, López S, Arias |

| | | | |
|---|---|---|---|
| | | | C, Muñoz-Medina JE |
| EPI_ISL_960308 | NLZOH, Laboratory for Virology | NLZOH, Laboratory for Virology | Katarina Prosenc (Laboratory for Virology), Cesare Camma (IZSAM), Erik Alm (ECDC) |
| EPI_ISL_961800 | Laboratorio de Infectología, Servicio de Infectología, Hospital Universitario Dr. José Eleuterio González - Universidad Autónoma de Nuevo León | Laboratorio de Infectología Molecular, Departamento de Bioquímica y Medicina Molecular,Facultad de Medicina - Universidad Autónoma de Nuevo León | Kame A. Galán-Huerta, María F. Herrera-Saldivar, Natalia Martínez-Acuña, Sonia A. Lozano-Sepúlveda, Daniel Arellanos-Soto, Ana M. Rivas-Estilla, Paola Bocanegra-Ibarias, Samantha M. Flores-Treviño, Elvira Garza-González, Eduardo Perez-Alba, Laura Nuzzolo-Shihadeh, Adrian Camacho-Ortiz |
| EPI_ISL_961902, EPI_ISL_961951, EPI_ISL_961998, EPI_ISL_961999, EPI_ISL_962001, EPI_ISL_962016, EPI_ISL_962017, EPI_ISL_962018 | Illinois Department of Public Health | Gagnon Lab, Southern Illinois University | Keith Gagnon |
| EPI_ISL_965183 | NLZOH, Laboratory for Virology | NLZOH, Laboratory for Virology | Katarina Prosenc (Laboratory for Virology), Cesare Camma (IZSAM), Erik Alm (ECDC) |
| EPI_ISL_965562, EPI_ISL_965587, EPI_ISL_965592, EPI_ISL_965666, EPI_ISL_965694, EPI_ISL_965715, EPI_ISL_965716 | Dutch COVID-19 response team | Medical Microbiology, Maastricht University Medical Centre | Jozef Dingemans*, Brian van der Veer*, Erik Beuken, Carmen Reumkens, Lieke van Alphen, Christian Hoebe, Paul Savelkoul |
| EPI_ISL_966360, EPI_ISL_966361, EPI_ISL_966362, EPI_ISL_966363, EPI_ISL_966364 | Kentucky State Public Health Lab | Kentucky State Public Health Lab | Stephanie Lunn, Karim George, Joshua Tobias, William Grooms, Vaneet Arora, Matthew Johnson, Rachel Zinner, Rhonda Lucas |
| EPI_ISL_968140, EPI_ISL_968144 | Clinical Molecular Microbiology Laboratory, UNC Hospital | Dirk Dittmer | Justin T. Landis , Razia Moorad , Brent A. Eason, Melissa B. Miller, Linda Pluta, Dirk Dittmer, Angelica Juarez, Cecilia Thompson, Shawn Hawken, Cameroon Grant, Evelyn Hoffman, Patricio Cano, Jason Wong, Carolina Caro-Vegas, Ryan McNamara, Blossom Damania. |
| EPI_ISL_976906 | Thai Red Cross Emerging Infectious Diseases Health Science Centre, Chulalongkorn Hospital, Faculty of Medicine, Chulalongkorn University | Thai Red Cross Emerging Infectious Diseases Center and Faculty of Medicine, Chulalongkorn University | Opass Putcharoen, Watsamon Jantarabenjakul, Pattama Torvorapanit, Gompol Suwanpimolkul, Leilani Paitoonpong, Sopon Iamsirithaworn, Sininat Petcharat, Yutthana Joyjinda, Weenassarin Ampoot, Apaporn Rodpan, Thiravat Hemachudha, Supaporn Wacharapluesadee |
| EPI_ISL_977357 | University of Zambia, School of Veterinary Medicine | UNZAVET and PATH | Mulenga Mwenda-Chimfwembe, Ngonda Saasa, Daniel Bridges |
| EPI_ISL_979152, EPI_ISL_979153, EPI_ISL_979154, EPI_ISL_979155, EPI_ISL_979156, EPI_ISL_979157, EPI_ISL_979158 | Santa Clara County Public Health Laboratory | Chan-Zuckerberg Biohub | CZB Cliahub Consortium |
