## Supplementary material for "Detection and characterization of the SARS-CoV-2 lineage B.1.526 in New York": Supp. Table 3 GISAID Acknowledgment part 2: gisaid_hcov-19_acknowledgement_table_2021_02_12_16-11.pdf

We gratefully acknowledge the following Authors from the Originating laboratories responsible for obtaining the specimens, as well as the Submitting laboratories where the genome data were generated and shared via GISAID, on which this research is based.

All Submitters of data may be contacted directly via www.gisaid.org

Authors are sorted alphabetically.

| Accession ID | Originating Laboratory | Submitting Laboratory | Authors |
|---|---|---|---|
| EPI_ISL_428857 | MRCG at LSHTM Genomics lab | MRCG at LSHTM Genomics lab | Sesay et al |
| EPI_ISL_431778 | Virology, Wageningen Bioveterinary Research | Virology, Wageningen Bioveterinary Research | Oreshkova,N., Vreman,S., Molenaar,R.J., Harders,F., Hakze van der Honing,R.W., Gerhards,N., Bouwstra,R., Hissink,H., Smit,L., Tacken,M., Weesendorp,E., Stegeman,A. and van der Poel,W.H.M. |
| EPI_ISL_434488, EPI_ISL_434511 | Laboratoire National de Sante, Microbiology, Virology | Laboratoire National de Sante, Microbiology, Epidemiology and Microbial Genomics | Anke Wienecke-Baldacchino, Ardashel Latsuzbaia, Jessica Tapp, Catherine Ragimbeau, Guillaume Fournier, Tamir Abdelrahman, Trung Nguyen Nguyen, Joel Mossong |
| EPI_ISL_434674 | Narhalsan Backa vardcentral | The Public Health Agency of Sweden | Mats Olsson, Oskar Karlsson Lindsjo, Maria Lind Karlberg, Anna-Malin Linde, Olov Svartstrom, Anna Risberg, Theresa Enkirch, Mia Brytting, Karin Tegmark-Wisell |
| EPI_ISL_434675, EPI_ISL_434676 | Surbrunns VC | The Public Health Agency of Sweden | Erik Embring, Oskar Karlsson Lindsjo, Maria Lind Karlberg, Anna-Malin Linde, Olov Svartstrom, Anna Risberg, Theresa Enkirch, Mia Brytting, Karin Tegmark-Wisell |
| EPI_ISL_435048 | Laboratory of Applied Genetics | RSE "National Center for Biotechnology" | Alexandr Shevtsov, Ilyas Akhmetollayev, Viktoriya Lutsay, Asylulan Amirgazin, Ruslan Kalendar, Yerlan Ramanculov |
| EPI_ISL_435055 | Gujarat Biotechnology Research Centre | Gujarat Biotechnology Research Centre | Tejas Shah, Ankit Hinsu, Pritesh Sabara, Apurvasinh Puvar, Janvi Raval, Monika Gandhi, Pinal Trivedi, Maharshi Pandya, Amit Kanani, Akanksha Verma, Nitin Savaliya, Raghawendra Kumar, Dinesh Kumar, Zuber Saiyed, Dipa Kinariwala, Disha Patel, Binita Aring, Geeta Vaghela, Sonia Barve, Bhavesh Modi, Kairavi Joshi, Gaurishankar Shrimali, Nidhi Sood, Pranay Shah, R D Dixit, Snehal Bagatharia, Kamlesh J Upadhyay, Ramesh Pandit, Anjali Rajwal, Chaitanya Joshi, Madhvi Joshi |
| EPI_ISL_435056 | Gujarat Biotechnology Research Centre | Gujarat Biotechnology Research Centre | Maharshi Pandya, Amit Kanani, Akanksha Verma, Nitin Savaliya, Raghawendra Kumar, Dinesh Kumar, Zuber Saiyed, Dipa Kinariwala, Disha Patel, Binita Aring, Geeta Vaghela, Sonia Barve, Bhavesh Modi, Kairavi Joshi, Gaurishankar Shrimali, Nidhi Sood, Pranay Shah, R D Dixit, Snehal Bagatharia, Kamlesh J Upadhyay, Ramesh Pandit, Tejas Shah, Ankit Hinsu, Pritesh Sabara, Apurvasinh Puvar, Janvi Raval, Monika Gandhi, Pinal Trivedi, Afzal Ansari, Chaitanya Joshi, Madhvi Joshi |
| EPI_ISL_435394, EPI_ISL_435395, EPI_ISL_435396, EPI_ISL_435397, EPI_ISL_435398, EPI_ISL_435399, EPI_ISL_435400, EPI_ISL_435401, EPI_ISL_435402 | Gundersen Molecular Diagnostics Laboratory | Kabara Cancer Research Institute | Craig S. Richmond, Paraic A. Kenny |
| EPI_ISL_435487, EPI_ISL_435488, EPI_ISL_435490, EPI_ISL_435491, EPI_ISL_435492, EPI_ISL_435497, EPI_ISL_435499, EPI_ISL_435500, EPI_ISL_435502, EPI_ISL_435503, EPI_ISL_435515, EPI_ISL_435516, EPI_ISL_435517, EPI_ISL_435518, EPI_ISL_435519, EPI_ISL_435520, EPI_ISL_435521, EPI_ISL_435522, EPI_ISL_435523, EPI_ISL_435524, EPI_ISL_435525, EPI_ISL_435526, EPI_ISL_435527, EPI_ISL_435528, EPI_ISL_435534, EPI_ISL_435536, EPI_ISL_435537, EPI_ISL_435538, EPI_ISL_435539, EPI_ISL_435540, EPI_ISL_435541, EPI_ISL_435542, EPI_ISL_435543, EPI_ISL_435544, EPI_ISL_435545, EPI_ISL_435546, EPI_ISL_435547, EPI_ISL_435548, EPI_ISL_435549 | | | |
| see above | NYU Langone Health | Departments of Pathology and Medicine, New York University School of Medicine | Maria Aguero-Rosenfeld, Brendan Belovarac, Margaret Black, Ludovic Boytard, John Cadley, Paolo Cotzia, John Chen, Dacia Dimartino, Xiaojun Feng, Tatyana Gindin, Emily Guzman, Adriana Heguy, Megan Hogan, Emily Huang, George Jour, Lawrence H. Lin, Raven Luther, Andrew Lytle, Christian Marier, Matthew T. Maurano, Mark J. Mulligan, Peter Meyn, Raquel Ordonez Ciriza, Iman Osman, Jared Pinnell, Vanessa Raabe, Sitharam Ramaswami, Amy Rapkiewicz, Andre M. Ribeiro-dos-Santos, Marie Samanovic-Golden, Antonio Serrano, Guomiao Shen, Matija Snuderl, Theodore Vougiouklakis, Nick Vulpescu, Gael Westby, Paul Zappile, Yutong Zhang |
| EPI_ISL_435550, EPI_ISL_435551, EPI_ISL_435552, EPI_ISL_435553, EPI_ISL_435554 | LSUHS Emerging Viral Threat Laboratory | Microbial Genome Sequencing Center | Rona S. Scott, Jeremy P. Kamil, John A. Vanchiere, Camille F. Abshire, Abida Siddiqa, Byeong-Jae Lee, Chan-ki Min, Md Maksudul Alam, Monica Gestal-Cartele, Edna Ondari, Adam Greer, Malgorzata Bienkowska-Haba, Katarzyna Zwolinska, Jason M. Bodily, Andrew D. Yurochko, Paul M. Weinberger, Christopher G. Kevil, Martin J. Sapp, Daniel J. Snyder, Vaughn S. Cooper |
| EPI_ISL_435555, EPI_ISL_435556, EPI_ISL_435557, EPI_ISL_435558, EPI_ISL_435559, EPI_ISL_435560, EPI_ISL_435561, EPI_ISL_435562, EPI_ISL_435563, EPI_ISL_435564, EPI_ISL_435565, EPI_ISL_435566, EPI_ISL_435567, EPI_ISL_435568 | | | |
| see above | LSUHS Emerging Viral Threat Laboratory | Microbial Genome Sequencing Center | John A. Vanchiere, Jeremy P. Kamil, Rona S. Scott, Camille F. Abshire, Abida Siddiqa, Byeong-Jae Lee, Chan-ki Min, Md Maksudul Alam, Monica Gestal-Cartele, Edna Ondari, Adam Greer, Malgorzata Bienkowska-Haba, Katarzyna Zwolinska, Jason M. Bodily, Andrew D. Yurochko, Paul M. Weinberger, Christopher G. Kevil, Martin J. Sapp, Daniel J. Snyder, Vaughn S. Cooper |
| EPI_ISL_435569, EPI_ISL_435570, EPI_ISL_435571, EPI_ISL_435572, EPI_ISL_435573, EPI_ISL_435574, EPI_ISL_435575, EPI_ISL_435576, EPI_ISL_435577, EPI_ISL_435578, EPI_ISL_435579 | | | |
| see above | LSUHS Emerging Viral Threat Laboratory | Microbial Genome Sequencing Center | Jeremy P. Kamil, John A. Vanchiere, Rona S. Scott, Camille F. Abshire, Abida Siddiqa, Byeong-Jae Lee, Chan-ki Min, Md Maksudul Alam, Monica Gestal-Cartele, Edna Ondari, Adam Greer, Malgorzata Bienkowska-Haba, Katarzyna Zwolinska, Jason M. Bodily, Andrew D. Yurochko, Paul M. Weinberger, Christopher G. Kevil, Martin J. Sapp, Daniel J. Snyder, Vaughn S. Cooper |
| EPI_ISL_435691, EPI_ISL_435692, EPI_ISL_435693, EPI_ISL_435694, EPI_ISL_435695, EPI_ISL_435696 | National Public Health Laboratory, National Centre for Infectious Diseases | National Public Health Laboratory, National Centre for Infectious Diseases | Mak Tze Minn, Octavia Sophie, Chavatte Jean-Marc, Cui Lin, Lin Raymond Tzer Pin |
| EPI_ISL_435721 | NYU Langone Health | Departments of Pathology and Medicine, New York University School of Medicine | Maria Aguero-Rosenfeld, Brendan Belovarac, Margaret Black, Ludovic Boytard, John Cadley, Paolo Cotzia, John Chen, Dacia Dimartino, Xiaojun Feng, Tatyana Gindin, Emily Guzman, Adriana Heguy, Megan Hogan, Emily Huang, George Jour, Lawrence H. Lin, Raven Luther, Andrew Lytle, Christian Marier, Matthew T. Maurano, Mark J. Mulligan, Peter Meyn, Raquel Ordonez Ciriza, Iman Osman, Jared Pinnell, Vanessa Raabe, Sitharam Ramaswami, Amy Rapkiewicz, Andre M. Ribeiro-dos-Santos, Marie Samanovic-Golden, Antonio Serrano, Guomiao Shen, Matija Snuderl, Theodore Vougiouklakis, Nick Vulpescu, Gael Westby, Paul Zappile, Yutong Zhang |
| EPI_ISL_436137, EPI_ISL_436138, EPI_ISL_436139, EPI_ISL_436140, EPI_ISL_436141 | District Surveillance Unit | Department of Neurovirology, National Institute of Mental Health and Neuroscience (NIMHANS) | Chitra Pattabiraman, Vijayalakshmi Reddy, Harsha PK, Risha Rasheed, Shafeeq S Hameed, Manjunatha Venkataswamy, Anita Desai, Ravi Vasanthapuram |
| EPI_ISL_436456, EPI_ISL_436457, EPI_ISL_436459, EPI_ISL_436460, EPI_ISL_436461 | National Centre for Disease control (NCDC) | NCDC/CSIR-IGIB | Pramod Kumar#, Rajesh Pandey#, Pooja Sharma, Mahesh S Dhar, Vivekanand A, Bharathram Uppili, Himanshu Vashisht, Saruchi Wadhwa, Nishu Tyagi, Uma Sharma, Priyanka Singh, Hemlata Lall, Meena Datta, Poonam Gupta, Nidhi Saini, Aarti Tewari, Bibhash Nandi, Dhirendra Kumar, Satyabrata Bag, Varun Jaiswal, Hema Gogia, Preeti Madan, Simrita Singh, Prateek Singh, Debasis Dash, Mitali Mukerji, Manju Bala, Sandhya Kabra, Sujeet Singh, Mohammed Faruq, Anurag Agrawal*, Partha Rakshit* |
| EPI_ISL_436559, EPI_ISL_436560, EPI_ISL_436561, EPI_ISL_436562, EPI_ISL_436563 | Florida Bureau of Public Health Laboratories | Florida Bureau of Public Health Laboratories | Sarah Schmedes, Jason Blanton |
| EPI_ISL_436565, EPI_ISL_436613, EPI_ISL_436614, EPI_ISL_436615, EPI_ISL_436616, EPI_ISL_436617, EPI_ISL_436618, EPI_ISL_436619, EPI_ISL_436620, EPI_ISL_436621, EPI_ISL_436622, EPI_ISL_436623, EPI_ISL_436624, EPI_ISL_436625, EPI_ISL_436626, EPI_ISL_436640 | | | |
| see above | University of Wisconsin-Madison AIDS Vaccine Research Laboratories | University of Wisconsin-Madison AIDS Vaccine Research Laboratories | Gage Moreno, Katarina Braun, et al. AIDS Vaccine Research Laboratories |
| EPI_ISL_436683 | County of Santa Clara Public Health Department | Chan-Zuckerberg Biohub | CZB Cliahub Consortium |

| | | | |
|---|---|---|---|
| EPI_ISL_436731 | Ospedale Civile S. Liberatore di Atri | Istituto Zooprofilattico Sperimentale dell'Abruzzo e Molise "G.Caporale" | Lorusso A, Marcacci M, Di Domenico M, Ancora M, Curini V, Mangone I, Rinaldi A, Di Pasquale A, Cammà C, Puglia I, Savini G |
| EPI_ISL_437068, EPI_ISL_437069, EPI_ISL_437070, EPI_ISL_437071, EPI_ISL_437072, EPI_ISL_437073, EPI_ISL_437074, EPI_ISL_437075, EPI_ISL_437079, EPI_ISL_437081, EPI_ISL_437082 | | | |
| see above | County of Santa Clara Public Health | Chan-Zuckerberg Biohub | CZB Cliahub Consortium |
| EPI_ISL_437438 | Department of MicroBiology, Government Medical College, Surat | Gujarat Biotechnology Research Centre | Amit Kanani, Akanksha Verma, Nitin Savaliya, Raghawendra Kumar, Dinesh Kumar, Zuber Saiyed, Dipa Kinariwala, Disha Patel, Binita Aring, Neeta Khandelwal, Geeta Vaghela, Sonia Barve, Bhavesh Modi, Kairavi Joshi, Gaurishankar Shrimali, Nidhi Sood, Pranay Shah, R D Dixit, Snehal Bagatharia, Kamlesh J Upadhyay, Ramesh Pandit, Tejas Shah, Ankit Hinsu, Pritesh Sabara, Apurvasinh Puvar, Janvi Raval, Monika Gandhi, Pinal Trivedi, Maharshi Pandya, Neelam Nathani, Chaitanya Joshi, Madhvi Joshi |
| EPI_ISL_437439 | Department of MicroBiology, Government Medical College, Surat | Gujarat Biotechnology Research Centre | Akanksha Verma, Nitin Savaliya, Raghawendra Kumar, Dinesh Kumar, Zuber Saiyed, Dipa Kinariwala, Disha Patel, Binita Aring, Neeta Khandelwal, Geeta Vaghela, Sonia Barve, Bhavesh Modi, Kairavi Joshi, Gaurishankar Shrimali, Nidhi Sood, Pranay Shah, R D Dixit, Snehal Bagatharia, Kamlesh J Upadhyay, Ramesh Pandit, Tejas Shah, Ankit Hinsu, Pritesh Sabara, Apurvasinh Puvar, Janvi Raval, Monika Gandhi, Pinal Trivedi, Maharshi Pandya, Amit Kanani, Armi Chaudhari, Chaitanya Joshi, Madhvi Joshi |
| EPI_ISL_437440 | Department of MicroBiology, Government Medical College, Surat | Gujarat Biotechnology Research Centre | Nitin Savaliya, Raghawendra Kumar, Dinesh Kumar, Zuber Saiyed, Dipa Kinariwala, Disha Patel, Binita Aring, Neeta Khandelwal, Geeta Vaghela, Sonia Barve, Bhavesh Modi, Kairavi Joshi, Gaurishankar Shrimali, Nidhi Sood, Pranay Shah, R D Dixit, Snehal Bagatharia, Kamlesh J Upadhyay, Ramesh Pandit, Tejas Shah, Ankit Hinsu, Pritesh Sabara, Apurvasinh Puvar, Janvi Raval, Monika Gandhi, Pinal Trivedi, Maharshi Pandya, Amit Kanani, Akanksha Verma, Bhavya Jindal, Chaitanya Joshi, Madhvi Joshi |
| EPI_ISL_437441 | Department of MicroBiology, Government Medical College, Surat | Gujarat Biotechnology Research Centre | Raghawendra Kumar, Dinesh Kumar, Zuber Saiyed, Dipa Kinariwala, Disha Patel, Binita Aring, Neeta Khandelwal, Geeta Vaghela, Sonia Barve, Bhavesh Modi, Kairavi Joshi, Gaurishankar Shrimali, Nidhi Sood, Pranay Shah, R D Dixit, Snehal Bagatharia, Kamlesh J Upadhyay, Ramesh Pandit, Tejas Shah, Ankit Hinsu, Pritesh Sabara, Apurvasinh Puvar, Janvi Raval, Monika Gandhi, Pinal Trivedi, Maharshi Pandya, Amit Kanani, Akanksha Verma, Nitin Savaliya, Anjali Rajwar, Chaitanya Joshi, Madhvi Joshi |
| EPI_ISL_437444 | Department of MicroBiology, Government Medical College, Surat | Gujarat Biotechnology Research Centre | Dipa Kinariwala, Disha Patel, Binita Aring, Neeta Khandelwal, Geeta Vaghela, Sonia Barve, Bhavesh Modi, Kairavi Joshi, Gaurishankar Shrimali, Nidhi Sood, Pranay Shah, R D Dixit, Snehal Bagatharia, Kamlesh J Upadhyay, Ramesh Pandit, Tejas Shah, Ankit Hinsu, Pritesh Sabara, Apurvasinh Puvar, Janvi Raval, Monika Gandhi, Pinal Trivedi, Maharshi Pandya, Amit Kanani, Akanksha Verma, Nitin Savaliya, Raghawendra Kumar, Dinesh Kumar, Zuber Saiyed, Pooja P Doshi, Chaitanya Joshi, Madhvi Joshi |
| EPI_ISL_437445 | B.J. Medical College and Civil hospital | Gujarat Biotechnology Research Centre | Disha Patel, Binita Aring, Neeta Khandelwal, Geeta Vaghela, Sonia Barve, Bhavesh Modi, Kairavi Joshi, Gaurishankar Shrimali, Nidhi Sood, Pranay Shah, R D Dixit, Snehal Bagatharia, Kamlesh J Upadhyay, Ramesh Pandit, Tejas Shah, Ankit Hinsu, Pritesh Sabara, Apurvasinh Puvar, Janvi Raval, Monika Gandhi, Pinal Trivedi, Maharshi Pandya, Amit Kanani, Akanksha Verma, Nitin Savaliya, Raghawendra Kumar, Dinesh Kumar, Zuber Saiyed, Dipa Kinariwala, Nidhi Patel, Chaitanya Joshi, Madhvi Joshi |
| EPI_ISL_437446 | B.J. Medical College and Civil hospital | Gujarat Biotechnology Research Centre | Binita Aring, Neeta Khandelwal, Geeta Vaghela, Sonia Barve, Bhavesh Modi, Kairavi Joshi, Gaurishankar Shrimali, Nidhi Sood, Pranay Shah, R D Dixit, Snehal Bagatharia, Kamlesh J Upadhyay, Ramesh Pandit, Tejas Shah, Ankit Hinsu, Pritesh Sabara, Apurvasinh Puvar, Janvi Raval, Monika Gandhi, Pinal Trivedi, Maharshi Pandya, Amit Kanani, Akanksha Verma, Nitin Savaliya, Raghawendra Kumar, Dinesh Kumar, Zuber Saiyed, Dipa Kinariwala, Disha Patel, Priti Pandita, Chaitanya Joshi, Madhvi Joshi |
| EPI_ISL_437447 | B.J. Medical College and Civil hospital | Gujarat Biotechnology Research Centre | Neeta Khandelwal, Geeta Vaghela, Sonia Barve, Bhavesh Modi, Kairavi Joshi, Gaurishankar Shrimali, Nidhi Sood, Pranay Shah, R D Dixit, Snehal Bagatharia, Kamlesh J Upadhyay, Ramesh Pandit, Tejas Shah, Ankit Hinsu, Pritesh Sabara, Apurvasinh Puvar, Janvi Raval, Monika Gandhi, Pinal Trivedi, Maharshi Pandya, Amit Kanani, Akanksha Verma, Nitin Savaliya, Raghawendra Kumar, Dinesh Kumar, Zuber Saiyed, Dipa Kinariwala, Disha Patel, Binita Aring, Neha Rajpara, Chaitanya Joshi, Madhvi Joshi |
| EPI_ISL_437448 | B.J. Medical College and Civil hospital | Gujarat Biotechnology Research Centre | Geeta Vaghela, Sonia Barve, Bhavesh Modi, Kairavi Joshi, Gaurishankar Shrimali, Nidhi Sood, Pranay Shah, R D Dixit, Snehal Bagatharia, Kamlesh J Upadhyay, Ramesh Pandit, Tejas Shah, Ankit Hinsu, Pritesh Sabara, Apurvasinh Puvar, Janvi Raval, Monika Gandhi, Pinal Trivedi, Maharshi Pandya, Amit Kanani, Akanksha Verma, Nitin Savaliya, Raghawendra Kumar, Dinesh Kumar, Zuber Saiyed, Dipa Kinariwala, Disha Patel, Binita Aring, Neeta Khandelwal, Afzal Ansari, Chaitanya Joshi, Madhvi Joshi |
| EPI_ISL_437449 | B.J. Medical College and Civil hospital | Gujarat Biotechnology Research Centre | Sonia Barve, Bhavesh Modi, Kairavi Joshi, Gaurishankar Shrimali, Nidhi Sood, Pranay Shah, R D Dixit, Snehal Bagatharia, Kamlesh J Upadhyay, Ramesh Pandit, Tejas Shah, Ankit Hinsu, Pritesh Sabara, Apurvasinh Puvar, Janvi Raval, Monika Gandhi, Pinal Trivedi, Maharshi Pandya, Amit Kanani, Akanksha Verma, Nitin Savaliya, Raghawendra Kumar, Dinesh Kumar, Zuber Saiyed, Dipa Kinariwala, Disha Patel, Binita Aring, Neeta Khandelwal, Geeta Vaghela, Neelam Nathani, Chaitanya Joshi, Madhvi Joshi |
| EPI_ISL_437450 | B.J. Medical College and Civil hospital | Gujarat Biotechnology Research Centre | Bhavesh Modi, Kairavi Joshi, Gaurishankar Shrimali, Nidhi Sood, Pranay Shah, R D Dixit, Snehal Bagatharia, Kamlesh J Upadhyay, Ramesh Pandit, Tejas Shah, Ankit Hinsu, Pritesh Sabara, Apurvasinh Puvar, Janvi Raval, Monika Gandhi, Pinal Trivedi, Maharshi Pandya, Amit Kanani, Akanksha Verma, Nitin Savaliya, Raghawendra Kumar, Dinesh Kumar, Zuber Saiyed, Dipa Kinariwala, Disha Patel, Binita Aring, Neeta Khandelwal, Geeta Vaghela, Sonia Barve, Armi Chaudhari, Chaitanya Joshi, Madhvi Joshi |
| EPI_ISL_437451 | B.J. Medical College and Civil hospital | Gujarat Biotechnology Research Centre | Kairavi Joshi, Gaurishankar Shrimali, Nidhi Sood, Pranay Shah, R D Dixit, Snehal Bagatharia, Kamlesh J Upadhyay, Ramesh Pandit, Tejas Shah, Ankit Hinsu, Pritesh Sabara, Apurvasinh Puvar, Janvi Raval, Monika Gandhi, Pinal Trivedi, Maharshi Pandya, Amit Kanani, Akanksha Verma, Nitin Savaliya, Raghawendra Kumar, Dinesh Kumar, Zuber Saiyed, Dipa Kinariwala, Disha Patel, Binita Aring, Neeta Khandelwal, Geeta Vaghela, Sonia Barve, Bhavesh Modi, Bhavya Jindal, Chaitanya Joshi, Madhvi Joshi |
| EPI_ISL_437452 | B.J. Medical College and Civil hospital | Gujarat Biotechnology Research Centre | Gaurishankar Shrimali, Nidhi Sood, Pranay Shah, R D Dixit, Snehal Bagatharia, Kamlesh J Upadhyay, Ramesh Pandit, Tejas Shah, Ankit Hinsu, Pritesh Sabara, Apurvasinh Puvar, Janvi Raval, Monika Gandhi, Pinal Trivedi, Maharshi Pandya, Amit Kanani, Akanksha Verma, Nitin Savaliya, Raghawendra Kumar, Dinesh Kumar, Zuber Saiyed, Dipa Kinariwala, Disha Patel, Binita Aring, Neeta Khandelwal, Geeta Vaghela, Sonia Barve, Bhavesh Modi, Kairavi Joshi, Chaitanya Joshi, Anjali Rajwar, Madhvi Joshi |
| EPI_ISL_437453 | B.J. Medical College and Civil hospital | Gujarat Biotechnology Research Centre | Nidhi Sood, Pranay Shah, R D Dixit, Snehal Bagatharia, Kamlesh J Upadhyay, Ramesh Pandit, Tejas Shah, Ankit Hinsu, Pritesh Sabara, Apurvasinh Puvar, Janvi Raval, Monika Gandhi, Pinal Trivedi, Maharshi Pandya, Amit Kanani, Akanksha Verma, Nitin Savaliya, Raghawendra Kumar, Dinesh Kumar, Zuber Saiyed, Dipa Kinariwala, Disha Patel, Binita Aring, Neeta Khandelwal, Geeta Vaghela, Sonia Barve, Bhavesh Modi, Kairavi Joshi, Gaurishankar Shrimali, Chaitanya Joshi, Dipeshwari Shewale, Madhvi Joshi |
| EPI_ISL_437454 | B.J. Medical College and Civil hospital | Gujarat Biotechnology Research Centre | Pranay Shah, R D Dixit, Snehal Bagatharia, Kamlesh J Upadhyay, Ramesh Pandit, Tejas Shah, Ankit Hinsu, Pritesh Sabara, Apurvasinh Puvar, Janvi Raval, Monika Gandhi, Pinal Trivedi, Maharshi Pandya, Amit Kanani, Akanksha Verma, Nitin Savaliya, Raghawendra Kumar, Dinesh Kumar, Zuber Saiyed, Dipa Kinariwala, Disha Patel, Binita Aring, Neeta Khandelwal, Geeta Vaghela, Sonia Barve, Bhavesh Modi, Kairavi Joshi, Gaurishankar Shrimali, Nidhi Sood, Chaitanya Joshi, Sharmistha Majumdar, Madhvi Joshi |
| EPI_ISL_437518 | Alaska State Virology Laboratory | Alaska State Virology Laboratory | Jack Chen, Ph.D. |
| EPI_ISL_437525, EPI_ISL_437526, EPI_ISL_437527 | OHSU Lab Services Molecular Microbiology Lab | Oregon SARS-CoV-2 Genome Sequencing Center | Brendan L. O'Connell, Ruth V. Nichols, Alec J. Hirsch, Guang Fan, Daniel N. Streblow, William B. Messer, Andrew C. Adey, Benjamin N. Bimber, Brian J. O'Roak |
| EPI_ISL_437549, EPI_ISL_437567, EPI_ISL_437568, EPI_ISL_437569, EPI_ISL_437570, EPI_ISL_437571, EPI_ISL_437572, EPI_ISL_437573, EPI_ISL_437577 | Scripps Medical Laboratory | Andersen lab at Scripps Research | SEARCH Alliance San Diego with Michael Quigley, Ellen Stefanski, Ian Mchardy |
| EPI_ISL_438159, EPI_ISL_438160, EPI_ISL_438162, EPI_ISL_438163, EPI_ISL_438164, EPI_ISL_438165, EPI_ISL_438166, EPI_ISL_438167, EPI_ISL_438168, EPI_ISL_438169, EPI_ISL_438170, EPI_ISL_438171, EPI_ISL_438172, EPI_ISL_438173 | | | |
| see above | Seattle Flu Study | Seattle Flu Study | Chu et al |
| EPI_ISL_438203, EPI_ISL_438204, EPI_ISL_438205, EPI_ISL_438207, EPI_ISL_438209, EPI_ISL_438210, EPI_ISL_438211, EPI_ISL_438212, EPI_ISL_438213, EPI_ISL_438214, EPI_ISL_438215, EPI_ISL_438216, EPI_ISL_438217, EPI_ISL_438218, EPI_ISL_438219, EPI_ISL_438220, EPI_ISL_438221 | | | |

| | | | |
|---|---|---|---|
| see above | Washington State Department of Health | Seattle Flu Study | Chu et al |
| EPI_ISL_438553, EPI_ISL_438554, EPI_ISL_438555, EPI_ISL_438556, EPI_ISL_438557, EPI_ISL_438558, EPI_ISL_438559, EPI_ISL_438561, EPI_ISL_438562, EPI_ISL_438563, EPI_ISL_438564, EPI_ISL_438566, EPI_ISL_438568, EPI_ISL_438570, EPI_ISL_438574, EPI_ISL_438575, EPI_ISL_438576, EPI_ISL_438577, EPI_ISL_438578, EPI_ISL_438580, EPI_ISL_438581, EPI_ISL_438582, EPI_ISL_438583, EPI_ISL_438584, EPI_ISL_438585, EPI_ISL_438586, EPI_ISL_438587, EPI_ISL_438588, EPI_ISL_438589, EPI_ISL_438590, EPI_ISL_438591, EPI_ISL_438592, EPI_ISL_438593, EPI_ISL_438594, EPI_ISL_438595, EPI_ISL_438596, EPI_ISL_438597, EPI_ISL_438598, EPI_ISL_438599, EPI_ISL_438600, EPI_ISL_438601, EPI_ISL_438602, EPI_ISL_438603, EPI_ISL_438604, EPI_ISL_438605, EPI_ISL_438606, EPI_ISL_438607, EPI_ISL_438608, EPI_ISL_438609, EPI_ISL_438610, EPI_ISL_438612, EPI_ISL_438613, EPI_ISL_438614, EPI_ISL_438615, EPI_ISL_438616, EPI_ISL_438617, EPI_ISL_438618, EPI_ISL_438619, EPI_ISL_438620, EPI_ISL_438621, EPI_ISL_438622, EPI_ISL_438623, EPI_ISL_438624, EPI_ISL_438625, EPI_ISL_438626, EPI_ISL_438627, EPI_ISL_438628, EPI_ISL_438629, EPI_ISL_438630, EPI_ISL_438631, EPI_ISL_438632, EPI_ISL_438633, EPI_ISL_438634, EPI_ISL_438635, EPI_ISL_438636, EPI_ISL_438637, EPI_ISL_438638, EPI_ISL_438639, EPI_ISL_438640, EPI_ISL_438641, EPI_ISL_438642, EPI_ISL_438643, EPI_ISL_438644, EPI_ISL_438645, EPI_ISL_438646, EPI_ISL_438647, EPI_ISL_438648, EPI_ISL_438649, EPI_ISL_438650, EPI_ISL_438651, EPI_ISL_438652, EPI_ISL_438653, EPI_ISL_438654, EPI_ISL_438656, EPI_ISL_438657, EPI_ISL_438658, EPI_ISL_438659, EPI_ISL_438660, EPI_ISL_438661, EPI_ISL_438662, EPI_ISL_438663, EPI_ISL_438664, EPI_ISL_438665, EPI_ISL_438666, EPI_ISL_438667, EPI_ISL_438668, EPI_ISL_438669, EPI_ISL_438670, EPI_ISL_438671, EPI_ISL_438672, EPI_ISL_438673, EPI_ISL_438674, EPI_ISL_438675, EPI_ISL_438676, EPI_ISL_438677, EPI_ISL_438678, EPI_ISL_438679, EPI_ISL_438680, EPI_ISL_438681, EPI_ISL_438682, EPI_ISL_438683, EPI_ISL_438684, EPI_ISL_438685, EPI_ISL_438686, EPI_ISL_438687, EPI_ISL_438689, EPI_ISL_438690, EPI_ISL_438691, EPI_ISL_438692, EPI_ISL_438693, EPI_ISL_438694, EPI_ISL_438695, EPI_ISL_438696, EPI_ISL_438697, EPI_ISL_438698, EPI_ISL_438700, EPI_ISL_438702, EPI_ISL_438703, EPI_ISL_438704, EPI_ISL_438705, EPI_ISL_438706, EPI_ISL_438707, EPI_ISL_438708, EPI_ISL_438709, EPI_ISL_438710, EPI_ISL_438712, EPI_ISL_438713, EPI_ISL_438714, EPI_ISL_438715, EPI_ISL_438716, EPI_ISL_438717, EPI_ISL_438718, EPI_ISL_438719, EPI_ISL_438720, EPI_ISL_438722, EPI_ISL_438724, EPI_ISL_438726, EPI_ISL_438727, EPI_ISL_438728, EPI_ISL_438729, EPI_ISL_438730, EPI_ISL_438731, EPI_ISL_438732, EPI_ISL_438733, EPI_ISL_438734, EPI_ISL_438735, EPI_ISL_438736, EPI_ISL_438737, EPI_ISL_438738, EPI_ISL_438740, EPI_ISL_438741, EPI_ISL_438742, EPI_ISL_438745 | | | |
| see above | Department of Pathology, University of Cambridge | COVID-19 Genomics UK (COG-UK) Consortium | Luke W Meredith, M. Est©e Trk , Myra Hosmillo, William L. Hamilton, Martin D. Curran, Theresa Feltwell, Grant Hall, Anna Yakovleva, Fahad A Khokhar, Charlotte J. Houldcroft, Laura G Caller, Aminu S. Jahun, Sarah L. Caddy, Ian Goodfellow |
| EPI_ISL_438968, EPI_ISL_438969, EPI_ISL_438971 | Keio University School of Medicine | Keio University School of Medicine | Kenjiro Kosaki |
| EPI_ISL_438976, EPI_ISL_438977, EPI_ISL_438978, EPI_ISL_438980, EPI_ISL_438981, EPI_ISL_438982, EPI_ISL_438983, EPI_ISL_438984, EPI_ISL_438985, EPI_ISL_438986, EPI_ISL_438987, EPI_ISL_438988, EPI_ISL_438989, EPI_ISL_438990, EPI_ISL_438991, EPI_ISL_438994, EPI_ISL_438995, EPI_ISL_438996, EPI_ISL_438997, EPI_ISL_438998, EPI_ISL_438999, EPI_ISL_439000, EPI_ISL_439001, EPI_ISL_439002, EPI_ISL_439003, EPI_ISL_439004, EPI_ISL_439005, EPI_ISL_439006, EPI_ISL_439007, EPI_ISL_439008, EPI_ISL_439009, EPI_ISL_439010, EPI_ISL_439011, EPI_ISL_439012, EPI_ISL_439013, EPI_ISL_439014, EPI_ISL_439015, EPI_ISL_439016, EPI_ISL_439017, EPI_ISL_439018, EPI_ISL_439019, EPI_ISL_439020, EPI_ISL_439021, EPI_ISL_439022, EPI_ISL_439026, EPI_ISL_439027, EPI_ISL_439028, EPI_ISL_439029, EPI_ISL_439030, EPI_ISL_439031, EPI_ISL_439032, EPI_ISL_439033, EPI_ISL_439034, EPI_ISL_439035, EPI_ISL_439036, EPI_ISL_439037, EPI_ISL_439039, EPI_ISL_439042, EPI_ISL_439043, EPI_ISL_439044 | | | |
| see above | West of Scotland Specialist Virology Centre, NHSGGC / MRC-University of Glasgow Centre for Virus Research | COVID-19 Genomics UK (COG-UK) Consortium | Ana da Silva Filipe, Natasha Johnson, Kathy Smollett, Daniel Mair, Stephen Carmichael, Lily Tong, Jenna Nichols, Elihu Aranday-Cortes, Kirstyn Brunker, Yasmin Parr, Kyriaki Nomikou; Sarah McDonald, Marc Niebel, Patawee Asamaphan; Richard Orton, Joseph Hughes, Sreenu Vattipally, David L Robertson; Alasdair MacLean, Rory Gunson; Kathy Li, Natasha Jesudason, Rajiv Shah, James Shepherd, Antonia Ho, Emma Thomson |
| EPI_ISL_439153, EPI_ISL_439154, EPI_ISL_439155, EPI_ISL_439156, EPI_ISL_439157, EPI_ISL_439158, EPI_ISL_439159, EPI_ISL_439161, EPI_ISL_439162, EPI_ISL_439163, EPI_ISL_439164, EPI_ISL_439165, EPI_ISL_439166, EPI_ISL_439167, EPI_ISL_439168, EPI_ISL_439169, EPI_ISL_439170, EPI_ISL_439171, EPI_ISL_439172, EPI_ISL_439173, EPI_ISL_439174, EPI_ISL_439175, EPI_ISL_439176, EPI_ISL_439177, EPI_ISL_439178, EPI_ISL_439179, EPI_ISL_439180, EPI_ISL_439181, EPI_ISL_439182, EPI_ISL_439183, EPI_ISL_439184, EPI_ISL_439185, EPI_ISL_439186, EPI_ISL_439187, EPI_ISL_439188, EPI_ISL_439189, EPI_ISL_439190, EPI_ISL_439191, EPI_ISL_439192, EPI_ISL_439193, EPI_ISL_439194, EPI_ISL_439195, EPI_ISL_439196, EPI_ISL_439197, EPI_ISL_439198, EPI_ISL_439199, EPI_ISL_439200, EPI_ISL_439201, EPI_ISL_439202, EPI_ISL_439203, EPI_ISL_439204, EPI_ISL_439205, EPI_ISL_439206, EPI_ISL_439207, EPI_ISL_439208, EPI_ISL_439209, EPI_ISL_439210, EPI_ISL_439211, EPI_ISL_439212, EPI_ISL_439213, EPI_ISL_439214, EPI_ISL_439215, EPI_ISL_439216, EPI_ISL_439217, EPI_ISL_439218, EPI_ISL_439219, EPI_ISL_439220, EPI_ISL_439221, EPI_ISL_439222, EPI_ISL_439223, EPI_ISL_439224, EPI_ISL_439225, EPI_ISL_439226, EPI_ISL_439227, EPI_ISL_439228, EPI_ISL_439229, EPI_ISL_439230, EPI_ISL_439231, EPI_ISL_439232, EPI_ISL_439233, EPI_ISL_439234, EPI_ISL_439235, EPI_ISL_439236, EPI_ISL_439237, EPI_ISL_439238, EPI_ISL_439239, EPI_ISL_439240, EPI_ISL_439241, EPI_ISL_439242, EPI_ISL_439243, EPI_ISL_439244, EPI_ISL_439245, EPI_ISL_439246, EPI_ISL_439247, EPI_ISL_439248, EPI_ISL_439249, EPI_ISL_439251, EPI_ISL_439252, EPI_ISL_439253, EPI_ISL_439254, EPI_ISL_439255, EPI_ISL_439256, EPI_ISL_439257, EPI_ISL_439258, EPI_ISL_439259, EPI_ISL_439260, EPI_ISL_439261, EPI_ISL_439262, EPI_ISL_439263, EPI_ISL_439264 | | | |
| see above | Virology Department, Royal Infirmary of Edinburgh, NHS Lothian / School of Biological Sciences, University of Edinburgh / Institute of Genetics and Molecular Medicine, University of Edinburgh | COVID-19 Genomics UK (COG-UK) Consortium | McHugh M, Dewar R, Rooke S, Gallagher M, Balcaza C, O'ÄôToole Å, Scher E, Hill V, McCrone JT, Colquhoun R, Yu X, Jackson B, Rambaut A, Williams TC, Templeton K |
| EPI_ISL_439821, EPI_ISL_440864, EPI_ISL_440865, EPI_ISL_440866, EPI_ISL_440867, EPI_ISL_440869, EPI_ISL_440870, EPI_ISL_440871, EPI_ISL_440872, EPI_ISL_440873, EPI_ISL_440874, EPI_ISL_440875, EPI_ISL_440876, EPI_ISL_440877, EPI_ISL_440878, EPI_ISL_440879, EPI_ISL_440880, EPI_ISL_440881, EPI_ISL_440882, EPI_ISL_440883, EPI_ISL_440891, EPI_ISL_440892, EPI_ISL_440897, EPI_ISL_440898, EPI_ISL_440899, EPI_ISL_440900, EPI_ISL_440901, EPI_ISL_440902, EPI_ISL_440903, EPI_ISL_440904, EPI_ISL_440905, EPI_ISL_440906, EPI_ISL_440907, EPI_ISL_440908, EPI_ISL_440909, EPI_ISL_440910, EPI_ISL_440911, EPI_ISL_440913, EPI_ISL_440916, EPI_ISL_440917, EPI_ISL_440922, EPI_ISL_440923, EPI_ISL_440924, EPI_ISL_440925, EPI_ISL_440926, EPI_ISL_440927, EPI_ISL_440928, EPI_ISL_440929, EPI_ISL_440930, EPI_ISL_440931, EPI_ISL_440932, EPI_ISL_440933, EPI_ISL_440934, EPI_ISL_440935, EPI_ISL_440936, EPI_ISL_440937, EPI_ISL_440938, EPI_ISL_440939, EPI_ISL_440940, EPI_ISL_440941, EPI_ISL_440942, EPI_ISL_440943, EPI_ISL_440944, EPI_ISL_440946, EPI_ISL_440947, EPI_ISL_440948, EPI_ISL_440949 | | | |
| see above | Liverpool Clinical Laboratories | COVID-19 Genomics UK (COG-UK) Consortium | Sam Haldenby, Anita Lucaci, Steve Paterson, Julian Hiscox, Alistair Darby, M Almsaud, A Alrezaihi, Muhannad Alruwaili, Stuart D Armstrong, Jones Benjamin , Eleanor G Bentley, Anu Chawla, Jordan J Clark, Angela Cowell, Richard Eccles, Isabel Garca-Dorival, Matthew Gemmell, Alessandro Gerada, PKF Gilmore, Richard Gregory, Ximeng Han, Catherine Hartley, Margaret Hughes, Miren Iturriza-Gomara, James Johnson, L Luu, Jenifer Manson , Charlotte Nelson, Elaine O'ÄôToole, Cassie Olateju, Rebekah Penrice-Randal¬†, Lucille Rainbow, N.P Randle, Trevor Ian Robinson, Parul Sharma, Ghada T Shawli, James P Stewart , Neil Swainston, Ecaterina Vamos, Joanne Watts, Mark Whitehead |
| EPI_ISL_441350, EPI_ISL_441351, EPI_ISL_441352, EPI_ISL_441353, EPI_ISL_441354 | University College London, Great Ormond Street Hospital for Children NHS Foundation Trust, Imperial College Healthcare NHS Trust | COVID-19 Genomics UK (COG-UK) Consortium | Sergi Castellano, Rachel Williams, Mark Kristiansen, Paola Resende Silva, Sunando Roy, Tony Brooks, Helena Tutill, Paola Niola, Patricia Dyal, Charlotte Williams, Leysa Forrest, Yasmin Panchbhaya, Jacqueline Findlay, Sam Weeks, Julianne Brown, Kathryn Harris, Paul Randell, James Price, Alison Holmes, Judith Breuer |
| EPI_ISL_441516, EPI_ISL_441517, EPI_ISL_441518, EPI_ISL_441519, EPI_ISL_441520, EPI_ISL_441522, EPI_ISL_441523, EPI_ISL_441524, EPI_ISL_441525, EPI_ISL_441526, EPI_ISL_441527, EPI_ISL_441528, EPI_ISL_441529, EPI_ISL_441530, EPI_ISL_441531, EPI_ISL_441532, EPI_ISL_441533, EPI_ISL_441534, EPI_ISL_441535, EPI_ISL_441544, EPI_ISL_441545, EPI_ISL_441546, EPI_ISL_441851, EPI_ISL_441852, EPI_ISL_441853, EPI_ISL_441854, EPI_ISL_441855, EPI_ISL_441856, EPI_ISL_441857, EPI_ISL_441858, EPI_ISL_441859, EPI_ISL_441860, EPI_ISL_441861, EPI_ISL_441862 | | | |
| see above | Queens Medical Centre, Clinical Microbiology Department / DeepSeq Nottingham | COVID-19 Genomics UK (COG-UK) Consortium | Gemma Clark, Wendy Smith, Manjinder Khakh, Hannah Howson-Wells, Jonathan Ball, Patrick McClure, Joseph Chappell, Theocharis Tsoleridis, Nadine Holmes, Matthew Carlisle, Christopher Moore, Fei Sang, Johnny Debebe, Victoria Wright, Matthew Loose |
| EPI_ISL_442369, EPI_ISL_442420, EPI_ISL_442444, EPI_ISL_442496 | Virology Department, Sheffield Teaching Hospitals NHS Foundation Trust/Department of Infection, Immunity and Cardiovascular Disease, The Medical School, University of Sheffield | COVID-19 Genomics UK (COG-UK) Consortium | Thushan de Silva, Matthew Parker, Nikki Smith, Adri Angyal, Rebecca Brown, Luke Green, Rachel Tucker, Paul Parsons, Danielle Groves, Katie Johnson, Laura Carrilero, Alex Keeley, Dave Partridge, Matthew Wyles, Benjamin Lindsey, Mehmet Yavuz, Mohammad Raza, Cariad Evans |
| EPI_ISL_443192 | National Public Health Laboratory, National Centre for Infectious Diseases | National Public Health Laboratory, National Centre for Infectious Diseases | Mak Tze Minn, Octavia Sophie, Chavatte Jean-Marc, Cui Lin, Lin Raymond Tzer Pin |
| EPI_ISL_444055, EPI_ISL_444056 | UCSF Clinical Microbiology Laboratory | Chan-Zuckerberg Biohub | CZB Cliahub Consortium |
| EPI_ISL_444267, EPI_ISL_444268, EPI_ISL_444269, EPI_ISL_444270, EPI_ISL_444271, EPI_ISL_444272 | University College London, Great Ormond Street Hospital for Children NHS Foundation Trust, Imperial College Healthcare NHS Trust | COVID-19 Genomics UK (COG-UK) Consortium | Sergi Castellano, Rachel Williams, Mark Kristiansen, Paola Resende Silva, Sunando Roy, Tony Brooks, Helena Tutill, Paola Niola, Patricia Dyal, Charlotte Williams, Leysa Forrest, Yasmin Panchbhaya, Jacqueline Findlay, Sam Weeks, Julianne Brown, Kathryn Harris, Paul Randell, James Price, Alison Holmes, Judith Breuer |
| EPI_ISL_444320, EPI_ISL_444332, EPI_ISL_444333, EPI_ISL_444334, EPI_ISL_444335, EPI_ISL_444336, EPI_ISL_444348, EPI_ISL_444352, EPI_ISL_444362, EPI_ISL_444392, EPI_ISL_444393, EPI_ISL_444407, EPI_ISL_444408, EPI_ISL_444410, EPI_ISL_444413, EPI_ISL_444414, EPI_ISL_444415, EPI_ISL_444416, EPI_ISL_444417, EPI_ISL_444419, EPI_ISL_444432 | | | |
| see above | Department of Pathology, University of Cambridge | COVID-19 Genomics UK (COG-UK) Consortium | Luke W Meredith, M. Estée Török , Myra Hosmillo, William L. Hamilton, Martin D. Curran, Theresa Feltwell, Grant Hall, Anna Yakovleva, Fahad A Khokhar, Charlotte J. Houldcroft, Laura G Caller, Aminu S. Jahun, Sarah L. Caddy, Ian Goodfellow |
| EPI_ISL_444456 | B.J. Medical College and Civil hospital | Gujarat Biotechnology Research Centre | R D Dixit, Snehal Bagatharia, Kamlesh J Upadhyay, Ramesh Pandit, Tejas Shah, Ankit Hinsu, Pritesh Sabara, Apurvasinh Puvar, Janvi Raval, Monika Gandhi, Pinal Trivedi, Maharshi Pandya, Amit Kanani, Akanksha Verma, Nitin Savaliya, Raghawendra Kumar, Dinesh Kumar, Zuber Saiyed, Dipa Kinariwala, Disha Patel, Binita Aring, Neeta Khandelwal, Geeta Vaghela, Sonia Barve, Bhavesh Modi, Kairavi Joshi, Gaurishankar Shrimali, Nidhi Sood, Pranay Shah, Pooja P Doshi, Chaitanya Joshi, Madhvi Joshi |
| EPI_ISL_444457 | B.J. Medical College and Civil hospital | Gujarat Biotechnology Research Centre | Snehal Bagatharia, Kamlesh J Upadhyay, Ramesh Pandit, Tejas Shah, Ankit Hinsu, Pritesh Sabara, Apurvasinh Puvar, Janvi Raval, Monika Gandhi, Pinal Trivedi, Maharshi Pandya, Amit Kanani, Akanksha Verma, Nitin Savaliya, Raghawendra Kumar, Dinesh Kumar, Zuber Saiyed, Dipa Kinariwala, Disha Patel, Binita Aring, Neeta Khandelwal, Geeta Vaghela, Sonia Barve, Bhavesh Modi, Kairavi Joshi, Gaurishankar Shrimali, Nidhi Sood, Pranay Shah, R D Dixit, Nidhi Patel, Chaitanya Joshi, Madhvi Joshi |
| EPI_ISL_444458 | B.J. Medical College and Civil hospital | Gujarat Biotechnology Research Centre | Kamlesh J Upadhyay, Ramesh Pandit, Tejas Shah, Ankit Hinsu, Pritesh Sabara, Apurvasinh Puvar, Janvi Raval, Monika Gandhi, Pinal Trivedi, Maharshi Pandya, Amit Kanani, Akanksha Verma, Nitin Savaliya, Raghawendra Kumar, Dinesh Kumar, Zuber Saiyed, Dipa Kinariwala, Disha Patel, Binita Aring, Neeta Khandelwal, Geeta Vaghela, Sonia Barve, Bhavesh Modi, Kairavi Joshi, Gaurishankar Shrimali, Nidhi Sood, Pranay Shah, R D Dixit, Snehal Bagatharia, Priti Pandita, Chaitanya Joshi, Madhvi Joshi |
| EPI_ISL_445015, EPI_ISL_445016, | Florida Bureau of Public Health Laboratories | Florida Bureau of Public Health Laboratories | Sarah Schmedes, Jason Blanton |

| | | | |
|---|---|---|---|
| EPI_ISL_445017, EPI_ISL_445018, EPI_ISL_445029, EPI_ISL_445030, EPI_ISL_445045 | | | |
| EPI_ISL_445054, EPI_ISL_445055, EPI_ISL_445056, EPI_ISL_445057, EPI_ISL_445058, EPI_ISL_445059, EPI_ISL_445060, EPI_ISL_445061, EPI_ISL_445063, EPI_ISL_445066, EPI_ISL_445067, EPI_ISL_445069, EPI_ISL_445070, EPI_ISL_445071, EPI_ISL_445072, EPI_ISL_445073, EPI_ISL_445074, EPI_ISL_445076 | | | |
| see above | Laboratoire National de Sante, Microbiology, Virology | Laboratoire National de Sante, Microbiology, Epidemiology and Microbial Genomics | Anke Wienecke-Baldacchino, Ardashel Latsuzbaia, Jessica Tapp, Catherine Ragimbeau, Guillaume Fournier, Tamir Abdelrahman, Trung Nguyen Nguyen, Joel Mossong |
| EPI_ISL_445234, EPI_ISL_445235 | Wasterlakarna | The Public Health Agency of Sweden | Frida Ahlfors, Oskar Karlsson Lindsjo, Maria Lind Karlberg, Anna-Malin Linde, Olov Svartstrom, Anna Risberg, Theresa Enkirch, Mia Brytting, Karin Tegmark-Wisell |
| EPI_ISL_445236 | Narhalsan Backa vardcentral | The Public Health Agency of Sweden | Mats Olsson, Oskar Karlsson Lindsjo, Maria Lind Karlberg, Anna-Malin Linde, Olov Svartstrom, Anna Risberg, Theresa Enkirch, Mia Brytting, Karin Tegmark-Wisell |
| EPI_ISL_445237 | Narhalsan Molnlycke, Barn och ungdomsmedicin | The Public Health Agency of Sweden | Mats Reimer, Oskar Karlsson Lindsjo, Maria Lind Karlberg, Anna-Malin Linde, Olov Svartstrom, Anna Risberg, Theresa Enkirch, Mia Brytting, Karin Tegmark-Wisell |
| EPI_ISL_445238 | Ã-resundslakarna | The Public Health Agency of Sweden | Del Akrawi, Oskar Karlsson Lindsjo, Maria Lind Karlberg, Anna-Malin Linde, Olov Svartstrom, Anna Risberg, Theresa Enkirch, Mia Brytting, Karin Tegmark-Wisell |
| EPI_ISL_445240 | Ulltuna Vardcentral | The Public Health Agency of Sweden | Heidi Lindback, Oskar Karlsson Lindsjo, Maria Lind Karlberg, Anna-Malin Linde, Olov Svartstrom, Anna Risberg, Theresa Enkirch, Mia Brytting, Karin Tegmark-Wisell |
| EPI_ISL_445241 | Ã-restadsklinikens VC | The Public Health Agency of Sweden | Lisa Kjellberg / Laura Plavitu, Oskar Karlsson Lindsjo, Maria Lind Karlberg, Anna-Malin Linde, Olov Svartstrom, Anna Risberg, Theresa Enkirch, Mia Brytting, Karin Tegmark-Wisell |
| EPI_ISL_445242 | Jokkmokks Halsocentral | The Public Health Agency of Sweden | Markus Beland, Oskar Karlsson Lindsjo, Maria Lind Karlberg, Anna-Malin Linde, Olov Svartstrom, Anna Risberg, Theresa Enkirch, Mia Brytting, Karin Tegmark-Wisell |
| EPI_ISL_445244 | Akbiomed lab | Tejgaon College bmb lab | Md.Abdul kaium,Md.Easin Arafat |
| EPI_ISL_446758, EPI_ISL_446759, EPI_ISL_446760, EPI_ISL_446761, EPI_ISL_446762, EPI_ISL_446763, EPI_ISL_446768, EPI_ISL_446770, EPI_ISL_446771, EPI_ISL_446772, EPI_ISL_446773, EPI_ISL_446774, EPI_ISL_446775, EPI_ISL_446776, EPI_ISL_446782, EPI_ISL_446783, EPI_ISL_446784, EPI_ISL_446785, EPI_ISL_446789, EPI_ISL_446790, EPI_ISL_446791, EPI_ISL_446792, EPI_ISL_446793, EPI_ISL_446794, EPI_ISL_446795, EPI_ISL_446796, EPI_ISL_446797, EPI_ISL_446798, EPI_ISL_446851, EPI_ISL_446865, EPI_ISL_446866, EPI_ISL_446867, EPI_ISL_446870, EPI_ISL_446871, EPI_ISL_446896, EPI_ISL_446897, EPI_ISL_446946, EPI_ISL_446947, EPI_ISL_446948, EPI_ISL_446949, EPI_ISL_446950, EPI_ISL_446951, EPI_ISL_446952, EPI_ISL_446953, EPI_ISL_446954, EPI_ISL_446955, EPI_ISL_446956, EPI_ISL_446957, EPI_ISL_446958, EPI_ISL_446959, EPI_ISL_446960, EPI_ISL_446961, EPI_ISL_446962, EPI_ISL_446967, EPI_ISL_446968, EPI_ISL_446969, EPI_ISL_446970, EPI_ISL_446971, EPI_ISL_446972, EPI_ISL_446973, EPI_ISL_446974, EPI_ISL_446986, EPI_ISL_446987, EPI_ISL_446988, EPI_ISL_446989, EPI_ISL_446990 | | | |
| see above | Wales Specialist Virology Centre | Public Health Wales Microbiology Cardiff | Catherine Moore, Johnathan Evans, Laura Gifford, Malorie Perry, Simon Cottrell, Alec Birchley, Alexander Adams, Amy Gaskin, Bree Gatica-Wilcox, Jason Coombes, Lauren Gilbert, Lee Graham, Nicole Pacchiarini, Sara Kumziene-Summerhayes, Sarah Taylor, Sophie Jones, Sara Rey, Matthew Bull, Joanne Watkins, Sally Corden, Tom Connor |
| EPI_ISL_447065, EPI_ISL_447077, EPI_ISL_447080, EPI_ISL_447176, EPI_ISL_447185, EPI_ISL_447205, EPI_ISL_447210, EPI_ISL_447219, EPI_ISL_447226 | Michigan Department of Health and Human Services, Bureau of Laboratories | Michigan Department of Health and Human Services, Bureau of Laboratories | Blankenship HM, Riner D, Soehnlen MK |
| EPI_ISL_447230, EPI_ISL_447231, EPI_ISL_447232, EPI_ISL_447233, EPI_ISL_447234, EPI_ISL_447235, EPI_ISL_447236, EPI_ISL_447237, EPI_ISL_447238, EPI_ISL_447239, EPI_ISL_447240, EPI_ISL_447241, EPI_ISL_447242, EPI_ISL_447243, EPI_ISL_447244, EPI_ISL_447245, EPI_ISL_447246, EPI_ISL_447247, EPI_ISL_447248, EPI_ISL_447249 | | | |
| see above | Viral Respiratory Lab, National Institute for Biomedical Research (INRB) | Pathogen Sequencing Lab, National Institute for Biomedical Research (INRB) | Placide Mbala-Kingebeni, Edith Nkwembe, Eddy Kinganda-Lusamaki, Amuri Aziza, Francisca Muyembe Mawete, Catherine Pratt, Matthias Pauthner, Josh Quick, Allison Black, James Hadfield, Trevor Bedford, Ian Goodfellow, Andrew Rambaut, Nick Loman, Kristian Andersen, Michael Wiley, Steve Ahuka-Mundeke, Jean-Jacques Muyembe Tamfum |
| EPI_ISL_447331, EPI_ISL_447332, EPI_ISL_447333 | Clinical Virology Unit, Hadassah Hebrew University Medical Center | Stern Lab | Stern Lab |
| EPI_ISL_447451, EPI_ISL_447452, EPI_ISL_447453, EPI_ISL_447454, EPI_ISL_447455, EPI_ISL_447456 | Clinical Microbiology Laboratory, Sheba Medical Center | Stern Lab | Stern Lab |
| EPI_ISL_447551 | GMERS Medical College and Hospital, Gandhinagar | Gujarat Biotechnology Research Centre | Nitin Savaliya, Raghawendra Kumar, Dinesh Kumar, Zuber Saiyed, Dipa Kinariwala, Disha Patel, Binita Aring, Neeta Khandelwal, Geeta Vaghela, Sonia Barve, Bhavesh Modi, Kairavi Joshi, Gaurishankar Shrimali, Nidhi Sood, Pranay Shah, R D Dixit, Snehal Bagatharia, Kamlesh J Upadhyay, Ramesh Pandit, Tejas Shah, Ankit Hinsu, Pritesh Sabara, Apurvasinh Puvar, Janvi Raval, Monika Gandhi, Pinal Trivedi, Maharshi Pandya, Amit Kanani, Akanksha Verma, Bhavya Jindal, Chaitanya Joshi, Madhvi Joshi |
| EPI_ISL_447554 | GMERS Medical College and Hospital, Gandhinagar | Gujarat Biotechnology Research Centre | Zuber Saiyed, Dipa Kinariwala, Disha Patel, Binita Aring, Neeta Khandelwal, Geeta Vaghela, Sonia Barve, Bhavesh Modi, Kairavi Joshi, Gaurishankar Shrimali, Nidhi Sood, Pranay Shah, R D Dixit, Snehal Bagatharia, Kamlesh J Upadhyay, Ramesh Pandit, Tejas Shah, Ankit Hinsu, Pritesh Sabara, Apurvasinh Puvar, Janvi Raval, Monika Gandhi, Pinal Trivedi, Maharshi Pandya, Amit Kanani, Akanksha Verma, Nitin Savaliya, Raghawendra Kumar, Dinesh Kumar, Sharmistha Majumdar, Chaitanya Joshi, Madhvi Joshi |
| EPI_ISL_447587 | Tamil Nadu Veterinary and Animal Sciences University | CSIR-Centre for Cellular and Molecular Biology | K Kaveri, S Sivasubramanian, S Vennila, P Padmapriya, R Kiruba, S Magesh, G Dhinakar Raj, G Ravi Kumar, Payel Mukherjee, Tulasi Nagabandi, Namami Gaur, Sakshi Shambhavi, Lamuk Zaveri, Shagufta Khan, Purushotham Vodnala, Sofia Banu, Priya Singh, Dhiviya Vedagiri, Divya Gupta, Vishal Sah, Santosh Kumar Kuncha, Krishnan Harinivas Harshan, Archana Bharadwaj Siva, Karthik Bharadwaj Tallapaka, Kumarasamy Thangaraj, Rakesh K Mishra, Divya Tej Sowpati |
| EPI_ISL_447597, EPI_ISL_447598, EPI_ISL_447599, EPI_ISL_447600, EPI_ISL_447601, EPI_ISL_447602, EPI_ISL_447603, EPI_ISL_447604, EPI_ISL_447605, EPI_ISL_447606, EPI_ISL_447607 | | | |
| see above | Viral Respiratory Lab, National Institute for Biomedical Research (INRB) | Pathogen Sequencing Lab, National Institute for Biomedical Research (INRB) | Placide Mbala-Kingebeni, Edith Nkwembe, Eddy Kinganda-Lusamaki, Amuri Aziza, Francisca Muyembe Mawete, Catherine Pratt, Matthias Pauthner, Josh Quick, Allison Black, James Hadfield, Trevor Bedford, Ian Goodfellow, Andrew Rambaut, Nick Loman, Kristian Andersen, Michael Wiley, Steve Ahuka-Mundeke, Jean-Jacques Muyembe Tamfum |
| EPI_ISL_447631 | Virology, Wageningen Bioveterinary Research | Virology, Wageningen Bioveterinary Research | Oreshkova,N., Vreman,S., Molenaar,R.J., Harders,F., Hakze van der Honing,R.W., Gerhards,N., Bouwstra,R., Hissink,H., Smit,L., Tacken,M., Weesendorp,E., Stegeman,A., van der Poel,W.H.M., Engelsma,M.Y. |
| EPI_ISL_447897 | Genome Centre | Genome Centre | A. S. M. Rubayet Ul Alam, M. Rafiul Islam, M. Shaminur Rahman, Md. Tanvir Islam, Md. Shazid Hasan, Pravas Chandra Roy, Habiba Ibnat, MD. Ali Ahasan Setu, Tanay Chakrovarty, Sourav Dutta Dip, Ruhul Amin, Md. Nur Kabidul Islam, Ovinu Kibria Islam, Hassan Md. Al-Emran, Shireen Nigar, Selina Akter, Md. Nazmul Hasan, Iqbal Kabir Jahid, Md. Anwar Hossain |
| EPI_ISL_447922, EPI_ISL_447923, EPI_ISL_447924, EPI_ISL_447925, EPI_ISL_447927, EPI_ISL_447928, EPI_ISL_447929 | University of Birmingham | COVID-19 Genomics UK (COG-UK) Consortium | Claire McMurray, Joanne Stockton, Samuel Nicholls, Radoslaw Poplawski, Will Rowe, Josh Quick, Nicholas Loman, Celina M Whalley, Andrew Bosworth, Charlotte Poxon, Kasun Wanigasooriya, Oliver Pickles, Mike Kidd, Alex Richter, Andrew D Beggs, Husam Osman, Andrew Bosworth |
| EPI_ISL_448192, EPI_ISL_448193, EPI_ISL_448194, EPI_ISL_448195, EPI_ISL_448196, EPI_ISL_448197, EPI_ISL_448198, EPI_ISL_448199, EPI_ISL_448200, EPI_ISL_448201, EPI_ISL_448202, EPI_ISL_448203, EPI_ISL_448204, EPI_ISL_448205, EPI_ISL_448206, EPI_ISL_448207, EPI_ISL_448208, EPI_ISL_448209, EPI_ISL_448210, EPI_ISL_448211, EPI_ISL_448212, EPI_ISL_448213, EPI_ISL_448214, EPI_ISL_448215, EPI_ISL_448216, EPI_ISL_448217, EPI_ISL_448218, EPI_ISL_448219, EPI_ISL_448220, EPI_ISL_448221 | | | |
| see above | West of Scotland Specialist Virology Centre, NHSGGC / MRC-University of Glasgow Centre for Virus Research | COVID-19 Genomics UK (COG-UK) Consortium | Ana da Silva Filipe, Natasha Johnson, Kathy Smollett, Daniel Mair, Stephen Carmichael, Lily Tong, Jenna Nichols, Elihu Aranday-Cortes, Kirstyn Brunker, Yasmin Parr, Kyriaki Nomikou, Sarah McDonald, Marc Niebel, Patawee Asamaphan, Richard Orton, Joseph Hughes, Sreenu Vattipally, David L Robertson, Alasdair MacLean, Rory Gunson, Kathy Li, Natasha Jesudason, Rajiv Shah, James Shepherd, Antonia Ho, Emma Thomson |

| | | | |
|---|---|---|---|
| EPI_ISL_448309, EPI_ISL_448310, EPI_ISL_448311, EPI_ISL_448312, EPI_ISL_448313, EPI_ISL_448314, EPI_ISL_448315, EPI_ISL_448316, EPI_ISL_448317, EPI_ISL_448318, EPI_ISL_448319, EPI_ISL_448320, EPI_ISL_448321, EPI_ISL_448322, EPI_ISL_448323, EPI_ISL_448324, EPI_ISL_448325, EPI_ISL_448326, EPI_ISL_448327, EPI_ISL_448328, EPI_ISL_448329, EPI_ISL_448330, EPI_ISL_448331, EPI_ISL_448332, EPI_ISL_448333, EPI_ISL_448334, EPI_ISL_448335, EPI_ISL_448336, EPI_ISL_448337, EPI_ISL_448338, EPI_ISL_448339, EPI_ISL_448340, EPI_ISL_448341, EPI_ISL_448342, EPI_ISL_448343, EPI_ISL_448344, EPI_ISL_448345, EPI_ISL_448346, EPI_ISL_448347, EPI_ISL_448348 | | | |
| see above | Quadram Institute Bioscience | COVID-19 Genomics UK (COG-UK) Consortium | Dave J. Baker, Gemma L. Kay, Alp Aydin, Thanh Le-Viet, Steven Rudder, Ana P. Tedim, Anastasia Kolyva, Maria Diaz, Leonardo de Oliveira Martins, Nabil-Fareed Alikhan, Lizzie Meadows, Rachael Stanley, Ngozi Elumogo, Muhammed Yasir, Nicholas M. Thomson, Alexander J Trotter, Rachel Gilroy, Samuel Bloomfield, Claire Stuart, Andrew Bell, Reenesh Prakash, Samir Dervisevic, Alison E. Mather, John Wain, Mark Webber, Andrew J. Page, Justin O'Grady |
| EPI_ISL_448438, EPI_ISL_448439, EPI_ISL_448440, EPI_ISL_448441, EPI_ISL_448442, EPI_ISL_448443, EPI_ISL_448444, EPI_ISL_448445, EPI_ISL_448446 | Queens Medical Centre, Clinical Microbiology Department / DeepSeq Nottingham | COVID-19 Genomics UK (COG-UK) Consortium | Gemma Clark, Wendy Smith, Manjinder Khakh, Hannah Howson-Wells, Jonathan Ball, Patrick McClure, Joseph Chappell, Theocharis Tsoleridis, Nadine Holmes, Matthew Carlisle, Christopher Moore, Fei Sang, Johnny Debebe, Victoria Wright, Matthew Loose |
| EPI_ISL_448978, EPI_ISL_448983, EPI_ISL_448988, EPI_ISL_448989, EPI_ISL_448991, EPI_ISL_448997, EPI_ISL_449000, EPI_ISL_449006, EPI_ISL_449009, EPI_ISL_449011, EPI_ISL_449014, EPI_ISL_449017, EPI_ISL_449018, EPI_ISL_449019, EPI_ISL_449021, EPI_ISL_449023, EPI_ISL_449025, EPI_ISL_449027, EPI_ISL_449029, EPI_ISL_449031, EPI_ISL_449032 | | | |
| see above | Quadram Institute Bioscience | COVID-19 Genomics UK (COG-UK) Consortium | Dave J. Baker, Gemma L. Kay, Alp Aydin, Thanh Le-Viet, Steven Rudder, Ana P. Tedim, Anastasia Kolyva, Maria Diaz, Leonardo de Oliveira Martins, Nabil-Fareed Alikhan, Lizzie Meadows, Rachael Stanley, Ngozi Elumogo, Muhammed Yasir, Nicholas M. Thomson, Alexander J Trotter, Rachel Gilroy, Samuel Bloomfield, Claire Stuart, Andrew Bell, Reenesh Prakash, Samir Dervisevic, Alison E. Mather, John Wain, Mark Webber, Andrew J. Page, Justin O'Grady |
| EPI_ISL_449176, EPI_ISL_449177, EPI_ISL_449178, EPI_ISL_449179, EPI_ISL_449180, EPI_ISL_449181, EPI_ISL_449182, EPI_ISL_449183, EPI_ISL_449184, EPI_ISL_449185, EPI_ISL_449189, EPI_ISL_449190, EPI_ISL_449191, EPI_ISL_449192, EPI_ISL_449193, EPI_ISL_449194, EPI_ISL_449195, EPI_ISL_449196, EPI_ISL_449197, EPI_ISL_449198, EPI_ISL_449202, EPI_ISL_449203, EPI_ISL_449204, EPI_ISL_449205, EPI_ISL_449232, EPI_ISL_449233, EPI_ISL_449234, EPI_ISL_449235, EPI_ISL_449236, EPI_ISL_449237, EPI_ISL_449238, EPI_ISL_449239, EPI_ISL_449240, EPI_ISL_449241, EPI_ISL_449242, EPI_ISL_449243, EPI_ISL_449244, EPI_ISL_449245, EPI_ISL_449246, EPI_ISL_449247, EPI_ISL_449248 | | | |
| see above | West of Scotland Specialist Virology Centre, NHSGGC / MRC-University of Glasgow Centre for Virus Research | COVID-19 Genomics UK (COG-UK) Consortium | Ana da Silva Filipe, Natasha Johnson, Kathy Smollett, Daniel Mair, Stephen Carmichael, Lily Tong, Jenna Nichols, Elihu Aranday-Cortes, Kirstyn Brunker, Yasmin Parr, Kyriaki Nomikou, Sarah McDonald, Marc Niebel, Patawee Asamaphan, Richard Orton, Joseph Hughes, Sreenu Vattipally, David L Robertson, Alasdair MacLean, Rory Gunson, Kathy Li, Natasha Jesudason, Rajiv Shah, James Shepherd, Antonia Ho, Emma Thomson |
| EPI_ISL_449278, EPI_ISL_449279, EPI_ISL_449280, EPI_ISL_449281, EPI_ISL_449282, EPI_ISL_449283, EPI_ISL_449284, EPI_ISL_449285, EPI_ISL_449286, EPI_ISL_449287, EPI_ISL_449288, EPI_ISL_449289, EPI_ISL_449290, EPI_ISL_449291, EPI_ISL_449292, EPI_ISL_449293, EPI_ISL_449294, EPI_ISL_449295, EPI_ISL_449296, EPI_ISL_449297, EPI_ISL_449298, EPI_ISL_449299, EPI_ISL_449300, EPI_ISL_449301, EPI_ISL_449302 | | | |
| see above | Virology Department, Royal Infirmary of Edinburgh, NHS Lothian / School of Biological Sciences, University of Edinburgh / Institute of Genetics and Molecular Medicine, University of Edinburgh | COVID-19 Genomics UK (COG-UK) Consortium | McHugh M, Dewar R, Rooke S, Gallagher M, Balcaza C, O'Toole Á, Scher E, Hill V, McCrone JT, Colquhoun R, Yu X, Jackson B, Rambaut A, Williams TC, Templeton K |
| EPI_ISL_450192 | Scripps Medical Laboratory | Andersen lab at Scripps Research | SEARCH Alliance San Diego with Michael Quigley, Ellen Stefanski, Ian Mchardy |
| EPI_ISL_450248, EPI_ISL_450249, EPI_ISL_450250, EPI_ISL_450251, EPI_ISL_450252, EPI_ISL_450253, EPI_ISL_450254, EPI_ISL_450255, EPI_ISL_450256, EPI_ISL_450257, EPI_ISL_450258, EPI_ISL_450259, EPI_ISL_450260, EPI_ISL_450261, EPI_ISL_450262, EPI_ISL_450263, EPI_ISL_450264, EPI_ISL_450265, EPI_ISL_450266, EPI_ISL_450267, EPI_ISL_450268, EPI_ISL_450269, EPI_ISL_450270, EPI_ISL_450271, EPI_ISL_450272, EPI_ISL_450273, EPI_ISL_450274, EPI_ISL_450275, EPI_ISL_450276, EPI_ISL_450277, EPI_ISL_450278, EPI_ISL_450279, EPI_ISL_450280, EPI_ISL_450281, EPI_ISL_450282, EPI_ISL_450283, EPI_ISL_450284, EPI_ISL_450285, EPI_ISL_450286, EPI_ISL_450287, EPI_ISL_450288, EPI_ISL_450289, EPI_ISL_450290, EPI_ISL_450291, EPI_ISL_450292, EPI_ISL_450293 | | | |
| see above | WHO National Influenza Centre Russian Federation | WHO National Influenza Centre Russian Federation | Andrey Komissarov, Artem Fadeev, Mariia Sergeeva, Anna Ivanova, Tamila Musaeva, Ksenia Komissarova, Mariia Timofeeva, Veronica Eder, Mariia Pisareva, Daria Danilenko |
| EPI_ISL_450499 | Molecular Pathology, Mehr Pathobiology Lab | Molecular Pathology, Mehr Pathobiology Lab | Shabadori,R., Soleimani Dodaran,M., Mirzapour,Z., Kamali,M. and Hamedi,D. |
| EPI_ISL_450505 | Molecular Pathology, Mehr Pathobiology Lab | Molecular Pathology, Mehr Pathobiology Lab | Soleimani Dodaran,M., Soleimani Dodaran,M., Mirzapour,Z., Shabadori,R., Kamali,M. and Hamedi,D. |
| EPI_ISL_450814 | Huslakarna Varmbadhuset Varberg | The Public Health Agency of Sweden | Johanna Hilmersson, Anna-Malin Linde, Maria Lind Karlberg, Oskar Karlsson Lindsjo, Olov Svartstrom, Anna Risberg, Theresa Enkirch, Mia Brytting, Karin Tegmark-Wisell |
| EPI_ISL_450815 | Narhalsan Molnlycke, Barn och ungdomsmedicin | The Public Health Agency of Sweden | Mats Reimer, Anna-Malin Linde, Maria Lind Karlberg, Oskar Karlsson Lindsjo, Olov Svartstrom, Anna Risberg, Theresa Enkirch, Mia Brytting, Karin Tegmark-Wisell |
| EPI_ISL_450816 | orestadsklinikens VC | The Public Health Agency of Sweden | Lisa Kjellberg / Laura Plavitu, Anna-Malin Linde, Maria Lind Karlberg, Oskar Karlsson Lindsjo, Olov Svartstrom, Anna Risberg, Theresa Enkirch, Mia Brytting, Karin Tegmark-Wisell |
| EPI_ISL_450817 | Narhalsan Backa vardcentral | The Public Health Agency of Sweden | Mats Olsson, Anna-Malin Linde, Maria Lind Karlberg, Oskar Karlsson Lindsjo, Olov Svartstrom, Anna Risberg, Theresa Enkirch, Mia Brytting, Karin Tegmark-Wisell |
| EPI_ISL_450818 | Aneby VC | The Public Health Agency of Sweden | Ken Granath, Anna-Malin Linde, Maria Lind Karlberg, Oskar Karlsson Lindsjo, Olov Svartstrom, Anna Risberg, Theresa Enkirch, Mia Brytting, Karin Tegmark-Wisell |
| EPI_ISL_450819 | Hovas Askim Familjelakare och BVC | The Public Health Agency of Sweden | Anna Wendel, Anna-Malin Linde, Maria Lind Karlberg, Oskar Karlsson Lindsjo, Olov Svartstrom, Anna Risberg, Theresa Enkirch, Mia Brytting, Karin Tegmark-Wisell |
| EPI_ISL_450820 | Narhalsan Molnlycke, Barn och ungdomsmedicin | The Public Health Agency of Sweden | Mats Reimer, Anna-Malin Linde, Maria Lind Karlberg, Oskar Karlsson Lindsjo, Olov Svartstrom, Anna Risberg, Theresa Enkirch, Mia Brytting, Karin Tegmark-Wisell |
| EPI_ISL_450821, EPI_ISL_450822 | Jarpens HC | The Public Health Agency of Sweden | Gunilla Johansson, Anna-Malin Linde, Maria Lind Karlberg, Oskar Karlsson Lindsjo, Olov Svartstrom, Anna Risberg, Theresa Enkirch, Mia Brytting, Karin Tegmark-Wisell |
| EPI_ISL_450823 | Narhalsan Backa vardcentral | The Public Health Agency of Sweden | Mats Olsson, Anna-Malin Linde, Maria Lind Karlberg, Oskar Karlsson Lindsjo, Olov Svartstrom, Anna Risberg, Theresa Enkirch, Mia Brytting, Karin Tegmark-Wisell |
| EPI_ISL_450824 | Ulltuna Vardcentral | The Public Health Agency of Sweden | Heidi Lindback, Anna-Malin Linde, Maria Lind Karlberg, Oskar Karlsson Lindsjo, Olov Svartstrom, Anna Risberg, Theresa Enkirch, Mia Brytting, Karin Tegmark-Wisell |
| EPI_ISL_451199 | Uganda Virus Research Institute | MRC/UVRI & LSHTM Uganda Research Unit | Dan Lule Bugembe, John Kayiwa, My V.T Phan, Phionah Tushabe, Stephen Balinandi, Beatrice Dhaala, Deogratius Ssemwanga, Jonas Lexow, Henry Mwebesa, Jane Aceng, Henry Kyobe, Julius Lutwama, Pontiano Kaleebu, Matthew Cotten |
| EPI_ISL_451204, EPI_ISL_451212, EPI_ISL_451215 | LSUHS Emerging Viral Threat Laboratory | Microbial Genome Sequencing Center | Jeremy P. Kamil, John A. Vanchiere, Rona S. Scott, Camille F. Abshire, Abida Siddiqa, Byeong-Jae Lee, Chan-ki Min, Md Maksudul Alam, Monica Gestal-Cartele, Edna Ondari, Adam Greer, Malgorzata Bienkowska-Haba, Katarzyna Zwolinska, Jason M. Bodily, Andrew D. Yurochko, Paul M. Weinberger, Christopher G. Kevil, Martin J. Sapp, Daniel J. Snyder, Vaughn S. Cooper |
| EPI_ISL_451221 | LSUHS Emerging Viral Threat Laboratory | Microbial Genome Sequencing Center | Rona S. Scott, Jeremy P. Kamil, John A. Vanchiere, Camille F. Abshire, Abida Siddiqa, Byeong-Jae Lee, Chan-ki Min, Md Maksudul Alam, Monica Gestal-Cartele, Edna Ondari, Adam Greer, Malgorzata Bienkowska-Haba, Katarzyna Zwolinska, Jason M. Bodily, Andrew D. Yurochko, Paul M. Weinberger, Christopher G. Kevil, Martin J. Sapp, Daniel J. Snyder, Vaughn S. Cooper |
| EPI_ISL_451400 | Laboratoire de Recherche et d'Analyse Médicale de la Gendarmerie Royale | Laboratoire de Recherche et d'Analyse Médicale de la Gendarmerie Royale | Sanaâ LEMRISS, Amal SOUIRI, Saâd EL KABBAJ |
| EPI_ISL_451630 | Pathology West - NSW Health Pathology | NSW Health Pathology - Institute of Clinical Pathology and Medical Research; Westmead Hospital; University of Sydney | CIDM-PH et al. |
| EPI_ISL_451642 | Pathology Sydney South West - NSW Health Pathology | NSW Health Pathology - Institute of Clinical Pathology and Medical Research; Westmead Hospital; University of Sydney | CIDM-PH et al. |
| EPI_ISL_451646, EPI_ISL_451647 | Laboratory of Molecular Biology, Diagnostyka sp. z o.o. | Laboratory of Recombinant Vaccines | Lukasz Rabalski, Anna Piotrowska-Mietelska, Maciej Kosinski, Boguslaw Szewczyk, Krystyna Bienkowska-Szewczyk |

| | | | |
|---|---|---|---|
| EPI_ISL_451648 | Hematology Laboratory, Section of Molecular Diagnostics, University Clinical Centre, Medical University of Gdansk | Laboratory of Recombinant Vaccines | Lukasz Rabalski, Adam Sodol, Aneta Szulc, Krzysztof Lewandowski, Ewa Milosz, Marlena Robakowska, Boguslaw Szewczyk, Krystyna Bienkowska-Szewczyk |
| EPI_ISL_452059, EPI_ISL_452060, EPI_ISL_452083, EPI_ISL_452084, EPI_ISL_452085, EPI_ISL_452086, EPI_ISL_452087, EPI_ISL_452088, EPI_ISL_452089, EPI_ISL_452090, EPI_ISL_452091 | | | |
| see above | Department of Clinical Microbiology, Copenhagen University Hospital, Hvidovre, Kettegaard Alle 30, 2650 Hvidovre. | Albertsen lab, Department of Chemistry and Bioscience, Aalborg University, Denmark | Rasmus Kirkegaard |
| EPI_ISL_452143 | Yale COVID-19 Biorepository | Grubaugh Lab - Yale School of Public Health | Joseph Fauver, Tara Alpert, Anderson Brito, Anne Wyllie, Chantal Vogels, Mary Petrone, Cole Jensen, Chaney Kalinich, Isabel Ott, Arnau Casanovas, Catherine Muenker, Adam Moore, Alice Lu, Maria Tokuyama, Patrick Wong, Peiwen Lu, Saad Omer, Richard Martinello, Allison Nelson, Shelli Farhadian, Akiko Iwasaki, Charlese Dela Cruz, Albert Ko, Nathan Grubaugh |
| EPI_ISL_452152 | CUB Hopital Erasme Laboratoire d'Anatomie Pathologique | CUB Hopital Erasme Laboratoire d'Anatomie Pathologique | Prof. Isabelle Salmon, Dr Nicky D'Haene |
| EPI_ISL_452198, EPI_ISL_452199, EPI_ISL_452200, EPI_ISL_452201, EPI_ISL_452215, EPI_ISL_452217 | NIV Influenza | NIV Influenza | Potdar V |
| EPI_ISL_452237 | Huddinge VC | The Public Health Agency of Sweden | Anders Johansson, Anna-Malin Linde, Maria Lind Karlberg, Oskar Karlsson Lindsjo, Olov Svartstrom, Anna Risberg, Theresa Enkirch, Mia Brytting, Karin Tegmark-Wisell |
| EPI_ISL_452239 | Narhalsan Backa vardcentral | The Public Health Agency of Sweden | Mats Olsson, Anna-Malin Linde, Maria Lind Karlberg, Oskar Karlsson Lindsjo, Olov Svartstrom, Anna Risberg, Theresa Enkirch, Mia Brytting, Karin Tegmark-Wisell |
| EPI_ISL_452240 | Huddinge VC | The Public Health Agency of Sweden | Anders Johansson, Anna-Malin Linde, Maria Lind Karlberg, Oskar Karlsson Lindsjo, Olov Svartstrom, Anna Risberg, Theresa Enkirch, Mia Brytting, Karin Tegmark-Wisell |
| EPI_ISL_452241 | Huslakarna Varmbadhuset Varberg | The Public Health Agency of Sweden | Johanna Hilmersson, Anna-Malin Linde, Maria Lind Karlberg, Oskar Karlsson Lindsjo, Olov Svartstrom, Anna Risberg, Theresa Enkirch, Mia Brytting, Karin Tegmark-Wisell |
| EPI_ISL_452789, EPI_ISL_452790 | ICAR-National Institute of High Security Animal Diseases | ICAR-National Institute of High Security Animal Diseases | Anamika Mishra, Ashutosh Aasdev, Sandeep Bhatia, Harshad Murugkar, Chakradhar Tosh, Niranjan Mishra, Shanmugasundaram Nagarajan, Katherukamem Rajukumar, Richa Sood, G Venkatesh, Atul Kumar Pateriya, Manoj Kumar, Shashi Bhushan Sudhakar, Fateh Singh, Sethil Kumar D, Senmannan Kalaiyarasu, Pradeep Gandhale, Naveen Kumar, Chandan Kumar Dubey, Sushil Tripathi, Sandeep Kumar Jhade, Meghna Tripathi, Suman Kumari Shah, Pushpendra Singh, Pushpendra Namdeo, Suman Mishra, Rupal Singh, Vishnupriya Patil, Dipesh Kumar Nayak, Vijendra Pal Singh, Ashwin Ashok Raut |
| EPI_ISL_452824, EPI_ISL_452825, EPI_ISL_452826, EPI_ISL_452827, EPI_ISL_452828, EPI_ISL_452829, EPI_ISL_452830, EPI_ISL_452831, EPI_ISL_452832, EPI_ISL_452836, EPI_ISL_452837, EPI_ISL_452838, EPI_ISL_452839, EPI_ISL_452840, EPI_ISL_452841, EPI_ISL_452842, EPI_ISL_452843, EPI_ISL_452844, EPI_ISL_452845, EPI_ISL_452846, EPI_ISL_452847, EPI_ISL_452848, EPI_ISL_452849, EPI_ISL_452850, EPI_ISL_452851, EPI_ISL_452852, EPI_ISL_452853, EPI_ISL_452854, EPI_ISL_452855, EPI_ISL_452856 | | | |
| see above | Virginia DCLS | Virginia DCLS | Virginia DCLS |
| EPI_ISL_453075, EPI_ISL_453076, EPI_ISL_453077, EPI_ISL_453078, EPI_ISL_453079, EPI_ISL_453080, EPI_ISL_453081, EPI_ISL_453082, EPI_ISL_453083, EPI_ISL_453084, EPI_ISL_453085, EPI_ISL_453086, EPI_ISL_453087, EPI_ISL_453088, EPI_ISL_453089, EPI_ISL_453090, EPI_ISL_453092, EPI_ISL_453095, EPI_ISL_453096 | | | |
| see above | West of Scotland Specialist Virology Centre, NHSGGC / MRC-University of Glasgow Centre for Virus Research | COVID-19 Genomics UK (COG-UK) Consortium | Ana da Silva Filipe, Natasha Johnson, Kathy Smollett, Daniel Mair, Stephen Carmichael, Lily Tong, Jenna Nichols, Elihu Aranday-Cortes, Kirstyn Brunker, Yasmin Parr, Kyriaki Nomikou; Sarah McDonald, Marc Niebel, Patawee Asamaphan; Richard Orton, Joseph Hughes, Sreenu Vattipally, David L Robertson; Alasdair MacLean, Rory Gunson; Kathy Li, Natasha Jesudason, Rajiv Shah, James Shepherd, Antonia Ho, Emma Thomson |
| EPI_ISL_453099, EPI_ISL_453110, EPI_ISL_453111, EPI_ISL_453112, EPI_ISL_453113, EPI_ISL_453114, EPI_ISL_453115, EPI_ISL_453116, EPI_ISL_453117, EPI_ISL_453118, EPI_ISL_453119, EPI_ISL_453120, EPI_ISL_453121, EPI_ISL_453122, EPI_ISL_453123, EPI_ISL_453124, EPI_ISL_453125, EPI_ISL_453126, EPI_ISL_453127, EPI_ISL_453128, EPI_ISL_453191, EPI_ISL_453192 | | | |
| see above | Virology Department, Royal Infirmary of Edinburgh, NHS Lothian / School of Biological Sciences, University of Edinburgh / Institute of Genetics and Molecular Medicine, University of Edinburgh | COVID-19 Genomics UK (COG-UK) Consortium | McHugh M, Dewar R, Rooke S, Gallagher M, Balcaza C, O'Toole Á, Scher E, Hill V, McCrone JT, Colquhoun R, Yu X, Jackson B, Rambaut A, Williams TC, Templeton K |
| EPI_ISL_453209, EPI_ISL_453334, EPI_ISL_453393, EPI_ISL_453458 | Liverpool Clinical Laboratories | COVID-19 Genomics UK (COG-UK) Consortium | Sam Haldenby, Anita Lucaci, Steve Paterson, Julian Hiscox, Alistair Darby, M Almsaud, A Alrezaihi, Muhannad Alruwaili, Stuart D Armstrong, Jones Benjamin , Eleanor G Bentley, Anu Chawla, Jordan J Clark, Angela Cowell, Richard Eccles, Isabel García-Dorival, Matthew Gemmell, Alessandro Gerada, PKF Gilmore, Richard Gregory, Ximeng Han, Catherine Hartley, Margaret Hughes, Miren Iturriza-Gomara, James Johnson, L Luu, Jenifer Manson , Charlotte Nelson, Elaine O'Toole, Cassie Olateju, Rebekah Penrice-Randal , Lucille Rainbow, N.P Randle, Trevor Ian Robinson, Parul Sharma, Ghada T Shawli, James P Stewart , Neil Swainston, Ecaterina Vamos, Joanne Watts, Mark Whitehead |
| EPI_ISL_453463 | University College London, Great Ormond Street Hospital for Children NHS Foundation Trust, Imperial College Healthcare NHS Trust | COVID-19 Genomics UK (COG-UK) Consortium | Sergi Castellano, Rachel Williams, Mark Kristiansen, Paola Resende Silva, Sunando Roy, Tony Brooks, Helena Tutill, Paola Niola, Patricia Dyal, Charlotte Williams, Leysa Forrest, Yasmin Panchbhaya, Jacqueline Findlay, Sam Weeks, Julianne Brown, Kathryn Harris, Paul Randell, James Price, Alison Holmes, Judith Breuer |
| EPI_ISL_453538, EPI_ISL_453539, EPI_ISL_453542 | Northumbria University / South Tees Hospitals NHS Foundation Trust / North Cumbria Integrated Care NHS Foundation Trust / North Tees and Hartlepool NHS Foundation Trust / Newcastle Hospitals NHS Foundation Trust | COVID-19 Genomics UK (COG-UK) Consortium | Darren L Smith,Andrew Nelson,Matthew Bashton,Greg R Young,Joshua Loh,John Allan,Mohammad A Tariq,Giles S Holt,Gary Black,Wen C Yew,Lynn Dover ,Paul Baker,Steve Liggett,Sarah Essex,Jane Greenaway ,Debra Padgett,Clive Graham,Garren Scott,Edward Barton ,Emma Swindells ,Brendan Payne,Jennifer Collins,Yusri Taha,Gary Eltringham |
| EPI_ISL_453543 | Northumbria University / South Tees Hospitals NHS Foundation Trust / North Cumbria Integrated Care NHS Foundation Trust / North Tees and Hartlepool NHS Foundation Trust / Newcastle Hospitals NHS Foundation Trust | Northumbria University | Darren L Smith,Andrew Nelson,Matthew Bashton,Greg R Young,Joshua Loh,John Allan,Mohammad A Tariq,Giles S Holt,Gary Black,Wen C Yew,Lynn Dover ,Paul Baker,Steve Liggett,Sarah Essex,Jane Greenaway ,Debra Padgett,Clive Graham,Garren Scott,Edward Barton ,Emma Swindells ,Brendan Payne,Jennifer Collins,Yusri Taha,Gary Eltringham |
| EPI_ISL_453544 | Northumbria University / South Tees Hospitals NHS Foundation Trust / North Cumbria Integrated Care NHS Foundation Trust / North Tees and Hartlepool NHS Foundation Trust / Newcastle Hospitals NHS Foundation Trust | COVID-19 Genomics UK (COG-UK) Consortium | Darren L Smith,Andrew Nelson,Matthew Bashton,Greg R Young,Joshua Loh,John Allan,Mohammad A Tariq,Giles S Holt,Gary Black,Wen C Yew,Lynn Dover ,Paul Baker,Steve Liggett,Sarah Essex,Jane Greenaway ,Debra Padgett,Clive Graham,Garren Scott,Edward Barton ,Emma Swindells ,Brendan Payne,Jennifer Collins,Yusri Taha,Gary Eltringham |
| EPI_ISL_453545, EPI_ISL_453546, EPI_ISL_453547, EPI_ISL_453548, EPI_ISL_453549, EPI_ISL_453550, EPI_ISL_453551, EPI_ISL_453552, EPI_ISL_453553, EPI_ISL_453554, EPI_ISL_453555, EPI_ISL_453556, EPI_ISL_453557, EPI_ISL_453558, EPI_ISL_453559, EPI_ISL_453560, EPI_ISL_453561, EPI_ISL_453562, EPI_ISL_453563, EPI_ISL_453564, EPI_ISL_453565, EPI_ISL_453566, EPI_ISL_453567, EPI_ISL_453568, EPI_ISL_453569, EPI_ISL_453570, EPI_ISL_453571, EPI_ISL_453572, EPI_ISL_453573, EPI_ISL_453574, EPI_ISL_453575, EPI_ISL_453576, EPI_ISL_453577, EPI_ISL_453578, EPI_ISL_453579, EPI_ISL_453580, EPI_ISL_453583, EPI_ISL_453584, EPI_ISL_453592, EPI_ISL_453595, EPI_ISL_453597, EPI_ISL_453598, EPI_ISL_453599, EPI_ISL_453600, EPI_ISL_453601, EPI_ISL_453602, EPI_ISL_453603, EPI_ISL_453604, EPI_ISL_453605, EPI_ISL_453606, EPI_ISL_453607, EPI_ISL_453608, EPI_ISL_453609, EPI_ISL_453610, EPI_ISL_453611, EPI_ISL_453612, EPI_ISL_453613 | | | |
| see above | Quadram Institute Bioscience | COVID-19 Genomics UK (COG-UK) Consortium | Dave J. Baker, Gemma L. Kay, Alp Aydin, Thanh Le-Viet, Steven Rudder, Ana P. Tedim, Anastasia Kolyva, Maria Diaz, Leonardo de Oliveira Martins, Nabil-Fareed Alikhan, Lizzie Meadows, Rachael Stanley, Ngozi Elumogo, Muhammed Yasir, Nicholas M. Thomson, Alexander J Trotter, Rachel Gilroy, Samuel Bloomfield, Claire Stuart, Andrew Bell, Reenesh Prakash, Samir Dervisevic, Alison E. Mather, John Wain, Mark Webber, Andrew J. Page, Justin O'Grady |
| EPI_ISL_453636, EPI_ISL_453638, EPI_ISL_453639, EPI_ISL_453640, EPI_ISL_453641, EPI_ISL_453642 | Queens Medical Centre, Clinical Microbiology Department / DeepSeq Nottingham | COVID-19 Genomics UK (COG-UK) Consortium | Gemma Clark, Wendy Smith, Manjinder Khakh, Hannah Howson-Wells, Jonathan Ball, Patrick McClure, Joseph Chappell, Theocharis Tsoleridis, Nadine Holmes, Matthew Carlisle, Christopher Moore, Fei Sang, Johnny Debebe, Victoria Wright, Matthew Loose |
| EPI_ISL_453665 | Centre for Enzyme Innovation, University of Portsmouth / Translational Research Laboratory, Portsmouth Hospitals NHS Trust | University of Portsmouth | Angela Beckett,,Yann Bourgeois,,Garry Scarlett,,Sharon Glaysher,,Scott Elliott,,Kelly Bicknell,,Robert Impey,,Allyson Lloyd,,Sarah Wyllie,,Ethan Butcher,,Anoop Chauhan,,Samuel Robson |
| EPI_ISL_453666, EPI_ISL_453667, | Centre for Enzyme Innovation, University of Portsmouth / | COVID-19 Genomics UK (COG-UK) Consortium | Angela Beckett,,Yann Bourgeois,,Garry Scarlett,,Sharon Glaysher,,Scott Elliott,,Kelly Bicknell,,Robert Impey,,Allyson Lloyd,,Sarah Wyllie,,Ethan |

| | | | |
|---|---|---|---|
| EPI_ISL_453668, EPI_ISL_453669, EPI_ISL_453670, EPI_ISL_453671 | Translational Research Laboratory, Portsmouth Hospitals NHS Trust | | Butcher,,Anoop Chauhan,,Samuel Robson |
| EPI_ISL_453696, EPI_ISL_453697, EPI_ISL_453698, EPI_ISL_453699, EPI_ISL_453700, EPI_ISL_453701, EPI_ISL_453702, EPI_ISL_453703, EPI_ISL_453704, EPI_ISL_453705, EPI_ISL_453706, EPI_ISL_453707, EPI_ISL_453708, EPI_ISL_453709, EPI_ISL_453710, EPI_ISL_453711, EPI_ISL_453712, EPI_ISL_453713, EPI_ISL_453714, EPI_ISL_453715, EPI_ISL_453716, EPI_ISL_453717, EPI_ISL_453718, EPI_ISL_453720, EPI_ISL_453721, EPI_ISL_453722, EPI_ISL_453723, EPI_ISL_453724, EPI_ISL_453725, EPI_ISL_453726, EPI_ISL_453727, EPI_ISL_453728, EPI_ISL_453729, EPI_ISL_453730, EPI_ISL_453731, EPI_ISL_453732, EPI_ISL_453733, EPI_ISL_453735, EPI_ISL_453736, EPI_ISL_453737, EPI_ISL_453738, EPI_ISL_453739, EPI_ISL_453740, EPI_ISL_453741, EPI_ISL_453742, EPI_ISL_453743, EPI_ISL_453744, EPI_ISL_453745, EPI_ISL_453746, EPI_ISL_453747, EPI_ISL_453748, EPI_ISL_453749, EPI_ISL_453750, EPI_ISL_453751, EPI_ISL_453752, EPI_ISL_453753, EPI_ISL_453754, EPI_ISL_453755, EPI_ISL_453756, EPI_ISL_453757, EPI_ISL_453758, EPI_ISL_453759, EPI_ISL_453760, EPI_ISL_453761, EPI_ISL_453762, EPI_ISL_453763, EPI_ISL_453764, EPI_ISL_453765, EPI_ISL_453766, EPI_ISL_453768, EPI_ISL_453769, EPI_ISL_453770, EPI_ISL_453773, EPI_ISL_453774, EPI_ISL_453775, EPI_ISL_453776, EPI_ISL_453777, EPI_ISL_453778 | | | |
| see above | Virology Department, Sheffield Teaching Hospitals NHS Foundation Trust/Department of Infection, Immunity and Cardiovascular Disease, The Medical School, University of Sheffield | COVID-19 Genomics UK (COG-UK) Consortium | Thushan de Silva, Matthew Parker, Nikki Smith, Adri Angyal, Rebecca Brown, Luke Green, Rachel Tucker, Paul Parsons, Danielle Groves, Katie Johnson, Laura Carrilero, Alex Keeley, Dave Partridge, Matthew Wyles, Benjamin Lindsey, Mehmet Yavuz, Mohammad Raza, Cariad Evans |
| EPI_ISL_453998, EPI_ISL_453999, EPI_ISL_454107, EPI_ISL_454116, EPI_ISL_454118, EPI_ISL_454120, EPI_ISL_454123, EPI_ISL_454124, EPI_ISL_454126, EPI_ISL_454197, EPI_ISL_454198, EPI_ISL_454199, EPI_ISL_454200, EPI_ISL_454201, EPI_ISL_454279, EPI_ISL_454280, EPI_ISL_454281, EPI_ISL_454282, EPI_ISL_454283, EPI_ISL_454304 | | | |
| see above | unknown | Instituto Nacional de Saude (INSA) | Borges et al |
| EPI_ISL_454501, EPI_ISL_454502, EPI_ISL_454503, EPI_ISL_454504, EPI_ISL_454505, EPI_ISL_454506, EPI_ISL_454507, EPI_ISL_454509, EPI_ISL_454510, EPI_ISL_454511, EPI_ISL_454512, EPI_ISL_454513, EPI_ISL_454514, EPI_ISL_454515, EPI_ISL_454516, EPI_ISL_454517, EPI_ISL_454518, EPI_ISL_454519, EPI_ISL_454520 | | | |
| see above | RSE "National Center for Biotechnology" | RSE "National Center for Biotechnology" | Alexandr Shevtsov, Ilyas Akhmetollayev, Viktoriya Lutsay, Asylulan Amirgazin, Askar Abdaliyev, Akbota Rakhmetova, Zabira Aushakhmetova, Ruslan Kalendar, Yerlan Ramankulov |
| EPI_ISL_454524, EPI_ISL_454566, EPI_ISL_454567, EPI_ISL_454568, EPI_ISL_454569, EPI_ISL_454570 | NIV Influenza | NIV Influenza | Potdar V |
| EPI_ISL_454582 | Laboratory of virology, National Center of Expertise | Laboratory of molecular-genetic research, National Center of Expertise, Kazakhstan National Center for Biotechnology, Kazakhstan | Abdaliyev Askar, Shevtsov Alexandr, Akhmetollayev Ilyas, Kalendar Ruslan, Rakhmetova Akbota, , Lutsay Viktoriya, Amirgazin Asylulan, Aushakhmetova Zabira, Ramankulov Yerlan |
| EPI_ISL_454584, EPI_ISL_454586 | Laboratory of virology, National Center of Expertise | Laboratory of molecular-genetic research, National Center for Expertise, Kazakhstan National Center for Biotechnology, Kazakhstan | Abdaliyev Askar, Shevtsov Alexandr, Akhmetollayev Ilyas, Kalendar Ruslan, Rakhmetova Akbota, , Lutsay Viktoriya, Amirgazin Asylulan, Aushakhmetova Zabira, Ramankulov Yerlan |
| EPI_ISL_454587 | Laboratory of virology, National Center of Expertise | Laboratory of molecular-genetic research, National Center of Expertise, Kazakhstan National Center for Biotechnology, Kazakhstan | Abdaliyev Askar, Shevtsov Alexandr, Akhmetollayev Ilyas, Kalendar Ruslan, Rakhmetova Akbota, , Lutsay Viktoriya, Amirgazin Asylulan, Aushakhmetova Zabira, Ramankulov Yerlan |
| EPI_ISL_454589 | Laboratory of virology, National Center of Expertise | Laboratory of molecular-genetic research, National Center for Expertise, Kazakhstan National Center for Biotechnology, Kazakhstan | Abdaliyev Askar, Shevtsov Alexandr, Akhmetollayev Ilyas, Kalendar Ruslan, Rakhmetova Akbota, , Lutsay Viktoriya, Amirgazin Asylulan, Aushakhmetova Zabira, Ramankulov Yerlan |
| EPI_ISL_454590, EPI_ISL_454591, EPI_ISL_454593, EPI_ISL_454594 | Laboratory of virology, National Center of Expertise | Laboratory of molecular-genetic research, National Center of Expertise, Kazakhstan National Center for Biotechnology, Kazakhstan | Abdaliyev Askar, Shevtsov Alexandr, Akhmetollayev Ilyas, Kalendar Ruslan, Rakhmetova Akbota, , Lutsay Viktoriya, Amirgazin Asylulan, Aushakhmetova Zabira, Ramankulov Yerlan |
| EPI_ISL_454596, EPI_ISL_454597 | Laboratory of virology, National Center of Expertise | Laboratory of molecular-genetic research, National Center for Expertise, Kazakhstan National Center for Biotechnology, Kazakhstan | Abdaliyev Askar, Shevtsov Alexandr, Akhmetollayev Ilyas, Kalendar Ruslan, Rakhmetova Akbota, , Lutsay Viktoriya, Amirgazin Asylulan, Aushakhmetova Zabira, Ramankulov Yerlan |
| EPI_ISL_454598 | Laboratory of virology, National Center of Expertise | Laboratory of molecular-genetic research, National Center of Expertise, Kazakhstan National Center for Biotechnology, Kazakhstan | Abdaliyev Askar, Shevtsov Alexandr, Akhmetollayev Ilyas, Kalendar Ruslan, Rakhmetova Akbota, , Lutsay Viktoriya, Amirgazin Asylulan, Aushakhmetova Zabira, Ramankulov Yerlan |
| EPI_ISL_454599, EPI_ISL_454600 | Laboratory of virology, National Center of Expertise | Laboratory of molecular-genetic research, National Center for Expertise, Kazakhstan National Center for Biotechnology, Kazakhstan | Abdaliyev Askar, Shevtsov Alexandr, Akhmetollayev Ilyas, Kalendar Ruslan, Rakhmetova Akbota, , Lutsay Viktoriya, Amirgazin Asylulan, Aushakhmetova Zabira, Ramankulov Yerlan |
| EPI_ISL_454601 | Laboratory of virology, National Center of Expertise | Laboratory of molecular-genetic research, National Center of Expertise, Kazakhstan National Center for Biotechnology, Kazakhstan | Abdaliyev Askar, Shevtsov Alexandr, Akhmetollayev Ilyas, Kalendar Ruslan, Rakhmetova Akbota, , Lutsay Viktoriya, Amirgazin Asylulan, Aushakhmetova Zabira, Ramankulov Yerlan |
| EPI_ISL_454602 | Croatian Institute of Public Health | University of Zagreb, Centre for research and knowledge transfer in biotechnology | Irena Tabain, Tatjana Vilibic-Cavlek, Jelena Ivancic Jelecki, Anamarija Slovic |
| EPI_ISL_454603 | Laboratory of virology, National Center of Expertise | Laboratory of molecular-genetic research, National Center of Expertise, Kazakhstan National Center for Biotechnology, Kazakhstan | Abdaliyev Askar, Shevtsov Alexandr, Akhmetollayev Ilyas, Kalendar Ruslan, Rakhmetova Akbota, , Lutsay Viktoriya, Amirgazin Asylulan, Aushakhmetova Zabira, Ramankulov Yerlan |
| EPI_ISL_454604 | Laboratory of virology, National Center of Expertise | Laboratory of molecular-genetic research, National Center for Expertise, Kazakhstan National Center for Biotechnology, Kazakhstan | Abdaliyev Askar, Shevtsov Alexandr, Akhmetollayev Ilyas, Kalendar Ruslan, Rakhmetova Akbota, , Lutsay Viktoriya, Amirgazin Asylulan, Aushakhmetova Zabira, Ramankulov Yerlan |
| EPI_ISL_454615, EPI_ISL_454616, EPI_ISL_454617, EPI_ISL_454618, EPI_ISL_454619 | UCSF Clinical Microbiology Laboratory | Chan-Zuckerberg Biohub | CZB Cliahub Consortium |
| EPI_ISL_454645 | CT-Dr. Katherine A. Kelley State Public Health Lab | Pathogen Discovery, Respiratory Viruses Branch, Division of Viral Diseases, Centers for Disease Control and Prevention | Jing Zhang, Ying Tao, Clinton R. Paden, Anna Uehara, Krista Queen, Yan Li, Haibin Wang, Zachary Weiner, Bettina Bankamp, Suxiang Tong |
| EPI_ISL_454649, EPI_ISL_454650 | VI-US Virgin Islands Department of Health | Pathogen Discovery, Respiratory Viruses Branch, Division of Viral Diseases, Centers for Disease Control and Prevention | Ying Tao, Clinton R. Paden, Jing Zhang, Anna Uehara, Krista Queen, Yan Li, Haibin Wang, Zachary Weiner, Bettina Bankamp, Suxiang Tong |
| EPI_ISL_454783, EPI_ISL_454784 | Dutch COVID-19 response team | National Institute for Public Health and the Environment (RIVM) | Adam Meijer, Harry Vennema, Jeroen Cremer, Sharon van den Brink, Pieter Overduin, Florian Zwagemaker, Dennis Schmitz, Chantal Reusken, on behalf of the national COVID-19 response team |
| EPI_ISL_454795 | Veterinary Specialized Institute Kraljevo | Veterinary Specialized Institute Kraljevo | Vidanovic,D., Tesovic,B., Sekler,M., Dmitric,M., Debeljak,Z., Matovic,K., Vaskovic,N., Petrovic,T., Volkening,J. and Alfonso,C.L. |
| EPI_ISL_454830, EPI_ISL_454832 | SMS Medical College, Jaipur | CSIR Institute of Genomics and Integrative Biology | Sudhir Bhandari, Rahul Bhoyar, Mohammed Imran, Mohit Divakar, Disha Sharma, Anshul Kumar, Bani Jolly, Rahul Sahlot, Abhinav Jain, Paras Sehgal, Gyan Ranjan, Vinod Scaria, Sridhar Sivasubbu, Sandeep K Mathur |
| EPI_ISL_455028, EPI_ISL_455029, EPI_ISL_455030, EPI_ISL_455031, EPI_ISL_455032, EPI_ISL_455033, EPI_ISL_455035 | Pathology West - NSW Health Pathology | NSW Health Pathology - Institute of Clinical Pathology and Medical Research; Westmead Hospital; University of Sydney | CIDM-PH et al. |
| EPI_ISL_455359 | Emory Molecular Diagnostics Laboratory, Emory Healthcare | Piantadosi Lab, Emory Department of Pathology | Ahmed Babiker, Anne Piantadosi |

| | | | |
|---|---|---|---|
| EPI_ISL_455648, EPI_ISL_455649, EPI_ISL_455650, EPI_ISL_455651, EPI_ISL_455652, EPI_ISL_455653, EPI_ISL_455654 | ICMR-National Institute of Cholera and Enteric Diseases | National Institute of Biomedical Genomics | Arindam Maitra, Mamta Chawla Sarkar, Sreedhar Chinnaswamy, Hasina Banu, Ananya Chatterjee, Shanta Dutta, Saumitra Das |
| EPI_ISL_455682 | University of Florida | University of Florida | Lednicky,J.A., Wu,C.-Y., Lauzardo,M. and Morris,J.G. |
| EPI_ISL_456107, EPI_ISL_456109, EPI_ISL_456110, EPI_ISL_456111, EPI_ISL_456112, EPI_ISL_456113 | NYU Langone Health | Departments of Pathology and Medicine, New York University School of Medicine | Maria Aguero-Rosenfeld, Brendan Belovarac, Margaret Black, Ludovic Boytard, John Cadley, Paolo Cotzia, John Chen, Dacia Dimartino, Xiaojun Feng, Tatyana Gindin, Emily Guzman, Adriana Heguy, Megan Hogan, Emily Huang, George Jour, Alireza Khodadadi-Jamayran, Lawrence H. Lin, Raven Luther, Andrew Lytle, Christian Marier, Matthew T. Maurano, Mark J. Mulligan, Peter Meyn, Raquel Ordonez Ciriza, Iman Osman, Jared Pinnell, Vanessa Raabe, Sitharam Ramaswami, Amy Rapkiewicz, Andre M. Ribeiro-dos-Santos, Marie Samanovic-Golden, Antonio Serrano, Guomiao Shen, Matija Snuderl, Theodore Vougiouklakis, Nick Vulpescu, Gael Westby, Paul Zappile, Yutong Zhang |
| EPI_ISL_456144, EPI_ISL_456153, EPI_ISL_456154 | Instituto Nacional de Salud - Unidad de Secuenciación y Análisis Genómico | Instituto Nacional de Salud, Universidad Cooperativa de Colombia, Instituto Alexander von Humboldt, Imperial College-London, London School of Hygiene & Tropical Medicine | Katherine Laiton-Donato, Diego A. Álvarez-Díaz, Carlos Franco-Muñoz, Jose A. Usme-Ciro, Gloria Puerto, Nicolas D. Franco-Sierra, Mailyn A.Gonzalez, Zulma M. Cucunubá, Christian Julian VillabonaArenas, Liz Villabona-Arenas, Sussy Echeverria, Astrid C. Flórez, Sergio Gomez-Rangel, Luz Dary Rodriguez, Juliana Barbosa, Erika Ospitia, Diana Marcela Walteros-Acero, Martha Lucia Ospina Martinez, Marcela Mercado-Reyes. |
| EPI_ISL_456388, EPI_ISL_456389, EPI_ISL_456390, EPI_ISL_456391, EPI_ISL_456392, EPI_ISL_456393, EPI_ISL_456394 | LabPLUS | Institute of Environmental Science and Research (ESR) | Matt Storey, Xiaoyun Ren, Anja Werno, Antje van der Linden, Arlo Upton, Chris Mansell, David Hammer, Dragana Drinkovic, Erasmus Smit, Gary McAuliffe, Hana Sofia Andersson, James Ussher, Jill Sherwood, Josh Freeman, Julia Howard, Juliet Elvy, Mary DeAlmeida, Matt Blakiston, Matthew Rogers, Max Bloomfield, Michael Addidle, Michelle Balm, Sally Roberts, Sarah Jefferies, Sharmini Muttaiyah, Susan Morpeth, Susan Taylor, Timothy Blackmore, Vani Sathyendran, Veronica Playle, Virginia Hope, Erasmus Smit, Lauren Jelly, Joep de Ligt |
| EPI_ISL_456395, EPI_ISL_456396, EPI_ISL_456397 | North Shore Hospital | Institute of Environmental Science and Research (ESR) | Matt Storey, Xiaoyun Ren, Anja Werno, Antje van der Linden, Arlo Upton, Chris Mansell, David Hammer, Dragana Drinkovic, Erasmus Smit, Gary McAuliffe, Hana Sofia Andersson, James Ussher, Jill Sherwood, Josh Freeman, Julia Howard, Juliet Elvy, Mary DeAlmeida, Matt Blakiston, Matthew Rogers, Max Bloomfield, Michael Addidle, Michelle Balm, Sally Roberts, Sarah Jefferies, Sharmini Muttaiyah, Susan Morpeth, Susan Taylor, Timothy Blackmore, Vani Sathyendran, Veronica Playle, Virginia Hope, Erasmus Smit, Lauren Jelly, Joep de Ligt |
| EPI_ISL_456398, EPI_ISL_456399, EPI_ISL_456400, EPI_ISL_456401, EPI_ISL_456402 | Wellington SCL | Institute of Environmental Science and Research (ESR) | Matt Storey, Xiaoyun Ren, Anja Werno, Antje van der Linden, Arlo Upton, Chris Mansell, David Hammer, Dragana Drinkovic, Erasmus Smit, Gary McAuliffe, Hana Sofia Andersson, James Ussher, Jill Sherwood, Josh Freeman, Julia Howard, Juliet Elvy, Mary DeAlmeida, Matt Blakiston, Matthew Rogers, Max Bloomfield, Michael Addidle, Michelle Balm, Sally Roberts, Sarah Jefferies, Sharmini Muttaiyah, Susan Morpeth, Susan Taylor, Timothy Blackmore, Vani Sathyendran, Veronica Playle, Virginia Hope, Erasmus Smit, Lauren Jelly, Joep de Ligt |
| EPI_ISL_456444, EPI_ISL_456445, EPI_ISL_456446, EPI_ISL_456447, EPI_ISL_456448, EPI_ISL_456449, EPI_ISL_456450, EPI_ISL_456451, EPI_ISL_456452, EPI_ISL_456453, EPI_ISL_456454, EPI_ISL_456455 | | | |
| see above | Victorian Infectious Diseases Reference Laboratory (VIDRL) | Microbiological Diagnostic Unit Public Health Laboratory and Victorian Infectious Diseases Reference Laboratory, Doherty Institute | Caly L., Seemann T., Sait, M., Schultz M., Druce J., Sherry, N. |
| EPI_ISL_456767, EPI_ISL_456769, EPI_ISL_456770 | West of Scotland Specialist Virology Centre, NHSGGC / MRC-University of Glasgow Centre for Virus Research | COVID-19 Genomics UK (COG-UK) Consortium | Ana da Silva Filipe, Natasha Johnson, Kathy Smollett, Daniel Mair, Stephen Carmichael, Lily Tong, Jenna Nichols, Elihu Aranday-Cortes, Kirstyn Brunker, Yasmin Parr, Kyriaki Nomikou; Sarah McDonald, Marc Niebel, Patawee Asamaphan; Richard Orton, Joseph Hughes, Sreenu Vattipally, David L Robertson; Alasdair MacLean, Rory Gunson; Kathy Li, Natasha Jesudason, Rajiv Shah, James Shepherd, Antonia Ho, Emma Thomson |
| EPI_ISL_456929, EPI_ISL_456930, EPI_ISL_456931, EPI_ISL_456932, EPI_ISL_456933, EPI_ISL_456934, EPI_ISL_456935, EPI_ISL_456936, EPI_ISL_456937, EPI_ISL_456938, EPI_ISL_456940, EPI_ISL_456957, EPI_ISL_456958, EPI_ISL_456959, EPI_ISL_456960, EPI_ISL_456961, EPI_ISL_456962, EPI_ISL_456963, EPI_ISL_456964, EPI_ISL_456965 | | | |
| see above | Virology Department, Royal Infirmary of Edinburgh, NHS Lothian / School of Biological Sciences, University of Edinburgh / Institute of Genetics and Molecular Medicine, University of Edinburgh | COVID-19 Genomics UK (COG-UK) Consortium | McHugh M, Dewar R, Rooke S, Gallagher M, Balcaza C, O'Toole Á, Scher E, Hill V, McCrone JT, Colquhoun R, Yu X, Jackson B, Rambaut A, Williams TC, Templeton K |
| EPI_ISL_457119, EPI_ISL_457120, EPI_ISL_457121, EPI_ISL_457125, EPI_ISL_457226, EPI_ISL_457233, EPI_ISL_457239, EPI_ISL_457248, EPI_ISL_457250, EPI_ISL_457255, EPI_ISL_457256, EPI_ISL_457258 | | | |
| see above | University of Exeter | COVID-19 Genomics UK (COG-UK) Consortium | Ben Temperton,Aaron Jeffries,Michelle Michelsen,Joanna Warwick-Dugdale,Audrey Farbos,Robyn Manley,Stephen Michell,Jane Masoli |
| EPI_ISL_457303 | Northumbria University / South Tees Hospitals NHS Foundation Trust / North Cumbria Integrated Care NHS Foundation Trust / North Tees and Hartlepool NHS Foundation Trust / Newcastle Hospitals NHS Foundation Trust | COVID-19 Genomics UK (COG-UK) Consortium | Darren L Smith,Andrew Nelson,Matthew Bashton,Greg R Young,Joshua Loh,John Allan,Mohammad A Tariq,Giles S Holt,Gary Black,Wen C Yew,Lynn Dover,Paul Baker,Steve Liggett,Sarah Essex,Jane Greenaway,Debra Padgett,Clive Graham,Garren Scott,Edward Barton,Emma Swindells,Brendan Payne,Jennifer Collins,Yusri Taha,Gary Eltringham |
| EPI_ISL_457582, EPI_ISL_457583, EPI_ISL_457586, EPI_ISL_457589, EPI_ISL_457597, EPI_ISL_457602, EPI_ISL_457615, EPI_ISL_457622, EPI_ISL_457634, EPI_ISL_457644, EPI_ISL_457652, EPI_ISL_457666 | | | |
| see above | Virology Department, Sheffield Teaching Hospitals NHS Foundation Trust/Department of Infection, Immunity and Cardiovascular Disease, The Medical School, University of Sheffield | COVID-19 Genomics UK (COG-UK) Consortium | Thushan de Silva, Matthew Parker, Nikki Smith, Adri Angyal, Rebecca Brown, Luke Green, Rachel Tucker, Paul Parsons, Danielle Groves, Katie Johnson, Laura Carrilero, Alex Keeley, Dave Partridge, Matthew Wyles, Benjamin Lindsey, Mehmet Yavuz, Mohammad Raza, Cariad Evans |
| EPI_ISL_457856, EPI_ISL_457857, EPI_ISL_457858, EPI_ISL_457859, EPI_ISL_457860, EPI_ISL_457861, EPI_ISL_457862, EPI_ISL_457863, EPI_ISL_457864, EPI_ISL_457865, EPI_ISL_457866, EPI_ISL_457867, EPI_ISL_457869, EPI_ISL_457870, EPI_ISL_457872, EPI_ISL_457877, EPI_ISL_457878, EPI_ISL_457879, EPI_ISL_457880, EPI_ISL_457881, EPI_ISL_457882, EPI_ISL_457883, EPI_ISL_457884, EPI_ISL_457885, EPI_ISL_457887, EPI_ISL_457888, EPI_ISL_457889 | | | |
| see above | KEMRI-CGMR-C | KEMRI-Wellcome Trust Research Programme/KEMRI-CGMR-C Kilifi | Githinji G. et al 2020 |
| EPI_ISL_457972, EPI_ISL_457973 | Laboratorio de Biología Molecular Asociación Española Primera en Salud | Departments of Pathology and Medicine, New York University School of Medicine | Maria Victoria Elizondo, Maria Noel Zubillaga, Gonzalo Manrique, Paul Zappile, Gael Westby, Matthew T Maurano, Christian Marier, Adriana Heguy |
| EPI_ISL_458030 | King Institute of Preventive Medicine & Research | CSIR-Centre for Cellular and Molecular Biology | K.Kaveri,S.Sivasubramanian,S.Vennila,P.Padmapriya,R.Kiruba,S.Magesh,G. Dhinakar Raj, G. Ravikumar, P. Azhahianambi,K Thangaraj,Payel Mukherjee, Sofia Banu, Priya Singh, Dhiviya Vedagiri, Divya Gupta, Vishal Sah, Santosh Kumar Kuncha, Krishnan Harinivas Harshan, Archana Bharadwaj Siva, Karthik Bharadwaj Tallapaka, Shagufta Khan, Lamuk Zaveri, Namami Gaur, Sakshi Shambhavi, Tulasi Nagabandi, Purushotham Vodnala, Rakesh K Mishra, Divya Tej Sowpati |
| EPI_ISL_458031 | King Institute of Preventive Medicine & Research | CSIR-Centre for Cellular and Molecular Biology | K.Kaveri,S.Sivasubramanian,S.Vennila,P.Padmapriya,R.Kiruba,S.Magesh,G. Dhinakar Raj, G. Ravikumar, P. Azhahianambi, K Thangaraj,Sofia Banu, Payel Mukherjee, Priya Singh, Dhiviya Vedagiri, Divya Gupta, Vishal Sah, Santosh Kumar Kuncha, Krishnan Harinivas Harshan, Archana Bharadwaj Siva, Karthik Bharadwaj Tallapaka, Shagufta Khan, Lamuk Zaveri, Namami Gaur, Sakshi Shambhavi, Tulasi Nagabandi, Purushotham Vodnala, Rakesh K Mishra, Divya Tej Sowpati |
| EPI_ISL_458062 | CSIR-Centre for Cellular and Molecular Biology | CSIR-Centre for Cellular and Molecular Biology | Payel Mukherjee, Sofia Banu, Priya Singh, Dhiviya Vedagiri, Divya Gupta, Vishal Sah, Santosh Kumar Kuncha, Krishnan Harinivas Harshan, Archana Bharadwaj Siva, Karthik Bharadwaj Tallapaka, Shagufta Khan, Lamuk Zaveri, Namami Gaur, Sakshi Shambhavi, Tulasi Nagabandi, Purushotham Vodnala, Rakesh K Mishra, Sonu Uday, Sudipta Mondal, Annapoorna P Karthyayani, Debabrata Jana, Debrya Saha, Divya Tej Sowpati |
| EPI_ISL_458063 | CSIR-Centre for Cellular and Molecular Biology | CSIR-Centre for Cellular and Molecular Biology | Sofia Banu, Payel Mukherjee, Priya Singh, Dhiviya Vedagiri, Divya Gupta, Vishal Sah, Santosh Kumar Kuncha, Krishnan Harinivas Harshan, Archana Bharadwaj Siva, Karthik Bharadwaj Tallapaka, Shagufta Khan, Lamuk Zaveri, Namami Gaur, Sakshi Shambhavi, Tulasi Nagabandi, Purushotham Vodnala, Gokulan C G, Gunjan Purohit, Hanuman Tulashiram Kale, Pankaj Kumar, Prachand Issarapu, Rakesh K Mishra, Divya Tej Sowpati |
| EPI_ISL_458073, EPI_ISL_458074, EPI_ISL_458076 | CSIR-Centre for Cellular and Molecular Biology | CSIR-Centre for Cellular and Molecular Biology | Dhiviya Vedagiri, Divya Gupta, Vishal Sah, Payel Mukherjee, Sofia Banu, Priya Singh, Santosh Kumar Kuncha, Archana Bharadwaj Siva, Karthik Bharadwaj Tallapaka, Shagufta Khan, Lamuk Zaveri, Namami Gaur, Sakshi Shambhavi, Tulasi Nagabandi, Purushotham Vodnala, Rakesh K Mishra, Divya Tej Sowpati, Krishnan Harinivas Harshan |
| EPI_ISL_458121, EPI_ISL_458126 | Oman National Influenza Centre | Department of Microbiology and Immunology-SQUH | Fahad Zadjali, Samira Al-Maruqi, Amina Al Jardani, Khulood Al-Mammary, Hanan Al-kindi, Fatma BaAlawi, Hamida AL Barwani, Zeyana AL-Dahmani, Intisar Al-Shukri, Aisha Al-Busaidi, Aisha Al-Amri, Ahlam Al-Amri, Mohammed Al-Tobi, Samiha Al Kharusi, Abdulla Balkhair |

| | | | |
|---|---|---|---|
| EPI_ISL_458141, EPI_ISL_458145, EPI_ISL_458146, EPI_ISL_458147 | Evandro Chagas Institute | Evandro Chagas Institute | Santos, M.C.; Silva, A.M.; Junior, W.D.C.; Barbagelata, L.S.; Ferreira, J.A.; Sousa, E.M.A.; da Silva, P.S.; Resque, H.R; Martins, L.C.; Sousa Junior, E.C.;Viana, G.M.R |
| EPI_ISL_458242, EPI_ISL_458255, EPI_ISL_458266, EPI_ISL_458272, EPI_ISL_458277, EPI_ISL_458278, EPI_ISL_458280, EPI_ISL_458282 | Scripps Medical Laboratory | Andersen lab at Scripps Research | SEARCH Alliance San Diego with Michael Quigley, Ellen Stefanski, Ian Mchardy |
| EPI_ISL_458287 | Biosafety Department PCL3 | Biosafety Department PCL3 | Lemriss,S., Souiri,A. and El Kabbaj,S. |
| EPI_ISL_458722, EPI_ISL_458738, EPI_ISL_458742, EPI_ISL_458762, EPI_ISL_458764, EPI_ISL_458765, EPI_ISL_458766, EPI_ISL_458769, EPI_ISL_458788 | PHE South West Regional Laboratory, National Infection Service | Wellcome Sanger Institute for the COVID-19 Genomics UK (COG-UK) consortium | Stephanie Hutchings, Hannah Pymont, Dr Peter Muir, Barry Vipond, Rich Hopes; and Alex Alderton, Roberto Amato, Sonia Goncalves, Ewan Harrison, David K. Jackson, Ian Johnston, Dominic Kwiatkowski, Cordelia Langford, John Sillitoe on behalf of the Wellcome Sanger Institute COVID-19 Surveillance Team (http://www.sanger.ac.uk/covid-team) |
| EPI_ISL_458800 | PHE South West Regional Laboratory, National Infection Service | Wellcome Sanger Institute for the COVID-19 Genomics UK (COG-UK) Consortium | Stephanie Hutchings, Hannah Pymont, Dr Peter Muir, Barry Vipond, Rich Hopes; and Alex Alderton, Roberto Amato, Sonia Goncalves, Ewan Harrison, David K. Jackson, Ian Johnston, Dominic Kwiatkowski, Cordelia Langford, John Sillitoe on behalf of the Wellcome Sanger Institute COVID-19 Surveillance Team |
| EPI_ISL_458808, EPI_ISL_458809, EPI_ISL_458810, EPI_ISL_458820, EPI_ISL_458834, EPI_ISL_458846, EPI_ISL_458849, EPI_ISL_458860, EPI_ISL_458864, EPI_ISL_458869, EPI_ISL_458894, EPI_ISL_458906, EPI_ISL_458908 | | | |
| see above | PHE South West Regional Laboratory, National Infection Service | Wellcome Sanger Institute for the COVID-19 Genomics UK (COG-UK) consortium | Stephanie Hutchings, Hannah Pymont, Dr Peter Muir, Barry Vipond, Rich Hopes; and Alex Alderton, Roberto Amato, Sonia Goncalves, Ewan Harrison, David K. Jackson, Ian Johnston, Dominic Kwiatkowski, Cordelia Langford, John Sillitoe on behalf of the Wellcome Sanger Institute COVID-19 Surveillance Team (http://www.sanger.ac.uk/covid-team) |
| EPI_ISL_459330, EPI_ISL_459332, EPI_ISL_459337, EPI_ISL_459338, EPI_ISL_459340, EPI_ISL_459342, EPI_ISL_459347, EPI_ISL_459349, EPI_ISL_459352, EPI_ISL_459354, EPI_ISL_459356, EPI_ISL_459357, EPI_ISL_459365, EPI_ISL_459366, EPI_ISL_459367, EPI_ISL_459368, EPI_ISL_459369, EPI_ISL_459374, EPI_ISL_459376, EPI_ISL_459378, EPI_ISL_459379, EPI_ISL_459387, EPI_ISL_459394, EPI_ISL_459397, EPI_ISL_459399, EPI_ISL_459401, EPI_ISL_459404, EPI_ISL_459406, EPI_ISL_459407, EPI_ISL_459408, EPI_ISL_459409 | | | |
| see above | Regional Virus Laboratory, Belfast Health and Social Care Trust | Wellcome Sanger Institute for the COVID-19 Genomics UK (COG-UK) consortium | Conall McCaughey, James McKenna, Tanya Curran, Susan Feeney, Alison Watt, Ciara Cox, Mairead Connor, Zoltan Molnar, David Simpson, Derek Fairley; and Alex Alderton, Roberto Amato, Sonia Goncalves, Ewan Harrison, David K. Jackson, Ian Johnston, Dominic Kwiatkowski, Cordelia Langford, John Sillitoe on behalf of the Wellcome Sanger Institute COVID-19 Surveillance Team (http://www.sanger.ac.uk/covid-team) |
| EPI_ISL_459411, EPI_ISL_459412, EPI_ISL_459413, EPI_ISL_459414, EPI_ISL_459421, EPI_ISL_459423, EPI_ISL_459425, EPI_ISL_459435, EPI_ISL_459437, EPI_ISL_459440, EPI_ISL_459441, EPI_ISL_459444, EPI_ISL_459448, EPI_ISL_459453, EPI_ISL_459455, EPI_ISL_459457, EPI_ISL_459459, EPI_ISL_459462, EPI_ISL_459463, EPI_ISL_459465, EPI_ISL_459468, EPI_ISL_459471, EPI_ISL_459475, EPI_ISL_459477, EPI_ISL_459478, EPI_ISL_459480, EPI_ISL_459496, EPI_ISL_459498, EPI_ISL_459501, EPI_ISL_459503 | | | |
| see above | Department of Pathology, University of Cambridge | Wellcome Sanger Institute for the COVID-19 Genomics UK (COG-UK) consortium | Luke W Meredith, M. Estée Török , Myra Hosmillo, William L. Hamilton, Martin D. Curran, Theresa Feltwell, Grant Hall, Anna Yakovleva, Fahad A Khokhar, Charlotte J. Houldcroft, Laura G Caller, Aminu S. Jahun, Sarah L. Caddy, Ian Goodfellow; and Alex Alderton, Roberto Amato, Sonia Goncalves, Ewan Harrison, David K. Jackson, Ian Johnston, Dominic Kwiatkowski, Cordelia Langford, John Sillitoe on behalf of the Wellcome Sanger Institute COVID-19 Surveillance Team (http://www.sanger.ac.uk/covid-team) |
| EPI_ISL_459977, EPI_ISL_459978, EPI_ISL_459983 | Institut Pasteur du Maroc | Institut Pasteur du Maroc | Marion Barbet, Sylvie Behillil, Méline Bizard, Angela Brisebarre, Camille Capel, Etienne Simon-Lorière, Vincent Enouf, Maud Vanpeene, Sylvie van der Werf, Latifa Anga, Abdellah Faouzi, Anass Abbad, Mjid Eloualid, Jalal Nourlil, Anderrahmane Maaroufi |
| EPI_ISL_460603 | NYU Langone Health | Departments of Pathology and Medicine, New York University School of Medicine | Maria Aguero-Rosenfeld, Brendan Belovarac, Margaret Black, Ludovic Boytard, John Cadley, Paolo Cotzia, John Chen, Dacia Dimartino, Xiaojun Feng, Tatyana Gindin, Emily Guzman, Adriana Heguy, Megan Hogan, Emily Huang, George Jour, Alireza Khodadadi-Jamayran, Lawrence H. Lin, Raven Luther, Andrew Lytle, Christian Marier, Matthew T. Maurano, Mark J. Mulligan, Peter Meyn, Raquel Ordonez Ciriza, Iman Osman, Jared Pinnell, Vanessa Raabe, Sitharam Ramaswami, Amy Rapkiewicz, Andre M. Ribeiro-dos-Santos, Marie Samanovic-Golden, Antonio Serrano, Guomiao Shen, Matija Snuderl, Theodore Vougiouklakis, Nick Vulpescu, Gael Westby, Paul Zappile, Yutong Zhang |
| EPI_ISL_460770, EPI_ISL_460771, EPI_ISL_460773, EPI_ISL_460774, EPI_ISL_460775, EPI_ISL_460776, EPI_ISL_460777, EPI_ISL_460785, EPI_ISL_460786, EPI_ISL_460787, EPI_ISL_460833, EPI_ISL_460834, EPI_ISL_460843, EPI_ISL_460849, EPI_ISL_460924, EPI_ISL_460925, EPI_ISL_460926, EPI_ISL_460939, EPI_ISL_460946, EPI_ISL_460947, EPI_ISL_461010, EPI_ISL_461012, EPI_ISL_461016, EPI_ISL_461017, EPI_ISL_461019, EPI_ISL_461020, EPI_ISL_461021, EPI_ISL_461023, EPI_ISL_461024, EPI_ISL_461025, EPI_ISL_461026, EPI_ISL_461042, EPI_ISL_461043, EPI_ISL_461044, EPI_ISL_461045, EPI_ISL_461047, EPI_ISL_461139, EPI_ISL_461140, EPI_ISL_461177, EPI_ISL_461183, EPI_ISL_461193, EPI_ISL_461194, EPI_ISL_461195, EPI_ISL_461196, EPI_ISL_461197, EPI_ISL_461204, EPI_ISL_461205, EPI_ISL_461207, EPI_ISL_461209, EPI_ISL_461210, EPI_ISL_461211, EPI_ISL_461228, EPI_ISL_461229, EPI_ISL_461230, EPI_ISL_461231, EPI_ISL_461267, EPI_ISL_461269, EPI_ISL_461270, EPI_ISL_461271, EPI_ISL_461272, EPI_ISL_461273, EPI_ISL_461293, EPI_ISL_461294, EPI_ISL_461295, EPI_ISL_461296, EPI_ISL_461297, EPI_ISL_461298, EPI_ISL_461299, EPI_ISL_461302, EPI_ISL_461340, EPI_ISL_461341, EPI_ISL_461342, EPI_ISL_461350, EPI_ISL_461351, EPI_ISL_461352, EPI_ISL_461353, EPI_ISL_461355, EPI_ISL_461356, EPI_ISL_461357, EPI_ISL_461358, EPI_ISL_461359, EPI_ISL_461360, EPI_ISL_461361, EPI_ISL_461362, EPI_ISL_461363, EPI_ISL_461364 | | | |
| see above | Dutch COVID-19 response team | Erasmus Medical Center | Bas Oude Munnink, David Nieuwenhuijse, Reina Sikkema, Claudia Schapendonk, Irina Chestakova, Anne van der Linden, Theo Bestebroer, Stefan van Nieuwkoop, Mark Pronk, Pascal Lexmond, Corien Swaan, Manon Haverkate, Madelief Mollers, Mart Stein, Sandra Kengne Kamga Mobou, Jeroen van Kampen, Jolanda Voermans, Aura Timen, Corine GeurtsvanKessel, Annemiek van der Eijk, Richard Molenkamp, Marion Koopmans, on behalf of the Dutch national COVID-19 response team. |
| EPI_ISL_461589, EPI_ISL_461590, EPI_ISL_461591, EPI_ISL_461592, EPI_ISL_461593, EPI_ISL_461594, EPI_ISL_461595, EPI_ISL_461596, EPI_ISL_461597, EPI_ISL_461598, EPI_ISL_461599, EPI_ISL_461600, EPI_ISL_461601, EPI_ISL_461602, EPI_ISL_461603, EPI_ISL_461604, EPI_ISL_461605, EPI_ISL_461606, EPI_ISL_461607, EPI_ISL_461608, EPI_ISL_461609, EPI_ISL_461610, EPI_ISL_461611, EPI_ISL_461612, EPI_ISL_461613, EPI_ISL_461614, EPI_ISL_461615, EPI_ISL_461616, EPI_ISL_461617, EPI_ISL_461618, EPI_ISL_461619, EPI_ISL_461620, EPI_ISL_461621, EPI_ISL_461622, EPI_ISL_461623, EPI_ISL_461624, EPI_ISL_461625, EPI_ISL_461626, EPI_ISL_461627, EPI_ISL_461628, EPI_ISL_461629, EPI_ISL_461630, EPI_ISL_461631, EPI_ISL_461632, EPI_ISL_461633, EPI_ISL_461634, EPI_ISL_461635, EPI_ISL_461636, EPI_ISL_461637, EPI_ISL_461638, EPI_ISL_461639, EPI_ISL_461640, EPI_ISL_461641, EPI_ISL_461642, EPI_ISL_461643, EPI_ISL_461644, EPI_ISL_461645, EPI_ISL_461646, EPI_ISL_461647, EPI_ISL_461648, EPI_ISL_461649, EPI_ISL_461650, EPI_ISL_461651, EPI_ISL_461652, EPI_ISL_461653, EPI_ISL_461654, EPI_ISL_461655, EPI_ISL_461656, EPI_ISL_461657, EPI_ISL_461658, EPI_ISL_461659, EPI_ISL_461660, EPI_ISL_461661, EPI_ISL_461662, EPI_ISL_461663, EPI_ISL_461664, EPI_ISL_461665, EPI_ISL_461666, EPI_ISL_461667, EPI_ISL_461668, EPI_ISL_461669, EPI_ISL_461670, EPI_ISL_461671, EPI_ISL_461672, EPI_ISL_461673, EPI_ISL_461674, EPI_ISL_461675, EPI_ISL_461676, EPI_ISL_461677, EPI_ISL_461678, EPI_ISL_461679, EPI_ISL_461680, EPI_ISL_461681, EPI_ISL_461682, EPI_ISL_461683, EPI_ISL_461684, EPI_ISL_461685, EPI_ISL_461686, EPI_ISL_461687, EPI_ISL_461688, EPI_ISL_461689, EPI_ISL_461690, EPI_ISL_461691, EPI_ISL_461692, EPI_ISL_461693, EPI_ISL_461694, EPI_ISL_461695, EPI_ISL_461696, EPI_ISL_461697, EPI_ISL_461698, EPI_ISL_461699, EPI_ISL_461700 | | | |
| see above | West of Scotland Specialist Virology Centre, NHSGGC / MRC-University of Glasgow Centre for Virus Research | COVID-19 Genomics UK (COG-UK) Consortium | Ana da Silva Filipe, Natasha Johnson, Kathy Smollett, Daniel Mair, Stephen Carmichael, Lily Tong, Jenna Nichols, Elihu Aranday-Cortes, Kirstyn Brunker, Yasmin Parr, Kyriaki Nomikou; Sarah McDonald, Marc Niebel, Patawee Asamaphan; Richard Orton, Joseph Hughes, Sreenu Vattipally, David L Robertson; Alasdair MacLean, Rory Gunson; Kathy Li, Natasha Jesudason, Rajiv Shah, James Shepherd, Antonia Ho, Emma Thomson |
| EPI_ISL_461740, EPI_ISL_461741, EPI_ISL_461742, EPI_ISL_461743, EPI_ISL_461744, EPI_ISL_461745, EPI_ISL_461746, EPI_ISL_461747, EPI_ISL_461748, EPI_ISL_461749, EPI_ISL_461750, EPI_ISL_461751, EPI_ISL_461752, EPI_ISL_461753, EPI_ISL_461754, EPI_ISL_461755, EPI_ISL_461756, EPI_ISL_461757, EPI_ISL_461758 | | | |
| see above | Virology Department, Royal Infirmary of Edinburgh, NHS Lothian / School of Biological Sciences, University of Edinburgh / Institute of Genetics and Molecular Medicine, University of Edinburgh | COVID-19 Genomics UK (COG-UK) Consortium | McHugh M, Dewar R, Rooke S, Gallagher M, Balcaza C, O'Toole Á, Scher E, Hill V, McCrone JT, Colquhoun R, Yu X, Jackson B, Rambaut A, Williams TC, Templeton K |
| EPI_ISL_461763, EPI_ISL_461764, EPI_ISL_461765, EPI_ISL_461766, EPI_ISL_461767, EPI_ISL_461768 | University College London, Great Ormond Street Hospital for Children NHS Foundation Trust, Imperial College Healthcare NHS Trust | COVID-19 Genomics UK (COG-UK) Consortium | Sergi Castellano, Rachel Williams, Mark Kristiansen, Paola Resende Silva, Sunando Roy, Tony Brooks, Helena Tutill, Paola Niola, Patricia Dyal, Charlotte Williams, Leysa Forrest, Yasmin Panchbhaya, Jacqueline Findlay, Sam Weeks, Julianne Brown, Kathryn Harris, Paul Randell, James Price, Alison Holmes, Judith Breuer |
| EPI_ISL_461810, EPI_ISL_461827, EPI_ISL_461828 | Quadram Institute Bioscience | COVID-19 Genomics UK (COG-UK) Consortium | Dave J. Baker, Gemma L. Kay, Alp Aydin, Thanh Le-Viet, Steven Rudder, Ana P. Tedim, Anastasia Kolyva, Maria Diaz, Leonardo de Oliveira Martins, Nabil-Fareed Alikhan, Lizzie Meadows, Rachael Stanley, Ngozi Elumogo, Muhammed Yasir, Nicholas M. Thomson, Alexander J Trotter, Rachel Gilroy, Samuel Bloomfield, Claire Stuart, Andrew Bell, Reenesh Prakash, Samir Dervisevic, Alison E. Mather, John Wain, Mark Webber, Andrew J. Page, Justin O'Grady |
| EPI_ISL_462035, EPI_ISL_462038 | Virology Department, Sheffield Teaching Hospitals NHS Foundation Trust/Department of Infection, Immunity and Cardiovascular Disease, The Medical School, University of Sheffield | COVID-19 Genomics UK (COG-UK) Consortium | Thushan de Silva, Matthew Parker, Nikki Smith, Adri Angyal, Rebecca Brown, Luke Green, Rachel Tucker, Paul Parsons, Danielle Groves, Katie Johnson, Laura Carrilero, Alex Keeley, Dave Partridge, Matthew Wyles, Benjamin Lindsey, Mehmet Yavuz, Mohammad Raza, Cariad Evans |
| EPI_ISL_462151, EPI_ISL_462152, EPI_ISL_462153, EPI_ISL_462154, EPI_ISL_462155, EPI_ISL_462156, | KU Leuven, Rega Institute, Clinical and Epidemiological Virology | KU Leuven, Rega Institute, Clinical and Epidemiological Virology | Tony Wawina-Bokalanga, Bert Vanmechelen, Joan Marti-Carerras, Piet Maes |

| | | | |
|---|---|---|---|
| EPI_ISL_462157 | | | |
| EPI_ISL_462307, EPI_ISL_462308, EPI_ISL_462309, EPI_ISL_462310, EPI_ISL_462311, EPI_ISL_462312, EPI_ISL_462313, EPI_ISL_462314, EPI_ISL_462315, EPI_ISL_462316, EPI_ISL_462317, EPI_ISL_462318, EPI_ISL_462319, EPI_ISL_462320, EPI_ISL_462343 | | | |
| see above | National Public Health Laboratory, National Centre for Infectious Diseases | National Public Health Laboratory, National Centre for Infectious Diseases | Mak TM, Octavia S, Chavatte JM, Cui L, Lin RTP |
| EPI_ISL_462437, EPI_ISL_462438 | unknown | Laboratory Diagnostic | Vidanovic,D., Tesovic,B., Banovic Djeri,B., Knezevic,A., Jankovic,M., Sekler,M., Dmitric,M., Petrovic,T., Volkening,J., Afonso,C.L. |
| EPI_ISL_462466, EPI_ISL_462470 | Clinical Center, University of Sarajevo | Charite Universitatsmedizin Berlin, Institute of Virology | Victor M Corman, Jorn Beheim-Schwarzbach, Barbara Muehlemann, Talitha Veith, Julia Schneider, Terry Jones, Amela Dedeic-Ljubovic, Irma Salimovic-Besic, Suzana Arapcic, Almedina Hadzihasanovic-Moro, Selma Mutevelic, Christian Drosten |
| EPI_ISL_462718, EPI_ISL_462719, EPI_ISL_462720, EPI_ISL_462721, EPI_ISL_462722, EPI_ISL_462723, EPI_ISL_462724, EPI_ISL_462725, EPI_ISL_462726, EPI_ISL_462727, EPI_ISL_462728, EPI_ISL_462729, EPI_ISL_462730, EPI_ISL_462731, EPI_ISL_462732, EPI_ISL_462733, EPI_ISL_462734 | | | |
| see above | Michigan Department of Health and Human Services, Bureau of Laboratories | Michigan Department of Health and Human Services, Bureau of Laboratories | Blankenship HM, Riner D, Soehnlen MK |
| EPI_ISL_463094, EPI_ISL_463095, EPI_ISL_463096, EPI_ISL_463097 | Virginia DCLS | Virginia DCLS | Virginia DCLS |
| EPI_ISL_463303 | United Christian Hospital | Hong Kong Department of Health | Mak Gannon C.K., Cheng Peter K.C., Lam Edman T.K., Chan Rickjason C.W., Tsang Dominic N.C. |
| EPI_ISL_463324, EPI_ISL_463325, EPI_ISL_463326, EPI_ISL_463327, EPI_ISL_463328, EPI_ISL_463329, EPI_ISL_463331, EPI_ISL_463332, EPI_ISL_463333, EPI_ISL_463334, EPI_ISL_463335, EPI_ISL_463336, EPI_ISL_463337, EPI_ISL_463338, EPI_ISL_463339, EPI_ISL_463340, EPI_ISL_463341, EPI_ISL_463342, EPI_ISL_463343, EPI_ISL_463344, EPI_ISL_463345, EPI_ISL_463346, EPI_ISL_463347, EPI_ISL_463348, EPI_ISL_463349, EPI_ISL_463350, EPI_ISL_463351, EPI_ISL_463352, EPI_ISL_463353, EPI_ISL_463354, EPI_ISL_463355, EPI_ISL_463356, EPI_ISL_463357, EPI_ISL_463358, EPI_ISL_463359, EPI_ISL_463360, EPI_ISL_463361, EPI_ISL_463362, EPI_ISL_463369, EPI_ISL_463370, EPI_ISL_463371, EPI_ISL_463375, EPI_ISL_463378, EPI_ISL_463385, EPI_ISL_463392, EPI_ISL_463408, EPI_ISL_463409, EPI_ISL_463410, EPI_ISL_463411, EPI_ISL_463412, EPI_ISL_463413, EPI_ISL_463414, EPI_ISL_463415, EPI_ISL_463416, EPI_ISL_463417, EPI_ISL_463418, EPI_ISL_463419, EPI_ISL_463420, EPI_ISL_463421, EPI_ISL_463422, EPI_ISL_463423, EPI_ISL_463424, EPI_ISL_463425, EPI_ISL_463426, EPI_ISL_463427, EPI_ISL_463428, EPI_ISL_463429, EPI_ISL_463430, EPI_ISL_463431, EPI_ISL_463437, EPI_ISL_463438 | | | |
| see above | Washington State Department of Health | Seattle Flu Study | Chu et al |
| EPI_ISL_463996, EPI_ISL_464021, EPI_ISL_464030, EPI_ISL_464038, EPI_ISL_464041, EPI_ISL_464054, EPI_ISL_464061, EPI_ISL_464062 | Unity Health Toronto | Ontario Institute for Cancer Research | Ramzi Fattouh,Larissa M. Matukas,Mark Downing,Annette Gower,Karel Boissinot,Samira Mubareka,TIBDN,Ilinca Lungu,Bernard Lam,Jeremy Johns,Paul Krzyzanowski,Richard de Borja,Philip Zuzarte,Jared Simpson |
| EPI_ISL_465262, EPI_ISL_465263, EPI_ISL_465283, EPI_ISL_465298, EPI_ISL_465299, EPI_ISL_465300, EPI_ISL_465301, EPI_ISL_465302, EPI_ISL_465303, EPI_ISL_465304, EPI_ISL_465305, EPI_ISL_465306, EPI_ISL_465307, EPI_ISL_465311, EPI_ISL_465312, EPI_ISL_465315, EPI_ISL_465316, EPI_ISL_465317, EPI_ISL_465318, EPI_ISL_465319, EPI_ISL_465325, EPI_ISL_465326, EPI_ISL_465327, EPI_ISL_465328, EPI_ISL_465329, EPI_ISL_465330, EPI_ISL_465331, EPI_ISL_465332, EPI_ISL_465333, EPI_ISL_465334, EPI_ISL_465335, EPI_ISL_465337, EPI_ISL_465338, EPI_ISL_465339, EPI_ISL_465340, EPI_ISL_465341, EPI_ISL_465342, EPI_ISL_465344, EPI_ISL_465350, EPI_ISL_465355, EPI_ISL_465357, EPI_ISL_465358, EPI_ISL_465359, EPI_ISL_465360, EPI_ISL_465361, EPI_ISL_465362, EPI_ISL_465363, EPI_ISL_465364, EPI_ISL_465365, EPI_ISL_465366, EPI_ISL_465367, EPI_ISL_465368, EPI_ISL_465369, EPI_ISL_465370, EPI_ISL_465371, EPI_ISL_465372, EPI_ISL_465373, EPI_ISL_465374, EPI_ISL_465375, EPI_ISL_465376, EPI_ISL_465377, EPI_ISL_465378, EPI_ISL_465379, EPI_ISL_465380, EPI_ISL_465381, EPI_ISL_465382, EPI_ISL_465383, EPI_ISL_465384, EPI_ISL_465385, EPI_ISL_465386, EPI_ISL_465387, EPI_ISL_465388, EPI_ISL_465389, EPI_ISL_465390, EPI_ISL_465391, EPI_ISL_465392, EPI_ISL_465393, EPI_ISL_465394, EPI_ISL_465395, EPI_ISL_465399, EPI_ISL_465401, EPI_ISL_465402, EPI_ISL_465403, EPI_ISL_465404, EPI_ISL_465405, EPI_ISL_465407, EPI_ISL_465408, EPI_ISL_465409, EPI_ISL_465411, EPI_ISL_465412, EPI_ISL_465413, EPI_ISL_465414, EPI_ISL_465415, EPI_ISL_465416, EPI_ISL_465417, EPI_ISL_465418, EPI_ISL_465419, EPI_ISL_465420, EPI_ISL_465421, EPI_ISL_465422, EPI_ISL_465423, EPI_ISL_465424, EPI_ISL_465425, EPI_ISL_465426, EPI_ISL_465427, EPI_ISL_465428, EPI_ISL_465429, EPI_ISL_465430, EPI_ISL_465432, EPI_ISL_465434, EPI_ISL_465435, EPI_ISL_465436, EPI_ISL_465439, EPI_ISL_465440, EPI_ISL_465441, EPI_ISL_465443, EPI_ISL_465444, EPI_ISL_465449, EPI_ISL_465450, EPI_ISL_465453, EPI_ISL_465456, EPI_ISL_465523, EPI_ISL_465715, EPI_ISL_465716 | | | |
| see above | Respiratory Virus Unit, Microbiology Services Colindale, Public Health England | Respiratory Virus Unit, Microbiology Services Colindale, Public Health England | PHE Covid Sequencing Team |
| EPI_ISL_466888, EPI_ISL_466890, EPI_ISL_466891, EPI_ISL_466892, EPI_ISL_466893, EPI_ISL_466894, EPI_ISL_466895, EPI_ISL_466896, EPI_ISL_466897 | Max von Pettenkofer Institute, Virology, National Reference Center for Retroviruses, LMU München | Laboratory for Functional Genome Analysis, Dept. Genomics, Gene Center of the LMU Munich | Max Muenchhoff, Stefan Krebs, Alexander Graf, Oliver Keppler, Helmut Blum |
| EPI_ISL_467352, EPI_ISL_467353, EPI_ISL_467354, EPI_ISL_467355, EPI_ISL_467356, EPI_ISL_467357, EPI_ISL_467358, EPI_ISL_467359, EPI_ISL_467360, EPI_ISL_467364 | Laboratory of Respiratory Viruses and Measles, Oswaldo Cruz Institute, FIOCRUZ | Laboratory of Respiratory Viruses and Measles, Oswaldo Cruz Institute, FIOCRUZ | Paola Resende, Luciana Appolinario, Fernando Motta, Anna Carolina Paixão, Ana Carolina Mendonça, Aline Mattos, Milene Miranda, Cristiana Garcia, Braulia Caetano, Maria Ogrzewalska, Jonathan Lopes, Marilda Siqueira |
| EPI_ISL_467376 | RSUP Fatmawati | Eijkman Institute for Molecular Biology, Ministry of Research and Technology/National Agency for Research and Innovation | Edison Johar, Frilasita A Yudhaputri, Hidayat Trimarsanto, David H Muljono, Safarina G Malik, Khin Saw Myint, Amin Soebandrio |
| EPI_ISL_467378, EPI_ISL_467388, EPI_ISL_467391, EPI_ISL_467392, EPI_ISL_467393, EPI_ISL_467394, EPI_ISL_467395, EPI_ISL_467396 | NYU Langone Health | Departments of Pathology and Medicine, New York University School of Medicine | Maria Aguero-Rosenfeld, Brendan Belovarac, Margaret Black, Ludovic Boytard, John Cadley, Paolo Cotzia, John Chen, Dacia Dimartino, Xiaojun Feng, Tatyana Gindin, Emily Guzman, Adriana Heguy, Megan Hogan, Emily Huang, George Jour, Alireza Khodadadi-Jamayran, Lawrence H. Lin, Raven Luther, Andrew Lytle, Christian Marier, Matthew T. Maurano, Mark J. Mulligan, Peter Meyn, Raquel Ordonez Ciriza, Iman Osman, Jared Pinnell, Vanessa Raabe, Sitharam Ramaswami, Amy Rapkiewicz, Andre M. Ribeiro-dos-Santos, Marie Samanovic-Golden, Antonio Serrano, Guomiao Shen, Matija Snuderl, Theodore Vougiouklakis, Nick Vulpescu, Gael Westby, Paul Zappile, Yutong Zhang |
| EPI_ISL_467780 | National Influenza Centre Romania | Charite Universitatsmedizin Berlin, Institute of Virology | Victor M Corman, Jorn Beheim-Schwarzbach, Barbara Muehlemann, Talitha Veith, Julia Schneider, Terry Jones, L. Ustea, N. Paraschiv, M. Lazar, Christian Drosten |
| EPI_ISL_467793, EPI_ISL_467794, EPI_ISL_467795, EPI_ISL_467796, EPI_ISL_467797, EPI_ISL_467798, EPI_ISL_467799, EPI_ISL_467800, EPI_ISL_467801, EPI_ISL_467802, EPI_ISL_467803, EPI_ISL_467804, EPI_ISL_467805, EPI_ISL_467806, EPI_ISL_467807, EPI_ISL_467808 | | | |
| see above | Virginia DCLS | Virginia DCLS | Virginia DCLS |
| EPI_ISL_467945 | Montefiore Medical Center, Dept. of Pathology, Clinical Virology | Albert Einstein College of Medicine, Dept. of Microbiology & Immunology, Chandran lab | J. Maximilian Fels, Saad Khan, Ryan Forster, Karin A. Skalina, Ariel S. Wirchnianski, Denise Haslwanter, Catalina Florez, Robert H. Bortz III, M. Eugenia Dieterle, Ethan Laudermilch, Rohit K. Jangra, Amanda Mengotto, Duncan Kimmel, Shahina B. Maqbool, John M. Greally, Wendy A. Szymczak, Amy S. Fox, Michael B. Prystowsky, D. Yitzchak Goldstein, Johanna P. Daily, Libusha Kelly, Kartik Chandran |
| EPI_ISL_467949 | Innovative Genomics Institute, UC Berkeley | Innovative Genomics Institute, UC Berkeley | Stacia Wyman, Haridha Shivram, Liana Lareau, Shana McDevitt, Justin Choi |
| EPI_ISL_468070 | Child Health Research Foundation | Child Health Research Foundation | Senjuti Saha, Roly Malaker, Md Saiful Islam Sajib, Hafizur Rahman, Maksuda Islam, Samir K Saha |
| EPI_ISL_468085, EPI_ISL_468087, EPI_ISL_468088, EPI_ISL_468089, EPI_ISL_468090, EPI_ISL_468091, EPI_ISL_468092, EPI_ISL_468093, EPI_ISL_468094, EPI_ISL_468095, EPI_ISL_468096, EPI_ISL_468097, EPI_ISL_468098, EPI_ISL_468099, EPI_ISL_468100, EPI_ISL_468101, EPI_ISL_468102 | | | |
| see above | OHSU Lab Services Molecular Microbiology Lab | Oregon SARS-CoV-2 Genome Sequencing Center | Brendan L. O'Connell, Ruth V. Nichols, Alec J. Hirsch, Guang Fan, Daniel N. Streblow, William B. Messer, Andrew C. Adey, Benjamin N. Bimber, Brian J. O'Roak |
| EPI_ISL_468226, EPI_ISL_468227, EPI_ISL_468228, EPI_ISL_468229, EPI_ISL_468230, EPI_ISL_468231, EPI_ISL_468232, EPI_ISL_468233, EPI_ISL_468234, EPI_ISL_468235, EPI_ISL_468236, EPI_ISL_468237, EPI_ISL_468238, EPI_ISL_468239, EPI_ISL_468240, EPI_ISL_468241, EPI_ISL_468242, EPI_ISL_468243, EPI_ISL_468244, EPI_ISL_468245, EPI_ISL_468246, EPI_ISL_468247, EPI_ISL_468248, EPI_ISL_468249 | | | |
| see above | Viollier AG | Department of Biosystems Science and Engineering, ETH Zürich | Christian Beisel, Sarah Nadeau, Ivan Topolsky, Pedro Ferreira, Philipp Jablonski, Susana Posada-Céspedes, Tobias Schär, Ina Nissen, Natascha Santacroce, Elodie Burcklen, Christiane Beckmann, Maurice Redondo, Olivier Kobel, Christoph Noppen, Sophie Seidel, Noemie Santamaria de Souza, Niko Beerenwinkel, Tanja Stadler |
| EPI_ISL_468321 | Hospital Universitario da USP | Instituto Adolfo Lutz, Interdiciplinary Procedures Center, Strategic Laboratory | Claudio Tavares Sacchi, Claudia Regina Gonçalves, Erica Valessa Ramos Gomes |
| EPI_ISL_468339, EPI_ISL_468340 | Microbiology Service, University Hospital of A Coruna-Biomedical Research Institute | Genomes & Disease, Center for Research in Molecular Medicine and Chronic Diseases, University of Santiago de Compostela | Kelly Conde, Jorge Arca, Soraya Rumbo, Juan A. Vallejo, M Poza, G Bou, Ana Pequeno-Valtierra, Jorge Rodriguez-Castro, Javier Temes, Daniel Garcia-Souto, Martin Santamarina, Cristina Gomez, Jose M. C. Tubio |
| EPI_ISL_468345, EPI_ISL_468346, EPI_ISL_468347, EPI_ISL_468348 | County of Santa Clara Public Health Department | Chan-Zuckerberg Biohub | CZB Cliahub Consortium |

| | | | |
|---|---|---|---|
| EPI_ISL_468383, EPI_ISL_468384, EPI_ISL_468385, EPI_ISL_468386, EPI_ISL_468387 | Alameda County Public Health Lab | Chan-Zuckerberg Biohub | CZB Cliahub Consortium |
| EPI_ISL_468421, EPI_ISL_468422, EPI_ISL_468423, EPI_ISL_468424 | County of San Luis Obispo Public Health Laboratory | Chan-Zuckerberg Biohub | CZB Cliahub Consortium |
| EPI_ISL_468482, EPI_ISL_468483, EPI_ISL_468484, EPI_ISL_468485, EPI_ISL_468486, EPI_ISL_468487 | Ventura County Public Health Lab | Chan-Zuckerberg Biohub | CZB Cliahub Consortium |
| EPI_ISL_468548, EPI_ISL_468549 | San Joaquin County Public Health Lab | Chan-Zuckerberg Biohub | CZB Cliahub Consortium |
| EPI_ISL_468600, EPI_ISL_468601, EPI_ISL_468602, EPI_ISL_468603, EPI_ISL_468604, EPI_ISL_468605, EPI_ISL_468606, EPI_ISL_468607 | Orange County Public Health Lab | Chan-Zuckerberg Biohub | CZB Cliahub Consortium |
| EPI_ISL_468719 | University of Florida | University of Florida | Elbadry,M.A., Subramaniam,K., Waltzek,T.B., Stephenson,C.J., Gibson,J.C., Alam,M.M., Lauzardo,M., Morris,J.G., Lednicky,J.A. |
| EPI_ISL_468720, EPI_ISL_468723 | University of Florida | University of Florida | Stephenson,C.J., Subramaniam,K., Waltzek,T.B., Lauzardo,M., Morris,J.G., Lednicky,J.A. |
| EPI_ISL_468981 | Servicio de Microbiología, Hospital Universitario Son Espases | SeqCOVID-SPAIN consortium/IBV(CSIC) | Carla López-Causapé, Jordi Reina, Antonio Oliver and SeqCOVID-SPAIN consortium |
| EPI_ISL_469119 | National Public Health Laboratory, National Centre for Infectious Diseases | National Public Health Laboratory, National Centre for Infectious Diseases | Mak TM, Octavia S, Chavatte JM, Cui L, Lin RTP |
| EPI_ISL_469529, EPI_ISL_469530, EPI_ISL_469532, EPI_ISL_469533, EPI_ISL_469535, EPI_ISL_469536, EPI_ISL_469540, EPI_ISL_469543, EPI_ISL_469547, EPI_ISL_469549, EPI_ISL_469554, EPI_ISL_469558, EPI_ISL_469560, EPI_ISL_469563, EPI_ISL_469567, EPI_ISL_469574, EPI_ISL_469579, EPI_ISL_469580, EPI_ISL_469583, EPI_ISL_469584, EPI_ISL_469586, EPI_ISL_469588, EPI_ISL_469589, EPI_ISL_469593, EPI_ISL_469595, EPI_ISL_469596, EPI_ISL_469610, EPI_ISL_469611, EPI_ISL_469614, EPI_ISL_469619, EPI_ISL_469624, EPI_ISL_469626, EPI_ISL_469632, EPI_ISL_469633, EPI_ISL_469637, EPI_ISL_469640, EPI_ISL_469641, EPI_ISL_469643, EPI_ISL_469649, EPI_ISL_469651, EPI_ISL_469655, EPI_ISL_469657, EPI_ISL_469660, EPI_ISL_469663, EPI_ISL_469665, EPI_ISL_469667, EPI_ISL_469672, EPI_ISL_469673, EPI_ISL_469678, EPI_ISL_469679, EPI_ISL_469684, EPI_ISL_469685, EPI_ISL_469686, EPI_ISL_469692, EPI_ISL_469696, EPI_ISL_469697, EPI_ISL_469701, EPI_ISL_469705, EPI_ISL_469707, EPI_ISL_469710, EPI_ISL_469713, EPI_ISL_469718, EPI_ISL_469723, EPI_ISL_469725, EPI_ISL_469726, EPI_ISL_469733, EPI_ISL_469735, EPI_ISL_469739, EPI_ISL_469741, EPI_ISL_469742, EPI_ISL_469743, EPI_ISL_469745, EPI_ISL_469750, EPI_ISL_469755, EPI_ISL_469758, EPI_ISL_469761, EPI_ISL_469764, EPI_ISL_469765, EPI_ISL_469767, EPI_ISL_469769, EPI_ISL_469777, EPI_ISL_469779, EPI_ISL_469785, EPI_ISL_469788, EPI_ISL_469792, EPI_ISL_469804 | | | |
| see above | PHE South West Regional Laboratory, National Infection Service | Wellcome Sanger Institute for the COVID-19 Genomics UK (COG-UK) consortium | Stephanie Hutchings, Hannah Pymont, Dr Peter Muir, Barry Vipond, Rich Hopes; and Alex Alderton, Roberto Amato, Sonia Goncalves, Ewan Harrison, David K. Jackson, Ian Johnston, Dominic Kwiatkowski, Cordelia Langford, John Sillitoe on behalf of the Wellcome Sanger Institute COVID-19 Surveillance Team (http://www.sanger.ac.uk/covid-team) |
| EPI_ISL_469805 | NU-OMICS DNA Sequencing research facility, Northumbria University | Wellcome Sanger Institute for the COVID-19 Genomics UK (COG-UK) consortium | Chris Duncan, Sheia Waugh, Shirelle Burton-Fanning, Gary Eltringham, Jennifer Collins, Brendan Payne, Yusri Taha, Emma Swindells, Jane Greenaway, Edward Barton, Garren Scott, Debra Padgett, Clive Graham, Sarah Essex, Steve Ligget, Paul Baker, Lynn Dover, Wen Yew, Gary Black, John Allan, Joshua Loh, Greg Young, Matthew Bashton, Andrew Nelson, Darren Smith and Alex Alderton, Roberto Amato, Sonia Goncalves, Ewan Harrison, David K. Jackson, Ian Johnston, Dominic Kwiatkowski, Cordelia Langford, John Sillitoe on behalf of the Wellcome Sanger Institute COVID-19 Surveillance Team (http://www.sanger.ac.uk/covid-team) |
| EPI_ISL_469808, EPI_ISL_469821, EPI_ISL_469839 | PHE South West Regional Laboratory, National Infection Service | Wellcome Sanger Institute for the COVID-19 Genomics UK (COG-UK) consortium | Stephanie Hutchings, Hannah Pymont, Dr Peter Muir, Barry Vipond, Rich Hopes; and Alex Alderton, Roberto Amato, Sonia Goncalves, Ewan Harrison, David K. Jackson, Ian Johnston, Dominic Kwiatkowski, Cordelia Langford, John Sillitoe on behalf of the Wellcome Sanger Institute COVID-19 Surveillance Team (http://www.sanger.ac.uk/covid-team) |
| EPI_ISL_469844, EPI_ISL_469849, EPI_ISL_469851, EPI_ISL_469852, EPI_ISL_469853, EPI_ISL_469854, EPI_ISL_469855, EPI_ISL_469858, EPI_ISL_469860, EPI_ISL_469869, EPI_ISL_469870, EPI_ISL_469878, EPI_ISL_469880, EPI_ISL_469882, EPI_ISL_469889 | | | |
| see above | Regional Virus Laboratory, Belfast Health and Social Care Trust | Wellcome Sanger Institute for the COVID-19 Genomics UK (COG-UK) consortium | Conall McCaughey, James McKenna, Tanya Curran, Susan Feeney, Alison Watt, Ciara Cox, Mairead Connor, Zoltan Molnar, David Simpson, Derek Fairley; and Alex Alderton, Roberto Amato, Sonia Goncalves, Ewan Harrison, David K. Jackson, Ian Johnston, Dominic Kwiatkowski, Cordelia Langford, John Sillitoe on behalf of the Wellcome Sanger Institute COVID-19 Surveillance Team (http://www.sanger.ac.uk/covid-team) |
| EPI_ISL_469921, EPI_ISL_469926 | Virology Department, Sheffield Teaching Hospitals NHS Foundation Trust / Department of Infection, Immunity and Cardiovascular Disease, The Medical School, University of Sheffield | Wellcome Sanger Institute for the COVID-19 Genomics UK (COG-UK) consortium | Thushan de Silva, Matthew Parker,Adri Angyal, Rebecca Brown, Luke Green, Rachel Tucker, Paul Parsons, Danielle Groves, Alex Keeley, Dave Partridge, Matthew Wyles, Benjamin Lindsey, Mehmet Yavuz, Mohammad Raza, Cariad Evans and Alex Alderton, Roberto Amato, Sonia Goncalves, Ewan Harrison, David K. Jackson, Ian Johnston, Dominic Kwiatkowski, Cordelia Langford, John Sillitoe on behalf of the Wellcome Sanger Institute COVID-19 Surveillance Team (http://www.sanger.ac.uk/covid-team) |
| EPI_ISL_470013, EPI_ISL_470015, EPI_ISL_470027, EPI_ISL_470029, EPI_ISL_470033, EPI_ISL_470048, EPI_ISL_470060, EPI_ISL_470062, EPI_ISL_470067, EPI_ISL_470077, EPI_ISL_470078 | | | |
| see above | Regional Virus Laboratory, Belfast Health and Social Care Trust | Wellcome Sanger Institute for the COVID-19 Genomics UK (COG-UK) consortium | Conall McCaughey, James McKenna, Tanya Curran, Susan Feeney, Alison Watt, Ciara Cox, Mairead Connor, Zoltan Molnar, David Simpson, Derek Fairley; and Alex Alderton, Roberto Amato, Sonia Goncalves, Ewan Harrison, David K. Jackson, Ian Johnston, Dominic Kwiatkowski, Cordelia Langford, John Sillitoe on behalf of the Wellcome Sanger Institute COVID-19 Surveillance Team (http://www.sanger.ac.uk/covid-team) |
| EPI_ISL_470096, EPI_ISL_470097, EPI_ISL_470098, EPI_ISL_470099, EPI_ISL_470100, EPI_ISL_470107, EPI_ISL_470110, EPI_ISL_470113, EPI_ISL_470117, EPI_ISL_470118, EPI_ISL_470120, EPI_ISL_470122, EPI_ISL_470123, EPI_ISL_470131, EPI_ISL_470132, EPI_ISL_470135, EPI_ISL_470136, EPI_ISL_470137, EPI_ISL_470138, EPI_ISL_470140, EPI_ISL_470142, EPI_ISL_470143, EPI_ISL_470145, EPI_ISL_470146, EPI_ISL_470148, EPI_ISL_470150, EPI_ISL_470151, EPI_ISL_470152, EPI_ISL_470154, EPI_ISL_470157, EPI_ISL_470160, EPI_ISL_470163, EPI_ISL_470164, EPI_ISL_470169, EPI_ISL_470170, EPI_ISL_470172, EPI_ISL_470173, EPI_ISL_470174, EPI_ISL_470175, EPI_ISL_470178, EPI_ISL_470181, EPI_ISL_470183, EPI_ISL_470184, EPI_ISL_470186, EPI_ISL_470189, EPI_ISL_470192, EPI_ISL_470193, EPI_ISL_470194, EPI_ISL_470195, EPI_ISL_470196, EPI_ISL_470197, EPI_ISL_470198, EPI_ISL_470201, EPI_ISL_470203, EPI_ISL_470204, EPI_ISL_470205, EPI_ISL_470207, EPI_ISL_470208, EPI_ISL_470211, EPI_ISL_470213, EPI_ISL_470214, EPI_ISL_470216, EPI_ISL_470219, EPI_ISL_470220, EPI_ISL_470222, EPI_ISL_470225, EPI_ISL_470226, EPI_ISL_470227, EPI_ISL_470228, EPI_ISL_470230, EPI_ISL_470231, EPI_ISL_470240, EPI_ISL_470242, EPI_ISL_470245, EPI_ISL_470246, EPI_ISL_470247, EPI_ISL_470248, EPI_ISL_470249, EPI_ISL_470252, EPI_ISL_470258, EPI_ISL_470259, EPI_ISL_470262, EPI_ISL_470264, EPI_ISL_470265, EPI_ISL_470266, EPI_ISL_470267, EPI_ISL_470268, EPI_ISL_470270, EPI_ISL_470271, EPI_ISL_470274, EPI_ISL_470275, EPI_ISL_470276, EPI_ISL_470281, EPI_ISL_470285, EPI_ISL_470290, EPI_ISL_470291, EPI_ISL_470295, EPI_ISL_470296, EPI_ISL_470297, EPI_ISL_470300, EPI_ISL_470301, EPI_ISL_470302, EPI_ISL_470306, EPI_ISL_470309, EPI_ISL_470312, EPI_ISL_470315, EPI_ISL_470316, EPI_ISL_470317, EPI_ISL_470318, EPI_ISL_470319, EPI_ISL_470320, EPI_ISL_470321, EPI_ISL_470322, EPI_ISL_470323, EPI_ISL_470324, EPI_ISL_470327, EPI_ISL_470328, EPI_ISL_470329, EPI_ISL_470332, EPI_ISL_470335, EPI_ISL_470337, EPI_ISL_470341, EPI_ISL_470342, EPI_ISL_470343, EPI_ISL_470347, EPI_ISL_470348, EPI_ISL_470349, EPI_ISL_470352 | | | |
| see above | Department of Pathology, University of Cambridge | Wellcome Sanger Institute for the COVID-19 Genomics UK (COG-UK) consortium | Luke W Meredith, M. Estée Török , Myra Hosmillo, William L. Hamilton, Martin D. Curran, Theresa Feltwell, Grant Hall, Anna Yakovleva, Fahad A Khokhar, Charlotte J. Houldcroft, Laura G Caller, Aminu S. Jahun, Sarah L. Caddy, Ian Goodfellow; and Alex Alderton, Roberto Amato, Sonia Goncalves, Ewan Harrison, David K. Jackson, Ian Johnston, Dominic Kwiatkowski, Cordelia Langford, John Sillitoe on behalf of the Wellcome Sanger Institute COVID-19 Surveillance Team (http://www.sanger.ac.uk/covid-team) |
| EPI_ISL_470545, EPI_ISL_470546, EPI_ISL_470547, EPI_ISL_470548, EPI_ISL_470549, EPI_ISL_470550, EPI_ISL_470551, EPI_ISL_470552, EPI_ISL_470553, EPI_ISL_470565, EPI_ISL_470567 | | | |
| see above | Utah Public Health Laboratory | Utah Public Health Laboratory | Erin Young, Kelly Oakeson |
| EPI_ISL_470639, EPI_ISL_470640, EPI_ISL_470641, EPI_ISL_470642, EPI_ISL_470643, EPI_ISL_470644, EPI_ISL_470645, EPI_ISL_470646, EPI_ISL_470647, EPI_ISL_470648, EPI_ISL_470649, EPI_ISL_470650 | | | |
| see above | Laboratorio de Virologia Molecular / UFRJ | Bioinformatics Laboratory / LNCC | Alexandra Gerber, Ana Paula Guimarães, Luiz Gonzaga Paula de Almeida, Ronaldo da Silva Francisco Junior, Mariane Talon, Filipe Romero, Átila Duque Rossi, Terezinha Marta Pereira, working group UFRJ, Jaqueline Goes de Jesus, Ingra Morales Claro, Ester Cerdeira Sabino, Nuno Rodrigues Faria, CADDE-group, Laboratorio Hermes Pardini, Laboratorio Simile, working group UFMG, Amilcar Tanuri, Carolina Voloch, Renato Santana Aguiar e Ana Tereza Vasconcelos |
| EPI_ISL_470668, EPI_ISL_470669, EPI_ISL_470670, EPI_ISL_470671, EPI_ISL_470672, EPI_ISL_470673, EPI_ISL_470674, EPI_ISL_470675, EPI_ISL_470676, EPI_ISL_470677 | Utah Public Health Laboratory | Utah Public Health Laboratory | Erin Young, Kelly Oakeson |
| EPI_ISL_470737, EPI_ISL_470738, EPI_ISL_470739, EPI_ISL_470740, EPI_ISL_470741 | Utah Public Health Laboratory | Utah Public Health Laboratory | Heidi Butz, Erin Young, Kelly Oakeson |
| EPI_ISL_470802 | State Key Laboratory of Agriculture Microbiology | State Key Laboratory of Agriculture Microbiology, Huazhong | Zhong Zou |

| | | | |
|---|---|---|---|
| Agric | | | |
| EPI_ISL_470854, EPI_ISL_470875 | PathWest Laboratory Medicine WA | PathWest Laboratory Medicine WA | Chisha Sikazwe, Jurissa Lang, Avram Levy, David Smith and David Speers |
| EPI_ISL_471173 | Hospital of Southern Norway - Kristiansand, Department of Medical Microbiology | Norwegian Institute of Public Health, Department of Virology | Kathrine Stene-Johansen, Kamilla Heddeland Instefjord, Hilde Elshaug, Rasmus Riis Kopperud, Karoline Bragstad, Olav Hungnes |
| EPI_ISL_471174 | Ostfold Hospital Trust - Kalnes, Centre for Laboratory Medicine, Section for gene technology and infection serology | Norwegian Institute of Public Health, Department of Virology | Kathrine Stene-Johansen, Kamilla Heddeland Instefjord, Hilde Elshaug, Rasmus Riis Kopperud, Karoline Bragstad, Olav Hungnes |
| EPI_ISL_471194, EPI_ISL_471195, EPI_ISL_471196, EPI_ISL_471197, EPI_ISL_471198, EPI_ISL_471200, EPI_ISL_471215, EPI_ISL_471216, EPI_ISL_471218, EPI_ISL_471228, EPI_ISL_471229, EPI_ISL_471242 | | | |
| see above | Wisconsin State Laboratory of Hygiene Communicable Disease Division | Wisconsin State Laboratory of Hygiene Communicable Disease Division | Kelsey R. Florek, Abigail C. Shockey |
| EPI_ISL_471396, EPI_ISL_471397, EPI_ISL_471398, EPI_ISL_471399, EPI_ISL_471400, EPI_ISL_471401, EPI_ISL_471402, EPI_ISL_471403, EPI_ISL_471411, EPI_ISL_471412, EPI_ISL_471413 | | | |
| see above | Viral Respiratory Lab, National Institute for Biomedical Research (INRB) | Pathogen Sequencing Lab, National Institute for Biomedical Research (INRB) | Placide Mbala-Kingebeni, Edith Nkwembe, Eddy Kinganda-Lusamaki, Amuri Aziza, Francisca Muyembe Mawete, Catherine Pratt, Matthias Pauthner, Josh Quick, Allison Black, James Hadfield, Trevor Bedford, Ian Goodfellow, Andrew Rambaut, Nick Loman, Kristian Andersen, Michael Wiley, Steve Ahuka-Mundeke, Jean-Jacques Muyembe Tamfum |
| EPI_ISL_471641, EPI_ISL_471642 | CSIR-Centre for Cellular and Molecular Biology | CSIR-Centre for Cellular and Molecular Biology | Dhiviya Vedagiri, Divya Gupta, Vishal Sah, Payel Mukherjee, Sofia Banu, Priya Singh, Santosh Kumar Kuncha, Archana Bharadwaj Siva, Karthik Bharadwaj Tallapaka, Shagufta Khan, Lamuk Zaveri, Namami Gaur, Sakshi Shambhavi, Tulasi Nagabandi, Purushotham Vodnala, Rakesh K Mishra, Divya Tej Sowpati, Krishnan Harinivas Harshan |
| EPI_ISL_471643 | CSIR-Centre for Cellular and Molecular Biology | CSIR-Centre for Cellular and Molecular Biology | Tulasi Nagabandi, Namami Gaur, Sakshi Shambhavi, Lamuk Zaveri, Shagufta Khan, Purushotham Vodnala, Payel Mukherjee, Sofia Banu, Priya Singh, Dhiviya Vedagiri, Divya Gupta, Vishal Sah, Santosh Kumar Kuncha, Krishnan Harinivas Harshan, Archana Bharadwaj Siva, Karthik Bharadwaj Tallapaka,G. Aditya Kumar, Koushick Sivakumar, Pooja Ramesh Gupta, Rajan Kumar Jha, Shraddha Vijay Lahoti, Rakesh K Mishra, Divya Tej Sowpati |
| EPI_ISL_471729, EPI_ISL_471743, EPI_ISL_471744, EPI_ISL_471745, EPI_ISL_471746, EPI_ISL_471747, EPI_ISL_471753, EPI_ISL_471754, EPI_ISL_471774, EPI_ISL_471776, EPI_ISL_471777, EPI_ISL_471778, EPI_ISL_471779, EPI_ISL_471780, EPI_ISL_471781, EPI_ISL_471782, EPI_ISL_471783, EPI_ISL_471784, EPI_ISL_471785, EPI_ISL_471786, EPI_ISL_471787, EPI_ISL_471789, EPI_ISL_471790, EPI_ISL_471791, EPI_ISL_471792, EPI_ISL_471793, EPI_ISL_471794, EPI_ISL_471797, EPI_ISL_471798, EPI_ISL_471800, EPI_ISL_471801, EPI_ISL_471802, EPI_ISL_471803, EPI_ISL_471804, EPI_ISL_471805, EPI_ISL_471806, EPI_ISL_471807, EPI_ISL_471808, EPI_ISL_471809, EPI_ISL_471810, EPI_ISL_471846, EPI_ISL_471847, EPI_ISL_471850, EPI_ISL_471857, EPI_ISL_471859, EPI_ISL_471870, EPI_ISL_471891 | | | |
| see above | Michigan Department of Health and Human Services, Bureau of Laboratories | Michigan Department of Health and Human Services, Bureau of Laboratories | Blankenship HM, Riner D, Soehnlen MK |
| EPI_ISL_471911, EPI_ISL_471913, EPI_ISL_471914, EPI_ISL_471915, EPI_ISL_471917, EPI_ISL_471918, EPI_ISL_471919, EPI_ISL_471920, EPI_ISL_471921, EPI_ISL_471922, EPI_ISL_471925, EPI_ISL_471926, EPI_ISL_471927, EPI_ISL_471928, EPI_ISL_471930, EPI_ISL_471931, EPI_ISL_471933, EPI_ISL_471934, EPI_ISL_471935, EPI_ISL_471937, EPI_ISL_471938, EPI_ISL_471939, EPI_ISL_471940, EPI_ISL_471941, EPI_ISL_471942, EPI_ISL_471943, EPI_ISL_471944, EPI_ISL_471945, EPI_ISL_471946, EPI_ISL_471947, EPI_ISL_471948, EPI_ISL_471949, EPI_ISL_471950, EPI_ISL_471952, EPI_ISL_471953 | | | |
| see above | University of Exeter | COVID-19 Genomics UK (COG-UK) Consortium | Ben Temperton,Aaron Jeffries,Michelle Michelsen,Joanna Warwick-Dugdale,Audrey Farbos,Robyn Manley,Stephen Michell,Jane Masoli |
| EPI_ISL_471993, EPI_ISL_471994, EPI_ISL_471995, EPI_ISL_471997, EPI_ISL_471998, EPI_ISL_471999, EPI_ISL_472002, EPI_ISL_472003, EPI_ISL_472004, EPI_ISL_472005, EPI_ISL_472006, EPI_ISL_472007, EPI_ISL_472008, EPI_ISL_472009, EPI_ISL_472010, EPI_ISL_472011, EPI_ISL_472012, EPI_ISL_472013, EPI_ISL_472014, EPI_ISL_472015, EPI_ISL_472016 | | | |
| see above | Liverpool Clinical Laboratories | COVID-19 Genomics UK (COG-UK) Consortium | Sam Haldenby, Anita Lucaci, Steve Paterson, Julian Hiscox, Alistair Darby, M Almsaud, A Alrezaihi, Muhannad Alruwaili, Stuart D Armstrong, Jones Benjamin, Eleanor G Bentley, Anu Chawla, Jordan J Clark, Angela Cowell, Richard Eccles, Isabel García-Dorival, Matthew Gemmell, Alessandro Gerada, PKF Gilmore, Richard Gregory, Ximeng Han, Catherine Hartley, Margaret Hughes, Miren Iturriza-Gomara, James Johnson, L Luu, Jenifer Manson, Charlotte Nelson, Elaine O'Toole, Cassie Olateju, Rebekah Penrice-Randal , Lucille Rainbow, N.P Randle, Trevor Ian Robinson, Parul Sharma, Ghada T Shawli, James P Stewart, Neil Swainston, Ecaterina Vamos, Joanne Watts, Mark Whitehead |
| EPI_ISL_472418 | Queens Medical Centre, Clinical Microbiology Department / DeepSeq Nottingham | COVID-19 Genomics UK (COG-UK) Consortium | Gemma Clark, Wendy Smith, Manjinder Khakh, Vicki M Fleming, Michelle M Lister, Hannah Howson-Wells, Jonathan Ball, Patrick McClure, Joseph Chappell, Theocharis Tsoleridis, Nadine Holmes, Matthew Carlisle, Christopher Moore, Fei Sang, Johnny Debebe, Victoria Wright, Matthew Loose |
| EPI_ISL_472433, EPI_ISL_472439, EPI_ISL_472447, EPI_ISL_472457, EPI_ISL_472459, EPI_ISL_472461, EPI_ISL_472463, EPI_ISL_472467, EPI_ISL_472475, EPI_ISL_472484, EPI_ISL_472489, EPI_ISL_472491, EPI_ISL_472502, EPI_ISL_472503, EPI_ISL_472504, EPI_ISL_472551, EPI_ISL_472554, EPI_ISL_472562, EPI_ISL_472570, EPI_ISL_472589, EPI_ISL_472594, EPI_ISL_472596, EPI_ISL_472614, EPI_ISL_472618, EPI_ISL_472624, EPI_ISL_472626, EPI_ISL_472637, EPI_ISL_472641, EPI_ISL_472651, EPI_ISL_472660, EPI_ISL_472661, EPI_ISL_472686, EPI_ISL_472688, EPI_ISL_472707, EPI_ISL_472848, EPI_ISL_472851, EPI_ISL_472858, EPI_ISL_472859, EPI_ISL_472871, EPI_ISL_472872, EPI_ISL_472886, EPI_ISL_472898, EPI_ISL_472935, EPI_ISL_472967, EPI_ISL_472978, EPI_ISL_472982, EPI_ISL_472988, EPI_ISL_473006, EPI_ISL_473037, EPI_ISL_473059, EPI_ISL_473065, EPI_ISL_473070, EPI_ISL_473284, EPI_ISL_473285, EPI_ISL_473300 | | | |
| see above | Wales Specialist Virology Centre Sequencing lab: Pathogen Genomics Unit | COVID-19 Genomics UK (COG-UK) Consortium | Catherine Moore, Johnathan Evans, Laura Gifford, Malorie Perry, Simon Cottrell, Angela Marchbank, Alec Birchley, Alexander Adams, Amy Gaskin, Bree Gatica-Wilcox, Jason Coombes, Joel Southgate, Lauren Gilbert, Lee Graham, Nicole Pacchiarini, Sara Kumziene-Summerhayes, Sarah Taylor, Sophie Jones, Sara Rey, Matthew Bull, Joanne Watkins, Sally Corden, Tom Connor |
| EPI_ISL_473552, EPI_ISL_473553, EPI_ISL_473554, EPI_ISL_473555, EPI_ISL_473556, EPI_ISL_473557, EPI_ISL_473558, EPI_ISL_473559, EPI_ISL_473560, EPI_ISL_473561, EPI_ISL_473562, EPI_ISL_473563, EPI_ISL_473564, EPI_ISL_473565, EPI_ISL_473566, EPI_ISL_473567, EPI_ISL_473568, EPI_ISL_473570, EPI_ISL_473571, EPI_ISL_473572, EPI_ISL_473573, EPI_ISL_473574, EPI_ISL_473575, EPI_ISL_473576, EPI_ISL_473577, EPI_ISL_473578, EPI_ISL_473579, EPI_ISL_473580, EPI_ISL_473581, EPI_ISL_473582, EPI_ISL_473583, EPI_ISL_473584, EPI_ISL_473585, EPI_ISL_473586, EPI_ISL_473587, EPI_ISL_473588, EPI_ISL_473589, EPI_ISL_473590, EPI_ISL_473591, EPI_ISL_473592, EPI_ISL_473593, EPI_ISL_473594, EPI_ISL_473595, EPI_ISL_473596, EPI_ISL_473597, EPI_ISL_473598, EPI_ISL_473599, EPI_ISL_473600, EPI_ISL_473601, EPI_ISL_473602, EPI_ISL_473603, EPI_ISL_473604, EPI_ISL_473605, EPI_ISL_473606, EPI_ISL_473607, EPI_ISL_473608, EPI_ISL_473609, EPI_ISL_473610, EPI_ISL_473611, EPI_ISL_473612, EPI_ISL_473613, EPI_ISL_473614, EPI_ISL_473615, EPI_ISL_473616, EPI_ISL_473617, EPI_ISL_473618, EPI_ISL_473619, EPI_ISL_473620, EPI_ISL_473621, EPI_ISL_473622, EPI_ISL_473623, EPI_ISL_473624, EPI_ISL_473625, EPI_ISL_473626, EPI_ISL_473627, EPI_ISL_473628, EPI_ISL_473629, EPI_ISL_473630, EPI_ISL_473631, EPI_ISL_473632, EPI_ISL_473633, EPI_ISL_473634, EPI_ISL_473635, EPI_ISL_473636, EPI_ISL_473637, EPI_ISL_473638, EPI_ISL_473639, EPI_ISL_473640, EPI_ISL_473641, EPI_ISL_473642, EPI_ISL_473643, EPI_ISL_473644, EPI_ISL_473645 | | | |
| see above | West of Scotland Specialist Virology Centre, NHSGGC / MRC-University of Glasgow Centre for Virus Research | COVID-19 Genomics UK (COG-UK) Consortium | Ana da Silva Filipe, Natasha Johnson, Kathy Smollett, Daniel Mair, Stephen Carmichael, Lily Tong, Jenna Nichols, Elihu Aranday-Cortes, Kirstyn Brunker, Yasmin Parr, Alice Broos, Kyriaki Nomikou; Sarah McDonald, Marc Niebel, Patawee Asamaphan; Richard Orton, Joseph Hughes, Sreenu Vattipally, David L Robertson; Alasdair MacLean, Rory Gunson; Kathy Li, Natasha Jesudason, Rajiv Shah, James Shepherd, Antonia Ho, Emma Thomson |
| EPI_ISL_473784, EPI_ISL_473785, EPI_ISL_473786, EPI_ISL_473787, EPI_ISL_473788 | Virology Department, Royal Infirmary of Edinburgh, NHS Lothian / School of Biological Sciences, University of Edinburgh / Institute of Genetics and Molecular Medicine, University of Edinburgh | COVID-19 Genomics UK (COG-UK) Consortium | McHugh M, Dewar R, Rooke S, Gallagher M, Balcaza C, O'Toole Á, Scher E, Hill V, McCrone JT, Colquhoun R, Yu X, Jackson B, Rambaut A, Williams TC, Templeton K |
| EPI_ISL_473959, EPI_ISL_473961, EPI_ISL_473963, EPI_ISL_473964, EPI_ISL_473965, EPI_ISL_473966, EPI_ISL_473970, EPI_ISL_473974, EPI_ISL_473988, EPI_ISL_473989, EPI_ISL_473991, EPI_ISL_474023, EPI_ISL_474027, EPI_ISL_474032, EPI_ISL_474034, EPI_ISL_474037, EPI_ISL_474044, EPI_ISL_474052, EPI_ISL_474053, EPI_ISL_474054, EPI_ISL_474057, EPI_ISL_474063, EPI_ISL_474065, EPI_ISL_474069, EPI_ISL_474087, EPI_ISL_474096, EPI_ISL_474098, EPI_ISL_474102, EPI_ISL_474103, EPI_ISL_474106, EPI_ISL_474107, EPI_ISL_474119, EPI_ISL_474124, EPI_ISL_474137, EPI_ISL_474138, EPI_ISL_474147, EPI_ISL_474150, EPI_ISL_474154, EPI_ISL_474169, EPI_ISL_474192, EPI_ISL_474195, EPI_ISL_474200, EPI_ISL_474222, EPI_ISL_474403, EPI_ISL_474404, EPI_ISL_474406, EPI_ISL_474411, EPI_ISL_474412, EPI_ISL_474420, EPI_ISL_474421, EPI_ISL_474422, EPI_ISL_474423, EPI_ISL_474424, EPI_ISL_474425, EPI_ISL_474426, EPI_ISL_474427, EPI_ISL_474428, EPI_ISL_474429, EPI_ISL_474430, EPI_ISL_474431, EPI_ISL_474432, EPI_ISL_474433, EPI_ISL_474434, EPI_ISL_474435, EPI_ISL_474436, EPI_ISL_474437, EPI_ISL_474438, EPI_ISL_474439, EPI_ISL_474440, EPI_ISL_474441, EPI_ISL_474442, EPI_ISL_474443, EPI_ISL_474444, EPI_ISL_474445, EPI_ISL_474446, EPI_ISL_474447, EPI_ISL_474448, EPI_ISL_474449, EPI_ISL_474450, EPI_ISL_474451, EPI_ISL_474452, EPI_ISL_474453, EPI_ISL_474454, EPI_ISL_474455, EPI_ISL_474456, EPI_ISL_474457, EPI_ISL_474458, EPI_ISL_474459, EPI_ISL_474460, EPI_ISL_474461, EPI_ISL_474462, EPI_ISL_474463, EPI_ISL_474464, EPI_ISL_474465, EPI_ISL_474466, EPI_ISL_474467, EPI_ISL_474468, EPI_ISL_474469, EPI_ISL_474470, EPI_ISL_474471, EPI_ISL_474472, EPI_ISL_474473, EPI_ISL_474474, EPI_ISL_474475, EPI_ISL_474476, EPI_ISL_474477, EPI_ISL_474478, EPI_ISL_474479, EPI_ISL_474480, EPI_ISL_474481, EPI_ISL_474482, EPI_ISL_474483, EPI_ISL_474484, EPI_ISL_474485, EPI_ISL_474486, EPI_ISL_474487, EPI_ISL_474488, EPI_ISL_474489, EPI_ISL_474490, EPI_ISL_474491, EPI_ISL_474492, EPI_ISL_474493, EPI_ISL_474494, EPI_ISL_474495, EPI_ISL_474496, EPI_ISL_474497, EPI_ISL_474498, EPI_ISL_474499, EPI_ISL_474500, EPI_ISL_474501, EPI_ISL_474502, EPI_ISL_474503, EPI_ISL_474504, EPI_ISL_474505, EPI_ISL_474506, EPI_ISL_474507, EPI_ISL_474508, EPI_ISL_474509, EPI_ISL_474510, EPI_ISL_474511, EPI_ISL_474512, EPI_ISL_474513, EPI_ISL_474514, EPI_ISL_474515, EPI_ISL_474516, EPI_ISL_474517, EPI_ISL_474518, EPI_ISL_474519, EPI_ISL_474520, EPI_ISL_474521, EPI_ISL_474522, EPI_ISL_474523, EPI_ISL_474524, EPI_ISL_474525, EPI_ISL_474526, EPI_ISL_474527, EPI_ISL_474528, EPI_ISL_474529, EPI_ISL_474530, EPI_ISL_474535, EPI_ISL_474539, EPI_ISL_474542, EPI_ISL_474543, EPI_ISL_474555, EPI_ISL_474556, EPI_ISL_474560, EPI_ISL_474564, EPI_ISL_474565, EPI_ISL_474576, EPI_ISL_474620, EPI_ISL_474622, EPI_ISL_474736, EPI_ISL_474737, EPI_ISL_474738, EPI_ISL_474739, EPI_ISL_474740, EPI_ISL_474741, EPI_ISL_474742, EPI_ISL_474743, EPI_ISL_474744, EPI_ISL_474745 | | | |
| see above | Wales Specialist Virology Centre Sequencing lab: Pathogen Genomics Unit | COVID-19 Genomics UK (COG-UK) Consortium | Catherine Moore, Johnathan Evans, Laura Gifford, Malorie Perry, Simon Cottrell, Angela Marchbank, Alec Birchley, Alexander Adams, Amy Gaskin, Bree Gatica-Wilcox, Jason Coombes, Joel Southgate, Lauren Gilbert, Lee Graham, Nicole Pacchiarini, Sara Kumziene-Summerhayes, Sarah Taylor, Sophie Jones, Sara Rey, Matthew Bull, Joanne Watkins, Sally Corden, Tom Connor |
| EPI_ISL_474830, EPI_ISL_474849 | Complejo Hospitalario Universitario de Albacete | SeqCOVID-SPAIN consortium/IBV(CSIC) | Encarnacion Simarro Córdoba, Julia Lozano Serra, Lorena Robles Fonseca , Monica Parra Grandes, Caridad Sainz de Baranda Camino and SeqCOVID-SPAIN consortium |
| EPI_ISL_474966, EPI_ISL_474967, EPI_ISL_474972, EPI_ISL_474984, EPI_ISL_474985, EPI_ISL_474991, | Israel Central Virology laboratory | Israel Central Virology laboratory | Neta Zuckerman, Efrat Dahan Bucris, Oran Erster, Ella Mendelson, Michal Mandelboim |

| | | | |
|---|---|---|---|
| EPI_ISL_475016, EPI_ISL_475017, EPI_ISL_475020 | | | |
| EPI_ISL_475127 | Umea klinisk mikrobiologi | The Public Health Agency of Sweden | Oskar Karlsson Lindsjo, Maria Lind Karlberg, Mattias Haukland, Reza Advani, Olov Svartstrom, Anna-Malin Linde, Sandra Broddesson, Petra Edquist, Shamam Muradrasoli, Anna Risberg, Karin Tegmark-Wisell |
| EPI_ISL_475160, EPI_ISL_475161, EPI_ISL_475162, EPI_ISL_475163 | Klinisk mikrobiologi Vasternorrland | The Public Health Agency of Sweden | Oskar Karlsson Lindsjo, Maria Lind Karlberg, Mattias Haukland, Reza Advani, Olov Svartstrom, Anna-Malin Linde, Sandra Broddesson, Petra Edquist, Shamam Muradrasoli, Anna Risberg, Karin Tegmark-Wisell |
| EPI_ISL_475164 | Halmstad klinisk mikrobiologi | The Public Health Agency of Sweden | Oskar Karlsson Lindsjo, Maria Lind Karlberg, Mattias Haukland, Reza Advani, Olov Svartstrom, Anna-Malin Linde, Sandra Broddesson, Petra Edquist, Shamam Muradrasoli, Anna Risberg, Karin Tegmark-Wisell |
| EPI_ISL_475261, EPI_ISL_475269, EPI_ISL_475270, EPI_ISL_475271, EPI_ISL_475272, EPI_ISL_475273, EPI_ISL_475274, EPI_ISL_475301, EPI_ISL_475302, EPI_ISL_475303, EPI_ISL_475306, EPI_ISL_475307, EPI_ISL_475308, EPI_ISL_475309, EPI_ISL_475310, EPI_ISL_475311, EPI_ISL_475312, EPI_ISL_475313, EPI_ISL_475314, EPI_ISL_475315, EPI_ISL_475316, EPI_ISL_475317, EPI_ISL_475318, EPI_ISL_475319, EPI_ISL_475320, EPI_ISL_475331, EPI_ISL_475332, EPI_ISL_475334, EPI_ISL_475335, EPI_ISL_475336, EPI_ISL_475337, EPI_ISL_475338, EPI_ISL_475339 | | | |
| see above | Centre for Enzyme Innovation, University of Portsmouth / Translational Research Laboratory, Portsmouth Hospitals NHS Trust | COVID-19 Genomics UK (COG-UK) Consortium | Angela Beckett,Yann Bourgeois,Garry Scarlett,Sharon Glaysher,Scott Elliott,Kelly Bicknell,Robert Impey,Allyson Lloyd,Sarah Wyllie,Ethan Butcher,Anoop Chauhan,Samuel Robson |
| EPI_ISL_475384, EPI_ISL_475386, EPI_ISL_475392, EPI_ISL_475404, EPI_ISL_475483 | Virology Department, Sheffield Teaching Hospitals NHS Foundation Trust/Department of Infection, Immunity and Cardiovascular Disease, The Medical School, University of Sheffield | COVID-19 Genomics UK (COG-UK) Consortium | Thushan de Silva, Matthew Parker, Nikki Smith, Adri Angyal, Rebecca Brown, Luke Green, Rachel Tucker, Paul Parsons, Danielle Groves, Katie Johnson, Laura Carrilero, Alex Keeley, Dave Partridge, Matthew Wyles, Benjamin Lindsey, Mehmet Yavuz, Mohammad Raza, Cariad Evans |
| EPI_ISL_475835, EPI_ISL_475842, EPI_ISL_475852, EPI_ISL_475861, EPI_ISL_475862, EPI_ISL_475865, EPI_ISL_475885 | Austrian Agency for Health and Food Safety (AGES) | Bergthaler laboratory, CeMM Research Center for Molecular Medicine of the Austrian Academy of Sciences | Alexandra Popa, Benedikt Agerer, Henrique Colaco, Lukas Endler, Jakob-Wendelin Genger, Alexander Lercher, Mark Smyth, Thomas Penz, Michael Schuster, Jan Laine, Martin Senekowitsch, Judith Aberle, Stephan Aberle, Peter Hufnagl, Daniela Schmid, Franz Allerberger, Elisabeth Puchhammer-Stoeckl, Manfred Nairz, Guenter Weiss, Gregor Hörmann, Kinga Rigler-Hohenwarter, Rainer Gattringer, Wegene Borena, Dorothee von Laer, Christoph Bock, Andreas Bergthaler |
| EPI_ISL_475939, EPI_ISL_475940, EPI_ISL_475941, EPI_ISL_475944, EPI_ISL_475945, EPI_ISL_475946, EPI_ISL_475947, EPI_ISL_475989, EPI_ISL_475990, EPI_ISL_475991, EPI_ISL_475992 | | | |
| see above | National Public Health Laboratory, National Centre for Infectious Diseases | National Public Health Laboratory, National Centre for Infectious Diseases | Mak TM, Octavia S, Chavatte JM, Cui L, Lin RTP |
| EPI_ISL_476031, EPI_ISL_476032, EPI_ISL_476033, EPI_ISL_476034, EPI_ISL_476035, EPI_ISL_476036, EPI_ISL_476037, EPI_ISL_476038, EPI_ISL_476039, EPI_ISL_476040 | Michigan Department of Health and Human Services, Bureau of Laboratories | Michigan Department of Health and Human Services, Bureau of Laboratories | Blankenship HM, Riner D, Soehnlen MK |
| EPI_ISL_476068 | University of Debrecen, Department of Medical Microbiology | National Laboratory of Virology, Szentágothai Research Centre | Endre Gábor Tóth, Balázs Somogyi, Brigitta Zana, Eszter Csoma, Ferenc Jakab, Gábor Kemenesi |
| EPI_ISL_476120, EPI_ISL_476121, EPI_ISL_476122, EPI_ISL_476123, EPI_ISL_476124, EPI_ISL_476125, EPI_ISL_476126, EPI_ISL_476127, EPI_ISL_476128, EPI_ISL_476129, EPI_ISL_476130 | | | |
| see above | Viollier AG | Department of Biosystems Science and Engineering, ETH Zürich | Christian Beisel, Sarah Nadeau, Ivan Topolsky, Pedro Ferreira, Philipp Jablonski, Susana Posada-Céspedes, Tobias Schär, Ina Nissen, Natascha Santacroce, Elodie Burcklen, Christiane Beckmann, Maurice Redondo, Olivier Kobel, Christoph Noppen, Sophie Seidel, Noemie Santamaria de Souza, Niko Beerenwinkel, Tanja Stadler |
| EPI_ISL_476346, EPI_ISL_476347, EPI_ISL_476348, EPI_ISL_476349, EPI_ISL_476400, EPI_ISL_476401, EPI_ISL_476408, EPI_ISL_476409, EPI_ISL_476416, EPI_ISL_476417, EPI_ISL_476418, EPI_ISL_476419, EPI_ISL_476420, EPI_ISL_476421, EPI_ISL_476422, EPI_ISL_476423, EPI_ISL_476424, EPI_ISL_476425 | | | |
| see above | Laboratório de Patologia Clínica - UNICAMP | Laboratório de Estudos de Vírus Emergentes - UNICAMP | José Luiz Proença-Modena, Magnun Nueldo Nunes dos Santos, Angelica Schreiber, Julia Forato,Camila Simeoni, Marcilio Jorge Fumagalli, Mariene Ribeiro Amorim, Darlan da Silva Candido, Nuno Rodrigues Faria, Julien Theze, Luiz Gonzaga,Jaqueline Goes Jesus e William Marciel de Souza |
| EPI_ISL_476496 | Hospital Garrahan | Héritas | Dalmacio Pereyra, Roberta Crespo, Mauricio Grisolia, Cristian Rohr, Andrea Mangano, Maria Florencia Fernandez, Fabian Fay, Martin Vazquez |
| EPI_ISL_476706, EPI_ISL_476707, EPI_ISL_476708, EPI_ISL_476709, EPI_ISL_476710, EPI_ISL_476711, EPI_ISL_476712, EPI_ISL_476713, EPI_ISL_476714, EPI_ISL_476715, EPI_ISL_476716, EPI_ISL_476717, EPI_ISL_476718, EPI_ISL_476719 | | | |
| see above | Minnesota Department of Health, Public Health Laboratory | Minnesota Department of Health, Public Health Laboratory | Matt Plumb, Jacob Garfin, and Xiong Wang |
| EPI_ISL_476841 | National Influenza Centre for Northern Greece | National Influenza Centre for Northern Greece | Maria Christoforidi |
| EPI_ISL_476991 | KU Leuven, Rega Institute, Clinical and Epidemiological Virology | KU Leuven, Rega Institute, Clinical and Epidemiological Virology | Tony Wawina-Bokalanga, Joan Marti-Carerras, Bert Vanmechelen, Piet Maes |
| EPI_ISL_477008, EPI_ISL_477009, EPI_ISL_477013 | University of Debrecen, Department of Medical Microbiology | National Laboratory of Virology, Szentágothai Research Centre | Endre Gábor Tóth, Balázs Somogyi, Brigitta Zana, Eszter Csoma, Ferenc Jakab, Gábor Kemenesi |
| EPI_ISL_477187, EPI_ISL_477188, EPI_ISL_477189, EPI_ISL_477190, EPI_ISL_477191 | Department of Laboratory Medicine Tan Tock Seng Hospital | Department of Laboratory Medicine Tan Tock Seng Hospital | Chen YYC, Zair X, Li C, Tang WY, Maurer-Stroh S, Barkham TMS, Nagarajan N, Sessions OM |
| EPI_ISL_477276, EPI_ISL_477293 | Mayo Clinic & Mayo Clinic Laboratories | Minnesota Department of Health, Public Health Laboratory | Matt Plumb, Jacob Garfin, Kelly Pung, and Xiong Wang |
| EPI_ISL_477631, EPI_ISL_477632, EPI_ISL_477634 | Virginia DCLS | Virginia DCLS | Virginia DCLS |
| EPI_ISL_477744, EPI_ISL_477745, EPI_ISL_477746, EPI_ISL_477747, EPI_ISL_477748, EPI_ISL_477749, EPI_ISL_477750, EPI_ISL_477751, EPI_ISL_477752, EPI_ISL_477753, EPI_ISL_477754 | | | |
| see above | University of Birmingham | COVID-19 Genomics UK (COG-UK) Consortium | Institute of Microbiology, University of Birmingham: Claire McMurray, Joanne Stockton, Samuel Nicholls, Radoslaw Poplawski, Will Rowe, Josh Quick, Nicholas Loman. University of Birmingham Testing Laboratory: Celina M Whalley, Andrew Bosworth, Charlotte Poxon, Kasun Wanigasooriya, Oliver Pickles, Mike Kidd, Alex Richter, Andrew D Beggs PHE Heartlands Lab: Husam Osman, Andrew Bosworth. Queen Elizabeth Hospital: Anna Casey |
| EPI_ISL_477819 | Department of Pathology, University of Cambridge | COVID-19 Genomics UK (COG-UK) Consortium | Luke W Meredith, M. Estée Török, Myra Hosmillo, William L. Hamilton, Martin D. Curran, Theresa Feltwell, Grant Hall, Anna Yakovleva, Fahad A Khokhar, Charlotte J. Houldcroft, Laura G Caller, Aminu S. Jahun, Sarah L. Caddy, Yasmin Chaudhry, Malte Pinckert, Ian Goodfellow |
| EPI_ISL_478029, EPI_ISL_478030, EPI_ISL_478031, EPI_ISL_478032, EPI_ISL_478033, EPI_ISL_478034, EPI_ISL_478035, EPI_ISL_478036, EPI_ISL_478037, EPI_ISL_478038, EPI_ISL_478039, EPI_ISL_478040, EPI_ISL_478041, EPI_ISL_478042, EPI_ISL_478043, EPI_ISL_478044, EPI_ISL_478045, EPI_ISL_478046, EPI_ISL_478047, EPI_ISL_478048, EPI_ISL_478049, EPI_ISL_478050, EPI_ISL_478051, EPI_ISL_478054, EPI_ISL_478055 | | | |
| see above | West of Scotland Specialist Virology Centre, NHSGGC / MRC-University of Glasgow Centre for Virus Research | COVID-19 Genomics UK (COG-UK) Consortium | Ana da Silva Filipe, Natasha Johnson, Kathy Smollett, Daniel Mair, Stephen Carmichael, Lily Tong, Jenna Nichols, Elihu Aranday-Cortes, Kirstyn Brunker, Yasmin Parr, Alice Broos, Kyriaki Nomikou; Sarah McDonald, Marc Niebel, Patawee Asamaphan; Richard Orton, Joseph Hughes, Sreenu Vattipally, David L Robertson; Alasdair MacLean, Rory Gunson; Kathy Li, Natasha Jesudason, Rajiv Shah, James Shepherd, Antonia Ho, Emma Thomson |
| EPI_ISL_478277, EPI_ISL_478288 | University of Exeter | COVID-19 Genomics UK (COG-UK) Consortium | Ben Temperton,Aaron Jeffries,Michelle Michelsen,Joanna Warwick-Dugdale,Audrey Farbos,Robyn Manley,Stephen Michell,Jane Masoli |
| EPI_ISL_478705 | South Eastern Area Laboratory Services (SEALS) | NSW Health Pathology - Institute of Clinical Pathology and Medical Research; Westmead Hospital; University of Sydney | CIDM-PH et al. |
| EPI_ISL_478728 | Queens Medical Centre, Clinical Microbiology Department / DeepSeq Nottingham | COVID-19 Genomics UK (COG-UK) Consortium | Gemma Clark, Wendy Smith, Manjinder Khakh, Vicki M Fleming, Michelle M Lister, Hannah Howson-Wells, Jonathan Ball, Patrick McClure, Joseph Chappell, Theocharis Tsoleridis, Nadine Holmes, Matthew Carlisle, Christopher Moore, Fei Sang, Johnny Debebe, Victoria Wright, Matthew Loose |
| EPI_ISL_478851, EPI_ISL_478866, EPI_ISL_478875, EPI_ISL_478876, EPI_ISL_478877, EPI_ISL_478879, EPI_ISL_478880, EPI_ISL_478882, EPI_ISL_478886, EPI_ISL_478887, EPI_ISL_478888, EPI_ISL_478889, EPI_ISL_478890, EPI_ISL_478893, EPI_ISL_478894, EPI_ISL_478895, EPI_ISL_478897, EPI_ISL_478898, EPI_ISL_478900, EPI_ISL_478901, EPI_ISL_478903, EPI_ISL_478905, EPI_ISL_478906, EPI_ISL_478907, EPI_ISL_478908, EPI_ISL_478909, EPI_ISL_478911, EPI_ISL_478916, EPI_ISL_478917, EPI_ISL_478920, EPI_ISL_478922, EPI_ISL_478924, EPI_ISL_478939, EPI_ISL_478946, EPI_ISL_478957, EPI_ISL_478963, EPI_ISL_478972, EPI_ISL_478991, EPI_ISL_478995, EPI_ISL_478998, EPI_ISL_479001, EPI_ISL_479003, EPI_ISL_479006, EPI_ISL_479008, EPI_ISL_479010, EPI_ISL_479012, EPI_ISL_479014, EPI_ISL_479017, EPI_ISL_479020, EPI_ISL_479022, EPI_ISL_479027, EPI_ISL_479042, EPI_ISL_479046, EPI_ISL_479047, | | | |

| | | | |
|---|---|---|---|
| EPI_ISL_479048, EPI_ISL_479056, EPI_ISL_479064, EPI_ISL_479071, EPI_ISL_479082, EPI_ISL_479087, EPI_ISL_479091, EPI_ISL_479095, EPI_ISL_479097, EPI_ISL_479102, EPI_ISL_479113, EPI_ISL_479114, EPI_ISL_479116, EPI_ISL_479120, EPI_ISL_479122, EPI_ISL_479126, EPI_ISL_479128, EPI_ISL_479129, EPI_ISL_479133, EPI_ISL_479138, EPI_ISL_479144, EPI_ISL_479146, EPI_ISL_479154, EPI_ISL_479159, EPI_ISL_479161, EPI_ISL_479164, EPI_ISL_479167, EPI_ISL_479171 | | | |
| see above | Oxford Viromics, NDM, University of Oxford; Oxford University Hospitals; Basingstoke and North Hampshire Hospital | COVID-19 Genomics UK (COG-UK) Consortium | Tanya Golubchik, David Bonsall, George Macintyre, Amy Trebes, Mariateresa de Cesare, Catrin Moore, Alex Mobbs, Anita Justice, Robert Shaw, Monique Andersson, Timothy Peto, Emma Wise, Nathan Moore, Jessica Lynch, Nick Cortes, Matilde Mori, Stephen Kidd, David Buck, John Todd, Christophe Fraser |
| EPI_ISL_479200, EPI_ISL_479201, EPI_ISL_479212, EPI_ISL_479219, EPI_ISL_479222, EPI_ISL_479224, EPI_ISL_479227, EPI_ISL_479240, EPI_ISL_479248, EPI_ISL_479249, EPI_ISL_479253, EPI_ISL_479258, EPI_ISL_479265, EPI_ISL_479267, EPI_ISL_479268, EPI_ISL_479274, EPI_ISL_479278, EPI_ISL_479279, EPI_ISL_479280 | | | |
| see above | Virology Department, Sheffield Teaching Hospitals NHS Foundation Trust/Department of Infection, Immunity and Cardiovascular Disease, The Medical School, University of Sheffield | COVID-19 Genomics UK (COG-UK) Consortium | Thushan de Silva, Matthew Parker, Nikki Smith, Adri Angyal, Rebecca Brown, Luke Green, Rachel Tucker, Paul Parsons, Danielle Groves, Katie Johnson, Laura Carrilero, Alex Keeley, Dave Partridge, Matthew Wyles, Benjamin Lindsey, Mehmet Yavuz, Mohammad Raza, Cariad Evans |
| EPI_ISL_479309, EPI_ISL_479312, EPI_ISL_479331, EPI_ISL_479348, EPI_ISL_479356, EPI_ISL_479360, EPI_ISL_479376, EPI_ISL_479393, EPI_ISL_479401, EPI_ISL_479412, EPI_ISL_479413, EPI_ISL_479415, EPI_ISL_479425, EPI_ISL_479439, EPI_ISL_479445, EPI_ISL_479454, EPI_ISL_479462, EPI_ISL_479464 | | | |
| see above | Wales Specialist Virology Centre Sequencing lab: Pathogen Genomics Unit | COVID-19 Genomics UK (COG-UK) Consortium | Catherine Moore, Johnathan Evans, Laura Gifford, Malorie Perry, Simon Cottrell, Angela Marchbank, Alec Birchley, Alexander Adams, Amy Gaskin, Bree Gatica-Wilcox, Jason Coombes, Joel Southgate, Lauren Gilbert, Lee Graham, Nicole Pacchiarini, Sara Kumziene-Summerhayes, Sarah Taylor, Sophie Jones, Sara Rey, Matthew Bull, Joanne Watkins, Sally Corden, Tom Connor |
| EPI_ISL_479482, EPI_ISL_479483, EPI_ISL_479485, EPI_ISL_479486, EPI_ISL_479487, EPI_ISL_479488, EPI_ISL_479489, EPI_ISL_479491, EPI_ISL_479492 | Department of Laboratory Medicine Tan Tock Seng Hospital | Department of Laboratory Medicine Tan Tock Seng Hospital | Chen YYC, Zair X, Li C, Tang WY, Maurer-Stroh S, Barkham TMS, Nagarajan N, Sessions OM |
| EPI_ISL_479496, EPI_ISL_479497, EPI_ISL_479498, EPI_ISL_479499, EPI_ISL_479500, EPI_ISL_479501 | NIV Influenza | NIV Influenza | Potdar V |
| EPI_ISL_479574, EPI_ISL_479582 | National Public Health Laboratory, National Centre for Infectious Diseases | National Public Health Laboratory, National Centre for Infectious Diseases | Mak TM, Octavia S, Zhou Z, Chavatte JM, Cui L, Lin RTP |
| EPI_ISL_480235, EPI_ISL_480236, EPI_ISL_480237, EPI_ISL_480238, EPI_ISL_480239, EPI_ISL_480240, EPI_ISL_480241, EPI_ISL_480243, EPI_ISL_480244, EPI_ISL_480245, EPI_ISL_480246, EPI_ISL_480247, EPI_ISL_480248, EPI_ISL_480264 | | | |
| see above | Genomic Laboratory (GLAB) (Conjoint lab of Health Directorate of Istanbul and Istanbul Technical University) | Genomic Laboratory (GLAB), Istanbul Technical University | Ilker Karacan, Tugba Kizilboga Akgun, Bugra Agaoglu, Gizem Alkurt, Jale Yildiz, Betsi Köse, Elifnaz Çelik, Arzu Irvem, Yasemin Kendir Demirkol, Ozlem Akgun Dogan, Mehtap Aydn, Levent Doganay, Gizem Dinler Doganay |
| EPI_ISL_480299, EPI_ISL_480304, EPI_ISL_480306 | National Reference Laboratory "Influenza and acute respiratory diseases" | NRL-HIV | Ivan Ivanov, Ivailo Alexiev, Ivva Philipova |
| EPI_ISL_480330 | Quadram Institute Bioscience | COVID-19 Genomics UK (COG-UK) Consortium | Dave J. Baker, Gemma L. Kay, Alp Aydin, Thanh Le-Viet, Steven Rudder, Ana P. Tedim, Anastasia Kolyva, Maria Diaz, Leonardo de Oliveira Martins, Nabil-Fareed Alikhan, Lizzie Meadows, Rachael Stanley, Ngozi Elumogo, Muhammed Yasir, Nicholas M. Thomson, Alexander J Trotter, Rachel Gilroy, Samuel Bloomfield, Claire Stuart, Andrew Bell, Reenesh Prakash, Samir Dervisevic, Alison E. Mather, John Wain, Mark Webber, Andrew J. Page, Justin O'Grady |
| EPI_ISL_480331, EPI_ISL_480336, EPI_ISL_480337, EPI_ISL_480342, EPI_ISL_480343 | Microbial Genomics Laboratory, Institut Pasteur de Montevideo | Microbial Genomics Laboratory, Institut Pasteur de Montevideo | Cecilia Salazar, Marianoel Pereira, Ignacio Ferrés, Gonzalo Moratorio, Pilar Moreno, Gregorio Iraola |
| EPI_ISL_480402, EPI_ISL_480403 | University of Wisconsin-Madison AIDS Vaccine Research Laboratories | University of Wisconsin-Madison AIDS Vaccine Research Laboratories | Gage Moreno, Katarina Braun, et al. AIDS Vaccine Research Laboratories |
| EPI_ISL_480431 | Laboratorio de Biología Molecular Asociación Española Primera en Salud | Departments of Pathology and Medicine, New York University School of Medicine | Maria Victoria Elizondo, Maria Noel Zubillaga, Gonzalo Manrique, Paul Zappile, Gael Westby, Matthew T Maurano, Christian Marier, Adriana Heguy |
| EPI_ISL_480594, EPI_ISL_480595 | Victorian Infectious Diseases Reference Laboratory (VIDRL) | VIDRL and MDU-PHL | Caly L., Seemann T., Sait, M., Schultz M., Druce J., Sherry, N. |
| EPI_ISL_480620 | Microbiological Diagnostic Unit - Public Health Laboratory (MDU-PHL) | MDU-PHL | Seemann T., Schultz M., Sait, M., Sherry, N. |
| EPI_ISL_480864, EPI_ISL_480866, EPI_ISL_480867, EPI_ISL_480868, EPI_ISL_480869, EPI_ISL_480870, EPI_ISL_480874, EPI_ISL_480875, EPI_ISL_480876, EPI_ISL_480877, EPI_ISL_480878, EPI_ISL_480879, EPI_ISL_480880, EPI_ISL_480881, EPI_ISL_480882, EPI_ISL_480883, EPI_ISL_480884, EPI_ISL_480885, EPI_ISL_480886, EPI_ISL_480887, EPI_ISL_480888, EPI_ISL_480889 | | | |
| see above | Florida Bureau of Public Health Laboratories | Florida Bureau of Public Health Laboratories | Sarah Schmedes, Jason Blanton |
| EPI_ISL_481069, EPI_ISL_481084 | Hospital General Universitario Gregorio Marañón | SeqCOVID-SPAIN consortium/IBV(CSIC) | Laura Pérez-Lago, Marta Herranz, Jon Sicilia, Julia Suárez, Pilar Catalán, Patricia Muñoz, Darío García de Viedma and SeqCOVID-SPAIN consortium |
| EPI_ISL_481208 | Furst Medical Laboratory | Norwegian Institute of Public Health, Department of Virology | Kathrine Stene-Johansen, Kamilla Heddeland Instefjord, Hilde Elshaug, Rasmus Riis Kopperud, Karoline Bragstad, Olav Hungnes |
| EPI_ISL_481209, EPI_ISL_481212 | Ostfold Hospital Trust - Kalnes, Centre for Laboratory Medicine, Section for gene technology and infection serology | Norwegian Institute of Public Health, Department of Virology | Kathrine Stene-Johansen, Kamilla Heddeland Instefjord, Hilde Elshaug, Rasmus Riis Kopperud, Karoline Bragstad, Olav Hungnes |
| EPI_ISL_481221, EPI_ISL_481222 | Lab voor klinische biologie | Onderzoeksgroep Virologie | Laurens Lambrechts, Nick Vereecke, Marthe Pauwels, Bruno Verhasselt, Linos Vandekerckhove, Hans Nauwynck, Sebastiaan Theuns |
| EPI_ISL_481255, EPI_ISL_481257, EPI_ISL_481258, EPI_ISL_481259, EPI_ISL_481260, EPI_ISL_481261, EPI_ISL_481263 | Department of Emerging Infectious Diseases, Institute of Tropical Medicine, Nagasaki University | Department of Emerging Infectious Diseases, Institute of Tropical Medicine, Nagasaki University | Jiro Yasuda, Rokusuke Yoshikawa, Yuichiro Furusato, Haruka Abe |
| EPI_ISL_481483 | Department for Virology, Molecular Biology and Genome Research, R. G. Lugar Center for Public Health Research, National Center for Disease Control and Public Health (NCDC) of Georgia. | Department for Virology, Molecular Biology and Genome Research, R. G. Lugar Center for Public Health Research, National Center for Disease Control and Public Health (NCDC) of Georgia. | Nino Berishvili, Tata Imnadze, Giorgi Tomashvili, Ana Papkiauri, Meri Pantsulaia, Gvantsa Brachveli, Gvantsa Chanturia, Ann Machablishvili, Nato Kotaria, Marine Murtskhvaladze, Lela Sabadze, Mari Gavashelidze, Tamar Jashiashvili, Tea Tevdoradze, Ketevan Sidamonidze, Ekaterine Khmaladze, Ekaterine Zhghenti, Roena Sukhiashvili, Mariam Zakalashvili, Lela Urushadze, Magda Dgebuadze, Davit Tsaguria, Ekaterine Zangaladze, Adam Kotorashvili, Maia Alkhazashvili, Irma Burjanadze, Anna Kasradze, Khatuna Zakhashvili, Paata Imnadze, Amiran Gamkrelidze. |
| EPI_ISL_481510, EPI_ISL_481511, EPI_ISL_481512 | Prof. Massimo Zollo CEINGE TASK-FORCE COVID19 - Regione Campania | Prof. Massimo Zollo CEINGE TASK-FORCE COVID19 - Regione Campania | Veronica Ferrucci1,2, Dae young Kong8, Fatemeh asadzadeh1,2, Laura Marrone1,2, Roberto Siciliano1,2, Rino Cerino3, Giovanna Fusco3, Marika Comegna1,2, Angelo Boccia2, Maurizio Viscardi3, Giorgia Borriello3, Sergio Brandi3, Claudia Tiberio4, Luigi Atripaldi4, Giovanni Paolella1,2, Giuseppe Castaldo1,2, Stefano Pascarella4, Martina Bianchi4, Lorenzo Chiariotti1,2, Jae Myun Lee5, Jae Ho Jung6, Kyong Seop Yun7, Hong Yeoul Kim 7,8* and Massimo Zollo1,2* 1 CEINGE Biotecnologie Avanzate, Naples, Italia 2 Dipartimento di Medicina Molecolare e Biotecnologie Mediche DMMBM University of Naples Federico II, Italia 3 Istituto Zooprofilattico Sperimentale del Mezzogiorno, Naples, Italia 4 -U.O.C. di Patologia Clinica Ospedale D. Cotugno, Azienda Sanitaria Ospedali dei Colli, Naples, Italy. 5 Università La Sapienza di Roma, Italia 6 Department of Microbiology, Yonsei University College of Medicine, Seoul, Korea 7 Department of Surgery, Yonsei University College of Medicine, Seoul, Korea 8 Haim bio co., Ltd, , Indust |
| EPI_ISL_481595, EPI_ISL_481596, EPI_ISL_481597, EPI_ISL_481598, EPI_ISL_481599, EPI_ISL_481600, EPI_ISL_481601, EPI_ISL_481602, EPI_ISL_481603, EPI_ISL_481604, EPI_ISL_481605, EPI_ISL_481606, EPI_ISL_481607, EPI_ISL_481608, EPI_ISL_481609, EPI_ISL_481610, EPI_ISL_481611, EPI_ISL_481612, EPI_ISL_481613, EPI_ISL_481614, EPI_ISL_481615, EPI_ISL_481616, EPI_ISL_481617, EPI_ISL_481618, EPI_ISL_481619, EPI_ISL_481620, EPI_ISL_481621 | | | |
| see above | Department of Virology and Immunology, University of Helsinki and Helsinki University Hospital, Huslab Finland | Department of Virology, Faculty of Medicine, University of Helsinki, Helsinki, Finland | Teemu Smura, Hannimari Kallio-Kokko, Jenni Virtanen, Maija Suvanto, Sari Hannula, Harri Kangas, Pekka Ellonen, Olli Vapalahti |
| EPI_ISL_481716, EPI_ISL_481741, EPI_ISL_481759, EPI_ISL_481760, EPI_ISL_481761, EPI_ISL_481762 | Prof. Massimo Zollo CEINGE TASK-FORCE COVID19 - Regione Campania | Prof. Massimo Zollo CEINGE TASK-FORCE COVID19 - Regione Campania | Veronica Ferrucci1,2, Dae young Kong8, Fatemeh asadzadeh1,2, Laura Marrone1,2, Roberto Siciliano1,2, Rino Cerino3, Giovanna Fusco3, Marika Comegna1,2, Angelo Boccia2, Maurizio Viscardi3, Giorgia Borriello3, Sergio Brandi3, Claudio Tiberio4, Luigi Atripaldi4, Giovanni Paolella1,2, Giuseppe Castaldo1,2, Stefano Pascarella4, Martina Bianchi4, Lorenzo Chiariotti1,2, Jae Myun Lee5, Jae Ho Jung6, Kyong Seop Yun7, Hong Yeoul Kim 7,8* and |

| | | | |
|---|---|---|---|
| | | | Massimo Zollo1,2* 1 CEINGE Biotecnologie Avanzate, Naples, Italia 2 Dipartimento di Medicina Molecolare e Biotecnologie Mediche DMMBM University of Naples Federico II, Italia 3 Istituto Zooprofilattico Sperimentale del Mezzogiorno, Naples, Italia 4 -U.O.C. di Patologia Clinica Ospedale D. Cotugno, Azienda Sanitaria Ospedali dei Colli, Naples, Italy. 5 Università La Sapienza di Roma, Italia 6 Department of Microbiology, Yonsei University College of Medicine, Seoul, Korea 7 Department of Surgery, Yonsei University College of Medicine, Seoul, Korea 8 Haim bio co., Ltd, , Indust |
| EPI_ISL_481763, EPI_ISL_481764, EPI_ISL_481765, EPI_ISL_481772, EPI_ISL_481779, EPI_ISL_481780, EPI_ISL_481782, EPI_ISL_481793, EPI_ISL_481799, EPI_ISL_481801, EPI_ISL_481804, EPI_ISL_481807, EPI_ISL_481812, EPI_ISL_481815, EPI_ISL_481822, EPI_ISL_481824, EPI_ISL_481829, EPI_ISL_481832, EPI_ISL_481833, EPI_ISL_481839, EPI_ISL_481843, EPI_ISL_481845, EPI_ISL_481853, EPI_ISL_481860, EPI_ISL_481864, EPI_ISL_481872, EPI_ISL_481876, EPI_ISL_481886, EPI_ISL_481910, EPI_ISL_481911, EPI_ISL_481914, EPI_ISL_481918, EPI_ISL_481919, EPI_ISL_481920, EPI_ISL_481921, EPI_ISL_481922, EPI_ISL_481929, EPI_ISL_481933, EPI_ISL_481938, EPI_ISL_481939, EPI_ISL_481942, EPI_ISL_481943, EPI_ISL_481945, EPI_ISL_481947, EPI_ISL_481948, EPI_ISL_481956, EPI_ISL_481959, EPI_ISL_481962, EPI_ISL_481964, EPI_ISL_481971, EPI_ISL_481972, EPI_ISL_481973, EPI_ISL_481974, EPI_ISL_481978, EPI_ISL_481980, EPI_ISL_481982, EPI_ISL_481991, EPI_ISL_481994, EPI_ISL_481997, EPI_ISL_481999, EPI_ISL_482000, EPI_ISL_482004, EPI_ISL_482014, EPI_ISL_482016, EPI_ISL_482017, EPI_ISL_482018, EPI_ISL_482021, EPI_ISL_482023, EPI_ISL_482024, EPI_ISL_482025, EPI_ISL_482027, EPI_ISL_482030 | | | |
| see above | PHE South West Regional Laboratory, National Infection Service | Wellcome Sanger Institute for the COVID-19 Genomics UK (COG-UK) consortium | Stephanie Hutchings, Hannah Pymont, Dr Peter Muir, Barry Vipond, Rich Hopes; and Alex Alderton, Roberto Amato, Sonia Goncalves, Ewan Harrison, David K. Jackson, Ian Johnston, Dominic Kwiatkowski, Cordelia Langford, John Sillitoe on behalf of the Wellcome Sanger Institute COVID-19 Surveillance Team (http://www.sanger.ac.uk/covid-team) |
| EPI_ISL_482036, EPI_ISL_482038, EPI_ISL_482044, EPI_ISL_482054, EPI_ISL_482055 | Regional Virus Laboratory, Belfast Health and Social Care Trust | Wellcome Sanger Institute for the COVID-19 Genomics UK (COG-UK) consortium | Conall McCaughey, James McKenna, Tanya Curran, Susan Feeney, Alison Watt, Ciara Cox, Mairead Connor, Zoltan Molnar, David Simpson, Derek Fairley; and Alex Alderton, Roberto Amato, Sonia Goncalves, Ewan Harrison, David K. Jackson, Ian Johnston, Dominic Kwiatkowski, Cordelia Langford, John Sillitoe on behalf of the Wellcome Sanger Institute COVID-19 Surveillance Team (http://www.sanger.ac.uk/covid-team) |
| EPI_ISL_482060, EPI_ISL_482062, EPI_ISL_482063, EPI_ISL_482064, EPI_ISL_482067, EPI_ISL_482068 | The Department of Microbiology, Torbay and South Devon NHS Foundation Trust | Wellcome Sanger Institute for the COVID-19 Genomics UK (COG-UK) consortium | Amy Hurd, Sophie Lloyd, Anthony Mogridge, Jack Howe, Helen Brown, Gary Booth, Mel Brown, Cheryl Bailiss, Michelle Harrison and Alex Alderton, Roberto Amato, Sonia Goncalves, Ewan Harrison, David K. Jackson, Ian Johnston, Dominic Kwiatkowski, Cordelia Langford, John Sillitoe on behalf of the Wellcome Sanger Institute COVID-19 Surveillance Team (http://www.sanger.ac.uk/covid-team) |
| EPI_ISL_482077, EPI_ISL_482105 | Regional Virus Laboratory, Belfast Health and Social Care Trust | Wellcome Sanger Institute for the COVID-19 Genomics UK (COG-UK) consortium | Conall McCaughey, James McKenna, Tanya Curran, Susan Feeney, Alison Watt, Ciara Cox, Mairead Connor, Zoltan Molnar, David Simpson, Derek Fairley; and Alex Alderton, Roberto Amato, Sonia Goncalves, Ewan Harrison, David K. Jackson, Ian Johnston, Dominic Kwiatkowski, Cordelia Langford, John Sillitoe on behalf of the Wellcome Sanger Institute COVID-19 Surveillance Team (http://www.sanger.ac.uk/covid-team) |
| EPI_ISL_482120, EPI_ISL_482121, EPI_ISL_482122, EPI_ISL_482123, EPI_ISL_482127, EPI_ISL_482129, EPI_ISL_482130 | The Department of Microbiology, Torbay and South Devon NHS Foundation Trust | Wellcome Sanger Institute for the COVID-19 Genomics UK (COG-UK) consortium | Amy Hurd, Sophie Lloyd, Anthony Mogridge, Jack Howe, Helen Brown, Gary Booth, Mel Brown, Cheryl Bailiss, Michelle Harrison and Alex Alderton, Roberto Amato, Sonia Goncalves, Ewan Harrison, David K. Jackson, Ian Johnston, Dominic Kwiatkowski, Cordelia Langford, John Sillitoe on behalf of the Wellcome Sanger Institute COVID-19 Surveillance Team (http://www.sanger.ac.uk/covid-team) |
| EPI_ISL_482160, EPI_ISL_482161, EPI_ISL_482162, EPI_ISL_482163, EPI_ISL_482164, EPI_ISL_482165, EPI_ISL_482166, EPI_ISL_482167, EPI_ISL_482168, EPI_ISL_482169, EPI_ISL_482170, EPI_ISL_482171, EPI_ISL_482172, EPI_ISL_482173, EPI_ISL_482174, EPI_ISL_482175, EPI_ISL_482176, EPI_ISL_482177, EPI_ISL_482178, EPI_ISL_482179, EPI_ISL_482180, EPI_ISL_482181, EPI_ISL_482182, EPI_ISL_482183, EPI_ISL_482184, EPI_ISL_482185, EPI_ISL_482186, EPI_ISL_482187, EPI_ISL_482188, EPI_ISL_482189, EPI_ISL_482190, EPI_ISL_482191, EPI_ISL_482192, EPI_ISL_482193, EPI_ISL_482194, EPI_ISL_482195, EPI_ISL_482196, EPI_ISL_482197, EPI_ISL_482198, EPI_ISL_482199, EPI_ISL_482200, EPI_ISL_482201, EPI_ISL_482202, EPI_ISL_482203, EPI_ISL_482204, EPI_ISL_482205, EPI_ISL_482206, EPI_ISL_482207, EPI_ISL_482208, EPI_ISL_482209, EPI_ISL_482210, EPI_ISL_482211, EPI_ISL_482212, EPI_ISL_482213, EPI_ISL_482214, EPI_ISL_482215, EPI_ISL_482216, EPI_ISL_482217, EPI_ISL_482218 | | | |
| see above | University College London, Great Ormond Street Hospital for Children NHS Foundation Trust, Imperial College Healthcare NHS Trust | Wellcome Sanger Institute for the COVID-19 Genomics UK (COG-UK) consortium | Sergi Castellano, Rachel Williams, Mark Kristiansen, Paola Resende Silva, Sunando Roy, Tony Brooks, Helena Tutill, Paola Niola, Patricia Dyal, Charlotte Williams, Leysa Forrest, Yasmin Panchbhaya, Jacqueline Findlay, Sam Weeks, Julianne Brown, Kathryn Harris, Paul Randell, James Price, Alison Holmes, Judith Breuer and Alex Alderton, Roberto Amato, Sonia Goncalves, Ewan Harrison, David K. Jackson, Ian Johnston, Dominic Kwiatkowski, Cordelia Langford, John Sillitoe on behalf of the Wellcome Sanger Institute COVID-19 Surveillance Team (http://www.sanger.ac.uk/covid-team) |
| EPI_ISL_482358, EPI_ISL_482359, EPI_ISL_482360, EPI_ISL_482361, EPI_ISL_482362, EPI_ISL_482363, EPI_ISL_482364, EPI_ISL_482365, EPI_ISL_482366, EPI_ISL_482367, EPI_ISL_482368, EPI_ISL_482369, EPI_ISL_482370, EPI_ISL_482371, EPI_ISL_482372, EPI_ISL_482373, EPI_ISL_482374, EPI_ISL_482375, EPI_ISL_482376, EPI_ISL_482377, EPI_ISL_482378, EPI_ISL_482379, EPI_ISL_482380, EPI_ISL_482381, EPI_ISL_482382 | | | |
| see above | Providence St. Joseph Health Molecular Genomics Laboratory | Providence St. Joseph Health Molecular Genomics Laboratory | Alexa K Dowdell, Brian D Piening, Fred L Robinson, Carlo B Bifulco, Mary Campbell |
| EPI_ISL_482735, EPI_ISL_482736, EPI_ISL_482739 | LNR National Reference Laboratory, Mohammed VI University of Health Sciences | Medical Biotechnology Laboratory, Rabat Medical and Pharmacy School, Mohammed The Vth University in Rabat | Meriem LAAMARTI, Souad KARTTI, Rokia LAAMARTI , M.W. CHEMAO-ELFIHRI, Loubna ALLAM, Mouna OUADGHIRI, Imane SMYEJ, Jalila RAHOUI, Houda BENRAHMA, Jalil El ATAR, Idrissa DIAWARA, Rachid EL JAOUDI, Laila SBABOU, Chakib NEJJARI, Saaid AMZAZI, Rachid MENTAG, Lahcen BELYAMANI and Azeddine IBRAHIMI |
| EPI_ISL_482820 | Centre de Recerca en Sanitat Animal (IRTA-CReSA) | IrsiCaixa AIDS Research Lab | J. Segalés, M. Puig, J. Rodon, C. Avila-Nieto, J. Carrillo, G. Cantero, M.T. Terrón, S. Cruz, M. Parera ,M. Noguera-Julián, N. Izquierdo-Useros, V. Guallar, E. Vidal, A. Valencia, I. Blanco, J. Blanco, B. Clotet, J. Vergara-Alert |
| EPI_ISL_482968, EPI_ISL_482969 | Mayo Clinic & Mayo Clinic Laboratories | Minnesota Department of Health, Public Health Laboratory | Matt Plumb, Jacob Garfin, and Xiong Wang |
| EPI_ISL_483141, EPI_ISL_483142, EPI_ISL_483143, EPI_ISL_483144, EPI_ISL_483145, EPI_ISL_483146, EPI_ISL_483147, EPI_ISL_483152, EPI_ISL_483153 | Robert Koch Institute, ZBS1 Highly Pathogenic Viruses, Berlin, Germany | Robert Koch Institute, Bioinformatics MF1, Berlin, Germany | Janine Michel, Andrea Thuermer, Oliver Drechsel, Rene Kmiecinski, Stephan Fuchs, Max v. Kleist, Andreas Nitsche |
| EPI_ISL_483212, EPI_ISL_483252, EPI_ISL_483253, EPI_ISL_483255, EPI_ISL_483257, EPI_ISL_483262, EPI_ISL_483263, EPI_ISL_483264, EPI_ISL_483265, EPI_ISL_483266, EPI_ISL_483267, EPI_ISL_483268, EPI_ISL_483269, EPI_ISL_483270, EPI_ISL_483271, EPI_ISL_483273, EPI_ISL_483275, EPI_ISL_483277, EPI_ISL_483280, EPI_ISL_483291, EPI_ISL_483296, EPI_ISL_483393, EPI_ISL_483397, EPI_ISL_483412, EPI_ISL_483422, EPI_ISL_483449, EPI_ISL_483450, EPI_ISL_483451, EPI_ISL_483452, EPI_ISL_483453, EPI_ISL_483454, EPI_ISL_483455, EPI_ISL_483456, EPI_ISL_483463, EPI_ISL_483464, EPI_ISL_483466, EPI_ISL_483467, EPI_ISL_483469, EPI_ISL_483470, EPI_ISL_483472, EPI_ISL_483473, EPI_ISL_483474, EPI_ISL_483475 | | | |
| see above | UC San Diego Center for Advanced Laboratory Medicine | Andersen lab at Scripps Research | SEARCH Alliance San Diego with David Pride, Ji H Shin |
| EPI_ISL_483601 | National Public Health Laboratory, National Centre for Infectious Diseases | National Public Health Laboratory, National Centre for Infectious Diseases | Mak TM, Octavia S, Zhou Z, Chavatte JM, Cui L, Lin RTP |
| EPI_ISL_483677, EPI_ISL_483682, EPI_ISL_483685 | University Hospital Zurich | Department of Biosystems Science and Engineering, ETH Zürich | Christian Beisel, Sarah Nadeau, Ivan Topolsky, Pedro Ferreira, Philipp Jablonski, Susana Posada-Céspedes, Tobias Schär, Ina Nissen, Natascha Santacroce, Elodie Burcklen, Julia Martinez-Gomez, Phil Cheng, Mitch Levesque, Philipp Bosshard, Niko Beerenwinkel, Tanja Stadler |
| EPI_ISL_484355, EPI_ISL_484363, EPI_ISL_484364, EPI_ISL_484365, EPI_ISL_484368, EPI_ISL_484369 | Lincolnshire Hospitals and DeepSeq Nottingham | COVID-19 Genomics UK (COG-UK) Consortium | Nichola Duckworth, Tim Sloan, Sarah Walsh, Jonathan Ball, Patrick McClure, Joeseph Chappell, Nadine Holmes, Matthew Carlisle, Christopher Moore, Fei Sang, Johnny Debebe, Victoria Wright, Matthew Loose |
| EPI_ISL_484374 | Queens Medical Centre, Clinical Microbiology Department / DeepSeq Nottingham | COVID-19 Genomics UK (COG-UK) Consortium | Gemma Clark, Wendy Smith, Manjinder Khakh, Vicki M Fleming, Michelle M Lister, Hannah Howson-Wells, Jonathan Ball, Patrick McClure, Joseph Chappell, Theocharis Tsoleridis, Nadine Holmes, Matthew Carlisle, Christopher Moore, Fei Sang, Johnny Debebe, Victoria Wright, Matthew Loose |
| EPI_ISL_484392, EPI_ISL_484394, EPI_ISL_484395, EPI_ISL_484398, EPI_ISL_484401, EPI_ISL_484403 | Lincolnshire Hospitals and DeepSeq Nottingham | COVID-19 Genomics UK (COG-UK) Consortium | Nichola Duckworth, Tim Sloan, Sarah Walsh, Jonathan Ball, Patrick McClure, Joeseph Chappell, Nadine Holmes, Matthew Carlisle, Christopher Moore, Fei Sang, Johnny Debebe, Victoria Wright, Matthew Loose |
| EPI_ISL_484408, EPI_ISL_484410, EPI_ISL_484411, EPI_ISL_484412, EPI_ISL_484413, EPI_ISL_484414 | Centre for Enzyme Innovation, University of Portsmouth / Translational Research Laboratory, Portsmouth Hospitals NHS Trust | COVID-19 Genomics UK (COG-UK) Consortium | Angela Beckett,Yann Bourgeois,Garry Scarlett,Sharon Glaysher,Scott Elliott,Kelly Bicknell,Robert Impey,Allyson Lloyd,Sarah Wyllie,Ethan Butcher,Anoop Chauhan,Samuel Robson |
| EPI_ISL_484463, EPI_ISL_484472, EPI_ISL_484477, EPI_ISL_484514, EPI_ISL_484518 | Virology Department, Sheffield Teaching Hospitals NHS Foundation Trust/Department of Infection, Immunity and Cardiovascular Disease, The Medical School, University of Sheffield | COVID-19 Genomics UK (COG-UK) Consortium | Thushan de Silva, Matthew Parker, Nikki Smith, Adri Angyal, Rebecca Brown, Luke Green, Rachel Tucker, Paul Parsons, Danielle Groves, Katie Johnson, Laura Carrilero, Alex Keeley, Dave Partridge, Matthew Wyles, Benjamin Lindsey, Mehmet Yavuz, Mohammad Raza, Cariad Evans |
| EPI_ISL_484652, EPI_ISL_484653, EPI_ISL_484654, EPI_ISL_484655, EPI_ISL_484656 | West of Scotland Specialist Virology Centre, NHSGGC / MRC-University of Glasgow Centre for Virus Research | COVID-19 Genomics UK (COG-UK) Consortium | Ana da Silva Filipe, Natasha Johnson, Kathy Smollett, Daniel Mair, Stephen Carmichael, Lily Tong, Jenna Nichols, Elihu Aranday-Cortes, Kirstyn Brunker, Yasmin Parr, Alice Broos, Kyriaki Nomikou; Sarah McDonald, Marc Niebel, Patawee Asamaphan; Richard Orton, Joseph Hughes, Sreenu Vattipally, David L Robertson; Alasdair MacLean, Rory Gunson; Kathy Li, Natasha Jesudason, Rajiv Shah, James Shepherd, Antonia Ho, Emma Thomson |
| EPI_ISL_485043, EPI_ISL_485044, EPI_ISL_485045, EPI_ISL_485046, EPI_ISL_485047, EPI_ISL_485048, EPI_ISL_485049, EPI_ISL_485050, EPI_ISL_485051, EPI_ISL_485052, EPI_ISL_485053, EPI_ISL_485054, EPI_ISL_485055, EPI_ISL_485056, EPI_ISL_485057, EPI_ISL_485058, EPI_ISL_485059, EPI_ISL_485060, EPI_ISL_485061, EPI_ISL_485062, EPI_ISL_485063, EPI_ISL_485064, EPI_ISL_485065, EPI_ISL_485066, EPI_ISL_485067, EPI_ISL_485068, EPI_ISL_485069, EPI_ISL_485070, EPI_ISL_485071, EPI_ISL_485072, EPI_ISL_485073, EPI_ISL_485074, EPI_ISL_485075, EPI_ISL_485076, EPI_ISL_485077, EPI_ISL_485078, EPI_ISL_485079, EPI_ISL_485080, EPI_ISL_485081, EPI_ISL_485082, EPI_ISL_485083, EPI_ISL_485084, EPI_ISL_485085, EPI_ISL_485086, EPI_ISL_485087, EPI_ISL_485088, EPI_ISL_485089, EPI_ISL_485090, EPI_ISL_485091, EPI_ISL_485092, EPI_ISL_485093, EPI_ISL_485094, EPI_ISL_485095, EPI_ISL_485096, | | | |

| | | | |
|---|---|---|---|
| EPI_ISL_485097, EPI_ISL_485098, EPI_ISL_485099, EPI_ISL_485100, EPI_ISL_485105, EPI_ISL_485106, EPI_ISL_485107, EPI_ISL_485108, EPI_ISL_485109, EPI_ISL_485110, EPI_ISL_485111, EPI_ISL_485112, EPI_ISL_485113, EPI_ISL_485114, EPI_ISL_485115, EPI_ISL_485116, EPI_ISL_485211, EPI_ISL_485213, EPI_ISL_485214, EPI_ISL_485215, EPI_ISL_485216, EPI_ISL_485217, EPI_ISL_485218, EPI_ISL_485219, EPI_ISL_485220, EPI_ISL_485221, EPI_ISL_485222, EPI_ISL_485223, EPI_ISL_485224, EPI_ISL_485225, EPI_ISL_485226, EPI_ISL_485227, EPI_ISL_485228, EPI_ISL_485229, EPI_ISL_485230, EPI_ISL_485231, EPI_ISL_485232, EPI_ISL_485233, EPI_ISL_485234, EPI_ISL_485235, EPI_ISL_485236, EPI_ISL_485237, EPI_ISL_485238, EPI_ISL_485239, EPI_ISL_485240, EPI_ISL_485241, EPI_ISL_485242, EPI_ISL_485243, EPI_ISL_485244, EPI_ISL_485245, EPI_ISL_485246, EPI_ISL_485247 | | | |
| see above | River Road Testing Lab | Ginkgo Bioworks Clinical Laboratory | Rebecca C. Christofferson, Stephania A. Cormier, Luan V. Dinh, E. Handly Mayton, Hollis R. O'Neil, Thaya Stoufflet, Malaika Mckenzie-Bennett, James McGann, Jim Griffin, Keith Robison, Alex Plocik, Becky Schilling, Rebecca Littlefield, Michelle Spencer, Birgitte Simen |
| EPI_ISL_485584, EPI_ISL_485596, EPI_ISL_485598, EPI_ISL_485599, EPI_ISL_485600, EPI_ISL_485601 | Instituto de diagnóstico y Referencia Epidemiologicos (INDRE) | Instituto de diagnóstico y Referencia Epidemiologicos (INDRE) | Barrera-Badillo,G., Ramirez-Gonzalez,E. |
| EPI_ISL_485874, EPI_ISL_485875, EPI_ISL_485876, EPI_ISL_485877, EPI_ISL_485878, EPI_ISL_485879, EPI_ISL_485880, EPI_ISL_485881, EPI_ISL_485882, EPI_ISL_485883, EPI_ISL_485884, EPI_ISL_485885, EPI_ISL_485892, EPI_ISL_485893, EPI_ISL_485894, EPI_ISL_485895, EPI_ISL_485896 | | | |
| see above | River Road Testing Lab | Ginkgo Bioworks Clinical Laboratory | Rebecca C. Christofferson, Stephania A. Cormier, Luan V. Dinh, E. Handly Mayton, Hollis R. O'Neil, Thaya Stoufflet, Malaika Mckenzie-Bennett, James McGann, Jim Griffin, Keith Robison, Alex Plocik, Becky Schilling, Rebecca Littlefield, Michelle Spencer, Birgitte Simen |
| EPI_ISL_486014, EPI_ISL_486015, EPI_ISL_486016, EPI_ISL_486019, EPI_ISL_486022, EPI_ISL_486023, EPI_ISL_486024, EPI_ISL_486112 | UW Virology Lab | UW Virology Lab | Pavitra Roychoudhury, Hong Xie, Lasata Shrestha, Amin Addetia, Truong Nguyen, Victoria M Rachleff, Meei-Li Huang, Keith R Jerome, Alexander Greninger |
| EPI_ISL_486133, EPI_ISL_486134, EPI_ISL_486135, EPI_ISL_486136, EPI_ISL_486137, EPI_ISL_486138, EPI_ISL_486139, EPI_ISL_486140, EPI_ISL_486141, EPI_ISL_486142, EPI_ISL_486143, EPI_ISL_486144, EPI_ISL_486145, EPI_ISL_486146, EPI_ISL_486147, EPI_ISL_486148, EPI_ISL_486149, EPI_ISL_486155, EPI_ISL_486156, EPI_ISL_486157, EPI_ISL_486158, EPI_ISL_486159, EPI_ISL_486191, EPI_ISL_486192, EPI_ISL_486193, EPI_ISL_486194, EPI_ISL_486195, EPI_ISL_486196, EPI_ISL_486197, EPI_ISL_486198, EPI_ISL_486199, EPI_ISL_486200, EPI_ISL_486201, EPI_ISL_486202, EPI_ISL_486203, EPI_ISL_486204, EPI_ISL_486205, EPI_ISL_486206, EPI_ISL_486207, EPI_ISL_486208, EPI_ISL_486209, EPI_ISL_486210, EPI_ISL_486211, EPI_ISL_486212, EPI_ISL_486213, EPI_ISL_486214, EPI_ISL_486215, EPI_ISL_486216, EPI_ISL_486217 | | | |
| see above | Orange County Public Health Laboratory | Chan-Zuckerberg Biohub | CZB Cliahub Consortium |
| EPI_ISL_486293, EPI_ISL_486294, EPI_ISL_486295, EPI_ISL_486296, EPI_ISL_486297, EPI_ISL_486298, EPI_ISL_486299, EPI_ISL_486300, EPI_ISL_486301, EPI_ISL_486302, EPI_ISL_486303, EPI_ISL_486304, EPI_ISL_486305, EPI_ISL_486306, EPI_ISL_486307, EPI_ISL_486308, EPI_ISL_486309, EPI_ISL_486336, EPI_ISL_486337, EPI_ISL_486338 | | | |
| see above | San Joaquin County Public Health Lab | Chan-Zuckerberg Biohub | CZB Cliahub Consortium |
| EPI_ISL_486383 | CV Raman Hospital | Department of Neurovirology, National Institute of Mental Health and Neuroscience (NIMHANS) | Chitra Pattabiraman, Vijayalakshmi Reddy, Harsha PK, Risha Rasheed, Shafeeq S Hameed, Manjunatha Venkataswamy, Anita Desai, Ravi Vasanthapuram |
| EPI_ISL_486394 | MIMS | Department of Neurovirology, National Institute of Mental Health and Neuroscience (NIMHANS) | Chitra Pattabiraman, Vijayalakshmi Reddy, Harsha PK, Risha Rasheed, Shafeeq S Hameed, Manjunatha Venkataswamy, Anita Desai, Ravi Vasanthapuram |
| EPI_ISL_486395 | BIMS | Department of Neurovirology, National Institute of Mental Health and Neuroscience (NIMHANS) | Chitra Pattabiraman, Vijayalakshmi Reddy, Harsha PK, Risha Rasheed, Shafeeq S Hameed, Manjunatha Venkataswamy, Anita Desai, Ravi Vasanthapuram |
| EPI_ISL_486396 | Jayanagar General Hospital to Victoria Hospital | Department of Neurovirology, National Institute of Mental Health and Neuroscience (NIMHANS) | Chitra Pattabiraman, Vijayalakshmi Reddy, Harsha PK, Risha Rasheed, Shafeeq S Hameed, Manjunatha Venkataswamy, Anita Desai, Ravi Vasanthapuram |
| EPI_ISL_486397 | KC General Hospital | Department of Neurovirology, National Institute of Mental Health and Neuroscience (NIMHANS) | Chitra Pattabiraman, Vijayalakshmi Reddy, Harsha PK, Risha Rasheed, Shafeeq S Hameed, Manjunatha Venkataswamy, Anita Desai, Ravi Vasanthapuram |
| EPI_ISL_486398 | MIMS | Department of Neurovirology, National Institute of Mental Health and Neuroscience (NIMHANS) | Chitra Pattabiraman, Vijayalakshmi Reddy, Harsha PK, Risha Rasheed, Shafeeq S Hameed, Manjunatha Venkataswamy, Anita Desai, Ravi Vasanthapuram |
| EPI_ISL_486857, EPI_ISL_486858, EPI_ISL_486859, EPI_ISL_486860, EPI_ISL_486861, EPI_ISL_486862, EPI_ISL_486863, EPI_ISL_486864, EPI_ISL_486866, EPI_ISL_486867, EPI_ISL_486868, EPI_ISL_486869, EPI_ISL_486870, EPI_ISL_486871, EPI_ISL_486873 | | | |
| see above | Institut Pasteur Dakar | Institut Pasteur de Dakar | Ndongo Dia, Moussa Moise Diagne, Mamadou Diop, Marie Henriette Dior Ndione, Mamadou Malado Jallow, Safietou Sanke, Ousmane Faye, Amadou Alpha Sall. |
| EPI_ISL_486881 | CV Raman Hospital | Department of Neurovirology, National Institute of Mental Health and Neuroscience (NIMHANS) | Chitra Pattabiraman, Vijayalakshmi Reddy, Harsha PK, Risha Rasheed, Shafeeq S Hameed, Manjunatha Venkataswamy, Anita Desai, Ravi Vasanthapuram |
| EPI_ISL_487090, EPI_ISL_487099, EPI_ISL_487105, EPI_ISL_487108 | Nigeria Centre for Disease Control (NCDC) | African Centre of Excellence for Genomics of Infectious Diseases (ACEGID), Redeemer's University, Ede, Osun State, Nigeria | Oluniyi P.E., Ajogbasile F.V., Kayode A., Oguzie J., Olawoye I., Uwanibe J., Olumade T., Folarin O.A., Ihekweazu C., Happi C.T. |
| EPI_ISL_487113 | Nigeria Centre for Disease Control (NCDC) | Redeemer's University, ACEGID | Oluniyi P.E., Ajogbasile F.V., Kayode A., Oguzie J., Olawoye I., Uwanibe J., Olumade T., Folarin O.A., Ihekweazu C., Happi C.T. |
| EPI_ISL_487193, EPI_ISL_487194, EPI_ISL_487195, EPI_ISL_487196, EPI_ISL_487197, EPI_ISL_487198, EPI_ISL_487199, EPI_ISL_487200, EPI_ISL_487201, EPI_ISL_487202, EPI_ISL_487203, EPI_ISL_487204, EPI_ISL_487205, EPI_ISL_487206, EPI_ISL_487207, EPI_ISL_487208, EPI_ISL_487209, EPI_ISL_487210, EPI_ISL_487211, EPI_ISL_487212, EPI_ISL_487213, EPI_ISL_487214, EPI_ISL_487215, EPI_ISL_487216, EPI_ISL_487217, EPI_ISL_487218, EPI_ISL_487219, EPI_ISL_487220, EPI_ISL_487221, EPI_ISL_487222, EPI_ISL_487223, EPI_ISL_487224, EPI_ISL_487225, EPI_ISL_487226, EPI_ISL_487244 | | | |
| see above | Utah Public Health Laboratory | Utah Public Health Laboratory | Heidi Butz, Erin Young, Kelly Oakeson |
| EPI_ISL_487525, EPI_ISL_487529, EPI_ISL_487534, EPI_ISL_487537, EPI_ISL_487540, EPI_ISL_487546, EPI_ISL_487550, EPI_ISL_487556, EPI_ISL_487557, EPI_ISL_487558, EPI_ISL_487559, EPI_ISL_487562, EPI_ISL_487564, EPI_ISL_487568, EPI_ISL_487572, EPI_ISL_487578, EPI_ISL_487581, EPI_ISL_487582, EPI_ISL_487584, EPI_ISL_487585, EPI_ISL_487589, EPI_ISL_487595, EPI_ISL_487597, EPI_ISL_487600, EPI_ISL_487606, EPI_ISL_487607, EPI_ISL_487614, EPI_ISL_487615, EPI_ISL_487616, EPI_ISL_487621, EPI_ISL_487625, EPI_ISL_487626, EPI_ISL_487627, EPI_ISL_487628, EPI_ISL_487630, EPI_ISL_487631, EPI_ISL_487632, EPI_ISL_487633 | | | |
| see above | University College London, Great Ormond Street Hospital for Children NHS Foundation Trust, Imperial College Healthcare NHS Trust | Wellcome Sanger Institute for the COVID-19 Genomics UK (COG-UK) consortium | Sergi Castellano, Rachel Williams, Mark Kristiansen, Paola Resende Silva, Sunando Roy, Tony Brooks, Helena Tutill, Paola Niola, Patricia Dyal, Charlotte Williams, Leysa Forrest, Yasmin Panchbhaya, Jacqueline Findlay, Sam Weeks, Julianne Brown, Kathryn Harris, Paul Randell, James Price, Alison Holmes, Judith Breuer and Alex Alderton, Roberto Amato, Sonia Goncalves, Ewan Harrison, David K. Jackson, Ian Johnston, Dominic Kwiatkowski, Cordelia Langford, John Sillitoe on behalf of the Wellcome Sanger Institute COVID-19 Surveillance Team (http://www.sanger.ac.uk/covid-team) |
| EPI_ISL_487649, EPI_ISL_487677, EPI_ISL_487693, EPI_ISL_487701, EPI_ISL_487705, EPI_ISL_487723, EPI_ISL_487739, EPI_ISL_487749, EPI_ISL_487752, EPI_ISL_487754, EPI_ISL_487755, EPI_ISL_487756, EPI_ISL_487760, EPI_ISL_487761 | | | |
| see above | Virology Department, Royal Infirmary of Edinburgh, NHS Lothian / School of Biological Sciences, University of Edinburgh | Wellcome Sanger Institute for the COVID-19 Genomics UK (COG-UK) consortium | McHugh M, Dewar R, Rooke S, O'Toole Á, Scher E, Hill V, McCrone JT, Colquhoun R, Yu X, Jackson B, Rambaut A, Templeton K and Alex Alderton, Roberto Amato, Sonia Goncalves, Ewan Harrison, David K. Jackson, Ian Johnston, Dominic Kwiatkowski, Cordelia Langford, John Sillitoe on behalf of the Wellcome Sanger Institute COVID-19 Surveillance Team (http://www.sanger.ac.uk/covid-team) |
| EPI_ISL_487767 | Virology Department, Royal Infirmary of Edinburgh, NHS Lothian / School of Biological Sciences, University of Edinburgh | Wellcome Sanger Institute for the COVID-19 Genomics UK (COG-UK) Consortium | McHugh M, Dewar R, Rooke S, O'Toole Á, Scher E, Hill V, McCrone JT, Colquhoun R, Yu X, Jackson B, Rambaut A, Templeton K and Alex Alderton, Roberto Amato, Sonia Goncalves, Ewan Harrison, David K. Jackson, Ian Johnston, Dominic Kwiatkowski, Cordelia Langford, John Sillitoe on behalf of the Wellcome Sanger Institute COVID-19 Surveillance Team |
| EPI_ISL_487778, EPI_ISL_487779, EPI_ISL_487781, EPI_ISL_487797, EPI_ISL_487799, EPI_ISL_487801, EPI_ISL_487818, EPI_ISL_487820, EPI_ISL_487835, EPI_ISL_487837, EPI_ISL_487839, EPI_ISL_487841, EPI_ISL_487844, EPI_ISL_487857, EPI_ISL_487859, EPI_ISL_487878, EPI_ISL_487879, EPI_ISL_487895, EPI_ISL_487900, EPI_ISL_487910, EPI_ISL_487917, EPI_ISL_487932, EPI_ISL_487938, EPI_ISL_487941, EPI_ISL_487944, EPI_ISL_487946, EPI_ISL_487953, EPI_ISL_487954, EPI_ISL_487956, EPI_ISL_487978, EPI_ISL_487979, EPI_ISL_487984, EPI_ISL_487997, EPI_ISL_488000, EPI_ISL_488001 | | | |
| see above | Virology Department, Royal Infirmary of Edinburgh, NHS Lothian / School of Biological Sciences, University of Edinburgh | Wellcome Sanger Institute for the COVID-19 Genomics UK (COG-UK) consortium | McHugh M, Dewar R, Rooke S, O'Toole Á, Scher E, Hill V, McCrone JT, Colquhoun R, Yu X, Jackson B, Rambaut A, Templeton K and Alex Alderton, Roberto Amato, Sonia Goncalves, Ewan Harrison, David K. Jackson, Ian Johnston, Dominic Kwiatkowski, Cordelia Langford, John Sillitoe on behalf of the Wellcome Sanger Institute COVID-19 Surveillance Team (http://www.sanger.ac.uk/covid-team) |
| EPI_ISL_488003, EPI_ISL_488004, EPI_ISL_488006, EPI_ISL_488007, EPI_ISL_488008, EPI_ISL_488011, EPI_ISL_488012, EPI_ISL_488013, EPI_ISL_488018, EPI_ISL_488019, EPI_ISL_488021, EPI_ISL_488022, EPI_ISL_488023, EPI_ISL_488024, EPI_ISL_488026, EPI_ISL_488027, EPI_ISL_488028, EPI_ISL_488029, EPI_ISL_488030, EPI_ISL_488031, EPI_ISL_488034, EPI_ISL_488036, EPI_ISL_488046, EPI_ISL_488048, EPI_ISL_488050, EPI_ISL_488052, EPI_ISL_488056, EPI_ISL_488060, EPI_ISL_488062, EPI_ISL_488063, EPI_ISL_488068, EPI_ISL_488070, EPI_ISL_488078, EPI_ISL_488085, EPI_ISL_488086, EPI_ISL_488090, EPI_ISL_488093, EPI_ISL_488099, EPI_ISL_488100, EPI_ISL_488101, EPI_ISL_488102, EPI_ISL_488106, EPI_ISL_488107, EPI_ISL_488114, EPI_ISL_488117, EPI_ISL_488123, EPI_ISL_488129, EPI_ISL_488130, EPI_ISL_488132, EPI_ISL_488136, EPI_ISL_488140, EPI_ISL_488141, EPI_ISL_488144, EPI_ISL_488145, EPI_ISL_488149, EPI_ISL_488151, EPI_ISL_488154, EPI_ISL_488156, EPI_ISL_488159, EPI_ISL_488170, EPI_ISL_488174, EPI_ISL_488175 | | | |
| see above | NU-OMICS DNA Sequencing research facility, Northumbria University | Wellcome Sanger Institute for the COVID-19 Genomics UK (COG-UK) consortium | Chris Duncan, Sheia Waugh, Shirelle Burton-Fanning, Gary Eltringham, Jennifer Collins, Brendan Payne, Yusri Taha, Emma Swindells, Jane Greenaway, Edward Barton, Garren Scott, Debra Padgett, Clive Graham, Sarah Essex, Steve Ligget, Paul Baker, Lynn Dover, Wen Yew, Gary Black, John Allan, Joshua Loh, Greg Young, Matthew Bashton, Andrew Nelson, Darren Smith and Alex Alderton, Roberto Amato, Sonia Goncalves, Ewan Harrison, David K. |

| | | | |
|---|---|---|---|
| | | | Jackson, Ian Johnston, Dominic Kwiatkowski, Cordelia Langford, John Sillitoe on behalf of the Wellcome Sanger Institute COVID-19 Surveillance Team (http://www.sanger.ac.uk/covid-team) |
| EPI_ISL_488188, EPI_ISL_488191, EPI_ISL_488197, EPI_ISL_488198, EPI_ISL_488203, EPI_ISL_488205, EPI_ISL_488206, EPI_ISL_488207, EPI_ISL_488208, EPI_ISL_488209, EPI_ISL_488210, EPI_ISL_488212, EPI_ISL_488213, EPI_ISL_488217, EPI_ISL_488220, EPI_ISL_488222, EPI_ISL_488225, EPI_ISL_488227, EPI_ISL_488229, EPI_ISL_488231, EPI_ISL_488234, EPI_ISL_488235, EPI_ISL_488236, EPI_ISL_488243, EPI_ISL_488245, EPI_ISL_488246, EPI_ISL_488247, EPI_ISL_488248, EPI_ISL_488249, EPI_ISL_488252, EPI_ISL_488255, EPI_ISL_488256, EPI_ISL_488257, EPI_ISL_488259, EPI_ISL_488260, EPI_ISL_488263, EPI_ISL_488264, EPI_ISL_488271, EPI_ISL_488274, EPI_ISL_488275, EPI_ISL_488276, EPI_ISL_488283, EPI_ISL_488284, EPI_ISL_488285, EPI_ISL_488286, EPI_ISL_488289, EPI_ISL_488290, EPI_ISL_488292, EPI_ISL_488294, EPI_ISL_488297, EPI_ISL_488299, EPI_ISL_488301, EPI_ISL_488302, EPI_ISL_488304, EPI_ISL_488307, EPI_ISL_488308, EPI_ISL_488313, EPI_ISL_488314, EPI_ISL_488316, EPI_ISL_488317, EPI_ISL_488318, EPI_ISL_488319, EPI_ISL_488320, EPI_ISL_488322, EPI_ISL_488323, EPI_ISL_488327, EPI_ISL_488330, EPI_ISL_488332, EPI_ISL_488333, EPI_ISL_488336, EPI_ISL_488338, EPI_ISL_488339, EPI_ISL_488343, EPI_ISL_488344, EPI_ISL_488346, EPI_ISL_488348, EPI_ISL_488352, EPI_ISL_488353, EPI_ISL_488354, EPI_ISL_488357, EPI_ISL_488360, EPI_ISL_488366, EPI_ISL_488368, EPI_ISL_488369, EPI_ISL_488370, EPI_ISL_488375, EPI_ISL_488376, EPI_ISL_488377, EPI_ISL_488378, EPI_ISL_488379, EPI_ISL_488380, EPI_ISL_488381, EPI_ISL_488383, EPI_ISL_488384, EPI_ISL_488387, EPI_ISL_488391, EPI_ISL_488393, EPI_ISL_488395, EPI_ISL_488396, EPI_ISL_488397, EPI_ISL_488398, EPI_ISL_488401, EPI_ISL_488402, EPI_ISL_488403, EPI_ISL_488405, EPI_ISL_488406, EPI_ISL_488407 | | | |
| see above | PHE South West Regional Laboratory, National Infection Service | Wellcome Sanger Institute for the COVID-19 Genomics UK (COG-UK) consortium | Stephanie Hutchings, Hannah Pymont, Dr Peter Muir, Barry Vipond, Rich Hopes; and Alex Alderton, Roberto Amato, Sonia Goncalves, Ewan Harrison, David K. Jackson, Ian Johnston, Dominic Kwiatkowski, Cordelia Langford, John Sillitoe on behalf of the Wellcome Sanger Institute COVID-19 Surveillance Team (http://www.sanger.ac.uk/covid-team) |
| EPI_ISL_488409 | PHE South West Regional Laboratory, National Infection Service | Wellcome Sanger Institute for the COVID-19 Genomics UK (COG-UK) Consortium | Stephanie Hutchings, Hannah Pymont, Dr Peter Muir, Barry Vipond, Rich Hopes; and Alex Alderton, Roberto Amato, Sonia Goncalves, Ewan Harrison, David K. Jackson, Ian Johnston, Dominic Kwiatkowski, Cordelia Langford, John Sillitoe on behalf of the Wellcome Sanger Institute COVID-19 Surveillance Team |
| EPI_ISL_488412, EPI_ISL_488415, EPI_ISL_488418, EPI_ISL_488421, EPI_ISL_488426, EPI_ISL_488429, EPI_ISL_488430, EPI_ISL_488431, EPI_ISL_488433, EPI_ISL_488436, EPI_ISL_488437, EPI_ISL_488438, EPI_ISL_488440, EPI_ISL_488441, EPI_ISL_488442, EPI_ISL_488443, EPI_ISL_488448, EPI_ISL_488449, EPI_ISL_488452, EPI_ISL_488454 | | | |
| see above | PHE South West Regional Laboratory, National Infection Service | Wellcome Sanger Institute for the COVID-19 Genomics UK (COG-UK) consortium | Stephanie Hutchings, Hannah Pymont, Dr Peter Muir, Barry Vipond, Rich Hopes; and Alex Alderton, Roberto Amato, Sonia Goncalves, Ewan Harrison, David K. Jackson, Ian Johnston, Dominic Kwiatkowski, Cordelia Langford, John Sillitoe on behalf of the Wellcome Sanger Institute COVID-19 Surveillance Team (http://www.sanger.ac.uk/covid-team) |
| EPI_ISL_488562, EPI_ISL_488632, EPI_ISL_488636 | NU-OMICS DNA Sequencing research facility, Northumbria University | Wellcome Sanger Institute for the COVID-19 Genomics UK (COG-UK) consortium | Chris Duncan, Sheia Waugh, Shirelle Burton-Fanning, Gary Eltringham, Jennifer Collins, Brendan Payne, Yusri Taha, Emma Swindells, Jane Greenaway, Edward Barton, Garren Scott, Debra Padgett, Clive Graham, Sarah Essex, Steve Ligget, Paul Baker, Lynn Dover, Wen Yew, Gary Black, John Allan, Joshua Loh, Greg Young, Matthew Bashton, Andrew Nelson, Darren Smith and Alex Alderton, Roberto Amato, Sonia Goncalves, Ewan Harrison, David K. Jackson, Ian Johnston, Dominic Kwiatkowski, Cordelia Langford, John Sillitoe on behalf of the Wellcome Sanger Institute COVID-19 Surveillance Team (http://www.sanger.ac.uk/covid-team) |
| EPI_ISL_488855, EPI_ISL_488863 | Department of Medical Microbiology, Western Sussex Hospitals NHS Foundation Trust, St Richard's Hospital | Wellcome Sanger Institute for the COVID-19 Genomics UK (COG-UK) consortium | Manasa Mutingwende, Sarah Lowdon, Olga Podplomyk, Michelle Erkiert, Jonathan Lewis, Paul Randell and Alex Alderton, Roberto Amato, Sonia Goncalves, Ewan Harrison, David K. Jackson, Ian Johnston, Dominic Kwiatkowski, Cordelia Langford, John Sillitoe on behalf of the Wellcome Sanger Institute COVID-19 Surveillance Team (http://www.sanger.ac.uk/covid-team) |
| EPI_ISL_489162, EPI_ISL_489165, EPI_ISL_489166, EPI_ISL_489168, EPI_ISL_489172, EPI_ISL_489173, EPI_ISL_489176, EPI_ISL_489182, EPI_ISL_489183, EPI_ISL_489184, EPI_ISL_489193, EPI_ISL_489200, EPI_ISL_489203, EPI_ISL_489208, EPI_ISL_489209, EPI_ISL_489212 | | | |
| see above | Regional Virus Laboratory, Belfast Health and Social Care Trust | Wellcome Sanger Institute for the COVID-19 Genomics UK (COG-UK) consortium | Conall McCaughey, James McKenna, Tanya Curran, Susan Feeney, Alison Watt, Ciara Cox, Mairead Connor, Zoltan Molnar, David Simpson, Derek Fairley; and Alex Alderton, Roberto Amato, Sonia Goncalves, Ewan Harrison, David K. Jackson, Ian Johnston, Dominic Kwiatkowski, Cordelia Langford, John Sillitoe on behalf of the Wellcome Sanger Institute COVID-19 Surveillance Team (http://www.sanger.ac.uk/covid-team) |
| EPI_ISL_489213 | Regional Virus Laboratory, Belfast Health and Social Care Trust | Wellcome Sanger Institute for the COVID-19 Genomics UK (COG-UK) Consortium | Conall McCaughey, James McKenna, Tanya Curran, Susan Feeney, Alison Watt, Ciara Cox, Mairead Connor, Zoltan Molnar, David Simpson, Derek Fairley; and Alex Alderton, Roberto Amato, Sonia Goncalves, Ewan Harrison, David K. Jackson, Ian Johnston, Dominic Kwiatkowski, Cordelia Langford, John Sillitoe on behalf of the Wellcome Sanger Institute COVID-19 Surveillance Team |
| EPI_ISL_489218, EPI_ISL_489220, EPI_ISL_489226, EPI_ISL_489227, EPI_ISL_489230, EPI_ISL_489236, EPI_ISL_489240, EPI_ISL_489244, EPI_ISL_489245, EPI_ISL_489246, EPI_ISL_489247, EPI_ISL_489307, EPI_ISL_489311, EPI_ISL_489316, EPI_ISL_489319, EPI_ISL_489320, EPI_ISL_489321, EPI_ISL_489324, EPI_ISL_489334, EPI_ISL_489335, EPI_ISL_489346, EPI_ISL_489347, EPI_ISL_489350, EPI_ISL_489351, EPI_ISL_489352, EPI_ISL_489356 | | | |
| see above | Regional Virus Laboratory, Belfast Health and Social Care Trust | Wellcome Sanger Institute for the COVID-19 Genomics UK (COG-UK) consortium | Conall McCaughey, James McKenna, Tanya Curran, Susan Feeney, Alison Watt, Ciara Cox, Mairead Connor, Zoltan Molnar, David Simpson, Derek Fairley; and Alex Alderton, Roberto Amato, Sonia Goncalves, Ewan Harrison, David K. Jackson, Ian Johnston, Dominic Kwiatkowski, Cordelia Langford, John Sillitoe on behalf of the Wellcome Sanger Institute COVID-19 Surveillance Team (http://www.sanger.ac.uk/covid-team) |
| EPI_ISL_489358 | Regional Virus Laboratory, Belfast Health and Social Care Trust | Wellcome Sanger Institute for the COVID-19 Genomics UK (COG-UK) Consortium | Conall McCaughey, James McKenna, Tanya Curran, Susan Feeney, Alison Watt, Ciara Cox, Mairead Connor, Zoltan Molnar, David Simpson, Derek Fairley; and Alex Alderton, Roberto Amato, Sonia Goncalves, Ewan Harrison, David K. Jackson, Ian Johnston, Dominic Kwiatkowski, Cordelia Langford, John Sillitoe on behalf of the Wellcome Sanger Institute COVID-19 Surveillance Team |
| EPI_ISL_489361, EPI_ISL_489362, EPI_ISL_489365, EPI_ISL_489367, EPI_ISL_489368, EPI_ISL_489371, EPI_ISL_489372, EPI_ISL_489374, EPI_ISL_489376 | Regional Virus Laboratory, Belfast Health and Social Care Trust | Wellcome Sanger Institute for the COVID-19 Genomics UK (COG-UK) consortium | Conall McCaughey, James McKenna, Tanya Curran, Susan Feeney, Alison Watt, Ciara Cox, Mairead Connor, Zoltan Molnar, David Simpson, Derek Fairley; and Alex Alderton, Roberto Amato, Sonia Goncalves, Ewan Harrison, David K. Jackson, Ian Johnston, Dominic Kwiatkowski, Cordelia Langford, John Sillitoe on behalf of the Wellcome Sanger Institute COVID-19 Surveillance Team (http://www.sanger.ac.uk/covid-team) |
| EPI_ISL_489381, EPI_ISL_489382, EPI_ISL_489383, EPI_ISL_489384, EPI_ISL_489385, EPI_ISL_489386, EPI_ISL_489387, EPI_ISL_489388, EPI_ISL_489389, EPI_ISL_489390, EPI_ISL_489394, EPI_ISL_489395, EPI_ISL_489397, EPI_ISL_489398, EPI_ISL_489399, EPI_ISL_489400, EPI_ISL_489402, EPI_ISL_489403, EPI_ISL_489404, EPI_ISL_489405, EPI_ISL_489406, EPI_ISL_489409, EPI_ISL_489410, EPI_ISL_489411, EPI_ISL_489412, EPI_ISL_489414, EPI_ISL_489415, EPI_ISL_489416, EPI_ISL_489417, EPI_ISL_489419, EPI_ISL_489420, EPI_ISL_489421, EPI_ISL_489422, EPI_ISL_489423, EPI_ISL_489424, EPI_ISL_489425, EPI_ISL_489426, EPI_ISL_489427, EPI_ISL_489428, EPI_ISL_489430, EPI_ISL_489431, EPI_ISL_489432, EPI_ISL_489433, EPI_ISL_489434, EPI_ISL_489435, EPI_ISL_489436, EPI_ISL_489437, EPI_ISL_489438, EPI_ISL_489439, EPI_ISL_489440, EPI_ISL_489441, EPI_ISL_489443, EPI_ISL_489444, EPI_ISL_489445, EPI_ISL_489446, EPI_ISL_489447, EPI_ISL_489448, EPI_ISL_489450, EPI_ISL_489451, EPI_ISL_489452, EPI_ISL_489453, EPI_ISL_489454, EPI_ISL_489456, EPI_ISL_489457, EPI_ISL_489458, EPI_ISL_489459, EPI_ISL_489460, EPI_ISL_489461, EPI_ISL_489462, EPI_ISL_489463, EPI_ISL_489464, EPI_ISL_489465, EPI_ISL_489466, EPI_ISL_489467, EPI_ISL_489468, EPI_ISL_489469, EPI_ISL_489471, EPI_ISL_489472, EPI_ISL_489473, EPI_ISL_489474, EPI_ISL_489475, EPI_ISL_489476, EPI_ISL_489477, EPI_ISL_489478, EPI_ISL_489479, EPI_ISL_489481, EPI_ISL_489482, EPI_ISL_489483, EPI_ISL_489484, EPI_ISL_489485, EPI_ISL_489486, EPI_ISL_489487, EPI_ISL_489488, EPI_ISL_489490, EPI_ISL_489491, EPI_ISL_489492, EPI_ISL_489493, EPI_ISL_489494, EPI_ISL_489495, EPI_ISL_489496, EPI_ISL_489497, EPI_ISL_489498, EPI_ISL_489499, EPI_ISL_489500, EPI_ISL_489501, EPI_ISL_489502, EPI_ISL_489503, EPI_ISL_489504, EPI_ISL_489505, EPI_ISL_489506, EPI_ISL_489507, EPI_ISL_489508, EPI_ISL_489509, EPI_ISL_489510, EPI_ISL_489511, EPI_ISL_489512, EPI_ISL_489513, EPI_ISL_489514, EPI_ISL_489515, EPI_ISL_489516, EPI_ISL_489517, EPI_ISL_489518, EPI_ISL_489519, EPI_ISL_489520, EPI_ISL_489521, EPI_ISL_489522, EPI_ISL_489524, EPI_ISL_489525, EPI_ISL_489526, EPI_ISL_489527, EPI_ISL_489528, EPI_ISL_489529, EPI_ISL_489531, EPI_ISL_489532, EPI_ISL_489533, EPI_ISL_489534, EPI_ISL_489535, EPI_ISL_489538, EPI_ISL_489539, EPI_ISL_489540, EPI_ISL_489541, EPI_ISL_489542, EPI_ISL_489544, EPI_ISL_489545, EPI_ISL_489546, EPI_ISL_489548, EPI_ISL_489549, EPI_ISL_489550, EPI_ISL_489551, EPI_ISL_489552, EPI_ISL_489553, EPI_ISL_489554, EPI_ISL_489555, EPI_ISL_489556, EPI_ISL_489557, EPI_ISL_489558, EPI_ISL_489559, EPI_ISL_489560, EPI_ISL_489561, EPI_ISL_489562, EPI_ISL_489563, EPI_ISL_489564, EPI_ISL_489565, EPI_ISL_489566, EPI_ISL_489568, EPI_ISL_489569, EPI_ISL_489570, EPI_ISL_489571, EPI_ISL_489572, EPI_ISL_489573, EPI_ISL_489574, EPI_ISL_489575, EPI_ISL_489576, EPI_ISL_489577 | | | |
| see above | Department of Pathology, University of Cambridge | Wellcome Sanger Institute for the COVID-19 Genomics UK (COG-UK) consortium | Luke W Meredith, M. Estée Török , Myra Hosmillo, William L. Hamilton, Martin D. Curran, Theresa Feltwell, Grant Hall, Anna Yakovleva, Fahad A Khokhar, Charlotte J. Houldcroft, Laura G Caller, Aminu S. Jahun, Sarah L. Caddy, Ian Goodfellow; and Alex Alderton, Roberto Amato, Sonia Goncalves, Ewan Harrison, David K. Jackson, Ian Johnston, Dominic Kwiatkowski, Cordelia Langford, John Sillitoe on behalf of the Wellcome Sanger Institute COVID-19 Surveillance Team (http://www.sanger.ac.uk/covid-team) |
| EPI_ISL_489744, EPI_ISL_489751, EPI_ISL_489777, EPI_ISL_489778, EPI_ISL_489779, EPI_ISL_489780, EPI_ISL_489781, EPI_ISL_489782, EPI_ISL_489783, EPI_ISL_489784, EPI_ISL_489785, EPI_ISL_489786, EPI_ISL_489787, EPI_ISL_489788, EPI_ISL_489789, EPI_ISL_489790, EPI_ISL_489791, EPI_ISL_489792, EPI_ISL_489793 | | | |
| see above | Florida Bureau of Public Health Laboratories | Florida Bureau of Public Health Laboratories | Sarah Schmedes, Jason Blanton |
| EPI_ISL_490206 | München Klinik Schwabing | MGZ Medical Genetics Center | Dieter A. Wolf, Elke Holinski-Feder |
| EPI_ISL_490255, EPI_ISL_490256 | National Institute for Communicable Diseases of the National Health Laboratory Service | National Institute for Communicable Diseases of the National Health Laboratory Service | Allam M, Ismail A, Khumalo Z, Kwenda S, Mtshali P, Mnyameni F, Mohale T, Subramoney K, Bhiman JN |
| EPI_ISL_490468 | Northumbria University / South Tees Hospitals NHS Foundation Trust / North Cumbria Integrated Care NHS Foundation Trust / North Tees and Hartlepool NHS Foundation Trust / Newcastle Hospitals NHS Foundation Trust | COVID-19 Genomics UK (COG-UK) Consortium | Darren L Smith,Andrew Nelson,Matthew Bashton,Greg R Young,Joshua Loh,John Allan,Mohammad A Tariq,Giles S Holt,Gary Black,Wen C Yew,Lynn Dover,Paul Baker,Steve Liggett,Sarah Essex,Jane Greenaway,Debra Padgett,Clive Graham,Garren Scott,Edward Barton,Emma Swindells,Brendan Payne,Jennifer Collins,Yusri Taha,Gary Eltringham |
| EPI_ISL_490496, EPI_ISL_490517, EPI_ISL_490518, EPI_ISL_490519, EPI_ISL_490520, EPI_ISL_490521, EPI_ISL_490522, EPI_ISL_490523, EPI_ISL_490525, EPI_ISL_490528, EPI_ISL_490529, EPI_ISL_490532, EPI_ISL_490533, EPI_ISL_490535, EPI_ISL_490539, EPI_ISL_490540, EPI_ISL_490541, EPI_ISL_490542, EPI_ISL_490545, EPI_ISL_490547, EPI_ISL_490549, EPI_ISL_490550, EPI_ISL_490553, EPI_ISL_490554, EPI_ISL_490555, EPI_ISL_490556 | | | |
| see above | Quadram Institute Bioscience | COVID-19 Genomics UK (COG-UK) Consortium | Dave J. Baker, Gemma L. Kay, Alp Aydin, Thanh Le-Viet, Steven Rudder, Ana P. Tedim, Anastasia Kolyva, Maria Diaz, Leonardo de Oliveira Martins, Nabil-Fareed Alikhan, Lizzie Meadows, Rachael Stanley, Ngozi Elumogo, Muhammed Yasir, Nicholas M. Thomson, Alexander J Trotter, Rachel Gilroy, Samuel Bloomfield, Claire Stuart, Andrew Bell, Reenesh Prakash, Samir Dervisevic, Alison E. Mather, John Wain, Mark Webber, Andrew J. Page, Justin |

| | | | |
|---|---|---|---|
| | | | O'Grady |
| EPI_ISL_490585, EPI_ISL_490591, EPI_ISL_490608, EPI_ISL_490609, EPI_ISL_490611, EPI_ISL_490612, EPI_ISL_490615, EPI_ISL_490622, EPI_ISL_490625, EPI_ISL_490626, EPI_ISL_490627, EPI_ISL_490629, EPI_ISL_490630, EPI_ISL_490637, EPI_ISL_490642 | | | |
| see above | Virology Department, Sheffield Teaching Hospitals NHS Foundation Trust/Department of Infection, Immunity and Cardiovascular Disease, The Medical School, University of Sheffield | COVID-19 Genomics UK (COG-UK) Consortium | Thushan de Silva, Matthew Parker, Nikki Smith, Adri Angyal, Rebecca Brown, Luke Green, Rachel Tucker, Paul Parsons, Danielle Groves, Katie Johnson, Laura Carrilero, Alex Keeley, Dave Partridge, Matthew Wyles, Benjamin Lindsey, Mehmet Yavuz, Mohammad Raza, Cariad Evans |
| EPI_ISL_491068, EPI_ISL_491069, EPI_ISL_491070, EPI_ISL_491071, EPI_ISL_491072 | Suceava County Emergency Hospital | "Stefan cel Mare" University Metagenomics Lab | Lobiuc Andrei et al. |
| EPI_ISL_491073 | Suceava County Emergency Hospital | "Stefan cel Mare" University Metagenomics Lab | Lobiuc Andrei, Antoniadis Panagiotis et al. |
| EPI_ISL_491074 | Suceava County Emergency Hospital | "Stefan cel Mare" University Metagenomics Lab | Lobiuc Andrei et al. |
| EPI_ISL_491075 | Suceava County Emergency Hospital | "Stefan cel Mare" University Metagenomics Lab | Lobiuc Andrei, Antoniadis Panagiotis et al. |
| EPI_ISL_491076 | Suceava County Emergency Hospital | "Stefan cel Mare" University Metagenomics Lab | Lobiuc Andrei et al. |
| EPI_ISL_491077 | Suceava County Emergency Hospital | "Stefan cel Mare" University Metagenomics Lab | Lobiuc Andrei, Antoniadis Panagiotis et al. |
| EPI_ISL_491078 | Suceava County Emergency Hospital | "Stefan cel Mare" University Metagenomics Lab | Lobiuc Andrei et al. |
| EPI_ISL_491079 | Suceava County Emergency Hospital | "Stefan cel Mare" University Metagenomics Lab | Lobiuc Andrei, Antoniadis Panagiotis et al. |
| EPI_ISL_491107, EPI_ISL_491108, EPI_ISL_491109 | SC Department of Health and Environmental Control | SC Department of Health and Environmental Control | Flores,H. |
| EPI_ISL_491125, EPI_ISL_491126, EPI_ISL_491127, EPI_ISL_491130, EPI_ISL_491131, EPI_ISL_491132 | Oman-National Influenza Center | Biotechnology & OMICs Laboratory | Samira Al-Mahruqi, Abdul Latif Khan, Samiha Al-Kharusi, Adil Khan , Ahmed Al-Rawahi, Sajjad Asaf, Amina Al-Jardani, Hanan Al-Kindi, Intisar Al-Shukri, Ahlam Al-Amri, Aisha Al-Amri, Aisha Al-Busaidi, Adil Al-Wahaibi, Seif Al-Abri, Ahmed Al-Harrasi |
| EPI_ISL_491140, EPI_ISL_491142, EPI_ISL_491143 | Oman-National Influenza Center | Biotechnology & OMICs Laboratory | Samiha Al-Kharusi, Sajjad Asaf, Abdul Latif Khan, Samira Al-Mahruqi, Adil Khan, Ahmed Al-Rawahi, Amina Al-Jardani, Hanan Al-Kindi, Intisar Al-Shukri, Ahlam Al-Amri, Aisha Al-Amri, Aisha Al-Busaidi, Adil Al-Wahaibi, Seif Al-Abri, Ahmed Al-Harrasi |
| EPI_ISL_491146 | Oman-National Influenza Center | Biotechnology & OMICs Laboratory | Abdul Latif Khan, Samira Al-Mahruqi, Ahmed Al-Harrasi, Samiha Al-Kharusi, Adil Khan, Ahmed Al-Rawahi, Sajjad Asaf, Amina Al-Jardani, Hanan Al-Kindi, Intisar Al-Shukri, Ahlam Al-Amri, Aisha Al-Amri, Aisha Al-Busaidi, Adil Al-Wahaibi, Seif Al-Abri. |
| EPI_ISL_491162, EPI_ISL_491163, EPI_ISL_491164, EPI_ISL_491165, EPI_ISL_491166, EPI_ISL_491168, EPI_ISL_491171 | Oman-National Influenza Center | Biotechnology & OMICs Laboratory | Sajjad Asaf, Samiha Al-Kharusi, Ahmed Al-Harrasi, Samira Al-Mahruqi, Adil Khan, Ahmed Al-Rawahi, Abdul Latif Khan, Amina Al-Jardani, Hanan Al-Kindi, Intisar Al-Shukri, Ahlam Al-Amri, Aisha Al-Amri, Aisha Al-Busaidi, Adil Al-Wahaibi, Seif Al-Abri. |
| EPI_ISL_491432 | Laboratorio de Referencia Nacional de Virus Respiratorio. Instituto Nacional de Salud.Perú | Laboratorio de Referencia Nacional de Biotecnología y Biología Molecular. Instituto Nacional de Salud.Perú | Carlos Padilla Rojas, Karolyn Vega Chozo, Priscila Lope Pari, Omar Caceres Rey, Marco Galarza Perez, Maribel Huaringa Nuñez, Johanna Balbuena Torres, Henri Bailon Calderon, Nancy Rojas Serrano. |
| EPI_ISL_491433, EPI_ISL_491434, EPI_ISL_491435 | Laboratorio de Referencia Nacional de Virus Respiratorio. Instituto Nacional de Salud Perú | Laboratorio de Referencia Nacional de Biotecnología y Biología Molecular. Instituto Nacional de Salud Perú | Carlos Padilla Rojas, Karolyn Vega Chozo, Priscila Lope Pari, Omar Caceres Rey, Marco Galarza Perez, Maribel Huaringa Nuñez, Johanna Balbuena Torres, Henri Bailon Calderon, Nancy Rojas Serrano. |
| EPI_ISL_491436 | Laboratorio de Referencia Nacional de Virus Respiratorio. Instituto Nacional de Salud Perú | Laboratorio de Referencia Nacional de Biotecnología y Biología Molecular. Instituto Nacional de Salud Perú | Carlos Padilla Rojas, Karolyn Vega Chozo, Priscila Lope Pari, Omar Caceres Rey, Marco Galarza Perez, Maribel Huaringa Nuñez, Johanna Balbuena Torres, Henri Bailon Calderon, Nancy Rojas Serrano |
| EPI_ISL_491496, EPI_ISL_491497, EPI_ISL_491498, EPI_ISL_491505, EPI_ISL_491506, EPI_ISL_491507, EPI_ISL_491509, EPI_ISL_491510, EPI_ISL_491511, EPI_ISL_491513, EPI_ISL_491514, EPI_ISL_491515, EPI_ISL_491517, EPI_ISL_491520, EPI_ISL_491521, EPI_ISL_491523, EPI_ISL_491527, EPI_ISL_491531, EPI_ISL_491532, EPI_ISL_491534, EPI_ISL_491535, EPI_ISL_491538, EPI_ISL_491542, EPI_ISL_491543, EPI_ISL_491547, EPI_ISL_491550, EPI_ISL_491567, EPI_ISL_491570, EPI_ISL_491571, EPI_ISL_491575, EPI_ISL_491577, EPI_ISL_491579, EPI_ISL_491580, EPI_ISL_491583, EPI_ISL_491594, EPI_ISL_491595, EPI_ISL_491597, EPI_ISL_491603, EPI_ISL_491609, EPI_ISL_491613, EPI_ISL_491617, EPI_ISL_491621, EPI_ISL_491622, EPI_ISL_491626, EPI_ISL_491627, EPI_ISL_491631, EPI_ISL_491632, EPI_ISL_491641, EPI_ISL_491647 | | | |
| see above | Virology Department, Royal Infirmary of Edinburgh, NHS Lothian / School of Biological Sciences, University of Edinburgh | Wellcome Sanger Institute for the COVID-19 Genomics UK (COG-UK) consortium | McHugh M, Dewar R, Rooke S, O'Toole Á, Scher E, Hill V, McCrone JT, Colquhoun R, Yu X, Jackson B, Rambaut A, Templeton K and Alex Alderton, Roberto Amato, Sonia Goncalves, Ewan Harrison, David K. Jackson, Ian Johnston, Dominic Kwiatkowski, Cordelia Langford, John Sillitoe on behalf of the Wellcome Sanger Institute COVID-19 Surveillance Team (http://www.sanger.ac.uk/covid-team) |
| EPI_ISL_491655 | Virology Department, Royal Infirmary of Edinburgh, NHS Lothian / School of Biological Sciences, University of Edinburgh | Wellcome Sanger Institute for the COVID-19 Genomics UK (COG-UK) Consortium | McHugh M, Dewar R, Rooke S, O'Toole Á, Scher E, Hill V, McCrone JT, Colquhoun R, Yu X, Jackson B, Rambaut A, Templeton K and Alex Alderton, Roberto Amato, Sonia Goncalves, Ewan Harrison, David K. Jackson, Ian Johnston, Dominic Kwiatkowski, Cordelia Langford, John Sillitoe on behalf of the Wellcome Sanger Institute COVID-19 Surveillance Team |
| EPI_ISL_491659, EPI_ISL_491660, EPI_ISL_491663, EPI_ISL_491665, EPI_ISL_491671, EPI_ISL_491673, EPI_ISL_491674, EPI_ISL_491677, EPI_ISL_491678, EPI_ISL_491679, EPI_ISL_491682, EPI_ISL_491684, EPI_ISL_491685, EPI_ISL_491688, EPI_ISL_491693, EPI_ISL_491695 | | | |
| see above | Virology Department, Royal Infirmary of Edinburgh, NHS Lothian / School of Biological Sciences, University of Edinburgh | Wellcome Sanger Institute for the COVID-19 Genomics UK (COG-UK) consortium | McHugh M, Dewar R, Rooke S, O'Toole Á, Scher E, Hill V, McCrone JT, Colquhoun R, Yu X, Jackson B, Rambaut A, Templeton K and Alex Alderton, Roberto Amato, Sonia Goncalves, Ewan Harrison, David K. Jackson, Ian Johnston, Dominic Kwiatkowski, Cordelia Langford, John Sillitoe on behalf of the Wellcome Sanger Institute COVID-19 Surveillance Team (http://www.sanger.ac.uk/covid-team) |
| EPI_ISL_491944, EPI_ISL_491945 | Instituto Nacional de Investigación en Salud Pública - INSPI | INSPI - Charité | Alfredo Bruno Caicedo, Domenica de Mora Coloma, Andres Moreira-Soto, Anna-Lena Sander, Nina Krause, Maritza Olmedo,Denisses Portugal, Manuel Gonzalez, Silvia Salgado, Alberto Orlando, Alexandra Usiña, Juan Carlos Zeballos, Jan Felix Drexler |
| EPI_ISL_492132, EPI_ISL_492180 | SA Pathology | SA Pathology | Lex Leong, Chuan Kok Lim, Mark Turra, Ivan Bastian, Geoff Higgins |
| EPI_ISL_492246, EPI_ISL_492254, EPI_ISL_492284, EPI_ISL_492316, EPI_ISL_492328, EPI_ISL_492329, EPI_ISL_492416 | PHE South West Regional Laboratory, National Infection Service | Wellcome Sanger Institute for the COVID-19 Genomics UK (COG-UK) consortium | Stephanie Hutchings, Hannah Pymont, Dr Peter Muir, Barry Vipond, Rich Hopes; and Alex Alderton, Roberto Amato, Sonia Goncalves, Ewan Harrison, David K. Jackson, Ian Johnston, Dominic Kwiatkowski, Cordelia Langford, John Sillitoe on behalf of the Wellcome Sanger Institute COVID-19 Surveillance Team (http://www.sanger.ac.uk/covid-team) |
| EPI_ISL_492450, EPI_ISL_492451, EPI_ISL_492455, EPI_ISL_492459, EPI_ISL_492463, EPI_ISL_492468, EPI_ISL_492470, EPI_ISL_492471, EPI_ISL_492473, EPI_ISL_492476, EPI_ISL_492479, EPI_ISL_492480, EPI_ISL_492481, EPI_ISL_492482, EPI_ISL_492485, EPI_ISL_492486, EPI_ISL_492487, EPI_ISL_492488, EPI_ISL_492489, EPI_ISL_492491, EPI_ISL_492494 | | | |
| see above | NU-OMICS DNA Sequencing research facility, Northumbria University | Wellcome Sanger Institute for the COVID-19 Genomics UK (COG-UK) consortium | Chris Duncan, Sheia Waugh, Shirelle Burton-Fanning, Gary Eltringham, Jennifer Collins, Brendan Payne, Yusri Taha, Emma Swindells, Jane Greenaway, Edward Barton, Garren Scott, Debra Padgett, Clive Graham, Sarah Essex, Steve Ligget, Paul Baker, Lynn Dover, Wen Yew, Gary Black, John Allan, Joshua Loh, Greg Young, Matthew Bashton, Andrew Nelson, Darren Smith and Alex Alderton, Roberto Amato, Sonia Goncalves, Ewan Harrison, David K. Jackson, Ian Johnston, Dominic Kwiatkowski, Cordelia Langford, John Sillitoe on behalf of the Wellcome Sanger Institute COVID-19 Surveillance Team (http://www.sanger.ac.uk/covid-team) |
| EPI_ISL_492747, EPI_ISL_492756 | University College London, Great Ormond Street Hospital for Children NHS Foundation Trust, Imperial College Healthcare NHS Trust | Wellcome Sanger Institute for the COVID-19 Genomics UK (COG-UK) consortium | Sergi Castellano, Rachel Williams, Mark Kristiansen, Paola Resende Silva, Sunando Roy, Tony Brooks, Helena Tutill, Paola Niola, Patricia Dyal, Charlotte Williams, Leysa Forrest, Yasmin Panchbhaya, Jacqueline Findlay, Sam Weeks, Julianne Brown, Kathryn Harris, Paul Randell, James Price, Alison Holmes, Judith Breuer and Alex Alderton, Roberto Amato, Sonia Goncalves, Ewan Harrison, David K. Jackson, Ian Johnston, Dominic Kwiatkowski, Cordelia Langford, John Sillitoe on behalf of the Wellcome Sanger Institute COVID-19 Surveillance Team (http://www.sanger.ac.uk/covid-team) |
| EPI_ISL_492839 | Department of Medical Microbiology, Western Sussex Hospitals NHS Foundation Trust, St Richard's Hospital | Wellcome Sanger Institute for the COVID-19 Genomics UK (COG-UK) consortium | Manasa Mutingwende, Sarah Lowdon, Olga Podplomyk, Michelle Erkiert, Jonathan Lewis, Paul Randell and Alex Alderton, Roberto Amato, Sonia Goncalves, Ewan Harrison, David K. Jackson, Ian Johnston, Dominic Kwiatkowski, Cordelia Langford, John Sillitoe on behalf of the Wellcome Sanger Institute COVID-19 Surveillance Team (http://www.sanger.ac.uk/covid-team) |
| EPI_ISL_492844 | Royal Free Hospital / Health Services Laboratories | Wellcome Sanger Institute for the COVID-19 Genomics UK (COG-UK) consortium | Tanzina Haque, Tabitha Mahungu, Dianne Irish, Cate Goodlad, Jenny Cross, Judith Heaney and Alex Alderton, Roberto Amato, Sonia Goncalves, Ewan Harrison, David K. Jackson, Ian Johnston, Dominic Kwiatkowski, Cordelia Langford, John Sillitoe on behalf of the Wellcome Sanger Institute COVID-19 Surveillance Team (http://www.sanger.ac.uk/covid-team) |

| | | | |
|---|---|---|---|
| EPI_ISL_492870, EPI_ISL_492884, EPI_ISL_492912 | Department of Medical Microbiology, Western Sussex Hospitals NHS Foundation Trust, St Richard's Hospital | Wellcome Sanger Institute for the COVID-19 Genomics UK (COG-UK) consortium | Manasa Mutingwende, Sarah Lowdon, Olga Podplomyk, Michelle Erkiert, Jonathan Lewis, Paul Randell and Alex Alderton, Roberto Amato, Sonia Goncalves, Ewan Harrison, David K. Jackson, Ian Johnston, Dominic Kwiatkowski, Cordelia Langford, John Sillitoe on behalf of the Wellcome Sanger Institute COVID-19 Surveillance Team (http://www.sanger.ac.uk/covid-team) |
| EPI_ISL_493087, EPI_ISL_493088, EPI_ISL_493089, EPI_ISL_493090, EPI_ISL_493091, EPI_ISL_493092, EPI_ISL_493093, EPI_ISL_493094, EPI_ISL_493095, EPI_ISL_493096, EPI_ISL_493097, EPI_ISL_493098, EPI_ISL_493099, EPI_ISL_493100, EPI_ISL_493101, EPI_ISL_493102, EPI_ISL_493103, EPI_ISL_493104, EPI_ISL_493105, EPI_ISL_493106, EPI_ISL_493107, EPI_ISL_493108, EPI_ISL_493109, EPI_ISL_493110, EPI_ISL_493111, EPI_ISL_493112, EPI_ISL_493113, EPI_ISL_493114, EPI_ISL_493115, EPI_ISL_493116, EPI_ISL_493117, EPI_ISL_493118, EPI_ISL_493119, EPI_ISL_493120, EPI_ISL_493121, EPI_ISL_493122, EPI_ISL_493123, EPI_ISL_493124 | | | |
| see above | Utah Public Health Laboratory | Utah Public Health Laboratory | Heidi Butz, Erin Young, Kelly Oakeson |
| EPI_ISL_493337 | Instituto de Diagnostico y Referencia Epidemiologicos (INDRE) | Instituto de Diagnostico y Referencia Epidemiologicos (INDRE) | Gisela Barrera-Badillo , Abril Rodriguez-Maldonado, Claudia Wong-Arambula , Natividad Cruz-Ortiz, Tatiana Nunez-Garcia, Dayanira Arellano-Suarez, Fabiola Garces-Ayala, Edgar Mendieta-Condado, Lucia Hernandez-Rivas, Irma Lopez-Martinez, Ernesto Ramirez-Gonzalez. |
| EPI_ISL_493350 | Ostfold Hospital Trust - Kalnes, Centre for Laboratory Medicine, Section for gene technology and infection serology | Norwegian Institute of Public Health, Department of Virology | Kathrine Stene-Johansen, Kamilla Heddeland Instefjord, Hilde Elshaug, Rasmus Riis Kopperud, Karoline Bragstad, Olav Hungnes |
| EPI_ISL_493394, EPI_ISL_493395, EPI_ISL_493396, EPI_ISL_493397, EPI_ISL_493401, EPI_ISL_493402, EPI_ISL_493403, EPI_ISL_493408, EPI_ISL_493412, EPI_ISL_493420 | National Public Health Laboratory, National Centre for Infectious Diseases | National Public Health Laboratory, National Centre for Infectious Diseases | Mak TM, Octavia S, Zhou Z, Chavatte JM, Cui L, Lin RTP |
| EPI_ISL_493472, EPI_ISL_493473, EPI_ISL_493474, EPI_ISL_493475, EPI_ISL_493476, EPI_ISL_493477, EPI_ISL_493478, EPI_ISL_493479, EPI_ISL_493480, EPI_ISL_493481, EPI_ISL_493482, EPI_ISL_493483, EPI_ISL_493484, EPI_ISL_493485, EPI_ISL_493486, EPI_ISL_493487 | | | |
| see above | Northumbria University / South Tees Hospitals NHS Foundation Trust / North Cumbria Integrated Care NHS Foundation Trust / North Tees and Hartlepool NHS Foundation Trust / Newcastle Hospitals NHS Foundation Trust | COVID-19 Genomics UK (COG-UK) Consortium | Darren L Smith,Andrew Nelson,Matthew Bashton,Greg R Young,Joshua Loh,John Allan,Mohammad A Tariq,Giles S Holt,Gary Black,Wen C Yew,Lynn Dover,Paul Baker,Steve Liggett,Sarah Essex,Jane Greenaway,Debra Padgett,Clive Graham,Garren Scott,Edward Barton,Emma Swindells,Brendan Payne,Jennifer Collins,Yusri Taha,Gary Eltringham |
| EPI_ISL_493563, EPI_ISL_493564, EPI_ISL_493565, EPI_ISL_493566, EPI_ISL_493567, EPI_ISL_493568, EPI_ISL_493569, EPI_ISL_493570, EPI_ISL_493571, EPI_ISL_493572, EPI_ISL_493573, EPI_ISL_493574, EPI_ISL_493575, EPI_ISL_493576, EPI_ISL_493577, EPI_ISL_493578, EPI_ISL_493579, EPI_ISL_493580, EPI_ISL_493581, EPI_ISL_493582, EPI_ISL_493583, EPI_ISL_493584, EPI_ISL_493585, EPI_ISL_493586, EPI_ISL_493587, EPI_ISL_493588, EPI_ISL_493589, EPI_ISL_493590, EPI_ISL_493591, EPI_ISL_493592, EPI_ISL_493593, EPI_ISL_493594 | | | |
| see above | Lincolnshire Hospitals and DeepSeq Nottingham | COVID-19 Genomics UK (COG-UK) Consortium | Nichola Duckworth, Tim Sloan, Sarah Walsh, Jonathan Ball, Patrick McClure, Joeseph Chappell, Nadine Holmes, Matthew Carlisle, Christopher Moore, Fei Sang, Johnny Debebe, Victoria Wright, Matthew Loose |
| EPI_ISL_493685, EPI_ISL_493686, EPI_ISL_493689, EPI_ISL_493698, EPI_ISL_493711, EPI_ISL_493718, EPI_ISL_493722, EPI_ISL_493733, EPI_ISL_493734, EPI_ISL_493740 | Virology Department, Sheffield Teaching Hospitals NHS Foundation Trust/Department of Infection, Immunity and Cardiovascular Disease, The Medical School, University of Sheffield | COVID-19 Genomics UK (COG-UK) Consortium | Thushan de Silva, Matthew Parker, Nikki Smith, Adri Angyal, Rebecca Brown, Luke Green, Rachel Tucker, Paul Parsons, Danielle Groves, Katie Johnson, Laura Carrilero, Alex Keeley, Dave Partridge, Matthew Wyles, Benjamin Lindsey, Mehmet Yavuz, Mohammad Raza, Cariad Evans |
| EPI_ISL_493825, EPI_ISL_493826, EPI_ISL_493827, EPI_ISL_493828, EPI_ISL_493829, EPI_ISL_493830, EPI_ISL_493832, EPI_ISL_493833 | West of Scotland Specialist Virology Centre, NHSGGC / MRC-University of Glasgow Centre for Virus Research | COVID-19 Genomics UK (COG-UK) Consortium | Ana da Silva Filipe, Natasha Johnson, Kathy Smollett, Daniel Mair, Stephen Carmichael, Lily Tong, Jenna Nichols, Elihu Aranday-Cortes, Kirstyn Brunker, Yasmin Parr, Alice Broos, Kyriaki Nomikou; Sarah McDonald, Marc Niebel, Patawee Asamaphan; Richard Orton, Joseph Hughes, Sreenu Vattipally, David L Robertson; Alasdair MacLean, Rory Gunson; Kathy Li, Natasha Jesudason, Rajiv Shah, James Shepherd, Antonia Ho, Emma Thomson |
| EPI_ISL_494101, EPI_ISL_494155, EPI_ISL_494156, EPI_ISL_494169, EPI_ISL_494173, EPI_ISL_494191, EPI_ISL_494204, EPI_ISL_494210, EPI_ISL_494212, EPI_ISL_494214, EPI_ISL_494220, EPI_ISL_494223, EPI_ISL_494224, EPI_ISL_494226, EPI_ISL_494232, EPI_ISL_494241, EPI_ISL_494248, EPI_ISL_494266, EPI_ISL_494267, EPI_ISL_494269, EPI_ISL_494272, EPI_ISL_494273, EPI_ISL_494274, EPI_ISL_494278, EPI_ISL_494281, EPI_ISL_494283, EPI_ISL_494284, EPI_ISL_494288, EPI_ISL_494289, EPI_ISL_494290, EPI_ISL_494292, EPI_ISL_494293, EPI_ISL_494295, EPI_ISL_494301, EPI_ISL_494306, EPI_ISL_494308, EPI_ISL_494309, EPI_ISL_494312, EPI_ISL_494315, EPI_ISL_494316, EPI_ISL_494318, EPI_ISL_494320, EPI_ISL_494324, EPI_ISL_494325, EPI_ISL_494329, EPI_ISL_494338, EPI_ISL_494339, EPI_ISL_494347, EPI_ISL_494349, EPI_ISL_494363, EPI_ISL_494365, EPI_ISL_494368, EPI_ISL_494369 | | | |
| see above | Wales Specialist Virology Centre Sequencing lab: Pathogen Genomics Unit | COVID-19 Genomics UK (COG-UK) Consortium | Catherine Moore, Johnathan Evans, Laura Gifford, Malorie Perry, Simon Cottrell, Angela Marchbank, Alec Birchley, Alexander Adams, Amy Gaskin, Bree Gatica-Wilcox, Jason Coombes, Joel Southgate, Lauren Gilbert, Lee Graham, Nicole Pacchiarini, Sara Kumziene-Summerhayes, Sarah Taylor, Sophie Jones, Sara Rey, Matthew Bull, Joanne Watkins, Sally Corden, Tom Connor |
| EPI_ISL_494430, EPI_ISL_494431, EPI_ISL_494432, EPI_ISL_494433, EPI_ISL_494434, EPI_ISL_494435, EPI_ISL_494436, EPI_ISL_494437, EPI_ISL_494442, EPI_ISL_494443, EPI_ISL_494445, EPI_ISL_494568, EPI_ISL_494569, EPI_ISL_494589, EPI_ISL_494590, EPI_ISL_494593, EPI_ISL_494594, EPI_ISL_494595, EPI_ISL_494620, EPI_ISL_494625, EPI_ISL_494626, EPI_ISL_494633, EPI_ISL_494634 | | | |
| see above | San Diego County Public Health Laboratory | Andersen lab at Scripps Research | SEARCH Alliance San Diego with Tracy Basler, Jovan Shephard, Brett Austin |
| EPI_ISL_494646, EPI_ISL_494673, EPI_ISL_494676, EPI_ISL_494677, EPI_ISL_494678, EPI_ISL_494679, EPI_ISL_494680, EPI_ISL_494681, EPI_ISL_494683, EPI_ISL_494684, EPI_ISL_494691, EPI_ISL_494692, EPI_ISL_494695, EPI_ISL_494696, EPI_ISL_494698, EPI_ISL_494699, EPI_ISL_494700, EPI_ISL_494704, EPI_ISL_494705, EPI_ISL_494707, EPI_ISL_494709, EPI_ISL_494710, EPI_ISL_494713 | | | |
| see above | Scripps Medical Laboratory | Andersen lab at Scripps Research | SEARCH Alliance San Diego with Michael Quigley, Ellen Stefanski, Ian Mchardy |
| EPI_ISL_494745, EPI_ISL_494746 | San Diego County Public Health Laboratory | Andersen lab at Scripps Research | SEARCH Alliance San Diego with Tracy Basler, Jovan Shephard, Brett Austin |
| EPI_ISL_495081, EPI_ISL_495082, EPI_ISL_495083, EPI_ISL_495084, EPI_ISL_495085, EPI_ISL_495087, EPI_ISL_495089, EPI_ISL_495094, EPI_ISL_495095, EPI_ISL_495096 | Department of Medical Microbiology, Western Sussex Hospitals NHS Foundation Trust, St Richard's Hospital | Wellcome Sanger Institute for the COVID-19 Genomics UK (COG-UK) consortium | Manasa Mutingwende, Sarah Lowdon, Olga Podplomyk, Michelle Erkiert, Jonathan Lewis, Paul Randell and Alex Alderton, Roberto Amato, Sonia Goncalves, Ewan Harrison, David K. Jackson, Ian Johnston, Dominic Kwiatkowski, Cordelia Langford, John Sillitoe on behalf of the Wellcome Sanger Institute COVID-19 Surveillance Team (http://www.sanger.ac.uk/covid-team) |
| EPI_ISL_495098, EPI_ISL_495102, EPI_ISL_495105, EPI_ISL_495109 | PHE South West Regional Laboratory, National Infection Service | Wellcome Sanger Institute for the COVID-19 Genomics UK (COG-UK) consortium | Stephanie Hutchings, Hannah Pymont, Dr Peter Muir, Barry Vipond, Rich Hopes; and Alex Alderton, Roberto Amato, Sonia Goncalves, Ewan Harrison, David K. Jackson, Ian Johnston, Dominic Kwiatkowski, Cordelia Langford, John Sillitoe on behalf of the Wellcome Sanger Institute COVID-19 Surveillance Team (http://www.sanger.ac.uk/covid-team) |
| EPI_ISL_495448 | Kafkas University, Faculty of Medicine, Department of Medical Microbiology | Kafkas University, Faculty of Medicine, Department of Medical Microbiology | Murat Karamese, Didem Ozgur, E. Ediz Tutuncu |
| EPI_ISL_495584, EPI_ISL_495585, EPI_ISL_495586, EPI_ISL_495587, EPI_ISL_495589, EPI_ISL_495590, EPI_ISL_495591 | University of Michigan Clinical Microbiology Laboratory | Lauring Lab, University of Michigan, Department of Microbiology and Immunology | Valesano et al. |
| EPI_ISL_496356, EPI_ISL_496357, EPI_ISL_496358, EPI_ISL_496359, EPI_ISL_496360, EPI_ISL_496361, EPI_ISL_496362, EPI_ISL_496363, EPI_ISL_496364, EPI_ISL_496365, EPI_ISL_496366, EPI_ISL_496367, EPI_ISL_496368, EPI_ISL_496369 | | | |
| see above | Infectolab | Andersen lab at Scripps Research | SEARCH Alliance San Diego with Samuel Navarro Alvarez, Carlos A. Cota Haros, Octavio Renteria Pacheco |
| EPI_ISL_496522, EPI_ISL_496527, EPI_ISL_496534, EPI_ISL_496549, EPI_ISL_496550, EPI_ISL_496551, EPI_ISL_497888 | B.J. Govt. Medical College | National Centre For Cell Science | Dhiraj Paul, Kunal Jani, Radha Chauhan, Janesh Kumar, Vasudevan Seshadri, Girdhari Lal, Rajesh Karyakarte, Suvarna Joshi, Murlidhar Tambe, Sourav Sen, Santosh Karade, Kavita Bala Anand, Shelinder Pal Singh Shergill, Rajiv Mohan Gupta, Manoj Kumar Bhat, Arvind Sahu, Maharashtra COVID-19 Study Group, DBT's PAN-INDIA 1000 SARS-CoV2 RNA genome sequencing consortium, Yogesh S Shouche |
| EPI_ISL_498229, EPI_ISL_498230, EPI_ISL_498231, EPI_ISL_498232, EPI_ISL_498233, EPI_ISL_498234, EPI_ISL_498235, EPI_ISL_498236, EPI_ISL_498237 | Institut Pasteur de Dakar | Institut Pasteur de Dakar | Ndongo Dia, Moussa Moise Diagne, Mamadou Diop, Marie Henriette Dior Ndione, Mamadou Malado Jallow, Safietou Sankhe Mbengue, Ousmane Faye, Amadou Alpha Sall. |
| EPI_ISL_498640, EPI_ISL_498641, EPI_ISL_498642, EPI_ISL_498643, EPI_ISL_498644, EPI_ISL_498645, EPI_ISL_498646, EPI_ISL_498647, EPI_ISL_498648, EPI_ISL_498649, EPI_ISL_498650, EPI_ISL_498651, EPI_ISL_498652, EPI_ISL_498653, EPI_ISL_498654, EPI_ISL_498655, EPI_ISL_498656, EPI_ISL_498657, EPI_ISL_498658, EPI_ISL_498659, EPI_ISL_498660, EPI_ISL_498661, EPI_ISL_498662, EPI_ISL_498663, EPI_ISL_498664, EPI_ISL_498665, EPI_ISL_498666, EPI_ISL_498667 | | | |

| | | | |
|---|---|---|---|
| see above | Utah Public Health Laboratory | Utah Public Health Laboratory | Heidi Butz, Erin Young, Kelly Oakeson |
| EPI_ISL_498713 | Quest Diagnostics | Quest Diagnostics | Rosenthal,S.H., Gerasimova,A., Kagan,R.M. and Owen, R. |
| EPI_ISL_499315, EPI_ISL_499316, EPI_ISL_499317, EPI_ISL_499318, EPI_ISL_499319, EPI_ISL_499320, EPI_ISL_499321, EPI_ISL_499322, EPI_ISL_499323, EPI_ISL_499324, EPI_ISL_499325, EPI_ISL_499326, EPI_ISL_499327, EPI_ISL_499328, EPI_ISL_499329 | | | |
| see above | Centre for Enzyme Innovation, University of Portsmouth / Translational Research Laboratory, Portsmouth Hospitals NHS Trust | COVID-19 Genomics UK (COG-UK) Consortium | Angela Beckett,Yann Bourgeois,Garry Scarlett,Sharon Glaysher,Scott Elliott,Kelly Bicknell,Robert Impey,Allyson Lloyd,Sarah Wyllie,Ethan Butcher,Anoop Chauhan,Samuel Robson |
| EPI_ISL_499506, EPI_ISL_499784, EPI_ISL_499785, EPI_ISL_499787, EPI_ISL_499788, EPI_ISL_499807 | Northumbria University / South Tees Hospitals NHS Foundation Trust / North Cumbria Integrated Care NHS Foundation Trust / North Tees and Hartlepool NHS Foundation Trust / Newcastle Hospitals NHS Foundation Trust | COVID-19 Genomics UK (COG-UK) Consortium | Darren L Smith,Andrew Nelson,Matthew Bashton,Greg R Young,Joshua Loh,John Allan,Mohammad A Tariq,Giles S Holt,Gary Black,Wen C Yew,Lynn Dover,Paul Baker,Steve Liggett,Sarah Essex,Jane Greenaway,Debra Padgett,Clive Graham,Garren Scott,Edward Barton,Emma Swindells,Brendan Payne,Jennifer Collins,Yusri Taha,Gary Eltringham |
| EPI_ISL_500160, EPI_ISL_500165, EPI_ISL_500173, EPI_ISL_500192, EPI_ISL_500193, EPI_ISL_500194, EPI_ISL_500195, EPI_ISL_500196, EPI_ISL_500197, EPI_ISL_500198, EPI_ISL_500199, EPI_ISL_500200, EPI_ISL_500205, EPI_ISL_500206, EPI_ISL_500208, EPI_ISL_500210, EPI_ISL_500217, EPI_ISL_500222, EPI_ISL_500223 | | | |
| see above | Hospital Universitario Virgen de las Nieves de Granada-SAS | SeqCOVID-SPAIN consortium/IBV(CSIC) | Mercedes Pérez Ruiz, Sara Sanbonmatsu Gámez, Irene Pedrosa Corral, José M. Navarro-Marí and SeqCOVID-SPAIN consortium |
| EPI_ISL_500287, EPI_ISL_500288, EPI_ISL_500336, EPI_ISL_500337, EPI_ISL_500338, EPI_ISL_500339, EPI_ISL_500340, EPI_ISL_500341, EPI_ISL_500343, EPI_ISL_500344, EPI_ISL_500345, EPI_ISL_500346, EPI_ISL_500347, EPI_ISL_500349, EPI_ISL_500350, EPI_ISL_500351, EPI_ISL_500352, EPI_ISL_500353, EPI_ISL_500354, EPI_ISL_500355, EPI_ISL_500356, EPI_ISL_500357, EPI_ISL_500359, EPI_ISL_500361, EPI_ISL_500362, EPI_ISL_500363, EPI_ISL_500364 | | | |
| see above | Servicio de Microbiología, Hospital Miguel Servet, Zaragoza | SeqCOVID-SPAIN consortium/IBV(CSIC) | Antonio Rezusta López, Alexander Tristancho Baró, Ana Milagro, Yolanda Gracia Grataloup, Nieves Martínez Cameo and SeqCOVID-SPAIN consortium |
| EPI_ISL_500621 | Area of Virology, Serology and Virology Division (SAViD), New South Wales Health Pathology Randwick | Area of Virology, Serology and Virology Division (SAViD), New South Wales Health Pathology Randwick | Rawlinson, W. |
| EPI_ISL_500859, EPI_ISL_500860, EPI_ISL_500861, EPI_ISL_500862, EPI_ISL_500863 | Virginia DCLS | Virginia DCLS | Virginia DCLS |
| EPI_ISL_500951, EPI_ISL_500952 | Respiratory Virus Unit, Microbiology Services Colindale, Public Health England | Respiratory Virus Unit, Microbiology Services Colindale, Public Health England | PHE Covid Sequencing Team |
| EPI_ISL_501152, EPI_ISL_501153, EPI_ISL_501154 | University of Washington Virology Lab | University of Washington Virology Lab | Pavitra Roychoudhury, Hong Xie, Lasata Shrestha, Amin Addetia, Truong Nguyen, Victoria M Rachleff, Meei-Li Huang, Keith R Jerome, Alexander Greninger |
| EPI_ISL_501174 | Baylor College of Medicine | Baylor College of Medicine: HGSC | Vasanthi Avadhanula, Erin Nicholson, David Henke, Pedro Piedra, Harsha Doddapaneni, Donna Muzny, Qingchang Meng, Hsu Chao, Zeineen Momin, Hua Shen, George Weissenberger, Kavya Kottapalli, Yimiti Meiheerguli, Sejal Salvi, Ginger Metcalf, Vipin Menon, Sara J.J. Cregeen, Matthew C. Ross, Tulin Ayvaz, Richard Sucgang, Kristi L. Hoffman, Matthew Wong, Joseph F. Petrosino |
| EPI_ISL_501192 | Department of Medical Microbiology, University Malaya Medical Centre | Department of Medical Microbiology, Faculty of Medicine, University of Malaya | Yoong Min CHONG, Jennifer Chong, I-Ching SAM, Yoke Fun CHAN, University Malaya Medical Centre COVID Team |
| EPI_ISL_501230 | Hellenic Pasteur Institute, Public Health Laboratories | Hellenic Pasteur Institute, National Influenza Reference laboratory of Southern Greece & Unit of Bioinformatics and Applied Genomics | Vasiliki Pogka, Timokratis Karamitros, Athanasios Kossyvakis, Antonios Kalliaropoulos, Horefti Elina, Evangelidou Maria, Androniki Voulgari-Kokota, Aspasia Kontou, Andreas Mentis |
| EPI_ISL_501231, EPI_ISL_501248, EPI_ISL_501249 | Hellenic Pasteur Institute, National Influenza Reference laboratory of Southern Greece & Unit of Bioinformatics and Applied Genomics | Hellenic Pasteur Institute, National Influenza Reference laboratory of Southern Greece & Unit of Bioinformatics and Applied Genomics | Vasiliki Pogka, Timokratis Karamitros, Athanasios Kossyvakis, Antonios Kalliaropoulos, Horefti Elina, Evangelidou Maria, Androniki Voulgari-Kokota, Aspasia Kontou, Andreas Mentis |
| EPI_ISL_501557, EPI_ISL_501562, EPI_ISL_501602, EPI_ISL_501613 | Department of Medical Microbiology, Western Sussex Hospitals NHS Foundation Trust, St Richard's Hospital | Wellcome Sanger Institute for the COVID-19 Genomics UK (COG-UK) consortium | Manasa Mutingwende, Sarah Lowdon, Olga Podplomyk, Michelle Erkiert, Jonathan Lewis, Paul Randell and Alex Alderton, Roberto Amato, Sonia Goncalves, Ewan Harrison, David K. Jackson, Ian Johnston, Dominic Kwiatkowski, Cordelia Langford, John Sillitoe on behalf of the Wellcome Sanger Institute COVID-19 Surveillance Team (http://www.sanger.ac.uk/covid-team) |
| EPI_ISL_501625 | Lab Microbiology, Pathology Department, William Harvey Hospital | Wellcome Sanger Institute for the COVID-19 Genomics UK (COG-UK) consortium | Samuel Moses, Hannah Lowe, Felicity Ryan and Alex Alderton, Roberto Amato, Sonia Goncalves, Ewan Harrison, David K. Jackson, Ian Johnston, Dominic Kwiatkowski, Cordelia Langford, John Sillitoe on behalf of the Wellcome Sanger Institute COVID-19 Surveillance Team (http://www.sanger.ac.uk/covid-team) |
| EPI_ISL_507005 | Department of Laboratory Medicine, Tan Tock Seng Hospital | Department of Laboratory Medicine, Tan Tock Seng Hospital | Chen YYC, Zair X, Li C, Tang WY, Maurer-Stroh S, Barkham TMS, Nagarajan N, Sessions OM |
| EPI_ISL_507041, EPI_ISL_507042, EPI_ISL_507109 | University College London Hospital | COVID-19 Genomics UK (COG-UK) Consortium | Judith Heaney, Matthew Byott, Catherine Houlihan, Dan Frampton, Stuart Kirk, Moira Spyer and Eleni Nastouli |
| EPI_ISL_507198, EPI_ISL_507199, EPI_ISL_507200 | Virology Department, Royal Infirmary of Edinburgh, NHS Lothian / School of Biological Sciences, University of Edinburgh / Institute of Genetics and Molecular Medicine, University of Edinburgh | COVID-19 Genomics UK (COG-UK) Consortium | McHugh M, Dewar R, Rooke S, Gallagher M, Balcaza C, O'Toole Á, Scher E, Hill V, McCrone JT, Colquhoun R, Yu X, Jackson B, Rambaut A, Williams TC, Templeton K |
| EPI_ISL_507225, EPI_ISL_507226, EPI_ISL_507227, EPI_ISL_507229, EPI_ISL_507230, EPI_ISL_507293, EPI_ISL_507294, EPI_ISL_507295, EPI_ISL_507296 | WHO National Influenza Centre Russian Federation | WHO National Influenza Centre Russian Federation | Andrey Komissarov, Artem Fadeev, Mariia Sergeeva, Anna Ivanova, Daria Danilenko |
| EPI_ISL_507937, EPI_ISL_507938, EPI_ISL_507939, EPI_ISL_507940, EPI_ISL_507941, EPI_ISL_507942, EPI_ISL_507943, EPI_ISL_507944, EPI_ISL_507945, EPI_ISL_507946, EPI_ISL_507947, EPI_ISL_507948, EPI_ISL_507949, EPI_ISL_507950, EPI_ISL_507951, EPI_ISL_507952, EPI_ISL_507953, EPI_ISL_507954, EPI_ISL_507955, EPI_ISL_507956 | | | |
| see above | Minnesota Department of Health, Public Health Laboratory | Minnesota Department of Health, Public Health Laboratory | Matt Plumb, Jacob Garfin, and Xiong Wang |
| EPI_ISL_507997, EPI_ISL_508020, EPI_ISL_508021, EPI_ISL_508066 | New Mexico Department of Health Scientific Laboratory Division | Center for Global Health, University of New Mexico Health Sciences Center | Daryl Domman, Kurt Schwalm, Twila Kunde, Joseph Hicks, Michael Edwards, Darrell Dinwiddie |
| EPI_ISL_508341, EPI_ISL_508342, EPI_ISL_508343, EPI_ISL_508344, EPI_ISL_508345, EPI_ISL_508346, EPI_ISL_508347, EPI_ISL_508348, EPI_ISL_508350, EPI_ISL_508351, EPI_ISL_508352, EPI_ISL_508353, EPI_ISL_508354, EPI_ISL_508355, EPI_ISL_508356 | | | |
| see above | Institute of Post Graduate Medical Education & Research | National Institute of Biomedical Genomics | Arindam Maitra, Aritra Biswas, Jayeeta Haldar, Raja Ray, Monimoy Banerjee, Saumitra Das |
| EPI_ISL_508656, EPI_ISL_508657, EPI_ISL_508662, EPI_ISL_508664, EPI_ISL_508665, EPI_ISL_508666, EPI_ISL_508667, EPI_ISL_508668, EPI_ISL_508669, EPI_ISL_508670, EPI_ISL_508671, EPI_ISL_508673, EPI_ISL_508685, EPI_ISL_508686 | | | |
| see above | Departamento de Microbiología, CDB, Hospital Clínic, Barcelona | SeqCOVID-SPAIN consortium/IBV(CSIC) | Andrea Vergara, Mikel Martínez, Elisa Rubio, Jéssica Navero, Aida Peiró and SeqCOVID-SPAIN consortium |
| EPI_ISL_509059, EPI_ISL_509063, EPI_ISL_509064, EPI_ISL_509065 | OHSU Lab Services Molecular Microbiology Lab | Oregon SARS-CoV-2 Genome Sequencing Center | Brendan L. O'Connell, Ruth V. Nichols, Sally B. Grindstaff, Alec J. Hirsch, Guang Fan, Daniel N. Streblow, William B. Messer, Andrew C. Adey, Benjamin N. Bimber, Brian J. O'Roak |
| EPI_ISL_509416 | Acibadem Labcell Cellular Therapy Laboratory | Acibadem Mehmet Ali Aydinlar University School of Medicine, Medical Genetics Department | Ozden Hatirnaz Ng, Sezer Akyoney, Ilayda Sahin, Gunseli Bayram Akcapinar, Ozkan Ozdemir, Derya Dilek Kancagi, Gozde Sir Karakus, Bulut Yurtsever, Cihan Tastan, Ercument Ovali, Ugur Ozbek |
| EPI_ISL_509437, EPI_ISL_509442 | The Princess Alexandra Hospital | Wellcome Sanger Institute for the COVID-19 Genomics UK (COG-UK) consortium | Nick Levene, Louise Lopez, Lynn Monaghan, Jessica Scott, Claudia McCrea and Alex Alderton, Roberto Amato, Sonia Goncalves, Ewan Harrison, David K. Jackson, Ian Johnston, Dominic Kwiatkowski, Cordelia Langford, John Sillitoe on behalf of the Wellcome Sanger Institute COVID-19 Surveillance Team (http://www.sanger.ac.uk/covid-team) |

| | | | |
|---|---|---|---|
| EPI_ISL_509453 | Maryland Department of Health | Maryland Department of Health | Keller,E. |
| EPI_ISL_509576, EPI_ISL_509578 | Utah Public Health Laboratory | Utah Public Health Laboratory | Heidi Butz, Erin Young, Kelly Oakeson |
| EPI_ISL_509690, EPI_ISL_509691 | Utah Public Health Laboratory | Pathogen Discovery, Respiratory Viruses Branch, Division of Viral Diseases, Centers for Disease Control and Prevention | Ying Tao, Jing Zhang, Krista Queen, Anna Uehara, Yan Li, Clinton Paden, Haibin Wang, Suxiang Tong |
| EPI_ISL_509705 | Wisconsin Department of Health Services | Pathogen Discovery, Respiratory Viruses Branch, Division of Viral Diseases, Centers for Disease Control and Prevention | Ying Tao, Jing Zhang, Krista Queen, Anna Uehara, Yan Li, Clinton Paden, Haibin Wang, Suxiang Tong |
| EPI_ISL_509708 | Utah Public Health Laboratory | Pathogen Discovery, Respiratory Viruses Branch, Division of Viral Diseases, Centers for Disease Control and Prevention | Jing Zhang, Ying Tao, Krista Queen, Anna Uehara, Yan Li, Clinton Paden, Haibin Wang, Suxiang Tong |
| EPI_ISL_510057 | Servicio de Microbiología. HRU de Málaga. Servicio Andaluz de Salud | SeqCOVID-SPAIN consortium/IBV(CSIC) | Inmaculada de Toro Peinado. MªConcepción Mediavilla Gradolph. Begoña Palop Borrás and SeqCOVID-SPAIN consortium |
| EPI_ISL_510065, EPI_ISL_510073, EPI_ISL_510074, EPI_ISL_510075 | Instituto de Investigaciones Biomédicas de Barcelona (CSIC), Hospital Clinic i Provincial de Barcelona, Instituto de Biomedicina de Valencia (CSIC), Hospital de Sant Pau | SeqCOVID-SPAIN consortium/IBV(CSIC) | Anna M. Planas, Mª Angeles Marcos, Miguel J. Martínez, Andrea Vergara, Alex Soriano, Jordi Pérez Tur, Israel Fernández Cadenas and SeqCOVID-SPAIN consortium |
| EPI_ISL_510118, EPI_ISL_510140, EPI_ISL_510143 | Hospital General Universitario Gregorio Marañón | SeqCOVID-SPAIN consortium/IBV(CSIC) | Laura Pérez-Lago, Marta Herranz, Jon Sicilia, Julia Suárez, Pilar Catalán, Patricia Muñoz, Darío García de Viedma and SeqCOVID-SPAIN consortium |
| EPI_ISL_510160 | Utah Public Health Laboratory | Pathogen Discovery, Respiratory Viruses Branch, Division of Viral Diseases, Centers for Disease Control and Prevention | Mary L. Killian, Kerrie Franzen, Krista Queen, Anna Uehara, Ying Tao, Yan Li, Jing Zhang, Clinton R. Paden, Haibin Wang, Suxiang Tong |
| EPI_ISL_510337, EPI_ISL_510338, EPI_ISL_510339, EPI_ISL_510340, EPI_ISL_510343, EPI_ISL_510344, EPI_ISL_510345, EPI_ISL_510346, EPI_ISL_510347, EPI_ISL_510348, EPI_ISL_510349, EPI_ISL_510350, EPI_ISL_510351, EPI_ISL_510352, EPI_ISL_510353, EPI_ISL_510354, EPI_ISL_510355, EPI_ISL_510356, EPI_ISL_510357, EPI_ISL_510358, EPI_ISL_510359, EPI_ISL_510360, EPI_ISL_510361, EPI_ISL_510362, EPI_ISL_510363, EPI_ISL_510364, EPI_ISL_510365, EPI_ISL_510366, EPI_ISL_510367, EPI_ISL_510368, EPI_ISL_510369, EPI_ISL_510370, EPI_ISL_510371, EPI_ISL_510372, EPI_ISL_510373, EPI_ISL_510374, EPI_ISL_510375, EPI_ISL_510376, EPI_ISL_510377, EPI_ISL_510378, EPI_ISL_510379, EPI_ISL_510380, EPI_ISL_510381, EPI_ISL_510382, EPI_ISL_510383, EPI_ISL_510384, EPI_ISL_510385 | | | |
| see above | Servicio de Microbiología, Hospital Miguel Servet, Zaragoza | SeqCOVID-SPAIN consortium/IBV(CSIC) | Antonio Rezusta López, Alexander Tristancho Baró, Ana Milagro, Yolanda Gracia Grataloup, Nieves Martínez Cameo and SeqCOVID-SPAIN consortium |
| EPI_ISL_510819 | Klinisk mikrobiologi Vasternorrland | The Public Health Agency of Sweden | Oskar Karlsson Lindsjo, Maria Lind Karlberg, Mattias Haukland, Reza Advani, Olov Svartstrom, Anna-Malin Linde, Sandra Broddesson, Petra Edquist, Mia Brytting, Anna Risberg, Karin Tegmark-Wisell |
| EPI_ISL_510934, EPI_ISL_510951, EPI_ISL_510952, EPI_ISL_511139, EPI_ISL_511140 | Instituto Nacional de Saude (INSA) | Instituto Nacional de Saude (INSA) | Borges et al |
| EPI_ISL_511182, EPI_ISL_511183, EPI_ISL_511184 | Instituto Nacional de Saude (INSA) and Instituto Gulbenkian de Ciencia (IGC) | Instituto Nacional de Saude (INSA) and Instituto Gulbenkian de Ciencia (IGC) | Borges et al |
| EPI_ISL_511505 | Instituto Nacional de Saude (INSA) | Instituto Nacional de Saude (INSA) and Instituto Gulbenkian de Ciencia (IGC) | Borges et al |
| EPI_ISL_511697, EPI_ISL_511698, EPI_ISL_511699, EPI_ISL_511700, EPI_ISL_511701, EPI_ISL_511702, EPI_ISL_511703, EPI_ISL_511704 | Instituto Nacional de Saude (INSA) | Instituto Nacional de Saude (INSA) | Borges et al |
| EPI_ISL_512221, EPI_ISL_512222, EPI_ISL_512223, EPI_ISL_512224, EPI_ISL_512225, EPI_ISL_512226, EPI_ISL_512228, EPI_ISL_512229, EPI_ISL_512230, EPI_ISL_512231, EPI_ISL_512233, EPI_ISL_512234, EPI_ISL_512271, EPI_ISL_512272, EPI_ISL_512273 | | | |
| see above | San Diego County Public Health Laboratory | Andersen lab at Scripps Research | SEARCH Alliance San Diego with Tracy Basler, Jovan Shephard, Brett Austin |
| EPI_ISL_512401, EPI_ISL_512402, EPI_ISL_512403, EPI_ISL_512404, EPI_ISL_512414, EPI_ISL_512415, EPI_ISL_512416, EPI_ISL_512417, EPI_ISL_512418, EPI_ISL_512425 | Centre for Enzyme Innovation, University of Portsmouth / Translational Research Laboratory, Portsmouth Hospitals NHS Trust | COVID-19 Genomics UK (COG-UK) Consortium | Angela Beckett,Yann Bourgeois,Garry Scarlett,Sharon Glaysher,Scott Elliott,Kelly Bicknell,Robert Impey,Allyson Lloyd,Sarah Wyllie,Ethan Butcher,Anoop Chauhan,Samuel Robson |
| EPI_ISL_512639, EPI_ISL_512640 | National Laboratory for Influenza/Virology reference laboratory, Public Health Center of the Ministry of Health of Ukraine | Respiratory Virus Unit, Microbiology Services Colindale, Public Health England | PHE Covid Sequencing Team, Dr. Iryna Demchyshyna |
| EPI_ISL_512780 | Utah Public Health Laboratory, Utah Public Health Laboratory Infectious Disease submission group | Utah Public Health Laboratory, Utah Public Health Laboratory Infectious Disease submission group | Butz,H.A., Young,E.L., Oakeson,K. |
| EPI_ISL_512813 | Kenema Government Hospital, Ministry of Health and Sanitation | Kenema Government Hospital, Ministry of Health and Sanitation | Goba,A., Momoh,M., Sandi,J., Tomkins-Tinch,C., Siddle,K., Mehta,S., Oluniyi,P., Jalloh,S., Park,D., Andersen,K., Garry,R., Happi,C., Grant,D., Olawoye,I. |
| EPI_ISL_512844 | Department of Medical Research | DMR_Myanmar | Myat Htut Nyunt, Hnin Ohnmar Soe, Kay Thi Aye, Wah Wah Aung,Yi Yi Kyaw, Aung Kyaw Kyaw, Theingi Win Myat, Phyu Win Ei, Aung Zaw Latt, Nan Aye Thida Oo, Lai Lai San, Su Mon Win, Ni Ni Zaw, Htin Lin, Hlaing Myat Thu, Zaw Than Htun |
| EPI_ISL_512944, EPI_ISL_512945 | Pathogen Genomics Lab King Abdullah University of Science and Technology(KAUST) | Pathogen Genomics Lab King Abdullah University of Science and Technology(KAUST) | Fadwa Alofi, Sharif Hala, Rahul P Salunke, Sara Mfarrej, Amit Kumar Subudhi, Fathia Ben Rached, Amanda, Luke, Afrah Alsomali, Asim Khogeer, Jumana Taha, Abdulaziz Alahmadi, Kahled Algithami, Raeece Naeem, Anwar Hashem, Naif Almontashiri, Arnab Pain |
| EPI_ISL_512946, EPI_ISL_512947, EPI_ISL_512948, EPI_ISL_512949, EPI_ISL_512950, EPI_ISL_512951, EPI_ISL_512952, EPI_ISL_512953, EPI_ISL_512956, EPI_ISL_512957, EPI_ISL_512958, EPI_ISL_512959, EPI_ISL_512960, EPI_ISL_512961, EPI_ISL_512962, EPI_ISL_512963, EPI_ISL_512964, EPI_ISL_512965, EPI_ISL_512973, EPI_ISL_512976, EPI_ISL_512977, EPI_ISL_512978, EPI_ISL_512979, EPI_ISL_512980, EPI_ISL_512981, EPI_ISL_512982, EPI_ISL_512983, EPI_ISL_512984 | | | |
| see above | Pathogen Genomics Lab King Abdullah University of Science and Technology(KAUST) | Pathogen Genomics Lab King Abdullah University of Science and Technology(KAUST) | Sara Mfarrej, Raeece Naeem, Rahul P Salunke, Sharif Hala, Fadwa Alofi, Amit Kumar Subudhi, Fathia Ben Rached, Afrah Alsomali, Jumana Taha, Abdulaziz Alahmadi, Asim Khogeer, Nashwa Al-khotani, Anwar Hashem, Naif Almontashiri, Arnab Pain |
| EPI_ISL_513127, EPI_ISL_513128, EPI_ISL_513129, EPI_ISL_513130, EPI_ISL_513131, EPI_ISL_513132, EPI_ISL_513133, EPI_ISL_513134, EPI_ISL_513135, EPI_ISL_513136, EPI_ISL_513137, EPI_ISL_513138, EPI_ISL_513139, EPI_ISL_513140, EPI_ISL_513141, EPI_ISL_513142, EPI_ISL_513143, EPI_ISL_513144, EPI_ISL_513145, EPI_ISL_513146 | | | |
| see above | Pathogen Genomics Lab King Abdullah University of Science and Technology(KAUST) | Pathogen Genomics Lab King Abdullah University of Science and Technology(KAUST) | Sharif Hala, Fadwa Alofi, Sara Mfarrej, Amit Kumar Subudhi, Rahul P Salunke, Fathia Ben Rached, Amanda Ooi, Luke Esau, Afrah Alsomali, Asim Khogeer, Jumana Taha, Abdulaziz Alahmadi, Kahled Algithami, Raeece Naeem, Anwar Hashem, Naif Almontashiri, Arnab Pain |
| EPI_ISL_513147, EPI_ISL_513148, EPI_ISL_513149, EPI_ISL_513150, EPI_ISL_513151, EPI_ISL_513152, EPI_ISL_513153, EPI_ISL_513154, EPI_ISL_513155, EPI_ISL_513156, EPI_ISL_513157, EPI_ISL_513158, EPI_ISL_513159, EPI_ISL_513160, EPI_ISL_513161 | | | |
| see above | Pathogen Genomics Lab King Abdullah University of Science and Technology(KAUST) | Pathogen Genomics Lab King Abdullah University of Science and Technology(KAUST) | Fadwa Alofi, Sharif Hala, Rahul P Salunke, Sara Mfarrej, Amit Kumar Subudhi, Fathia Ben Rached, Amanda, Luke, Afrah Alsomali, Asim Khogeer, Jumana Taha, Abdulaziz Alahmadi, Kahled Algithami, Raeece Naeem, Anwar Hashem, Naif Almontashiri, Arnab Pain |
| EPI_ISL_513169, EPI_ISL_513173 | Pathogen Genomics Lab King Abdullah University of Science and Technology(KAUST) | Pathogen Genomics Lab King Abdullah University of Science and Technology(KAUST) | Sara Mfarrej, Raeece Naeem, Rahul P Salunke, Sharif Hala, Fadwa Alofi, Amit Kumar Subudhi, Fathia Ben Rached, Afrah Alsomali, Jumana Taha, Abdulaziz Alahmadi, Asim Khogeer, Nashwa Al-khotani, Anwar Hashem, Naif Almontashiri, Arnab Pain |
| EPI_ISL_513214, EPI_ISL_513215, EPI_ISL_513216, EPI_ISL_513217, EPI_ISL_513218, EPI_ISL_513219, EPI_ISL_513220, EPI_ISL_513221, EPI_ISL_513222, EPI_ISL_513223, EPI_ISL_513224, EPI_ISL_513225, EPI_ISL_513226 | | | |
| see above | Pathogen Genomics Lab King Abdullah University of Science and Technology(KAUST) | Pathogen Genomics Lab King Abdullah University of Science and Technology(KAUST) | Sharif Hala, Fadwa Alofi, Sara Mfarrej, Amit Kumar Subudhi, Rahul P Salunke, Fathia Ben Rached, Amanda Ooi, Luke Esau, Afrah Alsomali, Asim Khogeer, Jumana Taha, Abdulaziz Alahmadi, Kahled Algithami, Raeece Naeem, Anwar Hashem, Naif Almontashiri, Arnab Pain |
| EPI_ISL_513239, EPI_ISL_513248 | Pathogen Genomics Lab King Abdullah University of Science and Technology(KAUST) | Pathogen Genomics Lab King Abdullah University of Science and Technology(KAUST) | Amit Kumar Subudhi, Rahul P Salunke, Sara Mfarrej, Sharif Hala, Fadwa Alofi, Fathia Ben Rached, Afrah Alsomali, Asim Khogeer, Nashwa Al-khotani, Raeece Naeem, Anwar Hashem, Naif Almontashiri, Arnab Pain |
| EPI_ISL_513249, EPI_ISL_513250, EPI_ISL_513251, EPI_ISL_513252, EPI_ISL_513253, EPI_ISL_513254, EPI_ISL_513255, EPI_ISL_513256, EPI_ISL_513257, EPI_ISL_513258, EPI_ISL_513259, EPI_ISL_513260, EPI_ISL_513261, EPI_ISL_513262, EPI_ISL_513263, EPI_ISL_513264 | | | |

| | | | |
|---|---|---|---|
| see above | Pathogen Genomics Lab King Abdullah University of Science and Technology(KAUST) | Pathogen Genomics Lab King Abdullah University of Science and Technology(KAUST) | Afrah Alsomali, Fathia Ben Rached, Raeece Naeem, Sharif Hala,Rahul P Salunke, Amanda Ooi, Luke Esau, Sara Mfarrej, Amit Kumar Subudhi, Fadwa Alofi, Asim Khogeer, Kahled Algithami, Anwar Hashem, Naif Almontashiri, Arnab Pain |
| EPI_ISL_513265 | University of Miami Immunology and Histocompatibility Laboratory | University of Miami Immunology and Histocompatibility Laboratory | Emilio Margolles-Clark, PhD and Phillip Ruiz, MD, PhD |
| EPI_ISL_513552, EPI_ISL_513553, EPI_ISL_513554, EPI_ISL_513555, EPI_ISL_513556, EPI_ISL_513557, EPI_ISL_513558, EPI_ISL_513559, EPI_ISL_513560, EPI_ISL_513561, EPI_ISL_513562, EPI_ISL_513563, EPI_ISL_513564, EPI_ISL_513565, EPI_ISL_513566, EPI_ISL_513567, EPI_ISL_513568, EPI_ISL_513569, EPI_ISL_513570, EPI_ISL_513571, EPI_ISL_513572 | | | |
| see above | Programa de Oncovirologia, Instituto Nacional de Câncer | Programa de Oncovirologia, Instituto Nacional de Câncer | Juliana D. Siqueira, Livia R. Goes, Brunna M. Alves, Claudia Cicala,James Arthos, João P.B. Viola, Andreia C. de Melo, Marcelo A. Soares |
| EPI_ISL_513877 | San Francisco Public Health Laboratory | Chan-Zuckerberg Biohub | CZB Cliahub Consortium |
| EPI_ISL_514062, EPI_ISL_514063, EPI_ISL_514064 | Navy and Marine Corps Public Health Center | Pathogen Discovery, Respiratory Viruses Branch, Division of Viral Diseases, Centers for Disease Control and Prevention | Yan Li, Anna Montmayeur, Krista Queen, Jing Zhang, Ying Tao, Anna Uehara, Rachel Marine, Clinton R. Paden, Haibin Wang, Suxiang Tong |
| EPI_ISL_514065 | Navy and Marine Corps Public Health Center | Pathogen Discovery, Respiratory Viruses Branch, Division of Viral Diseases, Centers for Disease Control and Prevention | Krista Queen, Yan Li, Anna Montmayeur, Jing Zhang, Ying Tao, Anna Uehara, Rachel Marine, Clinton R. Paden, Haibin Wang, Suxiang Tong |
| EPI_ISL_514066 | Navy and Marine Corps Public Health Center | Pathogen Discovery, Respiratory Viruses Branch, Division of Viral Diseases, Centers for Disease Control and Prevention | Yan Li, Anna Montmayeur, Krista Queen, Jing Zhang, Ying Tao, Anna Uehara, Rachel Marine, Clinton R. Paden, Haibin Wang, Suxiang Tong |
| EPI_ISL_514067, EPI_ISL_514068 | Navy and Marine Corps Public Health Center | Pathogen Discovery, Respiratory Viruses Branch, Division of Viral Diseases, Centers for Disease Control and Prevention | Krista Queen, Yan Li, Anna Montmayeur, Jing Zhang, Ying Tao, Anna Uehara, Rachel Marine, Clinton R. Paden, Haibin Wang, Suxiang Tong |
| EPI_ISL_514069, EPI_ISL_514070 | Navy and Marine Corps Public Health Center | Pathogen Discovery, Respiratory Viruses Branch, Division of Viral Diseases, Centers for Disease Control and Prevention | Yan Li, Anna Montmayeur, Krista Queen, Jing Zhang, Ying Tao, Anna Uehara, Rachel Marine, Clinton R. Paden, Haibin Wang, Suxiang Tong |
| EPI_ISL_514071, EPI_ISL_514072 | Navy and Marine Corps Public Health Center | Pathogen Discovery, Respiratory Viruses Branch, Division of Viral Diseases, Centers for Disease Control and Prevention | Krista Queen, Yan Li, Anna Montmayeur, Jing Zhang, Ying Tao, Anna Uehara, Rachel Marine, Clinton R. Paden, Haibin Wang, Suxiang Tong |
| EPI_ISL_514073, EPI_ISL_514074 | Navy and Marine Corps Public Health Center | Pathogen Discovery, Respiratory Viruses Branch, Division of Viral Diseases, Centers for Disease Control and Prevention | Yan Li, Anna Montmayeur, Krista Queen, Jing Zhang, Ying Tao, Anna Uehara, Rachel Marine, Clinton R. Paden, Haibin Wang, Suxiang Tong |
| EPI_ISL_514075, EPI_ISL_514076 | Navy and Marine Corps Public Health Center | Pathogen Discovery, Respiratory Viruses Branch, Division of Viral Diseases, Centers for Disease Control and Prevention | Krista Queen, Yan Li, Anna Montmayeur, Jing Zhang, Ying Tao, Anna Uehara, Rachel Marine, Clinton R. Paden, Haibin Wang, Suxiang Tong |
| EPI_ISL_514077, EPI_ISL_514078, EPI_ISL_514079 | Navy and Marine Corps Public Health Center | Pathogen Discovery, Respiratory Viruses Branch, Division of Viral Diseases, Centers for Disease Control and Prevention | Yan Li, Anna Montmayeur, Krista Queen, Jing Zhang, Ying Tao, Anna Uehara, Rachel Marine, Clinton R. Paden, Haibin Wang, Suxiang Tong |
| EPI_ISL_514080, EPI_ISL_514081, EPI_ISL_514082, EPI_ISL_514083 | Navy and Marine Corps Public Health Center | Pathogen Discovery, Respiratory Viruses Branch, Division of Viral Diseases, Centers for Disease Control and Prevention | Krista Queen, Yan Li, Anna Montmayeur, Jing Zhang, Ying Tao, Anna Uehara, Rachel Marine, Clinton R. Paden, Haibin Wang, Suxiang Tong |
| EPI_ISL_514084 | Navy and Marine Corps Public Health Center | Pathogen Discovery, Respiratory Viruses Branch, Division of Viral Diseases, Centers for Disease Control and Prevention | Yan Li, Anna Montmayeur, Krista Queen, Jing Zhang, Ying Tao, Anna Uehara, Rachel Marine, Clinton R. Paden, Haibin Wang, Suxiang Tong |
| EPI_ISL_514085, EPI_ISL_514086, EPI_ISL_514087 | Navy and Marine Corps Public Health Center | Pathogen Discovery, Respiratory Viruses Branch, Division of Viral Diseases, Centers for Disease Control and Prevention | Krista Queen, Yan Li, Anna Montmayeur, Jing Zhang, Ying Tao, Anna Uehara, Rachel Marine, Clinton R. Paden, Haibin Wang, Suxiang Tong |
| EPI_ISL_514088 | Navy and Marine Corps Public Health Center | Pathogen Discovery, Respiratory Viruses Branch, Division of Viral Diseases, Centers for Disease Control and Prevention | Yan Li, Anna Montmayeur, Krista Queen, Jing Zhang, Ying Tao, Anna Uehara, Rachel Marine, Clinton R. Paden, Haibin Wang, Suxiang Tong |
| EPI_ISL_514089 | Navy and Marine Corps Public Health Center | Pathogen Discovery, Respiratory Viruses Branch, Division of Viral Diseases, Centers for Disease Control and Prevention | Krista Queen, Yan Li, Anna Montmayeur, Jing Zhang, Ying Tao, Anna Uehara, Rachel Marine, Clinton R. Paden, Haibin Wang, Suxiang Tong |
| EPI_ISL_514090, EPI_ISL_514091, EPI_ISL_514092 | Navy and Marine Corps Public Health Center | Pathogen Discovery, Respiratory Viruses Branch, Division of Viral Diseases, Centers for Disease Control and Prevention | Yan Li, Anna Montmayeur, Krista Queen, Jing Zhang, Ying Tao, Anna Uehara, Rachel Marine, Clinton R. Paden, Haibin Wang, Suxiang Tong |
| EPI_ISL_514126 | National Scientific Center for Especially Dangerous Infections (NSCEDI) | Kazakh National Agrarian University (KazNAU) TreeGene LLP Genetic Laboratory | Tabynov Kaissar, Beloussov Vyacheslav, Strochkov Vitaliy, Sandybayev Nurlan, Tabynov Kairat, Turebekov Nurkeldy, Granica Joanna, Solomadin Maksim, Yerubayev Toktassyn, Yespolov Tlektes |
| EPI_ISL_514128 | Navy and Marine Corps Public Health Center | Centers for Disease Control and Prevention | Krista Queen, Yan Li, Anna Montmayeur, Jing Zhang, Ying Tao, Anna Uehara, Rachel Marine, Clinton R. Paden, Haibin Wang, Suxiang Tong |
| EPI_ISL_514339, EPI_ISL_514340, EPI_ISL_514341, EPI_ISL_514342 | Laboratorio de Referencia Nacional de Virus Respiratorio. Instituto Nacional de Salud Perú | Laboratorio de Referencia Nacional de Biotecnología y Biología Molecular. Instituto Nacional de Salud Perú | Carlos Padilla Rojas, Karolyn Vega Chozo, Priscila Lope Pari, Omar Caceres Rey, Marco Galarza Perez, Maribel Huaringa Nuñez, Johanna Balbuena Torres, Henri Bailon Calderon, Nancy Rojas Serrano. |
| EPI_ISL_514431 | Laboratory Diagnostic, Veterinary Specialized Institute Kraljevo | Laboratory Diagnostic, Veterinary Specialized Institute Kraljevo | Vidanovic,D., Tesovic,B., Knezevic,A., Jankovic,M., Sekler,M., Banovic Djeri,B., Volkening,J., Afonso,C., Petrovic,T. |
| EPI_ISL_514503, EPI_ISL_514504, EPI_ISL_514505, EPI_ISL_514506, EPI_ISL_514507, EPI_ISL_514508, EPI_ISL_514509, EPI_ISL_514510, EPI_ISL_514511, EPI_ISL_514512, EPI_ISL_514513 | | | |
| see above | Centre for Enzyme Innovation, University of Portsmouth / Translational Research Laboratory, Portsmouth Hospitals NHS Trust | COVID-19 Genomics UK (COG-UK) Consortium | Angela Beckett,Yann Bourgeois,Garry Scarlett,Sharon Glaysher,Scott Elliott,Kelly Bicknell,Robert Impey,Allyson Lloyd,Sarah Wyllie,Ethan Butcher,Anoop Chauhan,Samuel Robson |
| EPI_ISL_515103 | Department of Biochemistry, Cell and Molecular Biology | WACCBIP, University of Ghana | Ngoi,J.M., Quashie,P., Morang'a,C.M., Amuzu,D.S., Adu,B., Kumordjie,S., Eshun,M., Boatemaa,L., Magnussen,V., Kotey,E., Tei-Maya,F., Arjarquah,A., Mutungi,J.K., Bediako,Y., Asante,I., Bonney,E., Kyei,G.B., Bonney,K., Amenga-Etego,L.N., Anang,A.K., Awandare,G.A., Ampofo,W. |
| EPI_ISL_515262 | Minnesota Department of Health, Public Health Laboratory | Minnesota Department of Health, Public Health Laboratory | Matt Plumb, Jacob Garfin, and Xiong Wang |
| EPI_ISL_515386, EPI_ISL_515387, EPI_ISL_515388, EPI_ISL_515389, EPI_ISL_515390, EPI_ISL_515391, EPI_ISL_515392, EPI_ISL_515393, EPI_ISL_515394 | Nevada State Public Health Laboratory | Nevada State Public Health Laboratory | Richard Tillett, Joel R. Sevinsky, Paul Hartley, Heather Kerwin, David Jackson, Subhash C. Verma, Cyprian Rossetto, Andrew Gorzalski, Chris Laverdure, Natalie Crawford, Stephanie Van Hooser, and Mark Pandori |
| EPI_ISL_515542 | Vigilância Epidemiológica de Leme | Instituto Adolfo Lutz, Interdiciplinary Procedures Center, Strategic Laboratory | Claudio Tavares Sacchi, Claudia Regina Gonçalves, Erica Valessa Ramos Gomes |
| EPI_ISL_515566 | PS Municipal Dr Lauro Ribas Braga | Instituto Adolfo Lutz, Interdiciplinary Procedures Center, Strategic Laboratory | Claudio Tavares Sacchi, Claudia Regina Gonçalves, Erica Valessa Ramos Gomes |
| EPI_ISL_515916 | California Department of Public Health | California Department of Public Health | CDPH IDLB COVIDNet |
| EPI_ISL_515938, EPI_ISL_515939, EPI_ISL_515940, EPI_ISL_515941, EPI_ISL_515942 | CV RAMAN HOSPITAL | Department of Neurovirology, National Institute of Mental Health and Neuroscience (NIMHANS) | Chitra Pattabiraman,Vijayalakshmi Reddy, Harsha PK, Risha Rasheed, Pramada Prasad, Shafeeq S Hameed, Manjunatha Venkataswamy, Anita Desai, Ravi Vasanthapuram |
| EPI_ISL_516491 | University of Wisconsin-Madison AIDS Vaccine Research Laboratories | University of Wisconsin-Madison AIDS Vaccine Research Laboratories | Gage Moreno, Katarina Braun, et al. AIDS Vaccine Research Laboratories |
| EPI_ISL_516712 | Virginia DCLS | Virginia DCLS | Virginia DCLS |
| EPI_ISL_517782 | Florida Bureau of Public Health Laboratories | Florida Bureau of Public Health Laboratories | Sarah Schmedes, Jason Blanton |

| | | | |
|---|---|---|---|
| EPI_ISL_518843, EPI_ISL_518844, EPI_ISL_518845 | Oman-National Influenza Center | Biotechnology & OMICs Laboratory, Natural & Medical Sciences Research Center, University of Nizwa | Samiha Al-Kharusi, Sajjad Asaf, Abdul Latif Khan, Samira Al-Mahruqi, Adil Khan, Ahmed Al-Rawahi, Amina Al-Jardani, Hanan Al-Kindi, Intisar Al-Shukri, Ahlam Al-Amri, Aisha Al-Amri, Aisha Al-Busaidi, Adil Al-Wahaibi, Seif Al-Abri, Ahmed Al-Harrasi |
| EPI_ISL_518846, EPI_ISL_518847, EPI_ISL_518848, EPI_ISL_518849, EPI_ISL_518850 | Oman-National Influenza Center | Biotechnology & OMICs Laboratory, Natural & Medical Sciences Research Center, University of Nizwa | Abdul Latif Khan, Samira Al-Mahruqi, Ahmed Al-Harrasi, Samiha Al-Kharusi, Adil Khan, Ahmed Al-Rawahi, Sajjad Asaf, Amina Al-Jardani, Hanan Al-Kindi, Intisar Al-Shukri, Ahlam Al-Amri, Aisha Al-Amri, Aisha Al-Busaidi, Adil Al-Wahaibi, Seif Al-Abri. |
| EPI_ISL_520666, EPI_ISL_520667, EPI_ISL_520672, EPI_ISL_520739, EPI_ISL_520744 | Mohammed Bin Rashid University of Medicine and Health Sciences | Al Jalila Genomics Center | Ahmad Abou Tayoun, Tom Loney, Hamda Khansaheb, Sathishkumar Ramaswamy, Divinlal Harilal, Zulfa Omar Deesi, Rupa Murthy Varghese, Hanan Al Suwaidi, Abdulmajeed Alkhaja, Mohammed Uddin, Rifat Hamoudi, Rabih Halwani, Abiola Catherine Senok, Qutayba Hamid, Norbert Nowotny, Alawi Alsheikh-Ali |
| EPI_ISL_522254, EPI_ISL_522255, EPI_ISL_522256, EPI_ISL_522257, EPI_ISL_522258, EPI_ISL_522259, EPI_ISL_522260, EPI_ISL_522261, EPI_ISL_522262, EPI_ISL_522263, EPI_ISL_522271, EPI_ISL_522272 | | | |
| see above | Utah Public Health Laboratory | Utah Public Health Laboratory | Erin Young, Kelly Oakeson |
| EPI_ISL_522591, EPI_ISL_522597, EPI_ISL_522624, EPI_ISL_522631, EPI_ISL_522637, EPI_ISL_522639, EPI_ISL_522642, EPI_ISL_522645, EPI_ISL_522674, EPI_ISL_522675, EPI_ISL_522676, EPI_ISL_522678, EPI_ISL_522679, EPI_ISL_522680, EPI_ISL_522681 | | | |
| see above | Royal Hobart Hospital Microbiology Department | MDU-PHL | Cooley L., van Haeften R., Seemann T., Sait M., Schultz, M.B., Sherry N. |
| EPI_ISL_522855 | ULSS9 Distretto di Bussolengo | Istituto Zooprofilattico Sperimentale delle Venezie | Adelaide Milani, Alessia Schivo, Annalisa Salviato, Erika Giorgia Quaranta, Ambra Pastori, Bianca Zecchin, Alice Fusaro, Isabella Monne, Calogero Terregino, Antonia Ricci |
| EPI_ISL_522857 | ULSS9 Scaligera | Istituto Zooprofilattico Sperimentale delle Venezie | Adelaide Milani, Alessia Schivo, Annalisa Salviato, Erika Giorgia Quaranta, Ambra Pastori, Bianca Zecchin, Alice Fusaro, Isabella Monne, Calogero Terregino, Antonia Ricci |
| EPI_ISL_523135, EPI_ISL_523170, EPI_ISL_523171, EPI_ISL_523174, EPI_ISL_523205, EPI_ISL_523237, EPI_ISL_523238, EPI_ISL_523425, EPI_ISL_523426, EPI_ISL_523429, EPI_ISL_523483, EPI_ISL_523484, EPI_ISL_523485, EPI_ISL_523529, EPI_ISL_523603, EPI_ISL_523692 | | | |
| see above | Dutch COVID-19 response team | Erasmus Medical Center | Bas Oude Munnink, David Nieuwenhuijse, Reina Sikkema, Claudia Schapendonk, Irina Chestakova, Anne van der Linden, Theo Bestebroer, Stefan van Nieuwkoop, Mark Pronk, Pascal Lexmond, Corien Swaan, Manon Haverkate, Madelief Mollers, Mart Stein, Sandra Kengne Kamga Mobou, Jeroen van Kampen, Jolanda Voermans, Aura Timen, Corine GeurtsvanKessel, Annemiek van der Eijk, Richard Molenkamp, Marion Koopmans, on behalf of the Dutch national COVID-19 response team. |
| EPI_ISL_523928 | Center of Medical Microbiology, Virology, and Hospital Hygiene, University of Duesseldorf | Center of Medical Microbiology, Virology, and Hospital Hygiene, University of Duesseldorf | Maximilian Damagnez, Alexander Dilthey, Torsten Houwaart, Malte Kohns Vasconcelos, Marek Korencak, Jessica Nicolai, Klaus Pfeffer, Hendrik Streeck, Daniel Strelow, Jörg Timm, Andreas Walker, Tobias Wienemann |
| EPI_ISL_523986 | Ama Dr Jose Soares Hungria | Instituto Adolfo Lutz, Interdiciplinary Procedures Center, Strategic Laboratory | Claudio Tavares Sacchi, Claudia Regina Gonçalves, Erica Valessa Ramos Gomes |
| EPI_ISL_523988 | Hospital Sao Paulo de Ensino da Unifesp | Instituto Adolfo Lutz, Interdiciplinary Procedures Center, Strategic Laboratory | Claudio Tavares Sacchi, Claudia Regina Gonçalves, Erica Valessa Ramos Gomes |
| EPI_ISL_523989 | AMA Jardim Joamar | Instituto Adolfo Lutz, Interdiciplinary Procedures Center, Strategic Laboratory | Claudio Tavares Sacchi, Claudia Regina Gonçalves, Erica Valessa Ramos Gomes |
| EPI_ISL_523991, EPI_ISL_523992 | Hospital Municipal Carmen Prudente | Instituto Adolfo Lutz, Interdiciplinary Procedures Center, Strategic Laboratory | Claudio Tavares Sacchi, Claudia Regina Gonçalves, Erica Valessa Ramos Gomes |
| EPI_ISL_523995, EPI_ISL_523996, EPI_ISL_524049, EPI_ISL_524050, EPI_ISL_524051 | WHO National Influenza Centre Russian Federation | WHO National Influenza Centre Russian Federation | Andrey Komissarov, Artem Fadeev, Mariia Sergeeva, Anna Ivanova, Daria Danilenko |
| EPI_ISL_524070 | Texas Department of State Health Services | Texas Department of State Health Services | Rashmi Tuladhar, Bonnie Oh, Cara Akrout, Jenny Zhang, Maliha Rahman, Anita Pokharel, Myong Koag, Chun Wang, Rachel Lee, Grace Kubin |
| EPI_ISL_524435 | Department of Immunology, The Scripps Research Institute | Department of Immunology, The Scripps Research Institute | SEARCH Alliance,S.D., Pride,D., Shin,J.H. |
| EPI_ISL_524466 | PS Municipal Dr Lauro Ribas Braga | Instituto Adolfo Lutz, Interdiciplinary Procedures Center, Strategic Laboratory | Claudio Tavares Sacchi, Claudia Regina Gonçalves, Erica Valessa Ramos Gomes |
| EPI_ISL_524467 | Hospital Municipal Dr. Moysés Deutsch | Instituto Adolfo Lutz, Interdiciplinary Procedures Center, Strategic Laboratory | Claudio Tavares Sacchi, Claudia Regina Gonçalves, Erica Valessa Ramos Gomes |
| EPI_ISL_524578, EPI_ISL_524608 | Department of Pathology, University of Cambridge | Wellcome Sanger Institute for the COVID-19 Genomics UK (COG-UK) consortium | Luke W Meredith, M. Estée Török , Myra Hosmillo, William L. Hamilton, Martin D. Curran, Theresa Feltwell, Grant Hall, Anna Yakovleva, Fahad A Khokhar, Charlotte J. Houldcroft, Laura G Caller, Aminu S. Jahun, Sarah L. Caddy, Ian Goodfellow; and Alex Alderton, Roberto Amato, Sonia Goncalves, Ewan Harrison, David K. Jackson, Ian Johnston, Dominic Kwiatkowski, Cordelia Langford, John Sillitoe on behalf of the Wellcome Sanger Institute COVID-19 Surveillance Team (http://www.sanger.ac.uk/covid-team) |
| EPI_ISL_524788, EPI_ISL_524794, EPI_ISL_524796 | Evandro Chagas Institute | Evandro Chagas Institute | Santos, M.C.; Silva, A.M.; Junior, W.D.C.; Barbagelata, L.S.; Ferreira, J.A.; Sousa, E.M.A.; da Silva, P.S.; Resque, H.R; Martins, L.C.; Sousa Junior, E.C.;Viana, G.M.R |
| EPI_ISL_524879 | MD PHL | MD PHL | Maryland Department of Health Laboratories Administration |
| EPI_ISL_525424 | Oman-National Influenza Center | Biotechnology & OMICs Laboratory | Samira Al-Mahruqi, Abdul Latif Khan, Samiha Al-Kharusi, Adil Khan , Ahmed Al-Rawahi, Sajjad Asaf, Amina Al-Jardani, Hanan Al-Kindi, Intisar Al-Shukri, Ahlam Al-Amri, Aisha Al-Amri, Aisha Al-Busaidi, Adil Al-Wahaibi, Seif Al-Abri, Ahmed Al-Harrasi |
| EPI_ISL_525686 | Wadsworth Center, New York State Department of Health | Wadsworth Center, New York State Department of Health | Kirsten St. George, Daryl M. Lamson, Sara Griesemer, Jonathan Plitnick, Navjot Singh, Matthew D. Shudt, Erica Lasek-Nesselquist |
| EPI_ISL_525786, EPI_ISL_525787, EPI_ISL_525788, EPI_ISL_525789 | Texas Department of State Health Services | Texas Department of State Health Services | Jenny Zhang, Rashmi Tuladhar, Bonnie Oh, Maliha Rahman, Anita Pokharel, Myong Koag, Chun Wang, Rachel Lee, Grace Kubin |
| EPI_ISL_526218, EPI_ISL_526226, EPI_ISL_526228, EPI_ISL_526231, EPI_ISL_526235, EPI_ISL_526238 | Hungarian Defence Forces Military Medical Centre | National Laboratory of Virology, Szentágothai Research Centre | Endre Gábor Tóth, Balázs Somogyi, Bálint Eszenyi, Ferenc Jakab, Gábor Kemenesi |
| EPI_ISL_526479 | Virology Department, Royal Infirmary of Edinburgh, NHS Lothian / School of Biological Sciences, University of Edinburgh / Institute of Genetics and Molecular Medicine, University of Edinburgh | COVID-19 Genomics UK (COG-UK) Consortium | McHugh M, Dewar R, Rooke S, Gallagher M, Balcaza C, O'Toole Á, Scher E, Hill V, McCrone JT, Colquhoun R, Yu X, Jackson B, Rambaut A, Williams TC, Templeton K |
| EPI_ISL_526554, EPI_ISL_526555, EPI_ISL_526556, EPI_ISL_526559, EPI_ISL_526560, EPI_ISL_526561, EPI_ISL_526562, EPI_ISL_526563 | Florida Bureau of Public Health Laboratories | Florida Bureau of Public Health Laboratories | Sarah Schmedes, Jason Blanton |
| EPI_ISL_526755, EPI_ISL_526756, EPI_ISL_526757, EPI_ISL_526758, EPI_ISL_526759, EPI_ISL_526760, EPI_ISL_526761, EPI_ISL_526762, EPI_ISL_526763, EPI_ISL_526764, EPI_ISL_526765, EPI_ISL_526766, EPI_ISL_526767, EPI_ISL_526768, EPI_ISL_526769, EPI_ISL_526770 | | | |
| see above | Respiratory Virus Unit, Microbiology Services Colindale, Public Health England | Respiratory Virus Unit, Microbiology Services Colindale, Public Health England | PHE Covid Sequencing Team |
| EPI_ISL_527060 | Area of Virology, Serology and Virology Division (SAViD), New South Wales Health Pathology Randwick | Area of Virology, Serology and Virology Division (SAViD), New South Wales Health Pathology Randwick | Rawlinson, W. |
| EPI_ISL_527257, EPI_ISL_527340, EPI_ISL_527341, EPI_ISL_527342, EPI_ISL_527343, EPI_ISL_527344, EPI_ISL_527345, EPI_ISL_527346, EPI_ISL_527347, EPI_ISL_527348, EPI_ISL_527349, EPI_ISL_527350, EPI_ISL_527351, EPI_ISL_527352, EPI_ISL_527353, EPI_ISL_527354, EPI_ISL_527355, EPI_ISL_527356, EPI_ISL_527357 | | | |
| see above | Respiratory Virus Unit, Microbiology Services Colindale, | Respiratory Virus Unit, Microbiology Services Colindale, | PHE Covid Sequencing Team |

| | | | |
|---|---|---|---|
| | Public Health England | Public Health England | |
| EPI_ISL_527376, EPI_ISL_527377 | National Public Health Laboratory, National Centre for Infectious Diseases | National Public Health Laboratory, National Centre for Infectious Diseases | Mak TM, Octavia S, Zhou Z, Cui L, Lin RTP |
| EPI_ISL_527423, EPI_ISL_527424, EPI_ISL_527430, EPI_ISL_527448, EPI_ISL_527464, EPI_ISL_527471, EPI_ISL_527481, EPI_ISL_527482, EPI_ISL_527485, EPI_ISL_527486, EPI_ISL_527488 | | | |
| see above | Colorado State University - Ebel Lab | Colorado State University - Ebel Lab | Greg Ebel et al. |
| EPI_ISL_527573 | Minnesota Department of Health, Public Health Laboratory | Minnesota Department of Health, Public Health Laboratory | Matt Plumb, Jacob Garfin, and Xiong Wang |
| EPI_ISL_527667, EPI_ISL_527668, EPI_ISL_527669, EPI_ISL_527670, EPI_ISL_527671, EPI_ISL_527672, EPI_ISL_527673, EPI_ISL_527674, EPI_ISL_527675, EPI_ISL_527677, EPI_ISL_527723, EPI_ISL_527733 | | | |
| see above | MN PHL Division, Minnesota Department of Health | Pathogen Discovery, Respiratory Viruses Branch, Division of Viral Diseases, Centers for Disease Control and Prevention | Yan Li, Anna Montmayeur, Jing Zhang, Krista Queen, Ying Tao, Anna Uehara, Rachel Marine, Clinton R. Paden, Haibin Wang, Suxiang Tong |
| EPI_ISL_527820, EPI_ISL_527821, EPI_ISL_527822, EPI_ISL_527823, EPI_ISL_527824, EPI_ISL_527825, EPI_ISL_527826, EPI_ISL_527827, EPI_ISL_527828, EPI_ISL_527829, EPI_ISL_527830 | | | |
| see above | Texas Department of State Health Services | Texas Department of State Health Services | Bonnie Oh, Rashmi Tuladhar, Jenny Zhang, Maliha Rahman, Anita Pokharel, Myong Koag, Chun Wang, Rachel Lee, Grace Kubin |
| EPI_ISL_527859 | Hospital Municipal Vereador Jose Storopolli | Instituto Adolfo Lutz, Interdiciplinary Procedures Center, Strategic Laboratory | Claudio Tavares Sacchi, Claudia Regina Gonçalves, Erica Valessa Ramos Gomes |
| EPI_ISL_527861 | Hospital e Maternidade Celso Pierro | Instituto Adolfo Lutz, Interdiciplinary Procedures Center, Strategic Laboratory | Av. Dr. Arnaldo, 355 - Brazil, Cerqueira Cesar, São Paulo - SP, 01246-1301 |
| EPI_ISL_527864 | Hospital e Pronto Socorro Comunitário Vila Iolanda | Instituto Adolfo Lutz, Interdiciplinary Procedures Center, Strategic Laboratory | Claudio Tavares Sacchi, Claudia Regina Gonçalves, Erica Valessa Ramos Gomes |
| EPI_ISL_527865 | Hospital e Maternidade São Cristóvão | Instituto Adolfo Lutz, Interdiciplinary Procedures Center, Strategic Laboratory | Claudio Tavares Sacchi, Claudia Regina Gonçalves, Erica Valessa Ramos Gomes |
| EPI_ISL_527866 | PS Municipal Dr Lauro Ribas Braga | Instituto Adolfo Lutz, Interdiciplinary Procedures Center, Strategic Laboratory | Av. Dr. Arnaldo, 355 - Brazil, Cerqueira Cesar, São Paulo - SP, 01246-1301 |
| EPI_ISL_527867 | Pronto Socorro Municipal - Balneario São José | Instituto Adolfo Lutz, Interdiciplinary Procedures Center, Strategic Laboratory | Claudio Tavares Sacchi, Claudia Regina Gonçalves, Erica Valessa Ramos Gomes |
| EPI_ISL_527869 | Hospital Municipal Carmen Prudente | Instituto Adolfo Lutz, Interdiciplinary Procedures Center, Strategic Laboratory | Claudio Tavares Sacchi, Claudia Regina Gonçalves, Erica Valessa Ramos Gomes |
| EPI_ISL_527870 | Hospital Municipal Mário Gatti | Instituto Adolfo Lutz, Interdiciplinary Procedures Center, Strategic Laboratory | Claudio Tavares Sacchi, Claudia Regina Gonçalves, Erica Valessa Ramos Gomes |
| EPI_ISL_527884, EPI_ISL_527885, EPI_ISL_527886 | Nigeria Centre for Disease Control (NCDC) | African Centre of Excellence for Genomics of Infectious Diseases (ACEGID), Redeemer's University, Ede, Osun State, Nigeria | Oluniyi P.E. et al |
| EPI_ISL_528390, EPI_ISL_528405, EPI_ISL_528407, EPI_ISL_528431, EPI_ISL_528432, EPI_ISL_528433, EPI_ISL_528434, EPI_ISL_528435 | Respiratory Virus Unit, Microbiology Services Colindale, Public Health England | Respiratory Virus Unit, Microbiology Services Colindale, Public Health England | PHE Covid Sequencing Team |
| EPI_ISL_528710, EPI_ISL_528711, EPI_ISL_528712, EPI_ISL_528713, EPI_ISL_528714, EPI_ISL_528715, EPI_ISL_528716, EPI_ISL_528717, EPI_ISL_528719, EPI_ISL_528720, EPI_ISL_528721 | | | |
| see above | Alsafar - Khalifa University Abu Dhabi | Alsafar - Khalifa University Abu Dhabi | Andreas Henschel, Gihan Daw Elbait, Samuel Feng, Rifat Hamoudi, Ernesto Damiani, Guan Tay, Habiba Alsafar |
| EPI_ISL_529153 | Department of Immunology, The Scripps Research Institute | Andersen lab at Scripps Research | Quigley, M., Stefanski, E., Mchardy, I. with SEARCH Alliance San Diego |
| EPI_ISL_529178 | South Carolina Department of Health and Environmental Control | South Carolina Department of Health and Environmental Control | Haley V. Flores |
| EPI_ISL_529922, EPI_ISL_529923, EPI_ISL_529924, EPI_ISL_529925, EPI_ISL_529926, EPI_ISL_529927, EPI_ISL_529928, EPI_ISL_529929, EPI_ISL_529930, EPI_ISL_529931, EPI_ISL_529932, EPI_ISL_529933, EPI_ISL_529934, EPI_ISL_529935 | | | |
| see above | Virginia Division of Consolidated Laboratory Services | Virginia Division of Consolidated Laboratory Services | Virginia DCLS |
| EPI_ISL_529965 | RSUD Dr. Soetomo | Institute of Tropical Disease, Universitas Airlangga | Krisnoadi Rahardjo, Aldise M Nastri, Jezzy R Dewantari, Rima R Prasetya, Joni Wahyuhadi, Gatot Soegiarto, Laksmi Wulandari, Retno A Setyoningrum, Resti Yudhawati, Yohko K Shimizu, Mitsuhiro Nishimura, Yasuko Mori, Soetjipto, Kazufumi Shimizu, Maria I Lusida |
| EPI_ISL_530024 | Hospital Universitario 12 de Octubre | Hospital Universitario 12 de Octubre | Esther Viedma, Raúl Recio, Sara González, Elias Dahdouh, Fernando Lázaro, Natalia Stella, Julio García, Juan Carlos Galán, Rafael Cantón, Mª Dolores Folgueira, Rafael Delgado, Jesús Mingorance |
| EPI_ISL_530025, EPI_ISL_530026, EPI_ISL_530027, EPI_ISL_530030, EPI_ISL_530031, EPI_ISL_530032, EPI_ISL_530033, EPI_ISL_530036, EPI_ISL_530037, EPI_ISL_530038 | Hospital Universitario La Paz | Hospital Universitario La Paz | María Rodríguez, Elias Dahdouh, Sara González, Raúl Recio, Fernando Lázaro, Esther Viedma, Natalia Stella, Julio García, Juan Carlos Galán, Rafael Cantón, Mª Dolores Folgueira, Rafael Delgado, Jesús Mingorance |
| EPI_ISL_530052, EPI_ISL_530053, EPI_ISL_530054, EPI_ISL_530055, EPI_ISL_530056, EPI_ISL_530057, EPI_ISL_530058, EPI_ISL_530059, EPI_ISL_530060, EPI_ISL_530061, EPI_ISL_530062, EPI_ISL_530063, EPI_ISL_530064, EPI_ISL_530065 | | | |
| see above | Hospital Universitario La Paz | Hospital Universitario La Paz | Elias Dahdouh, Sara González, Raúl Recio, Fernando Lázaro, Esther Viedma, Natalia Stella, Julio García, Juan Carlos Galán, Rafael Cantón, Mª Dolores Folgueira, Rafael Delgado, Jesús Mingorance |
| EPI_ISL_530070, EPI_ISL_530073, EPI_ISL_530085, EPI_ISL_530092 | Hospital Universitario La Paz | Hospital Universitario La Paz | María Rodríguez, Elias Dahdouh, Sara González, Raúl Recio, Fernando Lázaro, Esther Viedma, Natalia Stella, Julio García, Juan Carlos Galán, Rafael Cantón, Mª Dolores Folgueira, Rafael Delgado, Jesús Mingorance |
| EPI_ISL_530100, EPI_ISL_530101, EPI_ISL_530102, EPI_ISL_530103, EPI_ISL_530104, EPI_ISL_530108 | Hospital Universitario Ramón y Cajal | Hospital Universitario La Paz | Raúl Recio, Sara González, Elias Dahdouh, Fernando Lázaro, Esther Viedma, Natalia Stella, Julio García, Juan Carlos Galán, Rafael Cantón, Mª Dolores Folgueira, Rafael Delgado, Jesús Mingorance |
| EPI_ISL_530208, EPI_ISL_530209, EPI_ISL_530210, EPI_ISL_530211, EPI_ISL_530212, EPI_ISL_530213, EPI_ISL_530214, EPI_ISL_530215 | Minnesota Department of Health, Public Health Laboratory | Minnesota Department of Health, Public Health Laboratory | Matt Plumb, Jacob Garfin, and Xiong Wang |
| EPI_ISL_534322 | PS Mun Julio Tupy | Instituto Adolfo Lutz, Interdiciplinary Procedures Center, Strategic Laboratory | Claudio Tavares Sacchi, Claudia Regina Gonçalves, Erica Valessa Ramos Gomes |
| EPI_ISL_534323 | Hospital e Pronto Socorro Comunitario Vila Yolanda | Instituto Adolfo Lutz, Interdiciplinary Procedures Center, Strategic Laboratory | Claudio Tavares Sacchi, Claudia Regina Gonçalves, Erica Valessa Ramos Gomes |
| EPI_ISL_534324 | Hospital Mun Ver Jose Storopolli | Instituto Adolfo Lutz, Interdiciplinary Procedures Center, Strategic Laboratory | Claudio Tavares Sacchi, Claudia Regina Gonçalves, Erica Valessa Ramos Gomes |
| EPI_ISL_534325 | Unidade de Vigilancia em Saude de Guarulhos | Instituto Adolfo Lutz, Interdiciplinary Procedures Center, Strategic Laboratory | Claudio Tavares Sacchi, Claudia Regina Gonçalves, Erica Valessa Ramos Gomes |

| | | | |
|---|---|---|---|
| EPI_ISL_534333 | Hospital Universitario La Paz | Hospital Universitario La Paz | María Rodríguez, Elias Dahdouh, Sara González, Raúl Recio, Fernando Lázaro, Esther Viedma, Natalia Stella, Julio García, Juan Carlos Galán, Rafael Cantón, Ma Dolores Folgueira, Rafael Delgado, Jesús Mingorance |
| EPI_ISL_534823, EPI_ISL_534825, EPI_ISL_534829, EPI_ISL_534843, EPI_ISL_534847, EPI_ISL_534848, EPI_ISL_534866, EPI_ISL_534881, EPI_ISL_534909, EPI_ISL_534912, EPI_ISL_534922, EPI_ISL_534927, EPI_ISL_534942, EPI_ISL_534951, EPI_ISL_534962, EPI_ISL_534965, EPI_ISL_534967, EPI_ISL_534969, EPI_ISL_534972, EPI_ISL_534973, EPI_ISL_534974, EPI_ISL_534984, EPI_ISL_534988, EPI_ISL_534990, EPI_ISL_534993, EPI_ISL_534994, EPI_ISL_534995, EPI_ISL_534997, EPI_ISL_534998, EPI_ISL_535004, EPI_ISL_535012, EPI_ISL_535013, EPI_ISL_535014, EPI_ISL_535019 | | | |
| see above | Oxford Viromics, NDM, University of Oxford; Oxford University Hospitals; Basingstoke and North Hampshire Hospital | COVID-19 Genomics UK (COG-UK) Consortium | Tanya Golubchik, David Bonsall, George Macintyre, Amy Trebes, Mariateresa de Cesare, Catrin Moore, Alex Mobbs, Anita Justice, Robert Shaw, Monique Andersson, Timothy Peto, Emma Wise, Nathan Moore, Jessica Lynch, Nick Cortes, Matilde Mori, Stephen Kidd, David Buck, John Todd, Christophe Fraser |
| EPI_ISL_537474, EPI_ISL_537478, EPI_ISL_537484, EPI_ISL_537491, EPI_ISL_537499, EPI_ISL_537534, EPI_ISL_537549 | UCLA Pathology Clinical Microbiology Lab | Kruglyak Lab | Guo et al. |
| EPI_ISL_537618, EPI_ISL_537619, EPI_ISL_537620, EPI_ISL_537621, EPI_ISL_537622, EPI_ISL_537623, EPI_ISL_537624, EPI_ISL_537625, EPI_ISL_537626, EPI_ISL_537627, EPI_ISL_537628, EPI_ISL_537629, EPI_ISL_537630, EPI_ISL_537631, EPI_ISL_537632, EPI_ISL_537633, EPI_ISL_537634, EPI_ISL_537635, EPI_ISL_537636, EPI_ISL_537637, EPI_ISL_537638, EPI_ISL_537639, EPI_ISL_537640, EPI_ISL_537641, EPI_ISL_537642, EPI_ISL_537643, EPI_ISL_537644, EPI_ISL_537645, EPI_ISL_537646, EPI_ISL_537647, EPI_ISL_537648, EPI_ISL_537649, EPI_ISL_537650, EPI_ISL_537651, EPI_ISL_537652, EPI_ISL_537653, EPI_ISL_537654, EPI_ISL_537655, EPI_ISL_537656, EPI_ISL_537657, EPI_ISL_537658, EPI_ISL_537659, EPI_ISL_537660, EPI_ISL_537661, EPI_ISL_537662, EPI_ISL_537663, EPI_ISL_537664, EPI_ISL_537665, EPI_ISL_537666, EPI_ISL_537667, EPI_ISL_537668, EPI_ISL_537669, EPI_ISL_537670, EPI_ISL_537671, EPI_ISL_537672, EPI_ISL_537673, EPI_ISL_537674, EPI_ISL_537675, EPI_ISL_537676 | | | |
| see above | Universidad de León | SeqCOVID-SPAIN consortium/IBV(CSIC) | Ana Carvajal, Vicente Martín, Héctor Argüello, Juan M. Fregeneda, Tania Fernández-Villa, Antonio J. Molina and SeqCOVID-SPAIN consortium |
| EPI_ISL_538002 | Servicio de Microbiologia. Hospital Arnau de Vilanova | SeqCOVID-SPAIN consortium/IBV(CSIC) | Victoria Dominguez, Maria Alma Bracho, Griselda De Marco, Lidia Ruiz Roldan, Neris Garcia-Gonzalez, Inma Galán Vendrell, Sandra Carbo, Loreto Ferrús Abad, Paula Ruiz-Hueso, Mariana Reyes-Prieto, Vicente Soriano Chirona, Ivan Ansari, Llúcia Martínez-Priego, Giuseppe D'Auria, Fernando Gonzalez-Candelas and SeqCOVID-SPAIN consortium |
| EPI_ISL_538808, EPI_ISL_538868, EPI_ISL_538920, EPI_ISL_538927, EPI_ISL_539029, EPI_ISL_539039, EPI_ISL_539048 | Leeds Teaching Hospitals NHS Trust and Public Health England, National Infection Service (Leeds laboratory) | Wellcome Sanger Institute for the COVID-19 Genomics UK (COG-UK) consortium | Louissa Macfarlane-Smith, Holli Carden, Katherine L. Harper, Antony Hale and Alex Alderton, Roberto Amato, Sonia Goncalves, Ewan Harrison, David K. Jackson, Ian Johnston, Dominic Kwiatkowski, Cordelia Langford, John Sillitoe on behalf of the Wellcome Sanger Institute COVID-19 Surveillance Team |
| EPI_ISL_539321 | KWR Watercycle Research Institute | Erasmus Medical Center | Ray Izquierdo-Lara, Goffe Elsinga, Leo Heijnen, Bas B. Oude Munnink, Claudia M. E. Schapendonk, David Nieuwenhuijse, Matthijs Kon, Lu Lu, Frank M. Aarestrup, Samantha Lycett, Gertjan Medema, Marion P.G. Koopmans, Miranda de Graaf |
| EPI_ISL_539780 | National Institute of Public Health (Czech Republic) | State Veterinary Institute Prague | Nagy, A; Jirincova, H; Novakova, L; Trnka, D; Vecerova, J. |
| EPI_ISL_539797, EPI_ISL_539798, EPI_ISL_539799, EPI_ISL_539800 | Wyoming Public Health Laboratory | Wyoming Public Health Laboratory | Noah Hull, Rob Christensen, Jim Mildenberger, Joel Sevinsky, Cari Sloma, and Wanda Manley |
| EPI_ISL_540436 | WI State Laboratory of Hygiene | Pathogen Discovery, Respiratory Viruses Branch, Division of Viral Diseases, Centers for Disease Control and Prevention | Yan Li, Jing Zhang, Anna Montmayeur, Krista Queen,Ying Tao, Anna Uehara, Clinton R. Paden, Rachel Marine, Haibin Wang, Suxiang Tong |
| EPI_ISL_541056, EPI_ISL_541057 | Hospital Clínico Universitario de Santiago de Compostela | SeqCOVID-SPAIN consortium/Institute of Biomedicine of Valencia, IBV-CSIC | José Javier Costa Alcalde, Antonio Aguilera Guirao, Mª Luisa Pérez del Molino Bernal, Amparo Coira Nieto, Gema Barbeito Castiñeiras, Rocio Trastoy Pena and SeqCOVID-SPAIN consortium |
| EPI_ISL_541152, EPI_ISL_541155, EPI_ISL_541156, EPI_ISL_541211 | Florida Bureau of Public Health Laboratories, Florida Department of Health | Florida Bureau of Public Health Laboratories, Florida Department of Health | Schmedes,S., Blanton,J. |
| EPI_ISL_541357, EPI_ISL_541358, EPI_ISL_541359, EPI_ISL_541360, EPI_ISL_541361 | Laboratory of Respiratory Viruses and Measles, Oswaldo Cruz Institute, FIOCRUZ | Laboratory of Respiratory Viruses and Measles, Oswaldo Cruz Institute, FIOCRUZ | Paola Resende, Luciana Appolinario, Fernando Motta, Anna Carolina Paixão, Ana Carolina Mendonça, Jonathan Lopes, Marilda Siqueira |
| EPI_ISL_541377, EPI_ISL_541378, EPI_ISL_541379, EPI_ISL_541385, EPI_ISL_541386, EPI_ISL_541387, EPI_ISL_541388, EPI_ISL_541389, EPI_ISL_541392, EPI_ISL_541393, EPI_ISL_541394, EPI_ISL_541395, EPI_ISL_541396 | | | |
| see above | LACEN/SE | Laboratory of Respiratory Viruses and Measles, Oswaldo Cruz Institute, FIOCRUZ | Paola Resende, Luciana Appolinario, Fernando Motta, Anna Carolina Paixão, Ana Carolina Mendonça, Jonathan Lopes, Clioma Santos, Marilda Siqueira |
| EPI_ISL_541397 | Laboratório de Virologia Comparada e Ambiental- LVCA-IOC | Laboratory of Respiratory Viruses and Measles, Oswaldo Cruz Institute, FIOCRUZ | Paola Resende, Luciana Appolinario, Tulio Machado Fumian, Tatiana Prado, Camille Ferreira Mannarino, Fernando Motta, Ana Carolina Mendonça, Marilda Siqueira, Marize Pereira Miagostovich |
| EPI_ISL_541697, EPI_ISL_541698, EPI_ISL_541700 | National Institute of Virology, NIV Influenza | National Institute of Virology, NIV Influenza | Potdar V |
| EPI_ISL_541763, EPI_ISL_541765 | Barts Health NHS Trust | Wellcome Sanger Institute for the COVID-19 Genomics UK (COG-UK) consortium | Teresa Cutino-Moguel, Mark Hopkins, Beatrix Kele, David Harrington and Alex Alderton, Roberto Amato, Sonia Goncalves, Ewan Harrison, David K. Jackson, Ian Johnston, Dominic Kwiatkowski, Cordelia Langford, John Sillitoe on behalf of the Wellcome Sanger Institute COVID-19 Surveillance Team |
| EPI_ISL_541882 | Hospital General Universitario Gregorio Marañón | SeqCOVID-SPAIN consortium/IBV(CSIC) | Laura Pérez-Lago, Marta Herranz, Jon Sicilia, Julia Suárez, Pilar Catalán, Patricia Muñoz, Darío García de Viedma and SeqCOVID-SPAIN consortium |
| EPI_ISL_542524, EPI_ISL_542527, EPI_ISL_542528, EPI_ISL_542529, EPI_ISL_542530, EPI_ISL_542531, EPI_ISL_542533, EPI_ISL_542534, EPI_ISL_542536, EPI_ISL_542538, EPI_ISL_542539, EPI_ISL_542540, EPI_ISL_542542, EPI_ISL_542543, EPI_ISL_542545, EPI_ISL_542546, EPI_ISL_542547, EPI_ISL_542549, EPI_ISL_542554, EPI_ISL_542562, EPI_ISL_542563, EPI_ISL_542567, EPI_ISL_542568, EPI_ISL_542569, EPI_ISL_542570, EPI_ISL_542575, EPI_ISL_542576, EPI_ISL_542577, EPI_ISL_542583, EPI_ISL_542588, EPI_ISL_542592, EPI_ISL_542593, EPI_ISL_542597, EPI_ISL_542598, EPI_ISL_542601, EPI_ISL_542603, EPI_ISL_542604, EPI_ISL_542608, EPI_ISL_542609, EPI_ISL_542613, EPI_ISL_542631, EPI_ISL_542632, EPI_ISL_542636, EPI_ISL_542639, EPI_ISL_542640, EPI_ISL_542641, EPI_ISL_542646, EPI_ISL_542652, EPI_ISL_542657, EPI_ISL_542658, EPI_ISL_542661, EPI_ISL_542666, EPI_ISL_542687, EPI_ISL_542689, EPI_ISL_542693, EPI_ISL_542694, EPI_ISL_542695, EPI_ISL_542704, EPI_ISL_542711, EPI_ISL_542713, EPI_ISL_542714, EPI_ISL_542721, EPI_ISL_542731, EPI_ISL_542733, EPI_ISL_542744, EPI_ISL_542752, EPI_ISL_542761, EPI_ISL_542769, EPI_ISL_542770, EPI_ISL_542773, EPI_ISL_542791, EPI_ISL_542792, EPI_ISL_542797, EPI_ISL_542798, EPI_ISL_542855, EPI_ISL_542876, EPI_ISL_542879, EPI_ISL_542905, EPI_ISL_542907 | | | |
| see above | Houston Methodist Hospital | Houston Methodist Hospital | S. Wesley Long, Randall J. Olsen, Paul A. Christensen, David W. Bernard, James J. Davis, Maulik Shukla, Marcus Nguyen, Matthew Ojeda Saavedra, Concepcion C. Cantu, Prasanti Yerramilli, Layne Pruitt, Sishir Subedi, Hung-Che Kuo, Heather Hendrickson, Ghazaleh Eskandari, Hoang A. T. Nguyen, J. Hunter Long, Muthiah Kumaraswami, Jule Goike, Daniel Boutz, Jimmy Gollihar, Jason S. McLellan, Chia-Wei Chou, Kamyab Javanmardi, Ilya J. Finkelstein, and James M. Musser |
| EPI_ISL_542939, EPI_ISL_542940, EPI_ISL_542976 | TriCore Reference Laboratories | Center for Global Health, University of New Mexico Health Sciences Center | Daryl Domman, Kurt Schwalm, Twila Kunde, Joseph Hicks, Michael Edwards, Darrell Dinwiddie |
| EPI_ISL_545239, EPI_ISL_545279, EPI_ISL_545280, EPI_ISL_545299, EPI_ISL_545845, EPI_ISL_545894, EPI_ISL_545895, EPI_ISL_545896 | Houston Methodist Hospital | Houston Methodist Hospital | S. Wesley Long, Randall J. Olsen, Paul A. Christensen, David W. Bernard, James J. Davis, Maulik Shukla, Marcus Nguyen, Matthew Ojeda Saavedra, Concepcion C. Cantu, Prasanti Yerramilli, Layne Pruitt, Sishir Subedi, Hung-Che Kuo, Heather Hendrickson, Ghazaleh Eskandari, Hoang A. T. Nguyen, J. Hunter Long, Muthiah Kumaraswami, Jule Goike, Daniel Boutz, Jimmy Gollihar, Jason S. McLellan, Chia-Wei Chou, Kamyab Javanmardi, Ilya J. Finkelstein, and James M. Musser |
| EPI_ISL_547479, EPI_ISL_547486, EPI_ISL_547496, EPI_ISL_547506, EPI_ISL_547507, EPI_ISL_547508, EPI_ISL_547522, EPI_ISL_547538 | Dutch COVID-19 response team | National Institute for Public Health and the Environment (RIVM) | Adam Meijer, Harry Vennema, Jeroen Cremer, Sharon van den Brink, Bas van der Veer, AnneMarie van den Brandt, Florian Zwagemaker, Dennis Schmitz, Chantal Reusken, on behalf of the national COVID-19 response team |
| EPI_ISL_548955 | Max von Pettenkofer Institute, Virology, National Reference Center for Retroviruses, LMU München | Laboratory for Functional Genome Analysis, Dept. Genomics, Gene Center of the LMU Munich | Max Muenchhoff, Stefan Krebs, Alexander Graf, Oliver Keppler, Helmut Blum |
| EPI_ISL_548966, EPI_ISL_548967, EPI_ISL_548968, EPI_ISL_548969, EPI_ISL_548970, EPI_ISL_548971 | Expo2020 Emergency Center | Agiomix | Walaa Allam, Cherif Ben Hamada, Cengiz Yakicier, Walid Dridi, Rashid Mohammed, Tamer Degheidy |
| EPI_ISL_549020 | KWR Watercycle Research Institute | Erasmus Medical Center | Ray Izquierdo-Lara, Goffe Elsinga, Leo Heijnen, Bas B. Oude Munnink, Claudia M. E. Schapendonk, David Nieuwenhuijse, Matthijs Kon, Lu Lu, Frank M. Aarestrup, Samantha Lycett, Gertjan Medema, Marion P.G. Koopmans, Miranda de Graaf |
| EPI_ISL_549181, EPI_ISL_549244, | Florida Bureau of Public Health Laboratories | Florida Bureau of Public Health Laboratories | Sarah Schmedes, Jason Blanton |

| | | | |
|---|---|---|---|
| EPI_ISL_549245 | | | |
| EPI_ISL_559696 | Lighthouse Lab in Milton Keynes | Wellcome Sanger Institute for the COVID-19 Genomics UK (COG-UK) consortium | The Lighthouse Lab in Milton Keynes and Alex Alderton, Roberto Amato, Sonia Goncalves, Ewan Harrison, David K. Jackson, Ian Johnston, Dominic Kwiatkowski, Cordelia Langford, John Sillitoe on behalf of the Wellcome Sanger Institute COVID-19 Surveillance Team (http://www.sanger.ac.uk/covid-team) |
| EPI_ISL_560572, EPI_ISL_560580 | hôpital | National Reference Center for Viruses of Respiratory Infections, Institut Pasteur, Paris | Sylvie Behillil, Fabiana Gambaro, Etienne Simon-Lorière, Vincent Enouf, Maud Vanpeene, Sylvie van der Werf |
| EPI_ISL_560584, EPI_ISL_560586 | Hopital | National Reference Center for Viruses of Respiratory Infections, Institut Pasteur, Paris | Sylvie Behillil, Fabiana Gambaro, Etienne Simon-Lorière, Vincent Enouf, Maud Vanpeene, Sylvie van der Werf |
| EPI_ISL_560633 | Hospital | National Reference Center for Viruses of Respiratory Infections, Institut Pasteur, Paris | Sylvie Behillil, Fabiana Gambaro, Etienne Simon-Lorière, Vincent Enouf, Maud Vanpeene, Sylvie van der Werf |
| EPI_ISL_560806 | Maryland Public Health Laboratory | Maryland Public Health Laboratory | Maryland Department of Health Laboratories Administration |
| EPI_ISL_561523, EPI_ISL_562066, EPI_ISL_562539 | Victorian Infectious Diseases Reference Laboratory (VIDRL) | VIDRL and MDU-PHL | Caly, L., Seemann, T., Sait, M., Schultz, M. B., Druce J., Sherry, N. |
| EPI_ISL_565890 | Michigan Department of Health and Human Services, Bureau of Laboratories | Michigan Department of Health and Human Services, Bureau of Laboratories | Blankenship HM, Riner D, Soehnlen MK |
| EPI_ISL_568480 | Virology, Iran University of Medical Sciences | Virology, Iran University of Medical Sciences | Keyvani,H., Ranjbar,Mm., Keyvani,F., Soleimani,S. |
| EPI_ISL_568481, EPI_ISL_568489 | Virology, Iran University of Medical Sciences | Virology, Iran University of Medical Sciences | Keyvani,H., Ranjbar,M.M., Keyvani,F., Soleimani,S. |
| EPI_ISL_568490, EPI_ISL_568499, EPI_ISL_568502 | Virology, Iran University of Medical Sciences | Virology, Iran University of Medical Sciences | Keyvani,H., Ranjbar,Mm., Soleimani,S., Keyvani,F. |
| EPI_ISL_568563 | Department of Infectious Diseases and Immunology, National Hospital Organization Nagoya Medical Center | Clinical Research Center, National Hospital Organization Nagoya Medical Center | Yoshihiro Nakata, Hirotaka Ode, Mai Kubota, Masakazu Matsuda, Kazuhiro Matsuoka, Nakasuji Miho, Mikiko Mori, Mayumi Imahashi, Yoshiyuki Yokomaku, Yasumasa Iwatani |
| EPI_ISL_568688 | RS Pelni | Eijkman Institute for Molecular Biology, Ministry of Research and Technology/National Agency for Research and Innovation | Frilasita A Yudhaputri, Edison Johar, Hidayat Trimarsanto, Iskandar A Adnan, Willy Agustine, David H Muljono, Safarina G Malik, Herawati Sudoyo, Khin Saw Myint, Amin Soebandrio |
| EPI_ISL_568709, EPI_ISL_568710, EPI_ISL_568711, EPI_ISL_568712 | KEMRI-Wellcome Trust Research Programme/KEMRI-CGMR-C Kilifi | KEMRI-Wellcome Trust Research Programme/KEMRI-CGMR-C Kilifi | Githinji et al 2020 |
| EPI_ISL_568876 | Florida Bureau of Public Health Laboratories | Florida Bureau of Public Health Laboratories | Sarah Schmedes, Jason Blanton |
| EPI_ISL_568947, EPI_ISL_568948, EPI_ISL_568949, EPI_ISL_568950, EPI_ISL_568951, EPI_ISL_568952, EPI_ISL_568953, EPI_ISL_568954, EPI_ISL_568955 | MEPHI, Aix Marseille University | MEPHI, Aix Marseille University | Anthony LEVASSEUR |
| EPI_ISL_569849 | Omsk Research Institute of Natural Focal Infections | WHO National Influenza Centre Russian Federation | Artem Fadeev, Ekaterina Gradoboeva, Ekaterina Savkina, Daria Nashatyreva, Elena Poleshchuk, Aleksei Vasilenko, Valery Yakimenko, Andrey Komissarov |
| EPI_ISL_570192, EPI_ISL_570195, EPI_ISL_570196, EPI_ISL_570197, EPI_ISL_570198, EPI_ISL_570199, EPI_ISL_570200 | UW Virology Lab | UW Virology Lab | Pavitra Roychoudhury, Hong Xie, Lasata Shrestha, Amin Addetia, Victoria M Rachleff, Meei-Li Huang, Keith R Jerome, Alexander Greninger |
| EPI_ISL_572335, EPI_ISL_572336, EPI_ISL_572337, EPI_ISL_572338, EPI_ISL_572340, EPI_ISL_572341, EPI_ISL_572342, EPI_ISL_572343, EPI_ISL_572344, EPI_ISL_572346, EPI_ISL_572347, EPI_ISL_572348, EPI_ISL_572349, EPI_ISL_572350, EPI_ISL_572351, EPI_ISL_572352, EPI_ISL_572354, EPI_ISL_572355 | | | |
| see above | LACEN/PE | WallauLab, Aggeu Magalhaes Institute | Marcelo Henrique Santos Paiva, Duschinka Ribeiro Duarte Guedes, Cássia Docena, Matheus Filgueira Bezerra, Filipe Zimmer Dezordi, Laís Ceschini Machado, Larissa Krokovsky, Elisama Helvecio, Alexandre Freitas da Silva, Luydson Richardson Silva Vasconcelos, Antonio Mauro Rezende, Severino Jefferson Ribeiro da Silva, Kamila Gaudêncio da Silva Sales, Bruna Santos Lima Figueiredo de Sá, Derciliano Lopes da Cruz, Claudio Eduardo Cavalcanti, Armando de Menezes Neto, Caroline Targino Alves da Silva, Renata Pessôa Germano Mendes, Maria Almerice Lopes da Silva, Tiago Gräf, Paola Cristina Resende, Gonzalo Bello0, Michelle da Silva Barros, Wheverton Ricardo Correia do Nascimento,, Rodrigo Moraes Loyo Arcoverde, Luciane Caroline Albuquerque Bezerra, Sinval Pinto Brandão Filho, Constância Flávia Junqueira Ayres, Gabriel Luz Wallau |
| EPI_ISL_574578 | Hospital Municipal Mário Gatti | Instituto Adolfo Lutz, Interdiciplinary Procedures Center, Strategic Laboratory | Claudio Tavares Sacchi, Claudia Regina Gonçalves, Erica Valessa Ramos Gomes, Karoline Rodrigues Campos |
| EPI_ISL_574579 | Hospital Municipal Dr. Ignacio Proença de Gouvea | Instituto Adolfo Lutz, Interdiciplinary Procedures Center, Strategic Laboratory | Claudio Tavares Sacchi, Claudia Regina Gonçalves, Erica Valessa Ramos Gomes, Karoline Rodrigues Campos |
| EPI_ISL_574580 | Hospital Cidade Tiradentes Carmen Prudente | Instituto Adolfo Lutz, Interdiciplinary Procedures Center, Strategic Laboratory | Claudio Tavares Sacchi, Claudia Regina Gonçalves, Erica Valessa Ramos Gomes, Karoline Rodrigues Campos |
| EPI_ISL_574582, EPI_ISL_574589 | Hospital Municipal Dr. Jose Soares Hungria | Instituto Adolfo Lutz, Interdiciplinary Procedures Center, Strategic Laboratory | Claudio Tavares Sacchi, Claudia Regina Gonçalves, Erica Valessa Ramos Gomes, Karoline Rodrigues Campos |
| EPI_ISL_574590 | Unidade de Printo Atendimento UPA I Santa Isabel | Instituto Adolfo Lutz, Interdiciplinary Procedures Center, Strategic Laboratory | Claudio Tavares Sacchi, Claudia Regina Gonçalves, Erica Valessa Ramos Gomes, Karoline Rodrigues Campos |
| EPI_ISL_574610 | Puskesmas Tambora | Eijkman Institute for Molecular Biology, Ministry of Research and Technology/National Agency for Research and Innovation | Frilasita A Yudhaputri, Edison Johar, Hidayat Trimarsanto, Iskandar A Adnan, Willy Agustine, David H Muljono, Safarina G Malik, Herawati Sudoyo, Khin Saw Myint, Amin Soebandrio |
| EPI_ISL_574862 | Institute for Infectious Diseases, University of Bern | Institute for Infectious Diseases, University of Bern | Michel C Koch, Christian Baumann, Miguel A Terrazos Miani, Cora Sägesser, Stephen L Leib, Peter Keller, Franziska Suter-Riniker, Alban Ramette |
| EPI_ISL_575074, EPI_ISL_575075, EPI_ISL_575076, EPI_ISL_575077, EPI_ISL_575078, EPI_ISL_575079, EPI_ISL_575080, EPI_ISL_575081, EPI_ISL_575082, EPI_ISL_575083, EPI_ISL_575084, EPI_ISL_575085, EPI_ISL_575086, EPI_ISL_575087, EPI_ISL_575088, EPI_ISL_575089, EPI_ISL_575090, EPI_ISL_575091, EPI_ISL_575092, EPI_ISL_575093, EPI_ISL_575096 | | | |
| see above | Utah Public Health Laboratory | Utah Public Health Laboratory | Erin Young, Kelly Oakeson |
| EPI_ISL_576178 | DC Public Health Lab/ Dept. of Forensic Sciences | Pathogen Discovery, Respiratory Viruses Branch, Division of Viral Diseases, Centers for Disease Control and Prevention | Ying Tao, Jing Zhang, Brian Lynch, Yan Li, Krista Queen, Anna Uehara, Clinton R. Paden, Peter Cook, Haibin Wang, Suxiang Tong |
| EPI_ISL_576514, EPI_ISL_576515, EPI_ISL_576516, EPI_ISL_576517 | UW Virology Lab | UW Virology Lab | Pavitra Roychoudhury, Hong Xie, Lasata Shrestha, Amin Addetia, Victoria M Rachleff, Meei-Li Huang, Keith R Jerome, Alexander Greninger |
| EPI_ISL_577193, EPI_ISL_577194, EPI_ISL_577195 | Centre for Enzyme Innovation, University of Portsmouth / Translational Research Laboratory, Portsmouth Hospitals NHS Trust | COVID-19 Genomics UK (COG-UK) Consortium | Angela Beckett,Yann Bourgeois,Garry Scarlett,Sharon Glaysher,Scott Elliott,Kelly Bicknell,Robert Impey,Allyson Lloyd,Sarah Wyllie,Ethan Butcher,Anoop Chauhan,Samuel Robson |
| EPI_ISL_578160, EPI_ISL_578161, EPI_ISL_578162, EPI_ISL_578163, EPI_ISL_578164, EPI_ISL_578165, EPI_ISL_578166, EPI_ISL_578167, EPI_ISL_578185 | University of Michigan Clinical Microbiology Laboratory | Lauring Lab, University of Michigan, Department of Microbiology and Immunology | Valesano |

| | | | |
|---|---|---|---|
| EPI_ISL_578356, EPI_ISL_578357, EPI_ISL_578358, EPI_ISL_578359, EPI_ISL_578360, EPI_ISL_578361, EPI_ISL_578362, EPI_ISL_578363, EPI_ISL_578364, EPI_ISL_578365, EPI_ISL_578485, EPI_ISL_578488, EPI_ISL_578489, EPI_ISL_578490, EPI_ISL_578491, EPI_ISL_578494, EPI_ISL_578495, EPI_ISL_578496, EPI_ISL_578497, EPI_ISL_578498, EPI_ISL_578501, EPI_ISL_578502, EPI_ISL_578503, EPI_ISL_578506, EPI_ISL_578508, EPI_ISL_578509, EPI_ISL_578510, EPI_ISL_578511, EPI_ISL_578512, EPI_ISL_578513, EPI_ISL_578514, EPI_ISL_578515, EPI_ISL_578516, EPI_ISL_578517, EPI_ISL_578518, EPI_ISL_578519, EPI_ISL_578520, EPI_ISL_578521, EPI_ISL_578522, EPI_ISL_578523, EPI_ISL_578524, EPI_ISL_578525, EPI_ISL_578526, EPI_ISL_578527, EPI_ISL_578528, EPI_ISL_578529, EPI_ISL_578530, EPI_ISL_578531, EPI_ISL_578532, EPI_ISL_578533, EPI_ISL_578534, EPI_ISL_578535, EPI_ISL_578536, EPI_ISL_578537, EPI_ISL_578538, EPI_ISL_578539, EPI_ISL_578540, EPI_ISL_578541, EPI_ISL_578542, EPI_ISL_578543, EPI_ISL_578544, EPI_ISL_578545, EPI_ISL_578547, EPI_ISL_578548, EPI_ISL_578549, EPI_ISL_578550, EPI_ISL_578551, EPI_ISL_578552, EPI_ISL_578553, EPI_ISL_578554, EPI_ISL_578555, EPI_ISL_578556, EPI_ISL_578557, EPI_ISL_578558, EPI_ISL_578559, EPI_ISL_578560, EPI_ISL_578561, EPI_ISL_578562, EPI_ISL_578563, EPI_ISL_578564, EPI_ISL_578565, EPI_ISL_578566, EPI_ISL_578567, EPI_ISL_578568, EPI_ISL_578585, EPI_ISL_578586, EPI_ISL_578587 | | | |
| see above | Wisconsin State Laboratory of Hygiene Communicable Disease Division | Wisconsin State Laboratory of Hygiene Communicable Disease Division | Kelsey R. Florek, Abigail C. Shockey |
| EPI_ISL_579404 | Middlemore Hospital | Institute of Environmental Science and Research (ESR) | Xiaoyun Ren, Matt Storey, Nikki Freed, Muhammad Faisal, Jing Wang, Hermes Perez, Anja Werno, Antje van der Linden, Arlo Upton, Chris Mansell, David Hammer, Dragana Drinkovic, Gary McAuliffe, Hana Sofia Andersson, James Ussher, Jill Sherwood, Josh Freeman, Julia Howard, Juliet Elvy, Mary DeAlmeida, Matt Blakiston, Matthew Rogers, Max Bloomfield, Michael Addidle, Michelle Balm, Sally Roberts, Sarah Jefferies, Sharmini Muttaiyah, Susan Morpeth, Susan Taylor, Timothy Blackmore, Vani Sathyendran, Veronica Playle, Virginia Hope, Erasmus Smit, Lauren Jelly, Olin Silander, Joep de Ligt |
| EPI_ISL_579421 | LabTests | Institute of Environmental Science and Research (ESR) | Xiaoyun Ren, Matt Storey, Nikki Freed, Muhammad Faisal, Jing Wang, Hermes Perez, Anja Werno, Antje van der Linden, Arlo Upton, Chris Mansell, David Hammer, Dragana Drinkovic, Gary McAuliffe, Hana Sofia Andersson, James Ussher, Jill Sherwood, Josh Freeman, Julia Howard, Juliet Elvy, Mary DeAlmeida, Matt Blakiston, Matthew Rogers, Max Bloomfield, Michael Addidle, Michelle Balm, Sally Roberts, Sarah Jefferies, Sharmini Muttaiyah, Susan Morpeth, Susan Taylor, Timothy Blackmore, Vani Sathyendran, Veronica Playle, Virginia Hope, Erasmus Smit, Lauren Jelly, Olin Silander, Joep de Ligt |
| EPI_ISL_579505, EPI_ISL_579506 | Canterbury Health Laboratories | Institute of Environmental Science and Research (ESR) | Xiaoyun Ren, Matt Storey, Nikki Freed, Muhammad Faisal, Jing Wang, Hermes Perez, Anja Werno, Antje van der Linden, Arlo Upton, Chris Mansell, David Hammer, Dragana Drinkovic, Gary McAuliffe, Hana Sofia Andersson, James Ussher, Jill Sherwood, Josh Freeman, Julia Howard, Juliet Elvy, Mary DeAlmeida, Matt Blakiston, Matthew Rogers, Max Bloomfield, Michael Addidle, Michelle Balm, Sally Roberts, Sarah Jefferies, Sharmini Muttaiyah, Susan Morpeth, Susan Taylor, Timothy Blackmore, Vani Sathyendran, Veronica Playle, Virginia Hope, Erasmus Smit, Lauren Jelly, Olin Silander, Joep de Ligt |
| EPI_ISL_579528, EPI_ISL_579532, EPI_ISL_579533, EPI_ISL_579534, EPI_ISL_579535 | QEII Health Sciences Centre | National Microbiology Laboratory (NML) | Anna Majer, Shari Tyson, Grace Seo, Philip Mabon, Darian Hole, Elsie Grudeski, Rhiannon Huzarewich, Russell Mandes, Anneliese Landgraff, Jennifer Tanner, Natalie Knox, Morag Graham, Gary Van Domselaar, Todd Hatchette, Jason LeBlanc, Nathalie Bastien, Yan Li, Timothy Booth, CanCOGeN's metadata curation team, Public Health Agency of Canada's CanCOGeN team |
| EPI_ISL_581453, EPI_ISL_581467, EPI_ISL_581468, EPI_ISL_581485 | Medizinische Klinik Innere Medizin I, Universitätsklinikum Tübingen | NGS Competence Center Tübingen, Institut für Medizinische Mikrobiologie und Hygiene, Universitätsklinikum Tübingen | Angel Angelov |
| EPI_ISL_581715, EPI_ISL_581916, EPI_ISL_581917, EPI_ISL_581918, EPI_ISL_581919, EPI_ISL_581920 | University Hospital Basel, Clinical Virology | University Hospital Basel, Clinical Bacteriology | Madlen Stange, Alfredo Mari, Tim Roloff, Helena MB Seth-Smith, Michael Schweitzer, Myrta Brunner, Karoline Leuzinger, Kirstine K. Soegaard, Alexander Gensch, Sarah Tschudin-Sutter, Simon Fuchs, Julia Bielicki, Hans Pargger, Martin Siegemund, Christian Nickel, Roland Bingisser, Michael Osthoff, Stefano Bassetti, Rita Schneider-Sliwa, Manuel Battegay, Hans Hirsch, Adrian Egli |
| EPI_ISL_582180, EPI_ISL_582197, EPI_ISL_582201, EPI_ISL_582203, EPI_ISL_582212, EPI_ISL_582215, EPI_ISL_582217 | TriCore Reference Laboratories | Center for Global Health, University of New Mexico Health Sciences Center | Daryl Domman, Kurt Schwalm, Twila Kunde, Joseph Hicks, Michael Edwards, Darrell Dinwiddie |
| EPI_ISL_582325, EPI_ISL_582408, EPI_ISL_582507 | Cadham Provincial Laboratory | National Microbiology Laboratory (NML) | Anna Majer, Shari Tyson, Grace Seo, Philip Mabon, Elsie Grudeski, Rhiannon Huzarewich, Russell Mandes, Anneliese Landgraff, Jennifer Tanner, Natalie Knox, Morag Graham, Gary Van Domselaar, Paul Van Caeseele, Jared Bullard, David Alexander, Kerry Dust, Nathalie Bastien, Yan Li, Timothy Booth, Darian Hole, Madison Chapel, CanCOGeN's metadata curation team, Public Health Agency of Canada CanCOGeN team |
| EPI_ISL_582635, EPI_ISL_582636 | Sheikh Khalifa Medical City | Molecular/Surveillance lab Sheikh Khalifa Medical City | Amirtharaj Francis, Sajeed Abdul, Hala Imambaccus, Sahar Almarzooqi, Hiba Saud, Stefan Weber |
| EPI_ISL_583430, EPI_ISL_583437, EPI_ISL_583456 | Memorial Sloan Kettering Cancer Center | van Bakel Laboratory, Genetics and Genomics Sciences, Icahn School of Medicine at Mount Sinai | Teresa Aydillo, Ana S. Gonzalez-Reiche, Sadaf Aslam, Adriana van de Guchte, Zenab Khan, Ajay Obla, Jayeeta Dutta, Harm van Bakel, Judith Aberg, Adolfo García-Sastre, Gunjan Shah, Tobias Hohl, Genovefa Papanicolaou, Miguel-Angel Perales, Kent Sepkowitz, Ngolela Esther Babady, and Mini Kamboj |
| EPI_ISL_583458 | Garcia-Sastre Laboratory, Department of Microbiology, Icahn School of Medicine at Mount Sinai | van Bakel Laboratory, Genetics and Genomics Sciences, Icahn School of Medicine at Mount Sinai | Teresa Aydillo, Ana S. Gonzalez-Reiche, Sadaf Aslam, Adriana van de Guchte, Zenab Khan, Ajay Obla, Jayeeta Dutta, Harm van Bakel, Judith Aberg, Adolfo García-Sastre, Gunjan Shah, Tobias Hohl, Genovefa Papanicolaou, Miguel-Angel Perales, Kent Sepkowitz, Ngolela Esther Babady, and Mini Kamboj |
| EPI_ISL_583494 | CS II Dr. Antonio Vicoso Moreira de Rezende Sumare | Instituto Adolfo Lutz, Interdiciplinary Procedures Center, Strategic Laboratory | Claudio Tavares Sacchi, Claudia Regina Gonçalves, Erica Valessa Ramos Gomes, Karoline Rodrigues Campos |
| EPI_ISL_583615, EPI_ISL_583616 | Institut für Virologie am Department für Hygiene, Mikrobiologie und Public Health | Bergthaler laboratory, CeMM Research Center for Molecular Medicine of the Austrian Academy of Sciences | Alexandra Popa, Benedikt Agerer, Henrique Colaco, Lukas Endler, Jakob-Wendelin Genger, Alexander Lercher, Mark Smyth, Thomas Penz, Michael Schuster, Jan Laine, Martin Senekowitsch, Judith Aberle, Stephan Aberle, Peter Hufnagl, Daniela Schmid, Franz Allerberger, Elisabeth Puchhammer-Stoeckl, Manfred Nairz, Guenter Weiss, Gregor Hörmann, Kinga Rigler-Hohenwarter, Rainer Gattringer, Wegene Borena, Dorothee von Laer, Gernot Walder, Peter Obrist, Christian Paar, Sabine Sussitz-Rack, Gunther Vogl, Adi Steinrigl, Christoph Bock, Andreas Bergthaler |
| EPI_ISL_583635, EPI_ISL_583644, EPI_ISL_583652, EPI_ISL_583674, EPI_ISL_583681, EPI_ISL_583689, EPI_ISL_583690 | Austrian Agency for Health and Food Safety (AGES) | Bergthaler laboratory, CeMM Research Center for Molecular Medicine of the Austrian Academy of Sciences | Alexandra Popa, Benedikt Agerer, Henrique Colaco, Lukas Endler, Jakob-Wendelin Genger, Alexander Lercher, Mark Smyth, Thomas Penz, Michael Schuster, Jan Laine, Martin Senekowitsch, Judith Aberle, Stephan Aberle, Peter Hufnagl, Daniela Schmid, Franz Allerberger, Elisabeth Puchhammer-Stoeckl, Manfred Nairz, Guenter Weiss, Gregor Hörmann, Kinga Rigler-Hohenwarter, Rainer Gattringer, Wegene Borena, Dorothee von Laer, Gernot Walder, Peter Obrist, Christian Paar, Sabine Sussitz-Rack, Gunther Vogl, Adi Steinrigl, Christoph Bock, Andreas Bergthaler |
| EPI_ISL_583696, EPI_ISL_583701, EPI_ISL_583721, EPI_ISL_583725 | Center for Virology, Medical University of Vienna | Bergthaler laboratory, CeMM Research Center for Molecular Medicine of the Austrian Academy of Sciences | Alexandra Popa, Benedikt Agerer, Henrique Colaco, Lukas Endler, Jakob-Wendelin Genger, Alexander Lercher, Mark Smyth, Thomas Penz, Michael Schuster, Jan Laine, Martin Senekowitsch, Judith Aberle, Stephan Aberle, Peter Hufnagl, Daniela Schmid, Franz Allerberger, Elisabeth Puchhammer-Stoeckl, Manfred Nairz, Guenter Weiss, Gregor Hörmann, Kinga Rigler-Hohenwarter, Rainer Gattringer, Wegene Borena, Dorothee von Laer, Gernot Walder, Peter Obrist, Christian Paar, Sabine Sussitz-Rack, Gunther Vogl, Adi Steinrigl, Christoph Bock, Andreas Bergthaler |
| EPI_ISL_583732, EPI_ISL_583845, EPI_ISL_583846 | Dr. Gernot Walder GmbH | Bergthaler laboratory, CeMM Research Center for Molecular Medicine of the Austrian Academy of Sciences | Alexandra Popa, Benedikt Agerer, Henrique Colaco, Lukas Endler, Jakob-Wendelin Genger, Alexander Lercher, Mark Smyth, Thomas Penz, Michael Schuster, Jan Laine, Martin Senekowitsch, Judith Aberle, Stephan Aberle, Peter Hufnagl, Daniela Schmid, Franz Allerberger, Elisabeth Puchhammer-Stoeckl, Manfred Nairz, Guenter Weiss, Gregor Hörmann, Kinga Rigler-Hohenwarter, Rainer Gattringer, Wegene Borena, Dorothee von Laer, Gernot Walder, Peter Obrist, Christian Paar, Sabine Sussitz-Rack, Gunther Vogl, Adi Steinrigl, Christoph Bock, Andreas Bergthaler |
| EPI_ISL_583867 | Institute for Laboratory Diagnostics and Microbiology, Klinikum Klagenfurt am Worthersee | Bergthaler laboratory, CeMM Research Center for Molecular Medicine of the Austrian Academy of Sciences | Alexandra Popa, Benedikt Agerer, Henrique Colaco, Lukas Endler, Jakob-Wendelin Genger, Alexander Lercher, Mark Smyth, Thomas Penz, Michael Schuster, Jan Laine, Martin Senekowitsch, Judith Aberle, Stephan Aberle, Peter Hufnagl, Daniela Schmid, Franz Allerberger, Elisabeth Puchhammer-Stoeckl, Manfred Nairz, Guenter Weiss, Gregor Hörmann, Kinga Rigler-Hohenwarter, Rainer Gattringer, Wegene Borena, Dorothee von Laer, Gernot Walder, Peter Obrist, Christian Paar, Sabine Sussitz-Rack, Gunther Vogl, Adi Steinrigl, Christoph Bock, Andreas Bergthaler |
| EPI_ISL_584447, EPI_ISL_584448, EPI_ISL_584449, EPI_ISL_584450, EPI_ISL_584451, EPI_ISL_584452, EPI_ISL_584453, EPI_ISL_584454, EPI_ISL_584455, EPI_ISL_584456, EPI_ISL_584457, EPI_ISL_584458 | | | |
| see above | UHCW / University of Warwick | COVID-19 Genomics UK (COG-UK) Consortium | Richard Stark, Chrystala Constantinidou, Meera Unnikrishnan, Laura Baxter, Jeff Cheng, Grace Taylor-Joyce, Hannah Elizabeth Bridgewater, Lucy Frost, Sarojini Pandey, Paul Brown, Tauqeer Alam, Sascha Ott, Dimitris Grammatopoulos |
| EPI_ISL_584609, EPI_ISL_584610, EPI_ISL_584611, EPI_ISL_584612, EPI_ISL_584613 | Liverpool Clinical Laboratories | COVID-19 Genomics UK (COG-UK) Consortium | Sam Haldenby, Anita Lucaci, Steve Paterson, Julian Hiscox, Alistair Darby, M Almsaud, A Alrezaihi, Muhannad Alruwaili, Stuart D Armstrong, Jones Benjamin, Eleanor G Bentley, Anu Chawla, Jordan J Clark, Angela Cowell, Richard Eccles, Isabel García-Dorival, Matthew Gemmell, Alessandro Gerada, PKF Gilmore, Richard Gregory, Ximeng Han, Catherine Hartley, Margaret Hughes, Miren Iturriza-Gomara, James Johnson, L Luu, Jenifer Manson, Charlotte Nelson, Elaine O'Toole, Cassie Olateju, Rebekah Penrice-Randal , Lucille Rainbow, N.P Randle, Trevor Ian Robinson, Parul Sharma, Ghada T Shawli, James P Stewart, Neil Swainston, Ecaterina Vamos, Joanne Watts, Mark Whitehead |
| EPI_ISL_586274, EPI_ISL_586275, EPI_ISL_586309 | Toronto Invasive Bacterial Diseases Network | McMaster University | Allison McGeer, Patryk Aftanas, Hooman Derakhshani, Angel Li, Kuganya Nirmalarajah, Emily Panousis, Ahmed Draia, Jalees Nasir, Michael Surette, Samira Mubareka, Andrew G. McArthur |
| EPI_ISL_589297, EPI_ISL_589423, EPI_ISL_589440, EPI_ISL_589519, EPI_ISL_589521 | Lighthouse Lab in Milton Keynes | Wellcome Sanger Institute for the COVID-19 Genomics UK (COG-UK) consortium | The Lighthouse Lab in Milton Keynes and Alex Alderton, Roberto Amato, Sonia Goncalves, Ewan Harrison, David K. Jackson, Ian Johnston, Dominic Kwiatkowski, Cordelia Langford, John Sillitoe on behalf of the Wellcome Sanger Institute COVID-19 Surveillance Team |
| EPI_ISL_590750, EPI_ISL_590751 | University of Michigan Clinical Microbiology Laboratory | Lauring Lab, University of Michigan, Department of | Valesano |

| | | | |
|---|---|---|---|
| | | Microbiology and Immunology | |
| EPI_ISL_593483, EPI_ISL_593486, EPI_ISL_593487, EPI_ISL_593489 | Eastern Ontario Regional Laboratory Association | McMaster University | Leanne Mortimer, Hooman Derakhshani, Emily Panousis, Ahmed Draia, Jalees Nasir, Robert Slinger, Andrew G. McArthur |
| EPI_ISL_594119, EPI_ISL_594120, EPI_ISL_594121 | Yale COVID-19 Biorepository | Grubaugh Lab - Yale School of Public Health | Joseph Fauver, Tara Alpert, Anderson Brito, Anne Wyllie, Chantal Vogels, Mary Petrone, Chaney Kalinich, Isabel Ott, Arnau Casanovas, Catherine Muenker, Adam Moore, Alice Lu, Maria Tokuyama, Patrick Wong, Peiwen Lu, Saad Omer, Richard Martinello, Allison Nelson, Shelli Farhadian, Akiko Iwasaki, Charlese Dela Cruz, Albert Ko, Nathan Grubaugh |
| EPI_ISL_594206, EPI_ISL_594208, EPI_ISL_594209 | Michigan Department of Health and Human Services, Bureau of Laboratories | Michigan Department of Health and Human Services, Bureau of Laboratories | Blankenship HM, Riner D, Soehnlen MK |
| EPI_ISL_594418, EPI_ISL_594419, EPI_ISL_594420, EPI_ISL_594421, EPI_ISL_594422, EPI_ISL_594423, EPI_ISL_594424, EPI_ISL_594425, EPI_ISL_594426, EPI_ISL_594427, EPI_ISL_594428, EPI_ISL_594429, EPI_ISL_594430, EPI_ISL_594431, EPI_ISL_594432, EPI_ISL_594433 | | | |
| see above | Utah Public Health Laboratory | Utah Public Health Laboratory | Erin Young, Kelly Oakeson |
| EPI_ISL_594463 | MN Department of Health | Pathogen Discovery, Respiratory Viruses Branch, Division of Viral Diseases, Centers for Disease Control and Prevention | Ying Tao, Yan Li, Clinton Paden, Jing Zhang, Krista Queen, Anna Uehara, Haibin Wang, Julu Bhatnagar, Suxiang Tong |
| EPI_ISL_596638 | St.Vincent's University Hospital | St.Vincent's University Hospital | Mary Lucey, Guerrino Macori, Niamh Mullane, Una Sutton-Fitzpatrick, Gabriel Gonzalez, Suzie Coughlan, Aisling Purcell, Lynda Fenelon, Séamus Fanning, Kirsten Schaffer |
| EPI_ISL_600430 | Institute of Epidemiology Disease Control And Research | Institute for Developing Science and Health Initiatives | Lauren Cowley, Mokibul Hassan Afrad, Sadia Isfat Ara Rahman, Md. Mahfuz-Al-mamun, Firadausi Qadri, Tahmina Shirin |
| EPI_ISL_602550 | Center of Medical Microbiology, Virology, and Hospital Hygiene, University of Duesseldorf | Center of Medical Microbiology, Virology, and Hospital Hygiene, University of Duesseldorf | Maximilian Damagnez, Alexander Dilthey, Torsten Houwaart, Malte Kohns Vasconcelos, Marek Korencak, Nadine Lübke, Jessica Nicolai, Klaus Pfeffer, Hendrik Streeck, Daniel Strelow, Jörg Timm, Andreas Walker, Tobias Wienemann |
| EPI_ISL_602562, EPI_ISL_602563 | Department of Microbiology and Immunology, Stony Brook University | Department of Microbiology and Immunology, Stony Brook University | Nascimento Conde,J., Luperchio,A.M., Gorbunova,E.E., Mackow,E.R. |
| EPI_ISL_603112, EPI_ISL_603113, EPI_ISL_603114, EPI_ISL_603115, EPI_ISL_603116, EPI_ISL_603117 | Lithuanian University of Health Sciences Hospital, Department of Laboratory Medicine | Lithuanian University of Health Sciences, Molecular cardiology lab. | Lukas Zemaitis, Ingrida Olendrait, Arnoldas Pautienius, Kamile Tamusauskaite, Dovydas Gecys, Laura Pareckaite, Vaiva Lesauskaite, Astra Vitkauskiene |
| EPI_ISL_613837, EPI_ISL_613839 | Florida Bureau of Public Health Laboratories | Florida Bureau of Public Health Laboratories | Sarah Schmedes, Jason Blanton |
| EPI_ISL_614115 | Virginia DCLS | Virginia DCLS | Virginia DCLS |
| EPI_ISL_614396 | Laboratorio Biologia Molecolare Sars Cov2 - UOC Laboratorio Analisi - Servizio Medicina di Laboratorio, Ospedale "San Francesco" - ATS-ASSL Nuoro | Laboratorio specialistico UOC Ematologia - Ospedale "San Francesco" - ATS-ASSL Nuoro | Piras Giovanna, Fancello Tatiana, Asproni Rosanna, Fiamma Maura, Monne Maria Itria, Toja Alessandro, Sanna Filomena, Floris Anna Rita, Sulis Vincenzo, Palmas Angelo Domenico, Casu Gavino, Lo Maglio Iana, Mameli Giuseppe |
| EPI_ISL_614419, EPI_ISL_614433, EPI_ISL_614434, EPI_ISL_614447, EPI_ISL_614484, EPI_ISL_614485, EPI_ISL_614697, EPI_ISL_614698, EPI_ISL_614699, EPI_ISL_614700, EPI_ISL_614701, EPI_ISL_614702, EPI_ISL_614703, EPI_ISL_614704, EPI_ISL_614705, EPI_ISL_614706, EPI_ISL_614707, EPI_ISL_614708, EPI_ISL_614709, EPI_ISL_614710, EPI_ISL_614724, EPI_ISL_614726, EPI_ISL_614727, EPI_ISL_614728, EPI_ISL_614729, EPI_ISL_614730, EPI_ISL_614731, EPI_ISL_614732, EPI_ISL_614733, EPI_ISL_614734, EPI_ISL_614735, EPI_ISL_614736, EPI_ISL_614737, EPI_ISL_614738, EPI_ISL_614739, EPI_ISL_614740, EPI_ISL_614741, EPI_ISL_614742, EPI_ISL_614743, EPI_ISL_614744, EPI_ISL_614745, EPI_ISL_614746, EPI_ISL_614747, EPI_ISL_614748, EPI_ISL_614777 | | | |
| see above | Department of Virus and Microbiological Special Diagnostics, Statens Serum Institut, Denmark | Albertsen lab, Department of Chemistry and Bioscience, Aalborg University, Denmark | Danish Covid-19 Genome Consortia |
| EPI_ISL_615122, EPI_ISL_615123 | Texas Department of State Health Services | Texas Department of State Health Services | Rashmi Tuladhar, Bonnie Oh, Jenny Zhang, Maliha Rahman, Anita Pokharel, Myong Koag, Chung Wang, Rachel Lee, Grace Kubin, Mayela Pedrueza |
| EPI_ISL_618662, EPI_ISL_618720, EPI_ISL_618730, EPI_ISL_618740, EPI_ISL_618768, EPI_ISL_618800, EPI_ISL_618801, EPI_ISL_618804, EPI_ISL_622342, EPI_ISL_622343, EPI_ISL_622344, EPI_ISL_622345, EPI_ISL_622346, EPI_ISL_622347, EPI_ISL_622348, EPI_ISL_622349, EPI_ISL_622350, EPI_ISL_622351, EPI_ISL_622355, EPI_ISL_622356, EPI_ISL_622357, EPI_ISL_622358, EPI_ISL_622359, EPI_ISL_622360, EPI_ISL_622361, EPI_ISL_622362, EPI_ISL_622363, EPI_ISL_622364, EPI_ISL_622369, EPI_ISL_622370, EPI_ISL_622371, EPI_ISL_622372, EPI_ISL_622373, EPI_ISL_622374, EPI_ISL_622375, EPI_ISL_622376, EPI_ISL_622377, EPI_ISL_622378, EPI_ISL_622379, EPI_ISL_622380, EPI_ISL_622381, EPI_ISL_622382, EPI_ISL_622383, EPI_ISL_622384, EPI_ISL_622385, EPI_ISL_622386, EPI_ISL_622387, EPI_ISL_622388, EPI_ISL_622389, EPI_ISL_622390, EPI_ISL_622391, EPI_ISL_622392, EPI_ISL_622393, EPI_ISL_622394, EPI_ISL_622395, EPI_ISL_622396, EPI_ISL_622399, EPI_ISL_622400, EPI_ISL_622401, EPI_ISL_622402, EPI_ISL_622404, EPI_ISL_622405, EPI_ISL_622432, EPI_ISL_622478, EPI_ISL_622501, EPI_ISL_622512, EPI_ISL_622513, EPI_ISL_622531, EPI_ISL_622538, EPI_ISL_622545, EPI_ISL_622547, EPI_ISL_622548, EPI_ISL_622584, EPI_ISL_622627, EPI_ISL_622641, EPI_ISL_622642, EPI_ISL_622643, EPI_ISL_622645, EPI_ISL_622646, EPI_ISL_622647, EPI_ISL_622648, EPI_ISL_622649, EPI_ISL_622650, EPI_ISL_622651, EPI_ISL_622652, EPI_ISL_622653, EPI_ISL_622654, EPI_ISL_622655, EPI_ISL_622656, EPI_ISL_622657, EPI_ISL_622658, EPI_ISL_622659, EPI_ISL_622660, EPI_ISL_622661, EPI_ISL_622662, EPI_ISL_622663, EPI_ISL_622664, EPI_ISL_622665 | | | |
| see above | Department of Virus and Microbiological Special Diagnostics, Statens Serum Institut, Denmark | Albertsen lab, Department of Chemistry and Bioscience, Aalborg University, Denmark | Danish Covid-19 Genome Consortia |
| EPI_ISL_623106, EPI_ISL_623110, EPI_ISL_623112, EPI_ISL_623114, EPI_ISL_623118, EPI_ISL_623120, EPI_ISL_623122, EPI_ISL_623124, EPI_ISL_623127, EPI_ISL_623130, EPI_ISL_623139, EPI_ISL_623142, EPI_ISL_623150, EPI_ISL_623160, EPI_ISL_623166, EPI_ISL_623169 | | | |
| see above | Laboratorio de Virologia Molecular / UFRJ | Bioinformatics Laboratory / LNCC | Carolina M Voloch, Ronaldo S Francisco Jr, Luiz G P de Almeida, Otavio J. Brustolini, Cynthia C Cardoso, Alexandra L Gerber, Ana Paula de C Guimarães, Diana Mariani, Covid19-UFRJ Workgroup, Luís Cristóvão Pôrto, Renato S Aguiar, Terezinha M P P Castiñeiras, Orlando C. Ferreira, Amilcar Tanuri, Ana Tereza R de Vasconcelos |
| EPI_ISL_623206, EPI_ISL_623207, EPI_ISL_623208, EPI_ISL_623209, EPI_ISL_623210 | Utah Public Health Laboratory | Utah Public Health Laboratory | Erin Young, Kelly Oakeson |
| EPI_ISL_626216, EPI_ISL_626217, EPI_ISL_626218, EPI_ISL_626223 | Institute for Virology, University Hospital Essen | Center of Medical Microbiology, Virology, and Hospital Hygiene, University of Duesseldorf | Olympia E. Anastasiou, Ulf Dittmer, Maximilian Damagnez, Alexander Dilthey, Torsten Houwaart, Lisanna Hülse, Malte Kohns Vasconcelos, Nadine Lübke, Jessica Nicolai, Klaus Pfeffer, Daniel Strelow, Jörg Timm, Andreas Walker, Tobias Wienemann |
| EPI_ISL_626412, EPI_ISL_626413, EPI_ISL_626414, EPI_ISL_626415, EPI_ISL_626416, EPI_ISL_626417, EPI_ISL_626418, EPI_ISL_626419, EPI_ISL_626420, EPI_ISL_626421, EPI_ISL_626422, EPI_ISL_626423, EPI_ISL_626424, EPI_ISL_626425, EPI_ISL_626426, EPI_ISL_626427, EPI_ISL_626428, EPI_ISL_626429, EPI_ISL_626430, EPI_ISL_626431, EPI_ISL_626432, EPI_ISL_626433, EPI_ISL_626434, EPI_ISL_626435, EPI_ISL_626436, EPI_ISL_626437, EPI_ISL_626438, EPI_ISL_626439, EPI_ISL_626440 | | | |
| see above | Northwestern Memorial Hospital | Ozer Lab | Ramon Lorenzo-Redondo, Hannah H. Nam, Scott C. Roberts, Lacy M. Simons, Chad J. Achenbach, Lawrence J. Jennings, Chao Qi, Alan R. Hauser, Michael G. Ison, Judd F. Hultquist, Egon A. Ozer |
| EPI_ISL_629017 | Centro de Biotecnología Vegetal, Universidad Andrés Bello, Center for Genome Regulation | Center for Mathematical Modeling and Center for Genome Regulation. Santiago, Chile | Bastias M, Sanhueza D, Travisany D, Allende ML, Maass A, González M, Bustos F, Arriagada G, Montecino, M, Orellana A, Castro E, Meneses C. |
| EPI_ISL_629106, EPI_ISL_629107 | Laboratoire du Centre Hospitalier Annecy Genevois | CNR Virus des Infections Respiratoires - France SUD | Antonin Bal, Gregory Destras, Gwendolyne Burfin, Hadrien Règue, Quentin Semanas, Martine Valette, Bruno Lina, Hélène Petitprez, Bruno Chanzy, Laurence Josset |
| EPI_ISL_631396 | Wisconsin State Laboratory of Hygiene Communicable Disease Division | Wisconsin State Laboratory of Hygiene Communicable Disease Division | Kelsey R. Florek, Abigail C. Shockey |
| EPI_ISL_631719, EPI_ISL_631720, EPI_ISL_631721, EPI_ISL_631743, EPI_ISL_631744, EPI_ISL_631745, EPI_ISL_631746, EPI_ISL_631747 | Jamaica Hospital Medical Center | New York City Public Health Laboratory | Jade Wang, et al. |
| EPI_ISL_631748 | St Barnabas Hospital | New York City Public Health Laboratory | Jade Wang, et al. |
| EPI_ISL_631777 | Jamaica Hospital Medical Center | New York City Public Health Laboratory | Jade Wang, et al. |
| EPI_ISL_631778, EPI_ISL_631779, EPI_ISL_631780, EPI_ISL_631781, EPI_ISL_631782, EPI_ISL_631783 | St Barnabas Hospital | New York City Public Health Laboratory | Jade Wang, et al. |
| EPI_ISL_631841 | OCME Office Of Chief Medical Examiner | New York City Public Health Laboratory | Jade Wang, et al. |
| EPI_ISL_631856, EPI_ISL_631857, | St Barnabas Hospital | New York City Public Health Laboratory | Jade Wang, et al. |

| | | | |
|---|---|---|---|
| EPI_ISL_631858, EPI_ISL_631859, EPI_ISL_631911, EPI_ISL_631912 | | | |
| EPI_ISL_634896, EPI_ISL_634897, EPI_ISL_634898, EPI_ISL_634899, EPI_ISL_634900, EPI_ISL_634901, EPI_ISL_634902 | Utah Public Health Laboratory | Utah Public Health Laboratory | Erin L. Young, Kelly F. Oakeson |
| EPI_ISL_635271, EPI_ISL_635272, EPI_ISL_635273, EPI_ISL_635274, EPI_ISL_635275 | Institute of Microbiology and Immunology, Faculty of Medicine, University of Ljubljana | Institute of Microbiology and Immunology, Faculty of Medicine, University of Ljubljana | Tomaž Mark Zorec, Samo Zakotnik, Miša Korva, Tatjana Avši - Županc, Mario Poljak |
| EPI_ISL_635322, EPI_ISL_635325, EPI_ISL_635326, EPI_ISL_635327, EPI_ISL_635329, EPI_ISL_635330, EPI_ISL_635331 | San Diego County Public Health Laboratory | Andersen lab at Scripps Research | SEARCH Alliance San Diego with Tracy Basler, Jovan Shephard, Brett Austin |
| EPI_ISL_636526, EPI_ISL_636555, EPI_ISL_636561 | Dutch COVID-19 response team | National Institute for Public Health and the Environment (RIVM) | Adam Meijer, Harry Vennema, Jeroen Cremer, Sharon van den Brink, Bas van der Veer, AnneMarie van den Brandt, Florian Zwagemaker, Dennis Schmitz, Chantal Reusken, on behalf of the national COVID-19 response team |
| EPI_ISL_636960 | Pathogen Genomics Lab King Abdullah University of Science and Technology(KAUST) | Pathogen Genomics Lab King Abdullah University of Science and Technology(KAUST) | Sharif Hala, Fadwa Alofi, Sara Mfarrej, Amit Kumar Subudhi, Rahul P Salunke, Fathia Ben Rached, Amanda Ooi, Luke Esau, Afrah Alsomali, Asim Khogeer, Jumana Taha, Abdulaziz Alahmadi, Kahled Algithami, Raeece Naeem, Anwar Hashem, Naif Almontashiri, Arnab Pain |
| EPI_ISL_637109 | Laboratorio Biologia Molecolare Sars Cov2 - UOC Laboratorio Analisi - Servizio Medicina di Laboratorio, Ospedale "San Francesco" - ATS-ASSL Nuoro | Laboratorio specialistico UOC Ematologia - Ospedale "San Francesco" - ATS-ASSL Nuoro | Piras Giovanna, Fancello Tatiana, Asproni Rosanna, Fiamma Maura, Monne Maria Itria, Toja Alessandro, Sanna Filomena, Floris Anna Rita, Sulis Vincenzo, Palmas Angelo Domenico, Casu Gavino, Lo Maglio Iana, Mameli Giuseppe |
| EPI_ISL_640080 | Mfuleni CDC wc MFU | NHLS/UCT | Arash Iranzadeh, Deelan Doolabh, Lynn Tyers, Bruna Galvao, Innocent Mudau, Marvin Hsiao, Kruger Marais, Diana Hardie, Stephen Korsman, Carolyn Williamson |
| EPI_ISL_640082 | Heideveld Emergency Centre | NHLS/UCT | Arash Iranzadeh, Deelan Doolabh, Lynn Tyers, Bruna Galvao, Innocent Mudau, Marvin Hsiao, Kruger Marais, Diana Hardie, Stephen Korsman, Carolyn Williamson |
| EPI_ISL_640100 | Dr Abdurahman CDC wc DAC | NHLS/UCT | Arash Iranzadeh, Deelan Doolabh, Lynn Tyers, Bruna Galvao, Innocent Mudau, Marvin Hsiao, Kruger Marais, Diana Hardie, Stephen Korsman, Carolyn Williamson |
| EPI_ISL_640244, EPI_ISL_640245, EPI_ISL_640246, EPI_ISL_640255, EPI_ISL_640257, EPI_ISL_640265 | MVZ Laborärzte Singen | MVZ Laborärzte Singen | Jonas Schmidt, Frithjof Blessing, Sandro Berghaus, Folker Wenzel |
| EPI_ISL_641312, EPI_ISL_641314 | Rocky Mountain Laboratories, RTS Genomics Unit, National Institute of Allergy and Infectious Diseases, National Institutes of Health | Rocky Mountain Laboratories, RTS Genomics Unit, National Institute of Allergy and Infectious Diseases, National Institutes of Health | Avanzato,V.A., Matson,M.J., Seifert,S.N., Pryce,R., Williamson,B.N., Anzick,S.L., Barbian,K., Judson,S.D., Fischer,E.R., Martens,C., Bowden,T.A., de Wit,E., Riedo,F.X., Munster,V.J., Siefert,S.N., WIlliamson,B.N., Anzick,S., Martens,C.A. |
| EPI_ISL_641531 | CHU de Nice - Hôpital Archet 13 | CNR Virus des Infections Respiratoires - France SUD | Antonin Bal, Géraldine Gonfrier, Gregory Destras, Gwendolyne Burfin, Hadrien Règue, Quentin Semanas, Martine Valette, Bruno Lina, Valérie Giordanengo, Laurence Josset |
| EPI_ISL_641532 | CHU de Nice - Hôpital Archet 14 | CNR Virus des Infections Respiratoires - France SUD | Antonin Bal, Géraldine Gonfrier, Gregory Destras, Gwendolyne Burfin, Hadrien Règue, Quentin Semanas, Martine Valette, Bruno Lina, Valérie Giordanengo, Laurence Josset |
| EPI_ISL_641548, EPI_ISL_641549 | CHU Clermont-Ferrand | CNR Virus des Infections Respiratoires - France SUD | Antonin Bal, Gregory Destras, Gwendolyne Burfin, Hadrien Règue, Quentin Semanas, Martine Valette, Bruno Lina, Christine Archimbaud, Amélie Brebion, Hélène Chabrolles, Martine Chambon, Audrey Mirand, Christel Regagnon, Maxime Bisseux, Patricia Combes, Cécile Henquell, Laurence Josset |
| EPI_ISL_644170, EPI_ISL_644185 | Texas Department of State Health Services | Texas Department of State Health Services | Rashmi Tuladhar, Bonnie Oh, Jenny Zhang, Maliha Rahman, Anita Pokharel, Myong Koag, Chung Wang, Rachel Lee, Grace Kubin, Mayela Pedrueza |
| EPI_ISL_644209, EPI_ISL_644215, EPI_ISL_644221, EPI_ISL_644222, EPI_ISL_644231, EPI_ISL_644240, EPI_ISL_644241, EPI_ISL_644242, EPI_ISL_644246, EPI_ISL_644247, EPI_ISL_644249, EPI_ISL_644264, EPI_ISL_644265, EPI_ISL_644266, EPI_ISL_644267, EPI_ISL_644268, EPI_ISL_644269, EPI_ISL_644270, EPI_ISL_644271, EPI_ISL_644272, EPI_ISL_644275, EPI_ISL_644281, EPI_ISL_644294, EPI_ISL_644295, EPI_ISL_644298, EPI_ISL_644304, EPI_ISL_644325 | | | |
| see above | CEPHR / Vincent's Hospital | Irish Coronavirus Sequencing Consortium - National Virus Reference Laboratory | Michael Carr, Gabriel Gonzalez, Alejandro Abner Garcia Leon, Patrick Mallon |
| EPI_ISL_644694, EPI_ISL_644695 | CHU Montpellier | CNR Virus des Infections Respiratoires - France SUD | Antonin Bal, Gregory Destras, Gwendolyne Burfin, Hadrien Règue, Quentin Semanas, Martine Valette, Bruno Lina, Michel Segondy, Vincent Foulongne, Laurence Josset |
| EPI_ISL_644703, EPI_ISL_644704, EPI_ISL_644705, EPI_ISL_644706, EPI_ISL_644708, EPI_ISL_644709 | Unité des Virus Émergents | CNR Virus des Infections Respiratoires - France SUD | Antonin Bal, Gregory Destras, Gwendolyne Burfin, Hadrien Règue, Quentin Semanas, Martine Valette, Bruno Lina, Laetitia Ninove, Léa Luciani, Antoine Nougairède, Laurence Josset |
| EPI_ISL_645000, EPI_ISL_645001, EPI_ISL_645002, EPI_ISL_645003, EPI_ISL_645010, EPI_ISL_645031, EPI_ISL_645035, EPI_ISL_645038, EPI_ISL_645044, EPI_ISL_645045, EPI_ISL_645046, EPI_ISL_645100, EPI_ISL_645102, EPI_ISL_645103, EPI_ISL_645104, EPI_ISL_645108 | | | |
| see above | Human Genome Variation Research Group, Malopolska Centre of Biotechnology | Human Genome Variation Research Group, Malopolska Centre of Biotechnology | Kowalski,M., Pospiech,E., Klajmon,A., Gromowski,T., Pisarek,A., Marszalek,K., Kopera,K., Foremny,J., Swadzba,J., Sanak,M., Owczarek,K., Dabrowska,A., Szczepanski,A., Botwina,P., Labaj,P.P., Pyrc,K., Branicki,W. |
| EPI_ISL_645207, EPI_ISL_645208 | CHU Nîmes | CNR Virus des Infections Respiratoires - France SUD | Antonin Bal, Gregory Destras, Gwendolyne Burfin, Hadrien Règue, Quentin Semanas, Martine Valette, Bruno Lina, Jean-Philippe Lavigne, Stephan Robin, Maxence Lotellier, Marie-Josée Carles, Laurence Josset |
| EPI_ISL_647985 | RI State Health Laboratories | Pathogen Discovery, Respiratory Viruses Branch, Division of Viral Diseases, Centers for Disease Control and Prevention | Yan Li, Jing Zhang, Ying Tao, Brian Lynch, Krista Queen, Anna Montmayeur, Anna Uehara, Clinton R. Paden, Rachel Marine, Haibin Wang, Suxiang Tong |
| EPI_ISL_647986 | Indiana State Department of Health | Pathogen Discovery, Respiratory Viruses Branch, Division of Viral Diseases, Centers for Disease Control and Prevention | Yan Li, Jing Zhang, Ying Tao, Brian Lynch, Krista Queen, Anna Montmayeur, Anna Uehara, Clinton R. Paden, Rachel Marine, Haibin Wang, Suxiang Tong |
| EPI_ISL_648265, EPI_ISL_648266 | Brigham and Women's Hospital | Jonathan Li laboratory | Jonathan Z. Li, Manish C. Choudhary, Upasana D. Adhikari, George Eng, Douglas S Kwon |
| EPI_ISL_648315, EPI_ISL_648316, EPI_ISL_648317, EPI_ISL_648319, EPI_ISL_648321, EPI_ISL_648323, EPI_ISL_648325, EPI_ISL_648326, EPI_ISL_648331, EPI_ISL_648332, EPI_ISL_648333 | | | |
| see above | Laboratorio de Investigaciones de Baney | University Hospital Basel, Clinical Bacteriology | Carlos Cortes, Claudia Daubenberger, Adrian Egli, Guillermo Garcia, Salome Hosch, Bonifacio Manguire Nlavo, Alfredo Mari, Maximilian Mpina, Elizabeth Nyakarungu, Diosdado Odjama Nseng Ada, Mitoha Ondo O Ayekaba, Tim Roloff, Tobias Schindler, Helena Seth-Smith, Madlen Stange, Philip Wonder Phiri |
| EPI_ISL_648578, EPI_ISL_648579 | Utah Public Health Laboratory | Utah Public Health Laboratory | Erin Young, Kelly Oakeson |
| EPI_ISL_648614, EPI_ISL_648615, EPI_ISL_648616, EPI_ISL_648617, EPI_ISL_648618, EPI_ISL_648619, EPI_ISL_648620, EPI_ISL_648621, EPI_ISL_648622, EPI_ISL_648623, EPI_ISL_648624, EPI_ISL_648625, EPI_ISL_648626, EPI_ISL_648627, EPI_ISL_648628, EPI_ISL_648629, EPI_ISL_648630, EPI_ISL_648631, EPI_ISL_648632, EPI_ISL_648633, EPI_ISL_648634, EPI_ISL_648635, EPI_ISL_648636, EPI_ISL_648637, EPI_ISL_648638, EPI_ISL_648639, EPI_ISL_648640, EPI_ISL_648641, EPI_ISL_648642, EPI_ISL_648643, EPI_ISL_648644, EPI_ISL_648645, EPI_ISL_648646, EPI_ISL_648647, EPI_ISL_648648, EPI_ISL_648649, EPI_ISL_648650, EPI_ISL_648651, EPI_ISL_648652, EPI_ISL_648653, EPI_ISL_648742 | | | |
| see above | Department of Laboratory Medicine, Tan Tock Seng Hospital | Department of Laboratory Medicine, Tan Tock Seng Hospital | Chen YYC, Zair X, Lim JX, Li C, Tang WY, Maurer-Stroh S, Barkham TMS, Nagarajan N, Sessions OM |
| EPI_ISL_649186, EPI_ISL_649187 | CHU Nantes | CNR Virus des Infections Respiratoires - France SUD | Antonin Bal, Louise Castain, Gregory Destras, Gwendolyne Burfin, Hadrien Règue, Quentin Semanas, Martine Valette, Bruno Lina, Celine Bressollette, Laurence Josset |
| EPI_ISL_649951, EPI_ISL_649952 | CHU de Saint-Étienne Hôpital Nord | CNR Virus des Infections Respiratoires - France SUD | Antonin Bal, Gregory Destras, Gwendolyne Burfin, Hadrien Règue, Quentin Semanas, Martine Valette, Bruno Lina, Issam Bechri, Manon Vogrig, Marine Delorme, Bruno Pozzetto, Thomas Bourlet, Sylvie Gonzalo, Sylvie Pillet, Laurence Josset |
| EPI_ISL_653149, EPI_ISL_653310, | Florida Bureau of Public Health Laboratories | Florida Bureau of Public Health Laboratories | Sarah Schmedes, Jason Blanton |

| | | | |
|---|---|---|---|
| EPI_ISL_653311, EPI_ISL_653312 | | | |
| EPI_ISL_654619 | Servicio de Microbiología, Hospital Universitario Central de Asturias | SeqCOVID-SPAIN consortium/IBV(CSIC) | Cristián Castelló Abietar, Jose A. Boga, Susana Rojo-Alba, Marta Elena Álvarez-Argüelles, Santiago Melón and SeqCOVID-SPAIN consortium |
| EPI_ISL_654626 | Hospital Universitario Virgen de las Nieves de Granada-SAS | SeqCOVID-SPAIN consortium/IBV(CSIC) | Sara Sanbonmatsu Gámez, Mercedes Pérez Ruiz, Irene Pedrosa Corral, José M. Navarro-Marí and SeqCOVID-SPAIN consortium |
| EPI_ISL_654629, EPI_ISL_654671, EPI_ISL_654680 | Servicio de Microbiología, Hospital Universitario Central de Asturias | SeqCOVID-SPAIN consortium/IBV(CSIC) | Cristián Castelló Abietar, Jose A. Boga, Susana Rojo-Alba, Marta Elena Álvarez-Argüelles, Santiago Melón and SeqCOVID-SPAIN consortium |
| EPI_ISL_654883 | Pasteur Institute in Ho Chi Minh city | Department of Microbiology and Immunology - Pasteur Institute in Ho Chi Minh city | Nguyn Thanh Long, ào Huy Mnh, Phm Th Thu Hng, V Phm Hng Nhung, Cao Minh Thng, Hunh Th Kim Loan, Nguyn Hoàng Quân, Hunh Phng Tho, Hoàng Nh ào, Nguyn Trung Hiu, Nguyn Hoàng Anh, Nguyn Thu Ngc, Lê Hoàng Chng, Nguyn Th Ngc Tho, ng Thanh Giang, Nguyn Th Thanh Thng, Hoàng Minh, Trn Th Hng Kim, Phm Duy Quang, Lng Chn Quang, Hoàng Quc Cng, Phan Trng Lân, Nguyn V Thng |
| EPI_ISL_654937 | Texas Department of State Health Services | Texas Department of State Health Services | Rashmi Tuladhar, Bonnie Oh, Jenny Zhang, Maliha Rahman, Anita Pokharel, Myong Koag, Chung Wang, Rachel Lee, Grace Kubin, Mayela Pedrueza, James Daniel Bonser |
| EPI_ISL_660365, EPI_ISL_660366, EPI_ISL_660367 | CHU Clermont-Ferrand | CNR Virus des Infections Respiratoires - France SUD | Antonin Bal, Gregory Destras, Gwendolyne Burfin, Hadrien Règue, Quentin Semanas, Martine Valette, Bruno Lina, Christine Archimbaud, Amélie Brebion, Hélène Chabrolles, Martine Chambon, Audrey Mirand, Christel Regagnon, Maxime Bisseux, Patricia Combes, Cécile Henquell, Laurence Josset |
| EPI_ISL_660432 | CHU de Saint-Étienne Hôpital Nord | CNR Virus des Infections Respiratoires - France SUD | Antonin Bal, Gregory Destras, Gwendolyne Burfin, Hadrien Règue, Quentin Semanas, Martine Valette, Bruno Lina, Issam Bechri, Manon Vogrig, Marine Delorme, Bruno Pozzetto, Thomas Bourlet, Sylvie Gonzalo, Sylvie Pillet, Laurence Josset |
| EPI_ISL_660696, EPI_ISL_660697, EPI_ISL_660698, EPI_ISL_660699, EPI_ISL_660700, EPI_ISL_660701, EPI_ISL_660702 | CHU Montpellier | CNR Virus des Infections Respiratoires - France SUD | Antonin Bal, Gregory Destras, Gwendolyne Burfin, Hadrien Règue, Quentin Semanas, Martine Valette, Bruno Lina, Michel Segondy, Vincent Foulongne, Laurence Josset |
| EPI_ISL_660726, EPI_ISL_660727, EPI_ISL_660728, EPI_ISL_660729, EPI_ISL_660730 | CHU Nîmes | CNR Virus des Infections Respiratoires - France SUD | Antonin Bal, Gregory Destras, Gwendolyne Burfin, Hadrien Règue, Quentin Semanas, Martine Valette, Bruno Lina, Jean-Philippe Lavigne, Stephan Robin, Maxence Lotellier, Marie-Josée Carles, Laurence Josset |
| EPI_ISL_660735 | Unité des Virus Émergents | CNR Virus des Infections Respiratoires - France SUD | Antonin Bal, Gregory Destras, Gwendolyne Burfin, Hadrien Règue, Quentin Semanas, Martine Valette, Bruno Lina, Laetitia Ninove, Léa Luciani, Antoine Nougairède, Laurence Josset |
| EPI_ISL_664471, EPI_ISL_664521, EPI_ISL_664540, EPI_ISL_665149, EPI_ISL_665150, EPI_ISL_665151, EPI_ISL_665161, EPI_ISL_665208 | University College London Hospital | COVID-19 Genomics UK (COG-UK) Consortium | Judith Heaney, Matthew Byott, Catherine Houlihan, Dan Frampton, Stuart Kirk, Moira Spyer and Eleni Nastouli |
| EPI_ISL_665256, EPI_ISL_665257, EPI_ISL_666608, EPI_ISL_666610, EPI_ISL_666611, EPI_ISL_666612, EPI_ISL_666614, EPI_ISL_666615 | Dept. of Microbiology and Infection Control, Akershus University Hospital HF | Dept. of Microbiology and Infection Control, Akershus University Hospital HF | Hege Vangstein Aamot, Alexander Hesselberg Løvestad, Silje Bakken Jørgensen, Nina Handal, Ole Herman Ambur |
| EPI_ISL_666678, EPI_ISL_666679 | Laboratoire du Centre Hospitalier Annecy Genevois | CNR Virus des Infections Respiratoires - France SUD | Antonin Bal, Gregory Destras, Gwendolyne Burfin, Hadrien Règue, Quentin Semanas, Martine Valette, Bruno Lina, Hélène Petitprez, Bruno Chanzy, Laurence Josset |
| EPI_ISL_666711 | CHU de Limoges | CNR Virus des Infections Respiratoires - France SUD | Antonin Bal, Gregory Destras, Gwendolyne Burfin, Hadrien Règue, Quentin Semanas, Martine Valette, Bruno Lina, Sylvie Rogez, Laurence Josset |
| EPI_ISL_666721, EPI_ISL_666722, EPI_ISL_666723 | CHU Nantes | CNR Virus des Infections Respiratoires - France SUD | Antonin Bal, Louise Castain, Gregory Destras, Gwendolyne Burfin, Hadrien Règue, Quentin Semanas, Martine Valette, Bruno Lina, Virginie Ferré, Celine Bressollette, Laurence Josset |
| EPI_ISL_667079 | OHSU Lab Services Molecular Microbiology Lab | Oregon SARS-CoV-2 Genome Sequencing Center | Brendan L. O'Connell, Ruth V. Nichols, Sally Grindstaff, Alec J. Hirsch, Donna Hansel, Guang Fan, Daniel N. Streblow, William B. Messer, Andrew C. Adey, Benjamin N. Bimber, Brian J. O'Roak |
| EPI_ISL_671341, EPI_ISL_671342, EPI_ISL_671356, EPI_ISL_671357, EPI_ISL_671358 | National Virus Reference Laboratory | Irish Coronavirus Sequencing Consortium - Teagasc Moorepark | Calm Walsh, Genuity Ireland |
| EPI_ISL_672030, EPI_ISL_672144, EPI_ISL_672145, EPI_ISL_672146, EPI_ISL_672149, EPI_ISL_672150, EPI_ISL_672151, EPI_ISL_672152 | The Ashley Laboratory, Stanford University | Chan-Zuckerberg Biohub | CZB Cliahub Consortium |
| EPI_ISL_672669, EPI_ISL_672733, EPI_ISL_672734, EPI_ISL_672735, EPI_ISL_672736, EPI_ISL_672737, EPI_ISL_672738, EPI_ISL_672739, EPI_ISL_672740, EPI_ISL_672741, EPI_ISL_672742, EPI_ISL_672743, EPI_ISL_672744, EPI_ISL_672745 | | | |
| see above | Hospital das Clínicas da Faculdade de Medicina da Universidade de São Paulo (HC-FMUSP) | Laboratório de Parasitologia Médica - Instituto de Medicina Tropical - Universidade de São Paulo | Brazil-UK Centre for Arbovirus Discovery Diagnosis Genomics and Epidemiology (CADDE) Genomic Network - Instituto de Medicina Tropical |
| EPI_ISL_676522, EPI_ISL_676523 | Uppsala klinisk mikrobiologi | The Public Health Agency of Sweden | Department of Microbiology, The Public Health Agency of Sweden |
| EPI_ISL_676541 | TNMC & Nair ch. Hospital | CSIR-Institute of Genomics and Integrative Biology | Rajesh Pandey, Jayanthi Shastri, Akshay Kanakan, Janani Srinivasa Vasudevan, Ranjeet Maurya, Sachee Agrawal, Nirjhar Chatterjee, Swapneil Parikh, Manish Pathak, Subrat Thanapati, Jasmina Savak, Suresh Poojari, Mahesh Sangar, Amol Borse, Shweta Kawankar, Vasil Nachan, Mayuresh Vishwanathan, Shruthi Sachidanandan, Shrutika Pophale, Utkarsha Yelve |
| EPI_ISL_676597, EPI_ISL_676598, EPI_ISL_676599 | Scientific Veterinary Institute Novi Sad | Veterinary Specialized Institute "Kraljevo", Serbia | Vidanovic,D., Tesovic,B., Knezevic,A., Jovanovic,T., Jankovic,M., Sekler,M., Banovic Djeri,B., Petrovic,T., Volkening,J., Afonso,C. |
| EPI_ISL_676684, EPI_ISL_676866, EPI_ISL_676867, EPI_ISL_676868, EPI_ISL_676869, EPI_ISL_676870, EPI_ISL_676871, EPI_ISL_676872, EPI_ISL_676873, EPI_ISL_676874, EPI_ISL_676875, EPI_ISL_676876, EPI_ISL_676877, EPI_ISL_676878, EPI_ISL_676879, EPI_ISL_676880, EPI_ISL_676881, EPI_ISL_676882, EPI_ISL_676883, EPI_ISL_676884, EPI_ISL_676886, EPI_ISL_676892, EPI_ISL_676965 | | | |
| see above | Wadsworth Center, New York State Department.of Health | Wadsworth Center, New York State Department.of Health | Kirsten St. George, Daryl M. Lamson, Alexis Russel, Jonathan Plitnick, Navjot Singh, John Kelly, Sara Griesemer, Erasmus Schneider, Erica Lasek-Nesselquist |
| EPI_ISL_677252, EPI_ISL_677255, EPI_ISL_677264, EPI_ISL_677288, EPI_ISL_677289, EPI_ISL_677290 | Colorado Department of Public Health and Environment | Colorado Department of Puplic Health and Environment | Laura Bankers, Molly Hetherington-Rauth, Shannon Ely, Shannon R. Matzinger, Sarah Elizabeth Totten, Emily A. Travanty |
| EPI_ISL_677637 | Colorado Department of Public Health and Environment | Colorado Department of Puplic Health and Environment | Laura Bankers, Molly C. Hetherington-Rauth, Shannon Ely, Shannon R. Matzinger, Sarah Elizabeth Totten, Emily A. Travanty |
| EPI_ISL_677679, EPI_ISL_677683 | Vanda Pharmaceuticals Clinical Site | Vanda Pharmaceuticals | Vanda Pharmaceuticals |
| EPI_ISL_677938, EPI_ISL_677939 | Pathogen Genomics Lab King Abdullah University of Science and Technology(KAUST) | Pathogen Genomics Lab King Abdullah University of Science and Technology(KAUST) | Sara Mfarrej, Raushan Nugmanova, Olga Douvropoulou, Sharif Hala, Raeece Naeem, Afrah Alsomali, Fadwa Alofi, Asim Khogeer, Jumana Taha, Abdulaziz Alahmadi, Kahled Algithami, Anwar Hashem, Naif Almontashiri, Arnab Pain |
| EPI_ISL_678030, EPI_ISL_678031 | Pathogen Genomics Lab King Abdullah University of Science and Technology(KAUST) | Pathogen Genomics Lab King Abdullah University of Science and Technology(KAUST) | Amit Kumar Subudhi, Sara Mfarrej, Amanda Ooi, Luke Esau, Sharif Hala, Raeece Naeem, Fadwa Alofi, Afrah Alsomali, Asim Khogeer, Jumana Taha, Abdulaziz Alahmadi, Kahled Algithami, Anwar Hashem, Naif Almontashiri, Arnab Pain |
| EPI_ISL_678032 | Pathogen Genomics Lab King Abdullah University of Science and Technology(KAUST) | Pathogen Genomics Lab King Abdullah University of Science and Technology(KAUST) | Sara Mfarrej, Raushan Nugmanova, Olga Douvropoulou, Sharif Hala, Raeece Naeem, Fadwa Alofi, Asim Khogeer, Afrah Alsomali, Jumana Taha, Abdulaziz Alahmadi, Kahled Algithami, Anwar Hashem, Naif Almontashiri, Arnab Pain |
| EPI_ISL_678050, EPI_ISL_678067, EPI_ISL_678124, EPI_ISL_678125, EPI_ISL_678126, EPI_ISL_678128, EPI_ISL_678129, EPI_ISL_678130, EPI_ISL_678131, EPI_ISL_678132, EPI_ISL_678133, EPI_ISL_678134 | | | |

| | | | |
|---|---|---|---|
| see above | Pathogen Genomics Lab King Abdullah University of Science and Technology(KAUST) | Pathogen Genomics Lab King Abdullah University of Science and Technology(KAUST) | Luke Esau, Amanda Ooi, Sharif Hala, Raeece Naeem, Sara Mfarrej, Asim Khogeer, Fadwa Alofi, Afrah Alsomali, Jumana Taha, Abdulaziz Alahmadi, Kahled Algithami, Anwar Hashem, Naif Almontashiri, Arnab Pain |
| EPI_ISL_678145, EPI_ISL_678146 | Pathogen Genomics Lab King Abdullah University of Science and Technology(KAUST) | Pathogen Genomics Lab King Abdullah University of Science and Technology(KAUST) | Sara Mfarrej, Luke Esau, Amanda Ooi, Sharif Hala, Raeece Naeem, Asim Khogeer, Fadwa Alofi, Afrah Alsomali, Jumana Taha, Abdulaziz Alahmadi, Kahled Algithami, Anwar Hashem, Naif Almontashiri, Arnab Pain |
| EPI_ISL_678246 | Pathogen Genomics Lab King Abdullah University of Science and Technology(KAUST) | Pathogen Genomics Lab King Abdullah University of Science and Technology(KAUST) | Luke Esau, Amanda Ooi, Sharif Hala, Raeece Naeem, Sara Mfarrej, Asim Khogeer, Fadwa Alofi, Afrah Alsomali, Jumana Taha, Abdulaziz Alahmadi, Kahled Algithami, Anwar Hashem, Naif Almontashiri, Arnab Pain |
| EPI_ISL_678397 | Suceava County Emergency Hospital | "Stefan cel Mare" University Metagenomics Lab | Lobiuc Andrei, Puscaselu Roxana |
| EPI_ISL_678487 | Veterinary Specialized Institute "Sabac", Serbia | Veterinary Specialized Institute "Kraljevo", Serbia | Vidanovic,D., Tesovic,B., Knezevic,A., Jovanovic,T., Jankovic,M., Sekler,M., Banovic Djeri,B., Petrovic,T., Mrkovacki, S., Volkening,J., Afonso,C. |
| EPI_ISL_678507, EPI_ISL_678534, EPI_ISL_678535, EPI_ISL_678536 | CNR Virus des Infections Respiratoires - France SUD | CNR Virus des Infections Respiratoires - France SUD | Antonin Bal, Gregory Destras, Gwendolyne Burfin, Solenne Brun, Martine Valette, Bruno Lina, Laurence Josset |
| EPI_ISL_679968 | Lincolnshire Hospitals and DeepSeq Nottingham | COVID-19 Genomics UK (COG-UK) Consortium | Nichola Duckworth, Tim Sloan, Sarah Walsh, Jonathan Ball, Patrick McClure, Joeseph Chappell, Nadine Holmes, Matthew Carlisle, Christopher Moore, Fei Sang, Johnny Debebe, Victoria Wright, Matthew Loose |
| EPI_ISL_680288 | Regional Virus Laboratory, Belfast Health and Social Care Trust | COVID-19 Genomics UK (COG-UK) Consortium | Conall McCaughey, James McKenna, Tanya Curran, Susan Feeney, Alison Watt, Ciara Cox, Mairead Connor, Zoltan Molnar, David Simpson, Derek Fairley |
| EPI_ISL_681712, EPI_ISL_681713, EPI_ISL_681740, EPI_ISL_681741, EPI_ISL_681770 | University Hospital Limerick | Irish Coronavirus Sequencing Consortium - Teagasc Moorepark | Paul Cotter, Fiona Crispie, Amy Fitzpatrick, John Kenny, Elaine Lawton, Carolyn Meaney, Patrick Stapleton, Calum Walsh |
| EPI_ISL_681987, EPI_ISL_681989, EPI_ISL_681993, EPI_ISL_681994, EPI_ISL_681996, EPI_ISL_681997 | UPMC Clinical Microbiology Laboratory | Microbial Genomic Epidemiology Laboratory, University of Pittsburgh | Mustapha M. Mustapha, Jane W. Marsh, Dan Snyder, Marissa P. Griffith, Stephanie L. Mitchell, Vatsala R. Srinivasa, Kady D. Waggle, Chinelo Ezeonwuku, Vaughn S. Cooper, Lee H. Harrison |
| EPI_ISL_684107 | CNR Virus des Infections Respiratoires - France SUD | CNR Virus des Infections Respiratoires - France SUD | Antonin Bal, Gregory Destras, Gwendolyne Burfin, Quentin Semanas, Martine Valette, Bruno Lina, Laurence Josset |
| EPI_ISL_691619 | Servicio de Microbiología, Hospital Universitario Son Espases | SeqCOVID-SPAIN consortium/IBV(CSIC) | Carla López-Causapé, Jordi Reina, Antonio Oliver and SeqCOVID-SPAIN consortium |
| EPI_ISL_692950, EPI_ISL_692951, EPI_ISL_692952, EPI_ISL_692953, EPI_ISL_692954, EPI_ISL_692955, EPI_ISL_692956, EPI_ISL_692957, EPI_ISL_692958, EPI_ISL_693115, EPI_ISL_693136, EPI_ISL_693137, EPI_ISL_693138, EPI_ISL_693139, EPI_ISL_693140, EPI_ISL_693141, EPI_ISL_693142, EPI_ISL_693143, EPI_ISL_693144, EPI_ISL_693145, EPI_ISL_693146, EPI_ISL_693147, EPI_ISL_693148, EPI_ISL_693149, EPI_ISL_693158, EPI_ISL_693185, EPI_ISL_693186 | | | |
| see above | Massachusetts State Public Health Laboratory | Massachusetts State Public Health Laboratory | Andrew Lang, Timelia Fink, Glen Gallagher, Sandra Smole |
| EPI_ISL_693203 | Hospital Municipal Doutor Arthur Ribeiro de Saboya | Instituto Adolfo Lutz, Interdiciplinary Procedures Center, Strategic Laboratory | Claudio Tavares Sacchi, Claudia Regina Gonçalves, Erica Valessa Ramos Gomes, Karoline Rodrigues Campos |
| EPI_ISL_694139 | TGen North | TGen North | Jolene Bowers, Megan Folkerts, Chris French, Hayley Yaglom, Ashlyn Pfeiffer, Darrin Lemmer, Dave Engelthaler, The Arizona COVID Genomics Union (ACGU) |
| EPI_ISL_694156, EPI_ISL_694157, EPI_ISL_694158, EPI_ISL_694159, EPI_ISL_694160, EPI_ISL_694161, EPI_ISL_694162, EPI_ISL_694163, EPI_ISL_694164, EPI_ISL_694165, EPI_ISL_694166, EPI_ISL_694167, EPI_ISL_694168, EPI_ISL_694169, EPI_ISL_694170, EPI_ISL_694171, EPI_ISL_694172, EPI_ISL_694173, EPI_ISL_694174, EPI_ISL_694175, EPI_ISL_694176, EPI_ISL_694177, EPI_ISL_694178, EPI_ISL_694179, EPI_ISL_694180, EPI_ISL_694181, EPI_ISL_694182, EPI_ISL_694183, EPI_ISL_694184, EPI_ISL_694185, EPI_ISL_694186, EPI_ISL_694187, EPI_ISL_694188, EPI_ISL_694189, EPI_ISL_694190, EPI_ISL_694191, EPI_ISL_694192, EPI_ISL_694193, EPI_ISL_694194, EPI_ISL_694195, EPI_ISL_694196, EPI_ISL_694197, EPI_ISL_694198, EPI_ISL_694199, EPI_ISL_694200, EPI_ISL_694201, EPI_ISL_694202, EPI_ISL_694203, EPI_ISL_694204, EPI_ISL_694205, EPI_ISL_694206, EPI_ISL_694207, EPI_ISL_694208, EPI_ISL_694209, EPI_ISL_694210, EPI_ISL_694211, EPI_ISL_694212, EPI_ISL_694213, EPI_ISL_694214, EPI_ISL_694215, EPI_ISL_694216, EPI_ISL_694217, EPI_ISL_694218, EPI_ISL_694248, EPI_ISL_694249, EPI_ISL_694250, EPI_ISL_694251, EPI_ISL_694252, EPI_ISL_694253, EPI_ISL_694254, EPI_ISL_694255, EPI_ISL_694256, EPI_ISL_694257, EPI_ISL_694258, EPI_ISL_694268, EPI_ISL_694269, EPI_ISL_694270, EPI_ISL_694271, EPI_ISL_694272 | | | |
| see above | AZ SPHL, Arizona Department of Health Services | TGen North | Jolene Bowers, Megan Folkerts, Chris French, Hayley Yaglom, Ashlyn Pfeiffer, Darrin Lemmer, Dave Engelthaler, The Arizona COVID Genomics Union (ACGU) |
| EPI_ISL_695770, EPI_ISL_695786, EPI_ISL_695829, EPI_ISL_696102, EPI_ISL_696103, EPI_ISL_696104, EPI_ISL_696105, EPI_ISL_696106, EPI_ISL_696109, EPI_ISL_696110, EPI_ISL_696111, EPI_ISL_696114, EPI_ISL_696115, EPI_ISL_696116, EPI_ISL_696117, EPI_ISL_696119, EPI_ISL_696122, EPI_ISL_696127, EPI_ISL_696131, EPI_ISL_696133, EPI_ISL_696134, EPI_ISL_696136, EPI_ISL_696137, EPI_ISL_696139, EPI_ISL_696140, EPI_ISL_696141, EPI_ISL_696142, EPI_ISL_696143, EPI_ISL_696145, EPI_ISL_696146, EPI_ISL_696147, EPI_ISL_696148, EPI_ISL_696149, EPI_ISL_696150, EPI_ISL_696151, EPI_ISL_696152, EPI_ISL_696153, EPI_ISL_696154, EPI_ISL_696156, EPI_ISL_696159, EPI_ISL_696160, EPI_ISL_696161, EPI_ISL_696162, EPI_ISL_696163, EPI_ISL_696164, EPI_ISL_696165, EPI_ISL_696166, EPI_ISL_696169, EPI_ISL_696170, EPI_ISL_696171, EPI_ISL_696172, EPI_ISL_696174, EPI_ISL_696175, EPI_ISL_696176, EPI_ISL_696177, EPI_ISL_696261, EPI_ISL_696269, EPI_ISL_696273, EPI_ISL_696282, EPI_ISL_696286, EPI_ISL_696296, EPI_ISL_696299, EPI_ISL_696302, EPI_ISL_696304, EPI_ISL_696307, EPI_ISL_696310, EPI_ISL_696314 | | | |
| see above | Sonora Quest Laboratories, Laboratory Sciences of Arizona | TGen North | Jolene Bowers, Megan Folkerts, Chris French, Hayley Yaglom, Ashlyn Pfeiffer, Darrin Lemmer, Dave Engelthaler, The Arizona COVID Genomics Union (ACGU) |
| EPI_ISL_699658 | Hematopathology Laboratory, ACTREC, TMC | Hematopathology Laboratory, ACTREC, TMC | Hematopathology Laboratory, ACTREC |
| EPI_ISL_700392, EPI_ISL_700393, EPI_ISL_700394, EPI_ISL_700395 | Centre hospitalier Métropole Savoie | CNR Virus des Infections Respiratoires - France SUD | Antonin Bal, Carine Dumollard, Gregory Destras, Gwendolyne Burfin, Hadrien Règue, Quentin Semanas, Martine Valette, Bruno Lina, Jérôme Grosjean, Laurence Josset |
| EPI_ISL_700412, EPI_ISL_700454, EPI_ISL_700459 | Heideveld CDC wc HVP | NHLS/UCT | Arash Iranzadeh, Deelan Doolabh, Lynn Tyers, Bruna Galvao, Innocent Mudau, Marvin Hsiao, Kruger Marais, Diana Hardie, Stephen Korsman, Carolyn Williamson |
| EPI_ISL_700480 | Hanover Park CHC wc HPH | NHLS/UCT | Arash Iranzadeh, Deelan Doolabh, Lynn Tyers, Bruna Galvao, Innocent Mudau, Marvin Hsiao, Kruger Marais, Diana Hardie, Stephen Korsman, Carolyn Williamson |
| EPI_ISL_700545 | Heideveld Emergency Centre | NHLS/UCT | Arash Iranzadeh, Deelan Doolabh, Lynn Tyers, Bruna Galvao, Innocent Mudau, Marvin Hsiao, Kruger Marais, Diana Hardie, Stephen Korsman, Carolyn Williamson |
| EPI_ISL_700567, EPI_ISL_700591 | Dr Abdurahman CDC wc DAC | NHLS/UCT | Arash Iranzadeh, Deelan Doolabh, Lynn Tyers, Bruna Galvao, Innocent Mudau, Marvin Hsiao, Kruger Marais, Diana Hardie, Stephen Korsman, Carolyn Williamson |
| EPI_ISL_710131, EPI_ISL_710134, EPI_ISL_710135, EPI_ISL_710137, EPI_ISL_710139, EPI_ISL_710140, EPI_ISL_710143, EPI_ISL_710144, EPI_ISL_710150, EPI_ISL_710151, EPI_ISL_710153, EPI_ISL_710159, EPI_ISL_710160, EPI_ISL_710161, EPI_ISL_710164, EPI_ISL_710166, EPI_ISL_710171, EPI_ISL_710172, EPI_ISL_710175, EPI_ISL_710176, EPI_ISL_710178, EPI_ISL_710179, EPI_ISL_710180, EPI_ISL_710181, EPI_ISL_710182, EPI_ISL_710183, EPI_ISL_710184, EPI_ISL_710185, EPI_ISL_710186, EPI_ISL_710187, EPI_ISL_710188, EPI_ISL_710189, EPI_ISL_710190, EPI_ISL_710191, EPI_ISL_710192, EPI_ISL_710193, EPI_ISL_710194, EPI_ISL_710195, EPI_ISL_710196, EPI_ISL_710197, EPI_ISL_710198, EPI_ISL_710199, EPI_ISL_710200, EPI_ISL_710201, EPI_ISL_710202, EPI_ISL_710203, EPI_ISL_710204, EPI_ISL_710205, EPI_ISL_710206, EPI_ISL_710207, EPI_ISL_710208, EPI_ISL_710259, EPI_ISL_710290, EPI_ISL_710301 | | | |
| see above | Colorado Department of Public Health and Environment | Colorado Department of Puplic Health and Environment | Laura Bankers, Molly C. Hetherington-Rauth, Shannon Ely, Shannon R. Matzinger, Sarah Elizabeth Totten, Emily A. Travanty |
| EPI_ISL_714253 | Department of Virus and Microbiological Special Diagnostics, Statens Serum Institut, Copenhagen, Denmark | Albertsen Lab, Department of Chemistry and Bioscience, Aalborg University, Denmark | Danish Covid-19 Genome Consortium |
| EPI_ISL_717723, EPI_ISL_717725, EPI_ISL_717730, EPI_ISL_717733, EPI_ISL_717736, EPI_ISL_717738, EPI_ISL_717743, EPI_ISL_717748, EPI_ISL_717753, EPI_ISL_717755, EPI_ISL_717758 | | | |
| see above | Kingston Health Sciences Centre and Queen's University | Ontario Institute for Cancer Research | Prameet M. Sheth,Calvin Sjaarda,Robert Colautti,Katya Douchant,Ilinca Lungu,Bernard Lam,Paul Krzyzanowski,Michael Laszloffy,Lawrence E. Heisler,Richard de Borja,Jared T. Simpson |
| EPI_ISL_717843, EPI_ISL_717844 | Laboratorio de Virologia Molecular / UFRJ | Bioinformatics Laboratory / LNCC | Carolina M Voloch, Ronaldo da Silva F Jr, Luiz G P de Almeida, Cynthia C Cardoso, Otavio Bustrolini, Alexandra L Gerber, Ana Paula de C Guimarães, Diana Mariani, Andréa Cony Cavalcanti, Claudia dos Santos Rodrigues, Terezinha M P P Castiñeira, Amilcar Tanuri, Ana Tereza R de Vasconcelos |
| EPI_ISL_718139, EPI_ISL_718144, EPI_ISL_718158 | Ministry of Health Hospitals | Institute of Health and Community Medicine | David Perera, Ooi Mong How, Chua Hock Hin, Tonnii Sia Loong Loong, Wong Jyn Shan, Wong Kiing Aik, Chan Chia Jui |
| EPI_ISL_721989, EPI_ISL_721996, EPI_ISL_721997, EPI_ISL_721999, EPI_ISL_722001, EPI_ISL_722003, EPI_ISL_722008, EPI_ISL_722009, EPI_ISL_722015, EPI_ISL_722019, EPI_ISL_722021, EPI_ISL_722024, EPI_ISL_722025, EPI_ISL_722026, EPI_ISL_722027, EPI_ISL_722028, EPI_ISL_722036, EPI_ISL_722037, EPI_ISL_722038, EPI_ISL_722039, EPI_ISL_722044, EPI_ISL_722046, EPI_ISL_722047, EPI_ISL_722048, EPI_ISL_722049, EPI_ISL_722050, EPI_ISL_722051, EPI_ISL_722052, EPI_ISL_722053, EPI_ISL_722056, EPI_ISL_722061, EPI_ISL_722072, EPI_ISL_722073, EPI_ISL_722076, EPI_ISL_722087, EPI_ISL_722088, EPI_ISL_722091, EPI_ISL_722102, EPI_ISL_722110, EPI_ISL_722112, EPI_ISL_722128, EPI_ISL_722129 | | | |
| see above | Hospital das Clínicas Universidade de São Paulo Medical School | Laboratório de Parasitologia Médica - Instituto de Medicina Tropical - Universidade de São Paulo | Brazil-UK Centre for Arbovirus Discovery Diagnosis Genomics and Epidemiology (CADDE) Genomic Network - Instituto de Medicina Tropical |

| | | | |
|---|---|---|---|
| EPI_ISL_728361 | Hospital Universitari Vall d'Hebron (HUVH) - Vall d'Hebron Research Institute (VHIR) | Hospital Universitari Vall d'Hebron - Vall d'hebron Research Institut (VHIR) | Maria Piñana, Josep F Abril, Cristina Andrés, Damir Garcia-Cehic, Aroa Silgado, Maria Carmen Martín, Carla Castillo, Ariadna Rando, Maria Gema Codina, Juliana Esperalba, Elena Sulleiro, Tomàs Pumarola, Josep Quer, Andrés Antón |
| EPI_ISL_728362 | Hospital Universitari Vall d'Hebron (HUVH) - Vall d'Hebron Research Institute (VHIR) | Hospital Universitari Vall d'Hebron - Vall d'Hebron Research Institute (VHIR) | Maria Piñana, Josep F Abril, Cristina Andrés, Damir Garcia-Cehic, Aroa Silgado, Maria Carmen Martín, Carla Castillo, Ariadna Rando, Maria Gema Codina, Juliana Esperalba, Elena Sulleiro, Tomàs Pumarola, Josep Quer, Andrés Antón |
| EPI_ISL_728364, EPI_ISL_728365 | Hospital Universitari Vall d'Hebron (HUVH) - Vall d'Hebron Research Institute (VHIR) | Hospital Universitari Vall d'Hebron (HUVH) - Vall d'Hebron Research Institute (VHIR) | Maria Piñana, Josep F Abril, Cristina Andrés, Damir Garcia-Cehic, Aroa Silgado, Maria Carmen Martín, Carla Castillo, Ariadna Rando, Maria Gema Codina, Juliana Esperalba, Elena Sulleiro, Tomàs Pumarola, Josep Quer, Andrés Antón |
| EPI_ISL_729834 | Laboratorio Central de Saude Publica do Estado do Rio Grande do Sul (LACEN-RS) | Laboratory of Respiratory Viruses and Measles, Oswaldo Cruz Institute, FIOCRUZ | Paola Resende, Luciana Appolinario, Fernando Motta, Anna Carolina Paixão, Ana Carolina Mendonça, Tatiana Schaffer Gregianini, Marilda Tereza Mar da Rosa, Marilda Siqueira |
| EPI_ISL_729920, EPI_ISL_729937, EPI_ISL_729938, EPI_ISL_729939, EPI_ISL_729940, EPI_ISL_729942, EPI_ISL_729943, EPI_ISL_729959, EPI_ISL_729996, EPI_ISL_729997, EPI_ISL_729998, EPI_ISL_729999, EPI_ISL_730000, EPI_ISL_730028, EPI_ISL_730029, EPI_ISL_730030 | | | |
| see above | Nigeria Centre for Disease Control (NCDC) | African Centre of Excellence for Genomics of Infectious Diseases (ACEGID), Redeemer's University, Ede, Osun State, Nigeria | Oluniyi P.E. et al |
| EPI_ISL_730145 | Scripps Medical Laboratory | Andersen lab at Scripps Research | SEARCH Alliance San Diego with Michael Quigley, Ellen Stefanski, Ian Mchardy |
| EPI_ISL_732749, EPI_ISL_732759 | Centro de Investigación Biomédica de La Rioja - Hospital San Pedro Logroño | SeqCOVID-SPAIN consortium/IBV(CSIC) | María de Toro, José Manuel Azcona Gutiérrez, María Pilar Bea Escudero, Miriam Blasco Alberdi and SeqCOVID-SPAIN consortium |
| EPI_ISL_734632, EPI_ISL_734633, EPI_ISL_734634, EPI_ISL_734635, EPI_ISL_734636, EPI_ISL_734637, EPI_ISL_734638, EPI_ISL_734639, EPI_ISL_734640, EPI_ISL_734641, EPI_ISL_734642, EPI_ISL_734643, EPI_ISL_734644, EPI_ISL_734645, EPI_ISL_734646, EPI_ISL_734647, EPI_ISL_734648, EPI_ISL_734649, EPI_ISL_734650, EPI_ISL_734651, EPI_ISL_734652, EPI_ISL_734653, EPI_ISL_734654, EPI_ISL_734655, EPI_ISL_734656, EPI_ISL_735244, EPI_ISL_735248 | | | |
| see above | UZ Leuven, National Reference Laboratory for Coronaviruses, Laboratory Medicine, Leuven, Belgium | KU Leuven, Rega Institute, Clinical and Epidemiological Virology | Tony Wawina-Bokalanga, Joan Marti-Carerras, Bert Vanmechelen, Piet Maes |
| EPI_ISL_735263, EPI_ISL_735264, EPI_ISL_735265, EPI_ISL_735266, EPI_ISL_735281 | Genomic Laboratory (GLAB) (Conjoint lab of Health Directorate of Istanbul and Istanbul Technical University) | Genomic Laboratory (GLAB), Istanbul Technical University | Ilker Karacan, Tugba Kizilboga Akgun, Nihat Bugra Agaoglu, Payam Zolfagharian, Mehtap Aydin, Gizem Alkurt, Jale Yildiz, Betsi Kose, Nisan Denizce Can, Ayse Serra Ozel, Nilsun Altunal, Arzu Irvem, Yasemin Kendir Demirkol, Ozlem Akgun Dogan, Levent Doganay, Gizem Dinler Doganay |
| EPI_ISL_738247, EPI_ISL_738275, EPI_ISL_738276, EPI_ISL_738277, EPI_ISL_738278, EPI_ISL_738279, EPI_ISL_738280, EPI_ISL_738281 | Connecticut Veterans' Affairs Hospital | Grubaugh Lab - Yale School of Public Health | Joseph Fauver, Tara Alpert, Chantal Vogels, Mary Petrone, Isabel Ott, Ellen Foxman, Shaili Gupta, Danielle Plank, Nathan Grubaugh |
| EPI_ISL_738820 | Napa-Solano-Yolo- Marin County (NSYM) Public Health Laboratories | Chan-Zuckerberg Biohub | CZB Cliahub Consortium |
| EPI_ISL_740182 | Laboratoire national de santé, Microbiology, Virology | Laboratoire national de santé, Microbiology, Microbial Genomics Platform | Anke Wienecke-Baldacchino, Catherine Ragimbeau, Tamir Abdelrahman, Jessica Tapp, Fatu Djabi |
| EPI_ISL_751202 | Pathogen Genomics Lab King Abdullah University of Science and Technology(KAUST) | Pathogen Genomics Lab King Abdullah University of Science and Technology(KAUST) | Sara Mfarrej, Raushan Nugmanova, Olga Douvropoulou, Sharif Hala, Raeece Naeem, Asim Khogeer, Fadwa Alofi, Afrah Alsomali, Jumana Taha, Abdulaziz Alahmadi, Kahled Algithami, Anwar Hashem, Naif Almontashiri, Arnab Pain |
| EPI_ISL_751203 | Pathogen Genomics Lab King Abdullah University of Science and Technology(KAUST) | Pathogen Genomics Lab King Abdullah University of Science and Technology(KAUST) | Sharif Hala, Raeece Naeem, Raushan Nugmanova, Olga Douvropoulou, Rahul P Salunke, Fadwa Alofi, Asim Khogeer, Afrah Alsomali, Jumana Taha, Abdulaziz Alahmadi, Kahled Algithami, Anwar Hashem, Naif Almontashiri, Arnab Pain |
| EPI_ISL_751218 | Pathogen Genomics Lab King Abdullah University of Science and Technology(KAUST) | Pathogen Genomics Lab King Abdullah University of Science and Technology(KAUST) | Sharif Hala, Fathia Ben Rached, Sara Mfarrej, Raushan Nugmanova, Olga Douvropoulou, Rahul P Salunke, Raeece Naeem, Fadwa Alofi, Asim Khogeer, Afrah Alsomali, Jumana Taha, Abdulaziz Alahmadi, Kahled Algithami, Anwar Hashem, Naif Almontashiri, Arnab Pain |
| EPI_ISL_751336, EPI_ISL_751338 | IRCCS Sacro Cuore Don Calabria Hospital, Department of Infectious, Tropical Diseases & Microbiology | University of Verona, Department of Biotechnology | Antonio Mori, Michela Deiana, Elena Pomari, Chiara Piubelli; Giulia Lopatriello, Luca Marcolungo, Cristina Beltrami, Chiara Degli Esposti, Emanuela Cosentino, Massimo Delledonne |
| EPI_ISL_752613, EPI_ISL_752620, EPI_ISL_752634, EPI_ISL_752635, EPI_ISL_752639, EPI_ISL_752662, EPI_ISL_752669 | State Laboratories Division, Hawaii State Department of Health | State Laboratories Division, Hawaii State Department of Health | Pamela O'Brien, Sabrina Diemert, Drew Kuwazaki, Razvan Sultana, Edward Desmond |
| EPI_ISL_753702, EPI_ISL_753746, EPI_ISL_753841, EPI_ISL_753949 | Charité Universitätsmedizin Berlin, Institut für Virologie/Labor Berlin | Charité Universitätsmedizin Berlin, Institut für Virologie | Victor M Corman, Jörn Beheim-Schwarzbach, Barbara Mühlemann, Julia Schneider, Talitha Veith, Terry Jones, Christian Drosten |
| EPI_ISL_754229 | The Republican Research and Practical Center for Epidemiology and Microbiology (RRPCEM) | WHO National Influenza Centre Russian Federation | Elena Gasich, Kirill Bulda, Anatoly Krasko, Andrey Komissarov, Artem Fadeev, Anna Ivanova, Kseniya Komissarova, Dmitry Bazhenov, Daria Danilenko, Ksenia Safina, Elena Nabieva, Georgii Bazykin, Dmitry Lioznov |
| EPI_ISL_759865 | Molecular, ProgenaBiome | Molecular, ProgenaBiome | Papoutsis,A.J. |
| EPI_ISL_759956 | Husada Utama Hospital | Institute of Tropical Disease, Universitas Airlangga | Jezzy R Dewantari, Rima R Prasetya, Krisnoadi Rahardjo, Aldise M Nastri, Didi Dewanto, Gatot Soegiarto, Laksmi Wulandari, Resti Yudhawati, Soetjipto, Yasuko Mori, Maria I Lusida, Kazufumi Shimizu |
| EPI_ISL_763076 | Ohio Department Of Health | Ohio Department of Health | Jennifer Holmes, Eric Brandt, Keoni Omura, Glen McGillivary, Caitlin McDonnell, Kirtana Ramadugu, Kelsey Florek, Heather Blankenship, Quanta Brown, and Tammy Bannerman |
| EPI_ISL_763319 | Ohio Department of Health Laboratory | Ohio Department of Health laboratory | Jennifer Holmes, Eric Brandt, Keoni Omura, Glen McGillivary, Caitlin McDonnell, Kirtana Ramadugu, Kelsey Florek, Heather Blankenship, Quanta Brown, and Tammy Bannerman |
| EPI_ISL_763321 | Ohio Department of Health Laboratory | Ohio Depart of Health Laboratory | Jennifer Holmes, Eric Brandt, Keoni Omura, Glen McGillivary, Caitlin McDonnell, Kirtana Ramadugu, Kelsey Florek, Heather Blankenship, Quanta Brown, and Tammy Bannerman |
| EPI_ISL_763364 | AIID | Irish Coronavirus Sequencing Consortium - National Virus Reference Laboratory | Michael Carr, Gabriel Gonzalez, Alejandro Abner Garcia Leon, Patrick Mallon |
| EPI_ISL_765209 | Ohio Department of Health Laboratory | Ohio Department of Health Laboratory | Holmes, Jennifer; Eric Brandt, Glen McGillivary, Keoni Omura, Caitlin McDonnell, Heather Blankenship, Kelsey Florek, Kirtana Ramadugu, Erica Leasure, Quanta Brown and Tammy Bannerman |
| EPI_ISL_765222 | Ohio Department of Health Laboratory | Ohio Department of Health Laboratory | Holmes, Jennifer; Eric Brandt, Keoni Omura, Glen McGillivary, Caitiln McDonnell, Kirtana Ramadugu, Erica Leasure, Kelsey Florek, Heather Blankenship, Quanta Brown, and Tammy Bannerman |
| EPI_ISL_766043 | RS Pelni | Eijkman Institute for Molecular Biology, Ministry of Research and Technology/National Agency for Research and Innovation | Frilasita A Yudhaputri, Edison Johar, Hidayat Trimarsanto, David H Muljono, Safarina G Malik, Khin Saw Myint, Amin Soebandrio |
| EPI_ISL_766044 | RSUP Fatmawati | Eijkman Institute for Molecular Biology, Ministry of Research and Technology/National Agency for Research and Innovation | Frilasita A Yudhaputri, Edison Johar, Hidayat Trimarsanto, David H Muljono, Safarina G Malik, Khin Saw Myint, Amin Soebandrio |
| EPI_ISL_768735, EPI_ISL_768738 | Child Health Research Foundation | Child Health Research Foundation | Senjuti Saha, Afroza Akter Tanni, Roly Malaker, Sharmistha Goswami, Syed Muktadir Al Sium, Arif Mohammad Tanmoy, Md Hafizur Rahman, Samir K Saha |
| EPI_ISL_770632, EPI_ISL_770775, EPI_ISL_770776, EPI_ISL_770777 | Ohio Department of Health Laboratory | Ohio Department of Health Laboratory | Holmes, Jennifer; Eric Brandt, Keoni Omura, Glen McGillivary, Caitiln McDonnell, Kirtana Ramadugu, Erica Leasure, Kelsey Florek, Heather Blankenship, Quanta Brown, and Tammy Bannerman |
| EPI_ISL_771178, EPI_ISL_771225 | Colorado Department of Public Health and Environment | Colorado Department of Puplic Health and Environment | Laura Bankers, Molly C. Hetherington-Rauth, Diana Ir, Shannon Ely, Shannon R. Matzinger, Sarah Elizabeth Totten, Emily A. Travanty |

| | | | |
|---|---|---|---|
| EPI_ISL_775261 | INT Fondazione Pascale | INT Fondazione Pascale | INT Fondazione Pascale |
| EPI_ISL_775600, EPI_ISL_775601, EPI_ISL_775602 | Ohio Department of Health Laboratory | Ohio Department of Health Laboratory | Holmes, Jennifer; Eric Brandt, Keoni Omura, Glen McGillivary, Caitiln McDonnell, Kirtana Ramadugu, Erica Leasure, Kelsey Florek, Heather Blankenship, Quanta Brown, and Tammy Bannerman |
| EPI_ISL_775603, EPI_ISL_775605, EPI_ISL_775606 | Ohio Department of Health Laboratory | Ohio Department of Health Laboratory | Holmes, Jennifer; Eric Brandt, Keoni Omura, Glen McGillivary, Caitiln McDonnell, Kirtana Ramadugu, Erica Leasure, Kelsey Florek, Heather Blankenship, Quanta Brown, and Tammy Bannerman |
| EPI_ISL_775621, EPI_ISL_775653, EPI_ISL_775696, EPI_ISL_775723, EPI_ISL_775724, EPI_ISL_775737, EPI_ISL_775758, EPI_ISL_775908, EPI_ISL_775939, EPI_ISL_776470, EPI_ISL_776471, EPI_ISL_776472, EPI_ISL_776474, EPI_ISL_776475, EPI_ISL_776476, EPI_ISL_776477, EPI_ISL_776478, EPI_ISL_776479, EPI_ISL_776480, EPI_ISL_776481, EPI_ISL_776482, EPI_ISL_776483, EPI_ISL_776486, EPI_ISL_776487, EPI_ISL_776488, EPI_ISL_776489, EPI_ISL_776490, EPI_ISL_776491, EPI_ISL_776492, EPI_ISL_776493, EPI_ISL_776494, EPI_ISL_776498, EPI_ISL_776499, EPI_ISL_776500, EPI_ISL_776505, EPI_ISL_776506, EPI_ISL_776507, EPI_ISL_776509, EPI_ISL_776510, EPI_ISL_776515, EPI_ISL_776527, EPI_ISL_776554 | | | |
| see above | University Medical Center Hamburg Eppendorf | Heinrich Pette Institute, Leibniz Institute for Experimental Virology | Alexis Robitaille, Thomas Günther, Johannes Knobloch, Martin Aepfelbacher, Nicole Fischer, Adam Grundhoff |
| EPI_ISL_776567, EPI_ISL_776568 | Ohio Department of Health Laboratory | Ohio Department of Health Laboratory | Holmes, Jennifer; Eric Brandt, Keoni Omura, Glen McGillivary, Caitlin McDonnell, Kirtana Ramadugu, Erica Leasure, Kelsey Florek, Heather Blankenship, Quanta Brown, and Tammy Bannerman |
| EPI_ISL_776734, EPI_ISL_776749 | UW Virology Lab | UW Virology Lab | Pavitra Roychoudhury, Hong Xie, Lasata Shrestha, Meei-Li Huang, Keith R Jerome, Alexander Greninger |
| EPI_ISL_776755, EPI_ISL_776756 | Instituto Adolfo Lutz - Central | Instituto Adolfo Lutz, Interdiciplinary Procedures Center, Strategic Laboratory | Claudio Tavares Sacchi, Claudia Regina Gonçalves, Erica Valessa Ramos Gomes, Karoline Rodrigues Campos |
| EPI_ISL_776773, EPI_ISL_776774, EPI_ISL_776869, EPI_ISL_776872, EPI_ISL_776874, EPI_ISL_776876, EPI_ISL_776994 | Ohio Department of Health Laboratory | Ohio Department of Health Laboratory | Holmes, Jennifer; Eric Brandt, Keoni Omura, Glen McGillivary, Caitlin McDonnell, Kirtana Ramadugu, Erica Leasure, Kelsey Florek, Heather Blankenship, Quanta Brown, and Tammy Bannerman |
| EPI_ISL_778673, EPI_ISL_778674, EPI_ISL_778675, EPI_ISL_778676, EPI_ISL_778677, EPI_ISL_778678, EPI_ISL_778679, EPI_ISL_778680, EPI_ISL_778681, EPI_ISL_778682, EPI_ISL_778735 | | | |
| see above | Istituto Zooprofilattico Sperimentale del Mezzogiorno | TIGEM | Patrizia Annunziata, Andrea Ballabio, Valentina Bouche, Davide Cacchiarelli (CorrespAuthor), Pellegrino Cerino, Chiara Colantuono, Lucio Di Filippo, Antonio Grimaldi, Antonio Limone, Gabriella Loconte, Anna Manfredi, Francesco Panariello, Biancamaria Pierri, Marcello Salvi, Lucia Vassallo |
| EPI_ISL_778855, EPI_ISL_778856, EPI_ISL_778857, EPI_ISL_778858, EPI_ISL_778859, EPI_ISL_778861, EPI_ISL_778862, EPI_ISL_778864 | Ohio Department of Health Laboratory | Ohio Department of Health Laboratory | Holmes, Jennifer; Eric Brandt, Keoni Omura, Glen McGillivary, Caitlin McDonnell, Kirtana Ramadugu, Erica Leasure, Kelsey Florek, Heather Blankenship, Quanta Brown, and Tammy Bannerman |
| EPI_ISL_779734, EPI_ISL_779737 | Servicio de Microbiología, Hospital Miguel Servet, Zaragoza | SeqCOVID-SPAIN consortium/IBV(CSIC) | Antonio Rezusta López, Alexander Tristancho Baró, Ana Milagro, Yolanda Gracia Grataloup, Nieves Martínez Cameo and SeqCOVID-SPAIN consortium |
| EPI_ISL_780377, EPI_ISL_780379, EPI_ISL_780380, EPI_ISL_780417, EPI_ISL_780419, EPI_ISL_780421, EPI_ISL_780422, EPI_ISL_780423, EPI_ISL_780424, EPI_ISL_780425, EPI_ISL_780426, EPI_ISL_780670, EPI_ISL_781113 | | | |
| see above | Ohio Department of Health Laboratory | Ohio Department of Health Laboratory | Holmes, Jennifer; Eric Brandt, Keoni Omura, Glen McGillivary, Caitlin McDonnell, Kirtana Ramadugu, Erica Leasure, Kelsey Florek, Heather Blankenship, Quanta Brown, and Tammy Bannerman |
| EPI_ISL_788945, EPI_ISL_788947 | Ospedale "Di Venere" | Beaconlab (Bioinformatics, Evolution and Comparative Genomics lab), Dept of Biosciences, University on Milan | Iacobellis M, d'Avenia M, Piluscio R, Parisi A, Chiara M, Manzari C, Pesole G |
| EPI_ISL_789104 | Ohio Department of Health Laboratory | Ohio Department of Health Laboratory | Holmes, Jennifer; Eric Brandt, Keoni Omura, Glen McGillivary, Caitlin McDonnell, Kirtana Ramadugu, Erica Leasure, Kelsey Florek, Heather Blankenship, Quanta Brown, and Tammy Bannerman |
| EPI_ISL_792109, EPI_ISL_792110, EPI_ISL_792111, EPI_ISL_792112 | Instituto Adolfo Lutz - Central | Instituto Adolfo Lutz, Interdiciplinary Procedures Center, Strategic Laboratory | Claudio Tavares Sacchi, Claudia Regina Gonçalves, Erica Valessa Ramos Gomes, Karoline Rodrigues Campos |
| EPI_ISL_792117, EPI_ISL_792118, EPI_ISL_792119, EPI_ISL_792120, EPI_ISL_792121, EPI_ISL_792122, EPI_ISL_792123, EPI_ISL_792124, EPI_ISL_792125, EPI_ISL_792126, EPI_ISL_792127, EPI_ISL_792128 | | | |
| see above | Laboratorio de Virología del Hospital de Niños Dr. Ricardo Gutierrez | Área de Secuenciación del Laboratorio de Virología del Hospital de Niños Dr. Ricardo Gutierrez on behalf of 'Proyecto Argentino Interinstitucional de genomica de SARS-CoV-2' (PAIS Consortium) | Nabaes Jodar, MS; Goya, S; Natale, MI; Lusso, S; Gravis, E; Acevedo, ME; Alvarez Lopez, C; Alexay, S; Jacques, O; Mistchenko, AS; Valinotto, LE; Viegas, M. |
| EPI_ISL_792291, EPI_ISL_792293, EPI_ISL_792294, EPI_ISL_792295, EPI_ISL_792298 | Laboratorio de genética y biología molecular del Hospital de trauma y emergencia Dr Federico Abete | Área de Secuenciación del Laboratorio de Virología del Hospital de Niños Dr. Ricardo Gutierrez on behalf of 'Proyecto Argentino Interinstitucional de genomica de SARS-CoV-2' (PAIS Consortium) | Nabaes Jodar, MS; Goya, S; Natale, MI; Lusso, S; Primost, I; Gallino, MI; Valinotto, LE; Viegas, M. |
| EPI_ISL_792306, EPI_ISL_792315 | Laboratorio del Hospital El Cruce Dr. Néstor C. Kirchner | Área de Secuenciación del Laboratorio de Virología del Hospital de Niños Dr. Ricardo Gutierrez on behalf of 'Proyecto Argentino Interinstitucional de genomica de SARS-CoV-2' (PAIS Consortium) | Nabaes Jodar, MS; Goya, S; Natale, MI; Lusso, S; Zubieta, M; Rahhal, M; Valinotto, LE; Viegas, M. |
| EPI_ISL_792324, EPI_ISL_792325, EPI_ISL_792330, EPI_ISL_792331 | Instituto de Investigaciones Biomédicas en Retrovirus y SIDA | Área de Secuenciación del Laboratorio de Virología del Hospital de Niños Dr. Ricardo Gutierrez on behalf of 'Proyecto Argentino Interinstitucional de genomica de SARS-CoV-2' (PAIS Consortium) | Nabaes Jodar, MS; Goya, S; Natale, MI; Lusso, S; Seery, V; Remes Lenicov, F; Salomón, H; Valinotto, LE; Viegas, M. |
| EPI_ISL_792334, EPI_ISL_792335, EPI_ISL_792347, EPI_ISL_792349, EPI_ISL_792350 | Laboratorio del Hospital Interzonal General de Agudos Evita | Área de Secuenciación del Laboratorio de Virología del Hospital de Niños Dr. Ricardo Gutierrez on behalf of 'Proyecto Argentino Interinstitucional de genomica de SARS-CoV-2' (PAIS Consortium) | Nabaes Jodar, MS; Goya, S; Natale, MI; Lusso, S; Desimone, I; Luczac, E; Serrano, L; Grossi, O; Musto, A; Valinotto, LE; Viegas, M. |
| EPI_ISL_792368 | Laboratorio de Virologia - HIEAyC San Juan de Dios | Área de Secuenciación del Laboratorio de Virología del Hospital de Niños Dr. Ricardo Gutierrez on behalf of 'Proyecto Argentino Interinstitucional de genomica de SARS-CoV-2' (PAIS Consortium) | Nabaes Jodar, MS; Goya, S; Natale, MI; Lusso, S; Ferioli, M; Colmeiro, M; Gatelli, A; Ercole, R; Valinotto, LE; Viegas, M. |
| EPI_ISL_792448, EPI_ISL_792466 | Laboratorio del Hospital Regional Ushuaia Gdor. Ernesto Campos | Hospital Regional Ushuaia - Centro Austral De Investigaciones Científicas - Universidad Nacional De Tierra Del Fuego on behalf of 'Proyecto Argentino Interinstitucional de genomica de SARS-CoV-2' (PAIS Consortium) | Ceballos, SG; Nardi, CF; Gramundi, ID; Gallego, F; De Roccis, CA; Castro, G; Cáceres, SB; Yulan, CB; Boutureira, MF. |
| EPI_ISL_792496, EPI_ISL_792499 | Laboratorio Central Mg. Luis Alfredo Pianciola | Hospital Regional Ushuaia - Centro Austral De Investigaciones Científicas - Universidad Nacional De Tierra Del Fuego on behalf of 'Proyecto Argentino Interinstitucional de genomica de SARS-CoV-2' (PAIS Consortium) | Ceballos, SG; Nardi, CF; Gramundi, ID; Gallego, F; Pinto, C; Ziehm, MC; Mazzeo, M; Pianciola, L. |
| EPI_ISL_792507, EPI_ISL_792508, EPI_ISL_792509 | Laboratorio de Inmunología del Hospital Perrando e Instituto de Medicina Regional de la UNNE | Grupo de Genómica y Bioinformática del Instituto de Investigación de la Cadena Láctea CONICET-INTA on behalf of 'Proyecto Argentino Interinstitucional de genomica de SARS-CoV-2' (PAIS Consortium) | Eberhardt, MF; Irazoqui, JM; Foussal, MD; Deluca, G; Ayala, NA; Gómez, MV; Giusiano, G; Lucero, H; Marin, M; Lescano, L; Cayré, A; Amadio, AF |

| | | | |
|---|---|---|---|
| EPI_ISL_792570, EPI_ISL_792573, EPI_ISL_792574, EPI_ISL_792575, EPI_ISL_792578, EPI_ISL_792579, EPI_ISL_792580, EPI_ISL_792581, EPI_ISL_792582, EPI_ISL_792583, EPI_ISL_792584, EPI_ISL_792585, EPI_ISL_792586, EPI_ISL_792587, EPI_ISL_792588, EPI_ISL_792589, EPI_ISL_792590 | | | |
| see above | LACEN-PB | Laboratory of Respiratory Viruses and Measles, Oswaldo Cruz Institute, FIOCRUZ | Paola Resende, Luciana Appolinario, Fernando Motta, Anna Carolina Paixao, Ana Carolina Mendonca, João Felipe Bezerra, Romero Henrique Teixeira de Vasconcelos, Dalane Loudal Florentino Teixeira, Thiago Franco de Oliveira Carneiro, Marilda Siqueira |
| EPI_ISL_796128 | Laboratorio de Microbiología. Hospital General Universitario de Elda, Alicante | SeqCOVID-SPAIN consortium/IBV(CSIC) | Mª Isabel Gascón Ros, Cristina Torregrosa Hetland, Eva Pastor Boix, Paloma Cascales Ramos and SeqCOVID-SPAIN consortium |
| EPI_ISL_801538 | TNMC & BYL NAIR CH. HOSPITAL | CSIR-IGIB | Rajesh Pandey, Jayanthi Shastri, Akshay Kanakan, Janani Srinivasa Vasudevan, Ranjeet Maurya, Sachee Agrawal, Nirjhar Chatterjee, Swapneil Parikh, Manish Pathak, Subrat Thanapati, Jasmina Savak, Suresh Poojari, Mahesh Sangar, Amol Borse, Shweta Kawankar, Vasil Nachan, Mayuresh Vishwanathan, Shruthi Sachidanandan, Shrutika Pophale, Utkarsha Yelve |
| EPI_ISL_801539, EPI_ISL_801554 | Laboratory of Molecular Virology, Pontificia Universidad Católica de Chile | MSHS Pathogen Surveillance Program | Leonardo I. Almonacid, Ana S. Gonzalez-Reiche, Matthew M. Hernandez, Jorge Levican, Ana Maria Contreras, Carlos Palma, Tamara García-Salum, Zenab Khan, Adriana van De Guchte, Ajay Obla, Jayeeta Dutta, Bremy Alburquerque, Eileen Serrano, Constanza Maldonado, M. Belen Leyton, Erick Salinas, Hala Alshammary, Juan Soto, Shwetha Hara Sridhar, Ying-Chih Wang, Melissa Smith, Robert Sebra, Marcela Ferres, Adolfo Garcia-Sastre, Edward C. Holmes, Viviana Simon, Harm van Bakel, Rafael A. Medina. |
| EPI_ISL_801561, EPI_ISL_801569, EPI_ISL_801570, EPI_ISL_801575, EPI_ISL_801577 | Laboratory of Molecular Virology, Pontificia Universidad Católica de Chile | MSHS Pathogen Surveillance Program | Leonardo I. Almonacid, Ana S. Gonzalez-Reiche, Matthew M. Hernandez, Jorge Levican, Tamara García-Salum, Zenab Khan, Adriana van De Guchte, Ajay Obla, Jayeeta Dutta, Bremy Alburquerque, Eileen Serrano, Erick Salinas, Manuel Ampuero, Hala Alshammary, Juan Soto, Shwetha Hara Sridhar, Ying-Chih Wang, Melissa Smith, Robert Sebra, Fernando Valiente, Aldo Gaggero, Adolfo Garcia-Sastre, Edward C. Holmes, Viviana Simon, Harm van Bakel, Rafael A. Medina. |
| EPI_ISL_801585, EPI_ISL_801592, EPI_ISL_801595 | Laboratory of Molecular Virology, Pontificia Universidad Católica de Chile | MSHS Pathogen Surveillance Program | Leonardo I. Almonacid, Ana S. Gonzalez-Reiche, Matthew M. Hernandez, Jorge Levican, Tamara García-Salum, Zenab Khan, Adriana van De Guchte, Ajay Obla, Jayeeta Dutta, Bremy Alburquerque, Eileen Serrano, Erick Salinas, Hala Alshammary, Juan Soto, Shwetha Hara Sridhar, Ying-Chih Wang, Melissa Smith, Robert Sebra, Adolfo Garcia-Sastre, Edward C. Holmes, Viviana Simon, Harm van Bakel, Rafael A. Medina |
| EPI_ISL_801598 | Laboratory of Molecular Virology, Pontificia Universidad Católica de Chile | MSHS Pathogen Surveillance Program | Leonardo I. Almonacid, Ana S. Gonzalez-Reiche, Matthew M. Hernandez, Jorge Levican, Tamara García-Salum, Zenab Khan, Adriana van De Guchte, Ajay Obla, Jayeeta Dutta, Bremy Alburquerque, Eileen Serrano, Erick Salinas, Manuel Ampuero, Hala Alshammary, Juan Soto, Shwetha Hara Sridhar, Ying-Chih Wang, Melissa Smith, Robert Sebra, Fernando Valiente, Aldo Gaggero, Adolfo Garcia-Sastre, Edward C. Holmes, Viviana Simon, Harm van Bakel, Rafael A. Medina. |
| EPI_ISL_801713, EPI_ISL_801714, EPI_ISL_801715, EPI_ISL_801716, EPI_ISL_801717, EPI_ISL_801718, EPI_ISL_801719, EPI_ISL_801720, EPI_ISL_801721, EPI_ISL_801722, EPI_ISL_801723, EPI_ISL_801724, EPI_ISL_801725, EPI_ISL_801726, EPI_ISL_801727, EPI_ISL_801730, EPI_ISL_801731, EPI_ISL_801733, EPI_ISL_801740 | | | |
| see above | Laboratory of Molecular Virology, Pontificia Universidad Católica de Chile | MSHS Pathogen Surveillance Program | Leonardo I. Almonacid, Ana S. Gonzalez-Reiche, Matthew M. Hernandez, Jorge Levican, Ana Maria Contreras, Carlos Palma, Tamara García-Salum, Zenab Khan, Adriana van De Guchte, Ajay Obla, Jayeeta Dutta, Bremy Alburquerque, Eileen Serrano, Constanza Maldonado, M. Belen Leyton, Erick Salinas, Hala Alshammary, Juan Soto, Shwetha Hara Sridhar, Ying-Chih Wang, Melissa Smith, Robert Sebra, Marcela Ferres, Adolfo Garcia-Sastre, Edward C. Holmes, Viviana Simon, Harm van Bakel, Rafael A. Medina. |
| EPI_ISL_801853, EPI_ISL_801854, EPI_ISL_801856, EPI_ISL_801858 | Laboratory of Molecular Virology, Pontificia Universidad Católica de Chile | MSHS Pathogen Surveillance Program | Leonardo I. Almonacid, Ana S. Gonzalez-Reiche, Matthew M. Hernandez, Jorge Levican, Tamara García-Salum, Zenab Khan, Adriana van De Guchte, Ajay Obla, Jayeeta Dutta, Bremy Alburquerque, Eileen Serrano, Erick Salinas, Manuel Ampuero, Hala Alshammary, Juan Soto, Shwetha Hara Sridhar, Ying-Chih Wang, Melissa Smith, Robert Sebra, Fernando Valiente, Aldo Gaggero, Adolfo Garcia-Sastre, Edward C. Holmes, Viviana Simon, Harm van Bakel, Rafael A. Medina. |
| EPI_ISL_801896, EPI_ISL_801929, EPI_ISL_802031, EPI_ISL_802032, EPI_ISL_802033, EPI_ISL_802034, EPI_ISL_802035, EPI_ISL_802036, EPI_ISL_802037, EPI_ISL_802038 | MSHS Clinical Microbiology Laboratories | MSHS Pathogen Surveillance Program | Ana S. Gonzalez-Reiche, Hala Alshammary, Mitchell J. Sullivan, Brianne Ciferri, Ajay Obla, Angela Amoako, Mahmoud Awawda, Elena Hirsch, Ashley S. Salimbangon, Levy Sominsky, Katherine Beach, Kayla Russo, Charles Gleason, Shelcie Fabre, Giulio Kleiner, Zenab Khan, Bremy Alburquerque, Adriana van de Guchte, Komal Srivastava, Matthew M. Hernandez, Jayeeta Dutta, Denise Jurczyszak, Emily Ferreri, Rachel Chernet, Nancy Francoeur, Betsaida Salom Melo, Irina Oussenko, Gintaras Deikus, Juan Soto, Shwetha Hara Sridhar, Ying-Chih Wang, Kathryn Twyman, Andrew Kasarskis, Deena R. Altman, Robert Sebra, Adolfo García-Sastre, Marta Luksza, Gopi Patel, Sarah Schaefer, Melissa Gitman, Michael D. Nowak, Alberto Paniz-Mondolfi, Emilia Mia Sordillo, Viviana Simon, Harm van Bakel |
| EPI_ISL_804017 | Facultad de Medicina, Universidad de Atacama | Facultad de Ciencias de la Vida, UNAB | Dayán Sanhueza, Macarena Bastías, César Echeverría, Sebastián Wolter, Waldo Díaz, Jorge Olivares, Eduardo Castro, Claudio Meneses |
| EPI_ISL_804436, EPI_ISL_804439, EPI_ISL_804446 | Dutch COVID-19 response team | National Institute for Public Health and the Environment (RIVM) | Adam Meijer, Harry Vennema, Dirk Eggink, Matthijs Welkers, Jeroen Cremer, Sharon van den Brink, Bas van der Veer, AnneMarie van den Brandt, Florian Zwagemaker, Dennis Schmitz, Chantal Reusken, on behalf of the national COVID-19 response team |
| EPI_ISL_804985, EPI_ISL_804987, EPI_ISL_804988, EPI_ISL_804989, EPI_ISL_804992, EPI_ISL_804993, EPI_ISL_804998, EPI_ISL_805008, EPI_ISL_805009, EPI_ISL_805010, EPI_ISL_805014, EPI_ISL_805015, EPI_ISL_805017, EPI_ISL_805018, EPI_ISL_805019, EPI_ISL_805021, EPI_ISL_805022, EPI_ISL_805024, EPI_ISL_805025, EPI_ISL_805026, EPI_ISL_805027, EPI_ISL_805028, EPI_ISL_805029, EPI_ISL_805030, EPI_ISL_805031, EPI_ISL_805032, EPI_ISL_805033, EPI_ISL_805034, EPI_ISL_805035, EPI_ISL_805036, EPI_ISL_805037, EPI_ISL_805038, EPI_ISL_805039, EPI_ISL_805040, EPI_ISL_805041, EPI_ISL_805042, EPI_ISL_805043, EPI_ISL_805044, EPI_ISL_805045, EPI_ISL_805046, EPI_ISL_805047, EPI_ISL_805048, EPI_ISL_805049, EPI_ISL_805050, EPI_ISL_805051, EPI_ISL_805052, EPI_ISL_805053, EPI_ISL_805054, EPI_ISL_805055, EPI_ISL_805056, EPI_ISL_805057, EPI_ISL_805058, EPI_ISL_805059, EPI_ISL_805060, EPI_ISL_805061, EPI_ISL_805062, EPI_ISL_805063, EPI_ISL_805064, EPI_ISL_805065, EPI_ISL_805066, EPI_ISL_805067, EPI_ISL_805068, EPI_ISL_805069, EPI_ISL_805070, EPI_ISL_805071, EPI_ISL_805072, EPI_ISL_805073, EPI_ISL_805074, EPI_ISL_805076, EPI_ISL_805077, EPI_ISL_805079, EPI_ISL_805080, EPI_ISL_805081, EPI_ISL_805082, EPI_ISL_805083, EPI_ISL_805084, EPI_ISL_805085, EPI_ISL_805086, EPI_ISL_805087, EPI_ISL_805088, EPI_ISL_805089, EPI_ISL_805090, EPI_ISL_805091, EPI_ISL_805092, EPI_ISL_805093, EPI_ISL_805094, EPI_ISL_805095, EPI_ISL_805096, EPI_ISL_805097, EPI_ISL_805098, EPI_ISL_805099, EPI_ISL_805100, EPI_ISL_805101, EPI_ISL_805102, EPI_ISL_805103, EPI_ISL_805104, EPI_ISL_805105, EPI_ISL_805106, EPI_ISL_805107, EPI_ISL_805108, EPI_ISL_805109, EPI_ISL_805110, EPI_ISL_805111, EPI_ISL_805112, EPI_ISL_805113, EPI_ISL_805114, EPI_ISL_805115, EPI_ISL_805116, EPI_ISL_805117, EPI_ISL_805118, EPI_ISL_805119, EPI_ISL_805120, EPI_ISL_805121, EPI_ISL_805123, EPI_ISL_805124, EPI_ISL_805125, EPI_ISL_805126, EPI_ISL_805128, EPI_ISL_805129, EPI_ISL_805130, EPI_ISL_805131, EPI_ISL_805132, EPI_ISL_805134, EPI_ISL_805135, EPI_ISL_805136, EPI_ISL_805137, EPI_ISL_805138, EPI_ISL_805139, EPI_ISL_805141, EPI_ISL_805142, EPI_ISL_805143, EPI_ISL_805144, EPI_ISL_805145, EPI_ISL_805146, EPI_ISL_805148, EPI_ISL_805150, EPI_ISL_805151, EPI_ISL_805152, EPI_ISL_805153, EPI_ISL_805154, EPI_ISL_805155, EPI_ISL_805156, EPI_ISL_805157, EPI_ISL_805158, EPI_ISL_805159, EPI_ISL_805160, EPI_ISL_805161, EPI_ISL_805162, EPI_ISL_805163, EPI_ISL_805164, EPI_ISL_805165, EPI_ISL_805167, EPI_ISL_805168, EPI_ISL_805169, EPI_ISL_805170, EPI_ISL_805171, EPI_ISL_805172, EPI_ISL_805173, EPI_ISL_805174, EPI_ISL_805175, EPI_ISL_805176, EPI_ISL_805177, EPI_ISL_805178, EPI_ISL_805179, EPI_ISL_805180, EPI_ISL_805182, EPI_ISL_805183, EPI_ISL_805184, EPI_ISL_805187, EPI_ISL_805189, EPI_ISL_805190, EPI_ISL_805192, EPI_ISL_805194, EPI_ISL_805195, EPI_ISL_805196, EPI_ISL_805197, EPI_ISL_805198, EPI_ISL_805199, EPI_ISL_805200, EPI_ISL_805201, EPI_ISL_805203, EPI_ISL_805205, EPI_ISL_805206, EPI_ISL_805207, EPI_ISL_805209, EPI_ISL_805210, EPI_ISL_805211, EPI_ISL_805212, EPI_ISL_805213, EPI_ISL_805216, EPI_ISL_805217, EPI_ISL_805218, EPI_ISL_805219, EPI_ISL_805221, EPI_ISL_805222, EPI_ISL_805223, EPI_ISL_805224, EPI_ISL_805227, EPI_ISL_805230, EPI_ISL_805232, EPI_ISL_805233, EPI_ISL_805236, EPI_ISL_805237, EPI_ISL_805239, EPI_ISL_805243, EPI_ISL_805245, EPI_ISL_805246, EPI_ISL_805248, EPI_ISL_805250, EPI_ISL_805251, EPI_ISL_805252, EPI_ISL_805254, EPI_ISL_805255, EPI_ISL_805256, EPI_ISL_805258, EPI_ISL_805266, EPI_ISL_805273, EPI_ISL_805281, EPI_ISL_805301, EPI_ISL_805303, EPI_ISL_805311, EPI_ISL_805315, EPI_ISL_805325, EPI_ISL_805331, EPI_ISL_805344, EPI_ISL_805359, EPI_ISL_805377, EPI_ISL_805379, EPI_ISL_805385, EPI_ISL_805386, EPI_ISL_805390, EPI_ISL_805394, EPI_ISL_805417, EPI_ISL_805420, EPI_ISL_805423, EPI_ISL_805430, EPI_ISL_805431, EPI_ISL_805438, EPI_ISL_805450, EPI_ISL_805452, EPI_ISL_805454, EPI_ISL_805456, EPI_ISL_805459, EPI_ISL_805464, EPI_ISL_805475, EPI_ISL_805477, EPI_ISL_805478, EPI_ISL_805479, EPI_ISL_805480, EPI_ISL_805489, EPI_ISL_805492, EPI_ISL_805497, EPI_ISL_805499, EPI_ISL_805500, EPI_ISL_805505, EPI_ISL_805506, EPI_ISL_805507, EPI_ISL_805510, EPI_ISL_805511, EPI_ISL_805524, EPI_ISL_805535, EPI_ISL_805538, EPI_ISL_805562, EPI_ISL_805568, EPI_ISL_805578, EPI_ISL_805580, EPI_ISL_805589, EPI_ISL_805591, EPI_ISL_805592, EPI_ISL_805593, EPI_ISL_805594, EPI_ISL_805597, EPI_ISL_805603, EPI_ISL_805609, EPI_ISL_805613, EPI_ISL_805614, EPI_ISL_805620, EPI_ISL_805622, EPI_ISL_805625, EPI_ISL_805629, EPI_ISL_805630, EPI_ISL_805631, EPI_ISL_805633, EPI_ISL_805634, EPI_ISL_805636, EPI_ISL_805638, EPI_ISL_805639, EPI_ISL_805641, EPI_ISL_805642, EPI_ISL_805643, EPI_ISL_805644, EPI_ISL_805647, EPI_ISL_805652, EPI_ISL_805655, EPI_ISL_805657, EPI_ISL_805660, EPI_ISL_805668, EPI_ISL_805672, EPI_ISL_805675, EPI_ISL_805676, EPI_ISL_805680, EPI_ISL_805686, EPI_ISL_805687, EPI_ISL_805688, EPI_ISL_805689, EPI_ISL_805694, EPI_ISL_805709, EPI_ISL_805718, EPI_ISL_805721, EPI_ISL_805722, EPI_ISL_805724, EPI_ISL_805729, EPI_ISL_805734, EPI_ISL_805737, EPI_ISL_805739, EPI_ISL_805742, EPI_ISL_805748, EPI_ISL_805757, EPI_ISL_805760, EPI_ISL_805762, EPI_ISL_805765, EPI_ISL_805769, EPI_ISL_805776, EPI_ISL_805778, EPI_ISL_805783, EPI_ISL_805790, EPI_ISL_805794, EPI_ISL_805799, EPI_ISL_805803, EPI_ISL_805808, EPI_ISL_805810, EPI_ISL_805812, EPI_ISL_805817, EPI_ISL_805818, EPI_ISL_805819, EPI_ISL_805828, EPI_ISL_805829, EPI_ISL_805832, EPI_ISL_805834, EPI_ISL_805839, EPI_ISL_805841, EPI_ISL_805851, EPI_ISL_805856, EPI_ISL_805858, EPI_ISL_805867, EPI_ISL_805876, EPI_ISL_805877, EPI_ISL_805878, EPI_ISL_805883, EPI_ISL_805890, EPI_ISL_805894, EPI_ISL_805896, EPI_ISL_805909, EPI_ISL_805910, EPI_ISL_805912, EPI_ISL_805915, EPI_ISL_805924, EPI_ISL_805925, EPI_ISL_805928, EPI_ISL_805933, EPI_ISL_805940, EPI_ISL_805945, EPI_ISL_805947, EPI_ISL_805961, EPI_ISL_805966, EPI_ISL_805969, EPI_ISL_805970, EPI_ISL_805972, EPI_ISL_805973, EPI_ISL_805982, EPI_ISL_805983, EPI_ISL_805984, EPI_ISL_805990, EPI_ISL_805993, EPI_ISL_805996, EPI_ISL_806001, EPI_ISL_806008, EPI_ISL_806016, EPI_ISL_806024, EPI_ISL_806038, EPI_ISL_806051, EPI_ISL_806052, EPI_ISL_806053, EPI_ISL_806060, EPI_ISL_806071, EPI_ISL_806075, EPI_ISL_806084, EPI_ISL_806085, EPI_ISL_806091, EPI_ISL_806097, EPI_ISL_806108, EPI_ISL_806109, EPI_ISL_806120, EPI_ISL_806128, EPI_ISL_806138, EPI_ISL_806170, EPI_ISL_806177, EPI_ISL_806185, EPI_ISL_806220, EPI_ISL_806242, EPI_ISL_806262, EPI_ISL_806315, EPI_ISL_806322, EPI_ISL_806336, EPI_ISL_806395, EPI_ISL_806415, EPI_ISL_806438, EPI_ISL_806443, EPI_ISL_806475, EPI_ISL_806479, EPI_ISL_806485, EPI_ISL_806487, EPI_ISL_806489, EPI_ISL_806490, EPI_ISL_806502, EPI_ISL_806503, EPI_ISL_806511, EPI_ISL_806512, EPI_ISL_806517, EPI_ISL_806518, EPI_ISL_806523, EPI_ISL_806525 | | | |
| see above | Alberta Precision Labs (APL) | Alberta Precision Labs (APL) | Gordon P, Lam LG, Pabbaraju K, Wong A, Ma R, Li V, Melin A, Tipples G, Berenger B, Zelyas N, Kellner J, Bernier F, Chui L, Croxen M |
| EPI_ISL_807150 | Makmal Kesihatan Awam Kebangsaan, Kementerian Kesihatan Malaysia. | Integrative Pharmacogenomics Institute (iPROMISE) | Mohd Zaki Salleh, Teh Lay Kek, Ariza Adnan, Sazzli Shahlan Kassim, Siti Hamimah Sheikh Abdul Kadir, Fadzilah Mohd Nor, Mohd Nur Fakhruzzaman Noorizhab, Siti Farah Alwani Mohd Nawi, Wang Seok Mui, Hani Mat Hussin, Nor Zahrin Hasran, Lim Wai Feng |
| EPI_ISL_807151, EPI_ISL_807153 | Makmal Kesihatan Awam Kebangsaan, Kementerian Kesihatan Malaysia | Integrative Pharmacogenomics Institute (iPROMISE) | Mohd Zaki Salleh, Teh Lay Kek, Ariza Adnan, Sazzli Shahlan Kassim, Siti Hamimah Sheikh Abdul Kadir, Fadzilah Mohd Nor, Mohd Nur Fakhruzzaman Noorizhab, Siti Farah Alwani Mohd Nawi, Wang Seok Mui, Hani Mat Hussin, Nor Zahrin Hasran, Lim Wai Feng |
| EPI_ISL_812248 | North Dakota Department of Health, Public Health Laboratory | North Dakota Department of Health, Public Health Laboratory | Lisa Wingerter |
| EPI_ISL_812326, EPI_ISL_812336 | United States Air Force School of Aerospace Medicine | United States Air Force School of Aerospace Medicine | Anthony Fries, Jennifer Meyer, Amanda Javorina, Sarah Purves, William Gruner, Clarise Starr, Elizabeth Macias |
| EPI_ISL_814070, EPI_ISL_814072, EPI_ISL_814077 | Ministry of Health Turkey | Ministry of Health Turkey | Fatma Bayrakdar, Yasemin Cogun, Süleyman Yalcin, Aye Baak Alta, Gülay Korukluolu |
| EPI_ISL_816775, EPI_ISL_816776, EPI_ISL_816777, EPI_ISL_816778, EPI_ISL_816779, EPI_ISL_816780, EPI_ISL_816781, EPI_ISL_816782, EPI_ISL_816783, EPI_ISL_816784, EPI_ISL_816785, EPI_ISL_816786, EPI_ISL_816787, EPI_ISL_816788, EPI_ISL_816790, EPI_ISL_816791, EPI_ISL_816792, EPI_ISL_816793, EPI_ISL_816794, EPI_ISL_816795 | | | |

| | | | |
|---|---|---|---|
| see above | Bioinformatics and Biostatistics Lab, Advanced Sequencing Facility | COVID-19 Genomics UK (COG-UK) Consortium | Aengus Stewart,Jerome Nicod,Chelsea Sawyer,Laura Cubitt,Harshil Patel,Margaret Crawford |
| EPI_ISL_819331, EPI_ISL_819332, EPI_ISL_819333, EPI_ISL_819334 | Hospital Universitari Vall d'Hebron - Vall d'Hebron Institut de Rercerca | Hospital Universitari Vall d'Hebron | Cristina Andrés, Maria Piñana, Josep F Abril, Damir Garcia-Cehic, Ariadna Rando, Juliana Esperalba, Maria Gema Codina, Carla Castillo, Maria Carmen Martin, Tomàs Pumarola, Josep Quer, Andrés Antón |
| EPI_ISL_825838, EPI_ISL_825858, EPI_ISL_825859, EPI_ISL_825860, EPI_ISL_825861, EPI_ISL_825862, EPI_ISL_825867, EPI_ISL_825875, EPI_ISL_825876, EPI_ISL_825877, EPI_ISL_825878, EPI_ISL_825880, EPI_ISL_825881, EPI_ISL_825882, EPI_ISL_825883, EPI_ISL_825884, EPI_ISL_825885, EPI_ISL_825886, EPI_ISL_825887, EPI_ISL_825888, EPI_ISL_825889, EPI_ISL_826263, EPI_ISL_826264, EPI_ISL_826265, EPI_ISL_826266, EPI_ISL_826267, EPI_ISL_826268 | | | |
| see above | Laboratoire de santé publique du Québec | Laboratoire de santé publique du Québec | Sandrine Moreira, Ioannis Ragoussis, Guillaume Bourque, Jesse Shapiro, Mark Lathrop and Michel Roger on behalf of the CoVSeQ research group (http://covseq.ca/researchgroup) |
| EPI_ISL_831065, EPI_ISL_831066, EPI_ISL_831067, EPI_ISL_831068, EPI_ISL_831069, EPI_ISL_831070, EPI_ISL_831071, EPI_ISL_831072, EPI_ISL_831073, EPI_ISL_831074, EPI_ISL_831075, EPI_ISL_831076, EPI_ISL_831077, EPI_ISL_831079, EPI_ISL_831080 | | | |
| see above | Hospital Universitario La Paz (Madrid) | SeqCOVID-SPAIN consortium/IBV(CSIC) | María Rodríguez-Tejedor, Elias Dahdouh, Fernando Lázaro-Perona, Jesús Mingorance and SeqCOVID-SPAIN consortium |
| EPI_ISL_833154 | Instituto Adolfo Lutz - Central | Instituto Adolfo Lutz, Interdiciplinary Procedures Center, Strategic Laboratory | Claudio Tavares Sacchi, Claudia Regina Gonçalves, Erica Valessa Ramos Gomes, Karoline Rodrigues Campos |
| EPI_ISL_833523, EPI_ISL_833525, EPI_ISL_833531, EPI_ISL_833535, EPI_ISL_833544, EPI_ISL_833545, EPI_ISL_833546, EPI_ISL_833552, EPI_ISL_833553, EPI_ISL_833554, EPI_ISL_833556, EPI_ISL_833558, EPI_ISL_833559, EPI_ISL_833560, EPI_ISL_833561, EPI_ISL_833562, EPI_ISL_833563 | | | |
| see above | Istituto Zooprofilattico Sperimentale del Mezzogiorno | TIGEM | Patrizia Annunziata, Andrea Ballabio, Valentina Bouche, Davide Cacchiarelli (CorrespAuthor), Pellegrino Cerino, Chiara Colantuono, Lucio Di Filippo, Antonio Grimaldi, Antonio Limone, Gabriella Loconte, Anna Manfredi, Francesco Panariello, Biancamaria Pierri, Marcello Salvi, Lucia Vassallo |
| EPI_ISL_833573 | Veterinary Specialized Instute "Nis" | Veterinary Specialized Institute "Sabac", Serbia | Vidanovic,D., Tesovic,B., Mrkovacki, S., Vujinovic,S., .,Knezevic,A., Jovanovic,T., Jankovic,M., Sekler,M., Banovic Djeri,B., Petrovic,T., Volkening,J., Afonso,C. |
| EPI_ISL_837573, EPI_ISL_837578 | Centro Nacional de Enfermedades Tropicales (CENETROP) | Laboratory of Respiratory Viruses and Measles, Oswaldo Cruz Institute, FIOCRUZ | Paola Resende, Roxana Loayza, Cinthia Avila, Luciana Appolinario, Fernando Motta, Anna Carolina Paixao, Ana Carolina Mendonca, Marilda Siqueira |
| EPI_ISL_837605, EPI_ISL_837668, EPI_ISL_837669, EPI_ISL_837670, EPI_ISL_837671, EPI_ISL_837672, EPI_ISL_837673, EPI_ISL_837674, EPI_ISL_837675, EPI_ISL_837676, EPI_ISL_837677, EPI_ISL_837678 | | | |
| see above | Instituto Nacional de Enfermedades Respiratorias (INER) | Instituto Nacional de Enfermedades Respiratorias (INER) | Celia Boukadida, Margarita Matías-Florentino, Alma Rincón-Rubio, Hector Esteban Paz-Juárez, Olivia Briceño, Edgar Sevilla-Reyes, Fidencio Mejía-Nepomuceno, Mario Mújica-Sánchez, Eduardo Becerril-Vargas, José Arturo Martínez-Orozco, Alejandra Hernández-Terán, Jorge Salas-Hernández, Santiago Ávila-Ríos, Joel Armando Vázquez-Pérez |
| EPI_ISL_842793, EPI_ISL_842794, EPI_ISL_842795, EPI_ISL_842809, EPI_ISL_842817, EPI_ISL_842824, EPI_ISL_842827, EPI_ISL_842833, EPI_ISL_842836, EPI_ISL_842837 | Barts Health NHS Trust | COVID-19 Genomics UK (COG-UK) Consortium | CUTINO-MOGUEL, Maria-Teresa; HARRINGTON, David; OWOYEMI, Dola; SHYLINI, Raghavendran; BROAD, Claire; KELE, Beatrix |
| EPI_ISL_848212, EPI_ISL_848226, EPI_ISL_848284, EPI_ISL_848285, EPI_ISL_848468 | Illinois Department of Public Health | Gagnon Lab, Southern Illinois University | Keith Gagnon |
| EPI_ISL_848587, EPI_ISL_848588, EPI_ISL_848589, EPI_ISL_848617, EPI_ISL_848618, EPI_ISL_848619, EPI_ISL_848620 | Evandro Chagas Institute | Evandro Chagas Institute | Santos, M.C.; Silva, A.M.; Junior, W.D.C.; Barbagelata, L.S.; Ferreira, J.A.; Sousa, E.M.A.; da Silva, P.S.; Pinheiro, K.C.; L.C.; Sousa Junior, E.C. |
| EPI_ISL_850183, EPI_ISL_850184 | Division of Emerging Infectious Diseases, Bureau of Infectious Diseases Diagnosis Control, Korea Disease Control and Prevention Agency | Division of Emerging Infectious Diseases, Bureau of Infectious Diseases Diagnosis Control, Korea Disease Control and Prevention Agency | Ae Kyung Park, Il-Hwan Kim, Heui Man Kim, Jeong-Min Kim, Namjoo Lee, Chaeyoung Lee, Sang Hee Woo, Eun-Jin Kim |
| EPI_ISL_854515, EPI_ISL_854516, EPI_ISL_854517, EPI_ISL_854518, EPI_ISL_854519, EPI_ISL_854520, EPI_ISL_854521, EPI_ISL_854522, EPI_ISL_854523, EPI_ISL_854524, EPI_ISL_854525, EPI_ISL_854526, EPI_ISL_854527, EPI_ISL_854528, EPI_ISL_854529, EPI_ISL_854530, EPI_ISL_854531, EPI_ISL_854532, EPI_ISL_854533, EPI_ISL_854534, EPI_ISL_854535, EPI_ISL_854536, EPI_ISL_854537, EPI_ISL_854538, EPI_ISL_854539, EPI_ISL_854540, EPI_ISL_854541, EPI_ISL_854542, EPI_ISL_854543, EPI_ISL_854544, EPI_ISL_854545, EPI_ISL_854546, EPI_ISL_854547, EPI_ISL_854548, EPI_ISL_854549, EPI_ISL_854550, EPI_ISL_854551, EPI_ISL_854552, EPI_ISL_854553, EPI_ISL_854554, EPI_ISL_854555 | | | |
| see above | Alberta Precision Labs (APL) | Alberta Precision Labs (APL) | Gordon P, Lam LG, Pabbaraju K, Wong A, Ma R, Li V, Melin A, Tipples G, Berenger B, Zelyas N, Kellner J, Bernier F, Chui L, Croxen M |
| EPI_ISL_855348 | Department of Epidemiology, Infectious Disease Control and Prevention, Hiroshima University, Japan | Department of Epidemiology, Infectious Disease Control and Prevention, Hiroshima University, Japan | Junko Tanaka, Kazuaki Takahashi, Masao Kuwabara, Eisaku Kishita, Shintaro Nagashima, Ko Ko |
| EPI_ISL_857037, EPI_ISL_857038 | The Ashley Laboratory, Stanford University | Chan-Zuckerberg Biohub | CZB Cliahub Consortium |
| EPI_ISL_861637 | Hospital Nipo Brasileiro | Instituto Adolfo Lutz, Interdiciplinary Procedures Center, Strategic Laboratory | Claudio Tavares Sacchi, Claudia Regina Gonçalves, Erica Valessa Ramos Gomes, Karoline Rodrigues Campos |
| EPI_ISL_861638 | Hospital Municipal Dr. Moyses Deutsch | Instituto Adolfo Lutz, Interdiciplinary Procedures Center, Strategic Laboratory | Claudio Tavares Sacchi, Claudia Regina Gonçalves, Erica Valessa Ramos Gomes, Karoline Rodrigues Campos |
| EPI_ISL_861639 | Hospital Sao Paulo de Ensino da Unifesp | Instituto Adolfo Lutz, Interdiciplinary Procedures Center, Strategic Laboratory | Claudio Tavares Sacchi, Claudia Regina Gonçalves, Erica Valessa Ramos Gomes, Karoline Rodrigues Campos |
| EPI_ISL_861640, EPI_ISL_861641 | Hospital Municipal Dr. Moyses Deutsch | Instituto Adolfo Lutz, Interdiciplinary Procedures Center, Strategic Laboratory | Claudio Tavares Sacchi, Claudia Regina Gonçalves, Erica Valessa Ramos Gomes, Karoline Rodrigues Campos |
| EPI_ISL_861642, EPI_ISL_861663 | Instituto Adolfo Lutz - Central | Instituto Adolfo Lutz, Interdiciplinary Procedures Center, Strategic Laboratory | Claudio Tavares Sacchi, Claudia Regina Gonçalves, Erica Valessa Ramos Gomes, Karoline Rodrigues Campos |
| EPI_ISL_861867 | LATE - Laboratório de Técnicas Especiais - Hospital Israelita Albert Einstein | LATE - Laboratório de Técnicas Especiais - Hospital Israelita Albert Einstein | Deyvid Amgarten, Fernanda de Mello Malta, Raquel Riyuzo, Ana Paula Moreira Salles, Pedro Henrique Sebe Rodrigues, João Renato Rebello Pinho |
| EPI_ISL_862863, EPI_ISL_862864, EPI_ISL_862865 | University Hospital Kyoto Prefectural University of Medicine | Fujita health University, School of Medicine, Department of Microbiology | Masahiro suzuki, Yohei Doi |
| EPI_ISL_862867 | Yokohama City University Hospital | Fujita health University, School of Medicine, Department of Microbiology | Masahiro suzuki, Yohei Doi |
| EPI_ISL_862871 | Nara Medical University Hospital | Fujita health University, School of Medicine, Department of Microbiology | Masahiro suzuki, Yohei Doi |
| EPI_ISL_862873 | Japanese Red Cross Narita Hospital | Fujita health University, School of Medicine, Department of Microbiology | Masahiro suzuki, Yohei Doi |
| EPI_ISL_862874 | Japanese Red Cross Nagoya Daiichi Hospital | Fujita health University, School of Medicine, Department of Microbiology | Masahiro suzuki, Yohei Doi |
| EPI_ISL_862877, EPI_ISL_862878, EPI_ISL_862880 | Nara Medical University Hospital | Fujita health University, School of Medicine, Department of Microbiology | Masahiro suzuki, Yohei Doi |
| EPI_ISL_869144, EPI_ISL_869145, EPI_ISL_869146, EPI_ISL_869147, | National Virus Reference Laboratory | Irish Coronavirus Sequencing Consortium - Teagasc Moorepark | Calum Walsh, Genuity Ireland |

| | | | |
|---|---|---|---|
| EPI_ISL_871815 | | | |
| EPI_ISL_872027 | Ospedale "Di Venere" | Beaconlab (Bioinformatics, Evolution and Comparative Genomics lab), Dept of Biosciences, University on Milan | Iacobellis M, d'Avenia M, Piluscio R, Parisi A, Chiara M, Manzari C, Pesole G |
| EPI_ISL_872097 | Instituto de Diagnostico y Referencia Epidemiologicos (INDRE) | Instituto de Diagnostico y Referencia Epidemiologicos (INDRE) | Abril Rodriguez-Maldonado, Gisela Barrera-Badillo ,Claudia Wong-Arambula , Natividad Cruz-Ortiz, Tatiana Nunez-Garcia, Dayanira Arellano-Suarez, Fabiola Garces-Ayala, Adnan-Araiza Rodríguez, Edgar Mendieta-Condado, Lucia Hernandez-Rivas, Irma Lopez-Martinez, Ernesto Ramirez-Gonzalez. |
| EPI_ISL_872098 | Instituto de Diagnostico y Referencia Epidemiologicos (INDRE) | Instituto de Diagnostico y Referencia Epidemiologicos (INDRE) | Claudia Wong-Arambula, Gisela Barrera-Badillo , Abril Rodriguez-Maldonado, Natividad Cruz-Ortiz, Tatiana Nunez-Garcia, Dayanira Arellano-Suarez, Fabiola Garces-Ayala, Adnan-Araiza Rodríguez, Edgar Mendieta-Condado, Lucia Hernandez-Rivas, Irma Lopez-Martinez, Ernesto Ramirez-Gonzalez. |
| EPI_ISL_876811, EPI_ISL_876812 | Istituto Zooprofilattico Sperimentale della Puglia e della Basilicata | Istituto Zooprofilattico Sperimentale della Puglia e della Basilicata | Parisi A., Bianco A., Capozzi L., Del Sambro L., Manzulli V, Rondinone V., Pace L., Cipolletta D., Galante D. |
| EPI_ISL_877032, EPI_ISL_877033 | 1.AO Universitaria 'S. Giovanni di Dio e Ruggi D'Aragona, Scuola Medica Salernitana' Hospital / 2.UOC di Virologia e Microbiologia, Università della Campania 'L. Vanvitelli' / 3.AO Universitaria 'Federico II' Napoli Hospital / 4.AORN 'San Giuseppe Moscati' Avellino Hospital / 5.AO 'San Pio - presidio G. Rummo' Benevento Hospital / 6.AO 'Sant'Anna e San Sebastiano' Caserta Hospital / 7.PO 'Maria Santissima Addolorata' Eboli Hospital / 8.Biogem Istituto di Ricerche Genetiche | 1. Genome Research Center for Health (CRGS) / 2. Laboratory of Molecular Medicine and Genomics(LMMGe) / 3. Center for Research in Pure and Applied Mathematics (CRMPA) | Giorgio Giurato (Corresponding Author), Francesca Rizzo (Corresponding Author), Alessandro Weisz (Corresponding Author), Gianluigi Franci, Giovanni Nassa, Pasquale Pagliano, Roberta Tarallo, Elena Alexandrova, Ylenia D'Agostino, Carlo Ferravante, Jessica Lamberti, Viola Melone, Domenico Memoli, Valeria Mirici Cappa, Domenico Palumbo, Giovanni Pecoraro, Assunta Sellitto, Oriana Strianese, Ilaria Terenzi, Giuseppe Fenza, Aniello Gentile, Antonello Saccomanno, Sonia Amabile, Teresa Rocco, Annamaria Salvati, Emilia Vaccaro, Massimiliano Galdiero, Michele Cennamo, Giuseppe Portella, Maria Grazia Foti, Mariarosaria Ingino, Maria Landi, Maurizio Fumi, Vincenzo Rocco, Rita Greco, Vittoria Letizia, Arnolfo Petruzziello, Maddalena Schioppa, Gregorio Goffredi, Francesca Marciano, Michele Caraglia, Alessia Cossu, Marianna Scrima |
| EPI_ISL_877600, EPI_ISL_877601, EPI_ISL_877602, EPI_ISL_877603, EPI_ISL_877604, EPI_ISL_877605, EPI_ISL_877762, EPI_ISL_877763 | Clinical Molecular Microbiology Laboratory, UNC Hospital | Dirk Dittmer | Razia Moorad , Justin T. Landis , Brent A. Eason, Melissa B. Miller, Linda Pluta, Dirk Dittmer, Angelica Juarez, Cecilia Thompson , Cameroon Grant, Evelyn Hoffman, Patricio Cano, Jason Wong, Carolina Caro-Vegas, Blossom Damania. |
| EPI_ISL_883171 | Hannover Medical School, Institute of Virology | Hannover Medical School, Institute of Virology | Lars Steinbrück, Jasper Götting |
| EPI_ISL_883191 | UK Tübingen, Medical Microbiology | Hannover Medical School, Institute of Virology | Lars Steinbrück, Jasper Götting |
| EPI_ISL_887111 | Institute of Medical Microbiology and Virology, University Hospital Carl Gustav Carus, TU Dresden | DRESDEN-concept Genome Center, CMCB, TU Dresden | Beil J., Brown T., Büttner L., Gscheidel N., Hochauf-Stange K., Klemroth S., Lindemann D., Mehnert G., Petzold A., Reinhardt S., Rost F., Sameith K., Winkler S. |
| EPI_ISL_887170, EPI_ISL_887174, EPI_ISL_887264, EPI_ISL_887292, EPI_ISL_887293, EPI_ISL_887298, EPI_ISL_887302, EPI_ISL_887320 | Protzer Lab | Protzer Lab, Gagneur Lab, Robert Koch Institut | Ulrike Protzer, Dieter Hoffmann, Eva Schulte, Andrea Theumer, Oliver Drechsel, Max von Kleist,Aleksandar Radonic,Stephan Fuchs, Alexander Karollus, Julien Gagneur |
| EPI_ISL_889394 | Olomouc University Hospital | Institute of Applied Biotechnologies a.s. | Petr Klempt, Ondej Brzo, Martin Kašný, Kateina Kvapilová, Marián Hajdúch, Petr Kvapil |
| EPI_ISL_891155 | DPH, Massachusetts State Public Health Lab | DPH, Massachusetts State Public Health Lab | Lang,A.S., Fink,T., Gallagher,G.R., Smole,S.C. |
| EPI_ISL_891266 | Institute of Biocides and Medical Ecology, Belgrade, Serbia | Virology department Institute of microbiology and immunology Faculty of Medicine University of Belgrade | Banko Ana, Miljanovic Danijela, Milicevic Ognjen, Loncar Ana, Abazovic Dzihan, Despot Dragana |
| EPI_ISL_895765, EPI_ISL_895769, EPI_ISL_895788, EPI_ISL_895816, EPI_ISL_895818, EPI_ISL_895819 | Molecular biology division, Institute of Clinical Biochemistry and Diagnostics, Charles University, Faculty of Medicine in Hradec Králové and University Hospital Hradec Králové | Molecular biology division, Institute of Clinical Biochemistry and Diagnostics, Charles University, Faculty of Medicine in Hradec Králové and University Hospital Hradec Králové | Helena Kovaíková, Petr Brož, Ivana Baranová, Kateina Hrochová, Tereza Baková, Jitka Novotná, Kateina Pehlíková, Vladimír Palika. Cooperation project with BioVendor-R&D and bioinformatics company BIOXSYS s r.o. |
| EPI_ISL_896125, EPI_ISL_900068, EPI_ISL_900084, EPI_ISL_900114, EPI_ISL_900140, EPI_ISL_900334, EPI_ISL_900344, EPI_ISL_900426, EPI_ISL_900455 | MEPHI, Aix Marseille University | MEPHI, Aix Marseille University | Anthony LEVASSEUR |
| EPI_ISL_903929 | VT Dept. of Health Laboratory | Genomics and Discovery, Respiratory Viruses Branch, Division of Viral Diseases, Centers for Disease Control and Prevention | Krista Queen, Yan Li, Ying Tao, Jing Zhang, Anna Uehara, Anna Montmayeur, Clinton R. Paden, Peter W. Cook, Rachel Marine, Mili Sheth, Jasmine Padilla, Sarah Nobles, Mark Burroughs, Lori Rowe, Haibin Wang, Ben L. Rambo-Martin, Dhwani Batra, Justin Lee, Suxiang Tong |
| EPI_ISL_906066 | Hospital Nipo Brasileiro | Instituto Adolfo Lutz, Interdiciplinary Procedures Center, Strategic Laboratory | Claudio Tavares Sacchi, Claudia Regina Gonçalves, Erica Valessa Ramos Gomes, Karoline Rodrigues Campos |
| EPI_ISL_913473 | Klinisk mikrobiologi | The Public Health Agency of Sweden | Anna-Malin Linde, Maria Lind Karlberg, Carlo Berg, Oskar Karlsson Lindsjo, Sofia Stamouli, Reza Advani, Mattias Haukland, Petra Holmstrom, Noura Walai, Petra Edquist, Mia Brytting, Anna Risberg, Karin Tegmark-Wisell |
| EPI_ISL_914047, EPI_ISL_914052, EPI_ISL_914053, EPI_ISL_914054, EPI_ISL_914055, EPI_ISL_914056, EPI_ISL_914057, EPI_ISL_914058, EPI_ISL_914059, EPI_ISL_914060, EPI_ISL_914061, EPI_ISL_914062, EPI_ISL_914063, EPI_ISL_914064, EPI_ISL_914065, EPI_ISL_914066 | | | |
| see above | TGen North | TGen North | "Jolene Bowers, Megan Folkerts, Chris French, Hayley Yaglom, Ashlyn Pfeiffer, Darrin Lemmer, Dave Engelthaler, The Arizona COVID Genomics Union (ACGU)" |
| EPI_ISL_914973, EPI_ISL_915080, EPI_ISL_915081, EPI_ISL_915082, EPI_ISL_915083, EPI_ISL_915084, EPI_ISL_915085, EPI_ISL_915086, EPI_ISL_915087, EPI_ISL_915088, EPI_ISL_915089, EPI_ISL_915090, EPI_ISL_915091, EPI_ISL_915092, EPI_ISL_915093, EPI_ISL_915094, EPI_ISL_915095, EPI_ISL_915096, EPI_ISL_915097, EPI_ISL_915098, EPI_ISL_915099, EPI_ISL_915100, EPI_ISL_915101, EPI_ISL_915102, EPI_ISL_915103, EPI_ISL_915104, EPI_ISL_915105, EPI_ISL_915106, EPI_ISL_915107, EPI_ISL_915108, EPI_ISL_915109, EPI_ISL_915110, EPI_ISL_915111, EPI_ISL_915112, EPI_ISL_915113, EPI_ISL_915114, EPI_ISL_915115, EPI_ISL_915116, EPI_ISL_915117, EPI_ISL_915118, EPI_ISL_915119, EPI_ISL_915120, EPI_ISL_915121, EPI_ISL_915122, EPI_ISL_915123, EPI_ISL_915124, EPI_ISL_915125, EPI_ISL_915126, EPI_ISL_915127, EPI_ISL_915128, EPI_ISL_915129, EPI_ISL_915130, EPI_ISL_915131, EPI_ISL_915132, EPI_ISL_915133, EPI_ISL_915134, EPI_ISL_915135, EPI_ISL_915136, EPI_ISL_915137, EPI_ISL_915138, EPI_ISL_915139, EPI_ISL_915140, EPI_ISL_915141, EPI_ISL_915142, EPI_ISL_915143, EPI_ISL_915144, EPI_ISL_915145, EPI_ISL_915146, EPI_ISL_915147, EPI_ISL_915148 | | | |
| see above | QEII Health Sciences Centre | National Microbiology Laboratory (NML) | Anna Majer, Shari Tyson, Grace Seo, Philip Mabon, Elsie Grudeski, Rhiannon Huzarewich, Russell Mandes, Anneliese Landgraff, Jennifer Tanner, Natalie Knox, Morag Graham, Gary Van Domselaar, Todd Hatchette, Jason LeBlanc, Janice Pettipas, Dan Gaston, Nathalie Bastien, Yan Li, Timothy Booth, Darian Hole, Madison Chapel, CanCOGeN's metadata curation team, Public Health Agency of Canada CanCOGeN team |
| EPI_ISL_918414 | 1.AO Universitaria 'S. Giovanni di Dio e Ruggi D'Aragona, Scuola Medica Salernitana' Hospital / 2.UOC di Virologia e Microbiologia, Università della Campania 'L. Vanvitelli' / 3.AO Universitaria 'Federico II' Napoli Hospital / 4.AORN 'San Giuseppe Moscati' Avellino Hospital / 5.AO 'San Pio - presidio G. Rummo' Benevento Hospital / 6.AO 'Sant'Anna e San Sebastiano' Caserta Hospital / 7.PO 'Maria Santissima Addolorata' Eboli Hospital / 8.Biogem Istituto di Ricerche Genetiche | 1. Genome Research Center for Health (CRGS) / 2. Laboratory of Molecular Medicine and Genomics(LMMGe) / 3. Center for Research in Pure and Applied Mathematics (CRMPA) | Giorgio Giurato (Corresponding Author), Francesca Rizzo (Corresponding Author), Alessandro Weisz (Corresponding Author), Gianluigi Franci, Giovanni Nassa, Pasquale Pagliano, Roberta Tarallo, Elena Alexandrova, Ylenia D'Agostino, Carlo Ferravante, Jessica Lamberti, Viola Melone, Domenico Memoli, Valeria Mirici Cappa, Domenico Palumbo, Giovanni Pecoraro, Assunta Sellitto, Oriana Strianese, Ilaria Terenzi, Giuseppe Fenza, Aniello Gentile, Antonello Saccomanno, Sonia Amabile, Teresa Rocco, Annamaria Salvati, Emilia Vaccaro, Massimiliano Galdiero, Michele Cennamo, Giuseppe Portella, Maria Grazia Foti, Mariarosaria Ingino, Maria Landi, Maurizio Fumi, Vincenzo Rocco, Rita Greco, Vittoria Letizia, Arnolfo Petruzziello, Maddalena Schioppa, Gregorio Goffredi, Francesca Marciano, Michele Caraglia, Alessia Cossu, Marianna Scrima |
| EPI_ISL_933631, EPI_ISL_933632 | Toronto Invasive Bacterial Diseases Network | McMaster University | Allison McGeer, Patryk Aftanas, Hooman Derakhshani, Angel Li, Kuganya Nirmalarajah, Emily Panousis, Ahmed Draia, Jalees Nasir, Michael Surette, Samira Mubareka, Andrew G. McArthur |
| EPI_ISL_934419, EPI_ISL_935103 | 1.AO Universitaria 'S. Giovanni di Dio e Ruggi D'Aragona, Scuola Medica Salernitana' Hospital / 2.UOC di Virologia e Microbiologia, Università della Campania 'L. Vanvitelli' / 3.AO Universitaria 'Federico II' Napoli Hospital / 4.AORN 'San Giuseppe Moscati' Avellino Hospital / 5.AO 'San Pio - presidio | 1. Genome Research Center for Health (CRGS) / 2. Laboratory of Molecular Medicine and Genomics(LMMGe) / 3. Center for Research in Pure and Applied Mathematics (CRMPA) | Giorgio Giurato (Corresponding Author), Francesca Rizzo (Corresponding Author), Alessandro Weisz (Corresponding Author), Gianluigi Franci, Giovanni Nassa, Pasquale Pagliano, Roberta Tarallo, Elena Alexandrova, Ylenia D'Agostino, Carlo Ferravante, Jessica Lamberti, Viola Melone, Domenico Memoli, Valeria Mirici Cappa, Domenico Palumbo, Giovanni Pecoraro, Assunta Sellitto, Oriana Strianese, Ilaria Terenzi, Giuseppe Fenza, Aniello Gentile, Antonello Saccomanno, Sonia Amabile, Teresa Rocco, Annamaria Salvati, Emilia Vaccaro, Massimiliano Galdiero, Michele Cennamo, Giuseppe Portella, Maria Grazia Foti, Mariarosaria Ingino, Maria Landi, Maurizio Fumi, Vincenzo Rocco, Rita Greco, Vittoria Letizia, Arnolfo Petruzziello, Maddalena Schioppa, |

| | | | |
|---|---|---|---|
| | G. Rummo' Benevento Hospital / 6.AO 'Sant'Anna e San Sebastiano' Caserta Hospital / 7.PO 'Maria Santissima Addolorata' Eboli Hospital / 8.Biogem Istituto di Ricerche Genetiche | | Gregorio Goffredi, Francesca Marciano, Michele Caraglia, Alessia Cossu, Marianna Scrima, Edmondo Adorisio, Morena D'Avenia, Michela Iacobellis, Rosanna Piluscio, Giorgio Dirani, Vittorio Sambri, Simona Semprini, Silvia Zanoli, Francesco Curcio, Stefania Marzinotto, Andreina Baj, Fausto Sessa. |
| EPI_ISL_940452, EPI_ISL_940453, EPI_ISL_940460, EPI_ISL_940471, EPI_ISL_940498, EPI_ISL_940502 | Hôpital Bichat Claude Bernard, Laboratoire de Virologie | IAME UMR1137 Inserm, Université de Paris, Hôpital Bichat | Antoine Bridier-Nahmias, Amélie Recoing, Quentin Le Hingrat, Lena Daniel, Siham Hamri, Gilles Collin, Alexandre Storto, Mélanie Bertine, Charlotte Charpentier, Nadhira Houhou-Fidouh, Diane Descamps, Benoit Visseaux |
| EPI_ISL_941141 | Centro de Investigaciones en Microbiología y Biotecnología-UR (CIMBIUR), Facultad de Ciencias Naturales, Universidad del Rosario, Bogotá, Colombia | Centro de Investigaciones en Microbiología y Biotecnología-UR (CIMBIUR), Facultad de Ciencias Naturales, Universidad del Rosario, Bogotá, Colombia Icahn School of Medicine at Mount Sinai, New York, USA | Nathalia Ballesteros, Marina Muñoz, Luz Helena Patiño, Carolina Hernández, Felipe González-Casabianca, Iván Carroll, Mauricio Santos-Vega, Jaime Cascante, Andrés Angel, Alejandro Feged-Rivadeneira, Mónica Palma-Cuero, Carolina Flórez, Sergio Gomez, Adriana van de Guchte, Zenab Khan, Jayeeta Dutta, Hala Alejel Alshammary, Ana S. Gonzalez-Reiche, Matthew M. Hernandez, Emilia Mia Sordillo, Viviana Simon, Harm van Bakel, Alberto Paniz-Mondolfi, Juan David Ramírez |
| EPI_ISL_941948, EPI_ISL_941950, EPI_ISL_941961, EPI_ISL_941968, EPI_ISL_941969, EPI_ISL_941970, EPI_ISL_941971, EPI_ISL_941972, EPI_ISL_941973, EPI_ISL_941974, EPI_ISL_941975, EPI_ISL_941976, EPI_ISL_941983 | | | |
| see above | Centro de Investigaciones en Microbiología y Biotecnología-UR (CIMBIUR), Facultad de Ciencias Naturales, Universidad del Rosario, Bogotá, Colombia | Centro de Investigaciones en Microbiología y Biotecnología-UR (CIMBIUR), Facultad de Ciencias Naturales, Universidad del Rosario, Bogotá, Colombia Instituto Nacional de Salud, Bogotá, Colombia Icahn School of Medicine at Mount Sinai, New York, USA | Luz Helena Patiño, Marina Muñoz, Nathalia Ballesteros, Carolina Hernández, Carolina Flórez, Sergio Gomez, Adriana van de Guchte, Zenab Khan, Jayeeta Dutta, Hala Alejel Alshammary, Ana S. Gonzalez-Reiche, Matthew M. Hernandez, Emilia Mia Sordillo, Viviana Simon, Harm van Bakel, Alberto Paniz-Mondolfi, Juan David Ramírez |
| EPI_ISL_943624, EPI_ISL_943626, EPI_ISL_943627, EPI_ISL_943628, EPI_ISL_943631, EPI_ISL_943632, EPI_ISL_943634 | Utah Public Health Laboratory | Utah Public Health Laboratory | Erin L. Young, Kelly F. Oakeson, Tara Gallagher |
| EPI_ISL_944636 | Centro de Investigaciones en Microbiología y Biotecnología-UR (CIMBIUR), Facultad de Ciencias Naturales, Universidad del Rosario, Bogotá, Colombia | Centro de Investigaciones en Microbiología y Biotecnología-UR (CIMBIUR), Facultad de Ciencias Naturales, Universidad del Rosario, Bogotá, Colombia Instituto Nacional de Salud, Bogotá, Colombia Icahn School of Medicine at Mount Sinai, New York, USA | Luz Helena Patiño, Marina Muñoz, Nathalia Ballesteros, Carolina Hernández, Carolina Flórez, Sergio Gomez, Adriana van de Guchte, Zenab Khan, Jayeeta Dutta, Hala Alejel Alshammary, Ana S. Gonzalez-Reiche, Matthew M. Hernandez, Emilia Mia Sordillo, Viviana Simon, Harm van Bakel, Alberto Paniz-Mondolfi, Juan David Ramírez |
| EPI_ISL_953295, EPI_ISL_953307, EPI_ISL_953308, EPI_ISL_953317, EPI_ISL_953318, EPI_ISL_953323, EPI_ISL_953334, EPI_ISL_953347, EPI_ISL_953348, EPI_ISL_953349, EPI_ISL_953367 | | | |
| see above | Bioinformatics and Biostatistics Lab, Advanced Sequencing Facility | COVID-19 Genomics UK (COG-UK) Consortium | Aengus Stewart,Jerome Nicod,Chelsea Sawyer,Laura Cubitt,Harshil Patel,Margaret Crawford |
| EPI_ISL_953407, EPI_ISL_953419 | Laboratorio de Investigaciones de Baney | "Swiss Tropical and Public Health Institute" | "Carlos Cortes, Claudia Daubenberger, Guillermo Garcia, Salome Hosch, Bonifacio Manguire Nlavo, Maximilian Mpina, Elizabeth Nyakarungu, Diosdado Odjama Nseng Ada, Mitoha Ondo O Ayekaba, Tobias Schindler, Philip Wonder Phiri" |
| EPI_ISL_961013 | Viral Respiratory Lab, National Institute for Biomedical Research (INRB) | Pathogen Sequencing Lab, National Institute for Biomedical Research (INRB) | Placide Mbala-Kingebeni, Edith Nkwembe, Eddy Kinganda-Lusamaki, Amuri Aziza, Francisca Muyembe Mawete, Emmanuel Lokilo Lofiko, Jean-Claude Makangara Cigolo, Catherine Pratt, Matthias Pauthner, Josh Quick, Allison Black, James Hadfield, Trevor Bedford, Ian Goodfellow, Andrew Rambaut, Nick Loman, Kristian Andersen, Michael Wiley, Steve Ahuka-Mundeke, Jean-Jacques Muyembe Tamfum |
| EPI_ISL_961797 | Laboratorio de Infectología, Servicio de Infectología, Hospital Universitario Dr. José Eleuterio González - Universidad Autónoma de Nuevo León | Laboratorio de Infectología Molecular, Departamento de Bioquímica y Medicina Molecular,Facultad de Medicina - Universidad Autónoma de Nuevo León | Kame A. Galán-Huerta, María F. Herrera-Saldivar, Natalia Martínez-Acuña, Sonia A. Lozano-Sepúlveda, Daniel Arellanos-Soto, Ana M. Rivas-Estilla, Paola Bocanegra-Ibarias, Samantha M. Flores-Treviño, Elvira Garza-González, Eduardo Perez-Alba, Laura Nuzzolo-Shihadeh, Adrian Camacho-Ortiz |
| EPI_ISL_961904, EPI_ISL_961952, EPI_ISL_961953, EPI_ISL_961954, EPI_ISL_962045, EPI_ISL_962046, EPI_ISL_962047, EPI_ISL_962048, EPI_ISL_962049, EPI_ISL_962050, EPI_ISL_962051, EPI_ISL_962052, EPI_ISL_962054 | | | |
| see above | Illinois Department of Public Health | Gagnon Lab, Southern Illinois University | Keith Gagnon |
| EPI_ISL_968151 | Clinical Molecular Microbiology Laboratory, UNC Hospital | Dirk Dittmer | Justin T. Landis , Razia Moorad , Brent A. Eason, Melissa B. Miller, Linda Pluta, Dirk Dittmer, Angelica Juarez, Cecilia Thompson, Shawn Hawken, Cameroon Grant, Evelyn Hoffman, Patricio Cano, Jason Wong, Carolina Caro-Vegas, Ryan McNamara, Blossom Damania. |
| EPI_ISL_977093, EPI_ISL_977096, EPI_ISL_977097, EPI_ISL_977098 | Massachusetts General Hospital | Infectious Disease Program, Broad Institute of Harvard and MIT | Lemieux,J.E., Siddle,K.J., Shaw,B., Adams,G., Pierce,V., Turbett,S., Anahtar,M., Branda,J., Slater,D., Harris,J., Lin,A.E., Gladden-Young,A., Lagerborg,K., Rudy,M., DeRuff,K., Carter,A., Normandin,E., Bauer,M., Reilly,S., Tomkins-Tinch,C., Loreth,C., Chaluvadi,S., Neumann,A., Cusick,C., Chapman,S.B., Gnirke,A., Flowers,K., Cerrato,F., Birren,B.W., Gallagher,G., Smole,S., Park,D.J., MacInnis,B.L., Ryan,E., LaRocque,R., Rosenberg,E. and Sabeti,P.C. |
| EPI_ISL_978492, EPI_ISL_978514, EPI_ISL_978521, EPI_ISL_978528 | Central Public Health Laboratory - LACEN -Bahia, Salvador, Brazil | Central Public Health Laboratory - LACEN -Bahia, Salvador, Brazil | Stephane Tosta, Luciana Oliveira, Vanessa Nardy,Patrícia Cajado,Marcela Gómez, Breno Dominguez, Jaqueline Gomes, Vagner Fonseca,Marta Giovanetti,Luiz Alcantara, Felicidade Pereira, Arabela Leal |
| EPI_ISL_979291 | Cadham Provincial laboratory | National Microbiology Laboratory (NML) | Anna Majer, Shari Tyson, Grace Seo, Philip Mabon, Elsie Grudeski, Rhiannon Huzarewich, Russell Mandes, Anneliese Landgraff, Jennifer Tanner, Natalie Knox, Morag Graham, Gary Van Domselaar, Paul Van Caeseele, Jared Bullard, David Alexander, Kerry Dust, Nathalie Bastien, Yan Li, Timothy Booth, Darian Hole, Madison Chapel, Kirsten Biggar, CanCOGeN's metadata curation team, Public Health Agency of Canada CanCOGeN team |
| EPI_ISL_981216, EPI_ISL_981217, EPI_ISL_981218, EPI_ISL_981219, EPI_ISL_981220, EPI_ISL_981221, EPI_ISL_981222, EPI_ISL_981223, EPI_ISL_981224 | National Virus Reference Laboratory | Irish Coronavirus Sequencing Consortium - Helixworks | Sachin Chalapati, Conor Crosbie, Nimesh Pinnamaneni |
