## Supplementary material for "Detection and characterization of the SARS-CoV-2 lineage B.1.526 in New York": Supp. Table 3 GISAID Acknowledgment part 2: gisaid_hcov-19_acknowledgement_table_2021_02_12_16-12.pdf

We gratefully acknowledge the following Authors from the Originating laboratories responsible for obtaining the specimens, as well as the Submitting laboratories where the genome data were generated and shared via GISAID, on which this research is based.

All Submitters of data may be contacted directly via www.gisaid.org

Authors are sorted alphabetically.

| Accession ID | Originating Laboratory | Submitting Laboratory | Authors |
|---|---|---|---|
| EPI_ISL_435473, EPI_ISL_435474 | Rady's Childrens Hospital | Andersen lab at Scripps Research | SEARCH Alliance San Diego |
| EPI_ISL_436606 | University of Wisconsin-Madison AIDS Vaccine Research Laboratories | University of Wisconsin-Madison AIDS Vaccine Research Laboratories | Gage Moreno, Katarina Braun, et al. AIDS Vaccine Research Laboratories |
| EPI_ISL_436725 | RSA/RP Villa San Giovanni - Gruppo Edos | Istituto Zooprofilattico Sperimentale dell'Abruzzo e Molise "G.Caporale" | Lorusso A, Marcacci M, Di Domenico M, Ancora M, Curini V, Mangone I, Rinaldi A, Di Pasquale A, Cammà C, Puglia I, Savini G |
| EPI_ISL_436726, EPI_ISL_436727, EPI_ISL_436728, EPI_ISL_436729 | SERVIZIO DI IGIENE E SANITÀ PUBBLICA ASL Teramo | Istituto Zooprofilattico Sperimentale dell'Abruzzo e Molise "G.Caporale" | Lorusso A, Marcacci M, Di Domenico M, Ancora M, Curini V, Mangone I, Rinaldi A, Di Pasquale A, Cammà C, Puglia I, Savini G |
| EPI_ISL_436730 | Servizio di igiene epidemiologia e sanità pubblica (Siesp) Chieti | Istituto Zooprofilattico Sperimentale dell'Abruzzo e Molise "G.Caporale" | Lorusso A, Marcacci M, Di Domenico M, Ancora M, Curini V, Mangone I, Rinaldi A, Di Pasquale A, Cammà C, Puglia I, Savini G |
| EPI_ISL_436732 | Ospedale Civile S. Liberatore di Atri | Istituto Zooprofilattico Sperimentale dell'Abruzzo e Molise "G.Caporale" | Lorusso A, Marcacci M, Di Domenico M, Ancora M, Curini V, Mangone I, Rinaldi A, Di Pasquale A, Cammà C, Puglia I, Savini G |
| EPI_ISL_436891, EPI_ISL_436892, EPI_ISL_436893, EPI_ISL_436894, EPI_ISL_436895, EPI_ISL_436896, EPI_ISL_436897, EPI_ISL_436898, EPI_ISL_436899, EPI_ISL_436900 | Gundersen Molecular Diagnostics Laboratory | Kabara Cancer Research Institute | Craig S. Richmond, Paraic A. Kenny |
| EPI_ISL_437442 | Department of MicroBiology, Government Medical College, Surat | Gujarat Biotechnology Research Centre | Dinesh Kumar, Zuber Saiyed, Dipa Kinariwala, Disha Patel, Binita Aring, Neeta Khandelwal, Geeta Vaghela, Sonia Barve, Bhavesh Modi, Kairavi Joshi, Gaurishankar Shrimali, Nidhi Sood, Pranay Shah, R D Dixit, Snehal Bagatharia, Kamlesh J Upadhyay, Ramesh Pandit, Tejas Shah, Ankit Hinsu, Pritesh Sabara, Apurvasinh Puvar, Janvi Raval, Monika Gandhi, Pinal Trivedi, Maharshi Pandya, Amit Kanani, Akanksha Verma, Nitin Savaliya, Raghawendra Kumar, Dipeshwari Shewale, Chaitanya Joshi, Madhvi Joshi |
| EPI_ISL_437443 | Department of MicroBiology, Government Medical College, Surat | Gujarat Biotechnology Research Centre | Zuber Saiyed, Dipa Kinariwala, Disha Patel, Binita Aring, Neeta Khandelwal, Geeta Vaghela, Sonia Barve, Bhavesh Modi, Kairavi Joshi, Gaurishankar Shrimali, Nidhi Sood, Pranay Shah, R D Dixit, Snehal Bagatharia, Kamlesh J Upadhyay, Ramesh Pandit, Tejas Shah, Ankit Hinsu, Pritesh Sabara, Apurvasinh Puvar, Janvi Raval, Monika Gandhi, Pinal Trivedi, Maharshi Pandya, Amit Kanani, Akanksha Verma, Nitin Savaliya, Raghawendra Kumar, Dinesh Kumar, Sharmistha Majumdar, Chaitanya Joshi, Madhvi Joshi |
| EPI_ISL_437550, EPI_ISL_437551 | Scripps Medical Laboratory | Andersen lab at Scripps Research | SEARCH Alliance San Diego with Michael Quigley, Ellen Stefanski, Ian Mchardy |
| EPI_ISL_438174, EPI_ISL_438175 | Seattle Flu Study | Seattle Flu Study | Chu et al |
| EPI_ISL_438699, EPI_ISL_438701, EPI_ISL_438711, EPI_ISL_438721, EPI_ISL_438723, EPI_ISL_438725, EPI_ISL_438739, EPI_ISL_438743, EPI_ISL_438744, EPI_ISL_438746, EPI_ISL_438747, EPI_ISL_438748 | | | |
| see above | Department of Pathology, University of Cambridge | COVID-19 Genomics UK (COG-UK) Consortium | Luke W Meredith, M. Est©e Trk , Myra Hosmillo, William L. Hamilton, Martin D. Curran, Theresa Feltwell, Grant Hall, Anna Yakovleva, Fahad A Khokhar, Charlotte J. Houldcroft, Laura G Caller, Aminu S. Jahun, Sarah L. Caddy, Ian Goodfellow |
| EPI_ISL_439038, EPI_ISL_439040, EPI_ISL_439041 | West of Scotland Specialist Virology Centre, NHSGGC / MRC-University of Glasgow Centre for Virus Research | COVID-19 Genomics UK (COG-UK) Consortium | Ana da Silva Filipe, Natasha Johnson, Kathy Smollett, Daniel Mair, Stephen Carmichael, Lily Tong, Jenna Nichols, Elihu Aranday-Cortes, Kirstyn Brunker, Yasmin Parr, Kyriaki Nomikou; Sarah McDonald, Marc Niebel, Patawee Asamaphan; Richard Orton, Joseph Hughes, Sreenu Vattipally, David L Robertson; Alasdair MacLean, Rory Gunson; Kathy Li, Natasha Jesudason, Rajiv Shah, James Shepherd, Antonia Ho, Emma Thomson |
| EPI_ISL_439250, EPI_ISL_439288, EPI_ISL_439289, EPI_ISL_439290, EPI_ISL_439291, EPI_ISL_439292, EPI_ISL_439293, EPI_ISL_439294, EPI_ISL_439295, EPI_ISL_439296, EPI_ISL_439297, EPI_ISL_439298, EPI_ISL_439299, EPI_ISL_439300, EPI_ISL_439301, EPI_ISL_439302, EPI_ISL_439303, EPI_ISL_439304, EPI_ISL_439305, EPI_ISL_439306, EPI_ISL_439307, EPI_ISL_439308, EPI_ISL_439309, EPI_ISL_439310, EPI_ISL_439311, EPI_ISL_439312, EPI_ISL_439313, EPI_ISL_439314, EPI_ISL_439315, EPI_ISL_439316, EPI_ISL_439317, EPI_ISL_439318, EPI_ISL_439319, EPI_ISL_439320, EPI_ISL_439321, EPI_ISL_439322, EPI_ISL_439323, EPI_ISL_439324, EPI_ISL_439325, EPI_ISL_439326, EPI_ISL_439327, EPI_ISL_439328, EPI_ISL_439329, EPI_ISL_439330, EPI_ISL_439331, EPI_ISL_439332, EPI_ISL_439333, EPI_ISL_439334, EPI_ISL_439335, EPI_ISL_439336, EPI_ISL_439337, EPI_ISL_439338, EPI_ISL_439339, EPI_ISL_439340, EPI_ISL_439341, EPI_ISL_439342, EPI_ISL_439343, EPI_ISL_439344, EPI_ISL_439345, EPI_ISL_439346, EPI_ISL_439347, EPI_ISL_439348, EPI_ISL_439349, EPI_ISL_439350, EPI_ISL_439351, EPI_ISL_439352, EPI_ISL_439353, EPI_ISL_439354, EPI_ISL_439355, EPI_ISL_439356, EPI_ISL_439357, EPI_ISL_439358, EPI_ISL_439359, EPI_ISL_439360, EPI_ISL_439361, EPI_ISL_439362, EPI_ISL_439363, EPI_ISL_439364, EPI_ISL_439365, EPI_ISL_439366, EPI_ISL_439367, EPI_ISL_439665, EPI_ISL_439666, EPI_ISL_439667, EPI_ISL_439668, EPI_ISL_439669, EPI_ISL_439670 | | | |
| see above | Virology Department, Royal Infirmary of Edinburgh, NHS Lothian / School of Biological Sciences, University of Edinburgh / Institute of Genetics and Molecular Medicine, University of Edinburgh | COVID-19 Genomics UK (COG-UK) Consortium | McHugh M, Dewar R, Rooke S, Gallagher M, Balcaza C, O'ÄôToole Å, Scher E, Hill V, McCrone JT, Colquhoun R, Yu X, Jackson B, Rambaut A, Williams TC, Templeton K |
| EPI_ISL_440945, EPI_ISL_440950 | Liverpool Clinical Laboratories | COVID-19 Genomics UK (COG-UK) Consortium | Sam Haldenby, Anita Lucaci, Steve Paterson, Julian Hiscox, Alistair Darby, M Almsaud, A Alrezaihi, Muhannad Alruwaili, Stuart D Armstrong, Jones Benjamin , Eleanor G Bentley, Anu Chawla, Jordan J Clark, Angela Cowell, Richard Eccles, Isabel Garca-Dorival, Matthew Gemmell, Alessandro Gerada, PKF Gilmore, Richard Gregory, Ximeng Han, Catherine Hartley, Margaret Hughes, Miren Iturriza-Gomara, James Johnson, L Luu, Jenifer Manson , Charlotte Nelson, Elaine O'ÄôToole, Cassie Olateju, Rebekah Penrice-Randal¬†, Lucille Rainbow, N.P Randle, Trevor Ian Robinson, Parul Sharma, Ghada T Shawli, James P Stewart , Neil Swainston, Ecaterina Vamos, Joanne Watts, Mark Whitehead |
| EPI_ISL_441536, EPI_ISL_441537, EPI_ISL_441538, EPI_ISL_441539, EPI_ISL_441863, EPI_ISL_441864, EPI_ISL_441865, EPI_ISL_441866, EPI_ISL_441867, EPI_ISL_441868, EPI_ISL_441869, EPI_ISL_441870, EPI_ISL_441871, EPI_ISL_441872, EPI_ISL_441873, EPI_ISL_441874, EPI_ISL_441875, EPI_ISL_441876, EPI_ISL_441877, EPI_ISL_441878, EPI_ISL_441879, EPI_ISL_441880, EPI_ISL_441881, EPI_ISL_441882, EPI_ISL_441883, EPI_ISL_441884, EPI_ISL_441885, EPI_ISL_441886, EPI_ISL_441887, EPI_ISL_441888, EPI_ISL_441889, EPI_ISL_441890, EPI_ISL_441891, EPI_ISL_441892, EPI_ISL_441893, EPI_ISL_441894, EPI_ISL_441895, EPI_ISL_441896, EPI_ISL_441897 | | | |
| see above | Queens Medical Centre, Clinical Microbiology Department / DeepSeq Nottingham | COVID-19 Genomics UK (COG-UK) Consortium | Gemma Clark, Wendy Smith, Manjinder Khakh, Hannah Howson-Wells, Jonathan Ball, Patrick McClure, Joseph Chappell, Theocharis Tsoleridis, Nadine Holmes, Matthew Carlisle, Christopher Moore, Fei Sang, Johnny Debebe, Victoria Wright, Matthew Loose |
| EPI_ISL_444023, EPI_ISL_444024, EPI_ISL_444025, EPI_ISL_444026 | County of Santa Clara Public Health | Chan-Zuckerberg Biohub | CZB Cliahub Consortium |
| EPI_ISL_444051, EPI_ISL_444052, EPI_ISL_444053, EPI_ISL_444054, EPI_ISL_444057, EPI_ISL_444058, EPI_ISL_444059, EPI_ISL_444060, EPI_ISL_444061, EPI_ISL_444062, EPI_ISL_444063, EPI_ISL_444064, EPI_ISL_444065, EPI_ISL_444066, EPI_ISL_444067, EPI_ISL_444068, EPI_ISL_444069, EPI_ISL_444070, EPI_ISL_444071, EPI_ISL_444072, EPI_ISL_444073, EPI_ISL_444074, EPI_ISL_444075, EPI_ISL_444076, EPI_ISL_444077, EPI_ISL_444078 | | | |
| see above | UCSF Clinical Microbiology Laboratory | Chan-Zuckerberg Biohub | CZB Cliahub Consortium |
| EPI_ISL_444313, EPI_ISL_444314, EPI_ISL_444315, EPI_ISL_444316, EPI_ISL_444317, EPI_ISL_444323, EPI_ISL_444324, EPI_ISL_444325, EPI_ISL_444326, EPI_ISL_444327, EPI_ISL_444328, EPI_ISL_444329, EPI_ISL_444330, EPI_ISL_444331, EPI_ISL_444337, EPI_ISL_444338, EPI_ISL_444339, EPI_ISL_444340, EPI_ISL_444341, EPI_ISL_444342, EPI_ISL_444344, EPI_ISL_444346, EPI_ISL_444347, EPI_ISL_444350, EPI_ISL_444351, EPI_ISL_444353, EPI_ISL_444354, EPI_ISL_444356, EPI_ISL_444357, EPI_ISL_444358, EPI_ISL_444359, EPI_ISL_444360, EPI_ISL_444361, EPI_ISL_444363, EPI_ISL_444364, EPI_ISL_444365, EPI_ISL_444366, EPI_ISL_444367, EPI_ISL_444368, EPI_ISL_444369, EPI_ISL_444370, EPI_ISL_444371, EPI_ISL_444372, EPI_ISL_444373, EPI_ISL_444374, EPI_ISL_444376, EPI_ISL_444377, EPI_ISL_444378, EPI_ISL_444379, EPI_ISL_444380, EPI_ISL_444381, EPI_ISL_444382, EPI_ISL_444384, EPI_ISL_444385, EPI_ISL_444386, EPI_ISL_444387, EPI_ISL_444388, EPI_ISL_444389, EPI_ISL_444390, EPI_ISL_444391 | | | |
| see above | Department of Pathology, University of Cambridge | COVID-19 Genomics UK (COG-UK) Consortium | Luke W Meredith, M. Estée Török , Myra Hosmillo, William L. Hamilton, Martin D. Curran, Theresa Feltwell, Grant Hall, Anna Yakovleva, Fahad A Khokhar, Charlotte J. Houldcroft, Laura G Caller, Aminu S. Jahun, Sarah L. Caddy, Ian Goodfellow |
| EPI_ISL_444459 | B.J. Medical College and Civil hospital | Gujarat Biotechnology Research Centre | Ramesh Pandit, Tejas Shah, Ankit Hinsu, Pritesh Sabara, Apurvasinh Puvar, Janvi Raval, Monika Gandhi, Pinal Trivedi, Maharshi Pandya, Amit Kanani, |

| | | | |
|---|---|---|---|
| | | | Akanksha Verma, Nitin Savaliya, Raghawendra Kumar, Dinesh Kumar, Zuber Saiyed, Dipa Kinariwala, Disha Patel, Binita Aring, Neeta Khandelwal, Geeta Vaghela, Sonia Barve, Bhavesh Modi, Kairavi Joshi, Gaurishankar Shrimali, Nidhi Sood, Pranay Shah, R D Dixit, Snehal Bagatharia, Kamlesh J Upadhyay, Neha Rajpara, Chaitanya Joshi, Madhvi Joshi |
| EPI_ISL_444460 | B.J. Medical College and Civil hospital | Gujarat Biotechnology Research Centre | Tejas Shah, Ankit Hinsu, Pritesh Sabara, Apurvasinh Puvar, Janvi Raval, Monika Gandhi, Pinal Trivedi, Maharshi Pandya, Amit Kanani, Akanksha Verma, Nitin Savaliya, Raghawendra Kumar, Dinesh Kumar, Zuber Saiyed, Dipa Kinariwala, Disha Patel, Binita Aring, Neeta Khandelwal, Geeta Vaghela, Sonia Barve, Bhavesh Modi, Kairavi Joshi, Gaurishankar Shrimali, Nidhi Sood, Pranay Shah, R D Dixit, Snehal Bagatharia, Kamlesh J Upadhyay, Ramesh Pandit, Afzal Ansari, Chaitanya Joshi, Madhvi Joshi |
| EPI_ISL_444461 | B.J. Medical College and Civil hospital | Gujarat Biotechnology Research Centre | Ankit Hinsu, Pritesh Sabara, Apurvasinh Puvar, Janvi Raval, Monika Gandhi, Pinal Trivedi, Maharshi Pandya, Amit Kanani, Akanksha Verma, Nitin Savaliya, Raghawendra Kumar, Dinesh Kumar, Zuber Saiyed, Dipa Kinariwala, Disha Patel, Binita Aring, Neeta Khandelwal, Geeta Vaghela, Sonia Barve, Bhavesh Modi, Kairavi Joshi, Gaurishankar Shrimali, Nidhi Sood, Pranay Shah, R D Dixit, Snehal Bagatharia, Kamlesh J Upadhyay, Ramesh Pandit, Neelam Nathani, Chaitanya Joshi, Madhvi Joshi, Tejas Shah |
| EPI_ISL_444462 | B.J. Medical College and Civil hospital | Gujarat Biotechnology Research Centre | Pritesh Sabara, Apurvasinh Puvar, Janvi Raval, Monika Gandhi, Pinal Trivedi, Maharshi Pandya, Amit Kanani, Akanksha Verma, Nitin Savaliya, Raghawendra Kumar, Dinesh Kumar, Zuber Saiyed, Dipa Kinariwala, Disha Patel, Binita Aring, Neeta Khandelwal, Geeta Vaghela, Sonia Barve, Bhavesh Modi, Kairavi Joshi, Gaurishankar Shrimali, Nidhi Sood, Pranay Shah, R D Dixit, Snehal Bagatharia, Kamlesh J Upadhyay, Ramesh Pandit, Tejas Shah, Ankit Hinsu, Armi Chaudhari, Chaitanya Joshi, Madhvi Joshi |
| EPI_ISL_444463 | B.J. Medical College and Civil hospital | Gujarat Biotechnology Research Centre | Apurvasinh Puvar, Janvi Raval, Monika Gandhi, Pinal Trivedi, Maharshi Pandya, Amit Kanani, Akanksha Verma, Nitin Savaliya, Raghawendra Kumar, Dinesh Kumar, Zuber Saiyed, Dipa Kinariwala, Disha Patel, Binita Aring, Neeta Khandelwal, Geeta Vaghela, Sonia Barve, Bhavesh Modi, Kairavi Joshi, Gaurishankar Shrimali, Nidhi Sood, Pranay Shah, R D Dixit, Snehal Bagatharia, Kamlesh J Upadhyay, Ramesh Pandit, Tejas Shah, Ankit Hinsu, Pritesh Sabara, Bhavya Jindal, Chaitanya Joshi, Madhvi Joshi |
| EPI_ISL_444464 | B.J. Medical College and Civil hospital | Gujarat Biotechnology Research Centre | Janvi Raval, Monika Gandhi, Pinal Trivedi, Maharshi Pandya, Amit Kanani, Akanksha Verma, Nitin Savaliya, Raghawendra Kumar, Dinesh Kumar, Zuber Saiyed, Dipa Kinariwala, Disha Patel, Binita Aring, Neeta Khandelwal, Geeta Vaghela, Sonia Barve, Bhavesh Modi, Kairavi Joshi, Gaurishankar Shrimali, Nidhi Sood, Pranay Shah, R D Dixit, Snehal Bagatharia, Kamlesh J Upadhyay, Ramesh Pandit, Tejas Shah, Ankit Hinsu, Pritesh Sabara, Apurvasinh Puvar, Dipeshwari Shewale, Chaitanya Joshi, Madhvi Joshi |
| EPI_ISL_444465 | B.J. Medical College and Civil hospital | Gujarat Biotechnology Research Centre | Monika Gandhi, Pinal Trivedi, Maharshi Pandya, Amit Kanani, Akanksha Verma, Nitin Savaliya, Raghawendra Kumar, Dinesh Kumar, Zuber Saiyed, Dipa Kinariwala, Disha Patel, Binita Aring, Neeta Khandelwal, Geeta Vaghela, Sonia Barve, Bhavesh Modi, Kairavi Joshi, Gaurishankar Shrimali, Nidhi Sood, Pranay Shah, R D Dixit, Snehal Bagatharia, Kamlesh J Upadhyay, Ramesh Pandit, Tejas Shah, Ankit Hinsu, Pritesh Sabara, Apurvasinh Puvar, Janvi Raval, Anjali Rajwar, Chaitanya Joshi, Madhvi Joshi |
| EPI_ISL_444466 | B.J. Medical College and Civil hospital | Gujarat Biotechnology Research Centre | Pinal Trivedi, Maharshi Pandya, Amit Kanani, Akanksha Verma, Nitin Savaliya, Raghawendra Kumar, Dinesh Kumar, Zuber Saiyed, Dipa Kinariwala, Disha Patel, Binita Aring, Neeta Khandelwal, Geeta Vaghela, Sonia Barve, Bhavesh Modi, Kairavi Joshi, Gaurishankar Shrimali, Nidhi Sood, Pranay Shah, R D Dixit, Snehal Bagatharia, Kamlesh J Upadhyay, Ramesh Pandit, Tejas Shah, Ankit Hinsu, Pritesh Sabara, Apurvasinh Puvar, Janvi Raval, Monika Gandhi, Sharmistha Majumdar, Chaitanya Joshi, Madhvi Joshi |
| EPI_ISL_444467 | B.J. Medical College and Civil hospital | Gujarat Biotechnology Research Centre | Maharshi Pandya, Amit Kanani, Akanksha Verma, Nitin Savaliya, Raghawendra Kumar, Dinesh Kumar, Zuber Saiyed, Dipa Kinariwala, Disha Patel, Binita Aring, Neeta Khandelwal, Geeta Vaghela, Sonia Barve, Bhavesh Modi, Kairavi Joshi, Gaurishankar Shrimali, Nidhi Sood, Pranay Shah, R D Dixit, Snehal Bagatharia, Kamlesh J Upadhyay, Ramesh Pandit, Tejas Shah, Ankit Hinsu, Pritesh Sabara, Apurvasinh Puvar, Janvi Raval, Monika Gandhi, Pinal Trivedi, Pooja P Doshi, Chaitanya Joshi, Madhvi Joshi |
| EPI_ISL_444468 | B.J. Medical College and Civil hospital | Gujarat Biotechnology Research Centre | Amit Kanani, Akanksha Verma, Nitin Savaliya, Raghawendra Kumar, Dinesh Kumar, Zuber Saiyed, Dipa Kinariwala, Disha Patel, Binita Aring, Neeta Khandelwal, Geeta Vaghela, Sonia Barve, Bhavesh Modi, Kairavi Joshi, Gaurishankar Shrimali, Nidhi Sood, Pranay Shah, R D Dixit, Snehal Bagatharia, Kamlesh J Upadhyay, Ramesh Pandit, Tejas Shah, Ankit Hinsu, Pritesh Sabara, Apurvasinh Puvar, Janvi Raval, Monika Gandhi, Pinal Trivedi, Maharshi Pandya, Nidhi Patel, Chaitanya Joshi, Madhvi Joshi |
| EPI_ISL_444469 | B.J. Medical College and Civil hospital | Gujarat Biotechnology Research Centre | Akanksha Verma, Nitin Savaliya, Raghawendra Kumar, Dinesh Kumar, Zuber Saiyed, Dipa Kinariwala, Disha Patel, Binita Aring, Neeta Khandelwal, Geeta Vaghela, Sonia Barve, Bhavesh Modi, Kairavi Joshi, Gaurishankar Shrimali, Nidhi Sood, Pranay Shah, R D Dixit, Snehal Bagatharia, Kamlesh J Upadhyay, Ramesh Pandit, Tejas Shah, Ankit Hinsu, Pritesh Sabara, Apurvasinh Puvar, Janvi Raval, Monika Gandhi, Pinal Trivedi, Maharshi Pandya, Amit Kanani, Priti Pandita, Chaitanya Joshi, Madhvi Joshi |
| EPI_ISL_444470 | B.J. Medical College and Civil hospital | Gujarat Biotechnology Research Centre | Nitin Savaliya, Raghawendra Kumar, Dinesh Kumar, Zuber Saiyed, Dipa Kinariwala, Disha Patel, Binita Aring, Neeta Khandelwal, Geeta Vaghela, Sonia Barve, Bhavesh Modi, Kairavi Joshi, Gaurishankar Shrimali, Nidhi Sood, Pranay Shah, R D Dixit, Snehal Bagatharia, Kamlesh J Upadhyay, Ramesh Pandit, Tejas Shah, Ankit Hinsu, Pritesh Sabara, Apurvasinh Puvar, Janvi Raval, Monika Gandhi, Pinal Trivedi, Maharshi Pandya, Amit Kanani, Akanksha Verma, Neha Rajpara, Chaitanya Joshi, Madhvi Joshi |
| EPI_ISL_444471 | B.J. Medical College and Civil hospital | Gujarat Biotechnology Research Centre | Raghawendra Kumar, Dinesh Kumar, Zuber Saiyed, Dipa Kinariwala, Disha Patel, Binita Aring, Neeta Khandelwal, Geeta Vaghela, Sonia Barve, Bhavesh Modi, Kairavi Joshi, Gaurishankar Shrimali, Nidhi Sood, Pranay Shah, R D Dixit, Snehal Bagatharia, Kamlesh J Upadhyay, Ramesh Pandit, Tejas Shah, Ankit Hinsu, Pritesh Sabara, Apurvasinh Puvar, Janvi Raval, Monika Gandhi, Pinal Trivedi, Maharshi Pandya, Amit Kanani, Akanksha Verma, Nitin Savaliya, Afzal Ansari, Chaitanya Joshi, Madhvi Joshi |
| EPI_ISL_444472 | B.J. Medical College and Civil hospital | Gujarat Biotechnology Research Centre | Dinesh Kumar, Zuber Saiyed, Dipa Kinariwala, Disha Patel, Binita Aring, Neeta Khandelwal, Geeta Vaghela, Sonia Barve, Bhavesh Modi, Kairavi Joshi, Gaurishankar Shrimali, Nidhi Sood, Pranay Shah, R D Dixit, Snehal Bagatharia, Kamlesh J Upadhyay, Ramesh Pandit, Tejas Shah, Ankit Hinsu, Pritesh Sabara, Apurvasinh Puvar, Janvi Raval, Monika Gandhi, Pinal Trivedi, Maharshi Pandya, Amit Kanani, Akanksha Verma, Nitin Savaliya, Raghawendra Kumar, Neelam Nathani, Chaitanya Joshi, Madhvi Joshi |
| EPI_ISL_444473 | B.J. Medical College and Civil hospital | Gujarat Biotechnology Research Centre | Zuber Saiyed, Dipa Kinariwala, Disha Patel, Binita Aring, Neeta Khandelwal, Geeta Vaghela, Sonia Barve, Bhavesh Modi, Kairavi Joshi, Gaurishankar Shrimali, Nidhi Sood, Pranay Shah, R D Dixit, Snehal Bagatharia, Kamlesh J Upadhyay, Ramesh Pandit, Tejas Shah, Ankit Hinsu, Pritesh Sabara, Apurvasinh Puvar, Janvi Raval, Monika Gandhi, Pinal Trivedi, Maharshi Pandya, Amit Kanani, Akanksha Verma, Nitin Savaliya, Raghawendra Kumar, Dinesh Kumar, Armi Chaudhari, Chaitanya Joshi, Madhvi Joshi |
| EPI_ISL_444474 | B.J. Medical College and Civil hospital | Gujarat Biotechnology Research Centre | Dipa Kinariwala, Disha Patel, Binita Aring, Neeta Khandelwal, Geeta Vaghela, Sonia Barve, Bhavesh Modi, Kairavi Joshi, Gaurishankar Shrimali, Nidhi Sood, Pranay Shah, R D Dixit, Snehal Bagatharia, Kamlesh J Upadhyay, Ramesh Pandit, Tejas Shah, Ankit Hinsu, Pritesh Sabara, Apurvasinh Puvar, Janvi Raval, Monika Gandhi, Pinal Trivedi, Maharshi Pandya, Amit Kanani, Akanksha Verma, Nitin Savaliya, Raghawendra Kumar, Dinesh Kumar, Zuber Saiyed, Bhavya Jindal, Chaitanya Joshi, Madhvi Joshi |
| EPI_ISL_444475 | B.J. Medical College and Civil hospital | Gujarat Biotechnology Research Centre | Disha Patel, Binita Aring, Neeta Khandelwal, Geeta Vaghela, Sonia Barve, Bhavesh Modi, Kairavi Joshi, Gaurishankar Shrimali, Nidhi Sood, Pranay Shah, R D Dixit, Snehal Bagatharia, Kamlesh J Upadhyay, Ramesh Pandit, Tejas Shah, Ankit Hinsu, Pritesh Sabara, Apurvasinh Puvar, Janvi Raval, Monika Gandhi, Pinal Trivedi, Maharshi Pandya, Amit Kanani, Akanksha Verma, Nitin Savaliya, Raghawendra Kumar, Dinesh Kumar, Zuber Saiyed, Dipa Kinariwala, Dipeshwari Shewale, Chaitanya Joshi, Madhvi Joshi |
| EPI_ISL_444476 | B.J. Medical College and Civil hospital | Gujarat Biotechnology Research Centre | Binita Aring, Neeta Khandelwal, Geeta Vaghela, Sonia Barve, Bhavesh Modi, Kairavi Joshi, Gaurishankar Shrimali, Nidhi Sood, Pranay Shah, R D Dixit, Snehal Bagatharia, Kamlesh J Upadhyay, Ramesh Pandit, Tejas Shah, Ankit Hinsu, Pritesh Sabara, Apurvasinh Puvar, Janvi Raval, Monika Gandhi, Pinal Trivedi, Maharshi Pandya, Amit Kanani, Akanksha Verma, Nitin Savaliya, Raghawendra Kumar, Dinesh Kumar, Zuber Saiyed, Dipa Kinariwala, Disha Patel, Chaitanya Joshi, Madhvi Joshi, Dipeshwari Shewale |
| EPI_ISL_444477 | B.J. Medical College and Civil hospital | Gujarat Biotechnology Research Centre | Neeta Khandelwal, Geeta Vaghela, Sonia Barve, Bhavesh Modi, Kairavi Joshi, Gaurishankar Shrimali, Nidhi Sood, Pranay Shah, R D Dixit, Snehal Bagatharia, Kamlesh J Upadhyay, Ramesh Pandit, Tejas Shah, Ankit Hinsu, Pritesh Sabara, Apurvasinh Puvar, Janvi Raval, Monika Gandhi, Pinal Trivedi, Maharshi Pandya, Amit Kanani, Akanksha Verma, Nitin Savaliya, Raghawendra Kumar, Dinesh Kumar, Zuber Saiyed, Dipa Kinariwala, Disha Patel, Binita Aring, Sharmistha Majumdar, Chaitanya Joshi, Madhvi Joshi |
| EPI_ISL_444478 | B.J. Medical College and Civil hospital | Gujarat Biotechnology Research Centre | Geeta Vaghela, Sonia Barve, Bhavesh Modi, Kairavi Joshi, Gaurishankar Shrimali, Nidhi Sood, Pranay Shah, R D Dixit, Snehal Bagatharia, Kamlesh J Upadhyay, Ramesh Pandit, Tejas Shah, Ankit Hinsu, Pritesh Sabara, Apurvasinh Puvar, Janvi Raval, Monika Gandhi, Pinal Trivedi, Maharshi Pandya, Amit Kanani, Akanksha Verma, Nitin Savaliya, Raghawendra Kumar, Dinesh Kumar, Zuber Saiyed, Dipa Kinariwala, Disha Patel, Binita Aring, Neeta Khandelwal, |

| | | | |
|---|---|---|---|
| | | | Pooja P Doshi, Chaitanya Joshi, Madhvi Joshi |
| EPI_ISL_444479 | B.J. Medical College and Civil hospital | Gujarat Biotechnology Research Centre | Sonia Barve, Bhavesh Modi, Kairavi Joshi, Gaurishankar Shrimali, Nidhi Sood, Pranay Shah, R D Dixit, Snehal Bagatharia, Kamlesh J Upadhyay, Ramesh Pandit, Tejas Shah, Ankit Hinsu, Pritesh Sabara, Apurvasinh Puvar, Janvi Raval, Monika Gandhi, Pinal Trivedi, Maharshi Pandya, Amit Kanani, Akanksha Verma, Nitin Savaliya, Raghawendra Kumar, Dinesh Kumar, Zuber Saiyed, Dipa Kinariwala, Disha Patel, Binita Aring, Neeta Khandelwal, Geeta Vaghela, Nidhi Patel, Chaitanya Joshi, Madhvi Joshi |
| EPI_ISL_444480 | B.J. Medical College and Civil hospital | Gujarat Biotechnology Research Centre | Bhavesh Modi, Kairavi Joshi, Gaurishankar Shrimali, Nidhi Sood, Pranay Shah, R D Dixit, Snehal Bagatharia, Kamlesh J Upadhyay, Ramesh Pandit, Tejas Shah, Ankit Hinsu, Pritesh Sabara, Apurvasinh Puvar, Janvi Raval, Monika Gandhi, Pinal Trivedi, Maharshi Pandya, Amit Kanani, Akanksha Verma, Nitin Savaliya, Raghawendra Kumar, Dinesh Kumar, Zuber Saiyed, Dipa Kinariwala, Disha Patel, Binita Aring, Neeta Khandelwal, Geeta Vaghela, Sonia Barve, Priti Pandita, Chaitanya Joshi, Madhvi Joshi |
| EPI_ISL_444481 | B.J. Medical College and Civil hospital | Gujarat Biotechnology Research Centre | Kairavi Joshi, Gaurishankar Shrimali, Nidhi Sood, Pranay Shah, R D Dixit, Snehal Bagatharia, Kamlesh J Upadhyay, Ramesh Pandit, Tejas Shah, Ankit Hinsu, Pritesh Sabara, Apurvasinh Puvar, Janvi Raval, Monika Gandhi, Pinal Trivedi, Maharshi Pandya, Amit Kanani, Akanksha Verma, Nitin Savaliya, Raghawendra Kumar, Dinesh Kumar, Zuber Saiyed, Dipa Kinariwala, Disha Patel, Binita Aring, Neeta Khandelwal, Geeta Vaghela, Sonia Barve, Bhavesh Modi, Neha Rajpara, Chaitanya Joshi, Madhvi Joshi |
| EPI_ISL_444482 | Gujarat Biotechnology Research Centre | Gujarat Biotechnology Research Centre | Gaurishankar Shrimali, Nidhi Sood, Pranay Shah, R D Dixit, Snehal Bagatharia, Kamlesh J Upadhyay, Ramesh Pandit, Tejas Shah, Ankit Hinsu, Pritesh Sabara, Apurvasinh Puvar, Janvi Raval, Monika Gandhi, Pinal Trivedi, Maharshi Pandya, Amit Kanani, Akanksha Verma, Nitin Savaliya, Raghawendra Kumar, Dinesh Kumar, Zuber Saiyed, Dipa Kinariwala, Disha Patel, Binita Aring, Neeta Khandelwal, Geeta Vaghela, Sonia Barve, Bhavesh Modi, Kairavi Joshi, Afzal Ansari, Chaitanya Joshi, Madhvi Joshi |
| EPI_ISL_444483 | Gujarat Biotechnology Research Centre | Gujarat Biotechnology Research Centre | Nidhi Sood, Pranay Shah, R D Dixit, Snehal Bagatharia, Kamlesh J Upadhyay, Ramesh Pandit, Tejas Shah, Ankit Hinsu, Pritesh Sabara, Apurvasinh Puvar, Janvi Raval, Monika Gandhi, Pinal Trivedi, Maharshi Pandya, Amit Kanani, Akanksha Verma, Nitin Savaliya, Raghawendra Kumar, Dinesh Kumar, Zuber Saiyed, Dipa Kinariwala, Disha Patel, Binita Aring, Neeta Khandelwal, Geeta Vaghela, Sonia Barve, Bhavesh Modi, Kairavi Joshi, Gaurishankar Shrimali, Neelam Nathani, Chaitanya Joshi, Madhvi Joshi |
| EPI_ISL_444484 | Gujarat Biotechnology Research Centre | Gujarat Biotechnology Research Centre | Pranay Shah, R D Dixit, Snehal Bagatharia, Kamlesh J Upadhyay, Ramesh Pandit, Tejas Shah, Ankit Hinsu, Pritesh Sabara, Apurvasinh Puvar, Janvi Raval, Monika Gandhi, Pinal Trivedi, Maharshi Pandya, Amit Kanani, Akanksha Verma, Nitin Savaliya, Raghawendra Kumar, Dinesh Kumar, Zuber Saiyed, Dipa Kinariwala, Disha Patel, Binita Aring, Neeta Khandelwal, Geeta Vaghela, Sonia Barve, Bhavesh Modi, Kairavi Joshi, Gaurishankar Shrimali, Nidhi Sood, Armi Chaudhari, Chaitanya Joshi, Madhvi Joshi |
| EPI_ISL_444485 | Gujarat Biotechnology Research Centre | Gujarat Biotechnology Research Centre | R D Dixit, Snehal Bagatharia, Kamlesh J Upadhyay, Ramesh Pandit, Tejas Shah, Ankit Hinsu, Pritesh Sabara, Apurvasinh Puvar, Janvi Raval, Monika Gandhi, Pinal Trivedi, Maharshi Pandya, Amit Kanani, Akanksha Verma, Nitin Savaliya, Raghawendra Kumar, Dinesh Kumar, Zuber Saiyed, Dipa Kinariwala, Disha Patel, Binita Aring, Neeta Khandelwal, Geeta Vaghela, Sonia Barve, Bhavesh Modi, Kairavi Joshi, Gaurishankar Shrimali, Nidhi Sood, Pranay Shah, Bhavya Jindal, Chaitanya Joshi, Madhvi Joshi |
| EPI_ISL_444486 | Gujarat Biotechnology Research Centre | Gujarat Biotechnology Research Centre | Snehal Bagatharia, Kamlesh J Upadhyay, Ramesh Pandit, Tejas Shah, Ankit Hinsu, Pritesh Sabara, Apurvasinh Puvar, Janvi Raval, Monika Gandhi, Pinal Trivedi, Maharshi Pandya, Amit Kanani, Akanksha Verma, Nitin Savaliya, Raghawendra Kumar, Dinesh Kumar, Zuber Saiyed, Dipa Kinariwala, Disha Patel, Binita Aring, Neeta Khandelwal, Geeta Vaghela, Sonia Barve, Bhavesh Modi, Kairavi Joshi, Gaurishankar Shrimali, Nidhi Sood, Pranay Shah, R D Dixit, Dipeshwari Shewale, Chaitanya Joshi, Madhvi Joshi |
| EPI_ISL_445019, EPI_ISL_445020, EPI_ISL_445021, EPI_ISL_445022, EPI_ISL_445023, EPI_ISL_445024, EPI_ISL_445025, EPI_ISL_445026, EPI_ISL_445027, EPI_ISL_445028, EPI_ISL_445031, EPI_ISL_445032, EPI_ISL_445033, EPI_ISL_445034, EPI_ISL_445035, EPI_ISL_445036, EPI_ISL_445037, EPI_ISL_445038, EPI_ISL_445039, EPI_ISL_445040, EPI_ISL_445041, EPI_ISL_445042, EPI_ISL_445043, EPI_ISL_445044, EPI_ISL_445046, EPI_ISL_445047, EPI_ISL_445048, EPI_ISL_445049, EPI_ISL_445050 | | | |
| see above | Florida Bureau of Public Health Laboratories | Florida Bureau of Public Health Laboratories | Sarah Schmedes, Jason Blanton |
| EPI_ISL_445053, EPI_ISL_445062, EPI_ISL_445064, EPI_ISL_445068 | Laboratoire National de Sante, Microbiology, Virology | Laboratoire National de Sante, Microbiology, Epidemiology and Microbial Genomics | Anke Wienecke-Baldacchino, Ardashel Latsuzbaia, Jessica Tapp, Catherine Ragimbeau, Guillaume Fournier, Tamir Abdelrahman, Trung Nguyen Nguyen, Joel Mossong |
| EPI_ISL_445118 | Rady's Childrens Hospital | Andersen lab at Scripps Research | SEARCH Alliance San Diego |
| EPI_ISL_445213 | DNA Solution Ltd | DNA Solution Ltd | Md. Imran Khan, Kazi Nadim Hasan, Abu Sufian, Mohammed Nafiz Imtiaz Polol, Abdul Khaleque, Mizanur Rahman, MSM Chowdhury, Hasan Ul Haider, Mamudul Hasan Razu, Mala Khan, Mohammad Fazle Alam Rabbi |
| EPI_ISL_445214, EPI_ISL_445215, EPI_ISL_445216, EPI_ISL_445217 | DNA Solution Ltd. | DNA Solution Ltd. | Md. Imran Khan, Kazi Nadim Hasan, Abu Sufian, Mohammed Nafiz Imtiaz Polol, Abdul Khaleque, Mizanur Rahman, MSM Chowdhury, Hasan Ul Haider, Mamudul Hasan Razu, Mala Khan, Mohammad Fazle Alam Rabbi |
| EPI_ISL_445220 | Laborarory for Respiratory Viruses, "Cantacuzino" National Military-Medical Institute for Resararch and Development | Cantacuzino Institute | M.Lazar, L.Ustea, A.Cretu |
| EPI_ISL_445243 | Laboratory for Respiratory Viruses, Cantacuzino National Military-Medical Institute for Research and Development | Cantacuzino Institute | M.Lazar, L.Ustea, A.Cretu |
| EPI_ISL_447030 | B.J. Medical College and Civil hospital | Gujarat Biotechnology Research Centre | Kamlesh J Upadhyay, Ramesh Pandit, Tejas Shah, Ankit Hinsu, Pritesh Sabara, Apurvasinh Puvar, Janvi Raval, Monika Gandhi, Pinal Trivedi, Maharshi Pandya, Amit Kanani, Akanksha Verma, Nitin Savaliya, Raghawendra Kumar, Dinesh Kumar, Zuber Saiyed, Dipa Kinariwala, Disha Patel, Binita Aring, Neeta Khandelwal, Geeta Vaghela, Sonia Barve, Bhavesh Modi, Kairavi Joshi, Gaurishankar Shrimali, Nidhi Sood, Pranay Shah, R D Dixit, Snehal Bagatharia, Anjali Rajwar, Chaitanya Joshi, Madhvi Joshi |
| EPI_ISL_447031 | B.J. Medical College and Civil hospital | Gujarat Biotechnology Research Centre | Ramesh Pandit, Tejas Shah, Ankit Hinsu, Pritesh Sabara, Apurvasinh Puvar, Janvi Raval, Monika Gandhi, Pinal Trivedi, Maharshi Pandya, Amit Kanani, Akanksha Verma, Nitin Savaliya, Raghawendra Kumar, Dinesh Kumar, Zuber Saiyed, Dipa Kinariwala, Disha Patel, Binita Aring, Neeta Khandelwal, Geeta Vaghela, Sonia Barve, Bhavesh Modi, Kairavi Joshi, Gaurishankar Shrimali, Nidhi Sood, Pranay Shah, R D Dixit, Snehal Bagatharia, Kamlesh J Upadhyay, Sharmistha Majumdar, Chaitanya Joshi, Madhvi Joshi |
| EPI_ISL_447032 | B.J. Medical College and Civil hospital | Gujarat Biotechnology Research Centre | Tejas Shah, Ankit Hinsu, Pritesh Sabara, Apurvasinh Puvar, Janvi Raval, Monika Gandhi, Pinal Trivedi, Maharshi Pandya, Amit Kanani, Akanksha Verma, Nitin Savaliya, Raghawendra Kumar, Dinesh Kumar, Zuber Saiyed, Dipa Kinariwala, Disha Patel, Binita Aring, Neeta Khandelwal, Geeta Vaghela, Sonia Barve, Bhavesh Modi, Kairavi Joshi, Gaurishankar Shrimali, Nidhi Sood, Pranay Shah, R D Dixit, Snehal Bagatharia, Ramesh Pandit, Kamlesh J Upadhyay, Pooja P Doshi, Chaitanya Joshi, Madhvi Joshi |
| EPI_ISL_447033 | B.J. Medical College and Civil hospital | Gujarat Biotechnology Research Centre | Ankit Hinsu, Pritesh Sabara, Apurvasinh Puvar, Janvi Raval, Monika Gandhi, Pinal Trivedi, Maharshi Pandya, Amit Kanani, Akanksha Verma, Nitin Savaliya, Raghawendra Kumar, Dinesh Kumar, Zuber Saiyed, Dipa Kinariwala, Disha Patel, Binita Aring, Neeta Khandelwal, Geeta Vaghela, Sonia Barve, Bhavesh Modi, Kairavi Joshi, Gaurishankar Shrimali, Nidhi Sood, Pranay Shah, R D Dixit, Snehal Bagatharia, Kamlesh J Upadhyay, Ramesh Pandit, Tejas Shah, Nidhi Patel, Chaitanya Joshi, Madhvi Joshi |
| EPI_ISL_447034 | B.J. Medical College and Civil hospital | Gujarat Biotechnology Research Centre | Pritesh Sabara, Apurvasinh Puvar, Janvi Raval, Monika Gandhi, Pinal Trivedi, Maharshi Pandya, Amit Kanani, Akanksha Verma, Nitin Savaliya, Raghawendra Kumar, Dinesh Kumar, Zuber Saiyed, Dipa Kinariwala, Disha Patel, Binita Aring, Neeta Khandelwal, Geeta Vaghela, Sonia Barve, Bhavesh Modi, Kairavi Joshi, Gaurishankar Shrimali, Nidhi Sood, Pranay Shah, R D Dixit, Snehal Bagatharia, Kamlesh J Upadhyay, Ramesh Pandit, Tejas Shah, Ankit Hinsu, Priti Pandita, Chaitanya Joshi, Madhvi Joshi |
| EPI_ISL_447035 | B.J. Medical College and Civil hospital | Gujarat Biotechnology Research Centre | Apurvasinh Puvar, Janvi Raval, Monika Gandhi, Pinal Trivedi, Maharshi Pandya, Amit Kanani, Akanksha Verma, Nitin Savaliya, Raghawendra Kumar, Dinesh Kumar, Zuber Saiyed, Dipa Kinariwala, Disha Patel, Binita Aring, Neeta Khandelwal, Geeta Vaghela, Sonia Barve, Bhavesh Modi, Kairavi Joshi, Gaurishankar Shrimali, Nidhi Sood, Pranay Shah, R D Dixit, Snehal Bagatharia, Kamlesh J Upadhyay, Ramesh Pandit, Tejas Shah, Ankit Hinsu, Pritesh Sabara, Neha Rajpara, Chaitanya Joshi, Madhvi Joshi |
| EPI_ISL_447036 | B.J. Medical College and Civil hospital | Gujarat Biotechnology Research Centre | Janvi Raval, Monika Gandhi, Pinal Trivedi, Maharshi Pandya, Amit Kanani, Akanksha Verma, Nitin Savaliya, Raghawendra Kumar, Dinesh Kumar, Zuber Saiyed, Dipa Kinariwala, Disha Patel, Binita Aring, Neeta Khandelwal, Geeta Vaghela, Sonia Barve, Bhavesh Modi, Kairavi Joshi, Gaurishankar Shrimali, Nidhi Sood, Pranay Shah, R D Dixit, Snehal Bagatharia, Kamlesh J Upadhyay, Ramesh Pandit, Tejas Shah, Ankit Hinsu, Pritesh Sabara, Apurvasinh Puvar, Afzal Ansari, Chaitanya Joshi, Madhvi Joshi |

| | | | |
|---|---|---|---|
| EPI_ISL_447037 | B.J. Medical College and Civil hospital | Gujarat Biotechnology Research Centre | Monika Gandhi, Pinal Trivedi, Maharshi Pandya, Amit Kanani, Akanksha Verma, Nitin Savaliya, Raghawendra Kumar, Dinesh Kumar, Zuber Saiyed, Dipa Kinariwala, Disha Patel, Binita Aring, Neeta Khandelwal, Geeta Vaghela, Sonia Barve, Bhavesh Modi, Kairavi Joshi, Gaurishankar Shrimali, Nidhi Sood, Pranay Shah, R D Dixit, Snehal Bagatharia, Kamlesh J Upadhyay, Ramesh Pandit, Tejas Shah, Ankit Hinsu, Pritesh Sabara, Apurvasinh Puvar, Janvi Raval, Neelam Nathani, Chaitanya Joshi, Madhvi Joshi |
| EPI_ISL_447038 | B.J. Medical College and Civil hospital | Gujarat Biotechnology Research Centre | Pinal Trivedi, Maharshi Pandya, Amit Kanani, Akanksha Verma, Nitin Savaliya, Raghawendra Kumar, Dinesh Kumar, Zuber Saiyed, Dipa Kinariwala, Disha Patel, Binita Aring, Neeta Khandelwal, Geeta Vaghela, Sonia Barve, Bhavesh Modi, Kairavi Joshi, Gaurishankar Shrimali, Nidhi Sood, Pranay Shah, R D Dixit, Snehal Bagatharia, Kamlesh J Upadhyay, Ramesh Pandit, Tejas Shah, Ankit Hinsu, Pritesh Sabara, Apurvasinh Puvar, Janvi Raval, Monika Gandhi, Armi Chaudhari, Chaitanya Joshi, Madhvi Joshi |
| EPI_ISL_447039 | B.J. Medical College and Civil hospital | Gujarat Biotechnology Research Centre | Maharshi Pandya, Amit Kanani, Akanksha Verma, Nitin Savaliya, Raghawendra Kumar, Dinesh Kumar, Zuber Saiyed, Dipa Kinariwala, Disha Patel, Binita Aring, Neeta Khandelwal, Geeta Vaghela, Sonia Barve, Bhavesh Modi, Kairavi Joshi, Gaurishankar Shrimali, Nidhi Sood, Pranay Shah, R D Dixit, Snehal Bagatharia, Kamlesh J Upadhyay, Ramesh Pandit, Tejas Shah, Ankit Hinsu, Pritesh Sabara, Apurvasinh Puvar, Janvi Raval, Monika Gandhi, Pinal Trivedi, Bhavya Jindal, Chaitanya Joshi, Madhvi Joshi |
| EPI_ISL_447040 | B.J. Medical College and Civil hospital | Gujarat Biotechnology Research Centre | Amit Kanani, Akanksha Verma, Nitin Savaliya, Raghawendra Kumar, Dinesh Kumar, Zuber Saiyed, Dipa Kinariwala, Disha Patel, Binita Aring, Neeta Khandelwal, Geeta Vaghela, Sonia Barve, Bhavesh Modi, Kairavi Joshi, Gaurishankar Shrimali, Nidhi Sood, Pranay Shah, R D Dixit, Snehal Bagatharia, Kamlesh J Upadhyay, Ramesh Pandit, Tejas Shah, Ankit Hinsu, Pritesh Sabara, Apurvasinh Puvar, Janvi Raval, Monika Gandhi, Pinal Trivedi, Maharshi Pandya, Anjali Rajwar, Chaitanya Joshi, Madhvi Joshi |
| EPI_ISL_447041 | B.J. Medical College and Civil hospital | Gujarat Biotechnology Research Centre | Akanksha Verma, Nitin Savaliya, Raghawendra Kumar, Dinesh Kumar, Zuber Saiyed, Dipa Kinariwala, Disha Patel, Binita Aring, Neeta Khandelwal, Geeta Vaghela, Sonia Barve, Bhavesh Modi, Kairavi Joshi, Gaurishankar Shrimali, Nidhi Sood, Pranay Shah, R D Dixit, Snehal Bagatharia, Kamlesh J Upadhyay, Ramesh Pandit, Tejas Shah, Ankit Hinsu, Pritesh Sabara, Apurvasinh Puvar, Janvi Raval, Monika Gandhi, Pinal Trivedi, Maharshi Pandya, Amit Kanani, Sharmistha Majumdar, Chaitanya Joshi, Madhvi Joshi |
| EPI_ISL_447042 | B.J. Medical College and Civil hospital | Gujarat Biotechnology Research Centre | Nitin Savaliya, Raghawendra Kumar, Dinesh Kumar, Zuber Saiyed, Dipa Kinariwala, Disha Patel, Binita Aring, Neeta Khandelwal, Geeta Vaghela, Sonia Barve, Bhavesh Modi, Kairavi Joshi, Gaurishankar Shrimali, Nidhi Sood, Pranay Shah, R D Dixit, Snehal Bagatharia, Kamlesh J Upadhyay, Ramesh Pandit, Tejas Shah, Ankit Hinsu, Pritesh Sabara, Apurvasinh Puvar, Janvi Raval, Monika Gandhi, Pinal Trivedi, Maharshi Pandya, Amit Kanani, Akanksha Verma, Pooja P Doshi, Chaitanya Joshi, Madhvi Joshi |
| EPI_ISL_447043 | B.J. Medical College and Civil hospital | Gujarat Biotechnology Research Centre | Raghawendra Kumar, Dinesh Kumar, Zuber Saiyed, Dipa Kinariwala, Disha Patel, Binita Aring, Neeta Khandelwal, Geeta Vaghela, Sonia Barve, Bhavesh Modi, Kairavi Joshi, Gaurishankar Shrimali, Nidhi Sood, Pranay Shah, R D Dixit, Snehal Bagatharia, Kamlesh J Upadhyay, Ramesh Pandit, Tejas Shah, Ankit Hinsu, Pritesh Sabara, Apurvasinh Puvar, Janvi Raval, Monika Gandhi, Pinal Trivedi, Maharshi Pandya, Amit Kanani, Akanksha Verma, Nitin Savaliya, Nidhi Patel, Chaitanya Joshi, Madhvi Joshi |
| EPI_ISL_447044 | B.J. Medical College and Civil hospital | Gujarat Biotechnology Research Centre | Dinesh Kumar, Zuber Saiyed, Dipa Kinariwala, Disha Patel, Binita Aring, Neeta Khandelwal, Geeta Vaghela, Sonia Barve, Bhavesh Modi, Kairavi Joshi, Gaurishankar Shrimali, Nidhi Sood, Pranay Shah, R D Dixit, Snehal Bagatharia, Kamlesh J Upadhyay, Ramesh Pandit, Tejas Shah, Ankit Hinsu, Pritesh Sabara, Apurvasinh Puvar, Janvi Raval, Monika Gandhi, Pinal Trivedi, Maharshi Pandya, Amit Kanani, Akanksha Verma, Nitin Savaliya, Raghawendra Kumar, Priti Pandita, Chaitanya Joshi, Madhvi Joshi |
| EPI_ISL_447045 | B.J. Medical College and Civil hospital | Gujarat Biotechnology Research Centre | Zuber Saiyed, Dipa Kinariwala, Disha Patel, Binita Aring, Neeta Khandelwal, Geeta Vaghela, Sonia Barve, Bhavesh Modi, Kairavi Joshi, Gaurishankar Shrimali, Nidhi Sood, Pranay Shah, R D Dixit, Snehal Bagatharia, Kamlesh J Upadhyay, Ramesh Pandit, Tejas Shah, Ankit Hinsu, Pritesh Sabara, Apurvasinh Puvar, Janvi Raval, Monika Gandhi, Pinal Trivedi, Maharshi Pandya, Amit Kanani, Akanksha Verma, Nitin Savaliya, Raghawendra Kumar, Dinesh Kumar, Neha Rajpara, Chaitanya Joshi, Madhvi Joshi |
| EPI_ISL_447046 | B.J. Medical College and Civil hospital | Gujarat Biotechnology Research Centre | Dipa Kinariwala, Disha Patel, Binita Aring, Neeta Khandelwal, Geeta Vaghela, Sonia Barve, Bhavesh Modi, Kairavi Joshi, Gaurishankar Shrimali, Nidhi Sood, Pranay Shah, R D Dixit, Snehal Bagatharia, Kamlesh J Upadhyay, Ramesh Pandit, Tejas Shah, Ankit Hinsu, Pritesh Sabara, Apurvasinh Puvar, Janvi Raval, Monika Gandhi, Pinal Trivedi, Maharshi Pandya, Amit Kanani, Akanksha Verma, Nitin Savaliya, Raghawendra Kumar, Dinesh Kumar, Zuber Saiyed, Afzal Ansari, Chaitanya Joshi, Madhvi Joshi |
| EPI_ISL_447047 | GMERS Medical College and Hospital, Gandhinagar | Gujarat Biotechnology Research Centre | Disha Patel, Binita Aring, Neeta Khandelwal, Geeta Vaghela, Sonia Barve, Bhavesh Modi, Kairavi Joshi, Gaurishankar Shrimali, Nidhi Sood, Pranay Shah, R D Dixit, Snehal Bagatharia, Kamlesh J Upadhyay, Ramesh Pandit, Tejas Shah, Ankit Hinsu, Pritesh Sabara, Apurvasinh Puvar, Janvi Raval, Monika Gandhi, Pinal Trivedi, Maharshi Pandya, Amit Kanani, Akanksha Verma, Nitin Savaliya, Raghawendra Kumar, Dinesh Kumar, Zuber Saiyed, Dipa Kinariwala, Neelam Nathani, Chaitanya Joshi, Madhvi Joshi |
| EPI_ISL_447048 | GMERS Medical College and Hospital, Gandhinagar | Gujarat Biotechnology Research Centre | Binita Aring, Neeta Khandelwal, Geeta Vaghela, Sonia Barve, Bhavesh Modi, Kairavi Joshi, Gaurishankar Shrimali, Nidhi Sood, Pranay Shah, R D Dixit, Snehal Bagatharia, Kamlesh J Upadhyay, Ramesh Pandit, Tejas Shah, Ankit Hinsu, Pritesh Sabara, Apurvasinh Puvar, Janvi Raval, Monika Gandhi, Pinal Trivedi, Maharshi Pandya, Amit Kanani, Akanksha Verma, Nitin Savaliya, Raghawendra Kumar, Dinesh Kumar, Zuber Saiyed, Dipa Kinariwala, Disha Patel, Armi Chaudhari, Chaitanya Joshi, Madhvi Joshi |
| EPI_ISL_447049 | GMERS Medical College and Hospital, Gandhinagar | Gujarat Biotechnology Research Centre | Neeta Khandelwal, Geeta Vaghela, Sonia Barve, Bhavesh Modi, Kairavi Joshi, Gaurishankar Shrimali, Nidhi Sood, Pranay Shah, R D Dixit, Snehal Bagatharia, Kamlesh J Upadhyay, Ramesh Pandit, Tejas Shah, Ankit Hinsu, Pritesh Sabara, Apurvasinh Puvar, Janvi Raval, Monika Gandhi, Pinal Trivedi, Maharshi Pandya, Amit Kanani, Akanksha Verma, Nitin Savaliya, Raghawendra Kumar, Dinesh Kumar, Zuber Saiyed, Dipa Kinariwala, Disha Patel, Binita Aring, Bhavya Jindal, Chaitanya Joshi, Madhvi Joshi |
| EPI_ISL_447050 | GMERS Medical College and Hospital, Gandhinagar | Gujarat Biotechnology Research Centre | Geeta Vaghela, Sonia Barve, Bhavesh Modi, Kairavi Joshi, Gaurishankar Shrimali, Nidhi Sood, Pranay Shah, R D Dixit, Snehal Bagatharia, Kamlesh J Upadhyay, Ramesh Pandit, Tejas Shah, Ankit Hinsu, Pritesh Sabara, Apurvasinh Puvar, Janvi Raval, Monika Gandhi, Pinal Trivedi, Maharshi Pandya, Amit Kanani, Akanksha Verma, Nitin Savaliya, Raghawendra Kumar, Dinesh Kumar, Zuber Saiyed, Dipa Kinariwala, Disha Patel, Binita Aring, Neeta Khandelwal, Dipeshwari Shewale, Chaitanya Joshi, Madhvi Joshi |
| EPI_ISL_447051 | GMERS Medical College and Hospital, Gandhinagar | Gujarat Biotechnology Research Centre | Sonia Barve, Bhavesh Modi, Kairavi Joshi, Gaurishankar Shrimali, Nidhi Sood, Pranay Shah, R D Dixit, Snehal Bagatharia, Kamlesh J Upadhyay, Ramesh Pandit, Tejas Shah, Ankit Hinsu, Pritesh Sabara, Apurvasinh Puvar, Janvi Raval, Monika Gandhi, Pinal Trivedi, Maharshi Pandya, Amit Kanani, Akanksha Verma, Nitin Savaliya, Raghawendra Kumar, Dinesh Kumar, Zuber Saiyed, Dipa Kinariwala, Disha Patel, Binita Aring, Neeta Khandelwal, Geeta Vaghela, Anjali Rajwar, Chaitanya Joshi, Madhvi Joshi |
| EPI_ISL_447052 | GMERS Medical College and Hospital, Gandhinagar | Gujarat Biotechnology Research Centre | Bhavesh Modi, Kairavi Joshi, Gaurishankar Shrimali, Nidhi Sood, Pranay Shah, R D Dixit, Snehal Bagatharia, Kamlesh J Upadhyay, Ramesh Pandit, Tejas Shah, Ankit Hinsu, Pritesh Sabara, Apurvasinh Puvar, Janvi Raval, Monika Gandhi, Pinal Trivedi, Maharshi Pandya, Amit Kanani, Akanksha Verma, Nitin Savaliya, Raghawendra Kumar, Dinesh Kumar, Zuber Saiyed, Dipa Kinariwala, Disha Patel, Binita Aring, Neeta Khandelwal, Geeta Vaghela, Sonia Barve, Sharmistha Majumdar, Chaitanya Joshi, Madhvi Joshi |
| EPI_ISL_447053 | GMERS Medical College and Hospital, Gandhinagar | Gujarat Biotechnology Research Centre | Kairavi Joshi, Gaurishankar Shrimali, Nidhi Sood, Pranay Shah, R D Dixit, Snehal Bagatharia, Kamlesh J Upadhyay, Ramesh Pandit, Tejas Shah, Ankit Hinsu, Pritesh Sabara, Apurvasinh Puvar, Janvi Raval, Monika Gandhi, Pinal Trivedi, Maharshi Pandya, Amit Kanani, Akanksha Verma, Nitin Savaliya, Raghawendra Kumar, Dinesh Kumar, Zuber Saiyed, Dipa Kinariwala, Disha Patel, Binita Aring, Neeta Khandelwal, Geeta Vaghela, Sonia Barve, Bhavesh Modi, Pooja P Doshi, Chaitanya Joshi, Madhvi Joshi |
| EPI_ISL_447054 | Cantacuzino National Military-Medical Institute for Research and Development | Cantacuzino Institute | M.Lazar, L.Ustea, A.Cretu |
| EPI_ISL_447055 | Department for Virology, Molecular Biology and Genome Research, R. G. Lugar Center for Public Health Research, National Center for Disease Control and Public Health (NCDC) of Georgia. | Department for Virology, Molecular Biology and Genome Research, R. G. Lugar Center for Public Health Research, National Center for Disease Control and Public Health (NCDC) of Georgia. | Meri Pantsulaia, Gvantsa Brachveli, Giorgi Tomashvili, Gvantsa Chanturia, Ann Machablishvili, Nato Kotaria, Marine Murtskhvaladze, Lela Sabadze, Mari Gavashelidze, Ana Papkiauri, Tata Imnadze, Tamar Jashiashvili, Tea Tevdoradze, Ketevan Sidamonidze, Ekaterine Khmaladze, Ekaterine Zhghenti, Roena Sukhiashvili, Mariam Zakalashvili, Lela Urushadze, Magda Dgebuadze, Davit Tsaguria, Ekaterine Zangaladze, Nino Berishvili, Adam Kotorashvili, Maia Alkhazashvili, Irma Burjanadze, Anna Kasradze, Khatuna Zakhashvili, Paata Imnadze, Amiran Gamkrelidze. |
| EPI_ISL_447056 | Department for Virology, Molecular Biology and Genome Research, R. G. Lugar Center for Public Health Research, National Center for Disease Control and Public Health (NCDC) of Georgia. | Department for Virology, Molecular Biology and Genome Research, R. G. Lugar Center for Public Health Research, National Center for Disease Control and Public Health (NCDC) of Georgia. | Gvantsa Brachveli, Meri Pantsulaia, Giorgi Tomashvili, Gvantsa Chanturia, Ann Machablishvili, Nato Kotaria, Marine Murtskhvaladze, Lela Sabadze, Mari Gavashelidze, Ana Papkiauri, Gvantsa Brachveli, Tata Imnadze, Tamar Jashiashvili, Tea Tevdoradze, Ketevan Sidamonidze, Ekaterine Khmaladze, Ekaterine Zhghenti, Roena Sukhiashvili, Mariam Zakalashvili, Lela Urushadze, Magda Dgebuadze, Davit Tsaguria, Ekaterine Zangaladze, Nino Berishvili, Adam Kotorashvili, Maia Alkhazashvili, Irma Burjanadze, Anna Kasradze, Khatuna Zakhashvili, Paata Imnadze, Amiran Gamkrelidze. |

| | | | |
|---|---|---|---|
| EPI_ISL_447070, EPI_ISL_447073 | Michigan Department of Health and Human Services, Bureau of Laboratories | Michigan Department of Health and Human Services, Bureau of Laboratories | Blankenship HM, Riner D, Soehnlen MK |
| EPI_ISL_447547 | GMERS Medical College and Hospital, Gandhinagar | Gujarat Biotechnology Research Centre | Pinal Trivedi, Maharshi Pandya, Amit Kanani, Akanksha Verma, Nitin Savaliya, Raghawendra Kumar, Dinesh Kumar, Zuber Saiyed, Dipa Kinariwala, Disha Patel, Binita Aring, Neeta Khandelwal, Geeta Vaghela, Sonia Barve, Bhavesh Modi, Kairavi Joshi, Gaurishankar Shrimali, Nidhi Sood, Pranay Shah, R D Dixit, Snehal Bagatharia, Kamlesh J Upadhyay, Ramesh Pandit, Tejas Shah, Ankit Hinsu, Pritesh Sabara, Apurvasinh Puvar, Janvi Raval, Monika Gandhi, Neha Rajpara, Chaitanya Joshi, Madhvi Joshi |
| EPI_ISL_447548 | GMERS Medical College and Hospital, Gandhinagar | Gujarat Biotechnology Research Centre | Maharshi Pandya, Amit Kanani, Akanksha Verma, Nitin Savaliya, Raghawendra Kumar, Dinesh Kumar, Zuber Saiyed, Dipa Kinariwala, Disha Patel, Binita Aring, Neeta Khandelwal, Geeta Vaghela, Sonia Barve, Bhavesh Modi, Kairavi Joshi, Gaurishankar Shrimali, Nidhi Sood, Pranay Shah, R D Dixit, Snehal Bagatharia, Kamlesh J Upadhyay, Ramesh Pandit, Tejas Shah, Ankit Hinsu, Pritesh Sabara, Apurvasinh Puvar, Janvi Raval, Monika Gandhi, Pinal Trivedi, Afzal Ansari, Chaitanya Joshi, Madhvi Joshi |
| EPI_ISL_447549 | GMERS Medical College and Hospital, Gandhinagar | Gujarat Biotechnology Research Centre | Amit Kanani, Akanksha Verma, Nitin Savaliya, Raghawendra Kumar, Dinesh Kumar, Zuber Saiyed, Dipa Kinariwala, Disha Patel, Binita Aring, Neeta Khandelwal, Geeta Vaghela, Sonia Barve, Bhavesh Modi, Kairavi Joshi, Gaurishankar Shrimali, Nidhi Sood, Pranay Shah, R D Dixit, Snehal Bagatharia, Kamlesh J Upadhyay, Ramesh Pandit, Tejas Shah, Ankit Hinsu, Pritesh Sabara, Apurvasinh Puvar, Janvi Raval, Monika Gandhi, Pinal Trivedi, Maharshi Pandya, Neelam Nathani, Chaitanya Joshi, Madhvi Joshi |
| EPI_ISL_447550 | GMERS Medical College and Hospital, Gandhinagar | Gujarat Biotechnology Research Centre | Akanksha Verma, Nitin Savaliya, Raghawendra Kumar, Dinesh Kumar, Zuber Saiyed, Dipa Kinariwala, Disha Patel, Binita Aring, Neeta Khandelwal, Geeta Vaghela, Sonia Barve, Bhavesh Modi, Kairavi Joshi, Gaurishankar Shrimali, Nidhi Sood, Pranay Shah, R D Dixit, Snehal Bagatharia, Kamlesh J Upadhyay, Ramesh Pandit, Tejas Shah, Ankit Hinsu, Pritesh Sabara, Apurvasinh Puvar, Janvi Raval, Monika Gandhi, Pinal Trivedi, Maharshi Pandya, Amit Kanani, Armi Chaudhari, Chaitanya Joshi, Madhvi Joshi |
| EPI_ISL_447552 | GMERS Medical College and Hospital, Gandhinagar | Gujarat Biotechnology Research Centre | Raghawendra Kumar, Dinesh Kumar, Zuber Saiyed, Dipa Kinariwala, Disha Patel, Binita Aring, Neeta Khandelwal, Geeta Vaghela, Sonia Barve, Bhavesh Modi, Kairavi Joshi, Gaurishankar Shrimali, Nidhi Sood, Pranay Shah, R D Dixit, Snehal Bagatharia, Kamlesh J Upadhyay, Ramesh Pandit, Tejas Shah, Ankit Hinsu, Pritesh Sabara, Apurvasinh Puvar, Janvi Raval, Monika Gandhi, Pinal Trivedi, Maharshi Pandya, Amit Kanani, Akanksha Verma, Nitin Savaliya, Anjali Rajwar, Chaitanya Joshi, Madhvi Joshi |
| EPI_ISL_447553 | GMERS Medical College and Hospital, Gandhinagar | Gujarat Biotechnology Research Centre | Dinesh Kumar, Zuber Saiyed, Dipa Kinariwala, Disha Patel, Binita Aring, Neeta Khandelwal, Geeta Vaghela, Sonia Barve, Bhavesh Modi, Kairavi Joshi, Gaurishankar Shrimali, Nidhi Sood, Pranay Shah, R D Dixit, Snehal Bagatharia, Kamlesh J Upadhyay, Ramesh Pandit, Tejas Shah, Ankit Hinsu, Pritesh Sabara, Apurvasinh Puvar, Janvi Raval, Monika Gandhi, Pinal Trivedi, Maharshi Pandya, Amit Kanani, Akanksha Verma, Nitin Savaliya, Raghawendra Kumar, Dipeshwari Shewale, Chaitanya Joshi, Madhvi Joshi |
| EPI_ISL_447555 | GMERS Medical College and Hospital, Gandhinagar | Gujarat Biotechnology Research Centre | Dipa Kinariwala, Disha Patel, Binita Aring, Neeta Khandelwal, Geeta Vaghela, Sonia Barve, Bhavesh Modi, Kairavi Joshi, Gaurishankar Shrimali, Nidhi Sood, Pranay Shah, R D Dixit, Snehal Bagatharia, Kamlesh J Upadhyay, Ramesh Pandit, Tejas Shah, Ankit Hinsu, Pritesh Sabara, Apurvasinh Puvar, Janvi Raval, Monika Gandhi, Pinal Trivedi, Maharshi Pandya, Amit Kanani, Akanksha Verma, Nitin Savaliya, Raghawendra Kumar, Dinesh Kumar, Zuber Saiyed, Pooja P Doshi, Chaitanya Joshi, Madhvi Joshi |
| EPI_ISL_447622 | Department of Laboratory Medicine, National Taiwan University Hospital | Microbial Genomics Core Lab, National Taiwan University Centers of Genomic and Precision Medicine | Shiou-Hwei Yeh, You-Yu Lin, Ya-Yun Lai, Chiao-Ling Li, Shan-Chwen Chang, Pei-Jer Chen, Sui-Yuan Chang |
| EPI_ISL_447623, EPI_ISL_447624, EPI_ISL_447625, EPI_ISL_447626, EPI_ISL_447627, EPI_ISL_447628, EPI_ISL_447629, EPI_ISL_447630 | Virology, Wageningen Bioveterinary Research | Virology, Wageningen Bioveterinary Research | van der Poel,W.H.M., Hakze van der Honing,R.W., Harders,F. |
| EPI_ISL_447632 | Virology, Wageningen Bioveterinary Research | Virology, Wageningen Bioveterinary Research | Oreshkova,N., Vreman,S., Molenaar,R.J., Harders,F., Hakze van der Honing,R.W., Gerhards,N., Bouwstra,R., Hissink,H., Smit,L., Tacken,M., Weesendorp,E., Stegeman,A., van der Poel,W., Engelsma,M.Y. |
| EPI_ISL_447905 | University of Florida | University of Florida | Elbadry,M.A., Subramaniam,K., Waltzek,T.B., Gibson,J.C., Stephenson,C.J., Alam,M.M., Morris,J.G. Jr. and Lednicky,J.A. |
| EPI_ISL_447926, EPI_ISL_447930, EPI_ISL_447931, EPI_ISL_447932, EPI_ISL_447933, EPI_ISL_447934, EPI_ISL_447935, EPI_ISL_447936, EPI_ISL_447937, EPI_ISL_447938, EPI_ISL_447939, EPI_ISL_447940, EPI_ISL_447941, EPI_ISL_447942, EPI_ISL_447943, EPI_ISL_447944 | | | |
| see above | University of Birmingham | COVID-19 Genomics UK (COG-UK) Consortium | Claire McMurray, Joanne Stockton, Samuel Nicholls, Radoslaw Poplawski, Will Rowe, Josh Quick, Nicholas Loman, Celina M Whalley, Andrew Bosworth, Charlotte Poxon, Kasun Wanigasooriya, Oliver Pickles, Mike Kidd, Alex Richter, Andrew D Beggs, Husam Osman, Andrew Bosworth |
| EPI_ISL_447945, EPI_ISL_447946, EPI_ISL_447947, EPI_ISL_447948, EPI_ISL_447949, EPI_ISL_447950, EPI_ISL_447951, EPI_ISL_447952, EPI_ISL_447953, EPI_ISL_447954, EPI_ISL_447955, EPI_ISL_447956, EPI_ISL_447957, EPI_ISL_447958, EPI_ISL_447959, EPI_ISL_447960, EPI_ISL_447961, EPI_ISL_447962, EPI_ISL_447963, EPI_ISL_447964, EPI_ISL_447965, EPI_ISL_447966, EPI_ISL_447967, EPI_ISL_447968, EPI_ISL_447969, EPI_ISL_447970, EPI_ISL_447971, EPI_ISL_447972, EPI_ISL_447973, EPI_ISL_447974, EPI_ISL_447975, EPI_ISL_447976, EPI_ISL_447977, EPI_ISL_447978, EPI_ISL_447979, EPI_ISL_447980, EPI_ISL_447981, EPI_ISL_447982, EPI_ISL_447983, EPI_ISL_447987, EPI_ISL_447988, EPI_ISL_447989, EPI_ISL_447990, EPI_ISL_447991, EPI_ISL_447992, EPI_ISL_447993, EPI_ISL_447994, EPI_ISL_447995, EPI_ISL_447996, EPI_ISL_447997, EPI_ISL_447998, EPI_ISL_447999, EPI_ISL_448000, EPI_ISL_448001, EPI_ISL_448002, EPI_ISL_448003, EPI_ISL_448004, EPI_ISL_448005, EPI_ISL_448006, EPI_ISL_448007, EPI_ISL_448008, EPI_ISL_448009, EPI_ISL_448010, EPI_ISL_448011, EPI_ISL_448012, EPI_ISL_448013, EPI_ISL_448015, EPI_ISL_448016, EPI_ISL_448017, EPI_ISL_448018, EPI_ISL_448019, EPI_ISL_448020, EPI_ISL_448021, EPI_ISL_448022, EPI_ISL_448023, EPI_ISL_448024, EPI_ISL_448025, EPI_ISL_448026, EPI_ISL_448027, EPI_ISL_448028, EPI_ISL_448029, EPI_ISL_448031, EPI_ISL_448032, EPI_ISL_448033, EPI_ISL_448034, EPI_ISL_448035, EPI_ISL_448036, EPI_ISL_448037, EPI_ISL_448039, EPI_ISL_448040, EPI_ISL_448041, EPI_ISL_448042, EPI_ISL_448043, EPI_ISL_448044, EPI_ISL_448045, EPI_ISL_448046, EPI_ISL_448047, EPI_ISL_448051, EPI_ISL_448052, EPI_ISL_448054, EPI_ISL_448055, EPI_ISL_448056, EPI_ISL_448057, EPI_ISL_448058, EPI_ISL_448059, EPI_ISL_448060, EPI_ISL_448063, EPI_ISL_448064, EPI_ISL_448104 | | | |
| see above | Department of Pathology, University of Cambridge | COVID-19 Genomics UK (COG-UK) Consortium | Luke W Meredith, M. Estée Török , Myra Hosmillo, William L. Hamilton, Martin D. Curran, Theresa Feltwell, Grant Hall, Anna Yakovleva, Fahad A Khokhar, Charlotte J. Houldcroft, Laura G Caller, Aminu S. Jahun, Sarah L. Caddy, Ian Goodfellow |
| EPI_ISL_448349, EPI_ISL_448350, EPI_ISL_448351, EPI_ISL_448352, EPI_ISL_448353, EPI_ISL_448354, EPI_ISL_448355, EPI_ISL_448356, EPI_ISL_448357, EPI_ISL_448358, EPI_ISL_448359, EPI_ISL_448360, EPI_ISL_448361, EPI_ISL_448362, EPI_ISL_448363, EPI_ISL_448364, EPI_ISL_448365, EPI_ISL_448366, EPI_ISL_448367, EPI_ISL_448368, EPI_ISL_448369, EPI_ISL_448370, EPI_ISL_448371, EPI_ISL_448372, EPI_ISL_448373, EPI_ISL_448374, EPI_ISL_448375, EPI_ISL_448376, EPI_ISL_448377, EPI_ISL_448378, EPI_ISL_448379, EPI_ISL_448380, EPI_ISL_448381, EPI_ISL_448382, EPI_ISL_448383, EPI_ISL_448384, EPI_ISL_448385, EPI_ISL_448386, EPI_ISL_448387, EPI_ISL_448388 | | | |
| see above | Quadram Institute Bioscience | COVID-19 Genomics UK (COG-UK) Consortium | Dave J. Baker, Gemma L. Kay, Alp Aydin, Thanh Le-Viet, Steven Rudder, Ana P. Tedim, Anastasia Kolyva, Maria Diaz, Leonardo de Oliveira Martins, Nabil-Fareed Alikhan, Lizzie Meadows, Rachael Stanley, Ngozi Elumogo, Muhammed Yasir, Nicholas M. Thomson, Alexander J Trotter, Rachel Gilroy, Samuel Bloomfield, Claire Stuart, Andrew Bell, Reenesh Prakash, Samir Dervisevic, Alison E. Mather, John Wain, Mark Webber, Andrew J. Page, Justin O'Grady |
| EPI_ISL_448389, EPI_ISL_448390, EPI_ISL_448391, EPI_ISL_448392, EPI_ISL_448393, EPI_ISL_448394, EPI_ISL_448395, EPI_ISL_448396, EPI_ISL_448397, EPI_ISL_448398, EPI_ISL_448399, EPI_ISL_448400, EPI_ISL_448401, EPI_ISL_448402, EPI_ISL_448403, EPI_ISL_448404, EPI_ISL_448405, EPI_ISL_448406, EPI_ISL_448407, EPI_ISL_448408, EPI_ISL_448409, EPI_ISL_448410, EPI_ISL_448411, EPI_ISL_448412, EPI_ISL_448413, EPI_ISL_448414, EPI_ISL_448419, EPI_ISL_448420, EPI_ISL_448421, EPI_ISL_448422, EPI_ISL_448423, EPI_ISL_448424, EPI_ISL_448425, EPI_ISL_448426, EPI_ISL_448427, EPI_ISL_448428, EPI_ISL_448429, EPI_ISL_448430, EPI_ISL_448431, EPI_ISL_448432, EPI_ISL_448433, EPI_ISL_448447, EPI_ISL_448448 | | | |
| see above | Queens Medical Centre, Clinical Microbiology Department / DeepSeq Nottingham | COVID-19 Genomics UK (COG-UK) Consortium | Gemma Clark, Wendy Smith, Manjinder Khakh, Hannah Howson-Wells, Jonathan Ball, Patrick McClure, Joseph Chappell, Theocharis Tsoleridis, Nadine Holmes, Matthew Carlisle, Christopher Moore, Fei Sang, Johnny Debebe, Victoria Wright, Matthew Loose |
| EPI_ISL_449033, EPI_ISL_449034, EPI_ISL_449035, EPI_ISL_449036, EPI_ISL_449037, EPI_ISL_449038, EPI_ISL_449039, EPI_ISL_449040, EPI_ISL_449041, EPI_ISL_449042, EPI_ISL_449043, EPI_ISL_449044, EPI_ISL_449045, EPI_ISL_449046, EPI_ISL_449047, EPI_ISL_449048, EPI_ISL_449049, EPI_ISL_449050, EPI_ISL_449051, EPI_ISL_449052, EPI_ISL_449053, EPI_ISL_449054, EPI_ISL_449055, EPI_ISL_449056, EPI_ISL_449057, EPI_ISL_449058, EPI_ISL_449059, EPI_ISL_449060, EPI_ISL_449061, EPI_ISL_449062, EPI_ISL_449063, EPI_ISL_449064, EPI_ISL_449065, EPI_ISL_449066, EPI_ISL_449067, EPI_ISL_449068, EPI_ISL_449069, EPI_ISL_449070, EPI_ISL_449072, EPI_ISL_449086, EPI_ISL_449088 | | | |
| see above | Quadram Institute Bioscience | COVID-19 Genomics UK (COG-UK) Consortium | Dave J. Baker, Gemma L. Kay, Alp Aydin, Thanh Le-Viet, Steven Rudder, Ana P. Tedim, Anastasia Kolyva, Maria Diaz, Leonardo de Oliveira Martins, Nabil-Fareed Alikhan, Lizzie Meadows, Rachael Stanley, Ngozi Elumogo, Muhammed Yasir, Nicholas M. Thomson, Alexander J Trotter, Rachel Gilroy, Samuel Bloomfield, Claire Stuart, Andrew Bell, Reenesh Prakash, Samir Dervisevic, Alison E. Mather, John Wain, Mark Webber, Andrew J. Page, Justin O'Grady |
| EPI_ISL_449186, EPI_ISL_449187, EPI_ISL_449199, EPI_ISL_449200, EPI_ISL_449201, EPI_ISL_449206, EPI_ISL_449207, EPI_ISL_449208, EPI_ISL_449209, EPI_ISL_449210, EPI_ISL_449211, EPI_ISL_449212, EPI_ISL_449213, EPI_ISL_449214, EPI_ISL_449215, EPI_ISL_449216, EPI_ISL_449217, EPI_ISL_449218, EPI_ISL_449219, EPI_ISL_449220, EPI_ISL_449221, EPI_ISL_449222, EPI_ISL_449223, EPI_ISL_449249, EPI_ISL_449250, EPI_ISL_449251, EPI_ISL_449252, EPI_ISL_449253, EPI_ISL_449254, EPI_ISL_449255, EPI_ISL_449256, EPI_ISL_449257, EPI_ISL_449258 | | | |
| see above | West of Scotland Specialist Virology Centre, NHSGGC / MRC-University of Glasgow Centre for Virus Research | COVID-19 Genomics UK (COG-UK) Consortium | Ana da Silva Filipe, Natasha Johnson, Kathy Smollett, Daniel Mair, Stephen Carmichael, Lily Tong, Jenna Nichols, Elihu Aranday-Cortes, Kirstyn Brunker, Yasmin Parr, Kyriaki Nomikou, Sarah McDonald, Marc Niebel, Patawee Asamaphan, Richard Orton, Joseph Hughes, Sreenu Vattipally, David L Robertson, Alasdair MacLean, Rory Gunson, Kathy Li, Natasha Jesudason, Rajiv Shah, James Shepherd, Antonia Ho, Emma Thomson |

| | | | |
|---|---|---|---|
| EPI_ISL_449259, EPI_ISL_449260, EPI_ISL_449261, EPI_ISL_449262, EPI_ISL_449263, EPI_ISL_449264, EPI_ISL_449265, EPI_ISL_449266, EPI_ISL_449267, EPI_ISL_449268, EPI_ISL_449269, EPI_ISL_449270, EPI_ISL_449271, EPI_ISL_449272, EPI_ISL_449303, EPI_ISL_449304, EPI_ISL_449305, EPI_ISL_449306, EPI_ISL_449307, EPI_ISL_449308, EPI_ISL_449309, EPI_ISL_449310, EPI_ISL_449324, EPI_ISL_449325, EPI_ISL_449326, EPI_ISL_449327, EPI_ISL_449328, EPI_ISL_449329 | | | |
| see above | Virology Department, Royal Infirmary of Edinburgh, NHS Lothian / School of Biological Sciences, University of Edinburgh / Institute of Genetics and Molecular Medicine, University of Edinburgh | COVID-19 Genomics UK (COG-UK) Consortium | McHugh M, Dewar R, Rooke S, Gallagher M, Balcaza C, O'Toole Á, Scher E, Hill V, McCrone JT, Colquhoun R, Yu X, Jackson B, Rambaut A, Williams TC, Templeton K |
| EPI_ISL_449604, EPI_ISL_449605, EPI_ISL_449606, EPI_ISL_449607, EPI_ISL_449608, EPI_ISL_449609, EPI_ISL_449610, EPI_ISL_449611, EPI_ISL_449612, EPI_ISL_449613, EPI_ISL_449614, EPI_ISL_449615, EPI_ISL_449616, EPI_ISL_449617, EPI_ISL_449618, EPI_ISL_449619, EPI_ISL_449620, EPI_ISL_449621, EPI_ISL_449622, EPI_ISL_449623, EPI_ISL_449624 | | | |
| see above | Liverpool Clinical Laboratories | COVID-19 Genomics UK (COG-UK) Consortium | Sam Haldenby, Anita Lucaci, Steve Paterson, Julian Hiscox, Alistair Darby, M Almsaud, A Alrezaihi, Muhannad Alruwaili, Stuart D Armstrong, Jones Benjamin , Eleanor G Bentley, Anu Chawla, Jordan J Clark, Angela Cowell, Richard Eccles, Isabel García-Dorival, Matthew Gemmell, Alessandro Gerada, PKF Gilmore, Richard Gregory, Ximeng Han, Catherine Hartley, Margaret Hughes, Miren Iturriza-Gomara, James Johnson, L Luu, Jenifer Manson , Charlotte Nelson, Elaine O'Toole, Cassie Olateju, Rebekah Penrice-Randal , Lucille Rainbow, N.P Randle, Trevor Ian Robinson, Parul Sharma, Ghada T Shawli, James P Stewart , Neil Swainston, Ecaterina Vamos, Joanne Watts, Mark Whitehead |
| EPI_ISL_450344 | Bangladesh Institute of Tropical & Infectious Diseases, COVID-19 Testing Laboratory | Basic and Applied Research on Jute Project | Rasel Ahmed, Md. Sabbir Hossain, Shah Md Tamim Kabir, Emdadul Mannan Emdad, Md. Nazmul Haq Rony, Eaftekhar Ahmed Rana, Paritous Kumar Biswas, M A Hassan Chowdhury, Md. Shakeel Ahmed, Md. Samiul Haque, Md. Monjurul Alam, Md. Sharifur Rahman, A S M Anwarul Huq, Md. Shahidul Islam, Goutam Buddha Das, AMAM Zonaed Siddiki |
| EPI_ISL_450520, EPI_ISL_450521, EPI_ISL_450522, EPI_ISL_450523 | Centrl Laboratorija | Latvian Biomedical Research and Study Centre | Ivars Silamielis, Kaspars Megnis, Monta Ustinova, ikita Zrelovs, Vita Rovte, Stella Lapia, Jana Oste, Marta Priedte, Uga Dumpis, Jnis Kloviš |
| EPI_ISL_450600 | Michigan Department of Health and Human Services, Bureau of Laboratories | Michigan Department of Health and Human Services, Bureau of Laboratories | Blankenship HM; Riner D; Soehnlen MK |
| EPI_ISL_450790 | Pandit Deendayal Upadhyay Government Medical College, Rajkot | Gujarat Biotechnology Research Centre | Zarna Patel, Pritesh Sabara, Apurvasinh Puvar, Janvi Raval, Monika Gandhi, Pinal Trivedi, Maharshi Pandya, Amit Kanani, Nidhi Patel, Nitin Savaliya, Raghawendra Kumar, Dinesh Kumar, Zuber Saiyed, Komal Patel, Labdhi Pandya, Snehal Bagatharia, Prakash Modi, Sejul Antala, Manish Pattani, Ramesh Pandit, Tejas Shah, Ankit Hinsu, Bhavesh Modi, Gaurishankar Shrimali, R D Dixit, A M Kadri, Sharmistha Majumdar, Chaitanya Joshi, Madhvi Joshi |
| EPI_ISL_450791 | Pandit Deendayal Upadhyay Government Medical College, Rajkot | Gujarat Biotechnology Research Centre | Zarna Patel, Apurvasinh Puvar, Janvi Raval, Monika Gandhi, Pinal Trivedi, Maharshi Pandya, Amit Kanani, Nidhi Patel, Nitin Savaliya, Raghawendra Kumar, Dinesh Kumar, Zuber Saiyed, Komal Patel, Labdhi Pandya, Snehal Bagatharia, Prakash Modi, Sejul Antala, Manish Pattani, Ramesh Pandit, Tejas Shah, Ankit Hinsu, Pritesh Sabara, Bhavesh Modi, Gaurishankar Shrimali, R D Dixit, A M Kadri, Pooja P Doshi, Chaitanya Joshi, Madhvi Joshi |
| EPI_ISL_450825 | Narhalsan Backa vardcentral | The Public Health Agency of Sweden | Mats Olsson, Anna-Malin Linde, Maria Lind Karlberg, Oskar Karlsson Lindsjo, Olov Svartstrom, Anna Risberg, Theresa Enkirch, Mia Brytting, Karin Tegmark-Wisell |
| EPI_ISL_450826, EPI_ISL_450827, EPI_ISL_450828 | Uppsala Narakut Aleris | The Public Health Agency of Sweden | Annika Nilsson, Anna-Malin Linde, Maria Lind Karlberg, Oskar Karlsson Lindsjo, Olov Svartstrom, Anna Risberg, Theresa Enkirch, Mia Brytting, Karin Tegmark-Wisell |
| EPI_ISL_450829 | Narhalsan Sjobo vardcentral | The Public Health Agency of Sweden | Lovisa Hjerten, Anna-Malin Linde, Maria Lind Karlberg, Oskar Karlsson Lindsjo, Olov Svartstrom, Anna Risberg, Theresa Enkirch, Mia Brytting, Karin Tegmark-Wisell |
| EPI_ISL_450830 | Narhalsan Olskroken VC | The Public Health Agency of Sweden | Mahin Ghoroghi, Anna-Malin Linde, Maria Lind Karlberg, Oskar Karlsson Lindsjo, Olov Svartstrom, Anna Risberg, Theresa Enkirch, Mia Brytting, Karin Tegmark-Wisell |
| EPI_ISL_450831 | Wetterhalsan | The Public Health Agency of Sweden | Anders Tengblad, Anna-Malin Linde, Maria Lind Karlberg, Oskar Karlsson Lindsjo, Olov Svartstrom, Anna Risberg, Theresa Enkirch, Mia Brytting, Karin Tegmark-Wisell |
| EPI_ISL_450832 | Byjorden vardcentral | The Public Health Agency of Sweden | Pernilla Brunman, Anna-Malin Linde, Maria Lind Karlberg, Oskar Karlsson Lindsjo, Olov Svartstrom, Anna Risberg, Theresa Enkirch, Mia Brytting, Karin Tegmark-Wisell |
| EPI_ISL_451154 | Government Medical College, Vadodara | Gujarat Biotechnology Research Centre | Manish Pattani, Tanuja Javadekar , Maharshi Pandya, Amit Kanani, Nidhi Patel, Nitin Savaliya, Raghawendra Kumar, Dinesh Kumar, Zuber Saiyed, Komal Patel, Labdhi Pandya, Snehal Bagatharia, Ramesh Pandit, Tejas Shah, Ankit Hinsu, Pritesh Sabara, Apurvasinh Puvar, Janvi Raval, Zarna Patel, Monika Gandhi, Pinal Trivedi, Bhavesh Modi, Gaurishankar Shrimali, R D Dixit, A M Kadri, Neelam Nathani, Chaitanya Joshi, Madhvi Joshi |
| EPI_ISL_451155 | Government Medical College, Vadodara | Gujarat Biotechnology Research Centre | Tanuja Javadekar , Manish Pattani, Amit Kanani, Nidhi Patel, Nitin Savaliya, Raghawendra Kumar, Dinesh Kumar, Zuber Saiyed, Komal Patel, Labdhi Pandya, Snehal Bagatharia, Ramesh Pandit, Tejas Shah, Ankit Hinsu, Pritesh Sabara, Apurvasinh Puvar, Janvi Raval, Zarna Patel, Monika Gandhi, Pinal Trivedi, Maharshi Pandya, Bhavesh Modi, Gaurishankar Shrimali, R D Dixit, A M Kadri, Armi Chaudhari, Chaitanya Joshi, Madhvi Joshi |
| EPI_ISL_451156 | Government Medical College, Vadodara | Gujarat Biotechnology Research Centre | Amit Kanani, Nidhi Patel, Nitin Savaliya, Raghawendra Kumar, Dinesh Kumar, Zuber Saiyed, Komal Patel, Labdhi Pandya, Snehal Bagatharia, Ramesh Pandit, Tejas Shah, Ankit Hinsu, Pritesh Sabara, Apurvasinh Puvar, Janvi Raval, Zarna Patel, Monika Gandhi, Pinal Trivedi, Maharshi Pandya, Manish Pattani, Tanuja Javadekar , Bhavesh Modi, Gaurishankar Shrimali, R D Dixit, A M Kadri, Bhavya Jindal, Chaitanya Joshi, Madhvi Joshi |
| EPI_ISL_451157 | Government Medical College, Vadodara | Gujarat Biotechnology Research Centre | Nidhi Patel, Nitin Savaliya, Raghawendra Kumar, Dinesh Kumar, Zuber Saiyed, Komal Patel, Labdhi Pandya, Snehal Bagatharia, Ramesh Pandit, Tejas Shah, Ankit Hinsu, Pritesh Sabara, Apurvasinh Puvar, Janvi Raval, Zarna Patel, Monika Gandhi, Pinal Trivedi, Maharshi Pandya, Manish Pattani, Tanuja Javadekar , Amit Kanani, Bhavesh Modi, Gaurishankar Shrimali, R D Dixit, A M Kadri, Camellia Chakraborty, Chaitanya Joshi, Madhvi Joshi |
| EPI_ISL_451158 | Government Medical College, Vadodara | Gujarat Biotechnology Research Centre | Nitin Savaliya, Raghawendra Kumar, Dinesh Kumar, Zuber Saiyed, Komal Patel, Labdhi Pandya, Snehal Bagatharia, Ramesh Pandit, Tejas Shah, Ankit Hinsu, Pritesh Sabara, Apurvasinh Puvar, Janvi Raval, Zarna Patel, Monika Gandhi, Pinal Trivedi, Maharshi Pandya, Manish Pattani, Tanuja Javadekar , Amit Kanani, Nidhi Patel, Bhavesh Modi, Gaurishankar Shrimali, R D Dixit, A M Kadri, Siddhant Kumar, Chaitanya Joshi, Madhvi Joshi |
| EPI_ISL_451159 | Government Medical College, Vadodara | Gujarat Biotechnology Research Centre | Raghawendra Kumar, Dinesh Kumar, Zuber Saiyed, Komal Patel, Labdhi Pandya, Snehal Bagatharia, Ramesh Pandit, Tejas Shah, Ankit Hinsu, Pritesh Sabara, Apurvasinh Puvar, Janvi Raval, Zarna Patel, Monika Gandhi, Pinal Trivedi, Maharshi Pandya, Manish Pattani, Tanuja Javadekar , Amit Kanani, Nidhi Patel, Nitin Savaliya, Bhavesh Modi, Gaurishankar Shrimali, R D Dixit, A M Kadri, Sharmistha Majumdar, Chaitanya Joshi, Madhvi Joshi |
| EPI_ISL_451160 | Government Medical College, Vadodara | Gujarat Biotechnology Research Centre | Dinesh Kumar, Zuber Saiyed, Komal Patel, Labdhi Pandya, Snehal Bagatharia, Ramesh Pandit, Tejas Shah, Ankit Hinsu, Pritesh Sabara, Apurvasinh Puvar, Janvi Raval, Zarna Patel, Monika Gandhi, Pinal Trivedi, Maharshi Pandya, Manish Pattani, Tanuja Javadekar , Amit Kanani, Nidhi Patel, Nitin Savaliya, Raghawendra Kumar, Bhavesh Modi, Gaurishankar Shrimali, R D Dixit, A M Kadri, Pooja P Doshi, Chaitanya Joshi, Madhvi Joshi |
| EPI_ISL_451161 | Government Medical College, Vadodara | Gujarat Biotechnology Research Centre | Zuber Saiyed, Komal Patel, Labdhi Pandya, Snehal Bagatharia, Ramesh Pandit, Tejas Shah, Ankit Hinsu, Pritesh Sabara, Apurvasinh Puvar, Janvi Raval, Zarna Patel, Monika Gandhi, Pinal Trivedi, Maharshi Pandya, Manish Pattani, Tanuja Javadekar , Amit Kanani, Nidhi Patel, Nitin Savaliya, Raghawendra Kumar, Dinesh Kumar, Bhavesh Modi, Gaurishankar Shrimali, R D Dixit, A M Kadri, Akanksha Verma, Chaitanya Joshi, Madhvi Joshi |
| EPI_ISL_451162 | Government Medical College, Vadodara | Gujarat Biotechnology Research Centre | Komal Patel, Labdhi Pandya, Snehal Bagatharia, Ramesh Pandit, Tejas Shah, Ankit Hinsu, Pritesh Sabara, Apurvasinh Puvar, Janvi Raval, Zarna Patel, Monika Gandhi, Pinal Trivedi, Maharshi Pandya, Manish Pattani, Tanuja Javadekar , Amit Kanani, Nidhi Patel, Nitin Savaliya, Raghawendra Kumar, Dinesh Kumar, Zuber Saiyed, Bhavesh Modi, Gaurishankar Shrimali, R D Dixit, A M Kadri, Priti Pandita, Chaitanya Joshi, Madhvi Joshi |
| EPI_ISL_451163 | Government Medical College, Vadodara | Gujarat Biotechnology Research Centre | Labdhi Pandya, Snehal Bagatharia, Ramesh Pandit, Tejas Shah, Ankit Hinsu, Pritesh Sabara, Apurvasinh Puvar, Janvi Raval, Zarna Patel, Monika Gandhi, Pinal Trivedi, Maharshi Pandya, Manish Pattani, Tanuja Javadekar , Amit Kanani, Nidhi Patel, Nitin Savaliya, Raghawendra Kumar, Dinesh Kumar, Zuber Saiyed, Komal Patel, Bhavesh Modi, Gaurishankar Shrimali, R D Dixit, A M Kadri, Pragya Sharma, Chaitanya Joshi, Madhvi Joshi |
| EPI_ISL_451197, EPI_ISL_451198, EPI_ISL_451200, EPI_ISL_451201, EPI_ISL_451202 | Uganda Virus Research Institute | MRC/UVRI & LSHTM Uganda Research Unit | Dan Lule Bugembe, John Kayiwa, My V.T Phan, Phionah Tushabe, Stephen Balinandi, Beatrice Dhaala, Deogratius Ssemwanga, Jonas Lexow, Henry Mwebesa, Jane Aceng, Henry Kyobe, Julius Lutwama, Pontiano Kaleebu, Matthew Cotten |
| EPI_ISL_451203, EPI_ISL_451205, EPI_ISL_451206, EPI_ISL_451207, EPI_ISL_451208, EPI_ISL_451209, EPI_ISL_451210, EPI_ISL_451213, EPI_ISL_451214, EPI_ISL_451216, EPI_ISL_451217, EPI_ISL_451218 | | | |
| see above | LSUHS Emerging Viral Threat Laboratory | Microbial Genome Sequencing Center | Jeremy P. Kamil, John A. Vanchiere, Rona S. Scott, Camille F. Abshire, Abida Siddiqa, Byeong-Jae Lee, Chan-ki Min, Md Maksudul Alam, Monica Gestal-Cartele, Edna Ondari, Adam Greer, Malgorzata Bienkowska-Haba, Katarzyna Zwolinska, Jason M. Bodily, Andrew D. Yurochko, Paul M. |

| | | | |
|---|---|---|---|
| | | | Weinberger, Christopher G. Kevil, Martin J. Sapp, Daniel J. Snyder, Vaughn S. Cooper |
| EPI_ISL_451219, EPI_ISL_451220, EPI_ISL_451222, EPI_ISL_451223, EPI_ISL_451224, EPI_ISL_451225, EPI_ISL_451226, EPI_ISL_451227, EPI_ISL_451228, EPI_ISL_451229 | LSUHS Emerging Viral Threat Laboratory | Microbial Genome Sequencing Center | Rona S. Scott, Jeremy P. Kamil, John A. Vanchiere, Camille F. Abshire, Abida Siddiqa, Byeong-Jae Lee, Chan-ki Min, Md Maksudul Alam, Monica Gestal-Cartele, Edna Ondari, Adam Greer, Malgorzata Bienkowska-Haba, Katarzyna Zwolinska, Jason M. Bodily, Andrew D. Yurochko, Paul M. Weinberger, Christopher G. Kevil, Martin J. Sapp, Daniel J. Snyder, Vaughn S. Cooper |
| EPI_ISL_451230, EPI_ISL_451231, EPI_ISL_451232, EPI_ISL_451233, EPI_ISL_451234, EPI_ISL_451235, EPI_ISL_451236, EPI_ISL_451239, EPI_ISL_451240, EPI_ISL_451241, EPI_ISL_451246, EPI_ISL_451247 | | | |
| see above | LSUHS Emerging Viral Threat Laboratory | Microbial Genome Sequencing Center | John A. Vanchiere, Jeremy P. Kamil, Rona S. Scott, Camille F. Abshire, Abida Siddiqa, Byeong-Jae Lee, Chan-ki Min, Md Maksudul Alam, Monica Gestal-Cartele, Edna Ondari, Adam Greer, Malgorzata Bienkowska-Haba, Katarzyna Zwolinska, Jason M. Bodily, Andrew D. Yurochko, Paul M. Weinberger, Christopher G. Kevil, Martin J. Sapp, Daniel J. Snyder, Vaughn S. Cooper |
| EPI_ISL_451644, EPI_ISL_451645 | Laboratory of Molecular Biology, Diagnostyka sp. z o.o. | Laboratory of Recombinant Vaccines | Lukasz Rabalski, Anna Piotrowska-Mietelska, Maciej Kosinski, Boguslaw Szewczyk, Krystyna Bienkowska-Szewczyk |
| EPI_ISL_451649, EPI_ISL_451650, EPI_ISL_451651, EPI_ISL_451652, EPI_ISL_451653, EPI_ISL_451654 | Hematology Laboratory, Section of Molecular Diagnostics, University Clinical Centre, Medical University of Gdansk | Laboratory of Recombinant Vaccines | Lukasz Rabalski, Adam Sodol, Aneta Szulc, Krzysztof Lewandowski, Ewa Milosz, Marlena Robakowska, Boguslaw Szewczyk, Krystyna Bienkowska-Szewczyk |
| EPI_ISL_451666 | M.P Shah Government Medocal college Jamnagar | Gujarat Biotechnology Research Centre | Binita Aring, Janvi Raval, Zarna Patel, Monika Gandhi, Pinal Trivedi, Maharshi Pandya, Amit Kanani, Nidhi Patel, Nitin Savaliya, Raghawendra Kumar, Dinesh Kumar, Zuber Saiyed, Komal Patel, Labdhi Pandya, Snehal Bagatharia, Ramesh Pandit, Tejas Shah, Ankit Hinsu, Pritesh Sabara, Apurvasinh Puvar, Bhavesh Modi, Gaurishankar Shrimali, R D Dixit, A M Kadri, Akanksha Verma, Chaitanya Joshi, Madhvi Joshi, |
| EPI_ISL_452061, EPI_ISL_452062, EPI_ISL_452063, EPI_ISL_452064, EPI_ISL_452065, EPI_ISL_452066, EPI_ISL_452067, EPI_ISL_452068, EPI_ISL_452069, EPI_ISL_452070, EPI_ISL_452071, EPI_ISL_452072, EPI_ISL_452073, EPI_ISL_452074, EPI_ISL_452075, EPI_ISL_452076, EPI_ISL_452077, EPI_ISL_452078, EPI_ISL_452081 | | | |
| see above | Department of Clinical Microbiology, Copenhagen University Hospital, Hvidovre, Kettegaard Alle 30, 2650 Hvidovre. | Albertsen lab, Department of Chemistry and Bioscience, Aalborg University, Denmark | Rasmus Kirkegaard |
| EPI_ISL_452137 | VI-US Virgin Islands Department of Health | Pathogen Discovery, Respiratory Viruses Branch, Division of Viral Diseases, Centers for Disease Control and Prevention | Anna Uehara, Yan Li, Anna Montmayeur, Ying Tao, Krista Queen, Jing Zhang, Clinton R. Paden, Rachel Marine, Haibin Wang, Bettina Bankamp, Zachary Weiner, Suxiang Tong |
| EPI_ISL_452144, EPI_ISL_452146, EPI_ISL_452147 | Yale COVID-19 Biorepository | Grubaugh Lab - Yale School of Public Health | Joseph Fauver, Tara Alpert, Anderson Brito, Anne Wyllie, Chantal Vogels, Mary Petrone, Cole Jensen, Chaney Kalinich, Isabel Ott, Arnau Casanovas, Catherine Muenker, Adam Moore, Alice Lu, Maria Tokuyama, Patrick Wong, Peiwen Lu, Saad Omer, Richard Martinello, Allison Nelson, Shelli Farhadian, Akiko Iwasaki, Charlese Dela Cruz, Albert Ko, Nathan Grubaugh |
| EPI_ISL_452216 | NIV Influenza | NIV Influenza | Potdar V |
| EPI_ISL_452238 | Narhalsan Sjobo vardcentral | The Public Health Agency of Sweden | Lovisa Hjerten, Anna-Malin Linde, Maria Lind Karlberg, Oskar Karlsson Lindsjo, Olov Svartstrom, Anna Risberg, Theresa Enkirch, Mia Brytting, Karin Tegmark-Wisell |
| EPI_ISL_452242 | Wernstedt Medical AB | The Public Health Agency of Sweden | Eva Sandberg, Anna-Malin Linde, Maria Lind Karlberg, Oskar Karlsson Lindsjo, Olov Svartstrom, Anna Risberg, Theresa Enkirch, Mia Brytting, Karin Tegmark-Wisell |
| EPI_ISL_452262, EPI_ISL_452263, EPI_ISL_452276, EPI_ISL_452277, EPI_ISL_452278, EPI_ISL_452279, EPI_ISL_452280, EPI_ISL_452281, EPI_ISL_452282, EPI_ISL_452283, EPI_ISL_452284 | | | |
| see above | Michigan Department of Health and Human Services, Bureau of Laboratories | Michigan Department of Health and Human Services, Bureau of Laboratories | Blankenship HM, Riner D, Soehnlen MK |
| EPI_ISL_452791, EPI_ISL_452792, EPI_ISL_452793 | ICAR-National Institute of High Security Animal Diseases | ICAR-National Institute of High Security Animal Diseases | Anamika Mishra, Ashutosh Aasdev, Sandeep Bhatia, Harshad Murugkar, Chakradhar Tosh, Niranjan Mishra, Shanmugasundaram Nagarajan, Katherukamem Rajukumar, Richa Sood, G Venkatesh, Atul Kumar Pateriya, Manoj Kumar, Shashi Bhushan Sudhakar, Fateh Singh, Sethil Kumar D, Senmannan Kalaiyarasu, Pradeep Gandhale, Naveen Kumar, Chandan Kumar Dubey, Sushil Tripathi, Sandeep Kumar Jhade, Meghna Tripathi, Suman Kumari Shah, Pushpendra Singh, Pushpendra Namdeo, Suman Mishra, Rupal Singh, Vishnupriya Patil, Dipesh Kumar Nayak, Vijendra Pal Singh, Ashwin Ashok Raut |
| EPI_ISL_453097 | West of Scotland Specialist Virology Centre, NHSGGC / MRC-University of Glasgow Centre for Virus Research | COVID-19 Genomics UK (COG-UK) Consortium | Ana da Silva Filipe, Natasha Johnson, Kathy Smollett, Daniel Mair, Stephen Carmichael, Lily Tong, Jenna Nichols, Elihu Aranday-Cortes, Kirstyn Brunker, Yasmin Parr, Kyriaki Nomikou; Sarah McDonald, Marc Niebel, Patawee Asamaphan; Richard Orton, Joseph Hughes, Sreenu Vattipally, David L Robertson; Alasdair MacLean, Rory Gunson; Kathy Li, Natasha Jesudason, Rajiv Shah, James Shepherd, Antonia Ho, Emma Thomson |
| EPI_ISL_453101, EPI_ISL_453133, EPI_ISL_453134, EPI_ISL_453135, EPI_ISL_453136, EPI_ISL_453137, EPI_ISL_453138, EPI_ISL_453139, EPI_ISL_453140 | Virology Department, Royal Infirmary of Edinburgh, NHS Lothian / School of Biological Sciences, University of Edinburgh / Institute of Genetics and Molecular Medicine, University of Edinburgh | COVID-19 Genomics UK (COG-UK) Consortium | McHugh M, Dewar R, Rooke S, Gallagher M, Balcaza C, O'Toole Á, Scher E, Hill V, McCrone JT, Colquhoun R, Yu X, Jackson B, Rambaut A, Williams TC, Templeton K |
| EPI_ISL_453195, EPI_ISL_453433, EPI_ISL_453434, EPI_ISL_453435, EPI_ISL_453436, EPI_ISL_453437, EPI_ISL_453438, EPI_ISL_453439, EPI_ISL_453440, EPI_ISL_453441, EPI_ISL_453442, EPI_ISL_453443, EPI_ISL_453444, EPI_ISL_453445, EPI_ISL_453446, EPI_ISL_453447, EPI_ISL_453448, EPI_ISL_453449, EPI_ISL_453450, EPI_ISL_453451, EPI_ISL_453452, EPI_ISL_453453, EPI_ISL_453454, EPI_ISL_453455, EPI_ISL_453456, EPI_ISL_453457, EPI_ISL_453459 | | | |
| see above | Liverpool Clinical Laboratories | COVID-19 Genomics UK (COG-UK) Consortium | Sam Haldenby, Anita Lucaci, Steve Paterson, Julian Hiscox, Alistair Darby, M Almsaud, A Alrezaihi, Muhannad Alruwaili, Stuart D Armstrong, Jones Benjamin , Eleanor G Bentley, Anu Chawla, Jordan J Clark, Angela Cowell, Richard Eccles, Isabel García-Dorival, Matthew Gemmell, Alessandro Gerada, PKF Gilmore, Richard Gregory, Ximeng Han, Catherine Hartley, Matthew Hughes, Miren Iturriza-Gomara, James Johnson, L Luu, Jenifer Manson , Charlotte Nelson, Elaine O'Toole, Cassie Olateju, Rebekah Penrice-Randal , Lucille Rainbow, N.P Randle, Trevor Ian Robinson, Parul Sharma, Ghada T Shawli, James P Stewart , Neil Swainston, Ecaterina Vamos, Joanne Watts, Mark Whitehead |
| EPI_ISL_453533, EPI_ISL_453534, EPI_ISL_453535, EPI_ISL_453536, EPI_ISL_453537, EPI_ISL_453540, EPI_ISL_453541 | Northumbria University / South Tees Hospitals NHS Foundation Trust / North Cumbria Integrated Care NHS Foundation Trust / North Tees and Hartlepool NHS Foundation Trust / Newcastle Hospitals NHS Foundation Trust | COVID-19 Genomics UK (COG-UK) Consortium | Darren L Smith,Andrew Nelson,Matthew Bashton,Greg R Young,Joshua Loh,John Allan,Mohammad A Tariq,Giles S Holt,Gary Black,Wen C Yew,Lynn Dover ,Paul Baker,Steve Liggett,Sarah Essex,Jane Greenaway ,Debra Padgett,Clive Graham,Garren Scott,Edward Barton ,Emma Swindells ,Brendan Payne,Jennifer Collins,Yusri Taha,Gary Eltringham |
| EPI_ISL_453581, EPI_ISL_453582, EPI_ISL_453585, EPI_ISL_453586, EPI_ISL_453587, EPI_ISL_453588, EPI_ISL_453589, EPI_ISL_453590, EPI_ISL_453591, EPI_ISL_453593, EPI_ISL_453594, EPI_ISL_453596 | | | |
| see above | Quadram Institute Bioscience | COVID-19 Genomics UK (COG-UK) Consortium | Dave J. Baker, Gemma L. Kay, Alp Aydin, Thanh Le-Viet, Steven Rudder, Ana P. Tedim, Anastasia Kolyva, Maria Diaz, Leonardo de Oliveira Martins, Nabil-Fareed Alikhan, Lizzie Meadows, Rachael Stanley, Ngozi Elumogo, Muhammed Yasir, Nicholas M. Thomson, Alexander J Trotter, Rachel Gilroy, Samuel Bloomfield, Claire Stuart, Andrew Bell, Reenesh Prakash, Samir Dervisevic, Alison E. Mather, John Wain, Mark Webber, Andrew J. Page, Justin O'Grady |
| EPI_ISL_453614, EPI_ISL_453615, EPI_ISL_453626, EPI_ISL_453627, EPI_ISL_453628, EPI_ISL_453629, EPI_ISL_453630, EPI_ISL_453631, EPI_ISL_453632, EPI_ISL_453643 | Queens Medical Centre, Clinical Microbiology Department / DeepSeq Nottingham | COVID-19 Genomics UK (COG-UK) Consortium | Gemma Clark, Wendy Smith, Manjinder Khakh, Hannah Howson-Wells, Jonathan Ball, Patrick McClure, Joseph Chappell, Theocharis Tsoleridis, Nadine Holmes, Matthew Carlisle, Christopher Moore, Fei Sang, Johnny Debebe, Victoria Wright, Matthew Loose |
| EPI_ISL_453672, EPI_ISL_453673, EPI_ISL_453674, EPI_ISL_453675, EPI_ISL_453676, EPI_ISL_453677, EPI_ISL_453678, EPI_ISL_453679, EPI_ISL_453680, EPI_ISL_453681, EPI_ISL_453682, EPI_ISL_453683, EPI_ISL_453684, EPI_ISL_453685, EPI_ISL_453686, EPI_ISL_453687, EPI_ISL_453688, EPI_ISL_453689, EPI_ISL_453690, EPI_ISL_453691, EPI_ISL_453692, EPI_ISL_453693, EPI_ISL_453694, EPI_ISL_453695 | | | |
| see above | Centre for Enzyme Innovation, University of Portsmouth / Translational Research Laboratory, Portsmouth Hospitals NHS Trust | COVID-19 Genomics UK (COG-UK) Consortium | Angela Beckett,,Yann Bourgeois,,Garry Scarlett,,Sharon Glaysher,,Scott Elliott,,Kelly Bicknell,,Robert Impey,,Allyson Lloyd,,Sarah Wyllie,,Ethan Butcher,,Anoop Chauhan,,Samuel Robson |
| EPI_ISL_453734 | Virology Department, Sheffield Teaching Hospitals NHS | COVID-19 Genomics UK (COG-UK) Consortium | Thushan de Silva, Matthew Parker, Nikki Smith, Adri Angyal, Rebecca Brown, Luke Green, Rachel Tucker, Paul Parsons, Danielle Groves, Katie Johnson, |

| | | | |
|---|---|---|---|
| | Foundation Trust/Department of Infection, Immunity and Cardiovascular Disease, The Medical School, University of Sheffield | | Laura Carrilero, Alex Keeley, Dave Partridge, Matthew Wyles, Benjamin Lindsey, Mehmet Yavuz, Mohammad Raza, Cariad Evans |
| EPI_ISL_454213, EPI_ISL_454214, EPI_ISL_454215, EPI_ISL_454216, EPI_ISL_454217, EPI_ISL_454218, EPI_ISL_454219, EPI_ISL_454220, EPI_ISL_454221, EPI_ISL_454272, EPI_ISL_454284, EPI_ISL_454285, EPI_ISL_454286, EPI_ISL_454316 | | | |
| see above | unknown | Instituto Nacional de Saude (INSA) | Borges et al |
| EPI_ISL_454585 | Laboratory of virology, National Center of Expertise | Laboratory of molecular-genetic research, National Center for Expertise, Kazakhstan National Center for Biotechnology, Kazakhstan | Abdaliyev Askar, Shevtsov Alexandr, Akhmetollayev Ilyas, Kalendar Ruslan, Rakhmetova Akbota, , Lutsay Viktoriya, Amirgazin Asylulan, Aushakhmetova Zabira, Ramankulov Yerlan |
| EPI_ISL_454614, EPI_ISL_454620, EPI_ISL_454621, EPI_ISL_454622, EPI_ISL_454623, EPI_ISL_454624, EPI_ISL_454625, EPI_ISL_454626, EPI_ISL_454627, EPI_ISL_454628, EPI_ISL_454629, EPI_ISL_454630, EPI_ISL_454631, EPI_ISL_454632, EPI_ISL_454633, EPI_ISL_454634 | | | |
| see above | UCSF Clinical Microbiology Laboratory | Chan-Zuckerberg Biohub | CZB Cliahub Consortium |
| EPI_ISL_454644 | VI-US Virgin Islands Department of Health | Pathogen Discovery, Respiratory Viruses Branch, Division of Viral Diseases, Centers for Disease Control and Prevention | Jing Zhang, Ying Tao, Clinton R. Paden, Anna Uehara, Krista Queen, Yan Li, Haibin Wang, Zachary Weiner, Bettina Bankamp, Suxiang Tong |
| EPI_ISL_454647, EPI_ISL_454648, EPI_ISL_454651, EPI_ISL_454652 | VI-US Virgin Islands Department of Health | Pathogen Discovery, Respiratory Viruses Branch, Division of Viral Diseases, Centers for Disease Control and Prevention | Ying Tao, Clinton R. Paden, Jing Zhang, Anna Uehara, Krista Queen, Yan Li, Haibin Wang, Zachary Weiner, Bettina Bankamp, Suxiang Tong |
| EPI_ISL_454688, EPI_ISL_454689 | County of Santa Clara Public Health Department | Chan-Zuckerberg Biohub | CZB Cliahub Consortium |
| EPI_ISL_454831, EPI_ISL_454833 | SMS Medical College, Jaipur | CSIR Institute of Genomics and Integrative Biology | Sudhir Bhandari, Rahul Bhoyar, Mohammed Imran, Mohit Divakar, Disha Sharma, Anshul Kumar, Bani Jolly, Rahul Sahlot, Abhinav Jain, Paras Sehgal, Gyan Ranjan, Vinod Scaria, Sridhar Sivasubbu, Sandeep K Mathur |
| EPI_ISL_455015 | Pandit Deendayal Upadhyay Government Medical College, Rajkot | Gujarat Biotechnology Research Centre | Snehal Bagatharia, Prakash Modi, Sejul Antala, Manish Pattani, Ramesh Pandit, Tejas Shah, Ankit Hinsu, Pritesh Sabara, Apurvasinh Puvar, Janvi Raval, Zarna Patel, Monika Gandhi, Pinal Trivedi, Maharshi Pandya, Amit Kanani, Nidhi Patel, Nitin Savaliya, Raghawendra Kumar, Dinesh Kumar, Zuber Saiyed, Komal Patel, Labdhi Pandya, Neha Rajpara, Bhavesh Modi, Gaurishankar Shrimali, R D Dixit, A M Kadri, Umang Mishra, Chaitanya Joshi, Madhvi Joshi |
| EPI_ISL_455016 | Pandit Deendayal Upadhyay Government Medical College, Rajkot | Gujarat Biotechnology Research Centre | Prakash Modi, Sejul Antala, Manish Pattani, Ramesh Pandit, Tejas Shah, Ankit Hinsu, Pritesh Sabara, Apurvasinh Puvar, Janvi Raval, Zarna Patel, Monika Gandhi, Pinal Trivedi, Maharshi Pandya, Amit Kanani, Nidhi Patel, Nitin Savaliya, Raghawendra Kumar, Dinesh Kumar, Zuber Saiyed, Komal Patel, Labdhi Pandya, Snehal Bagatharia, Afzal Ansari, Bhavesh Modi, Gaurishankar Shrimali, R D Dixit, A M Kadri, Umang Mishra, Chaitanya Joshi, Madhvi Joshi |
| EPI_ISL_455017 | Government Medical College, Vadodara | Gujarat Biotechnology Research Centre | Tanuja Javadekar , R N Daveshwar, Ramesh Pandit, Tejas Shah, Ankit Hinsu, Pritesh Sabara, Apurvasinh Puvar, Janvi Raval, Zarna Patel, Monika Gandhi, Pinal Trivedi, Maharshi Pandya, Amit Kanani, Nidhi Patel, Nitin Savaliya, Raghawendra Kumar, Dinesh Kumar, Zuber Saiyed, Komal Patel, Labdhi Pandya, Snehal Bagatharia, Fenil Patel, Bhavesh Modi, Gaurishankar Shrimali, R D Dixit, A M Kadri, Umang Mishra, Chaitanya Joshi, Madhvi Joshi, |
| EPI_ISL_455018 | Government Medical College, Vadodara | Gujarat Biotechnology Research Centre | R N Daveshwar, Ramesh Pandit, Tejas Shah, Ankit Hinsu, Pritesh Sabara, Apurvasinh Puvar, Janvi Raval, Zarna Patel, Monika Gandhi, Pinal Trivedi, Maharshi Pandya, Amit Kanani, Nidhi Patel, Nitin Savaliya, Raghawendra Kumar, Dinesh Kumar, Zuber Saiyed, Komal Patel, Labdhi Pandya, Snehal Bagatharia, Tanuja Javadekar , Neelam Nathani, Bhavesh Modi, Gaurishankar Shrimali, R D Dixit, A M Kadri, Umang Mishra, Chaitanya Joshi, Madhvi Joshi, |
| EPI_ISL_455019 | Government Medical College, Vadodara | Gujarat Biotechnology Research Centre | Ramesh Pandit, Tejas Shah, Ankit Hinsu, Pritesh Sabara, Apurvasinh Puvar, Janvi Raval, Zarna Patel, Monika Gandhi, Pinal Trivedi, Maharshi Pandya, Amit Kanani, Nidhi Patel, Nitin Savaliya, Raghawendra Kumar, Dinesh Kumar, Zuber Saiyed, Komal Patel, Labdhi Pandya, Snehal Bagatharia, Tanuja Javadekar , R N Daveshwar, Armi Chaudhari, Bhavesh Modi, Gaurishankar Shrimali, R D Dixit, A M Kadri, Umang Mishra, Chaitanya Joshi, Madhvi Joshi, |
| EPI_ISL_455020 | Government Medical College, Vadodara | Gujarat Biotechnology Research Centre | Tejas Shah, Ankit Hinsu, Pritesh Sabara, Apurvasinh Puvar, Janvi Raval, Zarna Patel, Monika Gandhi, Pinal Trivedi, Maharshi Pandya, Amit Kanani, Nidhi Patel, Nitin Savaliya, Raghawendra Kumar, Dinesh Kumar, Zuber Saiyed, Komal Patel, Labdhi Pandya, Snehal Bagatharia, Tanuja Javadekar , R N Daveshwar, Ramesh Pandit, Bhavya Jindal, Bhavesh Modi, Gaurishankar Shrimali, R D Dixit, A M Kadri, Umang Mishra, Chaitanya Joshi, Madhvi Joshi, |
| EPI_ISL_455021 | Government Medical College, Vadodara | Gujarat Biotechnology Research Centre | Ankit Hinsu, Pritesh Sabara, Apurvasinh Puvar, Janvi Raval, Zarna Patel, Monika Gandhi, Pinal Trivedi, Maharshi Pandya, Amit Kanani, Nidhi Patel, Nitin Savaliya, Raghawendra Kumar, Dinesh Kumar, Zuber Saiyed, Komal Patel, Labdhi Pandya, Snehal Bagatharia, Tanuja Javadekar , R N Daveshwar, Ramesh Pandit, Tejas Shah, Camellia Chakraborty, Bhavesh Modi, Gaurishankar Shrimali, R D Dixit, A M Kadri, Umang Mishra, Chaitanya Joshi, Madhvi Joshi, |
| EPI_ISL_455022 | Government Medical College, Vadodara | Gujarat Biotechnology Research Centre | Pritesh Sabara, Apurvasinh Puvar, Janvi Raval, Zarna Patel, Monika Gandhi, Pinal Trivedi, Maharshi Pandya, Amit Kanani, Nidhi Patel, Nitin Savaliya, Raghawendra Kumar, Dinesh Kumar, Zuber Saiyed, Komal Patel, Labdhi Pandya, Snehal Bagatharia, Tanuja Javadekar , R N Daveshwar, Ramesh Pandit, Tejas Shah, Ankit Hinsu, Siddhant Kumar, Bhavesh Modi, Gaurishankar Shrimali, R D Dixit, A M Kadri, Umang Mishra, Chaitanya Joshi, Madhvi Joshi, |
| EPI_ISL_455023 | Government Medical College, Vadodara | Gujarat Biotechnology Research Centre | Apurvasinh Puvar, Janvi Raval, Zarna Patel, Monika Gandhi, Pinal Trivedi, Maharshi Pandya, Amit Kanani, Nidhi Patel, Nitin Savaliya, Raghawendra Kumar, Dinesh Kumar, Zuber Saiyed, Komal Patel, Labdhi Pandya, Snehal Bagatharia, Tanuja Javadekar , R N Daveshwar, Ramesh Pandit, Tejas Shah, Ankit Hinsu, Pritesh Sabara, Priyanka P Vatsa, Bhavesh Modi, Gaurishankar Shrimali, R D Dixit, A M Kadri, Umang Mishra, Chaitanya Joshi, Madhvi Joshi, |
| EPI_ISL_455024 | Government Medical College, Vadodara | Gujarat Biotechnology Research Centre | Janvi Raval, Zarna Patel, Monika Gandhi, Pinal Trivedi, Maharshi Pandya, Amit Kanani, Nidhi Patel, Nitin Savaliya, Raghawendra Kumar, Dinesh Kumar, Zuber Saiyed, Komal Patel, Labdhi Pandya, Snehal Bagatharia, Tanuja Javadekar , R N Daveshwar, Ramesh Pandit, Tejas Shah, Ankit Hinsu, Pritesh Sabara, Apurvasinh Puvar, Pooja P Doshi, Bhavesh Modi, Gaurishankar Shrimali, R D Dixit, A M Kadri, Umang Mishra, Chaitanya Joshi, Madhvi Joshi, |
| EPI_ISL_455025 | Government Medical College, Vadodara | Gujarat Biotechnology Research Centre | Zarna Patel, Monika Gandhi, Pinal Trivedi, Maharshi Pandya, Amit Kanani, Nidhi Patel, Nitin Savaliya, Raghawendra Kumar, Dinesh Kumar, Zuber Saiyed, Komal Patel, Labdhi Pandya, Snehal Bagatharia, Tanuja Javadekar , R N Daveshwar, Ramesh Pandit, Tejas Shah, Ankit Hinsu, Pritesh Sabara, Apurvasinh Puvar, Janvi Raval, Akanksha Verma, Bhavesh Modi, Gaurishankar Shrimali, R D Dixit, A M Kadri, Umang Mishra, Chaitanya Joshi, Madhvi Joshi, |
| EPI_ISL_455026 | Government Medical College, Vadodara | Gujarat Biotechnology Research Centre | Monika Gandhi, Pinal Trivedi, Maharshi Pandya, Amit Kanani, Nidhi Patel, Nitin Savaliya, Raghawendra Kumar, Dinesh Kumar, Zuber Saiyed, Komal Patel, Labdhi Pandya, Snehal Bagatharia, Tanuja Javadekar , R N Daveshwar, Ramesh Pandit, Tejas Shah, Ankit Hinsu, Pritesh Sabara, Apurvasinh Puvar, Janvi Raval, Zarna Patel, Priti Pandita, Bhavesh Modi, Gaurishankar Shrimali, R D Dixit, A M Kadri, Umang Mishra, Chaitanya Joshi, Madhvi Joshi, |
| EPI_ISL_455027 | Government Medical College, Vadodara | Gujarat Biotechnology Research Centre | Pinal Trivedi, Maharshi Pandya, Amit Kanani, Nidhi Patel, Nitin Savaliya, Raghawendra Kumar, Dinesh Kumar, Zuber Saiyed, Komal Patel, Labdhi Pandya, Snehal Bagatharia, Tanuja Javadekar , R N Daveshwar, Ramesh Pandit, Tejas Shah, Ankit Hinsu, Pritesh Sabara, Apurvasinh Puvar, Janvi Raval, Zarna Patel, Monika Gandhi, Pragya Sharma, Bhavesh Modi, Gaurishankar Shrimali, R D Dixit, A M Kadri, Umang Mishra, Chaitanya Joshi, Madhvi Joshi, |
| EPI_ISL_455055, EPI_ISL_455057, EPI_ISL_455060, EPI_ISL_455061, EPI_ISL_455062, EPI_ISL_455066 | Pathology West - NSW Health Pathology | NSW Health Pathology - Institute of Clinical Pathology and Medical Research; Westmead Hospital; University of Sydney | CIDM-PH et al. |
| EPI_ISL_455102 | Kungsors VC | The Public Health Agency of Sweden | Jessica Karlsson, Anna-Malin Linde, Maria Lind Karlberg, Oskar Karlsson Lindsjo, Olov Svartstrom, Anna Risberg, Theresa Enkirch, Mia Brytting, Karin Tegmark-Wisell |
| EPI_ISL_455103 | Huslakarna Varmbadhuset Varberg | The Public Health Agency of Sweden | Johanna Hilmersson, Anna-Malin Linde, Maria Lind Karlberg, Oskar Karlsson Lindsjo, Olov Svartstrom, Anna Risberg, Theresa Enkirch, Mia Brytting, Karin Tegmark-Wisell |
| EPI_ISL_455629, EPI_ISL_455630, EPI_ISL_455631, EPI_ISL_455632, EPI_ISL_455633, EPI_ISL_455634, EPI_ISL_455637, EPI_ISL_455638 | KRISP, KZN Research Innovation and Sequencing Platform | KRISP, KZN Research Innovation and Sequencing Platform | Giandhari J, Pillay S, Lessells R, Chimukangara B, Deforche K, Tegally H, Wilkinson E, de Oliveira T |
| EPI_ISL_455655, EPI_ISL_455656, EPI_ISL_455657, EPI_ISL_455658, EPI_ISL_455659, EPI_ISL_455660, EPI_ISL_455661, EPI_ISL_455662, EPI_ISL_455663, EPI_ISL_455664, EPI_ISL_455665, EPI_ISL_455666, EPI_ISL_455667, EPI_ISL_455669, EPI_ISL_455670, EPI_ISL_455671, EPI_ISL_455672, EPI_ISL_455673, EPI_ISL_455674, EPI_ISL_455675, EPI_ISL_455676, EPI_ISL_455677, EPI_ISL_455678, EPI_ISL_455679 | | | |
| see above | ICMR-National Institute of Cholera and Enteric Diseases | National Institute of Biomedical Genomics | Arindam Maitra, Mamta Chawla Sarkar, Sreedhar Chinnaswamy, Hasina Banu, Ananya Chatterjee, Shanta Dutta, Saumitra Das |
| EPI_ISL_455785, EPI_ISL_455786, | REGIONAL VRDL,ICMR-RMRC BBSR | Immunogenomics lab, Institute of Life Sciences, Bhubaneswar | Sunil Raghav, Jyotirmayee Turuk, Arup Ghosh, Atimukta Jha, Viplov K. Biswas, Swati Madhulika, Manasi Priyadarshini, Shuchi Smita, Jaya Singh Khastri, |

| | | | |
|---|---|---|---|
| EPI_ISL_455787, EPI_ISL_455788 | | | Rupesh Dash, Soma Chattopadhyay, Ghulam Hussain Syed, Shanti Senapati, Tushar K. Beuria, Debdutta Bhattacharya, Rajeeb Swain, Punit Prasad, COVID-19 team of ILS & RMRC, Orissa COVID-19 study group, DBT's PAN-INDIA 1000 SARS-CoV2 RNA genome sequencing consortium, Sanghamitra Pati, Ajay Parida |
| EPI_ISL_456114, EPI_ISL_456115 | NYU Langone Health | Departments of Pathology and Medicine, New York University School of Medicine | Maria Aguero-Rosenfeld, Brendan Belovarac, Margaret Black, Ludovic Boytard, John Cadley, Paolo Cotzia, John Chen, Dacia Dimartino, Xiaojun Feng, Tatyana Gindin, Emily Guzman, Adriana Heguy, Megan Hogan, Emily Huang, George Jour, Alireza Khodadadi-Jamayran, Lawrence H. Lin, Raven Luther, Andrew Lytle, Christian Marier, Matthew T. Maurano, Mark J. Mulligan, Peter Meyn, Raquel Ordonez Ciriza, Iman Osman, Jared Pinnell, Vanessa Raabe, Sitharam Ramaswami, Amy Rapkiewicz, Andre M. Ribeiro-dos-Santos, Marie Samanovic-Golden, Antonio Serrano, Guomiao Shen, Matija Snuderl, Theodore Vougiouklakis, Nick Vulpescu, Gael Westby, Paul Zappile, Yutong Zhang |
| EPI_ISL_456403 | Middlemore Hospital | Institute of Environmental Science and Research (ESR) | Matt Storey, Xiaoyun Ren, Anja Werno, Antje van der Linden, Arlo Upton, Chris Mansell, David Hammer, Dragana Drinkovic, Erasmus Smit, Gary McAuliffe, Hana Sofia Andersson, James Ussher, Jill Sherwood, Josh Freeman, Julia Howard, Juliet Elvy, Mary DeAlmeida, Matt Blakiston, Matthew Rogers, Max Bloomfield, Michael Addidle, Michelle Balm, Sally Roberts, Sarah Jefferies, Sharmini Muttaiyah, Susan Morpeth, Susan Taylor, Timothy Blackmore, Vani Sathyendran, Veronica Playle, Virginia Hope, Erasmus Smit, Lauren Jelly, Joep de Ligt |
| EPI_ISL_456456, EPI_ISL_456457, EPI_ISL_456458, EPI_ISL_456459, EPI_ISL_456460, EPI_ISL_456461, EPI_ISL_456462, EPI_ISL_456463, EPI_ISL_456464, EPI_ISL_456465, EPI_ISL_456466, EPI_ISL_456467, EPI_ISL_456468, EPI_ISL_456469, EPI_ISL_456470, EPI_ISL_456471, EPI_ISL_456472, EPI_ISL_456473, EPI_ISL_456474, EPI_ISL_456475, EPI_ISL_456476, EPI_ISL_456477 | | | |
| see above | Victorian Infectious Diseases Reference Laboratory (VIDRL) | Microbiological Diagnostic Unit Public Health Laboratory and Victorian Infectious Diseases Reference Laboratory, Doherty Institute | Caly L., Seemann T., Sait, M., Schultz M., Druce J., Sherry, N. |
| EPI_ISL_456479 | Microbiological Diagnostic Unit Public Health Laboratory | Microbiological Diagnostic Unit Public Health Laboratory, The Peter Doherty Institute for Infection and Immunity | Seemann T., Schultz M., Sait, M., Sherry, N. |
| EPI_ISL_456480, EPI_ISL_456481, EPI_ISL_456482, EPI_ISL_456483, EPI_ISL_456492, EPI_ISL_456538, EPI_ISL_456539, EPI_ISL_456540, EPI_ISL_456541, EPI_ISL_456542, EPI_ISL_456544, EPI_ISL_456546, EPI_ISL_456557, EPI_ISL_456629, EPI_ISL_456630, EPI_ISL_456632, EPI_ISL_456633, EPI_ISL_456634, EPI_ISL_456635, EPI_ISL_456636 | | | |
| see above | Victorian Infectious Diseases Reference Laboratory (VIDRL) | Microbiological Diagnostic Unit Public Health Laboratory and Victorian Infectious Diseases Reference Laboratory, Doherty Institute | Caly L., Seemann T., Sait, M., Schultz M., Druce J., Sherry, N. |
| EPI_ISL_456670, EPI_ISL_456671, EPI_ISL_456672, EPI_ISL_456673 | University of Birmingham | COVID-19 Genomics UK (COG-UK) Consortium | Loman Lab: Claire McMurray, Joanne Stockton, Samuel Nicholls, Radoslaw Poplawski, Will Rowe, Josh Quick, Nicholas Loman // UHB Lab: Celina M Whalley, Andrew Bosworth, Charlotte Poxon, Kasun Wanigasooriya, Oliver Pickles, Mike Kidd, Alex Richter, Andrew D Beggs // PHE Heartlands Lab: Husam Osman, Andrew Bosworth |
| EPI_ISL_456768, EPI_ISL_456771, EPI_ISL_456775, EPI_ISL_456776, EPI_ISL_456777, EPI_ISL_456778, EPI_ISL_456779, EPI_ISL_456780, EPI_ISL_456781, EPI_ISL_456785, EPI_ISL_456786, EPI_ISL_456787, EPI_ISL_456788, EPI_ISL_456789, EPI_ISL_456790, EPI_ISL_456791, EPI_ISL_456792, EPI_ISL_456793, EPI_ISL_456794, EPI_ISL_456795, EPI_ISL_456796, EPI_ISL_456797, EPI_ISL_456798 | | | |
| see above | West of Scotland Specialist Virology Centre, NHSGGC / MRC-University of Glasgow Centre for Virus Research | COVID-19 Genomics UK (COG-UK) Consortium | Ana da Silva Filipe, Natasha Johnson, Kathy Smollett, Daniel Mair, Stephen Carmichael, Lily Tong, Jenna Nichols, Elihu Aranday-Cortes, Kirstyn Brunker, Yasmin Parr, Kyriaki Nomikou; Sarah McDonald, Marc Niebel, Patawee Asamaphan; Richard Orton, Joseph Hughes, Sreenu Vattipally, David L Robertson; Alasdair MacLean, Rory Gunson; Kathy Li, Natasha Jesudason, Rajiv Shah, James Shepherd, Antonia Ho, Emma Thomson |
| EPI_ISL_457227, EPI_ISL_457228, EPI_ISL_457229, EPI_ISL_457230, EPI_ISL_457232, EPI_ISL_457236, EPI_ISL_457238, EPI_ISL_457240, EPI_ISL_457242, EPI_ISL_457245, EPI_ISL_457246, EPI_ISL_457251, EPI_ISL_457252, EPI_ISL_457253, EPI_ISL_457254, EPI_ISL_457257 | | | |
| see above | University of Exeter | COVID-19 Genomics UK (COG-UK) Consortium | Ben Temperton,Aaron Jeffries,Michelle Michelsen,Joanna Warwick-Dugdale,Audrey Farbos,Robyn Manley,Stephen Michell,Jane Masoli |
| EPI_ISL_457311, EPI_ISL_457312, EPI_ISL_457322, EPI_ISL_457323 | Northumbria University / South Tees Hospitals NHS Foundation Trust / North Cumbria Integrated Care NHS Foundation Trust / North Tees and Hartlepool NHS Foundation Trust / Newcastle Hospitals NHS Foundation Trust | COVID-19 Genomics UK (COG-UK) Consortium | Darren L Smith,Andrew Nelson,Matthew Bashton,Greg R Young,Joshua Loh,John Allan,Mohammad A Tariq,Giles S Holt,Gary Black,Wen C Yew,Lynn Dover,Paul Baker,Steve Liggett,Sarah Essex,Jane Greenaway,Debra Padgett,Clive Graham,Garren Scott,Edward Barton,Emma Swindells,Brendan Payne,Jennifer Collins,Yusri Taha,Gary Eltringham |
| EPI_ISL_457326, EPI_ISL_457327, EPI_ISL_457371, EPI_ISL_457372, EPI_ISL_457373, EPI_ISL_457374, EPI_ISL_457375, EPI_ISL_457376, EPI_ISL_457377, EPI_ISL_457378, EPI_ISL_457379, EPI_ISL_457380, EPI_ISL_457381, EPI_ISL_457382, EPI_ISL_457383, EPI_ISL_457384, EPI_ISL_457385, EPI_ISL_457386, EPI_ISL_457387, EPI_ISL_457388, EPI_ISL_457389, EPI_ISL_457390, EPI_ISL_457391, EPI_ISL_457392, EPI_ISL_457393, EPI_ISL_457394, EPI_ISL_457395, EPI_ISL_457396, EPI_ISL_457397, EPI_ISL_457398, EPI_ISL_457399, EPI_ISL_457400, EPI_ISL_457401, EPI_ISL_457402, EPI_ISL_457403, EPI_ISL_457404, EPI_ISL_457405, EPI_ISL_457406, EPI_ISL_457407, EPI_ISL_457408, EPI_ISL_457409, EPI_ISL_457410, EPI_ISL_457411, EPI_ISL_457412, EPI_ISL_457413, EPI_ISL_457414, EPI_ISL_457415, EPI_ISL_457416, EPI_ISL_457417, EPI_ISL_457418, EPI_ISL_457419, EPI_ISL_457420, EPI_ISL_457421, EPI_ISL_457422, EPI_ISL_457423, EPI_ISL_457424, EPI_ISL_457425, EPI_ISL_457426, EPI_ISL_457427, EPI_ISL_457428, EPI_ISL_457429, EPI_ISL_457430, EPI_ISL_457431, EPI_ISL_457432, EPI_ISL_457433, EPI_ISL_457434, EPI_ISL_457435, EPI_ISL_457436, EPI_ISL_457437, EPI_ISL_457438, EPI_ISL_457439, EPI_ISL_457440, EPI_ISL_457441, EPI_ISL_457442, EPI_ISL_457443, EPI_ISL_457444, EPI_ISL_457445, EPI_ISL_457446, EPI_ISL_457447, EPI_ISL_457448, EPI_ISL_457449, EPI_ISL_457450, EPI_ISL_457451, EPI_ISL_457452, EPI_ISL_457471, EPI_ISL_457502 | | | |
| see above | Quadram Institute Bioscience | COVID-19 Genomics UK (COG-UK) Consortium | Dave J. Baker, Gemma L. Kay, Alp Aydin, Thanh Le-Viet, Steven Rudder, Ana P. Tedim, Anastasia Kolyva, Maria Diaz, Leonardo de Oliveira Martins, Nabil-Fareed Alikhan, Lizzie Meadows, Rachael Stanley, Ngozi Elumogo, Muhammed Yasir, Nicholas M. Thomson, Alexander J Trotter, Rachel Gilroy, Samuel Bloomfield, Claire Stuart, Andrew Bell, Reenesh Prakash, Samir Dervisevic, Alison E. Mather, John Wain, Mark Webber, Andrew J. Page, Justin O'Grady |
| EPI_ISL_457573, EPI_ISL_457575, EPI_ISL_457576, EPI_ISL_457577, EPI_ISL_457578 | Queens Medical Centre, Clinical Microbiology Department / DeepSeq Nottingham | COVID-19 Genomics UK (COG-UK) Consortium | Gemma Clark, Wendy Smith, Manjinder Khakh, Hannah Howson-Wells, Jonathan Ball, Patrick McClure, Joseph Chappell, Theocharis Tsoleridis, Nadine Holmes, Matthew Carlisle, Christopher Moore, Fei Sang, Johnny Debebe, Victoria Wright, Matthew Loose |
| EPI_ISL_457581, EPI_ISL_457585, EPI_ISL_457588, EPI_ISL_457591, EPI_ISL_457592, EPI_ISL_457593, EPI_ISL_457594, EPI_ISL_457596, EPI_ISL_457599, EPI_ISL_457600, EPI_ISL_457605, EPI_ISL_457606, EPI_ISL_457607, EPI_ISL_457609, EPI_ISL_457616, EPI_ISL_457617, EPI_ISL_457618, EPI_ISL_457620, EPI_ISL_457621, EPI_ISL_457623, EPI_ISL_457628, EPI_ISL_457631, EPI_ISL_457633, EPI_ISL_457636, EPI_ISL_457637, EPI_ISL_457640, EPI_ISL_457645, EPI_ISL_457647, EPI_ISL_457651, EPI_ISL_457655, EPI_ISL_457657, EPI_ISL_457660, EPI_ISL_457662, EPI_ISL_457664, EPI_ISL_457667, EPI_ISL_457669, EPI_ISL_457670, EPI_ISL_457672, EPI_ISL_457674, EPI_ISL_457677, EPI_ISL_457678, EPI_ISL_457686 | | | |
| see above | Virology Department, Sheffield Teaching Hospitals NHS Foundation Trust/Department of Infection, Immunity and Cardiovascular Disease, The Medical School, University of Sheffield | COVID-19 Genomics UK (COG-UK) Consortium | Thushan de Silva, Matthew Parker, Nikki Smith, Adri Angyal, Rebecca Brown, Luke Green, Rachel Tucker, Paul Parsons, Danielle Groves, Katie Johnson, Laura Carrilero, Alex Keeley, Dave Partridge, Matthew Wyles, Benjamin Lindsey, Mehmet Yavuz, Mohammad Raza, Cariad Evans |
| EPI_ISL_457891, EPI_ISL_457892, EPI_ISL_457893, EPI_ISL_457894, EPI_ISL_457896, EPI_ISL_457897, EPI_ISL_457898, EPI_ISL_457899, EPI_ISL_457900, EPI_ISL_457901, EPI_ISL_457902, EPI_ISL_457903, EPI_ISL_457904, EPI_ISL_457906, EPI_ISL_457907 | | | |
| see above | KEMRI-CGMR-C | KEMRI-Wellcome Trust Research Programme/KEMRI-CGMR-C Kilifi | Githinji G. et al 2020 |
| EPI_ISL_457983, EPI_ISL_457984 | Oman-NIC | Department of Microbiology and Immunology-SQUH | Fahad Zadjali, Samira Al-Maruqi, Amina Al Jardani, Khulood Al-Mammary, Hanan Al-kindi, Fatma BaAlawi, Hamida AL Barwani, Zeyana AL-Dahmani, Intisar Al-Shukri, Aisha Al-Busaidi, Aisha Al-Amri, Ahlam Al-Amri, Mohammed Al-Tobi, Samiha Al Kharusi, Abdulla Balkhair |
| EPI_ISL_458032 | King Institute of Preventive Medicine & Research | CSIR-Centre for Cellular and Molecular Biology | K.Kaveri,S.Sivasubramanian,S.Vennila,P.Padmapriya,R.Kiruba,S.Magesh,G. Dhinakar Raj, G. Ravikumar, P. Azhahianambi, K Thangaraj,Shagufta Khan, Lamuk Zaveri, Namami Gaur, Sakshi Shambhavi, Tulasi Nagabandi, Purushotham Vodnala, Payel Mukherjee, Sofia Banu, Priya Singh, Dhiviya Vedagiri, Divya Gupta, Vishal Sah, Santosh Kumar Kuncha, Krishnan Harinivas Harshan, Archana Bharadwaj Siva, Karthik Bharadwaj Tallapaka, Rakesh K Mishra, Divya Tej Sowpati |
| EPI_ISL_458033 | King Institute of Preventive Medicine & Research | CSIR-Centre for Cellular and Molecular Biology | K.Kaveri,S.Sivasubramanian,S.Vennila,P.Padmapriya,R.Kiruba,S.Magesh,G. Dhinakar Raj, G. Ravikumar, P. Azhahianambi, K Thangaraj,Lamuk Zaveri, Shagufta Khan, Namami Gaur, Sakshi Shambhavi, Tulasi Nagabandi, Purushotham Vodnala, Payel Mukherjee, Sofia Banu, Priya Singh, Dhiviya Vedagiri, Divya Gupta, Vishal Sah, Santosh Kumar Kuncha, Krishnan Harinivas Harshan, Archana Bharadwaj Siva, Karthik Bharadwaj Tallapaka, Rakesh K Mishra, Divya Tej Sowpati |
| EPI_ISL_458034 | King Institute of Preventive Medicine & Research | CSIR-Centre for Cellular and Molecular Biology | K.Kaveri,S.Sivasubramanian,S.Vennila,P.Padmapriya,R.Kiruba,S.Magesh,G. Dhinakar Raj, G. Ravikumar, P. Azhahianambi, K Thangaraj, Namami Gaur, Sakshi Shambhavi, Lamuk Zaveri, Shagufta Khan, Tulasi Nagabandi, Purushotham Vodnala, Payel Mukherjee, Sofia Banu, Priya Singh, Dhiviya Vedagiri, Divya Gupta, Vishal Sah, Santosh Kumar Kuncha, Krishnan Harinivas Harshan, Archana Bharadwaj Siva, Karthik Bharadwaj Tallapaka, Rakesh K Mishra, Divya Tej Sowpati |
| EPI_ISL_458035 | King Institute of Preventive Medicine & Research | CSIR-Centre for Cellular and Molecular Biology | K.Kaveri,S.Sivasubramanian,S.Vennila,P.Padmapriya,R.Kiruba,S.Magesh,G. Dhinakar Raj, G. Ravikumar, P. Azhahianambi, K Thangaraj, Tulasi |

| | | | |
|---|---|---|---|
| | | | Nagabandi, Namami Gaur, Sakshi Shambhavi, Lamuk Zaveri, Shagufta Khan, Purushotham Vodnala, Payel Mukherjee, Sofia Banu, Priya Singh, Dhiviya Vedagiri, Divya Gupta, Vishal Sah, Santosh Kumar Kuncha, Krishnan Harinivas Harshan, Archana Bharadwaj Siva, Karthik Bharadwaj Tallapaka, Rakesh K Mishra, Divya Tej Sowpati |
| EPI_ISL_458036 | King Institute of Preventive Medicine & Research | CSIR-Centre for Cellular and Molecular Biology | K.Kaveri,S.Sivasubramanian,S.Vennila,P.Padmapriya,R.Kiruba,S.Magesh,G. Dhinakar Raj, G. Ravikumar, R. P. Aravindh Babu, K Thangaraj, Payel Mukherjee, Sofia Banu, Priya Singh, Dhiviya Vedagiri, Divya Gupta, Vishal Sah, Santosh Kumar Kuncha, Krishnan Harinivas Harshan, Archana Bharadwaj Siva, Karthik Bharadwaj Tallapaka, Shagufta Khan, Lamuk Zaveri, Namami Gaur, Sakshi Shambhavi, Tulasi Nagabandi, Purushotham Vodnala, Rakesh K Mishra, Divya Tej Sowpati |
| EPI_ISL_458037 | King Institute of Preventive Medicine & Research | CSIR-Centre for Cellular and Molecular Biology | K.Kaveri,S.Sivasubramanian,S.Vennila,P.Padmapriya,R.Kiruba,S.Magesh,G. Dhinakar Raj, G. Ravikumar, R. P. Aravindh Babu, K Thangaraj, Sofia Banu, Payel Mukherjee, Priya Singh, Dhiviya Vedagiri, Divya Gupta, Vishal Sah, Santosh Kumar Kuncha, Krishnan Harinivas Harshan, Archana Bharadwaj Siva, Karthik Bharadwaj Tallapaka, Shagufta Khan, Lamuk Zaveri, Namami Gaur, Sakshi Shambhavi, Tulasi Nagabandi, Purushotham Vodnala, Rakesh K Mishra, Divya Tej Sowpati |
| EPI_ISL_458038 | King Institute of Preventive Medicine & Research | CSIR-Centre for Cellular and Molecular Biology | K.Kaveri,S.Sivasubramanian,S.Vennila,P.Padmapriya,R.Kiruba,S.Magesh,G. Dhinakar Raj, G. Ravikumar, R. P. Aravindh Babu, K Thangaraj, Shagufta Khan, Lamuk Zaveri, Namami Gaur, Sakshi Shambhavi, Tulasi Nagabandi, Purushotham Vodnala, Payel Mukherjee, Sofia Banu, Priya Singh, Dhiviya Vedagiri, Divya Gupta, Vishal Sah, Santosh Kumar Kuncha, Krishnan Harinivas Harshan, Archana Bharadwaj Siva, Karthik Bharadwaj Tallapaka, Rakesh K Mishra, Divya Tej Sowpati |
| EPI_ISL_458116, EPI_ISL_458118, EPI_ISL_458124, EPI_ISL_458128 | Oman National Influenza Centre | Department of Microbiology and Immunology-SQUH | Fahad Zadjali, Samira Al-Maruqi, Amina Al Jardani, Khulood Al-Mammary, Hanan Al-kindi, Fatma BaAlawi, Hamida AL Barwani, Zeyana AL-Dahmani, Intisar Al-Shukri, Aisha Al-Busaidi, Aisha Al-Amri, Ahlam Al-Amri, Mohammed Al-Tobi, Samiha Al Kharusi, Abdulla Balkhair |
| EPI_ISL_458148 | Evandro Chagas Institute | Evandro Chagas Institute | Santos, M.C.; Silva, A.M.; Junior, W.D.C.; Barbagelata, L.S.; Ferreira, J.A.; Sousa, E.M.A.; da Silva, P.S.; Resque, H.R; Martins, L.C.; Sousa Junior, E.C.;Viana, G.M.R |
| EPI_ISL_458237, EPI_ISL_458238, EPI_ISL_458244, EPI_ISL_458249, EPI_ISL_458253, EPI_ISL_458254, EPI_ISL_458263, EPI_ISL_458264, EPI_ISL_458269, EPI_ISL_458271 | Scripps Medical Laboratory | Andersen lab at Scripps Research | SEARCH Alliance San Diego with Michael Quigley, Ellen Stefanski, Ian Mchardy |
| EPI_ISL_458292, EPI_ISL_458293 | Dirk Dittmer | Dirk Dittmer | Aubrey,B.G., Caro-Vegas,C.P., Dittmer,D., Eason,A.B., Juarez,A., Landis,J.T., Mcnamara,R.P., Miller,M.B., Moorad,R., Pluta,L.J., Seltzer,T.A., Thompson,C., Vahrson,W., Villamor,F. |
| EPI_ISL_458299, EPI_ISL_458300, EPI_ISL_458301, EPI_ISL_458304, EPI_ISL_458307, EPI_ISL_458308, EPI_ISL_458310, EPI_ISL_458312, EPI_ISL_458313, EPI_ISL_458314, EPI_ISL_458318, EPI_ISL_458319, EPI_ISL_458321, EPI_ISL_458322, EPI_ISL_458323, EPI_ISL_458324, EPI_ISL_458325, EPI_ISL_458327, EPI_ISL_458328, EPI_ISL_458329, EPI_ISL_458330, EPI_ISL_458331, EPI_ISL_458333, EPI_ISL_458334, EPI_ISL_458335, EPI_ISL_458337, EPI_ISL_458338, EPI_ISL_458339, EPI_ISL_458342, EPI_ISL_458344, EPI_ISL_458345, EPI_ISL_458346, EPI_ISL_458347, EPI_ISL_458350, EPI_ISL_458351, EPI_ISL_458352, EPI_ISL_458354, EPI_ISL_458355, EPI_ISL_458356, EPI_ISL_458358, EPI_ISL_458360, EPI_ISL_458362, EPI_ISL_458363, EPI_ISL_458364, EPI_ISL_458366, EPI_ISL_458367, EPI_ISL_458369, EPI_ISL_458376, EPI_ISL_458379, EPI_ISL_458382, EPI_ISL_458383, EPI_ISL_458385, EPI_ISL_458388, EPI_ISL_458389, EPI_ISL_458390, EPI_ISL_458392, EPI_ISL_458393, EPI_ISL_458394, EPI_ISL_458397, EPI_ISL_458399, EPI_ISL_458400, EPI_ISL_458402, EPI_ISL_458403, EPI_ISL_458404, EPI_ISL_458406, EPI_ISL_458408, EPI_ISL_458410, EPI_ISL_458411, EPI_ISL_458413, EPI_ISL_458414, EPI_ISL_458418, EPI_ISL_458420, EPI_ISL_458421, EPI_ISL_458423, EPI_ISL_458424, EPI_ISL_458427, EPI_ISL_458428, EPI_ISL_458429, EPI_ISL_458431, EPI_ISL_458432, EPI_ISL_458434, EPI_ISL_458437, EPI_ISL_458438, EPI_ISL_458441, EPI_ISL_458442, EPI_ISL_458445, EPI_ISL_458446, EPI_ISL_458447, EPI_ISL_458452, EPI_ISL_458453, EPI_ISL_458454, EPI_ISL_458458, EPI_ISL_458460, EPI_ISL_458461, EPI_ISL_458462, EPI_ISL_458464, EPI_ISL_458465, EPI_ISL_458469, EPI_ISL_458470, EPI_ISL_458471, EPI_ISL_458472, EPI_ISL_458474, EPI_ISL_458476, EPI_ISL_458479, EPI_ISL_458481, EPI_ISL_458483, EPI_ISL_458484, EPI_ISL_458485, EPI_ISL_458486, EPI_ISL_458487, EPI_ISL_458488, EPI_ISL_458489, EPI_ISL_458490 | | | |
| see above | PHE South West Regional Laboratory, National Infection Service | Wellcome Sanger Institute for the COVID-19 Genomics UK (COG-UK) consortium | Stephanie Hutchings, Hannah Pymont, Dr Peter Muir, Barry Vipond, Rich Hopes; and Alex Alderton, Roberto Amato, Sonia Goncalves, Ewan Harrison, David K. Jackson, Ian Johnston, Dominic Kwiatkowski, Cordelia Langford, John Sillitoe on behalf of the Wellcome Sanger Institute COVID-19 Surveillance Team (http://www.sanger.ac.uk/covid-team) |
| EPI_ISL_458491 | PHE South West Regional Laboratory, National Infection Service | Wellcome Sanger Institute for the COVID-19 Genomics UK (COG-UK) Consortium | Stephanie Hutchings, Hannah Pymont, Dr Peter Muir, Barry Vipond, Rich Hopes; and Alex Alderton, Roberto Amato, Sonia Goncalves, Ewan Harrison, David K. Jackson, Ian Johnston, Dominic Kwiatkowski, Cordelia Langford, John Sillitoe on behalf of the Wellcome Sanger Institute COVID-19 Surveillance Team |
| EPI_ISL_458492, EPI_ISL_458493, EPI_ISL_458494, EPI_ISL_458495, EPI_ISL_458496, EPI_ISL_458497, EPI_ISL_458502, EPI_ISL_458504, EPI_ISL_458505, EPI_ISL_458506, EPI_ISL_458508, EPI_ISL_458510, EPI_ISL_458513, EPI_ISL_458514, EPI_ISL_458515 | | | |
| see above | PHE South West Regional Laboratory, National Infection Service | Wellcome Sanger Institute for the COVID-19 Genomics UK (COG-UK) consortium | Stephanie Hutchings, Hannah Pymont, Dr Peter Muir, Barry Vipond, Rich Hopes; and Alex Alderton, Roberto Amato, Sonia Goncalves, Ewan Harrison, David K. Jackson, Ian Johnston, Dominic Kwiatkowski, Cordelia Langford, John Sillitoe on behalf of the Wellcome Sanger Institute COVID-19 Surveillance Team (http://www.sanger.ac.uk/covid-team) |
| EPI_ISL_459167, EPI_ISL_459168, EPI_ISL_459169, EPI_ISL_459170, EPI_ISL_459171, EPI_ISL_459172, EPI_ISL_459174, EPI_ISL_459177, EPI_ISL_459179, EPI_ISL_459180, EPI_ISL_459181, EPI_ISL_459183, EPI_ISL_459184, EPI_ISL_459186, EPI_ISL_459187, EPI_ISL_459189, EPI_ISL_459190, EPI_ISL_459192, EPI_ISL_459194, EPI_ISL_459196, EPI_ISL_459198, EPI_ISL_459199, EPI_ISL_459201, EPI_ISL_459202, EPI_ISL_459203, EPI_ISL_459205, EPI_ISL_459210, EPI_ISL_459211, EPI_ISL_459212, EPI_ISL_459213, EPI_ISL_459214, EPI_ISL_459216, EPI_ISL_459218, EPI_ISL_459219, EPI_ISL_459220, EPI_ISL_459221, EPI_ISL_459223, EPI_ISL_459224, EPI_ISL_459225, EPI_ISL_459226, EPI_ISL_459227, EPI_ISL_459230, EPI_ISL_459232, EPI_ISL_459233, EPI_ISL_459235, EPI_ISL_459236, EPI_ISL_459237, EPI_ISL_459238, EPI_ISL_459239, EPI_ISL_459240 | | | |
| see above | Department of Pathology, University of Cambridge | Wellcome Sanger Institute for the COVID-19 Genomics UK (COG-UK) consortium | Luke W Meredith, M. Estée Török , Myra Hosmillo, William L. Hamilton, Martin D. Curran, Theresa Feltwell, Grant Hall, Anna Yakovleva, Fahad A Khokhar, Charlotte J. Houldcroft, Laura G Caller, Aminu S. Jahun, Sarah L. Caddy, Ian Goodfellow; and Alex Alderton, Roberto Amato, Sonia Goncalves, Ewan Harrison, David K. Jackson, Ian Johnston, Dominic Kwiatkowski, Cordelia Langford, John Sillitoe on behalf of the Wellcome Sanger Institute COVID-19 Surveillance Team (http://www.sanger.ac.uk/covid-team) |
| EPI_ISL_459241 | Department of Pathology, University of Cambridge | Wellcome Sanger Institute for the COVID-19 Genomics UK (COG-UK) Consortium | Luke W Meredith, M. Estée Török , Myra Hosmillo, William L. Hamilton, Martin D. Curran, Theresa Feltwell, Grant Hall, Anna Yakovleva, Fahad A Khokhar, Charlotte J. Houldcroft, Laura G Caller, Aminu S. Jahun, Sarah L. Caddy, Ian Goodfellow; and Alex Alderton, Roberto Amato, Sonia Goncalves, Ewan Harrison, David K. Jackson, Ian Johnston, Dominic Kwiatkowski, Cordelia Langford, John Sillitoe on behalf of the Wellcome Sanger Institute COVID-19 Surveillance Team |
| EPI_ISL_459242 | Department of Pathology, University of Cambridge | Wellcome Sanger Institute for the COVID-19 Genomics UK (COG-UK) consortium | Luke W Meredith, M. Estée Török , Myra Hosmillo, William L. Hamilton, Martin D. Curran, Theresa Feltwell, Grant Hall, Anna Yakovleva, Fahad A Khokhar, Charlotte J. Houldcroft, Laura G Caller, Aminu S. Jahun, Sarah L. Caddy, Ian Goodfellow; and Alex Alderton, Roberto Amato, Sonia Goncalves, Ewan Harrison, David K. Jackson, Ian Johnston, Dominic Kwiatkowski, Cordelia Langford, John Sillitoe on behalf of the Wellcome Sanger Institute COVID-19 Surveillance Team (http://www.sanger.ac.uk/covid-team) |
| EPI_ISL_459243 | Department of Pathology, University of Cambridge | Wellcome Sanger Institute for the COVID-19 Genomics UK (COG-UK) Consortium | Luke W Meredith, M. Estée Török , Myra Hosmillo, William L. Hamilton, Martin D. Curran, Theresa Feltwell, Grant Hall, Anna Yakovleva, Fahad A Khokhar, Charlotte J. Houldcroft, Laura G Caller, Aminu S. Jahun, Sarah L. Caddy, Ian Goodfellow; and Alex Alderton, Roberto Amato, Sonia Goncalves, Ewan Harrison, David K. Jackson, Ian Johnston, Dominic Kwiatkowski, Cordelia Langford, John Sillitoe on behalf of the Wellcome Sanger Institute COVID-19 Surveillance Team |
| EPI_ISL_459244, EPI_ISL_459245, EPI_ISL_459246, EPI_ISL_459247, EPI_ISL_459248, EPI_ISL_459249, EPI_ISL_459250, EPI_ISL_459251, EPI_ISL_459252, EPI_ISL_459253, EPI_ISL_459254, EPI_ISL_459255, EPI_ISL_459256, EPI_ISL_459257 | | | |
| see above | Department of Pathology, University of Cambridge | Wellcome Sanger Institute for the COVID-19 Genomics UK (COG-UK) consortium | Luke W Meredith, M. Estée Török , Myra Hosmillo, William L. Hamilton, Martin D. Curran, Theresa Feltwell, Grant Hall, Anna Yakovleva, Fahad A Khokhar, Charlotte J. Houldcroft, Laura G Caller, Aminu S. Jahun, Sarah L. Caddy, Ian Goodfellow; and Alex Alderton, Roberto Amato, Sonia Goncalves, Ewan Harrison, David K. Jackson, Ian Johnston, Dominic Kwiatkowski, Cordelia Langford, John Sillitoe on behalf of the Wellcome Sanger Institute COVID-19 Surveillance Team (http://www.sanger.ac.uk/covid-team) |
| EPI_ISL_459258 | Department of Pathology, University of Cambridge | Wellcome Sanger Institute for the COVID-19 Genomics UK (COG-UK) Consortium | Luke W Meredith, M. Estée Török , Myra Hosmillo, William L. Hamilton, Martin D. Curran, Theresa Feltwell, Grant Hall, Anna Yakovleva, Fahad A Khokhar, Charlotte J. Houldcroft, Laura G Caller, Aminu S. Jahun, Sarah L. Caddy, Ian Goodfellow; and Alex Alderton, Roberto Amato, Sonia Goncalves, Ewan Harrison, David K. Jackson, Ian Johnston, Dominic Kwiatkowski, Cordelia Langford, John Sillitoe on behalf of the Wellcome Sanger Institute COVID-19 Surveillance Team |
| EPI_ISL_459259, EPI_ISL_459260, EPI_ISL_459261, EPI_ISL_459262, EPI_ISL_459263, EPI_ISL_459264, EPI_ISL_459265, EPI_ISL_459266, EPI_ISL_459267, EPI_ISL_459268, EPI_ISL_459269, EPI_ISL_459270, EPI_ISL_459271, EPI_ISL_459272, EPI_ISL_459273, EPI_ISL_459274, EPI_ISL_459275, EPI_ISL_459276, EPI_ISL_459277, EPI_ISL_459278, EPI_ISL_459279, EPI_ISL_459280, EPI_ISL_459281, EPI_ISL_459282, EPI_ISL_459283, EPI_ISL_459284, EPI_ISL_459285, EPI_ISL_459286, EPI_ISL_459287, EPI_ISL_459288, EPI_ISL_459289, EPI_ISL_459290, EPI_ISL_459291, EPI_ISL_459292, EPI_ISL_459293, EPI_ISL_459294, EPI_ISL_459295, EPI_ISL_459296, EPI_ISL_459297, EPI_ISL_459304, EPI_ISL_459305, EPI_ISL_459306, EPI_ISL_459307, EPI_ISL_459308, EPI_ISL_459309, EPI_ISL_459310, EPI_ISL_459311, EPI_ISL_459312, EPI_ISL_459313, EPI_ISL_459314, EPI_ISL_459315, EPI_ISL_459316, EPI_ISL_459317, EPI_ISL_459318, EPI_ISL_459319, EPI_ISL_459320, EPI_ISL_459321, EPI_ISL_459322, EPI_ISL_459323, EPI_ISL_459410, EPI_ISL_459415, EPI_ISL_459416, EPI_ISL_459417, EPI_ISL_459418, EPI_ISL_459419, EPI_ISL_459420, EPI_ISL_459422, EPI_ISL_459424, EPI_ISL_459426, EPI_ISL_459427, EPI_ISL_459428, EPI_ISL_459429, | | | |

| | | | |
|---|---|---|---|
| EPI_ISL_459430, EPI_ISL_459431, EPI_ISL_459432, EPI_ISL_459433, EPI_ISL_459434, EPI_ISL_459436, EPI_ISL_459438, EPI_ISL_459439, EPI_ISL_459442, EPI_ISL_459443, EPI_ISL_459445, EPI_ISL_459446, EPI_ISL_459447, EPI_ISL_459449, EPI_ISL_459450, EPI_ISL_459451, EPI_ISL_459452, EPI_ISL_459454, EPI_ISL_459456, EPI_ISL_459458, EPI_ISL_459460, EPI_ISL_459461, EPI_ISL_459464, EPI_ISL_459466, EPI_ISL_459467, EPI_ISL_459469, EPI_ISL_459470, EPI_ISL_459472, EPI_ISL_459473, EPI_ISL_459474, EPI_ISL_459476, EPI_ISL_459479, EPI_ISL_459481, EPI_ISL_459482, EPI_ISL_459483, EPI_ISL_459484, EPI_ISL_459485, EPI_ISL_459486, EPI_ISL_459487, EPI_ISL_459488, EPI_ISL_459489, EPI_ISL_459490, EPI_ISL_459491, EPI_ISL_459492, EPI_ISL_459493, EPI_ISL_459494, EPI_ISL_459495, EPI_ISL_459497, EPI_ISL_459499, EPI_ISL_459500, EPI_ISL_459502, EPI_ISL_459504 | | | |
| see above | Department of Pathology, University of Cambridge | Wellcome Sanger Institute for the COVID-19 Genomics UK (COG-UK) consortium | Luke W Meredith, M. Estée Török , Myra Hosmillo, William L. Hamilton, Martin D. Curran, Theresa Feltwell, Grant Hall, Anna Yakovleva, Fahad A Khokhar, Charlotte J. Houldcroft, Laura G Caller, Aminu S. Jahun, Sarah L. Caddy, Ian Goodfellow; and Alex Alderton, Roberto Amato, Sonia Goncalves, Ewan Harrison, David K. Jackson, Ian Johnston, Dominic Kwiatkowski, Cordelia Langford, John Sillitoe on behalf of the Wellcome Sanger Institute COVID-19 Surveillance Team (http://www.sanger.ac.uk/covid-team) |
| EPI_ISL_459913, EPI_ISL_459914, EPI_ISL_459942, EPI_ISL_459943, EPI_ISL_459944, EPI_ISL_459945, EPI_ISL_459946, EPI_ISL_459947, EPI_ISL_459949, EPI_ISL_459950, EPI_ISL_459951 | | | |
| see above | Devki Devi Foundation, a unit of Max Healthcare | CSIR-IGIB/Max | Rajesh Pandey#, Samreen Siddiqui, Pooja Sharma, Bansidhar Tarai, Vivekanand A, Bharathram Uppili, Saruchi Wadhwa, Nishu Tyagi, Mitali Mukerji, Bansidhar Tarai, Poonam Das, Sujeet Jha, Mohammed Faruq, Vinita Jha, Anurag Agrawal |
| EPI_ISL_460032, EPI_ISL_460561, EPI_ISL_460566, EPI_ISL_460572, EPI_ISL_460589, EPI_ISL_460590, EPI_ISL_460592, EPI_ISL_460594, EPI_ISL_460596, EPI_ISL_460597, EPI_ISL_460599, EPI_ISL_460600, EPI_ISL_460601 | | | |
| see above | Michigan Department of Health and Human Services, Bureau of Laboratories | Michigan Department of Health and Human Services, Bureau of Laboratories | Blankenship HM, Riner D, Soehnlen MK |
| EPI_ISL_460788, EPI_ISL_460835, EPI_ISL_460838, EPI_ISL_460854, EPI_ISL_460855, EPI_ISL_460856, EPI_ISL_460857, EPI_ISL_460858, EPI_ISL_460859, EPI_ISL_460860, EPI_ISL_460935, EPI_ISL_460938, EPI_ISL_460942, EPI_ISL_460943, EPI_ISL_460944, EPI_ISL_460945, EPI_ISL_461041, EPI_ISL_461074, EPI_ISL_461075, EPI_ISL_461141, EPI_ISL_461142, EPI_ISL_461179, EPI_ISL_461184, EPI_ISL_461185, EPI_ISL_461186, EPI_ISL_461187, EPI_ISL_461188, EPI_ISL_461189, EPI_ISL_461198, EPI_ISL_461199, EPI_ISL_461201, EPI_ISL_461206, EPI_ISL_461208, EPI_ISL_461218, EPI_ISL_461219, EPI_ISL_461225, EPI_ISL_461232, EPI_ISL_461237, EPI_ISL_461238, EPI_ISL_461239, EPI_ISL_461240, EPI_ISL_461274, EPI_ISL_461275, EPI_ISL_461276, EPI_ISL_461277, EPI_ISL_461300, EPI_ISL_461301, EPI_ISL_461303, EPI_ISL_461304, EPI_ISL_461305, EPI_ISL_461306, EPI_ISL_461310, EPI_ISL_461312, EPI_ISL_461313, EPI_ISL_461315, EPI_ISL_461325, EPI_ISL_461326, EPI_ISL_461327, EPI_ISL_461328, EPI_ISL_461329, EPI_ISL_461330, EPI_ISL_461331, EPI_ISL_461332, EPI_ISL_461333, EPI_ISL_461343, EPI_ISL_461354, EPI_ISL_461365, EPI_ISL_461368, EPI_ISL_461369, EPI_ISL_461370, EPI_ISL_461371, EPI_ISL_461372, EPI_ISL_461373, EPI_ISL_461375, EPI_ISL_461376, EPI_ISL_461377, EPI_ISL_461398 | | | |
| see above | Dutch COVID-19 response team | Erasmus Medical Center | Bas Oude Munnink, David Nieuwenhuijse, Reina Sikkema, Claudia Schapendonk, Irina Chestakova, Anne van der Linden, Theo Bestebroer, Stefan van Nieuwkoop, Mark Pronk, Pascal Lexmond, Corien Swaan, Manon Haverkate, Madelief Mollers, Mart Stein, Sandra Kengne Kamga Mobou, Jeroen van Kampen, Jolanda Voermans, Aura Timen, Corine GeurtsvanKessel, Annemiek van der Eijk, Richard Molenkamp, Marion Koopmans, on behalf of the Dutch national COVID-19 response team. |
| EPI_ISL_461406, EPI_ISL_461407 | UW Virology Lab | UW Virology Lab | Pavitra Roychoudhury, Amin Addetia, Hong Xie, Lasata Shrestha, Truong Nguyen, Meei-Li Huang, Keith Jerome, Alexander Greninger |
| EPI_ISL_461478 | Government Medical College, Vadodara | Gujarat Biotechnology Research Centre | Fenil Patel, Nidhi Patel, Nitin Savaliya, Raghawendra Kumar, Dinesh Kumar, Zuber Saiyed, Komal Patel, Labdhi Pandya, Snehal Bagatharia, Tanuja Javadekar , R N Daveshwar, Tejas Shah, Ankit Hinsu, Pritesh Sabara, Apurvasinh Puvar, Janvi Raval, Zarna Patel, Monika Gandhi, Pinal Trivedi, Maharshi Pandya, R D Dixit, A M Kadri, Harsh Bakshi, Chaitanya Joshi, Madhvi Joshi, |
| EPI_ISL_461479 | Government Medical College, Vadodara | Gujarat Biotechnology Research Centre | Neelam Nathani, Nitin Savaliya, Raghawendra Kumar, Dinesh Kumar, Zuber Saiyed, Komal Patel, Labdhi Pandya, Snehal Bagatharia, Tanuja Javadekar , R N Daveshwar, Tejas Shah, Ankit Hinsu, Pritesh Sabara, Apurvasinh Puvar, Janvi Raval, Zarna Patel, Monika Gandhi, Pinal Trivedi, Maharshi Pandya, Nidhi Patel, R D Dixit, A M Kadri, Harsh Bakshi, Chaitanya Joshi, Madhvi Joshi, |
| EPI_ISL_461480 | Government Medical College, Vadodara | Gujarat Biotechnology Research Centre | Armi Chaudhari, Raghawendra Kumar, Dinesh Kumar, Zuber Saiyed, Komal Patel, Labdhi Pandya, Snehal Bagatharia, Tanuja Javadekar , R N Daveshwar, Tejas Shah, Ankit Hinsu, Pritesh Sabara, Apurvasinh Puvar, Janvi Raval, Zarna Patel, Monika Gandhi, Pinal Trivedi, Maharshi Pandya, Nidhi Patel, Nitin Savaliya, R D Dixit, A M Kadri, Harsh Bakshi, Chaitanya Joshi, Madhvi Joshi, |
| EPI_ISL_461481 | Pandit Deendayal Upadhyay Government Medical College, Rajkot | Gujarat Biotechnology Research Centre | Bhavya Jindal, Dinesh Kumar, Zuber Saiyed, Komal Patel, Labdhi Pandya, Snehal Bagatharia, Prakash Modi, Sejul Antala, Manish Pattani, Tejas Shah, Ankit Hinsu, Pritesh Sabara, Apurvasinh Puvar, Janvi Raval, Zarna Patel, Monika Gandhi, Pinal Trivedi, Maharshi Pandya, Nidhi Patel, Nitin Savaliya, Raghawendra Kumar, R D Dixit, A M Kadri, Harsh Bakshi, Chaitanya Joshi, Madhvi Joshi |
| EPI_ISL_461482 | Pandit Deendayal Upadhyay Government Medical College, Rajkot | Gujarat Biotechnology Research Centre | Anjali Rajwar, Zuber Saiyed, Komal Patel, Labdhi Pandya, Snehal Bagatharia, Prakash Modi, Sejul Antala, Manish Pattani, Tejas Shah, Ankit Hinsu, Pritesh Sabara, Apurvasinh Puvar, Janvi Raval, Zarna Patel, Monika Gandhi, Pinal Trivedi, Maharshi Pandya, Nidhi Patel, Nitin Savaliya, Raghawendra Kumar, Dinesh Kumar, R D Dixit, A M Kadri, Harsh Bakshi, Chaitanya Joshi, Madhvi Joshi |
| EPI_ISL_461769 | University College London, Great Ormond Street Hospital for Children NHS Foundation Trust, Imperial College Healthcare NHS Trust | COVID-19 Genomics UK (COG-UK) Consortium | Sergi Castellano, Rachel Williams, Mark Kristiansen, Paola Resende Silva, Sunando Roy, Tony Brooks, Helena Tutill, Paola Niola, Patricia Dyal, Charlotte Williams, Leysa Forrest, Yasmin Panchbhaya, Jacqueline Findlay, Sam Weeks, Julianne Brown, Kathryn Harris, Paul Randell, James Price, Alison Holmes, Judith Breuer |
| EPI_ISL_461791, EPI_ISL_461792 | Northumbria University / South Tees Hospitals NHS Foundation Trust / North Cumbria Integrated Care NHS Foundation Trust / North Tees and Hartlepool NHS Foundation Trust / Newcastle Hospitals NHS Foundation Trust | COVID-19 Genomics UK (COG-UK) Consortium | Darren L Smith,Andrew Nelson,Matthew Bashton,Greg R Young,Joshua Loh,John Allan,Mohammad A Tariq,Giles S Holt,Gary Black,Wen C Yew,Lynn Dover,Paul Baker,Steve Liggett,Sarah Essex,Jane Greenaway,Debra Padgett,Clive Graham,Garren Scott,Edward Barton,Emma Swindells,Brendan Payne,Jennifer Collins,Yusri Taha,Gary Eltringham |
| EPI_ISL_461805, EPI_ISL_461807, EPI_ISL_461808, EPI_ISL_461809, EPI_ISL_461811, EPI_ISL_461812, EPI_ISL_461813, EPI_ISL_461814, EPI_ISL_461815, EPI_ISL_461816, EPI_ISL_461817, EPI_ISL_461818, EPI_ISL_461819, EPI_ISL_461820, EPI_ISL_461821, EPI_ISL_461822, EPI_ISL_461823, EPI_ISL_461824, EPI_ISL_461825, EPI_ISL_461826, EPI_ISL_461829, EPI_ISL_461830, EPI_ISL_461831, EPI_ISL_461832, EPI_ISL_461833, EPI_ISL_461834, EPI_ISL_461835, EPI_ISL_461836, EPI_ISL_461837, EPI_ISL_461838, EPI_ISL_461839, EPI_ISL_461840, EPI_ISL_461841, EPI_ISL_461842, EPI_ISL_461843, EPI_ISL_461844, EPI_ISL_461845, EPI_ISL_461846, EPI_ISL_461847, EPI_ISL_461848, EPI_ISL_461849, EPI_ISL_461850, EPI_ISL_461851, EPI_ISL_461852, EPI_ISL_461853, EPI_ISL_461884, EPI_ISL_461885, EPI_ISL_461886, EPI_ISL_461887, EPI_ISL_461888, EPI_ISL_461889, EPI_ISL_461890, EPI_ISL_461891, EPI_ISL_461892, EPI_ISL_461893 | | | |
| see above | Quadram Institute Bioscience | COVID-19 Genomics UK (COG-UK) Consortium | Dave J. Baker, Gemma L. Kay, Alp Aydin, Thanh Le-Viet, Steven Rudder, Ana P. Tedim, Anastasia Kolyva, Maria Diaz, Leonardo de Oliveira Martins, Nabil-Fareed Alikhan, Lizzie Meadows, Rachael Stanley, Ngozi Elumogo, Muhammed Yasir, Nicholas M. Thomson, Alexander J Trotter, Rachel Gilroy, Samuel Bloomfield, Claire Stuart, Andrew Bell, Reenesh Prakash, Samir Dervisevic, Alison E. Mather, John Wain, Mark Webber, Andrew J. Page, Justin O'Grady |
| EPI_ISL_461999, EPI_ISL_462006, EPI_ISL_462009, EPI_ISL_462015, EPI_ISL_462021, EPI_ISL_462025, EPI_ISL_462036, EPI_ISL_462047, EPI_ISL_462055, EPI_ISL_462059, EPI_ISL_462069 | | | |
| see above | Virology Department, Sheffield Teaching Hospitals NHS Foundation Trust/Department of Infection, Immunity and Cardiovascular Disease, The Medical School, University of Sheffield | COVID-19 Genomics UK (COG-UK) Consortium | Thushan de Silva, Matthew Parker, Nikki Smith, Adri Angyal, Rebecca Brown, Luke Green, Rachel Tucker, Paul Parsons, Danielle Groves, Katie Johnson, Laura Carrilero, Alex Keeley, Dave Partridge, Matthew Wyles, Benjamin Lindsey, Mehmet Yavuz, Mohammad Raza, Cariad Evans |
| EPI_ISL_462302, EPI_ISL_462303, EPI_ISL_462321, EPI_ISL_462322, EPI_ISL_462323, EPI_ISL_462324, EPI_ISL_462325, EPI_ISL_462326, EPI_ISL_462327, EPI_ISL_462361, EPI_ISL_462406, EPI_ISL_462411, EPI_ISL_462412 | | | |
| see above | National Public Health Laboratory, National Centre for Infectious Diseases | National Public Health Laboratory, National Centre for Infectious Diseases | Mak TM, Octavia S, Chavatte JM, Cui L, Lin RTP |
| EPI_ISL_462436 | unknown | Laboratory Diagnostic | Vidanovic,D., Tesovic,B., Banovic Djeri,B., Knezevic,A., Jankovic,M., Sekler,M., Dmitric,M., Petrovic,T., Volkening,J., Afonso,C.L. |
| EPI_ISL_462636, EPI_ISL_462637, EPI_ISL_462638, EPI_ISL_462639, EPI_ISL_462640, EPI_ISL_462641, EPI_ISL_462642, EPI_ISL_462643, EPI_ISL_462644, EPI_ISL_462645, EPI_ISL_462646, EPI_ISL_462647, EPI_ISL_462648, EPI_ISL_462649, EPI_ISL_462650, EPI_ISL_462651, EPI_ISL_462652, EPI_ISL_462653, EPI_ISL_462654, EPI_ISL_462655, EPI_ISL_462656, EPI_ISL_462657, EPI_ISL_462658, EPI_ISL_462659, EPI_ISL_462660, EPI_ISL_462661, EPI_ISL_462662, EPI_ISL_462663, EPI_ISL_462664, EPI_ISL_462665, EPI_ISL_462666, EPI_ISL_462667, EPI_ISL_462668, EPI_ISL_462669, EPI_ISL_462670, EPI_ISL_462671, EPI_ISL_462672, EPI_ISL_462673, EPI_ISL_462674, EPI_ISL_462675, EPI_ISL_462676, EPI_ISL_462677, EPI_ISL_462678, EPI_ISL_462679, EPI_ISL_462680, EPI_ISL_462681, EPI_ISL_462682, EPI_ISL_462683, EPI_ISL_462684, EPI_ISL_462685, EPI_ISL_462686, EPI_ISL_462687, EPI_ISL_462688, EPI_ISL_462689, EPI_ISL_462690, EPI_ISL_462691, EPI_ISL_462692, EPI_ISL_462693, EPI_ISL_462694, EPI_ISL_462695, EPI_ISL_462696, EPI_ISL_462697, EPI_ISL_462698, EPI_ISL_462699, EPI_ISL_462700, EPI_ISL_462701, EPI_ISL_462735, EPI_ISL_462736 | | | |
| see above | Michigan Department of Health and Human Services, Bureau of Laboratories | Michigan Department of Health and Human Services, Bureau of Laboratories | Blankenship HM, Riner D, Soehnlen MK |
| EPI_ISL_462845, EPI_ISL_462846, EPI_ISL_462847, EPI_ISL_462848, EPI_ISL_462849, EPI_ISL_462850, EPI_ISL_462851, EPI_ISL_462852, EPI_ISL_462853, EPI_ISL_462854, EPI_ISL_462855, EPI_ISL_462856, EPI_ISL_462857, EPI_ISL_462858, EPI_ISL_462859, EPI_ISL_462860, EPI_ISL_462861, EPI_ISL_462862, EPI_ISL_462863, EPI_ISL_462864 | | | |
| see above | Minnesota Department of Health, Public Health Laboratory | Minnesota Department of Health, Public Health Laboratory | Matt Plumb, Jacob Garfin, and Xiong Wang |
| EPI_ISL_462990 | University Clinical Centre of the Republic of Srpska | University of Sarajevo, Veterinary Faculty | Teufik, G., Šejla, G., Toni, E., Maja, T., Mirsada, H., Aida, K., Alma, Š.A. |
| EPI_ISL_463098, EPI_ISL_463099, | Virginia DCLS | Virginia DCLS | Virginia DCLS |

| | | | |
|---|---|---|---|
| EPI_ISL_463100, EPI_ISL_463101, EPI_ISL_463124, EPI_ISL_463126 | | | |
| EPI_ISL_463379, EPI_ISL_463380, EPI_ISL_463381, EPI_ISL_463382, EPI_ISL_463383, EPI_ISL_463384, EPI_ISL_463386, EPI_ISL_463387, EPI_ISL_463388, EPI_ISL_463389, EPI_ISL_463390, EPI_ISL_463391, EPI_ISL_463393, EPI_ISL_463394, EPI_ISL_463395, EPI_ISL_463396, EPI_ISL_463397, EPI_ISL_463398, EPI_ISL_463399, EPI_ISL_463400, EPI_ISL_463401, EPI_ISL_463402, EPI_ISL_463403, EPI_ISL_463404, EPI_ISL_463405, EPI_ISL_463406, EPI_ISL_463407, EPI_ISL_463432, EPI_ISL_463433, EPI_ISL_463434, EPI_ISL_463435, EPI_ISL_463436, EPI_ISL_463439, EPI_ISL_463440, EPI_ISL_463441, EPI_ISL_463442, EPI_ISL_463443, EPI_ISL_463444, EPI_ISL_463445, EPI_ISL_463446, EPI_ISL_463449, EPI_ISL_463450, EPI_ISL_463451, EPI_ISL_463452, EPI_ISL_463453, EPI_ISL_463454, EPI_ISL_463455, EPI_ISL_463456, EPI_ISL_463457, EPI_ISL_463458, EPI_ISL_463459, EPI_ISL_463460, EPI_ISL_463461, EPI_ISL_463462, EPI_ISL_463463, EPI_ISL_463464, EPI_ISL_463465, EPI_ISL_463466, EPI_ISL_463467, EPI_ISL_463472, EPI_ISL_463478, EPI_ISL_463479, EPI_ISL_463480, EPI_ISL_463481, EPI_ISL_463482, EPI_ISL_463483, EPI_ISL_463484, EPI_ISL_463485, EPI_ISL_463486, EPI_ISL_463487, EPI_ISL_463488, EPI_ISL_463489, EPI_ISL_463490, EPI_ISL_463491, EPI_ISL_463492, EPI_ISL_463493, EPI_ISL_463494, EPI_ISL_463495, EPI_ISL_463496, EPI_ISL_463497, EPI_ISL_463498, EPI_ISL_463499, EPI_ISL_463500, EPI_ISL_463501, EPI_ISL_463502, EPI_ISL_463503, EPI_ISL_463504, EPI_ISL_463505, EPI_ISL_463506, EPI_ISL_463507, EPI_ISL_463508, EPI_ISL_463509, EPI_ISL_463510, EPI_ISL_463511, EPI_ISL_463512, EPI_ISL_463513, EPI_ISL_463514, EPI_ISL_463515, EPI_ISL_463516, EPI_ISL_463517, EPI_ISL_463518, EPI_ISL_463519, EPI_ISL_463520, EPI_ISL_463521, EPI_ISL_463522, EPI_ISL_463523, EPI_ISL_463524, EPI_ISL_463525, EPI_ISL_463526, EPI_ISL_463527, EPI_ISL_463528, EPI_ISL_463529, EPI_ISL_463530, EPI_ISL_463531, EPI_ISL_463532, EPI_ISL_463533, EPI_ISL_463534, EPI_ISL_463535, EPI_ISL_463536, EPI_ISL_463537, EPI_ISL_463538, EPI_ISL_463539, EPI_ISL_463540, EPI_ISL_463541, EPI_ISL_463542, EPI_ISL_463543, EPI_ISL_463544, EPI_ISL_463545, EPI_ISL_463546, EPI_ISL_463547, EPI_ISL_463548, EPI_ISL_463549, EPI_ISL_463550, EPI_ISL_463551, EPI_ISL_463552, EPI_ISL_463553, EPI_ISL_463554, EPI_ISL_463555, EPI_ISL_463556, EPI_ISL_463557, EPI_ISL_463558, EPI_ISL_463559, EPI_ISL_463560, EPI_ISL_463561, EPI_ISL_463562, EPI_ISL_463563, EPI_ISL_463564, EPI_ISL_463565, EPI_ISL_463566, EPI_ISL_463567, EPI_ISL_463568, EPI_ISL_463569, EPI_ISL_463570, EPI_ISL_463571, EPI_ISL_463572, EPI_ISL_463573, EPI_ISL_463574, EPI_ISL_463575, EPI_ISL_463576, EPI_ISL_463577, EPI_ISL_463578, EPI_ISL_463579, EPI_ISL_463580, EPI_ISL_463581, EPI_ISL_463582, EPI_ISL_463583, EPI_ISL_463584, EPI_ISL_463585, EPI_ISL_463586, EPI_ISL_463587, EPI_ISL_463588, EPI_ISL_463589, EPI_ISL_463590, EPI_ISL_463591, EPI_ISL_463592, EPI_ISL_463593, EPI_ISL_463594, EPI_ISL_463595, EPI_ISL_463596, EPI_ISL_463597, EPI_ISL_463598, EPI_ISL_463599, EPI_ISL_463600, EPI_ISL_463601, EPI_ISL_463602, EPI_ISL_463603, EPI_ISL_463604, EPI_ISL_463605, EPI_ISL_463606, EPI_ISL_463607, EPI_ISL_463608, EPI_ISL_463609, EPI_ISL_463610, EPI_ISL_463611, EPI_ISL_463612, EPI_ISL_463613, EPI_ISL_463614, EPI_ISL_463615, EPI_ISL_463616, EPI_ISL_463617, EPI_ISL_463618, EPI_ISL_463619, EPI_ISL_463620, EPI_ISL_463621, EPI_ISL_463622, EPI_ISL_463623, EPI_ISL_463624, EPI_ISL_463625, EPI_ISL_463626, EPI_ISL_463627, EPI_ISL_463628, EPI_ISL_463629, EPI_ISL_463630, EPI_ISL_463634, EPI_ISL_463635, EPI_ISL_463641, EPI_ISL_463642, EPI_ISL_463644, EPI_ISL_463645, EPI_ISL_463646, EPI_ISL_463647, EPI_ISL_463648, EPI_ISL_463649, EPI_ISL_463656, EPI_ISL_463660, EPI_ISL_463662, EPI_ISL_463663 | | | |
| see above | Washington State Department of Health | Seattle Flu Study | Chu et al |
| EPI_ISL_463748 | Department of Molecular Virology, Cyprus Institute of Neurology and Genetics | Department of Molecular Virology, Cyprus Institute of Neurology and Genetics | Jan Richter, George Krashias, Christina Tryfonos, Stavros Bashiardes, Dana Koptides, Christina Christodoulou |
| EPI_ISL_464008, EPI_ISL_464029, EPI_ISL_464064 | Unity Health Toronto | Ontario Institute for Cancer Research | Ramzi Fattouh,Larissa M. Matukas,Mark Downing,Annette Gower,Karel Boissinot,Samira Mubareka,TIBDN,Ilinca Lungu,Bernard Lam,Jeremy Johns,Paul Krzyzanowski,Richard de Borja,Philip Zuzarte,Jared Simpson |
| EPI_ISL_465433, EPI_ISL_465437, EPI_ISL_465438, EPI_ISL_465442, EPI_ISL_465445, EPI_ISL_465446, EPI_ISL_465448, EPI_ISL_465451, EPI_ISL_465452, EPI_ISL_465454, EPI_ISL_465455, EPI_ISL_465457, EPI_ISL_465458, EPI_ISL_465459, EPI_ISL_465460, EPI_ISL_465461, EPI_ISL_465462, EPI_ISL_465463, EPI_ISL_465464, EPI_ISL_465465, EPI_ISL_465466, EPI_ISL_465467, EPI_ISL_465468, EPI_ISL_465469, EPI_ISL_465470, EPI_ISL_465471, EPI_ISL_465472, EPI_ISL_465474, EPI_ISL_465475, EPI_ISL_465476, EPI_ISL_465477, EPI_ISL_465478, EPI_ISL_465479, EPI_ISL_465480, EPI_ISL_465481, EPI_ISL_465482, EPI_ISL_465483, EPI_ISL_465484, EPI_ISL_465485, EPI_ISL_465486, EPI_ISL_465487, EPI_ISL_465489, EPI_ISL_465490, EPI_ISL_465491, EPI_ISL_465492, EPI_ISL_465493, EPI_ISL_465494, EPI_ISL_465495, EPI_ISL_465496, EPI_ISL_465497, EPI_ISL_465498, EPI_ISL_465500, EPI_ISL_465501, EPI_ISL_465502, EPI_ISL_465503, EPI_ISL_465504, EPI_ISL_465505, EPI_ISL_465506, EPI_ISL_465507, EPI_ISL_465508, EPI_ISL_465509, EPI_ISL_465510, EPI_ISL_465511, EPI_ISL_465512, EPI_ISL_465513, EPI_ISL_465514, EPI_ISL_465515, EPI_ISL_465516, EPI_ISL_465517, EPI_ISL_465518, EPI_ISL_465519, EPI_ISL_465520, EPI_ISL_465521, EPI_ISL_465522, EPI_ISL_465524, EPI_ISL_465525, EPI_ISL_465526, EPI_ISL_465527, EPI_ISL_465528, EPI_ISL_465529, EPI_ISL_465530, EPI_ISL_465531, EPI_ISL_465532, EPI_ISL_465533, EPI_ISL_465534, EPI_ISL_465535, EPI_ISL_465536, EPI_ISL_465537, EPI_ISL_465538, EPI_ISL_465539, EPI_ISL_465540, EPI_ISL_465541, EPI_ISL_465542, EPI_ISL_465543, EPI_ISL_465544, EPI_ISL_465545, EPI_ISL_465546, EPI_ISL_465547, EPI_ISL_465548, EPI_ISL_465549, EPI_ISL_465550, EPI_ISL_465551, EPI_ISL_465552, EPI_ISL_465553, EPI_ISL_465554, EPI_ISL_465555, EPI_ISL_465556, EPI_ISL_465557, EPI_ISL_465558, EPI_ISL_465559, EPI_ISL_465560, EPI_ISL_465561, EPI_ISL_465562, EPI_ISL_465563, EPI_ISL_465564, EPI_ISL_465565, EPI_ISL_465566, EPI_ISL_465567, EPI_ISL_465568, EPI_ISL_465569, EPI_ISL_465570, EPI_ISL_465571, EPI_ISL_465572, EPI_ISL_465573, EPI_ISL_465574, EPI_ISL_465575, EPI_ISL_465576, EPI_ISL_465577, EPI_ISL_465578, EPI_ISL_465579, EPI_ISL_465580, EPI_ISL_465581, EPI_ISL_465582, EPI_ISL_465583, EPI_ISL_465584, EPI_ISL_465585, EPI_ISL_465586, EPI_ISL_465587, EPI_ISL_465588, EPI_ISL_465589, EPI_ISL_465590, EPI_ISL_465591, EPI_ISL_465592, EPI_ISL_465593, EPI_ISL_465594, EPI_ISL_465595, EPI_ISL_465596, EPI_ISL_465597, EPI_ISL_465598, EPI_ISL_465599, EPI_ISL_465600, EPI_ISL_465602, EPI_ISL_465603, EPI_ISL_465604, EPI_ISL_465605, EPI_ISL_465607, EPI_ISL_465609, EPI_ISL_465610, EPI_ISL_465612, EPI_ISL_465614, EPI_ISL_465620, EPI_ISL_465624, EPI_ISL_465626, EPI_ISL_465627, EPI_ISL_465629, EPI_ISL_465704, EPI_ISL_465705, EPI_ISL_465706, EPI_ISL_465707, EPI_ISL_465708, EPI_ISL_465709, EPI_ISL_465710, EPI_ISL_465711, EPI_ISL_465712, EPI_ISL_465713, EPI_ISL_465714, EPI_ISL_465717, EPI_ISL_465718 | | | |
| see above | Respiratory Virus Unit, Microbiology Services Colindale, Public Health England | Respiratory Virus Unit, Microbiology Services Colindale, Public Health England | PHE Covid Sequencing Team |
| EPI_ISL_466648 | Innovative Genomics Institute, UCB | Innovative Genomics Institute, UCB | Stacia Wyman, Haridha Shivram, Liana Lareau, Shana McDevitt, Justin Choi |
| EPI_ISL_466654, EPI_ISL_466655, EPI_ISL_466656, EPI_ISL_466657, EPI_ISL_466658, EPI_ISL_466659, EPI_ISL_466660, EPI_ISL_466661, EPI_ISL_466662, EPI_ISL_466665, EPI_ISL_466666, EPI_ISL_466667, EPI_ISL_466668, EPI_ISL_466669, EPI_ISL_466670, EPI_ISL_466671, EPI_ISL_466672, EPI_ISL_466673, EPI_ISL_466674, EPI_ISL_466675, EPI_ISL_466676, EPI_ISL_466677 | | | |
| see above | Nebraska Public Health Laboratory | UNMC COVID-19 Response Team | UNMC COVID-19 Response Team |
| EPI_ISL_466898, EPI_ISL_466899, EPI_ISL_466900, EPI_ISL_466901, EPI_ISL_466902, EPI_ISL_466903, EPI_ISL_466904, EPI_ISL_466905, EPI_ISL_466906, EPI_ISL_466907 | Max von Pettenkofer Institute, Virology, National Reference Center for Retroviruses, LMU München | Laboratory for Functional Genome Analysis, Dept. Genomics, Gene Center of the LMU Munich | Max Muenchhoff, Stefan Krebs, Alexander Graf, Oliver Keppler, Helmut Blum |
| EPI_ISL_467361, EPI_ISL_467362, EPI_ISL_467363, EPI_ISL_467365, EPI_ISL_467366, EPI_ISL_467367, EPI_ISL_467368, EPI_ISL_467369, EPI_ISL_467370, EPI_ISL_467371 | Laboratory of Respiratory Viruses and Measles, Oswaldo Cruz Institute, FIOCRUZ | Laboratory of Respiratory Viruses and Measles, Oswaldo Cruz Institute, FIOCRUZ | Paola Resende, Luciana Appolinario, Fernando Motta, Anna Carolina Paixão, Ana Carolina Mendonça, Aline Mattos, Milene Miranda, Cristiana Garcia, Braulia Caetano, Maria Ogrzewalska, Jonathan Lopes, Marilda Siqueira |
| EPI_ISL_467377, EPI_ISL_467380, EPI_ISL_467387, EPI_ISL_467389, EPI_ISL_467397, EPI_ISL_467398, EPI_ISL_467399, EPI_ISL_467401, EPI_ISL_467402 | NYU Langone Health | Departments of Pathology and Medicine, New York University School of Medicine | Maria Aguero-Rosenfeld, Brendan Belovarac, Margaret Black, Ludovic Boytard, John Cadley, Paolo Cotzia, John Chen, Dacia Dimartino, Xiaojun Feng, Tatyana Gindin, Emily Guzman, Adriana Heguy, Megan Hogan, Emily Huang, George Jour, Alireza Khodadadi-Jamayran, Lawrence H. Lin, Raven Luther, Andrew Lytle, Christian Marier, Matthew T. Maurano, Mark J. Mulligan, Peter Meyn, Raquel Ordonez Ciriza, Iman Osman, Jared Pinnell, Vanessa Raabe, Sitharam Ramaswami, Amy Rapkiewicz, Andre M. Ribeiro-dos-Santos, Marie Samanovic-Golden, Antonio Serrano, Guomiao Shen, Matija Snuderl, Theodore Vougiouklakis, Nick Vulpescu, Gael Westby, Paul Zappile, Yutong Zhang |
| EPI_ISL_467667 | Florida Bureau of Public Health Laboratories | Florida Bureau of Public Health Laboratories | Schmedes,S. and Blanton,J. |
| EPI_ISL_467778, EPI_ISL_467779, EPI_ISL_467781 | National Influenza Centre Romania | Charite Universitatsmedizin Berlin, Institute of Virology | Victor M Corman, Jorn Beheim-Schwarzbach, Barbara Muehlemann, Talitha Veith, Julia Schneider, Terry Jones, L. Ustea, N. Paraschiv, M. Lazar, Christian Drosten |
| EPI_ISL_467928, EPI_ISL_467929, EPI_ISL_467930, EPI_ISL_467931, EPI_ISL_467932, EPI_ISL_467933, EPI_ISL_467934, EPI_ISL_467941 | Virginia DCLS | Virginia DCLS | Virginia DCLS |
| EPI_ISL_468044, EPI_ISL_468045, EPI_ISL_468046, EPI_ISL_468047, EPI_ISL_468048, EPI_ISL_468049, EPI_ISL_468050, EPI_ISL_468051, EPI_ISL_468052, EPI_ISL_468053, EPI_ISL_468054, EPI_ISL_468055 | | | |
| see above | Egyptian National Cancer Institute (ENCI) | Egyptian National Cancer Institute (ENCI) | Zekri, Abdel Rahman N, Amer,K.E., Ahmed,O.S., Soliman,H.K., Hafez,M.M., Bahnassy,A.A., Abdelhamid,W., Gad,A., Ali,M., Hassan,W., Samir,M., Raouf,A., Hamdy,M.S., Soliman,M.S., Elsissy,M.H., Elkhateeb,S.M., Ezzelarab,M.H., Abouelhoda, Mohamed |
| EPI_ISL_468056 | Egyptian National Cancer Institute (ENCI) | Egyptian National Cancer Institute (ENCI) | Zekri, Abdel Rahman N., Amer,K.E., Ahmed,O.S., Soliman,H.K., Ali,M.A., Hassan,W.A., Mahmoud,A.A., Khattab,A.A., Hafez,M.M., Abouelhoda, Mohamed |
| EPI_ISL_468057, EPI_ISL_468058, EPI_ISL_468059 | Egyptian National Cancer Institute (ENCI) | Egyptian National Cancer Institute (ENCI) | Zekri, Abdel Rahman N, Amer,K.E., Ahmed,O.S., Soliman,H.K., Hafez,M.M., Bahnassy,A.A., Abdelhamid,W., Gad,A., Ali,M., Hassan,W., Samir,M., Raouf,A., Hamdy,M.S., Soliman,M.S., Elsissy,M.H., Elkhateeb,S.M., Ezzelarab,M.H., Abouelhoda, Mohamed |
| EPI_ISL_468060, EPI_ISL_468061, EPI_ISL_468062 | Egyptian National Cancer Institute (ENCI) | Egyptian National Cancer Institute (ENCI) | Zekri, Abdel Rahman N., Amer,K.E., Ahmed,O.S., Soliman,H.K., Ali,M.A., Hassan,W.A., Mahmoud,A.A., Khattab,A.A., Hafez,M.M., Abouelhoda, Mohamed |
| EPI_ISL_468078 | Child Health Research Foundation | Child Health Research Foundation | Senjuti Saha, Roly Malaker, Md Saiful Islam Sajib, Hafizur Rahman, Afroza Akter Tanni, Syed Muktadir Al Sium, Maksuda Islam, Samir K Saha |
| EPI_ISL_468103, EPI_ISL_468104, EPI_ISL_468105, EPI_ISL_468106, EPI_ISL_468107, EPI_ISL_468108, EPI_ISL_468109, EPI_ISL_468115, EPI_ISL_468116 | OHSU Lab Services Molecular Microbiology Lab | Oregon SARS-CoV-2 Genome Sequencing Center | Brendan L. O'Connell, Ruth V. Nichols, Alec J. Hirsch, Guang Fan, Daniel N. Streblow, William B. Messer, Andrew C. Adey, Benjamin N. Bimber, Brian J. O'Roak |
| EPI_ISL_468250, EPI_ISL_468251, EPI_ISL_468252, EPI_ISL_468253, EPI_ISL_468254, EPI_ISL_468255, EPI_ISL_468256, EPI_ISL_468257, EPI_ISL_468258, EPI_ISL_468259, EPI_ISL_468260, EPI_ISL_468261, EPI_ISL_468262, EPI_ISL_468263, EPI_ISL_468264, EPI_ISL_468265, EPI_ISL_468266, EPI_ISL_468267, EPI_ISL_468268, EPI_ISL_468269, EPI_ISL_468270, EPI_ISL_468271, EPI_ISL_468272, EPI_ISL_468273, EPI_ISL_468274, EPI_ISL_468275, EPI_ISL_468276, EPI_ISL_468277, EPI_ISL_468278, EPI_ISL_468279, EPI_ISL_468280, EPI_ISL_468281, EPI_ISL_468282, EPI_ISL_468283, EPI_ISL_468284, EPI_ISL_468285, EPI_ISL_468286, EPI_ISL_468287, EPI_ISL_468288, EPI_ISL_468289, EPI_ISL_468290, EPI_ISL_468291, EPI_ISL_468292, EPI_ISL_468293, EPI_ISL_468294, EPI_ISL_468295, EPI_ISL_468296, EPI_ISL_468298, EPI_ISL_468299, EPI_ISL_468300, EPI_ISL_468301, EPI_ISL_468302, EPI_ISL_468303, EPI_ISL_468304 | | | |

| | | | |
|---|---|---|---|
| see above | Viollier AG | Department of Biosystems Science and Engineering, ETH Zürich | Christian Beisel, Sarah Nadeau, Ivan Topolsky, Pedro Ferreira, Philipp Jablonski, Susana Posada-Céspedes, Tobias Schär, Ina Nissen, Natascha Santacroce, Elodie Burcklen, Christiane Beckmann, Maurice Redondo, Olivier Kobel, Christoph Noppen, Sophie Seidel, Noemie Santamaria de Souza, Niko Beerenwinkel, Tanja Stadler |
| EPI_ISL_468333, EPI_ISL_468334, EPI_ISL_468335, EPI_ISL_468337 | Microbiology Service, University Hospital of A Coruna-Biomedical Research Institute | Genomes & Disease, Center for Research in Molecular Medicine and Chronic Diseases, University of Santiago de Compostela | Kelly Conde, Jorge Arca, Soraya Rumbo, Juan A. Vallejo, M Poza, G Bou, Ana Pequeno-Valtierra, Jorge Rodriguez-Castro, Javier Temes, Daniel Garcia-Souto, Martin Santamarina, Cristina Gomez, Jose M. C. Tubio |
| EPI_ISL_468425, EPI_ISL_468426, EPI_ISL_468427, EPI_ISL_468428, EPI_ISL_468429, EPI_ISL_468430, EPI_ISL_468431, EPI_ISL_468432, EPI_ISL_468433, EPI_ISL_468434, EPI_ISL_468435 | | | |
| see above | County of San Luis Obispo Public Health Laboratory | Chan-Zuckerberg Biohub | CZB Cliahub Consortium |
| EPI_ISL_468473, EPI_ISL_468474, EPI_ISL_468475, EPI_ISL_468476, EPI_ISL_468477, EPI_ISL_468478, EPI_ISL_468479, EPI_ISL_468480, EPI_ISL_468481, EPI_ISL_468488, EPI_ISL_468489, EPI_ISL_468494 | | | |
| see above | Ventura County Public Health Lab | Chan-Zuckerberg Biohub | CZB Cliahub Consortium |
| EPI_ISL_468550 | San Joaquin County Public Health Lab | Chan-Zuckerberg Biohub | CZB Cliahub Consortium |
| EPI_ISL_468747, EPI_ISL_468748, EPI_ISL_468749, EPI_ISL_468750, EPI_ISL_468751 | Facultad de Medicina UC | Center for Mathematical Modeling and Center for Genome Regulation. Santiago, Chile | Gaete A, Travisany D, Palma R, Urra C, Varas M, Allende ML, Maass A, González M, Ferres M. |
| EPI_ISL_468753, EPI_ISL_468754, EPI_ISL_468755, EPI_ISL_468756, EPI_ISL_468757, EPI_ISL_468758, EPI_ISL_468759 | Laboratorio de Biologia Molecular, Facultad de Medicina, Universidad de Atacama | Center for Mathematical Modeling and Center for Genome Regulation. Santiago, Chile | Gaete A, Travisany D, Palma R, Urra C, Varas M, Allende ML, Maass A, González M, C Echeverria |
| EPI_ISL_469055 | Jourcentralen | The Public Health Agency of Sweden | Oskar Karlsson Lindsjo, Maria Lind Karlberg, Mattias Haukland, Reza Advani, Olov Svartstrom, Anna-Malin Linde, Sandra Broddesson, Petra Edquist, Shamam Muradrasoli, Anna Risberg, Karin Tegmark-Wisell |
| EPI_ISL_469275 | Egyptian National Cancer Institute (ENCI) | Human Genome Center | Zekri, Abdel Rahman N, Amer,K.E., Ahmed,O.S., Soliman,H.K., Hafez,M.M., Bahnassy,A.A., Abdelhamid,W., Gad,A., Ali,M., Hassan,W., Samir,M., Raouf,A., Hamdy,M.S., Soliman,M.S., Elsissy,M.H., Elkhateeb,S.M., Ezzelarab,M.H., Abouelhoda, Mohamed |
| EPI_ISL_469277 | Mohammed Bin Rashid University of Medicine and Health Sciences | Al Jalila Genomics Center | Ahmad Abou Tayoun, Tom Loney, Hamda Khansaheb, Sathishkumar Ramaswamy, Divinlal Harilal, Zulfa Omar Deesi, Rupa Murthy Varghese, Hanan Al Suwaidi, Abdulmajeed Alkhaja, Mohammed Uddin, Rifat Hamoudi, Rabih Halwani, Abiola Catherine Senok, Qutayba Hamid, Norbert Nowotny, Alawi Alsheikh-Ali |
| EPI_ISL_469287, EPI_ISL_469288, EPI_ISL_469290 | Keio University Hospital | Keio University Hospital | Kenjiro Kosaki |
| EPI_ISL_469383, EPI_ISL_469464, EPI_ISL_469465, EPI_ISL_469471, EPI_ISL_469477, EPI_ISL_469478, EPI_ISL_469482, EPI_ISL_469485, EPI_ISL_469486, EPI_ISL_469489, EPI_ISL_469499, EPI_ISL_469527, EPI_ISL_469814, EPI_ISL_469817, EPI_ISL_469819, EPI_ISL_469820, EPI_ISL_469823, EPI_ISL_469826, EPI_ISL_469831, EPI_ISL_469836, EPI_ISL_469841 | | | |
| see above | PHE South West Regional Laboratory, National Infection Service | Wellcome Sanger Institute for the COVID-19 Genomics UK (COG-UK) consortium | Stephanie Hutchings, Hannah Pymont, Dr Peter Muir, Barry Vipond, Rich Hopes; and Alex Alderton, Roberto Amato, Sonia Goncalves, Ewan Harrison, David K. Jackson, Ian Johnston, Dominic Kwiatkowski, Cordelia Langford, John Sillitoe on behalf of the Wellcome Sanger Institute COVID-19 Surveillance Team (http://www.sanger.ac.uk/covid-team) |
| EPI_ISL_469845, EPI_ISL_469846, EPI_ISL_469847, EPI_ISL_469848, EPI_ISL_469856, EPI_ISL_469861, EPI_ISL_469863, EPI_ISL_469866, EPI_ISL_469867, EPI_ISL_469871, EPI_ISL_469873, EPI_ISL_469874, EPI_ISL_469875, EPI_ISL_469877, EPI_ISL_469883, EPI_ISL_469884, EPI_ISL_469886, EPI_ISL_469887, EPI_ISL_469888, EPI_ISL_469890 | | | |
| see above | Regional Virus Laboratory, Belfast Health and Social Care Trust | Wellcome Sanger Institute for the COVID-19 Genomics UK (COG-UK) consortium | Conall McCaughey, James McKenna, Tanya Curran, Susan Feeney, Alison Watt, Ciara Cox, Mairead Connor, Zoltan Molnar, David Simpson, Derek Fairley; and Alex Alderton, Roberto Amato, Sonia Goncalves, Ewan Harrison, David K. Jackson, Ian Johnston, Dominic Kwiatkowski, Cordelia Langford, John Sillitoe on behalf of the Wellcome Sanger Institute COVID-19 Surveillance Team (http://www.sanger.ac.uk/covid-team) |
| EPI_ISL_469914 | PHE South West Regional Laboratory, National Infection Service | Wellcome Sanger Institute for the COVID-19 Genomics UK (COG-UK) consortium | Stephanie Hutchings, Hannah Pymont, Dr Peter Muir, Barry Vipond, Rich Hopes; and Alex Alderton, Roberto Amato, Sonia Goncalves, Ewan Harrison, David K. Jackson, Ian Johnston, Dominic Kwiatkowski, Cordelia Langford, John Sillitoe on behalf of the Wellcome Sanger Institute COVID-19 Surveillance Team (http://www.sanger.ac.uk/covid-team) |
| EPI_ISL_470790, EPI_ISL_470791, EPI_ISL_470792, EPI_ISL_470793, EPI_ISL_470794, EPI_ISL_470795, EPI_ISL_470796, EPI_ISL_470797, EPI_ISL_470798 | M Health Fairview | Minnesota Department of Health, Public Health Laboratory | Matt Plumb, Jacob Garfin, and Xiong Wang |
| EPI_ISL_470839 | PathWest Laboratory Medicine WA | PathWest Laboratory Medicine WA | Chisha Sikazwe, Jurissa Lang, Avram Levy, David Smith and David Speers |
| EPI_ISL_470878, EPI_ISL_470879, EPI_ISL_470880 | National Institute for Communicable Diseases of the National Health Laboratory Service | National Institute for Communicable Diseases of the National Health Laboratory Service | Allam M, Ismail A, Khumalo Z, Kwenda S, van Heusden P, Mtshali P, Mnyameni F, Mohale T, Subramoney K, Bhiman JN |
| EPI_ISL_471161, EPI_ISL_471162, EPI_ISL_471163, EPI_ISL_471164, EPI_ISL_471165, EPI_ISL_471166, EPI_ISL_471167 | MRCG at LSHTM Genomics lab | MRCG at LSHTM Genomics lab | Sesay et al |
| EPI_ISL_471175 | Oslo University Hospital, Department of Medical Microbiology | Norwegian Institute of Public Health, Department of Virology | Kathrine Stene-Johansen, Kamilla Heddeland Instefjord, Hilde Elshaug, Rasmus Riis Kopperud, Karoline Bragstad, Olav Hungnes |
| EPI_ISL_471176 | Hospital of Southern Norway - Kristiansand, Department of Medical Microbiology | Norwegian Institute of Public Health, Department of Virology | Kathrine Stene-Johansen, Kamilla Heddeland Instefjord, Hilde Elshaug, Rasmus Riis Kopperud, Karoline Bragstad, Olav Hungnes |
| EPI_ISL_471177 | Oslo University Hospital, Department of Medical Microbiology | Norwegian Institute of Public Health, Department of Virology | Kathrine Stene-Johansen, Kamilla Heddeland Instefjord, Hilde Elshaug, Rasmus Riis Kopperud, Karoline Bragstad, Olav Hungnes |
| EPI_ISL_471404, EPI_ISL_471405, EPI_ISL_471406, EPI_ISL_471407, EPI_ISL_471408, EPI_ISL_471409, EPI_ISL_471410, EPI_ISL_471414, EPI_ISL_471415 | Viral Respiratory Lab, National Institute for Biomedical Research (INRB) | Pathogen Sequencing Lab, National Institute for Biomedical Research (INRB) | Placide Mbala-Kingebeni, Edith Nkwembe, Eddy Kinganda-Lusamaki, Amuri Aziza, Francisca Muyembe Mawete, Catherine Pratt, Matthias Pauthner, Josh Quick, Allison Black, James Hadfield, Trevor Bedford, Ian Goodfellow, Andrew Rambaut, Nick Loman, Kristian Andersen, Michael Wiley, Steve Ahuka-Mundeke, Jean-Jacques Muyembe Tamfum |
| EPI_ISL_471562, EPI_ISL_471581, EPI_ISL_471582 | Hosp. Municipal Prof. Dr. Alípio Corrêa Netto | Instituto Adolfo Lutz, Interdiciplinary Procedures Center, Strategic Laboratory | Claudio Tavares Sacchi, Claudia Regina Gonçalves, Erica Valessa Ramos Gomes |
| EPI_ISL_471583 | King Institute of Preventive Medicine & Research | CSIR-Centre for Cellular and Molecular Biology | K.Kaveri,S.Sivasubramanian,S.Vennila,P.Padmapriya,R.Kiruba,S.Magesh,G. Dhinakar Raj, G. Ravikumar, P. Azhahianambi,K Thangaraj,Payel Mukherjee, Sofia Banu, Priya Singh, Dhiviya Vedagiri, Divya Gupta, Vishal Sah, Santosh Kumar Kuncha, Krishnan Harinivas Harshan, Archana Bharadwaj Siva, Karthik Bharadwaj Tallapaka, Shagufta Khan, Lamuk Zaveri, Namami Gaur, Sakshi Shambhavi, Tulasi Nagabandi, Purushotham Vodnala, Rakesh K Mishra, Divya Tej Sowpati |
| EPI_ISL_471647 | Hospital Municipal de Barueri Dr. Francisco Moran | Instituto Adolfo Lutz, Interdiciplinary Procedures Center, Strategic Laboratory | Claudio Tavares Sacchi, Claudia Regina Gonçalves, Erica Valessa Ramos Gomes |
| EPI_ISL_471648 | UBS e Pronto Socorro Jd. Jacira | Instituto Adolfo Lutz, Interdiciplinary Procedures Center, Strategic Laboratory | Claudio Tavares Sacchi, Claudia Regina Gonçalves, Erica Valessa Ramos Gomes |
| EPI_ISL_471679, EPI_ISL_471680, EPI_ISL_471681, EPI_ISL_471682, EPI_ISL_471683, EPI_ISL_471684, EPI_ISL_471685, EPI_ISL_471686, EPI_ISL_471687, EPI_ISL_471688, EPI_ISL_471689, EPI_ISL_471690, EPI_ISL_471691, EPI_ISL_471692, EPI_ISL_471693, EPI_ISL_471720, EPI_ISL_471721, EPI_ISL_471722, EPI_ISL_471723, EPI_ISL_471724, EPI_ISL_471725, EPI_ISL_471726, EPI_ISL_471727, EPI_ISL_471728, EPI_ISL_471730, EPI_ISL_471731, EPI_ISL_471732, EPI_ISL_471733, EPI_ISL_471734, EPI_ISL_471735, EPI_ISL_471736, EPI_ISL_471737, EPI_ISL_471738, EPI_ISL_471755, EPI_ISL_471756, EPI_ISL_471757, | | | |

| | | | |
|---|---|---|---|
| EPI_ISL_471758, EPI_ISL_471759, EPI_ISL_471760, EPI_ISL_471761, EPI_ISL_471762, EPI_ISL_471763, EPI_ISL_471764, EPI_ISL_471765, EPI_ISL_471766, EPI_ISL_471767, EPI_ISL_471768, EPI_ISL_471769, EPI_ISL_471770, EPI_ISL_471771, EPI_ISL_471772, EPI_ISL_471773, EPI_ISL_471795, EPI_ISL_471811, EPI_ISL_471812, EPI_ISL_471813, EPI_ISL_471814, EPI_ISL_471815, EPI_ISL_471816, EPI_ISL_471817, EPI_ISL_471818, EPI_ISL_471835, EPI_ISL_471860, EPI_ISL_471861, EPI_ISL_471863, EPI_ISL_471864, EPI_ISL_471865, EPI_ISL_471866, EPI_ISL_471867, EPI_ISL_471868, EPI_ISL_471869, EPI_ISL_471871, EPI_ISL_471872, EPI_ISL_471874, EPI_ISL_471875, EPI_ISL_471877, EPI_ISL_471878, EPI_ISL_471879, EPI_ISL_471880, EPI_ISL_471881, EPI_ISL_471882, EPI_ISL_471883, EPI_ISL_471884, EPI_ISL_471886, EPI_ISL_471889, EPI_ISL_471890, EPI_ISL_471910 | | | |
| see above | Michigan Department of Health and Human Services, Bureau of Laboratories | Michigan Department of Health and Human Services, Bureau of Laboratories | Blankenship HM, Riner D, Soehnlen MK |
| EPI_ISL_471912, EPI_ISL_471932, EPI_ISL_471936, EPI_ISL_471951, EPI_ISL_471986, EPI_ISL_471987, EPI_ISL_471988, EPI_ISL_471989, EPI_ISL_471990, EPI_ISL_471991 | University of Exeter | COVID-19 Genomics UK (COG-UK) Consortium | Ben Temperton,Aaron Jeffries,Michelle Michelsen,Joanna Warwick-Dugdale,Audrey Farbos,Robyn Manley,Stephen Michell,Jane Masoli |
| EPI_ISL_472259, EPI_ISL_472260, EPI_ISL_472265, EPI_ISL_472266 | Northumbria University / South Tees Hospitals NHS Foundation Trust / North Cumbria Integrated Care NHS Foundation Trust / North Tees and Hartlepool NHS Foundation Trust / Newcastle Hospitals NHS Foundation Trust | COVID-19 Genomics UK (COG-UK) Consortium | Darren L Smith,Andrew Nelson,Matthew Bashton,Greg R Young,Joshua Loh,John Allan,Mohammad A Tariq,Giles S Holt,Gary Black,Wen C Yew,Lynn Dover,Paul Baker,Steve Liggett,Sarah Essex,Jane Greenaway,Debra Padgett,Clive Graham,Garren Scott,Edward Barton,Emma Swindells,Brendan Payne,Jennifer Collins,Yusri Taha,Gary Eltringham |
| EPI_ISL_472296, EPI_ISL_472297, EPI_ISL_472298, EPI_ISL_472299, EPI_ISL_472300, EPI_ISL_472301, EPI_ISL_472302, EPI_ISL_472303, EPI_ISL_472304, EPI_ISL_472305, EPI_ISL_472306, EPI_ISL_472307, EPI_ISL_472310, EPI_ISL_472312, EPI_ISL_472314 | | | |
| see above | Quadram Institute Bioscience | COVID-19 Genomics UK (COG-UK) Consortium | Dave J. Baker, Gemma L. Kay, Alp Aydin, Thanh Le-Viet, Steven Rudder, Ana P. Tedim, Anastasia Kolyva, Maria Diaz, Leonardo de Oliveira Martins, Nabil-Fareed Alikhan, Lizzie Meadows, Rachael Stanley, Ngozi Elumogo, Muhammed Yasir, Nicholas M. Thomson, Alexander J Trotter, Rachel Gilroy, Samuel Bloomfield, Claire Stuart, Andrew Bell, Reenesh Prakash, Samir Dervisevic, Alison E. Mather, John Wain, Mark Webber, Andrew J. Page, Justin O'Grady |
| EPI_ISL_472407, EPI_ISL_472408, EPI_ISL_472411, EPI_ISL_472419, EPI_ISL_472420, EPI_ISL_472421, EPI_ISL_472422, EPI_ISL_472423, EPI_ISL_472425, EPI_ISL_472426, EPI_ISL_472427, EPI_ISL_472428, EPI_ISL_472429 | | | |
| see above | Queens Medical Centre, Clinical Microbiology Department / DeepSeq Nottingham | COVID-19 Genomics UK (COG-UK) Consortium | Gemma Clark, Wendy Smith, Manjinder Khakh, Vicki M Fleming, Michelle M Lister, Hannah Howson-Wells, Jonathan Ball, Patrick McClure, Joseph Chappell, Theocharis Tsoleridis, Nadine Holmes, Matthew Carlisle, Christopher Moore, Fei Sang, Johnny Debebe, Victoria Wright, Matthew Loose |
| EPI_ISL_472430, EPI_ISL_472431 | Queens Medical Centre, Clinical Microbiology Department / DeepSeq Nottingham | COVID-19 Genomics UK (COG-UK) Consortium | Nichola Duckworth, Tim Sloan, Sarah Walsh, Jonathan Ball, Patrick McClure, Joeseph Chappell, Nadine Holmes, Matthew Carlisle, Christopher Moore, Fei Sang, Johnny Debebe, Victoria Wright, Matthew Loose |
| EPI_ISL_472437, EPI_ISL_472443, EPI_ISL_472446, EPI_ISL_472448, EPI_ISL_472453, EPI_ISL_472465, EPI_ISL_472474, EPI_ISL_472476, EPI_ISL_472483, EPI_ISL_472485, EPI_ISL_472492, EPI_ISL_472494, EPI_ISL_472496, EPI_ISL_472497, EPI_ISL_472505, EPI_ISL_472510, EPI_ISL_472512, EPI_ISL_472513, EPI_ISL_472515, EPI_ISL_472516, EPI_ISL_472536, EPI_ISL_472537, EPI_ISL_472544, EPI_ISL_472548, EPI_ISL_472555, EPI_ISL_472564, EPI_ISL_472566, EPI_ISL_472571, EPI_ISL_472582, EPI_ISL_472584, EPI_ISL_472587, EPI_ISL_472590, EPI_ISL_472598, EPI_ISL_472611, EPI_ISL_472615, EPI_ISL_472616, EPI_ISL_472617, EPI_ISL_472622, EPI_ISL_472627, EPI_ISL_472629, EPI_ISL_472633, EPI_ISL_472643, EPI_ISL_472645, EPI_ISL_472662, EPI_ISL_472663, EPI_ISL_472664, EPI_ISL_472665, EPI_ISL_472668, EPI_ISL_472669, EPI_ISL_472672, EPI_ISL_472683, EPI_ISL_472702, EPI_ISL_472705, EPI_ISL_472710, EPI_ISL_472711, EPI_ISL_472713, EPI_ISL_472716, EPI_ISL_472718, EPI_ISL_472719, EPI_ISL_472724, EPI_ISL_472727, EPI_ISL_472730, EPI_ISL_472732, EPI_ISL_472836, EPI_ISL_472908, EPI_ISL_472948, EPI_ISL_472990, EPI_ISL_473283, EPI_ISL_473287, EPI_ISL_473291, EPI_ISL_473294, EPI_ISL_473299, EPI_ISL_473304 | | | |
| see above | Wales Specialist Virology Centre Sequencing lab: Pathogen Genomics Unit | COVID-19 Genomics UK (COG-UK) Consortium | Catherine Moore, Johnathan Evans, Laura Gifford, Malorie Perry, Simon Cottrell, Angela Marchbank, Alec Birchley, Alexander Adams, Amy Gaskin, Bree Gatica-Wilcox, Jason Coombes, Joel Southgate, Lauren Gilbert, Lee Graham, Nicole Pacchiarini, Sara Kumziene-Summerhayes, Sarah Taylor, Sophie Jones, Sara Rey, Matthew Bull, Joanne Watkins, Sally Corden, Tom Connor |
| EPI_ISL_473313, EPI_ISL_473314, EPI_ISL_473315, EPI_ISL_473316, EPI_ISL_473317, EPI_ISL_473321, EPI_ISL_473322, EPI_ISL_473323, EPI_ISL_473324, EPI_ISL_473325 | University of Birmingham | COVID-19 Genomics UK (COG-UK) Consortium | Institute of Microbiology, University of Birmingham: Claire McMurray, Joanne Stockton, Samuel Nicholls, Radoslaw Poplawski, Will Rowe, Josh Quick, Nicholas Loman, University of Birmingham Testing Laboratory: Celina M Whalley, Andrew Bosworth, Charlotte Poxon, Kasun Wanigasooriya, Oliver Pickles, Mike Kidd, Alex Richter, Andrew D Beggs PHE Heartlands Lab: Husam Osman, Andrew Bosworth. Queen Elizabeth Hospital: Anna Casey |
| EPI_ISL_473510, EPI_ISL_473646 | West of Scotland Specialist Virology Centre, NHSGGC / MRC-University of Glasgow Centre for Virus Research | COVID-19 Genomics UK (COG-UK) Consortium | Ana da Silva Filipe, Natasha Johnson, Kathy Smollett, Daniel Mair, Stephen Carmichael, Lily Tong, Jenna Nichols, Elihu Aranday-Cortes, Kirstyn Brunker, Yasmin Parr, Alice Broos, Kyriaki Nomikou; Sarah McDonald, Marc Niebel, Patawee Asamaphan; Richard Orton, Joseph Hughes, Sreenu Vattipally, David L Robertson; Alasdair MacLean, Rory Gunson; Kathy Li, Natasha Jesudason, Rajiv Shah, James Shepherd, Antonia Ho, Emma Thomson |
| EPI_ISL_473790, EPI_ISL_473855, EPI_ISL_473856, EPI_ISL_473857, EPI_ISL_473858, EPI_ISL_473859, EPI_ISL_473860, EPI_ISL_473861, EPI_ISL_473862, EPI_ISL_473863, EPI_ISL_473864, EPI_ISL_473865, EPI_ISL_473866, EPI_ISL_473867, EPI_ISL_473868, EPI_ISL_473869, EPI_ISL_473870, EPI_ISL_473871, EPI_ISL_473949, EPI_ISL_473950 | | | |
| see above | Virology Department, Royal Infirmary of Edinburgh, NHS Lothian / School of Biological Sciences, University of Edinburgh / Institute of Genetics and Molecular Medicine, University of Edinburgh | COVID-19 Genomics UK (COG-UK) Consortium | McHugh M, Dewar R, Rooke S, Gallagher M, Balcaza C, O'Toole Á, Scher E, Hill V, McCrone JT, Colquhoun R, Yu X, Jackson B, Rambaut A, Williams TC, Templeton K |
| EPI_ISL_473956, EPI_ISL_473962, EPI_ISL_473976, EPI_ISL_473978, EPI_ISL_473979, EPI_ISL_473980, EPI_ISL_473981, EPI_ISL_473984, EPI_ISL_473986, EPI_ISL_473993, EPI_ISL_473996, EPI_ISL_474002, EPI_ISL_474004, EPI_ISL_474005, EPI_ISL_474007, EPI_ISL_474011, EPI_ISL_474012, EPI_ISL_474017, EPI_ISL_474026, EPI_ISL_474028, EPI_ISL_474046, EPI_ISL_474049, EPI_ISL_474059, EPI_ISL_474060, EPI_ISL_474061, EPI_ISL_474062, EPI_ISL_474067, EPI_ISL_474068, EPI_ISL_474070, EPI_ISL_474082, EPI_ISL_474083, EPI_ISL_474084, EPI_ISL_474086, EPI_ISL_474088, EPI_ISL_474089, EPI_ISL_474092, EPI_ISL_474093, EPI_ISL_474100, EPI_ISL_474101, EPI_ISL_474104, EPI_ISL_474109, EPI_ISL_474110, EPI_ISL_474111, EPI_ISL_474118, EPI_ISL_474122, EPI_ISL_474127, EPI_ISL_474129, EPI_ISL_474139, EPI_ISL_474141, EPI_ISL_474145, EPI_ISL_474146, EPI_ISL_474148, EPI_ISL_474156, EPI_ISL_474161, EPI_ISL_474162, EPI_ISL_474164, EPI_ISL_474166, EPI_ISL_474174, EPI_ISL_474175, EPI_ISL_474176, EPI_ISL_474179, EPI_ISL_474180, EPI_ISL_474181, EPI_ISL_474182, EPI_ISL_474184, EPI_ISL_474186, EPI_ISL_474189, EPI_ISL_474191, EPI_ISL_474196, EPI_ISL_474205, EPI_ISL_474211, EPI_ISL_474215, EPI_ISL_474217, EPI_ISL_474219, EPI_ISL_474531, EPI_ISL_474532, EPI_ISL_474533, EPI_ISL_474534, EPI_ISL_474536, EPI_ISL_474537, EPI_ISL_474538, EPI_ISL_474540, EPI_ISL_474541, EPI_ISL_474544, EPI_ISL_474545, EPI_ISL_474546, EPI_ISL_474547, EPI_ISL_474548, EPI_ISL_474549, EPI_ISL_474550, EPI_ISL_474551, EPI_ISL_474552, EPI_ISL_474553, EPI_ISL_474554, EPI_ISL_474557, EPI_ISL_474558, EPI_ISL_474559, EPI_ISL_474561, EPI_ISL_474562, EPI_ISL_474563, EPI_ISL_474566, EPI_ISL_474567, EPI_ISL_474568, EPI_ISL_474569, EPI_ISL_474570, EPI_ISL_474571, EPI_ISL_474572, EPI_ISL_474573, EPI_ISL_474574, EPI_ISL_474575, EPI_ISL_474577, EPI_ISL_474578, EPI_ISL_474579, EPI_ISL_474580, EPI_ISL_474581, EPI_ISL_474582, EPI_ISL_474583, EPI_ISL_474584, EPI_ISL_474585, EPI_ISL_474586, EPI_ISL_474587, EPI_ISL_474588, EPI_ISL_474589, EPI_ISL_474590, EPI_ISL_474591, EPI_ISL_474592, EPI_ISL_474593, EPI_ISL_474594, EPI_ISL_474595, EPI_ISL_474596, EPI_ISL_474597, EPI_ISL_474598, EPI_ISL_474599, EPI_ISL_474600, EPI_ISL_474601, EPI_ISL_474602, EPI_ISL_474603, EPI_ISL_474604, EPI_ISL_474605, EPI_ISL_474606, EPI_ISL_474607, EPI_ISL_474608, EPI_ISL_474609, EPI_ISL_474610, EPI_ISL_474611, EPI_ISL_474612, EPI_ISL_474613, EPI_ISL_474614, EPI_ISL_474615, EPI_ISL_474616, EPI_ISL_474617, EPI_ISL_474618, EPI_ISL_474619, EPI_ISL_474621, EPI_ISL_474623, EPI_ISL_474624, EPI_ISL_474625, EPI_ISL_474626, EPI_ISL_474627, EPI_ISL_474628, EPI_ISL_474629, EPI_ISL_474630, EPI_ISL_474631, EPI_ISL_474632, EPI_ISL_474633, EPI_ISL_474634, EPI_ISL_474635, EPI_ISL_474636, EPI_ISL_474637, EPI_ISL_474638, EPI_ISL_474639, EPI_ISL_474640, EPI_ISL_474641, EPI_ISL_474642, EPI_ISL_474643, EPI_ISL_474644, EPI_ISL_474645, EPI_ISL_474646, EPI_ISL_474647, EPI_ISL_474648, EPI_ISL_474649, EPI_ISL_474650, EPI_ISL_474651, EPI_ISL_474652, EPI_ISL_474653, EPI_ISL_474654, EPI_ISL_474655, EPI_ISL_474656, EPI_ISL_474657, EPI_ISL_474658, EPI_ISL_474659, EPI_ISL_474660, EPI_ISL_474661, EPI_ISL_474662, EPI_ISL_474663, EPI_ISL_474664, EPI_ISL_474665, EPI_ISL_474666, EPI_ISL_474667, EPI_ISL_474668, EPI_ISL_474669, EPI_ISL_474670, EPI_ISL_474671, EPI_ISL_474672, EPI_ISL_474673, EPI_ISL_474674, EPI_ISL_474675, EPI_ISL_474676, EPI_ISL_474677, EPI_ISL_474678, EPI_ISL_474679, EPI_ISL_474680, EPI_ISL_474681, EPI_ISL_474682, EPI_ISL_474683, EPI_ISL_474684, EPI_ISL_474685, EPI_ISL_474686, EPI_ISL_474687, EPI_ISL_474688, EPI_ISL_474689, EPI_ISL_474690, EPI_ISL_474691, EPI_ISL_474692, EPI_ISL_474693, EPI_ISL_474694, EPI_ISL_474695, EPI_ISL_474696, EPI_ISL_474697, EPI_ISL_474698, EPI_ISL_474699, EPI_ISL_474700, EPI_ISL_474701, EPI_ISL_474702, EPI_ISL_474703, EPI_ISL_474704, EPI_ISL_474705, EPI_ISL_474706, EPI_ISL_474707, EPI_ISL_474708, EPI_ISL_474709, EPI_ISL_474710, EPI_ISL_474711, EPI_ISL_474712, EPI_ISL_474713, EPI_ISL_474714, EPI_ISL_474715, EPI_ISL_474716, EPI_ISL_474717, EPI_ISL_474718, EPI_ISL_474719, EPI_ISL_474720, EPI_ISL_474721, EPI_ISL_474722, EPI_ISL_474723, EPI_ISL_474724, EPI_ISL_474725, EPI_ISL_474726, EPI_ISL_474727, EPI_ISL_474728, EPI_ISL_474729, EPI_ISL_474730, EPI_ISL_474731, EPI_ISL_474732, EPI_ISL_474733, EPI_ISL_474734, EPI_ISL_474735, EPI_ISL_474746, EPI_ISL_474747, EPI_ISL_474748, EPI_ISL_474749, EPI_ISL_474750, EPI_ISL_474751, EPI_ISL_474752, EPI_ISL_474753, EPI_ISL_474754, EPI_ISL_474755, EPI_ISL_474756, EPI_ISL_474757, EPI_ISL_474758, EPI_ISL_474759, EPI_ISL_474760, EPI_ISL_474761, EPI_ISL_474762, EPI_ISL_474763, EPI_ISL_474764, EPI_ISL_474765, EPI_ISL_474766, EPI_ISL_474767, EPI_ISL_474768, EPI_ISL_474769, EPI_ISL_474770, EPI_ISL_474771, EPI_ISL_474772, EPI_ISL_474773, EPI_ISL_474774, EPI_ISL_474775, EPI_ISL_474776, EPI_ISL_474777, EPI_ISL_474778, EPI_ISL_474779, EPI_ISL_474780, EPI_ISL_474781, EPI_ISL_474782, EPI_ISL_474783, EPI_ISL_474784, EPI_ISL_474785, EPI_ISL_474786, EPI_ISL_474787, EPI_ISL_474788, EPI_ISL_474789, EPI_ISL_474790, EPI_ISL_474791, EPI_ISL_474792, EPI_ISL_474793, EPI_ISL_474794, EPI_ISL_474795, EPI_ISL_474796 | | | |
| see above | Wales Specialist Virology Centre Sequencing lab: Pathogen Genomics Unit | COVID-19 Genomics UK (COG-UK) Consortium | Catherine Moore, Johnathan Evans, Laura Gifford, Malorie Perry, Simon Cottrell, Angela Marchbank, Alec Birchley, Alexander Adams, Amy Gaskin, Bree Gatica-Wilcox, Jason Coombes, Joel Southgate, Lauren Gilbert, Lee Graham, Nicole Pacchiarini, Sara Kumziene-Summerhayes, Sarah Taylor, Sophie Jones, Sara Rey, Matthew Bull, Joanne Watkins, Sally Corden, Tom Connor |
| EPI_ISL_475009 | Israel Central Virology laboratory | Israel Central Virology laboratory | Neta Zuckerman, Efrat Dahan Bucris, Oran Erster, Ella Mendelson, Michal Mandelboim |
| EPI_ISL_475176, EPI_ISL_475177, EPI_ISL_475178, EPI_ISL_475179, EPI_ISL_475180, EPI_ISL_475181, EPI_ISL_475182, EPI_ISL_475183, EPI_ISL_475184, EPI_ISL_475185, EPI_ISL_475186, EPI_ISL_475187, EPI_ISL_475188, EPI_ISL_475189, EPI_ISL_475190, EPI_ISL_475191, EPI_ISL_475192, EPI_ISL_475193, EPI_ISL_475194, EPI_ISL_475195, EPI_ISL_475196, EPI_ISL_475197, EPI_ISL_475198, EPI_ISL_475199, EPI_ISL_475200, EPI_ISL_475201, EPI_ISL_475202, EPI_ISL_475203, EPI_ISL_475204, EPI_ISL_475205, EPI_ISL_475206, EPI_ISL_475207, EPI_ISL_475208, EPI_ISL_475209, EPI_ISL_475211, EPI_ISL_475216 | | | |
| see above | Nebraska Public Health Laboratory | UNMC COVID-19 Response Team | UNMC COVID-19 Response Team |
| EPI_ISL_475246, EPI_ISL_475247, EPI_ISL_475249, EPI_ISL_475250, EPI_ISL_475251, EPI_ISL_475252, EPI_ISL_475253, EPI_ISL_475254, EPI_ISL_475255, EPI_ISL_475256, EPI_ISL_475257, EPI_ISL_475258, EPI_ISL_475262, EPI_ISL_475263, EPI_ISL_475264, EPI_ISL_475265, EPI_ISL_475267, EPI_ISL_475268, EPI_ISL_475275, EPI_ISL_475276, EPI_ISL_475277, EPI_ISL_475278 | | | |
| see above | Centre for Enzyme Innovation, University of Portsmouth / Translational Research Laboratory, Portsmouth Hospitals NHS Trust | COVID-19 Genomics UK (COG-UK) Consortium | Angela Beckett,Yann Bourgeois,Garry Scarlett,Sharon Glaysher,Scott Elliott,Kelly Bicknell,Robert Impey,Allyson Lloyd,Sarah Wyllie,Ethan Butcher,Anoop Chauhan,Samuel Robson |

| | | | |
|---|---|---|---|
| EPI_ISL_475342, EPI_ISL_475343, EPI_ISL_475345, EPI_ISL_475346, EPI_ISL_475350, EPI_ISL_475351, EPI_ISL_475353, EPI_ISL_475354, EPI_ISL_475355, EPI_ISL_475360, EPI_ISL_475361, EPI_ISL_475362, EPI_ISL_475363, EPI_ISL_475365, EPI_ISL_475366, EPI_ISL_475368, EPI_ISL_475369, EPI_ISL_475370, EPI_ISL_475371, EPI_ISL_475372, EPI_ISL_475373, EPI_ISL_475380, EPI_ISL_475381, EPI_ISL_475382, EPI_ISL_475383, EPI_ISL_475385, EPI_ISL_475387, EPI_ISL_475388, EPI_ISL_475395, EPI_ISL_475396, EPI_ISL_475402, EPI_ISL_475405, EPI_ISL_475407, EPI_ISL_475409, EPI_ISL_475410, EPI_ISL_475413, EPI_ISL_475414, EPI_ISL_475415, EPI_ISL_475416, EPI_ISL_475419, EPI_ISL_475421, EPI_ISL_475424, EPI_ISL_475425, EPI_ISL_475427, EPI_ISL_475428, EPI_ISL_475429, EPI_ISL_475433, EPI_ISL_475436, EPI_ISL_475438, EPI_ISL_475443, EPI_ISL_475444, EPI_ISL_475445, EPI_ISL_475447, EPI_ISL_475448, EPI_ISL_475450, EPI_ISL_475452, EPI_ISL_475454, EPI_ISL_475455, EPI_ISL_475459, EPI_ISL_475463, EPI_ISL_475464, EPI_ISL_475470, EPI_ISL_475471, EPI_ISL_475473, EPI_ISL_475475, EPI_ISL_475477, EPI_ISL_475479, EPI_ISL_475480, EPI_ISL_475484, EPI_ISL_475485, EPI_ISL_475486, EPI_ISL_475489, EPI_ISL_475491, EPI_ISL_475494, EPI_ISL_475496, EPI_ISL_475499, EPI_ISL_475503, EPI_ISL_475504, EPI_ISL_475505, EPI_ISL_475506, EPI_ISL_475508 | | | |
| see above | Virology Department, Sheffield Teaching Hospitals NHS Foundation Trust/Department of Infection, Immunity and Cardiovascular Disease, The Medical School, University of Sheffield | COVID-19 Genomics UK (COG-UK) Consortium | Thushan de Silva, Matthew Parker, Nikki Smith, Adri Angyal, Rebecca Brown, Luke Green, Rachel Tucker, Paul Parsons, Danielle Groves, Katie Johnson, Laura Carrilero, Alex Keeley, Dave Partridge, Matthew Wyles, Benjamin Lindsey, Mehmet Yavuz, Mohammad Raza, Cariad Evans |
| EPI_ISL_475511 | Orestadsklinikens VC | The Public Health Agency of Sweden | Oskar Karlsson Lindsjo, Maria Lind Karlberg, Mattias Haukland, Reza Advani, Olov Svartstrom, Anna-Malin Linde, Sandra Broddesson, Mia Brytting, Anna Risberg, Karin Tegmark-Wisell |
| EPI_ISL_475512 | Din Klinik | The Public Health Agency of Sweden | Oskar Karlsson Lindsjo, Maria Lind Karlberg, Mattias Haukland, Reza Advani, Olov Svartstrom, Anna-Malin Linde, Sandra Broddesson, Mia Brytting, Anna Risberg, Karin Tegmark-Wisell |
| EPI_ISL_475513 | Huddinge VC | The Public Health Agency of Sweden | Oskar Karlsson Lindsjo, Maria Lind Karlberg, Mattias Haukland, Reza Advani, Olov Svartstrom, Anna-Malin Linde, Sandra Broddesson, Mia Brytting, Anna Risberg, Karin Tegmark-Wisell |
| EPI_ISL_475514 | Uppsala Narakut Aleris | The Public Health Agency of Sweden | Oskar Karlsson Lindsjo, Maria Lind Karlberg, Mattias Haukland, Reza Advani, Olov Svartstrom, Anna-Malin Linde, Sandra Broddesson, Mia Brytting, Anna Risberg, Karin Tegmark-Wisell |
| EPI_ISL_475541 | Follinge Halsocentral | The Public Health Agency of Sweden | Oskar Karlsson Lindsjo, Maria Lind Karlberg, Mattias Haukland, Reza Advani, Olov Svartstrom, Anna-Malin Linde, Sandra Broddesson, Mia Brytting, Anna Risberg, Karin Tegmark-Wisell |
| EPI_ISL_475722 | Egyptian National Cancer Institute (ENCI) | Egyptian National Cancer Institute (ENCI) | Zekri, Abdel Rahman N, Amer,K.E., Ahmed,O.S., Soliman,H.K., Hafez,M.M., Bahnassy,A.A., Abdelhamid,W., Gad,A., Ali,M., Hassan,W., Samir,M., Raouf,A., Hamdy,M.S., Soliman,M.S., Elsissy,M.H., Elkhateeb,S.M., Ezzelarab,M.H., Abouelhoda, Mohamed |
| EPI_ISL_475745, EPI_ISL_475746, EPI_ISL_475747, EPI_ISL_475748, EPI_ISL_475749, EPI_ISL_475750, EPI_ISL_475751, EPI_ISL_475752, EPI_ISL_475753 | Medical Ain Shams Research Institute (MASRI), Ain Shams University | Medical Ain Shams Research Institute (MASRI), Ain Shams University | Hesham Elghazaly , Sara Hassan Agwa, Mahmoud Elmeteini , Ahmad Moustafa , Ashraf Omar, Osama Mansour, Samia Abdo, Hala Hafez, Ghada Ismael , Shaimaa Moustafa , Aya Mohamed, Reham Mamdouh , Hoda Abd Elsatar, Manal Hamdy Elsaid, Fatma Ebied |
| EPI_ISL_475903, EPI_ISL_475904 | Zentralinstitut für medizinische und chemische Labordiagnostik, Universitätskliniken Insbruck | Bergthaler laboratory, CeMM Research Center for Molecular Medicine of the Austrian Academy of Sciences | Alexandra Popa, Benedikt Agerer, Henrique Colaco, Lukas Endler, Jakob-Wendelin Genger, Alexander Lercher, Mark Smyth, Thomas Penz, Michael Schuster, Jan Laine, Martin Senekowitsch, Judith Aberle, Stephan Aberle, Peter Hufnagl, Daniela Schmid, Franz Allerberger, Elisabeth Puchhammer-Stoeckl, Manfred Nairz, Guenter Weiss, Gregor Hörmann, Kinga Rigler-Hohenwarter, Rainer Gattringer, Wegene Borena, Dorothee von Laer, Christoph Bock, Andreas Bergthaler |
| EPI_ISL_475938, EPI_ISL_475942, EPI_ISL_475993, EPI_ISL_475994, EPI_ISL_475995 | National Public Health Laboratory, National Centre for Infectious Diseases | National Public Health Laboratory, National Centre for Infectious Diseases | Mak TM, Octavia S, Chavatte JM, Cui L, Lin RTP |
| EPI_ISL_476025 | Laboratoire de Recherche et d'Analyses Médicales de la Gendarmerie Royale | Laboratoire de Recherche et d'Analyses Médicales de la Gendarmerie Royale | Sanaâ LEMRISS, Amal Souiri, Saâd EL KABBAJ |
| EPI_ISL_476041, EPI_ISL_476042, EPI_ISL_476043, EPI_ISL_476044, EPI_ISL_476045, EPI_ISL_476046, EPI_ISL_476047, EPI_ISL_476048, EPI_ISL_476049, EPI_ISL_476050 | Michigan Department of Health and Human Services, Bureau of Laboratories | Michigan Department of Health and Human Services, Bureau of Laboratories | Blankenship HM, Riner D, Soehnlen MK |
| EPI_ISL_476101, EPI_ISL_476102, EPI_ISL_476103, EPI_ISL_476104, EPI_ISL_476105, EPI_ISL_476106, EPI_ISL_476107, EPI_ISL_476108, EPI_ISL_476109, EPI_ISL_476110, EPI_ISL_476112, EPI_ISL_476113, EPI_ISL_476114, EPI_ISL_476115, EPI_ISL_476116, EPI_ISL_476117, EPI_ISL_476118, EPI_ISL_476119 | | | |
| see above | Viollier AG | Department of Biosystems Science and Engineering, ETH Zürich | Christian Beisel, Sarah Nadeau, Ivan Topolsky, Pedro Ferreira, Philipp Jablonski, Susana Posada-Céspedes, Tobias Schär, Ina Nissen, Natascha Santacroce, Elodie Burcklen, Christiane Beckmann, Maurice Redondo, Olivier Kobel, Christoph Noppen, Sophie Seidel, Noemie Santamaria de Souza, Niko Beerenwinkel, Tanja Stadler |
| EPI_ISL_476410, EPI_ISL_476411, EPI_ISL_476412, EPI_ISL_476413, EPI_ISL_476414, EPI_ISL_476415 | Laboratório de Patologia Clínica - UNICAMP | Laboratório de Estudos de Vírus Emergentes - UNICAMP | José Luiz Proença-Modena, Magnun Nueldo Nunes dos Santos, Angelica Schreiber, Julia Forato,Camila Simeoni, Marcilio Jorge Fumagalli, Mariene Ribeiro Amorim, Darlan da Silva Candido, Nuno Rodrigues Faria, Julien Theze, Luiz Gonzaga,Jaqueline Goes Jesus e William Marciel de Souza |
| EPI_ISL_476720, EPI_ISL_476721, EPI_ISL_476722, EPI_ISL_476723, EPI_ISL_476724, EPI_ISL_476725, EPI_ISL_476726, EPI_ISL_476727, EPI_ISL_476728, EPI_ISL_476729, EPI_ISL_476730, EPI_ISL_476731, EPI_ISL_476732, EPI_ISL_476733, EPI_ISL_476734, EPI_ISL_476735, EPI_ISL_476736, EPI_ISL_476737, EPI_ISL_476738, EPI_ISL_476739, EPI_ISL_476740, EPI_ISL_476741, EPI_ISL_476742, EPI_ISL_476743, EPI_ISL_476744, EPI_ISL_476745, EPI_ISL_476746, EPI_ISL_476747, EPI_ISL_476748 | | | |
| see above | Minnesota Department of Health, Public Health Laboratory | Minnesota Department of Health, Public Health Laboratory | Matt Plumb, Jacob Garfin, and Xiong Wang |
| EPI_ISL_476832 | Medical Biology Department, Kocaeli University | Medical Genetics Department, Kocaeli University | Savli H, Cine N, Sunnetci-Akkoyunlu D, Eren-Keskin S, Ilgazli A, Akhan S, Karadenizli A, Kasap M, Sayan M, Akpinar G, Canturk NZ. |
| EPI_ISL_476849, EPI_ISL_476850, EPI_ISL_476852, EPI_ISL_476853 | Defence Research & Development Establishment (DRDE) | Defence Research & Development Establishment (DRDE) | Shashi Sharma, Paban Kumar Dash, Sushil Kumar Sharma, Ambuj Shrivastava, Jyoti S. Kumar |
| EPI_ISL_476992, EPI_ISL_476993, EPI_ISL_476994, EPI_ISL_476995, EPI_ISL_476996, EPI_ISL_476997, EPI_ISL_476998, EPI_ISL_476999, EPI_ISL_477000 | KU Leuven, Rega Institute, Clinical and Epidemiological Virology | KU Leuven, Rega Institute, Clinical and Epidemiological Virology | Tony Wawina-Bokalanga, Joan Marti-Carerras, Bert Vanmechelen, Piet Maes |
| EPI_ISL_477039, EPI_ISL_477040, EPI_ISL_477041, EPI_ISL_477042, EPI_ISL_477043, EPI_ISL_477044, EPI_ISL_477045, EPI_ISL_477046, EPI_ISL_477047, EPI_ISL_477048, EPI_ISL_477049, EPI_ISL_477050, EPI_ISL_477051, EPI_ISL_477052, EPI_ISL_477053, EPI_ISL_477054, EPI_ISL_477055, EPI_ISL_477056, EPI_ISL_477057, EPI_ISL_477058, EPI_ISL_477059, EPI_ISL_477060, EPI_ISL_477061, EPI_ISL_477062, EPI_ISL_477063, EPI_ISL_477065, EPI_ISL_477066, EPI_ISL_477067, EPI_ISL_477068, EPI_ISL_477069, EPI_ISL_477070, EPI_ISL_477071, EPI_ISL_477072, EPI_ISL_477073, EPI_ISL_477074, EPI_ISL_477075 | | | |
| see above | BCCDC Public Health Laboratory | BCCDC Public Health Laboratory | Richard Harrigan, Hope Lapointe, Jinny Choi, Kimia Kamelian, John Tyson,Terry Snutch, Linda Hoang, Inna Sekirov, Paul Levett, Mel Krajden, Natalie Prystajecky |
| EPI_ISL_477128 | Child Health Research Foundation | Child Health Research Foundation | Senjuti Saha, Md Saiful Islam Sajib, Roly Malaker, Md Hafizur Rahman, Afroza Akter Tanni, Syed Muktadir Al Sium, Maksuda Islam, Samir K Saha |
| EPI_ISL_477169 | Department for Virology, Molecular Biology and Genome Research, R. G. Lugar Center for Public Health Research, National Center for Disease Control and Public Health (NCDC) of Georgia. | Department for Virology, Molecular Biology and Genome Research, R. G. Lugar Center for Public Health Research, National Center for Disease Control and Public Health (NCDC) of Georgia. | Tata Imnadze, Giorgi Tomashvili, Meri Pantsulaia, Gvantsa Brachveli, Gvantsa Chanturia, Ann Machablishvili, Nato Kotaria, Marine Murtskhvaladze, Lela Sabadze, Mari Gavashelidze, Ana Papkiauri, Tamar Jashiashvili, Tea Tevdoradze, Ketevan Sidamonidze, Ekaterine Khmaladze, Ekaterine Zhghenti, Roena Sukhiashvili, Mariam Zakalashvili, Lela Urushadze, Magda Dgebuadze, Davit Tsaguria, Ekaterine Zangaladze, Nino Berishvili, Adam Kotorashvili, Maia Alkhazashvili, Irma Burjanadze, Anna Kasradze, Khatuna Zakhashvili, Paata Imnadze, Amiran Gamkrelidze. |
| EPI_ISL_477192 | Department of Laboratory Medicine Tan Tock Seng Hospital | Department of Laboratory Medicine Tan Tock Seng Hospital | Chen YYC, Zair X, Li C, Tang WY, Maurer-Stroh S, Barkham TMS, Nagarajan N, Sessions OM |
| EPI_ISL_477244, EPI_ISL_477245, EPI_ISL_477246 | Institute for Stem Cell Science and Regenerative Medicine | National Centre for Biological Sciences | Farhan Ali, Vanessa Molin Paynter, Srikar Krishna, Mohak Sharda, Shah-e-Jahan Gulzar, Awadhesh Pandit, Varadha Sundarmurthy, Uma Ramakrishnan, Dasaradhi Palakodeti, Aswin Seshasayee |
| EPI_ISL_477277, EPI_ISL_477278, EPI_ISL_477279 | M Health Fairview | Minnesota Department of Health, Public Health Laboratory | Matt Plumb, Jacob Garfin, Kelly Pung, and Xiong Wang |
| EPI_ISL_477294 | Mayo Clinic & Mayo Clinic Laboratories | Minnesota Department of Health, Public Health Laboratory | Matt Plumb, Jacob Garfin, Kelly Pung, and Xiong Wang |

| | | | |
|---|---|---|---|
| EPI_ISL_477633, EPI_ISL_477635, EPI_ISL_477636, EPI_ISL_477637, EPI_ISL_477638, EPI_ISL_477639, EPI_ISL_477640, EPI_ISL_477641, EPI_ISL_477642, EPI_ISL_477643, EPI_ISL_477644, EPI_ISL_477645, EPI_ISL_477646, EPI_ISL_477647, EPI_ISL_477648, EPI_ISL_477649, EPI_ISL_477650, EPI_ISL_477651, EPI_ISL_477652, EPI_ISL_477653, EPI_ISL_477654, EPI_ISL_477655, EPI_ISL_477656, EPI_ISL_477657, EPI_ISL_477658, EPI_ISL_477659, EPI_ISL_477660, EPI_ISL_477661, EPI_ISL_477662, EPI_ISL_477663 | | | |
| see above | Virginia DCLS | Virginia DCLS | Virginia DCLS |
| EPI_ISL_477763, EPI_ISL_477764, EPI_ISL_477765, EPI_ISL_477766, EPI_ISL_477767, EPI_ISL_477768, EPI_ISL_477769, EPI_ISL_477770, EPI_ISL_477771, EPI_ISL_477772, EPI_ISL_477773, EPI_ISL_477774, EPI_ISL_477775, EPI_ISL_477776, EPI_ISL_477777, EPI_ISL_477778, EPI_ISL_477779, EPI_ISL_477780, EPI_ISL_477781, EPI_ISL_477782 | | | |
| see above | University of Birmingham | COVID-19 Genomics UK (COG-UK) Consortium | Institute of Microbiology, University of Birmingham: Claire McMurray, Joanne Stockton, Samuel Nicholls, Radoslaw Poplawski, Will Rowe, Josh Quick, Nicholas Loman. University of Birmingham Testing Laboratory: Celina M Whalley, Andrew Bosworth, Charlotte Poxon, Kasun Wanigasooriya, Oliver Pickles, Mike Kidd, Alex Richter, Andrew D Beggs PHE Heartlands Lab: Husam Osman, Andrew Bosworth. Queen Elizabeth Hospital: Anna Casey |
| EPI_ISL_477783, EPI_ISL_477784 | Department of Pathology, University of Cambridge | COVID-19 Genomics UK (COG-UK) Consortium | Luke W Meredith, M. Estée Török, Myra Hosmillo, William L. Hamilton, Martin D. Curran, Theresa Feltwell, Grant Hall, Anna Yakovleva, Fahad A Khokhar, Charlotte J. Houldcroft, Laura G Caller, Aminu S. Jahun, Sarah L. Caddy, Yasmin Chaudhry, Malte Pinckert, Ian Goodfellow |
| EPI_ISL_478053 | West of Scotland Specialist Virology Centre, NHSGGC / MRC-University of Glasgow Centre for Virus Research | COVID-19 Genomics UK (COG-UK) Consortium | Ana da Silva Filipe, Natasha Johnson, Kathy Smollett, Daniel Mair, Stephen Carmichael, Lily Tong, Jenna Nichols, Elihu Aranday-Cortes, Kirstyn Brunker, Yasmin Parr, Alice Broos, Kyriaki Nomikou; Sarah McDonald, Marc Niebel, Patawee Asamaphan; Richard Orton, Joseph Hughes, Sreenu Vattipally, David L Robertson; Alasdair MacLean, Rory Gunson; Kathy Li, Natasha Jesudason, Rajiv Shah, James Shepherd, Antonia Ho, Emma Thomson |
| EPI_ISL_478253 | Virology Department, Royal Infirmary of Edinburgh, NHS Lothian / School of Biological Sciences, University of Edinburgh / Institute of Genetics and Molecular Medicine, University of Edinburgh | COVID-19 Genomics UK (COG-UK) Consortium | McHugh M, Dewar R, Rooke S, Gallagher M, Balcaza C, O'Toole Á, Scher E, Hill V, McCrone JT, Colquhoun R, Yu X, Jackson B, Rambaut A, Williams TC, Templeton K |
| EPI_ISL_478531, EPI_ISL_478532 | Northumbria University / South Tees Hospitals NHS Foundation Trust / North Cumbria Integrated Care NHS Foundation Trust / North Tees and Hartlepool NHS Foundation Trust / Newcastle Hospitals NHS Foundation Trust | COVID-19 Genomics UK (COG-UK) Consortium | Darren L Smith,Andrew Nelson,Matthew Bashton,Greg R Young,Joshua Loh,John Allan,Mohammad A Tariq,Giles S Holt,Gary Black,Wen C Yew,Lynn Dover,Paul Baker,Steve Liggett,Sarah Essex,Jane Greenaway,Debra Padgett,Clive Graham,Garren Scott,Edward Barton,Emma Swindells,Brendan Payne,Jennifer Collins,Yusri Taha,Gary Eltringham |
| EPI_ISL_478912, EPI_ISL_478923, EPI_ISL_478937, EPI_ISL_478953, EPI_ISL_478967, EPI_ISL_478983, EPI_ISL_478997, EPI_ISL_479004, EPI_ISL_479026, EPI_ISL_479028, EPI_ISL_479030, EPI_ISL_479031, EPI_ISL_479034, EPI_ISL_479036, EPI_ISL_479044, EPI_ISL_479050, EPI_ISL_479057, EPI_ISL_479066, EPI_ISL_479069, EPI_ISL_479074, EPI_ISL_479076, EPI_ISL_479080, EPI_ISL_479093, EPI_ISL_479099, EPI_ISL_479100, EPI_ISL_479104, EPI_ISL_479108, EPI_ISL_479111, EPI_ISL_479125, EPI_ISL_479132 | | | |
| see above | Oxford Viromics, NDM, University of Oxford; Oxford University Hospitals; Basingstoke and North Hampshire Hospital | COVID-19 Genomics UK (COG-UK) Consortium | Tanya Golubchik, David Bonsall, George Macintyre, Amy Trebes, Mariateresa de Cesare, Catrin Moore, Alex Mobbs, Anita Justice, Robert Shaw, Monique Andersson, Timothy Peto, Emma Wise, Nathan Moore, Jessica Lynch, Nick Cortes, Matilde Mori, Stephen Kidd, David Buck, John Todd, Christophe Fraser |
| EPI_ISL_479225 | Virology Department, Sheffield Teaching Hospitals NHS Foundation Trust/Department of Infection, Immunity and Cardiovascular Disease, The Medical School, University of Sheffield | COVID-19 Genomics UK (COG-UK) Consortium | Thushan de Silva, Matthew Parker, Nikki Smith, Adri Angyal, Rebecca Brown, Luke Green, Rachel Tucker, Paul Parsons, Danielle Groves, Katie Johnson, Laura Carrilero, Alex Keeley, Dave Partridge, Matthew Wyles, Benjamin Lindsey, Mehmet Yavuz, Mohammad Raza, Cariad Evans |
| EPI_ISL_479311, EPI_ISL_479320, EPI_ISL_479321, EPI_ISL_479328, EPI_ISL_479358, EPI_ISL_479377, EPI_ISL_479381, EPI_ISL_479406, EPI_ISL_479414, EPI_ISL_479453, EPI_ISL_479460, EPI_ISL_479481 | | | |
| see above | Wales Specialist Virology Centre Sequencing lab: Pathogen Genomics Unit | COVID-19 Genomics UK (COG-UK) Consortium | Catherine Moore, Johnathan Evans, Laura Gifford, Malorie Perry, Simon Cottrell, Angela Marchbank, Alec Birchley, Alexander Adams, Amy Gaskin, Bree Gatica-Wilcox, Jason Coombes, Joel Southgate, Lauren Gilbert, Lee Graham, Nicole Pacchiarini, Sara Kumziene-Summerhayes, Sarah Taylor, Sophie Jones, Sara Rey, Matthew Bull, Joanne Watkins, Sally Corden, Tom Connor |
| EPI_ISL_479484, EPI_ISL_479490 | Department of Laboratory Medicine Tan Tock Seng Hospital | Department of Laboratory Medicine Tan Tock Seng Hospital | Chen YYC, Zair X, Li C, Tang WY, Maurer-Stroh S, Barkham TMS, Nagarajan N, Sessions OM |
| EPI_ISL_479502, EPI_ISL_479503, EPI_ISL_479504, EPI_ISL_479505, EPI_ISL_479506, EPI_ISL_479507, EPI_ISL_479508, EPI_ISL_479509, EPI_ISL_479510, EPI_ISL_479511, EPI_ISL_479512, EPI_ISL_479513, EPI_ISL_479514 | | | |
| see above | NIV Influenza | NIV Influenza | Potdar V |
| EPI_ISL_480224 | National Reference Laboratory "Influenza and acute respiratory diseases" | NRL-HIV | Ivan Ivanov, Ivailo Alexiev, Ivva Philipova |
| EPI_ISL_480242, EPI_ISL_480249, EPI_ISL_480250, EPI_ISL_480251, EPI_ISL_480252, EPI_ISL_480253, EPI_ISL_480254, EPI_ISL_480255, EPI_ISL_480256, EPI_ISL_480257, EPI_ISL_480258, EPI_ISL_480259, EPI_ISL_480260, EPI_ISL_480261, EPI_ISL_480262, EPI_ISL_480263, EPI_ISL_480265, EPI_ISL_480266, EPI_ISL_480267, EPI_ISL_480268, EPI_ISL_480269, EPI_ISL_480277 | | | |
| see above | Genomic Laboratory (GLAB) (Conjoint lab of Health Directorate of Istanbul and Istanbul Technical University) | Genomic Laboratory (GLAB), Istanbul Technical University | Ilker Karacan, Tugba Kizilboga Akgun, Bugra Agaoglu, Gizem Alkurt, Jale Yildiz, Betsi Köse, Elifnaz Çelik, Arzu Irvem, Yasemin Kendir Demirkol, Ozlem Akgun Dogan, Mehtap Aydn, Levent Doganay, Gizem Dinler Doganay |
| EPI_ISL_480295 | Institute for Stem Cell Science and Regenerative Medicine | National Centre for Biological Sciences | Farhan Ali, Vanessa Molin Paynter, Srikar Krishna, Mohak Sharda, Shah-e-Jahan Gulzar, Awadhesh Pandit, Varadha Sundarmurthy, Uma Ramakrishnan, Dasaradhi Palakodeti, Aswin Seshasayee |
| EPI_ISL_480301 | National Reference Laboratory "Influenza and acute respiratory diseases" | NRL-HIV | Ivan Ivanov, Ivailo Alexiev, Ivva Philipova |
| EPI_ISL_480404, EPI_ISL_480405, EPI_ISL_480406, EPI_ISL_480407, EPI_ISL_480408, EPI_ISL_480409, EPI_ISL_480410, EPI_ISL_480411, EPI_ISL_480412, EPI_ISL_480413 | University of Wisconsin-Madison AIDS Vaccine Research Laboratories | University of Wisconsin-Madison AIDS Vaccine Research Laboratories | Gage Moreno, Katarina Braun, et al. AIDS Vaccine Research Laboratories |
| EPI_ISL_480598, EPI_ISL_480599, EPI_ISL_480608, EPI_ISL_480609 | Victorian Infectious Diseases Reference Laboratory (VIDRL) | VIDRL and MDU-PHL | Caly L., Seemann T., Sait, M., Schultz M., Druce J., Sherry, N. |
| EPI_ISL_480890, EPI_ISL_480891, EPI_ISL_480892, EPI_ISL_480893, EPI_ISL_480894, EPI_ISL_480895, EPI_ISL_480896, EPI_ISL_480897, EPI_ISL_480898, EPI_ISL_480899, EPI_ISL_480900, EPI_ISL_480901, EPI_ISL_480902, EPI_ISL_480903, EPI_ISL_480904, EPI_ISL_480905, EPI_ISL_480906, EPI_ISL_480907, EPI_ISL_480908, EPI_ISL_480909, EPI_ISL_480910, EPI_ISL_480911, EPI_ISL_480912, EPI_ISL_480913, EPI_ISL_480914, EPI_ISL_480915, EPI_ISL_480916, EPI_ISL_480917 | | | |
| see above | Florida Bureau of Public Health Laboratories | Florida Bureau of Public Health Laboratories | Sarah Schmedes, Jason Blanton |
| EPI_ISL_481042, EPI_ISL_481046, EPI_ISL_481047, EPI_ISL_481049, EPI_ISL_481071, EPI_ISL_481079, EPI_ISL_481105, EPI_ISL_481107 | Hospital General Universitario Gregorio Marañón | SeqCOVID-SPAIN consortium/IBV(CSIC) | Laura Pérez-Lago, Marta Herranz, Jon Sicilia, Julia Suárez, Pilar Catalán, Patricia Muñoz, Darío García de Viedma and SeqCOVID-SPAIN consortium |
| EPI_ISL_481206 | Hospital of Southern Norway - Kristiansand, Department of Medical Microbiology | Norwegian Institute of Public Health, Department of Virology | Kathrine Stene-Johansen, Kamilla Heddeland Instefjord, Hilde Elshaug, Rasmus Riis Kopperud, Karoline Bragstad, Olav Hungnes |
| EPI_ISL_481210, EPI_ISL_481211 | Ostfold Hospital Trust - Kalnes, Centre for Laboratory Medicine, Section for gene technology and infection serology | Norwegian Institute of Public Health, Department of Virology | Kathrine Stene-Johansen, Kamilla Heddeland Instefjord, Hilde Elshaug, Rasmus Riis Kopperud, Karoline Bragstad, Olav Hungnes |
| EPI_ISL_481227, EPI_ISL_481230, EPI_ISL_481231, EPI_ISL_481232 | Lab voor klinische biologie | Onderzoeksgroep Virologie | Nick Vereecke, Laurens Lambrechts, Marthe Pauwels, Bruno Verhasselt, Linos Vandekerckhove, Hans Nauwynck, Sebastiaan Theuns |
| EPI_ISL_481380 | Department for Virology, Molecular Biology and Genome Research, R. G. Lugar Center for Public Health Research, National Center for Disease Control and Public Health (NCDC) of Georgia. | Department for Virology, Molecular Biology and Genome Research, R. G. Lugar Center for Public Health Research, National Center for Disease Control and Public Health (NCDC) of Georgia. | Ana Papkiauri, Tata Imnadze, Giorgi Tomashvili, Meri Pantsulaia, Gvantsa Brachveli, Gvantsa Chanturia, Ann Machablishvili, Nato Kotaria, Marine Murtskhvaladze, Lela Sabadze, Mari Gavashelidze, Tamar Jashiashvili, Tea Tevdoradze, Ketevan Sidamonidze, Ekaterine Khmaladze, Ekaterine Zhghenti, Roena Sukhiashvili, Mariam Zakalashvili, Lela Urushadze, Magda Dgebuadze, Davit Tsaguria, Ekaterine Zangaladze, Nino Berishvili, Adam Kotorashvili, Maia Alkhazashvili, Irma Burjanadze, Anna Kasradze, Khatuna Zakhashvili, Paata Imnadze, Amiran Gamkrelidze. |
| EPI_ISL_481624, EPI_ISL_481625, EPI_ISL_481626, EPI_ISL_481627, EPI_ISL_481628, EPI_ISL_481629, EPI_ISL_481630, EPI_ISL_481631, EPI_ISL_481632, EPI_ISL_481633, EPI_ISL_481634, EPI_ISL_481635, EPI_ISL_481636, EPI_ISL_481637, EPI_ISL_481638, EPI_ISL_481674, EPI_ISL_481675, EPI_ISL_481676, EPI_ISL_481677, EPI_ISL_481678, EPI_ISL_481679, EPI_ISL_481680, EPI_ISL_481681 | | | |

| | | | |
|---|---|---|---|
| see above | Department of Virology and Immunology, University of Helsinki and Helsinki University Hospital, Huslab Finland | Department of Virology, Faculty of Medicine, University of Helsinki, Helsinki, Finland | Teemu Smura, Hannimari Kallio-Kokko, Jenni Virtanen, Maija Suvanto, Sari Hannula, Harri Kangas, Pekka Ellonen, Olli Vapalahti |
| EPI_ISL_481766, EPI_ISL_481770, EPI_ISL_481781, EPI_ISL_481786, EPI_ISL_481788, EPI_ISL_481789, EPI_ISL_481798, EPI_ISL_481802, EPI_ISL_481806, EPI_ISL_481808, EPI_ISL_481811, EPI_ISL_481835, EPI_ISL_481840, EPI_ISL_481848, EPI_ISL_481849, EPI_ISL_481855, EPI_ISL_481859, EPI_ISL_481861, EPI_ISL_481873, EPI_ISL_481877, EPI_ISL_481878, EPI_ISL_481881, EPI_ISL_481882, EPI_ISL_481893, EPI_ISL_481895, EPI_ISL_481897, EPI_ISL_481899, EPI_ISL_481907, EPI_ISL_481913, EPI_ISL_481915, EPI_ISL_481925, EPI_ISL_481926, EPI_ISL_481934, EPI_ISL_481941, EPI_ISL_481950, EPI_ISL_481952, EPI_ISL_481969, EPI_ISL_481987, EPI_ISL_481990, EPI_ISL_481993, EPI_ISL_482001, EPI_ISL_482007, EPI_ISL_482012 | | | |
| see above | PHE South West Regional Laboratory, National Infection Service | Wellcome Sanger Institute for the COVID-19 Genomics UK (COG-UK) consortium | Stephanie Hutchings, Hannah Pymont, Dr Peter Muir, Barry Vipond, Rich Hopes; and Alex Alderton, Roberto Amato, Sonia Goncalves, Ewan Harrison, David K. Jackson, Ian Johnston, Dominic Kwiatkowski, Cordelia Langford, John Sillitoe on behalf of the Wellcome Sanger Institute COVID-19 Surveillance Team (http://www.sanger.ac.uk/covid-team) |
| EPI_ISL_482028 | PHE South West Regional Laboratory, National Infection Service | Wellcome Sanger Institute for the COVID-19 Genomics UK (COG-UK) Consortium | Stephanie Hutchings, Hannah Pymont, Dr Peter Muir, Barry Vipond, Rich Hopes; and Alex Alderton, Roberto Amato, Sonia Goncalves, Ewan Harrison, David K. Jackson, Ian Johnston, Dominic Kwiatkowski, Cordelia Langford, John Sillitoe on behalf of the Wellcome Sanger Institute COVID-19 Surveillance Team |
| EPI_ISL_482033, EPI_ISL_482047, EPI_ISL_482050 | Regional Virus Laboratory, Belfast Health and Social Care Trust | Wellcome Sanger Institute for the COVID-19 Genomics UK (COG-UK) consortium | Conall McCaughey, James McKenna, Tanya Curran, Susan Feeney, Alison Watt, Ciara Cox, Mairead Connor, Zoltan Molnar, David Simpson, Derek Fairley; and Alex Alderton, Roberto Amato, Sonia Goncalves, Ewan Harrison, David K. Jackson, Ian Johnston, Dominic Kwiatkowski, Cordelia Langford, John Sillitoe on behalf of the Wellcome Sanger Institute COVID-19 Surveillance Team (http://www.sanger.ac.uk/covid-team) |
| EPI_ISL_482058, EPI_ISL_482059, EPI_ISL_482061, EPI_ISL_482065, EPI_ISL_482066 | The Department of Microbiology, Torbay and South Devon NHS Foundation Trust | Wellcome Sanger Institute for the COVID-19 Genomics UK (COG-UK) consortium | Amy Hurd, Sophie Lloyd, Anthony Mogridge, Jack Howe, Helen Brown, Gary Booth, Mel Brown, Cheryl Bailiss, Michelle Harrison and Alex Alderton, Roberto Amato, Sonia Goncalves, Ewan Harrison, David K. Jackson, Ian Johnston, Dominic Kwiatkowski, Cordelia Langford, John Sillitoe on behalf of the Wellcome Sanger Institute COVID-19 Surveillance Team (http://www.sanger.ac.uk/covid-team) |
| EPI_ISL_482069 | Microbiology Department, Hereford County Hospital | Wellcome Sanger Institute for the COVID-19 Genomics UK (COG-UK) consortium | Alison Johnson, Venkat Sivaprakasam, Fenella Halstead, Jane Thomas, Wendy Hogsden, Samantha Lamb and Alex Alderton, Roberto Amato, Sonia Goncalves, Ewan Harrison, David K. Jackson, Ian Johnston, Dominic Kwiatkowski, Cordelia Langford, John Sillitoe on behalf of the Wellcome Sanger Institute COVID-19 Surveillance Team (http://www.sanger.ac.uk/covid-team) |
| EPI_ISL_482075 | Regional Virus Laboratory, Belfast Health and Social Care Trust | Wellcome Sanger Institute for the COVID-19 Genomics UK (COG-UK) consortium | Conall McCaughey, James McKenna, Tanya Curran, Susan Feeney, Alison Watt, Ciara Cox, Mairead Connor, Zoltan Molnar, David Simpson, Derek Fairley; and Alex Alderton, Roberto Amato, Sonia Goncalves, Ewan Harrison, David K. Jackson, Ian Johnston, Dominic Kwiatkowski, Cordelia Langford, John Sillitoe on behalf of the Wellcome Sanger Institute COVID-19 Surveillance Team (http://www.sanger.ac.uk/covid-team) |
| EPI_ISL_482076, EPI_ISL_482078, EPI_ISL_482081, EPI_ISL_482084, EPI_ISL_482085, EPI_ISL_482095 | Microbiology Department, Hereford County Hospital | Wellcome Sanger Institute for the COVID-19 Genomics UK (COG-UK) consortium | Alison Johnson, Venkat Sivaprakasam, Fenella Halstead, Jane Thomas, Wendy Hogsden, Samantha Lamb and Alex Alderton, Roberto Amato, Sonia Goncalves, Ewan Harrison, David K. Jackson, Ian Johnston, Dominic Kwiatkowski, Cordelia Langford, John Sillitoe on behalf of the Wellcome Sanger Institute COVID-19 Surveillance Team (http://www.sanger.ac.uk/covid-team) |
| EPI_ISL_482096, EPI_ISL_482097 | Regional Virus Laboratory, Belfast Health and Social Care Trust | Wellcome Sanger Institute for the COVID-19 Genomics UK (COG-UK) consortium | Conall McCaughey, James McKenna, Tanya Curran, Susan Feeney, Alison Watt, Ciara Cox, Mairead Connor, Zoltan Molnar, David Simpson, Derek Fairley; and Alex Alderton, Roberto Amato, Sonia Goncalves, Ewan Harrison, David K. Jackson, Ian Johnston, Dominic Kwiatkowski, Cordelia Langford, John Sillitoe on behalf of the Wellcome Sanger Institute COVID-19 Surveillance Team (http://www.sanger.ac.uk/covid-team) |
| EPI_ISL_482100 | Microbiology Department, Hereford County Hospital | Wellcome Sanger Institute for the COVID-19 Genomics UK (COG-UK) consortium | Alison Johnson, Venkat Sivaprakasam, Fenella Halstead, Jane Thomas, Wendy Hogsden, Samantha Lamb and Alex Alderton, Roberto Amato, Sonia Goncalves, Ewan Harrison, David K. Jackson, Ian Johnston, Dominic Kwiatkowski, Cordelia Langford, John Sillitoe on behalf of the Wellcome Sanger Institute COVID-19 Surveillance Team (http://www.sanger.ac.uk/covid-team) |
| EPI_ISL_482101 | Regional Virus Laboratory, Belfast Health and Social Care Trust | Wellcome Sanger Institute for the COVID-19 Genomics UK (COG-UK) consortium | Conall McCaughey, James McKenna, Tanya Curran, Susan Feeney, Alison Watt, Ciara Cox, Mairead Connor, Zoltan Molnar, David Simpson, Derek Fairley; and Alex Alderton, Roberto Amato, Sonia Goncalves, Ewan Harrison, David K. Jackson, Ian Johnston, Dominic Kwiatkowski, Cordelia Langford, John Sillitoe on behalf of the Wellcome Sanger Institute COVID-19 Surveillance Team (http://www.sanger.ac.uk/covid-team) |
| EPI_ISL_482103, EPI_ISL_482114 | Microbiology Department, Hereford County Hospital | Wellcome Sanger Institute for the COVID-19 Genomics UK (COG-UK) consortium | Alison Johnson, Venkat Sivaprakasam, Fenella Halstead, Jane Thomas, Wendy Hogsden, Samantha Lamb and Alex Alderton, Roberto Amato, Sonia Goncalves, Ewan Harrison, David K. Jackson, Ian Johnston, Dominic Kwiatkowski, Cordelia Langford, John Sillitoe on behalf of the Wellcome Sanger Institute COVID-19 Surveillance Team (http://www.sanger.ac.uk/covid-team) |
| EPI_ISL_482115 | Regional Virus Laboratory, Belfast Health and Social Care Trust | Wellcome Sanger Institute for the COVID-19 Genomics UK (COG-UK) consortium | Conall McCaughey, James McKenna, Tanya Curran, Susan Feeney, Alison Watt, Ciara Cox, Mairead Connor, Zoltan Molnar, David Simpson, Derek Fairley; and Alex Alderton, Roberto Amato, Sonia Goncalves, Ewan Harrison, David K. Jackson, Ian Johnston, Dominic Kwiatkowski, Cordelia Langford, John Sillitoe on behalf of the Wellcome Sanger Institute COVID-19 Surveillance Team (http://www.sanger.ac.uk/covid-team) |
| EPI_ISL_482117, EPI_ISL_482119 | Microbiology Department, Hereford County Hospital | Wellcome Sanger Institute for the COVID-19 Genomics UK (COG-UK) consortium | Alison Johnson, Venkat Sivaprakasam, Fenella Halstead, Jane Thomas, Wendy Hogsden, Samantha Lamb and Alex Alderton, Roberto Amato, Sonia Goncalves, Ewan Harrison, David K. Jackson, Ian Johnston, Dominic Kwiatkowski, Cordelia Langford, John Sillitoe on behalf of the Wellcome Sanger Institute COVID-19 Surveillance Team (http://www.sanger.ac.uk/covid-team) |
| EPI_ISL_482124, EPI_ISL_482131 | North West London Pathology, Imperial College Healthcare NHS Trust | Wellcome Sanger Institute for the COVID-19 Genomics UK (COG-UK) consortium | Ling Li, Paul Randell, David Muir, Frankie Bolt, Alison Holmes, James Price, Aileen Rowan, Graham Taylor, Anjna Badhan, Carolina Herrera and Alex Alderton, Roberto Amato, Sonia Goncalves, Ewan Harrison, David K. Jackson, Ian Johnston, Dominic Kwiatkowski, Cordelia Langford, John Sillitoe on behalf of the Wellcome Sanger Institute COVID-19 Surveillance Team (http://www.sanger.ac.uk/covid-team) |
| EPI_ISL_482138 | North West London Pathology, Imperial College Healthcare NHS Trust | Wellcome Sanger Institute for the COVID-19 Genomics UK (COG-UK) Consortium | Ling Li, Paul Randell, David Muir, Frankie Bolt, Alison Holmes, James Price, Aileen Rowan, Graham Taylor, Anjna Badhan, Carolina Herrera and Alex Alderton, Roberto Amato, Sonia Goncalves, Ewan Harrison, David K. Jackson, Ian Johnston, Dominic Kwiatkowski, Cordelia Langford, John Sillitoe on behalf of the Wellcome Sanger Institute COVID-19 Surveillance Team |
| EPI_ISL_482140, EPI_ISL_482141, EPI_ISL_482142, EPI_ISL_482143, EPI_ISL_482144, EPI_ISL_482145, EPI_ISL_482146, EPI_ISL_482147, EPI_ISL_482148, EPI_ISL_482149, EPI_ISL_482150, EPI_ISL_482151, EPI_ISL_482152, EPI_ISL_482153, EPI_ISL_482154, EPI_ISL_482155, EPI_ISL_482157, EPI_ISL_482158, EPI_ISL_482159 | | | |
| see above | Regional Virus Laboratory, Belfast Health and Social Care Trust | Wellcome Sanger Institute for the COVID-19 Genomics UK (COG-UK) consortium | Conall McCaughey, James McKenna, Tanya Curran, Susan Feeney, Alison Watt, Ciara Cox, Mairead Connor, Zoltan Molnar, David Simpson, Derek Fairley; and Alex Alderton, Roberto Amato, Sonia Goncalves, Ewan Harrison, David K. Jackson, Ian Johnston, Dominic Kwiatkowski, Cordelia Langford, John Sillitoe on behalf of the Wellcome Sanger Institute COVID-19 Surveillance Team (http://www.sanger.ac.uk/covid-team) |
| EPI_ISL_482383, EPI_ISL_482384, EPI_ISL_482385, EPI_ISL_482386, EPI_ISL_482387, EPI_ISL_482388, EPI_ISL_482389, EPI_ISL_482390, EPI_ISL_482391, EPI_ISL_482392, EPI_ISL_482393, EPI_ISL_482394, EPI_ISL_482395, EPI_ISL_482396, EPI_ISL_482397, EPI_ISL_482398, EPI_ISL_482399, EPI_ISL_482400, EPI_ISL_482401, EPI_ISL_482402, EPI_ISL_482403, EPI_ISL_482404, EPI_ISL_482405, EPI_ISL_482406, EPI_ISL_482407, EPI_ISL_482408, EPI_ISL_482409, EPI_ISL_482440 | | | |
| see above | Providence St. Joseph Health Molecular Genomics Laboratory | Providence St. Joseph Health Molecular Genomics Laboratory | Alexa K Dowdell, Brian D Piening, Fred L Robinson, Carlo B Bifulco, Mary Campbell |
| EPI_ISL_482759, EPI_ISL_482760, EPI_ISL_482761, EPI_ISL_482765, EPI_ISL_483035 | Medical Ain Shams Research Institute (MASRI), Ain Shams University | Medical Ain Shams Research Institute (MASRI), Ain Shams University | Hesham Elghazaly, Sara Hassan Agwa, Ahmad Moustafa, Hala Hafez, Sara Elnakeep, Shaimaa Moustafa, Aya Mohamed, Reham Mamdouh, Ghada Ismael, Ashraf Omar, Osama Mansour, Mahmoud Elmeitini |
| EPI_ISL_483148, EPI_ISL_483149, EPI_ISL_483150, EPI_ISL_483151, EPI_ISL_483154, EPI_ISL_483155, EPI_ISL_483156, EPI_ISL_483157 | Robert Koch Institute, ZBS1 Highly Pathogenic Viruses, Berlin, Germany | Robert Koch Institute, Bioinformatics MF1, Berlin, Germany | Janine Michel, Andrea Thuermer, Oliver Drechsel, Rene Kmiecinski, Stephan Fuchs, Max v. Kleist, Andreas Nitsche |
| EPI_ISL_483204, EPI_ISL_483205, EPI_ISL_483206, EPI_ISL_483207, EPI_ISL_483210, EPI_ISL_483213, EPI_ISL_483214, EPI_ISL_483215, EPI_ISL_483219, EPI_ISL_483220, EPI_ISL_483224, EPI_ISL_483272, EPI_ISL_483274, EPI_ISL_483276, EPI_ISL_483278, EPI_ISL_483281, EPI_ISL_483284, EPI_ISL_483285, EPI_ISL_483287, EPI_ISL_483289, EPI_ISL_483290, EPI_ISL_483293, EPI_ISL_483294, EPI_ISL_483295, EPI_ISL_483297, EPI_ISL_483324, EPI_ISL_483325, EPI_ISL_483326, EPI_ISL_483329, EPI_ISL_483330, EPI_ISL_483332, EPI_ISL_483333, EPI_ISL_483334, EPI_ISL_483394, EPI_ISL_483395, EPI_ISL_483396, EPI_ISL_483403, EPI_ISL_483404, EPI_ISL_483405, EPI_ISL_483406, EPI_ISL_483407, EPI_ISL_483408, EPI_ISL_483409, EPI_ISL_483410, EPI_ISL_483411, EPI_ISL_483413, EPI_ISL_483414, EPI_ISL_483415, EPI_ISL_483416, EPI_ISL_483417, EPI_ISL_483418, EPI_ISL_483419, EPI_ISL_483420, EPI_ISL_483421, EPI_ISL_483423 | | | |
| see above | UC San Diego Center for Advanced Laboratory Medicine | Andersen lab at Scripps Research | SEARCH Alliance San Diego with David Pride, Ji H Shin |
| EPI_ISL_483604 | National Public Health Laboratory, National Centre for Infectious Diseases | National Public Health Laboratory, National Centre for Infectious Diseases | Mak TM, Octavia S, Zhou Z, Chavatte JM, Cui L, Lin RTP |
| EPI_ISL_484270, EPI_ISL_484271, EPI_ISL_484272, EPI_ISL_484273, EPI_ISL_484274, EPI_ISL_484275, EPI_ISL_484276, EPI_ISL_484277, EPI_ISL_484278, EPI_ISL_484279, EPI_ISL_484280, EPI_ISL_484281, EPI_ISL_484282, EPI_ISL_484283, EPI_ISL_484284, EPI_ISL_484285, EPI_ISL_484286, EPI_ISL_484287, EPI_ISL_484288, EPI_ISL_484289, EPI_ISL_484290, EPI_ISL_484291, EPI_ISL_484292, EPI_ISL_484293, EPI_ISL_484294, EPI_ISL_484295, EPI_ISL_484296, EPI_ISL_484325, EPI_ISL_484326, EPI_ISL_484327 | | | |

| | | | |
|---|---|---|---|
| see above | Northumbria University / South Tees Hospitals NHS Foundation Trust / North Cumbria Integrated Care NHS Foundation Trust / North Tees and Hartlepool NHS Foundation Trust / Newcastle Hospitals NHS Foundation Trust | COVID-19 Genomics UK (COG-UK) Consortium | Darren L Smith,Andrew Nelson,Matthew Bashton,Greg R Young,Joshua Loh,John Allan,Mohammad A Tariq,Giles S Holt,Gary Black,Wen C Yew,Lynn Dover,Paul Baker,Steve Liggett,Sarah Essex,Jane Greenaway,Debra Padgett,Clive Graham,Garren Scott,Edward Barton,Emma Swindells,Brendan Payne,Jennifer Collins,Yusri Taha,Gary Eltringham |
| EPI_ISL_484354, EPI_ISL_484356, EPI_ISL_484357, EPI_ISL_484358, EPI_ISL_484359, EPI_ISL_484360, EPI_ISL_484361, EPI_ISL_484362, EPI_ISL_484366, EPI_ISL_484367, EPI_ISL_484370, EPI_ISL_484371, EPI_ISL_484372, EPI_ISL_484373, EPI_ISL_484393, EPI_ISL_484396, EPI_ISL_484397, EPI_ISL_484399, EPI_ISL_484400, EPI_ISL_484402, EPI_ISL_484404 | | | |
| see above | Lincolnshire Hospitals and DeepSeq Nottingham | COVID-19 Genomics UK (COG-UK) Consortium | Nichola Duckworth, Tim Sloan, Sarah Walsh, Jonathan Ball, Patrick McClure, Joeseph Chappell, Nadine Holmes, Matthew Carlisle, Christopher Moore, Fei Sang, Johnny Debebe, Victoria Wright, Matthew Loose |
| EPI_ISL_484512 | Virology Department, Sheffield Teaching Hospitals NHS Foundation Trust/Department of Infection, Immunity and Cardiovascular Disease, The Medical School, University of Sheffield | COVID-19 Genomics UK (COG-UK) Consortium | Thushan de Silva, Matthew Parker, Nikki Smith, Adri Angyal, Rebecca Brown, Luke Green, Rachel Tucker, Paul Parsons, Danielle Groves, Katie Johnson, Laura Carrilero, Alex Keeley, Dave Partridge, Matthew Wyles, Benjamin Lindsey, Mehmet Yavuz, Mohammad Raza, Cariad Evans |
| EPI_ISL_485019, EPI_ISL_485020, EPI_ISL_485021, EPI_ISL_485022, EPI_ISL_485023, EPI_ISL_485024, EPI_ISL_485025, EPI_ISL_485026, EPI_ISL_485027, EPI_ISL_485028, EPI_ISL_485029, EPI_ISL_485030, EPI_ISL_485031, EPI_ISL_485032, EPI_ISL_485033, EPI_ISL_485034, EPI_ISL_485035, EPI_ISL_485036, EPI_ISL_485037, EPI_ISL_485038, EPI_ISL_485039, EPI_ISL_485040, EPI_ISL_485041, EPI_ISL_485042, EPI_ISL_485101, EPI_ISL_485102, EPI_ISL_485103, EPI_ISL_485104 | | | |
| see above | River Road Testing Lab | Ginkgo Bioworks Clinical Laboratory | Rebecca C. Christofferson, Stephania A. Cormier, Luan V. Dinh, E. Handly Mayton, Hollis R. O'Neil, Thaya Stoufflet, Malaika Mckenzie-Bennett, James McGann, Jim Griffin, Keith Robison, Alex Plocik, Becky Schilling, Rebecca Littlefield, Michelle Spencer, Birgitte Simen |
| EPI_ISL_485575, EPI_ISL_485576, EPI_ISL_485577 | Instituto de diagnóstico y Referencia Epidemiologicos (INDRE) | Instituto de diagnóstico y Referencia Epidemiologicos (INDRE) | Barrera-Badillo,G., Ramirez-Gonzalez,E. |
| EPI_ISL_485873 | River Road Testing Lab | Ginkgo Bioworks Clinical Laboratory | Rebecca C. Christofferson, Stephania A. Cormier, Luan V. Dinh, E. Handly Mayton, Hollis R. O'Neil, Thaya Stoufflet, Malaika Mckenzie-Bennett, James McGann, Jim Griffin, Keith Robison, Alex Plocik, Becky Schilling, Rebecca Littlefield, Michelle Spencer, Birgitte Simen |
| EPI_ISL_486150, EPI_ISL_486151, EPI_ISL_486152, EPI_ISL_486153, EPI_ISL_486154, EPI_ISL_486160, EPI_ISL_486161, EPI_ISL_486162, EPI_ISL_486163, EPI_ISL_486164, EPI_ISL_486165, EPI_ISL_486166, EPI_ISL_486167, EPI_ISL_486168, EPI_ISL_486169, EPI_ISL_486170, EPI_ISL_486171, EPI_ISL_486172, EPI_ISL_486173, EPI_ISL_486174, EPI_ISL_486175, EPI_ISL_486176, EPI_ISL_486177, EPI_ISL_486178, EPI_ISL_486179, EPI_ISL_486180, EPI_ISL_486181, EPI_ISL_486182, EPI_ISL_486183, EPI_ISL_486222, EPI_ISL_486223, EPI_ISL_486225, EPI_ISL_486252, EPI_ISL_486253, EPI_ISL_486254, EPI_ISL_486255, EPI_ISL_486256, EPI_ISL_486257, EPI_ISL_486258, EPI_ISL_486259, EPI_ISL_486260, EPI_ISL_486261, EPI_ISL_486262, EPI_ISL_486263, EPI_ISL_486264, EPI_ISL_486265, EPI_ISL_486266, EPI_ISL_486267, EPI_ISL_486268, EPI_ISL_486269, EPI_ISL_486270, EPI_ISL_486271, EPI_ISL_486272, EPI_ISL_486273, EPI_ISL_486274, EPI_ISL_486275 | | | |
| see above | Orange County Public Health Laboratory | Chan-Zuckerberg Biohub | CZB Cliahub Consortium |
| EPI_ISL_486310, EPI_ISL_486311, EPI_ISL_486312, EPI_ISL_486313, EPI_ISL_486314, EPI_ISL_486315, EPI_ISL_486316, EPI_ISL_486317, EPI_ISL_486318, EPI_ISL_486319, EPI_ISL_486320, EPI_ISL_486321, EPI_ISL_486322, EPI_ISL_486323, EPI_ISL_486324, EPI_ISL_486325, EPI_ISL_486326, EPI_ISL_486327, EPI_ISL_486328, EPI_ISL_486329, EPI_ISL_486330 | | | |
| see above | San Joaquin County Public Health Lab | Chan-Zuckerberg Biohub | CZB Cliahub Consortium |
| EPI_ISL_486384 | DH | Department of Neurovirology, National Institute of Mental Health and Neuroscience (NIMHANS) | Chitra Pattabiraman, Vijayalakshmi Reddy, Harsha PK, Risha Rasheed, Shafeeq S Hameed, Manjunatha Venkataswamy, Anita Desai, Ravi Vasanthapuram |
| EPI_ISL_486385, EPI_ISL_486386 | Victoria Hospital | Department of Neurovirology, National Institute of Mental Health and Neuroscience (NIMHANS) | Chitra Pattabiraman, Vijayalakshmi Reddy, Harsha PK, Risha Rasheed, Shafeeq S Hameed, Manjunatha Venkataswamy, Anita Desai, Ravi Vasanthapuram |
| EPI_ISL_486387 | DH | Department of Neurovirology, National Institute of Mental Health and Neuroscience (NIMHANS) | Chitra Pattabiraman, Vijayalakshmi Reddy, Harsha PK, Risha Rasheed, Shafeeq S Hameed, Manjunatha Venkataswamy, Anita Desai, Ravi Vasanthapuram |
| EPI_ISL_486391 | Centrl laboratorija | Latvian Biomedical Research and Study Centre | Ivars Silamielis, Kaspars Megnis, Monta Ustinova, ikita Zrelovs, Vita Rovte, Stella Lapia, Jana Oste, Marta Priedte, Uga Dumpis, Jnis Kloviš |
| EPI_ISL_486399 | MIMS | Department of Neurovirology, National Institute of Mental Health and Neuroscience (NIMHANS) | Chitra Pattabiraman, Vijayalakshmi Reddy, Harsha PK, Risha Rasheed, Shafeeq S Hameed, Manjunatha Venkataswamy, Anita Desai, Ravi Vasanthapuram |
| EPI_ISL_486400 | Victoria Hospital | Department of Neurovirology, National Institute of Mental Health and Neuroscience (NIMHANS) | Chitra Pattabiraman, Vijayalakshmi Reddy, Harsha PK, Risha Rasheed, Shafeeq S Hameed, Manjunatha Venkataswamy, Anita Desai, Ravi Vasanthapuram |
| EPI_ISL_486401, EPI_ISL_486402, EPI_ISL_486403 | DH | Department of Neurovirology, National Institute of Mental Health and Neuroscience (NIMHANS) | Chitra Pattabiraman, Vijayalakshmi Reddy, Harsha PK, Risha Rasheed, Shafeeq S Hameed, Manjunatha Venkataswamy, Anita Desai, Ravi Vasanthapuram |
| EPI_ISL_486412, EPI_ISL_486415, EPI_ISL_486416, EPI_ISL_486419, EPI_ISL_486420, EPI_ISL_486421, EPI_ISL_486437 | Centrl laboratorija | Latvian Biomedical Research and Study Centre | Ivars Silamielis, Kaspars Megnis, Monta Ustinova, ikita Zrelovs, Vita Rovte, Stella Lapia, Jana Oste, Marta Priedte, Uga Dumpis, Jnis Kloviš |
| EPI_ISL_486673 | Institute for Stem Cell Science and Regenerative Medicine | National Centre for Biological Sciences | Farhan Ali, Vanessa Molin Paynter, Srikar Krishna, Mohak Sharda, Shah-e-Jahan Gulzar, Awadhesh Pandit, Varadha Sundarmurthy, Uma Ramakrishnan, Dasaradhi Palakodeti, Aswin Seshasayee |
| EPI_ISL_486897, EPI_ISL_486898, EPI_ISL_486899, EPI_ISL_486900, EPI_ISL_486901, EPI_ISL_486902, EPI_ISL_486903, EPI_ISL_486904, EPI_ISL_486905, EPI_ISL_486906, EPI_ISL_486907, EPI_ISL_486908, EPI_ISL_486909, EPI_ISL_486910, EPI_ISL_486911 | | | |
| see above | Tokyo Metoropolitan Institute of Public Health | Tokyo Metoropolitan Institute of Public Health | Asakura,H., Yoshida,I., Kumagai,R., Chiba,T., Sadamasu,K., Nagashima,M. |
| EPI_ISL_487095, EPI_ISL_487104 | Nigeria Centre for Disease Control (NCDC) | African Centre of Excellence for Genomics of Infectious Diseases (ACEGID), Redeemer's University, Ede, Osun State, Nigeria | Oluniyi P.E., Ajogbasile F.V., Kayode A., Oguzie J., Olawoye I., Uwanibe J., Olumade T., Folarin O.A., Ihekweazu C., Happi C.T. |
| EPI_ISL_487192 | Viral Respiratory Lab, National institute for Biomedical Research (INRB) | Pathogen Sequencing Lab, National Institute for Biomedical Research (INRB) | Placide Mbala-Kingebeni, Edith Nkwembe, Eddy Kinganda-Lusamaki, Amuri Aziza, Francisca Muyembe-Mawete, Emmanuel Lokilo-Lofiko, Catherine Pratt, Matthias Pauthner, Josh Quick, Allison Black, James Hadfield, Trevor Bedford, Ian Goodfellow, Andrew Rambault, Nick Loman, Kristian Andersen, Michael Wiley, Steve Ahuka-Mundeke, Jean-Jacques Muyembe Tamfum. |
| EPI_ISL_487365, EPI_ISL_487369 | Viral Respiratory Lab, National Institute for Biomedical Research (INRB) | Pathogen Sequencing Lab, National Institute for Biomedical Research (INRB) | Placide Mbala-Kingebeni, Edith Nkwembe, Eddy Kinganda-Lusamaki, Amuri Aziza, Francisca Muyembe-Mawete, Emmanuel Lokilo-Lofiko, Catherine Pratt, Matthias Pauthner, Josh Quick, Allison Black, James Hadfield, Trevor Bedford, Ian Goodfellow, Andrew Rambault, Nick Loman, Kristian Andersen, Michael Wiley, Steve Ahuka-Mundeke, Jean-Jacques Muyembe Tamfum |
| EPI_ISL_487527, EPI_ISL_487531, EPI_ISL_487532, EPI_ISL_487533, EPI_ISL_487538, EPI_ISL_487545, EPI_ISL_487547, EPI_ISL_487555, EPI_ISL_487560, EPI_ISL_487570, EPI_ISL_487575, EPI_ISL_487577, EPI_ISL_487592, EPI_ISL_487602, EPI_ISL_487608, EPI_ISL_487611, EPI_ISL_487620, EPI_ISL_487624 | | | |
| see above | PHE South West Regional Laboratory, National Infection Service | Wellcome Sanger Institute for the COVID-19 Genomics UK (COG-UK) consortium | Stephanie Hutchings, Hannah Pymont, Dr Peter Muir, Barry Vipond, Rich Hopes; and Alex Alderton, Roberto Amato, Sonia Goncalves, Ewan Harrison, David K. Jackson, Ian Johnston, Dominic Kwiatkowski, Cordelia Langford, John Sillitoe on behalf of the Wellcome Sanger Institute COVID-19 Surveillance Team (http://www.sanger.ac.uk/covid-team) |
| EPI_ISL_487651, EPI_ISL_487658, EPI_ISL_487659, EPI_ISL_487664, EPI_ISL_487665, EPI_ISL_487666, EPI_ISL_487668, EPI_ISL_487669, EPI_ISL_487674, EPI_ISL_487680, EPI_ISL_487683, EPI_ISL_487690, EPI_ISL_487691, EPI_ISL_487734, EPI_ISL_487735, EPI_ISL_487736, EPI_ISL_487737, EPI_ISL_487751, EPI_ISL_487758, EPI_ISL_487763, EPI_ISL_487766, EPI_ISL_487768, EPI_ISL_487772, EPI_ISL_487774, EPI_ISL_487775, EPI_ISL_487777, EPI_ISL_487780, EPI_ISL_487786, EPI_ISL_487793, EPI_ISL_487798, EPI_ISL_487800, EPI_ISL_487810, EPI_ISL_487827, EPI_ISL_487830, EPI_ISL_487843, EPI_ISL_487849, EPI_ISL_487854, EPI_ISL_487860, EPI_ISL_487862, EPI_ISL_487873, EPI_ISL_487881, EPI_ISL_487884, EPI_ISL_487885, EPI_ISL_487890, EPI_ISL_487891, EPI_ISL_487896, EPI_ISL_487903, EPI_ISL_487904, EPI_ISL_487905, EPI_ISL_487906, EPI_ISL_487920, EPI_ISL_487921, EPI_ISL_487926, EPI_ISL_487931, EPI_ISL_487939, EPI_ISL_487945, EPI_ISL_487947, EPI_ISL_487948, EPI_ISL_487951, EPI_ISL_487964, EPI_ISL_487976, EPI_ISL_487986, EPI_ISL_487990, EPI_ISL_487992, EPI_ISL_487994 | | | |
| see above | Virology Department, Royal Infirmary of Edinburgh, NHS Lothian / School of Biological Sciences, University of Edinburgh | Wellcome Sanger Institute for the COVID-19 Genomics UK (COG-UK) consortium | McHugh M, Dewar R, Rooke S, O'Toole Á, Scher E, Hill V, McCrone JT, Colquhoun R, Yu X, Jackson B, Rambaut A, Templeton K and Alex Alderton, Roberto Amato, Sonia Goncalves, Ewan Harrison, David K. Jackson, Ian Johnston, Dominic Kwiatkowski, Cordelia Langford, John Sillitoe on behalf of the Wellcome Sanger Institute COVID-19 Surveillance Team (http://www.sanger.ac.uk/covid-team) |
| EPI_ISL_488005, EPI_ISL_488009, EPI_ISL_488014, EPI_ISL_488015, EPI_ISL_488016, EPI_ISL_488020, EPI_ISL_488025, EPI_ISL_488032, EPI_ISL_488033, EPI_ISL_488041, EPI_ISL_488051, EPI_ISL_488053, EPI_ISL_488066, EPI_ISL_488069, EPI_ISL_488072, EPI_ISL_488075, EPI_ISL_488077, EPI_ISL_488079, EPI_ISL_488087, EPI_ISL_488089, EPI_ISL_488096, EPI_ISL_488097, EPI_ISL_488108, EPI_ISL_488109, EPI_ISL_488110, EPI_ISL_488116, EPI_ISL_488119, EPI_ISL_488120, EPI_ISL_488124, EPI_ISL_488127, EPI_ISL_488128, EPI_ISL_488137, EPI_ISL_488138, EPI_ISL_488142, EPI_ISL_488143, EPI_ISL_488146, EPI_ISL_488150, EPI_ISL_488152, EPI_ISL_488153, EPI_ISL_488157, EPI_ISL_488160, EPI_ISL_488161, EPI_ISL_488164, EPI_ISL_488165, EPI_ISL_488166, EPI_ISL_488168, EPI_ISL_488169, EPI_ISL_488172, EPI_ISL_488622, EPI_ISL_488776 | | | |

| | | | |
|---|---|---|---|
| see above | NU-OMICS DNA Sequencing research facility, Northumbria University | Wellcome Sanger Institute for the COVID-19 Genomics UK (COG-UK) consortium | Chris Duncan, Sheia Waugh, Shirelle Burton-Fanning, Gary Eltringham, Jennifer Collins, Brendan Payne, Yusri Taha, Emma Swindells, Jane Greenaway, Edward Barton, Garren Scott, Debra Padgett, Clive Graham, Sarah Essex, Steve Ligget, Paul Baker, Lynn Dover, Wen Yew, Gary Black, John Allan, Joshua Loh, Greg Young, Matthew Bashton, Andrew Nelson, Darren Smith and Alex Alderton, Roberto Amato, Sonia Goncalves, Ewan Harrison, David K. Jackson, Ian Johnston, Dominic Kwiatkowski, Cordelia Langford, John Sillitoe on behalf of the Wellcome Sanger Institute COVID-19 Surveillance Team (http://www.sanger.ac.uk/covid-team) |
| EPI_ISL_488872 | Department of Medical Microbiology, Western Sussex Hospitals NHS Foundation Trust, St Richard's Hospital | Wellcome Sanger Institute for the COVID-19 Genomics UK (COG-UK) consortium | Manasa Mutingwende, Sarah Lowdon, Olga Podplomyk, Michelle Erkiert, Jonathan Lewis, Paul Randell and Alex Alderton, Roberto Amato, Sonia Goncalves, Ewan Harrison, David K. Jackson, Ian Johnston, Dominic Kwiatkowski, Cordelia Langford, John Sillitoe on behalf of the Wellcome Sanger Institute COVID-19 Surveillance Team (http://www.sanger.ac.uk/covid-team) |
| EPI_ISL_489070, EPI_ISL_489112, EPI_ISL_489143, EPI_ISL_489144 | NU-OMICS DNA Sequencing research facility, Northumbria University | Wellcome Sanger Institute for the COVID-19 Genomics UK (COG-UK) consortium | Chris Duncan, Sheia Waugh, Shirelle Burton-Fanning, Gary Eltringham, Jennifer Collins, Brendan Payne, Yusri Taha, Emma Swindells, Jane Greenaway, Edward Barton, Garren Scott, Debra Padgett, Clive Graham, Sarah Essex, Steve Ligget, Paul Baker, Lynn Dover, Wen Yew, Gary Black, John Allan, Joshua Loh, Greg Young, Matthew Bashton, Andrew Nelson, Darren Smith and Alex Alderton, Roberto Amato, Sonia Goncalves, Ewan Harrison, David K. Jackson, Ian Johnston, Dominic Kwiatkowski, Cordelia Langford, John Sillitoe on behalf of the Wellcome Sanger Institute COVID-19 Surveillance Team (http://www.sanger.ac.uk/covid-team) |
| EPI_ISL_489157, EPI_ISL_489167, EPI_ISL_489174, EPI_ISL_489175, EPI_ISL_489181, EPI_ISL_489187, EPI_ISL_489194, EPI_ISL_489222, EPI_ISL_489225, EPI_ISL_489231, EPI_ISL_489233, EPI_ISL_489235, EPI_ISL_489239, EPI_ISL_489250, EPI_ISL_489257, EPI_ISL_489260, EPI_ISL_489261, EPI_ISL_489263, EPI_ISL_489264, EPI_ISL_489267, EPI_ISL_489271, EPI_ISL_489273, EPI_ISL_489274, EPI_ISL_489279, EPI_ISL_489284, EPI_ISL_489287, EPI_ISL_489294, EPI_ISL_489296, EPI_ISL_489297, EPI_ISL_489298, EPI_ISL_489302, EPI_ISL_489303, EPI_ISL_489304, EPI_ISL_489305, EPI_ISL_489308, EPI_ISL_489322, EPI_ISL_489323, EPI_ISL_489325, EPI_ISL_489326, EPI_ISL_489327, EPI_ISL_489329, EPI_ISL_489330, EPI_ISL_489331, EPI_ISL_489336, EPI_ISL_489337, EPI_ISL_489338, EPI_ISL_489339, EPI_ISL_489340, EPI_ISL_489343, EPI_ISL_489353, EPI_ISL_489355, EPI_ISL_489359, EPI_ISL_489363, EPI_ISL_489364, EPI_ISL_489370, EPI_ISL_489373, EPI_ISL_489378, EPI_ISL_489379 | | | |
| see above | Regional Virus Laboratory, Belfast Health and Social Care Trust | Wellcome Sanger Institute for the COVID-19 Genomics UK (COG-UK) consortium | Conall McCaughey, James McKenna, Tanya Curran, Susan Feeney, Alison Watt, Ciara Cox, Mairead Connor, Zoltan Molnar, David Simpson, Derek Fairley; and Alex Alderton, Roberto Amato, Sonia Goncalves, Ewan Harrison, David K. Jackson, Ian Johnston, Dominic Kwiatkowski, Cordelia Langford, John Sillitoe on behalf of the Wellcome Sanger Institute COVID-19 Surveillance Team (http://www.sanger.ac.uk/covid-team) |
| EPI_ISL_489794, EPI_ISL_489795, EPI_ISL_489796, EPI_ISL_489797, EPI_ISL_489798, EPI_ISL_489799 | Florida Bureau of Public Health Laboratories | Florida Bureau of Public Health Laboratories | Sarah Schmedes, Jason Blanton |
| EPI_ISL_490201 | National Influenza Centre of INSPI-Ecuador | INSPI - Charité | Alfredo Bruno Caicedo, Domenica de Mora Coloma, Andres Moreira-Soto, Anna-Lena Sander, Nina Krause, Maritza Olmedo,Denisses Portugal, Manuel Gonzalez, Silvia Salgado, Alberto Orlando, Alexandra Usiña, Juan Carlos Zeballos,Jan Felix Drexler |
| EPI_ISL_490257 | National Institute for Communicable Diseases of the National Health Laboratory Service | National Institute for Communicable Diseases of the National Health Laboratory Service | Allam M, Ismail A, Khumalo Z, Kwenda S, Mtshali P, Mnyameni F, Mohale T, Subramoney K, Bhiman JN |
| EPI_ISL_490491, EPI_ISL_490492, EPI_ISL_490493, EPI_ISL_490494, EPI_ISL_490495 | Northumbria University / South Tees Hospitals NHS Foundation Trust / North Cumbria Integrated Care NHS Foundation Trust / North Tees and Hartlepool NHS Foundation Trust / Newcastle Hospitals NHS Foundation Trust | COVID-19 Genomics UK (COG-UK) Consortium | Darren L Smith,Andrew Nelson,Matthew Bashton,Greg R Young,Joshua Loh,John Allan,Mohammad A Tariq,Giles S Holt,Gary Black,Wen C Yew,Lynn Dover,Paul Baker,Steve Liggett,Sarah Essex,Jane Greenaway,Debra Padgett,Clive Graham,Garren Scott,Edward Barton,Emma Swindells,Brendan Payne,Jennifer Collins,Yusri Taha,Gary Eltringham |
| EPI_ISL_490581, EPI_ISL_490587, EPI_ISL_490588, EPI_ISL_490589, EPI_ISL_490593, EPI_ISL_490595, EPI_ISL_490596, EPI_ISL_490599, EPI_ISL_490601, EPI_ISL_490602, EPI_ISL_490605, EPI_ISL_490616, EPI_ISL_490618, EPI_ISL_490620, EPI_ISL_490623, EPI_ISL_490624, EPI_ISL_490628, EPI_ISL_490633, EPI_ISL_490636, EPI_ISL_490639, EPI_ISL_490641, EPI_ISL_490644 | | | |
| see above | Virology Department, Sheffield Teaching Hospitals NHS Foundation Trust/Department of Infection, Immunity and Cardiovascular Disease, The Medical School, University of Sheffield | COVID-19 Genomics UK (COG-UK) Consortium | Thushan de Silva, Matthew Parker, Nikki Smith, Adri Angyal, Rebecca Brown, Luke Green, Rachel Tucker, Paul Parsons, Danielle Groves, Katie Johnson, Laura Carrilero, Alex Keeley, Dave Partridge, Matthew Wyles, Benjamin Lindsey, Mehmet Yavuz, Mohammad Raza, Cariad Evans |
| EPI_ISL_490693 | West of Scotland Specialist Virology Centre, NHSGGC / MRC-University of Glasgow Centre for Virus Research | COVID-19 Genomics UK (COG-UK) Consortium | Ana da Silva Filipe, Natasha Johnson, Kathy Smollett, Daniel Mair, Stephen Carmichael, Lily Tong, Jenna Nichols, Elihu Aranday-Cortes, Kirstyn Brunker, Yasmin Parr, Alice Broos, Kyriaki Nomikou; Sarah McDonald, Marc Niebel, Patawee Asamaphan; Richard Orton, Joseph Hughes, Sreenu Vattipally, David L Robertson; Alasdair MacLean, Rory Gunson; Kathy Li, Natasha Jesudason, Rajiv Shah, James Shepherd, Antonia Ho, Emma Thomson |
| EPI_ISL_491080 | Suceava County Emergency Hospital | "Stefan cel Mare" University Metagenomics Lab | Lobiuc Andrei, Antoniadis Panagiotis et al. |
| EPI_ISL_491081 | Suceava County Emergency Hospital | "Stefan cel Mare" University Metagenomics Lab | Lobiuc Andrei et al. |
| EPI_ISL_491082 | Suceava County Emergency Hospital | "Stefan cel Mare" University Metagenomics Lab | Lobiuc Andrei, Antoniadis Panagiotis et al. |
| EPI_ISL_491083, EPI_ISL_491084, EPI_ISL_491086 | Suceava County Emergency Hospital | "Stefan cel Mare" University Metagenomics Lab | Lobiuc Andrei et al. |
| EPI_ISL_491087 | Suceava County Emergency Hospital | "Stefan cel Mare" University Metagenomics Lab | Lobiuc Andrei, Antoniadis Panagiotis et al. |
| EPI_ISL_491110 | SC Department of Health and Environmental Control | SC Department of Health and Environmental Control | Flores,H. |
| EPI_ISL_491115 | Cicin-Sain Lab | Cicin-Sain Lab | M. Zeeshan Chaudhry, Kathrin Eschke, Yeonsu Kim, Luka Cicin-Sain |
| EPI_ISL_491122, EPI_ISL_491124 | Oman-National Influenza Center | Biotechnology & OMICs Laboratory | Samira Al-Mahruqi, Abdul Latif Khan, Samiha Al-Kharusi, Adil Khan , Ahmed Al-Rawahi, Sajjad Asaf, Amina Al-Jardani, Hanan Al-Kindi, Intisar Al-Shukri, Ahlam Al-Amri, Aisha Al-Amri, Aisha Al-Busaidi, Adil Al-Wahaibi, Seif Al-Abri, Ahmed Al-Harrasi |
| EPI_ISL_491133, EPI_ISL_491136 | Oman-National Influenza Center | Biotechnology & OMICs Laboratory | Samiha Al-Kharusi, Sajjad Asaf, Abdul Latif Khan, Samira Al-Mahruqi, Adil Khan, Ahmed Al-Rawahi, Amina Al-Jardani, Hanan Al-Kindi, Intisar Al-Shukri, Ahlam Al-Amri, Aisha Al-Amri, Aisha Al-Busaidi, Adil Al-Wahaibi, Seif Al-Abri, Ahmed Al-Harrasi |
| EPI_ISL_491169, EPI_ISL_491170 | Oman-National Influenza Center | Biotechnology & OMICs Laboratory | Sajjad Asaf, Samiha Al-Kharusi, Ahmed Al-Harrasi, Samira Al-Mahruqi, Adil Khan, Ahmed Al-Rawahi, Abdul Latif Khan, Amina Al-Jardani, Hanan Al-Kindi, Intisar Al-Shukri, Ahlam Al-Amri, Aisha Al-Amri, Aisha Al-Busaidi, Adil Al-Wahaibi, Seif Al-Abri. |
| EPI_ISL_491494, EPI_ISL_491503, EPI_ISL_491533, EPI_ISL_491556, EPI_ISL_491573, EPI_ISL_491586, EPI_ISL_491593, EPI_ISL_491598, EPI_ISL_491614, EPI_ISL_491619, EPI_ISL_491651, EPI_ISL_491652, EPI_ISL_491656, EPI_ISL_491676, EPI_ISL_491680, EPI_ISL_491683, EPI_ISL_491690, EPI_ISL_491700 | | | |
| see above | Virology Department, Royal Infirmary of Edinburgh, NHS Lothian / School of Biological Sciences, University of Edinburgh | Wellcome Sanger Institute for the COVID-19 Genomics UK (COG-UK) consortium | McHugh M, Dewar R, Rooke S, O'Toole Á, Scher E, Hill V, McCrone JT, Colquhoun R, Yu X, Jackson B, Rambaut A, Templeton K and Alex Alderton, Roberto Amato, Sonia Goncalves, Ewan Harrison, David K. Jackson, Ian Johnston, Dominic Kwiatkowski, Cordelia Langford, John Sillitoe on behalf of the Wellcome Sanger Institute COVID-19 Surveillance Team (http://www.sanger.ac.uk/covid-team) |
| EPI_ISL_491941 | Centro de Investigaciones, Universidad de Especialidades Espíritu Santo | Institute of Microbiology, Universidad San Francisco de Quito | Derly Andrade, Juan Carlos Fernandez, Belén Prado-Vivar, Sully Márquez, Juan José Guadalupe, Monica Becerra-Wong, Bernardo Gutiérrez, Gabriel Morey, Ruben Armas, Jose Pedro Barberan, Fernando Espinoza, Edith Lopez, Verónica Barragán, Patricio Rojas-Silva, Gabriel Trueba, Michelle Grunauer, Paúl Cárdenas |
| EPI_ISL_491948 | Instituto Nacional de Investigación en Salud Pública - INSPI | INSPI - Charité | Alfredo Bruno Caicedo, Domenica de Mora Coloma, Andres Moreira-Soto, Anna-Lena Sander, Nina Krause, Maritza Olmedo,Denisses Portugal, Manuel Gonzalez, Silvia Salgado, Alberto Orlando, Alexandra Usiña, Juan Carlos Zeballos, Jan Felix Drexler |
| EPI_ISL_492033, EPI_ISL_492037 | Instituto de Biologia do Exército | Laboratório Metabolismo Macromolecular FirminoTorres de Castro, Instituto de Biofísica Carlos Chagas Filho, Universidade Federal do Rio de Janeiro | Bianca Catarina Azevedo Cabral, Aline Rosa Vianna de Souza, Caleb GM Santos, Marcos Dornelas-Ribeiro, Tatiana LS Nogueira, Nádia Vaez Gonçalves da Cruz, Elizabeth Valentin, Marcio da Costa Cipitelli, Virginia Sara Grancieri do Amaral, Rodrigo Soares de Moura Neto, Clarissa Damaso, Rosane Silva |
| EPI_ISL_492038 | Instituto de Biologia do Exército | Laboratório Metabolismo Macromolecular FirminoTorres de Castro, Instituto de Biofísica Carlos Chagas Filho, Universidade Federal do Rio de Janeiro | Bianca Catarina Azevedo Cabral, Aline Rosa Vianna de Souza, Nádia Vaez Gonçalves da Cruz, Caleb GM Santos, Marcos Dornelas-Ribeiro, Tatiana LS Nogueira, Elizabeth Valentin, Marcio da Costa Cipitelli, Virginia Sara Grancieri do Amaral, Rodrigo Soares de Moura Neto, Clarissa Damaso, Rosane Silva |
| EPI_ISL_492039 | Instituto de Biologia do Exército | Laboratório Metabolismo Macromolecular FirminoTorres de Castro, Instituto de Biofísica Carlos Chagas Filho, Universidade Federal do Rio de Janeiro | Bianca Catarina Azevedo Cabral, Aline Rosa Vianna de Souza, Tatiana LS Nogueira, Nádia Vaez Gonçalves da Cruz, Caleb GM Santos, Marcos Dornelas-Ribeiro, Elizabeth Valentin, Marcio da Costa Cipitelli, Virginia Sara Grancieri do Amaral, Rodrigo Soares de Moura Neto, Clarissa Damaso, Rosane Silva |
| EPI_ISL_492040 | Instituto de Biologia do Exército | Laboratório Metabolismo Macromolecular FirminoTorres de | Bianca Catarina Azevedo Cabral, Aline Rosa Vianna de Souza , Marcos Dornelas-Ribeiro, Tatiana LS Nogueira, Nádia Vaez Gonçalves da Cruz, Caleb GM |

| | | | |
|---|---|---|---|
| | | Castro, Instituto de Biofísica Carlos Chagas Filho, Universidade Federal do Rio de Janeiro | Santos, Elizabeth Valentin, Marcio da Costa Cipitelli, Virginia Sara Grancieri do Amaral, Rodrigo Soares de Moura Neto, Clarissa Damaso, Rosane Silva |
| EPI_ISL_492041 | Instituto de Biologia do Exército | Laboratório Metabolismo Macromolecular FirminoTorres de Castro, Instituto de Biofísica Carlos Chagas Filho, Universidade Federal do Rio de Janeiro | Bianca Catarina Azevedo Cabral, Aline Rosa Vianna de Souza, Caleb GM Santos, Marcos Dornelas-Ribeiro, Tatiana LS Nogueira, Nádia Vaez Gonçalves da Cruz, Elizabeth Valentin, Marcio da Costa Cipitelli, Virginia Sara Grancieri do Amaral, Rodrigo Soares de Moura Neto, Clarissa Damaso, Rosane Silva |
| EPI_ISL_492212, EPI_ISL_492221, EPI_ISL_492225, EPI_ISL_492226, EPI_ISL_492227, EPI_ISL_492229, EPI_ISL_492230, EPI_ISL_492232, EPI_ISL_492233, EPI_ISL_492234, EPI_ISL_492235, EPI_ISL_492238, EPI_ISL_492239, EPI_ISL_492244, EPI_ISL_492245, EPI_ISL_492248, EPI_ISL_492249, EPI_ISL_492250, EPI_ISL_492252, EPI_ISL_492253, EPI_ISL_492255, EPI_ISL_492258, EPI_ISL_492260, EPI_ISL_492264, EPI_ISL_492265, EPI_ISL_492270, EPI_ISL_492271, EPI_ISL_492273, EPI_ISL_492274, EPI_ISL_492276, EPI_ISL_492277, EPI_ISL_492278, EPI_ISL_492283, EPI_ISL_492289, EPI_ISL_492292, EPI_ISL_492293, EPI_ISL_492294, EPI_ISL_492297, EPI_ISL_492299, EPI_ISL_492300, EPI_ISL_492303, EPI_ISL_492305, EPI_ISL_492306, EPI_ISL_492310, EPI_ISL_492314, EPI_ISL_492319, EPI_ISL_492320, EPI_ISL_492322, EPI_ISL_492325, EPI_ISL_492327, EPI_ISL_492330, EPI_ISL_492331, EPI_ISL_492335, EPI_ISL_492336, EPI_ISL_492338, EPI_ISL_492339, EPI_ISL_492342, EPI_ISL_492348, EPI_ISL_492349, EPI_ISL_492350, EPI_ISL_492352, EPI_ISL_492355, EPI_ISL_492360, EPI_ISL_492362, EPI_ISL_492363, EPI_ISL_492365, EPI_ISL_492367, EPI_ISL_492370, EPI_ISL_492372, EPI_ISL_492373, EPI_ISL_492375, EPI_ISL_492378, EPI_ISL_492379, EPI_ISL_492381, EPI_ISL_492383, EPI_ISL_492384, EPI_ISL_492385, EPI_ISL_492386, EPI_ISL_492390, EPI_ISL_492391, EPI_ISL_492392, EPI_ISL_492395, EPI_ISL_492396, EPI_ISL_492399, EPI_ISL_492400, EPI_ISL_492401, EPI_ISL_492404, EPI_ISL_492405, EPI_ISL_492406, EPI_ISL_492408, EPI_ISL_492410, EPI_ISL_492411, EPI_ISL_492417, EPI_ISL_492421, EPI_ISL_492423, EPI_ISL_492427, EPI_ISL_492428, EPI_ISL_492433, EPI_ISL_492434, EPI_ISL_492437, EPI_ISL_492440, EPI_ISL_492442, EPI_ISL_492445, EPI_ISL_492446, EPI_ISL_492447, EPI_ISL_492448, EPI_ISL_492449 | | | |
| see above | PHE South West Regional Laboratory, National Infection Service | Wellcome Sanger Institute for the COVID-19 Genomics UK (COG-UK) consortium | Stephanie Hutchings, Hannah Pymont, Dr Peter Muir, Barry Vipond, Rich Hopes; and Alex Alderton, Roberto Amato, Sonia Goncalves, Ewan Harrison, David K. Jackson, Ian Johnston, Dominic Kwiatkowski, Cordelia Langford, John Sillitoe on behalf of the Wellcome Sanger Institute COVID-19 Surveillance Team (http://www.sanger.ac.uk/covid-team) |
| EPI_ISL_492460, EPI_ISL_492465, EPI_ISL_492466, EPI_ISL_492475, EPI_ISL_492477, EPI_ISL_492478 | NU-OMICS DNA Sequencing research facility, Northumbria University | Wellcome Sanger Institute for the COVID-19 Genomics UK (COG-UK) consortium | Chris Duncan, Sheia Waugh, Shirelle Burton-Fanning, Gary Eltringham, Jennifer Collins, Brendan Payne, Yusri Taha, Emma Swindells, Jane Greenaway, Edward Barton, Garren Scott, Debra Padgett, Clive Graham, Sarah Essex, Steve Ligget, Paul Baker, Lynn Dover, Wen Yew, Gary Black, John Allan, Joshua Loh, Greg Young, Matthew Bashton, Andrew Nelson, Darren Smith and Alex Alderton, Roberto Amato, Sonia Goncalves, Ewan Harrison, David K. Jackson, Ian Johnston, Dominic Kwiatkowski, Cordelia Langford, John Sillitoe on behalf of the Wellcome Sanger Institute COVID-19 Surveillance Team (http://www.sanger.ac.uk/covid-team) |
| EPI_ISL_492497 | University College London, Great Ormond Street Hospital for Children NHS Foundation Trust, Imperial College Healthcare NHS Trust | Wellcome Sanger Institute for the COVID-19 Genomics UK (COG-UK) Consortium | Sergi Castellano, Rachel Williams, Mark Kristiansen, Paola Resende Silva, Sunando Roy, Tony Brooks, Helena Tutill, Paola Niola, Patricia Dyal, Charlotte Williams, Leysa Forrest, Yasmin Panchbhaya, Jacqueline Findlay, Sam Weeks, Julianne Brown, Kathryn Harris, Paul Randell, James Price, Alison Holmes, Judith Breuer and Alex Alderton, Roberto Amato, Sonia Goncalves, Ewan Harrison, David K. Jackson, Ian Johnston, Dominic Kwiatkowski, Cordelia Langford, John Sillitoe on behalf of the Wellcome Sanger Institute COVID-19 Surveillance Team |
| EPI_ISL_492836, EPI_ISL_492848, EPI_ISL_492853, EPI_ISL_492855, EPI_ISL_492858, EPI_ISL_492859, EPI_ISL_492866, EPI_ISL_492871 | Department of Medical Microbiology, Western Sussex Hospitals NHS Foundation Trust, St Richard's Hospital | Wellcome Sanger Institute for the COVID-19 Genomics UK (COG-UK) consortium | Manasa Mutingwende, Sarah Lowdon, Olga Podplomyk, Michelle Erkiert, Jonathan Lewis, Paul Randell and Alex Alderton, Roberto Amato, Sonia Goncalves, Ewan Harrison, David K. Jackson, Ian Johnston, Dominic Kwiatkowski, Cordelia Langford, John Sillitoe on behalf of the Wellcome Sanger Institute COVID-19 Surveillance Team (http://www.sanger.ac.uk/covid-team) |
| EPI_ISL_492877 | Royal Free Hospital / Health Services Laboratories | Wellcome Sanger Institute for the COVID-19 Genomics UK (COG-UK) consortium | Tanzina Haque, Tabitha Mahungu, Dianne Irish, Cate Goodlad, Jenny Cross, Judith Heaney and Alex Alderton, Roberto Amato, Sonia Goncalves, Ewan Harrison, David K. Jackson, Ian Johnston, Dominic Kwiatkowski, Cordelia Langford, John Sillitoe on behalf of the Wellcome Sanger Institute COVID-19 Surveillance Team (http://www.sanger.ac.uk/covid-team) |
| EPI_ISL_492878, EPI_ISL_492881, EPI_ISL_492889, EPI_ISL_492892, EPI_ISL_492896, EPI_ISL_492898, EPI_ISL_492900, EPI_ISL_492904, EPI_ISL_492907, EPI_ISL_492909 | Department of Medical Microbiology, Western Sussex Hospitals NHS Foundation Trust, St Richard's Hospital | Wellcome Sanger Institute for the COVID-19 Genomics UK (COG-UK) consortium | Manasa Mutingwende, Sarah Lowdon, Olga Podplomyk, Michelle Erkiert, Jonathan Lewis, Paul Randell and Alex Alderton, Roberto Amato, Sonia Goncalves, Ewan Harrison, David K. Jackson, Ian Johnston, Dominic Kwiatkowski, Cordelia Langford, John Sillitoe on behalf of the Wellcome Sanger Institute COVID-19 Surveillance Team (http://www.sanger.ac.uk/covid-team) |
| EPI_ISL_493130 | Functional Genomics Core University of South Carolina / Prisma Health-Midlands | Functional Genomics Core, University of South Carolina, | Hao Ji, Diego Altomare, B.Celia Cui, Mengqian Chen, Alyssa Clay-Glimour, Michael Wyatt, Phillip Buckhaults, Helmut Albrecht, Michael Shtutman |
| EPI_ISL_493209, EPI_ISL_493210, EPI_ISL_493211 | Functional Genomics Core University of South Carolina / Prisma Health-Midlands | Functional Genomics Core, University of South Carolina | Hao Ji, Diego Altomare, B.Celia Cui, Mengqian Chen, Alyssa Clay-Glimour, Michael Wyatt, Phillip Buckhaults, Helmut Albrecht, Michael Shtutman |
| EPI_ISL_493338 | Instituto de Diagnostico y Referencia Epidemiologicos (INDRE) | Instituto de Diagnostico y Referencia Epidemiologicos (INDRE) | Gisela Barrera-Badillo , Abril Rodriguez-Maldonado, Claudia Wong-Arambula , Natividad Cruz-Ortiz, Tatiana Nunez-Garcia, Dayanira Arellano-Suarez, Fabiola Garces-Ayala, Edgar Mendieta-Condado, Lucia Hernandez-Rivas, Irma Lopez-Martinez, Ernesto Ramirez-Gonzalez. |
| EPI_ISL_493339 | Instituto de Diagnostico y Referencia Epidemiologicos (INDRE) | Instituto de Diagnostico y Referencia Epidemiologicos (INDRE) | Gisela Barrera-Badillo , Abril Rodriguez-Maldonado, Claudia Wong-Arambula , Natividad Cruz-Ortiz, Tatiana Nunez-Garcia, Dayanira Arellano-Suarez, Adnan Araiza-Rodriguez, Edgar Mendieta-Condado, Lucia Hernandez-Rivas, Irma Lopez-Martinez, Ernesto Ramirez-Gonzalez. |
| EPI_ISL_493398, EPI_ISL_493405 | National Public Health Laboratory, National Centre for Infectious Diseases | National Public Health Laboratory, National Centre for Infectious Diseases | Mak TM, Octavia S, Zhou Z, Chavatte JM, Cui L, Lin RTP |
| EPI_ISL_493443, EPI_ISL_493444, EPI_ISL_493445, EPI_ISL_493446, EPI_ISL_493447, EPI_ISL_493448, EPI_ISL_493449, EPI_ISL_493450, EPI_ISL_493451 | University of Birmingham | COVID-19 Genomics UK (COG-UK) Consortium | Institute of Microbiology, University of Birmingham: Claire McMurray, Joanne Stockton, Samuel Nicholls, Radoslaw Poplawski, Will Rowe, Josh Quick, Nicholas Loman. University of Birmingham Testing Laboratory: Celina M Whalley, Andrew Bosworth, Charlotte Poxon, Kasun Wanigasooriya, Oliver Pickles, Mike Kidd, Alex Richter, Andrew D Beggs PHE Heartlands Lab: Husam Osman, Andrew Bosworth. Queen Elizabeth Hospital: Anna Casey |
| EPI_ISL_493509, EPI_ISL_493515, EPI_ISL_493516, EPI_ISL_493517, EPI_ISL_493518, EPI_ISL_493519, EPI_ISL_493520, EPI_ISL_493521, EPI_ISL_493522, EPI_ISL_493524, EPI_ISL_493525, EPI_ISL_493526, EPI_ISL_493527, EPI_ISL_493528, EPI_ISL_493529, EPI_ISL_493530, EPI_ISL_493531, EPI_ISL_493532, EPI_ISL_493533 | | | |
| see above | Northumbria University / South Tees Hospitals NHS Foundation Trust / North Cumbria Integrated Care NHS Foundation Trust / North Tees and Hartlepool NHS Foundation Trust / Newcastle Hospitals NHS Foundation Trust | COVID-19 Genomics UK (COG-UK) Consortium | Darren L Smith,Andrew Nelson,Matthew Bashton,Greg R Young,Joshua Loh,John Allan,Mohammad A Tariq,Giles S Holt,Gary Black,Wen C Yew,Lynn Dover,Paul Baker,Steve Liggett,Sarah Essex,Jane Greenaway,Debra Padgett,Clive Graham,Garren Scott,Edward Barton,Emma Swindells,Brendan Payne,Jennifer Collins,Yusri Taha,Gary Eltringham |
| EPI_ISL_493595, EPI_ISL_493596, EPI_ISL_493597, EPI_ISL_493598, EPI_ISL_493599, EPI_ISL_493600 | Lincolnshire Hospitals and DeepSeq Nottingham | COVID-19 Genomics UK (COG-UK) Consortium | Nichola Duckworth, Tim Sloan, Sarah Walsh, Jonathan Ball, Patrick McClure, Joeseph Chappell, Nadine Holmes, Matthew Carlisle, Christopher Moore, Fei Sang, Johnny Debebe, Victoria Wright, Matthew Loose |
| EPI_ISL_493631, EPI_ISL_493632, EPI_ISL_493635, EPI_ISL_493636 | Centre for Enzyme Innovation, University of Portsmouth / Translational Research Laboratory, Portsmouth Hospitals NHS Trust | COVID-19 Genomics UK (COG-UK) Consortium | Angela Beckett,Yann Bourgeois,Garry Scarlett,Sharon Glaysher,Scott Elliott,Kelly Bicknell,Robert Impey,Allyson Lloyd,Sarah Wyllie,Ethan Butcher,Anoop Chauhan,Samuel Robson |
| EPI_ISL_493726, EPI_ISL_493727, EPI_ISL_493728 | Virology Department, Sheffield Teaching Hospitals NHS Foundation Trust/Department of Infection, Immunity and Cardiovascular Disease, The Medical School, University of Sheffield | COVID-19 Genomics UK (COG-UK) Consortium | Thushan de Silva, Matthew Parker, Nikki Smith, Adri Angyal, Rebecca Brown, Luke Green, Rachel Tucker, Paul Parsons, Danielle Groves, Katie Johnson, Laura Carrilero, Alex Keeley, Dave Partridge, Matthew Wyles, Benjamin Lindsey, Mehmet Yavuz, Mohammad Raza, Cariad Evans |
| EPI_ISL_493831, EPI_ISL_493834, EPI_ISL_493835, EPI_ISL_493836, EPI_ISL_493837, EPI_ISL_493838, EPI_ISL_493839, EPI_ISL_493844, EPI_ISL_493845, EPI_ISL_493846, EPI_ISL_493847, EPI_ISL_493848, EPI_ISL_493849, EPI_ISL_493850, EPI_ISL_493851, EPI_ISL_493853, EPI_ISL_493854, EPI_ISL_493855, EPI_ISL_493856, EPI_ISL_493857, EPI_ISL_493858, EPI_ISL_493859, EPI_ISL_493860, EPI_ISL_493861, EPI_ISL_493862, EPI_ISL_493863, EPI_ISL_493864, EPI_ISL_493865, EPI_ISL_493866, EPI_ISL_493867, EPI_ISL_493873, EPI_ISL_493874, EPI_ISL_493875, EPI_ISL_493876, EPI_ISL_493877, EPI_ISL_493878, EPI_ISL_493879, EPI_ISL_493880, EPI_ISL_493881, EPI_ISL_493882, EPI_ISL_493883, EPI_ISL_493884, EPI_ISL_493885, EPI_ISL_493886, EPI_ISL_493887, EPI_ISL_493888, EPI_ISL_493889, EPI_ISL_493890 | | | |
| see above | West of Scotland Specialist Virology Centre, NHSGGC / MRC-University of Glasgow Centre for Virus Research | COVID-19 Genomics UK (COG-UK) Consortium | Ana da Silva Filipe, Natasha Johnson, Kathy Smollett, Daniel Mair, Stephen Carmichael, Lily Tong, Jenna Nichols, Elihu Aranday-Cortes, Kirstyn Brunker, Yasmin Parr, Alice Broos, Kyriaki Nomikou; Sarah McDonald, Marc Niebel, Patawee Asamaphan; Richard Orton, Joseph Hughes, Sreenu Vattipally, David L Robertson; Alasdair MacLean, Rory Gunson; Kathy Li, Natasha Jesudason, Rajiv Shah, James Shepherd, Antonia Ho, Emma Thomson |
| EPI_ISL_493898, EPI_ISL_493899, EPI_ISL_493900, EPI_ISL_493901, EPI_ISL_493902, EPI_ISL_493903, EPI_ISL_493904 | Virology Department, Royal Infirmary of Edinburgh, NHS Lothian / School of Biological Sciences, University of Edinburgh / Institute of Genetics and Molecular Medicine, University of Edinburgh | COVID-19 Genomics UK (COG-UK) Consortium | McHugh M, Dewar R, Rooke S, Gallagher M, Balcaza C, O'Toole Á, Scher E, Hill V, McCrone JT, Colquhoun R, Yu X, Jackson B, Rambaut A, Williams TC, Templeton K |

| | | | |
|---|---|---|---|
| EPI_ISL_493979, EPI_ISL_493999, EPI_ISL_494165, EPI_ISL_494166, EPI_ISL_494182, EPI_ISL_494265 | Wales Specialist Virology Centre Sequencing lab: Pathogen Genomics Unit | COVID-19 Genomics UK (COG-UK) Consortium | Catherine Moore, Johnathan Evans, Laura Gifford, Malorie Perry, Simon Cottrell, Angela Marchbank, Alec Birchley, Alexander Adams, Amy Gaskin, Bree Gatica-Wilcox, Jason Coombes, Joel Southgate, Lauren Gilbert, Lee Graham, Nicole Pacchiarini, Sara Kumziene-Summerhayes, Sarah Taylor, Sophie Jones, Sara Rey, Matthew Bull, Joanne Watkins, Sally Corden, Tom Connor |
| EPI_ISL_494375, EPI_ISL_494376, EPI_ISL_494382, EPI_ISL_494384, EPI_ISL_494387, EPI_ISL_494388, EPI_ISL_494389, EPI_ISL_494393, EPI_ISL_494396, EPI_ISL_494397, EPI_ISL_494399, EPI_ISL_494401, EPI_ISL_494403, EPI_ISL_494414, EPI_ISL_494415, EPI_ISL_494422, EPI_ISL_494423, EPI_ISL_494424, EPI_ISL_494425, EPI_ISL_494426, EPI_ISL_494428, EPI_ISL_494429, EPI_ISL_494441, EPI_ISL_494454, EPI_ISL_494456, EPI_ISL_494463, EPI_ISL_494466, EPI_ISL_494467, EPI_ISL_494468, EPI_ISL_494469, EPI_ISL_494475, EPI_ISL_494476, EPI_ISL_494477, EPI_ISL_494478, EPI_ISL_494480, EPI_ISL_494481, EPI_ISL_494482, EPI_ISL_494483, EPI_ISL_494484, EPI_ISL_494485, EPI_ISL_494486, EPI_ISL_494487, EPI_ISL_494489, EPI_ISL_494490, EPI_ISL_494491, EPI_ISL_494492, EPI_ISL_494493, EPI_ISL_494494, EPI_ISL_494495, EPI_ISL_494496, EPI_ISL_494497, EPI_ISL_494498, EPI_ISL_494499, EPI_ISL_494500, EPI_ISL_494501, EPI_ISL_494502 | | | |
| see above | San Diego County Public Health Laboratory | Andersen lab at Scripps Research | SEARCH Alliance San Diego with Tracy Basler, Jovan Shephard, Brett Austin |
| EPI_ISL_494504, EPI_ISL_494505, EPI_ISL_494506, EPI_ISL_494507, EPI_ISL_494508, EPI_ISL_494509, EPI_ISL_494510, EPI_ISL_494511, EPI_ISL_494512, EPI_ISL_494513, EPI_ISL_494514, EPI_ISL_494515, EPI_ISL_494516, EPI_ISL_494517, EPI_ISL_494518, EPI_ISL_494519, EPI_ISL_494520, EPI_ISL_494522, EPI_ISL_494523, EPI_ISL_494524, EPI_ISL_494525, EPI_ISL_494526, EPI_ISL_494527, EPI_ISL_494528, EPI_ISL_494529, EPI_ISL_494530, EPI_ISL_494531, EPI_ISL_494532, EPI_ISL_494533, EPI_ISL_494534, EPI_ISL_494535, EPI_ISL_494536, EPI_ISL_494537, EPI_ISL_494542, EPI_ISL_494543, EPI_ISL_494544, EPI_ISL_494545, EPI_ISL_494546, EPI_ISL_494547, EPI_ISL_494548, EPI_ISL_494549, EPI_ISL_494550, EPI_ISL_494551, EPI_ISL_494552 | | | |
| see above | Quest Diagnostics | Quest Diagnostics | Anderson,B.P., Rosenthal,S.H., Gerasimova,A., Kagan,R.M. and Owen, R. |
| EPI_ISL_494588, EPI_ISL_494591, EPI_ISL_494621 | San Diego County Public Health Laboratory | Andersen lab at Scripps Research | SEARCH Alliance San Diego with Tracy Basler, Jovan Shephard, Brett Austin |
| EPI_ISL_494639, EPI_ISL_494641, EPI_ISL_494645, EPI_ISL_494649, EPI_ISL_494651, EPI_ISL_494654, EPI_ISL_494656, EPI_ISL_494663, EPI_ISL_494664, EPI_ISL_494669, EPI_ISL_494670, EPI_ISL_494674, EPI_ISL_494675, EPI_ISL_494682, EPI_ISL_494689, EPI_ISL_494690, EPI_ISL_494697, EPI_ISL_494701, EPI_ISL_494702, EPI_ISL_494712 | | | |
| see above | Scripps Medical Laboratory | Andersen lab at Scripps Research | SEARCH Alliance San Diego with Michael Quigley, Ellen Stefanski, Ian Mchardy |
| EPI_ISL_495092 | Department of Medical Microbiology, Western Sussex Hospitals NHS Foundation Trust, St Richard's Hospital | Wellcome Sanger Institute for the COVID-19 Genomics UK (COG-UK) consortium | Manasa Mutingwende, Sarah Lowdon, Olga Podplomyk, Michelle Erkiert, Jonathan Lewis, Paul Randell and Alex Alderton, Roberto Amato, Sonia Goncalves, Ewan Harrison, David K. Jackson, Ian Johnston, Dominic Kwiatkowski, Cordelia Langford, John Sillitoe on behalf of the Wellcome Sanger Institute COVID-19 Surveillance Team (http://www.sanger.ac.uk/covid-team) |
| EPI_ISL_495097, EPI_ISL_495101, EPI_ISL_495104, EPI_ISL_495106, EPI_ISL_495107, EPI_ISL_495110, EPI_ISL_495112, EPI_ISL_495113, EPI_ISL_495114, EPI_ISL_495119, EPI_ISL_495120 | | | |
| see above | PHE South West Regional Laboratory, National Infection Service | Wellcome Sanger Institute for the COVID-19 Genomics UK (COG-UK) consortium | Stephanie Hutchings, Hannah Pymont, Dr Peter Muir, Barry Vipond, Rich Hopes; and Alex Alderton, Roberto Amato, Sonia Goncalves, Ewan Harrison, David K. Jackson, Ian Johnston, Dominic Kwiatkowski, Cordelia Langford, John Sillitoe on behalf of the Wellcome Sanger Institute COVID-19 Surveillance Team (http://www.sanger.ac.uk/covid-team) |
| EPI_ISL_495383, EPI_ISL_495384, EPI_ISL_495385 | Florida Bureau of Public Health Laboratories | Florida Bureau of Public Health Laboratories | Sarah Schmedes, Jason Blanton |
| EPI_ISL_495414, EPI_ISL_495417, EPI_ISL_495419, EPI_ISL_495428, EPI_ISL_495432, EPI_ISL_495434, EPI_ISL_495441, EPI_ISL_495442, EPI_ISL_495443 | Kafkas University, Faculty of Medicine, Department of Medical Microbiology | Kafkas University, Faculty of Medicine, Department of Medical Microbiology | Murat Karamese, Didem Ozgur, E. Ediz Tutuncu |
| EPI_ISL_495565, EPI_ISL_495567 | University of Michigan Clinical Microbiology Laboratory | Lauring Lab, University of Michigan, Department of Microbiology and Immunology | Valesano et al. |
| EPI_ISL_495654, EPI_ISL_495655, EPI_ISL_495659, EPI_ISL_495660, EPI_ISL_495662, EPI_ISL_495664, EPI_ISL_495665 | Seattle Flu Study | Seattle Flu Study | Deborah A. Nickerson, Chris D. Frazar, Jover Lee, Benjamin Pelle, Matthew Richardson, Amanda Adler, Elisabeth Brandstetter, Peter D. Han, Kairsten Fay, Misja Ilcisin, Kirsten Lacombe, Thomas R. Sibley, Melissa Truong, Caitlin R. Wolf, Karen Cowgill, Stephanie Schrag, Jeff Duchin, Michael Boeckh, Janet A. Englund, Michael Famulare, Barry R. Lutz, Mark J. Rieder, Lea M. Starita, Matthew Thompson, Helen Y. Chu, Trevor Bedford, Jay Shendure |
| EPI_ISL_496339, EPI_ISL_496340, EPI_ISL_496341, EPI_ISL_496342, EPI_ISL_496343, EPI_ISL_496344, EPI_ISL_496345, EPI_ISL_496346, EPI_ISL_496347, EPI_ISL_496348, EPI_ISL_496349, EPI_ISL_496350, EPI_ISL_496351, EPI_ISL_496352, EPI_ISL_496353, EPI_ISL_496354, EPI_ISL_496355, EPI_ISL_496374, EPI_ISL_496375, EPI_ISL_496376 | | | |
| see above | Infectolab | Andersen lab at Scripps Research | SEARCH Alliance San Diego with Samuel Navarro Alvarez, Carlos A. Cota Haros, Octavio Renteria Pacheco |
| EPI_ISL_496491, EPI_ISL_496492, EPI_ISL_496506, EPI_ISL_496507, EPI_ISL_496508, EPI_ISL_496509, EPI_ISL_496510, EPI_ISL_496511, EPI_ISL_496512, EPI_ISL_496513 | Viral Respiratory Lab, National Institute for Biomedical Research (INRB) | Pathogen Sequencing Lab, National Institute for Biomedical Research (INRB) | Placide Mbala-Kingebeni, Edith Nkwembe, Eddy Kinganda-Lusamaki, Amuri Aziza, Francisca Muyembe Mawete, Emmanuel Lokilo Lofiko, Catherine Pratt, Matthias Pauthner, Josh Quick, Allison Black, James Hadfield, Trevor Bedford, Ian Goodfellow, Andrew Rambaut, Nick Loman, Kristian Andersen, Michael Wiley, Steve Ahuka-Mundeke, Jean-Jacques Muyembe Tamfum |
| EPI_ISL_496555, EPI_ISL_496556, EPI_ISL_496557, EPI_ISL_496586, EPI_ISL_497879 | National Centre For Cell Science | National Centre For Cell Science | Dhiraj Paul, Kunal Jani, Radha Chauhan, Janesh Kumar, Vasudevan Seshadri, Girdhari Lal, Rajesh Karyakarte, Suvarna Joshi, Murlidhar Tambe, Sourav Sen, Santosh Karade, Kavita Bala Anand, Shelinder Pal Singh Shergill, Rajiv Mohan Gupta, Manoj Kumar Bhat, Arvind Sahu, Maharashtra COVID-19 Study Group, DBT's PAN-INDIA 1000 SARS-CoV2 RNA genome sequencing consortium, Yogesh S Shouche |
| EPI_ISL_498668, EPI_ISL_498669, EPI_ISL_498670, EPI_ISL_498671, EPI_ISL_498672, EPI_ISL_498673, EPI_ISL_498674, EPI_ISL_498675, EPI_ISL_498676, EPI_ISL_498677 | Utah Public Health Laboratory | Utah Public Health Laboratory | Heidi Butz, Erin Young, Kelly Oakeson |
| EPI_ISL_498695, EPI_ISL_498696, EPI_ISL_498700, EPI_ISL_498701, EPI_ISL_498702, EPI_ISL_498703, EPI_ISL_498704, EPI_ISL_498714, EPI_ISL_498715, EPI_ISL_498716, EPI_ISL_498717, EPI_ISL_498718, EPI_ISL_498719, EPI_ISL_498720, EPI_ISL_498721, EPI_ISL_498722, EPI_ISL_498723, EPI_ISL_498724, EPI_ISL_498725, EPI_ISL_498726, EPI_ISL_498727, EPI_ISL_498729, EPI_ISL_498730, EPI_ISL_498731, EPI_ISL_498732, EPI_ISL_498733, EPI_ISL_498734, EPI_ISL_498735, EPI_ISL_498737, EPI_ISL_498740, EPI_ISL_498741, EPI_ISL_498742, EPI_ISL_498743, EPI_ISL_498744, EPI_ISL_498745, EPI_ISL_498746, EPI_ISL_498747 | | | |
| see above | Quest Diagnostics | Quest Diagnostics | Rosenthal,S.H., Gerasimova,A., Kagan,R.M. and Owen, R. |
| EPI_ISL_499365, EPI_ISL_499372, EPI_ISL_499382 | Wales Specialist Virology Centre Sequencing lab: Pathogen Genomics Unit | COVID-19 Genomics UK (COG-UK) Consortium | Catherine Moore, Johnathan Evans, Laura Gifford, Malorie Perry, Simon Cottrell, Angela Marchbank, Alec Birchley, Alexander Adams, Amy Gaskin, Bree Gatica-Wilcox, Jason Coombes, Joel Southgate, Lauren Gilbert, Lee Graham, Nicole Pacchiarini, Sara Kumziene-Summerhayes, Sarah Taylor, Sophie Jones, Sara Rey, Matthew Bull, Joanne Watkins, Sally Corden, Tom Connor |
| EPI_ISL_499507, EPI_ISL_499508, EPI_ISL_499558, EPI_ISL_499789, EPI_ISL_499790, EPI_ISL_499791, EPI_ISL_499792, EPI_ISL_499793, EPI_ISL_499794, EPI_ISL_499795, EPI_ISL_499797, EPI_ISL_499798, EPI_ISL_499799, EPI_ISL_499800, EPI_ISL_499801, EPI_ISL_499802, EPI_ISL_499803, EPI_ISL_499804, EPI_ISL_499805 | | | |
| see above | Northumbria University / South Tees Hospitals NHS Foundation Trust / North Cumbria Integrated Care NHS Foundation Trust / North Tees and Hartlepool NHS Foundation Trust / Newcastle Hospitals NHS Foundation Trust | COVID-19 Genomics UK (COG-UK) Consortium | Darren L Smith,Andrew Nelson,Matthew Bashton,Greg R Young,Joshua Loh,John Allan,Mohammad A Tariq,Giles S Holt,Gary Black,Wen C Yew,Lynn Dover,Paul Baker,Steve Liggett,Sarah Essex,Jane Greenaway,Debra Padgett,Clive Graham,Garren Scott,Edward Barton,Emma Swindells,Brendan Payne,Jennifer Collins,Yusri Taha,Gary Eltringham |
| EPI_ISL_500158, EPI_ISL_500159 | Complejo Hospitalario Universitario de Albacete | SeqCOVID-SPAIN consortium/IBV(CSIC) | Encarnacion Simarro Córdoba, Julia Lozano Serra, Lorena Robles Fonseca , Monica Parra Grandes, Caridad Sainz de Baranda Camino and SeqCOVID-SPAIN consortium |
| EPI_ISL_500161 | Hospital Universitario Virgen de las Nieves de Granada-SAS | SeqCOVID-SPAIN consortium/IBV(CSIC) | Mercedes Pérez Ruiz, Sara Sanbonmatsu Gámez, Irene Pedrosa Corral, José M. Navarro-Marí and SeqCOVID-SPAIN consortium |
| EPI_ISL_500166 | Complejo Hospitalario Universitario de Albacete | SeqCOVID-SPAIN consortium/IBV(CSIC) | Encarnacion Simarro Córdoba, Julia Lozano Serra, Lorena Robles Fonseca , Monica Parra Grandes, Caridad Sainz de Baranda Camino and SeqCOVID-SPAIN consortium |
| EPI_ISL_500201 | Hospital Universitario Virgen de las Nieves de Granada-SAS | SeqCOVID-SPAIN consortium/IBV(CSIC) | Mercedes Pérez Ruiz, Sara Sanbonmatsu Gámez, Irene Pedrosa Corral, José M. Navarro-Marí and SeqCOVID-SPAIN consortium |
| EPI_ISL_500207, EPI_ISL_500218 | Complejo Hospitalario Universitario de Albacete | SeqCOVID-SPAIN consortium/IBV(CSIC) | Encarnacion Simarro Córdoba, Julia Lozano Serra, Lorena Robles Fonseca , Monica Parra Grandes, Caridad Sainz de Baranda Camino and |

| | | | |
|---|---|---|---|
| | | | SeqCOVID-SPAIN consortium |
| EPI_ISL_500290, EPI_ISL_500291, EPI_ISL_500360, EPI_ISL_500365, EPI_ISL_500366 | Servicio de Microbiología, Hospital Miguel Servet, Zaragoza | SeqCOVID-SPAIN consortium/IBV(CSIC) | Antonio Rezusta López, Alexander Tristancho Baró, Ana Milagro, Yolanda Gracia Grataloup, Nieves Martínez Cameo and SeqCOVID-SPAIN consortium |
| EPI_ISL_500622, EPI_ISL_500623 | Area of Virology, Serology and Virology Division (SAViD), New South Wales Health Pathology Randwick | Area of Virology, Serology and Virology Division (SAViD), New South Wales Health Pathology Randwick | Rawlinson, W. |
| EPI_ISL_500953 | Respiratory Virus Unit, Microbiology Services Colindale, Public Health England | Respiratory Virus Unit, Microbiology Services Colindale, Public Health England | PHE Covid Sequencing Team |
| EPI_ISL_500956, EPI_ISL_500962, EPI_ISL_500965, EPI_ISL_500972, EPI_ISL_500975, EPI_ISL_500985, EPI_ISL_500993, EPI_ISL_500999, EPI_ISL_501018, EPI_ISL_501020, EPI_ISL_501027, EPI_ISL_501028, EPI_ISL_501030, EPI_ISL_501033, EPI_ISL_501050, EPI_ISL_501051, EPI_ISL_501069, EPI_ISL_501071 | | | |
| see above | Regional Virus Laboratory, Belfast Health and Social Care Trust | Wellcome Sanger Institute for the COVID-19 Genomics UK (COG-UK) consortium | Conall McCaughey, James McKenna, Tanya Curran, Susan Feeney, Alison Watt, Ciara Cox, Mairead Connor, Zoltan Molnar, David Simpson, Derek Fairley; and Alex Alderton, Roberto Amato, Sonia Goncalves, Ewan Harrison, David K. Jackson, Ian Johnston, Dominic Kwiatkowski, Cordelia Langford, John Sillitoe on behalf of the Wellcome Sanger Institute COVID-19 Surveillance Team (http://www.sanger.ac.uk/covid-team) |
| EPI_ISL_501200 | Department of Medical Microbiology, University Malaya Medical Centre | Department of Medical Microbiology, Faculty of Medicine, University of Malaya | Yoong Min CHONG, Jennifer Chong, I-Ching SAM, Yoke Fun CHAN, University Malaya Medical Centre COVID Team |
| EPI_ISL_501232, EPI_ISL_501233 | Hellenic Pasteur Institute, National Influenza Reference laboratory of Southern Greece & Unit of Bioinformatics and Applied Genomics | Hellenic Pasteur Institute, National Influenza Reference laboratory of Southern Greece & Unit of Bioinformatics and Applied Genomics | Vasiliki Pogka, Timokratis Karamitros, Athanasios Kossyvakis, Antonios Kalliaropoulos, Horefti Elina, Evangelidou Maria, Androniki Voulgari-Kokota, Aspasia Kontou, Andreas Mentis |
| EPI_ISL_507006 | Department of Laboratory Medicine, Tan Tock Seng Hospital | Department of Laboratory Medicine, Tan Tock Seng Hospital | Chen YYC, Zair X, Li C, Tang WY, Maurer-Stroh S, Barkham TMS, Nagarajan N, Sessions OM |
| EPI_ISL_507007 | Yaftabad Hospital, Covid Lab Center | Tehran University of Medical Sciences | Shahabzadeh,Z., Hosseinzadeh Gharajeh,N., Hashemian,S.M. and Barati,O. |
| EPI_ISL_507097, EPI_ISL_507098, EPI_ISL_507099, EPI_ISL_507100 | University College London Hospital | COVID-19 Genomics UK (COG-UK) Consortium | Judith Heaney, Matthew Byott, Catherine Houlihan, Dan Frampton, Stuart Kirk, Moira Spyer and Eleni Nastouli |
| EPI_ISL_507110, EPI_ISL_507111, EPI_ISL_507112, EPI_ISL_507113, EPI_ISL_507114, EPI_ISL_507115, EPI_ISL_507116, EPI_ISL_507117, EPI_ISL_507118 | Northumbria University / South Tees Hospitals NHS Foundation Trust / North Cumbria Integrated Care NHS Foundation Trust / North Tees and Hartlepool NHS Foundation Trust / Newcastle Hospitals NHS Foundation Trust | COVID-19 Genomics UK (COG-UK) Consortium | Darren L Smith,Andrew Nelson,Matthew Bashton,Greg R Young,Joshua Loh,John Allan,Mohammad A Tariq,Giles S Holt,Gary Black,Wen C Yew,Lynn Dover,Paul Baker,Steve Liggett,Sarah Essex,Jane Greenaway,Debra Padgett,Clive Graham,Garren Scott,Edward Barton,Emma Swindells,Brendan Payne,Jennifer Collins,Yusri Taha,Gary Eltringham |
| EPI_ISL_507181, EPI_ISL_507197 | Virology Department, Royal Infirmary of Edinburgh, NHS Lothian / School of Biological Sciences, University of Edinburgh / Institute of Genetics and Molecular Medicine, University of Edinburgh | COVID-19 Genomics UK (COG-UK) Consortium | McHugh M, Dewar R, Rooke S, Gallagher M, Balcaza C, O'Toole Á, Scher E, Hill V, McCrone JT, Colquhoun R, Yu X, Jackson B, Rambaut A, Williams TC, Templeton K |
| EPI_ISL_507205 | WHO National Influenza Centre Russian Federation | WHO National Influenza Centre Russian Federation | Andrey Komissarov, Artem Fadeev, Mariia Sergeeva, Anna Ivanova, Daria Danilenko |
| EPI_ISL_507206, EPI_ISL_507207, EPI_ISL_507208, EPI_ISL_507209, EPI_ISL_507210, EPI_ISL_507211, EPI_ISL_507212 | Department of Experimental Modeling and Pathogenesis of Infectious Diseases | WHO National Influenza Centre Russian Federation | Andrey Komissarov, Artem Fadeev, Mariia Sergeeva, Anna Ivanova, Daria Danilenko |
| EPI_ISL_507216, EPI_ISL_507217, EPI_ISL_507218, EPI_ISL_507219, EPI_ISL_507220, EPI_ISL_507221, EPI_ISL_507222, EPI_ISL_507223, EPI_ISL_507224, EPI_ISL_507231, EPI_ISL_507232, EPI_ISL_507233, EPI_ISL_507234, EPI_ISL_507235, EPI_ISL_507236, EPI_ISL_507237, EPI_ISL_507238, EPI_ISL_507239, EPI_ISL_507240, EPI_ISL_507241, EPI_ISL_507242, EPI_ISL_507243, EPI_ISL_507244, EPI_ISL_507245, EPI_ISL_507246, EPI_ISL_507247, EPI_ISL_507248, EPI_ISL_507249, EPI_ISL_507250, EPI_ISL_507251, EPI_ISL_507252, EPI_ISL_507253, EPI_ISL_507254, EPI_ISL_507255, EPI_ISL_507300 | | | |
| see above | WHO National Influenza Centre Russian Federation | WHO National Influenza Centre Russian Federation | Andrey Komissarov, Artem Fadeev, Mariia Sergeeva, Anna Ivanova, Daria Danilenko |
| EPI_ISL_507984, EPI_ISL_508001, EPI_ISL_508009, EPI_ISL_508010, EPI_ISL_508012, EPI_ISL_508015, EPI_ISL_508018, EPI_ISL_508022, EPI_ISL_508023, EPI_ISL_508038, EPI_ISL_508039, EPI_ISL_508040, EPI_ISL_508041, EPI_ISL_508042, EPI_ISL_508043, EPI_ISL_508061, EPI_ISL_508064, EPI_ISL_508065, EPI_ISL_508067, EPI_ISL_508080, EPI_ISL_508082, EPI_ISL_508083, EPI_ISL_508084, EPI_ISL_508085, EPI_ISL_508086, EPI_ISL_508087, EPI_ISL_508089, EPI_ISL_508090, EPI_ISL_508091 | | | |
| see above | New Mexico Department of Health Scientific Laboratory Division | Center for Global Health, University of New Mexico Health Sciences Center | Daryl Domman, Kurt Schwalm, Twila Kunde, Joseph Hicks, Michael Edwards, Darrell Dinwiddie |
| EPI_ISL_508357, EPI_ISL_508358, EPI_ISL_508359, EPI_ISL_508360, EPI_ISL_508361, EPI_ISL_508362, EPI_ISL_508363, EPI_ISL_508364, EPI_ISL_508365, EPI_ISL_508366, EPI_ISL_508367, EPI_ISL_508369, EPI_ISL_508370, EPI_ISL_508371, EPI_ISL_508372, EPI_ISL_508373, EPI_ISL_508374, EPI_ISL_508375 | | | |
| see above | Institute of Post Graduate Medical Education & Research | National Institute of Biomedical Genomics | Arindam Maitra, Aritra Biswas, Jayeeta Haldar, Raja Ray, Monimoy Banerjee, Saumitra Das |
| EPI_ISL_508423, EPI_ISL_508424, EPI_ISL_508433, EPI_ISL_508439, EPI_ISL_508440 | Mahatma Gandhi Institute of Medical Sciences | National Institute of Biomedical Genomics | Arindam Maitra, Vijayshri Deotale, Rahul Narang, Deepashri Maraskolhe, Saumitra Das |
| EPI_ISL_508444, EPI_ISL_508445, EPI_ISL_508446, EPI_ISL_508447, EPI_ISL_508448, EPI_ISL_508449, EPI_ISL_508450, EPI_ISL_508451, EPI_ISL_508452 | ICMR-National Institute of Cholera and Enteric Diseases | National Institute of Biomedical Genomics | Arindam Maitra, Mamta Chawla Sarkar, Sreedhar Chinnaswamy, Hasina Banu, Ananya Chatterjee, Shanta Dutta, Saumitra Das |
| EPI_ISL_508658, EPI_ISL_508660, EPI_ISL_508672, EPI_ISL_508674, EPI_ISL_508677 | Departamento de Microbiología, CDB, Hospital Clínic, Barcelona | SeqCOVID-SPAIN consortium/IBV(CSIC) | Andrea Vergara, Mikel Martínez, Elisa Rubio, Jéssica Navero, Aida Peiró and SeqCOVID-SPAIN consortium |
| EPI_ISL_509060, EPI_ISL_509061, EPI_ISL_509062, EPI_ISL_509066, EPI_ISL_509067, EPI_ISL_509068 | OHSU Lab Services Molecular Microbiology Lab | Oregon SARS-CoV-2 Genome Sequencing Center | Brendan L. O'Connell, Ruth V. Nichols, Sally B. Grindstaff, Alec J. Hirsch, Guang Fan, Daniel N. Streblow, William B. Messer, Andrew C. Adey, Benjamin N. Bimber, Brian J. O'Roak |
| EPI_ISL_509436, EPI_ISL_509438, EPI_ISL_509439, EPI_ISL_509440, EPI_ISL_509441 | The Princess Alexandra Hospital | Wellcome Sanger Institute for the COVID-19 Genomics UK (COG-UK) consortium | Nick Levene, Louise Lopez, Lynn Monaghan, Jessica Scott, Claudia McCrea and Alex Alderton, Roberto Amato, Sonia Goncalves, Ewan Harrison, David K. Jackson, Ian Johnston, Dominic Kwiatkowski, Cordelia Langford, John Sillitoe on behalf of the Wellcome Sanger Institute COVID-19 Surveillance Team (http://www.sanger.ac.uk/covid-team) |
| EPI_ISL_509499, EPI_ISL_509500 | Area of Virology, Serology and Virology Division (SAViD), New South Wales Health Pathology Randwick | Area of Virology, Serology and Virology Division (SAViD), New South Wales Health Pathology Randwick | Rawlinson, W. |
| EPI_ISL_509527, EPI_ISL_509528, EPI_ISL_509529, EPI_ISL_509530, EPI_ISL_509531, EPI_ISL_509532, EPI_ISL_509533, EPI_ISL_509534, EPI_ISL_509535, EPI_ISL_509536, EPI_ISL_509537, EPI_ISL_509538, EPI_ISL_509539, EPI_ISL_509540, EPI_ISL_509541, EPI_ISL_509542, EPI_ISL_509543, EPI_ISL_509544, EPI_ISL_509545, EPI_ISL_509546, EPI_ISL_509547, EPI_ISL_509548, EPI_ISL_509549, EPI_ISL_509550, EPI_ISL_509551, EPI_ISL_509552, EPI_ISL_509553, EPI_ISL_509554, EPI_ISL_509555, EPI_ISL_509556, EPI_ISL_509557, EPI_ISL_509558, EPI_ISL_509559, EPI_ISL_509560, EPI_ISL_509561, EPI_ISL_509562, EPI_ISL_509563 | | | |
| see above | Utah Public Health Laboratory | Utah Public Health Laboratory | Heidi Butz, Erin Young, Kelly Oakeson |
| EPI_ISL_509692, EPI_ISL_509693, EPI_ISL_509694 | Utah Public Health Laboratory | Pathogen Discovery, Respiratory Viruses Branch, Division of Viral Diseases, Centers for Disease Control and Prevention | Ying Tao, Jing Zhang, Krista Queen, Anna Uehara, Yan Li, Clinton Paden, Haibin Wang, Suxiang Tong |
| EPI_ISL_509844, EPI_ISL_509845, EPI_ISL_509939 | University of Wisconsin-Madison AIDS Vaccine Research Laboratories | University of Wisconsin-Madison AIDS Vaccine Research Laboratories | Gage Moreno, Katarina Braun, et al. AIDS Vaccine Research Laboratories |
| EPI_ISL_510059, EPI_ISL_510060, | Servicio de Microbiología. HRU de Málaga. Servicio Andaluz | SeqCOVID-SPAIN consortium/IBV(CSIC) | Inmaculada de Toro Peinado. MªConcepción Mediavilla Gradolph. Begoña Palop Borrás and SeqCOVID-SPAIN consortium |

| | | | |
|---|---|---|---|
| EPI_ISL_510061, EPI_ISL_510062 | de Salud | | |
| EPI_ISL_510064, EPI_ISL_510076 | Instituto de Investigaciones Biomédicas de Barcelona (CSIC), Hospital Clinic i Provincial de Barcelona, Instituto de Biomedicina de Valencia (CSIC), Hospital de Sant Pau | SeqCOVID-SPAIN consortium/IBV(CSIC) | Anna M. Planas, Mª Angeles Marcos, Miguel J. Martínez, Andrea Vergara, Alex Soriano, Jordi Pérez Tur, Israel Fernández Cadenas and SeqCOVID-SPAIN consortium |
| EPI_ISL_510106, EPI_ISL_510116, EPI_ISL_510126, EPI_ISL_510127, EPI_ISL_510133 | Hospital General Universitario Gregorio Marañón | SeqCOVID-SPAIN consortium/IBV(CSIC) | Laura Pérez-Lago, Marta Herranz, Jon Sicilia, Julia Suárez, Pilar Catalán, Patricia Muñoz, Darío García de Viedma and SeqCOVID-SPAIN consortium |
| EPI_ISL_510246 | Hospital de la Santa Creu i Sant Pau. Servicio de Microbiología | SeqCOVID-SPAIN consortium/IBV(CSIC) | Ferran Navarro, Núria Rabella, Elisenda Miró and SeqCOVID-SPAIN consortium |
| EPI_ISL_510386, EPI_ISL_510387, EPI_ISL_510388, EPI_ISL_510389 | Servicio de Microbiología, Hospital Miguel Servet, Zaragoza | SeqCOVID-SPAIN consortium/IBV(CSIC) | Antonio Rezusta López, Alexander Tristancho Baró, Ana Milagro, Yolanda Gracia Grataloup, Nieves Martínez Cameo and SeqCOVID-SPAIN consortium |
| EPI_ISL_510425, EPI_ISL_510426, EPI_ISL_510448, EPI_ISL_510449 | Hospital Universitario Virgen de las Nieves de Granada-SAS | SeqCOVID-SPAIN consortium/IBV(CSIC) | Mercedes Pérez Ruiz, Sara Sanbonmatsu Gámez, Irene Pedrosa Corral, José M. Navarro-Marí and SeqCOVID-SPAIN consortium |
| EPI_ISL_510527 | Unite des virus emergents, UMR190 | Unite des virus emergents, UMR190 | Baronti, C., Piorkowski,G., Coutard,B., Charrel,R. and de Lamballerie,X. |
| EPI_ISL_510549 | Division of Viral Diseases, Center for Laboratory Control of Infectious Diseases, Korea Centers for Diseases Control and Prevention | Division of Viral Diseases, Center for Laboratory Control of Infectious Diseases, Korea Centers for Diseases Control and Prevention | Jeong-Min Kim, Yoon-Seok Chung, Namjoo Lee, Sang Hee Woo, Hye-Jun Jo, Heui Man Kim, Jun-Sub Kim, Myung Guk Han |
| EPI_ISL_510820, EPI_ISL_510821, EPI_ISL_510822 | Klinisk mikrobiologi centralsjukhuset Karlstad | The Public Health Agency of Sweden | Oskar Karlsson Lindsjo, Maria Lind Karlberg, Mattias Haukland, Reza Advani, Olov Svartstrom, Anna-Malin Linde, Sandra Broddesson, Petra Edquist, Mia Brytting, Anna Risberg, Karin Tegmark-Wisell |
| EPI_ISL_511103, EPI_ISL_511106, EPI_ISL_511141 | Instituto Nacional de Saude (INSA) | Instituto Nacional de Saude (INSA) | Borges et al |
| EPI_ISL_511508 | Instituto Nacional de Saude (INSA) | Instituto Nacional de Saude (INSA) and Instituto Gulbenkian de Ciencia (IGC) | Borges et al |
| EPI_ISL_511877 | Johns Hopkins Hospital Department of Pathology | Johns Hopkins Hospital Department of Pathology | Peter M. Thielen, Thomas Mehoke, Shirlee Wohl, Srividya Ramakrishnan, Melanie Kirsche, Amanda Ernlund, Craig Howser, Kristina Zudock, Oluwaseun Falade-Nwulia, Norah Sadowski, Paul Morris, Mark Hopkins, Yunfan Fan, Nidia Trovao, Victoria Gniazdowski, Michael C. Schatz, Stuart C. Ray, Winston Timp, Heba H. Mostafa |
| EPI_ISL_511923 | Mahatma Gandhi Institute of Medical Sciences | National Institute of Biomedical Genomics - DBT's PAN-INDIA 1000 SARS--CoV-2 RNA Genome Sequencing Consortium | Arindam Maitra, Vijayshri Deotale, Rahul Narang, Deepashri Maraskolhe, Saumitra Das |
| EPI_ISL_512107 | Florida Bureau of Public Health Laboratories | Florida Bureau of Public Health Laboratories | Sarah Schmedes, Jason Blanton |
| EPI_ISL_512427, EPI_ISL_512435 | Centre for Enzyme Innovation, University of Portsmouth / Translational Research Laboratory, Portsmouth Hospitals NHS Trust | COVID-19 Genomics UK (COG-UK) Consortium | Angela Beckett,Yann Bourgeois,Garry Scarlett,Sharon Glaysher,Scott Elliott,Kelly Bicknell,Robert Impey,Allyson Lloyd,Sarah Wyllie,Ethan Butcher,Anoop Chauhan,Samuel Robson |
| EPI_ISL_512775, EPI_ISL_512776 | Utah Public Health Laboratory, Utah Public Health Laboratory Infectious Disease submission group | Utah Public Health Laboratory, Utah Public Health Laboratory Infectious Disease submission group | Butz,H.A., Young,E.L., Oakeson,K. |
| EPI_ISL_512821 | Kenema Government Hospital, Ministry of Health and Sanitation | Kenema Government Hospital, Ministry of Health and Sanitation | Goba,A., Momoh,M., Sandi,J., Tomkins-Tinch,C., Siddle,K., Mehta,S., Oluniyi,P., Jalloh,S., Park,D., Andersen,K., Garry,R., Happi,C., Grant,D., Olawoye,I. |
| EPI_ISL_512954, EPI_ISL_512955, EPI_ISL_512966, EPI_ISL_512967, EPI_ISL_512968, EPI_ISL_512969, EPI_ISL_512970, EPI_ISL_512971, EPI_ISL_512972, EPI_ISL_512974, EPI_ISL_512975, EPI_ISL_512985, EPI_ISL_512986, EPI_ISL_512987, EPI_ISL_512988 | | | |
| see above | Pathogen Genomics Lab King Abdullah University of Science and Technology(KAUST) | Pathogen Genomics Lab King Abdullah University of Science and Technology(KAUST) | Sara Mfarrej, Raeece Naeem, Rahul P Salunke, Sharif Hala, Fadwa Alofi, Amit Kumar Subudhi, Fathia Ben Rached, Afrah Alsomali, Jumana Taha, Abdulaziz Alahmadi, Asim Khogeer, Nashwa Al-khotani, Anwar Hashem, Naif Almontashiri, Arnab Pain |
| EPI_ISL_512990, EPI_ISL_512991, EPI_ISL_512992 | Pathogen Genomics Lab King Abdullah University of Science and Technology(KAUST) | Pathogen Genomics Lab King Abdullah University of Science and Technology(KAUST) | Amit Kumar Subudhi, Rahul P Salunke, Sara Mfarrej, Sharif Hala, Fadwa Alofi, Fathia Ben Rached, Afrah Alsomali, Asim Khogeer, Nashwa Al-khotani, Raeece Naeem, Anwar Hashem, Naif Almontashiri, Arnab Pain |
| EPI_ISL_513168 | Pathogen Genomics Lab King Abdullah University of Science and Technology(KAUST) | Pathogen Genomics Lab King Abdullah University of Science and Technology(KAUST) | Sara Mfarrej, Raeece Naeem, Rahul P Salunke, Sharif Hala, Fadwa Alofi, Amit Kumar Subudhi, Fathia Ben Rached, Afrah Alsomali, Jumana Taha, Abdulaziz Alahmadi, Asim Khogeer, Nashwa Al-khotani, Anwar Hashem, Naif Almontashiri, Arnab Pain |
| EPI_ISL_513175, EPI_ISL_513176, EPI_ISL_513181 | Pathogen Genomics Lab King Abdullah University of Science and Technology(KAUST) | Pathogen Genomics Lab King Abdullah University of Science and Technology(KAUST) | Amit Kumar Subudhi, Rahul P Salunke, Sara Mfarrej, Sharif Hala, Fadwa Alofi, Fathia Ben Rached, Afrah Alsomali, Asim Khogeer, Nashwa Al-khotani, Raeece Naeem, Anwar Hashem, Naif Almontashiri, Arnab Pain |
| EPI_ISL_513519, EPI_ISL_513520, EPI_ISL_513521, EPI_ISL_513522, EPI_ISL_513523, EPI_ISL_513524, EPI_ISL_513525, EPI_ISL_513526, EPI_ISL_513527, EPI_ISL_513528, EPI_ISL_513529, EPI_ISL_513573, EPI_ISL_513574, EPI_ISL_513575, EPI_ISL_513576, EPI_ISL_513577, EPI_ISL_513578, EPI_ISL_513579, EPI_ISL_513580 | | | |
| see above | Programa de Oncovirologia, Instituto Nacional de Câncer | Programa de Oncovirologia, Instituto Nacional de Câncer | Juliana D. Siqueira, Livia R. Goes, Brunna M. Alves, Claudia Cicala,James Arthos, João P.B. Viola, Andreia C. de Melo, Marcelo A. Soares |
| EPI_ISL_513589, EPI_ISL_513590, EPI_ISL_513597, EPI_ISL_513598, EPI_ISL_513599, EPI_ISL_513600, EPI_ISL_513601, EPI_ISL_513602, EPI_ISL_513603, EPI_ISL_513604, EPI_ISL_513605, EPI_ISL_513606, EPI_ISL_513607, EPI_ISL_513608, EPI_ISL_513609, EPI_ISL_513610, EPI_ISL_513611, EPI_ISL_513615, EPI_ISL_513616, EPI_ISL_513619 | | | |
| see above | Viral Respiratory Lab, National Institute for Biomedical Research (INRB) | Pathogen Sequencing Lab, National Institute for Biomedical Research (INRB) | Placide Mbala-Kingebeni, Edith Nkwembe, Eddy Kinganda-Lusamaki, Amuri Aziza, Francisca Muyembe Mawete, Emmanuel Lokilo Lofiko, Catherine Pratt, Matthias Pauthner, Josh Quick, Allison Black, James Hadfield, Trevor Bedford, Ian Goodfellow, Andrew Rambaut, Nick Loman, Kristian Andersen, Michael Wiley, Steve Ahuka-Mundeke, Jean-Jacques Muyembe Tamfum |
| EPI_ISL_513640, EPI_ISL_513641, EPI_ISL_513642, EPI_ISL_513643, EPI_ISL_513644, EPI_ISL_513645, EPI_ISL_513646, EPI_ISL_513647, EPI_ISL_513648, EPI_ISL_513649, EPI_ISL_513650, EPI_ISL_513651, EPI_ISL_513652, EPI_ISL_513653, EPI_ISL_513654, EPI_ISL_513655, EPI_ISL_513656, EPI_ISL_513657, EPI_ISL_513658, EPI_ISL_513659, EPI_ISL_513660, EPI_ISL_513661, EPI_ISL_513662, EPI_ISL_513663, EPI_ISL_513664, EPI_ISL_513665, EPI_ISL_513666, EPI_ISL_513667, EPI_ISL_513668, EPI_ISL_513669, EPI_ISL_513670, EPI_ISL_513671, EPI_ISL_513672, EPI_ISL_513673, EPI_ISL_513674, EPI_ISL_513675, EPI_ISL_513676, EPI_ISL_513677, EPI_ISL_513678, EPI_ISL_513679, EPI_ISL_513680, EPI_ISL_513681, EPI_ISL_513682, EPI_ISL_513683, EPI_ISL_513684, EPI_ISL_513685, EPI_ISL_513686, EPI_ISL_513687, EPI_ISL_513688, EPI_ISL_513689, EPI_ISL_513690, EPI_ISL_513691, EPI_ISL_513692, EPI_ISL_513693, EPI_ISL_513694, EPI_ISL_513695, EPI_ISL_513696, EPI_ISL_513697, EPI_ISL_513698, EPI_ISL_513699, EPI_ISL_513700, EPI_ISL_513701, EPI_ISL_513702, EPI_ISL_513703, EPI_ISL_513704, EPI_ISL_513705, EPI_ISL_513706 | | | |
| see above | Utah Public Health Laboratory | Utah Public Health Laboratory | Heidi Butz, Erin Young, Kelly Oakeson |
| EPI_ISL_513896, EPI_ISL_513897, EPI_ISL_513898, EPI_ISL_513899, EPI_ISL_513900, EPI_ISL_513901 | UCSF Clinical Microbiology Laboratory | Chan-Zuckerberg Biohub | CZB Cliahub Consortium |
| EPI_ISL_513941, EPI_ISL_513942 | Utah Public Health Laboratory, Utah Public Health Laboratory Infectious Disease submission group | Utah Public Health Laboratory, Utah Public Health Laboratory Infectious Disease submission group | Butz,H.A., Young,E.L. and Oakeson,K. |
| EPI_ISL_514104, EPI_ISL_514105, EPI_ISL_514106, EPI_ISL_514107, EPI_ISL_514108, EPI_ISL_514114, EPI_ISL_514115, EPI_ISL_514116, EPI_ISL_514117, EPI_ISL_514118, EPI_ISL_514119 | | | |
| see above | Viral Respiratory Lab, National Institute for Biomedical Research (INRB) | Pathogen Sequencing Lab, National Institute for Biomedical Research (INRB) | Placide Mbala-Kingebeni, Edith Nkwembe, Eddy Kinganda-Lusamaki, Amuri Aziza, Francisca Muyembe Mawete, Emmanuel Lokilo Lofiko, Catherine Pratt, Matthias Pauthner, Josh Quick, Allison Black, James Hadfield, Trevor Bedford, Ian Goodfellow, Andrew Rambaut, Nick Loman, Kristian Andersen, Michael Wiley, Steve Ahuka-Mundeke, Jean-Jacques Muyembe Tamfum |
| EPI_ISL_514430 | Laboratory Diagnostic, Veterinary Specialized Institute Kraljevo | Laboratory Diagnostic, Veterinary Specialized Institute Kraljevo | Vidanovic,D., Tesovic,B., Knezevic,A., Jankovic,M., Sekler,M., Banovic Djeri,B., Volkening,J., Afonso,C., Petrovic,T. |
| EPI_ISL_514616, EPI_ISL_514617 | Minnesota Department of Health, Public Health Laboratory | Minnesota Department of Health, Public Health Laboratory | Matt Plumb, Jacob Garfin, and Xiong Wang |
| EPI_ISL_514753 | Yaftabad Hospital, COVID Lab Center | University of Tabriz | Shahabzadeh,Z., Hosseinzadeh Gharajeh,N., Hashemian,S.M. and Barati,O. |

| | | | |
|---|---|---|---|
| EPI_ISL_515395, EPI_ISL_515396, EPI_ISL_515397, EPI_ISL_515398, EPI_ISL_515399, EPI_ISL_515400, EPI_ISL_515401, EPI_ISL_515402, EPI_ISL_515403, EPI_ISL_515404, EPI_ISL_515405, EPI_ISL_515406, EPI_ISL_515407, EPI_ISL_515408, EPI_ISL_515409, EPI_ISL_515410, EPI_ISL_515411, EPI_ISL_515412, EPI_ISL_515413, EPI_ISL_515414, EPI_ISL_515462 | | | |
| see above | Nevada State Public Health Laboratory | Nevada State Public Health Laboratory | Richard Tillett, Joel R. Sevinsky, Paul Hartley, Heather Kerwin, David Jackson, Subhash C. Verma, Cyprian Rossetto, Andrew Gorzalski, Chris Laverdure, Natalie Crawford, Stephanie Van Hooser, and Mark Pandori |
| EPI_ISL_515943, EPI_ISL_515944 | ESIC | Department of Neurovirology, National Institute of Mental Health and Neuroscience (NIMHANS) | Chitra Pattabiraman,Vijayalakshmi Reddy, Harsha PK, Risha Rasheed, Pramada Prasad, Shafeeq S Hameed, Manjunatha Venkataswamy, Anita Desai, Ravi Vasanthapuram |
| EPI_ISL_515945, EPI_ISL_515946, EPI_ISL_515947, EPI_ISL_515948, EPI_ISL_515949 | DH | Department of Neurovirology, National Institute of Mental Health and Neuroscience (NIMHANS) | Chitra Pattabiraman,Vijayalakshmi Reddy, Harsha PK, Risha Rasheed, Pramada Prasad, Shafeeq S Hameed, Manjunatha Venkataswamy, Anita Desai, Ravi Vasanthapuram |
| EPI_ISL_516387 | Michigan Department of Health and Human Services, Bureau of Laboratories | Michigan Department of Health and Human Services, Bureau of Laboratories | Blankenship HM, Riner D, Soehnlen MK |
| EPI_ISL_516655, EPI_ISL_516656, EPI_ISL_516657, EPI_ISL_516658, EPI_ISL_516659, EPI_ISL_516660, EPI_ISL_516661, EPI_ISL_516662, EPI_ISL_516663, EPI_ISL_516664, EPI_ISL_516665, EPI_ISL_516666, EPI_ISL_516667, EPI_ISL_516668, EPI_ISL_516669, EPI_ISL_516670, EPI_ISL_516671, EPI_ISL_516672, EPI_ISL_516673, EPI_ISL_516674, EPI_ISL_516675, EPI_ISL_516676, EPI_ISL_516687, EPI_ISL_516688, EPI_ISL_516689, EPI_ISL_516690, EPI_ISL_516691, EPI_ISL_516713, EPI_ISL_516714, EPI_ISL_516715, EPI_ISL_516716, EPI_ISL_516717, EPI_ISL_516718, EPI_ISL_516719, EPI_ISL_516720 | | | |
| see above | Virginia DCLS | Virginia DCLS | Virginia DCLS |
| EPI_ISL_517583, EPI_ISL_517584, EPI_ISL_517585, EPI_ISL_517586, EPI_ISL_517587 | Virology Department, Royal Infirmary of Edinburgh, NHS Lothian / School of Biological Sciences, University of Edinburgh / Institute of Genetics and Molecular Medicine, University of Edinburgh | COVID-19 Genomics UK (COG-UK) Consortium | McHugh M, Dewar R, Rooke S, Gallagher M, Balcaza C, O'Toole Á, Scher E, Hill V, McCrone JT, Colquhoun R, Yu X, Jackson B, Rambaut A, Williams TC, Templeton K |
| EPI_ISL_517612, EPI_ISL_517613, EPI_ISL_517614, EPI_ISL_517615 | Academic Hospital Paramaribo | Erasmus Medical Center | Bas Oude Munnink, Dion Gajadin, Ed Ijzerman, Emmanuelle Munger, Gary Gummels, Ingrid Krishnadath, Lycke Woittiez, Marion Koopmans, Mireille Van de Veer, Princes Wongsowidjojo, Radjesh Ori, Rohma Banwari, Stephen Vreden |
| EPI_ISL_517776, EPI_ISL_517777 | Florida Bureau of Public Health Laboratories | Florida Bureau of Public Health Laboratories | Sarah Schmedes, Jason Blanton |
| EPI_ISL_518851, EPI_ISL_518852, EPI_ISL_518853, EPI_ISL_518854 | Oman-National Influenza Center | Biotechnology & OMICs Laboratory, Natural & Medical Sciences Research Center, University of Nizwa | Abdul Latif Khan, Samira Al-Mahruqi, Ahmed Al-Harrasi, Samiha Al-Kharusi, Adil Khan, Ahmed Al-Rawahi, Sajjad Asaf, Amina Al-Jardani, Hanan Al-Kindi, Intisar Al-Shukri, Ahlam Al-Amri, Aisha Al-Amri, Aisha Al-Busaidi, Adil Al-Wahaibi, Seif Al-Abri. |
| EPI_ISL_520663, EPI_ISL_520673, EPI_ISL_520716, EPI_ISL_520738, EPI_ISL_520740, EPI_ISL_520741, EPI_ISL_520742 | Mohammed Bin Rashid University of Medicine and Health Sciences | Al Jalila Genomics Center | Ahmad Abou Tayoun, Tom Loney, Hamda Khansaheb, Sathishkumar Ramaswamy, Divinlal Harilal, Zulfa Omar Deesi, Rupa Murthy Varghese, Hanan Al Suwaidi, Abdulmajeed Alkhaja, Mohammed Uddin, Rifat Hamoudi, Rabih Halwani, Abiola Catherine Senok, Qutayba Hamid, Norbert Nowotny, Alawi Alsheikh-Ali |
| EPI_ISL_521903 | Victorian Infectious Diseases Reference Laboratory (VIDRL) | VIDRL and MDU-PHL | Caly L., Seemann T., Sait, M., Schultz M., Druce J., Sherry, N. |
| EPI_ISL_522264, EPI_ISL_522265, EPI_ISL_522266, EPI_ISL_522267, EPI_ISL_522268, EPI_ISL_522269, EPI_ISL_522270, EPI_ISL_522273, EPI_ISL_522274, EPI_ISL_522275, EPI_ISL_522276, EPI_ISL_522277, EPI_ISL_522278, EPI_ISL_522279, EPI_ISL_522280, EPI_ISL_522281, EPI_ISL_522282, EPI_ISL_522283, EPI_ISL_522284, EPI_ISL_522285, EPI_ISL_522286, EPI_ISL_522287, EPI_ISL_522288, EPI_ISL_522289, EPI_ISL_522290, EPI_ISL_522291, EPI_ISL_522292, EPI_ISL_522293, EPI_ISL_522294, EPI_ISL_522295, EPI_ISL_522296, EPI_ISL_522297, EPI_ISL_522298, EPI_ISL_522299, EPI_ISL_522300, EPI_ISL_522301, EPI_ISL_522302, EPI_ISL_522303, EPI_ISL_522304, EPI_ISL_522305, EPI_ISL_522306, EPI_ISL_522307, EPI_ISL_522308, EPI_ISL_522309, EPI_ISL_522310, EPI_ISL_522311, EPI_ISL_522312, EPI_ISL_522313, EPI_ISL_522314, EPI_ISL_522315, EPI_ISL_522316, EPI_ISL_522317 | | | |
| see above | Utah Public Health Laboratory | Utah Public Health Laboratory | Erin Young, Kelly Oakeson |
| EPI_ISL_522439, EPI_ISL_522440 | Center for Laboratory Control of Infectious Diseases, Korea Centers for Diseases Control and Prevention | Center for Laboratory Control of Infectious Diseases, Korea Centers for Diseases Control and Prevention | Junyoung Kim, Ae Kyung Park, Eunkyung Shin, Jin Sun No, Jeong-Min Kim, Yoon-Seok Chung, Heui Man Kim, Myung Guk Han |
| EPI_ISL_522596, EPI_ISL_522615, EPI_ISL_522619, EPI_ISL_522621, EPI_ISL_522625, EPI_ISL_522627, EPI_ISL_522629, EPI_ISL_522632, EPI_ISL_522682, EPI_ISL_522683, EPI_ISL_522684 | | | |
| see above | Royal Hobart Hospital Microbiology Department | MDU-PHL | Cooley L., van Haeften R., Seemann T., Sait M., Schultz, M.B., Sherry N. |
| EPI_ISL_522859 | ULSS9 Scaligera | Istituto Zooprofilattico Sperimentale delle Venezie | Adelaide Milani, Alessia Schivo, Annalisa Salviato, Erika Giorgia Quaranta, Ambra Pastori, Bianca Zecchin, Alice Fusaro, Isabella Monne, Calogero Terregino, Antonia Ricci |
| EPI_ISL_522991, EPI_ISL_522992, EPI_ISL_522994, EPI_ISL_522997, EPI_ISL_522999, EPI_ISL_523000, EPI_ISL_523001, EPI_ISL_523002, EPI_ISL_523003, EPI_ISL_523004, EPI_ISL_523010, EPI_ISL_523016, EPI_ISL_523022, EPI_ISL_523028, EPI_ISL_523034, EPI_ISL_523040, EPI_ISL_523046, EPI_ISL_523068, EPI_ISL_523070, EPI_ISL_523072, EPI_ISL_523073, EPI_ISL_523075, EPI_ISL_523085, EPI_ISL_523089, EPI_ISL_523090 | | | |
| see above | Dutch COVID-19 response team | Erasmus Medical Center | OH consortium |
| EPI_ISL_523123, EPI_ISL_523172, EPI_ISL_523175, EPI_ISL_523180, EPI_ISL_523196, EPI_ISL_523197, EPI_ISL_523198, EPI_ISL_523354, EPI_ISL_523355, EPI_ISL_523402, EPI_ISL_523427, EPI_ISL_523430, EPI_ISL_523431, EPI_ISL_523432, EPI_ISL_523433, EPI_ISL_523434, EPI_ISL_523435, EPI_ISL_523436, EPI_ISL_523437, EPI_ISL_523468, EPI_ISL_523495, EPI_ISL_523496, EPI_ISL_523548, EPI_ISL_523606 | | | |
| see above | Dutch COVID-19 response team | Erasmus Medical Center | Bas Oude Munnink, David Nieuwenhuijse, Reina Sikkema, Claudia Schapendonk, Irina Chestakova, Anne van der Linden, Theo Bestebroer, Stefan van Nieuwkoop, Mark Pronk, Pascal Lexmond, Corien Swaan, Manon Haverkate, Madelief Mollers, Mart Stein, Sandra Kengne Kamga Mobou, Jeroen van Kampen, Jolanda Voermans, Aura Timen, Corine GeurtsvanKessel, Annemiek van der Eijk, Richard Molenkamp, Marion Koopmans, on behalf of the Dutch national COVID-19 response team. |
| EPI_ISL_523997, EPI_ISL_523998, EPI_ISL_524052 | WHO National Influenza Centre Russian Federation | WHO National Influenza Centre Russian Federation | Andrey Komissarov, Artem Fadeev, Mariia Sergeeva, Anna Ivanova, Daria Danilenko |
| EPI_ISL_524468 | Hospital Municipal Vereador Jose Storopolli | Instituto Adolfo Lutz, Interdiciplinary Procedures Center, Strategic Laboratory | Claudio Tavares Sacchi, Claudia Regina Gonçalves, Erica Valessa Ramos Gomes |
| EPI_ISL_524573, EPI_ISL_524574, EPI_ISL_524575, EPI_ISL_524576, EPI_ISL_524577, EPI_ISL_524579, EPI_ISL_524580, EPI_ISL_524581, EPI_ISL_524582, EPI_ISL_524583, EPI_ISL_524584, EPI_ISL_524585, EPI_ISL_524586, EPI_ISL_524587, EPI_ISL_524588, EPI_ISL_524589 | | | |
| see above | Department of Pathology, University of Cambridge | Wellcome Sanger Institute for the COVID-19 Genomics UK (COG-UK) consortium | Luke W Meredith, M. Estée Török , Myra Hosmillo, William L. Hamilton, Martin D. Curran, Theresa Feltwell, Grant Hall, Anna Yakovleva, Fahad A Khokhar, Charlotte J. Houldcroft, Laura G Caller, Aminu S. Jahun, Sarah L. Caddy, Ian Goodfellow; and Alex Alderton, Roberto Amato, Sonia Goncalves, Ewan Harrison, David K. Jackson, Ian Johnston, Dominic Kwiatkowski, Cordelia Langford, John Sillitoe on behalf of the Wellcome Sanger Institute COVID-19 Surveillance Team (http://www.sanger.ac.uk/covid-team) |
| EPI_ISL_524590 | Department of Pathology, University of Cambridge | Wellcome Sanger Institute for the COVID-19 Genomics UK (COG-UK) Consortium | Luke W Meredith, M. Estée Török , Myra Hosmillo, William L. Hamilton, Martin D. Curran, Theresa Feltwell, Grant Hall, Anna Yakovleva, Fahad A Khokhar, Charlotte J. Houldcroft, Laura G Caller, Aminu S. Jahun, Sarah L. Caddy, Ian Goodfellow; and Alex Alderton, Roberto Amato, Sonia Goncalves, Ewan Harrison, David K. Jackson, Ian Johnston, Dominic Kwiatkowski, Cordelia Langford, John Sillitoe on behalf of the Wellcome Sanger Institute COVID-19 Surveillance Team |
| EPI_ISL_524591, EPI_ISL_524592, EPI_ISL_524593, EPI_ISL_524594 | Department of Pathology, University of Cambridge | Wellcome Sanger Institute for the COVID-19 Genomics UK (COG-UK) consortium | Luke W Meredith, M. Estée Török , Myra Hosmillo, William L. Hamilton, Martin D. Curran, Theresa Feltwell, Grant Hall, Anna Yakovleva, Fahad A Khokhar, Charlotte J. Houldcroft, Laura G Caller, Aminu S. Jahun, Sarah L. Caddy, Ian Goodfellow; and Alex Alderton, Roberto Amato, Sonia Goncalves, Ewan Harrison, David K. Jackson, Ian Johnston, Dominic Kwiatkowski, Cordelia Langford, John Sillitoe on behalf of the Wellcome Sanger Institute COVID-19 Surveillance Team (http://www.sanger.ac.uk/covid-team) |
| EPI_ISL_524595 | Department of Pathology, University of Cambridge | Wellcome Sanger Institute for the COVID-19 Genomics UK (COG-UK) Consortium | Luke W Meredith, M. Estée Török , Myra Hosmillo, William L. Hamilton, Martin D. Curran, Theresa Feltwell, Grant Hall, Anna Yakovleva, Fahad A Khokhar, Charlotte J. Houldcroft, Laura G Caller, Aminu S. Jahun, Sarah L. Caddy, Ian Goodfellow; and Alex Alderton, Roberto Amato, Sonia Goncalves, Ewan Harrison, David K. Jackson, Ian Johnston, Dominic Kwiatkowski, Cordelia Langford, John Sillitoe on behalf of the Wellcome Sanger Institute COVID-19 Surveillance Team |
| EPI_ISL_524596, EPI_ISL_524597, EPI_ISL_524598, EPI_ISL_524599, EPI_ISL_524600, EPI_ISL_524601, EPI_ISL_524602, EPI_ISL_524603, EPI_ISL_524604, EPI_ISL_524605, EPI_ISL_524606, EPI_ISL_524607, EPI_ISL_524609, EPI_ISL_524610, EPI_ISL_524611, EPI_ISL_524612, EPI_ISL_524613, EPI_ISL_524614, EPI_ISL_524615, EPI_ISL_524616, EPI_ISL_524617, EPI_ISL_524618 | | | |
| see above | Department of Pathology, University of Cambridge | Wellcome Sanger Institute for the COVID-19 Genomics UK (COG-UK) consortium | Luke W Meredith, M. Estée Török , Myra Hosmillo, William L. Hamilton, Martin D. Curran, Theresa Feltwell, Grant Hall, Anna Yakovleva, Fahad A Khokhar, Charlotte J. Houldcroft, Laura G Caller, Aminu S. Jahun, Sarah L. Caddy, Ian Goodfellow; and Alex Alderton, Roberto Amato, Sonia Goncalves, Ewan |

| | | | |
|---|---|---|---|
| | | | Harrison, David K. Jackson, Ian Johnston, Dominic Kwiatkowski, Cordelia Langford, John Sillitoe on behalf of the Wellcome Sanger Institute COVID-19 Surveillance Team (http://www.sanger.ac.uk/covid-team) |
| EPI_ISL_524783, EPI_ISL_524784, EPI_ISL_524786, EPI_ISL_524790, EPI_ISL_524791, EPI_ISL_524793 | Evandro Chagas Institute | Evandro Chagas Institute | Santos, M.C.; Silva, A.M.; Junior, W.D.C.; Barbagelata, L.S.; Ferreira, J.A.; Sousa, E.M.A.; da Silva, P.S.; Resque, H.R; Martins, L.C.; Sousa Junior, E.C.;Viana, G.M.R |
| EPI_ISL_525476 | Centre for Dengue Research | Centre for Dengue Research | Chandima Jeewandara, Deshni Jayathilaka, Dinuka Ariyaratne, Laksiri Gomes, Diyanath Ranasinghe, Dinuka Guruge, Ruwan Wijayamuni, Gathsaurie Neelika Malavige |
| EPI_ISL_526219, EPI_ISL_526227, EPI_ISL_526237 | Hungarian Defence Forces Military Medical Centre | National Laboratory of Virology, Szentágothai Research Centre | Endre Gábor Tóth, Balázs Somogyi, Bálint Eszenyi, Ferenc Jakab, Gábor Kemenesi |
| EPI_ISL_526472, EPI_ISL_526473, EPI_ISL_526474, EPI_ISL_526480, EPI_ISL_526481, EPI_ISL_526482, EPI_ISL_526483, EPI_ISL_526485, EPI_ISL_526486, EPI_ISL_526487, EPI_ISL_526488, EPI_ISL_526489, EPI_ISL_526495, EPI_ISL_526496, EPI_ISL_526497, EPI_ISL_526498, EPI_ISL_526499, EPI_ISL_526500, EPI_ISL_526501, EPI_ISL_526502, EPI_ISL_526503, EPI_ISL_526504, EPI_ISL_526506, EPI_ISL_526507, EPI_ISL_526508, EPI_ISL_526509, EPI_ISL_526510, EPI_ISL_526511, EPI_ISL_526512, EPI_ISL_526514, EPI_ISL_526515, EPI_ISL_526516, EPI_ISL_526517 | | | |
| see above | Virology Department, Royal Infirmary of Edinburgh, NHS Lothian / School of Biological Sciences, University of Edinburgh / Institute of Genetics and Molecular Medicine, University of Edinburgh | COVID-19 Genomics UK (COG-UK) Consortium | McHugh M, Dewar R, Rooke S, Gallagher M, Balcaza C, O'Toole Á, Scher E, Hill V, McCrone JT, Colquhoun R, Yu X, Jackson B, Rambaut A, Williams TC, Templeton K |
| EPI_ISL_526542, EPI_ISL_526543, EPI_ISL_526544, EPI_ISL_526545 | Respiratory Virus Unit, Microbiology Services Colindale, Public Health England | Respiratory Virus Unit, Microbiology Services Colindale, Public Health England | PHE Covid Sequencing Team |
| EPI_ISL_526553, EPI_ISL_526564 | Florida Bureau of Public Health Laboratories | Florida Bureau of Public Health Laboratories | Sarah Schmedes, Jason Blanton |
| EPI_ISL_526771, EPI_ISL_526772, EPI_ISL_526773 | Respiratory Virus Unit, Microbiology Services Colindale, Public Health England | Respiratory Virus Unit, Microbiology Services Colindale, Public Health England | PHE Covid Sequencing Team |
| EPI_ISL_526933, EPI_ISL_526934, EPI_ISL_526950, EPI_ISL_526951, EPI_ISL_526953, EPI_ISL_526955, EPI_ISL_526956 | Instituto Nacional de Salud, Bogotá, Colombia | Instituto Nacional de Salud, Bogotá, Colombia | Katherine Laiton-Donato, Diego A. Álvarez-Díaz, Carlos Franco-Muñoz, Mauricio Pacheco-Montealegre, Jonathan Reales, Diego Andrés Prada, Jose A. Usme-Ciro, Zulma M. Cucunubá, Christian Julian VillabonaArenas, Liz Villabona-Arenas, Sussy Echeverria, Astrid C. Flórez, Carolina Ferro, Diana Marcela Walteros-Acero, Franklin Prieto, Carlos Andrés Durán, Martha Lucia Ospina Martinez, Marcela Mercado-Reyes |
| EPI_ISL_527358 | Respiratory Virus Unit, Microbiology Services Colindale, Public Health England | Respiratory Virus Unit, Microbiology Services Colindale, Public Health England | PHE Covid Sequencing Team |
| EPI_ISL_527375 | National Public Health Laboratory, National Centre for Infectious Diseases | National Public Health Laboratory, National Centre for Infectious Diseases | Mak TM, Octavia S, Zhou Z, Cui L, Lin RTP |
| EPI_ISL_527420, EPI_ISL_527425, EPI_ISL_527426, EPI_ISL_527461, EPI_ISL_527487 | Colorado State University - Ebel Lab | Colorado State University - Ebel Lab | Greg Ebel et al. |
| EPI_ISL_527678, EPI_ISL_527679, EPI_ISL_527680, EPI_ISL_527681, EPI_ISL_527682, EPI_ISL_527683, EPI_ISL_527684, EPI_ISL_527685, EPI_ISL_527686, EPI_ISL_527687, EPI_ISL_527688, EPI_ISL_527689, EPI_ISL_527690, EPI_ISL_527691, EPI_ISL_527694, EPI_ISL_527695 | | | |
| see above | MN PHL Division, Minnesota Department of Health | Pathogen Discovery, Respiratory Viruses Branch, Division of Viral Diseases, Centers for Disease Control and Prevention | Yan Li, Anna Montmayeur, Jing Zhang, Krista Queen, Ying Tao, Anna Uehara, Rachel Marine, Clinton R. Paden, Haibin Wang, Suxiang Tong |
| EPI_ISL_527714, EPI_ISL_527717, EPI_ISL_527720, EPI_ISL_527731 | MN PHL Division, Minnesota Department of Health | Pathogen Discovery, Respiratory Viruses Branch, Division of Viral Diseases, Centers for Disease Control and Prevention | Krista Queen, Brian Lynch, Yan Li, Anna Montmayeur, Jing Zhang, Ying Tao, Anna Uehara, Rachel Marine, Clinton R. Paden, Haibin Wang, Suxiang Tong |
| EPI_ISL_527879 | Nigeria Centre for Disease Control (NCDC) | African Centre of Excellence for Genomics of Infectious Diseases (ACEGID), Redeemer's University, Ede, Osun State, Nigeria | Oluniyi P.E. et al |
| EPI_ISL_528382, EPI_ISL_528383, EPI_ISL_528384, EPI_ISL_528385 | Translational Health Science and Technology Institute -ESIC medical college and hospital, Faridabad | THSTI Bioassay laboratory | Saurabh Kumar, Jigme Wangchuk, Anil Kumar Pandey, Asim Das, Guruprasad R. Medigeshi |
| EPI_ISL_528391, EPI_ISL_528408, EPI_ISL_528409, EPI_ISL_528410, EPI_ISL_528411, EPI_ISL_528412, EPI_ISL_528413, EPI_ISL_528414, EPI_ISL_528415, EPI_ISL_528416, EPI_ISL_528417, EPI_ISL_528418 | | | |
| see above | Respiratory Virus Unit, Microbiology Services Colindale, Public Health England | Respiratory Virus Unit, Microbiology Services Colindale, Public Health England | PHE Covid Sequencing Team |
| EPI_ISL_528611 | National Genomics Core-Center for DNA Fingerprinting and Diagnostics | National Genomics Core- Center for DNA Fingerprinting and Diagnostics (NGC-CDFD)- DBT's PAN-INDIA-1000 Genome consortium | Divya Vashisht, Bala Pratyusha, Heena Shah, G Shashikanth, Vinay Donipadi, K.Manohar, Madhumohan Rao, SPR Prasad, Yogesh Patidar, Arjita Jaiswal, Arpita Singh, Devanshi Gupta, Romila Moirangthem, Sanjana Sarkar, Shivani Yadav, R Harinarayanan, Rashna Bhandari, Murali Dharan Bashyam, Debashish Mitra, Ashwin Dalal |
| EPI_ISL_528686, EPI_ISL_528688, EPI_ISL_528690, EPI_ISL_528692, EPI_ISL_528697, EPI_ISL_528709, EPI_ISL_528718 | Alsafar - Khalifa University Abu Dhabi | Alsafar - Khalifa University Abu Dhabi | Andreas Henschel, Gihan Daw Elbait, Samuel Feng, Rifat Hamoudi, Ernesto Damiani, Guan Tay, Habiba Alsafar |
| EPI_ISL_528934, EPI_ISL_528935, EPI_ISL_528936, EPI_ISL_528937, EPI_ISL_528938, EPI_ISL_528939, EPI_ISL_528940 | Agenzia di Tutela della Salute di Bergamo | Istituto Zooprofilattico Sperimentale dell'Abruzzo e Molise "G.Caporale" | Lorusso A, Marcacci M, Di Domenico M, Curini V, Ancora M, Cammà C, Rinaldi A, Mangone I, Di Pasquale A, Puglia I, Savini G. |
| EPI_ISL_528992 | Ospedale SS Annunziata-Sulmona | Istituto Zooprofilattico Sperimentale dell'Abruzzo e Molise "G.Caporale" | Lorusso A, Marcacci M, Di Domenico M, Curini V, Ancora M, Cammà C, Rinaldi A, Mangone I, Di Pasquale A, Puglia I, Savini G. |
| EPI_ISL_528993 | Ospedale Civile S. Liberatore-Atri | Istituto Zooprofilattico Sperimentale dell'Abruzzo e Molise "G.Caporale" | Lorusso A, Marcacci M, Di Domenico M, Curini V, Ancora M, Cammà C, Rinaldi A, Mangone I, Di Pasquale A, Puglia I, Savini G. |
| EPI_ISL_529019 | RSA/RP Villa San Giovanni - Gruppo Edos | Istituto Zooprofilattico Sperimentale dell'Abruzzo e Molise "G.Caporale" | Lorusso A, Marcacci M, Di Domenico M, Curini V, Ancora M, Cammà C, Rinaldi A, Mangone I, Di Pasquale A, Puglia I, Savini G. |
| EPI_ISL_529020, EPI_ISL_529021 | Ospedale Civile S. Liberatore-Atri | Istituto Zooprofilattico Sperimentale dell'Abruzzo e Molise "G.Caporale" | Lorusso A, Marcacci M, Di Domenico M, Curini V, Ancora M, Cammà C, Rinaldi A, Mangone I, Di Pasquale A, Puglia I, Savini G. |
| EPI_ISL_529154 | Department of Immunology, The Scripps Research Institute | Andersen lab at Scripps Research | Basler, T., Shephard, J., Austin, B. with SEARCH Alliance San Diego |
| EPI_ISL_529180, EPI_ISL_529181, EPI_ISL_529182, EPI_ISL_529183, EPI_ISL_529184 | South Carolina Department of Health and Environmental Control | South Carolina Department of Health and Environmental Control | Haley V. Flores |
| EPI_ISL_529438 | University of Birmingham | COVID-19 Genomics UK (COG-UK) Consortium | Institute of Microbiology, University of Birmingham: Claire McMurray, Joanne Stockton, Samuel Nicholls, Radoslaw Poplawski, Will Rowe, Josh Quick, Nicholas Loman. University of Birmingham Testing Laboratory: Celina M Whalley, Andrew Bosworth, Charlotte Poxon, Kasun Wanigasooriya, Oliver Pickles, Mike Kidd, Alex Richter, Andrew D Beggs PHE Heartlands Lab: Husam Osman, Andrew Bosworth. Queen Elizabeth Hospital: Anna Casey |
| EPI_ISL_529689 | Virology Department, Royal Infirmary of Edinburgh, NHS Lothian / School of Biological Sciences, University of Edinburgh / Institute of Genetics and Molecular Medicine, | COVID-19 Genomics UK (COG-UK) Consortium | McHugh M, Dewar R, Rooke S, Gallagher M, Balcaza C, O'Toole Á, Scher E, Hill V, McCrone JT, Colquhoun R, Yu X, Jackson B, Rambaut A, Williams TC, Templeton K |

| | | | |
|---|---|---|---|
| | University of Edinburgh | | |
| EPI_ISL_530034, EPI_ISL_530035, EPI_ISL_530074, EPI_ISL_530082, EPI_ISL_530083, EPI_ISL_530084, EPI_ISL_530086, EPI_ISL_530087, EPI_ISL_530088, EPI_ISL_530093 | Hospital Universitario La Paz | Hospital Universitario La Paz | María Rodríguez, Elias Dahdouh, Sara González, Raúl Recio, Fernando Lázaro, Esther Viedma, Natalia Stella, Julio García, Juan Carlos Galán, Rafael Cantón, Mª Dolores Folgueira, Rafael Delgado, Jesús Mingorance |
| EPI_ISL_530171 | Hennepin County Medical Center | Minnesota Department of Health, Public Health Laboratory | Matt Plumb, Jacob Garfin, and Xiong Wang |
| EPI_ISL_530199, EPI_ISL_530200, EPI_ISL_530201, EPI_ISL_530202, EPI_ISL_530203, EPI_ISL_530204, EPI_ISL_530205, EPI_ISL_530206, EPI_ISL_530207 | Minnesota Department of Health, Public Health Laboratory | Minnesota Department of Health, Public Health Laboratory | Matt Plumb, Jacob Garfin, and Xiong Wang |
| EPI_ISL_534311 | UPA III 26 de Agosto | Instituto Adolfo Lutz, Interdiciplinary Procedures Center, Strategic Laboratory | Claudio Tavares Sacchi, Claudia Regina Gonçalves, Erica Valessa Ramos Gomes |
| EPI_ISL_534312 | Distrito Sanitario Sul | Instituto Adolfo Lutz, Interdiciplinary Procedures Center, Strategic Laboratory | Claudio Tavares Sacchi, Claudia Regina Gonçalves, Erica Valessa Ramos Gomes |
| EPI_ISL_534314 | Hospital Universitario da USP de SP | Instituto Adolfo Lutz, Interdiciplinary Procedures Center, Strategic Laboratory | Claudio Tavares Sacchi, Claudia Regina Gonçalves, Erica Valessa Ramos Gomes |
| EPI_ISL_534315 | Serviço de Verificação de Óbitos SVO Guaruulhos | Instituto Adolfo Lutz, Interdiciplinary Procedures Center, Strategic Laboratory | Claudio Tavares Sacchi, Claudia Regina Gonçalves, Erica Valessa Ramos Gomes |
| EPI_ISL_534316 | OS Mun Santana Lauro Ribas Braga | Instituto Adolfo Lutz, Interdiciplinary Procedures Center, Strategic Laboratory | Claudio Tavares Sacchi, Claudia Regina Gonçalves, Erica Valessa Ramos Gomes |
| EPI_ISL_534318 | Hospital Municipal Antonio Giglio | Instituto Adolfo Lutz, Interdiciplinary Procedures Center, Strategic Laboratory | Claudio Tavares Sacchi, Claudia Regina Gonçalves, Erica Valessa Ramos Gomes |
| EPI_ISL_534326 | Notre Dame Intermedica Saude AS | Instituto Adolfo Lutz, Interdiciplinary Procedures Center, Strategic Laboratory | Claudio Tavares Sacchi, Claudia Regina Gonçalves, Erica Valessa Ramos Gomes |
| EPI_ISL_534331, EPI_ISL_534332 | Hospital Universitario La Paz | Hospital Universitario La Paz | María Rodríguez, Elias Dahdouh, Sara González, Raúl Recio, Fernando Lázaro, Esther Viedma, Natalia Stella, Julio García, Juan Carlos Galán, Rafael Cantón, Ma Dolores Folgueira, Rafael Delgado, Jesús Mingorance |
| EPI_ISL_534770, EPI_ISL_534774, EPI_ISL_534801, EPI_ISL_534805, EPI_ISL_534807, EPI_ISL_534913, EPI_ISL_534958, EPI_ISL_534970, EPI_ISL_534977, EPI_ISL_534992, EPI_ISL_534996, EPI_ISL_535003, EPI_ISL_535005, EPI_ISL_535008, EPI_ISL_535009, EPI_ISL_535011, EPI_ISL_535015, EPI_ISL_535017 | | | |
| see above | Oxford Viromics, NDM, University of Oxford; Oxford University Hospitals; Basingstoke and North Hampshire Hospital | COVID-19 Genomics UK (COG-UK) Consortium | Tanya Golubchik, David Bonsall, George Macintyre, Amy Trebes, Mariateresa de Cesare, Catrin Moore, Alex Mobbs, Anita Justice, Robert Shaw, Monique Andersson, Timothy Peto, Emma Wise, Nathan Moore, Jessica Lynch, Nick Cortes, Matilde Mori, Stephen Kidd, David Buck, John Todd, Christophe Fraser |
| EPI_ISL_536400 | Fareham Creek Veterinary Surgery | MRC-University of Glasgow Centre for Virus Research | Margaret J Hosie, Ilaria Epifano, Vanessa Herder, Richard J Orton, Andrew Stevenson, Natasha Johnson, Daniel Goldfarb, Emma MacDonald, Lynn Stevenson, Frazer Bell, Dawn Dunbar, Michael McDonald, Fiona Howie, Bryn Tennant, Darcy Herrity, Ana Da Silva Filipe, Daniel G Streicker, Brian J Willett, Pablo R Murcia, Ruth F Jarrett, David L Robertson and William Weir |
| EPI_ISL_536501, EPI_ISL_536502, EPI_ISL_536503, EPI_ISL_536504, EPI_ISL_536505, EPI_ISL_536506, EPI_ISL_536507, EPI_ISL_536508, EPI_ISL_536509, EPI_ISL_536510, EPI_ISL_536511, EPI_ISL_536512, EPI_ISL_536513, EPI_ISL_536515, EPI_ISL_536516, EPI_ISL_536519 | | | |
| see above | Instituto Nacional de Salud | Laboratorio de Infecciones Respiratorias Agudas | Eduardo Juscamayta Lopez, David Tarazona, Faviola Valdivia Guerrero, Nancy Rojas Serrano, Dennis Carhuaricra, Lenin Maturrano Hernandez, Ronnie Gavilan Chavez |
| EPI_ISL_537208, EPI_ISL_537212, EPI_ISL_537223, EPI_ISL_537227, EPI_ISL_537229, EPI_ISL_537231, EPI_ISL_537242, EPI_ISL_537256, EPI_ISL_537273, EPI_ISL_537274, EPI_ISL_537275, EPI_ISL_537276, EPI_ISL_537277, EPI_ISL_537278 | | | |
| see above | Virology Department, Sheffield Teaching Hospitals NHS Foundation Trust / Department of Infection, Immunity and Cardiovascular Disease, The Medical School, University of Sheffield | Wellcome Sanger Institute for the COVID-19 Genomics UK (COG-UK) consortium | Thushan de Silva, Matthew Parker,Adri Angyal, Rebecca Brown, Luke Green, Rachel Tucker, Paul Parsons, Danielle Groves, Alex Keeley, Dave Partridge, Matthew Wyles, Benjamin Lindsey, Mehmet Yavuz, Mohammad Raza, Cariad Evans and Alex Alderton, Roberto Amato, Sonia Goncalves, Ewan Harrison, David K. Jackson, Ian Johnston, Dominic Kwiatkowski, Cordelia Langford, John Sillitoe on behalf of the Wellcome Sanger Institute COVID-19 Surveillance Team |
| EPI_ISL_537279 | Virology Department, Sheffield Teaching Hospitals NHS Foundation Trust / Department of Infection, Immunity and Cardiovascular Disease, The Medical School, University of Sheffield | Wellcome Sanger Institute for the COVID-19 Genomics UK (COG-UK) Consortium | Thushan de Silva, Matthew Parker,Adri Angyal, Rebecca Brown, Luke Green, Rachel Tucker, Paul Parsons, Danielle Groves, Alex Keeley, Dave Partridge, Matthew Wyles, Benjamin Lindsey, Mehmet Yavuz, Mohammad Raza, Cariad Evans and Alex Alderton, Roberto Amato, Sonia Goncalves, Ewan Harrison, David K. Jackson, Ian Johnston, Dominic Kwiatkowski, Cordelia Langford, John Sillitoe on behalf of the Wellcome Sanger Institute COVID-19 Surveillance Team |
| EPI_ISL_537280, EPI_ISL_537281, EPI_ISL_537282, EPI_ISL_537283, EPI_ISL_537284 | Virology Department, Sheffield Teaching Hospitals NHS Foundation Trust / Department of Infection, Immunity and Cardiovascular Disease, The Medical School, University of Sheffield | Wellcome Sanger Institute for the COVID-19 Genomics UK (COG-UK) consortium | Thushan de Silva, Matthew Parker,Adri Angyal, Rebecca Brown, Luke Green, Rachel Tucker, Paul Parsons, Danielle Groves, Alex Keeley, Dave Partridge, Matthew Wyles, Benjamin Lindsey, Mehmet Yavuz, Mohammad Raza, Cariad Evans and Alex Alderton, Roberto Amato, Sonia Goncalves, Ewan Harrison, David K. Jackson, Ian Johnston, Dominic Kwiatkowski, Cordelia Langford, John Sillitoe on behalf of the Wellcome Sanger Institute COVID-19 Surveillance Team |
| EPI_ISL_537285 | Virology Department, Sheffield Teaching Hospitals NHS Foundation Trust / Department of Infection, Immunity and Cardiovascular Disease, The Medical School, University of Sheffield | Wellcome Sanger Institute for the COVID-19 Genomics UK (COG-UK) Consortium | Thushan de Silva, Matthew Parker,Adri Angyal, Rebecca Brown, Luke Green, Rachel Tucker, Paul Parsons, Danielle Groves, Alex Keeley, Dave Partridge, Matthew Wyles, Benjamin Lindsey, Mehmet Yavuz, Mohammad Raza, Cariad Evans and Alex Alderton, Roberto Amato, Sonia Goncalves, Ewan Harrison, David K. Jackson, Ian Johnston, Dominic Kwiatkowski, Cordelia Langford, John Sillitoe on behalf of the Wellcome Sanger Institute COVID-19 Surveillance Team |
| EPI_ISL_537505, EPI_ISL_537506, EPI_ISL_537516, EPI_ISL_537518, EPI_ISL_537519, EPI_ISL_537536, EPI_ISL_537545 | UCLA Pathology Clinical Microbiology Lab | Kruglyak Lab | Guo et al. |
| EPI_ISL_537677, EPI_ISL_537678, EPI_ISL_537679 | Universidad de León | SeqCOVID-SPAIN consortium/IBV(CSIC) | Ana Carvajal, Vicente Martín, Héctor Argüello, Juan M. Fregeneda, Tania Fernández-Villa, Antonio J. Molina and SeqCOVID-SPAIN consortium |
| EPI_ISL_537773, EPI_ISL_537774, EPI_ISL_537775, EPI_ISL_537776, EPI_ISL_537777, EPI_ISL_537778, EPI_ISL_537779, EPI_ISL_537780, EPI_ISL_537781 | Servicio de Microbiología, Hospital Miguel Servet, Zaragoza | SeqCOVID-SPAIN consortium/IBV(CSIC) | Antonio Rezusta López, Alexander Tristancho Baró, Ana Milagro, Yolanda Gracia Grataloup, Nieves Martínez Cameo and SeqCOVID-SPAIN consortium |
| EPI_ISL_538245, EPI_ISL_538246, EPI_ISL_538247, EPI_ISL_538248, EPI_ISL_538249, EPI_ISL_538253 | TriCore Reference Laboratories | Center for Global Health, University of New Mexico Health Sciences Center | Daryl Domman, Kurt Schwalm, Twila Kunde, Joseph Hicks, Michael Edwards, Darrell Dinwiddie |
| EPI_ISL_538757 | Leeds Teaching Hospitals NHS Trust and Public Health England, National Infection Service (Leeds laboratory) | Wellcome Sanger Institute for the COVID-19 Genomics UK (COG-UK) consortium | Louissa Macfarlane-Smith, Holli Carden, Katherine L. Harper, Antony Hale and Alex Alderton, Roberto Amato, Sonia Goncalves, Ewan Harrison, David K. Jackson, Ian Johnston, Dominic Kwiatkowski, Cordelia Langford, John Sillitoe on behalf of the Wellcome Sanger Institute COVID-19 Surveillance Team |
| EPI_ISL_538818 | Leeds Teaching Hospitals NHS Trust and Public Health England, National Infection Service (Leeds laboratory) | Wellcome Sanger Institute for the COVID-19 Genomics UK (COG-UK) Consortium | Louissa Macfarlane-Smith, Holli Carden, Katherine L. Harper, Antony Hale and Alex Alderton, Roberto Amato, Sonia Goncalves, Ewan Harrison, David K. Jackson, Ian Johnston, Dominic Kwiatkowski, Cordelia Langford, John Sillitoe on behalf of the Wellcome Sanger Institute COVID-19 Surveillance Team |
| EPI_ISL_538835, EPI_ISL_538849, | Leeds Teaching Hospitals NHS Trust and Public Health | Wellcome Sanger Institute for the COVID-19 Genomics UK | Louissa Macfarlane-Smith, Holli Carden, Katherine L. Harper, Antony Hale and Alex Alderton, Roberto Amato, Sonia Goncalves, Ewan Harrison, David K. |

| | | | |
|---|---|---|---|
| EPI_ISL_538916 | England, National Infection Service (Leeds laboratory) | (COG-UK) consortium | Jackson, Ian Johnston, Dominic Kwiatkowski, Cordelia Langford, John Sillitoe on behalf of the Wellcome Sanger Institute COVID-19 Surveillance Team |
| EPI_ISL_538917 | Leeds Teaching Hospitals NHS Trust and Public Health England, National Infection Service (Leeds laboratory) | Wellcome Sanger Institute for the COVID-19 Genomics UK (COG-UK) Consortium | Louissa Macfarlane-Smith, Holli Carden, Katherine L. Harper, Antony Hale and Alex Alderton, Roberto Amato, Sonia Goncalves, Ewan Harrison, David K. Jackson, Ian Johnston, Dominic Kwiatkowski, Cordelia Langford, John Sillitoe on behalf of the Wellcome Sanger Institute COVID-19 Surveillance Team |
| EPI_ISL_538992, EPI_ISL_538999 | Leeds Teaching Hospitals NHS Trust and Public Health England, National Infection Service (Leeds laboratory) | Wellcome Sanger Institute for the COVID-19 Genomics UK (COG-UK) consortium | Louissa Macfarlane-Smith, Holli Carden, Katherine L. Harper, Antony Hale and Alex Alderton, Roberto Amato, Sonia Goncalves, Ewan Harrison, David K. Jackson, Ian Johnston, Dominic Kwiatkowski, Cordelia Langford, John Sillitoe on behalf of the Wellcome Sanger Institute COVID-19 Surveillance Team |
| EPI_ISL_539006 | Leeds Teaching Hospitals NHS Trust and Public Health England, National Infection Service (Leeds laboratory) | Wellcome Sanger Institute for the COVID-19 Genomics UK (COG-UK) Consortium | Louissa Macfarlane-Smith, Holli Carden, Katherine L. Harper, Antony Hale and Alex Alderton, Roberto Amato, Sonia Goncalves, Ewan Harrison, David K. Jackson, Ian Johnston, Dominic Kwiatkowski, Cordelia Langford, John Sillitoe on behalf of the Wellcome Sanger Institute COVID-19 Surveillance Team |
| EPI_ISL_539154 | Leeds Teaching Hospitals NHS Trust and Public Health England, National Infection Service (Leeds laboratory) | Wellcome Sanger Institute for the COVID-19 Genomics UK (COG-UK) consortium | Louissa Macfarlane-Smith, Holli Carden, Katherine L. Harper, Antony Hale and Alex Alderton, Roberto Amato, Sonia Goncalves, Ewan Harrison, David K. Jackson, Ian Johnston, Dominic Kwiatkowski, Cordelia Langford, John Sillitoe on behalf of the Wellcome Sanger Institute COVID-19 Surveillance Team |
| EPI_ISL_539483, EPI_ISL_539484, EPI_ISL_539486, EPI_ISL_539488 | Civil Hospital, Rupnagar | CSIR-Institute of Microbial Technology | Kanika Bansal, Sanjeet Kumar, Anu Singh, Debarghya Ghose, Amandeep Kaur, Rajesh Kumar Mishra, Poushali Chakraborty, Harsh Goar, Navin Baid, Ashwani Kumar, Dipak Dutta, Sanjeev Khosla, Prabhu B. Patil |
| EPI_ISL_539801, EPI_ISL_539802 | Wyoming Public Health Laboratory | Wyoming Public Health Laboratory | Noah Hull, Rob Christensen, Jim Mildenberger, Joel Sevinsky, Cari Sloma, and Wanda Manley |
| EPI_ISL_539824, EPI_ISL_539825, EPI_ISL_539826, EPI_ISL_539827, EPI_ISL_539828 | Minnesota Department of Health, Public Health Laboratory | Minnesota Department of Health, Public Health Laboratory | Matt Plumb, Jacob Garfin, and Xiong Wang |
| EPI_ISL_540421, EPI_ISL_540422 | Wyoming Public Health Laboratory | Wyoming Public Health Laboratory | Noah Hull, Rob Christensen, Jim Mildenberger, Joel Sevinsky, Cari Sloma, and Wanda Manley |
| EPI_ISL_540435 | VA-Division of Consolidated Laboratory Services | Pathogen Discovery, Respiratory Viruses Branch, Division of Viral Diseases, Centers for Disease Control and Prevention | Krista Queen, Yan Li, Jing Zhang, Anna Montmayeur, Krista Queen,Ying Tao, Anna Uehara, Clinton R. Paden, Rachel Marine, Haibin Wang, Suxiang Tong |
| EPI_ISL_540874, EPI_ISL_540875, EPI_ISL_540876, EPI_ISL_540877 | Virology Department, Royal Infirmary of Edinburgh, NHS Lothian / School of Biological Sciences, University of Edinburgh / Institute of Genetics and Molecular Medicine, University of Edinburgh | COVID-19 Genomics UK (COG-UK) Consortium | McHugh M, Dewar R, Rooke S, Gallagher M, Balcaza C, O'Toole Á, Scher E, Hill V, McCrone JT, Colquhoun R, Yu X, Jackson B, Rambaut A, Williams TC, Templeton K |
| EPI_ISL_541020, EPI_ISL_541021, EPI_ISL_541022, EPI_ISL_541023, EPI_ISL_541024, EPI_ISL_541025, EPI_ISL_541026, EPI_ISL_541027, EPI_ISL_541028, EPI_ISL_541029, EPI_ISL_541030, EPI_ISL_541031, EPI_ISL_541032, EPI_ISL_541033 | | | |
| see above | Servicio de Microbiología, Hospital Miguel Servet, Zaragoza | SeqCOVID-SPAIN consortium/Institute of Biomedicine of Valencia, IBV-CSIC | Antonio Rezusta López, Alexander Tristancho Baró, Ana Milagro, Yolanda Gracia Grataloup, Nieves Martínez Cameo and SeqCOVID-SPAIN consortium |
| EPI_ISL_541058, EPI_ISL_541059, EPI_ISL_541060, EPI_ISL_541061, EPI_ISL_541062, EPI_ISL_541063, EPI_ISL_541064, EPI_ISL_541065 | Hospital Clínico Universitario de Santiago de Compostela | SeqCOVID-SPAIN consortium/Institute of Biomedicine of Valencia, IBV-CSIC | José Javier Costa Alcalde, Antonio Aguilera Guirao, Mª Luisa Pérez del Molino Bernal, Amparo Coira Nieto, Gema Barbeito Castiñeiras, Rocio Trastoy Pena and SeqCOVID-SPAIN consortium |
| EPI_ISL_541067, EPI_ISL_541068, EPI_ISL_541069 | Hospital de la Santa Creu i Sant Pau. Servicio de Microbiología | SeqCOVID-SPAIN consortium/Institute of Biomedicine of Valencia, IBV-CSIC | Ferran Navarro, Núria Rabella, Elisenda Miró and SeqCOVID-SPAIN consortium |
| EPI_ISL_541362, EPI_ISL_541363, EPI_ISL_541364, EPI_ISL_541365, EPI_ISL_541366, EPI_ISL_541367, EPI_ISL_541368, EPI_ISL_541369 | Laboratory of Respiratory Viruses and Measles, Oswaldo Cruz Institute, FIOCRUZ | Laboratory of Respiratory Viruses and Measles, Oswaldo Cruz Institute, FIOCRUZ | Paola Resende, Luciana Appolinario, Fernando Motta, Anna Carolina Paixão, Ana Carolina Mendonça, Jonathan Lopes, Marilda Siqueira |
| EPI_ISL_541380, EPI_ISL_541381, EPI_ISL_541382, EPI_ISL_541383, EPI_ISL_541384, EPI_ISL_541390, EPI_ISL_541391 | LACEN/SE | Laboratory of Respiratory Viruses and Measles, Oswaldo Cruz Institute, FIOCRUZ | Paola Resende, Luciana Appolinario, Fernando Motta, Anna Carolina Paixão, Ana Carolina Mendonça, Jonathan Lopes, Clioma Santos, Marilda Siqueira |
| EPI_ISL_541722 | National Institute of Virology, NIV Influenza | National Institute of Virology, NIV Influenza | Potdar V |
| EPI_ISL_542535, EPI_ISL_542548, EPI_ISL_542552, EPI_ISL_542553, EPI_ISL_542556, EPI_ISL_542557, EPI_ISL_542589, EPI_ISL_542596, EPI_ISL_542635, EPI_ISL_542670, EPI_ISL_542671, EPI_ISL_542673, EPI_ISL_542674, EPI_ISL_542676, EPI_ISL_542677, EPI_ISL_542678, EPI_ISL_542679, EPI_ISL_542682, EPI_ISL_542683, EPI_ISL_542684, EPI_ISL_542685, EPI_ISL_542688, EPI_ISL_542690, EPI_ISL_542691, EPI_ISL_542699, EPI_ISL_542700, EPI_ISL_542703, EPI_ISL_542707, EPI_ISL_542708, EPI_ISL_542710, EPI_ISL_542712, EPI_ISL_542715, EPI_ISL_542716, EPI_ISL_542717, EPI_ISL_542718, EPI_ISL_542719, EPI_ISL_542720, EPI_ISL_542722, EPI_ISL_542723, EPI_ISL_542725, EPI_ISL_542726, EPI_ISL_542729, EPI_ISL_542730, EPI_ISL_542734, EPI_ISL_542735, EPI_ISL_542736, EPI_ISL_542737, EPI_ISL_542738, EPI_ISL_542739, EPI_ISL_542740, EPI_ISL_542741, EPI_ISL_542743, EPI_ISL_542745, EPI_ISL_542746, EPI_ISL_542747, EPI_ISL_542748, EPI_ISL_542749, EPI_ISL_542750, EPI_ISL_542751, EPI_ISL_542753, EPI_ISL_542754, EPI_ISL_542755, EPI_ISL_542756, EPI_ISL_542759, EPI_ISL_542760, EPI_ISL_542762, EPI_ISL_542763, EPI_ISL_542764, EPI_ISL_542765, EPI_ISL_542766, EPI_ISL_542767, EPI_ISL_542768, EPI_ISL_542771, EPI_ISL_542772, EPI_ISL_542774, EPI_ISL_542820, EPI_ISL_542875, EPI_ISL_542877, EPI_ISL_542882, EPI_ISL_542888, EPI_ISL_542891, EPI_ISL_542893, EPI_ISL_542908, EPI_ISL_542911, EPI_ISL_542914, EPI_ISL_542916, EPI_ISL_542931 | | | |
| see above | Houston Methodist Hospital | Houston Methodist Hospital | S. Wesley Long, Randall J. Olsen, Paul A. Christensen, David W. Bernard, James J. Davis, Maulik Shukla, Marcus Nguyen, Matthew Ojeda Saavedra, Concepcion C. Cantu, Prasanti Yerramilli, Layne Pruitt, Sishir Subedi, Hung-Che Kuo, Heather Hendrickson, Ghazaleh Eskandari, Hoang A. T. Nguyen, J. Hunter Long, Muthiah Kumaraswami, Jule Goike, Daniel Boutz, Jimmy Gollihar, Jason S. McLellan, Chia-Wei Chou, Kamyab Javanmardi, Ilya J. Finkelstein, and James M. Musser |
| EPI_ISL_542943, EPI_ISL_542945, EPI_ISL_542947, EPI_ISL_542949, EPI_ISL_542952, EPI_ISL_542953, EPI_ISL_542954, EPI_ISL_542963, EPI_ISL_542968, EPI_ISL_542970 | TriCore Reference Laboratories | Center for Global Health, University of New Mexico Health Sciences Center | Daryl Domman, Kurt Schwalm, Twila Kunde, Joseph Hicks, Michael Edwards, Darrell Dinwiddie |
| EPI_ISL_545061, EPI_ISL_545080, EPI_ISL_545081, EPI_ISL_545085, EPI_ISL_545089, EPI_ISL_545090, EPI_ISL_545092, EPI_ISL_545093, EPI_ISL_545094, EPI_ISL_545104, EPI_ISL_545223, EPI_ISL_545229, EPI_ISL_545233, EPI_ISL_545235, EPI_ISL_545287, EPI_ISL_545310, EPI_ISL_545313 | | | |
| see above | Houston Methodist Hospital | Houston Methodist Hospital | S. Wesley Long, Randall J. Olsen, Paul A. Christensen, David W. Bernard, James J. Davis, Maulik Shukla, Marcus Nguyen, Matthew Ojeda Saavedra, Concepcion C. Cantu, Prasanti Yerramilli, Layne Pruitt, Sishir Subedi, Hung-Che Kuo, Heather Hendrickson, Ghazaleh Eskandari, Hoang A. T. Nguyen, J. Hunter Long, Muthiah Kumaraswami, Jule Goike, Daniel Boutz, Jimmy Gollihar, Jason S. McLellan, Chia-Wei Chou, Kamyab Javanmardi, Ilya J. Finkelstein, and James M. Musser |
| EPI_ISL_547455, EPI_ISL_547456, EPI_ISL_547476, EPI_ISL_547477, EPI_ISL_547497, EPI_ISL_547498, EPI_ISL_547509, EPI_ISL_547511 | Dutch COVID-19 response team | National Institute for Public Health and the Environment (RIVM) | Adam Meijer, Harry Vennema, Jeroen Cremer, Sharon van den Brink, Bas van der Veer, AnneMarie van den Brandt, Florian Zwagemaker, Dennis Schmitz, Chantal Reusken, on behalf of the national COVID-19 response team |
| EPI_ISL_547570 | PS Municipal Dr Augusto Gomes de Mattos | Instituto Adolfo Lutz, Interdiciplinary Procedures Center, Strategic Laboratory | Claudio Tavares Sacchi, Claudia Regina Gonçalves, Erica Valessa Ramos Gomes, Karoline Rodrigues Campos |
| EPI_ISL_548962, EPI_ISL_548963 | Klinisk mikrobiologi, Region Västerbotten | Unit for Biological Agents, Department for CBRN Defence and Security, Swedish Defence Research Agency | FOI Bioinformatics team |
| EPI_ISL_559663, EPI_ISL_559667, EPI_ISL_559669 | Lighthouse Lab in Milton Keynes | Wellcome Sanger Institute for the COVID-19 Genomics UK (COG-UK) consortium | The Lighthouse Lab in Milton Keynes and Alex Alderton, Roberto Amato, Sonia Goncalves, Ewan Harrison, David K. Jackson, Ian Johnston, Dominic Kwiatkowski, Cordelia Langford, John Sillitoe on behalf of the Wellcome Sanger Institute COVID-19 Surveillance Team (http://www.sanger.ac.uk/covid-team) |
| EPI_ISL_559670 | Lighthouse Lab in Milton Keynes | Wellcome Sanger Institute for the COVID-19 Genomics UK (COG-UK) Consortium | The Lighthouse Lab in Milton Keynes and Alex Alderton, Roberto Amato, Sonia Goncalves, Ewan Harrison, David K. Jackson, Ian Johnston, Dominic Kwiatkowski, Cordelia Langford, John Sillitoe on behalf of the Wellcome Sanger Institute COVID-19 Surveillance Team |
| EPI_ISL_559672, EPI_ISL_559674, EPI_ISL_559675, EPI_ISL_559676, EPI_ISL_559679, EPI_ISL_559683, EPI_ISL_559687, EPI_ISL_559690, EPI_ISL_559694, EPI_ISL_559695, EPI_ISL_559697, EPI_ISL_559700 | | | |

| | | | |
|---|---|---|---|
| see above | Lighthouse Lab in Milton Keynes | Wellcome Sanger Institute for the COVID-19 Genomics UK (COG-UK) consortium | The Lighthouse Lab in Milton Keynes and Alex Alderton, Roberto Amato, Sonia Goncalves, Ewan Harrison, David K. Jackson, Ian Johnston, Dominic Kwiatkowski, Cordelia Langford, John Sillitoe on behalf of the Wellcome Sanger Institute COVID-19 Surveillance Team (http://www.sanger.ac.uk/covid-team) |
| EPI_ISL_559974, EPI_ISL_559975, EPI_ISL_559976, EPI_ISL_559977 | Virology Department, Royal Infirmary of Edinburgh, NHS Lothian / School of Biological Sciences, University of Edinburgh / Institute of Genetics and Molecular Medicine, University of Edinburgh | COVID-19 Genomics UK (COG-UK) Consortium | McHugh M, Dewar R, Rooke S, Gallagher M, Balcaza C, O'Toole Á, Scher E, Hill V, McCrone JT, Colquhoun R, Yu X, Jackson B, Rambaut A, Williams TC, Templeton K |
| EPI_ISL_560362 | TriCore Reference Laboratories | Center for Global Health, University of New Mexico Health Sciences Center | Daryl Domman, Kurt Schwalm, Twila Kunde, Joseph Hicks, Michael Edwards, Darrell Dinwiddie |
| EPI_ISL_560634, EPI_ISL_560635 | Hospital | National Reference Center for Viruses of Respiratory Infections, Institut Pasteur, Paris | Sylvie Behillil, Fabiana Gambaro, Etienne Simon-Lorière, Vincent Enouf, Maud Vanpeene, Sylvie van der Werf |
| EPI_ISL_560895, EPI_ISL_560896, EPI_ISL_560897, EPI_ISL_560898, EPI_ISL_560899, EPI_ISL_560900, EPI_ISL_560901, EPI_ISL_560902, EPI_ISL_560903, EPI_ISL_560904, EPI_ISL_560905, EPI_ISL_560906, EPI_ISL_560907, EPI_ISL_560908, EPI_ISL_560909, EPI_ISL_560910, EPI_ISL_560911, EPI_ISL_560912, EPI_ISL_560913, EPI_ISL_560914, EPI_ISL_560915, EPI_ISL_560916, EPI_ISL_560917, EPI_ISL_560918, EPI_ISL_560919 | | | |
| see above | Utah Public Health Laboratory | Utah Public Health Laboratory | Erin Young, Kelly Oakeson |
| EPI_ISL_560926 | Texas Department of State Health Services | Texas Department of State Health Services | Rashmi Tuladhar, Bonnie Oh, Jenny Zhang, Maliha Rahman, Anita Pokharel, Myong Koag, Chun Wang, Rachel Lee, Grace Kubin |
| EPI_ISL_561015, EPI_ISL_561016 | MRCG at LSHTM Genomics lab | MRCG at LSHTM Genomics lab | Abdul Karim sesay, Abdoulie Kanteh, Jarra Manneh, Mariama Kujabi, Bakary Sanyang |
| EPI_ISL_561344 | Civil Hospital, Rupnagar | CSIR-Institute of Microbial Technology | Kanika Bansal, Sanjeet Kumar, Anu Singh, Debarghya Ghose, Amandeep Kaur, Rajesh Kumar Mishra, Poushali Chakraborty, Harsh Goar, Navin Baid, Ashwani Kumar, Dipak Dutta, Sanjeev Khosla, Prabhu B. Patil |
| EPI_ISL_565891, EPI_ISL_565892, EPI_ISL_565893, EPI_ISL_565894, EPI_ISL_565895, EPI_ISL_565896, EPI_ISL_565899, EPI_ISL_565900, EPI_ISL_565901, EPI_ISL_565902 | Michigan Department of Health and Human Services, Bureau of Laboratories | Michigan Department of Health and Human Services, Bureau of Laboratories | Blankenship HM, Riner D, Soehnlen MK |
| EPI_ISL_568478 | Virology, Iran University of Medical Sciences | Virology, Iran University of Medical Sciences | Keyvani,H., Ranjbar,M.M., Keyvani,F., Soleimani,S. |
| EPI_ISL_568479 | Virology, Iran University of Medical Sciences | Virology, Iran University of Medical Sciences | Keyvani,H., Ranjbar,Mm., Keyvani,F., soleimani,s. |
| EPI_ISL_568494 | Virology, Iran University of Medical Sciences | Virology, Iran University of Medical Sciences | Keyvani,H., Ranjbar,M.M., Keyvani,F., Soleimani,S. |
| EPI_ISL_568501, EPI_ISL_568503 | Virology, Iran University of Medical Sciences | Virology, Iran University of Medical Sciences | Keyvani,H., Ranjbar,Mm., Soleimani,S., Keyvani,F. |
| EPI_ISL_568504, EPI_ISL_568505, EPI_ISL_568506 | Virology, Iran University of Medical Sciences | Virology, Iran University of Medical Sciences | Keyvani,H., Ranjbar,Mm., Keyvani,F., Soleimani,S. |
| EPI_ISL_568509 | Virology, Iran University of Medical Sciences | Virology, Iran University of Medical Sciences | Keyvani,H., Ranjbar,Mm., Soleimani,S., Keyvani,F. |
| EPI_ISL_568713, EPI_ISL_568714, EPI_ISL_568715, EPI_ISL_568717 | KEMRI-Wellcome Trust Research Programme/KEMRI-CGMR-C Kilifi | KEMRI-Wellcome Trust Research Programme/KEMRI-CGMR-C Kilifi | Githinji et al 2020 |
| EPI_ISL_568875 | Malaysia Genome Institute | Malaysia Genome Institute | Mohd Noor Mat Isa, Irni Suhayu Sapian, Yusuf Muhammad Noor, Nurhezreen Md Iqbal, Mohd Faizal Abu Bakar, Enizza Kasim, Shamsidar Sopie, Siti Noraini Othman, Azrin Ahmad, Nor Azfa Johari, Shahrul Hisham Zainal Ariffin |
| EPI_ISL_568958, EPI_ISL_568959, EPI_ISL_568960, EPI_ISL_568961, EPI_ISL_568962, EPI_ISL_568963, EPI_ISL_568964, EPI_ISL_568965, EPI_ISL_568966, EPI_ISL_568967, EPI_ISL_568968, EPI_ISL_568969, EPI_ISL_568970 | | | |
| see above | MEPHI, Aix Marseille University | MEPHI, Aix Marseille University | Anthony LEVASSEUR |
| EPI_ISL_569610 | Avera McKennan Hospital | South Dakota Public Health Laboratory | Matt Plumb, Jacob Garfin, Xiong Wang, and Chris Carlson |
| EPI_ISL_569796, EPI_ISL_569797, EPI_ISL_569811, EPI_ISL_569854, EPI_ISL_569857 | Omsk Research Institute of Natural Focal Infections | WHO National Influenza Centre Russian Federation | Artem Fadeev, Ekaterina Gradoboeva, Ekaterina Savkina, Daria Nashatyreva, Elena Poleshchuk, Aleksei Vasilenko, Valery Yakimenko, Andrey Komissarov |
| EPI_ISL_569974, EPI_ISL_570010 | Unity Health Toronto | Ontario Institute for Cancer Research | Ramzi Fattouh, Larissa M. Matukas, Yan Chen,Mark Downing, Trina Otterman, Karel Boissinot, Wai Sum Siu, Zhi Cui, Le Luu, Samira Mubareka, TIBDN, Ilinca Lungu, Bernard Lam, Jeremy Johns, Paul Krzyzanowski, Richard de Borja, Felicia Vincelli, Philip Zuzarte, Jared T. Simpson |
| EPI_ISL_570036, EPI_ISL_570037, EPI_ISL_570038, EPI_ISL_570043, EPI_ISL_570049, EPI_ISL_570202, EPI_ISL_570203, EPI_ISL_570205, EPI_ISL_570206, EPI_ISL_570207, EPI_ISL_570208, EPI_ISL_570210, EPI_ISL_570211, EPI_ISL_570212, EPI_ISL_570213, EPI_ISL_570216 | | | |
| see above | UW Virology Lab | UW Virology Lab | Pavitra Roychoudhury, Hong Xie, Lasata Shrestha, Amin Addetia, Victoria M Rachleff, Meei-Li Huang, Keith R Jerome, Alexander Greninger |
| EPI_ISL_572330 | Institute for Virology, University Hospital Duesseldorf, Medical Faculty, Heinrich-Heine-University Duesseldorf | Institute for Virology, University Hospital Duesseldorf, Medical Faculty, Heinrich-Heine-University Duesseldorf | Maximilian Damagnez, Verena Keitel, Björn Jensen, Nadine Lübke, Lisa Müller, Philipp Ostermann, Tina Senff, Ortwin Adams, Philipp Albrecht, Gerald Antoch, Johannes Bode, Edwin Bölke, Saskia Elben, Torsten Feldt, Johannes C. Fischer, , Anselm Kunstein, Caroline Klindt, Alexander Killer, Tom Lüdde, Annemarie Mohring, Jennifer Neubert, Heiner Schaal, Ansgar Schulz, Jörg Timm, Andreas Walker |
| EPI_ISL_572353, EPI_ISL_572356, EPI_ISL_572357, EPI_ISL_572358, EPI_ISL_572359, EPI_ISL_572360, EPI_ISL_572361, EPI_ISL_572362, EPI_ISL_572363, EPI_ISL_572364, EPI_ISL_572365, EPI_ISL_572366, EPI_ISL_572367, EPI_ISL_572368, EPI_ISL_572375 | | | |
| see above | LACEN/PE | WallauLab, Aggeu Magalhaes Institute | Marcelo Henrique Santos Paiva, Duschinka Ribeiro Duarte Guedes, Cássia Docena, Matheus Filgueira Bezerra, Filipe Zimmer Dezordi, Laís Ceschini Machado, Larissa Krokovsky, Elisama Helvecio, Alexandre Freitas da Silva, Luydson Richardson Silva Vasconcelos, Antonio Mauro Rezende, Severino Jefferson Ribeiro da Silva, Kamila Gaudêncio da Silva Sales, Bruna Santos Lima Figueiredo de Sá, Derciliano Lopes da Cruz, Claudio Eduardo Cavalcanti, Armando de Menezes Neto, Caroline Targino Alves da Silva, Renata Pessôa Germano Mendes, Maria Almerice Lopes da Silva, Tiago Gräf, Paola Cristina Resende, Gonzalo Bello0, Michelle da Silva Barros, Wheverton Ricardo Correia do Nascimento,, Rodrigo Moraes Loyo Arcoverde, Luciane Caroline Albuquerque Bezerra, Sinval Pinto Brandão Filho, Constância Flávia Junqueira Ayres, Gabriel Luz Wallau |
| EPI_ISL_572595, EPI_ISL_572648, EPI_ISL_572663, EPI_ISL_573358, EPI_ISL_573359, EPI_ISL_573360, EPI_ISL_573361, EPI_ISL_573362, EPI_ISL_573363, EPI_ISL_573364, EPI_ISL_573365, EPI_ISL_573366, EPI_ISL_573367, EPI_ISL_573368, EPI_ISL_573369, EPI_ISL_573370, EPI_ISL_573371 | | | |
| see above | Northumbria University / South Tees Hospitals NHS Foundation Trust / North Cumbria Integrated Care NHS Foundation Trust / North Tees and Hartlepool NHS Foundation Trust / Newcastle Hospitals NHS Foundation Trust | COVID-19 Genomics UK (COG-UK) Consortium | Darren L Smith,Andrew Nelson,Matthew Bashton,Greg R Young,Joshua Loh,John Allan,Mohammad A Tariq,Giles S Holt,Gary Black,Wen C Yew,Lynn Dover,Paul Baker,Steve Liggett,Sarah Essex,Jane Greenaway,Debra Padgett,Clive Graham,Garren Scott,Edward Barton,Emma Swindells,Brendan Payne,Jennifer Collins,Yusri Taha,Gary Eltringham |
| EPI_ISL_574583 | Secretaria Municipal de Saude de Jandira | Instituto Adolfo Lutz, Interdiciplinary Procedures Center, Strategic Laboratory | Claudio Tavares Sacchi, Claudia Regina Gonçalves, Erica Valessa Ramos Gomes, Karoline Rodrigues Campos |
| EPI_ISL_574588 | Hospital Estadual Sumare | Instituto Adolfo Lutz, Interdiciplinary Procedures Center, Strategic Laboratory | Claudio Tavares Sacchi, Claudia Regina Gonçalves, Erica Valessa Ramos Gomes, Karoline Rodrigues Campos |
| EPI_ISL_574593 | CS II Dr. Antonio Vicoso Moreira de Rezende Sumare | Instituto Adolfo Lutz, Interdiciplinary Procedures Center, Strategic Laboratory | Claudio Tavares Sacchi, Claudia Regina Gonçalves, Erica Valessa Ramos Gomes, Karoline Rodrigues Campos |
| EPI_ISL_575094, EPI_ISL_575095, EPI_ISL_575246, EPI_ISL_575249, EPI_ISL_575250, EPI_ISL_575251, EPI_ISL_575252, EPI_ISL_575256, EPI_ISL_575257, EPI_ISL_575258, EPI_ISL_575259 | | | |
| see above | Utah Public Health Laboratory | Utah Public Health Laboratory | Erin Young, Kelly Oakeson |
| EPI_ISL_577623 | The National Institute of Public Health | State Veterinary Institute Prague | Nagy,A;Jirincova,H;Novakova,L;Trnka,D;Vecerova,J |
| EPI_ISL_577738 | Institute of Virology, Biomedical Research Center of the Slovak Academy of Sciences, Bratislava | Faculty of Natural Sciences, Comenius University, Bratislava | Viktória Hodorová, Kristína Boršová, Broa Brejová, Viktória abanová, Dominika Friová, Sabina Fumaová Havlíková, Juraj Kopáek, Martina Liková, ubomíra Lukáiková, Martina Neboháová, Monika Sláviková, Edita Staroová, Elena Tichá, Tomáš Vina, Jozef Nosek, Boris Klempa |

| | | | |
|---|---|---|---|
| EPI_ISL_577743, EPI_ISL_577744, EPI_ISL_577745, EPI_ISL_577746 | Dutch COVID-19 response team | Erasmus Medical Center | Bas Oude Munnink, Reina Sikkema, David Nieuwenhuijse, Irina Chestakova, Anne van der Linden, Marjan Boter, Emmanuelle Munger, Corine GeurtsvanKessel, Annemiek van der Eijk, Richard Molenkamp, Marion Koopmans, on behalf of the Dutch national COVID-19 response team. |
| EPI_ISL_578169 | University of Michigan Clinical Microbiology Laboratory | Lauring Lab, University of Michigan, Department of Microbiology and Immunology | Valesano |
| EPI_ISL_579403 | LabPLUS | Institute of Environmental Science and Research (ESR) | Xiaoyun Ren, Matt Storey, Nikki Freed, Muhammad Faisal, Jing Wang, Hermes Perez, Anja Werno, Antje van der Linden, Arlo Upton, Chris Mansell, David Hammer, Dragana Drinkovic, Gary McAuliffe, Hana Sofia Andersson, James Ussher, Jill Sherwood, Josh Freeman, Julia Howard, Juliet Elvy, Mary DeAlmeida, Matt Blakiston, Matthew Rogers, Max Bloomfield, Michael Addidle, Michelle Balm, Sally Roberts, Sarah Jefferies, Sharmini Muttaiyah, Susan Morpeth, Susan Taylor, Timothy Blackmore, Vani Sathyendran, Veronica Playle, Virginia Hope, Erasmus Smit, Lauren Jelly, Olin Silander, Joep de Ligt |
| EPI_ISL_579423, EPI_ISL_579424 | LabTests | Institute of Environmental Science and Research (ESR) | Xiaoyun Ren, Matt Storey, Nikki Freed, Muhammad Faisal, Jing Wang, Hermes Perez, Anja Werno, Antje van der Linden, Arlo Upton, Chris Mansell, David Hammer, Dragana Drinkovic, Gary McAuliffe, Hana Sofia Andersson, James Ussher, Jill Sherwood, Josh Freeman, Julia Howard, Juliet Elvy, Mary DeAlmeida, Matt Blakiston, Matthew Rogers, Max Bloomfield, Michael Addidle, Michelle Balm, Sally Roberts, Sarah Jefferies, Sharmini Muttaiyah, Susan Morpeth, Susan Taylor, Timothy Blackmore, Vani Sathyendran, Veronica Playle, Virginia Hope, Erasmus Smit, Lauren Jelly, Olin Silander, Joep de Ligt |
| EPI_ISL_579426 | Canterbury Health Laboratories | Institute of Environmental Science and Research (ESR) | Xiaoyun Ren, Matt Storey, Nikki Freed, Muhammad Faisal, Jing Wang, Hermes Perez, Anja Werno, Antje van der Linden, Arlo Upton, Chris Mansell, David Hammer, Dragana Drinkovic, Gary McAuliffe, Hana Sofia Andersson, James Ussher, Jill Sherwood, Josh Freeman, Julia Howard, Juliet Elvy, Mary DeAlmeida, Matt Blakiston, Matthew Rogers, Max Bloomfield, Michael Addidle, Michelle Balm, Sally Roberts, Sarah Jefferies, Sharmini Muttaiyah, Susan Morpeth, Susan Taylor, Timothy Blackmore, Vani Sathyendran, Veronica Playle, Virginia Hope, Erasmus Smit, Lauren Jelly, Olin Silander, Joep de Ligt |
| EPI_ISL_579536 | QEII Health Sciences Centre | National Microbiology Laboratory (NML) | Anna Majer, Shari Tyson, Grace Seo, Philip Mabon, Darian Hole, Elsie Grudeski, Rhiannon Huzarewich, Russell Mandes, Anneliese Landgraff, Jennifer Tanner, Natalie Knox, Morag Graham, Gary Van Domselaar, Todd Hatchette, Jason LeBlanc, Nathalie Bastien, Yan Li, Timothy Booth, CanCOGeN's metadata curation team, Public Health Agency of Canada's CanCOGeN team |
| EPI_ISL_581460 | Medizinische Klinik Innere Medizin I, Universitätsklinikum Tübingen | NGS Competence Center Tübingen, Institut für Medizinische Mikrobiologie und Hygiene, Universitätsklinikum Tübingen | Angel Angelov |
| EPI_ISL_581488 | Fondation Congolaise pour la recherche medicale (FCRM) | NGS Competence Center Tübingen, Institut für Medizinische Mikrobiologie und Hygiene, Universitätsklinikum Tübingen | Angel Angelov |
| EPI_ISL_581921, EPI_ISL_581922, EPI_ISL_581923, EPI_ISL_581924, EPI_ISL_581925, EPI_ISL_581959, EPI_ISL_581960 | University Hospital Basel, Clinical Virology | University Hospital Basel, Clinical Bacteriology | Madlen Stange, Alfredo Mari, Tim Roloff, Helena MB Seth-Smith, Michael Schweitzer, Myrta Brunner, Karoline Leuzinger, Kirstine K. Soegaard, Alexander Gensch, Sarah Tschudin-Sutter, Simon Fuchs, Julia Bielicki, Hans Pargger, Martin Siegemund, Christian Nickel, Roland Bingisser, Michael Osthoff, Stefano Bassetti, Rita Schneider-Sliwa, Manuel Battegay, Hans Hirsch, Adrian Egli |
| EPI_ISL_582153, EPI_ISL_582155, EPI_ISL_582161, EPI_ISL_582164, EPI_ISL_582172, EPI_ISL_582178, EPI_ISL_582182, EPI_ISL_582191, EPI_ISL_582196, EPI_ISL_582199, EPI_ISL_582205, EPI_ISL_582207, EPI_ISL_582218 | | | |
| see above | TriCore Reference Laboratories | Center for Global Health, University of New Mexico Health Sciences Center | Daryl Domman, Kurt Schwalm, Twila Kunde, Joseph Hicks, Michael Edwards, Darrell Dinwiddie |
| EPI_ISL_582262, EPI_ISL_582263, EPI_ISL_582401 | Cadham Provincial Laboratory | National Microbiology Laboratory (NML) | Anna Majer, Shari Tyson, Grace Seo, Philip Mabon, Elsie Grudeski, Rhiannon Huzarewich, Russell Mandes, Anneliese Landgraff, Jennifer Tanner, Natalie Knox, Morag Graham, Gary Van Domselaar, Paul Van Caeseele, Jared Bullard, David Alexander, Kerry Dust, Nathalie Bastien, Yan Li, Timothy Booth, Darian Hole, Madison Chapel, CanCOGeN's metadata curation team, Public Health Agency of Canada CanCOGeN team |
| EPI_ISL_582637, EPI_ISL_582638, EPI_ISL_582639, EPI_ISL_582640, EPI_ISL_582641 | Sheikh Khalifa Medical City | Molecular/Surveillance lab Sheikh Khalifa Medical City | Amirtharaj Francis, Sajeed Abdul, Hala Imambaccus, Sahar Almarzooqi, Hiba Saud, Stefan Weber |
| EPI_ISL_582792 | Orebro klinisk mikrobiologi | The Public Health Agency of Sweden | Anna-Malin Linde, Maria Lind Karlberg, Mattias Haukland, Reza Advani, Olov Svartstrom, Oskar Karlsson Lindsjo, Sandra Broddesson, Petra Edquist, Mia Brytting, Anna Risberg, Karin Tegmark-Wisell |
| EPI_ISL_583187, EPI_ISL_583188, EPI_ISL_583189 | San Bernardino County Public Health Lab | Chan-Zuckerberg Biohub | CZB Cliahub Consortium |
| EPI_ISL_583445, EPI_ISL_583446, EPI_ISL_583455, EPI_ISL_583465, EPI_ISL_583470, EPI_ISL_583478 | Memorial Sloan Kettering Cancer Center | van Bakel Laboratory, Genetics and Genomics Sciences, Icahn School of Medicine at Mount Sinai | Teresa Aydillo, Ana S. Gonzalez-Reiche, Sadaf Aslam, Adriana van de Guchte, Zenab Khan, Ajay Obla, Jayeeta Dutta, Harm van Bakel, Judith Aberg, Adolfo García-Sastre, Gunjan Shah, Tobias Hohl, Genovefa Papanicolaou, Miguel-Angel Perales, Kent Sepkowitz, Ngolela Esther Babady, and Mini Kamboj |
| EPI_ISL_583493 | Vigilância em Saúde de Cajamar | Instituto Adolfo Lutz, Interdiciplinary Procedures Center, Strategic Laboratory | Claudio Tavares Sacchi, Claudia Regina Gonçalves, Erica Valessa Ramos Gomes, Karoline Rodrigues Campos |
| EPI_ISL_583694, EPI_ISL_583697, EPI_ISL_583710, EPI_ISL_583711, EPI_ISL_583716, EPI_ISL_583718, EPI_ISL_583720, EPI_ISL_583726 | Center for Virology, Medical University of Vienna | Bergthaler laboratory, CeMM Research Center for Molecular Medicine of the Austrian Academy of Sciences | Alexandra Popa, Benedikt Agerer, Henrique Colaco, Lukas Endler, Jakob-Wendelin Genger, Alexander Lercher, Mark Smyth, Thomas Penz, Michael Schuster, Jan Laine, Martin Senekowitsch, Judith Aberle, Stephan Aberle, Peter Hufnagl, Daniela Schmid, Franz Allerberger, Elisabeth Puchhammer-Stoeckl, Manfred Nairz, Guenter Weiss, Gregor Hörmann, Kinga Rigler-Hohenwarter, Rainer Gattringer, Wegene Borena, Dorothee von Laer, Gernot Walder, Peter Obrist, Christian Paar, Sabine Sussitz-Rack, Gunther Vogl, Adi Steinrigl, Christoph Bock, Andreas Bergthaler |
| EPI_ISL_583733, EPI_ISL_583734, EPI_ISL_583735, EPI_ISL_583736, EPI_ISL_583737, EPI_ISL_583738, EPI_ISL_583741, EPI_ISL_583742, EPI_ISL_583743, EPI_ISL_583847 | Dr. Gernot Walder GmbH | Bergthaler laboratory, CeMM Research Center for Molecular Medicine of the Austrian Academy of Sciences | Alexandra Popa, Benedikt Agerer, Henrique Colaco, Lukas Endler, Jakob-Wendelin Genger, Alexander Lercher, Mark Smyth, Thomas Penz, Michael Schuster, Jan Laine, Martin Senekowitsch, Judith Aberle, Stephan Aberle, Peter Hufnagl, Daniela Schmid, Franz Allerberger, Elisabeth Puchhammer-Stoeckl, Manfred Nairz, Guenter Weiss, Gregor Hörmann, Kinga Rigler-Hohenwarter, Rainer Gattringer, Wegene Borena, Dorothee von Laer, Gernot Walder, Peter Obrist, Christian Paar, Sabine Sussitz-Rack, Gunther Vogl, Adi Steinrigl, Christoph Bock, Andreas Bergthaler |
| EPI_ISL_583914 | DH | Department of Neurovirology, National Institute of Mental Health and Neuroscience (NIMHANS) | Chitra Pattabiraman,Vijayalakshmi Reddy, Harsha PK, Risha Rasheed, Pramada Prasad, Shafeeq S Hameed, Manjunatha Venkataswamy, Anita Desai, Ravi Vasanthapuram |
| EPI_ISL_583958 | UOC Microbiologia e Virologia, Azienda Ospedaliera Universitaria Senese, Siena, Italy | Dipartimento di Biotecnologie Mediche | Maria Grazia Cusi, David Pinzauti, Claudia Gandolfo, Gabriele Anichini, Gianni Pozzi, Francesco Santoro |
| EPI_ISL_584283 | Department of Pathology, University of Cambridge | COVID-19 Genomics UK (COG-UK) Consortium | Aminu S. Jahun, Yasmin Chaudhry, Grant Hall, Iliana Georgana, Myra Hosmillo, Martin D. Curran, Malte Pinckert, Surendra Parmar, Ian Goodfellow |
| EPI_ISL_584459, EPI_ISL_584460, EPI_ISL_584461, EPI_ISL_584462, EPI_ISL_584463, EPI_ISL_584464, EPI_ISL_584465, EPI_ISL_584466, EPI_ISL_584467, EPI_ISL_584468, EPI_ISL_584469, EPI_ISL_584470, EPI_ISL_584471, EPI_ISL_584472, EPI_ISL_584473, EPI_ISL_584474, EPI_ISL_584475, EPI_ISL_584476, EPI_ISL_584477, EPI_ISL_584478, EPI_ISL_584479, EPI_ISL_584480, EPI_ISL_584481, EPI_ISL_584482, EPI_ISL_584483, EPI_ISL_584484, EPI_ISL_584485, EPI_ISL_584486, EPI_ISL_584487, EPI_ISL_584488, EPI_ISL_584489, EPI_ISL_584490, EPI_ISL_584491, EPI_ISL_584492, EPI_ISL_584493, EPI_ISL_584494, EPI_ISL_584495, EPI_ISL_584496, EPI_ISL_584497, EPI_ISL_584498, EPI_ISL_584499, EPI_ISL_584500, EPI_ISL_584502, EPI_ISL_584503, EPI_ISL_584569, EPI_ISL_584570, EPI_ISL_584571, EPI_ISL_584572, EPI_ISL_584573, EPI_ISL_584574, EPI_ISL_584575, EPI_ISL_584576, EPI_ISL_584578, EPI_ISL_584579, EPI_ISL_584580 | | | |
| see above | UHCW / University of Warwick | COVID-19 Genomics UK (COG-UK) Consortium | Richard Stark, Chrystala Constantinidou, Meera Unnikrishnan, Laura Baxter, Jeff Cheng, Grace Taylor-Joyce, Hannah Elizabeth Bridgewater, Lucy Frost, Sarojini Pandey, Paul Brown, Tauqeer Alam, Sascha Ott, Dimitris Grammatopoulos |
| EPI_ISL_584681 | Northumbria University / South Tees Hospitals NHS Foundation Trust / North Cumbria Integrated Care NHS Foundation Trust / North Tees and Hartlepool NHS Foundation Trust / Newcastle Hospitals NHS Foundation Trust | COVID-19 Genomics UK (COG-UK) Consortium | Darren L Smith,Andrew Nelson,Matthew Bashton,Greg R Young,Joshua Loh,John Allan,Mohammad A Tariq,Giles S Holt,Gary Black,Wen C Yew,Lynn Dover,Paul Baker,Steve Liggett,Sarah Essex,Jane Greenaway,Debra Padgett,Clive Graham,Garren Scott,Edward Barton,Emma Swindells,Brendan Payne,Jennifer Collins,Yusri Taha,Gary Eltringham |
| EPI_ISL_586276, EPI_ISL_586277, EPI_ISL_586310, EPI_ISL_586312, EPI_ISL_586319, EPI_ISL_586320, EPI_ISL_586322, EPI_ISL_586323, EPI_ISL_586324, EPI_ISL_586325, EPI_ISL_586326, EPI_ISL_586327, EPI_ISL_586332 | | | |
| see above | Toronto Invasive Bacterial Diseases Network | McMaster University | Allison McGeer, Patryk Aftanas, Hooman Derakhshani, Angel Li, Kuganya Nirmalarajah, Emily Panousis, Ahmed Draia, Jalees Nasir, Michael Surette, Samira Mubareka, Andrew G. McArthur |
| EPI_ISL_589271, EPI_ISL_589273 | Lighthouse Lab in Alderley Park | Wellcome Sanger Institute for the COVID-19 Genomics UK (COG-UK) consortium | Jacquelyn Wynn, Mairead Hyland, The Lighthouse Lab in Alderley Park and Alex Alderton, Roberto Amato, Sonia Goncalves, Ewan Harrison, David K. Jackson, Ian Johnston, Dominic Kwiatkowski, Cordelia Langford, John Sillitoe on behalf of the Wellcome Sanger Institute COVID-19 Surveillance Team |

| | | | |
|---|---|---|---|
| EPI_ISL_589304 | Lighthouse Lab in Milton Keynes | Wellcome Sanger Institute for the COVID-19 Genomics UK (COG-UK) consortium | The Lighthouse Lab in Milton Keynes and Alex Alderton, Roberto Amato, Sonia Goncalves, Ewan Harrison, David K. Jackson, Ian Johnston, Dominic Kwiatkowski, Cordelia Langford, John Sillitoe on behalf of the Wellcome Sanger Institute COVID-19 Surveillance Team |
| EPI_ISL_589305 | Lighthouse Lab in Alderley Park | Wellcome Sanger Institute for the COVID-19 Genomics UK (COG-UK) consortium | Jacquelyn Wynn, Mairead Hyland, The Lighthouse Lab in Alderley Park and Alex Alderton, Roberto Amato, Sonia Goncalves, Ewan Harrison, David K. Jackson, Ian Johnston, Dominic Kwiatkowski, Cordelia Langford, John Sillitoe on behalf of the Wellcome Sanger Institute COVID-19 Surveillance Team |
| EPI_ISL_589334 | Lighthouse Lab in Milton Keynes | Wellcome Sanger Institute for the COVID-19 Genomics UK (COG-UK) Consortium | The Lighthouse Lab in Milton Keynes and Alex Alderton, Roberto Amato, Sonia Goncalves, Ewan Harrison, David K. Jackson, Ian Johnston, Dominic Kwiatkowski, Cordelia Langford, John Sillitoe on behalf of the Wellcome Sanger Institute COVID-19 Surveillance Team |
| EPI_ISL_589335 | Lighthouse Lab in Milton Keynes | Wellcome Sanger Institute for the COVID-19 Genomics UK (COG-UK) consortium | The Lighthouse Lab in Milton Keynes and Alex Alderton, Roberto Amato, Sonia Goncalves, Ewan Harrison, David K. Jackson, Ian Johnston, Dominic Kwiatkowski, Cordelia Langford, John Sillitoe on behalf of the Wellcome Sanger Institute COVID-19 Surveillance Team |
| EPI_ISL_589343 | Lighthouse Lab in Alderley Park | Wellcome Sanger Institute for the COVID-19 Genomics UK (COG-UK) consortium | Jacquelyn Wynn, Mairead Hyland, The Lighthouse Lab in Alderley Park and Alex Alderton, Roberto Amato, Sonia Goncalves, Ewan Harrison, David K. Jackson, Ian Johnston, Dominic Kwiatkowski, Cordelia Langford, John Sillitoe on behalf of the Wellcome Sanger Institute COVID-19 Surveillance Team |
| EPI_ISL_589355 | Lighthouse Lab in Milton Keynes | Wellcome Sanger Institute for the COVID-19 Genomics UK (COG-UK) consortium | The Lighthouse Lab in Milton Keynes and Alex Alderton, Roberto Amato, Sonia Goncalves, Ewan Harrison, David K. Jackson, Ian Johnston, Dominic Kwiatkowski, Cordelia Langford, John Sillitoe on behalf of the Wellcome Sanger Institute COVID-19 Surveillance Team |
| EPI_ISL_589371 | Lighthouse Lab in Alderley Park | Wellcome Sanger Institute for the COVID-19 Genomics UK (COG-UK) consortium | Jacquelyn Wynn, Mairead Hyland, The Lighthouse Lab in Alderley Park and Alex Alderton, Roberto Amato, Sonia Goncalves, Ewan Harrison, David K. Jackson, Ian Johnston, Dominic Kwiatkowski, Cordelia Langford, John Sillitoe on behalf of the Wellcome Sanger Institute COVID-19 Surveillance Team |
| EPI_ISL_589377 | Lighthouse Lab in Milton Keynes | Wellcome Sanger Institute for the COVID-19 Genomics UK (COG-UK) consortium | The Lighthouse Lab in Milton Keynes and Alex Alderton, Roberto Amato, Sonia Goncalves, Ewan Harrison, David K. Jackson, Ian Johnston, Dominic Kwiatkowski, Cordelia Langford, John Sillitoe on behalf of the Wellcome Sanger Institute COVID-19 Surveillance Team |
| EPI_ISL_589401, EPI_ISL_589405, EPI_ISL_589406 | Lighthouse Lab in Alderley Park | Wellcome Sanger Institute for the COVID-19 Genomics UK (COG-UK) consortium | Jacquelyn Wynn, Mairead Hyland, The Lighthouse Lab in Alderley Park and Alex Alderton, Roberto Amato, Sonia Goncalves, Ewan Harrison, David K. Jackson, Ian Johnston, Dominic Kwiatkowski, Cordelia Langford, John Sillitoe on behalf of the Wellcome Sanger Institute COVID-19 Surveillance Team |
| EPI_ISL_589407, EPI_ISL_589417 | Lighthouse Lab in Milton Keynes | Wellcome Sanger Institute for the COVID-19 Genomics UK (COG-UK) consortium | The Lighthouse Lab in Milton Keynes and Alex Alderton, Roberto Amato, Sonia Goncalves, Ewan Harrison, David K. Jackson, Ian Johnston, Dominic Kwiatkowski, Cordelia Langford, John Sillitoe on behalf of the Wellcome Sanger Institute COVID-19 Surveillance Team |
| EPI_ISL_589427, EPI_ISL_589428 | Lighthouse Lab in Alderley Park | Wellcome Sanger Institute for the COVID-19 Genomics UK (COG-UK) consortium | Jacquelyn Wynn, Mairead Hyland, The Lighthouse Lab in Alderley Park and Alex Alderton, Roberto Amato, Sonia Goncalves, Ewan Harrison, David K. Jackson, Ian Johnston, Dominic Kwiatkowski, Cordelia Langford, John Sillitoe on behalf of the Wellcome Sanger Institute COVID-19 Surveillance Team |
| EPI_ISL_589430 | Lighthouse Lab in Milton Keynes | Wellcome Sanger Institute for the COVID-19 Genomics UK (COG-UK) consortium | The Lighthouse Lab in Milton Keynes and Alex Alderton, Roberto Amato, Sonia Goncalves, Ewan Harrison, David K. Jackson, Ian Johnston, Dominic Kwiatkowski, Cordelia Langford, John Sillitoe on behalf of the Wellcome Sanger Institute COVID-19 Surveillance Team |
| EPI_ISL_589450, EPI_ISL_589470, EPI_ISL_589487, EPI_ISL_589489 | Lighthouse Lab in Alderley Park | Wellcome Sanger Institute for the COVID-19 Genomics UK (COG-UK) consortium | Jacquelyn Wynn, Mairead Hyland, The Lighthouse Lab in Alderley Park and Alex Alderton, Roberto Amato, Sonia Goncalves, Ewan Harrison, David K. Jackson, Ian Johnston, Dominic Kwiatkowski, Cordelia Langford, John Sillitoe on behalf of the Wellcome Sanger Institute COVID-19 Surveillance Team |
| EPI_ISL_589493 | Lighthouse Lab in Milton Keynes | Wellcome Sanger Institute for the COVID-19 Genomics UK (COG-UK) consortium | The Lighthouse Lab in Milton Keynes and Alex Alderton, Roberto Amato, Sonia Goncalves, Ewan Harrison, David K. Jackson, Ian Johnston, Dominic Kwiatkowski, Cordelia Langford, John Sillitoe on behalf of the Wellcome Sanger Institute COVID-19 Surveillance Team |
| EPI_ISL_589504, EPI_ISL_589507 | Lighthouse Lab in Alderley Park | Wellcome Sanger Institute for the COVID-19 Genomics UK (COG-UK) consortium | Jacquelyn Wynn, Mairead Hyland, The Lighthouse Lab in Alderley Park and Alex Alderton, Roberto Amato, Sonia Goncalves, Ewan Harrison, David K. Jackson, Ian Johnston, Dominic Kwiatkowski, Cordelia Langford, John Sillitoe on behalf of the Wellcome Sanger Institute COVID-19 Surveillance Team |
| EPI_ISL_589520 | Lighthouse Lab in Milton Keynes | Wellcome Sanger Institute for the COVID-19 Genomics UK (COG-UK) consortium | The Lighthouse Lab in Milton Keynes and Alex Alderton, Roberto Amato, Sonia Goncalves, Ewan Harrison, David K. Jackson, Ian Johnston, Dominic Kwiatkowski, Cordelia Langford, John Sillitoe on behalf of the Wellcome Sanger Institute COVID-19 Surveillance Team |
| EPI_ISL_589524 | Lighthouse Lab in Alderley Park | Wellcome Sanger Institute for the COVID-19 Genomics UK (COG-UK) consortium | Jacquelyn Wynn, Mairead Hyland, The Lighthouse Lab in Alderley Park and Alex Alderton, Roberto Amato, Sonia Goncalves, Ewan Harrison, David K. Jackson, Ian Johnston, Dominic Kwiatkowski, Cordelia Langford, John Sillitoe on behalf of the Wellcome Sanger Institute COVID-19 Surveillance Team |
| EPI_ISL_589526 | Lighthouse Lab in Milton Keynes | Wellcome Sanger Institute for the COVID-19 Genomics UK (COG-UK) consortium | The Lighthouse Lab in Milton Keynes and Alex Alderton, Roberto Amato, Sonia Goncalves, Ewan Harrison, David K. Jackson, Ian Johnston, Dominic Kwiatkowski, Cordelia Langford, John Sillitoe on behalf of the Wellcome Sanger Institute COVID-19 Surveillance Team |
| EPI_ISL_589535, EPI_ISL_589550, EPI_ISL_589554, EPI_ISL_589561 | Lighthouse Lab in Alderley Park | Wellcome Sanger Institute for the COVID-19 Genomics UK (COG-UK) consortium | Jacquelyn Wynn, Mairead Hyland, The Lighthouse Lab in Alderley Park and Alex Alderton, Roberto Amato, Sonia Goncalves, Ewan Harrison, David K. Jackson, Ian Johnston, Dominic Kwiatkowski, Cordelia Langford, John Sillitoe on behalf of the Wellcome Sanger Institute COVID-19 Surveillance Team |
| EPI_ISL_589567 | Lighthouse Lab in Milton Keynes | Wellcome Sanger Institute for the COVID-19 Genomics UK (COG-UK) Consortium | The Lighthouse Lab in Milton Keynes and Alex Alderton, Roberto Amato, Sonia Goncalves, Ewan Harrison, David K. Jackson, Ian Johnston, Dominic Kwiatkowski, Cordelia Langford, John Sillitoe on behalf of the Wellcome Sanger Institute COVID-19 Surveillance Team |
| EPI_ISL_589568 | Lighthouse Lab in Alderley Park | Wellcome Sanger Institute for the COVID-19 Genomics UK (COG-UK) consortium | Jacquelyn Wynn, Mairead Hyland, The Lighthouse Lab in Alderley Park and Alex Alderton, Roberto Amato, Sonia Goncalves, Ewan Harrison, David K. Jackson, Ian Johnston, Dominic Kwiatkowski, Cordelia Langford, John Sillitoe on behalf of the Wellcome Sanger Institute COVID-19 Surveillance Team |
| EPI_ISL_590833, EPI_ISL_590834, EPI_ISL_590835, EPI_ISL_590836 | Institute of Medical Virology, University of Zurich | Institute of Medical Virology, University of Zurich | Marie O. Pohl, Idoia Busnadiego, Verena Kufner, Stefan Schmutz, Maryam Zaheri, Irene Abela, Alexandra Trkola, Michael Huber, Silke Stertz, Benjamin G. Hale |
| EPI_ISL_591320, EPI_ISL_591322 | Virology, Iran University of Medical Sciences | Virology, Iran University of Medical Sciences | Keyvani,H., Ranjbar,Mm., Keyvani,F., Soleimani,S. |
| EPI_ISL_591323 | Virology, Iran University of Medical Sciences | Virology, Iran University of Medical Sciences | Keyvani,H., Ranjbar,Mm., Soleimani,S., Keyvani,F. |
| EPI_ISL_591325 | Virology, Iran University of Medical Sciences | Virology, Iran University of Medical Sciences | Keyvani,H., Ranjbar,Mm., Keyvani,F. and Soleimani,S. |
| EPI_ISL_591331 | Dipartimento di Biotecnologie Mediche, University of Siena | Dipartimento di Biotecnologie Mediche, University of Siena | Cusi,M.G., Pinzauti,D., Gandolfo,C., Anichini,G., Pozzi,G., Santoro,F. |
| EPI_ISL_591370, EPI_ISL_591371, EPI_ISL_591372, EPI_ISL_591373, EPI_ISL_591374, EPI_ISL_591379, EPI_ISL_591380, EPI_ISL_591381, EPI_ISL_591382, EPI_ISL_591383, EPI_ISL_591384, EPI_ISL_591385, EPI_ISL_591386, EPI_ISL_591387, EPI_ISL_591388, EPI_ISL_591389, EPI_ISL_591390 | | | |
| see above | Pathogen Genomics Center, National Institute of Infectious Diseases | Pathogen Genomics Center, National Institute of Infectious Diseases | Tsuyoshi Sekizuka, Kentaro Itokawa, Rina Tanaka, Masanori Hashino, Makoto Kuroda |
| EPI_ISL_593478 | Brigham and Women's Hospital | Jonathan Li Laboratory | Manish C. Choudhary, James Regan, Jonathan Z. Li |
| EPI_ISL_593490, EPI_ISL_593491, EPI_ISL_593492 | Eastern Ontario Regional Laboratory Association | McMaster University | Leanne Mortimer, Hooman Derakhshani, Emily Panousis, Ahmed Draia, Jalees Nasir, Robert Slinger, Andrew G. McArthur |
| EPI_ISL_593667 | Pathology West - NSW Health Pathology | NSW Health Pathology - Institute of Clinical Pathology and Medical Research; Westmead Hospital; University of Sydney | CIDM-PH et al. |
| EPI_ISL_593746, EPI_ISL_593747 | South Eastern Area Laboratory Services (SEALS) | NSW Health Pathology - Institute of Clinical Pathology and Medical Research; Westmead Hospital; University of Sydney | CIDM-PH et al. |
| EPI_ISL_594122, EPI_ISL_594124, EPI_ISL_594125 | Yale COVID-19 Biorepository | Grubaugh Lab - Yale School of Public Health | Joseph Fauver, Tara Alpert, Anderson Brito, Anne Wyllie, Chantal Vogels, Mary Petrone, Chaney Kalinich, Isabel Ott, Arnau Casanovas, Catherine Muenker, Adam Moore, Alice Lu, Maria Tokuyama, Patrick Wong, Peiwen Lu, Saad Omer, Richard Martinello, Allison Nelson, Shelli Farhadian, Akiko Iwasaki, Charlese Dela Cruz, Albert Ko, Nathan Grubaugh |
| EPI_ISL_594210, EPI_ISL_594211, EPI_ISL_594212, EPI_ISL_594213, EPI_ISL_594214, EPI_ISL_594215 | Michigan Department of Health and Human Services, Bureau of Laboratories | Michigan Department of Health and Human Services, Bureau of Laboratories | Blankenship HM, Riner D, Soehnlen MK |
| EPI_ISL_596382 | Virology, Iran University of Medical Sciences | Virology, Iran University of Medical Sciences | Keyvani,H., Ranjbar,Mm., Keyvani,F., Soleimani,S. |
| EPI_ISL_596384 | Virology, Iran University of Medical Sciences | Virology, Iran University of Medical Sciences | Keyvani,H., Ranjbar,M.M., Keyvani,F., Soleimani,S. |
| EPI_ISL_596627, EPI_ISL_596628, EPI_ISL_596629, EPI_ISL_596630, EPI_ISL_596631, EPI_ISL_596632, EPI_ISL_596633, EPI_ISL_596634, EPI_ISL_596635, EPI_ISL_596636, EPI_ISL_596637, EPI_ISL_596639, EPI_ISL_596640, EPI_ISL_596667, EPI_ISL_596676 | | | |
| see above | St.Vincent's University Hospital | St.Vincent's University Hospital | Mary Lucey, Guerrino Macori, Niamh Mullane, Una Sutton-Fitzpatrick, Gabriel Gonzalez, Suzie Coughlan, Aisling Purcell, Lynda Fenelon, Séamus Fanning, Kirsten Schaffer |

| | | | |
|---|---|---|---|
| EPI_ISL_600437 | Institute of Epidemiology Disease Control And Research | Institute for Developing Science and Health Initiatives | Lauren Cowley, Mokibul Hassan Afrad, Sadia Isfat Ara Rahman, Md. Mahfuz-Al-mamun, Firadausi Qadri, Tahmina Shirin |
| EPI_ISL_605799, EPI_ISL_605801, EPI_ISL_605802, EPI_ISL_605804, EPI_ISL_605805, EPI_ISL_605806, EPI_ISL_605807, EPI_ISL_605808, EPI_ISL_605809 | Clinical Virology Laboratory, Institute of Liver and Biliary Sciences | ILBS - IGIB | Ekta Gupta, Sheetalnath Rooge, Abhishek Padhi, Reshu Agarwal, Jaswinder Singh Maras, Shridhar Sivasubbu, Vinod Scaria, Shvetank Sharma |
| EPI_ISL_605882, EPI_ISL_605883, EPI_ISL_605884, EPI_ISL_605885, EPI_ISL_605886, EPI_ISL_605887, EPI_ISL_605888, EPI_ISL_605889, EPI_ISL_605890, EPI_ISL_605891, EPI_ISL_605892, EPI_ISL_605893, EPI_ISL_605894, EPI_ISL_605895, EPI_ISL_605896, EPI_ISL_605897, EPI_ISL_605898, EPI_ISL_605899, EPI_ISL_605900, EPI_ISL_605901, EPI_ISL_605902, EPI_ISL_605903, EPI_ISL_605904, EPI_ISL_605905, EPI_ISL_605906, EPI_ISL_605907, EPI_ISL_605908 | | | |
| see above | NGS Lab, DNA SOLUTION LTD. | NGS Lab, DNA SOLUTION LTD. | Khan,M.I., Hasan,K.N., Sufian,A., Polol,M.N.I., Khaleque,A., Rahman,M., Chowdhury,M., Haider,H.U., Razu,M.H., Khan,M., Rabbi,M.F.A. |
| EPI_ISL_610250, EPI_ISL_610253 | New Mexico Department of Health Scientific Laboratory Division | Center for Global Health, University of New Mexico Health Sciences Center | Daryl Domman, Kurt Schwalm, Twila Kunde, Joseph Hicks, Michael Edwards, Darrell Dinwiddie |
| EPI_ISL_610263, EPI_ISL_610264 | TriCore Reference Laboratories | Center for Global Health, University of New Mexico Health Sciences Center | Daryl Domman, Kurt Schwalm, Twila Kunde, Joseph Hicks, Michael Edwards, Darrell Dinwiddie |
| EPI_ISL_610268, EPI_ISL_610270, EPI_ISL_610272, EPI_ISL_610274, EPI_ISL_610275, EPI_ISL_610276, EPI_ISL_610277, EPI_ISL_610278, EPI_ISL_610279, EPI_ISL_610280, EPI_ISL_610281 | | | |
| see above | New Mexico Department of Health Scientific Laboratory Division | Center for Global Health, University of New Mexico Health Sciences Center | Daryl Domman, Kurt Schwalm, Twila Kunde, Joseph Hicks, Michael Edwards, Darrell Dinwiddie |
| EPI_ISL_610282 | TriCore Reference Laboratories | Center for Global Health, University of New Mexico Health Sciences Center | Daryl Domman, Kurt Schwalm, Twila Kunde, Joseph Hicks, Michael Edwards, Darrell Dinwiddie |
| EPI_ISL_611751 | Department of Pathology, University of Cambridge | COVID-19 Genomics UK (COG-UK) Consortium | Aminu S. Jahun, Yasmin Chaudhry, Grant Hall, Iliana Georgana, Myra Hosmillo, Martin D. Curran, Malte Pinckert, Surendra Parmar, Ian Goodfellow |
| EPI_ISL_614104, EPI_ISL_614105, EPI_ISL_614106, EPI_ISL_614109, EPI_ISL_614110, EPI_ISL_614116, EPI_ISL_614117, EPI_ISL_614118, EPI_ISL_614119 | Virginia DCLS | Virginia DCLS | Virginia DCLS |
| EPI_ISL_614397 | Laboratorio Biologia Molecolare Sars Cov2 - UOC Laboratorio Analisi - Servizio Medicina di Laboratorio, Ospedale "San Francesco" - ATS-ASSL Nuoro | Laboratorio specialistico UOC Ematologia - Ospedale "San Francesco" - ATS-ASSL Nuoro | Piras Giovanna, Fancello Tatiana, Asproni Rosanna, Fiamma Maura, Monne Maria Itria, Toja Alessandro, Sanna Filomena, Floris Anna Rita, Sulis Vincenzo, Palmas Angelo Domenico, Casu Gavino, Lo Maglio Iana, Mameli Giuseppe |
| EPI_ISL_614410, EPI_ISL_614413, EPI_ISL_614418, EPI_ISL_614432, EPI_ISL_614448, EPI_ISL_614470, EPI_ISL_614471, EPI_ISL_614749, EPI_ISL_614750, EPI_ISL_614751, EPI_ISL_614753, EPI_ISL_614754, EPI_ISL_614759, EPI_ISL_614760, EPI_ISL_614762, EPI_ISL_614765, EPI_ISL_614766, EPI_ISL_614767, EPI_ISL_614769, EPI_ISL_614771, EPI_ISL_614773, EPI_ISL_614776, EPI_ISL_614778, EPI_ISL_614786, EPI_ISL_614787, EPI_ISL_614788, EPI_ISL_614789, EPI_ISL_614791, EPI_ISL_614792, EPI_ISL_614793, EPI_ISL_614794, EPI_ISL_614795, EPI_ISL_614814, EPI_ISL_614815 | | | |
| see above | Department of Virus and Microbiological Special Diagnostics, Statens Serum Institut, Denmark | Albertsen lab, Department of Chemistry and Bioscience, Aalborg University, Denmark | Danish Covid-19 Genome Consortia |
| EPI_ISL_614889 | Laboratorio Biologia Molecolare Sars Cov2 - UOC Laboratorio Analisi - Servizio Medicina di Laboratorio, Ospedale "San Francesco" - ATS-ASSL Nuoro | Laboratorio specialistico UOC Ematologia - Ospedale "San Francesco" - ATS-ASSL Nuoro | Piras Giovanna, Fancello Tatiana, Asproni Rosanna, Fiamma Maura, Monne Maria Itria, Toja Alessandro, Sanna Filomena, Floris Anna Rita, Sulis Vincenzo, Palmas Angelo Domenico, Casu Gavino, Lo Maglio Iana, Mameli Giuseppe |
| EPI_ISL_615079 | Orebro klinisk mikrobiologi | The Public Health Agency of Sweden | Anna-Malin Linde, Maria Lind Karlberg, Mattias Haukland, Reza Advani, Olov Svartstrom, Oskar Karlsson Lindsjo, Sandra Broddesson, Petra Edquist, Mia Brytting, Anna Risberg, Karin Tegmark-Wisell |
| EPI_ISL_615124 | Texas Department of State Health Services | Texas Department of State Health Services | Rashmi Tuladhar, Bonnie Oh, Jenny Zhang, Maliha Rahman, Anita Pokharel, Myong Koag, Chung Wang, Rachel Lee, Grace Kubin, Mayela Pedrueza |
| EPI_ISL_616923, EPI_ISL_617382, EPI_ISL_617384, EPI_ISL_617941, EPI_ISL_618802, EPI_ISL_618803, EPI_ISL_618807, EPI_ISL_622352, EPI_ISL_622353, EPI_ISL_622354, EPI_ISL_622406, EPI_ISL_622407, EPI_ISL_622421, EPI_ISL_622427, EPI_ISL_622428, EPI_ISL_622431, EPI_ISL_622435, EPI_ISL_622463, EPI_ISL_622464, EPI_ISL_622465, EPI_ISL_622466, EPI_ISL_622467, EPI_ISL_622468, EPI_ISL_622469, EPI_ISL_622470, EPI_ISL_622471, EPI_ISL_622472, EPI_ISL_622473, EPI_ISL_622474, EPI_ISL_622475, EPI_ISL_622476, EPI_ISL_622477, EPI_ISL_622492, EPI_ISL_622514, EPI_ISL_622522, EPI_ISL_622523, EPI_ISL_622525, EPI_ISL_622526, EPI_ISL_622527, EPI_ISL_622528, EPI_ISL_622529, EPI_ISL_622530, EPI_ISL_622532, EPI_ISL_622533, EPI_ISL_622534, EPI_ISL_622535, EPI_ISL_622536, EPI_ISL_622537, EPI_ISL_622539, EPI_ISL_622540, EPI_ISL_622541, EPI_ISL_622542, EPI_ISL_622543, EPI_ISL_622544, EPI_ISL_622546, EPI_ISL_622575, EPI_ISL_622576, EPI_ISL_622577, EPI_ISL_622578, EPI_ISL_622579, EPI_ISL_622581, EPI_ISL_622582, EPI_ISL_622583, EPI_ISL_622585, EPI_ISL_622586, EPI_ISL_622588, EPI_ISL_622589, EPI_ISL_622590, EPI_ISL_622591, EPI_ISL_622592, EPI_ISL_622593, EPI_ISL_622594, EPI_ISL_622595, EPI_ISL_622615, EPI_ISL_622616, EPI_ISL_622617, EPI_ISL_622618, EPI_ISL_622619, EPI_ISL_622620, EPI_ISL_622621, EPI_ISL_622622, EPI_ISL_622623, EPI_ISL_622624, EPI_ISL_622625, EPI_ISL_622626, EPI_ISL_622628, EPI_ISL_622629, EPI_ISL_622630, EPI_ISL_622631, EPI_ISL_622632, EPI_ISL_622633, EPI_ISL_622634, EPI_ISL_622635, EPI_ISL_622636, EPI_ISL_622637, EPI_ISL_622638, EPI_ISL_622639, EPI_ISL_622640 | | | |
| see above | Department of Virus and Microbiological Special Diagnostics, Statens Serum Institut, Denmark | Albertsen lab, Department of Chemistry and Bioscience, Aalborg University, Denmark | Danish Covid-19 Genome Consortia |
| EPI_ISL_623109, EPI_ISL_623117, EPI_ISL_623119, EPI_ISL_623125, EPI_ISL_623129, EPI_ISL_623133, EPI_ISL_623134, EPI_ISL_623136, EPI_ISL_623141, EPI_ISL_623144, EPI_ISL_623146, EPI_ISL_623148, EPI_ISL_623153, EPI_ISL_623154, EPI_ISL_623157, EPI_ISL_623161, EPI_ISL_623162, EPI_ISL_623163, EPI_ISL_623164, EPI_ISL_623167 | | | |
| see above | Laboratorio de Virologia Molecular / UFRJ | Bioinformatics Laboratory / LNCC | Carolina M Voloch, Ronaldo S Francisco Jr, Luiz G P de Almeida, Otavio J. Brustolini, Cynthia C Cardoso, Alexandra L Gerber, Ana Paula de C Guimarães, Diana Mariani, Covid19-UFRJ Workgroup, Luís Cristóvão Pôrto, Renato S Aguiar, Terezinha M P P Castiñeiras, Orlando C. Ferreira, Amilcar Tanuri, Ana Tereza R de Vasconcelos |
| EPI_ISL_625456 | Virology Unit, Institut Pasteur de Madagascar | Virology Unit, Institut Pasteur de Madagascar | Christian Ranaivoson, Cara Brook, Norosoa Razanajatovo, Vida Ahyong, Tsiry Randriambolamanantsoa, Michelle Tan, Vololoniaina Raharinosy, Helisoa Razafimanjato, Cristina M. Tato, Joseph L. DeRisi, Soa Fy Andriamandimby, Jean-Michel Heraud, Philippe Dussart |
| EPI_ISL_626219, EPI_ISL_626220, EPI_ISL_626221 | Institute for Virology, University Hospital Essen | Center of Medical Microbiology, Virology, and Hospital Hygiene, University of Duesseldorf | Olympia E. Anastasiou, Ulf Dittmer, Maximilian Damagnez, Alexander Dilthey, Torsten Houwaart, Lisanna Hülse, Malte Kohns Vasconcelos, Nadine Lübke, Jessica Nicolai, Klaus Pfeffer, Daniel Strelow, Jörg Timm, Andreas Walker, Tobias Wienemann |
| EPI_ISL_626441, EPI_ISL_626442, EPI_ISL_626443, EPI_ISL_626444, EPI_ISL_626445, EPI_ISL_626446, EPI_ISL_626447, EPI_ISL_626448, EPI_ISL_626449, EPI_ISL_626450, EPI_ISL_626451, EPI_ISL_626452, EPI_ISL_626453, EPI_ISL_626454, EPI_ISL_626455, EPI_ISL_626456, EPI_ISL_626457, EPI_ISL_626458, EPI_ISL_626459 | | | |
| see above | Northwestern Memorial Hospital | Ozer Lab | Ramon Lorenzo-Redondo, Hannah H. Nam, Scott C. Roberts, Lacy M. Simons, Chad J. Achenbach, Lawrence J. Jennings, Chao Qi, Alan R. Hauser, Michael G. Ison, Judd F. Hultquist, Egon A. Ozer |
| EPI_ISL_629011, EPI_ISL_629012, EPI_ISL_629014, EPI_ISL_629015, EPI_ISL_629016, EPI_ISL_629018, EPI_ISL_629019, EPI_ISL_629020, EPI_ISL_629021, EPI_ISL_629022 | Centro de Biotecnología Vegetal, Universidad Andrés Bello, Center for Genome Regulation | Center for Mathematical Modeling and Center for Genome Regulation. Santiago, Chile | Bastias M, Sanhueza D, Travisany D, Allende ML, Maass A, González M, Bustos F, Arriagada G, Montecino, M, Orellana A, Castro E, Meneses C. |
| EPI_ISL_629031, EPI_ISL_629032, EPI_ISL_629033, EPI_ISL_629034 | Omsk Research Institute of Natural Focal Infections | WHO National Influenza Centre Russian Federation | Artem Fadeev, Ekaterina Gradoboeva, Ekaterina Savkina, Daria Nashatyreva, Elena Poleshchuk, Aleksei Vasilenko, Valery Yakimenko, Andrey Komissarov |
| EPI_ISL_629108 | Laboratoire du Centre Hospitalier Annecy Genevois | CNR Virus des Infections Respiratoires - France SUD | Antonin Bal, Gregory Destras, Gwendolyne Burfin, Hadrien Règue, Quentin Semanas, Martine Valette, Bruno Lina, Hélène Petitprez, Bruno Chanzy, Laurence Josset |
| EPI_ISL_631749, EPI_ISL_631750, EPI_ISL_631751, EPI_ISL_631784 | St Barnabas Hospital | New York City Public Health Laboratory | Jade Wang, et al. |
| EPI_ISL_631840, EPI_ISL_631842, EPI_ISL_631843, EPI_ISL_631844, EPI_ISL_631845, EPI_ISL_631846 | OCME Office Of Chief Medical Examiner | New York City Public Health Laboratory | Jade Wang, et al. |

| | | | |
|---|---|---|---|
| EPI_ISL_631939, EPI_ISL_631940, EPI_ISL_631941, EPI_ISL_631942, EPI_ISL_631943 | St Barnabas Hospital | New York City Public Health Laboratory | Jade Wang, et al. |
| EPI_ISL_632313 | NU-sjukvården | Clinical Microbiology, Sahlgrenska University Hospital, | Johan Ringlander, Josefin Olausson, Hedvig Engström Jakobsson, Magnus Lindh |
| EPI_ISL_632909 | Cadham Provincial laboratory | Cadham Provincial laboratory | Anna Majer, Shari Tyson, Grace Seo, Philip Mabon, Elsie Grudeski, Rhiannon Huzarewich, Russell Mandes, Anneliese Landgraff, Jennifer Tanner, Natalie Knox, Morag Graham, Gary Van Domselaar, Paul Van Caeseele, Jared Bullard, David Alexander, Kerry Dust, Nathalie Bastien, Yan Li, Timothy Booth, Darian Hole, Madison Chapel, CanCOGeN's metadata curation team, Public Health Agency of Canada CanCOGeN team |
| EPI_ISL_634837 | Laboratoire de virologie, CHU de Grenoble - CS 10217 - 38043 Grenoble cedex 29 | CNR Virus des Infections Respiratoires - France SUD | Antonin Bal, Gregory Destras, Gwendolyne Burfin, Hadrien Règue, Quentin Semanas, Martine Valette, Bruno Lina, Sylvie Larrat, Laurence Josset |
| EPI_ISL_634903, EPI_ISL_634904, EPI_ISL_634905, EPI_ISL_634906, EPI_ISL_634907, EPI_ISL_634908, EPI_ISL_634909, EPI_ISL_634910, EPI_ISL_634911, EPI_ISL_634912, EPI_ISL_634913, EPI_ISL_634914, EPI_ISL_634915, EPI_ISL_634916, EPI_ISL_634917, EPI_ISL_634918, EPI_ISL_634919, EPI_ISL_634920, EPI_ISL_634921, EPI_ISL_634922 | | | |
| see above | Utah Public Health Laboratory | Utah Public Health Laboratory | Erin L. Young, Kelly F. Oakeson |
| EPI_ISL_635277 | Institute of Microbiology and Immunology, Faculty of Medicine, University of Ljubljana | Institute of Microbiology and Immunology, Faculty of Medicine, University of Ljubljana | Tomaž Mark Zorec, Samo Zakotnik, Miša Korva, Tatjana Avši - Županc, Mario Poljak |
| EPI_ISL_636476, EPI_ISL_636477 | CNR Virus des Infections Respiratoires - France SUD | CNR Virus des Infections Respiratoires - France SUD | Antonin Bal, Gregory Destras, Gwendolyne Burfin, Hadrien Règue, Alexandre Gaymard, Maude Bouscambert-Duchamp, Florence Morfin-Sherpa, Martine Valette, Bruno Lina, Laurence Josset |
| EPI_ISL_636478 | Centre Hospitalier Pierre Oudot | CNR Virus des Infections Respiratoires - France SUD | Antonin Bal, Gregory Destras, Gwendolyne Burfin, Hadrien Règue, Alexandre Gaymard, Maude Bouscambert-Duchamp, Florence Morfin-Sherpa, Martine Valette, Bruno Lina, Laurence Josset |
| EPI_ISL_636479, EPI_ISL_636487, EPI_ISL_636489, EPI_ISL_636490 | CNR Virus des Infections Respiratoires - France SUD | CNR Virus des Infections Respiratoires - France SUD | Antonin Bal, Gregory Destras, Gwendolyne Burfin, Hadrien Règue, Alexandre Gaymard, Maude Bouscambert-Duchamp, Florence Morfin-Sherpa, Martine Valette, Bruno Lina, Laurence Josset |
| EPI_ISL_636525, EPI_ISL_636543, EPI_ISL_636598 | Dutch COVID-19 response team | National Institute for Public Health and the Environment (RIVM) | Adam Meijer, Harry Vennema, Jeroen Cremer, Sharon van den Brink, Bas van der Veer, AnneMarie van den Brandt, Florian Zwagemaker, Dennis Schmitz, Chantal Reusken, on behalf of the national COVID-19 response team |
| EPI_ISL_636948, EPI_ISL_636959 | Public Health Ontario Laboratory | Public Health Ontario Laboratory | Vanessa G Allen, Philip Banh, Richard de Borja, Yao Chen, Alireza Eshaghi, Nahuel Fittipaldi, Christine Frantz, Jonathan B Gubbay, Jennifer L Guthrie, Lawrence Heisler, Esha Joshi, Michael Laszloffy, Aimin Li, Michael CY Li, Dean Maxwell, Sandeep Nagra, Samir N Patel, Heather Rilkoff, Jared Simpson, Karthikeyan Sivaraman, Yogi Sundaravadanam, Sarah Teatero, Andre Villegas, Sandra Zittermann |
| EPI_ISL_637075 | Department of Infectious Diseases and Immunology, National Hospital Organization Nagoya Medical Center | Clinical Research Center, National Hospital Organization Nagoya Medical Center | Yoshihiro Nakata, Hirotaka Ode, Mai Kubota, Masakazu Matsuda, Kazuhiro Matsuoka, Miho Nakasuji, Mikiko Mori, Mayumi Imahashi, Yoshiyuki Yokomaku, Yasumasa Iwatani |
| EPI_ISL_637107 | Laboratorio Biologia Molecolare Sars Cov2 - UOC Laboratorio Analisi - Servizio Medicina di Laboratorio, Ospedale "San Francesco" - ATS-ASSL Nuoro | Laboratorio specialistico UOC Ematologia - Ospedale "San Francesco" - ATS-ASSL Nuoro | Piras Giovanna, Fancello Tatiana, Asproni Rosanna, Fiamma Maura, Monne Maria Itria, Toja Alessandro, Sanna Filomena, Floris Anna Rita, Sulis Vincenzo, Palmas Angelo Domenico, Casu Gavino, Lo Maglio Iana, Mameli Giuseppe |
| EPI_ISL_637153 | Respiratory Virus Unit, Microbiology Services Colindale, Public Health England | COVID-19 Genomics UK (COG-UK) Consortium | PHE Covid Sequencing Team |
| EPI_ISL_640083 | Khayelitsha (Site B) CHC wc KHC | NHLS/UCT | Arash Iranzadeh, Deelan Doolabh, Lynn Tyers, Bruna Galvao, Innocent Mudau, Marvin Hsiao, Kruger Marais, Diana Hardie, Stephen Korsman, Carolyn Williamson |
| EPI_ISL_640101 | Heideveld CDC wc HVP | NHLS/UCT | Arash Iranzadeh, Deelan Doolabh, Lynn Tyers, Bruna Galvao, Innocent Mudau, Marvin Hsiao, Kruger Marais, Diana Hardie, Stephen Korsman, Carolyn Williamson |
| EPI_ISL_640102 | Vanguard CHC wc VGC | NHLS/UCT | Arash Iranzadeh, Deelan Doolabh, Lynn Tyers, Bruna Galvao, Innocent Mudau, Marvin Hsiao, Kruger Marais, Diana Hardie, Stephen Korsman, Carolyn Williamson |
| EPI_ISL_640103 | Heideveld CDC wc HVP | NHLS/UCT | Arash Iranzadeh, Deelan Doolabh, Lynn Tyers, Bruna Galvao, Innocent Mudau, Marvin Hsiao, Kruger Marais, Diana Hardie, Stephen Korsman, Carolyn Williamson |
| EPI_ISL_640104 | Guguletu CHC wc GDH | NHLS/UCT | Arash Iranzadeh, Deelan Doolabh, Lynn Tyers, Bruna Galvao, Innocent Mudau, Marvin Hsiao, Kruger Marais, Diana Hardie, Stephen Korsman, Carolyn Williamson |
| EPI_ISL_640105 | Nyanga CDC wc NGC | NHLS/UCT | Arash Iranzadeh, Deelan Doolabh, Lynn Tyers, Bruna Galvao, Innocent Mudau, Marvin Hsiao, Kruger Marais, Diana Hardie, Stephen Korsman, Carolyn Williamson |
| EPI_ISL_640106 | Heideveld CDC wc HVP | NHLS/UCT | Arash Iranzadeh, Deelan Doolabh, Lynn Tyers, Bruna Galvao, Innocent Mudau, Marvin Hsiao, Kruger Marais, Diana Hardie, Stephen Korsman, Carolyn Williamson |
| EPI_ISL_640219, EPI_ISL_640221, EPI_ISL_640233, EPI_ISL_640252, EPI_ISL_640261, EPI_ISL_640267 | MVZ Laborärzte Singen | MVZ Laborärzte Singen | Jonas Schmidt, Frithjof Blessing, Sandro Berghaus, Folker Wenzel |
| EPI_ISL_641320, EPI_ISL_641321, EPI_ISL_641322, EPI_ISL_641323, EPI_ISL_641325, EPI_ISL_641326, EPI_ISL_641331, EPI_ISL_641359 | Devki Devi Foundation, a unit of Max Healthcare | CSIR Institute of Genomics and Integrative Biology (CSIR-IGIB) / Max | Rajesh Pandey#, Samreen Siddiqui, Pooja Sharma, Bansidhar Tarai, Vivekanand A, Bharathram Uppili, Saruchi Wadhwa, Nishu Tyagi, Mitali Mukerji, Poonam Das, Sujeet Jha, Mohammed Faruq, Vinitaa Jha, Anurag Agrawal |
| EPI_ISL_641533 | CHU de Nice - Hôpital Archet 15 | CNR Virus des Infections Respiratoires - France SUD | Antonin Bal, Géraldine Gonfrier, Gregory Destras, Gwendolyne Burfin, Hadrien Règue, Quentin Semanas, Martine Valette, Bruno Lina, Valérie Giordanengo, Laurence Josset |
| EPI_ISL_644211, EPI_ISL_644216 | CEPHR / Vincent's Hospital | Irish Coronavirus Sequencing Consortium - National Virus Reference Laboratory | Michael Carr, Gabriel Gonzalez, Alejandro Abner Garcia Leon, Patrick Mallon |
| EPI_ISL_644229 | CEPHR / Mater Hospital | Irish Coronavirus Sequencing Consortium - National Virus Reference Laboratory | Michael Carr, Gabriel Gonzalez, Alejandro Abner Garcia Leon, Patrick Mallon |
| EPI_ISL_644233, EPI_ISL_644237 | CEPHR / Vincent's Hospital | Irish Coronavirus Sequencing Consortium - National Virus Reference Laboratory | Michael Carr, Gabriel Gonzalez, Alejandro Abner Garcia Leon, Patrick Mallon |
| EPI_ISL_644250, EPI_ISL_644251, EPI_ISL_644253, EPI_ISL_644254 | CEPHR / Mater Hospital | Irish Coronavirus Sequencing Consortium - National Virus Reference Laboratory | Michael Carr, Gabriel Gonzalez, Alejandro Abner Garcia Leon, Patrick Mallon |
| EPI_ISL_644277, EPI_ISL_644280, EPI_ISL_644284, EPI_ISL_644297, EPI_ISL_644299, EPI_ISL_644301, EPI_ISL_644302, EPI_ISL_644315, EPI_ISL_644317, EPI_ISL_644329, EPI_ISL_644336, EPI_ISL_644338, EPI_ISL_644341, EPI_ISL_644344 | | | |
| see above | CEPHR / Vincent's Hospital | Irish Coronavirus Sequencing Consortium - National Virus Reference Laboratory | Michael Carr, Gabriel Gonzalez, Alejandro Abner Garcia Leon, Patrick Mallon |
| EPI_ISL_644680 | CHU de Limoges | CNR Virus des Infections Respiratoires - France SUD | Antonin Bal, Gregory Destras, Gwendolyne Burfin, Hadrien Règue, Quentin Semanas, Martine Valette, Bruno Lina, Sylvie Rogez, Laurence Josset |
| EPI_ISL_644696 | CHU Montpellier | CNR Virus des Infections Respiratoires - France SUD | Antonin Bal, Gregory Destras, Gwendolyne Burfin, Hadrien Règue, Quentin Semanas, Martine Valette, Bruno Lina, Michel Segondy, Vincent Foulongne, Laurence Josset |
| EPI_ISL_644707 | Unité des Virus Émergents | CNR Virus des Infections Respiratoires - France SUD | Antonin Bal, Gregory Destras, Gwendolyne Burfin, Hadrien Règue, Quentin Semanas, Martine Valette, Bruno Lina, Laetitia Ninove, Léa Luciani, Antoine Nougairède, Laurence Josset |

| | | | |
|---|---|---|---|
| EPI_ISL_644738, EPI_ISL_644740, EPI_ISL_644751, EPI_ISL_644752, EPI_ISL_644756, EPI_ISL_644758, EPI_ISL_644762, EPI_ISL_644763, EPI_ISL_644765, EPI_ISL_644770, EPI_ISL_644780, EPI_ISL_644791, EPI_ISL_644792, EPI_ISL_644793, EPI_ISL_644794, EPI_ISL_644795, EPI_ISL_644796, EPI_ISL_644797, EPI_ISL_644798, EPI_ISL_644800, EPI_ISL_644801, EPI_ISL_644802, EPI_ISL_644803, EPI_ISL_644804, EPI_ISL_644805, EPI_ISL_644806, EPI_ISL_644807, EPI_ISL_644808, EPI_ISL_644809, EPI_ISL_644810, EPI_ISL_644811 | | | |
| see above | National Microbiology Reference Laboratory | Quadram Institute Bioscience | Thanh Le Viet, Andrew J. Page, Justin O'Grady, Gemma Kay, David Baker, Gaetan Thilliez, Ana-Victoria Gutierrez, Robert Kingsley, Leonardo de Oliveira Martins, Sekesai Zinyowera, Tatenda Takawira, Muchaneta Mugabe, Gibson Mhlanga, Portia Manangazira, Andrew Tarupiwa, Hlanai Gumbo, Agnes Juru, Charles Nyagupe, Alexander Goredema, Isaac Phiri, Barbra Murwira, Beuty Makamure, Tapfumanei Mashe |
| EPI_ISL_645039, EPI_ISL_645040, EPI_ISL_645041, EPI_ISL_645075, EPI_ISL_645076, EPI_ISL_645101 | Human Genome Variation Research Group, Malopolska Centre of Biotechnology | Human Genome Variation Research Group, Malopolska Centre of Biotechnology | Kowalski,M., Pospiech,E., Klajmon,A., Gromowski,T., Pisarek,A., Marszalek,K., Kopera,K., Foremny,J., Swadzba,J., Sanak,M., Owczarek,K., Dabrowska,A., Szczepanski,A., Botwina,P., Labaj,P.P., Pyrc,K., Branicki,W. |
| EPI_ISL_645172 | CHU Bordeaux | CNR Virus des Infections Respiratoires - France SUD | Antonin Bal, Gregory Destras, Gwendolyne Burfin, Hadrien Règue, Quentin Semanas, Martine Valette, Bruno Lina, Pantxika Bellecave, Camille Ciccone, Isabelle Garrigue, Marie-Edith Lafon, Pascale Trimoulet, Laurence Josset |
| EPI_ISL_645185 | CHU de Saint-Étienne Hôpital Nord | CNR Virus des Infections Respiratoires - France SUD | Antonin Bal, Gregory Destras, Gwendolyne Burfin, Hadrien Règue, Quentin Semanas, Martine Valette, Bruno Lina, Issam Bechri, Manon Vogrig, Marine Delorme, Bruno Pozzetto, Thomas Bourlet, Sylvie Gonzalo, Sylvie Pillet, Laurence Josset |
| EPI_ISL_647979, EPI_ISL_647980, EPI_ISL_647981, EPI_ISL_647982 | National Microbiology Reference Laboratory | Quadram Institute Bioscience | Thanh Le Viet, Andrew J. Page, Justin O'Grady, Gemma Kay, David Baker, Gaetan Thilliez, Ana-Victoria Gutierrez, Robert Kingsley, Leonardo de Oliveira Martins, Sekesai Zinyowera, Tatenda Takawira, Muchaneta Mugabe, Gibson Mhlanga, Portia Manangazira, Andrew Tarupiwa, Hlanai Gumbo, Agnes Juru, Charles Nyagupe, Alexander Goredema, Isaac Phiri, Barbra Murwira, Beuty Makamure, Tapfumanei Mashe |
| EPI_ISL_648334, EPI_ISL_648335, EPI_ISL_648336, EPI_ISL_648340, EPI_ISL_648341, EPI_ISL_648342, EPI_ISL_648343 | Laboratorio de Investigaciones de Baney | University Hospital Basel, Clinical Bacteriology | Carlos Cortes, Claudia Daubenberger, Adrian Egli, Guillermo Garcia, Salome Hosch, Bonifacio Manguire Nlavo, Alfredo Mari, Maximilian Mpina, Elizabeth Nyakarungu, Diosdado Odjama Nseng Ada, Mitoha Ondo O Ayekaba, Tim Roloff, Tobias Schindler, Helena Seth-Smith, Madlen Stange, Philip Wonder Phiri |
| EPI_ISL_648600 | Laboratorio de Infectología Servicio de Infectología Hospital Universitario Dr. José Eleuterio González - Universidad Autónoma de Nuevo León | Laboratorio de Infectología Molecular Departamento de Bioquímica y Medicina Molecular Facultad de Medicina - Universidad Autónoma de Nuevo León | Kame A. Galán-Huerta, María F. Herrera-Saldivar, Natalia Martínez-Acuña, Sonia A. Lozano-Sepúlveda, Daniel Arellanos-Soto, Ana M. Rivas-Estilla, Paola Bocanegra-Ibarias, Samantha M. Flores-Treviño, Elvira Garza-González, Eduardo Perez-Alba, Laura Nuzzolo-Shihadeh, Adrian Camacho-Ortiz, Roberto Montes-de-Oca, Consuelo Treviño-Garza, Manuel E. de-la-O-Cavazos |
| EPI_ISL_648654, EPI_ISL_648655, EPI_ISL_648656, EPI_ISL_648657, EPI_ISL_648658, EPI_ISL_648659, EPI_ISL_648660, EPI_ISL_648661, EPI_ISL_648662, EPI_ISL_648663, EPI_ISL_648664, EPI_ISL_648665, EPI_ISL_648666, EPI_ISL_648667, EPI_ISL_648668, EPI_ISL_648669, EPI_ISL_648670, EPI_ISL_648671, EPI_ISL_648672, EPI_ISL_648673, EPI_ISL_648674, EPI_ISL_648675, EPI_ISL_648676, EPI_ISL_648743, EPI_ISL_648744, EPI_ISL_648745, EPI_ISL_648746, EPI_ISL_648747 | | | |
| see above | Department of Laboratory Medicine, Tan Tock Seng Hospital | Department of Laboratory Medicine, Tan Tock Seng Hospital | Chen YYC, Zair X, Lim JX, Li C, Tang WY, Maurer-Stroh S, Barkham TMS, Nagarajan N, Sessions OM |
| EPI_ISL_649971 | CHU Bordeaux | CNR Virus des Infections Respiratoires - France SUD | Antonin Bal, Gregory Destras, Gwendolyne Burfin, Hadrien Règue, Quentin Semanas, Martine Valette, Bruno Lina, Pantxika Bellecave, Camille Ciccone, Isabelle Garrigue, Marie-Edith Lafon, Pascale Trimoulet, Laurence Josset |
| EPI_ISL_653150 | Florida Bureau of Public Health Laboratories | Florida Bureau of Public Health Laboratories | Sarah Schmedes, Jason Blanton |
| EPI_ISL_653869 | Translational Health Science and Technology Institute | National Institute of Biomedical Genomics | Arindam Maitra, Guruprasad Medigeshi, Sharanabasava Patil, Anbalagan Ananthraj, Madhu Pareek, Imran Khan, Gagandeep Kang, Saumitra Das |
| EPI_ISL_654681 | Servicio de Microbiología, Hospital Universitario Central de Asturias | SeqCOVID-SPAIN consortium/IBV(CSIC) | Cristián Castelló Abietar, Jose A. Boga, Susana Rojo-Alba, Marta Elena Álvarez-Argüelles, Santiago Melón and SeqCOVID-SPAIN consortium |
| EPI_ISL_654695 | Minnesota Department of Health, Public Health Laboratory | Minnesota Department of Health, Public Health Laboratory | Matt Plumb, Jacob Garfin, Alexandra Lorentz, and Xiong Wang |
| EPI_ISL_654735, EPI_ISL_654747, EPI_ISL_654768, EPI_ISL_654769, EPI_ISL_654938, EPI_ISL_654939, EPI_ISL_654940, EPI_ISL_654941 | Texas Department of State Health Services | Texas Department of State Health Services | Rashmi Tuladhar, Bonnie Oh, Jenny Zhang, Maliha Rahman, Anita Pokharel, Myong Koag, Chung Wang, Rachel Lee, Grace Kubin, Mayela Pedrueza, James Daniel Bonser |
| EPI_ISL_660350 | CHU de Saint-Étienne Hôpital Nord | CNR Virus des Infections Respiratoires - France SUD | Antonin Bal, Gregory Destras, Gwendolyne Burfin, Hadrien Règue, Quentin Semanas, Martine Valette, Bruno Lina, Issam Bechri, Manon Vogrig, Marine Delorme, Bruno Pozzetto, Thomas Bourlet, Sylvie Gonzalo, Sylvie Pillet, Laurence Josset |
| EPI_ISL_660368 | CHU Clermont-Ferrand | CNR Virus des Infections Respiratoires - France SUD | Antonin Bal, Gregory Destras, Gwendolyne Burfin, Hadrien Règue, Quentin Semanas, Martine Valette, Bruno Lina, Christine Archimbaud, Amélie Brebion, Hélène Chabrolles, Martine Chambon, Audrey Mirand, Christel Regagnon, Maxime Bisseux, Patricia Combes, Cécile Henquell, Laurence Josset |
| EPI_ISL_660372 | CHU Bordeaux | CNR Virus des Infections Respiratoires - France SUD | Antonin Bal, Gregory Destras, Gwendolyne Burfin, Hadrien Règue, Quentin Semanas, Martine Valette, Bruno Lina, Pantxika Bellecave, Camille Ciccone, Isabelle Garrigue, Marie-Edith Lafon, Pascale Trimoulet, Laurence Josset |
| EPI_ISL_660703, EPI_ISL_660704, EPI_ISL_660705, EPI_ISL_660706, EPI_ISL_660707 | CHU Montpellier | CNR Virus des Infections Respiratoires - France SUD | Antonin Bal, Gregory Destras, Gwendolyne Burfin, Hadrien Règue, Quentin Semanas, Martine Valette, Bruno Lina, Michel Segondy, Vincent Foulongne, Laurence Josset |
| EPI_ISL_660732 | Unité des Virus Émergents | CNR Virus des Infections Respiratoires - France SUD | Antonin Bal, Gregory Destras, Gwendolyne Burfin, Hadrien Règue, Quentin Semanas, Martine Valette, Bruno Lina, Laetitia Ninove, Léa Luciani, Antoine Nougairède, Laurence Josset |
| EPI_ISL_663239 | CHU Poitiers | CNR Virus des Infections Respiratoires - France SUD | Antonin Bal, Gregory Destras, Gwendolyne Burfin, Hadrien Règue, Quentin Semanas, Martine Valette, Bruno Lina, Agnès Beby-Defaux, Magali Garcia, Clément Jousselin, Nicolas Lévêque, Laurence Josset |
| EPI_ISL_665255, EPI_ISL_666603 | Dept. of Microbiology and Infection Control, Akershus University Hospital HF | Dept. of Microbiology and Infection Control, Akershus University Hospital HF | Hege Vangstein Aamot, Alexander Hesselberg Løvestad, Silje Bakken Jørgensen, Nina Handal, Ole Herman Ambur |
| EPI_ISL_666680, EPI_ISL_666681 | Laboratoire du Centre Hospitalier Annecy Genevois | CNR Virus des Infections Respiratoires - France SUD | Antonin Bal, Gregory Destras, Gwendolyne Burfin, Hadrien Règue, Quentin Semanas, Martine Valette, Bruno Lina, Hélène Petitprez, Bruno Chanzy, Laurence Josset |
| EPI_ISL_666718, EPI_ISL_666719, EPI_ISL_666720 | CHU de Limoges | CNR Virus des Infections Respiratoires - France SUD | Antonin Bal, Gregory Destras, Gwendolyne Burfin, Hadrien Règue, Quentin Semanas, Martine Valette, Bruno Lina, Sylvie Rogez, Laurence Josset |
| EPI_ISL_666724 | CHU Nantes | CNR Virus des Infections Respiratoires - France SUD | Antonin Bal, Louise Castain, Gregory Destras, Gwendolyne Burfin, Hadrien Règue, Quentin Semanas, Martine Valette, Bruno Lina, Virginie Ferré, Celine Bressollette, Laurence Josset |
| EPI_ISL_666820 | Florida Bureau of Public Health Laboratories | Florida Bureau of Public Health Laboratories | Sarah Schmedes, Jason Blanton |
| EPI_ISL_667776 | Texas Department of State Health Services | Texas Department of State Health Services | Pedrueza,M., Oh,B., Zhang,J., Rahman,M., Pokharel,A., Tuladhar,R., Koag,M., Wang,C., Lee,R., Kubin,G. and Bonser,J. |
| EPI_ISL_671337, EPI_ISL_671338, EPI_ISL_671339, EPI_ISL_671340, EPI_ISL_671343, EPI_ISL_671344, EPI_ISL_671345, EPI_ISL_671346, EPI_ISL_671355 | National Virus Reference Laboratory | Irish Coronavirus Sequencing Consortium - Teagasc Moorepark | Calm Walsh, Genuity Ireland |
| EPI_ISL_672050, EPI_ISL_672054, EPI_ISL_672060, EPI_ISL_672064 | Alameda County Public Health Lab | Chan-Zuckerberg Biohub | CZB Cliahub Consortium |
| EPI_ISL_672071, EPI_ISL_672105, EPI_ISL_672147, EPI_ISL_672148, EPI_ISL_672254, EPI_ISL_672255 | The Ashley Laboratory, Stanford University | Chan-Zuckerberg Biohub | CZB Cliahub Consortium |
| EPI_ISL_672336, EPI_ISL_672337, EPI_ISL_672338, EPI_ISL_672339, EPI_ISL_672340, EPI_ISL_672341, EPI_ISL_672439, EPI_ISL_672440, EPI_ISL_672441, EPI_ISL_672442, EPI_ISL_672443, EPI_ISL_672444, EPI_ISL_672445, EPI_ISL_672446, EPI_ISL_672447 | | | |
| see above | Alameda County Public Health Lab | Chan-Zuckerberg Biohub | CZB Cliahub Consortium |

| | | | |
|---|---|---|---|
| EPI_ISL_676521, EPI_ISL_676525 | Uppsala klinisk mikrobiologi | The Public Health Agency of Sweden | Department of Microbiology, The Public Health Agency of Sweden |
| EPI_ISL_676578, EPI_ISL_676600 | Scientific Veterinary Institute Novi Sad | Veterinary Specialized Institute "Kraljevo", Serbia | Vidanovic,D., Tesovic,B., Knezevic,A., Jovanovic,T., Jankovic,M., Sekler,M., Banovic Djeri,B., Petrovic,T., Volkening,J., Afonso,C. |
| EPI_ISL_676647 | Texas Department of State Health Services | Texas Department of State Health Services | Rashmi Tuladhar, Bonnie Oh, Jenny Zhang, Maliha Rahman, Anita Pokharel, Myong Koag, Chung Wang, Rachel Lee, Grace Kubin, Mayela Pedrueza, James Daniel Bonser |
| EPI_ISL_676664, EPI_ISL_676670, EPI_ISL_676678, EPI_ISL_676719, EPI_ISL_676885, EPI_ISL_676887, EPI_ISL_676888, EPI_ISL_676889, EPI_ISL_676890, EPI_ISL_676891, EPI_ISL_676894, EPI_ISL_676895, EPI_ISL_676896, EPI_ISL_676897, EPI_ISL_676898, EPI_ISL_676899, EPI_ISL_676900, EPI_ISL_676901, EPI_ISL_676902, EPI_ISL_676903, EPI_ISL_676904, EPI_ISL_676905, EPI_ISL_676906 | | | |
| see above | Wadsworth Center, New York State Department.of Health | Wadsworth Center, New York State Department.of Health | Kirsten St. George, Daryl M. Lamson, Alexis Russel, Jonathan Plitnick, Navjot Singh, John Kelly, Sara Griesemer, Erasmus Schneider, Erica Lasek-Nesselquist |
| EPI_ISL_677684, EPI_ISL_677692 | Vanda Pharmaceuticals Clinical Site | Vanda Pharmaceuticals | Vanda Pharmaceuticals |
| EPI_ISL_677941, EPI_ISL_677942 | Pathogen Genomics Lab King Abdullah University of Science and Technology(KAUST) | Pathogen Genomics Lab King Abdullah University of Science and Technology(KAUST) | Sara Mfarrej, Amanda Ooi, Luke Esau, Sharif Hala, Raeece Naeem, Afrah Alsomali, Fadwa Alofi, Asim Khogeer, Jumana Taha, Abdulaziz Alahmadi, Kahled Algithami, Anwar Hashem, Naif Almontashiri, Arnab Pain |
| EPI_ISL_678033 | Pathogen Genomics Lab King Abdullah University of Science and Technology(KAUST) | Pathogen Genomics Lab King Abdullah University of Science and Technology(KAUST) | Sara Mfarrej, Olga Douvropoulou, Raushan Nugmanova, Sharif Hala, Raeece Naeem, Amanda Ooi, Luke Esau, Fadwa Alofi, Afrah Alsomali, Asim Khogeer, Jumana Taha, Abdulaziz Alahmadi, Kahled Algithami, Anwar Hashem, Naif Almontashiri, Arnab Pain |
| EPI_ISL_678044 | Pathogen Genomics Lab King Abdullah University of Science and Technology(KAUST) | Pathogen Genomics Lab King Abdullah University of Science and Technology(KAUST) | Sara Mfarrej, Luke Esau, Amanda Ooi, Sharif Hala, Raeece Naeem, Asim Khogeer, Fadwa Alofi, Afrah Alsomali, Jumana Taha, Abdulaziz Alahmadi, Kahled Algithami, Anwar Hashem, Naif Almontashiri, Arnab Pain |
| EPI_ISL_678277, EPI_ISL_678280, EPI_ISL_678283 | Mikrobiologie, RARI | Mikrobiologie, RARI | Krasnov,Y.M., Naryshkina,E.A., Guseva,N.P., Sosedova,E.A., Fedorov,A.V., Badanin,D.V., Sharapova,N.A., Portenko,S.A., Shcherbakova,S.A., Kutyrev,V.V. |
| EPI_ISL_678488, EPI_ISL_678490 | Veterinary Specialized Institute "Sabac", Serbia | Veterinary Specialized Institute "Kraljevo", Serbia | Vidanovic,D., Tesovic,B., Knezevic,A., Jovanovic,T., Jankovic,M., Sekler,M., Banovic Djeri,B., Petrovic,T., Mrkovacki, S., Volkening,J., Afonso,C. |
| EPI_ISL_678537, EPI_ISL_678538, EPI_ISL_678539, EPI_ISL_678540, EPI_ISL_678541, EPI_ISL_678542 | CNR Virus des Infections Respiratoires - France SUD | CNR Virus des Infections Respiratoires - France SUD | Antonin Bal, Gregory Destras, Gwendolyne Burfin, Solenne Brun, Martine Valette, Bruno Lina, Laurence Josset |
| EPI_ISL_681679, EPI_ISL_681680, EPI_ISL_681681, EPI_ISL_681682, EPI_ISL_681683 | Molecular Medicine Laboratory, University of Magallanes | Centro Asistencial Docente y de Investigacion, Universidad de Magallanes | Jorge González, Jacqueline Aldridge, Diego Alvarez, Marco Montes de Oca, Hermy Alvarez, Roberto Uribe-Paredes, Marcelo Navarrete |
| EPI_ISL_681703 | Zurita & Zurita Laboratorios | Zurita & Zurita Laboratorios | Gabriela Sevillano, Camilo Zurita-Salinas, Jeannete Zurita |
| EPI_ISL_681714, EPI_ISL_681715 | University Hospital Limerick | Irish Coronavirus Sequencing Consortium - Teagasc Moorepark | Paul Cotter, Fiona Crispie, Amy Fitzpatrick, John Kenny, Carolyn Meaney, Patrick Stapleton, Calum Walsh |
| EPI_ISL_681716, EPI_ISL_681717, EPI_ISL_681742, EPI_ISL_681743 | University Hospital Limerick | Irish Coronavirus Sequencing Consortium - Teagasc Moorepark | Paul Cotter, Fiona Crispie, Amy Fitzpatrick, John Kenny, Elaine Lawton, Carolyn Meaney, Patrick Stapleton, Calum Walsh |
| EPI_ISL_681745 | University Hospital Limerick | Irish Coronavirus Sequencing Consortium - Teagasc Moorepark | Paul Cotter, Fiona Crispie, Amy Fitzpatrick, John Kenny, Carolyn Meaney, Patrick Stapleton, Calum Walsh |
| EPI_ISL_681746 | University Hospital Limerick | Irish Coronavirus Sequencing Consortium - Teagasc Moorepark | Paul Cotter, Fiona Crispie, Amy Fitzpatrick, John Kenny, Elaine Lawton, Carolyn Meaney, Patrick Stapleton, Calum Walsh |
| EPI_ISL_681747, EPI_ISL_681748 | University Hospital Limerick | Irish Coronavirus Sequencing Consortium - Teagasc Moorepark | Paul Cotter, Fiona Crispie, Amy Fitzpatrick, John Kenny, Carolyn Meaney, Patrick Stapleton, Calum Walsh |
| EPI_ISL_683596 | Hospital Clínico Universitario Lozano Blesa de Zaragoza (España) | SeqCOVID-SPAIN consortium/IBV(CSIC) | Rafael Benito, Sonia Algarate, Jessica Bueno and SeqCOVID-SPAIN consortium |
| EPI_ISL_685281 | Gunma Prefectural Institute of Public Health and Environmental Sciences | Pathogen Genomics Center, National Institute of Infectious Diseases | Tsuyoshi Sekizuka, Kentaro Itokawa, Rina Tanaka, Masanori Hashino, Makoto Kuroda |
| EPI_ISL_685283, EPI_ISL_685284, EPI_ISL_685285, EPI_ISL_685286, EPI_ISL_685287, EPI_ISL_685288, EPI_ISL_685289, EPI_ISL_685290, EPI_ISL_685291, EPI_ISL_685292, EPI_ISL_685293, EPI_ISL_685294, EPI_ISL_685295, EPI_ISL_685296, EPI_ISL_685311, EPI_ISL_685312, EPI_ISL_685344, EPI_ISL_685345, EPI_ISL_685346, EPI_ISL_685347, EPI_ISL_685348, EPI_ISL_685349, EPI_ISL_685350, EPI_ISL_685351, EPI_ISL_685352, EPI_ISL_685353, EPI_ISL_685390, EPI_ISL_685391, EPI_ISL_685392, EPI_ISL_685393, EPI_ISL_685394, EPI_ISL_685395, EPI_ISL_685396, EPI_ISL_685397, EPI_ISL_685398, EPI_ISL_685399, EPI_ISL_685400, EPI_ISL_685401, EPI_ISL_685402, EPI_ISL_685403, EPI_ISL_685404, EPI_ISL_685405, EPI_ISL_685406 | | | |
| see above | Pathogen Genomics Center, National Institute of Infectious Diseases | Pathogen Genomics Center, National Institute of Infectious Diseases | Tsuyoshi Sekizuka, Kentaro Itokawa, Rina Tanaka, Masanori Hashino, Makoto Kuroda |
| EPI_ISL_685454, EPI_ISL_685455, EPI_ISL_685456, EPI_ISL_685457, EPI_ISL_685458, EPI_ISL_685459, EPI_ISL_685460, EPI_ISL_685461, EPI_ISL_685462, EPI_ISL_685463, EPI_ISL_685464, EPI_ISL_685465, EPI_ISL_685466, EPI_ISL_685467, EPI_ISL_685468, EPI_ISL_685469, EPI_ISL_685470, EPI_ISL_685471, EPI_ISL_685472, EPI_ISL_685473, EPI_ISL_685474, EPI_ISL_685475, EPI_ISL_685476, EPI_ISL_685477, EPI_ISL_685478, EPI_ISL_685479, EPI_ISL_685480, EPI_ISL_685481, EPI_ISL_685482, EPI_ISL_685483, EPI_ISL_685484, EPI_ISL_685485, EPI_ISL_685486, EPI_ISL_685487, EPI_ISL_685488, EPI_ISL_685489, EPI_ISL_685490, EPI_ISL_685491, EPI_ISL_685492, EPI_ISL_685493, EPI_ISL_685494, EPI_ISL_685495, EPI_ISL_685496, EPI_ISL_685497 | | | |
| see above | Tokyo Metropolitan Institute of Public Health | Pathogen Genomics Center, National Institute of Infectious Diseases | Tsuyoshi Sekizuka, Kentaro Itokawa, Rina Tanaka, Masanori Hashino, Makoto Kuroda |
| EPI_ISL_685502 | Yamagata Prefectural Institute of Public Health | Pathogen Genomics Center, National Institute of Infectious Diseases | Tsuyoshi Sekizuka, Kentaro Itokawa, Rina Tanaka, Masanori Hashino, Makoto Kuroda |
| EPI_ISL_685505, EPI_ISL_685521, EPI_ISL_685522, EPI_ISL_685523, EPI_ISL_685574, EPI_ISL_685575, EPI_ISL_685576, EPI_ISL_685577, EPI_ISL_685578, EPI_ISL_685579, EPI_ISL_685580, EPI_ISL_685581, EPI_ISL_685582, EPI_ISL_685583, EPI_ISL_685604, EPI_ISL_685605, EPI_ISL_685638, EPI_ISL_685639, EPI_ISL_685640, EPI_ISL_685641, EPI_ISL_685642, EPI_ISL_685643, EPI_ISL_685644, EPI_ISL_685645, EPI_ISL_685646, EPI_ISL_685712, EPI_ISL_685713, EPI_ISL_685714, EPI_ISL_685715, EPI_ISL_685716, EPI_ISL_685717, EPI_ISL_685718, EPI_ISL_685719, EPI_ISL_685720, EPI_ISL_685721, EPI_ISL_685722, EPI_ISL_685723, EPI_ISL_685724, EPI_ISL_685725, EPI_ISL_685726, EPI_ISL_685727, EPI_ISL_685728, EPI_ISL_685729, EPI_ISL_685730, EPI_ISL_685731, EPI_ISL_685732, EPI_ISL_685733, EPI_ISL_685744, EPI_ISL_685745, EPI_ISL_685775, EPI_ISL_685776, EPI_ISL_685777, EPI_ISL_685778, EPI_ISL_685779, EPI_ISL_685904, EPI_ISL_685909, EPI_ISL_685910, EPI_ISL_685911, EPI_ISL_685912, EPI_ISL_685913, EPI_ISL_685914, EPI_ISL_685915, EPI_ISL_685916, EPI_ISL_685917 | | | |
| see above | Pathogen Genomics Center, National Institute of Infectious Diseases | Pathogen Genomics Center, National Institute of Infectious Diseases | Tsuyoshi Sekizuka, Kentaro Itokawa, Rina Tanaka, Masanori Hashino, Makoto Kuroda |
| EPI_ISL_685922, EPI_ISL_685923, EPI_ISL_685924 | Fukuoka Institute of Health and Environmental Sciences | Pathogen Genomics Center, National Institute of Infectious Diseases | Tsuyoshi Sekizuka, Kentaro Itokawa, Rina Tanaka, Masanori Hashino, Makoto Kuroda |
| EPI_ISL_685928, EPI_ISL_685937, EPI_ISL_685938, EPI_ISL_685939, EPI_ISL_685940, EPI_ISL_685941, EPI_ISL_685942, EPI_ISL_685943, EPI_ISL_685944, EPI_ISL_685945, EPI_ISL_685946, EPI_ISL_685948, EPI_ISL_685949, EPI_ISL_685950, EPI_ISL_685951, EPI_ISL_685952, EPI_ISL_685956 | | | |
| see above | Pathogen Genomics Center, National Institute of Infectious Diseases | Pathogen Genomics Center, National Institute of Infectious Diseases | Tsuyoshi Sekizuka, Kentaro Itokawa, Rina Tanaka, Masanori Hashino, Makoto Kuroda |
| EPI_ISL_685963, EPI_ISL_685964, EPI_ISL_685965, EPI_ISL_685966, EPI_ISL_685967, EPI_ISL_685968, EPI_ISL_685969, EPI_ISL_685970, EPI_ISL_685971, EPI_ISL_685972, EPI_ISL_685973, EPI_ISL_685974, EPI_ISL_685975, EPI_ISL_685976, EPI_ISL_685977, EPI_ISL_685978, EPI_ISL_685979, EPI_ISL_685980, EPI_ISL_685981, EPI_ISL_685982, EPI_ISL_685983, EPI_ISL_685984, EPI_ISL_685985, EPI_ISL_685986 | | | |
| see above | Tokyo Metropolitan Institute of Public Health | Pathogen Genomics Center, National Institute of Infectious Diseases | Tsuyoshi Sekizuka, Kentaro Itokawa, Rina Tanaka, Masanori Hashino, Makoto Kuroda |
| EPI_ISL_686074, EPI_ISL_686075, EPI_ISL_686076, EPI_ISL_686077, EPI_ISL_686078, EPI_ISL_686079, EPI_ISL_686080, EPI_ISL_686081, EPI_ISL_686082 | Pathogen Genomics Center, National Institute of Infectious Diseases | Pathogen Genomics Center, National Institute of Infectious Diseases | Tsuyoshi Sekizuka, Kentaro Itokawa, Rina Tanaka, Masanori Hashino, Makoto Kuroda |

| | | | |
|---|---|---|---|
| EPI_ISL_687043, EPI_ISL_687044, EPI_ISL_687045, EPI_ISL_687046, EPI_ISL_687047, EPI_ISL_687048, EPI_ISL_687049, EPI_ISL_687050, EPI_ISL_687051, EPI_ISL_687052, EPI_ISL_687053, EPI_ISL_687054, EPI_ISL_687055, EPI_ISL_687056, EPI_ISL_687057, EPI_ISL_687058, EPI_ISL_687059, EPI_ISL_687060, EPI_ISL_687061, EPI_ISL_687062, EPI_ISL_687063, EPI_ISL_687064, EPI_ISL_687065, EPI_ISL_687066, EPI_ISL_687067, EPI_ISL_687068, EPI_ISL_687069, EPI_ISL_687070, EPI_ISL_687071, EPI_ISL_687072, EPI_ISL_687073 | | | |
| see above | Kanagawa Prefectural Institute of Public Health | Pathogen Genomics Center, National Institute of Infectious Diseases | Tsuyoshi Sekizuka, Kentaro Itokawa, Rina Tanaka, Masanori Hashino, Makoto Kuroda |
| EPI_ISL_687923, EPI_ISL_687931, EPI_ISL_688812, EPI_ISL_688813, EPI_ISL_688814 | Pathogen Genomics Center, National Institute of Infectious Diseases | Pathogen Genomics Center, National Institute of Infectious Diseases | Tsuyoshi Sekizuka, Kentaro Itokawa, Rina Tanaka, Masanori Hashino, Makoto Kuroda |
| EPI_ISL_690619 | Saitama Prefectural Institute of Public Health | Pathogen Genomics Center, National Institute of Infectious Diseases | Tsuyoshi Sekizuka, Kentaro Itokawa, Rina Tanaka, Masanori Hashino, Makoto Kuroda |
| EPI_ISL_690624, EPI_ISL_690634 | Pathogen Genomics Center, National Institute of Infectious Diseases | Pathogen Genomics Center, National Institute of Infectious Diseases | Tsuyoshi Sekizuka, Kentaro Itokawa, Rina Tanaka, Masanori Hashino, Makoto Kuroda |
| EPI_ISL_690656, EPI_ISL_690658, EPI_ISL_690659, EPI_ISL_690660, EPI_ISL_690661, EPI_ISL_690662, EPI_ISL_690663, EPI_ISL_690664, EPI_ISL_690665, EPI_ISL_690666, EPI_ISL_690667, EPI_ISL_690668, EPI_ISL_690669, EPI_ISL_690670, EPI_ISL_690671, EPI_ISL_690672, EPI_ISL_690673, EPI_ISL_690674, EPI_ISL_690675, EPI_ISL_690676, EPI_ISL_690677, EPI_ISL_690678, EPI_ISL_690679, EPI_ISL_690680, EPI_ISL_690681, EPI_ISL_690682, EPI_ISL_690683 | | | |
| see above | Saitama Prefectural Institute of Public Health | Pathogen Genomics Center, National Institute of Infectious Diseases | Tsuyoshi Sekizuka, Kentaro Itokawa, Rina Tanaka, Masanori Hashino, Makoto Kuroda |
| EPI_ISL_690687, EPI_ISL_690688, EPI_ISL_690741, EPI_ISL_690742, EPI_ISL_690743 | Pathogen Genomics Center, National Institute of Infectious Diseases | Pathogen Genomics Center, National Institute of Infectious Diseases | Tsuyoshi Sekizuka, Kentaro Itokawa, Rina Tanaka, Masanori Hashino, Makoto Kuroda |
| EPI_ISL_690835, EPI_ISL_690836 | Kanagawa Prefectural Institute of Public Health | Pathogen Genomics Center, National Institute of Infectious Diseases | Tsuyoshi Sekizuka, Kentaro Itokawa, Rina Tanaka, Masanori Hashino, Makoto Kuroda |
| EPI_ISL_690842, EPI_ISL_690843 | Sakai City Institute of Public Health | Pathogen Genomics Center, National Institute of Infectious Diseases | Tsuyoshi Sekizuka, Kentaro Itokawa, Rina Tanaka, Masanori Hashino, Makoto Kuroda |
| EPI_ISL_690973, EPI_ISL_690974, EPI_ISL_690975, EPI_ISL_690976, EPI_ISL_690977, EPI_ISL_691012, EPI_ISL_691013, EPI_ISL_691014, EPI_ISL_691105, EPI_ISL_691106, EPI_ISL_691107, EPI_ISL_691108, EPI_ISL_691226, EPI_ISL_691227, EPI_ISL_691232, EPI_ISL_691233, EPI_ISL_691234, EPI_ISL_691235, EPI_ISL_691236, EPI_ISL_691237, EPI_ISL_691238, EPI_ISL_691239, EPI_ISL_691240, EPI_ISL_691241, EPI_ISL_691248, EPI_ISL_691259, EPI_ISL_691273, EPI_ISL_691274, EPI_ISL_691500, EPI_ISL_691501, EPI_ISL_691502, EPI_ISL_691503, EPI_ISL_691504, EPI_ISL_691505, EPI_ISL_691506, EPI_ISL_691861, EPI_ISL_692044, EPI_ISL_692045, EPI_ISL_692046, EPI_ISL_692059, EPI_ISL_692060, EPI_ISL_692527, EPI_ISL_692597, EPI_ISL_692598, EPI_ISL_692599 | | | |
| see above | Pathogen Genomics Center, National Institute of Infectious Diseases | Pathogen Genomics Center, National Institute of Infectious Diseases | Tsuyoshi Sekizuka, Kentaro Itokawa, Rina Tanaka, Masanori Hashino, Makoto Kuroda |
| EPI_ISL_692832, EPI_ISL_692833, EPI_ISL_693062, EPI_ISL_693063, EPI_ISL_693064, EPI_ISL_693065, EPI_ISL_693066, EPI_ISL_693067, EPI_ISL_693068, EPI_ISL_693069, EPI_ISL_693070, EPI_ISL_693071, EPI_ISL_693072, EPI_ISL_693073, EPI_ISL_693074, EPI_ISL_693075, EPI_ISL_693076, EPI_ISL_693077, EPI_ISL_693078, EPI_ISL_693079, EPI_ISL_693080, EPI_ISL_693081, EPI_ISL_693082, EPI_ISL_693083, EPI_ISL_693084, EPI_ISL_693085, EPI_ISL_693086, EPI_ISL_693087, EPI_ISL_693088, EPI_ISL_693089, EPI_ISL_693090, EPI_ISL_693091, EPI_ISL_693092, EPI_ISL_693093, EPI_ISL_693094, EPI_ISL_693095, EPI_ISL_693096, EPI_ISL_693097, EPI_ISL_693098, EPI_ISL_693099, EPI_ISL_693100, EPI_ISL_693101, EPI_ISL_693102, EPI_ISL_693103, EPI_ISL_693104, EPI_ISL_693105, EPI_ISL_693106, EPI_ISL_693107, EPI_ISL_693108, EPI_ISL_693109, EPI_ISL_693110, EPI_ISL_693111, EPI_ISL_693112, EPI_ISL_693113, EPI_ISL_693114, EPI_ISL_693116, EPI_ISL_693117, EPI_ISL_693118, EPI_ISL_693119, EPI_ISL_693120, EPI_ISL_693121, EPI_ISL_693122, EPI_ISL_693123, EPI_ISL_693124, EPI_ISL_693125, EPI_ISL_693126, EPI_ISL_693127, EPI_ISL_693128, EPI_ISL_693129, EPI_ISL_693130, EPI_ISL_693131, EPI_ISL_693132, EPI_ISL_693133, EPI_ISL_693134, EPI_ISL_693135, EPI_ISL_693151, EPI_ISL_693152, EPI_ISL_693178, EPI_ISL_693179, EPI_ISL_693180, EPI_ISL_693181, EPI_ISL_693182, EPI_ISL_693183, EPI_ISL_693184, EPI_ISL_693187, EPI_ISL_693188, EPI_ISL_693189, EPI_ISL_693190, EPI_ISL_693191 | | | |
| see above | Massachusetts State Public Health Laboratory | Massachusetts State Public Health Laboratory | Andrew Lang, Timelia Fink, Glen Gallagher, Sandra Smole |
| EPI_ISL_693196 | Hospital Santa Clara | Instituto Adolfo Lutz, Interdiciplinary Procedures Center, Strategic Laboratory | Claudio Tavares Sacchi, Claudia Regina Gonçalves, Erica Valessa Ramos Gomes, Karoline Rodrigues Campos |
| EPI_ISL_693197 | Central de Rede de Frio Municipal | Instituto Adolfo Lutz, Interdiciplinary Procedures Center, Strategic Laboratory | Claudio Tavares Sacchi, Claudia Regina Gonçalves, Erica Valessa Ramos Gomes, Karoline Rodrigues Campos |
| EPI_ISL_693198 | Santa Casa de Misericordia de Sao Paulo - Hospital Central | Instituto Adolfo Lutz, Interdiciplinary Procedures Center, Strategic Laboratory | Claudio Tavares Sacchi, Claudia Regina Gonçalves, Erica Valessa Ramos Gomes, Karoline Rodrigues Campos |
| EPI_ISL_693200 | Hospital e Maternidade Mairipora | Instituto Adolfo Lutz, Interdiciplinary Procedures Center, Strategic Laboratory | Claudio Tavares Sacchi, Claudia Regina Gonçalves, Erica Valessa Ramos Gomes, Karoline Rodrigues Campos |
| EPI_ISL_694140, EPI_ISL_694141, EPI_ISL_694142, EPI_ISL_694143, EPI_ISL_694144, EPI_ISL_694145, EPI_ISL_694146, EPI_ISL_694147, EPI_ISL_694219, EPI_ISL_694220, EPI_ISL_694221, EPI_ISL_694222, EPI_ISL_694223, EPI_ISL_694224, EPI_ISL_694225, EPI_ISL_694226, EPI_ISL_694227, EPI_ISL_694228, EPI_ISL_694229, EPI_ISL_694230, EPI_ISL_694231, EPI_ISL_694232, EPI_ISL_694233 | | | |
| see above | TGen North | TGen North | Jolene Bowers, Megan Folkerts, Chris French, Hayley Yaglom, Ashlyn Pfeiffer, Darrin Lemmer, Dave Engelthaler, The Arizona COVID Genomics Union (ACGU) |
| EPI_ISL_694259, EPI_ISL_694260, EPI_ISL_694261, EPI_ISL_694262, EPI_ISL_694263, EPI_ISL_694264, EPI_ISL_694265, EPI_ISL_694266, EPI_ISL_694267, EPI_ISL_694273, EPI_ISL_694274, EPI_ISL_694275, EPI_ISL_694276, EPI_ISL_694277, EPI_ISL_694278, EPI_ISL_694279, EPI_ISL_694280, EPI_ISL_694281, EPI_ISL_694282, EPI_ISL_694283, EPI_ISL_694284, EPI_ISL_694285, EPI_ISL_694286, EPI_ISL_694287, EPI_ISL_694288, EPI_ISL_694289, EPI_ISL_694290, EPI_ISL_694291, EPI_ISL_694292, EPI_ISL_694293, EPI_ISL_694294, EPI_ISL_694295, EPI_ISL_694296, EPI_ISL_694297, EPI_ISL_694298, EPI_ISL_694299, EPI_ISL_694300, EPI_ISL_694301, EPI_ISL_694302, EPI_ISL_694303, EPI_ISL_694304, EPI_ISL_694305, EPI_ISL_694306, EPI_ISL_694307, EPI_ISL_694308, EPI_ISL_694309, EPI_ISL_694310, EPI_ISL_694311, EPI_ISL_694312, EPI_ISL_694313, EPI_ISL_694314, EPI_ISL_694315 | | | |
| see above | AZ SPHL, Arizona Department of Health Services | TGen North | Jolene Bowers, Megan Folkerts, Chris French, Hayley Yaglom, Ashlyn Pfeiffer, Darrin Lemmer, Dave Engelthaler, The Arizona COVID Genomics Union (ACGU) |
| EPI_ISL_695755, EPI_ISL_695756, EPI_ISL_695757, EPI_ISL_695758, EPI_ISL_695759, EPI_ISL_695760, EPI_ISL_695761, EPI_ISL_695762, EPI_ISL_695763, EPI_ISL_695764, EPI_ISL_695765, EPI_ISL_695766, EPI_ISL_695767, EPI_ISL_695768, EPI_ISL_695769, EPI_ISL_695771, EPI_ISL_695772, EPI_ISL_695773, EPI_ISL_695774, EPI_ISL_695775, EPI_ISL_695776, EPI_ISL_695777, EPI_ISL_695778, EPI_ISL_695779, EPI_ISL_695780, EPI_ISL_695781, EPI_ISL_695782, EPI_ISL_695783, EPI_ISL_695784, EPI_ISL_695785, EPI_ISL_695787, EPI_ISL_695788, EPI_ISL_695789, EPI_ISL_695790, EPI_ISL_695791, EPI_ISL_695792, EPI_ISL_695793, EPI_ISL_695794, EPI_ISL_695795, EPI_ISL_695796, EPI_ISL_695797, EPI_ISL_695798, EPI_ISL_695799, EPI_ISL_695800, EPI_ISL_695801, EPI_ISL_695802, EPI_ISL_695803, EPI_ISL_695804, EPI_ISL_695805, EPI_ISL_695811, EPI_ISL_695816, EPI_ISL_695817, EPI_ISL_695822, EPI_ISL_695824, EPI_ISL_695825, EPI_ISL_695826, EPI_ISL_695827, EPI_ISL_695840, EPI_ISL_695842, EPI_ISL_695843, EPI_ISL_695844, EPI_ISL_695845, EPI_ISL_695846, EPI_ISL_695847, EPI_ISL_695848, EPI_ISL_695849, EPI_ISL_695850, EPI_ISL_695851, EPI_ISL_695852, EPI_ISL_695853, EPI_ISL_695854, EPI_ISL_695855, EPI_ISL_695856, EPI_ISL_695857, EPI_ISL_695858, EPI_ISL_695859, EPI_ISL_695860, EPI_ISL_695861, EPI_ISL_695862, EPI_ISL_695863, EPI_ISL_695864, EPI_ISL_695865, EPI_ISL_695866, EPI_ISL_695867, EPI_ISL_695868, EPI_ISL_695869, EPI_ISL_695870, EPI_ISL_695871, EPI_ISL_695872, EPI_ISL_695873, EPI_ISL_695874, EPI_ISL_695875, EPI_ISL_695877, EPI_ISL_695878, EPI_ISL_695879, EPI_ISL_695880, EPI_ISL_695881, EPI_ISL_695882, EPI_ISL_695883, EPI_ISL_695884, EPI_ISL_695885, EPI_ISL_695886, EPI_ISL_695887, EPI_ISL_695888, EPI_ISL_695889, EPI_ISL_695890, EPI_ISL_695891, EPI_ISL_695892, EPI_ISL_695902, EPI_ISL_695907, EPI_ISL_695912, EPI_ISL_695922, EPI_ISL_695928, EPI_ISL_695933, EPI_ISL_695973, EPI_ISL_695975, EPI_ISL_695994, EPI_ISL_695996, EPI_ISL_696018, EPI_ISL_696020, EPI_ISL_696025, EPI_ISL_696027, EPI_ISL_696028, EPI_ISL_696029, EPI_ISL_696030, EPI_ISL_696031, EPI_ISL_696032, EPI_ISL_696033, EPI_ISL_696034, EPI_ISL_696035, EPI_ISL_696036, EPI_ISL_696037, EPI_ISL_696038, EPI_ISL_696039, EPI_ISL_696040, EPI_ISL_696041, EPI_ISL_696042, EPI_ISL_696043, EPI_ISL_696044, EPI_ISL_696045, EPI_ISL_696046, EPI_ISL_696047, EPI_ISL_696048, EPI_ISL_696049, EPI_ISL_696050, EPI_ISL_696051, EPI_ISL_696052, EPI_ISL_696053, EPI_ISL_696054, EPI_ISL_696055, EPI_ISL_696056, EPI_ISL_696057, EPI_ISL_696058, EPI_ISL_696059, EPI_ISL_696060, EPI_ISL_696061, EPI_ISL_696062, EPI_ISL_696063, EPI_ISL_696064, EPI_ISL_696065, EPI_ISL_696066, EPI_ISL_696067, EPI_ISL_696068, EPI_ISL_696069, EPI_ISL_696070, EPI_ISL_696071, EPI_ISL_696072, EPI_ISL_696073, EPI_ISL_696074, EPI_ISL_696075, EPI_ISL_696076, EPI_ISL_696077, EPI_ISL_696078, EPI_ISL_696079, EPI_ISL_696080, EPI_ISL_696081, EPI_ISL_696082, EPI_ISL_696083, EPI_ISL_696084, EPI_ISL_696085, EPI_ISL_696086, EPI_ISL_696087, EPI_ISL_696088, EPI_ISL_696089, EPI_ISL_696090, EPI_ISL_696091, EPI_ISL_696092, EPI_ISL_696093, EPI_ISL_696094, EPI_ISL_696095, EPI_ISL_696096, EPI_ISL_696097, EPI_ISL_696098, EPI_ISL_696099, EPI_ISL_696100, EPI_ISL_696101, EPI_ISL_696107, EPI_ISL_696112, EPI_ISL_696113, EPI_ISL_696118, EPI_ISL_696120, EPI_ISL_696121, EPI_ISL_696132, EPI_ISL_696155, EPI_ISL_696157, EPI_ISL_696179, EPI_ISL_696180, EPI_ISL_696181, EPI_ISL_696182, EPI_ISL_696183, EPI_ISL_696184, EPI_ISL_696185, EPI_ISL_696186, EPI_ISL_696187, EPI_ISL_696188, EPI_ISL_696189, EPI_ISL_696190, EPI_ISL_696191, EPI_ISL_696193, EPI_ISL_696194, EPI_ISL_696195, EPI_ISL_696196, EPI_ISL_696198, EPI_ISL_696199, EPI_ISL_696200, EPI_ISL_696201, EPI_ISL_696202, EPI_ISL_696203, EPI_ISL_696204, EPI_ISL_696205, EPI_ISL_696206, EPI_ISL_696207, EPI_ISL_696209, EPI_ISL_696210, EPI_ISL_696211, EPI_ISL_696212, EPI_ISL_696213, EPI_ISL_696214, EPI_ISL_696215, EPI_ISL_696216, EPI_ISL_696217, EPI_ISL_696219, EPI_ISL_696220, EPI_ISL_696222, EPI_ISL_696224, EPI_ISL_696225, EPI_ISL_696227, EPI_ISL_696228, EPI_ISL_696229, EPI_ISL_696230, EPI_ISL_696231, EPI_ISL_696232, EPI_ISL_696233, EPI_ISL_696234, EPI_ISL_696235, EPI_ISL_696236, EPI_ISL_696237, EPI_ISL_696238, EPI_ISL_696239, EPI_ISL_696240, EPI_ISL_696241, EPI_ISL_696242, EPI_ISL_696243, EPI_ISL_696244, EPI_ISL_696245, EPI_ISL_696247, EPI_ISL_696248, EPI_ISL_696250, EPI_ISL_696251, EPI_ISL_696252, EPI_ISL_696253, EPI_ISL_696254, EPI_ISL_696255, EPI_ISL_696316, EPI_ISL_696317, EPI_ISL_696318, EPI_ISL_696319, EPI_ISL_696320, EPI_ISL_696321, EPI_ISL_696322, EPI_ISL_696323, EPI_ISL_696324, EPI_ISL_696325, EPI_ISL_696326, EPI_ISL_696327, EPI_ISL_696328, EPI_ISL_696329, EPI_ISL_696330, EPI_ISL_696331, EPI_ISL_696332, EPI_ISL_696333, EPI_ISL_696334, EPI_ISL_696335, EPI_ISL_696336, EPI_ISL_696337, EPI_ISL_696338, EPI_ISL_696339, EPI_ISL_696340, EPI_ISL_696341, EPI_ISL_696342, EPI_ISL_696343, EPI_ISL_696344, EPI_ISL_696345, EPI_ISL_696346, EPI_ISL_696347, EPI_ISL_696348, EPI_ISL_696349, EPI_ISL_696351, EPI_ISL_696352, EPI_ISL_696353, EPI_ISL_696354, EPI_ISL_696355, EPI_ISL_696356, EPI_ISL_696357, EPI_ISL_696358, EPI_ISL_696361, EPI_ISL_696363, EPI_ISL_696365, EPI_ISL_696366, EPI_ISL_696368, EPI_ISL_696369, EPI_ISL_696370, EPI_ISL_696371, EPI_ISL_696372, EPI_ISL_696374, EPI_ISL_696375, EPI_ISL_696376, EPI_ISL_696377, EPI_ISL_696378, EPI_ISL_696380, EPI_ISL_696381, EPI_ISL_696382, EPI_ISL_696383, EPI_ISL_696384, EPI_ISL_696385, EPI_ISL_696386, EPI_ISL_696387, EPI_ISL_696388, EPI_ISL_696389, EPI_ISL_696390, EPI_ISL_696391, EPI_ISL_696392, EPI_ISL_696393, EPI_ISL_696394, EPI_ISL_696395, EPI_ISL_696396, EPI_ISL_696397, EPI_ISL_696398, EPI_ISL_696399, EPI_ISL_696400, EPI_ISL_696401, EPI_ISL_696402, EPI_ISL_696403, EPI_ISL_696404, EPI_ISL_696405, EPI_ISL_696406, EPI_ISL_696407, EPI_ISL_696408, EPI_ISL_696409, EPI_ISL_696410, EPI_ISL_696411, EPI_ISL_696412, EPI_ISL_696413, EPI_ISL_696414, EPI_ISL_696421, EPI_ISL_696438, EPI_ISL_696439, EPI_ISL_696440, EPI_ISL_696441, EPI_ISL_696443, EPI_ISL_696444, EPI_ISL_696445, EPI_ISL_696447, EPI_ISL_696448, EPI_ISL_696449, EPI_ISL_696450 | | | |
| see above | Sonora Quest Laboratories, Laboratory Sciences of Arizona | TGen North | Jolene Bowers, Megan Folkerts, Chris French, Hayley Yaglom, Ashlyn Pfeiffer, Darrin Lemmer, Dave Engelthaler, The Arizona COVID Genomics Union (ACGU) |
| EPI_ISL_699659, EPI_ISL_699660, EPI_ISL_699661, EPI_ISL_699662, EPI_ISL_699663, EPI_ISL_699664, EPI_ISL_699665, EPI_ISL_699666, EPI_ISL_699667, EPI_ISL_699668, EPI_ISL_699669, EPI_ISL_699670, EPI_ISL_699671, EPI_ISL_699672, EPI_ISL_699673, EPI_ISL_699674, EPI_ISL_699675, EPI_ISL_699676, EPI_ISL_699677, EPI_ISL_699678, EPI_ISL_699679, EPI_ISL_699680, EPI_ISL_699681, EPI_ISL_699682, EPI_ISL_699683, EPI_ISL_699684 | | | |

| | | | |
|---|---|---|---|
| see above | Hematopathology Laboratory, ACTREC, TMC | Hematopathology Laboratory, ACTREC, TMC | Hematopathology Laboratory, ACTREC |
| EPI_ISL_700362 | Laboratoire de virologie, CHU de Grenoble - CS 10217 - 38043 Grenoble cedex 23 | CNR Virus des Infections Respiratoires - France SUD | Antonin Bal, Gregory Destras, Gwendolyne Burfin, Hadrien Règue, Quentin Semanas, Martine Valette, Bruno Lina, Sylvie Larrat, Laurence Josset |
| EPI_ISL_700363 | Laboratoire de virologie, CHU de Grenoble - CS 10217 - 38043 Grenoble cedex 24 | CNR Virus des Infections Respiratoires - France SUD | Antonin Bal, Gregory Destras, Gwendolyne Burfin, Hadrien Règue, Quentin Semanas, Martine Valette, Bruno Lina, Sylvie Larrat, Laurence Josset |
| EPI_ISL_700364 | Laboratoire de virologie, CHU de Grenoble - CS 10217 - 38043 Grenoble cedex 25 | CNR Virus des Infections Respiratoires - France SUD | Antonin Bal, Gregory Destras, Gwendolyne Burfin, Hadrien Règue, Quentin Semanas, Martine Valette, Bruno Lina, Sylvie Larrat, Laurence Josset |
| EPI_ISL_700365 | Laboratoire de virologie, CHU de Grenoble - CS 10217 - 38043 Grenoble cedex 26 | CNR Virus des Infections Respiratoires - France SUD | Antonin Bal, Gregory Destras, Gwendolyne Burfin, Hadrien Règue, Quentin Semanas, Martine Valette, Bruno Lina, Sylvie Larrat, Laurence Josset |
| EPI_ISL_700366 | Laboratoire de virologie, CHU de Grenoble - CS 10217 - 38043 Grenoble cedex 27 | CNR Virus des Infections Respiratoires - France SUD | Antonin Bal, Gregory Destras, Gwendolyne Burfin, Hadrien Règue, Quentin Semanas, Martine Valette, Bruno Lina, Sylvie Larrat, Laurence Josset |
| EPI_ISL_700367 | Laboratoire de virologie, CHU de Grenoble - CS 10217 - 38043 Grenoble cedex 28 | CNR Virus des Infections Respiratoires - France SUD | Antonin Bal, Gregory Destras, Gwendolyne Burfin, Hadrien Règue, Quentin Semanas, Martine Valette, Bruno Lina, Sylvie Larrat, Laurence Josset |
| EPI_ISL_700368 | Laboratoire de virologie, CHU de Grenoble - CS 10217 - 38043 Grenoble cedex 29 | CNR Virus des Infections Respiratoires - France SUD | Antonin Bal, Gregory Destras, Gwendolyne Burfin, Hadrien Règue, Quentin Semanas, Martine Valette, Bruno Lina, Sylvie Larrat, Laurence Josset |
| EPI_ISL_700369 | Laboratoire de virologie, CHU de Grenoble - CS 10217 - 38043 Grenoble cedex 30 | CNR Virus des Infections Respiratoires - France SUD | Antonin Bal, Gregory Destras, Gwendolyne Burfin, Hadrien Règue, Quentin Semanas, Martine Valette, Bruno Lina, Sylvie Larrat, Laurence Josset |
| EPI_ISL_700370 | Laboratoire de virologie, CHU de Grenoble - CS 10217 - 38043 Grenoble cedex 31 | CNR Virus des Infections Respiratoires - France SUD | Antonin Bal, Gregory Destras, Gwendolyne Burfin, Hadrien Règue, Quentin Semanas, Martine Valette, Bruno Lina, Sylvie Larrat, Laurence Josset |
| EPI_ISL_700397, EPI_ISL_700402, EPI_ISL_700403, EPI_ISL_700405, EPI_ISL_700406, EPI_ISL_700407, EPI_ISL_700409 | Centre hospitalier Métropole Savoie | CNR Virus des Infections Respiratoires - France SUD | Antonin Bal, Carine Dumollard, Gregory Destras, Gwendolyne Burfin, Hadrien Règue, Quentin Semanas, Martine Valette, Bruno Lina, Jérôme Grosjean, Laurence Josset |
| EPI_ISL_700411 | Hanover Park CHC wc HPH | NHLS/UCT | Arash Iranzadeh, Deelan Doolabh, Lynn Tyers, Bruna Galvao, Innocent Mudau, Marvin Hsiao, Kruger Marais, Diana Hardie, Stephen Korsman, Carolyn Williamson |
| EPI_ISL_700415 | Guguletu CHC wc GDH | NHLS/UCT | Arash Iranzadeh, Deelan Doolabh, Lynn Tyers, Bruna Galvao, Innocent Mudau, Marvin Hsiao, Kruger Marais, Diana Hardie, Stephen Korsman, Carolyn Williamson |
| EPI_ISL_700418, EPI_ISL_700463 | Hanover Park CHC wc HPH | NHLS/UCT | Arash Iranzadeh, Deelan Doolabh, Lynn Tyers, Bruna Galvao, Innocent Mudau, Marvin Hsiao, Kruger Marais, Diana Hardie, Stephen Korsman, Carolyn Williamson |
| EPI_ISL_700471, EPI_ISL_700486 | Nyanga CDC wc NGC | NHLS/UCT | Arash Iranzadeh, Deelan Doolabh, Lynn Tyers, Bruna Galvao, Innocent Mudau, Marvin Hsiao, Kruger Marais, Diana Hardie, Stephen Korsman, Carolyn Williamson |
| EPI_ISL_700491 | Guguletu CHC wc GDH | NHLS/UCT | Arash Iranzadeh, Deelan Doolabh, Lynn Tyers, Bruna Galvao, Innocent Mudau, Marvin Hsiao, Kruger Marais, Diana Hardie, Stephen Korsman, Carolyn Williamson |
| EPI_ISL_700508 | Heideveld CDC wc HVP | NHLS/UCT | Arash Iranzadeh, Deelan Doolabh, Lynn Tyers, Bruna Galvao, Innocent Mudau, Marvin Hsiao, Kruger Marais, Diana Hardie, Stephen Korsman, Carolyn Williamson |
| EPI_ISL_700511 | Nyanga CDC wc NGC | NHLS/UCT | Arash Iranzadeh, Deelan Doolabh, Lynn Tyers, Bruna Galvao, Innocent Mudau, Marvin Hsiao, Kruger Marais, Diana Hardie, Stephen Korsman, Carolyn Williamson |
| EPI_ISL_700512, EPI_ISL_700514 | Guguletu CHC wc GDH | NHLS/UCT | Arash Iranzadeh, Deelan Doolabh, Lynn Tyers, Bruna Galvao, Innocent Mudau, Marvin Hsiao, Kruger Marais, Diana Hardie, Stephen Korsman, Carolyn Williamson |
| EPI_ISL_700522 | Heideveld CDC wc HVP | NHLS/UCT | Arash Iranzadeh, Deelan Doolabh, Lynn Tyers, Bruna Galvao, Innocent Mudau, Marvin Hsiao, Kruger Marais, Diana Hardie, Stephen Korsman, Carolyn Williamson |
| EPI_ISL_700525 | Dr Abdurahman CDC wc DAC | NHLS/UCT | Arash Iranzadeh, Deelan Doolabh, Lynn Tyers, Bruna Galvao, Innocent Mudau, Marvin Hsiao, Kruger Marais, Diana Hardie, Stephen Korsman, Carolyn Williamson |
| EPI_ISL_700544 | Masincedane Clinic wc MAS | NHLS/UCT | Arash Iranzadeh, Deelan Doolabh, Lynn Tyers, Bruna Galvao, Innocent Mudau, Marvin Hsiao, Kruger Marais, Diana Hardie, Stephen Korsman, Carolyn Williamson |
| EPI_ISL_700582 | Dr Abdurahman CDC wc DAC | NHLS/UCT | Arash Iranzadeh, Deelan Doolabh, Lynn Tyers, Bruna Galvao, Innocent Mudau, Marvin Hsiao, Kruger Marais, Diana Hardie, Stephen Korsman, Carolyn Williamson |
| EPI_ISL_700584 | Heideveld Emergency Centre | NHLS/UCT | Arash Iranzadeh, Deelan Doolabh, Lynn Tyers, Bruna Galvao, Innocent Mudau, Marvin Hsiao, Kruger Marais, Diana Hardie, Stephen Korsman, Carolyn Williamson |
| EPI_ISL_708020 | Virology, Universitätsklinikum des Saarlandes | Epigenetics, Saarland University | Kathrin Kattler, Markus Vogelgesang, Stefan Lohse, Sascha Tierling, Sigrun Smola, Jörn Walter |
| EPI_ISL_710209, EPI_ISL_710210 | Colorado Department of Public Health and Environment | Colorado Department of Puplic Health and Environment | Laura Bankers, Molly C. Hetherington-Rauth, Shannon Ely, Shannon R. Matzinger, Sarah Elizabeth Totten, Emily A. Travanty |
| EPI_ISL_714191 | Department of Virus and Microbiological Special Diagnostics, Statens Serum Institut, Copenhagen, Denmark | Albertsen Lab, Department of Chemistry and Bioscience, Aalborg University, Denmark | Danish Covid-19 Genome Consortium |
| EPI_ISL_717724, EPI_ISL_717732, EPI_ISL_717739, EPI_ISL_717761, EPI_ISL_717763 | Kingston Health Sciences Centre and Queen's University | Ontario Institute for Cancer Research | Prameet M. Sheth,Calvin Sjaarda,Robert Colautti,Katya Douchant,Ilinca Lungu,Bernard Lam,Paul Krzyzanowski,Michael Laszloffy,Lawrence E. Heisler,Richard de Borja,Jared T. Simpson |
| EPI_ISL_718159 | Ministry of Health Hospitals | Institute of Health and Community Medicine | David Perera, Ooi Mong How, Chua Hock Hin, Tonnii Sia Loong Loong, Wong Jyn Shan, Wong Kiing Aik, Chan Chia Jui |
| EPI_ISL_721993, EPI_ISL_721994, EPI_ISL_721998, EPI_ISL_722016, EPI_ISL_722020, EPI_ISL_722022, EPI_ISL_722040, EPI_ISL_722041, EPI_ISL_722060, EPI_ISL_722062, EPI_ISL_722064, EPI_ISL_722071, EPI_ISL_722081, EPI_ISL_722086, EPI_ISL_722089, EPI_ISL_722092, EPI_ISL_722094, EPI_ISL_722098, EPI_ISL_722103, EPI_ISL_722107, EPI_ISL_722109, EPI_ISL_722116, EPI_ISL_722117, EPI_ISL_722124, EPI_ISL_722125, EPI_ISL_722126 | | | |
| see above | Hospital das Clínicas Universidade de São Paulo Medical School | Laboratório de Parasitologia Médica - Instituto de Medicina Tropical - Universidade de São Paulo | Brazil-UK Centre for Arbovirus Discovery Diagnosis Genomics and Epidemiology (CADDE) Genomic Network - Instituto de Medicina Tropical |
| EPI_ISL_723045 | Hematopathology Laboratory, ACTREC, TMC | Hematopathology Laboratory, ACTREC, TMC | Hematopathology Laboratory, ACTREC |
| EPI_ISL_729921, EPI_ISL_729936, EPI_ISL_729960, EPI_ISL_729961, EPI_ISL_729962, EPI_ISL_729963, EPI_ISL_730001, EPI_ISL_730002, EPI_ISL_730003, EPI_ISL_730004, EPI_ISL_730005, EPI_ISL_730006, EPI_ISL_730007, EPI_ISL_730008, EPI_ISL_730023 | | | |
| see above | Nigeria Centre for Disease Control (NCDC) | African Centre of Excellence for Genomics of Infectious Diseases (ACEGID), Redeemer's University, Ede, Osun State, Nigeria | Oluniyi P.E. et al |
| EPI_ISL_730062, EPI_ISL_730063, EPI_ISL_730064, EPI_ISL_730065 | Yale Clinical Virology Laboratory | Grubaugh Lab - Yale School of Public Health | Joseph Fauver, Tara Alpert, Anderson Brito, Annie Watkins, Anne Wyllie, Chantal Vogels, Mary Petrone, Chaney Kalinich, Isabel Ott, Arnau Casanovas, Catherine Muenker, Adam Moore, Alice Lu, Maria Tokuyama, Patrick Wong, Peiwen Lu, Saad Omer, Richard Martinello, Allison Nelson, Shelli Farhadian, Akiko Iwasaki, Charlese Dela Cruz, Albert Ko, Nathan Grubaugh |
| EPI_ISL_730137, EPI_ISL_730139, EPI_ISL_730141, EPI_ISL_730142, EPI_ISL_730143, EPI_ISL_730144, EPI_ISL_730146, EPI_ISL_730147, EPI_ISL_730148, EPI_ISL_730150, EPI_ISL_730151, EPI_ISL_730155, EPI_ISL_730159 | | | |

| | | | |
|---|---|---|---|
| see above | Scripps Medical Laboratory | Andersen lab at Scripps Research | SEARCH Alliance San Diego with Michael Quigley, Ellen Stefanski, Ian Mchardy |
| EPI_ISL_734500, EPI_ISL_734657, EPI_ISL_734658, EPI_ISL_734659, EPI_ISL_734660, EPI_ISL_734661, EPI_ISL_734662, EPI_ISL_734663, EPI_ISL_734664, EPI_ISL_734665, EPI_ISL_734666, EPI_ISL_734667, EPI_ISL_734668, EPI_ISL_734669, EPI_ISL_734670, EPI_ISL_734671, EPI_ISL_734672, EPI_ISL_734673, EPI_ISL_734674, EPI_ISL_734675, EPI_ISL_734676, EPI_ISL_734677, EPI_ISL_734678, EPI_ISL_734679, EPI_ISL_734680, EPI_ISL_734681, EPI_ISL_734682, EPI_ISL_734683, EPI_ISL_734684, EPI_ISL_734685, EPI_ISL_734686, EPI_ISL_734687, EPI_ISL_734688, EPI_ISL_734689, EPI_ISL_734690, EPI_ISL_734691, EPI_ISL_734692, EPI_ISL_734693, EPI_ISL_734694, EPI_ISL_734695, EPI_ISL_734696, EPI_ISL_734697, EPI_ISL_734698, EPI_ISL_734699, EPI_ISL_734700, EPI_ISL_734701, EPI_ISL_734702, EPI_ISL_734703, EPI_ISL_735249 | | | |
| see above | UZ Leuven, National Reference Laboratory for Coronaviruses, Laboratory Medicine, Leuven, Belgium | KU Leuven, Rega Institute, Clinical and Epidemiological Virology | Tony Wawina-Bokalanga, Joan Marti-Carerras, Bert Vanmechelen, Piet Maes |
| EPI_ISL_735267, EPI_ISL_735268, EPI_ISL_735269, EPI_ISL_735270, EPI_ISL_735271, EPI_ISL_735272, EPI_ISL_735273, EPI_ISL_735274, EPI_ISL_735275, EPI_ISL_735276, EPI_ISL_735277, EPI_ISL_735278, EPI_ISL_735279, EPI_ISL_735280, EPI_ISL_735282, EPI_ISL_735283, EPI_ISL_735284, EPI_ISL_735285, EPI_ISL_735286 | | | |
| see above | Genomic Laboratory (GLAB) (Conjoint lab of Health Directorate of Istanbul and Istanbul Technical University) | Genomic Laboratory (GLAB), Istanbul Technical University | Ilker Karacan, Tugba Kizilboga Akgun, Nihat Bugra Agaoglu, Payam Zolfagharian, Mehtap Aydin, Gizem Alkurt, Jale Yildiz, Betsi Kose, Nisan Denizce Can, Ayse Serra Ozel, Nilsun Altunal, Arzu Irvem, Yasemin Kendir Demirkol, Ozlem Akgun Dogan, Levent Doganay, Gizem Dinler Doganay |
| EPI_ISL_737934, EPI_ISL_737936, EPI_ISL_737946, EPI_ISL_737948, EPI_ISL_737949, EPI_ISL_737950, EPI_ISL_737962, EPI_ISL_737963 | Uganda Central Public Health Lab and Uganda Virus Research Institute | MRC/UVRI & LSHTM Uganda Research Unit | Matthew Cotten, Dan Lule Bugembe, My V.T. Phan, Pontiano Kaleebu et al. |
| EPI_ISL_738271, EPI_ISL_738272, EPI_ISL_738273, EPI_ISL_738274, EPI_ISL_738298, EPI_ISL_738309 | Connecticut Veterans' Affairs Hospital | Grubaugh Lab - Yale School of Public Health | Joseph Fauver, Tara Alpert, Chantal Vogels, Mary Petrone, Isabel Ott, Ellen Foxman, Shaili Gupta, Danielle Plank, Nathan Grubaugh |
| EPI_ISL_738777, EPI_ISL_738897, EPI_ISL_739301, EPI_ISL_739514, EPI_ISL_739635 | Napa-Solano-Yolo- Marin County (NSYM) Public Health Laboratories | Chan-Zuckerberg Biohub | CZB Cliahub Consortium |
| EPI_ISL_751252 | Devki Devi Foundation, a unit of Max Healthcare | CSIR-IGIB/Max | Rajesh Pandey#, Samreen Siddiqui, Janani Srinivasa Vasudevan, Akshay Kanakan, Ranjeet Maurya, Uzma Shamim, Bansidhar Tarai, Akansha Tyagi, Mitali Mukerji, Poonam Das, Sujeet Jha, Mohammed Faruq, Anurag Agrawal |
| EPI_ISL_751333 | IRCCS Sacro Cuore Don Calabria Hospital, Department of Infectious, Tropical Diseases & Microbiology | University of Verona, Department of Biotechnology | Antonio Mori, Michela Deiana, Elena Pomari, Chiara Piubelli; Giulia Lopatriello, Luca Marcolungo, Cristina Beltrami, Chiara Degli Esposti, Emanuela Cosentino, Massimo Delledonne |
| EPI_ISL_752645 | State Laboratories Division, Hawaii State Department of Health | State Laboratories Division, Hawaii State Department of Health | Pamela O'Brien, Sabrina Diemert, Drew Kuwazaki, Razvan Sultana, Edward Desmond |
| EPI_ISL_753738, EPI_ISL_753767, EPI_ISL_753772, EPI_ISL_753800, EPI_ISL_753805, EPI_ISL_753806, EPI_ISL_753823, EPI_ISL_753834, EPI_ISL_753950, EPI_ISL_753951, EPI_ISL_753952, EPI_ISL_754041 | | | |
| see above | Charité Universitätsmedizin Berlin, Institut für Virologie/Labor Berlin | Charité Universitätsmedizin Berlin, Institut für Virologie | Victor M Corman, Jörn Beheim-Schwarzbach, Barbara Mühlemann, Julia Schneider, Talitha Veith, Terry Jones, Christian Drosten |
| EPI_ISL_754927, EPI_ISL_754930, EPI_ISL_754946 | California Department of Public Health | California Department of Public Health | CDPH IDLB COVIDNet |
| EPI_ISL_755587 | Infection and Immunology, Translational Health Science and Technology Institute | Infection and Immunology, Translational Health Science and Technology Institute | Kumar,S., Wangchuk,J., Pandey,A.K., Das,A. and Medigeshi,G.R. |
| EPI_ISL_759860, EPI_ISL_759861 | Molecular, ProgenaBiome | Molecular, ProgenaBiome | Papoutsis,A.J. |
| EPI_ISL_759868 | Virology, Iran University of Medical Sciences | Razi Vaccine and Serum Research Institute, Co. | Keyvani,H., Ranjbar,Mm., Keyvani,F. and Soleimani,S. |
| EPI_ISL_759957 | Siti Khodijah Hospital | Institute of Tropical Disease, Universitas Airlangga | Rima R Prasetya, Krisnoadi Rahardjo, Aldise M Nastri, Jezzy R Dewantari, Muhammad Hamdan, Gatot Soegiarto, Laksmi Wulandari, Resti Yudhawati, Soetjipto, Yasuko Mori, Maria I Lusida, Kazufumi Shimizu |
| EPI_ISL_759958 | Universitas Airlangga Hospital | Institute of Tropical Disease, Universitas Airlangga | Maria I Lusida, Krisnoadi Rahardjo, Aldise M Nastri, Jezzy R Dewantari, Rima R Prasetya, Nasronudin, Gatot Soegiarto, Laksmi Wulandari, Resti Yudhawati, Soetjipto, Yasuko Mori, Kazufumi Shimizu |
| EPI_ISL_759964 | Mitra Keluarga Hospital Waru | Institute of Tropical Disease, Universitas Airlangga | Jezzy R Dewantari, Rima R Prasetya, Krisnoadi Rahardjo, Aldise M Nastri, Christina Dian Anggraeni, Gatot Soegiarto, Laksmi Wulandari, Resti Yudhawati, Yasuko Mori, Soetjipto, Kazufumi Shimizu ,Maria I Lusida |
| EPI_ISL_763362 | AIID | Irish Coronavirus Sequencing Consortium - National Virus Reference Laboratory | Michael Carr, Gabriel Gonzalez, Alejandro Abner Garcia Leon, Patrick Mallon |
| EPI_ISL_766045 | RS Pelni | Eijkman Institute for Molecular Biology, Ministry of Research and Technology/National Agency for Research and Innovation | Frilasita A Yudhaputri, Edison Johar, Hidayat Trimarsanto, David H Muljono, Safarina G Malik, Khin Saw Myint, Amin Soebandrio |
| EPI_ISL_767728, EPI_ISL_767752, EPI_ISL_767753, EPI_ISL_767754 | National Virus Reference Laboratory | Irish Coronavirus Sequencing Consortium - Teagasc Moorepark | Calum Walsh, Genuity Ireland |
| EPI_ISL_774872, EPI_ISL_774873, EPI_ISL_774874, EPI_ISL_774875, EPI_ISL_774876, EPI_ISL_774877, EPI_ISL_774878, EPI_ISL_774879, EPI_ISL_774880, EPI_ISL_774881, EPI_ISL_774882, EPI_ISL_774883, EPI_ISL_774884, EPI_ISL_774885, EPI_ISL_774886, EPI_ISL_774887, EPI_ISL_774888, EPI_ISL_774889, EPI_ISL_774890, EPI_ISL_774891, EPI_ISL_774892, EPI_ISL_774893, EPI_ISL_774894, EPI_ISL_774895, EPI_ISL_774896, EPI_ISL_774897, EPI_ISL_774898, EPI_ISL_774899, EPI_ISL_774900, EPI_ISL_774901, EPI_ISL_774902, EPI_ISL_774903, EPI_ISL_774904, EPI_ISL_774905, EPI_ISL_774906, EPI_ISL_774907, EPI_ISL_774908, EPI_ISL_774909, EPI_ISL_774910, EPI_ISL_774911, EPI_ISL_774912, EPI_ISL_774913 | | | |
| see above | Designated Reference Institute for Chemical Measurements (DRICM) | DNA SOLUTION LTD. | Md. Imran Khan, Kazi Nadim Hasan, Abu Sufian, Jannatun Naima, Abdul Khaleque, Mizanur Rahman, MSM Chowdhury, Hasan Ul Haider, Mamudul Hasan Razu, Mala Khan, Mohammad Fazle Alam Rabbi |
| EPI_ISL_775612, EPI_ISL_775641, EPI_ISL_775655, EPI_ISL_775834, EPI_ISL_776467, EPI_ISL_776468, EPI_ISL_776469, EPI_ISL_776473, EPI_ISL_776484, EPI_ISL_776485, EPI_ISL_776495, EPI_ISL_776496, EPI_ISL_776501, EPI_ISL_776502, EPI_ISL_776503, EPI_ISL_776522, EPI_ISL_776526 | | | |
| see above | University Medical Center Hamburg Eppendorf | Heinrich Pette Institute, Leibniz Institute for Experimental Virology | Alexis Robitaille, Thomas Günther, Johannes Knobloch, Martin Aepfelbacher, Nicole Fischer, Adam Grundhoff |
| EPI_ISL_776657, EPI_ISL_776732, EPI_ISL_776747, EPI_ISL_776748 | UW Virology Lab | UW Virology Lab | Pavitra Roychoudhury, Hong Xie, Lasata Shrestha, Meei-Li Huang, Keith R Jerome, Alexander Greninger |
| EPI_ISL_776757, EPI_ISL_776758 | Instituto Adolfo Lutz - Regional de Marilia | Instituto Adolfo Lutz, Interdiciplinary Procedures Center, Strategic Laboratory | Claudio Tavares Sacchi, Claudia Regina Gonçalves, Erica Valessa Ramos Gomes, Karoline Rodrigues Campos |
| EPI_ISL_776759 | Instituto Adolfo Lutz - Central | Instituto Adolfo Lutz, Interdiciplinary Procedures Center, Strategic Laboratory | Claudio Tavares Sacchi, Claudia Regina Gonçalves, Erica Valessa Ramos Gomes, Karoline Rodrigues Campos |
| EPI_ISL_777490, EPI_ISL_777902, EPI_ISL_778644 | Hospital General Universitario Gregorio Marañón | Hospital General Universitario Gregorio Marañón | Sergio Buenestado Serrano, Pedro Sola Campoy, Laura Perez-Lago, Pilar Catalán, Patricia Muñoz, Darío García de Viedma |
| EPI_ISL_778683, EPI_ISL_778684, EPI_ISL_778685, EPI_ISL_778736 | Istituto Zooprofilattico Sperimentale del Mezzogiorno | TIGEM | Patrizia Annunziata, Andrea Ballabio, Valentina Bouche, Davide Cacchiarelli (CorrespAuthor), Pellegrino Cerino, Chiara Colantuono, Lucio Di Filippo, Antonio Grimaldi, Antonio Limone, Gabriella Loconte, Anna Manfredi, Francesco Panariello, Biancamaria Pierri, Marcello Salvi, Lucia Vassallo |
| EPI_ISL_779172 | Laboratorio de Infectología, Servicio de Infectología, Hospital Universitario Dr. José Eleuterio González - Universidad Autónoma de Nuevo León | Laboratorio de Infectología Molecular, Departamento de Bioquímica y Medicina Molecular,Facultad de Medicina - Universidad Autónoma de Nuevo León | Kame A. Galán-Huerta, María F. Herrera-Saldivar, Natalia Martínez-Acuña, Sonia A. Lozano-Sepúlveda, Daniel Arellanos-Soto, Ana M. Rivas-Estilla, Paola Bocanegra-Ibarias, Samantha M. Flores-Treviño, Elvira Garza-González, Eduardo Perez-Alba, Laura Nuzzolo-Shihadeh, Adrian Camacho-Ortiz |
| EPI_ISL_779257, EPI_ISL_779263 | Jamil-ur-Rahman Center for Genome Research, Dr. Panjwani Center for Molecular Medicine and Drug Research | Jamil-ur-Rahman Center for Genome Research, Dr. Panjwani Center for Molecular Medicine and Drug Research | Shakeel,M., Irfan,M., Nisa,Z., Rashid,M., Ansari,S., Khan,I. |
| EPI_ISL_779735 | Servicio de Microbiología, Hospital Miguel Servet, Zaragoza | SeqCOVID-SPAIN consortium/IBV(CSIC) | Antonio Rezusta López, Alexander Tristancho Baró, Ana Milagro, Yolanda Gracia Grataloup, Nieves Martínez Cameo and SeqCOVID-SPAIN consortium |

| | | | |
|---|---|---|---|
| EPI_ISL_788946, EPI_ISL_788965 | Ospedale "Di Venere" | Beaconlab (Bioinformatics, Evolution and Comparative Genomics lab), Dept of Biosciences, University on Milan | Iacobellis M, d'Avenia M, Piluscio R, Parisi A, Chiara M, Manzari C, Pesole G |
| EPI_ISL_791819, EPI_ISL_791820, EPI_ISL_791821, EPI_ISL_791822, EPI_ISL_791823, EPI_ISL_791824, EPI_ISL_791825, EPI_ISL_791826, EPI_ISL_791827, EPI_ISL_791828, EPI_ISL_791829, EPI_ISL_791830, EPI_ISL_791831, EPI_ISL_791832, EPI_ISL_791833, EPI_ISL_791834, EPI_ISL_791835, EPI_ISL_791836, EPI_ISL_791837, EPI_ISL_791838, EPI_ISL_791839, EPI_ISL_791840, EPI_ISL_791841, EPI_ISL_791842, EPI_ISL_791843, EPI_ISL_791844, EPI_ISL_791845, EPI_ISL_791846, EPI_ISL_791847, EPI_ISL_791848, EPI_ISL_791849, EPI_ISL_791850, EPI_ISL_791851, EPI_ISL_791852, EPI_ISL_791853, EPI_ISL_791854, EPI_ISL_791855, EPI_ISL_791856, EPI_ISL_791857, EPI_ISL_791858, EPI_ISL_791859, EPI_ISL_791860, EPI_ISL_791861, EPI_ISL_791862, EPI_ISL_791863, EPI_ISL_791864, EPI_ISL_791865, EPI_ISL_791866, EPI_ISL_791867, EPI_ISL_791868, EPI_ISL_791869, EPI_ISL_791870, EPI_ISL_791871, EPI_ISL_791872, EPI_ISL_791873, EPI_ISL_791874, EPI_ISL_791875, EPI_ISL_791876, EPI_ISL_791877, EPI_ISL_791878, EPI_ISL_791879, EPI_ISL_791880, EPI_ISL_791881, EPI_ISL_791882, EPI_ISL_791883, EPI_ISL_791884, EPI_ISL_791885, EPI_ISL_791886, EPI_ISL_791887, EPI_ISL_791888, EPI_ISL_791889, EPI_ISL_791890, EPI_ISL_791891, EPI_ISL_791892, EPI_ISL_791893, EPI_ISL_791894, EPI_ISL_791895, EPI_ISL_791896, EPI_ISL_791897, EPI_ISL_791898, EPI_ISL_791899, EPI_ISL_791900, EPI_ISL_791901, EPI_ISL_791902, EPI_ISL_791903, EPI_ISL_791904, EPI_ISL_791905, EPI_ISL_791906, EPI_ISL_791907, EPI_ISL_791908, EPI_ISL_791909, EPI_ISL_791910, EPI_ISL_791911, EPI_ISL_791912, EPI_ISL_791913, EPI_ISL_791914, EPI_ISL_791915, EPI_ISL_791948, EPI_ISL_791949, EPI_ISL_791950, EPI_ISL_791951, EPI_ISL_791952, EPI_ISL_791963, EPI_ISL_791964, EPI_ISL_791965, EPI_ISL_791966, EPI_ISL_791976 | | | |
| see above | Massachusetts General Hospital | Infectious Disease Program, Broad Institute of Harvard and MIT | Lemieux,J.E., Siddle,K.J., Shaw,B., Adams,G., Pierce,V., Turbett,S., Anahtar,M., Branda,J., Slater,D., Harris,J., Lin,A.E., Gladden-Young,A., Lagerborg,K., Rudy,M., DeRuff,K., Carter,A., Normandin,E., Bauer,M., Reilly,S., Tomkins-Tinch,C., Loreth,C., Chaluvadi,S., Neumann,A., Cusick,C., Chapman,S.B., Gnirke,A., Flowers,K., Cerrato,F., Birren,B.W., Gallagher,G., Smole,S., Park,D.J., MacInnis,B.L., Ryan,E., LaRocque,R., Rosenberg,E. and Sabeti,P.C. |
| EPI_ISL_792113, EPI_ISL_792114 | Instituto Adolfo Lutz - Central | Instituto Adolfo Lutz, Interdiciplinary Procedures Center, Strategic Laboratory | Claudio Tavares Sacchi, Claudia Regina Gonçalves, Erica Valessa Ramos Gomes, Karoline Rodrigues Campos |
| EPI_ISL_792129, EPI_ISL_792130, EPI_ISL_792131, EPI_ISL_792132, EPI_ISL_792133, EPI_ISL_792134, EPI_ISL_792135, EPI_ISL_792136, EPI_ISL_792137, EPI_ISL_792138, EPI_ISL_792139, EPI_ISL_792140, EPI_ISL_792141, EPI_ISL_792142, EPI_ISL_792143, EPI_ISL_792144, EPI_ISL_792145, EPI_ISL_792146, EPI_ISL_792147, EPI_ISL_792148, EPI_ISL_792149, EPI_ISL_792150, EPI_ISL_792151, EPI_ISL_792152, EPI_ISL_792153, EPI_ISL_792154, EPI_ISL_792155, EPI_ISL_792156, EPI_ISL_792157, EPI_ISL_792158, EPI_ISL_792159, EPI_ISL_792160, EPI_ISL_792161, EPI_ISL_792162, EPI_ISL_792163, EPI_ISL_792164, EPI_ISL_792165, EPI_ISL_792166, EPI_ISL_792167, EPI_ISL_792168, EPI_ISL_792169, EPI_ISL_792170, EPI_ISL_792171, EPI_ISL_792172, EPI_ISL_792173, EPI_ISL_792174, EPI_ISL_792286 | | | |
| see above | Laboratorio de Virología del Hospital de Niños Dr. Ricardo Gutierrez | Área de Secuenciación del Laboratorio de Virología del Hospital de Niños Dr. Ricardo Gutierrez on behalf of 'Proyecto Argentino Interinstitucional de genomica de SARS-CoV-2' (PAIS Consortium) | Nabaes Jodar, MS; Goya, S; Natale, MI; Lusso, S; Gravis, E; Acevedo, ME; Alvarez Lopez, C; Alexay, S; Jacques, O; Mistchenko, AS; Valinotto, LE; Viegas, M. |
| EPI_ISL_792292, EPI_ISL_792296, EPI_ISL_792297 | Laboratorio de genética y biología molecular del Hospital de trauma y emergencia Dr Federico Abete | Área de Secuenciación del Laboratorio de Virología del Hospital de Niños Dr. Ricardo Gutierrez on behalf of 'Proyecto Argentino Interinstitucional de genomica de SARS-CoV-2' (PAIS Consortium) | Nabaes Jodar, MS; Goya, S; Natale, MI; Lusso, S; Primost, I; Gallino, MI; Valinotto, LE; Viegas, M. |
| EPI_ISL_792301 | Departamento de Biología y genética molecular, IACA Laboratorios. | Área de Secuenciación del Laboratorio de Virología del Hospital de Niños Dr. Ricardo Gutierrez on behalf of 'Proyecto Argentino Interinstitucional de genomica de SARS-CoV-2' (PAIS Consortium) | Nabaes Jodar, MS; Goya, S; Natale, MI; Lusso, S; Tittarelli, E; Suárez, A; Masciovecchio MV; Streitenberger ER; Mistchenko, AS; Valinotto, LE; Viegas, M. |
| EPI_ISL_792312, EPI_ISL_792313, EPI_ISL_792314 | Laboratorio del Hospital El Cruce Dr. Néstor C. Kirchner | Área de Secuenciación del Laboratorio de Virología del Hospital de Niños Dr. Ricardo Gutierrez on behalf of 'Proyecto Argentino Interinstitucional de genomica de SARS-CoV-2' (PAIS Consortium) | Nabaes Jodar, MS; Goya, S; Natale, MI; Lusso, S; Zubieta, M; Rahhal, M; Valinotto, LE; Viegas, M. |
| EPI_ISL_792332 | Instituto de Investigaciones Biomédicas en Retrovirus y SIDA | Área de Secuenciación del Laboratorio de Virología del Hospital de Niños Dr. Ricardo Gutierrez on behalf of 'Proyecto Argentino Interinstitucional de genomica de SARS-CoV-2' (PAIS Consortium) | Nabaes Jodar, MS; Goya, S; Natale, MI; Lusso, S; Seery, V; Remes Lenicov, F; Salomón, H; Valinotto, LE; Viegas, M. |
| EPI_ISL_792336, EPI_ISL_792337, EPI_ISL_792338, EPI_ISL_792339, EPI_ISL_792340, EPI_ISL_792341, EPI_ISL_792342 | Laboratorio del Hospital Interzonal General de Agudos Evita | Área de Secuenciación del Laboratorio de Virología del Hospital de Niños Dr. Ricardo Gutierrez on behalf of 'Proyecto Argentino Interinstitucional de genomica de SARS-CoV-2' (PAIS Consortium) | Nabaes Jodar, MS; Goya, S; Natale, MI; Lusso, S; Desimone, I; Luczac, E; Serrano, L; Grossi, O; Musto, A; Valinotto, LE; Viegas, M. |
| EPI_ISL_792367, EPI_ISL_792369 | Laboratorio de Virologia - HIEAyC San Juan de Dios | Área de Secuenciación del Laboratorio de Virología del Hospital de Niños Dr. Ricardo Gutierrez on behalf of 'Proyecto Argentino Interinstitucional de genomica de SARS-CoV-2' (PAIS Consortium) | Nabaes Jodar, MS; Goya, S; Natale, MI; Lusso, S; Ferioli, M; Colmeiro, M; Gatelli, A; Ercole, R; Valinotto, LE; Viegas, M. |
| EPI_ISL_792386, EPI_ISL_792389, EPI_ISL_792390, EPI_ISL_792391 | Plataforma de Servicios Biotecnólogicos: UTTIPP/PSB , Universidad Nacional de Quilmes. | Área de Secuenciación del Laboratorio de Virología del Hospital de Niños Dr. Ricardo Gutierrez on behalf of 'Proyecto Argentino Interinstitucional de genomica de SARS-CoV-2' (PAIS Consortium) | Nabaes Jodar, MS; Goya, S; Natale, MI; Lusso, S; Farina, H; Goñi, S; Cardama, G; Castello, A; Valinotto, LE; Viegas, M. |
| EPI_ISL_792402, EPI_ISL_792403, EPI_ISL_792404, EPI_ISL_792409 | LABORATORIO DE CAMPAÑA COVID 19 INTA | Instituto de Biotecnología, IABIMO (CONICET), Instituto de Virología, IVIT(CONICET), Instituto de Patobiología, IPVET(CONICET), CICVyA, INTA on behalf of 'Proyecto Argentino Interinstitucional de genomica de SARS-CoV-2' (PAIS Consortium) | König, GA; Peralta, AV; Distéfano, AJ; Zavallo, D; Muñoz Hidalgo, MG; Vera, PA; Fass, M; Farber, MD; Cacciabué, MPD; Pedroarias, VC; Lozano Calderón, LC; Bengoa Luoni, S; Asurmendi, S; Gioffré, A; Ferández PC; Zumárraga, M; Delgado, F; Díaz Carrasco, J; Garbaccio, S; Álvarez, I; Diagiacomo, S; Franco, L; Schammas, J; Rimondi, A; Soria, I;Craig, MI; Chimeno, S; Gómez, E; Klepp, L; Lucero, MS; Sioya, B; Del Médico, P; Rodríguez, A; Dus Santos, MJ; Olivera, V; Trono, K; Rosende, J; Fusco, W; Huertas, P; Vissani, A; Perea, A; Pereda, A; Paniego, NB; Rivarola, M; Puebla, AF; Viegas, M. |
| EPI_ISL_792425 | Laboratorio de Inmunología del Hospital Perrando e Instituto de Medicina Regional de la UNNE | Instituto de Biotecnología, IABIMO (CONICET), Instituto de Virología, IVIT(CONICET), Instituto de Patobiología, IPVET(CONICET), CICVyA, INTA on behalf of 'Proyecto Argentino Interinstitucional de genomica de SARS-CoV-2' (PAIS Consortium) | König, GA; Peralta, AV; Distéfano, AJ; Zavallo, D; Muñoz Hidalgo, MG; Vera, PA; Fass, M; Farber, MD; Cacciabué, MPD; Pedroarias, VC; Lozano Calderón, LC; Bengoa Luoni, S; Asurmendi, S; Foussal, MD; Deluca, G; Ayala, NA; Gómez, MV; Giusiano, G; Lucero, H; Marin, M; Lescano, L; Cayré, A; Paniego, NB; Rivarola, M; Puebla, AF; Viegas, M. |
| EPI_ISL_792444, EPI_ISL_792446, EPI_ISL_792450, EPI_ISL_792457 | Laboratorio del Hospital Regional Ushuaia Gdor. Ernesto Campos | Hospital Regional Ushuaia - Centro Austral De Investigaciones Científicas - Universidad Nacional De Tierra Del Fuego on behalf of 'Proyecto Argentino Interinstitucional de genomica de SARS-CoV-2' (PAIS Consortium) | Ceballos, SG; Nardi, CF; Gramundi, ID; Gallego, F; De Roccis, CA; Castro, G; Cáceres, SB; Yulan, CB; Boutureira, MF. |
| EPI_ISL_792498 | Laboratorio Central Mg. Luis Alfredo Pianciola | Hospital Regional Ushuaia - Centro Austral De Investigaciones Científicas - Universidad Nacional De Tierra Del Fuego on behalf of 'Proyecto Argentino Interinstitucional de genomica de SARS-CoV-2' (PAIS Consortium) | Ceballos, SG; Nardi, CF; Gramundi, ID; Gallego, F; Pinto, C; Ziehm, MC; Mazzeo, M; Pianciola, L. |
| EPI_ISL_792510 | Laboratorio de Inmunología del Hospital Perrando e Instituto de Medicina Regional de la UNNE | Grupo de Genómica y Bioinformática del Instituto de Investigación de la Cadena Láctea CONICET-INTA on behalf of 'Proyecto Argentino Interinstitucional de genomica de SARS-CoV-2' (PAIS Consortium) | Eberhardt, MF; Irazoqui, JM; Foussal, MD; Deluca, G; Ayala, NA; Gómez, MV; Giusiano, G; Lucero, H; Marin, M; Lescano, L; Cayré, A; Amadio, AF |
| EPI_ISL_792571, EPI_ISL_792598 | LACEN-PB | Laboratory of Respiratory Viruses and Measles, Oswaldo Cruz Institute, FIOCRUZ | Paola Resende, Luciana Appolinario, Fernando Motta, Anna Carolina Paixao, Ana Carolina Mendonca, João Felipe Bezerra, Romero Henrique Teixeira de Vasconcelos, Dalane Loudal Florentino Teixeira, Thiago Franco de Oliveira Carneiro, Marilda Siqueira |
| EPI_ISL_792643 | LACEN-AL | Laboratory of Respiratory Viruses and Measles, Oswaldo Cruz Institute, FIOCRUZ | Paola Resende, Luciana Appolinario, Fernando Motta, Anna Carolina Paixao, Ana Carolina Mendonca, Anderson Brandao Leite, Marilda Siqueira |
| EPI_ISL_796129 | Laboratorio de Microbiología. Hospital General Universitario de Elda, Alicante | SeqCOVID-SPAIN consortium/IBV(CSIC) | Mª Isabel Gascón Ros, Cristina Torregrosa Hetland, Eva Pastor Boix, Paloma Cascales Ramos and SeqCOVID-SPAIN consortium |

| | | | |
|---|---|---|---|
| EPI_ISL_801402 | Laboratório Central de Saúde Pública do Amazonas - LACEN-AM | Laboratorio de Ecologia de Doencas Transmissiveis na Amazonia, Instituto Leonidas e Maria Deane - Fiocruz Amazonia | Valdinete Nascimento, Victor Souza, André Corado, Fernanda Nascimento, George Silva, Ágatha Costa, Debora Duarte, Luciana Gonçalves, Maria Júlia Brandão, Michele Jesus, Felipe Naveca |
| EPI_ISL_801544, EPI_ISL_801548, EPI_ISL_801551, EPI_ISL_801552 | Laboratory of Molecular Virology, Pontificia Universidad Católica de Chile | MSHS Pathogen Surveillance Program | Leonardo I. Almonacid, Ana S. Gonzalez-Reiche, Matthew M. Hernandez, Jorge Levican, Ana Maria Contreras, Carlos Palma, Tamara García-Salum, Zenab Khan, Adriana van De Guchte, Ajay Obla, Jayeeta Dutta, Bremy Alburquerque, Eileen Serrano, Constanza Maldonado, M. Belen Leyton, Erick Salinas, Hala Alshammary, Juan Soto, Shwetha Hara Sridhar, Ying-Chih Wang, Melissa Smith, Robert Sebra, Marcela Ferres, Adolfo Garcia-Sastre, Edward C. Holmes, Viviana Simon, Harm van Bakel, Rafael A. Medina. |
| EPI_ISL_801555, EPI_ISL_801556, EPI_ISL_801558, EPI_ISL_801559, EPI_ISL_801560, EPI_ISL_801562, EPI_ISL_801563, EPI_ISL_801564, EPI_ISL_801565, EPI_ISL_801566, EPI_ISL_801567, EPI_ISL_801572, EPI_ISL_801573, EPI_ISL_801574, EPI_ISL_801579, EPI_ISL_801580 | | | |
| see above | Laboratory of Molecular Virology, Pontificia Universidad Católica de Chile | MSHS Pathogen Surveillance Program | Leonardo I. Almonacid, Ana S. Gonzalez-Reiche, Matthew M. Hernandez, Jorge Levican, Tamara García-Salum, Zenab Khan, Adriana van De Guchte, Ajay Obla, Jayeeta Dutta, Bremy Alburquerque, Eileen Serrano, Erick Salinas, Manuel Ampuero, Hala Alshammary, Juan Soto, Shwetha Hara Sridhar, Ying-Chih Wang, Melissa Smith, Robert Sebra, Fernando Valiente, Aldo Gaggero, Adolfo Garcia-Sastre, Edward C. Holmes, Viviana Simon, Harm van Bakel, Rafael A. Medina. |
| EPI_ISL_801593, EPI_ISL_801594, EPI_ISL_801597 | Laboratory of Molecular Virology, Pontificia Universidad Católica de Chile | MSHS Pathogen Surveillance Program | Leonardo I. Almonacid, Ana S. Gonzalez-Reiche, Matthew M. Hernandez, Jorge Levican, Tamara García-Salum, Zenab Khan, Adriana van De Guchte, Ajay Obla, Jayeeta Dutta, Bremy Alburquerque, Eileen Serrano, Erick Salinas, Hala Alshammary, Juan Soto, Shwetha Hara Sridhar, Ying-Chih Wang, Melissa Smith, Robert Sebra, Adolfo Garcia-Sastre, Edward C. Holmes, Viviana Simon, Harm van Bakel, Rafael A. Medina |
| EPI_ISL_801728, EPI_ISL_801729, EPI_ISL_801732, EPI_ISL_801734, EPI_ISL_801735, EPI_ISL_801737, EPI_ISL_801738, EPI_ISL_801739, EPI_ISL_801741, EPI_ISL_801742, EPI_ISL_801743, EPI_ISL_801744, EPI_ISL_801745, EPI_ISL_801746, EPI_ISL_801747, EPI_ISL_801748, EPI_ISL_801749, EPI_ISL_801750, EPI_ISL_801751, EPI_ISL_801752, EPI_ISL_801753, EPI_ISL_801754, EPI_ISL_801755, EPI_ISL_801756, EPI_ISL_801757, EPI_ISL_801758, EPI_ISL_801759, EPI_ISL_801760, EPI_ISL_801761, EPI_ISL_801762, EPI_ISL_801764 | | | |
| see above | Laboratory of Molecular Virology, Pontificia Universidad Católica de Chile | MSHS Pathogen Surveillance Program | Leonardo I. Almonacid, Ana S. Gonzalez-Reiche, Matthew M. Hernandez, Jorge Levican, Ana Maria Contreras, Carlos Palma, Tamara García-Salum, Zenab Khan, Adriana van De Guchte, Ajay Obla, Jayeeta Dutta, Bremy Alburquerque, Eileen Serrano, Constanza Maldonado, M. Belen Leyton, Erick Salinas, Hala Alshammary, Juan Soto, Shwetha Hara Sridhar, Ying-Chih Wang, Melissa Smith, Robert Sebra, Marcela Ferres, Adolfo Garcia-Sastre, Edward C. Holmes, Viviana Simon, Harm van Bakel, Rafael A. Medina. |
| EPI_ISL_801860 | Laboratory of Molecular Virology, Pontificia Universidad Católica de Chile | MSHS Pathogen Surveillance Program | Leonardo I. Almonacid, Ana S. Gonzalez-Reiche, Matthew M. Hernandez, Jorge Levican, Tamara García-Salum, Zenab Khan, Adriana van De Guchte, Ajay Obla, Jayeeta Dutta, Bremy Alburquerque, Eileen Serrano, Erick Salinas, Manuel Ampuero, Hala Alshammary, Juan Soto, Shwetha Hara Sridhar, Ying-Chih Wang, Melissa Smith, Robert Sebra, Fernando Valiente, Aldo Gaggero, Adolfo Garcia-Sastre, Edward C. Holmes, Viviana Simon, Harm van Bakel, Rafael A. Medina. |
| EPI_ISL_801861, EPI_ISL_801867, EPI_ISL_801891, EPI_ISL_801944, EPI_ISL_801945, EPI_ISL_801950, EPI_ISL_801951, EPI_ISL_801961, EPI_ISL_801962, EPI_ISL_801977, EPI_ISL_801978, EPI_ISL_802039, EPI_ISL_802040, EPI_ISL_802041, EPI_ISL_802042, EPI_ISL_802043, EPI_ISL_802044, EPI_ISL_802045, EPI_ISL_802046, EPI_ISL_802047, EPI_ISL_802048, EPI_ISL_802049, EPI_ISL_802050, EPI_ISL_802051, EPI_ISL_802052, EPI_ISL_802053, EPI_ISL_802054, EPI_ISL_802055, EPI_ISL_802056, EPI_ISL_802057, EPI_ISL_802058, EPI_ISL_802059, EPI_ISL_802060, EPI_ISL_802061, EPI_ISL_802062, EPI_ISL_802063, EPI_ISL_802064, EPI_ISL_802065, EPI_ISL_802066, EPI_ISL_802067, EPI_ISL_802068, EPI_ISL_802069, EPI_ISL_802070, EPI_ISL_802071, EPI_ISL_802072, EPI_ISL_802073, EPI_ISL_802074, EPI_ISL_802075, EPI_ISL_802076, EPI_ISL_802077, EPI_ISL_802078, EPI_ISL_802079, EPI_ISL_802080, EPI_ISL_802081, EPI_ISL_802082, EPI_ISL_802083, EPI_ISL_802084, EPI_ISL_802085, EPI_ISL_802086 | | | |
| see above | MSHS Clinical Microbiology Laboratories | MSHS Pathogen Surveillance Program | Ana S. Gonzalez-Reiche, Hala Alshammary, Mitchell J. Sullivan, Brianne Ciferri, Ajay Obla, Angela Amoako, Mahmoud Awawda, Elena Hirsch, Ashley S. Salimbangon, Levy Sominsky, Katherine Beach, Kayla Russo, Charles Gleason, Shelcie Fabre, Giulio Kleiner, Zenab Khan, Bremy Alburquerque, Adriana van de Guchte, Komal Srivastava, Matthew M. Hernandez, Jayeeta Dutta, Denise Jurczyszak, Emily Ferreri, Rachel Chernet, Nancy Francoeur, Betsaida Salom Melo, Irina Oussenko, Gintaras Deikus, Juan Soto, Shwetha Hara Sridhar, Ying-Chih Wang, Kathryn Twyman, Andrew Kasarskis, Deena R. Altman, Robert Sebra, Adolfo García-Sastre, Marta Luksza, Gopi Patel, Sarah Schaefer, Melissa Gitman, Michael D. Nowak, Alberto Paniz-Mondolfi, Emilia Mia Sordillo, Viviana Simon, Harm van Bakel |
| EPI_ISL_802501 | Ohio Department of Health Laboratory | Ohio Department of Health Laboratory | Holmes, Jennifer; Eric Brandt, Keoni Omura, Glen McGillivary, Caitlin McDonnell, Kirtana Ramadugu, Erica Leasure, Kelsey Florek, Heather Blankenship, Quanta Brown, and Tammy Bannerman |
| EPI_ISL_804387 | Dutch COVID-19 response team | National Institute for Public Health and the Environment (RIVM) | Adam Meijer, Harry Vennema, Dirk Eggink, Matthijs Welkers, Jeroen Cremer, Sharon van den Brink, Bas van der Veer, AnneMarie van den Brandt, Florian Zwagemaker, Dennis Schmitz, Chantal Reusken, on behalf of the national COVID-19 response team |
| EPI_ISL_804990, EPI_ISL_804991, EPI_ISL_804995, EPI_ISL_805002, EPI_ISL_805004, EPI_ISL_805013, EPI_ISL_805016, EPI_ISL_805147, EPI_ISL_805204, EPI_ISL_805259, EPI_ISL_805278, EPI_ISL_805292, EPI_ISL_805316, EPI_ISL_805324, EPI_ISL_805329, EPI_ISL_805339, EPI_ISL_805372, EPI_ISL_805409, EPI_ISL_805410, EPI_ISL_805426, EPI_ISL_805429, EPI_ISL_805436, EPI_ISL_805444, EPI_ISL_805449, EPI_ISL_805457, EPI_ISL_805460, EPI_ISL_805469, EPI_ISL_805472, EPI_ISL_805476, EPI_ISL_805482, EPI_ISL_805484, EPI_ISL_805485, EPI_ISL_805487, EPI_ISL_805488, EPI_ISL_805491, EPI_ISL_805494, EPI_ISL_805498, EPI_ISL_805501, EPI_ISL_805504, EPI_ISL_805514, EPI_ISL_805516, EPI_ISL_805518, EPI_ISL_805519, EPI_ISL_805526, EPI_ISL_805528, EPI_ISL_805532, EPI_ISL_805537, EPI_ISL_805540, EPI_ISL_805552, EPI_ISL_805553, EPI_ISL_805555, EPI_ISL_805557, EPI_ISL_805561, EPI_ISL_805564, EPI_ISL_805565, EPI_ISL_805567, EPI_ISL_805569, EPI_ISL_805570, EPI_ISL_805572, EPI_ISL_805575, EPI_ISL_805583, EPI_ISL_805584, EPI_ISL_805585, EPI_ISL_805586, EPI_ISL_805596, EPI_ISL_805598, EPI_ISL_805604, EPI_ISL_805605, EPI_ISL_805606, EPI_ISL_805607, EPI_ISL_805608, EPI_ISL_805612, EPI_ISL_805617, EPI_ISL_805621, EPI_ISL_805626, EPI_ISL_805627, EPI_ISL_805640, EPI_ISL_805650, EPI_ISL_805662, EPI_ISL_805665, EPI_ISL_805667, EPI_ISL_805677, EPI_ISL_805681, EPI_ISL_805682, EPI_ISL_805684, EPI_ISL_805690, EPI_ISL_805691, EPI_ISL_805696, EPI_ISL_805697, EPI_ISL_805699, EPI_ISL_805700, EPI_ISL_805704, EPI_ISL_805706, EPI_ISL_805707, EPI_ISL_805717, EPI_ISL_805719, EPI_ISL_805720, EPI_ISL_805725, EPI_ISL_805731, EPI_ISL_805732, EPI_ISL_805733, EPI_ISL_805740, EPI_ISL_805741, EPI_ISL_805747, EPI_ISL_805750, EPI_ISL_805751, EPI_ISL_805754, EPI_ISL_805756, EPI_ISL_805767, EPI_ISL_805768, EPI_ISL_805771, EPI_ISL_805773, EPI_ISL_805774, EPI_ISL_805777, EPI_ISL_805780, EPI_ISL_805781, EPI_ISL_805785, EPI_ISL_805786, EPI_ISL_805791, EPI_ISL_805793, EPI_ISL_805796, EPI_ISL_805797, EPI_ISL_805798, EPI_ISL_805800, EPI_ISL_805804, EPI_ISL_805807, EPI_ISL_805811, EPI_ISL_805816, EPI_ISL_805827, EPI_ISL_805835, EPI_ISL_805837, EPI_ISL_805838, EPI_ISL_805842, EPI_ISL_805847, EPI_ISL_805848, EPI_ISL_805849, EPI_ISL_805862, EPI_ISL_805870, EPI_ISL_805884, EPI_ISL_805887, EPI_ISL_805888, EPI_ISL_805889, EPI_ISL_805892, EPI_ISL_805902, EPI_ISL_805905, EPI_ISL_805907, EPI_ISL_805914, EPI_ISL_805916, EPI_ISL_805919, EPI_ISL_805927, EPI_ISL_805931, EPI_ISL_805934, EPI_ISL_805936, EPI_ISL_805944, EPI_ISL_805950, EPI_ISL_805952, EPI_ISL_805959, EPI_ISL_805967, EPI_ISL_805968, EPI_ISL_805987, EPI_ISL_805989, EPI_ISL_805992, EPI_ISL_805997, EPI_ISL_806011, EPI_ISL_806020, EPI_ISL_806026, EPI_ISL_806031, EPI_ISL_806036, EPI_ISL_806044, EPI_ISL_806046, EPI_ISL_806049, EPI_ISL_806059, EPI_ISL_806065, EPI_ISL_806069, EPI_ISL_806074, EPI_ISL_806077, EPI_ISL_806099, EPI_ISL_806105, EPI_ISL_806106, EPI_ISL_806123, EPI_ISL_806141, EPI_ISL_806142, EPI_ISL_806145, EPI_ISL_806155, EPI_ISL_806167, EPI_ISL_806188, EPI_ISL_806214, EPI_ISL_806218, EPI_ISL_806225, EPI_ISL_806238, EPI_ISL_806249, EPI_ISL_806252, EPI_ISL_806265, EPI_ISL_806269, EPI_ISL_806277, EPI_ISL_806281, EPI_ISL_806324, EPI_ISL_806337, EPI_ISL_806339, EPI_ISL_806345, EPI_ISL_806358, EPI_ISL_806362, EPI_ISL_806364, EPI_ISL_806393, EPI_ISL_806426, EPI_ISL_806466, EPI_ISL_806478, EPI_ISL_806481, EPI_ISL_806484, EPI_ISL_806492, EPI_ISL_806494, EPI_ISL_806497, EPI_ISL_806504, EPI_ISL_806505 | | | |
| see above | Alberta Precision Labs (APL) | Alberta Precision Labs (APL) | Gordon P, Lam LG, Pabbaraju K, Wong A, Ma R, Li V, Melin A, Tipples G, Berenger B, Zelyas N, Kellner J, Bernier F, Chui L, Croxen M |
| EPI_ISL_812245, EPI_ISL_812247 | North Dakota Department of Health, Public Health Laboratory | North Dakota Department of Health, Public Health Laboratory | Lisa Wingerter |
| EPI_ISL_812793 | Genomics Program, Children Cancer Hospital | Genomics Program, Children Cancer Hospital | Hatem,A., Hadad,A., Abouelnaga,S., Amer,K., Salah,H., Farawyla,H., Halafawy,A., Mansour,T., shalaby,L., Hassan,W., Soliman,M., Gomaa,C., Hassan,R., Soliman,S., Monuir,G., Hammad,M., Hussein,S., Abdo,I., Jalal,D., El-Zayat,M., El-Shaqnqery,H., Diab,A., Bakry,U., Samir,O., Magdeldin,S., Sayed,A. |
| EPI_ISL_814071 | Ministry of Health Turkey | Ministry of Health Turkey | Fatma Bayrakdar, Yasemin Cogun, Süleyman Yalcin, Aye Baak Alta, Gülay Korukluolu |
| EPI_ISL_816135, EPI_ISL_816136, EPI_ISL_816432 | Ohio Department of Health Laboratory | Ohio Department of Health Laboratory | Holmes, Jennifer; Eric Brandt, Keoni Omura, Glen McGillivary, Caitlin McDonnell, Kirtana Ramadugu, Erica Leasure, Kelsey Florek, Heather Blankenship, Quanta Brown, and Tammy Bannerman |
| EPI_ISL_816723, EPI_ISL_816724, EPI_ISL_816725, EPI_ISL_816726, EPI_ISL_816727, EPI_ISL_816728, EPI_ISL_816729, EPI_ISL_816730, EPI_ISL_816731, EPI_ISL_816732, EPI_ISL_816733, EPI_ISL_816734, EPI_ISL_816796, EPI_ISL_816797, EPI_ISL_816798, EPI_ISL_816799, EPI_ISL_816800, EPI_ISL_816801, EPI_ISL_816819, EPI_ISL_816820 | | | |
| see above | Bioinformatics and Biostatistics Lab, Advanced Sequencing Facility | COVID-19 Genomics UK (COG-UK) Consortium | Aengus Stewart,Jerome Nicod,Chelsea Sawyer,Laura Cubitt,Harshil Patel,Margaret Crawford |
| EPI_ISL_819127 | Ohio Department of Health Laboratory | Ohio Department of Health Laboratory | Holmes, Jennifer; Eric Brandt, Keoni Omura, Glen McGillivary, Caitlin McDonnell, Kirtana Ramadugu, Erica Leasure, Kelsey Florek, Heather Blankenship, Quanta Brown, and Tammy Bannerman |
| EPI_ISL_819295, EPI_ISL_819297, EPI_ISL_819335, EPI_ISL_819336, EPI_ISL_819337 | Hospital Universitari Vall d'Hebron - Vall d'Hebron Institut de Rercerca | Hospital Universitari Vall d'Hebron | Cristina Andrés, Maria Piñana, Josep F Abril, Damir Garcia-Cehic, Ariadna Rando, Juliana Esperalba, Maria Gema Codina, Carla Castillo, Maria Carmen Martin, Tomàs Pumarola, Josep Quer, Andrés Antón |
| EPI_ISL_824332, EPI_ISL_824333, EPI_ISL_824384, EPI_ISL_824385 | Ohio Department of Health Laboratory | Ohio Department of Health Laboratory | Holmes, Jennifer; Eric Brandt, Keoni Omura, Glen McGillivary, Caitlin McDonnell, Kirtana Ramadugu, Erica Leasure, Kelsey Florek, Heather Blankenship, Quanta Brown, and Tammy Bannerman |
| EPI_ISL_825947, EPI_ISL_825951, EPI_ISL_825956, EPI_ISL_825959, EPI_ISL_825960, EPI_ISL_825962, EPI_ISL_825966, EPI_ISL_825977, EPI_ISL_825979, EPI_ISL_825980, EPI_ISL_825981, EPI_ISL_825982, EPI_ISL_825983, EPI_ISL_825984, EPI_ISL_825985, EPI_ISL_825986, EPI_ISL_825987, EPI_ISL_825988, EPI_ISL_825989, EPI_ISL_825990, EPI_ISL_825991, EPI_ISL_825992, EPI_ISL_825993, EPI_ISL_825994, EPI_ISL_825995, EPI_ISL_825996, EPI_ISL_825997, EPI_ISL_825998, EPI_ISL_825999, EPI_ISL_826000, EPI_ISL_826001, EPI_ISL_826002, EPI_ISL_826003, EPI_ISL_826004, EPI_ISL_826005, EPI_ISL_826006, EPI_ISL_826007, EPI_ISL_826008, EPI_ISL_826009, EPI_ISL_826010, EPI_ISL_826011, EPI_ISL_826012, EPI_ISL_826013, EPI_ISL_826014, EPI_ISL_826015, EPI_ISL_826016, EPI_ISL_826017, EPI_ISL_826018, EPI_ISL_826019, EPI_ISL_826020, EPI_ISL_826021, EPI_ISL_826022, EPI_ISL_826023, EPI_ISL_826024, EPI_ISL_826025, EPI_ISL_826026, EPI_ISL_826027 | | | |
| see above | Laboratoire de santé publique du Québec | Laboratoire de santé publique du Québec | Sandrine Moreira, Ioannis Ragoussis, Guillaume Bourque, Jesse Shapiro, Mark Lathrop and Michel Roger on behalf of the CoVSeQ research group (http://covseq.ca/researchgroup) |

| | | | |
|---|---|---|---|
| EPI_ISL_831078, EPI_ISL_831081, EPI_ISL_831082, EPI_ISL_831083, EPI_ISL_831084 | Hospital Universitario La Paz (Madrid) | SeqCOVID-SPAIN consortium/IBV(CSIC) | María Rodríguez-Tejedor, Elias Dahdouh, Fernando Lázaro-Perona, Jesús Mingorance and SeqCOVID-SPAIN consortium |
| EPI_ISL_831904 | New Mexico Department of Health Scientific Laboratory | New Mexico Department of Health Scientific Laboratory | Ellie Johnson, Anastacia Griego-Fisher, D'eldra Malone |
| EPI_ISL_833155 | Instituto Adolfo Lutz - Central | Instituto Adolfo Lutz, Interdiciplinary Procedures Center, Strategic Laboratory | Claudio Tavares Sacchi, Claudia Regina Gonçalves, Erica Valessa Ramos Gomes, Karoline Rodrigues Campos |
| EPI_ISL_833200 | Hôpital Bichat Claude Bernard, Laboratoire de Virologie | IAME UMR1137 Inserm, Université de Paris, Hôpital Bichat | Antoine Bridier, Amélie Recoing, Quentin Le Hingrat, Lena Daniel, Siham Hamri, Gilles Collin, Alexandre Storto, Mélanie Bertine, Charlotte Charpentier, Nadhira Houhou-Fidouh, Diane Descamps, Benoit Visseaux |
| EPI_ISL_833521 | Veterinary Specialized Instute "Nis" | Veterinary Specialized Institute "Kraljevo", Serbia | Vidanovic,D., Tesovic,B., Manic,M., Petrovic,M.,Knezevic,A., Jovanovic,T., Jankovic,M., Sekler,M., Banovic Djeri,B., Petrovic,T., Volkening,J., Afonso,C. |
| EPI_ISL_833522, EPI_ISL_833524, EPI_ISL_833526, EPI_ISL_833528, EPI_ISL_833529, EPI_ISL_833530, EPI_ISL_833533, EPI_ISL_833534, EPI_ISL_833536, EPI_ISL_833542, EPI_ISL_833543, EPI_ISL_833547, EPI_ISL_833548, EPI_ISL_833549, EPI_ISL_833550, EPI_ISL_833551, EPI_ISL_833555, EPI_ISL_833564, EPI_ISL_833565, EPI_ISL_833566, EPI_ISL_833567, EPI_ISL_833568, EPI_ISL_833569, EPI_ISL_833570 | | | |
| see above | Istituto Zooprofilattico Sperimentale del Mezzogiorno | TIGEM | Patrizia Annunziata, Andrea Ballabio, Valentina Bouche, Davide Cacchiarelli (CorrespAuthor), Pellegrino Cerino, Chiara Colantuono, Lucio Di Filippo, Antonio Grimaldi, Antonio Limone, Gabriella Loconte, Anna Manfredi, Francesco Panariello, Biancamaria Pierri, Marcello Salvi, Lucia Vassallo |
| EPI_ISL_837550, EPI_ISL_837571, EPI_ISL_837574, EPI_ISL_837575, EPI_ISL_837577 | Centro Nacional de Enfermedades Tropicales (CENETROP) | Laboratory of Respiratory Viruses and Measles, Oswaldo Cruz Institute, FIOCRUZ | Paola Resende, Roxana Loayza, Cinthia Avila, Luciana Appolinario, Fernando Motta, Anna Carolina Paixao, Ana Carolina Mendonca, Marilda Siqueira |
| EPI_ISL_837679, EPI_ISL_837680, EPI_ISL_837681, EPI_ISL_837682, EPI_ISL_837683, EPI_ISL_837684 | Instituto Nacional de Enfermedades Respiratorias (INER) | Instituto Nacional de Enfermedades Respiratorias (INER) | Celia Boukadida, Margarita Matías-Florentino, Alma Rincón-Rubio, Hector Esteban Paz-Juárez, Olivia Briceño, Edgar Sevilla-Reyes, Fidencio Mejía-Nepomuceno, Mario Mújica-Sánchez, Eduardo Becerril-Vargas, José Arturo Martínez-Orozco, Alejandra Hernández-Terán, Jorge Salas-Hernández, Santiago Ávila-Ríos, Joel Armando Vázquez-Pérez |
| EPI_ISL_842655, EPI_ISL_842658 | University College London Hospital | COVID-19 Genomics UK (COG-UK) Consortium | Judith Heaney, Matthew Byott, Catherine Houlihan, Dan Frampton, Stuart Kirk, Moira Spyer and Eleni Nastouli |
| EPI_ISL_842829 | Barts Health NHS Trust | COVID-19 Genomics UK (COG-UK) Consortium | CUTINO-MOGUEL, Maria-Teresa; HARRINGTON, David; OWOYEMI, Dola; SHYLINI, Raghavendran; BROAD, Claire; KELE, Beatrix |
| EPI_ISL_848208, EPI_ISL_848215, EPI_ISL_848222, EPI_ISL_848261, EPI_ISL_848262, EPI_ISL_848269, EPI_ISL_848286, EPI_ISL_848287, EPI_ISL_848288, EPI_ISL_848289, EPI_ISL_848290, EPI_ISL_848291, EPI_ISL_848292, EPI_ISL_848293, EPI_ISL_848294, EPI_ISL_848295, EPI_ISL_848450, EPI_ISL_848460, EPI_ISL_848469, EPI_ISL_848470, EPI_ISL_848471, EPI_ISL_848472, EPI_ISL_848473, EPI_ISL_848474, EPI_ISL_848475, EPI_ISL_848476, EPI_ISL_848477, EPI_ISL_848478, EPI_ISL_848479, EPI_ISL_848480 | | | |
| see above | Illinois Department of Public Health | Gagnon Lab, Southern Illinois University | Keith Gagnon |
| EPI_ISL_848583, EPI_ISL_848584, EPI_ISL_848585, EPI_ISL_848586, EPI_ISL_848616 | Evandro Chagas Institute | Evandro Chagas Institute | Santos, M.C.; Silva, A.M.; Junior, W.D.C.; Barbagelata, L.S.; Ferreira, J.A.; Sousa, E.M.A.; da Silva, P.S.; Pinheiro, K.C.; L.C.; Sousa Junior, E.C. |
| EPI_ISL_850185, EPI_ISL_850186 | Division of Emerging Infectious Diseases, Bureau of Infectious Diseases Diagnosis Control, Korea Disease Control and Prevention Agency | Division of Emerging Infectious Diseases, Bureau of Infectious Diseases Diagnosis Control, Korea Disease Control and Prevention Agency | Ae Kyung Park, Il-Hwan Kim, Heui Man Kim, Jeong-Min Kim, Namjoo Lee, Chaeyoung Lee, Sang Hee Woo, Eun-Jin Kim |
| EPI_ISL_852659, EPI_ISL_852660 | Institute of Virology, Medical Center, University of Freiburg, Freiburg, Germany | Institute of Virology, Clinial Virus Genomics, Medical Center, University of Freiburg, Freiburg, Germany | Jonas Fuchs, Lisa Kern, Sandra Reuter, Hajo Grundmann, Marcus Panning |
| EPI_ISL_852834, EPI_ISL_852835, EPI_ISL_852836, EPI_ISL_852837, EPI_ISL_852838, EPI_ISL_852839, EPI_ISL_852840, EPI_ISL_852841, EPI_ISL_853407, EPI_ISL_853408, EPI_ISL_853409, EPI_ISL_853410, EPI_ISL_853411 | | | |
| see above | Laboratoire de santé publique du Québec | Laboratoire de santé publique du Québec | Sandrine Moreira, Ioannis Ragoussis, Guillaume Bourque, Jesse Shapiro, Mark Lathrop and Michel Roger on behalf of the CoVSeQ research group (http://covseq.ca/researchgroup) |
| EPI_ISL_853820, EPI_ISL_853821 | Department of Pathology, Landeskrankenhaus Graz II, Medical University Graz | Bergthaler laboratory, CeMM Research Center for Molecular Medicine of the Austrian Academy of Sciences | Lukas Endler, Alexandra Popa, Benedikt Agerer, Jakob-Wendelin Genger, Alexander Lercher, Anna Schedl, Thomas Penz, Michael Schuster, Jan Laine, Martin Senekowitsch, Christoph Bock, Andreas Bergthaler |
| EPI_ISL_853884 | Center for Virology, Medical University of Vienna | Bergthaler laboratory, CeMM Research Center for Molecular Medicine of the Austrian Academy of Sciences | Lukas Endler, Alexandra Popa, Benedikt Agerer, Jakob-Wendelin Genger, Alexander Lercher, Anna Schedl, Thomas Penz, Michael Schuster, Jan Laine, Martin Senekowitsch, Christoph Bock, Andreas Bergthaler |
| EPI_ISL_854556, EPI_ISL_854557, EPI_ISL_854558, EPI_ISL_854559, EPI_ISL_854560, EPI_ISL_854561, EPI_ISL_854562, EPI_ISL_854563, EPI_ISL_854564, EPI_ISL_854565, EPI_ISL_854566, EPI_ISL_854567, EPI_ISL_854568, EPI_ISL_854569, EPI_ISL_854570, EPI_ISL_854571, EPI_ISL_854572, EPI_ISL_854573, EPI_ISL_854574, EPI_ISL_854575, EPI_ISL_854576 | | | |
| see above | Alberta Precision Labs (APL) | Alberta Precision Labs (APL) | Gordon P, Lam LG, Pabbaraju K, Wong A, Ma R, Li V, Melin A, Tipples G, Berenger B, Zelyas N, Kellner J, Bernier F, Chui L, Croxen M |
| EPI_ISL_859936, EPI_ISL_859937, EPI_ISL_859938, EPI_ISL_859939, EPI_ISL_859940, EPI_ISL_859941, EPI_ISL_859942, EPI_ISL_859943, EPI_ISL_859944, EPI_ISL_859945, EPI_ISL_859946, EPI_ISL_859947, EPI_ISL_859948, EPI_ISL_859949, EPI_ISL_859950, EPI_ISL_859951, EPI_ISL_859952, EPI_ISL_859953, EPI_ISL_859954, EPI_ISL_859955, EPI_ISL_859956, EPI_ISL_859957, EPI_ISL_859958, EPI_ISL_859959, EPI_ISL_859960, EPI_ISL_859961, EPI_ISL_859962, EPI_ISL_859963, EPI_ISL_859964, EPI_ISL_859965, EPI_ISL_859966, EPI_ISL_859967, EPI_ISL_859968, EPI_ISL_859969, EPI_ISL_859970, EPI_ISL_859971, EPI_ISL_859972, EPI_ISL_859973, EPI_ISL_859974, EPI_ISL_859975, EPI_ISL_859976, EPI_ISL_859977, EPI_ISL_859978, EPI_ISL_859979, EPI_ISL_859980, EPI_ISL_859981, EPI_ISL_859982, EPI_ISL_859983, EPI_ISL_859993, EPI_ISL_859994, EPI_ISL_859995, EPI_ISL_859996, EPI_ISL_859997, EPI_ISL_859998, EPI_ISL_860000, EPI_ISL_860001, EPI_ISL_860002, EPI_ISL_860003, EPI_ISL_860004, EPI_ISL_860005, EPI_ISL_860006, EPI_ISL_860007, EPI_ISL_860008, EPI_ISL_860009, EPI_ISL_860010, EPI_ISL_860011, EPI_ISL_860012, EPI_ISL_860013, EPI_ISL_860014, EPI_ISL_860015, EPI_ISL_860016, EPI_ISL_860017, EPI_ISL_860018, EPI_ISL_860019, EPI_ISL_860020, EPI_ISL_860021, EPI_ISL_860022, EPI_ISL_860023, EPI_ISL_860024, EPI_ISL_860025 | | | |
| see above | BTC, Khalifa University | BTC, Khalifa University | Al Safar et al |
| EPI_ISL_861421 | Instituto de Diagnostico y Referencia Epidemiologicos (INDRE) | Instituto de Diagnostico y Referencia Epidemiologicos (INDRE) | Claudia Wong-Arambula, Gisela Barrera-Badillo , Abril Rodriguez-Maldonado, Natividad Cruz-Ortiz, Tatiana Nunez-Garcia, Dayanira Arellano-Suarez, Fabiola Garces-Ayala, Adnan-Araiza Rodríguez, Edgar Mendieta-Condado, Lucia Hernandez-Rivas, Irma Lopez-Martinez, Ernesto Ramirez-Gonzalez. |
| EPI_ISL_861643 | Instituto Adolfo Lutz - Central | Instituto Adolfo Lutz, Interdiciplinary Procedures Center, Strategic Laboratory | Claudio Tavares Sacchi, Claudia Regina Gonçalves, Erica Valessa Ramos Gomes, Karoline Rodrigues Campos |
| EPI_ISL_862868 | Fujita Health University Bantane Hospital | Fujita health University, School of Medicine, Department of Microbiology | Masahiro suzuki, Yohei Doi |
| EPI_ISL_862875 | Komatsu Municipal Hospital | Fujita health University, School of Medicine, Department of Microbiology | Masahiro suzuki, Yohei Doi |
| EPI_ISL_862876 | NTT East Sapporo Hospital | Fujita health University, School of Medicine, Department of Microbiology | Masahiro suzuki, Yohei Doi |
| EPI_ISL_862881 | Kitakyushu Municipal Medical Center | Fujita health University, School of Medicine, Department of Microbiology | Masahiro suzuki, Yohei Doi |
| EPI_ISL_862882, EPI_ISL_862883 | University Hospital Kyoto Prefectural University of Medicine | Fujita health University, School of Medicine, Department of Microbiology | Masahiro suzuki, Yohei Doi |
| EPI_ISL_872099 | Instituto de Diagnostico y Referencia Epidemiologicos (INDRE) | Instituto de Diagnostico y Referencia Epidemiologicos (INDRE) | Fabiola Garces-Ayala, Gisela Barrera-Badillo , Abril Rodriguez-Maldonado, Claudia Wong-Arambula , Natividad Cruz-Ortiz, Tatiana Nunez-Garcia, Dayanira Arellano-Suarez, Adnan-Araiza Rodríguez, Edgar Mendieta-Condado, Lucia Hernandez-Rivas, Irma Lopez-Martinez, Ernesto Ramirez-Gonzalez. |
| EPI_ISL_872897 | Stem Cell Research and Development Center | Stem Cell Research and Development Center | Igo Syaiful Ihsan, Eryk Hendrianto, Naimah Putri, Deya Karsari, Nora Ertanti, Aristika Dinaryanti, Purwati |
| EPI_ISL_876969, EPI_ISL_876970, EPI_ISL_877034 | 1.AO Universitaria 'S. Giovanni di Dio e Ruggi D'Aragona, Scuola Medica Salernitana' Hospital / 2.UOC di Virologia e Microbiologia, Università della Campania 'L. Vanvitelli' / 3.AO Universitaria 'Federico II' Napoli Hospital / 4.AORN 'San Giuseppe Moscati' Avellino Hospital / 5.AO 'San Pio - presidio G. Rummo' Benevento Hospital / 6.AO 'Sant'Anna e San | 1. Genome Research Center for Health (CRGS) / 2. Laboratory of Molecular Medicine and Genomics(LMMGe) / 3. Center for Research in Pure and Applied Mathematics (CRMPA) | Giorgio Giurato (Corresponding Author), Francesca Rizzo (Corresponding Author), Alessandro Weisz (Corresponding Author), Gianluigi Franci, Giovanni Nassa, Pasquale Pagliano, Roberta Tarallo, Elena Alexandrova, Ylenia D'Agostino, Carlo Ferravante, Jessica Lamberti, Viola Melone, Domenico Memoli, Valeria Mirici Cappa, Domenico Palumbo, Giovanni Pecoraro, Assunta Sellitto, Oriana Strianese, Ilaria Terenzi, Giuseppe Fenza, Aniello Gentile, Antonello Saccomanno, Sonia Amabile, Teresa Rocco, Annamaria Salvati, Emilia Vaccaro, Massimiliano Galdiero, Michele Cennamo, Giuseppe Portella, Maria Grazia Foti, Mariarosaria Ingino, Maria Landi, Maurizio Fumi, Vincenzo Rocco, Rita Greco, Vittoria Letizia, Arnolfo Petruzziello, Maddalena Schioppa, Gregorio Goffredi, Francesca Marciano, Michele Caraglia, Alessia Cossu, Marianna Scrima |

| | | | |
|---|---|---|---|
| | Sebastiano' Caserta Hospital / 7.PO 'Maria Santissima Addolorata' Eboli Hospital / 8.Biogem Istituto di Ricerche Genetiche | | |
| EPI_ISL_877606, EPI_ISL_877607, EPI_ISL_877608 | Clinical Molecular Microbiology Laboratory, UNC Hospital | Dirk Dittmer | Razia Moorad , Justin T. Landis , Brent A. Eason, Melissa B. Miller, Linda Pluta, Dirk Dittmer, Angelica Juarez, Cecilia Thompson , Cameroon Grant, Evelyn Hoffman, Patricio Cano, Jason Wong, Carolina Caro-Vegas, Blossom Damania. |
| EPI_ISL_883192 | UK Tübingen, Medical Microbiology | Hannover Medical School, Institute of Virology | Lars Steinbrück, Jasper Götting |
| EPI_ISL_887186, EPI_ISL_887188, EPI_ISL_887196, EPI_ISL_887200, EPI_ISL_887206, EPI_ISL_887218, EPI_ISL_887237, EPI_ISL_887239, EPI_ISL_887290, EPI_ISL_887309 | Protzer Lab | Protzer Lab, Gagneur Lab, Robert Koch Institut | Ulrike Protzer, Dieter Hoffmann, Eva Schulte, Andrea Theumer, Oliver Drechsel, Max von Kleist,Aleksandar Radonic,Stephan Fuchs, Alexander Karollus, Julien Gagneur |
| EPI_ISL_890033 | Laboratoire de santé publique du Québec | Laboratoire de santé publique du Québec | Sandrine Moreira, Ioannis Ragoussis, Guillaume Bourque, Jesse Shapiro, Mark Lathrop and Michel Roger on behalf of the CoVSeQ research group |
| EPI_ISL_891156, EPI_ISL_891157, EPI_ISL_891158, EPI_ISL_891159 | DPH, Massachusetts State Public Health Lab | DPH, Massachusetts State Public Health Lab | Lang,A.S., Fink,T., Gallagher,G.R., Smole,S.C. |
| EPI_ISL_895754, EPI_ISL_895789 | Molecular biology division, Institute of Clinical Biochemistry and Diagnostics, Charles University, Faculty of Medicine in Hradec Králové and University Hospital Hradec Králové | Molecular biology division, Institute of Clinical Biochemistry and Diagnostics, Charles University, Faculty of Medicine in Hradec Králové and University Hospital Hradec Králové | Helena Kovaíková, Petr Brož, Ivana Baranová, Kateina Hrochová, Tereza Baková, Jitka Novotná, Kateina Pehlíková, Vladimír Palika. Cooperation project with BioVendor-R&D and bioinformatics company BIOXSYS s r.o. |
| EPI_ISL_896576, EPI_ISL_896577, EPI_ISL_896578, EPI_ISL_896579, EPI_ISL_896580, EPI_ISL_896581 | Department of Medical Research | DMR | Myat Htut Nyunt |
| EPI_ISL_897537, EPI_ISL_897538, EPI_ISL_897539, EPI_ISL_897540, EPI_ISL_897541, EPI_ISL_897542, EPI_ISL_897543 | Pathogen Genomics Center, National Institute of Infectious Diseases | Pathogen Genomics Center, National Institute of Infectious Diseases | Tsuyoshi Sekizuka, Kentaro Itokawa, Rina Tanaka, Masanori Hashino, Makoto Kuroda |
| EPI_ISL_898325 | Toyama Institute of Health | Pathogen Genomics Center, National Institute of Infectious Diseases | Tsuyoshi Sekizuka, Kentaro Itokawa, Rina Tanaka, Masanori Hashino, Makoto Kuroda |
| EPI_ISL_900057, EPI_ISL_900148, EPI_ISL_900165, EPI_ISL_900457 | MEPHI, Aix Marseille University | MEPHI, Aix Marseille University | Anthony LEVASSEUR |
| EPI_ISL_901251, EPI_ISL_901391, EPI_ISL_902209, EPI_ISL_902210, EPI_ISL_902211, EPI_ISL_902212, EPI_ISL_902309, EPI_ISL_902351, EPI_ISL_902352, EPI_ISL_902353, EPI_ISL_902354, EPI_ISL_902355, EPI_ISL_902356, EPI_ISL_902357, EPI_ISL_902358, EPI_ISL_902359, EPI_ISL_902360, EPI_ISL_902361, EPI_ISL_902362, EPI_ISL_902363, EPI_ISL_902364, EPI_ISL_902365, EPI_ISL_902366, EPI_ISL_906466 | | | |
| see above | Pathogen Genomics Center, National Institute of Infectious Diseases | Pathogen Genomics Center, National Institute of Infectious Diseases | Tsuyoshi Sekizuka, Kentaro Itokawa, Rina Tanaka, Masanori Hashino, Makoto Kuroda |
| EPI_ISL_913474 | Klinisk mikrobiologi | The Public Health Agency of Sweden | Anna-Malin Linde, Maria Lind Karlberg, Carlo Berg, Oskar Karlsson Lindsjo, Sofia Stamouli, Reza Advani, Mattias Haukland, Petra Holmstrom, Noura Walai, Petra Edquist, Mia Brytting, Anna Risberg, Karin Tegmark-Wisell |
| EPI_ISL_914067, EPI_ISL_914068, EPI_ISL_914069, EPI_ISL_914070, EPI_ISL_914073, EPI_ISL_914074, EPI_ISL_914075, EPI_ISL_914076, EPI_ISL_914077, EPI_ISL_914078, EPI_ISL_914079, EPI_ISL_914080, EPI_ISL_914081, EPI_ISL_914082, EPI_ISL_914083, EPI_ISL_914084, EPI_ISL_914085, EPI_ISL_914086, EPI_ISL_914087, EPI_ISL_914088, EPI_ISL_914089, EPI_ISL_914090, EPI_ISL_914091, EPI_ISL_914092, EPI_ISL_914093, EPI_ISL_914094, EPI_ISL_914095, EPI_ISL_914242 | | | |
| see above | TGen North | TGen North | "Jolene Bowers, Megan Folkerts, Chris French, Hayley Yaglom, Ashlyn Pfeiffer, Darrin Lemmer, Dave Engelthaler, The Arizona COVID Genomics Union (ACGU)" |
| EPI_ISL_914974, EPI_ISL_914975, EPI_ISL_915149, EPI_ISL_915150, EPI_ISL_915151, EPI_ISL_915152, EPI_ISL_915153, EPI_ISL_915154, EPI_ISL_915155, EPI_ISL_915156, EPI_ISL_915157, EPI_ISL_915158, EPI_ISL_915159, EPI_ISL_915160, EPI_ISL_915161, EPI_ISL_915162, EPI_ISL_915163, EPI_ISL_915164, EPI_ISL_915165, EPI_ISL_915166, EPI_ISL_915167, EPI_ISL_915168, EPI_ISL_915169, EPI_ISL_915170, EPI_ISL_915171, EPI_ISL_915172, EPI_ISL_915173 | | | |
| see above | QEII Health Sciences Centre | National Microbiology Laboratory (NML) | Anna Majer, Shari Tyson, Grace Seo, Philip Mabon, Elsie Grudeski, Rhiannon Huzarewich, Russell Mandes, Anneliese Landgraff, Jennifer Tanner, Natalie Knox, Morag Graham, Gary Van Domselaar, Todd Hatchette, Jason LeBlanc, Janice Pettipas, Dan Gaston, Nathalie Bastien, Yan Li, Timothy Booth, Darian Hole, Madison Chapel, CanCOGeN's metadata curation team, Public Health Agency of Canada CanCOGeN team |
| EPI_ISL_933633, EPI_ISL_933634, EPI_ISL_933635, EPI_ISL_933636 | Toronto Invasive Bacterial Diseases Network | McMaster University | Allison McGeer, Patryk Aftanas, Hooman Derakhshani, Angel Li, Kuganya Nirmalarajah, Emily Panousis, Ahmed Draia, Jalees Nasir, Michael Surette, Samira Mubareka, Andrew G. McArthur |
| EPI_ISL_934323 | Klinisk mikrobiologi | The Public Health Agency of Sweden | Anna-Malin Linde, Maria Lind Karlberg, Carlo Berg, Oskar Karlsson Lindsjo, Sofia Stamouli, Reza Advani, Mattias Haukland, Petra Holmstrom, Noura Walai, Petra Edquist, Mia Brytting, Anna Risberg, Karin Tegmark-Wisell |
| EPI_ISL_939671, EPI_ISL_939672 | Division of Infectious Diseases, University Hospital Zürich | Institute of Medical Virology, University of Zurich | Stefan Schmutz, Maryam Zaheri, Verena Kufner, Annette Audigé, Maria Grünberg, Kevin Steiner, Jon Huder, Cyril Shah, Riccarda Capaul, Guido Bloemberg, Jürg Böni, Michael Huber, Alexandra Trkola |
| EPI_ISL_940210, EPI_ISL_940505 | Hôpital Bichat Claude Bernard, Laboratoire de Virologie | IAME UMR1137 Inserm, Université de Paris, Hôpital Bichat | Antoine Bridier-Nahmias, Amélie Recoing, Quentin Le Hingrat, Lena Daniel, Siham Hamri, Gilles Collin, Alexandre Storto, Mélanie Bertine, Charlotte Charpentier, Nadhira Houhou-Fidouh, Diane Descamps, Benoit Visseaux |
| EPI_ISL_941104, EPI_ISL_941105, EPI_ISL_941106, EPI_ISL_941107, EPI_ISL_941108, EPI_ISL_941109, EPI_ISL_941110, EPI_ISL_941111, EPI_ISL_941112, EPI_ISL_941113, EPI_ISL_941114, EPI_ISL_941115, EPI_ISL_941116, EPI_ISL_941117, EPI_ISL_941118, EPI_ISL_941119, EPI_ISL_941120, EPI_ISL_941121, EPI_ISL_941122, EPI_ISL_941123, EPI_ISL_941124, EPI_ISL_941125, EPI_ISL_941126, EPI_ISL_941127, EPI_ISL_941128, EPI_ISL_941129, EPI_ISL_941130, EPI_ISL_941131, EPI_ISL_941132, EPI_ISL_941133, EPI_ISL_941135, EPI_ISL_941136, EPI_ISL_941137, EPI_ISL_941138, EPI_ISL_941139, EPI_ISL_941140, EPI_ISL_941142, EPI_ISL_941143, EPI_ISL_941144, EPI_ISL_941145, EPI_ISL_941146, EPI_ISL_941147, EPI_ISL_941148, EPI_ISL_941149, EPI_ISL_941150, EPI_ISL_941151, EPI_ISL_941152, EPI_ISL_941153, EPI_ISL_941154, EPI_ISL_941155, EPI_ISL_941156, EPI_ISL_941157, EPI_ISL_941158, EPI_ISL_941159, EPI_ISL_941160, EPI_ISL_941162 | | | |
| see above | Centro de Investigaciones en Microbiología y Biotecnología-UR (CIMBIUR), Facultad de Ciencias Naturales, Universidad del Rosario, Bogotá, Colombia | Centro de Investigaciones en Microbiología y Biotecnología-UR (CIMBIUR), Facultad de Ciencias Naturales, Universidad del Rosario, Bogotá, Colombia Icahn School of Medicine at Mount Sinai, New York, USA | Nathalia Ballesteros, Marina Muñoz, Luz Helena Patiño, Carolina Hernández, Felipe González-Casabianca, Iván Carroll, Mauricio Santos-Vega, Jaime Cascante, Andrés Angel, Alejandro Feged-Rivadeneira, Mónica Palma-Cuero, Carolina Flórez, Sergio Gomez, Adriana van de Guchte, Zenab Khan, Jayeeta Dutta, Hala Alejel Alshammary, Ana S. Gonzalez-Reiche, Matthew M. Hernandez, Emilia Mia Sordillo, Viviana Simon, Harm van Bakel, Alberto Paniz-Mondolfi, Juan David Ramírez |
| EPI_ISL_941944, EPI_ISL_941945, EPI_ISL_941962, EPI_ISL_941964, EPI_ISL_941965, EPI_ISL_941977, EPI_ISL_941978, EPI_ISL_941979, EPI_ISL_941980, EPI_ISL_941981, EPI_ISL_941982, EPI_ISL_942000, EPI_ISL_942001, EPI_ISL_942002 | | | |
| see above | Centro de Investigaciones en Microbiología y Biotecnología-UR (CIMBIUR), Facultad de Ciencias Naturales, Universidad del Rosario, Bogotá, Colombia | Centro de Investigaciones en Microbiología y Biotecnología-UR (CIMBIUR), Facultad de Ciencias Naturales, Universidad del Rosario, Bogotá, Colombia Instituto Nacional de Salud, Bogotá, Colombia Icahn School of Medicine at Mount Sinai, New York, USA | Luz Helena Patiño, Marina Muñoz, Nathalia Ballesteros, Carolina Hernández, Carolina Flórez, Sergio Gomez, Adriana van de Guchte, Zenab Khan, Jayeeta Dutta, Hala Alejel Alshammary, Ana S. Gonzalez-Reiche, Matthew M. Hernandez, Emilia Mia Sordillo, Viviana Simon, Harm van Bakel, Alberto Paniz-Mondolfi, Juan David Ramírez |
| EPI_ISL_944645 | Medical Microbiology, Radboud University Medical Center | Medical Microbiology, Radboud University Medical Center | Varghese,F.S., Overheul,G.J., Coolen,J.P. and van Rij,R.P. |
| EPI_ISL_953358, EPI_ISL_953373, EPI_ISL_953374 | Bioinformatics and Biostatistics Lab, Advanced Sequencing Facility | COVID-19 Genomics UK (COG-UK) Consortium | Aengus Stewart,Jerome Nicod,Chelsea Sawyer,Laura Cubitt,Harshil Patel,Margaret Crawford |
| EPI_ISL_953409, EPI_ISL_953421 | Laboratorio de Investigaciones de Baney | "Swiss Tropical and Public Health Institute" | "Carlos Cortes, Claudia Daubenberger, Guillermo Garcia, Salome Hosch, Bonifacio Manguire Nlavo, Maximilian Mpina, Elizabeth Nyakarungu, Diosdado Odjama Nseng Ada, Mitoha Ondo O Ayekaba, Tobias Schindler, Philip Wonder Phiri" |
| EPI_ISL_956307 | Primasatya Husada Citra Hospital | Institute of Tropical Disease, Universitas Airlangga | Krisnoadi Rahardjo, Aldise M Nastri, Jezzy R Dewantari, Rima R Prasetya, Pudji Djanuartono, Gatot Soegiarto, Laksmi Wulandari, Resti Yudhawati, Yasuko Mori, Soetjipto, Kazufumi Shimizu, Maria I Lusida |
| EPI_ISL_960174 | Guguletu CHC wc GDH | National Health Laboratory Service/UCT | Arash Iranzadeh, Deelan Doolabh, Lynn Tyers, Bruna Galvao, Innocent Mudau, Marvin Hsiao, Kruger Marais, Diana Hardie, Stephen Korsman, Carolyn |

| | | | |
|---|---|---|---|
| | | | Williamson |
| EPI_ISL_960306 | NLZOH, Laboratory for Virology | NLZOH, Laboratory for Virology | Katarina Prosenc (Laboratory for Virology), Cesare Camma (IZSAM), Erik Alm (ECDC) |
| EPI_ISL_961014 | Viral Respiratory Lab, National Institute for Biomedical Research (INRB) | Pathogen Sequencing Lab, National Institute for Biomedical Research (INRB) | Placide Mbala-Kingebeni, Edith Nkwembe, Eddy Kinganda-Lusamaki, Amuri Aziza, Francisca Muyembe Mawete, Emmanuel Lokilo Lofiko, Jean-Claude Makangara Cigolo, Catherine Pratt, Matthias Pauthner, Josh Quick, Allison Black, James Hadfield, Trevor Bedford, Ian Goodfellow, Andrew Rambaut, Nick Loman, Kristian Andersen, Michael Wiley, Steve Ahuka-Mundeke, Jean-Jacques Muyembe Tamfum |
| EPI_ISL_961883, EPI_ISL_961884, EPI_ISL_961885, EPI_ISL_961887, EPI_ISL_961894, EPI_ISL_961895, EPI_ISL_961903, EPI_ISL_961908, EPI_ISL_961911, EPI_ISL_961912, EPI_ISL_961913, EPI_ISL_961915, EPI_ISL_961916, EPI_ISL_961917, EPI_ISL_961919, EPI_ISL_961922, EPI_ISL_961923, EPI_ISL_961943, EPI_ISL_961955, EPI_ISL_961956, EPI_ISL_961961, EPI_ISL_961963, EPI_ISL_961964, EPI_ISL_961965, EPI_ISL_961979, EPI_ISL_961986, EPI_ISL_961987, EPI_ISL_961989, EPI_ISL_961995, EPI_ISL_961997, EPI_ISL_962000, EPI_ISL_962002, EPI_ISL_962003, EPI_ISL_962005, EPI_ISL_962006, EPI_ISL_962007, EPI_ISL_962008, EPI_ISL_962009, EPI_ISL_962019, EPI_ISL_962020, EPI_ISL_962021, EPI_ISL_962022, EPI_ISL_962023, EPI_ISL_962024, EPI_ISL_962025, EPI_ISL_962026, EPI_ISL_962027, EPI_ISL_962028, EPI_ISL_962029, EPI_ISL_962030, EPI_ISL_962031, EPI_ISL_962032, EPI_ISL_962033, EPI_ISL_962034, EPI_ISL_962035, EPI_ISL_962036, EPI_ISL_962037, EPI_ISL_962038, EPI_ISL_962039, EPI_ISL_962040, EPI_ISL_962041, EPI_ISL_962042, EPI_ISL_962043, EPI_ISL_962044, EPI_ISL_962053, EPI_ISL_962055, EPI_ISL_962056, EPI_ISL_962057, EPI_ISL_962058, EPI_ISL_962059, EPI_ISL_962060, EPI_ISL_962061, EPI_ISL_962062, EPI_ISL_962063, EPI_ISL_962064, EPI_ISL_962065, EPI_ISL_962066 | | | |
| see above | Illinois Department of Public Health | Gagnon Lab, Southern Illinois University | Keith Gagnon |
| EPI_ISL_968177, EPI_ISL_968178, EPI_ISL_968179 | Clinical Molecular Microbiology Laboratory, UNC Hospital | Dirk Dittmer | Justin T. Landis , Razia Moorad , Brent A. Eason, Melissa B. Miller, Linda Pluta, Dirk Dittmer, Angelica Juarez, Cecilia Thompson, Shawn Hawken, Cameroon Grant, Evelyn Hoffman, Patricio Cano, Jason Wong, Carolina Caro-Vegas, Ryan McNamara, Blossom Damania. |
| EPI_ISL_978491 | Central Public Health Laboratory - LACEN -Bahia, Salvador, Brazil | Central Public Health Laboratory - LACEN -Bahia, Salvador, Brazil | Stephane Tosta, Luciana Oliveira, Vanessa Nardy,Patrícia Cajado,Marcela Gómez, Breno Dominguez, Jaqueline Gomes, Vagner Fonseca,Marta Giovanetti,Luiz Alcantara, Felicidade Pereira, Arabela Leal |
| EPI_ISL_981225, EPI_ISL_981226, EPI_ISL_981227, EPI_ISL_981228, EPI_ISL_981229, EPI_ISL_981230, EPI_ISL_981231, EPI_ISL_981232, EPI_ISL_981233, EPI_ISL_981234, EPI_ISL_981235, EPI_ISL_981236, EPI_ISL_981237, EPI_ISL_981238, EPI_ISL_981239, EPI_ISL_981240, EPI_ISL_981241, EPI_ISL_981242 | | | |
| see above | National Virus Reference Laboratory | Irish Coronavirus Sequencing Consortium - Helixworks | Sachin Chalapati, Conor Crosbie, Nimesh Pinnamaneni |
