## Supplementary material for "Detection and characterization of the SARS-CoV-2 lineage B.1.526 in New York": Supp. Table 3 GISAID Acknowledgment part 2: gisaid_hcov-19_acknowledgement_table_2021_02_12_16-13.pdf

We gratefully acknowledge the following Authors from the Originating laboratories responsible for obtaining the specimens, as well as the Submitting laboratories where the genome data were generated and shared via GISAID, on which this research is based.

All Submitters of data may be contacted directly via www.gisaid.org

Authors are sorted alphabetically.

| Accession ID | Originating Laboratory | Submitting Laboratory | Authors |
|---|---|---|---|
| EPI_ISL_445075 | Laboratoire National de Sante, Microbiology, Virology | Laboratoire National de Sante, Microbiology, Epidemiology and Microbial Genomics | Anke Wienecke-Baldacchino, Ardashel Latsuzbaia, Jessica Tapp, Catherine Ragimbeau, Guillaume Fournier, Tamir Abdelrahman, Trung Nguyen Nguyen, Joel Mossong |
| EPI_ISL_447534 | Gujarat Biotechnology Research Centre | Gujarat Biotechnology Research Centre | Gaurishankar Shrimali, Nidhi Sood, Pranay Shah, R D Dixit, Snehal Bagatharia, Kamlesh J Upadhyay, Ramesh Pandit, Tejas Shah, Ankit Hinsu, Pritesh Sabara, Apurvasinh Puvar, Janvi Raval, Monika Gandhi, Pinal Trivedi, Maharshi Pandya, Amit Kanani, Akanksha Verma, Nitin Savaliya, Raghawendra Kumar, Dinesh Kumar, Zuber Saiyed, Dipa Kinariwala, Disha Patel, Binita Aring, Neeta Khandelwal, Geeta Vaghela, Sonia Barve, Bhavesh Modi, Kairavi Joshi, Nidhi Patel, Chaitanya Joshi, Madhvi Joshi |
| EPI_ISL_447535 | Gujarat Biotechnology Research Centre | Gujarat Biotechnology Research Centre | Nidhi Sood, Pranay Shah, R D Dixit, Snehal Bagatharia, Kamlesh J Upadhyay, Ramesh Pandit, Tejas Shah, Ankit Hinsu, Pritesh Sabara, Apurvasinh Puvar, Janvi Raval, Monika Gandhi, Pinal Trivedi, Maharshi Pandya, Amit Kanani, Akanksha Verma, Nitin Savaliya, Raghawendra Kumar, Dinesh Kumar, Zuber Saiyed, Dipa Kinariwala, Disha Patel, Binita Aring, Neeta Khandelwal, Geeta Vaghela, Sonia Barve, Bhavesh Modi, Kairavi Joshi, Gaurishankar Shrimali, Priti Pandita, Chaitanya Joshi, Madhvi Joshi |
| EPI_ISL_447536 | Gujarat Biotechnology Research Centre | Gujarat Biotechnology Research Centre | Pranay Shah, R D Dixit, Snehal Bagatharia, Kamlesh J Upadhyay, Ramesh Pandit, Tejas Shah, Ankit Hinsu, Pritesh Sabara, Apurvasinh Puvar, Janvi Raval, Monika Gandhi, Pinal Trivedi, Maharshi Pandya, Amit Kanani, Akanksha Verma, Nitin Savaliya, Raghawendra Kumar, Dinesh Kumar, Zuber Saiyed, Dipa Kinariwala, Disha Patel, Binita Aring, Neeta Khandelwal, Geeta Vaghela, Sonia Barve, Bhavesh Modi, Kairavi Joshi, Gaurishankar Shrimali, Nidhi Sood, Neha Rajpara, Chaitanya Joshi, Madhvi Joshi |
| EPI_ISL_447537 | Gujarat Biotechnology Research Centre | Gujarat Biotechnology Research Centre | R D Dixit, Snehal Bagatharia, Kamlesh J Upadhyay, Ramesh Pandit, Tejas Shah, Ankit Hinsu, Pritesh Sabara, Apurvasinh Puvar, Janvi Raval, Monika Gandhi, Pinal Trivedi, Maharshi Pandya, Amit Kanani, Akanksha Verma, Nitin Savaliya, Raghawendra Kumar, Dinesh Kumar, Zuber Saiyed, Dipa Kinariwala, Disha Patel, Binita Aring, Neeta Khandelwal, Geeta Vaghela, Sonia Barve, Bhavesh Modi, Kairavi Joshi, Gaurishankar Shrimali, Nidhi Sood, Pranay Shah, Afzal Ansari, Chaitanya Joshi, Madhvi Joshi |
| EPI_ISL_447538 | Gujarat Biotechnology Research Centre | Gujarat Biotechnology Research Centre | Snehal Bagatharia, Kamlesh J Upadhyay, Ramesh Pandit, Tejas Shah, Ankit Hinsu, Pritesh Sabara, Apurvasinh Puvar, Janvi Raval, Monika Gandhi, Pinal Trivedi, Maharshi Pandya, Amit Kanani, Akanksha Verma, Nitin Savaliya, Raghawendra Kumar, Dinesh Kumar, Zuber Saiyed, Dipa Kinariwala, Disha Patel, Binita Aring, Neeta Khandelwal, Geeta Vaghela, Sonia Barve, Bhavesh Modi, Kairavi Joshi, Gaurishankar Shrimali, Nidhi Sood, Pranay Shah, R D Dixit, Neelam Nathani, Chaitanya Joshi, Madhvi Joshi |
| EPI_ISL_447539 | Gujarat Biotechnology Research Centre | Gujarat Biotechnology Research Centre | Kamlesh J Upadhyay, Ramesh Pandit, Tejas Shah, Ankit Hinsu, Pritesh Sabara, Apurvasinh Puvar, Janvi Raval, Monika Gandhi, Pinal Trivedi, Maharshi Pandya, Amit Kanani, Akanksha Verma, Nitin Savaliya, Raghawendra Kumar, Dinesh Kumar, Zuber Saiyed, Dipa Kinariwala, Disha Patel, Binita Aring, Neeta Khandelwal, Geeta Vaghela, Sonia Barve, Bhavesh Modi, Kairavi Joshi, Gaurishankar Shrimali, Nidhi Sood, Pranay Shah, R D Dixit, Snehal Bagatharia, Armi Chaudhari, Chaitanya Joshi, Madhvi Joshi |
| EPI_ISL_447540 | Gujarat Biotechnology Research Centre | Gujarat Biotechnology Research Centre | Ramesh Pandit, Tejas Shah, Ankit Hinsu, Pritesh Sabara, Apurvasinh Puvar, Janvi Raval, Monika Gandhi, Pinal Trivedi, Maharshi Pandya, Amit Kanani, Akanksha Verma, Nitin Savaliya, Raghawendra Kumar, Dinesh Kumar, Zuber Saiyed, Dipa Kinariwala, Disha Patel, Binita Aring, Neeta Khandelwal, Geeta Vaghela, Sonia Barve, Bhavesh Modi, Kairavi Joshi, Gaurishankar Shrimali, Nidhi Sood, Pranay Shah, R D Dixit, Snehal Bagatharia, Kamlesh J Upadhyay, Bhavya Jindal, Chaitanya Joshi, Madhvi Joshi |
| EPI_ISL_447541 | Gujarat Biotechnology Research Centre | Gujarat Biotechnology Research Centre | Tejas Shah, Ankit Hinsu, Pritesh Sabara, Apurvasinh Puvar, Janvi Raval, Monika Gandhi, Pinal Trivedi, Maharshi Pandya, Amit Kanani, Akanksha Verma, Nitin Savaliya, Raghawendra Kumar, Dinesh Kumar, Zuber Saiyed, Dipa Kinariwala, Disha Patel, Binita Aring, Neeta Khandelwal, Geeta Vaghela, Sonia Barve, Bhavesh Modi, Kairavi Joshi, Gaurishankar Shrimali, Nidhi Sood, Pranay Shah, R D Dixit, Snehal Bagatharia, Kamlesh J Upadhyay, Ramesh Pandit, Anjali Rajwar, Chaitanya Joshi, Madhvi Joshi |
| EPI_ISL_447542 | Gujarat Biotechnology Research Centre | Gujarat Biotechnology Research Centre | Ankit Hinsu, Pritesh Sabara, Apurvasinh Puvar, Janvi Raval, Monika Gandhi, Pinal Trivedi, Maharshi Pandya, Amit Kanani, Akanksha Verma, Nitin Savaliya, Raghawendra Kumar, Dinesh Kumar, Zuber Saiyed, Dipa Kinariwala, Disha Patel, Binita Aring, Neeta Khandelwal, Geeta Vaghela, Sonia Barve, Bhavesh Modi, Kairavi Joshi, Gaurishankar Shrimali, Nidhi Sood, Pranay Shah, R D Dixit, Snehal Bagatharia, Kamlesh J Upadhyay, Ramesh Pandit, Tejas Shah, Dipeshwari Shewale, Chaitanya Joshi, Madhvi Joshi |
| EPI_ISL_447543 | Gujarat Biotechnology Research Centre | Gujarat Biotechnology Research Centre | Pritesh Sabara, Apurvasinh Puvar, Janvi Raval, Monika Gandhi, Pinal Trivedi, Maharshi Pandya, Amit Kanani, Akanksha Verma, Nitin Savaliya, Raghawendra Kumar, Dinesh Kumar, Zuber Saiyed, Dipa Kinariwala, Disha Patel, Binita Aring, Neeta Khandelwal, Geeta Vaghela, Sonia Barve, Bhavesh Modi, Kairavi Joshi, Gaurishankar Shrimali, Nidhi Sood, Pranay Shah, R D Dixit, Snehal Bagatharia, Kamlesh J Upadhyay, Ramesh Pandit, Tejas Shah, Ankit Hinsu, Sharmistha Majumdar, Chaitanya Joshi, Madhvi Joshi |
| EPI_ISL_447544 | Gujarat Biotechnology Research Centre | Gujarat Biotechnology Research Centre | Apurvasinh Puvar, Janvi Raval, Monika Gandhi, Pinal Trivedi, Maharshi Pandya, Amit Kanani, Akanksha Verma, Nitin Savaliya, Raghawendra Kumar, Dinesh Kumar, Zuber Saiyed, Dipa Kinariwala, Disha Patel, Binita Aring, Neeta Khandelwal, Geeta Vaghela, Sonia Barve, Bhavesh Modi, Kairavi Joshi, Gaurishankar Shrimali, Nidhi Sood, Pranay Shah, R D Dixit, Snehal Bagatharia, Kamlesh J Upadhyay, Ramesh Pandit, Tejas Shah, Ankit Hinsu, Pritesh Sabara, Pooja P Doshi, Chaitanya Joshi, Madhvi Joshi |
| EPI_ISL_447545 | Gujarat Biotechnology Research Centre | Gujarat Biotechnology Research Centre | Janvi Raval, Monika Gandhi, Pinal Trivedi, Maharshi Pandya, Amit Kanani, Akanksha Verma, Nitin Savaliya, Raghawendra Kumar, Dinesh Kumar, Zuber Saiyed, Dipa Kinariwala, Disha Patel, Binita Aring, Neeta Khandelwal, Geeta Vaghela, Sonia Barve, Bhavesh Modi, Kairavi Joshi, Gaurishankar Shrimali, Nidhi Sood, Pranay Shah, R D Dixit, Snehal Bagatharia, Kamlesh J Upadhyay, Ramesh Pandit, Tejas Shah, Ankit Hinsu, Pritesh Sabara, Apurvasinh Puvar, Nidhi Patel, Chaitanya Joshi, Madhvi Joshi |
| EPI_ISL_447546 | Gujarat Biotechnology Research Centre | Gujarat Biotechnology Research Centre | Monika Gandhi, Pinal Trivedi, Maharshi Pandya, Amit Kanani, Akanksha Verma, Nitin Savaliya, Raghawendra Kumar, Dinesh Kumar, Zuber Saiyed, Dipa Kinariwala, Disha Patel, Binita Aring, Neeta Khandelwal, Geeta Vaghela, Sonia Barve, Bhavesh Modi, Kairavi Joshi, Gaurishankar Shrimali, Nidhi Sood, Pranay Shah, R D Dixit, Snehal Bagatharia, Kamlesh J Upadhyay, Ramesh Pandit, Tejas Shah, Ankit Hinsu, Pritesh Sabara, Apurvasinh Puvar, Janvi Raval, Priti Pandita, Chaitanya Joshi, Madhvi Joshi |
| EPI_ISL_447590 | Genome Centre | Genome Centre | A. S. M. Rubayet Ul Alam, M. Rafiul Islam, M. Shaminur Rahman, Md. Tanvir Islam, Md. Shazid Hasan, Pravas Chandra Roy, Habiba Ibnat, MD. Ali Ahasan Setu, Tanay Chakrovarty, Sourav Dutta Dip, Ruhul Amin, Md Nur Kabidul Azam, Ovinu Kibria Islam, Hassan M. Al-Emran, Shireen Nigar, Selina Akter, Md. Nazmul Hasan, Iqbal Kabir Jahid, M. Anwar Hossain |
| EPI_ISL_447633, EPI_ISL_447634 | Virology, Wageningen Bioveterinary Research | Virology, Wageningen Bioveterinary Research | Oreshkova,N., Vreman,S., Molenaar,R.J., Harders,F., Hakze van der Honing,R.W., Gerhards,N., Bouwstra,R., Hissink,H., Smit,L., Tacken,M., Weesendorp,E., Stegeman,A., van der Poel,W., Engelsma,M.Y. |
| EPI_ISL_447904 | National Institute of Biotechnology | National Institute of Biotechnology | Md. Moniruzzaman, Mohammad Uzzal Hossain, Md. Nazrul Islam, Md. Hadisur Rahman, Irfan Ahmed, Tahia Anan Rahman, Arittra Bhattacharjee, Md. Ruhul Amin, Asif Rashid, Chaman Ara Keya, Keshob Chandra Das, Md. Salimullah |
| EPI_ISL_448048, EPI_ISL_448049, EPI_ISL_448050, EPI_ISL_448053, EPI_ISL_448061, EPI_ISL_448062, EPI_ISL_448065, EPI_ISL_448066, EPI_ISL_448067, EPI_ISL_448068, EPI_ISL_448069, EPI_ISL_448070, EPI_ISL_448071, EPI_ISL_448074, EPI_ISL_448075, EPI_ISL_448076, EPI_ISL_448077, EPI_ISL_448078, EPI_ISL_448079, EPI_ISL_448080, EPI_ISL_448081, EPI_ISL_448082, EPI_ISL_448083, EPI_ISL_448084, EPI_ISL_448086, EPI_ISL_448087, EPI_ISL_448088, EPI_ISL_448089, EPI_ISL_448090, EPI_ISL_448092, EPI_ISL_448096, EPI_ISL_448097, EPI_ISL_448098, EPI_ISL_448099, EPI_ISL_448100, EPI_ISL_448101, EPI_ISL_448102, EPI_ISL_448103, EPI_ISL_448106, EPI_ISL_448107, EPI_ISL_448108, EPI_ISL_448109, EPI_ISL_448110, EPI_ISL_448111, EPI_ISL_448112, EPI_ISL_448113, EPI_ISL_448114, EPI_ISL_448115 | | | |
| see above | Department of Pathology, University of Cambridge | COVID-19 Genomics UK (COG-UK) Consortium | Luke W Meredith, M. Estée Török , Myra Hosmillo, William L. Hamilton, Martin D. Curran, Theresa Feltwell, Grant Hall, Anna Yakovleva, Fahad A Khokhar, Charlotte J. Houldcroft, Laura G Caller, Aminu S. Jahun, Sarah L. Caddy, Ian Goodfellow |

| | | | |
|---|---|---|---|
| EPI_ISL_448417, EPI_ISL_448418 | Queens Medical Centre, Clinical Microbiology Department / DeepSeq Nottingham | COVID-19 Genomics UK (COG-UK) Consortium | Gemma Clark, Wendy Smith, Manjinder Khakh, Hannah Howson-Wells, Jonathan Ball, Patrick McClure, Joseph Chappell, Theocharis Tsoleridis, Nadine Holmes, Matthew Carlisle, Christopher Moore, Fei Sang, Johnny Debebe, Victoria Wright, Matthew Loose |
| EPI_ISL_449071, EPI_ISL_449073, EPI_ISL_449074, EPI_ISL_449075, EPI_ISL_449076, EPI_ISL_449077, EPI_ISL_449078, EPI_ISL_449079, EPI_ISL_449080, EPI_ISL_449081, EPI_ISL_449082, EPI_ISL_449083, EPI_ISL_449084, EPI_ISL_449085, EPI_ISL_449087, EPI_ISL_449089, EPI_ISL_449090, EPI_ISL_449091, EPI_ISL_449092, EPI_ISL_449093, EPI_ISL_449094, EPI_ISL_449095, EPI_ISL_449096, EPI_ISL_449097, EPI_ISL_449098 | | | |
| see above | Quadram Institute Bioscience | COVID-19 Genomics UK (COG-UK) Consortium | Dave J. Baker, Gemma L. Kay, Alp Aydin, Thanh Le-Viet, Steven Rudder, Ana P. Tedim, Anastasia Kolyva, Maria Diaz, Leonardo de Oliveira Martins, Nabil-Fareed Alikhan, Lizzie Meadows, Rachael Stanley, Ngozi Elumogo, Muhammed Yasir, Nicholas M. Thomson, Alexander J Trotter, Rachel Gilroy, Samuel Bloomfield, Claire Stuart, Andrew Bell, Reenesh Prakash, Samir Dervisevic, Alison E. Mather, John Wain, Mark Webber, Andrew J. Page, Justin O'Grady |
| EPI_ISL_449311, EPI_ISL_449312, EPI_ISL_449313, EPI_ISL_449314, EPI_ISL_449315, EPI_ISL_449316, EPI_ISL_449317, EPI_ISL_449318, EPI_ISL_449319, EPI_ISL_449320, EPI_ISL_449321, EPI_ISL_449322, EPI_ISL_449323 | | | |
| see above | Virology Department, Royal Infirmary of Edinburgh, NHS Lothian / School of Biological Sciences, University of Edinburgh / Institute of Genetics and Molecular Medicine, University of Edinburgh | COVID-19 Genomics UK (COG-UK) Consortium | McHugh M, Dewar R, Rooke S, Gallagher M, Balcaza C, O'Toole Á, Scher E, Hill V, McCrone JT, Colquhoun R, Yu X, Jackson B, Rambaut A, Williams TC, Templeton K |
| EPI_ISL_450190 | Rady's Childrens Hospital | Andersen lab at Scripps Research | SEARCH Alliance San Diego |
| EPI_ISL_450339, EPI_ISL_450340 | Bangladesh Institute of Tropical & Infectious Diseases, COVID-19 Testing Laboratory | Basic and Applied Research on Jute Project | Rasel Ahmed, Md. Sabbir Hossain, Shah Md Tamim Kabir, Emdadul Mannan Emdad, Md. Nazmul Haq Rony, Eaftekhar Ahmed Rana, Paritous Kumar Biswas, M A Hassan Chowdhury, Md. Shakeel Ahmed, Md. Samiul Haque, Md. Monjurul Alam, Md. Sharifur Rahman, A S M Anwarul Huq, Md. Shahidul Islam, Goutam Buddha Das, AMAM Zonaed Siddiki |
| EPI_ISL_450341 | Bangladesh Institute of Tropical & Infectious Diseases, COVID-19 Testing Laboratory | Basic and Applied Research on Jute Project | Md. Sabbir Hossain, Rasel Ahmed, Shah Md Tamim Kabir, Emdadul Mannan Emdad, Md. Nazmul Haq Rony, Eaftekhar Ahmed Rana, Paritous Kumar Biswas, M A Hassan Chowdhury, Md. Shakeel Ahmed, Md. Samiul Haque, Md. Monjurul Alam, Md. Sharifur Rahman, A S M Anwarul Huq, Md. Shahidul Islam, Goutam Buddha Das, AMAM Zonaed Siddiki |
| EPI_ISL_450342 | Bangladesh Institute of Tropical & Infectious Diseases, COVID-19 Testing Laboratory | Basic and Applied Research on Jute Project | Rasel Ahmed, Md. Sabbir Hossain, Shah Md Tamim Kabir, Emdadul Mannan Emdad, Md. Nazmul Haq Rony, Eaftekhar Ahmed Rana, Paritous Kumar Biswas, M A Hassan Chowdhury, Md. Shakeel Ahmed, Md. Samiul Haque, Md. Monjurul Alam, Md. Sharifur Rahman, A S M Anwarul Huq, Md. Shahidul Islam, Goutam Buddha Das, AMAM Zonaed Siddiki |
| EPI_ISL_450343, EPI_ISL_450345 | Bangladesh Institute of Tropical & Infectious Diseases, COVID-19 Testing Laboratory | Basic and Applied Research on Jute Project | Md. Sabbir Hossain, Rasel Ahmed, Shah Md Tamim Kabir, Emdadul Mannan Emdad, Md. Nazmul Haq Rony, Eaftekhar Ahmed Rana, Paritous Kumar Biswas, M A Hassan Chowdhury, Md. Shakeel Ahmed, Md. Samiul Haque, Md. Monjurul Alam, Md. Sharifur Rahman, A S M Anwarul Huq, Md. Shahidul Islam, Goutam Buddha Das, AMAM Zonaed Siddiki |
| EPI_ISL_450746 | Laboratory of Molecular Biology, Diagnostyka sp. z o.o. | Laboratory of Recombinant Vaccines | Lukasz Rabalski, Anna Piotrowska-Mietelska, Maciej Kosinski, Boguslaw Szewczyk, Krystyna Bienkowska-Szewczyk |
| EPI_ISL_450781 | Government Medical College-Bhavnagar | Gujarat Biotechnology Research Centre | Kairavi Desai, Saklain Malek, Shirish Patel, Ramesh Pandit, Tejas Shah, Ankit Hinsu, Pritesh Sabara, Apurvasinh Puvar, Janvi Raval, Zarna Patel, Monika Gandhi, Pinal Trivedi, Maharshi Pandya, Amit Kanani, Nidhi Patel, Nitin Savaliya, Raghawendra Kumar, Dinesh Kumar, Zuber Saiyed, Komal Patel, Labdhi Pandya, Snehal Bagatharia, Bhavesh Modi, Gaurishankar Shrimali, R D Dixit, A M Kadri, Akanksha Verma, Chaitanya Joshi, Madhvi Joshi |
| EPI_ISL_450784 | Government Medical College-Bhavnagar | Gujarat Biotechnology Research Centre | Zarna Patel, Ramesh Pandit, Tejas Shah, Ankit Hinsu, Pritesh Sabara, Apurvasinh Puvar, Janvi Raval, Monika Gandhi, Pinal Trivedi, Maharshi Pandya, Amit Kanani, Nidhi Patel, Nitin Savaliya, Raghawendra Kumar, Dinesh Kumar, Zuber Saiyed, Komal Patel, Labdhi Pandya, Snehal Bagatharia, Kairavi Desai, Saklain Malek, Shirish Patel, Bhavesh Modi, Gaurishankar Shrimali, R D Dixit, A M Kadri, Afzal Ansari, Chaitanya Joshi, Madhvi Joshi |
| EPI_ISL_450785 | Pandit Deendayal Upadhyay Government Medical College, Rajkot | Gujarat Biotechnology Research Centre | Prakash Modi, Sejul Antala, Manish Pattani, Apurvasinh Puvar, Janvi Raval, Zarna Patel, Monika Gandhi, Pinal Trivedi, Maharshi Pandya, Amit Kanani, Nidhi Patel, Nitin Savaliya, Raghawendra Kumar, Dinesh Kumar, Zuber Saiyed, Komal Patel, Labdhi Pandya, Snehal Bagatharia, Ramesh Pandit, Tejas Shah, Ankit Hinsu, Pritesh Sabara, Bhavesh Modi, Gaurishankar Shrimali, R D Dixit, A M Kadri, Neelam Nathani, Chaitanya Joshi, Madhvi Joshi |
| EPI_ISL_450786 | Pandit Deendayal Upadhyay Government Medical College, Rajkot | Gujarat Biotechnology Research Centre | Sejul Antala, Manish Pattani, Prakash Modi, Janvi Raval, Zarna Patel, Monika Gandhi, Pinal Trivedi, Maharshi Pandya, Amit Kanani, Nidhi Patel, Nitin Savaliya, Raghawendra Kumar, Dinesh Kumar, Zuber Saiyed, Komal Patel, Labdhi Pandya, Snehal Bagatharia, Ramesh Pandit, Tejas Shah, Ankit Hinsu, Pritesh Sabara, Apurvasinh Puvar, Bhavesh Modi, Gaurishankar Shrimali, R D Dixit, A M Kadri, Armi Chaudhari, Chaitanya Joshi, Madhvi Joshi |
| EPI_ISL_450787 | Pandit Deendayal Upadhyay Government Medical College, Rajkot | Gujarat Biotechnology Research Centre | Manish Pattani, Prakash Modi, Sejul Antala, Zarna Patel, Monika Gandhi, Pinal Trivedi, Maharshi Pandya, Amit Kanani, Nidhi Patel, Nitin Savaliya, Raghawendra Kumar, Dinesh Kumar, Zuber Saiyed, Komal Patel, Labdhi Pandya, Snehal Bagatharia, Ramesh Pandit, Tejas Shah, Ankit Hinsu, Pritesh Sabara, Apurvasinh Puvar, Janvi Raval, Bhavesh Modi, Gaurishankar Shrimali, R D Dixit, A M Kadri, Bhavya Jindal, Chaitanya Joshi, Madhvi Joshi |
| EPI_ISL_450788 | Pandit Deendayal Upadhyay Government Medical College, Rajkot | Gujarat Biotechnology Research Centre | Zarna Patel, Tejas Shah, Ankit Hinsu, Pritesh Sabara, Apurvasinh Puvar, Janvi Raval, Monika Gandhi, Pinal Trivedi, Maharshi Pandya, Amit Kanani, Nidhi Patel, Nitin Savaliya, Raghawendra Kumar, Dinesh Kumar, Zuber Saiyed, Komal Patel, Labdhi Pandya, Snehal Bagatharia, Prakash Modi, Sejul Antala, Manish Pattani, Ramesh Pandit, Bhavesh Modi, Gaurishankar Shrimali, R D Dixit, A M Kadri, Camellia Chakraborty, Chaitanya Joshi, Madhvi Joshi |
| EPI_ISL_450789 | Pandit Deendayal Upadhyay Government Medical College, Rajkot | Gujarat Biotechnology Research Centre | Ankit Hinsu, Pritesh Sabara, Apurvasinh Puvar, Janvi Raval, Zarna Patel, Monika Gandhi, Pinal Trivedi, Maharshi Pandya, Amit Kanani, Nidhi Patel, Nitin Savaliya, Raghawendra Kumar, Dinesh Kumar, Zuber Saiyed, Komal Patel, Labdhi Pandya, Snehal Bagatharia, Prakash Modi, Sejul Antala, Manish Pattani, Ramesh Pandit, Tejas Shah, Bhavesh Modi, Gaurishankar Shrimali, R D Dixit, A M Kadri, Siddhant Kumar, Chaitanya Joshi, Madhvi Joshi |
| EPI_ISL_450833 | Wetterhalsan | The Public Health Agency of Sweden | Anders Tengblad, Anna-Malin Linde, Maria Lind Karlberg, Oskar Karlsson Lindsjo, Olov Svartstrom, Anna Risberg, Theresa Enkirch, Mia Brytting, Karin Tegmark-Wisell |
| EPI_ISL_450839 | COVID-19 Laboratory Centre for Advanced Research in Sciences (CARS), University of Dhaka, Dhaka-1000, Bangladesh | DNA Solution Ltd | Sharif Akhteruzzaman, Zeba Islam Seraj, Nazmul Ahsan, Md Imdadul Hoque, MA Malek, Shahryar Nabi, Sabrina Moriom Elius, ABM Khademul Islam, Richard Malo, Imran Khan, Abu Sufian, Sabita Rezwana Rahman, Habibul Bari Shozib, Mamun Ahmed, AHM Nurun Nabi, Mohammad Riazul Islam, Md Mizanur Rahman, Md Ismail Hosen, Latiful Bari, Gazi Nurun Nahar, Haseena Khan, M Anwar Hossain. |
| EPI_ISL_450840 | COVID-19 Laboratory | DNA Solution Ltd. L-5 | Sharif Akhteruzzaman, Zeba Islam Seraj, Nazmul Ahsan, Md Imdadul Hoque, MA Malek, Shahryar Nabi, Sabrina Moriom Elius, ABM Khademul Islam, Richard Malo, Imran Khan, Abu Sufian, Sabita Rezwana Rahman, Habibul Bari Shozib, Mamun Ahmed, AHM Nurun Nabi, Mohammad Riazul Islam, Md Mizanur Rahman, Md Ismail Hosen, Latiful Bari, Gazi Nurun Nahar, Haseena Khan, M Anwar Hossain. |
| EPI_ISL_450841 | COVID-19 Laboratory | DNA Solution Ltd | Sharif Akhteruzzaman, Zeba Islam Seraj, Nazmul Ahsan, Md Imdadul Hoque, MA Malek, Shahryar Nabi, Sabrina Moriom Elius, ABM Khademul Islam, Richard Malo, Imran Khan, Abu Sufian, Sabita Rezwana Rahman, Habibul Bari Shozib, Mamun Ahmed, AHM Nurun Nabi, Mohammad Riazul Islam, Md Mizanur Rahman, Md Ismail Hosen, Latiful Bari, Gazi Nurun Nahar, Haseena Khan, M Anwar Hossain. |
| EPI_ISL_450842, EPI_ISL_450843 | COVID-19 Laboratory | DNA Solution Ltd. | Sharif Akhteruzzaman, Zeba Islam Seraj, Nazmul Ahsan, Md Imdadul Hoque, MA Malek, Shahryar Nabi, Sabrina Moriom Elius, ABM Khademul Islam, Richard Malo, Imran Khan, Abu Sufian, Sabita Rezwana Rahman, Habibul Bari Shozib, Mamun Ahmed, AHM Nurun Nabi, Mohammad Riazul Islam, Md Mizanur Rahman, Md Ismail Hosen, Latiful Bari, Gazi Nurun Nahar, Haseena Khan, M Anwar Hossain. |
| EPI_ISL_450913, EPI_ISL_450953, EPI_ISL_450993, EPI_ISL_451033, EPI_ISL_451073 | Center of Excellence in Clinical Virology | Center of Excellence in Clinical Virology | Puenpa,J., Chansaenroj,J., Nilyanimit,P., Auphimai,C., Yorsaeng,R., Suwannakarn,K., Poovorawan,Y. |
| EPI_ISL_451149 | M.P Shah Government Medocal college Jamnagar | Gujarat Biotechnology Research Centre | Janvi Raval, Zarna Patel, Monika Gandhi, Pinal Trivedi, Maharshi Pandya, Amit Kanani, Nidhi Patel, Nitin Savaliya, Raghawendra Kumar, Dinesh Kumar, Zuber Saiyed, Komal Patel, Labdhi Pandya, Snehal Bagatharia, Ramesh Pandit, Tejas Shah, Ankit Hinsu, Pritesh Sabara, Apurvasinh Puvar, Binita Aring, Bhavesh Modi, Gaurishankar Shrimali, R D Dixit, A M Kadri, Priti Pandita, Chaitanya Joshi, Madhvi Joshi, |
| EPI_ISL_451150 | M.P Shah Government Medocal college Jamnagar | Gujarat Biotechnology Research Centre | Zarna Patel, Monika Gandhi, Pinal Trivedi, Maharshi Pandya, Amit Kanani, Nidhi Patel, Nitin Savaliya, Raghawendra Kumar, Dinesh Kumar, Zuber Saiyed, Komal Patel, Labdhi Pandya, Snehal Bagatharia, Ramesh Pandit, Tejas Shah, Ankit Hinsu, Pritesh Sabara, Apurvasinh Puvar, Binita Aring, Janvi Raval, Bhavesh Modi, Gaurishankar Shrimali, R D Dixit, A M Kadri, Pragya Sharma, Chaitanya Joshi, Madhvi Joshi, |
| EPI_ISL_451151 | M.P Shah Government Medocal college Jamnagar | Gujarat Biotechnology Research Centre | Monika Gandhi, Pinal Trivedi, Maharshi Pandya, Amit Kanani, Nidhi Patel, Nitin Savaliya, Raghawendra Kumar, Dinesh Kumar, Zuber Saiyed, Komal Patel, Labdhi Pandya, Snehal Bagatharia, Ramesh Pandit, Tejas Shah, Ankit Hinsu, Pritesh Sabara, Apurvasinh Puvar, Binita Aring, Janvi Raval, Zarna Patel, Bhavesh Modi, Gaurishankar Shrimali, R D Dixit, A M Kadri, Neha Rajpara, Chaitanya Joshi, Madhvi Joshi, |

| | | | |
|---|---|---|---|
| EPI_ISL_451152 | M.P Shah Government Medocal college Jamnagar | Gujarat Biotechnology Research Centre | Pinal Trivedi, Maharshi Pandya, Amit Kanani, Nidhi Patel, Nitin Savaliya, Raghawendra Kumar, Dinesh Kumar, Zuber Saiyed, Komal Patel, Labdhi Pandya, Snehal Bagatharia, Ramesh Pandit, Tejas Shah, Ankit Hinsu, Pritesh Sabara, Apurvasinh Puvar, Binita Aring, Janvi Raval, Zarna Patel, Monika Gandhi, Bhavesh Modi, Gaurishankar Shrimali, R D Dixit, A M Kadri, Afzal Ansari, Chaitanya Joshi, Madhvi Joshi, |
| EPI_ISL_451153 | M.P Shah Government Medocal college Jamnagar | Gujarat Biotechnology Research Centre | Maharshi Pandya, Amit Kanani, Nidhi Patel, Nitin Savaliya, Raghawendra Kumar, Dinesh Kumar, Zuber Saiyed, Komal Patel, Labdhi Pandya, Snehal Bagatharia, Ramesh Pandit, Tejas Shah, Ankit Hinsu, Pritesh Sabara, Apurvasinh Puvar, Binita Aring, Janvi Raval, Zarna Patel, Monika Gandhi, Pinal Trivedi, Bhavesh Modi, Gaurishankar Shrimali, R D Dixit, A M Kadri, Fenil Patel, Chaitanya Joshi, Madhvi Joshi, |
| EPI_ISL_451211 | LSUHS Emerging Viral Threat Laboratory | Microbial Genome Sequencing Center | Jeremy P. Kamil, John A. Vanchiere, Rona S. Scott, Camille F. Abshire, Abida Siddiqa, Byeong-Jae Lee, Chan-ki Min, Md Maksudul Alam, Monica Gestal-Cartele, Edna Ondari, Adam Greer, Malgorzata Bienkowska-Haba, Katarzyna Zwolinska, Jason M. Bodily, Andrew D. Yurochko, Paul M. Weinberger, Christopher G. Kevil, Martin J. Sapp, Daniel J. Snyder, Vaughn S. Cooper |
| EPI_ISL_451237, EPI_ISL_451238, EPI_ISL_451242, EPI_ISL_451243, EPI_ISL_451244, EPI_ISL_451245 | LSUHS Emerging Viral Threat Laboratory | Microbial Genome Sequencing Center | John A. Vanchiere, Jeremy P. Kamil, Rona S. Scott, Camille F. Abshire, Abida Siddiqa, Byeong-Jae Lee, Chan-ki Min, Md Maksudul Alam, Monica Gestal-Cartele, Edna Ondari, Adam Greer, Malgorzata Bienkowska-Haba, Katarzyna Zwolinska, Jason M. Bodily, Andrew D. Yurochko, Paul M. Weinberger, Christopher G. Kevil, Martin J. Sapp, Daniel J. Snyder, Vaughn S. Cooper |
| EPI_ISL_451248, EPI_ISL_451249, EPI_ISL_451250, EPI_ISL_451251, EPI_ISL_451252, EPI_ISL_451253, EPI_ISL_451254, EPI_ISL_451255, EPI_ISL_451256, EPI_ISL_451257, EPI_ISL_451258, EPI_ISL_451259, EPI_ISL_451260, EPI_ISL_451261, EPI_ISL_451262, EPI_ISL_451263, EPI_ISL_451264, EPI_ISL_451265, EPI_ISL_451266, EPI_ISL_451267, EPI_ISL_451268, EPI_ISL_451269, EPI_ISL_451270, EPI_ISL_451271, EPI_ISL_451272, EPI_ISL_451273, EPI_ISL_451274, EPI_ISL_451275, EPI_ISL_451276, EPI_ISL_451277, EPI_ISL_451278, EPI_ISL_451279, EPI_ISL_451280, EPI_ISL_451281, EPI_ISL_451282, EPI_ISL_451283, EPI_ISL_451284, EPI_ISL_451285, EPI_ISL_451286, EPI_ISL_451287, EPI_ISL_451288, EPI_ISL_451289, EPI_ISL_451290, EPI_ISL_451291, EPI_ISL_451292, EPI_ISL_451293, EPI_ISL_451294, EPI_ISL_451295, EPI_ISL_451296, EPI_ISL_451297 | | | |
| see above | LSUHS Emerging Viral Threat Laboratory | Microbial Genome Sequencing Center | Jeremy P. Kamil, John A. Vanchiere, Rona S. Scott, Camille F. Abshire, Abida Siddiqa, Byeong-Jae Lee, Chan-ki Min, Md Maksudul Alam, Monica Gestal-Cartele, Edna Ondari, Adam Greer, Malgorzata Bienkowska-Haba, Katarzyna Zwolinska, Jason M. Bodily, Andrew D. Yurochko, Paul M. Weinberger, Christopher G. Kevil, Martin J. Sapp, Daniel J. Snyder, Vaughn S. Cooper |
| EPI_ISL_451655, EPI_ISL_451656, EPI_ISL_451657, EPI_ISL_451658, EPI_ISL_451659, EPI_ISL_451660, EPI_ISL_451661, EPI_ISL_451662, EPI_ISL_451663, EPI_ISL_451664 | State Sanitary Inspectorate | Laboratory of Recombinant Vaccines | Lukasz Rabalski, Boguslaw Szewczyk, Krystyna Bienkowska-Szewczyk, Jaroslaw Pinkas |
| EPI_ISL_452097, EPI_ISL_452098 | Department of Clinical Microbiology, Copenhagen University Hospital, Hvidovre, Kettegaard Alle 30, 2650 Hvidovre. | Albertsen lab, Department of Chemistry and Bioscience, Aalborg University, Denmark | Rasmus Kirkegaard |
| EPI_ISL_452138 | VI-US Virgin Islands Department of Health | Pathogen Discovery, Respiratory Viruses Branch, Division of Viral Diseases, Centers for Disease Control and Prevention | Anna Uehara, Yan Li, Anna Montmayeur, Ying Tao, Krista Queen, Jing Zhang, Clinton R. Paden, Rachel Marine, Haibin Wang, Bettina Bankamp, Zachary Weiner, Suxiang Tong |
| EPI_ISL_452145 | Yale COVID-19 Biorepository | Grubaugh Lab - Yale School of Public Health | Joseph Fauver, Tara Alpert, Anderson Brito, Anne Wyllie, Chantal Vogels, Mary Petrone, Cole Jensen, Chaney Kalinich, Isabel Ott, Arnau Casanovas, Catherine Muenker, Adam Moore, Alice Lu, Maria Tokuyama, Patrick Wong, Peiwen Lu, Saad Omer, Richard Martinello, Allison Nelson, Shelli Farhadian, Akiko Iwasaki, Charlese Dela Cruz, Albert Ko, Nathan Grubaugh |
| EPI_ISL_452794, EPI_ISL_452795 | ICAR-National Institute of High Security Animal Diseases | ICAR-National Institute of High Security Animal Diseases | Anamika Mishra, Ashutosh Aasdev, Sandeep Bhatia, Harshad Murugkar, Chakradhar Tosh, Niranjan Mishra, Shanmugasundaram Nagarajan, Katherukamem Rajukumar, Richa Sood, G Venkatesh, Atul Kumar Pateriya, Manoj Kumar, Shashi Bhushan Sudhakar, Fateh Singh, Sethil Kumar D, Senmannan Kalaiyarasu, Pradeep Gandhale, Naveen Kumar, Chandan Kumar Dubey, Sushil Tripathi, Sandeep Kumar Jhade, Meghna Tripathi, Suman Kumari Shah, Pushpendra Singh, Pushpendra Namdeo, Suman Mishra, Rupal Singh, Vishnupriya Patil, Dipesh Kumar Nayak, Vijendra Pal Singh, Ashwin Ashok Raut |
| EPI_ISL_452857, EPI_ISL_452858, EPI_ISL_452859, EPI_ISL_452860, EPI_ISL_452861, EPI_ISL_452862, EPI_ISL_452863, EPI_ISL_452864, EPI_ISL_452865, EPI_ISL_452866, EPI_ISL_452867, EPI_ISL_452868, EPI_ISL_452869, EPI_ISL_452870, EPI_ISL_452871, EPI_ISL_452872, EPI_ISL_452873, EPI_ISL_452874, EPI_ISL_452875, EPI_ISL_452876, EPI_ISL_452877, EPI_ISL_452878, EPI_ISL_452879, EPI_ISL_452880, EPI_ISL_452881, EPI_ISL_452882, EPI_ISL_452883, EPI_ISL_452884, EPI_ISL_452885, EPI_ISL_452886, EPI_ISL_452887, EPI_ISL_452888, EPI_ISL_452889, EPI_ISL_452890, EPI_ISL_452891, EPI_ISL_452892, EPI_ISL_452893, EPI_ISL_452894, EPI_ISL_452895, EPI_ISL_452896, EPI_ISL_452897, EPI_ISL_452898, EPI_ISL_452899, EPI_ISL_452900, EPI_ISL_452901, EPI_ISL_452903, EPI_ISL_452904, EPI_ISL_452905, EPI_ISL_452906, EPI_ISL_452910, EPI_ISL_452911, EPI_ISL_452912, EPI_ISL_452913, EPI_ISL_452914, EPI_ISL_452915, EPI_ISL_452916, EPI_ISL_452917, EPI_ISL_452918, EPI_ISL_452919, EPI_ISL_452920, EPI_ISL_452921, EPI_ISL_452922, EPI_ISL_452923, EPI_ISL_452924, EPI_ISL_452925, EPI_ISL_452926, EPI_ISL_452927, EPI_ISL_452928, EPI_ISL_452929, EPI_ISL_452930, EPI_ISL_452932, EPI_ISL_452933, EPI_ISL_452934, EPI_ISL_452935, EPI_ISL_452936, EPI_ISL_452937, EPI_ISL_452938, EPI_ISL_452939, EPI_ISL_452940, EPI_ISL_452941, EPI_ISL_452942, EPI_ISL_452943, EPI_ISL_452944, EPI_ISL_452945, EPI_ISL_452946, EPI_ISL_452947, EPI_ISL_452948, EPI_ISL_452949, EPI_ISL_452951, EPI_ISL_452952, EPI_ISL_452953, EPI_ISL_452954, EPI_ISL_452956, EPI_ISL_452957, EPI_ISL_452958, EPI_ISL_452959, EPI_ISL_452960, EPI_ISL_452961, EPI_ISL_452962, EPI_ISL_452963, EPI_ISL_452964, EPI_ISL_452965, EPI_ISL_452966, EPI_ISL_452967, EPI_ISL_452968, EPI_ISL_452969, EPI_ISL_452970, EPI_ISL_452971, EPI_ISL_452972, EPI_ISL_452973, EPI_ISL_452974, EPI_ISL_452975, EPI_ISL_452976, EPI_ISL_452977, EPI_ISL_452978, EPI_ISL_452979, EPI_ISL_452980, EPI_ISL_452981, EPI_ISL_452982, EPI_ISL_452983, EPI_ISL_452984, EPI_ISL_452985, EPI_ISL_452986, EPI_ISL_452987, EPI_ISL_452988, EPI_ISL_452989, EPI_ISL_452990, EPI_ISL_452991, EPI_ISL_452992, EPI_ISL_452993, EPI_ISL_452994, EPI_ISL_452995, EPI_ISL_452996, EPI_ISL_452997, EPI_ISL_452998, EPI_ISL_452999, EPI_ISL_453000, EPI_ISL_453005 | | | |
| see above | Department of Pathology, University of Cambridge | COVID-19 Genomics UK (COG-UK) Consortium | Luke W Meredith, M. Estée Török , Myra Hosmillo, William L. Hamilton, Martin D. Curran, Theresa Feltwell, Grant Hall, Anna Yakovleva, Fahad A Khokhar, Charlotte J. Houldcroft, Laura G Caller, Aminu S. Jahun, Sarah L. Caddy, Ian Goodfellow |
| EPI_ISL_453098 | West of Scotland Specialist Virology Centre, NHSGGC / MRC-University of Glasgow Centre for Virus Research | COVID-19 Genomics UK (COG-UK) Consortium | Ana da Silva Filipe, Natasha Johnson, Kathy Smollett, Daniel Mair, Stephen Carmichael, Lily Tong, Jenna Nichols, Elihu Aranday-Cortes, Kirstyn Brunker, Yasmin Parr, Kyriaki Nomikou; Sarah McDonald, Marc Niebel, Patawee Asamaphan; Richard Orton, Joseph Hughes, Sreenu Vattipally, David L Robertson; Alasdair MacLean, Rory Gunson; Kathy Li, Natasha Jesudason, Rajiv Shah, James Shepherd, Antonia Ho, Emma Thomson |
| EPI_ISL_453129, EPI_ISL_453130, EPI_ISL_453131, EPI_ISL_453132, EPI_ISL_453141, EPI_ISL_453142, EPI_ISL_453143, EPI_ISL_453144, EPI_ISL_453145, EPI_ISL_453146, EPI_ISL_453147, EPI_ISL_453148, EPI_ISL_453149, EPI_ISL_453150, EPI_ISL_453151, EPI_ISL_453152, EPI_ISL_453153, EPI_ISL_453154, EPI_ISL_453155, EPI_ISL_453167, EPI_ISL_453168, EPI_ISL_453169, EPI_ISL_453170, EPI_ISL_453171, EPI_ISL_453172, EPI_ISL_453173, EPI_ISL_453174, EPI_ISL_453175, EPI_ISL_453176, EPI_ISL_453177, EPI_ISL_453178, EPI_ISL_453179, EPI_ISL_453180, EPI_ISL_453181, EPI_ISL_453182, EPI_ISL_453183 | | | |
| see above | Virology Department, Royal Infirmary of Edinburgh, NHS Lothian / School of Biological Sciences, University of Edinburgh / Institute of Genetics and Molecular Medicine, University of Edinburgh | COVID-19 Genomics UK (COG-UK) Consortium | McHugh M, Dewar R, Rooke S, Gallagher M, Balcaza C, O'Toole Á, Scher E, Hill V, McCrone JT, Colquhoun R, Yu X, Jackson B, Rambaut A, Williams TC, Templeton K |
| EPI_ISL_453431, EPI_ISL_453432 | Liverpool Clinical Laboratories | COVID-19 Genomics UK (COG-UK) Consortium | Sam Haldenby, Anita Lucaci, Steve Paterson, Julian Hiscox, Alistair Darby, M Almsaud, A Alrezaihi, Muhannad Alruwaili, Stuart D Armstrong, Jones Benjamin , Eleanor G Bentley, Anu Chawla, Jordan J Clark, Angela Cowell, Richard Eccles, Isabel García-Dorival, Matthew Gemmell, Alessandro Gerada, PKF Gilmore, Richard Gregory, Ximeng Han, Catherine Hartley, Margaret Hughes, Miren Iturriza-Gomara, James Johnson, L Luu, Jenifer Manson , Charlotte Nelson, Elaine O'Toole, Cassie Olateju, Rebekah Penrice-Randal , Lucille Rainbow, N.P Randle, Trevor Ian Robinson, Parul Sharma, Ghada T Shawli, James P Stewart , Neil Swainston, Ecaterina Vamos, Joanne Watts, Mark Whitehead |
| EPI_ISL_453616, EPI_ISL_453617, EPI_ISL_453618, EPI_ISL_453619, EPI_ISL_453620, EPI_ISL_453621, EPI_ISL_453622, EPI_ISL_453623, EPI_ISL_453624, EPI_ISL_453625, EPI_ISL_453633, EPI_ISL_453634, EPI_ISL_453635, EPI_ISL_453645, EPI_ISL_453646, EPI_ISL_453647, EPI_ISL_453648, EPI_ISL_453649, EPI_ISL_453650, EPI_ISL_453651, EPI_ISL_453652, EPI_ISL_453653, EPI_ISL_453654, EPI_ISL_453655, EPI_ISL_453656, EPI_ISL_453657, EPI_ISL_453658, EPI_ISL_453659, EPI_ISL_453660, EPI_ISL_453661 | | | |
| see above | Queens Medical Centre, Clinical Microbiology Department / DeepSeq Nottingham | COVID-19 Genomics UK (COG-UK) Consortium | Gemma Clark, Wendy Smith, Manjinder Khakh, Hannah Howson-Wells, Jonathan Ball, Patrick McClure, Joseph Chappell, Theocharis Tsoleridis, Nadine Holmes, Matthew Carlisle, Christopher Moore, Fei Sang, Johnny Debebe, Victoria Wright, Matthew Loose |
| EPI_ISL_454273 | unknown | Instituto Nacional de Saude (INSA) | Borges et al |
| EPI_ISL_454497 | RSE "National Center for Biotechnology" | RSE "National Center for Biotechnology" | Alexandr Shevtsov, Ilyas Akhmetollayev, Viktoriya Lutsay, Asylulan Amirgazin, Askar Abdaliyev, Akbota Rakhmetova, Zabira Aushakhmetova, Ruslan Kalendar, Yerlan Ramankulov |
| EPI_ISL_454575 | Laboratory of virology, National Center of Expertise | Laboratory of molecular-genetic research, National Center for Expertise, Kazakhstan National Center for Biotechnology, Kazakhstan | Abdaliyev Askar, Shevtsov Alexandr, Akhmetollayev Ilyas, Kalendar Ruslan, Rakhmetova Akbota, , Lutsay Viktoriya, Amirgazin Asylulan, Aushakhmetova Zabira, Ramankulov Yerlan |
| EPI_ISL_454656, EPI_ISL_454657, EPI_ISL_454659, EPI_ISL_454669, EPI_ISL_454672, EPI_ISL_454675, EPI_ISL_454676, EPI_ISL_454677, EPI_ISL_454678, EPI_ISL_454680, EPI_ISL_454681, EPI_ISL_454682, EPI_ISL_454683, EPI_ISL_454685, EPI_ISL_454687 | | | |
| see above | County of Santa Clara Public Health Department | Chan-Zuckerberg Biohub | CZB Cliahub Consortium |
| EPI_ISL_455058 | Pathology West - NSW Health Pathology | NSW Health Pathology - Institute of Clinical Pathology and Medical Research; Westmead Hospital; University of Sydney | CIDM-PH et al. |
| EPI_ISL_455068 | Childrens Hospital Westmead | NSW Health Pathology - Institute of Clinical Pathology and Medical Research; Westmead Hospital; University of Sydney | CIDM-PH et al. |

| | | | |
|---|---|---|---|
| EPI_ISL_455069, EPI_ISL_455070 | Pathology West - NSW Health Pathology | NSW Health Pathology - Institute of Clinical Pathology and Medical Research; Westmead Hospital; University of Sydney | CIDM-PH et al. |
| EPI_ISL_455071 | Pathology Sydney South West - NSW Health Pathology | NSW Health Pathology - Institute of Clinical Pathology and Medical Research; Westmead Hospital; University of Sydney | CIDM-PH et al. |
| EPI_ISL_455072, EPI_ISL_455073 | Sullivan Nicolaides Pathology | NSW Health Pathology - Institute of Clinical Pathology and Medical Research; Westmead Hospital; University of Sydney | CIDM-PH et al. |
| EPI_ISL_455074 | ACT Pathology | NSW Health Pathology - Institute of Clinical Pathology and Medical Research; Westmead Hospital; University of Sydney | CIDM-PH et al. |
| EPI_ISL_455090, EPI_ISL_455091 | South Eastern Area Laboratory Services | NSW Health Pathology - Institute of Clinical Pathology and Medical Research; Westmead Hospital; University of Sydney | CIDM-PH et al. |
| EPI_ISL_455104 | Ulltuna Vardcentral | The Public Health Agency of Sweden | Heidi Lindback, Anna-Malin Linde, Maria Lind Karlberg, Oskar Karlsson Lindsjo, Olov Svartstrom, Anna Risberg, Theresa Enkirch, Mia Brytting, Karin Tegmark-Wisell |
| EPI_ISL_455105 | Wetterhalsan | The Public Health Agency of Sweden | Anders Tengblad, Anna-Malin Linde, Maria Lind Karlberg, Oskar Karlsson Lindsjo, Olov Svartstrom, Anna Risberg, Theresa Enkirch, Mia Brytting, Karin Tegmark-Wisell |
| EPI_ISL_455106 | Surbrunns VC | The Public Health Agency of Sweden | Erik Embring, Anna-Malin Linde, Maria Lind Karlberg, Oskar Karlsson Lindsjo, Olov Svartstrom, Anna Risberg, Theresa Enkirch, Mia Brytting, Karin Tegmark-Wisell |
| EPI_ISL_455107 | Narhalsan Mellerud | The Public Health Agency of Sweden | Maria Nykvist, Anna-Malin Linde, Maria Lind Karlberg, Oskar Karlsson Lindsjo, Olov Svartstrom, Anna Risberg, Theresa Enkirch, Mia Brytting, Karin Tegmark-Wisell |
| EPI_ISL_455108 | Smedby HC | The Public Health Agency of Sweden | Susanne Brunby, Anna-Malin Linde, Maria Lind Karlberg, Oskar Karlsson Lindsjo, Olov Svartstrom, Anna Risberg, Theresa Enkirch, Mia Brytting, Karin Tegmark-Wisell |
| EPI_ISL_455109 | Ulltuna Vardcentral | The Public Health Agency of Sweden | Heidi Lindback, Anna-Malin Linde, Maria Lind Karlberg, Oskar Karlsson Lindsjo, Olov Svartstrom, Anna Risberg, Theresa Enkirch, Mia Brytting, Karin Tegmark-Wisell |
| EPI_ISL_455110 | Scania Halsocenter, B288 | The Public Health Agency of Sweden | Christina Lergin, Anna-Malin Linde, Maria Lind Karlberg, Oskar Karlsson Lindsjo, Olov Svartstrom, Anna Risberg, Theresa Enkirch, Mia Brytting, Karin Tegmark-Wisell |
| EPI_ISL_455111 | Olof Norrby | The Public Health Agency of Sweden | Bla Kustens halsocentral, Anna-Malin Linde, Maria Lind Karlberg, Oskar Karlsson Lindsjo, Olov Svartstrom, Anna Risberg, Theresa Enkirch, Mia Brytting, Karin Tegmark-Wisell |
| EPI_ISL_455311 | REGIONAL VRDL,ICMR-RMRC BBSR | Immunogenomics group, Institute of Life Sciences, Bhubaneswar | Sunil Raghav, Jyotirmayee Turuk, Arup Ghosh, Atimukta Jha, Viplov K. Biswas, Swati Madhulika, Manasi Priyadarshini, Shuchi Smita, Jaya Singh Khastri, Rupesh Dash, Soma Chattopadhyay, Ghulam Hussain Syed, Shanti Senapati, Tushar K. Beuria, Debdutta Bhattacharya, Rajeeb Swain, Punit Prasad, COVID-19 team of ILS & RMRC, Orissa COVID-19 study group, DBT's PAN-INDIA 1000 SARS-CoV2 RNA genome sequencing consortium, Sanghamitra Pati, Ajay Parida |
| EPI_ISL_455468, EPI_ISL_455469, EPI_ISL_455470, EPI_ISL_455471, EPI_ISL_455472, EPI_ISL_455473, EPI_ISL_455474 | Laboratory for Respiratory Viruses, Cantacuzino National Military-Medical Institute for Research and Development | Cantacuzino Institute | M.Lazar, L.Ustea, A.Cretu, Tim Durfee |
| EPI_ISL_455475, EPI_ISL_455477, EPI_ISL_455479 | Laboratory for Respiratory Viruses, Cantacuzino National Military-Medical Institute for Research and Development | Cantacuzino Institute | M.Lazar, L.Ustea, A.Cretu, T.Durfee |
| EPI_ISL_455568, EPI_ISL_455569, EPI_ISL_455570, EPI_ISL_455571, EPI_ISL_455572 | Gundersen Molecular Diagnostics Laboratory | Kabara Cancer Research Institute | Craig S. Richmond, Paraic A. Kenny |
| EPI_ISL_455573, EPI_ISL_455574, EPI_ISL_455580 | Gundersen Clinical Microbiology Laboratory | Kabara Cancer Research Institute | Craig S. Richmond, Paraic A. Kenny |
| EPI_ISL_455635 | KRISP, KZN Research Innovation and Sequencing Platform | KRISP, KZN Research Innovation and Sequencing Platform | Giandhari J, Pillay S, Lessells R, Chimukangara B, Deforche K, Tegally H, Wilkinson E, de Oliveira T |
| EPI_ISL_455668 | ICMR-National Institute of Cholera and Enteric Diseases | National Institute of Biomedical Genomics | Arindam Maitra, Mamta Chawla Sarkar, Sreedhar Chinnaswamy, Hasina Banu, Ananya Chatterjee, Shanta Dutta, Saumitra Das |
| EPI_ISL_455749, EPI_ISL_455750, EPI_ISL_455751, EPI_ISL_455752, EPI_ISL_455753, EPI_ISL_455754, EPI_ISL_455755, EPI_ISL_455756, EPI_ISL_455757, EPI_ISL_455758, EPI_ISL_455759, EPI_ISL_455760, EPI_ISL_455761, EPI_ISL_455762, EPI_ISL_455763, EPI_ISL_455764, EPI_ISL_455765, EPI_ISL_455766, EPI_ISL_455767, EPI_ISL_455768, EPI_ISL_455769, EPI_ISL_455770 | | | |
| see above | REGIONAL VRDL,ICMR-RMRC BBSR | Immunogenomics lab, Institute of Life Sciences, Bhubaneswar | Sunil Raghav, Jyotirmayee Turuk, Arup Ghosh, Atimukta Jha, Viplov K. Biswas, Swati Madhulika, Manasi Priyadarshini, Shuchi Smita, Jaya Singh Khastri, Rupesh Dash, Soma Chattopadhyay, Ghulam Hussain Syed, Shanti Senapati, Tushar K. Beuria, Debdutta Bhattacharya, Rajeeb Swain, Punit Prasad, COVID-19 team of ILS & RMRC, Orissa COVID-19 study group, DBT's PAN-INDIA 1000 SARS-CoV2 RNA genome sequencing consortium, Sanghamitra Pati, Ajay Parida |
| EPI_ISL_455958, EPI_ISL_455959, EPI_ISL_455960, EPI_ISL_455961, EPI_ISL_455962, EPI_ISL_455963, EPI_ISL_455964, EPI_ISL_455965, EPI_ISL_455966, EPI_ISL_455967, EPI_ISL_455968, EPI_ISL_455969, EPI_ISL_455970, EPI_ISL_455971, EPI_ISL_455972, EPI_ISL_455973, EPI_ISL_455974, EPI_ISL_455975, EPI_ISL_455976, EPI_ISL_455977, EPI_ISL_455978, EPI_ISL_455979 | | | |
| see above | Department of Clinical Microbiology | GIGA Medical Genomics | Keith Durkin, Maria Artesi, Sébastien Bontems, Raphaël Boreux, Cécile Meex, Pierrette Melin, Marie-Pierre Hayette, Vincent Bours. |
| EPI_ISL_456478 | Microbiological Diagnostic Unit Public Health Laboratory | Microbiological Diagnostic Unit Public Health Laboratory, The Peter Doherty Institute for Infection and Immunity | Seemann T., Schultz M., Sait, M., Sherry, N. |
| EPI_ISL_456484, EPI_ISL_456485, EPI_ISL_456486, EPI_ISL_456487, EPI_ISL_456488, EPI_ISL_456489, EPI_ISL_456490, EPI_ISL_456491, EPI_ISL_456499, EPI_ISL_456500, EPI_ISL_456502, EPI_ISL_456503, EPI_ISL_456504, EPI_ISL_456505, EPI_ISL_456508, EPI_ISL_456509, EPI_ISL_456510, EPI_ISL_456511, EPI_ISL_456512, EPI_ISL_456513, EPI_ISL_456514, EPI_ISL_456515, EPI_ISL_456516, EPI_ISL_456517, EPI_ISL_456519, EPI_ISL_456520, EPI_ISL_456521, EPI_ISL_456522, EPI_ISL_456523, EPI_ISL_456524, EPI_ISL_456525, EPI_ISL_456526, EPI_ISL_456527, EPI_ISL_456528, EPI_ISL_456529, EPI_ISL_456530, EPI_ISL_456531, EPI_ISL_456532, EPI_ISL_456533, EPI_ISL_456534, EPI_ISL_456535, EPI_ISL_456536, EPI_ISL_456537, EPI_ISL_456543, EPI_ISL_456545, EPI_ISL_456547, EPI_ISL_456548, EPI_ISL_456549, EPI_ISL_456550, EPI_ISL_456551, EPI_ISL_456552, EPI_ISL_456553, EPI_ISL_456554, EPI_ISL_456555, EPI_ISL_456556, EPI_ISL_456558, EPI_ISL_456559, EPI_ISL_456560, EPI_ISL_456561, EPI_ISL_456562, EPI_ISL_456563, EPI_ISL_456564, EPI_ISL_456565, EPI_ISL_456566, EPI_ISL_456567, EPI_ISL_456568, EPI_ISL_456569, EPI_ISL_456570, EPI_ISL_456571, EPI_ISL_456572, EPI_ISL_456573, EPI_ISL_456574, EPI_ISL_456575, EPI_ISL_456576, EPI_ISL_456577, EPI_ISL_456579, EPI_ISL_456580, EPI_ISL_456581 | | | |
| see above | Victorian Infectious Diseases Reference Laboratory (VIDRL) | Microbiological Diagnostic Unit Public Health Laboratory and Victorian Infectious Diseases Reference Laboratory, Doherty Institute | Caly L., Seemann T., Sait, M., Schultz M., Druce J., Sherry, N. |
| EPI_ISL_456594, EPI_ISL_456595 | Microbiological Diagnostic Unit Public Health Laboratory | Microbiological Diagnostic Unit Public Health Laboratory, The Peter Doherty Institute for Infection and Immunity | Seemann T., Schultz M., Sait, M., Sherry, N. |
| EPI_ISL_456613, EPI_ISL_456631, EPI_ISL_456645 | Victorian Infectious Diseases Reference Laboratory (VIDRL) | Microbiological Diagnostic Unit Public Health Laboratory and Victorian Infectious Diseases Reference Laboratory, Doherty Institute | Caly L., Seemann T., Sait, M., Schultz M., Druce J., Sherry, N. |
| EPI_ISL_456666, EPI_ISL_456667, EPI_ISL_456668, EPI_ISL_456669, EPI_ISL_456674, EPI_ISL_456675, EPI_ISL_456676 | University of Birmingham | COVID-19 Genomics UK (COG-UK) Consortium | Loman Lab: Claire McMurray, Joanne Stockton, Samuel Nicholls, Radoslaw Poplawski, Will Rowe, Josh Quick, Nicholas Loman // UHB Lab: Celina M Whalley, Andrew Bosworth, Charlotte Poxon, Kasun Wanigasooriya, Oliver Pickles, Mike Kidd, Alex Richter, Andrew D Beggs // PHE Heartlands Lab: Husam Osman, Andrew Bosworth |
| EPI_ISL_456681, EPI_ISL_456682, | Department of Pathology, University of Cambridge | COVID-19 Genomics UK (COG-UK) Consortium | Luke W Meredith, M. Estée Török, Myra Hosmillo, William L. Hamilton, Martin D. Curran, Theresa Feltwell, Grant Hall, Anna Yakovleva, Fahad A Khokhar, |

| | | | |
|---|---|---|---|
| EPI_ISL_456695, EPI_ISL_456696, EPI_ISL_456697, EPI_ISL_456737 | | | Charlotte J. Houldcroft, Laura G Caller, Aminu S. Jahun, Sarah L. Caddy, Ian Goodfellow |
| EPI_ISL_456772, EPI_ISL_456773, EPI_ISL_456782, EPI_ISL_456783, EPI_ISL_456784, EPI_ISL_456799, EPI_ISL_456800, EPI_ISL_456801, EPI_ISL_456802, EPI_ISL_456803, EPI_ISL_456804, EPI_ISL_456805, EPI_ISL_456806, EPI_ISL_456807, EPI_ISL_456808, EPI_ISL_456809, EPI_ISL_456810, EPI_ISL_456811, EPI_ISL_456812, EPI_ISL_456813, EPI_ISL_456814, EPI_ISL_456815, EPI_ISL_456816, EPI_ISL_456817, EPI_ISL_456818, EPI_ISL_456819, EPI_ISL_456820, EPI_ISL_456821, EPI_ISL_456822, EPI_ISL_456823, EPI_ISL_456824, EPI_ISL_456825, EPI_ISL_456826, EPI_ISL_456827, EPI_ISL_456828, EPI_ISL_456829, EPI_ISL_456830, EPI_ISL_456831, EPI_ISL_456832, EPI_ISL_456833, EPI_ISL_456834, EPI_ISL_456835, EPI_ISL_456836, EPI_ISL_456837, EPI_ISL_456838, EPI_ISL_456839, EPI_ISL_456840, EPI_ISL_456841, EPI_ISL_456842, EPI_ISL_456843, EPI_ISL_456844, EPI_ISL_456845, EPI_ISL_456846, EPI_ISL_456847, EPI_ISL_456848, EPI_ISL_456849, EPI_ISL_456850, EPI_ISL_456851, EPI_ISL_456852, EPI_ISL_456853, EPI_ISL_456854, EPI_ISL_456855, EPI_ISL_456856, EPI_ISL_456857, EPI_ISL_456858, EPI_ISL_456859, EPI_ISL_456860, EPI_ISL_456861, EPI_ISL_456862, EPI_ISL_456863, EPI_ISL_456864, EPI_ISL_456865, EPI_ISL_456866, EPI_ISL_456867, EPI_ISL_456868, EPI_ISL_456869, EPI_ISL_456870, EPI_ISL_456871, EPI_ISL_456872, EPI_ISL_456873, EPI_ISL_456874, EPI_ISL_456875, EPI_ISL_456876, EPI_ISL_456877, EPI_ISL_456878, EPI_ISL_456879, EPI_ISL_456882, EPI_ISL_456883, EPI_ISL_456884, EPI_ISL_456885, EPI_ISL_456886, EPI_ISL_456887, EPI_ISL_456888 | | | |
| see above | West of Scotland Specialist Virology Centre, NHSGGC / MRC-University of Glasgow Centre for Virus Research | COVID-19 Genomics UK (COG-UK) Consortium | Ana da Silva Filipe, Natasha Johnson, Kathy Smollett, Daniel Mair, Stephen Carmichael, Lily Tong, Jenna Nichols, Elihu Aranday-Cortes, Kirstyn Brunker, Yasmin Parr, Kyriaki Nomikou; Sarah McDonald, Marc Niebel, Patawee Asamaphan; Richard Orton, Joseph Hughes, Sreenu Vattipally, David L Robertson; Alasdair MacLean, Rory Gunson; Kathy Li, Natasha Jesudason, Rajiv Shah, James Shepherd, Antonia Ho, Emma Thomson |
| EPI_ISL_456941, EPI_ISL_456942, EPI_ISL_456943, EPI_ISL_456944, EPI_ISL_456945, EPI_ISL_456946, EPI_ISL_456947, EPI_ISL_456948, EPI_ISL_456949 | Virology Department, Royal Infirmary of Edinburgh, NHS Lothian / School of Biological Sciences, University of Edinburgh / Institute of Genetics and Molecular Medicine, University of Edinburgh | COVID-19 Genomics UK (COG-UK) Consortium | McHugh M, Dewar R, Rooke S, Gallagher M, Balcaza C, O'Toole Á, Scher E, Hill V, McCrone JT, Colquhoun R, Yu X, Jackson B, Rambaut A, Williams TC, Templeton K |
| EPI_ISL_457231, EPI_ISL_457234, EPI_ISL_457235, EPI_ISL_457237, EPI_ISL_457241, EPI_ISL_457243, EPI_ISL_457244, EPI_ISL_457247, EPI_ISL_457249 | University of Exeter | COVID-19 Genomics UK (COG-UK) Consortium | Ben Temperton,Aaron Jeffries,Michelle Michelsen,Joanna Warwick-Dugdale,Audrey Farbos,Robyn Manley,Stephen Michell,Jane Masoli |
| EPI_ISL_457259, EPI_ISL_457260, EPI_ISL_457261, EPI_ISL_457262, EPI_ISL_457263, EPI_ISL_457264, EPI_ISL_457265, EPI_ISL_457266, EPI_ISL_457267, EPI_ISL_457268, EPI_ISL_457269 | | | |
| see above | Liverpool Clinical Laboratories | COVID-19 Genomics UK (COG-UK) Consortium | Sam Haldenby, Anita Lucaci, Steve Paterson, Julian Hiscox, Alistair Darby, M Almsaud, A Alrezaihi, Muhannad Alruwaili, Stuart D Armstrong, Jones Benjamin, Eleanor G Bentley, Anu Chawla, Jordan J Clark, Angela Cowell, Richard Eccles, Isabel García-Dorival, Matthew Gemmell, Alessandro Gerada, PKF Gilmore, Richard Gregory, Ximeng Han, Catherine Hartley, Margaret Hughes, Miren Iturriza-Gomara, James Johnson, L Luu, Jenifer Manson, Charlotte Nelson, Elaine O'Toole, Cassie Olateju, Rebekah Penrice-Randal , Lucille Rainbow, N.P Randle, Trevor Ian Robinson, Parul Sharma, Ghada T Shawli, James P Stewart, Neil Swainston, Ecaterina Vamos, Joanne Watts, Mark Whitehead |
| EPI_ISL_457313, EPI_ISL_457314, EPI_ISL_457315, EPI_ISL_457316, EPI_ISL_457317, EPI_ISL_457318, EPI_ISL_457319, EPI_ISL_457320, EPI_ISL_457321, EPI_ISL_457324, EPI_ISL_457325 | | | |
| see above | Northumbria University / South Tees Hospitals NHS Foundation Trust / North Cumbria Integrated Care NHS Foundation Trust / North Tees and Hartlepool NHS Foundation Trust / Newcastle Hospitals NHS Foundation Trust | COVID-19 Genomics UK (COG-UK) Consortium | Darren L Smith,Andrew Nelson,Matthew Bashton,Greg R Young,Joshua Loh,John Allan,Mohammad A Tariq,Giles S Holt,Gary Black,Wen C Yew,Lynn Dover,Paul Baker,Steve Liggett,Sarah Essex,Jane Greenaway,Debra Padgett,Clive Graham,Garren Scott,Edward Barton,Emma Swindells,Brendan Payne,Jennifer Collins,Yusri Taha,Gary Eltringham |
| EPI_ISL_457328, EPI_ISL_457329, EPI_ISL_457330, EPI_ISL_457331, EPI_ISL_457332, EPI_ISL_457333, EPI_ISL_457334, EPI_ISL_457335, EPI_ISL_457336, EPI_ISL_457337, EPI_ISL_457338, EPI_ISL_457339, EPI_ISL_457340, EPI_ISL_457341, EPI_ISL_457342, EPI_ISL_457343, EPI_ISL_457344, EPI_ISL_457345, EPI_ISL_457346, EPI_ISL_457347, EPI_ISL_457348, EPI_ISL_457349, EPI_ISL_457350, EPI_ISL_457351, EPI_ISL_457352, EPI_ISL_457353, EPI_ISL_457354, EPI_ISL_457355, EPI_ISL_457356, EPI_ISL_457357, EPI_ISL_457358, EPI_ISL_457359, EPI_ISL_457360, EPI_ISL_457361, EPI_ISL_457362, EPI_ISL_457363, EPI_ISL_457364, EPI_ISL_457365, EPI_ISL_457366, EPI_ISL_457367, EPI_ISL_457368, EPI_ISL_457369, EPI_ISL_457370, EPI_ISL_457453, EPI_ISL_457454, EPI_ISL_457455, EPI_ISL_457456, EPI_ISL_457457, EPI_ISL_457458, EPI_ISL_457459, EPI_ISL_457460, EPI_ISL_457461, EPI_ISL_457462, EPI_ISL_457463, EPI_ISL_457464, EPI_ISL_457465, EPI_ISL_457466, EPI_ISL_457467, EPI_ISL_457468, EPI_ISL_457469, EPI_ISL_457470, EPI_ISL_457472, EPI_ISL_457473, EPI_ISL_457474, EPI_ISL_457475, EPI_ISL_457476, EPI_ISL_457477, EPI_ISL_457478, EPI_ISL_457479, EPI_ISL_457480, EPI_ISL_457481, EPI_ISL_457482, EPI_ISL_457483, EPI_ISL_457484, EPI_ISL_457485, EPI_ISL_457486, EPI_ISL_457487, EPI_ISL_457488, EPI_ISL_457489, EPI_ISL_457490, EPI_ISL_457491, EPI_ISL_457492, EPI_ISL_457493, EPI_ISL_457494, EPI_ISL_457495, EPI_ISL_457496, EPI_ISL_457497, EPI_ISL_457498, EPI_ISL_457499, EPI_ISL_457500, EPI_ISL_457501, EPI_ISL_457503, EPI_ISL_457504, EPI_ISL_457505, EPI_ISL_457506, EPI_ISL_457507, EPI_ISL_457508, EPI_ISL_457509, EPI_ISL_457510, EPI_ISL_457511, EPI_ISL_457512, EPI_ISL_457513, EPI_ISL_457514, EPI_ISL_457515, EPI_ISL_457516, EPI_ISL_457517, EPI_ISL_457518, EPI_ISL_457519, EPI_ISL_457520, EPI_ISL_457521, EPI_ISL_457522, EPI_ISL_457523, EPI_ISL_457524, EPI_ISL_457525, EPI_ISL_457526, EPI_ISL_457528, EPI_ISL_457529, EPI_ISL_457530, EPI_ISL_457531, EPI_ISL_457532, EPI_ISL_457533, EPI_ISL_457534, EPI_ISL_457535, EPI_ISL_457536, EPI_ISL_457537, EPI_ISL_457538, EPI_ISL_457539, EPI_ISL_457540, EPI_ISL_457541, EPI_ISL_457542, EPI_ISL_457543, EPI_ISL_457544, EPI_ISL_457545, EPI_ISL_457546, EPI_ISL_457547, EPI_ISL_457548, EPI_ISL_457549, EPI_ISL_457550, EPI_ISL_457551, EPI_ISL_457552, EPI_ISL_457553, EPI_ISL_457554, EPI_ISL_457555, EPI_ISL_457556, EPI_ISL_457557, EPI_ISL_457558, EPI_ISL_457559, EPI_ISL_457560, EPI_ISL_457561, EPI_ISL_457562, EPI_ISL_457563, EPI_ISL_457564, EPI_ISL_457565, EPI_ISL_457566, EPI_ISL_457567, EPI_ISL_457572 | | | |
| see above | Quadram Institute Bioscience | COVID-19 Genomics UK (COG-UK) Consortium | Dave J. Baker, Gemma L. Kay, Alp Aydin, Thanh Le-Viet, Steven Rudder, Ana P. Tedim, Anastasia Kolyva, Maria Diaz, Leonardo de Oliveira Martins, Nabil-Fareed Alikhan, Lizzie Meadows, Rachael Stanley, Ngozi Elumogo, Muhammed Yasir, Nicholas M. Thomson, Alexander J Trotter, Rachel Gilroy, Samuel Bloomfield, Claire Stuart, Andrew Bell, Reenesh Prakash, Samir Dervisevic, Alison E. Mather, John Wain, Mark Webber, Andrew J. Page, Justin O'Grady |
| EPI_ISL_457574, EPI_ISL_457579, EPI_ISL_457580 | Queens Medical Centre, Clinical Microbiology Department / DeepSeq Nottingham | COVID-19 Genomics UK (COG-UK) Consortium | Gemma Clark, Wendy Smith, Manjinder Khakh, Hannah Howson-Wells, Jonathan Ball, Patrick McClure, Joseph Chappell, Theocharis Tsoleridis, Nadine Holmes, Matthew Carlisle, Christopher Moore, Fei Sang, Johnny Debebe, Victoria Wright, Matthew Loose |
| EPI_ISL_457613, EPI_ISL_457635 | Virology Department, Sheffield Teaching Hospitals NHS Foundation Trust/Department of Infection, Immunity and Cardiovascular Disease, The Medical School, University of Sheffield | COVID-19 Genomics UK (COG-UK) Consortium | Thushan de Silva, Matthew Parker, Nikki Smith, Adri Angyal, Rebecca Brown, Luke Green, Rachel Tucker, Paul Parsons, Danielle Groves, Katie Johnson, Laura Carrilero, Alex Keeley, Dave Partridge, Matthew Wyles, Benjamin Lindsey, Mehmet Yavuz, Mohammad Raza, Cariad Evans |
| EPI_ISL_457705 | OMAN-NIC | Department of Microbiology and Immunology- SQUH | Fahad Zadjali, Samira Al-Maruqi, Amina Al Jardani, Khulood Al-Mammary, Hanan Al-kindi, Fatma BaAlawi, Hamida AL Barwani, Zeyana AL-Dahmani, Intisar Al-Shukri, Aisha Al-Busaidi, Aisha Al-Amri, Ahlam Al-Amri, Mohammed Al-Tobi, Samiha Al Kharusi, Abdulla Balkhair |
| EPI_ISL_458039 | King Institute of Preventive Medicine & Research | CSIR-Centre for Cellular and Molecular Biology | K.Kaveri,S.Sivasubramanian,S.Vennila,P.Padmapriya,R.Kiruba,S.Magesh,G. Dhinakar Raj, G. Ravikumar, R. P. Aravindh Babu, K Thangaraj, Lamuk Zaveri, Shagufta Khan, Namami Gaur, Sakshi Shambhavi, Tulasi Nagabandi, Purushotham Vodnala, Payel Mukherjee, Sofia Banu, Priya Singh, Dhiviya Vedagiri, Divya Gupta, Vishal Sah, Santosh Kumar Kuncha, Krishnan Harinivas Harshan, Archana Bharadwaj Siva, Karthik Bharadwaj Tallapaka, Rakesh K Mishra, Divya Tej Sowpati |
| EPI_ISL_458040 | King Institute of Preventive Medicine & Research | CSIR-Centre for Cellular and Molecular Biology | K.Kaveri,S.Sivasubramanian,S.Vennila,P.Padmapriya,R.Kiruba,S.Magesh,G. Dhinakar Raj, G. Ravikumar, R. P. Aravindh Babu, K Thangaraj, Namami Gaur, Sakshi Shambhavi, Lamuk Zaveri, Shagufta Khan, Tulasi Nagabandi, Purushotham Vodnala, Payel Mukherjee, Sofia Banu, Priya Singh, Dhiviya Vedagiri, Divya Gupta, Vishal Sah, Santosh Kumar Kuncha, Krishnan Harinivas Harshan, Archana Bharadwaj Siva, Karthik Bharadwaj Tallapaka, Rakesh K Mishra, Divya Tej Sowpati |
| EPI_ISL_458041 | King Institute of Preventive Medicine & Research | CSIR-Centre for Cellular and Molecular Biology | K.Kaveri,S.Sivasubramanian,S.Vennila,P.Padmapriya,R.Kiruba,S.Magesh,G. Dhinakar Raj, G. Ravikumar, R. P. Aravindh Babu, K Thangaraj, Tulasi Nagabandi, Namami Gaur, Sakshi Shambhavi, Lamuk Zaveri, Shagufta Khan, Purushotham Vodnala, Payel Mukherjee, Sofia Banu, Priya Singh, Dhiviya Vedagiri, Divya Gupta, Vishal Sah, Santosh Kumar Kuncha, Krishnan Harinivas Harshan, Archana Bharadwaj Siva, Karthik Bharadwaj Tallapaka, Rakesh K Mishra, Divya Tej Sowpati |
| EPI_ISL_458042 | King Institute of Preventive Medicine & Research | CSIR-Centre for Cellular and Molecular Biology | K.Kaveri,S.Sivasubramanian,S.Vennila,P.Padmapriya,R.Kiruba,S.Magesh,G. Dhinakar Raj, G. Ravikumar, M. Sekar, K Thangaraj, Payel Mukherjee, Sofia Banu, Priya Singh, Dhiviya Vedagiri, Divya Gupta, Vishal Sah, Santosh Kumar Kuncha, Krishnan Harinivas Harshan, Archana Bharadwaj Siva, Karthik Bharadwaj Tallapaka, Shagufta Khan, Lamuk Zaveri, Namami Gaur, Sakshi Shambhavi, Tulasi Nagabandi, Purushotham Vodnala, Rakesh K Mishra, Divya Tej Sowpati |
| EPI_ISL_458043 | King Institute of Preventive Medicine & Research | CSIR-Centre for Cellular and Molecular Biology | K.Kaveri,S.Sivasubramanian,S.Vennila,P.Padmapriya,R.Kiruba,S.Magesh,G. Dhinakar Raj, G. Ravikumar, M. Sekar, K Thangaraj,Sofia Banu, Payel Mukherjee, Priya Singh, Dhiviya Vedagiri, Divya Gupta, Vishal Sah, Santosh Kumar Kuncha, Krishnan Harinivas Harshan, Archana Bharadwaj Siva, Karthik Bharadwaj Tallapaka, Shagufta Khan, Lamuk Zaveri, Namami Gaur, Sakshi Shambhavi, Tulasi Nagabandi, Purushotham Vodnala, Rakesh K Mishra, Divya Tej Sowpati |
| EPI_ISL_458044 | King Institute of Preventive Medicine & Research | CSIR-Centre for Cellular and Molecular Biology | K.Kaveri,S.Sivasubramanian,S.Vennila,P.Padmapriya,R.Kiruba,S.Magesh,G. Dhinakar Raj, G. Ravikumar, M. Sekar, K Thangaraj,Shagufta Khan, Lamuk Zaveri, Namami Gaur, Sakshi Shambhavi, Tulasi Nagabandi, Purushotham Vodnala, Payel Mukherjee, Sofia Banu, Priya Singh, Dhiviya Vedagiri, Divya Gupta, Vishal Sah, Santosh Kumar Kuncha, Krishnan Harinivas Harshan, Archana Bharadwaj Siva, Karthik Bharadwaj Tallapaka, Rakesh K Mishra, Divya Tej Sowpati |

| | | | |
|---|---|---|---|
| EPI_ISL_458045 | CSIR-Centre for Cellular and Molecular Biology | CSIR-Centre for Cellular and Molecular Biology | Payel Mukherjee, Sofia Banu, Priya Singh, Dhiviya Vedagiri, Divya Gupta, Vishal Sah, Santosh Kumar Kuncha, Krishnan Harinivas Harshan, Archana Bharadwaj Siva, Karthik Bharadwaj Tallapaka, Shagufta Khan, Lamuk Zaveri, Namami Gaur, Sakshi Shambhavi, Tulasi Nagabandi, Purushotham Vodnala, G. Aditya Kumar, Koushick Sivakumar, Pooja Ramesh Gupta, Rajan Kumar Jha, Shraddha Vijay Lahoti, Rakesh K Mishra, Divya Tej Sowpati |
| EPI_ISL_458046 | CSIR-Centre for Cellular and Molecular Biology | CSIR-Centre for Cellular and Molecular Biology | Sofia Banu, Payel Mukherjee, Priya Singh, Dhiviya Vedagiri, Divya Gupta, Vishal Sah, Santosh Kumar Kuncha, Krishnan Harinivas Harshan, Archana Bharadwaj Siva, Karthik Bharadwaj Tallapaka, Shagufta Khan, Lamuk Zaveri, Namami Gaur, Sakshi Shambhavi, Tulasi Nagabandi, Purushotham Vodnala, Deepak Kumar, Devi Prasad Vijayashankar, Disha Nanda, Divya Das, Jotin Gogoi, Manish Bhattacharjee, Rakesh K Mishra, Divya Tej Sowpati |
| EPI_ISL_458047 | CSIR-Centre for Cellular and Molecular Biology | CSIR-Centre for Cellular and Molecular Biology | Shagufta Khan, Lamuk Zaveri, Namami Gaur, Sakshi Shambhavi, Tulasi Nagabandi, Purushotham Vodnala, Payel Mukherjee, Sofia Banu, Priya Singh, Dhiviya Vedagiri, Divya Gupta, Vishal Sah, Santosh Kumar Kuncha, Krishnan Harinivas Harshan, Archana Bharadwaj Siva, Karthik Bharadwaj Tallapaka, Disha Nanda, Divya Das, Jotin Gogoi, Manish Bhattacharjee, Ravi Prasad Mukku, Rakesh K Mishra, Divya Tej Sowpati |
| EPI_ISL_458048 | CSIR-Centre for Cellular and Molecular Biology | CSIR-Centre for Cellular and Molecular Biology | Lamuk Zaveri, Shagufta Khan, Namami Gaur, Sakshi Shambhavi, Tulasi Nagabandi, Purushotham Vodnala, Payel Mukherjee, Sofia Banu, Priya Singh, Dhiviya Vedagiri, Divya Gupta, Vishal Sah, Santosh Kumar Kuncha, Krishnan Harinivas Harshan, Archana Bharadwaj Siva, Karthik Bharadwaj Tallapaka, Renu Sudhakar, Somesh Gorde, Gangumala Srinivas Reddy, Sujoy Deb, Swati Bayyana, Rakesh K Mishra, Divya Tej Sowpati |
| EPI_ISL_458049 | CSIR-Centre for Cellular and Molecular Biology | CSIR-Centre for Cellular and Molecular Biology | Namami Gaur, Sakshi Shambhavi, Lamuk Zaveri, Shagufta Khan, Tulasi Nagabandi, Purushotham Vodnala, Payel Mukherjee, Sofia Banu, Priya Singh, Dhiviya Vedagiri, Divya Gupta, Vishal Sah, Santosh Kumar Kuncha, Krishnan Harinivas Harshan, Archana Bharadwaj Siva, Karthik Bharadwaj Tallapaka, Zeba Rizvi, Zuberwasim Sayyad, Kakade Aishwarya Arun, Amrutha H C, Ananga Ghosh, Rakesh K Mishra, Divya Tej Sowpati |
| EPI_ISL_458050 | CSIR-Centre for Cellular and Molecular Biology | CSIR-Centre for Cellular and Molecular Biology | Tulasi Nagabandi, Namami Gaur, Sakshi Shambhavi, Lamuk Zaveri, Shagufta Khan, Purushotham Vodnala, Payel Mukherjee, Sofia Banu, Priya Singh, Dhiviya Vedagiri, Divya Gupta, Vishal Sah, Santosh Kumar Kuncha, Krishnan Harinivas Harshan, Archana Bharadwaj Siva, Karthik Bharadwaj Tallapaka,Kezia J Ann, Radhika Khandelwal, Roshan Maku Venkata, Shemin Mansuri, Sonu Uday, Rakesh K Mishra, Divya Tej Sowpati |
| EPI_ISL_458051 | CSIR-Centre for Cellular and Molecular Biology | CSIR-Centre for Cellular and Molecular Biology | Payel Mukherjee, Sofia Banu, Priya Singh, Dhiviya Vedagiri, Divya Gupta, Vishal Sah, Santosh Kumar Kuncha, Krishnan Harinivas Harshan, Archana Bharadwaj Siva, Karthik Bharadwaj Tallapaka, Shagufta Khan, Lamuk Zaveri, Namami Gaur, Sakshi Shambhavi, Tulasi Nagabandi, Purushotham Vodnala, Gokulan C G, Gunjan Purohit, Hanuman Tulashiram Kale, Pankaj Kumar, Prachand Issarapu, Rakesh K Mishra, Divya Tej Sowpati |
| EPI_ISL_458052 | CSIR-Centre for Cellular and Molecular Biology | CSIR-Centre for Cellular and Molecular Biology | Sofia Banu, Payel Mukherjee, Priya Singh, Dhiviya Vedagiri, Divya Gupta, Vishal Sah, Santosh Kumar Kuncha, Krishnan Harinivas Harshan, Archana Bharadwaj Siva, Karthik Bharadwaj Tallapaka, Shagufta Khan, Lamuk Zaveri, Namami Gaur, Sakshi Shambhavi, Tulasi Nagabandi, Purushotham Vodnala,Preethi Jampala, Sharada Ravi Iyer, Sulagana Mukherjee, Swetha Sundar, Peddapuvala Sai Uday Kiran, Rakesh K Mishra, Divya Tej Sowpati |
| EPI_ISL_458053 | CSIR-Centre for Cellular and Molecular Biology | CSIR-Centre for Cellular and Molecular Biology | Shagufta Khan, Lamuk Zaveri, Namami Gaur, Sakshi Shambhavi, Tulasi Nagabandi, Purushotham Vodnala, Payel Mukherjee, Sofia Banu, Priya Singh, Dhiviya Vedagiri, Divya Gupta, Vishal Sah, Santosh Kumar Kuncha, Krishnan Harinivas Harshan, Archana Bharadwaj Siva, Karthik Bharadwaj Tallapaka,Umesh Kumar, Unis Ahmad Bhat, Ajay Sarawagi, Priyanka Pant, Rajkanwar Nathawat, Rakesh K Mishra, Divya Tej Sowpati |
| EPI_ISL_458054 | CSIR-Centre for Cellular and Molecular Biology | CSIR-Centre for Cellular and Molecular Biology | Lamuk Zaveri, Shagufta Khan, Namami Gaur, Sakshi Shambhavi, Tulasi Nagabandi, Purushotham Vodnala, Payel Mukherjee, Sofia Banu, Priya Singh, Dhiviya Vedagiri, Divya Gupta, Vishal Sah, Santosh Kumar Kuncha, Krishnan Harinivas Harshan, Archana Bharadwaj Siva, Karthik Bharadwaj Tallapaka,Umesh Kumar, Unis Ahmad Bhat, Ajay Sarawagi, Priyanka Pant, Rajkanwar Nathawat, Rakesh K Mishra, Divya Tej Sowpati |
| EPI_ISL_458055 | CSIR-Centre for Cellular and Molecular Biology | CSIR-Centre for Cellular and Molecular Biology | Namami Gaur, Sakshi Shambhavi, Lamuk Zaveri, Shagufta Khan, Tulasi Nagabandi, Purushotham Vodnala, Payel Mukherjee, Sofia Banu, Priya Singh, Dhiviya Vedagiri, Divya Gupta, Vishal Sah, Santosh Kumar Kuncha, Krishnan Harinivas Harshan, Archana Bharadwaj Siva, Karthik Bharadwaj Tallapaka, Nikhil Hajirnis, Pratheusa Maccha, M Soujanya Reddy,G. Aditya Kumar, Koushick Sivakumar, Rakesh K Mishra, Divya Tej Sowpati |
| EPI_ISL_458056 | CSIR-Centre for Cellular and Molecular Biology | CSIR-Centre for Cellular and Molecular Biology | Tulasi Nagabandi, Namami Gaur, Sakshi Shambhavi, Lamuk Zaveri, Shagufta Khan, Purushotham Vodnala, Payel Mukherjee, Sofia Banu, Priya Singh, Dhiviya Vedagiri, Divya Gupta, Vishal Sah, Santosh Kumar Kuncha, Krishnan Harinivas Harshan, Archana Bharadwaj Siva, Karthik Bharadwaj Tallapaka,G. Aditya Kumar, Koushick Sivakumar, Pooja Ramesh Gupta, Rajan Kumar Jha, Shraddha Vijay Lahoti, Rakesh K Mishra, Divya Tej Sowpati |
| EPI_ISL_458057 | CSIR-Centre for Cellular and Molecular Biology | CSIR-Centre for Cellular and Molecular Biology | Payel Mukherjee, Sofia Banu, Priya Singh, Dhiviya Vedagiri, Divya Gupta, Vishal Sah, Santosh Kumar Kuncha, Krishnan Harinivas Harshan, Archana Bharadwaj Siva, Karthik Bharadwaj Tallapaka, Shagufta Khan, Lamuk Zaveri, Namami Gaur, Sakshi Shambhavi, Tulasi Nagabandi, Purushotham Vodnala,Deepak Kumar, Devi Prasad Vijayashankar, Disha Nanda, Divya Das, Jotin Gogoi, Manish Bhattacharjee, Rakesh K Mishra, Divya Tej Sowpati |
| EPI_ISL_458058 | CSIR-Centre for Cellular and Molecular Biology | CSIR-Centre for Cellular and Molecular Biology | Sofia Banu, Payel Mukherjee, Priya Singh, Dhiviya Vedagiri, Divya Gupta, Vishal Sah, Santosh Kumar Kuncha, Krishnan Harinivas Harshan, Archana Bharadwaj Siva, Karthik Bharadwaj Tallapaka, Shagufta Khan, Lamuk Zaveri, Namami Gaur, Sakshi Shambhavi, Tulasi Nagabandi, Purushotham Vodnala, Disha Nanda, Divya Das, Jotin Gogoi, Manish Bhattacharjee, Ravi Prasad Mukku, Rakesh K Mishra, Divya Tej Sowpati |
| EPI_ISL_458059 | CSIR-Centre for Cellular and Molecular Biology | CSIR-Centre for Cellular and Molecular Biology | Shagufta Khan, Lamuk Zaveri, Namami Gaur, Sakshi Shambhavi, Tulasi Nagabandi, Purushotham Vodnala, Payel Mukherjee, Sofia Banu, Priya Singh, Dhiviya Vedagiri, Divya Gupta, Vishal Sah, Santosh Kumar Kuncha, Krishnan Harinivas Harshan, Archana Bharadwaj Siva, Karthik Bharadwaj Tallapaka, Renu Sudhakar, Somesh Gorde, Gangumala Srinivas Reddy, Sujoy Deb, Swati Bayyana, Rakesh K Mishra, Divya Tej Sowpati |
| EPI_ISL_458060 | CSIR-Centre for Cellular and Molecular Biology | CSIR-Centre for Cellular and Molecular Biology | Lamuk Zaveri, Shagufta Khan, Namami Gaur, Sakshi Shambhavi, Tulasi Nagabandi, Purushotham Vodnala, Payel Mukherjee, Sofia Banu, Priya Singh, Dhiviya Vedagiri, Divya Gupta, Vishal Sah, Santosh Kumar Kuncha, Krishnan Harinivas Harshan, Archana Bharadwaj Siva, Karthik Bharadwaj Tallapaka,Zeba Rizvi, Zuberwasim Sayyad, Kakade Aishwarya Arun, Amrutha H C, Ananga Ghosh, Rakesh K Mishra, Divya Tej Sowpati |
| EPI_ISL_458061 | CSIR-Centre for Cellular and Molecular Biology | CSIR-Centre for Cellular and Molecular Biology | Namami Gaur, Sakshi Shambhavi, Lamuk Zaveri, Shagufta Khan, Tulasi Nagabandi, Purushotham Vodnala, Payel Mukherjee, Sofia Banu, Priya Singh, Dhiviya Vedagiri, Divya Gupta, Vishal Sah, Santosh Kumar Kuncha, Krishnan Harinivas Harshan, Archana Bharadwaj Siva, Karthik Bharadwaj Tallapaka,Kezia J Ann, Radhika Khandelwal, Roshan Maku Venkata, Shemin Mansuri, Sonu Uday, Rakesh K Mishra, Divya Tej Sowpati |
| EPI_ISL_458064 | CSIR-Centre for Cellular and Molecular Biology | CSIR-Centre for Cellular and Molecular Biology | Shagufta Khan, Lamuk Zaveri, Namami Gaur, Sakshi Shambhavi, Tulasi Nagabandi, Purushotham Vodnala, Payel Mukherjee, Sofia Banu, Priya Singh, Dhiviya Vedagiri, Divya Gupta, Vishal Sah, Santosh Kumar Kuncha, Krishnan Harinivas Harshan, Archana Bharadwaj Siva, Karthik Bharadwaj Tallapaka,Preethi Jampala, Sharada Ravi Iyer, Sulagana Mukherjee, Swetha Sundar, Peddapuvala Sai Uday Kiran Rakesh K Mishra, Divya Tej Sowpati |
| EPI_ISL_458065 | CSIR-Centre for Cellular and Molecular Biology | CSIR-Centre for Cellular and Molecular Biology | Lamuk Zaveri, Shagufta Khan, Namami Gaur, Sakshi Shambhavi, Tulasi Nagabandi, Purushotham Vodnala, Payel Mukherjee, Sofia Banu, Priya Singh, Dhiviya Vedagiri, Divya Gupta, Vishal Sah, Santosh Kumar Kuncha, Krishnan Harinivas Harshan, Archana Bharadwaj Siva, Karthik Bharadwaj Tallapaka,Umesh Kumar, Unis Ahmad Bhat, Ajay Sarawagi, Priyanka Pant, Rajkanwar Nathawat, Rakesh K Mishra, Divya Tej Sowpati |
| EPI_ISL_458103 | Gujarat Biotechnology Research Centre | Gujarat Biotechnology Research Centre | Ramesh Pandit, Tejas Shah, Ankit Hinsu, Pritesh Sabara, Apurvasinh Puvar, Janvi Raval, Zarna Patel, Monika Gandhi, Pinal Trivedi, Maharshi Pandya, Amit Kanani, Nidhi Patel, Nitin Savaliya, Raghawendra Kumar, Dinesh Kumar, Zuber Saiyed, Komal Patel, Labdhi Pandya, Snehal Bagatharia, Armi Chaudhari, Bhavesh Modi, Gaurishankar Shrimali, R D Dixit, A M Kadri, Umang Mishra, Chaitanya Joshi, Madhvi Joshi, , , , , |
| EPI_ISL_458104 | Gujarat Biotechnology Research Centre | Gujarat Biotechnology Research Centre | Tejas Shah, Ankit Hinsu, Pritesh Sabara, Apurvasinh Puvar, Janvi Raval, Zarna Patel, Monika Gandhi, Pinal Trivedi, Maharshi Pandya, Amit Kanani, Nidhi Patel, Nitin Savaliya, Raghawendra Kumar, Dinesh Kumar, Zuber Saiyed, Komal Patel, Labdhi Pandya, Snehal Bagatharia, Ramesh Pandit, Bhavya Jindal, Bhavesh Modi, Gaurishankar Shrimali, R D Dixit, A M Kadri, Umang Mishra, Chaitanya Joshi, Madhvi Joshi, , , , , |
| EPI_ISL_458105 | Gujarat Biotechnology Research Centre | Gujarat Biotechnology Research Centre | Ankit Hinsu, Pritesh Sabara, Apurvasinh Puvar, Janvi Raval, Zarna Patel, Monika Gandhi, Pinal Trivedi, Maharshi Pandya, Amit Kanani, Nidhi Patel, Nitin Savaliya, Raghawendra Kumar, Dinesh Kumar, Zuber Saiyed, Komal Patel, Labdhi Pandya, Snehal Bagatharia, Ramesh Pandit, Tejas Shah, Camellia Chakraborty, Bhavesh Modi, Gaurishankar Shrimali, R D Dixit, A M Kadri, Umang Mishra, Chaitanya Joshi, Madhvi Joshi, , , , , |
| EPI_ISL_458106 | Gujarat Biotechnology Research Centre | Gujarat Biotechnology Research Centre | Pritesh Sabara, Apurvasinh Puvar, Janvi Raval, Zarna Patel, Monika Gandhi, Pinal Trivedi, Maharshi Pandya, Amit Kanani, Nidhi Patel, Nitin Savaliya, Raghawendra Kumar, Dinesh Kumar, Zuber Saiyed, Komal Patel, Labdhi Pandya, Snehal Bagatharia, Ramesh Pandit, Tejas Shah, Ankit Hinsu, Siddhant Kumar, Bhavesh Modi, Gaurishankar Shrimali, R D Dixit, A M Kadri, Umang Mishra, Chaitanya Joshi, Madhvi Joshi, , , , , |
| EPI_ISL_458107 | Gujarat Biotechnology Research Centre | Gujarat Biotechnology Research Centre | Apurvasinh Puvar, Janvi Raval, Zarna Patel, Monika Gandhi, Pinal Trivedi, Maharshi Pandya, Amit Kanani, Nidhi Patel, Nitin Savaliya, Raghawendra Kumar, Dinesh Kumar, Zuber Saiyed, Komal Patel, Labdhi Pandya, Snehal Bagatharia, Ramesh Pandit, Tejas Shah, Ankit Hinsu, Pritesh Sabara, Priyanka P Vatsa, Bhavesh Modi, Gaurishankar Shrimali, R D Dixit, A M Kadri, Umang Mishra, Chaitanya Joshi, Madhvi Joshi, , , , , |
| EPI_ISL_458108 | Gujarat Biotechnology Research Centre | Gujarat Biotechnology Research Centre | Janvi Raval, Zarna Patel, Monika Gandhi, Pinal Trivedi, Maharshi Pandya, Amit Kanani, Nidhi Patel, Nitin Savaliya, Raghawendra Kumar, Dinesh Kumar, Zuber Saiyed, Komal Patel, Labdhi Pandya, Snehal Bagatharia, Ramesh Pandit, Tejas Shah, Ankit Hinsu, Pritesh Sabara, Apurvasinh Puvar, Pooja P Doshi, Bhavesh Modi, Gaurishankar Shrimali, R D Dixit, A M Kadri, Umang Mishra, Chaitanya Joshi, Madhvi Joshi, , , , , |

| | | | |
|---|---|---|---|
| EPI_ISL_458117 | Oman National Influenza Centre | Department of Microbiology and Immunology-SQUH | Fahad Zadjali, Samira Al-Maruqi, Amina Al Jardani, Khulood Al-Mammary, Hanan Al-kindi, Fatma BaAlawi, Hamida AL Barwani, Zeyana AL-Dahmani, Intisar Al-Shukri, Aisha Al-Busaidi, Aisha Al-Amri, Ahlam Al-Amri, Mohammed Al-Tobi, Samiha Al Kharusi, Abdulla Balkhair |
| EPI_ISL_458133 | National Institute of Biotechnology | Bioinformatics Division, National Institute of Biotechnology | Mohammad Uzzal Hossain, Md. Moniruzzaman, Md. Salim Khan, Md. Nazrul Islam, Md. Hadisur Rahman, Arittra Bhattacharjee, Md. Ruhul Amin, Asif Rashid, Chaman Ara Keya, Keshob Chandra Das, Md. Salimullah |
| EPI_ISL_458150 | ANOUAL | ANOUAL | Jouali Farah, El Ansari Fatima Zahra, Marchoudi Nabila, Kasmi Yassine, Chenaoui Mohamed, El Aliani Aissam, Benhida Rachid, Azami Nawfel, Kitane Driss Lahlou, Loukman Salma, Fekkak Jamal |
| EPI_ISL_458516, EPI_ISL_458517, EPI_ISL_458518, EPI_ISL_458519, EPI_ISL_458520, EPI_ISL_458521, EPI_ISL_458522, EPI_ISL_458523, EPI_ISL_458524, EPI_ISL_458525, EPI_ISL_458526, EPI_ISL_458527, EPI_ISL_458528, EPI_ISL_458529, EPI_ISL_458530, EPI_ISL_458531, EPI_ISL_458532, EPI_ISL_458533, EPI_ISL_458534, EPI_ISL_458535, EPI_ISL_458536, EPI_ISL_458537, EPI_ISL_458538, EPI_ISL_458539, EPI_ISL_458540, EPI_ISL_458541, EPI_ISL_458542, EPI_ISL_458543, EPI_ISL_458544, EPI_ISL_458545, EPI_ISL_458546, EPI_ISL_458547, EPI_ISL_458548, EPI_ISL_458549, EPI_ISL_458550, EPI_ISL_458551, EPI_ISL_458552, EPI_ISL_458553, EPI_ISL_458554, EPI_ISL_458555, EPI_ISL_458556, EPI_ISL_458557, EPI_ISL_458558, EPI_ISL_458559, EPI_ISL_458560, EPI_ISL_458561, EPI_ISL_458562, EPI_ISL_458563, EPI_ISL_458564, EPI_ISL_458565, EPI_ISL_458566, EPI_ISL_458567, EPI_ISL_458568, EPI_ISL_458569, EPI_ISL_458570 | | | |
| see above | Department of Pathology, University of Cambridge | Wellcome Sanger Institute for the COVID-19 Genomics UK (COG-UK) consortium | Luke W Meredith, M. Estée Török , Myra Hosmillo, William L. Hamilton, Martin D. Curran, Theresa Feltwell, Grant Hall, Anna Yakovleva, Fahad A Khokhar, Charlotte J. Houldcroft, Laura G Caller, Aminu S. Jahun, Sarah L. Caddy, Ian Goodfellow; and Alex Alderton, Roberto Amato, Sonia Goncalves, Ewan Harrison, David K. Jackson, Ian Johnston, Dominic Kwiatkowski, Cordelia Langford, John Sillitoe on behalf of the Wellcome Sanger Institute COVID-19 Surveillance Team (http://www.sanger.ac.uk/covid-team) |
| EPI_ISL_458571 | Department of Pathology, University of Cambridge | Wellcome Sanger Institute for the COVID-19 Genomics UK (COG-UK) Consortium | Luke W Meredith, M. Estée Török , Myra Hosmillo, William L. Hamilton, Martin D. Curran, Theresa Feltwell, Grant Hall, Anna Yakovleva, Fahad A Khokhar, Charlotte J. Houldcroft, Laura G Caller, Aminu S. Jahun, Sarah L. Caddy, Ian Goodfellow; and Alex Alderton, Roberto Amato, Sonia Goncalves, Ewan Harrison, David K. Jackson, Ian Johnston, Dominic Kwiatkowski, Cordelia Langford, John Sillitoe on behalf of the Wellcome Sanger Institute COVID-19 Surveillance Team |
| EPI_ISL_458572, EPI_ISL_458573, EPI_ISL_458574, EPI_ISL_458575, EPI_ISL_458576, EPI_ISL_458577, EPI_ISL_459166 | Department of Pathology, University of Cambridge | Wellcome Sanger Institute for the COVID-19 Genomics UK (COG-UK) consortium | Luke W Meredith, M. Estée Török , Myra Hosmillo, William L. Hamilton, Martin D. Curran, Theresa Feltwell, Grant Hall, Anna Yakovleva, Fahad A Khokhar, Charlotte J. Houldcroft, Laura G Caller, Aminu S. Jahun, Sarah L. Caddy, Ian Goodfellow; and Alex Alderton, Roberto Amato, Sonia Goncalves, Ewan Harrison, David K. Jackson, Ian Johnston, Dominic Kwiatkowski, Cordelia Langford, John Sillitoe on behalf of the Wellcome Sanger Institute COVID-19 Surveillance Team (http://www.sanger.ac.uk/covid-team) |
| EPI_ISL_459173 | Department of Pathology, University of Cambridge | Wellcome Sanger Institute for the COVID-19 Genomics UK (COG-UK) Consortium | Luke W Meredith, M. Estée Török , Myra Hosmillo, William L. Hamilton, Martin D. Curran, Theresa Feltwell, Grant Hall, Anna Yakovleva, Fahad A Khokhar, Charlotte J. Houldcroft, Laura G Caller, Aminu S. Jahun, Sarah L. Caddy, Ian Goodfellow; and Alex Alderton, Roberto Amato, Sonia Goncalves, Ewan Harrison, David K. Jackson, Ian Johnston, Dominic Kwiatkowski, Cordelia Langford, John Sillitoe on behalf of the Wellcome Sanger Institute COVID-19 Surveillance Team |
| EPI_ISL_459175, EPI_ISL_459176, EPI_ISL_459178, EPI_ISL_459182, EPI_ISL_459185, EPI_ISL_459188, EPI_ISL_459191, EPI_ISL_459193, EPI_ISL_459195, EPI_ISL_459197, EPI_ISL_459200, EPI_ISL_459204, EPI_ISL_459206, EPI_ISL_459207, EPI_ISL_459208, EPI_ISL_459209, EPI_ISL_459215, EPI_ISL_459217, EPI_ISL_459222, EPI_ISL_459228, EPI_ISL_459229, EPI_ISL_459231, EPI_ISL_459234 | | | |
| see above | Department of Pathology, University of Cambridge | Wellcome Sanger Institute for the COVID-19 Genomics UK (COG-UK) consortium | Luke W Meredith, M. Estée Török , Myra Hosmillo, William L. Hamilton, Martin D. Curran, Theresa Feltwell, Grant Hall, Anna Yakovleva, Fahad A Khokhar, Charlotte J. Houldcroft, Laura G Caller, Aminu S. Jahun, Sarah L. Caddy, Ian Goodfellow; and Alex Alderton, Roberto Amato, Sonia Goncalves, Ewan Harrison, David K. Jackson, Ian Johnston, Dominic Kwiatkowski, Cordelia Langford, John Sillitoe on behalf of the Wellcome Sanger Institute COVID-19 Surveillance Team (http://www.sanger.ac.uk/covid-team) |
| EPI_ISL_459893, EPI_ISL_459894, EPI_ISL_459896, EPI_ISL_459898, EPI_ISL_459899, EPI_ISL_459900, EPI_ISL_459905 | Laboratoire National de Sante, Microbiology, Virology | Laboratoire National de Sante, Microbiology, Epidemiology and Microbial Genomics | Anke Wienecke-Baldacchino, Jessica Tapp, Guillaume Fournier, Tamir Abdelrahman, Trung Nguyen Nguyen, Catherine Ragimbeau |
| EPI_ISL_459911 | Devki Devi Foundation, a unit of Max Healthcare | CSIR-IGIB/Max | Rajesh Pandey#, Samreen Siddiqui, Pooja Sharma, Bansidhar Tarai, Vivekanand A, Bharathram Uppili, Saruchi Wadhwa, Nishu Tyagi, Mitali Mukerji, Poonam Das, Sujeet Jha, Mohammed Faruq, Vinita Jha, Anurag Agrawal |
| EPI_ISL_459912, EPI_ISL_459915, EPI_ISL_459916, EPI_ISL_459917, EPI_ISL_459918, EPI_ISL_459921, EPI_ISL_459922, EPI_ISL_459923, EPI_ISL_459924, EPI_ISL_459925, EPI_ISL_459931, EPI_ISL_459932, EPI_ISL_459933, EPI_ISL_459934, EPI_ISL_459935, EPI_ISL_459936, EPI_ISL_459937, EPI_ISL_459938, EPI_ISL_459939, EPI_ISL_459940, EPI_ISL_459941, EPI_ISL_459948 | | | |
| see above | Devki Devi Foundation, a unit of Max Healthcare | CSIR-IGIB/Max | Rajesh Pandey#, Samreen Siddiqui, Pooja Sharma, Bansidhar Tarai, Vivekanand A, Bharathram Uppili, Saruchi Wadhwa, Nishu Tyagi, Mitali Mukerji, Bansidhar Tarai, Poonam Das, Sujeet Jha, Mohammed Faruq, Vinita Jha, Anurag Agrawal |
| EPI_ISL_459992, EPI_ISL_459993, EPI_ISL_459994, EPI_ISL_459995, EPI_ISL_459996, EPI_ISL_459997, EPI_ISL_459998, EPI_ISL_460031, EPI_ISL_460033, EPI_ISL_460034, EPI_ISL_460035, EPI_ISL_460036, EPI_ISL_460037, EPI_ISL_460038, EPI_ISL_460039, EPI_ISL_460040, EPI_ISL_460041, EPI_ISL_460042, EPI_ISL_460043, EPI_ISL_460554, EPI_ISL_460555, EPI_ISL_460556, EPI_ISL_460557, EPI_ISL_460558, EPI_ISL_460559, EPI_ISL_460560, EPI_ISL_460563, EPI_ISL_460565, EPI_ISL_460567, EPI_ISL_460568, EPI_ISL_460569, EPI_ISL_460570, EPI_ISL_460571, EPI_ISL_460573, EPI_ISL_460574, EPI_ISL_460575, EPI_ISL_460576, EPI_ISL_460577, EPI_ISL_460578, EPI_ISL_460579, EPI_ISL_460580, EPI_ISL_460581, EPI_ISL_460582, EPI_ISL_460583, EPI_ISL_460585, EPI_ISL_460586, EPI_ISL_460587, EPI_ISL_460588, EPI_ISL_460591, EPI_ISL_460595 | | | |
| see above | Michigan Department of Health and Human Services, Bureau of Laboratories | Michigan Department of Health and Human Services, Bureau of Laboratories | Blankenship HM, Riner D, Soehnlen MK |
| EPI_ISL_460949, EPI_ISL_460951, EPI_ISL_460999, EPI_ISL_461000, EPI_ISL_461004, EPI_ISL_461007, EPI_ISL_461008, EPI_ISL_461083, EPI_ISL_461084, EPI_ISL_461085, EPI_ISL_461086, EPI_ISL_461087, EPI_ISL_461143, EPI_ISL_461144, EPI_ISL_461145, EPI_ISL_461212, EPI_ISL_461213, EPI_ISL_461214, EPI_ISL_461215, EPI_ISL_461216, EPI_ISL_461217, EPI_ISL_461224, EPI_ISL_461307, EPI_ISL_461308, EPI_ISL_461309, EPI_ISL_461311, EPI_ISL_461334, EPI_ISL_461335, EPI_ISL_461336, EPI_ISL_461366, EPI_ISL_461367, EPI_ISL_461378, EPI_ISL_461379, EPI_ISL_461380, EPI_ISL_461381, EPI_ISL_461383, EPI_ISL_461384, EPI_ISL_461385, EPI_ISL_461386, EPI_ISL_461387, EPI_ISL_461388, EPI_ISL_461389, EPI_ISL_461390, EPI_ISL_461391, EPI_ISL_461394, EPI_ISL_461396, EPI_ISL_461397 | | | |
| see above | Dutch COVID-19 response team | Erasmus Medical Center | Bas Oude Munnink, David Nieuwenhuijse, Reina Sikkema, Claudia Schapendonk, Irina Chestakova, Anne van der Linden, Theo Bestebroer, Stefan van Nieuwkoop, Mark Pronk, Pascal Lexmond, Corien Swaan, Manon Haverkate, Madelief Mollers, Mart Stein, Sandra Kengne Kamga Mobou, Jeroen van Kampen, Jolanda Voermans, Aura Timen, Corine GeurtsvanKessel, Annemiek van der Eijk, Richard Molenkamp, Marion Koopmans, on behalf of the Dutch national COVID-19 response team. |
| EPI_ISL_461701, EPI_ISL_461702, EPI_ISL_461703, EPI_ISL_461704 | West of Scotland Specialist Virology Centre, NHSGGC / MRC-University of Glasgow Centre for Virus Research | COVID-19 Genomics UK (COG-UK) Consortium | Ana da Silva Filipe, Natasha Johnson, Kathy Smollett, Daniel Mair, Stephen Carmichael, Lily Tong, Jenna Nichols, Elihu Aranday-Cortes, Kirstyn Brunker, Yasmin Parr, Kyriaki Nomikou; Sarah McDonald, Marc Niebel, Patawee Asamaphan; Richard Orton, Joseph Hughes, Sreenu Vattipally, David L Robertson; Alasdair MacLean, Rory Gunson; Kathy Li, Natasha Jesudason, Rajiv Shah, James Shepherd, Antonia Ho, Emma Thomson |
| EPI_ISL_461711, EPI_ISL_461712, EPI_ISL_461713, EPI_ISL_461714, EPI_ISL_461715, EPI_ISL_461716, EPI_ISL_461717, EPI_ISL_461718, EPI_ISL_461719, EPI_ISL_461720, EPI_ISL_461721, EPI_ISL_461722, EPI_ISL_461723, EPI_ISL_461724, EPI_ISL_461725, EPI_ISL_461726, EPI_ISL_461727, EPI_ISL_461728, EPI_ISL_461729, EPI_ISL_461730, EPI_ISL_461731 | | | |
| see above | Virology Department, Royal Infirmary of Edinburgh, NHS Lothian / School of Biological Sciences, University of Edinburgh / Institute of Genetics and Molecular Medicine, University of Edinburgh | COVID-19 Genomics UK (COG-UK) Consortium | McHugh M, Dewar R, Rooke S, Gallagher M, Balcaza C, O'Toole Á, Scher E, Hill V, McCrone JT, Colquhoun R, Yu X, Jackson B, Rambaut A, Williams TC, Templeton K |
| EPI_ISL_461770, EPI_ISL_461771, EPI_ISL_461772 | University College London, Great Ormond Street Hospital for Children NHS Foundation Trust, Imperial College Healthcare NHS Trust | COVID-19 Genomics UK (COG-UK) Consortium | Sergi Castellano, Rachel Williams, Mark Kristiansen, Paola Resende Silva, Sunando Roy, Tony Brooks, Helena Tutill, Paola Niola, Patricia Dyal, Charlotte Williams, Leysa Forrest, Yasmin Panchbhaya, Jacqueline Findlay, Sam Weeks, Julianne Brown, Kathryn Harris, Paul Randell, James Price, Alison Holmes, Judith Breuer |
| EPI_ISL_461793, EPI_ISL_461794, EPI_ISL_461800, EPI_ISL_461801, EPI_ISL_461802, EPI_ISL_461803, EPI_ISL_461804 | Northumbria University / South Tees Hospitals NHS Foundation Trust / North Cumbria Integrated Care NHS Foundation Trust / North Tees and Hartlepool NHS Foundation Trust / Newcastle Hospitals NHS Foundation Trust | COVID-19 Genomics UK (COG-UK) Consortium | Darren L Smith,Andrew Nelson,Matthew Bashton,Greg R Young,Joshua Loh,John Allan,Mohammad A Tariq,Giles S Holt,Gary Black,Wen C Yew,Lynn Dover,Paul Baker,Steve Liggett,Sarah Essex,Jane Greenaway,Debra Padgett,Clive Graham,Garren Scott,Edward Barton,Emma Swindells,Brendan Payne,Jennifer Collins,Yusri Taha,Gary Eltringham |
| EPI_ISL_461806, EPI_ISL_461868, EPI_ISL_461881, EPI_ISL_461882, EPI_ISL_461883 | Quadram Institute Bioscience | COVID-19 Genomics UK (COG-UK) Consortium | Dave J. Baker, Gemma L. Kay, Alp Aydin, Thanh Le-Viet, Steven Rudder, Ana P. Tedim, Anastasia Kolyva, Maria Diaz, Leonardo de Oliveira Martins, Nabil-Fareed Alikhan, Lizzie Meadows, Rachael Stanley, Ngozi Elumogo, Muhammed Yasir, Nicholas M. Thomson, Alexander J Trotter, Rachel Gilroy, Samuel Bloomfield, Claire Stuart, Andrew Bell, Reenesh Prakash, Samir Dervisevic, Alison E. Mather, John Wain, Mark Webber, Andrew J. Page, Justin O'Grady |
| EPI_ISL_461894 | Queens Medical Centre, Clinical Microbiology Department / | COVID-19 Genomics UK (COG-UK) Consortium | Gemma Clark, Wendy Smith, Manjinder Khakh, Hannah Howson-Wells, Jonathan Ball, Patrick McClure, Joseph Chappell, Theocharis Tsoleridis, Nadine |

| | | | |
|---|---|---|---|
| | DeepSeq Nottingham | | Holmes, Matthew Carlisle, Christopher Moore, Fei Sang, Johnny Debebe, Victoria Wright, Matthew Loose |
| EPI_ISL_461970, EPI_ISL_461971, EPI_ISL_461972, EPI_ISL_461973, EPI_ISL_461974, EPI_ISL_461975, EPI_ISL_461976, EPI_ISL_461977, EPI_ISL_461978, EPI_ISL_461979, EPI_ISL_461983, EPI_ISL_461984, EPI_ISL_461985, EPI_ISL_461986, EPI_ISL_461987, EPI_ISL_461988, EPI_ISL_461989, EPI_ISL_461990, EPI_ISL_461991, EPI_ISL_461992, EPI_ISL_461993, EPI_ISL_461994, EPI_ISL_461995, EPI_ISL_461996, EPI_ISL_461997 | | | |
| see above | Centre for Enzyme Innovation, University of Portsmouth / Translational Research Laboratory, Portsmouth Hospitals NHS Trust | COVID-19 Genomics UK (COG-UK) Consortium | Angela Beckett,Yann Bourgeois,Garry Scarlett,Sharon Glaysher,Scott Elliott,Kelly Bicknell,Robert Impey,Allyson Lloyd,Sarah Wyllie,Ethan Butcher,Anoop Chauhan,Samuel Robson |
| EPI_ISL_462003, EPI_ISL_462020, EPI_ISL_462024, EPI_ISL_462028, EPI_ISL_462030, EPI_ISL_462034, EPI_ISL_462039, EPI_ISL_462043, EPI_ISL_462044, EPI_ISL_462052, EPI_ISL_462056, EPI_ISL_462057, EPI_ISL_462058, EPI_ISL_462063, EPI_ISL_462065, EPI_ISL_462076, EPI_ISL_462078, EPI_ISL_462084 | | | |
| see above | Virology Department, Sheffield Teaching Hospitals NHS Foundation Trust/Department of Infection, Immunity and Cardiovascular Disease, The Medical School, University of Sheffield | COVID-19 Genomics UK (COG-UK) Consortium | Thushan de Silva, Matthew Parker, Nikki Smith, Adri Angyal, Rebecca Brown, Luke Green, Rachel Tucker, Paul Parsons, Danielle Groves, Katie Johnson, Laura Carrilero, Alex Keeley, Dave Partridge, Matthew Wyles, Benjamin Lindsey, Mehmet Yavuz, Mohammad Raza, Cariad Evans |
| EPI_ISL_462270, EPI_ISL_462271, EPI_ISL_462272, EPI_ISL_462273, EPI_ISL_462274, EPI_ISL_462275 | KU Leuven, Rega Institute, Clinical and Epidemiological Virology | KU Leuven, Rega Institute, Clinical and Epidemiological Virology | Tony Wawina-Bokalanga, Bert Vanmechelen, Joan Marti-Carerras, Piet Maes |
| EPI_ISL_462362, EPI_ISL_462372, EPI_ISL_462382, EPI_ISL_462393, EPI_ISL_462427 | National Public Health Laboratory, National Centre for Infectious Diseases | National Public Health Laboratory, National Centre for Infectious Diseases | Mak TM, Octavia S, Chavatte JM, Cui L, Lin RTP |
| EPI_ISL_462435 | unknown | Laboratory Diagnostic | Vidanovic,D., Tesovic,B., Banovic Djeri,B., Knezevic,A., Jankovic,M., Sekler,M., Dmitric,M., Petrovic,T., Volkening,J., Afonso,C.L. |
| EPI_ISL_462480 | Institute of Human Genetics, Polish Academy of Sciences | Institute of Human Genetics, Polish Academy of Sciences | Szymon Hryhorowicz, Adam Ustaszewski, Emilia Lis, Marta Kaczmarek-Ry, Micha Witt, Andrzej Pawski |
| EPI_ISL_462865, EPI_ISL_462866, EPI_ISL_462867, EPI_ISL_462868, EPI_ISL_462869, EPI_ISL_462870, EPI_ISL_462871, EPI_ISL_462872, EPI_ISL_462873, EPI_ISL_462874, EPI_ISL_462875, EPI_ISL_462876, EPI_ISL_462877, EPI_ISL_462878, EPI_ISL_462879, EPI_ISL_462880, EPI_ISL_462881, EPI_ISL_462882, EPI_ISL_462883, EPI_ISL_462884, EPI_ISL_462885, EPI_ISL_462886, EPI_ISL_462887, EPI_ISL_462888, EPI_ISL_462889, EPI_ISL_462890, EPI_ISL_462891, EPI_ISL_462892, EPI_ISL_462893, EPI_ISL_462894, EPI_ISL_462895, EPI_ISL_462896, EPI_ISL_462897, EPI_ISL_462898, EPI_ISL_462899, EPI_ISL_462900, EPI_ISL_462901, EPI_ISL_462902, EPI_ISL_462903, EPI_ISL_462904, EPI_ISL_462905, EPI_ISL_462906, EPI_ISL_462907, EPI_ISL_462908, EPI_ISL_462909, EPI_ISL_462910, EPI_ISL_462911 | | | |
| see above | Minnesota Department of Health, Public Health Laboratory | Minnesota Department of Health, Public Health Laboratory | Matt Plumb, Jacob Garfin, and Xiong Wang |
| EPI_ISL_462991 | Microbiology Division | Microbiology Division | Flores,H. |
| EPI_ISL_463010, EPI_ISL_463011, EPI_ISL_463012, EPI_ISL_463013, EPI_ISL_463014, EPI_ISL_463015, EPI_ISL_463016, EPI_ISL_463017, EPI_ISL_463018, EPI_ISL_463019, EPI_ISL_463020, EPI_ISL_463021, EPI_ISL_463022, EPI_ISL_463023, EPI_ISL_463024, EPI_ISL_463025, EPI_ISL_463026, EPI_ISL_463027, EPI_ISL_463028, EPI_ISL_463029, EPI_ISL_463030 | | | |
| see above | Institute of Life Sciences, Bhubaneswar | Immunogenomics lab, Institute of Life Sciences, Bhubaneswar | Sunil Raghav, Arup Ghosh, Atimukta Jha, Viplov K. Biswas, Swati Madhulika, Manasi Priyadarshini, Shuchi Smita, Kaushik Sen, Hiren G. Dodia, Deepak Singh, Jeky Chawla, Shamima Ansari, Rupesh Dash, Soma Chattopadhyay, Ghulam Hussain Syed, Shanti Senapati, Tushar K. Beuria, Rajeeb Swain, Punit Prasad, ILS COVID-19 TEAM, Orissa COVID-19 Study Group, DBT's PAN-INDIA 1000 SARS-CoV2 RNA genome sequencing consortium, Ajay Parida |
| EPI_ISL_463031, EPI_ISL_463032, EPI_ISL_463033, EPI_ISL_463034, EPI_ISL_463035, EPI_ISL_463036, EPI_ISL_463037, EPI_ISL_463038, EPI_ISL_463039, EPI_ISL_463040, EPI_ISL_463041, EPI_ISL_463042, EPI_ISL_463043, EPI_ISL_463044, EPI_ISL_463045, EPI_ISL_463046, EPI_ISL_463047, EPI_ISL_463048, EPI_ISL_463049, EPI_ISL_463050, EPI_ISL_463051 | | | |
| see above | Institute of Life Sciences, Bhubaneswar | Immunogenomics lab, Institute of Life Sciences, Bhubaneswar | Sunil Raghav, Arup Ghosh, Atimukta Jha, Viplov K. Biswas, Swati Madhulika, Manasi Priyadarshini, Shuchi Smita, O. P. Shriwas, Priyanka Mohapatra, Satya Ranjan Sahu, Aliva Minz, Debyashrita Barik, Rupesh Dash, Soma Chattopadhyay, Ghulam Hussain Syed, Shanti Senapati, Tushar K. Beuria, Rajeeb Swain, Punit Prasad, ILS COVID-19 TEAM, Orissa COVID-19 Study Group, DBT's PAN-INDIA 1000 SARS-CoV2 RNA genome sequencing consortium, Ajay Parida |
| EPI_ISL_463052, EPI_ISL_463053, EPI_ISL_463054, EPI_ISL_463055, EPI_ISL_463056, EPI_ISL_463057, EPI_ISL_463058, EPI_ISL_463059, EPI_ISL_463060, EPI_ISL_463061, EPI_ISL_463062, EPI_ISL_463063, EPI_ISL_463064, EPI_ISL_463065, EPI_ISL_463066, EPI_ISL_463067, EPI_ISL_463068, EPI_ISL_463069, EPI_ISL_463070, EPI_ISL_463071, EPI_ISL_463072 | | | |
| see above | Institute of Life Sciences, Bhubaneswar | Immunogenomics lab, Institute of Life Sciences, Bhubaneswar | Sunil Raghav, Arup Ghosh, Atimukta Jha, Viplov K. Biswas, Swati Madhulika, Manasi Priyadarshini, Shuchi Smita, Sifu Agarwal, Sanchari Chatterjee, Avula Kiran, Parej Nath, Supriya Suman, Rina Yadav, Rupesh Dash, Soma Chattopadhyay, Ghulam Hussain Syed, Shanti Senapati, Tushar K. Beuria, Rajeeb Swain, Punit Prasad, ILS COVID-19 TEAM, Orissa COVID-19 Study Group, DBT's PAN-INDIA 1000 SARS-CoV2 RNA genome sequencing consortium, Ajay Parida |
| EPI_ISL_463073, EPI_ISL_463074, EPI_ISL_463075, EPI_ISL_463076, EPI_ISL_463077, EPI_ISL_463078, EPI_ISL_463079, EPI_ISL_463080, EPI_ISL_463081, EPI_ISL_463082, EPI_ISL_463083, EPI_ISL_463084, EPI_ISL_463085 | | | |
| see above | Institute of Life Sciences, Bhubaneswar | Immunogenomics lab, Institute of Life Sciences, Bhubaneswar | Sunil Raghav, Arup Ghosh, Atimukta Jha, Viplov K. Biswas, Swati Madhulika, Manasi Priyadarshini, Shuchi Smita, Kautilya Kumar Jena, Sandhya Suranjika, Neha Singh, Eshna Laha, Saiket De, Rupesh Dash, Soma Chattopadhyay, Ghulam Hussain Syed, Shanti Senapati, Tushar K. Beuria, Rajeeb Swain, Punit Prasad, ILS COVID-19 TEAM, Orissa COVID-19 Study Group, DBT's PAN-INDIA 1000 SARS-CoV2 RNA genome sequencing consortium, Ajay Parida |
| EPI_ISL_463114, EPI_ISL_463115, EPI_ISL_463116, EPI_ISL_463117, EPI_ISL_463118, EPI_ISL_463119, EPI_ISL_463120, EPI_ISL_463121, EPI_ISL_463122, EPI_ISL_463123, EPI_ISL_463125, EPI_ISL_463127, EPI_ISL_463128, EPI_ISL_463129, EPI_ISL_463130, EPI_ISL_463131, EPI_ISL_463132, EPI_ISL_463133, EPI_ISL_463134, EPI_ISL_463135, EPI_ISL_463136, EPI_ISL_463137 | | | |
| see above | Virginia DCLS | Virginia DCLS | Virginia DCLS |
| EPI_ISL_463304 | Mrs Wu York Yu GOPC | Hong Kong Department of Health | Mak Gannon C.K., Cheng Peter K.C., Lam Edman T.K., Chan Rickjason C.W., Tsang Dominic N.C. |
| EPI_ISL_463305 | Queen Elizabeth Hospital | Hong Kong Department of Health | Mak Gannon C.K., Cheng Peter K.C., Lam Edman T.K., Chan Rickjason C.W., Tsang Dominic N.C. |
| EPI_ISL_463631, EPI_ISL_463632, EPI_ISL_463633, EPI_ISL_463636, EPI_ISL_463637, EPI_ISL_463638, EPI_ISL_463639, EPI_ISL_463640, EPI_ISL_463643, EPI_ISL_463650, EPI_ISL_463651, EPI_ISL_463652, EPI_ISL_463653, EPI_ISL_463654, EPI_ISL_463655, EPI_ISL_463657, EPI_ISL_463658, EPI_ISL_463659, EPI_ISL_463661, EPI_ISL_463664, EPI_ISL_463665, EPI_ISL_463666, EPI_ISL_463667, EPI_ISL_463668, EPI_ISL_463669, EPI_ISL_463670, EPI_ISL_463671, EPI_ISL_463672, EPI_ISL_463673, EPI_ISL_463674, EPI_ISL_463675, EPI_ISL_463676, EPI_ISL_463677, EPI_ISL_463678, EPI_ISL_463679, EPI_ISL_463680, EPI_ISL_463681, EPI_ISL_463682, EPI_ISL_463683, EPI_ISL_463684, EPI_ISL_463685, EPI_ISL_463686, EPI_ISL_463687, EPI_ISL_463688, EPI_ISL_463689, EPI_ISL_463690, EPI_ISL_463691, EPI_ISL_463692, EPI_ISL_463693, EPI_ISL_463694, EPI_ISL_463695, EPI_ISL_463696, EPI_ISL_463697, EPI_ISL_463698, EPI_ISL_463699, EPI_ISL_463700, EPI_ISL_463701, EPI_ISL_463702, EPI_ISL_463703, EPI_ISL_463704, EPI_ISL_463705 | | | |
| see above | Washington State Department of Health | Seattle Flu Study | Chu et al |
| EPI_ISL_464159 | National Institute of Laboratory Medicine and Referral Center | Genomic Research Lab, BCSIR | Shahina Akter, Abu Sayeed Mohammad Mahmud, Mohammad Samir Uzzaman, Eshrar Osman, Md. Ahasan Habib, Tanjina Akhter Banu, Md. Murshed Hasan Sarker, Barna Goswami, Iffat Jahan, Md. Saddam Hossain, Tasnim Nafisa, Md. Maruf Ahmed Molla, Mahmuda Yeasmin, Asish Kumar Ghosh, Arifa Akram, A. K. M. Shamsuzzaman, Sheikh Md. Selim Al Din, Utpal Chandra Ray, Salek Ahmed Sajib, Md. Salim Khan |
| EPI_ISL_464165 | National Institute of Laboratory Medicine and Referral Center | Genomic Research Lab, BCSIR | Barna Goswami, Abu Sayeed Mohammad Mahmud, Mohammad Samir Uzzaman, Eshrar Osman, Md. Ahasan Habib, Shahina Akter, Tanjina Akhter Banu, Md. Murshed Hasan Sarker, Iffat Jahan, Md. Saddam Hossain, Tasnim Nafisa, Md. Maruf Ahmed Molla, Mahmuda Yeasmin, Asish Kumar Ghosh, Arifa Akram, A. K. M. Shamsuzzaman, Sheikh Md. Selim Al Din, Utpal Chandra Ray, Salek Ahmed Sajib, Md. Salim Khan |
| EPI_ISL_465163, EPI_ISL_465164 | National Institute of Laboratory Medicine and Referral Center | Genomic Research Lab, BCSIR | Iffat Jahan, Abu Sayeed Mohammad Mahmud, Mohammad Samir Uzzaman, Eshrar Osman, Md. Ahasan Habib, Shahina Akter, Tanjina Akhter Banu, Md. Murshed Hasan Sarker, Barna Goswami, Md. Saddam Hossain, Tasnim Nafisa, Md. Maruf Ahmed Molla, Mahmuda Yeasmin, Asish Kumar Ghosh, Arifa Akram, A. K. M. Shamsuzzaman, Sheikh Md. Selim Al Din, Utpal Chandra Ray, Salek Ahmed Sajib, Md. Salim Khan |
| EPI_ISL_465601, EPI_ISL_465606, EPI_ISL_465608, EPI_ISL_465611, EPI_ISL_465613, EPI_ISL_465615, EPI_ISL_465616, EPI_ISL_465617, EPI_ISL_465618, EPI_ISL_465619, EPI_ISL_465621, EPI_ISL_465622, EPI_ISL_465623, EPI_ISL_465625, EPI_ISL_465628, EPI_ISL_465630, EPI_ISL_465631, EPI_ISL_465632, EPI_ISL_465633, EPI_ISL_465634, EPI_ISL_465635, EPI_ISL_465636, EPI_ISL_465637, EPI_ISL_465638, EPI_ISL_465639, EPI_ISL_465640, EPI_ISL_465641, EPI_ISL_465642, EPI_ISL_465643, EPI_ISL_465644, EPI_ISL_465645, EPI_ISL_465646, EPI_ISL_465647, EPI_ISL_465648, EPI_ISL_465649, EPI_ISL_465650, EPI_ISL_465651, EPI_ISL_465652, EPI_ISL_465653, EPI_ISL_465654, EPI_ISL_465655, EPI_ISL_465656, EPI_ISL_465657, EPI_ISL_465658, EPI_ISL_465659, EPI_ISL_465660, EPI_ISL_465661, EPI_ISL_465662, EPI_ISL_465663, EPI_ISL_465664, EPI_ISL_465665, EPI_ISL_465666, EPI_ISL_465667, EPI_ISL_465668, EPI_ISL_465669, EPI_ISL_465670, EPI_ISL_465671, EPI_ISL_465672 | | | |
| see above | Respiratory Virus Unit, Microbiology Services Colindale, Public Health England | Respiratory Virus Unit, Microbiology Services Colindale, Public Health England | PHE Covid Sequencing Team |

| | | | |
|---|---|---|---|
| EPI_ISL_466626, EPI_ISL_466629, EPI_ISL_466630, EPI_ISL_466636 | National Institute of Laboratory Medicine and Referral Center | Genomic Research Lab, BCSIR | Abu Sayeed Mohammad Mahmud, Mohammad Samir Uzzaman, Eshrar Osman, Md. Ahasan Habib, Shahina Akter, Tanjina Akhter Banu, Md. Murshed Hasan Sarker, Iffat Jahan, Barna Goswami, Md. Saddam Hossain, Tasnim Nafisa, Md. Maruf Ahmed Molla, Mahmuda Yeasmin, Asish Kumar Ghosh, Arifa Akram, A. K. M. Shamsuzzaman, Sheikh Md. Selim Al Din, Utpal Chandra Ray, Salek Ahmed Sajib, Md. Salim Khan |
| EPI_ISL_466652, EPI_ISL_466653, EPI_ISL_466663, EPI_ISL_466664, EPI_ISL_466678, EPI_ISL_466679, EPI_ISL_466680, EPI_ISL_466681, EPI_ISL_466682, EPI_ISL_466683 | Nebraska Public Health Laboratory | UNMC COVID-19 Response Team | UNMC COVID-19 Response Team |
| EPI_ISL_466686 | National Institute of Laboratory Medicine and Referral Center | Genomic Research Lab, BCSIR | Abu Sayeed Mohammad Mahmud, Mohammad Samir Uzzaman, Eshrar Osman, Md. Ahasan Habib, Shahina Akter, Tanjina Akhter Banu, Md. Murshed Hasan Sarker, Iffat Jahan, Barna Goswami, Md. Saddam Hossain, Tasnim Nafisa, Md. Maruf Ahmed Molla, Mahmuda Yeasmin, Asish Kumar Ghosh, Arifa Akram, A. K. M. Shamsuzzaman, Sheikh Md. Selim Al Din, Utpal Chandra Ray, Salek Ahmed Sajib, Md. Salim Khan |
| EPI_ISL_466839 | National Genomics Core-Center for DNA Fingerprinting and Diagnostics | National Genomics Core- Center for DNA Fingerprinting and Diagnostics (NGC-CDFD)- DBT's PAN-INDIA-1000 Genome consortium | Bala Pratyusha, Vinay Donipadi, G Shashikanth, Amrita Bhattacherjee, Rajeshree Sanyal, Raju Kumar, Ajay Kumar Chaudhary, Akash Chinchole, Brahmaji Sontyana, C. Arun Kumar, R HARINARAYANAN, RASHNA BHANDARI, MURALI DHARAN BASHYAM, DEBASHIS MITRA, DIVYA VASHISHT, ASHWIN DALAL |
| EPI_ISL_466840, EPI_ISL_466841, EPI_ISL_466842, EPI_ISL_466843, EPI_ISL_466844 | National Genomics Core-Center for DNA Fingerprinting and Diagnostics | National Genomics Core- Center for DNA Fingerprinting and Diagnostics (NGC-CDFD)- DBT's PAN-INDIA-1000 Genome consortium | Bala Pratyusha, Vinay Donipadi, G Shashikanth, Amrita Bhattacherjee, Nalini Raghunathan, Rajeshree Sanyal, Raju Kumar, Ajay Kumar Chaudhary, Akash Chinchole, Brahmaji Sontyana, C. Arun Kumar, R Harinarayanan, Rashna Bhandari, Murali Dharan Bashyam, Debashis Mitra, Divya Vashisht, Ashwin Dalal |
| EPI_ISL_466845, EPI_ISL_466846, EPI_ISL_466847 | National Genomics Core-Center for DNA Fingerprinting and Diagnostics | National Genomics Core- Center for DNA Fingerprinting and Diagnostics (NGC-CDFD)- DBT's PAN-INDIA-1000 Genome consortium | Bala Pratyusha, Vinay Donipadi, G Shashikanth, Amrita Bhattacherjee, Chandra Shekhar V, Chilakala Gangi Reddy, Chinthakindi KrishnaPrasad, Edurugatla Dinesh, Guru Raja, Hilal Ahmad Reshi, R HARINARAYANAN, RASHNA BHANDARI, MURALI DHARAN BASHYAM, DEBASHIS MITRA, DIVYA VASHISHT, ASHWIN DALAL |
| EPI_ISL_466848 | National Genomics Core-Center for DNA Fingerprinting and Diagnostics | National Genomics Core- Center for DNA Fingerprinting and Diagnostics (NGC-CDFD)- DBT's PAN-INDIA-1000 Genome consortium | Bala Pratyusha, Vinay Donipadi, G Shashikanth, Amrita Bhattacherjee, J. Mallikarjun, K. Viswakalyan, Kaisar Ahmad Lone, Kausika Kumar Malik, N. Sudheer, Neeraj Kumar, R HARINARAYANAN, RASHNA BHANDARI, MURALI DHARAN BASHYAM, DEBASHIS MITRA, DIVYA VASHISHT, ASHWIN DALAL |
| EPI_ISL_466908, EPI_ISL_466909 | Max von Pettenkofer Institute, Virology, National Reference Center for Retroviruses, LMU München | Laboratory for Functional Genome Analysis, Dept. Genomics, Gene Center of the LMU Munich | Max Muenchhoff, Stefan Krebs, Alexander Graf, Oliver Keppler, Helmut Blum |
| EPI_ISL_466997, EPI_ISL_466998, EPI_ISL_466999, EPI_ISL_467000, EPI_ISL_467001, EPI_ISL_467002, EPI_ISL_467003, EPI_ISL_467004, EPI_ISL_467005, EPI_ISL_467006, EPI_ISL_467007, EPI_ISL_467008, EPI_ISL_467009, EPI_ISL_467010, EPI_ISL_467011, EPI_ISL_467012, EPI_ISL_467013, EPI_ISL_467014, EPI_ISL_467015, EPI_ISL_467016 | | | |
| see above | Viollier AG | Department of Biosystems Science and Engineering, ETH Zürich | Christian Beisel, Sarah Nadeau, Ivan Topolsky, Pedro Ferreira, Philipp Jablonski, Susana Posada-Céspedes, Tobias Schär, Ina Nissen, Natascha Santacroce, Elodie Burcklen, Christiane Beckmann, Maurice Redondo, Olivier Kobel, Christoph Noppen, Sophie Seidel, Noemie Santamaria de Souza, Niko Beerenwinkel, Tanja Stadler |
| EPI_ISL_467190, EPI_ISL_467195, EPI_ISL_467234, EPI_ISL_467246 | Hospital General Universitario Gregorio Marañón | SeqCOVID-SPAIN consortium/IBV(CSIC) | Laura Pérez-Lago, Marta Herranz, Jon Sicilia, Julia Suárez, Pilar Catalán, Patricia Muñoz, Darío García de Viedma and SeqCOVID-SPAIN consortium |
| EPI_ISL_467298 | Nebraska Public Health Laboratory | UNMC COVID-19 Response Team | UNMC COVID-19 Response Team |
| EPI_ISL_467379, EPI_ISL_467384, EPI_ISL_467386, EPI_ISL_467390, EPI_ISL_467400, EPI_ISL_467403, EPI_ISL_467404, EPI_ISL_467405, EPI_ISL_467406, EPI_ISL_467407, EPI_ISL_467408 | | | |
| see above | NYU Langone Health | Departments of Pathology and Medicine, New York University School of Medicine | Maria Aguero-Rosenfeld, Brendan Belovarac, Margaret Black, Ludovic Boytard, John Cadley, Paolo Cotzia, John Chen, Dacia Dimartino, Xiaojun Feng, Tatyana Gindin, Emily Guzman, Adriana Heguy, Megan Hogan, Emily Huang, George Jour, Alireza Khodadadi-Jamayran, Lawrence H. Lin, Raven Luther, Andrew Lytle, Christian Marier, Matthew T. Maurano, Mark J. Mulligan, Peter Meyn, Raquel Ordonez Ciriza, Iman Osman, Jared Pinnell, Vanessa Raabe, Sitharam Ramaswami, Amy Rapkiewicz, Andre M. Ribeiro-dos-Santos, Marie Samanovic-Golden, Antonio Serrano, Guomiao Shen, Matija Snuderl, Theodore Vougiouklakis, Nick Vulpescu, Gael Westby, Paul Zappile, Yutong Zhang |
| EPI_ISL_467475, EPI_ISL_467476, EPI_ISL_467477, EPI_ISL_467478, EPI_ISL_467479, EPI_ISL_467480, EPI_ISL_467481, EPI_ISL_467482, EPI_ISL_467483, EPI_ISL_467484, EPI_ISL_467485, EPI_ISL_467486, EPI_ISL_467487, EPI_ISL_467488, EPI_ISL_467489, EPI_ISL_467490, EPI_ISL_467491 | | | |
| see above | Molecular Diagnostics Services (MDS) | KRISP, KZN Research Innovation and Sequencing Platform | Giandhari J, Pillay S, Lessells R, Chimukangara B, Mdlalose K, York D, Khan S, Tegally H, Wilkinson E, de Oliveira T |
| EPI_ISL_467935, EPI_ISL_467936, EPI_ISL_467937, EPI_ISL_467938, EPI_ISL_467942 | Virginia DCLS | Virginia DCLS | Virginia DCLS |
| EPI_ISL_467946, EPI_ISL_467947, EPI_ISL_467948 | Innovative Genomics Institute, UC Berkeley | Innovative Genomics Institute, UC Berkeley | Stacia Wyman, Haridha Shivram, Liana Lareau, Shana McDevitt, Justin Choi |
| EPI_ISL_467989 | SA Pathology | SA Pathology | Lex Leong, Chuan Kok Lim, Mark Turra, Ivan Bastian, Geoff Higgins |
| EPI_ISL_468073 | Child Health Research Foundation | Child Health Research Foundation | Senjuti Saha, Roly Malaker, Md Saiful Islam Sajib, Hafizur Rahman, Maksuda Islam, Samir K Saha |
| EPI_ISL_468075, EPI_ISL_468076 | Child Health Research Foundation | Child Health Research Foundation | Senjuti Saha, Roly Malaker, Md Saiful Islam Sajib, Hafizur Rahman, Afroza Akter Tanni, Syed Muktadir Al Sium, Maksuda Islam, Samir K Saha |
| EPI_ISL_468110, EPI_ISL_468111, EPI_ISL_468112, EPI_ISL_468113, EPI_ISL_468114, EPI_ISL_468117, EPI_ISL_468118, EPI_ISL_468119, EPI_ISL_468120, EPI_ISL_468121, EPI_ISL_468122, EPI_ISL_468123, EPI_ISL_468124, EPI_ISL_468125, EPI_ISL_468126, EPI_ISL_468127, EPI_ISL_468128, EPI_ISL_468129 | | | |
| see above | OHSU Lab Services Molecular Microbiology Lab | Oregon SARS-CoV-2 Genome Sequencing Center | Brendan L. O'Connell, Ruth V. Nichols, Alec J. Hirsch, Guang Fan, Daniel N. Streblow, William B. Messer, Andrew C. Adey, Benjamin N. Bimber, Brian J. O'Roak |
| EPI_ISL_468156 | [Romania, Bucharest] National Institute for Infectious Diseases "Prof. Dr. Matei Bal" | [Romania, Bucharest] National Institute for Infectious Diseases "Prof. Dr. Matei Bal" | Leontina Banica, Marius Cotic, Corina Casangiu, Marius Surleac, Simona Paraschiv |
| EPI_ISL_468331, EPI_ISL_468332, EPI_ISL_468336, EPI_ISL_468338, EPI_ISL_468341, EPI_ISL_468342, EPI_ISL_468344 | Microbiology Service, University Hospital of A Coruna-Biomedical Research Institute | Genomes & Disease, Center for Research in Molecular Medicine and Chronic Diseases, University of Santiago de Compostela | Kelly Conde, Jorge Arca, Soraya Rumbo, Juan A. Vallejo, M Poza, G Bou, Ana Pequeno-Valtierra, Jorge Rodriguez-Castro, Javier Temes, Daniel Garcia-Souto, Martin Santamarina, Cristina Gomez, Jose M. C. Tubio |
| EPI_ISL_468349, EPI_ISL_468350, EPI_ISL_468351, EPI_ISL_468352, EPI_ISL_468353 | County of Santa Clara Public Health Department | Chan-Zuckerberg Biohub | CZB Cliahub Consortium |
| EPI_ISL_468436, EPI_ISL_468437 | County of San Luis Obispo Public Health Laboratory | Chan-Zuckerberg Biohub | CZB Cliahub Consortium |
| EPI_ISL_468448, EPI_ISL_468450, EPI_ISL_468451 | Humboldt County Public Health Laboratory | Chan-Zuckerberg Biohub | CZB Cliahub Consortium |
| EPI_ISL_468462, EPI_ISL_468463, EPI_ISL_468464, EPI_ISL_468465, EPI_ISL_468466, EPI_ISL_468467, EPI_ISL_468468, EPI_ISL_468469, EPI_ISL_468470, EPI_ISL_468471, EPI_ISL_468472, EPI_ISL_468490, EPI_ISL_468491, EPI_ISL_468492, EPI_ISL_468493 | | | |
| see above | Ventura County Public Health Lab | Chan-Zuckerberg Biohub | CZB Cliahub Consortium |
| EPI_ISL_468551, EPI_ISL_468552, EPI_ISL_468553, EPI_ISL_468554, EPI_ISL_468555, EPI_ISL_468556, EPI_ISL_468557, EPI_ISL_468558 | San Joaquin County Public Health Lab | Chan-Zuckerberg Biohub | CZB Cliahub Consortium |
| EPI_ISL_468608, EPI_ISL_468609, EPI_ISL_468610, EPI_ISL_468611, | Orange County Public Health Lab | Chan-Zuckerberg Biohub | CZB Cliahub Consortium |

| | | | |
|---|---|---|---|
| EPI_ISL_468612, EPI_ISL_468613, EPI_ISL_468614 | | | |
| EPI_ISL_468654, EPI_ISL_468655 | Contra Costa Public Health Lab | Chan-Zuckerberg Biohub | CZB Cliahub Consortium |
| EPI_ISL_468718 | Environmental and Global Health | Environmental and Global Health | Stephenson,C.J., Subramaniam,K., Waltzek,T.B., Merck,L.H., Gibson,J.C., Morris,J.G. |
| EPI_ISL_469056 | Jourcentralen | The Public Health Agency of Sweden | Oskar Karlsson Lindsjo, Maria Lind Karlberg, Mattias Haukland, Reza Advani, Olov Svartstrom, Anna-Malin Linde, Sandra Broddesson, Petra Edquist, Shamam Muradrasoli, Anna Risberg, Karin Tegmark-Wisell |
| EPI_ISL_469057 | Inger Landgren | The Public Health Agency of Sweden | Oskar Karlsson Lindsjo, Maria Lind Karlberg, Mattias Haukland, Reza Advani, Olov Svartstrom, Anna-Malin Linde, Sandra Broddesson, Petra Edquist, Shamam Muradrasoli, Anna Risberg, Karin Tegmark-Wisell |
| EPI_ISL_469058 | Narhalsan Sjobo vardcentral | The Public Health Agency of Sweden | Oskar Karlsson Lindsjo, Maria Lind Karlberg, Mattias Haukland, Reza Advani, Olov Svartstrom, Anna-Malin Linde, Sandra Broddesson, Petra Edquist, Shamam Muradrasoli, Anna Risberg, Karin Tegmark-Wisell |
| EPI_ISL_469059 | Hovas Askim Familjelakare och BVC | The Public Health Agency of Sweden | Oskar Karlsson Lindsjo, Maria Lind Karlberg, Mattias Haukland, Reza Advani, Olov Svartstrom, Anna-Malin Linde, Sandra Broddesson, Petra Edquist, Shamam Muradrasoli, Anna Risberg, Karin Tegmark-Wisell |
| EPI_ISL_469060, EPI_ISL_469061 | Narhalsan Sjobo vardcentral | The Public Health Agency of Sweden | Oskar Karlsson Lindsjo, Maria Lind Karlberg, Mattias Haukland, Reza Advani, Olov Svartstrom, Anna-Malin Linde, Sandra Broddesson, Petra Edquist, Shamam Muradrasoli, Anna Risberg, Karin Tegmark-Wisell |
| EPI_ISL_469062 | Huddinge VC | The Public Health Agency of Sweden | Oskar Karlsson Lindsjo, Maria Lind Karlberg, Mattias Haukland, Reza Advani, Olov Svartstrom, Anna-Malin Linde, Sandra Broddesson, Petra Edquist, Shamam Muradrasoli, Anna Risberg, Karin Tegmark-Wisell |
| EPI_ISL_469066 | Surbrunns VC | The Public Health Agency of Sweden | Oskar Karlsson Lindsjo, Maria Lind Karlberg, Mattias Haukland, Reza Advani, Olov Svartstrom, Anna-Malin Linde, Sandra Broddesson, Petra Edquist, Shamam Muradrasoli, Anna Risberg, Karin Tegmark-Wisell |
| EPI_ISL_469096, EPI_ISL_469108 | National Public Health Laboratory, National Centre for Infectious Diseases | National Public Health Laboratory, National Centre for Infectious Diseases | Mak TM, Octavia S, Chavatte JM, Cui L, Lin RTP |
| EPI_ISL_469291, EPI_ISL_469292, EPI_ISL_469294, EPI_ISL_469295 | Keio University Hospital | Keio University Hospital | Kenjiro Kosaki |
| EPI_ISL_469310, EPI_ISL_469313, EPI_ISL_469315, EPI_ISL_469318, EPI_ISL_469319, EPI_ISL_469321, EPI_ISL_469323, EPI_ISL_469325, EPI_ISL_469326, EPI_ISL_469327, EPI_ISL_469334, EPI_ISL_469335, EPI_ISL_469336, EPI_ISL_469337, EPI_ISL_469338, EPI_ISL_469340, EPI_ISL_469341, EPI_ISL_469343, EPI_ISL_469344 | | | |
| see above | NU-OMICS DNA Sequencing research facility, Northumbria University | Wellcome Sanger Institute for the COVID-19 Genomics UK (COG-UK) consortium | Chris Duncan, Sheia Waugh, Shirelle Burton-Fanning, Gary Eltringham, Jennifer Collins, Brendan Payne, Yusri Taha, Emma Swindells, Jane Greenaway, Edward Barton, Garren Scott, Debra Padgett, Clive Graham, Sarah Essex, Steve Ligget, Paul Baker, Lynn Dover, Wen Yew, Gary Black, John Allan, Joshua Loh, Greg Young, Matthew Bashton, Andrew Nelson, Darren Smith and Alex Alderton, Roberto Amato, Sonia Goncalves, Ewan Harrison, David K. Jackson, Ian Johnston, Dominic Kwiatkowski, Cordelia Langford, John Sillitoe on behalf of the Wellcome Sanger Institute COVID-19 Surveillance Team (http://www.sanger.ac.uk/covid-team) |
| EPI_ISL_469345, EPI_ISL_469346, EPI_ISL_469348, EPI_ISL_469349, EPI_ISL_469350, EPI_ISL_469351, EPI_ISL_469352, EPI_ISL_469353, EPI_ISL_469354, EPI_ISL_469355, EPI_ISL_469356, EPI_ISL_469358, EPI_ISL_469360, EPI_ISL_469361, EPI_ISL_469362, EPI_ISL_469363, EPI_ISL_469364, EPI_ISL_469365, EPI_ISL_469366, EPI_ISL_469367, EPI_ISL_469368, EPI_ISL_469370, EPI_ISL_469371, EPI_ISL_469373, EPI_ISL_469375, EPI_ISL_469376, EPI_ISL_469377, EPI_ISL_469378, EPI_ISL_469379, EPI_ISL_469380, EPI_ISL_469381, EPI_ISL_469382, EPI_ISL_469384, EPI_ISL_469385, EPI_ISL_469388, EPI_ISL_469389, EPI_ISL_469390, EPI_ISL_469391, EPI_ISL_469393, EPI_ISL_469396, EPI_ISL_469397, EPI_ISL_469399, EPI_ISL_469401, EPI_ISL_469402, EPI_ISL_469403, EPI_ISL_469405, EPI_ISL_469407, EPI_ISL_469408, EPI_ISL_469409, EPI_ISL_469410, EPI_ISL_469411, EPI_ISL_469412, EPI_ISL_469414, EPI_ISL_469417, EPI_ISL_469420, EPI_ISL_469421, EPI_ISL_469422, EPI_ISL_469423, EPI_ISL_469424, EPI_ISL_469425, EPI_ISL_469426, EPI_ISL_469427, EPI_ISL_469428, EPI_ISL_469429, EPI_ISL_469430, EPI_ISL_469433, EPI_ISL_469434, EPI_ISL_469436, EPI_ISL_469437, EPI_ISL_469439, EPI_ISL_469440, EPI_ISL_469441, EPI_ISL_469442, EPI_ISL_469443, EPI_ISL_469444, EPI_ISL_469445, EPI_ISL_469446, EPI_ISL_469447, EPI_ISL_469449, EPI_ISL_469450, EPI_ISL_469451, EPI_ISL_469453, EPI_ISL_469454, EPI_ISL_469455, EPI_ISL_469456, EPI_ISL_469457, EPI_ISL_469458, EPI_ISL_469459, EPI_ISL_469460, EPI_ISL_469461, EPI_ISL_469462, EPI_ISL_469463, EPI_ISL_469466, EPI_ISL_469467, EPI_ISL_469468, EPI_ISL_469469, EPI_ISL_469470, EPI_ISL_469472, EPI_ISL_469473, EPI_ISL_469474, EPI_ISL_469475, EPI_ISL_469476, EPI_ISL_469479, EPI_ISL_469480, EPI_ISL_469481, EPI_ISL_469483, EPI_ISL_469484, EPI_ISL_469487, EPI_ISL_469488, EPI_ISL_469490, EPI_ISL_469491, EPI_ISL_469492, EPI_ISL_469493, EPI_ISL_469494, EPI_ISL_469495, EPI_ISL_469496, EPI_ISL_469497, EPI_ISL_469498, EPI_ISL_469500, EPI_ISL_469501, EPI_ISL_469502, EPI_ISL_469503, EPI_ISL_469504, EPI_ISL_469505, EPI_ISL_469506, EPI_ISL_469507, EPI_ISL_469508, EPI_ISL_469509, EPI_ISL_469510, EPI_ISL_469511, EPI_ISL_469512, EPI_ISL_469513, EPI_ISL_469514, EPI_ISL_469515, EPI_ISL_469516, EPI_ISL_469517, EPI_ISL_469518, EPI_ISL_469519, EPI_ISL_469520, EPI_ISL_469521, EPI_ISL_469522, EPI_ISL_469523, EPI_ISL_469524, EPI_ISL_469525, EPI_ISL_469807, EPI_ISL_469809, EPI_ISL_469811, EPI_ISL_469815, EPI_ISL_469816, EPI_ISL_469824, EPI_ISL_469827, EPI_ISL_469828, EPI_ISL_469829, EPI_ISL_469830, EPI_ISL_469832, EPI_ISL_469833, EPI_ISL_469835, EPI_ISL_469837, EPI_ISL_469838, EPI_ISL_469840, EPI_ISL_469842, EPI_ISL_469843 | | | |
| see above | PHE South West Regional Laboratory, National Infection Service | Wellcome Sanger Institute for the COVID-19 Genomics UK (COG-UK) consortium | Stephanie Hutchings, Hannah Pymont, Dr Peter Muir, Barry Vipond, Rich Hopes; and Alex Alderton, Roberto Amato, Sonia Goncalves, Ewan Harrison, David K. Jackson, Ian Johnston, Dominic Kwiatkowski, Cordelia Langford, John Sillitoe on behalf of the Wellcome Sanger Institute COVID-19 Surveillance Team (http://www.sanger.ac.uk/covid-team) |
| EPI_ISL_469891, EPI_ISL_469892, EPI_ISL_469893, EPI_ISL_469894, EPI_ISL_469895, EPI_ISL_469896, EPI_ISL_469897, EPI_ISL_469898, EPI_ISL_469899, EPI_ISL_469900, EPI_ISL_469901, EPI_ISL_469902, EPI_ISL_469903, EPI_ISL_469904, EPI_ISL_469905, EPI_ISL_469906, EPI_ISL_469907, EPI_ISL_469908, EPI_ISL_469909, EPI_ISL_469910, EPI_ISL_469911, EPI_ISL_469912, EPI_ISL_469913 | | | |
| see above | Department of Pathology, University of Cambridge | Wellcome Sanger Institute for the COVID-19 Genomics UK (COG-UK) consortium | Luke W Meredith, M. Estée Török , Myra Hosmillo, William L. Hamilton, Martin D. Curran, Theresa Feltwell, Grant Hall, Anna Yakovleva, Fahad A Khokhar, Charlotte J. Houldcroft, Laura G Caller, Aminu S. Jahun, Sarah L. Caddy, Ian Goodfellow; and Alex Alderton, Roberto Amato, Sonia Goncalves, Ewan Harrison, David K. Jackson, Ian Johnston, Dominic Kwiatkowski, Cordelia Langford, John Sillitoe on behalf of the Wellcome Sanger Institute COVID-19 Surveillance Team (http://www.sanger.ac.uk/covid-team) |
| EPI_ISL_470799, EPI_ISL_470800 | M Health Fairview | Minnesota Department of Health, Public Health Laboratory | Matt Plumb, Jacob Garfin, and Xiong Wang |
| EPI_ISL_470897, EPI_ISL_470899 | Pathogenic Microorganisms Variability Laboratory | Pathogenic Microorganisms Variability Laboratory | Alexey Shchetinin, Maria Nikiforova, Elena Shidlovskaya, Nadezhda Kuznetsova, Andrey Botikov, Alexander Gintsburg, Vladimir Gushchin |
| EPI_ISL_471168, EPI_ISL_471169, EPI_ISL_471170, EPI_ISL_471171 | MRCG at LSHTM Genomics lab | MRCG at LSHTM Genomics lab | Sesay et al |
| EPI_ISL_471584 | King Institute of Preventive Medicine & Research | CSIR-Centre for Cellular and Molecular Biology | K.Kaveri,S.Sivasubramanian,S.Vennila,P.Padmapriya,R.Kiruba,S.Magesh,G. Dhinakar Raj, G. Ravikumar, P. Azhahianambi,K Thangaraj,Payel Mukherjee, Sofia Banu, Priya Singh, Dhiviya Vedagiri, Divya Gupta, Vishal Sah, Santosh Kumar Kuncha, Krishnan Harinivas Harshan, Archana Bharadwaj Siva, Karthik Bharadwaj Tallapaka, Shagufta Khan, Lamuk Zaveri, Namami Gaur, Sakshi Shambhavi, Tulasi Nagabandi, Purushotham Vodnala, Rakesh K Mishra, Divya Tej Sowpati |
| EPI_ISL_471644 | CSIR-Centre for Cellular and Molecular Biology | CSIR-Centre for Cellular and Molecular Biology | Tulasi Nagabandi, Namami Gaur, Sakshi Shambhavi, Lamuk Zaveri, Shagufta Khan, Purushotham Vodnala, Payel Mukherjee, Sofia Banu, Priya Singh, Dhiviya Vedagiri, Divya Gupta, Vishal Sah, Santosh Kumar Kuncha, Krishnan Harinivas Harshan, Archana Bharadwaj Siva, Karthik Bharadwaj Tallapaka,Kezia J Ann, Radhika Khandelwal, Roshan Maku Venkata, Shemin Mansuri, Sonu Uday, Rakesh K Mishra, Divya Tej Sowpati |
| EPI_ISL_471694, EPI_ISL_471695, EPI_ISL_471696, EPI_ISL_471697, EPI_ISL_471698, EPI_ISL_471699, EPI_ISL_471700, EPI_ISL_471701, EPI_ISL_471702, EPI_ISL_471703, EPI_ISL_471704, EPI_ISL_471705, EPI_ISL_471706, EPI_ISL_471707, EPI_ISL_471708, EPI_ISL_471709, EPI_ISL_471710, EPI_ISL_471711, EPI_ISL_471712, EPI_ISL_471836, EPI_ISL_471837, EPI_ISL_471838, EPI_ISL_471839, EPI_ISL_471840, EPI_ISL_471841, EPI_ISL_471842, EPI_ISL_471843, EPI_ISL_471844, EPI_ISL_471873, EPI_ISL_471876, EPI_ISL_471887, EPI_ISL_471888, EPI_ISL_471892, EPI_ISL_471893, EPI_ISL_471894, EPI_ISL_471895, EPI_ISL_471896, EPI_ISL_471899, EPI_ISL_471900, EPI_ISL_471901, EPI_ISL_471902, EPI_ISL_471903, EPI_ISL_471905 | | | |
| see above | Michigan Department of Health and Human Services, Bureau of Laboratories | Michigan Department of Health and Human Services, Bureau of Laboratories | Blankenship HM, Riner D, Soehnlen MK |
| EPI_ISL_471977, EPI_ISL_471981, EPI_ISL_471982, EPI_ISL_471992 | University of Exeter | COVID-19 Genomics UK (COG-UK) Consortium | Ben Temperton,Aaron Jeffries,Michelle Michelsen,Joanna Warwick-Dugdale,Audrey Farbos,Robyn Manley,Stephen Michell,Jane Masoli |
| EPI_ISL_472022 | Liverpool Clinical Laboratories | COVID-19 Genomics UK (COG-UK) Consortium | Sam Haldenby, Anita Lucaci, Steve Paterson, Julian Hiscox, Alistair Darby, M Almsaud, A Alrezaihi, Muhannad Alruwaili, Stuart D Armstrong, Jones Benjamin, Eleanor G Bentley, Anu Chawla, Jordan J Clark, Angela Cowell, Richard Eccles, Isabel García-Dorival, Matthew Gemmell, Alessandro Gerada, PKF Gilmore, Richard Gregory, Ximeng Han, Catherine Hartley, Margaret Hughes, Miren Iturriza-Gomara, James Johnson, L Luu, Jenifer Manson, Charlotte Nelson, Elaine O'Toole, Cassie Olateju, Rebekah Penrice-Randal , Lucille Rainbow, N.P Randle, Trevor Ian Robinson, Parul Sharma, Ghada T Shawli, James P Stewart, Neil Swainston, Ecaterina Vamos, Joanne Watts, Mark Whitehead |
| EPI_ISL_472267, EPI_ISL_472268, EPI_ISL_472271, EPI_ISL_472284, | Northumbria University / South Tees Hospitals NHS Foundation Trust / North Cumbria Integrated Care NHS Foundation Trust / North Tees and Hartlepool NHS Foundation Trust / Newcastle Hospitals NHS Foundation Trust | COVID-19 Genomics UK (COG-UK) Consortium | Darren L Smith,Andrew Nelson,Matthew Bashton,Greg R Young,Joshua Loh,John Allan,Mohammad A Tariq,Giles S Holt,Gary Black,Wen C Yew,Lynn Dover,Paul Baker,Steve Liggett,Sarah Essex,Jane Greenaway,Debra Padgett,Clive Graham,Garren Scott,Edward Barton,Emma Swindells,Brendan Payne,Jennifer Collins,Yusri Taha,Gary Eltringham |

| | | | |
|---|---|---|---|
| EPI_ISL_472285, EPI_ISL_472286, EPI_ISL_472287, EPI_ISL_472288, EPI_ISL_472289 | | | |
| EPI_ISL_472308, EPI_ISL_472309, EPI_ISL_472311, EPI_ISL_472313, EPI_ISL_472315, EPI_ISL_472316, EPI_ISL_472317, EPI_ISL_472318, EPI_ISL_472319, EPI_ISL_472320, EPI_ISL_472321, EPI_ISL_472322, EPI_ISL_472371 | | | |
| see above | Quadram Institute Bioscience | COVID-19 Genomics UK (COG-UK) Consortium | Dave J. Baker, Gemma L. Kay, Alp Aydin, Thanh Le-Viet, Steven Rudder, Ana P. Tedim, Anastasia Kolyva, Maria Diaz, Leonardo de Oliveira Martins, Nabil-Fareed Alikhan, Lizzie Meadows, Rachael Stanley, Ngozi Elumogo, Muhammed Yasir, Nicholas M. Thomson, Alexander J Trotter, Rachel Gilroy, Samuel Bloomfield, Claire Stuart, Andrew Bell, Reenesh Prakash, Samir Dervisevic, Alison E. Mather, John Wain, Mark Webber, Andrew J. Page, Justin O'Grady |
| EPI_ISL_472424 | Queens Medical Centre, Clinical Microbiology Department / DeepSeq Nottingham | COVID-19 Genomics UK (COG-UK) Consortium | Gemma Clark, Wendy Smith, Manjinder Khakh, Vicki M Fleming, Michelle M Lister, Hannah Howson-Wells, Jonathan Ball, Patrick McClure, Joseph Chappell, Theocharis Tsoleridis, Nadine Holmes, Matthew Carlisle, Christopher Moore, Fei Sang, Johnny Debebe, Victoria Wright, Matthew Loose |
| EPI_ISL_472436, EPI_ISL_472438, EPI_ISL_472445, EPI_ISL_472454, EPI_ISL_472455, EPI_ISL_472460, EPI_ISL_472469, EPI_ISL_472470, EPI_ISL_472472, EPI_ISL_472473, EPI_ISL_472477, EPI_ISL_472480, EPI_ISL_472482, EPI_ISL_472498, EPI_ISL_472508, EPI_ISL_472514, EPI_ISL_472518, EPI_ISL_472520, EPI_ISL_472522, EPI_ISL_472523, EPI_ISL_472531, EPI_ISL_472532, EPI_ISL_472540, EPI_ISL_472542, EPI_ISL_472545, EPI_ISL_472550, EPI_ISL_472556, EPI_ISL_472560, EPI_ISL_472561, EPI_ISL_472563, EPI_ISL_472574, EPI_ISL_472576, EPI_ISL_472580, EPI_ISL_472583, EPI_ISL_472586, EPI_ISL_472597, EPI_ISL_472601, EPI_ISL_472602, EPI_ISL_472603, EPI_ISL_472606, EPI_ISL_472607, EPI_ISL_472608, EPI_ISL_472609, EPI_ISL_472610, EPI_ISL_472621, EPI_ISL_472623, EPI_ISL_472625, EPI_ISL_472628, EPI_ISL_472630, EPI_ISL_472631, EPI_ISL_472634, EPI_ISL_472635, EPI_ISL_472636, EPI_ISL_472638, EPI_ISL_472640, EPI_ISL_472644, EPI_ISL_472650, EPI_ISL_472653, EPI_ISL_472655, EPI_ISL_472656, EPI_ISL_472659, EPI_ISL_472666, EPI_ISL_472671, EPI_ISL_472673, EPI_ISL_472675, EPI_ISL_472678, EPI_ISL_472682, EPI_ISL_472684, EPI_ISL_472685, EPI_ISL_472687, EPI_ISL_472689, EPI_ISL_472691, EPI_ISL_472692, EPI_ISL_472694, EPI_ISL_472695, EPI_ISL_472696, EPI_ISL_472697, EPI_ISL_472699, EPI_ISL_472708, EPI_ISL_472712, EPI_ISL_472720, EPI_ISL_472721, EPI_ISL_472723, EPI_ISL_472729, EPI_ISL_472733, EPI_ISL_472735, EPI_ISL_472737, EPI_ISL_472738, EPI_ISL_472739, EPI_ISL_472740, EPI_ISL_472742, EPI_ISL_472743, EPI_ISL_472744, EPI_ISL_472745, EPI_ISL_472746, EPI_ISL_472747, EPI_ISL_472748, EPI_ISL_472749, EPI_ISL_472750, EPI_ISL_472751, EPI_ISL_472752, EPI_ISL_472753, EPI_ISL_472754, EPI_ISL_472755, EPI_ISL_472757, EPI_ISL_472758, EPI_ISL_472759, EPI_ISL_472760, EPI_ISL_472761, EPI_ISL_472762, EPI_ISL_472763, EPI_ISL_472764, EPI_ISL_472765, EPI_ISL_472766, EPI_ISL_472767, EPI_ISL_472768, EPI_ISL_472769, EPI_ISL_472770, EPI_ISL_472771, EPI_ISL_472772, EPI_ISL_472773, EPI_ISL_472774, EPI_ISL_472775, EPI_ISL_472776, EPI_ISL_472777, EPI_ISL_472778, EPI_ISL_472779, EPI_ISL_472780, EPI_ISL_472782, EPI_ISL_472783, EPI_ISL_472784, EPI_ISL_472785, EPI_ISL_472786, EPI_ISL_472787, EPI_ISL_472789, EPI_ISL_472790, EPI_ISL_472791, EPI_ISL_472792, EPI_ISL_472793, EPI_ISL_472794, EPI_ISL_472795, EPI_ISL_472796, EPI_ISL_472797, EPI_ISL_472798, EPI_ISL_472799, EPI_ISL_472800, EPI_ISL_472801, EPI_ISL_472803, EPI_ISL_472804, EPI_ISL_472805, EPI_ISL_472806, EPI_ISL_472807, EPI_ISL_472808, EPI_ISL_472809, EPI_ISL_472810, EPI_ISL_472811, EPI_ISL_472812, EPI_ISL_472813, EPI_ISL_472814, EPI_ISL_472815, EPI_ISL_472816, EPI_ISL_472817, EPI_ISL_472818, EPI_ISL_472819, EPI_ISL_472820, EPI_ISL_472821, EPI_ISL_472822, EPI_ISL_472823, EPI_ISL_472824, EPI_ISL_472825, EPI_ISL_472826, EPI_ISL_472827, EPI_ISL_472830, EPI_ISL_472831, EPI_ISL_472832, EPI_ISL_472833, EPI_ISL_472834, EPI_ISL_472835, EPI_ISL_472837, EPI_ISL_472838, EPI_ISL_472839, EPI_ISL_472840, EPI_ISL_472841, EPI_ISL_472842, EPI_ISL_472843, EPI_ISL_472870, EPI_ISL_472876, EPI_ISL_472895, EPI_ISL_472904, EPI_ISL_472909, EPI_ISL_472927, EPI_ISL_472938, EPI_ISL_472945, EPI_ISL_472953, EPI_ISL_472974, EPI_ISL_473015, EPI_ISL_473028, EPI_ISL_473029, EPI_ISL_473031, EPI_ISL_473038, EPI_ISL_473039, EPI_ISL_473073, EPI_ISL_473081, EPI_ISL_473092, EPI_ISL_473096, EPI_ISL_473108, EPI_ISL_473110, EPI_ISL_473118, EPI_ISL_473128, EPI_ISL_473130, EPI_ISL_473131, EPI_ISL_473133, EPI_ISL_473145, EPI_ISL_473148, EPI_ISL_473152, EPI_ISL_473155, EPI_ISL_473160, EPI_ISL_473164, EPI_ISL_473169, EPI_ISL_473181, EPI_ISL_473182, EPI_ISL_473197, EPI_ISL_473210, EPI_ISL_473222, EPI_ISL_473224, EPI_ISL_473232, EPI_ISL_473237, EPI_ISL_473243, EPI_ISL_473258, EPI_ISL_473263, EPI_ISL_473266, EPI_ISL_473267, EPI_ISL_473271, EPI_ISL_473276, EPI_ISL_473286, EPI_ISL_473289, EPI_ISL_473290, EPI_ISL_473292, EPI_ISL_473293, EPI_ISL_473295, EPI_ISL_473296, EPI_ISL_473297, EPI_ISL_473298, EPI_ISL_473301, EPI_ISL_473302, EPI_ISL_473303, EPI_ISL_473305 | | | |
| see above | Wales Specialist Virology Centre Sequencing lab: Pathogen Genomics Unit | COVID-19 Genomics UK (COG-UK) Consortium | Catherine Moore, Johnathan Evans, Laura Gifford, Malorie Perry, Simon Cottrell, Angela Marchbank, Alec Birchley, Alexander Adams, Amy Gaskin, Bree Gatica-Wilcox, Jason Coombes, Joel Southgate, Lauren Gilbert, Lee Graham, Nicole Pacchiarini, Sara Kumziene-Summerhayes, Sarah Taylor, Sophie Jones, Sara Rey, Matthew Bull, Joanne Watkins, Sally Corden, Tom Connor |
| EPI_ISL_473306, EPI_ISL_473307, EPI_ISL_473308, EPI_ISL_473309, EPI_ISL_473310, EPI_ISL_473311, EPI_ISL_473312, EPI_ISL_473318, EPI_ISL_473319, EPI_ISL_473320, EPI_ISL_473326, EPI_ISL_473327, EPI_ISL_473328, EPI_ISL_473329, EPI_ISL_473330, EPI_ISL_473331, EPI_ISL_473332, EPI_ISL_473333, EPI_ISL_473334, EPI_ISL_473335, EPI_ISL_473336, EPI_ISL_473337, EPI_ISL_473338, EPI_ISL_473339, EPI_ISL_473340, EPI_ISL_473341, EPI_ISL_473342, EPI_ISL_473343, EPI_ISL_473344, EPI_ISL_473354, EPI_ISL_473355, EPI_ISL_473356, EPI_ISL_473357, EPI_ISL_473358, EPI_ISL_473359, EPI_ISL_473360, EPI_ISL_473361, EPI_ISL_473362, EPI_ISL_473363, EPI_ISL_473364, EPI_ISL_473365, EPI_ISL_473366, EPI_ISL_473367, EPI_ISL_473368, EPI_ISL_473369, EPI_ISL_473394, EPI_ISL_473395, EPI_ISL_473396, EPI_ISL_473397, EPI_ISL_473426, EPI_ISL_473427, EPI_ISL_473428, EPI_ISL_473429 | | | |
| see above | University of Birmingham | COVID-19 Genomics UK (COG-UK) Consortium | Institute of Microbiology, University of Birmingham: Claire McMurray, Joanne Stockton, Samuel Nicholls, Radoslaw Poplawski, Will Rowe, Josh Quick, Nicholas Loman, University of Birmingham Testing Laboratory: Celina M Whalley, Andrew Bosworth, Charlotte Poxon, Kasun Wanigasooriya, Oliver Pickles, Mike Kidd, Alex Richter, Andrew D Beggs PHE Heartlands Lab: Husam Osman, Andrew Bosworth. Queen Elizabeth Hospital: Anna Casey |
| EPI_ISL_473647, EPI_ISL_473648, EPI_ISL_473649, EPI_ISL_473650, EPI_ISL_473651, EPI_ISL_473652, EPI_ISL_473653, EPI_ISL_473654, EPI_ISL_473655, EPI_ISL_473656, EPI_ISL_473657, EPI_ISL_473658, EPI_ISL_473659, EPI_ISL_473660, EPI_ISL_473661, EPI_ISL_473662, EPI_ISL_473663, EPI_ISL_473664, EPI_ISL_473665, EPI_ISL_473666, EPI_ISL_473667, EPI_ISL_473668, EPI_ISL_473669, EPI_ISL_473670, EPI_ISL_473671, EPI_ISL_473672, EPI_ISL_473673, EPI_ISL_473674, EPI_ISL_473675, EPI_ISL_473676, EPI_ISL_473677, EPI_ISL_473678, EPI_ISL_473679, EPI_ISL_473680, EPI_ISL_473681, EPI_ISL_473682, EPI_ISL_473683, EPI_ISL_473684, EPI_ISL_473685, EPI_ISL_473686, EPI_ISL_473687, EPI_ISL_473688, EPI_ISL_473689, EPI_ISL_473690, EPI_ISL_473691, EPI_ISL_473692, EPI_ISL_473693, EPI_ISL_473694, EPI_ISL_473695, EPI_ISL_473696, EPI_ISL_473697, EPI_ISL_473698, EPI_ISL_473699, EPI_ISL_473700, EPI_ISL_473701, EPI_ISL_473702, EPI_ISL_473726, EPI_ISL_473727, EPI_ISL_473729, EPI_ISL_473730, EPI_ISL_473770, EPI_ISL_473771, EPI_ISL_473772, EPI_ISL_473773, EPI_ISL_473774, EPI_ISL_473775, EPI_ISL_473776, EPI_ISL_473777 | | | |
| see above | West of Scotland Specialist Virology Centre, NHSGGC / MRC-University of Glasgow Centre for Virus Research | COVID-19 Genomics UK (COG-UK) Consortium | Ana da Silva Filipe, Natasha Johnson, Kathy Smollett, Daniel Mair, Stephen Carmichael, Lily Tong, Jenna Nichols, Elihu Aranday-Cortes, Kirstyn Brunker, Yasmin Parr, Alice Broos, Kyriaki Nomikou; Sarah McDonald, Marc Niebel, Patawee Asamaphan; Richard Orton, Joseph Hughes, Sreenu Vattipally, David L Robertson; Alasdair MacLean, Rory Gunson; Kathy Li, Natasha Jesudason, Rajiv Shah, James Shepherd, Antonia Ho, Emma Thomson |
| EPI_ISL_473795, EPI_ISL_473796, EPI_ISL_473797, EPI_ISL_473798, EPI_ISL_473799, EPI_ISL_473800, EPI_ISL_473801, EPI_ISL_473802, EPI_ISL_473803, EPI_ISL_473804, EPI_ISL_473805, EPI_ISL_473806, EPI_ISL_473807, EPI_ISL_473808, EPI_ISL_473809, EPI_ISL_473810, EPI_ISL_473811, EPI_ISL_473812, EPI_ISL_473872, EPI_ISL_473873, EPI_ISL_473874, EPI_ISL_473875, EPI_ISL_473876, EPI_ISL_473877, EPI_ISL_473878, EPI_ISL_473879, EPI_ISL_473880, EPI_ISL_473881, EPI_ISL_473882, EPI_ISL_473883, EPI_ISL_473884, EPI_ISL_473885, EPI_ISL_473886, EPI_ISL_473887, EPI_ISL_473888, EPI_ISL_473889, EPI_ISL_473890, EPI_ISL_473891, EPI_ISL_473892, EPI_ISL_473893, EPI_ISL_473894, EPI_ISL_473895, EPI_ISL_473896, EPI_ISL_473897, EPI_ISL_473898, EPI_ISL_473899, EPI_ISL_473900, EPI_ISL_473901, EPI_ISL_473902, EPI_ISL_473903, EPI_ISL_473904, EPI_ISL_473905, EPI_ISL_473951, EPI_ISL_473952, EPI_ISL_473953, EPI_ISL_473954 | | | |
| see above | Virology Department, Royal Infirmary of Edinburgh, NHS Lothian / School of Biological Sciences, University of Edinburgh / Institute of Genetics and Molecular Medicine, University of Edinburgh | COVID-19 Genomics UK (COG-UK) Consortium | McHugh M, Dewar R, Rooke S, Gallagher M, Balcaza C, O'Toole Á, Scher E, Hill V, McCrone JT, Colquhoun R, Yu X, Jackson B, Rambaut A, Williams TC, Templeton K |
| EPI_ISL_473957, EPI_ISL_473958, EPI_ISL_473960, EPI_ISL_473967, EPI_ISL_473968, EPI_ISL_473969, EPI_ISL_473971, EPI_ISL_473972, EPI_ISL_473973, EPI_ISL_473975, EPI_ISL_473977, EPI_ISL_473982, EPI_ISL_473983, EPI_ISL_473985, EPI_ISL_473987, EPI_ISL_473990, EPI_ISL_473992, EPI_ISL_473994, EPI_ISL_473995, EPI_ISL_473997, EPI_ISL_473998, EPI_ISL_473999, EPI_ISL_474000, EPI_ISL_474001, EPI_ISL_474003, EPI_ISL_474006, EPI_ISL_474008, EPI_ISL_474009, EPI_ISL_474010, EPI_ISL_474013, EPI_ISL_474014, EPI_ISL_474015, EPI_ISL_474016, EPI_ISL_474018, EPI_ISL_474019, EPI_ISL_474020, EPI_ISL_474021, EPI_ISL_474022, EPI_ISL_474024, EPI_ISL_474025, EPI_ISL_474029, EPI_ISL_474030, EPI_ISL_474031, EPI_ISL_474033, EPI_ISL_474035, EPI_ISL_474036, EPI_ISL_474038, EPI_ISL_474039, EPI_ISL_474040, EPI_ISL_474041, EPI_ISL_474042, EPI_ISL_474043, EPI_ISL_474045, EPI_ISL_474047, EPI_ISL_474048, EPI_ISL_474050, EPI_ISL_474051, EPI_ISL_474055, EPI_ISL_474056, EPI_ISL_474058, EPI_ISL_474064, EPI_ISL_474066, EPI_ISL_474071, EPI_ISL_474072, EPI_ISL_474073, EPI_ISL_474074, EPI_ISL_474075, EPI_ISL_474076, EPI_ISL_474077, EPI_ISL_474078, EPI_ISL_474079, EPI_ISL_474080, EPI_ISL_474081, EPI_ISL_474085, EPI_ISL_474090, EPI_ISL_474091, EPI_ISL_474094, EPI_ISL_474095, EPI_ISL_474097, EPI_ISL_474099, EPI_ISL_474105, EPI_ISL_474108, EPI_ISL_474112, EPI_ISL_474113, EPI_ISL_474114, EPI_ISL_474115, EPI_ISL_474116, EPI_ISL_474117, EPI_ISL_474120, EPI_ISL_474121, EPI_ISL_474123, EPI_ISL_474125, EPI_ISL_474126, EPI_ISL_474128, EPI_ISL_474130, EPI_ISL_474131, EPI_ISL_474132, EPI_ISL_474133, EPI_ISL_474134, EPI_ISL_474135, EPI_ISL_474136, EPI_ISL_474140, EPI_ISL_474142, EPI_ISL_474143, EPI_ISL_474144, EPI_ISL_474149, EPI_ISL_474151, EPI_ISL_474152, EPI_ISL_474153, EPI_ISL_474155, EPI_ISL_474157, EPI_ISL_474158, EPI_ISL_474159, EPI_ISL_474160, EPI_ISL_474163, EPI_ISL_474165, EPI_ISL_474167, EPI_ISL_474168, EPI_ISL_474170, EPI_ISL_474171, EPI_ISL_474172, EPI_ISL_474173, EPI_ISL_474177, EPI_ISL_474178, EPI_ISL_474183, EPI_ISL_474185, EPI_ISL_474187, EPI_ISL_474188, EPI_ISL_474190, EPI_ISL_474193, EPI_ISL_474194, EPI_ISL_474197, EPI_ISL_474198, EPI_ISL_474199, EPI_ISL_474201, EPI_ISL_474202, EPI_ISL_474203, EPI_ISL_474204, EPI_ISL_474206, EPI_ISL_474207, EPI_ISL_474208, EPI_ISL_474209, EPI_ISL_474210, EPI_ISL_474212, EPI_ISL_474213, EPI_ISL_474214, EPI_ISL_474216, EPI_ISL_474218, EPI_ISL_474220, EPI_ISL_474221 | | | |
| see above | Wales Specialist Virology Centre Sequencing lab: Pathogen Genomics Unit | COVID-19 Genomics UK (COG-UK) Consortium | Catherine Moore, Johnathan Evans, Laura Gifford, Malorie Perry, Simon Cottrell, Angela Marchbank, Alec Birchley, Alexander Adams, Amy Gaskin, Bree Gatica-Wilcox, Jason Coombes, Joel Southgate, Lauren Gilbert, Lee Graham, Nicole Pacchiarini, Sara Kumziene-Summerhayes, Sarah Taylor, Sophie Jones, Sara Rey, Matthew Bull, Joanne Watkins, Sally Corden, Tom Connor |
| EPI_ISL_474814, EPI_ISL_474840, EPI_ISL_474906 | Complejo Hospitalario Universitario de Albacete | SeqCOVID-SPAIN consortium/IBV(CSIC) | Encarnacion Simarro Córdoba, Julia Lozano Serra, Lorena Robles Fonseca , Monica Parra Grandes, Caridad Sainz de Baranda Camino and SeqCOVID-SPAIN consortium |
| EPI_ISL_474950 | Hospital Universitario Virgen de las Nieves de Granada-SAS | SeqCOVID-SPAIN consortium/IBV(CSIC) | Mercedes Pérez Ruiz, Sara Sanbonmatsu Gámez, Irene Pedrosa Corral, José M. Navarro-Marí and SeqCOVID-SPAIN consortium |
| EPI_ISL_475174, EPI_ISL_475175, EPI_ISL_475210, EPI_ISL_475212, EPI_ISL_475213, EPI_ISL_475214, EPI_ISL_475215, EPI_ISL_475217, EPI_ISL_475218, EPI_ISL_475219, EPI_ISL_475220, EPI_ISL_475221, EPI_ISL_475222, EPI_ISL_475223, EPI_ISL_475224, EPI_ISL_475225, EPI_ISL_475226, EPI_ISL_475227, EPI_ISL_475228, EPI_ISL_475229, EPI_ISL_475230, EPI_ISL_475231, EPI_ISL_475232, EPI_ISL_475233, EPI_ISL_475234, EPI_ISL_475235, EPI_ISL_475236, EPI_ISL_475237 | | | |
| see above | Nebraska Public Health Laboratory | UNMC COVID-19 Response Team | UNMC COVID-19 Response Team |
| EPI_ISL_475240, EPI_ISL_475241, EPI_ISL_475242, EPI_ISL_475243, EPI_ISL_475244, EPI_ISL_475245 | Centre for Enzyme Innovation, University of Portsmouth / Translational Research Laboratory, Portsmouth Hospitals NHS Trust | COVID-19 Genomics UK (COG-UK) Consortium | Angela Beckett,Yann Bourgeois,Garry Scarlett,Sharon Glaysher,Scott Elliott,Kelly Bicknell,Robert Impey,Allyson Lloyd,Sarah Wyllie,Ethan Butcher,Anoop Chauhan,Samuel Robson |
| EPI_ISL_475349, EPI_ISL_475352, EPI_ISL_475356, EPI_ISL_475357, EPI_ISL_475359, EPI_ISL_475364, EPI_ISL_475367, EPI_ISL_475374, EPI_ISL_475375, EPI_ISL_475379, EPI_ISL_475394, EPI_ISL_475397, EPI_ISL_475398, EPI_ISL_475406, EPI_ISL_475408, EPI_ISL_475411, EPI_ISL_475412, EPI_ISL_475422, EPI_ISL_475426, EPI_ISL_475430, EPI_ISL_475431, EPI_ISL_475432, EPI_ISL_475435, EPI_ISL_475437, EPI_ISL_475440, EPI_ISL_475441, EPI_ISL_475446, EPI_ISL_475449, EPI_ISL_475451, EPI_ISL_475458, EPI_ISL_475460, EPI_ISL_475462, EPI_ISL_475465, EPI_ISL_475467, EPI_ISL_475468, EPI_ISL_475472, EPI_ISL_475476, EPI_ISL_475482, EPI_ISL_475487, EPI_ISL_475488, EPI_ISL_475490, EPI_ISL_475493, EPI_ISL_475498, EPI_ISL_475501, EPI_ISL_475510 | | | |
| see above | Virology Department, Sheffield Teaching Hospitals NHS Foundation Trust/Department of Infection, Immunity and Cardiovascular Disease, The Medical School, University of Sheffield | COVID-19 Genomics UK (COG-UK) Consortium | Thushan de Silva, Matthew Parker, Nikki Smith, Adri Angyal, Rebecca Brown, Luke Green, Rachel Tucker, Paul Parsons, Danielle Groves, Katie Johnson, Laura Carrilero, Alex Keeley, Dave Partridge, Matthew Wyles, Benjamin Lindsey, Mehmet Yavuz, Mohammad Raza, Cariad Evans |
| EPI_ISL_475515 | Lakargruppen | The Public Health Agency of Sweden | Oskar Karlsson Lindsjo, Maria Lind Karlberg, Mattias Haukland, Reza Advani, Olov Svartstrom, Anna-Malin Linde, Sandra Broddesson, Mia Brytting, Anna |

| | | | |
|---|---|---|---|
| | | | Risberg, Karin Tegmark-Wisell |
| EPI_ISL_475516, EPI_ISL_475517 | Uppsala Narakut Aleris | The Public Health Agency of Sweden | Oskar Karlsson Lindsjo, Maria Lind Karlberg, Mattias Haukland, Reza Advani, Olov Svartstrom, Anna-Malin Linde, Sandra Broddesson, Mia Brytting, Anna Risberg, Karin Tegmark-Wisell |
| EPI_ISL_475518 | Trollbackens VC | The Public Health Agency of Sweden | Oskar Karlsson Lindsjo, Maria Lind Karlberg, Mattias Haukland, Reza Advani, Olov Svartstrom, Anna-Malin Linde, Sandra Broddesson, Mia Brytting, Anna Risberg, Karin Tegmark-Wisell |
| EPI_ISL_475519 | Orsa VC | The Public Health Agency of Sweden | Oskar Karlsson Lindsjo, Maria Lind Karlberg, Mattias Haukland, Reza Advani, Olov Svartstrom, Anna-Malin Linde, Sandra Broddesson, Mia Brytting, Anna Risberg, Karin Tegmark-Wisell |
| EPI_ISL_475542 | Kungsholmsdoktorn | The Public Health Agency of Sweden | Oskar Karlsson Lindsjo, Maria Lind Karlberg, Mattias Haukland, Reza Advani, Olov Svartstrom, Anna-Malin Linde, Sandra Broddesson, Mia Brytting, Anna Risberg, Karin Tegmark-Wisell |
| EPI_ISL_475543 | Surbrunns VC | The Public Health Agency of Sweden | Oskar Karlsson Lindsjo, Maria Lind Karlberg, Mattias Haukland, Reza Advani, Olov Svartstrom, Anna-Malin Linde, Sandra Broddesson, Mia Brytting, Anna Risberg, Karin Tegmark-Wisell |
| EPI_ISL_475566 | Vardcentralen Brinken | The Public Health Agency of Sweden | Oskar Karlsson Lindsjo, Maria Lind Karlberg, Mattias Haukland, Reza Advani, Olov Svartstrom, Anna-Malin Linde, Sandra Broddesson, Mia Brytting, Anna Risberg, Karin Tegmark-Wisell |
| EPI_ISL_475570 | Genome Center | Genome Center | A. S. M. Rubayet- Ul- Alam, Ovinu Kibria Islam, Md. Shazid Hasan, Hassan M. Al-Emran, Shireen Nigar, Selina Akter, Pravas Chandra Roy, Md. Tanvir Islam, Shovon Lal Sarkar, M. Shaminur Rahman, M. Rafiul Islam, Habiba Ibnat, Md Nur Kabidul Azam, Chakrabortty Atonu, Proshanto Kumar Das, Md. Hasan al Pramanik, Md. Zannat Ali, Shohanur Rahaman, Md. Aminul Islam, Ashok Kumar, Md. Nazmul Hasan, Md. Iqbal Kabir Jahid, Md. Anwar Hossain |
| EPI_ISL_475725, EPI_ISL_475726, EPI_ISL_475727, EPI_ISL_475728, EPI_ISL_475729, EPI_ISL_475730, EPI_ISL_475731, EPI_ISL_475732, EPI_ISL_475733, EPI_ISL_475734, EPI_ISL_475735, EPI_ISL_475736, EPI_ISL_475737, EPI_ISL_475738, EPI_ISL_475739, EPI_ISL_475740, EPI_ISL_475741, EPI_ISL_475742, EPI_ISL_475743, EPI_ISL_475744 | | | |
| see above | Utah Public Health Laboratory | Utah Public Health Laboratory | Erin Young, Kelly Oakeson |
| EPI_ISL_475936 | Universitaetsklinik für Innere Medizin II Innsbruck | Bergthaler laboratory, CeMM Research Center for Molecular Medicine of the Austrian Academy of Sciences | Alexandra Popa, Benedikt Agerer, Henrique Colaco, Lukas Endler, Jakob-Wendelin Genger, Alexander Lercher, Mark Smyth, Thomas Penz, Michael Schuster, Jan Laine, Martin Senekowitsch, Judith Aberle, Stephan Aberle, Peter Hufnagl, Daniela Schmid, Franz Allerberger, Elisabeth Puchhammer-Stoeckl, Manfred Nairz, Guenter Weiss, Gregor Hörmann, Kinga Rigler-Hohenwarter, Rainer Gattringer, Wegene Borena, Dorothee von Laer, Christoph Bock, Andreas Bergthaler |
| EPI_ISL_476024 | Laboratoire de Recherche et d'Analyses Médicales de la Gendarmerie Royale | Laboratoire de Recherche et d'Analyses Médicales de la Gendarmerie Royale | Sanaâ Lemriss, Amal SOUIRI, Nabil Lemzaoui, Omar Mestoui, Mohamed Labioui, Nabil Ouaariba, Ayoub Jibjibe, Mahmoud Yartaoui, Mohamed Chahmi, Marouane El Rhouila, Samiha Sellak, Nadia Kandoussi, Saâd El Kabbaj. |
| EPI_ISL_476026 | Laboratoire de Recherche et d'Analyses Médicales de la Gendarmerie Royale | Laboratoire de Recherche et d'Analyses Médicales de la Gendarmerie Royale | Sanaâ Lemriss, Amal SOUIRI, Saâd EL KABBAJ |
| EPI_ISL_476079, EPI_ISL_476080, EPI_ISL_476081, EPI_ISL_476082, EPI_ISL_476083, EPI_ISL_476084, EPI_ISL_476085 | Viollier AG | Department of Biosystems Science and Engineering, ETH Zürich | Christian Beisel, Sarah Nadeau, Ivan Topolsky, Pedro Ferreira, Philipp Jablonski, Susana Posada-Céspedes, Tobias Schär, Ina Nissen, Natascha Santacroce, Elodie Burcklen, Christiane Beckmann, Maurice Redondo, Olivier Kobel, Christoph Noppen, Sophie Seidel, Noemie Santamaria de Souza, Niko Beerenwinkel, Tanja Stadler |
| EPI_ISL_476561 | Hospital Garrahan | Héritas | Roberta Crespo, Dalmacio Pereyra, Mauricio Grisolia, Cristian Rohr, Andrea Mangano, Maria Florencia Fernandez, Fabian Fay, Martin Vazquez |
| EPI_ISL_476563 | Hospital de Pediatría "Prof. Dr. Juan P Garrahan" | Héritas | Dalmacio Pereyra, Roberta Crespo, Mauricio Grisolia, Cristian Rohr, Andrea Mangano, Maria Florencia Fernandez, Fabian Fay, Martin Vazquez |
| EPI_ISL_476565 | Hospital de Pediatría "Prof. Dr. Juan P Garrahan" | Héritas | Andrea Mangano, Maria Florencia Fernandez, Dalmacio Pereyra, Roberta Crespo, Mauricio Grisolia, Cristian Rohr, Fabian Fay, Martin Vazquez |
| EPI_ISL_476568 | Hospital de Pediatría "Prof. Dr. Juan P Garrahan" | Héritas | Cristian Rohr, Andrea Mangano, Maria Florencia Fernandez, Dalmacio Pereyra, Roberta Crespo, Mauricio Grisolia, Fabian Fay, Martin Vazquez |
| EPI_ISL_476749, EPI_ISL_476750, EPI_ISL_476751, EPI_ISL_476752, EPI_ISL_476753, EPI_ISL_476754, EPI_ISL_476755 | Minnesota Department of Health, Public Health Laboratory | Minnesota Department of Health, Public Health Laboratory | Matt Plumb, Jacob Garfin, and Xiong Wang |
| EPI_ISL_476844, EPI_ISL_476854, EPI_ISL_476883, EPI_ISL_476884, EPI_ISL_476888, EPI_ISL_476889, EPI_ISL_476890 | Defence Research & Development Establishment (DRDE) | Defence Research & Development Establishment (DRDE) | Shashi Sharma, Paban Kumar Dash, Sushil Kumar Sharma, Ambuj Shrivastava, Jyoti S. Kumar |
| EPI_ISL_477001, EPI_ISL_477002, EPI_ISL_477003, EPI_ISL_477004, EPI_ISL_477005, EPI_ISL_477006, EPI_ISL_477007 | KU Leuven, Rega Institute, Clinical and Epidemiological Virology | KU Leuven, Rega Institute, Clinical and Epidemiological Virology | Tony Wawina-Bokalanga, Joan Marti-Carerras, Bert Vanmechelen, Piet Maes |
| EPI_ISL_477125, EPI_ISL_477126 | Child Health Research Foundation | Child Health Research Foundation | Senjuti Saha, Md Saiful Islam Sajib, Roly Malaker, Md Hafizur Rahman, Afroza Akter Tanni, Syed Muktadir Al Sium, Maksuda Islam, Samir K Saha |
| EPI_ISL_477248, EPI_ISL_477249, EPI_ISL_477250, EPI_ISL_477251, EPI_ISL_477255, EPI_ISL_477256, EPI_ISL_477257, EPI_ISL_477258, EPI_ISL_477259, EPI_ISL_477260 | Institute for Stem Cell Science and Regenerative Medicine | National Centre for Biological Sciences | Farhan Ali, Vanessa Molin Paynter, Srikar Krishna, Mohak Sharda, Shah-e-Jahan Gulzar, Awadhesh Pandit, Varadha Sundarmurthy, Uma Ramakrishnan, Dasaradhi Palakodeti, Aswin Seshasayee |
| EPI_ISL_477280, EPI_ISL_477281, EPI_ISL_477282, EPI_ISL_477283, EPI_ISL_477284, EPI_ISL_477285, EPI_ISL_477286, EPI_ISL_477287 | M Health Fairview | Minnesota Department of Health, Public Health Laboratory | Matt Plumb, Jacob Garfin, Kelly Pung, and Xiong Wang |
| EPI_ISL_477295, EPI_ISL_477296 | Mayo Clinic & Mayo Clinic Laboratories | Minnesota Department of Health, Public Health Laboratory | Matt Plumb, Jacob Garfin, Kelly Pung, and Xiong Wang |
| EPI_ISL_477664, EPI_ISL_477665, EPI_ISL_477666, EPI_ISL_477667, EPI_ISL_477668, EPI_ISL_477669, EPI_ISL_477670, EPI_ISL_477671, EPI_ISL_477672 | Virginia DCLS | Virginia DCLS | Virginia DCLS |
| EPI_ISL_477785 | Department of Pathology, University of Cambridge | COVID-19 Genomics UK (COG-UK) Consortium | Luke W Meredith, M. Estée Török, Myra Hosmillo, William L. Hamilton, Martin D. Curran, Theresa Feltwell, Grant Hall, Anna Yakovleva, Fahad A Khokhar, Charlotte J. Houldcroft, Laura G Caller, Aminu S. Jahun, Sarah L. Caddy, Yasmin Chaudhry, Malte Pinckert, Ian Goodfellow |
| EPI_ISL_478056, EPI_ISL_478057, EPI_ISL_478058, EPI_ISL_478059, EPI_ISL_478060, EPI_ISL_478061, EPI_ISL_478062, EPI_ISL_478063, EPI_ISL_478064, EPI_ISL_478065, EPI_ISL_478066, EPI_ISL_478067, EPI_ISL_478068, EPI_ISL_478069, EPI_ISL_478070, EPI_ISL_478071, EPI_ISL_478072, EPI_ISL_478073, EPI_ISL_478074, EPI_ISL_478075, EPI_ISL_478076, EPI_ISL_478121, EPI_ISL_478130 | | | |
| see above | West of Scotland Specialist Virology Centre, NHSGGC / MRC-University of Glasgow Centre for Virus Research | COVID-19 Genomics UK (COG-UK) Consortium | Ana da Silva Filipe, Natasha Johnson, Kathy Smollett, Daniel Mair, Stephen Carmichael, Lily Tong, Jenna Nichols, Elihu Aranday-Cortes, Kirstyn Brunker, Yasmin Parr, Alice Broos, Kyriaki Nomikou; Sarah McDonald, Marc Niebel, Patawee Asamaphan; Richard Orton, Joseph Hughes, Sreenu Vattipally, David L Robertson; Alasdair MacLean, Rory Gunson; Kathy Li, Natasha Jesudason, Rajiv Shah, James Shepherd, Antonia Ho, Emma Thomson |
| EPI_ISL_478163, EPI_ISL_478164, EPI_ISL_478165, EPI_ISL_478166, EPI_ISL_478167, EPI_ISL_478168, EPI_ISL_478169, EPI_ISL_478170, EPI_ISL_478171, EPI_ISL_478172, EPI_ISL_478173, EPI_ISL_478174, EPI_ISL_478175, EPI_ISL_478176, EPI_ISL_478177, EPI_ISL_478178, EPI_ISL_478179, EPI_ISL_478180, EPI_ISL_478181, EPI_ISL_478182, EPI_ISL_478183, EPI_ISL_478184, EPI_ISL_478185, EPI_ISL_478186, EPI_ISL_478187, EPI_ISL_478188, EPI_ISL_478189, EPI_ISL_478190, EPI_ISL_478191, EPI_ISL_478192, EPI_ISL_478193, EPI_ISL_478194, EPI_ISL_478195, EPI_ISL_478199, EPI_ISL_478200, EPI_ISL_478201, EPI_ISL_478202, EPI_ISL_478203, EPI_ISL_478204, EPI_ISL_478205, EPI_ISL_478206, EPI_ISL_478207, EPI_ISL_478208, EPI_ISL_478209, EPI_ISL_478210, EPI_ISL_478211, EPI_ISL_478212, EPI_ISL_478213, EPI_ISL_478214, EPI_ISL_478215, EPI_ISL_478216, EPI_ISL_478217, EPI_ISL_478218, EPI_ISL_478219, | | | |

| | | | |
|---|---|---|---|
| EPI_ISL_478220, EPI_ISL_478221, EPI_ISL_478222, EPI_ISL_478223, EPI_ISL_478224, EPI_ISL_478225, EPI_ISL_478226, EPI_ISL_478227, EPI_ISL_478228, EPI_ISL_478229, EPI_ISL_478230, EPI_ISL_478231, EPI_ISL_478232, EPI_ISL_478233, EPI_ISL_478234, EPI_ISL_478235, EPI_ISL_478236, EPI_ISL_478251, EPI_ISL_478254, EPI_ISL_478255, EPI_ISL_478257, EPI_ISL_478258, EPI_ISL_478259, EPI_ISL_478260, EPI_ISL_478261, EPI_ISL_478262, EPI_ISL_478263, EPI_ISL_478264, EPI_ISL_478265, EPI_ISL_478266, EPI_ISL_478267, EPI_ISL_478268, EPI_ISL_478269 | | | |
| see above | Virology Department, Royal Infirmary of Edinburgh, NHS Lothian / School of Biological Sciences, University of Edinburgh / Institute of Genetics and Molecular Medicine, University of Edinburgh | COVID-19 Genomics UK (COG-UK) Consortium | McHugh M, Dewar R, Rooke S, Gallagher M, Balcaza C, O'Toole Á, Scher E, Hill V, McCrone JT, Colquhoun R, Yu X, Jackson B, Rambaut A, Williams TC, Templeton K |
| EPI_ISL_478533, EPI_ISL_478534, EPI_ISL_478597, EPI_ISL_478598, EPI_ISL_478599, EPI_ISL_478600, EPI_ISL_478601, EPI_ISL_478602, EPI_ISL_478603, EPI_ISL_478604, EPI_ISL_478605, EPI_ISL_478606, EPI_ISL_478607, EPI_ISL_478662 | | | |
| see above | Northumbria University / South Tees Hospitals NHS Foundation Trust / North Cumbria Integrated Care NHS Foundation Trust / North Tees and Hartlepool NHS Foundation Trust / Newcastle Hospitals NHS Foundation Trust | COVID-19 Genomics UK (COG-UK) Consortium | Darren L Smith,Andrew Nelson,Matthew Bashton,Greg R Young,Joshua Loh,John Allan,Mohammad A Tariq,Giles S Holt,Gary Black,Wen C Yew,Lynn Dover,Paul Baker,Steve Liggett,Sarah Essex,Jane Greenaway,Debra Padgett,Clive Graham,Garren Scott,Edward Barton,Emma Swindells,Brendan Payne,Jennifer Collins,Yusri Taha,Gary Eltringham |
| EPI_ISL_479060, EPI_ISL_479085, EPI_ISL_479118 | Oxford Viromics, NDM, University of Oxford; Oxford University Hospitals; Basingstoke and North Hampshire Hospital | COVID-19 Genomics UK (COG-UK) Consortium | Tanya Golubchik, David Bonsall, George Macintyre, Amy Trebes, Mariateresa de Cesare, Catrin Moore, Alex Mobbs, Anita Justice, Robert Shaw, Monique Andersson, Timothy Peto, Emma Wise, Nathan Moore, Jessica Lynch, Nick Cortes, Matilde Mori, Stephen Kidd, David Buck, John Todd, Christophe Fraser |
| EPI_ISL_479198, EPI_ISL_479204, EPI_ISL_479210, EPI_ISL_479228, EPI_ISL_479229, EPI_ISL_479230, EPI_ISL_479231, EPI_ISL_479232, EPI_ISL_479236, EPI_ISL_479239, EPI_ISL_479244, EPI_ISL_479250, EPI_ISL_479256, EPI_ISL_479259, EPI_ISL_479261, EPI_ISL_479263, EPI_ISL_479264, EPI_ISL_479266, EPI_ISL_479270, EPI_ISL_479282, EPI_ISL_479283 | | | |
| see above | Virology Department, Sheffield Teaching Hospitals NHS Foundation Trust/Department of Infection, Immunity and Cardiovascular Disease, The Medical School, University of Sheffield | COVID-19 Genomics UK (COG-UK) Consortium | Thushan de Silva, Matthew Parker, Nikki Smith, Adri Angyal, Rebecca Brown, Luke Green, Rachel Tucker, Paul Parsons, Danielle Groves, Katie Johnson, Laura Carrilero, Alex Keeley, Dave Partridge, Matthew Wyles, Benjamin Lindsey, Mehmet Yavuz, Mohammad Raza, Cariad Evans |
| EPI_ISL_479318, EPI_ISL_479324, EPI_ISL_479326, EPI_ISL_479334, EPI_ISL_479341, EPI_ISL_479342, EPI_ISL_479344, EPI_ISL_479370, EPI_ISL_479371, EPI_ISL_479384, EPI_ISL_479438, EPI_ISL_479442, EPI_ISL_479457, EPI_ISL_479472, EPI_ISL_479478 | | | |
| see above | Wales Specialist Virology Centre Sequencing lab: Pathogen Genomics Unit | COVID-19 Genomics UK (COG-UK) Consortium | Catherine Moore, Johnathan Evans, Laura Gifford, Malorie Perry, Simon Cottrell, Angela Marchbank, Alec Birchley, Alexander Adams, Amy Gaskin, Bree Gatica-Wilcox, Jason Coombes, Joel Southgate, Lauren Gilbert, Lee Graham, Nicole Pacchiarini, Sara Kumziene-Summerhayes, Sarah Taylor, Sophie Jones, Sara Rey, Matthew Bull, Joanne Watkins, Sally Corden, Tom Connor |
| EPI_ISL_479515, EPI_ISL_479516, EPI_ISL_479517, EPI_ISL_479518, EPI_ISL_479519, EPI_ISL_479520, EPI_ISL_479521, EPI_ISL_479522, EPI_ISL_479523, EPI_ISL_479524, EPI_ISL_479525, EPI_ISL_479526, EPI_ISL_479527, EPI_ISL_479528, EPI_ISL_479529, EPI_ISL_479530, EPI_ISL_479531, EPI_ISL_479532, EPI_ISL_479533 | | | |
| see above | NIV Influenza | NIV Influenza | Potdar V |
| EPI_ISL_479584, EPI_ISL_479585, EPI_ISL_479586 | National Public Health Laboratory, National Centre for Infectious Diseases | National Public Health Laboratory, National Centre for Infectious Diseases | Mak TM, Octavia S, Zhou Z, Chavatte JM, Cui L, Lin RTP |
| EPI_ISL_479657, EPI_ISL_479658 | NIV Influenza | NIV Influenza | Potdar V |
| EPI_ISL_479755 | Institute for Stem Cell Science and Regenerative Medicine | National Centre for Biological Sciences | Farhan Ali, Vanessa Molin Paynter, Srikar Krishna, Mohak Sharda, Shah-e-Jahan Gulzar, Awadhesh Pandit, Varadha Sundarmurthy, Uma Ramakrishnan, Dasaradhi Palakodeti, Aswin Seshasayee |
| EPI_ISL_480270, EPI_ISL_480271, EPI_ISL_480272, EPI_ISL_480273, EPI_ISL_480274, EPI_ISL_480275, EPI_ISL_480276, EPI_ISL_480278, EPI_ISL_480279, EPI_ISL_480280, EPI_ISL_480281, EPI_ISL_480282, EPI_ISL_480283, EPI_ISL_480284, EPI_ISL_480285, EPI_ISL_480286, EPI_ISL_480287, EPI_ISL_480288, EPI_ISL_480289, EPI_ISL_480290, EPI_ISL_480291, EPI_ISL_480292 | | | |
| see above | Genomic Laboratory (GLAB) (Conjoint lab of Health Directorate of Istanbul and Istanbul Technical University) | Genomic Laboratory (GLAB), Istanbul Technical University | Ilker Karacan, Tugba Kizilboga Akgun, Bugra Agaoglu, Gizem Alkurt, Jale Yildiz, Betsi Köse, Elifnaz Çelik, Arzu Irvem, Yasemin Kendir Demirkol, Ozlem Akgun Dogan, Mehtap Aydn, Levent Doganay, Gizem Dinler Doganay |
| EPI_ISL_480298, EPI_ISL_480302, EPI_ISL_480308 | National Reference Laboratory "Influenza and acute respiratory diseases" | NRL-HIV | Ivan Ivanov, Ivailo Alexiev, Ivva Philipova |
| EPI_ISL_480375, EPI_ISL_480376 | University of Wisconsin-Madison AIDS Vaccine Research Laboratories | University of Wisconsin-Madison AIDS Vaccine Research Laboratories | Gage Moreno, Katarina Braun, et al. AIDS Vaccine Research Laboratories |
| EPI_ISL_480432, EPI_ISL_480433 | Laboratorio de Biología Molecular Asociación Española Primera en Salud | Departments of Pathology and Medicine, New York University School of Medicine | Maria Victoria Elizondo, Maria Noel Zubillaga, Gonzalo Manrique, Paul Zappile, Gael Westby, Matthew T Maurano, Christian Marier, Adriana Heguy |
| EPI_ISL_480597, EPI_ISL_480604, EPI_ISL_480605, EPI_ISL_480607, EPI_ISL_480610, EPI_ISL_480658, EPI_ISL_480659, EPI_ISL_480660, EPI_ISL_480661 | Victorian Infectious Diseases Reference Laboratory (VIDRL) | VIDRL and MDU-PHL | Caly L., Seemann T., Sait, M., Schultz M., Druce J., Sherry, N. |
| EPI_ISL_480745, EPI_ISL_480746 | Microbiological Diagnostic Unit - Public Health Laboratory (MDU-PHL) | MDU-PHL | Seemann T., Schultz M., Sait, M., Sherry, N. |
| EPI_ISL_480918, EPI_ISL_480919, EPI_ISL_480920, EPI_ISL_480921, EPI_ISL_480922, EPI_ISL_480923, EPI_ISL_480924, EPI_ISL_480925, EPI_ISL_480926, EPI_ISL_480927, EPI_ISL_480928, EPI_ISL_480929, EPI_ISL_480930, EPI_ISL_480931, EPI_ISL_480932, EPI_ISL_480933, EPI_ISL_480934, EPI_ISL_480935, EPI_ISL_480936, EPI_ISL_480937, EPI_ISL_480938, EPI_ISL_480939, EPI_ISL_480940, EPI_ISL_480941, EPI_ISL_480942, EPI_ISL_480943, EPI_ISL_480944, EPI_ISL_480945, EPI_ISL_480946, EPI_ISL_480947, EPI_ISL_480948, EPI_ISL_480949, EPI_ISL_480950 | | | |
| see above | Florida Bureau of Public Health Laboratories | Florida Bureau of Public Health Laboratories | Sarah Schmedes, Jason Blanton |
| EPI_ISL_481041, EPI_ISL_481045, EPI_ISL_481050, EPI_ISL_481053, EPI_ISL_481067, EPI_ISL_481068, EPI_ISL_481073, EPI_ISL_481095, EPI_ISL_481098, EPI_ISL_481099, EPI_ISL_481108 | | | |
| see above | Hospital General Universitario Gregorio Marañón | SeqCOVID-SPAIN consortium/IBV(CSIC) | Laura Pérez-Lago, Marta Herranz, Jon Sicilia, Julia Suárez, Pilar Catalán, Patricia Muñoz, Darío García de Viedma and SeqCOVID-SPAIN consortium |
| EPI_ISL_481207, EPI_ISL_481213 | Ostfold Hospital Trust - Kalnes, Centre for Laboratory Medicine, Section for gene technology and infection serology | Norwegian Institute of Public Health, Department of Virology | Kathrine Stene-Johansen, Kamilla Heddeland Instefjord, Hilde Elshaug, Rasmus Riis Kopperud, Karoline Bragstad, Olav Hungnes |
| EPI_ISL_481224, EPI_ISL_481226 | Lab voor klinische biologie | Onderzoeksgroep Virologie | Laurens Lambrechts, Nick Vereecke, Marthe Pauwels, Bruno Verhasselt, Linos Vandekerckhove, Hans Nauwynck, Sebastiaan Theuns |
| EPI_ISL_481233 | Lab voor klinische biologie | Onderzoeksgroep Virologie | Nick Vereecke, Laurens Lambrechts, Marthe Pauwels, Bruno Verhasselt, Linos Vandekerckhove, Hans Nauwynck, Sebastiaan Theuns |
| EPI_ISL_481536, EPI_ISL_481537, EPI_ISL_481550, EPI_ISL_481565, EPI_ISL_481571, EPI_ISL_481694, EPI_ISL_481695, EPI_ISL_481696, EPI_ISL_481697, EPI_ISL_481698, EPI_ISL_481699, EPI_ISL_481700, EPI_ISL_481701, EPI_ISL_481702, EPI_ISL_481703, EPI_ISL_481704, EPI_ISL_481705, EPI_ISL_481706, EPI_ISL_481714, EPI_ISL_481715, EPI_ISL_481717, EPI_ISL_481718, EPI_ISL_481719, EPI_ISL_481720, EPI_ISL_481721, EPI_ISL_481722, EPI_ISL_481723, EPI_ISL_481724, EPI_ISL_481725, EPI_ISL_481726, EPI_ISL_481727, EPI_ISL_481728, EPI_ISL_481729 | | | |
| see above | Department of Virology and Immunology, University of Helsinki and Helsinki University Hospital, Huslab Finland | Department of Virology, Faculty of Medicine, University of Helsinki, Helsinki, Finland | Teemu Smura, Hannimari Kallio-Kokko, Jenni Virtanen, Maija Suvanto, Sari Hannula, Harri Kangas, Pekka Ellonen, Olli Vapalahti |
| EPI_ISL_481769, EPI_ISL_481773, EPI_ISL_481774, EPI_ISL_481775, EPI_ISL_481776, EPI_ISL_481777, EPI_ISL_481778, EPI_ISL_481783, EPI_ISL_481785, EPI_ISL_481787, EPI_ISL_481790, EPI_ISL_481792, EPI_ISL_481795, EPI_ISL_481800, EPI_ISL_481809, EPI_ISL_481810, EPI_ISL_481814, EPI_ISL_481816, EPI_ISL_481819, EPI_ISL_481821, EPI_ISL_481823, EPI_ISL_481825, EPI_ISL_481826, EPI_ISL_481827, EPI_ISL_481828, EPI_ISL_481830, EPI_ISL_481836, EPI_ISL_481841, EPI_ISL_481844, EPI_ISL_481846, EPI_ISL_481850, EPI_ISL_481852, EPI_ISL_481856, EPI_ISL_481857, EPI_ISL_481862, EPI_ISL_481863, EPI_ISL_481866, EPI_ISL_481867, EPI_ISL_481868, EPI_ISL_481870, EPI_ISL_481871, EPI_ISL_481874, EPI_ISL_481875, EPI_ISL_481880, EPI_ISL_481888, EPI_ISL_481889, EPI_ISL_481890, EPI_ISL_481891, EPI_ISL_481894, EPI_ISL_481896, EPI_ISL_481900, EPI_ISL_481901, EPI_ISL_481903, EPI_ISL_481904, EPI_ISL_481905, EPI_ISL_481908, EPI_ISL_481912, EPI_ISL_481916, EPI_ISL_481917, EPI_ISL_481923, EPI_ISL_481927, EPI_ISL_481928, EPI_ISL_481930, EPI_ISL_481936, EPI_ISL_481937, EPI_ISL_481940, EPI_ISL_481944, EPI_ISL_481946, EPI_ISL_481951, EPI_ISL_481961, EPI_ISL_481965, EPI_ISL_482029 | | | |
| see above | PHE South West Regional Laboratory, National Infection Service | Wellcome Sanger Institute for the COVID-19 Genomics UK (COG-UK) consortium | Stephanie Hutchings, Hannah Pymont, Dr Peter Muir, Barry Vipond, Rich Hopes; and Alex Alderton, Roberto Amato, Sonia Goncalves, Ewan Harrison, David K. Jackson, Ian Johnston, Dominic Kwiatkowski, Cordelia Langford, John Sillitoe on behalf of the Wellcome Sanger Institute COVID-19 Surveillance Team (http://www.sanger.ac.uk/covid-team) |
| EPI_ISL_482071, EPI_ISL_482072, EPI_ISL_482073, EPI_ISL_482074, EPI_ISL_482079, EPI_ISL_482080, EPI_ISL_482083, EPI_ISL_482086, EPI_ISL_482087, EPI_ISL_482088, EPI_ISL_482089, EPI_ISL_482090, EPI_ISL_482092, EPI_ISL_482094, EPI_ISL_482098, EPI_ISL_482099, EPI_ISL_482104, EPI_ISL_482106, EPI_ISL_482107, EPI_ISL_482108, EPI_ISL_482109, EPI_ISL_482110, EPI_ISL_482111, EPI_ISL_482112, EPI_ISL_482113, EPI_ISL_482118 | | | |
| see above | Microbiology Department, Hereford County Hospital | Wellcome Sanger Institute for the COVID-19 Genomics UK | Alison Johnson, Venkat Sivaprakasam, Fenella Halstead, Jane Thomas, Wendy Hogsden, Samantha Lamb and Alex Alderton, Roberto Amato, Sonia |

| | | | |
|---|---|---|---|
| | | (COG-UK) consortium | Goncalves, Ewan Harrison, David K. Jackson, Ian Johnston, Dominic Kwiatkowski, Cordelia Langford, John Sillitoe on behalf of the Wellcome Sanger Institute COVID-19 Surveillance Team (http://www.sanger.ac.uk/covid-team) |
| EPI_ISL_482410, EPI_ISL_482411, EPI_ISL_482412, EPI_ISL_482413, EPI_ISL_482414, EPI_ISL_482415, EPI_ISL_482416, EPI_ISL_482417, EPI_ISL_482418, EPI_ISL_482419, EPI_ISL_482420, EPI_ISL_482421, EPI_ISL_482422, EPI_ISL_482423, EPI_ISL_482424, EPI_ISL_482425, EPI_ISL_482426, EPI_ISL_482427, EPI_ISL_482428, EPI_ISL_482429, EPI_ISL_482430, EPI_ISL_482431, EPI_ISL_482432, EPI_ISL_482433, EPI_ISL_482434, EPI_ISL_482435, EPI_ISL_482436, EPI_ISL_482437, EPI_ISL_482441, EPI_ISL_482442, EPI_ISL_482443, EPI_ISL_482444, EPI_ISL_482445 | | | |
| see above | Providence St. Joseph Health Molecular Genomics Laboratory | Providence St. Joseph Health Molecular Genomics Laboratory | Alexa K Dowdell, Brian D Piening, Fred L Robinson, Carlo B Bifulco, Mary Campbell |
| EPI_ISL_482494, EPI_ISL_482495, EPI_ISL_482496, EPI_ISL_482497, EPI_ISL_482498, EPI_ISL_482525 | National Centre for Disease control (NCDC) | NCDC/CSIR-IGIB | Pramod Kumar#, Rajesh Pandey#, Pooja Sharma, Mahesh S Dhar, Vivekanand A, Bharathram Uppili, Robin Marwal, Radhakrishanan VS, Saruchi Wadhwa, Nishu Tyagi, Uma Sharma, Priyanka Singh, Hemlata Lall, Meena Datta, Varun Jaiswal, Hema Gogia, Preeti Madan, Prateek Singh, Debasis Dash, Mitali Mukerji, Sandhya Kabra, Sujeet Singh, Mohammed Faruq, Anurag Agrawal*, Partha Rakshit* |
| EPI_ISL_482737 | LNR National Reference Laboratory, Mohammed VI University of Health Sciences | Medical Biotechnology Laboratory, Rabat Medical and Pharmacy School, Mohammed The Vth University in Rabat | Meriem LAAMARTI, Souad KARTTI, Rokia LAAMARTI , M.W. CHEMAO-ELFIHRI, Loubna ALLAM, Mouna OUADGHIRI, Imane SMYEJ, Jalila RAHOUI, Houda BENRAHMA, Jalil El ATAR, Idrissa DIAWARA, Rachid EL JAOUDI, Laila SBABOU, Chakib NEJJARI, Saaid AMZAZI, Rachid MENTAG, Lahcen BELYAMANI and Azeddine IBRAHIMI |
| EPI_ISL_482762, EPI_ISL_482763, EPI_ISL_482764, EPI_ISL_482766, EPI_ISL_482767, EPI_ISL_482768, EPI_ISL_482769, EPI_ISL_482770, EPI_ISL_482771, EPI_ISL_482772, EPI_ISL_482773, EPI_ISL_482774 | | | |
| see above | Medical Ain Shams Research Institute (MASRI), Ain Shams University | Medical Ain Shams Research Institute (MASRI), Ain Shams University | Hesham Elghazaly, Sara Hassan Agwa, Ahmad Moustafa, Hala Hafez, Sara Elnakeep, Shaimaa Moustafa, Aya Mohamed, Reham Mamdouh, Ghada Ismael, Ashraf Omar, Osama Mansour, Mahmoud Elmeitini |
| EPI_ISL_482946 | Minnesota Department of Health, Public Health Laboratory | Minnesota Department of Health, Public Health Laboratory | Matt Plumb, Jacob Garfin, and Xiong Wang |
| EPI_ISL_483018, EPI_ISL_483019, EPI_ISL_483020, EPI_ISL_483021, EPI_ISL_483022, EPI_ISL_483023, EPI_ISL_483024, EPI_ISL_483025, EPI_ISL_483026, EPI_ISL_483027, EPI_ISL_483028, EPI_ISL_483029, EPI_ISL_483030, EPI_ISL_483031, EPI_ISL_483032, EPI_ISL_483033 | | | |
| see above | Utah Public Health Laboratory | Utah Public Health Laboratory | Heidi Butz, Erin Young, Kelly Oakeson |
| EPI_ISL_483036, EPI_ISL_483037, EPI_ISL_483038 | Medical Ain Shams Research Institute (MASRI), Ain Shams University | Medical Ain Shams Research Institute (MASRI), Ain Shams University | Hesham Elghazaly, Sara Hassan Agwa, Ahmad Moustafa, Hala Hafez, Sara Elnakeep, Shaimaa Moustafa, Aya Mohamed, Reham Mamdouh, Ghada Ismael, Ashraf Omar, Osama Mansour, Mahmoud Elmeitini |
| EPI_ISL_483063, EPI_ISL_483064 | unknown | Virology, Ecole Nationale Veterinaire de Toulouse | Bessiere,P., Cadiergues,M.-C., Croville,G., Walch,M., Dubois,M., Izopet,J., Guerin,J.-L. |
| EPI_ISL_483208, EPI_ISL_483209, EPI_ISL_483211, EPI_ISL_483216, EPI_ISL_483217, EPI_ISL_483218, EPI_ISL_483221, EPI_ISL_483222, EPI_ISL_483223, EPI_ISL_483225, EPI_ISL_483226, EPI_ISL_483227, EPI_ISL_483228, EPI_ISL_483229, EPI_ISL_483230, EPI_ISL_483231, EPI_ISL_483232, EPI_ISL_483233, EPI_ISL_483234, EPI_ISL_483235, EPI_ISL_483236, EPI_ISL_483237, EPI_ISL_483279, EPI_ISL_483292, EPI_ISL_483298, EPI_ISL_483327, EPI_ISL_483328, EPI_ISL_483331, EPI_ISL_483335, EPI_ISL_483336, EPI_ISL_483337 | | | |
| see above | UC San Diego Center for Advanced Laboratory Medicine | Andersen lab at Scripps Research | SEARCH Alliance San Diego with David Pride, Ji H Shin |
| EPI_ISL_483398, EPI_ISL_483399, EPI_ISL_483400, EPI_ISL_483401, EPI_ISL_483402 | UC San Diego Center for Advanced Laboratory Medicine | Andersen lab at Scripps Research | Allison Smither, Gilberto Sabino-Santos, Patricia Snarski, Lilia Melnik, Antoinette Bell, Kaylynn Genemaras, Arnaud Drouin, Dahlene Fusco, Robert Garry with SEARCH Alliance San Diego |
| EPI_ISL_483580, EPI_ISL_483584, EPI_ISL_483586, EPI_ISL_483588, EPI_ISL_483607 | National Public Health Laboratory, National Centre for Infectious Diseases | National Public Health Laboratory, National Centre for Infectious Diseases | Mak TM, Octavia S, Zhou Z, Chavatte JM, Cui L, Lin RTP |
| EPI_ISL_484267, EPI_ISL_484268, EPI_ISL_484269, EPI_ISL_484297, EPI_ISL_484298, EPI_ISL_484299, EPI_ISL_484300, EPI_ISL_484301, EPI_ISL_484302, EPI_ISL_484303, EPI_ISL_484304, EPI_ISL_484305, EPI_ISL_484306, EPI_ISL_484307, EPI_ISL_484308, EPI_ISL_484309, EPI_ISL_484310, EPI_ISL_484311, EPI_ISL_484312, EPI_ISL_484313, EPI_ISL_484314, EPI_ISL_484315, EPI_ISL_484316, EPI_ISL_484317, EPI_ISL_484318, EPI_ISL_484319, EPI_ISL_484320, EPI_ISL_484321, EPI_ISL_484322, EPI_ISL_484323, EPI_ISL_484324 | | | |
| see above | Northumbria University / South Tees Hospitals NHS Foundation Trust / North Cumbria Integrated Care NHS Foundation Trust / North Tees and Hartlepool NHS Foundation Trust / Newcastle Hospitals NHS Foundation Trust | COVID-19 Genomics UK (COG-UK) Consortium | Darren L Smith,Andrew Nelson,Matthew Bashton,Greg R Young,Joshua Loh,John Allan,Mohammad A Tariq,Giles S Holt,Gary Black,Wen C Yew,Lynn Dover,Paul Baker,Steve Liggett,Sarah Essex,Jane Greenaway,Debra Padgett,Clive Graham,Garren Scott,Edward Barton,Emma Swindells,Brendan Payne,Jennifer Collins,Yusri Taha,Gary Eltringham |
| EPI_ISL_484436, EPI_ISL_484438, EPI_ISL_484443, EPI_ISL_484444, EPI_ISL_484449, EPI_ISL_484452, EPI_ISL_484454, EPI_ISL_484457, EPI_ISL_484458, EPI_ISL_484462, EPI_ISL_484465, EPI_ISL_484466, EPI_ISL_484468, EPI_ISL_484483, EPI_ISL_484490, EPI_ISL_484496, EPI_ISL_484500, EPI_ISL_484503, EPI_ISL_484507 | | | |
| see above | Virology Department, Sheffield Teaching Hospitals NHS Foundation Trust/Department of Infection, Immunity and Cardiovascular Disease, The Medical School, University of Sheffield | COVID-19 Genomics UK (COG-UK) Consortium | Thushan de Silva, Matthew Parker, Nikki Smith, Adri Angyal, Rebecca Brown, Luke Green, Rachel Tucker, Paul Parsons, Danielle Groves, Katie Johnson, Laura Carrilero, Alex Keeley, Dave Partridge, Matthew Wyles, Benjamin Lindsey, Mehmet Yavuz, Mohammad Raza, Cariad Evans |
| EPI_ISL_484657 | West of Scotland Specialist Virology Centre, NHSGGC / MRC-University of Glasgow Centre for Virus Research | COVID-19 Genomics UK (COG-UK) Consortium | Ana da Silva Filipe, Natasha Johnson, Kathy Smollett, Daniel Mair, Stephen Carmichael, Lily Tong, Jenna Nichols, Elihu Aranday-Cortes, Kirstyn Brunker, Yasmin Parr, Alice Broos, Kyriaki Nomikou; Sarah McDonald, Marc Niebel, Patawee Asamaphan; Richard Orton, Joseph Hughes, Sreenu Vattipally, David L Robertson; Alasdair MacLean, Rory Gunson; Kathy Li, Natasha Jesudason, Rajiv Shah, James Shepherd, Antonia Ho, Emma Thomson |
| EPI_ISL_484681, EPI_ISL_484682 | Virology Department, Royal Infirmary of Edinburgh, NHS Lothian / School of Biological Sciences, University of Edinburgh / Institute of Genetics and Molecular Medicine, University of Edinburgh | COVID-19 Genomics UK (COG-UK) Consortium | McHugh M, Dewar R, Rooke S, Gallagher M, Balcaza C, O'Toole Á, Scher E, Hill V, McCrone JT, Colquhoun R, Yu X, Jackson B, Rambaut A, Williams TC, Templeton K |
| EPI_ISL_484693, EPI_ISL_484694, EPI_ISL_484695, EPI_ISL_484696, EPI_ISL_484697, EPI_ISL_484698, EPI_ISL_484706, EPI_ISL_484707 | Department of Clinical Microbiology | GIGA Medical Genomics | Keith Durkin, Maria Artesi, Sébastien Bontems, Raphaël Boreux, Cécile Meex, Axelle Chaslain, Céline Fombellida-Lopez, Pierrette Melin, Marie-Pierre Hayette, Vincent Bours. |
| EPI_ISL_485809 | Institut für Virologie und Epidemiologie der Viruskrankheiten, Universitätsklinikum Tübingen | NGS Competence Center Tübingen, Institut für Medizinische Mikrobiologie und Hygiene, Universitätsklinikum Tübingen | Angelov at al. |
| EPI_ISL_485810, EPI_ISL_485811, EPI_ISL_485812, EPI_ISL_485813 | Institut für Virologie und Epidemiologie der Viruskrankheiten, Universitätsklinikum Tübingen | NGS Competence Center Tübingen, Institut für Medizinische Mikrobiologie und Hygiene, Universitätsklinikum Tübingen | Angelov et al. |
| EPI_ISL_486224, EPI_ISL_486226, EPI_ISL_486227, EPI_ISL_486228, EPI_ISL_486229, EPI_ISL_486230, EPI_ISL_486231, EPI_ISL_486232, EPI_ISL_486233, EPI_ISL_486234, EPI_ISL_486235, EPI_ISL_486236, EPI_ISL_486237, EPI_ISL_486238, EPI_ISL_486239, EPI_ISL_486240, EPI_ISL_486241, EPI_ISL_486242, EPI_ISL_486243, EPI_ISL_486244, EPI_ISL_486245, EPI_ISL_486246, EPI_ISL_486247, EPI_ISL_486248, EPI_ISL_486249, EPI_ISL_486250, EPI_ISL_486251, EPI_ISL_486276, EPI_ISL_486277, EPI_ISL_486278, EPI_ISL_486279 | | | |
| see above | Orange County Public Health Laboratory | Chan-Zuckerberg Biohub | CZB Cliahub Consortium |
| EPI_ISL_486280, EPI_ISL_486281, EPI_ISL_486282, EPI_ISL_486283, EPI_ISL_486284, EPI_ISL_486285, EPI_ISL_486286 | Humboldt County Public Health Laboratory | Chan-Zuckerberg Biohub | CZB Cliahub Consortium |
| EPI_ISL_486331, EPI_ISL_486332, EPI_ISL_486333, EPI_ISL_486334, EPI_ISL_486335 | San Joaquin County Public Health Lab | Chan-Zuckerberg Biohub | CZB Cliahub Consortium |
| EPI_ISL_486339, EPI_ISL_486340, EPI_ISL_486341, EPI_ISL_486342, EPI_ISL_486345, EPI_ISL_486346, EPI_ISL_486347, EPI_ISL_486348, EPI_ISL_486349 | UCSF Clinical Microbiology Laboratory | Chan-Zuckerberg Biohub | CZB Cliahub Consortium |
| EPI_ISL_486388, EPI_ISL_486389 | DH | Department of Neurovirology, National Institute of Mental Health and Neuroscience (NIMHANS) | Chitra Pattabiraman, Vijayalakshmi Reddy, Harsha PK, Risha Rasheed, Shafeeq S Hameed, Manjunatha Venkataswamy, Anita Desai, Ravi Vasanthapuram |

| | | | |
|---|---|---|---|
| EPI_ISL_486390 | Centrl laboratorija | Latvian Biomedical Research and Study Centre | Ivars Silamielis, Kaspars Megnis, Monta Ustinova, ikita Zrelovs, Vita Rovte, Stella Lapia, Jana Oste, Marta Priedte, Uga Dumpis, Jnis Kloviš |
| EPI_ISL_486404 | Victoria Hospital | Department of Neurovirology, National Institute of Mental Health and Neuroscience (NIMHANS) | Chitra Pattabiraman, Vijayalakshmi Reddy, Harsha PK, Risha Rasheed, Shafeeq S Hameed, Manjunatha Venkataswamy, Anita Desai, Ravi Vasanthapuram |
| EPI_ISL_486405, EPI_ISL_486406, EPI_ISL_486407, EPI_ISL_486408, EPI_ISL_486409 | DH | Department of Neurovirology, National Institute of Mental Health and Neuroscience (NIMHANS) | Chitra Pattabiraman, Vijayalakshmi Reddy, Harsha PK, Risha Rasheed, Shafeeq S Hameed, Manjunatha Venkataswamy, Anita Desai, Ravi Vasanthapuram |
| EPI_ISL_486410 | Centrala laboratorija | Latvian Biomedical Research and Study Centre | Ivars Silamielis, Kaspars Megnis, Monta Ustinova, ikita Zrelovs, Vita Rovte, Stella Lapia, Jana Oste, Marta Priedte, Uga Dumpis, Jnis Kloviš |
| EPI_ISL_486411, EPI_ISL_486413, EPI_ISL_486414 | Centrl laboratorija | Latvian Biomedical Research and Study Centre | Ivars Silamielis, Kaspars Megnis, Monta Ustinova, ikita Zrelovs, Vita Rovte, Stella Lapia, Jana Oste, Marta Priedte, Uga Dumpis, Jnis Kloviš |
| EPI_ISL_486417 | Centrala laboratorija | Latvian Biomedical Research and Study Centre | Ivars Silamielis, Kaspars Megnis, Monta Ustinova, ikita Zrelovs, Vita Rovte, Stella Lapia, Jana Oste, Marta Priedte, Uga Dumpis, Jnis Kloviš |
| EPI_ISL_486418 | Centrl laboratorija | Latvian Biomedical Research and Study Centre | Ivars Silamielis, Kaspars Megnis, Monta Ustinova, ikita Zrelovs, Vita Rovte, Stella Lapia, Jana Oste, Marta Priedte, Uga Dumpis, Jnis Kloviš |
| EPI_ISL_487089, EPI_ISL_487091, EPI_ISL_487092, EPI_ISL_487101, EPI_ISL_487102, EPI_ISL_487103, EPI_ISL_487107, EPI_ISL_487110 | Nigeria Centre for Disease Control (NCDC) | African Centre of Excellence for Genomics of Infectious Diseases (ACEGID), Redeemer's University, Ede, Osun State, Nigeria | Oluniyi P.E., Ajogbasile F.V., Kayode A., Oguzie J., Olawoye I., Uwanibe J., Olumade T., Folarin O.A., Ihekweazu C., Happi C.T. |
| EPI_ISL_487227, EPI_ISL_487228, EPI_ISL_487245, EPI_ISL_487246, EPI_ISL_487247, EPI_ISL_487248, EPI_ISL_487249, EPI_ISL_487250, EPI_ISL_487251, EPI_ISL_487252, EPI_ISL_487253, EPI_ISL_487254, EPI_ISL_487255, EPI_ISL_487256, EPI_ISL_487257, EPI_ISL_487258, EPI_ISL_487259 | | | |
| see above | Utah Public Health Laboratory | Utah Public Health Laboratory | Heidi Butz, Erin Young, Kelly Oakeson |
| EPI_ISL_487435 | Queen Astrid Military Hospital | Institute of Tropical Medicine | Philippe Selhorst, Colin Anthony |
| EPI_ISL_487657, EPI_ISL_487667, EPI_ISL_487682, EPI_ISL_487731, EPI_ISL_487732, EPI_ISL_487776, EPI_ISL_487783, EPI_ISL_487787, EPI_ISL_487795, EPI_ISL_487861, EPI_ISL_487866, EPI_ISL_487868, EPI_ISL_487872, EPI_ISL_487877, EPI_ISL_487899, EPI_ISL_487913, EPI_ISL_487977, EPI_ISL_487981, EPI_ISL_487989 | | | |
| see above | Virology Department, Royal Infirmary of Edinburgh, NHS Lothian / School of Biological Sciences, University of Edinburgh | Wellcome Sanger Institute for the COVID-19 Genomics UK (COG-UK) consortium | McHugh M, Dewar R, Rooke S, O'Toole Á, Scher E, Hill V, McCrone JT, Colquhoun R, Yu X, Jackson B, Rambaut A, Templeton K and Alex Alderton, Roberto Amato, Sonia Goncalves, Ewan Harrison, David K. Jackson, Ian Johnston, Dominic Kwiatkowski, Cordelia Langford, John Sillitoe on behalf of the Wellcome Sanger Institute COVID-19 Surveillance Team (http://www.sanger.ac.uk/covid-team) |
| EPI_ISL_488017, EPI_ISL_488039, EPI_ISL_488040, EPI_ISL_488042, EPI_ISL_488044, EPI_ISL_488045, EPI_ISL_488047, EPI_ISL_488049, EPI_ISL_488054, EPI_ISL_488057, EPI_ISL_488058, EPI_ISL_488061, EPI_ISL_488064, EPI_ISL_488065, EPI_ISL_488067, EPI_ISL_488071, EPI_ISL_488073, EPI_ISL_488074, EPI_ISL_488076, EPI_ISL_488080, EPI_ISL_488081, EPI_ISL_488082, EPI_ISL_488083, EPI_ISL_488084, EPI_ISL_488088, EPI_ISL_488091, EPI_ISL_488092, EPI_ISL_488095, EPI_ISL_488098, EPI_ISL_488103, EPI_ISL_488104, EPI_ISL_488105, EPI_ISL_488111, EPI_ISL_488112, EPI_ISL_488113, EPI_ISL_488115, EPI_ISL_488118, EPI_ISL_488121, EPI_ISL_488122, EPI_ISL_488125, EPI_ISL_488126, EPI_ISL_488131, EPI_ISL_488133, EPI_ISL_488134, EPI_ISL_488135, EPI_ISL_488139, EPI_ISL_488147, EPI_ISL_488148, EPI_ISL_488155, EPI_ISL_488158, EPI_ISL_488162, EPI_ISL_488163, EPI_ISL_488167, EPI_ISL_488171, EPI_ISL_488173, EPI_ISL_488176, EPI_ISL_488557 | | | |
| see above | NU-OMICS DNA Sequencing research facility, Northumbria University | Wellcome Sanger Institute for the COVID-19 Genomics UK (COG-UK) consortium | Chris Duncan, Sheia Waugh, Shirelle Burton-Fanning, Gary Eltringham, Jennifer Collins, Brendan Payne, Yusri Taha, Emma Swindells, Jane Greenaway, Edward Barton, Garren Scott, Debra Padgett, Clive Graham, Sarah Essex, Steve Ligget, Paul Baker, Lynn Dover, Wen Yew, Gary Black, John Allan, Joshua Loh, Greg Young, Matthew Bashton, Andrew Nelson, Darren Smith and Alex Alderton, Roberto Amato, Sonia Goncalves, Ewan Harrison, David K. Jackson, Ian Johnston, Dominic Kwiatkowski, Cordelia Langford, John Sillitoe on behalf of the Wellcome Sanger Institute COVID-19 Surveillance Team (http://www.sanger.ac.uk/covid-team) |
| EPI_ISL_488840, EPI_ISL_488847, EPI_ISL_488854, EPI_ISL_488856, EPI_ISL_488861 | Department of Medical Microbiology, Western Sussex Hospitals NHS Foundation Trust, St Richard's Hospital | Wellcome Sanger Institute for the COVID-19 Genomics UK (COG-UK) consortium | Manasa Mutingwende, Sarah Lowdon, Olga Podplomyk, Michelle Erkiert, Jonathan Lewis, Paul Randell and Alex Alderton, Roberto Amato, Sonia Goncalves, Ewan Harrison, David K. Jackson, Ian Johnston, Dominic Kwiatkowski, Cordelia Langford, John Sillitoe on behalf of the Wellcome Sanger Institute COVID-19 Surveillance Team (http://www.sanger.ac.uk/covid-team) |
| EPI_ISL_488873 | Microbiology Department, Hereford County Hospital | Wellcome Sanger Institute for the COVID-19 Genomics UK (COG-UK) consortium | Alison Johnson, Venkat Sivaprakasam, Fenella Halstead, Jane Thomas, Wendy Hogsden, Samantha Lamb and Alex Alderton, Roberto Amato, Sonia Goncalves, Ewan Harrison, David K. Jackson, Ian Johnston, Dominic Kwiatkowski, Cordelia Langford, John Sillitoe on behalf of the Wellcome Sanger Institute COVID-19 Surveillance Team (http://www.sanger.ac.uk/covid-team) |
| EPI_ISL_489064, EPI_ISL_489065, EPI_ISL_489066, EPI_ISL_489068, EPI_ISL_489069, EPI_ISL_489071, EPI_ISL_489072, EPI_ISL_489074, EPI_ISL_489075, EPI_ISL_489077, EPI_ISL_489078, EPI_ISL_489079, EPI_ISL_489081, EPI_ISL_489082, EPI_ISL_489083, EPI_ISL_489084, EPI_ISL_489086, EPI_ISL_489087, EPI_ISL_489088, EPI_ISL_489090, EPI_ISL_489092, EPI_ISL_489096, EPI_ISL_489097, EPI_ISL_489098, EPI_ISL_489100, EPI_ISL_489101, EPI_ISL_489103, EPI_ISL_489104, EPI_ISL_489106, EPI_ISL_489107, EPI_ISL_489108, EPI_ISL_489109, EPI_ISL_489110, EPI_ISL_489111, EPI_ISL_489114, EPI_ISL_489116, EPI_ISL_489117, EPI_ISL_489118, EPI_ISL_489119, EPI_ISL_489122, EPI_ISL_489123, EPI_ISL_489124, EPI_ISL_489125, EPI_ISL_489126, EPI_ISL_489127, EPI_ISL_489128, EPI_ISL_489130, EPI_ISL_489131, EPI_ISL_489132, EPI_ISL_489133, EPI_ISL_489134, EPI_ISL_489135, EPI_ISL_489136, EPI_ISL_489138, EPI_ISL_489139, EPI_ISL_489145, EPI_ISL_489146, EPI_ISL_489147, EPI_ISL_489148, EPI_ISL_489149, EPI_ISL_489150, EPI_ISL_489152, EPI_ISL_489153, EPI_ISL_489154, EPI_ISL_489155 | | | |
| see above | NU-OMICS DNA Sequencing research facility, Northumbria University | Wellcome Sanger Institute for the COVID-19 Genomics UK (COG-UK) consortium | Chris Duncan, Sheia Waugh, Shirelle Burton-Fanning, Gary Eltringham, Jennifer Collins, Brendan Payne, Yusri Taha, Emma Swindells, Jane Greenaway, Edward Barton, Garren Scott, Debra Padgett, Clive Graham, Sarah Essex, Steve Ligget, Paul Baker, Lynn Dover, Wen Yew, Gary Black, John Allan, Joshua Loh, Greg Young, Matthew Bashton, Andrew Nelson, Darren Smith and Alex Alderton, Roberto Amato, Sonia Goncalves, Ewan Harrison, David K. Jackson, Ian Johnston, Dominic Kwiatkowski, Cordelia Langford, John Sillitoe on behalf of the Wellcome Sanger Institute COVID-19 Surveillance Team (http://www.sanger.ac.uk/covid-team) |
| EPI_ISL_489159, EPI_ISL_489160, EPI_ISL_489180, EPI_ISL_489199, EPI_ISL_489229, EPI_ISL_489238, EPI_ISL_489242, EPI_ISL_489251, EPI_ISL_489253, EPI_ISL_489258, EPI_ISL_489259, EPI_ISL_489265, EPI_ISL_489268, EPI_ISL_489270, EPI_ISL_489272, EPI_ISL_489276, EPI_ISL_489277, EPI_ISL_489278, EPI_ISL_489280, EPI_ISL_489281, EPI_ISL_489282, EPI_ISL_489283, EPI_ISL_489285, EPI_ISL_489286, EPI_ISL_489288, EPI_ISL_489289, EPI_ISL_489290, EPI_ISL_489291, EPI_ISL_489292, EPI_ISL_489293, EPI_ISL_489295, EPI_ISL_489299, EPI_ISL_489301, EPI_ISL_489310, EPI_ISL_489313, EPI_ISL_489332 | | | |
| see above | Regional Virus Laboratory, Belfast Health and Social Care Trust | Wellcome Sanger Institute for the COVID-19 Genomics UK (COG-UK) consortium | Conall McCaughey, James McKenna, Tanya Curran, Susan Feeney, Alison Watt, Ciara Cox, Mairead Connor, Zoltan Molnar, David Simpson, Derek Fairley; and Alex Alderton, Roberto Amato, Sonia Goncalves, Ewan Harrison, David K. Jackson, Ian Johnston, Dominic Kwiatkowski, Cordelia Langford, John Sillitoe on behalf of the Wellcome Sanger Institute COVID-19 Surveillance Team (http://www.sanger.ac.uk/covid-team) |
| EPI_ISL_489809, EPI_ISL_489810, EPI_ISL_489811, EPI_ISL_489812, EPI_ISL_489813, EPI_ISL_489814, EPI_ISL_489815, EPI_ISL_489816, EPI_ISL_489817, EPI_ISL_489818, EPI_ISL_489819, EPI_ISL_489820, EPI_ISL_489821, EPI_ISL_489822, EPI_ISL_489823, EPI_ISL_489824, EPI_ISL_489825, EPI_ISL_489826, EPI_ISL_489827, EPI_ISL_489828, EPI_ISL_489829, EPI_ISL_489830, EPI_ISL_489831, EPI_ISL_489832 | | | |
| see above | Florida Bureau of Public Health Laboratories | Florida Bureau of Public Health Laboratories | Sarah Schmedes, Jason Blanton |
| EPI_ISL_490052, EPI_ISL_490059, EPI_ISL_490060, EPI_ISL_490061, EPI_ISL_490062, EPI_ISL_490063, EPI_ISL_490064, EPI_ISL_490065, EPI_ISL_490066, EPI_ISL_490067, EPI_ISL_490068, EPI_ISL_490069, EPI_ISL_490070, EPI_ISL_490072, EPI_ISL_490073, EPI_ISL_490075, EPI_ISL_490076, EPI_ISL_490077, EPI_ISL_490078 | | | |
| see above | National Public Health Laboratory, National Centre for Infectious Diseases | National Public Health Laboratory, National Centre for Infectious Diseases | Mak TM, Octavia S, Zhou Z, Chavatte JM, Cui L, Lin RTP |
| EPI_ISL_490089 | Institute for Medical Research, Infectious Disease Research Centre, National Institutes of Health, Ministry of Health Malaysia | Institute for Medical Research, Infectious Disease Research Centre, National Institutes of Health, Ministry of Health Malaysia | Suppiah J, Mohd-Zawawi Z, Kamel K, Kalyanasundram J, Thayan R |
| EPI_ISL_490208 | München Klinik Schwabing | MGZ Medical Genetics Center | Dieter A. Wolf, Elke Holinski-Feder |
| EPI_ISL_490258 | National Institute for Communicable Diseases of the National Health Laboratory Service | National Institute for Communicable Diseases of the National Health Laboratory Service | Allam M, Ismail A, Khumalo Z, Kwenda S, Mtshali P, Mnyameni F, Mohale T, Subramoney K, Bhiman JN |
| EPI_ISL_490471, EPI_ISL_490473, EPI_ISL_490475, EPI_ISL_490476, EPI_ISL_490477, EPI_ISL_490478, EPI_ISL_490479, EPI_ISL_490480, EPI_ISL_490481, EPI_ISL_490482, EPI_ISL_490483, EPI_ISL_490484, EPI_ISL_490485, EPI_ISL_490486, EPI_ISL_490487, EPI_ISL_490488, EPI_ISL_490489, EPI_ISL_490490 | | | |
| see above | Northumbria University / South Tees Hospitals NHS Foundation Trust / North Cumbria Integrated Care NHS Foundation Trust / North Tees and Hartlepool NHS Foundation Trust / Newcastle Hospitals NHS Foundation Trust | COVID-19 Genomics UK (COG-UK) Consortium | Darren L Smith,Andrew Nelson,Matthew Bashton,Greg R Young,Joshua Loh,John Allan,Mohammad A Tariq,Giles S Holt,Gary Black,Wen C Yew,Lynn Dover,Paul Baker,Steve Liggett,Sarah Essex,Jane Greenaway,Debra Padgett,Clive Graham,Garren Scott,Edward Barton,Emma Swindells,Brendan Payne,Jennifer Collins,Yusri Taha,Gary Eltringham |
| EPI_ISL_490557, EPI_ISL_490558, EPI_ISL_490559 | Quadram Institute Bioscience | COVID-19 Genomics UK (COG-UK) Consortium | Dave J. Baker, Gemma L. Kay, Alp Aydin, Thanh Le-Viet, Steven Rudder, Ana P. Tedim, Anastasia Kolyva, Maria Diaz, Leonardo de Oliveira Martins, Nabil-Fareed Alikhan, Lizzie Meadows, Rachael Stanley, Ngozi Elumogo, Muhammed Yasir, Nicholas M. Thomson, Alexander J Trotter, Rachel Gilroy, |

| | | | |
|---|---|---|---|
| | | | Samuel Bloomfield, Claire Stuart, Andrew Bell, Reenesh Prakash, Samir Dervisevic, Alison E. Mather, John Wain, Mark Webber, Andrew J. Page, Justin O'Grady |
| EPI_ISL_490582, EPI_ISL_490583, EPI_ISL_490584, EPI_ISL_490586, EPI_ISL_490590, EPI_ISL_490592, EPI_ISL_490594, EPI_ISL_490597, EPI_ISL_490598, EPI_ISL_490600, EPI_ISL_490603, EPI_ISL_490604, EPI_ISL_490606, EPI_ISL_490607, EPI_ISL_490610, EPI_ISL_490614, EPI_ISL_490617, EPI_ISL_490619, EPI_ISL_490631, EPI_ISL_490632, EPI_ISL_490634, EPI_ISL_490638, EPI_ISL_490640, EPI_ISL_490643, EPI_ISL_490645 | | | |
| see above | Virology Department, Sheffield Teaching Hospitals NHS Foundation Trust/Department of Infection, Immunity and Cardiovascular Disease, The Medical School, University of Sheffield | COVID-19 Genomics UK (COG-UK) Consortium | Thushan de Silva, Matthew Parker, Nikki Smith, Adri Angyal, Rebecca Brown, Luke Green, Rachel Tucker, Paul Parsons, Danielle Groves, Katie Johnson, Laura Carrilero, Alex Keeley, Dave Partridge, Matthew Wyles, Benjamin Lindsey, Mehmet Yavuz, Mohammad Raza, Cariad Evans |
| EPI_ISL_490694 | West of Scotland Specialist Virology Centre, NHSGGC / MRC-University of Glasgow Centre for Virus Research | COVID-19 Genomics UK (COG-UK) Consortium | Ana da Silva Filipe, Natasha Johnson, Kathy Smollett, Daniel Mair, Stephen Carmichael, Lily Tong, Jenna Nichols, Elihu Aranday-Cortes, Kirstyn Brunker, Yasmin Parr, Alice Broos, Kyriaki Nomikou; Sarah McDonald, Marc Niebel, Patawee Asamaphan; Richard Orton, Joseph Hughes, Sreenu Vattipally, David L Robertson; Alasdair MacLean, Rory Gunson; Kathy Li, Natasha Jesudason, Rajiv Shah, James Shepherd, Antonia Ho, Emma Thomson |
| EPI_ISL_490710, EPI_ISL_490712, EPI_ISL_490717, EPI_ISL_490725, EPI_ISL_490729, EPI_ISL_490730, EPI_ISL_490737, EPI_ISL_490738, EPI_ISL_490744, EPI_ISL_490745, EPI_ISL_490760, EPI_ISL_490776, EPI_ISL_490777, EPI_ISL_490779, EPI_ISL_490784, EPI_ISL_490789, EPI_ISL_490792, EPI_ISL_490801, EPI_ISL_490804, EPI_ISL_490805, EPI_ISL_490807, EPI_ISL_490810, EPI_ISL_490813, EPI_ISL_490816, EPI_ISL_490819, EPI_ISL_490824, EPI_ISL_490826, EPI_ISL_490831, EPI_ISL_490843, EPI_ISL_490844, EPI_ISL_490860 | | | |
| see above | Wales Specialist Virology Centre Sequencing lab: Pathogen Genomics Unit | COVID-19 Genomics UK (COG-UK) Consortium | Catherine Moore, Johnathan Evans, Laura Gifford, Malorie Perry, Simon Cottrell, Angela Marchbank, Alec Birchley, Alexander Adams, Amy Gaskin, Bree Gatica-Wilcox, Jason Coombes, Joel Southgate, Lauren Gilbert, Lee Graham, Nicole Pacchiarini, Sara Kumziene-Summerhayes, Sarah Taylor, Sophie Jones, Sara Rey, Matthew Bull, Joanne Watkins, Sally Corden, Tom Connor |
| EPI_ISL_491088 | Suceava County Emergency Hospital | "Stefan cel Mare" University Metagenomics Lab | Lobiuc Andrei, Antoniadis Panagiotis et al. |
| EPI_ISL_491089 | Suceava County Emergency Hospital | "Stefan cel Mare" University Metagenomics Lab | Lobiuc Andrei et al. |
| EPI_ISL_491138, EPI_ISL_491141, EPI_ISL_491145 | Oman-National Influenza Center | Biotechnology & OMICs Laboratory | Samiha Al-Kharusi, Sajjad Asaf, Abdul Latif Khan, Samira Al-Mahruqi, Adil Khan, Ahmed Al-Rawahi, Amina Al-Jardani, Hanan Al-Kindi, Intisar Al-Shukri, Ahlam Al-Amri, Aisha Al-Amri, Aisha Al-Busaidi, Adil Al-Wahaibi, Seif Al-Abri, Ahmed Al-Harrasi |
| EPI_ISL_491153 | Oman-National Influenza Center | Biotechnology & OMICs Laboratory | Abdul Latif Khan, Samira Al-Mahruqi, Ahmed Al-Harrasi, Samiha Al-Kharusi, Adil Khan, Ahmed Al-Rawahi, Sajjad Asaf, Amina Al-Jardani, Hanan Al-Kindi, Intisar Al-Shukri, Ahlam Al-Amri, Aisha Al-Amri, Aisha Al-Busaidi, Adil Al-Wahaibi, Seif Al-Abri. |
| EPI_ISL_491947, EPI_ISL_491950, EPI_ISL_491953 | Instituto Nacional de Investigación en Salud Pública - INSPI | INSPI - Charité | Alfredo Bruno Caicedo, Domenica de Mora Coloma, Andres Moreira-Soto, Anna-Lena Sander, Nina Krause, Maritza Olmedo,Denisses Portugal, Manuel Gonzalez, Silvia Salgado, Alberto Orlando, Alexandra Usiña, Juan Carlos Zeballos, Jan Felix Drexler |
| EPI_ISL_491968, EPI_ISL_491969, EPI_ISL_491970, EPI_ISL_491971, EPI_ISL_491972, EPI_ISL_491973, EPI_ISL_491974, EPI_ISL_491975, EPI_ISL_491976, EPI_ISL_491977 | Oman-NIC | Department of Microbiology and Immunology-SQUH | Fahad Zadjali, Samira Al-Maruqi, Amina Al Jardani, Khulood Al-Mammary, Hanan Al-kindi, Fatma BaAlawi, Hamida AL Barwani, Zeyana AL-Dahmani, Intisar Al-Shukri, Aisha Al-Busaidi, Aisha Al-Amri, Ahlam Al-Amri, Mohammed Al-Tobi, Samiha Al Kharusi, Abdulla Balkhair |
| EPI_ISL_492029, EPI_ISL_492030, EPI_ISL_492031 | Child Health Research Foundation | Child Health Research Foundation | Senjuti Saha, Md Saiful Islam Sajib, Roly Malaker, Md Hafizur Rahman, Afroza Akter Tanni, Syed Muktadir Al Sium, Maksuda Islam, Samir K Saha |
| EPI_ISL_492838, EPI_ISL_492842 | Department of Medical Microbiology, Western Sussex Hospitals NHS Foundation Trust, St Richard's Hospital | Wellcome Sanger Institute for the COVID-19 Genomics UK (COG-UK) consortium | Manasa Mutingwende, Sarah Lowdon, Olga Podplomyk, Michelle Erkiert, Jonathan Lewis, Paul Randell and Alex Alderton, Roberto Amato, Sonia Goncalves, Ewan Harrison, David K. Jackson, Ian Johnston, Dominic Kwiatkowski, Cordelia Langford, John Sillitoe on behalf of the Wellcome Sanger Institute COVID-19 Surveillance Team (http://www.sanger.ac.uk/covid-team) |
| EPI_ISL_492843 | Royal Free Hospital / Health Services Laboratories | Wellcome Sanger Institute for the COVID-19 Genomics UK (COG-UK) consortium | Tanzina Haque, Tabitha Mahungu, Dianne Irish, Cate Goodlad, Jenny Cross, Judith Heaney and Alex Alderton, Roberto Amato, Sonia Goncalves, Ewan Harrison, David K. Jackson, Ian Johnston, Dominic Kwiatkowski, Cordelia Langford, John Sillitoe on behalf of the Wellcome Sanger Institute COVID-19 Surveillance Team (http://www.sanger.ac.uk/covid-team) |
| EPI_ISL_492854, EPI_ISL_492857, EPI_ISL_492860, EPI_ISL_492861, EPI_ISL_492867, EPI_ISL_492873, EPI_ISL_492891, EPI_ISL_492893, EPI_ISL_492895, EPI_ISL_492899, EPI_ISL_492910, EPI_ISL_492913 | | | |
| see above | Department of Medical Microbiology, Western Sussex Hospitals NHS Foundation Trust, St Richard's Hospital | Wellcome Sanger Institute for the COVID-19 Genomics UK (COG-UK) consortium | Manasa Mutingwende, Sarah Lowdon, Olga Podplomyk, Michelle Erkiert, Jonathan Lewis, Paul Randell and Alex Alderton, Roberto Amato, Sonia Goncalves, Ewan Harrison, David K. Jackson, Ian Johnston, Dominic Kwiatkowski, Cordelia Langford, John Sillitoe on behalf of the Wellcome Sanger Institute COVID-19 Surveillance Team (http://www.sanger.ac.uk/covid-team) |
| EPI_ISL_492924, EPI_ISL_492926 | NU-OMICS DNA Sequencing research facility, Northumbria University | Wellcome Sanger Institute for the COVID-19 Genomics UK (COG-UK) consortium | Chris Duncan, Sheia Waugh, Shirelle Burton-Fanning, Gary Eltringham, Jennifer Collins, Brendan Payne, Yusri Taha, Emma Swindells, Jane Greenaway, Edward Barton, Garren Scott, Debra Padgett, Clive Graham, Sarah Essex, Steve Ligget, Paul Baker, Lynn Dover, Wen Yew, Gary Black, John Allan, Joshua Loh, Greg Young, Matthew Bashton, Andrew Nelson, Darren Smith and Alex Alderton, Roberto Amato, Sonia Goncalves, Ewan Harrison, David K. Jackson, Ian Johnston, Dominic Kwiatkowski, Cordelia Langford, John Sillitoe on behalf of the Wellcome Sanger Institute COVID-19 Surveillance Team (http://www.sanger.ac.uk/covid-team) |
| EPI_ISL_492929 | NU-OMICS DNA Sequencing research facility, Northumbria University | Wellcome Sanger Institute for the COVID-19 Genomics UK (COG-UK) Consortium | Chris Duncan, Sheia Waugh, Shirelle Burton-Fanning, Gary Eltringham, Jennifer Collins, Brendan Payne, Yusri Taha, Emma Swindells, Jane Greenaway, Edward Barton, Garren Scott, Debra Padgett, Clive Graham, Sarah Essex, Steve Ligget, Paul Baker, Lynn Dover, Wen Yew, Gary Black, John Allan, Joshua Loh, Greg Young, Matthew Bashton, Andrew Nelson, Darren Smith and Alex Alderton, Roberto Amato, Sonia Goncalves, Ewan Harrison, David K. Jackson, Ian Johnston, Dominic Kwiatkowski, Cordelia Langford, John Sillitoe on behalf of the Wellcome Sanger Institute COVID-19 Surveillance Team |
| EPI_ISL_492930 | NU-OMICS DNA Sequencing research facility, Northumbria University | Wellcome Sanger Institute for the COVID-19 Genomics UK (COG-UK) consortium | Chris Duncan, Sheia Waugh, Shirelle Burton-Fanning, Gary Eltringham, Jennifer Collins, Brendan Payne, Yusri Taha, Emma Swindells, Jane Greenaway, Edward Barton, Garren Scott, Debra Padgett, Clive Graham, Sarah Essex, Steve Ligget, Paul Baker, Lynn Dover, Wen Yew, Gary Black, John Allan, Joshua Loh, Greg Young, Matthew Bashton, Andrew Nelson, Darren Smith and Alex Alderton, Roberto Amato, Sonia Goncalves, Ewan Harrison, David K. Jackson, Ian Johnston, Dominic Kwiatkowski, Cordelia Langford, John Sillitoe on behalf of the Wellcome Sanger Institute COVID-19 Surveillance Team (http://www.sanger.ac.uk/covid-team) |
| EPI_ISL_492931 | NU-OMICS DNA Sequencing research facility, Northumbria University | Wellcome Sanger Institute for the COVID-19 Genomics UK (COG-UK) Consortium | Chris Duncan, Sheia Waugh, Shirelle Burton-Fanning, Gary Eltringham, Jennifer Collins, Brendan Payne, Yusri Taha, Emma Swindells, Jane Greenaway, Edward Barton, Garren Scott, Debra Padgett, Clive Graham, Sarah Essex, Steve Ligget, Paul Baker, Lynn Dover, Wen Yew, Gary Black, John Allan, Joshua Loh, Greg Young, Matthew Bashton, Andrew Nelson, Darren Smith and Alex Alderton, Roberto Amato, Sonia Goncalves, Ewan Harrison, David K. Jackson, Ian Johnston, Dominic Kwiatkowski, Cordelia Langford, John Sillitoe on behalf of the Wellcome Sanger Institute COVID-19 Surveillance Team |
| EPI_ISL_492933, EPI_ISL_492941, EPI_ISL_492944, EPI_ISL_492949, EPI_ISL_492951, EPI_ISL_492952, EPI_ISL_492954, EPI_ISL_492957, EPI_ISL_492958, EPI_ISL_492960, EPI_ISL_492967, EPI_ISL_492975 | | | |
| see above | NU-OMICS DNA Sequencing research facility, Northumbria University | Wellcome Sanger Institute for the COVID-19 Genomics UK (COG-UK) consortium | Chris Duncan, Sheia Waugh, Shirelle Burton-Fanning, Gary Eltringham, Jennifer Collins, Brendan Payne, Yusri Taha, Emma Swindells, Jane Greenaway, Edward Barton, Garren Scott, Debra Padgett, Clive Graham, Sarah Essex, Steve Ligget, Paul Baker, Lynn Dover, Wen Yew, Gary Black, John Allan, Joshua Loh, Greg Young, Matthew Bashton, Andrew Nelson, Darren Smith and Alex Alderton, Roberto Amato, Sonia Goncalves, Ewan Harrison, David K. Jackson, Ian Johnston, Dominic Kwiatkowski, Cordelia Langford, John Sillitoe on behalf of the Wellcome Sanger Institute COVID-19 Surveillance Team (http://www.sanger.ac.uk/covid-team) |
| EPI_ISL_493004, EPI_ISL_493005, EPI_ISL_493006, EPI_ISL_493007, EPI_ISL_493008, EPI_ISL_493009, EPI_ISL_493010, EPI_ISL_493011, EPI_ISL_493012, EPI_ISL_493013, EPI_ISL_493014, EPI_ISL_493015, EPI_ISL_493016, EPI_ISL_493017, EPI_ISL_493018, EPI_ISL_493019, EPI_ISL_493020, EPI_ISL_493021, EPI_ISL_493022, EPI_ISL_493023, EPI_ISL_493024, EPI_ISL_493025, EPI_ISL_493026, EPI_ISL_493027, EPI_ISL_493028, EPI_ISL_493029, EPI_ISL_493030, EPI_ISL_493031, EPI_ISL_493032, EPI_ISL_493033, EPI_ISL_493034, EPI_ISL_493035, EPI_ISL_493036, EPI_ISL_493037, EPI_ISL_493038, EPI_ISL_493039, EPI_ISL_493040, EPI_ISL_493041, EPI_ISL_493042, EPI_ISL_493043, EPI_ISL_493044, EPI_ISL_493045, EPI_ISL_493046, EPI_ISL_493047, EPI_ISL_493048, EPI_ISL_493049, EPI_ISL_493050, EPI_ISL_493051, EPI_ISL_493052, EPI_ISL_493054, EPI_ISL_493055, EPI_ISL_493056, EPI_ISL_493057, EPI_ISL_493058, EPI_ISL_493059 | | | |
| see above | Utah Public Health Laboratory | Utah Public Health Laboratory | Heidi Butz, Erin Young, Kelly Oakeson |
| EPI_ISL_493212 | Functional Genomics Core University of South Carolina / Prisma Health-Midlands | Functional Genomics Core, University of South Carolina | Hao Ji, Diego Altomare, B.Celia Cui, Mengqian Chen, Alyssa Clay-Glimour, Michael Wyatt, Phillip Buckhaults, Helmut Albrecht, Michael Shtutman |
| EPI_ISL_493330 | INMI Lazzaro Spallanzani IRCCS | INMI Lazzaro Spallanzani IRCCS | Cesare E.M. Gruber, Martina Rueca, Barbara Bartolini, Francesco Messina, Maria R. Capobianchi, Antonino Di Caro |
| EPI_ISL_493331 | INMI Lazzaro Spallanzani IRCCS | INMI Lazzaro Spallanzani IRCCS | Martina Rueca, Cesare E.M. Gruber, Barbara Bartolini, Francesco Messina, Maria R. Capobianchi, Antonino Di Caro |

| | | | |
|---|---|---|---|
| EPI_ISL_493344 | Instituto de Diagnostico y Referencia Epidemiologicos (INDRE) | Instituto de Diagnostico y Referencia Epidemiologicos (INDRE) | Ernesto Ramirez-Gonzalez, Abril Rodriguez-Maldonado, Claudia Wong-Arambula , Natividad Cruz-Ortiz, Tatiana Nunez-Garcia, Dayanira Arellano-Suarez, Adnan Araiza-Rodriguez, Fabiola Garces-Ayala, Lucia Hernandez-Rivas, Irma Lopez-Martinez, Gisela Barrera-Badillo. |
| EPI_ISL_493399, EPI_ISL_493400, EPI_ISL_493404, EPI_ISL_493407, EPI_ISL_493409, EPI_ISL_493411 | National Public Health Laboratory, National Centre for Infectious Diseases | National Public Health Laboratory, National Centre for Infectious Diseases | Mak TM, Octavia S, Zhou Z, Chavatte JM, Cui L, Lin RTP |
| EPI_ISL_493426 | Functional Genomics Core University of South Carolina / Prisma Health-Midlands | Functional Genomics Core, University of South Carolina, | Hao Ji, Diego Altomare, B.Celia Cui, Mengqian Chen, Alyssa Clay-Glimour, Michael Wyatt, Phillip Buckhaults, Helmut Albrecht, Michael Shtutman |
| EPI_ISL_493427 | Functional Genomics Core University of South Carolina / Prisma Health-Midlands | Functional Genomics Core, University of South Carolina | Hao Ji, Diego Altomare, B.Celia Cui, Mengqian Chen, Alyssa Clay-Glimour, Michael Wyatt, Phillip Buckhaults, Helmut Albrecht, Michael Shtutman |
| EPI_ISL_493452, EPI_ISL_493453, EPI_ISL_493454, EPI_ISL_493459, EPI_ISL_493498, EPI_ISL_493499, EPI_ISL_493500, EPI_ISL_493501, EPI_ISL_493502, EPI_ISL_493503, EPI_ISL_493504, EPI_ISL_493505, EPI_ISL_493506, EPI_ISL_493507, EPI_ISL_493508, EPI_ISL_493510, EPI_ISL_493511, EPI_ISL_493512, EPI_ISL_493513, EPI_ISL_493514, EPI_ISL_493523, EPI_ISL_493534, EPI_ISL_493535, EPI_ISL_493536, EPI_ISL_493537, EPI_ISL_493538, EPI_ISL_493539, EPI_ISL_493540 | | | |
| see above | Northumbria University / South Tees Hospitals NHS Foundation Trust / North Cumbria Integrated Care NHS Foundation Trust / North Tees and Hartlepool NHS Foundation Trust / Newcastle Hospitals NHS Foundation Trust | COVID-19 Genomics UK (COG-UK) Consortium | Darren L Smith,Andrew Nelson,Matthew Bashton,Greg R Young,Joshua Loh,John Allan,Mohammad A Tariq,Giles S Holt,Gary Black,Wen C Yew,Lynn Dover,Paul Baker,Steve Liggett,Sarah Essex,Jane Greenaway,Debra Padgett,Clive Graham,Garren Scott,Edward Barton,Emma Swindells,Brendan Payne,Jennifer Collins,Yusri Taha,Gary Eltringham |
| EPI_ISL_493616, EPI_ISL_493617, EPI_ISL_493618, EPI_ISL_493619, EPI_ISL_493620, EPI_ISL_493621, EPI_ISL_493622, EPI_ISL_493623, EPI_ISL_493633, EPI_ISL_493634 | Centre for Enzyme Innovation, University of Portsmouth / Translational Research Laboratory, Portsmouth Hospitals NHS Trust | COVID-19 Genomics UK (COG-UK) Consortium | Angela Beckett,Yann Bourgeois,Garry Scarlett,Sharon Glaysher,Scott Elliott,Kelly Bicknell,Robert Impey,Allyson Lloyd,Sarah Wyllie,Ethan Butcher,Anoop Chauhan,Samuel Robson |
| EPI_ISL_493667, EPI_ISL_493679, EPI_ISL_493680, EPI_ISL_493692, EPI_ISL_493694, EPI_ISL_493695, EPI_ISL_493699, EPI_ISL_493701, EPI_ISL_493703, EPI_ISL_493704, EPI_ISL_493706, EPI_ISL_493707, EPI_ISL_493708, EPI_ISL_493710, EPI_ISL_493712, EPI_ISL_493713, EPI_ISL_493715, EPI_ISL_493720, EPI_ISL_493721, EPI_ISL_493723, EPI_ISL_493725, EPI_ISL_493729, EPI_ISL_493731, EPI_ISL_493732, EPI_ISL_493737, EPI_ISL_493738 | | | |
| see above | Virology Department, Sheffield Teaching Hospitals NHS Foundation Trust/Department of Infection, Immunity and Cardiovascular Disease, The Medical School, University of Sheffield | COVID-19 Genomics UK (COG-UK) Consortium | Thushan de Silva, Matthew Parker, Nikki Smith, Adri Angyal, Rebecca Brown, Luke Green, Rachel Tucker, Paul Parsons, Danielle Groves, Katie Johnson, Laura Carrilero, Alex Keeley, Dave Partridge, Matthew Wyles, Benjamin Lindsey, Mehmet Yavuz, Mohammad Raza, Cariad Evans |
| EPI_ISL_493840, EPI_ISL_493841, EPI_ISL_493842, EPI_ISL_493843, EPI_ISL_493852, EPI_ISL_493868, EPI_ISL_493869, EPI_ISL_493870, EPI_ISL_493871, EPI_ISL_493872 | West of Scotland Specialist Virology Centre, NHSGGC / MRC-University of Glasgow Centre for Virus Research | COVID-19 Genomics UK (COG-UK) Consortium | Ana da Silva Filipe, Natasha Johnson, Kathy Smollett, Daniel Mair, Stephen Carmichael, Lily Tong, Jenna Nichols, Elihu Aranday-Cortes, Kirstyn Brunker, Yasmin Parr, Alice Broos, Kyriaki Nomikou; Sarah McDonald, Marc Niebel, Patawee Asamaphan; Richard Orton, Joseph Hughes, Sreenu Vattipally, David L Robertson; Alasdair MacLean, Rory Gunson; Kathy Li, Natasha Jesudason, Rajiv Shah, James Shepherd, Antonia Ho, Emma Thomson |
| EPI_ISL_493905, EPI_ISL_493906, EPI_ISL_493907, EPI_ISL_493908, EPI_ISL_493909, EPI_ISL_493910, EPI_ISL_493911, EPI_ISL_493912, EPI_ISL_493913, EPI_ISL_493914, EPI_ISL_493915, EPI_ISL_493916, EPI_ISL_493917, EPI_ISL_493918, EPI_ISL_493919, EPI_ISL_493920, EPI_ISL_493921, EPI_ISL_493922, EPI_ISL_493923, EPI_ISL_493924, EPI_ISL_493925, EPI_ISL_493926, EPI_ISL_493927, EPI_ISL_493928, EPI_ISL_493929, EPI_ISL_493930, EPI_ISL_493931, EPI_ISL_493932, EPI_ISL_493933, EPI_ISL_493934, EPI_ISL_493935, EPI_ISL_493936, EPI_ISL_493937, EPI_ISL_493938, EPI_ISL_493939, EPI_ISL_493940, EPI_ISL_493941, EPI_ISL_493942, EPI_ISL_493950, EPI_ISL_493962 | | | |
| see above | Virology Department, Royal Infirmary of Edinburgh, NHS Lothian / School of Biological Sciences, University of Edinburgh / Institute of Genetics and Molecular Medicine, University of Edinburgh | COVID-19 Genomics UK (COG-UK) Consortium | McHugh M, Dewar R, Rooke S, Gallagher M, Balcaza C, O'Toole Á, Scher E, Hill V, McCrone JT, Colquhoun R, Yu X, Jackson B, Rambaut A, Williams TC, Templeton K |
| EPI_ISL_493978, EPI_ISL_493980, EPI_ISL_493981, EPI_ISL_494152 | Wales Specialist Virology Centre Sequencing lab: Pathogen Genomics Unit | COVID-19 Genomics UK (COG-UK) Consortium | Catherine Moore, Johnathan Evans, Laura Gifford, Malorie Perry, Simon Cottrell, Angela Marchbank, Alec Birchley, Alexander Adams, Amy Gaskin, Bree Gatica-Wilcox, Jason Coombes, Joel Southgate, Lauren Gilbert, Lee Graham, Nicole Pacchiarini, Sara Kumziene-Summerhayes, Sarah Taylor, Sophie Jones, Sara Rey, Matthew Bull, Joanne Watkins, Sally Corden, Tom Connor |
| EPI_ISL_494450, EPI_ISL_494451, EPI_ISL_494452, EPI_ISL_494453, EPI_ISL_494455, EPI_ISL_494457, EPI_ISL_494458, EPI_ISL_494459, EPI_ISL_494460, EPI_ISL_494461, EPI_ISL_494462, EPI_ISL_494464, EPI_ISL_494465, EPI_ISL_494470, EPI_ISL_494471, EPI_ISL_494472, EPI_ISL_494473, EPI_ISL_494474, EPI_ISL_494479, EPI_ISL_494488, EPI_ISL_494503 | | | |
| see above | San Diego County Public Health Laboratory | Andersen lab at Scripps Research | SEARCH Alliance San Diego with Tracy Basler, Jovan Shephard, Brett Austin |
| EPI_ISL_494553 | Functional Genomics Core University of South Carolina / Prisma Health-Midlands | Functional Genomics Core, University of South Carolina | Hao Ji, Diego Altomare, B.Celia Cui, Mengqian Chen, Alyssa Clay-Glimour, Michael Wyatt, Phillip Buckhaults, Helmut Albrecht, Michael Shtutman |
| EPI_ISL_494577, EPI_ISL_494578, EPI_ISL_494579, EPI_ISL_494580, EPI_ISL_494581, EPI_ISL_494582, EPI_ISL_494583, EPI_ISL_494584, EPI_ISL_494585, EPI_ISL_494586, EPI_ISL_494587 | | | |
| see above | San Diego County Public Health Laboratory | Andersen lab at Scripps Research | SEARCH Alliance San Diego with Tracy Basler, Jovan Shephard, Brett Austin |
| EPI_ISL_494635, EPI_ISL_494636, EPI_ISL_494637, EPI_ISL_494638, EPI_ISL_494640, EPI_ISL_494642, EPI_ISL_494643, EPI_ISL_494644, EPI_ISL_494647, EPI_ISL_494648, EPI_ISL_494650, EPI_ISL_494652, EPI_ISL_494653, EPI_ISL_494655, EPI_ISL_494657, EPI_ISL_494658, EPI_ISL_494659, EPI_ISL_494660, EPI_ISL_494661, EPI_ISL_494662, EPI_ISL_494666, EPI_ISL_494671 | | | |
| see above | Scripps Medical Laboratory | Andersen lab at Scripps Research | SEARCH Alliance San Diego with Michael Quigley, Ellen Stefanski, Ian Mchardy |
| EPI_ISL_495088, EPI_ISL_495093 | Department of Medical Microbiology, Western Sussex Hospitals NHS Foundation Trust, St Richard's Hospital | Wellcome Sanger Institute for the COVID-19 Genomics UK (COG-UK) consortium | Manasa Mutingwende, Sarah Lowdon, Olga Podplomyk, Michelle Erkiert, Jonathan Lewis, Paul Randell and Alex Alderton, Roberto Amato, Sonia Goncalves, Ewan Harrison, David K. Jackson, Ian Johnston, Dominic Kwiatkowski, Cordelia Langford, John Sillitoe on behalf of the Wellcome Sanger Institute COVID-19 Surveillance Team (http://www.sanger.ac.uk/covid-team) |
| EPI_ISL_495115 | PHE South West Regional Laboratory, National Infection Service | Wellcome Sanger Institute for the COVID-19 Genomics UK (COG-UK) consortium | Stephanie Hutchings, Hannah Pymont, Dr Peter Muir, Barry Vipond, Rich Hopes; and Alex Alderton, Roberto Amato, Sonia Goncalves, Ewan Harrison, David K. Jackson, Ian Johnston, Dominic Kwiatkowski, Cordelia Langford, John Sillitoe on behalf of the Wellcome Sanger Institute COVID-19 Surveillance Team (http://www.sanger.ac.uk/covid-team) |
| EPI_ISL_495161 | CSIR-Centre for Cellular and Molecular Biology | CSIR-Centre for Cellular and Molecular Biology | Onkar Kulkarni,Sofia Banu, Payel Mukherjee, Priya Singh, Dhiviya Vedagiri, Divya Gupta, Vishal Sah, Santosh Kumar Kuncha, Krishnan Harinivas Harshan, Archana Bharadwaj Siva, Karthik Bharadwaj Tallapaka, Shagufta Khan, Lamuk Zaveri,Nikhil Hajirnis, M Soujanya Reddy, Pratheusa Maccha, Namami Gaur, Sakshi Shambhavi, Tulasi Nagabandi, Purushotham Vodnala, Deepak Kumar, Devi Prasad Vijayashankar, Disha Nanda, Divya Das, Jotin Gogoi, Manish Bhattacharjee, Rakesh K Mishra, Divya Tej Sowpati |
| EPI_ISL_495162 | CSIR-Centre for Cellular and Molecular Biology | CSIR-Centre for Cellular and Molecular Biology | Onkar Kulkarni, Payel Mukherjee, Sofia Banu, Priya Singh, Dhiviya Vedagiri, Divya Gupta, Vishal Sah, Santosh Kumar Kuncha, Krishnan Harinivas Harshan, Archana Bharadwaj Siva, Karthik Bharadwaj Tallapaka, Shagufta Khan, Lamuk Zaveri,Nikhil Hajirnis, M Soujanya Reddy, Pratheusa Maccha, Namami Gaur, Sakshi Shambhavi, Tulasi Nagabandi, Purushotham Vodnala, Deepak Kumar, Devi Prasad Vijayashankar, Disha Nanda, Divya Das, Jotin Gogoi, Manish Bhattacharjee, Rakesh K Mishra, Divya Tej Sowpati |
| EPI_ISL_495274 | Osmania Medical College | CSIR-Centre for Cellular and Molecular Biology | Shashikala Reddy, Mahboob Khan, Lamuk Zaveri, Sofia Banu,Payel Mukherjee, Shagufta Khan,Priya Singh, Onkar Kulkarni, Dhiviya Vedagiri, Divya Gupta, Vishal Sah, Santosh Kumar Kuncha, Krishnan Harinivas Harshan, Archana Bharadwaj Siva, Karthik Bharadwaj Tallapaka,Nikhil Hajirnis, M Soujanya Reddy, Pratheusa Maccha, Namami Gaur, Sakshi Shambhavi, Tulasi Nagabandi, Purushotham Vodnala, Rakesh K Mishra, Divya Tej Sowpati |
| EPI_ISL_495275 | Osmania Medical College | CSIR-Centre for Cellular and Molecular Biology | Shashikala Reddy, Mahboob Khan, Onkar Kulkarni, Payel Mukherjee, Sofia Banu, Priya Singh, Dhiviya Vedagiri, Divya Gupta, Vishal Sah, Santosh Kumar Kuncha, Krishnan Harinivas Harshan, Archana Bharadwaj Siva, Karthik Bharadwaj Tallapaka, Shagufta Khan, Lamuk Zaveri, Nikhil Hajirnis, M Soujanya Reddy, Pratheusa Maccha,Namami Gaur, Sakshi Shambhavi, Tulasi Nagabandi, Purushotham Vodnala, Rakesh K Mishra, Divya Tej Sowpati |
| EPI_ISL_495276 | Osmania Medical College | CSIR-Centre for Cellular and Molecular Biology | Shashikala Reddy, Mahboob Khan,Payel Mukherjee, Sofia Banu, Priya Singh, Onkar Kulkarni, Dhiviya Vedagiri, Divya Gupta, Vishal Sah, Santosh Kumar Kuncha, Krishnan Harinivas Harshan, Archana Bharadwaj Siva, Karthik Bharadwaj Tallapaka, Shagufta Khan, Lamuk Zaveri, Namami Gaur, Sakshi Shambhavi, Nikhil Hajirnis, M Soujanya Reddy, Pratheusa Maccha,Tulasi Nagabandi, Purushotham Vodnala, Rakesh K Mishra, Divya Tej Sowpati |

| | | | |
|---|---|---|---|
| EPI_ISL_495277 | Osmania Medical College | CSIR-Centre for Cellular and Molecular Biology | Shashikala Reddy, Mahboob Khan, Namami Gaur, Lamuk Zaveri,Shagufta Khan, Nikhil Hajirnis, M Soujanya Reddy, Pratheusa Maccha, Sofia Banu, Onkar Kulkarni, Payel Mukherjee, Priya Singh, Dhiviya Vedagiri, Divya Gupta, Vishal Sah, Santosh Kumar Kuncha, Krishnan Harinivas Harshan, Archana Bharadwaj Siva, Karthik Bharadwaj Tallapaka,Sakshi Shambhavi, Tulasi Nagabandi, Purushotham Vodnala, Rakesh K Mishra, Divya Tej Sowpati |
| EPI_ISL_495278 | Osmania Medical College | CSIR-Centre for Cellular and Molecular Biology | Shashikala Reddy, Mahboob Khan,Tulasi Nagabandi, Namami Gaur, Lamuk Zaveri,Shagufta Khan,Nikhil Hajirnis, M Soujanya Reddy, Pratheusa Maccha, Sofia Banu,Payel Mukherjee, Priya Singh, Onkar Kulkarni, Dhiviya Vedagiri, Divya Gupta, Vishal Sah, Santosh Kumar Kuncha, Krishnan Harinivas Harshan, Archana Bharadwaj Siva, Karthik Bharadwaj Tallapaka,Sakshi Shambhavi, Purushotham Vodnala, Rakesh K Mishra, Divya Tej Sowpati |
| EPI_ISL_495279 | Osmania Medical College | CSIR-Centre for Cellular and Molecular Biology | Shashikala Reddy, Mahboob Khan,Sakshi Shambhavi,Tulasi Nagabandi, Namami Gaur, Lamuk Zaveri,Shagufta Khan, Nikhil Hajirnis, M Soujanya Reddy, Pratheusa Maccha,Sofia Banu,Payel Mukherjee, Priya Singh,Onkar Kulkarni, Dhiviya Vedagiri, Divya Gupta, Vishal Sah, Santosh Kumar Kuncha, Krishnan Harinivas Harshan, Archana Bharadwaj Siva, Karthik Bharadwaj Tallapaka,Sakshi Shambhavi, Purushotham Vodnala, Rakesh K Mishra, Divya Tej Sowpati |
| EPI_ISL_495280 | Osmania Medical College | CSIR-Centre for Cellular and Molecular Biology | Shashikala Reddy, Mahboob Khan,Payel Mukherjee, Sofia Banu, Priya Singh, Onkar Kulkarni, Dhiviya Vedagiri, Divya Gupta, Vishal Sah, Santosh Kumar Kuncha, Krishnan Harinivas Harshan, Archana Bharadwaj Siva, Karthik Bharadwaj Tallapaka, Shagufta Khan, Lamuk Zaveri, Nikhil Hajirnis, M Soujanya Reddy, Pratheusa Maccha,Namami Gaur, Sakshi Shambhavi, Tulasi Nagabandi, Purushotham Vodnala, Rakesh K Mishra, Divya Tej Sowpati |
| EPI_ISL_495286 | Osmania Medical College | CSIR-Centre for Cellular and Molecular Biology | Shashikala Reddy, Mahboob Khan, Sofia Banu,Payel Mukherjee, Priya Singh,Onkar Kulkarni, Dhiviya Vedagiri, Divya Gupta, Vishal Sah, Santosh Kumar Kuncha, Krishnan Harinivas Harshan, Archana Bharadwaj Siva, Karthik Bharadwaj Tallapaka, Shagufta Khan, Lamuk Zaveri, Namami Gaur, Sakshi Shambhavi, Nikhil Hajirnis, M Soujanya Reddy, Pratheusa Maccha,Tulasi Nagabandi, Purushotham Vodnala, Rakesh K Mishra, Divya Tej Sowpati |
| EPI_ISL_495287 | Osmania Medical College | CSIR-Centre for Cellular and Molecular Biology | Shashikala Reddy, Mahboob Khan,Onkar Kulkarni, Sofia Banu,Payel Mukherjee, Priya Singh, Dhiviya Vedagiri, Divya Gupta, Vishal Sah, Santosh Kumar Kuncha, Krishnan Harinivas Harshan, Archana Bharadwaj Siva, Karthik Bharadwaj Tallapaka, Shagufta Khan, Nikhil Hajirnis, M Soujanya Reddy, Pratheusa Maccha, Lamuk Zaveri, Namami Gaur, Sakshi Shambhavi, Tulasi Nagabandi, Purushotham Vodnala, Rakesh K Mishra, Divya Tej Sowpati |
| EPI_ISL_495289 | Osmania Medical College | CSIR-Centre for Cellular and Molecular Biology | Shashikala Reddy, Mahboob Khan, Namami Gaur, Lamuk Zaveri,Shagufta Khan, Sofia Banu, Onkar Kulkarni, Payel Mukherjee, Priya Singh, Dhiviya Vedagiri, Divya Gupta, Vishal Sah, Santosh Kumar Kuncha, Krishnan Harinivas Harshan, Archana Bharadwaj Siva, Karthik Bharadwaj Tallapaka,Sakshi Shambhavi, Nikhil Hajirnis, M Soujanya Reddy, Pratheusa Maccha, Tulasi Nagabandi, Purushotham Vodnala, Rakesh K Mishra, Divya Tej Sowpati |
| EPI_ISL_495290 | Osmania Medical College | CSIR-Centre for Cellular and Molecular Biology | Shashikala Reddy, Mahboob Khan,Tulasi Nagabandi, Namami Gaur, Lamuk Zaveri,Shagufta Khan, Sofia Banu,Payel Mukherjee, Priya Singh, Onkar Kulkarni, Dhiviya Vedagiri, Divya Gupta, Vishal Sah, Santosh Kumar Kuncha, Krishnan Harinivas Harshan, Archana Bharadwaj Siva, Karthik Bharadwaj Tallapaka,Sakshi Shambhavi, Nikhil Hajirnis, M Soujanya Reddy, Pratheusa Maccha, Purushotham Vodnala, Rakesh K Mishra, Divya Tej Sowpati |
| EPI_ISL_495291 | Osmania Medical College | CSIR-Centre for Cellular and Molecular Biology | Shashikala Reddy, Mahboob Khan,Onkar Kulkarni,Shagufta Khan, Sofia Banu,Payel Mukherjee, Priya Singh, Dhiviya Vedagiri, Divya Gupta, Vishal Sah, Santosh Kumar Kuncha, Krishnan Harinivas Harshan, Archana Bharadwaj Siva, Karthik Bharadwaj Tallapaka, Lamuk Zaveri, Nikhil Hajirnis, M Soujanya Reddy, Pratheusa Maccha, Namami Gaur, Sakshi Shambhavi, Tulasi Nagabandi, Purushotham Vodnala, Rakesh K Mishra, Divya Tej Sowpati |
| EPI_ISL_495292 | Osmania Medical College | CSIR-Centre for Cellular and Molecular Biology | Shashikala Reddy, Mahboob Khan,Shagufta Khan, Sofia Banu,Payel Mukherjee, Priya Singh,Onkar Kulkarni, Dhiviya Vedagiri, Divya Gupta, Vishal Sah, Santosh Kumar Kuncha, Krishnan Harinivas Harshan, Archana Bharadwaj Siva, Karthik Bharadwaj Tallapaka, Lamuk Zaveri, Namami Gaur, Sakshi Shambhavi,Nikhil Hajirnis, M Soujanya Reddy, Pratheusa Maccha, Tulasi Nagabandi, Purushotham Vodnala, Rakesh K Mishra, Divya Tej Sowpati |
| EPI_ISL_495293 | Osmania Medical College | CSIR-Centre for Cellular and Molecular Biology | Shashikala Reddy, Mahboob Khan, Lamuk Zaveri, Sofia Banu,Payel Mukherjee, Shagufta Khan,Priya Singh, Onkar Kulkarni, Dhiviya Vedagiri, Divya Gupta, Vishal Sah, Santosh Kumar Kuncha, Krishnan Harinivas Harshan, Archana Bharadwaj Siva, Karthik Bharadwaj Tallapaka, Namami Gaur, Sakshi Shambhavi, Nikhil Hajirnis, M Soujanya Reddy, Pratheusa Maccha, Tulasi Nagabandi, Purushotham Vodnala, Rakesh K Mishra, Divya Tej Sowpati |
| EPI_ISL_495294 | Osmania Medical College | CSIR-Centre for Cellular and Molecular Biology | Shashikala Reddy, Mahboob Khan,Onkar Kulkarni, Lamuk Zaveri, Sofia Banu,Payel Mukherjee, Shagufta Khan,Priya Singh, Dhiviya Vedagiri, Divya Gupta, Vishal Sah, Santosh Kumar Kuncha, Krishnan Harinivas Harshan, Archana Bharadwaj Siva, Karthik Bharadwaj Tallapaka, Namami Gaur, Sakshi Shambhavi, Nikhil Hajirnis, M Soujanya Reddy, Pratheusa Maccha, Tulasi Nagabandi, Purushotham Vodnala, Rakesh K Mishra, Divya Tej Sowpati |
| EPI_ISL_495295 | Osmania Medical College | CSIR-Centre for Cellular and Molecular Biology | Shashikala Reddy, Mahboob Khan, Sofia Banu,Payel Mukherjee, Priya Singh,Onkar Kulkarni, Dhiviya Vedagiri, Divya Gupta, Vishal Sah, Santosh Kumar Kuncha, Krishnan Harinivas Harshan, Archana Bharadwaj Siva, Karthik Bharadwaj Tallapaka, Shagufta Khan, Lamuk Zaveri,Nikhil Hajirnis, M Soujanya Reddy, Pratheusa Maccha, Namami Gaur, Sakshi Shambhavi, Tulasi Nagabandi, Purushotham Vodnala, Rakesh K Mishra, Divya Tej Sowpati |
| EPI_ISL_495296 | Osmania Medical College | CSIR-Centre for Cellular and Molecular Biology | Shashikala Reddy, Mahboob Khan,Shagufta Khan, Sofia Banu,Payel Mukherjee, Priya Singh,Onkar Kulkarni, Dhiviya Vedagiri, Divya Gupta, Vishal Sah, Santosh Kumar Kuncha, Krishnan Harinivas Harshan, Archana Bharadwaj Siva, Karthik Bharadwaj Tallapaka, Lamuk Zaveri, Nikhil Hajirnis, M Soujanya Reddy, Pratheusa Maccha, Namami Gaur, Sakshi Shambhavi, Tulasi Nagabandi, Purushotham Vodnala, Rakesh K Mishra, Divya Tej Sowpati |
| EPI_ISL_495297 | Osmania Medical College | CSIR-Centre for Cellular and Molecular Biology | Shashikala Reddy, Mahboob Khan,Sakshi Shambhavi,Tulasi Nagabandi, Namami Gaur, Lamuk Zaveri,Shagufta Khan, Sofia Banu,Payel Mukherjee, Priya Singh,Onkar Kulkarni, Dhiviya Vedagiri, Divya Gupta, Vishal Sah, Santosh Kumar Kuncha, Krishnan Harinivas Harshan, Archana Bharadwaj Siva, Karthik Bharadwaj Tallapaka,Sakshi Shambhavi, Nikhil Hajirnis, M Soujanya Reddy, Pratheusa Maccha,Purushotham Vodnala, Rakesh K Mishra, Divya Tej Sowpati |
| EPI_ISL_495418, EPI_ISL_495423, EPI_ISL_495426, EPI_ISL_495430, EPI_ISL_495435, EPI_ISL_495437, EPI_ISL_495439, EPI_ISL_495455, EPI_ISL_495457 | Kafkas University, Faculty of Medicine, Department of Medical Microbiology | Kafkas University, Faculty of Medicine, Department of Medical Microbiology | Murat Karamese, Didem Ozgur, E. Ediz Tutuncu |
| EPI_ISL_495566, EPI_ISL_495568, EPI_ISL_495569, EPI_ISL_495592 | University of Michigan Clinical Microbiology Laboratory | Lauring Lab, University of Michigan, Department of Microbiology and Immunology | Valesano et al. |
| EPI_ISL_495629, EPI_ISL_495630, EPI_ISL_495631, EPI_ISL_495632, EPI_ISL_495633, EPI_ISL_495634, EPI_ISL_495635, EPI_ISL_495636, EPI_ISL_495637, EPI_ISL_495638, EPI_ISL_495639, EPI_ISL_495640, EPI_ISL_495641 | | | |
| see above | Viral Respiratory Lab, National Institute for Biomedical Research (INRB) | Pathogen Sequencing Lab, National Institute for Biomedical Research (INRB) | Placide Mbala-Kingebeni, Edith Nkwembe, Eddy Kinganda-Lusamaki, Amuri Aziza, Francisca Muyembe Mawete, Emmanuel Lokilo Lofiko, Catherine Pratt, Matthias Pauthner, Josh Quick, Allison Black, James Hadfield, Trevor Bedford, Ian Goodfellow, Andrew Rambaut, Nick Loman, Kristian Andersen, Michael Wiley, Steve Ahuka-Mundeke, Jean-Jacques Muyembe Tamfum |
| EPI_ISL_495642, EPI_ISL_495643, EPI_ISL_495644, EPI_ISL_495645, EPI_ISL_495646, EPI_ISL_495647, EPI_ISL_495648, EPI_ISL_495649, EPI_ISL_495650, EPI_ISL_495651, EPI_ISL_495652, EPI_ISL_495653 | | | |
| see above | Washington State Department of Health | Seattle Flu Study | Deborah A. Nickerson, Chris D. Frazar, Jover Lee, Benjamin Pelle, Matthew Richardson, Amanda Adler, Elisabeth Brandstetter, Peter D. Han, Kairsten Fay, Misja Ilcisin, Kirsten Lacombe, Thomas R. Sibley, Melissa Truong, Caitlin R. Wolf, Romesh Gautom, Geoff Melly, Brian Hiatt, Philip Dykema, Scott Lindquist, Michael Boeckh, Janet A. Englund, Michael Famulare, Barry R. Lutz, Mark J. Rieder, Lea M. Starita, Matthew Thompson, Helen Y. Chu, Jay Shendure, Trevor Bedford |
| EPI_ISL_495666, EPI_ISL_495667, EPI_ISL_495668, EPI_ISL_495669, EPI_ISL_495670, EPI_ISL_495671, EPI_ISL_495672, EPI_ISL_495673, EPI_ISL_495674, EPI_ISL_495675, EPI_ISL_495676, EPI_ISL_495677, EPI_ISL_495678, EPI_ISL_495679, EPI_ISL_495680, EPI_ISL_495681, EPI_ISL_495682, EPI_ISL_495683, EPI_ISL_495684, EPI_ISL_495685, EPI_ISL_495686, EPI_ISL_495687, EPI_ISL_495688, EPI_ISL_495689, EPI_ISL_495690, EPI_ISL_495691, EPI_ISL_495692, EPI_ISL_495693, EPI_ISL_495694, EPI_ISL_495695, EPI_ISL_495696, EPI_ISL_495697, EPI_ISL_495698, EPI_ISL_495699, EPI_ISL_495700, EPI_ISL_495701, EPI_ISL_495702, EPI_ISL_495703, EPI_ISL_495704, EPI_ISL_495705, EPI_ISL_495706, EPI_ISL_495707, EPI_ISL_495708, EPI_ISL_495709, EPI_ISL_495710, EPI_ISL_495711, EPI_ISL_495712, EPI_ISL_495713, EPI_ISL_495714, EPI_ISL_495715, EPI_ISL_495716, EPI_ISL_495717, EPI_ISL_495718, EPI_ISL_495719, EPI_ISL_495720, EPI_ISL_495721, EPI_ISL_495722, EPI_ISL_495723, EPI_ISL_495724, EPI_ISL_495725, EPI_ISL_495726, EPI_ISL_495727, EPI_ISL_495728, EPI_ISL_495729, EPI_ISL_495730, EPI_ISL_495731, EPI_ISL_495732, EPI_ISL_495733, EPI_ISL_495734, EPI_ISL_495735, EPI_ISL_495736, EPI_ISL_495737, EPI_ISL_495738, EPI_ISL_495739, EPI_ISL_495740, EPI_ISL_495741, EPI_ISL_495742, EPI_ISL_495743, EPI_ISL_495744, EPI_ISL_495745, EPI_ISL_495746, EPI_ISL_495747, EPI_ISL_495748, EPI_ISL_495749, EPI_ISL_495750, EPI_ISL_495751, EPI_ISL_495752, EPI_ISL_495753, EPI_ISL_495754, EPI_ISL_495755, EPI_ISL_495756, EPI_ISL_495757, EPI_ISL_495758, EPI_ISL_495759, EPI_ISL_495760, EPI_ISL_495761, EPI_ISL_495762, EPI_ISL_495763, EPI_ISL_495764, EPI_ISL_495765, EPI_ISL_495766, EPI_ISL_495767, EPI_ISL_495768, EPI_ISL_495769, EPI_ISL_495770, EPI_ISL_495771, EPI_ISL_495772, EPI_ISL_495773, EPI_ISL_495774, EPI_ISL_495775, EPI_ISL_495776, EPI_ISL_495777, EPI_ISL_495778, EPI_ISL_495779, EPI_ISL_495780, EPI_ISL_495781, EPI_ISL_495782, EPI_ISL_495783, EPI_ISL_495784, EPI_ISL_495785, EPI_ISL_495786, EPI_ISL_495787, EPI_ISL_495788, EPI_ISL_495789, EPI_ISL_495790, EPI_ISL_495791, EPI_ISL_495792, EPI_ISL_495793, EPI_ISL_495794, EPI_ISL_495795, EPI_ISL_495796, EPI_ISL_495797, EPI_ISL_495798, EPI_ISL_495799, EPI_ISL_495802, EPI_ISL_495808, EPI_ISL_495811, EPI_ISL_495818, EPI_ISL_495821 | | | |
| see above | Washington State Department of Health | Seattle Flu Study | Deborah A. Nickerson, Chris D. Frazar, Jover Lee, Benjamin Pelle, Matthew Richardson, Amanda Adler, Elisabeth Brandstetter, Peter D. Han, Kairsten Fay, Misja Ilcisin, Kirsten Lacombe, Thomas R. Sibley, Melissa Truong, Caitlin R. Wolf, Romesh Gautom, Geoff |
| EPI_ISL_496493, EPI_ISL_496494, EPI_ISL_496495, EPI_ISL_496496, EPI_ISL_496497, EPI_ISL_496498, EPI_ISL_496499, EPI_ISL_496500, EPI_ISL_496501, EPI_ISL_496502, EPI_ISL_496503, EPI_ISL_496504, EPI_ISL_496505, EPI_ISL_496514, EPI_ISL_496515, EPI_ISL_496516, EPI_ISL_496517 | | | |
| see above | Viral Respiratory Lab, National Institute for Biomedical Research (INRB) | Pathogen Sequencing Lab, National Institute for Biomedical Research (INRB) | Placide Mbala-Kingebeni, Edith Nkwembe, Eddy Kinganda-Lusamaki, Amuri Aziza, Francisca Muyembe Mawete, Emmanuel Lokilo Lofiko, Catherine Pratt, Matthias Pauthner, Josh Quick, Allison Black, James Hadfield, Trevor Bedford, Ian Goodfellow, Andrew Rambaut, Nick Loman, Kristian Andersen, Michael |

| | | | |
|---|---|---|---|
| | | | Wiley, Steve Ahuka-Mundeke, Jean-Jacques Muyembe Tamfum |
| EPI_ISL_496524, EPI_ISL_496536, EPI_ISL_496558, EPI_ISL_496559, EPI_ISL_496560, EPI_ISL_496587, EPI_ISL_497877, EPI_ISL_497878 | National Centre For Cell Science | National Centre For Cell Science | Dhiraj Paul, Kunal Jani, Radha Chauhan, Janesh Kumar, Vasudevan Seshadri, Girdhari Lal, Rajesh Karyakarte, Suvarna Joshi, Murlidhar Tambe, Sourav Sen, Santosh Karade, Kavita Bala Anand, Shelinder Pal Singh Shergill, Rajiv Mohan Gupta, Manoj Kumar Bhat, Arvind Sahu, Maharashtra COVID-19 Study Group, DBT's PAN-INDIA 1000 SARS-CoV2 RNA genome sequencing consortium, Yogesh S Shouche |
| EPI_ISL_498127, EPI_ISL_498128, EPI_ISL_498129, EPI_ISL_498130, EPI_ISL_498131, EPI_ISL_498132, EPI_ISL_498133, EPI_ISL_498134, EPI_ISL_498135, EPI_ISL_498136, EPI_ISL_498137, EPI_ISL_498138, EPI_ISL_498139 | | | |
| see above | Department of Clinical Microbiology | GIGA Medical Genomics | Keith Durkin, Maria Artesi, Sébastien Bontems, Raphaël Boreux, Cécile Meex, Axelle Chaslain, Céline Fombellida-Lopez, Pierrette Melin, Marie-Pierre Hayette, Vincent Bours. |
| EPI_ISL_498270, EPI_ISL_498271 | Department of Microbiology, The University of Hong Kong | Department of Microbiology, The University of Hong Kong | Kelvin K.W. To, Kwok-Yung Yuen |
| EPI_ISL_498534 | ACT Pathology | Schwessinger Lab | Ashley Jones, Benjamin Schwessinger, Robert Lanfear, Robyn N Hall, Megan McDonald, Ming-Dao Chia, Kevin Murray, Craig Kennedy, Karina Kennedy |
| EPI_ISL_498566 | National Public Health Laboratory, National Centre for Infectious Diseases | National Public Health Laboratory, National Centre for Infectious Diseases | Mak TM, Octavia S, Zhou Z, Chavatte JM, Cui L, Lin RTP |
| EPI_ISL_498678, EPI_ISL_498679, EPI_ISL_498680, EPI_ISL_498681, EPI_ISL_498682, EPI_ISL_498683, EPI_ISL_498684, EPI_ISL_498685, EPI_ISL_498686, EPI_ISL_498687, EPI_ISL_498688, EPI_ISL_498689, EPI_ISL_498690 | | | |
| see above | Utah Public Health Laboratory | Utah Public Health Laboratory | Heidi Butz, Erin Young, Kelly Oakeson |
| EPI_ISL_499363, EPI_ISL_499394, EPI_ISL_499396, EPI_ISL_499399, EPI_ISL_499405, EPI_ISL_499412, EPI_ISL_499444, EPI_ISL_499452, EPI_ISL_499453, EPI_ISL_499454, EPI_ISL_499456 | | | |
| see above | Wales Specialist Virology Centre Sequencing lab: Pathogen Genomics Unit | COVID-19 Genomics UK (COG-UK) Consortium | Catherine Moore, Johnathan Evans, Laura Gifford, Malorie Perry, Simon Cottrell, Angela Marchbank, Alec Birchley, Alexander Adams, Amy Gaskin, Bree Gatica-Wilcox, Jason Coombes, Joel Southgate, Lauren Gilbert, Lee Graham, Nicole Pacchiarini, Sara Kumziene-Summerhayes, Sarah Taylor, Sophie Jones, Sara Rey, Matthew Bull, Joanne Watkins, Sally Corden, Tom Connor |
| EPI_ISL_499796, EPI_ISL_499806 | Northumbria University / South Tees Hospitals NHS Foundation Trust / North Cumbria Integrated Care NHS Foundation Trust / North Tees and Hartlepool NHS Foundation Trust / Newcastle Hospitals NHS Foundation Trust | COVID-19 Genomics UK (COG-UK) Consortium | Darren L Smith,Andrew Nelson,Matthew Bashton,Greg R Young,Joshua Loh,John Allan,Mohammad A Tariq,Giles S Holt,Gary Black,Wen C Yew,Lynn Dover,Paul Baker,Steve Liggett,Sarah Essex,Jane Greenaway,Debra Padgett,Clive Graham,Garren Scott,Edward Barton,Emma Swindells,Brendan Payne,Jennifer Collins,Yusri Taha,Gary Eltringham |
| EPI_ISL_500367 | Servicio de Microbiología, Hospital Miguel Servet, Zaragoza | SeqCOVID-SPAIN consortium/IBV(CSIC) | Antonio Rezusta López, Alexander Tristancho Baró, Ana Milagro, Yolanda Gracia Grataloup, Nieves Martínez Cameo and SeqCOVID-SPAIN consortium |
| EPI_ISL_500703 | Area of Virology, Serology and Virology Division (SAViD), New South Wales Health Pathology Randwick | Area of Virology, Serology and Virology Division (SAViD), New South Wales Health Pathology Randwick | Rawlinson, W. |
| EPI_ISL_500821, EPI_ISL_500822, EPI_ISL_500823, EPI_ISL_500824, EPI_ISL_500825, EPI_ISL_500826, EPI_ISL_500827, EPI_ISL_500828, EPI_ISL_500829 | National Institute for Biological Standards and Control | National Institute for Biological Standards and Control | Javier Martin, Dimitra Klapsa, Thomas Wilton |
| EPI_ISL_500960, EPI_ISL_500961, EPI_ISL_500964, EPI_ISL_500970, EPI_ISL_500971, EPI_ISL_500973, EPI_ISL_500974, EPI_ISL_500977, EPI_ISL_500983, EPI_ISL_500988, EPI_ISL_500990, EPI_ISL_500991, EPI_ISL_500992, EPI_ISL_500995, EPI_ISL_500996, EPI_ISL_501002, EPI_ISL_501006, EPI_ISL_501007, EPI_ISL_501008, EPI_ISL_501011, EPI_ISL_501012, EPI_ISL_501014, EPI_ISL_501016, EPI_ISL_501017, EPI_ISL_501019, EPI_ISL_501031, EPI_ISL_501042, EPI_ISL_501043, EPI_ISL_501044, EPI_ISL_501047, EPI_ISL_501049, EPI_ISL_501052, EPI_ISL_501054, EPI_ISL_501055, EPI_ISL_501057, EPI_ISL_501060, EPI_ISL_501062, EPI_ISL_501066 | | | |
| see above | Regional Virus Laboratory, Belfast Health and Social Care Trust | Wellcome Sanger Institute for the COVID-19 Genomics UK (COG-UK) consortium | Conall McCaughey, James McKenna, Tanya Curran, Susan Feeney, Alison Watt, Ciara Cox, Mairead Connor, Zoltan Molnar, David Simpson, Derek Fairley; and Alex Alderton, Roberto Amato, Sonia Goncalves, Ewan Harrison, David K. Jackson, Ian Johnston, Dominic Kwiatkowski, Cordelia Langford, John Sillitoe on behalf of the Wellcome Sanger Institute COVID-19 Surveillance Team (http://www.sanger.ac.uk/covid-team) |
| EPI_ISL_501155, EPI_ISL_501156, EPI_ISL_501157, EPI_ISL_501158, EPI_ISL_501159, EPI_ISL_501160 | University of Washington Virology Lab | University of Washington Virology Lab | Pavitra Roychoudhury, Hong Xie, Lasata Shrestha, Amin Addetia, Truong Nguyen, Victoria M Rachleff, Meei-Li Huang, Keith R Jerome, Alexander Greninger |
| EPI_ISL_501234, EPI_ISL_501235, EPI_ISL_501236, EPI_ISL_501237, EPI_ISL_501238, EPI_ISL_501250 | Hellenic Pasteur Institute, National Influenza Reference laboratory of Southern Greece & Unit of Bioinformatics and Applied Genomics | Hellenic Pasteur Institute, National Influenza Reference laboratory of Southern Greece & Unit of Bioinformatics and Applied Genomics | Vasiliki Pogka, Timokratis Karamitros, Athanasios Kossyvakis, Antonios Kalliaropoulos, Horefti Elina, Evangelidou Maria, Androniki Voulgari-Kokota, Aspasia Kontou, Andreas Mentis |
| EPI_ISL_501617, EPI_ISL_501627, EPI_ISL_501630 | Lab Microbiology, Pathology Department, William Harvey Hospital | Wellcome Sanger Institute for the COVID-19 Genomics UK (COG-UK) consortium | Samuel Moses, Hannah Lowe, Felicity Ryan and Alex Alderton, Roberto Amato, Sonia Goncalves, Ewan Harrison, David K. Jackson, Ian Johnston, Dominic Kwiatkowski, Cordelia Langford, John Sillitoe on behalf of the Wellcome Sanger Institute COVID-19 Surveillance Team (http://www.sanger.ac.uk/covid-team) |
| EPI_ISL_504185 | Discovery DNA | Discovery DNA | Dustin Hittel, Marina Kerr, Leo Dimnik, Desmond Koo, Alice Li, Aneal Khan |
| EPI_ISL_507046, EPI_ISL_507101, EPI_ISL_507102, EPI_ISL_507103 | University College London Hospital | COVID-19 Genomics UK (COG-UK) Consortium | Judith Heaney, Matthew Byott, Catherine Houlihan, Dan Frampton, Stuart Kirk, Moira Spyer and Eleni Nastouli |
| EPI_ISL_507184, EPI_ISL_507185, EPI_ISL_507186, EPI_ISL_507187, EPI_ISL_507188, EPI_ISL_507189, EPI_ISL_507190, EPI_ISL_507191, EPI_ISL_507192, EPI_ISL_507193, EPI_ISL_507194, EPI_ISL_507195, EPI_ISL_507196 | | | |
| see above | Virology Department, Royal Infirmary of Edinburgh, NHS Lothian / School of Biological Sciences, University of Edinburgh / Institute of Genetics and Molecular Medicine, University of Edinburgh | COVID-19 Genomics UK (COG-UK) Consortium | McHugh M, Dewar R, Rooke S, Gallagher M, Balcaza C, O'Toole Á, Scher E, Hill V, McCrone JT, Colquhoun R, Yu X, Jackson B, Rambaut A, Williams TC, Templeton K |
| EPI_ISL_507213, EPI_ISL_507214, EPI_ISL_507215 | Department of Experimental Modeling and Pathogenesis of Infectious Diseases | WHO National Influenza Centre Russian Federation | Andrey Komissarov, Artem Fadeev, Mariia Sergeeva, Anna Ivanova, Daria Danilenko |
| EPI_ISL_507256, EPI_ISL_507257, EPI_ISL_507258, EPI_ISL_507259, EPI_ISL_507260, EPI_ISL_507261, EPI_ISL_507262, EPI_ISL_507263, EPI_ISL_507264, EPI_ISL_507265, EPI_ISL_507266, EPI_ISL_507267, EPI_ISL_507268, EPI_ISL_507269, EPI_ISL_507270, EPI_ISL_507271, EPI_ISL_507272, EPI_ISL_507274, EPI_ISL_507275 | | | |
| see above | WHO National Influenza Centre Russian Federation | WHO National Influenza Centre Russian Federation | Andrey Komissarov, Artem Fadeev, Mariia Sergeeva, Anna Ivanova, Daria Danilenko |
| EPI_ISL_507765, EPI_ISL_507769, EPI_ISL_507772 | Michigan Department of Health and Human Services, Bureau of Laboratories | Michigan Department of Health and Human Services, Bureau of Laboratories | Blankenship HM, Riner D, Soehnlen MK |
| EPI_ISL_507985, EPI_ISL_507986, EPI_ISL_507987, EPI_ISL_507988, EPI_ISL_507989, EPI_ISL_507990, EPI_ISL_507991, EPI_ISL_507992, EPI_ISL_507993, EPI_ISL_507994, EPI_ISL_507999, EPI_ISL_508000, EPI_ISL_508002, EPI_ISL_508003, EPI_ISL_508004, EPI_ISL_508006, EPI_ISL_508007, EPI_ISL_508008, EPI_ISL_508011, EPI_ISL_508013, EPI_ISL_508014, EPI_ISL_508016, EPI_ISL_508019, EPI_ISL_508024, EPI_ISL_508025, EPI_ISL_508026, EPI_ISL_508027, EPI_ISL_508028, EPI_ISL_508029, EPI_ISL_508030, EPI_ISL_508032, EPI_ISL_508033, EPI_ISL_508037, EPI_ISL_508044, EPI_ISL_508045, EPI_ISL_508046, EPI_ISL_508047, EPI_ISL_508048, EPI_ISL_508050, EPI_ISL_508051, EPI_ISL_508052, EPI_ISL_508054, EPI_ISL_508055, EPI_ISL_508056, EPI_ISL_508057, EPI_ISL_508058, EPI_ISL_508059, EPI_ISL_508062, EPI_ISL_508063, EPI_ISL_508068, EPI_ISL_508069, EPI_ISL_508071, EPI_ISL_508073, EPI_ISL_508078, EPI_ISL_508079, EPI_ISL_508081, EPI_ISL_508088, EPI_ISL_508092, EPI_ISL_508093, EPI_ISL_508094, EPI_ISL_508095, EPI_ISL_508096, EPI_ISL_508097, EPI_ISL_508098, EPI_ISL_508099, EPI_ISL_508100, EPI_ISL_508101, EPI_ISL_508102, EPI_ISL_508103, EPI_ISL_508104, EPI_ISL_508105, EPI_ISL_508106, EPI_ISL_508107, EPI_ISL_508108, EPI_ISL_508109, EPI_ISL_508110, EPI_ISL_508111, EPI_ISL_508112, EPI_ISL_508113, EPI_ISL_508114, EPI_ISL_508115, EPI_ISL_508116, EPI_ISL_508117, EPI_ISL_508118, EPI_ISL_508119, EPI_ISL_508120 | | | |
| see above | New Mexico Department of Health Scientific Laboratory Division | Center for Global Health, University of New Mexico Health Sciences Center | Daryl Domman, Kurt Schwalm, Twila Kunde, Joseph Hicks, Michael Edwards, Darrell Dinwiddie |
| EPI_ISL_508314, EPI_ISL_508324, EPI_ISL_508328 | Indian Institute of Science | National Institute of Biomedical Genomics | Arindam Maitra, Bharath K Sundararaj, Harsha Raheja, N. Srinivasan, Deepak K Saini, Amit Singh, Saumitra Das |
| EPI_ISL_508377, EPI_ISL_508378, EPI_ISL_508379, EPI_ISL_508380, EPI_ISL_508381, EPI_ISL_508382, EPI_ISL_508383, EPI_ISL_508384, EPI_ISL_508386, EPI_ISL_508387, EPI_ISL_508388, EPI_ISL_508389, EPI_ISL_508390, EPI_ISL_508391, EPI_ISL_508392, EPI_ISL_508393, EPI_ISL_508395, EPI_ISL_508396, EPI_ISL_508397 | | | |
| see above | Institute of Post Graduate Medical Education & Research | National Institute of Biomedical Genomics | Arindam Maitra, Aritra Biswas, Jayeeta Haldar, Raja Ray, Monimoy Banerjee, Saumitra Das |

| | | | |
|---|---|---|---|
| EPI_ISL_508425, EPI_ISL_508426, EPI_ISL_508427 | Mahatma Gandhi Institute of Medical Sciences | National Institute of Biomedical Genomics | Arindam Maitra, Vijayshri Deotale, Rahul Narang, Deepashri Maraskolhe, Saumitra Das |
| EPI_ISL_508504, EPI_ISL_508506 | Translational Health Science and Technology Institute | National Institute of Biomedical Genomics | Arindam Maitra, Guruprasad Medigeshi, Sharanabasava Patil, Anbalagan Ananthraj, Madhu Pareek, Imran Khan, Gagandeep Kang, Saumitra Das |
| EPI_ISL_509095, EPI_ISL_509096, EPI_ISL_509097, EPI_ISL_509098, EPI_ISL_509099, EPI_ISL_509100, EPI_ISL_509101, EPI_ISL_509102, EPI_ISL_509103, EPI_ISL_509104, EPI_ISL_509105, EPI_ISL_509106, EPI_ISL_509107, EPI_ISL_509108, EPI_ISL_509109, EPI_ISL_509110, EPI_ISL_509111, EPI_ISL_509121, EPI_ISL_509122, EPI_ISL_509123, EPI_ISL_509124, EPI_ISL_509125, EPI_ISL_509126, EPI_ISL_509127, EPI_ISL_509128, EPI_ISL_509129, EPI_ISL_509130, EPI_ISL_509131 | | | |
| see above | OHSU Lab Services Molecular Microbiology Lab | Oregon SARS-CoV-2 Genome Sequencing Center | Brendan L. O'Connell, Ruth V. Nichols, Sally B. Grindstaff, Alec J. Hirsch, Guang Fan, Daniel N. Streblow, William B. Messer, Andrew C. Adey, Benjamin N. Bimber, Brian J. O'Roak |
| EPI_ISL_509424, EPI_ISL_509425, EPI_ISL_509427, EPI_ISL_509428 | Microbiology and Immunology, University of South Alabama | Microbiology and Immunology, University of South Alabama | Wood,R.R., Roberts,R.A., Houserova,D., Borchert,G.M., Fouty,B., Rayner,J.O. |
| EPI_ISL_509443 | The Princess Alexandra Hospital | Wellcome Sanger Institute for the COVID-19 Genomics UK (COG-UK) consortium | Nick Levene, Louise Lopez, Lynn Monaghan, Jessica Scott, Claudia McCrea and Alex Alderton, Roberto Amato, Sonia Goncalves, Ewan Harrison, David K. Jackson, Ian Johnston, Dominic Kwiatkowski, Cordelia Langford, John Sillitoe on behalf of the Wellcome Sanger Institute COVID-19 Surveillance Team (http://www.sanger.ac.uk/covid-team) |
| EPI_ISL_509686 | M Health Fairview | Minnesota Department of Health, Public Health Laboratory | Matt Plumb, Jacob Garfin, and Xiong Wang |
| EPI_ISL_509709 | Utah Public Health Laboratory | Pathogen Discovery, Respiratory Viruses Branch, Division of Viral Diseases, Centers for Disease Control and Prevention | Jing Zhang, Ying Tao, Krista Queen, Anna Uehara, Yan Li, Clinton Paden, Haibin Wang, Suxiang Tong |
| EPI_ISL_509834, EPI_ISL_509835, EPI_ISL_509836, EPI_ISL_509837, EPI_ISL_509838, EPI_ISL_509846, EPI_ISL_509847, EPI_ISL_509848, EPI_ISL_509849, EPI_ISL_509850, EPI_ISL_509851, EPI_ISL_509933, EPI_ISL_509934, EPI_ISL_509935, EPI_ISL_509936, EPI_ISL_509937, EPI_ISL_509938, EPI_ISL_509940, EPI_ISL_509941, EPI_ISL_509942, EPI_ISL_509943, EPI_ISL_509944, EPI_ISL_509945, EPI_ISL_509946, EPI_ISL_509947, EPI_ISL_509948, EPI_ISL_509949, EPI_ISL_509950, EPI_ISL_509958, EPI_ISL_509959, EPI_ISL_509998 | | | |
| see above | University of Wisconsin-Madison AIDS Vaccine Research Laboratories | University of Wisconsin-Madison AIDS Vaccine Research Laboratories | Gage Moreno, Katarina Braun, et al. AIDS Vaccine Research Laboratories |
| EPI_ISL_510063 | Servicio de Microbiología. HRU de Málaga. Servicio Andaluz de Salud | SeqCOVID-SPAIN consortium/IBV(CSIC) | Inmaculada de Toro Peinado. MªConcepción Mediavilla Gradolph. Begoña Palop Borrás and SeqCOVID-SPAIN consortium |
| EPI_ISL_510103, EPI_ISL_510115, EPI_ISL_510125, EPI_ISL_510130, EPI_ISL_510136 | Hospital General Universitario Gregorio Marañón | SeqCOVID-SPAIN consortium/IBV(CSIC) | Laura Pérez-Lago, Marta Herranz, Jon Sicilia, Julia Suárez, Pilar Catalán, Patricia Muñoz, Darío García de Viedma and SeqCOVID-SPAIN consortium |
| EPI_ISL_510305, EPI_ISL_510306, EPI_ISL_510307, EPI_ISL_510308 | Hospital San Pedro de Alcántara (Cáceres) | SeqCOVID-SPAIN consortium/IBV(CSIC) | Cristina Muñoz Cuevas, Guadalupe Rodriguez Rodriguez and SeqCOVID-SPAIN consortium |
| EPI_ISL_510390 | Servicio de Microbiología, Hospital Miguel Servet, Zaragoza | SeqCOVID-SPAIN consortium/IBV(CSIC) | Antonio Rezusta López, Alexander Tristancho Baró, Ana Milagro, Yolanda Gracia Grataloup, Nieves Martínez Cameo and SeqCOVID-SPAIN consortium |
| EPI_ISL_510427 | Hospital Universitario Virgen de las Nieves de Granada-SAS | SeqCOVID-SPAIN consortium/IBV(CSIC) | Mercedes Pérez Ruiz, Sara Sanbonmatsu Gámez, Irene Pedrosa Corral, José M. Navarro-Marí and SeqCOVID-SPAIN consortium |
| EPI_ISL_510510, EPI_ISL_510511, EPI_ISL_510512, EPI_ISL_510513, EPI_ISL_510514, EPI_ISL_510515, EPI_ISL_510522 | Servicio de Microbiología, Laboratori Clínic Metropolitana Nord. Hospital Universitari Germans Trias i Pujol. Institut d'Investigació en Ciències de la Salut Germans Trias i Pujol (IGTP) | SeqCOVID-SPAIN consortium/IBV(CSIC) | Elisa Martró, Antoni E. Bordoy, Anna Not, Adrián Antuori, Anabel Fernández, Nona Romaní and SeqCOVID-SPAIN consortium |
| EPI_ISL_510535 | Molecular Virology, Instituto Carlos Chagas / Fiocruz Paraná | Universidade Federal do Parana (UFPR) | Suzukawa,A., Tscha,M., Zanluca,C., Raboni,S., Duarte dos Santos,C. |
| EPI_ISL_510550, EPI_ISL_510551, EPI_ISL_510552, EPI_ISL_510553, EPI_ISL_510554, EPI_ISL_510555, EPI_ISL_510556, EPI_ISL_510557, EPI_ISL_510558, EPI_ISL_510559, EPI_ISL_510560, EPI_ISL_510561, EPI_ISL_510562, EPI_ISL_510563, EPI_ISL_510564, EPI_ISL_510565, EPI_ISL_510566, EPI_ISL_510567, EPI_ISL_510568, EPI_ISL_510569, EPI_ISL_510570, EPI_ISL_510571, EPI_ISL_510572, EPI_ISL_510573, EPI_ISL_510574, EPI_ISL_510575, EPI_ISL_510576, EPI_ISL_510577, EPI_ISL_510578, EPI_ISL_510579, EPI_ISL_510580, EPI_ISL_510581, EPI_ISL_510582, EPI_ISL_510583, EPI_ISL_510584, EPI_ISL_510585, EPI_ISL_510586, EPI_ISL_510587, EPI_ISL_510588, EPI_ISL_510589, EPI_ISL_510590, EPI_ISL_510591, EPI_ISL_510592, EPI_ISL_510593, EPI_ISL_510594, EPI_ISL_510595, EPI_ISL_510596, EPI_ISL_510597, EPI_ISL_510598, EPI_ISL_510599, EPI_ISL_510600, EPI_ISL_510601, EPI_ISL_510602, EPI_ISL_510603, EPI_ISL_510604, EPI_ISL_510605, EPI_ISL_510606, EPI_ISL_510607, EPI_ISL_510608, EPI_ISL_510614 | | | |
| see above | Division of Viral Diseases, Center for Laboratory Control of Infectious Diseases, Korea Centers for Diseases Control and Prevention | Division of Viral Diseases, Center for Laboratory Control of Infectious Diseases, Korea Centers for Diseases Control and Prevention | Jeong-Min Kim, Yoon-Seok Chung, Namjoo Lee, Sang Hee Woo, Hye-Jun Jo, Heui Man Kim, Jun-Sub Kim, Myung Guk Han |
| EPI_ISL_510823, EPI_ISL_510824 | Klinisk mikrobiologi centralsjukhuset Karlstad | The Public Health Agency of Sweden | Oskar Karlsson Lindsjo, Maria Lind Karlberg, Mattias Haukland, Reza Advani, Olov Svartstrom, Anna-Malin Linde, Sandra Broddesson, Petra Edquist, Mia Brytting, Anna Risberg, Karin Tegmark-Wisell |
| EPI_ISL_510825, EPI_ISL_510826 | Kalmar klinisk mikrobiologi | The Public Health Agency of Sweden | Oskar Karlsson Lindsjo, Maria Lind Karlberg, Mattias Haukland, Reza Advani, Olov Svartstrom, Anna-Malin Linde, Sandra Broddesson, Petra Edquist, Mia Brytting, Anna Risberg, Karin Tegmark-Wisell |
| EPI_ISL_510829 | Unilabs Eskilstuna | The Public Health Agency of Sweden | Oskar Karlsson Lindsjo, Maria Lind Karlberg, Mattias Haukland, Reza Advani, Olov Svartstrom, Anna-Malin Linde, Sandra Broddesson, Petra Edquist, Mia Brytting, Anna Risberg, Karin Tegmark-Wisell |
| EPI_ISL_510832, EPI_ISL_510833 | Kalmar klinisk mikrobiologi | The Public Health Agency of Sweden | Oskar Karlsson Lindsjo, Maria Lind Karlberg, Mattias Haukland, Reza Advani, Olov Svartstrom, Anna-Malin Linde, Sandra Broddesson, Petra Edquist, Mia Brytting, Anna Risberg, Karin Tegmark-Wisell |
| EPI_ISL_511030, EPI_ISL_511104, EPI_ISL_511105, EPI_ISL_511142, EPI_ISL_511143, EPI_ISL_511683, EPI_ISL_511684 | Instituto Nacional de Saude (INSA) | Instituto Nacional de Saude (INSA) | Borges et al |
| EPI_ISL_511902, EPI_ISL_511904 | Institute of Post Graduate Medical Education & Research | National Institute of Biomedical Genomics - DBT's PAN-INDIA 1000 SARS--CoV-2 RNA Genome Sequencing Consortium | Arindam Maitra, Aritra Biswas, Jayeeta Haldar, Raja Ray, Monimoy Banerjee, Saumitra Das |
| EPI_ISL_511924, EPI_ISL_511925 | Mahatma Gandhi Institute of Medical Sciences | National Institute of Biomedical Genomics - DBT's PAN-INDIA 1000 SARS--CoV-2 RNA Genome Sequencing Consortium | Arindam Maitra, Vijayshri Deotale, Rahul Narang, Deepashri Maraskolhe, Saumitra Das |
| EPI_ISL_511948 | Indian Institute of Science | National Institute of Biomedical Genomics - DBT's PAN-INDIA 1000 SARS--CoV-2 RNA Genome Sequencing Consortium | Arindam Maitra, Bharath K Sundararaj, Harsha Raheja, N. Srinivasan, Deepak K Saini, Amit Singh, Saumitra Das |
| EPI_ISL_512162, EPI_ISL_512165, EPI_ISL_512168, EPI_ISL_512169, EPI_ISL_512170, EPI_ISL_512172, EPI_ISL_512174, EPI_ISL_512175, EPI_ISL_512178, EPI_ISL_512180, EPI_ISL_512181, EPI_ISL_512182, EPI_ISL_512183, EPI_ISL_512184, EPI_ISL_512185, EPI_ISL_512186, EPI_ISL_512187, EPI_ISL_512189, EPI_ISL_512190, EPI_ISL_512191, EPI_ISL_512192, EPI_ISL_512193, EPI_ISL_512194, EPI_ISL_512195, EPI_ISL_512198, EPI_ISL_512201, EPI_ISL_512232 | | | |
| see above | San Diego County Public Health Laboratory | Andersen lab at Scripps Research | SEARCH Alliance San Diego with Tracy Basler, Jovan Shephard, Brett Austin |
| EPI_ISL_512296, EPI_ISL_512297 | Hematology Laboratory, Section of Molecular Diagnostics, University Clinical Centre, Medical University of Gdansk | Department of Virology, Faculty of Medicine, University of Helsinki, Helsinki, Finland | Maciej Grzybek, Marlena Robakowska, Aneta Szulc, Ewa Miosz, Olli Vapalahti, Teemu Smura |
| EPI_ISL_512336, EPI_ISL_512337, EPI_ISL_512338, EPI_ISL_512340 | University of Exeter | COVID-19 Genomics UK (COG-UK) Consortium | Ben Temperton,Aaron Jeffries,Michelle Michelsen,Joanna Warwick-Dugdale,Audrey Farbos,Robyn Manley,Stephen Michell,Jane Masoli |
| EPI_ISL_512366, EPI_ISL_512368, EPI_ISL_512369, EPI_ISL_512370, EPI_ISL_512371, EPI_ISL_512374, EPI_ISL_512375, EPI_ISL_512376, EPI_ISL_512378 | Quadram Institute Bioscience | COVID-19 Genomics UK (COG-UK) Consortium | Dave J. Baker, Gemma L. Kay, Alp Aydin, Thanh Le-Viet, Steven Rudder, Ana P. Tedim, Anastasia Kolyva, Maria Diaz, Leonardo de Oliveira Martins, Nabil-Fareed Alikhan, Lizzie Meadows, Rachael Stanley, Ngozi Elumogo, Muhammed Yasir, Nicholas M. Thomson, Alexander J Trotter, Rachel Gilroy, Samuel Bloomfield, Claire Stuart, Andrew Bell, Reenesh Prakash, Samir Dervisevic, Alison E. Mather, John Wain, Mark Webber, Andrew J. Page, Justin O'Grady |
| EPI_ISL_512529 | Wales Specialist Virology Centre Sequencing lab: Pathogen | COVID-19 Genomics UK (COG-UK) Consortium | Catherine Moore, Johnathan Evans, Laura Gifford, Malorie Perry, Simon Cottrell, Angela Marchbank, Alec Birchley, Alexander Adams, Amy Gaskin, Bree |

| | | | |
|---|---|---|---|
| | Genomics Unit | | Gatica-Wilcox, Jason Coombes, Joel Southgate, Lauren Gilbert, Lee Graham, Nicole Pacchiarini, Sara Kumziene-Summerhayes, Sarah Taylor, Sophie Jones, Sara Rey, Matthew Bull, Joanne Watkins, Sally Corden, Tom Connor |
| EPI_ISL_512619, EPI_ISL_512626, EPI_ISL_512627, EPI_ISL_512636, EPI_ISL_512637, EPI_ISL_512641, EPI_ISL_512642, EPI_ISL_512643, EPI_ISL_512644 | National Laboratory for Influenza/Virology reference laboratory, Public Health Center of the Ministry of Health of Ukraine | Respiratory Virus Unit, Microbiology Services Colindale, Public Health England | PHE Covid Sequencing Team, Dr. Iryna Demchyshyna |
| EPI_ISL_512715, EPI_ISL_512716, EPI_ISL_512737 | PathWest Laboratory Medicine WA | PathWest Laboratory Medicine WA Microbial Surveillance Unit | PathWest Laboratory Medicine WA Microbial Surveillance Unit |
| EPI_ISL_512810 | National Laboratory for Influenza/Virology reference laboratory, Public Health Center of the Ministry of Health of Ukraine | Respiratory Virus Unit, Microbiology Services Colindale, Public Health England | PHE Covid Sequencing Team, Dr. Iryna Demchyshyna |
| EPI_ISL_513170, EPI_ISL_513171, EPI_ISL_513172, EPI_ISL_513174 | Pathogen Genomics Lab King Abdullah University of Science and Technology(KAUST) | Pathogen Genomics Lab King Abdullah University of Science and Technology(KAUST) | Sara Mfarrej, Raeece Naeem, Rahul P Salunke, Sharif Hala, Fadwa Alofi, Amit Kumar Subudhi, Fathia Ben Rached, Afrah Alsomali, Jumana Taha, Abdulaziz Alahmadi, Asim Khogeer, Nashwa Al-khotani, Anwar Hashem, Naif Almontashiri, Arnab Pain |
| EPI_ISL_513177, EPI_ISL_513178, EPI_ISL_513179, EPI_ISL_513180 | Pathogen Genomics Lab King Abdullah University of Science and Technology(KAUST) | Pathogen Genomics Lab King Abdullah University of Science and Technology(KAUST) | Amit Kumar Subudhi, Rahul P Salunke, Sara Mfarrej, Sharif Hala, Fadwa Alofi, Fathia Ben Rached, Afrah Alsomali, Asim Khogeer, Nashwa Al-khotani, Raeece Naeem, Anwar Hashem, Naif Almontashiri, Arnab Pain |
| EPI_ISL_513195 | Pathogen Genomics Lab King Abdullah University of Science and Technology(KAUST) | Pathogen Genomics Lab King Abdullah University of Science and Technology(KAUST) | Afrah Alsomali, Fathia Ben Rached, Raeece Naeem, Sharif Hala,Rahul P Salunke, Amanda Ooi, Luke Esau, Sara Mfarrej, Amit Kumar Subudhi, Fadwa Alofi, Asim Khogeer, Kahled Algithami, Anwar Hashem, Naif Almontashiri, Arnab Pain |
| EPI_ISL_513198, EPI_ISL_513205, EPI_ISL_513206 | Pathogen Genomics Lab King Abdullah University of Science and Technology(KAUST) | Pathogen Genomics Lab King Abdullah University of Science and Technology(KAUST) | Rahul P Salunke, Sharif Hala, Raeece Naeem, Sara Mfarrej, Amit Kumar Subudhi, Amanda Ooi, Luke Esau, Fadwa Alofi, Fathia Ben Rached, Afrah Alsomali, Asim Khogeer, Ahmad Bakur Mahmoud, Anwar Hashem, Naif Almontashiri, Arnab Pain |
| EPI_ISL_513208, EPI_ISL_513209, EPI_ISL_513211 | Pathogen Genomics Lab King Abdullah University of Science and Technology(KAUST) | Pathogen Genomics Lab King Abdullah University of Science and Technology(KAUST) | Raeece Naeem, Rahul P Salunke, Sharif Hala, Sara Mfarrej, Amit Kumar Subudhi, Fadwa Alofi, Fathia Ben Rached, Afrah Alsomali, Asim Khogeer, Ahmad Bakur Mahmoud, Anwar Hashem, Naif Almontashiri, Arnab Pain |
| EPI_ISL_513581, EPI_ISL_513582, EPI_ISL_513583 | Programa de Oncovirologia, Instituto Nacional de Câncer | Programa de Oncovirologia, Instituto Nacional de Câncer | Juliana D. Siqueira, Livia R. Goes, Brunna M. Alves, Claudia Cicala,James Arthos, João P.B. Viola, Andreia C. de Melo, Marcelo A. Soares |
| EPI_ISL_513592, EPI_ISL_513593, EPI_ISL_513594, EPI_ISL_513595, EPI_ISL_513596, EPI_ISL_513612, EPI_ISL_513613, EPI_ISL_513614, EPI_ISL_513617, EPI_ISL_513618, EPI_ISL_513620, EPI_ISL_513621, EPI_ISL_513622, EPI_ISL_513623 | | | |
| see above | Viral Respiratory Lab, National Institute for Biomedical Research (INRB) | Pathogen Sequencing Lab, National Institute for Biomedical Research (INRB) | Placide Mbala-Kingebeni, Edith Nkwembe, Eddy Kinganda-Lusamaki, Amuri Aziza, Francisca Muyembe Mawete, Emmanuel Lokilo Lofiko, Catherine Pratt, Matthias Pauthner, Josh Quick, Allison Black, James Hadfield, Trevor Bedford, Ian Goodfellow, Andrew Rambaut, Nick Loman, Kristian Andersen, Michael Wiley, Steve Ahuka-Mundeke, Jean-Jacques Muyembe Tamfum |
| EPI_ISL_513788 | County of Santa Clara Public Health Department | Chan-Zuckerberg Biohub | CZB Cliahub Consortium |
| EPI_ISL_513789, EPI_ISL_513790, EPI_ISL_513791, EPI_ISL_513792, EPI_ISL_513793, EPI_ISL_513794, EPI_ISL_513795, EPI_ISL_513796, EPI_ISL_513797, EPI_ISL_513798, EPI_ISL_513799, EPI_ISL_513800, EPI_ISL_513801, EPI_ISL_513802, EPI_ISL_513803, EPI_ISL_513804, EPI_ISL_513805, EPI_ISL_513806, EPI_ISL_513807, EPI_ISL_513808, EPI_ISL_513809, EPI_ISL_513810, EPI_ISL_513811, EPI_ISL_513812, EPI_ISL_513813, EPI_ISL_513814, EPI_ISL_513815, EPI_ISL_513816, EPI_ISL_513817, EPI_ISL_513818 | | | |
| see above | Orange County Public Health Laboratory | Chan-Zuckerberg Biohub | CZB Cliahub Consortium |
| EPI_ISL_513840, EPI_ISL_513841, EPI_ISL_513845, EPI_ISL_513855 | Humboldt County Public Health Laboratory | Chan-Zuckerberg Biohub | CZB Cliahub Consortium |
| EPI_ISL_513889, EPI_ISL_513890, EPI_ISL_513891, EPI_ISL_513903, EPI_ISL_513908, EPI_ISL_513909, EPI_ISL_513910 | UCSF Clinical Microbiology Laboratory | Chan-Zuckerberg Biohub | CZB Cliahub Consortium |
| EPI_ISL_513925 | Microbiology & Bioinformatics and Biostatistics, Kohat University of Science and Technology (Pakistan) & Shanghai Jiao Tong University (China) | Microbiology & Bioinformatics and Biostatistics, Kohat University of Science and Technology (Pakistan) & Shanghai Jiao Tong University (China) | Khan,M.T., Khan,T.A., Ali,S., Khan,A.S., Muhammad,N. and Wei,D.Q. |
| EPI_ISL_514109, EPI_ISL_514110, EPI_ISL_514111, EPI_ISL_514112, EPI_ISL_514113, EPI_ISL_514120, EPI_ISL_514121, EPI_ISL_514122, EPI_ISL_514123, EPI_ISL_514124, EPI_ISL_514125 | | | |
| see above | Viral Respiratory Lab, National Institute for Biomedical Research (INRB) | Pathogen Sequencing Lab, National Institute for Biomedical Research (INRB) | Placide Mbala-Kingebeni, Edith Nkwembe, Eddy Kinganda-Lusamaki, Amuri Aziza, Francisca Muyembe Mawete, Emmanuel Lokilo Lofiko, Catherine Pratt, Matthias Pauthner, Josh Quick, Allison Black, James Hadfield, Trevor Bedford, Ian Goodfellow, Andrew Rambaut, Nick Loman, Kristian Andersen, Michael Wiley, Steve Ahuka-Mundeke, Jean-Jacques Muyembe Tamfum |
| EPI_ISL_514131 | Rondônia Central Public Health Laboratory (LACEN/RO), vinculated to State Health Secretariat of Rondônia (SESAU/RO) | Molecular Virology Laboratory of Oswaldo Cruz Foundation of Rondônia | Luan Felipo Botelho-Souza, Felipe Souza Nogueira-Lima, Tárcio Peixoto Roca, Alcione de Oliveira dos Santos, Felipe Gomes Naveca, Adriana Cristina Salvador Maia, Cicileia Correia da Silva, Aline Linhares Ferreira de Melo Mendonça, Celina Aparecida Bertoni Lugtenburg, Camila Flávia Gomes Azzi, Juliana Loca Furtado, Suelen Cavalcante, Rita de Cássia Pontello Rampazzo, Caio Henrique Nemeth Santos, Alice Paula Di Sabatino Guimarães, Jansen Fernandes de Medeiros, Fernando Rodrigues Máximo, Juan Miguel Vilallobos-Salcedo and Deusilene Souza Vieira1 |
| EPI_ISL_514132 | Rondônia Central Public Health Laboratory (LACEN/RO), vinculated to State Health Secretariat of Rondônia (SESAU/RO) | Molecular Virology Laboratory of Oswaldo Cruz Foundation of Rondônia | Luan Felipo Botelho-Souza, Felipe Souza Nogueira-Lima, Tárcio Peixoto Roca, Alcione de Oliveira dos Santos, Felipe Gomes Naveca, Adriana Cristina Salvador Maia, Cicileia Correia da Silva, Aline Linhares Ferreira de Melo Mendonça, Celina Aparecida Bertoni Lugtenburg, Camila Flávia Gomes Azzi, Juliana Loca Furtado, Suelen Cavalcante, Rita de Cássia Pontello Rampazzo, Caio Henrique Nemeth Santos, Alice Paula Di Sabatino Guimarães, Jansen Fernandes de Medeiros, Fernando Rodrigues Máximo, Juan Miguel Vilallobos-Salcedo and Deusilene Souza Vieira. |
| EPI_ISL_514133, EPI_ISL_514134, EPI_ISL_514135, EPI_ISL_514136, EPI_ISL_514137 | Rondônia Central Public Health Laboratory (LACEN/RO), vinculated to State Health Secretariat of Rondônia (SESAU/RO) | Molecular Virology Laboratory of Oswaldo Cruz Foundation of Rondônia | Luan Felipo Botelho-Souza, Felipe Souza Nogueira-Lima, Tárcio Peixoto Roca, Alcione de Oliveira dos Santos, Felipe Gomes Naveca, Adriana Cristina Salvador Maia, Cicileia Correia da Silva, Aline Linhares Ferreira de Melo Mendonça, Celina Aparecida Bertoni Lugtenburg, Camila Flávia Gomes Azzi, Juliana Loca Furtado, Suelen Cavalcante, Rita de Cássia Pontello Rampazzo, Caio Henrique Nemeth Santos, Alice Paula Di Sabatino Guimarães, Jansen Fernandes de Medeiros, Fernando Rodrigues Máximo, Juan Miguel Vilallobos-Salcedo and Deusilene Souza Vieira |
| EPI_ISL_514149, EPI_ISL_514150, EPI_ISL_514151 | Florida Bureau of Public Health Laboratories | Florida Bureau of Public Health Laboratories | Sarah Schmedes, Jason Blanton |
| EPI_ISL_514426 | Laboratory Diagnostic, Veterinary Specialized Institute Kraljevo | Laboratory Diagnostic, Veterinary Specialized Institute Kraljevo | Vidanovic,D., Tesovic,B., Knezevic,A., Jankovic,M., Sekler,M., Banovic Djeri,B., Volkening,J., Afonso,C., Petrovic,T. |
| EPI_ISL_514618 | Minnesota Department of Health, Public Health Laboratory | Minnesota Department of Health, Public Health Laboratory | Matt Plumb, Jacob Garfin, and Xiong Wang |
| EPI_ISL_515250, EPI_ISL_515251, EPI_ISL_515252, EPI_ISL_515253, EPI_ISL_515254, EPI_ISL_515255, EPI_ISL_515256, EPI_ISL_515257, EPI_ISL_515258, EPI_ISL_515259, EPI_ISL_515260, EPI_ISL_515261 | | | |
| see above | Texas Department of State Health Services | Texas Department of State Health Services | Rashmi Tuladhar, Bonnie Oh, Cara Akrout, Jenny Zhang, Maliha Rahman, Anita Pokharel, Myong Koag, Chun Wang, Rachel Lee, Grace Kubin |
| EPI_ISL_515415, EPI_ISL_515416, EPI_ISL_515417, EPI_ISL_515418, EPI_ISL_515419, EPI_ISL_515420, EPI_ISL_515421, EPI_ISL_515422, EPI_ISL_515423, EPI_ISL_515424, EPI_ISL_515425, EPI_ISL_515426, EPI_ISL_515427, EPI_ISL_515461 | | | |
| see above | Nevada State Public Health Laboratory | Nevada State Public Health Laboratory | Richard Tillett, Joel R. Sevinsky, Paul Hartley, Heather Kerwin, David Jackson, Subhash C. Verma, Cyprian Rossetto, Andrew Gorzalski, Chris Laverdure, Natalie Crawford, Stephanie Van Hooser, and Mark Pandori |
| EPI_ISL_515950, EPI_ISL_515951, EPI_ISL_515952, EPI_ISL_515953 | VICTORIA HOSPITAL | Department of Neurovirology, National Institute of Mental Health and Neuroscience (NIMHANS) | Chitra Pattabiraman,Vijayalakshmi Reddy, Harsha PK, Risha Rasheed, Pramada Prasad, Shafeeq S Hameed, Manjunatha Venkataswamy, Anita Desai, Ravi Vasanthapuram |
| EPI_ISL_515954, EPI_ISL_515955 | DH | Department of Neurovirology, National Institute of Mental Health and Neuroscience (NIMHANS) | Chitra Pattabiraman,Vijayalakshmi Reddy, Harsha PK, Risha Rasheed, Pramada Prasad, Shafeeq S Hameed, Manjunatha Venkataswamy, Anita Desai, Ravi Vasanthapuram |
| EPI_ISL_515956 | SHEKAR HOSPITAL | Department of Neurovirology, National Institute of Mental | Chitra Pattabiraman,Vijayalakshmi Reddy, Harsha PK, Risha Rasheed, Pramada Prasad, Shafeeq S Hameed, Manjunatha Venkataswamy, Anita Desai, |

| | | | |
|---|---|---|---|
| | | Health and Neuroscience (NIMHANS) | Ravi Vasanthapuram |
| EPI_ISL_515957, EPI_ISL_515958, EPI_ISL_515959 | DH | Department of Neurovirology, National Institute of Mental Health and Neuroscience (NIMHANS) | Chitra Pattabiraman,Vijayalakshmi Reddy, Harsha PK, Risha Rasheed, Pramada Prasad, Shafeeq S Hameed, Manjunatha Venkataswamy, Anita Desai, Ravi Vasanthapuram |
| EPI_ISL_515960 | JGH | Department of Neurovirology, National Institute of Mental Health and Neuroscience (NIMHANS) | Chitra Pattabiraman,Vijayalakshmi Reddy, Harsha PK, Risha Rasheed, Pramada Prasad, Shafeeq S Hameed, Manjunatha Venkataswamy, Anita Desai, Ravi Vasanthapuram |
| EPI_ISL_515961, EPI_ISL_515962 | DH | Department of Neurovirology, National Institute of Mental Health and Neuroscience (NIMHANS) | Chitra Pattabiraman,Vijayalakshmi Reddy, Harsha PK, Risha Rasheed, Pramada Prasad, Shafeeq S Hameed, Manjunatha Venkataswamy, Anita Desai, Ravi Vasanthapuram |
| EPI_ISL_515963, EPI_ISL_515964 | VICTORIA HOSPITAL | Department of Neurovirology, National Institute of Mental Health and Neuroscience (NIMHANS) | Chitra Pattabiraman,Vijayalakshmi Reddy, Harsha PK, Risha Rasheed, Pramada Prasad, Shafeeq S Hameed, Manjunatha Venkataswamy, Anita Desai, Ravi Vasanthapuram |
| EPI_ISL_515965 | DH | Department of Neurovirology, National Institute of Mental Health and Neuroscience (NIMHANS) | Chitra Pattabiraman,Vijayalakshmi Reddy, Harsha PK, Risha Rasheed, Pramada Prasad, Shafeeq S Hameed, Manjunatha Venkataswamy, Anita Desai, Ravi Vasanthapuram |
| EPI_ISL_515966, EPI_ISL_515967 | VICTORIA HOSPITAL | Department of Neurovirology, National Institute of Mental Health and Neuroscience (NIMHANS) | Chitra Pattabiraman,Vijayalakshmi Reddy, Harsha PK, Risha Rasheed, Pramada Prasad, Shafeeq S Hameed, Manjunatha Venkataswamy, Anita Desai, Ravi Vasanthapuram |
| EPI_ISL_515968, EPI_ISL_515969, EPI_ISL_515970, EPI_ISL_515971, EPI_ISL_515972, EPI_ISL_515973 | MIMS | Department of Neurovirology, National Institute of Mental Health and Neuroscience (NIMHANS) | Chitra Pattabiraman,Vijayalakshmi Reddy, Harsha PK, Risha Rasheed, Pramada Prasad, Shafeeq S Hameed, Manjunatha Venkataswamy, Anita Desai, Ravi Vasanthapuram |
| EPI_ISL_516622 | Instituto de Diagnostico y Referencia Epidemiologicos (INDRE) | Instituto de Diagnostico y Referencia Epidemiologicos (INDRE) | Gisela Barrera-Badillo , Abril Rodriguez-Maldonado, Claudia Wong-Arambula , Natividad Cruz-Ortiz, Tatiana Nunez-Garcia, Dayanira Arellano-Suarez, Fabiola Garces-Ayala, Edgar Mendieta-Condado, Lucia Hernandez-Rivas, Irma Lopez-Martinez, Ernesto Ramirez-Gonzalez. |
| EPI_ISL_516677, EPI_ISL_516678, EPI_ISL_516679, EPI_ISL_516680, EPI_ISL_516681, EPI_ISL_516682, EPI_ISL_516683, EPI_ISL_516684, EPI_ISL_516692, EPI_ISL_516693, EPI_ISL_516694, EPI_ISL_516695, EPI_ISL_516696, EPI_ISL_516697, EPI_ISL_516698, EPI_ISL_516699, EPI_ISL_516700, EPI_ISL_516701, EPI_ISL_516702, EPI_ISL_516703, EPI_ISL_516704, EPI_ISL_516705, EPI_ISL_516706, EPI_ISL_516707, EPI_ISL_516708, EPI_ISL_516709, EPI_ISL_516710, EPI_ISL_516711 | | | |
| see above | Virginia DCLS | Virginia DCLS | Virginia DCLS |
| EPI_ISL_516806 | Rumah Sakit PKU Gamping | Genetics Working Group (Pokja Genetik) Faculty of Medicine, Public Health and Nursing Universitas Gadjah Mada (FK-KMK UGM); Disease Investigation Center Wates Ministry of Agriculture Indonesia; Department of Microbiology FK-KMK UGM; Laboratorium Diagnostik Yayasan Tahija World Mosquito Program (WMP) Yogyakarta Center for Tropical Medicine FK-KMK UGM; Integrated Research center FK-KMK UGM; Department of Computer Science and Electronics FMIPA UGM | Gunadi, Hendra Wibawa, . Marcellus, Mohamad S. Hakim, Edwin W. Daniwijaya, Ludhang P. Rizki, Endah Supriyati, Eggi Arguni, Titik Nuryastuti, Tri Wibawa, Dwi AA Nugrahaningsih, . Afiahayati , . Siswanto, Ardorisye Saptaty Fornia, Kemala Athollah |
| EPI_ISL_516947, EPI_ISL_516949, EPI_ISL_516950, EPI_ISL_516951, EPI_ISL_516952, EPI_ISL_516953, EPI_ISL_516954, EPI_ISL_516955, EPI_ISL_516956, EPI_ISL_516957, EPI_ISL_516958, EPI_ISL_516959, EPI_ISL_516963, EPI_ISL_516964, EPI_ISL_516965, EPI_ISL_516966, EPI_ISL_516967, EPI_ISL_516968 | | | |
| see above | King Georges Medical University | CSIR-National Botanical Research Institute | Priti Prasad, Shantanu Prakash, Kishan Sahu, Babita Singh, Suruchi Shukla, Hricha Mishra, Danish Nasar Khan , Om Prakash, MLB Bhatt, SK Barik, Mehar H.Asif,Samir V. Sawant,Amita Jain, Sumit Kr. Bag |
| EPI_ISL_517460 | Liverpool Clinical Laboratories | COVID-19 Genomics UK (COG-UK) Consortium | Sam Haldenby, Anita Lucaci, Steve Paterson, Julian Hiscox, Alistair Darby, M Almsaud, A Alrezaihi, Muhannad Alruwaili, Stuart D Armstrong, Jones Benjamin, Eleanor G Bentley, Anu Chawla, Jordan J Clark, Angela Cowell, Richard Eccles, Isabel García-Dorival, Matthew Gemmell, Alessandro Gerada, PKF Gilmore, Richard Gregory, Ximeng Han, Catherine Hartley, Margaret Hughes, Miren Iturriza-Gomara, James Johnson, L Luu, Jenifer Manson, Charlotte Nelson, Elaine O'Toole, Cassie Olateju, Rebekah Penrice-Randal , Lucille Rainbow, N.P Randle, Trevor Ian Robinson, Parul Sharma, Ghada T Shawli, James P Stewart, Neil Swainston, Ecaterina Vamos, Joanne Watts, Mark Whitehead |
| EPI_ISL_517616 | Academic Hospital Paramaribo | Erasmus Medical Center | Bas Oude Munnink, Dion Gajadin, Ed Ijzerman, Emmanuelle Munger, Gary Gummels, Ingrid Krishnadath, Lycke Woittiez, Marion Koopmans, Mireille Van de Veer, Princes Wongsowidjojo, Radjesh Ori, Rohma Banwari, Stephen Vreden |
| EPI_ISL_517775, EPI_ISL_517778, EPI_ISL_517940, EPI_ISL_517941, EPI_ISL_517942 | Florida Bureau of Public Health Laboratories | Florida Bureau of Public Health Laboratories | Sarah Schmedes, Jason Blanton |
| EPI_ISL_517960, EPI_ISL_517961, EPI_ISL_517962, EPI_ISL_517963, EPI_ISL_517964, EPI_ISL_517965, EPI_ISL_517966, EPI_ISL_517967, EPI_ISL_517968, EPI_ISL_517969, EPI_ISL_517970, EPI_ISL_517971, EPI_ISL_517972, EPI_ISL_517973, EPI_ISL_517974, EPI_ISL_517975, EPI_ISL_517976, EPI_ISL_517977, EPI_ISL_517978, EPI_ISL_517979, EPI_ISL_517980, EPI_ISL_517981, EPI_ISL_517982, EPI_ISL_517983, EPI_ISL_517984, EPI_ISL_517985, EPI_ISL_517986, EPI_ISL_517987, EPI_ISL_517988, EPI_ISL_517989, EPI_ISL_517990, EPI_ISL_517991, EPI_ISL_517992, EPI_ISL_517993, EPI_ISL_517994, EPI_ISL_517995 | | | |
| see above | Texas Department of State Health Services | Texas Department of State Health Services | Rashmi Tuladhar, Bonnie Oh, Cara Akrout, Jenny Zhang, Maliha Rahman, Anita Pokharel, Myong Koag, Chun Wang, Rachel Lee, Grace Kubin |
| EPI_ISL_518855 | Minnesota Department of Health, Public Health Laboratory | Minnesota Department of Health, Public Health Laboratory | Matt Plumb, Jacob Garfin, and Xiong Wang |
| EPI_ISL_518900 | Mayo Clinic & Mayo Clinic Laboratories | Minnesota Department of Health, Public Health Laboratory | Matt Plumb, Jacob Garfin, and Xiong Wang |
| EPI_ISL_520674, EPI_ISL_520675, EPI_ISL_520681, EPI_ISL_520682, EPI_ISL_520683, EPI_ISL_520684, EPI_ISL_520685, EPI_ISL_520686 | Mohammed Bin Rashid University of Medicine and Health Sciences | Al Jalila Genomics Center | Ahmad Abou Tayoun, Tom Loney, Hamda Khansaheb, Sathishkumar Ramaswamy, Divinlal Harilal, Zulfa Omar Deesi, Rupa Murthy Varghese, Hanan Al Suwaidi, Abdulmajeed Alkhaja, Mohammed Uddin, Rifat Hamoudi, Rabih Halwani, Abiola Catherine Senok, Qutayba Hamid, Norbert Nowotny, Alawi Alsheikh-Ali |
| EPI_ISL_521904 | Victorian Infectious Diseases Reference Laboratory (VIDRL) | VIDRL and MDU-PHL | Caly L., Seemann T., Sait, M., Schultz M., Druce J., Sherry, N. |
| EPI_ISL_522318, EPI_ISL_522319, EPI_ISL_522320, EPI_ISL_522321, EPI_ISL_522322, EPI_ISL_522323, EPI_ISL_522324, EPI_ISL_522325, EPI_ISL_522326, EPI_ISL_522327, EPI_ISL_522328, EPI_ISL_522329, EPI_ISL_522330, EPI_ISL_522331, EPI_ISL_522332, EPI_ISL_522333, EPI_ISL_522334, EPI_ISL_522335, EPI_ISL_522336, EPI_ISL_522337, EPI_ISL_522338, EPI_ISL_522339, EPI_ISL_522340 | | | |
| see above | Utah Public Health Laboratory | Utah Public Health Laboratory | Erin Young, Kelly Oakeson |
| EPI_ISL_522351, EPI_ISL_522352, EPI_ISL_522353, EPI_ISL_522354, EPI_ISL_522355, EPI_ISL_522356, EPI_ISL_522357, EPI_ISL_522358, EPI_ISL_522359, EPI_ISL_522360, EPI_ISL_522361, EPI_ISL_522362, EPI_ISL_522363, EPI_ISL_522364, EPI_ISL_522365, EPI_ISL_522366, EPI_ISL_522367, EPI_ISL_522368, EPI_ISL_522369, EPI_ISL_522370, EPI_ISL_522371 | | | |
| see above | Texas Department of State Health Services | Texas Department of State Health Services | Rashmi Tuladhar, Bonnie Oh, Cara Akrout, Jenny Zhang, Maliha Rahman, Anita Pokharel, Myong Koag, Chun Wang, Rachel Lee, Grace Kubin |
| EPI_ISL_522441, EPI_ISL_522442, EPI_ISL_522443 | Center for Laboratory Control of Infectious Diseases, Korea Centers for Diseases Control and Prevention | Center for Laboratory Control of Infectious Diseases, Korea Centers for Diseases Control and Prevention | Junyoung Kim, Ae Kyung Park, Eunkyung Shin, Jin Sun No, Jeong-Min Kim, Yoon-Seok Chung, Heui Man Kim, Myung Guk Han |
| EPI_ISL_522686, EPI_ISL_522688 | Royal Hobart Hospital Microbiology Department | MDU-PHL | Cooley L., van Haeften R., Seemann T., Sait M., Schultz, M.B., Sherry N. |
| EPI_ISL_522777, EPI_ISL_522778, EPI_ISL_522779, EPI_ISL_522780, EPI_ISL_522781, EPI_ISL_522782, EPI_ISL_522783, EPI_ISL_522784, EPI_ISL_522785, EPI_ISL_522786, EPI_ISL_522787, EPI_ISL_522788, EPI_ISL_522789, EPI_ISL_522790, EPI_ISL_522791, EPI_ISL_522792, EPI_ISL_522793, EPI_ISL_522794, EPI_ISL_522795 | | | |
| see above | Virginia DCLS | Virginia DCLS | Virginia DCLS |
| EPI_ISL_522967, EPI_ISL_522968, EPI_ISL_522969 | Texas Department of State Health Services | Texas Department of State Health Services | Rashmi Tuladhar, Bonnie Oh, Cara Akrout, Jenny Zhang, Maliha Rahman, Anita Pokharel, Myong Koag, Chun Wang, Rachel Lee, Grace Kubin |
| EPI_ISL_523091, EPI_ISL_523092, EPI_ISL_523093, EPI_ISL_523094, EPI_ISL_523095, EPI_ISL_523096 | Dutch COVID-19 response team | Erasmus Medical Center | OH consortium |
| EPI_ISL_523181, EPI_ISL_523182, EPI_ISL_523183, EPI_ISL_523184, EPI_ISL_523203, EPI_ISL_523204, EPI_ISL_523239, EPI_ISL_523303, EPI_ISL_523304, EPI_ISL_523305, EPI_ISL_523306, EPI_ISL_523307, EPI_ISL_523308, EPI_ISL_523352, EPI_ISL_523356, EPI_ISL_523357, EPI_ISL_523358, EPI_ISL_523359, | | | |

| | | | |
|---|---|---|---|
| EPI_ISL_523367, EPI_ISL_523368, EPI_ISL_523369, EPI_ISL_523370, EPI_ISL_523371, EPI_ISL_523401, EPI_ISL_523403, EPI_ISL_523406, EPI_ISL_523408, EPI_ISL_523438, EPI_ISL_523439, EPI_ISL_523440, EPI_ISL_523441, EPI_ISL_523442, EPI_ISL_523467, EPI_ISL_523469, EPI_ISL_523497, EPI_ISL_523498, EPI_ISL_523499, EPI_ISL_523547, EPI_ISL_523549, EPI_ISL_523550, EPI_ISL_523557, EPI_ISL_523569, EPI_ISL_523572, EPI_ISL_523573, EPI_ISL_523614 | | | |
| see above | Dutch COVID-19 response team | Erasmus Medical Center | Bas Oude Munnink, David Nieuwenhuijse, Reina Sikkema, Claudia Schapendonk, Irina Chestakova, Anne van der Linden, Theo Bestebroer, Stefan van Nieuwkoop, Mark Pronk, Pascal Lexmond, Corien Swaan, Manon Haverkate, Madelief Mollers, Mart Stein, Sandra Kengne Kamga Mobou, Jeroen van Kampen, Jolanda Voermans, Aura Timen, Corine GeurtsvanKessel, Annemiek van der Eijk, Richard Molenkamp, Marion Koopmans, on behalf of the Dutch national COVID-19 response team. |
| EPI_ISL_523999, EPI_ISL_524000, EPI_ISL_524001, EPI_ISL_524002, EPI_ISL_524003, EPI_ISL_524004, EPI_ISL_524005, EPI_ISL_524006, EPI_ISL_524007, EPI_ISL_524008, EPI_ISL_524009 | | | |
| see above | WHO National Influenza Centre Russian Federation | WHO National Influenza Centre Russian Federation | Andrey Komissarov, Artem Fadeev, Mariia Sergeeva, Anna Ivanova, Daria Danilenko |
| EPI_ISL_524071, EPI_ISL_524072 | Texas Department of State Health Services | Texas Department of State Health Services | Rashmi Tuladhar, Bonnie Oh, Cara Akrout, Jenny Zhang, Maliha Rahman, Anita Pokharel, Myong Koag, Chun Wang, Rachel Lee, Grace Kubin |
| EPI_ISL_524108, EPI_ISL_524109, EPI_ISL_524110, EPI_ISL_524111, EPI_ISL_524112, EPI_ISL_524113, EPI_ISL_524114, EPI_ISL_524115, EPI_ISL_524116, EPI_ISL_524117, EPI_ISL_524118, EPI_ISL_524119, EPI_ISL_524120, EPI_ISL_524121, EPI_ISL_524122, EPI_ISL_524125, EPI_ISL_524126, EPI_ISL_524127, EPI_ISL_524128, EPI_ISL_524129, EPI_ISL_524130, EPI_ISL_524132, EPI_ISL_524133, EPI_ISL_524134, EPI_ISL_524135, EPI_ISL_524136, EPI_ISL_524137, EPI_ISL_524138, EPI_ISL_524140, EPI_ISL_524141, EPI_ISL_524142, EPI_ISL_524143, EPI_ISL_524144, EPI_ISL_524151, EPI_ISL_524152, EPI_ISL_524153, EPI_ISL_524154, EPI_ISL_524155, EPI_ISL_524162, EPI_ISL_524163, EPI_ISL_524164, EPI_ISL_524165, EPI_ISL_524166, EPI_ISL_524172, EPI_ISL_524173, EPI_ISL_524174, EPI_ISL_524181, EPI_ISL_524182, EPI_ISL_524183, EPI_ISL_524190, EPI_ISL_524191, EPI_ISL_524192, EPI_ISL_524193, EPI_ISL_524194, EPI_ISL_524201, EPI_ISL_524202, EPI_ISL_524203, EPI_ISL_524210, EPI_ISL_524211, EPI_ISL_524212, EPI_ISL_524213, EPI_ISL_524303, EPI_ISL_524312, EPI_ISL_524321, EPI_ISL_524330, EPI_ISL_524337, EPI_ISL_524344, EPI_ISL_524352, EPI_ISL_524359, EPI_ISL_524363, EPI_ISL_524364, EPI_ISL_524373, EPI_ISL_524374, EPI_ISL_524375, EPI_ISL_524376, EPI_ISL_524379, EPI_ISL_524380, EPI_ISL_524381, EPI_ISL_524382, EPI_ISL_524384, EPI_ISL_524385, EPI_ISL_524386, EPI_ISL_524388, EPI_ISL_524389, EPI_ISL_524390, EPI_ISL_524391, EPI_ISL_524392, EPI_ISL_524393, EPI_ISL_524395, EPI_ISL_524396, EPI_ISL_524397, EPI_ISL_524398, EPI_ISL_524399, EPI_ISL_524402, EPI_ISL_524403, EPI_ISL_524404, EPI_ISL_524405, EPI_ISL_524406, EPI_ISL_524408, EPI_ISL_524409, EPI_ISL_524410, EPI_ISL_524411, EPI_ISL_524412, EPI_ISL_524415, EPI_ISL_524416, EPI_ISL_524417, EPI_ISL_524418, EPI_ISL_524419, EPI_ISL_524420, EPI_ISL_524423 | | | |
| see above | Utah Public Health Laboratory | Utah Public Health Laboratory | Erin L. Young, Kelly Oakeson, Tara Gallagher, Michael T. Pyne, E. Susan Slechta, Melanie A. Mallory, Jeffrey B. Stevenson, Salika M. Shakir, David R. Hillyard |
| EPI_ISL_524436 | Department of Immunology, The Scripps Research Institute | Department of Immunology, The Scripps Research Institute | Smither,A., Sabino-Santos,G., Snarski,P., Melnik,L., Bell,A., Genemaras,K., Drouin,A., Fusco,D., Garry,R., SEARCH Alliance,S.D. |
| EPI_ISL_524469 | Santa Casa de Misericordia de Sao Paulo | Instituto Adolfo Lutz, Interdiciplinary Procedures Center, Strategic Laboratory | Claudio Tavares Sacchi, Claudia Regina Gonçalves, Erica Valessa Ramos Gomes |
| EPI_ISL_524470 | Hospital do Servidor Público Estadual Francisco Morato de Oliveira | Instituto Adolfo Lutz, Interdiciplinary Procedures Center, Strategic Laboratory | Claudio Tavares Sacchi, Claudia Regina Gonçalves, Erica Valessa Ramos Gomes |
| EPI_ISL_524785, EPI_ISL_524789, EPI_ISL_524799, EPI_ISL_524800 | Evandro Chagas Institute | Evandro Chagas Institute | Santos, M.C.; Silva, A.M.; Junior, W.D.C.; Barbagelata, L.S.; Ferreira, J.A.; Sousa, E.M.A.; da Silva, P.S.; Resque, H.R; Martins, L.C.; Sousa Junior, E.C.;Viana, G.M.R |
| EPI_ISL_524904, EPI_ISL_524905, EPI_ISL_524906, EPI_ISL_524907, EPI_ISL_524908, EPI_ISL_524912, EPI_ISL_524913, EPI_ISL_524914, EPI_ISL_524915, EPI_ISL_524916, EPI_ISL_524917, EPI_ISL_524923, EPI_ISL_524924, EPI_ISL_524925, EPI_ISL_524926, EPI_ISL_524927, EPI_ISL_524932, EPI_ISL_524933, EPI_ISL_524936, EPI_ISL_524937, EPI_ISL_524939, EPI_ISL_524940, EPI_ISL_524941, EPI_ISL_524942, EPI_ISL_524947, EPI_ISL_524948, EPI_ISL_524949, EPI_ISL_524951, EPI_ISL_524952, EPI_ISL_524953, EPI_ISL_524958, EPI_ISL_524959, EPI_ISL_524961, EPI_ISL_524962, EPI_ISL_524965, EPI_ISL_524966, EPI_ISL_524967, EPI_ISL_524971, EPI_ISL_524972, EPI_ISL_524973, EPI_ISL_524993, EPI_ISL_525048, EPI_ISL_525049, EPI_ISL_525050, EPI_ISL_525051, EPI_ISL_525052, EPI_ISL_525053, EPI_ISL_525054, EPI_ISL_525055, EPI_ISL_525056, EPI_ISL_525057, EPI_ISL_525058, EPI_ISL_525059, EPI_ISL_525060, EPI_ISL_525061, EPI_ISL_525062, EPI_ISL_525063, EPI_ISL_525064, EPI_ISL_525065, EPI_ISL_525066, EPI_ISL_525067, EPI_ISL_525068, EPI_ISL_525069, EPI_ISL_525070, EPI_ISL_525071, EPI_ISL_525072, EPI_ISL_525073, EPI_ISL_525074, EPI_ISL_525075, EPI_ISL_525076, EPI_ISL_525077, EPI_ISL_525078, EPI_ISL_525079, EPI_ISL_525080, EPI_ISL_525081, EPI_ISL_525082, EPI_ISL_525083, EPI_ISL_525084, EPI_ISL_525085, EPI_ISL_525086, EPI_ISL_525087, EPI_ISL_525088, EPI_ISL_525089, EPI_ISL_525090, EPI_ISL_525091, EPI_ISL_525092, EPI_ISL_525093, EPI_ISL_525094, EPI_ISL_525095, EPI_ISL_525096, EPI_ISL_525097, EPI_ISL_525098, EPI_ISL_525099, EPI_ISL_525100, EPI_ISL_525101, EPI_ISL_525102, EPI_ISL_525103, EPI_ISL_525104, EPI_ISL_525105, EPI_ISL_525106, EPI_ISL_525107, EPI_ISL_525108, EPI_ISL_525109, EPI_ISL_525110, EPI_ISL_525111, EPI_ISL_525112, EPI_ISL_525113, EPI_ISL_525114, EPI_ISL_525115, EPI_ISL_525116, EPI_ISL_525117, EPI_ISL_525183, EPI_ISL_525207, EPI_ISL_525208, EPI_ISL_525209, EPI_ISL_525210, EPI_ISL_525211, EPI_ISL_525212, EPI_ISL_525213, EPI_ISL_525214, EPI_ISL_525215, EPI_ISL_525216, EPI_ISL_525217, EPI_ISL_525218, EPI_ISL_525219, EPI_ISL_525220, EPI_ISL_525221, EPI_ISL_525222, EPI_ISL_525223, EPI_ISL_525224, EPI_ISL_525225, EPI_ISL_525226, EPI_ISL_525227, EPI_ISL_525228, EPI_ISL_525229, EPI_ISL_525230, EPI_ISL_525231, EPI_ISL_525233, EPI_ISL_525235, EPI_ISL_525236, EPI_ISL_525237, EPI_ISL_525238, EPI_ISL_525239, EPI_ISL_525240, EPI_ISL_525241, EPI_ISL_525243, EPI_ISL_525244, EPI_ISL_525245, EPI_ISL_525246, EPI_ISL_525247, EPI_ISL_525248, EPI_ISL_525249, EPI_ISL_525250, EPI_ISL_525251, EPI_ISL_525252, EPI_ISL_525253, EPI_ISL_525254, EPI_ISL_525255, EPI_ISL_525256, EPI_ISL_525257, EPI_ISL_525258, EPI_ISL_525259, EPI_ISL_525260, EPI_ISL_525261, EPI_ISL_525262, EPI_ISL_525263, EPI_ISL_525264, EPI_ISL_525265, EPI_ISL_525267, EPI_ISL_525268, EPI_ISL_525269, EPI_ISL_525270, EPI_ISL_525271, EPI_ISL_525272, EPI_ISL_525273, EPI_ISL_525274, EPI_ISL_525276, EPI_ISL_525277, EPI_ISL_525278, EPI_ISL_525279, EPI_ISL_525280, EPI_ISL_525281, EPI_ISL_525282, EPI_ISL_525283, EPI_ISL_525284, EPI_ISL_525285, EPI_ISL_525286, EPI_ISL_525287, EPI_ISL_525288, EPI_ISL_525289, EPI_ISL_525290, EPI_ISL_525291, EPI_ISL_525292, EPI_ISL_525293, EPI_ISL_525295, EPI_ISL_525296, EPI_ISL_525297, EPI_ISL_525298, EPI_ISL_525299, EPI_ISL_525300, EPI_ISL_525301, EPI_ISL_525302, EPI_ISL_525303, EPI_ISL_525304, EPI_ISL_525305, EPI_ISL_525306, EPI_ISL_525307, EPI_ISL_525308, EPI_ISL_525309, EPI_ISL_525310, EPI_ISL_525311, EPI_ISL_525312, EPI_ISL_525314, EPI_ISL_525315, EPI_ISL_525316, EPI_ISL_525317, EPI_ISL_525318, EPI_ISL_525319, EPI_ISL_525320, EPI_ISL_525321, EPI_ISL_525322, EPI_ISL_525323, EPI_ISL_525324, EPI_ISL_525325, EPI_ISL_525326, EPI_ISL_525327, EPI_ISL_525328, EPI_ISL_525329, EPI_ISL_525330, EPI_ISL_525332, EPI_ISL_525333, EPI_ISL_525334, EPI_ISL_525335, EPI_ISL_525336 | | | |
| see above | Utah Public Health Laboratory | Utah Public Health Laboratory | Erin L. Young, Kelly Oakeson, Tara Gallagher, Michael T. Pyne, E. Susan Slechta, Melanie A. Mallory, Jeffrey B. Stevenson, Salika M. Shakir, David R. Hillyard |
| EPI_ISL_525467 | Universidad Iberoamericana | International Centre for Genetic Engineering and Biotechnology (ICGEB) and ARGO Open Lab Platform | Robert Paulino-Ramirez, Eileen Riego, Alejandro Vallejo Degaudenzi, Victor Virgilio Calderon, Leandro Tapia, Patricia Leon, Danilo Licastro, Simeone Dal Monego, Sreejith Rajasekharan and Alessandro Marcello. |
| EPI_ISL_525468 | Universidad Iberoamericana | International Centre for Genetic Engineering and Biotechnology (ICGEB) and ARGO Open Lab Platform | Robert Paulino-Ramirez, Eileen Riego, Alejandro Vallejo Degaudenzi, Victor Virgilio Calderon, Leandro Tapia,Patricia Leon,Danilo Licastro, Simeone Dal Monego, Sreejith Rajasekharan and Alessandro Marcello. |
| EPI_ISL_525469, EPI_ISL_525470, EPI_ISL_525471 | Universidad Iberoamericana | International Centre for Genetic Engineering and Biotechnology (ICGEB) and ARGO Open Lab Platform | Robert Paulino-Ramirez, Eileen Riego, Alejandro Vallejo Degaudenzi, Victor Virgilio Calderon, Leandro Tapia, Patricia Leon, Danilo Licastro, Simeone Dal Monego, Sreejith Rajasekharan and Alessandro Marcello. |
| EPI_ISL_525807 | OHSU Lab Services Molecular Microbiology Lab | Oregon SARS-CoV-2 Genome Sequencing Center | Brendan L. O'Connell, Ruth V. Nichols, Alec J. Hirsch, Guang Fan, Daniel N. Streblow, William B. Messer, Andrew C. Adey, Benjamin N. Bimber, Brian J. O'Roak |
| EPI_ISL_526220, EPI_ISL_526221, EPI_ISL_526222, EPI_ISL_526236 | Hungarian Defence Forces Military Medical Centre | National Laboratory of Virology, Szentágothai Research Centre | Endre Gábor Tóth, Balázs Somogyi, Bálint Eszenyi, Ferenc Jakab, Gábor Kemenesi |
| EPI_ISL_526253, EPI_ISL_526254, EPI_ISL_526255, EPI_ISL_526256, EPI_ISL_526257, EPI_ISL_526258, EPI_ISL_526259, EPI_ISL_526260 | Unity Health Toronto | Ontario Institute for Cancer Research | Ramzi Fattouh, Larissa M. Matukas, Mark Downing, Annette Gower, Karel Boissinot, Samira Mubareka, TIBDN, Ilinca Lungu, Bernard Lam, Jeremy Johns, Paul Krzyzanowski, Richard de Borja, Felicia Vincelli, Philip Zuzarte, Jared Simpson |
| EPI_ISL_526484, EPI_ISL_526505, EPI_ISL_526513 | Virology Department, Royal Infirmary of Edinburgh, NHS Lothian / School of Biological Sciences, University of Edinburgh / Institute of Genetics and Molecular Medicine, University of Edinburgh | COVID-19 Genomics UK (COG-UK) Consortium | McHugh M, Dewar R, Rooke S, Gallagher M, Balcaza C, O'Toole Á, Scher E, Hill V, McCrone JT, Colquhoun R, Yu X, Jackson B, Rambaut A, Williams TC, Templeton K |
| EPI_ISL_526546, EPI_ISL_526547, EPI_ISL_526548, EPI_ISL_526549, EPI_ISL_526550 | Respiratory Virus Unit, Microbiology Services Colindale, Public Health England | Respiratory Virus Unit, Microbiology Services Colindale, Public Health England | PHE Covid Sequencing Team |
| EPI_ISL_526565, EPI_ISL_526566 | Florida Bureau of Public Health Laboratories | Florida Bureau of Public Health Laboratories | Sarah Schmedes, Jason Blanton |
| EPI_ISL_526686, EPI_ISL_526688 | Faith Laboratory, Immunology Institute, Icahn School of Medicine at Mount Sinai | van Bakel Laboratory, Genetics and Genomics Sciences, Icahn School of Medicine at Mount Sinai | Graham J. Britton, Alice Chen-Liaw, Francesca Cossarini, Alexandra Livanos, Matthew P. Spindler, Tamar Plitt, Joseph Eggers, Ilaria Mogno, Ana S. Gonzalez-Reiche, Sophia Sui, Michael Tankelevich, Lauren Tal Grinspan, Rebekah E. Dixon, Divya Jha, Gustavo Martinez-Delgado, Fatima Amanat, Daisy Hoagland, Benjamin R. tenOever, Marla C. Dubinsky, Miriam Merad, Harm Van Bakel, Florian Krammer, Gerold Bongers, Saurabh Mehandru and Jeremiah J. Faith |
| EPI_ISL_526775, EPI_ISL_526776, EPI_ISL_526777, EPI_ISL_526778, EPI_ISL_526779, EPI_ISL_526780, EPI_ISL_526781, EPI_ISL_526782, EPI_ISL_526783, EPI_ISL_526784, EPI_ISL_526785, EPI_ISL_526786, EPI_ISL_526787, EPI_ISL_526788, EPI_ISL_526789, EPI_ISL_526790, EPI_ISL_526791, EPI_ISL_526792, EPI_ISL_526793, EPI_ISL_526794, EPI_ISL_526795, EPI_ISL_526796 | | | |
| see above | Virginia DCLS | Virginia DCLS | Virginia DCLS |
| EPI_ISL_526932, EPI_ISL_526949 | Instituto Nacional de Salud, Bogotá, Colombia | Instituto Nacional de Salud, Bogotá, Colombia | Katherine Laiton-Donato, Diego A. Álvarez-Díaz, Carlos Franco-Muñoz, Mauricio Pacheco-Montealegre, Jonathan Reales, Diego Andrés Prada, Jose A. Usme-Ciro, Zulma M. Cucunubá, Christian Julian VillabonaArenas, Liz Villabona-Arenas, Sussy Echeverria, Astrid C. Flórez, Carolina Ferro, Diana Marcela Walteros-Acero, Franklin Prieto, Carlos Andrés Durán, Martha Lucia Ospina Martinez, Marcela Mercado-Reyes |
| EPI_ISL_527401, EPI_ISL_527427, EPI_ISL_527467, EPI_ISL_527479, | Colorado State University - Ebel Lab | Colorado State University - Ebel Lab | Greg Ebel et al. |

| | | | |
|---|---|---|---|
| EPI_ISL_527480 | | | |
| EPI_ISL_527676, EPI_ISL_527696, EPI_ISL_527697, EPI_ISL_527698 | MN PHL Division, Minnesota Department of Health | Pathogen Discovery, Respiratory Viruses Branch, Division of Viral Diseases, Centers for Disease Control and Prevention | Yan Li, Anna Montmayeur, Jing Zhang, Krista Queen, Ying Tao, Anna Uehara, Rachel Marine, Clinton R. Paden, Haibin Wang, Suxiang Tong |
| EPI_ISL_527699 | MN PHL Division, Minnesota Department of Health | Pathogen Discovery, Respiratory Viruses Branch, Division of Viral Diseases, Centers for Disease Control and Prevention | Krista Queen, Brian Lynch, Yan Li, Anna Montmayeur, Jing Zhang, Ying Tao, Anna Uehara, Rachel Marine, Clinton R. Paden, Haibin Wang, Suxiang Tong |
| EPI_ISL_527700 | MN PHL Division, Minnesota Department of Health | Pathogen Discovery, Respiratory Viruses Branch, Division of Viral Diseases, Centers for Disease Control and Prevention | Yan Li, Anna Montmayeur, Jing Zhang, Krista Queen, Ying Tao, Anna Uehara, Rachel Marine, Clinton R. Paden, Haibin Wang, Suxiang Tong |
| EPI_ISL_527701 | MN PHL Division, Minnesota Department of Health | Pathogen Discovery, Respiratory Viruses Branch, Division of Viral Diseases, Centers for Disease Control and Prevention | Krista Queen, Brian Lynch, Yan Li, Anna Montmayeur, Jing Zhang, Ying Tao, Anna Uehara, Rachel Marine, Clinton R. Paden, Haibin Wang, Suxiang Tong |
| EPI_ISL_527703, EPI_ISL_527704 | MN PHL Division, Minnesota Department of Health | Pathogen Discovery, Respiratory Viruses Branch, Division of Viral Diseases, Centers for Disease Control and Prevention | Yan Li, Anna Montmayeur, Jing Zhang, Krista Queen, Ying Tao, Anna Uehara, Rachel Marine, Clinton R. Paden, Haibin Wang, Suxiang Tong |
| EPI_ISL_527734 | MN PHL Division, Minnesota Department of Health | Pathogen Discovery, Respiratory Viruses Branch, Division of Viral Diseases, Centers for Disease Control and Prevention | Krista Queen, Brian Lynch, Yan Li, Anna Montmayeur, Jing Zhang, Ying Tao, Anna Uehara, Rachel Marine, Clinton R. Paden, Haibin Wang, Suxiang Tong |
| EPI_ISL_527872, EPI_ISL_527882, EPI_ISL_527893 | Nigeria Centre for Disease Control (NCDC) | African Centre of Excellence for Genomics of Infectious Diseases (ACEGID), Redeemer's University, Ede, Osun State, Nigeria | Oluniyi P.E. et al |
| EPI_ISL_528392, EPI_ISL_528393, EPI_ISL_528394 | Respiratory Virus Unit, Microbiology Services Colindale, Public Health England | Respiratory Virus Unit, Microbiology Services Colindale, Public Health England | PHE Covid Sequencing Team |
| EPI_ISL_528419, EPI_ISL_528421 | TNMC & BYL NAIR CH. HOSPITAL | Institute of Genomics and Integrative Biology - Council of Scientific and Industrial Research | Rajesh Pandey, Jayanthi Shastri, Akshay Kanakan, Vivekanand A, Janani Srinivasa Vasudevan, Ranjeet Maurya, Sachee Agrawal, Nirjhar Chatterjee, Swapneil Parikh, Manish Pathak, Subrat Thanapati, Jasmina Savak, Suresh Poojari, Mahesh Sangar, Amol Borse, Shweta Kawankar, Vasil Nachan, Mayuresh Vishwanathan, Shruthi Sachidanandan, Shrutika Pophale, Utkarsha Yelve |
| EPI_ISL_528425 | P. D. Hinduja Hospital and Medical Research Centre | Institute of Genomics and Integrative Biology - Council of Scientific and Industrial Research | Rajesh Pandey, Jayanthi Shastri, Akshay Kanakan, Vivekanand A, Janani Srinivasa Vasudevan, Ranjeet Maurya, Sachee Agrawal, Nirjhar Chatterjee, Swapneil Parikh, Manish Pathak, Subrat Thanapati, Jasmina Savak, Suresh Poojari, Mahesh Sangar, Amol Borse, Shweta Kawankar, Vasil Nachan, Mayuresh Vishwanathan, Shruthi Sachidanandan, Shrutika Pophale, Utkarsha Yelve |
| EPI_ISL_528427 | National Genomics Core-Center for DNA Fingerprinting and Diagnostics | National Genomics Core- Center for DNA Fingerprinting and Diagnostics (NGC-CDFD)- DBT's PAN-INDIA-1000 Genome consortium | Bala Pratyusha, Heena Shah, G Shashikanth, Vinay Donipadi, Edurugatla Dinesh, Guru Raja, Hilal Ahmad Reshi, J. Mallikarjun, K. Viswakalyan, Kaisar Ahmad Lone,Kausika Kumar Malik, N. Sudheer, R Harinarayanan, Rashna Bhandari, Murali Dharan Bashyam, Debashish Mitra, Divya Vashisht, Ashwin Dalal |
| EPI_ISL_528436, EPI_ISL_528437, EPI_ISL_528439 | Respiratory Virus Unit, Microbiology Services Colindale, Public Health England | Respiratory Virus Unit, Microbiology Services Colindale, Public Health England | PHE Covid Sequencing Team |
| EPI_ISL_528601, EPI_ISL_528602 | National Genomics Core-Center for DNA Fingerprinting and Diagnostics | National Genomics Core- Center for DNA Fingerprinting and Diagnostics (NGC-CDFD)- DBT's PAN-INDIA-1000 Genome consortium | Bala Pratyusha, Heena Shah, G Shashikanth, Vinay Donipadi, Edurugatla Dinesh, Guru Raja, Hilal Ahmad Reshi, J. Mallikarjun, K. Viswakalyan, Kaisar Ahmad Lone,Kausika Kumar Malik, N. Sudheer, R Harinarayanan, Rashna Bhandari, Murali Dharan Bashyam, Debashish Mitra, Divya Vashisht, Ashwin Dalal |
| EPI_ISL_528603 | National Genomics Core-Center for DNA Fingerprinting and Diagnostics | National Genomics Core- Center for DNA Fingerprinting and Diagnostics (NGC-CDFD)- DBT's PAN-INDIA-1000 Genome consortium | Ashwin Dalal, Bala Pratyusha, Heena Shah, G Shashikanth, Vinay Donipadi, Neeraj Kumar, Niteen Pathak, Pradipta Hore, Rahul Baroi, Sayantan Goswami, Shaffiqu T S, Shalini Aricthota, Sobhan Babu, R Harinarayanan, Rashna Bhandari, Murali Dharan Bashyam, Debashish Mitra, Divya Vashisht |
| EPI_ISL_528612, EPI_ISL_528613 | National Genomics Core-Center for DNA Fingerprinting and Diagnostics | National Genomics Core- Center for DNA Fingerprinting and Diagnostics (NGC-CDFD)- DBT's PAN-INDIA-1000 Genome consortium | Divya Vashisht, Bala Pratyusha, Heena Shah, G Shashikanth, Vinay Donipadi, K.Manohar, Madhumohan Rao, SPR Prasad, Yogesh Patidar, Arjita Jaiswal, Arpita Singh, Devanshi Gupta, Romila Moirangthem, Sanjana Sarkar, Shivani Yadav, R Harinarayanan, Rashna Bhandari, Murali Dharan Bashyam, Debashish Mitra, Ashwin Dalal |
| EPI_ISL_528687, EPI_ISL_528689, EPI_ISL_528693, EPI_ISL_528698, EPI_ISL_528699, EPI_ISL_528701, EPI_ISL_528704, EPI_ISL_528706 | Alsafar - Khalifa University Abu Dhabi | Alsafar - Khalifa University Abu Dhabi | Andreas Henschel, Gihan Daw Elbait, Samuel Feng, Rifat Hamoudi, Ernesto Damiani, Guan Tay, Habiba Alsafar |
| EPI_ISL_528749 | Santosa Hospital Bandung Central | School of Life Sciences and Technology & School of Pharmacy-Institut Teknologi Bandung; Molecular Genetics Laboratory-Faculty of Medicine-Universitas Padjadjaran; Laboratorium Kesehatan Provinsi Jawa Barat | Husna Nugrahapraja, Azzania Fibriani, Catur Riani, Marselina Irasonia Tan, Yunia Sribudiani, Tarwadi, Ema Rahmawati, Savira Ekawardhani, Hesti Lina Wiraswati, Ryan Bayusantika Ristandi, Rifky Waluyajati Rachman, Cut Nur Cinthia Alamanda, Lia Faridah, Davin H. E. Setiamarga, Rizki Mardian, Hammam Riza, Sony Solistia Wirawan, Agung Eru Wibowo, Irvan Faizal |
| EPI_ISL_528788, EPI_ISL_528792, EPI_ISL_528793, EPI_ISL_528794, EPI_ISL_528799, EPI_ISL_528801, EPI_ISL_528802, EPI_ISL_528803, EPI_ISL_528804, EPI_ISL_528805, EPI_ISL_528806 | | | |
| see above | Microbiology Department, Barking Havering and Redbridge University Hospitals NHS trust | Wellcome Sanger Institute for the COVID-19 Genomics UK (COG-UK) consortium | Amy Ash, Fatima Ali, Cherian Koshy and Alex Alderton, Roberto Amato, Sonia Goncalves, Ewan Harrison, David K. Jackson, Ian Johnston, Dominic Kwiatkowski, Cordelia Langford, John Sillitoe on behalf of the Wellcome Sanger Institute COVID-19 Surveillance Team (http://www.sanger.ac.uk/covid-team) |
| EPI_ISL_528930, EPI_ISL_528931 | Respiratory Virus Unit, Microbiology Services Colindale, Public Health England | Respiratory Virus Unit, Microbiology Services Colindale, Public Health England | PHE Covid Sequencing Team |
| EPI_ISL_528941, EPI_ISL_528942, EPI_ISL_528943, EPI_ISL_528944, EPI_ISL_528945, EPI_ISL_528946, EPI_ISL_528947, EPI_ISL_528948, EPI_ISL_528949 | Agenzia di Tutela della Salute di Bergamo | Istituto Zooprofilattico Sperimentale dell'Abruzzo e Molise "G.Caporale" | Lorusso A, Marcacci M, Di Domenico M, Curini V, Ancora M, Cammà C, Rinaldi A, Mangone I, Di Pasquale A, Puglia I, Savini G. |
| EPI_ISL_528994, EPI_ISL_528995, EPI_ISL_528996, EPI_ISL_528997, EPI_ISL_528998, EPI_ISL_528999, EPI_ISL_529000, EPI_ISL_529001, EPI_ISL_529002, EPI_ISL_529003, EPI_ISL_529004, EPI_ISL_529005 | | | |
| see above | Servizio di igiene epidemiologia e sanità pubblica (SIESP)-Chieti | Istituto Zooprofilattico Sperimentale dell'Abruzzo e Molise "G.Caporale" | Lorusso A, Marcacci M, Di Domenico M, Curini V, Ancora M, Cammà C, Rinaldi A, Mangone I, Di Pasquale A, Puglia I, Savini G. |
| EPI_ISL_529022 | Ospedale "Ss. Annunziata" | Istituto Zooprofilattico Sperimentale dell'Abruzzo e Molise "G.Caporale" | Lorusso A, Marcacci M, Di Domenico M, Curini V, Ancora M, Cammà C, Rinaldi A, Mangone I, Di Pasquale A, Puglia I, Savini G. |
| EPI_ISL_529023, EPI_ISL_529024, EPI_ISL_529025 | Servizio di igiene epidemiologia e sanità pubblica (SIESP)-Chieti | Istituto Zooprofilattico Sperimentale dell'Abruzzo e Molise "G.Caporale" | Lorusso A, Marcacci M, Di Domenico M, Curini V, Ancora M, Cammà C, Rinaldi A, Mangone I, Di Pasquale A, Puglia I, Savini G. |
| EPI_ISL_529138 | RSAL Dr. Ramelan Surabaya | Institute of Tropical Disease, Universitas Airlangga | Jezzy R Dewantari, Rima R Prasetya, Krisnoadi Rahardjo, Aldise M Nastri, Radito Soesanto, Gatot Soegiarto, Laksmi Wulandari, Retno A Setyoningrum, Resti Yudhawati, Yohko K Shimizu, Mitsuhiro Nishimura, Yasuko Mori, Soetjipto, Kazufumi Shimizu, Maria I Lusida |
| EPI_ISL_529185, EPI_ISL_529186, EPI_ISL_529187, EPI_ISL_529188, EPI_ISL_529189 | South Carolina Department of Health and Environmental Control | South Carolina Department of Health and Environmental Control | Haley V. Flores |
| EPI_ISL_529384, EPI_ISL_529518 | University of Birmingham | COVID-19 Genomics UK (COG-UK) Consortium | Institute of Microbiology, University of Birmingham: Claire McMurray, Joanne Stockton, Samuel Nicholls, Radoslaw Poplawski, Will Rowe, Josh Quick, Nicholas Loman. University of Birmingham Testing Laboratory: Celina M Whalley, Andrew Bosworth, Charlotte Poxon, Kasun Wanigasooriya, Oliver Pickles, Mike Kidd, Alex Richter, Andrew D Beggs PHE Heartlands Lab: Husam Osman, Andrew Bosworth. Queen Elizabeth Hospital: Anna Casey |
| EPI_ISL_529966 | RSUD Sidoarjo | Institute of Tropical Disease, Universitas Airlangga | Kazufumi Shimizu, Krisnoadi Rahardjo, Aldise M Nastri, Jezzy R Dewantari, Rima R Prasetya, Atok Irawan, Gatot Soegiarto, Laksmi Wulandari, Retno A |

| | | | |
|---|---|---|---|
| | | | Setyoningrum, Resti Yudhawati, Yohko K Shimizu, Mitsuhiro Nishimura, Yasuko Mori, Soetjipto, Maria I Lusida |
| EPI_ISL_530028, EPI_ISL_530029, EPI_ISL_530071, EPI_ISL_530075, EPI_ISL_530076, EPI_ISL_530077, EPI_ISL_530078, EPI_ISL_530079, EPI_ISL_530089, EPI_ISL_530090 | Hospital Universitario La Paz | Hospital Universitario La Paz | María Rodríguez, Elias Dahdouh, Sara González, Raúl Recio, Fernando Lázaro, Esther Viedma, Natalia Stella, Julio García, Juan Carlos Galán, Rafael Cantón, Mª Dolores Folgueira, Rafael Delgado, Jesús Mingorance |
| EPI_ISL_530114, EPI_ISL_530115, EPI_ISL_530116, EPI_ISL_530117, EPI_ISL_530118 | Hospital Universitario Ramón y Cajal | Hospital Universitario La Paz | María Rodríguez, Elias Dahdouh, Sara González, Raúl Recio, Fernando Lázaro, Esther Viedma, Natalia Stella, Julio García, Juan Carlos Galán, Rafael Cantón, Mª Dolores Folgueira, Rafael Delgado, Jesús Mingorance |
| EPI_ISL_530124 | Seattle Flu Study | Seattle Flu Study | Deborah A. Nickerson, Chris D. Frazar, Jover Lee, Benjamin Pelle, Matthew Richardson, Amanda Adler, Elisabeth Brandstetter, Peter D. Han, Kairsten Fay, Misja Ilcisin, Kirsten Lacombe, Thomas R. Sibley, Melissa Truong, Caitlin R. Wolf, Michael Boeckh, Janet A. Englund, Michael Famulare, Barry R. Lutz, Mark J. Rieder, Lea M. Starita, Matthew Thompson, Jay Shendure, Trevor Bedford, Helen Y. Chu |
| EPI_ISL_530190, EPI_ISL_530191, EPI_ISL_530192, EPI_ISL_530193, EPI_ISL_530194, EPI_ISL_530195, EPI_ISL_530196, EPI_ISL_530197, EPI_ISL_530198 | Minnesota Department of Health, Public Health Laboratory | Minnesota Department of Health, Public Health Laboratory | Matt Plumb, Jacob Garfin, and Xiong Wang |
| EPI_ISL_532143, EPI_ISL_532144, EPI_ISL_532145, EPI_ISL_532148, EPI_ISL_532149 | Lighthouse Lab in Glasgow | Wellcome Sanger Institute for the COVID-19 Genomics UK (COG-UK) consortium | Harper VanSteenhouse, Yumi Kasai, David Gray, Carol Clugston, Anna Dominiczak and Alex Alderton, Roberto Amato, Sonia Goncalves, Ewan Harrison, David K. Jackson, Ian Johnston, Dominic Kwiatkowski, Cordelia Langford, John Sillitoe |
| EPI_ISL_532153 | NHSGGC West of Scotland Specialist Virology Centre / MRC-University of Glasgow Centre for Virus Research | Wellcome Sanger Institute for the COVID-19 Genomics UK (COG-UK) consortium | Ana da Silva Filipe, Natasha Johnson, Kathy Smollett, Daniel Mair, Stephen Carmichael, Lily Tong, Jenna Nichols, Elihu Aranday-Cortes, Kirstyn Brunker, Yasmin Parr, Kyriaki Nomikou; Sarah McDonald, Marc Niebel, Patawee Asamaphan; Richard Orton, Joseph Hughes, Sreenu Vattipally, David L Robertson; Alasdair MacLean, Rory Gunson; Kathy Li, Natasha Jesudason, Rajiv Shah, James Shepherd, Antonia Ho, Alice Broos, Emma Thomson and Alex Alderton, Roberto Amato, Sonia Goncalves, Ewan Harrison, David K. Jackson, Ian Johnston, Dominic Kwiatkowski, Cordelia Langford, John Sillitoe |
| EPI_ISL_532156, EPI_ISL_532157, EPI_ISL_532161, EPI_ISL_532162, EPI_ISL_532164, EPI_ISL_532165, EPI_ISL_532167, EPI_ISL_532169, EPI_ISL_532170, EPI_ISL_532172, EPI_ISL_532174, EPI_ISL_532175, EPI_ISL_532176, EPI_ISL_532177, EPI_ISL_532179, EPI_ISL_532181, EPI_ISL_532182, EPI_ISL_532183, EPI_ISL_532184, EPI_ISL_532187, EPI_ISL_532188, EPI_ISL_532190 | | | |
| see above | Lighthouse Lab in Glasgow | Wellcome Sanger Institute for the COVID-19 Genomics UK (COG-UK) consortium | Harper VanSteenhouse, Yumi Kasai, David Gray, Carol Clugston, Anna Dominiczak and Alex Alderton, Roberto Amato, Sonia Goncalves, Ewan Harrison, David K. Jackson, Ian Johnston, Dominic Kwiatkowski, Cordelia Langford, John Sillitoe |
| EPI_ISL_532192, EPI_ISL_532194, EPI_ISL_532196, EPI_ISL_532197 | NHSGGC West of Scotland Specialist Virology Centre / MRC-University of Glasgow Centre for Virus Research | Wellcome Sanger Institute for the COVID-19 Genomics UK (COG-UK) consortium | Ana da Silva Filipe, Natasha Johnson, Kathy Smollett, Daniel Mair, Stephen Carmichael, Lily Tong, Jenna Nichols, Elihu Aranday-Cortes, Kirstyn Brunker, Yasmin Parr, Kyriaki Nomikou; Sarah McDonald, Marc Niebel, Patawee Asamaphan; Richard Orton, Joseph Hughes, Sreenu Vattipally, David L Robertson; Alasdair MacLean, Rory Gunson; Kathy Li, Natasha Jesudason, Rajiv Shah, James Shepherd, Antonia Ho, Alice Broos, Emma Thomson and Alex Alderton, Roberto Amato, Sonia Goncalves, Ewan Harrison, David K. Jackson, Ian Johnston, Dominic Kwiatkowski, Cordelia Langford, John Sillitoe |
| EPI_ISL_532199, EPI_ISL_532201 | Lighthouse Lab in Glasgow | Wellcome Sanger Institute for the COVID-19 Genomics UK (COG-UK) consortium | Harper VanSteenhouse, Yumi Kasai, David Gray, Carol Clugston, Anna Dominiczak and Alex Alderton, Roberto Amato, Sonia Goncalves, Ewan Harrison, David K. Jackson, Ian Johnston, Dominic Kwiatkowski, Cordelia Langford, John Sillitoe |
| EPI_ISL_532202, EPI_ISL_532203 | NHSGGC West of Scotland Specialist Virology Centre / MRC-University of Glasgow Centre for Virus Research | Wellcome Sanger Institute for the COVID-19 Genomics UK (COG-UK) consortium | Ana da Silva Filipe, Natasha Johnson, Kathy Smollett, Daniel Mair, Stephen Carmichael, Lily Tong, Jenna Nichols, Elihu Aranday-Cortes, Kirstyn Brunker, Yasmin Parr, Kyriaki Nomikou; Sarah McDonald, Marc Niebel, Patawee Asamaphan; Richard Orton, Joseph Hughes, Sreenu Vattipally, David L Robertson; Alasdair MacLean, Rory Gunson; Kathy Li, Natasha Jesudason, Rajiv Shah, James Shepherd, Antonia Ho, Alice Broos, Emma Thomson and Alex Alderton, Roberto Amato, Sonia Goncalves, Ewan Harrison, David K. Jackson, Ian Johnston, Dominic Kwiatkowski, Cordelia Langford, John Sillitoe |
| EPI_ISL_532204, EPI_ISL_532205, EPI_ISL_532206, EPI_ISL_532207, EPI_ISL_532211, EPI_ISL_532213 | Lighthouse Lab in Glasgow | Wellcome Sanger Institute for the COVID-19 Genomics UK (COG-UK) consortium | Harper VanSteenhouse, Yumi Kasai, David Gray, Carol Clugston, Anna Dominiczak and Alex Alderton, Roberto Amato, Sonia Goncalves, Ewan Harrison, David K. Jackson, Ian Johnston, Dominic Kwiatkowski, Cordelia Langford, John Sillitoe |
| EPI_ISL_532214 | NHSGGC West of Scotland Specialist Virology Centre / MRC-University of Glasgow Centre for Virus Research | Wellcome Sanger Institute for the COVID-19 Genomics UK (COG-UK) consortium | Ana da Silva Filipe, Natasha Johnson, Kathy Smollett, Daniel Mair, Stephen Carmichael, Lily Tong, Jenna Nichols, Elihu Aranday-Cortes, Kirstyn Brunker, Yasmin Parr, Kyriaki Nomikou; Sarah McDonald, Marc Niebel, Patawee Asamaphan; Richard Orton, Joseph Hughes, Sreenu Vattipally, David L Robertson; Alasdair MacLean, Rory Gunson; Kathy Li, Natasha Jesudason, Rajiv Shah, James Shepherd, Antonia Ho, Alice Broos, Emma Thomson and Alex Alderton, Roberto Amato, Sonia Goncalves, Ewan Harrison, David K. Jackson, Ian Johnston, Dominic Kwiatkowski, Cordelia Langford, John Sillitoe |
| EPI_ISL_532215, EPI_ISL_532216 | Lighthouse Lab in Glasgow | Wellcome Sanger Institute for the COVID-19 Genomics UK (COG-UK) consortium | Harper VanSteenhouse, Yumi Kasai, David Gray, Carol Clugston, Anna Dominiczak and Alex Alderton, Roberto Amato, Sonia Goncalves, Ewan Harrison, David K. Jackson, Ian Johnston, Dominic Kwiatkowski, Cordelia Langford, John Sillitoe |
| EPI_ISL_532219 | NHSGGC West of Scotland Specialist Virology Centre / MRC-University of Glasgow Centre for Virus Research | Wellcome Sanger Institute for the COVID-19 Genomics UK (COG-UK) consortium | Ana da Silva Filipe, Natasha Johnson, Kathy Smollett, Daniel Mair, Stephen Carmichael, Lily Tong, Jenna Nichols, Elihu Aranday-Cortes, Kirstyn Brunker, Yasmin Parr, Kyriaki Nomikou; Sarah McDonald, Marc Niebel, Patawee Asamaphan; Richard Orton, Joseph Hughes, Sreenu Vattipally, David L Robertson; Alasdair MacLean, Rory Gunson; Kathy Li, Natasha Jesudason, Rajiv Shah, James Shepherd, Antonia Ho, Alice Broos, Emma Thomson and Alex Alderton, Roberto Amato, Sonia Goncalves, Ewan Harrison, David K. Jackson, Ian Johnston, Dominic Kwiatkowski, Cordelia Langford, John Sillitoe |
| EPI_ISL_532222, EPI_ISL_532223, EPI_ISL_532224, EPI_ISL_532225, EPI_ISL_532226, EPI_ISL_532227, EPI_ISL_532236, EPI_ISL_532240 | Lighthouse Lab in Glasgow | Wellcome Sanger Institute for the COVID-19 Genomics UK (COG-UK) consortium | Harper VanSteenhouse, Yumi Kasai, David Gray, Carol Clugston, Anna Dominiczak and Alex Alderton, Roberto Amato, Sonia Goncalves, Ewan Harrison, David K. Jackson, Ian Johnston, Dominic Kwiatkowski, Cordelia Langford, John Sillitoe |
| EPI_ISL_532241 | NHSGGC West of Scotland Specialist Virology Centre / MRC-University of Glasgow Centre for Virus Research | Wellcome Sanger Institute for the COVID-19 Genomics UK (COG-UK) consortium | Ana da Silva Filipe, Natasha Johnson, Kathy Smollett, Daniel Mair, Stephen Carmichael, Lily Tong, Jenna Nichols, Elihu Aranday-Cortes, Kirstyn Brunker, Yasmin Parr, Kyriaki Nomikou; Sarah McDonald, Marc Niebel, Patawee Asamaphan; Richard Orton, Joseph Hughes, Sreenu Vattipally, David L Robertson; Alasdair MacLean, Rory Gunson; Kathy Li, Natasha Jesudason, Rajiv Shah, James Shepherd, Antonia Ho, Alice Broos, Emma Thomson and Alex Alderton, Roberto Amato, Sonia Goncalves, Ewan Harrison, David K. Jackson, Ian Johnston, Dominic Kwiatkowski, Cordelia Langford, John Sillitoe |
| EPI_ISL_532242, EPI_ISL_532243 | Lighthouse Lab in Glasgow | Wellcome Sanger Institute for the COVID-19 Genomics UK (COG-UK) consortium | Harper VanSteenhouse, Yumi Kasai, David Gray, Carol Clugston, Anna Dominiczak and Alex Alderton, Roberto Amato, Sonia Goncalves, Ewan Harrison, David K. Jackson, Ian Johnston, Dominic Kwiatkowski, Cordelia Langford, John Sillitoe |
| EPI_ISL_532244 | NHSGGC West of Scotland Specialist Virology Centre / MRC-University of Glasgow Centre for Virus Research | Wellcome Sanger Institute for the COVID-19 Genomics UK (COG-UK) consortium | Ana da Silva Filipe, Natasha Johnson, Kathy Smollett, Daniel Mair, Stephen Carmichael, Lily Tong, Jenna Nichols, Elihu Aranday-Cortes, Kirstyn Brunker, Yasmin Parr, Kyriaki Nomikou; Sarah McDonald, Marc Niebel, Patawee Asamaphan; Richard Orton, Joseph Hughes, Sreenu Vattipally, David L Robertson; Alasdair MacLean, Rory Gunson; Kathy Li, Natasha Jesudason, Rajiv Shah, James Shepherd, Antonia Ho, Alice Broos, Emma Thomson and Alex Alderton, Roberto Amato, Sonia Goncalves, Ewan Harrison, David K. Jackson, Ian Johnston, Dominic Kwiatkowski, Cordelia Langford, John Sillitoe |
| EPI_ISL_532245, EPI_ISL_532246, EPI_ISL_532247 | Lighthouse Lab in Glasgow | Wellcome Sanger Institute for the COVID-19 Genomics UK (COG-UK) consortium | Harper VanSteenhouse, Yumi Kasai, David Gray, Carol Clugston, Anna Dominiczak and Alex Alderton, Roberto Amato, Sonia Goncalves, Ewan Harrison, David K. Jackson, Ian Johnston, Dominic Kwiatkowski, Cordelia Langford, John Sillitoe |
| EPI_ISL_532249 | NHSGGC West of Scotland Specialist Virology Centre / MRC-University of Glasgow Centre for Virus Research | Wellcome Sanger Institute for the COVID-19 Genomics UK (COG-UK) consortium | Ana da Silva Filipe, Natasha Johnson, Kathy Smollett, Daniel Mair, Stephen Carmichael, Lily Tong, Jenna Nichols, Elihu Aranday-Cortes, Kirstyn Brunker, Yasmin Parr, Kyriaki Nomikou; Sarah McDonald, Marc Niebel, Patawee Asamaphan; Richard Orton, Joseph Hughes, Sreenu Vattipally, David L Robertson; Alasdair MacLean, Rory Gunson; Kathy Li, Natasha Jesudason, Rajiv Shah, James Shepherd, Antonia Ho, Alice Broos, Emma Thomson and Alex Alderton, Roberto Amato, Sonia Goncalves, Ewan Harrison, David K. Jackson, Ian Johnston, Dominic Kwiatkowski, Cordelia Langford, John Sillitoe |
| EPI_ISL_532250, EPI_ISL_532252, EPI_ISL_532253, EPI_ISL_532256, EPI_ISL_532260, EPI_ISL_532264, EPI_ISL_532267, EPI_ISL_532270 | Lighthouse Lab in Glasgow | Wellcome Sanger Institute for the COVID-19 Genomics UK (COG-UK) consortium | Harper VanSteenhouse, Yumi Kasai, David Gray, Carol Clugston, Anna Dominiczak and Alex Alderton, Roberto Amato, Sonia Goncalves, Ewan Harrison, David K. Jackson, Ian Johnston, Dominic Kwiatkowski, Cordelia Langford, John Sillitoe |
| EPI_ISL_532271, EPI_ISL_532276 | NHSGGC West of Scotland Specialist Virology Centre / | Wellcome Sanger Institute for the COVID-19 Genomics UK | Ana da Silva Filipe, Natasha Johnson, Kathy Smollett, Daniel Mair, Stephen Carmichael, Lily Tong, Jenna Nichols, Elihu Aranday-Cortes, Kirstyn Brunker, |

| | | | |
|---|---|---|---|
| | MRC-University of Glasgow Centre for Virus Research | (COG-UK) consortium | Yasmin Parr, Kyriaki Nomikou; Sarah McDonald, Marc Niebel, Patawee Asamaphan; Richard Orton, Joseph Hughes, Sreenu Vattipally, David L Robertson; Alasdair MacLean, Rory Gunson; Kathy Li, Natasha Jesudason, Rajiv Shah, James Shepherd, Antonia Ho, Alice Broos, Emma Thomson and Alex Alderton, Roberto Amato, Sonia Goncalves, Ewan Harrison, David K. Jackson, Ian Johnston, Dominic Kwiatkowski, Cordelia Langford, John Sillitoe |
| EPI_ISL_532280, EPI_ISL_532281, EPI_ISL_532282, EPI_ISL_533345 | Lighthouse Lab in Glasgow | Wellcome Sanger Institute for the COVID-19 Genomics UK (COG-UK) consortium | Harper VanSteenhouse, Yumi Kasai, David Gray, Carol Clugston, Anna Dominiczak and Alex Alderton, Roberto Amato, Sonia Goncalves, Ewan Harrison, David K. Jackson, Ian Johnston, Dominic Kwiatkowski, Cordelia Langford, John Sillitoe |
| EPI_ISL_533346 | NHSGGC West of Scotland Specialist Virology Centre / MRC-University of Glasgow Centre for Virus Research | Wellcome Sanger Institute for the COVID-19 Genomics UK (COG-UK) consortium | Ana da Silva Filipe, Natasha Johnson, Kathy Smollett, Daniel Mair, Stephen Carmichael, Lily Tong, Jenna Nichols, Elihu Aranday-Cortes, Kirstyn Brunker, Yasmin Parr, Kyriaki Nomikou; Sarah McDonald, Marc Niebel, Patawee Asamaphan; Richard Orton, Joseph Hughes, Sreenu Vattipally, David L Robertson; Alasdair MacLean, Rory Gunson; Kathy Li, Natasha Jesudason, Rajiv Shah, James Shepherd, Antonia Ho, Alice Broos, Emma Thomson and Alex Alderton, Roberto Amato, Sonia Goncalves, Ewan Harrison, David K. Jackson, Ian Johnston, Dominic Kwiatkowski, Cordelia Langford, John Sillitoe |
| EPI_ISL_533347, EPI_ISL_533348, EPI_ISL_533349 | Lighthouse Lab in Glasgow | Wellcome Sanger Institute for the COVID-19 Genomics UK (COG-UK) consortium | Harper VanSteenhouse, Yumi Kasai, David Gray, Carol Clugston, Anna Dominiczak and Alex Alderton, Roberto Amato, Sonia Goncalves, Ewan Harrison, David K. Jackson, Ian Johnston, Dominic Kwiatkowski, Cordelia Langford, John Sillitoe |
| EPI_ISL_533350, EPI_ISL_533351 | NHSGGC West of Scotland Specialist Virology Centre / MRC-University of Glasgow Centre for Virus Research | Wellcome Sanger Institute for the COVID-19 Genomics UK (COG-UK) consortium | Ana da Silva Filipe, Natasha Johnson, Kathy Smollett, Daniel Mair, Stephen Carmichael, Lily Tong, Jenna Nichols, Elihu Aranday-Cortes, Kirstyn Brunker, Yasmin Parr, Kyriaki Nomikou; Sarah McDonald, Marc Niebel, Patawee Asamaphan; Richard Orton, Joseph Hughes, Sreenu Vattipally, David L Robertson; Alasdair MacLean, Rory Gunson; Kathy Li, Natasha Jesudason, Rajiv Shah, James Shepherd, Antonia Ho, Alice Broos, Emma Thomson and Alex Alderton, Roberto Amato, Sonia Goncalves, Ewan Harrison, David K. Jackson, Ian Johnston, Dominic Kwiatkowski, Cordelia Langford, John Sillitoe |
| EPI_ISL_533352, EPI_ISL_533353, EPI_ISL_533354 | Lighthouse Lab in Glasgow | Wellcome Sanger Institute for the COVID-19 Genomics UK (COG-UK) consortium | Harper VanSteenhouse, Yumi Kasai, David Gray, Carol Clugston, Anna Dominiczak and Alex Alderton, Roberto Amato, Sonia Goncalves, Ewan Harrison, David K. Jackson, Ian Johnston, Dominic Kwiatkowski, Cordelia Langford, John Sillitoe |
| EPI_ISL_533355 | NHSGGC West of Scotland Specialist Virology Centre / MRC-University of Glasgow Centre for Virus Research | Wellcome Sanger Institute for the COVID-19 Genomics UK (COG-UK) consortium | Ana da Silva Filipe, Natasha Johnson, Kathy Smollett, Daniel Mair, Stephen Carmichael, Lily Tong, Jenna Nichols, Elihu Aranday-Cortes, Kirstyn Brunker, Yasmin Parr, Kyriaki Nomikou; Sarah McDonald, Marc Niebel, Patawee Asamaphan; Richard Orton, Joseph Hughes, Sreenu Vattipally, David L Robertson; Alasdair MacLean, Rory Gunson; Kathy Li, Natasha Jesudason, Rajiv Shah, James Shepherd, Antonia Ho, Alice Broos, Emma Thomson and Alex Alderton, Roberto Amato, Sonia Goncalves, Ewan Harrison, David K. Jackson, Ian Johnston, Dominic Kwiatkowski, Cordelia Langford, John Sillitoe |
| EPI_ISL_533356, EPI_ISL_533357 | Lighthouse Lab in Glasgow | Wellcome Sanger Institute for the COVID-19 Genomics UK (COG-UK) consortium | Harper VanSteenhouse, Yumi Kasai, David Gray, Carol Clugston, Anna Dominiczak and Alex Alderton, Roberto Amato, Sonia Goncalves, Ewan Harrison, David K. Jackson, Ian Johnston, Dominic Kwiatkowski, Cordelia Langford, John Sillitoe |
| EPI_ISL_533358 | NHSGGC West of Scotland Specialist Virology Centre / MRC-University of Glasgow Centre for Virus Research | Wellcome Sanger Institute for the COVID-19 Genomics UK (COG-UK) consortium | Ana da Silva Filipe, Natasha Johnson, Kathy Smollett, Daniel Mair, Stephen Carmichael, Lily Tong, Jenna Nichols, Elihu Aranday-Cortes, Kirstyn Brunker, Yasmin Parr, Kyriaki Nomikou; Sarah McDonald, Marc Niebel, Patawee Asamaphan; Richard Orton, Joseph Hughes, Sreenu Vattipally, David L Robertson; Alasdair MacLean, Rory Gunson; Kathy Li, Natasha Jesudason, Rajiv Shah, James Shepherd, Antonia Ho, Alice Broos, Emma Thomson and Alex Alderton, Roberto Amato, Sonia Goncalves, Ewan Harrison, David K. Jackson, Ian Johnston, Dominic Kwiatkowski, Cordelia Langford, John Sillitoe |
| EPI_ISL_533359, EPI_ISL_533360, EPI_ISL_533361, EPI_ISL_533362, EPI_ISL_533363, EPI_ISL_533364 | Lighthouse Lab in Glasgow | Wellcome Sanger Institute for the COVID-19 Genomics UK (COG-UK) consortium | Harper VanSteenhouse, Yumi Kasai, David Gray, Carol Clugston, Anna Dominiczak and Alex Alderton, Roberto Amato, Sonia Goncalves, Ewan Harrison, David K. Jackson, Ian Johnston, Dominic Kwiatkowski, Cordelia Langford, John Sillitoe |
| EPI_ISL_533365 | NHSGGC West of Scotland Specialist Virology Centre / MRC-University of Glasgow Centre for Virus Research | Wellcome Sanger Institute for the COVID-19 Genomics UK (COG-UK) consortium | Ana da Silva Filipe, Natasha Johnson, Kathy Smollett, Daniel Mair, Stephen Carmichael, Lily Tong, Jenna Nichols, Elihu Aranday-Cortes, Kirstyn Brunker, Yasmin Parr, Kyriaki Nomikou; Sarah McDonald, Marc Niebel, Patawee Asamaphan; Richard Orton, Joseph Hughes, Sreenu Vattipally, David L Robertson; Alasdair MacLean, Rory Gunson; Kathy Li, Natasha Jesudason, Rajiv Shah, James Shepherd, Antonia Ho, Alice Broos, Emma Thomson and Alex Alderton, Roberto Amato, Sonia Goncalves, Ewan Harrison, David K. Jackson, Ian Johnston, Dominic Kwiatkowski, Cordelia Langford, John Sillitoe |
| EPI_ISL_533366, EPI_ISL_533367, EPI_ISL_533368, EPI_ISL_533369, EPI_ISL_533370, EPI_ISL_533371, EPI_ISL_533372, EPI_ISL_533373, EPI_ISL_533374, EPI_ISL_533375 | Lighthouse Lab in Glasgow | Wellcome Sanger Institute for the COVID-19 Genomics UK (COG-UK) consortium | Harper VanSteenhouse, Yumi Kasai, David Gray, Carol Clugston, Anna Dominiczak and Alex Alderton, Roberto Amato, Sonia Goncalves, Ewan Harrison, David K. Jackson, Ian Johnston, Dominic Kwiatkowski, Cordelia Langford, John Sillitoe |
| EPI_ISL_533376 | NHSGGC West of Scotland Specialist Virology Centre / MRC-University of Glasgow Centre for Virus Research | Wellcome Sanger Institute for the COVID-19 Genomics UK (COG-UK) consortium | Ana da Silva Filipe, Natasha Johnson, Kathy Smollett, Daniel Mair, Stephen Carmichael, Lily Tong, Jenna Nichols, Elihu Aranday-Cortes, Kirstyn Brunker, Yasmin Parr, Kyriaki Nomikou; Sarah McDonald, Marc Niebel, Patawee Asamaphan; Richard Orton, Joseph Hughes, Sreenu Vattipally, David L Robertson; Alasdair MacLean, Rory Gunson; Kathy Li, Natasha Jesudason, Rajiv Shah, James Shepherd, Antonia Ho, Alice Broos, Emma Thomson and Alex Alderton, Roberto Amato, Sonia Goncalves, Ewan Harrison, David K. Jackson, Ian Johnston, Dominic Kwiatkowski, Cordelia Langford, John Sillitoe |
| EPI_ISL_533377, EPI_ISL_533378, EPI_ISL_533379, EPI_ISL_533380, EPI_ISL_533381, EPI_ISL_533382, EPI_ISL_533383, EPI_ISL_533384, EPI_ISL_533385, EPI_ISL_533386, EPI_ISL_533387, EPI_ISL_533388, EPI_ISL_533389, EPI_ISL_533390, EPI_ISL_533391, EPI_ISL_533392, EPI_ISL_533393 | | | |
| see above | Lighthouse Lab in Glasgow | Wellcome Sanger Institute for the COVID-19 Genomics UK (COG-UK) consortium | Harper VanSteenhouse, Yumi Kasai, David Gray, Carol Clugston, Anna Dominiczak and Alex Alderton, Roberto Amato, Sonia Goncalves, Ewan Harrison, David K. Jackson, Ian Johnston, Dominic Kwiatkowski, Cordelia Langford, John Sillitoe |
| EPI_ISL_533394 | NHSGGC West of Scotland Specialist Virology Centre / MRC-University of Glasgow Centre for Virus Research | Wellcome Sanger Institute for the COVID-19 Genomics UK (COG-UK) consortium | Ana da Silva Filipe, Natasha Johnson, Kathy Smollett, Daniel Mair, Stephen Carmichael, Lily Tong, Jenna Nichols, Elihu Aranday-Cortes, Kirstyn Brunker, Yasmin Parr, Kyriaki Nomikou; Sarah McDonald, Marc Niebel, Patawee Asamaphan; Richard Orton, Joseph Hughes, Sreenu Vattipally, David L Robertson; Alasdair MacLean, Rory Gunson; Kathy Li, Natasha Jesudason, Rajiv Shah, James Shepherd, Antonia Ho, Alice Broos, Emma Thomson and Alex Alderton, Roberto Amato, Sonia Goncalves, Ewan Harrison, David K. Jackson, Ian Johnston, Dominic Kwiatkowski, Cordelia Langford, John Sillitoe |
| EPI_ISL_533395 | Lighthouse Lab in Glasgow | Wellcome Sanger Institute for the COVID-19 Genomics UK (COG-UK) consortium | Harper VanSteenhouse, Yumi Kasai, David Gray, Carol Clugston, Anna Dominiczak and Alex Alderton, Roberto Amato, Sonia Goncalves, Ewan Harrison, David K. Jackson, Ian Johnston, Dominic Kwiatkowski, Cordelia Langford, John Sillitoe |
| EPI_ISL_533396 | NHSGGC West of Scotland Specialist Virology Centre / MRC-University of Glasgow Centre for Virus Research | Wellcome Sanger Institute for the COVID-19 Genomics UK (COG-UK) consortium | Ana da Silva Filipe, Natasha Johnson, Kathy Smollett, Daniel Mair, Stephen Carmichael, Lily Tong, Jenna Nichols, Elihu Aranday-Cortes, Kirstyn Brunker, Yasmin Parr, Kyriaki Nomikou; Sarah McDonald, Marc Niebel, Patawee Asamaphan; Richard Orton, Joseph Hughes, Sreenu Vattipally, David L Robertson; Alasdair MacLean, Rory Gunson; Kathy Li, Natasha Jesudason, Rajiv Shah, James Shepherd, Antonia Ho, Alice Broos, Emma Thomson and Alex Alderton, Roberto Amato, Sonia Goncalves, Ewan Harrison, David K. Jackson, Ian Johnston, Dominic Kwiatkowski, Cordelia Langford, John Sillitoe |
| EPI_ISL_533397, EPI_ISL_533398, EPI_ISL_533399 | Lighthouse Lab in Glasgow | Wellcome Sanger Institute for the COVID-19 Genomics UK (COG-UK) consortium | Harper VanSteenhouse, Yumi Kasai, David Gray, Carol Clugston, Anna Dominiczak and Alex Alderton, Roberto Amato, Sonia Goncalves, Ewan Harrison, David K. Jackson, Ian Johnston, Dominic Kwiatkowski, Cordelia Langford, John Sillitoe |
| EPI_ISL_533400 | Lighthouse Lab in Glasgow | Wellcome Sanger Institute for the COVID-19 Genomics UK (COG-UK) consortium | Harper VanSteenhouse, Yumi Kasai, David Gray, Carol Clugston, Anna Dominiczak and Alex Alderton, Roberto Amato, Sonia Goncalves, Ewan Harrison, David K. Jackson, Ian Johnston, Dominic Kwiatkowski, Cordelia Langford, John Sillitoe on behalf of the Wellcome Sanger Institute COVID-19 Surveillance Team |
| EPI_ISL_533401, EPI_ISL_533402 | Lighthouse Lab in Glasgow | Wellcome Sanger Institute for the COVID-19 Genomics UK (COG-UK) consortium | Harper VanSteenhouse, Yumi Kasai, David Gray, Carol Clugston, Anna Dominiczak and Alex Alderton, Roberto Amato, Sonia Goncalves, Ewan Harrison, David K. Jackson, Ian Johnston, Dominic Kwiatkowski, Cordelia Langford, John Sillitoe |
| EPI_ISL_533403 | Lighthouse Lab in Glasgow | Wellcome Sanger Institute for the COVID-19 Genomics UK (COG-UK) Consortium | Harper VanSteenhouse, Yumi Kasai, David Gray, Carol Clugston, Anna Dominiczak and Alex Alderton, Roberto Amato, Sonia Goncalves, Ewan Harrison, David K. Jackson, Ian Johnston, Dominic Kwiatkowski, Cordelia Langford, John Sillitoe on behalf of the Wellcome Sanger Institute COVID-19 Surveillance Team |
| EPI_ISL_533404, EPI_ISL_533405 | Lighthouse Lab in Glasgow | Wellcome Sanger Institute for the COVID-19 Genomics UK (COG-UK) consortium | Harper VanSteenhouse, Yumi Kasai, David Gray, Carol Clugston, Anna Dominiczak and Alex Alderton, Roberto Amato, Sonia Goncalves, Ewan Harrison, David K. Jackson, Ian Johnston, Dominic Kwiatkowski, Cordelia Langford, John Sillitoe |
| EPI_ISL_533406 | NHSGGC West of Scotland Specialist Virology Centre / MRC-University of Glasgow Centre for Virus Research | Wellcome Sanger Institute for the COVID-19 Genomics UK (COG-UK) consortium | Ana da Silva Filipe, Natasha Johnson, Kathy Smollett, Daniel Mair, Stephen Carmichael, Lily Tong, Jenna Nichols, Elihu Aranday-Cortes, Kirstyn Brunker, Yasmin Parr, Kyriaki Nomikou; Sarah McDonald, Marc Niebel, Patawee Asamaphan; Richard Orton, Joseph Hughes, Sreenu Vattipally, David L Robertson; Alasdair MacLean, Rory Gunson; Kathy Li, Natasha Jesudason, Rajiv Shah, James Shepherd, Antonia Ho, Alice Broos, Emma Thomson and Alex Alderton, Roberto Amato, Sonia Goncalves, Ewan Harrison, David K. Jackson, Ian Johnston, Dominic Kwiatkowski, Cordelia Langford, John Sillitoe |
| EPI_ISL_533407, EPI_ISL_533408, EPI_ISL_533409, EPI_ISL_533410, EPI_ISL_533411, EPI_ISL_533412, EPI_ISL_533413, EPI_ISL_533414, EPI_ISL_533415, EPI_ISL_533416, EPI_ISL_533417, EPI_ISL_533418 | | | |
| see above | Lighthouse Lab in Glasgow | Wellcome Sanger Institute for the COVID-19 Genomics UK (COG-UK) consortium | Harper VanSteenhouse, Yumi Kasai, David Gray, Carol Clugston, Anna Dominiczak and Alex Alderton, Roberto Amato, Sonia Goncalves, Ewan Harrison, David K. Jackson, Ian Johnston, Dominic Kwiatkowski, Cordelia Langford, John Sillitoe |

| | | | |
|---|---|---|---|
| EPI_ISL_533419 | NHSGGC West of Scotland Specialist Virology Centre / MRC-University of Glasgow Centre for Virus Research | Wellcome Sanger Institute for the COVID-19 Genomics UK (COG-UK) consortium | Ana da Silva Filipe, Natasha Johnson, Kathy Smollett, Daniel Mair, Stephen Carmichael, Lily Tong, Jenna Nichols, Elihu Aranday-Cortes, Kirstyn Brunker, Yasmin Parr, Kyriaki Nomikou; Sarah McDonald, Marc Niebel, Patawee Asamaphan; Richard Orton, Joseph Hughes, Sreenu Vattipally, David L Robertson; Alasdair MacLean, Rory Gunson; Kathy Li, Natasha Jesudason, Rajiv Shah, James Shepherd, Antonia Ho, Alice Broos, Emma Thomson and Alex Alderton, Roberto Amato, Sonia Goncalves, Ewan Harrison, David K. Jackson, Ian Johnston, Dominic Kwiatkowski, Cordelia Langford, John Sillitoe |
| EPI_ISL_533420 | Lighthouse Lab in Glasgow | Wellcome Sanger Institute for the COVID-19 Genomics UK (COG-UK) consortium | Harper VanSteenhouse, Yumi Kasai, David Gray, Carol Clugston, Anna Dominiczak and Alex Alderton, Roberto Amato, Sonia Goncalves, Ewan Harrison, David K. Jackson, Ian Johnston, Dominic Kwiatkowski, Cordelia Langford, John Sillitoe |
| EPI_ISL_533421 | NHSGGC West of Scotland Specialist Virology Centre / MRC-University of Glasgow Centre for Virus Research | Wellcome Sanger Institute for the COVID-19 Genomics UK (COG-UK) consortium | Ana da Silva Filipe, Natasha Johnson, Kathy Smollett, Daniel Mair, Stephen Carmichael, Lily Tong, Jenna Nichols, Elihu Aranday-Cortes, Kirstyn Brunker, Yasmin Parr, Kyriaki Nomikou; Sarah McDonald, Marc Niebel, Patawee Asamaphan; Richard Orton, Joseph Hughes, Sreenu Vattipally, David L Robertson; Alasdair MacLean, Rory Gunson; Kathy Li, Natasha Jesudason, Rajiv Shah, James Shepherd, Antonia Ho, Alice Broos, Emma Thomson and Alex Alderton, Roberto Amato, Sonia Goncalves, Ewan Harrison, David K. Jackson, Ian Johnston, Dominic Kwiatkowski, Cordelia Langford, John Sillitoe |
| EPI_ISL_533422, EPI_ISL_533423, EPI_ISL_533424, EPI_ISL_533425, EPI_ISL_533426, EPI_ISL_533427, EPI_ISL_533428, EPI_ISL_533429, EPI_ISL_533430, EPI_ISL_533431 | Lighthouse Lab in Glasgow | Wellcome Sanger Institute for the COVID-19 Genomics UK (COG-UK) consortium | Harper VanSteenhouse, Yumi Kasai, David Gray, Carol Clugston, Anna Dominiczak and Alex Alderton, Roberto Amato, Sonia Goncalves, Ewan Harrison, David K. Jackson, Ian Johnston, Dominic Kwiatkowski, Cordelia Langford, John Sillitoe |
| EPI_ISL_533432, EPI_ISL_533433, EPI_ISL_533434 | Lighthouse Lab in Glasgow | Wellcome Sanger Institute for the COVID-19 Genomics UK (COG-UK) consortium | Harper VanSteenhouse, Yumi Kasai, David Gray, Carol Clugston, Anna Dominiczak and Alex Alderton, Roberto Amato, Sonia Goncalves, Ewan Harrison, David K. Jackson, Ian Johnston, Dominic Kwiatkowski, Cordelia Langford, John Sillitoe on behalf of the Wellcome Sanger Institute COVID-19 Surveillance Team |
| EPI_ISL_534317 | Hospital Geral de Itapevi | Instituto Adolfo Lutz, Interdiciplinary Procedures Center, Strategic Laboratory | Claudio Tavares Sacchi, Claudia Regina Gonçalves, Erica Valessa Ramos Gomes |
| EPI_ISL_534319, EPI_ISL_534320 | Hospital do Serv Pub ESTAFCO Morato de Oliveira | Instituto Adolfo Lutz, Interdiciplinary Procedures Center, Strategic Laboratory | Claudio Tavares Sacchi, Claudia Regina Gonçalves, Erica Valessa Ramos Gomes |
| EPI_ISL_534321 | PS e Maternidade Nair Fonseca Leitao Arantes | Instituto Adolfo Lutz, Interdiciplinary Procedures Center, Strategic Laboratory | Claudio Tavares Sacchi, Claudia Regina Gonçalves, Erica Valessa Ramos Gomes |
| EPI_ISL_534330 | Hospital Universitario La Paz | Hospital Universitario La Paz | María Rodríguez, Elias Dahdouh, Sara González, Raúl Recio, Fernando Lázaro, Esther Viedma, Natalia Stella, Julio García, Juan Carlos Galán, Rafael Cantón, Ma Dolores Folgueira, Rafael Delgado, Jesús Mingorance |
| EPI_ISL_534776, EPI_ISL_534836, EPI_ISL_534908, EPI_ISL_534920, EPI_ISL_534930, EPI_ISL_534960, EPI_ISL_534963, EPI_ISL_534964, EPI_ISL_534968, EPI_ISL_535000 | Oxford Viromics, NDM, University of Oxford; Oxford University Hospitals; Basingstoke and North Hampshire Hospital | COVID-19 Genomics UK (COG-UK) Consortium | Tanya Golubchik, David Bonsall, George Macintyre, Amy Trebes, Mariateresa de Cesare, Catrin Moore, Alex Mobbs, Anita Justice, Robert Shaw, Monique Andersson, Timothy Peto, Emma Wise, Nathan Moore, Jessica Lynch, Nick Cortes, Matilde Mori, Stephen Kidd, David Buck, John Todd, Christophe Fraser |
| EPI_ISL_535341, EPI_ISL_535342 | LA Office of Public Health Laboratories | Pathogen Discovery, Respiratory Viruses Branch, Division of Viral Diseases, Centers for Disease Control and Prevention | Jing Zhang, Ying Tao, Yan Li, Krista Queen, Anna Uehara, Clinton Paden, Haibin Wang, Suxiang Tong |
| EPI_ISL_535343 | LA Office of Public Health Laboratories | Pathogen Discovery, Respiratory Viruses Branch, Division of Viral Diseases, Centers for Disease Control and Prevention | Ying Tao, Jing Zhang, Yan Li, Krista Queen, Anna Uehara, Clinton Paden, Haibin Wang, Suxiang Tong |
| EPI_ISL_535344, EPI_ISL_535345 | LA Office of Public Health Laboratories | Pathogen Discovery, Respiratory Viruses Branch, Division of Viral Diseases, Centers for Disease Control and Prevention | Jing Zhang, Ying Tao, Yan Li, Krista Queen, Anna Uehara, Clinton Paden, Haibin Wang, Suxiang Tong |
| EPI_ISL_536426 | National Public Health Laboratory, National Centre for Infectious Diseases | National Public Health Laboratory, National Centre for Infectious Diseases | Mak TM, Octavia S, Zhou Z, Cui L, Lin RTP |
| EPI_ISL_536514, EPI_ISL_536517, EPI_ISL_536518, EPI_ISL_536520, EPI_ISL_536521 | Instituto Nacional de Salud | Laboratorio de Infecciones Respiratorias Agudas | Eduardo Juscamayta Lopez, David Tarazona, Faviola Valdivia Guerrero, Nancy Rojas Serrano, Dennis Carhuaricra, Lenin Maturrano Hernandez, Ronnie Gavilan Chavez |
| EPI_ISL_537207, EPI_ISL_537209, EPI_ISL_537214, EPI_ISL_537215, EPI_ISL_537216, EPI_ISL_537218, EPI_ISL_537220, EPI_ISL_537221, EPI_ISL_537222, EPI_ISL_537224, EPI_ISL_537225, EPI_ISL_537228, EPI_ISL_537230, EPI_ISL_537232, EPI_ISL_537235, EPI_ISL_537237, EPI_ISL_537238, EPI_ISL_537239, EPI_ISL_537240, EPI_ISL_537241, EPI_ISL_537244, EPI_ISL_537246, EPI_ISL_537248, EPI_ISL_537249, EPI_ISL_537250, EPI_ISL_537251, EPI_ISL_537253 | | | |
| see above | Virology Department, Sheffield Teaching Hospitals NHS Foundation Trust / Department of Infection, Immunity and Cardiovascular Disease, The Medical School, University of Sheffield | Wellcome Sanger Institute for the COVID-19 Genomics UK (COG-UK) consortium | Thushan de Silva, Matthew Parker,Adri Angyal, Rebecca Brown, Luke Green, Rachel Tucker, Paul Parsons, Danielle Groves, Alex Keeley, Dave Partridge, Matthew Wyles, Benjamin Lindsey, Mehmet Yavuz, Mohammad Raza, Cariad Evans and Alex Alderton, Roberto Amato, Sonia Goncalves, Ewan Harrison, David K. Jackson, Ian Johnston, Dominic Kwiatkowski, Cordelia Langford, John Sillitoe on behalf of the Wellcome Sanger Institute COVID-19 Surveillance Team |
| EPI_ISL_537254 | Virology Department, Sheffield Teaching Hospitals NHS Foundation Trust / Department of Infection, Immunity and Cardiovascular Disease, The Medical School, University of Sheffield | Wellcome Sanger Institute for the COVID-19 Genomics UK (COG-UK) Consortium | Thushan de Silva, Matthew Parker,Adri Angyal, Rebecca Brown, Luke Green, Rachel Tucker, Paul Parsons, Danielle Groves, Alex Keeley, Dave Partridge, Matthew Wyles, Benjamin Lindsey, Mehmet Yavuz, Mohammad Raza, Cariad Evans and Alex Alderton, Roberto Amato, Sonia Goncalves, Ewan Harrison, David K. Jackson, Ian Johnston, Dominic Kwiatkowski, Cordelia Langford, John Sillitoe on behalf of the Wellcome Sanger Institute COVID-19 Surveillance Team |
| EPI_ISL_537255, EPI_ISL_537260, EPI_ISL_537261, EPI_ISL_537262, EPI_ISL_537263, EPI_ISL_537264, EPI_ISL_537265, EPI_ISL_537269, EPI_ISL_537271, EPI_ISL_537272 | Virology Department, Sheffield Teaching Hospitals NHS Foundation Trust / Department of Infection, Immunity and Cardiovascular Disease, The Medical School, University of Sheffield | Wellcome Sanger Institute for the COVID-19 Genomics UK (COG-UK) consortium | Thushan de Silva, Matthew Parker,Adri Angyal, Rebecca Brown, Luke Green, Rachel Tucker, Paul Parsons, Danielle Groves, Alex Keeley, Dave Partridge, Matthew Wyles, Benjamin Lindsey, Mehmet Yavuz, Mohammad Raza, Cariad Evans and Alex Alderton, Roberto Amato, Sonia Goncalves, Ewan Harrison, David K. Jackson, Ian Johnston, Dominic Kwiatkowski, Cordelia Langford, John Sillitoe on behalf of the Wellcome Sanger Institute COVID-19 Surveillance Team |
| EPI_ISL_537508, EPI_ISL_537510, EPI_ISL_537517, EPI_ISL_537526, EPI_ISL_537532, EPI_ISL_537548 | UCLA Pathology Clinical Microbiology Lab | Kruglyak Lab | Guo et al. |
| EPI_ISL_537680, EPI_ISL_537681, EPI_ISL_537682, EPI_ISL_537683, EPI_ISL_537684, EPI_ISL_537685, EPI_ISL_537686, EPI_ISL_537687 | Universidad de León | SeqCOVID-SPAIN consortium/IBV(CSIC) | Ana Carvajal, Vicente Martín, Héctor Argüello, Juan M. Fregeneda, Tania Fernández-Villa, Antonio J. Molina and SeqCOVID-SPAIN consortium |
| EPI_ISL_537782, EPI_ISL_537783, EPI_ISL_537784, EPI_ISL_537785 | Servicio de Microbiología, Hospital Miguel Servet, Zaragoza | SeqCOVID-SPAIN consortium/IBV(CSIC) | Antonio Rezusta López, Alexander Tristancho Baró, Ana Milagro, Yolanda Gracia Grataloup, Nieves Martínez Cameo and SeqCOVID-SPAIN consortium |
| EPI_ISL_538251, EPI_ISL_538252, EPI_ISL_538254, EPI_ISL_538255, EPI_ISL_538256, EPI_ISL_538257, EPI_ISL_538258, EPI_ISL_538259, EPI_ISL_538260 | TriCore Reference Laboratories | Center for Global Health, University of New Mexico Health Sciences Center | Daryl Domman, Kurt Schwalm, Twila Kunde, Joseph Hicks, Michael Edwards, Darrell Dinwiddie |
| EPI_ISL_539489, EPI_ISL_539490 | Civil Hospital, Rupnagar | CSIR-Institute of Microbial Technology | Kanika Bansal, Sanjeet Kumar, Anu Singh, Debarghya Ghose, Amandeep Kaur, Rajesh Kumar Mishra, Poushali Chakraborty, Harsh Goar, Navin Baid, Ashwani Kumar, Dipak Dutta, Sanjeev Khosla, Prabhu B. Patil |
| EPI_ISL_539533, EPI_ISL_539534, EPI_ISL_539535, EPI_ISL_539536, | Hospital Clínic | Instituto de Salud Carlos III | Iglesias-Caballero, M. Molinero Calamita, M. González-Esguevillas, M. Camarero, S. Pozo, F. Casas, I. Jiménez, P. Jiménez, M. Zaballos, A. Monzón, S. Varona, S. Juliá, M. Cuesta, I, M.A Marcos |

| | | | |
|---|---|---|---|
| EPI_ISL_539537 | | | |
| EPI_ISL_539829, EPI_ISL_539830, EPI_ISL_539831 | Minnesota Department of Health, Public Health Laboratory | Minnesota Department of Health, Public Health Laboratory | Matt Plumb, Jacob Garfin, and Xiong Wang |
| EPI_ISL_540430 | Microbial Genomics Laboratory, Institut Pasteur de Montevideo, Montevideo, Uruguay | Microbial Genomics Laboratory, Institut Pasteur de Montevideo, Montevideo, Uruguay | Cecilia Salazar, Marianoel Pereira, Gonzalo Moratorio, Pilar Moreno, Gregorio Iraola |
| EPI_ISL_541034, EPI_ISL_541035, EPI_ISL_541036 | Servicio de Microbiología, Hospital Miguel Servet, Zaragoza | SeqCOVID-SPAIN consortium/Institute of Biomedicine of Valencia, IBV-CSIC | Antonio Rezusta López, Alexander Tristancho Baró, Ana Milagro, Yolanda Gracia Grataloup, Nieves Martínez Cameo and SeqCOVID-SPAIN consortium |
| EPI_ISL_541039, EPI_ISL_541041 | Hospital Clínico Universitario de Santiago de Compostela | SeqCOVID-SPAIN consortium/Institute of Biomedicine of Valencia, IBV-CSIC | José Javier Costa Alcalde, Antonio Aguilera Guirao, Mª Luisa Pérez del Molino Bernal, Amparo Coira Nieto, Gema Barbeito Castiñeiras, Rocio Trastoy Pena and SeqCOVID-SPAIN consortium |
| EPI_ISL_541157 | Florida Bureau of Public Health Laboratories, Florida Department of Health | Florida Bureau of Public Health Laboratories, Florida Department of Health | Schmedes,S., Blanton,J. |
| EPI_ISL_541335 | The National Institute of Public Health | Sídlištní 136/24 165 03, Prague Czech Republic | Nagy,A; Jirincova,H; Novakova,L; Trnka,D; Vecerova,J |
| EPI_ISL_541398 | Laboratório de Virologia Comparada e Ambiental- LVCA-IOC | Laboratory of Respiratory Viruses and Measles, Oswaldo Cruz Institute, FIOCRUZ | Paola Resende, Luciana Appolinario, Tulio Machado Fumian, Tatiana Prado, Camille Ferreira Mannarino, Fernando Motta, Ana Carolina Mendonça, Marilda Siqueira, Marize Pereira Miagostovich |
| EPI_ISL_541764, EPI_ISL_541781 | Barts Health NHS Trust | Wellcome Sanger Institute for the COVID-19 Genomics UK (COG-UK) consortium | Teresa Cutino-Moguel, Mark Hopkins, Beatrix Kele, David Harrington and Alex Alderton, Roberto Amato, Sonia Goncalves, Ewan Harrison, David K. Jackson, Ian Johnston, Dominic Kwiatkowski, Cordelia Langford, John Sillitoe on behalf of the Wellcome Sanger Institute COVID-19 Surveillance Team |
| EPI_ISL_542804, EPI_ISL_542809, EPI_ISL_542810, EPI_ISL_542811, EPI_ISL_542812, EPI_ISL_542813, EPI_ISL_542814, EPI_ISL_542815, EPI_ISL_542816, EPI_ISL_542817, EPI_ISL_542818, EPI_ISL_542819, EPI_ISL_542822, EPI_ISL_542823, EPI_ISL_542824, EPI_ISL_542825, EPI_ISL_542826, EPI_ISL_542827, EPI_ISL_542828, EPI_ISL_542829, EPI_ISL_542830, EPI_ISL_542831, EPI_ISL_542834, EPI_ISL_542837, EPI_ISL_542839, EPI_ISL_542840, EPI_ISL_542841, EPI_ISL_542845, EPI_ISL_542848, EPI_ISL_542850, EPI_ISL_542851, EPI_ISL_542853, EPI_ISL_542854, EPI_ISL_542856, EPI_ISL_542857, EPI_ISL_542858, EPI_ISL_542859, EPI_ISL_542860, EPI_ISL_542861, EPI_ISL_542862, EPI_ISL_542863, EPI_ISL_542864, EPI_ISL_542865, EPI_ISL_542866, EPI_ISL_542867, EPI_ISL_542868, EPI_ISL_542869, EPI_ISL_542870, EPI_ISL_542871, EPI_ISL_542872, EPI_ISL_542873, EPI_ISL_542874, EPI_ISL_542881, EPI_ISL_542883, EPI_ISL_542884, EPI_ISL_542885, EPI_ISL_542886, EPI_ISL_542887, EPI_ISL_542889, EPI_ISL_542890, EPI_ISL_542892, EPI_ISL_542894, EPI_ISL_542895, EPI_ISL_542896, EPI_ISL_542897, EPI_ISL_542898, EPI_ISL_542899, EPI_ISL_542910, EPI_ISL_542915, EPI_ISL_542920, EPI_ISL_542923, EPI_ISL_542927 | | | |
| see above | Houston Methodist Hospital | Houston Methodist Hospital | S. Wesley Long, Randall J. Olsen, Paul A. Christensen, David W. Bernard, James J. Davis, Maulik Shukla, Marcus Nguyen, Matthew Ojeda Saavedra, Concepcion C. Cantu, Prasanti Yerramilli, Layne Pruitt, Sishir Subedi, Hung-Che Kuo, Heather Hendrickson, Ghazaleh Eskandari, Hoang A. T. Nguyen, J. Hunter Long, Muthiah Kumaraswami, Jule Goike, Daniel Boutz, Jimmy Gollihar, Jason S. McLellan, Chia-Wei Chou, Kamyab Javanmardi, Ilya J. Finkelstein, and James M. Musser |
| EPI_ISL_542936, EPI_ISL_542937, EPI_ISL_542941, EPI_ISL_542942, EPI_ISL_542944, EPI_ISL_542946, EPI_ISL_542951, EPI_ISL_542955, EPI_ISL_542956, EPI_ISL_542957, EPI_ISL_542958, EPI_ISL_542959, EPI_ISL_542960, EPI_ISL_542961, EPI_ISL_542962, EPI_ISL_542964, EPI_ISL_542965, EPI_ISL_542966, EPI_ISL_542967, EPI_ISL_542969, EPI_ISL_542972, EPI_ISL_542973, EPI_ISL_542974, EPI_ISL_542975, EPI_ISL_542977, EPI_ISL_542978, EPI_ISL_542980, EPI_ISL_542981, EPI_ISL_542982 | | | |
| see above | TriCore Reference Laboratories | Center for Global Health, University of New Mexico Health Sciences Center | Daryl Domman, Kurt Schwalm, Twila Kunde, Joseph Hicks, Michael Edwards, Darrell Dinwiddie |
| EPI_ISL_545069, EPI_ISL_545070, EPI_ISL_545071, EPI_ISL_545073, EPI_ISL_545074, EPI_ISL_545075, EPI_ISL_545076, EPI_ISL_545077, EPI_ISL_545078, EPI_ISL_545079, EPI_ISL_545082, EPI_ISL_545083, EPI_ISL_545084, EPI_ISL_545086, EPI_ISL_545087, EPI_ISL_545088, EPI_ISL_545091, EPI_ISL_545095, EPI_ISL_545096, EPI_ISL_545097, EPI_ISL_545098, EPI_ISL_545099, EPI_ISL_545100, EPI_ISL_545101, EPI_ISL_545102, EPI_ISL_545103, EPI_ISL_545105, EPI_ISL_545106, EPI_ISL_545107, EPI_ISL_545108, EPI_ISL_545109, EPI_ISL_545110, EPI_ISL_545111, EPI_ISL_545134, EPI_ISL_545188, EPI_ISL_545189, EPI_ISL_545190, EPI_ISL_545191, EPI_ISL_545192, EPI_ISL_545193, EPI_ISL_545194, EPI_ISL_545199, EPI_ISL_545200, EPI_ISL_545201, EPI_ISL_545218, EPI_ISL_545221, EPI_ISL_545222, EPI_ISL_545224, EPI_ISL_545225, EPI_ISL_545226, EPI_ISL_545227, EPI_ISL_545228, EPI_ISL_545230, EPI_ISL_545231, EPI_ISL_545232, EPI_ISL_545234, EPI_ISL_545236, EPI_ISL_545237, EPI_ISL_545297, EPI_ISL_545298, EPI_ISL_545309, EPI_ISL_545343, EPI_ISL_545344 | | | |
| see above | Houston Methodist Hospital | Houston Methodist Hospital | S. Wesley Long, Randall J. Olsen, Paul A. Christensen, David W. Bernard, James J. Davis, Maulik Shukla, Marcus Nguyen, Matthew Ojeda Saavedra, Concepcion C. Cantu, Prasanti Yerramilli, Layne Pruitt, Sishir Subedi, Hung-Che Kuo, Heather Hendrickson, Ghazaleh Eskandari, Hoang A. T. Nguyen, J. Hunter Long, Muthiah Kumaraswami, Jule Goike, Daniel Boutz, Jimmy Gollihar, Jason S. McLellan, Chia-Wei Chou, Kamyab Javanmardi, Ilya J. Finkelstein, and James M. Musser |
| EPI_ISL_548442 | Ventura County Public Health Lab | Chan-Zuckerberg Biohub | CZB Cliahub Consortium |
| EPI_ISL_548942, EPI_ISL_548943, EPI_ISL_548944, EPI_ISL_548945, EPI_ISL_548946 | Institute of Microbiology, University of Veterinary and Animal sciences | Institute of Microbiology, University of Veterinary and Animal sciences | Yaqub,T., Nawaz,M., Ali,M.A., Altaf,I., Raza,S., Shabbir,M.A., Ashraf,M.A., Aziz,S.Z., Cheema,S.Q., Shah,M.B., Hassan,S., Rafique,S., Sardar,N., Mehmood,A., Aziz,M.W., Fazal,S., Khan,N., Khan,M.T., Attique,M.M., Asif,A., Anwar,M., Awan,N.A., Younis,M.U., Bhatti,M.A., Tahir,Z., Mukhtar,N., Sarwar,H., Rana,M.S., Shabbir,M.Z. |
| EPI_ISL_548956 | Max von Pettenkofer Institute, Virology, National Reference Center for Retroviruses, LMU München | Laboratory for Functional Genome Analysis, Dept. Genomics, Gene Center of the LMU Munich | Max Muenchhoff, Stefan Krebs, Alexander Graf, Oliver Keppler, Helmut Blum |
| EPI_ISL_549185, EPI_ISL_549186, EPI_ISL_549187, EPI_ISL_549188, EPI_ISL_549246, EPI_ISL_549247, EPI_ISL_549248, EPI_ISL_549249, EPI_ISL_549250, EPI_ISL_549251, EPI_ISL_549252, EPI_ISL_549253, EPI_ISL_549254, EPI_ISL_549255, EPI_ISL_549256, EPI_ISL_549257 | | | |
| see above | Florida Bureau of Public Health Laboratories | Florida Bureau of Public Health Laboratories | Sarah Schmedes, Jason Blanton |
| EPI_ISL_559662, EPI_ISL_559664, EPI_ISL_559665, EPI_ISL_559666, EPI_ISL_559668, EPI_ISL_559673, EPI_ISL_559677, EPI_ISL_559678, EPI_ISL_559680, EPI_ISL_559682, EPI_ISL_559684, EPI_ISL_559686, EPI_ISL_559688, EPI_ISL_559689, EPI_ISL_559691, EPI_ISL_559692, EPI_ISL_559693, EPI_ISL_559698, EPI_ISL_559699, EPI_ISL_559701, EPI_ISL_559702, EPI_ISL_559704 | | | |
| see above | Lighthouse Lab in Milton Keynes | Wellcome Sanger Institute for the COVID-19 Genomics UK (COG-UK) consortium | The Lighthouse Lab in Milton Keynes and Alex Alderton, Roberto Amato, Sonia Goncalves, Ewan Harrison, David K. Jackson, Ian Johnston, Dominic Kwiatkowski, Cordelia Langford, John Sillitoe on behalf of the Wellcome Sanger Institute COVID-19 Surveillance Team (http://www.sanger.ac.uk/covid-team) |
| EPI_ISL_559705, EPI_ISL_559707, EPI_ISL_559709 | Lighthouse Lab in Milton Keynes | Wellcome Sanger Institute for the COVID-19 Genomics UK (COG-UK) consortium | The Lighthouse Lab in Milton Keynes and Alex Alderton, Roberto Amato, Sonia Goncalves, Ewan Harrison, David K. Jackson, Ian Johnston, Dominic Kwiatkowski, Cordelia Langford, John Sillitoe on behalf of the Wellcome Sanger Institute COVID-19 Surveillance Team |
| EPI_ISL_559713, EPI_ISL_559716, EPI_ISL_559717 | Lighthouse Lab in Milton Keynes | Wellcome Sanger Institute for the COVID-19 Genomics UK (COG-UK) consortium | The Lighthouse Lab in Milton Keynes and Alex Alderton, Roberto Amato, Sonia Goncalves, Ewan Harrison, David K. Jackson, Ian Johnston, Dominic Kwiatkowski, Cordelia Langford, John Sillitoe on behalf of the Wellcome Sanger Institute COVID-19 Surveillance Team (http://www.sanger.ac.uk/covid-team) |
| EPI_ISL_559728 | Lighthouse Lab in Milton Keynes | Wellcome Sanger Institute for the COVID-19 Genomics UK (COG-UK) consortium | The Lighthouse Lab in Milton Keynes and Alex Alderton, Roberto Amato, Sonia Goncalves, Ewan Harrison, David K. Jackson, Ian Johnston, Dominic Kwiatkowski, Cordelia Langford, John Sillitoe on behalf of the Wellcome Sanger Institute COVID-19 Surveillance Team |
| EPI_ISL_559730, EPI_ISL_559732, EPI_ISL_559736 | Lighthouse Lab in Milton Keynes | Wellcome Sanger Institute for the COVID-19 Genomics UK (COG-UK) consortium | The Lighthouse Lab in Milton Keynes and Alex Alderton, Roberto Amato, Sonia Goncalves, Ewan Harrison, David K. Jackson, Ian Johnston, Dominic Kwiatkowski, Cordelia Langford, John Sillitoe on behalf of the Wellcome Sanger Institute COVID-19 Surveillance Team (http://www.sanger.ac.uk/covid-team) |
| EPI_ISL_559741, EPI_ISL_559742 | Lighthouse Lab in Milton Keynes | Wellcome Sanger Institute for the COVID-19 Genomics UK (COG-UK) consortium | The Lighthouse Lab in Milton Keynes and Alex Alderton, Roberto Amato, Sonia Goncalves, Ewan Harrison, David K. Jackson, Ian Johnston, Dominic Kwiatkowski, Cordelia Langford, John Sillitoe on behalf of the Wellcome Sanger Institute COVID-19 Surveillance Team |
| EPI_ISL_559743 | Lighthouse Lab in Milton Keynes | Wellcome Sanger Institute for the COVID-19 Genomics UK (COG-UK) consortium | The Lighthouse Lab in Alderley Park and Alex Alderton, Roberto Amato, Sonia Goncalves, Ewan Harrison, David K. Jackson, Ian Johnston, Dominic Kwiatkowski, Cordelia Langford, John Sillitoe on behalf of the Wellcome Sanger Institute COVID-19 Surveillance Team |
| EPI_ISL_559744, EPI_ISL_559745 | Lighthouse Lab in Milton Keynes | Wellcome Sanger Institute for the COVID-19 Genomics UK (COG-UK) consortium | The Lighthouse Lab in Milton Keynes and Alex Alderton, Roberto Amato, Sonia Goncalves, Ewan Harrison, David K. Jackson, Ian Johnston, Dominic Kwiatkowski, Cordelia Langford, John Sillitoe on behalf of the Wellcome Sanger Institute COVID-19 Surveillance Team (http://www.sanger.ac.uk/covid-team) |
| EPI_ISL_559751 | Lighthouse Lab in Milton Keynes | Wellcome Sanger Institute for the COVID-19 Genomics UK (COG-UK) consortium | The Lighthouse Lab in Milton Keynes and Alex Alderton, Roberto Amato, Sonia Goncalves, Ewan Harrison, David K. Jackson, Ian Johnston, Dominic Kwiatkowski, Cordelia Langford, John Sillitoe on behalf of the Wellcome Sanger Institute COVID-19 Surveillance Team |
| EPI_ISL_560325, EPI_ISL_560326, EPI_ISL_560331, EPI_ISL_560332, EPI_ISL_560334, EPI_ISL_560336, EPI_ISL_560337, EPI_ISL_560338, EPI_ISL_560339, EPI_ISL_560340, EPI_ISL_560342, EPI_ISL_560344, EPI_ISL_560346, EPI_ISL_560347, EPI_ISL_560348, EPI_ISL_560349, EPI_ISL_560350, EPI_ISL_560351, EPI_ISL_560352, EPI_ISL_560353, EPI_ISL_560355, EPI_ISL_560356, EPI_ISL_560359, EPI_ISL_560360, EPI_ISL_560361 | | | |
| see above | TriCore Reference Laboratories | Center for Global Health, University of New Mexico Health Sciences Center | Daryl Domman, Kurt Schwalm, Twila Kunde, Joseph Hicks, Michael Edwards, Darrell Dinwiddie |

| | | | |
|---|---|---|---|
| EPI_ISL_560407 | Istituto Zooprofilattico Sperimentale del Mezzogiorno | INMI Lazzaro Spallanzani IRCCS | Barbara Bartolini, Cesare E.M. Gruber, Martina Rueca, Francesco Messina, Antonino Di Caro, Giovanna Fusco, Maurizio Viscardi, Giorgia Borriello, Sergio Brandi, Maria R. Capobianchi |
| EPI_ISL_560644 | Hospital | National Reference Center for Viruses of Respiratory Infections, Institut Pasteur, Paris | Sylvie Behillil, Fabiana Gambaro, Etienne Simon-Lorière, Vincent Enouf, Maud Vanpeene, Sylvie van der Werf |
| EPI_ISL_560927, EPI_ISL_560928 | Texas Department of State Health Services | Texas Department of State Health Services | Rashmi Tuladhar, Bonnie Oh, Jenny Zhang, Maliha Rahman, Anita Pokharel, Myong Koag, Chun Wang, Rachel Lee, Grace Kubin |
| EPI_ISL_561017, EPI_ISL_561018, EPI_ISL_561019, EPI_ISL_561020, EPI_ISL_561021, EPI_ISL_561022, EPI_ISL_561023 | MRCG at LSHTM Genomics lab | MRCG at LSHTM Genomics lab | Abdul Karim sesay, Abdoulie Kanteh, Jarra Manneh, Mariama Kujabi, Bakary Sanyang |
| EPI_ISL_561799 | Victorian Infectious Diseases Reference Laboratory (VIDRL) | VIDRL and MDU-PHL | Caly, L., Seemann, T., Sait, M., Schultz, M. B., Druce J., Sherry, N. |
| EPI_ISL_565897, EPI_ISL_565898, EPI_ISL_565903, EPI_ISL_565904, EPI_ISL_565905 | Michigan Department of Health and Human Services, Bureau of Laboratories | Michigan Department of Health and Human Services, Bureau of Laboratories | Blankenship HM, Riner D, Soehnlen MK |
| EPI_ISL_568629, EPI_ISL_568630, EPI_ISL_568631, EPI_ISL_568632, EPI_ISL_568633, EPI_ISL_568634, EPI_ISL_568635, EPI_ISL_568636, EPI_ISL_568637 | Florida Bureau of Public Health Laboratories | Florida Bureau of Public Health Laboratories | Sarah Schmedes, Jason Blanton |
| EPI_ISL_568716, EPI_ISL_568718, EPI_ISL_568719, EPI_ISL_568720, EPI_ISL_568721, EPI_ISL_568724, EPI_ISL_568726, EPI_ISL_568727, EPI_ISL_568728, EPI_ISL_568729, EPI_ISL_568730, EPI_ISL_568731, EPI_ISL_568732, EPI_ISL_568733, EPI_ISL_568734, EPI_ISL_568735, EPI_ISL_568736, EPI_ISL_568737, EPI_ISL_568738, EPI_ISL_568739, EPI_ISL_568740, EPI_ISL_568741, EPI_ISL_568742, EPI_ISL_568743, EPI_ISL_568744, EPI_ISL_568745, EPI_ISL_568746, EPI_ISL_568747, EPI_ISL_568748, EPI_ISL_568749, EPI_ISL_568750, EPI_ISL_568751, EPI_ISL_568752, EPI_ISL_568753, EPI_ISL_568754, EPI_ISL_568755, EPI_ISL_568756, EPI_ISL_568757, EPI_ISL_568758, EPI_ISL_568759, EPI_ISL_568760, EPI_ISL_568761, EPI_ISL_568762, EPI_ISL_568763, EPI_ISL_568764, EPI_ISL_568765, EPI_ISL_568766, EPI_ISL_568767, EPI_ISL_568768, EPI_ISL_568769, EPI_ISL_568770, EPI_ISL_568771, EPI_ISL_568772, EPI_ISL_568773, EPI_ISL_568774, EPI_ISL_568775, EPI_ISL_568776, EPI_ISL_568777, EPI_ISL_568778, EPI_ISL_568779, EPI_ISL_568780, EPI_ISL_568781, EPI_ISL_568782, EPI_ISL_568783, EPI_ISL_568784, EPI_ISL_568785, EPI_ISL_568786, EPI_ISL_568787, EPI_ISL_568788, EPI_ISL_568789, EPI_ISL_568790, EPI_ISL_568791, EPI_ISL_568792, EPI_ISL_568793, EPI_ISL_568794, EPI_ISL_568795, EPI_ISL_568796, EPI_ISL_568797, EPI_ISL_568798, EPI_ISL_568799, EPI_ISL_568800, EPI_ISL_568801, EPI_ISL_568802, EPI_ISL_568803, EPI_ISL_568804, EPI_ISL_568805, EPI_ISL_568806, EPI_ISL_568807, EPI_ISL_568808, EPI_ISL_568809, EPI_ISL_568810, EPI_ISL_568811, EPI_ISL_568812, EPI_ISL_568813, EPI_ISL_568814, EPI_ISL_568815, EPI_ISL_568816, EPI_ISL_568817, EPI_ISL_568818, EPI_ISL_568819, EPI_ISL_568820, EPI_ISL_568821, EPI_ISL_568822, EPI_ISL_568823, EPI_ISL_568824, EPI_ISL_568825, EPI_ISL_568827, EPI_ISL_568828, EPI_ISL_568829, EPI_ISL_568830, EPI_ISL_568832, EPI_ISL_568833, EPI_ISL_568841, EPI_ISL_568842, EPI_ISL_568843, EPI_ISL_568844, EPI_ISL_568845, EPI_ISL_568846, EPI_ISL_568847 | | | |
| see above | KEMRI-Wellcome Trust Research Programme/KEMRI-CGMR-C Kilifi | KEMRI-Wellcome Trust Research Programme/KEMRI-CGMR-C Kilifi | Githinji et al 2020 |
| EPI_ISL_568971, EPI_ISL_568972, EPI_ISL_568973, EPI_ISL_568974, EPI_ISL_568975, EPI_ISL_568976, EPI_ISL_568977, EPI_ISL_568978, EPI_ISL_568979, EPI_ISL_568980, EPI_ISL_568981 | | | |
| see above | MEPHI, Aix Marseille University | MEPHI, Aix Marseille University | Anthony LEVASSEUR |
| EPI_ISL_569611 | Sioux Falls Urgent Care | South Dakota Public Health Laboratory | Matt Plumb, Jacob Garfin, Xiong Wang, and Chris Carlson |
| EPI_ISL_569667, EPI_ISL_569668, EPI_ISL_569669, EPI_ISL_569670 | Lee Lab | Lee Lab | Sung Yong Park, Gina Faraci, Pamela M. Ward, Jane F. Emerson, and Ha Youn Lee |
| EPI_ISL_569735, EPI_ISL_569736, EPI_ISL_569737, EPI_ISL_569738, EPI_ISL_569739, EPI_ISL_569740, EPI_ISL_569741, EPI_ISL_569742, EPI_ISL_569743, EPI_ISL_569744, EPI_ISL_569745, EPI_ISL_569763, EPI_ISL_569802, EPI_ISL_569809, EPI_ISL_569812, EPI_ISL_569817, EPI_ISL_569818, EPI_ISL_569821, EPI_ISL_569822, EPI_ISL_569825, EPI_ISL_569832, EPI_ISL_569837, EPI_ISL_569840, EPI_ISL_569848, EPI_ISL_569855 | | | |
| see above | Omsk Research Institute of Natural Focal Infections | WHO National Influenza Centre Russian Federation | Artem Fadeev, Ekaterina Gradoboeva, Ekaterina Savkina, Daria Nashatyreva, Elena Poleshchuk, Aleksei Vasilenko, Valery Yakimenko, Andrey Komissarov |
| EPI_ISL_569967, EPI_ISL_570013, EPI_ISL_570026 | Unity Health Toronto | Ontario Institute for Cancer Research | Ramzi Fattouh, Larissa M. Matukas, Yan Chen,Mark Downing, Trina Otterman, Karel Boissinot, Wai Sum Siu, Zhi Cui, Le Luu, Samira Mubareka, TIBDN, Ilinca Lungu, Bernard Lam, Jeremy Johns, Paul Krzyzanowski, Richard de Borja, Felicia Vincelli, Philip Zuzarte, Jared T. Simpson |
| EPI_ISL_570214, EPI_ISL_570215, EPI_ISL_570217, EPI_ISL_570218, EPI_ISL_570219, EPI_ISL_570220, EPI_ISL_570221, EPI_ISL_570222, EPI_ISL_570223, EPI_ISL_570224, EPI_ISL_570225, EPI_ISL_570226, EPI_ISL_570227, EPI_ISL_570228, EPI_ISL_570229, EPI_ISL_570230, EPI_ISL_570231, EPI_ISL_570232, EPI_ISL_570233, EPI_ISL_570234, EPI_ISL_570235, EPI_ISL_570236, EPI_ISL_570237, EPI_ISL_570238, EPI_ISL_570239, EPI_ISL_570240, EPI_ISL_570241, EPI_ISL_570242, EPI_ISL_570243, EPI_ISL_570244, EPI_ISL_570245, EPI_ISL_570246, EPI_ISL_570247, EPI_ISL_570248, EPI_ISL_570249, EPI_ISL_570250, EPI_ISL_570251 | | | |
| see above | UW Virology Lab | UW Virology Lab | Pavitra Roychoudhury, Hong Xie, Lasata Shrestha, Amin Addetia, Victoria M Rachleff, Meei-Li Huang, Keith R Jerome, Alexander Greninger |
| EPI_ISL_572339, EPI_ISL_572369, EPI_ISL_572370, EPI_ISL_572371, EPI_ISL_572372, EPI_ISL_572373, EPI_ISL_572374, EPI_ISL_572376, EPI_ISL_572377, EPI_ISL_572378, EPI_ISL_572379, EPI_ISL_572380, EPI_ISL_572381, EPI_ISL_572382, EPI_ISL_572383, EPI_ISL_572384, EPI_ISL_572385, EPI_ISL_572386, EPI_ISL_572387, EPI_ISL_572388, EPI_ISL_572389, EPI_ISL_572390, EPI_ISL_572391, EPI_ISL_572392, EPI_ISL_572393, EPI_ISL_572394, EPI_ISL_572395, EPI_ISL_572396 | | | |
| see above | LACEN/PE | WallauLab, Aggeu Magalhaes Institute | Marcelo Henrique Santos Paiva, Duschinka Ribeiro Duarte Guedes, Cássia Docena, Matheus Filgueira Bezerra, Filipe Zimmer Dezordi, Laís Ceschini Machado, Larissa Krokovsky, Elisama Helvecio, Alexandre Freitas da Silva, Luydson Richardson Silva Vasconcelos, Antonio Mauro Rezende, Severino Jefferson Ribeiro da Silva, Kamila Gaudêncio da Silva Sales, Bruna Santos Lima Figueiredo de Sá, Derciliano Lopes da Cruz, Claudio Eduardo Cavalcanti, Armando de Menezes Neto, Caroline Targino Alves da Silva, Renata Pessôa Germano Mendes, Maria Almerice Lopes da Silva, Tiago Gräf, Paola Cristina Resende, Gonzalo Bello0, Michelle da Silva Barros, Wheverton Ricardo Correia do Nascimento,, Rodrigo Moraes Loyo Arcoverde, Luciane Caroline Albuquerque Bezerra, Sinval Pinto Brandão Filho, Constância Flávia Junqueira Ayres, Gabriel Luz Wallau |
| EPI_ISL_572402, EPI_ISL_572404, EPI_ISL_572429, EPI_ISL_572431, EPI_ISL_572443, EPI_ISL_572445, EPI_ISL_572451, EPI_ISL_572473, EPI_ISL_572500, EPI_ISL_572502, EPI_ISL_572552, EPI_ISL_572554, EPI_ISL_572557, EPI_ISL_572562, EPI_ISL_572578, EPI_ISL_572588, EPI_ISL_572601, EPI_ISL_572650, EPI_ISL_572669, EPI_ISL_572677, EPI_ISL_572679, EPI_ISL_572682, EPI_ISL_572687, EPI_ISL_572688, EPI_ISL_572691, EPI_ISL_572696, EPI_ISL_572710, EPI_ISL_572736, EPI_ISL_572739, EPI_ISL_572748, EPI_ISL_572750, EPI_ISL_572769, EPI_ISL_572771, EPI_ISL_572773, EPI_ISL_572813, EPI_ISL_572815, EPI_ISL_572843, EPI_ISL_573236, EPI_ISL_573237, EPI_ISL_573238, EPI_ISL_573239, EPI_ISL_573240, EPI_ISL_573241, EPI_ISL_573242, EPI_ISL_573243, EPI_ISL_573244, EPI_ISL_573245 | | | |
| see above | Liverpool Clinical Laboratories | COVID-19 Genomics UK (COG-UK) Consortium | Sam Haldenby, Anita Lucaci, Steve Paterson, Julian Hiscox, Alistair Darby, M Almsaud, A Alrezaihi, Muhannad Alruwaili, Stuart D Armstrong, Jones Benjamin, Eleanor G Bentley, Anu Chawla, Jordan J Clark, Angela Cowell, Richard Eccles, Isabel García-Dorival, Matthew Gemmell, Alessandro Gerada, PKF Gilmore, Richard Gregory, Ximeng Han, Catherine Hartley, Margaret Hughes, Miren Iturriza-Gomara, James Johnson, L Luu, Jenifer Manson, Charlotte Nelson, Elaine O'Toole, Cassie Olateju, Rebekah Penrice-Randal , Lucille Rainbow, N.P Randle, Trevor Ian Robinson, Parul Sharma, Ghada T Shawli, James P Stewart, Neil Swainston, Ecaterina Vamos, Joanne Watts, Mark Whitehead |
| EPI_ISL_574295, EPI_ISL_574296, EPI_ISL_574297, EPI_ISL_574298, EPI_ISL_574299, EPI_ISL_574300 | LSUHS Emerging Viral Threat Laboratory | Microbial Genome Sequencing Center | Jeremy P. Kamil, Rona S. Scott, Maarten Van Diest, Malgorzata Bienkowska-Haba, Katarzyna Zwolinska, Andrew D. Yurochko, Christopher G. Kevil, Martin J. Sapp, Daniel J. Snyder, Vaughn S. Cooper, John A. Vanchiere |
| EPI_ISL_574301, EPI_ISL_574302, EPI_ISL_574303, EPI_ISL_574304, EPI_ISL_574305, EPI_ISL_574306, EPI_ISL_574307, EPI_ISL_574308, EPI_ISL_574309, EPI_ISL_574310, EPI_ISL_574311, EPI_ISL_574312, EPI_ISL_574313, EPI_ISL_574314, EPI_ISL_574315, EPI_ISL_574316 | | | |
| see above | LSUHS Emerging Viral Threat Laboratory | Microbial Genome Sequencing Center | Rona S. Scott, Jeremy P. Kamil, Maarten Van Diest, Malgorzata Bienkowska-Haba, Katarzyna Zwolinska, Andrew D. Yurochko, Christopher G. Kevil, Martin J. Sapp, Daniel J. Snyder, Vaughn S. Cooper, John A. Vanchiere |
| EPI_ISL_574434, EPI_ISL_574435, EPI_ISL_574436, EPI_ISL_574437, EPI_ISL_574438, EPI_ISL_574439, EPI_ISL_574440, EPI_ISL_574441, EPI_ISL_574442, EPI_ISL_574443, EPI_ISL_574444, EPI_ISL_574445, EPI_ISL_574446, EPI_ISL_574447, EPI_ISL_574448, EPI_ISL_574449, EPI_ISL_574450, EPI_ISL_574451, EPI_ISL_574452, EPI_ISL_574453, EPI_ISL_574454, EPI_ISL_574455, EPI_ISL_574456, EPI_ISL_574457, EPI_ISL_574458, EPI_ISL_574459, EPI_ISL_574460, EPI_ISL_574461, EPI_ISL_574462, EPI_ISL_574463, EPI_ISL_574464, EPI_ISL_574465, EPI_ISL_574466, EPI_ISL_574467, EPI_ISL_574468, EPI_ISL_574469, EPI_ISL_574470, EPI_ISL_574471, EPI_ISL_574472, EPI_ISL_574473 | | | |
| see above | Texas Department of State Health Services | Texas Department of State Health Services | Rashmi Tuladhar, Bonnie Oh, Jenny Zhang, Maliha Rahman, Anita Pokharel, Myong Koag, Chun Wang, Rachel Lee, Grace Kubin |
| EPI_ISL_574611 | RS Premier Bintaro | Eijkman Institute for Molecular Biology, Ministry of Research and Technology/National Agency for Research and Innovation | Frilasita A Yudhaputri, Edison Johar, Hidayat Trimarsanto, Iskandar A Adnan, Willy Agustine, David H Muljono, Safarina G Malik, Herawati Sudoyo, Khin Saw Myint, Amin Soebandrio |
| EPI_ISL_574648 | Seattle Flu Study | Seattle Flu Study | Deborah A. Nickerson, Chris D. Frazar, Jover Lee, Benjamin Pelle, Matthew Richardson, Amanda Adler, Elisabeth Brandstetter, Peter D. Han, Kairsten Fay, Misja Ilcisin, Kirsten Lacombe, Thomas R. Sibley, Melissa Truong, Caitlin R. Wolf, Michael Boeckh, Janet A. Englund, Michael Famulare, Barry R. Lutz, Mark J. Rieder, Lea M. Starita, Matthew Thompson, Jay Shendure, Trevor Bedford, Helen Y. Chu |
| EPI_ISL_575117, EPI_ISL_575118, EPI_ISL_575119, EPI_ISL_575120, EPI_ISL_575121, EPI_ISL_575122, EPI_ISL_575123, EPI_ISL_575124, EPI_ISL_575125, EPI_ISL_575126, EPI_ISL_575127, EPI_ISL_575128, EPI_ISL_575129, EPI_ISL_575130, EPI_ISL_575131, EPI_ISL_575132, EPI_ISL_575133, EPI_ISL_575134, | | | |

| | | | |
|---|---|---|---|
| EPI_ISL_575135, EPI_ISL_575136, EPI_ISL_575137, EPI_ISL_575138, EPI_ISL_575139, EPI_ISL_575140, EPI_ISL_575141, EPI_ISL_575142, EPI_ISL_575143, EPI_ISL_575144, EPI_ISL_575148, EPI_ISL_575247, EPI_ISL_575248, EPI_ISL_575253, EPI_ISL_575254, EPI_ISL_575255, EPI_ISL_575260, EPI_ISL_575261, EPI_ISL_575262, EPI_ISL_575263, EPI_ISL_575264, EPI_ISL_575265, EPI_ISL_575266, EPI_ISL_575267, EPI_ISL_575268, EPI_ISL_575269, EPI_ISL_575270, EPI_ISL_575271, EPI_ISL_575272, EPI_ISL_575273, EPI_ISL_575274, EPI_ISL_575275, EPI_ISL_575276, EPI_ISL_575277, EPI_ISL_575278, EPI_ISL_575279, EPI_ISL_575280, EPI_ISL_575281, EPI_ISL_575282, EPI_ISL_575283, EPI_ISL_575284, EPI_ISL_575285, EPI_ISL_575286, EPI_ISL_575287, EPI_ISL_575288 | | | |
| see above | Utah Public Health Laboratory | Utah Public Health Laboratory | Erin Young, Kelly Oakeson |
| EPI_ISL_576281 | Texas Department of State Health Services | Texas Department of State Health Services | Rashmi Tuladhar, Bonnie Oh, Jenny Zhang, Maliha Rahman, Anita Pokharel, Myong Koag, Chun Wang, Rachel Lee, Grace Kubin |
| EPI_ISL_576366, EPI_ISL_576367, EPI_ISL_576368, EPI_ISL_576369, EPI_ISL_576370 | Utah Public Health Laboratory, Utah Public Health Laboratory Infectious Disease submission group | Utah Public Health Laboratory, Utah Public Health Laboratory Infectious Disease submission group | Young,E.L., Oakeson,K., Sangster,A., Hirschi,B., Butz,H. |
| EPI_ISL_577606 | Infectious Diseases and Tropical Medicine Research Center, Infectious Diseases and Tropical Medicine Research Center | Infectious Diseases and Tropical Medicine Research Center, Infectious Diseases and Tropical Medicine Research Center | Ahangarzadeh,S. |
| EPI_ISL_577631, EPI_ISL_577632 | The National Institute of Public Health | State Veterinary Institute Prague | Nagy,A;Jirincova,H;Novakova,L;Trnka,D;Vecerova,J |
| EPI_ISL_577739 | Institute of Virology, Biomedical Research Center of the Slovak Academy of Sciences, Bratislava | Faculty of Natural Sciences, Comenius University, Bratislava | Kristína Boršová, Viktória Hodorová, Broa Brejová, Viktória abanová, Dominika Friová, Sabina Fumaová Havlíková, Juraj Kopáek, Martina Liková, ubomíra Lukáiková, Martina Neboháová, Monika Sláviková, Edita Staroová, Elena Tichá, Tomáš Vina, Boris Klempa, Jozef Nosek |
| EPI_ISL_577872, EPI_ISL_577873, EPI_ISL_577874 | Dutch COVID-19 response team | Erasmus Medical Center | Bas Oude Munnink, Reina Sikkema, David Nieuwenhuijse, Irina Chestakova, Anne van der Linden, Marjan Boter, Emmanuelle Munger, Corine GeurtsvanKessel, Annemiek van der Eijk, Richard Molenkamp, Marion Koopmans, on behalf of the Dutch national COVID-19 response team. |
| EPI_ISL_578170, EPI_ISL_578171, EPI_ISL_578172 | University of Michigan Clinical Microbiology Laboratory | Lauring Lab, University of Michigan, Department of Microbiology and Immunology | Valesano |
| EPI_ISL_578186 | Hospital General Juan Ramón Jiménez | Instituto de Salud Carlos III | Iglesias-Caballero, M. Molinero Calamita, M. González-Esguevillas, M. Camarero, S. Pozo, F. Casas, I. Jiménez, P. Jiménez, M. Zaballos, A. Monzón, S. Varona, S. Juliá, M. Cuesta, I, J. Saavedra |
| EPI_ISL_578579, EPI_ISL_578588 | Wisconsin State Laboratory of Hygiene Communicable Disease Division | Wisconsin State Laboratory of Hygiene Communicable Disease Division | Kelsey R. Florek, Abigail C. Shockey |
| EPI_ISL_578704, EPI_ISL_578705, EPI_ISL_578706, EPI_ISL_578707, EPI_ISL_578708, EPI_ISL_578709, EPI_ISL_578710, EPI_ISL_578711, EPI_ISL_578712, EPI_ISL_578713, EPI_ISL_578714, EPI_ISL_578715, EPI_ISL_578716, EPI_ISL_578717, EPI_ISL_578718, EPI_ISL_578719, EPI_ISL_578720, EPI_ISL_578721, EPI_ISL_578722, EPI_ISL_578723, EPI_ISL_578724, EPI_ISL_578725, EPI_ISL_578726 | | | |
| see above | LSUHS Emerging Viral Threat Laboratory | Microbial Genome Sequencing Center | Maarten Van Diest, Jeremy P. Kamil, Rona S. Scott, Malgorzata Bienkowska-Haba, Katarzyna Zwolinska, Andrew D. Yurochko, Christopher G. Kevil, Martin J. Sapp, Daniel J. Snyder, Vaughn S. Cooper, John A. Vanchiere |
| EPI_ISL_579406, EPI_ISL_579407 | North Shore Hospital | Institute of Environmental Science and Research (ESR) | Xiaoyun Ren, Matt Storey, Nikki Freed, Muhammad Faisal, Jing Wang, Hermes Perez, Anja Werno, Antje van der Linden, Arlo Upton, Chris Mansell, David Hammer, Dragana Drinkovic, Gary McAuliffe, Hana Sofia Andersson, James Ussher, Jill Sherwood, Josh Freeman, Julia Howard, Juliet Elvy, Mary DeAlmeida, Matt Blakiston, Matthew Rogers, Max Bloomfield, Michael Addidle, Michelle Balm, Sally Roberts, Sarah Jefferies, Sharmini Muttaiyah, Susan Morpeth, Susan Taylor, Timothy Blackmore, Vani Sathyendran, Veronica Playle, Virginia Hope, Erasmus Smit, Lauren Jelly, Olin Silander, Joep de Ligt |
| EPI_ISL_579422 | LabTests | Institute of Environmental Science and Research (ESR) | Xiaoyun Ren, Matt Storey, Nikki Freed, Muhammad Faisal, Jing Wang, Hermes Perez, Anja Werno, Antje van der Linden, Arlo Upton, Chris Mansell, David Hammer, Dragana Drinkovic, Gary McAuliffe, Hana Sofia Andersson, James Ussher, Jill Sherwood, Josh Freeman, Julia Howard, Juliet Elvy, Mary DeAlmeida, Matt Blakiston, Matthew Rogers, Max Bloomfield, Michael Addidle, Michelle Balm, Sally Roberts, Sarah Jefferies, Sharmini Muttaiyah, Susan Morpeth, Susan Taylor, Timothy Blackmore, Vani Sathyendran, Veronica Playle, Virginia Hope, Erasmus Smit, Lauren Jelly, Olin Silander, Joep de Ligt |
| EPI_ISL_580633, EPI_ISL_580635, EPI_ISL_580636, EPI_ISL_580640, EPI_ISL_580645, EPI_ISL_580648, EPI_ISL_580650, EPI_ISL_580651, EPI_ISL_580655, EPI_ISL_580660, EPI_ISL_580663, EPI_ISL_580674, EPI_ISL_580675, EPI_ISL_580677, EPI_ISL_580678, EPI_ISL_580679, EPI_ISL_580680, EPI_ISL_580681, EPI_ISL_580682 | | | |
| see above | Lighthouse Lab in Alderley Park | Wellcome Sanger Institute for the COVID-19 Genomics UK (COG-UK) consortium | Jacquelyn Wynn, Mairead Hyland, The Lighthouse Lab in Alderley Park and Alex Alderton, Roberto Amato, Sonia Goncalves, Ewan Harrison, David K. Jackson, Ian Johnston, Dominic Kwiatkowski, Cordelia Langford, John Sillitoe on behalf of the Wellcome Sanger Institute COVID-19 Surveillance Team |
| EPI_ISL_580686 | Lighthouse Lab in Alderley Park | Wellcome Sanger Institute for the COVID-19 Genomics UK (COG-UK) Consortium | Jacquelyn Wynn, Mairead Hyland, The Lighthouse Lab in Alderley Park and Alex Alderton, Roberto Amato, Sonia Goncalves, Ewan Harrison, David K. Jackson, Ian Johnston, Dominic Kwiatkowski, Cordelia Langford, John Sillitoe on behalf of the Wellcome Sanger Institute COVID-19 Surveillance Team |
| EPI_ISL_580689, EPI_ISL_580690, EPI_ISL_580691, EPI_ISL_580692, EPI_ISL_580695, EPI_ISL_580696, EPI_ISL_580697, EPI_ISL_580699, EPI_ISL_580706 | Lighthouse Lab in Alderley Park | Wellcome Sanger Institute for the COVID-19 Genomics UK (COG-UK) consortium | Jacquelyn Wynn, Mairead Hyland, The Lighthouse Lab in Alderley Park and Alex Alderton, Roberto Amato, Sonia Goncalves, Ewan Harrison, David K. Jackson, Ian Johnston, Dominic Kwiatkowski, Cordelia Langford, John Sillitoe on behalf of the Wellcome Sanger Institute COVID-19 Surveillance Team |
| EPI_ISL_580708 | Lighthouse Lab in Alderley Park | Wellcome Sanger Institute for the COVID-19 Genomics UK (COG-UK) Consortium | Jacquelyn Wynn, Mairead Hyland, The Lighthouse Lab in Alderley Park and Alex Alderton, Roberto Amato, Sonia Goncalves, Ewan Harrison, David K. Jackson, Ian Johnston, Dominic Kwiatkowski, Cordelia Langford, John Sillitoe on behalf of the Wellcome Sanger Institute COVID-19 Surveillance Team |
| EPI_ISL_580710, EPI_ISL_580720, EPI_ISL_580722, EPI_ISL_580723, EPI_ISL_580730, EPI_ISL_580734, EPI_ISL_580737, EPI_ISL_580738, EPI_ISL_580740, EPI_ISL_580754, EPI_ISL_580763, EPI_ISL_580764, EPI_ISL_580769, EPI_ISL_580771 | | | |
| see above | Lighthouse Lab in Alderley Park | Wellcome Sanger Institute for the COVID-19 Genomics UK (COG-UK) consortium | Jacquelyn Wynn, Mairead Hyland, The Lighthouse Lab in Alderley Park and Alex Alderton, Roberto Amato, Sonia Goncalves, Ewan Harrison, David K. Jackson, Ian Johnston, Dominic Kwiatkowski, Cordelia Langford, John Sillitoe on behalf of the Wellcome Sanger Institute COVID-19 Surveillance Team |
| EPI_ISL_580780 | Lighthouse Lab in Alderley Park | Wellcome Sanger Institute for the COVID-19 Genomics UK (COG-UK) Consortium | Jacquelyn Wynn, Mairead Hyland, The Lighthouse Lab in Alderley Park and Alex Alderton, Roberto Amato, Sonia Goncalves, Ewan Harrison, David K. Jackson, Ian Johnston, Dominic Kwiatkowski, Cordelia Langford, John Sillitoe on behalf of the Wellcome Sanger Institute COVID-19 Surveillance Team |
| EPI_ISL_580784, EPI_ISL_580787, EPI_ISL_580788, EPI_ISL_580789, EPI_ISL_580791, EPI_ISL_580821, EPI_ISL_580822, EPI_ISL_580823 | Lighthouse Lab in Alderley Park | Wellcome Sanger Institute for the COVID-19 Genomics UK (COG-UK) consortium | Jacquelyn Wynn, Mairead Hyland, The Lighthouse Lab in Alderley Park and Alex Alderton, Roberto Amato, Sonia Goncalves, Ewan Harrison, David K. Jackson, Ian Johnston, Dominic Kwiatkowski, Cordelia Langford, John Sillitoe on behalf of the Wellcome Sanger Institute COVID-19 Surveillance Team |
| EPI_ISL_580824 | Lighthouse Lab in Alderley Park | Wellcome Sanger Institute for the COVID-19 Genomics UK (COG-UK) Consortium | Jacquelyn Wynn, Mairead Hyland, The Lighthouse Lab in Alderley Park and Alex Alderton, Roberto Amato, Sonia Goncalves, Ewan Harrison, David K. Jackson, Ian Johnston, Dominic Kwiatkowski, Cordelia Langford, John Sillitoe on behalf of the Wellcome Sanger Institute COVID-19 Surveillance Team |
| EPI_ISL_580825, EPI_ISL_580828, EPI_ISL_580833, EPI_ISL_580835, EPI_ISL_580849, EPI_ISL_580851, EPI_ISL_580853 | Lighthouse Lab in Alderley Park | Wellcome Sanger Institute for the COVID-19 Genomics UK (COG-UK) consortium | Jacquelyn Wynn, Mairead Hyland, The Lighthouse Lab in Alderley Park and Alex Alderton, Roberto Amato, Sonia Goncalves, Ewan Harrison, David K. Jackson, Ian Johnston, Dominic Kwiatkowski, Cordelia Langford, John Sillitoe on behalf of the Wellcome Sanger Institute COVID-19 Surveillance Team |
| EPI_ISL_581458, EPI_ISL_581459, EPI_ISL_581475 | Medizinische Klinik Innere Medizin I, Universitätsklinikum Tübingen | NGS Competence Center Tübingen, Institut für Medizinische Mikrobiologie und Hygiene, Universitätsklinikum Tübingen | Angel Angelov |
| EPI_ISL_581926, EPI_ISL_581927 | University Hospital Basel, Clinical Virology | University Hospital Basel, Clinical Bacteriology | Madlen Stange, Alfredo Mari, Tim Roloff, Helena MB Seth-Smith, Michael Schweitzer, Myrta Brunner, Karoline Leuzinger, Kirstine K. Soegaard, Alexander Gensch, Sarah Tschudin-Sutter, Simon Fuchs, Julia Bielicki, Hans Pargger, Martin Siegemund, Christian Nickel, Roland Bingisser, Michael Osthoff, Stefano Bassetti, Rita Schneider-Sliwa, Manuel Battegay, Hans Hirsch, Adrian Egli |
| EPI_ISL_582003 | Infectious Diseases and Tropical Medicine Research Center, Infectious Diseases and Tropical Medicine Research Center | Infectious Diseases and Tropical Medicine Research Center, Infectious Diseases and Tropical Medicine Research Center | Haghjoo javanmard,S., Ahangarzadeh,S., Shariati,L., Ataei,B., Ranjbar,M.M., Shoaei,P. |
| EPI_ISL_582005 | Infectious Diseases and Tropical Medicine Research Center, Infectious Diseases and Tropical Medicine Research Center | Infectious Diseases and Tropical Medicine Research Center, Infectious Diseases and Tropical Medicine Research Center | Haghjooy Javanmard,S., Shariati,L., Ahangarzadeh,S., Ataei,B., Shoaei,P., Ranjbar,M.M. |
| EPI_ISL_582006 | Infectious Diseases and Tropical Medicine Research Center, Infectious Diseases and Tropical Medicine Research Center | Infectious Diseases and Tropical Medicine Research Center, Infectious Diseases and Tropical Medicine Research Center | Shariati,L., Haghjooy Javanmard,S., Ahangarzadeh,S., Ataei,B., Shoaei,P., Ranjbar,M.M. |

| | | | |
|---|---|---|---|
| EPI_ISL_582007 | Infectious Diseases and Tropical Medicine Research Center, Infectious Diseases and Tropical Medicine Research Center | Infectious Diseases and Tropical Medicine Research Center, Infectious Diseases and Tropical Medicine Research Center | Ahangarzadeh,S., Haghjooy Javanmard,S., Ataei,B., Shariati,L., Shoaei,P., Ranjbar,M.M. |
| EPI_ISL_582008 | Infectious Diseases and Tropical Medicine Research Center, Infectious Diseases and Tropical Medicine Research Center | Infectious Diseases and Tropical Medicine Research Center, Infectious Diseases and Tropical Medicine Research Center | Haghjooy Javanmard,S., Ahangarzadeh,S., Shariati,L., Shoaei,P., Ranjbar,M.M., Ataei,B. |
| EPI_ISL_582009 | Infectious Diseases and Tropical Medicine Research Center, Infectious Diseases and Tropical Medicine Research Center | Infectious Diseases and Tropical Medicine Research Center, Infectious Diseases and Tropical Medicine Research Center | Haghjooy Javanmard,S., Ahangarzadeh,S., Ataei,B., Shariati,L., Shoaei,P., Ranjbar,M.M. |
| EPI_ISL_582013 | Infectious Diseases and Tropical Medicine Research Center, Infectious Diseases and Tropical Medicine Research Center | Infectious Diseases and Tropical Medicine Research Center, Infectious Diseases and Tropical Medicine Research Center | Shariati,L., Haghjooy Javanmard,S., Ahangarzadeh,S., Ataei,B., Shoaei,P. |
| EPI_ISL_582014 | Infectious Diseases and Tropical Medicine Research Center, Infectious Diseases and Tropical Medicine Research Center | Infectious Diseases and Tropical Medicine Research Center, Infectious Diseases and Tropical Medicine Research Center | Ahangarzadeh,S., Haghjoo Javanmard,S., Shariati,L., Ataei,B., Ranjbar,M.M., Shoaei,P. |
| EPI_ISL_582015 | Infectious Diseases and Tropical Medicine Research Center, Infectious Diseases and Tropical Medicine Research Center | Infectious Diseases and Tropical Medicine Research Center, Infectious Diseases and Tropical Medicine Research Center | Shariati,L., Ahangarzadeh,S., Haghjooy Javanmard,S., Ataei,B., Ranjbar,M.M., Shoaei,P. |
| EPI_ISL_582148, EPI_ISL_582149, EPI_ISL_582150, EPI_ISL_582151, EPI_ISL_582152, EPI_ISL_582154, EPI_ISL_582156, EPI_ISL_582157, EPI_ISL_582158, EPI_ISL_582159, EPI_ISL_582160, EPI_ISL_582162, EPI_ISL_582163, EPI_ISL_582165, EPI_ISL_582166, EPI_ISL_582167, EPI_ISL_582168, EPI_ISL_582169, EPI_ISL_582170, EPI_ISL_582171, EPI_ISL_582173, EPI_ISL_582174, EPI_ISL_582175, EPI_ISL_582176, EPI_ISL_582177, EPI_ISL_582179, EPI_ISL_582181, EPI_ISL_582183, EPI_ISL_582184, EPI_ISL_582185, EPI_ISL_582186, EPI_ISL_582187, EPI_ISL_582188, EPI_ISL_582189, EPI_ISL_582190, EPI_ISL_582192, EPI_ISL_582193, EPI_ISL_582194, EPI_ISL_582195, EPI_ISL_582198, EPI_ISL_582200, EPI_ISL_582202, EPI_ISL_582208, EPI_ISL_582210, EPI_ISL_582211, EPI_ISL_582213, EPI_ISL_582214, EPI_ISL_582216, EPI_ISL_582219 | | | |
| see above | TriCore Reference Laboratories | Center for Global Health, University of New Mexico Health Sciences Center | Daryl Domman, Kurt Schwalm, Twila Kunde, Joseph Hicks, Michael Edwards, Darrell Dinwiddie |
| EPI_ISL_582283 | Cadham Provincial Laboratory | National Microbiology Laboratory (NML) | Anna Majer, Shari Tyson, Grace Seo, Philip Mabon, Elsie Grudeski, Rhiannon Huzarewich, Russell Mandes, Anneliese Landgraff, Jennifer Tanner, Natalie Knox, Morag Graham, Gary Van Domselaar, Paul Van Caeseele, Jared Bullard, David Alexander, Kerry Dust, Nathalie Bastien, Yan Li, Timothy Booth, Darian Hole, Madison Chapel, CanCOGeN's metadata curation team, Public Health Agency of Canada CanCOGeN team |
| EPI_ISL_583173, EPI_ISL_583174, EPI_ISL_583175, EPI_ISL_583176, EPI_ISL_583177, EPI_ISL_583178, EPI_ISL_583179, EPI_ISL_583180, EPI_ISL_583181, EPI_ISL_583182, EPI_ISL_583183, EPI_ISL_583184, EPI_ISL_583185, EPI_ISL_583186 | | | |
| see above | San Bernardino County Public Health Lab | Chan-Zuckerberg Biohub | CZB Cliahub Consortium |
| EPI_ISL_583438, EPI_ISL_583440 | Memorial Sloan Kettering Cancer Center | van Bakel Laboratory, Genetics and Genomics Sciences, Icahn School of Medicine at Mount Sinai | Teresa Aydillo, Ana S. Gonzalez-Reiche, Sadaf Aslam, Adriana van de Guchte, Zenab Khan, Ajay Obla, Jayeeta Dutta, Harm van Bakel, Judith Aberg, Adolfo García-Sastre, Gunjan Shah, Tobias Hohl, Genovefa Papanicolaou, Miguel-Angel Perales, Kent Sepkowitz, Ngolela Esther Babady, and Mini Kamboj |
| EPI_ISL_583442 | Garcia-Sastre Laboratory, Department of Microbiology, Icahn School of Medicine at Mount Sinai | van Bakel Laboratory, Genetics and Genomics Sciences, Icahn School of Medicine at Mount Sinai | Teresa Aydillo, Ana S. Gonzalez-Reiche, Sadaf Aslam, Adriana van de Guchte, Zenab Khan, Ajay Obla, Jayeeta Dutta, Harm van Bakel, Judith Aberg, Adolfo García-Sastre, Gunjan Shah, Tobias Hohl, Genovefa Papanicolaou, Miguel-Angel Perales, Kent Sepkowitz, Ngolela Esther Babady, and Mini Kamboj |
| EPI_ISL_583447, EPI_ISL_583448, EPI_ISL_583457 | Memorial Sloan Kettering Cancer Center | van Bakel Laboratory, Genetics and Genomics Sciences, Icahn School of Medicine at Mount Sinai | Teresa Aydillo, Ana S. Gonzalez-Reiche, Sadaf Aslam, Adriana van de Guchte, Zenab Khan, Ajay Obla, Jayeeta Dutta, Harm van Bakel, Judith Aberg, Adolfo García-Sastre, Gunjan Shah, Tobias Hohl, Genovefa Papanicolaou, Miguel-Angel Perales, Kent Sepkowitz, Ngolela Esther Babady, and Mini Kamboj |
| EPI_ISL_583459 | Garcia-Sastre Laboratory, Department of Microbiology, Icahn School of Medicine at Mount Sinai | van Bakel Laboratory, Genetics and Genomics Sciences, Icahn School of Medicine at Mount Sinai | Teresa Aydillo, Ana S. Gonzalez-Reiche, Sadaf Aslam, Adriana van de Guchte, Zenab Khan, Ajay Obla, Jayeeta Dutta, Harm van Bakel, Judith Aberg, Adolfo García-Sastre, Gunjan Shah, Tobias Hohl, Genovefa Papanicolaou, Miguel-Angel Perales, Kent Sepkowitz, Ngolela Esther Babady, and Mini Kamboj |
| EPI_ISL_583460, EPI_ISL_583480 | Memorial Sloan Kettering Cancer Center | van Bakel Laboratory, Genetics and Genomics Sciences, Icahn School of Medicine at Mount Sinai | Teresa Aydillo, Ana S. Gonzalez-Reiche, Sadaf Aslam, Adriana van de Guchte, Zenab Khan, Ajay Obla, Jayeeta Dutta, Harm van Bakel, Judith Aberg, Adolfo García-Sastre, Gunjan Shah, Tobias Hohl, Genovefa Papanicolaou, Miguel-Angel Perales, Kent Sepkowitz, Ngolela Esther Babady, and Mini Kamboj |
| EPI_ISL_583630, EPI_ISL_583631, EPI_ISL_583632, EPI_ISL_583637, EPI_ISL_583638, EPI_ISL_583639, EPI_ISL_583646, EPI_ISL_583647, EPI_ISL_583654, EPI_ISL_583655, EPI_ISL_583656, EPI_ISL_583663, EPI_ISL_583664, EPI_ISL_583669, EPI_ISL_583676, EPI_ISL_583677, EPI_ISL_583683, EPI_ISL_583684, EPI_ISL_583685 | | | |
| see above | Austrian Agency for Health and Food Safety (AGES) | Bergthaler laboratory, CeMM Research Center for Molecular Medicine of the Austrian Academy of Sciences | Alexandra Popa, Benedikt Agerer, Henrique Colaco, Lukas Endler, Jakob-Wendelin Genger, Alexander Lercher, Mark Smyth, Thomas Penz, Michael Schuster, Jan Laine, Martin Senekowitsch, Judith Aberle, Stephan Aberle, Peter Hufnagl, Daniela Schmid, Franz Allerberger, Elisabeth Puchhammer-Stoeckl, Manfred Nairz, Guenter Weiss, Gregor Hörmann, Kinga Rigler-Hohenwarter, Rainer Gattringer, Wegene Borena, Dorothee von Laer, Gernot Walder, Peter Obrist, Christian Paar, Sabine Sussitz-Rack, Gunther Vogl, Adi Steinrigl, Christoph Bock, Andreas Bergthaler |
| EPI_ISL_583693, EPI_ISL_583702, EPI_ISL_583706, EPI_ISL_583707, EPI_ISL_583708, EPI_ISL_583717, EPI_ISL_583719, EPI_ISL_583723 | Center for Virology, Medical University of Vienna | Bergthaler laboratory, CeMM Research Center for Molecular Medicine of the Austrian Academy of Sciences | Alexandra Popa, Benedikt Agerer, Henrique Colaco, Lukas Endler, Jakob-Wendelin Genger, Alexander Lercher, Mark Smyth, Thomas Penz, Michael Schuster, Jan Laine, Martin Senekowitsch, Judith Aberle, Stephan Aberle, Peter Hufnagl, Daniela Schmid, Franz Allerberger, Elisabeth Puchhammer-Stoeckl, Manfred Nairz, Guenter Weiss, Gregor Hörmann, Kinga Rigler-Hohenwarter, Rainer Gattringer, Wegene Borena, Dorothee von Laer, Gernot Walder, Peter Obrist, Christian Paar, Sabine Sussitz-Rack, Gunther Vogl, Adi Steinrigl, Christoph Bock, Andreas Bergthaler |
| EPI_ISL_583739, EPI_ISL_583740 | Dr. Gernot Walder GmbH | Bergthaler laboratory, CeMM Research Center for Molecular Medicine of the Austrian Academy of Sciences | Alexandra Popa, Benedikt Agerer, Henrique Colaco, Lukas Endler, Jakob-Wendelin Genger, Alexander Lercher, Mark Smyth, Thomas Penz, Michael Schuster, Jan Laine, Martin Senekowitsch, Judith Aberle, Stephan Aberle, Peter Hufnagl, Daniela Schmid, Franz Allerberger, Elisabeth Puchhammer-Stoeckl, Manfred Nairz, Guenter Weiss, Gregor Hörmann, Kinga Rigler-Hohenwarter, Rainer Gattringer, Wegene Borena, Dorothee von Laer, Gernot Walder, Peter Obrist, Christian Paar, Sabine Sussitz-Rack, Gunther Vogl, Adi Steinrigl, Christoph Bock, Andreas Bergthaler |
| EPI_ISL_583856 | Institute for Laboratory Diagnostics and Microbiology, Klinikum Klagenfurt am Worthersee | Bergthaler laboratory, CeMM Research Center for Molecular Medicine of the Austrian Academy of Sciences | Alexandra Popa, Benedikt Agerer, Henrique Colaco, Lukas Endler, Jakob-Wendelin Genger, Alexander Lercher, Mark Smyth, Thomas Penz, Michael Schuster, Jan Laine, Martin Senekowitsch, Judith Aberle, Stephan Aberle, Peter Hufnagl, Daniela Schmid, Franz Allerberger, Elisabeth Puchhammer-Stoeckl, Manfred Nairz, Guenter Weiss, Gregor Hörmann, Kinga Rigler-Hohenwarter, Rainer Gattringer, Wegene Borena, Dorothee von Laer, Gernot Walder, Peter Obrist, Christian Paar, Sabine Sussitz-Rack, Gunther Vogl, Adi Steinrigl, Christoph Bock, Andreas Bergthaler |
| EPI_ISL_583965, EPI_ISL_583966 | UOC Microbiologia e Virologia, Azienda Ospedaliera Universitaria Senese, Siena, Italy | Dipartimento di Biotecnologie Mediche | Maria Grazia Cusi, David Pinzauti, Claudia Gandolfo, Gabriele Anichini, Gianni Pozzi, Francesco Santoro |
| EPI_ISL_584073 | The National Institute of Public Health | State Veterinary Institute Prague | Nagy,A;Jirincova,H;Novakova,L;Trnka,D;Vecerova,J |
| EPI_ISL_584123, EPI_ISL_584124, EPI_ISL_584125, EPI_ISL_584126, EPI_ISL_584127, EPI_ISL_584128, EPI_ISL_584129, EPI_ISL_584130, EPI_ISL_584131, EPI_ISL_584132, EPI_ISL_584133, EPI_ISL_584134, EPI_ISL_584135, EPI_ISL_584136, EPI_ISL_584137, EPI_ISL_584138, EPI_ISL_584139, EPI_ISL_584140, EPI_ISL_584141, EPI_ISL_584142, EPI_ISL_584143, EPI_ISL_584144, EPI_ISL_584145, EPI_ISL_584146, EPI_ISL_584147, EPI_ISL_584148, EPI_ISL_584149, EPI_ISL_584150, EPI_ISL_584151, EPI_ISL_584152, EPI_ISL_584153, EPI_ISL_584154, EPI_ISL_584155, EPI_ISL_584156 | | | |
| see above | Texas Department of State Health Services | Texas Department of State Health Services | Rashmi Tuladhar, Bonnie Oh, Jenny Zhang, Maliha Rahman, Anita Pokharel, Mayela Pedrueza, Myong Koag, Chun Wang, Rachel Lee, Grace Kubin |
| EPI_ISL_584501, EPI_ISL_584504, EPI_ISL_584505, EPI_ISL_584506, EPI_ISL_584507, EPI_ISL_584508, EPI_ISL_584509, EPI_ISL_584510, EPI_ISL_584511, EPI_ISL_584512, EPI_ISL_584513, EPI_ISL_584514, EPI_ISL_584515, EPI_ISL_584516, EPI_ISL_584517, EPI_ISL_584518, EPI_ISL_584519 | | | |
| see above | UHCW / University of Warwick | COVID-19 Genomics UK (COG-UK) Consortium | Richard Stark, Chrystala Constantinidou, Meera Unnikrishnan, Laura Baxter, Jeff Cheng, Grace Taylor-Joyce, Hannah Elizabeth Bridgewater, Lucy Frost, Sarojini Pandey, Paul Brown, Tauqeer Alam, Sascha Ott, Dimitris Grammatopoulos |
| EPI_ISL_586313, EPI_ISL_586314, EPI_ISL_586315, EPI_ISL_586316, EPI_ISL_586317, EPI_ISL_586318, EPI_ISL_586333, EPI_ISL_586335, EPI_ISL_586338, EPI_ISL_586343 | Toronto Invasive Bacterial Diseases Network | McMaster University | Allison McGeer, Patryk Aftanas, Hooman Derakhshani, Angel Li, Kuganya Nirmalarajah, Emily Panousis, Ahmed Draia, Jalees Nasir, Michael Surette, Samira Mubareka, Andrew G. McArthur |
| EPI_ISL_589268, EPI_ISL_589272, EPI_ISL_589274, EPI_ISL_589275, EPI_ISL_589277, EPI_ISL_589279, EPI_ISL_589280, EPI_ISL_589281, EPI_ISL_589283, EPI_ISL_589284, EPI_ISL_589285, EPI_ISL_589286, EPI_ISL_589287, EPI_ISL_589288, EPI_ISL_589290, EPI_ISL_589296, EPI_ISL_589299, EPI_ISL_589301 | | | |
| see above | Lighthouse Lab in Alderley Park | Wellcome Sanger Institute for the COVID-19 Genomics UK (COG-UK) consortium | Jacquelyn Wynn, Mairead Hyland, The Lighthouse Lab in Alderley Park and Alex Alderton, Roberto Amato, Sonia Goncalves, Ewan Harrison, David K. Jackson, Ian Johnston, Dominic Kwiatkowski, Cordelia Langford, John Sillitoe on behalf of the Wellcome Sanger Institute COVID-19 Surveillance Team |
| EPI_ISL_589302 | Lighthouse Lab in Alderley Park | Wellcome Sanger Institute for the COVID-19 Genomics UK (COG-UK) Consortium | Jacquelyn Wynn, Mairead Hyland, The Lighthouse Lab in Alderley Park and Alex Alderton, Roberto Amato, Sonia Goncalves, Ewan Harrison, David K. Jackson, Ian Johnston, Dominic Kwiatkowski, Cordelia Langford, John Sillitoe on behalf of the Wellcome Sanger Institute COVID-19 Surveillance Team |
| EPI_ISL_589303, EPI_ISL_589306, EPI_ISL_589307, EPI_ISL_589308, EPI_ISL_589309, EPI_ISL_589310, EPI_ISL_589311, EPI_ISL_589312, EPI_ISL_589313, EPI_ISL_589314, EPI_ISL_589316, EPI_ISL_589319, EPI_ISL_589320, EPI_ISL_589321, EPI_ISL_589322, EPI_ISL_589323, EPI_ISL_589326, EPI_ISL_589327 | | | |
| see above | Lighthouse Lab in Alderley Park | Wellcome Sanger Institute for the COVID-19 Genomics UK (COG-UK) consortium | Jacquelyn Wynn, Mairead Hyland, The Lighthouse Lab in Alderley Park and Alex Alderton, Roberto Amato, Sonia Goncalves, Ewan Harrison, David K. Jackson, Ian Johnston, Dominic Kwiatkowski, Cordelia Langford, John Sillitoe on behalf of the Wellcome Sanger Institute COVID-19 Surveillance Team |

| | | | |
|---|---|---|---|
| EPI_ISL_589329 | Lighthouse Lab in Alderley Park | Wellcome Sanger Institute for the COVID-19 Genomics UK (COG-UK) Consortium | Jacquelyn Wynn, Mairead Hyland, The Lighthouse Lab in Alderley Park and Alex Alderton, Roberto Amato, Sonia Goncalves, Ewan Harrison, David K. Jackson, Ian Johnston, Dominic Kwiatkowski, Cordelia Langford, John Sillitoe on behalf of the Wellcome Sanger Institute COVID-19 Surveillance Team |
| EPI_ISL_589331, EPI_ISL_589332, EPI_ISL_589333, EPI_ISL_589336, EPI_ISL_589338, EPI_ISL_589339, EPI_ISL_589340 | Lighthouse Lab in Alderley Park | Wellcome Sanger Institute for the COVID-19 Genomics UK (COG-UK) consortium | Jacquelyn Wynn, Mairead Hyland, The Lighthouse Lab in Alderley Park and Alex Alderton, Roberto Amato, Sonia Goncalves, Ewan Harrison, David K. Jackson, Ian Johnston, Dominic Kwiatkowski, Cordelia Langford, John Sillitoe on behalf of the Wellcome Sanger Institute COVID-19 Surveillance Team |
| EPI_ISL_589342, EPI_ISL_589344 | Lighthouse Lab in Alderley Park | Wellcome Sanger Institute for the COVID-19 Genomics UK (COG-UK) Consortium | Jacquelyn Wynn, Mairead Hyland, The Lighthouse Lab in Alderley Park and Alex Alderton, Roberto Amato, Sonia Goncalves, Ewan Harrison, David K. Jackson, Ian Johnston, Dominic Kwiatkowski, Cordelia Langford, John Sillitoe on behalf of the Wellcome Sanger Institute COVID-19 Surveillance Team |
| EPI_ISL_589345, EPI_ISL_589346, EPI_ISL_589347, EPI_ISL_589350, EPI_ISL_589351, EPI_ISL_589352, EPI_ISL_589353, EPI_ISL_589356, EPI_ISL_589358, EPI_ISL_589360, EPI_ISL_589361, EPI_ISL_589362, EPI_ISL_589363, EPI_ISL_589364, EPI_ISL_589365, EPI_ISL_589366, EPI_ISL_589368, EPI_ISL_589369, EPI_ISL_589374, EPI_ISL_589375 | | | |
| see above | Lighthouse Lab in Alderley Park | Wellcome Sanger Institute for the COVID-19 Genomics UK (COG-UK) consortium | Jacquelyn Wynn, Mairead Hyland, The Lighthouse Lab in Alderley Park and Alex Alderton, Roberto Amato, Sonia Goncalves, Ewan Harrison, David K. Jackson, Ian Johnston, Dominic Kwiatkowski, Cordelia Langford, John Sillitoe on behalf of the Wellcome Sanger Institute COVID-19 Surveillance Team |
| EPI_ISL_589376 | Lighthouse Lab in Alderley Park | Wellcome Sanger Institute for the COVID-19 Genomics UK (COG-UK) Consortium | Jacquelyn Wynn, Mairead Hyland, The Lighthouse Lab in Alderley Park and Alex Alderton, Roberto Amato, Sonia Goncalves, Ewan Harrison, David K. Jackson, Ian Johnston, Dominic Kwiatkowski, Cordelia Langford, John Sillitoe on behalf of the Wellcome Sanger Institute COVID-19 Surveillance Team |
| EPI_ISL_589379, EPI_ISL_589380, EPI_ISL_589382, EPI_ISL_589383, EPI_ISL_589384, EPI_ISL_589386, EPI_ISL_589387, EPI_ISL_589388, EPI_ISL_589389, EPI_ISL_589390, EPI_ISL_589391, EPI_ISL_589392, EPI_ISL_589393, EPI_ISL_589395, EPI_ISL_589396, EPI_ISL_589397, EPI_ISL_589399, EPI_ISL_589400, EPI_ISL_589402, EPI_ISL_589403, EPI_ISL_589408, EPI_ISL_589409, EPI_ISL_589410, EPI_ISL_589411, EPI_ISL_589415, EPI_ISL_589418, EPI_ISL_589419, EPI_ISL_589421, EPI_ISL_589422, EPI_ISL_589424, EPI_ISL_589425, EPI_ISL_589433, EPI_ISL_589435, EPI_ISL_589438, EPI_ISL_589439, EPI_ISL_589441, EPI_ISL_589442, EPI_ISL_589443, EPI_ISL_589444, EPI_ISL_589445, EPI_ISL_589446, EPI_ISL_589447, EPI_ISL_589449, EPI_ISL_589451, EPI_ISL_589453, EPI_ISL_589454, EPI_ISL_589455, EPI_ISL_589457, EPI_ISL_589460, EPI_ISL_589461, EPI_ISL_589462, EPI_ISL_589463, EPI_ISL_589464, EPI_ISL_589465, EPI_ISL_589468, EPI_ISL_589469, EPI_ISL_589471, EPI_ISL_589475, EPI_ISL_589476, EPI_ISL_589477, EPI_ISL_589480, EPI_ISL_589483, EPI_ISL_589486, EPI_ISL_589490, EPI_ISL_589492, EPI_ISL_589495, EPI_ISL_589496, EPI_ISL_589498, EPI_ISL_589500, EPI_ISL_589501, EPI_ISL_589502, EPI_ISL_589506, EPI_ISL_589510, EPI_ISL_589512, EPI_ISL_589514, EPI_ISL_589515, EPI_ISL_589516, EPI_ISL_589517, EPI_ISL_589518, EPI_ISL_589525, EPI_ISL_589527, EPI_ISL_589528, EPI_ISL_589529, EPI_ISL_589534, EPI_ISL_589536, EPI_ISL_589537, EPI_ISL_589538, EPI_ISL_589541, EPI_ISL_589543, EPI_ISL_589544, EPI_ISL_589545, EPI_ISL_589546, EPI_ISL_589548, EPI_ISL_589549, EPI_ISL_589552, EPI_ISL_589555, EPI_ISL_589557, EPI_ISL_589558, EPI_ISL_589559, EPI_ISL_589560, EPI_ISL_589562, EPI_ISL_589563, EPI_ISL_589565, EPI_ISL_589566, EPI_ISL_589569, EPI_ISL_589570 | | | |
| see above | Lighthouse Lab in Alderley Park | Wellcome Sanger Institute for the COVID-19 Genomics UK (COG-UK) consortium | Jacquelyn Wynn, Mairead Hyland, The Lighthouse Lab in Alderley Park and Alex Alderton, Roberto Amato, Sonia Goncalves, Ewan Harrison, David K. Jackson, Ian Johnston, Dominic Kwiatkowski, Cordelia Langford, John Sillitoe on behalf of the Wellcome Sanger Institute COVID-19 Surveillance Team |
| EPI_ISL_589765 | Lighthouse Lab in Alderley Park | Wellcome Sanger Institute for the COVID-19 Genomics UK (COG-UK) consortium | Jacquelyn Wynn, Mairead Hyland, The Lighthouse Lab in Alderley Park and Alex Alderton, Roberto Amato, Sonia Goncalves, Ewan Harrison, David K. Jackson, Ian Johnston, Dominic Kwiatkowski, Cordelia Langford, John Sillitoe on behalf of the Wellcome Sanger Institute COVID-19 Surveillance Team (http://www.sanger.ac.uk/covid-team) |
| EPI_ISL_591062 | Infectious Diseases and Tropical Medicine Research Center, Infectious Diseases and Tropical Medicine Research Center | Infectious Diseases and Tropical Medicine Research Center, Infectious Diseases and Tropical Medicine Research Center | Haghjooy Javanmard,S., Ahangarzadeh,S., Shariati,L., Ataei,B., Shoaei,P., Ranjbar,M.M. |
| EPI_ISL_591063 | Infectious Diseases and Tropical Medicine Research Center, Infectious Diseases and Tropical Medicine Research Center | Infectious Diseases and Tropical Medicine Research Center, Infectious Diseases and Tropical Medicine Research Center | Ahangarzadeh,S., Haghjooy Javanmard,S., Shariati,L. |
| EPI_ISL_591064 | Infectious Diseases and Tropical Medicine Research Center, Infectious Diseases and Tropical Medicine Research Center | Infectious Diseases and Tropical Medicine Research Center, Infectious Diseases and Tropical Medicine Research Center | Haghjooy Javanmard,S., Shariati,l., Ahangarzadeh,S., Ataei,B., Shoaei,P. |
| EPI_ISL_591065 | Infectious Diseases and Tropical Medicine Research Center, Infectious Diseases and Tropical Medicine Research Center | Infectious Diseases and Tropical Medicine Research Center, Infectious Diseases and Tropical Medicine Research Center | Haghjooy Javanmard,S., Ahangarzadeh,S., Shariati,L., Ataei,B., Shoaei,P., Ranjbar,M.M. |
| EPI_ISL_591281, EPI_ISL_591282, EPI_ISL_591283, EPI_ISL_591284, EPI_ISL_591285, EPI_ISL_591286, EPI_ISL_591287, EPI_ISL_591288, EPI_ISL_591289, EPI_ISL_591290, EPI_ISL_591291, EPI_ISL_591292, EPI_ISL_591293, EPI_ISL_591294, EPI_ISL_591295, EPI_ISL_591296, EPI_ISL_591297, EPI_ISL_591298, EPI_ISL_591299, EPI_ISL_591300, EPI_ISL_591301, EPI_ISL_591302, EPI_ISL_591303, EPI_ISL_591304, EPI_ISL_591305, EPI_ISL_591309 | | | |
| see above | Utah Public Health Laboratory | Utah Public Health Laboratory | Erin Young, Kelly Oakeson |
| EPI_ISL_591338, EPI_ISL_591339 | Dipartimento di Biotecnologie Mediche, University of Siena | Dipartimento di Biotecnologie Mediche, University of Siena | Cusi,M.G., Pinzauti,D., Gandolfo,C., Anichini,G., Pozzi,G., Santoro,F. |
| EPI_ISL_591341 | Infectious Diseases and Tropical Medicine Research Center, Infectious Diseases and Tropical Medicine Research Center | Infectious Diseases and Tropical Medicine Research Center, Infectious Diseases and Tropical Medicine Research Center | Haghjooy Javanmard,S., Ahangarzadeh,S., Shariati,L., Ataei,B., Shoaei,P., Ranjbar,M.M., Abutalebian,S. |
| EPI_ISL_591342 | Infectious Diseases and Tropical Medicine Research Center, Infectious Diseases and Tropical Medicine Research Center | Infectious Diseases and Tropical Medicine Research Center, Infectious Diseases and Tropical Medicine Research Center | Haghjooy Javanmard,S., Ahangarzadeh,S., Shariati,L., Abutalebian,S., Ataei,B., Ranjbar,M.M., Shoaei,P. |
| EPI_ISL_591344, EPI_ISL_591345 | Infectious Diseases and Tropical Medicine Research Center, Infectious Diseases and Tropical Medicine Research Center | Infectious Diseases and Tropical Medicine Research Center, Infectious Diseases and Tropical Medicine Research Center | Ahangarzadeh,S., Haghjooy Javanmard,S., Abutalebian,S., Shoaei,P., Ataei,B., Shariati,L. |
| EPI_ISL_591521 | Programa de Virología, Facultad de Medicina UC | Center for Mathematical Modeling and Center for Genome Regulation. Santiago, Chile | Gaggero A, Valiente F, Gaete A, Travisany D, Palma R, Urra C, Varas M, Allende ML, Maass A, González M, Ferres M. |
| EPI_ISL_593479 | Brigham and Women's Hospital | Jonathan Li Laboratory | Manish C. Choudhary, James Regan, Jonathan Z. Li |
| EPI_ISL_593493, EPI_ISL_593494, EPI_ISL_593495, EPI_ISL_593496, EPI_ISL_593497, EPI_ISL_593498, EPI_ISL_593499, EPI_ISL_593501, EPI_ISL_593502, EPI_ISL_593503, EPI_ISL_593549, EPI_ISL_593550 | | | |
| see above | Eastern Ontario Regional Laboratory Association | McMaster University | Leanne Mortimer, Hooman Derakhshani, Emily Panousis, Ahmed Draia, Jalees Nasir, Robert Slinger, Andrew G. McArthur |
| EPI_ISL_593999, EPI_ISL_594000, EPI_ISL_594001, EPI_ISL_594002, EPI_ISL_594003, EPI_ISL_594004, EPI_ISL_594005, EPI_ISL_594006 | Utah Public Health Laboratory | Utah Public Health Laboratory | Erin Young, Kelly Oakeson |
| EPI_ISL_594123 | Yale COVID-19 Biorepository | Grubaugh Lab - Yale School of Public Health | Joseph Fauver, Tara Alpert, Anderson Brito, Anne Wyllie, Chantal Vogels, Mary Petrone, Chaney Kalinich, Isabel Ott, Arnau Casanovas, Catherine Muenker, Adam Moore, Alice Lu, Maria Tokuyama, Patrick Wong, Peiwen Lu, Saad Omer, Richard Martinello, Allison Nelson, Shelli Farhadian, Akiko Iwasaki, Charlese Dela Cruz, Albert Ko, Nathan Grubaugh |
| EPI_ISL_594216 | Michigan Department of Health and Human Services, Bureau of Laboratories | Michigan Department of Health and Human Services, Bureau of Laboratories | Blankenship HM, Riner D, Soehnlen MK |
| EPI_ISL_596318, EPI_ISL_596349, EPI_ISL_596352 | Pathogenic Microorganisms Variability Laboratory | WHO National Influenza Centre Russian Federation | Andrey Komissarov, Artem Fadeev, Anna Ivanova, Kseniya Komissarova, Dmitry Bazhenov, Daria Danilenko, Dmitry Lioznov, Nadezhda Kuznetsova, Elena Shidlovskaya, Elizaveta Divisenko, Ekaterina Milashenko, Kirill Krasnoslobotsev, Evgeniya Mukasheva, Anna Ignatieva, Svetlana Trushakova, Alexey Shchetinin, Maria Nikiforova, Andrey Pochtovyy, Valeria Bacalin, Evgeny Usachev, Olga Burgasova, Ludmila Kolobukhina, Svetlana Smetanina, Elena Burtseva, Artem Tkachuk, Vladimir Gushchin, Alexander Gintsburg |
| EPI_ISL_600535, EPI_ISL_600543, EPI_ISL_600570 | Institute of Epidemiology Disease Control And Research | Institute for Developing Science and Health Initiatives | Lauren Cowley, Mokibul Hassan Afrad, Sadia Isfat Ara Rahman, Md. Mahfuz-Al-mamun, Firadausi Qadri, Tahmina Shirin |
| EPI_ISL_603053, EPI_ISL_603054, EPI_ISL_603055, EPI_ISL_603059, EPI_ISL_603061, EPI_ISL_603065, EPI_ISL_603066, EPI_ISL_603068, EPI_ISL_603069, EPI_ISL_603070, EPI_ISL_603072, EPI_ISL_603074, EPI_ISL_603078, EPI_ISL_603079 | | | |
| see above | Utah Public Health Laboratory, Utah Public Health Laboratory Infectious Disease submission group | Utah Public Health Laboratory, Utah Public Health Laboratory Infectious Disease submission group | Young,E.L., Oakeson,K. |
| EPI_ISL_603178 | INMI Lazzaro Spallanzani IRCCS | INMI Lazzaro Spallanzani IRCCS | Martina Rueca, Francesco Messina, Cesare E.M. Gruber, Barbara Bartolini, Simone Lanini, Emanuela Giombini, Antonino Di Caro, Maria R. Capobianchi |
| EPI_ISL_603187 | INMI Lazzaro Spallanzani IRCCS | INMI Lazzaro Spallanzani IRCCS | Cesare E.M. Gruber, Martina Rueca, Francesco Messina, Barbara Bartolini, Simone Lanini, Emanuela Giombini, Antonino Di Caro, Maria R. Capobianchi |
| EPI_ISL_610251, EPI_ISL_610252, EPI_ISL_610254, EPI_ISL_610255, EPI_ISL_610256, EPI_ISL_610257, EPI_ISL_610258, EPI_ISL_610259, EPI_ISL_610260, EPI_ISL_610261, EPI_ISL_610262, EPI_ISL_610265, EPI_ISL_610266, EPI_ISL_610267 | | | |
| see above | TriCore Reference Laboratories | Center for Global Health, University of New Mexico Health Sciences Center | Daryl Domman, Kurt Schwalm, Twila Kunde, Joseph Hicks, Michael Edwards, Darrell Dinwiddie |

| | | | |
|---|---|---|---|
| EPI_ISL_610269, EPI_ISL_610271, EPI_ISL_610273 | New Mexico Department of Health Scientific Laboratory Division | Center for Global Health, University of New Mexico Health Sciences Center | Daryl Domman, Kurt Schwalm, Twila Kunde, Joseph Hicks, Michael Edwards, Darrell Dinwiddie |
| EPI_ISL_611565, EPI_ISL_612042, EPI_ISL_612046, EPI_ISL_612443 | Liverpool Clinical Laboratories | COVID-19 Genomics UK (COG-UK) Consortium | Sam Haldenby, Anita Lucaci, Steve Paterson, Julian Hiscox, Alistair Darby, M Almsaud, A Alrezaihi, Muhannad Alruwaili, Stuart D Armstrong, Jones Benjamin, Eleanor G Bentley, Anu Chawla, Jordan J Clark, Angela Cowell, Richard Eccles, Isabel García-Dorival, Matthew Gemmell, Alessandro Gerada, PKF Gilmore, Richard Gregory, Ximeng Han, Catherine Hartley, Margaret Hughes, Miren Iturriza-Gomara, James Johnson, L Luu, Jenifer Manson, Charlotte Nelson, Elaine O'Toole, Cassie Olateju, Rebekah Penrice-Randal , Lucille Rainbow, N.P Randle, Trevor Ian Robinson, Parul Sharma, Ghada T Shawli, James P Stewart, Neil Swainston, Ecaterina Vamos, Joanne Watts, Mark Whitehead |
| EPI_ISL_614100, EPI_ISL_614101, EPI_ISL_614102, EPI_ISL_614103, EPI_ISL_614107, EPI_ISL_614108 | Virginia DCLS | Virginia DCLS | Virginia DCLS |
| EPI_ISL_614424, EPI_ISL_614425, EPI_ISL_614426, EPI_ISL_614449, EPI_ISL_614452, EPI_ISL_614460, EPI_ISL_614465, EPI_ISL_614466, EPI_ISL_614467, EPI_ISL_614468, EPI_ISL_614498, EPI_ISL_614499, EPI_ISL_614501, EPI_ISL_614502, EPI_ISL_614503, EPI_ISL_614504, EPI_ISL_614505, EPI_ISL_614506, EPI_ISL_614507, EPI_ISL_614508, EPI_ISL_614509, EPI_ISL_614510, EPI_ISL_614511, EPI_ISL_614512, EPI_ISL_614513, EPI_ISL_614785, EPI_ISL_614796, EPI_ISL_614797, EPI_ISL_614798, EPI_ISL_614799, EPI_ISL_614800, EPI_ISL_614801, EPI_ISL_614802, EPI_ISL_614803, EPI_ISL_614804, EPI_ISL_614805, EPI_ISL_614806, EPI_ISL_614807, EPI_ISL_614808, EPI_ISL_614809, EPI_ISL_614810, EPI_ISL_614812, EPI_ISL_614813, EPI_ISL_614816, EPI_ISL_614829 | | | |
| see above | Department of Virus and Microbiological Special Diagnostics, Statens Serum Institut, Denmark | Albertsen lab, Department of Chemistry and Bioscience, Aalborg University, Denmark | Danish Covid-19 Genome Consortia |
| EPI_ISL_615076, EPI_ISL_615078 | Uppsala klinisk mikrobiologi | The Public Health Agency of Sweden | Anna-Malin Linde, Maria Lind Karlberg, Mattias Haukland, Reza Advani, Olov Svartstrom, Oskar Karlsson Lindsjo, Sandra Broddesson, Petra Edquist, Mia Brytting, Anna Risberg, Karin Tegmark-Wisell |
| EPI_ISL_615086 | Karolinska Universitetslaboratoriet | The Public Health Agency of Sweden | Anna-Malin Linde, Maria Lind Karlberg, Mattias Haukland, Reza Advani, Olov Svartstrom, Oskar Karlsson Lindsjo, Sandra Broddesson, Petra Edquist, Mia Brytting, Anna Risberg, Karin Tegmark-Wisell |
| EPI_ISL_615094, EPI_ISL_615095, EPI_ISL_615096 | Umea klinisk mikrobiologi | The Public Health Agency of Sweden | Anna-Malin Linde, Maria Lind Karlberg, Mattias Haukland, Reza Advani, Olov Svartstrom, Oskar Karlsson Lindsjo, Sandra Broddesson, Petra Edquist, Mia Brytting, Anna Risberg, Karin Tegmark-Wisell |
| EPI_ISL_615097, EPI_ISL_615098 | Klinisk mikrobiologi Lanssjukhuset Ryhov, Jonkoping | The Public Health Agency of Sweden | Anna-Malin Linde, Maria Lind Karlberg, Mattias Haukland, Reza Advani, Olov Svartstrom, Oskar Karlsson Lindsjo, Sandra Broddesson, Petra Edquist, Mia Brytting, Anna Risberg, Karin Tegmark-Wisell |
| EPI_ISL_615125, EPI_ISL_615126, EPI_ISL_615127, EPI_ISL_615128, EPI_ISL_615129, EPI_ISL_615130 | Texas Department of State Health Services | Texas Department of State Health Services | Rashmi Tuladhar, Bonnie Oh, Jenny Zhang, Maliha Rahman, Anita Pokharel, Myong Koag, Chung Wang, Rachel Lee, Grace Kubin, Mayela Pedrueza |
| EPI_ISL_617939, EPI_ISL_617940, EPI_ISL_617942, EPI_ISL_617943, EPI_ISL_617944, EPI_ISL_617945, EPI_ISL_617946, EPI_ISL_617947, EPI_ISL_617948, EPI_ISL_617949, EPI_ISL_617950, EPI_ISL_617951, EPI_ISL_617952, EPI_ISL_618027, EPI_ISL_618098, EPI_ISL_622410, EPI_ISL_622411, EPI_ISL_622420, EPI_ISL_622423, EPI_ISL_622424, EPI_ISL_622425, EPI_ISL_622426, EPI_ISL_622430, EPI_ISL_622433, EPI_ISL_622434, EPI_ISL_622436, EPI_ISL_622497, EPI_ISL_622498, EPI_ISL_622517, EPI_ISL_622587 | | | |
| see above | Department of Virus and Microbiological Special Diagnostics, Statens Serum Institut, Denmark | Albertsen lab, Department of Chemistry and Bioscience, Aalborg University, Denmark | Danish Covid-19 Genome Consortia |
| EPI_ISL_623113, EPI_ISL_623121, EPI_ISL_623123, EPI_ISL_623131, EPI_ISL_623143, EPI_ISL_623149, EPI_ISL_623151, EPI_ISL_623155, EPI_ISL_623158, EPI_ISL_623165 | Laboratorio de Virologia Molecular / UFRJ | Bioinformatics Laboratory / LNCC | Carolina M Voloch, Ronaldo S Francisco Jr, Luiz G P de Almeida, Otavio J. Brustolini, Cynthia C Cardoso, Alexandra L Gerber, Ana Paula de C Guimarães, Diana Mariani, Covid19-UFRJ Workgroup, Luís Cristóvão Pôrto, Renato S Aguiar, Terezinha M P P Castiñeiras, Orlando C. Ferreira, Amilcar Tanuri, Ana Tereza R de Vasconcelos |
| EPI_ISL_625459 | Child Health Research Foundation | Child Health Research Foundation | Senjuti Saha, Md Saiful Islam Sajib, Nikkon Sarkar, Syed Muktadir Al Sium, Afroza Akter Tanni, Roly Malaker, Arif Mohammad Tanmoy, Md Hafizur Rahman, Samir K Saha |
| EPI_ISL_626222 | Institute for Virology, University Hospital Essen | Center of Medical Microbiology, Virology, and Hospital Hygiene, University of Duesseldorf | Olympia E. Anastasiou, Ulf Dittmer, Maximilian Damagnez, Alexander Dilthey, Torsten Houwaart, Lisanna Hülse, Malte Kohns Vasconcelos, Nadine Lübke, Jessica Nicolai, Klaus Pfeffer, Daniel Strelow, Jörg Timm, Andreas Walker, Tobias Wienemann |
| EPI_ISL_626460, EPI_ISL_626461, EPI_ISL_626462, EPI_ISL_626463, EPI_ISL_626469, EPI_ISL_626470, EPI_ISL_626471, EPI_ISL_626472, EPI_ISL_626473, EPI_ISL_626474, EPI_ISL_626475, EPI_ISL_626476 | | | |
| see above | Northwestern Memorial Hospital | Ozer Lab | Ramon Lorenzo-Redondo, Hannah H. Nam, Scott C. Roberts, Lacy M. Simons, Chad J. Achenbach, Lawrence J. Jennings, Chao Qi, Alan R. Hauser, Michael G. Ison, Judd F. Hultquist, Egon A. Ozer |
| EPI_ISL_628757 | UHAS COVID-19 Lab | UHAS COVID-19 Lab | Kwabena O. Duedu, Jones Gyamfi, Reuben Ayivor-Djanie, John O. Gyapong and the UHAS COVID-19 Lab Team |
| EPI_ISL_628964, EPI_ISL_628965, EPI_ISL_628966, EPI_ISL_628967, EPI_ISL_628968, EPI_ISL_628969, EPI_ISL_628970, EPI_ISL_628971, EPI_ISL_628972, EPI_ISL_628973, EPI_ISL_628974, EPI_ISL_628975, EPI_ISL_628976, EPI_ISL_628977, EPI_ISL_628978, EPI_ISL_628979, EPI_ISL_628980, EPI_ISL_628981, EPI_ISL_628982, EPI_ISL_628983, EPI_ISL_628984, EPI_ISL_628985, EPI_ISL_628986, EPI_ISL_628987, EPI_ISL_628988, EPI_ISL_628989, EPI_ISL_628990, EPI_ISL_628991, EPI_ISL_628992, EPI_ISL_628993, EPI_ISL_628994, EPI_ISL_628995, EPI_ISL_628996, EPI_ISL_628997, EPI_ISL_628998, EPI_ISL_628999 | | | |
| see above | Utah Public Health Laboratory | Utah Public Health Laboratory | Erin Young, Kelly Oakeson |
| EPI_ISL_631932 | OCME Office Of Chief Medical Examiner | New York City Public Health Laboratory | Jade Wang, et al. |
| EPI_ISL_631933 | St Barnabas Hospital | New York City Public Health Laboratory | Jade Wang, et al. |
| EPI_ISL_631934 | OCME Office Of Chief Medical Examiner | New York City Public Health Laboratory | Jade Wang, et al. |
| EPI_ISL_631935 | St Barnabas Hospital | New York City Public Health Laboratory | Jade Wang, et al. |
| EPI_ISL_631936, EPI_ISL_631937, EPI_ISL_631938 | OCME Office Of Chief Medical Examiner | New York City Public Health Laboratory | Jade Wang, et al. |
| EPI_ISL_631944, EPI_ISL_631945, EPI_ISL_631946, EPI_ISL_631947, EPI_ISL_631948, EPI_ISL_631949 | St Barnabas Hospital | New York City Public Health Laboratory | Jade Wang, et al. |
| EPI_ISL_631956 | OCME Office Of Chief Medical Examiner | New York City Public Health Laboratory | Jade Wang, et al. |
| EPI_ISL_631957 | St Barnabas Hospital | New York City Public Health Laboratory | Jade Wang, et al. |
| EPI_ISL_635279 | Institute of Microbiology and Immunology, Faculty of Medicine, University of Ljubljana | Institute of Microbiology and Immunology, Faculty of Medicine, University of Ljubljana | Tomaž Mark Zorec, Samo Zakotnik, Miša Korva, Tatjana Avši - Županc, Mario Poljak |
| EPI_ISL_635300, EPI_ISL_635301, EPI_ISL_635302, EPI_ISL_635303, EPI_ISL_635304, EPI_ISL_635305, EPI_ISL_635306, EPI_ISL_635307, EPI_ISL_635308, EPI_ISL_635309, EPI_ISL_635310, EPI_ISL_635311, EPI_ISL_635312, EPI_ISL_635313, EPI_ISL_635314, EPI_ISL_635315, EPI_ISL_635316, EPI_ISL_635317, EPI_ISL_635318, EPI_ISL_635319, EPI_ISL_635320, EPI_ISL_635321, EPI_ISL_635323, EPI_ISL_635324, EPI_ISL_635328, EPI_ISL_635374, EPI_ISL_635375 | | | |
| see above | San Diego County Public Health Laboratory | Andersen lab at Scripps Research | SEARCH Alliance San Diego with Tracy Basler, Jovan Shephard, Brett Austin |
| EPI_ISL_635479, EPI_ISL_635480, EPI_ISL_635481, EPI_ISL_635482, EPI_ISL_635483, EPI_ISL_635484, EPI_ISL_635485 | Centro de Diagnstico COVID-19 UABC Tijuana | Andersen lab at Scripps Research | SEARCH Alliance San Diego with Idanya Rubí Serafín Higuera, Manuel Sánchez Alavez, Jorge Luis Jiménez Niebla, Germán Ibarra, Jonathan Vincent Baena, Oscar Efrén Zazueta Fierro |
| EPI_ISL_636480 | Centre Hospitalier Pierre Oudot | CNR Virus des Infections Respiratoires - France SUD | Antonin Bal, Gregory Destras, Gwendolyne Burfin, Hadrien Règue, Alexandre Gaymard, Maude Bouscambert-Duchamp, Florence Morfin-Sherpa, Martine Valette, Bruno Lina, Laurence Josset |
| EPI_ISL_636481 | Centre Hospitalier de Bourg en Bresse | CNR Virus des Infections Respiratoires - France SUD | Antonin Bal, Gregory Destras, Gwendolyne Burfin, Hadrien Règue, Alexandre Gaymard, Maude Bouscambert-Duchamp, Florence Morfin-Sherpa, Martine Valette, Bruno Lina, Laurence Josset |

| | | | |
|---|---|---|---|
| EPI_ISL_636482, EPI_ISL_636483 | Centre Hospitalier Pierre Oudot | CNR Virus des Infections Respiratoires - France SUD | Antonin Bal, Gregory Destras, Gwendolyne Burfin, Hadrien Règue, Alexandre Gaymard, Maude Bouscambert-Duchamp, Florence Morfin-Sherpa, Martine Valette, Bruno Lina, Laurence Josset |
| EPI_ISL_636484, EPI_ISL_636485 | CNR Virus des Infections Respiratoires - France SUD | CNR Virus des Infections Respiratoires - France SUD | Antonin Bal, Gregory Destras, Gwendolyne Burfin, Hadrien Règue, Alexandre Gaymard, Maude Bouscambert-Duchamp, Florence Morfin-Sherpa, Martine Valette, Bruno Lina, Laurence Josset |
| EPI_ISL_636486 | Centre Hospitalier Pierre Oudot | CNR Virus des Infections Respiratoires - France SUD | Antonin Bal, Gregory Destras, Gwendolyne Burfin, Hadrien Règue, Alexandre Gaymard, Maude Bouscambert-Duchamp, Florence Morfin-Sherpa, Martine Valette, Bruno Lina, Laurence Josset |
| EPI_ISL_636573 | Dutch COVID-19 response team | National Institute for Public Health and the Environment (RIVM) | Adam Meijer, Harry Vennema, Jeroen Cremer, Sharon van den Brink, Bas van der Veer, AnneMarie van den Brandt, Florian Zwagemaker, Dennis Schmitz, Chantal Reusken, on behalf of the national COVID-19 response team |
| EPI_ISL_636964 | Pathogen Genomics Lab King Abdullah University of Science and Technology(KAUST) | Pathogen Genomics Lab King Abdullah University of Science and Technology(KAUST) | Rahul P Salunke, Sharif Hala, Raeece Naeem, Sara Mfarrej, Amit Kumar Subudhi, Amanda Ooi, Luke Esau, Fadwa Alofi, Fathia Ben Rached, Afrah Alsomali, Asim Khogeer, Ahmad Bakur Mahmoud, Anwar Hashem, Naif Almontashiri, Arnab Pain |
| EPI_ISL_636983, EPI_ISL_636984, EPI_ISL_636985 | Virology Lab, National Institute for Biomedical Research (INRB) | Project group Epidemology of Highly Pathogenic Microorganisms, Robert Koch-Institute | Jean-Jacques Muyembe Tamfum, Steve Ahuka-Mundeke, Eddy Kinganda-Lusamaki, Gabriel Mbunsu, Sheila Makiala, Essia Belarbi, Jasmin Schlotterbeck, Grit Schubert, Fabian Leendertz |
| EPI_ISL_639928, EPI_ISL_639929, EPI_ISL_639933, EPI_ISL_639935, EPI_ISL_639936, EPI_ISL_639937 | Omsk Research Institute of Natural Focal Infections | WHO National Influenza Centre Russian Federation | Artem Fadeev, Ekaterina Gradoboeva, Ekaterina Savkina, Daria Nashatyreva, Elena Poleshchuk, Aleksei Vasilenko, Valery Yakimenko, Andrey Komissarov |
| EPI_ISL_640081 | Vanguard CHC wc VGC | NHLS/UCT | Arash Iranzadeh, Deelan Doolabh, Lynn Tyers, Bruna Galvao, Innocent Mudau, Marvin Hsiao, Kruger Marais, Diana Hardie, Stephen Korsman, Carolyn Williamson |
| EPI_ISL_640231, EPI_ISL_640270 | MVZ Laborärzte Singen | MVZ Laborärzte Singen | Jonas Schmidt, Frithjof Blessing, Sandro Berghaus, Folker Wenzel |
| EPI_ISL_641313, EPI_ISL_641315 | Rocky Mountain Laboratories, RTS Genomics Unit, National Institute of Allergy and Infectious Diseases, National Institutes of Health | Rocky Mountain Laboratories, RTS Genomics Unit, National Institute of Allergy and Infectious Diseases, National Institutes of Health | Avanzato,V.A., Matson,M.J., Seifert,S.N., Pryce,R., Williamson,B.N., Anzick,S.L., Barbian,K., Judson,S.D., Fischer,E.R., Martens,C., Bowden,T.A., de Wit,E., Riedo,F.X., Munster,V.J., Siefert,S.N., WIlliamson,B.N., Anzick,S., Martens,C.A. |
| EPI_ISL_641534 | CHU de Nice - Hôpital Archet 16 | CNR Virus des Infections Respiratoires - France SUD | Antonin Bal, Géraldine Gonfrier, Gregory Destras, Gwendolyne Burfin, Hadrien Règue, Quentin Semanas, Martine Valette, Bruno Lina, Valérie Giordanengo, Laurence Josset |
| EPI_ISL_644218, EPI_ISL_644224 | CEPHR / Vincent's Hospital | Irish Coronavirus Sequencing Consortium - National Virus Reference Laboratory | Michael Carr, Gabriel Gonzalez, Alejandro Abner Garcia Leon, Patrick Mallon |
| EPI_ISL_644230 | CEPHR / Mater Hospital | Irish Coronavirus Sequencing Consortium - National Virus Reference Laboratory | Michael Carr, Gabriel Gonzalez, Alejandro Abner Garcia Leon, Patrick Mallon |
| EPI_ISL_644236, EPI_ISL_644255, EPI_ISL_644307, EPI_ISL_644308, EPI_ISL_644323, EPI_ISL_644330, EPI_ISL_644337, EPI_ISL_644345 | CEPHR / Vincent's Hospital | Irish Coronavirus Sequencing Consortium - National Virus Reference Laboratory | Michael Carr, Gabriel Gonzalez, Alejandro Abner Garcia Leon, Patrick Mallon |
| EPI_ISL_644572, EPI_ISL_644580, EPI_ISL_644581 | Veterinary Specialized Institute "Kraljevo", Serbia | Veterinary Specialized Institute "Kraljevo", Serbia | Vidanovic,D., Tesovic,B., Knezevic,A., Jovanovic,T., Jankovic,M., Sekler,M., Banovic Djeri,B., Petrovic,T., Volkening,J., Afonso,C. |
| EPI_ISL_645025, EPI_ISL_645026, EPI_ISL_645027, EPI_ISL_645028, EPI_ISL_645032, EPI_ISL_645033, EPI_ISL_645034, EPI_ISL_645043, EPI_ISL_645047, EPI_ISL_645048, EPI_ISL_645049, EPI_ISL_645050, EPI_ISL_645051, EPI_ISL_645052, EPI_ISL_645053, EPI_ISL_645054, EPI_ISL_645068, EPI_ISL_645069, EPI_ISL_645070, EPI_ISL_645071, EPI_ISL_645083, EPI_ISL_645084, EPI_ISL_645085, EPI_ISL_645086, EPI_ISL_645087, EPI_ISL_645088, EPI_ISL_645089, EPI_ISL_645090, EPI_ISL_645091, EPI_ISL_645092, EPI_ISL_645093, EPI_ISL_645094, EPI_ISL_645095, EPI_ISL_645096, EPI_ISL_645097, EPI_ISL_645098, EPI_ISL_645099, EPI_ISL_645105, EPI_ISL_645106, EPI_ISL_645107 | | | |
| see above | Human Genome Variation Research Group, Malopolska Centre of Biotechnology | Human Genome Variation Research Group, Malopolska Centre of Biotechnology | Kowalski,M., Pospiech,E., Klajmon,A., Gromowski,T., Pisarek,A., Marszalek,K., Kopera,K., Foremny,J., Swadzba,J., Sanak,M., Owczarek,K., Dabrowska,A., Szczepanski,A., Botwina,P., Labaj,P.P., Pyrc,K., Branicki,W. |
| EPI_ISL_645186, EPI_ISL_645187 | CHU de Saint-Étienne Hôpital Nord | CNR Virus des Infections Respiratoires - France SUD | Antonin Bal, Gregory Destras, Gwendolyne Burfin, Hadrien Règue, Quentin Semanas, Martine Valette, Bruno Lina, Issam Bechri, Manon Vogrig, Marine Delorme, Bruno Pozzetto, Thomas Bourlet, Sylvie Gonzalo, Sylvie Pillet, Laurence Josset |
| EPI_ISL_645212 | Unité des Virus Émergents | CNR Virus des Infections Respiratoires - France SUD | Antonin Bal, Gregory Destras, Gwendolyne Burfin, Hadrien Règue, Quentin Semanas, Martine Valette, Bruno Lina, Laetitia Ninove, Léa Luciani, Antoine Nougairède, Laurence Josset |
| EPI_ISL_648267, EPI_ISL_648268, EPI_ISL_648269, EPI_ISL_648270 | Brigham and Women's Hospital | Jonathan Li laboratory | Jonathan Z. Li, Manish C. Choudhary, Upasana D. Adhikari, George Eng, Douglas S Kwon |
| EPI_ISL_648344, EPI_ISL_648345, EPI_ISL_648346, EPI_ISL_648347, EPI_ISL_648348, EPI_ISL_648349 | Laboratorio de Investigaciones de Baney | University Hospital Basel, Clinical Bacteriology | Carlos Cortes, Claudia Daubenberger, Adrian Egli, Guillermo Garcia, Salome Hosch, Bonifacio Manguire Nlavo, Alfredo Mari, Maximilian Mpina, Elizabeth Nyakarungu, Diosdado Odjama Nseng Ada, Mitoha Ondo O Ayekaba, Tim Roloff, Tobias Schindler, Helena Seth-Smith, Madlen Stange, Philip Wonder Phiri |
| EPI_ISL_648531, EPI_ISL_648532, EPI_ISL_648533, EPI_ISL_648534 | Tulare County Public Health Lab | Chan-Zuckerberg Biohub | CZB Cliahub Consortium |
| EPI_ISL_648580, EPI_ISL_648581, EPI_ISL_648582, EPI_ISL_648583, EPI_ISL_648584, EPI_ISL_648585, EPI_ISL_648586, EPI_ISL_648587, EPI_ISL_648588, EPI_ISL_648589, EPI_ISL_648590, EPI_ISL_648591, EPI_ISL_648592, EPI_ISL_648593, EPI_ISL_648594 | | | |
| see above | Utah Public Health Laboratory | Utah Public Health Laboratory | Erin Young, Kelly Oakeson |
| EPI_ISL_648748, EPI_ISL_648749, EPI_ISL_648750, EPI_ISL_648751, EPI_ISL_648752, EPI_ISL_648753, EPI_ISL_648754, EPI_ISL_648755, EPI_ISL_648756, EPI_ISL_648757, EPI_ISL_648758, EPI_ISL_648759, EPI_ISL_648760, EPI_ISL_648761, EPI_ISL_648762, EPI_ISL_648763, EPI_ISL_648764, EPI_ISL_648765, EPI_ISL_648766, EPI_ISL_648767, EPI_ISL_648768, EPI_ISL_648769, EPI_ISL_648770, EPI_ISL_648771, EPI_ISL_648772, EPI_ISL_648773, EPI_ISL_648774, EPI_ISL_648775, EPI_ISL_648776, EPI_ISL_648777, EPI_ISL_648778, EPI_ISL_648779, EPI_ISL_648780, EPI_ISL_648781, EPI_ISL_648782, EPI_ISL_648783, EPI_ISL_648784, EPI_ISL_648785, EPI_ISL_648786, EPI_ISL_648787, EPI_ISL_648788, EPI_ISL_648789, EPI_ISL_648790, EPI_ISL_648791, EPI_ISL_648792, EPI_ISL_648793, EPI_ISL_648794 | | | |
| see above | Department of Laboratory Medicine, Tan Tock Seng Hospital | Department of Laboratory Medicine, Tan Tock Seng Hospital | Chen YYC, Zair X, Lim JX, Li C, Tang WY, Maurer-Stroh S, Barkham TMS, Nagarajan N, Sessions OM |
| EPI_ISL_649188 | Unité des Virus Émergents | CNR Virus des Infections Respiratoires - France SUD | Antonin Bal, Gregory Destras, Gwendolyne Burfin, Hadrien Règue, Quentin Semanas, Martine Valette, Bruno Lina, Laetitia Ninove, Léa Luciani, Antoine Nougairède, Laurence Josset |
| EPI_ISL_653151, EPI_ISL_653152, EPI_ISL_653211, EPI_ISL_653284, EPI_ISL_653285, EPI_ISL_653286, EPI_ISL_653287, EPI_ISL_653288, EPI_ISL_653289 | Florida Bureau of Public Health Laboratories | Florida Bureau of Public Health Laboratories | Sarah Schmedes, Jason Blanton |
| EPI_ISL_653826 | Institute of Post Graduate Medical Education & Research | National Institute of Biomedical Genomics | Arindam Maitra, Aritra Biswas, Jayeeta Haldar, Raja Ray, Monimoy Banerjee, Saumitra Das |
| EPI_ISL_653870, EPI_ISL_653903 | Translational Health Science and Technology Institute | National Institute of Biomedical Genomics | Arindam Maitra, Guruprasad Medigeshi, Sharanabasava Patil, Anbalagan Ananthraj, Madhu Pareek, Imran Khan, Gagandeep Kang, Saumitra Das |
| EPI_ISL_654628, EPI_ISL_654630 | Servicio de Microbiología, Hospital Universitario Central de Asturias | SeqCOVID-SPAIN consortium/IBV(CSIC) | Cristián Castelló Abietar, Jose A. Boga, Susana Rojo-Alba, Marta Elena Álvarez-Argüelles, Santiago Melón and SeqCOVID-SPAIN consortium |
| EPI_ISL_654734, EPI_ISL_654736, EPI_ISL_654738, EPI_ISL_654740, EPI_ISL_654741, EPI_ISL_654743, EPI_ISL_654744, EPI_ISL_654745, EPI_ISL_654746, EPI_ISL_654748, EPI_ISL_654749, EPI_ISL_654750, EPI_ISL_654751, EPI_ISL_654752, EPI_ISL_654756, EPI_ISL_654757, EPI_ISL_654758, EPI_ISL_654760, EPI_ISL_654761, EPI_ISL_654762, EPI_ISL_654763, EPI_ISL_654766, EPI_ISL_654767, EPI_ISL_654770, EPI_ISL_654771, EPI_ISL_654772, EPI_ISL_654773 | | | |
| see above | Texas Department of State Health Services | Texas Department of State Health Services | Rashmi Tuladhar, Bonnie Oh, Jenny Zhang, Maliha Rahman, Anita Pokharel, Myong Koag, Chung Wang, Rachel Lee, Grace Kubin, Mayela Pedrueza, James Daniel Bonser |
| EPI_ISL_660351, EPI_ISL_660352 | CHU de Saint-Étienne Hôpital Nord | CNR Virus des Infections Respiratoires - France SUD | Antonin Bal, Gregory Destras, Gwendolyne Burfin, Hadrien Règue, Quentin Semanas, Martine Valette, Bruno Lina, Issam Bechri, Manon Vogrig, Marine |

| | | | |
|---|---|---|---|
| | | | Delorme, Bruno Pozzetto, Thomas Bourlet, Sylvie Gonzalo, Sylvie Pillet, Laurence Josset |
| EPI_ISL_660369, EPI_ISL_660370 | CHU Clermont-Ferrand | CNR Virus des Infections Respiratoires - France SUD | Antonin Bal, Gregory Destras, Gwendolyne Burfin, Hadrien Règue, Quentin Semanas, Martine Valette, Bruno Lina, Christine Archimbaud, Amélie Brebion, Hélène Chabrolles, Martine Chambon, Audrey Mirand, Christel Regagnon, Maxime Bisseux, Patricia Combes, Cécile Henquell, Laurence Josset |
| EPI_ISL_660444 | Hospital de Braga | Instituto Nacional de Saude (INSA) | Borges et al |
| EPI_ISL_660450, EPI_ISL_660451 | Laboratoire de Microbiologie CHU Souro Sanou | Centre Muraz | Abdoul-Salam Ouedraogo, Yacouba Sawadogo, Essia Belarbi, Grit Schubert, Fabian Leendertz, Arsène Zongo, Soumeya Ouangraoua, Zekiba Tarnagda, Lassana Sangaré, Halidou Tinto |
| EPI_ISL_660708, EPI_ISL_660709 | CHU Montpellier | CNR Virus des Infections Respiratoires - France SUD | Antonin Bal, Gregory Destras, Gwendolyne Burfin, Hadrien Règue, Quentin Semanas, Martine Valette, Bruno Lina, Michel Segondy, Vincent Foulongne, Laurence Josset |
| EPI_ISL_660738, EPI_ISL_660739, EPI_ISL_660740, EPI_ISL_660741 | Unité des Virus Émergents | CNR Virus des Infections Respiratoires - France SUD | Antonin Bal, Gregory Destras, Gwendolyne Burfin, Hadrien Règue, Quentin Semanas, Martine Valette, Bruno Lina, Laetitia Ninove, Léa Luciani, Antoine Nougairède, Laurence Josset |
| EPI_ISL_663236, EPI_ISL_663237, EPI_ISL_663240, EPI_ISL_663241 | CHU Poitiers | CNR Virus des Infections Respiratoires - France SUD | Antonin Bal, Gregory Destras, Gwendolyne Burfin, Hadrien Règue, Quentin Semanas, Martine Valette, Bruno Lina, Agnès Beby-Defaux, Magali Garcia, Clément Jousselin, Nicolas Lévêque, Laurence Josset |
| EPI_ISL_665258, EPI_ISL_665259 | Dept. of Microbiology and Infection Control, Akershus University Hospital HF | Dept. of Microbiology and Infection Control, Akershus University Hospital HF | Hege Vangstein Aamot, Alexander Hesselberg Løvestad, Silje Bakken Jørgensen, Nina Handal, Ole Herman Ambur |
| EPI_ISL_666712, EPI_ISL_666713, EPI_ISL_666714, EPI_ISL_666715, EPI_ISL_666716, EPI_ISL_666717 | CHU de Limoges | CNR Virus des Infections Respiratoires - France SUD | Antonin Bal, Gregory Destras, Gwendolyne Burfin, Hadrien Règue, Quentin Semanas, Martine Valette, Bruno Lina, Sylvie Rogez, Laurence Josset |
| EPI_ISL_666832, EPI_ISL_666833, EPI_ISL_666834, EPI_ISL_666857 | Florida Bureau of Public Health Laboratories | Florida Bureau of Public Health Laboratories | Sarah Schmedes, Jason Blanton |
| EPI_ISL_667083 | OHSU Lab Services Molecular Microbiology Lab | Oregon SARS-CoV-2 Genome Sequencing Center | Brendan L. O'Connell, Ruth V. Nichols, Sally Grindstaff, Alec J. Hirsch, Donna Hansel, Guang Fan, Daniel N. Streblow, William B. Messer, Andrew C. Adey, Benjamin N. Bimber, Brian J. O'Roak |
| EPI_ISL_671347, EPI_ISL_671348, EPI_ISL_671349, EPI_ISL_671350, EPI_ISL_671351, EPI_ISL_671352, EPI_ISL_671353 | National Virus Reference Laboratory | Irish Coronavirus Sequencing Consortium - Teagasc Moorepark | Calm Walsh, Genuity Ireland |
| EPI_ISL_672014, EPI_ISL_672032, EPI_ISL_672038, EPI_ISL_672040, EPI_ISL_672053, EPI_ISL_672066, EPI_ISL_672078, EPI_ISL_672082, EPI_ISL_672092, EPI_ISL_672095 | Alameda County Public Health Lab | Chan-Zuckerberg Biohub | CZB Cliahub Consortium |
| EPI_ISL_672134, EPI_ISL_672135, EPI_ISL_672136, EPI_ISL_672137, EPI_ISL_672138, EPI_ISL_672139, EPI_ISL_672142, EPI_ISL_672143, EPI_ISL_672193, EPI_ISL_672194, EPI_ISL_672239, EPI_ISL_672242, EPI_ISL_672244, EPI_ISL_672245, EPI_ISL_672246, EPI_ISL_672247, EPI_ISL_672249, EPI_ISL_672251, EPI_ISL_672252, EPI_ISL_672256, EPI_ISL_672258, EPI_ISL_672259, EPI_ISL_672260, EPI_ISL_672261, EPI_ISL_672262, EPI_ISL_672263, EPI_ISL_672264, EPI_ISL_672265, EPI_ISL_672267, EPI_ISL_672270 | | | |
| see above | The Ashley Laboratory, Stanford University | Chan-Zuckerberg Biohub | CZB Cliahub Consortium |
| EPI_ISL_672305, EPI_ISL_672306, EPI_ISL_672307, EPI_ISL_672308, EPI_ISL_672309, EPI_ISL_672310, EPI_ISL_672311, EPI_ISL_672312, EPI_ISL_672313, EPI_ISL_672314, EPI_ISL_672315, EPI_ISL_672316, EPI_ISL_672317, EPI_ISL_672318, EPI_ISL_672319, EPI_ISL_672320, EPI_ISL_672321, EPI_ISL_672322, EPI_ISL_672323, EPI_ISL_672324, EPI_ISL_672325, EPI_ISL_672327, EPI_ISL_672328, EPI_ISL_672329, EPI_ISL_672335, EPI_ISL_672342, EPI_ISL_672343, EPI_ISL_672344, EPI_ISL_672345, EPI_ISL_672346, EPI_ISL_672347, EPI_ISL_672348, EPI_ISL_672349, EPI_ISL_672350, EPI_ISL_672351, EPI_ISL_672352, EPI_ISL_672353, EPI_ISL_672450, EPI_ISL_672451, EPI_ISL_672452, EPI_ISL_672453 | | | |
| see above | Alameda County Public Health Lab | Chan-Zuckerberg Biohub | CZB Cliahub Consortium |
| EPI_ISL_672568, EPI_ISL_672569 | Utah Public Health Laboratory | Utah Public Health Laboratory | Erin Young, Kelly Oakeson |
| EPI_ISL_672678 | DB Diagnosticos do Brasil | Laboratório de Parasitologia Médica - Instituto de Medicina Tropical - Universidade de São Paulo | Brazil-UK Centre for Arbovirus Discovery Diagnosis Genomics and Epidemiology (CADDE) Genomic Network - Instituto de Medicina Tropical |
| EPI_ISL_676585, EPI_ISL_676586, EPI_ISL_676587, EPI_ISL_676588, EPI_ISL_676589 | Scientific Veterinary Institute Novi Sad | Veterinary Specialized Institute "Kraljevo", Serbia | Vidanovic,D., Tesovic,B., Knezevic,A., Jovanovic,T., Jankovic,M., Sekler,M., Banovic Djeri,B., Petrovic,T., Volkening,J., Afonso,C. |
| EPI_ISL_676613 | Texas Department of State Health Services | Texas Department of State Health Services | Rashmi Tuladhar, Bonnie Oh, Jenny Zhang, Maliha Rahman, Anita Pokharel, Myong Koag, Chung Wang, Rachel Lee, Grace Kubin, Mayela Pedrueza, James Daniel Bonser |
| EPI_ISL_676654, EPI_ISL_676672, EPI_ISL_676674, EPI_ISL_676675, EPI_ISL_676677, EPI_ISL_676682, EPI_ISL_676683, EPI_ISL_676685, EPI_ISL_676692, EPI_ISL_676693, EPI_ISL_676696, EPI_ISL_676698, EPI_ISL_676705, EPI_ISL_676712, EPI_ISL_676717, EPI_ISL_676720, EPI_ISL_676721, EPI_ISL_676722, EPI_ISL_676724, EPI_ISL_676725, EPI_ISL_676893, EPI_ISL_676907, EPI_ISL_676908, EPI_ISL_676909, EPI_ISL_676910, EPI_ISL_676911, EPI_ISL_676912, EPI_ISL_676913, EPI_ISL_676914, EPI_ISL_676915, EPI_ISL_676916, EPI_ISL_676917, EPI_ISL_676918, EPI_ISL_676919, EPI_ISL_676920, EPI_ISL_676921, EPI_ISL_676922, EPI_ISL_676924, EPI_ISL_676925, EPI_ISL_676926, EPI_ISL_676927, EPI_ISL_676928, EPI_ISL_676929, EPI_ISL_676930, EPI_ISL_676931, EPI_ISL_676932, EPI_ISL_676933, EPI_ISL_676934, EPI_ISL_676935, EPI_ISL_676936, EPI_ISL_676937, EPI_ISL_676938, EPI_ISL_676939, EPI_ISL_676940, EPI_ISL_676941, EPI_ISL_677666, EPI_ISL_677667 | | | |
| see above | Wadsworth Center, New York State Department.of Health | Wadsworth Center, New York State Department.of Health | Kirsten St. George, Daryl M. Lamson, Alexis Russel, Jonathan Plitnick, Navjot Singh, John Kelly, Sara Griesemer, Erasmus Schneider, Erica Lasek-Nesselquist |
| EPI_ISL_677685, EPI_ISL_677686, EPI_ISL_677687, EPI_ISL_677688 | Vanda Pharmaceuticals Clinical Site | Vanda Pharmaceuticals | Vanda Pharmaceuticals |
| EPI_ISL_677945 | Pathogen Genomics Lab King Abdullah University of Science and Technology(KAUST) | Pathogen Genomics Lab King Abdullah University of Science and Technology(KAUST) | Sara Mfarrej, Luke Esau, Amanda Ooi, Sharif Hala, Raeece Naeem, Fadwa Alofi, Afrah Alsomali, Asim Khogeer, Jumana Taha, Abdulaziz Alahmadi, Kahled Algithami, Anwar Hashem, Naif Almontashiri, Arnab Pain |
| EPI_ISL_677965 | Pathogen Genomics Lab King Abdullah University of Science and Technology(KAUST) | Pathogen Genomics Lab King Abdullah University of Science and Technology(KAUST) | Sara Mfarrej, Olga Douvropoulou, Raushan Nugmanova, Sharif Hala, Raeece Naeem, Amanda Ooi, Luke Esau, Fadwa Alofi, Afrah Alsomali, Asim Khogeer, Jumana Taha, Abdulaziz Alahmadi, Kahled Algithami, Anwar Hashem, Naif Almontashiri, Arnab Pain |
| EPI_ISL_678034 | Pathogen Genomics Lab King Abdullah University of Science and Technology(KAUST) | Pathogen Genomics Lab King Abdullah University of Science and Technology(KAUST) | Sara Mfarrej, Olga Douvropoulou, Raushan Nugmanova, Sharif Hala, Raeece Naeem, Fadwa Alofi, Asim Khogeer, Afrah Alsomali, Jumana Taha, Abdulaziz Alahmadi, Kahled Algithami, Anwar Hashem, Naif Almontashiri, Arnab Pain |
| EPI_ISL_678508, EPI_ISL_678543, EPI_ISL_678544 | CNR Virus des Infections Respiratoires - France SUD | CNR Virus des Infections Respiratoires - France SUD | Antonin Bal, Gregory Destras, Gwendolyne Burfin, Solenne Brun, Martine Valette, Bruno Lina, Laurence Josset |
| EPI_ISL_679967 | Lincolnshire Hospitals and DeepSeq Nottingham | COVID-19 Genomics UK (COG-UK) Consortium | Nichola Duckworth, Tim Sloan, Sarah Walsh, Jonathan Ball, Patrick McClure, Joeseph Chappell, Nadine Holmes, Matthew Carlisle, Christopher Moore, Fei Sang, Johnny Debebe, Victoria Wright, Matthew Loose |
| EPI_ISL_681758, EPI_ISL_681761, EPI_ISL_681762, EPI_ISL_681763, EPI_ISL_681774, EPI_ISL_681775 | University Hospital Limerick | Irish Coronavirus Sequencing Consortium - Teagasc Moorepark | Paul Cotter, Fiona Crispie, Amy Fitzpatrick, John Kenny, Elaine Lawton, Carolyn Meaney, Patrick Stapleton, Calum Walsh |
| EPI_ISL_681998, EPI_ISL_682003 | UPMC Clinical Microbiology Laboratory | Microbial Genomic Epidemiology Laboratory, University of Pittsburgh | Mustapha M. Mustapha, Jane W. Marsh, Dan Snyder, Marissa P. Griffith, Stephanie L. Mitchell, Vatsala R. Srinivasa, Kady D. Waggle, Chinelo Ezeonwuku, Vaughn S. Cooper, Lee H. Harrison |
| EPI_ISL_691682 | Hospital General Universitario de Ciudad Real | Instituto de Salud Carlos III | Iglesias-Caballero, M. Camarero, S. Molinero Calamita, M. González-Esguevillas, M. Pozo, F. Casas, I. Jiménez, P. Jiménez, M. Zaballos, A. Monzón, S. Varona, S. Juliá, M. Cuesta, I. Jarilla, M. |
| EPI_ISL_691683 | Hospital General Universitario de Ciudad Real | Instituto de Salud Carlos III | Iglesias-Caballero, M. Camarero, S. Molinero Calamita, M. González-Esguevillas, M. Pozo, F. Casas, I. Jiménez, P. Jiménez, M. Zaballos, A. Monzón, S. Varona, S. Juliá, M. Cuesta, I. Madrigal, M. |

| | | | |
|---|---|---|---|
| EPI_ISL_691684 | Complejo Hospitalario Universitario de Albacete | Instituto de Salud Carlos III | Iglesias-Caballero, M. Camarero, S. Molinero Calamita, M. González-Esguevillas, M. Pozo, F. Casas, I. Jiménez, P. Jiménez, M. Zaballos, A. Monzón, S. Varona, S. Juliá, M. Cuesta, I. Martínez, E. |
| EPI_ISL_692831, EPI_ISL_692949, EPI_ISL_692970, EPI_ISL_692971, EPI_ISL_692976, EPI_ISL_692977, EPI_ISL_692978, EPI_ISL_692979, EPI_ISL_692980, EPI_ISL_692981, EPI_ISL_692982, EPI_ISL_692983, EPI_ISL_692984, EPI_ISL_692985, EPI_ISL_692986, EPI_ISL_692987, EPI_ISL_692988, EPI_ISL_692989, EPI_ISL_692990, EPI_ISL_692991, EPI_ISL_692992, EPI_ISL_692993, EPI_ISL_692994, EPI_ISL_692995, EPI_ISL_692996, EPI_ISL_692997, EPI_ISL_692998, EPI_ISL_692999, EPI_ISL_693000, EPI_ISL_693001, EPI_ISL_693002, EPI_ISL_693003, EPI_ISL_693004, EPI_ISL_693005, EPI_ISL_693006, EPI_ISL_693007, EPI_ISL_693008, EPI_ISL_693009, EPI_ISL_693010, EPI_ISL_693011, EPI_ISL_693012, EPI_ISL_693013, EPI_ISL_693014, EPI_ISL_693015, EPI_ISL_693016, EPI_ISL_693017, EPI_ISL_693018, EPI_ISL_693019, EPI_ISL_693020, EPI_ISL_693021, EPI_ISL_693022, EPI_ISL_693023, EPI_ISL_693024, EPI_ISL_693025, EPI_ISL_693026, EPI_ISL_693027, EPI_ISL_693028, EPI_ISL_693029, EPI_ISL_693030, EPI_ISL_693031, EPI_ISL_693032, EPI_ISL_693033, EPI_ISL_693034, EPI_ISL_693035, EPI_ISL_693036, EPI_ISL_693037, EPI_ISL_693038, EPI_ISL_693039, EPI_ISL_693040, EPI_ISL_693041, EPI_ISL_693042, EPI_ISL_693043, EPI_ISL_693044, EPI_ISL_693045, EPI_ISL_693046, EPI_ISL_693047, EPI_ISL_693048, EPI_ISL_693049, EPI_ISL_693050, EPI_ISL_693051, EPI_ISL_693052, EPI_ISL_693053, EPI_ISL_693054, EPI_ISL_693055, EPI_ISL_693056, EPI_ISL_693057, EPI_ISL_693058, EPI_ISL_693059, EPI_ISL_693060, EPI_ISL_693061, EPI_ISL_693150, EPI_ISL_693153, EPI_ISL_693155, EPI_ISL_693156, EPI_ISL_693157, EPI_ISL_693176, EPI_ISL_693177 | | | |
| see above | Massachusetts State Public Health Laboratory | Massachusetts State Public Health Laboratory | Andrew Lang, Timelia Fink, Glen Gallagher, Sandra Smole |
| EPI_ISL_693545, EPI_ISL_693546, EPI_ISL_693547 | Instituto Nacional de Saude (INSA) | Instituto Nacional de Saude (INSA) | Borges et al |
| EPI_ISL_694009, EPI_ISL_694010, EPI_ISL_694011, EPI_ISL_694012, EPI_ISL_694013, EPI_ISL_694014 | TGen North | TGen North | Hayley D Yaglom, Marette Gebhardt, Ashlyn Pfeiffer, Mary Ellen Ormsby, Daniel Jasso-Selles, Darrin Lemmer, Megan Folkerts, Chris French, Matthew Maurer, Jolene R Bowers, David M Engelthaler |
| EPI_ISL_694234, EPI_ISL_694235, EPI_ISL_694236, EPI_ISL_694237, EPI_ISL_694238, EPI_ISL_694239, EPI_ISL_694240, EPI_ISL_694241, EPI_ISL_694242, EPI_ISL_694243, EPI_ISL_694244, EPI_ISL_694245, EPI_ISL_694246, EPI_ISL_694247, EPI_ISL_694316, EPI_ISL_694317, EPI_ISL_694318, EPI_ISL_694319, EPI_ISL_694320, EPI_ISL_694321, EPI_ISL_694322, EPI_ISL_694323, EPI_ISL_694324, EPI_ISL_694325, EPI_ISL_694331 | | | |
| see above | TGen North | TGen North | Jolene Bowers, Megan Folkerts, Chris French, Hayley Yaglom, Ashlyn Pfeiffer, Darrin Lemmer, Dave Engelthaler, The Arizona COVID Genomics Union (ACGU) |
| EPI_ISL_694524, EPI_ISL_694525, EPI_ISL_694526, EPI_ISL_694527, EPI_ISL_694528, EPI_ISL_694529, EPI_ISL_694530, EPI_ISL_694531, EPI_ISL_694532, EPI_ISL_694533, EPI_ISL_694534, EPI_ISL_694535, EPI_ISL_694536, EPI_ISL_694537, EPI_ISL_694538, EPI_ISL_694539, EPI_ISL_694540, EPI_ISL_694541, EPI_ISL_694542, EPI_ISL_694543, EPI_ISL_694544, EPI_ISL_694545 | | | |
| see above | AZ SPHL, Arizona Department of Health Services | TGen North | Jolene Bowers, Megan Folkerts, Chris French, Hayley Yaglom, Ashlyn Pfeiffer, Darrin Lemmer, Dave Engelthaler, The Arizona COVID Genomics Union (ACGU) |
| EPI_ISL_696256, EPI_ISL_696257, EPI_ISL_696258, EPI_ISL_696259, EPI_ISL_696260, EPI_ISL_696262, EPI_ISL_696263, EPI_ISL_696264, EPI_ISL_696265, EPI_ISL_696266, EPI_ISL_696267, EPI_ISL_696268, EPI_ISL_696270, EPI_ISL_696271, EPI_ISL_696272, EPI_ISL_696275, EPI_ISL_696276, EPI_ISL_696277, EPI_ISL_696278, EPI_ISL_696279, EPI_ISL_696280, EPI_ISL_696281, EPI_ISL_696283, EPI_ISL_696284, EPI_ISL_696285, EPI_ISL_696287, EPI_ISL_696288, EPI_ISL_696289, EPI_ISL_696290, EPI_ISL_696291, EPI_ISL_696295, EPI_ISL_696297, EPI_ISL_696298, EPI_ISL_696300, EPI_ISL_696301, EPI_ISL_696303, EPI_ISL_696305, EPI_ISL_696308, EPI_ISL_696311, EPI_ISL_696312, EPI_ISL_696313, EPI_ISL_696315 | | | |
| see above | Sonora Quest Laboratories, Laboratory Sciences of Arizona | TGen North | Jolene Bowers, Megan Folkerts, Chris French, Hayley Yaglom, Ashlyn Pfeiffer, Darrin Lemmer, Dave Engelthaler, The Arizona COVID Genomics Union (ACGU) |
| EPI_ISL_698155, EPI_ISL_698156, EPI_ISL_698157, EPI_ISL_698158, EPI_ISL_698159, EPI_ISL_698160, EPI_ISL_698161, EPI_ISL_698162, EPI_ISL_698163, EPI_ISL_698164, EPI_ISL_698165, EPI_ISL_698166, EPI_ISL_698167, EPI_ISL_698168, EPI_ISL_698169, EPI_ISL_698178, EPI_ISL_698179, EPI_ISL_698180, EPI_ISL_698181, EPI_ISL_698182, EPI_ISL_698189, EPI_ISL_698198, EPI_ISL_698199, EPI_ISL_698200, EPI_ISL_698211, EPI_ISL_698213, EPI_ISL_698214, EPI_ISL_698215, EPI_ISL_698216, EPI_ISL_698217, EPI_ISL_698218, EPI_ISL_698219, EPI_ISL_698220, EPI_ISL_698221, EPI_ISL_698222, EPI_ISL_698223, EPI_ISL_698224, EPI_ISL_698225, EPI_ISL_698226, EPI_ISL_698227, EPI_ISL_698228, EPI_ISL_698229, EPI_ISL_698230, EPI_ISL_698231, EPI_ISL_698232, EPI_ISL_698233, EPI_ISL_698265, EPI_ISL_698266, EPI_ISL_698267, EPI_ISL_698281, EPI_ISL_698282, EPI_ISL_698283, EPI_ISL_698284, EPI_ISL_698285, EPI_ISL_698295, EPI_ISL_698296, EPI_ISL_698297, EPI_ISL_698298, EPI_ISL_698299, EPI_ISL_698300, EPI_ISL_698301, EPI_ISL_698302, EPI_ISL_698303, EPI_ISL_698304, EPI_ISL_698305, EPI_ISL_698306, EPI_ISL_698307, EPI_ISL_698308, EPI_ISL_698309, EPI_ISL_698310, EPI_ISL_698460, EPI_ISL_698515, EPI_ISL_698516, EPI_ISL_698517, EPI_ISL_698518, EPI_ISL_698519, EPI_ISL_698520, EPI_ISL_698521, EPI_ISL_698522, EPI_ISL_698523, EPI_ISL_698524, EPI_ISL_698525, EPI_ISL_698526, EPI_ISL_698527, EPI_ISL_698528, EPI_ISL_698529, EPI_ISL_698530, EPI_ISL_698531, EPI_ISL_698532, EPI_ISL_698533, EPI_ISL_698534, EPI_ISL_698535, EPI_ISL_698536, EPI_ISL_698537, EPI_ISL_698538, EPI_ISL_698539, EPI_ISL_698540, EPI_ISL_698541, EPI_ISL_698542, EPI_ISL_698543, EPI_ISL_698544, EPI_ISL_698545, EPI_ISL_698546, EPI_ISL_698547, EPI_ISL_698548, EPI_ISL_698549, EPI_ISL_698550, EPI_ISL_698551, EPI_ISL_698552, EPI_ISL_698556, EPI_ISL_698557, EPI_ISL_698558, EPI_ISL_698559, EPI_ISL_698560, EPI_ISL_698561, EPI_ISL_698562, EPI_ISL_698563, EPI_ISL_698564, EPI_ISL_698565, EPI_ISL_698567, EPI_ISL_698568, EPI_ISL_698578, EPI_ISL_698579, EPI_ISL_698580, EPI_ISL_698581, EPI_ISL_698582, EPI_ISL_698583, EPI_ISL_698584, EPI_ISL_698585, EPI_ISL_698586, EPI_ISL_698587, EPI_ISL_698588, EPI_ISL_698589, EPI_ISL_698590, EPI_ISL_698591, EPI_ISL_698592, EPI_ISL_698593, EPI_ISL_698594, EPI_ISL_698595, EPI_ISL_698596, EPI_ISL_698597, EPI_ISL_698598, EPI_ISL_698599, EPI_ISL_698600, EPI_ISL_698601, EPI_ISL_698602, EPI_ISL_698603, EPI_ISL_698604, EPI_ISL_698605, EPI_ISL_698606, EPI_ISL_698607, EPI_ISL_698608, EPI_ISL_698609, EPI_ISL_698610, EPI_ISL_698611, EPI_ISL_698612, EPI_ISL_698613, EPI_ISL_698614, EPI_ISL_698615, EPI_ISL_698708, EPI_ISL_698709, EPI_ISL_698710, EPI_ISL_698711, EPI_ISL_698712, EPI_ISL_698713, EPI_ISL_698714, EPI_ISL_698715, EPI_ISL_698723, EPI_ISL_698724, EPI_ISL_698725, EPI_ISL_698726, EPI_ISL_698727, EPI_ISL_698728, EPI_ISL_698729, EPI_ISL_698730, EPI_ISL_698731, EPI_ISL_698732, EPI_ISL_698733, EPI_ISL_698734, EPI_ISL_698735, EPI_ISL_698736, EPI_ISL_698737, EPI_ISL_698738, EPI_ISL_698739, EPI_ISL_698740, EPI_ISL_698741, EPI_ISL_698742, EPI_ISL_698743, EPI_ISL_698744, EPI_ISL_698745, EPI_ISL_698746, EPI_ISL_698747, EPI_ISL_698749, EPI_ISL_698750, EPI_ISL_698751, EPI_ISL_698752, EPI_ISL_698753, EPI_ISL_698754, EPI_ISL_698755, EPI_ISL_698756, EPI_ISL_698757, EPI_ISL_698758, EPI_ISL_698759, EPI_ISL_698760, EPI_ISL_698761, EPI_ISL_698762, EPI_ISL_698763, EPI_ISL_698764, EPI_ISL_698765, EPI_ISL_698766, EPI_ISL_698767, EPI_ISL_698768, EPI_ISL_698769, EPI_ISL_698770, EPI_ISL_698771, EPI_ISL_698772, EPI_ISL_698773, EPI_ISL_698774, EPI_ISL_698775, EPI_ISL_698776, EPI_ISL_698777, EPI_ISL_698778, EPI_ISL_698779, EPI_ISL_698780, EPI_ISL_698781, EPI_ISL_698782, EPI_ISL_698783, EPI_ISL_698784, EPI_ISL_698785, EPI_ISL_698786, EPI_ISL_698787, EPI_ISL_698788, EPI_ISL_698789, EPI_ISL_698790, EPI_ISL_698791, EPI_ISL_698792, EPI_ISL_698793, EPI_ISL_698794, EPI_ISL_698795, EPI_ISL_698796, EPI_ISL_698797, EPI_ISL_698798, EPI_ISL_698799, EPI_ISL_698800, EPI_ISL_698801, EPI_ISL_698802, EPI_ISL_698803, EPI_ISL_698804, EPI_ISL_698805, EPI_ISL_698806, EPI_ISL_698807, EPI_ISL_698808, EPI_ISL_698809, EPI_ISL_698810, EPI_ISL_698811, EPI_ISL_698812, EPI_ISL_698813, EPI_ISL_698814, EPI_ISL_698815, EPI_ISL_698816, EPI_ISL_698817, EPI_ISL_698818, EPI_ISL_698819, EPI_ISL_698820, EPI_ISL_698821, EPI_ISL_698822, EPI_ISL_698823, EPI_ISL_698824, EPI_ISL_698825, EPI_ISL_698826, EPI_ISL_698827, EPI_ISL_698828, EPI_ISL_698829, EPI_ISL_698830, EPI_ISL_698831, EPI_ISL_698832, EPI_ISL_698833, EPI_ISL_698834, EPI_ISL_698835, EPI_ISL_698836, EPI_ISL_698837, EPI_ISL_698838, EPI_ISL_698839, EPI_ISL_698840, EPI_ISL_698841, EPI_ISL_698842, EPI_ISL_698843, EPI_ISL_698844, EPI_ISL_698845, EPI_ISL_698846, EPI_ISL_698847, EPI_ISL_698848, EPI_ISL_698849, EPI_ISL_698850, EPI_ISL_698851, EPI_ISL_698852, EPI_ISL_698853, EPI_ISL_698854, EPI_ISL_698855, EPI_ISL_698856, EPI_ISL_698857, EPI_ISL_698858, EPI_ISL_698859, EPI_ISL_698879, EPI_ISL_698880, EPI_ISL_698881, EPI_ISL_698882, EPI_ISL_698883, EPI_ISL_698884, EPI_ISL_698885, EPI_ISL_698886, EPI_ISL_698887, EPI_ISL_698888, EPI_ISL_698889, EPI_ISL_698890, EPI_ISL_698891, EPI_ISL_698892, EPI_ISL_698893, EPI_ISL_698894, EPI_ISL_698895, EPI_ISL_698896, EPI_ISL_698897, EPI_ISL_698898, EPI_ISL_698899, EPI_ISL_698900, EPI_ISL_698901, EPI_ISL_698902, EPI_ISL_698903, EPI_ISL_698904, EPI_ISL_698905, EPI_ISL_698912, EPI_ISL_698913, EPI_ISL_698914, EPI_ISL_698915, EPI_ISL_698916, EPI_ISL_698917, EPI_ISL_698926 | | | |
| see above | Group 42 (G42) Healthcare, Abu Dhabi, United Arab Emirates; Department of Health, The United Arab Emirates | G42 Healthcare | Rong Liu, Pei Wu, Sally Mahmoud, Ke Liang, Pauline Ogrodzki, Pengjuan Liu, Stephen S. Francis, Tao Ma, Hanif Khalak, Fang Chen, Denghui Liu, Junhua Li, Weibin Liu, Wenjun He, Xinyu Huang, Zhaorong Yuan, Long Lin, Nan Qiao, Xin Meng, Budoor Alqarni, Javier Quilez, Vinay Kusuma, Xin Jin, Xavier Anton, Ashish Koshy, Huanming Yang, Xun Xu, Jian Wang, Peng Xiao, Nawal Ahmed Mohamed Al Kaabi, Mohammed Saifuddin Fasihuddin, Siyang Liu, Walid Abbas Zaher |
| EPI_ISL_699685, EPI_ISL_699686, EPI_ISL_699687, EPI_ISL_699688, EPI_ISL_699689, EPI_ISL_699690, EPI_ISL_699691, EPI_ISL_699692, EPI_ISL_699693, EPI_ISL_699694, EPI_ISL_699695, EPI_ISL_699696, EPI_ISL_699697, EPI_ISL_699698, EPI_ISL_699699, EPI_ISL_699700, EPI_ISL_699701, EPI_ISL_699702, EPI_ISL_699703, EPI_ISL_699704, EPI_ISL_699705, EPI_ISL_699706, EPI_ISL_699707, EPI_ISL_699708, EPI_ISL_699709, EPI_ISL_699710, EPI_ISL_699711, EPI_ISL_699712, EPI_ISL_699713, EPI_ISL_699714, EPI_ISL_699715, EPI_ISL_699716, EPI_ISL_699717, EPI_ISL_699718, EPI_ISL_699723 | | | |
| see above | Hematopathology Laboratory, ACTREC, TMC | Hematopathology Laboratory, ACTREC, TMC | Hematopathology Laboratory, ACTREC |
| EPI_ISL_700398, EPI_ISL_700399, EPI_ISL_700400, EPI_ISL_700401, EPI_ISL_700404, EPI_ISL_700408, EPI_ISL_700410 | Centre hospitalier Métropole Savoie | CNR Virus des Infections Respiratoires - France SUD | Antonin Bal, Carine Dumollard, Gregory Destras, Gwendolyne Burfin, Hadrien Règue, Quentin Semanas, Martine Valette, Bruno Lina, Jérôme Grosjean, Laurence Josset |
| EPI_ISL_700455 | Guguletu CHC wc GDH | NHLS/UCT | Arash Iranzadeh, Deelan Doolabh, Lynn Tyers, Bruna Galvao, Innocent Mudau, Marvin Hsiao, Kruger Marais, Diana Hardie, Stephen Korsman, Carolyn Williamson |
| EPI_ISL_700457 | Hanover Park CHC wc HPH | NHLS/UCT | Arash Iranzadeh, Deelan Doolabh, Lynn Tyers, Bruna Galvao, Innocent Mudau, Marvin Hsiao, Kruger Marais, Diana Hardie, Stephen Korsman, Carolyn Williamson |
| EPI_ISL_700462 | Vanguard CHC wc VGC | NHLS/UCT | Arash Iranzadeh, Deelan Doolabh, Lynn Tyers, Bruna Galvao, Innocent Mudau, Marvin Hsiao, Kruger Marais, Diana Hardie, Stephen Korsman, Carolyn Williamson |
| EPI_ISL_700472 | Nyanga CDC wc NGC | NHLS/UCT | Arash Iranzadeh, Deelan Doolabh, Lynn Tyers, Bruna Galvao, Innocent Mudau, Marvin Hsiao, Kruger Marais, Diana Hardie, Stephen Korsman, Carolyn Williamson |
| EPI_ISL_700481 | Guguletu CHC wc GDH | NHLS/UCT | Arash Iranzadeh, Deelan Doolabh, Lynn Tyers, Bruna Galvao, Innocent Mudau, Marvin Hsiao, Kruger Marais, Diana Hardie, Stephen Korsman, Carolyn Williamson |
| EPI_ISL_700494 | Hanover Park CHC wc HPH | NHLS/UCT | Arash Iranzadeh, Deelan Doolabh, Lynn Tyers, Bruna Galvao, Innocent Mudau, Marvin Hsiao, Kruger Marais, Diana Hardie, Stephen Korsman, Carolyn Williamson |
| EPI_ISL_700503 | Guguletu CHC wc GDH | NHLS/UCT | Arash Iranzadeh, Deelan Doolabh, Lynn Tyers, Bruna Galvao, Innocent Mudau, Marvin Hsiao, Kruger Marais, Diana Hardie, Stephen Korsman, Carolyn Williamson |
| EPI_ISL_700513 | Dr Abdurahman CDC wc DAC | NHLS/UCT | Arash Iranzadeh, Deelan Doolabh, Lynn Tyers, Bruna Galvao, Innocent Mudau, Marvin Hsiao, Kruger Marais, Diana Hardie, Stephen Korsman, Carolyn Williamson |
| EPI_ISL_700515, EPI_ISL_700516 | Hanover Park CHC wc HPH | NHLS/UCT | Arash Iranzadeh, Deelan Doolabh, Lynn Tyers, Bruna Galvao, Innocent Mudau, Marvin Hsiao, Kruger Marais, Diana Hardie, Stephen Korsman, Carolyn Williamson |

| | | | |
|---|---|---|---|
| EPI_ISL_700523 | Nyanga CDC wc NGC | NHLS/UCT | Arash Iranzadeh, Deelan Doolabh, Lynn Tyers, Bruna Galvao, Innocent Mudau, Marvin Hsiao, Kruger Marais, Diana Hardie, Stephen Korsman, Carolyn Williamson |
| EPI_ISL_700530 | Guguletu CHC wc GDH | NHLS/UCT | Arash Iranzadeh, Deelan Doolabh, Lynn Tyers, Bruna Galvao, Innocent Mudau, Marvin Hsiao, Kruger Marais, Diana Hardie, Stephen Korsman, Carolyn Williamson |
| EPI_ISL_700561, EPI_ISL_700566 | Hanover Park CHC wc HPH | NHLS/UCT | Arash Iranzadeh, Deelan Doolabh, Lynn Tyers, Bruna Galvao, Innocent Mudau, Marvin Hsiao, Kruger Marais, Diana Hardie, Stephen Korsman, Carolyn Williamson |
| EPI_ISL_700574 | Heideveld Emergency Centre | NHLS/UCT | Arash Iranzadeh, Deelan Doolabh, Lynn Tyers, Bruna Galvao, Innocent Mudau, Marvin Hsiao, Kruger Marais, Diana Hardie, Stephen Korsman, Carolyn Williamson |
| EPI_ISL_700576, EPI_ISL_700578 | Guguletu CHC wc GDH | NHLS/UCT | Arash Iranzadeh, Deelan Doolabh, Lynn Tyers, Bruna Galvao, Innocent Mudau, Marvin Hsiao, Kruger Marais, Diana Hardie, Stephen Korsman, Carolyn Williamson |
| EPI_ISL_700580 | Nyanga CDC wc NGC | NHLS/UCT | Arash Iranzadeh, Deelan Doolabh, Lynn Tyers, Bruna Galvao, Innocent Mudau, Marvin Hsiao, Kruger Marais, Diana Hardie, Stephen Korsman, Carolyn Williamson |
| EPI_ISL_700585 | Hanover Park CHC wc HPH | NHLS/UCT | Arash Iranzadeh, Deelan Doolabh, Lynn Tyers, Bruna Galvao, Innocent Mudau, Marvin Hsiao, Kruger Marais, Diana Hardie, Stephen Korsman, Carolyn Williamson |
| EPI_ISL_700595 | Heideveld Emergency Centre | NHLS/UCT | Arash Iranzadeh, Deelan Doolabh, Lynn Tyers, Bruna Galvao, Innocent Mudau, Marvin Hsiao, Kruger Marais, Diana Hardie, Stephen Korsman, Carolyn Williamson |
| EPI_ISL_708021 | Virology, Universitätsklinikum des Saarlandes | Epigenetics, Saarland University | Kathrin Kattler, Markus Vogelgesang, Stefan Lohse, Sascha Tierling, Sigrun Smola, Jörn Walter |
| EPI_ISL_708505 | Los Angeles County Public Health Laboratory | Los Angeles County Public Health Laboratory | P. Hemarajata et al. |
| EPI_ISL_708608, EPI_ISL_708618, EPI_ISL_708622, EPI_ISL_708634, EPI_ISL_708635, EPI_ISL_708642, EPI_ISL_708646, EPI_ISL_708647, EPI_ISL_708651 | Texas Department of State Health Services | Texas Department of State Health Services | Rashmi Tuladhar, Bonnie Oh, Jenny Zhang, Maliha Rahman, Anita Pokharel, Myong Koag, Chung Wang, Rachel Lee, Grace Kubin, Mayela Pedrueza, James Daniel Bonser |
| EPI_ISL_708725 | Los Angeles County Public Health Laboratory | Los Angeles County Public Health Laboratory | P. Hemarajata et al. |
| EPI_ISL_708828, EPI_ISL_708832, EPI_ISL_708833, EPI_ISL_708834, EPI_ISL_708837, EPI_ISL_708838 | Group 42 (G42) Healthcare, Abu Dhabi, United Arab Emirates; Department of Health, The United Arab Emirates | G42 Healthcare | Rong Liu, Pei Wu, Sally Mahmoud, Ke Liang, Pauline Ogrodzki, Pengjuan Liu, Stephen S. Francis, Tao Ma, Hanif Khalak, Fang Chen, Denghui Liu, Junhua Li, Weibin Liu, Wenjun He, Xinyu Huang, Zhaorong Yuan, Long Lin, Nan Qiao, Xin Meng, Budoor Alqarni, Javier Quilez, Vinay Kusuma, Xin Jin, Xavier Anton, Ashish Koshy, Huanming Yang, Xun Xu, Jian Wang, Peng Xiao, Nawal Ahmed Mohamed Al Kaabi, Mohammed Saifuddin Fasihuddin, Siyang Liu, Walid Abbas Zaher |
| EPI_ISL_710146, EPI_ISL_710148, EPI_ISL_710156, EPI_ISL_710211 | Colorado Department of Public Health and Environment | Colorado Department of Puplic Health and Environment | Laura Bankers, Molly C. Hetherington-Rauth, Shannon Ely, Shannon R. Matzinger, Sarah Elizabeth Totten, Emily A. Travanty |
| EPI_ISL_717731, EPI_ISL_717744 | Kingston Health Sciences Centre and Queen's University | Ontario Institute for Cancer Research | Prameet M. Sheth,Calvin Sjaarda,Robert Colautti,Katya Douchant,Ilinca Lungu,Bernard Lam,Paul Krzyzanowski,Michael Laszloffy,Lawrence E. Heisler,Richard de Borja,Jared T. Simpson |
| EPI_ISL_718133, EPI_ISL_718154, EPI_ISL_718165, EPI_ISL_718166 | Ministry of Health Hospitals | Institute of Health and Community Medicine | David Perera, Ooi Mong How, Chua Hock Hin, Tonnii Sia Loong Loong, Wong Jyn Shan, Wong Kiing Aik, Chan Chia Jui |
| EPI_ISL_721988, EPI_ISL_721990, EPI_ISL_721991, EPI_ISL_721995, EPI_ISL_722006, EPI_ISL_722007, EPI_ISL_722010, EPI_ISL_722011, EPI_ISL_722017, EPI_ISL_722058, EPI_ISL_722066, EPI_ISL_722069, EPI_ISL_722070, EPI_ISL_722074, EPI_ISL_722077, EPI_ISL_722093, EPI_ISL_722095, EPI_ISL_722096, EPI_ISL_722097, EPI_ISL_722100, EPI_ISL_722104, EPI_ISL_722105, EPI_ISL_722106, EPI_ISL_722111, EPI_ISL_722113, EPI_ISL_722115, EPI_ISL_722120, EPI_ISL_722121, EPI_ISL_722123 | | | |
| see above | Hospital das Clínicas Universidade de São Paulo Medical School | Laboratório de Parasitologia Médica - Instituto de Medicina Tropical - Universidade de São Paulo | Brazil-UK Centre for Arbovirus Discovery Diagnosis Genomics and Epidemiology (CADDE) Genomic Network - Instituto de Medicina Tropical |
| EPI_ISL_723082, EPI_ISL_723083 | Institute of Medical Genetics and Applied Genomics | Institute of Medical Genetics and Applied Genomics | Caspar Gross, Tina Ganzenmüller, Siri Göpel, Michaela Pogoda, Daniela Bezdan, Michael Sonnabend, Angel Angelov, Nicolas Casadei, Stephan Ossowski, Thomas Iftner, Michael Bitzer |
| EPI_ISL_723118 | Minnesota Department of Health, Public Health Laboratory | Minnesota Department of Health, Public Health Laboratory | Alexandra Lorentz, Jacob Garfin, Matt Plumb, and Xiong Wang |
| EPI_ISL_729794, EPI_ISL_729797, EPI_ISL_729798, EPI_ISL_729820, EPI_ISL_729821, EPI_ISL_729822, EPI_ISL_729823, EPI_ISL_729824, EPI_ISL_729825, EPI_ISL_729826, EPI_ISL_729827, EPI_ISL_729828, EPI_ISL_729829, EPI_ISL_729830, EPI_ISL_729831 | | | |
| see above | Laboratorio Central de Saude Publica do Estado do Rio Grande do Sul (LACEN-RS) | Laboratory of Respiratory Viruses and Measles, Oswaldo Cruz Institute, FIOCRUZ | Paola Resende, Luciana Appolinario, Fernando Motta, Anna Carolina Paixão, Ana Carolina Mendonça, Tatiana Schaffer Gregianini, Marilda Tereza Mar da Rosa, Marilda Siqueira |
| EPI_ISL_729944, EPI_ISL_729945, EPI_ISL_729947, EPI_ISL_729948, EPI_ISL_729949, EPI_ISL_729950, EPI_ISL_729964, EPI_ISL_730009, EPI_ISL_730010, EPI_ISL_730011, EPI_ISL_730032, EPI_ISL_730033, EPI_ISL_730036 | | | |
| see above | Nigeria Centre for Disease Control (NCDC) | African Centre of Excellence for Genomics of Infectious Diseases (ACEGID), Redeemer's University, Ede, Osun State, Nigeria | Oluniyi P.E. et al |
| EPI_ISL_730066, EPI_ISL_730067, EPI_ISL_730068, EPI_ISL_730069, EPI_ISL_730070, EPI_ISL_730071, EPI_ISL_730072, EPI_ISL_730073, EPI_ISL_730074, EPI_ISL_730075, EPI_ISL_730076, EPI_ISL_730077, EPI_ISL_730078 | | | |
| see above | Yale Clinical Virology Laboratory | Grubaugh Lab - Yale School of Public Health | Joseph Fauver, Tara Alpert, Anderson Brito, Annie Watkins, Anne Wyllie, Chantal Vogels, Mary Petrone, Chaney Kalinich, Isabel Ott, Arnau Casanovas, Catherine Muenker, Adam Moore, Alice Lu, Maria Tokuyama, Patrick Wong, Peiwen Lu, Saad Omer, Richard Martinello, Allison Nelson, Shelli Farhadian, Akiko Iwasaki, Charlese Dela Cruz, Albert Ko, Nathan Grubaugh |
| EPI_ISL_730140, EPI_ISL_730149, EPI_ISL_730152, EPI_ISL_730154 | Scripps Medical Laboratory | Andersen lab at Scripps Research | SEARCH Alliance San Diego with Michael Quigley, Ellen Stefanski, Ian Mchardy |
| EPI_ISL_731946, EPI_ISL_731947 | Instituto Nacional de Saude (INSA) | Instituto Nacional de Saude (INSA) | Borges et al |
| EPI_ISL_732556, EPI_ISL_732557, EPI_ISL_732558 | Bundeswehr Institute of Microbiology | Bundeswehr Institute of Microbiology | Elham Khatamzas, Markus Antwerpen, Mathias Walter, Alexandra Rehn, Sabine Zange, Enrico Georgi, Michael von Bergwelt-Baildon, Roman Wölfel |
| EPI_ISL_733163 | Pathogenic Microorganisms Variability Laboratory | WHO National Influenza Centre Russian Federation | Andrey Komissarov, Artem Fadeev, Anna Ivanova, Kseniya Komissarova, Dmitry Bazhenov, Daria Danilenko, Ksenia Safina, Elena Nabieva, Georgii Bazykin, Nadezhda Kuznetsova, Elena Shidlovskaya, Sergey Alkhovsky, Tatyana Vishnevskaya, Elizaveta Divisenko, Alexey Shchetinin, Maria Nikiforova, Andrey Pochtovyy, Evgeny Usachev, Elena Vokalova, Maxim Rubalsky, Oleg Rubalsky, Artem Tkachuk, Vladimir Gushchin, Alexander Gintsburg, Dmitry Lioznov |
| EPI_ISL_734492, EPI_ISL_734704, EPI_ISL_734705, EPI_ISL_734706, EPI_ISL_734707, EPI_ISL_734708, EPI_ISL_734709, EPI_ISL_734710, EPI_ISL_734711, EPI_ISL_734712, EPI_ISL_734713, EPI_ISL_734714, EPI_ISL_734715, EPI_ISL_734716, EPI_ISL_734717, EPI_ISL_734718, EPI_ISL_734719, EPI_ISL_734720, EPI_ISL_734721, EPI_ISL_734722, EPI_ISL_734723, EPI_ISL_734724, EPI_ISL_734725, EPI_ISL_734726, EPI_ISL_734727, EPI_ISL_734728, EPI_ISL_734729, EPI_ISL_734730, EPI_ISL_734731, EPI_ISL_734732, EPI_ISL_734733, EPI_ISL_734734, EPI_ISL_734735, EPI_ISL_735245, EPI_ISL_735246 | | | |
| see above | UZ Leuven, National Reference Laboratory for Coronaviruses, Laboratory Medicine, Leuven, Belgium | KU Leuven, Rega Institute, Clinical and Epidemiological Virology | Tony Wawina-Bokalanga, Joan Marti-Carerras, Bert Vanmechelen, Piet Maes |
| EPI_ISL_735287, EPI_ISL_735288, EPI_ISL_735289, EPI_ISL_735290, EPI_ISL_735291, EPI_ISL_735292, EPI_ISL_735293, EPI_ISL_735294, EPI_ISL_735295, EPI_ISL_735296, EPI_ISL_735297, EPI_ISL_735298, EPI_ISL_735299, EPI_ISL_735300, EPI_ISL_735301, EPI_ISL_735302, EPI_ISL_735303, EPI_ISL_735304, EPI_ISL_735305, EPI_ISL_735306, EPI_ISL_735307, EPI_ISL_735308, EPI_ISL_735309, EPI_ISL_735310, EPI_ISL_735311, EPI_ISL_735434 | | | |
| see above | Genomic Laboratory (GLAB) (Conjoint lab of Health Directorate of Istanbul and Istanbul Technical University) | Genomic Laboratory (GLAB), Istanbul Technical University | Ilker Karacan, Tugba Kizilboga Akgun, Nihat Bugra Agaoglu, Payam Zolfagharian, Mehtap Aydin, Gizem Alkurt, Jale Yildiz, Betsi Kose, Nisan Denizce Can, Ayse Serra Ozel, Nilsun Altunal, Arzu Irvem, Yasemin Kendir Demirkol, Ozlem Akgun Dogan, Levent Doganay, Gizem Dinler Doganay |
| EPI_ISL_737932, EPI_ISL_737933, EPI_ISL_737937, EPI_ISL_737938, EPI_ISL_737947, EPI_ISL_737952, EPI_ISL_737953, EPI_ISL_737960, EPI_ISL_737961, EPI_ISL_737964, EPI_ISL_737966, EPI_ISL_737967, EPI_ISL_737968, EPI_ISL_737969, EPI_ISL_737970 | | | |

| | | | |
|---|---|---|---|
| see above | Uganda Central Public Health Lab and Uganda Virus Research Institute | MRC/UVRI & LSHTM Uganda Research Unit | Matthew Cotten, Dan Lule Bugembe, My V.T. Phan, Pontiano Kaleebu et al. |
| EPI_ISL_738197, EPI_ISL_738214, EPI_ISL_738215, EPI_ISL_738216 | UZ Leuven, National Reference Laboratory for Coronaviruses, Laboratory Medicine, Leuven, Belgium | KU Leuven, Rega Institute, Clinical and Epidemiological Virology | Tony Wawina-Bokalanga, Joan Marti-Carerras, Bert Vanmechelen, Piet Maes |
| EPI_ISL_738289 | Connecticut Veterans' Affairs Hospital | Grubaugh Lab - Yale School of Public Health | Joseph Fauver, Tara Alpert, Chantal Vogels, Mary Petrone, Isabel Ott, Ellen Foxman, Shaili Gupta, Danielle Plank, Nathan Grubaugh |
| EPI_ISL_738296, EPI_ISL_738297 | Yale Pathology Lab | Grubaugh Lab - Yale School of Public Health | Joseph Fauver, Tara Alpert, Chantal Vogels, Mary Petrone, Isabel Ott, Ellen Foxman, Shaili Gupta, Danielle Plank, Nathan Grubaugh |
| EPI_ISL_738307, EPI_ISL_738308, EPI_ISL_738310, EPI_ISL_738311, EPI_ISL_738312 | Connecticut Veterans' Affairs Hospital | Grubaugh Lab - Yale School of Public Health | Joseph Fauver, Tara Alpert, Chantal Vogels, Mary Petrone, Isabel Ott, Ellen Foxman, Shaili Gupta, Danielle Plank, Nathan Grubaugh |
| EPI_ISL_739249 | Santa Clara County Public Health Laboratory | Chan-Zuckerberg Biohub | CZB Cliahub Consortium |
| EPI_ISL_739251 | Napa-Solano-Yolo- Marin County (NSYM) Public Health Laboratories | Chan-Zuckerberg Biohub | CZB Cliahub Consortium |
| EPI_ISL_739615 | Santa Clara County Public Health Laboratory | Chan-Zuckerberg Biohub | CZB Cliahub Consortium |
| EPI_ISL_744819 | Laboratoire national de santé, Microbiology, Virology | Laboratoire national de santé, Microbiology, Epidemiology and Microbial Genomics | Anke Wienecke-Baldacchino, Catherine Ragimbeau, Tamir Abdelrahman, Jessica Tapp, Fatu Djabi, Trung Nguyen Nguyen |
| EPI_ISL_746320 | Mubarak Al-Kabeer Hospital | Virology Unit, Microbiology Department, Faculty of Medicine, Kuwait University | Nada Madi |
| EPI_ISL_746322 | Mubarak Al-Kabeer Hospital | Virology Unit, Department of Microbiology, Faculty of Medicine, Kuwait University | Nada Madi |
| EPI_ISL_747615 | Instituto de Investigaciones Biologicas Clemente Estable | Institut Pasteur de Montevideo | Daiana Mir, Natalia Rego, Paola Cristina Resende, Fernando Lopez-Tort, Tamara Fernandez-Calero, Veronica Noya, Mariana Brandes, Tania Possi, Mailen Arleo, Natalia Reyes, Matias Victoria, Andres Lizasoain, Matias Castells, Leticia Maya, Matías Salvo, Tatiana Schäffer Gregianini, Marilda Tereza Mar da Rosa, Letícia Garay Martins, Cecilia Alonso, Yasser Vega, Cecilia Salazar, Ignacio Ferrés, Pablo Smircich, Jose Sotelo, Ighor Arantes, Luciana Appolinario, Ana Carolina Mendonça, Maria Jose Benitez-Galeano, Martín Graña, Camila Simoes, Fernando Motta, Marilda Mendonça Siqueira, Gonzalo Bello, Rodney Colina, Lucia Spangenberg |
| EPI_ISL_752663, EPI_ISL_752670, EPI_ISL_752671 | State Laboratories Division, Hawaii State Department of Health | State Laboratories Division, Hawaii State Department of Health | Pamela O'Brien, Sabrina Diemert, Drew Kuwazaki, Razvan Sultana, Edward Desmond |
| EPI_ISL_753712, EPI_ISL_753725, EPI_ISL_753760, EPI_ISL_753775, EPI_ISL_753812, EPI_ISL_753824, EPI_ISL_753953, EPI_ISL_753954, EPI_ISL_753955 | Charité Universitätsmedizin Berlin, Institut für Virologie/Labor Berlin | Charité Universitätsmedizin Berlin, Institut für Virologie | Victor M Corman, Jörn Beheim-Schwarzbach, Barbara Mühlemann, Julia Schneider, Talitha Veith, Terry Jones, Christian Drosten |
| EPI_ISL_754903 | Laboratory Diagnostics and Clinical Immunology of Developmental Age, Medical University of Warsaw | genXone SA, Research & Development Laboratory; The Faculty of Mathematics, Informatics and Mechanics of the University of Warsaw | Maciej Sykulski, Grzegorz Nowicki, Monika Makowska-Woniak, Jakub Grabowski, Natalia Drwska-Matelska, ukasz Krych, Micha Kaszuba, Anna Gambin, Urszula Demkow |
| EPI_ISL_755304 | Laboratorio de Virología, Centro de Biotecnología Acuícola, Universidad de Santiato de Chile | Center for Mathematical Modeling and Center for Genome Regulation. Santiago, Chile | A. Sandino, Reyes-López F., Bastias M, Sanhueza D, Travisany D, Allende ML, Maass A, González M, Bustos F, Arriagada G, Montecino, M, Orellana A, Castro E, Meneses C. |
| EPI_ISL_766046 | RSUP Fatmawati | Eijkman Institute for Molecular Biology, Ministry of Research and Technology/National Agency for Research and Innovation | Frilasita A Yudhaputri, Edison Johar, Hidayat Trimarsanto, David H Muljono, Safarina G Malik, Khin Saw Myint, Amin Soebandrio |
| EPI_ISL_768729, EPI_ISL_768732 | Child Health Research Foundation | Child Health Research Foundation | Senjuti Saha, Afroza Akter Tanni, Roly Malaker, Sharmistha Goswami, Syed Muktadir Al Sium, Arif Mohammad Tanmoy, Md Hafizur Rahman, Samir K Saha |
| EPI_ISL_768760, EPI_ISL_768762, EPI_ISL_768764 | AIID | Irish Coronavirus Sequencing Consortium - National Virus Reference Laboratory | Michael Carr, Gabriel Gonzalez, Alejandro Abner Garcia Leon, Patrick Mallon |
| EPI_ISL_770827 | Eastern Ontario Regional Laboratory Association | McMaster University | Leanne Mortimer, Hooman Derakhshani, Emily Panousis, Ahmed Draia, Jalees Nasir, Robert Slinger, Andrew G. McArthur |
| EPI_ISL_771202, EPI_ISL_771218, EPI_ISL_771226, EPI_ISL_771227, EPI_ISL_771228 | Colorado Department of Public Health and Environment | Colorado Department of Puplic Health and Environment | Laura Bankers, Molly C. Hetherington-Rauth, Diana Ir, Shannon Ely, Shannon R. Matzinger, Sarah Elizabeth Totten, Emily A. Travanty |
| EPI_ISL_775746, EPI_ISL_775757, EPI_ISL_775921, EPI_ISL_776455, EPI_ISL_776520, EPI_ISL_776558 | University Medical Center Hamburg Eppendorf | Heinrich Pette Institute, Leibniz Institute for Experimental Virology | Alexis Robitaille, Thomas Günther, Johannes Knobloch, Martin Aepfelbacher, Nicole Fischer, Adam Grundhoff |
| EPI_ISL_776659, EPI_ISL_776735, EPI_ISL_776736, EPI_ISL_776737, EPI_ISL_776738, EPI_ISL_776739 | UW Virology Lab | UW Virology Lab | Pavitra Roychoudhury, Hong Xie, Lasata Shrestha, Meei-Li Huang, Keith R Jerome, Alexander Greninger |
| EPI_ISL_780026 | Hospital General Universitario Gregorio Marañón | SeqCOVID-SPAIN consortium/IBV(CSIC) | Darío García de Viedma, Laura Pérez-Lago, Marta Herranz, Jon Sicilia, Julia Suárez, Pilar Catalán, Patricia Muñoz and SeqCOVID-SPAIN consortium |
| EPI_ISL_780661 | Houston Methodist Hospital | Houston Methodist Hospital | S. Wesley Long, Randall J. Olsen, Paul A. Christensen, David W. Bernard, James J. Davis, Maulik Shukla, Marcus Nguyen, Matthew Ojeda Saavedra, Prasanti Yerramilli, Layne Pruitt, Sishir Subedi, Heather Hendrickson, and James M. Musser |
| EPI_ISL_788967 | Ospedale "Di Venere" | Beaconlab (Bioinformatics, Evolution and Comparative Genomics lab), Dept of Biosciences, University on Milan | Iacobellis M, d'Avenia M, Piluscio R, Parisi A, Chiara M, Manzari C, Pesole G |
| EPI_ISL_791812, EPI_ISL_791813, EPI_ISL_791814, EPI_ISL_791916, EPI_ISL_791917, EPI_ISL_791918, EPI_ISL_791919, EPI_ISL_791920, EPI_ISL_791921, EPI_ISL_791922, EPI_ISL_791923, EPI_ISL_791924, EPI_ISL_791925, EPI_ISL_791926, EPI_ISL_791927, EPI_ISL_791928, EPI_ISL_791929, EPI_ISL_791930, EPI_ISL_791931, EPI_ISL_791932, EPI_ISL_791933, EPI_ISL_791934, EPI_ISL_791935, EPI_ISL_791936, EPI_ISL_791937, EPI_ISL_791938, EPI_ISL_791939, EPI_ISL_791940, EPI_ISL_791941, EPI_ISL_791942, EPI_ISL_791943, EPI_ISL_791944, EPI_ISL_791945, EPI_ISL_791946, EPI_ISL_791947, EPI_ISL_791953, EPI_ISL_791954, EPI_ISL_791955, EPI_ISL_791956, EPI_ISL_791957, EPI_ISL_791958, EPI_ISL_791959, EPI_ISL_791960, EPI_ISL_791961, EPI_ISL_791962, EPI_ISL_791967, EPI_ISL_791968, EPI_ISL_791969, EPI_ISL_791970, EPI_ISL_791971, EPI_ISL_791972, EPI_ISL_791973, EPI_ISL_791974, EPI_ISL_791975 | | | |
| see above | Massachusetts General Hospital | Infectious Disease Program, Broad Institute of Harvard and MIT | Lemieux,J.E., Siddle,K.J., Shaw,B., Adams,G., Pierce,V., Turbett,S., Anahtar,M., Branda,J., Slater,D., Harris,J., Lin,A.E., Gladden-Young,A., Lagerborg,K., Rudy,M., DeRuff,K., Carter,A., Normandin,E., Bauer,M., Reilly,S., Tomkins-Tinch,C., Loreth,C., Chaluvadi,S., Neumann,A., Cusick,C., Chapman,S.B., Gnirke,A., Flowers,K., Cerrato,F., Birren,B.W., Gallagher,G., Smole,S., Park,D.J., MacInnis,B.L., Ryan,E., LaRocque,R., Rosenberg,E. and Sabeti,P.C. |
| EPI_ISL_792175, EPI_ISL_792176, EPI_ISL_792177, EPI_ISL_792178, EPI_ISL_792179, EPI_ISL_792180, EPI_ISL_792181, EPI_ISL_792182, EPI_ISL_792183, EPI_ISL_792184, EPI_ISL_792185, EPI_ISL_792186, EPI_ISL_792187, EPI_ISL_792188, EPI_ISL_792189, EPI_ISL_792190, EPI_ISL_792191, EPI_ISL_792192, EPI_ISL_792193, EPI_ISL_792194, EPI_ISL_792195, EPI_ISL_792196, EPI_ISL_792197, EPI_ISL_792198, EPI_ISL_792199, EPI_ISL_792200, EPI_ISL_792201, EPI_ISL_792202, EPI_ISL_792203, EPI_ISL_792204, EPI_ISL_792205, EPI_ISL_792206, EPI_ISL_792207, EPI_ISL_792208, EPI_ISL_792209, EPI_ISL_792210, EPI_ISL_792211, EPI_ISL_792212, EPI_ISL_792213, EPI_ISL_792214, EPI_ISL_792215, EPI_ISL_792216, EPI_ISL_792217, EPI_ISL_792218, EPI_ISL_792219, EPI_ISL_792220, EPI_ISL_792221, EPI_ISL_792222, EPI_ISL_792223, EPI_ISL_792224, EPI_ISL_792225, EPI_ISL_792226, EPI_ISL_792227, EPI_ISL_792228, EPI_ISL_792229, EPI_ISL_792230, EPI_ISL_792231, EPI_ISL_792232, EPI_ISL_792233, EPI_ISL_792234, EPI_ISL_792235, EPI_ISL_792236, EPI_ISL_792237, EPI_ISL_792238, EPI_ISL_792239, EPI_ISL_792240, EPI_ISL_792241, EPI_ISL_792242, EPI_ISL_792243, EPI_ISL_792287, EPI_ISL_792288, EPI_ISL_792289 | | | |
| see above | Laboratorio de Virología del Hospital de Niños Dr. Ricardo Gutierrez | Área de Secuenciación del Laboratorio de Virología del Hospital de Niños Dr. Ricardo Gutierrez on behalf of 'Proyecto Argentino Interinstitucional de genomica de SARS-CoV-2' (PAIS Consortium) | Nabaes Jodar, MS; Goya, S; Natale, MI; Lusso, S; Gravis, E; Acevedo, ME; Alvarez Lopez, C; Alexay, S; Jacques, O; Mistchenko, AS; Valinotto, LE; Viegas, M. |
| EPI_ISL_792290 | Laboratorio de Virología del Hospital de Niños Dr. Ricardo Gutierrez | Área de Secuenciación del Laboratorio de Virología del Hospital de Niños Dr. Ricardo Gutierrez on behalf of 'Proyecto | Nabaes Jodar, MS; Goya, S; Natale, MI; Lusso, S; Primost, I; Gallino, MI; Valinotto, LE; Viegas, M. |

| | | | |
|---|---|---|---|
| | | Argentino Interinstitucional de genomica de SARS-CoV-2' (PAIS Consortium) | |
| EPI_ISL_792316, EPI_ISL_792317, EPI_ISL_792318, EPI_ISL_792319 | Laboratorio del Hospital El Cruce Dr. Néstor C. Kirchner | Área de Secuenciación del Laboratorio de Virología del Hospital de Niños Dr. Ricardo Gutierrez on behalf of 'Proyecto Argentino Interinstitucional de genomica de SARS-CoV-2' (PAIS Consortium) | Nabaes Jodar, MS; Goya, S; Natale, MI; Lusso, S; Zubieta, M; Rahhal, M; Valinotto, LE; Viegas, M. |
| EPI_ISL_792321, EPI_ISL_792322, EPI_ISL_792323 | Instituto de Investigaciones Biomédicas en Retrovirus y SIDA | Área de Secuenciación del Laboratorio de Virología del Hospital de Niños Dr. Ricardo Gutierrez on behalf of 'Proyecto Argentino Interinstitucional de genomica de SARS-CoV-2' (PAIS Consortium) | Nabaes Jodar, MS; Goya, S; Natale, MI; Lusso, S; Seery, V; Remes Lenicov, F; Salomón, H; Valinotto, LE; Viegas, M. |
| EPI_ISL_792344 | Laboratorio del Hospital Interzonal General de Agudos Evita | Área de Secuenciación del Laboratorio de Virología del Hospital de Niños Dr. Ricardo Gutierrez on behalf of 'Proyecto Argentino Interinstitucional de genomica de SARS-CoV-2' (PAIS Consortium) | Nabaes Jodar, MS; Goya, S; Natale, MI; Lusso, S; Desimone, I; Luczac, E; Serrano, L; Grossi, O; Musto, A; Valinotto, LE; Viegas, M. |
| EPI_ISL_792375 | Laboratorio de salud pública, Facultad de Ciencias Exactas, Universidad Nacional de La Plata | Área de Secuenciación del Laboratorio de Virología del Hospital de Niños Dr. Ricardo Gutierrez on behalf of 'Proyecto Argentino Interinstitucional de genomica de SARS-CoV-2' (PAIS Consortium) | Nabaes Jodar, MS; Goya, S; Natale, MI; Lusso, S; Angeletti, A; Nadalich, V; Cordero, A; Toro, R ; Valinotto, LE; Viegas, M. |
| EPI_ISL_792392, EPI_ISL_792393, EPI_ISL_792394, EPI_ISL_792395 | Plataforma de Servicios Biotecnólogicos: UTTIPP/PSB , Universidad Nacional de Quilmes. | Área de Secuenciación del Laboratorio de Virología del Hospital de Niños Dr. Ricardo Gutierrez on behalf of 'Proyecto Argentino Interinstitucional de genomica de SARS-CoV-2' (PAIS Consortium) | Nabaes Jodar, MS; Goya, S; Natale, MI; Lusso, S; Farina, H; Goñi, S; Cardama, G; Castello, A; Valinotto, LE; Viegas, M. |
| EPI_ISL_792405, EPI_ISL_792406, EPI_ISL_792407, EPI_ISL_792408, EPI_ISL_792410, EPI_ISL_792411 | LABORATORIO DE CAMPAÑA COVID 19 INTA | Instituto de Biotecnología, IABIMO (CONICET), Instituto de Virología, IVIT(CONICET), Instituto de Patobiología, IPVET(CONICET), CICVyA, INTA on behalf of 'Proyecto Argentino Interinstitucional de genomica de SARS-CoV-2' (PAIS Consortium) | König, GA; Peralta, AV; Distéfano, AJ; Zavallo, D; Muñoz Hidalgo, MG; Vera, PA; Fass, M; Farber, MD; Cacciabué, MPD; Pedroarias, VC; Lozano Calderón, LC; Bengoa Luoni, S; Asurmendi, S; Gioffré, A; Ferández PC; Zumárraga, M; Delgado, F; Díaz Carrasco, J; Garbaccio, S; Álvarez, I; Diagiacomo, S; Franco, L; Schammas, J; Rimondi, A; Soria, I;Craig, MI; Chimeno, S; Gómez, E; Klepp, L; Lucero, MS; Sioya, B; Del Médico, P; Rodríguez, A; Dus Santos, MJ; Olivera, V; Trono, K; Rosende, J; Fusco, W; Huertas, P; Vissani, A; Perea, A; Pereda, A; Paniego, NB; Rivarola, M; Puebla, AF; Viegas, M. |
| EPI_ISL_792426, EPI_ISL_792427, EPI_ISL_792428, EPI_ISL_792429, EPI_ISL_792430 | Laboratorio de Inmunología del Hospital Perrando e Instituto de Medicina Regional de la UNNE | Instituto de Biotecnología, IABIMO (CONICET), Instituto de Virología, IVIT(CONICET), Instituto de Patobiología, IPVET(CONICET), CICVyA, INTA on behalf of 'Proyecto Argentino Interinstitucional de genomica de SARS-CoV-2' (PAIS Consortium) | König, GA; Peralta, AV; Distéfano, AJ; Zavallo, D; Muñoz Hidalgo, MG; Vera, PA; Fass, M; Farber, MD; Cacciabué, MPD; Pedroarias, VC; Lozano Calderón, LC; Bengoa Luoni, S; Asurmendi, S; Foussal, MD; Deluca, G; Ayala, NA; Gómez, MV; Giusiano, G; Lucero, H; Marin, M; Lescano, L; Cayré, A; Paniego, NB; Rivarola, M; Puebla, AF; Viegas, M. |
| EPI_ISL_792640, EPI_ISL_792644 | LACEN-AL | Laboratory of Respiratory Viruses and Measles, Oswaldo Cruz Institute, FIOCRUZ | Paola Resende, Luciana Appolinario, Fernando Motta, Anna Carolina Paixao, Ana Carolina Mendonca, Anderson Brandao Leite, Marilda Siqueira |
| EPI_ISL_794596 | Central Laboratories, Egyptian Ministry of Health and Population | Central Laboratories, Egyptian Ministry of Health and Population | Roshdy,W.H., Kayed,A.E., Mostafa,A., El-Shesheny,R., Khalifa,M.K., Shehata,M., Shawky,S., Saleh,M., Gomaa,M., El Taweel,A., Mahmoud,S.H., Moatasim,Y., Kutkat,O., Kamel,M.N., Mahrous,N., El Sayes,M.A., El Guindy,N.M., Naguib,A., Kandeil,A., Kayali,G., Ali,M.A. |
| EPI_ISL_794607 | Central Public Health Laboratories, Egyptian Ministry of Health and Population | Central Public Health Laboratories, Egyptian Ministry of Health and Population | El-Shesheny,R., Roshdy,W.H., Kayed,A.E., Mostafa,A., Khalifa,M.K., Shehata,M., Shawky,S., Saleh,M., Gomaa,M., El Taweel,A., Mahmoud,S.H., Moatasim,Y., Kutkat,O., Kamel,M.N., Mahrous,N., El Sayes,M.A., El Guindy,N.M., Naguib,A., Kandeil,A., Kayali,G., Ali,M.A. |
| EPI_ISL_796130, EPI_ISL_796131, EPI_ISL_796132, EPI_ISL_796133 | Laboratorio de Microbiología. Hospital General Universitario de Elda, Alicante | SeqCOVID-SPAIN consortium/IBV(CSIC) | Mª Isabel Gascón Ros, Cristina Torregrosa Hetland, Eva Pastor Boix, Paloma Cascales Ramos and SeqCOVID-SPAIN consortium |
| EPI_ISL_801403 | Laboratório Central de Saúde Pública do Amazonas - LACEN-AM | Laboratorio de Ecologia de Doencas Transmissiveis na Amazonia, Instituto Leonidas e Maria Deane - Fiocruz Amazonia | Valdinete Nascimento, Victor Souza, André Corado, Fernanda Nascimento, George Silva, Ágatha Costa, Debora Duarte, Luciana Gonçalves, Maria Júlia Brandão, Michele Jesus, Felipe Naveca |
| EPI_ISL_801545 | Laboratory of Molecular Virology, Pontificia Universidad Católica de Chile | MSHS Pathogen Surveillance Program | Leonardo I. Almonacid, Ana S. Gonzalez-Reiche, Matthew M. Hernandez, Jorge Levican, Ana Maria Contreras, Carlos Palma, Tamara García-Salum, Zenab Khan, Adriana van De Guchte, Ajay Obla, Jayeeta Dutta, Bremy Alburquerque, Eileen Serrano, Constanza Maldonado, M. Belen Leyton, Erick Salinas, Hala Alshammary, Juan Soto, Shwetha Hara Sridhar, Ying-Chih Wang, Melissa Smith, Robert Sebra, Marcela Ferres, Adolfo Garcia-Sastre, Edward C. Holmes, Viviana Simon, Harm van Bakel, Rafael A. Medina. |
| EPI_ISL_801557, EPI_ISL_801568, EPI_ISL_801571, EPI_ISL_801576, EPI_ISL_801578 | Laboratory of Molecular Virology, Pontificia Universidad Católica de Chile | MSHS Pathogen Surveillance Program | Leonardo I. Almonacid, Ana S. Gonzalez-Reiche, Matthew M. Hernandez, Jorge Levican, Tamara García-Salum, Zenab Khan, Adriana van De Guchte, Ajay Obla, Jayeeta Dutta, Bremy Alburquerque, Eileen Serrano, Erick Salinas, Manuel Ampuero, Hala Alshammary, Juan Soto, Shwetha Hara Sridhar, Ying-Chih Wang, Melissa Smith, Robert Sebra, Fernando Valiente, Aldo Gaggero, Adolfo Garcia-Sastre, Edward C. Holmes, Viviana Simon, Harm van Bakel, Rafael A. Medina. |
| EPI_ISL_801763, EPI_ISL_801765, EPI_ISL_801766, EPI_ISL_801767, EPI_ISL_801768, EPI_ISL_801769, EPI_ISL_801770, EPI_ISL_801771, EPI_ISL_801772, EPI_ISL_801773, EPI_ISL_801774, EPI_ISL_801775, EPI_ISL_801776, EPI_ISL_801777, EPI_ISL_801778, EPI_ISL_801779, EPI_ISL_801780, EPI_ISL_801781, EPI_ISL_801782, EPI_ISL_801783, EPI_ISL_801784, EPI_ISL_801785, EPI_ISL_801786, EPI_ISL_801787, EPI_ISL_801788, EPI_ISL_801789, EPI_ISL_801790, EPI_ISL_801791, EPI_ISL_801792, EPI_ISL_801793, EPI_ISL_801794, EPI_ISL_801795, EPI_ISL_801796, EPI_ISL_801797, EPI_ISL_801798, EPI_ISL_801799, EPI_ISL_801800, EPI_ISL_801801, EPI_ISL_801802, EPI_ISL_801803, EPI_ISL_801804, EPI_ISL_801805, EPI_ISL_801806, EPI_ISL_801807, EPI_ISL_801808, EPI_ISL_801809, EPI_ISL_801810, EPI_ISL_801811, EPI_ISL_801812, EPI_ISL_801813, EPI_ISL_801814, EPI_ISL_801815, EPI_ISL_801816, EPI_ISL_801817, EPI_ISL_801818, EPI_ISL_801819, EPI_ISL_801820, EPI_ISL_801821, EPI_ISL_801822, EPI_ISL_801823, EPI_ISL_801824, EPI_ISL_801825, EPI_ISL_801826, EPI_ISL_801827, EPI_ISL_801828, EPI_ISL_801829, EPI_ISL_801830, EPI_ISL_801831, EPI_ISL_801832, EPI_ISL_801833, EPI_ISL_801834 | | | |
| see above | Laboratory of Molecular Virology, Pontificia Universidad Católica de Chile | MSHS Pathogen Surveillance Program | Leonardo I. Almonacid, Ana S. Gonzalez-Reiche, Matthew M. Hernandez, Jorge Levican, Ana Maria Contreras, Carlos Palma, Tamara García-Salum, Zenab Khan, Adriana van De Guchte, Ajay Obla, Jayeeta Dutta, Bremy Alburquerque, Eileen Serrano, Constanza Maldonado, M. Belen Leyton, Erick Salinas, Hala Alshammary, Juan Soto, Shwetha Hara Sridhar, Ying-Chih Wang, Melissa Smith, Robert Sebra, Marcela Ferres, Adolfo Garcia-Sastre, Edward C. Holmes, Viviana Simon, Harm van Bakel, Rafael A. Medina. |
| EPI_ISL_801855, EPI_ISL_801857, EPI_ISL_801859 | Laboratory of Molecular Virology, Pontificia Universidad Católica de Chile | MSHS Pathogen Surveillance Program | Leonardo I. Almonacid, Ana S. Gonzalez-Reiche, Matthew M. Hernandez, Jorge Levican, Tamara García-Salum, Zenab Khan, Adriana van De Guchte, Ajay Obla, Jayeeta Dutta, Bremy Alburquerque, Eileen Serrano, Erick Salinas, Manuel Ampuero, Hala Alshammary, Juan Soto, Shwetha Hara Sridhar, Ying-Chih Wang, Melissa Smith, Robert Sebra, Fernando Valiente, Aldo Gaggero, Adolfo Garcia-Sastre, Edward C. Holmes, Viviana Simon, Harm van Bakel, Rafael A. Medina. |
| EPI_ISL_801960, EPI_ISL_802087, EPI_ISL_802088, EPI_ISL_802089, EPI_ISL_802090, EPI_ISL_802091, EPI_ISL_802092, EPI_ISL_802093 | MSHS Clinical Microbiology Laboratories | MSHS Pathogen Surveillance Program | Ana S. Gonzalez-Reiche, Hala Alshammary, Mitchell J. Sullivan, Brianne Ciferri, Ajay Obla, Angela Amoako, Mahmoud Awawda, Elena Hirsch, Ashley S. Salimbangon, Levy Sominsky, Katherine Beach, Kayla Russo, Charles Gleason, Shelcie Fabre, Giulio Kleiner, Zenab Khan, Bremy Alburquerque, Adriana van de Guchte, Komal Srivastava, Matthew M. Hernandez, Jayeeta Dutta, Denise Jurczyszak, Emily Ferreri, Rachel Chernet, Nancy Francoeur, Betsaida Salom Melo, Irina Oussenko, Gintaras Deikus, Juan Soto, Shwetha Hara Sridhar, Ying-Chih Wang, Kathryn Twyman, Andrew Kasarskis, Deena R. Altman, Robert Sebra, Adolfo García-Sastre, Marta Luksza, Gopi Patel, Sarah Schaefer, Melissa Gitman, Michael D. Nowak, Alberto Paniz-Mondolfi, Emilia Mia Sordillo, Viviana Simon, Harm van Bakel |
| EPI_ISL_804868, EPI_ISL_804869, EPI_ISL_804886, EPI_ISL_804923 | DC Public Health Lab/ Dept. of Forensic Sciences | DC Public Health Lab/ Dept. of Forensic Sciences | Scott Nguyen, Elizabeth Zelaya, Connie Maza, Monica Mann, Brittany Hamilton, David Payne, Jocelyn Hauser |
| EPI_ISL_804999, EPI_ISL_805001, EPI_ISL_805003, EPI_ISL_805007, EPI_ISL_805023, EPI_ISL_805122, EPI_ISL_805283, EPI_ISL_805296, EPI_ISL_805310, EPI_ISL_805312, EPI_ISL_805330, EPI_ISL_805337, EPI_ISL_805341, EPI_ISL_805346, EPI_ISL_805351, EPI_ISL_805357, EPI_ISL_805361, EPI_ISL_805362, EPI_ISL_805363, EPI_ISL_805368, EPI_ISL_805376, EPI_ISL_805378, EPI_ISL_805381, EPI_ISL_805384, EPI_ISL_805387, EPI_ISL_805395, EPI_ISL_805396, EPI_ISL_805400, EPI_ISL_805405, EPI_ISL_805411, EPI_ISL_805418, EPI_ISL_805424, EPI_ISL_805427, EPI_ISL_805434, EPI_ISL_805435, EPI_ISL_805446, EPI_ISL_805453, EPI_ISL_805455, EPI_ISL_805458, EPI_ISL_805462, EPI_ISL_805465, EPI_ISL_805468, EPI_ISL_805490, EPI_ISL_805503, EPI_ISL_805515, EPI_ISL_805527, EPI_ISL_805529, EPI_ISL_805531, EPI_ISL_805533, EPI_ISL_805534, EPI_ISL_805539, EPI_ISL_805542, EPI_ISL_805543, EPI_ISL_805547, EPI_ISL_805548, EPI_ISL_805550, EPI_ISL_805554, EPI_ISL_805559, EPI_ISL_805576, EPI_ISL_805577, EPI_ISL_805579, EPI_ISL_805582, EPI_ISL_805590, EPI_ISL_805595, EPI_ISL_805600, EPI_ISL_805601, EPI_ISL_805610, EPI_ISL_805618, EPI_ISL_805645, EPI_ISL_805646, EPI_ISL_805649, EPI_ISL_805659, EPI_ISL_805664, EPI_ISL_805673, EPI_ISL_805678, EPI_ISL_805693, EPI_ISL_805708, EPI_ISL_805713, EPI_ISL_805715, EPI_ISL_805726, EPI_ISL_805735, EPI_ISL_805736, EPI_ISL_805738, EPI_ISL_805744, EPI_ISL_805745, EPI_ISL_805749, EPI_ISL_805759, EPI_ISL_805772, EPI_ISL_805775, EPI_ISL_805788, EPI_ISL_805792, EPI_ISL_805802, EPI_ISL_805806, EPI_ISL_805813, EPI_ISL_805824, EPI_ISL_805830, EPI_ISL_805831, EPI_ISL_805833, EPI_ISL_805840, EPI_ISL_805845, EPI_ISL_805853, EPI_ISL_805864, EPI_ISL_805865, EPI_ISL_805879, EPI_ISL_805882, EPI_ISL_805895, EPI_ISL_805900, EPI_ISL_805901, | | | |

| | | | |
|---|---|---|---|
| EPI_ISL_805920, EPI_ISL_805939, EPI_ISL_805956, EPI_ISL_805962, EPI_ISL_805986, EPI_ISL_805988, EPI_ISL_805998, EPI_ISL_806000, EPI_ISL_806004, EPI_ISL_806010, EPI_ISL_806014, EPI_ISL_806022, EPI_ISL_806028, EPI_ISL_806035, EPI_ISL_806039, EPI_ISL_806042, EPI_ISL_806045, EPI_ISL_806062, EPI_ISL_806076, EPI_ISL_806083, EPI_ISL_806087, EPI_ISL_806096, EPI_ISL_806118, EPI_ISL_806121, EPI_ISL_806125, EPI_ISL_806143, EPI_ISL_806144, EPI_ISL_806146, EPI_ISL_806147, EPI_ISL_806160, EPI_ISL_806173, EPI_ISL_806182, EPI_ISL_806183, EPI_ISL_806198, EPI_ISL_806202, EPI_ISL_806204, EPI_ISL_806208, EPI_ISL_806213, EPI_ISL_806217, EPI_ISL_806219, EPI_ISL_806227, EPI_ISL_806228, EPI_ISL_806229, EPI_ISL_806232, EPI_ISL_806236, EPI_ISL_806271, EPI_ISL_806273, EPI_ISL_806274, EPI_ISL_806275, EPI_ISL_806297, EPI_ISL_806299, EPI_ISL_806305, EPI_ISL_806306, EPI_ISL_806307, EPI_ISL_806308, EPI_ISL_806316, EPI_ISL_806326, EPI_ISL_806335, EPI_ISL_806338, EPI_ISL_806356, EPI_ISL_806357, EPI_ISL_806361, EPI_ISL_806369, EPI_ISL_806401, EPI_ISL_806414, EPI_ISL_806418, EPI_ISL_806433, EPI_ISL_806434, EPI_ISL_806441, EPI_ISL_806447, EPI_ISL_806452, EPI_ISL_806453, EPI_ISL_806456, EPI_ISL_806457, EPI_ISL_806467, EPI_ISL_806468, EPI_ISL_806496, EPI_ISL_806521 | | | |
| see above | Alberta Precision Labs (APL) | Alberta Precision Labs (APL) | Gordon P, Lam LG, Pabbaraju K, Wong A, Ma R, Li V, Melin A, Tipples G, Berenger B, Zelyas N, Kellner J, Bernier F, Chui L, Croxen M |
| EPI_ISL_812246 | North Dakota Department of Health, Public Health Laboratory | North Dakota Department of Health, Public Health Laboratory | Lisa Wingerter |
| EPI_ISL_812341 | United States Air Force School of Aerospace Medicine | United States Air Force School of Aerospace Medicine | Anthony Fries, Jennifer Meyer, Amanda Javorina, Sarah Purves, William Gruner, Clarise Starr, Elizabeth Macias |
| EPI_ISL_812784, EPI_ISL_812785, EPI_ISL_812789, EPI_ISL_812790, EPI_ISL_812791 | Genomics Program, Children Cancer Hospital | Genomics Program, Children Cancer Hospital | Hatem,A., Hadad,A., Abouelnaga,S., Amer,K., Salah,H., Farawyla,H., Halafawy,A., Mansour,T., shalaby,L., Hassan,W., Soliman,M., Gomaa,C., Hassan,R., Soliman,S., Monuir,G., Hammad,M., Hussein,S., Abdo,I., Jalal,D., El-Zayat,M., El-Shaqnqery,H., Diab,A., Bakry,U., Samir,O., Magdeldin,S., Sayed,A. |
| EPI_ISL_814073, EPI_ISL_814087, EPI_ISL_814088, EPI_ISL_814089 | Ministry of Health Turkey | Ministry of Health Turkey | Fatma Bayrakdar, Yasemin Cogun, Süleyman Yalcin, Aye Baak Alta, Gülay Korukluolu |
| EPI_ISL_816686, EPI_ISL_816687, EPI_ISL_816688, EPI_ISL_816689, EPI_ISL_816690, EPI_ISL_816691, EPI_ISL_816692, EPI_ISL_816695, EPI_ISL_816696, EPI_ISL_816697, EPI_ISL_816698, EPI_ISL_816699, EPI_ISL_816700, EPI_ISL_816701, EPI_ISL_816702, EPI_ISL_816703, EPI_ISL_816704, EPI_ISL_816705, EPI_ISL_816706, EPI_ISL_816707, EPI_ISL_816708, EPI_ISL_816709, EPI_ISL_816710, EPI_ISL_816711, EPI_ISL_816712, EPI_ISL_816735 | | | |
| see above | Bioinformatics and Biostatistics Lab, Advanced Sequencing Facility | COVID-19 Genomics UK (COG-UK) Consortium | Aengus Stewart,Jerome Nicod,Chelsea Sawyer,Laura Cubitt,Harshil Patel,Margaret Crawford |
| EPI_ISL_819294, EPI_ISL_819338, EPI_ISL_819339, EPI_ISL_819340, EPI_ISL_819341, EPI_ISL_819342 | Hospital Universitari Vall d'Hebron - Vall d'Hebron Institut de Rercerca | Hospital Universitari Vall d'Hebron | Cristina Andrés, Maria Piñana, Josep F Abril, Damir Garcia-Cehic, Ariadna Rando, Juliana Esperalba, Maria Gema Codina, Carla Castillo, Maria Carmen Martin, Tomàs Pumarola, Josep Quer, Andrés Antón |
| EPI_ISL_825890, EPI_ISL_825891, EPI_ISL_825892, EPI_ISL_825894, EPI_ISL_825895, EPI_ISL_825896, EPI_ISL_825898, EPI_ISL_825899, EPI_ISL_825900, EPI_ISL_825901, EPI_ISL_825903, EPI_ISL_825904, EPI_ISL_825905, EPI_ISL_825907, EPI_ISL_825920, EPI_ISL_825921, EPI_ISL_825922, EPI_ISL_825923, EPI_ISL_825924, EPI_ISL_825925, EPI_ISL_825942, EPI_ISL_825943, EPI_ISL_825944, EPI_ISL_825945, EPI_ISL_825946, EPI_ISL_825948, EPI_ISL_825950, EPI_ISL_825952, EPI_ISL_825953, EPI_ISL_825954, EPI_ISL_825955, EPI_ISL_825957, EPI_ISL_825958, EPI_ISL_825961, EPI_ISL_825963, EPI_ISL_825964, EPI_ISL_825965, EPI_ISL_825968, EPI_ISL_825969, EPI_ISL_825970, EPI_ISL_825971, EPI_ISL_825974, EPI_ISL_825978, EPI_ISL_826028, EPI_ISL_826029, EPI_ISL_826030, EPI_ISL_826031, EPI_ISL_826032, EPI_ISL_826033, EPI_ISL_826034, EPI_ISL_826035, EPI_ISL_826036, EPI_ISL_826037, EPI_ISL_826038, EPI_ISL_826039, EPI_ISL_826040, EPI_ISL_826041, EPI_ISL_826042, EPI_ISL_826043, EPI_ISL_826044, EPI_ISL_826045, EPI_ISL_826046, EPI_ISL_826047, EPI_ISL_826048, EPI_ISL_826049, EPI_ISL_826050, EPI_ISL_826051, EPI_ISL_826052, EPI_ISL_826053, EPI_ISL_826054, EPI_ISL_826055, EPI_ISL_826056, EPI_ISL_826057, EPI_ISL_826058, EPI_ISL_826059, EPI_ISL_826060, EPI_ISL_826061, EPI_ISL_826062, EPI_ISL_826063, EPI_ISL_826064, EPI_ISL_826065, EPI_ISL_826066, EPI_ISL_826067, EPI_ISL_826068, EPI_ISL_826069, EPI_ISL_826070, EPI_ISL_826071, EPI_ISL_826072, EPI_ISL_826073, EPI_ISL_826074, EPI_ISL_826075, EPI_ISL_826076, EPI_ISL_826078, EPI_ISL_826079, EPI_ISL_826080, EPI_ISL_826081, EPI_ISL_826082, EPI_ISL_826083, EPI_ISL_826084, EPI_ISL_826085, EPI_ISL_826086, EPI_ISL_826087, EPI_ISL_826088, EPI_ISL_826089, EPI_ISL_826090, EPI_ISL_826091, EPI_ISL_826092, EPI_ISL_826093, EPI_ISL_826094, EPI_ISL_826095, EPI_ISL_826133, EPI_ISL_826134, EPI_ISL_826135, EPI_ISL_826136, EPI_ISL_826137, EPI_ISL_826139, EPI_ISL_826140, EPI_ISL_826141, EPI_ISL_826142, EPI_ISL_826143, EPI_ISL_826145, EPI_ISL_826147, EPI_ISL_826149, EPI_ISL_826150, EPI_ISL_826151, EPI_ISL_826152, EPI_ISL_826153, EPI_ISL_826154, EPI_ISL_826155, EPI_ISL_826156, EPI_ISL_826157, EPI_ISL_826158, EPI_ISL_826159, EPI_ISL_826160, EPI_ISL_826161, EPI_ISL_826162, EPI_ISL_826163, EPI_ISL_826164, EPI_ISL_826165, EPI_ISL_826166, EPI_ISL_826167, EPI_ISL_826168, EPI_ISL_826169, EPI_ISL_826170, EPI_ISL_826171, EPI_ISL_826172, EPI_ISL_826173, EPI_ISL_826174, EPI_ISL_826175, EPI_ISL_826176, EPI_ISL_826177, EPI_ISL_826178, EPI_ISL_826179, EPI_ISL_826180, EPI_ISL_826181, EPI_ISL_826182, EPI_ISL_826183, EPI_ISL_826185, EPI_ISL_826186, EPI_ISL_826187, EPI_ISL_826188, EPI_ISL_826189, EPI_ISL_826190, EPI_ISL_826191, EPI_ISL_826192, EPI_ISL_826193, EPI_ISL_826194, EPI_ISL_826195, EPI_ISL_826196, EPI_ISL_826197, EPI_ISL_826198, EPI_ISL_826199, EPI_ISL_826200, EPI_ISL_826201, EPI_ISL_826202, EPI_ISL_826203, EPI_ISL_826204, EPI_ISL_826205, EPI_ISL_826206, EPI_ISL_826207, EPI_ISL_826208, EPI_ISL_826209, EPI_ISL_826210, EPI_ISL_826211, EPI_ISL_826212, EPI_ISL_826213, EPI_ISL_826214, EPI_ISL_826215, EPI_ISL_826216, EPI_ISL_826217, EPI_ISL_826218, EPI_ISL_826219, EPI_ISL_826220, EPI_ISL_826224, EPI_ISL_826225, EPI_ISL_826234 | | | |
| see above | Laboratoire de santé publique du Québec | Laboratoire de santé publique du Québec | Sandrine Moreira, Ioannis Ragoussis, Guillaume Bourque, Jesse Shapiro, Mark Lathrop and Michel Roger on behalf of the CoVSeQ research group (http://covseq.ca/researchgroup) |
| EPI_ISL_826592, EPI_ISL_826647, EPI_ISL_826648, EPI_ISL_826649, EPI_ISL_826650 | Montefiore Medical Center | Albert Einstein College of Medicine, Dept. of Microbiology & Immunology, Chandran lab | J. Maximilian Fels, Saad Khan, Ryan Forster, Karin A. Skalina, Surksha Sirichand, Amy S. Fox, Aviv Bergman, William B. Mitchell, Lucia R. Wolgast, Wendy Szymczak, Robert H. Bortz III, M. Eugenia Dieterle, Catalina Florez, Denise Haslwanter, Rohit K. Jangra, Ethan Laudermilch, Ariel S. Wirchnianski, Jason Barnhill, David L. Goldman, Hnin Khine, D. Yitzchak Goldstein, Johanna P. Daily, Kartik Chandran, Libusha Kelly |
| EPI_ISL_833518, EPI_ISL_833520 | Veterinary Specialized Instute "Nis" | Veterinary Specialized Institute "Kraljevo", Serbia | Vidanovic,D., Tesovic,B., Manic,M., Petrovic,M.,Knezevic,A., Jovanovic,T., Jankovic,M., Sekler,M., Banovic Djeri,B., Petrovic,T., Volkening,J., Afonso,C. |
| EPI_ISL_833527, EPI_ISL_833532, EPI_ISL_833557 | Istituto Zooprofilattico Sperimentale del Mezzogiorno | TIGEM | Patrizia Annunziata, Andrea Ballabio, Valentina Bouche, Davide Cacchiarelli (CorrespAuthor), Pellegrino Cerino, Chiara Colantuono, Lucio Di Filippo, Antonio Grimaldi, Antonio Limone, Gabriella Loconte, Anna Manfredi, Francesco Panariello, Biancamaria Pierri, Marcello Salvi, Lucia Vassallo |
| EPI_ISL_837553, EPI_ISL_837563, EPI_ISL_837564, EPI_ISL_837565, EPI_ISL_837566, EPI_ISL_837567, EPI_ISL_837568, EPI_ISL_837569, EPI_ISL_837570 | Centro Nacional de Enfermedades Tropicales (CENETROP) | Laboratory of Respiratory Viruses and Measles, Oswaldo Cruz Institute, FIOCRUZ | Paola Resende, Roxana Loayza, Cinthia Avila, Luciana Appolinario, Fernando Motta, Anna Carolina Paixao, Ana Carolina Mendonca, Marilda Siqueira |
| EPI_ISL_837685, EPI_ISL_837686, EPI_ISL_837687, EPI_ISL_837688, EPI_ISL_837689, EPI_ISL_837690, EPI_ISL_837691, EPI_ISL_837692, EPI_ISL_837693, EPI_ISL_837694, EPI_ISL_837695, EPI_ISL_837696, EPI_ISL_837697, EPI_ISL_837698 | | | |
| see above | Instituto Nacional de Enfermedades Respiratorias (INER) | Instituto Nacional de Enfermedades Respiratorias (INER) | Celia Boukadida, Margarita Matías-Florentino, Alma Rincón-Rubio, Hector Esteban Paz-Juárez, Olivia Briceño, Edgar Sevilla-Reyes, Fidencio Mejía-Nepomuceno, Mario Mújica-Sánchez, Eduardo Becerril-Vargas, José Arturo Martínez-Orozco, Alejandra Hernández-Terán, Jorge Salas-Hernández, Santiago Ávila-Ríos, Joel Armando Vázquez-Pérez |
| EPI_ISL_842792 | University College London Hospital | COVID-19 Genomics UK (COG-UK) Consortium | Judith Heaney, Matthew Byott, Catherine Houlihan, Dan Frampton, Stuart Kirk, Moira Spyer and Eleni Nastouli |
| EPI_ISL_842798, EPI_ISL_842799, EPI_ISL_842800, EPI_ISL_842801, EPI_ISL_842802 | Barts Health NHS Trust | COVID-19 Genomics UK (COG-UK) Consortium | CUTINO-MOGUEL, Maria-Teresa; HARRINGTON, David; OWOYEMI, Dola; SHYLINI, Raghavendran; BROAD, Claire; KELE, Beatrix |
| EPI_ISL_848580, EPI_ISL_848581, EPI_ISL_848582, EPI_ISL_848615 | Evandro Chagas Institute | Evandro Chagas Institute | Santos, M.C.; Silva, A.M.; Junior, W.D.C.; Barbagelata, L.S.; Ferreira, J.A.; Sousa, E.M.A.; da Silva, P.S.; Pinheiro, K.C.; L.C.; Sousa Junior, E.C. |
| EPI_ISL_849151, EPI_ISL_849152, EPI_ISL_849156, EPI_ISL_849157, EPI_ISL_849158, EPI_ISL_849159, EPI_ISL_849160, EPI_ISL_849161, EPI_ISL_849162, EPI_ISL_849163, EPI_ISL_849164, EPI_ISL_849165, EPI_ISL_849166, EPI_ISL_849167, EPI_ISL_849168, EPI_ISL_849169, EPI_ISL_849170, EPI_ISL_849171, EPI_ISL_849172, EPI_ISL_849173, EPI_ISL_849174, EPI_ISL_849186, EPI_ISL_849187, EPI_ISL_849188, EPI_ISL_849189, EPI_ISL_849190, EPI_ISL_849191, EPI_ISL_849192, EPI_ISL_849193, EPI_ISL_849194, EPI_ISL_849195, EPI_ISL_849196, EPI_ISL_849197, EPI_ISL_849198 | | | |
| see above | Servicio Virosis Respiratorias-Departamento Virología-INEI | Instituto Nacional Enfermedades Infecciosas C.G.Malbran | Baumeister E., Avaro M., Benedetti E., Russo M., Dattero ME, Pontoriero A., Cisterna D., Molina V., Perandones C., Tuduri E., Lorenzo F., Poklepovich T., Campos J. |
| EPI_ISL_850187, EPI_ISL_850188, EPI_ISL_850189, EPI_ISL_850190 | Division of Emerging Infectious Diseases, Bureau of Infectious Diseases Diagnosis Control, Korea Disease Control and Prevention Agency | Division of Emerging Infectious Diseases, Bureau of Infectious Diseases Diagnosis Control, Korea Disease Control and Prevention Agency | Ae Kyung Park, Il-Hwan Kim, Heui Man Kim, Jeong-Min Kim, Namjoo Lee, Chaeyoung Lee, Sang Hee Woo, Eun-Jin Kim |
| EPI_ISL_853286 | UPMC Clinical Microbiology Laboratory | Microbial Genome Sequencing Center; Microbial Genomic Epidemiology Laboratory | Mustapha M. Mustapha, Jane W. Marsh, Dan Snyder, Marissa P. Griffith, Stephanie L. Mitchell, Vatsala R. Srinivasa, Kady D. Waggle, Chinelo Ezeonwuku, Vaughn S. Cooper, Lee H. Harrison |
| EPI_ISL_853405, EPI_ISL_853412, EPI_ISL_853413, EPI_ISL_853414 | Laboratoire de santé publique du Québec | Laboratoire de santé publique du Québec | Sandrine Moreira, Ioannis Ragoussis, Guillaume Bourque, Jesse Shapiro, Mark Lathrop and Michel Roger on behalf of the CoVSeQ research group (http://covseq.ca/researchgroup) |
| EPI_ISL_853885, EPI_ISL_853960 | Center for Virology, Medical University of Vienna | Bergthaler laboratory, CeMM Research Center for Molecular Medicine of the Austrian Academy of Sciences | Lukas Endler, Alexandra Popa, Benedikt Agerer, Jakob-Wendelin Genger, Alexander Lercher, Anna Schedl, Thomas Penz, Michael Schuster, Jan Laine, Martin Senekowitsch, Christoph Bock, Andreas Bergthaler |
| EPI_ISL_854239 | Department of Pathology, Landeskrankenhaus Graz II, Medical University Graz | Bergthaler laboratory, CeMM Research Center for Molecular Medicine of the Austrian Academy of Sciences | Lukas Endler, Alexandra Popa, Benedikt Agerer, Jakob-Wendelin Genger, Alexander Lercher, Anna Schedl, Thomas Penz, Michael Schuster, Jan Laine, Martin Senekowitsch, Christoph Bock, Andreas Bergthaler |
| EPI_ISL_854577, EPI_ISL_854578, EPI_ISL_854579, EPI_ISL_854580, EPI_ISL_854581, EPI_ISL_854582, EPI_ISL_854583, EPI_ISL_854584, EPI_ISL_854585, EPI_ISL_854586, EPI_ISL_854587, EPI_ISL_854588 | | | |
| see above | Alberta Precision Labs (APL) | Alberta Precision Labs (APL) | Gordon P, Lam LG, Pabbaraju K, Wong A, Ma R, Li V, Melin A, Tipples G, Berenger B, Zelyas N, Kellner J, Bernier F, Chui L, Croxen M |
| EPI_ISL_856974, EPI_ISL_856985, EPI_ISL_857029, EPI_ISL_857030, | The Ashley Laboratory, Stanford University | Chan-Zuckerberg Biohub | CZB Cliahub Consortium |

| | | | |
|---|---|---|---|
| EPI_ISL_857031, EPI_ISL_857032, EPI_ISL_857033, EPI_ISL_857034, EPI_ISL_857035, EPI_ISL_857036 | | | |
| EPI_ISL_857050 | Colorado Department of Public Health and Environment | Colorado Department of Puplic Health and Environment | Laura Bankers, Molly C. Hetherington-Rauth, Diana Ir, Shannon Ely, Shannon R. Matzinger, Sarah Elizabeth Totten, Emily A. Travanty |
| EPI_ISL_859984, EPI_ISL_859985, EPI_ISL_859986, EPI_ISL_859987, EPI_ISL_859988, EPI_ISL_859989, EPI_ISL_859990, EPI_ISL_859991, EPI_ISL_859992, EPI_ISL_859999 | BTC, Khalifa University | BTC, Khalifa University | Al Safar et al |
| EPI_ISL_861644 | Hospital Santa Virginia | Instituto Adolfo Lutz, Interdiciplinary Procedures Center, Strategic Laboratory | Claudio Tavares Sacchi, Claudia Regina Gonçalves, Erica Valessa Ramos Gomes, Karoline Rodrigues Campos |
| EPI_ISL_861645 | Hospital e Pronto Socorro Comunitario Vila Iolanda | Instituto Adolfo Lutz, Interdiciplinary Procedures Center, Strategic Laboratory | Claudio Tavares Sacchi, Claudia Regina Gonçalves, Erica Valessa Ramos Gomes, Karoline Rodrigues Campos |
| EPI_ISL_861650 | Hospital Santa Marcelina Sao Paulo | Instituto Adolfo Lutz, Interdiciplinary Procedures Center, Strategic Laboratory | Claudio Tavares Sacchi, Claudia Regina Gonçalves, Erica Valessa Ramos Gomes, Karoline Rodrigues Campos |
| EPI_ISL_861651 | AMA Jardim Brasil | Instituto Adolfo Lutz, Interdiciplinary Procedures Center, Strategic Laboratory | Claudio Tavares Sacchi, Claudia Regina Gonçalves, Erica Valessa Ramos Gomes, Karoline Rodrigues Campos |
| EPI_ISL_861652 | AMA Wamberto Dias da Costa | Instituto Adolfo Lutz, Interdiciplinary Procedures Center, Strategic Laboratory | Claudio Tavares Sacchi, Claudia Regina Gonçalves, Erica Valessa Ramos Gomes, Karoline Rodrigues Campos |
| EPI_ISL_861653 | Hospital Santa Virgínia | Instituto Adolfo Lutz, Interdiciplinary Procedures Center, Strategic Laboratory | Claudio Tavares Sacchi, Claudia Regina Gonçalves, Erica Valessa Ramos Gomes, Karoline Rodrigues Campos |
| EPI_ISL_861654, EPI_ISL_861655 | Hospital Santa Marcelina Sao Paulo | Instituto Adolfo Lutz, Interdiciplinary Procedures Center, Strategic Laboratory | Claudio Tavares Sacchi, Claudia Regina Gonçalves, Erica Valessa Ramos Gomes, Karoline Rodrigues Campos |
| EPI_ISL_861656 | UPA de Jandira | Instituto Adolfo Lutz, Interdiciplinary Procedures Center, Strategic Laboratory | Claudio Tavares Sacchi, Claudia Regina Gonçalves, Erica Valessa Ramos Gomes, Karoline Rodrigues Campos |
| EPI_ISL_861659, EPI_ISL_861660, EPI_ISL_861661 | PS e Maternidade Nair Fonseca Leitao Arantes | Instituto Adolfo Lutz, Interdiciplinary Procedures Center, Strategic Laboratory | Claudio Tavares Sacchi, Claudia Regina Gonçalves, Erica Valessa Ramos Gomes, Karoline Rodrigues Campos |
| EPI_ISL_862879 | Ehime Prefectural Central Hospital | Fujita health University, School of Medicine, Department of Microbiology | Masahiro suzuki, Yohei Doi |
| EPI_ISL_862884 | University Hospital Kyoto Prefectural University of Medicine | Fujita health University, School of Medicine, Department of Microbiology | Masahiro suzuki, Yohei Doi |
| EPI_ISL_862885 | Japanese Red Cross Narita Hospital | Fujita health University, School of Medicine, Department of Microbiology | Masahiro suzuki, Yohei Doi |
| EPI_ISL_875543, EPI_ISL_875548, EPI_ISL_875549 | Instituto de Biotecnologia - UNESP-Botucatu-SP | Instituto de Biotecnologia - UNESP-Botucatu-SP | Leila Sabrina Ullmann; Fábio Sossai Possebon, Camila Dantas Malossi, Paula Rahal, Paulo Inacio da Costa, João Pessoa Araújo Jr. |
| EPI_ISL_883176, EPI_ISL_883177, EPI_ISL_883178 | UK Tübingen, Medical Microbiology | Hannover Medical School, Institute of Virology | Lars Steinbrück, Jasper Götting |
| EPI_ISL_884483 | Molecular Microbiology & Immunology, University of Missouri | Molecular Microbiology & Immunology, University of Missouri | Tang,C.Y., Li,T., Hang,J., Lidl,G.M., Wan,X.-F. |
| EPI_ISL_887201, EPI_ISL_887205, EPI_ISL_887286, EPI_ISL_887317, EPI_ISL_887328, EPI_ISL_887345 | Protzer Lab | Protzer Lab, Gagneur Lab, Robert Koch Institut | Ulrike Protzer, Dieter Hoffmann, Eva Schulte, Andrea Theumer, Oliver Drechsel, Max von Kleist,Aleksandar Radonic,Stephan Fuchs, Alexander Karollus, Julien Gagneur |
| EPI_ISL_890034, EPI_ISL_890035, EPI_ISL_890036, EPI_ISL_890037, EPI_ISL_890038, EPI_ISL_890039, EPI_ISL_890040, EPI_ISL_890041, EPI_ISL_890042, EPI_ISL_890043, EPI_ISL_890044, EPI_ISL_890045, EPI_ISL_890046, EPI_ISL_890047, EPI_ISL_890049 | | | |
| see above | Laboratoire de santé publique du Québec | Laboratoire de santé publique du Québec | Sandrine Moreira, Ioannis Ragoussis, Guillaume Bourque, Jesse Shapiro, Mark Lathrop and Michel Roger on behalf of the CoVSeQ research group |
| EPI_ISL_891160, EPI_ISL_891161, EPI_ISL_891162, EPI_ISL_891163, EPI_ISL_891164, EPI_ISL_891165, EPI_ISL_891166, EPI_ISL_891167, EPI_ISL_891168, EPI_ISL_891169, EPI_ISL_891170, EPI_ISL_891171, EPI_ISL_891172, EPI_ISL_891174, EPI_ISL_891176, EPI_ISL_891193 | | | |
| see above | DPH, Massachusetts State Public Health Lab | DPH, Massachusetts State Public Health Lab | Lang,A.S., Fink,T., Gallagher,G.R., Smole,S.C. |
| EPI_ISL_895729, EPI_ISL_895813 | Molecular biology division, Institute of Clinical Biochemistry and Diagnostics, Charles University, Faculty of Medicine in Hradec Králové and University Hospital Hradec Králové | Molecular biology division, Institute of Clinical Biochemistry and Diagnostics, Charles University, Faculty of Medicine in Hradec Králové and University Hospital Hradec Králové | Helena Kovaíková, Petr Brož, Ivana Baranová, Kateina Hrochová, Tereza Baková, Jitka Novotná, Kateina Pehlíková, Vladimír Palika. Cooperation project with BioVendor-R&D and bioinformatics company BIOXSYS s r.o. |
| EPI_ISL_900726 | Bozeman Health Deaconess Hospital | Wiedenheft lab, Montana State University | Artem Nemudryi, Anna Nemudraia, Tanner Wiegand, Joseph Nichols, Deann T. Snyder, Jodi F. Hedges, Calvin Cicha, Helen Lee, Karl K. Vanderwood, Diane Bimczok, Mark A. Jutila and Blake Wiedenheft |
| EPI_ISL_903572 | VT Dept. of Health Laboratory | Genomics and Discovery, Respiratory Viruses Branch, Division of Viral Diseases, Centers for Disease Control and Prevention | Krista Queen, Yan Li, Ying Tao, Jing Zhang, Anna Uehara, Anna Montmayeur, Clinton R. Paden, Peter W. Cook, Rachel Marine, Mili Sheth, Jasmine Padilla, Sarah Nobles, Mark Burroughs, Lori Rowe, Haibin Wang, Ben L. Rambo-Martin, Dhwani Batra, Justin Lee, Suxiang Tong |
| EPI_ISL_906067 | PS e Maternidade Nair Fonseca Leitao Arantes | Instituto Adolfo Lutz, Interdiciplinary Procedures Center, Strategic Laboratory | Claudio Tavares Sacchi, Claudia Regina Gonçalves, Erica Valessa Ramos Gomes, Karoline Rodrigues Campos |
| EPI_ISL_913325 | Klinisk mikrobiologi | The Public Health Agency of Sweden | Anna-Malin Linde, Maria Lind Karlberg, Carlo Berg, Oskar Karlsson Lindsjo, Sofia Stamouli, Reza Advani, Mattias Haukland, Petra Holmstrom, Noura Walai, Petra Edquist, Mia Brytting, Anna Risberg, Karin Tegmark-Wisell |
| EPI_ISL_914096, EPI_ISL_914097, EPI_ISL_914098, EPI_ISL_914099, EPI_ISL_914100, EPI_ISL_914101, EPI_ISL_914102, EPI_ISL_914103, EPI_ISL_914104, EPI_ISL_914105, EPI_ISL_914106, EPI_ISL_914107, EPI_ISL_914108, EPI_ISL_914109, EPI_ISL_914110, EPI_ISL_914111, EPI_ISL_914112, EPI_ISL_914113, EPI_ISL_914114, EPI_ISL_914115, EPI_ISL_914116, EPI_ISL_914117, EPI_ISL_914118, EPI_ISL_914119, EPI_ISL_914120, EPI_ISL_914121, EPI_ISL_914122, EPI_ISL_914123, EPI_ISL_914124, EPI_ISL_914125, EPI_ISL_914126, EPI_ISL_914127, EPI_ISL_914128, EPI_ISL_914129, EPI_ISL_914130, EPI_ISL_914131, EPI_ISL_914132, EPI_ISL_914133, EPI_ISL_914134, EPI_ISL_914135, EPI_ISL_914137, EPI_ISL_914138, EPI_ISL_914141 | | | |
| see above | TGen North | TGen North | "Jolene Bowers, Megan Folkerts, Chris French, Hayley Yaglom, Ashlyn Pfeiffer, Darrin Lemmer, Dave Engelthaler, The Arizona COVID Genomics Union (ACGU)" |
| EPI_ISL_915174, EPI_ISL_915175, EPI_ISL_915176, EPI_ISL_915177, EPI_ISL_915178, EPI_ISL_915179, EPI_ISL_915180, EPI_ISL_915181, EPI_ISL_915182, EPI_ISL_915183, EPI_ISL_915184, EPI_ISL_915185, EPI_ISL_915186, EPI_ISL_915187, EPI_ISL_915188, EPI_ISL_915189, EPI_ISL_915190, EPI_ISL_915191 | | | |
| see above | QEII Health Sciences Centre | National Microbiology Laboratory (NML) | Anna Majer, Shari Tyson, Grace Seo, Philip Mabon, Elsie Grudeski, Rhiannon Huzarewich, Russell Mandes, Anneliese Landgraff, Jennifer Tanner, Natalie Knox, Morag Graham, Gary Van Domselaar, Todd Hatchette, Jason LeBlanc, Janice Pettipas, Dan Gaston, Nathalie Bastien, Yan Li, Timothy Booth, Darian Hole, Madison Chapel, CanCOGeN's metadata curation team, Public Health Agency of Canada CanCOGeN team |
| EPI_ISL_933637, EPI_ISL_933638 | Toronto Invasive Bacterial Diseases Network | McMaster University | Allison McGeer, Patryk Aftanas, Hooman Derakhshani, Angel Li, Kuganya Nirmalarajah, Emily Panousis, Ahmed Draia, Jalees Nasir, Michael Surette, Samira Mubareka, Andrew G. McArthur |
| EPI_ISL_934341 | Klinisk mikrobiologi | The Public Health Agency of Sweden | Anna-Malin Linde, Maria Lind Karlberg, Carlo Berg, Oskar Karlsson Lindsjo, Sofia Stamouli, Reza Advani, Mattias Haukland, Petra Holmstrom, Noura Walai, Petra Edquist, Mia Brytting, Anna Risberg, Karin Tegmark-Wisell |
| EPI_ISL_935522 | University of Massachusetts Medical School | Infectious Disease Program, Broad Institute of Harvard and | Tomkins-Tinch, Christopher H.; Daly, Jennifer S.; Gladden-Young, Adrianne; Theodoropoulos, Nicole M.; Madaio, Michael; Yu, Neng; Vanguri, Vijay K; |

| | | | |
|---|---|---|---|
| | | MIT | Siddle, Katherine J.; Adams, Gordon; Krasilnikova, Lydia A.; Movahedi, Babak; Bozorgzadeh, Adel; Simin, Karl; Lemieux, Jacob E.; Luban, Jeremy; Park, Daniel J.; MacInnis, Bronwyn L.; Sabeti, Pardis C.; Levitz, Stuart M. |
| EPI_ISL_936487 | Infectious Disease Biology, Institute of Life Sciences | Infectious Disease Biology, Institute of Life Sciences | Syed,G.H., Singh,B., Avula,K., Ghosh,A., Jha,A., Laha,E.,Madhulika,S., Priyadarhini,M., Puwan,S.S., Singh,N., Singh,D.,Datey,A., Biswas,V.K., Das,R., Senapati,S., Beuria,T.K., Swain,R.,Prasad,P., Chattopadhyay,S., Raghav,S., Team,I.C. and Parida,A. |
| EPI_ISL_936498, EPI_ISL_936499, EPI_ISL_936500 | Northwestern Memorial Hospital | Ozer Lab | Ramon Lorenzo-Redondo, Lacy M. Simons, Chad J. Achenbach, Lawrence J. Jennings, Michael G. Ison, Judd F. Hultquist, Egon A. Ozer |
| EPI_ISL_940273, EPI_ISL_940350 | Hôpital Bichat Claude Bernard, Laboratoire de Virologie | IAME UMR1137 Inserm, Université de Paris, Hôpital Bichat | Antoine Bridier-Nahmias, Amélie Recoing, Quentin Le Hingrat, Lena Daniel, Siham Hamri, Gilles Collin, Alexandre Storto, Mélanie Bertine, Charlotte Charpentier, Nadhira Houhou-Fidouh, Diane Descamps, Benoit Visseaux |
| EPI_ISL_941161 | Centro de Investigaciones en Microbiología y Biotecnología-UR (CIMBIUR), Facultad de Ciencias Naturales, Universidad del Rosario, Bogotá, Colombia | Centro de Investigaciones en Microbiología y Biotecnología-UR (CIMBIUR), Facultad de Ciencias Naturales, Universidad del Rosario, Bogotá, Colombia Icahn School of Medicine at Mount Sinai, New York, USA | Nathalia Ballesteros, Marina Muñoz, Luz Helena Patiño, Carolina Hernández, Felipe González-Casabianca, Iván Carroll, Mauricio Santos-Vega, Jaime Cascante, Andrés Angel, Alejandro Feged-Rivadeneira, Mónica Palma-Cuero, Carolina Flórez, Sergio Gomez, Adriana van de Guchte, Zenab Khan, Jayeeta Dutta, Hala Alejel Alshammary, Ana S. Gonzalez-Reiche, Matthew M. Hernandez, Emilia Mia Sordillo, Viviana Simon, Harm van Bakel, Alberto Paniz-Mondolfi, Juan David Ramírez |
| EPI_ISL_941984, EPI_ISL_941985 | Instituto Nacional de Salud, Bogotá, Colombia | Centro de Investigaciones en Microbiología y Biotecnología-UR (CIMBIUR), Facultad de Ciencias Naturales, Universidad del Rosario, Bogotá, Colombia Instituto Nacional de Salud, Bogotá, Colombia Icahn School of Medicine at Mount Sinai, New York, USA | Luz Helena Patiño, Marina Muñoz, Nathalia Ballesteros, Carolina Hernández, Carolina Flórez, Sergio Gomez, Adriana van de Guchte, Zenab Khan, Jayeeta Dutta, Hala Alejel Alshammary, Ana S. Gonzalez-Reiche, Matthew M. Hernandez, Emilia Mia Sordillo, Viviana Simon, Harm van Bakel, Alberto Paniz-Mondolfi, Juan David Ramírez |
| EPI_ISL_953403, EPI_ISL_953423, EPI_ISL_953424 | Laboratorio de Investigaciones de Baney | "Swiss Tropical and Public Health Institute" | "Carlos Cortes, Claudia Daubenberger, Guillermo Garcia, Salome Hosch, Bonifacio Manguire Nlavo, Maximilian Mpina, Elizabeth Nyakarungu, Diosdado Odjama Nseng Ada, Mitoha Ondo O Ayekaba, Tobias Schindler, Philip Wonder Phiri" |
| EPI_ISL_954892 | Colorado Department of Public Health and Environment | Colorado Department of Puplic Health and Environment | Laura Bankers, Molly C. Hetherington-Rauth, Diana Ir, Shannon Ely, Shannon R. Matzinger, Sarah Elizabeth Totten, Emily A. Travanty |
| EPI_ISL_956270 | Universitas Airlangga Hospital | Institute of Tropical Disease, Universitas Airlangga | Aldise M Nastri, Jezzy R Dewantari, Rima R Prasetya, Krisnoadi Rahardjo, Nasronudin, Gatot Soegiarto, Laksmi Wulandari, Resti Yudhawati, Soetjipto, Yasuko Mori, Maria I Lusida, Kazufumi Shimizu |
| EPI_ISL_956308 | RSUD Dr. M. Soewandhie | Institute of Tropical Disease, Universitas Airlangga | Jezzy R Dewantari, Rima R Prasetya, Krisnoadi Rahardjo, Aldise M Nastri, Febria Rachmanita, Gatot Soegiarto, Laksmi Wulandari, Resti Yudhawati, Yasuko Mori, Soetjipto, Kazufumi Shimizu, Maria I Lusida |
| EPI_ISL_960304 | NLZOH, Laboratory for Virology | NLZOH, Laboratory for Virology | Katarina Prosenc (Laboratory for Virology), Cesare Camma (IZSAM), Erik Alm (ECDC) |
| EPI_ISL_961015, EPI_ISL_961016 | Viral Respiratory Lab, National Institute for Biomedical Research (INRB) | Pathogen Sequencing Lab, National Institute for Biomedical Research (INRB) | Placide Mbala-Kingebeni, Edith Nkwembe, Eddy Kinganda-Lusamaki, Amuri Aziza, Francisca Muyembe Mawete, Emmanuel Lokilo Lofiko, Jean-Claude Makangara Cigolo, Catherine Pratt, Matthias Pauthner, Josh Quick, Allison Black, James Hadfield, Trevor Bedford, Ian Goodfellow, Andrew Rambaut, Nick Loman, Kristian Andersen, Michael Wiley, Steve Ahuka-Mundeke, Jean-Jacques Muyembe Tamfum |
| EPI_ISL_961796 | Laboratorio de Infectología, Servicio de Infectología, Hospital Universitario Dr. José Eleuterio González - Universidad Autónoma de Nuevo León | Laboratorio de Infectología Molecular, Departamento de Bioquímica y Medicina Molecular,Facultad de Medicina - Universidad Autónoma de Nuevo León | Kame A. Galán-Huerta, María F. Herrera-Saldivar, Natalia Martínez-Acuña, Sonia A. Lozano-Sepúlveda, Daniel Arellanos-Soto, Ana M. Rivas-Estilla, Paola Bocanegra-Ibarias, Samantha M. Flores-Treviño, Elvira Garza-González, Eduardo Perez-Alba, Laura Nuzzolo-Shihadeh, Adrian Camacho-Ortiz |
| EPI_ISL_962378 | Washington State Department of Health | Seattle Flu Study | Deborah A. Nickerson, Chris D. Frazar, Jover Lee, Benjamin Pelle, Erica Ryke, Matthew Richardson, Amanda Adler, Elisabeth Brandstetter, Peter D. Han, Kairsten Fay, Misja Ilcisin, Kirsten Lacombe, Thomas R. Sibley, Melissa Truong, Caitlin R. Wolf, Romesh Gautom, Geoff Melly, Brian Hiatt, Philip Dykema, Scott Lindquist, Michael Boeckh, Janet A. Englund, Michael Famulare, Barry R. Lutz, Mark J. Rieder, Lea M. Starita, Matthew Thompson, Helen Y. Chu, Jay Shendure, Trevor Bedford |
| EPI_ISL_968180, EPI_ISL_968181, EPI_ISL_968182 | Clinical Molecular Microbiology Laboratory, UNC Hospital | Dirk Dittmer | Justin T. Landis , Razia Moorad , Brent A. Eason, Melissa B. Miller, Linda Pluta, Dirk Dittmer, Angelica Juarez, Cecilia Thompson, Shawn Hawken, Cameroon Grant, Evelyn Hoffman, Patricio Cano, Jason Wong, Carolina Caro-Vegas, Ryan McNamara, Blossom Damania. |
| EPI_ISL_977469 | University of Zambia, School of Veterinary Medicine | UNZAVET and PATH | Mulenga Mwenda-Chimfwembe, Ngonda Saasa, Daniel Bridges |
| EPI_ISL_978534 | Centre de Virologie des Maladies infectueuses Tropicales | Functional Genomic Platform UATRS-biology, CNRST | Tahar BAJJOU, Farida HILALI, Marouane MELLOUL, Mly Abdelaziz ELALAOUI, Sanaa ALAOUI-Amine, Hemlali Mouhssine, Abderrazzak Rfaki, Elmostafa BENAISSA, , Abdelilah LARAQUI, Nadia Touil, Yassine SEKHSOKH, Mostafa ELOUENNASS, Khalid ENNIBI, Elmostafa EL FAHIME |
| EPI_ISL_978535 | Centre de Virologie des Maladies infectueuses Tropicales | Functional Genomic Platform UATRS-biology, CNRST | Tahar BAJJOU, Marouane MELLOUL, Farida HILALI, , Mly Abdelaziz ELALAOUI, Sanaa ALAOUI-Amine, Hemlali Mouhssine, Abderrazzak Rfaki, Elmostafa BENAISSA, , Abdelilah LARAQUI, Nadia Touil, Yassine SEKHSOKH, Mostafa ELOUENNASS, Khalid ENNIBI, Elmostafa EL FAHIME |
| EPI_ISL_978536 | Centre de Virologie des Maladies infectueuses Tropicales | Functional Genomic Platform UATRS-biology, CNRST | Farida HILALI, Mly Abdelaziz ELALAOUI,Tahar BAJJOU, Marouane MELLOUL, Sanaa ALAOUI-Amine, Hemlali Mouhssine, Abderrazzak Rfaki, Elmostafa BENAISSA, , Abdelilah LARAQUI, Nadia Touil, Mostafa ELOUENNASS, Khalid ENNIBI, Elmostafa EL FAHIME, Yassine SEKHSOK |
| EPI_ISL_978537 | Centre de Virologie des Maladies infectueuses Tropicales | Functional Genomic Platform UATRS-biology, CNRST | Farida HILALI, Mly Abdelaziz ELALAOUI,Tahar BAJJOU, Marouane MELLOUL, Sanaa ALAOUI-Amine, Hemlali Mouhssine, Abderrazzak Rfaki, Elmostafa BENAISSA, , Abdelilah LARAQUI, Nadia Touil, Mostafa ELOUENNASS, Khalid ENNIBI, Elmostafa EL FAHIME, Yassine SEKHSOKH |
| EPI_ISL_978538 | Centre de Virologie des Maladies infectueuses Tropicales | Functional Genomic Platform UATRS-biology, CNRST | Mly Abdelaziz ELALAOUI, Tahar BAJJOU, Marouane MELLOUL, Farida HILALI, Sanaa ALAOUI-Amine, Hemlali Mouhssine, Abderrazzak Rfaki, Elmostafa BENAISSA, , Abdelilah LARAQUI, Nadia Touil, Yassine SEKHSOKH, Mostafa ELOUENNASS, Khalid ENNIBI, Elmostafa EL FAHIME |
| EPI_ISL_978539 | Centre de Virologie des Maladies infectueuses Tropicales | Functional Genomic Platform UATRS-biology, CNRST | Mly Abdelaziz ELALAOUI, Tahar BAJJOU, Marouane MELLOUL, Farida HILALI, Sanaa ALAOUI-Amine, Hemlali Mouhssine, Abderrazzak Rfaki, Elmostafa BENAISSA, , Abdelilah LARAQUI, Nadia Touil, Yassine SEKHSOKH, , Elmostafa EL FAHIME, Mostafa ELOUENNASS, Khalid ENNIBI, |
| EPI_ISL_978540 | Centre de Virologie des Maladies infectueuses Tropicales | Functional Genomic Platform UATRS-biology, CNRST | Abdelilah LARAQUI, Nadia Touil, Mly Abdelaziz ELALAOUI, Sanaa ALAOUI-Amine, Tahar BAJJOU, Marouane MELLOUL, Farida HILALI, Hemlali Mouhssine, Abderrazzak Rfaki, Elmostafa BENAISSA, Yassine SEKHSOKH, , Elmostafa EL FAHIME, Mostafa ELOUENNASS, Khalid ENNIBI. |
| EPI_ISL_978541 | Centre de Virologie des Maladies infectueuses Tropicales | Functional Genomic Platform UATRS-biology, CNRST | Marouane MELLOUL, Abdelilah LARAQUI, Nadia Touil, Mly Abdelaziz ELALAOUI, Sanaa ALAOUI-Amine, Tahar BAJJOU, Farida HILALI, Hemlali Mouhssine, Abderrazzak Rfaki, Elmostafa BENAISSA, Yassine SEKHSOKH, Elmostafa EL FAHIME, Mostafa ELOUENNASS, Khalid ENNIBI. |
| EPI_ISL_978542 | Centre de Virologie des Maladies infectueuses Tropicales | Functional Genomic Platform UATRS-biology, CNRST | Marouane MELLOUL, Abdelilah LARAQUI, Nadia Touil, Mly Abdelaziz ELALAOUI, Sanaa ALAOUI-Amine, Tahar BAJJOU, Farida HILALI, Hemlali Mouhssine, Abderrazzak Rfaki, Elmostafa BENAISSA, Elmostafa EL FAHIME, Mostafa ELOUENNASS, Khalid ENNIBI, Yassine SEKHSOKH |
| EPI_ISL_978543 | Centre de Virologie des Maladies infectueuses Tropicales | Functional Genomic Platform UATRS-biology, CNRST | Yassine SEKHSOKH, Mly Abdelaziz ELALAOUI, Marouane MELLOUL, Abdelilah LARAQUI, Nadia Touil, , Sanaa ALAOUI-Amine, Tahar BAJJOU, Farida HILALI, Hemlali Mouhssine, Abderrazzak Rfaki, Elmostafa BENAISSA, Elmostafa EL FAHIME, Mostafa ELOUENNASS, Khalid ENNIBI |
| EPI_ISL_978544 | Centre de Virologie des Maladies infectueuses Tropicales | Functional Genomic Platform UATRS-biology, CNRST | Elmostafa EL FAHIME, Elmostafa BENAISSA, Marouane MELLOUL, Abdelilah LARAQUI, Nadia Touil, Mly Abdelaziz ELALAOUI, Sanaa ALAOUI-Amine, Yassine SEKHSOKH, Farida HILALI, Hemlali Mouhssine, Abderrazzak Rfaki, , Mostafa ELOUENNASS, Khalid ENNIBI, Tahar BAJJOU |
| EPI_ISL_978545 | Centre de Virologie des Maladies infectueuses Tropicales | Functional Genomic Platform UATRS-biology, CNRST | Mly Abdelaziz ELALAOUI, Marouane MELLOUL, Sanaa ALAOUI-Amine, Elmostafa EL FAHIME, Elmostafa BENAISSA, Abdelilah LARAQUI, Nadia Touil, Yassine SEKHSOKH, Farida HILALI, Hemlali Mouhssine, Abderrazzak Rfaki, , Mostafa ELOUENNASS, Khalid ENNIBI, Tahar BAJJOU |
| EPI_ISL_978546 | Centre de Virologie des Maladies infectueuses Tropicales | Functional Genomic Platform UATRS-biology, CNRST | Farida HILALI, Mly Abdelaziz ELALAOUI,Tahar BAJJOU, Marouane MELLOUL, Sanaa ALAOUI-Amine, Hemlali Mouhssine, Abderrazzak Rfaki, Elmostafa BENAISSA, , Abdelilah LARAQUI, Nadia Touil, Mostafa ELOUENNASS, Khalid ENNIBI, Elmostafa EL FAHIME, Yassine SEKHSOKH |
| EPI_ISL_978547 | Centre de Virologie des Maladies infectueuses Tropicales | Functional Genomic Platform UATRS-biology, CNRST | Nadia Touil, Farida HILALI, Mly Abdelaziz ELALAOUI,Tahar BAJJOU, Marouane MELLOUL, Sanaa ALAOUI-Amine, Hemlali Mouhssine, Abderrazzak Rfaki, Elmostafa BENAISSA, , Abdelilah LARAQUI, Mostafa ELOUENNASS, Khalid ENNIBI, Elmostafa EL FAHIME, Yassine SEKHSOKH |
| EPI_ISL_978548 | Centre de Virologie des Maladies infectueuses Tropicales | Functional Genomic Platform UATRS-biology, CNRST | Nadia Touil, Farida HILALI, Mly Abdelaziz ELALAOUI, Marouane MELLOUL, Sanaa ALAOUI-Amine, Hemlali Mouhssine, Abderrazzak Rfaki, Elmostafa BENAISSA, , Abdelilah LARAQUI, Mostafa ELOUENNASS, Khalid ENNIBI, Elmostafa EL FAHIME, Yassine SEKHSOKH, Tahar BAJJOU |
| EPI_ISL_978549 | Centre de Virologie des Maladies infectueuses Tropicales | Functional Genomic Platform UATRS-biology, CNRST | Elmostafa EL FAHIME, Hemlali Mouhssine, Elmostafa BENAISSA, Marouane MELLOUL, Abdelilah LARAQUI, Nadia Touil, Mly Abdelaziz ELALAOUI, Sanaa ALAOUI-Amine, Safaa GHOULAM, Tahar BAJJOU, Farida HILALI, Abderrazzak Rfaki,Yassine SEKHSOKH, Mostafa ELOUENNASS, Khalid ENNIBI. |
| EPI_ISL_978550 | Centre de Virologie des Maladies infectueuses Tropicales | Functional Genomic Platform UATRS-biology, CNRST | Hemlali Mouhssine, Taha Chouati, Marouane MELLOUL, Abdelilah LARAQUI, Nadia Touil, Mly Abdelaziz ELALAOUI, Sanaa ALAOUI-Amine, Safaa |

| | | | |
|---|---|---|---|
| | | | Ghoulam, Tahar BAJJOU, Farida HILALI, Abderrazzak Rfaki, Elmostafa BENAISSA, Elmostafa EL FAHIME, Yassine SEKHSOKH, Mostafa ELOUENNASS, Khalid ENNIBI |
| EPI_ISL_978551, EPI_ISL_978552 | Centre de Virologie des Maladies infectueuses Tropicales | Functional Genomic Platform UATRS-biology, CNRST | Sanaa ALAOUI-Amine, Hemlali Mouhssine, Elmostafa BENAISSA, Marouane MELLOUL, Abdelilah LARAQUI, Nadia Touil, Mly Abdelaziz ELALAOUI, Safaa GHOULAM, Tahar BAJJOU, Farida HILALI, Abderrazzak Rfaki,Yassine SEKHSOKH, Mostafa ELOUENNASS, Khalid ENNIBI, Elmostafa EL FAHIME. |
| EPI_ISL_979150, EPI_ISL_979151 | Santa Clara County Public Health Laboratory | Chan-Zuckerberg Biohub | CZB Cliahub Consortium |
| EPI_ISL_981038 | Hospital Bariloche | Laboratorio Central Mg. Luis Alfredo Pianciola on behalf of 'Proyecto Argentino Interinstitucional de genomica de SARS-CoV-2' (PAIS Consortium) | L Pianciola, M Mazzeo, C Ziehm, C Pintos, M Fernandez, J Ousset, M Nabaes, M Viegas. |
| EPI_ISL_981243, EPI_ISL_981244, EPI_ISL_981245, EPI_ISL_981246, EPI_ISL_981247, EPI_ISL_981248, EPI_ISL_981249, EPI_ISL_981250, EPI_ISL_981251, EPI_ISL_981252, EPI_ISL_981253, EPI_ISL_981254, EPI_ISL_981255, EPI_ISL_981256, EPI_ISL_981257, EPI_ISL_981258, EPI_ISL_981259, EPI_ISL_981260, EPI_ISL_981261, EPI_ISL_981262, EPI_ISL_981263, EPI_ISL_981264, EPI_ISL_981265, EPI_ISL_981266, EPI_ISL_981268 | | | |
| see above | National Virus Reference Laboratory | Irish Coronavirus Sequencing Consortium - Helixworks | Sachin Chalapati, Conor Crosbie, Nimesh Pinnamaneni |
