## Supplementary material for "Detection and characterization of the SARS-CoV-2 lineage B.1.526 in New York": Supp. Table 3 GISAID Acknowledgment part 2: gisaid_hcov-19_acknowledgement_table_2021_02_12_22-4.pdf

We gratefully acknowledge the following Authors from the Originating laboratories responsible for obtaining the specimens, as well as the Submitting laboratories where the genome data were generated and shared via GISAID, on which this research is based.

All Submitters of data may be contacted directly via www.gisaid.org

Authors are sorted alphabetically.

| Accession ID | Originating Laboratory | Submitting Laboratory | Authors |
|---|---|---|---|
| EPI_ISL_483486, EPI_ISL_483501, EPI_ISL_483502, EPI_ISL_483505, EPI_ISL_483506, EPI_ISL_483507, EPI_ISL_483508, EPI_ISL_483510, EPI_ISL_483511, EPI_ISL_483512, EPI_ISL_483513 | | | |
| see above | UC San Diego Center for Advanced Laboratory Medicine | Andersen lab at Scripps Research | SEARCH Alliance San Diego with David Pride, Ji H Shin |
| EPI_ISL_483531, EPI_ISL_483532, EPI_ISL_483536, EPI_ISL_483537, EPI_ISL_483538, EPI_ISL_483540 | San Diego County Public Health Laboratory | Andersen lab at Scripps Research | SEARCH Alliance San Diego with Tracy Basler, Jovan Shephard, Brett Austin |
| EPI_ISL_483590, EPI_ISL_483608, EPI_ISL_483611, EPI_ISL_483614, EPI_ISL_483615, EPI_ISL_483618, EPI_ISL_483620 | National Public Health Laboratory, National Centre for Infectious Diseases | National Public Health Laboratory, National Centre for Infectious Diseases | Mak TM, Octavia S, Zhou Z, Chavatte JM, Cui L, Lin RTP |
| EPI_ISL_483686, EPI_ISL_483687 | National Institute of Laboratory Medicine and Referral Center | Genomic Research Lab, BCSIR | Md. Murshed Hasan Sarkar, Abu Sayeed Mohammad Mahmud, Mohammad Samir Uzzaman, Eshrar Osman, Md. Ahasan Habib, Shahina Akter, Tanjina Akhter Banu, Barna Goswami, Iffat Jahan, Md. Saddam Hossain, Tasnim Nafisa, Md. Maruf Ah [illegible] Shamsuzzaman, Sheikh Md. Selim Al Din, Utpal Chandra Ray, Salek Ahmed Sajib, Md. Salim Khan |
| EPI_ISL_483688 | Genomic Research Lab, BCSIR | Genomic Research Lab, BCSIR | Md. Murshed Hasan Sarkar, Abu Sayeed Mohammad Mahmud, Mohammad Samir Uzzaman, Eshrar Osman, Md. Ahasan Habib, Shahina Akter, Tanjina Akhter Banu, Barna Goswami, Iffat Jahan, Md. Saddam Hossain, Tasnim Nafisa, Md. Maruf Ah [illegible] Shamsuzzaman, Sheikh Md. Selim Al Din, Utpal Chandra Ray, Salek Ahmed Sajib, Md. Salim Khan |
| EPI_ISL_483689, EPI_ISL_483690, EPI_ISL_483692 | National Institute of Laboratory Medicine and Referral Center | Genomic Research Lab, BCSIR | Md. Murshed Hasan Sarkar, Abu Sayeed Mohammad Mahmud, Mohammad Samir Uzzaman, Eshrar Osman, Md. Ahasan Habib, Shahina Akter, Tanjina Akhter Banu, Barna Goswami, Iffat Jahan, Md. Saddam Hossain, Tasnim Nafisa, Md. Maruf Ah [illegible] Shamsuzzaman, Sheikh Md. Selim Al Din, Utpal Chandra Ray, Salek Ahmed Sajib, Md. Salim Khan |
| EPI_ISL_483703, EPI_ISL_483705, EPI_ISL_483707, EPI_ISL_483710 | National Institute of Laboratory Medicine and Referral Center | Genomic Research Lab, BCSIR | Abu Sayeed Mohammad Mahmud, Mohammad Samir Uzzaman, Eshrar Osman, Md. Ahasan Habib, Shahina Akter, Tanjina Akhter Banu, Md. Murshed Hasan Sarkar, Barna Goswami, Iffat Jahan, Md. Saddam Hossain, Tasnim Nafisa, Md. Maruf Ah [illegible] Shamsuzzaman, Sheikh Md. Selim Al Din, Utpal Chandra Ray, Salek Ahmed Sajib, Md. Salim Khan |
| EPI_ISL_483914, EPI_ISL_483915 | Department of Pathology, University of Cambridge | COVID-19 Genomics UK (COG-UK) Consortium | Luke W Meredith, M. Estée Török, Myra Hosmillo, William L. Hamilton, Martin D. Curran, Theresa Feltwell, Grant Hall, Anna Yakovleva, Fahad A Khokhar, Charlotte J. Houldcroft, Laura G Caller, Aminu S. Jahun, Sarah L. Caddy, Yas [illegible] |
| EPI_ISL_484220 | University of Birmingham | COVID-19 Genomics UK (COG-UK) Consortium | Institute of Microbiology, University of Birmingham: Claire McMurray, Joanne Stockton, Samuel Nicholls, Radoslaw Poplawski, Will Rowe, Josh Quick, Nicholas Loman. University of Birmingham Testing Laboratory: Celina M Whalley, Andrew Bosworth, Ch [illegible] Richter, Andrew D Beggs PHE Heartlands Lab: Husam Osman, Andrew Bosworth. Queen Elizabeth Hospital: Anna Casey |
| EPI_ISL_484350, EPI_ISL_484351 | Queens Medical Centre, Clinical Microbiology Department / DeepSeq Nottingham | COVID-19 Genomics UK (COG-UK) Consortium | Gemma Clark, Wendy Smith, Manjinder Khakh, Vicki M Fleming, Michelle M Lister, Hannah Howson-Wells, Jonathan Ball, Patrick McClure, Joseph Chappell, Theocharis Tsoleridis, Nadine Holmes, Matthew Carlisle, Christopher Moore, Fei S [illegible] |
| EPI_ISL_484376 | University Hospitals Of Leicester NHS Trust and DeepSeq Nottingham | COVID-19 Genomics UK (COG-UK) Consortium | Christopher Holmes, Paul Bird, Thomas Helmer, Karlie Fallon, Julian Tang, Jonathan Ball, Patrick McClure, Joeseph Chappell, Nadine Holmes, Matthew Carlisle, Christopher Moore, Fei Sang, Johnny Debebe, Vict [illegible] |
| EPI_ISL_484390, EPI_ISL_484391 | Queens Medical Centre, Clinical Microbiology Department / DeepSeq Nottingham | COVID-19 Genomics UK (COG-UK) Consortium | Gemma Clark, Wendy Smith, Manjinder Khakh, Vicki M Fleming, Michelle M Lister, Hannah Howson-Wells, Jonathan Ball, Patrick McClure, Joseph Chappell, Theocharis Tsoleridis, Nadine Holmes, Matthew Carlisle, Christopher Moore, Fei S [illegible] |
| EPI_ISL_484455 | Virology Department, Sheffield Teaching Hospitals NHS Foundation Trust/Department of Infection, Immunity and Cardiovascular Disease, The Medical School, University of Sheffield | COVID-19 Genomics UK (COG-UK) Consortium | Thushan de Silva, Matthew Parker, Nikki Smith, Adri Angyal, Rebecca Brown, Luke Green, Rachel Tucker, Paul Parsons, Danielle Groves, Katie Johnson, Laura Carrilero, Alex Keeley, Dave Partridge, Matthew Wyles, Benjamin Lindsey, [illegible] |
| EPI_ISL_484685, EPI_ISL_484686, EPI_ISL_484687, EPI_ISL_484688, EPI_ISL_484689, EPI_ISL_484690, EPI_ISL_484691 | Wales Specialist Virology Centre Sequencing lab: Pathogen Genomics Unit | COVID-19 Genomics UK (COG-UK) Consortium | Catherine Moore, Johnathan Evans, Laura Gifford, Malorie Perry, Simon Cottrell, Angela Marchbank, Alec Birchley, Alexander Adams, Amy Gaskin, Bree Gatica-Wilcox, Jason Coombes, Joel Southgate, Lauren Gilbert, Lee Graham, Nicole Pacchiarini, Sa [illegible] Matthew Bull, Joanne Watkins, Sally Corden, Tom Connor |
| EPI_ISL_484892, EPI_ISL_484893, EPI_ISL_484894, EPI_ISL_484895, EPI_ISL_484896, EPI_ISL_484897, EPI_ISL_484898, EPI_ISL_484899, EPI_ISL_484900, EPI_ISL_484901, EPI_ISL_484902, EPI_ISL_484903, EPI_ISL_484904, EPI_ISL_484905, EPI_ISL_484906, EPI_ISL_484907, EPI_ISL_484908, EPI_ISL_484909, EPI_I [illegible] EPI_ISL_484916, EPI_ISL_484917, EPI_ISL_484920, EPI_ISL_484924, EPI_ISL_484926, EPI_ISL_484927, EPI_ISL_484929, EPI_ISL_484930, EPI_ISL_484931, EPI_ISL_484932, EPI_ISL_484933, EPI_ISL_484934, EPI_ISL_484935, EPI_ISL_484936, EPI_ISL_484937, EPI_ISL_484938, EPI_ISL_484939, EPI_ISL_484940, EPI_I [illegible] EPI_ISL_484945, EPI_ISL_484946, EPI_ISL_484949, EPI_ISL_484950, EPI_ISL_484952, EPI_ISL_484953, EPI_ISL_484954, EPI_ISL_484959, EPI_ISL_484962, EPI_ISL_484967, EPI_ISL_484968, EPI_ISL_484969, EPI_ISL_484970, EPI_ISL_484971, EPI_ISL_484972, EPI_ISL_484973, EPI_ISL_484974, EPI_ISL_484975, EPI_I [illegible] EPI_ISL_484980, EPI_ISL_484981, EPI_ISL_484982, EPI_ISL_484983, EPI_ISL_484984, EPI_ISL_484985, EPI_ISL_484989, EPI_ISL_484991, EPI_ISL_484992, EPI_ISL_484993, EPI_ISL_484994, EPI_ISL_484995, EPI_ISL_484996, EPI_ISL_484997, EPI_ISL_484998, EPI_ISL_484999, EPI_ISL_485000 | | | |
| see above | University of Wisconsin-Madison AIDS Vaccine Research Laboratories | University of Wisconsin-Madison AIDS Vaccine Research Laboratories | Gage Moreno, Katarina Braun, et al. AIDS Vaccine Research Laboratories |

| | | | |
|---|---|---|---|
| EPI_ISL_485401 | Communicable Disease Laboratory, Public Health Directorate | Communicable Disease Laboratory, Public Health Directorate | Zaed,A., Al-Wasti,H., Al-Taif,Z. and Shehab,F. |

EPI_ISL_486515, EPI_ISL_486516, EPI_ISL_486517, EPI_ISL_486518, EPI_ISL_486519, EPI_ISL_486520, EPI_ISL_486521, EPI_ISL_486522, EPI_ISL_486523, EPI_ISL_486524, EPI_ISL_486525, EPI_ISL_486526, EPI_ISL_486527, EPI_ISL_486528, EPI_ISL_486529, EPI_ISL_486530, EPI_ISL_486531, EPI_ISL_486532, EPI_I[illegible]
EPI_ISL_486537, EPI_ISL_486538

| | | | |
|---|---|---|---|
| see above | Viollier AG | Department of Biosystems Science and Engineering, ETH Zürich | Christian Beisel, Sarah Nadeau, Ivan Topolsky, Pedro Ferreira, Philipp Jablonski, Susana Posada-Céspedes, Tobias Schär, Ina Nissen, Natascha Santacroce, Elodie Burcklen, Christiane Beckmann, Maurice Redondo, Olivier Kobel, Christoph Noppen, Sop[illegible] Stadler |
| EPI_ISL_486845, EPI_ISL_486846, EPI_ISL_486847, EPI_ISL_486848, EPI_ISL_486849, EPI_ISL_486850, EPI_ISL_486851 | Institute of Microbiology, Universidad San Francisco de Quito | Institute of Microbiology, Universidad San Francisco de Quito | Belén Prado-Vivar, Sully Márquez, Juan José Guadalupe, Monica Becerra-Wong, Carla Torres, Bernardo Gutiérrez, Jonathan Araujo, Verónica Barragán, Patricio Rojas-Silva, Gabriel Trueba, Michelle Gruna[illegible] |
| EPI_ISL_486887 | National Influenza Center, Bahrain | National Influenza Center, Bahrain | Zaed,A., Altaif,Z., Shehab,F., AlWasti,H. |
| EPI_ISL_486888 | National Influenza Center, Bahrain | National Influenza Center, Bahrain | AlWasti,H., Altaif,Z., Zaed,A., Shehab,F. |
| EPI_ISL_486889 | National Influenza Center, Bahrain | National Influenza Center, Bahrain | Altaif,Z., AlWasti,H., Shehab,F., Zaed,A. |
| EPI_ISL_487106 | Nigeria Centre for Disease Control (NCDC) | African Centre of Excellence for Genomics of Infectious Diseases (ACEGID), Redeemer's University, Ede, Osun State, Nigeria | Oluniyi P.E., Ajogbasile F.V., Kayode A., Oguzie J., Olawoye I., Uwanibe J., Olumade T., Folarin O.A., Ihekweazu C., Happi C.T. |
| EPI_ISL_487273 | unknown | Communicable Disease Laboratory, Public Health Directorate | Zaed,A., Shehab,F., AlWasti,H., Altaif,Z. |
| EPI_ISL_487274 | unknown | Communicable Disease Laboratory, Public Health Directorate | AlWasti,H., AlTaif,Z., Zaed,A., Shehab,F. |

EPI_ISL_487277, EPI_ISL_487278, EPI_ISL_487279, EPI_ISL_487280, EPI_ISL_487281, EPI_ISL_487282, EPI_ISL_487283, EPI_ISL_487284, EPI_ISL_487285, EPI_ISL_487286, EPI_ISL_487287, EPI_ISL_487288, EPI_ISL_487289, EPI_ISL_487290, EPI_ISL_487291, EPI_ISL_487292, EPI_ISL_487293, EPI_ISL_487294, EPI_I[illegible]
EPI_ISL_487299, EPI_ISL_487300, EPI_ISL_487301, EPI_ISL_487302, EPI_ISL_487303, EPI_ISL_487313, EPI_ISL_487316, EPI_ISL_487318, EPI_ISL_487320, EPI_ISL_487321, EPI_ISL_487325, EPI_ISL_487328, EPI_ISL_487348

| | | | |
|---|---|---|---|
| see above | NHLS-IALCH | KRISP, KZN Research Innovation and Sequencing Platform | Giandhari J, Pillay S, Lessells R, Chimukangara B, Mdlalose K, York D, Khan S, Tegally H, Wilkinson E, de Oliveira T |

EPI_ISL_489896, EPI_ISL_489900, EPI_ISL_489901, EPI_ISL_489921, EPI_ISL_489922, EPI_ISL_489923, EPI_ISL_489924, EPI_ISL_489925, EPI_ISL_489926, EPI_ISL_489927, EPI_ISL_489928, EPI_ISL_489929, EPI_ISL_489930, EPI_ISL_489931, EPI_ISL_489932, EPI_ISL_489933, EPI_ISL_489935

| | | | |
|---|---|---|---|
| see above | Gundersen Molecular Diagnostics Laboratory | Kabara Cancer Research Institute | Craig S. Richmond, Paraic A. Kenny |

EPI_ISL_489959, EPI_ISL_489960, EPI_ISL_489961, EPI_ISL_489962, EPI_ISL_489963, EPI_ISL_489964, EPI_ISL_489965, EPI_ISL_489966, EPI_ISL_489967, EPI_ISL_489968, EPI_ISL_489969, EPI_ISL_489970, EPI_ISL_489971, EPI_ISL_489972, EPI_ISL_489983, EPI_ISL_489984, EPI_ISL_489985, EPI_ISL_489986

| | | | |
|---|---|---|---|
| see above | Viollier AG | Department of Biosystems Science and Engineering, ETH Zürich | Christian Beisel, Sarah Nadeau, Ivan Topolsky, Pedro Ferreira, Philipp Jablonski, Susana Posada-Céspedes, Tobias Schär, Ina Nissen, Natascha Santacroce, Elodie Burcklen, Christiane Beckmann, Maurice Redondo, Olivier Kobel, Christoph Noppen, Sop[illegible] Stadler |
| EPI_ISL_490036 | Pathology West - NSW Health Pathology | NSW Health Pathology - Institute of Clinical Pathology and Medical Research; Westmead Hospital; University of Sydney | CIDM-PH et al. |
| EPI_ISL_490049, EPI_ISL_490050, EPI_ISL_490053, EPI_ISL_490054, EPI_ISL_490079 | National Public Health Laboratory, National Centre for Infectious Diseases | National Public Health Laboratory, National Centre for Infectious Diseases | Mak TM, Octavia S, Zhou Z, Chavatte JM, Cui L, Lin RTP |
| EPI_ISL_490317, EPI_ISL_490319, EPI_ISL_490320, EPI_ISL_490321, EPI_ISL_490322, EPI_ISL_490323, EPI_ISL_490324, EPI_ISL_490325, EPI_ISL_490326 | Department of Pathology, University of Cambridge | COVID-19 Genomics UK (COG-UK) Consortium | Luke W Meredith, M. Estée Török, Myra Hosmillo, William L. Hamilton, Martin D. Curran, Theresa Feltwell, Grant Hall, Anna Yakovleva, Fahad A Khokhar, Charlotte J. Houldcroft, Laura G Caller, Aminu S. Jahun, Sarah L. Caddy, Yas[illegible] |
| EPI_ISL_490330 | West of Scotland Specialist Virology Centre, NHSGGC / MRC-University of Glasgow Centre for Virus Research | COVID-19 Genomics UK (COG-UK) Consortium | Ana da Silva Filipe, Natasha Johnson, Kathy Smollett, Daniel Mair, Stephen Carmichael, Lily Tong, Jenna Nichols, Elihu Aranday-Cortes, Kirstyn Brunker, Yasmin Parr, Alice Broos, Kyriaki Nomikou; Sarah McDonald, Marc Niebel, Patawee Asamaphan; R[illegible] Alasdair MacLean, Rory Gunson; Kathy Li, Natasha Jesudason, Rajiv Shah, James Shepherd, Antonia Ho, Emma Thomson |

EPI_ISL_490395, EPI_ISL_490396, EPI_ISL_490397, EPI_ISL_490398, EPI_ISL_490399, EPI_ISL_490400, EPI_ISL_490401, EPI_ISL_490402, EPI_ISL_490403, EPI_ISL_490404, EPI_ISL_490405, EPI_ISL_490406

| | | | |
|---|---|---|---|
| see above | Liverpool Clinical Laboratories | COVID-19 Genomics UK (COG-UK) Consortium | Sam Haldenby, Anita Lucaci, Steve Paterson, Julian Hiscox, Alistair Darby, M Almsaud, A Alrezaihi, Muhannad Alruwaili, Stuart D Armstrong, Jones Benjamin, Eleanor G Bentley, Anu Chawla, Jordan J Clark, Angela Cowell, Richard Eccles, Isabel Garc[illegible] Richard Gregory, Ximeng Han, Catherine Hartley, Margaret Hughes, Miren Iturriza-Gomara, James Johnson, L Luu, Jenifer Manson, Charlotte Nelson, Elaine O'Toole, Cassie Olateju, Rebekah Penrice-Randal , Lucille Rainbow, N.P Randle, Trevor Ian [illegible] Swainston, Ecaterina Vamos, Joanne Watts, Mark Whitehead |
| EPI_ISL_490544, EPI_ISL_490548 | Quadram Institute Bioscience | COVID-19 Genomics UK (COG-UK) Consortium | Dave J. Baker, Gemma L. Kay, Alp Aydin, Thanh Le-Viet, Steven Rudder, Ana P. Tedim, Anastasia Kolyva, Maria Diaz, Leonardo de Oliveira Martins, Nabil-Fareed Alikhan, Lizzie Meadows, Rachael Stanley, Ngozi Elumogo, Muhammed Yasir, Nicholas M[illegible] Claire Stuart, Andrew Bell, Reenesh Prakash, Samir Dervisevic, Alison E. Mather, John Wain, Mark Webber, Andrew J. Page, Justin O'Grady |
| EPI_ISL_490698, EPI_ISL_490699, EPI_ISL_490700, | West of Scotland Specialist Virology Centre, NHSGGC / MRC-University of Glasgow | COVID-19 Genomics UK (COG-UK) Consortium | Ana da Silva Filipe, Natasha Johnson, Kathy Smollett, Daniel Mair, Stephen Carmichael, Lily Tong, Jenna Nichols, Elihu Aranday-Cortes, Kirstyn Brunker, Yasmin Parr, Alice Broos, Kyriaki Nomikou; Sarah McDonald, Marc Niebel, Patawee Asamaphan; R[illegible] Alasdair MacLean, Rory Gunson; Kathy Li, Natasha Jesudason, Rajiv Shah, James Shepherd, Antonia Ho, Emma Thomson |

| | | | |
|---|---|---|---|
| EPI_ISL_490701, EPI_ISL_490702, EPI_ISL_490703 | Centre for Virus Research | | |
| EPI_ISL_490763 | Wales Specialist Virology Centre Sequencing lab: Pathogen Genomics Unit | COVID-19 Genomics UK (COG-UK) Consortium | Catherine Moore, Johnathan Evans, Laura Gifford, Malorie Perry, Simon Cottrell, Angela Marchbank, Alec Birchley, Alexander Adams, Amy Gaskin, Bree Gatica-Wilcox, Jason Coombes, Joel Southgate, Lauren Gilbert, Lee Graham, Nicole Pacchiarini, Sa Matthew Bull, Joanne Watkins, Sally Corden, Tom Connor |
| EPI_ISL_491298 | Rural Health Unit - Calauan, Laguna | Research Institute for Tropical Medicine | Tujan, M.A.A., Onza,O.J.T., Polotan,F.G.M., Medado,I.A.P., Bautista,C.T., Brunker, K., Mercado,E.S., Manalo, D.L., Demetria, C.S. |
| EPI_ISL_491299, EPI_ISL_491300, EPI_ISL_491301, EPI_ISL_491302, EPI_ISL_491303, EPI_ISL_491304, EPI_ISL_491305, EPI_ISL_491306, EPI_ISL_491307, EPI_ISL_491308, EPI_ISL_491309, EPI_ISL_491310, EPI_ISL_491311, EPI_ISL_491312, EPI_ISL_491313, EPI_ISL_491314, EPI_ISL_491315, EPI_ISL_491316, EPI_I EPI_ISL_491321, EPI_ISL_491322, EPI_ISL_491323, EPI_ISL_491324, EPI_ISL_491325, EPI_ISL_491326, EPI_ISL_491327, EPI_ISL_491328, EPI_ISL_491329, EPI_ISL_491330, EPI_ISL_491331, EPI_ISL_491332, EPI_ISL_491333, EPI_ISL_491334, EPI_ISL_491335, EPI_ISL_491336, EPI_ISL_491337, EPI_ISL_491338, EPI_I EPI_ISL_491343, EPI_ISL_491344, EPI_ISL_491345, EPI_ISL_491346, EPI_ISL_491347, EPI_ISL_491348, EPI_ISL_491349, EPI_ISL_491350, EPI_ISL_491351, EPI_ISL_491352, EPI_ISL_491353, EPI_ISL_491354, EPI_ISL_491355, EPI_ISL_491356, EPI_ISL_491357, EPI_ISL_491358, EPI_ISL_491359, EPI_ISL_491360, EPI_I EPI_ISL_491365, EPI_ISL_491366, EPI_ISL_491367, EPI_ISL_491368, EPI_ISL_491369, EPI_ISL_491370, EPI_ISL_491371, EPI_ISL_491372, EPI_ISL_491373, EPI_ISL_491376, EPI_ISL_491377, EPI_ISL_491380, EPI_ISL_491381, EPI_ISL_491383, EPI_ISL_491384, EPI_ISL_491386, EPI_ISL_491387, EPI_ISL_491389, EPI_I EPI_ISL_491394, EPI_ISL_491395, EPI_ISL_491397, EPI_ISL_491398, EPI_ISL_491400, EPI_ISL_491401, EPI_ISL_491402, EPI_ISL_491405, EPI_ISL_491407, EPI_ISL_491412, EPI_ISL_491413, EPI_ISL_491414, EPI_ISL_491415, EPI_ISL_491416, EPI_ISL_491417, EPI_ISL_491421, EPI_ISL_491422 | | | |
| see above | University of Wisconsin-Madison AIDS Vaccine Research Laboratories | University of Wisconsin-Madison AIDS Vaccine Research Laboratories | Gage Moreno, Katarina Braun, et al. AIDS Vaccine Research Laboratories |
| EPI_ISL_491475 | Protacio Hospital | Research Institute for Tropical Medicine | Ma. Angelica Tujan, Othoniel Jan Onza, Francisco Gerardo Polotan, Inez Andrea Medado, Criselda Bautista, Kirstyn Brunker, Edelwisa Mercado, Daria Manalo, Catalino Demetria |
| EPI_ISL_491717, EPI_ISL_491718, EPI_ISL_491719, EPI_ISL_491720, EPI_ISL_491721 | Respiratory Virus Unit, Microbiology Services Colindale, Public Health England | Respiratory Virus Unit, Microbiology Services Colindale, Public Health England | PHE Covid Sequencing Team |
| EPI_ISL_491936 | Institute of Microbiology, Universidad San Francisco de Quito | Institute of Microbiology, Universidad San Francisco de Quito | Belén Prado-Vivar, Sully Márquez, Juan José Guadalupe, Monica Becerra-Wong, Bernardo Gutiérrez, Carlos Guerrero, Verónica Barragán, Patricio Rojas-Silva, Gabriel Trueba, Michelle Grunauer, Pa |
| EPI_ISL_491937, EPI_ISL_491938 | Institute of Microbiology, Universidad San Francisco de Quito | Institute of Microbiology, Universidad San Francisco de Quito | Belén Prado-Vivar, Sully Márquez, Juan José Guadalupe, Monica Becerra-Wong, Bernardo Gutiérrez, Rosario Erazo, Verónica Barragán, Patricio Rojas-Silva, Gabriel Trueba, Michelle Grunauer, Pa |
| EPI_ISL_492075, EPI_ISL_492076, EPI_ISL_492077, EPI_ISL_492078, EPI_ISL_492079, EPI_ISL_492080, EPI_ISL_492081, EPI_ISL_492082, EPI_ISL_492083, EPI_ISL_492084, EPI_ISL_492085, EPI_ISL_492086 | | | |
| see above | Institute for Public Health of the Republic of North Macedonia | Charite Universitatsmedizin Berlin, Institute of Virology | Victor M Corman, Joern Beheim-Schwarzbach, Barbara Muhlemann, Talitha Veith, Julia Schneider, Elizabeta Jancheska, Maja Kuzmanovska, Golubinka Bosevska, Terry Jones, Christian Dro |
| EPI_ISL_492988 | Centrl laboratorija | Latvian Biomedical Research and Study Centre | Ivars Silamielis, Kaspars Megnis, Monta Ustinova, ikita Zrelovs, Vita Rovte, Stella Lapia, Jana Oste, Marta Priedte, Uga Dumpis, Jnis Kloviš |
| EPI_ISL_493002 | Respiratory Virus Unit, Microbiology Services Colindale, Public Health England | Respiratory Virus Unit, Microbiology Services Colindale, Public Health England | PHE Covid Sequencing Team |
| EPI_ISL_493354, EPI_ISL_493355 | Oslo University Hospital, Department of Medical Microbiology | Norwegian Institute of Public Health, Department of Virology | Kathrine Stene-Johansen, Kamilla Heddeland Instefjord, Hilde Elshaug, Rasmus Riis Kopperud, Karoline Bragstad, Olav Hungnes |
| EPI_ISL_493356, EPI_ISL_493357, EPI_ISL_493358, EPI_ISL_493359, EPI_ISL_493360, EPI_ISL_493361, EPI_ISL_493362, EPI_ISL_493363, EPI_ISL_493364, EPI_ISL_493365, EPI_ISL_493366, EPI_ISL_493367, EPI_ISL_493368, EPI_ISL_493369, EPI_ISL_493370, EPI_ISL_493371, EPI_ISL_493372, EPI_ISL_493373, EPI_I EPI_ISL_493378, EPI_ISL_493379, EPI_ISL_493380 | | | |
| see above | Furst Medical Laboratory | Norwegian Institute of Public Health, Department of Virology | Kathrine Stene-Johansen, Kamilla Heddeland Instefjord, Hilde Elshaug, Rasmus Riis Kopperud, Karoline Bragstad, Olav Hungnes |
| EPI_ISL_493381, EPI_ISL_493382, EPI_ISL_493383 | Medical Microbiology Unit, Department for Laboratory Medicine, Drammen Hospital, Vestre Viken Health Trust, | Norwegian Institute of Public Health, Department of Virology | Kathrine Stene-Johansen, Kamilla Heddeland Instefjord, Hilde Elshaug, Rasmus Riis Kopperud, Karoline Bragstad, Olav Hungnes |
| EPI_ISL_493384 | Hospital of Southern Norway - Kristiansand, Department of Medical Microbiology | Norwegian Institute of Public Health, Department of Virology | Kathrine Stene-Johansen, Kamilla Heddeland Instefjord, Hilde Elshaug, Rasmus Riis Kopperud, Karoline Bragstad, Olav Hungnes |
| EPI_ISL_493385, EPI_ISL_493386, EPI_ISL_493387, EPI_ISL_493388, EPI_ISL_493389 | Oslo University Hospital, Department of Medical Microbiology | Norwegian Institute of Public Health, Department of Virology | Kathrine Stene-Johansen, Kamilla Heddeland Instefjord, Hilde Elshaug, Rasmus Riis Kopperud, Karoline Bragstad, Olav Hungnes |
| EPI_ISL_493668, EPI_ISL_493671, EPI_ISL_493674, EPI_ISL_493675, EPI_ISL_493676, EPI_ISL_493687, EPI_ISL_493688, EPI_ISL_493690, EPI_ISL_493730, EPI_ISL_493736 | Virology Department, Sheffield Teaching Hospitals NHS Foundation Trust/Department of Infection, Immunity and Cardiovascular Disease, The Medical School, University of Sheffield | COVID-19 Genomics UK (COG-UK) Consortium | Thushan de Silva, Matthew Parker, Nikki Smith, Adri Angyal, Rebecca Brown, Luke Green, Rachel Tucker, Paul Parsons, Danielle Groves, Katie Johnson, Laura Carrilero, Alex Keeley, Dave Partridge, Matthew Wyles, Benjamin Lindsey, |
| EPI_ISL_493974 | Virology Department, Royal Infirmary of Edinburgh, NHS Lothian / School of Biological Sciences, University of Edinburgh / Institute of Genetics and Molecular | COVID-19 Genomics UK (COG-UK) Consortium | McHugh M, Dewar R, Rooke S, Gallagher M, Balcaza C, O'Toole Á, Scher E, Hill V, McCrone JT, Colquhoun R, Yu X, Jackson B, Rambaut A, Williams TC, Templeton K |

| | | | |
|---|---|---|---|
| | Medicine, University of Edinburgh | | |
| EPI_ISL_493985, EPI_ISL_493990, EPI_ISL_493992, EPI_ISL_493993, EPI_ISL_494001, EPI_ISL_494005, EPI_ISL_494008, EPI_ISL_494010, EPI_ISL_494013, EPI_ISL_494020, EPI_ISL_494026, EPI_ISL_494027, EPI_ISL_494028, EPI_ISL_494032, EPI_ISL_494033, EPI_ISL_494036, EPI_ISL_494038, EPI_ISL_494043, EPI_I EPI_ISL_494050, EPI_ISL_494053, EPI_ISL_494054, EPI_ISL_494057, EPI_ISL_494058, EPI_ISL_494059, EPI_ISL_494060, EPI_ISL_494062, EPI_ISL_494066, EPI_ISL_494071, EPI_ISL_494072, EPI_ISL_494075, EPI_ISL_494077, EPI_ISL_494078, EPI_ISL_494080, EPI_ISL_494088, EPI_ISL_494091, EPI_ISL_494094, EPI_I EPI_ISL_494107, EPI_ISL_494109, EPI_ISL_494115, EPI_ISL_494118, EPI_ISL_494121, EPI_ISL_494122, EPI_ISL_494127, EPI_ISL_494130, EPI_ISL_494132, EPI_ISL_494133, EPI_ISL_494139, EPI_ISL_494140, EPI_ISL_494141, EPI_ISL_494146 | | | |
| see above | Wales Specialist Virology Centre Sequencing lab: Pathogen Genomics Unit | COVID-19 Genomics UK (COG-UK) Consortium | Catherine Moore, Johnathan Evans, Laura Gifford, Malorie Perry, Simon Cottrell, Angela Marchbank, Alec Birchley, Alexander Adams, Amy Gaskin, Bree Gatica-Wilcox, Jason Coombes, Joel Southgate, Lauren Gilbert, Lee Graham, Nicole Pacchiarini, Sa Matthew Bull, Joanne Watkins, Sally Corden, Tom Connor |
| EPI_ISL_494558 | Functional Genomics Core University of South Carolina / Prisma Health-Midlands | Functional Genomics Core, University of South Carolina | Hao Ji, Diego Altomare, B.Celia Cui, Mengqian Chen, Alyssa Clay-Glimour, Michael Wyatt, Phillip Buckhaults, Helmut Albrecht, Michael Shtutman |
| EPI_ISL_494570 | Rady's Childrens Hospital | Andersen lab at Scripps Research | SEARCH Alliance San Diego |
| EPI_ISL_495020 | Government Medical College, Bhavnagar | Gujarat Biotechnology Research Centre | Kairavi Desai, Saklin Malek, Shirish Patel, Nitin Savaliya, Raghawendra Kumar, Dinesh Kumar, Zuber Saiyed, Komal Patel, Labdhi Pandya, Afzal Ansari, Nikha Trivedi, Apurvasinh Puvar, Janvi Raval, Zarna Patel, Monika Gandhi, Pinal Trivedi, Maharshi Joshi, Madhvi Joshi |
| EPI_ISL_495021 | Government Medical College, Bhavnagar | Gujarat Biotechnology Research Centre | Saklin Malek, Shirish Patel, Kairavi Desai, Raghawendra Kumar, Dinesh Kumar, Zuber Saiyed, Komal Patel, Labdhi Pandya, Afzal Ansari, Nikha Trivedi, Apurvasinh Puvar, Janvi Raval, Zarna Patel, Monika Gandhi, Pinal Trivedi, Maharshi Pandya, Nidhi I Joshi, Madhvi Joshi |
| EPI_ISL_495022 | Government Medical College, Bhavnagar | Gujarat Biotechnology Research Centre | Shirish Patel, Kairavi Desai, Saklin Malek, Dinesh Kumar, Zuber Saiyed, Komal Patel, Labdhi Pandya, Afzal Ansari, Nikha Trivedi, Apurvasinh Puvar, Janvi Raval, Zarna Patel, Monika Gandhi, Pinal Trivedi, Maharshi Pandya, Nidhi Patel, Nitin Savaliya, R Joshi, Madhvi Joshi |
| EPI_ISL_495516, EPI_ISL_495523, EPI_ISL_495524, EPI_ISL_495525, EPI_ISL_495526, EPI_ISL_495527, EPI_ISL_495528, EPI_ISL_495529, EPI_ISL_495530, EPI_ISL_495532, EPI_ISL_495533, EPI_ISL_495534, EPI_ISL_495547, EPI_ISL_495553 | | | |
| see above | NHLS-IALCH | KRISP, KZN Research Innovation and Sequencing Platform | Giandhari J, Pillay S, Lessells R, Chimukangara B, Mdlalose K, York D, Khan S, Tegally H, Wilkinson E, de Oliveira T |
| EPI_ISL_497312, EPI_ISL_497333, EPI_ISL_497338 | Washington State Department of Health | Seattle Flu Study | Deborah A. Nickerson, Chris D. Frazar, Jover Lee, Benjamin Pelle, Matthew Richardson, Amanda Adler, Elisabeth Brandstetter, Peter D. Han, Kairsten Fay, Misja Ilcisin, Kirsten Lacombe, Thomas R. Sibley, Melissa Truong, Caitlin R. Wolf, Romesh Gautc Boeckh, Janet A. Englund, Michael Famulare, Barry R. Lutz, Mark J. Rieder, Lea M. Starita, Matthew Thompson, Helen Y. Chu, Jay Shendure, Trevor Bedford |
| EPI_ISL_497852 | Department of Microbiology, The University of Hong Kong | Department of Microbiology, The University of Hong Kong | Kelvin K.W. To, Kwok-Yung Yuen |
| EPI_ISL_498057, EPI_ISL_498058, EPI_ISL_498059 | NHLS-IALCH | KRISP, KZN Research Innovation and Sequencing Platform | Giandhari J, Pillay S, Lessells R, Chimukangara B, Mdlalose K, York D, Khan S, Tegally H, Wilkinson E, de Oliveira T |
| EPI_ISL_498141, EPI_ISL_498142, EPI_ISL_498143, EPI_ISL_498144, EPI_ISL_498145, EPI_ISL_498146, EPI_ISL_498147, EPI_ISL_498148 | Department of Clinical Microbiology | GIGA Medical Genomics | Keith Durkin, Maria Artesi, Sébastien Bontems, Raphaël Boreux, Cécile Meex, Axelle Chaslain, Céline Fombellida-Lopez, Pierrette Melin, Marie-Pierre Hayette, Vincent Bours. |
| EPI_ISL_498570, EPI_ISL_498573, EPI_ISL_498574, EPI_ISL_498575, EPI_ISL_498576 | National Public Health Laboratory, National Centre for Infectious Diseases | National Public Health Laboratory, National Centre for Infectious Diseases | Mak TM, Octavia S, Zhou Z, Chavatte JM, Cui L, Lin RTP |
| EPI_ISL_499356, EPI_ISL_499360, EPI_ISL_499362, EPI_ISL_499364, EPI_ISL_499367, EPI_ISL_499368, EPI_ISL_499371, EPI_ISL_499376, EPI_ISL_499377, EPI_ISL_499380, EPI_ISL_499385, EPI_ISL_499390, EPI_ISL_499395, EPI_ISL_499408, EPI_ISL_499414, EPI_ISL_499415, EPI_ISL_499417, EPI_ISL_499418, EPI_I EPI_ISL_499424, EPI_ISL_499425, EPI_ISL_499426, EPI_ISL_499430, EPI_ISL_499435, EPI_ISL_499436, EPI_ISL_499437, EPI_ISL_499438, EPI_ISL_499439, EPI_ISL_499443, EPI_ISL_499446, EPI_ISL_499449, EPI_ISL_499450, EPI_ISL_499455, EPI_ISL_499459 | | | |
| see above | Wales Specialist Virology Centre Sequencing lab: Pathogen Genomics Unit | COVID-19 Genomics UK (COG-UK) Consortium | Catherine Moore, Johnathan Evans, Laura Gifford, Malorie Perry, Simon Cottrell, Angela Marchbank, Alec Birchley, Alexander Adams, Amy Gaskin, Bree Gatica-Wilcox, Jason Coombes, Joel Southgate, Lauren Gilbert, Lee Graham, Nicole Pacchiarini, Sa Matthew Bull, Joanne Watkins, Sally Corden, Tom Connor |
| EPI_ISL_499494, EPI_ISL_499503, EPI_ISL_499504, EPI_ISL_499546, EPI_ISL_499547, EPI_ISL_499548, EPI_ISL_499549, EPI_ISL_499553, EPI_ISL_499556, EPI_ISL_499560, EPI_ISL_499561, EPI_ISL_499568, EPI_ISL_499602, EPI_ISL_499606, EPI_ISL_499626 | | | |
| see above | Northumbria University / South Tees Hospitals NHS Foundation Trust / North Cumbria Integrated Care NHS Foundation Trust / North Tees and Hartlepool NHS Foundation Trust / Newcastle Hospitals NHS Foundation Trust | COVID-19 Genomics UK (COG-UK) Consortium | Darren L Smith,Andrew Nelson,Matthew Bashton,Greg R Young,Joshua Loh,John Allan,Mohammad A Tariq,Giles S Holt,Gary Black,Wen C Yew,Lynn Dover,Paul Baker,Steve Liggett,Sarah Essex,Jane Greenaway,Debra Padgett,Clive Graham,Garre Collins,Yusri Taha,Gary Eltringham |
| EPI_ISL_499675 | Liverpool Clinical Laboratories | COVID-19 Genomics UK (COG-UK) Consortium | Sam Haldenby, Anita Lucaci, Steve Paterson, Julian Hiscox, Alistair Darby, M Almsaud, A Alrezaihi, Muhannad Alruwaili, Stuart D Armstrong, Jones Benjamin, Eleanor G Bentley, Anu Chawla, Jordan J Clark, Angela Cowell, Richard Eccles, Isabel Garc Richard Gregory, Ximeng Han, Catherine Hartley, Margaret Hughes, Miren Iturriza-Gomara, James Johnson, L Luu, Jenifer Manson, Charlotte Nelson, Elaine O'Toole, Cassie Olateju, Rebekah Penrice-Randal , Lucille Rainbow, N.P Randle, Trevor Ian Swainston, Ecaterina Vamos, Joanne Watts, Mark Whitehead |
| EPI_ISL_499778, EPI_ISL_499779, EPI_ISL_499780, EPI_ISL_499781, EPI_ISL_499786 | Northumbria University / South Tees Hospitals NHS Foundation Trust / North Cumbria Integrated Care NHS Foundation Trust / North Tees and Hartlepool NHS Foundation Trust / Newcastle Hospitals NHS Foundation Trust | COVID-19 Genomics UK (COG-UK) Consortium | Darren L Smith,Andrew Nelson,Matthew Bashton,Greg R Young,Joshua Loh,John Allan,Mohammad A Tariq,Giles S Holt,Gary Black,Wen C Yew,Lynn Dover,Paul Baker,Steve Liggett,Sarah Essex,Jane Greenaway,Debra Padgett,Clive Graham,Garre Collins,Yusri Taha,Gary Eltringham |
| EPI_ISL_499810, EPI_ISL_499829, EPI_ISL_499861, EPI_ISL_499869 | University Hospitals Of Leicester NHS Trust and DeepSeq Nottingham | COVID-19 Genomics UK (COG-UK) Consortium | Christopher Holmes, Paul Bird, Thomas Helmer, Karlie Fallon, Julian Tang, Jonathan Ball, Patrick McClure, Joeseph Chappell, Nadine Holmes, Matthew Carlisle, Christopher Moore, Fei Sang, Johnny Debebe, Vict |
| EPI_ISL_499877 | Liverpool Clinical | COVID-19 Genomics UK | Sam Haldenby, Anita Lucaci, Steve Paterson, Julian Hiscox, Alistair Darby, M Almsaud, A Alrezaihi, Muhannad Alruwaili, Stuart D Armstrong, Jones Benjamin, Eleanor G Bentley, Anu Chawla, Jordan J Clark, Angela Cowell, Richard Eccles, Isabel Garc |

| | | | |
|---|---|---|---|
| | Laboratories | (COG-UK) Consortium | Richard Gregory, Ximeng Han, Catherine Hartley, Margaret Hughes, Miren Iturriza-Gomara, James Johnson, L Luu, Jenifer Manson, Charlotte Nelson, Elaine O'Toole, Cassie Olateju, Rebekah Penrice-Randal , Lucille Rainbow, N.P Randle, Trevor Ian Swainston, Ecaterina Vamos, Joanne Watts, Mark Whitehead |
| EPI_ISL_499894, EPI_ISL_499895, EPI_ISL_499910, EPI_ISL_499912 | University Hospitals Of Leicester NHS Trust and DeepSeq Nottingham | COVID-19 Genomics UK (COG-UK) Consortium | Christopher Holmes, Paul Bird, Thomas Helmer, Karlie Fallon, Julian Tang, Jonathan Ball, Patrick McClure, Joeseph Chappell, Nadine Holmes, Matthew Carlisle, Christopher Moore, Fei Sang, Johnny Debebe, Vict |
| EPI_ISL_499927 | Liverpool Clinical Laboratories | COVID-19 Genomics UK (COG-UK) Consortium | Sam Haldenby, Anita Lucaci, Steve Paterson, Julian Hiscox, Alistair Darby, M Almsaud, A Alrezaihi, Muhannad Alruwaili, Stuart D Armstrong, Jones Benjamin, Eleanor G Bentley, Anu Chawla, Jordan J Clark, Angela Cowell, Richard Eccles, Isabel Garc Richard Gregory, Ximeng Han, Catherine Hartley, Margaret Hughes, Miren Iturriza-Gomara, James Johnson, L Luu, Jenifer Manson, Charlotte Nelson, Elaine O'Toole, Cassie Olateju, Rebekah Penrice-Randal , Lucille Rainbow, N.P Randle, Trevor Ian Swainston, Ecaterina Vamos, Joanne Watts, Mark Whitehead |
| EPI_ISL_499933, EPI_ISL_499950 | University Hospitals Of Leicester NHS Trust and DeepSeq Nottingham | COVID-19 Genomics UK (COG-UK) Consortium | Christopher Holmes, Paul Bird, Thomas Helmer, Karlie Fallon, Julian Tang, Jonathan Ball, Patrick McClure, Joeseph Chappell, Nadine Holmes, Matthew Carlisle, Christopher Moore, Fei Sang, Johnny Debebe, Vict |
| EPI_ISL_499953, EPI_ISL_499965 | Liverpool Clinical Laboratories | COVID-19 Genomics UK (COG-UK) Consortium | Sam Haldenby, Anita Lucaci, Steve Paterson, Julian Hiscox, Alistair Darby, M Almsaud, A Alrezaihi, Muhannad Alruwaili, Stuart D Armstrong, Jones Benjamin, Eleanor G Bentley, Anu Chawla, Jordan J Clark, Angela Cowell, Richard Eccles, Isabel Garc Richard Gregory, Ximeng Han, Catherine Hartley, Margaret Hughes, Miren Iturriza-Gomara, James Johnson, L Luu, Jenifer Manson, Charlotte Nelson, Elaine O'Toole, Cassie Olateju, Rebekah Penrice-Randal , Lucille Rainbow, N.P Randle, Trevor Ian Swainston, Ecaterina Vamos, Joanne Watts, Mark Whitehead |
| EPI_ISL_499976, EPI_ISL_499977, EPI_ISL_499978, EPI_ISL_499979, EPI_ISL_499980, EPI_ISL_499981, EPI_ISL_499982 | University Hospitals Of Leicester NHS Trust and DeepSeq Nottingham | COVID-19 Genomics UK (COG-UK) Consortium | Christopher Holmes, Paul Bird, Thomas Helmer, Karlie Fallon, Julian Tang, Jonathan Ball, Patrick McClure, Joeseph Chappell, Nadine Holmes, Matthew Carlisle, Christopher Moore, Fei Sang, Johnny Debebe, Vict |
| EPI_ISL_500029 | Liverpool Clinical Laboratories | COVID-19 Genomics UK (COG-UK) Consortium | Sam Haldenby, Anita Lucaci, Steve Paterson, Julian Hiscox, Alistair Darby, M Almsaud, A Alrezaihi, Muhannad Alruwaili, Stuart D Armstrong, Jones Benjamin, Eleanor G Bentley, Anu Chawla, Jordan J Clark, Angela Cowell, Richard Eccles, Isabel Garc Richard Gregory, Ximeng Han, Catherine Hartley, Margaret Hughes, Miren Iturriza-Gomara, James Johnson, L Luu, Jenifer Manson, Charlotte Nelson, Elaine O'Toole, Cassie Olateju, Rebekah Penrice-Randal , Lucille Rainbow, N.P Randle, Trevor Ian Swainston, Ecaterina Vamos, Joanne Watts, Mark Whitehead |
| EPI_ISL_500542, EPI_ISL_500543, EPI_ISL_500544, EPI_ISL_500545, EPI_ISL_500546, EPI_ISL_500547, EPI_ISL_500548, EPI_ISL_500549, EPI_ISL_500554, EPI_ISL_500555 | Singapore General Hospital | Department of Microbiology | Nurdyana Abdul Rahman, Kun Lee Lim, Chenhao Li, Kian Sing Chan, Lynette Oon, Kern Rei Chng, Niranjan Nagarajan, Karrie Ko |
| EPI_ISL_500768 | Furst Medical Laboratory | Norwegian Institute of Public Health, Department of Virology | Kathrine Stene-Johansen, Kamilla Heddeland Instefjord, Hilde Elshaug, Rasmus Riis Kopperud, Karoline Bragstad, Olav Hungnes |
| EPI_ISL_500776, EPI_ISL_500777, EPI_ISL_500778 | Foerde Hospital, Department of Microbiology | Norwegian Institute of Public Health, Department of Virology | Kathrine Stene-Johansen, Kamilla Heddeland Instefjord, Hilde Elshaug, Rasmus Riis Kopperud, Karoline Bragstad, Olav Hungnes |
| EPI_ISL_501084, EPI_ISL_501085, EPI_ISL_501098, EPI_ISL_501103, EPI_ISL_501104, EPI_ISL_501105, EPI_ISL_501106, EPI_ISL_501107, EPI_ISL_501108, EPI_ISL_501109, EPI_ISL_501110, EPI_ISL_501111, EPI_ISL_501112, EPI_ISL_501113, EPI_ISL_501114, EPI_ISL_501115, EPI_ISL_501116, EPI_ISL_501117, EPI_I EPI_ISL_501125, EPI_ISL_501126, EPI_ISL_501127, EPI_ISL_501128 | | | |
| see above | University of Washington Virology Lab | University of Washington Virology Lab | Pavitra Roychoudhury, Hong Xie, Lasata Shrestha, Amin Addetia, Truong Nguyen, Victoria M Rachleff, Meei-Li Huang, Keith R Jerome, Alexander Greninger |
| EPI_ISL_502779 | LACEN/PE | LABBE, Federal University of Pernambuco | WILSON JOSE DA SILVA JUNIOR, HEIDI LACERDA ALVES DA CRUZ, MARCOS DA SILVEIRA REGUEIRA NETO, BRUNO SAMPAIO, SERGIO DE SA LEITAO PAIVA JUNIOR, ZILDENE DE SOUSA SILVEIRA, MAIRA GALDINO DA ROCHA PITTA, JUNIOR, ANTONIO CARLOS DE FREITAS, VALDIR DE QUEIROZ BALBINO. |
| EPI_ISL_502875 | LACEN/PE | LABBE, Federal University of Pernambuco | WILSON JOSE DA SILVA JUNIOR, HEIDI LACERDA ALVES DA CRUZ, MARCOS DA SILVEIRA REGUEIRA NETO, BRUNO SAMPAIO, SERGIO DE SA LEITAO PAIVA JUNIOR, ZILDENE DE SOUSA SILVEIRA, MAIRA GALDINO DA ROCHA PITTA LIMA NETO, MARCOS ANTONIO DE MORAIS JUNIOR, ANTONIO CARLOS DE FREITAS, VALDIR DE QUEIROZ BALBINO. |
| EPI_ISL_507420, EPI_ISL_507432, EPI_ISL_507438 | Michigan Department of Health and Human Services, Bureau of Laboratories | Michigan Department of Health and Human Services, Bureau of Laboratories | Blankenship HM, Riner D, Soehnlen MK |
| EPI_ISL_507957, EPI_ISL_507960 | Mayo Clinic & Mayo Clinic Laboratories | Minnesota Department of Health, Public Health Laboratory | Matt Plumb, Jacob Garfin, and Xiong Wang |
| EPI_ISL_508175 | All india institute of Medical Sciences Rishikesh | National Institute of Biomedical Genomics | Arindam Maitra, Deepjyoti Kalita, Amit Mangla, Ravi Kant, Saumitra Das |
| EPI_ISL_508419 | Maulana Azad Medical College | National Institute of Biomedical Genomics | Arindam Maitra, Sonal Saxena, Vikas Manchanda, Oves Siddiqui, Saumitra Das |
| EPI_ISL_508756, EPI_ISL_508757, EPI_ISL_508758, EPI_ISL_508759, EPI_ISL_508760 | Florida Bureau of Public Health Laboratories | Florida Bureau of Public Health Laboratories | Sarah Schmedes, Jason Blanton |
| EPI_ISL_509612, EPI_ISL_509613, EPI_ISL_509614, EPI_ISL_509643, EPI_ISL_509644, EPI_ISL_509645, EPI_ISL_509646, EPI_ISL_509647, EPI_ISL_509648, EPI_ISL_509649, EPI_ISL_509650, EPI_ISL_509651, EPI_ISL_509652, EPI_ISL_509653, EPI_ISL_509654 | | | |
| see above | Servicio de Microbiología. Hospital Universitario Donostia. OSI Donostialdea. Área de Enfermedades Infecciosas, Grupo de Infección Respiratoria y Resistencia Antimicrobiana. Instituto de Investigación Sanitaria Biodonostia | SeqCOVID-SPAIN consortium/IBV(CSIC) | Gustavo Cilla, Milagrosa Montes, Luis Piñeiro, Jose Maria Marimón and SeqCOVID-SPAIN consortium |
| EPI_ISL_509821, EPI_ISL_509822, EPI_ISL_509854, | University of Wisconsin-Madison AIDS Vaccine Research | University of Wisconsin-Madison AIDS Vaccine Research | Gage Moreno, Katarina Braun, et al. AIDS Vaccine Research Laboratories |

| | | | |
|---|---|---|---|
| EPI_ISL_509997 | Laboratories | Laboratories | |
| EPI_ISL_510084 | Prince of Wales Hospital | Hong Kong Department of Health | Mak Gannon C.K., Lam Edman T.K., Chan Rickjason C.W., Tsang Dominic N.C. |
| EPI_ISL_510428, EPI_ISL_510430, EPI_ISL_510432, EPI_ISL_510435, EPI_ISL_510436, EPI_ISL_510453, EPI_ISL_510457 | Hospital Universitario Virgen de las Nieves de Granada-SAS | SeqCOVID-SPAIN consortium/IBV(CSIC) | Mercedes Pérez Ruiz, Sara Sanbonmatsu Gámez, Irene Pedrosa Corral, José M. Navarro-Marí and SeqCOVID-SPAIN consortium |
| EPI_ISL_510524 | University of Wisconsin-Madison AIDS Vaccine Research Laboratories | University of Wisconsin-Madison AIDS Vaccine Research Laboratories | Gage Moreno, Katarina Braun, et al. AIDS Vaccine Research Laboratories |
| EPI_ISL_510528 | Communicable Disease Laboratory, Public Health Directorate | Communicable Disease Laboratory, Public Health Directorate | Al Wasti,H. and AlTaif,Z. |
| EPI_ISL_510531 | Communicable Disease Laboratory, Public Health Directorate | Communicable Disease Laboratory, Public Health Directorate | Shehab,F., Altaif,Z., AlWasti,H., Zaed,A. |
| EPI_ISL_510533, EPI_ISL_510534 | Faculty of Medicine, Bursa Uludag University | Bursa Uludag University | Temel, S.G., Saglik, I., Özemri Sa, ., Alemdar, A. |
| EPI_ISL_511955, EPI_ISL_511957, EPI_ISL_511969, EPI_ISL_511971, EPI_ISL_511977 | PHE South West Regional Laboratory, National Infection Service | Wellcome Sanger Institute for the COVID-19 Genomics UK (COG-UK) consortium | Stephanie Hutchings, Hannah Pymont, Dr Peter Muir, Barry Vipond, Rich Hopes; and Alex Alderton, Roberto Amato, Sonia Goncalves, Ewan Harrison, David K. Jackson, Ian Johnston, Dominic Kwiatkowski, Cordelia Langford, John Sillitoe on behalf [illegible] (http://www.sanger.ac.uk/covid-team) |
| EPI_ISL_512060 | B.J. Medical College and Civil hospital, Ahmedabad | Gujarat Biotechnology Research Centre | Maharshi Pandya, Nidhi Patel, Nitin Savaliya, Raghawendra Kumar, Dinesh Kumar, Zuber Saiyed, Komal Patel, Labdhi Pandya, Afzal Ansari, Nikha Trivedi, Pranay Shah, Kamlesh J Upadhyay, Sanjay Kapadia, Apurvasinh Puvar, Janvi Raval, Zarna Pate[illegible] Chaitanya Joshi, Madhvi Joshi |
| EPI_ISL_512061 | B.J. Medical College and Civil hospital, Ahmedabad | Gujarat Biotechnology Research Centre | Nidhi Patel, Nitin Savaliya, Raghawendra Kumar, Dinesh Kumar, Zuber Saiyed, Komal Patel, Labdhi Pandya, Afzal Ansari, Nikha Trivedi, Pranay Shah, Kamlesh J Upadhyay, Sanjay Kapadia, Apurvasinh Puvar, Janvi Raval, Zarna Patel, Monika Gandhi, [illegible] Chaitanya Joshi, Madhvi Joshi |
| EPI_ISL_512062 | B.J. Medical College and Civil hospital, Ahmedabad | Gujarat Biotechnology Research Centre | Nitin Savaliya, Raghawendra Kumar, Dinesh Kumar, Zuber Saiyed, Komal Patel, Labdhi Pandya, Afzal Ansari, Nikha Trivedi, Pranay Shah, Kamlesh J Upadhyay, Sanjay Kapadia, Apurvasinh Puvar, Janvi Raval, Zarna Patel, Monika Gandhi, Pinal Trivedi[illegible] Chaitanya Joshi, Madhvi Joshi |
| EPI_ISL_512063 | B.J. Medical College and Civil hospital, Ahmedabad | Gujarat Biotechnology Research Centre | Raghawendra Kumar, Dinesh Kumar, Zuber Saiyed, Komal Patel, Labdhi Pandya, Afzal Ansari, Nikha Trivedi, Pranay Shah, Kamlesh J Upadhyay, Sanjay Kapadia, Apurvasinh Puvar, Janvi Raval, Zarna Patel, Monika Gandhi, Pinal Trivedi, Maharshi Pan[illegible] Chaitanya Joshi, Madhvi Joshi |
| EPI_ISL_512064 | B.J. Medical College and Civil hospital, Ahmedabad | Gujarat Biotechnology Research Centre | Dinesh Kumar, Zuber Saiyed, Komal Patel, Labdhi Pandya, Afzal Ansari, Nikha Trivedi, Pranay Shah, Kamlesh J Upadhyay, Sanjay Kapadia, Apurvasinh Puvar, Janvi Raval, Zarna Patel, Monika Gandhi, Pinal Trivedi, Maharshi Pandya, Nidhi Patel, Nitin [illegible] Chaitanya Joshi, Madhvi Joshi |
| EPI_ISL_512065 | B.J. Medical College and Civil hospital, Ahmedabad | Gujarat Biotechnology Research Centre | Zuber Saiyed, Komal Patel, Labdhi Pandya, Afzal Ansari, Nikha Trivedi, Pranay Shah, Kamlesh J Upadhyay, Sanjay Kapadia, Apurvasinh Puvar, Janvi Raval, Zarna Patel, Monika Gandhi, Pinal Trivedi, Maharshi Pandya, Nidhi Patel, Nitin Savaliya, Ragha[illegible] Chaitanya Joshi, Madhvi Joshi |
| EPI_ISL_512149, EPI_ISL_512153, EPI_ISL_512154, EPI_ISL_512155, EPI_ISL_512156, EPI_ISL_512157 | Alaska State Virology Laboratory | Alaska State Virology Laboratory | Chen J et al with Pathogenomics group Dagdag R, Redlinger M, Milton E, George W, Kovalenko A, Drown DM, Bortz E |
| EPI_ISL_512246, EPI_ISL_512247, EPI_ISL_512253, EPI_ISL_512254, EPI_ISL_512255, EPI_ISL_512256, EPI_ISL_512257, EPI_ISL_512258, EPI_ISL_512259, EPI_ISL_512260, EPI_ISL_512261, EPI_ISL_512262, EPI_ISL_512263, EPI_ISL_512264, EPI_ISL_512265, EPI_ISL_512266, EPI_ISL_512268, EPI_ISL_512269, EPI_I[illegible] | | | |
| see above | San Diego County Public Health Laboratory | Andersen lab at Scripps Research | SEARCH Alliance San Diego with Tracy Basler, Jovan Shephard, Brett Austin |
| EPI_ISL_512332 | Department of Pathology, University of Cambridge | COVID-19 Genomics UK (COG-UK) Consortium | Luke W Meredith, M. Estée Török, Myra Hosmillo, William L. Hamilton, Martin D. Curran, Theresa Feltwell, Grant Hall, Anna Yakovleva, Fahad A Khokhar, Charlotte J. Houldcroft, Laura G Caller, Aminu S. Jahun, Sarah L. Caddy, Yas[illegible] |
| EPI_ISL_512363, EPI_ISL_512364, EPI_ISL_512365 | Northumbria University / South Tees Hospitals NHS Foundation Trust / North Cumbria Integrated Care NHS Foundation Trust / North Tees and Hartlepool NHS Foundation Trust / Newcastle Hospitals NHS Foundation Trust | COVID-19 Genomics UK (COG-UK) Consortium | Darren L Smith,Andrew Nelson,Matthew Bashton,Greg R Young,Joshua Loh,John Allan,Mohammad A Tariq,Giles S Holt,Gary Black,Wen C Yew,Lynn Dover,Paul Baker,Steve Liggett,Sarah Essex,Jane Greenaway,Debra Padgett,Clive Graham,Garre[illegible] Collins,Yusri Taha,Gary Eltringham |
| EPI_ISL_512475 | West of Scotland Specialist Virology Centre, NHSGGC / MRC-University of Glasgow Centre for Virus Research | COVID-19 Genomics UK (COG-UK) Consortium | Ana da Silva Filipe, Natasha Johnson, Kathy Smollett, Daniel Mair, Stephen Carmichael, Lily Tong, Jenna Nichols, Elihu Aranday-Cortes, Kirstyn Brunker, Yasmin Parr, Alice Broos, Kyriaki Nomikou; Sarah McDonald, Marc Niebel, Patawee Asamaphan; R[illegible] Alasdair MacLean, Rory Gunson; Kathy Li, Natasha Jesudason, Rajiv Shah, James Shepherd, Antonia Ho, Emma Thomson |
| EPI_ISL_512508 | Wales Specialist Virology Centre Sequencing lab: Pathogen Genomics Unit | COVID-19 Genomics UK (COG-UK) Consortium | Catherine Moore, Johnathan Evans, Laura Gifford, Malorie Perry, Simon Cottrell, Angela Marchbank, Alec Birchley, Alexander Adams, Amy Gaskin, Bree Gatica-Wilcox, Jason Coombes, Joel Southgate, Lauren Gilbert, Lee Graham, Nicole Pacchiarini, Sa[illegible] Matthew Bull, Joanne Watkins, Sally Corden, Tom Connor |
| EPI_ISL_512557, EPI_ISL_512558, EPI_ISL_512560, EPI_ISL_512561, EPI_ISL_512562, EPI_ISL_512563, EPI_ISL_512564, EPI_ISL_512565, EPI_ISL_512566, EPI_ISL_512567, EPI_ISL_512568, EPI_ISL_512569, EPI_ISL_512570, EPI_ISL_512571, EPI_ISL_512572, EPI_ISL_512573, EPI_ISL_512574, EPI_ISL_512575, EPI_I[illegible] | | | |
| see above | Florida Bureau of Public Health Laboratories | Florida Bureau of Public Health Laboratories | Sarah Schmedes, Jason Blanton |
| EPI_ISL_512597, EPI_ISL_512598, EPI_ISL_512599, EPI_ISL_512600, EPI_ISL_512601, EPI_ISL_512612, EPI_ISL_512613, EPI_ISL_512614, EPI_ISL_512615, EPI_ISL_512628, EPI_ISL_512629, EPI_ISL_512630, EPI_ISL_512631, EPI_ISL_512632 | | | |
| see above | National Laboratory for Influenza/Virology reference laboratory, Public Health Center of the Ministry of Health of Ukraine | Respiratory Virus Unit, Microbiology Services Colindale, Public Health England | PHE Covid Sequencing Team, Dr. Iryna Demchyshyna |

| | | | |
|---|---|---|---|
| EPI_ISL_512647 | Latvijas Infektoloijas centrs | Latvian Biomedical Research and Study Centre | Ivars Silamielis, Kaspars Megnis, Monta Ustinova, ikita Zrelovs, Vita Rovte, Jeena Storoženko, Tatjana Kolupajeva, Oksana Savicka, Uga Dumpis, Jnis Kloviš |
| EPI_ISL_512676, EPI_ISL_512677 | Florida Bureau of Public Health Laboratories | Florida Bureau of Public Health Laboratories | Sarah Schmedes, Jason Blanton |
| EPI_ISL_512874, EPI_ISL_512875, EPI_ISL_512876, EPI_ISL_512877, EPI_ISL_512882, EPI_ISL_512883, EPI_ISL_512890, EPI_ISL_512891, EPI_ISL_512892, EPI_ISL_512893, EPI_ISL_512894, EPI_ISL_512895, EPI_ISL_512897, EPI_ISL_512898, EPI_ISL_512899, EPI_ISL_512900, EPI_ISL_512901, EPI_ISL_512902 | | | |
| see above | Pathogen Genomics Lab King Abdullah University of Science and Technology(KAUST) | Pathogen Genomics Lab King Abdullah University of Science and Technology(KAUST) | Raeece Naeem, Rahul P Salunke, Sharif Hala, Sara Mfarrej, Amit Kumar Subudhi, Fadwa Alofi, Fathia Ben Rached, Afrah Alsomali, Asim Khogeer, Ahmad Bakur Mahmoud, Anwar Hashem, Naif Almonta |
| EPI_ISL_512904, EPI_ISL_512906, EPI_ISL_512907 | Pathogen Genomics Lab King Abdullah University of Science and Technology(KAUST) | Pathogen Genomics Lab King Abdullah University of Science and Technology(KAUST) | Fathia Ben Rached, Raeece Naeem, Sharif Hala, Fadwa Alofi, Rahul P Salunke, Sara Mfarrej, Amit Kumar Subudhi, Afrah Alsomali, Asim Khogeer, Ahmad Bakur Mahmoud, Anwar Hashem, Naif Almonta |
| EPI_ISL_513064, EPI_ISL_513065, EPI_ISL_513071, EPI_ISL_513072 | Pathogen Genomics Lab King Abdullah University of Science and Technology(KAUST) | Pathogen Genomics Lab King Abdullah University of Science and Technology(KAUST) | Raeece Naeem, Rahul P Salunke, Sharif Hala, Sara Mfarrej, Amit Kumar Subudhi, Fadwa Alofi, Fathia Ben Rached, Afrah Alsomali, Asim Khogeer, Ahmad Bakur Mahmoud, Anwar Hashem, Naif Almonta |
| EPI_ISL_513267, EPI_ISL_513268, EPI_ISL_513269, EPI_ISL_513270, EPI_ISL_513271, EPI_ISL_513272, EPI_ISL_513273, EPI_ISL_513274, EPI_ISL_513275, EPI_ISL_513276, EPI_ISL_513277, EPI_ISL_513278, EPI_ISL_513279, EPI_ISL_513280, EPI_ISL_513281, EPI_ISL_513282, EPI_ISL_513283, EPI_ISL_513284, EPI_I | | | |
| see above | University of Miami Immunology and Histocompatibility Laboratory | University of Miami Immunology and Histocompatibility Laboratory | Emilio Margolles-Clark, PhD and Phillip Ruiz, MD, PhD |
| EPI_ISL_513785, EPI_ISL_513786 | County of Santa Clara Public Health Department | Chan-Zuckerberg Biohub | CZB Cliahub Consortium |
| EPI_ISL_514194, EPI_ISL_514195, EPI_ISL_514196, EPI_ISL_514197, EPI_ISL_514198, EPI_ISL_514199, EPI_ISL_514200, EPI_ISL_514201, EPI_ISL_514202, EPI_ISL_514203, EPI_ISL_514204, EPI_ISL_514205, EPI_ISL_514206, EPI_ISL_514207, EPI_ISL_514208, EPI_ISL_514209 | | | |
| see above | Florida Bureau of Public Health Laboratories | Florida Bureau of Public Health Laboratories | Sarah Schmedes, Jason Blanton |
| EPI_ISL_514600 | B.J. Medical College and Civil hospital, Ahmedabad | Gujarat Biotechnology Research Centre | Raghawendra Kumar, Dinesh Kumar, Zuber Saiyed, Komal Patel, Labdhi Pandya, Afzal Ansari, Nikha Trivedi, Pranay Shah, Kamlesh J Upadhyay, Sanjay Kapadia, Apurvasinh Puvar, Janvi Raval, Zarna Patel, Monika Gandhi, Pinal Trivedi, Maharshi Pan<br>Chaitanya Joshi, Madhvi Joshi |
| EPI_ISL_514601 | B.J. Medical College and Civil hospital, Ahmedabad | Gujarat Biotechnology Research Centre | Dinesh Kumar, Zuber Saiyed, Komal Patel, Labdhi Pandya, Afzal Ansari, Nikha Trivedi, Pranay Shah, Kamlesh J Upadhyay, Sanjay Kapadia, Apurvasinh Puvar, Janvi Raval, Zarna Patel, Monika Gandhi, Pinal Trivedi, Maharshi Pandya, Nidhi Patel, Nitin S<br>Chaitanya Joshi, Madhvi Joshi |
| EPI_ISL_514602 | B.J. Medical College and Civil hospital, Ahmedabad | Gujarat Biotechnology Research Centre | Zuber Saiyed, Komal Patel, Labdhi Pandya, Afzal Ansari, Nikha Trivedi, Pranay Shah, Kamlesh J Upadhyay, Sanjay Kapadia, Apurvasinh Puvar, Janvi Raval, Zarna Patel, Monika Gandhi, Pinal Trivedi, Maharshi Pandya, Nidhi Patel, Nitin Savaliya, Ragha<br>Chaitanya Joshi, Madhvi Joshi |
| EPI_ISL_514603 | B.J. Medical College and Civil hospital, Ahmedabad | Gujarat Biotechnology Research Centre | Komal Patel, Labdhi Pandya, Afzal Ansari, Nikha Trivedi, Pranay Shah, Kamlesh J Upadhyay, Sanjay Kapadia, Apurvasinh Puvar, Janvi Raval, Zarna Patel, Monika Gandhi, Pinal Trivedi, Maharshi Pandya, Nidhi Patel, Nitin Savaliya, Raghawendra Kuma<br>Chaitanya Joshi, Madhvi Joshi |
| EPI_ISL_514604 | B.J. Medical College and Civil hospital, Ahmedabad | Gujarat Biotechnology Research Centre | Labdhi Pandya, Afzal Ansari, Nikha Trivedi, Pranay Shah, Kamlesh J Upadhyay, Sanjay Kapadia, Apurvasinh Puvar, Janvi Raval, Zarna Patel, Monika Gandhi, Pinal Trivedi, Maharshi Pandya, Nidhi Patel, Nitin Savaliya, Raghawendra Kumar, Dinesh Kum<br>Chaitanya Joshi, Madhvi Joshi |
| EPI_ISL_514605 | B.J. Medical College and Civil hospital, Ahmedabad | Gujarat Biotechnology Research Centre | Afzal Ansari, Nikha Trivedi, Pranay Shah, Kamlesh J Upadhyay, Sanjay Kapadia, Apurvasinh Puvar, Janvi Raval, Zarna Patel, Monika Gandhi, Pinal Trivedi, Maharshi Pandya, Nidhi Patel, Nitin Savaliya, Raghawendra Kumar, Dinesh Kumar, Zuber Saiye<br>Chaitanya Joshi, Madhvi Joshi |
| EPI_ISL_514606 | B.J. Medical College and Civil hospital, Ahmedabad | Gujarat Biotechnology Research Centre | Nikha Trivedi, Pranay Shah, Kamlesh J Upadhyay, Sanjay Kapadia, Apurvasinh Puvar, Janvi Raval, Zarna Patel, Monika Gandhi, Pinal Trivedi, Maharshi Pandya, Nidhi Patel, Nitin Savaliya, Raghawendra Kumar, Dinesh Kumar, Zuber Saiyed, Komal Pate<br>Chaitanya Joshi, Madhvi Joshi |
| EPI_ISL_514607 | B.J. Medical College and Civil hospital, Ahmedabad | Gujarat Biotechnology Research Centre | Pranay Shah, Kamlesh J Upadhyay, Sanjay Kapadia, Apurvasinh Puvar, Janvi Raval, Zarna Patel, Monika Gandhi, Pinal Trivedi, Maharshi Pandya, Nidhi Patel, Nitin Savaliya, Raghawendra Kumar, Dinesh Kumar, Zuber Saiyed, Komal Patel, Labdhi Panc<br>Chaitanya Joshi, Madhvi Joshi |
| EPI_ISL_514608 | B.J. Medical College and Civil hospital, Ahmedabad | Gujarat Biotechnology Research Centre | Kamlesh J Upadhyay, Sanjay Kapadia, Apurvasinh Puvar, Janvi Raval, Zarna Patel, Monika Gandhi, Pinal Trivedi, Maharshi Pandya, Nidhi Patel, Nitin Savaliya, Raghawendra Kumar, Dinesh Kumar, Zuber Saiyed, Komal Patel, Labdhi Pandya, Afzal Ansa<br>Chaitanya Joshi, Madhvi Joshi |
| EPI_ISL_514609 | B.J. Medical College and Civil hospital, Ahmedabad | Gujarat Biotechnology Research Centre | Sanjay Kapadia, Apurvasinh Puvar, Janvi Raval, Zarna Patel, Monika Gandhi, Pinal Trivedi, Maharshi Pandya, Nidhi Patel, Nitin Savaliya, Raghawendra Kumar, Dinesh Kumar, Zuber Saiyed, Komal Patel, Labdhi Pandya, Afzal Ansari, Nikha Trivedi, Pran<br>Chaitanya Joshi, Madhvi Joshi |
| EPI_ISL_514620, EPI_ISL_514621, EPI_ISL_514622, EPI_ISL_514623, EPI_ISL_514624, EPI_ISL_514625, EPI_ISL_514626, EPI_ISL_514627, EPI_ISL_514628 | Mayo Clinic & Mayo Clinic Laboratories | Minnesota Department of Health, Public Health Laboratory | Matt Plumb, Jacob Garfin, and Xiong Wang |
| EPI_ISL_514630, EPI_ISL_514631, EPI_ISL_514632, EPI_ISL_514633, EPI_ISL_514634, EPI_ISL_514635 | M Health Fairview | Minnesota Department of Health, Public Health Laboratory | Matt Plumb, Jacob Garfin, and Xiong Wang |
| EPI_ISL_514957, EPI_ISL_514958, EPI_ISL_514959, EPI_ISL_514960, EPI_ISL_514961, EPI_ISL_514962, EPI_ISL_514963, EPI_ISL_514964, EPI_ISL_514965, EPI_ISL_514966, EPI_ISL_514967, EPI_ISL_514968, EPI_ISL_514969, EPI_ISL_514970, EPI_ISL_514971, EPI_ISL_514972, EPI_ISL_514973, EPI_ISL_514974, EPI_I<br>EPI_ISL_514979, EPI_ISL_514980, EPI_ISL_514981, EPI_ISL_514982, EPI_ISL_514983, EPI_ISL_514984, EPI_ISL_514985, EPI_ISL_514986, EPI_ISL_514987, EPI_ISL_514988, EPI_ISL_514989, EPI_ISL_514990, EPI_ISL_514991, EPI_ISL_514992, EPI_ISL_514993, EPI_ISL_514994, EPI_ISL_514995, EPI_ISL_514996, EPI_I<br>EPI_ISL_515001, EPI_ISL_515002, EPI_ISL_515003, EPI_ISL_515004, EPI_ISL_515012 | | | |
| see above | Division of Viral Diseases, Center for Laboratory Control of Infectious Diseases, Korea Centers for Diseases Control and Prevention | Division of Viral Diseases, Center for Laboratory Control of Infectious Diseases, Korea Centers for Diseases Control and Prevention | Jeong-Min Kim, Yoon-Seok Chung, Namjoo Lee, Sang Hee Woo, Hye-Jun Jo, Heui Man Kim, Jun-Sub Kim, Myung Guk Han |
| EPI_ISL_515138 | National Institute for Communicable Diseases of the National Health Laboratory Service | National Institute for Communicable Diseases of the National Health Laboratory Service | Allam M, Ismail A, Khumalo Z, Kwenda S, Mtshali P, Mnyameni F, Mohale T, Bhiman JN |
| EPI_ISL_515264, | Mayo Clinic & Mayo Clinic | Minnesota Department of | Matt Plumb, Jacob Garfin, and Xiong Wang |

| | | | |
|---|---|---|---|
| EPI_ISL_515265, EPI_ISL_515266 | Laboratories | Health, Public Health Laboratory | |
| EPI_ISL_515268 | M Health Fairview | Minnesota Department of Health, Public Health Laboratory | Matt Plumb, Jacob Garfin, and Xiong Wang |
| EPI_ISL_515525 | National Influenza Center - Instituto Adolfo Lutz | Instituto Adolfo Lutz, Interdiciplinary Procedures Center, Strategic Laboratory | Claudio Tavares Sacchi, Claudia Regina Gonçalves, Erica Valessa Ramos Gomes |
| EPI_ISL_515578, EPI_ISL_515583, EPI_ISL_515584, EPI_ISL_515585 | NHLS-IALCH | KRISP, KZN Research Innovation and Sequencing Platform | Giandhari J, Pillay S, Lessells R, Mdlalose K, York D, Khan S, Tegally H, Wilkinson E, de Oliveira T |
| EPI_ISL_515835, EPI_ISL_515837 | Medical Disagnostics Services (MDS) | KRISP, KZN Research Innovation and Sequencing Platform | Giandhari J, Pillay S, Lessells R, ChimukangaraB, Mdlalose K, York D, Khan S, Tegally H, Wilkinson E, de Oliveira T |
| EPI_ISL_515926, EPI_ISL_515927, EPI_ISL_515928, EPI_ISL_515929 | California Department of Public Health | California Department of Public Health | CDPH IDLB COVIDNet |
| EPI_ISL_516193 | Hospital Universitari Germans Trias i Pujol. | IrsiCaixa AIDS Research Lab | Marc Noguera-Julian, Mariona Parera, Maria Pilar Armengol, Marta Massanella, Ester Ballana, Lidia Ruiz, Nuria Izquierdo, Jorge Carrillo, Roger Paredes, Julia Blanco, Joaquim Segalés, Bonavent |
| EPI_ISL_516384, EPI_ISL_516405, EPI_ISL_516406, EPI_ISL_516407, EPI_ISL_516408, EPI_ISL_516410, EPI_ISL_516411 | Michigan Department of Health and Human Services, Bureau of Laboratories | Michigan Department of Health and Human Services, Bureau of Laboratories | Blankenship HM, Riner D, Soehnlen MK |
| EPI_ISL_516426, EPI_ISL_516427 | Clinical Hospital - Shtip | Research Center for Genetic Engineering and Biotechnology "Georgi D. Efremov" , Macedonian Academy of Sciences and Arts | RCGEB - MASA |
| EPI_ISL_516608 | Instituto de Diagnostico y Referencia Epidemiologicos (INDRE) | Instituto de Diagnostico y Referencia Epidemiologicos (INDRE) | Ernesto Ramirez-Gonzalez, Abril Rodriguez-Maldonado, Claudia Wong-Arambula , Natividad Cruz-Ortiz, Tatiana Nunez-Garcia, Dayanira Arellano-Suarez, Adnan Araiza-Rodriguez, Edgar Mendieta-Condado, Lucia Hernandez-Rivas |
| EPI_ISL_516609, EPI_ISL_516610 | Instituto de Diagnostico y Referencia Epidemiologicos (INDRE) | Instituto de Diagnostico y Referencia Epidemiologicos (INDRE) | Ernesto Ramirez-Gonzalez, Abril Rodriguez-Maldonado, Claudia Wong-Arambula , Natividad Cruz-Ortiz, Tatiana Nunez-Garcia, Dayanira Arellano-Suarez, Adnan Araiza-Rodriguez, Fabiola Garces-Ayala, Lucia Hernandez-Rivas, I |
| EPI_ISL_516613 | Instituto de Diagnostico y Referencia Epidemiologicos (INDRE) | Instituto de Diagnostico y Referencia Epidemiologicos (INDRE) | Ernesto Ramirez-Gonzalez, Abril Rodriguez-Maldonado, Claudia Wong-Arambula , Natividad Cruz-Ortiz, Tatiana Nunez-Garcia, Dayanira Arellano-Suarez, Adnan Araiza-Rodriguez, Edgar Mendieta-Condado, Lucia Hernandez-Rivas |
| EPI_ISL_516614, EPI_ISL_516615, EPI_ISL_516616 | Instituto de Diagnostico y Referencia Epidemiologicos (INDRE) | Instituto de Diagnostico y Referencia Epidemiologicos (INDRE) | Ernesto Ramirez-Gonzalez, Abril Rodriguez-Maldonado, Claudia Wong-Arambula , Natividad Cruz-Ortiz, Tatiana Nunez-Garcia, Dayanira Arellano-Suarez, Adnan Araiza-Rodriguez, Fabiola Garces-Ayala, Lucia Hernandez-Rivas, I |
| EPI_ISL_516621 | Instituto de Diagnostico y Referencia Epidemiologicos (INDRE) | Instituto de Diagnostico y Referencia Epidemiologicos (INDRE) | Gisela Barrera-Badillo , Abril Rodriguez-Maldonado, Claudia Wong-Arambula , Natividad Cruz-Ortiz, Tatiana Nunez-Garcia, Dayanira Arellano-Suarez, Adnan Araiza-Rodriguez, Edgar Mendieta-Condado, Lucia Hernandez-Rivas, Irm |
| EPI_ISL_516751, EPI_ISL_516752, EPI_ISL_516753, EPI_ISL_516754, EPI_ISL_516759 | van Bakel Laboratory, Genetics and Genomics Sciences, Icahn School of Medicine at Mount Sinai | van Bakel Laboratory, Genetics and Genomics Sciences, Icahn School of Medicine at Mount Sinai | Andrew G. Letizia, Irene Ramos, Ajay Obla, Carl Goforth, Dawn Weir, Yongchao Ge, Marcas M. Bamman, Jayeeta Dutta, Ethan Ellis, Luis Estrella, Mary-Catherine George, Ana S. Gonzalez-Reiche, Darnell Graham, Adriana van de Guchte, Ramiro Gutie Lizewski, Jan Marayag, Nada Marjanovic, Eugene V. Millar, Venugopalan Nair, German Nudelman, Edgar Nunez, Brian Pike, James Regeimbal, Stas Rirak , Ernesto Santa Ana, Rachel S. Gelernter Sealfon, Robert Sebra, Mark Simons, Alessandra Soares van Bakel, Stuart C. Sealfon |
| EPI_ISL_516901, EPI_ISL_516902, EPI_ISL_516903, EPI_ISL_516904, EPI_ISL_516905, EPI_ISL_516906, EPI_ISL_516907, EPI_ISL_516918, EPI_ISL_516919, EPI_ISL_516920, EPI_ISL_516921 | | | |
| see above | Israel Central Virology laboratory | Israel Central Virology laboratory | Neta Zuckerman, Efrat Dahan Bucris, Oran Erster, Ella Mendelson, Michal Mandelboim |
| EPI_ISL_517624, EPI_ISL_517625, EPI_ISL_517626, EPI_ISL_517627, EPI_ISL_517628, EPI_ISL_517629, EPI_ISL_517630, EPI_ISL_517631, EPI_ISL_517632, EPI_ISL_517633, EPI_ISL_517634, EPI_ISL_517635, EPI_ISL_517636, EPI_ISL_517637, EPI_ISL_517638, EPI_ISL_517639, EPI_ISL_517640, EPI_ISL_517641, EPI_I[illegible] EPI_ISL_517646, EPI_ISL_517647, EPI_ISL_517648, EPI_ISL_517649, EPI_ISL_517650, EPI_ISL_517651, EPI_ISL_517652, EPI_ISL_517653, EPI_ISL_517654, EPI_ISL_517655 | | | |
| see above | Academic Hospital Paramaribo | Erasmus Medical Center | Bas Oude Munnink, Dion Gajadin, Ed Ijzerman, Emmanuelle Munger, Gary Gummels, Ingrid Krishnadath, Lycke Woittiez, Marion Koopmans, Mireille Van de Veer, Princes Wongsowidjojo, Radjesh Ori, Rohma B |
| EPI_ISL_517792, EPI_ISL_517793, EPI_ISL_517794, EPI_ISL_517795, EPI_ISL_517796, EPI_ISL_517797, EPI_ISL_517798, EPI_ISL_517799, EPI_ISL_517800, EPI_ISL_517801, EPI_ISL_517802, EPI_ISL_517803 | | | |
| see above | Florida Bureau of Public Health Laboratories | Florida Bureau of Public Health Laboratories | Sarah Schmedes, Jason Blanton |
| EPI_ISL_518048, EPI_ISL_518049, EPI_ISL_518052, EPI_ISL_518053 | NHLS-IALCH | KRISP, KZN Research Innovation and Sequencing Platform | Giandhari J, Pillay S, Lessells R, Mdlalose K, York D, Khan S, Tegally H, Wilkinson E, de Oliveira T |
| EPI_ISL_518231, EPI_ISL_518414 | Microbiological Diagnostic Unit - Public Health Laboratory (MDU-PHL) | MDU-PHL | Seemann T., Schultz M., Sait, M., Sherry, N. |
| EPI_ISL_519322, EPI_ISL_519323, EPI_ISL_520278, EPI_ISL_520291, EPI_ISL_520371 | Victorian Infectious Diseases Reference Laboratory (VIDRL) | VIDRL and MDU-PHL | Caly L., Seemann T., Sait, M., Schultz M., Druce J., Sherry, N. |

| | | | |
|---|---|---|---|
| EPI_ISL_520616, EPI_ISL_520617, EPI_ISL_520618, EPI_ISL_520619, EPI_ISL_520646 | Microbiological Diagnostic Unit - Public Health Laboratory (MDU-PHL) | MDU-PHL | Seemann T., Schultz M., Sait, M., Sherry, N. |
| EPI_ISL_520749, EPI_ISL_520875, EPI_ISL_520905, EPI_ISL_520909, EPI_ISL_520910, EPI_ISL_520911, EPI_ISL_520912, EPI_ISL_520913 | Victorian Infectious Diseases Reference Laboratory (VIDRL) | VIDRL and MDU-PHL | Caly L., Seemann T., Sait, M., Schultz M., Druce J., Sherry, N. |
| EPI_ISL_521459, EPI_ISL_521460, EPI_ISL_521461 | Microbiological Diagnostic Unit - Public Health Laboratory (MDU-PHL) | MDU-PHL | Seemann T., Schultz M., Sait, M., Sherry, N. |
| EPI_ISL_521463, EPI_ISL_521464, EPI_ISL_521465, EPI_ISL_521466, EPI_ISL_521467, EPI_ISL_521468, EPI_ISL_521469, EPI_ISL_521470, EPI_ISL_521471, EPI_ISL_521472, EPI_ISL_521473, EPI_ISL_521474, EPI_ISL_521475, EPI_ISL_521477, EPI_ISL_521478, EPI_ISL_521479, EPI_ISL_521481, EPI_ISL_521482, EPI_I[illegible] EPI_ISL_521487, EPI_ISL_521488, EPI_ISL_521489, EPI_ISL_521490, EPI_ISL_521491, EPI_ISL_521492, EPI_ISL_521493, EPI_ISL_521502, EPI_ISL_521503, EPI_ISL_521504, EPI_ISL_521506, EPI_ISL_521508, EPI_ISL_521517, EPI_ISL_521519, EPI_ISL_521520, EPI_ISL_521521, EPI_ISL_521522, EPI_ISL_521523, EPI_I[illegible] | | | |
| see above | Victorian Infectious Diseases Reference Laboratory (VIDRL) | VIDRL and MDU-PHL | Caly L., Seemann T., Sait, M., Schultz M., Druce J., Sherry, N. |
| EPI_ISL_521633, EPI_ISL_521645, EPI_ISL_521653, EPI_ISL_521672, EPI_ISL_521711, EPI_ISL_521713, EPI_ISL_521714 | Microbiological Diagnostic Unit - Public Health Laboratory (MDU-PHL) | MDU-PHL | Seemann T., Schultz M., Sait, M., Sherry, N. |
| EPI_ISL_521853, EPI_ISL_521855 | Victorian Infectious Diseases Reference Laboratory (VIDRL) | VIDRL and MDU-PHL | Caly L., Seemann T., Sait, M., Schultz M., Druce J., Sherry, N. |
| EPI_ISL_521862 | Royal Darwin Hospital Pathology | MDU-PHL | Meumann, E., Caly L., Seemann T., Sait, M., Schultz M., Druce J., Sherry, N. |
| EPI_ISL_521919, EPI_ISL_521920, EPI_ISL_521921, EPI_ISL_521922 | Victorian Infectious Diseases Reference Laboratory (VIDRL) | VIDRL and MDU-PHL | Caly L., Seemann T., Sait, M., Schultz M., Druce J., Sherry, N. |
| EPI_ISL_521926, EPI_ISL_521928, EPI_ISL_521929, EPI_ISL_521930, EPI_ISL_521931, EPI_ISL_521932, EPI_ISL_521933, EPI_ISL_521934, EPI_ISL_521935, EPI_ISL_521939, EPI_ISL_521940, EPI_ISL_521941, EPI_ISL_521942, EPI_ISL_521943, EPI_ISL_521944 | | | |
| see above | Microbiological Diagnostic Unit - Public Health Laboratory (MDU-PHL) | MDU-PHL | Seemann T., Schultz M., Sait, M., Sherry, N. |
| EPI_ISL_521945, EPI_ISL_521946, EPI_ISL_521947, EPI_ISL_521948, EPI_ISL_521949, EPI_ISL_521950, EPI_ISL_521951, EPI_ISL_521952 | Victorian Infectious Diseases Reference Laboratory (VIDRL) | VIDRL and MDU-PHL | Caly L., Seemann T., Sait, M., Schultz M., Druce J., Sherry, N. |
| EPI_ISL_521972 | Microbiological Diagnostic Unit - Public Health Laboratory (MDU-PHL) | MDU-PHL | Seemann T., Schultz M., Sait, M., Sherry, N. |
| EPI_ISL_521975, EPI_ISL_521976, EPI_ISL_521978, EPI_ISL_521980, EPI_ISL_521981, EPI_ISL_521982, EPI_ISL_521983, EPI_ISL_521984, EPI_ISL_521985, EPI_ISL_521986, EPI_ISL_521987 | | | |
| see above | Victorian Infectious Diseases Reference Laboratory (VIDRL) | VIDRL and MDU-PHL | Caly L., Seemann T., Sait, M., Schultz M., Druce J., Sherry, N. |
| EPI_ISL_521990, EPI_ISL_521991, EPI_ISL_521992, EPI_ISL_521993, EPI_ISL_521994, EPI_ISL_521995, EPI_ISL_521996, EPI_ISL_521997, EPI_ISL_521998, EPI_ISL_521999, EPI_ISL_522000, EPI_ISL_522001, EPI_ISL_522002, EPI_ISL_522003, EPI_ISL_522004, EPI_ISL_522005, EPI_ISL_522006, EPI_ISL_522007, EPI_I[illegible] EPI_ISL_522012, EPI_ISL_522013, EPI_ISL_522014, EPI_ISL_522015, EPI_ISL_522016, EPI_ISL_522017, EPI_ISL_522020, EPI_ISL_522021, EPI_ISL_522022, EPI_ISL_522025, EPI_ISL_522029, EPI_ISL_522030, EPI_ISL_522031, EPI_ISL_522033, EPI_ISL_522034, EPI_ISL_522036, EPI_ISL_522037, EPI_ISL_522038, EPI_I[illegible] EPI_ISL_522044, EPI_ISL_522045, EPI_ISL_522046, EPI_ISL_522047, EPI_ISL_522048, EPI_ISL_522049, EPI_ISL_522050, EPI_ISL_522051, EPI_ISL_522052, EPI_ISL_522053, EPI_ISL_522054 | | | |
| see above | Microbiological Diagnostic Unit - Public Health Laboratory (MDU-PHL) | MDU-PHL | Seemann T., Schultz M., Sait, M., Sherry, N. |
| EPI_ISL_522056, EPI_ISL_522057, EPI_ISL_522058, EPI_ISL_522059, EPI_ISL_522060, EPI_ISL_522061, EPI_ISL_522062, EPI_ISL_522063, EPI_ISL_522064, EPI_ISL_522065, EPI_ISL_522066, EPI_ISL_522067, EPI_ISL_522068, EPI_ISL_522069, EPI_ISL_522070, EPI_ISL_522071, EPI_ISL_522072, EPI_ISL_522073, EPI_I[illegible] EPI_ISL_522078, EPI_ISL_522079, EPI_ISL_522080, EPI_ISL_522081, EPI_ISL_522082, EPI_ISL_522083, EPI_ISL_522084, EPI_ISL_522085, EPI_ISL_522086, EPI_ISL_522087, EPI_ISL_522088, EPI_ISL_522089, EPI_ISL_522091, EPI_ISL_522092, EPI_ISL_522093, EPI_ISL_522094, EPI_ISL_522095, EPI_ISL_522097, EPI_I[illegible] EPI_ISL_522102, EPI_ISL_522103, EPI_ISL_522104, EPI_ISL_522105, EPI_ISL_522106, EPI_ISL_522107, EPI_ISL_522108, EPI_ISL_522109, EPI_ISL_522110 | | | |
| see above | Victorian Infectious Diseases Reference Laboratory (VIDRL) | VIDRL and MDU-PHL | Caly L., Seemann T., Sait, M., Schultz M., Druce J., Sherry, N. |
| EPI_ISL_522111, EPI_ISL_522112, EPI_ISL_522113, EPI_ISL_522114, EPI_ISL_522115, EPI_ISL_522116, EPI_ISL_522117, EPI_ISL_522118 | Microbiological Diagnostic Unit - Public Health Laboratory (MDU-PHL) | MDU-PHL | Seemann T., Schultz M., Sait, M., Sherry, N. |
| EPI_ISL_522119, EPI_ISL_522120, EPI_ISL_522121, EPI_ISL_522122, EPI_ISL_522124, EPI_ISL_522125, EPI_ISL_522126, EPI_ISL_522127, EPI_ISL_522128, EPI_ISL_522129, EPI_ISL_522130, EPI_ISL_522131, EPI_ISL_522132, EPI_ISL_522133, EPI_ISL_522134, EPI_ISL_522135, EPI_ISL_522136, EPI_ISL_522137, EPI_I[illegible] | | | |

| | | | |
|---|---|---|---|
| EPI_ISL_522142, EPI_ISL_522143, EPI_ISL_522144, EPI_ISL_522145, EPI_ISL_522146, EPI_ISL_522147, EPI_ISL_522148, EPI_ISL_522149, EPI_ISL_522150, EPI_ISL_522151, EPI_ISL_522152, EPI_ISL_522153, EPI_ISL_522154, EPI_ISL_522155, EPI_ISL_522156, EPI_ISL_522157, EPI_ISL_522158, EPI_ISL_522159, EPI_I[illegible] EPI_ISL_522164, EPI_ISL_522165, EPI_ISL_522166, EPI_ISL_522167, EPI_ISL_522168, EPI_ISL_522169, EPI_ISL_522170, EPI_ISL_522171, EPI_ISL_522172, EPI_ISL_522173, EPI_ISL_522174, EPI_ISL_522175, EPI_ISL_522176, EPI_ISL_522177, EPI_ISL_522178, EPI_ISL_522179, EPI_ISL_522180, EPI_ISL_522181, EPI_I[illegible] EPI_ISL_522186, EPI_ISL_522187 | | | |
| see above | Victorian Infectious Diseases Reference Laboratory (VIDRL) | VIDRL and MDU-PHL | Caly L., Seemann T., Sait, M., Schultz M., Druce J., Sherry, N. |
| EPI_ISL_522188, EPI_ISL_522189, EPI_ISL_522190, EPI_ISL_522191, EPI_ISL_522192, EPI_ISL_522193, EPI_ISL_522195, EPI_ISL_522196, EPI_ISL_522197, EPI_ISL_522198, EPI_ISL_522199, EPI_ISL_522200, EPI_ISL_522201, EPI_ISL_522202, EPI_ISL_522203, EPI_ISL_522204, EPI_ISL_522205, EPI_ISL_522206, EPI_I[illegible] EPI_ISL_522211, EPI_ISL_522212, EPI_ISL_522213, EPI_ISL_522214, EPI_ISL_522215, EPI_ISL_522216, EPI_ISL_522217, EPI_ISL_522218, EPI_ISL_522219, EPI_ISL_522220, EPI_ISL_522221, EPI_ISL_522222, EPI_ISL_522223, EPI_ISL_522225, EPI_ISL_522226, EPI_ISL_522230, EPI_ISL_522231 | | | |
| see above | Microbiological Diagnostic Unit - Public Health Laboratory (MDU-PHL) | MDU-PHL | Seemann T., Schultz M., Sait, M., Sherry, N. |
| EPI_ISL_522233, EPI_ISL_522234, EPI_ISL_522235, EPI_ISL_522236, EPI_ISL_522237, EPI_ISL_522238, EPI_ISL_522239, EPI_ISL_522240, EPI_ISL_522241, EPI_ISL_522242 | Victorian Infectious Diseases Reference Laboratory (VIDRL) | VIDRL and MDU-PHL | Caly L., Seemann T., Sait, M., Schultz M., Druce J., Sherry, N. |
| EPI_ISL_522243, EPI_ISL_522244 | Microbiological Diagnostic Unit - Public Health Laboratory (MDU-PHL) | MDU-PHL | Seemann T., Schultz M., Sait, M., Sherry, N. |
| EPI_ISL_522461, EPI_ISL_522462, EPI_ISL_522463, EPI_ISL_522464 | Center for Laboratory Control of Infectious Diseases, Korea Centers for Diseases Control and Prevention | Center for Laboratory Control of Infectious Diseases, Korea Centers for Diseases Control and Prevention | Junyoung Kim, Ae Kyung Park, Eunkyung Shin, Jin Sun No, Jeong-Min Kim, Yoon-Seok Chung, Heui Man Kim, Myung Guk Han |
| EPI_ISL_522465, EPI_ISL_522466 | Division of Viral Diseases, Center for Laboratory Control of Infectious Diseases, Korea Centers for Diseases Control and Prevention | Division of Viral Diseases, Center for Laboratory Control of Infectious Diseases, Korea Centers for Diseases Control and Prevention | Jeong-Min Kim, Yoon-Seok Chung, Namjoo Lee, Sang Hee Woo, Hye-Jun Jo, Heui Man Kim, Jun-Sub Kim, Myung Guk Han |
| EPI_ISL_522467, EPI_ISL_522468 | Center for Laboratory Control of Infectious Diseases, Korea Centers for Diseases Control and Prevention | Center for Laboratory Control of Infectious Diseases, Korea Centers for Diseases Control and Prevention | Junyoung Kim, Ae Kyung Park, Eunkyung Shin, Jin Sun No, Jeong-Min Kim, Yoon-Seok Chung, Heui Man Kim, Myung Guk Han |
| EPI_ISL_523458, EPI_ISL_523459, EPI_ISL_523460, EPI_ISL_523461, EPI_ISL_523462, EPI_ISL_523463, EPI_ISL_523508, EPI_ISL_523516, EPI_ISL_523533, EPI_ISL_523595, EPI_ISL_523596, EPI_ISL_523597, EPI_ISL_523598, EPI_ISL_523602, EPI_ISL_523609, EPI_ISL_523627, EPI_ISL_523628, EPI_ISL_523629, EPI_I[illegible] EPI_ISL_523634, EPI_ISL_523662, EPI_ISL_523663, EPI_ISL_523665, EPI_ISL_523666, EPI_ISL_523667, EPI_ISL_523668, EPI_ISL_523669, EPI_ISL_523672, EPI_ISL_523673, EPI_ISL_523674, EPI_ISL_523675, EPI_ISL_523676, EPI_ISL_523677, EPI_ISL_523678, EPI_ISL_523679, EPI_ISL_523680, EPI_ISL_523681, EPI_I[illegible] | | | |
| see above | Dutch COVID-19 response team | Erasmus Medical Center | Bas Oude Munnink, David Nieuwenhuijse, Reina Sikkema, Claudia Schapendonk, Irina Chestakova, Anne van der Linden, Theo Bestebroer, Stefan van Nieuwkoop, Mark Pronk, Pascal Lexmond, Corien Swaan, Manon Haverkate, Madelief Mollers, Mart [illegible] Voermans, Aura Timen, Corine GeurtsvanKessel, Annemiek van der Eijk, Richard Molenkamp, Marion Koopmans, on behalf of the Dutch national COVID-19 response team. |
| EPI_ISL_525421 | B.J. Medical College and Civil hospital, Ahmedabad | Gujarat Biotechnology Research Centre | Zarna Patel, Monika Gandhi, Pinal Trivedi, Maharshi Pandya, Nidhi Patel, Nitin Savaliya, Raghawendra Kumar, Dinesh Kumar, Zuber Saiyed, Komal Patel, Labdhi Pandya, Afzal Ansari, Nikha Trivedi, Pranay Shah, Kamlesh J Upadhyay, Sanjay Kapadia, [illegible] Chaitanya Joshi, Madhvi Joshi |
| EPI_ISL_525422 | B.J. Medical College and Civil hospital, Ahmedabad | Gujarat Biotechnology Research Centre | Monika Gandhi, Pinal Trivedi, Maharshi Pandya, Nidhi Patel, Nitin Savaliya, Raghawendra Kumar, Dinesh Kumar, Zuber Saiyed, Komal Patel, Labdhi Pandya, Afzal Ansari, Nikha Trivedi, Pranay Shah, Kamlesh J Upadhyay, Sanjay Kapadia, Apurvasinh [illegible] Chaitanya Joshi, Madhvi Joshi |
| EPI_ISL_525492 | RSUP dr. SOERADJI TIRTONEGORO | Genetics Working Group (Pokja Genetik) Faculty of Medicine, Public Health and Nursing Universitas Gadjah Mada (FK-KMK UGM), Disease Investigation Center Wates Ministry of Agriculture Indonesia, Department of Microbiology FK-KMK UGM, Laboratorium Diagnostik Yayasan Tahija World Mosquito Program (WMP) Yogyakarta Center for Tropical Medicine FK-KMK UGM, Integrated Research Center FK-KMK UGM, Department of Computer Science and Electronics FMIPA UGM | Gunadi, Hendra Wibawa, . Marcellus, Mohamad S. Hakim, Edwin W. Daniwijaya, Ludhang P. Rizki, Endah Supriyati, Eggi Arguni, Titik Nuryastuti, Tri Wibawa, Dwi AA Nugrahaningsih, Afiahayati , . Siswanto, Kurniyanto, Indal[illegible] |
| EPI_ISL_525702, EPI_ISL_525703, EPI_ISL_525704, EPI_ISL_525705 | Seattle Flu Study | Seattle Flu Study | Deborah A. Nickerson, Chris D. Frazar, Jover Lee, Benjamin Pelle, Matthew Richardson, Amanda Adler, Elisabeth Brandstetter, Peter D. Han, Kairsten Fay, Misja Ilcisin, Kirsten Lacombe, Thomas R. Sibley, Melissa Truong, Caitlin R. Wolf, Karen Cowgill, [illegible] Michael Famulare, Barry R. Lutz, Mark J. Rieder, Lea M. Starita, Matthew Thompson, Helen Y. Chu, Trevor Bedford, Jay Shendure |
| EPI_ISL_525706 | Seattle Flu Study | Seattle Flu Study | Deborah A. Nickerson, Chris D. Frazar, Jover Lee, Benjamin Pelle, Matthew Richardson, Amanda Adler, Elisabeth Brandstetter, Peter D. Han, Kairsten Fay, Misja Ilcisin, Kirsten Lacombe, Thomas R. Sibley, Melissa Truong, Caitlin R. Wolf, Michael Boeckh[illegible] Lea M. Starita, Matthew Thompson, Jay Shendure, Trevor Bedford, Helen Y. Chu |
| EPI_ISL_525804, EPI_ISL_525805, EPI_ISL_525891, EPI_ISL_525894, EPI_ISL_525895, EPI_ISL_525896, EPI_ISL_525897, EPI_ISL_525898, EPI_ISL_525907, EPI_ISL_525908, EPI_ISL_525909, EPI_ISL_525910, EPI_ISL_525911, EPI_ISL_525912, EPI_ISL_525913, EPI_ISL_525914, EPI_ISL_525915, EPI_ISL_525916, EPI_I[illegible] EPI_ISL_525921, EPI_ISL_525922, EPI_ISL_525923, EPI_ISL_525930, EPI_ISL_525939, EPI_ISL_525968, EPI_ISL_525969, EPI_ISL_525970, EPI_ISL_526099 | | | |
| see above | OHSU Lab Services Molecular Microbiology Lab | Oregon SARS-CoV-2 Genome Sequencing Center | Brendan L. O'Connell, Ruth V. Nichols, Alec J. Hirsch, Guang Fan, Daniel N. Streblow, William B. Messer, Andrew C. Adey, Benjamin N. Bimber, Brian J. O'Roak |
| EPI_ISL_526246, EPI_ISL_526247, EPI_ISL_526248, EPI_ISL_526249 | Instituto Adolfo Lutz | Instituto Adolfo Lutz Laboratório de Vírus Respiratórios | Katia Corrêa de Oliveira Santos, Fabiana Cristina Pereira dos Santos, Maíra Marcele Birochi, Cecília Simões Santos, Ana Maria Sardinha Afonso, Maira do Carmo Sampaio Tavares Timenet[illegible] |

EPI_ISL_526868, EPI_ISL_526869, EPI_ISL_526870, EPI_ISL_526876, EPI_ISL_526877, EPI_ISL_526878, EPI_ISL_526879, EPI_ISL_526880, EPI_ISL_526881, EPI_ISL_526893, EPI_ISL_526894

| | | | |
|---|---|---|---|
| see above | Virginia DCLS | Virginia DCLS | Virginia DCLS |
| EPI_ISL_527018, EPI_ISL_527022, EPI_ISL_527023, EPI_ISL_527024, EPI_ISL_527025, EPI_ISL_527026 | Area of Virology, Serology and Virology Division (SAViD), New South Wales Health Pathology Randwick | Area of Virology, Serology and Virology Division (SAViD), New South Wales Health Pathology Randwick | Rawlinson, W. |
| EPI_ISL_527741 | Hospital Metropolitano | Inciensa, Instituto Costarricense de Investigación y Enseñanza en Nutrición y Salud | Francisco Duarte, Hebleen Porras, Claudio Soto-Garita, Estela Cordero, Adriana Godinez & Melany Calderon |
| EPI_ISL_527742 | Centro Nacional De Rehabilitacion Humberto Araya Rojas (Cenare) | Inciensa, Instituto Costarricense de Investigación y Enseñanza en Nutrición y Salud | Francisco Duarte, Hebleen Porras, Claudio Soto-Garita, Estela Cordero, Adriana Godinez & Melany Calderon |
| EPI_ISL_527743, EPI_ISL_527744 | Hospital Cima | Inciensa, Instituto Costarricense de Investigación y Enseñanza en Nutrición y Salud | Francisco Duarte, Hebleen Porras, Claudio Soto-Garita, Estela Cordero, Adriana Godinez & Melany Calderon |
| EPI_ISL_528423 | TNMC & BYL NAIR CH. HOSPITAL | Institute of Genomics and Integrative Biology - Council of Scientific and Industrial Research | Rajesh Pandey, Jayanthi Shastri, Akshay Kanakan, Vivekanand A, Janani Srinivasa Vasudevan, Ranjeet Maurya, Sachee Agrawal, Nirjhar Chatterjee, Swapneil Parikh, Manish Pathak, Subrat Thanapati, Jasmina Savak, Suresh Poojari, Mahesh Sar Vishwanathan, Shruthi Sachidanandan, Shrutika Pophale, Utkarsha Yelve |
| EPI_ISL_528522 | Alaska State Virology Laboratory | Alaska State Virology Laboratory | Chen J et al with Pathogenomics group Dagdag R, Redlinger M, Milton E, George W, Kovalenko A, Drown DM, Bortz E |
| EPI_ISL_528600 | National Genomics Core-Center for DNA Fingerprinting and Diagnostics | National Genomics Core-Center for DNA Fingerprinting and Diagnostics (NGC-CDFD)-DBT's PAN-INDIA-1000 Genome consortium | Bala Pratyusha, Heena Shah, G Shashikanth, Vinay Donipadi, Edurugatla Dinesh, Guru Raja, Hilal Ahmad Reshi, J. Mallikarjun, K. Viswakalyan, Kaisar Ahmad Lone,Kausika Kumar Malik, N. Sudheer, R Harinarayanan, Rashna Bhandari, Murali Dha |
| EPI_ISL_528703, EPI_ISL_528705 | Alsafar - Khalifa University Abu Dhabi | Alsafar - Khalifa University Abu Dhabi | Andreas Henschel, Gihan Daw Elbait, Samuel Feng, Rifat Hamoudi, Ernesto Damiani, Guan Tay, Habiba Alsafar |
| EPI_ISL_528747 | Santo Borromeus Hospital | School of Pharmacy & School of Life Sciences and Technology - Institut Teknologi Bandung; Molecular Genetics Laboratory-Faculty of Medicine-Universitas Padjadjaran; Laboratorium Kesehatan Provinsi Jawa Barat | Catur Riani, Marselina Irasonia Tan, Yunia Sribudiani, Azzania Fibriani, Husna Nugrahapraja, Tarwadi, Ema Rahmawati, Savira Ekawardhani, Hesti Lina Wiraswati, Ryan Bayusantika Ristandi, Rifky Waluyajati Rachman, Cut Nur Cinthia Alamanda, Lia Fa Solistia Wirawan, Agung Eru Wibowo, Irvan Faizal |
| EPI_ISL_529104, EPI_ISL_529105, EPI_ISL_529106, EPI_ISL_529107, EPI_ISL_529108 | Microbiology Division, SC DHEC | Microbiology Division, SC DHEC | Flores,H. |
| EPI_ISL_529596, EPI_ISL_529661, EPI_ISL_529662 | University of Birmingham | COVID-19 Genomics UK (COG-UK) Consortium | Institute of Microbiology, University of Birmingham: Claire McMurray, Joanne Stockton, Samuel Nicholls, Radoslaw Poplawski, Will Rowe, Josh Quick, Nicholas Loman. University of Birmingham Testing Laboratory: Celina M Whalley, Andrew Bosworth, Ch Richter, Andrew D Beggs PHE Heartlands Lab: Husam Osman, Andrew Bosworth. Queen Elizabeth Hospital: Anna Casey |
| EPI_ISL_529829, EPI_ISL_529830, EPI_ISL_529831, EPI_ISL_529832 | Michigan Department of Health and Human Services, Bureau of Laboratories | Michigan Department of Health and Human Services, Bureau of Laboratories | Blankenship HM, Riner D, Soehnlen MK |
| EPI_ISL_529936 | Virginia Division of Consolidated Laboratory Services | Virginia Division of Consolidated Laboratory Services | Virginia DCLS |
| EPI_ISL_530174 | Minnesota Department of Health, Public Health Laboratory | Minnesota Department of Health, Public Health Laboratory | Matt Plumb, Jacob Garfin, and Xiong Wang |
| EPI_ISL_533225 | Lighthouse Lab in Glasgow | Wellcome Sanger Institute for the COVID-19 Genomics UK (COG-UK) consortium | Harper VanSteenhouse, Yumi Kasai, David Gray, Carol Clugston, Anna Dominiczak and Alex Alderton, Roberto Amato, Sonia Goncalves, Ewan Harrison, David K. Jackson, Ian Johnston, Dominic Kwiatkowski, Cor |
| EPI_ISL_533228 | NHSGGC West of Scotland Specialist Virology Centre / MRC-University of Glasgow Centre for Virus Research | Wellcome Sanger Institute for the COVID-19 Genomics UK (COG-UK) consortium | Ana da Silva Filipe, Natasha Johnson, Kathy Smollett, Daniel Mair, Stephen Carmichael, Lily Tong, Jenna Nichols, Elihu Aranday-Cortes, Kirstyn Brunker, Yasmin Parr, Kyriaki Nomikou; Sarah McDonald, Marc Niebel, Patawee Asamaphan; Richard Orto MacLean, Rory Gunson; Kathy Li, Natasha Jesudason, Rajiv Shah, James Shepherd, Antonia Ho, Alice Broos, Emma Thomson and Alex Alderton, Roberto Amato, Sonia Goncalves, Ewan Harrison, David K. Jackson, Ian Johnston, Dom |
| EPI_ISL_533229 | Lighthouse Lab in Glasgow | Wellcome Sanger Institute for the COVID-19 Genomics UK (COG-UK) consortium | Harper VanSteenhouse, Yumi Kasai, David Gray, Carol Clugston, Anna Dominiczak and Alex Alderton, Roberto Amato, Sonia Goncalves, Ewan Harrison, David K. Jackson, Ian Johnston, Dominic Kwiatkowski, Cor |
| EPI_ISL_533230, EPI_ISL_533231, EPI_ISL_533232 | NHSGGC West of Scotland Specialist Virology Centre / MRC-University of Glasgow Centre for Virus Research | Wellcome Sanger Institute for the COVID-19 Genomics UK (COG-UK) consortium | Ana da Silva Filipe, Natasha Johnson, Kathy Smollett, Daniel Mair, Stephen Carmichael, Lily Tong, Jenna Nichols, Elihu Aranday-Cortes, Kirstyn Brunker, Yasmin Parr, Kyriaki Nomikou; Sarah McDonald, Marc Niebel, Patawee Asamaphan; Richard Orto MacLean, Rory Gunson; Kathy Li, Natasha Jesudason, Rajiv Shah, James Shepherd, Antonia Ho, Alice Broos, Emma Thomson and Alex Alderton, Roberto Amato, Sonia Goncalves, Ewan Harrison, David K. Jackson, Ian Johnston, Dom |
| EPI_ISL_533234 | Lighthouse Lab in Glasgow | Wellcome Sanger Institute | Harper VanSteenhouse, Yumi Kasai, David Gray, Carol Clugston, Anna Dominiczak and Alex Alderton, Roberto Amato, Sonia Goncalves, Ewan Harrison, David K. Jackson, Ian Johnston, Dominic Kwiatkowski, Cor |

| | | | |
|---|---|---|---|
| | | for the COVID-19 Genomics UK (COG-UK) consortium | |
| EPI_ISL_533236, EPI_ISL_533237 | NHSGGC West of Scotland Specialist Virology Centre / MRC-University of Glasgow Centre for Virus Research | Wellcome Sanger Institute for the COVID-19 Genomics UK (COG-UK) consortium | Ana da Silva Filipe, Natasha Johnson, Kathy Smollett, Daniel Mair, Stephen Carmichael, Lily Tong, Jenna Nichols, Elihu Aranday-Cortes, Kirstyn Brunker, Yasmin Parr, Kyriaki Nomikou; Sarah McDonald, Marc Niebel, Patawee Asamaphan; Richard Ortc MacLean, Rory Gunson; Kathy Li, Natasha Jesudason, Rajiv Shah, James Shepherd, Antonia Ho, Alice Broos, Emma Thomson and Alex Alderton, Roberto Amato, Sonia Goncalves, Ewan Harrison, David K. Jackson, Ian Johnston, Dom |
| EPI_ISL_533240 | Lighthouse Lab in Glasgow | Wellcome Sanger Institute for the COVID-19 Genomics UK (COG-UK) consortium | Harper VanSteenhouse, Yumi Kasai, David Gray, Carol Clugston, Anna Dominiczak and Alex Alderton, Roberto Amato, Sonia Goncalves, Ewan Harrison, David K. Jackson, Ian Johnston, Dominic Kwiatkowski, Cor |
| EPI_ISL_533241 | NHSGGC West of Scotland Specialist Virology Centre / MRC-University of Glasgow Centre for Virus Research | Wellcome Sanger Institute for the COVID-19 Genomics UK (COG-UK) consortium | Ana da Silva Filipe, Natasha Johnson, Kathy Smollett, Daniel Mair, Stephen Carmichael, Lily Tong, Jenna Nichols, Elihu Aranday-Cortes, Kirstyn Brunker, Yasmin Parr, Kyriaki Nomikou; Sarah McDonald, Marc Niebel, Patawee Asamaphan; Richard Ortc MacLean, Rory Gunson; Kathy Li, Natasha Jesudason, Rajiv Shah, James Shepherd, Antonia Ho, Alice Broos, Emma Thomson and Alex Alderton, Roberto Amato, Sonia Goncalves, Ewan Harrison, David K. Jackson, Ian Johnston, Dom |
| EPI_ISL_533245, EPI_ISL_533247, EPI_ISL_533248, EPI_ISL_533251 | Lighthouse Lab in Glasgow | Wellcome Sanger Institute for the COVID-19 Genomics UK (COG-UK) consortium | Harper VanSteenhouse, Yumi Kasai, David Gray, Carol Clugston, Anna Dominiczak and Alex Alderton, Roberto Amato, Sonia Goncalves, Ewan Harrison, David K. Jackson, Ian Johnston, Dominic Kwiatkowski, Cor |
| EPI_ISL_533252, EPI_ISL_533256 | NHSGGC West of Scotland Specialist Virology Centre / MRC-University of Glasgow Centre for Virus Research | Wellcome Sanger Institute for the COVID-19 Genomics UK (COG-UK) consortium | Ana da Silva Filipe, Natasha Johnson, Kathy Smollett, Daniel Mair, Stephen Carmichael, Lily Tong, Jenna Nichols, Elihu Aranday-Cortes, Kirstyn Brunker, Yasmin Parr, Kyriaki Nomikou; Sarah McDonald, Marc Niebel, Patawee Asamaphan; Richard Ortc MacLean, Rory Gunson; Kathy Li, Natasha Jesudason, Rajiv Shah, James Shepherd, Antonia Ho, Alice Broos, Emma Thomson and Alex Alderton, Roberto Amato, Sonia Goncalves, Ewan Harrison, David K. Jackson, Ian Johnston, Dom |
| EPI_ISL_533260 | Lighthouse Lab in Glasgow | Wellcome Sanger Institute for the COVID-19 Genomics UK (COG-UK) consortium | Harper VanSteenhouse, Yumi Kasai, David Gray, Carol Clugston, Anna Dominiczak and Alex Alderton, Roberto Amato, Sonia Goncalves, Ewan Harrison, David K. Jackson, Ian Johnston, Dominic Kwiatkowski, Cor |
| EPI_ISL_533262 | NHSGGC West of Scotland Specialist Virology Centre / MRC-University of Glasgow Centre for Virus Research | Wellcome Sanger Institute for the COVID-19 Genomics UK (COG-UK) consortium | Ana da Silva Filipe, Natasha Johnson, Kathy Smollett, Daniel Mair, Stephen Carmichael, Lily Tong, Jenna Nichols, Elihu Aranday-Cortes, Kirstyn Brunker, Yasmin Parr, Kyriaki Nomikou; Sarah McDonald, Marc Niebel, Patawee Asamaphan; Richard Ortc MacLean, Rory Gunson; Kathy Li, Natasha Jesudason, Rajiv Shah, James Shepherd, Antonia Ho, Alice Broos, Emma Thomson and Alex Alderton, Roberto Amato, Sonia Goncalves, Ewan Harrison, David K. Jackson, Ian Johnston, Dom |
| EPI_ISL_533263, EPI_ISL_533265, EPI_ISL_533266, EPI_ISL_533267, EPI_ISL_533268 | Lighthouse Lab in Glasgow | Wellcome Sanger Institute for the COVID-19 Genomics UK (COG-UK) consortium | Harper VanSteenhouse, Yumi Kasai, David Gray, Carol Clugston, Anna Dominiczak and Alex Alderton, Roberto Amato, Sonia Goncalves, Ewan Harrison, David K. Jackson, Ian Johnston, Dominic Kwiatkowski, Cor |
| EPI_ISL_533271 | NHSGGC West of Scotland Specialist Virology Centre / MRC-University of Glasgow Centre for Virus Research | Wellcome Sanger Institute for the COVID-19 Genomics UK (COG-UK) consortium | Ana da Silva Filipe, Natasha Johnson, Kathy Smollett, Daniel Mair, Stephen Carmichael, Lily Tong, Jenna Nichols, Elihu Aranday-Cortes, Kirstyn Brunker, Yasmin Parr, Kyriaki Nomikou; Sarah McDonald, Marc Niebel, Patawee Asamaphan; Richard Ortc MacLean, Rory Gunson; Kathy Li, Natasha Jesudason, Rajiv Shah, James Shepherd, Antonia Ho, Alice Broos, Emma Thomson and Alex Alderton, Roberto Amato, Sonia Goncalves, Ewan Harrison, David K. Jackson, Ian Johnston, Dom |
| EPI_ISL_534237, EPI_ISL_534238 | Lanssjukhuset Kalmar | The Public Health Agency of Sweden | Anna-Malin Linde, Maria Lind Karlberg, Mattias Haukland, Reza Advani, Olov Svartstrom, Oskar Karlsson Lindsjo, Sandra Broddesson, Petra Edquist, Mia Brytting, Anna Risberg, Karin Tegmark |
| EPI_ISL_534754 | Liverpool Clinical Laboratories | COVID-19 Genomics UK (COG-UK) Consortium | Sam Haldenby, Anita Lucaci, Steve Paterson, Julian Hiscox, Alistair Darby, M Almsaud, A Alrezaihi, Muhannad Alruwaili, Stuart D Armstrong, Jones Benjamin, Eleanor G Bentley, Anu Chawla, Jordan J Clark, Angela Cowell, Richard Eccles, Isabel Garc Richard Gregory, Ximeng Han, Catherine Hartley, Margaret Hughes, Miren Iturriza-Gomara, James Johnson, L Luu, Jenifer Manson, Charlotte Nelson, Elaine O'Toole, Cassie Olateju, Rebekah Penrice-Randal , Lucille Rainbow, N.P Randle, Trevor Ian Swainston, Ecaterina Vamos, Joanne Watts, Mark Whitehead |
| EPI_ISL_536482, EPI_ISL_536522 | Instituto Nacional de Salud | Laboratorio de Infecciones Respiratorias Agudas | Eduardo Juscamayta Lopez, David Tarazona, Faviola Valdivia Guerrero, Nancy Rojas Serrano, Dennis Carhuaricra, Lenin Maturrano Hernandez, Ronnie Gavilan Chavez |
| EPI_ISL_537737, EPI_ISL_537738, EPI_ISL_537739, EPI_ISL_537740, EPI_ISL_537741, EPI_ISL_537742, EPI_ISL_537743, EPI_ISL_537744, EPI_ISL_537745, EPI_ISL_537746, EPI_ISL_537747, EPI_ISL_537748, EPI_ISL_537749, EPI_ISL_537750, EPI_ISL_537751, EPI_ISL_537752, EPI_ISL_537753, EPI_ISL_537754, EPI_I: EPI_ISL_537759, EPI_ISL_537760, EPI_ISL_537761, EPI_ISL_537762, EPI_ISL_537763, EPI_ISL_537764, EPI_ISL_537765, EPI_ISL_537766, EPI_ISL_537767, EPI_ISL_537768, EPI_ISL_537769, EPI_ISL_537770, EPI_ISL_537771, EPI_ISL_537772 | | | |
| see above | Servicio de Microbiología, Hospital Miguel Servet, Zaragoza | SeqCOVID-SPAIN consortium/IBV(CSIC) | Antonio Rezusta López, Alexander Tristancho Baró, Ana Milagro, Yolanda Gracia Grataloup, Nieves Martínez Cameo and SeqCOVID-SPAIN consortium |
| EPI_ISL_538317, EPI_ISL_538318, EPI_ISL_538319, EPI_ISL_538320 | Texas Department of State Health Services | Texas Department of State Health Services | Bonnie Oh, Rashmi Tuladhar, Jenny Zhang, Maliha Rahman, Anita Pokharel, Myong Koag, Chun Wang, Rachel Lee, Grace Kubin |
| EPI_ISL_538390, EPI_ISL_538391, EPI_ISL_538392, EPI_ISL_538393, EPI_ISL_538394, EPI_ISL_538395, EPI_ISL_538396, EPI_ISL_538397, EPI_ISL_538398, EPI_ISL_538399, EPI_ISL_538400, EPI_ISL_538401, EPI_ISL_538402, EPI_ISL_538403, EPI_ISL_538404, EPI_ISL_538405, EPI_ISL_538406, EPI_ISL_538407, EPI_I: EPI_ISL_538412, EPI_ISL_538413, EPI_ISL_538414, EPI_ISL_538415, EPI_ISL_538416, EPI_ISL_538417, EPI_ISL_538418, EPI_ISL_538419, EPI_ISL_538420, EPI_ISL_538421, EPI_ISL_538432, EPI_ISL_538433, EPI_ISL_538434 | | | |
| see above | Microbiology Division, South Carolina Department of Health and Environmental Control | Microbiology Division, South Carolina Department of Health and Environmental Control | Flores,H. |
| EPI_ISL_538499 | RS Lavallete Malang East Java | National Institute of Health Research and Development | Pawestri, HA; Subangkit; Puspa, KD; Nugraha, AA; Ikawati, HD; Pangesti, KNA; Soekarso, T; Paisal; Setiawaty,V. |
| EPI_ISL_538654, EPI_ISL_538655, EPI_ISL_538656, EPI_ISL_538657, EPI_ISL_538658, EPI_ISL_538659, EPI_ISL_538660, EPI_ISL_538661, EPI_ISL_538662, EPI_ISL_538663, EPI_ISL_538664, EPI_ISL_538665, EPI_ISL_538666, EPI_ISL_538668, EPI_ISL_538669, EPI_ISL_538670 | | | |
| see above | Servicio de Microbiología. Hospital General Universitario de Castellón | SeqCOVID-SPAIN consortium/IBV(CSIC) | Rosario Moreno, María Dolores Tirado and SeqCOVID-SPAIN consortium |
| EPI_ISL_539541, EPI_ISL_539542, EPI_ISL_539547, EPI_ISL_539548 | Hospital Clínic | Instituto de Salud Carlos III | Iglesias-Caballero, M. Molinero Calamita, M. González-Esguevillas, M. Camarero, S. Pozo, F. Casas, I. Jiménez, P. Jiménez, M. Zaballos, A. Monzón, S. Varona, S. Juliá, M. Cuesta, I, M.A M |
| EPI_ISL_539881 | Kungsbacka Narakut | The Public Health Agency of Sweden | Anna-Malin Linde, Maria Lind Karlberg, Oskar Karlsson Lindsjo, Olov Svartstrom, Mattias Haukland, Reza Advani, Sandra Broddesson, Anna Risberg, Theresa Enkirch, Mia Brytting, Karin Tegma |
| EPI_ISL_539883 | Omtanken Grimmered | The Public Health Agency of Sweden | Anna-Malin Linde, Maria Lind Karlberg, Oskar Karlsson Lindsjo, Olov Svartstrom, Mattias Haukland, Reza Advani, Sandra Broddesson, Anna Risberg, Theresa Enkirch, Mia Brytting, Karin Tegma |
| EPI_ISL_539896 | PHE South West Regional Laboratory, National Infection | Wellcome Sanger Institute for the COVID-19 Genomics | Stephanie Hutchings, Hannah Pymont, Dr Peter Muir, Barry Vipond, Rich Hopes; and Alex Alderton, Roberto Amato, Sonia Goncalves, Ewan Harrison, David K. Jackson, Ian Johnston, Dominic Kwiatkowski, Cordelia Langford, John Sillitoe on behalf |

| | | | |
|---|---|---|---|
| | Service | UK (COG-UK) consortium | |
| EPI_ISL_541002, EPI_ISL_541003, EPI_ISL_541004 | Health and Environmental Research Institute of Gwangju Metropolitan city | Health and Environmental Research Institute of Gwangju Metropolitan city | Min Ji Kim, Ji-eun Lee |
| EPI_ISL_541009, EPI_ISL_541010, EPI_ISL_541011, EPI_ISL_541012, EPI_ISL_541013, EPI_ISL_541014, EPI_ISL_541015, EPI_ISL_541016, EPI_ISL_541017, EPI_ISL_541018, EPI_ISL_541019 | | | |
| see above | Servicio de Microbiología, Hospital Miguel Servet, Zaragoza | SeqCOVID-SPAIN consortium/Institute of Biomedicine of Valencia, IBV-CSIC | Antonio Rezusta López, Alexander Tristancho Baró, Ana Milagro, Yolanda Gracia Grataloup, Nieves Martínez Cameo and SeqCOVID-SPAIN consortium |
| EPI_ISL_541075, EPI_ISL_541076 | Hospital de la Santa Creu i Sant Pau. Servicio de Microbiología | SeqCOVID-SPAIN consortium/Institute of Biomedicine of Valencia, IBV-CSIC | Ferran Navarro, Núria Rabella, Elisenda Miró and SeqCOVID-SPAIN consortium |
| EPI_ISL_541243, EPI_ISL_541244, EPI_ISL_541245, EPI_ISL_541246, EPI_ISL_541247 | Florida Bureau of Public Health Laboratories, Florida Department of Health | Florida Bureau of Public Health Laboratories, Florida Department of Health | Schmedes,S., Blanton,J. |
| EPI_ISL_541663, EPI_ISL_541671, EPI_ISL_541672, EPI_ISL_541673, EPI_ISL_541674, EPI_ISL_541675, EPI_ISL_541676, EPI_ISL_541677, EPI_ISL_541678, EPI_ISL_541679 | Microbiology Division, South Carolina Department of Health and Environmental Control | Microbiology Division, South Carolina Department of Health and Environmental Control | Flores,H. |
| EPI_ISL_541741, EPI_ISL_541742, EPI_ISL_541743, EPI_ISL_541744, EPI_ISL_541745, EPI_ISL_541746, EPI_ISL_541747 | National Institute of Virology, NIV Influenza | National Institute of Virology, NIV Influenza | Potdar V |
| EPI_ISL_542023 | New Mexico Department of Health Scientific Laboratory | New Mexico Department of Health Scientific Laboratory | Ellie Johnson, Anastacia Griego-Fisher, D'Eldra Malone |
| EPI_ISL_542479, EPI_ISL_542480, EPI_ISL_542481, EPI_ISL_542482, EPI_ISL_542483, EPI_ISL_542484, EPI_ISL_542485 | Texas Department of State Health Services | Texas Department of State Health Services | Rashmi Tuladhar, Bonnie Oh,Jenny Zhang, Maliha Rahman, Anita Pokharel, Myong Koag, Chun Wang, Rachel Lee, Grace Kubin |
| EPI_ISL_543772, EPI_ISL_543784, EPI_ISL_544269, EPI_ISL_544305, EPI_ISL_544393, EPI_ISL_544454, EPI_ISL_544455, EPI_ISL_544458, EPI_ISL_544462, EPI_ISL_544464, EPI_ISL_544467, EPI_ISL_544468, EPI_ISL_544469, EPI_ISL_544470, EPI_ISL_544472, EPI_ISL_544474, EPI_ISL_544475, EPI_ISL_544476, EPI_I[illegible]<br>EPI_ISL_544482, EPI_ISL_544484, EPI_ISL_544486, EPI_ISL_544487, EPI_ISL_544488, EPI_ISL_544489, EPI_ISL_544491, EPI_ISL_544492, EPI_ISL_544494, EPI_ISL_544495, EPI_ISL_544497, EPI_ISL_544501, EPI_ISL_544507, EPI_ISL_544510, EPI_ISL_544511, EPI_ISL_544515, EPI_ISL_544516, EPI_ISL_544518, EPI_I[illegible]<br>EPI_ISL_544525, EPI_ISL_544526, EPI_ISL_544527, EPI_ISL_544530, EPI_ISL_544531, EPI_ISL_544533, EPI_ISL_544534, EPI_ISL_544535, EPI_ISL_544536, EPI_ISL_544537, EPI_ISL_544539, EPI_ISL_544543, EPI_ISL_544546, EPI_ISL_544552, EPI_ISL_544555, EPI_ISL_544557, EPI_ISL_544558, EPI_ISL_544560, EPI_I[illegible]<br>EPI_ISL_544574, EPI_ISL_544575, EPI_ISL_544577, EPI_ISL_544580, EPI_ISL_544583, EPI_ISL_544589, EPI_ISL_544590, EPI_ISL_544591, EPI_ISL_544593, EPI_ISL_544594, EPI_ISL_544600, EPI_ISL_544602, EPI_ISL_544603, EPI_ISL_544605, EPI_ISL_544607, EPI_ISL_544610, EPI_ISL_544612, EPI_ISL_544613, EPI_I[illegible]<br>EPI_ISL_544620, EPI_ISL_544621, EPI_ISL_544623, EPI_ISL_544624, EPI_ISL_544625, EPI_ISL_544626, EPI_ISL_544628, EPI_ISL_544631, EPI_ISL_544633, EPI_ISL_544634, EPI_ISL_544636, EPI_ISL_544637, EPI_ISL_544639, EPI_ISL_544641, EPI_ISL_544645, EPI_ISL_544647, EPI_ISL_544649, EPI_ISL_544651, EPI_I[illegible]<br>EPI_ISL_544661, EPI_ISL_544665, EPI_ISL_544667, EPI_ISL_544668, EPI_ISL_544670, EPI_ISL_544672, EPI_ISL_544673, EPI_ISL_544674, EPI_ISL_544675, EPI_ISL_544676, EPI_ISL_544677, EPI_ISL_544679, EPI_ISL_544680, EPI_ISL_544681, EPI_ISL_544682, EPI_ISL_544683, EPI_ISL_544684, EPI_ISL_544685, EPI_I[illegible]<br>EPI_ISL_544691, EPI_ISL_544692, EPI_ISL_544693, EPI_ISL_544694, EPI_ISL_544695, EPI_ISL_544697, EPI_ISL_544698, EPI_ISL_544699, EPI_ISL_544700, EPI_ISL_544701, EPI_ISL_544702, EPI_ISL_544703, EPI_ISL_544704, EPI_ISL_544706, EPI_ISL_544707, EPI_ISL_544708, EPI_ISL_544709, EPI_ISL_544710, EPI_I[illegible]<br>EPI_ISL_544716, EPI_ISL_544717, EPI_ISL_544718, EPI_ISL_544719, EPI_ISL_544720, EPI_ISL_544721, EPI_ISL_544722, EPI_ISL_544723, EPI_ISL_544724, EPI_ISL_544725, EPI_ISL_544726, EPI_ISL_544727, EPI_ISL_544728, EPI_ISL_544729, EPI_ISL_544730, EPI_ISL_544731, EPI_ISL_544732, EPI_ISL_544733, EPI_I[illegible]<br>EPI_ISL_544738, EPI_ISL_544739, EPI_ISL_544740, EPI_ISL_544741, EPI_ISL_544742, EPI_ISL_544743, EPI_ISL_544744, EPI_ISL_544745, EPI_ISL_544746, EPI_ISL_544747, EPI_ISL_544748, EPI_ISL_544749, EPI_ISL_544750, EPI_ISL_544751, EPI_ISL_544752, EPI_ISL_544753, EPI_ISL_544754, EPI_ISL_544755, EPI_I[illegible]<br>EPI_ISL_544760, EPI_ISL_544761, EPI_ISL_544762, EPI_ISL_544763, EPI_ISL_544764, EPI_ISL_544765, EPI_ISL_544766, EPI_ISL_544767, EPI_ISL_544768, EPI_ISL_544769, EPI_ISL_544770, EPI_ISL_544771, EPI_ISL_544772, EPI_ISL_544773, EPI_ISL_544774, EPI_ISL_544775, EPI_ISL_544776, EPI_ISL_544777, EPI_I[illegible]<br>EPI_ISL_544782, EPI_ISL_544783, EPI_ISL_544784, EPI_ISL_544785, EPI_ISL_544786, EPI_ISL_544787, EPI_ISL_544788, EPI_ISL_544789, EPI_ISL_544790, EPI_ISL_544791, EPI_ISL_544792, EPI_ISL_544793, EPI_ISL_544794, EPI_ISL_544795, EPI_ISL_544797, EPI_ISL_544798, EPI_ISL_544799, EPI_ISL_544800, EPI_I[illegible]<br>EPI_ISL_544806, EPI_ISL_544807, EPI_ISL_544808, EPI_ISL_544809, EPI_ISL_544810, EPI_ISL_544811, EPI_ISL_544813, EPI_ISL_544814, EPI_ISL_544815, EPI_ISL_544816, EPI_ISL_544817, EPI_ISL_544818, EPI_ISL_544819, EPI_ISL_544820, EPI_ISL_544821, EPI_ISL_544822, EPI_ISL_544824, EPI_ISL_544825, EPI_I[illegible]<br>EPI_ISL_544830, EPI_ISL_544831, EPI_ISL_544832, EPI_ISL_544834, EPI_ISL_544835, EPI_ISL_544836, EPI_ISL_544837, EPI_ISL_544838, EPI_ISL_544839, EPI_ISL_544840, EPI_ISL_544841, EPI_ISL_544842, EPI_ISL_544843, EPI_ISL_544844, EPI_ISL_544845, EPI_ISL_544846, EPI_ISL_544847, EPI_ISL_544848, EPI_I[illegible]<br>EPI_ISL_544853, EPI_ISL_544854, EPI_ISL_544855, EPI_ISL_544856, EPI_ISL_544857, EPI_ISL_544858, EPI_ISL_544859, EPI_ISL_544860, EPI_ISL_544861, EPI_ISL_544862, EPI_ISL_544863, EPI_ISL_544864, EPI_ISL_544865, EPI_ISL_544866, EPI_ISL_544867, EPI_ISL_544869, EPI_ISL_544870, EPI_ISL_544871, EPI_I[illegible]<br>EPI_ISL_544877, EPI_ISL_544878, EPI_ISL_544879, EPI_ISL_544880, EPI_ISL_544881, EPI_ISL_544884, EPI_ISL_544886, EPI_ISL_544888, EPI_ISL_544889, EPI_ISL_544891, EPI_ISL_544893, EPI_ISL_544895, EPI_ISL_544896, EPI_ISL_544899, EPI_ISL_544900, EPI_ISL_544902, EPI_ISL_544906, EPI_ISL_544907, EPI_I[illegible]<br>EPI_ISL_544923, EPI_ISL_544924, EPI_ISL_544929, EPI_ISL_544935, EPI_ISL_544936, EPI_ISL_544937, EPI_ISL_544939, EPI_ISL_544940, EPI_ISL_544941, EPI_ISL_544942, EPI_ISL_544943, EPI_ISL_544945, EPI_ISL_544946, EPI_ISL_544947, EPI_ISL_544949, EPI_ISL_544952, EPI_ISL_544953, EPI_ISL_544954, EPI_I[illegible]<br>EPI_ISL_545585, EPI_ISL_545586, EPI_ISL_545587, EPI_ISL_545588, EPI_ISL_545589, EPI_ISL_545591, EPI_ISL_545592, EPI_ISL_545593, EPI_ISL_545594, EPI_ISL_545595, EPI_ISL_545596, EPI_ISL_545597, EPI_ISL_545598, EPI_ISL_545599, EPI_ISL_545600, EPI_ISL_545602, EPI_ISL_545604, EPI_ISL_545605, EPI_I[illegible]<br>EPI_ISL_545614, EPI_ISL_545615, EPI_ISL_545616, EPI_ISL_545617, EPI_ISL_545618, EPI_ISL_545619, EPI_ISL_545620, EPI_ISL_545621, EPI_ISL_545622, EPI_ISL_545623, EPI_ISL_545624, EPI_ISL_545625, EPI_ISL_545626, EPI_ISL_545628, EPI_ISL_545629, EPI_ISL_545630, EPI_ISL_545631, EPI_ISL_545632, EPI_I[illegible]<br>EPI_ISL_545638, EPI_ISL_545642, EPI_ISL_545644, EPI_ISL_545646, EPI_ISL_545649, EPI_ISL_545650, EPI_ISL_545651, EPI_ISL_545654, EPI_ISL_545655, EPI_ISL_545657, EPI_ISL_545660, EPI_ISL_545661, EPI_ISL_545662, EPI_ISL_545663, EPI_ISL_545666, EPI_ISL_545669, EPI_ISL_545670, EPI_ISL_545671, EPI_I[illegible]<br>EPI_ISL_545678, EPI_ISL_545680, EPI_ISL_545683, EPI_ISL_545684, EPI_ISL_545685, EPI_ISL_545686, EPI_ISL_545687, EPI_ISL_545688, EPI_ISL_545689, EPI_ISL_545693, EPI_ISL_545694, EPI_ISL_545695, EPI_ISL_545696, EPI_ISL_545697, EPI_ISL_545698, EPI_ISL_545700, EPI_ISL_545701, EPI_ISL_545702, EPI_I[illegible]<br>EPI_ISL_545712, EPI_ISL_545713, EPI_ISL_545715, EPI_ISL_545716, EPI_ISL_545717, EPI_ISL_545718, EPI_ISL_545720, EPI_ISL_545721, EPI_ISL_545723, EPI_ISL_545725, EPI_ISL_545727, EPI_ISL_545728, EPI_ISL_545730, EPI_ISL_545731, EPI_ISL_545733, EPI_ISL_545734, EPI_ISL_545735, EPI_ISL_545736, EPI_I[illegible]<br>EPI_ISL_545745, EPI_ISL_545746, EPI_ISL_545747, EPI_ISL_545748, EPI_ISL_545749, EPI_ISL_545750, EPI_ISL_545751, EPI_ISL_545753, EPI_ISL_545756, EPI_ISL_545757, EPI_ISL_545758, EPI_ISL_545759, EPI_ISL_545760, EPI_ISL_545761, EPI_ISL_545762, EPI_ISL_545764, EPI_ISL_545768, EPI_ISL_545778, EPI_I[illegible]<br>EPI_ISL_546002, EPI_ISL_546003, EPI_ISL_546004, EPI_ISL_546005, EPI_ISL_546006, EPI_ISL_546007, EPI_ISL_546008, EPI_ISL_546009, EPI_ISL_546010, EPI_ISL_546011, EPI_ISL_546012, EPI_ISL_546013, EPI_ISL_546014, EPI_ISL_546015, EPI_ISL_546016, EPI_ISL_546017, EPI_ISL_546018, EPI_ISL_546020, EPI_I[illegible]<br>EPI_ISL_546025, EPI_ISL_546026, EPI_ISL_546027, EPI_ISL_546028, EPI_ISL_546029, EPI_ISL_546030, EPI_ISL_546031, EPI_ISL_546032, EPI_ISL_546033, EPI_ISL_546034, EPI_ISL_546035, EPI_ISL_546036, EPI_ISL_546037, EPI_ISL_546038, EPI_ISL_546039, EPI_ISL_546040, EPI_ISL_546041, EPI_ISL_546042, EPI_I[illegible]<br>EPI_ISL_546047, EPI_ISL_546048, EPI_ISL_546049, EPI_ISL_546050, EPI_ISL_546051, EPI_ISL_546052, EPI_ISL_546053, EPI_ISL_546054, EPI_ISL_546055, EPI_ISL_546056, EPI_ISL_546057, EPI_ISL_546058, EPI_ISL_546059, EPI_ISL_546060, EPI_ISL_546061, EPI_ISL_546062, EPI_ISL_546063, EPI_ISL_546064, EPI_I[illegible]<br>EPI_ISL_546071, EPI_ISL_546072, EPI_ISL_546073, EPI_ISL_546074, EPI_ISL_546077, EPI_ISL_546078, EPI_ISL_546079, EPI_ISL_546081, EPI_ISL_546082, EPI_ISL_546083, EPI_ISL_546084, EPI_ISL_546085, EPI_ISL_546086, EPI_ISL_546087, EPI_ISL_546088, EPI_ISL_546089, EPI_ISL_546090, EPI_ISL_546092, EPI_I[illegible]<br>EPI_ISL_546097, EPI_ISL_546098, EPI_ISL_546099, EPI_ISL_546100, EPI_ISL_546101, EPI_ISL_546102, EPI_ISL_546103, EPI_ISL_546104, EPI_ISL_546105, EPI_ISL_546106, EPI_ISL_546107, EPI_ISL_546108, EPI_ISL_546109, EPI_ISL_546110, EPI_ISL_546111, EPI_ISL_546112, EPI_ISL_546113, EPI_ISL_546114, EPI_I[illegible]<br>EPI_ISL_546119, EPI_ISL_546121, EPI_ISL_546122, EPI_ISL_546123, EPI_ISL_546124, EPI_ISL_546125, EPI_ISL_546126, EPI_ISL_546127, EPI_ISL_546128, EPI_ISL_546131, EPI_ISL_546132, EPI_ISL_546133, EPI_ISL_546134, EPI_ISL_546135, EPI_ISL_546136, EPI_ISL_546137, EPI_ISL_546138, EPI_ISL_546139, EPI_I[illegible]<br>EPI_ISL_546144, EPI_ISL_546145, EPI_ISL_546146, EPI_ISL_546147, EPI_ISL_546148, EPI_ISL_546149, EPI_ISL_546150, EPI_ISL_546151, EPI_ISL_546152, EPI_ISL_546153, EPI_ISL_546154, EPI_ISL_546155, EPI_ISL_546156, EPI_ISL_546157, EPI_ISL_546158, EPI_ISL_546159, EPI_ISL_546160, EPI_ISL_546161, EPI_I[illegible]<br>EPI_ISL_546166, EPI_ISL_546167, EPI_ISL_546168, EPI_ISL_546169, EPI_ISL_546170, EPI_ISL_546171, EPI_ISL_546172, EPI_ISL_546173, EPI_ISL_546174, EPI_ISL_546175, EPI_ISL_546176, EPI_ISL_546177, EPI_ISL_546178, EPI_ISL_546179, EPI_ISL_546180, EPI_ISL_546181, EPI_ISL_546182, EPI_ISL_546183, EPI_I[illegible]<br>EPI_ISL_546188, EPI_ISL_546189, EPI_ISL_546190, EPI_ISL_546191, EPI_ISL_546192, EPI_ISL_546193, EPI_ISL_546194, EPI_ISL_546195, EPI_ISL_546196, EPI_ISL_546197, EPI_ISL_546199, EPI_ISL_546200, EPI_ISL_546201, EPI_ISL_546202, EPI_ISL_546203, EPI_ISL_546204, EPI_ISL_546205, EPI_ISL_546206, EPI_I[illegible]<br>EPI_ISL_546211, EPI_ISL_546212, EPI_ISL_546213, EPI_ISL_546215, EPI_ISL_546216, EPI_ISL_546217, EPI_ISL_546218, EPI_ISL_546219, EPI_ISL_546220, EPI_ISL_546221, EPI_ISL_546222, EPI_ISL_546223, EPI_ISL_546224, EPI_ISL_546225, EPI_ISL_546226, EPI_ISL_546227, EPI_ISL_546228, EPI_ISL_546229, EPI_I[illegible]<br>EPI_ISL_546234, EPI_ISL_546235, EPI_ISL_546236, EPI_ISL_546237, EPI_ISL_546238, EPI_ISL_546239, EPI_ISL_546240, EPI_ISL_546241, EPI_ISL_546242, EPI_ISL_546243, EPI_ISL_546244, EPI_ISL_546245, EPI_ISL_546246, EPI_ISL_546247, EPI_ISL_546248, EPI_ISL_546249, EPI_ISL_546250, EPI_ISL_546251, EPI_I[illegible]<br>EPI_ISL_546256, EPI_ISL_546257, EPI_ISL_546258, EPI_ISL_546259, EPI_ISL_546260, EPI_ISL_546261, EPI_ISL_546262, EPI_ISL_546263, EPI_ISL_546264, EPI_ISL_546265, EPI_ISL_546266, EPI_ISL_546267, EPI_ISL_546268, EPI_ISL_546269, EPI_ISL_546270, EPI_ISL_546271, EPI_ISL_546272, EPI_ISL_546273, EPI_I[illegible]<br>EPI_ISL_546278, EPI_ISL_546279, EPI_ISL_546281, EPI_ISL_546282, EPI_ISL_546283, EPI_ISL_546284, EPI_ISL_546285, EPI_ISL_546286, EPI_ISL_546287, EPI_ISL_546289, EPI_ISL_546290, EPI_ISL_546291, EPI_ISL_546292, EPI_ISL_546293, EPI_ISL_546294, EPI_ISL_546296, EPI_ISL_546297, EPI_ISL_546298, EPI_I[illegible]<br>EPI_ISL_546303, EPI_ISL_546304, EPI_ISL_546306, EPI_ISL_546307, EPI_ISL_546308, EPI_ISL_546309, EPI_ISL_546310, EPI_ISL_546311, EPI_ISL_546312, EPI_ISL_546313, EPI_ISL_546314, EPI_ISL_546316, EPI_ISL_546317, EPI_ISL_546318, EPI_ISL_546319, EPI_ISL_546320, EPI_ISL_546321, EPI_ISL_546322, EPI_I[illegible]<br>EPI_ISL_546328, EPI_ISL_546329, EPI_ISL_546330, EPI_ISL_546331, EPI_ISL_546332, EPI_ISL_546333, EPI_ISL_546335, EPI_ISL_546336, EPI_ISL_546337, EPI_ISL_546338, EPI_ISL_546339, EPI_ISL_546340, EPI_ISL_546341, EPI_ISL_546343, EPI_ISL_546344, EPI_ISL_546345, EPI_ISL_546346, EPI_ISL_546347, EPI_I[illegible]<br>EPI_ISL_546353, EPI_ISL_546354, EPI_ISL_546355, EPI_ISL_546356, EPI_ISL_546357, EPI_ISL_546358, EPI_ISL_546359, EPI_ISL_546360, EPI_ISL_546361, EPI_ISL_546362, EPI_ISL_546363, EPI_ISL_546364, EPI_ISL_546365, EPI_ISL_546366, EPI_ISL_546367, EPI_ISL_546369, EPI_ISL_546370, EPI_ISL_546371, EPI_I[illegible] | | | |

EPI_ISL_546376, EPI_ISL_546377, EPI_ISL_546378, EPI_ISL_546379, EPI_ISL_546380, EPI_ISL_546381, EPI_ISL_546382, EPI_ISL_546383, EPI_ISL_546384, EPI_ISL_546385, EPI_ISL_546387, EPI_ISL_546388, EPI_ISL_546389, EPI_ISL_546390, EPI_ISL_546391, EPI_ISL_546392, EPI_ISL_546393, EPI_ISL_546394, EPI_I[illegible]
EPI_ISL_546399, EPI_ISL_546400, EPI_ISL_546401, EPI_ISL_546402, EPI_ISL_546403, EPI_ISL_546404, EPI_ISL_546405, EPI_ISL_546406, EPI_ISL_546407, EPI_ISL_546408, EPI_ISL_546409, EPI_ISL_546410, EPI_ISL_546411, EPI_ISL_546412, EPI_ISL_546414, EPI_ISL_546415, EPI_ISL_546416, EPI_ISL_546417, EPI_I[illegible]
EPI_ISL_546422, EPI_ISL_546423, EPI_ISL_546424, EPI_ISL_546425, EPI_ISL_546426, EPI_ISL_546427, EPI_ISL_546428, EPI_ISL_546429, EPI_ISL_546430, EPI_ISL_546431, EPI_ISL_546432, EPI_ISL_546433, EPI_ISL_546434, EPI_ISL_546435, EPI_ISL_546437, EPI_ISL_546438, EPI_ISL_546440, EPI_ISL_546441, EPI_I[illegible]
EPI_ISL_546446, EPI_ISL_546447, EPI_ISL_546448, EPI_ISL_546449, EPI_ISL_546450, EPI_ISL_546451, EPI_ISL_546452, EPI_ISL_546453, EPI_ISL_546454, EPI_ISL_546455, EPI_ISL_546456, EPI_ISL_546458, EPI_ISL_546459, EPI_ISL_546460, EPI_ISL_546461, EPI_ISL_546463, EPI_ISL_546464, EPI_ISL_546465, EPI_I[illegible]
EPI_ISL_546470, EPI_ISL_546472, EPI_ISL_546473, EPI_ISL_546474, EPI_ISL_546475, EPI_ISL_546476, EPI_ISL_546477, EPI_ISL_546478, EPI_ISL_546479, EPI_ISL_546480, EPI_ISL_546481, EPI_ISL_546482, EPI_ISL_546483, EPI_ISL_546484, EPI_ISL_546485, EPI_ISL_546486, EPI_ISL_546487, EPI_ISL_546488, EPI_I[illegible]
EPI_ISL_546493, EPI_ISL_546494, EPI_ISL_546496, EPI_ISL_546497, EPI_ISL_546498, EPI_ISL_546499, EPI_ISL_546500, EPI_ISL_546501, EPI_ISL_546502, EPI_ISL_546503, EPI_ISL_546504, EPI_ISL_546505, EPI_ISL_546506, EPI_ISL_546507, EPI_ISL_546508, EPI_ISL_546509, EPI_ISL_546510, EPI_ISL_546512, EPI_I[illegible]
EPI_ISL_546517, EPI_ISL_546518, EPI_ISL_546519, EPI_ISL_546520, EPI_ISL_546521, EPI_ISL_546522, EPI_ISL_546523, EPI_ISL_546524, EPI_ISL_546525, EPI_ISL_546526, EPI_ISL_546527, EPI_ISL_546528, EPI_ISL_546529, EPI_ISL_546530, EPI_ISL_546531, EPI_ISL_546532, EPI_ISL_546533, EPI_ISL_546534, EPI_I[illegible]
EPI_ISL_546539, EPI_ISL_546540, EPI_ISL_546541, EPI_ISL_546542, EPI_ISL_546543, EPI_ISL_546544, EPI_ISL_546545, EPI_ISL_546546, EPI_ISL_546547, EPI_ISL_546548, EPI_ISL_546549, EPI_ISL_546550, EPI_ISL_546551, EPI_ISL_546552, EPI_ISL_546553, EPI_ISL_546554, EPI_ISL_546556, EPI_ISL_546557, EPI_I[illegible]
EPI_ISL_546562, EPI_ISL_546563, EPI_ISL_546564, EPI_ISL_546566, EPI_ISL_546567, EPI_ISL_546568, EPI_ISL_546569, EPI_ISL_546570, EPI_ISL_546571, EPI_ISL_546572, EPI_ISL_546573, EPI_ISL_546574, EPI_ISL_546576, EPI_ISL_546577, EPI_ISL_546578, EPI_ISL_546579, EPI_ISL_546580, EPI_ISL_546581, EPI_I[illegible]
EPI_ISL_546586, EPI_ISL_546587, EPI_ISL_546588, EPI_ISL_546589, EPI_ISL_546590, EPI_ISL_546591, EPI_ISL_546592, EPI_ISL_546594, EPI_ISL_546595, EPI_ISL_546596, EPI_ISL_546597, EPI_ISL_546598, EPI_ISL_546599, EPI_ISL_546600, EPI_ISL_546601, EPI_ISL_546602, EPI_ISL_546603, EPI_ISL_546604, EPI_I[illegible]
EPI_ISL_546609, EPI_ISL_546610, EPI_ISL_546611, EPI_ISL_546612, EPI_ISL_546613, EPI_ISL_546614, EPI_ISL_546615, EPI_ISL_546616, EPI_ISL_546617, EPI_ISL_546618, EPI_ISL_546619, EPI_ISL_546620, EPI_ISL_546621, EPI_ISL_546622, EPI_ISL_546623, EPI_ISL_546624, EPI_ISL_546625, EPI_ISL_546626, EPI_I[illegible]
EPI_ISL_546631, EPI_ISL_546632, EPI_ISL_546633, EPI_ISL_546634, EPI_ISL_546635, EPI_ISL_546636, EPI_ISL_546637, EPI_ISL_546638, EPI_ISL_546639, EPI_ISL_546640, EPI_ISL_546641, EPI_ISL_546642, EPI_ISL_546643, EPI_ISL_546644, EPI_ISL_546645, EPI_ISL_546646, EPI_ISL_546647, EPI_ISL_546648, EPI_I[illegible]
EPI_ISL_546653, EPI_ISL_546654, EPI_ISL_546655, EPI_ISL_546657, EPI_ISL_546658, EPI_ISL_546659, EPI_ISL_546660, EPI_ISL_546661, EPI_ISL_546662, EPI_ISL_546663, EPI_ISL_546664, EPI_ISL_546665, EPI_ISL_546666, EPI_ISL_546667, EPI_ISL_546668, EPI_ISL_546669, EPI_ISL_546670, EPI_ISL_546671, EPI_I[illegible]
EPI_ISL_546676, EPI_ISL_546677, EPI_ISL_546679, EPI_ISL_546680, EPI_ISL_546681, EPI_ISL_546682, EPI_ISL_546683, EPI_ISL_546684, EPI_ISL_546685, EPI_ISL_546686, EPI_ISL_546687, EPI_ISL_546688, EPI_ISL_546689, EPI_ISL_546690, EPI_ISL_546691, EPI_ISL_546692, EPI_ISL_546693, EPI_ISL_546694, EPI_I[illegible]
EPI_ISL_546699, EPI_ISL_546700, EPI_ISL_546701, EPI_ISL_546702, EPI_ISL_546703, EPI_ISL_546704, EPI_ISL_546706, EPI_ISL_546707, EPI_ISL_546708, EPI_ISL_546709, EPI_ISL_546710, EPI_ISL_546711, EPI_ISL_546712, EPI_ISL_546713, EPI_ISL_546714, EPI_ISL_546715, EPI_ISL_546716, EPI_ISL_546717, EPI_I[illegible]
EPI_ISL_546723, EPI_ISL_546724, EPI_ISL_546725, EPI_ISL_546726, EPI_ISL_546727, EPI_ISL_546729, EPI_ISL_546730, EPI_ISL_546731, EPI_ISL_546732, EPI_ISL_546733, EPI_ISL_546734, EPI_ISL_546735, EPI_ISL_546736, EPI_ISL_546737, EPI_ISL_546739, EPI_ISL_546740, EPI_ISL_546741, EPI_ISL_546742, EPI_I[illegible]
EPI_ISL_546749, EPI_ISL_546750, EPI_ISL_546751, EPI_ISL_546752, EPI_ISL_546754, EPI_ISL_546755, EPI_ISL_546756, EPI_ISL_546757, EPI_ISL_546758, EPI_ISL_546759, EPI_ISL_546760, EPI_ISL_546761, EPI_ISL_546762, EPI_ISL_546763, EPI_ISL_546764, EPI_ISL_546765, EPI_ISL_546766, EPI_ISL_546767, EPI_I[illegible]
EPI_ISL_546773, EPI_ISL_546774, EPI_ISL_546776, EPI_ISL_546777, EPI_ISL_546778, EPI_ISL_546779, EPI_ISL_546780, EPI_ISL_546781, EPI_ISL_546782, EPI_ISL_546783, EPI_ISL_546784, EPI_ISL_546785, EPI_ISL_546787, EPI_ISL_546788, EPI_ISL_546789, EPI_ISL_546790, EPI_ISL_546791, EPI_ISL_546792, EPI_I[illegible]
EPI_ISL_546798, EPI_ISL_546799, EPI_ISL_546800, EPI_ISL_546801, EPI_ISL_546802, EPI_ISL_546803, EPI_ISL_546804, EPI_ISL_546805, EPI_ISL_546806, EPI_ISL_546807, EPI_ISL_546808, EPI_ISL_546809, EPI_ISL_546811, EPI_ISL_546812, EPI_ISL_546813, EPI_ISL_546814, EPI_ISL_546815, EPI_ISL_546816, EPI_I[illegible]
EPI_ISL_546821, EPI_ISL_546822, EPI_ISL_546823, EPI_ISL_546825, EPI_ISL_546826, EPI_ISL_546827, EPI_ISL_546828, EPI_ISL_546829, EPI_ISL_546830, EPI_ISL_546831, EPI_ISL_546832, EPI_ISL_546833, EPI_ISL_546835, EPI_ISL_546836, EPI_ISL_546837, EPI_ISL_546838, EPI_ISL_546839, EPI_ISL_546840, EPI_I[illegible]
EPI_ISL_546846, EPI_ISL_546847, EPI_ISL_546848, EPI_ISL_546849, EPI_ISL_546850, EPI_ISL_546851, EPI_ISL_546852, EPI_ISL_546853, EPI_ISL_546854, EPI_ISL_546855, EPI_ISL_546856, EPI_ISL_546857, EPI_ISL_546858, EPI_ISL_546859, EPI_ISL_546860, EPI_ISL_546861, EPI_ISL_546862, EPI_ISL_546863, EPI_I[illegible]
EPI_ISL_546869, EPI_ISL_546870, EPI_ISL_546871, EPI_ISL_546872, EPI_ISL_546873, EPI_ISL_546874, EPI_ISL_546875, EPI_ISL_546876, EPI_ISL_546877, EPI_ISL_546878, EPI_ISL_546879, EPI_ISL_546880, EPI_ISL_546881, EPI_ISL_546882, EPI_ISL_546883, EPI_ISL_546884, EPI_ISL_546885, EPI_ISL_546886, EPI_I[illegible]
EPI_ISL_546891, EPI_ISL_546892, EPI_ISL_546893, EPI_ISL_546894, EPI_ISL_546895, EPI_ISL_546896, EPI_ISL_546897, EPI_ISL_546898, EPI_ISL_546899, EPI_ISL_546900, EPI_ISL_546901, EPI_ISL_546902, EPI_ISL_546903, EPI_ISL_546904, EPI_ISL_546905, EPI_ISL_546906, EPI_ISL_546907, EPI_ISL_546908, EPI_I[illegible]
EPI_ISL_546914, EPI_ISL_546915, EPI_ISL_546916, EPI_ISL_546917, EPI_ISL_546918, EPI_ISL_546919, EPI_ISL_546920, EPI_ISL_546921, EPI_ISL_546922, EPI_ISL_546923, EPI_ISL_546924, EPI_ISL_546925, EPI_ISL_546926, EPI_ISL_546927, EPI_ISL_546928, EPI_ISL_546929, EPI_ISL_546930, EPI_ISL_546931, EPI_I[illegible]
EPI_ISL_546939, EPI_ISL_546941, EPI_ISL_546942, EPI_ISL_546943, EPI_ISL_546945, EPI_ISL_546946, EPI_ISL_546947, EPI_ISL_546948, EPI_ISL_546950, EPI_ISL_546951, EPI_ISL_546953, EPI_ISL_546954, EPI_ISL_546956, EPI_ISL_546957, EPI_ISL_546958, EPI_ISL_546961, EPI_ISL_546963, EPI_ISL_546968, EPI_I[illegible]
EPI_ISL_546975, EPI_ISL_546976, EPI_ISL_546977, EPI_ISL_546978, EPI_ISL_546979, EPI_ISL_546980, EPI_ISL_546981, EPI_ISL_546982, EPI_ISL_546984, EPI_ISL_546986, EPI_ISL_546987, EPI_ISL_546988, EPI_ISL_546990, EPI_ISL_546992, EPI_ISL_546993, EPI_ISL_546994, EPI_ISL_546995, EPI_ISL_546997, EPI_I[illegible]
EPI_ISL_547004, EPI_ISL_547005, EPI_ISL_547006, EPI_ISL_547007, EPI_ISL_547008, EPI_ISL_547009, EPI_ISL_547010, EPI_ISL_547011, EPI_ISL_547012, EPI_ISL_547013, EPI_ISL_547014, EPI_ISL_547016, EPI_ISL_547017, EPI_ISL_547018, EPI_ISL_547019, EPI_ISL_547020, EPI_ISL_547021, EPI_ISL_547022, EPI_I[illegible]
EPI_ISL_547027, EPI_ISL_547028, EPI_ISL_547029, EPI_ISL_547030, EPI_ISL_547031, EPI_ISL_547032, EPI_ISL_547033, EPI_ISL_547034, EPI_ISL_547035, EPI_ISL_547036, EPI_ISL_547038, EPI_ISL_547039, EPI_ISL_547040, EPI_ISL_547041, EPI_ISL_547042, EPI_ISL_547043, EPI_ISL_547044, EPI_ISL_547045, EPI_I[illegible]
EPI_ISL_547050, EPI_ISL_547051, EPI_ISL_547052, EPI_ISL_547053, EPI_ISL_547054, EPI_ISL_547055, EPI_ISL_547056, EPI_ISL_547057, EPI_ISL_547058, EPI_ISL_547059, EPI_ISL_547060, EPI_ISL_547061, EPI_ISL_547062, EPI_ISL_547063, EPI_ISL_547064, EPI_ISL_547065, EPI_ISL_547066, EPI_ISL_547067, EPI_I[illegible]
EPI_ISL_547076, EPI_ISL_547077, EPI_ISL_547078, EPI_ISL_547079, EPI_ISL_547080, EPI_ISL_547081, EPI_ISL_547082, EPI_ISL_547083, EPI_ISL_547084, EPI_ISL_547085, EPI_ISL_547086, EPI_ISL_547087, EPI_ISL_547088, EPI_ISL_547089, EPI_ISL_547090, EPI_ISL_547091, EPI_ISL_547092, EPI_ISL_547093, EPI_I[illegible]
EPI_ISL_547099, EPI_ISL_547100, EPI_ISL_547102, EPI_ISL_547103, EPI_ISL_547104, EPI_ISL_547105, EPI_ISL_547106, EPI_ISL_547107, EPI_ISL_547108, EPI_ISL_547110, EPI_ISL_547111, EPI_ISL_547112, EPI_ISL_547113, EPI_ISL_547114, EPI_ISL_547115, EPI_ISL_547116, EPI_ISL_547117, EPI_ISL_547118, EPI_I[illegible]
EPI_ISL_547124, EPI_ISL_547125, EPI_ISL_547126, EPI_ISL_547127, EPI_ISL_547128, EPI_ISL_547129, EPI_ISL_547130, EPI_ISL_547131, EPI_ISL_547133, EPI_ISL_547134, EPI_ISL_547135, EPI_ISL_547136, EPI_ISL_547137, EPI_ISL_547138, EPI_ISL_547139, EPI_ISL_547141, EPI_ISL_547142, EPI_ISL_547143, EPI_I[illegible]
EPI_ISL_547149, EPI_ISL_547150, EPI_ISL_547151, EPI_ISL_547153, EPI_ISL_547154, EPI_ISL_547155, EPI_ISL_547157, EPI_ISL_547158, EPI_ISL_547159, EPI_ISL_547160, EPI_ISL_547162, EPI_ISL_547163, EPI_ISL_547166, EPI_ISL_547167, EPI_ISL_547168, EPI_ISL_547169, EPI_ISL_547170, EPI_ISL_547171, EPI_I[illegible]
EPI_ISL_547177, EPI_ISL_547178, EPI_ISL_547180, EPI_ISL_547181, EPI_ISL_547183, EPI_ISL_547184, EPI_ISL_547186, EPI_ISL_547187, EPI_ISL_547188, EPI_ISL_547189, EPI_ISL_547190, EPI_ISL_547191, EPI_ISL_547192, EPI_ISL_547193, EPI_ISL_547194, EPI_ISL_547195, EPI_ISL_547196, EPI_ISL_547197, EPI_I[illegible]
EPI_ISL_547204, EPI_ISL_547215, EPI_ISL_547218, EPI_ISL_547222, EPI_ISL_547224, EPI_ISL_547229, EPI_ISL_547230, EPI_ISL_547231, EPI_ISL_547232, EPI_ISL_547239, EPI_ISL_547250, EPI_ISL_547256, EPI_ISL_547257, EPI_ISL_547259, EPI_ISL_547261, EPI_ISL_547265, EPI_ISL_547271, EPI_ISL_547273, EPI_I[illegible]
EPI_ISL_547283, EPI_ISL_547290, EPI_ISL_547293, EPI_ISL_547312, EPI_ISL_547317, EPI_ISL_547321, EPI_ISL_547327, EPI_ISL_547330, EPI_ISL_547332, EPI_ISL_547333, EPI_ISL_547338, EPI_ISL_547342, EPI_ISL_547347, EPI_ISL_547349, EPI_ISL_547353, EPI_ISL_547354, EPI_ISL_547355, EPI_ISL_547356, EPI_I[illegible]
EPI_ISL_547363, EPI_ISL_547364, EPI_ISL_547370, EPI_ISL_547372, EPI_ISL_547375, EPI_ISL_547376, EPI_ISL_547380, EPI_ISL_547383, EPI_ISL_547390, EPI_ISL_547393, EPI_ISL_547394, EPI_ISL_547395, EPI_ISL_547399, EPI_ISL_547402, EPI_ISL_547405, EPI_ISL_547408

| | | | |
|---|---|---|---|
| see above | Houston Methodist Hospital | Houston Methodist Hospital | S. Wesley Long, Randall J. Olsen, Paul A. Christensen, David W. Bernard, James J. Davis, Maulik Shukla, Marcus Nguyen, Matthew Ojeda Saavedra, Concepcion C. Cantu, Prasanti Yerramilli, Layne Pruitt, Sishir Subedi, Hung-Che Kuo, Heather Hendr[illegible] Muthiah Kumaraswami, Jule Goike, Daniel Boutz, Jimmy Gollihar, Jason S. McLellan, Chia-Wei Chou, Kamyab Javanmardi, Ilya J. Finkelstein, and James M. Musser |
| EPI_ISL_547576 | Secretaria Municipal de Saúde | Instituto Adolfo Lutz, Interdiciplinary Procedures Center, Strategic Laboratory | Claudio Tavares Sacchi, Claudia Regina Gonçalves, Erica Valessa Ramos Gomes, Karoline Rodrigues Campos |
| EPI_ISL_548133 | Middlemore Hospital | Institute of Environmental Science and Research (ESR) | Xiaoyun Ren, Matt Storey, Nikki Freed, Muhammad Faisal, Jing Wang, Hermes Perez, Anja Werno, Antje van der Linden, Arlo Upton, Chris Mansell, David Hammer, Dragana Drinkovic, Gary McAuliffe, Hana Sofia Andersson, James Ussher, Jill Sherwoo[illegible] Blakiston, Matthew Rogers, Max Bloomfield, Michael Addidle, Michelle Balm, Sally Roberts, Sarah Jefferies, Sharmini Muttaiyah, Susan Morpeth, Susan Taylor, Timothy Blackmore, Vani Sathyendran, Veronica Playle, Virginia Hope, Eras[illegible] |
| EPI_ISL_548455 | Ventura County Public Health Lab | Chan-Zuckerberg Biohub | CZB Cliahub Consortium |

EPI_ISL_549258, EPI_ISL_549259, EPI_ISL_549260, EPI_ISL_549261, EPI_ISL_549262, EPI_ISL_549263, EPI_ISL_549264, EPI_ISL_549265, EPI_ISL_549266, EPI_ISL_549267, EPI_ISL_549268, EPI_ISL_549269, EPI_ISL_549270

| | | | |
|---|---|---|---|
| see above | Florida Bureau of Public Health Laboratories | Florida Bureau of Public Health Laboratories | Sarah Schmedes, Jason Blanton |
| EPI_ISL_551356, EPI_ISL_551361 | Lighthouse Lab in Alderley Park | Wellcome Sanger Institute for the COVID-19 Genomics UK (COG-UK) consortium | The Lighthouse Lab in Alderley Park and Alex Alderton, Roberto Amato, Sonia Goncalves, Ewan Harrison, David K. Jackson, Ian Johnston, Dominic Kwiatkowski, Cordelia Langford, John Sillitoe on behalf of the Wellcome Sanger Institute COVID-[illegible] |
| EPI_ISL_551364 | Lighthouse Lab in Alderley Park | Wellcome Sanger Institute for the COVID-19 Genomics UK (COG-UK) consortium | The Lighthouse Lab in Alderley Park and Alex Alderton, Roberto Amato, Sonia Goncalves, Ewan Harrison, David K. Jackson, Ian Johnston, Dominic Kwiatkowski, Cordelia Langford, John Sillitoe on behalf of the Wellcome Sang[illegible] |
| EPI_ISL_551365, EPI_ISL_551366, EPI_ISL_551367, EPI_ISL_551368, EPI_ISL_551369, EPI_ISL_551375, EPI_ISL_551376 | Lighthouse Lab in Alderley Park | Wellcome Sanger Institute for the COVID-19 Genomics UK (COG-UK) consortium | The Lighthouse Lab in Alderley Park and Alex Alderton, Roberto Amato, Sonia Goncalves, Ewan Harrison, David K. Jackson, Ian Johnston, Dominic Kwiatkowski, Cordelia Langford, John Sillitoe on behalf of the Wellcome Sanger Institute COVID-[illegible] |
| EPI_ISL_551377 | Lighthouse Lab in Alderley Park | Wellcome Sanger Institute for the COVID-19 Genomics UK (COG-UK) consortium | The Lighthouse Lab in Alderley Park and Alex Alderton, Roberto Amato, Sonia Goncalves, Ewan Harrison, David K. Jackson, Ian Johnston, Dominic Kwiatkowski, Cordelia Langford, John Sillitoe on behalf of the Wellcome Sang[illegible] |

EPI_ISL_551378, EPI_ISL_551381, EPI_ISL_551382, EPI_ISL_551388, EPI_ISL_551394, EPI_ISL_551396, EPI_ISL_551398, EPI_ISL_551400, EPI_ISL_551401, EPI_ISL_551402, EPI_ISL_551404, EPI_ISL_551405, EPI_ISL_551407, EPI_ISL_551409, EPI_ISL_551410, EPI_ISL_551413

| | | | |
|---|---|---|---|
| see above | Lighthouse Lab in Alderley Park | Wellcome Sanger Institute for the COVID-19 Genomics UK (COG-UK) consortium | The Lighthouse Lab in Alderley Park and Alex Alderton, Roberto Amato, Sonia Goncalves, Ewan Harrison, David K. Jackson, Ian Johnston, Dominic Kwiatkowski, Cordelia Langford, John Sillitoe on behalf of the Wellcome Sanger Institute COVID-[illegible] |
| EPI_ISL_551415, EPI_ISL_551416 | Lighthouse Lab in Alderley Park | Wellcome Sanger Institute for the COVID-19 Genomics UK (COG-UK) consortium | The Lighthouse Lab in Alderley Park and Alex Alderton, Roberto Amato, Sonia Goncalves, Ewan Harrison, David K. Jackson, Ian Johnston, Dominic Kwiatkowski, Cordelia Langford, John Sillitoe on behalf of the Wellcome Sang[illegible] |
| EPI_ISL_551417, EPI_ISL_551418, EPI_ISL_551419, EPI_ISL_551428 | Lighthouse Lab in Alderley Park | Wellcome Sanger Institute for the COVID-19 Genomics UK (COG-UK) consortium | The Lighthouse Lab in Alderley Park and Alex Alderton, Roberto Amato, Sonia Goncalves, Ewan Harrison, David K. Jackson, Ian Johnston, Dominic Kwiatkowski, Cordelia Langford, John Sillitoe on behalf of the Wellcome Sanger Institute COVID-[illegible] |
| EPI_ISL_551429 | Lighthouse Lab in Alderley Park | Wellcome Sanger Institute for the COVID-19 Genomics UK (COG-UK) consortium | The Lighthouse Lab in Alderley Park and Alex Alderton, Roberto Amato, Sonia Goncalves, Ewan Harrison, David K. Jackson, Ian Johnston, Dominic Kwiatkowski, Cordelia Langford, John Sillitoe on behalf of the Wellcome Sang[illegible] |

| | | | |
|---|---|---|---|
| EPI_ISL_551430, EPI_ISL_551431, EPI_ISL_551432, EPI_ISL_551433, EPI_ISL_551434 | Lighthouse Lab in Alderley Park | Wellcome Sanger Institute for the COVID-19 Genomics UK (COG-UK) consortium | The Lighthouse Lab in Alderley Park and Alex Alderton, Roberto Amato, Sonia Goncalves, Ewan Harrison, David K. Jackson, Ian Johnston, Dominic Kwiatkowski, Cordelia Langford, John Sillitoe on behalf of the Wellcome Sanger Institute COVID- |
| EPI_ISL_551436, EPI_ISL_551437, EPI_ISL_551442, EPI_ISL_551444 | Lighthouse Lab in Alderley Park | Wellcome Sanger Institute for the COVID-19 Genomics UK (COG-UK) consortium | The Lighthouse Lab in Alderley Park and Alex Alderton, Roberto Amato, Sonia Goncalves, Ewan Harrison, David K. Jackson, Ian Johnston, Dominic Kwiatkowski, Cordelia Langford, John Sillitoe on behalf of the Wellcome Sang |
| EPI_ISL_551445, EPI_ISL_551446 | Lighthouse Lab in Alderley Park | Wellcome Sanger Institute for the COVID-19 Genomics UK (COG-UK) consortium | The Lighthouse Lab in Alderley Park and Alex Alderton, Roberto Amato, Sonia Goncalves, Ewan Harrison, David K. Jackson, Ian Johnston, Dominic Kwiatkowski, Cordelia Langford, John Sillitoe on behalf of the Wellcome Sanger Institute COVID- |
| EPI_ISL_551447, EPI_ISL_551451 | Lighthouse Lab in Alderley Park | Wellcome Sanger Institute for the COVID-19 Genomics UK (COG-UK) consortium | The Lighthouse Lab in Alderley Park and Alex Alderton, Roberto Amato, Sonia Goncalves, Ewan Harrison, David K. Jackson, Ian Johnston, Dominic Kwiatkowski, Cordelia Langford, John Sillitoe on behalf of the Wellcome Sang |
| EPI_ISL_551453 | Lighthouse Lab in Alderley Park | Wellcome Sanger Institute for the COVID-19 Genomics UK (COG-UK) Consortium | The Lighthouse Lab in Alderley Park and Alex Alderton, Roberto Amato, Sonia Goncalves, Ewan Harrison, David K. Jackson, Ian Johnston, Dominic Kwiatkowski, Cordelia Langford, John Sillitoe on behalf of the Wellcome Sang |
| EPI_ISL_551456 | Lighthouse Lab in Alderley Park | Wellcome Sanger Institute for the COVID-19 Genomics UK (COG-UK) consortium | The Lighthouse Lab in Alderley Park and Alex Alderton, Roberto Amato, Sonia Goncalves, Ewan Harrison, David K. Jackson, Ian Johnston, Dominic Kwiatkowski, Cordelia Langford, John Sillitoe on behalf of the Wellcome Sang |
| EPI_ISL_551459, EPI_ISL_551464, EPI_ISL_551466, EPI_ISL_551469 | Lighthouse Lab in Alderley Park | Wellcome Sanger Institute for the COVID-19 Genomics UK (COG-UK) consortium | The Lighthouse Lab in Alderley Park and Alex Alderton, Roberto Amato, Sonia Goncalves, Ewan Harrison, David K. Jackson, Ian Johnston, Dominic Kwiatkowski, Cordelia Langford, John Sillitoe on behalf of the Wellcome Sanger Institute COVID- |
| EPI_ISL_551471 | Lighthouse Lab in Alderley Park | Wellcome Sanger Institute for the COVID-19 Genomics UK (COG-UK) consortium | The Lighthouse Lab in Alderley Park and Alex Alderton, Roberto Amato, Sonia Goncalves, Ewan Harrison, David K. Jackson, Ian Johnston, Dominic Kwiatkowski, Cordelia Langford, John Sillitoe on behalf of the Wellcome Sang |
| EPI_ISL_551472, EPI_ISL_551473, EPI_ISL_551476, EPI_ISL_551477 | Lighthouse Lab in Alderley Park | Wellcome Sanger Institute for the COVID-19 Genomics UK (COG-UK) consortium | The Lighthouse Lab in Alderley Park and Alex Alderton, Roberto Amato, Sonia Goncalves, Ewan Harrison, David K. Jackson, Ian Johnston, Dominic Kwiatkowski, Cordelia Langford, John Sillitoe on behalf of the Wellcome Sanger Institute COVID- |
| EPI_ISL_551481, EPI_ISL_551482 | Lighthouse Lab in Alderley Park | Wellcome Sanger Institute for the COVID-19 Genomics UK (COG-UK) consortium | The Lighthouse Lab in Alderley Park and Alex Alderton, Roberto Amato, Sonia Goncalves, Ewan Harrison, David K. Jackson, Ian Johnston, Dominic Kwiatkowski, Cordelia Langford, John Sillitoe on behalf of the Wellcome Sang |
| EPI_ISL_551483 | Lighthouse Lab in Alderley Park | Wellcome Sanger Institute for the COVID-19 Genomics UK (COG-UK) consortium | The Lighthouse Lab in Alderley Park and Alex Alderton, Roberto Amato, Sonia Goncalves, Ewan Harrison, David K. Jackson, Ian Johnston, Dominic Kwiatkowski, Cordelia Langford, John Sillitoe on behalf of the Wellcome Sanger Institute COVID- |
| EPI_ISL_551484 | Lighthouse Lab in Alderley Park | Wellcome Sanger Institute for the COVID-19 Genomics UK (COG-UK) Consortium | The Lighthouse Lab in Alderley Park and Alex Alderton, Roberto Amato, Sonia Goncalves, Ewan Harrison, David K. Jackson, Ian Johnston, Dominic Kwiatkowski, Cordelia Langford, John Sillitoe on behalf of the Wellcome Sang |
| EPI_ISL_551485, EPI_ISL_551488, EPI_ISL_551489, EPI_ISL_551491, EPI_ISL_551493 | Lighthouse Lab in Alderley Park | Wellcome Sanger Institute for the COVID-19 Genomics UK (COG-UK) consortium | The Lighthouse Lab in Alderley Park and Alex Alderton, Roberto Amato, Sonia Goncalves, Ewan Harrison, David K. Jackson, Ian Johnston, Dominic Kwiatkowski, Cordelia Langford, John Sillitoe on behalf of the Wellcome Sanger Institute COVID- |
| EPI_ISL_551494 | Lighthouse Lab in Alderley Park | Wellcome Sanger Institute for the COVID-19 Genomics UK (COG-UK) consortium | The Lighthouse Lab in Alderley Park and Alex Alderton, Roberto Amato, Sonia Goncalves, Ewan Harrison, David K. Jackson, Ian Johnston, Dominic Kwiatkowski, Cordelia Langford, John Sillitoe on behalf of the Wellcome Sang |
| EPI_ISL_551495, EPI_ISL_551497, EPI_ISL_551499, EPI_ISL_551500, EPI_ISL_551502 | Lighthouse Lab in Alderley Park | Wellcome Sanger Institute for the COVID-19 Genomics UK (COG-UK) consortium | The Lighthouse Lab in Alderley Park and Alex Alderton, Roberto Amato, Sonia Goncalves, Ewan Harrison, David K. Jackson, Ian Johnston, Dominic Kwiatkowski, Cordelia Langford, John Sillitoe on behalf of the Wellcome Sanger Institute COVID- |
| EPI_ISL_551503 | Lighthouse Lab in Alderley Park | Wellcome Sanger Institute for the COVID-19 Genomics UK (COG-UK) consortium | The Lighthouse Lab in Alderley Park and Alex Alderton, Roberto Amato, Sonia Goncalves, Ewan Harrison, David K. Jackson, Ian Johnston, Dominic Kwiatkowski, Cordelia Langford, John Sillitoe on behalf of the Wellcome Sang |
| EPI_ISL_551504, EPI_ISL_551509, EPI_ISL_551510, EPI_ISL_551511, EPI_ISL_551515, EPI_ISL_551516 | Lighthouse Lab in Alderley Park | Wellcome Sanger Institute for the COVID-19 Genomics UK (COG-UK) consortium | The Lighthouse Lab in Alderley Park and Alex Alderton, Roberto Amato, Sonia Goncalves, Ewan Harrison, David K. Jackson, Ian Johnston, Dominic Kwiatkowski, Cordelia Langford, John Sillitoe on behalf of the Wellcome Sanger Institute COVID- |
| EPI_ISL_551518 | Lighthouse Lab in Alderley Park | Wellcome Sanger Institute for the COVID-19 Genomics UK (COG-UK) consortium | The Lighthouse Lab in Alderley Park and Alex Alderton, Roberto Amato, Sonia Goncalves, Ewan Harrison, David K. Jackson, Ian Johnston, Dominic Kwiatkowski, Cordelia Langford, John Sillitoe on behalf of the Wellcome Sang |
| EPI_ISL_551521, EPI_ISL_551523, EPI_ISL_551524, EPI_ISL_551526, EPI_ISL_551527, EPI_ISL_551528, EPI_ISL_551529, EPI_ISL_551531, EPI_ISL_551535, EPI_ISL_551537, EPI_ISL_551539 | | | |
| see above | Lighthouse Lab in Alderley Park | Wellcome Sanger Institute for the COVID-19 Genomics UK (COG-UK) consortium | The Lighthouse Lab in Alderley Park and Alex Alderton, Roberto Amato, Sonia Goncalves, Ewan Harrison, David K. Jackson, Ian Johnston, Dominic Kwiatkowski, Cordelia Langford, John Sillitoe on behalf of the Wellcome Sanger Institute COVID- |
| EPI_ISL_551540 | Lighthouse Lab in Alderley Park | Wellcome Sanger Institute for the COVID-19 Genomics UK (COG-UK) consortium | The Lighthouse Lab in Alderley Park and Alex Alderton, Roberto Amato, Sonia Goncalves, Ewan Harrison, David K. Jackson, Ian Johnston, Dominic Kwiatkowski, Cordelia Langford, John Sillitoe on behalf of the Wellcome Sang |
| EPI_ISL_551541 | Lighthouse Lab in Alderley Park | Wellcome Sanger Institute for the COVID-19 Genomics UK (COG-UK) consortium | The Lighthouse Lab in Alderley Park and Alex Alderton, Roberto Amato, Sonia Goncalves, Ewan Harrison, David K. Jackson, Ian Johnston, Dominic Kwiatkowski, Cordelia Langford, John Sillitoe on behalf of the Wellcome Sanger Institute COVID- |

| | | | |
|---|---|---|---|
| EPI_ISL_551542 | Lighthouse Lab in Alderley Park | Wellcome Sanger Institute for the COVID-19 Genomics UK (COG-UK) consortium | The Lighthouse Lab in Alderley Park and Alex Alderton, Roberto Amato, Sonia Goncalves, Ewan Harrison, David K. Jackson, Ian Johnston, Dominic Kwiatkowski, Cordelia Langford, John Sillitoe on behalf of the Wellcome Sang |
| EPI_ISL_551543 | Lighthouse Lab in Alderley Park | Wellcome Sanger Institute for the COVID-19 Genomics UK (COG-UK) consortium | The Lighthouse Lab in Alderley Park and Alex Alderton, Roberto Amato, Sonia Goncalves, Ewan Harrison, David K. Jackson, Ian Johnston, Dominic Kwiatkowski, Cordelia Langford, John Sillitoe on behalf of the Wellcome Sanger Institute COVID- |
| EPI_ISL_551544 | Lighthouse Lab in Alderley Park | Wellcome Sanger Institute for the COVID-19 Genomics UK (COG-UK) consortium | The Lighthouse Lab in Alderley Park and Alex Alderton, Roberto Amato, Sonia Goncalves, Ewan Harrison, David K. Jackson, Ian Johnston, Dominic Kwiatkowski, Cordelia Langford, John Sillitoe on behalf of the Wellcome Sang |
| EPI_ISL_551546, EPI_ISL_551547, EPI_ISL_551549, EPI_ISL_551550 | Lighthouse Lab in Alderley Park | Wellcome Sanger Institute for the COVID-19 Genomics UK (COG-UK) consortium | The Lighthouse Lab in Alderley Park and Alex Alderton, Roberto Amato, Sonia Goncalves, Ewan Harrison, David K. Jackson, Ian Johnston, Dominic Kwiatkowski, Cordelia Langford, John Sillitoe on behalf of the Wellcome Sanger Institute COVID- |
| EPI_ISL_551553 | Lighthouse Lab in Alderley Park | Wellcome Sanger Institute for the COVID-19 Genomics UK (COG-UK) consortium | The Lighthouse Lab in Alderley Park and Alex Alderton, Roberto Amato, Sonia Goncalves, Ewan Harrison, David K. Jackson, Ian Johnston, Dominic Kwiatkowski, Cordelia Langford, John Sillitoe on behalf of the Wellcome Sang |
| EPI_ISL_551555, EPI_ISL_551556, EPI_ISL_551557, EPI_ISL_551560, EPI_ISL_551562, EPI_ISL_551563, EPI_ISL_551564 | Lighthouse Lab in Alderley Park | Wellcome Sanger Institute for the COVID-19 Genomics UK (COG-UK) consortium | The Lighthouse Lab in Alderley Park and Alex Alderton, Roberto Amato, Sonia Goncalves, Ewan Harrison, David K. Jackson, Ian Johnston, Dominic Kwiatkowski, Cordelia Langford, John Sillitoe on behalf of the Wellcome Sanger Institute COVID- |
| EPI_ISL_551565, EPI_ISL_551566 | Lighthouse Lab in Alderley Park | Wellcome Sanger Institute for the COVID-19 Genomics UK (COG-UK) consortium | The Lighthouse Lab in Alderley Park and Alex Alderton, Roberto Amato, Sonia Goncalves, Ewan Harrison, David K. Jackson, Ian Johnston, Dominic Kwiatkowski, Cordelia Langford, John Sillitoe on behalf of the Wellcome Sang |
| EPI_ISL_551567, EPI_ISL_551568, EPI_ISL_551569 | Lighthouse Lab in Alderley Park | Wellcome Sanger Institute for the COVID-19 Genomics UK (COG-UK) consortium | The Lighthouse Lab in Alderley Park and Alex Alderton, Roberto Amato, Sonia Goncalves, Ewan Harrison, David K. Jackson, Ian Johnston, Dominic Kwiatkowski, Cordelia Langford, John Sillitoe on behalf of the Wellcome Sanger Institute COVID- |
| EPI_ISL_551570 | Lighthouse Lab in Alderley Park | Wellcome Sanger Institute for the COVID-19 Genomics UK (COG-UK) consortium | The Lighthouse Lab in Alderley Park and Alex Alderton, Roberto Amato, Sonia Goncalves, Ewan Harrison, David K. Jackson, Ian Johnston, Dominic Kwiatkowski, Cordelia Langford, John Sillitoe on behalf of the Wellcome Sang |
| EPI_ISL_551571, EPI_ISL_551573, EPI_ISL_551574, EPI_ISL_551576 | Lighthouse Lab in Alderley Park | Wellcome Sanger Institute for the COVID-19 Genomics UK (COG-UK) consortium | The Lighthouse Lab in Alderley Park and Alex Alderton, Roberto Amato, Sonia Goncalves, Ewan Harrison, David K. Jackson, Ian Johnston, Dominic Kwiatkowski, Cordelia Langford, John Sillitoe on behalf of the Wellcome Sanger Institute COVID- |
| EPI_ISL_551758, EPI_ISL_551786, EPI_ISL_551800 | Lighthouse Lab in Milton Keynes | Wellcome Sanger Institute for the COVID-19 Genomics UK (COG-UK) consortium | The Lighthouse Lab in Milton Keynes and Alex Alderton, Roberto Amato, Sonia Goncalves, Ewan Harrison, David K. Jackson, Ian Johnston, Dominic Kwiatkowski, Cordelia Langford, John Sillitoe on behalf of the Wellcome Sanger Institute COVID- |
| EPI_ISL_552798 | Lighthouse Lab in Alderley Park | Wellcome Sanger Institute for the COVID-19 Genomics UK (COG-UK) consortium | The Lighthouse Lab in Alderley Park and Alex Alderton, Roberto Amato, Sonia Goncalves, Ewan Harrison, David K. Jackson, Ian Johnston, Dominic Kwiatkowski, Cordelia Langford, John Sillitoe on behalf of the Wellcome Sanger Institute COVID- |
| EPI_ISL_552801, EPI_ISL_552805, EPI_ISL_552812, EPI_ISL_552817 | Lighthouse Lab in Alderley Park | Wellcome Sanger Institute for the COVID-19 Genomics UK (COG-UK) consortium | The Lighthouse Lab in Alderley Park and Alex Alderton, Roberto Amato, Sonia Goncalves, Ewan Harrison, David K. Jackson, Ian Johnston, Dominic Kwiatkowski, Cordelia Langford, John Sillitoe on behalf of the Wellcome Sang |
| EPI_ISL_552818 | Lighthouse Lab in Alderley Park | Wellcome Sanger Institute for the COVID-19 Genomics UK (COG-UK) Consortium | The Lighthouse Lab in Alderley Park and Alex Alderton, Roberto Amato, Sonia Goncalves, Ewan Harrison, David K. Jackson, Ian Johnston, Dominic Kwiatkowski, Cordelia Langford, John Sillitoe on behalf of the Wellcome Sang |
| EPI_ISL_552819, EPI_ISL_552827, EPI_ISL_552828, EPI_ISL_552830, EPI_ISL_552831 | Lighthouse Lab in Alderley Park | Wellcome Sanger Institute for the COVID-19 Genomics UK (COG-UK) consortium | The Lighthouse Lab in Alderley Park and Alex Alderton, Roberto Amato, Sonia Goncalves, Ewan Harrison, David K. Jackson, Ian Johnston, Dominic Kwiatkowski, Cordelia Langford, John Sillitoe on behalf of the Wellcome Sang |
| EPI_ISL_552836, EPI_ISL_552839 | Lighthouse Lab in Alderley Park | Wellcome Sanger Institute for the COVID-19 Genomics UK (COG-UK) consortium | The Lighthouse Lab in Alderley Park and Alex Alderton, Roberto Amato, Sonia Goncalves, Ewan Harrison, David K. Jackson, Ian Johnston, Dominic Kwiatkowski, Cordelia Langford, John Sillitoe on behalf of the Wellcome Sanger Institute COVID- |
| EPI_ISL_552840, EPI_ISL_552843, EPI_ISL_552844, EPI_ISL_552845, EPI_ISL_552847, EPI_ISL_552848, EPI_ISL_552851, EPI_ISL_552853 | Lighthouse Lab in Alderley Park | Wellcome Sanger Institute for the COVID-19 Genomics UK (COG-UK) consortium | The Lighthouse Lab in Alderley Park and Alex Alderton, Roberto Amato, Sonia Goncalves, Ewan Harrison, David K. Jackson, Ian Johnston, Dominic Kwiatkowski, Cordelia Langford, John Sillitoe on behalf of the Wellcome Sang |
| EPI_ISL_552854, EPI_ISL_552855, EPI_ISL_552856 | Lighthouse Lab in Alderley Park | Wellcome Sanger Institute for the COVID-19 Genomics UK (COG-UK) consortium | The Lighthouse Lab in Alderley Park and Alex Alderton, Roberto Amato, Sonia Goncalves, Ewan Harrison, David K. Jackson, Ian Johnston, Dominic Kwiatkowski, Cordelia Langford, John Sillitoe on behalf of the Wellcome Sanger Institute COVID- |
| EPI_ISL_552857 | Lighthouse Lab in Alderley Park | Wellcome Sanger Institute for the COVID-19 Genomics UK (COG-UK) consortium | The Lighthouse Lab in Alderley Park and Alex Alderton, Roberto Amato, Sonia Goncalves, Ewan Harrison, David K. Jackson, Ian Johnston, Dominic Kwiatkowski, Cordelia Langford, John Sillitoe on behalf of the Wellcome Sang |
| EPI_ISL_552858 | Lighthouse Lab in Alderley Park | Wellcome Sanger Institute for the COVID-19 Genomics UK (COG-UK) consortium | The Lighthouse Lab in Alderley Park and Alex Alderton, Roberto Amato, Sonia Goncalves, Ewan Harrison, David K. Jackson, Ian Johnston, Dominic Kwiatkowski, Cordelia Langford, John Sillitoe on behalf of the Wellcome Sanger Institute COVID- |
| EPI_ISL_552859, EPI_ISL_552861, EPI_ISL_552862 | Lighthouse Lab in Alderley Park | Wellcome Sanger Institute for the COVID-19 Genomics UK (COG-UK) consortium | The Lighthouse Lab in Alderley Park and Alex Alderton, Roberto Amato, Sonia Goncalves, Ewan Harrison, David K. Jackson, Ian Johnston, Dominic Kwiatkowski, Cordelia Langford, John Sillitoe on behalf of the Wellcome Sang |

| | | | |
|---|---|---|---|
| EPI_ISL_552863 | Lighthouse Lab in Alderley Park | Wellcome Sanger Institute for the COVID-19 Genomics UK (COG-UK) consortium | The Lighthouse Lab in Alderley Park and Alex Alderton, Roberto Amato, Sonia Goncalves, Ewan Harrison, David K. Jackson, Ian Johnston, Dominic Kwiatkowski, Cordelia Langford, John Sillitoe on behalf of the Wellcome Sanger Institute COVID- |
| EPI_ISL_552868 | Lighthouse Lab in Alderley Park | Wellcome Sanger Institute for the COVID-19 Genomics UK (COG-UK) consortium | The Lighthouse Lab in Alderley Park and Alex Alderton, Roberto Amato, Sonia Goncalves, Ewan Harrison, David K. Jackson, Ian Johnston, Dominic Kwiatkowski, Cordelia Langford, John Sillitoe on behalf of the Wellcome Sang |
| EPI_ISL_552869, EPI_ISL_552872, EPI_ISL_552873, EPI_ISL_552876 | Lighthouse Lab in Alderley Park | Wellcome Sanger Institute for the COVID-19 Genomics UK (COG-UK) consortium | The Lighthouse Lab in Alderley Park and Alex Alderton, Roberto Amato, Sonia Goncalves, Ewan Harrison, David K. Jackson, Ian Johnston, Dominic Kwiatkowski, Cordelia Langford, John Sillitoe on behalf of the Wellcome Sanger Institute COVID- |
| EPI_ISL_552877, EPI_ISL_552878, EPI_ISL_552883 | Lighthouse Lab in Alderley Park | Wellcome Sanger Institute for the COVID-19 Genomics UK (COG-UK) consortium | The Lighthouse Lab in Alderley Park and Alex Alderton, Roberto Amato, Sonia Goncalves, Ewan Harrison, David K. Jackson, Ian Johnston, Dominic Kwiatkowski, Cordelia Langford, John Sillitoe on behalf of the Wellcome Sang |
| EPI_ISL_552884 | Lighthouse Lab in Alderley Park | Wellcome Sanger Institute for the COVID-19 Genomics UK (COG-UK) consortium | The Lighthouse Lab in Alderley Park and Alex Alderton, Roberto Amato, Sonia Goncalves, Ewan Harrison, David K. Jackson, Ian Johnston, Dominic Kwiatkowski, Cordelia Langford, John Sillitoe on behalf of the Wellcome Sanger Institute COVID- |
| EPI_ISL_552886 | Lighthouse Lab in Alderley Park | Wellcome Sanger Institute for the COVID-19 Genomics UK (COG-UK) consortium | The Lighthouse Lab in Alderley Park and Alex Alderton, Roberto Amato, Sonia Goncalves, Ewan Harrison, David K. Jackson, Ian Johnston, Dominic Kwiatkowski, Cordelia Langford, John Sillitoe on behalf of the Wellcome Sang |
| EPI_ISL_552887, EPI_ISL_552889 | Lighthouse Lab in Alderley Park | Wellcome Sanger Institute for the COVID-19 Genomics UK (COG-UK) consortium | The Lighthouse Lab in Alderley Park and Alex Alderton, Roberto Amato, Sonia Goncalves, Ewan Harrison, David K. Jackson, Ian Johnston, Dominic Kwiatkowski, Cordelia Langford, John Sillitoe on behalf of the Wellcome Sanger Institute COVID- |
| EPI_ISL_552891, EPI_ISL_552892, EPI_ISL_552893 | Lighthouse Lab in Alderley Park | Wellcome Sanger Institute for the COVID-19 Genomics UK (COG-UK) consortium | The Lighthouse Lab in Alderley Park and Alex Alderton, Roberto Amato, Sonia Goncalves, Ewan Harrison, David K. Jackson, Ian Johnston, Dominic Kwiatkowski, Cordelia Langford, John Sillitoe on behalf of the Wellcome Sang |
| EPI_ISL_552894 | Lighthouse Lab in Alderley Park | Wellcome Sanger Institute for the COVID-19 Genomics UK (COG-UK) consortium | The Lighthouse Lab in Alderley Park and Alex Alderton, Roberto Amato, Sonia Goncalves, Ewan Harrison, David K. Jackson, Ian Johnston, Dominic Kwiatkowski, Cordelia Langford, John Sillitoe on behalf of the Wellcome Sanger Institute COVID- |
| EPI_ISL_552897 | Lighthouse Lab in Alderley Park | Wellcome Sanger Institute for the COVID-19 Genomics UK (COG-UK) consortium | The Lighthouse Lab in Alderley Park and Alex Alderton, Roberto Amato, Sonia Goncalves, Ewan Harrison, David K. Jackson, Ian Johnston, Dominic Kwiatkowski, Cordelia Langford, John Sillitoe on behalf of the Wellcome Sang |
| EPI_ISL_552900, EPI_ISL_552906, EPI_ISL_552908 | Lighthouse Lab in Alderley Park | Wellcome Sanger Institute for the COVID-19 Genomics UK (COG-UK) consortium | The Lighthouse Lab in Alderley Park and Alex Alderton, Roberto Amato, Sonia Goncalves, Ewan Harrison, David K. Jackson, Ian Johnston, Dominic Kwiatkowski, Cordelia Langford, John Sillitoe on behalf of the Wellcome Sanger Institute COVID- |
| EPI_ISL_552909 | Lighthouse Lab in Alderley Park | Wellcome Sanger Institute for the COVID-19 Genomics UK (COG-UK) consortium | The Lighthouse Lab in Alderley Park and Alex Alderton, Roberto Amato, Sonia Goncalves, Ewan Harrison, David K. Jackson, Ian Johnston, Dominic Kwiatkowski, Cordelia Langford, John Sillitoe on behalf of the Wellcome Sang |
| EPI_ISL_552911 | Lighthouse Lab in Alderley Park | Wellcome Sanger Institute for the COVID-19 Genomics UK (COG-UK) consortium | The Lighthouse Lab in Alderley Park and Alex Alderton, Roberto Amato, Sonia Goncalves, Ewan Harrison, David K. Jackson, Ian Johnston, Dominic Kwiatkowski, Cordelia Langford, John Sillitoe on behalf of the Wellcome Sanger Institute COVID- |
| EPI_ISL_552912, EPI_ISL_552917, EPI_ISL_552918, EPI_ISL_552923, EPI_ISL_552924, EPI_ISL_552929, EPI_ISL_552931, EPI_ISL_552935 | Lighthouse Lab in Alderley Park | Wellcome Sanger Institute for the COVID-19 Genomics UK (COG-UK) consortium | The Lighthouse Lab in Alderley Park and Alex Alderton, Roberto Amato, Sonia Goncalves, Ewan Harrison, David K. Jackson, Ian Johnston, Dominic Kwiatkowski, Cordelia Langford, John Sillitoe on behalf of the Wellcome Sang |
| EPI_ISL_552942 | Lighthouse Lab in Alderley Park | Wellcome Sanger Institute for the COVID-19 Genomics UK (COG-UK) consortium | The Lighthouse Lab in Alderley Park and Alex Alderton, Roberto Amato, Sonia Goncalves, Ewan Harrison, David K. Jackson, Ian Johnston, Dominic Kwiatkowski, Cordelia Langford, John Sillitoe on behalf of the Wellcome Sanger Institute COVID- |
| EPI_ISL_552944, EPI_ISL_552946, EPI_ISL_552960, EPI_ISL_552971 | Lighthouse Lab in Alderley Park | Wellcome Sanger Institute for the COVID-19 Genomics UK (COG-UK) consortium | The Lighthouse Lab in Alderley Park and Alex Alderton, Roberto Amato, Sonia Goncalves, Ewan Harrison, David K. Jackson, Ian Johnston, Dominic Kwiatkowski, Cordelia Langford, John Sillitoe on behalf of the Wellcome Sang |
| EPI_ISL_552972, EPI_ISL_552973 | Lighthouse Lab in Alderley Park | Wellcome Sanger Institute for the COVID-19 Genomics UK (COG-UK) Consortium | The Lighthouse Lab in Alderley Park and Alex Alderton, Roberto Amato, Sonia Goncalves, Ewan Harrison, David K. Jackson, Ian Johnston, Dominic Kwiatkowski, Cordelia Langford, John Sillitoe on behalf of the Wellcome Sang |
| EPI_ISL_552974, EPI_ISL_552976 | Lighthouse Lab in Alderley Park | Wellcome Sanger Institute for the COVID-19 Genomics UK (COG-UK) consortium | The Lighthouse Lab in Alderley Park and Alex Alderton, Roberto Amato, Sonia Goncalves, Ewan Harrison, David K. Jackson, Ian Johnston, Dominic Kwiatkowski, Cordelia Langford, John Sillitoe on behalf of the Wellcome Sanger Institute COVID- |
| EPI_ISL_552981 | Lighthouse Lab in Alderley Park | Wellcome Sanger Institute for the COVID-19 Genomics UK (COG-UK) consortium | The Lighthouse Lab in Alderley Park and Alex Alderton, Roberto Amato, Sonia Goncalves, Ewan Harrison, David K. Jackson, Ian Johnston, Dominic Kwiatkowski, Cordelia Langford, John Sillitoe on behalf of the Wellcome Sang |
| EPI_ISL_552983 | Lighthouse Lab in Alderley Park | Wellcome Sanger Institute for the COVID-19 Genomics UK (COG-UK) consortium | The Lighthouse Lab in Alderley Park and Alex Alderton, Roberto Amato, Sonia Goncalves, Ewan Harrison, David K. Jackson, Ian Johnston, Dominic Kwiatkowski, Cordelia Langford, John Sillitoe on behalf of the Wellcome Sanger Institute COVID- |
| EPI_ISL_552986 | Lighthouse Lab in Alderley Park | Wellcome Sanger Institute for the COVID-19 Genomics UK (COG-UK) consortium | The Lighthouse Lab in Alderley Park and Alex Alderton, Roberto Amato, Sonia Goncalves, Ewan Harrison, David K. Jackson, Ian Johnston, Dominic Kwiatkowski, Cordelia Langford, John Sillitoe on behalf of the Wellcome Sang |
| EPI_ISL_552996 | Lighthouse Lab in Alderley Park | Wellcome Sanger Institute for the COVID-19 Genomics UK (COG-UK) consortium | The Lighthouse Lab in Alderley Park and Alex Alderton, Roberto Amato, Sonia Goncalves, Ewan Harrison, David K. Jackson, Ian Johnston, Dominic Kwiatkowski, Cordelia Langford, John Sillitoe on behalf of the Wellcome Sanger Institute COVID- |
| EPI_ISL_552998, EPI_ISL_553001 | Lighthouse Lab in Alderley Park | Wellcome Sanger Institute for the COVID-19 Genomics UK (COG-UK) consortium | The Lighthouse Lab in Alderley Park and Alex Alderton, Roberto Amato, Sonia Goncalves, Ewan Harrison, David K. Jackson, Ian Johnston, Dominic Kwiatkowski, Cordelia Langford, John Sillitoe on behalf of the Wellcome Sang |

| | | | |
|---|---|---|---|
| EPI_ISL_553269, EPI_ISL_553271, EPI_ISL_553272 | Lighthouse Lab in Alderley Park | Wellcome Sanger Institute for the COVID-19 Genomics UK (COG-UK) consortium | The Lighthouse Lab in Alderley Park and Alex Alderton, Roberto Amato, Sonia Goncalves, Ewan Harrison, David K. Jackson, Ian Johnston, Dominic Kwiatkowski, Cordelia Langford, John Sillitoe on behalf of the Wellcome Sanger Institute COVID- |
| EPI_ISL_553273 | Lighthouse Lab in Alderley Park | Wellcome Sanger Institute for the COVID-19 Genomics UK (COG-UK) consortium | The Lighthouse Lab in Alderley Park and Alex Alderton, Roberto Amato, Sonia Goncalves, Ewan Harrison, David K. Jackson, Ian Johnston, Dominic Kwiatkowski, Cordelia Langford, John Sillitoe on behalf of the Wellcome Sang |
| EPI_ISL_553274, EPI_ISL_553277 | Lighthouse Lab in Alderley Park | Wellcome Sanger Institute for the COVID-19 Genomics UK (COG-UK) consortium | The Lighthouse Lab in Alderley Park and Alex Alderton, Roberto Amato, Sonia Goncalves, Ewan Harrison, David K. Jackson, Ian Johnston, Dominic Kwiatkowski, Cordelia Langford, John Sillitoe on behalf of the Wellcome Sanger Institute COVID- |
| EPI_ISL_553278 | Lighthouse Lab in Alderley Park | Wellcome Sanger Institute for the COVID-19 Genomics UK (COG-UK) consortium | The Lighthouse Lab in Alderley Park and Alex Alderton, Roberto Amato, Sonia Goncalves, Ewan Harrison, David K. Jackson, Ian Johnston, Dominic Kwiatkowski, Cordelia Langford, John Sillitoe on behalf of the Wellcome Sang |
| EPI_ISL_553279, EPI_ISL_553280 | Lighthouse Lab in Alderley Park | Wellcome Sanger Institute for the COVID-19 Genomics UK (COG-UK) consortium | The Lighthouse Lab in Alderley Park and Alex Alderton, Roberto Amato, Sonia Goncalves, Ewan Harrison, David K. Jackson, Ian Johnston, Dominic Kwiatkowski, Cordelia Langford, John Sillitoe on behalf of the Wellcome Sanger Institute COVID- |
| EPI_ISL_553281, EPI_ISL_553282, EPI_ISL_553283 | Lighthouse Lab in Alderley Park | Wellcome Sanger Institute for the COVID-19 Genomics UK (COG-UK) consortium | The Lighthouse Lab in Alderley Park and Alex Alderton, Roberto Amato, Sonia Goncalves, Ewan Harrison, David K. Jackson, Ian Johnston, Dominic Kwiatkowski, Cordelia Langford, John Sillitoe on behalf of the Wellcome Sang |
| EPI_ISL_553284, EPI_ISL_553285 | Lighthouse Lab in Alderley Park | Wellcome Sanger Institute for the COVID-19 Genomics UK (COG-UK) consortium | The Lighthouse Lab in Alderley Park and Alex Alderton, Roberto Amato, Sonia Goncalves, Ewan Harrison, David K. Jackson, Ian Johnston, Dominic Kwiatkowski, Cordelia Langford, John Sillitoe on behalf of the Wellcome Sanger Institute COVID- |
| EPI_ISL_553286, EPI_ISL_553288 | Lighthouse Lab in Alderley Park | Wellcome Sanger Institute for the COVID-19 Genomics UK (COG-UK) consortium | The Lighthouse Lab in Alderley Park and Alex Alderton, Roberto Amato, Sonia Goncalves, Ewan Harrison, David K. Jackson, Ian Johnston, Dominic Kwiatkowski, Cordelia Langford, John Sillitoe on behalf of the Wellcome Sang |
| EPI_ISL_553289, EPI_ISL_553290, EPI_ISL_553292, EPI_ISL_553293, EPI_ISL_553294 | Lighthouse Lab in Alderley Park | Wellcome Sanger Institute for the COVID-19 Genomics UK (COG-UK) consortium | The Lighthouse Lab in Alderley Park and Alex Alderton, Roberto Amato, Sonia Goncalves, Ewan Harrison, David K. Jackson, Ian Johnston, Dominic Kwiatkowski, Cordelia Langford, John Sillitoe on behalf of the Wellcome Sanger Institute COVID- |
| EPI_ISL_553296, EPI_ISL_553298, EPI_ISL_553299 | Lighthouse Lab in Alderley Park | Wellcome Sanger Institute for the COVID-19 Genomics UK (COG-UK) consortium | The Lighthouse Lab in Alderley Park and Alex Alderton, Roberto Amato, Sonia Goncalves, Ewan Harrison, David K. Jackson, Ian Johnston, Dominic Kwiatkowski, Cordelia Langford, John Sillitoe on behalf of the Wellcome Sang |
| EPI_ISL_553300 | Lighthouse Lab in Alderley Park | Wellcome Sanger Institute for the COVID-19 Genomics UK (COG-UK) consortium | The Lighthouse Lab in Alderley Park and Alex Alderton, Roberto Amato, Sonia Goncalves, Ewan Harrison, David K. Jackson, Ian Johnston, Dominic Kwiatkowski, Cordelia Langford, John Sillitoe on behalf of the Wellcome Sanger Institute COVID- |
| EPI_ISL_553304 | Lighthouse Lab in Alderley Park | Wellcome Sanger Institute for the COVID-19 Genomics UK (COG-UK) consortium | The Lighthouse Lab in Alderley Park and Alex Alderton, Roberto Amato, Sonia Goncalves, Ewan Harrison, David K. Jackson, Ian Johnston, Dominic Kwiatkowski, Cordelia Langford, John Sillitoe on behalf of the Wellcome Sang |
| EPI_ISL_553305 | Lighthouse Lab in Alderley Park | Wellcome Sanger Institute for the COVID-19 Genomics UK (COG-UK) consortium | The Lighthouse Lab in Alderley Park and Alex Alderton, Roberto Amato, Sonia Goncalves, Ewan Harrison, David K. Jackson, Ian Johnston, Dominic Kwiatkowski, Cordelia Langford, John Sillitoe on behalf of the Wellcome Sanger Institute COVID- |
| EPI_ISL_553306 | Lighthouse Lab in Alderley Park | Wellcome Sanger Institute for the COVID-19 Genomics UK (COG-UK) consortium | The Lighthouse Lab in Alderley Park and Alex Alderton, Roberto Amato, Sonia Goncalves, Ewan Harrison, David K. Jackson, Ian Johnston, Dominic Kwiatkowski, Cordelia Langford, John Sillitoe on behalf of the Wellcome Sang |
| EPI_ISL_553307 | Lighthouse Lab in Alderley Park | Wellcome Sanger Institute for the COVID-19 Genomics UK (COG-UK) consortium | The Lighthouse Lab in Alderley Park and Alex Alderton, Roberto Amato, Sonia Goncalves, Ewan Harrison, David K. Jackson, Ian Johnston, Dominic Kwiatkowski, Cordelia Langford, John Sillitoe on behalf of the Wellcome Sanger Institute COVID- |
| EPI_ISL_553308, EPI_ISL_553309 | Lighthouse Lab in Alderley Park | Wellcome Sanger Institute for the COVID-19 Genomics UK (COG-UK) consortium | The Lighthouse Lab in Alderley Park and Alex Alderton, Roberto Amato, Sonia Goncalves, Ewan Harrison, David K. Jackson, Ian Johnston, Dominic Kwiatkowski, Cordelia Langford, John Sillitoe on behalf of the Wellcome Sang |
| EPI_ISL_553310, EPI_ISL_553311 | Lighthouse Lab in Alderley Park | Wellcome Sanger Institute for the COVID-19 Genomics UK (COG-UK) consortium | The Lighthouse Lab in Alderley Park and Alex Alderton, Roberto Amato, Sonia Goncalves, Ewan Harrison, David K. Jackson, Ian Johnston, Dominic Kwiatkowski, Cordelia Langford, John Sillitoe on behalf of the Wellcome Sanger Institute COVID- |
| EPI_ISL_553312 | Lighthouse Lab in Alderley Park | Wellcome Sanger Institute for the COVID-19 Genomics UK (COG-UK) consortium | The Lighthouse Lab in Alderley Park and Alex Alderton, Roberto Amato, Sonia Goncalves, Ewan Harrison, David K. Jackson, Ian Johnston, Dominic Kwiatkowski, Cordelia Langford, John Sillitoe on behalf of the Wellcome Sang |
| EPI_ISL_553313 | Lighthouse Lab in Alderley Park | Wellcome Sanger Institute for the COVID-19 Genomics UK (COG-UK) consortium | The Lighthouse Lab in Alderley Park and Alex Alderton, Roberto Amato, Sonia Goncalves, Ewan Harrison, David K. Jackson, Ian Johnston, Dominic Kwiatkowski, Cordelia Langford, John Sillitoe on behalf of the Wellcome Sanger Institute COVID- |
| EPI_ISL_553315 | Lighthouse Lab in Alderley Park | Wellcome Sanger Institute for the COVID-19 Genomics UK (COG-UK) consortium | The Lighthouse Lab in Alderley Park and Alex Alderton, Roberto Amato, Sonia Goncalves, Ewan Harrison, David K. Jackson, Ian Johnston, Dominic Kwiatkowski, Cordelia Langford, John Sillitoe on behalf of the Wellcome Sang |
| EPI_ISL_553316, EPI_ISL_553317, EPI_ISL_553318 | Lighthouse Lab in Alderley Park | Wellcome Sanger Institute for the COVID-19 Genomics UK (COG-UK) consortium | The Lighthouse Lab in Alderley Park and Alex Alderton, Roberto Amato, Sonia Goncalves, Ewan Harrison, David K. Jackson, Ian Johnston, Dominic Kwiatkowski, Cordelia Langford, John Sillitoe on behalf of the Wellcome Sanger Institute COVID- |
| EPI_ISL_553319, EPI_ISL_553320 | Lighthouse Lab in Alderley Park | Wellcome Sanger Institute for the COVID-19 Genomics UK (COG-UK) consortium | The Lighthouse Lab in Alderley Park and Alex Alderton, Roberto Amato, Sonia Goncalves, Ewan Harrison, David K. Jackson, Ian Johnston, Dominic Kwiatkowski, Cordelia Langford, John Sillitoe on behalf of the Wellcome Sang |
| EPI_ISL_553321, EPI_ISL_553322, EPI_ISL_553324, EPI_ISL_553325, EPI_ISL_553327 | Lighthouse Lab in Alderley Park | Wellcome Sanger Institute for the COVID-19 Genomics UK (COG-UK) consortium | The Lighthouse Lab in Alderley Park and Alex Alderton, Roberto Amato, Sonia Goncalves, Ewan Harrison, David K. Jackson, Ian Johnston, Dominic Kwiatkowski, Cordelia Langford, John Sillitoe on behalf of the Wellcome Sanger Institute COVID- |
| EPI_ISL_553328 | Lighthouse Lab in Alderley Park | Wellcome Sanger Institute for the COVID-19 Genomics UK (COG-UK) consortium | The Lighthouse Lab in Alderley Park and Alex Alderton, Roberto Amato, Sonia Goncalves, Ewan Harrison, David K. Jackson, Ian Johnston, Dominic Kwiatkowski, Cordelia Langford, John Sillitoe on behalf of the Wellcome Sang |

| | | | |
|---|---|---|---|
| EPI_ISL_553329, EPI_ISL_553330 | Lighthouse Lab in Alderley Park | Wellcome Sanger Institute for the COVID-19 Genomics UK (COG-UK) consortium | The Lighthouse Lab in Alderley Park and Alex Alderton, Roberto Amato, Sonia Goncalves, Ewan Harrison, David K. Jackson, Ian Johnston, Dominic Kwiatkowski, Cordelia Langford, John Sillitoe on behalf of the Wellcome Sanger Institute COVID- |
| EPI_ISL_553331 | Lighthouse Lab in Alderley Park | Wellcome Sanger Institute for the COVID-19 Genomics UK (COG-UK) consortium | The Lighthouse Lab in Alderley Park and Alex Alderton, Roberto Amato, Sonia Goncalves, Ewan Harrison, David K. Jackson, Ian Johnston, Dominic Kwiatkowski, Cordelia Langford, John Sillitoe on behalf of the Wellcome Sang |
| EPI_ISL_553332, EPI_ISL_553333 | Lighthouse Lab in Alderley Park | Wellcome Sanger Institute for the COVID-19 Genomics UK (COG-UK) consortium | The Lighthouse Lab in Alderley Park and Alex Alderton, Roberto Amato, Sonia Goncalves, Ewan Harrison, David K. Jackson, Ian Johnston, Dominic Kwiatkowski, Cordelia Langford, John Sillitoe on behalf of the Wellcome Sanger Institute COVID- |
| EPI_ISL_553334 | Lighthouse Lab in Alderley Park | Wellcome Sanger Institute for the COVID-19 Genomics UK (COG-UK) consortium | The Lighthouse Lab in Alderley Park and Alex Alderton, Roberto Amato, Sonia Goncalves, Ewan Harrison, David K. Jackson, Ian Johnston, Dominic Kwiatkowski, Cordelia Langford, John Sillitoe on behalf of the Wellcome Sang |
| EPI_ISL_553335, EPI_ISL_553337 | Lighthouse Lab in Alderley Park | Wellcome Sanger Institute for the COVID-19 Genomics UK (COG-UK) consortium | The Lighthouse Lab in Alderley Park and Alex Alderton, Roberto Amato, Sonia Goncalves, Ewan Harrison, David K. Jackson, Ian Johnston, Dominic Kwiatkowski, Cordelia Langford, John Sillitoe on behalf of the Wellcome Sanger Institute COVID- |
| EPI_ISL_553338, EPI_ISL_553339 | Lighthouse Lab in Alderley Park | Wellcome Sanger Institute for the COVID-19 Genomics UK (COG-UK) consortium | The Lighthouse Lab in Alderley Park and Alex Alderton, Roberto Amato, Sonia Goncalves, Ewan Harrison, David K. Jackson, Ian Johnston, Dominic Kwiatkowski, Cordelia Langford, John Sillitoe on behalf of the Wellcome Sang |
| EPI_ISL_553340, EPI_ISL_553341, EPI_ISL_553342, EPI_ISL_553343 | Lighthouse Lab in Alderley Park | Wellcome Sanger Institute for the COVID-19 Genomics UK (COG-UK) consortium | The Lighthouse Lab in Alderley Park and Alex Alderton, Roberto Amato, Sonia Goncalves, Ewan Harrison, David K. Jackson, Ian Johnston, Dominic Kwiatkowski, Cordelia Langford, John Sillitoe on behalf of the Wellcome Sanger Institute COVID- |
| EPI_ISL_553344 | Lighthouse Lab in Alderley Park | Wellcome Sanger Institute for the COVID-19 Genomics UK (COG-UK) consortium | The Lighthouse Lab in Alderley Park and Alex Alderton, Roberto Amato, Sonia Goncalves, Ewan Harrison, David K. Jackson, Ian Johnston, Dominic Kwiatkowski, Cordelia Langford, John Sillitoe on behalf of the Wellcome Sang |
| EPI_ISL_553345, EPI_ISL_553610, EPI_ISL_553702, EPI_ISL_553708, EPI_ISL_553710, EPI_ISL_553713 | Lighthouse Lab in Alderley Park | Wellcome Sanger Institute for the COVID-19 Genomics UK (COG-UK) consortium | The Lighthouse Lab in Alderley Park and Alex Alderton, Roberto Amato, Sonia Goncalves, Ewan Harrison, David K. Jackson, Ian Johnston, Dominic Kwiatkowski, Cordelia Langford, John Sillitoe on behalf of the Wellcome Sanger Institute COVID- |
| EPI_ISL_553715, EPI_ISL_553717, EPI_ISL_553721 | Lighthouse Lab in Alderley Park | Wellcome Sanger Institute for the COVID-19 Genomics UK (COG-UK) consortium | The Lighthouse Lab in Alderley Park and Alex Alderton, Roberto Amato, Sonia Goncalves, Ewan Harrison, David K. Jackson, Ian Johnston, Dominic Kwiatkowski, Cordelia Langford, John Sillitoe on behalf of the Wellcome Sang |
| EPI_ISL_553723, EPI_ISL_553726, EPI_ISL_553727 | Lighthouse Lab in Alderley Park | Wellcome Sanger Institute for the COVID-19 Genomics UK (COG-UK) consortium | The Lighthouse Lab in Alderley Park and Alex Alderton, Roberto Amato, Sonia Goncalves, Ewan Harrison, David K. Jackson, Ian Johnston, Dominic Kwiatkowski, Cordelia Langford, John Sillitoe on behalf of the Wellcome Sanger Institute COVID- |
| EPI_ISL_553728 | Lighthouse Lab in Alderley Park | Wellcome Sanger Institute for the COVID-19 Genomics UK (COG-UK) consortium | The Lighthouse Lab in Alderley Park and Alex Alderton, Roberto Amato, Sonia Goncalves, Ewan Harrison, David K. Jackson, Ian Johnston, Dominic Kwiatkowski, Cordelia Langford, John Sillitoe on behalf of the Wellcome Sang |
| EPI_ISL_553729, EPI_ISL_553730 | Lighthouse Lab in Alderley Park | Wellcome Sanger Institute for the COVID-19 Genomics UK (COG-UK) consortium | The Lighthouse Lab in Alderley Park and Alex Alderton, Roberto Amato, Sonia Goncalves, Ewan Harrison, David K. Jackson, Ian Johnston, Dominic Kwiatkowski, Cordelia Langford, John Sillitoe on behalf of the Wellcome Sanger Institute COVID- |
| EPI_ISL_553731, EPI_ISL_553794 | Lighthouse Lab in Alderley Park | Wellcome Sanger Institute for the COVID-19 Genomics UK (COG-UK) consortium | The Lighthouse Lab in Alderley Park and Alex Alderton, Roberto Amato, Sonia Goncalves, Ewan Harrison, David K. Jackson, Ian Johnston, Dominic Kwiatkowski, Cordelia Langford, John Sillitoe on behalf of the Wellcome Sang |
| EPI_ISL_554563, EPI_ISL_555034 | Lighthouse Lab in Alderley Park | Wellcome Sanger Institute for the COVID-19 Genomics UK (COG-UK) consortium | The Lighthouse Lab in Alderley Park and Alex Alderton, Roberto Amato, Sonia Goncalves, Ewan Harrison, David K. Jackson, Ian Johnston, Dominic Kwiatkowski, Cordelia Langford, John Sillitoe on behalf of the Wellcome Sanger Institute COVID- |
| EPI_ISL_555181, EPI_ISL_556010, EPI_ISL_556018 | Lighthouse Lab in Alderley Park | Wellcome Sanger Institute for the COVID-19 Genomics UK (COG-UK) consortium | The Lighthouse Lab in Alderley Park and Alex Alderton, Roberto Amato, Sonia Goncalves, Ewan Harrison, David K. Jackson, Ian Johnston, Dominic Kwiatkowski, Cordelia Langford, John Sillitoe on behalf of the Wellcome Sang |
| EPI_ISL_556075 | Lighthouse Lab in Milton Keynes | Wellcome Sanger Institute for the COVID-19 Genomics UK (COG-UK) consortium | The Lighthouse Lab in Milton Keynes and Alex Alderton, Roberto Amato, Sonia Goncalves, Ewan Harrison, David K. Jackson, Ian Johnston, Dominic Kwiatkowski, Cordelia Langford, John Sillitoe on behalf of the Wellcome Sanger Institute COVID- |
| EPI_ISL_556081, EPI_ISL_556084, EPI_ISL_556089 | Lighthouse Lab in Alderley Park | Wellcome Sanger Institute for the COVID-19 Genomics UK (COG-UK) consortium | The Lighthouse Lab in Alderley Park and Alex Alderton, Roberto Amato, Sonia Goncalves, Ewan Harrison, David K. Jackson, Ian Johnston, Dominic Kwiatkowski, Cordelia Langford, John Sillitoe on behalf of the Wellcome Sanger Institute COVID- |
| EPI_ISL_556090 | Lighthouse Lab in Milton Keynes | Wellcome Sanger Institute for the COVID-19 Genomics UK (COG-UK) consortium | The Lighthouse Lab in Milton Keynes and Alex Alderton, Roberto Amato, Sonia Goncalves, Ewan Harrison, David K. Jackson, Ian Johnston, Dominic Kwiatkowski, Cordelia Langford, John Sillitoe on behalf of the Wellcome Sanger Institute COVID- |
| EPI_ISL_556101 | Lighthouse Lab in Alderley Park | Wellcome Sanger Institute for the COVID-19 Genomics UK (COG-UK) consortium | The Lighthouse Lab in Alderley Park and Alex Alderton, Roberto Amato, Sonia Goncalves, Ewan Harrison, David K. Jackson, Ian Johnston, Dominic Kwiatkowski, Cordelia Langford, John Sillitoe on behalf of the Wellcome Sang |
| EPI_ISL_556106, EPI_ISL_556107, EPI_ISL_556122 | Lighthouse Lab in Alderley Park | Wellcome Sanger Institute for the COVID-19 Genomics UK (COG-UK) consortium | The Lighthouse Lab in Alderley Park and Alex Alderton, Roberto Amato, Sonia Goncalves, Ewan Harrison, David K. Jackson, Ian Johnston, Dominic Kwiatkowski, Cordelia Langford, John Sillitoe on behalf of the Wellcome Sanger Institute COVID- |
| EPI_ISL_556130, EPI_ISL_556151, EPI_ISL_556184, EPI_ISL_556191 | Lighthouse Lab in Alderley Park | Wellcome Sanger Institute for the COVID-19 Genomics UK (COG-UK) consortium | The Lighthouse Lab in Alderley Park and Alex Alderton, Roberto Amato, Sonia Goncalves, Ewan Harrison, David K. Jackson, Ian Johnston, Dominic Kwiatkowski, Cordelia Langford, John Sillitoe on behalf of the Wellcome Sang |
| EPI_ISL_556202, EPI_ISL_556230, EPI_ISL_556258 | Lighthouse Lab in Alderley Park | Wellcome Sanger Institute for the COVID-19 Genomics UK (COG-UK) consortium | The Lighthouse Lab in Alderley Park and Alex Alderton, Roberto Amato, Sonia Goncalves, Ewan Harrison, David K. Jackson, Ian Johnston, Dominic Kwiatkowski, Cordelia Langford, John Sillitoe on behalf of the Wellcome Sanger Institute COVID- |
| EPI_ISL_556445, EPI_ISL_556458 | Lighthouse Lab in Alderley Park | Wellcome Sanger Institute for the COVID-19 Genomics | The Lighthouse Lab in Alderley Park and Alex Alderton, Roberto Amato, Sonia Goncalves, Ewan Harrison, David K. Jackson, Ian Johnston, Dominic Kwiatkowski, Cordelia Langford, John Sillitoe on behalf of the Wellcome Sang |

| | | | |
|---|---|---|---|
| | | UK (COG-UK) consortium | |
| EPI_ISL_556576, EPI_ISL_556577, EPI_ISL_556578, EPI_ISL_556579, EPI_ISL_556580, EPI_ISL_556581, EPI_ISL_556582, EPI_ISL_556583, EPI_ISL_556584, EPI_ISL_556587, EPI_ISL_556588, EPI_ISL_556589, EPI_ISL_556590, EPI_ISL_556591, EPI_ISL_556592, EPI_ISL_556593, EPI_ISL_556594, EPI_ISL_556595, EPI_I<br>EPI_ISL_556600, EPI_ISL_556601, EPI_ISL_556602, EPI_ISL_556603, EPI_ISL_556604, EPI_ISL_556605, EPI_ISL_556607, EPI_ISL_556608, EPI_ISL_556609, EPI_ISL_556610, EPI_ISL_556611, EPI_ISL_556614, EPI_ISL_556615, EPI_ISL_556616, EPI_ISL_556617, EPI_ISL_556618, EPI_ISL_556619, EPI_ISL_556620, EPI_I<br>EPI_ISL_556625, EPI_ISL_556626, EPI_ISL_556627, EPI_ISL_556628, EPI_ISL_556629, EPI_ISL_556630, EPI_ISL_556631, EPI_ISL_556632, EPI_ISL_556633, EPI_ISL_556634, EPI_ISL_556635, EPI_ISL_556636, EPI_ISL_556637, EPI_ISL_556638, EPI_ISL_556985, EPI_ISL_556986, EPI_ISL_556987, EPI_ISL_556988, EPI_I<br>EPI_ISL_556993, EPI_ISL_556994, EPI_ISL_556995, EPI_ISL_556996, EPI_ISL_556997, EPI_ISL_556998, EPI_ISL_557000, EPI_ISL_557001 | | | |
| see above | Lighthouse Lab in Alderley Park | Wellcome Sanger Institute for the COVID-19 Genomics UK (COG-UK) consortium | The Lighthouse Lab in Alderley Park and Alex Alderton, Roberto Amato, Sonia Goncalves, Ewan Harrison, David K. Jackson, Ian Johnston, Dominic Kwiatkowski, Cordelia Langford, John Sillitoe on behalf of the Wellcome Sanger Institute COVID- |
| EPI_ISL_557002 | Lighthouse Lab in Alderley Park | Wellcome Sanger Institute for the COVID-19 Genomics UK (COG-UK) Consortium | The Lighthouse Lab in Alderley Park and Alex Alderton, Roberto Amato, Sonia Goncalves, Ewan Harrison, David K. Jackson, Ian Johnston, Dominic Kwiatkowski, Cordelia Langford, John Sillitoe on behalf of the Wellcome Sang |
| EPI_ISL_557004, EPI_ISL_557005, EPI_ISL_557006, EPI_ISL_557007, EPI_ISL_557008, EPI_ISL_557009, EPI_ISL_557010, EPI_ISL_557011, EPI_ISL_557012, EPI_ISL_557013, EPI_ISL_557014, EPI_ISL_557015, EPI_ISL_557016, EPI_ISL_557017, EPI_ISL_557018, EPI_ISL_557019, EPI_ISL_557020, EPI_ISL_557021, EPI_I<br>EPI_ISL_557026, EPI_ISL_557027, EPI_ISL_557028, EPI_ISL_557029, EPI_ISL_557030, EPI_ISL_557031, EPI_ISL_557032, EPI_ISL_557033, EPI_ISL_557034, EPI_ISL_557035, EPI_ISL_557036, EPI_ISL_557037, EPI_ISL_557038, EPI_ISL_557039, EPI_ISL_557040, EPI_ISL_557041, EPI_ISL_557042 | | | |
| see above | Lighthouse Lab in Alderley Park | Wellcome Sanger Institute for the COVID-19 Genomics UK (COG-UK) consortium | The Lighthouse Lab in Alderley Park and Alex Alderton, Roberto Amato, Sonia Goncalves, Ewan Harrison, David K. Jackson, Ian Johnston, Dominic Kwiatkowski, Cordelia Langford, John Sillitoe on behalf of the Wellcome Sanger Institute COVID- |
| EPI_ISL_557043 | Lighthouse Lab in Alderley Park | Wellcome Sanger Institute for the COVID-19 Genomics UK (COG-UK) Consortium | The Lighthouse Lab in Alderley Park and Alex Alderton, Roberto Amato, Sonia Goncalves, Ewan Harrison, David K. Jackson, Ian Johnston, Dominic Kwiatkowski, Cordelia Langford, John Sillitoe on behalf of the Wellcome Sang |
| EPI_ISL_557044, EPI_ISL_557046, EPI_ISL_557047, EPI_ISL_557048, EPI_ISL_557049, EPI_ISL_557050, EPI_ISL_557051, EPI_ISL_557052, EPI_ISL_557053, EPI_ISL_557054, EPI_ISL_557055, EPI_ISL_557056, EPI_ISL_557057, EPI_ISL_557058, EPI_ISL_557059, EPI_ISL_557060, EPI_ISL_557061, EPI_ISL_557062, EPI_I<br>EPI_ISL_557067, EPI_ISL_557068, EPI_ISL_557069, EPI_ISL_557070, EPI_ISL_557071, EPI_ISL_557072, EPI_ISL_557073, EPI_ISL_557074, EPI_ISL_557075, EPI_ISL_557076, EPI_ISL_557077, EPI_ISL_557078, EPI_ISL_557079, EPI_ISL_557080, EPI_ISL_557081, EPI_ISL_557082, EPI_ISL_557083, EPI_ISL_557084, EPI_I<br>EPI_ISL_557089, EPI_ISL_557090, EPI_ISL_557091, EPI_ISL_557092, EPI_ISL_557093, EPI_ISL_557094, EPI_ISL_557095, EPI_ISL_557096, EPI_ISL_557097, EPI_ISL_557098, EPI_ISL_557099, EPI_ISL_557100, EPI_ISL_557101, EPI_ISL_557102, EPI_ISL_557103, EPI_ISL_557104, EPI_ISL_557105, EPI_ISL_557106, EPI_I | | | |
| see above | Lighthouse Lab in Alderley Park | Wellcome Sanger Institute for the COVID-19 Genomics UK (COG-UK) consortium | The Lighthouse Lab in Alderley Park and Alex Alderton, Roberto Amato, Sonia Goncalves, Ewan Harrison, David K. Jackson, Ian Johnston, Dominic Kwiatkowski, Cordelia Langford, John Sillitoe on behalf of the Wellcome Sanger Institute COVID- |
| EPI_ISL_557110 | Lighthouse Lab in Alderley Park | Wellcome Sanger Institute for the COVID-19 Genomics UK (COG-UK) Consortium | The Lighthouse Lab in Alderley Park and Alex Alderton, Roberto Amato, Sonia Goncalves, Ewan Harrison, David K. Jackson, Ian Johnston, Dominic Kwiatkowski, Cordelia Langford, John Sillitoe on behalf of the Wellcome Sang |
| EPI_ISL_557111, EPI_ISL_557112, EPI_ISL_557113, EPI_ISL_557114, EPI_ISL_557115, EPI_ISL_557116, EPI_ISL_557117, EPI_ISL_557118, EPI_ISL_557119, EPI_ISL_557120, EPI_ISL_557121, EPI_ISL_557122, EPI_ISL_557123, EPI_ISL_557124, EPI_ISL_557125, EPI_ISL_557126, EPI_ISL_557127, EPI_ISL_557128, EPI_I<br>EPI_ISL_557133, EPI_ISL_557134, EPI_ISL_557135, EPI_ISL_557136, EPI_ISL_557138, EPI_ISL_557139, EPI_ISL_557140, EPI_ISL_557141 | | | |
| see above | Lighthouse Lab in Alderley Park | Wellcome Sanger Institute for the COVID-19 Genomics UK (COG-UK) consortium | The Lighthouse Lab in Alderley Park and Alex Alderton, Roberto Amato, Sonia Goncalves, Ewan Harrison, David K. Jackson, Ian Johnston, Dominic Kwiatkowski, Cordelia Langford, John Sillitoe on behalf of the Wellcome Sanger Institute COVID- |
| EPI_ISL_557142 | Lighthouse Lab in Alderley Park | Wellcome Sanger Institute for the COVID-19 Genomics UK (COG-UK) Consortium | The Lighthouse Lab in Alderley Park and Alex Alderton, Roberto Amato, Sonia Goncalves, Ewan Harrison, David K. Jackson, Ian Johnston, Dominic Kwiatkowski, Cordelia Langford, John Sillitoe on behalf of the Wellcome Sang |
| EPI_ISL_557143, EPI_ISL_557144, EPI_ISL_557145, EPI_ISL_557146, EPI_ISL_557147, EPI_ISL_557148, EPI_ISL_557149, EPI_ISL_557150, EPI_ISL_557151, EPI_ISL_557152, EPI_ISL_557153, EPI_ISL_557154, EPI_ISL_557155, EPI_ISL_557156, EPI_ISL_557157, EPI_ISL_557158, EPI_ISL_557159, EPI_ISL_557160, EPI_I<br>EPI_ISL_557165, EPI_ISL_557351, EPI_ISL_557354 | | | |
| see above | Lighthouse Lab in Alderley Park | Wellcome Sanger Institute for the COVID-19 Genomics UK (COG-UK) consortium | The Lighthouse Lab in Alderley Park and Alex Alderton, Roberto Amato, Sonia Goncalves, Ewan Harrison, David K. Jackson, Ian Johnston, Dominic Kwiatkowski, Cordelia Langford, John Sillitoe on behalf of the Wellcome Sanger Institute COVID- |
| EPI_ISL_557355 | Lighthouse Lab in Milton Keynes | Wellcome Sanger Institute for the COVID-19 Genomics UK (COG-UK) Consortium | The Lighthouse Lab in Milton Keynes and Alex Alderton, Roberto Amato, Sonia Goncalves, Ewan Harrison, David K. Jackson, Ian Johnston, Dominic Kwiatkowski, Cordelia Langford, John Sillitoe on behalf of the Wellcome Sang |
| EPI_ISL_557356, EPI_ISL_557359, EPI_ISL_557361, EPI_ISL_557363, EPI_ISL_557366 | Lighthouse Lab in Alderley Park | Wellcome Sanger Institute for the COVID-19 Genomics UK (COG-UK) consortium | The Lighthouse Lab in Alderley Park and Alex Alderton, Roberto Amato, Sonia Goncalves, Ewan Harrison, David K. Jackson, Ian Johnston, Dominic Kwiatkowski, Cordelia Langford, John Sillitoe on behalf of the Wellcome Sanger Institute COVID- |
| EPI_ISL_557367, EPI_ISL_557368, EPI_ISL_557369, EPI_ISL_557370 | Lighthouse Lab in Milton Keynes | Wellcome Sanger Institute for the COVID-19 Genomics UK (COG-UK) consortium | The Lighthouse Lab in Milton Keynes and Alex Alderton, Roberto Amato, Sonia Goncalves, Ewan Harrison, David K. Jackson, Ian Johnston, Dominic Kwiatkowski, Cordelia Langford, John Sillitoe on behalf of the Wellcome Sanger Institute COVID- |
| EPI_ISL_557371, EPI_ISL_557373, EPI_ISL_557374 | Lighthouse Lab in Alderley Park | Wellcome Sanger Institute for the COVID-19 Genomics UK (COG-UK) consortium | The Lighthouse Lab in Alderley Park and Alex Alderton, Roberto Amato, Sonia Goncalves, Ewan Harrison, David K. Jackson, Ian Johnston, Dominic Kwiatkowski, Cordelia Langford, John Sillitoe on behalf of the Wellcome Sanger Institute COVID- |
| EPI_ISL_557375, EPI_ISL_557376 | Lighthouse Lab in Milton Keynes | Wellcome Sanger Institute for the COVID-19 Genomics UK (COG-UK) consortium | The Lighthouse Lab in Milton Keynes and Alex Alderton, Roberto Amato, Sonia Goncalves, Ewan Harrison, David K. Jackson, Ian Johnston, Dominic Kwiatkowski, Cordelia Langford, John Sillitoe on behalf of the Wellcome Sanger Institute COVID- |
| EPI_ISL_557378 | Lighthouse Lab in Alderley Park | Wellcome Sanger Institute for the COVID-19 Genomics UK (COG-UK) consortium | The Lighthouse Lab in Alderley Park and Alex Alderton, Roberto Amato, Sonia Goncalves, Ewan Harrison, David K. Jackson, Ian Johnston, Dominic Kwiatkowski, Cordelia Langford, John Sillitoe on behalf of the Wellcome Sanger Institute COVID- |
| EPI_ISL_557379 | Lighthouse Lab in Milton Keynes | Wellcome Sanger Institute for the COVID-19 Genomics UK (COG-UK) Consortium | The Lighthouse Lab in Milton Keynes and Alex Alderton, Roberto Amato, Sonia Goncalves, Ewan Harrison, David K. Jackson, Ian Johnston, Dominic Kwiatkowski, Cordelia Langford, John Sillitoe on behalf of the Wellcome Sang |
| EPI_ISL_557380, EPI_ISL_557381, EPI_ISL_557383 | Lighthouse Lab in Alderley Park | Wellcome Sanger Institute for the COVID-19 Genomics UK (COG-UK) consortium | The Lighthouse Lab in Alderley Park and Alex Alderton, Roberto Amato, Sonia Goncalves, Ewan Harrison, David K. Jackson, Ian Johnston, Dominic Kwiatkowski, Cordelia Langford, John Sillitoe on behalf of the Wellcome Sanger Institute COVID- |
| EPI_ISL_557385, EPI_ISL_557386 | Lighthouse Lab in Milton Keynes | Wellcome Sanger Institute for the COVID-19 Genomics UK (COG-UK) consortium | The Lighthouse Lab in Milton Keynes and Alex Alderton, Roberto Amato, Sonia Goncalves, Ewan Harrison, David K. Jackson, Ian Johnston, Dominic Kwiatkowski, Cordelia Langford, John Sillitoe on behalf of the Wellcome Sanger Institute COVID- |
| EPI_ISL_557388 | Lighthouse Lab in Alderley Park | Wellcome Sanger Institute for the COVID-19 Genomics UK (COG-UK) consortium | The Lighthouse Lab in Alderley Park and Alex Alderton, Roberto Amato, Sonia Goncalves, Ewan Harrison, David K. Jackson, Ian Johnston, Dominic Kwiatkowski, Cordelia Langford, John Sillitoe on behalf of the Wellcome Sanger Institute COVID- |
| EPI_ISL_557389 | Lighthouse Lab in Milton | Wellcome Sanger Institute | The Lighthouse Lab in Milton Keynes and Alex Alderton, Roberto Amato, Sonia Goncalves, Ewan Harrison, David K. Jackson, Ian Johnston, Dominic Kwiatkowski, Cordelia Langford, John Sillitoe on behalf of the Wellcome Sanger Institute COVID- |

| | | | |
|---|---|---|---|
| | Keynes | for the COVID-19 Genomics UK (COG-UK) consortium | |
| EPI_ISL_557390 | Lighthouse Lab in Alderley Park | Wellcome Sanger Institute for the COVID-19 Genomics UK (COG-UK) consortium | The Lighthouse Lab in Alderley Park and Alex Alderton, Roberto Amato, Sonia Goncalves, Ewan Harrison, David K. Jackson, Ian Johnston, Dominic Kwiatkowski, Cordelia Langford, John Sillitoe on behalf of the Wellcome Sanger Institute COVID- |
| EPI_ISL_557392, EPI_ISL_557395 | Lighthouse Lab in Milton Keynes | Wellcome Sanger Institute for the COVID-19 Genomics UK (COG-UK) consortium | The Lighthouse Lab in Milton Keynes and Alex Alderton, Roberto Amato, Sonia Goncalves, Ewan Harrison, David K. Jackson, Ian Johnston, Dominic Kwiatkowski, Cordelia Langford, John Sillitoe on behalf of the Wellcome Sanger Institute COVID- |
| EPI_ISL_557398, EPI_ISL_557399 | Lighthouse Lab in Alderley Park | Wellcome Sanger Institute for the COVID-19 Genomics UK (COG-UK) consortium | The Lighthouse Lab in Alderley Park and Alex Alderton, Roberto Amato, Sonia Goncalves, Ewan Harrison, David K. Jackson, Ian Johnston, Dominic Kwiatkowski, Cordelia Langford, John Sillitoe on behalf of the Wellcome Sanger Institute COVID- |
| EPI_ISL_557401, EPI_ISL_557403 | Lighthouse Lab in Milton Keynes | Wellcome Sanger Institute for the COVID-19 Genomics UK (COG-UK) consortium | The Lighthouse Lab in Milton Keynes and Alex Alderton, Roberto Amato, Sonia Goncalves, Ewan Harrison, David K. Jackson, Ian Johnston, Dominic Kwiatkowski, Cordelia Langford, John Sillitoe on behalf of the Wellcome Sanger Institute COVID- |
| EPI_ISL_557404, EPI_ISL_557405, EPI_ISL_557406 | Lighthouse Lab in Alderley Park | Wellcome Sanger Institute for the COVID-19 Genomics UK (COG-UK) consortium | The Lighthouse Lab in Alderley Park and Alex Alderton, Roberto Amato, Sonia Goncalves, Ewan Harrison, David K. Jackson, Ian Johnston, Dominic Kwiatkowski, Cordelia Langford, John Sillitoe on behalf of the Wellcome Sanger Institute COVID- |
| EPI_ISL_557407 | Lighthouse Lab in Milton Keynes | Wellcome Sanger Institute for the COVID-19 Genomics UK (COG-UK) consortium | The Lighthouse Lab in Milton Keynes and Alex Alderton, Roberto Amato, Sonia Goncalves, Ewan Harrison, David K. Jackson, Ian Johnston, Dominic Kwiatkowski, Cordelia Langford, John Sillitoe on behalf of the Wellcome Sanger Institute COVID- |
| EPI_ISL_557408 | Lighthouse Lab in Alderley Park | Wellcome Sanger Institute for the COVID-19 Genomics UK (COG-UK) consortium | The Lighthouse Lab in Alderley Park and Alex Alderton, Roberto Amato, Sonia Goncalves, Ewan Harrison, David K. Jackson, Ian Johnston, Dominic Kwiatkowski, Cordelia Langford, John Sillitoe on behalf of the Wellcome Sanger Institute COVID- |
| EPI_ISL_557409, EPI_ISL_557410 | Lighthouse Lab in Milton Keynes | Wellcome Sanger Institute for the COVID-19 Genomics UK (COG-UK) consortium | The Lighthouse Lab in Milton Keynes and Alex Alderton, Roberto Amato, Sonia Goncalves, Ewan Harrison, David K. Jackson, Ian Johnston, Dominic Kwiatkowski, Cordelia Langford, John Sillitoe on behalf of the Wellcome Sanger Institute COVID- |
| EPI_ISL_557411, EPI_ISL_557412, EPI_ISL_557414, EPI_ISL_557415, EPI_ISL_557417 | Lighthouse Lab in Alderley Park | Wellcome Sanger Institute for the COVID-19 Genomics UK (COG-UK) consortium | The Lighthouse Lab in Alderley Park and Alex Alderton, Roberto Amato, Sonia Goncalves, Ewan Harrison, David K. Jackson, Ian Johnston, Dominic Kwiatkowski, Cordelia Langford, John Sillitoe on behalf of the Wellcome Sanger Institute COVID- |
| EPI_ISL_557418 | Lighthouse Lab in Milton Keynes | Wellcome Sanger Institute for the COVID-19 Genomics UK (COG-UK) consortium | The Lighthouse Lab in Milton Keynes and Alex Alderton, Roberto Amato, Sonia Goncalves, Ewan Harrison, David K. Jackson, Ian Johnston, Dominic Kwiatkowski, Cordelia Langford, John Sillitoe on behalf of the Wellcome Sanger Institute COVID- |
| EPI_ISL_557419, EPI_ISL_557423, EPI_ISL_557425 | Lighthouse Lab in Alderley Park | Wellcome Sanger Institute for the COVID-19 Genomics UK (COG-UK) consortium | The Lighthouse Lab in Alderley Park and Alex Alderton, Roberto Amato, Sonia Goncalves, Ewan Harrison, David K. Jackson, Ian Johnston, Dominic Kwiatkowski, Cordelia Langford, John Sillitoe on behalf of the Wellcome Sanger Institute COVID- |
| EPI_ISL_557426 | Lighthouse Lab in Milton Keynes | Wellcome Sanger Institute for the COVID-19 Genomics UK (COG-UK) consortium | The Lighthouse Lab in Milton Keynes and Alex Alderton, Roberto Amato, Sonia Goncalves, Ewan Harrison, David K. Jackson, Ian Johnston, Dominic Kwiatkowski, Cordelia Langford, John Sillitoe on behalf of the Wellcome Sanger Institute COVID- |
| EPI_ISL_557428, EPI_ISL_557430, EPI_ISL_557433 | Lighthouse Lab in Alderley Park | Wellcome Sanger Institute for the COVID-19 Genomics UK (COG-UK) consortium | The Lighthouse Lab in Alderley Park and Alex Alderton, Roberto Amato, Sonia Goncalves, Ewan Harrison, David K. Jackson, Ian Johnston, Dominic Kwiatkowski, Cordelia Langford, John Sillitoe on behalf of the Wellcome Sanger Institute COVID- |
| EPI_ISL_557435 | Lighthouse Lab in Milton Keynes | Wellcome Sanger Institute for the COVID-19 Genomics UK (COG-UK) consortium | The Lighthouse Lab in Milton Keynes and Alex Alderton, Roberto Amato, Sonia Goncalves, Ewan Harrison, David K. Jackson, Ian Johnston, Dominic Kwiatkowski, Cordelia Langford, John Sillitoe on behalf of the Wellcome Sanger Institute COVID- |
| EPI_ISL_557436 | Lighthouse Lab in Alderley Park | Wellcome Sanger Institute for the COVID-19 Genomics UK (COG-UK) consortium | The Lighthouse Lab in Alderley Park and Alex Alderton, Roberto Amato, Sonia Goncalves, Ewan Harrison, David K. Jackson, Ian Johnston, Dominic Kwiatkowski, Cordelia Langford, John Sillitoe on behalf of the Wellcome Sanger Institute COVID- |
| EPI_ISL_557437, EPI_ISL_557438 | Lighthouse Lab in Milton Keynes | Wellcome Sanger Institute for the COVID-19 Genomics UK (COG-UK) consortium | The Lighthouse Lab in Milton Keynes and Alex Alderton, Roberto Amato, Sonia Goncalves, Ewan Harrison, David K. Jackson, Ian Johnston, Dominic Kwiatkowski, Cordelia Langford, John Sillitoe on behalf of the Wellcome Sanger Institute COVID- |
| EPI_ISL_557439, EPI_ISL_557440 | Lighthouse Lab in Alderley Park | Wellcome Sanger Institute for the COVID-19 Genomics UK (COG-UK) consortium | The Lighthouse Lab in Alderley Park and Alex Alderton, Roberto Amato, Sonia Goncalves, Ewan Harrison, David K. Jackson, Ian Johnston, Dominic Kwiatkowski, Cordelia Langford, John Sillitoe on behalf of the Wellcome Sanger Institute COVID- |
| EPI_ISL_557444, EPI_ISL_557445 | Lighthouse Lab in Milton Keynes | Wellcome Sanger Institute for the COVID-19 Genomics UK (COG-UK) consortium | The Lighthouse Lab in Milton Keynes and Alex Alderton, Roberto Amato, Sonia Goncalves, Ewan Harrison, David K. Jackson, Ian Johnston, Dominic Kwiatkowski, Cordelia Langford, John Sillitoe on behalf of the Wellcome Sanger Institute COVID- |
| EPI_ISL_557446, EPI_ISL_557447, EPI_ISL_557449, EPI_ISL_557453, EPI_ISL_557454 | Lighthouse Lab in Alderley Park | Wellcome Sanger Institute for the COVID-19 Genomics UK (COG-UK) consortium | The Lighthouse Lab in Alderley Park and Alex Alderton, Roberto Amato, Sonia Goncalves, Ewan Harrison, David K. Jackson, Ian Johnston, Dominic Kwiatkowski, Cordelia Langford, John Sillitoe on behalf of the Wellcome Sanger Institute COVID- |
| EPI_ISL_557456, EPI_ISL_557458, EPI_ISL_557461, EPI_ISL_557465 | Lighthouse Lab in Milton Keynes | Wellcome Sanger Institute for the COVID-19 Genomics UK (COG-UK) consortium | The Lighthouse Lab in Milton Keynes and Alex Alderton, Roberto Amato, Sonia Goncalves, Ewan Harrison, David K. Jackson, Ian Johnston, Dominic Kwiatkowski, Cordelia Langford, John Sillitoe on behalf of the Wellcome Sanger Institute COVID- |
| EPI_ISL_557466 | Lighthouse Lab in Alderley Park | Wellcome Sanger Institute for the COVID-19 Genomics UK (COG-UK) consortium | The Lighthouse Lab in Alderley Park and Alex Alderton, Roberto Amato, Sonia Goncalves, Ewan Harrison, David K. Jackson, Ian Johnston, Dominic Kwiatkowski, Cordelia Langford, John Sillitoe on behalf of the Wellcome Sanger Institute COVID- |
| EPI_ISL_557467 | Lighthouse Lab in Milton Keynes | Wellcome Sanger Institute for the COVID-19 Genomics UK (COG-UK) consortium | The Lighthouse Lab in Milton Keynes and Alex Alderton, Roberto Amato, Sonia Goncalves, Ewan Harrison, David K. Jackson, Ian Johnston, Dominic Kwiatkowski, Cordelia Langford, John Sillitoe on behalf of the Wellcome Sanger Institute COVID- |
| EPI_ISL_557468, EPI_ISL_557469, EPI_ISL_557471, | Lighthouse Lab in Alderley Park | Wellcome Sanger Institute for the COVID-19 Genomics UK (COG-UK) consortium | The Lighthouse Lab in Alderley Park and Alex Alderton, Roberto Amato, Sonia Goncalves, Ewan Harrison, David K. Jackson, Ian Johnston, Dominic Kwiatkowski, Cordelia Langford, John Sillitoe on behalf of the Wellcome Sanger Institute COVID- |

| | | | |
|---|---|---|---|
| EPI_ISL_557474, EPI_ISL_557476 | | | |
| EPI_ISL_557478 | Lighthouse Lab in Milton Keynes | Wellcome Sanger Institute for the COVID-19 Genomics UK (COG-UK) consortium | The Lighthouse Lab in Milton Keynes and Alex Alderton, Roberto Amato, Sonia Goncalves, Ewan Harrison, David K. Jackson, Ian Johnston, Dominic Kwiatkowski, Cordelia Langford, John Sillitoe on behalf of the Wellcome Sanger Institute COVID-[illegible] |
| EPI_ISL_557481, EPI_ISL_557482, EPI_ISL_557484, EPI_ISL_557485, EPI_ISL_557487, EPI_ISL_557488, EPI_ISL_557491 | Lighthouse Lab in Alderley Park | Wellcome Sanger Institute for the COVID-19 Genomics UK (COG-UK) consortium | The Lighthouse Lab in Alderley Park and Alex Alderton, Roberto Amato, Sonia Goncalves, Ewan Harrison, David K. Jackson, Ian Johnston, Dominic Kwiatkowski, Cordelia Langford, John Sillitoe on behalf of the Wellcome Sanger Institute COVID-[illegible] |
| EPI_ISL_557492 | Lighthouse Lab in Milton Keynes | Wellcome Sanger Institute for the COVID-19 Genomics UK (COG-UK) consortium | The Lighthouse Lab in Milton Keynes and Alex Alderton, Roberto Amato, Sonia Goncalves, Ewan Harrison, David K. Jackson, Ian Johnston, Dominic Kwiatkowski, Cordelia Langford, John Sillitoe on behalf of the Wellcome Sanger Institute COVID-[illegible] |
| EPI_ISL_557493 | Lighthouse Lab in Alderley Park | Wellcome Sanger Institute for the COVID-19 Genomics UK (COG-UK) consortium | The Lighthouse Lab in Alderley Park and Alex Alderton, Roberto Amato, Sonia Goncalves, Ewan Harrison, David K. Jackson, Ian Johnston, Dominic Kwiatkowski, Cordelia Langford, John Sillitoe on behalf of the Wellcome Sanger Institute COVID-[illegible] |
| EPI_ISL_557496 | Lighthouse Lab in Milton Keynes | Wellcome Sanger Institute for the COVID-19 Genomics UK (COG-UK) consortium | The Lighthouse Lab in Milton Keynes and Alex Alderton, Roberto Amato, Sonia Goncalves, Ewan Harrison, David K. Jackson, Ian Johnston, Dominic Kwiatkowski, Cordelia Langford, John Sillitoe on behalf of the Wellcome Sanger Institute COVID-[illegible] |
| EPI_ISL_557497, EPI_ISL_557499, EPI_ISL_557500 | Lighthouse Lab in Alderley Park | Wellcome Sanger Institute for the COVID-19 Genomics UK (COG-UK) consortium | The Lighthouse Lab in Alderley Park and Alex Alderton, Roberto Amato, Sonia Goncalves, Ewan Harrison, David K. Jackson, Ian Johnston, Dominic Kwiatkowski, Cordelia Langford, John Sillitoe on behalf of the Wellcome Sanger Institute COVID-[illegible] |
| EPI_ISL_557502, EPI_ISL_557503, EPI_ISL_557505 | Lighthouse Lab in Milton Keynes | Wellcome Sanger Institute for the COVID-19 Genomics UK (COG-UK) consortium | The Lighthouse Lab in Milton Keynes and Alex Alderton, Roberto Amato, Sonia Goncalves, Ewan Harrison, David K. Jackson, Ian Johnston, Dominic Kwiatkowski, Cordelia Langford, John Sillitoe on behalf of the Wellcome Sanger Institute COVID-[illegible] |
| EPI_ISL_557507, EPI_ISL_557511 | Lighthouse Lab in Alderley Park | Wellcome Sanger Institute for the COVID-19 Genomics UK (COG-UK) consortium | The Lighthouse Lab in Alderley Park and Alex Alderton, Roberto Amato, Sonia Goncalves, Ewan Harrison, David K. Jackson, Ian Johnston, Dominic Kwiatkowski, Cordelia Langford, John Sillitoe on behalf of the Wellcome Sanger Institute COVID-[illegible] |
| EPI_ISL_557512, EPI_ISL_557513, EPI_ISL_557516 | Lighthouse Lab in Milton Keynes | Wellcome Sanger Institute for the COVID-19 Genomics UK (COG-UK) consortium | The Lighthouse Lab in Milton Keynes and Alex Alderton, Roberto Amato, Sonia Goncalves, Ewan Harrison, David K. Jackson, Ian Johnston, Dominic Kwiatkowski, Cordelia Langford, John Sillitoe on behalf of the Wellcome Sanger Institute COVID-[illegible] |
| EPI_ISL_557518 | Lighthouse Lab in Alderley Park | Wellcome Sanger Institute for the COVID-19 Genomics UK (COG-UK) consortium | The Lighthouse Lab in Alderley Park and Alex Alderton, Roberto Amato, Sonia Goncalves, Ewan Harrison, David K. Jackson, Ian Johnston, Dominic Kwiatkowski, Cordelia Langford, John Sillitoe on behalf of the Wellcome Sanger Institute COVID-[illegible] |
| EPI_ISL_557521 | Lighthouse Lab in Milton Keynes | Wellcome Sanger Institute for the COVID-19 Genomics UK (COG-UK) consortium | The Lighthouse Lab in Milton Keynes and Alex Alderton, Roberto Amato, Sonia Goncalves, Ewan Harrison, David K. Jackson, Ian Johnston, Dominic Kwiatkowski, Cordelia Langford, John Sillitoe on behalf of the Wellcome Sanger Institute COVID-[illegible] |
| EPI_ISL_557522, EPI_ISL_557523, EPI_ISL_557526, EPI_ISL_557527, EPI_ISL_557528, EPI_ISL_557530, EPI_ISL_557531 | Lighthouse Lab in Alderley Park | Wellcome Sanger Institute for the COVID-19 Genomics UK (COG-UK) consortium | The Lighthouse Lab in Alderley Park and Alex Alderton, Roberto Amato, Sonia Goncalves, Ewan Harrison, David K. Jackson, Ian Johnston, Dominic Kwiatkowski, Cordelia Langford, John Sillitoe on behalf of the Wellcome Sanger Institute COVID-[illegible] |
| EPI_ISL_557532 | Lighthouse Lab in Milton Keynes | Wellcome Sanger Institute for the COVID-19 Genomics UK (COG-UK) consortium | The Lighthouse Lab in Milton Keynes and Alex Alderton, Roberto Amato, Sonia Goncalves, Ewan Harrison, David K. Jackson, Ian Johnston, Dominic Kwiatkowski, Cordelia Langford, John Sillitoe on behalf of the Wellcome Sanger Institute COVID-[illegible] |
| EPI_ISL_557533 | Lighthouse Lab in Alderley Park | Wellcome Sanger Institute for the COVID-19 Genomics UK (COG-UK) consortium | The Lighthouse Lab in Alderley Park and Alex Alderton, Roberto Amato, Sonia Goncalves, Ewan Harrison, David K. Jackson, Ian Johnston, Dominic Kwiatkowski, Cordelia Langford, John Sillitoe on behalf of the Wellcome Sanger Institute COVID-[illegible] |
| EPI_ISL_557535, EPI_ISL_557536 | Lighthouse Lab in Milton Keynes | Wellcome Sanger Institute for the COVID-19 Genomics UK (COG-UK) consortium | The Lighthouse Lab in Milton Keynes and Alex Alderton, Roberto Amato, Sonia Goncalves, Ewan Harrison, David K. Jackson, Ian Johnston, Dominic Kwiatkowski, Cordelia Langford, John Sillitoe on behalf of the Wellcome Sanger Institute COVID-[illegible] |
| EPI_ISL_557537, EPI_ISL_557539 | Lighthouse Lab in Alderley Park | Wellcome Sanger Institute for the COVID-19 Genomics UK (COG-UK) consortium | The Lighthouse Lab in Alderley Park and Alex Alderton, Roberto Amato, Sonia Goncalves, Ewan Harrison, David K. Jackson, Ian Johnston, Dominic Kwiatkowski, Cordelia Langford, John Sillitoe on behalf of the Wellcome Sanger Institute COVID-[illegible] |
| EPI_ISL_557540 | Lighthouse Lab in Milton Keynes | Wellcome Sanger Institute for the COVID-19 Genomics UK (COG-UK) consortium | The Lighthouse Lab in Milton Keynes and Alex Alderton, Roberto Amato, Sonia Goncalves, Ewan Harrison, David K. Jackson, Ian Johnston, Dominic Kwiatkowski, Cordelia Langford, John Sillitoe on behalf of the Wellcome Sanger Institute COVID-[illegible] |
| EPI_ISL_557541, EPI_ISL_557543, EPI_ISL_557544 | Lighthouse Lab in Alderley Park | Wellcome Sanger Institute for the COVID-19 Genomics UK (COG-UK) consortium | The Lighthouse Lab in Alderley Park and Alex Alderton, Roberto Amato, Sonia Goncalves, Ewan Harrison, David K. Jackson, Ian Johnston, Dominic Kwiatkowski, Cordelia Langford, John Sillitoe on behalf of the Wellcome Sanger Institute COVID-[illegible] |
| EPI_ISL_557545, EPI_ISL_557546, EPI_ISL_557549, EPI_ISL_557550 | Lighthouse Lab in Milton Keynes | Wellcome Sanger Institute for the COVID-19 Genomics UK (COG-UK) consortium | The Lighthouse Lab in Milton Keynes and Alex Alderton, Roberto Amato, Sonia Goncalves, Ewan Harrison, David K. Jackson, Ian Johnston, Dominic Kwiatkowski, Cordelia Langford, John Sillitoe on behalf of the Wellcome Sanger Institute COVID-[illegible] |
| EPI_ISL_557551, EPI_ISL_557553, EPI_ISL_557554, EPI_ISL_557556, EPI_ISL_557557, EPI_ISL_557558, EPI_ISL_557559, EPI_ISL_557560, EPI_ISL_557562, EPI_ISL_557563, EPI_ISL_557564, EPI_ISL_557565, EPI_ISL_557566, EPI_ISL_557568, EPI_ISL_557569, EPI_ISL_557571, EPI_ISL_557573, EPI_ISL_557574, EPI_I[illegible] EPI_ISL_557580, EPI_ISL_557584, EPI_ISL_557585, EPI_ISL_557586, EPI_ISL_557588, EPI_ISL_557589, EPI_ISL_557590, EPI_ISL_557591, EPI_ISL_557592, EPI_ISL_557594 | | | |
| see above | Lighthouse Lab in Alderley Park | Wellcome Sanger Institute for the COVID-19 Genomics UK (COG-UK) consortium | The Lighthouse Lab in Alderley Park and Alex Alderton, Roberto Amato, Sonia Goncalves, Ewan Harrison, David K. Jackson, Ian Johnston, Dominic Kwiatkowski, Cordelia Langford, John Sillitoe on behalf of the Wellcome Sang[illegible] |
| EPI_ISL_557596 | Lighthouse Lab in Alderley Park | Wellcome Sanger Institute for the COVID-19 Genomics UK (COG-UK) consortium | The Lighthouse Lab in Alderley Park and Alex Alderton, Roberto Amato, Sonia Goncalves, Ewan Harrison, David K. Jackson, Ian Johnston, Dominic Kwiatkowski, Cordelia Langford, John Sillitoe on behalf of the Wellcome Sanger Institute COVID-[illegible] |

| | | | |
|---|---|---|---|
| EPI_ISL_557599, EPI_ISL_557600, EPI_ISL_557601, EPI_ISL_557602, EPI_ISL_557604, EPI_ISL_557606 | Lighthouse Lab in Alderley Park | Wellcome Sanger Institute for the COVID-19 Genomics UK (COG-UK) consortium | The Lighthouse Lab in Alderley Park and Alex Alderton, Roberto Amato, Sonia Goncalves, Ewan Harrison, David K. Jackson, Ian Johnston, Dominic Kwiatkowski, Cordelia Langford, John Sillitoe on behalf of the Wellcome Sang |
| EPI_ISL_557607 | Lighthouse Lab in Alderley Park | Wellcome Sanger Institute for the COVID-19 Genomics UK (COG-UK) consortium | The Lighthouse Lab in Alderley Park and Alex Alderton, Roberto Amato, Sonia Goncalves, Ewan Harrison, David K. Jackson, Ian Johnston, Dominic Kwiatkowski, Cordelia Langford, John Sillitoe on behalf of the Wellcome Sanger Institute COVID- |
| EPI_ISL_557608, EPI_ISL_557610, EPI_ISL_557611 | Lighthouse Lab in Alderley Park | Wellcome Sanger Institute for the COVID-19 Genomics UK (COG-UK) consortium | The Lighthouse Lab in Alderley Park and Alex Alderton, Roberto Amato, Sonia Goncalves, Ewan Harrison, David K. Jackson, Ian Johnston, Dominic Kwiatkowski, Cordelia Langford, John Sillitoe on behalf of the Wellcome Sang |
| EPI_ISL_557612 | Lighthouse Lab in Alderley Park | Wellcome Sanger Institute for the COVID-19 Genomics UK (COG-UK) consortium | The Lighthouse Lab in Alderley Park and Alex Alderton, Roberto Amato, Sonia Goncalves, Ewan Harrison, David K. Jackson, Ian Johnston, Dominic Kwiatkowski, Cordelia Langford, John Sillitoe on behalf of the Wellcome Sanger Institute COVID- |
| EPI_ISL_557613, EPI_ISL_557614, EPI_ISL_557615, EPI_ISL_557616, EPI_ISL_557617, EPI_ISL_557619, EPI_ISL_557622, EPI_ISL_557623 | Lighthouse Lab in Alderley Park | Wellcome Sanger Institute for the COVID-19 Genomics UK (COG-UK) consortium | The Lighthouse Lab in Alderley Park and Alex Alderton, Roberto Amato, Sonia Goncalves, Ewan Harrison, David K. Jackson, Ian Johnston, Dominic Kwiatkowski, Cordelia Langford, John Sillitoe on behalf of the Wellcome Sang |
| EPI_ISL_557625 | Lighthouse Lab in Alderley Park | Wellcome Sanger Institute for the COVID-19 Genomics UK (COG-UK) Consortium | The Lighthouse Lab in Alderley Park and Alex Alderton, Roberto Amato, Sonia Goncalves, Ewan Harrison, David K. Jackson, Ian Johnston, Dominic Kwiatkowski, Cordelia Langford, John Sillitoe on behalf of the Wellcome Sang |
| EPI_ISL_557626, EPI_ISL_557628, EPI_ISL_557629, EPI_ISL_557630, EPI_ISL_557631, EPI_ISL_557633, EPI_ISL_557634, EPI_ISL_557635, EPI_ISL_557636, EPI_ISL_557637, EPI_ISL_557645, EPI_ISL_557648, EPI_ISL_557656, EPI_ISL_557659, EPI_ISL_557660, EPI_ISL_557661, EPI_ISL_557662, EPI_ISL_557664, EPI_I EPI_ISL_557671 | | | |
| see above | Lighthouse Lab in Alderley Park | Wellcome Sanger Institute for the COVID-19 Genomics UK (COG-UK) consortium | The Lighthouse Lab in Alderley Park and Alex Alderton, Roberto Amato, Sonia Goncalves, Ewan Harrison, David K. Jackson, Ian Johnston, Dominic Kwiatkowski, Cordelia Langford, John Sillitoe on behalf of the Wellcome Sang |
| EPI_ISL_557672 | Lighthouse Lab in Alderley Park | Wellcome Sanger Institute for the COVID-19 Genomics UK (COG-UK) consortium | The Lighthouse Lab in Alderley Park and Alex Alderton, Roberto Amato, Sonia Goncalves, Ewan Harrison, David K. Jackson, Ian Johnston, Dominic Kwiatkowski, Cordelia Langford, John Sillitoe on behalf of the Wellcome Sanger Institute COVID- |
| EPI_ISL_557674, EPI_ISL_557675, EPI_ISL_557676, EPI_ISL_557679, EPI_ISL_557680, EPI_ISL_557682, EPI_ISL_557688, EPI_ISL_557689, EPI_ISL_557693, EPI_ISL_557694, EPI_ISL_557696, EPI_ISL_557697, EPI_ISL_557699 | | | |
| see above | Lighthouse Lab in Alderley Park | Wellcome Sanger Institute for the COVID-19 Genomics UK (COG-UK) consortium | The Lighthouse Lab in Alderley Park and Alex Alderton, Roberto Amato, Sonia Goncalves, Ewan Harrison, David K. Jackson, Ian Johnston, Dominic Kwiatkowski, Cordelia Langford, John Sillitoe on behalf of the Wellcome Sang |
| EPI_ISL_557700 | Lighthouse Lab in Alderley Park | Wellcome Sanger Institute for the COVID-19 Genomics UK (COG-UK) consortium | The Lighthouse Lab in Alderley Park and Alex Alderton, Roberto Amato, Sonia Goncalves, Ewan Harrison, David K. Jackson, Ian Johnston, Dominic Kwiatkowski, Cordelia Langford, John Sillitoe on behalf of the Wellcome Sanger Institute COVID- |
| EPI_ISL_557702, EPI_ISL_557703, EPI_ISL_557706, EPI_ISL_557708, EPI_ISL_557710 | Lighthouse Lab in Alderley Park | Wellcome Sanger Institute for the COVID-19 Genomics UK (COG-UK) consortium | The Lighthouse Lab in Alderley Park and Alex Alderton, Roberto Amato, Sonia Goncalves, Ewan Harrison, David K. Jackson, Ian Johnston, Dominic Kwiatkowski, Cordelia Langford, John Sillitoe on behalf of the Wellcome Sang |
| EPI_ISL_557711 | Lighthouse Lab in Alderley Park | Wellcome Sanger Institute for the COVID-19 Genomics UK (COG-UK) Consortium | The Lighthouse Lab in Alderley Park and Alex Alderton, Roberto Amato, Sonia Goncalves, Ewan Harrison, David K. Jackson, Ian Johnston, Dominic Kwiatkowski, Cordelia Langford, John Sillitoe on behalf of the Wellcome Sang |
| EPI_ISL_557712 | Lighthouse Lab in Alderley Park | Wellcome Sanger Institute for the COVID-19 Genomics UK (COG-UK) consortium | The Lighthouse Lab in Alderley Park and Alex Alderton, Roberto Amato, Sonia Goncalves, Ewan Harrison, David K. Jackson, Ian Johnston, Dominic Kwiatkowski, Cordelia Langford, John Sillitoe on behalf of the Wellcome Sang |
| EPI_ISL_557713 | Lighthouse Lab in Alderley Park | Wellcome Sanger Institute for the COVID-19 Genomics UK (COG-UK) consortium | The Lighthouse Lab in Alderley Park and Alex Alderton, Roberto Amato, Sonia Goncalves, Ewan Harrison, David K. Jackson, Ian Johnston, Dominic Kwiatkowski, Cordelia Langford, John Sillitoe on behalf of the Wellcome Sanger Institute COVID- |
| EPI_ISL_557714, EPI_ISL_557717, EPI_ISL_557718, EPI_ISL_557719, EPI_ISL_557720, EPI_ISL_557726 | Lighthouse Lab in Alderley Park | Wellcome Sanger Institute for the COVID-19 Genomics UK (COG-UK) consortium | The Lighthouse Lab in Alderley Park and Alex Alderton, Roberto Amato, Sonia Goncalves, Ewan Harrison, David K. Jackson, Ian Johnston, Dominic Kwiatkowski, Cordelia Langford, John Sillitoe on behalf of the Wellcome Sang |
| EPI_ISL_557730 | Lighthouse Lab in Alderley Park | Wellcome Sanger Institute for the COVID-19 Genomics UK (COG-UK) Consortium | The Lighthouse Lab in Alderley Park and Alex Alderton, Roberto Amato, Sonia Goncalves, Ewan Harrison, David K. Jackson, Ian Johnston, Dominic Kwiatkowski, Cordelia Langford, John Sillitoe on behalf of the Wellcome Sang |
| EPI_ISL_557731, EPI_ISL_557733, EPI_ISL_557734, EPI_ISL_557736, EPI_ISL_557737, EPI_ISL_557762, EPI_ISL_557785, EPI_ISL_557835, EPI_ISL_557851, EPI_ISL_557887 | Lighthouse Lab in Alderley Park | Wellcome Sanger Institute for the COVID-19 Genomics UK (COG-UK) consortium | The Lighthouse Lab in Alderley Park and Alex Alderton, Roberto Amato, Sonia Goncalves, Ewan Harrison, David K. Jackson, Ian Johnston, Dominic Kwiatkowski, Cordelia Langford, John Sillitoe on behalf of the Wellcome Sang |
| EPI_ISL_557900 | Lighthouse Lab in Alderley Park | Wellcome Sanger Institute for the COVID-19 Genomics UK (COG-UK) consortium | The Lighthouse Lab in Alderley Park and Alex Alderton, Roberto Amato, Sonia Goncalves, Ewan Harrison, David K. Jackson, Ian Johnston, Dominic Kwiatkowski, Cordelia Langford, John Sillitoe on behalf of the Wellcome Sanger Institute COVID- |

| | | | |
|---|---|---|---|
| EPI_ISL_557914, EPI_ISL_557920, EPI_ISL_557929, EPI_ISL_557934, EPI_ISL_557935 | Lighthouse Lab in Alderley Park | Wellcome Sanger Institute for the COVID-19 Genomics UK (COG-UK) consortium | The Lighthouse Lab in Alderley Park and Alex Alderton, Roberto Amato, Sonia Goncalves, Ewan Harrison, David K. Jackson, Ian Johnston, Dominic Kwiatkowski, Cordelia Langford, John Sillitoe on behalf of the Wellcome Sang |
| EPI_ISL_557936 | Lighthouse Lab in Alderley Park | Wellcome Sanger Institute for the COVID-19 Genomics UK (COG-UK) consortium | The Lighthouse Lab in Alderley Park and Alex Alderton, Roberto Amato, Sonia Goncalves, Ewan Harrison, David K. Jackson, Ian Johnston, Dominic Kwiatkowski, Cordelia Langford, John Sillitoe on behalf of the Wellcome Sanger Institute COVID- |
| EPI_ISL_557939, EPI_ISL_557950, EPI_ISL_557951, EPI_ISL_557971, EPI_ISL_557978, EPI_ISL_558070, EPI_ISL_558458, EPI_ISL_558529, EPI_ISL_558608, EPI_ISL_559098, EPI_ISL_559099, EPI_ISL_559101, EPI_ISL_559102 | | | |
| see above | Lighthouse Lab in Alderley Park | Wellcome Sanger Institute for the COVID-19 Genomics UK (COG-UK) consortium | The Lighthouse Lab in Alderley Park and Alex Alderton, Roberto Amato, Sonia Goncalves, Ewan Harrison, David K. Jackson, Ian Johnston, Dominic Kwiatkowski, Cordelia Langford, John Sillitoe on behalf of the Wellcome Sang |
| EPI_ISL_559103 | Lighthouse Lab in Alderley Park | Wellcome Sanger Institute for the COVID-19 Genomics UK (COG-UK) consortium | The Lighthouse Lab in Alderley Park and Alex Alderton, Roberto Amato, Sonia Goncalves, Ewan Harrison, David K. Jackson, Ian Johnston, Dominic Kwiatkowski, Cordelia Langford, John Sillitoe on behalf of the Wellcome Sanger Institute COVID- |
| EPI_ISL_559105, EPI_ISL_559107, EPI_ISL_559108, EPI_ISL_559109, EPI_ISL_559110, EPI_ISL_559111, EPI_ISL_559112, EPI_ISL_559113, EPI_ISL_559114, EPI_ISL_559115, EPI_ISL_559116, EPI_ISL_559118 | | | |
| see above | Lighthouse Lab in Alderley Park | Wellcome Sanger Institute for the COVID-19 Genomics UK (COG-UK) consortium | The Lighthouse Lab in Alderley Park and Alex Alderton, Roberto Amato, Sonia Goncalves, Ewan Harrison, David K. Jackson, Ian Johnston, Dominic Kwiatkowski, Cordelia Langford, John Sillitoe on behalf of the Wellcome Sang |
| EPI_ISL_559120 | Lighthouse Lab in Alderley Park | Wellcome Sanger Institute for the COVID-19 Genomics UK (COG-UK) Consortium | The Lighthouse Lab in Alderley Park and Alex Alderton, Roberto Amato, Sonia Goncalves, Ewan Harrison, David K. Jackson, Ian Johnston, Dominic Kwiatkowski, Cordelia Langford, John Sillitoe on behalf of the Wellcome Sang |
| EPI_ISL_559121, EPI_ISL_559122, EPI_ISL_559123, EPI_ISL_559124, EPI_ISL_559127, EPI_ISL_559128 | Lighthouse Lab in Alderley Park | Wellcome Sanger Institute for the COVID-19 Genomics UK (COG-UK) consortium | The Lighthouse Lab in Alderley Park and Alex Alderton, Roberto Amato, Sonia Goncalves, Ewan Harrison, David K. Jackson, Ian Johnston, Dominic Kwiatkowski, Cordelia Langford, John Sillitoe on behalf of the Wellcome Sang |
| EPI_ISL_559129 | Lighthouse Lab in Alderley Park | Wellcome Sanger Institute for the COVID-19 Genomics UK (COG-UK) Consortium | The Lighthouse Lab in Alderley Park and Alex Alderton, Roberto Amato, Sonia Goncalves, Ewan Harrison, David K. Jackson, Ian Johnston, Dominic Kwiatkowski, Cordelia Langford, John Sillitoe on behalf of the Wellcome Sang |
| EPI_ISL_559134, EPI_ISL_559135, EPI_ISL_559137, EPI_ISL_559138, EPI_ISL_559139, EPI_ISL_559140, EPI_ISL_559141, EPI_ISL_559142, EPI_ISL_559143, EPI_ISL_559144, EPI_ISL_559145, EPI_ISL_559146, EPI_ISL_559148, EPI_ISL_559150, EPI_ISL_559151, EPI_ISL_559152, EPI_ISL_559153, EPI_ISL_559155, EPI_I[illegible] EPI_ISL_559162, EPI_ISL_559165, EPI_ISL_559168, EPI_ISL_559169, EPI_ISL_559170, EPI_ISL_559172, EPI_ISL_559173, EPI_ISL_559175 | | | |
| see above | Lighthouse Lab in Alderley Park | Wellcome Sanger Institute for the COVID-19 Genomics UK (COG-UK) consortium | The Lighthouse Lab in Alderley Park and Alex Alderton, Roberto Amato, Sonia Goncalves, Ewan Harrison, David K. Jackson, Ian Johnston, Dominic Kwiatkowski, Cordelia Langford, John Sillitoe on behalf of the Wellcome Sang |
| EPI_ISL_559176 | Lighthouse Lab in Alderley Park | Wellcome Sanger Institute for the COVID-19 Genomics UK (COG-UK) consortium | The Lighthouse Lab in Alderley Park and Alex Alderton, Roberto Amato, Sonia Goncalves, Ewan Harrison, David K. Jackson, Ian Johnston, Dominic Kwiatkowski, Cordelia Langford, John Sillitoe on behalf of the Wellcome Sanger Institute COVID- |
| EPI_ISL_559177, EPI_ISL_559178, EPI_ISL_559179, EPI_ISL_559180, EPI_ISL_559182, EPI_ISL_559183, EPI_ISL_559184, EPI_ISL_559185, EPI_ISL_559186, EPI_ISL_559188, EPI_ISL_559189 | | | |
| see above | Lighthouse Lab in Alderley Park | Wellcome Sanger Institute for the COVID-19 Genomics UK (COG-UK) consortium | The Lighthouse Lab in Alderley Park and Alex Alderton, Roberto Amato, Sonia Goncalves, Ewan Harrison, David K. Jackson, Ian Johnston, Dominic Kwiatkowski, Cordelia Langford, John Sillitoe on behalf of the Wellcome Sang |
| EPI_ISL_559190 | Lighthouse Lab in Alderley Park | Wellcome Sanger Institute for the COVID-19 Genomics UK (COG-UK) consortium | The Lighthouse Lab in Alderley Park and Alex Alderton, Roberto Amato, Sonia Goncalves, Ewan Harrison, David K. Jackson, Ian Johnston, Dominic Kwiatkowski, Cordelia Langford, John Sillitoe on behalf of the Wellcome Sanger Institute COVID- |
| EPI_ISL_559191 | Lighthouse Lab in Alderley Park | Wellcome Sanger Institute for the COVID-19 Genomics UK (COG-UK) consortium | The Lighthouse Lab in Alderley Park and Alex Alderton, Roberto Amato, Sonia Goncalves, Ewan Harrison, David K. Jackson, Ian Johnston, Dominic Kwiatkowski, Cordelia Langford, John Sillitoe on behalf of the Wellcome Sang |
| EPI_ISL_559192 | Lighthouse Lab in Alderley Park | Wellcome Sanger Institute for the COVID-19 Genomics UK (COG-UK) consortium | The Lighthouse Lab in Alderley Park and Alex Alderton, Roberto Amato, Sonia Goncalves, Ewan Harrison, David K. Jackson, Ian Johnston, Dominic Kwiatkowski, Cordelia Langford, John Sillitoe on behalf of the Wellcome Sanger Institute COVID- |
| EPI_ISL_559193, EPI_ISL_559196, EPI_ISL_559197, EPI_ISL_559198, EPI_ISL_559199, EPI_ISL_559200, EPI_ISL_559202, EPI_ISL_559203, EPI_ISL_559205 | Lighthouse Lab in Alderley Park | Wellcome Sanger Institute for the COVID-19 Genomics UK (COG-UK) consortium | The Lighthouse Lab in Alderley Park and Alex Alderton, Roberto Amato, Sonia Goncalves, Ewan Harrison, David K. Jackson, Ian Johnston, Dominic Kwiatkowski, Cordelia Langford, John Sillitoe on behalf of the Wellcome Sang |
| EPI_ISL_559206 | Lighthouse Lab in Alderley Park | Wellcome Sanger Institute for the COVID-19 Genomics UK (COG-UK) consortium | The Lighthouse Lab in Alderley Park and Alex Alderton, Roberto Amato, Sonia Goncalves, Ewan Harrison, David K. Jackson, Ian Johnston, Dominic Kwiatkowski, Cordelia Langford, John Sillitoe on behalf of the Wellcome Sanger Institute COVID- |
| EPI_ISL_559208, EPI_ISL_559209, EPI_ISL_559210, EPI_ISL_559211, EPI_ISL_559212 | Lighthouse Lab in Alderley Park | Wellcome Sanger Institute for the COVID-19 Genomics UK (COG-UK) consortium | The Lighthouse Lab in Alderley Park and Alex Alderton, Roberto Amato, Sonia Goncalves, Ewan Harrison, David K. Jackson, Ian Johnston, Dominic Kwiatkowski, Cordelia Langford, John Sillitoe on behalf of the Wellcome Sang |
| EPI_ISL_559213 | Lighthouse Lab in Alderley Park | Wellcome Sanger Institute for the COVID-19 Genomics UK (COG-UK) Consortium | The Lighthouse Lab in Alderley Park and Alex Alderton, Roberto Amato, Sonia Goncalves, Ewan Harrison, David K. Jackson, Ian Johnston, Dominic Kwiatkowski, Cordelia Langford, John Sillitoe on behalf of the Wellcome Sang |
| EPI_ISL_559214, EPI_ISL_559216, | Lighthouse Lab in Alderley Park | Wellcome Sanger Institute for the COVID-19 Genomics UK (COG-UK) consortium | The Lighthouse Lab in Alderley Park and Alex Alderton, Roberto Amato, Sonia Goncalves, Ewan Harrison, David K. Jackson, Ian Johnston, Dominic Kwiatkowski, Cordelia Langford, John Sillitoe on behalf of the Wellcome Sang |

| | | | |
|---|---|---|---|
| EPI_ISL_559217, EPI_ISL_559219, EPI_ISL_559220 | | | |
| EPI_ISL_559221 | Lighthouse Lab in Alderley Park | Wellcome Sanger Institute for the COVID-19 Genomics UK (COG-UK) consortium | The Lighthouse Lab in Alderley Park and Alex Alderton, Roberto Amato, Sonia Goncalves, Ewan Harrison, David K. Jackson, Ian Johnston, Dominic Kwiatkowski, Cordelia Langford, John Sillitoe on behalf of the Wellcome Sanger Institute COVID- |
| EPI_ISL_559222, EPI_ISL_559224, EPI_ISL_559225, EPI_ISL_559226, EPI_ISL_559227 | Lighthouse Lab in Alderley Park | Wellcome Sanger Institute for the COVID-19 Genomics UK (COG-UK) consortium | The Lighthouse Lab in Alderley Park and Alex Alderton, Roberto Amato, Sonia Goncalves, Ewan Harrison, David K. Jackson, Ian Johnston, Dominic Kwiatkowski, Cordelia Langford, John Sillitoe on behalf of the Wellcome Sang |
| EPI_ISL_559228 | Lighthouse Lab in Alderley Park | Wellcome Sanger Institute for the COVID-19 Genomics UK (COG-UK) Consortium | The Lighthouse Lab in Alderley Park and Alex Alderton, Roberto Amato, Sonia Goncalves, Ewan Harrison, David K. Jackson, Ian Johnston, Dominic Kwiatkowski, Cordelia Langford, John Sillitoe on behalf of the Wellcome Sang |
| EPI_ISL_559229, EPI_ISL_559230, EPI_ISL_559231, EPI_ISL_559233, EPI_ISL_559234, EPI_ISL_559237, EPI_ISL_559238, EPI_ISL_559239 | Lighthouse Lab in Alderley Park | Wellcome Sanger Institute for the COVID-19 Genomics UK (COG-UK) consortium | The Lighthouse Lab in Alderley Park and Alex Alderton, Roberto Amato, Sonia Goncalves, Ewan Harrison, David K. Jackson, Ian Johnston, Dominic Kwiatkowski, Cordelia Langford, John Sillitoe on behalf of the Wellcome Sang |
| EPI_ISL_559240 | Lighthouse Lab in Alderley Park | Wellcome Sanger Institute for the COVID-19 Genomics UK (COG-UK) consortium | The Lighthouse Lab in Alderley Park and Alex Alderton, Roberto Amato, Sonia Goncalves, Ewan Harrison, David K. Jackson, Ian Johnston, Dominic Kwiatkowski, Cordelia Langford, John Sillitoe on behalf of the Wellcome Sanger Institute COVID- |
| EPI_ISL_559242, EPI_ISL_559243, EPI_ISL_559245, EPI_ISL_559248, EPI_ISL_559249, EPI_ISL_559250, EPI_ISL_559251, EPI_ISL_559252, EPI_ISL_559255, EPI_ISL_559257, EPI_ISL_559258, EPI_ISL_559260 | | | |
| see above | Lighthouse Lab in Alderley Park | Wellcome Sanger Institute for the COVID-19 Genomics UK (COG-UK) consortium | The Lighthouse Lab in Alderley Park and Alex Alderton, Roberto Amato, Sonia Goncalves, Ewan Harrison, David K. Jackson, Ian Johnston, Dominic Kwiatkowski, Cordelia Langford, John Sillitoe on behalf of the Wellcome Sang |
| EPI_ISL_559261 | Lighthouse Lab in Alderley Park | Wellcome Sanger Institute for the COVID-19 Genomics UK (COG-UK) Consortium | The Lighthouse Lab in Alderley Park and Alex Alderton, Roberto Amato, Sonia Goncalves, Ewan Harrison, David K. Jackson, Ian Johnston, Dominic Kwiatkowski, Cordelia Langford, John Sillitoe on behalf of the Wellcome Sang |
| EPI_ISL_559262 | Lighthouse Lab in Alderley Park | Wellcome Sanger Institute for the COVID-19 Genomics UK (COG-UK) consortium | The Lighthouse Lab in Alderley Park and Alex Alderton, Roberto Amato, Sonia Goncalves, Ewan Harrison, David K. Jackson, Ian Johnston, Dominic Kwiatkowski, Cordelia Langford, John Sillitoe on behalf of the Wellcome Sang |
| EPI_ISL_559265 | Lighthouse Lab in Alderley Park | Wellcome Sanger Institute for the COVID-19 Genomics UK (COG-UK) consortium | The Lighthouse Lab in Alderley Park and Alex Alderton, Roberto Amato, Sonia Goncalves, Ewan Harrison, David K. Jackson, Ian Johnston, Dominic Kwiatkowski, Cordelia Langford, John Sillitoe on behalf of the Wellcome Sanger Institute COVID- |
| EPI_ISL_559266, EPI_ISL_559267, EPI_ISL_559268, EPI_ISL_559269, EPI_ISL_559270, EPI_ISL_559271, EPI_ISL_559272, EPI_ISL_559273, EPI_ISL_559274 | Lighthouse Lab in Alderley Park | Wellcome Sanger Institute for the COVID-19 Genomics UK (COG-UK) consortium | The Lighthouse Lab in Alderley Park and Alex Alderton, Roberto Amato, Sonia Goncalves, Ewan Harrison, David K. Jackson, Ian Johnston, Dominic Kwiatkowski, Cordelia Langford, John Sillitoe on behalf of the Wellcome Sang |
| EPI_ISL_559275 | Lighthouse Lab in Alderley Park | Wellcome Sanger Institute for the COVID-19 Genomics UK (COG-UK) Consortium | The Lighthouse Lab in Alderley Park and Alex Alderton, Roberto Amato, Sonia Goncalves, Ewan Harrison, David K. Jackson, Ian Johnston, Dominic Kwiatkowski, Cordelia Langford, John Sillitoe on behalf of the Wellcome Sang |
| EPI_ISL_559276, EPI_ISL_559279, EPI_ISL_559280, EPI_ISL_559281, EPI_ISL_559282, EPI_ISL_559285, EPI_ISL_559286, EPI_ISL_559288, EPI_ISL_559289, EPI_ISL_559291, EPI_ISL_559292, EPI_ISL_559293, EPI_ISL_559294, EPI_ISL_559295, EPI_ISL_559297, EPI_ISL_559298 | | | |
| see above | Lighthouse Lab in Alderley Park | Wellcome Sanger Institute for the COVID-19 Genomics UK (COG-UK) consortium | The Lighthouse Lab in Alderley Park and Alex Alderton, Roberto Amato, Sonia Goncalves, Ewan Harrison, David K. Jackson, Ian Johnston, Dominic Kwiatkowski, Cordelia Langford, John Sillitoe on behalf of the Wellcome Sang |
| EPI_ISL_560308 | UMMC-Health | WHO National Influenza Centre Russian Federation | Andrey Komissarov, Artem Fadeev, Anna Ivanova, Tatiana Platonova, Daria Danilenko |
| EPI_ISL_560327, EPI_ISL_560329, EPI_ISL_560341, EPI_ISL_560343 | TriCore Reference Laboratories | Center for Global Health, University of New Mexico Health Sciences Center | Daryl Domman, Kurt Schwalm, Twila Kunde, Joseph Hicks, Michael Edwards, Darrell Dinwiddie |
| EPI_ISL_560636 | Hospital | National Reference Center for Viruses of Respiratory Infections, Institut Pasteur, Paris | Sylvie Behillil, Fabiana Gambaro, Etienne Simon-Lorière, Vincent Enouf, Maud Vanpeene, Sylvie van der Werf |
| EPI_ISL_560637, EPI_ISL_560638, EPI_ISL_560639, EPI_ISL_560640, EPI_ISL_560641, EPI_ISL_560642 | Labo Analyses Med | National Reference Center for Viruses of Respiratory Infections, Institut Pasteur, Paris | Sylvie Behillil, Fabiana Gambaro, Etienne Simon-Lorière, Vincent Enouf, Maud Vanpeene, Sylvie van der Werf |
| EPI_ISL_560960, EPI_ISL_560961, EPI_ISL_560962, EPI_ISL_560963, EPI_ISL_560964, EPI_ISL_560965, EPI_ISL_560966, EPI_ISL_560967, EPI_ISL_560968, EPI_ISL_560969, EPI_ISL_560970 | | | |
| see above | Texas Department of State Health Services | Texas Department of State Health Services | Rashmi Tuladhar, Bonnie Oh, Jenny Zhang, Maliha Rahman, Anita Pokharel, Myong Koag, Chun Wang, Rachel Lee, Grace Kubin |
| EPI_ISL_561039, EPI_ISL_561040, EPI_ISL_561209, EPI_ISL_561212, EPI_ISL_561213, EPI_ISL_561214, EPI_ISL_561215, EPI_ISL_561216, EPI_ISL_561217, EPI_ISL_561218, EPI_ISL_561219, EPI_ISL_561328 | | | |
| see above | MRCG at LSHTM Genomics lab | MRCG at LSHTM Genomics lab | Abdul Karim sesay, Abdoulie Kanteh, Jarra Manneh, Mariama Kujabi, Bakary Sanyang |

| | | | |
|---|---|---|---|
| EPI_ISL_561749 | Microbiological Diagnostic Unit - Public Health Laboratory (MDU-PHL) | MDU-PHL | Seemann, T., Schultz M. B., Sait, M., Sherry, N. |
| EPI_ISL_561808 | Victorian Infectious Diseases Reference Laboratory (VIDRL) | VIDRL and MDU-PHL | Caly, L., Seemann, T., Sait, M., Schultz, M. B., Druce J., Sherry, N. |
| EPI_ISL_563981, EPI_ISL_563982, EPI_ISL_563983, EPI_ISL_563984, EPI_ISL_564268, EPI_ISL_564277, EPI_ISL_564279, EPI_ISL_564280, EPI_ISL_564284, EPI_ISL_564285, EPI_ISL_564288, EPI_ISL_564289, EPI_ISL_564292 | | | |
| see above | Microbiological Diagnostic Unit - Public Health Laboratory (MDU-PHL) | MDU-PHL | Seemann, T., Schultz M. B., Sait, M., Sherry, N. |
| EPI_ISL_565856 | Michigan Department of Health and Human Services, Bureau of Laboratories | Michigan Department of Health and Human Services, Bureau of Laboratories | Blankenship HM, Riner D, Soehnlen MK |
| EPI_ISL_565924, EPI_ISL_565925 | Servicio de Microbiología. Hospital Universitario Donostia. OSI Donostialdea. Área de Enfermedades Infecciosas, Grupo de Infección Respiratoria y Resistencia Antimicrobiana. Instituto de Investigación Sanitaria Biodonostia | SeqCOVID-SPAIN consortium/IBV(CSIC) | Gustavo Cilla, Milagrosa Montes, Luis Piñeiro, Jose Maria Marimón and SeqCOVID-SPAIN consortium |
| EPI_ISL_566013, EPI_ISL_566025 | Michigan Department of Health and Human Services, Bureau of Laboratories | Michigan Department of Health and Human Services, Bureau of Laboratories | Blankenship HM, Riner D, Soehnlen MK |
| EPI_ISL_566056, EPI_ISL_566057 | Respiratory Virus Unit, Microbiology Services Colindale, Public Health England | Respiratory Virus Unit, Microbiology Services Colindale, Public Health England | PHE Covid Sequencing Team |
| EPI_ISL_566080 | Pathogenic Microorganisms Variability Laboratory | WHO National Influenza Centre Russian Federation | Andrey Komissarov, Artem Fadeev, Anna Ivanova, Kseniya Komissarova, Daria Danilenko, Dmitry Lioznov, Nadezhda Kuznetsova, Elena Shidlovskaya, Elizaveta Divisenko, Ekaterina Milashenko, Kirill Krasnoslobotsev, Evgeniya Mukasheva, Anna Igna<br>Andrey Pochtovyy, Valeria Bacalin, Evgeny Usachev, Olga Burgasova, Ludmila Kolobukhina, Svetlana Smetanina, Elena Burtseva, Artem Tkachuk, Vladimir Gushchin, Alexander Gintsbur |
| EPI_ISL_566085, EPI_ISL_566086, EPI_ISL_566087, EPI_ISL_566088, EPI_ISL_566089, EPI_ISL_566090, EPI_ISL_566092, EPI_ISL_566093, EPI_ISL_566094, EPI_ISL_566096, EPI_ISL_566097, EPI_ISL_566098, EPI_ISL_566099, EPI_ISL_566100, EPI_ISL_566102, EPI_ISL_566104, EPI_ISL_566105 | | | |
| see above | GA Department of Public Health Laboratory | Pathogen Discovery, Respiratory Viruses Branch, Division of Viral Diseases, Centers for Disease Control and Prevention | Jing Zhang, Brian Lynch, Yan Li, Anna Montmayeur, Krista Queen, Ying Tao, Anna Uehara, Clinton R. Paden, Rachel Marine, Haibin Wang, Suxiang Tong |
| EPI_ISL_568467, EPI_ISL_568474 | Lighthouse Lab in Alderley Park | Wellcome Sanger Institute for the COVID-19 Genomics UK (COG-UK) consortium | Jacquelyn Wynn, Mairead Hyland, The Lighthouse Lab in Alderley Park and Alex Alderton, Roberto Amato, Sonia Goncalves, Ewan Harrison, David K. Jackson, Ian Johnston, Dominic Kwiatkowski, Cordelia Langford, John Sillitoe on behalf of the |
| EPI_ISL_568997 | MEPHI, Aix Marseille University | MEPHI, Aix Marseille University | Anthony LEVASSEUR |
| EPI_ISL_569776, EPI_ISL_569783, EPI_ISL_569819, EPI_ISL_569827, EPI_ISL_569833 | Omsk Research Institute of Natural Focal Infections | WHO National Influenza Centre Russian Federation | Artem Fadeev, Ekaterina Gradoboeva, Ekaterina Savkina, Daria Nashatyreva, Elena Poleshchuk, Aleksei Vasilenko, Valery Yakimenko, Andrey Komissarov |
| EPI_ISL_569893, EPI_ISL_569908, EPI_ISL_569915, EPI_ISL_569922, EPI_ISL_569929, EPI_ISL_569939, EPI_ISL_569942 | Innovative Genomics Institute, UC Berkeley | Innovative Genomics Institute, UC Berkeley | Stacia Wyman, Haridha Shivram, Phil Frankino, Liana Lareau, Shana McDevitt, Justin Choi |
| EPI_ISL_570003, EPI_ISL_570008 | Unity Health Toronto | Ontario Institute for Cancer Research | Ramzi Fattouh, Larissa M. Matukas, Yan Chen,Mark Downing, Trina Otterman, Karel Boissinot, Wai Sum Siu, Zhi Cui, Le Luu, Samira Mubareka, TIBDN, Ilinca Lungu, Bernard Lam, Jeremy Johns, Paul Krzyzanowski, Richard de Borja, |
| EPI_ISL_570401, EPI_ISL_570402, EPI_ISL_570403, EPI_ISL_570405, EPI_ISL_570406, EPI_ISL_570408, EPI_ISL_570409, EPI_ISL_570410, EPI_ISL_570411, EPI_ISL_570412, EPI_ISL_570413, EPI_ISL_570415, EPI_ISL_570416, EPI_ISL_570417, EPI_ISL_570418, EPI_ISL_570419, EPI_ISL_570420, EPI_ISL_570421, EPI_I<br>EPI_ISL_570443, EPI_ISL_570444, EPI_ISL_570466, EPI_ISL_570467, EPI_ISL_570468, EPI_ISL_570469, EPI_ISL_570470, EPI_ISL_570471, EPI_ISL_570472, EPI_ISL_570473, EPI_ISL_570474, EPI_ISL_570475, EPI_ISL_570476, EPI_ISL_570477, EPI_ISL_570478, EPI_ISL_570479, EPI_ISL_570480, EPI_ISL_570481, EPI_I<br>EPI_ISL_570490, EPI_ISL_570491, EPI_ISL_570492, EPI_ISL_570493, EPI_ISL_570494, EPI_ISL_570495, EPI_ISL_570496, EPI_ISL_570497, EPI_ISL_570498, EPI_ISL_570499, EPI_ISL_570500, EPI_ISL_570501, EPI_ISL_570502, EPI_ISL_570503, EPI_ISL_570504, EPI_ISL_570505, EPI_ISL_570506, EPI_ISL_570507, EPI_I<br>EPI_ISL_570512, EPI_ISL_570513, EPI_ISL_570514, EPI_ISL_570515, EPI_ISL_570516, EPI_ISL_570517, EPI_ISL_570518, EPI_ISL_570519, EPI_ISL_570520, EPI_ISL_570521, EPI_ISL_570522, EPI_ISL_570523, EPI_ISL_570524, EPI_ISL_570525, EPI_ISL_570526, EPI_ISL_570527, EPI_ISL_570528, EPI_ISL_570529, EPI_I<br>EPI_ISL_570534, EPI_ISL_570535, EPI_ISL_570538, EPI_ISL_570539, EPI_ISL_570540, EPI_ISL_570541, EPI_ISL_570542, EPI_ISL_570543, EPI_ISL_570544, EPI_ISL_570545, EPI_ISL_570546, EPI_ISL_570800, EPI_ISL_570801, EPI_ISL_570805, EPI_ISL_570809, EPI_ISL_570814, EPI_ISL_570818, EPI_ISL_570819, EPI_I<br>EPI_ISL_570836, EPI_ISL_570837, EPI_ISL_570841, EPI_ISL_570842, EPI_ISL_570844, EPI_ISL_570847, EPI_ISL_570849, EPI_ISL_570851, EPI_ISL_570858, EPI_ISL_570859, EPI_ISL_570862, EPI_ISL_570863, EPI_ISL_570864, EPI_ISL_570865, EPI_ISL_570866, EPI_ISL_570867, EPI_ISL_570868, EPI_ISL_570869, EPI_I<br>EPI_ISL_570874, EPI_ISL_570875, EPI_ISL_570876, EPI_ISL_570877, EPI_ISL_570878, EPI_ISL_570879, EPI_ISL_570880, EPI_ISL_570881, EPI_ISL_570882, EPI_ISL_570883, EPI_ISL_570884, EPI_ISL_570885, EPI_ISL_570886, EPI_ISL_570887, EPI_ISL_570888, EPI_ISL_570889, EPI_ISL_570890, EPI_ISL_570891, EPI_I<br>EPI_ISL_570897, EPI_ISL_570898, EPI_ISL_570899, EPI_ISL_570900, EPI_ISL_570902, EPI_ISL_570903, EPI_ISL_570904, EPI_ISL_570905, EPI_ISL_570906, EPI_ISL_570907, EPI_ISL_570908, EPI_ISL_570909, EPI_ISL_570910, EPI_ISL_570911, EPI_ISL_570913, EPI_ISL_570914, EPI_ISL_570915, EPI_ISL_570916, EPI_I<br>EPI_ISL_570921, EPI_ISL_570922, EPI_ISL_570923, EPI_ISL_570924, EPI_ISL_570925, EPI_ISL_570926, EPI_ISL_570927, EPI_ISL_570928, EPI_ISL_570929, EPI_ISL_570930, EPI_ISL_570931, EPI_ISL_570932, EPI_ISL_570933, EPI_ISL_570934, EPI_ISL_570935, EPI_ISL_570936, EPI_ISL_570937, EPI_ISL_570938, EPI_I<br>EPI_ISL_570946, EPI_ISL_570947, EPI_ISL_570948, EPI_ISL_570949, EPI_ISL_570950, EPI_ISL_570951, EPI_ISL_570952, EPI_ISL_570953, EPI_ISL_570955, EPI_ISL_570960, EPI_ISL_570961, EPI_ISL_570962, EPI_ISL_570965 | | | |
| see above | UW Virology Lab | UW Virology Lab | Pavitra Roychoudhury, Hong Xie, Lasata Shrestha, Amin Addetia, Victoria M Rachleff, Meei-Li Huang, Keith R Jerome, Alexander Greninger |
| EPI_ISL_572249, EPI_ISL_572251, EPI_ISL_572258, EPI_ISL_572259, EPI_ISL_572260, EPI_ISL_572261, EPI_ISL_572262, EPI_ISL_572263, EPI_ISL_572264, EPI_ISL_572265, EPI_ISL_572266, EPI_ISL_572272, EPI_ISL_572273, EPI_ISL_572277, EPI_ISL_572278, EPI_ISL_572279, EPI_ISL_572280, EPI_ISL_572282, EPI_I | | | |
| see above | Virginia DCLS | Virginia DCLS | Virginia DCLS |
| EPI_ISL_572397, EPI_ISL_573152 | Institute for Virology, University Hospital Duesseldorf, Medical Faculty, Heinrich-Heine-University Duesseldorf | Institute for Virology, University Hospital Duesseldorf, Medical Faculty, Heinrich-Heine-University Duesseldorf | Maximilian Damagnez, Verena Keitel, Björn Jensen, Nadine Lübke, Lisa Müller, Philipp Ostermann, Tina Senff, Ortwin Adams, Philipp Albrecht, Gerald Antoch, Johannes Bode, Edwin Bölke, Saskia Elben, Torsten Feldt, Johannes C. Fischer, , Anselm K<br>Mohring, Jennifer Neubert, Heiner Schaal, Ansgar Schulz, Jörg Timm, Andreas Walker |
| EPI_ISL_574120, EPI_ISL_574123, | Wales Specialist Virology Centre Sequencing lab: | COVID-19 Genomics UK (COG-UK) Consortium | Catherine Moore, Johnathan Evans, Laura Gifford, Malorie Perry, Simon Cottrell, Angela Marchbank, Alec Birchley, Alexander Adams, Amy Gaskin, Bree Gatica-Wilcox, Jason Coombes, Joel Southgate, Lauren Gilbert, Lee Graham, Nicole Pacchiarini, Sa<br>Matthew Bull, Joanne Watkins, Sally Corden, Tom Connor |

| | | | |
|---|---|---|---|
| EPI_ISL_574251 | Pathogen Genomics Unit | | |
| EPI_ISL_574259 | Institute for Virology, University Hospital Duesseldorf, Medical Faculty, Heinrich-Heine-University Duesseldorf | Institute for Virology, University Hospital Duesseldorf, Medical Faculty, Heinrich-Heine-University Duesseldorf | Maximilian Damagnez, Verena Keitel, Björn Jensen, Nadine Lübke, Lisa Müller, Philipp Ostermann, Tina Senff, Ortwin Adams, Philipp Albrecht, Gerald Antoch, Johannes Bode, Edwin Bölke, Saskia Elben, Torsten Feldt, Johannes C. Fischer, , Anselm K Mohring, Jennifer Neubert, Heiner Schaal, Ansgar Schulz, Jörg Timm, Andreas Walker |
| EPI_ISL_574476, EPI_ISL_574477, EPI_ISL_574478, EPI_ISL_576284, EPI_ISL_576285, EPI_ISL_576286, EPI_ISL_576287, EPI_ISL_576288, EPI_ISL_576289, EPI_ISL_576290, EPI_ISL_576291 | | | |
| see above | Texas Department of State Health Services | Texas Department of State Health Services | Rashmi Tuladhar, Bonnie Oh, Jenny Zhang, Maliha Rahman, Anita Pokharel, Myong Koag, Chun Wang, Rachel Lee, Grace Kubin |
| EPI_ISL_576326 | Texas Department of State Health Services | Texas Department of State Health Services | Rashmi Tuladhar, Bonnie Oh, Mayela Pedrueza, Jenny Zhang, Maliha Rahman, Anita Pokharel, Myong Koag, Chun Wang, Rachel Lee, Grace Kubin |
| EPI_ISL_576403, EPI_ISL_576404, EPI_ISL_576405 | UW Virology Lab | UW Virology Lab | Pavitra Roychoudhury, Hong Xie, Lasata Shrestha, Amin Addetia, Victoria M Rachleff, Meei-Li Huang, Keith R Jerome, Alexander Greninger |
| EPI_ISL_576520, EPI_ISL_576522, EPI_ISL_576524, EPI_ISL_576529, EPI_ISL_576530, EPI_ISL_576531, EPI_ISL_576534, EPI_ISL_576537, EPI_ISL_576539, EPI_ISL_576541, EPI_ISL_576544, EPI_ISL_576546, EPI_ISL_576547, EPI_ISL_576548, EPI_ISL_576549, EPI_ISL_576551, EPI_ISL_576553, EPI_ISL_576554, EPI_I[illegible] | | | |
| see above | Innovative Genomics Institute, UC Berkeley | Innovative Genomics Institute, UC Berkeley | Stacia Wyman, Haridha Shivram, Phil Frankino, Liana Lareau, Shana McDevitt, Justin Choi |
| EPI_ISL_577740, EPI_ISL_577741 | Institute of Virology, Biomedical Research Center of the Slovak Academy of Sciences, Bratislava | Faculty of Natural Sciences, Comenius University, Bratislava | Broa Brejová, Viktória Hodorová, Kristína Boršová, Viktória abanová, Dominika Friová, Sabina Fumaová Havlíková, Juraj Kopáek, Martina Liková, ubomíra Lukáiková, Martina Neboháová, Monika Sláviková, Edita Staroová, Elena Ti |
| EPI_ISL_578869, EPI_ISL_578870, EPI_ISL_578871, EPI_ISL_578872, EPI_ISL_578873, EPI_ISL_578874, EPI_ISL_578875, EPI_ISL_578876, EPI_ISL_578877, EPI_ISL_578878, EPI_ISL_578879, EPI_ISL_578880, EPI_ISL_578881, EPI_ISL_578882, EPI_ISL_578883, EPI_ISL_578884, EPI_ISL_578885, EPI_ISL_578886, EPI_I[illegible] EPI_ISL_578894, EPI_ISL_578899, EPI_ISL_578907, EPI_ISL_578908, EPI_ISL_578909, EPI_ISL_578910, EPI_ISL_578911, EPI_ISL_578914, EPI_ISL_578915, EPI_ISL_578916, EPI_ISL_578917, EPI_ISL_578918, EPI_ISL_578920, EPI_ISL_578921, EPI_ISL_578922, EPI_ISL_578924, EPI_ISL_578925, EPI_ISL_578926, EPI_I[illegible] EPI_ISL_578941, EPI_ISL_578942, EPI_ISL_578943, EPI_ISL_578944, EPI_ISL_578945, EPI_ISL_578946, EPI_ISL_578947, EPI_ISL_578948, EPI_ISL_578952, EPI_ISL_578954, EPI_ISL_578967, EPI_ISL_578968, EPI_ISL_578969, EPI_ISL_578970, EPI_ISL_578971, EPI_ISL_578972, EPI_ISL_578973, EPI_ISL_578974, EPI_I[illegible] EPI_ISL_578979, EPI_ISL_578980, EPI_ISL_578981, EPI_ISL_578982, EPI_ISL_578983, EPI_ISL_578984, EPI_ISL_578985, EPI_ISL_578986, EPI_ISL_578987, EPI_ISL_578988, EPI_ISL_578989, EPI_ISL_578990, EPI_ISL_578991, EPI_ISL_578992, EPI_ISL_578993, EPI_ISL_578994, EPI_ISL_578995, EPI_ISL_578996, EPI_I[illegible] EPI_ISL_579001, EPI_ISL_579002, EPI_ISL_579003, EPI_ISL_579004, EPI_ISL_579005, EPI_ISL_579006, EPI_ISL_579007 | | | |
| see above | LSUHS Emerging Viral Threat Laboratory | Microbial Genome Sequencing Center | Jeremy P. Kamil, Rona S. Scott, Maarten Van Diest, Malgorzata Bienkowska-Haba, Katarzyna Zwolinska, Andrew D. Yurochko, Christopher G. Kevil, Martin J. Sapp, Daniel J. Snyder, Vaughn S. Cooper, J |
| EPI_ISL_581391, EPI_ISL_581412, EPI_ISL_581424, EPI_ISL_581425, EPI_ISL_581426, EPI_ISL_581427, EPI_ISL_581429, EPI_ISL_581430, EPI_ISL_581431, EPI_ISL_581432, EPI_ISL_581446, EPI_ISL_581448 | | | |
| see above | Lighthouse Lab in Alderley Park | Wellcome Sanger Institute for the COVID-19 Genomics UK (COG-UK) consortium | Jacquelyn Wynn, Mairead Hyland, The Lighthouse Lab in Alderley Park and Alex Alderton, Roberto Amato, Sonia Goncalves, Ewan Harrison, David K. Jackson, Ian Johnston, Dominic Kwiatkowski, Cordelia Langford, John Sillitoe on behalf of the |
| EPI_ISL_581491, EPI_ISL_581493 | Fondation Congolaise pour la recherche medicale (FCRM) | NGS Competence Center Tübingen, Institut für Medizinische Mikrobiologie und Hygiene, Universitätsklinikum Tübingen | Angel Angelov |
| EPI_ISL_581875, EPI_ISL_581892, EPI_ISL_581908 | University Hospital Basel, Clinical Virology | University Hospital Basel, Clinical Bacteriology | Madlen Stange, Alfredo Mari, Tim Roloff, Helena MB Seth-Smith, Michael Schweitzer, Myrta Brunner, Karoline Leuzinger, Kirstine K. Soegaard, Alexander Gensch, Sarah Tschudin-Sutter, Simon Fuchs, Julia Bielicki, Hans Pargger, Martin Siegemund, Ch Rita Schneider-Sliwa, Manuel Battegay, Hans Hirsch, Adrian Egli |
| EPI_ISL_582029, EPI_ISL_582030 | Biology Department, College of Science, Al-Muthanna University | International Centre for Genetic Engineering and Biotechnology (ICGEB) and ARGO Open Lab Platform | Nihad Al-Rashedi, Danilo Licastro, Sreejith Rajasekharan, Simeone Dal Monego, Alessandro Marcello |
| EPI_ISL_582226, EPI_ISL_582232 | Wyoming Public Health Laboratory | Center for Global Health, University of New Mexico Health Sciences Center | Daryl Domman, Kurt Schwalm, Rob Christensen, Wanda Manley, Cari Sloma, Noah Hull, Darrell Dinwiddie |
| EPI_ISL_582809 | Gavle klinisk mikrobiologi | The Public Health Agency of Sweden | Anna-Malin Linde, Maria Lind Karlberg, Mattias Haukland, Reza Advani, Olov Svartstrom, Oskar Karlsson Lindsjo, Sandra Broddesson, Petra Edquist, Mia Brytting, Anna Risberg, Karin Tegmark |
| EPI_ISL_582836, EPI_ISL_582837, EPI_ISL_582838 | Klinisk mikrobiologi Vasternorrland | The Public Health Agency of Sweden | Anna-Malin Linde, Maria Lind Karlberg, Mattias Haukland, Reza Advani, Olov Svartstrom, Oskar Karlsson Lindsjo, Sandra Broddesson, Petra Edquist, Mia Brytting, Anna Risberg, Karin Tegmark |
| EPI_ISL_582851, EPI_ISL_582852, EPI_ISL_582853, EPI_ISL_582854, EPI_ISL_582855, EPI_ISL_582856, EPI_ISL_582857, EPI_ISL_582858, EPI_ISL_582859, EPI_ISL_582860, EPI_ISL_582861, EPI_ISL_582862, EPI_ISL_582863, EPI_ISL_582864, EPI_ISL_582865, EPI_ISL_582866, EPI_ISL_582867, EPI_ISL_582868, EPI_I[illegible] | | | |
| see above | County of Santa Clara Public Health Department | Chan-Zuckerberg Biohub | CZB Cliahub Consortium |
| EPI_ISL_583055, EPI_ISL_583056, EPI_ISL_583058, EPI_ISL_583059 | Humboldt County Public Health Laboratory | Chan-Zuckerberg Biohub | CZB Cliahub Consortium |
| EPI_ISL_583499 | Distrito Sanitario Sul Campinas | Instituto Adolfo Lutz, Interdiciplinary Procedures Center, Strategic Laboratory | Claudio Tavares Sacchi, Claudia Regina Gonçalves, Erica Valessa Ramos Gomes, Karoline Rodrigues Campos |
| EPI_ISL_583500 | Centro de Saude I Tacito Leite de Carvalho e Silva | Instituto Adolfo Lutz, Interdiciplinary Procedures Center, Strategic Laboratory | Claudio Tavares Sacchi, Claudia Regina Gonçalves, Erica Valessa Ramos Gomes, Karoline Rodrigues Campos |
| EPI_ISL_583505 | Casa de Saude Stella Maris | Instituto Adolfo Lutz, Interdiciplinary Procedures Center, Strategic Laboratory | Claudio Tavares Sacchi, Claudia Regina Gonçalves, Erica Valessa Ramos Gomes, Karoline Rodrigues Campos |
| EPI_ISL_583883, EPI_ISL_583884 | Austrian Agency for Health and Food Safety (AGES) | Bergthaler laboratory, CeMM Research Center for Molecular Medicine of the Austrian Academy of Sciences | Alexandra Popa, Benedikt Agerer, Henrique Colaco, Lukas Endler, Jakob-Wendelin Genger, Alexander Lercher, Mark Smyth, Thomas Penz, Michael Schuster, Jan Laine, Martin Senekowitsch, Judith Aberle, Stephan Aberle, Peter Hufnagl, Daniela Schmid Guenter Weiss, Gregor Hörmann, Kinga Rigler-Hohenwarter, Rainer Gattringer, Wegene Borena, Dorothee von Laer, Gernot Walder, Peter Obrist, Christian Paar, Sabine Sussitz-Rack, Gunther Vogl, Adi Steinrigl, Chris |

| | | | |
|---|---|---|---|
| EPI_ISL_584617, EPI_ISL_584618 | Liverpool Clinical Laboratories | COVID-19 Genomics UK (COG-UK) Consortium | Sam Haldenby, Anita Lucaci, Steve Paterson, Julian Hiscox, Alistair Darby, M Almsaud, A Alrezaihi, Muhannad Alruwaili, Stuart D Armstrong, Jones Benjamin, Eleanor G Bentley, Anu Chawla, Jordan J Clark, Angela Cowell, Richard Eccles, Isabel Garc [illegible] Richard Gregory, Ximeng Han, Catherine Hartley, Margaret Hughes, Miren Iturriza-Gomara, James Johnson, L Luu, Jenifer Manson, Charlotte Nelson, Elaine O'Toole, Cassie Olateju, Rebekah Penrice-Randal , Lucille Rainbow, N.P Randle, Trevor Ian [illegible] Swainston, Ecaterina Vamos, Joanne Watts, Mark Whitehead |
| EPI_ISL_586244, EPI_ISL_586245, EPI_ISL_586246, EPI_ISL_586247, EPI_ISL_586248, EPI_ISL_586249, EPI_ISL_586250, EPI_ISL_586251, EPI_ISL_586252, EPI_ISL_586253, EPI_ISL_586254, EPI_ISL_586255, EPI_ISL_586256, EPI_ISL_586257, EPI_ISL_586258, EPI_ISL_586259, EPI_ISL_586261, EPI_ISL_586262 | | | |
| see above | Alaska State Virology Laboratory | Alaska State Virology Laboratory | Jack Chen, Ph.D. |
| EPI_ISL_586513 | B.J. Medical College and Civil hospital, Ahmedabad | Gujarat Biotechnology Research Centre | Pinal Trivedi, Maharshi Pandya, Nidhi Patel, Nitin Savaliya, Raghawendra Kumar, Dinesh Kumar, Zuber Saiyed, Komal Patel, Labdhi Pandya, Afzal Ansari, Nikha Trivedi, Pranay Shah, Kamlesh J Upadhyay, Sanjay Kapadia, Apurvasinh Puvar, Janvi Rava [illegible] Chaitanya Joshi, Madhvi Joshi |
| EPI_ISL_586514 | B.J. Medical College and Civil hospital, Ahmedabad | Gujarat Biotechnology Research Centre | Maharshi Pandya, Nidhi Patel, Nitin Savaliya, Raghawendra Kumar, Dinesh Kumar, Zuber Saiyed, Komal Patel, Labdhi Pandya, Afzal Ansari, Nikha Trivedi, Pranay Shah, Kamlesh J Upadhyay, Sanjay Kapadia, Apurvasinh Puvar, Janvi Raval, Zarna Pate [illegible] Chaitanya Joshi, Madhvi Joshi |
| EPI_ISL_586515 | B.J. Medical College and Civil hospital, Ahmedabad | Gujarat Biotechnology Research Centre | Nidhi Patel, Nitin Savaliya, Raghawendra Kumar, Dinesh Kumar, Zuber Saiyed, Komal Patel, Labdhi Pandya, Afzal Ansari, Nikha Trivedi, Pranay Shah, Kamlesh J Upadhyay, Sanjay Kapadia, Apurvasinh Puvar, Janvi Raval, Zarna Patel, Monika Gandhi, [illegible] Chaitanya Joshi, Madhvi Joshi |
| EPI_ISL_586516 | B.J. Medical College and Civil hospital, Ahmedabad | Gujarat Biotechnology Research Centre | Nitin Savaliya, Raghawendra Kumar, Dinesh Kumar, Zuber Saiyed, Komal Patel, Labdhi Pandya, Afzal Ansari, Nikha Trivedi, Pranay Shah, Kamlesh J Upadhyay, Sanjay Kapadia, Apurvasinh Puvar, Janvi Raval, Zarna Patel, Monika Gandhi, Pinal Trivedi [illegible] Chaitanya Joshi, Madhvi Joshi |
| EPI_ISL_586517 | B.J. Medical College and Civil hospital, Ahmedabad | Gujarat Biotechnology Research Centre | Raghawendra Kumar, Dinesh Kumar, Zuber Saiyed, Komal Patel, Labdhi Pandya, Afzal Ansari, Nikha Trivedi, Pranay Shah, Kamlesh J Upadhyay, Sanjay Kapadia, Apurvasinh Puvar, Janvi Raval, Zarna Patel, Monika Gandhi, Pinal Trivedi, Maharshi Pan [illegible] Chaitanya Joshi, Madhvi Joshi |
| EPI_ISL_586518 | B.J. Medical College and Civil hospital, Ahmedabad | Gujarat Biotechnology Research Centre | Dinesh Kumar, Zuber Saiyed, Komal Patel, Labdhi Pandya, Afzal Ansari, Nikha Trivedi, Pranay Shah, Kamlesh J Upadhyay, Sanjay Kapadia, Apurvasinh Puvar, Janvi Raval, Zarna Patel, Monika Gandhi, Pinal Trivedi, Maharshi Pandya, Nidhi Patel, Nitin [illegible] Chaitanya Joshi, Madhvi Joshi |
| EPI_ISL_586519 | B.J. Medical College and Civil hospital, Ahmedabad | Gujarat Biotechnology Research Centre | Zuber Saiyed, Komal Patel, Labdhi Pandya, Afzal Ansari, Nikha Trivedi, Pranay Shah, Kamlesh J Upadhyay, Sanjay Kapadia, Apurvasinh Puvar, Janvi Raval, Zarna Patel, Monika Gandhi, Pinal Trivedi, Maharshi Pandya, Nidhi Patel, Nitin Savaliya, Ragha [illegible] Chaitanya Joshi, Madhvi Joshi |
| EPI_ISL_586520 | B.J. Medical College and Civil hospital, Ahmedabad | Gujarat Biotechnology Research Centre | Komal Patel, Labdhi Pandya, Afzal Ansari, Nikha Trivedi, Pranay Shah, Kamlesh J Upadhyay, Sanjay Kapadia, Apurvasinh Puvar, Janvi Raval, Zarna Patel, Monika Gandhi, Pinal Trivedi, Maharshi Pandya, Nidhi Patel, Nitin Savaliya, Raghawendra Kumar [illegible] Chaitanya Joshi, Madhvi Joshi |
| EPI_ISL_586521 | B.J. Medical College and Civil hospital, Ahmedabad | Gujarat Biotechnology Research Centre | Labdhi Pandya, Afzal Ansari, Nikha Trivedi, Pranay Shah, Kamlesh J Upadhyay, Sanjay Kapadia, Apurvasinh Puvar, Janvi Raval, Zarna Patel, Monika Gandhi, Pinal Trivedi, Maharshi Pandya, Nidhi Patel, Nitin Savaliya, Raghawendra Kumar, Dinesh Kum [illegible] Chaitanya Joshi, Madhvi Joshi |
| EPI_ISL_586522 | B.J. Medical College and Civil hospital, Ahmedabad | Gujarat Biotechnology Research Centre | Afzal Ansari, Nikha Trivedi, Pranay Shah, Kamlesh J Upadhyay, Sanjay Kapadia, Apurvasinh Puvar, Janvi Raval, Zarna Patel, Monika Gandhi, Pinal Trivedi, Maharshi Pandya, Nidhi Patel, Nitin Savaliya, Raghawendra Kumar, Dinesh Kumar, Zuber Saiye [illegible] Chaitanya Joshi, Madhvi Joshi |
| EPI_ISL_586523 | B.J. Medical College and Civil hospital, Ahmedabad | Gujarat Biotechnology Research Centre | Nikha Trivedi, Pranay Shah, Kamlesh J Upadhyay, Sanjay Kapadia, Apurvasinh Puvar, Janvi Raval, Zarna Patel, Monika Gandhi, Pinal Trivedi, Maharshi Pandya, Nidhi Patel, Nitin Savaliya, Raghawendra Kumar, Dinesh Kumar, Zuber Saiyed, Komal Pate [illegible] Chaitanya Joshi, Madhvi Joshi |
| EPI_ISL_586524 | B.J. Medical College and Civil hospital, Ahmedabad | Gujarat Biotechnology Research Centre | Pranay Shah, Kamlesh J Upadhyay, Sanjay Kapadia, Apurvasinh Puvar, Janvi Raval, Zarna Patel, Monika Gandhi, Pinal Trivedi, Maharshi Pandya, Nidhi Patel, Nitin Savaliya, Raghawendra Kumar, Dinesh Kumar, Zuber Saiyed, Komal Patel, Labdhi Panc [illegible] Chaitanya Joshi, Madhvi Joshi |
| EPI_ISL_586525 | B.J. Medical College and Civil hospital, Ahmedabad | Gujarat Biotechnology Research Centre | Kamlesh J Upadhyay, Sanjay Kapadia, Apurvasinh Puvar, Janvi Raval, Zarna Patel, Monika Gandhi, Pinal Trivedi, Maharshi Pandya, Nidhi Patel, Nitin Savaliya, Raghawendra Kumar, Dinesh Kumar, Zuber Saiyed, Komal Patel, Labdhi Pandya, Afzal Ansa [illegible] Chaitanya Joshi, Madhvi Joshi |
| EPI_ISL_586526 | B.J. Medical College and Civil hospital, Ahmedabad | Gujarat Biotechnology Research Centre | Sanjay Kapadia, Apurvasinh Puvar, Janvi Raval, Zarna Patel, Monika Gandhi, Pinal Trivedi, Maharshi Pandya, Nidhi Patel, Nitin Savaliya, Raghawendra Kumar, Dinesh Kumar, Zuber Saiyed, Komal Patel, Labdhi Pandya, Afzal Ansari, Nikha Trivedi, Pran [illegible] Chaitanya Joshi, Madhvi Joshi |
| EPI_ISL_586527 | B.J. Medical College and Civil hospital, Ahmedabad | Gujarat Biotechnology Research Centre | Apurvasinh Puvar, Janvi Raval, Zarna Patel, Monika Gandhi, Pinal Trivedi, Maharshi Pandya, Nidhi Patel, Nitin Savaliya, Raghawendra Kumar, Dinesh Kumar, Zuber Saiyed, Komal Patel, Labdhi Pandya, Afzal Ansari, Nikha Trivedi, Pranay Shah, Kamles [illegible] Chaitanya Joshi, Madhvi Joshi |
| EPI_ISL_586528 | B.J. Medical College and Civil hospital, Ahmedabad | Gujarat Biotechnology Research Centre | Zarna Patel, Monika Gandhi, Pinal Trivedi, Maharshi Pandya, Nidhi Patel, Nitin Savaliya, Raghawendra Kumar, Dinesh Kumar, Zuber Saiyed, Komal Patel, Labdhi Pandya, Afzal Ansari, Nikha Trivedi, Pranay Shah, Kamlesh J Upadhyay, Sanjay Kapadia, [illegible] Chaitanya Joshi, Madhvi Joshi |
| EPI_ISL_586529 | B.J. Medical College and Civil hospital, Ahmedabad | Gujarat Biotechnology Research Centre | Monika Gandhi, Pinal Trivedi, Maharshi Pandya, Nidhi Patel, Nitin Savaliya, Raghawendra Kumar, Dinesh Kumar, Zuber Saiyed, Komal Patel, Labdhi Pandya, Afzal Ansari, Nikha Trivedi, Pranay Shah, Kamlesh J Upadhyay, Sanjay Kapadia, Apurvasinh [illegible] Chaitanya Joshi, Madhvi Joshi |
| EPI_ISL_586530 | B.J. Medical College and Civil hospital, Ahmedabad | Gujarat Biotechnology Research Centre | Pinal Trivedi, Maharshi Pandya, Nidhi Patel, Nitin Savaliya, Raghawendra Kumar, Dinesh Kumar, Zuber Saiyed, Komal Patel, Labdhi Pandya, Afzal Ansari, Nikha Trivedi, Pranay Shah, Kamlesh J Upadhyay, Sanjay Kapadia, Apurvasinh Puvar, Janvi Rava [illegible] Chaitanya Joshi, Madhvi Joshi |
| EPI_ISL_590689 | B.J. Medical College and Civil hospital, Ahmedabad | Gujarat Biotechnology Research Centre | Janvi Raval, Zarna Patel, Monika Gandhi, Pinal Trivedi, Maharshi Pandya, Nidhi Patel, Nitin Savaliya, Raghawendra Kumar, Dinesh Kumar, Zuber Saiyed, Komal Patel, Labdhi Pandya, Afzal Ansari, Nikha Trivedi, Pranay Shah, Kamlesh J Upadhyay, San [illegible] Chaitanya Joshi, Madhvi Joshi |
| EPI_ISL_591545 | CHU Purpan - Laboratoire de Virologie - Institut Fédératif de Biologie | CHU Purpan - Laboratoire de Virologie - Institut Fédératif de Biologie | Latour J., Ranger N., Dubois M., Carcenac R., Harter A., Boyer P., Tremeaux P., Izopet J. |
| EPI_ISL_591999 | Microbiological Diagnostic Unit - Public Health Laboratory (MDU-PHL) | MDU-PHL | Seemann T., Schultz, M. B., Sait, M., Sherry, N. |
| EPI_ISL_593480, EPI_ISL_593553 | Brigham and Women's Hospital | Jonathan Li Laboratory | Manish C. Choudhary, James Regan, Jonathan Z. Li |
| EPI_ISL_593684, EPI_ISL_593685, EPI_ISL_593689, EPI_ISL_593690, EPI_ISL_593691, EPI_ISL_593692, EPI_ISL_593693 | South Eastern Area Laboratory Services (SEALS) | NSW Health Pathology - Institute of Clinical Pathology and Medical Research; Westmead Hospital; University of Sydney | CIDM-PH et al. |
| EPI_ISL_593865, EPI_ISL_593866 | CHU Purpan - Laboratoire de Virologie - Institut Fédératif de Biologie | CHU Purpan - Laboratoire de Virologie - Institut Fédératif de Biologie | Latour J., Ranger N., Dubois M., Carcenac R., Harter A., Boyer P., Tremeaux P., Izopet J. |
| EPI_ISL_594042 | Utah Public Health Laboratory | Utah Public Health Laboratory | Erin Young, Kelly Oakeson |
| EPI_ISL_594157 | Israel Institute for Biological Research | Israel Institute for Biological Research | Galia Zaide, Inbar Cohen-Gihon, Ofir Israeli, Dana Stein, Shay Weiss, Orly Laskar, Yoav Gal, Libby Weiss, Emanuelle Mamroud, Adi Beth-Din and Anat Zvi |
| EPI_ISL_594188 | Department of Pathology, School of Medicine, Imam Khomeini Hospital, Tehran University of Medical Sciences | Genetics Research Center, University of Social Welfare and Rehabilitation Sciences | Zohreh Fattahi, Marzieh Mohseni, Khadijeh Jalalvand, Azam Ghaziasadi, Seyedeh elham Mortazavi, Ali Jafarpour, Azar Hadadi, Alireza Abdollahi, Ali Jafarpour, Azam Ghaziasad, Seyedeh elham Mortazavi, Saber Soltani, Reza Najafipour, Kimia [illegible] |
| EPI_ISL_594320, | Florida Bureau of Public | Florida Bureau of Public | Sarah Schmedes, Jason Blanton |

| | | | |
|---|---|---|---|
| EPI_ISL_594327, EPI_ISL_594328, EPI_ISL_594329, EPI_ISL_594330, EPI_ISL_594331 | Health Laboratories | Health Laboratories | |
| EPI_ISL_594445, EPI_ISL_594446 | Utah Public Health Laboratory | Utah Public Health Laboratory | Erin Young, Kelly Oakeson |
| EPI_ISL_594465 | FL Bureau of Public Health Laboratories | Pathogen Discovery, Respiratory Viruses Branch, Division of Viral Diseases, Centers for Disease Control and Prevention | Ying Tao, Yan Li, Clinton Paden, Jing Zhang, Krista Queen, Anna Uehara, Haibin Wang, Julu Bhatnagar, Suxiang Tong |
| EPI_ISL_596542, EPI_ISL_596543, EPI_ISL_596545, EPI_ISL_596548, EPI_ISL_596549, EPI_ISL_596550, EPI_ISL_596552, EPI_ISL_596553, EPI_ISL_596554, EPI_ISL_596562 | Palestinian Ministry of Health | Molecular Genetics Lab | Nouar Qutob, Zaidoun Salah, Damien Richard, Hisham Darwish, Husam Sallam, Issa Shtayeh, Osama Najjar, Mahmoud Ruzayqat, Dana Najjar, Francois Balloux, Lucy van Dorp |
| EPI_ISL_596743 | PathWest Laboratory Medicine WA | PathWest Laboratory Medicine WA Microbial Surveillance Unit | PathWest Laboratory Medicine WA Microbial Surveillance Unit |
| EPI_ISL_602161 | Lighthouse Lab in Alderley Park | Wellcome Sanger Institute for the COVID-19 Genomics UK (COG-UK) consortium | Jacquelyn Wynn, Mairead Hyland, The Lighthouse Lab in Alderley Park and Alex Alderton, Roberto Amato, Sonia Goncalves, Ewan Harrison, David K. Jackson, Ian Johnston, Dominic Kwiatkowski, Cordelia Langford, John Sillitoe on behalf of the (http://www.sanger.ac.uk/covid-team) |
| EPI_ISL_602559, EPI_ISL_602560 | Department of Biology and Wildlife, Alaska State Virology Laboratory | Department of Biology and Wildlife, Alaska State Virology Laboratory | DeRonde,S., Deuling,H., Chen,J. |
| EPI_ISL_602627, EPI_ISL_602628 | AHRI-Sigal | KRISP, KZN Research Innovation and Sequencing Platform | Gazy I, Sigl A, Karim F, Cele S, Giandhari J, Pillay S, Tegally H, Wilkinson E, de Oliveira T |
| EPI_ISL_602948, EPI_ISL_602949, EPI_ISL_602950, EPI_ISL_602951, EPI_ISL_602952, EPI_ISL_602953, EPI_ISL_602954, EPI_ISL_602955 | Utah Public Health Laboratory | Utah Public Health Laboratory | Erin Young, Kelly Oakeson |
| EPI_ISL_603028 | Hospital Municipal Santa Ana | Instituto Adolfo Lutz, Interdiciplinary Procedures Center, Strategic Laboratory | Claudio Tavares Sacchi, Claudia Regina Gonçalves, Erica Valessa Ramos Gomes, Karoline Rodrigues Campos |
| EPI_ISL_605931 | Utah Public Health Laboratory, Utah Public Health Laboratory Infectious Disease submission group | Utah Public Health Laboratory, Utah Public Health Laboratory Infectious Disease submission group | Young,E.L. and Oakeson,K. |
| EPI_ISL_610040, EPI_ISL_610041 | Texas Department of State Health Services | Texas Department of State Health Services | Rashmi Tuladhar, Bonnie Oh, Jenny Zhang, Maliha Rahman, Anita Pokharel, Myong Koag, Chung Wang, Rachel Lee, Grace Kubin, Mayela Pedrueza |
| EPI_ISL_610211 | Department of Health Technology and Informatics, The Hong Kong Polytechnic University | Department of Health Technology and Informatics, The Hong Kong Polytechnic University | Siu,G.K.-H., Lee,L.-K., Leung,K.S.-S., Leung,J.S.-L., Ng,T.T.-L., Chan,C.T.-M., Tam,K.K.-G., Lao,H.-Y., Wu,A.K.-L., Yau,M.C.-Y., Lai,Y.W.-M., Fung,K.S.-C., Chau,S.K.-Y., Wong,B.K.-C., To,W.-K., Luk,K., Ho,A.Y.-M., Que,T.-L., Y |
| EPI_ISL_613783 | Florida Bureau of Public Health Laboratories | Florida Bureau of Public Health Laboratories | Sarah Schmedes, Jason Blanton |
| EPI_ISL_614370, EPI_ISL_614371, EPI_ISL_614372, EPI_ISL_614373, EPI_ISL_614374 | Molecular diagnostic unit for viral haemorrhagic fevers and emerging viruses, Bouaké CHU Laboratory | Project group Epidemology of Highly Pathogenic Microorganisms, Robert Koch-Institute | Chantal Akoua-Koffi, Diané Bamourou, Etilé Anoh, Essia Belarbi, Safiatou Karidioula, Grit Schubert, Adjaratou Traoré, Soundélé Maïté, Monemo Pacome, Coulibaly Mbegnan, Bamba Fatoumata Touré, Kra Ouf |
| EPI_ISL_615103 | Gavle klinisk mikrobiologi | The Public Health Agency of Sweden | Anna-Malin Linde, Maria Lind Karlberg, Mattias Haukland, Reza Advani, Olov Svartstrom, Oskar Karlsson Lindsjo, Sandra Broddesson, Petra Edquist, Mia Brytting, Anna Risberg, Karin Tegmark |
| EPI_ISL_618156, EPI_ISL_618158, EPI_ISL_618159, EPI_ISL_618165, EPI_ISL_618168, EPI_ISL_618169, EPI_ISL_618170, EPI_ISL_618171, EPI_ISL_618172, EPI_ISL_618173, EPI_ISL_618176, EPI_ISL_618177, EPI_ISL_618272, EPI_ISL_618294, EPI_ISL_618314, EPI_ISL_618375, EPI_ISL_622554, EPI_ISL_622555, EPI_I | | | |
| see above | Department of Virus and Microbiological Special Diagnostics, Statens Serum Institut, Denmark | Albertsen lab, Department of Chemistry and Bioscience, Aalborg University, Denmark | Danish Covid-19 Genome Consortia |
| EPI_ISL_622895, EPI_ISL_622896, EPI_ISL_622898, EPI_ISL_622899, EPI_ISL_622905, EPI_ISL_622909, EPI_ISL_622910, EPI_ISL_622913, EPI_ISL_622923, EPI_ISL_622924, EPI_ISL_622926, EPI_ISL_622934, EPI_ISL_622935, EPI_ISL_622936, EPI_ISL_622937, EPI_ISL_622938 | | | |
| see above | National Institute for Communicable Diseases of the National Health Laboratory Service | National Institute for Communicable Diseases of the National Health Laboratory Service | Allam M, Ismail A, Khumalo Z, Kwenda S, Mtshali P, Mnyameni F, Mohale T, Subramoney K, Bhiman JN |
| EPI_ISL_622974, EPI_ISL_622984, EPI_ISL_622991, | National Health Laboratory Service | National Institute for Communicable Diseases of the National Health | Allam M, Ismail A, Khumalo Z, Kwenda S, Mtshali P, Mnyameni F, Mohale T, Subramoney K, Bhiman JN |

| | | | |
|---|---|---|---|
| EPI_ISL_623043, EPI_ISL_623048 | | Laboratory Service | |
| EPI_ISL_623078 | Uppsala klinisk mikrobiologi | The Public Health Agency of Sweden | Anna-Malin Linde, Maria Lind Karlberg, Mattias Haukland, Reza Advani, Olov Svartstrom, Oskar Karlsson Lindsjo, Sandra Broddesson, Petra Edquist, Mia Brytting, Anna Risberg, Karin Tegmark |
| EPI_ISL_623080, EPI_ISL_623081, EPI_ISL_623082 | Klinisk mikrobiologi, Skanes universitetssjukhus, Lund | The Public Health Agency of Sweden | Anna-Malin Linde, Maria Lind Karlberg, Mattias Haukland, Reza Advani, Olov Svartstrom, Oskar Karlsson Lindsjo, Sandra Broddesson, Petra Edquist, Mia Brytting, Anna Risberg, Karin Tegmark |
| EPI_ISL_623094 | Klinisk mikrobiologi Lanssjukhuset Ryhov, Jonkoping | The Public Health Agency of Sweden | Anna-Malin Linde, Maria Lind Karlberg, Mattias Haukland, Reza Advani, Olov Svartstrom, Oskar Karlsson Lindsjo, Sandra Broddesson, Petra Edquist, Mia Brytting, Anna Risberg, Karin Tegmark |
| EPI_ISL_623095 | Gavle klinisk mikrobiologi | The Public Health Agency of Sweden | Anna-Malin Linde, Maria Lind Karlberg, Mattias Haukland, Reza Advani, Olov Svartstrom, Oskar Karlsson Lindsjo, Sandra Broddesson, Petra Edquist, Mia Brytting, Anna Risberg, Karin Tegmark |
| EPI_ISL_625468 | Child Health Research Foundation | Child Health Research Foundation | Senjuti Saha, Md Saiful Islam Sajib, Nikkon Sarkar, Syed Muktadir Al Sium, Afroza Akter Tanni, Roly Malaker, Arif Mohammad Tanmoy, Md Hafizur Rahman, Samir K Saha |
| EPI_ISL_626341, EPI_ISL_626346, EPI_ISL_626348 | Statens Serum Institute | Statens Serum Institute | Hammer, A.S., Quaade, M.L., Rasmussen, T.B., Fonager, J., Rasmussen, M., Mundbjerg, K., Lohse, L., Strandbygaard, B., Jorgensen, C.S., Afaro-Nunez, A., Rosenstierne, M.W., Halasa, T., Foomsgaard, A., Bel |
| EPI_ISL_626509, EPI_ISL_626510, EPI_ISL_626511, EPI_ISL_626512, EPI_ISL_626517, EPI_ISL_626518 | Northwestern Memorial Hospital | Ozer Lab | Ramon Lorenzo-Redondo, Hannah H. Nam, Scott C. Roberts, Lacy M. Simons, Chad J. Achenbach, Lawrence J. Jennings, Chao Qi, Alan R. Hauser, Michael G. Ison, Judd F. Hultquist, Egon A |
| EPI_ISL_631397, EPI_ISL_631431, EPI_ISL_631432, EPI_ISL_631433, EPI_ISL_631434, EPI_ISL_631458, EPI_ISL_631459, EPI_ISL_631460, EPI_ISL_631461, EPI_ISL_631462, EPI_ISL_631463, EPI_ISL_631464, EPI_ISL_631465, EPI_ISL_631466, EPI_ISL_631467, EPI_ISL_631468, EPI_ISL_631469, EPI_ISL_631470, EPI_I EPI_ISL_631475, EPI_ISL_631476, EPI_ISL_631498, EPI_ISL_631499, EPI_ISL_631500, EPI_ISL_631501 | | | |
| see above | Wisconsin State Laboratory of Hygiene Communicable Disease Division | Wisconsin State Laboratory of Hygiene Communicable Disease Division | Kelsey R. Florek, Abigail C. Shockey |
| EPI_ISL_631650 | Texas Department of State Health Services | Texas Department of State Health Services | Rashmi Tuladhar, Bonnie Oh, Jenny Zhang, Maliha Rahman, Anita Pokharel, Myong Koag, Chung Wang, Rachel Lee, Grace Kubin, Mayela Pedrueza |
| EPI_ISL_632771 | Dutch COVID-19 response team | Erasmus Medical Center | Bas Oude Munnink, David Nieuwenhuijse, Reina Sikkema, Claudia Schapendonk, Irina Chestakova, Anne van der Linden, Theo Bestebroer, Stefan van Nieuwkoop, Mark Pronk, Pascal Lexmond, Corien Swaan, Manon Haverkate, Madelief Mollers, Mart S Voermans, Aura Timen, Corine GeurtsvanKessel, Annemiek van der Eijk, Richard Molenkamp, Marion Koopmans, on behalf of the Dutch national COVID-19 response team. |
| EPI_ISL_635380, EPI_ISL_635385, EPI_ISL_635386, EPI_ISL_635388, EPI_ISL_635389, EPI_ISL_635390, EPI_ISL_635393, EPI_ISL_635394, EPI_ISL_635395, EPI_ISL_635396, EPI_ISL_635397, EPI_ISL_635399, EPI_ISL_635400, EPI_ISL_635401, EPI_ISL_635402, EPI_ISL_635454, EPI_ISL_635461, EPI_ISL_635462, EPI_I EPI_ISL_635474, EPI_ISL_635476 | | | |
| see above | San Diego County Public Health Laboratory | Andersen lab at Scripps Research | SEARCH Alliance San Diego with Tracy Basler, Jovan Shephard, Brett Austin |
| EPI_ISL_635536, EPI_ISL_635537, EPI_ISL_635538, EPI_ISL_635539, EPI_ISL_635540, EPI_ISL_635541, EPI_ISL_635542, EPI_ISL_635543, EPI_ISL_635544, EPI_ISL_635545, EPI_ISL_635546, EPI_ISL_635547, EPI_ISL_635548, EPI_ISL_635549, EPI_ISL_635550, EPI_ISL_635551 | | | |
| see above | Centro de Diagnstico COVID-19 UABC Tijuana | Andersen lab at Scripps Research | SEARCH Alliance San Diego with Idanya Rubí Serafín Higuera, Manuel Sánchez Alavez, Jorge Luis Jiménez Niebla, Germán Ibarra, Jonathan Vincent Baena, Oscar Efrén Zazueta Fierro |
| EPI_ISL_635777 | Biolab Diagnostic Laboratories | Andersen lab at Scripps Research | Issa Abu-Dayyeh, Ahmad Tibi, Lama Hussein, Lina Mohammad, Zein Naber, Amid Abdelnour with SEARCH Alliance San Diego |
| EPI_ISL_636090, EPI_ISL_636091, EPI_ISL_636092, EPI_ISL_636093, EPI_ISL_636094, EPI_ISL_636095, EPI_ISL_636096, EPI_ISL_636097, EPI_ISL_636099, EPI_ISL_636100, EPI_ISL_636101, EPI_ISL_636102, EPI_ISL_636103, EPI_ISL_636104, EPI_ISL_636105, EPI_ISL_636106, EPI_ISL_636107, EPI_ISL_636108, EPI_I EPI_ISL_636217, EPI_ISL_636238, EPI_ISL_636241, EPI_ISL_636242, EPI_ISL_636243, EPI_ISL_636244, EPI_ISL_636245, EPI_ISL_636246, EPI_ISL_636247, EPI_ISL_636251, EPI_ISL_636253, EPI_ISL_636257, EPI_ISL_636258, EPI_ISL_636259, EPI_ISL_636260, EPI_ISL_636261, EPI_ISL_636262, EPI_ISL_636263 | | | |
| see above | San Diego County Public Health Laboratory | Andersen lab at Scripps Research | SEARCH Alliance San Diego with Tracy Basler, Jovan Shephard, Brett Austin |
| EPI_ISL_636557 | Dutch COVID-19 response team | National Institute for Public Health and the Environment (RIVM) | Adam Meijer, Harry Vennema, Jeroen Cremer, Sharon van den Brink, Bas van der Veer, AnneMarie van den Brandt, Florian Zwagemaker, Dennis Schmitz, Chantal Reusken, on behalf of the national COVID |
| EPI_ISL_636966 | Pathogen Genomics Lab King Abdullah University of Science and Technology(KAUST) | Pathogen Genomics Lab King Abdullah University of Science and Technology(KAUST) | Fathia Ben Rached, Raeece Naeem, Sharif Hala, Fadwa Alofi, Rahul P Salunke, Sara Mfarrej, Amit Kumar Subudhi, Afrah Alsomali, Asim Khogeer, Ahmad Bakur Mahmoud, Anwar Hashem, Naif Almonta |
| EPI_ISL_639912, EPI_ISL_639913, EPI_ISL_639915, EPI_ISL_639921, EPI_ISL_639922, EPI_ISL_639941, EPI_ISL_639942, EPI_ISL_639943, EPI_ISL_639944, EPI_ISL_639945 | Omsk Research Institute of Natural Focal Infections | WHO National Influenza Centre Russian Federation | Artem Fadeev, Ekaterina Gradoboeva, Ekaterina Savkina, Daria Nashatyreva, Elena Poleshchuk, Aleksei Vasilenko, Valery Yakimenko, Andrey Komissarov |
| EPI_ISL_640065 | Mitchells Plain Hospital wc MPH | NHLS/UCT | Arash Iranzadeh, Deelan Doolabh, Lynn Tyers, Bruna Galvao, Innocent Mudau, Marvin Hsiao, Kruger Marais, Diana Hardie, Stephen Korsman, Carolyn Williamson |
| EPI_ISL_640066 | Groote Schuur Hospital wc GSH | NHLS/UCT | Arash Iranzadeh, Deelan Doolabh, Lynn Tyers, Bruna Galvao, Innocent Mudau, Marvin Hsiao, Kruger Marais, Diana Hardie, Stephen Korsman, Carolyn Williamson |
| EPI_ISL_644952, EPI_ISL_644953, EPI_ISL_644954 | Department of Infectious Diseases, Keio University School of Medicine, Tokyo, Japan | Center for Medical Genetics, Keio University School of Medicine, Tokyo, Japan | Kenjiro Kosaki, Yuka Iwasaki, Hirotsugu Ishizu, Haruhiko Siomi, Kodai Abe |
| EPI_ISL_648144 | Gavle klinisk mikrobiologi | The Public Health Agency of Sweden | Anna-Malin Linde, Maria Lind Karlberg, Mattias Haukland, Reza Advani, Olov Svartstrom, Oskar Karlsson Lindsjo, Sandra Broddesson, Petra Edquist, Mia Brytting, Anna Risberg, Karin Tegmark |
| EPI_ISL_648180 | The Public Health Agency of Sweden | The Public Health Agency of Sweden | Anna-Malin Linde, Maria Lind Karlberg, Mattias Haukland, Reza Advani, Olov Svartstrom, Oskar Karlsson Lindsjo, Sandra Broddesson, Petra Edquist, Mia Brytting, Anna Risberg, Karin Tegmark |

| | | | |
|---|---|---|---|
| EPI_ISL_648320, EPI_ISL_648369, EPI_ISL_648370, EPI_ISL_648371, EPI_ISL_648372 | Laboratorio de Investigaciones de Baney | University Hospital Basel, Clinical Bacteriology | Carlos Cortes, Claudia Daubenberger, Adrian Egli, Guillermo Garcia, Salome Hosch, Bonifacio Manguire Nlavo, Alfredo Mari, Maximilian Mpina, Elizabeth Nyakarungu, Diosdado Odjama Nseng Ada, Mitoha Ondo O Ayekaba, Tim Roloff, Tobias Schi |
| EPI_ISL_648714, EPI_ISL_648715, EPI_ISL_648716, EPI_ISL_648717, EPI_ISL_648718, EPI_ISL_648719, EPI_ISL_648720, EPI_ISL_648721, EPI_ISL_648722 | Department of Laboratory Medicine, Tan Tock Seng Hospital | Department of Laboratory Medicine, Tan Tock Seng Hospital | Chen YYC, Zair X, Lim JX, Li C, Tang WY, Maurer-Stroh S, Barkham TMS, Nagarajan N, Sessions OM |
| EPI_ISL_648845, EPI_ISL_648846, EPI_ISL_648847, EPI_ISL_648848, EPI_ISL_648849, EPI_ISL_648850, EPI_ISL_648851, EPI_ISL_648852, EPI_ISL_648853, EPI_ISL_648854, EPI_ISL_648855, EPI_ISL_648856, EPI_ISL_648857, EPI_ISL_648858, EPI_ISL_648859, EPI_ISL_648860, EPI_ISL_649008 | | | |
| see above | San Diego County Public Health Laboratory | Andersen lab at Scripps Research | SEARCH Alliance San Diego with Tracy Basler, Jovan Shephard, Brett Austin |
| EPI_ISL_649154 | Queen Astrid Military Hospital | Institute of Tropical Medicine | Philippe Selhorst, Colin Anthony |
| EPI_ISL_649164 | Laboratorio de Investigaciones de Baney | University Hospital Basel, Clinical Bacteriology | Carlos Cortes, Claudia Daubenberger, Adrian Egli, Guillermo Garcia, Salome Hosch, Bonifacio Manguire Nlavo, Alfredo Mari, Maximilian Mpina, Elizabeth Nyakarungu, Diosdado Odjama Nseng Ada, Mitoha Ondo O Ayekaba, Tim Roloff, Tobias Schi |
| EPI_ISL_653200, EPI_ISL_653201, EPI_ISL_653202, EPI_ISL_653203, EPI_ISL_653204, EPI_ISL_653205, EPI_ISL_653206, EPI_ISL_653207, EPI_ISL_653208, EPI_ISL_653218, EPI_ISL_653219, EPI_ISL_653226, EPI_ISL_653290, EPI_ISL_653291, EPI_ISL_653292, EPI_ISL_653293, EPI_ISL_653294, EPI_ISL_653295, EPI_I | | | |
| see above | Florida Bureau of Public Health Laboratories | Florida Bureau of Public Health Laboratories | Sarah Schmedes, Jason Blanton |
| EPI_ISL_653546, EPI_ISL_653547, EPI_ISL_653551, EPI_ISL_653554 | LSUHS Emerging Viral Threat Laboratory | Microbial Genome Sequencing Center | Jeremy P. Kamil, Rona S. Scott, Maarten Van Diest, Malgorzata Bienkowska-Haba, Katarzyna Zwolinska, Andrew D. Yurochko, Christopher G. Kevil, Martin J. Sapp, Daniel J. Snyder, Vaughn S. Cooper, J |
| EPI_ISL_653755, EPI_ISL_653757 | Instituto Nacional de Salud, Bogotá, Colombia | Instituto Nacional de Salud, Bogotá, Colombia | Katherine Laiton-Donato, Diego A. Álvarez-Díaz, Carlos Franco-Muñoz, Mauricio Pacheco-Montealegre, Jonathan Reales, Diego Andrés Prada, Jose A. Usme-Ciro, Zulma M. Cucunubá, Christian Julian VillabonaArenas, Liz Villabona-Arenas, Suss Walteros-Acero, Franklin Prieto, Carlos Andrés Durán, Martha Lucia Ospina Martinez, Marcela Mercado-Reyes |
| EPI_ISL_653854, EPI_ISL_653855, EPI_ISL_653856, EPI_ISL_653857, EPI_ISL_653858 | Maulana Azad Medical College | National Institute of Biomedical Genomics | Arindam Maitra, Sonal Saxena, Vikas Manchanda, Oves Siddiqui, Saumitra Das |
| EPI_ISL_653867, EPI_ISL_653868 | Translational Health Science and Technology Institute | National Institute of Biomedical Genomics | Arindam Maitra, Guruprasad Medigeshi, Sharanabasava Patil, Anbalagan Ananthraj, Madhu Pareek, Imran Khan, Gagandeep Kang, Saumitra Das |
| EPI_ISL_654166 | Hospital General Universitario Gregorio Marañón | SeqCOVID-SPAIN consortium/IBV(CSIC) | Darío García de Viedma, Laura Pérez-Lago, Marta Herranz, Jon Sicilia, Julia Suárez, Pilar Catalán, Patricia Muñoz and SeqCOVID-SPAIN consortium |
| EPI_ISL_657466, EPI_ISL_657467 | Servicio de Microbiología. Hospital General Universitario de Castellón | SeqCOVID-SPAIN consortium/IBV(CSIC) | Rosario Moreno Muñoz, María Dolores Tirado Balaguer and SeqCOVID-SPAIN consortium |
| EPI_ISL_660165 | NHLS-IALCH | KRISP, KZN Research Innovation and Sequencing Platform | Gazy I, Sigal A, Karim F, Cele S, Giandhari J, Pillay S, Tegally H, Wilkinson E, de Oliveira T |
| EPI_ISL_660445, EPI_ISL_660515 | Laboratoire de Microbiologie CHU Souro Sanou | Centre Muraz | Abdoul-Salam Ouedraogo, Yacouba Sawadogo, Essia Belarbi, Grit Schubert, Fabian Leendertz, Arsène Zongo, Soumeya Ouangraoua, Zekiba Tarnagda, Lassana Sangaré, Halidou Tint |
| EPI_ISL_661181, EPI_ISL_661184, EPI_ISL_661188, EPI_ISL_661191, EPI_ISL_661192, EPI_ISL_661195, EPI_ISL_661196 | Scientific Veterinary Institute Novi Sad | Veterinary Specialized Institute "Kraljevo", Serbia | Vidanovic,D., Tesovic,B., Knezevic,A., Jovanovic,T., Jankovic,M., Sekler,M., Banovic Djeri,B., Petrovic,T., Volkening,J., Afonso,C. |
| EPI_ISL_661283 | Gavle klinisk mikrobiologi | The Public Health Agency of Sweden | Department of Microbiology, The Public Health Agency of Sweden |
| EPI_ISL_666599 | Dept. of Microbiology and Infection Control, Akershus University Hospital HF | Dept. of Microbiology and Infection Control, Akershus University Hospital HF | Hege Vangstein Aamot, Alexander Hesselberg Løvestad, Silje Bakken Jørgensen, Nina Handal, Ole Herman Ambur |
| EPI_ISL_671607, EPI_ISL_671608, EPI_ISL_671610, EPI_ISL_671612, EPI_ISL_671613, EPI_ISL_671614, EPI_ISL_671617, EPI_ISL_671620, EPI_ISL_671624, EPI_ISL_671626, EPI_ISL_671628, EPI_ISL_671629, EPI_ISL_671632, EPI_ISL_671633, EPI_ISL_671634, EPI_ISL_671639, EPI_ISL_671641, EPI_ISL_671642, EPI_I | | | |
| see above | Texas Department of State Health Services | Texas Department of State Health Services | Rashmi Tuladhar, Bonnie Oh, Jenny Zhang, Maliha Rahman, Anita Pokharel, Myong Koag, Chung Wang, Rachel Lee, Grace Kubin, Mayela Pedrueza, James Daniel Bonser |
| EPI_ISL_672079, EPI_ISL_672173, EPI_ISL_672174, EPI_ISL_672183, EPI_ISL_672188, EPI_ISL_672206, EPI_ISL_672212, EPI_ISL_672213, EPI_ISL_672219, EPI_ISL_672220, EPI_ISL_672221, EPI_ISL_672224, EPI_ISL_672228, EPI_ISL_672243 | | | |
| see above | The Ashley Laboratory, Stanford University | Chan-Zuckerberg Biohub | CZB Cliahub Consortium |
| EPI_ISL_672598, EPI_ISL_672599, EPI_ISL_672600 | Infectious Diseases and Tropical Medicine Research Center, Infectious Diseases and Tropical Medicine Research Center | Infectious Diseases and Tropical Medicine Research Center, Infectious Diseases and Tropical Medicine Research Center | Ahangarzadeh,S., Haghjooy Javanmard,S., Shariati,L., Aboutalebian,S., Ataei,B. |
| EPI_ISL_672601 | Infectious Diseases and Tropical Medicine Research | Infectious Diseases and Tropical Medicine Research | Ahangarzadeh,S., Haghjooy Javanmard,S., Shariati,L., Aboutalebian,S., Ataei,B., Shoaei,P. |

| | | | |
|---|---|---|---|
| | Center, Infectious Diseases and Tropical Medicine Research Center | Center, Infectious Diseases and Tropical Medicine Research Center | |
| EPI_ISL_676582, EPI_ISL_676583, EPI_ISL_676584 | Scientific Veterinary Institute Novi Sad | Veterinary Specialized Institute "Kraljevo", Serbia | Vidanovic,D., Tesovic,B., Knezevic,A., Jovanovic,T., Jankovic,M., Sekler,M., Banovic Djeri,B., Petrovic,T., Volkening,J., Afonso,C. |
| EPI_ISL_676661, EPI_ISL_676668 | Masonic Medical Research Institute | Wadsworth Center, New York State Department.of Health | Nathan Tucker, Kirsten St. George, Daryl M. Lamson, Alexis Russel, Jonathan Plitnick, Navjot Singh, John Kelly, Sara Griesemer, Erasmus Schneider, Erica Lasek-Nesselquist |
| EPI_ISL_676983, EPI_ISL_676984, EPI_ISL_676985, EPI_ISL_676986 | Wadsworth Center, New York State Department.of Health | Wadsworth Center, New York State Department.of Health | Kirsten St. George, Daryl M. Lamson, Alexis Russel, Jonathan Plitnick, Navjot Singh, John Kelly, Sara Griesemer, Erasmus Schneider, Erica Lasek-Nesselquist |
| EPI_ISL_677004, EPI_ISL_677005, EPI_ISL_677006, EPI_ISL_677007, EPI_ISL_677008, EPI_ISL_677009, EPI_ISL_677052, EPI_ISL_677053, EPI_ISL_677054, EPI_ISL_677055, EPI_ISL_677056, EPI_ISL_677057, EPI_ISL_677058, EPI_ISL_677076 | | | |
| see above | Masonic Medical Research Institute | Wadsworth Center, New York State Department.of Health | Nathan Tucker, Kirsten St. George, Daryl M. Lamson, Alexis Russel, Jonathan Plitnick, Navjot Singh, John Kelly, Sara Griesemer, Erasmus Schneider, Erica Lasek-Nesselquist |
| EPI_ISL_677267, EPI_ISL_677268, EPI_ISL_677276, EPI_ISL_677293 | Colorado Department of Public Health and Environment | Colorado Department of Puplic Health and Environment | Laura Bankers, Molly Hetherington-Rauth, Shannon Ely, Shannon R. Matzinger, Sarah Elizabeth Totten, Emily A. Travanty |
| EPI_ISL_677711 | General Hospital - Ohrid | Research Center for Genetic Engineering and Biotechnology "Georgi D. Efremov" , Macedonian Academy of Sciences and Arts | RCGEB - MASA |
| EPI_ISL_677712, EPI_ISL_677713, EPI_ISL_677714 | General Hospital - Prilep | Research Center for Genetic Engineering and Biotechnology "Georgi D. Efremov" , Macedonian Academy of Sciences and Arts | RCGEB - MASA |
| EPI_ISL_677889, EPI_ISL_677891, EPI_ISL_677892, EPI_ISL_677894, EPI_ISL_677903, EPI_ISL_677905, EPI_ISL_677907 | Innovative Genomics Institute, UC Berkeley | Innovative Genomics Institute, UC Berkeley | Stacia Wyman, Haridha Shivram, Phil Frankino, Liana Lareau, Shana McDevitt, Justin Choi |
| EPI_ISL_677908, EPI_ISL_677909 | Pathogen Genomics Lab King Abdullah University of Science and Technology(KAUST) | Pathogen Genomics Lab King Abdullah University of Science and Technology(KAUST) | Sara Mfarrej, Raeece Naeem, Raushan Nugmanova, Olga Douvropoulou, Luke Esau, Amanda Ooi, Sharif Hala, Afrah Alsomali, Asim Khogeer, Fadwa Alofi,Jumana Taha, Abdulaziz Alahmadi, Kahled Algithami, Anwar Has[illegible] |
| EPI_ISL_677910 | Pathogen Genomics Lab King Abdullah University of Science and Technology(KAUST) | Pathogen Genomics Lab King Abdullah University of Science and Technology(KAUST) | Sara Mfarrej, Sharif Hala, Olga Douvropoulou, Raushan Nugmanova, Raeece Naeem, Afrah Alsomali, Asim Khogeer, Fadwa Alofi,Jumana Taha, Abdulaziz Alahmadi, Kahled Algithami, Anwar Hashem, Naif Al[illegible] |
| EPI_ISL_677911 | Pathogen Genomics Lab King Abdullah University of Science and Technology(KAUST) | Pathogen Genomics Lab King Abdullah University of Science and Technology(KAUST) | Muhammad Shuaib, Raeece Naeem, Sara Mfarrej, Raushan Nugmanova, Olga Douvropoulou, Luke Esau, Amanda Ooi, Sharif Hala, Afrah Alsomali, Asim Khogeer, Fadwa Alofi,Jumana Taha, Abdulaziz Alahmadi, Kahled Algithami, [illegible] |
| EPI_ISL_677912 | Pathogen Genomics Lab King Abdullah University of Science and Technology(KAUST) | Pathogen Genomics Lab King Abdullah University of Science and Technology(KAUST) | Sara Mfarrej, Raeece Naeem, Amanda Ooi, Luke Esau, Sharif Hala, Afrah Alsomali, Asim Khogeer, Fadwa Alofi,Jumana Taha, Abdulaziz Alahmadi, Kahled Algithami, Anwar Hashem, Naif Almontashi[illegible] |
| EPI_ISL_677913 | Pathogen Genomics Lab King Abdullah University of Science and Technology(KAUST) | Pathogen Genomics Lab King Abdullah University of Science and Technology(KAUST) | Sara Mfarrej, Raeece Naeem, Raushan Nugmanova, Olga Douvropoulou, Luke Esau, Amanda Ooi, Sharif Hala, Afrah Alsomali, Asim Khogeer, Fadwa Alofi,Jumana Taha, Abdulaziz Alahmadi, Kahled Algithami, Anwar Has[illegible] |
| EPI_ISL_677914 | Pathogen Genomics Lab King Abdullah University of Science and Technology(KAUST) | Pathogen Genomics Lab King Abdullah University of Science and Technology(KAUST) | Muhammad Shuaib, Raeece Naeem, Sara Mfarrej, Raushan Nugmanova, Olga Douvropoulou, Luke Esau, Amanda Ooi, Sharif Hala, Afrah Alsomali, Asim Khogeer, Fadwa Alofi,Jumana Taha, Abdulaziz Alahmadi, Kahled Algithami, [illegible] |
| EPI_ISL_677915 | Pathogen Genomics Lab King Abdullah University of Science and Technology(KAUST) | Pathogen Genomics Lab King Abdullah University of Science and Technology(KAUST) | Sara Mfarrej, Raeece Naeem, Amanda Ooi, Luke Esau, Sharif Hala, Afrah Alsomali, Asim Khogeer, Fadwa Alofi,Jumana Taha, Abdulaziz Alahmadi, Kahled Algithami, Anwar Hashem, Naif Almontashi[illegible] |
| EPI_ISL_677916 | Pathogen Genomics Lab King Abdullah University of Science and Technology(KAUST) | Pathogen Genomics Lab King Abdullah University of Science and Technology(KAUST) | Sara Mfarrej, Sharif Hala, Olga Douvropoulou, Raushan Nugmanova, Raeece Naeem, Afrah Alsomali, Asim Khogeer, Fadwa Alofi,Jumana Taha, Abdulaziz Alahmadi, Kahled Algithami, Anwar Hashem, Naif Al[illegible] |
| EPI_ISL_677917 | Pathogen Genomics Lab King Abdullah University of Science and Technology(KAUST) | Pathogen Genomics Lab King Abdullah University of Science and Technology(KAUST) | Sara Mfarrej, Raeece Naeem, Amanda Ooi, Luke Esau, Sharif Hala, Afrah Alsomali, Asim Khogeer, Fadwa Alofi,Jumana Taha, Abdulaziz Alahmadi, Kahled Algithami, Anwar Hashem, Naif Almontashi[illegible] |
| EPI_ISL_677918, EPI_ISL_677919 | Pathogen Genomics Lab King Abdullah University of | Pathogen Genomics Lab King Abdullah University of | Sara Mfarrej, Sharif Hala, Olga Douvropoulou, Raushan Nugmanova, Raeece Naeem, Afrah Alsomali, Asim Khogeer, Fadwa Alofi,Jumana Taha, Abdulaziz Alahmadi, Kahled Algithami, Anwar Hashem, Naif Al[illegible] |

| | | | |
|---|---|---|---|
| | Science and Technology(KAUST) | Science and Technology(KAUST) | |
| EPI_ISL_677920 | Pathogen Genomics Lab King Abdullah University of Science and Technology(KAUST) | Pathogen Genomics Lab King Abdullah University of Science and Technology(KAUST) | Sara Mfarrej, Raeece Naeem, Raushan Nugmanova, Olga Douvropoulou, Luke Esau, Amanda Ooi, Sharif Hala, Afrah Alsomali, Asim Khogeer, Fadwa Alofi,Jumana Taha, Abdulaziz Alahmadi, Kahled Algithami, Anwar Has |
| EPI_ISL_677921 | Pathogen Genomics Lab King Abdullah University of Science and Technology(KAUST) | Pathogen Genomics Lab King Abdullah University of Science and Technology(KAUST) | Sara Mfarrej, Sharif Hala, Olga Douvropoulou, Raushan Nugmanova, Raeece Naeem, Afrah Alsomali, Asim Khogeer, Fadwa Alofi,Jumana Taha, Abdulaziz Alahmadi, Kahled Algithami, Anwar Hashem, Naif Al |
| EPI_ISL_677922 | Pathogen Genomics Lab King Abdullah University of Science and Technology(KAUST) | Pathogen Genomics Lab King Abdullah University of Science and Technology(KAUST) | Muhammad Shuaib, Raeece Naeem, Raushan Nugmanova, Olga Douvropoulou, Sharif Hala, Sara Mfarrej, Afrah Alsomali, Asim Khogeer, Fadwa Alofi,Jumana Taha, Abdulaziz Alahmadi, Kahled Algithami, Anwar Hashe |
| EPI_ISL_677923 | Pathogen Genomics Lab King Abdullah University of Science and Technology(KAUST) | Pathogen Genomics Lab King Abdullah University of Science and Technology(KAUST) | Muhammad Shuaib, Sharif Hala, Sara Mfarrej, Luke Esau, Amanda Ooi, Raeece Naeem, Afrah Alsomali, Asim Khogeer, Fadwa Alofi,Jumana Taha, Abdulaziz Alahmadi, Kahled Algithami, Anwar Hashem, Naif A |
| EPI_ISL_677924 | Pathogen Genomics Lab King Abdullah University of Science and Technology(KAUST) | Pathogen Genomics Lab King Abdullah University of Science and Technology(KAUST) | Muhammad Shuaib, Raeece Naeem, Sara Mfarrej, Raushan Nugmanova, Olga Douvropoulou, Luke Esau, Amanda Ooi, Sharif Hala, Afrah Alsomali, Asim Khogeer, Fadwa Alofi,Jumana Taha, Abdulaziz Alahmadi, Kahled Algithami, |
| EPI_ISL_677925, EPI_ISL_677926 | Pathogen Genomics Lab King Abdullah University of Science and Technology(KAUST) | Pathogen Genomics Lab King Abdullah University of Science and Technology(KAUST) | Muhammad Shuaib, Sharif Hala, Sara Mfarrej, Luke Esau, Amanda Ooi, Raeece Naeem, Afrah Alsomali, Asim Khogeer, Fadwa Alofi,Jumana Taha, Abdulaziz Alahmadi, Kahled Algithami, Anwar Hashem, Naif A |
| EPI_ISL_677927 | Pathogen Genomics Lab King Abdullah University of Science and Technology(KAUST) | Pathogen Genomics Lab King Abdullah University of Science and Technology(KAUST) | Muhammad Shuaib, Raeece Naeem, Raushan Nugmanova, Olga Douvropoulou, Sharif Hala, Sara Mfarrej, Afrah Alsomali, Asim Khogeer, Fadwa Alofi,Jumana Taha, Abdulaziz Alahmadi, Kahled Algithami, Anwar Hashe |
| EPI_ISL_677928 | Pathogen Genomics Lab King Abdullah University of Science and Technology(KAUST) | Pathogen Genomics Lab King Abdullah University of Science and Technology(KAUST) | Sara Mfarrej, Sharif Hala, Olga Douvropoulou, Raushan Nugmanova, Raeece Naeem, Afrah Alsomali, Asim Khogeer, Fadwa Alofi,Jumana Taha, Abdulaziz Alahmadi, Kahled Algithami, Anwar Hashem, Naif Al |
| EPI_ISL_677929 | Pathogen Genomics Lab King Abdullah University of Science and Technology(KAUST) | Pathogen Genomics Lab King Abdullah University of Science and Technology(KAUST) | Sara Mfarrej, Raeece Naeem, Amanda Ooi, Luke Esau, Sharif Hala, Afrah Alsomali, Asim Khogeer, Fadwa Alofi,Jumana Taha, Abdulaziz Alahmadi, Kahled Algithami, Anwar Hashem, Naif Almontashi |
| EPI_ISL_677930, EPI_ISL_677931 | Pathogen Genomics Lab King Abdullah University of Science and Technology(KAUST) | Pathogen Genomics Lab King Abdullah University of Science and Technology(KAUST) | Sara Mfarrej, Sharif Hala, Olga Douvropoulou, Raushan Nugmanova, Raeece Naeem, Afrah Alsomali, Asim Khogeer, Fadwa Alofi,Jumana Taha, Abdulaziz Alahmadi, Kahled Algithami, Anwar Hashem, Naif Al |
| EPI_ISL_677932, EPI_ISL_677933, EPI_ISL_677934 | Pathogen Genomics Lab King Abdullah University of Science and Technology(KAUST) | Pathogen Genomics Lab King Abdullah University of Science and Technology(KAUST) | Sara Mfarrej, Raeece Naeem, Amanda Ooi, Luke Esau, Sharif Hala, Afrah Alsomali, Asim Khogeer, Fadwa Alofi,Jumana Taha, Abdulaziz Alahmadi, Kahled Algithami, Anwar Hashem, Naif Almontashi |
| EPI_ISL_677935, EPI_ISL_677936 | Pathogen Genomics Lab King Abdullah University of Science and Technology(KAUST) | Pathogen Genomics Lab King Abdullah University of Science and Technology(KAUST) | Sara Mfarrej, Sharif Hala, Olga Douvropoulou, Raushan Nugmanova, Raeece Naeem, Afrah Alsomali, Asim Khogeer, Fadwa Alofi,Jumana Taha, Abdulaziz Alahmadi, Kahled Algithami, Anwar Hashem, Naif Al |
| EPI_ISL_677950 | Pathogen Genomics Lab King Abdullah University of Science and Technology(KAUST) | Pathogen Genomics Lab King Abdullah University of Science and Technology(KAUST) | Sara Mfarrej, Sharif Hala, Luke Esau, Amanda Ooi, Raeece Naeem, Fadwa Alofi, Afrah Alsomali, Asim Khogeer, Jumana Taha, Abdulaziz Alahmadi, Kahled Algithami, Anwar Hashem, Naif Almontashi |
| EPI_ISL_677952 | Pathogen Genomics Lab King Abdullah University of Science and Technology(KAUST) | Pathogen Genomics Lab King Abdullah University of Science and Technology(KAUST) | Sara Mfarrej, Raushan Nugmanova, Olga Douvropoulou, Raeece Naeem, Fadwa Alofi, Afrah Alsomali, Asim Khogeer, Jumana Taha, Abdulaziz Alahmadi, Kahled Algithami, Anwar Hashem, Naif Almontashiri, |
| EPI_ISL_677971 | Pathogen Genomics Lab King Abdullah University of Science and Technology(KAUST) | Pathogen Genomics Lab King Abdullah University of Science and Technology(KAUST) | Sara Mfarrej, Olga Douvropoulou, Raushan Nugmanova, Sharif Hala, Raeece Naeem, Amanda Ooi, Luke Esau, Fadwa Alofi, Afrah Alsomali, Asim Khogeer, Jumana Taha, Abdulaziz Alahmadi, Kahled Algithami, Anwar Has |
| EPI_ISL_677991 | Pathogen Genomics Lab King Abdullah University of Science and Technology(KAUST) | Pathogen Genomics Lab King Abdullah University of Science and Technology(KAUST) | Olga Douvropoulou, Sara Mfarrej, Raushan Nugmanova, Sharif Hala, Raeece Naeem, Amanda Ooi, Luke Esau, Fadwa Alofi, Afrah Alsomali, Asim Khogeer, Jumana Taha, Abdulaziz Alahmadi, Kahled Algithami, Anwar Has |
| EPI_ISL_677992, EPI_ISL_677993, EPI_ISL_677994, EPI_ISL_677995, EPI_ISL_677996, EPI_ISL_677997, EPI_ISL_677998, EPI_ISL_677999 | Pathogen Genomics Lab King Abdullah University of Science and Technology(KAUST) | Pathogen Genomics Lab King Abdullah University of Science and Technology(KAUST) | Raeece Naeem, Sara Mfarrej, Sharif Hala, Olga Douvropoulou, Raushan Nugmanova, Fadwa Alofi, Asim Khogeer, Afrah Alsomali, Jumana Taha, Abdulaziz Alahmadi, Kahled Algithami, Anwar Hashem, Naif Al |
| EPI_ISL_678000, EPI_ISL_678001 | Pathogen Genomics Lab King Abdullah University of Science and Technology(KAUST) | Pathogen Genomics Lab King Abdullah University of Science and Technology(KAUST) | Raeece Naeem, Sara Mfarrej, Amanda Ooi, Luke Esau, Sharif Hala, Fadwa Alofi, Afrah Alsomali, Asim Khogeer, Jumana Taha, Abdulaziz Alahmadi, Kahled Algithami, Anwar Hashem, Naif Almontashi |

| | | | |
|---|---|---|---|
| EPI_ISL_678004, EPI_ISL_678005, EPI_ISL_678006, EPI_ISL_678007, EPI_ISL_678008, EPI_ISL_678009 | Pathogen Genomics Lab King Abdullah University of Science and Technology(KAUST) | Pathogen Genomics Lab King Abdullah University of Science and Technology(KAUST) | Sara Mfarrej, Raushan Nugmanova, Olga Douvropoulou, Raeece Naeem, Fadwa Alofi, Afrah Alsomali, Asim Khogeer, Jumana Taha, Abdulaziz Alahmadi, Kahled Algithami, Anwar Hashem, Naif Almontashiri, [illegible] |
| EPI_ISL_678049 | Pathogen Genomics Lab King Abdullah University of Science and Technology(KAUST) | Pathogen Genomics Lab King Abdullah University of Science and Technology(KAUST) | Muhammad Shuaib, Sara Mfarrej, Raushan Nugmanova, Olga Douvropoulou, Raeece Naeem, Sharif Hala, Luke Esau, Amanda Ooi, Asim Khogeer, Fadwa Alofi, Afrah Alsomali,Jumana Taha, Abdulaziz Alahmadi, Kahled Algithami, [illegible] |
| EPI_ISL_678064, EPI_ISL_678070, EPI_ISL_678083, EPI_ISL_678084, EPI_ISL_678085, EPI_ISL_678086, EPI_ISL_678087, EPI_ISL_678088 | Pathogen Genomics Lab King Abdullah University of Science and Technology(KAUST) | Pathogen Genomics Lab King Abdullah University of Science and Technology(KAUST) | Muhammad Shuaib, Raeece Naeem, Sharif Hala, Sara Mfarrej, Olga Douvropoulou, Raushan Nugmanova, Asim Khogeer, Fadwa Alofi, Afrah Alsomali, Jumana Taha, Abdulaziz Alahmadi, Kahled Algithami, Anwar Hashe[illegible] |
| EPI_ISL_678089 | Pathogen Genomics Lab King Abdullah University of Science and Technology(KAUST) | Pathogen Genomics Lab King Abdullah University of Science and Technology(KAUST) | Muhammad Shuaib, Amanda Ooi, Luke Esau, Sharif Hala, Raeece Naeem, Sara Mfarrej, Asim Khogeer, Fadwa Alofi, Afrah Alsomali,Jumana Taha, Abdulaziz Alahmadi, Kahled Algithami, Anwar Hashem, Naif [illegible] |
| EPI_ISL_678090, EPI_ISL_678091 | Pathogen Genomics Lab King Abdullah University of Science and Technology(KAUST) | Pathogen Genomics Lab King Abdullah University of Science and Technology(KAUST) | Muhammad Shuaib, Raeece Naeem, Sharif Hala, Sara Mfarrej, Olga Douvropoulou, Raushan Nugmanova, Asim Khogeer, Fadwa Alofi, Afrah Alsomali, Jumana Taha, Abdulaziz Alahmadi, Kahled Algithami, Anwar Hashe[illegible] |
| EPI_ISL_678092 | Pathogen Genomics Lab King Abdullah University of Science and Technology(KAUST) | Pathogen Genomics Lab King Abdullah University of Science and Technology(KAUST) | Muhammad Shuaib, Raeece Naeem, Raushan Nugmanova, Olga Douvropoulou, Sara Mfarrej, Sharif Hala, Asim Khogeer, Fadwa Alofi, Afrah Alsomali, Jumana Taha, Abdulaziz Alahmadi, Kahled Algithami, Anwar Hashe[illegible] |
| EPI_ISL_678093 | Pathogen Genomics Lab King Abdullah University of Science and Technology(KAUST) | Pathogen Genomics Lab King Abdullah University of Science and Technology(KAUST) | Muhammad Shuaib, Sara Mfarrej, Raeece Naeem,Raushan Nugmanova, Olga Douvropoulou, Sharif Hala, Asim Khogeer, Fadwa Alofi, Afrah Alsomali, Jumana Taha, Abdulaziz Alahmadi, Kahled Algithami, Anwar Hashe[illegible] |
| EPI_ISL_678149, EPI_ISL_678150 | Pathogen Genomics Lab King Abdullah University of Science and Technology(KAUST) | Pathogen Genomics Lab King Abdullah University of Science and Technology(KAUST) | Sara Mfarrej, Luke Esau, Amanda Ooi, Sharif Hala, Raeece Naeem, Awad Al-Omari, Samer Salih, Abbas Al Mutair, Arnab Pain |
| EPI_ISL_678151, EPI_ISL_678155, EPI_ISL_678156 | Pathogen Genomics Lab King Abdullah University of Science and Technology(KAUST) | Pathogen Genomics Lab King Abdullah University of Science and Technology(KAUST) | Sara Mfarrej, Raeece Naeem, Luke Esau, Amanda Ooi, Sharif Hala, Awad Al-Omari, Samer Salih, Abbas Al Mutair, Arnab Pain |
| EPI_ISL_678157, EPI_ISL_678158, EPI_ISL_678159 | Pathogen Genomics Lab King Abdullah University of Science and Technology(KAUST) | Pathogen Genomics Lab King Abdullah University of Science and Technology(KAUST) | Sara Mfarrej, Luke Esau, Amanda Ooi, Sharif Hala, Raeece Naeem, Awad Al-Omari, Samer Salih, Abbas Al Mutair, Arnab Pain |
| EPI_ISL_678172, EPI_ISL_678180, EPI_ISL_678226 | Pathogen Genomics Lab King Abdullah University of Science and Technology(KAUST) | Pathogen Genomics Lab King Abdullah University of Science and Technology(KAUST) | Olga Douvropoulou, Sara Mfarrej, Raushan Nugmanova, Sharif Hala, Raeece Naeem, Asim Khogeer, Fadwa Alofi, Afrah Alsomali, Jumana Taha, Abdulaziz Alahmadi, Kahled Algithami, Anwar Hashem, Naif Al[illegible] |
| EPI_ISL_678241 | Pathogen Genomics Lab King Abdullah University of Science and Technology(KAUST) | Pathogen Genomics Lab King Abdullah University of Science and Technology(KAUST) | Muhammad Shuaib, Raeece Naeem, Raushan Nugmanova, Olga Douvropoulou, Sharif Hala, Sara Mfarrej, Afrah Alsomali, Asim Khogeer, Fadwa Alofi,Jumana Taha, Abdulaziz Alahmadi, Kahled Algithami, Anwar Hashe[illegible] |
| EPI_ISL_678242 | Pathogen Genomics Lab King Abdullah University of Science and Technology(KAUST) | Pathogen Genomics Lab King Abdullah University of Science and Technology(KAUST) | Muhammad Shuaib, Sharif Hala, Sara Mfarrej, Luke Esau, Amanda Ooi, Raeece Naeem, Afrah Alsomali, Asim Khogeer, Fadwa Alofi,Jumana Taha, Abdulaziz Alahmadi, Kahled Algithami, Anwar Hashem, Naif [illegible] |
| EPI_ISL_678243 | Pathogen Genomics Lab King Abdullah University of Science and Technology(KAUST) | Pathogen Genomics Lab King Abdullah University of Science and Technology(KAUST) | Sara Mfarrej, Raeece Naeem, Raushan Nugmanova, Olga Douvropoulou, Luke Esau, Amanda Ooi, Sharif Hala, Afrah Alsomali, Asim Khogeer, Fadwa Alofi,Jumana Taha, Abdulaziz Alahmadi, Kahled Algithami, Anwar Has[illegible] |
| EPI_ISL_678245 | Pathogen Genomics Lab King Abdullah University of Science and Technology(KAUST) | Pathogen Genomics Lab King Abdullah University of Science and Technology(KAUST) | Sara Mfarrej, Luke Esau, Amanda Ooi, Sharif Hala, Raeece Naeem, Awad Al-Omari, Samer Salih, Abbas Al Mutair, Arnab Pain |
| EPI_ISL_678312, EPI_ISL_678319 | Area of Virology, Serology and Virology Division (SAViD), New South Wales Health Pathology Randwick | Virology Research Laboratory; Area of Virology, Serology and Virology Division (SAViD), New South Wales Health Pathology Randwick | Foster, C.; Au, J.; Ruiz Silva, M.; Deveson, I.; Bull, R.; Van Hal, S.; Rawlinson, W. |
| EPI_ISL_681322 | Environmental and Global Health, University of Florida | Environmental and Global Health, University of Florida | Loeb,J.C., Stephenson,C.J., Merck,L., Morris,J.G. and Lednicky,J.A. |
| EPI_ISL_681689, EPI_ISL_681690, EPI_ISL_681691 | Molecular Medicine Laboratory, University of Magallanes | Centro Asistencial Docente y de Investigacion, Universidad de Magallanes | Jorge González, Jacqueline Aldridge, Diego Alvarez, Marco Montes de Oca, Hermy Alvarez, Roberto Uribe-Paredes, Marcelo Navarrete |
| EPI_ISL_681832, | Molecular diagnostic unit for | Project group Epidemology | Chantal Akoua-Koffi, Diané Bamourou, Etilé Anoh, Essia Belarbi, Safiatou Karidioula, Grit Schubert, Adjaratou Traoré, Soundélé Maïté, Monemo Pacome, Coulibaly Mbegnan, Bamba Fatoumata Touré, Kra Ouf[illegible] |

| | | | |
|---|---|---|---|
| EPI_ISL_681833, EPI_ISL_681835 | viral haemorrhagic fevers and emerging viruses, Bouaké CHU Laboratory | of Highly Pathogenic Microorganisms, Robert Koch-Institute | |
| EPI_ISL_681843, EPI_ISL_681845, EPI_ISL_681849, EPI_ISL_681853, EPI_ISL_681854, EPI_ISL_681855, EPI_ISL_681856, EPI_ISL_681857, EPI_ISL_681860, EPI_ISL_681864, EPI_ISL_681865, EPI_ISL_681867, EPI_ISL_681868, EPI_ISL_681870, EPI_ISL_681873, EPI_ISL_681881, EPI_ISL_681882, EPI_ISL_681883 | | | |
| see above | Texas Department of State Health Services | Texas Department of State Health Services | Rashmi Tuladhar, Bonnie Oh, Jenny Zhang, Maliha Rahman, Anita Pokharel, Myong Koag, Chung Wang, Rachel Lee, Grace Kubin, Mayela Pedrueza, James Daniel Bonser |
| EPI_ISL_682005, EPI_ISL_682007, EPI_ISL_682008, EPI_ISL_682009, EPI_ISL_682010, EPI_ISL_682011, EPI_ISL_682012, EPI_ISL_682013, EPI_ISL_682014, EPI_ISL_682015, EPI_ISL_682016, EPI_ISL_682017, EPI_ISL_682018 | | | |
| see above | UPMC Clinical Microbiology Laboratory | Microbial Genomic Epidemiology Laboratory, University of Pittsburgh | Mustapha M. Mustapha, Jane W. Marsh, Dan Snyder, Marissa P. Griffith, Stephanie L. Mitchell, Vatsala R. Srinivasa, Kady D. Waggle, Chinelo Ezeonwuku, Vaughn S. Cooper, Lee H. Harris[illegible] |
| EPI_ISL_682235 | AREA DE SALUD ALAJUELA NORTE - CLINICA DR. MARCIAL RODRIGUEZ | Inciensa, Instituto Costarricense de Investigación y Enseñanza en Nutrición y Salud | Francisco Duarte, Hebleen Porras, Claudio Soto-Garita, Estela Cordero, Adriana Godinez & Melany Calderon |
| EPI_ISL_683433, EPI_ISL_683434, EPI_ISL_683435, EPI_ISL_683436, EPI_ISL_683437, EPI_ISL_683438, EPI_ISL_683439, EPI_ISL_683440, EPI_ISL_683441, EPI_ISL_683442, EPI_ISL_683443, EPI_ISL_683444, EPI_ISL_683445 | | | |
| see above | Texas Department of State Health Services | Texas Department of State Health Services | Rashmi Tuladhar, Bonnie Oh, Jenny Zhang, Maliha Rahman, Anita Pokharel, Myong Koag, Chung Wang, Rachel Lee, Grace Kubin, Mayela Pedrueza, James Daniel Bonser |
| EPI_ISL_683597 | Hospital Clínico Universitario Lozano Blesa de Zaragoza (España) | SeqCOVID-SPAIN consortium/IBV(CSIC) | Rafael Benito, Sonia Algarate, Jessica Bueno and SeqCOVID-SPAIN consortium |
| EPI_ISL_691621, EPI_ISL_691665 | Servicio de Microbiología, Hospital Universitario Son Espases | SeqCOVID-SPAIN consortium/IBV(CSIC) | Carla López-Causapé, Jordi Reina, Antonio Oliver and SeqCOVID-SPAIN consortium |
| EPI_ISL_693207 | Cs II Doutor Antonio Vicoso Moreira de Rezende | Instituto Adolfo Lutz, Interdiciplinary Procedures Center, Strategic Laboratory | Claudio Tavares Sacchi, Claudia Regina Gonçalves, Erica Valessa Ramos Gomes, Karoline Rodrigues Campos |
| EPI_ISL_693231 | Pronto Socorro Municipal de Santa Branca | Instituto Adolfo Lutz, Interdiciplinary Procedures Center, Strategic Laboratory | Claudio Tavares Sacchi, Claudia Regina Gonçalves, Erica Valessa Ramos Gomes, Karoline Rodrigues Campos |
| EPI_ISL_693234 | Upa Vereador Jose Da Rocha Goncalves | Instituto Adolfo Lutz, Interdiciplinary Procedures Center, Strategic Laboratory | Claudio Tavares Sacchi, Claudia Regina Gonçalves, Erica Valessa Ramos Gomes, Karoline Rodrigues Campos |
| EPI_ISL_693235 | Casmi Centro Atendimento Saude da Mulher e Infancia | Instituto Adolfo Lutz, Interdiciplinary Procedures Center, Strategic Laboratory | Claudio Tavares Sacchi, Claudia Regina Gonçalves, Erica Valessa Ramos Gomes, Karoline Rodrigues Campos |
| EPI_ISL_693238, EPI_ISL_693239 | Secao Centro de Diagnostico Secedi | Instituto Adolfo Lutz, Interdiciplinary Procedures Center, Strategic Laboratory | Claudio Tavares Sacchi, Claudia Regina Gonçalves, Erica Valessa Ramos Gomes, Karoline Rodrigues Campos |
| EPI_ISL_693240 | Centro de Vigilância a Saude de Diadema | Instituto Adolfo Lutz, Interdiciplinary Procedures Center, Strategic Laboratory | Claudio Tavares Sacchi, Claudia Regina Gonçalves, Erica Valessa Ramos Gomes, Karoline Rodrigues Campos |
| EPI_ISL_693241 | Hospital e Maternidade Sao Lucas | Instituto Adolfo Lutz, Interdiciplinary Procedures Center, Strategic Laboratory | Claudio Tavares Sacchi, Claudia Regina Gonçalves, Erica Valessa Ramos Gomes, Karoline Rodrigues Campos |
| EPI_ISL_693242 | Centro de Vigilância a Saude de Diadema | Instituto Adolfo Lutz, Interdiciplinary Procedures Center, Strategic Laboratory | Claudio Tavares Sacchi, Claudia Regina Gonçalves, Erica Valessa Ramos Gomes, Karoline Rodrigues Campos |
| EPI_ISL_693243 | Laboratório Municipal de Piracicaba | Instituto Adolfo Lutz, Interdiciplinary Procedures Center, Strategic Laboratory | Claudio Tavares Sacchi, Claudia Regina Gonçalves, Erica Valessa Ramos Gomes, Karoline Rodrigues Campos |
| EPI_ISL_693246 | Laboratorio Municipal de Rio Grande da Serra | Instituto Adolfo Lutz, Interdiciplinary Procedures Center, Strategic Laboratory | Claudio Tavares Sacchi, Claudia Regina Gonçalves, Erica Valessa Ramos Gomes, Karoline Rodrigues Campos |
| EPI_ISL_693247 | Secao Centro de Diagnostico Secedi | Instituto Adolfo Lutz, Interdiciplinary Procedures Center, Strategic Laboratory | Claudio Tavares Sacchi, Claudia Regina Gonçalves, Erica Valessa Ramos Gomes, Karoline Rodrigues Campos |
| EPI_ISL_693529, EPI_ISL_693540, EPI_ISL_693541, EPI_ISL_693564, EPI_ISL_693565, EPI_ISL_693566, EPI_ISL_693567, EPI_ISL_693568, EPI_ISL_693569, EPI_ISL_693570, EPI_ISL_693571, EPI_ISL_693572, EPI_ISL_693573, EPI_ISL_693574, EPI_ISL_693575, EPI_ISL_693576, EPI_ISL_693577, EPI_ISL_693578, EPI_I[illegible] EPI_ISL_693583, EPI_ISL_693584, EPI_ISL_693585, EPI_ISL_693586, EPI_ISL_693587, EPI_ISL_693588 | | | |
| see above | Instituto Nacional de Saude (INSA) | Instituto Nacional de Saude (INSA) | Borges et al |
| EPI_ISL_694853, EPI_ISL_694854, EPI_ISL_694855, EPI_ISL_694856, EPI_ISL_694857, EPI_ISL_694858, EPI_ISL_694859, EPI_ISL_694860, EPI_ISL_694861, EPI_ISL_694862, EPI_ISL_694863, EPI_ISL_694864, EPI_ISL_694865, EPI_ISL_694866, EPI_ISL_694867, EPI_ISL_694868, EPI_ISL_694869, EPI_ISL_694870, EPI_I[illegible] EPI_ISL_694877, EPI_ISL_694878, EPI_ISL_694879, EPI_ISL_694880, EPI_ISL_694881, EPI_ISL_694882, EPI_ISL_694883, EPI_ISL_694884, EPI_ISL_694951, EPI_ISL_694952, EPI_ISL_694953, EPI_ISL_694954, EPI_ISL_694955, EPI_ISL_694956, EPI_ISL_694957, EPI_ISL_694958, EPI_ISL_694959, EPI_ISL_694960, EPI_I[illegible] EPI_ISL_694965, EPI_ISL_694966, EPI_ISL_694967, EPI_ISL_694968, EPI_ISL_694969, EPI_ISL_694970, EPI_ISL_694971, EPI_ISL_694972, EPI_ISL_694973, EPI_ISL_694974, EPI_ISL_694975, EPI_ISL_694976, EPI_ISL_694977, EPI_ISL_694978, EPI_ISL_694979, EPI_ISL_694980, EPI_ISL_694981, EPI_ISL_694982, EPI_I[illegible] EPI_ISL_694987, EPI_ISL_694988, EPI_ISL_694989, EPI_ISL_694990, EPI_ISL_694991, EPI_ISL_694992, EPI_ISL_694993, EPI_ISL_694994, EPI_ISL_694995, EPI_ISL_694996, EPI_ISL_694997, EPI_ISL_694998, EPI_ISL_694999, EPI_ISL_695000, EPI_ISL_695001, EPI_ISL_695002, EPI_ISL_695003, EPI_ISL_695004, EPI_I[illegible] EPI_ISL_695009, EPI_ISL_695010, EPI_ISL_695011, EPI_ISL_695012, EPI_ISL_695013, EPI_ISL_695014, EPI_ISL_695015, EPI_ISL_695016, EPI_ISL_695017, EPI_ISL_695018, EPI_ISL_695019, EPI_ISL_695020, EPI_ISL_695021, EPI_ISL_695022, EPI_ISL_695023, EPI_ISL_695024, EPI_ISL_695025, EPI_ISL_695026, EPI_I[illegible] EPI_ISL_695031, EPI_ISL_695032, EPI_ISL_695033, EPI_ISL_695034, EPI_ISL_695035, EPI_ISL_695036, EPI_ISL_695037, EPI_ISL_695038, EPI_ISL_695039, EPI_ISL_695040, EPI_ISL_695041, EPI_ISL_695042, EPI_ISL_695043, EPI_ISL_695044, EPI_ISL_695045, EPI_ISL_695046, EPI_ISL_695047 | | | |
| see above | TGen North | TGen North | Jolene Bowers, Megan Folkerts, Chris French, Hayley Yaglom, Ashlyn Pfeiffer, Darrin Lemmer, Dave Engelthaler, The Arizona COVID Genomics Union (ACGU) |
| EPI_ISL_695166, EPI_ISL_695174, EPI_ISL_695175, EPI_ISL_695185, EPI_ISL_695186, EPI_ISL_695187, EPI_ISL_695188, EPI_ISL_695189, EPI_ISL_695190, EPI_ISL_695191, EPI_ISL_695192, EPI_ISL_695193, EPI_ISL_695194, EPI_ISL_695195, EPI_ISL_695196, EPI_ISL_695197, EPI_ISL_695198, EPI_ISL_695199, EPI_I[illegible] EPI_ISL_695204, EPI_ISL_695205, EPI_ISL_695206, EPI_ISL_695207, EPI_ISL_695208, EPI_ISL_695209, EPI_ISL_695210, EPI_ISL_695211, EPI_ISL_695212, EPI_ISL_695213, EPI_ISL_695214, EPI_ISL_695215, EPI_ISL_695216, EPI_ISL_695217, EPI_ISL_695218, EPI_ISL_695219, EPI_ISL_695220, EPI_ISL_695221, EPI_I[illegible] EPI_ISL_695226, EPI_ISL_695227, EPI_ISL_695228, EPI_ISL_695316, EPI_ISL_695317, EPI_ISL_695318, EPI_ISL_695319, EPI_ISL_695320, EPI_ISL_695321, EPI_ISL_695322, EPI_ISL_695323, EPI_ISL_695324, EPI_ISL_695325, EPI_ISL_695326, EPI_ISL_695327, EPI_ISL_695328, EPI_ISL_695329, EPI_ISL_695330, EPI_I[illegible] EPI_ISL_695335, EPI_ISL_695336, EPI_ISL_695337, EPI_ISL_695338, EPI_ISL_695339, EPI_ISL_695340, EPI_ISL_695341, EPI_ISL_695342, EPI_ISL_695343, EPI_ISL_695344, EPI_ISL_695345, EPI_ISL_695346, EPI_ISL_695347, EPI_ISL_695348, EPI_ISL_695349, EPI_ISL_695350, EPI_ISL_695351, EPI_ISL_695352, EPI_I[illegible] EPI_ISL_695359, EPI_ISL_695360, EPI_ISL_695361, EPI_ISL_695362, EPI_ISL_695363 | | | |
| see above | AZ SPHL, Arizona Department of Health | TGen North | Jolene Bowers, Megan Folkerts, Chris French, Hayley Yaglom, Ashlyn Pfeiffer, Darrin Lemmer, Dave Engelthaler, The Arizona COVID Genomics Union (ACGU) |

| | | | |
|---|---|---|---|
| | Services | | |
| EPI_ISL_698191, EPI_ISL_699027, EPI_ISL_699028, EPI_ISL_699029, EPI_ISL_699030, EPI_ISL_699031, EPI_ISL_699032, EPI_ISL_699033, EPI_ISL_699034, EPI_ISL_699035, EPI_ISL_699036, EPI_ISL_699037, EPI_ISL_699038, EPI_ISL_699039, EPI_ISL_699040, EPI_ISL_699042, EPI_ISL_699043, EPI_ISL_699044, EPI_I EPI_ISL_699049, EPI_ISL_699050, EPI_ISL_699051, EPI_ISL_699052, EPI_ISL_699053, EPI_ISL_699054, EPI_ISL_699055, EPI_ISL_699056, EPI_ISL_699057, EPI_ISL_699058, EPI_ISL_699059, EPI_ISL_699060, EPI_ISL_699061, EPI_ISL_699062, EPI_ISL_699063, EPI_ISL_699064, EPI_ISL_699065, EPI_ISL_699066, EPI_I EPI_ISL_699071, EPI_ISL_699072, EPI_ISL_699073, EPI_ISL_699074, EPI_ISL_699075, EPI_ISL_699076, EPI_ISL_699077, EPI_ISL_699078, EPI_ISL_699079, EPI_ISL_699080, EPI_ISL_699081, EPI_ISL_699082, EPI_ISL_699083, EPI_ISL_699084, EPI_ISL_699085, EPI_ISL_699086, EPI_ISL_699087, EPI_ISL_699088, EPI_I EPI_ISL_699094, EPI_ISL_699095, EPI_ISL_699096, EPI_ISL_699097, EPI_ISL_699098, EPI_ISL_699099, EPI_ISL_699100, EPI_ISL_699101, EPI_ISL_699102, EPI_ISL_699103, EPI_ISL_699104, EPI_ISL_699105, EPI_ISL_699106, EPI_ISL_699107, EPI_ISL_699108, EPI_ISL_699109, EPI_ISL_699110, EPI_ISL_699111, EPI_I | | | |
| see above | Group 42 (G42) Healthcare, Abu Dhabi, United Arab Emirates; Department of Health, The United Arab Emirates | G42 Healthcare | Rong Liu, Pei Wu, Sally Mahmoud, Ke Liang, Pauline Ogrodzki, Pengjuan Liu, Stephen S. Francis, Tao Ma, Hanif Khalak, Fang Chen, Denghui Liu, Junhua Li, Weibin Liu, Wenjun He, Xinyu Huang, Zhaorong Yuan, Long Lin, Nan Qiao, Xin Meng, Budoor Koshy, Huanming Yang, Xun Xu, Jian Wang, Peng Xiao, Nawal Ahmed Mohamed Al Kaabi, Mohammed Saifuddin Fasihuddin, Siyang Liu, Walid Abbas Zaher |
| EPI_ISL_699506 | Diagnostic Virology Laboratory, USDA National Veterinary Services Laboratories | Diagnostic Virology Laboratory, USDA National Veterinary Services Laboratories | Hamer,S.A., Pauvolid-Correa,A., Zecca,I.B., Davila,E., Auckland,L.D., Roundy,C.M., Tang,W., Torchetti,M., Killian,M.L., Jenkins-Moore,M., Akpalu,Y., Ghai,R.R., Spengler,J., Barton Behravesh,C., Fischer,R.S., Hamer |
| EPI_ISL_699852, EPI_ISL_699853, EPI_ISL_699854, EPI_ISL_699855, EPI_ISL_699856, EPI_ISL_699857, EPI_ISL_699858, EPI_ISL_699859, EPI_ISL_699860, EPI_ISL_699861, EPI_ISL_699862, EPI_ISL_699863, EPI_ISL_699864, EPI_ISL_699865, EPI_ISL_699866, EPI_ISL_699867, EPI_ISL_699868, EPI_ISL_699869, EPI_I EPI_ISL_699874, EPI_ISL_699875, EPI_ISL_699876, EPI_ISL_699877 | | | |
| see above | Hematopathology Laboratory, ACTREC, TMC | Hematopathology Laboratory, ACTREC, TMC | Hematopathology Laboratory, ACTREC |
| EPI_ISL_707906, EPI_ISL_707930 | Los Angeles County Public Health Laboratory | Los Angeles County Public Health Laboratory | P. Hemarajata et al. |
| EPI_ISL_708529 | Secretária Municipal de Saude de Fernandópolis | Instituto Adolfo Lutz, Interdiciplinary Procedures Center, Strategic Laboratory | Claudio Tavares Sacchi, Claudia Regina Gonçalves, Erica Valessa Ramos Gomes, Carlos Henrique Camargo, Karoline Rodrigues Campos, Fernanda Modesto Tolentino Binhardi, Maricelia Navarro Pinheiro Flores, Marcia Maria Costa |
| EPI_ISL_710043, EPI_ISL_710044, EPI_ISL_710045, EPI_ISL_710046, EPI_ISL_710047, EPI_ISL_710048, EPI_ISL_710049, EPI_ISL_710050, EPI_ISL_710051, EPI_ISL_710052, EPI_ISL_710053, EPI_ISL_710054, EPI_ISL_710055, EPI_ISL_710056, EPI_ISL_710057, EPI_ISL_710058, EPI_ISL_710059, EPI_ISL_710060, EPI_I EPI_ISL_710065, EPI_ISL_710066, EPI_ISL_710067, EPI_ISL_710068, EPI_ISL_710069, EPI_ISL_710070, EPI_ISL_710071, EPI_ISL_710072, EPI_ISL_710073, EPI_ISL_710074, EPI_ISL_710075, EPI_ISL_710076, EPI_ISL_710077, EPI_ISL_710078, EPI_ISL_710079, EPI_ISL_710080, EPI_ISL_710081, EPI_ISL_710082, EPI_I EPI_ISL_710087, EPI_ISL_710088, EPI_ISL_710089 | | | |
| see above | Texas Department of State Health Services | Texas Department of State Health Services | Anita Pokharel, Bonnie Oh, James Daniel Bonser, Rashmi Tuladhar, Mayela Pedrueza, Jenny Zhang, Maliha Rahman, Myong Koag, Chung Wang, Rachel Lee, Grace Kubin |
| EPI_ISL_710136, EPI_ISL_710213, EPI_ISL_710317 | Colorado Department of Public Health and Environment | Colorado Department of Puplic Health and Environment | Laura Bankers, Molly C. Hetherington-Rauth, Shannon Ely, Shannon R. Matzinger, Sarah Elizabeth Totten, Emily A. Travanty |
| EPI_ISL_710380, EPI_ISL_710401, EPI_ISL_710402, EPI_ISL_710403, EPI_ISL_710404, EPI_ISL_710407 | Texas Department of State Health Services | Texas Department of State Health Services | Rashmi Tuladhar, Bonnie Oh, Jenny Zhang, Maliha Rahman, Anita Pokharel, Myong Koag, Chung Wang, Rachel Lee, Grace Kubin, Mayela Pedrueza, James Daniel Bonser |
| EPI_ISL_710608 | Gavle klinisk mikrobiologi | The Public Health Agency of Sweden | Department of Microbiology, The Public Health Agency of Sweden |
| EPI_ISL_717787, EPI_ISL_717797 | LACEN RJ - Noel Nutels | Bioinformatics Laboratory / LNCC | Carolina M Voloch, Ronaldo da Silva F Jr, Luiz G P de Almeida, Cynthia C Cardoso, Otavio Bustrolini, Alexandra L Gerber, Ana Paula de C Guimarães, Diana Mariani, Andréa Cony Cavalcanti, Claudia dos Santos Rodrigues, Terezinha M P P |
| EPI_ISL_721642, EPI_ISL_721643, EPI_ISL_721644, EPI_ISL_721645 | Armed Forces Medical College | National Centre For Cell Science | Dhiraj Paul, Kunal Jani, Radha Chauhan, Janesh Kumar, Vasudevan Seshadri, Girdhari Lal, Rajesh Karyakarte, Suvarna Joshi, Murlidhar Tambe, Sourav Sen, Santosh Karade, Kavita Bala Anand, Shelinder Pal Singh Shergill, Rajiv Mohan Gu |
| EPI_ISL_721660, EPI_ISL_721661, EPI_ISL_721662, EPI_ISL_721663, EPI_ISL_722181, EPI_ISL_722183, EPI_ISL_722185, EPI_ISL_722186 | National Centre For Cell Science | National Centre For Cell Science | Dhiraj Paul, Kunal Jani, Radha Chauhan, Janesh Kumar, Vasudevan Seshadri, Girdhari Lal, Rajesh Karyakarte, Suvarna Joshi, Murlidhar Tambe, Sourav Sen, Santosh Karade, Kavita Bala Anand, Shelinder Pal Singh Shergill, Rajiv Mohan Gu |
| EPI_ISL_722195, EPI_ISL_722196 | Armed Forces Medical College | National Centre For Cell Science | Dhiraj Paul, Kunal Jani, Radha Chauhan, Janesh Kumar, Vasudevan Seshadri, Girdhari Lal, Rajesh Karyakarte, Suvarna Joshi, Murlidhar Tambe, Sourav Sen, Santosh Karade, Kavita Bala Anand, Shelinder Pal Singh Shergill, Rajiv Mohan Gu |
| EPI_ISL_722896 | Dipartimento di Scienze Biomediche e Oncologia Umana - Azienda Ospedaliero Universitaria Consorziale Policlinico | Istituto Zooprofilattico Sperimentale della Puglia e della Basilicata | Parisi A., Bianco A., Capozzi L., Del Sambro L., Chironna M., Loconsole D. |
| EPI_ISL_728257, EPI_ISL_728258, EPI_ISL_728263, EPI_ISL_728264, EPI_ISL_728265, EPI_ISL_728268 | B.J. Govt. Medical College | National Centre For Cell Science | Dhiraj Paul, Kunal Jani, Radha Chauhan, Janesh Kumar, Vasudevan Seshadri, Girdhari Lal, Rajesh Karyakarte, Suvarna Joshi, Murlidhar Tambe, Sourav Sen, Santosh Karade, Kavita Bala Anand, Shelinder Pal Singh Shergill, Rajiv Mohan Gu |
| EPI_ISL_729562 | A. Krumbholz, Labor Dr. Krause und Kollegen MVZ GmbH, Kiel | Charité Universitätsmedizin Berlin, Institut für Virologie | Victor M Corman, Barbara Mühlemann, Jörn Beheim-Schwarzbach, Talitha Veith, Julia Schneider, Terry Jones, Christian Drosten |
| EPI_ISL_729927, EPI_ISL_729951, EPI_ISL_729952, EPI_ISL_729953, EPI_ISL_729954, EPI_ISL_729966, EPI_ISL_729967, EPI_ISL_729968, EPI_ISL_729969, EPI_ISL_729987, EPI_ISL_729988, EPI_ISL_730018, EPI_ISL_730019, EPI_ISL_730020, EPI_ISL_730021, EPI_ISL_730022 | | | |
| see above | Nigeria Centre for Disease Control (NCDC) | African Centre of Excellence for Genomics of Infectious Diseases (ACEGID), Redeemer's University, Ede, Osun State, Nigeria | Oluniyi P.E. et al |
| EPI_ISL_730052 | Yale COVID-19 Biorepository | Grubaugh Lab - Yale School | Joseph Fauver, Tara Alpert, Anderson Brito, Annie Watkins, Anne Wyllie, Chantal Vogels, Mary Petrone, Chaney Kalinich, Isabel Ott, Arnau Casanovas, Catherine Muenker, Adam Moore, Alice Lu, Maria Tokuyama, Patrick Wong, Peiwen Lu, Saad Omer, |

| | | | |
|---|---|---|---|
| | | of Public Health | Charlese Dela Cruz, Albert Ko, Nathan Grubaugh |
| EPI_ISL_730088, EPI_ISL_730096, EPI_ISL_730103, EPI_ISL_730104, EPI_ISL_730118, EPI_ISL_730119, EPI_ISL_730321, EPI_ISL_730327, EPI_ISL_730331, EPI_ISL_730334 | San Diego County Public Health Laboratory | Andersen lab at Scripps Research | SEARCH Alliance San Diego with Tracy Basler, Jovan Shephard, Brett Austin |
| EPI_ISL_730576 | Gazi University Faculty of Medicine, Medical Virology Laboratory | Gazi University Faculty of Medicine, Medical Virology Laboratory | Erdem ahin, Gülendam Bozday, Hager Muftah, Selin Yiit, Shaknoza Sarzhanova, Özlem Güzel Tunçcan, Murat Dizbay, Il Fidan, Kayhan Çalar |
| EPI_ISL_732535, EPI_ISL_732536 | Bundeswehr Institute of Microbiology | Bundeswehr Institute of Microbiology | Elham Khatamzas, Markus Antwerpen, Mathias Walter, Alexandra Rehn, Sabine Zange, Enrico Georgi, Michael von Bergwelt-Baildon, Roman Wölfel |
| EPI_ISL_734777, EPI_ISL_734778, EPI_ISL_734779, EPI_ISL_734780, EPI_ISL_734781, EPI_ISL_734782, EPI_ISL_734783, EPI_ISL_734784, EPI_ISL_734785, EPI_ISL_734786, EPI_ISL_734787 | | | |
| see above | UZ Leuven, National Reference Laboratory for Coronaviruses, Laboratory Medicine, Leuven, Belgium | KU Leuven, Rega Institute, Clinical and Epidemiological Virology | Tony Wawina-Bokalanga, Joan Marti-Carerras, Bert Vanmechelen, Piet Maes |
| EPI_ISL_735350, EPI_ISL_735351, EPI_ISL_735352, EPI_ISL_735353, EPI_ISL_735354, EPI_ISL_735355, EPI_ISL_735356, EPI_ISL_735357, EPI_ISL_735358, EPI_ISL_735359, EPI_ISL_735360, EPI_ISL_735361, EPI_ISL_735362, EPI_ISL_735363, EPI_ISL_735364, EPI_ISL_735365 | | | |
| see above | Genomic Laboratory (GLAB) (Conjoint lab of Health Directorate of Istanbul and Istanbul Technical University) | Genomic Laboratory (GLAB), Istanbul Technical University | Ilker Karacan, Tugba Kizilboga Akgun, Nihat Bugra Agaoglu, Payam Zolfagharian, Mehtap Aydin, Gizem Alkurt, Jale Yildiz, Betsi Kose, Nisan Denizce Can, Ayse Serra Ozel, Nilsun Altunal, Arzu Irvem, Yasemin Kendir Demirkol, Ozlem Akg |
| EPI_ISL_735407 | Santa Casa de Marilia | Instituto Adolfo Lutz, Interdiciplinary Procedures Center, Strategic Laboratory | Claudio Tavares Sacchi, Claudia Regina Gonçalves, Erica Valessa Ramos Gomes, Karoline Rodrigues Campos |
| EPI_ISL_735408 | COVID 19 Centro de Combate ao Coronavirus CCC Jandira | Instituto Adolfo Lutz, Interdiciplinary Procedures Center, Strategic Laboratory | Claudio Tavares Sacchi, Claudia Regina Gonçalves, Erica Valessa Ramos Gomes, Karoline Rodrigues Campos |
| EPI_ISL_735414, EPI_ISL_735415 | Unidade de Pronto Atendimento de Agenor de Campos | Instituto Adolfo Lutz, Interdiciplinary Procedures Center, Strategic Laboratory | Claudio Tavares Sacchi, Claudia Regina Gonçalves, Erica Valessa Ramos Gomes, Karoline Rodrigues Campos |
| EPI_ISL_735416 | Centro de Saude II Dr Jose Paione Mococa | Instituto Adolfo Lutz, Interdiciplinary Procedures Center, Strategic Laboratory | Claudio Tavares Sacchi, Claudia Regina Gonçalves, Erica Valessa Ramos Gomes, Karoline Rodrigues Campos |
| EPI_ISL_735417 | Unidade de Pronto Atendimento de Agenor de Campos | Instituto Adolfo Lutz, Interdiciplinary Procedures Center, Strategic Laboratory | Claudio Tavares Sacchi, Claudia Regina Gonçalves, Erica Valessa Ramos Gomes, Karoline Rodrigues Campos |
| EPI_ISL_735418 | Hospital Regional do Vale do Paraiba | Instituto Adolfo Lutz, Interdiciplinary Procedures Center, Strategic Laboratory | Claudio Tavares Sacchi, Claudia Regina Gonçalves, Erica Valessa Ramos Gomes, Karoline Rodrigues Campos |
| EPI_ISL_738080 | Laboratory Services Section, Texas Department of State Health Services | Laboratory Services Section, Texas Department of State Health Services | Pokharel,A., Oh,B., Bonser,J.D., Tuladhar,R., Pedrueza,M., Zhang,J., Rahman,M., Koag,M., Wang,C., Lee,R. and Kubin,G. |
| EPI_ISL_738148, EPI_ISL_738153, EPI_ISL_738192 | Texas Department of State Health Services | Texas Department of State Health Services | Anita Pokharel, Bonnie Oh, James Daniel Bonser, Rashmi Tuladhar, Mayela Pedrueza, Jenny Zhang, Maliha Rahman, Myong Koag, Chung Wang, Rachel Lee, Grace Kubin |
| EPI_ISL_738514, EPI_ISL_738585, EPI_ISL_738616, EPI_ISL_738699, EPI_ISL_738804, EPI_ISL_738817, EPI_ISL_738822, EPI_ISL_738837, EPI_ISL_738876, EPI_ISL_738892, EPI_ISL_739018, EPI_ISL_739054, EPI_ISL_739056, EPI_ISL_739057, EPI_ISL_739111, EPI_ISL_739124, EPI_ISL_739140, EPI_ISL_739144, EPI_I<br>EPI_ISL_739270, EPI_ISL_739290, EPI_ISL_739309, EPI_ISL_739337, EPI_ISL_739391, EPI_ISL_739395, EPI_ISL_739434, EPI_ISL_739463, EPI_ISL_739484, EPI_ISL_739495, EPI_ISL_739522, EPI_ISL_739551, EPI_ISL_739552, EPI_ISL_739587, EPI_ISL_739590, EPI_ISL_739619 | | | |
| see above | Alameda County Public Health Lab | Chan-Zuckerberg Biohub | CZB Cliahub Consortium |
| EPI_ISL_739700 | Laboratoire national de santé, Microbiology, Virology | Laboratoire national de santé, Microbiology, Epidemiology and Microbial Genomics | Anke Wienecke-Baldacchino, Catherine Ragimbeau, Tamir Abdelrahman, Jessica Tapp, Fatu Djabi, Trung Nguyen Nguyen |
| EPI_ISL_739835, EPI_ISL_739870, EPI_ISL_739978 | Laboratoire national de santé, Microbiology, Virology | Laboratoire national de santé, Microbiology, Microbial Genomics Platform | Anke Wienecke-Baldacchino, Catherine Ragimbeau, Tamir Abdelrahman, Jessica Tapp, Fatu Djabi |
| EPI_ISL_740129 | Laboratoire national de santé, Microbiology, Virology | Laboratoire national de santé, Microbiology, Epidemiology and Microbial Genomics | Anke Wienecke-Baldacchino, Catherine Ragimbeau, Tamir Abdelrahman, Jessica Tapp, Fatu Djabi, Trung Nguyen Nguyen |
| EPI_ISL_740227, EPI_ISL_740259, EPI_ISL_740456, EPI_ISL_740486 | Laboratoire national de santé, Microbiology, Virology | Laboratoire national de santé, Microbiology, Microbial Genomics Platform | Anke Wienecke-Baldacchino, Catherine Ragimbeau, Tamir Abdelrahman, Jessica Tapp, Fatu Djabi |
| EPI_ISL_744295 | Laboratoire national de santé, Microbiology, Virology | Laboratoire national de santé, Microbiology, Epidemiology and Microbial Genomics | Anke Wienecke-Baldacchino, Catherine Ragimbeau, Tamir Abdelrahman, Jessica Tapp, Fatu Djabi, Trung Nguyen Nguyen |
| EPI_ISL_744367, EPI_ISL_744480, EPI_ISL_744631, | Laboratoire national de santé, Microbiology, Virology | Laboratoire national de santé, Microbiology, Microbial Genomics Platform | Anke Wienecke-Baldacchino, Catherine Ragimbeau, Tamir Abdelrahman, Jessica Tapp, Fatu Djabi |

| | | | |
|---|---|---|---|
| EPI_ISL_744724 | | | |
| EPI_ISL_745261, EPI_ISL_745262, EPI_ISL_745263, EPI_ISL_745264, EPI_ISL_745265, EPI_ISL_745266, EPI_ISL_745267, EPI_ISL_745268, EPI_ISL_745269, EPI_ISL_745270, EPI_ISL_745271, EPI_ISL_745272, EPI_ISL_745273, EPI_ISL_745274, EPI_ISL_745275, EPI_ISL_745276, EPI_ISL_745277, EPI_ISL_745278, EPI_I[illegible] EPI_ISL_745283, EPI_ISL_745284, EPI_ISL_745285, EPI_ISL_745286, EPI_ISL_745288, EPI_ISL_745289, EPI_ISL_745290, EPI_ISL_745291, EPI_ISL_745292, EPI_ISL_745293, EPI_ISL_745294, EPI_ISL_745295, EPI_ISL_745296, EPI_ISL_745297, EPI_ISL_745298, EPI_ISL_745299, EPI_ISL_745300, EPI_ISL_745301, EPI_I[illegible] EPI_ISL_745306 | | | |
| see above | Texas Department of State Health Services | Texas Department of State Health Services | Rashmi Tuladhar, Bonnie Oh, Jenny Zhang, Maliha Rahman, Anita Pokharel, Myong Koag, Chung Wang, Rachel Lee, Grace Kubin, Mayela Pedrueza, James Daniel Bonser |
| EPI_ISL_746319 | Genome Center | Genome Center | Md. Shazid Hasan, Hassan M. Al-Emran, Ovinu Kibria Islam, A. S. M. Rubayet- Ul- Alam, Selina Akter, Md. Tanvir Islam, Pravas Chandra Roy, Shovon Lal Sarkar, Najmuj Sakib, Nigar Sultana Meghla, S. M. Tanjil Shah, Shireen Nig[illegible] |
| EPI_ISL_746323 | Genome Center | Genome Center | Ovinu Kibria Islam, Hassan M. Al-Emran, A. S. M. Rubayet- Ul- Alam, Md. Shazid Hasan, Selina Akter, Md. Tanvir Islam, Pravas Chandra Roy, Shovon Lal Sarkar, Najmuj Sakib, Nigar Sultana Meghla, S. M. Tanjil Shah, Shireen Nig[illegible] |
| EPI_ISL_746324 | Genome Center | Genome Center | A. S. M. Rubayet- Ul- Alam, Ovinu Kibria Islam, Hassan M. Al-Emran, Md. Shazid Hasan, Selina Akter, Md. Tanvir Islam, Pravas Chandra Roy, Shovon Lal Sarkar, Najmuj Sakib, Nigar Sultana Meghla, S. M. Tanjil Shah, Shireen Nig[illegible] |
| EPI_ISL_746506, EPI_ISL_746545, EPI_ISL_746546, EPI_ISL_746547, EPI_ISL_746548, EPI_ISL_746549, EPI_ISL_746550, EPI_ISL_746551, EPI_ISL_746552, EPI_ISL_746553, EPI_ISL_746554, EPI_ISL_746555, EPI_ISL_746556, EPI_ISL_746557, EPI_ISL_746558, EPI_ISL_746559, EPI_ISL_746560, EPI_ISL_746561, EPI_I[illegible] EPI_ISL_746566, EPI_ISL_746567, EPI_ISL_746568, EPI_ISL_746569, EPI_ISL_746570, EPI_ISL_746571, EPI_ISL_746572, EPI_ISL_746573, EPI_ISL_746574, EPI_ISL_746575, EPI_ISL_746576, EPI_ISL_746577, EPI_ISL_746578, EPI_ISL_746579, EPI_ISL_746580, EPI_ISL_746581, EPI_ISL_746582, EPI_ISL_746583, EPI_I[illegible] EPI_ISL_746588, EPI_ISL_746589, EPI_ISL_746590, EPI_ISL_746591, EPI_ISL_746592, EPI_ISL_746593 | | | |
| see above | Genetica Molecular and Subdepartamento de Virologia ISP Chile | Instituto de Salud Publica de Chile | Javier Tognarelli, Barbara Parra, Loredana Arata, Jaime Lagos, Gisselle Barra, Patricia Bustos, Rodrigo Fasce, Andres Castillo, Jorge Fernandez |
| EPI_ISL_750168, EPI_ISL_750169, EPI_ISL_750170, EPI_ISL_750171, EPI_ISL_750172, EPI_ISL_750173, EPI_ISL_750174, EPI_ISL_750179 | Sanatorio Americano | Institut Pasteur de Montevideo | Daiana Mir, Natalia Rego, Paola Cristina Resende, Fernando Lopez-Tort, Tamara Fernandez-Calero, Veronica Noya, Mariana Brandes, Tania Possi, Mailen Arleo, Natalia Reyes, Matias Victoria, Andres Lizasoain, Matias Castells, Leticia Maya, Matías Sal[illegible] Garay Martins, Cecilia Alonso, Yasser Vega, Cecilia Salazar, Ignacio Ferrés, Pablo Smircich, Jose Sotelo, Ighor Arantes, Luciana Appolinario, Ana Carolina Mendonça, Maria Jose Benitez-Galeano, Martín Graña, Camila Simoes, Fernando Motta, Ma[illegible] Spangenberg |
| EPI_ISL_751680 | NJ Public Health and Environmental Laboratories | Genomics and Discovery, Respiratory Viruses Branch, Division of Viral Diseases, Centers for Disease Control and Prevention | Krista Queen, Yan Li, Ying Tao, Jing Zhang, Anna Uehara, Anna Montmayeur, Clinton R. Paden, Peter W. Cook,Rachel Marine, Mili Sheth, Haibin Wang, Justin Lee, Suxiang Tong |
| EPI_ISL_752666, EPI_ISL_752667, EPI_ISL_752673, EPI_ISL_752674, EPI_ISL_752675, EPI_ISL_752676, EPI_ISL_752677, EPI_ISL_752678, EPI_ISL_752679, EPI_ISL_752689, EPI_ISL_752690, EPI_ISL_752691, EPI_ISL_752704, EPI_ISL_752705, EPI_ISL_752707 | | | |
| see above | State Laboratories Division, Hawaii State Department of Health | State Laboratories Division, Hawaii State Department of Health | Pamela O'Brien, Sabrina Diemert, Drew Kuwazaki, Razvan Sultana, Edward Desmond |
| EPI_ISL_753713, EPI_ISL_753726, EPI_ISL_753733, EPI_ISL_753747, EPI_ISL_753757, EPI_ISL_753807, EPI_ISL_753808, EPI_ISL_753957, EPI_ISL_753960, EPI_ISL_753961, EPI_ISL_753962, EPI_ISL_753964, EPI_ISL_753969, EPI_ISL_753970, EPI_ISL_753971, EPI_ISL_753972, EPI_ISL_754022 | | | |
| see above | Charité Universitätsmedizin Berlin, Institut für Virologie/Labor Berlin | Charité Universitätsmedizin Berlin, Institut für Virologie | Victor M Corman, Jörn Beheim-Schwarzbach, Barbara Mühlemann, Julia Schneider, Talitha Veith, Terry Jones, Christian Drosten |
| EPI_ISL_754901 | Innovative Genomics Institute, UC Berkeley | Innovative Genomics Institute, UC Berkeley | Stacia Wyman, Haridha Shivram, Phil Frankino, Liana Lareau, Shana McDevitt, Justin Choi |
| EPI_ISL_754976, EPI_ISL_755006, EPI_ISL_755034, EPI_ISL_755060, EPI_ISL_755061 | California Department of Public Health | California Department of Public Health | CDPH IDLB COVIDNet |
| EPI_ISL_756341, EPI_ISL_756344, EPI_ISL_756346, EPI_ISL_756347, EPI_ISL_756351, EPI_ISL_756356 | Innovative Genomics Institute, UC Berkeley | Innovative Genomics Institute, UC Berkeley | Stacia Wyman, Haridha Shivram, Phil Frankino, Liana Lareau, Shana McDevitt, Justin Choi |
| EPI_ISL_765219, EPI_ISL_765220 | Instituto Nacional de Saude (INSA) | Instituto Nacional de Saude (INSA) | Borges et al |
| EPI_ISL_765896, EPI_ISL_765912, EPI_ISL_765916, EPI_ISL_765924, EPI_ISL_765929, EPI_ISL_765931, EPI_ISL_765933 | TXDSHS | TXDSHS | Rashmi Tuladhar, Bonnie Oh, Jenny Zhang, Maliha Rahman, Anita Pokharel, Myong Koag, Chung Wang, Rachel Lee, Grace Kubin, Mayela Pedrueza, James Daniel Bonser |
| EPI_ISL_766861, EPI_ISL_766862 | NIC Viral Respiratory Unit - Institut Pasteur of Algeria | National Reference Center for Viruses of Respiratory Infections, Institut Pasteur, Paris | Mélanie Albert, Marion Barbet, Sylvie Behillil, Méline Bizard, Angela Brisebarre, Flora Donati, Etienne Simon-Lorière, Vincent Enouf, Maud Vanpeene, Sylvie van der Werf, Fawzi Derrar |
| EPI_ISL_768730, EPI_ISL_768734, EPI_ISL_768739 | Child Health Research Foundation | Child Health Research Foundation | Senjuti Saha, Afroza Akter Tanni, Roly Malaker, Sharmistha Goswami, Syed Muktadir Al Sium, Arif Mohammad Tanmoy, Md Hafizur Rahman, Samir K Saha |
| EPI_ISL_774932, EPI_ISL_774934, EPI_ISL_774936, EPI_ISL_774938, EPI_ISL_774941, EPI_ISL_774943, EPI_ISL_774948, EPI_ISL_774949, EPI_ISL_774951, EPI_ISL_774952, EPI_ISL_774953, EPI_ISL_774954, EPI_ISL_774955, EPI_ISL_774956, EPI_ISL_774957, EPI_ISL_774958, EPI_ISL_774959, EPI_ISL_774960, EPI_I[illegible] EPI_ISL_774967, EPI_ISL_774968, EPI_ISL_774969, EPI_ISL_774970, EPI_ISL_774977, EPI_ISL_774979, EPI_ISL_774984, EPI_ISL_774987, EPI_ISL_774988, EPI_ISL_774990, EPI_ISL_774996, EPI_ISL_774997, EPI_ISL_774998, EPI_ISL_774999, EPI_ISL_775001, EPI_ISL_775002, EPI_ISL_775003, EPI_ISL_775007, EPI_I[illegible] | | | |
| see above | Designated Reference Institute for Chemical Measurements (DRICM) | DNA SOLUTION LTD. | Md. Imran Khan, Kazi Nadim Hasan, Abu Sufian, Jannatun Naima, Abdul Khaleque, Mizanur Rahman, MSM Chowdhury, Hasan Ul Haider, Mamudul Hasan Razu, Mala Khan, Mohammad Fazle Al[illegible] |
| EPI_ISL_776539, EPI_ISL_776546, EPI_ISL_776547 | University Medical Center Hamburg Eppendorf | Heinrich Pette Institute, Leibniz Institute for Experimental Virology | Alexis Robitaille, Thomas Günther, Johannes Knobloch, Martin Aepfelbacher, Nicole Fischer, Adam Grundhoff |
| EPI_ISL_776671, EPI_ISL_776677 | UW Virology Lab | UW Virology Lab | Pavitra Roychoudhury, Hong Xie, Lasata Shrestha, Meei-Li Huang, Keith R Jerome, Alexander Greninger |

| | | | |
|---|---|---|---|
| EPI_ISL_778737, EPI_ISL_778738, EPI_ISL_778739, EPI_ISL_778740, EPI_ISL_778741, EPI_ISL_778742, EPI_ISL_778743, EPI_ISL_778744, EPI_ISL_778745, EPI_ISL_778746, EPI_ISL_778747, EPI_ISL_778748, EPI_ISL_778749, EPI_ISL_778750, EPI_ISL_778751, EPI_ISL_778752, EPI_ISL_778753, EPI_ISL_778754, EPI_I EPI_ISL_778759, EPI_ISL_778760, EPI_ISL_778761, EPI_ISL_778762, EPI_ISL_778763, EPI_ISL_778764, EPI_ISL_778765, EPI_ISL_778766, EPI_ISL_778767, EPI_ISL_778768, EPI_ISL_778770, EPI_ISL_778771, EPI_ISL_778772, EPI_ISL_778779, EPI_ISL_778783, EPI_ISL_778784, EPI_ISL_778785, EPI_ISL_778786, EPI_I EPI_ISL_778791, EPI_ISL_778792, EPI_ISL_778793, EPI_ISL_778794, EPI_ISL_778795, EPI_ISL_778796, EPI_ISL_778797, EPI_ISL_778798, EPI_ISL_778799, EPI_ISL_778800, EPI_ISL_778801, EPI_ISL_778802, EPI_ISL_778803, EPI_ISL_778804, EPI_ISL_778805, EPI_ISL_778806, EPI_ISL_778807, EPI_ISL_778808, EPI_I EPI_ISL_778813, EPI_ISL_778814, EPI_ISL_778815, EPI_ISL_778816, EPI_ISL_778817 | | | |
| see above | Istituto Zooprofilattico Sperimentale del Mezzogiorno | TIGEM | Patrizia Annunziata, Andrea Ballabio, Valentina Bouche, Davide Cacchiarelli (CorrespAuthor), Pellegrino Cerino, Chiara Colantuono, Lucio Di Filippo, Antonio Grimaldi, Antonio Limone, Gabriella Loconte, Anna Manfredi, Francesco Panariel |
| EPI_ISL_779392 | Siloam Hospital Lippo Village | Mochtar Riady Institute for Nanotechnology-Universitas Pelita Harapan | Aksar C Lages, David Rustandi, Ivet M Suriapranata, Rinaldy Kusuma, Febi Andriani, Tri Shinta Kurniasih, Riska N Taufik, Young O Larasati, Rury M Wahyuni, Irawan Yusuf |
| EPI_ISL_779923, EPI_ISL_779926, EPI_ISL_779937 | Center of Medical Microbiology, Virology, and Hospital Hygiene, University of Duesseldorf | Center of Medical Microbiology, Virology, and Hospital Hygiene, University of Duesseldorf | Maximilian Damagnez, Alexander Dilthey, Ashley-Jane Duplessis, Torsten Houwaart, Lisanna Hülse, Malte Kohns Vasconcelos, Nadine Lübke, Jessica Nicolai, Klaus Pfeffer, Daniel Strelow, Teresa Tamayo, Jörg Timm, A |
| EPI_ISL_780023 | Hospital General Universitario Gregorio Marañón | SeqCOVID-SPAIN consortium/IBV(CSIC) | Darío García de Viedma, Laura Pérez-Lago, Marta Herranz, Jon Sicilia, Julia Suárez, Pilar Catalán, Patricia Muñoz and SeqCOVID-SPAIN consortium |
| EPI_ISL_780098, EPI_ISL_780162, EPI_ISL_780178, EPI_ISL_780183, EPI_ISL_780195, EPI_ISL_780209, EPI_ISL_780214, EPI_ISL_780244, EPI_ISL_780250, EPI_ISL_780253, EPI_ISL_780254, EPI_ISL_780486, EPI_ISL_780554, EPI_ISL_781072, EPI_ISL_783541, EPI_ISL_783558, EPI_ISL_783559, EPI_ISL_783567, EPI_I EPI_ISL_783577, EPI_ISL_783581, EPI_ISL_783582, EPI_ISL_783588, EPI_ISL_783590, EPI_ISL_783593, EPI_ISL_784184, EPI_ISL_784185, EPI_ISL_784186, EPI_ISL_784187, EPI_ISL_784188, EPI_ISL_784190, EPI_ISL_784191, EPI_ISL_784192, EPI_ISL_784193, EPI_ISL_784194, EPI_ISL_784195, EPI_ISL_784196, EPI_I EPI_ISL_784201, EPI_ISL_784202, EPI_ISL_784203, EPI_ISL_784204, EPI_ISL_784205, EPI_ISL_784206, EPI_ISL_784207, EPI_ISL_784208, EPI_ISL_784209, EPI_ISL_784210, EPI_ISL_784211, EPI_ISL_784213, EPI_ISL_784214, EPI_ISL_784215, EPI_ISL_784216, EPI_ISL_784217, EPI_ISL_784218, EPI_ISL_784219, EPI_I EPI_ISL_784224, EPI_ISL_784225, EPI_ISL_784226, EPI_ISL_784227, EPI_ISL_784228, EPI_ISL_784229, EPI_ISL_784230, EPI_ISL_784231, EPI_ISL_784232, EPI_ISL_784233, EPI_ISL_784234, EPI_ISL_784235, EPI_ISL_784236, EPI_ISL_784237, EPI_ISL_784238, EPI_ISL_784239, EPI_ISL_784240, EPI_ISL_784241, EPI_I EPI_ISL_784246, EPI_ISL_784248, EPI_ISL_784250, EPI_ISL_784280, EPI_ISL_784282, EPI_ISL_784285, EPI_ISL_784356, EPI_ISL_784357, EPI_ISL_784360, EPI_ISL_784361, EPI_ISL_784362, EPI_ISL_784364, EPI_ISL_784366, EPI_ISL_784367, EPI_ISL_784368, EPI_ISL_784369, EPI_ISL_784370, EPI_ISL_784371, EPI_I EPI_ISL_784379, EPI_ISL_784380, EPI_ISL_784393 | | | |
| see above | Houston Methodist Hospital | Houston Methodist Hospital | S. Wesley Long, Randall J. Olsen, Paul A. Christensen, David W. Bernard, James J. Davis, Maulik Shukla, Marcus Nguyen, Matthew Ojeda Saavedra, Prasanti Yerramilli, Layne Pruitt, Sishir Subedi, Heather Hendric |
| EPI_ISL_788938 | Centre de Recherche et de Formation en Infectiologie Guinée | TransVIHMI, IRD/INSERM/Monpellier University | Alpha Kabinet KEITA, Abdoul Karim SOUMAH, Abdoulaye TOURE, Moriba POVOGUI, Joel KOIVOGUI, Jean-louis MONEMOU, Mamadou Saliou SOW, Penda Malhado DIALLO, Mamadou Bhoye KEITA, Alimou CAMARA, Kaba KOUROUMA, Mandiou D Christelle BUTEL, Laetitia SERRANO, Ahidjo AYOUBA, Eric DELAPORTE, Martine PEETERS |
| EPI_ISL_792115 | Instituto Adolfo Lutz - Regional de Taubate | Instituto Adolfo Lutz, Interdiciplinary Procedures Center, Strategic Laboratory | Claudio Tavares Sacchi, Claudia Regina Gonçalves, Erica Valessa Ramos Gomes, Karoline Rodrigues Campos |
| EPI_ISL_792513, EPI_ISL_792517, EPI_ISL_792518, EPI_ISL_792519 | Laboratorio Central de la Ciudad de Santa Fe | Grupo de Genómica y Bioinformática del Instituto de Investigación de la Cadena Láctea CONICET-INTA on behalf of 'Proyecto Argentino Interinstitucional de genomica de SARS-CoV-2' (PAIS Consortium) | Eberhardt, MF; Irazoqui, JM; Ojeda, G; Rompato, G; Mugna, V; Pastor, C; Amadio, AF |
| EPI_ISL_792561, EPI_ISL_792568, EPI_ISL_792569, EPI_ISL_792597 | LACEN-PB | Laboratory of Respiratory Viruses and Measles, Oswaldo Cruz Institute, FIOCRUZ | Paola Resende, Luciana Appolinario, Fernando Motta, Anna Carolina Paixao, Ana Carolina Mendonca, João Felipe Bezerra, Romero Henrique Teixeira de Vasconcelos, Dalane Loudal Florentino Teixeira, Thiago Franco de |
| EPI_ISL_803867 | National Institute of Laboratory Medicine and Referral Center | Genomic Research Lab, BCSIR | Abu Sayeed Mohammad Mahmud, Mohammad Samir Uzzaman, Eshrar Osman, Md. Ahashan Habib, Shahina Akter, Tanjina Akhter Banu,Md. Murshed Hasan Sarkar,Barna Goswami, Iffat Jahan, Md. Saddam Hossain, Tasnim Nafisa, Md. Maruf Ah Shamsuzzaman, Monira Parveen, Md. Masum Hossain Arif, Md. Salim Khan |
| EPI_ISL_804863 | DC Public Health Lab/ Dept. of Forensic Sciences | DC Public Health Lab/ Dept. of Forensic Sciences | Scott Nguyen, Elizabeth Zelaya, Connie Maza, Monica Mann, Brittany Hamilton, David Payne, Jocelyn Hauser |
| EPI_ISL_806623, EPI_ISL_806624, EPI_ISL_806625, EPI_ISL_806638, EPI_ISL_806671, EPI_ISL_806672, EPI_ISL_806673, EPI_ISL_806674, EPI_ISL_806675, EPI_ISL_806676, EPI_ISL_806677, EPI_ISL_806678, EPI_ISL_806679, EPI_ISL_806680, EPI_ISL_806681, EPI_ISL_806682, EPI_ISL_806692, EPI_ISL_806693, EPI_I | | | |
| see above | KEMRI-Wellcome Trust Research Programme/KEMRI-CGMR-C Kilifi | KEMRI-Wellcome Trust Research Programme/KEMRI-CGMR-C Kilifi | Githinji et al |
| EPI_ISL_811154, EPI_ISL_811185 | Dharwad | CSIR Institute of Genomics and Integrative Biology | Dr. Shivarudrapp B Bhairappanavar, Rahul Bhoyar, Mohammed Imran, Mohit Divakar, Disha Sharma, Dr. Vijay A Yenagi, Dr. Suresh B Arakera, Dr. Amit Ugargol, Dr. Rgavendra B Nayak, Bani Jolly, Abhinav Jain, Paras Sehgal, Gy |
| EPI_ISL_812806, EPI_ISL_812813, EPI_ISL_812820, EPI_ISL_812829, EPI_ISL_812838, EPI_ISL_812839, EPI_ISL_812842, EPI_ISL_812847, EPI_ISL_812848, EPI_ISL_812856, EPI_ISL_812860, EPI_ISL_812863, EPI_ISL_812868, EPI_ISL_812872 | | | |
| see above | Genomics Program, Children Cancer Hospital | Genomics Program, Children Cancer Hospital | Hatem,A., Hadad,A., Abouelnaga,S., Amer,K., Salah,H., Farawyla,H., Halafawy,A., Mansour,T., shalaby,L., Hassan,W., Soliman,M., Gomaa,C., Hassan,R., Soliman,S., Monuir,G., Hammad,M., Hussein,S., Abdo,I., Jalal,D., El-Zayat,M., El-Shaqnq |
| EPI_ISL_816715, EPI_ISL_816716 | Bioinformatics and Biostatistics Lab, Advanced Sequencing Facility | COVID-19 Genomics UK (COG-UK) Consortium | Aengus Stewart,Jerome Nicod,Chelsea Sawyer,Laura Cubitt,Harshil Patel,Margaret Crawford |
| EPI_ISL_824386, EPI_ISL_824387 | Ohio Department of Health Laboratory | Ohio Department of Health Laboratory | Holmes, Jennifer; Eric Brandt, Keoni Omura, Glen McGillivary, Caitlin McDonnell, Kirtana Ramadugu, Erica Leasure, Kelsey Florek, Heather Blankenship, Quanta Brown, and Tammy Banner |
| EPI_ISL_824416, EPI_ISL_824417, EPI_ISL_824418, EPI_ISL_824419 | Hospital Universitari Vall d'Hebron - Vall d'Hebron Institut de Rercerca | Hospital Universitari Vall d'Hebron | Cristina Andrés, Maria Piñana, Josep F Abril, Damir Garcia-Cehic, Ariadna Rando, Juliana Esperalba, Maria Gema Codina, Carla Castillo, Maria Carmen Martin, Tomàs Pumarola, Josep Quer, And |
| EPI_ISL_825653, EPI_ISL_825654, EPI_ISL_825655, EPI_ISL_825656, EPI_ISL_825908, EPI_ISL_825909, EPI_ISL_825910, EPI_ISL_825911, EPI_ISL_825913 | Laboratoire de santé publique du Québec | Laboratoire de santé publique du Québec | Sandrine Moreira, Ioannis Ragoussis, Guillaume Bourque, Jesse Shapiro, Mark Lathrop and Michel Roger on behalf of the CoVSeQ research group (http://covseq.ca/researchgroup) |

| | | | |
|---|---|---|---|
| EPI_ISL_826823 | INSPI-CRN DE INFLUENZA Y OTROS VIRUS RESPIRATORIOS | Instituto de Salud Publica de Chile | Javier Tognarelli, Barbara Parra, Loredana Arata, Jaime Lagos, Gisselle Barra, Alfredo Bruno, Domenica de Mora, Solon Narvaez, Jimmy Garcez, Michelle Paez, Martiza Olmedo, Manuel Gonzalez, Patricia Bustos, Rodrigo Fa |
| EPI_ISL_831309, EPI_ISL_831314 | Texas Department of State Health Services | Texas Department of State Health Services | Anita Pokharel, Bonnie Oh, James Daniel Bonser, Rashmi Tuladhar, Mayela Pedrueza, Jenny Zhang, Maliha Rahman, Myong Koag, Chung Wang, Rachel Lee, Grace Kubin |
| EPI_ISL_831916 | New Mexico Department of Health Scientific Laboratory | New Mexico Department of Health Scientific Laboratory | Ellie Johnson, Anastacia Griego-Fisher, D'eldra Malone |
| EPI_ISL_832405 | OHSU Lab Services Molecular Microbiology Lab | Oregon SARS-CoV-2 Genome Sequencing Center | Brendan L. O'Connell, Ruth V. Nichols, Sally Grindstaff, Alec J. Hirsch, Donna Hansel, Guang Fan, Daniel N. Streblow, William B. Messer, Andrew C. Adey, Benjamin N. Bimber, Brian J. O'R |
| EPI_ISL_833130 | National Institute of Health Research and Development | National Institute of Health Research and Development | Agustiningsih;Adam,K;Wibowo,HA;Ramadhany,R;Rukminiati,Y;Pawestri,HA;Subangkit;Puspa,KD;Nugraha,AA;Ikawati,HD;Pangesti,KNA;Soekarso,T;Susilarini,NK;Hariastuti,NI;Nikmah,UA;Mursinah;Febriyani,A;Herman,R;Susanti,N;Herna;Febriyanti,T;Kurr |
| EPI_ISL_833192, EPI_ISL_833195 | Hôpital Bichat Claude Bernard, Laboratoire de Virologie | IAME UMR1137 Inserm, Université de Paris, Hôpital Bichat | Antoine Bridier, Amélie Recoing, Quentin Le Hingrat, Lena Daniel, Siham Hamri, Gilles Collin, Alexandre Storto, Mélanie Bertine, Charlotte Charpentier, Nadhira Houhou-Fidouh, Diane Descamps, Be |
| EPI_ISL_833393 | RS MMC Jakarta, Indonesia | Biosafety Level-3 Laboratory, Indonesian Institute of Sciences (LIPI) | Anggia Prasetyoputri, Isa Nuryana, Ade Andriani, Anik Budhi Dharmayanthi, Syam Budi Iryanto, Andri Wardiana, Ahmad Fathoni, Eva Erdayani, Linda Sukmarini, Ratih Asmana Ningrum |
| EPI_ISL_833503 | RS MMC, Jakarta, Indonesia | Biosafety Level-3 Laboratory, Indonesian Institute of Sciences (LIPI) | Anggia Prasetyoputri, Isa Nuryana, Anik Budhi Dharmayanthi, Syam Budi Iryanto, Andri Wardiana, Ade Andriani, Ahmad Fathoni, Idris, Ruby Setiawan, Ratih Asmana Ningrum |
| EPI_ISL_837551, EPI_ISL_837583 | Laboratorio Nacional de Salud | Laboratory of Respiratory Viruses and Measles, Oswaldo Cruz Institute, FIOCRUZ | Paola Resende, Cesar Roberto Conde Pereira, Claudia Estrada, Luciana Appolinario, Fernando Motta, Anna Carolina Paixao, Ana Carolina Mendonca, Marilda Siqueira |
| EPI_ISL_837743, EPI_ISL_837744, EPI_ISL_837745, EPI_ISL_837746, EPI_ISL_837747, EPI_ISL_837748, EPI_ISL_837749 | Instituto Nacional de Enfermedades Respiratorias (INER) | Instituto Nacional de Enfermedades Respiratorias (INER) | Celia Boukadida, Margarita Matías-Florentino, Alma Rincón-Rubio, Hector Esteban Paz-Juárez, Olivia Briceño, Edgar Sevilla-Reyes, Fidencio Mejía-Nepomuceno, Mario Mújica-Sánchez, Eduardo Becerril-Vargas, José Arturo Martínez-Orozco, Alejandra H Armando Vázquez-Pérez |
| EPI_ISL_848566, EPI_ISL_848610 | Evandro Chagas Institute | Evandro Chagas Institute | Santos, M.C.; Silva, A.M.; Junior, W.D.C.; Barbagelata, L.S.; Ferreira, J.A.; Sousa, E.M.A.; da Silva, P.S.; Pinheiro, K.C.; L.C.; Sousa Junior, E.C. |
| EPI_ISL_849279, EPI_ISL_849280, EPI_ISL_849282, EPI_ISL_849284, EPI_ISL_849285, EPI_ISL_849288, EPI_ISL_849297, EPI_ISL_849303, EPI_ISL_849304, EPI_ISL_849314, EPI_ISL_849315, EPI_ISL_849316 | | | |
| see above | Servicio Virosis Respiratorias-Departamento Virología-INEI | Instituto Nacional Enfermedades Infecciosas C.G.Malbran | Baumeister E., Avaro M., Benedetti E., Russo M., Dattero ME, Pontoriero A., Cisterna D., Molina V., Perandones C., Tuduri E., Lorenzo F., Poklepovich T., Campos J. |
| EPI_ISL_849692 | unknown | PHV-FSS | Son Nguyen et al. |
| EPI_ISL_850212, EPI_ISL_850213, EPI_ISL_850214, EPI_ISL_850215, EPI_ISL_850216, EPI_ISL_850217, EPI_ISL_850218, EPI_ISL_850219, EPI_ISL_850220, EPI_ISL_850221, EPI_ISL_850222, EPI_ISL_850223, EPI_ISL_850224, EPI_ISL_850225 | | | |
| see above | Division of Emerging Infectious Diseases, Bureau of Infectious Diseases Diagnosis Control, Korea Disease Control and Prevention Agency | Division of Emerging Infectious Diseases, Bureau of Infectious Diseases Diagnosis Control, Korea Disease Control and Prevention Agency | Ae Kyung Park, Il-Hwan Kim, Heui Man Kim, Jeong-Min Kim, Namjoo Lee, Chaeyoung Lee, Sang Hee Woo, Eun-Jin Kim |
| EPI_ISL_852564 | Max von Pettenkofer Institute, Virology, National Reference Center for Retroviruses, LMU München | Laboratory for Functional Genome Analysis, Dept. Genomics, Gene Center of the LMU Munich | Max Muenchhoff, Stefan Krebs, Alexander Graf, Oliver Keppler, Helmut Blum |
| EPI_ISL_852658 | Institute of Virology, Medical Center, University of Freiburg, Freiburg, Germany | Institute of Virology, Clinial Virus Genomics, Medical Center, University of Freiburg, Freiburg, Germany | Jonas Fuchs, Lisa Kern, Sandra Reuter, Hajo Grundmann, Marcus Panning |
| EPI_ISL_854977, EPI_ISL_854978, EPI_ISL_854979, EPI_ISL_854980, EPI_ISL_854981, EPI_ISL_854982, EPI_ISL_854983, EPI_ISL_854984, EPI_ISL_854985, EPI_ISL_854986, EPI_ISL_854987, EPI_ISL_854988, EPI_ISL_854989, EPI_ISL_854990, EPI_ISL_854991, EPI_ISL_854992, EPI_ISL_854993, EPI_ISL_854994, EPI_I: EPI_ISL_854999, EPI_ISL_855000, EPI_ISL_855001, EPI_ISL_855002, EPI_ISL_855003, EPI_ISL_855004, EPI_ISL_855005, EPI_ISL_855006, EPI_ISL_855007, EPI_ISL_855008, EPI_ISL_855009, EPI_ISL_855010, EPI_ISL_855011, EPI_ISL_855012, EPI_ISL_855013 | | | |
| see above | NGS Lab, DNA SOLUTION LTD. | NGS Lab, DNA SOLUTION LTD. | Khan,M.I., Hasan,K.N., Sufian,A., Hosen,M.B., Khaleque,A., Rahman,M., Chowdhury,M., Haider,H.U., Razu,M.H., Khan,M., Rabbi,M.F.A. |
| EPI_ISL_856987, EPI_ISL_856988, EPI_ISL_856989, EPI_ISL_856990, EPI_ISL_856991, EPI_ISL_856992 | The Ashley Laboratory, Stanford University | Chan-Zuckerberg Biohub | CZB Cliahub Consortium |
| EPI_ISL_859583, EPI_ISL_859584, EPI_ISL_859585, EPI_ISL_859586, EPI_ISL_859587, EPI_ISL_859588, EPI_ISL_859589, EPI_ISL_859590, EPI_ISL_859591, EPI_ISL_859592, EPI_ISL_859593, EPI_ISL_859594, EPI_ISL_859595, EPI_ISL_859596, EPI_ISL_859597, EPI_ISL_859598, EPI_ISL_859599, EPI_ISL_859600, EPI_I: EPI_ISL_859613, EPI_ISL_859614, EPI_ISL_859615, EPI_ISL_859616, EPI_ISL_859617, EPI_ISL_859618, EPI_ISL_859620, EPI_ISL_859624, EPI_ISL_859625, EPI_ISL_859626, EPI_ISL_859627, EPI_ISL_859628, EPI_ISL_859630, EPI_ISL_859631, EPI_ISL_859632, EPI_ISL_859633, EPI_ISL_859634, EPI_ISL_859635, EPI_I: EPI_ISL_859641, EPI_ISL_859645, EPI_ISL_859647, EPI_ISL_859648, EPI_ISL_859649, EPI_ISL_859650, EPI_ISL_859651 | | | |
| see above | BTC, Khalifa University | BTC, Khalifa University | Al Safar et al |
| EPI_ISL_860325, EPI_ISL_860329, EPI_ISL_860339, EPI_ISL_860341, EPI_ISL_860342, EPI_ISL_860343, EPI_ISL_860345, EPI_ISL_860350, EPI_ISL_860351, EPI_ISL_860364, EPI_ISL_860367, EPI_ISL_860368, EPI_ISL_860371, EPI_ISL_860388, EPI_ISL_860391, EPI_ISL_860393, EPI_ISL_860397, EPI_ISL_860398, EPI_I: EPI_ISL_860414, EPI_ISL_860417, EPI_ISL_860432, EPI_ISL_860436, EPI_ISL_860443, EPI_ISL_860447, EPI_ISL_860450, EPI_ISL_860455, EPI_ISL_860460, EPI_ISL_860461, EPI_ISL_860465, EPI_ISL_860466, EPI_ISL_860468, EPI_ISL_860469, EPI_ISL_860473, EPI_ISL_860485, EPI_ISL_860492, EPI_ISL_860495, EPI_I: EPI_ISL_860517, EPI_ISL_860518, EPI_ISL_860527, EPI_ISL_860531, EPI_ISL_860532, EPI_ISL_860537, EPI_ISL_860544 | | | |
| see above | MVZ Labor Krone GbR | Center of Medical Microbiology, Virology, and Hospital Hygiene, University of Duesseldorf | Dennis Deschka, Alexander Dilthey, Julia Fazaal, André Heimbach, Per Hoffmann, Torsten Houwaart, Malte Kohns Vasconcelos, Klaus Pfeffer, Bärbel Lippke, Kerstin Ludwig, Janine Silvery, Carsten Tiemann, Jörg Timm, A |
| EPI_ISL_861667 | Instituto Adolfo Lutz - | Instituto Adolfo Lutz, | Claudio Tavares Sacchi, Claudia Regina Gonçalves, Erica Valessa Ramos Gomes, Karoline Rodrigues Campos |

| | | | |
|---|---|---|---|
| | Regional de Rio Claro | Interdiciplinary Procedures Center, Strategic Laboratory | |
| EPI_ISL_862278, EPI_ISL_862279, EPI_ISL_862283, EPI_ISL_862297, EPI_ISL_862308, EPI_ISL_862437, EPI_ISL_862447, EPI_ISL_862458, EPI_ISL_862459, EPI_ISL_862460, EPI_ISL_862461, EPI_ISL_862465, EPI_ISL_862509, EPI_ISL_862510, EPI_ISL_862511, EPI_ISL_862513, EPI_ISL_862514, EPI_ISL_862515, EPI_I[illegible] EPI_ISL_862520, EPI_ISL_862534, EPI_ISL_862535, EPI_ISL_862536 | | | |
| see above | Kurnool Medical College (KMC) | CSIR Institute of Genomics and Integrative Biology | Pallavali Roja Rani, Mohamed Imran, J. Vijaya Lakshmi, Bani Jolly, S. Afsar, Abhinav Jain, Mohit Kumar Divakar, Panyam Suresh, Disha Sharma, Nambi Rajesh, Rahul C Bhoyar, Dasari Ankaiah, Sanaga Shanthi Kumari, Gyan Ranjan, Valluri Anitha Lavar[illegible] A. Surekha, Pulala Chandra, Rajamadugu Hymavathy, P R Vanaja, Vinod Scaria, Sridhar Sivasubbu |
| EPI_ISL_872640 | Pathogenic Microorganisms Variability Laboratory | Pathogenic Microorganisms Variability Laboratory | Alexey Shchetinin, Olesya Venchakova, Maria Nikiforova, Andrei Siniavin, Nadezhda Kuznetsova, Elena Shidlovskaya, Elizaveta Divisenko, Kirill Krasnoslobotsev, Evgeniya Mukasheva, Anna Ignatieva, Svetlana Trushakova, Andrey Pochtovyy, Valeria [illegible] Svetlana Smetanina, Elena Burtseva, Denis Logunov, Vladimir Gushchin, Alexander Gintsburg |
| EPI_ISL_876041 | Montefiore Medical Center | Albert Einstein College of Medicine, Dept. of Microbiology & Immunology, Chandran lab | J. Maximilian Fels, Saad Khan, Ryan Forster, Karin A. Skalina, Surksha Sirichand, Amy S. Fox, Aviv Bergman, William B. Mitchell, Lucia R. Wolgast, Wendy Szymczak, Robert H. Bortz III, M. Eugenia Dieterle, Catalina Florez, Denise Haslwanter, Rohit K[illegible] David L. Goldman, Hnin Khine, D. Yitzchak Goldstein, Johanna P. Daily, Kartik Chandran, Libusha Kelly |
| EPI_ISL_876824, EPI_ISL_876965, EPI_ISL_876966, EPI_ISL_876967 | Quest Diagnostics | Quest Diagnostics | Rosenthal,S.H., Gerasimova,A., Kagan,R.M., Anderson, B., Hua, M., Liu Y., Bernstein, L.E., Livingston, K.E., Perez, A., Shalhout, D.F., Shlyakhter, I.A., Owen, R., Tanpaiboon, P., Lacbawan[illegible] |
| EPI_ISL_877624, EPI_ISL_877625, EPI_ISL_877626, EPI_ISL_877627, EPI_ISL_877628, EPI_ISL_877629, EPI_ISL_877630, EPI_ISL_877631 | Clinical Molecular Microbiology Laboratory, UNC Hospital | Dirk Dittmer | Razia Moorad , Justin T. Landis , Brent A. Eason, Melissa B. Miller, Linda Pluta, Dirk Dittmer, Angelica Juarez, Cecilia Thompson , Cameroon Grant, Evelyn Hoffman, Patricio Cano, Jason Wong, Carolina Caro-V[illegible] |
| EPI_ISL_884461 | Molecular Microbiology & Immunology, University of Missouri | Molecular Microbiology & Immunology, University of Missouri | Tang,C.Y., Li,T., Hang,J., Lidl,G.M., Wan,X.-F. |
| EPI_ISL_890096, EPI_ISL_890097, EPI_ISL_890098 | Laboratoire de santé publique du Québec | Laboratoire de santé publique du Québec | Sandrine Moreira, Ioannis Ragoussis, Guillaume Bourque, Jesse Shapiro, Mark Lathrop and Michel Roger on behalf of the CoVSeQ research group |
| EPI_ISL_900249 | MEPHI, Aix Marseille University | MEPHI, Aix Marseille University | Anthony LEVASSEUR |
| EPI_ISL_900691, EPI_ISL_900692, EPI_ISL_900723, EPI_ISL_900724, EPI_ISL_900725 | Bozeman Health Deaconess Hospital | Wiedenheft lab, Montana State University | Artem Nemudryi, Anna Nemudraia, Tanner Wiegand, Joseph Nichols, Deann T. Snyder, Jodi F. Hedges, Calvin Cicha, Helen Lee, Karl K. Vanderwood, Diane Bimczok, Mark A. Jutila and Blake W[illegible] |
| EPI_ISL_903344 | Bozeman Health Deaconess Hospital | Wiedenheft lab, Montana State University | Artem Nemudryi, Anna Nemudraia, Tanner Wiegand, Joseph Nichols, Deann T. Snyder, Jodi F. Hedges, Calvin Cicha, Helen Lee, Karl K. Vanderwood, Diane Bimczok, Mark A. Jutila and Blake W[illegible] |
| EPI_ISL_914416, EPI_ISL_914420, EPI_ISL_914429, EPI_ISL_914430, EPI_ISL_914431, EPI_ISL_914432, EPI_ISL_914433, EPI_ISL_914434, EPI_ISL_914435, EPI_ISL_914436, EPI_ISL_914437, EPI_ISL_914438, EPI_ISL_914439, EPI_ISL_914440, EPI_ISL_914441, EPI_ISL_914442, EPI_ISL_914443, EPI_ISL_914444, EPI_I[illegible] EPI_ISL_914450, EPI_ISL_914451, EPI_ISL_914452, EPI_ISL_914453, EPI_ISL_914454, EPI_ISL_914455, EPI_ISL_914456, EPI_ISL_914457, EPI_ISL_914458, EPI_ISL_914459, EPI_ISL_914460, EPI_ISL_914461, EPI_ISL_914462, EPI_ISL_914463, EPI_ISL_914464, EPI_ISL_914465, EPI_ISL_914466, EPI_ISL_914468, EPI_I[illegible] EPI_ISL_914473, EPI_ISL_914474, EPI_ISL_914475, EPI_ISL_914476, EPI_ISL_914477, EPI_ISL_914478, EPI_ISL_914479, EPI_ISL_914480, EPI_ISL_914481, EPI_ISL_914482, EPI_ISL_914483, EPI_ISL_914484, EPI_ISL_914485, EPI_ISL_914486, EPI_ISL_914487 | | | |
| see above | TGen North | TGen North | "Jolene Bowers, Megan Folkerts, Chris French, Hayley Yaglom, Ashlyn Pfeiffer, Darrin Lemmer, Dave Engelthaler, The Arizona COVID Genomics Union (ACGU)" |
| EPI_ISL_930834, EPI_ISL_930835 | Virology, ICAR-National Research Centre on Equines | Virology, ICAR-National Research Centre on Equines | Kumar,N., Gulati,B.R., Barua,S., Riyesh,T., Shanmugasundram,K.,Khandelwal,N. and Kumar,R. |
| EPI_ISL_930836 | Virology, ICAR-National Research Centre on Equines | Virology, ICAR-National Research Centre on Equines | Gulati,B.R., Kumar,N., Barua,S., Riyesh,T., Kumar,R., Gupta,S.,Manuja,A., Kumar,B., Singha,H.S., Vaid,R.K., Anand,T., Bera,B.C., Virmani,N., Bhardwaj,A., Khandelwal,N., Kumar,R. and Pa[illegible] |
| EPI_ISL_930837 | Virology, ICAR-National Research Centre on Equines | Virology, ICAR-National Research Centre on Equines | Kumar,N., Gulati,B.R., Barua,S., Riyesh,T., Shanmugasundram,K.,Khandelwal,N. and Kumar,R. |
| EPI_ISL_930838, EPI_ISL_930839, EPI_ISL_930840, EPI_ISL_930841 | Virology, ICAR-National Research Centre on Equines | Virology, ICAR-National Research Centre on Equines | Gulati,B.R., Kumar,N., Barua,S., Riyesh,T., Kumar,R., Gupta,S.,Manuja,A., Kumar,B., Singha,H.S., Vaid,R.K., Anand,T., Bera,B.C., Virmani,N., Bhardwaj,A., Khandelwal,N., Kumar,R. and Pa[illegible] |
| EPI_ISL_930842, EPI_ISL_930843 | Virology, ICAR-National Research Centre on Equines | Virology, ICAR-National Research Centre on Equines | Kumar,N., Gulati,B.R., Barua,S., Riyesh,T., Shanmugasundram,K.,Khandelwal,N. and Kumar,R. |
| EPI_ISL_933645, EPI_ISL_933646, EPI_ISL_933647 | Toronto Invasive Bacterial Diseases Network | McMaster University | Allison McGeer, Patryk Aftanas, Hooman Derakhshani, Angel Li, Kuganya Nirmalarajah, Emily Panousis, Ahmed Draia, Jalees Nasir, Michael Surette, Samira Mubareka, Andrew G. McArth[illegible] |
| EPI_ISL_936551, EPI_ISL_936552, EPI_ISL_936553, EPI_ISL_936554, EPI_ISL_936555, EPI_ISL_936556, EPI_ISL_936557 | Northwestern Memorial Hospital | Ozer Lab | Ramon Lorenzo-Redondo, Lacy M. Simons, Chad J. Achenbach, Lawrence J. Jennings, Michael G. Ison, Judd F. Hultquist, Egon A. Ozer |
| EPI_ISL_940243, EPI_ISL_940539, EPI_ISL_940546 | Hôpital Bichat Claude Bernard, Laboratoire de Virologie | IAME UMR1137 Inserm, Université de Paris, Hôpital Bichat | Antoine Bridier-Nahmias, Amélie Recoing, Quentin Le Hingrat, Lena Daniel, Siham Hamri, Gilles Collin, Alexandre Storto, Mélanie Bertine, Charlotte Charpentier, Nadhira Houhou-Fidouh, Diane Descamps[illegible] |
| EPI_ISL_940608 | Laboratório Sao Lucas | Instituto Adolfo Lutz, Interdiciplinary Procedures Center, Strategic Laboratory | Claudio Tavares Sacchi, Claudia Regina Gonçalves, Erica Valessa Ramos Gomes, Karoline Rodrigues Campos |
| EPI_ISL_940900, EPI_ISL_940901 | Centers for Disease Control and Prevention, Dengue Branch | Centers for Disease Control and Prevention, Dengue Branch | Gilberto A. Santiago, Glenda Gonzalez, Betzabel Flores, Keyla Charriez, Gabriela Paz-Bailey, Jorge L. Munoz-Jordan |
| EPI_ISL_941951, | Instituto Nacional de Salud, | Centro de Investigaciones en | Luz Helena Patiño, Marina Muñoz, Nathalia Ballesteros, Carolina Hernández, Carolina Flórez, Sergio Gomez, Adriana van de Guchte, Zenab Khan, Jayeeta Dutta, Hala Alejel Alshammary, Ana S. Gonzalez-Reiche, Matthew M. Hernandez, Emilia Mia Sor[illegible] |

| | | | |
|---|---|---|---|
| EPI_ISL_941955, EPI_ISL_941991, EPI_ISL_941992, EPI_ISL_941993, EPI_ISL_941994 | Bogotá, Colombia | Microbiología y Biotecnología-UR (CIMBIUR), Facultad de Ciencias Naturales, Universidad del Rosario, Bogotá, Colombia Instituto Nacional de Salud, Bogotá, Colombia Icahn School of Medicine at Mount Sinai, New York, USA | David Ramírez |
| EPI_ISL_941995, EPI_ISL_941996 | Centro de Investigaciones en Microbiología y Biotecnología-UR (CIMBIUR), Facultad de Ciencias Naturales, Universidad del Rosario, Bogotá, Colombia | Centro de Investigaciones en Microbiología y Biotecnología-UR (CIMBIUR), Facultad de Ciencias Naturales, Universidad del Rosario, Bogotá, Colombia Instituto Nacional de Salud, Bogotá, Colombia Icahn School of Medicine at Mount Sinai, New York, USA | Luz Helena Patiño, Marina Muñoz, Nathalia Ballesteros, Carolina Hernández, Carolina Flórez, Sergio Gomez, Adriana van de Guchte, Zenab Khan, Jayeeta Dutta, Hala Alejel Alshammary, Ana S. Gonzalez-Reiche, Matthew M. Hernandez, Emilia Mia Sor David Ramírez |
| EPI_ISL_942003 | Instituto Nacional de Salud, Bogotá, Colombia | Centro de Investigaciones en Microbiología y Biotecnología-UR (CIMBIUR), Facultad de Ciencias Naturales, Universidad del Rosario, Bogotá, Colombia Instituto Nacional de Salud, Bogotá, Colombia Icahn School of Medicine at Mount Sinai, New York, USA | Luz Helena Patiño, Marina Muñoz, Nathalia Ballesteros, Carolina Hernández, Carolina Flórez, Sergio Gomez, Adriana van de Guchte, Zenab Khan, Jayeeta Dutta, Hala Alejel Alshammary, Ana S. Gonzalez-Reiche, Matthew M. Hernandez, Emilia Mia Sor David Ramírez |
| EPI_ISL_942004 | Centro de Investigaciones en Microbiología y Biotecnología-UR (CIMBIUR), Facultad de Ciencias Naturales, Universidad del Rosario, Bogotá, Colombia | Centro de Investigaciones en Microbiología y Biotecnología-UR (CIMBIUR), Facultad de Ciencias Naturales, Universidad del Rosario, Bogotá, Colombia Instituto Nacional de Salud, Bogotá, Colombia Icahn School of Medicine at Mount Sinai, New York, USA | Luz Helena Patiño, Marina Muñoz, Nathalia Ballesteros, Carolina Hernández, Carolina Flórez, Sergio Gomez, Adriana van de Guchte, Zenab Khan, Jayeeta Dutta, Hala Alejel Alshammary, Ana S. Gonzalez-Reiche, Matthew M. Hernandez, Emilia Mia Sor David Ramírez |
| EPI_ISL_943988 | LACEN do Estado de Goias | Instituto Adolfo Lutz, Interdiciplinary Procedures Center, Strategic Laboratory | Claudio Tavares Sacchi, Claudia Regina Gonçalves, Erica Valessa Ramos Gomes, Karoline Rodrigues Campos |
| EPI_ISL_956276 | Mitra Keluarga Hospital Waru | Institute of Tropical Disease, Universitas Airlangga | Maria I Lusida, Krisnoadi Rahardjo, Aldise M Nastri, Jezzy R Dewantari, Rima R Prasetya, Christina Dian Anggraeni, Gatot Soegiarto, Laksmi Wulandari, Resti Yudhawati, Soetjipto, Yasuko Mori, Kazu |
| EPI_ISL_960159 | Vanguard CHC wc VGC | National Health Laboratory Service/UCT | Arash Iranzadeh, Deelan Doolabh, Lynn Tyers, Bruna Galvao, Innocent Mudau, Marvin Hsiao, Kruger Marais, Diana Hardie, Stephen Korsman, Carolyn Williamson |
| EPI_ISL_960160, EPI_ISL_960161 | Dr Abdurahman CDC wc DAC | National Health Laboratory Service/UCT | Arash Iranzadeh, Deelan Doolabh, Lynn Tyers, Bruna Galvao, Innocent Mudau, Marvin Hsiao, Kruger Marais, Diana Hardie, Stephen Korsman, Carolyn Williamson |
| EPI_ISL_960305 | NLZOH, Laboratory for Virology | NLZOH, Laboratory for Virology | Katarina Prosenc (Laboratory for Virology), Cesare Camma (IZSAM), Erik Alm (ECDC) |
| EPI_ISL_961135, EPI_ISL_961136, EPI_ISL_961137, EPI_ISL_961140, EPI_ISL_961141, EPI_ISL_961142, EPI_ISL_961146, EPI_ISL_961147, EPI_ISL_961151, EPI_ISL_961153, EPI_ISL_961157, EPI_ISL_961160, EPI_ISL_961164, EPI_ISL_961167, EPI_ISL_961168, EPI_ISL_961169, EPI_ISL_961173, EPI_ISL_961174 | | | |
| see above | Texas Department of State Health Services | Texas Department of State Health Services | Bonnie Oh, Anita Pokharel, James Daniel Bonser, Myong Koag, Chung Wang, Rachel Lee, Grace Kubin, Rashmi Tuladhar, Mayela Pedrueza, Maliha Rahman, Jenny Zhang |
| EPI_ISL_961779, EPI_ISL_961780 | Laboratorio de Infectología, Servicio de Infectología, Hospital Universitario Dr. José Eleuterio González - Universidad Autónoma de Nuevo León | Laboratorio de Infectología Molecular, Departamento de Bioquímica y Medicina Molecular,Facultad de Medicina - Universidad Autónoma de Nuevo León | Kame A. Galán-Huerta, María F. Herrera-Saldivar, Natalia Martínez-Acuña, Sonia A. Lozano-Sepúlveda, Daniel Arellanos-Soto, Ana M. Rivas-Estilla, Paola Bocanegra-Ibarias, Samantha M. Flores-Treviño, Elvira Garza-González, Eduardo Pere |
| EPI_ISL_964901, EPI_ISL_964915 | Sistema de Emergencias | Laboratorio Central Mg. Luis Alfredo Pianciola on behalf of 'Proyecto Argentino Interinstitucional de genomica de SARS-CoV-2' (PAIS Consortium) | L Pianciola, M Mazzeo, C Ziehm, C Pintos, M Fernandez, J Ousset, M Nabaes, M Viegas. |
| EPI_ISL_965978 | Washington State Department of Health | Seattle Flu Study | Deborah A. Nickerson, Chris D. Frazar, Jover Lee, Benjamin Pelle, Erica Ryke, Matthew Richardson, Amanda Adler, Elisabeth Brandstetter, Peter D. Han, Kairsten Fay, Misja Ilcisin, Kirsten Lacombe, Thomas R. Sibley, Melissa Truong, Caitlin R. Wolf, Lindquist, Michael Boeckh, Janet A. Englund, Michael Famulare, Barry R. Lutz, Mark J. Rieder, Lea M. Starita, Matthew Thompson, Helen Y. Chu, Jay Shendure, Trevor Bedford |
| EPI_ISL_968156 | Clinical Molecular Microbiology Laboratory, UNC Hospital | Dirk Dittmer | Justin T. Landis , Razia Moorad , Brent A. Eason, Melissa B. Miller, Linda Pluta, Dirk Dittmer, Angelica Juarez, Cecilia Thompson, Shawn Hawken, Cameroon Grant, Evelyn Hoffman, Patricio Cano, Jason Wong, Carolina Caro-Ve |
| EPI_ISL_968250, EPI_ISL_968251, EPI_ISL_968253, EPI_ISL_968254, EPI_ISL_968255, EPI_ISL_968256, EPI_ISL_968257, EPI_ISL_968258, EPI_ISL_968259, EPI_ISL_968260, EPI_ISL_968261, EPI_ISL_968262, EPI_ISL_968263, EPI_ISL_968264, EPI_ISL_968265, EPI_ISL_968266, EPI_ISL_968267, EPI_ISL_968268, EPI_I: EPI_ISL_968273, EPI_ISL_968274, EPI_ISL_968275, EPI_ISL_968276, EPI_ISL_968277, EPI_ISL_968278, EPI_ISL_968279 | | | |
| see above | BCCDC Public Health Laboratory | BCCDC Public Health Laboratory | Prystajecky Natalie, Linda Hoang, Dan Fornika, John Tyson, Shannon Russell, Kim Macdonald, Kimia Kamelian, Ana Pacagnella, Corrinne Ng, Loretta Janz, Robert Azana Terry Snutch, Mel K |
| EPI_ISL_977086 | Massachusetts General | Infectious Disease Program, | Lemieux,J.E., Siddle,K.J., Shaw,B., Adams,G., Pierce,V., Turbett,S., Anahtar,M., Branda,J., Slater,D., Harris,J., Lin,A.E., Gladden-Young,A., Lagerborg,K., Rudy,M., DeRuff,K., Carter,A., Normandin,E., Bauer,M., Reilly,S., Tomkins-Tinch,C., Loreth,C., |

| | | | |
|---|---|---|---|
| | Hospital | Broad Institute of Harvard and MIT | Flowers,K., Cerrato,F., Birren,B.W., Gallagher,G., Smole,S., Park,D.J., MacInnis,B.L., Ryan,E., LaRocque,R., Rosenberg,E. and Sabeti,P.C. |
| EPI_ISL_977279, EPI_ISL_977325, EPI_ISL_977326, EPI_ISL_977335, EPI_ISL_977386, EPI_ISL_977387, EPI_ISL_977388, EPI_ISL_977389, EPI_ISL_977390, EPI_ISL_977391, EPI_ISL_977392, EPI_ISL_977394, EPI_ISL_977395, EPI_ISL_977396, EPI_ISL_977397, EPI_ISL_977398, EPI_ISL_977399 | | | |
| see above | University of Zambia, School of Veterinary Medicine | UNZAVET and PATH | Mulenga Mwenda-Chimfwembe, Ngonda Saasa, Daniel Bridges |
| EPI_ISL_978342, EPI_ISL_978343, EPI_ISL_978344, EPI_ISL_978345, EPI_ISL_978346 | Texas Department of State Health Services | Texas Department of State Health Services | Bonnie Oh, Anita Pokharel, James Daniel Bonser, Myong Koag, Chung Wang, Rachel Lee, Grace Kubin, Rashmi Tuladhar, Mayela Pedrueza, Maliha Rahman, Jenny Zhang |
| EPI_ISL_978488, EPI_ISL_978496 | Central Public Health Laboratory - LACEN -Bahia, Salvador, Brazil | Central Public Health Laboratory - LACEN -Bahia, Salvador, Brazil | Stephane Tosta, Luciana Oliveira, Vanessa Nardy,Patrícia Cajado,Marcela Gómez, Breno Dominguez, Jaqueline Gomes, Vagner Fonseca,Marta Giovanetti,Luiz Alcantara, Felicidade Pereira, Ara |
| EPI_ISL_979271, EPI_ISL_979272 | Cadham Provincial laboratory | National Microbiology Laboratory (NML) | Anna Majer, Shari Tyson, Grace Seo, Philip Mabon, Elsie Grudeski, Rhiannon Huzarewich, Russell Mandes, Anneliese Landgraff, Jennifer Tanner, Natalie Knox, Morag Graham, Gary Van Domselaar, Paul Van Caeseele, Jared Bullard, David Alexander, Madison Chapel, Kirsten Biggar, CanCOGeN's metadata curation team, Public Health Agency of Canada CanCOGeN team |
| EPI_ISL_983624, EPI_ISL_983625, EPI_ISL_983626, EPI_ISL_983627, EPI_ISL_983628, EPI_ISL_983629, EPI_ISL_983630, EPI_ISL_983631 | Texas Department of State Health Services | Texas Department of State Health Services | Bonnie Oh, Anita Pokharel, James Daniel Bonser, Myong Koag, Chung Wang, Rachel Lee, Grace Kubin, Rashmi Tuladhar, Mayela Pedrueza, Maliha Rahman, Jenny Zhang |
