## Supplementary material for "Detection and characterization of the SARS-CoV-2 lineage B.1.526 in New York": Supp. Table 3 GISAID Acknowledgment part 2: gisaid_hcov-19_acknowledgement_table_2021_02_12_22-5.pdf

We gratefully acknowledge the following Authors from the Originating laboratories responsible for obtaining the specimens, as well as the Submitting laboratories where the genome data were generated and shared via GISAID, on which this research is based.

All Submitters of data may be contacted directly via www.gisaid.org

Authors are sorted alphabetically.

| Accession ID | Originating Laboratory | Submitting Laboratory | Authors |
|---|---|---|---|
| EPI_ISL_486842, EPI_ISL_486843, EPI_ISL_486844 | Institute of Microbiology, Universidad San Francisco de Quito | Institute of Microbiology, Universidad San Francisco de Quito | Belén Prado-Vivar, Sully Márquez, Juan José Guadalupe, Monica Becerra-Wong, Carla Torres, Bernardo Gutiérrez, Fausto Maldonado, Geovanny Carzola, Verónica Barragán, Patricio Rojas-Silva, Gabriel Trueba, Michelle Grunauer, Paúl Cárdenas |
| EPI_ISL_487231 | Oklahoma State University Diagnostic Laboratory-Oklahoma Animal Disease Diagnostic Laboratory | Oklahoma Animal Disease Diagnostic Laboratory | Sai Narayanan, Teluguakula Narasaraju, John Ritchey, Girish Patil, Sunil More, Jerry Saliki, Anil Kaul, Akhilesh Ramachandran |
| EPI_ISL_487329, EPI_ISL_487330, EPI_ISL_487331, EPI_ISL_487332, EPI_ISL_487333, EPI_ISL_487334, EPI_ISL_487335, EPI_ISL_487336, EPI_ISL_487337, EPI_ISL_487338, EPI_ISL_487339, EPI_ISL_487340, EPI_ISL_487341 | | | |
| see above | Molecular Diagnostics Services (MDS) | KRISP, KZN Research Innovation and Sequencing Platform | Giandhari J, Pillay S, Lessells R, Chimukangara B, Mdlalose K, York D, Khan S, Tegally H, Wilkinson E, de Oliveira T |
| EPI_ISL_487362 | National Institute of Laboratory Medicine and Referral Center | Genomic Research Lab, BCSIR | Md. Ahasan Habib, Abu Sayeed Mohammad Mahmud, Mohammad Samir Uzzaman, Eshrar Osman, Shahina Akter, Tanjina Akhter Banu, Md. Murshed Hasan Sarkar, Barna Goswami, Iffat Jahan, Md. Saddam Hossain, Tasnim Nafisa, Md. Maruf Ahmed Molla, Mahmuda Yeasmin, Asish Kumar Ghosh, A. K. M. Shamsuzzaman, Sheikh Md. Selim Al Din, Utpal Chandra Ray, Salek Ahmed Sajib, Md. Salim Khan |
| EPI_ISL_487363, EPI_ISL_487364 | National Institute of Laboratory Medicine and Referral Center | Genomic Research Lab, BCSIR | Tanjina Akhter Banu, Abu Sayeed Mohammad Mahmud, Mohammad Samir Uzzaman, Eshrar Osman, Md. Ahasan Habib, Shahina Akter, Md. Murshed Hasan Sarkar, Barna Goswami, Iffat Jahan, Md. Saddam Hossain, Tasnim Nafisa, Md. Maruf Ahmed Molla, Mahmuda Yeasmin, Asish Kumar Ghosh, A. K. M. Shamsuzzaman, Sheikh Md. Selim Al Din, Utpal Chandra Ray, Salek Ahmed Sajib, Md. Salim Khan |
| EPI_ISL_487366, EPI_ISL_487367 | National Institute of Laboratory Medicine and Referral Center | Genomic Research Lab, BCSIR | Shahina Akter, Abu Sayeed Mohammad Mahmud, Mohammad Samir Uzzaman, Eshrar Osman, Md. Ahasan Habib, Tanjina Akhter Banu, Md. Murshed Hasan Sarkar, Barna Goswami, Iffat Jahan, Md. Saddam Hossain, Tasnim Nafisa, Md. Maruf Ahmed Molla, Mahmuda Yeasmin, Asish Kumar Ghosh, A. K. M. Shamsuzzaman, Sheikh Md. Selim Al Din, Utpal Chandra Ray, Salek Ahmed Sajib, Md. Salim Khan |
| EPI_ISL_487368, EPI_ISL_487371 | National Institute of Laboratory Medicine and Referral Center | Genomic Research Lab, BCSIR | Barna Goswami, Abu Sayeed Mohammad Mahmud, Mohammad Samir Uzzaman, Eshrar Osman, Md. Ahasan Habib, Shahina Akter, Tanjina Akhter Banu, Md. Murshed Hasan Sarkar, Iffat Jahan, Md. Saddam Hossain, Tasnim Nafisa, Md. Maruf Ahmed Molla, Mahmuda Yeasmin, Asish Kumar Ghosh, A. K. M. Shamsuzzaman, Sheikh Md. Selim Al Din, Utpal Chandra Ray, Salek Ahmed Sajib, Md. Salim Khan |
| EPI_ISL_487372, EPI_ISL_487373 | National Institute of Laboratory Medicine and Referral Center | Genomic Research Lab, BCSIR | Iffat Jahan, Abu Sayeed Mohammad Mahmud, Mohammad Samir Uzzaman, Eshrar Osman, Md. Ahasan Habib, Shahina Akter, Tanjina Akhter Banu, Md. Murshed Hasan Sarkar, Barna Goswami, Md. Saddam Hossain, Tasnim Nafisa, Md. Maruf Ahmed Molla, Mahmuda Yeasmin, Asish Kumar Ghosh, A. K. M. Shamsuzzaman, Sheikh Md. Selim Al Din, Utpal Chandra Ray, Salek Ahmed Sajib, Md. Salim Khan |
| EPI_ISL_487375, EPI_ISL_487376 | National Institute of Laboratory Medicine and Referral Center | Genomic Research Lab, BCSIR | Md. Saddam Hossain, Abu Sayeed Mohammad Mahmud, Mohammad Samir Uzzaman, Eshrar Osman, Md. Ahasan Habib, Shahina Akter, Tanjina Akhter Banu, Md. Murshed Hasan Sarkar, Barna Goswami, Iffat Jahan, Tasnim Nafisa, Md. Maruf Ahmed Molla, Mahmuda Yeasmin, Asish Kumar Ghosh, A. K. M. Shamsuzzaman, Sheikh Md. Selim Al Din, Utpal Chandra Ray, Salek Ahmed Sajib, Md. Salim Khan |
| EPI_ISL_487378, EPI_ISL_487380, EPI_ISL_487382, EPI_ISL_487383, EPI_ISL_487384, EPI_ISL_487385 | National Institute of Laboratory Medicine and Referral Center | Genomic Research Lab, BCSIR | Md. Murshed Hasan Sarkar, Abu Sayeed Mohammad Mahmud, Mohammad Samir Uzzaman, Eshrar Osman, Md. Ahasan Habib, Shahina Akter, Tanjina Akhter Banu, Barna Goswami, Iffat Jahan, Md. Saddam Hossain, Tasnim Nafisa, Md. Maruf Ahmed Molla, Mahmuda Yeasmin, Asish Kumar Ghosh, A. K. M. Shamsuzzaman, Sheikh Md. Selim Al Din, Utpal Chandra Ray, Salek Ahmed Sajib, Md. Salim Khan |
| EPI_ISL_487386, EPI_ISL_487392, EPI_ISL_487393, EPI_ISL_487394, EPI_ISL_487395, EPI_ISL_487396 | National Institute of Laboratory Medicine and Referral Center | Genomic Research Lab, BCSIR | Abu Sayeed Mohammad Mahmud, Mohammad Samir Uzzaman, Eshrar Osman, Md. Ahasan Habib, Shahina Akter, Tanjina Akhter Banu, Md. Murshed Hasan Sarkar, Barna Goswami, Iffat Jahan, Md. Saddam Hossain, Tasnim Nafisa, Md. Maruf Ahmed Molla, Mahmuda Yeasmin, Asish Kumar Ghosh, A. K. M. Shamsuzzaman, Sheikh Md. Selim Al Din, Utpal Chandra Ray, Salek Ahmed Sajib, Md. Salim Khan |
| EPI_ISL_489897, EPI_ISL_489898, EPI_ISL_489899, EPI_ISL_489902, EPI_ISL_489903, EPI_ISL_489934, EPI_ISL_489936, EPI_ISL_489937, EPI_ISL_489938, EPI_ISL_489939, EPI_ISL_489940, EPI_ISL_489941, EPI_ISL_489942, EPI_ISL_489943, EPI_ISL_489944, EPI_ISL_489945, EPI_ISL_489946, EPI_ISL_489947, EPI_ISL_489948, EPI_ISL_489949, EPI_ISL_489950, EPI_ISL_489951, EPI_ISL_489952, EPI_ISL_489953, EPI_ISL_489954, EPI_ISL_489955 | | | |
| see above | Gundersen Molecular Diagnostics Laboratory | Kabara Cancer Research Institute | Craig S. Richmond, Paraic A. Kenny |
| EPI_ISL_489973, EPI_ISL_489974, EPI_ISL_489975, EPI_ISL_489976, EPI_ISL_489977, EPI_ISL_489978, EPI_ISL_489979, EPI_ISL_489980, EPI_ISL_489981, EPI_ISL_489982 | Viollier AG | Department of Biosystems Science and Engineering, ETH Zürich | Christian Beisel, Sarah Nadeau, Ivan Topolsky, Pedro Ferreira, Philipp Jablonski, Susana Posada-Céspedes, Tobias Schär, Ina Nissen, Natascha Santacroce, Elodie Burcklen, Christiane Beckmann, Maurice Redondo, Olivier Kobel, Christoph Noppen, Sophie Seidel, Noemie Santamaria de Souza, Niko Beerenwinkel, Tanja Stadler |
| EPI_ISL_490038, EPI_ISL_490039 | Pathology West - NSW Health Pathology | NSW Health Pathology - Institute of Clinical Pathology and Medical Research; Westmead Hospital; University of Sydney | CIDM-PH et al. |
| EPI_ISL_490042 | South Eastern Area Laboratory Services (SEALS) | NSW Health Pathology - Institute of Clinical Pathology and Medical Research; Westmead Hospital; University of Sydney | CIDM-PH et al. |
| EPI_ISL_490043 | Pathology North Hunter- NSW Health Pathology | NSW Health Pathology - Institute of Clinical Pathology and Medical Research; Westmead Hospital; University of Sydney | CIDM-PH et al. |
| EPI_ISL_490044 | Pathology West - NSW Health Pathology | NSW Health Pathology - Institute of Clinical Pathology and Medical Research; Westmead Hospital; University of Sydney | CIDM-PH et al. |
| EPI_ISL_490045 | South Eastern Area Laboratory Services (SEALS) | NSW Health Pathology - Institute of Clinical Pathology and Medical Research; Westmead Hospital; University of Sydney | CIDM-PH et al. |
| EPI_ISL_490046 | Pathology West - NSW Health Pathology | NSW Health Pathology - Institute of Clinical Pathology and Medical Research; Westmead Hospital; University of Sydney | CIDM-PH et al. |
| EPI_ISL_490051, EPI_ISL_490055, EPI_ISL_490056, EPI_ISL_490057 | National Public Health Laboratory, National Centre for Infectious Diseases | National Public Health Laboratory, National Centre for Infectious Diseases | Mak TM, Octavia S, Zhou Z, Chavatte JM, Cui L, Lin RTP |
| EPI_ISL_490109, EPI_ISL_490110, EPI_ISL_490111 | National Institute of Laboratory Medicine and Referral Center | Genomic Research Lab, BCSIR | Md. Murshed Hasan Sarkar, Abu Sayeed Mohammad Mahmud, Mohammad Samir Uzzaman, Eshrar Osman, Md. Ahasan Habib, Shahina Akter, Tanjina Akhter Banu, Barna Goswami, Iffat Jahan, Md. Saddam Hossain, Tasnim Nafisa, Md. Maruf Ahmed Molla, Mahmuda Yeasmin, Asish Kumar Ghosh, A. K. M. Shamsuzzaman, Sheikh Md. Selim Al Din, Utpal Chandra Ray, Salek Ahmed Sajib, Md. Salim Khan |
| EPI_ISL_490113 | National Institute of Laboratory Medicine and Referral Center | Genomic Research Lab, BCSIR | Shahina Akter, Abu Sayeed Mohammad Mahmud, Mohammad Samir Uzzaman, Eshrar Osman, Md. Ahasan Habib, Tanjina Akhter Banu, Md. Murshed Hasan Sarkar, Barna Goswami, Iffat Jahan, Md. Saddam Hossain, Tasnim Nafisa, Md. Maruf Ahmed Molla, Mahmuda Yeasmin, Asish Kumar Ghosh, A. K. M. Shamsuzzaman, Sheikh Md. Selim Al Din, Utpal Chandra Ray, Salek Ahmed Sajib, Md. Salim Khan |
| EPI_ISL_490114 | National Institute of Laboratory Medicine and Referral Center | Genomic Research Lab, BCSIR | Tanjina Akhter Banu, Abu Sayeed Mohammad Mahmud, Mohammad Samir Uzzaman, Eshrar Osman, Md. Ahasan Habib, Shahina Akter, Md. Murshed Hasan Sarkar, Barna Goswami, Iffat Jahan, Md. Saddam Hossain, Tasnim Nafisa, Md. Maruf Ahmed Molla, Mahmuda Yeasmin, Asish Kumar Ghosh, A. K. M. Shamsuzzaman, Sheikh Md. Selim Al Din, Utpal Chandra Ray, Salek Ahmed Sajib, Md. Salim Khan |
| EPI_ISL_490144 | National Institute of Laboratory Medicine and Referral Center | Genomic Research Lab, BCSIR | Barna Goswami, Abu Sayeed Mohammad Mahmud, Mohammad Samir Uzzaman, Eshrar Osman, Md. Ahasan Habib, Shahina Akter, Tanjina Akhter Banu, Md. Murshed Hasan Sarkar, Iffat Jahan, Md. Saddam Hossain, Tasnim Nafisa, Md. Maruf Ahmed Molla, Mahmuda Yeasmin, Asish Kumar Ghosh, A. K. M. |

| | | | |
|---|---|---|---|
| | | | Shamsuzzaman, Sheikh Md. Selim Al Din, Utpal Chandra Ray, Salek Ahmed Sajib, Md. Salim Khan |
| EPI_ISL_490164 | National Institute of Laboratory Medicine and Referral Center | Genomic Research Lab, BCSIR | Iffat Jahan, Abu Sayeed Mohammad Mahmud, Mohammad Samir Uzzaman, Eshrar Osman, Md. Ahasan Habib, Shahina Akter, Tanjina Akhter Banu, Md. Murshed Hasan Sarkar, Barna Goswami, Md. Saddam Hossain, Tasnim Nafisa, Md. Maruf Ahmed Molla, Mahmuda Yeasmin, Asish Kumar Ghosh, A. K. M. Shamsuzzaman, Sheikh Md. Selim Al Din, Utpal Chandra Ray, Salek Ahmed Sajib, Md. Salim Khan |
| EPI_ISL_490165 | National Institute of Laboratory Medicine and Referral Center | Genomic Research Lab, BCSIR | Md. Saddam Hossain, Abu Sayeed Mohammad Mahmud, Mohammad Samir Uzzaman, Eshrar Osman, Md. Ahasan Habib, Shahina Akter, Tanjina Akhter Banu, Md. Murshed Hasan Sarkar, Barna Goswami, Iffat Jahan, Tasnim Nafisa, Md. Maruf Ahmed Molla, Mahmuda Yeasmin, Asish Kumar Ghosh, A. K. M. Shamsuzzaman, Sheikh Md. Selim Al Din, Utpal Chandra Ray, Salek Ahmed Sajib, Md. Salim Khan |
| EPI_ISL_490167, EPI_ISL_490168 | National Institute of Laboratory Medicine and Referral Center | Genomic Research Lab, BCSIR | Abu Sayeed Mohammad Mahmud, Mohammad Samir Uzzaman, Eshrar Osman, Md. Ahasan Habib, Shahina Akter, Tanjina Akhter Banu, Md. Murshed Hasan Sarkar, Barna Goswami, Iffat Jahan, Md. Saddam Hossain, Tasnim Nafisa, Md. Maruf Ahmed Molla, Mahmuda Yeasmin, Asish Kumar Ghosh, A. K. M. Shamsuzzaman, Sheikh Md. Selim Al Din, Utpal Chandra Ray, Salek Ahmed Sajib, Md. Salim Khan |
| EPI_ISL_490327, EPI_ISL_490328, EPI_ISL_490329 | Department of Pathology, University of Cambridge | COVID-19 Genomics UK (COG-UK) Consortium | Luke W Meredith, M. Estée Török, Myra Hosmillo, William L. Hamilton, Martin D. Curran, Theresa Feltwell, Grant Hall, Anna Yakovleva, Fahad A Khokhar, Charlotte J. Houldcroft, Laura G Caller, Aminu S. Jahun, Sarah L. Caddy, Yasmin Chaudhry, Malte Pinckert, Ian Goodfellow |
| EPI_ISL_490560, EPI_ISL_490561, EPI_ISL_490562 | Queens Medical Centre, Clinical Microbiology Department / DeepSeq Nottingham | COVID-19 Genomics UK (COG-UK) Consortium | Gemma Clark, Wendy Smith, Manjinder Khakh, Vicki M Fleming, Michelle M Lister, Hannah Howson-Wells, Jonathan Ball, Patrick McClure, Joseph Chappell, Theocharis Tsoleridis, Nadine Holmes, Matthew Carlisle, Christopher Moore, Fei Sang, Johnny Debebe, Victoria Wright, Matthew Loose |
| EPI_ISL_490704, EPI_ISL_490705, EPI_ISL_490706, EPI_ISL_490707, EPI_ISL_490708, EPI_ISL_490709 | West of Scotland Specialist Virology Centre, NHSGGC / MRC-University of Glasgow Centre for Virus Research | COVID-19 Genomics UK (COG-UK) Consortium | Ana da Silva Filipe, Natasha Johnson, Kathy Smollett, Daniel Mair, Stephen Carmichael, Lily Tong, Jenna Nichols, Elihu Aranday-Cortes, Kirstyn Brunker, Yasmin Parr, Alice Broos, Kyriaki Nomikou; Sarah McDonald, Marc Niebel, Patawee Asamaphan; Richard Orton, Joseph Hughes, Sreenu Vattipally, David L Robertson; Alasdair MacLean, Rory Gunson; Kathy Li, Natasha Jesudason, Rajiv Shah, James Shepherd, Antonia Ho, Emma Thomson |
| EPI_ISL_491374, EPI_ISL_491375, EPI_ISL_491378, EPI_ISL_491379, EPI_ISL_491382, EPI_ISL_491385, EPI_ISL_491388, EPI_ISL_491396, EPI_ISL_491399, EPI_ISL_491403, EPI_ISL_491404, EPI_ISL_491406, EPI_ISL_491408, EPI_ISL_491409, EPI_ISL_491410, EPI_ISL_491411, EPI_ISL_491418, EPI_ISL_491419, EPI_ISL_491420, EPI_ISL_491423, EPI_ISL_491424, EPI_ISL_491425, EPI_ISL_491426 | | | |
| see above | University of Wisconsin-Madison AIDS Vaccine Research Laboratories | University of Wisconsin-Madison AIDS Vaccine Research Laboratories | Gage Moreno, Katarina Braun, et al. AIDS Vaccine Research Laboratories |
| EPI_ISL_491722, EPI_ISL_491723, EPI_ISL_491724, EPI_ISL_491725, EPI_ISL_491726, EPI_ISL_491727, EPI_ISL_491728, EPI_ISL_491729, EPI_ISL_491730, EPI_ISL_491731, EPI_ISL_491732, EPI_ISL_491733, EPI_ISL_491734, EPI_ISL_491735 | | | |
| see above | Respiratory Virus Unit, Microbiology Services Colindale, Public Health England | Respiratory Virus Unit, Microbiology Services Colindale, Public Health England | PHE Covid Sequencing Team |
| EPI_ISL_491932, EPI_ISL_491939, EPI_ISL_491940 | Institute of Microbiology, Universidad San Francisco de Quito | Institute of Microbiology, Universidad San Francisco de Quito | Belén Prado-Vivar, Sully Márquez, Juan José Guadalupe, Monica Becerra-Wong, Bernardo Gutiérrez, Carlos Mena, Nabih Dahik, Verónica Barragán, Patricio Rojas-Silva, Gabriel Trueba, Michelle Grunauer, Paúl Cárdenas |
| EPI_ISL_492181, EPI_ISL_492182 | University of Arkansas for Medical Sciences (UAMS) | Department of Biomedical Informatics, University of Arkansas for Medical Sciences (UAMS) | Piroon Jenjaroenpun, David W Ussery, Thidathip Wongsurawat |
| EPI_ISL_492989, EPI_ISL_492990, EPI_ISL_492991, EPI_ISL_492992 | Centrl laboratorija | Latvian Biomedical Research and Study Centre | Ivars Silamielis, Kaspars Megnis, Monta Ustinova, ikita Zrelovs, Vita Rovte, Stella Lapia, Jana Oste, Marta Priedte, Uga Dumpis, Jnis Kloviš |
| EPI_ISL_492993, EPI_ISL_492994, EPI_ISL_492995, EPI_ISL_492996, EPI_ISL_492997, EPI_ISL_492998, EPI_ISL_492999, EPI_ISL_493000 | E. Gulbja Laboratorija | Latvian Biomedical Research and Study Centre | Ivars Silamielis, Kaspars Megnis, Monta Ustinova, ikita Zrelovs, Vita Rovte, Mikus Gavars, Dmitrijs Perminovs, Uga Dumpis, Jnis Kloviš |
| EPI_ISL_493541, EPI_ISL_493542, EPI_ISL_493543, EPI_ISL_493544, EPI_ISL_493545, EPI_ISL_493546 | Quadram Institute Bioscience | COVID-19 Genomics UK (COG-UK) Consortium | Dave J. Baker, Gemma L. Kay, Alp Aydin, Thanh Le-Viet, Steven Rudder, Ana P. Tedim, Anastasia Kolyva, Maria Diaz, Leonardo de Oliveira Martins, Nabil-Fareed Alikhan, Lizzie Meadows, Rachael Stanley, Ngozi Elumogo, Muhammed Yasir, Nicholas M. Thomson, Alexander J Trotter, Rachel Gilroy, Samuel Bloomfield, Claire Stuart, Andrew Bell, Reenesh Prakash, Samir Dervisevic, Alison E. Mather, John Wain, Mark Webber, Andrew J. Page, Justin O'Grady |
| EPI_ISL_493547, EPI_ISL_493548, EPI_ISL_493601, EPI_ISL_493602, EPI_ISL_493606, EPI_ISL_493608, EPI_ISL_493609 | Queens Medical Centre, Clinical Microbiology Department / DeepSeq Nottingham | COVID-19 Genomics UK (COG-UK) Consortium | Gemma Clark, Wendy Smith, Manjinder Khakh, Vicki M Fleming, Michelle M Lister, Hannah Howson-Wells, Jonathan Ball, Patrick McClure, Joseph Chappell, Theocharis Tsoleridis, Nadine Holmes, Matthew Carlisle, Christopher Moore, Fei Sang, Johnny Debebe, Victoria Wright, Matthew Loose |
| EPI_ISL_493666, EPI_ISL_493669, EPI_ISL_493683, EPI_ISL_493693, EPI_ISL_493697, EPI_ISL_493716, EPI_ISL_493735 | Virology Department, Sheffield Teaching Hospitals NHS Foundation Trust/Department of Infection, Immunity and Cardiovascular Disease, The Medical School, University of Sheffield | COVID-19 Genomics UK (COG-UK) Consortium | Thushan de Silva, Matthew Parker, Nikki Smith, Adri Angyal, Rebecca Brown, Luke Green, Rachel Tucker, Paul Parsons, Danielle Groves, Katie Johnson, Laura Carrilero, Alex Keeley, Dave Partridge, Matthew Wyles, Benjamin Lindsey, Mehmet Yavuz, Mohammad Raza, Cariad Evans |
| EPI_ISL_495023 | Government Medical College, Bhavnagar | Gujarat Biotechnology Research Centre | Nitin Savaliya, Raghawendra Kumar, Dinesh Kumar, Zuber Saiyed, Komal Patel, Labdhi Pandya, Afzal Ansari, Nikha Trivedi, Kairavi Desai, Saklin Malek, Shirish Patel, Apurvasinh Puvar, Janvi Raval, Zarna Patel, Monika Gandhi, Pinal Trivedi, Maharshi Pandya, Nidhi Patel, R D Dixit, A M Kadri, Harsh Bakshi, Chaitanya Joshi, Madhvi Joshi |
| EPI_ISL_495024 | Government Medical College, Bhavnagar | Gujarat Biotechnology Research Centre | Raghawendra Kumar, Dinesh Kumar, Zuber Saiyed, Komal Patel, Labdhi Pandya, Afzal Ansari, Nikha Trivedi, Kairavi Desai, Saklin Malek, Shirish Patel, Apurvasinh Puvar, Janvi Raval, Zarna Patel, Monika Gandhi, Pinal Trivedi, Maharshi Pandya, Nidhi Patel, Nitin Savaliya, R D Dixit, A M Kadri, Harsh Bakshi, Chaitanya Joshi, Madhvi Joshi |
| EPI_ISL_495025 | Government Medical College, Bhavnagar | Gujarat Biotechnology Research Centre | Dinesh Kumar, Zuber Saiyed, Komal Patel, Labdhi Pandya, Afzal Ansari, Nikha Trivedi, Kairavi Desai, Saklin Malek, Shirish Patel, Apurvasinh Puvar, Janvi Raval, Zarna Patel, Monika Gandhi, Pinal Trivedi, Maharshi Pandya, Nidhi Patel, Nitin Savaliya, Raghawendra Kumar, R D Dixit, A M Kadri, Harsh Bakshi, Chaitanya Joshi, Madhvi Joshi |
| EPI_ISL_495460, EPI_ISL_495461, EPI_ISL_495462, EPI_ISL_495463, EPI_ISL_495464, EPI_ISL_495465, EPI_ISL_495466, EPI_ISL_495467, EPI_ISL_495468, EPI_ISL_495469, EPI_ISL_495470, EPI_ISL_495471, EPI_ISL_495472, EPI_ISL_495473, EPI_ISL_495474, EPI_ISL_495475, EPI_ISL_495476, EPI_ISL_495477, EPI_ISL_495478, EPI_ISL_495479, EPI_ISL_495480, EPI_ISL_495481, EPI_ISL_495482, EPI_ISL_495483, EPI_ISL_495484, EPI_ISL_495485, EPI_ISL_495486, EPI_ISL_495487, EPI_ISL_495488, EPI_ISL_495489, EPI_ISL_495490, EPI_ISL_495491, EPI_ISL_495492, EPI_ISL_495493, EPI_ISL_495494, EPI_ISL_495495, EPI_ISL_495496, EPI_ISL_495497, EPI_ISL_495498, EPI_ISL_495499, EPI_ISL_495500, EPI_ISL_495501, EPI_ISL_495502, EPI_ISL_495503, EPI_ISL_495504, EPI_ISL_495505, EPI_ISL_495506, EPI_ISL_495507, EPI_ISL_495508, EPI_ISL_495509 | | | |
| see above | University of Wisconsin-Madison AIDS Vaccine Research Laboratories | University of Wisconsin-Madison AIDS Vaccine Research Laboratories | Gage Moreno, Katarina Braun, et al. AIDS Vaccine Research Laboratories |
| EPI_ISL_495535, EPI_ISL_495536, EPI_ISL_495537, EPI_ISL_495538, EPI_ISL_495539, EPI_ISL_495540, EPI_ISL_495541, EPI_ISL_495542 | Medical Disagnostics Services (MDS) | KRISP, KZN Research Innovation and Sequencing Platform | Giandhari J, Pillay S, Lessells R, Chimukangara B, Mdlalose K, York D, Khan S, Tegally H, Wilkinson E, de Oliveira T |
| EPI_ISL_495543, EPI_ISL_495544, EPI_ISL_495545, EPI_ISL_495546, EPI_ISL_495548, EPI_ISL_495549, EPI_ISL_495550, EPI_ISL_495551, EPI_ISL_495552, EPI_ISL_495554, EPI_ISL_495555, EPI_ISL_495556, EPI_ISL_495557, EPI_ISL_495558, EPI_ISL_495559, EPI_ISL_495560, EPI_ISL_495561, EPI_ISL_495562 | | | |
| see above | NHLS-IALCH | KRISP, KZN Research Innovation and Sequencing Platform | Giandhari J, Pillay S, Lessells R, Chimukangara B, Mdlalose K, York D, Khan S, Tegally H, Wilkinson E, de Oliveira T |
| EPI_ISL_496860, EPI_ISL_496861, EPI_ISL_496862, EPI_ISL_496863, EPI_ISL_496864 | Gorgas Memorial Laboratory of Health Studies | Gorgas Memorial Laboratory of Health Studies | Danilo Franco, Claudia Gonzalez Sandra Lopez-Verges, Alexander A Martinez |
| EPI_ISL_497768, EPI_ISL_497774, EPI_ISL_497776, EPI_ISL_497777, EPI_ISL_497779, EPI_ISL_497780, EPI_ISL_497782, EPI_ISL_497793, EPI_ISL_497795, EPI_ISL_497803, EPI_ISL_497807, EPI_ISL_497809, EPI_ISL_497814, EPI_ISL_497816, EPI_ISL_497817, EPI_ISL_497822, EPI_ISL_497825, EPI_ISL_497829, EPI_ISL_497830, EPI_ISL_497835, EPI_ISL_497836, EPI_ISL_497837, EPI_ISL_497841, EPI_ISL_497842, EPI_ISL_497843, EPI_ISL_497844, EPI_ISL_497851, EPI_ISL_497854, EPI_ISL_497855, EPI_ISL_497859, EPI_ISL_497861, EPI_ISL_497862, EPI_ISL_497863, EPI_ISL_497866 | | | |
| see above | Department of Microbiology, The University of Hong Kong | Department of Microbiology, The University of Hong Kong | Kelvin K.W. To, Kwok-Yung Yuen |
| EPI_ISL_498054, EPI_ISL_498055, EPI_ISL_498056, EPI_ISL_498060, EPI_ISL_498061, EPI_ISL_498062, EPI_ISL_498063, EPI_ISL_498064, EPI_ISL_498065, EPI_ISL_498066, EPI_ISL_498067, EPI_ISL_498068, EPI_ISL_498078, EPI_ISL_498081, EPI_ISL_498108, EPI_ISL_498114, EPI_ISL_498115, EPI_ISL_498116, | | | |

| | | | |
|---|---|---|---|
| EPI_ISL_498117, EPI_ISL_498118, EPI_ISL_498119, EPI_ISL_498120, EPI_ISL_498121, EPI_ISL_498122, EPI_ISL_498123, EPI_ISL_498124, EPI_ISL_498125, EPI_ISL_498126 | | | |
| see above | NHLS-IALCH | KRISP, KZN Research Innovation and Sequencing Platform | Giandhari J, Pillay S, Lessells R, Chimukangara B, Mdlalose K, York D, Khan S, Tegally H, Wilkinson E, de Oliveira T |
| EPI_ISL_498149, EPI_ISL_498150, EPI_ISL_498151 | Department of Clinical Microbiology | GIGA Medical Genomics | Keith Durkin, Maria Artesi, Sébastien Bontems, Raphaël Boreux, Cécile Meex, Axelle Chaslain, Céline Fombellida-Lopez, Pierrette Melin, Marie-Pierre Hayette, Vincent Bours. |
| EPI_ISL_498193 | Molecular Pathology Division, Department of Pathology, Hong Kong Sanatorium & Hospital | Molecular Pathology Division, Department of Pathology, Hong Kong Sanatorium & Hospital | Chun Hang AU, Wai Sing CHAN, Ho Yin LAM, Dona N. HO, Simon Y.M. LAM, Jonpaul S.T. ZEE, Tsun Leung CHAN, Edmond S.K. MA |
| EPI_ISL_498251, EPI_ISL_498252 | Institut Pasteur de Dakar | Institut Pasteur de Dakar | Ndongo Dia, Moussa Moise Diagne, Mamadou Diop, Marie Henriette Dior Ndione, Mamadou Malado Jallow, Safietou Sankhe Mbengue, Ousmane Faye, Amadou Alpha Sall. |
| EPI_ISL_498253, EPI_ISL_498254 | National Institute of Laboratory Medicine and Referral Center | Genomic Research Lab, BCSIR | Tanjina Akhter Banu, Abu Sayeed Mohammad Mahmud, Mohammad Samir Uzzaman, Eshrar Osman, Md. Ahasan Habib, Shahina Akter, Md. Murshed Hasan Sarkar, Barna Goswami, Iffat Jahan, Md. Saddam Hossain, Tasnim Nafisa, Md. Maruf Ahmed Molla, Mahmuda Yeasmin, Asish Kumar Ghosh, A. K. M. Shamsuzzaman, Sheikh Md. Selim Al Din, Utpal Chandra Ray, Salek Ahmed Sajib, Md. Salim Khan |
| EPI_ISL_498267 | National Institute of Laboratory Medicine and Referral Center | Genomic Research Lab, BCSIR | Barna Goswami, Abu Sayeed Mohammad Mahmud, Mohammad Samir Uzzaman, Eshrar Osman, Md. Ahasan Habib, Shahina Akter, Tanjina Akhter Banu, Md. Murshed Hasan Sarkar, Iffat Jahan, Md. Saddam Hossain, Tasnim Nafisa, Md. Maruf Ahmed Molla, Mahmuda Yeasmin, Asish Kumar Ghosh, A. K. M. Shamsuzzaman, Sheikh Md. Selim Al Din, Utpal Chandra Ray, Salek Ahmed Sajib, Md. Salim Khan |
| EPI_ISL_498544, EPI_ISL_498545, EPI_ISL_498546, EPI_ISL_498547, EPI_ISL_498548 | ACT Pathology | Schwessinger Lab | Ashley Jones, Benjamin Schwessinger, Robert Lanfear, Robyn N Hall, Megan McDonald, Ming-Dao Chia, Kevin Murray, Craig Kennedy, Karina Kennedy |
| EPI_ISL_498559, EPI_ISL_498560, EPI_ISL_498561, EPI_ISL_498562, EPI_ISL_498563 | Laboratory of Molecular Virology International Center for Genetic Engineering and Biotechnology (ICGEB) | ARGO Open Lab Platform for Genome Sequencing | Licastro D, Rajasekharan S, Dal Monego S, Segat L, D'Agaro P, Marcello A |
| EPI_ISL_498564, EPI_ISL_498565, EPI_ISL_498567, EPI_ISL_498568, EPI_ISL_498571, EPI_ISL_498577, EPI_ISL_498581, EPI_ISL_498582, EPI_ISL_498584, EPI_ISL_498585, EPI_ISL_498587, EPI_ISL_498588, EPI_ISL_498590, EPI_ISL_498591, EPI_ISL_498595, EPI_ISL_498597, EPI_ISL_498598, EPI_ISL_498600 | | | |
| see above | National Public Health Laboratory, National Centre for Infectious Diseases | National Public Health Laboratory, National Centre for Infectious Diseases | Mak TM, Octavia S, Zhou Z, Chavatte JM, Cui L, Lin RTP |
| EPI_ISL_498619, EPI_ISL_498620, EPI_ISL_498621, EPI_ISL_498622, EPI_ISL_498623, EPI_ISL_498624, EPI_ISL_498625, EPI_ISL_498626, EPI_ISL_498627 | Viollier AG | Department of Biosystems Science and Engineering, ETH Zürich | Christian Beisel, Sarah Nadeau, Ivan Topolsky, Pedro Ferreira, Philipp Jablonski, Susana Posada-Céspedes, Tobias Schär, Ina Nissen, Natascha Santacroce, Elodie Burcklen, Christiane Beckmann, Maurice Redondo, Olivier Kobel, Christoph Noppen, Sophie Seidel, Noemie Santamaria de Souza, Niko Beerenwinkel, Tanja Stadler |
| EPI_ISL_498761, EPI_ISL_498762 | Pathology West - NSW Health Pathology | NSW Health Pathology - Institute of Clinical Pathology and Medical Research; Westmead Hospital; University of Sydney | CIDM-PH et al. |
| EPI_ISL_498763 | Sydney South West Pathology Service (SSWPS) - Liverpool Hospital - NSW Health Pathology | NSW Health Pathology - Institute of Clinical Pathology and Medical Research; Westmead Hospital; University of Sydney | CIDM-PH et al. |
| EPI_ISL_498765 | Sydney South West Pathology Service (SSWPS) - Concord Repatriation General Hospital - NSW Health Pathology | NSW Health Pathology - Institute of Clinical Pathology and Medical Research; Westmead Hospital; University of Sydney | CIDM-PH et al. |
| EPI_ISL_498808 | National Institute of Laboratory Medicine and Referral Center | Genomic Research Lab, BCSIR | Abu Sayeed Mohammad Mahmud, Mohammad Samir Uzzaman, Eshrar Osman, Md. Ahasan Habib, Shahina Akter, Tanjina Akhter Banu, Md. Murshed Hasan Sarkar, Barna Goswami, Iffat Jahan, Md. Saddam Hossain, Tasnim Nafisa, Md. Maruf Ahmed Molla, Mahmuda Yeasmin, Asish Kumar Ghosh, A. K. M. Shamsuzzaman, Sheikh Md. Selim Al Din, Utpal Chandra Ray, Salek Ahmed Sajib, Md. Salim Khan |
| EPI_ISL_499357, EPI_ISL_499373, EPI_ISL_499379, EPI_ISL_499388, EPI_ISL_499393, EPI_ISL_499398, EPI_ISL_499404, EPI_ISL_499406, EPI_ISL_499432, EPI_ISL_499441, EPI_ISL_499458 | | | |
| see above | Wales Specialist Virology Centre Sequencing lab: Pathogen Genomics Unit | COVID-19 Genomics UK (COG-UK) Consortium | Catherine Moore, Johnathan Evans, Laura Gifford, Malorie Perry, Simon Cottrell, Angela Marchbank, Alec Birchley, Alexander Adams, Amy Gaskin, Bree Gatica-Wilcox, Jason Coombes, Joel Southgate, Lauren Gilbert, Lee Graham, Nicole Pacchiarini, Sara Kumziene-Summerhayes, Sarah Taylor, Sophie Jones, Sara Rey, Matthew Bull, Joanne Watkins, Sally Corden, Tom Connor |
| EPI_ISL_499493, EPI_ISL_499505, EPI_ISL_499550, EPI_ISL_499551, EPI_ISL_499554, EPI_ISL_499555, EPI_ISL_499559, EPI_ISL_499563, EPI_ISL_499587, EPI_ISL_499596, EPI_ISL_499597, EPI_ISL_499610 | | | |
| see above | Northumbria University / South Tees Hospitals NHS Foundation Trust / North Cumbria Integrated Care NHS Foundation Trust / North Tees and Hartlepool NHS Foundation Trust / Newcastle Hospitals NHS Foundation Trust | COVID-19 Genomics UK (COG-UK) Consortium | Darren L Smith,Andrew Nelson,Matthew Bashton,Greg R Young,Joshua Loh,John Allan,Mohammad A Tariq,Giles S Holt,Gary Black,Wen C Yew,Lynn Dover,Paul Baker,Steve Liggett,Sarah Essex,Jane Greenaway,Debra Padgett,Clive Graham,Garren Scott,Edward Barton,Emma Swindells,Brendan Payne,Jennifer Collins,Yusri Taha,Gary Eltringham |
| EPI_ISL_499691 | Liverpool Clinical Laboratories | COVID-19 Genomics UK (COG-UK) Consortium | Sam Haldenby, Anita Lucaci, Steve Paterson, Julian Hiscox, Alistair Darby, M Almsaud, A Alrezaihi, Muhannad Alruwaili, Stuart D Armstrong, Jones Benjamin, Eleanor G Bentley, Anu Chawla, Jordan J Clark, Angela Cowell, Richard Eccles, Isabel García-Dorival, Matthew Gemmell, Alessandro Gerada, PKF Gilmore, Richard Gregory, Ximeng Han, Catherine Hartley, Margaret Hughes, Miren Iturriza-Gomara, James Johnson, L Luu, Jenifer Manson, Charlotte Nelson, Elaine O'Toole, Cassie Olateju, Rebekah Penrice-Randal , Lucille Rainbow, N.P Randle, Trevor Ian Robinson, Parul Sharma, Ghada T Shawli, James P Stewart, Neil Swainston, Ecaterina Vamos, Joanne Watts, Mark Whitehead |
| EPI_ISL_499782, EPI_ISL_499783 | Northumbria University / South Tees Hospitals NHS Foundation Trust / North Cumbria Integrated Care NHS Foundation Trust / North Tees and Hartlepool NHS Foundation Trust / Newcastle Hospitals NHS Foundation Trust | COVID-19 Genomics UK (COG-UK) Consortium | Darren L Smith,Andrew Nelson,Matthew Bashton,Greg R Young,Joshua Loh,John Allan,Mohammad A Tariq,Giles S Holt,Gary Black,Wen C Yew,Lynn Dover,Paul Baker,Steve Liggett,Sarah Essex,Jane Greenaway,Debra Padgett,Clive Graham,Garren Scott,Edward Barton,Emma Swindells,Brendan Payne,Jennifer Collins,Yusri Taha,Gary Eltringham |
| EPI_ISL_499828 | University Hospitals Of Leicester NHS Trust and DeepSeq Nottingham | COVID-19 Genomics UK (COG-UK) Consortium | Christopher Holmes, Paul Bird, Thomas Helmer, Karlie Fallon, Julian Tang, Jonathan Ball, Patrick McClure, Joeseph Chappell, Nadine Holmes, Matthew Carlisle, Christopher Moore, Fei Sang, Johnny Debebe, Victoria Wright, Matthew Loose |
| EPI_ISL_499854 | Liverpool Clinical Laboratories | COVID-19 Genomics UK (COG-UK) Consortium | Sam Haldenby, Anita Lucaci, Steve Paterson, Julian Hiscox, Alistair Darby, M Almsaud, A Alrezaihi, Muhannad Alruwaili, Stuart D Armstrong, Jones Benjamin, Eleanor G Bentley, Anu Chawla, Jordan J Clark, Angela Cowell, Richard Eccles, Isabel García-Dorival, Matthew Gemmell, Alessandro Gerada, PKF Gilmore, Richard Gregory, Ximeng Han, Catherine Hartley, Margaret Hughes, Miren Iturriza-Gomara, James Johnson, L Luu, Jenifer Manson, Charlotte Nelson, Elaine O'Toole, Cassie Olateju, Rebekah Penrice-Randal , Lucille Rainbow, N.P Randle, Trevor Ian Robinson, Parul Sharma, Ghada T Shawli, James P Stewart, Neil Swainston, Ecaterina Vamos, Joanne Watts, Mark Whitehead |
| EPI_ISL_499855 | Department of Pathology, University of Cambridge | COVID-19 Genomics UK (COG-UK) Consortium | Luke W Meredith, M. Estée Török, Myra Hosmillo, William L. Hamilton, Martin D. Curran, Theresa Feltwell, Grant Hall, Anna Yakovleva, Fahad A Khokhar, Charlotte J. Houldcroft, Laura G Caller, Aminu S. Jahun, Sarah L. Caddy, Yasmin Chaudhry, Malte Pinckert, Ian Goodfellow |
| EPI_ISL_499858, EPI_ISL_499913 | University Hospitals Of Leicester NHS Trust and DeepSeq Nottingham | COVID-19 Genomics UK (COG-UK) Consortium | Christopher Holmes, Paul Bird, Thomas Helmer, Karlie Fallon, Julian Tang, Jonathan Ball, Patrick McClure, Joeseph Chappell, Nadine Holmes, Matthew Carlisle, Christopher Moore, Fei Sang, Johnny Debebe, Victoria Wright, Matthew Loose |
| EPI_ISL_499966, EPI_ISL_499968, EPI_ISL_499970, EPI_ISL_499971, EPI_ISL_499972, EPI_ISL_499973 | Department of Pathology, University of Cambridge | COVID-19 Genomics UK (COG-UK) Consortium | Luke W Meredith, M. Estée Török, Myra Hosmillo, William L. Hamilton, Martin D. Curran, Theresa Feltwell, Grant Hall, Anna Yakovleva, Fahad A Khokhar, Charlotte J. Houldcroft, Laura G Caller, Aminu S. Jahun, Sarah L. Caddy, Yasmin Chaudhry, Malte Pinckert, Ian Goodfellow |
| EPI_ISL_499983 | University Hospitals Of Leicester NHS Trust and DeepSeq Nottingham | COVID-19 Genomics UK (COG-UK) Consortium | Christopher Holmes, Paul Bird, Thomas Helmer, Karlie Fallon, Julian Tang, Jonathan Ball, Patrick McClure, Joeseph Chappell, Nadine Holmes, Matthew Carlisle, Christopher Moore, Fei Sang, Johnny Debebe, Victoria Wright, Matthew Loose |

| | | | |
|---|---|---|---|
| EPI_ISL_500030, EPI_ISL_500031, EPI_ISL_500032, EPI_ISL_500113 | Liverpool Clinical Laboratories | COVID-19 Genomics UK (COG-UK) Consortium | Sam Haldenby, Anita Lucaci, Steve Paterson, Julian Hiscox, Alistair Darby, M Almsaud, A Alrezaihi, Muhannad Alruwaili, Stuart D Armstrong, Jones Benjamin, Eleanor G Bentley, Anu Chawla, Jordan J Clark, Angela Cowell, Richard Eccles, Isabel García-Dorival, Matthew Gemmell, Alessandro Gerada, PKF Gilmore, Richard Gregory, Ximeng Han, Catherine Hartley, Margaret Hughes, Miren Iturriza-Gomara, James Johnson, L Luu, Jenifer Manson, Charlotte Nelson, Elaine O'Toole, Cassie Olateju, Rebekah Penrice-Randal , Lucille Rainbow, N.P Randle, Trevor Ian Robinson, Parul Sharma, Ghada T Shawli, James P Stewart, Neil Swainston, Ecaterina Vamos, Joanne Watts, Mark Whitehead |
| EPI_ISL_500556, EPI_ISL_500557, EPI_ISL_500558, EPI_ISL_500559, EPI_ISL_500560, EPI_ISL_500561 | Singapore General Hospital | Department of Microbiology | Nurdyana Abdul Rahman, Kun Lee Lim, Chenhao Li, Kian Sing Chan, Lynette Oon, Kern Rei Chng, Niranjan Nagarajan, Karrie Ko |
| EPI_ISL_500573, EPI_ISL_500574, EPI_ISL_500575, EPI_ISL_500576, EPI_ISL_500577, EPI_ISL_500578, EPI_ISL_500579, EPI_ISL_500580, EPI_ISL_500581, EPI_ISL_500582, EPI_ISL_500583, EPI_ISL_500584, EPI_ISL_500585, EPI_ISL_500586, EPI_ISL_500587, EPI_ISL_500588, EPI_ISL_500589, EPI_ISL_500590, EPI_ISL_500591, EPI_ISL_500592, EPI_ISL_500593, EPI_ISL_500594, EPI_ISL_500595 | | | |
| see above | National Virus Reference Laboratory | National Virus Reference Laboratory | Michael Carr, Gabriel Gonzalez, Jonathan Dean, Suzie Coughlan, Cillian F De Gascun |
| EPI_ISL_500707, EPI_ISL_500708, EPI_ISL_500709, EPI_ISL_500710, EPI_ISL_500711, EPI_ISL_500712, EPI_ISL_500719 | Respiratory Virus Unit, Microbiology Services Colindale, Public Health England | Respiratory Virus Unit, Microbiology Services Colindale, Public Health England | PHE Covid Sequencing Team |
| EPI_ISL_500769, EPI_ISL_500770, EPI_ISL_500771, EPI_ISL_500772, EPI_ISL_500773, EPI_ISL_500774 | Furst Medical Laboratory | Norwegian Institute of Public Health, Department of Virology | Kathrine Stene-Johansen, Kamilla Heddeland Instefjord, Hilde Elshaug, Rasmus Riis Kopperud, Karoline Bragstad, Olav Hungnes |
| EPI_ISL_500775 | Hospital of Southern Norway - Kristiansand, Department of Medical Microbiology | Norwegian Institute of Public Health, Department of Virology | Kathrine Stene-Johansen, Kamilla Heddeland Instefjord, Hilde Elshaug, Rasmus Riis Kopperud, Karoline Bragstad, Olav Hungnes |
| EPI_ISL_500779, EPI_ISL_500780, EPI_ISL_500781, EPI_ISL_500782, EPI_ISL_500783 | Akershus University Hospital, Department for Microbiology and Infectious Disease Control | Norwegian Institute of Public Health, Department of Virology | Kathrine Stene-Johansen, Kamilla Heddeland Instefjord, Hilde Elshaug, Rasmus Riis Kopperud, Karoline Bragstad, Olav Hungnes |
| EPI_ISL_500784, EPI_ISL_500785, EPI_ISL_500786, EPI_ISL_500787, EPI_ISL_500788, EPI_ISL_500789, EPI_ISL_500790 | Furst Medical Laboratory | Norwegian Institute of Public Health, Department of Virology | Kathrine Stene-Johansen, Kamilla Heddeland Instefjord, Hilde Elshaug, Rasmus Riis Kopperud, Karoline Bragstad, Olav Hungnes |
| EPI_ISL_500791, EPI_ISL_500792 | Hospital of Southern Norway - Kristiansand, Department of Medical Microbiology | Norwegian Institute of Public Health, Department of Virology | Kathrine Stene-Johansen, Kamilla Heddeland Instefjord, Hilde Elshaug, Rasmus Riis Kopperud, Karoline Bragstad, Olav Hungnes |
| EPI_ISL_500793, EPI_ISL_500794, EPI_ISL_500795 | Akershus University Hospital, Department for Microbiology and Infectious Disease Control | Norwegian Institute of Public Health, Department of Virology | Kathrine Stene-Johansen, Kamilla Heddeland Instefjord, Hilde Elshaug, Rasmus Riis Kopperud, Karoline Bragstad, Olav Hungnes |
| EPI_ISL_500800 | Oslo University Hospital, Department of Medical Microbiology | Norwegian Institute of Public Health, Department of Virology | Kathrine Stene-Johansen, Kamilla Heddeland Instefjord, Hilde Elshaug, Rasmus Riis Kopperud, Karoline Bragstad, Olav Hungnes |
| EPI_ISL_500877, EPI_ISL_500878, EPI_ISL_500879, EPI_ISL_500880, EPI_ISL_500881, EPI_ISL_500882, EPI_ISL_500883, EPI_ISL_500884, EPI_ISL_500885, EPI_ISL_500886, EPI_ISL_500887, EPI_ISL_500888, EPI_ISL_500889, EPI_ISL_500890, EPI_ISL_500891, EPI_ISL_500892, EPI_ISL_500893, EPI_ISL_500894, EPI_ISL_500895, EPI_ISL_500896, EPI_ISL_500897, EPI_ISL_500898, EPI_ISL_500899, EPI_ISL_500900, EPI_ISL_500901, EPI_ISL_500902, EPI_ISL_500903, EPI_ISL_500904, EPI_ISL_500905, EPI_ISL_500906, EPI_ISL_500907, EPI_ISL_500908, EPI_ISL_500909, EPI_ISL_500910, EPI_ISL_500911, EPI_ISL_500912, EPI_ISL_500913, EPI_ISL_500914, EPI_ISL_500915, EPI_ISL_500916, EPI_ISL_500917, EPI_ISL_500918, EPI_ISL_500919, EPI_ISL_500920, EPI_ISL_500921, EPI_ISL_500922, EPI_ISL_500923, EPI_ISL_500924, EPI_ISL_500925, EPI_ISL_500926, EPI_ISL_500927, EPI_ISL_500928, EPI_ISL_500929, EPI_ISL_500930, EPI_ISL_500931, EPI_ISL_500932, EPI_ISL_500933, EPI_ISL_500934, EPI_ISL_500935, EPI_ISL_500936, EPI_ISL_500937, EPI_ISL_500938, EPI_ISL_500939, EPI_ISL_500940, EPI_ISL_500941, EPI_ISL_500942, EPI_ISL_500943, EPI_ISL_500944, EPI_ISL_500945 | | | |
| see above | Viollier AG | Department of Biosystems Science and Engineering, ETH Zürich | Christian Beisel, Sarah Nadeau, Ivan Topolsky, Pedro Ferreira, Philipp Jablonski, Susana Posada-Céspedes, Tobias Schär, Ina Nissen, Natascha Santacroce, Elodie Burcklen, Christiane Beckmann, Maurice Redondo, Olivier Kobel, Christoph Noppen, Sophie Seidel, Noemie Santamaria de Souza, Niko Beerenwinkel, Tanja Stadler |
| EPI_ISL_501083, EPI_ISL_501118, EPI_ISL_501120, EPI_ISL_501124, EPI_ISL_501129, EPI_ISL_501130, EPI_ISL_501131, EPI_ISL_501132, EPI_ISL_501133, EPI_ISL_501134, EPI_ISL_501135, EPI_ISL_501136, EPI_ISL_501137, EPI_ISL_501138, EPI_ISL_501139, EPI_ISL_501140, EPI_ISL_501141, EPI_ISL_501142, EPI_ISL_501143, EPI_ISL_501144, EPI_ISL_501145, EPI_ISL_501146, EPI_ISL_501147, EPI_ISL_501148, EPI_ISL_501149, EPI_ISL_501150 | | | |
| see above | University of Washington Virology Lab | University of Washington Virology Lab | Pavitra Roychoudhury, Hong Xie, Lasata Shrestha, Amin Addetia, Truong Nguyen, Victoria M Rachleff, Meei-Li Huang, Keith R Jerome, Alexander Greninger |
| EPI_ISL_501262, EPI_ISL_501263, EPI_ISL_501264, EPI_ISL_501265, EPI_ISL_501266, EPI_ISL_501267, EPI_ISL_501268, EPI_ISL_501269, EPI_ISL_501270 | National Virus Reference Laboratory | National Virus Reference Laboratory | Michael Carr, Gabriel Gonzalez, Jonathan Dean, Suzie Coughlan, Cillian F De Gascun |
| EPI_ISL_501275, EPI_ISL_501284, EPI_ISL_501285 | E. Gulbja Laboratorija | Latvian Biomedical Research and Study Centre | Ivars Silamielis, Kaspars Megnis, Monta Ustinova, ikita Zrelovs, Vita Rovte, Mikus Gavars, Dmitrijs Perminovs, Uga Dumpis, Jnis Kloviš |
| EPI_ISL_501288, EPI_ISL_501289, EPI_ISL_501808, EPI_ISL_501817 | Centrl laboratorija | Latvian Biomedical Research and Study Centre | Ivars Silamielis, Kaspars Megnis, Monta Ustinova, ikita Zrelovs, Vita Rovte, Stella Lapia, Jana Oste, Marta Priedte, Uga Dumpis, Jnis Kloviš |
| EPI_ISL_501929, EPI_ISL_501936 | E. Gulbja Laboratorija | Latvian Biomedical Research and Study Centre | Ivars Silamielis, Kaspars Megnis, Monta Ustinova, ikita Zrelovs, Vita Rovte, Mikus Gavars, Dmitrijs Perminovs, Uga Dumpis, Jnis Kloviš |
| EPI_ISL_503958, EPI_ISL_504040 | National Institute of Laboratory Medicine and Referral Center | Genomic Research Lab, BCSIR | Md. Ahasan Habib, Abu Sayeed Mohammad Mahmud, Mohammad Samir Uzzaman, Eshrar Osman, Shahina Akter, Tanjina Akhter Banu, Md. Murshed Hasan Sarkar, Barna Goswami, Iffat Jahan, Md. Saddam Hossain, Tarannum Taznin, Tasnim Nafisa, Md. Maruf Ahmed Molla, Mahmuda Yeasmin, Asish Kumar Ghosh, A. K. M. Shamsuzzaman, Sheikh Md. Selim Al Din, Utpal Chandra Ray, Salek Ahmed Sajib, Md. Salim Khan |
| EPI_ISL_507418, EPI_ISL_507434, EPI_ISL_507435, EPI_ISL_507441, EPI_ISL_507442, EPI_ISL_507445 | Michigan Department of Health and Human Services, Bureau of Laboratories | Michigan Department of Health and Human Services, Bureau of Laboratories | Blankenship HM, Riner D, Soehnlen MK |
| EPI_ISL_507958, EPI_ISL_507961 | Mayo Clinic & Mayo Clinic Laboratories | Minnesota Department of Health, Public Health Laboratory | Matt Plumb, Jacob Garfin, and Xiong Wang |
| EPI_ISL_507963, EPI_ISL_507964, EPI_ISL_507965, EPI_ISL_507966, EPI_ISL_507967, EPI_ISL_507968, EPI_ISL_507969, EPI_ISL_507970, EPI_ISL_507971, EPI_ISL_507972 | Avera Mckennan Laboratory | Minnesota Department of Health, Public Health Laboratory | Matt Plumb, Jacob Garfin, and Xiong Wang |
| EPI_ISL_508738, EPI_ISL_508739, EPI_ISL_508761, EPI_ISL_508762 | Florida Bureau of Public Health Laboratories | Florida Bureau of Public Health Laboratories | Sarah Schmedes, Jason Blanton |
| EPI_ISL_509141, EPI_ISL_509142, EPI_ISL_509143, EPI_ISL_509144, EPI_ISL_509145, EPI_ISL_509146, EPI_ISL_509147, EPI_ISL_509148, EPI_ISL_509149, EPI_ISL_509150, EPI_ISL_509151, EPI_ISL_509152, EPI_ISL_509153 | | | |
| see above | OHSU Lab Services Molecular Microbiology Lab | Oregon SARS-CoV-2 Genome Sequencing Center | Brendan L. O'Connell, Ruth V. Nichols, Sally B. Grindstaff, Alec J. Hirsch, Guang Fan, Daniel N. Streblow, William B. Messer, Andrew C. Adey, Benjamin N. Bimber, Brian J. O'Roak |
| EPI_ISL_509229, EPI_ISL_509235, EPI_ISL_509247, EPI_ISL_509248, EPI_ISL_509249, EPI_ISL_509251, EPI_ISL_509253, EPI_ISL_509255, EPI_ISL_509256, EPI_ISL_509257, EPI_ISL_509258, EPI_ISL_509259, EPI_ISL_509261, EPI_ISL_509262, EPI_ISL_509263, EPI_ISL_509264, EPI_ISL_509266, EPI_ISL_509267, EPI_ISL_509268, EPI_ISL_509269, EPI_ISL_509270, EPI_ISL_509271, EPI_ISL_509272, EPI_ISL_509273, EPI_ISL_509274, EPI_ISL_509275, EPI_ISL_509276, EPI_ISL_509278, EPI_ISL_509280, EPI_ISL_509281, EPI_ISL_509282, EPI_ISL_509283, EPI_ISL_509284, EPI_ISL_509285, EPI_ISL_509292, EPI_ISL_509305, EPI_ISL_509311, EPI_ISL_509314, EPI_ISL_509315, EPI_ISL_509316, EPI_ISL_509317, EPI_ISL_509318, EPI_ISL_509319, EPI_ISL_509320, EPI_ISL_509321, EPI_ISL_509322, EPI_ISL_509323, EPI_ISL_509329, EPI_ISL_509330, EPI_ISL_509331, EPI_ISL_509332, EPI_ISL_509333, EPI_ISL_509334, EPI_ISL_509336, EPI_ISL_509337, EPI_ISL_509342, EPI_ISL_509344, EPI_ISL_509347, EPI_ISL_509349, EPI_ISL_509350, EPI_ISL_509351, EPI_ISL_509354 | | | |

| | | | |
|---|---|---|---|
| see above | NHLS-IALCH | KRISP, KZN Research Innovation and Sequencing Platform | Giandhari J, Pillay S, Lessells R, Mdlalose K, York D, Tegally H, Wilkinson E, de Oliveira T |
| EPI_ISL_509372, EPI_ISL_509376, EPI_ISL_509390 | Singapore General Hospital | Department of Microbiology | Nurdyana Abdul Rahman, Kun Lee Lim, Chenhao Li, Kian Sing Chan, Lynette Oon, Kern Rei Chng, Niranjan Nagarajan, Karrie Ko |
| EPI_ISL_509521 | Area of Virology, Serology and Virology Division (SAViD), New South Wales Health Pathology Randwick | Area of Virology, Serology and Virology Division (SAViD), New South Wales Health Pathology Randwick | Rawlinson, W. |
| EPI_ISL_509662 | Georgia Department of Health | Pathogen Discovery, Respiratory Viruses Branch, Division of Viral Diseases, Centers for Disease Control and Prevention | Yan Li, Anna Montmayeur, Jing Zhang, Krista Queen, Anna Uehara, Ying Tao, Rachel Marine, Clinton R. Paden, Haibin Wang, Suxiang Tong |
| EPI_ISL_509663, EPI_ISL_509664, EPI_ISL_509665, EPI_ISL_509666, EPI_ISL_509667, EPI_ISL_509668, EPI_ISL_509670, EPI_ISL_509671, EPI_ISL_509672, EPI_ISL_509673, EPI_ISL_509674, EPI_ISL_509675, EPI_ISL_509676, EPI_ISL_509677, EPI_ISL_509678, EPI_ISL_509679, EPI_ISL_509680, EPI_ISL_509681, EPI_ISL_509682 | | | |
| see above | AR Dept. of Health-Public Health Lab | Pathogen Discovery, Respiratory Viruses Branch, Division of Viral Diseases, Centers for Disease Control and Prevention | Yan Li, Anna Montmayeur, Jing Zhang, Krista Queen, Anna Uehara, Ying Tao, Rachel Marine, Clinton R. Paden, Haibin Wang, Suxiang Tong |
| EPI_ISL_509683, EPI_ISL_509684, EPI_ISL_509685 | AR Dept. of Health-Public Health Lab | Pathogen Discovery, Respiratory Viruses Branch, Division of Viral Diseases, Centers for Disease Control and Prevention | Jing Zhang, Yan Li, Anna Montmayeur, Krista Queen, Anna Uehara, Ying Tao, Rachel Marine, Clinton R. Paden, Haibin Wang, Suxiang Tong |
| EPI_ISL_509799, EPI_ISL_509800, EPI_ISL_509808, EPI_ISL_509815, EPI_ISL_509823, EPI_ISL_509852, EPI_ISL_509853, EPI_ISL_509855, EPI_ISL_509856, EPI_ISL_509857, EPI_ISL_509858, EPI_ISL_509859, EPI_ISL_509860, EPI_ISL_509861, EPI_ISL_509862, EPI_ISL_509863, EPI_ISL_509867, EPI_ISL_509868, EPI_ISL_509869, EPI_ISL_509870, EPI_ISL_509871, EPI_ISL_509872, EPI_ISL_509873, EPI_ISL_509874, EPI_ISL_509886, EPI_ISL_509894, EPI_ISL_509906, EPI_ISL_509908, EPI_ISL_509910, EPI_ISL_509911, EPI_ISL_509912, EPI_ISL_509914, EPI_ISL_509915, EPI_ISL_509916, EPI_ISL_509919, EPI_ISL_509924, EPI_ISL_509926, EPI_ISL_509928, EPI_ISL_509929, EPI_ISL_509964, EPI_ISL_509965, EPI_ISL_509966, EPI_ISL_509967, EPI_ISL_509968, EPI_ISL_509969, EPI_ISL_509970, EPI_ISL_509994, EPI_ISL_509995 | | | |
| see above | University of Wisconsin-Madison AIDS Vaccine Research Laboratories | University of Wisconsin-Madison AIDS Vaccine Research Laboratories | Gage Moreno, Katarina Braun, et al. AIDS Vaccine Research Laboratories |
| EPI_ISL_510081, EPI_ISL_510082 | Communicable Disease Branch | Hong Kong Department of Health | Mak Gannon C.K., Lam Edman T.K., Chan Rickjason C.W., Tsang Dominic N.C. |
| EPI_ISL_510085 | Yan Chai Hospital | Hong Kong Department of Health | Mak Gannon C.K., Lam Edman T.K., Chan Rickjason C.W., Tsang Dominic N.C. |
| EPI_ISL_510086 | Kwong Wah Hospital | Hong Kong Department of Health | Mak Gannon C.K., Lam Edman T.K., Chan Rickjason C.W., Tsang Dominic N.C. |
| EPI_ISL_510087 | Private medical practitioner | Hong Kong Department of Health | Mak Gannon C.K., Lam Edman T.K., Chan Rickjason C.W., Tsang Dominic N.C. |
| EPI_ISL_510088 | United Christian Hospital | Hong Kong Department of Health | Mak Gannon C.K., Lam Edman T.K., Chan Rickjason C.W., Tsang Dominic N.C. |
| EPI_ISL_510089 | Queen Elizabeth Hospital | Hong Kong Department of Health | Mak Gannon C.K., Lam Edman T.K., Chan Rickjason C.W., Tsang Dominic N.C. |
| EPI_ISL_510090 | Hong Kong Baptist Hospital | Hong Kong Department of Health | Mak Gannon C.K., Lam Edman T.K., Chan Rickjason C.W., Tsang Dominic N.C. |
| EPI_ISL_510091 | Kwong Wah Hospital | Hong Kong Department of Health | Mak Gannon C.K., Lam Edman T.K., Chan Rickjason C.W., Tsang Dominic N.C. |
| EPI_ISL_510092, EPI_ISL_510093, EPI_ISL_510094, EPI_ISL_510095, EPI_ISL_510096 | Queen Elizabeth Hospital | Hong Kong Department of Health | Mak Gannon C.K., Lam Edman T.K., Chan Rickjason C.W., Tsang Dominic N.C. |
| EPI_ISL_510097, EPI_ISL_510098 | United Christian Hospital | Hong Kong Department of Health | Mak Gannon C.K., Lam Edman T.K., Chan Rickjason C.W., Tsang Dominic N.C. |
| EPI_ISL_510099 | Queen Mary Hospital | Hong Kong Department of Health | Mak Gannon C.K., Lam Edman T.K., Chan Rickjason C.W., Tsang Dominic N.C. |
| EPI_ISL_510100 | Tuen Mun Hospital | Hong Kong Department of Health | Mak Gannon C.K., Lam Edman T.K., Chan Rickjason C.W., Tsang Dominic N.C. |
| EPI_ISL_510101 | Pamela Youde Nethersole Eastern Hospital | Hong Kong Department of Health | Mak Gannon C.K., Lam Edman T.K., Chan Rickjason C.W., Tsang Dominic N.C. |
| EPI_ISL_510439, EPI_ISL_510440, EPI_ISL_510446, EPI_ISL_510450, EPI_ISL_510452, EPI_ISL_510454, EPI_ISL_510455, EPI_ISL_510456, EPI_ISL_510458, EPI_ISL_510459 | Hospital Universitario Virgen de las Nieves de Granada-SAS | SeqCOVID-SPAIN consortium/IBV(CSIC) | Mercedes Pérez Ruiz, Sara Sanbonmatsu Gámez, Irene Pedrosa Corral, José M. Navarro-Marí and SeqCOVID-SPAIN consortium |
| EPI_ISL_510525 | University of Wisconsin-Madison AIDS Vaccine Research Laboratories | University of Wisconsin-Madison AIDS Vaccine Research Laboratories | Gage Moreno, Katarina Braun, et al. AIDS Vaccine Research Laboratories |
| EPI_ISL_510690, EPI_ISL_510691, EPI_ISL_510692, EPI_ISL_510693, EPI_ISL_510694, EPI_ISL_510695, EPI_ISL_510696, EPI_ISL_510697, EPI_ISL_510730, EPI_ISL_510731, EPI_ISL_510732, EPI_ISL_510733, EPI_ISL_510734 | | | |
| see above | Viollier AG | Department of Biosystems Science and Engineering, ETH Zürich | Christian Beisel, Sarah Nadeau, Ivan Topolsky, Pedro Ferreira, Philipp Jablonski, Susana Posada-Céspedes, Tobias Schär, Ina Nissen, Natascha Santacroce, Elodie Burcklen, Christiane Beckmann, Maurice Redondo, Olivier Kobel, Christoph Noppen, Sophie Seidel, Noemie Santamaria de Souza, Niko Beerenwinkel, Tanja Stadler |
| EPI_ISL_511879 | Laboratorium Kesehatan Provinsi Jawa Barat | Molecular Genetics Laboratory-Faculty of Medicine-Universitas Padjadjaran; School of Life Sciences and Technology & School of Pharmacy-Institut Teknologi Bandung; Laboratorium Kesehatan Provinsi Jawa Barat | Marselina Irasonia Tan, Yunia Sribudiani, Catur Riani, Azzania Fibriani, Husna Nugrahapraja, Tarwadi, Ema Rahmawati, Hesti Lina Wiraswati, Lia Faridah, Savira Ekawardhani, Ryan Bayusantika Ristandi, Rifky Waluyajati Rachman, Cut Nur Cinthia Alamanda, Hammam Riza, Soni Solistia Wirawan, Agung Eru Wibowo |
| EPI_ISL_512072 | Saikrishna Hospital, Mehsana | Gujarat Biotechnology Research Centre | Harshadbhai Parmar, Apurvasinh Puvar, Janvi Raval, Zarna Patel, Monika Gandhi, Pinal Trivedi, Maharshi Pandya, Nidhi Patel, Nitin Savaliya, Raghawendra Kumar, Dinesh Kumar, Zuber Saiyed, Komal Patel, Labdhi Pandya, Afzal Ansari, Nikha Trivedi, R D Dixit, A M Kadri, Harsh Bakshi, Chaitanya Joshi, Madhvi Joshi |
| EPI_ISL_512073 | Saikrishna Hospital, Mehsana | Gujarat Biotechnology Research Centre | Apurvasinh Puvar, Janvi Raval, Zarna Patel, Monika Gandhi, Pinal Trivedi, Maharshi Pandya, Nidhi Patel, Nitin Savaliya, Raghawendra Kumar, Dinesh Kumar, Zuber Saiyed, Komal Patel, Labdhi Pandya, Afzal Ansari, Nikha Trivedi, Harshadbhai Parmar, R D Dixit, A M Kadri, Harsh Bakshi, Chaitanya Joshi, Madhvi Joshi |
| EPI_ISL_512074 | Saikrishna Hospital, Mehsana | Gujarat Biotechnology Research Centre | Janvi Raval, Zarna Patel, Monika Gandhi, Pinal Trivedi, Maharshi Pandya, Nidhi Patel, Nitin Savaliya, Raghawendra Kumar, Dinesh Kumar, Zuber Saiyed, Komal Patel, Labdhi Pandya, Afzal Ansari, Nikha Trivedi, Harshadbhai Parmar, Apurvasinh Puvar, R D Dixit, A M Kadri, Harsh Bakshi, Chaitanya Joshi, Madhvi Joshi |
| EPI_ISL_512075 | Dr. RSS Hospital, Modasa | Gujarat Biotechnology Research Centre | Monika Gandhi, Pinal Trivedi, Maharshi Pandya, Nidhi Patel, Nitin Savaliya, Raghawendra Kumar, Dinesh Kumar, Zuber Saiyed, Komal Patel, Labdhi Pandya, Afzal Ansari, Nikha Trivedi, Harsh Chaudhari, Apurvasinh Puvar, Janvi Raval, Zarna Patel, R D Dixit, A M Kadri, Harsh Bakshi, Chaitanya Joshi, Madhvi Joshi |
| EPI_ISL_512076 | Dr. RSS Hospital, Modasa | Gujarat Biotechnology Research Centre | Pinal Trivedi, Maharshi Pandya, Nidhi Patel, Nitin Savaliya, Raghawendra Kumar, Dinesh Kumar, Zuber Saiyed, Komal Patel, Labdhi Pandya, Afzal Ansari, Nikha Trivedi, Harsh Chaudhari, Apurvasinh Puvar, Janvi Raval, Zarna Patel, Monika Gandhi, R D Dixit, A M Kadri, Harsh Bakshi, Chaitanya Joshi, Madhvi Joshi |
| EPI_ISL_512077 | Dr. RSS Hospital, Modasa | Gujarat Biotechnology Research Centre | Maharshi Pandya, Nidhi Patel, Nitin Savaliya, Raghawendra Kumar, Dinesh Kumar, Zuber Saiyed, Komal Patel, Labdhi Pandya, Afzal Ansari, Nikha Trivedi, Harsh Chaudhari, Apurvasinh Puvar, Janvi Raval, Zarna Patel, Monika Gandhi, Pinal Trivedi, R D Dixit, A M Kadri, Harsh Bakshi, Chaitanya Joshi, Madhvi Joshi |
| EPI_ISL_512235, EPI_ISL_512236, EPI_ISL_512237, EPI_ISL_512238, EPI_ISL_512239, EPI_ISL_512240, EPI_ISL_512241, EPI_ISL_512242, EPI_ISL_512243, EPI_ISL_512244, EPI_ISL_512245, EPI_ISL_512248, EPI_ISL_512249, EPI_ISL_512250, EPI_ISL_512251, EPI_ISL_512252, EPI_ISL_512267 | | | |
| see above | San Diego County Public Health Laboratory | Andersen lab at Scripps Research | SEARCH Alliance San Diego with Tracy Basler, Jovan Shephard, Brett Austin |
| EPI_ISL_512314 | E. Gulbja Laboratorija | Latvian Biomedical Research and Study Centre | Ivars Silamielis, Kaspars Megnis, Monta Ustinova, ikita Zrelovs, Vita Rovte, Mikus Gavars, Dmitrijs Perminovs, Uga Dumpis, Jnis Kloviš |
| EPI_ISL_512348, EPI_ISL_512349 | Northumbria University / South Tees Hospitals NHS Foundation Trust / North Cumbria Integrated Care NHS | COVID-19 Genomics UK (COG-UK) Consortium | Darren L Smith,Andrew Nelson,Matthew Bashton,Greg R Young,Joshua Loh,John Allan,Mohammad A Tariq,Giles S Holt,Gary Black,Wen C Yew,Lynn Dover,Paul Baker,Steve Liggett,Sarah Essex,Jane Greenaway,Debra Padgett,Clive Graham,Garren Scott,Edward Barton,Emma Swindells,Brendan |

| | | | |
|---|---|---|---|
| | Foundation Trust / North Tees and Hartlepool NHS Foundation Trust / Newcastle Hospitals NHS Foundation Trust | | Payne,Jennifer Collins,Yusri Taha,Gary Eltringham |
| EPI_ISL_512367 | Quadram Institute Bioscience | COVID-19 Genomics UK (COG-UK) Consortium | Dave J. Baker, Gemma L. Kay, Alp Aydin, Thanh Le-Viet, Steven Rudder, Ana P. Tedim, Anastasia Kolyva, Maria Diaz, Leonardo de Oliveira Martins, Nabil-Fareed Alikhan, Lizzie Meadows, Rachael Stanley, Ngozi Elumogo, Muhammed Yasir, Nicholas M. Thomson, Alexander J Trotter, Rachel Gilroy, Samuel Bloomfield, Claire Stuart, Andrew Bell, Reenesh Prakash, Samir Dervisevic, Alison E. Mather, John Wain, Mark Webber, Andrew J. Page, Justin O'Grady |
| EPI_ISL_512476 | West of Scotland Specialist Virology Centre, NHSGGC / MRC-University of Glasgow Centre for Virus Research | COVID-19 Genomics UK (COG-UK) Consortium | Ana da Silva Filipe, Natasha Johnson, Kathy Smollett, Daniel Mair, Stephen Carmichael, Lily Tong, Jenna Nichols, Elihu Aranday-Cortes, Kirstyn Brunker, Yasmin Parr, Alice Broos, Kyriaki Nomikou; Sarah McDonald, Marc Niebel, Patawee Asamaphan; Richard Orton, Joseph Hughes, Sreenu Vattipally, David L Robertson; Alasdair MacLean, Rory Gunson; Kathy Li, Natasha Jesudason, Rajiv Shah, James Shepherd, Antonia Ho, Emma Thomson |
| EPI_ISL_512487 | Wales Specialist Virology Centre Sequencing lab: Pathogen Genomics Unit | COVID-19 Genomics UK (COG-UK) Consortium | Catherine Moore, Johnathan Evans, Laura Gifford, Malorie Perry, Simon Cottrell, Angela Marchbank, Alec Birchley, Alexander Adams, Amy Gaskin, Bree Gatica-Wilcox, Jason Coombes, Joel Southgate, Lauren Gilbert, Lee Graham, Nicole Pacchiarini, Sara Kumziene-Summerhayes, Sarah Taylor, Sophie Jones, Sara Rey, Matthew Bull, Joanne Watkins, Sally Corden, Tom Connor |
| EPI_ISL_512616, EPI_ISL_512617 | National Laboratory for Influenza/Virology reference laboratory, Public Health Center of the Ministry of Health of Ukraine | Respiratory Virus Unit, Microbiology Services Colindale, Public Health England | PHE Covid Sequencing Team, Dr. Iryna Demchyshyna |
| EPI_ISL_512646 | E. Gulbja Laboratorija | Latvian Biomedical Research and Study Centre | Ivars Silamielis, Kaspars Megnis, Monta Ustinova, ikita Zrelovs, Vita Rovte, Mikus Gavars, Dmitrijs Perminovs, Uga Dumpis, Jnis Kloviš |
| EPI_ISL_512672, EPI_ISL_512673, EPI_ISL_512674, EPI_ISL_512675 | Hospital De Niños Dr. Carlos Saenz Herrera [San Jose/San Jose] | Inciensa, Instituto Costarricense de Investigación y Enseñanza en Nutrición y Salud | Francisco Duarte, Hebleen Porras, Claudio Soto-Garita, Estela Cordero, Adriana Godinez & Melany Calderon |
| EPI_ISL_512722, EPI_ISL_512723, EPI_ISL_512724, EPI_ISL_512725, EPI_ISL_512726, EPI_ISL_512727, EPI_ISL_512728, EPI_ISL_512729, EPI_ISL_512730 | PathWest Laboratory Medicine WA | PathWest Laboratory Medicine WA Microbial Surveillance Unit | PathWest Laboratory Medicine WA Microbial Surveillance Unit |
| EPI_ISL_512774 | Genomic Research Lab, BCSIR | Bangladesh Council of Scientific and Industrial Research | Abu Sayeed Mohammad Mahmud, Mohammad Samir Uzzaman, Eshrar Osman, Md. Ahasan Habib, Shahina Akter, Tanjina Akhter Banu, Md. Murshed Hasan Sarkar, Barna Goswami, Iffat Jahan, Md. Saddam Hossain, Tasnim Nafisa, Md. Maruf Ahmed Molla, Mahmuda Yeasmin, Asish Kumar Ghosh, A. K. M. Shamsuzzaman, Sheikh Md. Selim Al Din, Utpal Chandra Ray, Salek Ahmed Sajib, Md. Salim Khan |
| EPI_ISL_512846 | O.I.J. MORGUE JUDICIAL | Inciensa, Instituto Costarricense de Investigación y Enseñanza en Nutrición y Salud | Francisco Duarte, Hebleen Porras, Claudio Soto-Garita, Estela Cordero, Adriana Godinez & Melany Calderon |
| EPI_ISL_513287, EPI_ISL_513288, EPI_ISL_513289, EPI_ISL_513290, EPI_ISL_513291 | University of Miami Immunology and Histocompatibility Laboratory | University of Miami Immunology and Histocompatibility Laboratory | Emilio Margolles-Clark, PhD and Phillip Ruiz, MD, PhD |
| EPI_ISL_514145, EPI_ISL_514146, EPI_ISL_514210, EPI_ISL_514211, EPI_ISL_514212, EPI_ISL_514213, EPI_ISL_514214, EPI_ISL_514215, EPI_ISL_514216, EPI_ISL_514217, EPI_ISL_514218, EPI_ISL_514219, EPI_ISL_514220, EPI_ISL_514221, EPI_ISL_514222 | | | |
| see above | Florida Bureau of Public Health Laboratories | Florida Bureau of Public Health Laboratories | Sarah Schmedes, Jason Blanton |
| EPI_ISL_514435 | Dr. RSS Hospital, Modasa | Gujarat Biotechnology Research Centre | Harsh Chaudhari, Zarna Patel, Monika Gandhi, Pinal Trivedi, Maharshi Pandya, Nidhi Patel, Nitin Savaliya, Raghawendra Kumar, Dinesh Kumar, Zuber Saiyed, Komal Patel, Labdhi Pandya, Afzal Ansari, Nikha Trivedi, Apurvasinh Puvar, Janvi Raval, R D Dixit, A M Kadri, Harsh Bakshi, Chaitanya Joshi, Madhvi Joshi |
| EPI_ISL_514436 | Dr. RSS Hospital, Modasa | Gujarat Biotechnology Research Centre | Zarna Patel, Monika Gandhi, Pinal Trivedi, Maharshi Pandya, Nidhi Patel, Nitin Savaliya, Raghawendra Kumar, Dinesh Kumar, Zuber Saiyed, Komal Patel, Labdhi Pandya, Afzal Ansari, Nikha Trivedi, Harsh Chaudhari, Apurvasinh Puvar, Janvi Raval, R D Dixit, A M Kadri, Harsh Bakshi, Chaitanya Joshi, Madhvi Joshi |
| EPI_ISL_514437 | Dr. RSS Hospital, Modasa | Gujarat Biotechnology Research Centre | Nidhi Patel, Nitin Savaliya, Raghawendra Kumar, Dinesh Kumar, Zuber Saiyed, Komal Patel, Labdhi Pandya, Afzal Ansari, Nikha Trivedi, Harsh Chaudhari, Apurvasinh Puvar, Janvi Raval, Zarna Patel, Monika Gandhi, Pinal Trivedi, Maharshi Pandya, R D Dixit, A M Kadri, Harsh Bakshi, Chaitanya Joshi, Madhvi Joshi |
| EPI_ISL_514450 | Department of Pathology, University of Cambridge | COVID-19 Genomics UK (COG-UK) Consortium | Luke W Meredith, M. Estée Török, Myra Hosmillo, William L. Hamilton, Martin D. Curran, Theresa Feltwell, Grant Hall, Anna Yakovleva, Fahad A Khokhar, Charlotte J. Houldcroft, Laura G Caller, Aminu S. Jahun, Sarah L. Caddy, Yasmin Chaudhry, Malte Pinckert, Ian Goodfellow |
| EPI_ISL_514636 | Mayo Clinic & Mayo Clinic Laboratories | Minnesota Department of Health, Public Health Laboratory | Matt Plumb, Jacob Garfin, and Xiong Wang |
| EPI_ISL_515005, EPI_ISL_515006, EPI_ISL_515007, EPI_ISL_515008, EPI_ISL_515009, EPI_ISL_515010, EPI_ISL_515011, EPI_ISL_515013, EPI_ISL_515014, EPI_ISL_515015, EPI_ISL_515016, EPI_ISL_515017, EPI_ISL_515018, EPI_ISL_515019, EPI_ISL_515020, EPI_ISL_515021, EPI_ISL_515022, EPI_ISL_515023, EPI_ISL_515024, EPI_ISL_515025, EPI_ISL_515026, EPI_ISL_515027, EPI_ISL_515028, EPI_ISL_515029, EPI_ISL_515030, EPI_ISL_515031, EPI_ISL_515032, EPI_ISL_515033, EPI_ISL_515034, EPI_ISL_515035, EPI_ISL_515036, EPI_ISL_515037, EPI_ISL_515038, EPI_ISL_515039, EPI_ISL_515040, EPI_ISL_515041, EPI_ISL_515042, EPI_ISL_515043, EPI_ISL_515044, EPI_ISL_515045, EPI_ISL_515046, EPI_ISL_515047, EPI_ISL_515048 | | | |
| see above | Division of Viral Diseases, Center for Laboratory Control of Infectious Diseases, Korea Centers for Diseases Control and Prevention | Division of Viral Diseases, Center for Laboratory Control of Infectious Diseases, Korea Centers for Diseases Control and Prevention | Jeong-Min Kim, Yoon-Seok Chung, Namjoo Lee, Sang Hee Woo, Hye-Jun Jo, Heui Man Kim, Jun-Sub Kim, Myung Guk Han |
| EPI_ISL_515058, EPI_ISL_515065, EPI_ISL_515074, EPI_ISL_515080, EPI_ISL_515081 | Department of Clinical Microbiology | GIGA Medical Genomics | Keith Durkin, Maria Artesi, Sebastien Bontems, Raphael Boreux, Cecile Meex, Axelle Chaslain, Celine Fombellida-Lopez, Pierrette Melin, Marie-Pierre Hayette, Vincent Bours. |
| EPI_ISL_515133, EPI_ISL_515134, EPI_ISL_515135, EPI_ISL_515136, EPI_ISL_515137, EPI_ISL_515139, EPI_ISL_515140, EPI_ISL_515141, EPI_ISL_515142, EPI_ISL_515143, EPI_ISL_515144, EPI_ISL_515145, EPI_ISL_515146, EPI_ISL_515147, EPI_ISL_515148, EPI_ISL_515149, EPI_ISL_515150, EPI_ISL_515151, EPI_ISL_515152, EPI_ISL_515153, EPI_ISL_515154, EPI_ISL_515155, EPI_ISL_515156, EPI_ISL_515157, EPI_ISL_515158, EPI_ISL_515159, EPI_ISL_515160, EPI_ISL_515161, EPI_ISL_515162, EPI_ISL_515163, EPI_ISL_515164, EPI_ISL_515165, EPI_ISL_515166, EPI_ISL_515167, EPI_ISL_515168, EPI_ISL_515169, EPI_ISL_515170, EPI_ISL_515171, EPI_ISL_515172, EPI_ISL_515173, EPI_ISL_515174, EPI_ISL_515175, EPI_ISL_515176, EPI_ISL_515177, EPI_ISL_515178, EPI_ISL_515179, EPI_ISL_515180 | | | |
| see above | National Institute for Communicable Diseases of the National Health Laboratory Service | National Institute for Communicable Diseases of the National Health Laboratory Service | Allam M, Ismail A, Khumalo Z, Kwenda S, Mtshali P, Mnyameni F, Mohale T, Bhiman JN |
| EPI_ISL_515568, EPI_ISL_515572, EPI_ISL_515573, EPI_ISL_515574, EPI_ISL_515575, EPI_ISL_515576, EPI_ISL_515577, EPI_ISL_515579, EPI_ISL_515580, EPI_ISL_515581, EPI_ISL_515582, EPI_ISL_515586, EPI_ISL_515587, EPI_ISL_515588, EPI_ISL_515590, EPI_ISL_515591, EPI_ISL_515593, EPI_ISL_515594, EPI_ISL_515598, EPI_ISL_515599, EPI_ISL_515601, EPI_ISL_515608, EPI_ISL_515721 | | | |
| see above | NHLS-IALCH | KRISP, KZN Research Innovation and Sequencing Platform | Giandhari J, Pillay S, Lessells R, Mdlalose K, York D, Khan S, Tegally H, Wilkinson E, de Oliveira T |
| EPI_ISL_515824, EPI_ISL_515825, EPI_ISL_515826, EPI_ISL_515827, EPI_ISL_515828, EPI_ISL_515832, EPI_ISL_515833, EPI_ISL_515834, EPI_ISL_515836, EPI_ISL_515838, EPI_ISL_515839, EPI_ISL_515864, EPI_ISL_515869, EPI_ISL_515871, EPI_ISL_515873 | | | |
| see above | Medical Disagnostics Services (MDS) | KRISP, KZN Research Innovation and Sequencing Platform | Giandhari J, Pillay S, Lessells R, ChimukangaraB, Mdlalose K, York D, Khan S, Tegally H, Wilkinson E, de Oliveira T |
| EPI_ISL_516201, EPI_ISL_516202, EPI_ISL_516382, EPI_ISL_516383, EPI_ISL_516386, EPI_ISL_516389, EPI_ISL_516396, EPI_ISL_516398, EPI_ISL_516400, EPI_ISL_516401, EPI_ISL_516402, EPI_ISL_516404, EPI_ISL_516409, EPI_ISL_516412 | | | |
| see above | Michigan Department of Health and Human Services, Bureau of Laboratories | Michigan Department of Health and Human Services, Bureau of Laboratories | Blankenship HM, Riner D, Soehnlen MK |
| EPI_ISL_516428 | Clinical Hospital - Shtip | Research Center for Genetic Engineering and Biotechnology "Georgi D. Efremov" , Macedonian Academy of Sciences and Arts | RCGEB - MASA |
| EPI_ISL_516611 | Instituto de Diagnostico y Referencia Epidemiologicos (INDRE) | Instituto de Diagnostico y Referencia Epidemiologicos (INDRE) | Gisela Barrera-Badillo , Abril Rodriguez-Maldonado, Claudia Wong-Arambula , Natividad Cruz-Ortiz, Tatiana Nunez-Garcia, Dayanira Arellano-Suarez, Fabiola Garces-Ayala, Edgar Mendieta-Condado, Lucia Hernandez-Rivas, Irma Lopez-Martinez, Ernesto Ramirez-Gonzalez. |

| | | | |
|---|---|---|---|
| EPI_ISL_516612, EPI_ISL_516617 | Instituto de Diagnostico y Referencia Epidemiologicos (INDRE) | Instituto de Diagnostico y Referencia Epidemiologicos (INDRE) | Ernesto Ramirez-Gonzalez, Abril Rodriguez-Maldonado, Claudia Wong-Arambula , Natividad Cruz-Ortiz, Tatiana Nunez-Garcia, Dayanira Arellano-Suarez, Adnan Araiza-Rodriguez, Fabiola Garces-Ayala, Lucia Hernandez-Rivas, Irma Lopez-Martinez, Gisela Barrera-Badillo. |
| EPI_ISL_516723, EPI_ISL_516735, EPI_ISL_516736, EPI_ISL_516737, EPI_ISL_516738, EPI_ISL_516739, EPI_ISL_516740, EPI_ISL_516741, EPI_ISL_516742, EPI_ISL_516743, EPI_ISL_516744 | | | |
| see above | UCSF Clinical Microbiology Laboratory | Chan-Zuckerberg Biohub | CZB Cliahub Consortium |
| EPI_ISL_516755, EPI_ISL_516756, EPI_ISL_516757, EPI_ISL_516758, EPI_ISL_516760, EPI_ISL_516761, EPI_ISL_516762, EPI_ISL_516763, EPI_ISL_516764, EPI_ISL_516765, EPI_ISL_516766, EPI_ISL_516767, EPI_ISL_516768, EPI_ISL_516769, EPI_ISL_516770, EPI_ISL_516771, EPI_ISL_516773, EPI_ISL_516774, EPI_ISL_516776 | | | |
| see above | van Bakel Laboratory, Genetics and Genomics Sciences, Icahn School of Medicine at Mount Sinai | van Bakel Laboratory, Genetics and Genomics Sciences, Icahn School of Medicine at Mount Sinai | Andrew G. Letizia, Irene Ramos, Ajay Obla, Carl Goforth, Dawn Weir, Yongchao Ge, Marcas M. Bamman, Jayeeta Dutta, Ethan Ellis, Luis Estrella, Mary-Catherine George, Ana S. Gonzalez-Reiche, Darnell Graham, Adriana van de Guchte, Ramiro Gutierrez, Franca Jones, Aspasia Kalomoiri, Rhonda Lizewski, Stephen Lizewski, Jan Marayag, Nada Marjanovic, Eugene V. Millar, Venugopalan Nair, German Nudelman, Edgar Nunez, Brian Pike, James Regeimbal, Stas Rirak , Ernesto Santa Ana, Rachel S. Gelernter Sealfon, Robert Sebra, Mark Simons, Alessandra Soares-Schanoski, Michael Termini, Sindhu Vangeti, Carlos Williams, Harm van Bakel, Stuart C. Sealfon |
| EPI_ISL_516809, EPI_ISL_516818, EPI_ISL_516821, EPI_ISL_516822 | National Public Health Laboratory, National Centre for Infectious Diseases | National Public Health Laboratory, National Centre for Infectious Diseases | Mak TM, Octavia S, Zhou Z, Cui L, Lin RTP |
| EPI_ISL_516909, EPI_ISL_516910, EPI_ISL_516911, EPI_ISL_516912 | Israel Central Virology laboratory | Israel Central Virology laboratory | Neta Zuckerman, Efrat Dahan Bucris, Oran Erster, Ella Mendelson, Michal Mandelboim |
| EPI_ISL_517552 | Virology Department, Sheffield Teaching Hospitals NHS Foundation Trust/Department of Infection, Immunity and Cardiovascular Disease, The Medical School, University of Sheffield | COVID-19 Genomics UK (COG-UK) Consortium | Thushan de Silva, Matthew Parker, Nikki Smith, Adri Angyal, Rebecca Brown, Luke Green, Rachel Tucker, Paul Parsons, Danielle Groves, Katie Johnson, Laura Carrilero, Alex Keeley, Dave Partridge, Matthew Wyles, Benjamin Lindsey, Mehmet Yavuz, Mohammad Raza, Cariad Evans |
| EPI_ISL_517656, EPI_ISL_517657 | Academic Hospital Paramaribo | Erasmus Medical Center | Bas Oude Munnink, Dion Gadjadin, Ed Ijzerman, Emmanuelle Munger, Gary Gummels, Ingrid Krishnadath, Lycke Woittiez, Marion Koopmans, Mireille Van de Veer, Princes Wongsowidjojo, Radjesh Ori, Rohma Banwari, Stephen Vreden |
| EPI_ISL_517804, EPI_ISL_517805, EPI_ISL_517806, EPI_ISL_517807, EPI_ISL_517808, EPI_ISL_517809, EPI_ISL_517810, EPI_ISL_517811, EPI_ISL_517814, EPI_ISL_517815, EPI_ISL_517816, EPI_ISL_517817, EPI_ISL_517819, EPI_ISL_517839, EPI_ISL_517840, EPI_ISL_517841, EPI_ISL_517842, EPI_ISL_517843, EPI_ISL_517844, EPI_ISL_517845, EPI_ISL_517846, EPI_ISL_517847, EPI_ISL_517848, EPI_ISL_517849, EPI_ISL_517850, EPI_ISL_517851, EPI_ISL_517852, EPI_ISL_517853, EPI_ISL_517854, EPI_ISL_517855, EPI_ISL_517856, EPI_ISL_517857, EPI_ISL_517858, EPI_ISL_517859, EPI_ISL_517880, EPI_ISL_517881, EPI_ISL_517882, EPI_ISL_517883, EPI_ISL_517884, EPI_ISL_517885, EPI_ISL_517886, EPI_ISL_517887, EPI_ISL_517888, EPI_ISL_517889, EPI_ISL_517890, EPI_ISL_517891, EPI_ISL_517892, EPI_ISL_517895, EPI_ISL_517899 | | | |
| see above | Florida Bureau of Public Health Laboratories | Florida Bureau of Public Health Laboratories | Sarah Schmedes, Jason Blanton |
| EPI_ISL_517996, EPI_ISL_518000, EPI_ISL_518012, EPI_ISL_518017 | Singapore General Hospital | Department of Microbiology | Nurdyana Abdul Rahman, Kun Lee Lim, Chenhao Li, Kian Sing Chan, Lynette Oon, Kern Rei Chng, Niranjan Nagarajan, Karrie Ko |
| EPI_ISL_518033, EPI_ISL_518034, EPI_ISL_518035, EPI_ISL_518036, EPI_ISL_518042, EPI_ISL_518043, EPI_ISL_518044, EPI_ISL_518045, EPI_ISL_518050 | NHLS-IALCH | KRISP, KZN Research Innovation and Sequencing Platform | Giandhari J, Pillay S, Lessells R, Mdlalose K, York D, Khan S, Tegally H, Wilkinson E, de Oliveira T |
| EPI_ISL_518110, EPI_ISL_518121, EPI_ISL_518130 | Victorian Infectious Diseases Reference Laboratory (VIDRL) | VIDRL and MDU-PHL | Caly L., Seemann T., Sait, M., Schultz M., Druce J., Sherry, N. |
| EPI_ISL_518386, EPI_ISL_518472, EPI_ISL_518473, EPI_ISL_518480, EPI_ISL_518481, EPI_ISL_518482, EPI_ISL_518483, EPI_ISL_518484, EPI_ISL_518485, EPI_ISL_518486, EPI_ISL_518487, EPI_ISL_518488, EPI_ISL_518489, EPI_ISL_518490, EPI_ISL_518507, EPI_ISL_518508, EPI_ISL_518509, EPI_ISL_518788, EPI_ISL_518790 | | | |
| see above | Microbiological Diagnostic Unit - Public Health Laboratory (MDU-PHL) | MDU-PHL | Seemann T., Schultz M., Sait, M., Sherry, N. |
| EPI_ISL_518818 | Academic Hospital Paramaribo | Erasmus Medical Center | Bas Oude Munnink, Dion Gadjadin, Ed Ijzerman, Emmanuelle Munger, Gary Gummels, Ingrid Krishnadath, Lycke Woittiez, Marion Koopmans, Mireille Van de Veer, Princes Wongsowidjojo, Radjesh Ori, Rohma Banwari, Stephen Vreden |
| EPI_ISL_519017, EPI_ISL_519023, EPI_ISL_519025, EPI_ISL_519026, EPI_ISL_519027, EPI_ISL_519028 | Victorian Infectious Diseases Reference Laboratory (VIDRL) | VIDRL and MDU-PHL | Caly L., Seemann T., Sait, M., Schultz M., Druce J., Sherry, N. |
| EPI_ISL_519092, EPI_ISL_519337, EPI_ISL_519338, EPI_ISL_519358, EPI_ISL_519361, EPI_ISL_519362, EPI_ISL_519365, EPI_ISL_519367, EPI_ISL_519369 | Microbiological Diagnostic Unit - Public Health Laboratory (MDU-PHL) | MDU-PHL | Seemann T., Schultz M., Sait, M., Sherry, N. |
| EPI_ISL_519385 | Victorian Infectious Diseases Reference Laboratory (VIDRL) | VIDRL and MDU-PHL | Caly L., Seemann T., Sait, M., Schultz M., Druce J., Sherry, N. |
| EPI_ISL_519386, EPI_ISL_519388, EPI_ISL_519389, EPI_ISL_519390, EPI_ISL_519392, EPI_ISL_519394, EPI_ISL_519395, EPI_ISL_519397, EPI_ISL_519398, EPI_ISL_519399, EPI_ISL_519400, EPI_ISL_519401, EPI_ISL_519402, EPI_ISL_519403, EPI_ISL_519404, EPI_ISL_519405, EPI_ISL_519406, EPI_ISL_519407, EPI_ISL_519408, EPI_ISL_519409, EPI_ISL_519410, EPI_ISL_519411, EPI_ISL_519412, EPI_ISL_519417, EPI_ISL_519419, EPI_ISL_519420, EPI_ISL_519421, EPI_ISL_519423, EPI_ISL_519424, EPI_ISL_519425, EPI_ISL_519426, EPI_ISL_519428, EPI_ISL_519429, EPI_ISL_519430, EPI_ISL_519432, EPI_ISL_519433, EPI_ISL_519434, EPI_ISL_519436, EPI_ISL_519437, EPI_ISL_519438, EPI_ISL_519439, EPI_ISL_519440, EPI_ISL_519441, EPI_ISL_519443, EPI_ISL_519446, EPI_ISL_519449, EPI_ISL_519451, EPI_ISL_519453, EPI_ISL_519454, EPI_ISL_519459, EPI_ISL_519507, EPI_ISL_519508, EPI_ISL_519509, EPI_ISL_519512, EPI_ISL_519513 | | | |
| see above | Microbiological Diagnostic Unit - Public Health Laboratory (MDU-PHL) | MDU-PHL | Seemann T., Schultz M., Sait, M., Sherry, N. |
| EPI_ISL_519519 | Victorian Infectious Diseases Reference Laboratory (VIDRL) | VIDRL and MDU-PHL | Caly L., Seemann T., Sait, M., Schultz M., Druce J., Sherry, N. |
| EPI_ISL_519522, EPI_ISL_519564, EPI_ISL_519601, EPI_ISL_519650, EPI_ISL_519821, EPI_ISL_520011, EPI_ISL_520012, EPI_ISL_520013, EPI_ISL_520014, EPI_ISL_520015, EPI_ISL_520165, EPI_ISL_520169, EPI_ISL_520170, EPI_ISL_520171, EPI_ISL_520172, EPI_ISL_520173, EPI_ISL_520174, EPI_ISL_520175, EPI_ISL_520176, EPI_ISL_520177, EPI_ISL_520178, EPI_ISL_520179, EPI_ISL_520180 | | | |
| see above | Microbiological Diagnostic Unit - Public Health Laboratory (MDU-PHL) | MDU-PHL | Seemann T., Schultz M., Sait, M., Sherry, N. |
| EPI_ISL_520249, EPI_ISL_520250, EPI_ISL_520251, EPI_ISL_520252, EPI_ISL_520253, EPI_ISL_520254, EPI_ISL_520255, EPI_ISL_520256, EPI_ISL_520257, EPI_ISL_520258, EPI_ISL_520259, EPI_ISL_520260, EPI_ISL_520261, EPI_ISL_520262, EPI_ISL_520263, EPI_ISL_520264, EPI_ISL_520265, EPI_ISL_520266, EPI_ISL_520267, EPI_ISL_520268, EPI_ISL_520269, EPI_ISL_520270, EPI_ISL_520271, EPI_ISL_520272, EPI_ISL_520273, EPI_ISL_520274, EPI_ISL_520275, EPI_ISL_520276, EPI_ISL_520277, EPI_ISL_520279, EPI_ISL_520280, EPI_ISL_520281, EPI_ISL_520282, EPI_ISL_520283, EPI_ISL_520284, EPI_ISL_520285, EPI_ISL_520286, EPI_ISL_520287, EPI_ISL_520288, EPI_ISL_520289, EPI_ISL_520290, EPI_ISL_520292, EPI_ISL_520293, EPI_ISL_520294, EPI_ISL_520295, EPI_ISL_520296, EPI_ISL_520297, EPI_ISL_520298, EPI_ISL_520299, EPI_ISL_520300, EPI_ISL_520301, EPI_ISL_520302, EPI_ISL_520303, EPI_ISL_520304, EPI_ISL_520305, EPI_ISL_520306, EPI_ISL_520307, EPI_ISL_520309, EPI_ISL_520310, EPI_ISL_520311, EPI_ISL_520312, EPI_ISL_520313, EPI_ISL_520315, EPI_ISL_520316, EPI_ISL_520317, EPI_ISL_520318, EPI_ISL_520319, EPI_ISL_520320, EPI_ISL_520321, EPI_ISL_520322, EPI_ISL_520323, EPI_ISL_520324, EPI_ISL_520325, EPI_ISL_520326, EPI_ISL_520327, EPI_ISL_520328, EPI_ISL_520329, EPI_ISL_520331, EPI_ISL_520332, EPI_ISL_520333, EPI_ISL_520334, EPI_ISL_520335, EPI_ISL_520336, EPI_ISL_520337, EPI_ISL_520338, EPI_ISL_520339, EPI_ISL_520340, EPI_ISL_520341, EPI_ISL_520342, EPI_ISL_520343, EPI_ISL_520344, EPI_ISL_520345, EPI_ISL_520346, EPI_ISL_520347, EPI_ISL_520348, EPI_ISL_520349, EPI_ISL_520350, EPI_ISL_520351, EPI_ISL_520352, EPI_ISL_520353, EPI_ISL_520354, EPI_ISL_520355, EPI_ISL_520356, EPI_ISL_520357, EPI_ISL_520358, EPI_ISL_520359, EPI_ISL_520360, EPI_ISL_520361, EPI_ISL_520362, EPI_ISL_520363, EPI_ISL_520364, EPI_ISL_520365, EPI_ISL_520366, EPI_ISL_520367, EPI_ISL_520368, EPI_ISL_520369, EPI_ISL_520370, EPI_ISL_520372, EPI_ISL_520373, EPI_ISL_520374, EPI_ISL_520375, EPI_ISL_520376, EPI_ISL_520377, EPI_ISL_520378, EPI_ISL_520379, EPI_ISL_520380, EPI_ISL_520381, EPI_ISL_520382, EPI_ISL_520383, EPI_ISL_520384, EPI_ISL_520385, EPI_ISL_520386, EPI_ISL_520387, EPI_ISL_520388, EPI_ISL_520389, EPI_ISL_520390, EPI_ISL_520391 | | | |
| see above | Victorian Infectious Diseases Reference Laboratory (VIDRL) | VIDRL and MDU-PHL | Caly L., Seemann T., Sait, M., Schultz M., Druce J., Sherry, N. |
| EPI_ISL_520394, EPI_ISL_520395, EPI_ISL_520396, EPI_ISL_520397, EPI_ISL_520398, EPI_ISL_520399, EPI_ISL_520401, EPI_ISL_520402, EPI_ISL_520403, EPI_ISL_520404, EPI_ISL_520405, EPI_ISL_520406, EPI_ISL_520407, EPI_ISL_520408, EPI_ISL_520410, EPI_ISL_520411, EPI_ISL_520412, EPI_ISL_520416, EPI_ISL_520420, EPI_ISL_520421, EPI_ISL_520422, EPI_ISL_520434, EPI_ISL_520435, EPI_ISL_520436, EPI_ISL_520447, EPI_ISL_520448, EPI_ISL_520462, EPI_ISL_520463, EPI_ISL_520469, EPI_ISL_520476, EPI_ISL_520478, EPI_ISL_520479, EPI_ISL_520480, EPI_ISL_520483, EPI_ISL_520484, EPI_ISL_520486, EPI_ISL_520490, EPI_ISL_520491, EPI_ISL_520494, EPI_ISL_520499, EPI_ISL_520506, EPI_ISL_520515, EPI_ISL_520534, EPI_ISL_520535, EPI_ISL_520536, EPI_ISL_520537, EPI_ISL_520538, EPI_ISL_520539, EPI_ISL_520540, EPI_ISL_520541, EPI_ISL_520542, EPI_ISL_520550, EPI_ISL_520551, EPI_ISL_520566, EPI_ISL_520569, EPI_ISL_520570, EPI_ISL_520571, EPI_ISL_520572, EPI_ISL_520623, EPI_ISL_520631, EPI_ISL_520632, EPI_ISL_520633, EPI_ISL_520634 | | | |

| | | | |
|---|---|---|---|
| see above | Microbiological Diagnostic Unit - Public Health Laboratory (MDU-PHL) | MDU-PHL | Seemann T., Schultz M., Sait, M., Sherry, N. |
| EPI_ISL_520745, EPI_ISL_520746, EPI_ISL_520747, EPI_ISL_520748, EPI_ISL_520750, EPI_ISL_520751, EPI_ISL_520752, EPI_ISL_520753, EPI_ISL_520754, EPI_ISL_520755, EPI_ISL_520756, EPI_ISL_520757, EPI_ISL_520758, EPI_ISL_520759, EPI_ISL_520760, EPI_ISL_520761, EPI_ISL_520762, EPI_ISL_520763, EPI_ISL_520764 | | | |
| see above | Victorian Infectious Diseases Reference Laboratory (VIDRL) | VIDRL and MDU-PHL | Caly L., Seemann T., Sait, M., Schultz M., Druce J., Sherry, N. |
| EPI_ISL_520765, EPI_ISL_520767, EPI_ISL_520768, EPI_ISL_520769, EPI_ISL_520770, EPI_ISL_520771, EPI_ISL_520773, EPI_ISL_520774, EPI_ISL_520776, EPI_ISL_520777, EPI_ISL_520778, EPI_ISL_520779, EPI_ISL_520780, EPI_ISL_520781, EPI_ISL_520782, EPI_ISL_520783, EPI_ISL_520784, EPI_ISL_520785, EPI_ISL_520786, EPI_ISL_520787, EPI_ISL_520788 | | | |
| see above | Microbiological Diagnostic Unit - Public Health Laboratory (MDU-PHL) | MDU-PHL | Seemann T., Schultz M., Sait, M., Sherry, N. |
| EPI_ISL_520791, EPI_ISL_520792, EPI_ISL_520793, EPI_ISL_520794, EPI_ISL_520795, EPI_ISL_520796, EPI_ISL_520797, EPI_ISL_520798, EPI_ISL_520799, EPI_ISL_520800, EPI_ISL_520801, EPI_ISL_520802, EPI_ISL_520803, EPI_ISL_520804, EPI_ISL_520805, EPI_ISL_520806, EPI_ISL_520807, EPI_ISL_520808, EPI_ISL_520809, EPI_ISL_520810, EPI_ISL_520811, EPI_ISL_520812, EPI_ISL_520813, EPI_ISL_520814, EPI_ISL_520815, EPI_ISL_520816, EPI_ISL_520817, EPI_ISL_520818, EPI_ISL_520819, EPI_ISL_520820, EPI_ISL_520821, EPI_ISL_520822, EPI_ISL_520823, EPI_ISL_520824, EPI_ISL_520825, EPI_ISL_520826, EPI_ISL_520827, EPI_ISL_520828, EPI_ISL_520829, EPI_ISL_520830, EPI_ISL_520831, EPI_ISL_520832, EPI_ISL_520833, EPI_ISL_520834, EPI_ISL_520835, EPI_ISL_520836, EPI_ISL_520837, EPI_ISL_520838, EPI_ISL_520840, EPI_ISL_520841, EPI_ISL_520842, EPI_ISL_520843 | | | |
| see above | Victorian Infectious Diseases Reference Laboratory (VIDRL) | VIDRL and MDU-PHL | Caly L., Seemann T., Sait, M., Schultz M., Druce J., Sherry, N. |
| EPI_ISL_520844, EPI_ISL_520846, EPI_ISL_520848, EPI_ISL_520849, EPI_ISL_520851, EPI_ISL_520852, EPI_ISL_520855, EPI_ISL_520856, EPI_ISL_520858, EPI_ISL_520859, EPI_ISL_520861, EPI_ISL_520862 | | | |
| see above | Microbiological Diagnostic Unit - Public Health Laboratory (MDU-PHL) | MDU-PHL | Seemann T., Schultz M., Sait, M., Sherry, N. |
| EPI_ISL_520864, EPI_ISL_520865, EPI_ISL_520866, EPI_ISL_520867, EPI_ISL_520868, EPI_ISL_520869, EPI_ISL_520870, EPI_ISL_520871, EPI_ISL_520872, EPI_ISL_520873, EPI_ISL_520874, EPI_ISL_520876, EPI_ISL_520877, EPI_ISL_520878, EPI_ISL_520879, EPI_ISL_520880, EPI_ISL_520881, EPI_ISL_520882, EPI_ISL_520883, EPI_ISL_520884, EPI_ISL_520885, EPI_ISL_520886, EPI_ISL_520887, EPI_ISL_520888, EPI_ISL_520889, EPI_ISL_520890, EPI_ISL_520891, EPI_ISL_520892, EPI_ISL_520893, EPI_ISL_520894, EPI_ISL_520895, EPI_ISL_520896, EPI_ISL_520897, EPI_ISL_520898, EPI_ISL_520899, EPI_ISL_520900, EPI_ISL_520901, EPI_ISL_520902, EPI_ISL_520903, EPI_ISL_520904, EPI_ISL_520906, EPI_ISL_520907, EPI_ISL_520908, EPI_ISL_520914, EPI_ISL_520915, EPI_ISL_520916, EPI_ISL_520917, EPI_ISL_520918, EPI_ISL_520919, EPI_ISL_520920, EPI_ISL_520921, EPI_ISL_520922, EPI_ISL_520923, EPI_ISL_520924, EPI_ISL_520925, EPI_ISL_520926, EPI_ISL_520927, EPI_ISL_520928 | | | |
| see above | Victorian Infectious Diseases Reference Laboratory (VIDRL) | VIDRL and MDU-PHL | Caly L., Seemann T., Sait, M., Schultz M., Druce J., Sherry, N. |
| EPI_ISL_520930, EPI_ISL_520931, EPI_ISL_520932, EPI_ISL_520933, EPI_ISL_520934, EPI_ISL_520945, EPI_ISL_520946, EPI_ISL_520947, EPI_ISL_520948, EPI_ISL_520949, EPI_ISL_520950, EPI_ISL_520951, EPI_ISL_520952, EPI_ISL_520953, EPI_ISL_520954, EPI_ISL_520955, EPI_ISL_520956, EPI_ISL_520957, EPI_ISL_520958, EPI_ISL_520959, EPI_ISL_520960, EPI_ISL_520961, EPI_ISL_520962, EPI_ISL_520963, EPI_ISL_520964, EPI_ISL_520965, EPI_ISL_520966, EPI_ISL_520967, EPI_ISL_520968, EPI_ISL_520969, EPI_ISL_520970, EPI_ISL_520971, EPI_ISL_520972, EPI_ISL_520973, EPI_ISL_520974, EPI_ISL_520975, EPI_ISL_520976, EPI_ISL_520977, EPI_ISL_520978, EPI_ISL_520979, EPI_ISL_520980, EPI_ISL_520981, EPI_ISL_520982, EPI_ISL_520984, EPI_ISL_520985, EPI_ISL_520987, EPI_ISL_520988, EPI_ISL_520991, EPI_ISL_520992, EPI_ISL_520993, EPI_ISL_520994, EPI_ISL_520996, EPI_ISL_520997, EPI_ISL_520998, EPI_ISL_520999, EPI_ISL_521000, EPI_ISL_521001, EPI_ISL_521002, EPI_ISL_521003, EPI_ISL_521004, EPI_ISL_521005, EPI_ISL_521006, EPI_ISL_521007, EPI_ISL_521008, EPI_ISL_521009, EPI_ISL_521010, EPI_ISL_521011, EPI_ISL_521012, EPI_ISL_521013, EPI_ISL_521014, EPI_ISL_521015, EPI_ISL_521016, EPI_ISL_521018, EPI_ISL_521019, EPI_ISL_521020, EPI_ISL_521021, EPI_ISL_521022, EPI_ISL_521023, EPI_ISL_521024, EPI_ISL_521025, EPI_ISL_521026, EPI_ISL_521027, EPI_ISL_521028, EPI_ISL_521029, EPI_ISL_521030, EPI_ISL_521031, EPI_ISL_521032, EPI_ISL_521033, EPI_ISL_521034, EPI_ISL_521035, EPI_ISL_521036, EPI_ISL_521037, EPI_ISL_521038, EPI_ISL_521039, EPI_ISL_521040, EPI_ISL_521041, EPI_ISL_521042, EPI_ISL_521043, EPI_ISL_521044, EPI_ISL_521045, EPI_ISL_521046, EPI_ISL_521047, EPI_ISL_521048, EPI_ISL_521049, EPI_ISL_521050, EPI_ISL_521051, EPI_ISL_521052, EPI_ISL_521053, EPI_ISL_521054, EPI_ISL_521055, EPI_ISL_521056, EPI_ISL_521057, EPI_ISL_521058, EPI_ISL_521059, EPI_ISL_521060, EPI_ISL_521061, EPI_ISL_521062, EPI_ISL_521063, EPI_ISL_521064, EPI_ISL_521065, EPI_ISL_521066, EPI_ISL_521067, EPI_ISL_521068, EPI_ISL_521069, EPI_ISL_521070, EPI_ISL_521071, EPI_ISL_521072, EPI_ISL_521073, EPI_ISL_521074, EPI_ISL_521075, EPI_ISL_521076, EPI_ISL_521077, EPI_ISL_521081, EPI_ISL_521083, EPI_ISL_521084, EPI_ISL_521085, EPI_ISL_521086, EPI_ISL_521089, EPI_ISL_521091, EPI_ISL_521095, EPI_ISL_521096 | | | |
| see above | Microbiological Diagnostic Unit - Public Health Laboratory (MDU-PHL) | MDU-PHL | Seemann T., Schultz M., Sait, M., Sherry, N. |
| EPI_ISL_521102, EPI_ISL_521103, EPI_ISL_521104, EPI_ISL_521105, EPI_ISL_521106, EPI_ISL_521107, EPI_ISL_521108, EPI_ISL_521109, EPI_ISL_521110, EPI_ISL_521111, EPI_ISL_521112, EPI_ISL_521113, EPI_ISL_521114, EPI_ISL_521115, EPI_ISL_521116, EPI_ISL_521117, EPI_ISL_521118, EPI_ISL_521119, EPI_ISL_521120, EPI_ISL_521121, EPI_ISL_521122, EPI_ISL_521123, EPI_ISL_521124, EPI_ISL_521125, EPI_ISL_521126, EPI_ISL_521127, EPI_ISL_521128, EPI_ISL_521129, EPI_ISL_521130, EPI_ISL_521131, EPI_ISL_521132, EPI_ISL_521133, EPI_ISL_521134, EPI_ISL_521135, EPI_ISL_521136, EPI_ISL_521137, EPI_ISL_521138, EPI_ISL_521139, EPI_ISL_521140, EPI_ISL_521141, EPI_ISL_521142, EPI_ISL_521143, EPI_ISL_521144, EPI_ISL_521145, EPI_ISL_521146, EPI_ISL_521147, EPI_ISL_521148, EPI_ISL_521149, EPI_ISL_521150, EPI_ISL_521151, EPI_ISL_521152 | | | |
| see above | Victorian Infectious Diseases Reference Laboratory (VIDRL) | VIDRL and MDU-PHL | Caly L., Seemann T., Sait, M., Schultz M., Druce J., Sherry, N. |
| EPI_ISL_521451, EPI_ISL_521452, EPI_ISL_521453, EPI_ISL_521454, EPI_ISL_521455, EPI_ISL_521456, EPI_ISL_521457, EPI_ISL_521458 | Microbiological Diagnostic Unit - Public Health Laboratory (MDU-PHL) | MDU-PHL | Seemann T., Schultz M., Sait, M., Sherry, N. |
| EPI_ISL_521476, EPI_ISL_521480, EPI_ISL_521494, EPI_ISL_521495, EPI_ISL_521496, EPI_ISL_521497, EPI_ISL_521498, EPI_ISL_521499, EPI_ISL_521500, EPI_ISL_521501, EPI_ISL_521505, EPI_ISL_521507, EPI_ISL_521509, EPI_ISL_521510, EPI_ISL_521511, EPI_ISL_521512, EPI_ISL_521513, EPI_ISL_521514, EPI_ISL_521515, EPI_ISL_521516, EPI_ISL_521518, EPI_ISL_521525, EPI_ISL_521526, EPI_ISL_521527, EPI_ISL_521528, EPI_ISL_521529, EPI_ISL_521530, EPI_ISL_521531, EPI_ISL_521532, EPI_ISL_521533, EPI_ISL_521534, EPI_ISL_521535, EPI_ISL_521536, EPI_ISL_521537, EPI_ISL_521538, EPI_ISL_521539, EPI_ISL_521540, EPI_ISL_521541, EPI_ISL_521542, EPI_ISL_521543, EPI_ISL_521544, EPI_ISL_521545, EPI_ISL_521546, EPI_ISL_521547, EPI_ISL_521548, EPI_ISL_521549, EPI_ISL_521550, EPI_ISL_521551, EPI_ISL_521552, EPI_ISL_521553, EPI_ISL_521554, EPI_ISL_521555, EPI_ISL_521556, EPI_ISL_521557, EPI_ISL_521558, EPI_ISL_521559, EPI_ISL_521560, EPI_ISL_521561 | | | |
| see above | Victorian Infectious Diseases Reference Laboratory (VIDRL) | VIDRL and MDU-PHL | Caly L., Seemann T., Sait, M., Schultz M., Druce J., Sherry, N. |
| EPI_ISL_521562, EPI_ISL_521563, EPI_ISL_521564, EPI_ISL_521565, EPI_ISL_521566, EPI_ISL_521567, EPI_ISL_521568, EPI_ISL_521569, EPI_ISL_521570, EPI_ISL_521571, EPI_ISL_521572, EPI_ISL_521573, EPI_ISL_521574, EPI_ISL_521575, EPI_ISL_521576, EPI_ISL_521577, EPI_ISL_521578, EPI_ISL_521579, EPI_ISL_521580, EPI_ISL_521581, EPI_ISL_521582, EPI_ISL_521583, EPI_ISL_521585, EPI_ISL_521586, EPI_ISL_521587, EPI_ISL_521588, EPI_ISL_521589, EPI_ISL_521590, EPI_ISL_521591, EPI_ISL_521592, EPI_ISL_521593, EPI_ISL_521594, EPI_ISL_521595, EPI_ISL_521596, EPI_ISL_521597, EPI_ISL_521598, EPI_ISL_521599, EPI_ISL_521600, EPI_ISL_521601, EPI_ISL_521602, EPI_ISL_521603, EPI_ISL_521604, EPI_ISL_521605, EPI_ISL_521606, EPI_ISL_521607, EPI_ISL_521608, EPI_ISL_521609, EPI_ISL_521610, EPI_ISL_521611, EPI_ISL_521612, EPI_ISL_521613, EPI_ISL_521614, EPI_ISL_521615, EPI_ISL_521616, EPI_ISL_521617, EPI_ISL_521618, EPI_ISL_521619, EPI_ISL_521620, EPI_ISL_521621, EPI_ISL_521622, EPI_ISL_521623, EPI_ISL_521624, EPI_ISL_521625, EPI_ISL_521626, EPI_ISL_521627, EPI_ISL_521628, EPI_ISL_521629, EPI_ISL_521630, EPI_ISL_521631, EPI_ISL_521632, EPI_ISL_521634, EPI_ISL_521635, EPI_ISL_521636, EPI_ISL_521637, EPI_ISL_521638, EPI_ISL_521639, EPI_ISL_521640, EPI_ISL_521641, EPI_ISL_521642, EPI_ISL_521643, EPI_ISL_521644, EPI_ISL_521646, EPI_ISL_521647, EPI_ISL_521648, EPI_ISL_521649, EPI_ISL_521650, EPI_ISL_521651, EPI_ISL_521652, EPI_ISL_521654, EPI_ISL_521655, EPI_ISL_521656, EPI_ISL_521657, EPI_ISL_521658, EPI_ISL_521659, EPI_ISL_521660, EPI_ISL_521661, EPI_ISL_521662, EPI_ISL_521663, EPI_ISL_521664, EPI_ISL_521665, EPI_ISL_521666, EPI_ISL_521667, EPI_ISL_521668, EPI_ISL_521669, EPI_ISL_521670, EPI_ISL_521671, EPI_ISL_521673, EPI_ISL_521674, EPI_ISL_521675, EPI_ISL_521676, EPI_ISL_521677, EPI_ISL_521678, EPI_ISL_521679, EPI_ISL_521680, EPI_ISL_521681, EPI_ISL_521682, EPI_ISL_521683, EPI_ISL_521684, EPI_ISL_521685, EPI_ISL_521686, EPI_ISL_521687, EPI_ISL_521688, EPI_ISL_521689, EPI_ISL_521690, EPI_ISL_521691, EPI_ISL_521692, EPI_ISL_521693, EPI_ISL_521694, EPI_ISL_521695, EPI_ISL_521696, EPI_ISL_521697, EPI_ISL_521698, EPI_ISL_521699, EPI_ISL_521700, EPI_ISL_521701, EPI_ISL_521702, EPI_ISL_521703, EPI_ISL_521704, EPI_ISL_521705, EPI_ISL_521706, EPI_ISL_521707, EPI_ISL_521708, EPI_ISL_521709, EPI_ISL_521710, EPI_ISL_521712, EPI_ISL_521715, EPI_ISL_521716, EPI_ISL_521717, EPI_ISL_521718, EPI_ISL_521719, EPI_ISL_521720, EPI_ISL_521721, EPI_ISL_521722, EPI_ISL_521723, EPI_ISL_521724, EPI_ISL_521725, EPI_ISL_521726, EPI_ISL_521727, EPI_ISL_521728, EPI_ISL_521729, EPI_ISL_521730, EPI_ISL_521731, EPI_ISL_521732, EPI_ISL_521733, EPI_ISL_521734, EPI_ISL_521735, EPI_ISL_521736, EPI_ISL_521737, EPI_ISL_521738, EPI_ISL_521739, EPI_ISL_521740, EPI_ISL_521741, EPI_ISL_521742, EPI_ISL_521743, EPI_ISL_521744, EPI_ISL_521745, EPI_ISL_521746, EPI_ISL_521747, EPI_ISL_521748, EPI_ISL_521749, EPI_ISL_521750, EPI_ISL_521751, EPI_ISL_521752, EPI_ISL_521753, EPI_ISL_521754, EPI_ISL_521755, EPI_ISL_521756, EPI_ISL_521757, EPI_ISL_521758, EPI_ISL_521759, EPI_ISL_521760, EPI_ISL_521761, EPI_ISL_521762, EPI_ISL_521763, EPI_ISL_521764, EPI_ISL_521765, EPI_ISL_521766, EPI_ISL_521767, EPI_ISL_521768, EPI_ISL_521769, EPI_ISL_521770, EPI_ISL_521771, EPI_ISL_521772, EPI_ISL_521773, EPI_ISL_521775, EPI_ISL_521776, EPI_ISL_521777, EPI_ISL_521778, EPI_ISL_521779, EPI_ISL_521780, EPI_ISL_521781, EPI_ISL_521782, EPI_ISL_521783, EPI_ISL_521784, EPI_ISL_521785, EPI_ISL_521786, EPI_ISL_521787, EPI_ISL_521788, EPI_ISL_521790, EPI_ISL_521792, EPI_ISL_521793, EPI_ISL_521794, EPI_ISL_521795, EPI_ISL_521796, EPI_ISL_521798, EPI_ISL_521799, EPI_ISL_521800, EPI_ISL_521801, EPI_ISL_521802, EPI_ISL_521803, EPI_ISL_521804, EPI_ISL_521805, EPI_ISL_521806, EPI_ISL_521807, EPI_ISL_521808, EPI_ISL_521809, EPI_ISL_521810, EPI_ISL_521811, EPI_ISL_521812, EPI_ISL_521813, EPI_ISL_521814, EPI_ISL_521815, EPI_ISL_521816, EPI_ISL_521818, EPI_ISL_521819, EPI_ISL_521820, EPI_ISL_521822, EPI_ISL_521824, EPI_ISL_521826, EPI_ISL_521828, EPI_ISL_521829, EPI_ISL_521830, EPI_ISL_521831, EPI_ISL_521832, EPI_ISL_521833, EPI_ISL_521834, EPI_ISL_521835, EPI_ISL_521837, EPI_ISL_521838, EPI_ISL_521839 | | | |
| see above | Microbiological Diagnostic Unit - Public Health Laboratory (MDU-PHL) | MDU-PHL | Seemann T., Schultz M., Sait, M., Sherry, N. |
| EPI_ISL_521840, EPI_ISL_521841, EPI_ISL_521842, EPI_ISL_521843, EPI_ISL_521844, EPI_ISL_521845, EPI_ISL_521846, EPI_ISL_521847, EPI_ISL_521848, EPI_ISL_521849, EPI_ISL_521850, EPI_ISL_521851, EPI_ISL_521852, EPI_ISL_521854, EPI_ISL_521856, EPI_ISL_521857, EPI_ISL_521858, EPI_ISL_521859 | | | |
| see above | Victorian Infectious Diseases Reference Laboratory (VIDRL) | VIDRL and MDU-PHL | Caly L., Seemann T., Sait, M., Schultz M., Druce J., Sherry, N. |
| EPI_ISL_521860, EPI_ISL_521861 | Royal Darwin Hospital Pathology | MDU-PHL | Meumann, E., Caly L., Seemann T., Sait, M., Schultz M., Druce J., Sherry, N. |
| EPI_ISL_522224, EPI_ISL_522227, EPI_ISL_522228, EPI_ISL_522229, EPI_ISL_522232, EPI_ISL_522245, EPI_ISL_522246, EPI_ISL_522247, EPI_ISL_522248, EPI_ISL_522249, EPI_ISL_522250, EPI_ISL_522251, EPI_ISL_522252, EPI_ISL_522253 | | | |
| see above | Microbiological Diagnostic Unit - Public Health Laboratory (MDU-PHL) | MDU-PHL | Seemann T., Schultz M., Sait, M., Sherry, N. |
| EPI_ISL_522469, EPI_ISL_522470, EPI_ISL_522471, EPI_ISL_522472, EPI_ISL_522473 | Center for Laboratory Control of Infectious Diseases, Korea Centers for Diseases Control and Prevention | Center for Laboratory Control of Infectious Diseases, Korea Centers for Diseases Control and Prevention | Junyoung Kim, Ae Kyung Park, Eunkyung Shin, Jin Sun No, Jeong-Min Kim, Yoon-Seok Chung, Heui Man Kim, Myung Guk Han |
| EPI_ISL_522474, EPI_ISL_522475 | Division of Viral Diseases, Center for Laboratory Control of Infectious Diseases, Korea Centers for Diseases Control and Prevention | Division of Viral Diseases, Center for Laboratory Control of Infectious Diseases, Korea Centers for Diseases Control and Prevention | Jeong-Min Kim, Yoon-Seok Chung, Namjoo Lee, Sang Hee Woo, Hye-Jun Jo, Heui Man Kim, Jun-Sub Kim, Myung Guk Han |
| EPI_ISL_522476 | Center for Laboratory Control of Infectious Diseases, Korea | Center for Laboratory Control of Infectious Diseases, Korea | Junyoung Kim, Ae Kyung Park, Eunkyung Shin, Jin Sun No, Jeong-Min Kim, Yoon-Seok Chung, Heui Man Kim, Myung Guk Han |

| | | | |
|---|---|---|---|
| | Centers for Diseases Control and Prevention | Centers for Diseases Control and Prevention | |
| EPI_ISL_522477, EPI_ISL_522478 | Division of Viral Diseases, Center for Laboratory Control of Infectious Diseases, Korea Centers for Diseases Control and Prevention | Division of Viral Diseases, Center for Laboratory Control of Infectious Diseases, Korea Centers for Diseases Control and Prevention | Jeong-Min Kim, Yoon-Seok Chung, Namjoo Lee, Sang Hee Woo, Hye-Jun Jo, Heui Man Kim, Jun-Sub Kim, Myung Guk Han |
| EPI_ISL_522479, EPI_ISL_522480, EPI_ISL_522481, EPI_ISL_522482, EPI_ISL_522483 | Center for Laboratory Control of Infectious Diseases, Korea Centers for Diseases Control and Prevention | Center for Laboratory Control of Infectious Diseases, Korea Centers for Diseases Control and Prevention | Junyoung Kim, Ae Kyung Park, Eunkyung Shin, Jin Sun No, Jeong-Min Kim, Yoon-Seok Chung, Heui Man Kim, Myung Guk Han |
| EPI_ISL_522484 | Division of Viral Diseases, Center for Laboratory Control of Infectious Diseases, Korea Centers for Diseases Control and Prevention | Division of Viral Diseases, Center for Laboratory Control of Infectious Diseases, Korea Centers for Diseases Control and Prevention | Jeong-Min Kim, Yoon-Seok Chung, Namjoo Lee, Sang Hee Woo, Hye-Jun Jo, Heui Man Kim, Jun-Sub Kim, Myung Guk Han |
| EPI_ISL_522485 | Center for Laboratory Control of Infectious Diseases, Korea Centers for Diseases Control and Prevention | Center for Laboratory Control of Infectious Diseases, Korea Centers for Diseases Control and Prevention | Junyoung Kim, Ae Kyung Park, Eunkyung Shin, Jin Sun No, Jeong-Min Kim, Yoon-Seok Chung, Heui Man Kim, Myung Guk Han |
| EPI_ISL_522486, EPI_ISL_522487, EPI_ISL_522488, EPI_ISL_522489, EPI_ISL_522490 | Division of Viral Diseases, Center for Laboratory Control of Infectious Diseases, Korea Centers for Diseases Control and Prevention | Division of Viral Diseases, Center for Laboratory Control of Infectious Diseases, Korea Centers for Diseases Control and Prevention | Jeong-Min Kim, Yoon-Seok Chung, Namjoo Lee, Sang Hee Woo, Hye-Jun Jo, Heui Man Kim, Jun-Sub Kim, Myung Guk Han |
| EPI_ISL_522806, EPI_ISL_522807, EPI_ISL_522808, EPI_ISL_522809, EPI_ISL_522817, EPI_ISL_522818, EPI_ISL_522819 | Virginia DCLS | Virginia DCLS | Virginia DCLS |
| EPI_ISL_523035, EPI_ISL_523036, EPI_ISL_523037, EPI_ISL_523038, EPI_ISL_523039, EPI_ISL_523041, EPI_ISL_523042, EPI_ISL_523043, EPI_ISL_523044, EPI_ISL_523045, EPI_ISL_523047, EPI_ISL_523048, EPI_ISL_523049, EPI_ISL_523050, EPI_ISL_523051, EPI_ISL_523052, EPI_ISL_523053, EPI_ISL_523054, EPI_ISL_523055, EPI_ISL_523056, EPI_ISL_523057, EPI_ISL_523058, EPI_ISL_523059, EPI_ISL_523060, EPI_ISL_523061, EPI_ISL_523062, EPI_ISL_523063, EPI_ISL_523064, EPI_ISL_523065, EPI_ISL_523066, EPI_ISL_523067, EPI_ISL_523069, EPI_ISL_523071 | | | |
| see above | Dutch COVID-19 response team | Erasmus Medical Center | OH consortium |
| EPI_ISL_523121, EPI_ISL_523271, EPI_ISL_523272, EPI_ISL_523273, EPI_ISL_523345, EPI_ISL_523346, EPI_ISL_523347, EPI_ISL_523348, EPI_ISL_523351, EPI_ISL_523372, EPI_ISL_523380, EPI_ISL_523381, EPI_ISL_523416, EPI_ISL_523417, EPI_ISL_523418, EPI_ISL_523419, EPI_ISL_523464, EPI_ISL_523465, EPI_ISL_523511, EPI_ISL_523512, EPI_ISL_523513, EPI_ISL_523514, EPI_ISL_523515, EPI_ISL_523610, EPI_ISL_523611, EPI_ISL_523635, EPI_ISL_523636, EPI_ISL_523637, EPI_ISL_523638, EPI_ISL_523639, EPI_ISL_523640, EPI_ISL_523641, EPI_ISL_523642, EPI_ISL_523643, EPI_ISL_523644, EPI_ISL_523645, EPI_ISL_523646, EPI_ISL_523647, EPI_ISL_523648, EPI_ISL_523649, EPI_ISL_523659, EPI_ISL_523660, EPI_ISL_523661, EPI_ISL_523670, EPI_ISL_523671 | | | |
| see above | Dutch COVID-19 response team | Erasmus Medical Center | Bas Oude Munnink, David Nieuwenhuijse, Reina Sikkema, Claudia Schapendonk, Irina Chestakova, Anne van der Linden, Theo Bestebroer, Stefan van Nieuwkoop, Mark Pronk, Pascal Lexmond, Corien Swaan, Manon Haverkate, Madelief Mollers, Mart Stein, Sandra Kengne Kamga Mobou, Jeroen van Kampen, Jolanda Voermans, Aura Timen, Corine GeurtsvanKessel, Annemiek van der Eijk, Richard Molenkamp, Marion Koopmans, on behalf of the Dutch national COVID-19 response team. |
| EPI_ISL_523824 | Communicable Disease Branch | Hong Kong Department of Health | Mak Gannon C.K., Lam Edman T.K., Chan Rickjason C.W., Tsang Dominic N.C. |
| EPI_ISL_523825 | United Christian Hospital | Hong Kong Department of Health | Mak Gannon C.K., Lam Edman T.K., Chan Rickjason C.W., Tsang Dominic N.C. |
| EPI_ISL_523826, EPI_ISL_523827 | Queen Mary Hospital | Hong Kong Department of Health | Mak Gannon C.K., Lam Edman T.K., Chan Rickjason C.W., Tsang Dominic N.C. |
| EPI_ISL_523828, EPI_ISL_523829 | Tuen Mun Hospital | Hong Kong Department of Health | Mak Gannon C.K., Lam Edman T.K., Chan Rickjason C.W., Tsang Dominic N.C. |
| EPI_ISL_523830, EPI_ISL_523831 | Our Lady of Maryknoll Hospital | Hong Kong Department of Health | Mak Gannon C.K., Lam Edman T.K., Chan Rickjason C.W., Tsang Dominic N.C. |
| EPI_ISL_523835 | Queen Elizabeth Hospital | Hong Kong Department of Health | Mak Gannon C.K., Lam Edman T.K., Chan Rickjason C.W., Tsang Dominic N.C. |
| EPI_ISL_523837 | Yan Chai Hospital | Hong Kong Department of Health | Mak Gannon C.K., Lam Edman T.K., Chan Rickjason C.W., Tsang Dominic N.C. |
| EPI_ISL_524445, EPI_ISL_524446, EPI_ISL_524447, EPI_ISL_524448 | Singapore General Hospital | Department of Microbiology | Nurdyana Abdul Rahman, Kun Lee Lim, Chenhao Li, Kian Sing Chan, Lynette Oon, Kern Rei Chng, Niranjan Nagarajan, Karrie Ko |
| EPI_ISL_524732 | Dr. RSS Hospital, Modasa | Gujarat Biotechnology Research Centre | Harsh Chaudhari, Apurvasinh Puvar, Janvi Raval, Zarna Patel, Monika Gandhi, Pinal Trivedi, Maharshi Pandya, Nidhi Patel, Nitin Savaliya, Raghawendra Kumar, Dinesh Kumar, Zuber Saiyed, Komal Patel, Labdhi Pandya, Afzal Ansari, Nikha Trivedi, R D Dixit, A M Kadri, Harsh Bakshi, Chaitanya Joshi, Madhvi Joshi, Neena Doshi, Varsha Godbole |
| EPI_ISL_524733 | GMERS Medical College and Hospital, Dharpur, Patan | Gujarat Biotechnology Research Centre | A N Parmar, Apurvasinh Puvar, Janvi Raval, Zarna Patel, Monika Gandhi, Pinal Trivedi, Maharshi Pandya, Nidhi Patel, Nitin Savaliya, Raghawendra Kumar, Dinesh Kumar, Zuber Saiyed, Komal Patel, Labdhi Pandya, Afzal Ansari, Nikha Trivedi, R D Dixit, A M Kadri, Harsh Bakshi, Chaitanya Joshi, Madhvi Joshi |
| EPI_ISL_524734 | GMERS Medical College and Hospital, Dharpur, Patan | Gujarat Biotechnology Research Centre | Apurvasinh Puvar, Janvi Raval, Zarna Patel, Monika Gandhi, Pinal Trivedi, Maharshi Pandya, Nidhi Patel, Nitin Savaliya, Raghawendra Kumar, Dinesh Kumar, Zuber Saiyed, Komal Patel, Labdhi Pandya, Afzal Ansari, Nikha Trivedi, A N Parmar, R D Dixit, A M Kadri, Harsh Bakshi, Chaitanya Joshi, Madhvi Joshi |
| EPI_ISL_524752 | GMERS Medical College and Hospital, Dharpur, Patan | Gujarat Biotechnology Research Centre | Monika Gandhi, Pinal Trivedi, Maharshi Pandya, Nidhi Patel, Nitin Savaliya, Raghawendra Kumar, Dinesh Kumar, Zuber Saiyed, Komal Patel, Labdhi Pandya, Afzal Ansari, Nikha Trivedi, A N Parmar, Apurvasinh Puvar, Janvi Raval, Zarna Patel, R D Dixit, A M Kadri, Harsh Bakshi, Chaitanya Joshi, Madhvi Joshi |
| EPI_ISL_524753 | GMERS Medical College and Hospital, Dharpur, Patan | Gujarat Biotechnology Research Centre | Pinal Trivedi, Maharshi Pandya, Nidhi Patel, Nitin Savaliya, Raghawendra Kumar, Dinesh Kumar, Zuber Saiyed, Komal Patel, Labdhi Pandya, Afzal Ansari, Nikha Trivedi, A N Parmar, Apurvasinh Puvar, Janvi Raval, Zarna Patel, Monika Gandhi, R D Dixit, A M Kadri, Harsh Bakshi, Chaitanya Joshi, Madhvi Joshi |
| EPI_ISL_524754 | GMERS Medical College and Hospital, Dharpur, Patan | Gujarat Biotechnology Research Centre | Maharshi Pandya, Nidhi Patel, Nitin Savaliya, Raghawendra Kumar, Dinesh Kumar, Zuber Saiyed, Komal Patel, Labdhi Pandya, Afzal Ansari, Nikha Trivedi, A N Parmar, Apurvasinh Puvar, Janvi Raval, Zarna Patel, Monika Gandhi, Pinal Trivedi, R D Dixit, A M Kadri, Harsh Bakshi, Chaitanya Joshi, Madhvi Joshi |
| EPI_ISL_524755 | GMERS Medical College and Hospital, Dharpur, Patan | Gujarat Biotechnology Research Centre | Nidhi Patel, Nitin Savaliya, Raghawendra Kumar, Dinesh Kumar, Zuber Saiyed, Komal Patel, Labdhi Pandya, Afzal Ansari, Nikha Trivedi, A N Parmar, Apurvasinh Puvar, Janvi Raval, Zarna Patel, Monika Gandhi, Pinal Trivedi, Maharshi Pandya, R D Dixit, A M Kadri, Harsh Bakshi, Chaitanya Joshi, Madhvi Joshi |
| EPI_ISL_524756 | GMERS Medical College and Hospital, Dharpur, Patan | Gujarat Biotechnology Research Centre | Nitin Savaliya, Raghawendra Kumar, Dinesh Kumar, Zuber Saiyed, Komal Patel, Labdhi Pandya, Afzal Ansari, Nikha Trivedi, A N Parmar, Apurvasinh Puvar, Janvi Raval, Zarna Patel, Monika Gandhi, Pinal Trivedi, Maharshi Pandya, Nidhi Patel, R D Dixit, A M Kadri, Harsh Bakshi, Chaitanya Joshi, Madhvi Joshi |
| EPI_ISL_524757 | GMERS Medical College and Hospital, Dharpur, Patan | Gujarat Biotechnology Research Centre | Raghawendra Kumar, Dinesh Kumar, Zuber Saiyed, Komal Patel, Labdhi Pandya, Afzal Ansari, Nikha Trivedi, A N Parmar, Apurvasinh Puvar, Janvi Raval, Zarna Patel, Monika Gandhi, Pinal Trivedi, Maharshi Pandya, Nidhi Patel, Nitin Savaliya, R D Dixit, A M Kadri, Harsh Bakshi, Chaitanya Joshi, Madhvi Joshi |
| EPI_ISL_524758 | GMERS Medical College and Hospital, Dharpur, Patan | Gujarat Biotechnology Research Centre | Dinesh Kumar, Zuber Saiyed, Komal Patel, Labdhi Pandya, Afzal Ansari, Nikha Trivedi, A N Parmar, Apurvasinh Puvar, Janvi Raval, Zarna Patel, Monika Gandhi, Pinal Trivedi, Maharshi Pandya, Nidhi Patel, Nitin Savaliya, Raghawendra Kumar, R D Dixit, A M Kadri, Harsh Bakshi, Chaitanya Joshi, Madhvi Joshi |
| EPI_ISL_524759 | GMERS Medical College and Hospital, Dharpur, Patan | Gujarat Biotechnology Research Centre | Zuber Saiyed, Komal Patel, Labdhi Pandya, Afzal Ansari, Nikha Trivedi, A N Parmar, Apurvasinh Puvar, Janvi Raval, Zarna Patel, Monika Gandhi, Pinal Trivedi, Maharshi Pandya, Nidhi Patel, Nitin Savaliya, Raghawendra Kumar, Dinesh Kumar, R D Dixit, A M Kadri, Harsh Bakshi, Chaitanya Joshi, Madhvi Joshi |
| EPI_ISL_524760 | GMERS Medical College and Hospital, Dharpur, Patan | Gujarat Biotechnology Research Centre | Komal Patel, Labdhi Pandya, Afzal Ansari, Nikha Trivedi, A N Parmar, Apurvasinh Puvar, Janvi Raval, Zarna Patel, Monika Gandhi, Pinal Trivedi, Maharshi Pandya, Nidhi Patel, Nitin Savaliya, Raghawendra Kumar, Dinesh Kumar, Zuber Saiyed, R D Dixit, A M Kadri, Harsh Bakshi, Chaitanya Joshi, Madhvi Joshi |
| EPI_ISL_524761 | GMERS Medical College and Hospital, Dharpur, Patan | Gujarat Biotechnology Research Centre | Labdhi Pandya, Afzal Ansari, Nikha Trivedi, A N Parmar, Apurvasinh Puvar, Janvi Raval, Zarna Patel, Monika Gandhi, Pinal Trivedi, Maharshi Pandya, Nidhi Patel, Nitin Savaliya, Raghawendra Kumar, Dinesh Kumar, Zuber Saiyed, Komal Patel, R D Dixit, A M Kadri, Harsh Bakshi, Chaitanya Joshi, Madhvi Joshi |
| EPI_ISL_524762 | GMERS Medical College and Hospital, Dharpur, Patan | Gujarat Biotechnology Research Centre | Afzal Ansari, Nikha Trivedi, A N Parmar, Apurvasinh Puvar, Janvi Raval, Zarna Patel, Monika Gandhi, Pinal Trivedi, Maharshi Pandya, Nidhi Patel, Nitin Savaliya, Raghawendra Kumar, Dinesh Kumar, Zuber Saiyed, Komal Patel, Labdhi Pandya, R D Dixit, A M Kadri, Harsh Bakshi, Chaitanya Joshi, Madhvi Joshi |
| EPI_ISL_524763 | GMERS Medical College and Hospital, Dharpur, Patan | Gujarat Biotechnology Research Centre | Nikha Trivedi, A N Parmar, Apurvasinh Puvar, Janvi Raval, Zarna Patel, Monika Gandhi, Pinal Trivedi, Maharshi Pandya, Nidhi Patel, Nitin Savaliya, Raghawendra Kumar, Dinesh Kumar, Zuber Saiyed, Komal Patel, Labdhi Pandya, Afzal Ansari, R D Dixit, A M Kadri, Harsh Bakshi, Chaitanya Joshi, Madhvi |

| | | | |
|---|---|---|---|
| | | | Joshi |
| EPI_ISL_524764 | GMERS Medical College and Hospital, Dharpur, Patan | Gujarat Biotechnology Research Centre | A N Parmar, Apurvasinh Puvar, Janvi Raval, Zarna Patel, Monika Gandhi, Pinal Trivedi, Maharshi Pandya, Nidhi Patel, Nitin Savaliya, Raghawendra Kumar, Dinesh Kumar, Zuber Saiyed, Komal Patel, Labdhi Pandya, Afzal Ansari, Nikha Trivedi, R D Dixit, A M Kadri, Harsh Bakshi, Chaitanya Joshi, Madhvi Joshi |
| EPI_ISL_524765 | GMERS Medical College and Hospital, Dharpur, Patan | Gujarat Biotechnology Research Centre | Apurvasinh Puvar, Janvi Raval, Zarna Patel, Monika Gandhi, Pinal Trivedi, Maharshi Pandya, Nidhi Patel, Nitin Savaliya, Raghawendra Kumar, Dinesh Kumar, Zuber Saiyed, Komal Patel, Labdhi Pandya, Afzal Ansari, Nikha Trivedi, A N Parmar, R D Dixit, A M Kadri, Harsh Bakshi, Chaitanya Joshi, Madhvi Joshi |
| EPI_ISL_524766 | GMERS Medical College and Hospital, Dharpur, Patan | Gujarat Biotechnology Research Centre | Janvi Raval, Zarna Patel, Monika Gandhi, Pinal Trivedi, Maharshi Pandya, Nidhi Patel, Nitin Savaliya, Raghawendra Kumar, Dinesh Kumar, Zuber Saiyed, Komal Patel, Labdhi Pandya, Afzal Ansari, Nikha Trivedi, Apurvasinh Puvar, A N Parmar, R D Dixit, A M Kadri, Harsh Bakshi, Chaitanya Joshi, Madhvi Joshi |
| EPI_ISL_524880, EPI_ISL_524881, EPI_ISL_524882, EPI_ISL_524892, EPI_ISL_524893, EPI_ISL_524894, EPI_ISL_524895, EPI_ISL_524896, EPI_ISL_524897, EPI_ISL_524898, EPI_ISL_524899, EPI_ISL_524900, EPI_ISL_524901, EPI_ISL_524902, EPI_ISL_524903 | | | |
| see above | MD PHL | MD PHL | Maryland Department of Health Laboratories Administration |
| EPI_ISL_525189, EPI_ISL_525190, EPI_ISL_525191, EPI_ISL_525192, EPI_ISL_525200, EPI_ISL_525201, EPI_ISL_525202 | Virginia DCLS | Virginia DCLS | Virginia DCLS |
| EPI_ISL_525707, EPI_ISL_525708, EPI_ISL_525709, EPI_ISL_525710, EPI_ISL_525711, EPI_ISL_525712, EPI_ISL_525713, EPI_ISL_525714, EPI_ISL_525715, EPI_ISL_525716, EPI_ISL_525717, EPI_ISL_525718, EPI_ISL_525719 | | | |
| see above | Seattle Flu Study | Seattle Flu Study | Deborah A. Nickerson, Chris D. Frazar, Jover Lee, Benjamin Pelle, Matthew Richardson, Amanda Adler, Elisabeth Brandstetter, Peter D. Han, Kairsten Fay, Misja Ilcisin, Kirsten Lacombe, Thomas R. Sibley, Melissa Truong, Caitlin R. Wolf, Karen Cowgill, Stephanie Schrag, Jeff Duchin, Michael Boeckh, Janet A. Englund, Michael Famulare, Barry R. Lutz, Mark J. Rieder, Lea M. Starita, Matthew Thompson, Helen Y. Chu, Trevor Bedford, Jay Shendure |
| EPI_ISL_525901, EPI_ISL_525926, EPI_ISL_525927, EPI_ISL_525928, EPI_ISL_525929, EPI_ISL_525931, EPI_ISL_525932, EPI_ISL_525933, EPI_ISL_525934, EPI_ISL_525935, EPI_ISL_525936, EPI_ISL_525937, EPI_ISL_525938, EPI_ISL_525940, EPI_ISL_525941, EPI_ISL_525942, EPI_ISL_525943, EPI_ISL_525944, EPI_ISL_525945, EPI_ISL_525946, EPI_ISL_525947, EPI_ISL_525948, EPI_ISL_525949, EPI_ISL_525950, EPI_ISL_525951, EPI_ISL_525952, EPI_ISL_525953, EPI_ISL_525954, EPI_ISL_525955, EPI_ISL_525956, EPI_ISL_525957, EPI_ISL_525958, EPI_ISL_525959, EPI_ISL_525960, EPI_ISL_525961, EPI_ISL_525962, EPI_ISL_525963, EPI_ISL_525964, EPI_ISL_525965, EPI_ISL_525966, EPI_ISL_525967, EPI_ISL_526100, EPI_ISL_526101, EPI_ISL_526102 | | | |
| see above | OHSU Lab Services Molecular Microbiology Lab | Oregon SARS-CoV-2 Genome Sequencing Center | Brendan L. O'Connell, Ruth V. Nichols, Alec J. Hirsch, Guang Fan, Daniel N. Streblow, William B. Messer, Andrew C. Adey, Benjamin N. Bimber, Brian J. O'Roak |
| EPI_ISL_526225 | Hungarian Defence Forces Military Medical Centre | National Laboratory of Virology, Szentágothai Research Centre | Endre Gábor Tóth, Balázs Somogyi, Bálint Eszenyi, Ferenc Jakab, Gábor Kemenesi |
| EPI_ISL_526250, EPI_ISL_526251, EPI_ISL_526252 | Instituto Adolfo Lutz | Instituto Adolfo Lutz Laboratório de Vírus Respiratórios | Katia Corrêa de Oliveira Santos, Fabiana Cristina Pereira dos Santos, Maíra Marcele Birochi, Cecília Simões Santos, Ana Maria Sardinha Afonso, Maira do Carmo Sampaio Tavares Timenetsky |
| EPI_ISL_526389, EPI_ISL_526390 | Liverpool Clinical Laboratories | COVID-19 Genomics UK (COG-UK) Consortium | Sam Haldenby, Anita Lucaci, Steve Paterson, Julian Hiscox, Alistair Darby, M Almsaud, A Alrezaihi, Muhannad Alruwaili, Stuart D Armstrong, Jones Benjamin, Eleanor G Bentley, Anu Chawla, Jordan J Clark, Angela Cowell, Richard Eccles, Isabel García-Dorival, Matthew Gemmell, Alessandro Gerada, PKF Gilmore, Richard Gregory, Ximeng Han, Catherine Hartley, Margaret Hughes, Miren Iturriza-Gomara, James Johnson, L Luu, Jenifer Manson, Charlotte Nelson, Elaine O'Toole, Cassie Olateju, Rebekah Penrice-Randal , Lucille Rainbow, N.P Randle, Trevor Ian Robinson, Parul Sharma, Ghada T Shawli, James P Stewart, Neil Swainston, Ecaterina Vamos, Joanne Watts, Mark Whitehead |
| EPI_ISL_526437, EPI_ISL_526438, EPI_ISL_526439, EPI_ISL_526443, EPI_ISL_526444, EPI_ISL_526445, EPI_ISL_526446, EPI_ISL_526448, EPI_ISL_526449, EPI_ISL_526450 | Virology Department, Sheffield Teaching Hospitals NHS Foundation Trust/Department of Infection, Immunity and Cardiovascular Disease, The Medical School, University of Sheffield | COVID-19 Genomics UK (COG-UK) Consortium | Thushan de Silva, Matthew Parker, Nikki Smith, Adri Angyal, Rebecca Brown, Luke Green, Rachel Tucker, Paul Parsons, Danielle Groves, Katie Johnson, Laura Carrilero, Alex Keeley, Dave Partridge, Matthew Wyles, Benjamin Lindsey, Mehmet Yavuz, Mohammad Raza, Cariad Evans |
| EPI_ISL_526845, EPI_ISL_526846, EPI_ISL_526847, EPI_ISL_526848, EPI_ISL_526849, EPI_ISL_526850, EPI_ISL_526851, EPI_ISL_526852, EPI_ISL_526853, EPI_ISL_526854, EPI_ISL_526858, EPI_ISL_526859, EPI_ISL_526860, EPI_ISL_526861, EPI_ISL_526862, EPI_ISL_526871, EPI_ISL_526872, EPI_ISL_526873, EPI_ISL_526874, EPI_ISL_526875, EPI_ISL_526889, EPI_ISL_526890, EPI_ISL_526895, EPI_ISL_526896, EPI_ISL_526897, EPI_ISL_526898 | | | |
| see above | Virginia DCLS | Virginia DCLS | Virginia DCLS |
| EPI_ISL_526954, EPI_ISL_526967 | Instituto Nacional de Salud, Bogotá, Colombia | Instituto Nacional de Salud, Bogotá, Colombia | Katherine Laiton-Donato, Diego A. Álvarez-Díaz, Carlos Franco-Muñoz, Mauricio Pacheco-Montealegre, Jonathan Reales, Diego Andrés Prada, Jose A. Usme-Ciro, Zulma M. Cucunubá, Christian Julian VillabonaArenas, Liz Villabona-Arenas, Sussy Echeverria, Astrid C. Flórez, Carolina Ferro, Diana Marcela Walteros-Acero, Franklin Prieto, Carlos Andrés Durán, Martha Lucia Ospina Martinez, Marcela Mercado-Reyes |
| EPI_ISL_527019, EPI_ISL_527020, EPI_ISL_527021, EPI_ISL_527027, EPI_ISL_527028, EPI_ISL_527029, EPI_ISL_527030, EPI_ISL_527031, EPI_ISL_527032, EPI_ISL_527033, EPI_ISL_527034 | | | |
| see above | Area of Virology, Serology and Virology Division (SAViD), New South Wales Health Pathology Randwick | Area of Virology, Serology and Virology Division (SAViD), New South Wales Health Pathology Randwick | Rawlinson, W. |
| EPI_ISL_527575, EPI_ISL_527576, EPI_ISL_527577, EPI_ISL_527578, EPI_ISL_527579, EPI_ISL_527580, EPI_ISL_527581, EPI_ISL_527582, EPI_ISL_527583 | Minnesota Department of Health, Public Health Laboratory | Minnesota Department of Health, Public Health Laboratory | Matt Plumb, Jacob Garfin, and Xiong Wang |
| EPI_ISL_527651, EPI_ISL_527652, EPI_ISL_527653, EPI_ISL_527654, EPI_ISL_527655, EPI_ISL_527657 | AR Dept. of Health-Public Health Lab | Pathogen Discovery, Respiratory Viruses Branch, Division of Viral Diseases, Centers for Disease Control and Prevention | Ying Tao, Jing Zhang, Yan Li, Krista Queen, Anna Uehara, Clinton Paden, Haibin Wang, Suxiang Tong |
| EPI_ISL_527658, EPI_ISL_527659 | GA Department of Public Health Laboratory | Pathogen Discovery, Respiratory Viruses Branch, Division of Viral Diseases, Centers for Disease Control and Prevention | Yan Li, Anna Montmayeur, Jing Zhang, Krista Queen, Ying Tao, Anna Uehara, Rachel Marine, Clinton R. Paden, Haibin Wang, Suxiang Tong |
| EPI_ISL_527745 | Area De Salud Corredores | Inciensa, Instituto Costarricense de Investigación y Enseñanza en Nutrición y Salud | Francisco Duarte, Hebleen Porras, Claudio Soto-Garita, Estela Cordero, Adriana Godinez & Melany Calderon |
| EPI_ISL_527746, EPI_ISL_527747 | Area De Salud La Cruz | Inciensa, Instituto Costarricense de Investigación y Enseñanza en Nutrición y Salud | Francisco Duarte, Hebleen Porras, Claudio Soto-Garita, Estela Cordero, Adriana Godinez & Melany Calderon |
| EPI_ISL_527748 | Area De Salud Corredores | Inciensa, Instituto Costarricense de Investigación y Enseñanza en Nutrición y Salud | Francisco Duarte, Hebleen Porras, Claudio Soto-Garita, Estela Cordero, Adriana Godinez & Melany Calderon |
| EPI_ISL_527749 | Area De Salud Alajuela Norte - Clinica Dr. Marcial Rodriguez | Inciensa, Instituto Costarricense de Investigación y Enseñanza en Nutrición y Salud | Francisco Duarte, Hebleen Porras, Claudio Soto-Garita, Estela Cordero, Adriana Godinez & Melany Calderon |
| EPI_ISL_527750 | Hospital De Niños Dr. Carlos Saenz Herrera [San Jose/San Jose] | Inciensa, Instituto Costarricense de Investigación y Enseñanza en Nutrición y Salud | Francisco Duarte, Hebleen Porras, Claudio Soto-Garita, Estela Cordero, Adriana Godinez & Melany Calderon |
| EPI_ISL_527751 | Area De Salud Corredores | Inciensa, Instituto Costarricense de Investigación y Enseñanza en Nutrición y Salud | Francisco Duarte, Hebleen Porras, Claudio Soto-Garita, Estela Cordero, Adriana Godinez & Melany Calderon |
| EPI_ISL_527752 | Hospital Dr. Rafael A. Calderon Guardia | Inciensa, Instituto Costarricense de Investigación y Enseñanza en Nutrición y Salud | Francisco Duarte, Hebleen Porras, Claudio Soto-Garita, Estela Cordero, Adriana Godinez & Melany Calderon |
| EPI_ISL_527753, EPI_ISL_527754 | Hospital San Vicente De Paul | Inciensa, Instituto Costarricense de Investigación y Enseñanza en Nutrición y Salud | Francisco Duarte, Hebleen Porras, Claudio Soto-Garita, Estela Cordero, Adriana Godinez & Melany Calderon |
| EPI_ISL_527756 | Area De Salud Aserri | Inciensa, Instituto Costarricense de Investigación y | Francisco Duarte, Hebleen Porras, Claudio Soto-Garita, Estela Cordero, Adriana Godinez & Melany Calderon |

| | | | |
|---|---|---|---|
| | | Enseñanza en Nutrición y Salud | |
| EPI_ISL_527757 | Area De Salud Goicoechea 1 | Inciensa, Instituto Costarricense de Investigación y Enseñanza en Nutrición y Salud | Francisco Duarte, Hebleen Porras, Claudio Soto-Garita, Estela Cordero, Adriana Godinez & Melany Calderon |
| EPI_ISL_528426 | P. D. Hinduja Hospital and Medical Research Centre | Institute of Genomics and Integrative Biology - Council of Scientific and Industrial Research | Rajesh Pandey, Jayanthi Shastri, Akshay Kanakan, Vivekanand A, Janani Srinivasa Vasudevan, Ranjeet Maurya, Sachee Agrawal, Nirjhar Chatterjee, Swapneil Parikh, Manish Pathak, Subrat Thanapati, Jasmina Savak, Suresh Poojari, Mahesh Sangar, Amol Borse, Shweta Kawankar, Vasil Nachan, Mayuresh Vishwanathan, Shruthi Sachidanandan, Shrutika Pophale, Utkarsha Yelve |
| EPI_ISL_528485, EPI_ISL_528486, EPI_ISL_528487, EPI_ISL_528488, EPI_ISL_528521, EPI_ISL_528523, EPI_ISL_528524, EPI_ISL_528525, EPI_ISL_528526, EPI_ISL_528527, EPI_ISL_528528, EPI_ISL_528529, EPI_ISL_528530, EPI_ISL_528531, EPI_ISL_528532, EPI_ISL_528533, EPI_ISL_528534, EPI_ISL_528535, EPI_ISL_528536, EPI_ISL_528537 | | | |
| see above | Alaska State Virology Laboratory | Alaska State Virology Laboratory | Chen J et al with Pathogenomics group Dagdag R, Redlinger M, Milton E, George W, Kovalenko A, Drown DM, Bortz E |
| EPI_ISL_528707 | Alsafar - Khalifa University Abu Dhabi | Alsafar - Khalifa University Abu Dhabi | Andreas Henschel, Gihan Daw Elbait, Samuel Feng, Rifat Hamoudi, Ernesto Damiani, Guan Tay, Habiba Alsafar |
| EPI_ISL_529010, EPI_ISL_529011, EPI_ISL_529012 | Servizio Igiene Epidemiologia e Sanità Pubblica (SIESP)-L'Aquila | Istituto Zooprofilattico Sperimentale dell'Abruzzo e Molise "G.Caporale" | Lorusso A, Marcacci M, Di Domenico M, Curini V, Ancora M, Cammà C, Rinaldi A, Mangone I, Di Pasquale A, Puglia I, Savini G. |
| EPI_ISL_529013 | Presidio Ospedaliero "S.Filippo e Nicola"-Avezzano | Istituto Zooprofilattico Sperimentale dell'Abruzzo e Molise "G.Caporale" | Lorusso A, Marcacci M, Di Domenico M, Curini V, Ancora M, Cammà C, Rinaldi A, Mangone I, Di Pasquale A, Puglia I, Savini G. |
| EPI_ISL_529282 | Quadram Institute Bioscience | COVID-19 Genomics UK (COG-UK) Consortium | Dave J. Baker, Gemma L. Kay, Alp Aydin, Thanh Le-Viet, Steven Rudder, Ana P. Tedim, Anastasia Kolyva, Maria Diaz, Leonardo de Oliveira Martins, Nabil-Fareed Alikhan, Lizzie Meadows, Rachael Stanley, Ngozi Elumogo, Muhammed Yasir, Nicholas M. Thomson, Alexander J Trotter, Rachel Gilroy, Samuel Bloomfield, Claire Stuart, Andrew Bell, Reenesh Prakash, Samir Dervisevic, Alison E. Mather, John Wain, Mark Webber, Andrew J. Page, Justin O'Grady |
| EPI_ISL_529295 | Liverpool Clinical Laboratories | COVID-19 Genomics UK (COG-UK) Consortium | Sam Haldenby, Anita Lucaci, Steve Paterson, Julian Hiscox, Alistair Darby, M Almsaud, A Alrezaihi, Muhannad Alruwaili, Stuart D Armstrong, Jones Benjamin, Eleanor G Bentley, Anu Chawla, Jordan J Clark, Angela Cowell, Richard Eccles, Isabel García-Dorival, Matthew Gemmell, Alessandro Gerada, PKF Gilmore, Richard Gregory, Ximeng Han, Catherine Hartley, Margaret Hughes, Miren Iturriza-Gomara, James Johnson, L Luu, Jenifer Manson, Charlotte Nelson, Elaine O'Toole, Cassie Olateju, Rebekah Penrice-Randal , Lucille Rainbow, N.P Randle, Trevor Ian Robinson, Parul Sharma, Ghada T Shawli, James P Stewart, Neil Swainston, Ecaterina Vamos, Joanne Watts, Mark Whitehead |
| EPI_ISL_529321, EPI_ISL_529366, EPI_ISL_529408 | University of Birmingham | COVID-19 Genomics UK (COG-UK) Consortium | Institute of Microbiology, University of Birmingham: Claire McMurray, Joanne Stockton, Samuel Nicholls, Radoslaw Poplawski, Will Rowe, Josh Quick, Nicholas Loman. University of Birmingham Testing Laboratory: Celina M Whalley, Andrew Bosworth, Charlotte Poxon, Kasun Wanigasooriya, Oliver Pickles, Mike Kidd, Alex Richter, Andrew D Beggs PHE Heartlands Lab: Husam Osman, Andrew Bosworth. Queen Elizabeth Hospital: Anna Casey |
| EPI_ISL_529833, EPI_ISL_529852 | Michigan Department of Health and Human Services, Bureau of Laboratories | Michigan Department of Health and Human Services, Bureau of Laboratories | Blankenship HM, Riner D, Soehnlen MK |
| EPI_ISL_530173 | Minnesota Department of Health, Public Health Laboratory | Minnesota Department of Health, Public Health Laboratory | Matt Plumb, Jacob Garfin, and Xiong Wang |
| EPI_ISL_530250, EPI_ISL_530251 | Queensland Health Forensic and Scientific Services, Public Health Virology | Public Health Virology Laboratory, Forensic and Scientific Services, Queensland Health | Son Nguyen et al |
| EPI_ISL_532581, EPI_ISL_533010, EPI_ISL_533032, EPI_ISL_533042, EPI_ISL_533059, EPI_ISL_533084, EPI_ISL_533099, EPI_ISL_533104, EPI_ISL_533113, EPI_ISL_533150, EPI_ISL_533215 | | | |
| see above | Lighthouse Lab in Glasgow | Wellcome Sanger Institute for the COVID-19 Genomics UK (COG-UK) consortium | Harper VanSteenhouse, Yumi Kasai, David Gray, Carol Clugston, Anna Dominiczak and Alex Alderton, Roberto Amato, Sonia Goncalves, Ewan Harrison, David K. Jackson, Ian Johnston, Dominic Kwiatkowski, Cordelia Langford, John Sillitoe |
| EPI_ISL_534234, EPI_ISL_534235, EPI_ISL_534236 | Karolinska universitetslaboratoriet SOLNA | The Public Health Agency of Sweden | Anna-Malin Linde, Maria Lind Karlberg, Mattias Haukland, Reza Advani, Olov Svartstrom, Oskar Karlsson Lindsjo, Sandra Broddesson, Petra Edquist, Mia Brytting, Anna Risberg, Karin Tegmark-Wisell |
| EPI_ISL_534239, EPI_ISL_534241, EPI_ISL_534242, EPI_ISL_534243 | Norra Alvsborgs lanssjukhus | The Public Health Agency of Sweden | Anna-Malin Linde, Maria Lind Karlberg, Mattias Haukland, Reza Advani, Olov Svartstrom, Oskar Karlsson Lindsjo, Sandra Broddesson, Petra Edquist, Mia Brytting, Anna Risberg, Karin Tegmark-Wisell |
| EPI_ISL_534699, EPI_ISL_534700, EPI_ISL_534701, EPI_ISL_534702, EPI_ISL_534703 | MD PHL | MD PHL | Maryland Department of Health Laboratories Administration |
| EPI_ISL_534718 | Respiratory Virus Unit, Microbiology Services Colindale, Public Health England | Respiratory Virus Unit, Microbiology Services Colindale, Public Health England | PHE Covid Sequencing Team |
| EPI_ISL_534755 | Liverpool Clinical Laboratories | COVID-19 Genomics UK (COG-UK) Consortium | Sam Haldenby, Anita Lucaci, Steve Paterson, Julian Hiscox, Alistair Darby, M Almsaud, A Alrezaihi, Muhannad Alruwaili, Stuart D Armstrong, Jones Benjamin, Eleanor G Bentley, Anu Chawla, Jordan J Clark, Angela Cowell, Richard Eccles, Isabel García-Dorival, Matthew Gemmell, Alessandro Gerada, PKF Gilmore, Richard Gregory, Ximeng Han, Catherine Hartley, Margaret Hughes, Miren Iturriza-Gomara, James Johnson, L Luu, Jenifer Manson, Charlotte Nelson, Elaine O'Toole, Cassie Olateju, Rebekah Penrice-Randal , Lucille Rainbow, N.P Randle, Trevor Ian Robinson, Parul Sharma, Ghada T Shawli, James P Stewart, Neil Swainston, Ecaterina Vamos, Joanne Watts, Mark Whitehead |
| EPI_ISL_535296 | New Mexico Department of Health Scientific Laboratory | New Mexico Department of Health Scientific Laboratory | Ellie Johnson, Anastacia Griego-Fisher, D'Eldra Malone |
| EPI_ISL_535650 | AR Dept. of Health-Public Health Lab | Pathogen Discovery, Respiratory Viruses Branch, Division of Viral Diseases, Centers for Disease Control and Prevention | Brian Lynch, Yan Li, Jing Zhang, Ying Tao, Krista Queen, Anna Uehara, Clinton R. Paden, Rachel Marine, Haibin Wang, Suxiang Tong |
| EPI_ISL_535651 | AR Dept. of Health-Public Health Lab | Pathogen Discovery, Respiratory Viruses Branch, Division of Viral Diseases, Centers for Disease Control and Prevention | Yan Li, Jing Zhang, Ying Tao, Krista Queen, Brian Lynch, Anna Uehara, Clinton R. Paden, Rachel Marine, Haibin Wang, Suxiang Tong |
| EPI_ISL_535652 | AR Dept. of Health-Public Health Lab | Pathogen Discovery, Respiratory Viruses Branch, Division of Viral Diseases, Centers for Disease Control and Prevention | Brian Lynch, Yan Li, Jing Zhang, Ying Tao, Krista Queen, Anna Uehara, Clinton R. Paden, Rachel Marine, Haibin Wang, Suxiang Tong |
| EPI_ISL_535653 | AR Dept. of Health-Public Health Lab | Pathogen Discovery, Respiratory Viruses Branch, Division of Viral Diseases, Centers for Disease Control and Prevention | Yan Li, Jing Zhang, Ying Tao, Krista Queen, Brian Lynch, Anna Uehara, Clinton R. Paden, Rachel Marine, Haibin Wang, Suxiang Tong |
| EPI_ISL_535654 | AR Dept. of Health-Public Health Lab | Pathogen Discovery, Respiratory Viruses Branch, Division of Viral Diseases, Centers for Disease Control and Prevention | Brian Lynch, Yan Li, Jing Zhang, Ying Tao, Krista Queen, Anna Uehara, Clinton R. Paden, Rachel Marine, Haibin Wang, Suxiang Tong |
| EPI_ISL_535655, EPI_ISL_535656 | AR Dept. of Health-Public Health Lab | Pathogen Discovery, Respiratory Viruses Branch, Division of Viral Diseases, Centers for Disease Control and Prevention | Yan Li, Jing Zhang, Ying Tao, Krista Queen, Brian Lynch, Anna Uehara, Clinton R. Paden, Rachel Marine, Haibin Wang, Suxiang Tong |
| EPI_ISL_536477, EPI_ISL_536478, EPI_ISL_536479, EPI_ISL_536480, EPI_ISL_536481, EPI_ISL_536483, EPI_ISL_536484, EPI_ISL_536485, EPI_ISL_536486, EPI_ISL_536487, EPI_ISL_536488, EPI_ISL_536489, EPI_ISL_536490, EPI_ISL_536491, EPI_ISL_536492, EPI_ISL_536493, EPI_ISL_536494, EPI_ISL_536495, EPI_ISL_536496, EPI_ISL_536497, EPI_ISL_536498, EPI_ISL_536499, EPI_ISL_536500 | | | |
| see above | Instituto Nacional de Salud | Laboratorio de Infecciones Respiratorias Agudas | Eduardo Juscamayta Lopez, David Tarazona, Faviola Valdivia Guerrero, Nancy Rojas Serrano, Dennis Carhuaricra, Lenin Maturrano Hernandez, Ronnie Gavilan Chavez |
| EPI_ISL_537381 | Complejo Hospitalario Universitario de Vigo | SeqCOVID-SPAIN consortium/IBV(CSIC) | Benito Regueiro and SeqCOVID-SPAIN consortium |
| EPI_ISL_538651, EPI_ISL_538652, EPI_ISL_538653, EPI_ISL_538667, EPI_ISL_538671 | Servicio de Microbiología. Hospital General Universitario de Castellón | SeqCOVID-SPAIN consortium/IBV(CSIC) | Rosario Moreno, María Dolores Tirado and SeqCOVID-SPAIN consortium |
| EPI_ISL_539492 | CMS, Roorkee | CSIR-Institute of Microbial Technology | Kanika Bansal, Sanjeet Kumar, Anu Singh, Debarghya Ghose, Amandeep Kaur, Rajesh Kumar Mishra, Poushali Chakraborty, Harsh Goar, Navin Baid, Ashwani Kumar, Dipak Dutta, Sanjeev Khosla, Prabhu B. Patil |

| | | | |
|---|---|---|---|
| EPI_ISL_539495 | Centers for Disease Control and Prevention, Dengue Branch | Centers for Disease Control and Prevention, Dengue Branch | Gilberto A. Santiago, Glenda Gonzalez, Betzabel Flores, Keyla Charriez, Jorge L. Munoz-Jordan, Gabriela Paz-Bailey, Janice Perez, Vanessa Rivera-Amill, Diego Sainz de la Peña, Jorge Bertran |
| EPI_ISL_539804 | Yan Chai Hospital | Hong Kong Department of Health | Alan K.L. Tsang, Peter C.W. Yip, Edman T.K. Lam, Rickjason C.W. Chan, Dominic N.C. Tsang |
| EPI_ISL_539805 | Kwong Wah Hospital | Hong Kong Department of Health | Alan K.L. Tsang, Peter C.W. Yip, Edman T.K. Lam, Rickjason C.W. Chan, Dominic N.C. Tsang |
| EPI_ISL_539806, EPI_ISL_539807, EPI_ISL_539808 | Princess Margaret Hospital | Hong Kong Department of Health | Alan K.L. Tsang, Peter C.W. Yip, Edman T.K. Lam, Rickjason C.W. Chan, Dominic N.C. Tsang |
| EPI_ISL_539894 | PHE South West Regional Laboratory, National Infection Service | Wellcome Sanger Institute for the COVID-19 Genomics UK (COG-UK) consortium | Stephanie Hutchings, Hannah Pymont, Dr Peter Muir, Barry Vipond, Rich Hopes; and Alex Alderton, Roberto Amato, Sonia Goncalves, Ewan Harrison, David K. Jackson, Ian Johnston, Dominic Kwiatkowski, Cordelia Langford, John Sillitoe on behalf of the Wellcome Sanger Institute COVID-19 Surveillance Team |
| EPI_ISL_539895 | PHE South West Regional Laboratory, National Infection Service | Wellcome Sanger Institute for the COVID-19 Genomics UK (COG-UK) Consortium | Stephanie Hutchings, Hannah Pymont, Dr Peter Muir, Barry Vipond, Rich Hopes; and Alex Alderton, Roberto Amato, Sonia Goncalves, Ewan Harrison, David K. Jackson, Ian Johnston, Dominic Kwiatkowski, Cordelia Langford, John Sillitoe on behalf of the Wellcome Sanger Institute COVID-19 Surveillance Team |
| EPI_ISL_540444, EPI_ISL_540445, EPI_ISL_540446, EPI_ISL_540455 | University of Liège COVID-19 testing center | GIGA Medical Genomics | Keith Durkin, Maria Artesi, Emmanuel André, Marc Van Ranst, Fabrice Bureau, Laurent Gillet, Wouter Coppieters, Vincent Bours |
| EPI_ISL_541005, EPI_ISL_541006, EPI_ISL_541007 | Health and Environmental Research Institute of Gwangju Metropolitan city | Health and Environmental Research Institute of Gwangju Metropolitan city | Min Ji Kim, Ji-eun Lee |
| EPI_ISL_541038, EPI_ISL_541066 | Hospital Clínico Universitario de Santiago de Compostela | SeqCOVID-SPAIN consortium/Institute of Biomedicine of Valencia, IBV-CSIC | José Javier Costa Alcalde, Antonio Aguilera Guirao, Mª Luisa Pérez del Molino Bernal, Amparo Coira Nieto, Gema Barbeito Castiñeiras, Rocio Trastoy Pena and SeqCOVID-SPAIN consortium |
| EPI_ISL_541077, EPI_ISL_541078 | Hospital de la Santa Creu i Sant Pau. Servicio de Microbiología | SeqCOVID-SPAIN consortium/Institute of Biomedicine of Valencia, IBV-CSIC | Ferran Navarro, Núria Rabella, Elisenda Miró and SeqCOVID-SPAIN consortium |
| EPI_ISL_541083, EPI_ISL_541084, EPI_ISL_541085, EPI_ISL_541086, EPI_ISL_541087, EPI_ISL_541088, EPI_ISL_541089, EPI_ISL_541090, EPI_ISL_541091, EPI_ISL_541092, EPI_ISL_541093, EPI_ISL_541094, EPI_ISL_541095 | | | |
| see above | Servicio de Microbiología. Hospital Universitario Donostia. OSI Donostialdea. Área de Enfermedades Infecciosas, Grupo de Infección Respiratoria y Resistencia Antimicrobiana. Instituto de Investigación Sanitaria Biodonostia | SeqCOVID-SPAIN consortium/Institute of Biomedicine of Valencia, IBV-CSIC | Gustavo Cilla, Milagrosa Montes, Luis Piñeiro, Jose Maria Marimón and SeqCOVID-SPAIN consortium |
| EPI_ISL_541248, EPI_ISL_541249, EPI_ISL_541250 | Florida Bureau of Public Health Laboratories, Florida Department of Health | Florida Bureau of Public Health Laboratories, Florida Department of Health | Schmedes,S., Blanton,J. |
| EPI_ISL_541649, EPI_ISL_541651 | Laboratory Diagnostic, Veterinary Specialized Institute Kraljevo | Laboratory Diagnostic, Veterinary Specialized Institute Kraljevo | Vidanovic,D., Tesovic,B., Knezevic,A., Jovanovic,T., Jankovic,M., Sekler,M., Banovic Djeri,B., Volkening,J., Afonso,C., Petrovic,T. |
| EPI_ISL_541656 | Laboratory Diagnostic, Veterinary Specialized Institute Kraljevo | Laboratory Diagnostic, Veterinary Specialized Institute Kraljevo | Vidanovic,D., Tesovic,B., Knezevic,A., Jovanovic,T., Jankovic,M., Sekler,M., Banovic Djeri,B., Volkening,J., Afonso,C., Petrovic,T. |
| EPI_ISL_541657, EPI_ISL_541658, EPI_ISL_541659, EPI_ISL_541661 | Laboratory Diagnostic, Veterinary Specialized Institute Kraljevo | Laboratory Diagnostic, Veterinary Specialized Institute Kraljevo | Vidanovic,D., Tesovic,B., Knezevic,A., Jovanovic,T., Jankovic,M., Sekler,M., Banovic Djeri,B., Volkening,J., Afonso,C., Petrovic,T. |
| EPI_ISL_541664, EPI_ISL_541665, EPI_ISL_541666, EPI_ISL_541667, EPI_ISL_541668, EPI_ISL_541669, EPI_ISL_541670 | Microbiology Division, South Carolina Department of Health and Environmental Control | Microbiology Division, South Carolina Department of Health and Environmental Control | Flores,H. |
| EPI_ISL_541681, EPI_ISL_541740, EPI_ISL_541748, EPI_ISL_541749, EPI_ISL_541750, EPI_ISL_541751 | National Institute of Virology, NIV Influenza | National Institute of Virology, NIV Influenza | Potdar V |
| EPI_ISL_541773, EPI_ISL_541774 | Microbiology Department, Barking Havering and Redbridge University Hospitals NHS trust | Wellcome Sanger Institute for the COVID-19 Genomics UK (COG-UK) consortium | Amy Ash, Fatima Ali, Cherian Koshy and Alex Alderton, Roberto Amato, Sonia Goncalves, Ewan Harrison, David K. Jackson, Ian Johnston, Dominic Kwiatkowski, Cordelia Langford, John Sillitoe on behalf of the Wellcome Sanger Institute COVID-19 Surveillance Team |
| EPI_ISL_541885 | Hospital General Universitario Gregorio Marañón | SeqCOVID-SPAIN consortium/IBV(CSIC) | Laura Pérez-Lago, Marta Herranz, Jon Sicilia, Julia Suárez, Pilar Catalán, Patricia Muñoz, Darío García de Viedma and SeqCOVID-SPAIN consortium |
| EPI_ISL_542024, EPI_ISL_542025, EPI_ISL_542026, EPI_ISL_542027, EPI_ISL_542028, EPI_ISL_542029, EPI_ISL_542030, EPI_ISL_542031, EPI_ISL_542032, EPI_ISL_542033, EPI_ISL_542034, EPI_ISL_542035 | | | |
| see above | New Mexico Department of Health Scientific Laboratory | New Mexico Department of Health Scientific Laboratory | Ellie Johnson, Anastacia Griego-Fisher, D'Eldra Malone |
| EPI_ISL_542992, EPI_ISL_542993, EPI_ISL_542994, EPI_ISL_542995, EPI_ISL_542996, EPI_ISL_542997, EPI_ISL_542998 | TriCore Reference Laboratories | Center for Global Health, University of New Mexico Health Sciences Center | Daryl Domman, Kurt Schwalm, Twila Kunde, Joseph Hicks, Michael Edwards, Darrell Dinwiddie |
| EPI_ISL_545603, EPI_ISL_545607, EPI_ISL_545608, EPI_ISL_545611, EPI_ISL_545639, EPI_ISL_545641, EPI_ISL_545643, EPI_ISL_545647, EPI_ISL_545652, EPI_ISL_545656, EPI_ISL_545658, EPI_ISL_545664, EPI_ISL_545665, EPI_ISL_545667, EPI_ISL_545673, EPI_ISL_545674, EPI_ISL_545679, EPI_ISL_545681, EPI_ISL_545682, EPI_ISL_545690, EPI_ISL_545692, EPI_ISL_545710, EPI_ISL_545719, EPI_ISL_545729, EPI_ISL_545740, EPI_ISL_545741, EPI_ISL_545752, EPI_ISL_545754, EPI_ISL_545763, EPI_ISL_545765, EPI_ISL_545766, EPI_ISL_545767, EPI_ISL_545769, EPI_ISL_545771, EPI_ISL_545772, EPI_ISL_545773, EPI_ISL_545774, EPI_ISL_545775, EPI_ISL_545776, EPI_ISL_545777, EPI_ISL_545779, EPI_ISL_545781, EPI_ISL_545782, EPI_ISL_545783, EPI_ISL_545784, EPI_ISL_545785, EPI_ISL_545786, EPI_ISL_545787, EPI_ISL_545789, EPI_ISL_545790, EPI_ISL_545791, EPI_ISL_545792, EPI_ISL_545793, EPI_ISL_545794, EPI_ISL_545795, EPI_ISL_545796, EPI_ISL_545797, EPI_ISL_545798, EPI_ISL_545799, EPI_ISL_545800, EPI_ISL_545802, EPI_ISL_545803, EPI_ISL_545804, EPI_ISL_545805, EPI_ISL_545806, EPI_ISL_545807, EPI_ISL_545808, EPI_ISL_545809, EPI_ISL_545811, EPI_ISL_545812, EPI_ISL_545813, EPI_ISL_545815, EPI_ISL_545816, EPI_ISL_545818, EPI_ISL_545819, EPI_ISL_545820, EPI_ISL_545821, EPI_ISL_545822, EPI_ISL_545823, EPI_ISL_545824, EPI_ISL_545825, EPI_ISL_545827, EPI_ISL_545828, EPI_ISL_545829, EPI_ISL_545830, EPI_ISL_545831, EPI_ISL_545833, EPI_ISL_545834, EPI_ISL_545836, EPI_ISL_545837, EPI_ISL_545838, EPI_ISL_545839, EPI_ISL_545840, EPI_ISL_545841, EPI_ISL_545842, EPI_ISL_545843, EPI_ISL_545958, EPI_ISL_545959, EPI_ISL_545960, EPI_ISL_545961, EPI_ISL_545962, EPI_ISL_545963, EPI_ISL_545964, EPI_ISL_545965, EPI_ISL_545966, EPI_ISL_545967, EPI_ISL_545968, EPI_ISL_545969, EPI_ISL_545970, EPI_ISL_545971, EPI_ISL_545972, EPI_ISL_545973, EPI_ISL_545974, EPI_ISL_545975, EPI_ISL_545976, EPI_ISL_545977, EPI_ISL_545978, EPI_ISL_545979, EPI_ISL_545980, EPI_ISL_545981, EPI_ISL_545982, EPI_ISL_545983, EPI_ISL_545984, EPI_ISL_545985, EPI_ISL_545986, EPI_ISL_545987, EPI_ISL_545988, EPI_ISL_545989, EPI_ISL_545991, EPI_ISL_545992, EPI_ISL_545993, EPI_ISL_545994, EPI_ISL_545996, EPI_ISL_545998, EPI_ISL_545999, EPI_ISL_546000, EPI_ISL_546001, EPI_ISL_546748, EPI_ISL_546771, EPI_ISL_547015, EPI_ISL_547070, EPI_ISL_547101, EPI_ISL_547164, EPI_ISL_547185, EPI_ISL_547206, EPI_ISL_547207, EPI_ISL_547208, EPI_ISL_547209, EPI_ISL_547210, EPI_ISL_547211, EPI_ISL_547212, EPI_ISL_547213, EPI_ISL_547214, EPI_ISL_547216, EPI_ISL_547219, EPI_ISL_547221, EPI_ISL_547223, EPI_ISL_547225, EPI_ISL_547227, EPI_ISL_547233, EPI_ISL_547234, EPI_ISL_547235, EPI_ISL_547236, EPI_ISL_547237, EPI_ISL_547240, EPI_ISL_547241, EPI_ISL_547242, EPI_ISL_547243, EPI_ISL_547244, EPI_ISL_547246, EPI_ISL_547248, EPI_ISL_547252, EPI_ISL_547253, EPI_ISL_547258, EPI_ISL_547260, EPI_ISL_547262, EPI_ISL_547264, EPI_ISL_547266, EPI_ISL_547267, EPI_ISL_547268, EPI_ISL_547269, EPI_ISL_547270, EPI_ISL_547276, EPI_ISL_547277, EPI_ISL_547282, EPI_ISL_547284, EPI_ISL_547287, EPI_ISL_547289, EPI_ISL_547291, EPI_ISL_547292, EPI_ISL_547295, EPI_ISL_547296, EPI_ISL_547297, EPI_ISL_547298, EPI_ISL_547299, EPI_ISL_547301, EPI_ISL_547302, EPI_ISL_547303, EPI_ISL_547304, EPI_ISL_547305, EPI_ISL_547306, EPI_ISL_547307, EPI_ISL_547308, EPI_ISL_547309, EPI_ISL_547310, EPI_ISL_547311, EPI_ISL_547314, EPI_ISL_547316, EPI_ISL_547318, EPI_ISL_547320, EPI_ISL_547325, EPI_ISL_547326, EPI_ISL_547329, EPI_ISL_547331, EPI_ISL_547334, EPI_ISL_547336, EPI_ISL_547337, EPI_ISL_547339, EPI_ISL_547341, EPI_ISL_547344, EPI_ISL_547345, EPI_ISL_547351, EPI_ISL_547352, EPI_ISL_547366, EPI_ISL_547367, EPI_ISL_547368, EPI_ISL_547369, EPI_ISL_547373, EPI_ISL_547374, EPI_ISL_547377, EPI_ISL_547378, EPI_ISL_547379, EPI_ISL_547381, EPI_ISL_547382, EPI_ISL_547384, EPI_ISL_547385, EPI_ISL_547386, EPI_ISL_547387, EPI_ISL_547388, EPI_ISL_547389, EPI_ISL_547392, EPI_ISL_547396, EPI_ISL_547398, EPI_ISL_547400, EPI_ISL_547401, EPI_ISL_547403, EPI_ISL_547406, EPI_ISL_547407, EPI_ISL_547409, EPI_ISL_547410, EPI_ISL_547411, EPI_ISL_547412, EPI_ISL_547413, EPI_ISL_547414, EPI_ISL_547415, EPI_ISL_547417, EPI_ISL_547418, EPI_ISL_547419, EPI_ISL_547420, EPI_ISL_547421, EPI_ISL_547422, EPI_ISL_547423, EPI_ISL_547424, EPI_ISL_547425, EPI_ISL_547426, EPI_ISL_547427, EPI_ISL_547428, EPI_ISL_547429, EPI_ISL_547430, EPI_ISL_547431, EPI_ISL_547432 | | | |
| see above | Houston Methodist Hospital | Houston Methodist Hospital | S. Wesley Long, Randall J. Olsen, Paul A. Christensen, David W. Bernard, James J. Davis, Maulik Shukla, Marcus Nguyen, Matthew Ojeda Saavedra, Concepcion C. Cantu, Prasanti Yerramilli, Layne Pruitt, Sishir Subedi, Hung-Che Kuo, Heather Hendrickson, Ghazaleh Eskandari, Hoang A. T. Nguyen, J. Hunter Long, Muthiah Kumaraswami, Jule Goike, Daniel Boutz, Jimmy Gollihar, Jason S. McLellan, Chia-Wei Chou, Kamyab Javanmardi, Ilya J. Finkelstein, and James M. Musser |
| EPI_ISL_547577 | Hospital e Maternidade Nossa Senhora das Graças | Instituto Adolfo Lutz, Interdiciplinary Procedures Center, Strategic Laboratory | Claudio Tavares Sacchi, Claudia Regina Gonçalves, Erica Valessa Ramos Gomes, Karoline Rodrigues Campos |
| EPI_ISL_547578 | Hospital Doutor Domingos Leonardo Cerávolo | Instituto Adolfo Lutz, Interdiciplinary Procedures Center, Strategic Laboratory | Claudio Tavares Sacchi, Claudia Regina Gonçalves, Erica Valessa Ramos Gomes, Karoline Rodrigues Campos |
| EPI_ISL_547579 | Santa Casa de Misericórdia de Araçatuba | Instituto Adolfo Lutz, Interdiciplinary Procedures Center, Strategic Laboratory | Claudio Tavares Sacchi, Claudia Regina Gonçalves, Erica Valessa Ramos Gomes, Karoline Rodrigues Campos |

| | | | |
|---|---|---|---|
| EPI_ISL_547580 | Santa Casa da Misericórdia de Presidente Prudente | Instituto Adolfo Lutz, Interdiciplinary Procedures Center, Strategic Laboratory | Claudio Tavares Sacchi, Claudia Regina Gonçalves, Erica Valessa Ramos Gomes, Karoline Rodrigues Campos |
| EPI_ISL_547602, EPI_ISL_547603, EPI_ISL_547604, EPI_ISL_547605, EPI_ISL_547606, EPI_ISL_547607 | Gundersen Molecular Diagnostics Laboratory | Kabara Cancer Research Institute | Craig S. Richmond, Paraic A. Kenny |
| EPI_ISL_547668, EPI_ISL_547669, EPI_ISL_547670, EPI_ISL_547671, EPI_ISL_547673 | Gundersen Clinical Microbiology Laboratory | Kabara Cancer Research Institute | Craig S. Richmond, Paraic A. Kenny |
| EPI_ISL_548134, EPI_ISL_548138, EPI_ISL_548139 | Canterbury Health Laboratories | Institute of Environmental Science and Research (ESR) | Xiaoyun Ren, Matt Storey, Nikki Freed, Muhammad Faisal, Jing Wang, Hermes Perez, Anja Werno, Antje van der Linden, Arlo Upton, Chris Mansell, David Hammer, Dragana Drinkovic, Gary McAuliffe, Hana Sofia Andersson, James Ussher, Jill Sherwood, Josh Freeman, Julia Howard, Juliet Elvy, Mary DeAlmeida, Matt Blakiston, Matthew Rogers, Max Bloomfield, Michael Addidle, Michelle Balm, Sally Roberts, Sarah Jefferies, Sharmini Muttaiyah, Susan Morpeth, Susan Taylor, Timothy Blackmore, Vani Sathyendran, Veronica Playle, Virginia Hope, Erasmus Smit, Lauren Jelly, Olin Silander, Joep de Ligt |
| EPI_ISL_548255 | Lanssjukhuset Kalmar | The Public Health Agency of Sweden | Anna-Malin Linde, Maria Lind Karlberg, Mattias Haukland, Reza Advani, Olov Svartstrom, Oskar Karlsson Lindsjo, Sandra Broddesson, Petra Edquist, Mia Brytting, Anna Risberg, Karin Tegmark-Wisell |
| EPI_ISL_548256 | Centralsjukhuset | The Public Health Agency of Sweden | Anna-Malin Linde, Maria Lind Karlberg, Mattias Haukland, Reza Advani, Olov Svartstrom, Oskar Karlsson Lindsjo, Sandra Broddesson, Petra Edquist, Mia Brytting, Anna Risberg, Karin Tegmark-Wisell |
| EPI_ISL_548272, EPI_ISL_548276, EPI_ISL_548305, EPI_ISL_548313, EPI_ISL_548317 | Orange County Public Health Laboratory | Chan-Zuckerberg Biohub | CZB Cliahub Consortium |
| EPI_ISL_548382, EPI_ISL_548447, EPI_ISL_548452 | University of California, Davis | Chan-Zuckerberg Biohub | CZB Cliahub Consortium |
| EPI_ISL_549122 | Furst Medical Laboratory | Norwegian Institute of Public Health, Department of Virology | Kathrine Stene-Johansen, Kamilla Heddeland Instefjord, Hilde Elshaug, Rasmus Riis Kopperud, Hilde Synnøve Vollan, Karoline Bragstad, Olav Hungnes |
| EPI_ISL_549174 | Vestfold Hospital, Toensberg Department of Microbiology | Norwegian Institute of Public Health, Department of Virology | Kathrine Stene-Johansen, Kamilla Heddeland Instefjord, Hilde Elshaug, Rasmus Riis Kopperud, Hilde Synnøve Vollan, Karoline Bragstad, Olav Hungnes |
| EPI_ISL_551735, EPI_ISL_551736 | Lighthouse Lab in Alderley Park | Wellcome Sanger Institute for the COVID-19 Genomics UK (COG-UK) consortium | The Lighthouse Lab in Alderley Park and Alex Alderton, Roberto Amato, Sonia Goncalves, Ewan Harrison, David K. Jackson, Ian Johnston, Dominic Kwiatkowski, Cordelia Langford, John Sillitoe on behalf of the Wellcome Sanger Institute COVID-19 Surveillance Team |
| EPI_ISL_551737, EPI_ISL_551738, EPI_ISL_551739 | Lighthouse Lab in Milton Keynes | Wellcome Sanger Institute for the COVID-19 Genomics UK (COG-UK) consortium | The Lighthouse Lab in Milton Keynes and Alex Alderton, Roberto Amato, Sonia Goncalves, Ewan Harrison, David K. Jackson, Ian Johnston, Dominic Kwiatkowski, Cordelia Langford, John Sillitoe on behalf of the Wellcome Sanger Institute COVID-19 Surveillance Team (http://www.sanger.ac.uk/covid-team) |
| EPI_ISL_551740 | Lighthouse Lab in Alderley Park | Wellcome Sanger Institute for the COVID-19 Genomics UK (COG-UK) consortium | The Lighthouse Lab in Alderley Park and Alex Alderton, Roberto Amato, Sonia Goncalves, Ewan Harrison, David K. Jackson, Ian Johnston, Dominic Kwiatkowski, Cordelia Langford, John Sillitoe on behalf of the Wellcome Sanger Institute COVID-19 Surveillance Team (http://www.sanger.ac.uk/covid-team) |
| EPI_ISL_551741 | Lighthouse Lab in Milton Keynes | Wellcome Sanger Institute for the COVID-19 Genomics UK (COG-UK) consortium | The Lighthouse Lab in Milton Keynes and Alex Alderton, Roberto Amato, Sonia Goncalves, Ewan Harrison, David K. Jackson, Ian Johnston, Dominic Kwiatkowski, Cordelia Langford, John Sillitoe on behalf of the Wellcome Sanger Institute COVID-19 Surveillance Team (http://www.sanger.ac.uk/covid-team) |
| EPI_ISL_551742, EPI_ISL_551743 | Lighthouse Lab in Alderley Park | Wellcome Sanger Institute for the COVID-19 Genomics UK (COG-UK) consortium | The Lighthouse Lab in Alderley Park and Alex Alderton, Roberto Amato, Sonia Goncalves, Ewan Harrison, David K. Jackson, Ian Johnston, Dominic Kwiatkowski, Cordelia Langford, John Sillitoe on behalf of the Wellcome Sanger Institute COVID-19 Surveillance Team (http://www.sanger.ac.uk/covid-team) |
| EPI_ISL_551744, EPI_ISL_551745, EPI_ISL_551746, EPI_ISL_551747, EPI_ISL_551748 | Lighthouse Lab in Milton Keynes | Wellcome Sanger Institute for the COVID-19 Genomics UK (COG-UK) consortium | The Lighthouse Lab in Milton Keynes and Alex Alderton, Roberto Amato, Sonia Goncalves, Ewan Harrison, David K. Jackson, Ian Johnston, Dominic Kwiatkowski, Cordelia Langford, John Sillitoe on behalf of the Wellcome Sanger Institute COVID-19 Surveillance Team (http://www.sanger.ac.uk/covid-team) |
| EPI_ISL_551749 | Lighthouse Lab in Alderley Park | Wellcome Sanger Institute for the COVID-19 Genomics UK (COG-UK) consortium | The Lighthouse Lab in Alderley Park and Alex Alderton, Roberto Amato, Sonia Goncalves, Ewan Harrison, David K. Jackson, Ian Johnston, Dominic Kwiatkowski, Cordelia Langford, John Sillitoe on behalf of the Wellcome Sanger Institute COVID-19 Surveillance Team (http://www.sanger.ac.uk/covid-team) |
| EPI_ISL_551750 | Lighthouse Lab in Milton Keynes | Wellcome Sanger Institute for the COVID-19 Genomics UK (COG-UK) consortium | The Lighthouse Lab in Milton Keynes and Alex Alderton, Roberto Amato, Sonia Goncalves, Ewan Harrison, David K. Jackson, Ian Johnston, Dominic Kwiatkowski, Cordelia Langford, John Sillitoe on behalf of the Wellcome Sanger Institute COVID-19 Surveillance Team (http://www.sanger.ac.uk/covid-team) |
| EPI_ISL_551751 | Lighthouse Lab in Alderley Park | Wellcome Sanger Institute for the COVID-19 Genomics UK (COG-UK) consortium | The Lighthouse Lab in Alderley Park and Alex Alderton, Roberto Amato, Sonia Goncalves, Ewan Harrison, David K. Jackson, Ian Johnston, Dominic Kwiatkowski, Cordelia Langford, John Sillitoe on behalf of the Wellcome Sanger Institute COVID-19 Surveillance Team |
| EPI_ISL_551752, EPI_ISL_551753, EPI_ISL_551754 | Lighthouse Lab in Milton Keynes | Wellcome Sanger Institute for the COVID-19 Genomics UK (COG-UK) consortium | The Lighthouse Lab in Milton Keynes and Alex Alderton, Roberto Amato, Sonia Goncalves, Ewan Harrison, David K. Jackson, Ian Johnston, Dominic Kwiatkowski, Cordelia Langford, John Sillitoe on behalf of the Wellcome Sanger Institute COVID-19 Surveillance Team (http://www.sanger.ac.uk/covid-team) |
| EPI_ISL_551755, EPI_ISL_551756 | Lighthouse Lab in Alderley Park | Wellcome Sanger Institute for the COVID-19 Genomics UK (COG-UK) consortium | The Lighthouse Lab in Alderley Park and Alex Alderton, Roberto Amato, Sonia Goncalves, Ewan Harrison, David K. Jackson, Ian Johnston, Dominic Kwiatkowski, Cordelia Langford, John Sillitoe on behalf of the Wellcome Sanger Institute COVID-19 Surveillance Team |
| EPI_ISL_551757 | Lighthouse Lab in Milton Keynes | Wellcome Sanger Institute for the COVID-19 Genomics UK (COG-UK) consortium | The Lighthouse Lab in Milton Keynes and Alex Alderton, Roberto Amato, Sonia Goncalves, Ewan Harrison, David K. Jackson, Ian Johnston, Dominic Kwiatkowski, Cordelia Langford, John Sillitoe on behalf of the Wellcome Sanger Institute COVID-19 Surveillance Team (http://www.sanger.ac.uk/covid-team) |
| EPI_ISL_551759, EPI_ISL_551760, EPI_ISL_551761, EPI_ISL_551762 | Lighthouse Lab in Alderley Park | Wellcome Sanger Institute for the COVID-19 Genomics UK (COG-UK) consortium | The Lighthouse Lab in Alderley Park and Alex Alderton, Roberto Amato, Sonia Goncalves, Ewan Harrison, David K. Jackson, Ian Johnston, Dominic Kwiatkowski, Cordelia Langford, John Sillitoe on behalf of the Wellcome Sanger Institute COVID-19 Surveillance Team |
| EPI_ISL_551763 | Lighthouse Lab in Alderley Park | Wellcome Sanger Institute for the COVID-19 Genomics UK (COG-UK) consortium | The Lighthouse Lab in Alderley Park and Alex Alderton, Roberto Amato, Sonia Goncalves, Ewan Harrison, David K. Jackson, Ian Johnston, Dominic Kwiatkowski, Cordelia Langford, John Sillitoe on behalf of the Wellcome Sanger Institute COVID-19 Surveillance Team (http://www.sanger.ac.uk/covid-team) |
| EPI_ISL_551764 | Lighthouse Lab in Alderley Park | Wellcome Sanger Institute for the COVID-19 Genomics UK (COG-UK) consortium | The Lighthouse Lab in Alderley Park and Alex Alderton, Roberto Amato, Sonia Goncalves, Ewan Harrison, David K. Jackson, Ian Johnston, Dominic Kwiatkowski, Cordelia Langford, John Sillitoe on behalf of the Wellcome Sanger Institute COVID-19 Surveillance Team |
| EPI_ISL_551765, EPI_ISL_551766, EPI_ISL_551767, EPI_ISL_551768 | Lighthouse Lab in Milton Keynes | Wellcome Sanger Institute for the COVID-19 Genomics UK (COG-UK) consortium | The Lighthouse Lab in Milton Keynes and Alex Alderton, Roberto Amato, Sonia Goncalves, Ewan Harrison, David K. Jackson, Ian Johnston, Dominic Kwiatkowski, Cordelia Langford, John Sillitoe on behalf of the Wellcome Sanger Institute COVID-19 Surveillance Team (http://www.sanger.ac.uk/covid-team) |
| EPI_ISL_551769 | Lighthouse Lab in Alderley Park | Wellcome Sanger Institute for the COVID-19 Genomics UK (COG-UK) consortium | The Lighthouse Lab in Alderley Park and Alex Alderton, Roberto Amato, Sonia Goncalves, Ewan Harrison, David K. Jackson, Ian Johnston, Dominic Kwiatkowski, Cordelia Langford, John Sillitoe on behalf of the Wellcome Sanger Institute COVID-19 Surveillance Team |
| EPI_ISL_551770, EPI_ISL_551771 | Lighthouse Lab in Milton Keynes | Wellcome Sanger Institute for the COVID-19 Genomics UK (COG-UK) consortium | The Lighthouse Lab in Milton Keynes and Alex Alderton, Roberto Amato, Sonia Goncalves, Ewan Harrison, David K. Jackson, Ian Johnston, Dominic Kwiatkowski, Cordelia Langford, John Sillitoe on behalf of the Wellcome Sanger Institute COVID-19 Surveillance Team (http://www.sanger.ac.uk/covid-team) |
| EPI_ISL_551772 | Lighthouse Lab in Alderley Park | Wellcome Sanger Institute for the COVID-19 Genomics UK (COG-UK) consortium | The Lighthouse Lab in Alderley Park and Alex Alderton, Roberto Amato, Sonia Goncalves, Ewan Harrison, David K. Jackson, Ian Johnston, Dominic Kwiatkowski, Cordelia Langford, John Sillitoe on behalf of the Wellcome Sanger Institute COVID-19 Surveillance Team |
| EPI_ISL_551773, EPI_ISL_551774, EPI_ISL_551775 | Lighthouse Lab in Milton Keynes | Wellcome Sanger Institute for the COVID-19 Genomics UK (COG-UK) consortium | The Lighthouse Lab in Milton Keynes and Alex Alderton, Roberto Amato, Sonia Goncalves, Ewan Harrison, David K. Jackson, Ian Johnston, Dominic Kwiatkowski, Cordelia Langford, John Sillitoe on behalf of the Wellcome Sanger Institute COVID-19 Surveillance Team (http://www.sanger.ac.uk/covid-team) |
| EPI_ISL_551776 | Lighthouse Lab in Alderley Park | Wellcome Sanger Institute for the COVID-19 Genomics UK (COG-UK) consortium | The Lighthouse Lab in Alderley Park and Alex Alderton, Roberto Amato, Sonia Goncalves, Ewan Harrison, David K. Jackson, Ian Johnston, Dominic Kwiatkowski, Cordelia Langford, John Sillitoe on behalf of the Wellcome Sanger Institute COVID-19 Surveillance Team |
| EPI_ISL_551777 | Lighthouse Lab in Milton Keynes | Wellcome Sanger Institute for the COVID-19 Genomics UK (COG-UK) consortium | The Lighthouse Lab in Alderley Park and Alex Alderton, Roberto Amato, Sonia Goncalves, Ewan Harrison, David K. Jackson, Ian Johnston, Dominic Kwiatkowski, Cordelia Langford, John Sillitoe on behalf of the Wellcome Sanger Institute COVID-19 Surveillance Team |
| EPI_ISL_551778 | Lighthouse Lab in Milton Keynes | Wellcome Sanger Institute for the COVID-19 Genomics UK (COG-UK) consortium | The Lighthouse Lab in Milton Keynes and Alex Alderton, Roberto Amato, Sonia Goncalves, Ewan Harrison, David K. Jackson, Ian Johnston, Dominic Kwiatkowski, Cordelia Langford, John Sillitoe on behalf of the Wellcome Sanger Institute COVID-19 Surveillance Team (http://www.sanger.ac.uk/covid-team) |
| EPI_ISL_551779, EPI_ISL_551780 | Lighthouse Lab in Alderley Park | Wellcome Sanger Institute for the COVID-19 Genomics UK | The Lighthouse Lab in Alderley Park and Alex Alderton, Roberto Amato, Sonia Goncalves, Ewan Harrison, David K. Jackson, Ian Johnston, Dominic |

| | | | |
|---|---|---|---|
| | | (COG-UK) consortium | Kwiatkowski, Cordelia Langford, John Sillitoe on behalf of the Wellcome Sanger Institute COVID-19 Surveillance Team |
| EPI_ISL_551781, EPI_ISL_551782, EPI_ISL_551783 | Lighthouse Lab in Milton Keynes | Wellcome Sanger Institute for the COVID-19 Genomics UK (COG-UK) consortium | The Lighthouse Lab in Milton Keynes and Alex Alderton, Roberto Amato, Sonia Goncalves, Ewan Harrison, David K. Jackson, Ian Johnston, Dominic Kwiatkowski, Cordelia Langford, John Sillitoe on behalf of the Wellcome Sanger Institute COVID-19 Surveillance Team (http://www.sanger.ac.uk/covid-team) |
| EPI_ISL_551784, EPI_ISL_551785 | Lighthouse Lab in Alderley Park | Wellcome Sanger Institute for the COVID-19 Genomics UK (COG-UK) consortium | The Lighthouse Lab in Alderley Park and Alex Alderton, Roberto Amato, Sonia Goncalves, Ewan Harrison, David K. Jackson, Ian Johnston, Dominic Kwiatkowski, Cordelia Langford, John Sillitoe on behalf of the Wellcome Sanger Institute COVID-19 Surveillance Team |
| EPI_ISL_551787, EPI_ISL_551788 | Lighthouse Lab in Milton Keynes | Wellcome Sanger Institute for the COVID-19 Genomics UK (COG-UK) consortium | The Lighthouse Lab in Milton Keynes and Alex Alderton, Roberto Amato, Sonia Goncalves, Ewan Harrison, David K. Jackson, Ian Johnston, Dominic Kwiatkowski, Cordelia Langford, John Sillitoe on behalf of the Wellcome Sanger Institute COVID-19 Surveillance Team (http://www.sanger.ac.uk/covid-team) |
| EPI_ISL_551789 | Lighthouse Lab in Alderley Park | Wellcome Sanger Institute for the COVID-19 Genomics UK (COG-UK) consortium | The Lighthouse Lab in Alderley Park and Alex Alderton, Roberto Amato, Sonia Goncalves, Ewan Harrison, David K. Jackson, Ian Johnston, Dominic Kwiatkowski, Cordelia Langford, John Sillitoe on behalf of the Wellcome Sanger Institute COVID-19 Surveillance Team |
| EPI_ISL_551790, EPI_ISL_551791, EPI_ISL_551792, EPI_ISL_551793, EPI_ISL_551794 | Lighthouse Lab in Milton Keynes | Wellcome Sanger Institute for the COVID-19 Genomics UK (COG-UK) consortium | The Lighthouse Lab in Milton Keynes and Alex Alderton, Roberto Amato, Sonia Goncalves, Ewan Harrison, David K. Jackson, Ian Johnston, Dominic Kwiatkowski, Cordelia Langford, John Sillitoe on behalf of the Wellcome Sanger Institute COVID-19 Surveillance Team (http://www.sanger.ac.uk/covid-team) |
| EPI_ISL_551795, EPI_ISL_551796 | Lighthouse Lab in Alderley Park | Wellcome Sanger Institute for the COVID-19 Genomics UK (COG-UK) consortium | The Lighthouse Lab in Alderley Park and Alex Alderton, Roberto Amato, Sonia Goncalves, Ewan Harrison, David K. Jackson, Ian Johnston, Dominic Kwiatkowski, Cordelia Langford, John Sillitoe on behalf of the Wellcome Sanger Institute COVID-19 Surveillance Team |
| EPI_ISL_551797 | Lighthouse Lab in Milton Keynes | Wellcome Sanger Institute for the COVID-19 Genomics UK (COG-UK) consortium | The Lighthouse Lab in Milton Keynes and Alex Alderton, Roberto Amato, Sonia Goncalves, Ewan Harrison, David K. Jackson, Ian Johnston, Dominic Kwiatkowski, Cordelia Langford, John Sillitoe on behalf of the Wellcome Sanger Institute COVID-19 Surveillance Team (http://www.sanger.ac.uk/covid-team) |
| EPI_ISL_551798, EPI_ISL_551799 | Lighthouse Lab in Alderley Park | Wellcome Sanger Institute for the COVID-19 Genomics UK (COG-UK) consortium | The Lighthouse Lab in Alderley Park and Alex Alderton, Roberto Amato, Sonia Goncalves, Ewan Harrison, David K. Jackson, Ian Johnston, Dominic Kwiatkowski, Cordelia Langford, John Sillitoe on behalf of the Wellcome Sanger Institute COVID-19 Surveillance Team |
| EPI_ISL_551801 | Lighthouse Lab in Alderley Park | Wellcome Sanger Institute for the COVID-19 Genomics UK (COG-UK) consortium | The Lighthouse Lab in Alderley Park and Alex Alderton, Roberto Amato, Sonia Goncalves, Ewan Harrison, David K. Jackson, Ian Johnston, Dominic Kwiatkowski, Cordelia Langford, John Sillitoe on behalf of the Wellcome Sanger Institute COVID-19 Surveillance Team (http://www.sanger.ac.uk/covid-team) |
| EPI_ISL_551802 | Lighthouse Lab in Milton Keynes | Wellcome Sanger Institute for the COVID-19 Genomics UK (COG-UK) consortium | The Lighthouse Lab in Milton Keynes and Alex Alderton, Roberto Amato, Sonia Goncalves, Ewan Harrison, David K. Jackson, Ian Johnston, Dominic Kwiatkowski, Cordelia Langford, John Sillitoe on behalf of the Wellcome Sanger Institute COVID-19 Surveillance Team (http://www.sanger.ac.uk/covid-team) |
| EPI_ISL_551803, EPI_ISL_551804 | Lighthouse Lab in Alderley Park | Wellcome Sanger Institute for the COVID-19 Genomics UK (COG-UK) consortium | The Lighthouse Lab in Alderley Park and Alex Alderton, Roberto Amato, Sonia Goncalves, Ewan Harrison, David K. Jackson, Ian Johnston, Dominic Kwiatkowski, Cordelia Langford, John Sillitoe on behalf of the Wellcome Sanger Institute COVID-19 Surveillance Team |
| EPI_ISL_551805 | Lighthouse Lab in Milton Keynes | Wellcome Sanger Institute for the COVID-19 Genomics UK (COG-UK) consortium | The Lighthouse Lab in Milton Keynes and Alex Alderton, Roberto Amato, Sonia Goncalves, Ewan Harrison, David K. Jackson, Ian Johnston, Dominic Kwiatkowski, Cordelia Langford, John Sillitoe on behalf of the Wellcome Sanger Institute COVID-19 Surveillance Team (http://www.sanger.ac.uk/covid-team) |
| EPI_ISL_551806, EPI_ISL_551807, EPI_ISL_551808 | Lighthouse Lab in Alderley Park | Wellcome Sanger Institute for the COVID-19 Genomics UK (COG-UK) consortium | The Lighthouse Lab in Alderley Park and Alex Alderton, Roberto Amato, Sonia Goncalves, Ewan Harrison, David K. Jackson, Ian Johnston, Dominic Kwiatkowski, Cordelia Langford, John Sillitoe on behalf of the Wellcome Sanger Institute COVID-19 Surveillance Team |
| EPI_ISL_551809 | Lighthouse Lab in Milton Keynes | Wellcome Sanger Institute for the COVID-19 Genomics UK (COG-UK) consortium | The Lighthouse Lab in Milton Keynes and Alex Alderton, Roberto Amato, Sonia Goncalves, Ewan Harrison, David K. Jackson, Ian Johnston, Dominic Kwiatkowski, Cordelia Langford, John Sillitoe on behalf of the Wellcome Sanger Institute COVID-19 Surveillance Team (http://www.sanger.ac.uk/covid-team) |
| EPI_ISL_551810, EPI_ISL_551811, EPI_ISL_551812 | Lighthouse Lab in Alderley Park | Wellcome Sanger Institute for the COVID-19 Genomics UK (COG-UK) consortium | The Lighthouse Lab in Alderley Park and Alex Alderton, Roberto Amato, Sonia Goncalves, Ewan Harrison, David K. Jackson, Ian Johnston, Dominic Kwiatkowski, Cordelia Langford, John Sillitoe on behalf of the Wellcome Sanger Institute COVID-19 Surveillance Team |
| EPI_ISL_551813, EPI_ISL_551814 | Lighthouse Lab in Milton Keynes | Wellcome Sanger Institute for the COVID-19 Genomics UK (COG-UK) consortium | The Lighthouse Lab in Milton Keynes and Alex Alderton, Roberto Amato, Sonia Goncalves, Ewan Harrison, David K. Jackson, Ian Johnston, Dominic Kwiatkowski, Cordelia Langford, John Sillitoe on behalf of the Wellcome Sanger Institute COVID-19 Surveillance Team (http://www.sanger.ac.uk/covid-team) |
| EPI_ISL_551815 | Lighthouse Lab in Alderley Park | Wellcome Sanger Institute for the COVID-19 Genomics UK (COG-UK) consortium | The Lighthouse Lab in Alderley Park and Alex Alderton, Roberto Amato, Sonia Goncalves, Ewan Harrison, David K. Jackson, Ian Johnston, Dominic Kwiatkowski, Cordelia Langford, John Sillitoe on behalf of the Wellcome Sanger Institute COVID-19 Surveillance Team (http://www.sanger.ac.uk/covid-team) |
| EPI_ISL_551816, EPI_ISL_551817 | Lighthouse Lab in Milton Keynes | Wellcome Sanger Institute for the COVID-19 Genomics UK (COG-UK) consortium | The Lighthouse Lab in Milton Keynes and Alex Alderton, Roberto Amato, Sonia Goncalves, Ewan Harrison, David K. Jackson, Ian Johnston, Dominic Kwiatkowski, Cordelia Langford, John Sillitoe on behalf of the Wellcome Sanger Institute COVID-19 Surveillance Team (http://www.sanger.ac.uk/covid-team) |
| EPI_ISL_551818, EPI_ISL_551819 | Lighthouse Lab in Alderley Park | Wellcome Sanger Institute for the COVID-19 Genomics UK (COG-UK) consortium | The Lighthouse Lab in Alderley Park and Alex Alderton, Roberto Amato, Sonia Goncalves, Ewan Harrison, David K. Jackson, Ian Johnston, Dominic Kwiatkowski, Cordelia Langford, John Sillitoe on behalf of the Wellcome Sanger Institute COVID-19 Surveillance Team |
| EPI_ISL_551820, EPI_ISL_551821, EPI_ISL_551822 | Lighthouse Lab in Milton Keynes | Wellcome Sanger Institute for the COVID-19 Genomics UK (COG-UK) consortium | The Lighthouse Lab in Milton Keynes and Alex Alderton, Roberto Amato, Sonia Goncalves, Ewan Harrison, David K. Jackson, Ian Johnston, Dominic Kwiatkowski, Cordelia Langford, John Sillitoe on behalf of the Wellcome Sanger Institute COVID-19 Surveillance Team (http://www.sanger.ac.uk/covid-team) |
| EPI_ISL_551823, EPI_ISL_551824 | Lighthouse Lab in Alderley Park | Wellcome Sanger Institute for the COVID-19 Genomics UK (COG-UK) consortium | The Lighthouse Lab in Alderley Park and Alex Alderton, Roberto Amato, Sonia Goncalves, Ewan Harrison, David K. Jackson, Ian Johnston, Dominic Kwiatkowski, Cordelia Langford, John Sillitoe on behalf of the Wellcome Sanger Institute COVID-19 Surveillance Team |
| EPI_ISL_551825 | Lighthouse Lab in Alderley Park | Wellcome Sanger Institute for the COVID-19 Genomics UK (COG-UK) consortium | The Lighthouse Lab in Alderley Park and Alex Alderton, Roberto Amato, Sonia Goncalves, Ewan Harrison, David K. Jackson, Ian Johnston, Dominic Kwiatkowski, Cordelia Langford, John Sillitoe on behalf of the Wellcome Sanger Institute COVID-19 Surveillance Team (http://www.sanger.ac.uk/covid-team) |
| EPI_ISL_551826, EPI_ISL_551827 | Lighthouse Lab in Alderley Park | Wellcome Sanger Institute for the COVID-19 Genomics UK (COG-UK) consortium | The Lighthouse Lab in Alderley Park and Alex Alderton, Roberto Amato, Sonia Goncalves, Ewan Harrison, David K. Jackson, Ian Johnston, Dominic Kwiatkowski, Cordelia Langford, John Sillitoe on behalf of the Wellcome Sanger Institute COVID-19 Surveillance Team |
| EPI_ISL_551828, EPI_ISL_551829, EPI_ISL_551830, EPI_ISL_551831 | Lighthouse Lab in Milton Keynes | Wellcome Sanger Institute for the COVID-19 Genomics UK (COG-UK) consortium | The Lighthouse Lab in Milton Keynes and Alex Alderton, Roberto Amato, Sonia Goncalves, Ewan Harrison, David K. Jackson, Ian Johnston, Dominic Kwiatkowski, Cordelia Langford, John Sillitoe on behalf of the Wellcome Sanger Institute COVID-19 Surveillance Team (http://www.sanger.ac.uk/covid-team) |
| EPI_ISL_551832 | Lighthouse Lab in Alderley Park | Wellcome Sanger Institute for the COVID-19 Genomics UK (COG-UK) consortium | The Lighthouse Lab in Alderley Park and Alex Alderton, Roberto Amato, Sonia Goncalves, Ewan Harrison, David K. Jackson, Ian Johnston, Dominic Kwiatkowski, Cordelia Langford, John Sillitoe on behalf of the Wellcome Sanger Institute COVID-19 Surveillance Team (http://www.sanger.ac.uk/covid-team) |
| EPI_ISL_551833 | Lighthouse Lab in Milton Keynes | Wellcome Sanger Institute for the COVID-19 Genomics UK (COG-UK) consortium | The Lighthouse Lab in Milton Keynes and Alex Alderton, Roberto Amato, Sonia Goncalves, Ewan Harrison, David K. Jackson, Ian Johnston, Dominic Kwiatkowski, Cordelia Langford, John Sillitoe on behalf of the Wellcome Sanger Institute COVID-19 Surveillance Team (http://www.sanger.ac.uk/covid-team) |
| EPI_ISL_551834 | Lighthouse Lab in Alderley Park | Wellcome Sanger Institute for the COVID-19 Genomics UK (COG-UK) consortium | The Lighthouse Lab in Alderley Park and Alex Alderton, Roberto Amato, Sonia Goncalves, Ewan Harrison, David K. Jackson, Ian Johnston, Dominic Kwiatkowski, Cordelia Langford, John Sillitoe on behalf of the Wellcome Sanger Institute COVID-19 Surveillance Team |
| EPI_ISL_555509 | Lighthouse Lab in Alderley Park | Wellcome Sanger Institute for the COVID-19 Genomics UK (COG-UK) consortium | The Lighthouse Lab in Alderley Park and Alex Alderton, Roberto Amato, Sonia Goncalves, Ewan Harrison, David K. Jackson, Ian Johnston, Dominic Kwiatkowski, Cordelia Langford, John Sillitoe on behalf of the Wellcome Sanger Institute COVID-19 Surveillance Team (http://www.sanger.ac.uk/covid-team) |
| EPI_ISL_555515, EPI_ISL_555518, EPI_ISL_555521, EPI_ISL_555528, EPI_ISL_555530 | Lighthouse Lab in Alderley Park | Wellcome Sanger Institute for the COVID-19 Genomics UK (COG-UK) consortium | The Lighthouse Lab in Alderley Park and Alex Alderton, Roberto Amato, Sonia Goncalves, Ewan Harrison, David K. Jackson, Ian Johnston, Dominic Kwiatkowski, Cordelia Langford, John Sillitoe on behalf of the Wellcome Sanger Institute COVID-19 Surveillance Team |
| EPI_ISL_555536 | Lighthouse Lab in Alderley Park | Wellcome Sanger Institute for the COVID-19 Genomics UK (COG-UK) consortium | The Lighthouse Lab in Alderley Park and Alex Alderton, Roberto Amato, Sonia Goncalves, Ewan Harrison, David K. Jackson, Ian Johnston, Dominic Kwiatkowski, Cordelia Langford, John Sillitoe on behalf of the Wellcome Sanger Institute COVID-19 Surveillance Team (http://www.sanger.ac.uk/covid-team) |
| EPI_ISL_555538, EPI_ISL_555539, EPI_ISL_555554, EPI_ISL_555555, EPI_ISL_555562, EPI_ISL_555569, EPI_ISL_555571, EPI_ISL_555575, EPI_ISL_555578, EPI_ISL_555586, EPI_ISL_555591, EPI_ISL_555597, EPI_ISL_555603, EPI_ISL_555605, EPI_ISL_555609, EPI_ISL_555610, EPI_ISL_555612, EPI_ISL_555621, EPI_ISL_555624, EPI_ISL_555625 | | | |
| see above | Lighthouse Lab in Alderley Park | Wellcome Sanger Institute for the COVID-19 Genomics UK (COG-UK) consortium | The Lighthouse Lab in Alderley Park and Alex Alderton, Roberto Amato, Sonia Goncalves, Ewan Harrison, David K. Jackson, Ian Johnston, Dominic Kwiatkowski, Cordelia Langford, John Sillitoe on behalf of the Wellcome Sanger Institute COVID-19 Surveillance Team |
| EPI_ISL_555634, EPI_ISL_555637, EPI_ISL_555639, EPI_ISL_555641 | Lighthouse Lab in Alderley Park | Wellcome Sanger Institute for the COVID-19 Genomics UK (COG-UK) consortium | The Lighthouse Lab in Alderley Park and Alex Alderton, Roberto Amato, Sonia Goncalves, Ewan Harrison, David K. Jackson, Ian Johnston, Dominic Kwiatkowski, Cordelia Langford, John Sillitoe on behalf of the Wellcome Sanger Institute COVID-19 Surveillance Team (http://www.sanger.ac.uk/covid-team) |
| EPI_ISL_555644, EPI_ISL_555645, EPI_ISL_555646, EPI_ISL_555648, | Lighthouse Lab in Alderley Park | Wellcome Sanger Institute for the COVID-19 Genomics UK (COG-UK) consortium | The Lighthouse Lab in Alderley Park and Alex Alderton, Roberto Amato, Sonia Goncalves, Ewan Harrison, David K. Jackson, Ian Johnston, Dominic Kwiatkowski, Cordelia Langford, John Sillitoe on behalf of the Wellcome Sanger Institute COVID-19 Surveillance Team |

| | | | |
|---|---|---|---|
| EPI_ISL_555649, EPI_ISL_555652 | | | |
| EPI_ISL_555654 | Lighthouse Lab in Alderley Park | Wellcome Sanger Institute for the COVID-19 Genomics UK (COG-UK) consortium | The Lighthouse Lab in Alderley Park and Alex Alderton, Roberto Amato, Sonia Goncalves, Ewan Harrison, David K. Jackson, Ian Johnston, Dominic Kwiatkowski, Cordelia Langford, John Sillitoe on behalf of the Wellcome Sanger Institute COVID-19 Surveillance Team (http://www.sanger.ac.uk/covid-team) |
| EPI_ISL_555665, EPI_ISL_555675 | Lighthouse Lab in Alderley Park | Wellcome Sanger Institute for the COVID-19 Genomics UK (COG-UK) consortium | The Lighthouse Lab in Alderley Park and Alex Alderton, Roberto Amato, Sonia Goncalves, Ewan Harrison, David K. Jackson, Ian Johnston, Dominic Kwiatkowski, Cordelia Langford, John Sillitoe on behalf of the Wellcome Sanger Institute COVID-19 Surveillance Team |
| EPI_ISL_555676 | Lighthouse Lab in Alderley Park | Wellcome Sanger Institute for the COVID-19 Genomics UK (COG-UK) consortium | The Lighthouse Lab in Alderley Park and Alex Alderton, Roberto Amato, Sonia Goncalves, Ewan Harrison, David K. Jackson, Ian Johnston, Dominic Kwiatkowski, Cordelia Langford, John Sillitoe on behalf of the Wellcome Sanger Institute COVID-19 Surveillance Team (http://www.sanger.ac.uk/covid-team) |
| EPI_ISL_555684 | Lighthouse Lab in Alderley Park | Wellcome Sanger Institute for the COVID-19 Genomics UK (COG-UK) consortium | The Lighthouse Lab in Alderley Park and Alex Alderton, Roberto Amato, Sonia Goncalves, Ewan Harrison, David K. Jackson, Ian Johnston, Dominic Kwiatkowski, Cordelia Langford, John Sillitoe on behalf of the Wellcome Sanger Institute COVID-19 Surveillance Team |
| EPI_ISL_555686 | Lighthouse Lab in Alderley Park | Wellcome Sanger Institute for the COVID-19 Genomics UK (COG-UK) consortium | The Lighthouse Lab in Alderley Park and Alex Alderton, Roberto Amato, Sonia Goncalves, Ewan Harrison, David K. Jackson, Ian Johnston, Dominic Kwiatkowski, Cordelia Langford, John Sillitoe on behalf of the Wellcome Sanger Institute COVID-19 Surveillance Team (http://www.sanger.ac.uk/covid-team) |
| EPI_ISL_555687, EPI_ISL_555688, EPI_ISL_555689, EPI_ISL_555691, EPI_ISL_555692, EPI_ISL_555697, EPI_ISL_555701, EPI_ISL_555703, EPI_ISL_555704, EPI_ISL_555722, EPI_ISL_555723, EPI_ISL_555730, EPI_ISL_555733, EPI_ISL_555737 | | | |
| see above | Lighthouse Lab in Alderley Park | Wellcome Sanger Institute for the COVID-19 Genomics UK (COG-UK) consortium | The Lighthouse Lab in Alderley Park and Alex Alderton, Roberto Amato, Sonia Goncalves, Ewan Harrison, David K. Jackson, Ian Johnston, Dominic Kwiatkowski, Cordelia Langford, John Sillitoe on behalf of the Wellcome Sanger Institute COVID-19 Surveillance Team |
| EPI_ISL_555743, EPI_ISL_555747, EPI_ISL_555751 | Lighthouse Lab in Alderley Park | Wellcome Sanger Institute for the COVID-19 Genomics UK (COG-UK) consortium | The Lighthouse Lab in Alderley Park and Alex Alderton, Roberto Amato, Sonia Goncalves, Ewan Harrison, David K. Jackson, Ian Johnston, Dominic Kwiatkowski, Cordelia Langford, John Sillitoe on behalf of the Wellcome Sanger Institute COVID-19 Surveillance Team (http://www.sanger.ac.uk/covid-team) |
| EPI_ISL_555757, EPI_ISL_555760 | Lighthouse Lab in Alderley Park | Wellcome Sanger Institute for the COVID-19 Genomics UK (COG-UK) consortium | The Lighthouse Lab in Alderley Park and Alex Alderton, Roberto Amato, Sonia Goncalves, Ewan Harrison, David K. Jackson, Ian Johnston, Dominic Kwiatkowski, Cordelia Langford, John Sillitoe on behalf of the Wellcome Sanger Institute COVID-19 Surveillance Team |
| EPI_ISL_556074 | Lighthouse Lab in Alderley Park | Wellcome Sanger Institute for the COVID-19 Genomics UK (COG-UK) consortium | The Lighthouse Lab in Alderley Park and Alex Alderton, Roberto Amato, Sonia Goncalves, Ewan Harrison, David K. Jackson, Ian Johnston, Dominic Kwiatkowski, Cordelia Langford, John Sillitoe on behalf of the Wellcome Sanger Institute COVID-19 Surveillance Team (http://www.sanger.ac.uk/covid-team) |
| EPI_ISL_556078 | Lighthouse Lab in Alderley Park | Wellcome Sanger Institute for the COVID-19 Genomics UK (COG-UK) consortium | The Lighthouse Lab in Alderley Park and Alex Alderton, Roberto Amato, Sonia Goncalves, Ewan Harrison, David K. Jackson, Ian Johnston, Dominic Kwiatkowski, Cordelia Langford, John Sillitoe on behalf of the Wellcome Sanger Institute COVID-19 Surveillance Team |
| EPI_ISL_556080, EPI_ISL_556082 | Lighthouse Lab in Alderley Park | Wellcome Sanger Institute for the COVID-19 Genomics UK (COG-UK) consortium | The Lighthouse Lab in Alderley Park and Alex Alderton, Roberto Amato, Sonia Goncalves, Ewan Harrison, David K. Jackson, Ian Johnston, Dominic Kwiatkowski, Cordelia Langford, John Sillitoe on behalf of the Wellcome Sanger Institute COVID-19 Surveillance Team (http://www.sanger.ac.uk/covid-team) |
| EPI_ISL_556086 | Lighthouse Lab in Alderley Park | Wellcome Sanger Institute for the COVID-19 Genomics UK (COG-UK) consortium | The Lighthouse Lab in Alderley Park and Alex Alderton, Roberto Amato, Sonia Goncalves, Ewan Harrison, David K. Jackson, Ian Johnston, Dominic Kwiatkowski, Cordelia Langford, John Sillitoe on behalf of the Wellcome Sanger Institute COVID-19 Surveillance Team |
| EPI_ISL_556088 | Lighthouse Lab in Alderley Park | Wellcome Sanger Institute for the COVID-19 Genomics UK (COG-UK) consortium | The Lighthouse Lab in Alderley Park and Alex Alderton, Roberto Amato, Sonia Goncalves, Ewan Harrison, David K. Jackson, Ian Johnston, Dominic Kwiatkowski, Cordelia Langford, John Sillitoe on behalf of the Wellcome Sanger Institute COVID-19 Surveillance Team (http://www.sanger.ac.uk/covid-team) |
| EPI_ISL_556093, EPI_ISL_556094 | Lighthouse Lab in Milton Keynes | Wellcome Sanger Institute for the COVID-19 Genomics UK (COG-UK) consortium | The Lighthouse Lab in Milton Keynes and Alex Alderton, Roberto Amato, Sonia Goncalves, Ewan Harrison, David K. Jackson, Ian Johnston, Dominic Kwiatkowski, Cordelia Langford, John Sillitoe on behalf of the Wellcome Sanger Institute COVID-19 Surveillance Team (http://www.sanger.ac.uk/covid-team) |
| EPI_ISL_556096, EPI_ISL_556099, EPI_ISL_556100 | Lighthouse Lab in Alderley Park | Wellcome Sanger Institute for the COVID-19 Genomics UK (COG-UK) consortium | The Lighthouse Lab in Alderley Park and Alex Alderton, Roberto Amato, Sonia Goncalves, Ewan Harrison, David K. Jackson, Ian Johnston, Dominic Kwiatkowski, Cordelia Langford, John Sillitoe on behalf of the Wellcome Sanger Institute COVID-19 Surveillance Team |
| EPI_ISL_556102 | Lighthouse Lab in Milton Keynes | Wellcome Sanger Institute for the COVID-19 Genomics UK (COG-UK) consortium | The Lighthouse Lab in Milton Keynes and Alex Alderton, Roberto Amato, Sonia Goncalves, Ewan Harrison, David K. Jackson, Ian Johnston, Dominic Kwiatkowski, Cordelia Langford, John Sillitoe on behalf of the Wellcome Sanger Institute COVID-19 Surveillance Team (http://www.sanger.ac.uk/covid-team) |
| EPI_ISL_556103 | Lighthouse Lab in Alderley Park | Wellcome Sanger Institute for the COVID-19 Genomics UK (COG-UK) consortium | The Lighthouse Lab in Alderley Park and Alex Alderton, Roberto Amato, Sonia Goncalves, Ewan Harrison, David K. Jackson, Ian Johnston, Dominic Kwiatkowski, Cordelia Langford, John Sillitoe on behalf of the Wellcome Sanger Institute COVID-19 Surveillance Team |
| EPI_ISL_556105 | Lighthouse Lab in Alderley Park | Wellcome Sanger Institute for the COVID-19 Genomics UK (COG-UK) consortium | The Lighthouse Lab in Alderley Park and Alex Alderton, Roberto Amato, Sonia Goncalves, Ewan Harrison, David K. Jackson, Ian Johnston, Dominic Kwiatkowski, Cordelia Langford, John Sillitoe on behalf of the Wellcome Sanger Institute COVID-19 Surveillance Team (http://www.sanger.ac.uk/covid-team) |
| EPI_ISL_556108 | Lighthouse Lab in Alderley Park | Wellcome Sanger Institute for the COVID-19 Genomics UK (COG-UK) consortium | The Lighthouse Lab in Alderley Park and Alex Alderton, Roberto Amato, Sonia Goncalves, Ewan Harrison, David K. Jackson, Ian Johnston, Dominic Kwiatkowski, Cordelia Langford, John Sillitoe on behalf of the Wellcome Sanger Institute COVID-19 Surveillance Team |
| EPI_ISL_556109, EPI_ISL_556112 | Lighthouse Lab in Milton Keynes | Wellcome Sanger Institute for the COVID-19 Genomics UK (COG-UK) consortium | The Lighthouse Lab in Milton Keynes and Alex Alderton, Roberto Amato, Sonia Goncalves, Ewan Harrison, David K. Jackson, Ian Johnston, Dominic Kwiatkowski, Cordelia Langford, John Sillitoe on behalf of the Wellcome Sanger Institute COVID-19 Surveillance Team (http://www.sanger.ac.uk/covid-team) |
| EPI_ISL_556113, EPI_ISL_556114 | Lighthouse Lab in Alderley Park | Wellcome Sanger Institute for the COVID-19 Genomics UK (COG-UK) consortium | The Lighthouse Lab in Alderley Park and Alex Alderton, Roberto Amato, Sonia Goncalves, Ewan Harrison, David K. Jackson, Ian Johnston, Dominic Kwiatkowski, Cordelia Langford, John Sillitoe on behalf of the Wellcome Sanger Institute COVID-19 Surveillance Team (http://www.sanger.ac.uk/covid-team) |
| EPI_ISL_556115, EPI_ISL_556116 | Lighthouse Lab in Milton Keynes | Wellcome Sanger Institute for the COVID-19 Genomics UK (COG-UK) consortium | The Lighthouse Lab in Milton Keynes and Alex Alderton, Roberto Amato, Sonia Goncalves, Ewan Harrison, David K. Jackson, Ian Johnston, Dominic Kwiatkowski, Cordelia Langford, John Sillitoe on behalf of the Wellcome Sanger Institute COVID-19 Surveillance Team (http://www.sanger.ac.uk/covid-team) |
| EPI_ISL_556117, EPI_ISL_556119, EPI_ISL_556120, EPI_ISL_556121 | Lighthouse Lab in Alderley Park | Wellcome Sanger Institute for the COVID-19 Genomics UK (COG-UK) consortium | The Lighthouse Lab in Alderley Park and Alex Alderton, Roberto Amato, Sonia Goncalves, Ewan Harrison, David K. Jackson, Ian Johnston, Dominic Kwiatkowski, Cordelia Langford, John Sillitoe on behalf of the Wellcome Sanger Institute COVID-19 Surveillance Team (http://www.sanger.ac.uk/covid-team) |
| EPI_ISL_556123 | Lighthouse Lab in Milton Keynes | Wellcome Sanger Institute for the COVID-19 Genomics UK (COG-UK) consortium | The Lighthouse Lab in Milton Keynes and Alex Alderton, Roberto Amato, Sonia Goncalves, Ewan Harrison, David K. Jackson, Ian Johnston, Dominic Kwiatkowski, Cordelia Langford, John Sillitoe on behalf of the Wellcome Sanger Institute COVID-19 Surveillance Team (http://www.sanger.ac.uk/covid-team) |
| EPI_ISL_556124, EPI_ISL_556125 | Lighthouse Lab in Alderley Park | Wellcome Sanger Institute for the COVID-19 Genomics UK (COG-UK) consortium | The Lighthouse Lab in Alderley Park and Alex Alderton, Roberto Amato, Sonia Goncalves, Ewan Harrison, David K. Jackson, Ian Johnston, Dominic Kwiatkowski, Cordelia Langford, John Sillitoe on behalf of the Wellcome Sanger Institute COVID-19 Surveillance Team (http://www.sanger.ac.uk/covid-team) |
| EPI_ISL_556126 | Lighthouse Lab in Milton Keynes | Wellcome Sanger Institute for the COVID-19 Genomics UK (COG-UK) consortium | The Lighthouse Lab in Milton Keynes and Alex Alderton, Roberto Amato, Sonia Goncalves, Ewan Harrison, David K. Jackson, Ian Johnston, Dominic Kwiatkowski, Cordelia Langford, John Sillitoe on behalf of the Wellcome Sanger Institute COVID-19 Surveillance Team (http://www.sanger.ac.uk/covid-team) |
| EPI_ISL_556127, EPI_ISL_556128, EPI_ISL_556129, EPI_ISL_556132, EPI_ISL_556135 | Lighthouse Lab in Alderley Park | Wellcome Sanger Institute for the COVID-19 Genomics UK (COG-UK) consortium | The Lighthouse Lab in Alderley Park and Alex Alderton, Roberto Amato, Sonia Goncalves, Ewan Harrison, David K. Jackson, Ian Johnston, Dominic Kwiatkowski, Cordelia Langford, John Sillitoe on behalf of the Wellcome Sanger Institute COVID-19 Surveillance Team (http://www.sanger.ac.uk/covid-team) |
| EPI_ISL_556138, EPI_ISL_556139 | Lighthouse Lab in Milton Keynes | Wellcome Sanger Institute for the COVID-19 Genomics UK (COG-UK) consortium | The Lighthouse Lab in Milton Keynes and Alex Alderton, Roberto Amato, Sonia Goncalves, Ewan Harrison, David K. Jackson, Ian Johnston, Dominic Kwiatkowski, Cordelia Langford, John Sillitoe on behalf of the Wellcome Sanger Institute COVID-19 Surveillance Team (http://www.sanger.ac.uk/covid-team) |
| EPI_ISL_556140 | Lighthouse Lab in Alderley Park | Wellcome Sanger Institute for the COVID-19 Genomics UK (COG-UK) consortium | The Lighthouse Lab in Alderley Park and Alex Alderton, Roberto Amato, Sonia Goncalves, Ewan Harrison, David K. Jackson, Ian Johnston, Dominic Kwiatkowski, Cordelia Langford, John Sillitoe on behalf of the Wellcome Sanger Institute COVID-19 Surveillance Team (http://www.sanger.ac.uk/covid-team) |
| EPI_ISL_556141 | Lighthouse Lab in Alderley Park | Wellcome Sanger Institute for the COVID-19 Genomics UK (COG-UK) Consortium | The Lighthouse Lab in Alderley Park and Alex Alderton, Roberto Amato, Sonia Goncalves, Ewan Harrison, David K. Jackson, Ian Johnston, Dominic Kwiatkowski, Cordelia Langford, John Sillitoe on behalf of the Wellcome Sanger Institute COVID-19 Surveillance Team |
| EPI_ISL_556142, EPI_ISL_556143 | Lighthouse Lab in Alderley Park | Wellcome Sanger Institute for the COVID-19 Genomics UK (COG-UK) consortium | The Lighthouse Lab in Alderley Park and Alex Alderton, Roberto Amato, Sonia Goncalves, Ewan Harrison, David K. Jackson, Ian Johnston, Dominic Kwiatkowski, Cordelia Langford, John Sillitoe on behalf of the Wellcome Sanger Institute COVID-19 Surveillance Team |
| EPI_ISL_556144 | Lighthouse Lab in Alderley Park | Wellcome Sanger Institute for the COVID-19 Genomics UK (COG-UK) consortium | The Lighthouse Lab in Alderley Park and Alex Alderton, Roberto Amato, Sonia Goncalves, Ewan Harrison, David K. Jackson, Ian Johnston, Dominic Kwiatkowski, Cordelia Langford, John Sillitoe on behalf of the Wellcome Sanger Institute COVID-19 Surveillance Team (http://www.sanger.ac.uk/covid-team) |
| EPI_ISL_556145, EPI_ISL_556147 | Lighthouse Lab in Alderley Park | Wellcome Sanger Institute for the COVID-19 Genomics UK (COG-UK) consortium | The Lighthouse Lab in Alderley Park and Alex Alderton, Roberto Amato, Sonia Goncalves, Ewan Harrison, David K. Jackson, Ian Johnston, Dominic Kwiatkowski, Cordelia Langford, John Sillitoe on behalf of the Wellcome Sanger Institute COVID-19 Surveillance Team |
| EPI_ISL_556148 | Lighthouse Lab in Milton Keynes | Wellcome Sanger Institute for the COVID-19 Genomics UK | The Lighthouse Lab in Milton Keynes and Alex Alderton, Roberto Amato, Sonia Goncalves, Ewan Harrison, David K. Jackson, Ian Johnston, Dominic |

| | | | |
|---|---|---|---|
| | | (COG-UK) consortium | Kwiatkowski, Cordelia Langford, John Sillitoe on behalf of the Wellcome Sanger Institute COVID-19 Surveillance Team (http://www.sanger.ac.uk/covid-team) |
| EPI_ISL_556149, EPI_ISL_556152 | Lighthouse Lab in Alderley Park | Wellcome Sanger Institute for the COVID-19 Genomics UK (COG-UK) consortium | The Lighthouse Lab in Alderley Park and Alex Alderton, Roberto Amato, Sonia Goncalves, Ewan Harrison, David K. Jackson, Ian Johnston, Dominic Kwiatkowski, Cordelia Langford, John Sillitoe on behalf of the Wellcome Sanger Institute COVID-19 Surveillance Team (http://www.sanger.ac.uk/covid-team) |
| EPI_ISL_556153, EPI_ISL_556154 | Lighthouse Lab in Alderley Park | Wellcome Sanger Institute for the COVID-19 Genomics UK (COG-UK) consortium | The Lighthouse Lab in Alderley Park and Alex Alderton, Roberto Amato, Sonia Goncalves, Ewan Harrison, David K. Jackson, Ian Johnston, Dominic Kwiatkowski, Cordelia Langford, John Sillitoe on behalf of the Wellcome Sanger Institute COVID-19 Surveillance Team |
| EPI_ISL_556155, EPI_ISL_556156, EPI_ISL_556157, EPI_ISL_556159 | Lighthouse Lab in Alderley Park | Wellcome Sanger Institute for the COVID-19 Genomics UK (COG-UK) consortium | The Lighthouse Lab in Alderley Park and Alex Alderton, Roberto Amato, Sonia Goncalves, Ewan Harrison, David K. Jackson, Ian Johnston, Dominic Kwiatkowski, Cordelia Langford, John Sillitoe on behalf of the Wellcome Sanger Institute COVID-19 Surveillance Team (http://www.sanger.ac.uk/covid-team) |
| EPI_ISL_556161, EPI_ISL_556162 | Lighthouse Lab in Alderley Park | Wellcome Sanger Institute for the COVID-19 Genomics UK (COG-UK) consortium | The Lighthouse Lab in Alderley Park and Alex Alderton, Roberto Amato, Sonia Goncalves, Ewan Harrison, David K. Jackson, Ian Johnston, Dominic Kwiatkowski, Cordelia Langford, John Sillitoe on behalf of the Wellcome Sanger Institute COVID-19 Surveillance Team |
| EPI_ISL_556163, EPI_ISL_556164, EPI_ISL_556165, EPI_ISL_556166 | Lighthouse Lab in Milton Keynes | Wellcome Sanger Institute for the COVID-19 Genomics UK (COG-UK) consortium | The Lighthouse Lab in Milton Keynes and Alex Alderton, Roberto Amato, Sonia Goncalves, Ewan Harrison, David K. Jackson, Ian Johnston, Dominic Kwiatkowski, Cordelia Langford, John Sillitoe on behalf of the Wellcome Sanger Institute COVID-19 Surveillance Team (http://www.sanger.ac.uk/covid-team) |
| EPI_ISL_556167, EPI_ISL_556168 | Lighthouse Lab in Alderley Park | Wellcome Sanger Institute for the COVID-19 Genomics UK (COG-UK) consortium | The Lighthouse Lab in Alderley Park and Alex Alderton, Roberto Amato, Sonia Goncalves, Ewan Harrison, David K. Jackson, Ian Johnston, Dominic Kwiatkowski, Cordelia Langford, John Sillitoe on behalf of the Wellcome Sanger Institute COVID-19 Surveillance Team (http://www.sanger.ac.uk/covid-team) |
| EPI_ISL_556169 | Lighthouse Lab in Alderley Park | Wellcome Sanger Institute for the COVID-19 Genomics UK (COG-UK) consortium | The Lighthouse Lab in Alderley Park and Alex Alderton, Roberto Amato, Sonia Goncalves, Ewan Harrison, David K. Jackson, Ian Johnston, Dominic Kwiatkowski, Cordelia Langford, John Sillitoe on behalf of the Wellcome Sanger Institute COVID-19 Surveillance Team |
| EPI_ISL_556171 | Lighthouse Lab in Alderley Park | Wellcome Sanger Institute for the COVID-19 Genomics UK (COG-UK) consortium | The Lighthouse Lab in Alderley Park and Alex Alderton, Roberto Amato, Sonia Goncalves, Ewan Harrison, David K. Jackson, Ian Johnston, Dominic Kwiatkowski, Cordelia Langford, John Sillitoe on behalf of the Wellcome Sanger Institute COVID-19 Surveillance Team (http://www.sanger.ac.uk/covid-team) |
| EPI_ISL_556172 | Lighthouse Lab in Milton Keynes | Wellcome Sanger Institute for the COVID-19 Genomics UK (COG-UK) consortium | The Lighthouse Lab in Milton Keynes and Alex Alderton, Roberto Amato, Sonia Goncalves, Ewan Harrison, David K. Jackson, Ian Johnston, Dominic Kwiatkowski, Cordelia Langford, John Sillitoe on behalf of the Wellcome Sanger Institute COVID-19 Surveillance Team (http://www.sanger.ac.uk/covid-team) |
| EPI_ISL_556173, EPI_ISL_556174, EPI_ISL_556175, EPI_ISL_556177, EPI_ISL_556179, EPI_ISL_556180 | Lighthouse Lab in Alderley Park | Wellcome Sanger Institute for the COVID-19 Genomics UK (COG-UK) consortium | The Lighthouse Lab in Alderley Park and Alex Alderton, Roberto Amato, Sonia Goncalves, Ewan Harrison, David K. Jackson, Ian Johnston, Dominic Kwiatkowski, Cordelia Langford, John Sillitoe on behalf of the Wellcome Sanger Institute COVID-19 Surveillance Team (http://www.sanger.ac.uk/covid-team) |
| EPI_ISL_556181 | Lighthouse Lab in Alderley Park | Wellcome Sanger Institute for the COVID-19 Genomics UK (COG-UK) consortium | The Lighthouse Lab in Alderley Park and Alex Alderton, Roberto Amato, Sonia Goncalves, Ewan Harrison, David K. Jackson, Ian Johnston, Dominic Kwiatkowski, Cordelia Langford, John Sillitoe on behalf of the Wellcome Sanger Institute COVID-19 Surveillance Team |
| EPI_ISL_556182, EPI_ISL_556185, EPI_ISL_556188 | Lighthouse Lab in Alderley Park | Wellcome Sanger Institute for the COVID-19 Genomics UK (COG-UK) consortium | The Lighthouse Lab in Alderley Park and Alex Alderton, Roberto Amato, Sonia Goncalves, Ewan Harrison, David K. Jackson, Ian Johnston, Dominic Kwiatkowski, Cordelia Langford, John Sillitoe on behalf of the Wellcome Sanger Institute COVID-19 Surveillance Team (http://www.sanger.ac.uk/covid-team) |
| EPI_ISL_556189 | Lighthouse Lab in Alderley Park | Wellcome Sanger Institute for the COVID-19 Genomics UK (COG-UK) consortium | The Lighthouse Lab in Alderley Park and Alex Alderton, Roberto Amato, Sonia Goncalves, Ewan Harrison, David K. Jackson, Ian Johnston, Dominic Kwiatkowski, Cordelia Langford, John Sillitoe on behalf of the Wellcome Sanger Institute COVID-19 Surveillance Team |
| EPI_ISL_556190, EPI_ISL_556192, EPI_ISL_556193 | Lighthouse Lab in Alderley Park | Wellcome Sanger Institute for the COVID-19 Genomics UK (COG-UK) consortium | The Lighthouse Lab in Alderley Park and Alex Alderton, Roberto Amato, Sonia Goncalves, Ewan Harrison, David K. Jackson, Ian Johnston, Dominic Kwiatkowski, Cordelia Langford, John Sillitoe on behalf of the Wellcome Sanger Institute COVID-19 Surveillance Team (http://www.sanger.ac.uk/covid-team) |
| EPI_ISL_556194 | Lighthouse Lab in Alderley Park | Wellcome Sanger Institute for the COVID-19 Genomics UK (COG-UK) consortium | The Lighthouse Lab in Alderley Park and Alex Alderton, Roberto Amato, Sonia Goncalves, Ewan Harrison, David K. Jackson, Ian Johnston, Dominic Kwiatkowski, Cordelia Langford, John Sillitoe on behalf of the Wellcome Sanger Institute COVID-19 Surveillance Team |
| EPI_ISL_556195 | Lighthouse Lab in Milton Keynes | Wellcome Sanger Institute for the COVID-19 Genomics UK (COG-UK) consortium | The Lighthouse Lab in Milton Keynes and Alex Alderton, Roberto Amato, Sonia Goncalves, Ewan Harrison, David K. Jackson, Ian Johnston, Dominic Kwiatkowski, Cordelia Langford, John Sillitoe on behalf of the Wellcome Sanger Institute COVID-19 Surveillance Team (http://www.sanger.ac.uk/covid-team) |
| EPI_ISL_556196 | Lighthouse Lab in Alderley Park | Wellcome Sanger Institute for the COVID-19 Genomics UK (COG-UK) consortium | The Lighthouse Lab in Alderley Park and Alex Alderton, Roberto Amato, Sonia Goncalves, Ewan Harrison, David K. Jackson, Ian Johnston, Dominic Kwiatkowski, Cordelia Langford, John Sillitoe on behalf of the Wellcome Sanger Institute COVID-19 Surveillance Team (http://www.sanger.ac.uk/covid-team) |
| EPI_ISL_556197, EPI_ISL_556198 | Lighthouse Lab in Milton Keynes | Wellcome Sanger Institute for the COVID-19 Genomics UK (COG-UK) consortium | The Lighthouse Lab in Alderley Park and Alex Alderton, Roberto Amato, Sonia Goncalves, Ewan Harrison, David K. Jackson, Ian Johnston, Dominic Kwiatkowski, Cordelia Langford, John Sillitoe on behalf of the Wellcome Sanger Institute COVID-19 Surveillance Team |
| EPI_ISL_556200 | Lighthouse Lab in Alderley Park | Wellcome Sanger Institute for the COVID-19 Genomics UK (COG-UK) consortium | The Lighthouse Lab in Alderley Park and Alex Alderton, Roberto Amato, Sonia Goncalves, Ewan Harrison, David K. Jackson, Ian Johnston, Dominic Kwiatkowski, Cordelia Langford, John Sillitoe on behalf of the Wellcome Sanger Institute COVID-19 Surveillance Team (http://www.sanger.ac.uk/covid-team) |
| EPI_ISL_556201, EPI_ISL_556204 | Lighthouse Lab in Milton Keynes | Wellcome Sanger Institute for the COVID-19 Genomics UK (COG-UK) consortium | The Lighthouse Lab in Milton Keynes and Alex Alderton, Roberto Amato, Sonia Goncalves, Ewan Harrison, David K. Jackson, Ian Johnston, Dominic Kwiatkowski, Cordelia Langford, John Sillitoe on behalf of the Wellcome Sanger Institute COVID-19 Surveillance Team (http://www.sanger.ac.uk/covid-team) |
| EPI_ISL_556205, EPI_ISL_556206 | Lighthouse Lab in Alderley Park | Wellcome Sanger Institute for the COVID-19 Genomics UK (COG-UK) consortium | The Lighthouse Lab in Alderley Park and Alex Alderton, Roberto Amato, Sonia Goncalves, Ewan Harrison, David K. Jackson, Ian Johnston, Dominic Kwiatkowski, Cordelia Langford, John Sillitoe on behalf of the Wellcome Sanger Institute COVID-19 Surveillance Team (http://www.sanger.ac.uk/covid-team) |
| EPI_ISL_556207 | Lighthouse Lab in Milton Keynes | Wellcome Sanger Institute for the COVID-19 Genomics UK (COG-UK) Consortium | The Lighthouse Lab in Milton Keynes and Alex Alderton, Roberto Amato, Sonia Goncalves, Ewan Harrison, David K. Jackson, Ian Johnston, Dominic Kwiatkowski, Cordelia Langford, John Sillitoe on behalf of the Wellcome Sanger Institute COVID-19 Surveillance Team |
| EPI_ISL_556209, EPI_ISL_556210, EPI_ISL_556212, EPI_ISL_556215 | Lighthouse Lab in Alderley Park | Wellcome Sanger Institute for the COVID-19 Genomics UK (COG-UK) consortium | The Lighthouse Lab in Alderley Park and Alex Alderton, Roberto Amato, Sonia Goncalves, Ewan Harrison, David K. Jackson, Ian Johnston, Dominic Kwiatkowski, Cordelia Langford, John Sillitoe on behalf of the Wellcome Sanger Institute COVID-19 Surveillance Team (http://www.sanger.ac.uk/covid-team) |
| EPI_ISL_556216, EPI_ISL_556217, EPI_ISL_556219 | Lighthouse Lab in Alderley Park | Wellcome Sanger Institute for the COVID-19 Genomics UK (COG-UK) consortium | The Lighthouse Lab in Alderley Park and Alex Alderton, Roberto Amato, Sonia Goncalves, Ewan Harrison, David K. Jackson, Ian Johnston, Dominic Kwiatkowski, Cordelia Langford, John Sillitoe on behalf of the Wellcome Sanger Institute COVID-19 Surveillance Team |
| EPI_ISL_556220, EPI_ISL_556221 | Lighthouse Lab in Alderley Park | Wellcome Sanger Institute for the COVID-19 Genomics UK (COG-UK) consortium | The Lighthouse Lab in Alderley Park and Alex Alderton, Roberto Amato, Sonia Goncalves, Ewan Harrison, David K. Jackson, Ian Johnston, Dominic Kwiatkowski, Cordelia Langford, John Sillitoe on behalf of the Wellcome Sanger Institute COVID-19 Surveillance Team (http://www.sanger.ac.uk/covid-team) |
| EPI_ISL_556222 | Lighthouse Lab in Alderley Park | Wellcome Sanger Institute for the COVID-19 Genomics UK (COG-UK) consortium | The Lighthouse Lab in Alderley Park and Alex Alderton, Roberto Amato, Sonia Goncalves, Ewan Harrison, David K. Jackson, Ian Johnston, Dominic Kwiatkowski, Cordelia Langford, John Sillitoe on behalf of the Wellcome Sanger Institute COVID-19 Surveillance Team |
| EPI_ISL_556223 | Lighthouse Lab in Milton Keynes | Wellcome Sanger Institute for the COVID-19 Genomics UK (COG-UK) consortium | The Lighthouse Lab in Milton Keynes and Alex Alderton, Roberto Amato, Sonia Goncalves, Ewan Harrison, David K. Jackson, Ian Johnston, Dominic Kwiatkowski, Cordelia Langford, John Sillitoe on behalf of the Wellcome Sanger Institute COVID-19 Surveillance Team (http://www.sanger.ac.uk/covid-team) |
| EPI_ISL_556224, EPI_ISL_556225 | Lighthouse Lab in Alderley Park | Wellcome Sanger Institute for the COVID-19 Genomics UK (COG-UK) consortium | The Lighthouse Lab in Alderley Park and Alex Alderton, Roberto Amato, Sonia Goncalves, Ewan Harrison, David K. Jackson, Ian Johnston, Dominic Kwiatkowski, Cordelia Langford, John Sillitoe on behalf of the Wellcome Sanger Institute COVID-19 Surveillance Team (http://www.sanger.ac.uk/covid-team) |
| EPI_ISL_556226 | Lighthouse Lab in Alderley Park | Wellcome Sanger Institute for the COVID-19 Genomics UK (COG-UK) consortium | The Lighthouse Lab in Alderley Park and Alex Alderton, Roberto Amato, Sonia Goncalves, Ewan Harrison, David K. Jackson, Ian Johnston, Dominic Kwiatkowski, Cordelia Langford, John Sillitoe on behalf of the Wellcome Sanger Institute COVID-19 Surveillance Team |
| EPI_ISL_556227 | Lighthouse Lab in Alderley Park | Wellcome Sanger Institute for the COVID-19 Genomics UK (COG-UK) consortium | The Lighthouse Lab in Alderley Park and Alex Alderton, Roberto Amato, Sonia Goncalves, Ewan Harrison, David K. Jackson, Ian Johnston, Dominic Kwiatkowski, Cordelia Langford, John Sillitoe on behalf of the Wellcome Sanger Institute COVID-19 Surveillance Team (http://www.sanger.ac.uk/covid-team) |
| EPI_ISL_556228 | Lighthouse Lab in Alderley Park | Wellcome Sanger Institute for the COVID-19 Genomics UK (COG-UK) consortium | The Lighthouse Lab in Alderley Park and Alex Alderton, Roberto Amato, Sonia Goncalves, Ewan Harrison, David K. Jackson, Ian Johnston, Dominic Kwiatkowski, Cordelia Langford, John Sillitoe on behalf of the Wellcome Sanger Institute COVID-19 Surveillance Team |
| EPI_ISL_556229 | Lighthouse Lab in Alderley Park | Wellcome Sanger Institute for the COVID-19 Genomics UK (COG-UK) consortium | The Lighthouse Lab in Alderley Park and Alex Alderton, Roberto Amato, Sonia Goncalves, Ewan Harrison, David K. Jackson, Ian Johnston, Dominic Kwiatkowski, Cordelia Langford, John Sillitoe on behalf of the Wellcome Sanger Institute COVID-19 Surveillance Team (http://www.sanger.ac.uk/covid-team) |
| EPI_ISL_556236 | Lighthouse Lab in Alderley Park | Wellcome Sanger Institute for the COVID-19 Genomics UK (COG-UK) consortium | The Lighthouse Lab in Alderley Park and Alex Alderton, Roberto Amato, Sonia Goncalves, Ewan Harrison, David K. Jackson, Ian Johnston, Dominic Kwiatkowski, Cordelia Langford, John Sillitoe on behalf of the Wellcome Sanger Institute COVID-19 Surveillance Team |
| EPI_ISL_556237 | Lighthouse Lab in Milton Keynes | Wellcome Sanger Institute for the COVID-19 Genomics UK (COG-UK) consortium | The Lighthouse Lab in Milton Keynes and Alex Alderton, Roberto Amato, Sonia Goncalves, Ewan Harrison, David K. Jackson, Ian Johnston, Dominic Kwiatkowski, Cordelia Langford, John Sillitoe on behalf of the Wellcome Sanger Institute COVID-19 Surveillance Team (http://www.sanger.ac.uk/covid-team) |

| | | | |
|---|---|---|---|
| EPI_ISL_556238, EPI_ISL_556239, EPI_ISL_556240 | Lighthouse Lab in Alderley Park | Wellcome Sanger Institute for the COVID-19 Genomics UK (COG-UK) consortium | The Lighthouse Lab in Alderley Park and Alex Alderton, Roberto Amato, Sonia Goncalves, Ewan Harrison, David K. Jackson, Ian Johnston, Dominic Kwiatkowski, Cordelia Langford, John Sillitoe on behalf of the Wellcome Sanger Institute COVID-19 Surveillance Team (http://www.sanger.ac.uk/covid-team) |
| EPI_ISL_556241 | Lighthouse Lab in Alderley Park | Wellcome Sanger Institute for the COVID-19 Genomics UK (COG-UK) consortium | The Lighthouse Lab in Alderley Park and Alex Alderton, Roberto Amato, Sonia Goncalves, Ewan Harrison, David K. Jackson, Ian Johnston, Dominic Kwiatkowski, Cordelia Langford, John Sillitoe on behalf of the Wellcome Sanger Institute COVID-19 Surveillance Team |
| EPI_ISL_556243, EPI_ISL_556245, EPI_ISL_556246 | Lighthouse Lab in Alderley Park | Wellcome Sanger Institute for the COVID-19 Genomics UK (COG-UK) consortium | The Lighthouse Lab in Alderley Park and Alex Alderton, Roberto Amato, Sonia Goncalves, Ewan Harrison, David K. Jackson, Ian Johnston, Dominic Kwiatkowski, Cordelia Langford, John Sillitoe on behalf of the Wellcome Sanger Institute COVID-19 Surveillance Team (http://www.sanger.ac.uk/covid-team) |
| EPI_ISL_556247 | Lighthouse Lab in Alderley Park | Wellcome Sanger Institute for the COVID-19 Genomics UK (COG-UK) consortium | The Lighthouse Lab in Alderley Park and Alex Alderton, Roberto Amato, Sonia Goncalves, Ewan Harrison, David K. Jackson, Ian Johnston, Dominic Kwiatkowski, Cordelia Langford, John Sillitoe on behalf of the Wellcome Sanger Institute COVID-19 Surveillance Team |
| EPI_ISL_556248 | Lighthouse Lab in Milton Keynes | Wellcome Sanger Institute for the COVID-19 Genomics UK (COG-UK) consortium | The Lighthouse Lab in Milton Keynes and Alex Alderton, Roberto Amato, Sonia Goncalves, Ewan Harrison, David K. Jackson, Ian Johnston, Dominic Kwiatkowski, Cordelia Langford, John Sillitoe on behalf of the Wellcome Sanger Institute COVID-19 Surveillance Team (http://www.sanger.ac.uk/covid-team) |
| EPI_ISL_556249 | Lighthouse Lab in Alderley Park | Wellcome Sanger Institute for the COVID-19 Genomics UK (COG-UK) consortium | The Lighthouse Lab in Alderley Park and Alex Alderton, Roberto Amato, Sonia Goncalves, Ewan Harrison, David K. Jackson, Ian Johnston, Dominic Kwiatkowski, Cordelia Langford, John Sillitoe on behalf of the Wellcome Sanger Institute COVID-19 Surveillance Team |
| EPI_ISL_556250, EPI_ISL_556252 | Lighthouse Lab in Alderley Park | Wellcome Sanger Institute for the COVID-19 Genomics UK (COG-UK) consortium | The Lighthouse Lab in Alderley Park and Alex Alderton, Roberto Amato, Sonia Goncalves, Ewan Harrison, David K. Jackson, Ian Johnston, Dominic Kwiatkowski, Cordelia Langford, John Sillitoe on behalf of the Wellcome Sanger Institute COVID-19 Surveillance Team (http://www.sanger.ac.uk/covid-team) |
| EPI_ISL_556253 | Lighthouse Lab in Milton Keynes | Wellcome Sanger Institute for the COVID-19 Genomics UK (COG-UK) consortium | The Lighthouse Lab in Milton Keynes and Alex Alderton, Roberto Amato, Sonia Goncalves, Ewan Harrison, David K. Jackson, Ian Johnston, Dominic Kwiatkowski, Cordelia Langford, John Sillitoe on behalf of the Wellcome Sanger Institute COVID-19 Surveillance Team (http://www.sanger.ac.uk/covid-team) |
| EPI_ISL_556254 | Lighthouse Lab in Alderley Park | Wellcome Sanger Institute for the COVID-19 Genomics UK (COG-UK) consortium | The Lighthouse Lab in Alderley Park and Alex Alderton, Roberto Amato, Sonia Goncalves, Ewan Harrison, David K. Jackson, Ian Johnston, Dominic Kwiatkowski, Cordelia Langford, John Sillitoe on behalf of the Wellcome Sanger Institute COVID-19 Surveillance Team (http://www.sanger.ac.uk/covid-team) |
| EPI_ISL_556255, EPI_ISL_556257 | Lighthouse Lab in Alderley Park | Wellcome Sanger Institute for the COVID-19 Genomics UK (COG-UK) consortium | The Lighthouse Lab in Alderley Park and Alex Alderton, Roberto Amato, Sonia Goncalves, Ewan Harrison, David K. Jackson, Ian Johnston, Dominic Kwiatkowski, Cordelia Langford, John Sillitoe on behalf of the Wellcome Sanger Institute COVID-19 Surveillance Team |
| EPI_ISL_556260 | Lighthouse Lab in Alderley Park | Wellcome Sanger Institute for the COVID-19 Genomics UK (COG-UK) consortium | The Lighthouse Lab in Alderley Park and Alex Alderton, Roberto Amato, Sonia Goncalves, Ewan Harrison, David K. Jackson, Ian Johnston, Dominic Kwiatkowski, Cordelia Langford, John Sillitoe on behalf of the Wellcome Sanger Institute COVID-19 Surveillance Team (http://www.sanger.ac.uk/covid-team) |
| EPI_ISL_556261, EPI_ISL_556263 | Lighthouse Lab in Milton Keynes | Wellcome Sanger Institute for the COVID-19 Genomics UK (COG-UK) consortium | The Lighthouse Lab in Milton Keynes and Alex Alderton, Roberto Amato, Sonia Goncalves, Ewan Harrison, David K. Jackson, Ian Johnston, Dominic Kwiatkowski, Cordelia Langford, John Sillitoe on behalf of the Wellcome Sanger Institute COVID-19 Surveillance Team (http://www.sanger.ac.uk/covid-team) |
| EPI_ISL_556264, EPI_ISL_556265 | Lighthouse Lab in Alderley Park | Wellcome Sanger Institute for the COVID-19 Genomics UK (COG-UK) consortium | The Lighthouse Lab in Alderley Park and Alex Alderton, Roberto Amato, Sonia Goncalves, Ewan Harrison, David K. Jackson, Ian Johnston, Dominic Kwiatkowski, Cordelia Langford, John Sillitoe on behalf of the Wellcome Sanger Institute COVID-19 Surveillance Team (http://www.sanger.ac.uk/covid-team) |
| EPI_ISL_556266 | Lighthouse Lab in Milton Keynes | Wellcome Sanger Institute for the COVID-19 Genomics UK (COG-UK) consortium | The Lighthouse Lab in Milton Keynes and Alex Alderton, Roberto Amato, Sonia Goncalves, Ewan Harrison, David K. Jackson, Ian Johnston, Dominic Kwiatkowski, Cordelia Langford, John Sillitoe on behalf of the Wellcome Sanger Institute COVID-19 Surveillance Team (http://www.sanger.ac.uk/covid-team) |
| EPI_ISL_556268 | Lighthouse Lab in Alderley Park | Wellcome Sanger Institute for the COVID-19 Genomics UK (COG-UK) consortium | The Lighthouse Lab in Alderley Park and Alex Alderton, Roberto Amato, Sonia Goncalves, Ewan Harrison, David K. Jackson, Ian Johnston, Dominic Kwiatkowski, Cordelia Langford, John Sillitoe on behalf of the Wellcome Sanger Institute COVID-19 Surveillance Team |
| EPI_ISL_556269 | Lighthouse Lab in Milton Keynes | Wellcome Sanger Institute for the COVID-19 Genomics UK (COG-UK) consortium | The Lighthouse Lab in Milton Keynes and Alex Alderton, Roberto Amato, Sonia Goncalves, Ewan Harrison, David K. Jackson, Ian Johnston, Dominic Kwiatkowski, Cordelia Langford, John Sillitoe on behalf of the Wellcome Sanger Institute COVID-19 Surveillance Team (http://www.sanger.ac.uk/covid-team) |
| EPI_ISL_556270 | Lighthouse Lab in Alderley Park | Wellcome Sanger Institute for the COVID-19 Genomics UK (COG-UK) consortium | The Lighthouse Lab in Alderley Park and Alex Alderton, Roberto Amato, Sonia Goncalves, Ewan Harrison, David K. Jackson, Ian Johnston, Dominic Kwiatkowski, Cordelia Langford, John Sillitoe on behalf of the Wellcome Sanger Institute COVID-19 Surveillance Team (http://www.sanger.ac.uk/covid-team) |
| EPI_ISL_556271 | Lighthouse Lab in Alderley Park | Wellcome Sanger Institute for the COVID-19 Genomics UK (COG-UK) consortium | The Lighthouse Lab in Alderley Park and Alex Alderton, Roberto Amato, Sonia Goncalves, Ewan Harrison, David K. Jackson, Ian Johnston, Dominic Kwiatkowski, Cordelia Langford, John Sillitoe on behalf of the Wellcome Sanger Institute COVID-19 Surveillance Team |
| EPI_ISL_556272, EPI_ISL_556273, EPI_ISL_556274 | Lighthouse Lab in Alderley Park | Wellcome Sanger Institute for the COVID-19 Genomics UK (COG-UK) consortium | The Lighthouse Lab in Alderley Park and Alex Alderton, Roberto Amato, Sonia Goncalves, Ewan Harrison, David K. Jackson, Ian Johnston, Dominic Kwiatkowski, Cordelia Langford, John Sillitoe on behalf of the Wellcome Sanger Institute COVID-19 Surveillance Team (http://www.sanger.ac.uk/covid-team) |
| EPI_ISL_556275, EPI_ISL_556276 | Lighthouse Lab in Milton Keynes | Wellcome Sanger Institute for the COVID-19 Genomics UK (COG-UK) consortium | The Lighthouse Lab in Milton Keynes and Alex Alderton, Roberto Amato, Sonia Goncalves, Ewan Harrison, David K. Jackson, Ian Johnston, Dominic Kwiatkowski, Cordelia Langford, John Sillitoe on behalf of the Wellcome Sanger Institute COVID-19 Surveillance Team (http://www.sanger.ac.uk/covid-team) |
| EPI_ISL_556277, EPI_ISL_556278, EPI_ISL_556279, EPI_ISL_556281, EPI_ISL_556284 | Lighthouse Lab in Alderley Park | Wellcome Sanger Institute for the COVID-19 Genomics UK (COG-UK) consortium | The Lighthouse Lab in Alderley Park and Alex Alderton, Roberto Amato, Sonia Goncalves, Ewan Harrison, David K. Jackson, Ian Johnston, Dominic Kwiatkowski, Cordelia Langford, John Sillitoe on behalf of the Wellcome Sanger Institute COVID-19 Surveillance Team (http://www.sanger.ac.uk/covid-team) |
| EPI_ISL_556286 | Lighthouse Lab in Alderley Park | Wellcome Sanger Institute for the COVID-19 Genomics UK (COG-UK) consortium | The Lighthouse Lab in Alderley Park and Alex Alderton, Roberto Amato, Sonia Goncalves, Ewan Harrison, David K. Jackson, Ian Johnston, Dominic Kwiatkowski, Cordelia Langford, John Sillitoe on behalf of the Wellcome Sanger Institute COVID-19 Surveillance Team |
| EPI_ISL_556288 | Lighthouse Lab in Alderley Park | Wellcome Sanger Institute for the COVID-19 Genomics UK (COG-UK) consortium | The Lighthouse Lab in Alderley Park and Alex Alderton, Roberto Amato, Sonia Goncalves, Ewan Harrison, David K. Jackson, Ian Johnston, Dominic Kwiatkowski, Cordelia Langford, John Sillitoe on behalf of the Wellcome Sanger Institute COVID-19 Surveillance Team (http://www.sanger.ac.uk/covid-team) |
| EPI_ISL_556289 | Lighthouse Lab in Milton Keynes | Wellcome Sanger Institute for the COVID-19 Genomics UK (COG-UK) consortium | The Lighthouse Lab in Milton Keynes and Alex Alderton, Roberto Amato, Sonia Goncalves, Ewan Harrison, David K. Jackson, Ian Johnston, Dominic Kwiatkowski, Cordelia Langford, John Sillitoe on behalf of the Wellcome Sanger Institute COVID-19 Surveillance Team (http://www.sanger.ac.uk/covid-team) |
| EPI_ISL_556290, EPI_ISL_556292 | Lighthouse Lab in Alderley Park | Wellcome Sanger Institute for the COVID-19 Genomics UK (COG-UK) consortium | The Lighthouse Lab in Alderley Park and Alex Alderton, Roberto Amato, Sonia Goncalves, Ewan Harrison, David K. Jackson, Ian Johnston, Dominic Kwiatkowski, Cordelia Langford, John Sillitoe on behalf of the Wellcome Sanger Institute COVID-19 Surveillance Team (http://www.sanger.ac.uk/covid-team) |
| EPI_ISL_556294 | Lighthouse Lab in Milton Keynes | Wellcome Sanger Institute for the COVID-19 Genomics UK (COG-UK) consortium | The Lighthouse Lab in Milton Keynes and Alex Alderton, Roberto Amato, Sonia Goncalves, Ewan Harrison, David K. Jackson, Ian Johnston, Dominic Kwiatkowski, Cordelia Langford, John Sillitoe on behalf of the Wellcome Sanger Institute COVID-19 Surveillance Team (http://www.sanger.ac.uk/covid-team) |
| EPI_ISL_556295, EPI_ISL_556296, EPI_ISL_556299 | Lighthouse Lab in Alderley Park | Wellcome Sanger Institute for the COVID-19 Genomics UK (COG-UK) consortium | The Lighthouse Lab in Alderley Park and Alex Alderton, Roberto Amato, Sonia Goncalves, Ewan Harrison, David K. Jackson, Ian Johnston, Dominic Kwiatkowski, Cordelia Langford, John Sillitoe on behalf of the Wellcome Sanger Institute COVID-19 Surveillance Team (http://www.sanger.ac.uk/covid-team) |
| EPI_ISL_556300 | Lighthouse Lab in Milton Keynes | Wellcome Sanger Institute for the COVID-19 Genomics UK (COG-UK) consortium | The Lighthouse Lab in Milton Keynes and Alex Alderton, Roberto Amato, Sonia Goncalves, Ewan Harrison, David K. Jackson, Ian Johnston, Dominic Kwiatkowski, Cordelia Langford, John Sillitoe on behalf of the Wellcome Sanger Institute COVID-19 Surveillance Team (http://www.sanger.ac.uk/covid-team) |
| EPI_ISL_556301, EPI_ISL_556302, EPI_ISL_556303 | Lighthouse Lab in Alderley Park | Wellcome Sanger Institute for the COVID-19 Genomics UK (COG-UK) consortium | The Lighthouse Lab in Alderley Park and Alex Alderton, Roberto Amato, Sonia Goncalves, Ewan Harrison, David K. Jackson, Ian Johnston, Dominic Kwiatkowski, Cordelia Langford, John Sillitoe on behalf of the Wellcome Sanger Institute COVID-19 Surveillance Team (http://www.sanger.ac.uk/covid-team) |
| EPI_ISL_556516 | Lighthouse Lab in Cambridge | Wellcome Sanger Institute for the COVID-19 Genomics UK (COG-UK) consortium | Rob Howes, The Lighthouse Lab in Cambridge and Alex Alderton, Roberto Amato, Sonia Goncalves, Ewan Harrison, David K. Jackson, Ian Johnston, Dominic Kwiatkowski, Cordelia Langford, John Sillitoe on behalf of the Wellcome Sanger Institute COVID-19 Surveillance Team |
| EPI_ISL_556585, EPI_ISL_556586, EPI_ISL_557003, EPI_ISL_557045 | Lighthouse Lab in Alderley Park | Wellcome Sanger Institute for the COVID-19 Genomics UK (COG-UK) consortium | The Lighthouse Lab in Alderley Park and Alex Alderton, Roberto Amato, Sonia Goncalves, Ewan Harrison, David K. Jackson, Ian Johnston, Dominic Kwiatkowski, Cordelia Langford, John Sillitoe on behalf of the Wellcome Sanger Institute COVID-19 Surveillance Team (http://www.sanger.ac.uk/covid-team) |
| EPI_ISL_557357, EPI_ISL_557362 | Lighthouse Lab in Milton Keynes | Wellcome Sanger Institute for the COVID-19 Genomics UK (COG-UK) consortium | The Lighthouse Lab in Milton Keynes and Alex Alderton, Roberto Amato, Sonia Goncalves, Ewan Harrison, David K. Jackson, Ian Johnston, Dominic Kwiatkowski, Cordelia Langford, John Sillitoe on behalf of the Wellcome Sanger Institute COVID-19 Surveillance Team (http://www.sanger.ac.uk/covid-team) |
| EPI_ISL_557434, EPI_ISL_557452 | Lighthouse Lab in Alderley Park | Wellcome Sanger Institute for the COVID-19 Genomics UK (COG-UK) consortium | The Lighthouse Lab in Alderley Park and Alex Alderton, Roberto Amato, Sonia Goncalves, Ewan Harrison, David K. Jackson, Ian Johnston, Dominic Kwiatkowski, Cordelia Langford, John Sillitoe on behalf of the Wellcome Sanger Institute COVID-19 Surveillance Team (http://www.sanger.ac.uk/covid-team) |
| EPI_ISL_557555, EPI_ISL_557561, EPI_ISL_557567 | Lighthouse Lab in Alderley Park | Wellcome Sanger Institute for the COVID-19 Genomics UK (COG-UK) consortium | The Lighthouse Lab in Alderley Park and Alex Alderton, Roberto Amato, Sonia Goncalves, Ewan Harrison, David K. Jackson, Ian Johnston, Dominic Kwiatkowski, Cordelia Langford, John Sillitoe on behalf of the Wellcome Sanger Institute COVID-19 Surveillance Team |
| EPI_ISL_557570 | Lighthouse Lab in Alderley Park | Wellcome Sanger Institute for the COVID-19 Genomics UK (COG-UK) consortium | The Lighthouse Lab in Alderley Park and Alex Alderton, Roberto Amato, Sonia Goncalves, Ewan Harrison, David K. Jackson, Ian Johnston, Dominic Kwiatkowski, Cordelia Langford, John Sillitoe on behalf of the Wellcome Sanger Institute COVID-19 Surveillance Team (http://www.sanger.ac.uk/covid-team) |

| | | | |
|---|---|---|---|
| EPI_ISL_557572, EPI_ISL_557575, EPI_ISL_557581, EPI_ISL_557582, EPI_ISL_557583, EPI_ISL_557587, EPI_ISL_557593, EPI_ISL_557595, EPI_ISL_557597, EPI_ISL_557598, EPI_ISL_557603, EPI_ISL_557605, EPI_ISL_557609, EPI_ISL_557618, EPI_ISL_557620, EPI_ISL_557621, EPI_ISL_557624, EPI_ISL_557627, EPI_ISL_557632, EPI_ISL_557638, EPI_ISL_557639, EPI_ISL_557640, EPI_ISL_557641, EPI_ISL_557642, EPI_ISL_557643, EPI_ISL_557644, EPI_ISL_557646, EPI_ISL_557647, EPI_ISL_557649, EPI_ISL_557650, EPI_ISL_557651, EPI_ISL_557652, EPI_ISL_557653, EPI_ISL_557654, EPI_ISL_557655, EPI_ISL_557657 | | | |
| see above | Lighthouse Lab in Alderley Park | Wellcome Sanger Institute for the COVID-19 Genomics UK (COG-UK) consortium | The Lighthouse Lab in Alderley Park and Alex Alderton, Roberto Amato, Sonia Goncalves, Ewan Harrison, David K. Jackson, Ian Johnston, Dominic Kwiatkowski, Cordelia Langford, John Sillitoe on behalf of the Wellcome Sanger Institute COVID-19 Surveillance Team |
| EPI_ISL_557658 | Lighthouse Lab in Alderley Park | Wellcome Sanger Institute for the COVID-19 Genomics UK (COG-UK) consortium | The Lighthouse Lab in Alderley Park and Alex Alderton, Roberto Amato, Sonia Goncalves, Ewan Harrison, David K. Jackson, Ian Johnston, Dominic Kwiatkowski, Cordelia Langford, John Sillitoe on behalf of the Wellcome Sanger Institute COVID-19 Surveillance Team (http://www.sanger.ac.uk/covid-team) |
| EPI_ISL_557663, EPI_ISL_557666, EPI_ISL_557670, EPI_ISL_557673, EPI_ISL_557677, EPI_ISL_557678, EPI_ISL_557681, EPI_ISL_557683, EPI_ISL_557684 | Lighthouse Lab in Alderley Park | Wellcome Sanger Institute for the COVID-19 Genomics UK (COG-UK) consortium | The Lighthouse Lab in Alderley Park and Alex Alderton, Roberto Amato, Sonia Goncalves, Ewan Harrison, David K. Jackson, Ian Johnston, Dominic Kwiatkowski, Cordelia Langford, John Sillitoe on behalf of the Wellcome Sanger Institute COVID-19 Surveillance Team |
| EPI_ISL_557685 | Lighthouse Lab in Alderley Park | Wellcome Sanger Institute for the COVID-19 Genomics UK (COG-UK) consortium | The Lighthouse Lab in Alderley Park and Alex Alderton, Roberto Amato, Sonia Goncalves, Ewan Harrison, David K. Jackson, Ian Johnston, Dominic Kwiatkowski, Cordelia Langford, John Sillitoe on behalf of the Wellcome Sanger Institute COVID-19 Surveillance Team (http://www.sanger.ac.uk/covid-team) |
| EPI_ISL_557686, EPI_ISL_557687, EPI_ISL_557690, EPI_ISL_557691, EPI_ISL_557692, EPI_ISL_557695, EPI_ISL_557698 | Lighthouse Lab in Alderley Park | Wellcome Sanger Institute for the COVID-19 Genomics UK (COG-UK) consortium | The Lighthouse Lab in Alderley Park and Alex Alderton, Roberto Amato, Sonia Goncalves, Ewan Harrison, David K. Jackson, Ian Johnston, Dominic Kwiatkowski, Cordelia Langford, John Sillitoe on behalf of the Wellcome Sanger Institute COVID-19 Surveillance Team |
| EPI_ISL_557701 | Lighthouse Lab in Milton Keynes | Wellcome Sanger Institute for the COVID-19 Genomics UK (COG-UK) consortium | The Lighthouse Lab in Milton Keynes and Alex Alderton, Roberto Amato, Sonia Goncalves, Ewan Harrison, David K. Jackson, Ian Johnston, Dominic Kwiatkowski, Cordelia Langford, John Sillitoe on behalf of the Wellcome Sanger Institute COVID-19 Surveillance Team (http://www.sanger.ac.uk/covid-team) |
| EPI_ISL_557704, EPI_ISL_557705, EPI_ISL_557707 | Lighthouse Lab in Alderley Park | Wellcome Sanger Institute for the COVID-19 Genomics UK (COG-UK) consortium | The Lighthouse Lab in Alderley Park and Alex Alderton, Roberto Amato, Sonia Goncalves, Ewan Harrison, David K. Jackson, Ian Johnston, Dominic Kwiatkowski, Cordelia Langford, John Sillitoe on behalf of the Wellcome Sanger Institute COVID-19 Surveillance Team |
| EPI_ISL_557709 | Lighthouse Lab in Alderley Park | Wellcome Sanger Institute for the COVID-19 Genomics UK (COG-UK) consortium | The Lighthouse Lab in Alderley Park and Alex Alderton, Roberto Amato, Sonia Goncalves, Ewan Harrison, David K. Jackson, Ian Johnston, Dominic Kwiatkowski, Cordelia Langford, John Sillitoe on behalf of the Wellcome Sanger Institute COVID-19 Surveillance Team (http://www.sanger.ac.uk/covid-team) |
| EPI_ISL_557715, EPI_ISL_557716, EPI_ISL_557721, EPI_ISL_557722, EPI_ISL_557723, EPI_ISL_557724, EPI_ISL_557725, EPI_ISL_557727 | Lighthouse Lab in Alderley Park | Wellcome Sanger Institute for the COVID-19 Genomics UK (COG-UK) consortium | The Lighthouse Lab in Alderley Park and Alex Alderton, Roberto Amato, Sonia Goncalves, Ewan Harrison, David K. Jackson, Ian Johnston, Dominic Kwiatkowski, Cordelia Langford, John Sillitoe on behalf of the Wellcome Sanger Institute COVID-19 Surveillance Team |
| EPI_ISL_557728 | Lighthouse Lab in Alderley Park | Wellcome Sanger Institute for the COVID-19 Genomics UK (COG-UK) Consortium | The Lighthouse Lab in Alderley Park and Alex Alderton, Roberto Amato, Sonia Goncalves, Ewan Harrison, David K. Jackson, Ian Johnston, Dominic Kwiatkowski, Cordelia Langford, John Sillitoe on behalf of the Wellcome Sanger Institute COVID-19 Surveillance Team |
| EPI_ISL_557729, EPI_ISL_557732, EPI_ISL_557735, EPI_ISL_557738 | Lighthouse Lab in Alderley Park | Wellcome Sanger Institute for the COVID-19 Genomics UK (COG-UK) consortium | The Lighthouse Lab in Alderley Park and Alex Alderton, Roberto Amato, Sonia Goncalves, Ewan Harrison, David K. Jackson, Ian Johnston, Dominic Kwiatkowski, Cordelia Langford, John Sillitoe on behalf of the Wellcome Sanger Institute COVID-19 Surveillance Team |
| EPI_ISL_557739 | Lighthouse Lab in Alderley Park | Wellcome Sanger Institute for the COVID-19 Genomics UK (COG-UK) Consortium | The Lighthouse Lab in Alderley Park and Alex Alderton, Roberto Amato, Sonia Goncalves, Ewan Harrison, David K. Jackson, Ian Johnston, Dominic Kwiatkowski, Cordelia Langford, John Sillitoe on behalf of the Wellcome Sanger Institute COVID-19 Surveillance Team |
| EPI_ISL_557740, EPI_ISL_557741, EPI_ISL_557742 | Lighthouse Lab in Milton Keynes | Wellcome Sanger Institute for the COVID-19 Genomics UK (COG-UK) consortium | The Lighthouse Lab in Milton Keynes and Alex Alderton, Roberto Amato, Sonia Goncalves, Ewan Harrison, David K. Jackson, Ian Johnston, Dominic Kwiatkowski, Cordelia Langford, John Sillitoe on behalf of the Wellcome Sanger Institute COVID-19 Surveillance Team (http://www.sanger.ac.uk/covid-team) |
| EPI_ISL_557743, EPI_ISL_557744 | Lighthouse Lab in Alderley Park | Wellcome Sanger Institute for the COVID-19 Genomics UK (COG-UK) consortium | The Lighthouse Lab in Alderley Park and Alex Alderton, Roberto Amato, Sonia Goncalves, Ewan Harrison, David K. Jackson, Ian Johnston, Dominic Kwiatkowski, Cordelia Langford, John Sillitoe on behalf of the Wellcome Sanger Institute COVID-19 Surveillance Team |
| EPI_ISL_557745 | Lighthouse Lab in Milton Keynes | Wellcome Sanger Institute for the COVID-19 Genomics UK (COG-UK) consortium | The Lighthouse Lab in Milton Keynes and Alex Alderton, Roberto Amato, Sonia Goncalves, Ewan Harrison, David K. Jackson, Ian Johnston, Dominic Kwiatkowski, Cordelia Langford, John Sillitoe on behalf of the Wellcome Sanger Institute COVID-19 Surveillance Team (http://www.sanger.ac.uk/covid-team) |
| EPI_ISL_557746, EPI_ISL_557747 | Lighthouse Lab in Alderley Park | Wellcome Sanger Institute for the COVID-19 Genomics UK (COG-UK) consortium | The Lighthouse Lab in Alderley Park and Alex Alderton, Roberto Amato, Sonia Goncalves, Ewan Harrison, David K. Jackson, Ian Johnston, Dominic Kwiatkowski, Cordelia Langford, John Sillitoe on behalf of the Wellcome Sanger Institute COVID-19 Surveillance Team |
| EPI_ISL_557748 | Lighthouse Lab in Milton Keynes | Wellcome Sanger Institute for the COVID-19 Genomics UK (COG-UK) consortium | The Lighthouse Lab in Milton Keynes and Alex Alderton, Roberto Amato, Sonia Goncalves, Ewan Harrison, David K. Jackson, Ian Johnston, Dominic Kwiatkowski, Cordelia Langford, John Sillitoe on behalf of the Wellcome Sanger Institute COVID-19 Surveillance Team (http://www.sanger.ac.uk/covid-team) |
| EPI_ISL_557749 | Lighthouse Lab in Alderley Park | Wellcome Sanger Institute for the COVID-19 Genomics UK (COG-UK) consortium | The Lighthouse Lab in Alderley Park and Alex Alderton, Roberto Amato, Sonia Goncalves, Ewan Harrison, David K. Jackson, Ian Johnston, Dominic Kwiatkowski, Cordelia Langford, John Sillitoe on behalf of the Wellcome Sanger Institute COVID-19 Surveillance Team |
| EPI_ISL_557750 | Lighthouse Lab in Milton Keynes | Wellcome Sanger Institute for the COVID-19 Genomics UK (COG-UK) consortium | The Lighthouse Lab in Milton Keynes and Alex Alderton, Roberto Amato, Sonia Goncalves, Ewan Harrison, David K. Jackson, Ian Johnston, Dominic Kwiatkowski, Cordelia Langford, John Sillitoe on behalf of the Wellcome Sanger Institute COVID-19 Surveillance Team (http://www.sanger.ac.uk/covid-team) |
| EPI_ISL_557751, EPI_ISL_557752 | Lighthouse Lab in Alderley Park | Wellcome Sanger Institute for the COVID-19 Genomics UK (COG-UK) consortium | The Lighthouse Lab in Alderley Park and Alex Alderton, Roberto Amato, Sonia Goncalves, Ewan Harrison, David K. Jackson, Ian Johnston, Dominic Kwiatkowski, Cordelia Langford, John Sillitoe on behalf of the Wellcome Sanger Institute COVID-19 Surveillance Team |
| EPI_ISL_557753 | Lighthouse Lab in Milton Keynes | Wellcome Sanger Institute for the COVID-19 Genomics UK (COG-UK) consortium | The Lighthouse Lab in Milton Keynes and Alex Alderton, Roberto Amato, Sonia Goncalves, Ewan Harrison, David K. Jackson, Ian Johnston, Dominic Kwiatkowski, Cordelia Langford, John Sillitoe on behalf of the Wellcome Sanger Institute COVID-19 Surveillance Team (http://www.sanger.ac.uk/covid-team) |
| EPI_ISL_557754, EPI_ISL_557755, EPI_ISL_557756 | Lighthouse Lab in Alderley Park | Wellcome Sanger Institute for the COVID-19 Genomics UK (COG-UK) consortium | The Lighthouse Lab in Alderley Park and Alex Alderton, Roberto Amato, Sonia Goncalves, Ewan Harrison, David K. Jackson, Ian Johnston, Dominic Kwiatkowski, Cordelia Langford, John Sillitoe on behalf of the Wellcome Sanger Institute COVID-19 Surveillance Team |
| EPI_ISL_557757, EPI_ISL_557758 | Lighthouse Lab in Milton Keynes | Wellcome Sanger Institute for the COVID-19 Genomics UK (COG-UK) consortium | The Lighthouse Lab in Milton Keynes and Alex Alderton, Roberto Amato, Sonia Goncalves, Ewan Harrison, David K. Jackson, Ian Johnston, Dominic Kwiatkowski, Cordelia Langford, John Sillitoe on behalf of the Wellcome Sanger Institute COVID-19 Surveillance Team (http://www.sanger.ac.uk/covid-team) |
| EPI_ISL_557759 | Lighthouse Lab in Alderley Park | Wellcome Sanger Institute for the COVID-19 Genomics UK (COG-UK) consortium | The Lighthouse Lab in Alderley Park and Alex Alderton, Roberto Amato, Sonia Goncalves, Ewan Harrison, David K. Jackson, Ian Johnston, Dominic Kwiatkowski, Cordelia Langford, John Sillitoe on behalf of the Wellcome Sanger Institute COVID-19 Surveillance Team |
| EPI_ISL_557760 | Lighthouse Lab in Alderley Park | Wellcome Sanger Institute for the COVID-19 Genomics UK (COG-UK) Consortium | The Lighthouse Lab in Alderley Park and Alex Alderton, Roberto Amato, Sonia Goncalves, Ewan Harrison, David K. Jackson, Ian Johnston, Dominic Kwiatkowski, Cordelia Langford, John Sillitoe on behalf of the Wellcome Sanger Institute COVID-19 Surveillance Team |
| EPI_ISL_557761, EPI_ISL_557763 | Lighthouse Lab in Alderley Park | Wellcome Sanger Institute for the COVID-19 Genomics UK (COG-UK) consortium | The Lighthouse Lab in Alderley Park and Alex Alderton, Roberto Amato, Sonia Goncalves, Ewan Harrison, David K. Jackson, Ian Johnston, Dominic Kwiatkowski, Cordelia Langford, John Sillitoe on behalf of the Wellcome Sanger Institute COVID-19 Surveillance Team |
| EPI_ISL_557764, EPI_ISL_557765, EPI_ISL_557766 | Lighthouse Lab in Milton Keynes | Wellcome Sanger Institute for the COVID-19 Genomics UK (COG-UK) consortium | The Lighthouse Lab in Milton Keynes and Alex Alderton, Roberto Amato, Sonia Goncalves, Ewan Harrison, David K. Jackson, Ian Johnston, Dominic Kwiatkowski, Cordelia Langford, John Sillitoe on behalf of the Wellcome Sanger Institute COVID-19 Surveillance Team (http://www.sanger.ac.uk/covid-team) |
| EPI_ISL_557767 | Lighthouse Lab in Alderley Park | Wellcome Sanger Institute for the COVID-19 Genomics UK (COG-UK) consortium | The Lighthouse Lab in Alderley Park and Alex Alderton, Roberto Amato, Sonia Goncalves, Ewan Harrison, David K. Jackson, Ian Johnston, Dominic Kwiatkowski, Cordelia Langford, John Sillitoe on behalf of the Wellcome Sanger Institute COVID-19 Surveillance Team |
| EPI_ISL_557768 | Lighthouse Lab in Milton Keynes | Wellcome Sanger Institute for the COVID-19 Genomics UK (COG-UK) consortium | The Lighthouse Lab in Milton Keynes and Alex Alderton, Roberto Amato, Sonia Goncalves, Ewan Harrison, David K. Jackson, Ian Johnston, Dominic Kwiatkowski, Cordelia Langford, John Sillitoe on behalf of the Wellcome Sanger Institute COVID-19 Surveillance Team (http://www.sanger.ac.uk/covid-team) |
| EPI_ISL_557769, EPI_ISL_557770 | Lighthouse Lab in Alderley Park | Wellcome Sanger Institute for the COVID-19 Genomics UK (COG-UK) consortium | The Lighthouse Lab in Alderley Park and Alex Alderton, Roberto Amato, Sonia Goncalves, Ewan Harrison, David K. Jackson, Ian Johnston, Dominic Kwiatkowski, Cordelia Langford, John Sillitoe on behalf of the Wellcome Sanger Institute COVID-19 Surveillance Team |
| EPI_ISL_557771 | Lighthouse Lab in Milton Keynes | Wellcome Sanger Institute for the COVID-19 Genomics UK (COG-UK) consortium | The Lighthouse Lab in Milton Keynes and Alex Alderton, Roberto Amato, Sonia Goncalves, Ewan Harrison, David K. Jackson, Ian Johnston, Dominic Kwiatkowski, Cordelia Langford, John Sillitoe on behalf of the Wellcome Sanger Institute COVID-19 Surveillance Team (http://www.sanger.ac.uk/covid-team) |

| | | | |
|---|---|---|---|
| EPI_ISL_557772, EPI_ISL_557773 | Lighthouse Lab in Alderley Park | Wellcome Sanger Institute for the COVID-19 Genomics UK (COG-UK) consortium | The Lighthouse Lab in Alderley Park and Alex Alderton, Roberto Amato, Sonia Goncalves, Ewan Harrison, David K. Jackson, Ian Johnston, Dominic Kwiatkowski, Cordelia Langford, John Sillitoe on behalf of the Wellcome Sanger Institute COVID-19 Surveillance Team |
| EPI_ISL_557774 | Lighthouse Lab in Alderley Park | Wellcome Sanger Institute for the COVID-19 Genomics UK (COG-UK) consortium | The Lighthouse Lab in Alderley Park and Alex Alderton, Roberto Amato, Sonia Goncalves, Ewan Harrison, David K. Jackson, Ian Johnston, Dominic Kwiatkowski, Cordelia Langford, John Sillitoe on behalf of the Wellcome Sanger Institute COVID-19 Surveillance Team (http://www.sanger.ac.uk/covid-team) |
| EPI_ISL_557775 | Lighthouse Lab in Milton Keynes | Wellcome Sanger Institute for the COVID-19 Genomics UK (COG-UK) consortium | The Lighthouse Lab in Milton Keynes and Alex Alderton, Roberto Amato, Sonia Goncalves, Ewan Harrison, David K. Jackson, Ian Johnston, Dominic Kwiatkowski, Cordelia Langford, John Sillitoe on behalf of the Wellcome Sanger Institute COVID-19 Surveillance Team (http://www.sanger.ac.uk/covid-team) |
| EPI_ISL_557776, EPI_ISL_557777 | Lighthouse Lab in Alderley Park | Wellcome Sanger Institute for the COVID-19 Genomics UK (COG-UK) consortium | The Lighthouse Lab in Alderley Park and Alex Alderton, Roberto Amato, Sonia Goncalves, Ewan Harrison, David K. Jackson, Ian Johnston, Dominic Kwiatkowski, Cordelia Langford, John Sillitoe on behalf of the Wellcome Sanger Institute COVID-19 Surveillance Team |
| EPI_ISL_557778 | Lighthouse Lab in Milton Keynes | Wellcome Sanger Institute for the COVID-19 Genomics UK (COG-UK) consortium | The Lighthouse Lab in Milton Keynes and Alex Alderton, Roberto Amato, Sonia Goncalves, Ewan Harrison, David K. Jackson, Ian Johnston, Dominic Kwiatkowski, Cordelia Langford, John Sillitoe on behalf of the Wellcome Sanger Institute COVID-19 Surveillance Team (http://www.sanger.ac.uk/covid-team) |
| EPI_ISL_557779, EPI_ISL_557780 | Lighthouse Lab in Alderley Park | Wellcome Sanger Institute for the COVID-19 Genomics UK (COG-UK) consortium | The Lighthouse Lab in Alderley Park and Alex Alderton, Roberto Amato, Sonia Goncalves, Ewan Harrison, David K. Jackson, Ian Johnston, Dominic Kwiatkowski, Cordelia Langford, John Sillitoe on behalf of the Wellcome Sanger Institute COVID-19 Surveillance Team |
| EPI_ISL_557781 | Lighthouse Lab in Milton Keynes | Wellcome Sanger Institute for the COVID-19 Genomics UK (COG-UK) consortium | The Lighthouse Lab in Milton Keynes and Alex Alderton, Roberto Amato, Sonia Goncalves, Ewan Harrison, David K. Jackson, Ian Johnston, Dominic Kwiatkowski, Cordelia Langford, John Sillitoe on behalf of the Wellcome Sanger Institute COVID-19 Surveillance Team (http://www.sanger.ac.uk/covid-team) |
| EPI_ISL_557783 | Lighthouse Lab in Alderley Park | Wellcome Sanger Institute for the COVID-19 Genomics UK (COG-UK) consortium | The Lighthouse Lab in Alderley Park and Alex Alderton, Roberto Amato, Sonia Goncalves, Ewan Harrison, David K. Jackson, Ian Johnston, Dominic Kwiatkowski, Cordelia Langford, John Sillitoe on behalf of the Wellcome Sanger Institute COVID-19 Surveillance Team |
| EPI_ISL_557784, EPI_ISL_557786 | Lighthouse Lab in Milton Keynes | Wellcome Sanger Institute for the COVID-19 Genomics UK (COG-UK) consortium | The Lighthouse Lab in Milton Keynes and Alex Alderton, Roberto Amato, Sonia Goncalves, Ewan Harrison, David K. Jackson, Ian Johnston, Dominic Kwiatkowski, Cordelia Langford, John Sillitoe on behalf of the Wellcome Sanger Institute COVID-19 Surveillance Team (http://www.sanger.ac.uk/covid-team) |
| EPI_ISL_557787, EPI_ISL_557788 | Lighthouse Lab in Alderley Park | Wellcome Sanger Institute for the COVID-19 Genomics UK (COG-UK) consortium | The Lighthouse Lab in Alderley Park and Alex Alderton, Roberto Amato, Sonia Goncalves, Ewan Harrison, David K. Jackson, Ian Johnston, Dominic Kwiatkowski, Cordelia Langford, John Sillitoe on behalf of the Wellcome Sanger Institute COVID-19 Surveillance Team |
| EPI_ISL_557789 | Lighthouse Lab in Milton Keynes | Wellcome Sanger Institute for the COVID-19 Genomics UK (COG-UK) consortium | The Lighthouse Lab in Milton Keynes and Alex Alderton, Roberto Amato, Sonia Goncalves, Ewan Harrison, David K. Jackson, Ian Johnston, Dominic Kwiatkowski, Cordelia Langford, John Sillitoe on behalf of the Wellcome Sanger Institute COVID-19 Surveillance Team (http://www.sanger.ac.uk/covid-team) |
| EPI_ISL_557790 | Lighthouse Lab in Alderley Park | Wellcome Sanger Institute for the COVID-19 Genomics UK (COG-UK) consortium | The Lighthouse Lab in Alderley Park and Alex Alderton, Roberto Amato, Sonia Goncalves, Ewan Harrison, David K. Jackson, Ian Johnston, Dominic Kwiatkowski, Cordelia Langford, John Sillitoe on behalf of the Wellcome Sanger Institute COVID-19 Surveillance Team |
| EPI_ISL_557791 | Lighthouse Lab in Milton Keynes | Wellcome Sanger Institute for the COVID-19 Genomics UK (COG-UK) consortium | The Lighthouse Lab in Milton Keynes and Alex Alderton, Roberto Amato, Sonia Goncalves, Ewan Harrison, David K. Jackson, Ian Johnston, Dominic Kwiatkowski, Cordelia Langford, John Sillitoe on behalf of the Wellcome Sanger Institute COVID-19 Surveillance Team (http://www.sanger.ac.uk/covid-team) |
| EPI_ISL_557792 | Lighthouse Lab in Alderley Park | Wellcome Sanger Institute for the COVID-19 Genomics UK (COG-UK) consortium | The Lighthouse Lab in Alderley Park and Alex Alderton, Roberto Amato, Sonia Goncalves, Ewan Harrison, David K. Jackson, Ian Johnston, Dominic Kwiatkowski, Cordelia Langford, John Sillitoe on behalf of the Wellcome Sanger Institute COVID-19 Surveillance Team |
| EPI_ISL_557793 | Lighthouse Lab in Milton Keynes | Wellcome Sanger Institute for the COVID-19 Genomics UK (COG-UK) consortium | The Lighthouse Lab in Milton Keynes and Alex Alderton, Roberto Amato, Sonia Goncalves, Ewan Harrison, David K. Jackson, Ian Johnston, Dominic Kwiatkowski, Cordelia Langford, John Sillitoe on behalf of the Wellcome Sanger Institute COVID-19 Surveillance Team (http://www.sanger.ac.uk/covid-team) |
| EPI_ISL_557794 | Lighthouse Lab in Alderley Park | Wellcome Sanger Institute for the COVID-19 Genomics UK (COG-UK) consortium | The Lighthouse Lab in Alderley Park and Alex Alderton, Roberto Amato, Sonia Goncalves, Ewan Harrison, David K. Jackson, Ian Johnston, Dominic Kwiatkowski, Cordelia Langford, John Sillitoe on behalf of the Wellcome Sanger Institute COVID-19 Surveillance Team |
| EPI_ISL_557795 | Lighthouse Lab in Milton Keynes | Wellcome Sanger Institute for the COVID-19 Genomics UK (COG-UK) consortium | The Lighthouse Lab in Milton Keynes and Alex Alderton, Roberto Amato, Sonia Goncalves, Ewan Harrison, David K. Jackson, Ian Johnston, Dominic Kwiatkowski, Cordelia Langford, John Sillitoe on behalf of the Wellcome Sanger Institute COVID-19 Surveillance Team (http://www.sanger.ac.uk/covid-team) |
| EPI_ISL_557796 | Lighthouse Lab in Milton Keynes | Wellcome Sanger Institute for the COVID-19 Genomics UK (COG-UK) consortium | The Lighthouse Lab in Alderley Park and Alex Alderton, Roberto Amato, Sonia Goncalves, Ewan Harrison, David K. Jackson, Ian Johnston, Dominic Kwiatkowski, Cordelia Langford, John Sillitoe on behalf of the Wellcome Sanger Institute COVID-19 Surveillance Team |
| EPI_ISL_557797, EPI_ISL_557798, EPI_ISL_557799 | Lighthouse Lab in Alderley Park | Wellcome Sanger Institute for the COVID-19 Genomics UK (COG-UK) consortium | The Lighthouse Lab in Alderley Park and Alex Alderton, Roberto Amato, Sonia Goncalves, Ewan Harrison, David K. Jackson, Ian Johnston, Dominic Kwiatkowski, Cordelia Langford, John Sillitoe on behalf of the Wellcome Sanger Institute COVID-19 Surveillance Team |
| EPI_ISL_557800 | Lighthouse Lab in Milton Keynes | Wellcome Sanger Institute for the COVID-19 Genomics UK (COG-UK) consortium | The Lighthouse Lab in Milton Keynes and Alex Alderton, Roberto Amato, Sonia Goncalves, Ewan Harrison, David K. Jackson, Ian Johnston, Dominic Kwiatkowski, Cordelia Langford, John Sillitoe on behalf of the Wellcome Sanger Institute COVID-19 Surveillance Team (http://www.sanger.ac.uk/covid-team) |
| EPI_ISL_557801 | Lighthouse Lab in Alderley Park | Wellcome Sanger Institute for the COVID-19 Genomics UK (COG-UK) consortium | The Lighthouse Lab in Alderley Park and Alex Alderton, Roberto Amato, Sonia Goncalves, Ewan Harrison, David K. Jackson, Ian Johnston, Dominic Kwiatkowski, Cordelia Langford, John Sillitoe on behalf of the Wellcome Sanger Institute COVID-19 Surveillance Team (http://www.sanger.ac.uk/covid-team) |
| EPI_ISL_557802 | Lighthouse Lab in Milton Keynes | Wellcome Sanger Institute for the COVID-19 Genomics UK (COG-UK) consortium | The Lighthouse Lab in Milton Keynes and Alex Alderton, Roberto Amato, Sonia Goncalves, Ewan Harrison, David K. Jackson, Ian Johnston, Dominic Kwiatkowski, Cordelia Langford, John Sillitoe on behalf of the Wellcome Sanger Institute COVID-19 Surveillance Team (http://www.sanger.ac.uk/covid-team) |
| EPI_ISL_557803, EPI_ISL_557804, EPI_ISL_557805, EPI_ISL_557806 | Lighthouse Lab in Alderley Park | Wellcome Sanger Institute for the COVID-19 Genomics UK (COG-UK) consortium | The Lighthouse Lab in Alderley Park and Alex Alderton, Roberto Amato, Sonia Goncalves, Ewan Harrison, David K. Jackson, Ian Johnston, Dominic Kwiatkowski, Cordelia Langford, John Sillitoe on behalf of the Wellcome Sanger Institute COVID-19 Surveillance Team |
| EPI_ISL_557807, EPI_ISL_557808 | Lighthouse Lab in Milton Keynes | Wellcome Sanger Institute for the COVID-19 Genomics UK (COG-UK) consortium | The Lighthouse Lab in Milton Keynes and Alex Alderton, Roberto Amato, Sonia Goncalves, Ewan Harrison, David K. Jackson, Ian Johnston, Dominic Kwiatkowski, Cordelia Langford, John Sillitoe on behalf of the Wellcome Sanger Institute COVID-19 Surveillance Team (http://www.sanger.ac.uk/covid-team) |
| EPI_ISL_557809, EPI_ISL_557810 | Lighthouse Lab in Alderley Park | Wellcome Sanger Institute for the COVID-19 Genomics UK (COG-UK) consortium | The Lighthouse Lab in Alderley Park and Alex Alderton, Roberto Amato, Sonia Goncalves, Ewan Harrison, David K. Jackson, Ian Johnston, Dominic Kwiatkowski, Cordelia Langford, John Sillitoe on behalf of the Wellcome Sanger Institute COVID-19 Surveillance Team |
| EPI_ISL_557811, EPI_ISL_557812 | Lighthouse Lab in Milton Keynes | Wellcome Sanger Institute for the COVID-19 Genomics UK (COG-UK) consortium | The Lighthouse Lab in Milton Keynes and Alex Alderton, Roberto Amato, Sonia Goncalves, Ewan Harrison, David K. Jackson, Ian Johnston, Dominic Kwiatkowski, Cordelia Langford, John Sillitoe on behalf of the Wellcome Sanger Institute COVID-19 Surveillance Team (http://www.sanger.ac.uk/covid-team) |
| EPI_ISL_557813, EPI_ISL_557814, EPI_ISL_557815 | Lighthouse Lab in Alderley Park | Wellcome Sanger Institute for the COVID-19 Genomics UK (COG-UK) consortium | The Lighthouse Lab in Alderley Park and Alex Alderton, Roberto Amato, Sonia Goncalves, Ewan Harrison, David K. Jackson, Ian Johnston, Dominic Kwiatkowski, Cordelia Langford, John Sillitoe on behalf of the Wellcome Sanger Institute COVID-19 Surveillance Team |
| EPI_ISL_557816, EPI_ISL_557817 | Lighthouse Lab in Milton Keynes | Wellcome Sanger Institute for the COVID-19 Genomics UK (COG-UK) consortium | The Lighthouse Lab in Milton Keynes and Alex Alderton, Roberto Amato, Sonia Goncalves, Ewan Harrison, David K. Jackson, Ian Johnston, Dominic Kwiatkowski, Cordelia Langford, John Sillitoe on behalf of the Wellcome Sanger Institute COVID-19 Surveillance Team (http://www.sanger.ac.uk/covid-team) |
| EPI_ISL_557818, EPI_ISL_557819, EPI_ISL_557820 | Lighthouse Lab in Alderley Park | Wellcome Sanger Institute for the COVID-19 Genomics UK (COG-UK) consortium | The Lighthouse Lab in Alderley Park and Alex Alderton, Roberto Amato, Sonia Goncalves, Ewan Harrison, David K. Jackson, Ian Johnston, Dominic Kwiatkowski, Cordelia Langford, John Sillitoe on behalf of the Wellcome Sanger Institute COVID-19 Surveillance Team |
| EPI_ISL_557821, EPI_ISL_557822, EPI_ISL_557823, EPI_ISL_557824 | Lighthouse Lab in Milton Keynes | Wellcome Sanger Institute for the COVID-19 Genomics UK (COG-UK) consortium | The Lighthouse Lab in Milton Keynes and Alex Alderton, Roberto Amato, Sonia Goncalves, Ewan Harrison, David K. Jackson, Ian Johnston, Dominic Kwiatkowski, Cordelia Langford, John Sillitoe on behalf of the Wellcome Sanger Institute COVID-19 Surveillance Team (http://www.sanger.ac.uk/covid-team) |
| EPI_ISL_557825, EPI_ISL_557826 | Lighthouse Lab in Alderley Park | Wellcome Sanger Institute for the COVID-19 Genomics UK (COG-UK) consortium | The Lighthouse Lab in Alderley Park and Alex Alderton, Roberto Amato, Sonia Goncalves, Ewan Harrison, David K. Jackson, Ian Johnston, Dominic Kwiatkowski, Cordelia Langford, John Sillitoe on behalf of the Wellcome Sanger Institute COVID-19 Surveillance Team |
| EPI_ISL_557827, EPI_ISL_557828, EPI_ISL_557829, EPI_ISL_557830 | Lighthouse Lab in Milton Keynes | Wellcome Sanger Institute for the COVID-19 Genomics UK (COG-UK) consortium | The Lighthouse Lab in Milton Keynes and Alex Alderton, Roberto Amato, Sonia Goncalves, Ewan Harrison, David K. Jackson, Ian Johnston, Dominic Kwiatkowski, Cordelia Langford, John Sillitoe on behalf of the Wellcome Sanger Institute COVID-19 Surveillance Team (http://www.sanger.ac.uk/covid-team) |
| EPI_ISL_557831 | Lighthouse Lab in Alderley Park | Wellcome Sanger Institute for the COVID-19 Genomics UK (COG-UK) consortium | The Lighthouse Lab in Alderley Park and Alex Alderton, Roberto Amato, Sonia Goncalves, Ewan Harrison, David K. Jackson, Ian Johnston, Dominic Kwiatkowski, Cordelia Langford, John Sillitoe on behalf of the Wellcome Sanger Institute COVID-19 Surveillance Team |
| EPI_ISL_557832 | Lighthouse Lab in Milton Keynes | Wellcome Sanger Institute for the COVID-19 Genomics UK (COG-UK) consortium | The Lighthouse Lab in Milton Keynes and Alex Alderton, Roberto Amato, Sonia Goncalves, Ewan Harrison, David K. Jackson, Ian Johnston, Dominic Kwiatkowski, Cordelia Langford, John Sillitoe on behalf of the Wellcome Sanger Institute COVID-19 Surveillance Team (http://www.sanger.ac.uk/covid-team) |
| EPI_ISL_557833, EPI_ISL_557834, EPI_ISL_557836, EPI_ISL_557837 | Lighthouse Lab in Alderley Park | Wellcome Sanger Institute for the COVID-19 Genomics UK (COG-UK) consortium | The Lighthouse Lab in Alderley Park and Alex Alderton, Roberto Amato, Sonia Goncalves, Ewan Harrison, David K. Jackson, Ian Johnston, Dominic Kwiatkowski, Cordelia Langford, John Sillitoe on behalf of the Wellcome Sanger Institute COVID-19 Surveillance Team |

| | | | |
|---|---|---|---|
| EPI_ISL_557838 | Lighthouse Lab in Milton Keynes | Wellcome Sanger Institute for the COVID-19 Genomics UK (COG-UK) consortium | The Lighthouse Lab in Milton Keynes and Alex Alderton, Roberto Amato, Sonia Goncalves, Ewan Harrison, David K. Jackson, Ian Johnston, Dominic Kwiatkowski, Cordelia Langford, John Sillitoe on behalf of the Wellcome Sanger Institute COVID-19 Surveillance Team (http://www.sanger.ac.uk/covid-team) |
| EPI_ISL_557839 | Lighthouse Lab in Alderley Park | Wellcome Sanger Institute for the COVID-19 Genomics UK (COG-UK) consortium | The Lighthouse Lab in Alderley Park and Alex Alderton, Roberto Amato, Sonia Goncalves, Ewan Harrison, David K. Jackson, Ian Johnston, Dominic Kwiatkowski, Cordelia Langford, John Sillitoe on behalf of the Wellcome Sanger Institute COVID-19 Surveillance Team |
| EPI_ISL_557840 | Lighthouse Lab in Milton Keynes | Wellcome Sanger Institute for the COVID-19 Genomics UK (COG-UK) Consortium | The Lighthouse Lab in Milton Keynes and Alex Alderton, Roberto Amato, Sonia Goncalves, Ewan Harrison, David K. Jackson, Ian Johnston, Dominic Kwiatkowski, Cordelia Langford, John Sillitoe on behalf of the Wellcome Sanger Institute COVID-19 Surveillance Team |
| EPI_ISL_557841, EPI_ISL_557842 | Lighthouse Lab in Alderley Park | Wellcome Sanger Institute for the COVID-19 Genomics UK (COG-UK) consortium | The Lighthouse Lab in Alderley Park and Alex Alderton, Roberto Amato, Sonia Goncalves, Ewan Harrison, David K. Jackson, Ian Johnston, Dominic Kwiatkowski, Cordelia Langford, John Sillitoe on behalf of the Wellcome Sanger Institute COVID-19 Surveillance Team |
| EPI_ISL_557843 | Lighthouse Lab in Milton Keynes | Wellcome Sanger Institute for the COVID-19 Genomics UK (COG-UK) Consortium | The Lighthouse Lab in Milton Keynes and Alex Alderton, Roberto Amato, Sonia Goncalves, Ewan Harrison, David K. Jackson, Ian Johnston, Dominic Kwiatkowski, Cordelia Langford, John Sillitoe on behalf of the Wellcome Sanger Institute COVID-19 Surveillance Team |
| EPI_ISL_557844 | Lighthouse Lab in Alderley Park | Wellcome Sanger Institute for the COVID-19 Genomics UK (COG-UK) consortium | The Lighthouse Lab in Alderley Park and Alex Alderton, Roberto Amato, Sonia Goncalves, Ewan Harrison, David K. Jackson, Ian Johnston, Dominic Kwiatkowski, Cordelia Langford, John Sillitoe on behalf of the Wellcome Sanger Institute COVID-19 Surveillance Team |
| EPI_ISL_557845 | Lighthouse Lab in Milton Keynes | Wellcome Sanger Institute for the COVID-19 Genomics UK (COG-UK) consortium | The Lighthouse Lab in Milton Keynes and Alex Alderton, Roberto Amato, Sonia Goncalves, Ewan Harrison, David K. Jackson, Ian Johnston, Dominic Kwiatkowski, Cordelia Langford, John Sillitoe on behalf of the Wellcome Sanger Institute COVID-19 Surveillance Team (http://www.sanger.ac.uk/covid-team) |
| EPI_ISL_557846 | Lighthouse Lab in Alderley Park | Wellcome Sanger Institute for the COVID-19 Genomics UK (COG-UK) consortium | The Lighthouse Lab in Alderley Park and Alex Alderton, Roberto Amato, Sonia Goncalves, Ewan Harrison, David K. Jackson, Ian Johnston, Dominic Kwiatkowski, Cordelia Langford, John Sillitoe on behalf of the Wellcome Sanger Institute COVID-19 Surveillance Team |
| EPI_ISL_557847, EPI_ISL_557848, EPI_ISL_557849 | Lighthouse Lab in Milton Keynes | Wellcome Sanger Institute for the COVID-19 Genomics UK (COG-UK) consortium | The Lighthouse Lab in Milton Keynes and Alex Alderton, Roberto Amato, Sonia Goncalves, Ewan Harrison, David K. Jackson, Ian Johnston, Dominic Kwiatkowski, Cordelia Langford, John Sillitoe on behalf of the Wellcome Sanger Institute COVID-19 Surveillance Team (http://www.sanger.ac.uk/covid-team) |
| EPI_ISL_557850 | Lighthouse Lab in Alderley Park | Wellcome Sanger Institute for the COVID-19 Genomics UK (COG-UK) consortium | The Lighthouse Lab in Alderley Park and Alex Alderton, Roberto Amato, Sonia Goncalves, Ewan Harrison, David K. Jackson, Ian Johnston, Dominic Kwiatkowski, Cordelia Langford, John Sillitoe on behalf of the Wellcome Sanger Institute COVID-19 Surveillance Team |
| EPI_ISL_557852 | Lighthouse Lab in Alderley Park | Wellcome Sanger Institute for the COVID-19 Genomics UK (COG-UK) Consortium | The Lighthouse Lab in Alderley Park and Alex Alderton, Roberto Amato, Sonia Goncalves, Ewan Harrison, David K. Jackson, Ian Johnston, Dominic Kwiatkowski, Cordelia Langford, John Sillitoe on behalf of the Wellcome Sanger Institute COVID-19 Surveillance Team |
| EPI_ISL_557853, EPI_ISL_557854 | Lighthouse Lab in Milton Keynes | Wellcome Sanger Institute for the COVID-19 Genomics UK (COG-UK) consortium | The Lighthouse Lab in Milton Keynes and Alex Alderton, Roberto Amato, Sonia Goncalves, Ewan Harrison, David K. Jackson, Ian Johnston, Dominic Kwiatkowski, Cordelia Langford, John Sillitoe on behalf of the Wellcome Sanger Institute COVID-19 Surveillance Team (http://www.sanger.ac.uk/covid-team) |
| EPI_ISL_557855, EPI_ISL_557856, EPI_ISL_557857 | Lighthouse Lab in Alderley Park | Wellcome Sanger Institute for the COVID-19 Genomics UK (COG-UK) consortium | The Lighthouse Lab in Alderley Park and Alex Alderton, Roberto Amato, Sonia Goncalves, Ewan Harrison, David K. Jackson, Ian Johnston, Dominic Kwiatkowski, Cordelia Langford, John Sillitoe on behalf of the Wellcome Sanger Institute COVID-19 Surveillance Team |
| EPI_ISL_557858 | Lighthouse Lab in Milton Keynes | Wellcome Sanger Institute for the COVID-19 Genomics UK (COG-UK) consortium | The Lighthouse Lab in Milton Keynes and Alex Alderton, Roberto Amato, Sonia Goncalves, Ewan Harrison, David K. Jackson, Ian Johnston, Dominic Kwiatkowski, Cordelia Langford, John Sillitoe on behalf of the Wellcome Sanger Institute COVID-19 Surveillance Team (http://www.sanger.ac.uk/covid-team) |
| EPI_ISL_557859, EPI_ISL_557860, EPI_ISL_557861, EPI_ISL_557862, EPI_ISL_557863 | Lighthouse Lab in Alderley Park | Wellcome Sanger Institute for the COVID-19 Genomics UK (COG-UK) consortium | The Lighthouse Lab in Alderley Park and Alex Alderton, Roberto Amato, Sonia Goncalves, Ewan Harrison, David K. Jackson, Ian Johnston, Dominic Kwiatkowski, Cordelia Langford, John Sillitoe on behalf of the Wellcome Sanger Institute COVID-19 Surveillance Team |
| EPI_ISL_557864 | Lighthouse Lab in Alderley Park | Wellcome Sanger Institute for the COVID-19 Genomics UK (COG-UK) consortium | The Lighthouse Lab in Alderley Park and Alex Alderton, Roberto Amato, Sonia Goncalves, Ewan Harrison, David K. Jackson, Ian Johnston, Dominic Kwiatkowski, Cordelia Langford, John Sillitoe on behalf of the Wellcome Sanger Institute COVID-19 Surveillance Team (http://www.sanger.ac.uk/covid-team) |
| EPI_ISL_557865, EPI_ISL_557866, EPI_ISL_557867, EPI_ISL_557868, EPI_ISL_557869, EPI_ISL_557870 | Lighthouse Lab in Milton Keynes | Wellcome Sanger Institute for the COVID-19 Genomics UK (COG-UK) consortium | The Lighthouse Lab in Milton Keynes and Alex Alderton, Roberto Amato, Sonia Goncalves, Ewan Harrison, David K. Jackson, Ian Johnston, Dominic Kwiatkowski, Cordelia Langford, John Sillitoe on behalf of the Wellcome Sanger Institute COVID-19 Surveillance Team (http://www.sanger.ac.uk/covid-team) |
| EPI_ISL_557871, EPI_ISL_557872, EPI_ISL_557873, EPI_ISL_557874, EPI_ISL_557875 | Lighthouse Lab in Alderley Park | Wellcome Sanger Institute for the COVID-19 Genomics UK (COG-UK) consortium | The Lighthouse Lab in Alderley Park and Alex Alderton, Roberto Amato, Sonia Goncalves, Ewan Harrison, David K. Jackson, Ian Johnston, Dominic Kwiatkowski, Cordelia Langford, John Sillitoe on behalf of the Wellcome Sanger Institute COVID-19 Surveillance Team |
| EPI_ISL_557876, EPI_ISL_557877, EPI_ISL_557878, EPI_ISL_557879, EPI_ISL_557880 | Lighthouse Lab in Milton Keynes | Wellcome Sanger Institute for the COVID-19 Genomics UK (COG-UK) consortium | The Lighthouse Lab in Milton Keynes and Alex Alderton, Roberto Amato, Sonia Goncalves, Ewan Harrison, David K. Jackson, Ian Johnston, Dominic Kwiatkowski, Cordelia Langford, John Sillitoe on behalf of the Wellcome Sanger Institute COVID-19 Surveillance Team (http://www.sanger.ac.uk/covid-team) |
| EPI_ISL_557881, EPI_ISL_557882, EPI_ISL_557883 | Lighthouse Lab in Alderley Park | Wellcome Sanger Institute for the COVID-19 Genomics UK (COG-UK) consortium | The Lighthouse Lab in Alderley Park and Alex Alderton, Roberto Amato, Sonia Goncalves, Ewan Harrison, David K. Jackson, Ian Johnston, Dominic Kwiatkowski, Cordelia Langford, John Sillitoe on behalf of the Wellcome Sanger Institute COVID-19 Surveillance Team |
| EPI_ISL_557884 | Lighthouse Lab in Milton Keynes | Wellcome Sanger Institute for the COVID-19 Genomics UK (COG-UK) consortium | The Lighthouse Lab in Milton Keynes and Alex Alderton, Roberto Amato, Sonia Goncalves, Ewan Harrison, David K. Jackson, Ian Johnston, Dominic Kwiatkowski, Cordelia Langford, John Sillitoe on behalf of the Wellcome Sanger Institute COVID-19 Surveillance Team (http://www.sanger.ac.uk/covid-team) |
| EPI_ISL_557885 | Lighthouse Lab in Milton Keynes | Wellcome Sanger Institute for the COVID-19 Genomics UK (COG-UK) Consortium | The Lighthouse Lab in Milton Keynes and Alex Alderton, Roberto Amato, Sonia Goncalves, Ewan Harrison, David K. Jackson, Ian Johnston, Dominic Kwiatkowski, Cordelia Langford, John Sillitoe on behalf of the Wellcome Sanger Institute COVID-19 Surveillance Team |
| EPI_ISL_557886, EPI_ISL_557888 | Lighthouse Lab in Alderley Park | Wellcome Sanger Institute for the COVID-19 Genomics UK (COG-UK) consortium | The Lighthouse Lab in Alderley Park and Alex Alderton, Roberto Amato, Sonia Goncalves, Ewan Harrison, David K. Jackson, Ian Johnston, Dominic Kwiatkowski, Cordelia Langford, John Sillitoe on behalf of the Wellcome Sanger Institute COVID-19 Surveillance Team |
| EPI_ISL_557889, EPI_ISL_557890 | Lighthouse Lab in Milton Keynes | Wellcome Sanger Institute for the COVID-19 Genomics UK (COG-UK) consortium | The Lighthouse Lab in Milton Keynes and Alex Alderton, Roberto Amato, Sonia Goncalves, Ewan Harrison, David K. Jackson, Ian Johnston, Dominic Kwiatkowski, Cordelia Langford, John Sillitoe on behalf of the Wellcome Sanger Institute COVID-19 Surveillance Team (http://www.sanger.ac.uk/covid-team) |
| EPI_ISL_557891, EPI_ISL_557892 | Lighthouse Lab in Alderley Park | Wellcome Sanger Institute for the COVID-19 Genomics UK (COG-UK) consortium | The Lighthouse Lab in Alderley Park and Alex Alderton, Roberto Amato, Sonia Goncalves, Ewan Harrison, David K. Jackson, Ian Johnston, Dominic Kwiatkowski, Cordelia Langford, John Sillitoe on behalf of the Wellcome Sanger Institute COVID-19 Surveillance Team |
| EPI_ISL_557893, EPI_ISL_557894, EPI_ISL_557895 | Lighthouse Lab in Milton Keynes | Wellcome Sanger Institute for the COVID-19 Genomics UK (COG-UK) consortium | The Lighthouse Lab in Milton Keynes and Alex Alderton, Roberto Amato, Sonia Goncalves, Ewan Harrison, David K. Jackson, Ian Johnston, Dominic Kwiatkowski, Cordelia Langford, John Sillitoe on behalf of the Wellcome Sanger Institute COVID-19 Surveillance Team (http://www.sanger.ac.uk/covid-team) |
| EPI_ISL_557896 | Lighthouse Lab in Alderley Park | Wellcome Sanger Institute for the COVID-19 Genomics UK (COG-UK) Consortium | The Lighthouse Lab in Alderley Park and Alex Alderton, Roberto Amato, Sonia Goncalves, Ewan Harrison, David K. Jackson, Ian Johnston, Dominic Kwiatkowski, Cordelia Langford, John Sillitoe on behalf of the Wellcome Sanger Institute COVID-19 Surveillance Team |
| EPI_ISL_557897 | Lighthouse Lab in Alderley Park | Wellcome Sanger Institute for the COVID-19 Genomics UK (COG-UK) consortium | The Lighthouse Lab in Alderley Park and Alex Alderton, Roberto Amato, Sonia Goncalves, Ewan Harrison, David K. Jackson, Ian Johnston, Dominic Kwiatkowski, Cordelia Langford, John Sillitoe on behalf of the Wellcome Sanger Institute COVID-19 Surveillance Team |
| EPI_ISL_557898 | Lighthouse Lab in Milton Keynes | Wellcome Sanger Institute for the COVID-19 Genomics UK (COG-UK) consortium | The Lighthouse Lab in Milton Keynes and Alex Alderton, Roberto Amato, Sonia Goncalves, Ewan Harrison, David K. Jackson, Ian Johnston, Dominic Kwiatkowski, Cordelia Langford, John Sillitoe on behalf of the Wellcome Sanger Institute COVID-19 Surveillance Team (http://www.sanger.ac.uk/covid-team) |
| EPI_ISL_557899, EPI_ISL_557901, EPI_ISL_557902, EPI_ISL_557903, EPI_ISL_557904 | Lighthouse Lab in Alderley Park | Wellcome Sanger Institute for the COVID-19 Genomics UK (COG-UK) consortium | The Lighthouse Lab in Alderley Park and Alex Alderton, Roberto Amato, Sonia Goncalves, Ewan Harrison, David K. Jackson, Ian Johnston, Dominic Kwiatkowski, Cordelia Langford, John Sillitoe on behalf of the Wellcome Sanger Institute COVID-19 Surveillance Team |
| EPI_ISL_557905 | Lighthouse Lab in Milton Keynes | Wellcome Sanger Institute for the COVID-19 Genomics UK (COG-UK) consortium | The Lighthouse Lab in Milton Keynes and Alex Alderton, Roberto Amato, Sonia Goncalves, Ewan Harrison, David K. Jackson, Ian Johnston, Dominic Kwiatkowski, Cordelia Langford, John Sillitoe on behalf of the Wellcome Sanger Institute COVID-19 Surveillance Team (http://www.sanger.ac.uk/covid-team) |
| EPI_ISL_557906 | Lighthouse Lab in Alderley Park | Wellcome Sanger Institute for the COVID-19 Genomics UK (COG-UK) consortium | The Lighthouse Lab in Alderley Park and Alex Alderton, Roberto Amato, Sonia Goncalves, Ewan Harrison, David K. Jackson, Ian Johnston, Dominic Kwiatkowski, Cordelia Langford, John Sillitoe on behalf of the Wellcome Sanger Institute COVID-19 Surveillance Team |
| EPI_ISL_557907, EPI_ISL_557908, EPI_ISL_557909 | Lighthouse Lab in Milton Keynes | Wellcome Sanger Institute for the COVID-19 Genomics UK (COG-UK) consortium | The Lighthouse Lab in Milton Keynes and Alex Alderton, Roberto Amato, Sonia Goncalves, Ewan Harrison, David K. Jackson, Ian Johnston, Dominic Kwiatkowski, Cordelia Langford, John Sillitoe on behalf of the Wellcome Sanger Institute COVID-19 Surveillance Team (http://www.sanger.ac.uk/covid-team) |

| | | | |
|---|---|---|---|
| EPI_ISL_557910 | Lighthouse Lab in Alderley Park | Wellcome Sanger Institute for the COVID-19 Genomics UK (COG-UK) consortium | The Lighthouse Lab in Alderley Park and Alex Alderton, Roberto Amato, Sonia Goncalves, Ewan Harrison, David K. Jackson, Ian Johnston, Dominic Kwiatkowski, Cordelia Langford, John Sillitoe on behalf of the Wellcome Sanger Institute COVID-19 Surveillance Team (http://www.sanger.ac.uk/covid-team) |
| EPI_ISL_557911 | Lighthouse Lab in Milton Keynes | Wellcome Sanger Institute for the COVID-19 Genomics UK (COG-UK) consortium | The Lighthouse Lab in Milton Keynes and Alex Alderton, Roberto Amato, Sonia Goncalves, Ewan Harrison, David K. Jackson, Ian Johnston, Dominic Kwiatkowski, Cordelia Langford, John Sillitoe on behalf of the Wellcome Sanger Institute COVID-19 Surveillance Team (http://www.sanger.ac.uk/covid-team) |
| EPI_ISL_557912, EPI_ISL_557913 | Lighthouse Lab in Alderley Park | Wellcome Sanger Institute for the COVID-19 Genomics UK (COG-UK) consortium | The Lighthouse Lab in Alderley Park and Alex Alderton, Roberto Amato, Sonia Goncalves, Ewan Harrison, David K. Jackson, Ian Johnston, Dominic Kwiatkowski, Cordelia Langford, John Sillitoe on behalf of the Wellcome Sanger Institute COVID-19 Surveillance Team |
| EPI_ISL_557915 | Lighthouse Lab in Milton Keynes | Wellcome Sanger Institute for the COVID-19 Genomics UK (COG-UK) consortium | The Lighthouse Lab in Milton Keynes and Alex Alderton, Roberto Amato, Sonia Goncalves, Ewan Harrison, David K. Jackson, Ian Johnston, Dominic Kwiatkowski, Cordelia Langford, John Sillitoe on behalf of the Wellcome Sanger Institute COVID-19 Surveillance Team (http://www.sanger.ac.uk/covid-team) |
| EPI_ISL_557916 | Lighthouse Lab in Alderley Park | Wellcome Sanger Institute for the COVID-19 Genomics UK (COG-UK) consortium | The Lighthouse Lab in Alderley Park and Alex Alderton, Roberto Amato, Sonia Goncalves, Ewan Harrison, David K. Jackson, Ian Johnston, Dominic Kwiatkowski, Cordelia Langford, John Sillitoe on behalf of the Wellcome Sanger Institute COVID-19 Surveillance Team |
| EPI_ISL_557917 | Lighthouse Lab in Milton Keynes | Wellcome Sanger Institute for the COVID-19 Genomics UK (COG-UK) consortium | The Lighthouse Lab in Milton Keynes and Alex Alderton, Roberto Amato, Sonia Goncalves, Ewan Harrison, David K. Jackson, Ian Johnston, Dominic Kwiatkowski, Cordelia Langford, John Sillitoe on behalf of the Wellcome Sanger Institute COVID-19 Surveillance Team (http://www.sanger.ac.uk/covid-team) |
| EPI_ISL_557918 | Lighthouse Lab in Alderley Park | Wellcome Sanger Institute for the COVID-19 Genomics UK (COG-UK) consortium | The Lighthouse Lab in Alderley Park and Alex Alderton, Roberto Amato, Sonia Goncalves, Ewan Harrison, David K. Jackson, Ian Johnston, Dominic Kwiatkowski, Cordelia Langford, John Sillitoe on behalf of the Wellcome Sanger Institute COVID-19 Surveillance Team |
| EPI_ISL_557919, EPI_ISL_557921 | Lighthouse Lab in Milton Keynes | Wellcome Sanger Institute for the COVID-19 Genomics UK (COG-UK) consortium | The Lighthouse Lab in Milton Keynes and Alex Alderton, Roberto Amato, Sonia Goncalves, Ewan Harrison, David K. Jackson, Ian Johnston, Dominic Kwiatkowski, Cordelia Langford, John Sillitoe on behalf of the Wellcome Sanger Institute COVID-19 Surveillance Team (http://www.sanger.ac.uk/covid-team) |
| EPI_ISL_557922, EPI_ISL_557923, EPI_ISL_557924 | Lighthouse Lab in Alderley Park | Wellcome Sanger Institute for the COVID-19 Genomics UK (COG-UK) consortium | The Lighthouse Lab in Alderley Park and Alex Alderton, Roberto Amato, Sonia Goncalves, Ewan Harrison, David K. Jackson, Ian Johnston, Dominic Kwiatkowski, Cordelia Langford, John Sillitoe on behalf of the Wellcome Sanger Institute COVID-19 Surveillance Team |
| EPI_ISL_557925, EPI_ISL_557926, EPI_ISL_557927 | Lighthouse Lab in Milton Keynes | Wellcome Sanger Institute for the COVID-19 Genomics UK (COG-UK) consortium | The Lighthouse Lab in Milton Keynes and Alex Alderton, Roberto Amato, Sonia Goncalves, Ewan Harrison, David K. Jackson, Ian Johnston, Dominic Kwiatkowski, Cordelia Langford, John Sillitoe on behalf of the Wellcome Sanger Institute COVID-19 Surveillance Team (http://www.sanger.ac.uk/covid-team) |
| EPI_ISL_557928, EPI_ISL_557930, EPI_ISL_557931, EPI_ISL_557932 | Lighthouse Lab in Alderley Park | Wellcome Sanger Institute for the COVID-19 Genomics UK (COG-UK) consortium | The Lighthouse Lab in Alderley Park and Alex Alderton, Roberto Amato, Sonia Goncalves, Ewan Harrison, David K. Jackson, Ian Johnston, Dominic Kwiatkowski, Cordelia Langford, John Sillitoe on behalf of the Wellcome Sanger Institute COVID-19 Surveillance Team |
| EPI_ISL_557933, EPI_ISL_557937, EPI_ISL_557938, EPI_ISL_557940, EPI_ISL_557941, EPI_ISL_557942 | Lighthouse Lab in Milton Keynes | Wellcome Sanger Institute for the COVID-19 Genomics UK (COG-UK) consortium | The Lighthouse Lab in Milton Keynes and Alex Alderton, Roberto Amato, Sonia Goncalves, Ewan Harrison, David K. Jackson, Ian Johnston, Dominic Kwiatkowski, Cordelia Langford, John Sillitoe on behalf of the Wellcome Sanger Institute COVID-19 Surveillance Team (http://www.sanger.ac.uk/covid-team) |
| EPI_ISL_557943 | Lighthouse Lab in Alderley Park | Wellcome Sanger Institute for the COVID-19 Genomics UK (COG-UK) consortium | The Lighthouse Lab in Alderley Park and Alex Alderton, Roberto Amato, Sonia Goncalves, Ewan Harrison, David K. Jackson, Ian Johnston, Dominic Kwiatkowski, Cordelia Langford, John Sillitoe on behalf of the Wellcome Sanger Institute COVID-19 Surveillance Team |
| EPI_ISL_557944 | Lighthouse Lab in Milton Keynes | Wellcome Sanger Institute for the COVID-19 Genomics UK (COG-UK) consortium | The Lighthouse Lab in Milton Keynes and Alex Alderton, Roberto Amato, Sonia Goncalves, Ewan Harrison, David K. Jackson, Ian Johnston, Dominic Kwiatkowski, Cordelia Langford, John Sillitoe on behalf of the Wellcome Sanger Institute COVID-19 Surveillance Team (http://www.sanger.ac.uk/covid-team) |
| EPI_ISL_557945, EPI_ISL_557946 | Lighthouse Lab in Alderley Park | Wellcome Sanger Institute for the COVID-19 Genomics UK (COG-UK) consortium | The Lighthouse Lab in Alderley Park and Alex Alderton, Roberto Amato, Sonia Goncalves, Ewan Harrison, David K. Jackson, Ian Johnston, Dominic Kwiatkowski, Cordelia Langford, John Sillitoe on behalf of the Wellcome Sanger Institute COVID-19 Surveillance Team |
| EPI_ISL_557947, EPI_ISL_557948, EPI_ISL_557949 | Lighthouse Lab in Milton Keynes | Wellcome Sanger Institute for the COVID-19 Genomics UK (COG-UK) consortium | The Lighthouse Lab in Milton Keynes and Alex Alderton, Roberto Amato, Sonia Goncalves, Ewan Harrison, David K. Jackson, Ian Johnston, Dominic Kwiatkowski, Cordelia Langford, John Sillitoe on behalf of the Wellcome Sanger Institute COVID-19 Surveillance Team (http://www.sanger.ac.uk/covid-team) |
| EPI_ISL_557952 | Lighthouse Lab in Milton Keynes | Wellcome Sanger Institute for the COVID-19 Genomics UK (COG-UK) Consortium | The Lighthouse Lab in Milton Keynes and Alex Alderton, Roberto Amato, Sonia Goncalves, Ewan Harrison, David K. Jackson, Ian Johnston, Dominic Kwiatkowski, Cordelia Langford, John Sillitoe on behalf of the Wellcome Sanger Institute COVID-19 Surveillance Team |
| EPI_ISL_557953, EPI_ISL_557954 | Lighthouse Lab in Milton Keynes | Wellcome Sanger Institute for the COVID-19 Genomics UK (COG-UK) consortium | The Lighthouse Lab in Milton Keynes and Alex Alderton, Roberto Amato, Sonia Goncalves, Ewan Harrison, David K. Jackson, Ian Johnston, Dominic Kwiatkowski, Cordelia Langford, John Sillitoe on behalf of the Wellcome Sanger Institute COVID-19 Surveillance Team (http://www.sanger.ac.uk/covid-team) |
| EPI_ISL_557955 | Lighthouse Lab in Alderley Park | Wellcome Sanger Institute for the COVID-19 Genomics UK (COG-UK) consortium | The Lighthouse Lab in Alderley Park and Alex Alderton, Roberto Amato, Sonia Goncalves, Ewan Harrison, David K. Jackson, Ian Johnston, Dominic Kwiatkowski, Cordelia Langford, John Sillitoe on behalf of the Wellcome Sanger Institute COVID-19 Surveillance Team |
| EPI_ISL_557956 | Lighthouse Lab in Milton Keynes | Wellcome Sanger Institute for the COVID-19 Genomics UK (COG-UK) consortium | The Lighthouse Lab in Milton Keynes and Alex Alderton, Roberto Amato, Sonia Goncalves, Ewan Harrison, David K. Jackson, Ian Johnston, Dominic Kwiatkowski, Cordelia Langford, John Sillitoe on behalf of the Wellcome Sanger Institute COVID-19 Surveillance Team (http://www.sanger.ac.uk/covid-team) |
| EPI_ISL_557957 | Lighthouse Lab in Alderley Park | Wellcome Sanger Institute for the COVID-19 Genomics UK (COG-UK) consortium | The Lighthouse Lab in Alderley Park and Alex Alderton, Roberto Amato, Sonia Goncalves, Ewan Harrison, David K. Jackson, Ian Johnston, Dominic Kwiatkowski, Cordelia Langford, John Sillitoe on behalf of the Wellcome Sanger Institute COVID-19 Surveillance Team |
| EPI_ISL_557958, EPI_ISL_557959 | Lighthouse Lab in Milton Keynes | Wellcome Sanger Institute for the COVID-19 Genomics UK (COG-UK) consortium | The Lighthouse Lab in Milton Keynes and Alex Alderton, Roberto Amato, Sonia Goncalves, Ewan Harrison, David K. Jackson, Ian Johnston, Dominic Kwiatkowski, Cordelia Langford, John Sillitoe on behalf of the Wellcome Sanger Institute COVID-19 Surveillance Team (http://www.sanger.ac.uk/covid-team) |
| EPI_ISL_557960, EPI_ISL_557961, EPI_ISL_557962 | Lighthouse Lab in Alderley Park | Wellcome Sanger Institute for the COVID-19 Genomics UK (COG-UK) consortium | The Lighthouse Lab in Alderley Park and Alex Alderton, Roberto Amato, Sonia Goncalves, Ewan Harrison, David K. Jackson, Ian Johnston, Dominic Kwiatkowski, Cordelia Langford, John Sillitoe on behalf of the Wellcome Sanger Institute COVID-19 Surveillance Team |
| EPI_ISL_557963, EPI_ISL_557964, EPI_ISL_557965 | Lighthouse Lab in Milton Keynes | Wellcome Sanger Institute for the COVID-19 Genomics UK (COG-UK) consortium | The Lighthouse Lab in Milton Keynes and Alex Alderton, Roberto Amato, Sonia Goncalves, Ewan Harrison, David K. Jackson, Ian Johnston, Dominic Kwiatkowski, Cordelia Langford, John Sillitoe on behalf of the Wellcome Sanger Institute COVID-19 Surveillance Team (http://www.sanger.ac.uk/covid-team) |
| EPI_ISL_557966 | Lighthouse Lab in Alderley Park | Wellcome Sanger Institute for the COVID-19 Genomics UK (COG-UK) consortium | The Lighthouse Lab in Alderley Park and Alex Alderton, Roberto Amato, Sonia Goncalves, Ewan Harrison, David K. Jackson, Ian Johnston, Dominic Kwiatkowski, Cordelia Langford, John Sillitoe on behalf of the Wellcome Sanger Institute COVID-19 Surveillance Team (http://www.sanger.ac.uk/covid-team) |
| EPI_ISL_557967 | Lighthouse Lab in Milton Keynes | Wellcome Sanger Institute for the COVID-19 Genomics UK (COG-UK) consortium | The Lighthouse Lab in Milton Keynes and Alex Alderton, Roberto Amato, Sonia Goncalves, Ewan Harrison, David K. Jackson, Ian Johnston, Dominic Kwiatkowski, Cordelia Langford, John Sillitoe on behalf of the Wellcome Sanger Institute COVID-19 Surveillance Team (http://www.sanger.ac.uk/covid-team) |
| EPI_ISL_557968, EPI_ISL_557969 | Lighthouse Lab in Alderley Park | Wellcome Sanger Institute for the COVID-19 Genomics UK (COG-UK) consortium | The Lighthouse Lab in Alderley Park and Alex Alderton, Roberto Amato, Sonia Goncalves, Ewan Harrison, David K. Jackson, Ian Johnston, Dominic Kwiatkowski, Cordelia Langford, John Sillitoe on behalf of the Wellcome Sanger Institute COVID-19 Surveillance Team |
| EPI_ISL_557970, EPI_ISL_557972 | Lighthouse Lab in Milton Keynes | Wellcome Sanger Institute for the COVID-19 Genomics UK (COG-UK) consortium | The Lighthouse Lab in Milton Keynes and Alex Alderton, Roberto Amato, Sonia Goncalves, Ewan Harrison, David K. Jackson, Ian Johnston, Dominic Kwiatkowski, Cordelia Langford, John Sillitoe on behalf of the Wellcome Sanger Institute COVID-19 Surveillance Team (http://www.sanger.ac.uk/covid-team) |
| EPI_ISL_557973 | Lighthouse Lab in Alderley Park | Wellcome Sanger Institute for the COVID-19 Genomics UK (COG-UK) consortium | The Lighthouse Lab in Alderley Park and Alex Alderton, Roberto Amato, Sonia Goncalves, Ewan Harrison, David K. Jackson, Ian Johnston, Dominic Kwiatkowski, Cordelia Langford, John Sillitoe on behalf of the Wellcome Sanger Institute COVID-19 Surveillance Team |
| EPI_ISL_557974 | Lighthouse Lab in Milton Keynes | Wellcome Sanger Institute for the COVID-19 Genomics UK (COG-UK) consortium | The Lighthouse Lab in Milton Keynes and Alex Alderton, Roberto Amato, Sonia Goncalves, Ewan Harrison, David K. Jackson, Ian Johnston, Dominic Kwiatkowski, Cordelia Langford, John Sillitoe on behalf of the Wellcome Sanger Institute COVID-19 Surveillance Team (http://www.sanger.ac.uk/covid-team) |
| EPI_ISL_557975 | Lighthouse Lab in Alderley Park | Wellcome Sanger Institute for the COVID-19 Genomics UK (COG-UK) consortium | The Lighthouse Lab in Alderley Park and Alex Alderton, Roberto Amato, Sonia Goncalves, Ewan Harrison, David K. Jackson, Ian Johnston, Dominic Kwiatkowski, Cordelia Langford, John Sillitoe on behalf of the Wellcome Sanger Institute COVID-19 Surveillance Team |
| EPI_ISL_557976 | Lighthouse Lab in Milton Keynes | Wellcome Sanger Institute for the COVID-19 Genomics UK (COG-UK) consortium | The Lighthouse Lab in Milton Keynes and Alex Alderton, Roberto Amato, Sonia Goncalves, Ewan Harrison, David K. Jackson, Ian Johnston, Dominic Kwiatkowski, Cordelia Langford, John Sillitoe on behalf of the Wellcome Sanger Institute COVID-19 Surveillance Team (http://www.sanger.ac.uk/covid-team) |
| EPI_ISL_557977 | Lighthouse Lab in Milton Keynes | Wellcome Sanger Institute for the COVID-19 Genomics UK (COG-UK) consortium | The Lighthouse Lab in Alderley Park and Alex Alderton, Roberto Amato, Sonia Goncalves, Ewan Harrison, David K. Jackson, Ian Johnston, Dominic Kwiatkowski, Cordelia Langford, John Sillitoe on behalf of the Wellcome Sanger Institute COVID-19 Surveillance Team |
| EPI_ISL_557979 | Lighthouse Lab in Milton Keynes | Wellcome Sanger Institute for the COVID-19 Genomics UK (COG-UK) consortium | The Lighthouse Lab in Milton Keynes and Alex Alderton, Roberto Amato, Sonia Goncalves, Ewan Harrison, David K. Jackson, Ian Johnston, Dominic Kwiatkowski, Cordelia Langford, John Sillitoe on behalf of the Wellcome Sanger Institute COVID-19 Surveillance Team (http://www.sanger.ac.uk/covid-team) |
| EPI_ISL_557980, EPI_ISL_557981, EPI_ISL_557982, EPI_ISL_557983 | Lighthouse Lab in Alderley Park | Wellcome Sanger Institute for the COVID-19 Genomics UK (COG-UK) consortium | The Lighthouse Lab in Alderley Park and Alex Alderton, Roberto Amato, Sonia Goncalves, Ewan Harrison, David K. Jackson, Ian Johnston, Dominic Kwiatkowski, Cordelia Langford, John Sillitoe on behalf of the Wellcome Sanger Institute COVID-19 Surveillance Team |

| | | | |
|---|---|---|---|
| EPI_ISL_557984 | Lighthouse Lab in Milton Keynes | Wellcome Sanger Institute for the COVID-19 Genomics UK (COG-UK) consortium | The Lighthouse Lab in Milton Keynes and Alex Alderton, Roberto Amato, Sonia Goncalves, Ewan Harrison, David K. Jackson, Ian Johnston, Dominic Kwiatkowski, Cordelia Langford, John Sillitoe on behalf of the Wellcome Sanger Institute COVID-19 Surveillance Team (http://www.sanger.ac.uk/covid-team) |
| EPI_ISL_557985, EPI_ISL_557986, EPI_ISL_557987 | Lighthouse Lab in Alderley Park | Wellcome Sanger Institute for the COVID-19 Genomics UK (COG-UK) consortium | The Lighthouse Lab in Alderley Park and Alex Alderton, Roberto Amato, Sonia Goncalves, Ewan Harrison, David K. Jackson, Ian Johnston, Dominic Kwiatkowski, Cordelia Langford, John Sillitoe on behalf of the Wellcome Sanger Institute COVID-19 Surveillance Team |
| EPI_ISL_557988 | Lighthouse Lab in Alderley Park | Wellcome Sanger Institute for the COVID-19 Genomics UK (COG-UK) consortium | The Lighthouse Lab in Alderley Park and Alex Alderton, Roberto Amato, Sonia Goncalves, Ewan Harrison, David K. Jackson, Ian Johnston, Dominic Kwiatkowski, Cordelia Langford, John Sillitoe on behalf of the Wellcome Sanger Institute COVID-19 Surveillance Team (http://www.sanger.ac.uk/covid-team) |
| EPI_ISL_557989 | Lighthouse Lab in Milton Keynes | Wellcome Sanger Institute for the COVID-19 Genomics UK (COG-UK) consortium | The Lighthouse Lab in Milton Keynes and Alex Alderton, Roberto Amato, Sonia Goncalves, Ewan Harrison, David K. Jackson, Ian Johnston, Dominic Kwiatkowski, Cordelia Langford, John Sillitoe on behalf of the Wellcome Sanger Institute COVID-19 Surveillance Team (http://www.sanger.ac.uk/covid-team) |
| EPI_ISL_557990, EPI_ISL_557991, EPI_ISL_557992, EPI_ISL_557993, EPI_ISL_557994, EPI_ISL_557995, EPI_ISL_557996, EPI_ISL_557997, EPI_ISL_557998 | Lighthouse Lab in Alderley Park | Wellcome Sanger Institute for the COVID-19 Genomics UK (COG-UK) consortium | The Lighthouse Lab in Alderley Park and Alex Alderton, Roberto Amato, Sonia Goncalves, Ewan Harrison, David K. Jackson, Ian Johnston, Dominic Kwiatkowski, Cordelia Langford, John Sillitoe on behalf of the Wellcome Sanger Institute COVID-19 Surveillance Team |
| EPI_ISL_557999 | Lighthouse Lab in Alderley Park | Wellcome Sanger Institute for the COVID-19 Genomics UK (COG-UK) Consortium | The Lighthouse Lab in Alderley Park and Alex Alderton, Roberto Amato, Sonia Goncalves, Ewan Harrison, David K. Jackson, Ian Johnston, Dominic Kwiatkowski, Cordelia Langford, John Sillitoe on behalf of the Wellcome Sanger Institute COVID-19 Surveillance Team |
| EPI_ISL_558000 | Lighthouse Lab in Alderley Park | Wellcome Sanger Institute for the COVID-19 Genomics UK (COG-UK) consortium | The Lighthouse Lab in Alderley Park and Alex Alderton, Roberto Amato, Sonia Goncalves, Ewan Harrison, David K. Jackson, Ian Johnston, Dominic Kwiatkowski, Cordelia Langford, John Sillitoe on behalf of the Wellcome Sanger Institute COVID-19 Surveillance Team |
| EPI_ISL_558001 | Lighthouse Lab in Milton Keynes | Wellcome Sanger Institute for the COVID-19 Genomics UK (COG-UK) consortium | The Lighthouse Lab in Milton Keynes and Alex Alderton, Roberto Amato, Sonia Goncalves, Ewan Harrison, David K. Jackson, Ian Johnston, Dominic Kwiatkowski, Cordelia Langford, John Sillitoe on behalf of the Wellcome Sanger Institute COVID-19 Surveillance Team (http://www.sanger.ac.uk/covid-team) |
| EPI_ISL_558002, EPI_ISL_558003, EPI_ISL_558004, EPI_ISL_558005, EPI_ISL_558006, EPI_ISL_558007, EPI_ISL_558008, EPI_ISL_558009, EPI_ISL_558010, EPI_ISL_558011, EPI_ISL_558012, EPI_ISL_558013, EPI_ISL_558014, EPI_ISL_558015 | | | |
| see above | Lighthouse Lab in Alderley Park | Wellcome Sanger Institute for the COVID-19 Genomics UK (COG-UK) consortium | The Lighthouse Lab in Alderley Park and Alex Alderton, Roberto Amato, Sonia Goncalves, Ewan Harrison, David K. Jackson, Ian Johnston, Dominic Kwiatkowski, Cordelia Langford, John Sillitoe on behalf of the Wellcome Sanger Institute COVID-19 Surveillance Team |
| EPI_ISL_558016, EPI_ISL_558017 | Lighthouse Lab in Alderley Park | Wellcome Sanger Institute for the COVID-19 Genomics UK (COG-UK) consortium | The Lighthouse Lab in Alderley Park and Alex Alderton, Roberto Amato, Sonia Goncalves, Ewan Harrison, David K. Jackson, Ian Johnston, Dominic Kwiatkowski, Cordelia Langford, John Sillitoe on behalf of the Wellcome Sanger Institute COVID-19 Surveillance Team (http://www.sanger.ac.uk/covid-team) |
| EPI_ISL_558018, EPI_ISL_558019, EPI_ISL_558020 | Lighthouse Lab in Alderley Park | Wellcome Sanger Institute for the COVID-19 Genomics UK (COG-UK) consortium | The Lighthouse Lab in Alderley Park and Alex Alderton, Roberto Amato, Sonia Goncalves, Ewan Harrison, David K. Jackson, Ian Johnston, Dominic Kwiatkowski, Cordelia Langford, John Sillitoe on behalf of the Wellcome Sanger Institute COVID-19 Surveillance Team |
| EPI_ISL_558021 | Lighthouse Lab in Milton Keynes | Wellcome Sanger Institute for the COVID-19 Genomics UK (COG-UK) consortium | The Lighthouse Lab in Milton Keynes and Alex Alderton, Roberto Amato, Sonia Goncalves, Ewan Harrison, David K. Jackson, Ian Johnston, Dominic Kwiatkowski, Cordelia Langford, John Sillitoe on behalf of the Wellcome Sanger Institute COVID-19 Surveillance Team (http://www.sanger.ac.uk/covid-team) |
| EPI_ISL_558022 | Lighthouse Lab in Alderley Park | Wellcome Sanger Institute for the COVID-19 Genomics UK (COG-UK) consortium | The Lighthouse Lab in Alderley Park and Alex Alderton, Roberto Amato, Sonia Goncalves, Ewan Harrison, David K. Jackson, Ian Johnston, Dominic Kwiatkowski, Cordelia Langford, John Sillitoe on behalf of the Wellcome Sanger Institute COVID-19 Surveillance Team |
| EPI_ISL_558023 | Lighthouse Lab in Alderley Park | Wellcome Sanger Institute for the COVID-19 Genomics UK (COG-UK) consortium | The Lighthouse Lab in Alderley Park and Alex Alderton, Roberto Amato, Sonia Goncalves, Ewan Harrison, David K. Jackson, Ian Johnston, Dominic Kwiatkowski, Cordelia Langford, John Sillitoe on behalf of the Wellcome Sanger Institute COVID-19 Surveillance Team (http://www.sanger.ac.uk/covid-team) |
| EPI_ISL_558024 | Lighthouse Lab in Alderley Park | Wellcome Sanger Institute for the COVID-19 Genomics UK (COG-UK) Consortium | The Lighthouse Lab in Alderley Park and Alex Alderton, Roberto Amato, Sonia Goncalves, Ewan Harrison, David K. Jackson, Ian Johnston, Dominic Kwiatkowski, Cordelia Langford, John Sillitoe on behalf of the Wellcome Sanger Institute COVID-19 Surveillance Team |
| EPI_ISL_558025 | Lighthouse Lab in Alderley Park | Wellcome Sanger Institute for the COVID-19 Genomics UK (COG-UK) consortium | The Lighthouse Lab in Alderley Park and Alex Alderton, Roberto Amato, Sonia Goncalves, Ewan Harrison, David K. Jackson, Ian Johnston, Dominic Kwiatkowski, Cordelia Langford, John Sillitoe on behalf of the Wellcome Sanger Institute COVID-19 Surveillance Team |
| EPI_ISL_558026 | Lighthouse Lab in Milton Keynes | Wellcome Sanger Institute for the COVID-19 Genomics UK (COG-UK) consortium | The Lighthouse Lab in Milton Keynes and Alex Alderton, Roberto Amato, Sonia Goncalves, Ewan Harrison, David K. Jackson, Ian Johnston, Dominic Kwiatkowski, Cordelia Langford, John Sillitoe on behalf of the Wellcome Sanger Institute COVID-19 Surveillance Team (http://www.sanger.ac.uk/covid-team) |
| EPI_ISL_558027 | Lighthouse Lab in Alderley Park | Wellcome Sanger Institute for the COVID-19 Genomics UK (COG-UK) consortium | The Lighthouse Lab in Alderley Park and Alex Alderton, Roberto Amato, Sonia Goncalves, Ewan Harrison, David K. Jackson, Ian Johnston, Dominic Kwiatkowski, Cordelia Langford, John Sillitoe on behalf of the Wellcome Sanger Institute COVID-19 Surveillance Team |
| EPI_ISL_558028 | Lighthouse Lab in Milton Keynes | Wellcome Sanger Institute for the COVID-19 Genomics UK (COG-UK) consortium | The Lighthouse Lab in Milton Keynes and Alex Alderton, Roberto Amato, Sonia Goncalves, Ewan Harrison, David K. Jackson, Ian Johnston, Dominic Kwiatkowski, Cordelia Langford, John Sillitoe on behalf of the Wellcome Sanger Institute COVID-19 Surveillance Team (http://www.sanger.ac.uk/covid-team) |
| EPI_ISL_558029 | Lighthouse Lab in Alderley Park | Wellcome Sanger Institute for the COVID-19 Genomics UK (COG-UK) consortium | The Lighthouse Lab in Alderley Park and Alex Alderton, Roberto Amato, Sonia Goncalves, Ewan Harrison, David K. Jackson, Ian Johnston, Dominic Kwiatkowski, Cordelia Langford, John Sillitoe on behalf of the Wellcome Sanger Institute COVID-19 Surveillance Team |
| EPI_ISL_558030 | Lighthouse Lab in Milton Keynes | Wellcome Sanger Institute for the COVID-19 Genomics UK (COG-UK) consortium | The Lighthouse Lab in Milton Keynes and Alex Alderton, Roberto Amato, Sonia Goncalves, Ewan Harrison, David K. Jackson, Ian Johnston, Dominic Kwiatkowski, Cordelia Langford, John Sillitoe on behalf of the Wellcome Sanger Institute COVID-19 Surveillance Team (http://www.sanger.ac.uk/covid-team) |
| EPI_ISL_558031, EPI_ISL_558032, EPI_ISL_558033, EPI_ISL_558034, EPI_ISL_558035, EPI_ISL_558036 | Lighthouse Lab in Alderley Park | Wellcome Sanger Institute for the COVID-19 Genomics UK (COG-UK) consortium | The Lighthouse Lab in Alderley Park and Alex Alderton, Roberto Amato, Sonia Goncalves, Ewan Harrison, David K. Jackson, Ian Johnston, Dominic Kwiatkowski, Cordelia Langford, John Sillitoe on behalf of the Wellcome Sanger Institute COVID-19 Surveillance Team |
| EPI_ISL_558037 | Lighthouse Lab in Milton Keynes | Wellcome Sanger Institute for the COVID-19 Genomics UK (COG-UK) consortium | The Lighthouse Lab in Milton Keynes and Alex Alderton, Roberto Amato, Sonia Goncalves, Ewan Harrison, David K. Jackson, Ian Johnston, Dominic Kwiatkowski, Cordelia Langford, John Sillitoe on behalf of the Wellcome Sanger Institute COVID-19 Surveillance Team (http://www.sanger.ac.uk/covid-team) |
| EPI_ISL_558038, EPI_ISL_558039, EPI_ISL_558040 | Lighthouse Lab in Alderley Park | Wellcome Sanger Institute for the COVID-19 Genomics UK (COG-UK) consortium | The Lighthouse Lab in Alderley Park and Alex Alderton, Roberto Amato, Sonia Goncalves, Ewan Harrison, David K. Jackson, Ian Johnston, Dominic Kwiatkowski, Cordelia Langford, John Sillitoe on behalf of the Wellcome Sanger Institute COVID-19 Surveillance Team |
| EPI_ISL_558041 | Lighthouse Lab in Milton Keynes | Wellcome Sanger Institute for the COVID-19 Genomics UK (COG-UK) consortium | The Lighthouse Lab in Milton Keynes and Alex Alderton, Roberto Amato, Sonia Goncalves, Ewan Harrison, David K. Jackson, Ian Johnston, Dominic Kwiatkowski, Cordelia Langford, John Sillitoe on behalf of the Wellcome Sanger Institute COVID-19 Surveillance Team (http://www.sanger.ac.uk/covid-team) |
| EPI_ISL_558042, EPI_ISL_558043, EPI_ISL_558044, EPI_ISL_558045, EPI_ISL_558046, EPI_ISL_558047, EPI_ISL_558048, EPI_ISL_558049, EPI_ISL_558050, EPI_ISL_558051 | Lighthouse Lab in Alderley Park | Wellcome Sanger Institute for the COVID-19 Genomics UK (COG-UK) consortium | The Lighthouse Lab in Alderley Park and Alex Alderton, Roberto Amato, Sonia Goncalves, Ewan Harrison, David K. Jackson, Ian Johnston, Dominic Kwiatkowski, Cordelia Langford, John Sillitoe on behalf of the Wellcome Sanger Institute COVID-19 Surveillance Team |
| EPI_ISL_558052 | Lighthouse Lab in Alderley Park | Wellcome Sanger Institute for the COVID-19 Genomics UK (COG-UK) consortium | The Lighthouse Lab in Alderley Park and Alex Alderton, Roberto Amato, Sonia Goncalves, Ewan Harrison, David K. Jackson, Ian Johnston, Dominic Kwiatkowski, Cordelia Langford, John Sillitoe on behalf of the Wellcome Sanger Institute COVID-19 Surveillance Team (http://www.sanger.ac.uk/covid-team) |
| EPI_ISL_558053, EPI_ISL_558054, EPI_ISL_558055 | Lighthouse Lab in Alderley Park | Wellcome Sanger Institute for the COVID-19 Genomics UK (COG-UK) consortium | The Lighthouse Lab in Alderley Park and Alex Alderton, Roberto Amato, Sonia Goncalves, Ewan Harrison, David K. Jackson, Ian Johnston, Dominic Kwiatkowski, Cordelia Langford, John Sillitoe on behalf of the Wellcome Sanger Institute COVID-19 Surveillance Team |
| EPI_ISL_558056 | Lighthouse Lab in Alderley Park | Wellcome Sanger Institute for the COVID-19 Genomics UK (COG-UK) consortium | The Lighthouse Lab in Alderley Park and Alex Alderton, Roberto Amato, Sonia Goncalves, Ewan Harrison, David K. Jackson, Ian Johnston, Dominic Kwiatkowski, Cordelia Langford, John Sillitoe on behalf of the Wellcome Sanger Institute COVID-19 Surveillance Team (http://www.sanger.ac.uk/covid-team) |
| EPI_ISL_558057 | Lighthouse Lab in Alderley Park | Wellcome Sanger Institute for the COVID-19 Genomics UK (COG-UK) consortium | The Lighthouse Lab in Alderley Park and Alex Alderton, Roberto Amato, Sonia Goncalves, Ewan Harrison, David K. Jackson, Ian Johnston, Dominic Kwiatkowski, Cordelia Langford, John Sillitoe on behalf of the Wellcome Sanger Institute COVID-19 Surveillance Team |
| EPI_ISL_558058 | Lighthouse Lab in Milton Keynes | Wellcome Sanger Institute for the COVID-19 Genomics UK (COG-UK) consortium | The Lighthouse Lab in Milton Keynes and Alex Alderton, Roberto Amato, Sonia Goncalves, Ewan Harrison, David K. Jackson, Ian Johnston, Dominic Kwiatkowski, Cordelia Langford, John Sillitoe on behalf of the Wellcome Sanger Institute COVID-19 Surveillance Team (http://www.sanger.ac.uk/covid-team) |
| EPI_ISL_558059, EPI_ISL_558060, | Lighthouse Lab in Alderley Park | Wellcome Sanger Institute for the COVID-19 Genomics UK | The Lighthouse Lab in Alderley Park and Alex Alderton, Roberto Amato, Sonia Goncalves, Ewan Harrison, David K. Jackson, Ian Johnston, Dominic |

| | | | |
|---|---|---|---|
| EPI_ISL_558061, EPI_ISL_558062, EPI_ISL_558063 | | (COG-UK) consortium | Kwiatkowski, Cordelia Langford, John Sillitoe on behalf of the Wellcome Sanger Institute COVID-19 Surveillance Team |
| EPI_ISL_558064 | Lighthouse Lab in Alderley Park | Wellcome Sanger Institute for the COVID-19 Genomics UK (COG-UK) consortium | The Lighthouse Lab in Alderley Park and Alex Alderton, Roberto Amato, Sonia Goncalves, Ewan Harrison, David K. Jackson, Ian Johnston, Dominic Kwiatkowski, Cordelia Langford, John Sillitoe on behalf of the Wellcome Sanger Institute COVID-19 Surveillance Team (http://www.sanger.ac.uk/covid-team) |
| EPI_ISL_558065, EPI_ISL_558066, EPI_ISL_558067 | Lighthouse Lab in Alderley Park | Wellcome Sanger Institute for the COVID-19 Genomics UK (COG-UK) consortium | The Lighthouse Lab in Alderley Park and Alex Alderton, Roberto Amato, Sonia Goncalves, Ewan Harrison, David K. Jackson, Ian Johnston, Dominic Kwiatkowski, Cordelia Langford, John Sillitoe on behalf of the Wellcome Sanger Institute COVID-19 Surveillance Team |
| EPI_ISL_558068 | Lighthouse Lab in Milton Keynes | Wellcome Sanger Institute for the COVID-19 Genomics UK (COG-UK) consortium | The Lighthouse Lab in Milton Keynes and Alex Alderton, Roberto Amato, Sonia Goncalves, Ewan Harrison, David K. Jackson, Ian Johnston, Dominic Kwiatkowski, Cordelia Langford, John Sillitoe on behalf of the Wellcome Sanger Institute COVID-19 Surveillance Team (http://www.sanger.ac.uk/covid-team) |
| EPI_ISL_558069, EPI_ISL_558071, EPI_ISL_558072 | Lighthouse Lab in Alderley Park | Wellcome Sanger Institute for the COVID-19 Genomics UK (COG-UK) consortium | The Lighthouse Lab in Alderley Park and Alex Alderton, Roberto Amato, Sonia Goncalves, Ewan Harrison, David K. Jackson, Ian Johnston, Dominic Kwiatkowski, Cordelia Langford, John Sillitoe on behalf of the Wellcome Sanger Institute COVID-19 Surveillance Team |
| EPI_ISL_558073 | Lighthouse Lab in Alderley Park | Wellcome Sanger Institute for the COVID-19 Genomics UK (COG-UK) consortium | The Lighthouse Lab in Alderley Park and Alex Alderton, Roberto Amato, Sonia Goncalves, Ewan Harrison, David K. Jackson, Ian Johnston, Dominic Kwiatkowski, Cordelia Langford, John Sillitoe on behalf of the Wellcome Sanger Institute COVID-19 Surveillance Team (http://www.sanger.ac.uk/covid-team) |
| EPI_ISL_558074, EPI_ISL_558075, EPI_ISL_558077, EPI_ISL_558078, EPI_ISL_558079, EPI_ISL_558080, EPI_ISL_558081, EPI_ISL_558082, EPI_ISL_558083, EPI_ISL_558084, EPI_ISL_558085, EPI_ISL_558086, EPI_ISL_558087, EPI_ISL_558088 | | | |
| see above | Lighthouse Lab in Alderley Park | Wellcome Sanger Institute for the COVID-19 Genomics UK (COG-UK) consortium | The Lighthouse Lab in Alderley Park and Alex Alderton, Roberto Amato, Sonia Goncalves, Ewan Harrison, David K. Jackson, Ian Johnston, Dominic Kwiatkowski, Cordelia Langford, John Sillitoe on behalf of the Wellcome Sanger Institute COVID-19 Surveillance Team |
| EPI_ISL_558089 | Lighthouse Lab in Milton Keynes | Wellcome Sanger Institute for the COVID-19 Genomics UK (COG-UK) consortium | The Lighthouse Lab in Milton Keynes and Alex Alderton, Roberto Amato, Sonia Goncalves, Ewan Harrison, David K. Jackson, Ian Johnston, Dominic Kwiatkowski, Cordelia Langford, John Sillitoe on behalf of the Wellcome Sanger Institute COVID-19 Surveillance Team (http://www.sanger.ac.uk/covid-team) |
| EPI_ISL_558090, EPI_ISL_558091, EPI_ISL_558092, EPI_ISL_558093, EPI_ISL_558094, EPI_ISL_558095 | Lighthouse Lab in Alderley Park | Wellcome Sanger Institute for the COVID-19 Genomics UK (COG-UK) consortium | The Lighthouse Lab in Alderley Park and Alex Alderton, Roberto Amato, Sonia Goncalves, Ewan Harrison, David K. Jackson, Ian Johnston, Dominic Kwiatkowski, Cordelia Langford, John Sillitoe on behalf of the Wellcome Sanger Institute COVID-19 Surveillance Team |
| EPI_ISL_558096 | Lighthouse Lab in Milton Keynes | Wellcome Sanger Institute for the COVID-19 Genomics UK (COG-UK) consortium | The Lighthouse Lab in Milton Keynes and Alex Alderton, Roberto Amato, Sonia Goncalves, Ewan Harrison, David K. Jackson, Ian Johnston, Dominic Kwiatkowski, Cordelia Langford, John Sillitoe on behalf of the Wellcome Sanger Institute COVID-19 Surveillance Team (http://www.sanger.ac.uk/covid-team) |
| EPI_ISL_558097, EPI_ISL_558098, EPI_ISL_558099, EPI_ISL_558100, EPI_ISL_558101, EPI_ISL_558102 | Lighthouse Lab in Alderley Park | Wellcome Sanger Institute for the COVID-19 Genomics UK (COG-UK) consortium | The Lighthouse Lab in Alderley Park and Alex Alderton, Roberto Amato, Sonia Goncalves, Ewan Harrison, David K. Jackson, Ian Johnston, Dominic Kwiatkowski, Cordelia Langford, John Sillitoe on behalf of the Wellcome Sanger Institute COVID-19 Surveillance Team |
| EPI_ISL_558103 | Lighthouse Lab in Milton Keynes | Wellcome Sanger Institute for the COVID-19 Genomics UK (COG-UK) consortium | The Lighthouse Lab in Milton Keynes and Alex Alderton, Roberto Amato, Sonia Goncalves, Ewan Harrison, David K. Jackson, Ian Johnston, Dominic Kwiatkowski, Cordelia Langford, John Sillitoe on behalf of the Wellcome Sanger Institute COVID-19 Surveillance Team (http://www.sanger.ac.uk/covid-team) |
| EPI_ISL_558104, EPI_ISL_558105 | Lighthouse Lab in Alderley Park | Wellcome Sanger Institute for the COVID-19 Genomics UK (COG-UK) consortium | The Lighthouse Lab in Alderley Park and Alex Alderton, Roberto Amato, Sonia Goncalves, Ewan Harrison, David K. Jackson, Ian Johnston, Dominic Kwiatkowski, Cordelia Langford, John Sillitoe on behalf of the Wellcome Sanger Institute COVID-19 Surveillance Team |
| EPI_ISL_558106 | Lighthouse Lab in Milton Keynes | Wellcome Sanger Institute for the COVID-19 Genomics UK (COG-UK) consortium | The Lighthouse Lab in Milton Keynes and Alex Alderton, Roberto Amato, Sonia Goncalves, Ewan Harrison, David K. Jackson, Ian Johnston, Dominic Kwiatkowski, Cordelia Langford, John Sillitoe on behalf of the Wellcome Sanger Institute COVID-19 Surveillance Team (http://www.sanger.ac.uk/covid-team) |
| EPI_ISL_558107 | Lighthouse Lab in Alderley Park | Wellcome Sanger Institute for the COVID-19 Genomics UK (COG-UK) consortium | The Lighthouse Lab in Alderley Park and Alex Alderton, Roberto Amato, Sonia Goncalves, Ewan Harrison, David K. Jackson, Ian Johnston, Dominic Kwiatkowski, Cordelia Langford, John Sillitoe on behalf of the Wellcome Sanger Institute COVID-19 Surveillance Team (http://www.sanger.ac.uk/covid-team) |
| EPI_ISL_558108, EPI_ISL_558109, EPI_ISL_558110, EPI_ISL_558111, EPI_ISL_558112, EPI_ISL_558113, EPI_ISL_558114, EPI_ISL_558115, EPI_ISL_558116, EPI_ISL_558117, EPI_ISL_558118, EPI_ISL_558119, EPI_ISL_558120, EPI_ISL_558121, EPI_ISL_558122, EPI_ISL_558123, EPI_ISL_558124, EPI_ISL_558125 | | | |
| see above | Lighthouse Lab in Alderley Park | Wellcome Sanger Institute for the COVID-19 Genomics UK (COG-UK) consortium | The Lighthouse Lab in Alderley Park and Alex Alderton, Roberto Amato, Sonia Goncalves, Ewan Harrison, David K. Jackson, Ian Johnston, Dominic Kwiatkowski, Cordelia Langford, John Sillitoe on behalf of the Wellcome Sanger Institute COVID-19 Surveillance Team |
| EPI_ISL_558126 | Lighthouse Lab in Alderley Park | Wellcome Sanger Institute for the COVID-19 Genomics UK (COG-UK) Consortium | The Lighthouse Lab in Alderley Park and Alex Alderton, Roberto Amato, Sonia Goncalves, Ewan Harrison, David K. Jackson, Ian Johnston, Dominic Kwiatkowski, Cordelia Langford, John Sillitoe on behalf of the Wellcome Sanger Institute COVID-19 Surveillance Team |
| EPI_ISL_558127 | Lighthouse Lab in Milton Keynes | Wellcome Sanger Institute for the COVID-19 Genomics UK (COG-UK) consortium | The Lighthouse Lab in Milton Keynes and Alex Alderton, Roberto Amato, Sonia Goncalves, Ewan Harrison, David K. Jackson, Ian Johnston, Dominic Kwiatkowski, Cordelia Langford, John Sillitoe on behalf of the Wellcome Sanger Institute COVID-19 Surveillance Team (http://www.sanger.ac.uk/covid-team) |
| EPI_ISL_558128, EPI_ISL_558129, EPI_ISL_558130, EPI_ISL_558131, EPI_ISL_558132, EPI_ISL_558133, EPI_ISL_558134, EPI_ISL_558135, EPI_ISL_558136, EPI_ISL_558137, EPI_ISL_558138, EPI_ISL_558139, EPI_ISL_558140 | | | |
| see above | Lighthouse Lab in Alderley Park | Wellcome Sanger Institute for the COVID-19 Genomics UK (COG-UK) consortium | The Lighthouse Lab in Alderley Park and Alex Alderton, Roberto Amato, Sonia Goncalves, Ewan Harrison, David K. Jackson, Ian Johnston, Dominic Kwiatkowski, Cordelia Langford, John Sillitoe on behalf of the Wellcome Sanger Institute COVID-19 Surveillance Team |
| EPI_ISL_558141 | Lighthouse Lab in Alderley Park | Wellcome Sanger Institute for the COVID-19 Genomics UK (COG-UK) consortium | The Lighthouse Lab in Alderley Park and Alex Alderton, Roberto Amato, Sonia Goncalves, Ewan Harrison, David K. Jackson, Ian Johnston, Dominic Kwiatkowski, Cordelia Langford, John Sillitoe on behalf of the Wellcome Sanger Institute COVID-19 Surveillance Team (http://www.sanger.ac.uk/covid-team) |
| EPI_ISL_558142, EPI_ISL_558143, EPI_ISL_558144, EPI_ISL_558145, EPI_ISL_558146, EPI_ISL_558147, EPI_ISL_558148, EPI_ISL_558149, EPI_ISL_558150, EPI_ISL_558151, EPI_ISL_558152, EPI_ISL_558153, EPI_ISL_558154 | | | |
| see above | Lighthouse Lab in Alderley Park | Wellcome Sanger Institute for the COVID-19 Genomics UK (COG-UK) consortium | The Lighthouse Lab in Alderley Park and Alex Alderton, Roberto Amato, Sonia Goncalves, Ewan Harrison, David K. Jackson, Ian Johnston, Dominic Kwiatkowski, Cordelia Langford, John Sillitoe on behalf of the Wellcome Sanger Institute COVID-19 Surveillance Team |
| EPI_ISL_558155 | Lighthouse Lab in Milton Keynes | Wellcome Sanger Institute for the COVID-19 Genomics UK (COG-UK) consortium | The Lighthouse Lab in Milton Keynes and Alex Alderton, Roberto Amato, Sonia Goncalves, Ewan Harrison, David K. Jackson, Ian Johnston, Dominic Kwiatkowski, Cordelia Langford, John Sillitoe on behalf of the Wellcome Sanger Institute COVID-19 Surveillance Team (http://www.sanger.ac.uk/covid-team) |
| EPI_ISL_558156, EPI_ISL_558157, EPI_ISL_558158, EPI_ISL_558159, EPI_ISL_558160, EPI_ISL_558161, EPI_ISL_558162, EPI_ISL_558163, EPI_ISL_558164 | Lighthouse Lab in Alderley Park | Wellcome Sanger Institute for the COVID-19 Genomics UK (COG-UK) consortium | The Lighthouse Lab in Alderley Park and Alex Alderton, Roberto Amato, Sonia Goncalves, Ewan Harrison, David K. Jackson, Ian Johnston, Dominic Kwiatkowski, Cordelia Langford, John Sillitoe on behalf of the Wellcome Sanger Institute COVID-19 Surveillance Team |
| EPI_ISL_558165 | Lighthouse Lab in Alderley Park | Wellcome Sanger Institute for the COVID-19 Genomics UK (COG-UK) consortium | The Lighthouse Lab in Alderley Park and Alex Alderton, Roberto Amato, Sonia Goncalves, Ewan Harrison, David K. Jackson, Ian Johnston, Dominic Kwiatkowski, Cordelia Langford, John Sillitoe on behalf of the Wellcome Sanger Institute COVID-19 Surveillance Team (http://www.sanger.ac.uk/covid-team) |
| EPI_ISL_558166, EPI_ISL_558167, EPI_ISL_558168, EPI_ISL_558169, EPI_ISL_558170, EPI_ISL_558171, EPI_ISL_558172, EPI_ISL_558173, EPI_ISL_558174, EPI_ISL_558175, EPI_ISL_558176 | | | |
| see above | Lighthouse Lab in Alderley Park | Wellcome Sanger Institute for the COVID-19 Genomics UK (COG-UK) consortium | The Lighthouse Lab in Alderley Park and Alex Alderton, Roberto Amato, Sonia Goncalves, Ewan Harrison, David K. Jackson, Ian Johnston, Dominic Kwiatkowski, Cordelia Langford, John Sillitoe on behalf of the Wellcome Sanger Institute COVID-19 Surveillance Team |
| EPI_ISL_558177, EPI_ISL_558178 | Lighthouse Lab in Alderley Park | Wellcome Sanger Institute for the COVID-19 Genomics UK (COG-UK) consortium | The Lighthouse Lab in Alderley Park and Alex Alderton, Roberto Amato, Sonia Goncalves, Ewan Harrison, David K. Jackson, Ian Johnston, Dominic Kwiatkowski, Cordelia Langford, John Sillitoe on behalf of the Wellcome Sanger Institute COVID-19 Surveillance Team (http://www.sanger.ac.uk/covid-team) |
| EPI_ISL_558179, EPI_ISL_558180, EPI_ISL_558181, EPI_ISL_558182, EPI_ISL_558183, EPI_ISL_558184 | Lighthouse Lab in Alderley Park | Wellcome Sanger Institute for the COVID-19 Genomics UK (COG-UK) consortium | The Lighthouse Lab in Alderley Park and Alex Alderton, Roberto Amato, Sonia Goncalves, Ewan Harrison, David K. Jackson, Ian Johnston, Dominic Kwiatkowski, Cordelia Langford, John Sillitoe on behalf of the Wellcome Sanger Institute COVID-19 Surveillance Team |
| EPI_ISL_558185 | Lighthouse Lab in Alderley Park | Wellcome Sanger Institute for the COVID-19 Genomics UK (COG-UK) Consortium | The Lighthouse Lab in Alderley Park and Alex Alderton, Roberto Amato, Sonia Goncalves, Ewan Harrison, David K. Jackson, Ian Johnston, Dominic Kwiatkowski, Cordelia Langford, John Sillitoe on behalf of the Wellcome Sanger Institute COVID-19 Surveillance Team |
| EPI_ISL_558186, EPI_ISL_558187 | Lighthouse Lab in Alderley Park | Wellcome Sanger Institute for the COVID-19 Genomics UK (COG-UK) consortium | The Lighthouse Lab in Alderley Park and Alex Alderton, Roberto Amato, Sonia Goncalves, Ewan Harrison, David K. Jackson, Ian Johnston, Dominic Kwiatkowski, Cordelia Langford, John Sillitoe on behalf of the Wellcome Sanger Institute COVID-19 Surveillance Team |
| EPI_ISL_558188 | Lighthouse Lab in Alderley Park | Wellcome Sanger Institute for the COVID-19 Genomics UK (COG-UK) consortium | The Lighthouse Lab in Alderley Park and Alex Alderton, Roberto Amato, Sonia Goncalves, Ewan Harrison, David K. Jackson, Ian Johnston, Dominic Kwiatkowski, Cordelia Langford, John Sillitoe on behalf of the Wellcome Sanger Institute COVID-19 Surveillance Team (http://www.sanger.ac.uk/covid-team) |

| | | | |
|---|---|---|---|
| EPI_ISL_558189, EPI_ISL_558190, EPI_ISL_558191, EPI_ISL_558192, EPI_ISL_558193, EPI_ISL_558194 | Lighthouse Lab in Alderley Park | Wellcome Sanger Institute for the COVID-19 Genomics UK (COG-UK) consortium | The Lighthouse Lab in Alderley Park and Alex Alderton, Roberto Amato, Sonia Goncalves, Ewan Harrison, David K. Jackson, Ian Johnston, Dominic Kwiatkowski, Cordelia Langford, John Sillitoe on behalf of the Wellcome Sanger Institute COVID-19 Surveillance Team |
| EPI_ISL_558195 | Lighthouse Lab in Milton Keynes | Wellcome Sanger Institute for the COVID-19 Genomics UK (COG-UK) consortium | The Lighthouse Lab in Milton Keynes and Alex Alderton, Roberto Amato, Sonia Goncalves, Ewan Harrison, David K. Jackson, Ian Johnston, Dominic Kwiatkowski, Cordelia Langford, John Sillitoe on behalf of the Wellcome Sanger Institute COVID-19 Surveillance Team (http://www.sanger.ac.uk/covid-team) |
| EPI_ISL_558196, EPI_ISL_558197, EPI_ISL_558198, EPI_ISL_558199, EPI_ISL_558200, EPI_ISL_558201, EPI_ISL_558202, EPI_ISL_558203, EPI_ISL_558204, EPI_ISL_558205, EPI_ISL_558206, EPI_ISL_558207, EPI_ISL_558208, EPI_ISL_558209, EPI_ISL_558210, EPI_ISL_558211, EPI_ISL_558212, EPI_ISL_558213, EPI_ISL_558214 | | | |
| see above | Lighthouse Lab in Alderley Park | Wellcome Sanger Institute for the COVID-19 Genomics UK (COG-UK) consortium | The Lighthouse Lab in Alderley Park and Alex Alderton, Roberto Amato, Sonia Goncalves, Ewan Harrison, David K. Jackson, Ian Johnston, Dominic Kwiatkowski, Cordelia Langford, John Sillitoe on behalf of the Wellcome Sanger Institute COVID-19 Surveillance Team |
| EPI_ISL_558215 | Lighthouse Lab in Alderley Park | Wellcome Sanger Institute for the COVID-19 Genomics UK (COG-UK) consortium | The Lighthouse Lab in Alderley Park and Alex Alderton, Roberto Amato, Sonia Goncalves, Ewan Harrison, David K. Jackson, Ian Johnston, Dominic Kwiatkowski, Cordelia Langford, John Sillitoe on behalf of the Wellcome Sanger Institute COVID-19 Surveillance Team (http://www.sanger.ac.uk/covid-team) |
| EPI_ISL_558216, EPI_ISL_558217 | Lighthouse Lab in Alderley Park | Wellcome Sanger Institute for the COVID-19 Genomics UK (COG-UK) consortium | The Lighthouse Lab in Alderley Park and Alex Alderton, Roberto Amato, Sonia Goncalves, Ewan Harrison, David K. Jackson, Ian Johnston, Dominic Kwiatkowski, Cordelia Langford, John Sillitoe on behalf of the Wellcome Sanger Institute COVID-19 Surveillance Team |
| EPI_ISL_558218 | Lighthouse Lab in Alderley Park | Wellcome Sanger Institute for the COVID-19 Genomics UK (COG-UK) consortium | The Lighthouse Lab in Alderley Park and Alex Alderton, Roberto Amato, Sonia Goncalves, Ewan Harrison, David K. Jackson, Ian Johnston, Dominic Kwiatkowski, Cordelia Langford, John Sillitoe on behalf of the Wellcome Sanger Institute COVID-19 Surveillance Team (http://www.sanger.ac.uk/covid-team) |
| EPI_ISL_558219, EPI_ISL_558220, EPI_ISL_558221 | Lighthouse Lab in Alderley Park | Wellcome Sanger Institute for the COVID-19 Genomics UK (COG-UK) consortium | The Lighthouse Lab in Alderley Park and Alex Alderton, Roberto Amato, Sonia Goncalves, Ewan Harrison, David K. Jackson, Ian Johnston, Dominic Kwiatkowski, Cordelia Langford, John Sillitoe on behalf of the Wellcome Sanger Institute COVID-19 Surveillance Team |
| EPI_ISL_558222 | Lighthouse Lab in Alderley Park | Wellcome Sanger Institute for the COVID-19 Genomics UK (COG-UK) Consortium | The Lighthouse Lab in Alderley Park and Alex Alderton, Roberto Amato, Sonia Goncalves, Ewan Harrison, David K. Jackson, Ian Johnston, Dominic Kwiatkowski, Cordelia Langford, John Sillitoe on behalf of the Wellcome Sanger Institute COVID-19 Surveillance Team |
| EPI_ISL_558223, EPI_ISL_558224, EPI_ISL_558225, EPI_ISL_558226, EPI_ISL_558227, EPI_ISL_558228, EPI_ISL_558229, EPI_ISL_558230, EPI_ISL_558231, EPI_ISL_558232, EPI_ISL_558233 | | | |
| see above | Lighthouse Lab in Alderley Park | Wellcome Sanger Institute for the COVID-19 Genomics UK (COG-UK) consortium | The Lighthouse Lab in Alderley Park and Alex Alderton, Roberto Amato, Sonia Goncalves, Ewan Harrison, David K. Jackson, Ian Johnston, Dominic Kwiatkowski, Cordelia Langford, John Sillitoe on behalf of the Wellcome Sanger Institute COVID-19 Surveillance Team |
| EPI_ISL_558234 | Lighthouse Lab in Milton Keynes | Wellcome Sanger Institute for the COVID-19 Genomics UK (COG-UK) consortium | The Lighthouse Lab in Milton Keynes and Alex Alderton, Roberto Amato, Sonia Goncalves, Ewan Harrison, David K. Jackson, Ian Johnston, Dominic Kwiatkowski, Cordelia Langford, John Sillitoe on behalf of the Wellcome Sanger Institute COVID-19 Surveillance Team (http://www.sanger.ac.uk/covid-team) |
| EPI_ISL_558235, EPI_ISL_558236, EPI_ISL_559130 | Lighthouse Lab in Alderley Park | Wellcome Sanger Institute for the COVID-19 Genomics UK (COG-UK) consortium | The Lighthouse Lab in Alderley Park and Alex Alderton, Roberto Amato, Sonia Goncalves, Ewan Harrison, David K. Jackson, Ian Johnston, Dominic Kwiatkowski, Cordelia Langford, John Sillitoe on behalf of the Wellcome Sanger Institute COVID-19 Surveillance Team |
| EPI_ISL_560124 | Wales Specialist Virology Centre Sequencing lab: Pathogen Genomics Unit | COVID-19 Genomics UK (COG-UK) Consortium | Catherine Moore, Johnathan Evans, Laura Gifford, Malorie Perry, Simon Cottrell, Angela Marchbank, Alec Birchley, Alexander Adams, Amy Gaskin, Bree Gatica-Wilcox, Jason Coombes, Joel Southgate, Lauren Gilbert, Lee Graham, Nicole Pacchiarini, Sara Kumziene-Summerhayes, Sarah Taylor, Sophie Jones, Sara Rey, Matthew Bull, Joanne Watkins, Sally Corden, Tom Connor |
| EPI_ISL_560309, EPI_ISL_560310 | UMMC-Health | WHO National Influenza Centre Russian Federation | Andrey Komissarov, Artem Fadeev, Anna Ivanova, Tatiana Platonova, Daria Danilenko |
| EPI_ISL_560320 | CMS, Roorkee | CSIR-Institute of Microbial Technology | Kanika Bansal, Sanjeet Kumar, Anu Singh, Debarghya Ghose, Amandeep Kaur, Rajesh Kumar Mishra, Poushali Chakraborty, Harsh Goar, Navin Baid, Ashwani Kumar, Dipak Dutta, Sanjeev Khosla, Prabhu B. Patil |
| EPI_ISL_560330, EPI_ISL_560333 | TriCore Reference Laboratories | Center for Global Health, University of New Mexico Health Sciences Center | Daryl Domman, Kurt Schwalm, Twila Kunde, Joseph Hicks, Michael Edwards, Darrell Dinwiddie |
| EPI_ISL_560868, EPI_ISL_560873 | Utah Public Health Laboratory | Utah Public Health Laboratory | Erin Young, Kelly Oakeson |
| EPI_ISL_560982, EPI_ISL_560983, EPI_ISL_560984, EPI_ISL_560985 | Karolinska universitetslaboratoriet SOLNA | The Public Health Agency of Sweden | Anna-Malin Linde, Maria Lind Karlberg, Mattias Haukland, Reza Advani, Olov Svartstrom, Oskar Karlsson Lindsjo, Sandra Broddesson, Petra Edquist, Mia Brytting, Anna Risberg, Karin Tegmark-Wisell |
| EPI_ISL_561210, EPI_ISL_561221, EPI_ISL_561222, EPI_ISL_561223, EPI_ISL_561224, EPI_ISL_561225, EPI_ISL_561226, EPI_ISL_561227, EPI_ISL_561228, EPI_ISL_561229, EPI_ISL_561230 | | | |
| see above | MRCG at LSHTM Genomics lab | MRCG at LSHTM Genomics lab | Abdul Karim sesay, Abdoulie Kanteh, Jarra Manneh, Mariama Kujabi, Bakary Sanyang |
| EPI_ISL_561543, EPI_ISL_561706, EPI_ISL_561763, EPI_ISL_561851 | Microbiological Diagnostic Unit - Public Health Laboratory (MDU-PHL) | MDU-PHL | Seemann, T., Schultz M. B., Sait, M., Sherry, N. |
| EPI_ISL_562109 | Victorian Infectious Diseases Reference Laboratory (VIDRL) | VIDRL and MDU-PHL | Caly, L., Seemann, T., Sait, M., Schultz, M. B., Druce J., Sherry, N. |
| EPI_ISL_562137, EPI_ISL_562199, EPI_ISL_563985, EPI_ISL_563986 | Microbiological Diagnostic Unit - Public Health Laboratory (MDU-PHL) | MDU-PHL | Seemann, T., Schultz M. B., Sait, M., Sherry, N. |
| EPI_ISL_563987 | Victorian Infectious Diseases Reference Laboratory (VIDRL) | VIDRL and MDU-PHL | Caly, L., Seemann, T., Sait, M., Schultz, M. B., Druce J., Sherry, N. |
| EPI_ISL_564113, EPI_ISL_564114 | Microbiological Diagnostic Unit - Public Health Laboratory (MDU-PHL) | MDU-PHL | Seemann, T., Schultz M. B., Sait, M., Sherry, N. |
| EPI_ISL_564209 | Victorian Infectious Diseases Reference Laboratory (VIDRL) | VIDRL and MDU-PHL | Caly, L., Seemann, T., Sait, M., Schultz, M. B., Druce J., Sherry, N. |
| EPI_ISL_564269, EPI_ISL_564271, EPI_ISL_564274, EPI_ISL_564281, EPI_ISL_564286, EPI_ISL_564291, EPI_ISL_564294, EPI_ISL_564683, EPI_ISL_565376, EPI_ISL_565484, EPI_ISL_565498, EPI_ISL_565729 | | | |
| see above | Microbiological Diagnostic Unit - Public Health Laboratory (MDU-PHL) | MDU-PHL | Seemann, T., Schultz M. B., Sait, M., Sherry, N. |
| EPI_ISL_565843, EPI_ISL_565844, EPI_ISL_565845, EPI_ISL_565846, EPI_ISL_565847, EPI_ISL_565848, EPI_ISL_565849 | Michigan Department of Health and Human Services, Bureau of Laboratories | Michigan Department of Health and Human Services, Bureau of Laboratories | Blankenship HM, Riner D, Soehnlen MK |
| EPI_ISL_566059, EPI_ISL_566060, EPI_ISL_566061, EPI_ISL_566062 | Respiratory Virus Unit, Microbiology Services Colindale, Public Health England | Respiratory Virus Unit, Microbiology Services Colindale, Public Health England | PHE Covid Sequencing Team |
| EPI_ISL_566091, EPI_ISL_566095, EPI_ISL_566101, EPI_ISL_566103, EPI_ISL_566106, EPI_ISL_566107 | GA Department of Public Health Laboratory | Pathogen Discovery, Respiratory Viruses Branch, Division of Viral Diseases, Centers for Disease Control and Prevention | Jing Zhang, Brian Lynch, Yan Li, Anna Montmayeur, Krista Queen, Ying Tao, Anna Uehara, Clinton R. Paden, Rachel Marine, Haibin Wang, Suxiang Tong |
| EPI_ISL_568998, EPI_ISL_568999, EPI_ISL_569000, EPI_ISL_569001, EPI_ISL_569002, EPI_ISL_569003, EPI_ISL_569004, EPI_ISL_569005, EPI_ISL_569006 | MEPHI, Aix Marseille University | MEPHI, Aix Marseille University | Anthony LEVASSEUR |
| EPI_ISL_569612, EPI_ISL_569613 | Cheyenne River Health Center | South Dakota Public Health Laboratory | Matt Plumb, Jacob Garfin, Xiong Wang, and Chris Carlson |
| EPI_ISL_569614 | IHS Fort Yates Hospital | South Dakota Public Health Laboratory | Matt Plumb, Jacob Garfin, Xiong Wang, and Chris Carlson |
| EPI_ISL_569777, EPI_ISL_569778, EPI_ISL_569779, EPI_ISL_569780, EPI_ISL_569781, EPI_ISL_569782, EPI_ISL_569798, EPI_ISL_569800, EPI_ISL_569804, EPI_ISL_569805, EPI_ISL_569806, EPI_ISL_569814, EPI_ISL_569815, EPI_ISL_569816, EPI_ISL_569820, EPI_ISL_569828, EPI_ISL_569829, EPI_ISL_569834, EPI_ISL_569835, EPI_ISL_569845, EPI_ISL_569850, EPI_ISL_569853 | | | |

| | | | |
|---|---|---|---|
| see above | Omsk Research Institute of Natural Focal Infections | WHO National Influenza Centre Russian Federation | Artem Fadeev, Ekaterina Gradoboeva, Ekaterina Savkina, Daria Nashatyreva, Elena Poleshchuk, Aleksei Vasilenko, Valery Yakimenko, Andrey Komissarov |
| EPI_ISL_569887, EPI_ISL_569888, EPI_ISL_569889, EPI_ISL_569890, EPI_ISL_569891, EPI_ISL_569892, EPI_ISL_569894, EPI_ISL_569895, EPI_ISL_569896, EPI_ISL_569897, EPI_ISL_569898, EPI_ISL_569899, EPI_ISL_569900, EPI_ISL_569901, EPI_ISL_569902, EPI_ISL_569903, EPI_ISL_569904, EPI_ISL_569905, EPI_ISL_569906, EPI_ISL_569907, EPI_ISL_569909, EPI_ISL_569910, EPI_ISL_569911, EPI_ISL_569912, EPI_ISL_569913, EPI_ISL_569914, EPI_ISL_569916, EPI_ISL_569917, EPI_ISL_569918, EPI_ISL_569919, EPI_ISL_569921, EPI_ISL_569923, EPI_ISL_569924, EPI_ISL_569925, EPI_ISL_569926, EPI_ISL_569927, EPI_ISL_569928, EPI_ISL_569930, EPI_ISL_569931, EPI_ISL_569932, EPI_ISL_569933, EPI_ISL_569934, EPI_ISL_569935, EPI_ISL_569936, EPI_ISL_569937, EPI_ISL_569938, EPI_ISL_569940, EPI_ISL_569941, EPI_ISL_569943, EPI_ISL_569944, EPI_ISL_569945, EPI_ISL_569947, EPI_ISL_569948 | | | |
| see above | Innovative Genomics Institute, UC Berkeley | Innovative Genomics Institute, UC Berkeley | Stacia Wyman, Haridha Shivram, Phil Frankino, Liana Lareau, Shana McDevitt, Justin Choi |
| EPI_ISL_569981, EPI_ISL_569992 | Unity Health Toronto | Ontario Institute for Cancer Research | Ramzi Fattouh, Larissa M. Matukas, Yan Chen,Mark Downing, Trina Otterman, Karel Boissinot, Wai Sum Siu, Zhi Cui, Le Luu, Samira Mubareka, TIBDN, Ilinca Lungu, Bernard Lam, Jeremy Johns, Paul Krzyzanowski, Richard de Borja, Felicia Vincelli, Philip Zuzarte, Jared T. Simpson |
| EPI_ISL_570834, EPI_ISL_570850, EPI_ISL_570860, EPI_ISL_570942, EPI_ISL_570943, EPI_ISL_570944, EPI_ISL_570954, EPI_ISL_570956, EPI_ISL_570957, EPI_ISL_570958, EPI_ISL_570959, EPI_ISL_570963, EPI_ISL_570964, EPI_ISL_570966, EPI_ISL_570967, EPI_ISL_570968, EPI_ISL_570969 | | | |
| see above | UW Virology Lab | UW Virology Lab | Pavitra Roychoudhury, Hong Xie, Lasata Shrestha, Amin Addetia, Victoria M Rachleff, Meei-Li Huang, Keith R Jerome, Alexander Greninger |
| EPI_ISL_572220, EPI_ISL_572221, EPI_ISL_572222, EPI_ISL_572223, EPI_ISL_572224, EPI_ISL_572225, EPI_ISL_572226, EPI_ISL_572227, EPI_ISL_572228, EPI_ISL_572229, EPI_ISL_572230, EPI_ISL_572231, EPI_ISL_572232, EPI_ISL_572233, EPI_ISL_572234, EPI_ISL_572235, EPI_ISL_572236, EPI_ISL_572237, EPI_ISL_572257, EPI_ISL_572267, EPI_ISL_572268 | | | |
| see above | Virginia DCLS | Virginia DCLS | Virginia DCLS |
| EPI_ISL_572547 | Wales Specialist Virology Centre Sequencing lab: Pathogen Genomics Unit | COVID-19 Genomics UK (COG-UK) Consortium | Catherine Moore, Johnathan Evans, Laura Gifford, Malorie Perry, Simon Cottrell, Angela Marchbank, Alec Birchley, Alexander Adams, Amy Gaskin, Bree Gatica-Wilcox, Jason Coombes, Joel Southgate, Lauren Gilbert, Lee Graham, Nicole Pacchiarini, Sara Kumziene-Summerhayes, Sarah Taylor, Sophie Jones, Sara Rey, Matthew Bull, Joanne Watkins, Sally Corden, Tom Connor |
| EPI_ISL_573312, EPI_ISL_573357 | Northumbria University / South Tees Hospitals NHS Foundation Trust / North Cumbria Integrated Care NHS Foundation Trust / North Tees and Hartlepool NHS Foundation Trust / Newcastle Hospitals NHS Foundation Trust | COVID-19 Genomics UK (COG-UK) Consortium | Darren L Smith,Andrew Nelson,Matthew Bashton,Greg R Young,Joshua Loh,John Allan,Mohammad A Tariq,Giles S Holt,Gary Black,Wen C Yew,Lynn Dover,Paul Baker,Steve Liggett,Sarah Essex,Jane Greenaway,Debra Padgett,Clive Graham,Garren Scott,Edward Barton,Emma Swindells,Brendan Payne,Jennifer Collins,Yusri Taha,Gary Eltringham |
| EPI_ISL_574433 | National Institute of Health Research and Development | National Institute of Health Research and Development | Pawestri,HA;Subangkit;Puspa,KD;Nugraha,AA;Ikawati,HD;Pangesti, KNA;Soekarso,T;Susilarini,NK;Hariastuti,NI;Nikmah,UA;Mursinah;Febriyani,A;Herman,R;Susanti,N;Herna;Febriyanti,T; Nurhadi,M; Paisal;Ramadhany,R;Agustiningsih;Kurniawati,J;Kipuw,NL;Muna,F;Indalau,IL;Adam,K;Wibowo,HA;Rizki,A;Puspandari,N;Setiawaty,V |
| EPI_ISL_574543, EPI_ISL_574544, EPI_ISL_574545, EPI_ISL_574546, EPI_ISL_574554, EPI_ISL_574555, EPI_ISL_574556, EPI_ISL_574557, EPI_ISL_574558, EPI_ISL_574559, EPI_ISL_574560, EPI_ISL_574561, EPI_ISL_574562, EPI_ISL_574563, EPI_ISL_574564, EPI_ISL_574565, EPI_ISL_574566, EPI_ISL_574567, EPI_ISL_574568, EPI_ISL_574569, EPI_ISL_574570, EPI_ISL_574571, EPI_ISL_574572, EPI_ISL_574573, EPI_ISL_574574, EPI_ISL_574575 | | | |
| see above | Microbiology Division, South Carolina Department of Health and Environmental Control | Microbiology Division, South Carolina Department of Health and Environmental Control | Flores,H. |
| EPI_ISL_574594 | Hospital Escola da Universidade de Taubate | Instituto Adolfo Lutz, Interdiciplinary Procedures Center, Strategic Laboratory | Claudio Tavares Sacchi, Claudia Regina Gonçalves, Erica Valessa Ramos Gomes, Karoline Rodrigues Campos |
| EPI_ISL_575037, EPI_ISL_575239 | Utah Public Health Laboratory | Utah Public Health Laboratory | Erin Young, Kelly Oakeson |
| EPI_ISL_576127 | Oklahoma Animal Disease Diagnostic Laboratory, Oklahoma State University | Oklahoma Animal Disease Diagnostic Laboratory, Oklahoma State University | Narayanan,S., Ritchey,J.C., Patil,G., Narasaraju,T., More,S., Malayer,J.R., Saliki,J.T., Kaul,A., Ramachandran,A. |
| EPI_ISL_576526, EPI_ISL_576528, EPI_ISL_576533, EPI_ISL_576542, EPI_ISL_576543, EPI_ISL_576552 | Innovative Genomics Institute, UC Berkeley | Innovative Genomics Institute, UC Berkeley | Stacia Wyman, Haridha Shivram, Phil Frankino, Liana Lareau, Shana McDevitt, Justin Choi |
| EPI_ISL_577734 | Institute of Virology, Biomedical Research Center of the Slovak Academy of Sciences, Bratislava | Faculty of Natural Sciences, Comenius University, Bratislava | Viktória Hodorová, Kristína Boršová, Broa Brejová, Viktória abanová, Dominika Friová, Sabina Fumaová Havlíková, Juraj Kopáek, Martina Liková, ubomíra Lukáiková, Martina Neboháová, Monika Sláviková, Edita Staroová, Elena Tichá, Tomáš Vina, Jozef Nosek, Boris Klempa |
| EPI_ISL_578237, EPI_ISL_578238, EPI_ISL_578276 | National Virus Reference Laboratory | National Virus Reference Laboratory | Michael Carr, Gabriel Gonzalez, Jonathan Dean, Suzie Coughlan, Cillian F De Gascun |
| EPI_ISL_579010, EPI_ISL_579011, EPI_ISL_579013, EPI_ISL_579014, EPI_ISL_579015, EPI_ISL_579016, EPI_ISL_579017, EPI_ISL_579018, EPI_ISL_579020, EPI_ISL_579022, EPI_ISL_579024, EPI_ISL_579025, EPI_ISL_579027, EPI_ISL_579028, EPI_ISL_579030, EPI_ISL_579031, EPI_ISL_579034, EPI_ISL_579041 | | | |
| see above | LSUHS Emerging Viral Threat Laboratory | Microbial Genome Sequencing Center | Jeremy P. Kamil, Rona S. Scott, Maarten Van Diest, Malgorzata Bienkowska-Haba, Katarzyna Zwolinska, Andrew D. Yurochko, Christopher G. Kevil, Martin J. Sapp, Daniel J. Snyder, Vaughn S. Cooper, John A. Vanchiere |
| EPI_ISL_581393 | Lighthouse Lab in Milton Keynes | Wellcome Sanger Institute for the COVID-19 Genomics UK (COG-UK) consortium | The Lighthouse Lab in Milton Keynes and Alex Alderton, Roberto Amato, Sonia Goncalves, Ewan Harrison, David K. Jackson, Ian Johnston, Dominic Kwiatkowski, Cordelia Langford, John Sillitoe on behalf of the Wellcome Sanger Institute COVID-19 Surveillance Team |
| EPI_ISL_581433 | Lighthouse Lab in Alderley Park | Wellcome Sanger Institute for the COVID-19 Genomics UK (COG-UK) consortium | Jacquelyn Wynn, Mairead Hyland, The Lighthouse Lab in Alderley Park and Alex Alderton, Roberto Amato, Sonia Goncalves, Ewan Harrison, David K. Jackson, Ian Johnston, Dominic Kwiatkowski, Cordelia Langford, John Sillitoe on behalf of the Wellcome Sanger Institute COVID-19 Surveillance Team |
| EPI_ISL_581434 | Lighthouse Lab in Milton Keynes | Wellcome Sanger Institute for the COVID-19 Genomics UK (COG-UK) consortium | The Lighthouse Lab in Milton Keynes and Alex Alderton, Roberto Amato, Sonia Goncalves, Ewan Harrison, David K. Jackson, Ian Johnston, Dominic Kwiatkowski, Cordelia Langford, John Sillitoe on behalf of the Wellcome Sanger Institute COVID-19 Surveillance Team |
| EPI_ISL_581435 | Lighthouse Lab in Alderley Park | Wellcome Sanger Institute for the COVID-19 Genomics UK (COG-UK) consortium | Jacquelyn Wynn, Mairead Hyland, The Lighthouse Lab in Alderley Park and Alex Alderton, Roberto Amato, Sonia Goncalves, Ewan Harrison, David K. Jackson, Ian Johnston, Dominic Kwiatkowski, Cordelia Langford, John Sillitoe on behalf of the Wellcome Sanger Institute COVID-19 Surveillance Team |
| EPI_ISL_581487, EPI_ISL_581492 | Fondation Congolaise pour la recherche medicale (FCRM) | NGS Competence Center Tübingen, Institut für Medizinische Mikrobiologie und Hygiene, Universitätsklinikum Tübingen | Angel Angelov |
| EPI_ISL_581876, EPI_ISL_581877, EPI_ISL_581878, EPI_ISL_581879, EPI_ISL_581880, EPI_ISL_581881, EPI_ISL_581882, EPI_ISL_581883, EPI_ISL_581893, EPI_ISL_581894, EPI_ISL_581895, EPI_ISL_581896, EPI_ISL_581897, EPI_ISL_581898, EPI_ISL_581899 | | | |
| see above | University Hospital Basel, Clinical Virology | University Hospital Basel, Clinical Bacteriology | Madlen Stange, Alfredo Mari, Tim Roloff, Helena MB Seth-Smith, Michael Schweitzer, Myrta Brunner, Karoline Leuzinger, Kirstine K. Soegaard, Alexander Gensch, Sarah Tschudin-Sutter, Simon Fuchs, Julia Bielicki, Hans Pargger, Martin Siegemund, Christian Nickel, Roland Bingisser, Michael Osthoff, Stefano Bassetti, Rita Schneider-Sliwa, Manuel Battegay, Hans Hirsch, Adrian Egli |
| EPI_ISL_582097, EPI_ISL_582098, EPI_ISL_582099 | Hospital Universitario Marqués de Valdecilla - IDIVAL (Santander, Cantabria) | SeqCOVID-SPAIN consortium/IBV(CSIC) | María Eliecer Cano García, Mónica Gozalo Margüello, Jose Manuel Méndez Legaza, Daniel Pablo Marcos, Jesús Rodríguez Rodríguez, María Siller Ruiz and SeqCOVID-SPAIN consortium |
| EPI_ISL_582319, EPI_ISL_582494 | Cadham Provincial Laboratory | National Microbiology Laboratory (NML) | Anna Majer, Shari Tyson, Grace Seo, Philip Mabon, Elsie Grudeski, Rhiannon Huzarewich, Russell Mandes, Anneliese Landgraff, Jennifer Tanner, Natalie Knox, Morag Graham, Gary Van Domselaar, Paul Van Caeseele, Jared Bullard, David Alexander, Kerry Dust, Nathalie Bastien, Yan Li, Timothy Booth, Darian Hole, Madison Chapel, CanCOGeN's metadata curation team, Public Health Agency of Canada CanCOGeN team |
| EPI_ISL_582781, EPI_ISL_582782 | Uppsala klinisk mikrobiologi | The Public Health Agency of Sweden | Anna-Malin Linde, Maria Lind Karlberg, Mattias Haukland, Reza Advani, Olov Svartstrom, Oskar Karlsson Lindsjo, Sandra Broddesson, Petra Edquist, Mia Brytting, Anna Risberg, Karin Tegmark-Wisell |
| EPI_ISL_582791, EPI_ISL_582793 | Orebro klinisk mikrobiologi | The Public Health Agency of Sweden | Anna-Malin Linde, Maria Lind Karlberg, Mattias Haukland, Reza Advani, Olov Svartstrom, Oskar Karlsson Lindsjo, Sandra Broddesson, Petra Edquist, Mia Brytting, Anna Risberg, Karin Tegmark-Wisell |
| EPI_ISL_582797, EPI_ISL_582798 | Klinisk mikrobiologi Vasternorrland | The Public Health Agency of Sweden | Anna-Malin Linde, Maria Lind Karlberg, Mattias Haukland, Reza Advani, Olov Svartstrom, Oskar Karlsson Lindsjo, Sandra Broddesson, Petra Edquist, Mia Brytting, Anna Risberg, Karin Tegmark-Wisell |
| EPI_ISL_582799 | Stockholm_KUL Solna | The Public Health Agency of Sweden | Anna-Malin Linde, Maria Lind Karlberg, Mattias Haukland, Reza Advani, Olov Svartstrom, Oskar Karlsson Lindsjo, Sandra Broddesson, Petra Edquist, Mia Brytting, Anna Risberg, Karin Tegmark-Wisell |
| EPI_ISL_582800 | Klinisk mikrobiologi Vasternorrland | The Public Health Agency of Sweden | Anna-Malin Linde, Maria Lind Karlberg, Mattias Haukland, Reza Advani, Olov Svartstrom, Oskar Karlsson Lindsjo, Sandra Broddesson, Petra Edquist, Mia Brytting, Anna Risberg, Karin Tegmark-Wisell |

| | | | |
|---|---|---|---|
| EPI_ISL_582871, EPI_ISL_582872, EPI_ISL_582873, EPI_ISL_582874, EPI_ISL_582875, EPI_ISL_582876, EPI_ISL_582877, EPI_ISL_582878, EPI_ISL_582879 | County of Santa Clara Public Health Department | Chan-Zuckerberg Biohub | CZB Cliahub Consortium |
| EPI_ISL_583060, EPI_ISL_583061, EPI_ISL_583062, EPI_ISL_583063, EPI_ISL_583064, EPI_ISL_583065, EPI_ISL_583066, EPI_ISL_583067, EPI_ISL_583068, EPI_ISL_583069 | Humboldt County Public Health Laboratory | Chan-Zuckerberg Biohub | CZB Cliahub Consortium |
| EPI_ISL_583501 | Hospital Estadual de CampanhaCOVID 19 Barradas | Instituto Adolfo Lutz, Interdiciplinary Procedures Center, Strategic Laboratory | Claudio Tavares Sacchi, Claudia Regina Gonçalves, Erica Valessa Ramos Gomes, Karoline Rodrigues Campos |
| EPI_ISL_584619, EPI_ISL_584620 | Liverpool Clinical Laboratories | COVID-19 Genomics UK (COG-UK) Consortium | Sam Haldenby, Anita Lucaci, Steve Paterson, Julian Hiscox, Alistair Darby, M Almsaud, A Alrezaihi, Muhannad Alruwaili, Stuart D Armstrong, Jones Benjamin, Eleanor G Bentley, Anu Chawla, Jordan J Clark, Angela Cowell, Richard Eccles, Isabel García-Dorival, Matthew Gemmell, Alessandro Gerada, PKF Gilmore, Richard Gregory, Ximeng Han, Catherine Hartley, Margaret Hughes, Miren Iturriza-Gomara, James Johnson, L Luu, Jenifer Manson, Charlotte Nelson, Elaine O'Toole, Cassie Olateju, Rebekah Penrice-Randal , Lucille Rainbow, N.P Randle, Trevor Ian Robinson, Parul Sharma, Ghada T Shawli, James P Stewart, Neil Swainston, Ecaterina Vamos, Joanne Watts, Mark Whitehead |
| EPI_ISL_586263, EPI_ISL_586264, EPI_ISL_586265, EPI_ISL_586266 | Alaska State Virology Laboratory | Alaska State Virology Laboratory | Jack Chen, Ph.D. |
| EPI_ISL_586567, EPI_ISL_586568 | Saikrishna Hospital,Mehsana | Gujarat Biotechnology Research Centre | Zarna Patel, Monika Gandhi, Pinal Trivedi, Maharshi Pandya, Nidhi Patel, Nitin Savaliya, Raghawendra Kumar, Dinesh Kumar, Zuber Saiyed, Komal Patel, Labdhi Pandya, Afzal Ansari, Nikha Trivedi, Harshadbhai Parmar, Apurvasinh Puvar, Janvi Raval, R D Dixit, A M Kadri, Harsh Bakshi, Chaitanya Joshi, Madhvi Joshi |
| EPI_ISL_590693 | INMI Lazzaro Spallanzani IRCCS | INMI Lazzaro Spallanzani IRCCS | Martina Rueca, Barbara Bartolini, Cesare E.M. Gruber, Francesco Messina, Emanuela Giombini, Beatrice Valli, Eleonora Lalle, Simone Lanini, Francesco Vairo, Maria R. Capobianchi, Antonino Di Caro |
| EPI_ISL_590694 | INMI Lazzaro Spallanzani IRCCS | INMI Lazzaro Spallanzani IRCCS | Barbara Bartolini, Martina Rueca, Francesco Messina, Cesare E.M. Gruber, Emanuela Giombini, Beatrice Valli, Eleonora Lalle, Simone Lanini, Francesco Vairo, Maria R. Capobianchi, Antonino Di Caro |
| EPI_ISL_590695 | INMI Lazzaro Spallanzani IRCCS | INMI Lazzaro Spallanzani IRCCS | Cesare E.M. Gruber, Francesco Messina, Barbara Bartolini, Martina Rueca, Emanuela Giombini, Beatrice Valli, Eleonora Lalle, Simone Lanini, Francesco Vairo, Antonino Di Caro, Maria R. Capobianchi |
| EPI_ISL_590696 | INMI Lazzaro Spallanzani IRCCS | INMI Lazzaro Spallanzani IRCCS | Cesare E.M. Gruber, Barbara Bartolini, Francesco Messina, Martina Rueca, Emanuela Giombini, Beatrice Valli, Eleonora Lalle, Simone Lanini, Francesco Vairo, Antonino Di Caro, Maria R. Capobianchi |
| EPI_ISL_590697 | INMI Lazzaro Spallanzani IRCCS | INMI Lazzaro Spallanzani IRCCS | Martina Rueca, Cesare E.M. Gruber, Barbara Bartolini, Francesco Messina, Emanuela Giombini, Beatrice Valli, Eleonora Lalle, Simone Lanini, Francesco Vairo, Antonino Di Caro, Maria R. Capobianchi |
| EPI_ISL_590698 | INMI Lazzaro Spallanzani IRCCS | INMI Lazzaro Spallanzani IRCCS | Barbara Bartolini, Francesco Messina, Cesare E.M. Gruber, Martina Rueca, Emanuela Giombini, Beatrice Valli, Eleonora Lalle, Simone Lanini, Francesco Vairo, Maria R. Capobianchi, Antonino Di Caro |
| EPI_ISL_590752 | University of Michigan Clinical Microbiology Laboratory | Lauring Lab, University of Michigan, Department of Microbiology and Immunology | Valesano |
| EPI_ISL_593476, EPI_ISL_593477 | Victorian Infectious Diseases Reference Laboratory (VIDRL) | VIDRL and MDU-PHL | Caly L., Seemann T., Sait, M., Schultz, M. B., Druce J., Sherry, N. |
| EPI_ISL_593668 | Pathology West - NSW Health Pathology | NSW Health Pathology - Institute of Clinical Pathology and Medical Research; Westmead Hospital; University of Sydney | CIDM-PH et al. |
| EPI_ISL_593687, EPI_ISL_593688, EPI_ISL_593694, EPI_ISL_593695, EPI_ISL_593696, EPI_ISL_593697, EPI_ISL_593698, EPI_ISL_593699, EPI_ISL_593700, EPI_ISL_593701, EPI_ISL_593702, EPI_ISL_593703, EPI_ISL_593706, EPI_ISL_593707, EPI_ISL_593726 | | | |
| see above | South Eastern Area Laboratory Services (SEALS) | NSW Health Pathology - Institute of Clinical Pathology and Medical Research; Westmead Hospital; University of Sydney | CIDM-PH et al. |
| EPI_ISL_593867 | CHU Purpan - Laboratoire de Virologie - Institut Fédératif de Biologie | CHU Purpan - Laboratoire de Virologie - Institut Fédératif de Biologie | Latour J., Ranger N., Dubois M., Carcenac R., Harter A., Boyer P., Tremeaux P., Izopet J. |
| EPI_ISL_594043, EPI_ISL_594044, EPI_ISL_594045, EPI_ISL_594046, EPI_ISL_594047, EPI_ISL_594048, EPI_ISL_594049, EPI_ISL_594050, EPI_ISL_594051, EPI_ISL_594052, EPI_ISL_594053, EPI_ISL_594054, EPI_ISL_594055, EPI_ISL_594056, EPI_ISL_594057, EPI_ISL_594058, EPI_ISL_594059, EPI_ISL_594060, EPI_ISL_594061 | | | |
| see above | Utah Public Health Laboratory | Utah Public Health Laboratory | Erin Young, Kelly Oakeson |
| EPI_ISL_594288, EPI_ISL_594289, EPI_ISL_594290, EPI_ISL_594291, EPI_ISL_594292, EPI_ISL_594293, EPI_ISL_594294, EPI_ISL_594295, EPI_ISL_594296, EPI_ISL_594332, EPI_ISL_594333, EPI_ISL_594334, EPI_ISL_594335, EPI_ISL_594336, EPI_ISL_594337, EPI_ISL_594338, EPI_ISL_594339, EPI_ISL_594340, EPI_ISL_594341, EPI_ISL_594342, EPI_ISL_594343, EPI_ISL_594344, EPI_ISL_594345, EPI_ISL_594346, EPI_ISL_594347, EPI_ISL_594348, EPI_ISL_594349, EPI_ISL_594350, EPI_ISL_594351, EPI_ISL_594352, EPI_ISL_594353, EPI_ISL_594354, EPI_ISL_594355, EPI_ISL_594356, EPI_ISL_594357, EPI_ISL_594358, EPI_ISL_594359, EPI_ISL_594360, EPI_ISL_594361, EPI_ISL_594362, EPI_ISL_594363, EPI_ISL_594364, EPI_ISL_594365 | | | |
| see above | Florida Bureau of Public Health Laboratories | Florida Bureau of Public Health Laboratories | Sarah Schmedes, Jason Blanton |
| EPI_ISL_594464 | Texas Department of State Health Services Public Health Laboratory | Pathogen Discovery, Respiratory Viruses Branch, Division of Viral Diseases, Centers for Disease Control and Prevention | Ying Tao, Yan Li, Clinton Paden, Jing Zhang, Krista Queen, Anna Uehara, Haibin Wang, Julu Bhatnagar, Suxiang Tong |
| EPI_ISL_596544, EPI_ISL_596546, EPI_ISL_596547, EPI_ISL_596551, EPI_ISL_596556, EPI_ISL_596557 | Palestinian Ministry of Health | Molecular Genetics Lab | Nouar Qutob, Zaidoun Salah, Damien Richard, Hisham Darwish, Husam Sallam, Issa Shtayeh, Osama Najjar, Mahmoud Ruzayqat, Dana Najjar, Francois Balloux, Lucy van Dorp |
| EPI_ISL_596716, EPI_ISL_596726, EPI_ISL_596728, EPI_ISL_596737, EPI_ISL_596756, EPI_ISL_596775, EPI_ISL_596788, EPI_ISL_596803, EPI_ISL_596804 | PathWest Laboratory Medicine WA | PathWest Laboratory Medicine WA Microbial Surveillance Unit | PathWest Laboratory Medicine WA Microbial Surveillance Unit |
| EPI_ISL_600446, EPI_ISL_600466, EPI_ISL_600468, EPI_ISL_600471, EPI_ISL_600473, EPI_ISL_600476, EPI_ISL_600478, EPI_ISL_600489, EPI_ISL_600500, EPI_ISL_600502, EPI_ISL_600505, EPI_ISL_600507, EPI_ISL_600522, EPI_ISL_600524, EPI_ISL_600527 | | | |
| see above | Institute of Epidemiology Disease Control And Research | Institute for Developing Science and Health Initiatives | Lauren Cowley, Mokibul Hassan Afrad, Sadia Isfat Ara Rahman, Md. Mahfuz-Al-mamun, Firadausi Qadri, Tahmina Shirin |
| EPI_ISL_602242, EPI_ISL_602243, EPI_ISL_602244 | Texas Department of State Health Services | Texas Department of State Health Services | Rashmi Tuladhar, Bonnie Oh, Jenny Zhang, Maliha Rahman, Anita Pokharel, Myong Koag, Chung Wang, Rachel Lee, Grace Kubin, Mayela Pedrueza |
| EPI_ISL_602561 | Department of Biology and Wildlife, Alaska State Virology Laboratory | Department of Biology and Wildlife, Alaska State Virology Laboratory | DeRonde,S., Deuling,H., Chen,J. |
| EPI_ISL_602622 | AHRI-Sigal | KRISP, KZN Research Innovation and Sequencing Platform | Gazy I, Sigla, Karim F, Cele S, Giandhari J, Pillay S, Tegally H, Wilkinson E, de Oliveira T |
| EPI_ISL_602624, EPI_ISL_602626, EPI_ISL_602629, EPI_ISL_602631 | AHRI-Sigal | KRISP, KZN Research Innovation and Sequencing Platform | Gazy I, Sigl A, Karim F, Cele S, Giandhari J, Pillay S, Tegally H, Wilkinson E, de Oliveira T |
| EPI_ISL_603035 | Secretaria Municipal de Saúde | Instituto Adolfo Lutz, Interdiciplinary Procedures Center, Strategic Laboratory | Claudio Tavares Sacchi, Claudia Regina Gonçalves, Erica Valessa Ramos Gomes, Karoline Rodrigues Campos |

| | | | |
|---|---|---|---|
| EPI_ISL_603036 | Hospital Santa Ana | Instituto Adolfo Lutz, Interdiciplinary Procedures Center, Strategic Laboratory | Claudio Tavares Sacchi, Claudia Regina Gonçalves, Erica Valessa Ramos Gomes, Karoline Rodrigues Campos |
| EPI_ISL_605784 | AHRI-Sigal | KRISP, KZN Research Innovation and Sequencing Platform | Gazy I, Sigal A, Karim F, Cele S, Giandhari J, Pillay S, Tegally H, Wilkinson E, de Oliveira T |
| EPI_ISL_610042, EPI_ISL_610043, EPI_ISL_610044, EPI_ISL_610045, EPI_ISL_610046, EPI_ISL_610047 | Texas Department of State Health Services | Texas Department of State Health Services | Rashmi Tuladhar, Bonnie Oh, Jenny Zhang, Maliha Rahman, Anita Pokharel, Myong Koag, Chung Wang, Rachel Lee, Grace Kubin, Mayela Pedrueza |
| EPI_ISL_610178, EPI_ISL_610179, EPI_ISL_610180, EPI_ISL_610181, EPI_ISL_610185, EPI_ISL_610186, EPI_ISL_610187 | Department of Health Technology and Informatics, The Hong Kong Polytechnic University | Department of Health Technology and Informatics, The Hong Kong Polytechnic University | Siu,G.K.-H., Lee,L.-K., Leung,K.S.-S., Leung,J.S.-L., Ng,T.T.-L., Chan,C.T.-M., Tam,K.K.-G., Lao,H.-Y., Wu,A.K.-L., Yau,M.C.-Y., Lai,Y.W.-M., Fung,K.S.-C., Chau,S.K.-Y., Wong,B.K.-C., To,W.-K., Luk,K., Ho,A.Y.-M., Que,T.-L., Yip,K.-T., Yam,W.C., Shum,D.H.-K., Yip,S.P. |
| EPI_ISL_610283, EPI_ISL_610284, EPI_ISL_610285 | UW Virology Lab | UW Virology Lab | Pavitra Roychoudhury, Hong Xie, Lasata Shrestha, Meei-Li Huang, Keith R Jerome, Alexander Greninger |
| EPI_ISL_611627 | Liverpool Clinical Laboratories | COVID-19 Genomics UK (COG-UK) Consortium | Sam Haldenby, Anita Lucaci, Steve Paterson, Julian Hiscox, Alistair Darby, M Almsaud, A Alrezaihi, Muhannad Alruwaili, Stuart D Armstrong, Jones Benjamin, Eleanor G Bentley, Anu Chawla, Jordan J Clark, Angela Cowell, Richard Eccles, Isabel García-Dorival, Matthew Gemmell, Alessandro Gerada, PKF Gilmore, Richard Gregory, Ximeng Han, Catherine Hartley, Margaret Hughes, Miren Iturriza-Gomara, James Johnson, L Luu, Jenifer Manson, Charlotte Nelson, Elaine O'Toole, Cassie Olateju, Rebekah Penrice-Randal , Lucille Rainbow, N.P Randle, Trevor Ian Robinson, Parul Sharma, Ghada T Shawli, James P Stewart, Neil Swainston, Ecaterina Vamos, Joanne Watts, Mark Whitehead |
| EPI_ISL_613966, EPI_ISL_613967, EPI_ISL_613971, EPI_ISL_613976, EPI_ISL_613978, EPI_ISL_613979, EPI_ISL_613982, EPI_ISL_613983, EPI_ISL_613985, EPI_ISL_613990, EPI_ISL_613992, EPI_ISL_613998, EPI_ISL_614000, EPI_ISL_614001, EPI_ISL_614006, EPI_ISL_614008 | | | |
| see above | Texas Department of State Health Services | Texas Department of State Health Services | Rashmi Tuladhar, Bonnie Oh, Jenny Zhang, Maliha Rahman, Anita Pokharel, Myong Koag, Chung Wang, Rachel Lee, Grace Kubin, Mayela Pedrueza |
| EPI_ISL_614259, EPI_ISL_614260, EPI_ISL_614261, EPI_ISL_614262, EPI_ISL_614263, EPI_ISL_614264 | Wyoming Public Health Laboratory | Center for Global Health, University of New Mexico Health Sciences Center | Daryl Domman, Kurt Schwalm, Rob Christensen, Wanda Manley, Cari Sloma, Noah Hull, Darrell Dinwiddie |
| EPI_ISL_614375, EPI_ISL_614376 | Molecular diagnostic unit for viral haemorrhagic fevers and emerging viruses, Bouaké CHU Laboratory | Project group Epidemology of Highly Pathogenic Microorganisms, Robert Koch-Institute | Chantal Akoua-Koffi, Diané Bamourou, Etilé Anoh, Essia Belarbi, Safiatou Karidioula, Grit Schubert, Adjaratou Traoré, Soundélé Maïté, Monemo Pacome, Coulibaly Mbegnan, Bamba Fatoumata Touré, Kra Ouffoué, Fabian Leendertz |
| EPI_ISL_615165 | Texas Department of State Health Services | Texas Department of State Health Services | Rashmi Tuladhar, Bonnie Oh, Jenny Zhang, Maliha Rahman, Anita Pokharel, Myong Koag, Chung Wang, Rachel Lee, Grace Kubin, Mayela Pedrueza |
| EPI_ISL_618175, EPI_ISL_618185, EPI_ISL_618186, EPI_ISL_618187, EPI_ISL_618188, EPI_ISL_618189, EPI_ISL_618190, EPI_ISL_618196, EPI_ISL_618197, EPI_ISL_618198, EPI_ISL_618230, EPI_ISL_618273, EPI_ISL_618278, EPI_ISL_618280, EPI_ISL_618281, EPI_ISL_618289, EPI_ISL_618358 | | | |
| see above | Department of Virus and Microbiological Special Diagnostics, Statens Serum Institut, Denmark | Albertsen lab, Department of Chemistry and Bioscience, Aalborg University, Denmark | Danish Covid-19 Genome Consortia |
| EPI_ISL_620406 | Biomedical Science and Technology, Konkuk University, Laboratory of Cytokine Immunology (Room 601-1) IBST,Konkuk University | Biomedical Science and Technology, Konkuk University, Laboratory of Cytokine Immunology (Room 601-1) IBST,Konkuk University | Kim,S. |
| EPI_ISL_622560, EPI_ISL_622561, EPI_ISL_622562, EPI_ISL_622563, EPI_ISL_622564, EPI_ISL_622565, EPI_ISL_622566, EPI_ISL_622567, EPI_ISL_622568, EPI_ISL_622569, EPI_ISL_622570, EPI_ISL_622574, EPI_ISL_622601, EPI_ISL_622602, EPI_ISL_622613 | | | |
| see above | Department of Virus and Microbiological Special Diagnostics, Statens Serum Institut, Denmark | Albertsen lab, Department of Chemistry and Bioscience, Aalborg University, Denmark | Danish Covid-19 Genome Consortia |
| EPI_ISL_622894, EPI_ISL_622897, EPI_ISL_622900, EPI_ISL_622901, EPI_ISL_622902, EPI_ISL_622904, EPI_ISL_622906, EPI_ISL_622914, EPI_ISL_622916, EPI_ISL_622917, EPI_ISL_622918, EPI_ISL_622919, EPI_ISL_622920, EPI_ISL_622921, EPI_ISL_622922, EPI_ISL_622925, EPI_ISL_622927, EPI_ISL_622928, EPI_ISL_622929, EPI_ISL_622930, EPI_ISL_622931, EPI_ISL_622932, EPI_ISL_622933, EPI_ISL_622939, EPI_ISL_622940, EPI_ISL_622941 | | | |
| see above | National Institute for Communicable Diseases of the National Health Laboratory Service | National Institute for Communicable Diseases of the National Health Laboratory Service | Allam M, Ismail A, Khumalo Z, Kwenda S, Mtshali P, Mnyameni F, Mohale T, Subramoney K, Bhiman JN |
| EPI_ISL_622970, EPI_ISL_622987, EPI_ISL_623000, EPI_ISL_623019, EPI_ISL_623037, EPI_ISL_623052, EPI_ISL_623058, EPI_ISL_623072 | National Health Laboratory Service | National Institute for Communicable Diseases of the National Health Laboratory Service | Allam M, Ismail A, Khumalo Z, Kwenda S, Mtshali P, Mnyameni F, Mohale T, Subramoney K, Bhiman JN |
| EPI_ISL_625469, EPI_ISL_625470 | Child Health Research Foundation | Child Health Research Foundation | Senjuti Saha, Md Saiful Islam Sajib, Nikkon Sarkar, Syed Muktadir Al Sium, Afroza Akter Tanni, Roly Malaker, Arif Mohammad Tanmoy, Md Hafizur Rahman, Samir K Saha |
| EPI_ISL_626519, EPI_ISL_626520, EPI_ISL_626521, EPI_ISL_626522, EPI_ISL_626523, EPI_ISL_626524, EPI_ISL_626525, EPI_ISL_626526, EPI_ISL_626527 | Northwestern Memorial Hospital | Ozer Lab | Ramon Lorenzo-Redondo, Hannah H. Nam, Scott C. Roberts, Lacy M. Simons, Chad J. Achenbach, Lawrence J. Jennings, Chao Qi, Alan R. Hauser, Michael G. Ison, Judd F. Hultquist, Egon A. Ozer |
| EPI_ISL_628752, EPI_ISL_628754, EPI_ISL_628760 | UHAS COVID-19 Lab | UHAS COVID-19 Lab | Kwabena O. Duedu, Jones Gyamfi, Reuben Ayivor-Djanie, John O. Gyapong and the UHAS COVID-19 Lab Team |
| EPI_ISL_631480, EPI_ISL_631481, EPI_ISL_631482, EPI_ISL_631483, EPI_ISL_631484, EPI_ISL_631485, EPI_ISL_631486, EPI_ISL_631487, EPI_ISL_631488, EPI_ISL_631489 | Wisconsin State Laboratory of Hygiene Communicable Disease Division | Wisconsin State Laboratory of Hygiene Communicable Disease Division | Kelsey R. Florek, Abigail C. Shockey |
| EPI_ISL_632261 | Communicable Disease Laboratory, Public Health Directorate | Communicable Disease Laboratory, Public Health Directorate | AlWasti,H., AlTaif,Z., AlHujairi,Z., AlAbbas,Z. |
| EPI_ISL_632359, EPI_ISL_632366, EPI_ISL_632413, EPI_ISL_632772, EPI_ISL_632773, EPI_ISL_632774, EPI_ISL_632775 | Dutch COVID-19 response team | Erasmus Medical Center | Bas Oude Munnink, David Nieuwenhuijse, Reina Sikkema, Claudia Schapendonk, Irina Chestakova, Anne van der Linden, Theo Bestebroer, Stefan van Nieuwkoop, Mark Pronk, Pascal Lexmond, Corien Swaan, Manon Haverkate, Madelief Mollers, Mart Stein, Sandra Kengne Kamga Mobou, Jeroen van Kampen, Jolanda Voermans, Aura Timen, Corine GeurtsvanKessel, Annemiek van der Eijk, Richard Molenkamp, Marion Koopmans, on behalf of the Dutch national COVID-19 response team. |
| EPI_ISL_632964 | Department of Homeless Services | New York City Public Health Laboratory | Jade Wang, et al. |
| EPI_ISL_634951, EPI_ISL_634952, EPI_ISL_634953, EPI_ISL_634954, EPI_ISL_634955, EPI_ISL_634956, EPI_ISL_634957, EPI_ISL_634958, EPI_ISL_634959, EPI_ISL_634960, EPI_ISL_634961, EPI_ISL_634962, EPI_ISL_634963, EPI_ISL_634964, EPI_ISL_634965, EPI_ISL_634966, EPI_ISL_634967, EPI_ISL_634968, EPI_ISL_634969, EPI_ISL_634970, EPI_ISL_634971, EPI_ISL_634972, EPI_ISL_634973, EPI_ISL_634974, EPI_ISL_634975 | | | |
| see above | Texas Department of State Health Services | Texas Department of State Health Services | Rashmi Tuladhar, Bonnie Oh, Jenny Zhang, Maliha Rahman, Anita Pokharel, Myong Koag, Chung Wang, Rachel Lee, Grace Kubin, Mayela Pedrueza |
| EPI_ISL_635398, EPI_ISL_635403, EPI_ISL_635404, EPI_ISL_635405, EPI_ISL_635406, EPI_ISL_635407, EPI_ISL_635408, EPI_ISL_635409, EPI_ISL_635410, EPI_ISL_635411, EPI_ISL_635412, EPI_ISL_635413, EPI_ISL_635414, EPI_ISL_635415, EPI_ISL_635416, EPI_ISL_635417, EPI_ISL_635418, EPI_ISL_635419, EPI_ISL_635420, EPI_ISL_635421, EPI_ISL_635422, EPI_ISL_635423, EPI_ISL_635424, EPI_ISL_635425, EPI_ISL_635426, EPI_ISL_635427, EPI_ISL_635428, EPI_ISL_635429, EPI_ISL_635430, EPI_ISL_635431, EPI_ISL_635432, EPI_ISL_635433, EPI_ISL_635434, EPI_ISL_635435, EPI_ISL_635436, EPI_ISL_635437, EPI_ISL_635438, EPI_ISL_635439, EPI_ISL_635440, EPI_ISL_635441, EPI_ISL_635442, EPI_ISL_635443, EPI_ISL_635444, EPI_ISL_635445, EPI_ISL_635446, EPI_ISL_635447, EPI_ISL_635448, EPI_ISL_635449, EPI_ISL_635450, EPI_ISL_635451, EPI_ISL_635452, EPI_ISL_635453, EPI_ISL_635455, EPI_ISL_635456, EPI_ISL_635457, EPI_ISL_635458, EPI_ISL_635459, EPI_ISL_635460, EPI_ISL_635465, EPI_ISL_635467, EPI_ISL_635468, EPI_ISL_635470, EPI_ISL_635471, EPI_ISL_635472, EPI_ISL_635475, EPI_ISL_635477, EPI_ISL_635478 | | | |
| see above | San Diego County Public Health Laboratory | Andersen lab at Scripps Research | SEARCH Alliance San Diego with Tracy Basler, Jovan Shephard, Brett Austin |
| EPI_ISL_635552, EPI_ISL_635553, EPI_ISL_635554, EPI_ISL_635555, EPI_ISL_635556, EPI_ISL_635557, EPI_ISL_635558, EPI_ISL_635559, EPI_ISL_635560, EPI_ISL_635561, EPI_ISL_635562, EPI_ISL_635563, EPI_ISL_635564, EPI_ISL_635565, EPI_ISL_635566, EPI_ISL_635567, EPI_ISL_635568, EPI_ISL_635569, EPI_ISL_635570, EPI_ISL_635571, EPI_ISL_635572 | | | |

| | | | |
|---|---|---|---|
| see above | Centro de Diagnstico COVID-19 UABC Tijuana | Andersen lab at Scripps Research | SEARCH Alliance San Diego with Idanya Rubí Serafín Higuera, Manuel Sánchez Alavez, Jorge Luis Jiménez Niebla, Germán Ibarra, Jonathan Vincent Baena, Oscar Efrén Zazueta Fierro |
| EPI_ISL_635577, EPI_ISL_635578, EPI_ISL_635579, EPI_ISL_635581, EPI_ISL_635586, EPI_ISL_635590, EPI_ISL_635591, EPI_ISL_635593, EPI_ISL_635594, EPI_ISL_635598, EPI_ISL_635599, EPI_ISL_635600, EPI_ISL_635609, EPI_ISL_635613, EPI_ISL_635615, EPI_ISL_635617, EPI_ISL_635622, EPI_ISL_635623, EPI_ISL_635628, EPI_ISL_635629, EPI_ISL_635630, EPI_ISL_635631 | | | |
| see above | San Diego County Public Health Laboratory | Andersen lab at Scripps Research | SEARCH Alliance San Diego with Tracy Basler, Jovan Shephard, Brett Austin |
| EPI_ISL_635778, EPI_ISL_635782 | Biolab Diagnostic Laboratories | Andersen lab at Scripps Research | Issa Abu-Dayyeh, Ahmad Tibi, Lama Hussein, Lina Mohammad, Zein Naber, Amid Abdelnour with SEARCH Alliance San Diego |
| EPI_ISL_635988, EPI_ISL_635989, EPI_ISL_635995, EPI_ISL_635996, EPI_ISL_635997, EPI_ISL_635998, EPI_ISL_635999, EPI_ISL_636000, EPI_ISL_636001, EPI_ISL_636003, EPI_ISL_636004, EPI_ISL_636005, EPI_ISL_636006, EPI_ISL_636007, EPI_ISL_636008, EPI_ISL_636009, EPI_ISL_636010, EPI_ISL_636011, EPI_ISL_636012, EPI_ISL_636013, EPI_ISL_636014, EPI_ISL_636015, EPI_ISL_636016, EPI_ISL_636017, EPI_ISL_636018, EPI_ISL_636019, EPI_ISL_636020, EPI_ISL_636021, EPI_ISL_636180, EPI_ISL_636216, EPI_ISL_636248, EPI_ISL_636249, EPI_ISL_636250, EPI_ISL_636252, EPI_ISL_636264, EPI_ISL_636265, EPI_ISL_636266 | | | |
| see above | San Diego County Public Health Laboratory | Andersen lab at Scripps Research | SEARCH Alliance San Diego with Tracy Basler, Jovan Shephard, Brett Austin |
| EPI_ISL_636975, EPI_ISL_636979 | HP Pemba | KRISP, KZN Research Innovation and Sequencing Platform | Ismael N, Giandhari J, Pillay S, Tegally H, Wilkinson E, de Oliveira T, Nadia Sitoe, Paulo Arnaldo, Nedio Mabunda |
| EPI_ISL_636981 | City of Chimoio | KRISP, KZN Research Innovation and Sequencing Platform | Ismael N, Giandhari J, Pillay S, Tegally H, Wilkinson E, de Oliveira T, Nadia Sitoe, Paulo Arnaldo, Nedio Mabunda |
| EPI_ISL_639914, EPI_ISL_639916, EPI_ISL_639917, EPI_ISL_639918, EPI_ISL_639919, EPI_ISL_639920 | Omsk Research Institute of Natural Focal Infections | WHO National Influenza Centre Russian Federation | Artem Fadeev, Ekaterina Gradoboeva, Ekaterina Savkina, Daria Nashatyreva, Elena Poleshchuk, Aleksei Vasilenko, Valery Yakimenko, Andrey Komissarov |
| EPI_ISL_640067 | Groote Schuur Hospital wc GSH | NHLS/UCT | Arash Iranzadeh, Deelan Doolabh, Lynn Tyers, Bruna Galvao, Innocent Mudau, Marvin Hsiao, Kruger Marais, Diana Hardie, Stephen Korsman, Carolyn Williamson |
| EPI_ISL_640068 | Clinic-in-Asla | NHLS/UCT | Arash Iranzadeh, Deelan Doolabh, Lynn Tyers, Bruna Galvao, Innocent Mudau, Marvin Hsiao, Kruger Marais, Diana Hardie, Stephen Korsman, Carolyn Williamson |
| EPI_ISL_644208 | Texas Department of State Health Services | Texas Department of State Health Services | Rashmi Tuladhar, Bonnie Oh, Jenny Zhang, Maliha Rahman, Anita Pokharel, Myong Koag, Chung Wang, Rachel Lee, Grace Kubin, Mayela Pedrueza, James Daniel Bonser |
| EPI_ISL_644568, EPI_ISL_644570, EPI_ISL_644579 | Veterinary Specialized Institute "Kraljevo", Serbia | Veterinary Specialized Institute "Kraljevo", Serbia | Vidanovic,D., Tesovic,B., Knezevic,A., Jovanovic,T., Jankovic,M., Sekler,M., Banovic Djeri,B., Petrovic,T., Volkening,J., Afonso,C. |
| EPI_ISL_644955, EPI_ISL_644956, EPI_ISL_644957, EPI_ISL_644958, EPI_ISL_644959, EPI_ISL_644960 | Department of Infectious Diseases, Keio University School of Medicine, Tokyo, Japan | Center for Medical Genetics, Keio University School of Medicine, Tokyo, Japan | Kenjiro Kosaki, Yuka Iwasaki, Hirotsugu Ishizu, Haruhiko Siomi, Kodai Abe |
| EPI_ISL_648133 | Klinisk mikrobiologi centralsjukhuset Karlstad | The Public Health Agency of Sweden | Anna-Malin Linde, Maria Lind Karlberg, Mattias Haukland, Reza Advani, Olov Svartstrom, Oskar Karlsson Lindsjo, Sandra Broddesson, Petra Edquist, Mia Brytting, Anna Risberg, Karin Tegmark-Wisell |
| EPI_ISL_648145 | The Public Health Agency of Sweden | The Public Health Agency of Sweden | Anna-Malin Linde, Maria Lind Karlberg, Mattias Haukland, Reza Advani, Olov Svartstrom, Oskar Karlsson Lindsjo, Sandra Broddesson, Petra Edquist, Mia Brytting, Anna Risberg, Karin Tegmark-Wisell |
| EPI_ISL_648173 | Klinisk mikrobiologi NAL Trollhattan | The Public Health Agency of Sweden | Anna-Malin Linde, Maria Lind Karlberg, Mattias Haukland, Reza Advani, Olov Svartstrom, Oskar Karlsson Lindsjo, Sandra Broddesson, Petra Edquist, Mia Brytting, Anna Risberg, Karin Tegmark-Wisell |
| EPI_ISL_648322, EPI_ISL_648324, EPI_ISL_648373, EPI_ISL_648374, EPI_ISL_648375 | Laboratorio de Investigaciones de Baney | University Hospital Basel, Clinical Bacteriology | Carlos Cortes, Claudia Daubenberger, Adrian Egli, Guillermo Garcia, Salome Hosch, Bonifacio Manguire Nlavo, Alfredo Mari, Maximilian Mpina, Elizabeth Nyakarungu, Diosdado Odjama Nseng Ada, Mitoha Ondo O Ayekaba, Tim Roloff, Tobias Schindler, Helena Seth-Smith, Madlen Stange, Philip Wonder Phiri |
| EPI_ISL_648542 | Santa Clara County Public Health Laboratory | Chan-Zuckerberg Biohub | CZB Cliahub Consortium |
| EPI_ISL_648609 | Laboratorio de Infectología, Servicio de Infectología, Hospital Universitario Dr. José Eleuterio González - Universidad Autónoma de Nuevo León | Laboratorio de Infectología Molecular, Departamento de Bioquímica y Medicina Molecular,Facultad de Medicina - Universidad Autónoma de Nuevo León | Kame A. Galán-Huerta, María F. Herrera-Saldivar, Natalia Martínez-Acuña, Sonia A. Lozano-Sepúlveda, Daniel Arellanos-Soto, Ana M. Rivas-Estilla, Paola Bocanegra-Ibarias, Samantha M. Flores-Treviño, Elvira Garza-González, Eduardo Perez-Alba, Laura Nuzzolo-Shihadeh, Adrian Camacho-Ortiz, Roberto Montes-de-Oca, Consuelo Treviño-Garza, Manuel E. de-la-O-Cavazos |
| EPI_ISL_648678, EPI_ISL_648679, EPI_ISL_648723, EPI_ISL_648724, EPI_ISL_648725, EPI_ISL_648726, EPI_ISL_648727, EPI_ISL_648728, EPI_ISL_648729, EPI_ISL_648730, EPI_ISL_648731, EPI_ISL_648732, EPI_ISL_648733, EPI_ISL_648734 | | | |
| see above | Department of Laboratory Medicine, Tan Tock Seng Hospital | Department of Laboratory Medicine, Tan Tock Seng Hospital | Chen YYC, Zair X, Lim JX, Li C, Tang WY, Maurer-Stroh S, Barkham TMS, Nagarajan N, Sessions OM |
| EPI_ISL_648823, EPI_ISL_648824, EPI_ISL_648825, EPI_ISL_648826, EPI_ISL_648827, EPI_ISL_648828, EPI_ISL_648829, EPI_ISL_648831, EPI_ISL_648832, EPI_ISL_648833, EPI_ISL_648835, EPI_ISL_648836, EPI_ISL_648837, EPI_ISL_648838, EPI_ISL_648839, EPI_ISL_648840, EPI_ISL_648841, EPI_ISL_648842, EPI_ISL_648843, EPI_ISL_648844, EPI_ISL_648862, EPI_ISL_648863, EPI_ISL_648864, EPI_ISL_648865, EPI_ISL_648866, EPI_ISL_649033, EPI_ISL_649036, EPI_ISL_649037, EPI_ISL_649043, EPI_ISL_649047, EPI_ISL_649052, EPI_ISL_649055 | | | |
| see above | San Diego County Public Health Laboratory | Andersen lab at Scripps Research | SEARCH Alliance San Diego with Tracy Basler, Jovan Shephard, Brett Austin |
| EPI_ISL_649165, EPI_ISL_649166 | Laboratorio de Investigaciones de Baney | University Hospital Basel, Clinical Bacteriology | Carlos Cortes, Claudia Daubenberger, Adrian Egli, Guillermo Garcia, Salome Hosch, Bonifacio Manguire Nlavo, Alfredo Mari, Maximilian Mpina, Elizabeth Nyakarungu, Diosdado Odjama Nseng Ada, Mitoha Ondo O Ayekaba, Tim Roloff, Tobias Schindler, Helena Seth-Smith, Madlen Stange, Philip Wonder Phiri |
| EPI_ISL_653220, EPI_ISL_653221, EPI_ISL_653222, EPI_ISL_653227, EPI_ISL_653228, EPI_ISL_653229, EPI_ISL_653231, EPI_ISL_653232, EPI_ISL_653233, EPI_ISL_653300, EPI_ISL_653301, EPI_ISL_653302, EPI_ISL_653303, EPI_ISL_653304 | | | |
| see above | Florida Bureau of Public Health Laboratories | Florida Bureau of Public Health Laboratories | Sarah Schmedes, Jason Blanton |
| EPI_ISL_653548, EPI_ISL_653549, EPI_ISL_653550, EPI_ISL_653552, EPI_ISL_653553, EPI_ISL_653555, EPI_ISL_653556, EPI_ISL_653557, EPI_ISL_653558, EPI_ISL_653559, EPI_ISL_653560, EPI_ISL_653561, EPI_ISL_653562, EPI_ISL_653563, EPI_ISL_653564, EPI_ISL_653565, EPI_ISL_653566, EPI_ISL_653567, EPI_ISL_653568, EPI_ISL_653569, EPI_ISL_653570, EPI_ISL_653571, EPI_ISL_653572, EPI_ISL_653573, EPI_ISL_653574, EPI_ISL_653575, EPI_ISL_653576, EPI_ISL_653577, EPI_ISL_653578, EPI_ISL_653579, EPI_ISL_653580, EPI_ISL_653581, EPI_ISL_653582, EPI_ISL_653583, EPI_ISL_653584, EPI_ISL_653585, EPI_ISL_653586, EPI_ISL_653587, EPI_ISL_653588, EPI_ISL_653589, EPI_ISL_653591, EPI_ISL_653592, EPI_ISL_653593, EPI_ISL_653595, EPI_ISL_653596, EPI_ISL_653597, EPI_ISL_653598, EPI_ISL_653599, EPI_ISL_653601, EPI_ISL_653602, EPI_ISL_653603, EPI_ISL_653604, EPI_ISL_653605, EPI_ISL_653606, EPI_ISL_653608, EPI_ISL_653609 | | | |
| see above | LSUHS Emerging Viral Threat Laboratory | Microbial Genome Sequencing Center | Jeremy P. Kamil, Rona S. Scott, Maarten Van Diest, Malgorzata Bienkowska-Haba, Katarzyna Zwolinska, Andrew D. Yurochko, Christopher G. Kevil, Martin J. Sapp, Daniel J. Snyder, Vaughn S. Cooper, John A. Vanchiere |
| EPI_ISL_653756, EPI_ISL_653760 | Instituto Nacional de Salud, Bogotá, Colombia | Instituto Nacional de Salud, Bogotá, Colombia | Katherine Laiton-Donato, Diego A. Álvarez-Díaz, Carlos Franco-Muñoz, Mauricio Pacheco-Montealegre, Jonathan Reales, Diego Andrés Prada, Jose A. Usme-Ciro, Zulma M. Cucunubá, Christian Julian VillabonaArenas, Liz Villabona-Arenas, Sussy Echeverria, Astrid C. Flórez, Carolina Ferro, Diana Marcela Walteros-Acero, Franklin Prieto, Carlos Andrés Durán, Martha Lucia Ospina Martinez, Marcela Mercado-Reyes |
| EPI_ISL_654336, EPI_ISL_654338 | Hospital General Universitario Gregorio Marañón | SeqCOVID-SPAIN consortium/IBV(CSIC) | Darío García de Viedma, Laura Pérez-Lago, Marta Herranz, Jon Sicilia, Julia Suárez, Pilar Catalán, Patricia Muñoz and SeqCOVID-SPAIN consortium |
| EPI_ISL_654899 | Klinisk mikrobiologi | The Public Health Agency of Sweden | Anna-Malin Linde, Maria Lind Karlberg, Mattias Haukland, Reza Advani, Olov Svartstrom, Oskar Karlsson Lindsjo, Sandra Broddesson, Petra Edquist, Mia Brytting, Anna Risberg, Karin Tegmark-Wisell |
| EPI_ISL_660166, EPI_ISL_660167, EPI_ISL_660169 | NHLS-IALCH | KRISP, KZN Research Innovation and Sequencing Platform | Gazy I, Sigal A, Karim F, Cele S, Giandhari J, Pillay S, Tegally H, Wilkinson E, de Oliveira T |
| EPI_ISL_661182, EPI_ISL_661186, EPI_ISL_661187, EPI_ISL_661193, EPI_ISL_661197, EPI_ISL_661199 | Scientific Veterinary Institute Novi Sad | Veterinary Specialized Institute "Kraljevo", Serbia | Vidanovic,D., Tesovic,B., Knezevic,A., Jovanovic,T., Jankovic,M., Sekler,M., Banovic Djeri,B., Petrovic,T., Volkening,J., Afonso,C. |
| EPI_ISL_666600 | Dept. of Microbiology and Infection Control, Akershus University Hospital HF | Dept. of Microbiology and Infection Control, Akershus University Hospital HF | Hege Vangstein Aamot, Alexander Hesselberg Løvestad, Silje Bakken Jørgensen, Nina Handal, Ole Herman Ambur |
| EPI_ISL_666864 | Florida Bureau of Public Health Laboratories | Florida Bureau of Public Health Laboratories | Sarah Schmedes, Jason Blanton |
| EPI_ISL_666992, EPI_ISL_666999, EPI_ISL_667000, EPI_ISL_667001, EPI_ISL_667002, EPI_ISL_667005, EPI_ISL_667006, EPI_ISL_667007, EPI_ISL_667008, EPI_ISL_667009, EPI_ISL_667010, EPI_ISL_667011, EPI_ISL_667012, EPI_ISL_667013, EPI_ISL_667019, EPI_ISL_667020, EPI_ISL_667025, EPI_ISL_667026, EPI_ISL_667027, EPI_ISL_667028, EPI_ISL_667030, EPI_ISL_667031, EPI_ISL_667032, EPI_ISL_667034, EPI_ISL_667035, EPI_ISL_667036, EPI_ISL_667037, EPI_ISL_667038, EPI_ISL_667041 | | | |

| | | | |
|---|---|---|---|
| see above | San Diego County Public Health Laboratory | Andersen lab at Scripps Research | SEARCH Alliance San Diego with Tracy Basler, Jovan Shephard, Brett Austin |
| EPI_ISL_671844, EPI_ISL_671845, EPI_ISL_671846 | Servicio de Microbiología, Laboratori Clínic Metropolitana Nord. Hospital Universitari Germans Trias i Pujol. Institut d'Investigació en Ciències de la Salut Germans Trias i Pujol (IGTP) | SeqCOVID-SPAIN consortium/IBV(CSIC) | Elisa Martró, Antoni E. Bordoy, Anna Not, Adrián Antuori, Anabel Fernández, Nona Romaní, Verónica Saludes, Cristina Casañ and SeqCOVID-SPAIN consortium |
| EPI_ISL_671943 | CHU Purpan - Laboratoire de Virologie - Institut Fédératif de Biologie | CHU Purpan - Laboratoire de Virologie - Institut Fédératif de Biologie | Latour J., Ranger N., Dubois M., Carcenac R., Harter A., Boyer P., Tremeaux P., Izopet J. |
| EPI_ISL_672056, EPI_ISL_672190, EPI_ISL_672191, EPI_ISL_672274 | The Ashley Laboratory, Stanford University | Chan-Zuckerberg Biohub | CZB Cliahub Consortium |
| EPI_ISL_672383 | San Francisco Public Health Laboratory | Chan-Zuckerberg Biohub | CZB Cliahub Consortium |
| EPI_ISL_672580 | Infectious Diseases and Tropical Medicine Research Center, Infectious Diseases and Tropical Medicine Research Center | Infectious Diseases and Tropical Medicine Research Center, Infectious Diseases and Tropical Medicine Research Center | Ahangarzadeh,S., Shariati,L., Haghjooy Javanmard,S., Aboutalebian,S., Shoaei,P., Ataei,B. |
| EPI_ISL_672581, EPI_ISL_672583 | Infectious Diseases and Tropical Medicine Research Center, Infectious Diseases and Tropical Medicine Research Center | Infectious Diseases and Tropical Medicine Research Center, Infectious Diseases and Tropical Medicine Research Center | Ahangarzadeh,S., Ataei,B., Shariati,L., Haghjooy Javanmard,S., Shoaei,P., Aboutalebian,S. |
| EPI_ISL_672587 | Infectious Diseases and Tropical Medicine Research Center, Infectious Diseases and Tropical Medicine Research Center | Infectious Diseases and Tropical Medicine Research Center, Infectious Diseases and Tropical Medicine Research Center | Haghjooy Javanmard,S., Ahangarzadeh,S., Shariati,L., Shoaei,P. |
| EPI_ISL_672588, EPI_ISL_672589 | Infectious Diseases and Tropical Medicine Research Center, Infectious Diseases and Tropical Medicine Research Center | Infectious Diseases and Tropical Medicine Research Center, Infectious Diseases and Tropical Medicine Research Center | Ahangarzadeh,S., Haghjooy Javanmard,S., Shoaei,P., Ataei,B., Shariati,L. |
| EPI_ISL_672602 | Infectious Diseases and Tropical Medicine Research Center, Infectious Diseases and Tropical Medicine Research Center | Infectious Diseases and Tropical Medicine Research Center, Infectious Diseases and Tropical Medicine Research Center | Ahangarzadeh,S., Haghjooy Javanmard,S., Shariati,L., Aboutalebian,S., Ataei,B., Shoaei,P. |
| EPI_ISL_676495 | Klinisk mikrobiologi | The Public Health Agency of Sweden | Department of Microbiology, The Public Health Agency of Sweden |
| EPI_ISL_676527 | Uppsala klinisk mikrobiologi | The Public Health Agency of Sweden | Department of Microbiology, The Public Health Agency of Sweden |
| EPI_ISL_676579 | Scientific Veterinary Institute Novi Sad | Veterinary Specialized Institute "Kraljevo", Serbia | Vidanovic,D., Tesovic,B., Knezevic,A., Jovanovic,T., Jankovic,M., Sekler,M., Banovic Djeri,B., Petrovic,T., Volkening,J., Afonso,C. |
| EPI_ISL_676659, EPI_ISL_676680, EPI_ISL_676713 | Masonic Medical Research Institute | Wadsworth Center, New York State Department.of Health | Nathan Tucker, Kirsten St. George, Daryl M. Lamson, Alexis Russel, Jonathan Plitnick, Navjot Singh, John Kelly, Sara Griesemer, Erasmus Schneider, Erica Lasek-Nesselquist |
| EPI_ISL_676987, EPI_ISL_676988, EPI_ISL_676989, EPI_ISL_676990, EPI_ISL_676991, EPI_ISL_676992 | Wadsworth Center, New York State Department.of Health | Wadsworth Center, New York State Department.of Health | Kirsten St. George, Daryl M. Lamson, Alexis Russel, Jonathan Plitnick, Navjot Singh, John Kelly, Sara Griesemer, Erasmus Schneider, Erica Lasek-Nesselquist |
| EPI_ISL_677010, EPI_ISL_677011, EPI_ISL_677012, EPI_ISL_677013, EPI_ISL_677014, EPI_ISL_677015, EPI_ISL_677016, EPI_ISL_677017, EPI_ISL_677018, EPI_ISL_677019, EPI_ISL_677020, EPI_ISL_677021, EPI_ISL_677022, EPI_ISL_677023, EPI_ISL_677059, EPI_ISL_677060, EPI_ISL_677061, EPI_ISL_677062, EPI_ISL_677063, EPI_ISL_677064, EPI_ISL_677065, EPI_ISL_677066, EPI_ISL_677067, EPI_ISL_677068, EPI_ISL_677069, EPI_ISL_677070 | | | |
| see above | Masonic Medical Research Institute | Wadsworth Center, New York State Department.of Health | Nathan Tucker, Kirsten St. George, Daryl M. Lamson, Alexis Russel, Jonathan Plitnick, Navjot Singh, John Kelly, Sara Griesemer, Erasmus Schneider, Erica Lasek-Nesselquist |
| EPI_ISL_677251, EPI_ISL_677294 | Colorado Department of Public Health and Environment | Colorado Department of Puplic Health and Environment | Laura Bankers, Molly Hetherington-Rauth, Shannon Ely, Shannon R. Matzinger, Sarah Elizabeth Totten, Emily A. Travanty |
| EPI_ISL_677669 | Masonic Medical Research Institute | Wadsworth Center, New York State Department.of Health | Nathan Tucker, Kirsten St. George, Daryl M. Lamson, Alexis Russel, Jonathan Plitnick, Navjot Singh, John Kelly, Sara Griesemer, Erasmus Schneider, Erica Lasek-Nesselquist |
| EPI_ISL_677715 | Clinical Hospital - Bitola | Research Center for Genetic Engineering and Biotechnology "Georgi D. Efremov" , Macedonian Academy of Sciences and Arts | RCGEB - MASA |
| EPI_ISL_677716 | Clinical Hospital - Shtip | Research Center for Genetic Engineering and Biotechnology "Georgi D. Efremov" , Macedonian Academy of Sciences and Arts | RCGEB - MASA |
| EPI_ISL_677825, EPI_ISL_677893, EPI_ISL_677896, EPI_ISL_677898, EPI_ISL_677899, EPI_ISL_677901 | Innovative Genomics Institute, UC Berkeley | Innovative Genomics Institute, UC Berkeley | Stacia Wyman, Haridha Shivram, Phil Frankino, Liana Lareau, Shana McDevitt, Justin Choi |
| EPI_ISL_678094, EPI_ISL_678095 | Pathogen Genomics Lab King Abdullah University of Science and Technology(KAUST) | Pathogen Genomics Lab King Abdullah University of Science and Technology(KAUST) | Olga Douvropoulou, Sara Mfarrej, Raushan Nugmanova, Sharif Hala, Raeece Naeem, Asim Khogeer, Fadwa Alofi, Afrah Alsomali, Jumana Taha, Abdulaziz Alahmadi, Kahled Algithami, Anwar Hashem, Naif Almontashiri, Arnab Pain |
| EPI_ISL_678342 | Area of Virology, Serology and Virology Division (SAViD), New South Wales Health Pathology Randwick | Virology Research Laboratory; Area of Virology, Serology and Virology Division (SAViD), New South Wales Health Pathology Randwick | Foster, C.; Au, J.; Ruiz Silva, M.; Deveson, I.; Bull, R.; Van Hal, S.; Rawlinson, W. |
| EPI_ISL_678491, EPI_ISL_678492 | Veterinary Specialized Institute "Sabac", Serbia | Veterinary Specialized Institute "Kraljevo", Serbia | Vidanovic,D., Tesovic,B., Knezevic,A., Jovanovic,T., Jankovic,M., Sekler,M., Banovic Djeri,B., Petrovic,T., Mrkovacki, S., Volkening,J., Afonso,C. |
| EPI_ISL_681692, EPI_ISL_681693 | Molecular Medicine Laboratory, University of Magallanes | Centro Asistencial Docente y de Investigacion, Universidad de Magallanes | Jorge González, Jacqueline Aldridge, Diego Alvarez, Marco Montes de Oca, Hermy Alvarez, Roberto Uribe-Paredes, Marcelo Navarrete |
| EPI_ISL_681830, EPI_ISL_681836, EPI_ISL_681837 | Molecular diagnostic unit for viral haemorrhagic fevers and emerging viruses, Bouaké CHU Laboratory | Project group Epidemology of Highly Pathogenic Microorganisms, Robert Koch-Institute | Chantal Akoua-Koffi, Diané Bamourou, Etilé Anoh, Essia Belarbi, Safiatou Karidioula, Grit Schubert, Adjaratou Traoré, Soundélé Maïté, Monemo Pacome, Coulibaly Mbegnan, Bamba Fatoumata Touré, Kra Ouffoué, Fabian Leendertz |
| EPI_ISL_681874 | Texas Department of State Health Services | Texas Department of State Health Services | Rashmi Tuladhar, Bonnie Oh, Jenny Zhang, Maliha Rahman, Anita Pokharel, Myong Koag, Chung Wang, Rachel Lee, Grace Kubin, Mayela Pedrueza, James Daniel Bonser |
| EPI_ISL_682020 | UPMC Clinical Microbiology Laboratory | Microbial Genomic Epidemiology Laboratory, University of Pittsburgh | Mustapha M. Mustapha, Jane W. Marsh, Dan Snyder, Marissa P. Griffith, Stephanie L. Mitchell, Vatsala R. Srinivasa, Kady D. Waggle, Chinelo Ezeonwuku, Vaughn S. Cooper, Lee H. Harrison |
| EPI_ISL_682059 | Molecular diagnostic unit for viral haemorrhagic fevers and emerging viruses, Bouaké CHU Laboratory | Project group Epidemology of Highly Pathogenic Microorganisms, Robert Koch-Institute | Chantal Akoua-Koffi, Diané Bamourou, Etilé Anoh, Essia Belarbi, Safiatou Karidioula, Grit Schubert, Adjaratou Traoré, Soundélé Maïté, Monemo Pacome, Coulibaly Mbegnan, Bamba Fatoumata Touré, Kra Ouffoué, Fabian Leendertz |
| EPI_ISL_682236 | AREA DE SALUD LOS CHILES | Inciensa, Instituto Costarricense de Investigación y Enseñanza en Nutrición y Salud | Francisco Duarte, Hebleen Porras, Claudio Soto-Garita, Estela Cordero, Adriana Godinez, Melany Calderon & Mariel López |
| EPI_ISL_682237 | AREA DE SALUD CORREDORES | Inciensa, Instituto Costarricense de Investigación y Enseñanza en Nutrición y Salud | Francisco Duarte, Hebleen Porras, Claudio Soto-Garita, Estela Cordero, Adriana Godinez & Melany Calderon |
| EPI_ISL_682238 | AREA DE SALUD LA CRUZ | Inciensa, Instituto Costarricense de Investigación y Enseñanza en Nutrición y Salud | Francisco Duarte, Hebleen Porras, Claudio Soto-Garita, Estela Cordero, Adriana Godinez & Melany Calderon |
| EPI_ISL_682239 | HOSPITAL CIUDAD NEILY | Inciensa, Instituto Costarricense de Investigación y Enseñanza en Nutrición y Salud | Francisco Duarte, Hebleen Porras, Claudio Soto-Garita, Estela Cordero, Adriana Godinez & Melany Calderon |
| EPI_ISL_682240 | AREA DE SALUD GOICOCHEA 1 | Inciensa, Instituto Costarricense de Investigación y Enseñanza en Nutrición y Salud | Francisco Duarte, Hebleen Porras, Claudio Soto-Garita, Estela Cordero, Adriana Godinez & Melany Calderon |

| | | | |
|---|---|---|---|
| EPI_ISL_684000 | Utah Public Health Laboratory | Utah Public Health Laboratory | Erin Young, Kelly Oakeson |
| EPI_ISL_693208, EPI_ISL_693209 | Hospital Municipal Antonio Giglio | Instituto Adolfo Lutz, Interdiciplinary Procedures Center, Strategic Laboratory | Claudio Tavares Sacchi, Claudia Regina Gonçalves, Erica Valessa Ramos Gomes, Karoline Rodrigues Campos |
| EPI_ISL_693210 | Pronto-Socorro Dr. Osmar Mesquita | Instituto Adolfo Lutz, Interdiciplinary Procedures Center, Strategic Laboratory | Claudio Tavares Sacchi, Claudia Regina Gonçalves, Erica Valessa Ramos Gomes, Karoline Rodrigues Campos |
| EPI_ISL_693211 | Santa Casa de Misericordia e Maternidade | Instituto Adolfo Lutz, Interdiciplinary Procedures Center, Strategic Laboratory | Claudio Tavares Sacchi, Claudia Regina Gonçalves, Erica Valessa Ramos Gomes, Karoline Rodrigues Campos |
| EPI_ISL_693212 | Santa Casa de Misericordia de Braganca Paulista | Instituto Adolfo Lutz, Interdiciplinary Procedures Center, Strategic Laboratory | Claudio Tavares Sacchi, Claudia Regina Gonçalves, Erica Valessa Ramos Gomes, Karoline Rodrigues Campos |
| EPI_ISL_693215 | Secretaria Municipal de Saúde de Iracemapolis | Instituto Adolfo Lutz, Interdiciplinary Procedures Center, Strategic Laboratory | Claudio Tavares Sacchi, Claudia Regina Gonçalves, Erica Valessa Ramos Gomes, Karoline Rodrigues Campos |
| EPI_ISL_693216, EPI_ISL_693217 | Unidade de Vigilância Epiudemiológica de Araras | Instituto Adolfo Lutz, Interdiciplinary Procedures Center, Strategic Laboratory | Claudio Tavares Sacchi, Claudia Regina Gonçalves, Erica Valessa Ramos Gomes, Karoline Rodrigues Campos |
| EPI_ISL_693220, EPI_ISL_693223, EPI_ISL_693224 | Laboratório Municipal de Piracicaba | Instituto Adolfo Lutz, Interdiciplinary Procedures Center, Strategic Laboratory | Claudio Tavares Sacchi, Claudia Regina Gonçalves, Erica Valessa Ramos Gomes, Karoline Rodrigues Campos |
| EPI_ISL_693225 | Ubs Vila Rosa - Olímpia Gomes De Almeida | Instituto Adolfo Lutz, Interdiciplinary Procedures Center, Strategic Laboratory | Claudio Tavares Sacchi, Claudia Regina Gonçalves, Erica Valessa Ramos Gomes, Karoline Rodrigues Campos |
| EPI_ISL_693227 | UBS Vila Marchi | Instituto Adolfo Lutz, Interdiciplinary Procedures Center, Strategic Laboratory | Claudio Tavares Sacchi, Claudia Regina Gonçalves, Erica Valessa Ramos Gomes, Karoline Rodrigues Campos |
| EPI_ISL_693236 | Hospital Santa Marcelina Sao Paulo | Instituto Adolfo Lutz, Interdiciplinary Procedures Center, Strategic Laboratory | Claudio Tavares Sacchi, Claudia Regina Gonçalves, Erica Valessa Ramos Gomes, Karoline Rodrigues Campos |
| EPI_ISL_693237 | UPA Santa Isabel | Instituto Adolfo Lutz, Interdiciplinary Procedures Center, Strategic Laboratory | Claudio Tavares Sacchi, Claudia Regina Gonçalves, Erica Valessa Ramos Gomes, Karoline Rodrigues Campos |
| EPI_ISL_693244 | Centro Médico da Polícia Militar do Estado de Sao Paulo | Instituto Adolfo Lutz, Interdiciplinary Procedures Center, Strategic Laboratory | Claudio Tavares Sacchi, Claudia Regina Gonçalves, Erica Valessa Ramos Gomes, Karoline Rodrigues Campos |
| EPI_ISL_693245 | UPA Santa Isabel | Instituto Adolfo Lutz, Interdiciplinary Procedures Center, Strategic Laboratory | Claudio Tavares Sacchi, Claudia Regina Gonçalves, Erica Valessa Ramos Gomes, Karoline Rodrigues Campos |
| EPI_ISL_693248 | Centro Municipal de Epidemiologia e Imunizações | Instituto Adolfo Lutz, Interdiciplinary Procedures Center, Strategic Laboratory | Claudio Tavares Sacchi, Claudia Regina Gonçalves, Erica Valessa Ramos Gomes, Karoline Rodrigues Campos |
| EPI_ISL_693525, EPI_ISL_693530, EPI_ISL_693531, EPI_ISL_693533, EPI_ISL_693538, EPI_ISL_693542, EPI_ISL_693589, EPI_ISL_693590, EPI_ISL_693591, EPI_ISL_693592, EPI_ISL_693593, EPI_ISL_693594, EPI_ISL_693595, EPI_ISL_693596, EPI_ISL_693597, EPI_ISL_693598, EPI_ISL_693599, EPI_ISL_693600, EPI_ISL_693601, EPI_ISL_693602, EPI_ISL_693603, EPI_ISL_693604, EPI_ISL_693605, EPI_ISL_693606, EPI_ISL_693607, EPI_ISL_693608, EPI_ISL_693609, EPI_ISL_693610, EPI_ISL_693611, EPI_ISL_693612, EPI_ISL_693613, EPI_ISL_693614, EPI_ISL_693615, EPI_ISL_693616, EPI_ISL_693617, EPI_ISL_693618, EPI_ISL_693619, EPI_ISL_693620, EPI_ISL_693621, EPI_ISL_693622, EPI_ISL_693623, EPI_ISL_693624, EPI_ISL_693625 | | | |
| see above | Instituto Nacional de Saude (INSA) | Instituto Nacional de Saude (INSA) | Borges et al |
| EPI_ISL_693701, EPI_ISL_693707, EPI_ISL_693742 | Delaware Public Health Laboratory | Delaware Public Health Laboratory | Gregory Hovan |
| EPI_ISL_695048, EPI_ISL_695049, EPI_ISL_695050, EPI_ISL_695051, EPI_ISL_695052, EPI_ISL_695053, EPI_ISL_695054, EPI_ISL_695055, EPI_ISL_695056, EPI_ISL_695057, EPI_ISL_695058, EPI_ISL_695059, EPI_ISL_695060, EPI_ISL_695061, EPI_ISL_695062, EPI_ISL_695063, EPI_ISL_695064, EPI_ISL_695065, EPI_ISL_695066, EPI_ISL_695067, EPI_ISL_695068, EPI_ISL_695069, EPI_ISL_695070, EPI_ISL_695071, EPI_ISL_695072, EPI_ISL_695073, EPI_ISL_695074, EPI_ISL_695075, EPI_ISL_695076, EPI_ISL_695077, EPI_ISL_695078, EPI_ISL_695079, EPI_ISL_695080, EPI_ISL_695081, EPI_ISL_695082, EPI_ISL_695083, EPI_ISL_695084, EPI_ISL_695085, EPI_ISL_695086, EPI_ISL_695087, EPI_ISL_695088, EPI_ISL_695089, EPI_ISL_695090, EPI_ISL_695091, EPI_ISL_695092, EPI_ISL_695093, EPI_ISL_695094, EPI_ISL_695095, EPI_ISL_695096, EPI_ISL_695097, EPI_ISL_695098, EPI_ISL_695099, EPI_ISL_695100, EPI_ISL_695101, EPI_ISL_695102, EPI_ISL_695229, EPI_ISL_695230, EPI_ISL_695251, EPI_ISL_695252, EPI_ISL_695253, EPI_ISL_695254, EPI_ISL_695255, EPI_ISL_695256, EPI_ISL_695257, EPI_ISL_695258, EPI_ISL_695264 | | | |
| see above | TGen North | TGen North | Jolene Bowers, Megan Folkerts, Chris French, Hayley Yaglom, Ashlyn Pfeiffer, Darrin Lemmer, Dave Engelthaler, The Arizona COVID Genomics Union (ACGU) |
| EPI_ISL_695355, EPI_ISL_695356, EPI_ISL_695364, EPI_ISL_695365, EPI_ISL_695366, EPI_ISL_695367, EPI_ISL_695368, EPI_ISL_695369, EPI_ISL_695370, EPI_ISL_695371, EPI_ISL_695393, EPI_ISL_695394, EPI_ISL_695395, EPI_ISL_695396, EPI_ISL_695397, EPI_ISL_695398, EPI_ISL_695399, EPI_ISL_695400, EPI_ISL_695401, EPI_ISL_695402, EPI_ISL_695403, EPI_ISL_695404, EPI_ISL_695405, EPI_ISL_695406, EPI_ISL_695407, EPI_ISL_695408, EPI_ISL_695409, EPI_ISL_695410, EPI_ISL_695411, EPI_ISL_695412, EPI_ISL_695413, EPI_ISL_695414, EPI_ISL_695415, EPI_ISL_695416, EPI_ISL_695417, EPI_ISL_695418, EPI_ISL_695419, EPI_ISL_695420, EPI_ISL_695421, EPI_ISL_695422, EPI_ISL_695423, EPI_ISL_695424, EPI_ISL_695425, EPI_ISL_695426, EPI_ISL_695427, EPI_ISL_695428, EPI_ISL_695429, EPI_ISL_695430, EPI_ISL_695431, EPI_ISL_695432, EPI_ISL_695433, EPI_ISL_695434, EPI_ISL_695435, EPI_ISL_695436, EPI_ISL_695437, EPI_ISL_695438, EPI_ISL_695439, EPI_ISL_695440, EPI_ISL_695441, EPI_ISL_695442, EPI_ISL_695443, EPI_ISL_695444, EPI_ISL_695445, EPI_ISL_695446, EPI_ISL_695447, EPI_ISL_695448, EPI_ISL_695449, EPI_ISL_695450, EPI_ISL_695451, EPI_ISL_695452, EPI_ISL_695453, EPI_ISL_695454, EPI_ISL_695455, EPI_ISL_695456, EPI_ISL_695457, EPI_ISL_695458, EPI_ISL_695459, EPI_ISL_695460, EPI_ISL_695461, EPI_ISL_695462, EPI_ISL_695463, EPI_ISL_695464, EPI_ISL_695465, EPI_ISL_695466, EPI_ISL_695467, EPI_ISL_695468, EPI_ISL_695469, EPI_ISL_695470, EPI_ISL_695471, EPI_ISL_695472, EPI_ISL_695473, EPI_ISL_695474, EPI_ISL_695475, EPI_ISL_695476, EPI_ISL_695477, EPI_ISL_695478, EPI_ISL_695479, EPI_ISL_695480, EPI_ISL_695481, EPI_ISL_695482, EPI_ISL_695483, EPI_ISL_695484, EPI_ISL_695485, EPI_ISL_695486, EPI_ISL_695487, EPI_ISL_695488, EPI_ISL_695489, EPI_ISL_695490, EPI_ISL_695491, EPI_ISL_695492, EPI_ISL_695493, EPI_ISL_695494, EPI_ISL_695495, EPI_ISL_695496, EPI_ISL_695497, EPI_ISL_695498, EPI_ISL_695499, EPI_ISL_695500, EPI_ISL_695501, EPI_ISL_695502, EPI_ISL_695503, EPI_ISL_695504, EPI_ISL_695505, EPI_ISL_695506, EPI_ISL_695507, EPI_ISL_695508, EPI_ISL_695509, EPI_ISL_695510, EPI_ISL_695511, EPI_ISL_695512, EPI_ISL_695513, EPI_ISL_695514, EPI_ISL_695515, EPI_ISL_695516, EPI_ISL_695517, EPI_ISL_695518, EPI_ISL_695519, EPI_ISL_695520, EPI_ISL_695563, EPI_ISL_695564, EPI_ISL_695565, EPI_ISL_695566, EPI_ISL_695567, EPI_ISL_695568, EPI_ISL_695569, EPI_ISL_695570, EPI_ISL_695571, EPI_ISL_695572, EPI_ISL_695573, EPI_ISL_695574, EPI_ISL_695575, EPI_ISL_695576, EPI_ISL_695577, EPI_ISL_695578, EPI_ISL_695579, EPI_ISL_695580, EPI_ISL_695581, EPI_ISL_695582, EPI_ISL_695583, EPI_ISL_695584, EPI_ISL_695585, EPI_ISL_695586, EPI_ISL_695587, EPI_ISL_695588, EPI_ISL_695589, EPI_ISL_695590, EPI_ISL_695591, EPI_ISL_695592 | | | |
| see above | AZ SPHL, Arizona Department of Health Services | TGen North | Jolene Bowers, Megan Folkerts, Chris French, Hayley Yaglom, Ashlyn Pfeiffer, Darrin Lemmer, Dave Engelthaler, The Arizona COVID Genomics Union (ACGU) |
| EPI_ISL_699878, EPI_ISL_699879, EPI_ISL_699880, EPI_ISL_699881, EPI_ISL_699882, EPI_ISL_699883, EPI_ISL_699884, EPI_ISL_699885, EPI_ISL_699886, EPI_ISL_699887, EPI_ISL_699888, EPI_ISL_699889, EPI_ISL_699890, EPI_ISL_699891, EPI_ISL_699892, EPI_ISL_699893, EPI_ISL_699894, EPI_ISL_699895, EPI_ISL_699896, EPI_ISL_699897, EPI_ISL_699898, EPI_ISL_699899, EPI_ISL_699900, EPI_ISL_699901, EPI_ISL_699902, EPI_ISL_699903, EPI_ISL_699904, EPI_ISL_699905, EPI_ISL_699906, EPI_ISL_699907, EPI_ISL_699908, EPI_ISL_699909, EPI_ISL_699910, EPI_ISL_699911, EPI_ISL_699912, EPI_ISL_699913, EPI_ISL_699914, EPI_ISL_699915, EPI_ISL_699916, EPI_ISL_699917, EPI_ISL_699918, EPI_ISL_699919, EPI_ISL_699920, EPI_ISL_699921, EPI_ISL_699922, EPI_ISL_699923, EPI_ISL_699924, EPI_ISL_699925, EPI_ISL_699926, EPI_ISL_699927, EPI_ISL_699928, EPI_ISL_699929 | | | |
| see above | Hematopathology Laboratory, ACTREC, TMC | Hematopathology Laboratory, ACTREC, TMC | Hematopathology Laboratory, ACTREC |
| EPI_ISL_707776, EPI_ISL_707777 | Rwanda National Reference Laboratory | Rwanda National Reference Laboratory | Enatha Mukantwari,Jeanne d'Arc Umuringa |
| EPI_ISL_707907, EPI_ISL_707908, EPI_ISL_707909, EPI_ISL_707910, EPI_ISL_707911, EPI_ISL_707912, EPI_ISL_707913, EPI_ISL_707914, EPI_ISL_707915, EPI_ISL_707916, EPI_ISL_707917, EPI_ISL_707918, EPI_ISL_707919, EPI_ISL_707920, EPI_ISL_707921, EPI_ISL_707922, EPI_ISL_707923, EPI_ISL_707924, EPI_ISL_707925, EPI_ISL_707926, EPI_ISL_707931, EPI_ISL_707932 | | | |
| see above | Los Angeles County Public Health Laboratory | Los Angeles County Public Health Laboratory | P. Hemarajata et al. |
| EPI_ISL_710279 | Colorado Department of Public Health and Environment | Colorado Department of Puplic Health and Environment | Laura Bankers, Molly C. Hetherington-Rauth, Shannon Ely, Shannon R. Matzinger, Sarah Elizabeth Totten, Emily A. Travanty |
| EPI_ISL_710420 | Los Angeles County PHL | Los Angeles County PHL | P. Hemarajata et al. |
| EPI_ISL_714364 | Department of Virus and Microbiological Special Diagnostics, Statens Serum Institut, Copenhagen, Denmark | Albertsen Lab, Department of Chemistry and Bioscience, Aalborg University, Denmark | Danish Covid-19 Genome Consortium |
| EPI_ISL_717606 | Lab voor klinische biologie | Onderzoeksgroep Virologie | Laurens Lambrechts, Nick Vereecke, Marthe Pauwels, Bruno Verhasselt, Linos Vandekerckhove, Hans Nauwynck, Sebastiaan Theuns |
| EPI_ISL_717766 | UW Virology Lab | UW Virology Lab | Pavitra Roychoudhury, Hong Xie, Lasata Shrestha, Michelle Lin, Meei-Li Huang, Keith R Jerome, Alexander Greninger |
| EPI_ISL_717853, EPI_ISL_717854, EPI_ISL_717855, EPI_ISL_717856, EPI_ISL_717857, EPI_ISL_717858, EPI_ISL_717859, EPI_ISL_717860, EPI_ISL_717861, EPI_ISL_717862 | Laboratorio de Virologia Molecular / UFRJ | Bioinformatics Laboratory / LNCC | Carolina M Voloch, Ronaldo da Silva F Jr, Luiz G P de Almeida, Cynthia C Cardoso, Otavio Bustrolini, Alexandra L Gerber, Ana Paula de C Guimarães, Diana Mariani, Andréa Cony Cavalcanti, Claudia dos Santos Rodrigues, Terezinha M P P Castiñeira, Amilcar Tanuri, Ana Tereza R de Vasconcelos |

| | | | |
|---|---|---|---|
| EPI_ISL_718143 | Ministry of Health Hospitals | Institute of Health and Community Medicine | David Perera, Ooi Mong How, Chua Hock Hin, Tonnii Sia Loong Loong, Wong Jyn Shan, Wong Kiing Aik, Chan Chia Jui |
| EPI_ISL_721646, EPI_ISL_721664, EPI_ISL_721665, EPI_ISL_722178 | National Centre For Cell Science | National Centre For Cell Science | Dhiraj Paul, Kunal Jani, Radha Chauhan, Janesh Kumar, Vasudevan Seshadri, Girdhari Lal, Rajesh Karyakarte, Suvarna Joshi, Murlidhar Tambe, Sourav Sen, Santosh Karade, Kavita Bala Anand, Shelinder Pal Singh Shergill, Rajiv Mohan Gupta, Manoj Kumar Bhat, Arvind Sahu, Yogesh S Shouche |
| EPI_ISL_723048, EPI_ISL_723049 | Hematopathology Laboratory, ACTREC, TMC | Hematopathology Laboratory, ACTREC, TMC | Hematopathology Laboratory, ACTREC |
| EPI_ISL_729563 | A. Krumbholz, Labor Dr. Krause und Kollegen MVZ GmbH, Kiel | Charité Universitätsmedizin Berlin, Institut für Virologie | Victor M Corman, Barbara Mühlemann, Jörn Beheim-Schwarzbach, Talitha Veith, Julia Schneider, Terry Jones, Christian Drosten |
| EPI_ISL_729928, EPI_ISL_729929, EPI_ISL_729930, EPI_ISL_729931, EPI_ISL_729932, EPI_ISL_729933, EPI_ISL_729934, EPI_ISL_729935, EPI_ISL_729955, EPI_ISL_729970, EPI_ISL_729971, EPI_ISL_729972, EPI_ISL_729973, EPI_ISL_729974, EPI_ISL_729989, EPI_ISL_730024, EPI_ISL_730025, EPI_ISL_730026, EPI_ISL_730027, EPI_ISL_730044, EPI_ISL_730045 | | | |
| see above | Nigeria Centre for Disease Control (NCDC) | African Centre of Excellence for Genomics of Infectious Diseases (ACEGID), Redeemer's University, Ede, Osun State, Nigeria | Oluniyi P.E. et al |
| EPI_ISL_730082, EPI_ISL_730083 | Yale Clinical Virology Laboratory | Grubaugh Lab - Yale School of Public Health | Joseph Fauver, Tara Alpert, Anderson Brito, Annie Watkins, Anne Wyllie, Chantal Vogels, Mary Petrone, Chaney Kalinich, Isabel Ott, Arnau Casanovas, Catherine Muenker, Adam Moore, Alice Lu, Maria Tokuyama, Patrick Wong, Peiwen Lu, Saad Omer, Richard Martinello, Allison Nelson, Shelli Farhadian, Akiko Iwasaki, Charlese Dela Cruz, Albert Ko, Nathan Grubaugh |
| EPI_ISL_730126 | San Diego County Public Health Laboratory | Andersen lab at Scripps Research | SEARCH Alliance San Diego with Tracy Basler, Jovan Shephard, Brett Austin |
| EPI_ISL_730197, EPI_ISL_730198, EPI_ISL_730199, EPI_ISL_730200, EPI_ISL_730201, EPI_ISL_730202, EPI_ISL_730203, EPI_ISL_730204, EPI_ISL_730205 | Genomica Lab Molecular, M©xico | Andersen lab at Scripps Research | SEARCH Alliance San Diego with Jonathan Gonzalez Garcia, Jose Roman Chavez Mendez, Jose Horacio Reyna Verdugo, Martin Gonzalez Ibarra, Luis Alberto Rangel Gonzalez |
| EPI_ISL_730330, EPI_ISL_730343 | San Diego County Public Health Laboratory | Andersen lab at Scripps Research | SEARCH Alliance San Diego with Tracy Basler, Jovan Shephard, Brett Austin |
| EPI_ISL_732534 | Bundeswehr Institute of Microbiology | Bundeswehr Institute of Microbiology | Elham Khatamzas, Markus Antwerpen, Mathias Walter, Alexandra Rehn, Sabine Zange, Enrico Georgi, Michael von Bergwelt-Baildon, Roman Wölfel |
| EPI_ISL_732774 | Centro de Investigación Biomédica de La Rioja - Hospital San Pedro Logroño | SeqCOVID-SPAIN consortium/IBV(CSIC) | María de Toro, José Manuel Azcona Gutiérrez, María Pilar Bea Escudero, Miriam Blasco Alberdi and SeqCOVID-SPAIN consortium |
| EPI_ISL_732990, EPI_ISL_732991, EPI_ISL_732992 | UMMC-Health | WHO National Influenza Centre Russian Federation | Andrey Komissarov, Artem Fadeev, Anna Ivanova, Kseniya Komissarova, Dmitry Bazhenov, Tatiana Platonova, Daria Danilenko, Ksenia Safina, Elena Nabieva, Georgii Bazykin, Dmitry Lioznov |
| EPI_ISL_734788, EPI_ISL_734789, EPI_ISL_734790, EPI_ISL_734791, EPI_ISL_734792, EPI_ISL_734793, EPI_ISL_734794, EPI_ISL_734795, EPI_ISL_735247 | UZ Leuven, National Reference Laboratory for Coronaviruses, Laboratory Medicine, Leuven, Belgium | KU Leuven, Rega Institute, Clinical and Epidemiological Virology | Tony Wawina-Bokalanga, Joan Marti-Carerras, Bert Vanmechelen, Piet Maes |
| EPI_ISL_735374, EPI_ISL_735375, EPI_ISL_735376, EPI_ISL_735377 | Los Angeles County Public Health Laboratory | Los Angeles County Public Health Laboratory | P. Hemarajata et al. |
| EPI_ISL_735396 | Hospital de Camplanha COVID 19 SER | Instituto Adolfo Lutz, Interdiciplinary Procedures Center, Strategic Laboratory | Claudio Tavares Sacchi, Claudia Regina Gonçalves, Erica Valessa Ramos Gomes, Karoline Rodrigues Campos |
| EPI_ISL_735397 | Unidade Respiratória Nova Hortolandia | Instituto Adolfo Lutz, Interdiciplinary Procedures Center, Strategic Laboratory | Claudio Tavares Sacchi, Claudia Regina Gonçalves, Erica Valessa Ramos Gomes, Karoline Rodrigues Campos |
| EPI_ISL_735398 | Laboratorio Fleury | Instituto Adolfo Lutz, Interdiciplinary Procedures Center, Strategic Laboratory | Claudio Tavares Sacchi, Claudia Regina Gonçalves, Erica Valessa Ramos Gomes, Karoline Rodrigues Campos |
| EPI_ISL_735400 | Instituto Adolfo Lutz - Regional de Santos | Instituto Adolfo Lutz, Interdiciplinary Procedures Center, Strategic Laboratory | Claudio Tavares Sacchi, Claudia Regina Gonçalves, Erica Valessa Ramos Gomes, Karoline Rodrigues Campos |
| EPI_ISL_735401, EPI_ISL_735402, EPI_ISL_735403, EPI_ISL_735404 | Instituto Adolfo Lutz - Regional de Rio Claro | Instituto Adolfo Lutz, Interdiciplinary Procedures Center, Strategic Laboratory | Claudio Tavares Sacchi, Claudia Regina Gonçalves, Erica Valessa Ramos Gomes, Karoline Rodrigues Campos |
| EPI_ISL_735406 | Unidade de Pronto Atendimento UPA I Sta Isabel | Instituto Adolfo Lutz, Interdiciplinary Procedures Center, Strategic Laboratory | Claudio Tavares Sacchi, Claudia Regina Gonçalves, Erica Valessa Ramos Gomes, Karoline Rodrigues Campos |
| EPI_ISL_735410 | Instituto Adolfo Lutz - Regional de Rio Claro | Instituto Adolfo Lutz, Interdiciplinary Procedures Center, Strategic Laboratory | Claudio Tavares Sacchi, Claudia Regina Gonçalves, Erica Valessa Ramos Gomes, Karoline Rodrigues Campos |
| EPI_ISL_735411 | Centro de Vigilancia a Saude de Diadema | Instituto Adolfo Lutz, Interdiciplinary Procedures Center, Strategic Laboratory | Claudio Tavares Sacchi, Claudia Regina Gonçalves, Erica Valessa Ramos Gomes, Karoline Rodrigues Campos |
| EPI_ISL_735419 | UBS Alvarenga | Instituto Adolfo Lutz, Interdiciplinary Procedures Center, Strategic Laboratory | Claudio Tavares Sacchi, Claudia Regina Gonçalves, Erica Valessa Ramos Gomes, Karoline Rodrigues Campos |
| EPI_ISL_735420 | UBS Riacho Grande | Instituto Adolfo Lutz, Interdiciplinary Procedures Center, Strategic Laboratory | Claudio Tavares Sacchi, Claudia Regina Gonçalves, Erica Valessa Ramos Gomes, Karoline Rodrigues Campos |
| EPI_ISL_735421 | UBS Sta Terezinha | Instituto Adolfo Lutz, Interdiciplinary Procedures Center, Strategic Laboratory | Claudio Tavares Sacchi, Claudia Regina Gonçalves, Erica Valessa Ramos Gomes, Karoline Rodrigues Campos |
| EPI_ISL_735423, EPI_ISL_735424, EPI_ISL_735426 | Centro de Vigilancia a Saude de Diadema | Instituto Adolfo Lutz, Interdiciplinary Procedures Center, Strategic Laboratory | Claudio Tavares Sacchi, Claudia Regina Gonçalves, Erica Valessa Ramos Gomes, Karoline Rodrigues Campos |
| EPI_ISL_735427 | Instituto Adolfo Lutz - Regional de Santos | Instituto Adolfo Lutz, Interdiciplinary Procedures Center, Strategic Laboratory | Claudio Tavares Sacchi, Claudia Regina Gonçalves, Erica Valessa Ramos Gomes, Karoline Rodrigues Campos |
| EPI_ISL_735428, EPI_ISL_735429 | Hospital Nipo Brasileiro | Instituto Adolfo Lutz, Interdiciplinary Procedures Center, Strategic Laboratory | Claudio Tavares Sacchi, Claudia Regina Gonçalves, Erica Valessa Ramos Gomes, Karoline Rodrigues Campos |
| EPI_ISL_735430 | Instituto Adolfo Lutz - Regional de Santos | Instituto Adolfo Lutz, Interdiciplinary Procedures Center, Strategic Laboratory | Claudio Tavares Sacchi, Claudia Regina Gonçalves, Erica Valessa Ramos Gomes, Karoline Rodrigues Campos |
| EPI_ISL_735431, EPI_ISL_735432 | Hospital Nipo Brasileiro | Instituto Adolfo Lutz, Interdiciplinary Procedures Center, Strategic Laboratory | Claudio Tavares Sacchi, Claudia Regina Gonçalves, Erica Valessa Ramos Gomes, Karoline Rodrigues Campos |
| EPI_ISL_735433 | Posto de Atendimento Saude Cidade Pasc Cajati | Instituto Adolfo Lutz, Interdiciplinary Procedures Center, Strategic Laboratory | Claudio Tavares Sacchi, Claudia Regina Gonçalves, Erica Valessa Ramos Gomes, Karoline Rodrigues Campos |
| EPI_ISL_738009, EPI_ISL_738010, EPI_ISL_738011, EPI_ISL_738012 | Uganda Central Public Health Lab and Uganda Virus Research Institute | MRC/UVRI & LSHTM Uganda Research Unit | Matthew Cotten, Dan Lule Bugembe, My V.T. Phan, Pontiano Kaleebu et al. |
| EPI_ISL_738081, EPI_ISL_738082, EPI_ISL_738083 | Biology, College of Education | Biology, College of Education | Niranji,S.S., Al-Jaf,S.M., Mahmood,Z.H. |

| | | | |
|---|---|---|---|
| EPI_ISL_738221 | UZ Leuven, National Reference Laboratory for Coronaviruses, Laboratory Medicine, Leuven, Belgium | KU Leuven, Rega Institute, Clinical and Epidemiological Virology | Tony Wawina-Bokalanga, Joan Marti-Carerras, Bert Vanmechelen, Piet Maes |
| EPI_ISL_738565, EPI_ISL_738577, EPI_ISL_738584, EPI_ISL_738614, EPI_ISL_738648, EPI_ISL_738656, EPI_ISL_738659, EPI_ISL_738676, EPI_ISL_738677, EPI_ISL_738687, EPI_ISL_738715, EPI_ISL_738781, EPI_ISL_738847, EPI_ISL_738881, EPI_ISL_738891, EPI_ISL_738896, EPI_ISL_738992, EPI_ISL_739047, EPI_ISL_739077, EPI_ISL_739212, EPI_ISL_739233, EPI_ISL_739241, EPI_ISL_739243, EPI_ISL_739323, EPI_ISL_739324, EPI_ISL_739340, EPI_ISL_739459, EPI_ISL_739578, EPI_ISL_739600, EPI_ISL_739638, EPI_ISL_739648 | | | |
| see above | Alameda County Public Health Lab | Chan-Zuckerberg Biohub | CZB Cliahub Consortium |
| EPI_ISL_739753, EPI_ISL_739918, EPI_ISL_739977, EPI_ISL_740030, EPI_ISL_740062, EPI_ISL_740103, EPI_ISL_740343, EPI_ISL_740404, EPI_ISL_740469 | Laboratoire national de santé, Microbiology, Virology | Laboratoire national de santé, Microbiology, Microbial Genomics Platform | Anke Wienecke-Baldacchino, Catherine Ragimbeau, Tamir Abdelrahman, Jessica Tapp, Fatu Djabi |
| EPI_ISL_742530 | Wales Specialist Virology Centre Sequencing lab: Pathogen Genomics Unit | COVID-19 Genomics UK (COG-UK) Consortium | Catherine Moore, Johnathan Evans, Laura Gifford, Malorie Perry, Simon Cottrell, Angela Marchbank, Alec Birchley, Alexander Adams, Amy Gaskin, Bree Gatica-Wilcox, Jason Coombes, Joel Southgate, Lauren Gilbert, Lee Graham, Nicole Pacchiarini, Sara Kumziene-Summerhayes, Sarah Taylor, Sophie Jones, Sara Rey, Matthew Bull, Joanne Watkins, Sally Corden, Tom Connor |
| EPI_ISL_744249, EPI_ISL_744500, EPI_ISL_744586, EPI_ISL_744681, EPI_ISL_744968 | Laboratoire national de santé, Microbiology, Virology | Laboratoire national de santé, Microbiology, Microbial Genomics Platform | Anke Wienecke-Baldacchino, Catherine Ragimbeau, Tamir Abdelrahman, Jessica Tapp, Fatu Djabi |
| EPI_ISL_747414, EPI_ISL_747415, EPI_ISL_747416, EPI_ISL_747417, EPI_ISL_747418 | Division of Emerging Infectious Diseases, Bureau of Infectious Diseases Diagnosis Control, Korea Disease Control and Prevention Agency | Division of Emerging Infectious Diseases, Bureau of Infectious Diseases Diagnosis Control, Korea Disease Control and Prevention Agency | Ae Kyung Park, Il-Hwan Kim, Heui Man Kim, Jeong-Min Kim, Namjoo Lee, Chaeyoung Lee, Sang Hee Woo, Eun-Jin Kim |
| EPI_ISL_750176, EPI_ISL_750256, EPI_ISL_750430, EPI_ISL_750820, EPI_ISL_751011 | Sanatorio Americano | Institut Pasteur de Montevideo | Daiana Mir, Natalia Rego, Paola Cristina Resende, Fernando Lopez-Tort, Tamara Fernandez-Calero, Veronica Noya, Mariana Brandes, Tania Possi, Mailen Arleo, Natalia Reyes, Matias Victoria, Andres Lizasoain, Matias Castells, Leticia Maya, Matías Salvo, Tatiana Schäffer Gregianini, Marilda Tereza Mar da Rosa, Letícia Garay Martins, Cecilia Alonso, Yasser Vega, Cecilia Salazar, Ignacio Ferrés, Pablo Smircich, Jose Sotelo, Ighor Arantes, Luciana Appolinario, Ana Carolina Mendonça, Maria Jose Benitez-Galeano, Martín Graña, Camila Simoes, Fernando Motta, Marilda Mendonça Siqueira, Gonzalo Bello, Rodney Colina, Lucia Spangenberg |
| EPI_ISL_751184 | CENUR Litoral Norte - UdelaR, Salto, Uruguay | Institut Pasteur de Montevideo | Daiana Mir, Natalia Rego, Paola Cristina Resende, Fernando Lopez-Tort, Tamara Fernandez-Calero, Veronica Noya, Mariana Brandes, Tania Possi, Mailen Arleo, Natalia Reyes, Matias Victoria, Andres Lizasoain, Matias Castells, Leticia Maya, Matías Salvo, Tatiana Schäffer Gregianini, Marilda Tereza Mar da Rosa, Letícia Garay Martins, Cecilia Alonso, Yasser Vega, Cecilia Salazar, Ignacio Ferrés, Pablo Smircich, Jose Sotelo, Ighor Arantes, Luciana Appolinario, Ana Carolina Mendonça, Maria Jose Benitez-Galeano, Martín Graña, Camila Simoes, Fernando Motta, Marilda Mendonça Siqueira, Gonzalo Bello, Rodney Colina, Lucia Spangenberg |
| EPI_ISL_751504, EPI_ISL_751505, EPI_ISL_751506, EPI_ISL_751507, EPI_ISL_751508, EPI_ISL_751511, EPI_ISL_751512, EPI_ISL_751513, EPI_ISL_751514, EPI_ISL_751515, EPI_ISL_751516, EPI_ISL_751517, EPI_ISL_751518, EPI_ISL_751520, EPI_ISL_751521, EPI_ISL_751522, EPI_ISL_751523, EPI_ISL_751525, EPI_ISL_751528, EPI_ISL_751530, EPI_ISL_751531, EPI_ISL_751532, EPI_ISL_751535, EPI_ISL_751537, EPI_ISL_751538, EPI_ISL_751540 | | | |
| see above | Texas Department of State Health Services | Texas Department of State Health Services | Rashmi Tuladhar, Bonnie Oh, Jenny Zhang, Maliha Rahman, Anita Pokharel, Myong Koag, Chung Wang, Rachel Lee, Grace Kubin, Mayela Pedrueza, James Daniel Bonser |
| EPI_ISL_752692, EPI_ISL_752693, EPI_ISL_752694, EPI_ISL_752695, EPI_ISL_752696, EPI_ISL_752697, EPI_ISL_752698, EPI_ISL_752699, EPI_ISL_752708, EPI_ISL_752716, EPI_ISL_752844, EPI_ISL_752950, EPI_ISL_752951 | | | |
| see above | State Laboratories Division, Hawaii State Department of Health | State Laboratories Division, Hawaii State Department of Health | Pamela O'Brien, Sabrina Diemert, Drew Kuwazaki, Razvan Sultana, Edward Desmond |
| EPI_ISL_753809, EPI_ISL_753987, EPI_ISL_753990 | Charité Universitätsmedizin Berlin, Institut für Virologie/Labor Berlin | Charité Universitätsmedizin Berlin, Institut für Virologie | Victor M Corman, Jörn Beheim-Schwarzbach, Barbara Mühlemann, Julia Schneider, Talitha Veith, Terry Jones, Christian Drosten |
| EPI_ISL_754230 | The Republican Research and Practical Center for Epidemiology and Microbiology (RRPCEM) | WHO National Influenza Centre Russian Federation | Elena Gasich, Kirill Bulda, Anatoly Krasko, Andrey Komissarov, Artem Fadeev, Anna Ivanova, Kseniya Komissarova, Dmitry Bazhenov, Daria Danilenko, Ksenia Safina, Elena Nabieva, Georgii Bazykin, Dmitry Lioznov |
| EPI_ISL_754864, EPI_ISL_754878, EPI_ISL_754881, EPI_ISL_754885, EPI_ISL_754889, EPI_ISL_754893, EPI_ISL_754894, EPI_ISL_754895, EPI_ISL_754897, EPI_ISL_754899 | Innovative Genomics Institute, UC Berkeley | Innovative Genomics Institute, UC Berkeley | Stacia Wyman, Haridha Shivram, Phil Frankino, Liana Lareau, Shana McDevitt, Justin Choi |
| EPI_ISL_754932, EPI_ISL_754933, EPI_ISL_754956, EPI_ISL_754969, EPI_ISL_754977, EPI_ISL_754992, EPI_ISL_755018, EPI_ISL_755019, EPI_ISL_755042, EPI_ISL_755054, EPI_ISL_755055, EPI_ISL_755056 | | | |
| see above | California Department of Public Health | California Department of Public Health | CDPH IDLB COVIDNet |
| EPI_ISL_755305 | Laboratorio de Virología, Centro de Biotecnología Acuícola, Universidad de Santiato de Chile | Center for Mathematical Modeling and Center for Genome Regulation. Santiago, Chile | A. Sandino, Reyes-López F., Bastias M, Sanhueza D, Travisany D, Allende ML, Maass A, González M, Bustos F, Arriagada G, Montecino, M, Orellana A, Castro E, Meneses C. |
| EPI_ISL_756315, EPI_ISL_756323, EPI_ISL_756339, EPI_ISL_756340, EPI_ISL_756342, EPI_ISL_756343, EPI_ISL_756348, EPI_ISL_756349, EPI_ISL_756350, EPI_ISL_756352, EPI_ISL_756353, EPI_ISL_756354 | | | |
| see above | Innovative Genomics Institute, UC Berkeley | Innovative Genomics Institute, UC Berkeley | Stacia Wyman, Haridha Shivram, Phil Frankino, Liana Lareau, Shana McDevitt, Justin Choi |
| EPI_ISL_760150 | Division of Emerging Infectious Diseases, Bureau of Infectious Diseases Diagnosis Control, Korea Disease Control and Prevention Agency | Division of Emerging Infectious Diseases, Bureau of Infectious Diseases Diagnosis Control, Korea Disease Control and Prevention Agency | Ae Kyung Park, Il-Hwan Kim, Heui Man Kim, Jeong-Min Kim, Namjoo Lee, Chaeyoung Lee, Sang Hee Woo, Eun-Jin Kim |
| EPI_ISL_765221 | Instituto Nacional de Saude (INSA) | Instituto Nacional de Saude (INSA) | Borges et al |
| EPI_ISL_765661, EPI_ISL_765662, EPI_ISL_765663, EPI_ISL_765664, EPI_ISL_765665, EPI_ISL_765666, EPI_ISL_765667, EPI_ISL_765668, EPI_ISL_765669, EPI_ISL_765670, EPI_ISL_765671 | | | |
| see above | Massachusetts General Hospital | Infectious Disease Program, Broad Institute of Harvard and MIT | Lemieux,J.E., Siddle,K.J., Shaw,B., Adams,G., Pierce,V., Turbett,S., Anahtar,M., Branda,J., Slater,D., Harris,J., Lin,A.E., Gladden-Young,A., Lagerborg,K., Rudy,M., DeRuff,K., Carter,A., Normandin,E., Bauer,M., Reilly,S., Tomkins-Tinch,C., Loreth,C., Chaluvadi,S., Neumann,A., Cusick,C., Chapman,S.B., Gnirke,A., Flowers,K., Cerrato,F., Birren,B.W., Gallagher,G., Smole,S., Park,D.J., MacInnis,B.L., Ryan,E., LaRocque,R., Rosenberg,E. and Sabeti,P.C. |
| EPI_ISL_765911, EPI_ISL_765940 | TXDSHS | TXDSHS | Rashmi Tuladhar, Bonnie Oh, Jenny Zhang, Maliha Rahman, Anita Pokharel, Myong Koag, Chung Wang, Rachel Lee, Grace Kubin, Mayela Pedrueza, James Daniel Bonser |
| EPI_ISL_765994, EPI_ISL_765995 | USC Clinical Lab | Los Angeles County PHL | P. Hemarajata et al. |
| EPI_ISL_766015 | USC Clinical Lab | Los Angeles County PHL | P. Hemarajata el al. |
| EPI_ISL_771229 | Colorado Department of Public Health and Environment | Colorado Department of Puplic Health and Environment | Laura Bankers, Molly C. Hetherington-Rauth, Diana Ir, Shannon Ely, Shannon R. Matzinger, Sarah Elizabeth Totten, Emily A. Travanty |
| EPI_ISL_774914, EPI_ISL_774915, EPI_ISL_774916, EPI_ISL_774917, EPI_ISL_774918, EPI_ISL_774919, EPI_ISL_774920, EPI_ISL_774921, EPI_ISL_774922, EPI_ISL_774923, EPI_ISL_774924, EPI_ISL_774925, EPI_ISL_774926, EPI_ISL_774927, EPI_ISL_774928, EPI_ISL_774929, EPI_ISL_774930, EPI_ISL_774931, EPI_ISL_774933, EPI_ISL_774935, EPI_ISL_774937, EPI_ISL_774939, EPI_ISL_774940, EPI_ISL_774942, EPI_ISL_774944, EPI_ISL_774945, EPI_ISL_774946, EPI_ISL_774947, EPI_ISL_774950, EPI_ISL_774961, EPI_ISL_774966, EPI_ISL_774985, EPI_ISL_774992, EPI_ISL_774994, EPI_ISL_774995 | | | |
| see above | Designated Reference Institute for Chemical Measurements (DRICM) | DNA SOLUTION LTD. | Md. Imran Khan, Kazi Nadim Hasan, Abu Sufian, Jannatun Naima, Abdul Khaleque, Mizanur Rahman, MSM Chowdhury, Hasan Ul Haider, Mamudul Hasan Razu, Mala Khan, Mohammad Fazle Alam Rabbi |
| EPI_ISL_776664, EPI_ISL_776672, EPI_ISL_776674, EPI_ISL_776678, EPI_ISL_776679, EPI_ISL_776680, EPI_ISL_776683, EPI_ISL_776685 | UW Virology Lab | UW Virology Lab | Pavitra Roychoudhury, Hong Xie, Lasata Shrestha, Meei-Li Huang, Keith R Jerome, Alexander Greninger |

| | | | |
|---|---|---|---|
| EPI_ISL_776764, EPI_ISL_776765 | Instituto Adolfo Lutz - Regional de Santo Andre | Instituto Adolfo Lutz, Interdiciplinary Procedures Center, Strategic Laboratory | Claudio Tavares Sacchi, Claudia Regina Gonçalves, Erica Valessa Ramos Gomes, Karoline Rodrigues Campos |
| EPI_ISL_776871 | Hospital General Universitario Gregorio Marañón | Hospital General Universitario Gregorio Marañón | Sergio Buenestado Serrano, Pedro Sola Campoy, Laura Perez-Lago, Pilar Catalán, Patricia Muñoz, Darío García de Viedma |
| EPI_ISL_778769, EPI_ISL_778773, EPI_ISL_778774, EPI_ISL_778775, EPI_ISL_778776, EPI_ISL_778777, EPI_ISL_778778, EPI_ISL_778780, EPI_ISL_778781, EPI_ISL_778782 | Istituto Zooprofilattico Sperimentale del Mezzogiorno | TIGEM | Patrizia Annunziata, Andrea Ballabio, Valentina Bouche, Davide Cacchiarelli (CorrespAuthor), Pellegrino Cerino, Chiara Colantuono, Lucio Di Filippo, Antonio Grimaldi, Antonio Limone, Gabriella Loconte, Anna Manfredi, Francesco Panariello, Biancamaria Pierri, Marcello Salvi, Lucia Vassallo |
| EPI_ISL_779182, EPI_ISL_779183, EPI_ISL_779184 | Laboratorio de Infectología, Servicio de Infectología, Hospital Universitario Dr. José Eleuterio González - Universidad Autónoma de Nuevo León | Laboratorio de Infectología Molecular, Departamento de Bioquímica y Medicina Molecular,Facultad de Medicina - Universidad Autónoma de Nuevo León | Kame A. Galán-Huerta, María F. Herrera-Saldivar, Natalia Martínez-Acuña, Sonia A. Lozano-Sepúlveda, Daniel Arellanos-Soto, Ana M. Rivas-Estilla, Paola Bocanegra-Ibarias, Samantha M. Flores-Treviño, Elvira Garza-González, Eduardo Perez-Alba, Laura Nuzzolo-Shihadeh, Adrian Camacho-Ortiz |
| EPI_ISL_779272, EPI_ISL_779274, EPI_ISL_779276, EPI_ISL_779289, EPI_ISL_779290 | Jamil-ur-Rahman Center for Genome Research, Dr. Panjwani Center for Molecular Medicine and Drug Research | Jamil-ur-Rahman Center for Genome Research, Dr. Panjwani Center for Molecular Medicine and Drug Research | Shakeel,M., Irfan,M., Nisa,Z., Rashid,M., Ansari,S., Khan,I. |
| EPI_ISL_779711 | The University Hospital Brno | Institute of Applied Biotechnologies a.s. | Petr Klempt, Ondej Brzo, Martin Kašný, Petr Kvapil |
| EPI_ISL_779930, EPI_ISL_779936, EPI_ISL_779938 | Center of Medical Microbiology, Virology, and Hospital Hygiene, University of Duesseldorf | Center of Medical Microbiology, Virology, and Hospital Hygiene, University of Duesseldorf | Maximilian Damagnez, Alexander Dilthey, Ashley-Jane Duplessis, Torsten Houwaart, Lisanna Hülse, Malte Kohns Vasconcelos, Nadine Lübke, Jessica Nicolai, Klaus Pfeffer, Daniel Strelow, Teresa Tamayo, Jörg Timm, Andreas Walker, Tobias Wienemann |
| EPI_ISL_780097, EPI_ISL_780099, EPI_ISL_780100, EPI_ISL_780101, EPI_ISL_780102, EPI_ISL_780103, EPI_ISL_780104, EPI_ISL_780105, EPI_ISL_780106, EPI_ISL_780107, EPI_ISL_780108, EPI_ISL_780109, EPI_ISL_780110, EPI_ISL_780111, EPI_ISL_780112, EPI_ISL_780113, EPI_ISL_780115, EPI_ISL_780116, EPI_ISL_780117, EPI_ISL_780118, EPI_ISL_780119, EPI_ISL_780120, EPI_ISL_780121, EPI_ISL_780122, EPI_ISL_780123, EPI_ISL_780124, EPI_ISL_780126, EPI_ISL_780127, EPI_ISL_780128, EPI_ISL_780129, EPI_ISL_780130, EPI_ISL_780131, EPI_ISL_780132, EPI_ISL_780133, EPI_ISL_780134, EPI_ISL_780135, EPI_ISL_780136, EPI_ISL_780137, EPI_ISL_780138, EPI_ISL_780139, EPI_ISL_780140, EPI_ISL_780141, EPI_ISL_780142, EPI_ISL_780143, EPI_ISL_780144, EPI_ISL_780145, EPI_ISL_780146, EPI_ISL_780147, EPI_ISL_780148, EPI_ISL_780149, EPI_ISL_780150, EPI_ISL_780151, EPI_ISL_780152, EPI_ISL_780153, EPI_ISL_780154, EPI_ISL_780155, EPI_ISL_780156, EPI_ISL_780157, EPI_ISL_780158, EPI_ISL_780159, EPI_ISL_780160, EPI_ISL_780161, EPI_ISL_780163, EPI_ISL_780164, EPI_ISL_780165, EPI_ISL_780166, EPI_ISL_780167, EPI_ISL_780168, EPI_ISL_780169, EPI_ISL_780170, EPI_ISL_780171, EPI_ISL_780172, EPI_ISL_780173, EPI_ISL_780174, EPI_ISL_780175, EPI_ISL_780176, EPI_ISL_780177, EPI_ISL_780179, EPI_ISL_780180, EPI_ISL_780181, EPI_ISL_780182, EPI_ISL_780184, EPI_ISL_780185, EPI_ISL_780186, EPI_ISL_780187, EPI_ISL_780188, EPI_ISL_780189, EPI_ISL_780190, EPI_ISL_780191, EPI_ISL_780192, EPI_ISL_780193, EPI_ISL_780194, EPI_ISL_780196, EPI_ISL_780197, EPI_ISL_780198, EPI_ISL_780199, EPI_ISL_780200, EPI_ISL_780201, EPI_ISL_780202, EPI_ISL_780203, EPI_ISL_780204, EPI_ISL_780205, EPI_ISL_780206, EPI_ISL_780207, EPI_ISL_780208, EPI_ISL_780210, EPI_ISL_780211, EPI_ISL_780212, EPI_ISL_780213, EPI_ISL_780215, EPI_ISL_780216, EPI_ISL_780217, EPI_ISL_780218, EPI_ISL_780219, EPI_ISL_780220, EPI_ISL_780221, EPI_ISL_780222, EPI_ISL_780223, EPI_ISL_780224, EPI_ISL_780225, EPI_ISL_780226, EPI_ISL_780227, EPI_ISL_780228, EPI_ISL_780229, EPI_ISL_780230, EPI_ISL_780231, EPI_ISL_780232, EPI_ISL_780233, EPI_ISL_780234, EPI_ISL_780235, EPI_ISL_780236, EPI_ISL_780237, EPI_ISL_780238, EPI_ISL_780239, EPI_ISL_780240, EPI_ISL_780241, EPI_ISL_780242, EPI_ISL_780243, EPI_ISL_780245, EPI_ISL_780246, EPI_ISL_780247, EPI_ISL_780248, EPI_ISL_780249, EPI_ISL_780251, EPI_ISL_780252, EPI_ISL_780255, EPI_ISL_780256, EPI_ISL_780257, EPI_ISL_780258, EPI_ISL_780259, EPI_ISL_780260, EPI_ISL_780261, EPI_ISL_780262, EPI_ISL_780263, EPI_ISL_780264, EPI_ISL_780265, EPI_ISL_780267, EPI_ISL_780268, EPI_ISL_780269, EPI_ISL_780270, EPI_ISL_780271, EPI_ISL_780272, EPI_ISL_780273, EPI_ISL_780274, EPI_ISL_780276, EPI_ISL_780277, EPI_ISL_780278, EPI_ISL_780279, EPI_ISL_780280, EPI_ISL_780281, EPI_ISL_780282, EPI_ISL_780283, EPI_ISL_780284, EPI_ISL_780285, EPI_ISL_780286, EPI_ISL_780287, EPI_ISL_780288, EPI_ISL_780289, EPI_ISL_780290, EPI_ISL_780291, EPI_ISL_780292, EPI_ISL_780293, EPI_ISL_780294, EPI_ISL_780295, EPI_ISL_780296, EPI_ISL_780297, EPI_ISL_780298, EPI_ISL_780299, EPI_ISL_780300, EPI_ISL_780301, EPI_ISL_780302, EPI_ISL_780303, EPI_ISL_780304, EPI_ISL_780305, EPI_ISL_780306, EPI_ISL_780307, EPI_ISL_780308, EPI_ISL_780309, EPI_ISL_780310, EPI_ISL_780311, EPI_ISL_780312, EPI_ISL_780313, EPI_ISL_780314, EPI_ISL_780315, EPI_ISL_780316, EPI_ISL_780317, EPI_ISL_780318, EPI_ISL_780319, EPI_ISL_780320, EPI_ISL_780321, EPI_ISL_780322, EPI_ISL_780323, EPI_ISL_780324, EPI_ISL_780325, EPI_ISL_780326, EPI_ISL_780327, EPI_ISL_780328, EPI_ISL_780329, EPI_ISL_780330, EPI_ISL_780331, EPI_ISL_780332, EPI_ISL_780333, EPI_ISL_780334, EPI_ISL_780335, EPI_ISL_780336, EPI_ISL_780337, EPI_ISL_780338, EPI_ISL_780339, EPI_ISL_780340, EPI_ISL_780341, EPI_ISL_780342, EPI_ISL_780343, EPI_ISL_780344, EPI_ISL_780345, EPI_ISL_780346, EPI_ISL_780347, EPI_ISL_780348, EPI_ISL_780349, EPI_ISL_780350, EPI_ISL_780351, EPI_ISL_780352, EPI_ISL_780353, EPI_ISL_780354, EPI_ISL_780355, EPI_ISL_780356, EPI_ISL_780357, EPI_ISL_780358, EPI_ISL_780359, EPI_ISL_780360, EPI_ISL_780361, EPI_ISL_780362, EPI_ISL_780363, EPI_ISL_780364, EPI_ISL_780365, EPI_ISL_780366, EPI_ISL_780367, EPI_ISL_780368, EPI_ISL_780369, EPI_ISL_780370, EPI_ISL_780371, EPI_ISL_780372, EPI_ISL_780373, EPI_ISL_780374, EPI_ISL_780427, EPI_ISL_780428, EPI_ISL_780429, EPI_ISL_780430, EPI_ISL_780431, EPI_ISL_780432, EPI_ISL_780433, EPI_ISL_780434, EPI_ISL_780435, EPI_ISL_780436, EPI_ISL_780437, EPI_ISL_780438, EPI_ISL_780439, EPI_ISL_780440, EPI_ISL_780441, EPI_ISL_780442, EPI_ISL_780443, EPI_ISL_780444, EPI_ISL_780445, EPI_ISL_780446, EPI_ISL_780447, EPI_ISL_780448, EPI_ISL_780449, EPI_ISL_780450, EPI_ISL_780451, EPI_ISL_780452, EPI_ISL_780453, EPI_ISL_780454, EPI_ISL_780455, EPI_ISL_780456, EPI_ISL_780457, EPI_ISL_780458, EPI_ISL_780459, EPI_ISL_780460, EPI_ISL_780461, EPI_ISL_780462, EPI_ISL_780463, EPI_ISL_780464, EPI_ISL_780465, EPI_ISL_780466, EPI_ISL_780467, EPI_ISL_780468, EPI_ISL_780469, EPI_ISL_780470, EPI_ISL_780471, EPI_ISL_780472, EPI_ISL_780473, EPI_ISL_780474, EPI_ISL_780475, EPI_ISL_780476, EPI_ISL_780477, EPI_ISL_780478, EPI_ISL_780479, EPI_ISL_780480, EPI_ISL_780481, EPI_ISL_780482, EPI_ISL_780483, EPI_ISL_780484, EPI_ISL_780485, EPI_ISL_780487, EPI_ISL_780488, EPI_ISL_780489, EPI_ISL_780490, EPI_ISL_780491, EPI_ISL_780492, EPI_ISL_780493, EPI_ISL_780494, EPI_ISL_780495, EPI_ISL_780496, EPI_ISL_780497, EPI_ISL_780498, EPI_ISL_780499, EPI_ISL_780500, EPI_ISL_780501, EPI_ISL_780502, EPI_ISL_780503, EPI_ISL_780504, EPI_ISL_780505, EPI_ISL_780506, EPI_ISL_780507, EPI_ISL_780508, EPI_ISL_780509, EPI_ISL_780510, EPI_ISL_780511, EPI_ISL_780512, EPI_ISL_780513, EPI_ISL_780514, EPI_ISL_780515, EPI_ISL_780516, EPI_ISL_780517, EPI_ISL_780518, EPI_ISL_780519, EPI_ISL_780520, EPI_ISL_780521, EPI_ISL_780522, EPI_ISL_780523, EPI_ISL_780524, EPI_ISL_780525, EPI_ISL_780526, EPI_ISL_780527, EPI_ISL_780528, EPI_ISL_780529, EPI_ISL_780530, EPI_ISL_780531, EPI_ISL_780532, EPI_ISL_780533, EPI_ISL_780534, EPI_ISL_780535, EPI_ISL_780536, EPI_ISL_780537, EPI_ISL_780538, EPI_ISL_780539, EPI_ISL_780540, EPI_ISL_780541, EPI_ISL_780542, EPI_ISL_780543, EPI_ISL_780544, EPI_ISL_780545, EPI_ISL_780546, EPI_ISL_780547, EPI_ISL_780548, EPI_ISL_780549, EPI_ISL_780550, EPI_ISL_780551, EPI_ISL_780552, EPI_ISL_780553, EPI_ISL_780555, EPI_ISL_780556, EPI_ISL_780557, EPI_ISL_780558, EPI_ISL_780559, EPI_ISL_780560, EPI_ISL_780561, EPI_ISL_780562, EPI_ISL_780563, EPI_ISL_780564, EPI_ISL_780565, EPI_ISL_780566, EPI_ISL_780567, EPI_ISL_780568, EPI_ISL_780569, EPI_ISL_780570, EPI_ISL_780571, EPI_ISL_780572, EPI_ISL_780573, EPI_ISL_780574, EPI_ISL_780575, EPI_ISL_780576, EPI_ISL_780577, EPI_ISL_780578, EPI_ISL_780579, EPI_ISL_780580, EPI_ISL_780581, EPI_ISL_780582, EPI_ISL_780583, EPI_ISL_780584, EPI_ISL_780585, EPI_ISL_780586, EPI_ISL_780587, EPI_ISL_780588, EPI_ISL_780589, EPI_ISL_780590, EPI_ISL_780591, EPI_ISL_780592, EPI_ISL_780593, EPI_ISL_780594, EPI_ISL_780595, EPI_ISL_780596, EPI_ISL_780597, EPI_ISL_780598, EPI_ISL_780599, EPI_ISL_780600, EPI_ISL_780601, EPI_ISL_780602, EPI_ISL_780603, EPI_ISL_780604, EPI_ISL_780605, EPI_ISL_780606, EPI_ISL_780607, EPI_ISL_780608, EPI_ISL_780609, EPI_ISL_780610, EPI_ISL_780611, EPI_ISL_780612, EPI_ISL_780613, EPI_ISL_780614, EPI_ISL_780615, EPI_ISL_780616, EPI_ISL_780617, EPI_ISL_780618, EPI_ISL_780619, EPI_ISL_780620, EPI_ISL_780621, EPI_ISL_780622, EPI_ISL_780623, EPI_ISL_780624, EPI_ISL_780625, EPI_ISL_780626, EPI_ISL_780627, EPI_ISL_780628, EPI_ISL_780629, EPI_ISL_780630, EPI_ISL_780631, EPI_ISL_780632, EPI_ISL_780633, EPI_ISL_780634, EPI_ISL_780635, EPI_ISL_780636, EPI_ISL_780637, EPI_ISL_780638, EPI_ISL_780639, EPI_ISL_780640, EPI_ISL_780641, EPI_ISL_780642, EPI_ISL_780643, EPI_ISL_780644, EPI_ISL_780645, EPI_ISL_780646, EPI_ISL_780647, EPI_ISL_780648, EPI_ISL_780649, EPI_ISL_780650, EPI_ISL_780651, EPI_ISL_780652, EPI_ISL_780653, EPI_ISL_780654, EPI_ISL_780655, EPI_ISL_780656, EPI_ISL_780657, EPI_ISL_780658, EPI_ISL_780659, EPI_ISL_780660, EPI_ISL_780662, EPI_ISL_780663, EPI_ISL_780664, EPI_ISL_780665, EPI_ISL_780666, EPI_ISL_780667, EPI_ISL_780668, EPI_ISL_780669, EPI_ISL_780671, EPI_ISL_780672, EPI_ISL_780673, EPI_ISL_780674, EPI_ISL_780675, EPI_ISL_780676, EPI_ISL_780677, EPI_ISL_780678, EPI_ISL_780679, EPI_ISL_780680, EPI_ISL_780681, EPI_ISL_780682, EPI_ISL_780684, EPI_ISL_780685, EPI_ISL_780686, EPI_ISL_780687, EPI_ISL_780688, EPI_ISL_780689, EPI_ISL_780690, EPI_ISL_780691, EPI_ISL_780692, EPI_ISL_780693, EPI_ISL_780694, EPI_ISL_780695, EPI_ISL_780696, EPI_ISL_780697, EPI_ISL_780698, EPI_ISL_780699, EPI_ISL_780700, EPI_ISL_780701, EPI_ISL_780702, EPI_ISL_780703, EPI_ISL_780704, EPI_ISL_780705, EPI_ISL_780706, EPI_ISL_780707, EPI_ISL_780708, EPI_ISL_780709, EPI_ISL_780710, EPI_ISL_780711, EPI_ISL_780712, EPI_ISL_780713, EPI_ISL_780714, EPI_ISL_780715, EPI_ISL_780716, EPI_ISL_780717, EPI_ISL_780718, EPI_ISL_780719, EPI_ISL_780720, EPI_ISL_780721, EPI_ISL_780722, EPI_ISL_780723, EPI_ISL_780724, EPI_ISL_780725, EPI_ISL_780726, EPI_ISL_780727, EPI_ISL_780728, EPI_ISL_780729, EPI_ISL_780730, EPI_ISL_780731, EPI_ISL_780732, EPI_ISL_780733, EPI_ISL_780734, EPI_ISL_780735, EPI_ISL_780736, EPI_ISL_780737, EPI_ISL_780738, EPI_ISL_780739, EPI_ISL_780740, EPI_ISL_780741, EPI_ISL_780742, EPI_ISL_780743, EPI_ISL_780744, EPI_ISL_780745, EPI_ISL_780746, EPI_ISL_780747, EPI_ISL_780748, EPI_ISL_780749, EPI_ISL_780750, EPI_ISL_780751, EPI_ISL_780752, EPI_ISL_780753, EPI_ISL_780754, EPI_ISL_780755, EPI_ISL_780756, EPI_ISL_780757, EPI_ISL_780758, EPI_ISL_780759, EPI_ISL_780760, EPI_ISL_780761, EPI_ISL_780762, EPI_ISL_780763, EPI_ISL_780764, EPI_ISL_780765, EPI_ISL_780766, EPI_ISL_780767, EPI_ISL_780768, EPI_ISL_780769, EPI_ISL_780770, EPI_ISL_780771, EPI_ISL_780772, EPI_ISL_780773, EPI_ISL_780774, EPI_ISL_780775, EPI_ISL_780776, EPI_ISL_780777, EPI_ISL_780778, EPI_ISL_780779, EPI_ISL_780780, EPI_ISL_780781, EPI_ISL_780782, EPI_ISL_780783, EPI_ISL_780784, EPI_ISL_780785, EPI_ISL_780786, EPI_ISL_780787, EPI_ISL_780788, EPI_ISL_780789, EPI_ISL_780790, EPI_ISL_780791, EPI_ISL_780792, EPI_ISL_780793, EPI_ISL_780794, EPI_ISL_780795, EPI_ISL_780796, EPI_ISL_780797, EPI_ISL_780798, EPI_ISL_780799, EPI_ISL_780800, EPI_ISL_780801, EPI_ISL_780802, EPI_ISL_780803, EPI_ISL_780804, EPI_ISL_780805, EPI_ISL_780806, EPI_ISL_780807, EPI_ISL_780808, EPI_ISL_780809, EPI_ISL_780810, EPI_ISL_780811, EPI_ISL_780812, EPI_ISL_780813, EPI_ISL_780814, EPI_ISL_780815, EPI_ISL_780816, EPI_ISL_780817, EPI_ISL_780818, EPI_ISL_780819, EPI_ISL_780820, EPI_ISL_780821, EPI_ISL_780822, EPI_ISL_780823, EPI_ISL_780824, EPI_ISL_780825, EPI_ISL_780826, EPI_ISL_780827, EPI_ISL_780828, EPI_ISL_780829, EPI_ISL_780830, EPI_ISL_780831, EPI_ISL_780832, EPI_ISL_780833, EPI_ISL_780834, EPI_ISL_780835, EPI_ISL_780836, EPI_ISL_780837, EPI_ISL_780838, EPI_ISL_780839, EPI_ISL_780840, EPI_ISL_780841, EPI_ISL_780842, EPI_ISL_780843, EPI_ISL_780844, EPI_ISL_780845, EPI_ISL_780846, EPI_ISL_780847, EPI_ISL_780848, EPI_ISL_780849, EPI_ISL_780850, EPI_ISL_780851, EPI_ISL_780852, EPI_ISL_780853, EPI_ISL_780854, EPI_ISL_780855, EPI_ISL_780856, EPI_ISL_780857, EPI_ISL_780858, EPI_ISL_780859, EPI_ISL_780860, EPI_ISL_780861, EPI_ISL_780862, EPI_ISL_780863, EPI_ISL_780864, EPI_ISL_780865, EPI_ISL_780866, EPI_ISL_780867, EPI_ISL_780868, EPI_ISL_780869, EPI_ISL_780870, EPI_ISL_780871, EPI_ISL_780872, EPI_ISL_780873, EPI_ISL_780874, EPI_ISL_780875, EPI_ISL_780876, EPI_ISL_780877, EPI_ISL_780878, EPI_ISL_780879, EPI_ISL_780880, EPI_ISL_780881, EPI_ISL_780882, EPI_ISL_780883, EPI_ISL_780884, EPI_ISL_780885, EPI_ISL_780886, EPI_ISL_780887, EPI_ISL_780888, EPI_ISL_780889, EPI_ISL_780890, EPI_ISL_780891, EPI_ISL_780892, EPI_ISL_780893, EPI_ISL_780894, EPI_ISL_780895, EPI_ISL_780896, EPI_ISL_780897, EPI_ISL_780898, EPI_ISL_780899, EPI_ISL_780900, EPI_ISL_780901, EPI_ISL_780902, EPI_ISL_780903, EPI_ISL_780904, EPI_ISL_780905, EPI_ISL_780906, EPI_ISL_780907, EPI_ISL_780908, EPI_ISL_780909, EPI_ISL_780910, EPI_ISL_780911, EPI_ISL_780912, EPI_ISL_780913, EPI_ISL_780914, EPI_ISL_780915, EPI_ISL_780916, EPI_ISL_780917, EPI_ISL_780918, EPI_ISL_780919, EPI_ISL_780920, EPI_ISL_780921, EPI_ISL_780922, EPI_ISL_780923, EPI_ISL_780924, EPI_ISL_780925, EPI_ISL_780926, EPI_ISL_780927, EPI_ISL_780928, EPI_ISL_780929, EPI_ISL_780930, EPI_ISL_780931, EPI_ISL_780932, EPI_ISL_780933, EPI_ISL_780934, EPI_ISL_780935, EPI_ISL_780936, EPI_ISL_780937, EPI_ISL_780938, EPI_ISL_780939, EPI_ISL_780940, EPI_ISL_780941, EPI_ISL_780942, EPI_ISL_780943, EPI_ISL_780944, EPI_ISL_780945, EPI_ISL_780946, EPI_ISL_780947, EPI_ISL_780948, EPI_ISL_780949, EPI_ISL_780950, EPI_ISL_780951, EPI_ISL_780952, EPI_ISL_780953, EPI_ISL_780954, EPI_ISL_780955, EPI_ISL_780956, EPI_ISL_780957, EPI_ISL_780958, EPI_ISL_780959, EPI_ISL_780960, EPI_ISL_780961, EPI_ISL_780962, EPI_ISL_780963, EPI_ISL_780964, EPI_ISL_780965, EPI_ISL_780966, EPI_ISL_780967, EPI_ISL_780968, EPI_ISL_780969, EPI_ISL_780970, EPI_ISL_780971, EPI_ISL_780972, EPI_ISL_780973, EPI_ISL_780974, EPI_ISL_780975, EPI_ISL_780976, EPI_ISL_780977, EPI_ISL_780978, EPI_ISL_780979, EPI_ISL_780980, EPI_ISL_780981, EPI_ISL_780982, EPI_ISL_780983, EPI_ISL_780984, EPI_ISL_780985, EPI_ISL_780986, EPI_ISL_780987, EPI_ISL_780988, EPI_ISL_780989, EPI_ISL_780990, EPI_ISL_780991, EPI_ISL_780992, EPI_ISL_780993, EPI_ISL_780994, EPI_ISL_780995, EPI_ISL_780996, EPI_ISL_780997, EPI_ISL_780998, EPI_ISL_780999, EPI_ISL_781000, EPI_ISL_781001, EPI_ISL_781002, EPI_ISL_781003, EPI_ISL_781004, EPI_ISL_781005, EPI_ISL_781006, EPI_ISL_781007, EPI_ISL_781008, EPI_ISL_781009, EPI_ISL_781010, EPI_ISL_781011, EPI_ISL_781012, EPI_ISL_781013, EPI_ISL_781014, EPI_ISL_781015, EPI_ISL_781016, EPI_ISL_781017, EPI_ISL_781018, EPI_ISL_781019, EPI_ISL_781020, EPI_ISL_781021, EPI_ISL_781022, EPI_ISL_781023, EPI_ISL_781024, EPI_ISL_781025, EPI_ISL_781026, EPI_ISL_781027, EPI_ISL_781028, EPI_ISL_781029, EPI_ISL_781030, EPI_ISL_781031, EPI_ISL_781032, EPI_ISL_781033, EPI_ISL_781034, EPI_ISL_781035, EPI_ISL_781036, EPI_ISL_781037, EPI_ISL_781038, EPI_ISL_781039, EPI_ISL_781040, EPI_ISL_781041, EPI_ISL_781042, EPI_ISL_781043, EPI_ISL_781044, EPI_ISL_781045, EPI_ISL_781046, EPI_ISL_781047, EPI_ISL_781048, EPI_ISL_781049, EPI_ISL_781050, EPI_ISL_781051, EPI_ISL_781052, EPI_ISL_781053, EPI_ISL_781054, EPI_ISL_781055, EPI_ISL_781056, EPI_ISL_781057, EPI_ISL_781058, EPI_ISL_781059, EPI_ISL_781060, EPI_ISL_781061, EPI_ISL_781062, EPI_ISL_781063, EPI_ISL_781064, EPI_ISL_781065, EPI_ISL_781066, EPI_ISL_781067, EPI_ISL_781068, EPI_ISL_781069, EPI_ISL_781070, EPI_ISL_781071, EPI_ISL_781073, EPI_ISL_781074, EPI_ISL_781075, EPI_ISL_781076, EPI_ISL_781077, EPI_ISL_781078, EPI_ISL_781079, EPI_ISL_781080, EPI_ISL_781081, EPI_ISL_781082, EPI_ISL_781083, EPI_ISL_781084, EPI_ISL_781085, EPI_ISL_781086, EPI_ISL_781087, EPI_ISL_781088, EPI_ISL_781089, EPI_ISL_781090, EPI_ISL_781091, EPI_ISL_781092, EPI_ISL_781093, EPI_ISL_781094, EPI_ISL_781095, EPI_ISL_781096, EPI_ISL_781097, EPI_ISL_781098, EPI_ISL_781099, EPI_ISL_781100, EPI_ISL_781101, EPI_ISL_781102, EPI_ISL_781103, EPI_ISL_781104, EPI_ISL_781105, EPI_ISL_781106, EPI_ISL_781107, EPI_ISL_781108, EPI_ISL_781109, EPI_ISL_781110, EPI_ISL_781111, EPI_ISL_781112, EPI_ISL_781114, EPI_ISL_781115, EPI_ISL_781116, EPI_ISL_781117, EPI_ISL_781118, EPI_ISL_781119, EPI_ISL_781120, EPI_ISL_781121, EPI_ISL_781122, EPI_ISL_781123, EPI_ISL_781124, EPI_ISL_781125, EPI_ISL_781126, EPI_ISL_781127, EPI_ISL_781128, EPI_ISL_781129, EPI_ISL_781130, EPI_ISL_781131, EPI_ISL_781132, EPI_ISL_781133, EPI_ISL_781134, EPI_ISL_781135, EPI_ISL_781136, EPI_ISL_781137, EPI_ISL_781138, EPI_ISL_781139, EPI_ISL_781140, EPI_ISL_781141, EPI_ISL_781142, EPI_ISL_781143, EPI_ISL_781144, EPI_ISL_781145, EPI_ISL_781146, EPI_ISL_781147, EPI_ISL_781148, EPI_ISL_781149, EPI_ISL_781150, EPI_ISL_781151, EPI_ISL_781152, EPI_ISL_781153, EPI_ISL_781154, EPI_ISL_781155, EPI_ISL_781156, EPI_ISL_781157, EPI_ISL_781158, EPI_ISL_781159, EPI_ISL_781160, EPI_ISL_781161, EPI_ISL_781162, EPI_ISL_781163, EPI_ISL_781164, EPI_ISL_781165, EPI_ISL_781166, EPI_ISL_781167, EPI_ISL_781168, EPI_ISL_781169, EPI_ISL_781170, EPI_ISL_781171, EPI_ISL_781172, EPI_ISL_781173, EPI_ISL_781174, EPI_ISL_781175, EPI_ISL_781176, EPI_ISL_783536, EPI_ISL_783539, EPI_ISL_783542, EPI_ISL_783554, EPI_ISL_783555, EPI_ISL_783556, EPI_ISL_783557, EPI_ISL_783572, EPI_ISL_783573, EPI_ISL_783578, EPI_ISL_783583, EPI_ISL_783584, EPI_ISL_783585, EPI_ISL_783586, EPI_ISL_783587, EPI_ISL_783589, EPI_ISL_783591, EPI_ISL_783592, EPI_ISL_783788, EPI_ISL_783789, EPI_ISL_784189, EPI_ISL_784212, EPI_ISL_784247, EPI_ISL_784249, EPI_ISL_784251, EPI_ISL_784252, EPI_ISL_784253, EPI_ISL_784254, EPI_ISL_784255, EPI_ISL_784256, EPI_ISL_784257, EPI_ISL_784258, EPI_ISL_784259, EPI_ISL_784260, EPI_ISL_784261, EPI_ISL_784262, EPI_ISL_784263, EPI_ISL_784264, EPI_ISL_784265, EPI_ISL_784266, EPI_ISL_784267, EPI_ISL_784268, EPI_ISL_784269, EPI_ISL_784270, EPI_ISL_784271, EPI_ISL_784272, EPI_ISL_784273, EPI_ISL_784274, EPI_ISL_784275, EPI_ISL_784276, EPI_ISL_784277, EPI_ISL_784278, EPI_ISL_784279, EPI_ISL_784281, EPI_ISL_784283, EPI_ISL_784284, EPI_ISL_784286, EPI_ISL_784287, EPI_ISL_784288, EPI_ISL_784289, EPI_ISL_784290, EPI_ISL_784291, EPI_ISL_784292, EPI_ISL_784293, EPI_ISL_784294, EPI_ISL_784295, EPI_ISL_784296, EPI_ISL_784297, EPI_ISL_784298, EPI_ISL_784299, EPI_ISL_784300, EPI_ISL_784301, EPI_ISL_784302, EPI_ISL_784303, EPI_ISL_784304, EPI_ISL_784305, EPI_ISL_784306, EPI_ISL_784307, EPI_ISL_784308, EPI_ISL_784309, EPI_ISL_784310, EPI_ISL_784311, EPI_ISL_784312, EPI_ISL_784313, EPI_ISL_784314, EPI_ISL_784315, EPI_ISL_784316, EPI_ISL_784317, EPI_ISL_784318, EPI_ISL_784319, EPI_ISL_784320, EPI_ISL_784321, EPI_ISL_784322, EPI_ISL_784323, EPI_ISL_784324, EPI_ISL_784325, EPI_ISL_784326, EPI_ISL_784327, EPI_ISL_784328, EPI_ISL_784329, EPI_ISL_784330, EPI_ISL_784331, EPI_ISL_784332, EPI_ISL_784333, EPI_ISL_784334, EPI_ISL_784335, EPI_ISL_784336, EPI_ISL_784337, EPI_ISL_784338, EPI_ISL_784339, EPI_ISL_784340, EPI_ISL_784341, EPI_ISL_784342, EPI_ISL_784343, EPI_ISL_784344, EPI_ISL_784345, EPI_ISL_784346, EPI_ISL_784347, EPI_ISL_784348, EPI_ISL_784349, EPI_ISL_784350, EPI_ISL_784351, EPI_ISL_784352, EPI_ISL_784353, EPI_ISL_784354, EPI_ISL_784372, EPI_ISL_784378, EPI_ISL_784381, EPI_ISL_784382, EPI_ISL_784383, EPI_ISL_784384, EPI_ISL_784385, EPI_ISL_784386, EPI_ISL_784387, EPI_ISL_784388, EPI_ISL_784389, EPI_ISL_784390, EPI_ISL_784391, EPI_ISL_784392, EPI_ISL_784394, EPI_ISL_784395, EPI_ISL_784396, EPI_ISL_784397, EPI_ISL_784598, EPI_ISL_784599, EPI_ISL_785323, EPI_ISL_785324, EPI_ISL_785325, EPI_ISL_785326, EPI_ISL_785327, EPI_ISL_785328, EPI_ISL_785329, EPI_ISL_785330, EPI_ISL_785331, EPI_ISL_785332, EPI_ISL_785333, EPI_ISL_785334, EPI_ISL_785335, EPI_ISL_785336, EPI_ISL_785337, EPI_ISL_785338, EPI_ISL_785339, EPI_ISL_785340, EPI_ISL_785341, EPI_ISL_785342, EPI_ISL_785343, EPI_ISL_785344, EPI_ISL_785345, EPI_ISL_785346, EPI_ISL_785347, EPI_ISL_785348, EPI_ISL_785349, EPI_ISL_785350, EPI_ISL_785351, EPI_ISL_785352, EPI_ISL_785353, EPI_ISL_785354, EPI_ISL_785355, EPI_ISL_785356, EPI_ISL_785357, EPI_ISL_785358, EPI_ISL_785359, | | | |

EPI_ISL_785360, EPI_ISL_785361, EPI_ISL_785362, EPI_ISL_785363, EPI_ISL_785364, EPI_ISL_785365, EPI_ISL_785366, EPI_ISL_785367, EPI_ISL_785368, EPI_ISL_785369, EPI_ISL_785370, EPI_ISL_785371, EPI_ISL_785372, EPI_ISL_785373, EPI_ISL_785374, EPI_ISL_785375, EPI_ISL_785376, EPI_ISL_785377, EPI_ISL_785378, EPI_ISL_785379, EPI_ISL_785380, EPI_ISL_785381, EPI_ISL_785382, EPI_ISL_785383, EPI_ISL_785384, EPI_ISL_785385, EPI_ISL_785386, EPI_ISL_785387, EPI_ISL_785388, EPI_ISL_785389, EPI_ISL_785390, EPI_ISL_785391, EPI_ISL_785392, EPI_ISL_785393, EPI_ISL_785394, EPI_ISL_785395, EPI_ISL_785396, EPI_ISL_785397, EPI_ISL_785398, EPI_ISL_785399, EPI_ISL_785400, EPI_ISL_785401, EPI_ISL_785402, EPI_ISL_785403, EPI_ISL_785404, EPI_ISL_785405, EPI_ISL_785406, EPI_ISL_785407, EPI_ISL_785408, EPI_ISL_785409, EPI_ISL_785410, EPI_ISL_785411, EPI_ISL_785412, EPI_ISL_785413, EPI_ISL_785414, EPI_ISL_785415, EPI_ISL_785416, EPI_ISL_785417, EPI_ISL_785418, EPI_ISL_785419, EPI_ISL_785420, EPI_ISL_785421, EPI_ISL_785422, EPI_ISL_785423, EPI_ISL_785424, EPI_ISL_785425, EPI_ISL_785426, EPI_ISL_785427, EPI_ISL_785428, EPI_ISL_785429, EPI_ISL_785430, EPI_ISL_785431, EPI_ISL_785432, EPI_ISL_785433, EPI_ISL_785434, EPI_ISL_785435, EPI_ISL_785436, EPI_ISL_785437, EPI_ISL_785438, EPI_ISL_785439, EPI_ISL_785440, EPI_ISL_785441, EPI_ISL_785442, EPI_ISL_785443, EPI_ISL_785444, EPI_ISL_785445, EPI_ISL_785446, EPI_ISL_785447, EPI_ISL_785448, EPI_ISL_785449, EPI_ISL_785450, EPI_ISL_785451, EPI_ISL_785452, EPI_ISL_785453, EPI_ISL_785454, EPI_ISL_785455, EPI_ISL_785456, EPI_ISL_785457, EPI_ISL_785458, EPI_ISL_785459, EPI_ISL_785460, EPI_ISL_785461, EPI_ISL_785462, EPI_ISL_785463, EPI_ISL_785464, EPI_ISL_785465, EPI_ISL_785466, EPI_ISL_785467, EPI_ISL_785468, EPI_ISL_785469, EPI_ISL_785470, EPI_ISL_785471, EPI_ISL_785472, EPI_ISL_785473, EPI_ISL_785474, EPI_ISL_785475, EPI_ISL_785476, EPI_ISL_785477, EPI_ISL_785478, EPI_ISL_785479, EPI_ISL_785480, EPI_ISL_785481, EPI_ISL_785482, EPI_ISL_785483, EPI_ISL_785484, EPI_ISL_785486, EPI_ISL_785487, EPI_ISL_785488, EPI_ISL_785489, EPI_ISL_785490, EPI_ISL_785491, EPI_ISL_785492, EPI_ISL_785493, EPI_ISL_785494, EPI_ISL_785495, EPI_ISL_785496, EPI_ISL_785497, EPI_ISL_785498, EPI_ISL_785499, EPI_ISL_785500, EPI_ISL_785501, EPI_ISL_785502, EPI_ISL_785503, EPI_ISL_785504, EPI_ISL_785505, EPI_ISL_785506, EPI_ISL_785507, EPI_ISL_785508, EPI_ISL_785509, EPI_ISL_785510, EPI_ISL_785511, EPI_ISL_785512, EPI_ISL_785513, EPI_ISL_785514, EPI_ISL_785515, EPI_ISL_785516, EPI_ISL_785517, EPI_ISL_785518, EPI_ISL_785519, EPI_ISL_785520, EPI_ISL_785521, EPI_ISL_785522, EPI_ISL_785523, EPI_ISL_785524, EPI_ISL_785525, EPI_ISL_785526, EPI_ISL_785527, EPI_ISL_785528, EPI_ISL_785529, EPI_ISL_785530, EPI_ISL_785531, EPI_ISL_785532, EPI_ISL_785533, EPI_ISL_785534, EPI_ISL_785535, EPI_ISL_785536, EPI_ISL_785537, EPI_ISL_785538, EPI_ISL_785539, EPI_ISL_785540, EPI_ISL_785541, EPI_ISL_785542, EPI_ISL_785543, EPI_ISL_785544, EPI_ISL_785545, EPI_ISL_785546, EPI_ISL_785547, EPI_ISL_785548, EPI_ISL_785549, EPI_ISL_785550, EPI_ISL_785551, EPI_ISL_785552, EPI_ISL_785553, EPI_ISL_785554, EPI_ISL_785555, EPI_ISL_785556, EPI_ISL_785557, EPI_ISL_785558, EPI_ISL_785559, EPI_ISL_785560, EPI_ISL_785561, EPI_ISL_785562, EPI_ISL_785563, EPI_ISL_785564, EPI_ISL_785565, EPI_ISL_785566, EPI_ISL_785567, EPI_ISL_785568, EPI_ISL_785569, EPI_ISL_785570, EPI_ISL_785571, EPI_ISL_785572, EPI_ISL_785573, EPI_ISL_785574, EPI_ISL_785575, EPI_ISL_785576, EPI_ISL_785577, EPI_ISL_785578, EPI_ISL_785579, EPI_ISL_785580, EPI_ISL_785581, EPI_ISL_785582, EPI_ISL_785583, EPI_ISL_785584, EPI_ISL_785585, EPI_ISL_785586, EPI_ISL_785587, EPI_ISL_785588, EPI_ISL_785589, EPI_ISL_785590, EPI_ISL_785591, EPI_ISL_785592, EPI_ISL_785593, EPI_ISL_785594, EPI_ISL_785595, EPI_ISL_785596, EPI_ISL_785597, EPI_ISL_785598, EPI_ISL_785599, EPI_ISL_785600, EPI_ISL_785601, EPI_ISL_785602, EPI_ISL_785603, EPI_ISL_785604, EPI_ISL_785605, EPI_ISL_785606, EPI_ISL_785607, EPI_ISL_785608, EPI_ISL_785609, EPI_ISL_785610, EPI_ISL_785611, EPI_ISL_785612, EPI_ISL_785613, EPI_ISL_785614, EPI_ISL_785615, EPI_ISL_785616, EPI_ISL_785617, EPI_ISL_785618, EPI_ISL_785619, EPI_ISL_785620, EPI_ISL_785621, EPI_ISL_785622, EPI_ISL_785623, EPI_ISL_785624, EPI_ISL_785625, EPI_ISL_785626, EPI_ISL_785627, EPI_ISL_785628, EPI_ISL_785629, EPI_ISL_785630, EPI_ISL_785631, EPI_ISL_785632, EPI_ISL_785633, EPI_ISL_785634, EPI_ISL_785635, EPI_ISL_785636, EPI_ISL_785637, EPI_ISL_785638, EPI_ISL_785639, EPI_ISL_785640, EPI_ISL_785641, EPI_ISL_785642, EPI_ISL_785643, EPI_ISL_785644, EPI_ISL_785645, EPI_ISL_785646, EPI_ISL_785647, EPI_ISL_785648, EPI_ISL_785649, EPI_ISL_785650, EPI_ISL_785651, EPI_ISL_785652, EPI_ISL_785653, EPI_ISL_785654, EPI_ISL_785655, EPI_ISL_785656, EPI_ISL_785657, EPI_ISL_785658, EPI_ISL_785659, EPI_ISL_785660, EPI_ISL_785661, EPI_ISL_785662, EPI_ISL_785663, EPI_ISL_785664, EPI_ISL_785665, EPI_ISL_785666, EPI_ISL_785667, EPI_ISL_785668, EPI_ISL_785669, EPI_ISL_785670, EPI_ISL_785671, EPI_ISL_785672, EPI_ISL_785673, EPI_ISL_785674, EPI_ISL_785675, EPI_ISL_785676, EPI_ISL_785677, EPI_ISL_785678, EPI_ISL_785679, EPI_ISL_785680, EPI_ISL_785681, EPI_ISL_785682, EPI_ISL_785683, EPI_ISL_785684, EPI_ISL_785685, EPI_ISL_785686, EPI_ISL_785687, EPI_ISL_785688, EPI_ISL_785689, EPI_ISL_785690, EPI_ISL_785691, EPI_ISL_785692, EPI_ISL_785693, EPI_ISL_785694, EPI_ISL_785695, EPI_ISL_785696, EPI_ISL_785697, EPI_ISL_785698, EPI_ISL_785699, EPI_ISL_785700, EPI_ISL_785701, EPI_ISL_785702, EPI_ISL_785703, EPI_ISL_785704, EPI_ISL_785705, EPI_ISL_786398, EPI_ISL_786399, EPI_ISL_787298, EPI_ISL_787299, EPI_ISL_788198, EPI_ISL_788199, EPI_ISL_789106, EPI_ISL_789107, EPI_ISL_789109, EPI_ISL_789110, EPI_ISL_789111, EPI_ISL_789112, EPI_ISL_789113, EPI_ISL_789114, EPI_ISL_789116, EPI_ISL_789118, EPI_ISL_789119, EPI_ISL_789120, EPI_ISL_789122, EPI_ISL_789126, EPI_ISL_789127, EPI_ISL_789128, EPI_ISL_789130, EPI_ISL_789131, EPI_ISL_789132, EPI_ISL_789133, EPI_ISL_789136, EPI_ISL_789137, EPI_ISL_789138, EPI_ISL_789139, EPI_ISL_789142, EPI_ISL_789143, EPI_ISL_789146, EPI_ISL_789147, EPI_ISL_789150, EPI_ISL_789151, EPI_ISL_789152, EPI_ISL_789153, EPI_ISL_789154, EPI_ISL_789156, EPI_ISL_789157, EPI_ISL_789160, EPI_ISL_789161, EPI_ISL_789163, EPI_ISL_789165, EPI_ISL_789166, EPI_ISL_789167, EPI_ISL_789168, EPI_ISL_789169, EPI_ISL_789173, EPI_ISL_789174, EPI_ISL_789175, EPI_ISL_789176, EPI_ISL_789177, EPI_ISL_789178, EPI_ISL_789181, EPI_ISL_789182, EPI_ISL_789183, EPI_ISL_789184, EPI_ISL_789185, EPI_ISL_789186, EPI_ISL_789187, EPI_ISL_789188, EPI_ISL_789189, EPI_ISL_789190, EPI_ISL_789191, EPI_ISL_789192, EPI_ISL_789194, EPI_ISL_789195, EPI_ISL_789196, EPI_ISL_789197, EPI_ISL_789198, EPI_ISL_789199, EPI_ISL_789201, EPI_ISL_789202, EPI_ISL_789203, EPI_ISL_789204, EPI_ISL_789205, EPI_ISL_789206, EPI_ISL_789207, EPI_ISL_789208, EPI_ISL_789209, EPI_ISL_789210, EPI_ISL_789211, EPI_ISL_789212, EPI_ISL_789213, EPI_ISL_789214, EPI_ISL_789215, EPI_ISL_789216, EPI_ISL_789219, EPI_ISL_789220, EPI_ISL_789222, EPI_ISL_789223, EPI_ISL_789224, EPI_ISL_789225, EPI_ISL_789226, EPI_ISL_789227, EPI_ISL_789228, EPI_ISL_789229, EPI_ISL_789230, EPI_ISL_789234, EPI_ISL_789235, EPI_ISL_789236, EPI_ISL_789237, EPI_ISL_789238, EPI_ISL_789239, EPI_ISL_789240, EPI_ISL_789241, EPI_ISL_789242, EPI_ISL_789243, EPI_ISL_789244, EPI_ISL_789245, EPI_ISL_789246, EPI_ISL_789247, EPI_ISL_789248, EPI_ISL_789249, EPI_ISL_789250, EPI_ISL_789251, EPI_ISL_789252, EPI_ISL_789253, EPI_ISL_789254, EPI_ISL_789255, EPI_ISL_789256, EPI_ISL_789257, EPI_ISL_789258, EPI_ISL_789259, EPI_ISL_789261, EPI_ISL_789262, EPI_ISL_789263, EPI_ISL_789264, EPI_ISL_789265, EPI_ISL_789266, EPI_ISL_789267, EPI_ISL_789269, EPI_ISL_789270, EPI_ISL_789271, EPI_ISL_789274, EPI_ISL_789275, EPI_ISL_789276, EPI_ISL_789277, EPI_ISL_789278, EPI_ISL_789279, EPI_ISL_789280, EPI_ISL_789281, EPI_ISL_789282, EPI_ISL_789283, EPI_ISL_789284, EPI_ISL_789285, EPI_ISL_789286, EPI_ISL_789287, EPI_ISL_789288, EPI_ISL_789289, EPI_ISL_789290, EPI_ISL_789291, EPI_ISL_789292, EPI_ISL_789293, EPI_ISL_789294, EPI_ISL_789295, EPI_ISL_789296, EPI_ISL_789297, EPI_ISL_789298, EPI_ISL_789299, EPI_ISL_789300, EPI_ISL_789301, EPI_ISL_789302, EPI_ISL_789303, EPI_ISL_789304, EPI_ISL_789305, EPI_ISL_789306, EPI_ISL_789307, EPI_ISL_789308, EPI_ISL_789309, EPI_ISL_789310, EPI_ISL_789311, EPI_ISL_789312, EPI_ISL_789313, EPI_ISL_789314, EPI_ISL_789315, EPI_ISL_789316, EPI_ISL_789317, EPI_ISL_789318, EPI_ISL_789319, EPI_ISL_789320, EPI_ISL_789321, EPI_ISL_789322, EPI_ISL_789323, EPI_ISL_789324, EPI_ISL_789325, EPI_ISL_789326, EPI_ISL_789327, EPI_ISL_789328, EPI_ISL_789329, EPI_ISL_789330, EPI_ISL_789331, EPI_ISL_789332, EPI_ISL_789333, EPI_ISL_789334, EPI_ISL_789335, EPI_ISL_789336, EPI_ISL_789337, EPI_ISL_789338, EPI_ISL_789339, EPI_ISL_789340, EPI_ISL_789341, EPI_ISL_789342, EPI_ISL_789343, EPI_ISL_789344, EPI_ISL_789345, EPI_ISL_789346, EPI_ISL_789347, EPI_ISL_789348, EPI_ISL_789349, EPI_ISL_789350, EPI_ISL_789351, EPI_ISL_789352, EPI_ISL_789353, EPI_ISL_789354, EPI_ISL_789355, EPI_ISL_789356, EPI_ISL_789357, EPI_ISL_789358, EPI_ISL_789359, EPI_ISL_789360, EPI_ISL_789361, EPI_ISL_789362, EPI_ISL_789363, EPI_ISL_789364, EPI_ISL_789365, EPI_ISL_789366, EPI_ISL_789367, EPI_ISL_789368, EPI_ISL_789369, EPI_ISL_789370, EPI_ISL_789371, EPI_ISL_789372, EPI_ISL_789373, EPI_ISL_789374, EPI_ISL_789375, EPI_ISL_789376, EPI_ISL_789377, EPI_ISL_789379, EPI_ISL_789380, EPI_ISL_789381, EPI_ISL_789382, EPI_ISL_789383, EPI_ISL_789384, EPI_ISL_789385, EPI_ISL_789386, EPI_ISL_789387, EPI_ISL_789388, EPI_ISL_789389, EPI_ISL_789391, EPI_ISL_789392, EPI_ISL_789393, EPI_ISL_789394, EPI_ISL_789395, EPI_ISL_789396, EPI_ISL_789397, EPI_ISL_789398, EPI_ISL_789399, EPI_ISL_789400, EPI_ISL_789401, EPI_ISL_789402, EPI_ISL_789403, EPI_ISL_789404, EPI_ISL_789405, EPI_ISL_789406, EPI_ISL_789407, EPI_ISL_789408, EPI_ISL_789409, EPI_ISL_789410, EPI_ISL_789411, EPI_ISL_789412, EPI_ISL_789413, EPI_ISL_789414, EPI_ISL_789415, EPI_ISL_789416, EPI_ISL_789417, EPI_ISL_789418, EPI_ISL_789419, EPI_ISL_789420, EPI_ISL_789421, EPI_ISL_789422, EPI_ISL_789423, EPI_ISL_789424, EPI_ISL_789425, EPI_ISL_789426, EPI_ISL_789427, EPI_ISL_789428, EPI_ISL_789429, EPI_ISL_789430, EPI_ISL_789431, EPI_ISL_789432, EPI_ISL_789433, EPI_ISL_789434, EPI_ISL_789435, EPI_ISL_789436, EPI_ISL_789437, EPI_ISL_789438, EPI_ISL_789439, EPI_ISL_789440, EPI_ISL_789441, EPI_ISL_789442, EPI_ISL_789443, EPI_ISL_789444, EPI_ISL_789445, EPI_ISL_789446, EPI_ISL_789447, EPI_ISL_789448, EPI_ISL_789449, EPI_ISL_789450, EPI_ISL_789451, EPI_ISL_789452, EPI_ISL_789453, EPI_ISL_789454, EPI_ISL_789455, EPI_ISL_789456, EPI_ISL_789457, EPI_ISL_789458, EPI_ISL_789459, EPI_ISL_789460, EPI_ISL_789461, EPI_ISL_789462, EPI_ISL_789463, EPI_ISL_789464, EPI_ISL_789465, EPI_ISL_789466, EPI_ISL_789467, EPI_ISL_789468, EPI_ISL_789469, EPI_ISL_789470, EPI_ISL_789471, EPI_ISL_789472, EPI_ISL_789473, EPI_ISL_789474, EPI_ISL_789475, EPI_ISL_789476, EPI_ISL_789477, EPI_ISL_789478, EPI_ISL_789479, EPI_ISL_789480, EPI_ISL_789481, EPI_ISL_789482, EPI_ISL_789483, EPI_ISL_789484, EPI_ISL_789485, EPI_ISL_789486, EPI_ISL_789487, EPI_ISL_789488, EPI_ISL_789489, EPI_ISL_789490, EPI_ISL_789491, EPI_ISL_789492, EPI_ISL_789493, EPI_ISL_789494, EPI_ISL_789495, EPI_ISL_789496, EPI_ISL_789497, EPI_ISL_789498, EPI_ISL_789499, EPI_ISL_789500, EPI_ISL_789501, EPI_ISL_789502, EPI_ISL_789503, EPI_ISL_789504, EPI_ISL_789505, EPI_ISL_789506, EPI_ISL_789507, EPI_ISL_789508, EPI_ISL_789509, EPI_ISL_789510, EPI_ISL_789511, EPI_ISL_789512, EPI_ISL_789513, EPI_ISL_789514, EPI_ISL_789515, EPI_ISL_789516, EPI_ISL_789517, EPI_ISL_789518, EPI_ISL_789519, EPI_ISL_789520, EPI_ISL_789521, EPI_ISL_789522, EPI_ISL_789523, EPI_ISL_789524, EPI_ISL_789525, EPI_ISL_789526, EPI_ISL_789527, EPI_ISL_789528, EPI_ISL_789529, EPI_ISL_789530, EPI_ISL_789531, EPI_ISL_789532, EPI_ISL_789533, EPI_ISL_789534, EPI_ISL_789535, EPI_ISL_789536, EPI_ISL_789537, EPI_ISL_789538, EPI_ISL_789539, EPI_ISL_789540, EPI_ISL_789541, EPI_ISL_789542, EPI_ISL_789543, EPI_ISL_789544, EPI_ISL_789545, EPI_ISL_789546, EPI_ISL_789547, EPI_ISL_789548, EPI_ISL_789549, EPI_ISL_789550, EPI_ISL_789551, EPI_ISL_789552, EPI_ISL_789553, EPI_ISL_789554, EPI_ISL_789555, EPI_ISL_789556, EPI_ISL_789557, EPI_ISL_789558, EPI_ISL_789559, EPI_ISL_789560, EPI_ISL_789561, EPI_ISL_789562, EPI_ISL_789563, EPI_ISL_789564, EPI_ISL_789565, EPI_ISL_789566, EPI_ISL_789567, EPI_ISL_789568, EPI_ISL_789569, EPI_ISL_789570, EPI_ISL_789571, EPI_ISL_789572, EPI_ISL_789573, EPI_ISL_789574, EPI_ISL_789575, EPI_ISL_789576, EPI_ISL_789577, EPI_ISL_789578, EPI_ISL_789579, EPI_ISL_789580, EPI_ISL_789581, EPI_ISL_789582, EPI_ISL_789583, EPI_ISL_789584, EPI_ISL_789585, EPI_ISL_789586, EPI_ISL_789587, EPI_ISL_789588, EPI_ISL_789589, EPI_ISL_789590, EPI_ISL_789591, EPI_ISL_789592, EPI_ISL_789593, EPI_ISL_789594, EPI_ISL_789595, EPI_ISL_789596, EPI_ISL_789597, EPI_ISL_789598, EPI_ISL_789599, EPI_ISL_789600, EPI_ISL_789601, EPI_ISL_789602, EPI_ISL_789603, EPI_ISL_789604, EPI_ISL_789605, EPI_ISL_789606, EPI_ISL_789607, EPI_ISL_789608, EPI_ISL_789609, EPI_ISL_789610, EPI_ISL_789611, EPI_ISL_789612, EPI_ISL_789613, EPI_ISL_789614, EPI_ISL_789615, EPI_ISL_789616, EPI_ISL_789617, EPI_ISL_789618, EPI_ISL_789619, EPI_ISL_789620, EPI_ISL_789621, EPI_ISL_789622, EPI_ISL_789623, EPI_ISL_789624, EPI_ISL_789625, EPI_ISL_789626, EPI_ISL_789627, EPI_ISL_789628, EPI_ISL_789629, EPI_ISL_789630, EPI_ISL_789631, EPI_ISL_789632, EPI_ISL_789633, EPI_ISL_789634, EPI_ISL_789635, EPI_ISL_789636, EPI_ISL_789637, EPI_ISL_789638, EPI_ISL_789639, EPI_ISL_789640, EPI_ISL_789641, EPI_ISL_789642, EPI_ISL_789643, EPI_ISL_789644, EPI_ISL_789645, EPI_ISL_789646, EPI_ISL_789647, EPI_ISL_789648, EPI_ISL_789649, EPI_ISL_789650, EPI_ISL_789651, EPI_ISL_789652, EPI_ISL_789653, EPI_ISL_789654, EPI_ISL_789655, EPI_ISL_789656, EPI_ISL_789657, EPI_ISL_789658, EPI_ISL_789659, EPI_ISL_789660, EPI_ISL_789661, EPI_ISL_789662, EPI_ISL_789663, EPI_ISL_789664, EPI_ISL_789665, EPI_ISL_789666, EPI_ISL_789667, EPI_ISL_789668, EPI_ISL_789669, EPI_ISL_789670, EPI_ISL_789671, EPI_ISL_789672, EPI_ISL_789673, EPI_ISL_789674, EPI_ISL_789675, EPI_ISL_789676, EPI_ISL_789677, EPI_ISL_789678, EPI_ISL_789679, EPI_ISL_789680, EPI_ISL_789681, EPI_ISL_789682, EPI_ISL_789683, EPI_ISL_789684, EPI_ISL_789685, EPI_ISL_789686, EPI_ISL_789687, EPI_ISL_789688, EPI_ISL_789689, EPI_ISL_789690, EPI_ISL_789691, EPI_ISL_789692, EPI_ISL_789693, EPI_ISL_789694, EPI_ISL_789695, EPI_ISL_789696, EPI_ISL_789878

| | | | |
|---|---|---|---|
| see above | Houston Methodist Hospital | Houston Methodist Hospital | S. Wesley Long, Randall J. Olsen, Paul A. Christensen, David W. Bernard, James J. Davis, Maulik Shukla, Marcus Nguyen, Matthew Ojeda Saavedra, Prasanti Yerramilli, Layne Pruitt, Sishir Subedi, Heather Hendrickson, and James M. Musser |
| EPI_ISL_792116 | Instituto Adolfo Lutz - Regional de Taubate | Instituto Adolfo Lutz, Interdiciplinary Procedures Center, Strategic Laboratory | Claudio Tavares Sacchi, Claudia Regina Gonçalves, Erica Valessa Ramos Gomes, Karoline Rodrigues Campos |
| EPI_ISL_792544, EPI_ISL_792545, EPI_ISL_792546 | Laboratorio Central, Ministerio de Salud Córdoba | Instituto de Patologia Vegetal (CIAP-INTA) on behalf of 'Proyecto Argentino Interinstitucional de genomica de SARS-CoV-2' (PAIS Consortium) | Fernández, FD; Debat, HJ., Re, V; Pisano, MB; Castro, G.; Barbas, G. |
| EPI_ISL_792595, EPI_ISL_792596 | LACEN-PB | Laboratory of Respiratory Viruses and Measles, Oswaldo Cruz Institute, FIOCRUZ | Paola Resende, Luciana Appolinario, Fernando Motta, Anna Carolina Paixao, Ana Carolina Mendonca, João Felipe Bezerra, Romero Henrique Teixeira de Vasconcelos, Dalane Loudal Florentino Teixeira, Thiago Franco de Oliveira Carneiro, Marilda Siqueira |

EPI_ISL_801910, EPI_ISL_801963, EPI_ISL_802108, EPI_ISL_802109, EPI_ISL_802110, EPI_ISL_802111, EPI_ISL_802112, EPI_ISL_802113, EPI_ISL_802114, EPI_ISL_802115, EPI_ISL_802116, EPI_ISL_802117, EPI_ISL_802118, EPI_ISL_802119, EPI_ISL_802120, EPI_ISL_802121, EPI_ISL_802122, EPI_ISL_802123, EPI_ISL_802124, EPI_ISL_802125, EPI_ISL_802126, EPI_ISL_802127, EPI_ISL_802128, EPI_ISL_802129, EPI_ISL_802130, EPI_ISL_802131, EPI_ISL_802132, EPI_ISL_802133, EPI_ISL_802134, EPI_ISL_802135, EPI_ISL_802136

| | | | |
|---|---|---|---|
| see above | MSHS Clinical Microbiology Laboratories | MSHS Pathogen Surveillance Program | Ana S. Gonzalez-Reiche, Hala Alshammary, Mitchell J. Sullivan, Brianne Ciferri, Ajay Obla, Angela Amoako, Mahmoud Awawda, Elena Hirsch, Ashley S. Salimbangon, Levy Sominsky, Katherine Beach, Kayla Russo, Charles Gleason, Shelcie Fabre, Giulio Kleiner, Zenab Khan, Bremy Alburquerque, Adriana van de Guchte, Komal Srivastava, Matthew M. Hernandez, Jayeeta Dutta, Denise Jurczyszak, Emily Ferreri, Rachel Chernet, Nancy Francoeur, Betsaida Salom Melo, Irina Oussenko, Gintaras Deikus, Juan Soto, Shwetha Hara Sridhar, Ying-Chih Wang, Kathryn Twyman, Andrew Kasarskis, Deena R. Altman, Robert Sebra, Adolfo García-Sastre, Marta Luksza, Gopi Patel, Sarah Schaefer, Melissa Gitman, Michael D. Nowak, Alberto Paniz-Mondolfi, Emilia Mia Sordillo, Viviana Simon, Harm van Bakel |
| EPI_ISL_803138, EPI_ISL_803250, EPI_ISL_803300, EPI_ISL_803301, EPI_ISL_803302, EPI_ISL_803303, EPI_ISL_803304, EPI_ISL_803305 | Wisconsin State Laboratory of Hygiene Communicable Disease Division | Wisconsin State Laboratory of Hygiene Communicable Disease Division | Kelsey R. Florek, Abigail C. Shockey |
| EPI_ISL_804016 | Facultad de Medicina, Universidad de Atacama | Facultad de Ciencias de la Vida, UNAB | Dayán Sanhueza, Macarena Bastías, César Echeverría, Sebastián Wolter, Waldo Díaz, Jorge Olivares, Eduardo Castro, Claudio Meneses |
| EPI_ISL_804474, EPI_ISL_804583, EPI_ISL_804584 | MEPHI, Aix Marseille University | MEPHI, Aix Marseille University | Anthony LEVASSEUR |

EPI_ISL_806548, EPI_ISL_806549, EPI_ISL_806550, EPI_ISL_806551, EPI_ISL_806552, EPI_ISL_806553, EPI_ISL_806554, EPI_ISL_806555, EPI_ISL_806556, EPI_ISL_806557, EPI_ISL_806558, EPI_ISL_806559, EPI_ISL_806560, EPI_ISL_806561, EPI_ISL_806562, EPI_ISL_806563, EPI_ISL_806566, EPI_ISL_806626, EPI_ISL_806627, EPI_ISL_806628, EPI_ISL_806629, EPI_ISL_806630, EPI_ISL_806631, EPI_ISL_806632, EPI_ISL_806633, EPI_ISL_806634, EPI_ISL_806635, EPI_ISL_806639, EPI_ISL_806640

| | | | |
|---|---|---|---|
| see above | KEMRI-Wellcome Trust Research Programme/KEMRI-CGMR-C Kilifi | KEMRI-Wellcome Trust Research Programme/KEMRI-CGMR-C Kilifi | Githinji et al |

EPI_ISL_811151, EPI_ISL_811152, EPI_ISL_811155, EPI_ISL_811156, EPI_ISL_811161, EPI_ISL_811162, EPI_ISL_811165, EPI_ISL_811169, EPI_ISL_811180, EPI_ISL_811189, EPI_ISL_811190

| | | | |
|---|---|---|---|
| see above | Dharwad | CSIR Institute of Genomics and Integrative Biology | Dr. Shivarudrapp B Bhairappanavar, Rahul Bhoyar, Mohammed Imran, Mohit Divakar, Disha Sharma, Dr. Vijay A Yenagi, Dr. Suresh B Arakera, Dr. Amit |

| | | | |
|---|---|---|---|
| | | | Ugargol, Dr. Rgavendra B Nayak, Bani Jolly, Abhinav Jain, Paras Sehgal, Gyan Ranjan, Vinod Scaria, Sridhar Sivasubbu |
| EPI_ISL_812526 | United States Air Force School of Aerospace Medicine | United States Air Force School of Aerospace Medicine | Anthony Fries, Jennifer Meyer, Amanda Javorina, Sarah Purves, William Gruner, Clarise Starr, Elizabeth Macias |
| EPI_ISL_812796, EPI_ISL_812797, EPI_ISL_812798, EPI_ISL_812803, EPI_ISL_812804, EPI_ISL_812822, EPI_ISL_812837, EPI_ISL_812840, EPI_ISL_812855, EPI_ISL_812862, EPI_ISL_812866 | | | |
| see above | Genomics Program, Children Cancer Hospital | Genomics Program, Children Cancer Hospital | Hatem,A., Hadad,A., Abouelnaga,S., Amer,K., Salah,H., Farawyla,H., Halafawy,A., Mansour,T., shalaby,L., Hassan,W., Soliman,M., Gomaa,C., Hassan,R., Soliman,S., Monuir,G., Hammad,M., Hussein,S., Abdo,I., Jalal,D., El-Zayat,M., El-Shaqnqery,H., Diab,A., Bakry,U., Samir,O., Magdeldin,S., Sayed,A. |
| EPI_ISL_815265, EPI_ISL_815266, EPI_ISL_815268 | Centogene | Centogene | Peter Bauer, Krishna Kumar Kandaswamy, Vivi Hue-Trang Lieu |
| EPI_ISL_824389 | Ohio Department of Health Laboratory | Ohio Department of Health Laboratory | Holmes, Jennifer; Eric Brandt, Keoni Omura, Glen McGillivary, Caitlin McDonnell, Kirtana Ramadugu, Erica Leasure, Kelsey Florek, Heather Blankenship, Quanta Brown, and Tammy Bannerman |
| EPI_ISL_824420, EPI_ISL_824421, EPI_ISL_824422, EPI_ISL_824423, EPI_ISL_824424, EPI_ISL_824425, EPI_ISL_824426, EPI_ISL_824427, EPI_ISL_824428 | Hospital Universitari Vall d'Hebron - Vall d'Hebron Institut de Rercerca | Hospital Universitari Vall d'Hebron | Cristina Andrés, Maria Piñana, Josep F Abril, Damir Garcia-Cehic, Ariadna Rando, Juliana Esperalba, Maria Gema Codina, Carla Castillo, Maria Carmen Martin, Tomàs Pumarola, Josep Quer, Andrés Antón |
| EPI_ISL_825640, EPI_ISL_825643, EPI_ISL_825644, EPI_ISL_825645, EPI_ISL_825646, EPI_ISL_825647, EPI_ISL_825649, EPI_ISL_825657, EPI_ISL_825658, EPI_ISL_825659, EPI_ISL_825660, EPI_ISL_825661, EPI_ISL_825662, EPI_ISL_825663, EPI_ISL_825664, EPI_ISL_825912, EPI_ISL_825914, EPI_ISL_825915, EPI_ISL_825916, EPI_ISL_825917, EPI_ISL_825918, EPI_ISL_825919, EPI_ISL_825928, EPI_ISL_825929, EPI_ISL_825930, EPI_ISL_825931, EPI_ISL_825932, EPI_ISL_825933, EPI_ISL_825934, EPI_ISL_825935 | | | |
| see above | Laboratoire de santé publique du Québec | Laboratoire de santé publique du Québec | Sandrine Moreira, Ioannis Ragoussis, Guillaume Bourque, Jesse Shapiro, Mark Lathrop and Michel Roger on behalf of the CoVSeQ research group (http://covseq.ca/researchgroup) |
| EPI_ISL_826652, EPI_ISL_826653, EPI_ISL_826654 | Montefiore Medical Center | Albert Einstein College of Medicine, Dept. of Microbiology & Immunology, Chandran lab | J. Maximilian Fels, Saad Khan, Ryan Forster, Karin A. Skalina, Surksha Sirichand, Amy S. Fox, Aviv Bergman, William B. Mitchell, Lucia R. Wolgast, Wendy Szymczak, Robert H. Bortz III, M. Eugenia Dieterle, Catalina Florez, Denise Haslwanter, Rohit K. Jangra, Ethan Laudermilch, Ariel S. Wirchnianski, Jason Barnhill, David L. Goldman, Hnin Khine, D. Yitzchak Goldstein, Johanna P. Daily, Kartik Chandran, Libusha Kelly |
| EPI_ISL_831681, EPI_ISL_831683, EPI_ISL_831685 | Laboratório de Microbiologia Molecular - Universidade FEEVALE | Universidade Federal de Ciências da Saúde de Porto Alegre | Vinícius Bonetti Franceschi, Amanda de Menezes Mayer, Gabriel Dickin Caldana, Carla Andretta Moreira Neves, Patrícia Aline Gröhs Ferrareze, Gabriela Bettella Cybis, Ricardo Ariel Zimerman, Lívia Kmetzsch, Fernando Rosado Spilki, Claudia Elizabeth Thompson |
| EPI_ISL_831917 | New Mexico Department of Health Scientific Laboratory | New Mexico Department of Health Scientific Laboratory | Ellie Johnson, Anastacia Griego-Fisher, D'eldra Malone |
| EPI_ISL_833156 | Instituto Adolfo Lutz - Regional de Sorocaba | Instituto Adolfo Lutz, Interdiciplinary Procedures Center, Strategic Laboratory | Claudio Tavares Sacchi, Claudia Regina Gonçalves, Erica Valessa Ramos Gomes, Karoline Rodrigues Campos |
| EPI_ISL_833177 | National Institute of Laboratory Medicine and Referral Center | Genomic Research Lab, BCSIR | Shahina Akter, Mohammad Samir Uzzaman, Eshrar Osman, Md. Ahashan Habib,Tanjina Akhtar Banu, Abu Sayeed Mohammad Mahmud, Md. Murshed Hasan Sarkar,Barna Goswami, Iffat Jahan, Md. Saddam Hossain, Tasnim Nafisa, Md. Maruf Ahmed Molla, Mahmuda Yeasmin, Asish Kumar Ghosh, A. K. M. Shamsuzzaman, Monira Parveen, Md. Masum Hossain Arif, Md. Salim Khan |
| EPI_ISL_833179 | National Institute of Laboratory Medicine and Referral Center | Genomic Research Lab, BCSIR | Iffat Jahan, Mohammad Samir Uzzaman, Eshrar Osman, Md. Ahashan Habib, Shahina Akter, Tanjina Akhtar Banu, Abu Sayeed Mohammad Mahmud, Md. Murshed Hasan Sarkar, Barna Goswami, Md. Saddam Hossain, Tasnim Nafisa, Md. Maruf Ahmed Molla, Mahmuda Yeasmin, Asish Kumar Ghosh, A. K. M. Shamsuzzaman, Monira Parveen, Md. Masum Hossain Arif, Md. Salim Khan |
| EPI_ISL_833194, EPI_ISL_833197, EPI_ISL_833199 | Hôpital Bichat Claude Bernard, Laboratoire de Virologie | IAME UMR1137 Inserm, Université de Paris, Hôpital Bichat | Antoine Bridier, Amélie Recoing, Quentin Le Hingrat, Lena Daniel, Siham Hamri, Gilles Collin, Alexandre Storto, Mélanie Bertine, Charlotte Charpentier, Nadhira Houhou-Fidouh, Diane Descamps, Benoit Visseaux |
| EPI_ISL_833492 | RS MMC Jakarta, Indonesia | Biosafety Level-3 Laboratory, Indonesian Institute of Sciences (LIPI) | Isa Nuryana, Ade Andriani, Anik Budhi Dharmayanthi, Syam Budi Iryanto, Andri Wardiana, Anggia Prasetyoputri, Ahmad Fathoni, Nova Dilla Yanthi, Yuliawati, Ratih Asmana Ningrum |
| EPI_ISL_833494 | RS PMI Bogor, Indonesia | Biosafety Level-3 Laboratory, Indonesian Institute of Sciences (LIPI) | Ade Andriani, Anik Budhi Dharmayanthi, Syam Budi Iryanto, Andri Wardiana, Anggia Prasetyoputri, Isa Nuryana, Ahmad Fathoni, Maritsa Nurfatwa, Anky Zannati, Ratih Asmana Ningrum |
| EPI_ISL_833504 | RS MMC, Jakarta, Indonesia | Biosafety Level-3 Laboratory, Indonesian Institute of Sciences (LIPI) | Isa Nuryana, Anik Budhi Dharmayanthi, Syam Budi Iryanto, Andri Wardiana, Anggia Prasetyoputri, Ade Andriani, Ahmad Fathoni, Ahmad Randy, Listiana Oktavia, Ratih Asmana Ningrum |
| EPI_ISL_833505 | RS MMC, Jakarta, Indonesia | Biosafety Level-3 Laboratory, Indonesian Institute of Sciences (LIPI) | Ade Andriani, Anik Budhi Dharmayanthi, Syam Budi Iryanto, Andri Wardiana, Anggia Prasetyoputri, Isa Nuryana, Ahmad Fathoni, Muhammad Ridwan, Ratih Asmana Ningrum |
| EPI_ISL_833507 | RS MMC, Jakarta, Indonesia | Biosafety Level-3 Laboratory, Indonesian Institute of Sciences (LIPI) | Anggia Prasetyoputri, Anik Budhi Dharmayanthi, Syam Budi Iryanto, Andri Wardiana, Isa Nuryana, Ade Andriani, Ahmad Fathoni, Syaiful Rizal, Pangda Sopha, Ratih Asmana Ningrum |
| EPI_ISL_833575 | Veterinary Specialized Instute "Nis" | Veterinary Specialized Institute "Sabac", Serbia | Vidanovic,D., Tesovic,B., Mrkovacki, S., Vujinovic,S., .,Knezevic,A., Jovanovic,T., Jankovic,M., Sekler,M., Banovic Djeri,B., Petrovic,T., Volkening,J., Afonso,C. |
| EPI_ISL_837558, EPI_ISL_837560 | Centro Nacional de Enfermedades Tropicales (CENETROP) | Laboratory of Respiratory Viruses and Measles, Oswaldo Cruz Institute, FIOCRUZ | Paola Resende, Roxana Loayza, Cinthia Avila, Luciana Appolinario, Fernando Motta, Anna Carolina Paixao, Ana Carolina Mendonca, Marilda Siqueira |
| EPI_ISL_837584, EPI_ISL_837585 | Laboratorio Nacional de Salud | Laboratory of Respiratory Viruses and Measles, Oswaldo Cruz Institute, FIOCRUZ | Paola Resende, Cesar Roberto Conde Pereira, Claudia Estrada, Luciana Appolinario, Fernando Motta, Anna Carolina Paixao, Ana Carolina Mendonca, Marilda Siqueira |
| EPI_ISL_837750, EPI_ISL_837751, EPI_ISL_837752, EPI_ISL_837753, EPI_ISL_837754, EPI_ISL_837755, EPI_ISL_837756, EPI_ISL_837757, EPI_ISL_837758, EPI_ISL_837759, EPI_ISL_837760, EPI_ISL_837761, EPI_ISL_837762 | | | |
| see above | Instituto Nacional de Enfermedades Respiratorias (INER) | Instituto Nacional de Enfermedades Respiratorias (INER) | Celia Boukadida, Margarita Matías-Florentino, Alma Rincón-Rubio, Hector Esteban Paz-Juárez, Olivia Briceño, Edgar Sevilla-Reyes, Fidencio Mejía-Nepomuceno, Mario Mújica-Sánchez, Eduardo Becerril-Vargas, José Arturo Martínez-Orozco, Alejandra Hernández-Terán, Jorge Salas-Hernández, Santiago Ávila-Ríos, Joel Armando Vázquez-Pérez |
| EPI_ISL_842864 | Barts Health NHS Trust | COVID-19 Genomics UK (COG-UK) Consortium | CUTINO-MOGUEL, Maria-Teresa; HARRINGTON, David; OWOYEMI, Dola; SHYLINI, Raghavendran; BROAD, Claire; KELE, Beatrix |
| EPI_ISL_846599, EPI_ISL_846600, EPI_ISL_846628, EPI_ISL_846629, EPI_ISL_846630, EPI_ISL_846651, EPI_ISL_846655, EPI_ISL_847819, EPI_ISL_847823, EPI_ISL_847824 | Ohio Department of Health Laboratory | Ohio Department of Health Laboratory | Holmes, Jennifer; Eric Brandt, Keoni Omura, Glen McGillivary, Caitlin McDonnell, Kirtana Ramadugu, Erica Leasure, Kelsey Florek, Heather Blankenship, Quanta Brown, and Tammy Bannerman |
| EPI_ISL_847826 | COVID-19 National Reference Laboratory | COVID-19 National Reference Laboratory | Tahmineh Jalali, Mohammad Hassan Pouriayevali, Zahra Ahmadi, Marzieh Sadjadi, Mahsa Tavakoli, Zahra Fereydouni, Setareh Kashanian, Sanam Azad-Manjiri, Tahereh Mohammadi, Zabihollah Shoja, Parastoo Yekta, Farideh Niknam, Hessam Nemati, Ahmad Ghasemi, Sahar Khakifirouz, Sepideh Gerdooei, Maryam Rostamtabar,Sana Eybpoosh, Mohammad Mehdi Mortazavipour, Mohamad Sadegh Shams Nosrati, Zeynab VeisiZadeh, Amitis Ramezani, Kayhan Azadmanesh, Mostafa Salehi-Vaziri |
| EPI_ISL_847828, EPI_ISL_847842 | Ohio Department of Health Laboratory | Ohio Department of Health Laboratory | Holmes, Jennifer; Eric Brandt, Keoni Omura, Glen McGillivary, Caitlin McDonnell, Kirtana Ramadugu, Erica Leasure, Kelsey Florek, Heather Blankenship, Quanta Brown, and Tammy Bannerman |
| EPI_ISL_848076 | Laboratory of Clinical Research on Dermatozoonoses in Domestic Animals, Evandro Chagas National Institute of Infectious Diseases, Oswaldo Cruz Foundation (Fiocruz), Rio de Janeiro, RJ, Brazil. | Laboratory of Respiratory Viruses and Measles, Oswaldo Cruz Institute, FIOCRUZ | Guilherme Amaral Calvet, Michelle Fernanda Borges da Silva, Anielle de Pina Costa, Ezequias Batista Martins, Isabella Campos Vargas de Moraes, Lusiele Guaraldo, Patrícia Brasil, Sandro Antônio Pereira, Rodrigo Caldas Menezes, Isabella Dib Ferreira Gremião, Lucas Oliveira Keidel, Shanna Araujo dos Santos, Artur Augusto Velho Mendes Junior, Renato Orsini Ornellas, Maria Ogrzewalska, PaolaCristina Resende, Alex Pauvolid-Corrêa, Fernando do Couto Motta, Alice Sampaio Barreto da Rocha, Thiago C. Souza, Marilda Mendonça Siqueira |
| EPI_ISL_848210, EPI_ISL_848219, EPI_ISL_848296, EPI_ISL_848481, EPI_ISL_848482, EPI_ISL_848483 | Illinois Department of Public Health | Gagnon Lab, Southern Illinois University | Keith Gagnon |

| | | | |
|---|---|---|---|
| EPI_ISL_848562, EPI_ISL_848563, EPI_ISL_848564, EPI_ISL_848565, EPI_ISL_848605 | Evandro Chagas Institute | Evandro Chagas Institute | Santos, M.C.; Silva, A.M.; Junior, W.D.C.; Barbagelata, L.S.; Ferreira, J.A.; Sousa, E.M.A.; da Silva, P.S.; Pinheiro, K.C.; L.C.; Sousa Junior, E.C. |
| EPI_ISL_849120, EPI_ISL_849121, EPI_ISL_849122, EPI_ISL_849123, EPI_ISL_849124, EPI_ISL_849125, EPI_ISL_849126, EPI_ISL_849127, EPI_ISL_849128, EPI_ISL_849129, EPI_ISL_849130, EPI_ISL_849131, EPI_ISL_849132, EPI_ISL_849133, EPI_ISL_849134, EPI_ISL_849135, EPI_ISL_849136, EPI_ISL_849137, EPI_ISL_849138, EPI_ISL_849139, EPI_ISL_849140, EPI_ISL_849141, EPI_ISL_849142, EPI_ISL_849143, EPI_ISL_849144, EPI_ISL_849145, EPI_ISL_849146, EPI_ISL_849147 | | | |
| see above | Florida Bureau of Public Health Laboratories | Florida Bureau of Public Health Laboratories | Sarah Schmedes, Jason Blanton |
| EPI_ISL_849281, EPI_ISL_849289, EPI_ISL_849291, EPI_ISL_849292, EPI_ISL_849302, EPI_ISL_849306, EPI_ISL_849311, EPI_ISL_849312, EPI_ISL_849317, EPI_ISL_849320 | Servicio Virosis Respiratorias-Departamento Virología-INEI | Instituto Nacional Enfermedades Infecciosas C.G.Malbran | Baumeister E., Avaro M., Benedetti E., Russo M., Dattero ME, Pontoriero A., Cisterna D., Molina V., Perandones C., Tuduri E., Lorenzo F., Poklepovich T., Campos J. |
| EPI_ISL_849654 | Servizio Igiene Epidemiologia e Sanità Pubblica (SIESP)-L'Aquila | Istituto Zooprofilattico Sperimentale dell'Abruzzo e Molise "G.Caporale" | Lorusso A, Marcacci M, Di Domenico M, Curini V, Ancora M, Cammà C, Rinaldi A, Mangone I, Di Pasquale A, Puglia I, Savini G. |
| EPI_ISL_849937, EPI_ISL_849938, EPI_ISL_849949 | UC Davis- Department of Pathology and Laboratory Medicine | Chan-Zuckerberg Biohub | CZB Cliahub Consortium |
| EPI_ISL_850227, EPI_ISL_850228, EPI_ISL_850229 | Division of Emerging Infectious Diseases, Bureau of Infectious Diseases Diagnosis Control, Korea Disease Control and Prevention Agency | Division of Emerging Infectious Diseases, Bureau of Infectious Diseases Diagnosis Control, Korea Disease Control and Prevention Agency | Ae Kyung Park, Il-Hwan Kim, Heui Man Kim, Jeong-Min Kim, Namjoo Lee, Chaeyoung Lee, Sang Hee Woo, Eun-Jin Kim |
| EPI_ISL_852574 | Max von Pettenkofer Institute, Virology, National Reference Center for Retroviruses, LMU München | Laboratory for Functional Genome Analysis, Dept. Genomics, Gene Center of the LMU Munich | Max Muenchhoff, Stefan Krebs, Alexander Graf, Oliver Keppler, Helmut Blum |
| EPI_ISL_852617, EPI_ISL_852618, EPI_ISL_852620, EPI_ISL_852621, EPI_ISL_852623, EPI_ISL_852625, EPI_ISL_852626 | Ohio Department of Health Laboratory | Ohio Department of Health Laboratory | Holmes, Jennifer; Eric Brandt, Keoni Omura, Glen McGillivary, Caitlin McDonnell, Kirtana Ramadugu, Erica Leasure, Kelsey Florek, Heather Blankenship, Quanta Brown, and Tammy Bannerman |
| EPI_ISL_852807 | Institute of Virology, Medical Center, University of Freiburg, Freiburg, Germany | Institute of Virology, Clinial Virus Genomics, Medical Center, University of Freiburg, Freiburg, Germany | Jonas Fuchs, Lisa Kern, Sandra Reuter, Hajo Grundmann, Marcus Panning |
| EPI_ISL_853288, EPI_ISL_853290 | UPMC Clinical Microbiology Laboratory | Microbial Genome Sequencing Center; Microbial Genomic Epidemiology Laboratory | Mustapha M. Mustapha, Jane W. Marsh, Dan Snyder, Marissa P. Griffith, Stephanie L. Mitchell, Vatsala R. Srinivasa, Kady D. Waggle, Chinelo Ezeonwuku, Vaughn S. Cooper, Lee H. Harrison |
| EPI_ISL_859578, EPI_ISL_859579, EPI_ISL_859580, EPI_ISL_859581, EPI_ISL_859582, EPI_ISL_859621, EPI_ISL_859623, EPI_ISL_859629, EPI_ISL_859642, EPI_ISL_859643, EPI_ISL_859644, EPI_ISL_859646, EPI_ISL_859652, EPI_ISL_859653, EPI_ISL_859654, EPI_ISL_859655 | | | |
| see above | BTC, Khalifa University | BTC, Khalifa University | Al Safar et al |
| EPI_ISL_860256, EPI_ISL_860258, EPI_ISL_860296 | Ohio Department of Health Laboratory | Ohio Department of Health Laboratory | Holmes, Jennifer; Eric Brandt, Keoni Omura, Glen McGillivary, Caitlin McDonnell, Kirtana Ramadugu, Erica Leasure, Kelsey Florek, Heather Blankenship, Quanta Brown, and Tammy Bannerman |
| EPI_ISL_860336, EPI_ISL_860348, EPI_ISL_860355, EPI_ISL_860482, EPI_ISL_860516 | MVZ Labor Krone GbR | Center of Medical Microbiology, Virology, and Hospital Hygiene, University of Duesseldorf | Dennis Deschka, Alexander Dilthey, Julia Fazaal, André Heimbach, Per Hoffmann, Torsten Houwaart, Malte Kohns Vasconcelos, Klaus Pfeffer, Bärbel Lippke, Kerstin Ludwig, Janine Silvery, Carsten Tiemann, Jörg Timm, Andreas Walker, Tobias Wienemann |
| EPI_ISL_860549 | Ohio Department of Health Laboratory | Ohio Department of Health Laboratory | Holmes, Jennifer; Eric Brandt, Keoni Omura, Glen McGillivary, Caitlin McDonnell, Kirtana Ramadugu, Erica Leasure, Kelsey Florek, Heather Blankenship, Quanta Brown, and Tammy Bannerman |
| EPI_ISL_861665 | Instituto Adolfo Lutz - Regional de Taubate | Instituto Adolfo Lutz, Interdiciplinary Procedures Center, Strategic Laboratory | Claudio Tavares Sacchi, Claudia Regina Gonçalves, Erica Valessa Ramos Gomes, Karoline Rodrigues Campos |
| EPI_ISL_862268, EPI_ISL_862269, EPI_ISL_862273, EPI_ISL_862277, EPI_ISL_862281, EPI_ISL_862282, EPI_ISL_862287, EPI_ISL_862291, EPI_ISL_862292, EPI_ISL_862293, EPI_ISL_862294, EPI_ISL_862296, EPI_ISL_862302, EPI_ISL_862303, EPI_ISL_862305, EPI_ISL_862309, EPI_ISL_862310, EPI_ISL_862311, EPI_ISL_862312, EPI_ISL_862313, EPI_ISL_862438, EPI_ISL_862439, EPI_ISL_862440, EPI_ISL_862441, EPI_ISL_862442, EPI_ISL_862443, EPI_ISL_862444, EPI_ISL_862445, EPI_ISL_862446, EPI_ISL_862448, EPI_ISL_862449, EPI_ISL_862450, EPI_ISL_862451, EPI_ISL_862452, EPI_ISL_862453, EPI_ISL_862454, EPI_ISL_862455, EPI_ISL_862456, EPI_ISL_862457, EPI_ISL_862462, EPI_ISL_862463, EPI_ISL_862464, EPI_ISL_862466, EPI_ISL_862512, EPI_ISL_862521, EPI_ISL_862522, EPI_ISL_862523, EPI_ISL_862524, EPI_ISL_862525, EPI_ISL_862526, EPI_ISL_862527, EPI_ISL_862528, EPI_ISL_862529, EPI_ISL_862530, EPI_ISL_862531, EPI_ISL_862532, EPI_ISL_862533, EPI_ISL_862537, EPI_ISL_862538, EPI_ISL_862539, EPI_ISL_862540, EPI_ISL_862541, EPI_ISL_862542, EPI_ISL_862543 | | | |
| see above | Kurnool Medical College (KMC) | CSIR Institute of Genomics and Integrative Biology | Pallavali Roja Rani, Mohamed Imran, J. Vijaya Lakshmi, Bani Jolly, S. Afsar, Abhinav Jain, Mohit Kumar Divakar, Panyam Suresh, Disha Sharma, Nambi Rajesh, Rahul C Bhoyar, Dasari Ankaiah, Sanaga Shanthi Kumari, Gyan Ranjan, Valluri Anitha Lavanya, Mercy Rophina, S. Umadevi, Paras Sehgal, Avula Renuka Devi, A. Surekha, Pulala Chandra, Rajamadugu Hymavathy, P R Vanaja, Vinod Scaria, Sridhar Sivasubbu |
| EPI_ISL_875540 | Instituto de Biotecnologia - UNESP-Botucatu-SP | Instituto de Biotecnologia - UNESP-Botucatu-SP | Leila Sabrina Ullmann; Fábio Sossai Possebon, Camila Dantas Malossi, Paula Rahal, Paulo Inacio da Costa, João Pessoa Araújo Jr. |
| EPI_ISL_876035 | Montefiore Medical Center | Albert Einstein College of Medicine, Dept. of Microbiology & Immunology, Chandran lab | J. Maximilian Fels, Saad Khan, Ryan Forster, Karin A. Skalina, Surksha Sirichand, Amy S. Fox, Aviv Bergman, William B. Mitchell, Lucia R. Wolgast, Wendy Szymczak, Robert H. Bortz III, M. Eugenia Dieterle, Catalina Florez, Denise Haslwanter, Rohit K. Jangra, Ethan Laudermilch, Ariel S. Wirchnianski, Jason Barnhill, David L. Goldman, Hnin Khine, D. Yitzchak Goldstein, Johanna P. Daily, Kartik Chandran, Libusha Kelly |
| EPI_ISL_876958, EPI_ISL_876959, EPI_ISL_876960, EPI_ISL_876961, EPI_ISL_876962, EPI_ISL_876963, EPI_ISL_876964 | Quest Diagnostics | Quest Diagnostics | Rosenthal,S.H., Gerasimova,A., Kagan,R.M., Anderson, B., Hua, M., Liu Y., Bernstein, L.E., Livingston, K.E., Perez, A., Shalhout, D.F., Shlyakhter, I.A., Owen, R., Tanpaiboon, P., Lacbawan, F. |
| EPI_ISL_877632, EPI_ISL_877633, EPI_ISL_877634, EPI_ISL_877635, EPI_ISL_877636, EPI_ISL_877637, EPI_ISL_877638, EPI_ISL_877639, EPI_ISL_877640, EPI_ISL_877641, EPI_ISL_877642, EPI_ISL_877643, EPI_ISL_877644, EPI_ISL_877645, EPI_ISL_877646, EPI_ISL_877647 | | | |
| see above | Clinical Molecular Microbiology Laboratory, UNC Hospital | Dirk Dittmer | Razia Moorad , Justin T. Landis , Brent A. Eason, Melissa B. Miller, Linda Pluta, Dirk Dittmer, Angelica Juarez, Cecilia Thompson , Cameroon Grant, Evelyn Hoffman, Patricio Cano, Jason Wong, Carolina Caro-Vegas, Blossom Damania. |
| EPI_ISL_884344, EPI_ISL_884354, EPI_ISL_884425, EPI_ISL_884426, EPI_ISL_884427, EPI_ISL_884428, EPI_ISL_884431 | Infectious Diseases, Quest Diagnostics | Infectious Diseases, Quest Diagnostics | Rosenthal,S.H., Gerasimova,A., Kagan,R.M., Anderson,B., Bernstein,L.E., Livingston,K.E., Hua,M., Liu,Y., Shalhout,D.F., Owen,R., Lacbawan,F. |
| EPI_ISL_884445, EPI_ISL_884446, EPI_ISL_884447, EPI_ISL_884448, EPI_ISL_884449, EPI_ISL_884450, EPI_ISL_884451, EPI_ISL_884452, EPI_ISL_884453, EPI_ISL_884454, EPI_ISL_884455, EPI_ISL_884456, EPI_ISL_884457, EPI_ISL_884458, EPI_ISL_884459, EPI_ISL_884460, EPI_ISL_884462, EPI_ISL_884463, EPI_ISL_884464, EPI_ISL_884465, EPI_ISL_884466, EPI_ISL_884467, EPI_ISL_884468, EPI_ISL_884469, EPI_ISL_884470, EPI_ISL_884471, EPI_ISL_884472, EPI_ISL_884473, EPI_ISL_884474, EPI_ISL_884475, EPI_ISL_884476, EPI_ISL_884477, EPI_ISL_884478, EPI_ISL_884479, EPI_ISL_884480, EPI_ISL_884481, EPI_ISL_884482, EPI_ISL_884484, EPI_ISL_884485, EPI_ISL_884486, EPI_ISL_884487, EPI_ISL_884488, EPI_ISL_884489, EPI_ISL_884490, EPI_ISL_884491, EPI_ISL_884492, EPI_ISL_884493, EPI_ISL_884494, EPI_ISL_884495, EPI_ISL_884496, EPI_ISL_884497, EPI_ISL_884498, EPI_ISL_884499, EPI_ISL_884500, EPI_ISL_884501, EPI_ISL_884502, EPI_ISL_884503, EPI_ISL_884504, EPI_ISL_884505, EPI_ISL_884506, EPI_ISL_884507, EPI_ISL_884508, EPI_ISL_884509, EPI_ISL_884510, EPI_ISL_884511, EPI_ISL_884512, EPI_ISL_884513, EPI_ISL_884514, EPI_ISL_884515, EPI_ISL_884516, EPI_ISL_884517, EPI_ISL_884518, EPI_ISL_884519, EPI_ISL_884520, EPI_ISL_884521, EPI_ISL_884522, EPI_ISL_884523, EPI_ISL_884524, EPI_ISL_884525, EPI_ISL_884526, EPI_ISL_884527, EPI_ISL_884528, EPI_ISL_884529, EPI_ISL_884569, EPI_ISL_884579 | | | |
| see above | Molecular Microbiology & Immunology, University of Missouri | Molecular Microbiology & Immunology, University of Missouri | Tang,C.Y., Li,T., Hang,J., Lidl,G.M., Wan,X.-F. |
| EPI_ISL_888672 | Instituto de Biotecnologia - UNESP-Botucatu-SP | Instituto de Biotecnologia - UNESP-Botucatu-SP | Leila Sabrina Ullmann; Fábio Sossai Possebon, Camila Dantas Malossi, Paula Rahal, Paulo Inacio da Costa, João Pessoa Araújo Jr. |
| EPI_ISL_889331, EPI_ISL_889332, EPI_ISL_889333 | The University Hospital Brno | Institute of Applied Biotechnologies a.s. | Petr Klempt, Ondej Brzo, Martin Kašný, Kateina Kvapilová, Martina Lengerová, Petr Kvapil |
| EPI_ISL_890099, EPI_ISL_890100, EPI_ISL_890101, EPI_ISL_890102, EPI_ISL_890103, EPI_ISL_890104, EPI_ISL_890105, EPI_ISL_890106, EPI_ISL_890107, EPI_ISL_890108, EPI_ISL_890109 | | | |
| see above | Laboratoire de santé publique du Québec | Laboratoire de santé publique du Québec | Sandrine Moreira, Ioannis Ragoussis, Guillaume Bourque, Jesse Shapiro, Mark Lathrop and Michel Roger on behalf of the CoVSeQ research group |

| | | | |
|---|---|---|---|
| EPI_ISL_900128, EPI_ISL_900136, EPI_ISL_900187, EPI_ISL_900241, EPI_ISL_900371 | MEPHI, Aix Marseille University | MEPHI, Aix Marseille University | Anthony LEVASSEUR |
| EPI_ISL_900693, EPI_ISL_900694, EPI_ISL_900695, EPI_ISL_900696, EPI_ISL_900697, EPI_ISL_900698, EPI_ISL_900699, EPI_ISL_900700, EPI_ISL_900701, EPI_ISL_900702 | Bozeman Health Deaconess Hospital | Wiedenheft lab, Montana State University | Artem Nemudryi, Anna Nemudraia, Tanner Wiegand, Joseph Nichols, Deann T. Snyder, Jodi F. Hedges, Calvin Cicha, Helen Lee, Karl K. Vanderwood, Diane Bimczok, Mark A. Jutila and Blake Wiedenheft |
| EPI_ISL_903345 | Bozeman Health Deaconess Hospital | Wiedenheft lab, Montana State University | Artem Nemudryi, Anna Nemudraia, Tanner Wiegand, Joseph Nichols, Deann T. Snyder, Jodi F. Hedges, Calvin Cicha, Helen Lee, Karl K. Vanderwood, Diane Bimczok, Mark A. Jutila and Blake Wiedenheft |
| EPI_ISL_906101 | Child Health Research Foundation | Child Health Research Foundation | Senjuti Saha, Sharmistha Goswami, Afroza Akter Tanni, Syed Muktadir Al Sium, Arif Mohammad Tanmoy, Roly Malaker, Md Hafizur Rahman, Samir K Saha |
| EPI_ISL_914488, EPI_ISL_914489, EPI_ISL_914490, EPI_ISL_914491, EPI_ISL_914492, EPI_ISL_914494, EPI_ISL_914495, EPI_ISL_914496, EPI_ISL_914497, EPI_ISL_914498, EPI_ISL_914499, EPI_ISL_914500, EPI_ISL_914501, EPI_ISL_914502, EPI_ISL_914503, EPI_ISL_914504, EPI_ISL_914505, EPI_ISL_914506, EPI_ISL_914507, EPI_ISL_914508, EPI_ISL_914509, EPI_ISL_914510, EPI_ISL_914511, EPI_ISL_914512, EPI_ISL_914513, EPI_ISL_914514, EPI_ISL_914515, EPI_ISL_914516, EPI_ISL_914517, EPI_ISL_914518, EPI_ISL_914519, EPI_ISL_914520, EPI_ISL_914521, EPI_ISL_914522, EPI_ISL_914523, EPI_ISL_914524, EPI_ISL_914525, EPI_ISL_914526, EPI_ISL_914527, EPI_ISL_914528, EPI_ISL_914532 | | | |
| see above | TGen North | TGen North | "Jolene Bowers, Megan Folkerts, Chris French, Hayley Yaglom, Ashlyn Pfeiffer, Darrin Lemmer, Dave Engelthaler, The Arizona COVID Genomics Union (ACGU)" |
| EPI_ISL_915389, EPI_ISL_915390, EPI_ISL_915391 | Keio University School of Medicine | Keio University School of Medicine | Kenjiro Kosaki, Yuka Iwasaki, Hirotsugu Ishizu, Haruhiko Siomi, Kodai Abe |
| EPI_ISL_933648, EPI_ISL_933649 | Toronto Invasive Bacterial Diseases Network | McMaster University | Allison McGeer, Patryk Aftanas, Hooman Derakhshani, Angel Li, Kuganya Nirmalarajah, Emily Panousis, Ahmed Draia, Jalees Nasir, Michael Surette, Samira Mubareka, Andrew G. McArthur |
| EPI_ISL_936558, EPI_ISL_936559, EPI_ISL_936560, EPI_ISL_936561, EPI_ISL_936562, EPI_ISL_936563, EPI_ISL_936564, EPI_ISL_936565, EPI_ISL_936566, EPI_ISL_936567, EPI_ISL_936568, EPI_ISL_936569, EPI_ISL_936570, EPI_ISL_936571, EPI_ISL_936572, EPI_ISL_936573 | | | |
| see above | Northwestern Memorial Hospital | Ozer Lab | Ramon Lorenzo-Redondo, Lacy M. Simons, Chad J. Achenbach, Lawrence J. Jennings, Michael G. Ison, Judd F. Hultquist, Egon A. Ozer |
| EPI_ISL_940268, EPI_ISL_940278, EPI_ISL_940347, EPI_ISL_940540, EPI_ISL_940544, EPI_ISL_940549 | Hôpital Bichat Claude Bernard, Laboratoire de Virologie | IAME UMR1137 Inserm, Université de Paris, Hôpital Bichat | Antoine Bridier-Nahmias, Amélie Recoing, Quentin Le Hingrat, Lena Daniel, Siham Hamri, Gilles Collin, Alexandre Storto, Mélanie Bertine, Charlotte Charpentier, Nadhira Houhou-Fidouh, Diane Descamps, Benoit Visseaux |
| EPI_ISL_940905, EPI_ISL_940908 | Centers for Disease Control and Prevention, Dengue Branch | Centers for Disease Control and Prevention, Dengue Branch | Gilberto A. Santiago, Glenda Gonzalez, Betzabel Flores, Keyla Charriez, Gabriela Paz-Bailey, Jorge L. Munoz-Jordan |
| EPI_ISL_941947, EPI_ISL_941997, EPI_ISL_941998 | Centro de Investigaciones en Microbiología y Biotecnología-UR (CIMBIUR), Facultad de Ciencias Naturales, Universidad del Rosario, Bogotá, Colombia | Centro de Investigaciones en Microbiología y Biotecnología-UR (CIMBIUR), Facultad de Ciencias Naturales, Universidad del Rosario, Bogotá, Colombia Instituto Nacional de Salud, Bogotá, Colombia Icahn School of Medicine at Mount Sinai, New York, USA | Luz Helena Patiño, Marina Muñoz, Nathalia Ballesteros, Carolina Hernández, Carolina Flórez, Sergio Gomez, Adriana van de Guchte, Zenab Khan, Jayeeta Dutta, Hala Alejel Alshammary, Ana S. Gonzalez-Reiche, Matthew M. Hernandez, Emilia Mia Sordillo, Viviana Simon, Harm van Bakel, Alberto Paniz-Mondolfi, Juan David Ramírez |
| EPI_ISL_942929 | Instituto de Diagnostico y Referencia Epidemiologicos INDRE_RNLSP | Instituto de Diagnostico y Referencia Epidemiologicos (INDRE) | Claudia Wong-Arambula, Abril Rodriguez-Maldonado, Fabiola Garces-Ayala, Natividad Cruz-Ortiz, Tatiana Nunez-Garcia, Gisela Barrera-Badillo, Lucia Hernandez-Rivas, Irma Lopez-Martinez, Ernesto Ramirez-Gonzalez. |
| EPI_ISL_943600 | Lacen_RS | State Center for Health Surveillance. Rio Grande do Sul State Secretary of Health | Aline Campos, Amanda da Silva, Anelise Schaurich, Claudia Dornelles, Cynthia Molina, Fernanda Godinho, Lara Crescente, Leticia Garay, Regina Barcellos, Richard Salvato, Tatiana Gregianini, Vagner Fonseca |
| EPI_ISL_953408, EPI_ISL_953412, EPI_ISL_953414 | Laboratorio de Investigaciones de Baney | "Swiss Tropical and Public Health Institute" | "Carlos Cortes, Claudia Daubenberger, Guillermo Garcia, Salome Hosch, Bonifacio Manguire Nlavo, Maximilian Mpina, Elizabeth Nyakarungu, Diosdado Odjama Nseng Ada, Mitoha Ondo O Ayekaba, Tobias Schindler, Philip Wonder Phiri" |
| EPI_ISL_954302 | Laboratoire de santé publique du Québec | Laboratoire de santé publique du Québec | Sandrine Moreira, Ioannis Ragoussis, Guillaume Bourque, Jesse Shapiro, Mark Lathrop and Michel Roger on behalf of the CoVSeQ research group (http://covseq.ca/researchgroup) |
| EPI_ISL_959282 | National Influenza Center, Virology Department | National Influenza Center | J Yavarian, NZ Shafiei Jandaghi, V Salimi, A Nejati, K Sadeghi, F Ajaminejad,N Ghavvami and T Mokhtari Azad |
| EPI_ISL_960154, EPI_ISL_960155, EPI_ISL_960156, EPI_ISL_960157, EPI_ISL_960158 | Guguletu CHC wc GDH | National Health Laboratory Service/UCT | Arash Iranzadeh, Deelan Doolabh, Lynn Tyers, Bruna Galvao, Innocent Mudau, Marvin Hsiao, Kruger Marais, Diana Hardie, Stephen Korsman, Carolyn Williamson |
| EPI_ISL_961774, EPI_ISL_961775, EPI_ISL_961776, EPI_ISL_961777, EPI_ISL_961778 | Laboratorio de Infectología, Servicio de Infectología, Hospital Universitario Dr. José Eleuterio González - Universidad Autónoma de Nuevo León | Laboratorio de Infectología Molecular, Departamento de Bioquímica y Medicina Molecular,Facultad de Medicina - Universidad Autónoma de Nuevo León | Kame A. Galán-Huerta, María F. Herrera-Saldivar, Natalia Martínez-Acuña, Sonia A. Lozano-Sepúlveda, Daniel Arellanos-Soto, Ana M. Rivas-Estilla, Paola Bocanegra-Ibarias, Samantha M. Flores-Treviño, Elvira Garza-González, Eduardo Perez-Alba, Laura Nuzzolo-Shihadeh, Adrian Camacho-Ortiz |
| EPI_ISL_962916 | Servicio de Microbiología, Laboratori Clínic Metropolitana Nord. Hospital Universitari Germans Trias i Pujol. Institut d'Investigació en Ciències de la Salut Germans Trias i Pujol (IGTP) | SeqCOVID-SPAIN consortium/IBV(CSIC) | Elisa Martró, Antoni E. Bordoy, Anna Not, Adrián Antuori, Anabel Fernández, Nona Romaní, Verónica Saludes, Cristina Casañ and SeqCOVID-SPAIN consortium |
| EPI_ISL_964904 | Hospital Castro Rendon | Laboratorio Central Mg. Luis Alfredo Pianciola on behalf of 'Proyecto Argentino Interinstitucional de genomica de SARS-CoV-2' (PAIS Consortium) | L Pianciola, M Mazzeo, C Ziehm, C Pintos, M Fernandez, J Ousset, M Nabaes, M Viegas. |
| EPI_ISL_964911 | Hospital Cutral Co | Laboratorio Central Mg. Luis Alfredo Pianciola on behalf of 'Proyecto Argentino Interinstitucional de genomica de SARS-CoV-2' (PAIS Consortium) | L Pianciola, M Mazzeo, C Ziehm, C Pintos, M Fernandez, J Ousset, M Nabaes, M Viegas. |
| EPI_ISL_964912 | Hospital Bouquet Roldan | Laboratorio Central Mg. Luis Alfredo Pianciola on behalf of 'Proyecto Argentino Interinstitucional de genomica de SARS-CoV-2' (PAIS Consortium) | L Pianciola, M Mazzeo, C Ziehm, C Pintos, M Fernandez, J Ousset, M Nabaes, M Viegas. |
| EPI_ISL_965222 | Laboratorio de Infectología, Servicio de Infectología, Hospital Universitario Dr. José Eleuterio González - Universidad Autónoma de Nuevo León | Laboratorio de Infectología Molecular, Departamento de Bioquímica y Medicina Molecular,Facultad de Medicina - Universidad Autónoma de Nuevo León | Kame A. Galán-Huerta, María F. Herrera-Saldivar, Natalia Martínez-Acuña, Sonia A. Lozano-Sepúlveda, Daniel Arellanos-Soto, Ana M. Rivas-Estilla, Paola Bocanegra-Ibarias, Samantha M. Flores-Treviño, Elvira Garza-González, Eduardo Perez-Alba, Laura Nuzzolo-Shihadeh, Adrian Camacho-Ortiz |
| EPI_ISL_965979 | Washington State Department of Health | Seattle Flu Study | Deborah A. Nickerson, Chris D. Frazar, Jover Lee, Benjamin Pelle, Erica Ryke, Matthew Richardson, Amanda Adler, Elisabeth Brandstetter, Peter D. Han, Kairsten Fay, Misja Ilcisin, Kirsten Lacombe, Thomas R. Sibley, Melissa Truong, Caitlin R. Wolf, Romesh Gautom, Geoff Melly, Brian Hiatt, Philip Dykema, Scott Lindquist, Michael Boeckh, Janet A. Englund, Michael Famulare, Barry R. Lutz, Mark J. Rieder, Lea M. Starita, Matthew Thompson, Helen Y. Chu, Jay Shendure, Trevor Bedford |
| EPI_ISL_966940 | Technical Support Units for Scientific Research (UATRS), National Centre for Scientific and Technical Research (CNRST) | Technical Support Units for Scientific Research (UATRS), National Centre for Scientific and Technical Research (CNRST) | Touil,N., Rfaki,A., Hemlali,M., Alaoui,S.A., Melloul,M.,Elalaoui,M.A., Elannaz,H., Lahlou,a.I., Elouanass,M., Ennibi,H. and El Fahime,E. |
| EPI_ISL_968146, EPI_ISL_968157, EPI_ISL_968158, EPI_ISL_968184 | Clinical Molecular Microbiology Laboratory, UNC Hospital | Dirk Dittmer | Justin T. Landis , Razia Moorad , Brent A. Eason, Melissa B. Miller, Linda Pluta, Dirk Dittmer, Angelica Juarez, Cecilia Thompson, Shawn Hawken, Cameroon Grant, Evelyn Hoffman, Patricio Cano, Jason Wong, Carolina Caro-Vegas, Ryan McNamara, Blossom Damania. |
| EPI_ISL_968280, EPI_ISL_968281, EPI_ISL_968282, EPI_ISL_968283, EPI_ISL_968284, EPI_ISL_968285, EPI_ISL_968286, EPI_ISL_968287, EPI_ISL_968288, EPI_ISL_968289, EPI_ISL_968290, EPI_ISL_968291, EPI_ISL_968292, EPI_ISL_968293, EPI_ISL_968294, EPI_ISL_968295, EPI_ISL_968296, EPI_ISL_968297, EPI_ISL_968298, EPI_ISL_968299, EPI_ISL_968300, EPI_ISL_968301, EPI_ISL_968302, EPI_ISL_968303, EPI_ISL_968304, EPI_ISL_968305, EPI_ISL_968306, EPI_ISL_968307, EPI_ISL_968308, EPI_ISL_968309, EPI_ISL_968310, EPI_ISL_968311, EPI_ISL_968312, EPI_ISL_968313, EPI_ISL_968314, EPI_ISL_968315, | | | |

| | | | |
|---|---|---|---|
| EPI_ISL_968316 | | | |
| see above | BCCDC Public Health Laboratory | BCCDC Public Health Laboratory | Prystajecky Natalie, Linda Hoang, Dan Fornika, John Tyson, Shannon Russell, Kim Macdonald, Kimia Kamelian, Ana Pacagnella, Corrinne Ng, Loretta Janz, Robert Azana Terry Snutch, Mel Krajden |
| EPI_ISL_971451 | Cell culture Unit at CV-MIT belonging to HIMMV | Functional Genomic Platform UATRS-biology, CNRST | Nadia Touil, Abderrazzak Rfak, Mouhssine Hemlali, Sanaa ALAOUI-Amine, Marouane MELLOUL, Mly Abdelaziz ELALAOUI, Hicham Elannaz, Amine Idriss Lahlou, Mostafa ELOUENNASS, Khalid ENNIBI, Elmostafa EL FAHIME |
| EPI_ISL_977253, EPI_ISL_977265, EPI_ISL_977280, EPI_ISL_977281, EPI_ISL_977306, EPI_ISL_977311, EPI_ISL_977313, EPI_ISL_977314, EPI_ISL_977327, EPI_ISL_977328, EPI_ISL_977329, EPI_ISL_977330, EPI_ISL_977333, EPI_ISL_977393, EPI_ISL_977400, EPI_ISL_977401, EPI_ISL_977402, EPI_ISL_977403, EPI_ISL_977404, EPI_ISL_977405, EPI_ISL_977406, EPI_ISL_977407, EPI_ISL_977408, EPI_ISL_977409, EPI_ISL_977410, EPI_ISL_977412, EPI_ISL_977413, EPI_ISL_977414, EPI_ISL_977415 | | | |
| see above | University of Zambia, School of Veterinary Medicine | UNZAVET and PATH | Mulenga Mwenda-Chimfwembe, Ngonda Saasa, Daniel Bridges |
| EPI_ISL_981050 | Hospital Dr. Francisco López Lima | Laboratorio Central Mg. Luis Alfredo Pianciola on behalf of 'Proyecto Argentino Interinstitucional de genomica de SARS-CoV-2' (PAIS Consortium) | L Pianciola, M Mazzeo, C Ziehm, C Pintos, M Fernandez, J Ousset, M Nabaes, M Viegas. |
