## Supplementary material for "Detection and characterization of the SARS-CoV-2 lineage B.1.526 in New York": Supp. Table 3 GISAID Acknowledgment part 2: gisaid_hcov-19_acknowledgement_table_2021_02_12_23-2.pdf

We gratefully acknowledge the following Authors from the Originating laboratories responsible for obtaining the specimens, as well as the Submitting laboratories where the genome data were generated and shared via GISAID, on which this research is based.

All Submitters of data may be contacted directly via www.gisaid.org

Authors are sorted alphabetically.

| Accession ID | Originating Laboratory | Submitting Laboratory | Authors |
|---|---|---|---|
| EPI_ISL_500717 | Area of Virology, Serology and Virology Division (SAViD), New South Wales Health Pathology Randwick | Area of Virology, Serology and Virology Division (SAViD), New South Wales Health Pathology Randwick | Rawlinson, W |
| EPI_ISL_507160, EPI_ISL_507162, EPI_ISL_507170, EPI_ISL_507171 | Virology Department, Sheffield Teaching Hospitals NHS Foundation Trust/Department of Infection, Immunity and Cardiovascular Disease, The Medical School, University of Sheffield | COVID-19 Genomics UK (COG-UK) Consortium | Thushan de Silva, Matthew Parker, Nikki Smith, Adri Angyal, Rebecca Brown, Luke Green, Rachel Tucker, Paul Parsons, Danielle Groves, Katie Johnson, Laura Carrilero, Alex Keeley, Dave Partridge, Matthew Wyles, Benjamin Lindsey, Mehmet Yavuz, Mohammad Raza, Cariad Evans |
| EPI_ISL_508605, EPI_ISL_508606, EPI_ISL_508607, EPI_ISL_508608, EPI_ISL_510542, EPI_ISL_510543 | SA Pathology | SA Pathology | Lex Leong, Chuan Kok Lim, Mark Turra, Ivan Bastian, Geoff Higgins |
| EPI_ISL_510779, EPI_ISL_510780, EPI_ISL_510781, EPI_ISL_510782, EPI_ISL_510783, EPI_ISL_510784, EPI_ISL_510806, EPI_ISL_510807, EPI_ISL_510808, EPI_ISL_510809, EPI_ISL_511989, EPI_ISL_511993, EPI_ISL_511994, EPI_ISL_511995, EPI_ISL_511996, EPI_ISL_511997, EPI_ISL_511998, EPI_ISL_511999, EPI_ISL_512000, EPI_ISL_512001, EPI_ISL_512002, EPI_ISL_512003, EPI_ISL_512004, EPI_ISL_512005, EPI_ISL_512006, EPI_ISL_512007, EPI_ISL_512008, EPI_ISL_512009, EPI_ISL_512010, EPI_ISL_512011, EPI_ISL_512012, EPI_ISL_512013, EPI_ISL_512014, EPI_ISL_512015, EPI_ISL_512016, EPI_ISL_512017, EPI_ISL_512018, EPI_ISL_512019, EPI_ISL_512020, EPI_ISL_512021, EPI_ISL_512022, EPI_ISL_512023, EPI_ISL_512024, EPI_ISL_512025, EPI_ISL_512026, EPI_ISL_512027, EPI_ISL_512028, EPI_ISL_512029, EPI_ISL_512030, EPI_ISL_512031, EPI_ISL_512032, EPI_ISL_512033, EPI_ISL_512034, EPI_ISL_512035, EPI_ISL_512036, EPI_ISL_512037, EPI_ISL_512038, EPI_ISL_512039, EPI_ISL_512040, EPI_ISL_512041, EPI_ISL_512042, EPI_ISL_512043, EPI_ISL_512044, EPI_ISL_512045, EPI_ISL_512046, EPI_ISL_512047, EPI_ISL_512048, EPI_ISL_512049, EPI_ISL_512050, EPI_ISL_512051, EPI_ISL_512052, EPI_ISL_512053, EPI_ISL_512054 | | | |
| see above | Viollier AG | Department of Biosystems Science and Engineering, ETH Zürich | Christian Beisel, Sarah Nadeau, Ivan Topolsky, Pedro Ferreira, Philipp Jablonski, Susana Posada-Céspedes, Tobias Schär, Ina Nissen, Natascha Santacroce, Elodie Burcklen, Christiane Beckmann, Maurice Redondo, Olivier Kobel, Christoph Noppen, Sophie Seidel, Noemie Santamaria de Souza, Niko Beerenwinkel, Tanja Stadler |
| EPI_ISL_512066 | Sardar Vallabhbhai Patel Institute of Medical Sciences & Research | Gujarat Biotechnology Research Centre | Komal Patel, Labdhi Pandya, Afzal Ansari, Nikha Trivedi, Pranay Shah, Kamlesh J Upadhyay, Sanjay Kapadia, Apurvasinh Puvar, Janvi Raval, Zarna Patel, Monika Gandhi, Pinal Trivedi, Maharshi Pandya, Nidhi Patel, Nitin Savaliya, Raghawendra Kumar, Dinesh Kumar, Zuber Saiyed, R D Dixit, A M Kadri, Harsh Bakshi, Chaitanya Joshi, Madhvi Joshi |
| EPI_ISL_512067 | Sardar Vallabhbhai Patel Institute of Medical Sciences & Research | Gujarat Biotechnology Research Centre | Labdhi Pandya, Afzal Ansari, Nikha Trivedi, Pranay Shah, Kamlesh J Upadhyay, Sanjay Kapadia, Apurvasinh Puvar, Janvi Raval, Zarna Patel, Monika Gandhi, Pinal Trivedi, Maharshi Pandya, Nidhi Patel, Nitin Savaliya, Raghawendra Kumar, Dinesh Kumar, Zuber Saiyed, Komal Patel, R D Dixit, A M Kadri, Harsh Bakshi, Chaitanya Joshi, Madhvi Joshi |
| EPI_ISL_512068 | Sardar Vallabhbhai Patel Institute of Medical Sciences & Research | Gujarat Biotechnology Research Centre | Afzal Ansari, Nikha Trivedi, Pranay Shah, Kamlesh J Upadhyay, Sanjay Kapadia, Apurvasinh Puvar, Janvi Raval, Zarna Patel, Monika Gandhi, Pinal Trivedi, Maharshi Pandya, Nidhi Patel, Nitin Savaliya, Raghawendra Kumar, Dinesh Kumar, Zuber Saiyed, Komal Patel, Labdhi Pandya, R D Dixit, A M Kadri, Harsh Bakshi, Chaitanya Joshi, Madhvi Joshi |
| EPI_ISL_512069 | Sardar Vallabhbhai Patel Institute of Medical Sciences & Research | Gujarat Biotechnology Research Centre | Nikha Trivedi, Pranay Shah, Kamlesh J Upadhyay, Sanjay Kapadia, Apurvasinh Puvar, Janvi Raval, Zarna Patel, Monika Gandhi, Pinal Trivedi, Maharshi Pandya, Nidhi Patel, Nitin Savaliya, Raghawendra Kumar, Dinesh Kumar, Zuber Saiyed, Komal Patel, Labdhi Pandya, Afzal Ansari, R D Dixit, A M Kadri, Harsh Bakshi, Chaitanya Joshi, Madhvi Joshi |
| EPI_ISL_512089, EPI_ISL_512092, EPI_ISL_512093, EPI_ISL_512094, EPI_ISL_512095, EPI_ISL_512096, EPI_ISL_512097, EPI_ISL_512098, EPI_ISL_512099, EPI_ISL_512100, EPI_ISL_512103, EPI_ISL_512104, EPI_ISL_512105, EPI_ISL_512106 | | | |
| see above | National Virus Reference Laboratory | National Virus Reference Laboratory | Michael Carr, Gabriel Gonzalez, Jonathan Dean, Aditi Chaturvedi, Suzie Coughlan, Cillian F De Gascun |
| EPI_ISL_512330, EPI_ISL_512334 | Department of Pathology, University of Cambridge | COVID-19 Genomics UK (COG-UK) Consortium | Luke W Meredith, M. Estée Török, Myra Hosmillo, William L. Hamilton, Martin D. Curran, Theresa Feltwell, Grant Hall, Anna Yakovleva, Fahad A Khokhar, Charlotte J. Houldcroft, Laura G Caller, Aminu S. Jahun, Sarah L. Caddy, Yasmin Chaudhry, Malte Pinckert, Ian Goodfellow |
| EPI_ISL_512379, EPI_ISL_512381, EPI_ISL_512382, EPI_ISL_512383 | Queens Medical Centre, Clinical Microbiology Department / DeepSeq Nottingham | COVID-19 Genomics UK (COG-UK) Consortium | Gemma Clark, Wendy Smith, Manjinder Khakh, Vicki M Fleming, Michelle M Lister, Hannah Howson-Wells, Jonathan Ball, Patrick McClure, Joseph Chappell, Theocharis Tsoleridis, Nadine Holmes, Matthew Carlisle, Christopher Moore, Fei Sang, Johnny Debebe, Victoria Wright, Matthew Loose |
| EPI_ISL_512477, EPI_ISL_512478, EPI_ISL_512479, EPI_ISL_512480 | West of Scotland Specialist Virology Centre, NHSGGC / MRC-University of Glasgow Centre for Virus Research | COVID-19 Genomics UK (COG-UK) Consortium | Ana da Silva Filipe, Natasha Johnson, Kathy Smollett, Daniel Mair, Stephen Carmichael, Lily Tong, Jenna Nichols, Elihu Aranday-Cortes, Kirstyn Brunker, Yasmin Parr, Alice Broos, Kyriaki Nomikou; Sarah McDonald, Marc Niebel, Patawee Asamaphan; Richard Orton, Joseph Hughes, Sreenu Vattipally, David L Robertson; Alasdair MacLean, Rory Gunson; Kathy Li, Natasha Jesudason, Rajiv Shah, James Shepherd, Antonia Ho, Emma Thomson |
| EPI_ISL_512483, EPI_ISL_512492, EPI_ISL_512497, EPI_ISL_512498, EPI_ISL_512499, EPI_ISL_512509, EPI_ISL_512513, EPI_ISL_512514, EPI_ISL_512518, EPI_ISL_512519, EPI_ISL_512520, EPI_ISL_512521, EPI_ISL_512526, EPI_ISL_512528, EPI_ISL_512530, EPI_ISL_512536, EPI_ISL_512542, EPI_ISL_512544, EPI_ISL_512545 | | | |
| see above | Wales Specialist Virology Centre Sequencing lab: Pathogen Genomics Unit | COVID-19 Genomics UK (COG-UK) Consortium | Catherine Moore, Johnathan Evans, Laura Gifford, Malorie Perry, Simon Cottrell, Angela Marchbank, Alec Birchley, Alexander Adams, Amy Gaskin, Bree Gatica-Wilcox, Jason Coombes, Joel Southgate, Lauren Gilbert, Lee Graham, Nicole Pacchiarini, Sara Kumziene-Summerhayes, Sarah Taylor, Sophie Jones, Sara Rey, Matthew Bull, Joanne Watkins, Sally Corden, Tom Connor |
| EPI_ISL_512651, EPI_ISL_512652 | Latvijas Infektoloijas centrs | Latvian Biomedical Research and Study Centre | Ivars Silamielis, Kaspars Megnis, Monta Ustinova, ikita Zrelovs, Vita Rovte, Jeena Storoženko, Tatjana Kolupajeva, Oksana Savicka, Uga Dumpis, Jnis Kloviš |
| EPI_ISL_512831, EPI_ISL_512833, EPI_ISL_512837, EPI_ISL_512838, EPI_ISL_512839, EPI_ISL_512840, EPI_ISL_512841 | National Public Health Laboratory, National Centre for Infectious Diseases | National Public Health Laboratory, National Centre for Infectious Diseases | Mak TM, Octavia S, Zhou Z, Chavatte JM, Cui L, Lin RTP |
| EPI_ISL_513343, EPI_ISL_513344 | Children Westmead Hospital | NSW Health Pathology - Institute of Clinical Pathology and Medical Research; Westmead Hospital; University of Sydney | CIDM-PH et al. |
| EPI_ISL_513345, EPI_ISL_513346 | Pathology West - NSW Health Pathology | NSW Health Pathology - Institute of Clinical Pathology and Medical Research; Westmead Hospital; University of Sydney | CIDM-PH et al. |
| EPI_ISL_513348, EPI_ISL_513349 | 4Cyte Pathology | NSW Health Pathology - Institute of Clinical Pathology and Medical Research; Westmead Hospital; University of Sydney | CIDM-PH et al. |
| EPI_ISL_513350, EPI_ISL_513351 | Pathology West - NSW Health Pathology | NSW Health Pathology - Institute of Clinical Pathology and Medical Research; Westmead Hospital; University of Sydney | CIDM-PH et al. |
| EPI_ISL_513352 | South Eastern Area Laboratory Services (SEALS) | NSW Health Pathology - Institute of Clinical Pathology and Medical Research; Westmead Hospital; University of Sydney | CIDM-PH et al. |
| EPI_ISL_513353, EPI_ISL_513354, EPI_ISL_513355, EPI_ISL_513356 | St Vincent's Pathology (SydPath) | NSW Health Pathology - Institute of Clinical Pathology and Medical Research; Westmead Hospital; University of Sydney | CIDM-PH et al. |
| EPI_ISL_513357 | Douglas Hanly Moir | NSW Health Pathology - Institute of Clinical Pathology and Medical Research; Westmead Hospital; University of Sydney | CIDM-PH et al. |
| EPI_ISL_513358, EPI_ISL_513359 | Pathology West - NSW Health Pathology | NSW Health Pathology - Institute of Clinical Pathology and | CIDM-PH et al. |

| | | | |
|---|---|---|---|
| | | Medical Research; Westmead Hospital; University of Sydney | |
| EPI_ISL_513360 | Pathology North - Hunter - NSW Health Pathology | NSW Health Pathology - Institute of Clinical Pathology and Medical Research; Westmead Hospital; University of Sydney | CIDM-PH et al. |
| EPI_ISL_513361, EPI_ISL_513362, EPI_ISL_513363, EPI_ISL_513364, EPI_ISL_513365 | St Vincent's Pathology (SydPath) | NSW Health Pathology - Institute of Clinical Pathology and Medical Research; Westmead Hospital; University of Sydney | CIDM-PH et al. |
| EPI_ISL_513366, EPI_ISL_513367 | Sydney South West Pathology Service (SSWPS) - Liverpool Hospital - NSW Health Pathology | NSW Health Pathology - Institute of Clinical Pathology and Medical Research; Westmead Hospital; University of Sydney | CIDM-PH et al. |
| EPI_ISL_513368 | Pathology North - Hunter - NSW Health Pathology | NSW Health Pathology - Institute of Clinical Pathology and Medical Research; Westmead Hospital; University of Sydney | CIDM-PH et al. |
| EPI_ISL_513369, EPI_ISL_513370, EPI_ISL_513371 | Pathology West - NSW Health Pathology | NSW Health Pathology - Institute of Clinical Pathology and Medical Research; Westmead Hospital; University of Sydney | CIDM-PH et al. |
| EPI_ISL_513372 | St Vincent's Pathology (SydPath) | NSW Health Pathology - Institute of Clinical Pathology and Medical Research; Westmead Hospital; University of Sydney | CIDM-PH et al. |
| EPI_ISL_513373, EPI_ISL_513374 | Sydney South West Pathology Service (SSWPS) - Liverpool Hospital - NSW Health Pathology | NSW Health Pathology - Institute of Clinical Pathology and Medical Research; Westmead Hospital; University of Sydney | CIDM-PH et al. |
| EPI_ISL_513375, EPI_ISL_513376 | Pathology North - Royal North Shore Hospital - NSW Health Pathology | NSW Health Pathology - Institute of Clinical Pathology and Medical Research; Westmead Hospital; University of Sydney | CIDM-PH et al. |
| EPI_ISL_513377 | South Eastern Area Laboratory Services (SEALS) | NSW Health Pathology - Institute of Clinical Pathology and Medical Research; Westmead Hospital; University of Sydney | CIDM-PH et al. |
| EPI_ISL_513378 | Pathology West - NSW Health Pathology | NSW Health Pathology - Institute of Clinical Pathology and Medical Research; Westmead Hospital; University of Sydney | CIDM-PH et al. |
| EPI_ISL_513379 | St Vincent's Pathology (SydPath) | NSW Health Pathology - Institute of Clinical Pathology and Medical Research; Westmead Hospital; University of Sydney | CIDM-PH et al. |
| EPI_ISL_513380 | Pathology West - NSW Health Pathology | NSW Health Pathology - Institute of Clinical Pathology and Medical Research; Westmead Hospital; University of Sydney | CIDM-PH et al. |
| EPI_ISL_513381, EPI_ISL_513382, EPI_ISL_513383 | Sydney South West Pathology Service (SSWPS) - Liverpool Hospital - NSW Health Pathology | NSW Health Pathology - Institute of Clinical Pathology and Medical Research; Westmead Hospital; University of Sydney | CIDM-PH et al. |
| EPI_ISL_513384, EPI_ISL_513385 | St Vincent's Pathology (SydPath) | NSW Health Pathology - Institute of Clinical Pathology and Medical Research; Westmead Hospital; University of Sydney | CIDM-PH et al. |
| EPI_ISL_513386, EPI_ISL_513387, EPI_ISL_513388 | Pathology North - NSW Health Pathology | NSW Health Pathology - Institute of Clinical Pathology and Medical Research; Westmead Hospital; University of Sydney | CIDM-PH et al. |
| EPI_ISL_513390, EPI_ISL_513391, EPI_ISL_513392, EPI_ISL_513394, EPI_ISL_513395 | Pathology West - NSW Health Pathology | NSW Health Pathology - Institute of Clinical Pathology and Medical Research; Westmead Hospital; University of Sydney | CIDM-PH et al. |
| EPI_ISL_513400 | Sydney South West Pathology Service (SSWPS) - Royal Prince Alfred Hospital - NSW Health Pathology | NSW Health Pathology - Institute of Clinical Pathology and Medical Research; Westmead Hospital; University of Sydney | CIDM-PH et al. |
| EPI_ISL_513401 | Sydney South West Pathology Service (SSWPS) - Liverpool Hospital - NSW Health Pathology | NSW Health Pathology - Institute of Clinical Pathology and Medical Research; Westmead Hospital; University of Sydney | CIDM-PH et al. |
| EPI_ISL_513408 | Laverty Pathology | NSW Health Pathology - Institute of Clinical Pathology and Medical Research; Westmead Hospital; University of Sydney | CIDM-PH et al. |
| EPI_ISL_513409, EPI_ISL_513410 | Sydney South West Pathology Service (SSWPS) - Liverpool Hospital - NSW Health Pathology | NSW Health Pathology - Institute of Clinical Pathology and Medical Research; Westmead Hospital; University of Sydney | CIDM-PH et al. |
| EPI_ISL_513413 | 4Cyte Pathology | NSW Health Pathology - Institute of Clinical Pathology and Medical Research; Westmead Hospital; University of Sydney | CIDM-PH et al. |
| EPI_ISL_514129, EPI_ISL_514130 | National Institute of Laboratory Medicine and Referral Center | Genomic Research Lab, BCSIR | Md. Murshed Hasan Sarkar, Abu Sayeed Mohammad Mahmud, Mohammad Samir Uzzaman, Eshrar Osman, Md. Ahashan Habib, Shahina Akter, Tanjina Akhter Banu, Barna Goswami, Iffat Jahan, Md. Saddam Hossain, Tasnim Nafisa, Md. Maruf Ahmed Molla, Mahmuda Yeasmin, Asish Kumar Ghosh, A. K. M. Shamsuzzaman, Sheikh Md. Selim Al Din, Utpal Chandra Ray, Salek Ahmed Sajib, Md. Salim Khan |
| EPI_ISL_514233, EPI_ISL_514234, EPI_ISL_514235, EPI_ISL_514236 | National Institute of Laboratory Medicine and Referral Center | Genomic Research Lab, BCSIR | Md. Murshed Hasan Sarkar, Abu Sayeed Mohammad Mahmud, Mohammad Samir Uzzaman, Eshrar Osman, Md. Ahashan Habib, Shahina Akter, Tanjina Akhter Banu, Barna Goswami, Iffat Jahan, Md. Saddam Hossain, Tasnim Nafisa, Md. Maruf Ahmed Molla, Mahmuda Yeasmin, Asish Kumar Ghosh, A. K. M. Shamsuzzaman, Sheikh Md. Selim Al Din, Utpal Chandra Ray, Salek Ahmed Sajib, Md. Salim Khan |
| EPI_ISL_514237 | National Institute of Laboratory Medicine and Referral Center | Genomic Research Lab, BCSIR | Md. Saddam Hossain, Abu Sayeed Mohammad Mahmud, Mohammad Samir Uzzaman, Eshrar Osman, Md. Ahashan Habib, Shahina Akter, Tanjina Akhter Banu, Md. Murshed Hasan Sarkar, Barna Goswami, Iffat Jahan, Tasnim Nafisa, Md. Maruf Ahmed Molla, Mahmuda Yeasmin, Asish Kumar Ghosh, A. K. M. Shamsuzzaman, Sheikh Md. Selim Al Din, Utpal Chandra Ray, Salek Ahmed Sajib, Md. Salim Khan |
| EPI_ISL_514248, EPI_ISL_514249, EPI_ISL_514250 | National Institute of Laboratory Medicine and Referral Center | Genomic Research Lab, BCSIR | Abu Sayeed Mohammad Mahmud, Mohammad Samir Uzzaman, Eshrar Osman, Md. Ahashan Habib, Shahina Akter, Tanjina Akhter Banu, Md. Murshed Hasan Sarkar, Barna Goswami, Iffat Jahan, Md. Saddam Hossain, Tasnim Nafisa, Md. Maruf Ahmed Molla, Mahmuda Yeasmin, Asish Kumar Ghosh, A. K. M. Shamsuzzaman, Sheikh Md. Selim Al Din, Utpal Chandra Ray, Salek Ahmed Sajib, Md. Salim Khan |
| EPI_ISL_514305, EPI_ISL_514306 | Israel Central Virology laboratory | Israel Central Virology laboratory | Neta Zuckerman, Efrat Dahan Bucris, Oran Erster, Ella Mendelson, Michal Mandelboim |
| EPI_ISL_514343, EPI_ISL_514344, EPI_ISL_514350, EPI_ISL_514353 | Respiratory Virus Unit, Microbiology Services Colindale, Public Health England | Respiratory Virus Unit, Microbiology Services Colindale, Public Health England | PHE Covid Sequencing Team |
| EPI_ISL_514418, EPI_ISL_514420, EPI_ISL_514421, EPI_ISL_514422, EPI_ISL_514424 | National Institute for Communicable Diseases of the National Health Laboratory Service | National Institute for Communicable Diseases of the National Health Laboratory Service | Allam M, Ismail A, Khumalo Z, Kwenda S, Mtshali P, Mnyameni F, Mohale T, Bhiman JN |
| EPI_ISL_514441 | NSTU COVID-19 Diagnostic Center | NSU Genome Research Institute (NGRI), North South University | Dr. Muhammad Maqsud Hossain, Aura Rahman, Prof. Firoz Ahmed, Tahrima Huq, Abdus Sadique, Jahidul Alam, Md Aminul Islam, Prof. Md. Didar-Ul-Alam, Prof. Kazi Nadim Hasan, Prof. Abdul Khaleque, Prof, Hasan Mahmud Reza |
| EPI_ISL_514442 | NSTU COVID-19 Diagnostic Center | NSU Genome Research Institute (NGRI), North South University | Dr. Muhammad Maqsud Hossain, Aura Rahman, Prof. Firoz Ahmed, Tahrima Huq, Abdus Sadique, Jahidul Alam, Tamanna Afroze, Md Aminul Islam, Prof. Md. Didar-Ul-Alam, Prof. Kazi Nadim Hasan, Prof. Abdul Khaleque, Prof, Hasan Mahmud Reza |
| EPI_ISL_514444, EPI_ISL_514445, EPI_ISL_514446, EPI_ISL_514447 | Department of Pathology, University of Cambridge | COVID-19 Genomics UK (COG-UK) Consortium | Luke W Meredith, M. Estée Török, Myra Hosmillo, William L. Hamilton, Martin D. Curran, Theresa Feltwell, Grant Hall, Anna Yakovleva, Fahad A Khokhar, Charlotte J. Houldcroft, Laura G Caller, Aminu S. Jahun, Sarah L. Caddy, Yasmin Chaudhry, Malte Pinckert, Ian Goodfellow |
| EPI_ISL_514563, EPI_ISL_514564, EPI_ISL_514565, EPI_ISL_514567, EPI_ISL_514568, EPI_ISL_514569, EPI_ISL_514570, EPI_ISL_514571, EPI_ISL_514572, EPI_ISL_514573, EPI_ISL_514574, EPI_ISL_514575, EPI_ISL_514576, EPI_ISL_514578 | | | |
| see above | Wales Specialist Virology Centre Sequencing lab: Pathogen | COVID-19 Genomics UK (COG-UK) Consortium | Catherine Moore, Johnathan Evans, Laura Gifford, Malorie Perry, Simon Cottrell, Angela Marchbank, Alec Birchley, Alexander Adams, Amy Gaskin, Bree |

| | | | |
|---|---|---|---|
| | Genomics Unit | | Gatica-Wilcox, Jason Coombes, Joel Southgate, Lauren Gilbert, Lee Graham, Nicole Pacchiarini, Sara Kumziene-Summerhayes, Sarah Taylor, Sophie Jones, Sara Rey, Matthew Bull, Joanne Watkins, Sally Corden, Tom Connor |
| EPI_ISL_515185, EPI_ISL_515186 | Latvijas Infektoloijas centrs | Latvian Biomedical Research and Study Centre | Ivars Silamielis, Kaspars Megnis, Monta Ustinova, ikita Zrelovs, Vita Rovte, Jeena Storoženko, Tatjana Kolupajeva, Oksana Savicka, Uga Dumpis, Jnis Kloviš |
| EPI_ISL_515605, EPI_ISL_515606, EPI_ISL_515607, EPI_ISL_515609, EPI_ISL_515610, EPI_ISL_515611, EPI_ISL_515612, EPI_ISL_515668, EPI_ISL_515671, EPI_ISL_515672, EPI_ISL_515675, EPI_ISL_515679, EPI_ISL_515680, EPI_ISL_515681, EPI_ISL_515683, EPI_ISL_515685, EPI_ISL_515686, EPI_ISL_515699, EPI_ISL_515707, EPI_ISL_515709, EPI_ISL_515710, EPI_ISL_515716, EPI_ISL_515718, EPI_ISL_515719, EPI_ISL_515755, EPI_ISL_515757, EPI_ISL_515758, EPI_ISL_515759, EPI_ISL_515760, EPI_ISL_515761, EPI_ISL_515762, EPI_ISL_515763, EPI_ISL_515764, EPI_ISL_515765, EPI_ISL_515766, EPI_ISL_515767, EPI_ISL_515768, EPI_ISL_515769, EPI_ISL_515770, EPI_ISL_515771, EPI_ISL_515772, EPI_ISL_515773, EPI_ISL_515774, EPI_ISL_515775, EPI_ISL_515776, EPI_ISL_515777, EPI_ISL_515778, EPI_ISL_515779, EPI_ISL_515780, EPI_ISL_515781, EPI_ISL_515782, EPI_ISL_515783, EPI_ISL_515784, EPI_ISL_515785, EPI_ISL_515786, EPI_ISL_515787, EPI_ISL_515788, EPI_ISL_515789, EPI_ISL_515790, EPI_ISL_515791, EPI_ISL_515792, EPI_ISL_515793, EPI_ISL_515794, EPI_ISL_515795, EPI_ISL_515796, EPI_ISL_515797, EPI_ISL_515798, EPI_ISL_515799 | | | |
| see above | NHLS-IALCH | KRISP, KZN Research Innovation and Sequencing Platform | Giandhari J, Pillay S, Lessells R, Mdlalose K, York D, Khan S, Tegally H, Wilkinson E, de Oliveira T |
| EPI_ISL_516211, EPI_ISL_516212, EPI_ISL_516213, EPI_ISL_516214, EPI_ISL_516215, EPI_ISL_516216, EPI_ISL_516217, EPI_ISL_516218, EPI_ISL_516219, EPI_ISL_516220, EPI_ISL_516221, EPI_ISL_516222, EPI_ISL_516223 | | | |
| see above | Michigan Department of Health and Human Services, Bureau of Laboratories | Michigan Department of Health and Human Services, Bureau of Laboratories | Blankenship HM, Riner D, Soehnlen MK |
| EPI_ISL_516447, EPI_ISL_516448, EPI_ISL_516457, EPI_ISL_516458, EPI_ISL_516459, EPI_ISL_516464, EPI_ISL_516466, EPI_ISL_516471, EPI_ISL_516472, EPI_ISL_516474, EPI_ISL_516476, EPI_ISL_516479, EPI_ISL_516480, EPI_ISL_516481, EPI_ISL_516482, EPI_ISL_516483, EPI_ISL_516484, EPI_ISL_516485, EPI_ISL_516486, EPI_ISL_516487, EPI_ISL_516488, EPI_ISL_516489, EPI_ISL_516490, EPI_ISL_516492, EPI_ISL_516493, EPI_ISL_516494, EPI_ISL_516495, EPI_ISL_516497, EPI_ISL_516498, EPI_ISL_516499, EPI_ISL_516500, EPI_ISL_516501, EPI_ISL_516502, EPI_ISL_516503, EPI_ISL_516504, EPI_ISL_516505, EPI_ISL_516507, EPI_ISL_516508, EPI_ISL_516509, EPI_ISL_516510, EPI_ISL_516511, EPI_ISL_516512, EPI_ISL_516513, EPI_ISL_516514, EPI_ISL_516515, EPI_ISL_516516, EPI_ISL_516517, EPI_ISL_516518, EPI_ISL_516519, EPI_ISL_516521, EPI_ISL_516522, EPI_ISL_516523, EPI_ISL_516525, EPI_ISL_516526, EPI_ISL_516527, EPI_ISL_516528, EPI_ISL_516529, EPI_ISL_516530, EPI_ISL_516531, EPI_ISL_516532, EPI_ISL_516533, EPI_ISL_516534, EPI_ISL_516535, EPI_ISL_516536, EPI_ISL_516537, EPI_ISL_516538, EPI_ISL_516539, EPI_ISL_516540, EPI_ISL_516541, EPI_ISL_516542, EPI_ISL_516543, EPI_ISL_516544, EPI_ISL_516545, EPI_ISL_516546, EPI_ISL_516547, EPI_ISL_516548, EPI_ISL_516549 | | | |
| see above | University of Wisconsin-Madison AIDS Vaccine Research Laboratories | University of Wisconsin-Madison AIDS Vaccine Research Laboratories | Gage Moreno, Katarina Braun, et al. AIDS Vaccine Research Laboratories |
| EPI_ISL_516552, EPI_ISL_516553, EPI_ISL_516554, EPI_ISL_516555, EPI_ISL_516556 | Viollier AG | Department of Biosystems Science and Engineering, ETH Zürich | Christian Beisel, Sarah Nadeau, Ivan Topolsky, Pedro Ferreira, Philipp Jablonski, Susana Posada-Céspedes, Tobias Schär, Ina Nissen, Natascha Santacroce, Elodie Burcklen, Christiane Beckmann, Maurice Redondo, Olivier Kobel, Christoph Noppen, Sophie Seidel, Noemie Santamaria de Souza, Niko Beerenwinkel, Tanja Stadler |
| EPI_ISL_516651 | Institute of Microbiology, Universidad San Francisco de Quito | Institute of Microbiology, Universidad San Francisco de Quito | Prado-Vivar, Sully Márquez, Juan José Guadalupe, Monica Becerra-Wong, Bernardo Gutiérrez, Nabih Dahik, Carlos Mena, Eulalia Pazmiño, Carolina Pacheco, DamarisSandoya, Verónica Barragán, Patricio Rojas-Silva, Gabriel Trueba, Michelle Grunauer, Paúl Cárdenas |
| EPI_ISL_516652 | Institute of Microbiology, Universidad San Francisco de Quito | Institute of Microbiology, Universidad San Francisco de Quito | Prado-Vivar, Sully Márquez, Juan José Guadalupe, Monica Becerra-Wong, Bernardo Gutiérrez, Nabih Dahik, Carlos Mena, Edy Quizhpe, Yomara Napa, Verónica Barragán, Patricio Rojas-Silva, Gabriel Trueba, Michelle Grunauer, Paúl Cárdenas |
| EPI_ISL_517373 | Liverpool Clinical Laboratories | COVID-19 Genomics UK (COG-UK) Consortium | Sam Haldenby, Anita Lucaci, Steve Paterson, Julian Hiscox, Alistair Darby, M Almsaud, A Alrezaihi, Muhannad Alruwaili, Stuart D Armstrong, Jones Benjamin, Eleanor G Bentley, Anu Chawla, Jordan J Clark, Angela Cowell, Richard Eccles, Isabel García-Dorival, Matthew Gemmell, Alessandro Gerada, PKF Gilmore, Richard Gregory, Ximeng Han, Catherine Hartley, Margaret Hughes, Miren Iturriza-Gomara, James Johnson, L Luu, Jenifer Manson, Charlotte Nelson, Elaine O'Toole, Cassie Olateju, Rebekah Penrice-Randal , Lucille Rainbow, N.P Randle, Trevor Ian Robinson, Parul Sharma, Ghada T Shawli, James P Stewart, Neil Swainston, Ecaterina Vamos, Joanne Watts, Mark Whitehead |
| EPI_ISL_517544, EPI_ISL_517548, EPI_ISL_517553, EPI_ISL_517556, EPI_ISL_517566, EPI_ISL_517574 | Virology Department, Sheffield Teaching Hospitals NHS Foundation Trust/Department of Infection, Immunity and Cardiovascular Disease, The Medical School, University of Sheffield | COVID-19 Genomics UK (COG-UK) Consortium | Thushan de Silva, Matthew Parker, Nikki Smith, Adri Angyal, Rebecca Brown, Luke Green, Rachel Tucker, Paul Parsons, Danielle Groves, Katie Johnson, Laura Carrilero, Alex Keeley, Dave Partridge, Matthew Wyles, Benjamin Lindsey, Mehmet Yavuz, Mohammad Raza, Cariad Evans |
| EPI_ISL_517590, EPI_ISL_517593, EPI_ISL_517595, EPI_ISL_517597, EPI_ISL_517598, EPI_ISL_517599, EPI_ISL_517605 | Wales Specialist Virology Centre Sequencing lab: Pathogen Genomics Unit | COVID-19 Genomics UK (COG-UK) Consortium | Catherine Moore, Johnathan Evans, Laura Gifford, Malorie Perry, Simon Cottrell, Angela Marchbank, Alec Birchley, Alexander Adams, Amy Gaskin, Bree Gatica-Wilcox, Jason Coombes, Joel Southgate, Lauren Gilbert, Lee Graham, Nicole Pacchiarini, Sara Kumziene-Summerhayes, Sarah Taylor, Sophie Jones, Sara Rey, Matthew Bull, Joanne Watkins, Sally Corden, Tom Connor |
| EPI_ISL_517860, EPI_ISL_517861, EPI_ISL_517862, EPI_ISL_517863, EPI_ISL_517864, EPI_ISL_517865, EPI_ISL_517866, EPI_ISL_517867, EPI_ISL_517868, EPI_ISL_517869, EPI_ISL_517870, EPI_ISL_517922, EPI_ISL_517923, EPI_ISL_517933, EPI_ISL_517934, EPI_ISL_517935, EPI_ISL_517936, EPI_ISL_517937, EPI_ISL_517938, EPI_ISL_517939 | | | |
| see above | Florida Bureau of Public Health Laboratories | Florida Bureau of Public Health Laboratories | Sarah Schmedes, Jason Blanton |
| EPI_ISL_518001, EPI_ISL_518002, EPI_ISL_518003 | Singapore General Hospital | Department of Microbiology | Nurdyana Abdul Rahman, Kun Lee Lim, Chenhao Li, Kian Sing Chan, Lynette Oon, Kern Rei Chng, Niranjan Nagarajan, Karrie Ko |
| EPI_ISL_518063, EPI_ISL_518066, EPI_ISL_518067, EPI_ISL_518068, EPI_ISL_518069, EPI_ISL_518070, EPI_ISL_518071, EPI_ISL_518072, EPI_ISL_518073, EPI_ISL_518074, EPI_ISL_518075, EPI_ISL_518076, EPI_ISL_518079, EPI_ISL_518080, EPI_ISL_518093, EPI_ISL_518094, EPI_ISL_518095, EPI_ISL_518096, EPI_ISL_518097 | | | |
| see above | Microbiological Diagnostic Unit - Public Health Laboratory (MDU-PHL) | MDU-PHL | Seemann T., Schultz M., Sait, M., Sherry, N. |
| EPI_ISL_518148, EPI_ISL_518170, EPI_ISL_518171, EPI_ISL_518172, EPI_ISL_518173, EPI_ISL_518174, EPI_ISL_518175, EPI_ISL_518187, EPI_ISL_518195, EPI_ISL_518196, EPI_ISL_518197, EPI_ISL_518200, EPI_ISL_518201, EPI_ISL_518202, EPI_ISL_518203, EPI_ISL_518204, EPI_ISL_518205, EPI_ISL_518206, EPI_ISL_518207, EPI_ISL_518208, EPI_ISL_518209, EPI_ISL_518211, EPI_ISL_518213, EPI_ISL_518214, EPI_ISL_518215, EPI_ISL_518216, EPI_ISL_518217, EPI_ISL_518222, EPI_ISL_518224 | | | |
| see above | Victorian Infectious Diseases Reference Laboratory (VIDRL) | VIDRL and MDU-PHL | Caly L., Seemann T., Sait, M., Schultz M., Druce J., Sherry, N. |
| EPI_ISL_518232, EPI_ISL_518234, EPI_ISL_518236, EPI_ISL_518237, EPI_ISL_518241, EPI_ISL_518242, EPI_ISL_518243 | Microbiological Diagnostic Unit - Public Health Laboratory (MDU-PHL) | MDU-PHL | Seemann T., Schultz M., Sait, M., Sherry, N. |
| EPI_ISL_518244 | Victorian Infectious Diseases Reference Laboratory (VIDRL) | VIDRL and MDU-PHL | Caly L., Seemann T., Sait, M., Schultz M., Druce J., Sherry, N. |
| EPI_ISL_518250, EPI_ISL_518251, EPI_ISL_518252, EPI_ISL_518253, EPI_ISL_518254, EPI_ISL_518255, EPI_ISL_518256, EPI_ISL_518257, EPI_ISL_518258, EPI_ISL_518259, EPI_ISL_518260, EPI_ISL_518261, EPI_ISL_518262, EPI_ISL_518263, EPI_ISL_518264, EPI_ISL_518265, EPI_ISL_518266, EPI_ISL_518267, EPI_ISL_518268, EPI_ISL_518272, EPI_ISL_518273, EPI_ISL_518274, EPI_ISL_518285, EPI_ISL_518286, EPI_ISL_518287, EPI_ISL_518290, EPI_ISL_518294, EPI_ISL_518295, EPI_ISL_518296, EPI_ISL_518297, EPI_ISL_518298, EPI_ISL_518299, EPI_ISL_518300, EPI_ISL_518301, EPI_ISL_518302, EPI_ISL_518303, EPI_ISL_518304, EPI_ISL_518305, EPI_ISL_518306, EPI_ISL_518307, EPI_ISL_518308, EPI_ISL_518309, EPI_ISL_518310, EPI_ISL_518311, EPI_ISL_518312, EPI_ISL_518313, EPI_ISL_518314, EPI_ISL_518315, EPI_ISL_518317, EPI_ISL_518318, EPI_ISL_518319, EPI_ISL_518321, EPI_ISL_518324, EPI_ISL_518325, EPI_ISL_518326, EPI_ISL_518548, EPI_ISL_518621, EPI_ISL_518660, EPI_ISL_518668, EPI_ISL_518730, EPI_ISL_518731, EPI_ISL_518774, EPI_ISL_518775, EPI_ISL_518776, EPI_ISL_518777, EPI_ISL_518778, EPI_ISL_518779, EPI_ISL_518780, EPI_ISL_518781, EPI_ISL_518782, EPI_ISL_518783, EPI_ISL_518784, EPI_ISL_518785, EPI_ISL_518786, EPI_ISL_518787, EPI_ISL_518792, EPI_ISL_518793, EPI_ISL_518794, EPI_ISL_518795, EPI_ISL_518796, EPI_ISL_521164, EPI_ISL_521166, EPI_ISL_521167, EPI_ISL_521168, EPI_ISL_521170, EPI_ISL_521171, EPI_ISL_521172, EPI_ISL_521173, EPI_ISL_521174, EPI_ISL_521175, EPI_ISL_521176, EPI_ISL_521177, EPI_ISL_521178, EPI_ISL_521179, EPI_ISL_521180, EPI_ISL_521181, EPI_ISL_521182, EPI_ISL_521183, EPI_ISL_521184, EPI_ISL_521185, EPI_ISL_521186, EPI_ISL_521187, EPI_ISL_521188, EPI_ISL_521189, EPI_ISL_521190, EPI_ISL_521191, EPI_ISL_521192, EPI_ISL_521193, EPI_ISL_521194, EPI_ISL_521197, EPI_ISL_521198, EPI_ISL_521199, EPI_ISL_521200, EPI_ISL_521201, EPI_ISL_521207, EPI_ISL_521208, EPI_ISL_521210, EPI_ISL_521211, EPI_ISL_521213, EPI_ISL_521214, EPI_ISL_521215, EPI_ISL_521216, EPI_ISL_521217, EPI_ISL_521218, EPI_ISL_521219, EPI_ISL_521220, EPI_ISL_521221, EPI_ISL_521222, EPI_ISL_521223, EPI_ISL_521226, EPI_ISL_521227, EPI_ISL_521228, EPI_ISL_521229, EPI_ISL_521231, EPI_ISL_521232, EPI_ISL_521233, EPI_ISL_521234, EPI_ISL_521235, EPI_ISL_521236, EPI_ISL_521237, EPI_ISL_521238, EPI_ISL_521240, EPI_ISL_521241, EPI_ISL_521242, EPI_ISL_521243, EPI_ISL_521244, EPI_ISL_521247, EPI_ISL_521251, EPI_ISL_521252, EPI_ISL_521253, EPI_ISL_521254, EPI_ISL_521255, EPI_ISL_521256, EPI_ISL_521257, EPI_ISL_521258, EPI_ISL_521259 | | | |
| see above | Microbiological Diagnostic Unit - Public Health Laboratory (MDU-PHL) | MDU-PHL | Seemann T., Schultz M., Sait, M., Sherry, N. |
| EPI_ISL_521270, EPI_ISL_521271, EPI_ISL_521274, EPI_ISL_521276 | Victorian Infectious Diseases Reference Laboratory (VIDRL) | VIDRL and MDU-PHL | Caly L., Seemann T., Sait, M., Schultz M., Druce J., Sherry, N. |
| EPI_ISL_521278, EPI_ISL_521279, EPI_ISL_521280, EPI_ISL_521281, EPI_ISL_521282, EPI_ISL_521283, EPI_ISL_521284, EPI_ISL_521285, EPI_ISL_521286, EPI_ISL_521287, EPI_ISL_521288, EPI_ISL_521289, EPI_ISL_521290, EPI_ISL_521291, EPI_ISL_521292, EPI_ISL_521293, EPI_ISL_521294, EPI_ISL_521295, EPI_ISL_521296, EPI_ISL_521297, EPI_ISL_521298, EPI_ISL_521299, EPI_ISL_521300, EPI_ISL_521301, EPI_ISL_521303, EPI_ISL_521304, EPI_ISL_521305, EPI_ISL_521306, EPI_ISL_521307, EPI_ISL_521308, EPI_ISL_521309, EPI_ISL_521310, EPI_ISL_521311, EPI_ISL_521312, EPI_ISL_521313, EPI_ISL_521314 | | | |
| see above | Microbiological Diagnostic Unit - Public Health Laboratory (MDU-PHL) | MDU-PHL | Seemann T., Schultz M., Sait, M., Sherry, N. |
| EPI_ISL_522820, EPI_ISL_522821, | Virginia DCLS | Virginia DCLS | Virginia DCLS |

| | | | |
|---|---|---|---|
| EPI_ISL_522822, EPI_ISL_522823 | | | |
| EPI_ISL_522872 | Instituto Nacional de Medicina Genómica | Instituto Nacional de Medicina Genómica | Hidalgo-Miranda A, Mendoza-Vargas A, Reyes-Grajeda JP, Cisneros-Villanueva M, Cedro-Tanda A, Hurtado-Cordova E, Peñaloza-Figueroa F, Herrera-Montalvo LA |
| EPI_ISL_522873, EPI_ISL_522874, EPI_ISL_522875, EPI_ISL_522876 | Instituto Nacional de Medicina Genomica | Instituto Nacional de Medicina Genomica | Hidalgo-Miranda A, Mendoza-Vargas A, Reyes-Grajeda JP, Cisneros-Villanueva M, Cedro-Tanda A, Hurtado-Cordova E, Peñaloza-Figueroa F, Herrera-Montalvo LA |
| EPI_ISL_522879 | Instituto Nacional de Medicina Genómica | Instituto Nacional de Medicina Genómica | Hidalgo-Miranda A, Mendoza-Vargas A, Reyes-Grajeda JP, Cisneros-Villanueva M, Cedro-Tanda A, Hurtado-Cordova E, Peñaloza-Figueroa F, Herrera-Montalvo LA |
| EPI_ISL_522938 | University of Wisconsin-Madison AIDS Vaccine Research Laboratories | University of Wisconsin-Madison AIDS Vaccine Research Laboratories | Gage Moreno, Katarina Braun, et al. AIDS Vaccine Research Laboratories |
| EPI_ISL_522941 | Instituto Nacional de Medicina Genómica | Instituto Nacional de Medicina Genómica | Hidalgo-Miranda A, Mendoza-Vargas A, Reyes-Grajeda JP, Cisneros-Villanueva M, Cedro-Tanda A, Hurtado-Cordova E, Peñaloza-Figueroa F, Herrera-Montalvo LA |
| EPI_ISL_523083, EPI_ISL_523084, EPI_ISL_523086, EPI_ISL_523087, EPI_ISL_523088 | Dutch COVID-19 response team | Erasmus Medical Center | OH consortium |
| EPI_ISL_523137, EPI_ISL_523146, EPI_ISL_523517, EPI_ISL_523536, EPI_ISL_523537, EPI_ISL_523538, EPI_ISL_523687, EPI_ISL_523735, EPI_ISL_523736, EPI_ISL_523737, EPI_ISL_523738, EPI_ISL_523739, EPI_ISL_523741, EPI_ISL_523742, EPI_ISL_523743, EPI_ISL_523745 | | | |
| see above | Dutch COVID-19 response team | Erasmus Medical Center | Bas Oude Munnink, David Nieuwenhuijse, Reina Sikkema, Claudia Schapendonk, Irina Chestakova, Anne van der Linden, Theo Bestebroer, Stefan van Nieuwkoop, Mark Pronk, Pascal Lexmond, Corien Swaan, Manon Haverkate, Madelief Mollers, Mart Stein, Sandra Kengne Kamga Mobou, Jeroen van Kampen, Jolanda Voermans, Aura Timen, Corine GeurtsvanKessel, Annemiek van der Eijk, Richard Molenkamp, Marion Koopmans, on behalf of the Dutch national COVID-19 response team. |
| EPI_ISL_524486, EPI_ISL_524489 | PHE South West Regional Laboratory, National Infection Service | Wellcome Sanger Institute for the COVID-19 Genomics UK (COG-UK) consortium | Stephanie Hutchings, Hannah Pymont, Dr Peter Muir, Barry Vipond, Rich Hopes; and Alex Alderton, Roberto Amato, Sonia Goncalves, Ewan Harrison, David K. Jackson, Ian Johnston, Dominic Kwiatkowski, Cordelia Langford, John Sillitoe on behalf of the Wellcome Sanger Institute COVID-19 Surveillance Team (http://www.sanger.ac.uk/covid-team) |
| EPI_ISL_525203, EPI_ISL_525204, EPI_ISL_525205 | Virginia DCLS | Virginia DCLS | Virginia DCLS |
| EPI_ISL_525352, EPI_ISL_525353, EPI_ISL_525354, EPI_ISL_525355, EPI_ISL_525356, EPI_ISL_525357, EPI_ISL_525358, EPI_ISL_525359, EPI_ISL_525360, EPI_ISL_525361, EPI_ISL_525362, EPI_ISL_525363, EPI_ISL_525364, EPI_ISL_525365, EPI_ISL_525366, EPI_ISL_525367, EPI_ISL_525368, EPI_ISL_525369, EPI_ISL_525370 | | | |
| see above | National Virus Reference Laboratory | National Virus Reference Laboratory | Michael Carr, Gabriel Gonzalez, Jonathan Dean, Aditi Chaturvedi, Suzie Coughlan, Cillian F De Gascun |
| EPI_ISL_525741 | Seattle Flu Study | Seattle Flu Study | Deborah A. Nickerson, Chris D. Frazar, Jover Lee, Benjamin Pelle, Matthew Richardson, Amanda Adler, Elisabeth Brandstetter, Peter D. Han, Kairsten Fay, Misja Ilcisin, Kirsten Lacombe, Thomas R. Sibley, Melissa Truong, Caitlin R. Wolf, Karen Cowgill, Stephanie Schrag, Jeff Duchin, Michael Boeckh, Janet A. Englund, Michael Famulare, Barry R. Lutz, Mark J. Rieder, Lea M. Starita, Matthew Thompson, Helen Y. Chu, Trevor Bedford, Jay Shendure |
| EPI_ISL_525742 | Seattle Flu Study | Seattle Flu Study | Deborah A. Nickerson, Chris D. Frazar, Jover Lee, Benjamin Pelle, Matthew Richardson, Amanda Adler, Elisabeth Brandstetter, Peter D. Han, Kairsten Fay, Misja Ilcisin, Kirsten Lacombe, Thomas R. Sibley, Melissa Truong, Caitlin R. Wolf, Michael Boeckh, Janet A. Englund, Michael Famulare, Barry R. Lutz, Mark J. Rieder, Lea M. Starita, Matthew Thompson, Jay Shendure, Trevor Bedford, Helen Y. Chu |
| EPI_ISL_525743, EPI_ISL_525744, EPI_ISL_525745, EPI_ISL_525746, EPI_ISL_525747, EPI_ISL_525748, EPI_ISL_525749, EPI_ISL_525750, EPI_ISL_525751, EPI_ISL_525752, EPI_ISL_525753, EPI_ISL_525754, EPI_ISL_525756 | | | |
| see above | Seattle Flu Study | Seattle Flu Study | Deborah A. Nickerson, Chris D. Frazar, Jover Lee, Benjamin Pelle, Matthew Richardson, Amanda Adler, Elisabeth Brandstetter, Peter D. Han, Kairsten Fay, Misja Ilcisin, Kirsten Lacombe, Thomas R. Sibley, Melissa Truong, Caitlin R. Wolf, Karen Cowgill, Stephanie Schrag, Jeff Duchin, Michael Boeckh, Janet A. Englund, Michael Famulare, Barry R. Lutz, Mark J. Rieder, Lea M. Starita, Matthew Thompson, Helen Y. Chu, Trevor Bedford, Jay Shendure |
| EPI_ISL_525806, EPI_ISL_526004, EPI_ISL_526005, EPI_ISL_526006, EPI_ISL_526007, EPI_ISL_526008, EPI_ISL_526009, EPI_ISL_526010, EPI_ISL_526011, EPI_ISL_526012, EPI_ISL_526013, EPI_ISL_526014, EPI_ISL_526015, EPI_ISL_526016, EPI_ISL_526017, EPI_ISL_526018, EPI_ISL_526019, EPI_ISL_526020, EPI_ISL_526021, EPI_ISL_526022, EPI_ISL_526023, EPI_ISL_526024, EPI_ISL_526025, EPI_ISL_526026, EPI_ISL_526027, EPI_ISL_526028, EPI_ISL_526029, EPI_ISL_526030, EPI_ISL_526031, EPI_ISL_526032, EPI_ISL_526033, EPI_ISL_526034, EPI_ISL_526035, EPI_ISL_526036, EPI_ISL_526037, EPI_ISL_526038, EPI_ISL_526039, EPI_ISL_526040, EPI_ISL_526041, EPI_ISL_526042, EPI_ISL_526043, EPI_ISL_526044, EPI_ISL_526045, EPI_ISL_526046, EPI_ISL_526047, EPI_ISL_526048, EPI_ISL_526049, EPI_ISL_526050, EPI_ISL_526051, EPI_ISL_526052, EPI_ISL_526053, EPI_ISL_526054, EPI_ISL_526055, EPI_ISL_526056, EPI_ISL_526057, EPI_ISL_526058, EPI_ISL_526059, EPI_ISL_526060, EPI_ISL_526061, EPI_ISL_526062, EPI_ISL_526063, EPI_ISL_526064, EPI_ISL_526065, EPI_ISL_526066, EPI_ISL_526067, EPI_ISL_526068, EPI_ISL_526069, EPI_ISL_526070, EPI_ISL_526071, EPI_ISL_526072, EPI_ISL_526073, EPI_ISL_526074, EPI_ISL_526075, EPI_ISL_526076, EPI_ISL_526077, EPI_ISL_526078, EPI_ISL_526079, EPI_ISL_526080, EPI_ISL_526081, EPI_ISL_526082, EPI_ISL_526083, EPI_ISL_526084, EPI_ISL_526085, EPI_ISL_526107, EPI_ISL_526108, EPI_ISL_526109, EPI_ISL_526110, EPI_ISL_526111, EPI_ISL_526112, EPI_ISL_526113, EPI_ISL_526114 | | | |
| see above | OHSU Lab Services Molecular Microbiology Lab | Oregon SARS-CoV-2 Genome Sequencing Center | Brendan L. O'Connell, Ruth V. Nichols, Alec J. Hirsch, Guang Fan, Daniel N. Streblow, William B. Messer, Andrew C. Adey, Benjamin N. Bimber, Brian J. O'Roak |
| EPI_ISL_526115, EPI_ISL_526116, EPI_ISL_526117, EPI_ISL_526118 | Pathology West - NSW Health Pathology | NSW Health Pathology - Institute of Clinical Pathology and Medical Research; Westmead Hospital; University of Sydney | CIDM-PH et al. |
| EPI_ISL_526119, EPI_ISL_526120, EPI_ISL_526124, EPI_ISL_526126, EPI_ISL_526127 | Sydney South West Pathology Service (SSWPS) - Liverpool Hospital - NSW Health Pathology | NSW Health Pathology - Institute of Clinical Pathology and Medical Research; Westmead Hospital; University of Sydney | CIDM-PH et al. |
| EPI_ISL_526129 | The Children's Hospital at Westmead | NSW Health Pathology - Institute of Clinical Pathology and Medical Research; Westmead Hospital; University of Sydney | CIDM-PH et al. |
| EPI_ISL_526134, EPI_ISL_526135, EPI_ISL_526136, EPI_ISL_526137 | Sydney South West Pathology Service (SSWPS) - Royal Prince Alfred Hospital - NSW Health Pathology | NSW Health Pathology - Institute of Clinical Pathology and Medical Research; Westmead Hospital; University of Sydney | CIDM-PH et al. |
| EPI_ISL_526180, EPI_ISL_526181 | St Vincent's Pathology (SydPath) | NSW Health Pathology - Institute of Clinical Pathology and Medical Research; Westmead Hospital; University of Sydney | CIDM-PH et al. |
| EPI_ISL_526335, EPI_ISL_526336 | University of Birmingham | COVID-19 Genomics UK (COG-UK) Consortium | Institute of Microbiology, University of Birmingham: Claire McMurray, Joanne Stockton, Samuel Nicholls, Radoslaw Poplawski, Will Rowe, Josh Quick, Nicholas Loman. University of Birmingham Testing Laboratory: Celina M Whalley, Andrew Bosworth, Charlotte Poxon, Kasun Wanigasooriya, Oliver Pickles, Mike Kidd, Alex Richter, Andrew D Beggs PHE Heartlands Lab: Husam Osman, Andrew Bosworth. Queen Elizabeth Hospital: Anna Casey |
| EPI_ISL_526368, EPI_ISL_526370, EPI_ISL_526372, EPI_ISL_526374, EPI_ISL_526377, EPI_ISL_526379, EPI_ISL_526382, EPI_ISL_526383, EPI_ISL_526393 | Liverpool Clinical Laboratories | COVID-19 Genomics UK (COG-UK) Consortium | Sam Haldenby, Anita Lucaci, Steve Paterson, Julian Hiscox, Alistair Darby, M Almsaud, A Alrezaihi, Muhannad Alruwaili, Stuart D Armstrong, Jones Benjamin, Eleanor G Bentley, Anu Chawla, Jordan J Clark, Angela Cowell, Richard Eccles, Isabel García-Dorival, Matthew Gemmell, Alessandro Gerada, PKF Gilmore, Richard Gregory, Ximeng Han, Catherine Hartley, Margaret Hughes, Miren Iturriza-Gomara, James Johnson, L Luu, Jenifer Manson, Charlotte Nelson, Elaine O'Toole, Cassie Olateju, Rebekah Penrice-Randal , Lucille Rainbow, N.P Randle, Trevor Ian Robinson, Parul Sharma, Ghada T Shawli, James P Stewart, Neil Swainston, Ecaterina Vamos, Joanne Watts, Mark Whitehead |
| EPI_ISL_526395, EPI_ISL_526396, EPI_ISL_526397, EPI_ISL_526398, EPI_ISL_526399, EPI_ISL_526400, EPI_ISL_526401, EPI_ISL_526402, EPI_ISL_526403, EPI_ISL_526404, EPI_ISL_526405, EPI_ISL_526406, EPI_ISL_526407, EPI_ISL_526408, EPI_ISL_526409, EPI_ISL_526410, EPI_ISL_526411, EPI_ISL_526412, EPI_ISL_526413, EPI_ISL_526414, EPI_ISL_526415, EPI_ISL_526416, EPI_ISL_526417, EPI_ISL_526418, EPI_ISL_526419, EPI_ISL_526420, EPI_ISL_526421, EPI_ISL_526422 | | | |
| see above | Queens Medical Centre, Clinical Microbiology Department / DeepSeq Nottingham | COVID-19 Genomics UK (COG-UK) Consortium | Gemma Clark, Wendy Smith, Manjinder Khakh, Vicki M Fleming, Michelle M Lister, Hannah Howson-Wells, Jonathan Ball, Patrick McClure, Joseph Chappell, Theocharis Tsoleridis, Nadine Holmes, Matthew Carlisle, Christopher Moore, Fei Sang, Johnny Debebe, Victoria Wright, Matthew Loose |
| EPI_ISL_526431 | Lincolnshire Hospitals and DeepSeq Nottingham | COVID-19 Genomics UK (COG-UK) Consortium | Nichola Duckworth, Tim Sloan, Sarah Walsh, Jonathan Ball, Patrick McClure, Joeseph Chappell, Nadine Holmes, Matthew Carlisle, Christopher Moore, Fei Sang, Johnny Debebe, Victoria Wright, Matthew Loose |
| EPI_ISL_526552 | OHSU Lab Services Molecular Microbiology Lab | Oregon SARS-CoV-2 Genome Sequencing Center | Brendan L. O'Connell, Ruth V. Nichols, Alec J. Hirsch, Guang Fan, Daniel N. Streblow, William B. Messer, Andrew C. Adey, Benjamin N. Bimber, Brian J. O'Roak |
| EPI_ISL_526585, EPI_ISL_526586 | Florida Bureau of Public Health Laboratories | Florida Bureau of Public Health Laboratories | Sarah Schmedes, Jason Blanton |

| | | | |
|---|---|---|---|
| EPI_ISL_526689 | Pathology West - NSW Health Pathology | NSW Health Pathology - Institute of Clinical Pathology and Medical Research; Westmead Hospital; University of Sydney | CIDM-PH et al. |
| EPI_ISL_526707, EPI_ISL_526708, EPI_ISL_526709, EPI_ISL_526710, EPI_ISL_526711, EPI_ISL_526712, EPI_ISL_526713, EPI_ISL_526714, EPI_ISL_526715, EPI_ISL_526716, EPI_ISL_526717, EPI_ISL_526718, EPI_ISL_526719, EPI_ISL_526720, EPI_ISL_526721, EPI_ISL_526722, EPI_ISL_526723, EPI_ISL_526724, EPI_ISL_526725, EPI_ISL_526726, EPI_ISL_526727, EPI_ISL_526728, EPI_ISL_526729 | | | |
| see above | Division of Viral Diseases, Center for Laboratory Control of Infectious Diseases, Korea Centers for Diseases Control and Prevention | Division of Viral Diseases, Center for Laboratory Control of Infectious Diseases, Korea Centers for Diseases Control and Prevention | Jeong-Min Kim, Yoon-Seok Chung, Namjoo Lee, Sang Hee Woo, Hye-Jun Jo, Heui Man Kim, Jun-Sub Kim, Myung Guk Han |
| EPI_ISL_526730, EPI_ISL_526731 | Center for Laboratory Control of Infectious Diseases, Korea Centers for Diseases Control and Prevention | Center for Laboratory Control of Infectious Diseases, Korea Centers for Diseases Control and Prevention | Junyoung Kim, Ae Kyung Park, Eunkyung Shin, Jin Sun No, Jeong-Min Kim, Yoon-Seok Chung, Heui Man Kim, Myung Guk Han |
| EPI_ISL_526732, EPI_ISL_526733 | Division of Viral Diseases, Center for Laboratory Control of Infectious Diseases, Korea Centers for Diseases Control and Prevention | Division of Viral Diseases, Center for Laboratory Control of Infectious Diseases, Korea Centers for Diseases Control and Prevention | Jeong-Min Kim, Yoon-Seok Chung, Namjoo Lee, Sang Hee Woo, Hye-Jun Jo, Heui Man Kim, Jun-Sub Kim, Myung Guk Han |
| EPI_ISL_526734, EPI_ISL_526735, EPI_ISL_526736, EPI_ISL_526737, EPI_ISL_526738, EPI_ISL_526739, EPI_ISL_526740, EPI_ISL_526741, EPI_ISL_526742, EPI_ISL_526743, EPI_ISL_526744 | | | |
| see above | Center for Laboratory Control of Infectious Diseases, Korea Centers for Diseases Control and Prevention | Center for Laboratory Control of Infectious Diseases, Korea Centers for Diseases Control and Prevention | Junyoung Kim, Ae Kyung Park, Eunkyung Shin, Jin Sun No, Jeong-Min Kim, Yoon-Seok Chung, Heui Man Kim, Myung Guk Han |
| EPI_ISL_527063, EPI_ISL_527064 | Area of Virology, Serology and Virology Division (SAViD), New South Wales Health Pathology Randwick | Area of Virology, Serology and Virology Division (SAViD), New South Wales Health Pathology Randwick | Rawlinson, W. |
| EPI_ISL_527601 | Minnesota Department of Health, Public Health Laboratory | Minnesota Department of Health, Public Health Laboratory | Matt Plumb, Jacob Garfin, and Xiong Wang |
| EPI_ISL_527656 | AR Dept. of Health-Public Health Lab | Pathogen Discovery, Respiratory Viruses Branch, Division of Viral Diseases, Centers for Disease Control and Prevention | Ying Tao, Jing Zhang, Yan Li, Krista Queen, Anna Uehara, Clinton Paden, Haibin Wang, Suxiang Tong |
| EPI_ISL_527660, EPI_ISL_527661, EPI_ISL_527662, EPI_ISL_527663, EPI_ISL_527664, EPI_ISL_527665 | GA Department of Public Health Laboratory | Pathogen Discovery, Respiratory Viruses Branch, Division of Viral Diseases, Centers for Disease Control and Prevention | Yan Li, Anna Montmayeur, Jing Zhang, Krista Queen, Ying Tao, Anna Uehara, Rachel Marine, Clinton R. Paden, Haibin Wang, Suxiang Tong |
| EPI_ISL_527800, EPI_ISL_527807 | University of Wisconsin-Madison AIDS Vaccine Research Laboratories | University of Wisconsin-Madison AIDS Vaccine Research Laboratories | Gage Moreno, Katarina Braun, et al. AIDS Vaccine Research Laboratories |
| EPI_ISL_528509, EPI_ISL_528512, EPI_ISL_528516, EPI_ISL_528517, EPI_ISL_528518, EPI_ISL_528519, EPI_ISL_528520 | Alaska State Virology Laboratory | Alaska State Virology Laboratory | Chen J et al with Pathogenomics group Dagdag R, Redlinger M, Milton E, George W, Kovalenko A, Drown DM, Bortz E |
| EPI_ISL_528639, EPI_ISL_528640, EPI_ISL_528641, EPI_ISL_528642, EPI_ISL_528643, EPI_ISL_528644, EPI_ISL_528645, EPI_ISL_528646, EPI_ISL_528647, EPI_ISL_528648, EPI_ISL_528649, EPI_ISL_528650, EPI_ISL_528655, EPI_ISL_528656, EPI_ISL_528657, EPI_ISL_528658, EPI_ISL_528659, EPI_ISL_528660, EPI_ISL_528661, EPI_ISL_528662 | | | |
| see above | Virginia DCLS | Virginia DCLS | Virginia DCLS |
| EPI_ISL_528748 | Dinkes Provinsi Jawa Barat | School of Life Sciences and Technology & School of Pharmacy-Institut Teknologi Bandung; Molecular Genetics Laboratory-Faculty of Medicine-Universitas Padjadjaran; Laboratorium Kesehatan Provinsi Jawa Barat | Azzania Fibriani, Catur Riani, Marselina Irasonia Tan, Yunia Sribudiani, Husna Nugrahapraja, Tarwadi, Ema Rahmawati, Savira Ekawardhani, Hesti Lina Wiraswati, Ryan Bayusantika Ristandi, Rifky Waluyajati Rachman, Cut Nur Cinthia Alamanda, Lia Faridah, Gusti Ayu Prani Pradani, Adelina Khristiani Rahayu, Hammam Riza, Sony Solistia Wirawan, Agung Eru Wibowo, Irvan Faizal |
| EPI_ISL_528750 | Santo Borromeus Hospital | School of Life Sciences and Technology & School of Pharmacy-Institut Teknologi Bandung; Molecular Genetics Laboratory-Faculty of Medicine-Universitas Padjadjaran; Laboratorium Kesehatan Provinsi Jawa Barat | Marselina Irasonia Tan, Yunia Sribudiani, Catur Riani, Azzania Fibriani, Husna Nugrahapraja, Tarwadi, Ema Rahmawati, Savira Ekawardhani, Hesti Lina Wiraswati, Ryan Bayusantika Ristandi, Rifky Waluyajati Rachman, Cut Nur Cinthia Alamanda, Lia Faridah, Miftahul Faridl, Karimatu Khoirunnisa, Hammam Riza, Sony Solistia Wirawan, Agung Eru Wibowo, Irvan Faizal |
| EPI_ISL_528751 | Santo Borromeus Hospital | Molecular Genetics Laboratory-Faculty of Medicine-Universitas Padjadjaran; School of Life Sciences and Technology & School of Pharmacy-Institut Teknologi Bandung; Laboratorium Kesehatan Provinsi Jawa Barat | Yunia Sribudiani, Tri Hanggono Achmad, Mas Rizky A.A. Syamsunarno, Fensi Amalina, Catur Riani, Azzania Fibriani, Husna Nugrahapraja, Marselina Irasonia Tan, Tarwadi, Ema Rahmawati, Savira Ekawardhani, Hesti Lina Wiraswati, Ryan Bayusantika Ristandi, Rifky Waluyajati Rachman, Cut Nur Cinthia Alamanda, Lia Faridah, Hammam Riza, Sony Solistia Wirawan, Agung Eru Wibowo, Irvan Faizal |
| EPI_ISL_528752 | Dr. H. A. Rotinsulu Lung Hospital | School of Pharmacy & School of Life Sciences and Technology - Institut Teknologi Bandung; Molecular Genetics Laboratory-Faculty of Medicine-Universitas Padjadjaran; Laboratorium Kesehatan Provinsi Jawa Barat | Catur Riani, Marselina Irasonia Tan, Yunia Sribudiani, Azzania Fibriani, Husna Nugrahapraja, Tarwadi, Ema Rahmawati, Savira Ekawardhani, Hesti Lina Wiraswati, Ryan Bayusantika Ristandi, Rifky Waluyajati Rachman, Cut Nur Cinthia Alamanda, Lia Faridah, Gusti Ayu Prani Pradani, Adelina Khristiani Rahayu, Hammam Riza, Sony Solistia Wirawan, Agung Eru Wibowo, Irvan Faizal |
| EPI_ISL_528759 | Santo Borromeus Hospital | School of Life Sciences and Technology & School of Pharmacy-Institut Teknologi Bandung; Molecular Genetics Laboratory-Faculty of Medicine-Universitas Padjadjaran; Laboratorium Kesehatan Provinsi Jawa Barat | Husna Nugrahapraja, Azzania Fibriani, Catur Riani, Marselina Irasonia Tan, Yunia Sribudiani, Tarwadi, Ema Rahmawati, Savira Ekawardhani, Hesti Lina Wiraswati, Ryan Bayusantika Ristandi, Rifky Waluyajati Rachman, Cut Nur Cinthia Alamanda, Lia Faridah, Tri Hanggono Achmad, Mas Rizky A.A. Syamsunarno, Fensi Amalina, Hammam Riza, Sony Solistia Wirawan, Agung Eru Wibowo, Irvan Faizal |
| EPI_ISL_529027 | Respiratory Virus Unit, Microbiology Services Colindale, Public Health England | Respiratory Virus Unit, Microbiology Services Colindale, Public Health England | PHE Covid Sequencing Team |
| EPI_ISL_529234 | University of Birmingham | COVID-19 Genomics UK (COG-UK) Consortium | Institute of Microbiology, University of Birmingham: Claire McMurray, Joanne Stockton, Samuel Nicholls, Radoslaw Poplawski, Will Rowe, Josh Quick, Nicholas Loman. University of Birmingham Testing Laboratory: Celina M Whalley, Andrew Bosworth, Charlotte Poxon, Kasun Wanigasooriya, Oliver Pickles, Mike Kidd, Alex Richter, Andrew D Beggs PHE Heartlands Lab: Husam Osman, Andrew Bosworth. Queen Elizabeth Hospital: Anna Casey |
| EPI_ISL_529254, EPI_ISL_529259, EPI_ISL_529260 | Liverpool Clinical Laboratories | COVID-19 Genomics UK (COG-UK) Consortium | Sam Haldenby, Anita Lucaci, Steve Paterson, Julian Hiscox, Alistair Darby, M Almsaud, A Alrezaihi, Muhannad Alruwaili, Stuart D Armstrong, Jones Benjamin, Eleanor G Bentley, Anu Chawla, Jordan J Clark, Angela Cowell, Richard Eccles, Isabel García-Dorival, Matthew Gemmell, Alessandro Gerada, PKF Gilmore, Richard Gregory, Ximeng Han, Catherine Hartley, Margaret Hughes, Miren Iturriza-Gomara, James Johnson, L Luu, Jenifer Manson, Charlotte Nelson, Elaine O'Toole, Cassie Olateju, Rebekah Penrice-Randal , Lucille Rainbow, N.P Randle, Trevor Ian Robinson, Parul Sharma, Ghada T Shawli, James P Stewart, Neil Swainston, Ecaterina Vamos, Joanne Watts, Mark Whitehead |
| EPI_ISL_529262 | University of Birmingham | COVID-19 Genomics UK (COG-UK) Consortium | Institute of Microbiology, University of Birmingham: Claire McMurray, Joanne Stockton, Samuel Nicholls, Radoslaw Poplawski, Will Rowe, Josh Quick, Nicholas Loman. University of Birmingham Testing Laboratory: Celina M Whalley, Andrew Bosworth, Charlotte Poxon, Kasun Wanigasooriya, Oliver Pickles, Mike Kidd, Alex Richter, Andrew D Beggs PHE Heartlands Lab: Husam Osman, Andrew Bosworth. Queen Elizabeth Hospital: Anna Casey |
| EPI_ISL_529263, EPI_ISL_529274, EPI_ISL_529284, EPI_ISL_529287 | Liverpool Clinical Laboratories | COVID-19 Genomics UK (COG-UK) Consortium | Sam Haldenby, Anita Lucaci, Steve Paterson, Julian Hiscox, Alistair Darby, M Almsaud, A Alrezaihi, Muhannad Alruwaili, Stuart D Armstrong, Jones Benjamin, Eleanor G Bentley, Anu Chawla, Jordan J Clark, Angela Cowell, Richard Eccles, Isabel García-Dorival, Matthew Gemmell, Alessandro Gerada, PKF Gilmore, Richard Gregory, Ximeng Han, Catherine Hartley, Margaret Hughes, Miren Iturriza-Gomara, James Johnson, L Luu, Jenifer Manson, Charlotte Nelson, Elaine O'Toole, Cassie Olateju, Rebekah Penrice-Randal , Lucille Rainbow, N.P Randle, Trevor Ian Robinson, Parul Sharma, Ghada T Shawli, James P Stewart, Neil Swainston, Ecaterina Vamos, Joanne Watts, Mark Whitehead |
| EPI_ISL_529300 | University of Birmingham | COVID-19 Genomics UK (COG-UK) Consortium | Institute of Microbiology, University of Birmingham: Claire McMurray, Joanne Stockton, Samuel Nicholls, Radoslaw Poplawski, Will Rowe, Josh Quick, Nicholas Loman. University of Birmingham Testing Laboratory: Celina M Whalley, Andrew Bosworth, Charlotte Poxon, Kasun Wanigasooriya, Oliver Pickles, Mike Kidd, Alex Richter, Andrew D Beggs PHE Heartlands Lab: Husam Osman, Andrew Bosworth. Queen Elizabeth Hospital: Anna Casey |
| EPI_ISL_529305 | Liverpool Clinical Laboratories | COVID-19 Genomics UK (COG-UK) Consortium | Sam Haldenby, Anita Lucaci, Steve Paterson, Julian Hiscox, Alistair Darby, M Almsaud, A Alrezaihi, Muhannad Alruwaili, Stuart D Armstrong, Jones Benjamin, |

| | | | |
|---|---|---|---|
| | | | Eleanor G Bentley, Anu Chawla, Jordan J Clark, Angela Cowell, Richard Eccles, Isabel García-Dorival, Matthew Gemmell, Alessandro Gerada, PKF Gilmore, Richard Gregory, Ximeng Han, Catherine Hartley, Margaret Hughes, Miren Iturriza-Gomara, James Johnson, L Luu, Jenifer Manson, Charlotte Nelson, Elaine O'Toole, Cassie Olateju, Rebekah Penrice-Randal , Lucille Rainbow, N.P Randle, Trevor Ian Robinson, Parul Sharma, Ghada T Shawli, James P Stewart, Neil Swainston, Ecaterina Vamos, Joanne Watts, Mark Whitehead |
| EPI_ISL_529307, EPI_ISL_529310 | Queens Medical Centre, Clinical Microbiology Department / DeepSeq Nottingham | COVID-19 Genomics UK (COG-UK) Consortium | Gemma Clark, Wendy Smith, Manjinder Khakh, Vicki M Fleming, Michelle M Lister, Hannah Howson-Wells, Jonathan Ball, Patrick McClure, Joseph Chappell, Theocharis Tsoleridis, Nadine Holmes, Matthew Carlisle, Christopher Moore, Fei Sang, Johnny Debebe, Victoria Wright, Matthew Loose |
| EPI_ISL_529361 | University of Birmingham | COVID-19 Genomics UK (COG-UK) Consortium | Institute of Microbiology, University of Birmingham: Claire McMurray, Joanne Stockton, Samuel Nicholls, Radoslaw Poplawski, Will Rowe, Josh Quick, Nicholas Loman. University of Birmingham Testing Laboratory: Celina M Whalley, Andrew Bosworth, Charlotte Poxon, Kasun Wanigasooriya, Oliver Pickles, Mike Kidd, Alex Richter, Andrew D Beggs PHE Heartlands Lab: Husam Osman, Andrew Bosworth. Queen Elizabeth Hospital: Anna Casey |
| EPI_ISL_529383, EPI_ISL_529385, EPI_ISL_529410, EPI_ISL_529411, EPI_ISL_529414 | Queens Medical Centre, Clinical Microbiology Department / DeepSeq Nottingham | COVID-19 Genomics UK (COG-UK) Consortium | Gemma Clark, Wendy Smith, Manjinder Khakh, Vicki M Fleming, Michelle M Lister, Hannah Howson-Wells, Jonathan Ball, Patrick McClure, Joseph Chappell, Theocharis Tsoleridis, Nadine Holmes, Matthew Carlisle, Christopher Moore, Fei Sang, Johnny Debebe, Victoria Wright, Matthew Loose |
| EPI_ISL_529430 | University of Birmingham | COVID-19 Genomics UK (COG-UK) Consortium | Institute of Microbiology, University of Birmingham: Claire McMurray, Joanne Stockton, Samuel Nicholls, Radoslaw Poplawski, Will Rowe, Josh Quick, Nicholas Loman. University of Birmingham Testing Laboratory: Celina M Whalley, Andrew Bosworth, Charlotte Poxon, Kasun Wanigasooriya, Oliver Pickles, Mike Kidd, Alex Richter, Andrew D Beggs PHE Heartlands Lab: Husam Osman, Andrew Bosworth. Queen Elizabeth Hospital: Anna Casey |
| EPI_ISL_529436 | Queens Medical Centre, Clinical Microbiology Department / DeepSeq Nottingham | COVID-19 Genomics UK (COG-UK) Consortium | Gemma Clark, Wendy Smith, Manjinder Khakh, Vicki M Fleming, Michelle M Lister, Hannah Howson-Wells, Jonathan Ball, Patrick McClure, Joseph Chappell, Theocharis Tsoleridis, Nadine Holmes, Matthew Carlisle, Christopher Moore, Fei Sang, Johnny Debebe, Victoria Wright, Matthew Loose |
| EPI_ISL_529443 | University of Birmingham | COVID-19 Genomics UK (COG-UK) Consortium | Institute of Microbiology, University of Birmingham: Claire McMurray, Joanne Stockton, Samuel Nicholls, Radoslaw Poplawski, Will Rowe, Josh Quick, Nicholas Loman. University of Birmingham Testing Laboratory: Celina M Whalley, Andrew Bosworth, Charlotte Poxon, Kasun Wanigasooriya, Oliver Pickles, Mike Kidd, Alex Richter, Andrew D Beggs PHE Heartlands Lab: Husam Osman, Andrew Bosworth. Queen Elizabeth Hospital: Anna Casey |
| EPI_ISL_529452 | Quadram Institute Bioscience | COVID-19 Genomics UK (COG-UK) Consortium | Dave J. Baker, Gemma L. Kay, Alp Aydin, Thanh Le-Viet, Steven Rudder, Ana P. Tedim, Anastasia Kolyva, Maria Diaz, Leonardo de Oliveira Martins, Nabil-Fareed Alikhan, Lizzie Meadows, Rachael Stanley, Ngozi Elumogo, Muhammed Yasir, Nicholas M. Thomson, Alexander J Trotter, Rachel Gilroy, Samuel Bloomfield, Claire Stuart, Andrew Bell, Reenesh Prakash, Samir Dervisevic, Alison E. Mather, John Wain, Mark Webber, Andrew J. Page, Justin O'Grady |
| EPI_ISL_529454, EPI_ISL_529455, EPI_ISL_529456, EPI_ISL_529464 | Queens Medical Centre, Clinical Microbiology Department / DeepSeq Nottingham | COVID-19 Genomics UK (COG-UK) Consortium | Gemma Clark, Wendy Smith, Manjinder Khakh, Vicki M Fleming, Michelle M Lister, Hannah Howson-Wells, Jonathan Ball, Patrick McClure, Joseph Chappell, Theocharis Tsoleridis, Nadine Holmes, Matthew Carlisle, Christopher Moore, Fei Sang, Johnny Debebe, Victoria Wright, Matthew Loose |
| EPI_ISL_529466 | Liverpool Clinical Laboratories | COVID-19 Genomics UK (COG-UK) Consortium | Sam Haldenby, Anita Lucaci, Steve Paterson, Julian Hiscox, Alistair Darby, M Almsaud, A Alrezaihi, Muhannad Alruwaili, Stuart D Armstrong, Jones Benjamin, Eleanor G Bentley, Anu Chawla, Jordan J Clark, Angela Cowell, Richard Eccles, Isabel García-Dorival, Matthew Gemmell, Alessandro Gerada, PKF Gilmore, Richard Gregory, Ximeng Han, Catherine Hartley, Margaret Hughes, Miren Iturriza-Gomara, James Johnson, L Luu, Jenifer Manson, Charlotte Nelson, Elaine O'Toole, Cassie Olateju, Rebekah Penrice-Randal , Lucille Rainbow, N.P Randle, Trevor Ian Robinson, Parul Sharma, Ghada T Shawli, James P Stewart, Neil Swainston, Ecaterina Vamos, Joanne Watts, Mark Whitehead |
| EPI_ISL_529483, EPI_ISL_529486 | Queens Medical Centre, Clinical Microbiology Department / DeepSeq Nottingham | COVID-19 Genomics UK (COG-UK) Consortium | Gemma Clark, Wendy Smith, Manjinder Khakh, Vicki M Fleming, Michelle M Lister, Hannah Howson-Wells, Jonathan Ball, Patrick McClure, Joseph Chappell, Theocharis Tsoleridis, Nadine Holmes, Matthew Carlisle, Christopher Moore, Fei Sang, Johnny Debebe, Victoria Wright, Matthew Loose |
| EPI_ISL_529490 | Liverpool Clinical Laboratories | COVID-19 Genomics UK (COG-UK) Consortium | Sam Haldenby, Anita Lucaci, Steve Paterson, Julian Hiscox, Alistair Darby, M Almsaud, A Alrezaihi, Muhannad Alruwaili, Stuart D Armstrong, Jones Benjamin, Eleanor G Bentley, Anu Chawla, Jordan J Clark, Angela Cowell, Richard Eccles, Isabel García-Dorival, Matthew Gemmell, Alessandro Gerada, PKF Gilmore, Richard Gregory, Ximeng Han, Catherine Hartley, Margaret Hughes, Miren Iturriza-Gomara, James Johnson, L Luu, Jenifer Manson, Charlotte Nelson, Elaine O'Toole, Cassie Olateju, Rebekah Penrice-Randal , Lucille Rainbow, N.P Randle, Trevor Ian Robinson, Parul Sharma, Ghada T Shawli, James P Stewart, Neil Swainston, Ecaterina Vamos, Joanne Watts, Mark Whitehead |
| EPI_ISL_529493, EPI_ISL_529495 | University of Birmingham | COVID-19 Genomics UK (COG-UK) Consortium | Institute of Microbiology, University of Birmingham: Claire McMurray, Joanne Stockton, Samuel Nicholls, Radoslaw Poplawski, Will Rowe, Josh Quick, Nicholas Loman. University of Birmingham Testing Laboratory: Celina M Whalley, Andrew Bosworth, Charlotte Poxon, Kasun Wanigasooriya, Oliver Pickles, Mike Kidd, Alex Richter, Andrew D Beggs PHE Heartlands Lab: Husam Osman, Andrew Bosworth. Queen Elizabeth Hospital: Anna Casey |
| EPI_ISL_529510 | Liverpool Clinical Laboratories | COVID-19 Genomics UK (COG-UK) Consortium | Sam Haldenby, Anita Lucaci, Steve Paterson, Julian Hiscox, Alistair Darby, M Almsaud, A Alrezaihi, Muhannad Alruwaili, Stuart D Armstrong, Jones Benjamin, Eleanor G Bentley, Anu Chawla, Jordan J Clark, Angela Cowell, Richard Eccles, Isabel García-Dorival, Matthew Gemmell, Alessandro Gerada, PKF Gilmore, Richard Gregory, Ximeng Han, Catherine Hartley, Margaret Hughes, Miren Iturriza-Gomara, James Johnson, L Luu, Jenifer Manson, Charlotte Nelson, Elaine O'Toole, Cassie Olateju, Rebekah Penrice-Randal , Lucille Rainbow, N.P Randle, Trevor Ian Robinson, Parul Sharma, Ghada T Shawli, James P Stewart, Neil Swainston, Ecaterina Vamos, Joanne Watts, Mark Whitehead |
| EPI_ISL_529528, EPI_ISL_529529, EPI_ISL_529530, EPI_ISL_529531, EPI_ISL_529532 | Queens Medical Centre, Clinical Microbiology Department / DeepSeq Nottingham | COVID-19 Genomics UK (COG-UK) Consortium | Gemma Clark, Wendy Smith, Manjinder Khakh, Vicki M Fleming, Michelle M Lister, Hannah Howson-Wells, Jonathan Ball, Patrick McClure, Joseph Chappell, Theocharis Tsoleridis, Nadine Holmes, Matthew Carlisle, Christopher Moore, Fei Sang, Johnny Debebe, Victoria Wright, Matthew Loose |
| EPI_ISL_529560, EPI_ISL_529561, EPI_ISL_529562, EPI_ISL_529563, EPI_ISL_529564, EPI_ISL_529565, EPI_ISL_529566, EPI_ISL_529569, EPI_ISL_529582, EPI_ISL_529583, EPI_ISL_529584, EPI_ISL_529585, EPI_ISL_529586 | | | |
| see above | Quadram Institute Bioscience | COVID-19 Genomics UK (COG-UK) Consortium | Dave J. Baker, Gemma L. Kay, Alp Aydin, Thanh Le-Viet, Steven Rudder, Ana P. Tedim, Anastasia Kolyva, Maria Diaz, Leonardo de Oliveira Martins, Nabil-Fareed Alikhan, Lizzie Meadows, Rachael Stanley, Ngozi Elumogo, Muhammed Yasir, Nicholas M. Thomson, Alexander J Trotter, Rachel Gilroy, Samuel Bloomfield, Claire Stuart, Andrew Bell, Reenesh Prakash, Samir Dervisevic, Alison E. Mather, John Wain, Mark Webber, Andrew J. Page, Justin O'Grady |
| EPI_ISL_529618, EPI_ISL_529619, EPI_ISL_529620, EPI_ISL_529621 | University of Birmingham | COVID-19 Genomics UK (COG-UK) Consortium | Institute of Microbiology, University of Birmingham: Claire McMurray, Joanne Stockton, Samuel Nicholls, Radoslaw Poplawski, Will Rowe, Josh Quick, Nicholas Loman. University of Birmingham Testing Laboratory: Celina M Whalley, Andrew Bosworth, Charlotte Poxon, Kasun Wanigasooriya, Oliver Pickles, Mike Kidd, Alex Richter, Andrew D Beggs PHE Heartlands Lab: Husam Osman, Andrew Bosworth. Queen Elizabeth Hospital: Anna Casey |
| EPI_ISL_529697 | Wales Specialist Virology Centre Sequencing lab: Pathogen Genomics Unit | COVID-19 Genomics UK (COG-UK) Consortium | Catherine Moore, Johnathan Evans, Laura Gifford, Malorie Perry, Simon Cottrell, Angela Marchbank, Alec Birchley, Alexander Adams, Amy Gaskin, Bree Gatica-Wilcox, Jason Coombes, Joel Southgate, Lauren Gilbert, Lee Graham, Nicole Pacchiarini, Sara Kumziene-Summerhayes, Sarah Taylor, Sophie Jones, Sara Rey, Matthew Bull, Joanne Watkins, Sally Corden, Tom Connor |
| EPI_ISL_529843, EPI_ISL_529853, EPI_ISL_529854, EPI_ISL_529860, EPI_ISL_529861, EPI_ISL_529862, EPI_ISL_529863, EPI_ISL_529864, EPI_ISL_529865, EPI_ISL_529868, EPI_ISL_529872, EPI_ISL_529878, EPI_ISL_529879 | | | |
| see above | Michigan Department of Health and Human Services, Bureau of Laboratories | Michigan Department of Health and Human Services, Bureau of Laboratories | Blankenship HM, Riner D, Soehnlen MK |
| EPI_ISL_529937, EPI_ISL_529938, EPI_ISL_529939, EPI_ISL_529940, EPI_ISL_529941, EPI_ISL_529942 | Virginia DCLS | Virginia DCLS | Virginia DCLS |
| EPI_ISL_530240, EPI_ISL_530256, EPI_ISL_530257 | Queensland Health Forensic and Scientific Services, Public Health Virology | Public Health Virology Laboratory, Forensic and Scientific Services, Queensland Health | Son Nguyen et al |
| EPI_ISL_531944, EPI_ISL_531977, EPI_ISL_532065, EPI_ISL_532124, EPI_ISL_532541, EPI_ISL_532543, EPI_ISL_532544, EPI_ISL_532545, EPI_ISL_532547, EPI_ISL_532548, EPI_ISL_532549, EPI_ISL_532550, EPI_ISL_532552, EPI_ISL_532553, EPI_ISL_532554, EPI_ISL_532555, EPI_ISL_532556, EPI_ISL_532559, EPI_ISL_532565, EPI_ISL_532566, EPI_ISL_532570, EPI_ISL_532571, EPI_ISL_532573, EPI_ISL_532574, EPI_ISL_532575, EPI_ISL_532576, EPI_ISL_532577, EPI_ISL_532580 | | | |
| see above | Lighthouse Lab in Glasgow | Wellcome Sanger Institute for the COVID-19 Genomics UK (COG-UK) consortium | Harper VanSteenhouse, Yumi Kasai, David Gray, Carol Clugston, Anna Dominiczak and Alex Alderton, Roberto Amato, Sonia Goncalves, Ewan Harrison, David K. Jackson, Ian Johnston, Dominic Kwiatkowski, Cordelia Langford, John Sillitoe |
| EPI_ISL_532583 | Lighthouse Lab in Glasgow | Wellcome Sanger Institute for the COVID-19 Genomics UK (COG-UK) Consortium | Harper VanSteenhouse, Yumi Kasai, David Gray, Carol Clugston, Anna Dominiczak and Alex Alderton, Roberto Amato, Sonia Goncalves, Ewan Harrison, David K. Jackson, Ian Johnston, Dominic Kwiatkowski, Cordelia Langford, John Sillitoe on behalf of the Wellcome Sanger Institute COVID-19 Surveillance Team |

| | | | |
|---|---|---|---|
| EPI_ISL_532584, EPI_ISL_532585, EPI_ISL_532586, EPI_ISL_532590, EPI_ISL_532591, EPI_ISL_532592, EPI_ISL_532593 | Lighthouse Lab in Glasgow | Wellcome Sanger Institute for the COVID-19 Genomics UK (COG-UK) consortium | Harper VanSteenhouse, Yumi Kasai, David Gray, Carol Clugston, Anna Dominiczak and Alex Alderton, Roberto Amato, Sonia Goncalves, Ewan Harrison, David K. Jackson, Ian Johnston, Dominic Kwiatkowski, Cordelia Langford, John Sillitoe |
| EPI_ISL_532594 | NHSGGC West of Scotland Specialist Virology Centre / MRC-University of Glasgow Centre for Virus Research | Wellcome Sanger Institute for the COVID-19 Genomics UK (COG-UK) consortium | Ana da Silva Filipe, Natasha Johnson, Kathy Smollett, Daniel Mair, Stephen Carmichael, Lily Tong, Jenna Nichols, Elihu Aranday-Cortes, Kirstyn Brunker, Yasmin Parr, Kyriaki Nomikou; Sarah McDonald, Marc Niebel, Patawee Asamaphan; Richard Orton, Joseph Hughes, Sreenu Vattipally, David L Robertson; Alasdair MacLean, Rory Gunson; Kathy Li, Natasha Jesudason, Rajiv Shah, James Shepherd, Antonia Ho, Alice Broos, Emma Thomson and Alex Alderton, Roberto Amato, Sonia Goncalves, Ewan Harrison, David K. Jackson, Ian Johnston, Dominic Kwiatkowski, Cordelia Langford, John Sillitoe |
| EPI_ISL_532596, EPI_ISL_532597, EPI_ISL_532599, EPI_ISL_532601, EPI_ISL_532602, EPI_ISL_532603, EPI_ISL_532604, EPI_ISL_532605, EPI_ISL_532606, EPI_ISL_532607, EPI_ISL_532608, EPI_ISL_532609, EPI_ISL_532610, EPI_ISL_532611, EPI_ISL_532612, EPI_ISL_532613, EPI_ISL_532614, EPI_ISL_532616, EPI_ISL_532618, EPI_ISL_532619, EPI_ISL_532622, EPI_ISL_532623 | | | |
| see above | Lighthouse Lab in Glasgow | Wellcome Sanger Institute for the COVID-19 Genomics UK (COG-UK) consortium | Harper VanSteenhouse, Yumi Kasai, David Gray, Carol Clugston, Anna Dominiczak and Alex Alderton, Roberto Amato, Sonia Goncalves, Ewan Harrison, David K. Jackson, Ian Johnston, Dominic Kwiatkowski, Cordelia Langford, John Sillitoe |
| EPI_ISL_532625 | NHSGGC West of Scotland Specialist Virology Centre / MRC-University of Glasgow Centre for Virus Research | Wellcome Sanger Institute for the COVID-19 Genomics UK (COG-UK) consortium | Ana da Silva Filipe, Natasha Johnson, Kathy Smollett, Daniel Mair, Stephen Carmichael, Lily Tong, Jenna Nichols, Elihu Aranday-Cortes, Kirstyn Brunker, Yasmin Parr, Kyriaki Nomikou; Sarah McDonald, Marc Niebel, Patawee Asamaphan; Richard Orton, Joseph Hughes, Sreenu Vattipally, David L Robertson; Alasdair MacLean, Rory Gunson; Kathy Li, Natasha Jesudason, Rajiv Shah, James Shepherd, Antonia Ho, Alice Broos, Emma Thomson and Alex Alderton, Roberto Amato, Sonia Goncalves, Ewan Harrison, David K. Jackson, Ian Johnston, Dominic Kwiatkowski, Cordelia Langford, John Sillitoe |
| EPI_ISL_532628, EPI_ISL_532629, EPI_ISL_532630, EPI_ISL_532631, EPI_ISL_532632 | Lighthouse Lab in Glasgow | Wellcome Sanger Institute for the COVID-19 Genomics UK (COG-UK) consortium | Harper VanSteenhouse, Yumi Kasai, David Gray, Carol Clugston, Anna Dominiczak and Alex Alderton, Roberto Amato, Sonia Goncalves, Ewan Harrison, David K. Jackson, Ian Johnston, Dominic Kwiatkowski, Cordelia Langford, John Sillitoe |
| EPI_ISL_532633 | NHSGGC West of Scotland Specialist Virology Centre / MRC-University of Glasgow Centre for Virus Research | Wellcome Sanger Institute for the COVID-19 Genomics UK (COG-UK) consortium | Ana da Silva Filipe, Natasha Johnson, Kathy Smollett, Daniel Mair, Stephen Carmichael, Lily Tong, Jenna Nichols, Elihu Aranday-Cortes, Kirstyn Brunker, Yasmin Parr, Kyriaki Nomikou; Sarah McDonald, Marc Niebel, Patawee Asamaphan; Richard Orton, Joseph Hughes, Sreenu Vattipally, David L Robertson; Alasdair MacLean, Rory Gunson; Kathy Li, Natasha Jesudason, Rajiv Shah, James Shepherd, Antonia Ho, Alice Broos, Emma Thomson and Alex Alderton, Roberto Amato, Sonia Goncalves, Ewan Harrison, David K. Jackson, Ian Johnston, Dominic Kwiatkowski, Cordelia Langford, John Sillitoe |
| EPI_ISL_532634, EPI_ISL_532635, EPI_ISL_532636, EPI_ISL_532637, EPI_ISL_532638 | Lighthouse Lab in Glasgow | Wellcome Sanger Institute for the COVID-19 Genomics UK (COG-UK) consortium | Harper VanSteenhouse, Yumi Kasai, David Gray, Carol Clugston, Anna Dominiczak and Alex Alderton, Roberto Amato, Sonia Goncalves, Ewan Harrison, David K. Jackson, Ian Johnston, Dominic Kwiatkowski, Cordelia Langford, John Sillitoe |
| EPI_ISL_532639 | NHSGGC West of Scotland Specialist Virology Centre / MRC-University of Glasgow Centre for Virus Research | Wellcome Sanger Institute for the COVID-19 Genomics UK (COG-UK) consortium | Ana da Silva Filipe, Natasha Johnson, Kathy Smollett, Daniel Mair, Stephen Carmichael, Lily Tong, Jenna Nichols, Elihu Aranday-Cortes, Kirstyn Brunker, Yasmin Parr, Kyriaki Nomikou; Sarah McDonald, Marc Niebel, Patawee Asamaphan; Richard Orton, Joseph Hughes, Sreenu Vattipally, David L Robertson; Alasdair MacLean, Rory Gunson; Kathy Li, Natasha Jesudason, Rajiv Shah, James Shepherd, Antonia Ho, Alice Broos, Emma Thomson and Alex Alderton, Roberto Amato, Sonia Goncalves, Ewan Harrison, David K. Jackson, Ian Johnston, Dominic Kwiatkowski, Cordelia Langford, John Sillitoe |
| EPI_ISL_532640, EPI_ISL_532641, EPI_ISL_532642, EPI_ISL_532643, EPI_ISL_532644, EPI_ISL_532645, EPI_ISL_532646, EPI_ISL_532647 | Lighthouse Lab in Glasgow | Wellcome Sanger Institute for the COVID-19 Genomics UK (COG-UK) consortium | Harper VanSteenhouse, Yumi Kasai, David Gray, Carol Clugston, Anna Dominiczak and Alex Alderton, Roberto Amato, Sonia Goncalves, Ewan Harrison, David K. Jackson, Ian Johnston, Dominic Kwiatkowski, Cordelia Langford, John Sillitoe |
| EPI_ISL_532648 | NHSGGC West of Scotland Specialist Virology Centre / MRC-University of Glasgow Centre for Virus Research | Wellcome Sanger Institute for the COVID-19 Genomics UK (COG-UK) consortium | Ana da Silva Filipe, Natasha Johnson, Kathy Smollett, Daniel Mair, Stephen Carmichael, Lily Tong, Jenna Nichols, Elihu Aranday-Cortes, Kirstyn Brunker, Yasmin Parr, Kyriaki Nomikou; Sarah McDonald, Marc Niebel, Patawee Asamaphan; Richard Orton, Joseph Hughes, Sreenu Vattipally, David L Robertson; Alasdair MacLean, Rory Gunson; Kathy Li, Natasha Jesudason, Rajiv Shah, James Shepherd, Antonia Ho, Alice Broos, Emma Thomson and Alex Alderton, Roberto Amato, Sonia Goncalves, Ewan Harrison, David K. Jackson, Ian Johnston, Dominic Kwiatkowski, Cordelia Langford, John Sillitoe |
| EPI_ISL_532649, EPI_ISL_532653, EPI_ISL_532654, EPI_ISL_532655, EPI_ISL_532656, EPI_ISL_532657, EPI_ISL_532658, EPI_ISL_532660, EPI_ISL_532662, EPI_ISL_532663, EPI_ISL_532664, EPI_ISL_532665, EPI_ISL_532668 | | | |
| see above | Lighthouse Lab in Glasgow | Wellcome Sanger Institute for the COVID-19 Genomics UK (COG-UK) consortium | Harper VanSteenhouse, Yumi Kasai, David Gray, Carol Clugston, Anna Dominiczak and Alex Alderton, Roberto Amato, Sonia Goncalves, Ewan Harrison, David K. Jackson, Ian Johnston, Dominic Kwiatkowski, Cordelia Langford, John Sillitoe |
| EPI_ISL_532670 | NHSGGC West of Scotland Specialist Virology Centre / MRC-University of Glasgow Centre for Virus Research | Wellcome Sanger Institute for the COVID-19 Genomics UK (COG-UK) consortium | Ana da Silva Filipe, Natasha Johnson, Kathy Smollett, Daniel Mair, Stephen Carmichael, Lily Tong, Jenna Nichols, Elihu Aranday-Cortes, Kirstyn Brunker, Yasmin Parr, Kyriaki Nomikou; Sarah McDonald, Marc Niebel, Patawee Asamaphan; Richard Orton, Joseph Hughes, Sreenu Vattipally, David L Robertson; Alasdair MacLean, Rory Gunson; Kathy Li, Natasha Jesudason, Rajiv Shah, James Shepherd, Antonia Ho, Alice Broos, Emma Thomson and Alex Alderton, Roberto Amato, Sonia Goncalves, Ewan Harrison, David K. Jackson, Ian Johnston, Dominic Kwiatkowski, Cordelia Langford, John Sillitoe |
| EPI_ISL_532671, EPI_ISL_532672, EPI_ISL_532673, EPI_ISL_532674, EPI_ISL_532675, EPI_ISL_532676, EPI_ISL_532678, EPI_ISL_532679, EPI_ISL_532680, EPI_ISL_532681, EPI_ISL_532682, EPI_ISL_532684, EPI_ISL_532685, EPI_ISL_532686, EPI_ISL_532687, EPI_ISL_532688, EPI_ISL_532690, EPI_ISL_532692, EPI_ISL_532694, EPI_ISL_532695, EPI_ISL_532696 | | | |
| see above | Lighthouse Lab in Glasgow | Wellcome Sanger Institute for the COVID-19 Genomics UK (COG-UK) consortium | Harper VanSteenhouse, Yumi Kasai, David Gray, Carol Clugston, Anna Dominiczak and Alex Alderton, Roberto Amato, Sonia Goncalves, Ewan Harrison, David K. Jackson, Ian Johnston, Dominic Kwiatkowski, Cordelia Langford, John Sillitoe |
| EPI_ISL_532697 | NHSGGC West of Scotland Specialist Virology Centre / MRC-University of Glasgow Centre for Virus Research | Wellcome Sanger Institute for the COVID-19 Genomics UK (COG-UK) consortium | Ana da Silva Filipe, Natasha Johnson, Kathy Smollett, Daniel Mair, Stephen Carmichael, Lily Tong, Jenna Nichols, Elihu Aranday-Cortes, Kirstyn Brunker, Yasmin Parr, Kyriaki Nomikou; Sarah McDonald, Marc Niebel, Patawee Asamaphan; Richard Orton, Joseph Hughes, Sreenu Vattipally, David L Robertson; Alasdair MacLean, Rory Gunson; Kathy Li, Natasha Jesudason, Rajiv Shah, James Shepherd, Antonia Ho, Alice Broos, Emma Thomson and Alex Alderton, Roberto Amato, Sonia Goncalves, Ewan Harrison, David K. Jackson, Ian Johnston, Dominic Kwiatkowski, Cordelia Langford, John Sillitoe |
| EPI_ISL_532698, EPI_ISL_532699, EPI_ISL_532700, EPI_ISL_532701, EPI_ISL_532702, EPI_ISL_532703, EPI_ISL_532704, EPI_ISL_532707, EPI_ISL_532708, EPI_ISL_532709, EPI_ISL_532710, EPI_ISL_532711, EPI_ISL_532712, EPI_ISL_532713, EPI_ISL_532714 | | | |
| see above | Lighthouse Lab in Glasgow | Wellcome Sanger Institute for the COVID-19 Genomics UK (COG-UK) consortium | Harper VanSteenhouse, Yumi Kasai, David Gray, Carol Clugston, Anna Dominiczak and Alex Alderton, Roberto Amato, Sonia Goncalves, Ewan Harrison, David K. Jackson, Ian Johnston, Dominic Kwiatkowski, Cordelia Langford, John Sillitoe |
| EPI_ISL_532715 | NHSGGC West of Scotland Specialist Virology Centre / MRC-University of Glasgow Centre for Virus Research | Wellcome Sanger Institute for the COVID-19 Genomics UK (COG-UK) consortium | Ana da Silva Filipe, Natasha Johnson, Kathy Smollett, Daniel Mair, Stephen Carmichael, Lily Tong, Jenna Nichols, Elihu Aranday-Cortes, Kirstyn Brunker, Yasmin Parr, Kyriaki Nomikou; Sarah McDonald, Marc Niebel, Patawee Asamaphan; Richard Orton, Joseph Hughes, Sreenu Vattipally, David L Robertson; Alasdair MacLean, Rory Gunson; Kathy Li, Natasha Jesudason, Rajiv Shah, James Shepherd, Antonia Ho, Alice Broos, Emma Thomson and Alex Alderton, Roberto Amato, Sonia Goncalves, Ewan Harrison, David K. Jackson, Ian Johnston, Dominic Kwiatkowski, Cordelia Langford, John Sillitoe |
| EPI_ISL_532717, EPI_ISL_532718, EPI_ISL_532720, EPI_ISL_532721, EPI_ISL_532722, EPI_ISL_532723, EPI_ISL_532724, EPI_ISL_532725, EPI_ISL_532726, EPI_ISL_532727, EPI_ISL_532729, EPI_ISL_532731, EPI_ISL_532732 | | | |
| see above | Lighthouse Lab in Glasgow | Wellcome Sanger Institute for the COVID-19 Genomics UK (COG-UK) consortium | Harper VanSteenhouse, Yumi Kasai, David Gray, Carol Clugston, Anna Dominiczak and Alex Alderton, Roberto Amato, Sonia Goncalves, Ewan Harrison, David K. Jackson, Ian Johnston, Dominic Kwiatkowski, Cordelia Langford, John Sillitoe |
| EPI_ISL_532733 | NHSGGC West of Scotland Specialist Virology Centre / MRC-University of Glasgow Centre for Virus Research | Wellcome Sanger Institute for the COVID-19 Genomics UK (COG-UK) consortium | Ana da Silva Filipe, Natasha Johnson, Kathy Smollett, Daniel Mair, Stephen Carmichael, Lily Tong, Jenna Nichols, Elihu Aranday-Cortes, Kirstyn Brunker, Yasmin Parr, Kyriaki Nomikou; Sarah McDonald, Marc Niebel, Patawee Asamaphan; Richard Orton, Joseph Hughes, Sreenu Vattipally, David L Robertson; Alasdair MacLean, Rory Gunson; Kathy Li, Natasha Jesudason, Rajiv Shah, James Shepherd, Antonia Ho, Alice Broos, Emma Thomson and Alex Alderton, Roberto Amato, Sonia Goncalves, Ewan Harrison, David K. Jackson, Ian Johnston, Dominic Kwiatkowski, Cordelia Langford, John Sillitoe |
| EPI_ISL_532734, EPI_ISL_532735, EPI_ISL_532736, EPI_ISL_532737 | Lighthouse Lab in Glasgow | Wellcome Sanger Institute for the COVID-19 Genomics UK (COG-UK) consortium | Harper VanSteenhouse, Yumi Kasai, David Gray, Carol Clugston, Anna Dominiczak and Alex Alderton, Roberto Amato, Sonia Goncalves, Ewan Harrison, David K. Jackson, Ian Johnston, Dominic Kwiatkowski, Cordelia Langford, John Sillitoe |
| EPI_ISL_532738 | NHSGGC West of Scotland Specialist Virology Centre / MRC-University of Glasgow Centre for Virus Research | Wellcome Sanger Institute for the COVID-19 Genomics UK (COG-UK) consortium | Ana da Silva Filipe, Natasha Johnson, Kathy Smollett, Daniel Mair, Stephen Carmichael, Lily Tong, Jenna Nichols, Elihu Aranday-Cortes, Kirstyn Brunker, Yasmin Parr, Kyriaki Nomikou; Sarah McDonald, Marc Niebel, Patawee Asamaphan; Richard Orton, Joseph Hughes, Sreenu Vattipally, David L Robertson; Alasdair MacLean, Rory Gunson; Kathy Li, Natasha Jesudason, Rajiv Shah, James Shepherd, Antonia Ho, Alice Broos, Emma Thomson and Alex Alderton, Roberto Amato, Sonia Goncalves, Ewan Harrison, David K. Jackson, Ian Johnston, Dominic Kwiatkowski, Cordelia Langford, John Sillitoe |
| EPI_ISL_532739, EPI_ISL_532740, EPI_ISL_532741, EPI_ISL_532742, | Lighthouse Lab in Glasgow | Wellcome Sanger Institute for the COVID-19 Genomics UK (COG-UK) consortium | Harper VanSteenhouse, Yumi Kasai, David Gray, Carol Clugston, Anna Dominiczak and Alex Alderton, Roberto Amato, Sonia Goncalves, Ewan Harrison, David K. Jackson, Ian Johnston, Dominic Kwiatkowski, Cordelia Langford, John Sillitoe |

| | | | |
|---|---|---|---|
| EPI_ISL_532743 | | | |
| EPI_ISL_532744, EPI_ISL_532746 | NHSGGC West of Scotland Specialist Virology Centre / MRC-University of Glasgow Centre for Virus Research | Wellcome Sanger Institute for the COVID-19 Genomics UK (COG-UK) consortium | Ana da Silva Filipe, Natasha Johnson, Kathy Smollett, Daniel Mair, Stephen Carmichael, Lily Tong, Jenna Nichols, Elihu Aranday-Cortes, Kirstyn Brunker, Yasmin Parr, Kyriaki Nomikou; Sarah McDonald, Marc Niebel, Patawee Asamaphan; Richard Orton, Joseph Hughes, Sreenu Vattipally, David L Robertson; Alasdair MacLean, Rory Gunson; Kathy Li, Natasha Jesudason, Rajiv Shah, James Shepherd, Antonia Ho, Alice Broos, Emma Thomson and Alex Alderton, Roberto Amato, Sonia Goncalves, Ewan Harrison, David K. Jackson, Ian Johnston, Dominic Kwiatkowski, Cordelia Langford, John Sillitoe |
| EPI_ISL_532747, EPI_ISL_532748, EPI_ISL_532749 | Lighthouse Lab in Glasgow | Wellcome Sanger Institute for the COVID-19 Genomics UK (COG-UK) consortium | Harper VanSteenhouse, Yumi Kasai, David Gray, Carol Clugston, Anna Dominiczak and Alex Alderton, Roberto Amato, Sonia Goncalves, Ewan Harrison, David K. Jackson, Ian Johnston, Dominic Kwiatkowski, Cordelia Langford, John Sillitoe |
| EPI_ISL_532750 | NHSGGC West of Scotland Specialist Virology Centre / MRC-University of Glasgow Centre for Virus Research | Wellcome Sanger Institute for the COVID-19 Genomics UK (COG-UK) consortium | Ana da Silva Filipe, Natasha Johnson, Kathy Smollett, Daniel Mair, Stephen Carmichael, Lily Tong, Jenna Nichols, Elihu Aranday-Cortes, Kirstyn Brunker, Yasmin Parr, Kyriaki Nomikou; Sarah McDonald, Marc Niebel, Patawee Asamaphan; Richard Orton, Joseph Hughes, Sreenu Vattipally, David L Robertson; Alasdair MacLean, Rory Gunson; Kathy Li, Natasha Jesudason, Rajiv Shah, James Shepherd, Antonia Ho, Alice Broos, Emma Thomson and Alex Alderton, Roberto Amato, Sonia Goncalves, Ewan Harrison, David K. Jackson, Ian Johnston, Dominic Kwiatkowski, Cordelia Langford, John Sillitoe |
| EPI_ISL_532751, EPI_ISL_532752 | Lighthouse Lab in Glasgow | Wellcome Sanger Institute for the COVID-19 Genomics UK (COG-UK) consortium | Harper VanSteenhouse, Yumi Kasai, David Gray, Carol Clugston, Anna Dominiczak and Alex Alderton, Roberto Amato, Sonia Goncalves, Ewan Harrison, David K. Jackson, Ian Johnston, Dominic Kwiatkowski, Cordelia Langford, John Sillitoe |
| EPI_ISL_532753 | NHSGGC West of Scotland Specialist Virology Centre / MRC-University of Glasgow Centre for Virus Research | Wellcome Sanger Institute for the COVID-19 Genomics UK (COG-UK) consortium | Ana da Silva Filipe, Natasha Johnson, Kathy Smollett, Daniel Mair, Stephen Carmichael, Lily Tong, Jenna Nichols, Elihu Aranday-Cortes, Kirstyn Brunker, Yasmin Parr, Kyriaki Nomikou; Sarah McDonald, Marc Niebel, Patawee Asamaphan; Richard Orton, Joseph Hughes, Sreenu Vattipally, David L Robertson; Alasdair MacLean, Rory Gunson; Kathy Li, Natasha Jesudason, Rajiv Shah, James Shepherd, Antonia Ho, Alice Broos, Emma Thomson and Alex Alderton, Roberto Amato, Sonia Goncalves, Ewan Harrison, David K. Jackson, Ian Johnston, Dominic Kwiatkowski, Cordelia Langford, John Sillitoe |
| EPI_ISL_532754 | Lighthouse Lab in Glasgow | Wellcome Sanger Institute for the COVID-19 Genomics UK (COG-UK) consortium | Harper VanSteenhouse, Yumi Kasai, David Gray, Carol Clugston, Anna Dominiczak and Alex Alderton, Roberto Amato, Sonia Goncalves, Ewan Harrison, David K. Jackson, Ian Johnston, Dominic Kwiatkowski, Cordelia Langford, John Sillitoe |
| EPI_ISL_532755 | NHSGGC West of Scotland Specialist Virology Centre / MRC-University of Glasgow Centre for Virus Research | Wellcome Sanger Institute for the COVID-19 Genomics UK (COG-UK) consortium | Ana da Silva Filipe, Natasha Johnson, Kathy Smollett, Daniel Mair, Stephen Carmichael, Lily Tong, Jenna Nichols, Elihu Aranday-Cortes, Kirstyn Brunker, Yasmin Parr, Kyriaki Nomikou; Sarah McDonald, Marc Niebel, Patawee Asamaphan; Richard Orton, Joseph Hughes, Sreenu Vattipally, David L Robertson; Alasdair MacLean, Rory Gunson; Kathy Li, Natasha Jesudason, Rajiv Shah, James Shepherd, Antonia Ho, Alice Broos, Emma Thomson and Alex Alderton, Roberto Amato, Sonia Goncalves, Ewan Harrison, David K. Jackson, Ian Johnston, Dominic Kwiatkowski, Cordelia Langford, John Sillitoe |
| EPI_ISL_532756, EPI_ISL_532757, EPI_ISL_532758, EPI_ISL_532759, EPI_ISL_532760, EPI_ISL_532762, EPI_ISL_532763, EPI_ISL_532764 | Lighthouse Lab in Glasgow | Wellcome Sanger Institute for the COVID-19 Genomics UK (COG-UK) consortium | Harper VanSteenhouse, Yumi Kasai, David Gray, Carol Clugston, Anna Dominiczak and Alex Alderton, Roberto Amato, Sonia Goncalves, Ewan Harrison, David K. Jackson, Ian Johnston, Dominic Kwiatkowski, Cordelia Langford, John Sillitoe |
| EPI_ISL_532765 | NHSGGC West of Scotland Specialist Virology Centre / MRC-University of Glasgow Centre for Virus Research | Wellcome Sanger Institute for the COVID-19 Genomics UK (COG-UK) consortium | Ana da Silva Filipe, Natasha Johnson, Kathy Smollett, Daniel Mair, Stephen Carmichael, Lily Tong, Jenna Nichols, Elihu Aranday-Cortes, Kirstyn Brunker, Yasmin Parr, Kyriaki Nomikou; Sarah McDonald, Marc Niebel, Patawee Asamaphan; Richard Orton, Joseph Hughes, Sreenu Vattipally, David L Robertson; Alasdair MacLean, Rory Gunson; Kathy Li, Natasha Jesudason, Rajiv Shah, James Shepherd, Antonia Ho, Alice Broos, Emma Thomson and Alex Alderton, Roberto Amato, Sonia Goncalves, Ewan Harrison, David K. Jackson, Ian Johnston, Dominic Kwiatkowski, Cordelia Langford, John Sillitoe |
| EPI_ISL_532766 | Lighthouse Lab in Glasgow | Wellcome Sanger Institute for the COVID-19 Genomics UK (COG-UK) consortium | Harper VanSteenhouse, Yumi Kasai, David Gray, Carol Clugston, Anna Dominiczak and Alex Alderton, Roberto Amato, Sonia Goncalves, Ewan Harrison, David K. Jackson, Ian Johnston, Dominic Kwiatkowski, Cordelia Langford, John Sillitoe |
| EPI_ISL_532767 | NHSGGC West of Scotland Specialist Virology Centre / MRC-University of Glasgow Centre for Virus Research | Wellcome Sanger Institute for the COVID-19 Genomics UK (COG-UK) consortium | Ana da Silva Filipe, Natasha Johnson, Kathy Smollett, Daniel Mair, Stephen Carmichael, Lily Tong, Jenna Nichols, Elihu Aranday-Cortes, Kirstyn Brunker, Yasmin Parr, Kyriaki Nomikou; Sarah McDonald, Marc Niebel, Patawee Asamaphan; Richard Orton, Joseph Hughes, Sreenu Vattipally, David L Robertson; Alasdair MacLean, Rory Gunson; Kathy Li, Natasha Jesudason, Rajiv Shah, James Shepherd, Antonia Ho, Alice Broos, Emma Thomson and Alex Alderton, Roberto Amato, Sonia Goncalves, Ewan Harrison, David K. Jackson, Ian Johnston, Dominic Kwiatkowski, Cordelia Langford, John Sillitoe |
| EPI_ISL_532768, EPI_ISL_532769, EPI_ISL_532770, EPI_ISL_532771, EPI_ISL_532772, EPI_ISL_532773, EPI_ISL_532774, EPI_ISL_532775, EPI_ISL_532776, EPI_ISL_532777, EPI_ISL_532778, EPI_ISL_532779, EPI_ISL_532780, EPI_ISL_532781, EPI_ISL_532782, EPI_ISL_532783 | | | |
| see above | Lighthouse Lab in Glasgow | Wellcome Sanger Institute for the COVID-19 Genomics UK (COG-UK) consortium | Harper VanSteenhouse, Yumi Kasai, David Gray, Carol Clugston, Anna Dominiczak and Alex Alderton, Roberto Amato, Sonia Goncalves, Ewan Harrison, David K. Jackson, Ian Johnston, Dominic Kwiatkowski, Cordelia Langford, John Sillitoe |
| EPI_ISL_532784 | NHSGGC West of Scotland Specialist Virology Centre / MRC-University of Glasgow Centre for Virus Research | Wellcome Sanger Institute for the COVID-19 Genomics UK (COG-UK) consortium | Ana da Silva Filipe, Natasha Johnson, Kathy Smollett, Daniel Mair, Stephen Carmichael, Lily Tong, Jenna Nichols, Elihu Aranday-Cortes, Kirstyn Brunker, Yasmin Parr, Kyriaki Nomikou; Sarah McDonald, Marc Niebel, Patawee Asamaphan; Richard Orton, Joseph Hughes, Sreenu Vattipally, David L Robertson; Alasdair MacLean, Rory Gunson; Kathy Li, Natasha Jesudason, Rajiv Shah, James Shepherd, Antonia Ho, Alice Broos, Emma Thomson and Alex Alderton, Roberto Amato, Sonia Goncalves, Ewan Harrison, David K. Jackson, Ian Johnston, Dominic Kwiatkowski, Cordelia Langford, John Sillitoe |
| EPI_ISL_532785 | Lighthouse Lab in Glasgow | Wellcome Sanger Institute for the COVID-19 Genomics UK (COG-UK) consortium | Harper VanSteenhouse, Yumi Kasai, David Gray, Carol Clugston, Anna Dominiczak and Alex Alderton, Roberto Amato, Sonia Goncalves, Ewan Harrison, David K. Jackson, Ian Johnston, Dominic Kwiatkowski, Cordelia Langford, John Sillitoe |
| EPI_ISL_532786 | NHSGGC West of Scotland Specialist Virology Centre / MRC-University of Glasgow Centre for Virus Research | Wellcome Sanger Institute for the COVID-19 Genomics UK (COG-UK) consortium | Ana da Silva Filipe, Natasha Johnson, Kathy Smollett, Daniel Mair, Stephen Carmichael, Lily Tong, Jenna Nichols, Elihu Aranday-Cortes, Kirstyn Brunker, Yasmin Parr, Kyriaki Nomikou; Sarah McDonald, Marc Niebel, Patawee Asamaphan; Richard Orton, Joseph Hughes, Sreenu Vattipally, David L Robertson; Alasdair MacLean, Rory Gunson; Kathy Li, Natasha Jesudason, Rajiv Shah, James Shepherd, Antonia Ho, Alice Broos, Emma Thomson and Alex Alderton, Roberto Amato, Sonia Goncalves, Ewan Harrison, David K. Jackson, Ian Johnston, Dominic Kwiatkowski, Cordelia Langford, John Sillitoe |
| EPI_ISL_532787, EPI_ISL_532789, EPI_ISL_532790 | Lighthouse Lab in Glasgow | Wellcome Sanger Institute for the COVID-19 Genomics UK (COG-UK) consortium | Harper VanSteenhouse, Yumi Kasai, David Gray, Carol Clugston, Anna Dominiczak and Alex Alderton, Roberto Amato, Sonia Goncalves, Ewan Harrison, David K. Jackson, Ian Johnston, Dominic Kwiatkowski, Cordelia Langford, John Sillitoe |
| EPI_ISL_532791 | NHSGGC West of Scotland Specialist Virology Centre / MRC-University of Glasgow Centre for Virus Research | Wellcome Sanger Institute for the COVID-19 Genomics UK (COG-UK) consortium | Ana da Silva Filipe, Natasha Johnson, Kathy Smollett, Daniel Mair, Stephen Carmichael, Lily Tong, Jenna Nichols, Elihu Aranday-Cortes, Kirstyn Brunker, Yasmin Parr, Kyriaki Nomikou; Sarah McDonald, Marc Niebel, Patawee Asamaphan; Richard Orton, Joseph Hughes, Sreenu Vattipally, David L Robertson; Alasdair MacLean, Rory Gunson; Kathy Li, Natasha Jesudason, Rajiv Shah, James Shepherd, Antonia Ho, Alice Broos, Emma Thomson and Alex Alderton, Roberto Amato, Sonia Goncalves, Ewan Harrison, David K. Jackson, Ian Johnston, Dominic Kwiatkowski, Cordelia Langford, John Sillitoe |
| EPI_ISL_532792, EPI_ISL_532793, EPI_ISL_532794, EPI_ISL_532795, EPI_ISL_532796, EPI_ISL_532797, EPI_ISL_532798, EPI_ISL_532799, EPI_ISL_532800, EPI_ISL_532801, EPI_ISL_532802, EPI_ISL_532803, EPI_ISL_532804, EPI_ISL_532805, EPI_ISL_532807, EPI_ISL_532808, EPI_ISL_532809, EPI_ISL_532810 | | | |
| see above | Lighthouse Lab in Glasgow | Wellcome Sanger Institute for the COVID-19 Genomics UK (COG-UK) consortium | Harper VanSteenhouse, Yumi Kasai, David Gray, Carol Clugston, Anna Dominiczak and Alex Alderton, Roberto Amato, Sonia Goncalves, Ewan Harrison, David K. Jackson, Ian Johnston, Dominic Kwiatkowski, Cordelia Langford, John Sillitoe |
| EPI_ISL_532811 | NHSGGC West of Scotland Specialist Virology Centre / MRC-University of Glasgow Centre for Virus Research | Wellcome Sanger Institute for the COVID-19 Genomics UK (COG-UK) consortium | Ana da Silva Filipe, Natasha Johnson, Kathy Smollett, Daniel Mair, Stephen Carmichael, Lily Tong, Jenna Nichols, Elihu Aranday-Cortes, Kirstyn Brunker, Yasmin Parr, Kyriaki Nomikou; Sarah McDonald, Marc Niebel, Patawee Asamaphan; Richard Orton, Joseph Hughes, Sreenu Vattipally, David L Robertson; Alasdair MacLean, Rory Gunson; Kathy Li, Natasha Jesudason, Rajiv Shah, James Shepherd, Antonia Ho, Alice Broos, Emma Thomson and Alex Alderton, Roberto Amato, Sonia Goncalves, Ewan Harrison, David K. Jackson, Ian Johnston, Dominic Kwiatkowski, Cordelia Langford, John Sillitoe |
| EPI_ISL_532812, EPI_ISL_532813, EPI_ISL_532814, EPI_ISL_532815, EPI_ISL_532816, EPI_ISL_532817, EPI_ISL_532818, EPI_ISL_532819, EPI_ISL_532820, EPI_ISL_532822, EPI_ISL_532823, EPI_ISL_532824, EPI_ISL_532825, EPI_ISL_532826, EPI_ISL_532827, EPI_ISL_532828, EPI_ISL_532829, EPI_ISL_532830, EPI_ISL_532831, EPI_ISL_532832, EPI_ISL_532833, EPI_ISL_532834 | | | |
| see above | Lighthouse Lab in Glasgow | Wellcome Sanger Institute for the COVID-19 Genomics UK (COG-UK) consortium | Harper VanSteenhouse, Yumi Kasai, David Gray, Carol Clugston, Anna Dominiczak and Alex Alderton, Roberto Amato, Sonia Goncalves, Ewan Harrison, David K. Jackson, Ian Johnston, Dominic Kwiatkowski, Cordelia Langford, John Sillitoe |
| EPI_ISL_532835, EPI_ISL_532838, EPI_ISL_532844 | NHSGGC West of Scotland Specialist Virology Centre / MRC-University of Glasgow Centre for Virus Research | Wellcome Sanger Institute for the COVID-19 Genomics UK (COG-UK) consortium | Ana da Silva Filipe, Natasha Johnson, Kathy Smollett, Daniel Mair, Stephen Carmichael, Lily Tong, Jenna Nichols, Elihu Aranday-Cortes, Kirstyn Brunker, Yasmin Parr, Kyriaki Nomikou; Sarah McDonald, Marc Niebel, Patawee Asamaphan; Richard Orton, Joseph Hughes, Sreenu Vattipally, David L Robertson; Alasdair MacLean, Rory Gunson; Kathy Li, Natasha Jesudason, Rajiv Shah, James Shepherd, Antonia Ho, Alice Broos, Emma Thomson and Alex Alderton, Roberto Amato, Sonia Goncalves, Ewan Harrison, David K. Jackson, Ian Johnston, Dominic Kwiatkowski, Cordelia Langford, John Sillitoe |
| EPI_ISL_532849, EPI_ISL_532850, EPI_ISL_532854, EPI_ISL_532855 | Lighthouse Lab in Glasgow | Wellcome Sanger Institute for the COVID-19 Genomics UK (COG-UK) consortium | Harper VanSteenhouse, Yumi Kasai, David Gray, Carol Clugston, Anna Dominiczak and Alex Alderton, Roberto Amato, Sonia Goncalves, Ewan Harrison, David K. Jackson, Ian Johnston, Dominic Kwiatkowski, Cordelia Langford, John Sillitoe |
| EPI_ISL_532858 | NHSGGC West of Scotland Specialist Virology Centre / | Wellcome Sanger Institute for the COVID-19 Genomics UK | Ana da Silva Filipe, Natasha Johnson, Kathy Smollett, Daniel Mair, Stephen Carmichael, Lily Tong, Jenna Nichols, Elihu Aranday-Cortes, Kirstyn Brunker, |

| | | | |
|---|---|---|---|
| | MRC-University of Glasgow Centre for Virus Research | (COG-UK) consortium | Yasmin Parr, Kyriaki Nomikou; Sarah McDonald, Marc Niebel, Patawee Asamaphan; Richard Orton, Joseph Hughes, Sreenu Vattipally, David L Robertson; Alasdair MacLean, Rory Gunson; Kathy Li, Natasha Jesudason, Rajiv Shah, James Shepherd, Antonia Ho, Alice Broos, Emma Thomson and Alex Alderton, Roberto Amato, Sonia Goncalves, Ewan Harrison, David K. Jackson, Ian Johnston, Dominic Kwiatkowski, Cordelia Langford, John Sillitoe |
| EPI_ISL_532873 | Lighthouse Lab in Glasgow | Wellcome Sanger Institute for the COVID-19 Genomics UK (COG-UK) consortium | Harper VanSteenhouse, Yumi Kasai, David Gray, Carol Clugston, Anna Dominiczak and Alex Alderton, Roberto Amato, Sonia Goncalves, Ewan Harrison, David K. Jackson, Ian Johnston, Dominic Kwiatkowski, Cordelia Langford, John Sillitoe |
| EPI_ISL_532878, EPI_ISL_532885, EPI_ISL_532891, EPI_ISL_532896 | NHSGGC West of Scotland Specialist Virology Centre / MRC-University of Glasgow Centre for Virus Research | Wellcome Sanger Institute for the COVID-19 Genomics UK (COG-UK) consortium | Ana da Silva Filipe, Natasha Johnson, Kathy Smollett, Daniel Mair, Stephen Carmichael, Lily Tong, Jenna Nichols, Elihu Aranday-Cortes, Kirstyn Brunker, Yasmin Parr, Kyriaki Nomikou; Sarah McDonald, Marc Niebel, Patawee Asamaphan; Richard Orton, Joseph Hughes, Sreenu Vattipally, David L Robertson; Alasdair MacLean, Rory Gunson; Kathy Li, Natasha Jesudason, Rajiv Shah, James Shepherd, Antonia Ho, Alice Broos, Emma Thomson and Alex Alderton, Roberto Amato, Sonia Goncalves, Ewan Harrison, David K. Jackson, Ian Johnston, Dominic Kwiatkowski, Cordelia Langford, John Sillitoe |
| EPI_ISL_532903, EPI_ISL_532909, EPI_ISL_532912, EPI_ISL_532913, EPI_ISL_532924, EPI_ISL_532938, EPI_ISL_532945, EPI_ISL_532952, EPI_ISL_532953 | Lighthouse Lab in Glasgow | Wellcome Sanger Institute for the COVID-19 Genomics UK (COG-UK) consortium | Harper VanSteenhouse, Yumi Kasai, David Gray, Carol Clugston, Anna Dominiczak and Alex Alderton, Roberto Amato, Sonia Goncalves, Ewan Harrison, David K. Jackson, Ian Johnston, Dominic Kwiatkowski, Cordelia Langford, John Sillitoe |
| EPI_ISL_532962 | NHSGGC West of Scotland Specialist Virology Centre / MRC-University of Glasgow Centre for Virus Research | Wellcome Sanger Institute for the COVID-19 Genomics UK (COG-UK) consortium | Ana da Silva Filipe, Natasha Johnson, Kathy Smollett, Daniel Mair, Stephen Carmichael, Lily Tong, Jenna Nichols, Elihu Aranday-Cortes, Kirstyn Brunker, Yasmin Parr, Kyriaki Nomikou; Sarah McDonald, Marc Niebel, Patawee Asamaphan; Richard Orton, Joseph Hughes, Sreenu Vattipally, David L Robertson; Alasdair MacLean, Rory Gunson; Kathy Li, Natasha Jesudason, Rajiv Shah, James Shepherd, Antonia Ho, Alice Broos, Emma Thomson and Alex Alderton, Roberto Amato, Sonia Goncalves, Ewan Harrison, David K. Jackson, Ian Johnston, Dominic Kwiatkowski, Cordelia Langford, John Sillitoe |
| EPI_ISL_534207, EPI_ISL_534208 | Centrl laboratorija | Latvian Biomedical Research and Study Centre | Ivars Silamielis, Jnis Pjalkovskis, Kaspars Megnis, Monta Ustinova, ikita Zrelovs, Vita Rovte, Stella Lapia, Jana Oste, Marta Priedte, Uga Dumpis, Jnis Kloviš |
| EPI_ISL_534209 | Latvijas Infektoloijas centrs | Latvian Biomedical Research and Study Centre | Ivars Silamielis, Jnis Pjalkovskis, Kaspars Megnis, Monta Ustinova, ikita Zrelovs, Vita Rovte, Jeena Storoženko, Tatjana Kolupajeva, Oksana Savicka, Uga Dumpis, Jnis Kloviš |
| EPI_ISL_534233 | Capio S:t Gorans sjukhus | The Public Health Agency of Sweden | Anna-Malin Linde, Maria Lind Karlberg, Mattias Haukland, Reza Advani, Olov Svartstrom, Oskar Karlsson Lindsjo, Sandra Broddesson, Petra Edquist, Mia Brytting, Anna Risberg, Karin Tegmark-Wisell |
| EPI_ISL_534246, EPI_ISL_534247 | Universitetssjukhuset i Linkoping | The Public Health Agency of Sweden | Anna-Malin Linde, Maria Lind Karlberg, Mattias Haukland, Reza Advani, Olov Svartstrom, Oskar Karlsson Lindsjo, Sandra Broddesson, Petra Edquist, Mia Brytting, Anna Risberg, Karin Tegmark-Wisell |
| EPI_ISL_534248 | Laboratoriemedicin Vasternorrland | The Public Health Agency of Sweden | Anna-Malin Linde, Maria Lind Karlberg, Mattias Haukland, Reza Advani, Olov Svartstrom, Oskar Karlsson Lindsjo, Sandra Broddesson, Petra Edquist, Mia Brytting, Anna Risberg, Karin Tegmark-Wisell |
| EPI_ISL_534249, EPI_ISL_534251 | Gavle Sjukhus | The Public Health Agency of Sweden | Anna-Malin Linde, Maria Lind Karlberg, Mattias Haukland, Reza Advani, Olov Svartstrom, Oskar Karlsson Lindsjo, Sandra Broddesson, Petra Edquist, Mia Brytting, Anna Risberg, Karin Tegmark-Wisell |
| EPI_ISL_534732, EPI_ISL_534749, EPI_ISL_534750, EPI_ISL_534751 | Liverpool Clinical Laboratories | COVID-19 Genomics UK (COG-UK) Consortium | Sam Haldenby, Anita Lucaci, Steve Paterson, Julian Hiscox, Alistair Darby, M Almsaud, A Alrezaihi, Muhannad Alruwaili, Stuart D Armstrong, Jones Benjamin, Eleanor G Bentley, Anu Chawla, Jordan J Clark, Angela Cowell, Richard Eccles, Isabel García-Dorival, Matthew Gemmell, Alessandro Gerada, PKF Gilmore, Richard Gregory, Ximeng Han, Catherine Hartley, Margaret Hughes, Miren Iturriza-Gomara, James Johnson, L Luu, Jenifer Manson, Charlotte Nelson, Elaine O'Toole, Cassie Olateju, Rebekah Penrice-Randal , Lucille Rainbow, N.P Randle, Trevor Ian Robinson, Parul Sharma, Ghada T Shawli, James P Stewart, Neil Swainston, Ecaterina Vamos, Joanne Watts, Mark Whitehead |
| EPI_ISL_535039, EPI_ISL_535041, EPI_ISL_535042 | Centre for Enzyme Innovation, University of Portsmouth / Translational Research Laboratory, Portsmouth Hospitals NHS Trust | COVID-19 Genomics UK (COG-UK) Consortium | Angela Beckett,Yann Bourgeois,Garry Scarlett,Sharon Glaysher,Scott Elliott,Kelly Bicknell,Robert Impey,Allyson Lloyd,Sarah Wyllie,Ethan Butcher,Anoop Chauhan,Samuel Robson |
| EPI_ISL_535359, EPI_ISL_535360 | RI State Health Laboratories | Pathogen Discovery, Respiratory Viruses Branch, Division of Viral Diseases, Centers for Disease Control and Prevention | Jing Zhang, Ying Tao, Yan Li, Krista Queen, Anna Uehara, Clinton Paden, Haibin Wang, Suxiang Tong |
| EPI_ISL_535405, EPI_ISL_535406, EPI_ISL_535407, EPI_ISL_535408, EPI_ISL_535409, EPI_ISL_535410, EPI_ISL_535411, EPI_ISL_535412, EPI_ISL_535413, EPI_ISL_535414, EPI_ISL_535415, EPI_ISL_535416, EPI_ISL_535417, EPI_ISL_535418, EPI_ISL_535419, EPI_ISL_535420, EPI_ISL_535421, EPI_ISL_535422, EPI_ISL_535423, EPI_ISL_535424, EPI_ISL_535425, EPI_ISL_535426, EPI_ISL_535427, EPI_ISL_535428, EPI_ISL_535429, EPI_ISL_535430, EPI_ISL_535431, EPI_ISL_535432, EPI_ISL_535433, EPI_ISL_535434, EPI_ISL_535435, EPI_ISL_535436, EPI_ISL_535437, EPI_ISL_535438, EPI_ISL_535439, EPI_ISL_535440, EPI_ISL_535441, EPI_ISL_535442, EPI_ISL_535443, EPI_ISL_535444, EPI_ISL_535445, EPI_ISL_535446, EPI_ISL_535447, EPI_ISL_535450, EPI_ISL_535452, EPI_ISL_535453, EPI_ISL_535455, EPI_ISL_535456, EPI_ISL_535457, EPI_ISL_535459, EPI_ISL_535460, EPI_ISL_535461, EPI_ISL_535462, EPI_ISL_535463, EPI_ISL_535464, EPI_ISL_535465, EPI_ISL_535466, EPI_ISL_535467, EPI_ISL_535468, EPI_ISL_535469, EPI_ISL_535470, EPI_ISL_535471, EPI_ISL_535473, EPI_ISL_535474, EPI_ISL_535475, EPI_ISL_535476, EPI_ISL_535477, EPI_ISL_535478, EPI_ISL_535479, EPI_ISL_535481, EPI_ISL_535482, EPI_ISL_535483, EPI_ISL_535484, EPI_ISL_535485, EPI_ISL_535488, EPI_ISL_535497, EPI_ISL_535498, EPI_ISL_535499, EPI_ISL_535500, EPI_ISL_535502, EPI_ISL_535552, EPI_ISL_535553, EPI_ISL_535554, EPI_ISL_535555, EPI_ISL_535556, EPI_ISL_535557, EPI_ISL_535558, EPI_ISL_535559, EPI_ISL_535560, EPI_ISL_535561, EPI_ISL_535562, EPI_ISL_535563, EPI_ISL_535564, EPI_ISL_535565, EPI_ISL_535566, EPI_ISL_535567, EPI_ISL_535568, EPI_ISL_535569, EPI_ISL_535570, EPI_ISL_535571, EPI_ISL_535572 | | | |
| see above | NHLS-IALCH | KRISP, KZN Research Innovation and Sequencing Platform | Giandhari J, Pillay S, Lessells R, Mdlalose K, York D, Khan S, Tegally H, Wilkinson E, de Oliveira T |
| EPI_ISL_536574, EPI_ISL_536578 | University of Wisconsin-Madison AIDS Vaccine Research Laboratories | University of Wisconsin-Madison AIDS Vaccine Research Laboratories | Gage Moreno, Katarina Braun, et al. AIDS Vaccine Research Laboratories |
| EPI_ISL_536962, EPI_ISL_536967, EPI_ISL_536972, EPI_ISL_536978, EPI_ISL_536980, EPI_ISL_536989, EPI_ISL_536992, EPI_ISL_536997, EPI_ISL_536998, EPI_ISL_536999, EPI_ISL_537002, EPI_ISL_537007, EPI_ISL_537027, EPI_ISL_537033, EPI_ISL_537040, EPI_ISL_537059, EPI_ISL_537063, EPI_ISL_537070, EPI_ISL_537081, EPI_ISL_537089, EPI_ISL_537103, EPI_ISL_537104, EPI_ISL_537107, EPI_ISL_537108, EPI_ISL_537109, EPI_ISL_537115, EPI_ISL_537130, EPI_ISL_537132, EPI_ISL_537134, EPI_ISL_537142, EPI_ISL_537146 | | | |
| see above | Lighthouse Lab in Glasgow | Wellcome Sanger Institute for the COVID-19 Genomics UK (COG-UK) consortium | Harper VanSteenhouse, Yumi Kasai, David Gray, Carol Clugston, Anna Dominiczak and Alex Alderton, Roberto Amato, Sonia Goncalves, Ewan Harrison, David K. Jackson, Ian Johnston, Dominic Kwiatkowski, Cordelia Langford, John Sillitoe on behalf of the Wellcome Sanger Institute COVID-19 Surveillance Team |
| EPI_ISL_538238, EPI_ISL_538263, EPI_ISL_538264 | TriCore Reference Laboratories | Center for Global Health, University of New Mexico Health Sciences Center | Daryl Domman, Kurt Schwalm, Twila Kunde, Joseph Hicks, Michael Edwards, Darrell Dinwiddie |
| EPI_ISL_538502 | RSUD Sultan Imanudin Pangkalan Bun Center Kalimantan | National Institute of Health Research and Development | Pawestri, HA; Subangkit; Puspa, KD; Nugraha, AA; Ikawati, HD; Pangesti, KNA; Soekarso, T; Paisal; Setiawaty,V. |
| EPI_ISL_538504 | National Institute of Health Research and Development | National Institute of Health Research and Development | Pawestri, HA; Subangkit; Puspa, KD; Nugraha, AA; Ikawati, HD; Pangesti, KNA; Soekarso, T; Susilarini, NK; Hariastuti, NI; Nikmah, UA; Mursinah; Febriyani, A; Herman, R; Susanti, N; Herna; Febriyanti, T; Nurhadi, M; Paisal; Ramadhany, R; Agustiningsih; Kurniawati, J; Kipuw, NL; Muna, F; Indalau, IL; Adam, K; Wibowo, HA; Rizki, A; Puspandary, N; Setiawaty,V. |
| EPI_ISL_538512 | Balai Penelitian dan Pengembangan Biomedis Papua | National Institute of Health Research and Development | Pawestri, HA; Subangkit; Puspa, KD; Nugraha, AA; Ikawati, HD; Pangesti, KNA; Soekarso, T; Paisal; Pasaribu, M; Setiawaty,V. |
| EPI_ISL_538513 | Provincial Health Laboratory Bekasi West Java | National Institute of Health Research and Development | Pawestri, HA; Subangkit; Puspa, KD; Nugraha, AA; Ikawati, HD; Pangesti, KNA; Soekarso, T; Paisal; Setiawaty,V. |
| EPI_ISL_538517, EPI_ISL_538518, EPI_ISL_538519, EPI_ISL_538520 | Infectious Diseases, North Carolina State Laboratory of Public Health COVID-19 Response Team | Infectious Diseases, North Carolina State Laboratory of Public Health COVID-19 Response Team | Chase,K. |
| EPI_ISL_538523 | Infectious Diseases, North Carolina State Laboratory of Public Health COVID-19 Response Team | North Carolina State Laboratory of Public Health | Chase,K. |
| EPI_ISL_539885 | Narhalsan Fjallbacka VC | The Public Health Agency of Sweden | Anna-Malin Linde, Maria Lind Karlberg, Oskar Karlsson Lindsjo, Olov Svartstrom, Mattias Haukland, Reza Advani, Sandra Broddesson, Anna Risberg, Theresa Enkirch, Mia Brytting, Karin Tegmark-Wisell |
| EPI_ISL_540442, EPI_ISL_540443, EPI_ISL_540447, EPI_ISL_540448, EPI_ISL_540449, EPI_ISL_540450, EPI_ISL_540451, EPI_ISL_540452, EPI_ISL_540453, EPI_ISL_540454, EPI_ISL_540456, EPI_ISL_540457, EPI_ISL_540458, EPI_ISL_540459, EPI_ISL_540460, EPI_ISL_540461, EPI_ISL_540462, EPI_ISL_540463, EPI_ISL_540464, EPI_ISL_540465, EPI_ISL_540466, EPI_ISL_540467, EPI_ISL_540468 | | | |
| see above | University of Liège COVID-19 testing center | GIGA Medical Genomics | Keith Durkin, Maria Artesi, Emmanuel André, Marc Van Ranst, Fabrice Bureau, Laurent Gillet, Wouter Coppieters, Vincent Bours |
| EPI_ISL_540469, EPI_ISL_540470, EPI_ISL_540471, EPI_ISL_540472, EPI_ISL_540473, EPI_ISL_540474, EPI_ISL_540475, EPI_ISL_540476, EPI_ISL_540477, EPI_ISL_540478, EPI_ISL_540479, EPI_ISL_540480, EPI_ISL_540481, EPI_ISL_540482, EPI_ISL_540483, EPI_ISL_540484, EPI_ISL_540485, EPI_ISL_540486, EPI_ISL_540487, EPI_ISL_540488, EPI_ISL_540489, EPI_ISL_540490, EPI_ISL_540491, EPI_ISL_540492, EPI_ISL_540493, EPI_ISL_540494, EPI_ISL_540495, EPI_ISL_540496, EPI_ISL_540497, EPI_ISL_540498, EPI_ISL_540499, EPI_ISL_540500, EPI_ISL_540501, EPI_ISL_540502, EPI_ISL_540503, EPI_ISL_540504, EPI_ISL_540505, EPI_ISL_540506, EPI_ISL_540508 | | | |

| | | | |
|---|---|---|---|
| see above | Department of Clinical Microbiology | GIGA Medical Genomics | Keith Durkin, Maria Artesi, Sébastien Bontems, Raphaël Boreux, Bouchra Boujemla, Cécile Meex, Axelle Chaslain, Céline Fombellida-Lopez, Pierrette Melin, Marie-Pierre Hayette, Vincent Bours |
| EPI_ISL_540591, EPI_ISL_540592, EPI_ISL_540593, EPI_ISL_540594, EPI_ISL_540595, EPI_ISL_540596, EPI_ISL_540597, EPI_ISL_540598, EPI_ISL_540599, EPI_ISL_540600, EPI_ISL_540602, EPI_ISL_540603, EPI_ISL_540606, EPI_ISL_540607, EPI_ISL_540611, EPI_ISL_540612, EPI_ISL_540613, EPI_ISL_540614, EPI_ISL_540616, EPI_ISL_540617, EPI_ISL_540618, EPI_ISL_540619, EPI_ISL_540620, EPI_ISL_540621, EPI_ISL_540622, EPI_ISL_540623, EPI_ISL_540624, EPI_ISL_540625 | | | |
| see above | Liverpool Clinical Laboratories | COVID-19 Genomics UK (COG-UK) Consortium | Sam Haldenby, Anita Lucaci, Steve Paterson, Julian Hiscox, Alistair Darby, M Almsaud, A Alrezaihi, Muhannad Alruwaili, Stuart D Armstrong, Jones Benjamin, Eleanor G Bentley, Anu Chawla, Jordan J Clark, Angela Cowell, Richard Eccles, Isabel García-Dorival, Matthew Gemmell, Alessandro Gerada, PKF Gilmore, Richard Gregory, Ximeng Han, Catherine Hartley, Margaret Hughes, Miren Iturriza-Gomara, James Johnson, L Luu, Jenifer Manson, Charlotte Nelson, Elaine O'Toole, Cassie Olateju, Rebekah Penrice-Randal , Lucille Rainbow, N.P Randle, Trevor Ian Robinson, Parul Sharma, Ghada T Shawli, James P Stewart, Neil Swainston, Ecaterina Vamos, Joanne Watts, Mark Whitehead |
| EPI_ISL_541081 | Hospital de la Santa Creu i Sant Pau. Servicio de Microbiología | SeqCOVID-SPAIN consortium/Institute of Biomedicine of Valencia, IBV-CSIC | Ferran Navarro, Núria Rabella, Elisenda Miró and SeqCOVID-SPAIN consortium |
| EPI_ISL_541178, EPI_ISL_541180, EPI_ISL_541181, EPI_ISL_541182, EPI_ISL_541183, EPI_ISL_541184, EPI_ISL_541185, EPI_ISL_541186, EPI_ISL_541187, EPI_ISL_541188, EPI_ISL_541189, EPI_ISL_541190, EPI_ISL_541191, EPI_ISL_541192, EPI_ISL_541193, EPI_ISL_541194, EPI_ISL_541195, EPI_ISL_541196, EPI_ISL_541197, EPI_ISL_541198, EPI_ISL_541331 | | | |
| see above | Florida Bureau of Public Health Laboratories, Florida Department of Health | Florida Bureau of Public Health Laboratories, Florida Department of Health | Schmedes,S., Blanton,J. |
| EPI_ISL_541336, EPI_ISL_541337 | The National Institute of Public Health | State Veterinary Institute Prague | Nagy,A; Jirincova,H; Novakova,L; Trnka,D; Vecerova,J |
| EPI_ISL_541655 | Laboratory Diagnostic, Veterinary Specialized Institute Kraljevo | Laboratory Diagnostic, Veterinary Specialized Institute Kraljevo | Vidanovic,D., Tesovic,B., Knezevic,A., Jovanovic,T., Jankovic,M., Sekler,M., Banovic Djeri,B., Volkening,J., Afonso,C., Petrovic,T. |
| EPI_ISL_541695, EPI_ISL_541696, EPI_ISL_541699 | National Institute of Virology, NIV Influenza | National Institute of Virology, NIV Influenza | Potdar V |
| EPI_ISL_541891, EPI_ISL_541892, EPI_ISL_541893, EPI_ISL_541894, EPI_ISL_541895, EPI_ISL_541896, EPI_ISL_541897, EPI_ISL_541898, EPI_ISL_541899, EPI_ISL_541900, EPI_ISL_541901, EPI_ISL_541902, EPI_ISL_541903 | | | |
| see above | Hospital General Universitario Gregorio Marañón | SeqCOVID-SPAIN consortium/IBV(CSIC) | Laura Pérez-Lago, Marta Herranz, Jon Sicilia, Julia Suárez, Pilar Catalán, Patricia Muñoz, Darío García de Viedma and SeqCOVID-SPAIN consortium |
| EPI_ISL_542039, EPI_ISL_542040 | New Mexico Department of Health Scientific Laboratory | New Mexico Department of Health Scientific Laboratory | Ellie Johnson, Anastacia Griego-Fisher, D'Eldra Malone |
| EPI_ISL_542935 | TriCore Reference Laboratories | Center for Global Health, University of New Mexico Health Sciences Center | Daryl Domman, Kurt Schwalm, Twila Kunde, Joseph Hicks, Michael Edwards, Darrell Dinwiddie |
| EPI_ISL_544955 | Pathology West - NSW Health Pathology | NSW Health Pathology - Institute of Clinical Pathology and Medical Research; Westmead Hospital; University of Sydney | CIDM-PH et al. |
| EPI_ISL_544956 | Sydney South West Pathology Service (SSWPS) - Concord Repatriation General Hospital - NSW Health Pathology | NSW Health Pathology - Institute of Clinical Pathology and Medical Research; Westmead Hospital; University of Sydney | CIDM-PH et al. |
| EPI_ISL_545023 | Sydney South West Pathology Service (SSWPS) - Royal Prince Alfred Hospital - NSW Health Pathology | NSW Health Pathology - Institute of Clinical Pathology and Medical Research; Westmead Hospital; University of Sydney | CIDM-PH et al. |
| EPI_ISL_545580, EPI_ISL_545954, EPI_ISL_545956 | The National Institute of Public Health | State Veterinary Institute Prague | Nagy,A;Jirincova,H;Novakova,L;Trnka,D;Vecerova,J |
| EPI_ISL_545957 | The National Institute of Public Health | State Veterinary Institute Prague | Nagy,A; Jirincova,H; Novakova,L; Trnka,D; Vecerova,J |
| EPI_ISL_547433, EPI_ISL_547435 | Microbiology, Department of Pathology, St. Bernard's Hospital, Gibraltar Health Authority | Respiratory Virus Unit, Microbiology Services Colindale, Public Health England | PHE Covid Sequencing Team, Dr Nicholas Cortes (Gibraltar), Charlotte Gillborn-Jones (Gibraltar) |
| EPI_ISL_547541 | Dutch COVID-19 response team | National Institute for Public Health and the Environment (RIVM) | Adam Meijer, Harry Vennema, Jeroen Cremer, Sharon van den Brink, Bas van der Veer, AnneMarie van den Brandt, Florian Zwagemaker, Dennis Schmitz, Chantal Reusken, on behalf of the national COVID-19 response team |
| EPI_ISL_547638, EPI_ISL_547639, EPI_ISL_547640, EPI_ISL_547641, EPI_ISL_547642, EPI_ISL_547643, EPI_ISL_547644, EPI_ISL_547645, EPI_ISL_547646, EPI_ISL_547647, EPI_ISL_547648, EPI_ISL_547649, EPI_ISL_547650, EPI_ISL_547651, EPI_ISL_547652, EPI_ISL_547653, EPI_ISL_547654 | | | |
| see above | Gundersen Molecular Diagnostics Laboratory | Kabara Cancer Research Institute | Craig S. Richmond, Paraic A. Kenny |
| EPI_ISL_547674, EPI_ISL_547675 | Gundersen Clinical Microbiology Laboratory | Kabara Cancer Research Institute | Craig S. Richmond, Paraic A. Kenny |
| EPI_ISL_548145 | Middlemore Hospital | Institute of Environmental Science and Research (ESR) | Xiaoyun Ren, Matt Storey, Nikki Freed, Muhammad Faisal, Jing Wang, Hermes Perez, Anja Werno, Antje van der Linden, Arlo Upton, Chris Mansell, David Hammer, Dragana Drinkovic, Gary McAuliffe, Hana Sofia Andersson, James Ussher, Jill Sherwood, Josh Freeman, Julia Howard, Juliet Elvy, Mary DeAlmeida, Matt Blakiston, Matthew Rogers, Max Bloomfield, Michael Addidle, Michelle Balm, Sally Roberts, Sarah Jefferies, Sharmini Muttaiyah, Susan Morpeth, Susan Taylor, Timothy Blackmore, Vani Sathyendran, Veronica Playle, Virginia Hope, Erasmus Smit, Lauren Jelly, Olin Silander, Joep de Ligt |
| EPI_ISL_548337, EPI_ISL_548338, EPI_ISL_548339, EPI_ISL_548340, EPI_ISL_548341, EPI_ISL_548342, EPI_ISL_548343, EPI_ISL_548344 | County of Santa Clara Public Health Department | Chan-Zuckerberg Biohub | CZB Cliahub Consortium |
| EPI_ISL_548350, EPI_ISL_548351, EPI_ISL_548357, EPI_ISL_548359, EPI_ISL_548360, EPI_ISL_548361, EPI_ISL_548365, EPI_ISL_548366, EPI_ISL_548369, EPI_ISL_548370 | County of San Luis Obispo Public Health Laboratory | Chan-Zuckerberg Biohub | CZB Cliahub Consortium |
| EPI_ISL_548375, EPI_ISL_548378, EPI_ISL_548381, EPI_ISL_548383, EPI_ISL_548387, EPI_ISL_548389 | Orange County Public Health Laboratory | Chan-Zuckerberg Biohub | CZB Cliahub Consortium |
| EPI_ISL_548391 | County of San Luis Obispo Public Health Laboratory | Chan-Zuckerberg Biohub | CZB Cliahub Consortium |
| EPI_ISL_548393, EPI_ISL_548400, EPI_ISL_548405 | Orange County Public Health Laboratory | Chan-Zuckerberg Biohub | CZB Cliahub Consortium |
| EPI_ISL_548407 | County of Santa Clara Public Health Department | Chan-Zuckerberg Biohub | CZB Cliahub Consortium |
| EPI_ISL_548411, EPI_ISL_548412, EPI_ISL_548413 | Orange County Public Health Laboratory | Chan-Zuckerberg Biohub | CZB Cliahub Consortium |
| EPI_ISL_548417 | County of Santa Clara Public Health Department | Chan-Zuckerberg Biohub | CZB Cliahub Consortium |
| EPI_ISL_548424 | Orange County Public Health Laboratory | Chan-Zuckerberg Biohub | CZB Cliahub Consortium |
| EPI_ISL_548425, EPI_ISL_548428 | County of San Luis Obispo Public Health Laboratory | Chan-Zuckerberg Biohub | CZB Cliahub Consortium |
| EPI_ISL_548431 | Orange County Public Health Laboratory | Chan-Zuckerberg Biohub | CZB Cliahub Consortium |
| EPI_ISL_548432 | County of Santa Clara Public Health Department | Chan-Zuckerberg Biohub | CZB Cliahub Consortium |
| EPI_ISL_548436 | County of San Luis Obispo Public Health Laboratory | Chan-Zuckerberg Biohub | CZB Cliahub Consortium |

| | | | |
|---|---|---|---|
| EPI_ISL_548440, EPI_ISL_548441, EPI_ISL_548445 | Orange County Public Health Laboratory | Chan-Zuckerberg Biohub | CZB Cliahub Consortium |
| EPI_ISL_548453 | County of San Luis Obispo Public Health Laboratory | Chan-Zuckerberg Biohub | CZB Cliahub Consortium |
| EPI_ISL_548454, EPI_ISL_548456 | Orange County Public Health Laboratory | Chan-Zuckerberg Biohub | CZB Cliahub Consortium |
| EPI_ISL_548457 | County of San Luis Obispo Public Health Laboratory | Chan-Zuckerberg Biohub | CZB Cliahub Consortium |
| EPI_ISL_548458 | University of California, Davis | Chan-Zuckerberg Biohub | CZB Cliahub Consortium |
| EPI_ISL_548462, EPI_ISL_548463, EPI_ISL_548465, EPI_ISL_548467, EPI_ISL_548469, EPI_ISL_548484, EPI_ISL_548485, EPI_ISL_548488 | Orange County Public Health Laboratory | Chan-Zuckerberg Biohub | CZB Cliahub Consortium |
| EPI_ISL_548490 | University of California, Davis | Chan-Zuckerberg Biohub | CZB Cliahub Consortium |
| EPI_ISL_548491, EPI_ISL_548492 | Contra Costa Public Health Lab | Chan-Zuckerberg Biohub | CZB Cliahub Consortium |
| EPI_ISL_548493 | County of San Luis Obispo Public Health Laboratory | Chan-Zuckerberg Biohub | CZB Cliahub Consortium |
| EPI_ISL_548495, EPI_ISL_548499, EPI_ISL_548501 | Contra Costa Public Health Lab | Chan-Zuckerberg Biohub | CZB Cliahub Consortium |
| EPI_ISL_548502, EPI_ISL_548504 | County of San Luis Obispo Public Health Laboratory | Chan-Zuckerberg Biohub | CZB Cliahub Consortium |
| EPI_ISL_548505 | UCSF Clinical Microbiology Laboratory | Chan-Zuckerberg Biohub | CZB Cliahub Consortium |
| EPI_ISL_548506, EPI_ISL_548507, EPI_ISL_548508, EPI_ISL_548510 | Contra Costa Public Health Lab | Chan-Zuckerberg Biohub | CZB Cliahub Consortium |
| EPI_ISL_548513 | UCSF Clinical Microbiology Laboratory | Chan-Zuckerberg Biohub | CZB Cliahub Consortium |
| EPI_ISL_548515, EPI_ISL_548516 | Contra Costa Public Health Lab | Chan-Zuckerberg Biohub | CZB Cliahub Consortium |
| EPI_ISL_548517, EPI_ISL_548519, EPI_ISL_548521 | County of San Luis Obispo Public Health Laboratory | Chan-Zuckerberg Biohub | CZB Cliahub Consortium |
| EPI_ISL_548522 | UCSF Clinical Microbiology Laboratory | Chan-Zuckerberg Biohub | CZB Cliahub Consortium |
| EPI_ISL_548524, EPI_ISL_548527, EPI_ISL_548528, EPI_ISL_548531 | Contra Costa Public Health Lab | Chan-Zuckerberg Biohub | CZB Cliahub Consortium |
| EPI_ISL_548532 | County of San Luis Obispo Public Health Laboratory | Chan-Zuckerberg Biohub | CZB Cliahub Consortium |
| EPI_ISL_548534, EPI_ISL_548535, EPI_ISL_548537, EPI_ISL_548538, EPI_ISL_548539, EPI_ISL_548540 | Contra Costa Public Health Lab | Chan-Zuckerberg Biohub | CZB Cliahub Consortium |
| EPI_ISL_548541 | County of Santa Clara Public Health Department | Chan-Zuckerberg Biohub | CZB Cliahub Consortium |
| EPI_ISL_548542, EPI_ISL_548544 | County of San Luis Obispo Public Health Laboratory | Chan-Zuckerberg Biohub | CZB Cliahub Consortium |
| EPI_ISL_548548, EPI_ISL_548549, EPI_ISL_548552, EPI_ISL_548554 | Contra Costa Public Health Lab | Chan-Zuckerberg Biohub | CZB Cliahub Consortium |
| EPI_ISL_548555 | UCSF Clinical Microbiology Laboratory | Chan-Zuckerberg Biohub | CZB Cliahub Consortium |
| EPI_ISL_548556, EPI_ISL_548557 | Contra Costa Public Health Lab | Chan-Zuckerberg Biohub | CZB Cliahub Consortium |
| EPI_ISL_548560 | County of San Luis Obispo Public Health Laboratory | Chan-Zuckerberg Biohub | CZB Cliahub Consortium |
| EPI_ISL_548561, EPI_ISL_548564, EPI_ISL_548565, EPI_ISL_548566 | Contra Costa Public Health Lab | Chan-Zuckerberg Biohub | CZB Cliahub Consortium |
| EPI_ISL_548568 | County of Santa Clara Public Health Department | Chan-Zuckerberg Biohub | CZB Cliahub Consortium |
| EPI_ISL_548569 | Contra Costa Public Health Lab | Chan-Zuckerberg Biohub | CZB Cliahub Consortium |
| EPI_ISL_548571 | UCSF Clinical Microbiology Laboratory | Chan-Zuckerberg Biohub | CZB Cliahub Consortium |
| EPI_ISL_548572, EPI_ISL_548573, EPI_ISL_548574, EPI_ISL_548575, EPI_ISL_548579 | Contra Costa Public Health Lab | Chan-Zuckerberg Biohub | CZB Cliahub Consortium |
| EPI_ISL_548598, EPI_ISL_548600, EPI_ISL_548604, EPI_ISL_548605, EPI_ISL_548632, EPI_ISL_548642, EPI_ISL_548652, EPI_ISL_548657 | County of Santa Clara Public Health Department | Chan-Zuckerberg Biohub | CZB Cliahub Consortium |
| EPI_ISL_549027, EPI_ISL_549038, EPI_ISL_549048 | Furst Medical Laboratory | Norwegian Institute of Public Health, Department of Virology | Kathrine Stene-Johansen, Kamilla Heddeland Instefjord, Hilde Elshaug, Rasmus Riis Kopperud, Hilde Synnøve Vollan, Karoline Bragstad, Olav Hungnes |
| EPI_ISL_549051 | Vestfold Hospital, Toensberg Department of Microbiology | Norwegian Institute of Public Health, Department of Virology | Kathrine Stene-Johansen, Kamilla Heddeland Instefjord, Hilde Elshaug, Rasmus Riis Kopperud, Hilde Synnøve Vollan, Karoline Bragstad, Olav Hungnes |
| EPI_ISL_549069 | Furst Medical Laboratory | Norwegian Institute of Public Health, Department of Virology | Kathrine Stene-Johansen, Kamilla Heddeland Instefjord, Hilde Elshaug, Rasmus Riis Kopperud, Hilde Synnøve Vollan, Karoline Bragstad, Olav Hungnes |
| EPI_ISL_549070 | Medical Microbiology Unit, Department for Laboratory Medicine, Drammen Hospital, Vestre Viken Health Trust, | Norwegian Institute of Public Health, Department of Virology | Kathrine Stene-Johansen, Kamilla Heddeland Instefjord, Hilde Elshaug, Rasmus Riis Kopperud, Hilde Synnøve Vollan, Karoline Bragstad, Olav Hungnes |
| EPI_ISL_549071 | Furst Medical Laboratory | Norwegian Institute of Public Health, Department of Virology | Kathrine Stene-Johansen, Kamilla Heddeland Instefjord, Hilde Elshaug, Rasmus Riis Kopperud, Hilde Synnøve Vollan, Karoline Bragstad, Olav Hungnes |
| EPI_ISL_549081 | Medical Microbiology Unit, Department for Laboratory Medicine, Drammen Hospital, Vestre Viken Health Trust, | Norwegian Institute of Public Health, Department of Virology | Kathrine Stene-Johansen, Kamilla Heddeland Instefjord, Hilde Elshaug, Rasmus Riis Kopperud, Hilde Synnøve Vollan, Karoline Bragstad, Olav Hungnes |
| EPI_ISL_549089, EPI_ISL_549091 | Akershus University Hospital, Department for Microbiology and Infectious Disease Control | Norwegian Institute of Public Health, Department of Virology | Kathrine Stene-Johansen, Kamilla Heddeland Instefjord, Hilde Elshaug, Rasmus Riis Kopperud, Hilde Synnøve Vollan, Karoline Bragstad, Olav Hungnes |
| EPI_ISL_549092, EPI_ISL_549094, EPI_ISL_549095, EPI_ISL_549096, EPI_ISL_549098, EPI_ISL_549099, EPI_ISL_549100, EPI_ISL_549103, EPI_ISL_549104, EPI_ISL_549105, EPI_ISL_549106, EPI_ISL_549107, EPI_ISL_549108, EPI_ISL_549110, EPI_ISL_549111, EPI_ISL_549113, EPI_ISL_549115, EPI_ISL_549117, EPI_ISL_549118 | | | |
| see above | Ostfold Hospital Trust - Kalnes, Centre for Laboratory Medicine, Section for gene technology and infection serology | Norwegian Institute of Public Health, Department of Virology | Kathrine Stene-Johansen, Kamilla Heddeland Instefjord, Hilde Elshaug, Rasmus Riis Kopperud, Hilde Synnøve Vollan, Karoline Bragstad, Olav Hungnes |
| EPI_ISL_549124, EPI_ISL_549143, EPI_ISL_549154, EPI_ISL_549165 | Furst Medical Laboratory | Norwegian Institute of Public Health, Department of Virology | Kathrine Stene-Johansen, Kamilla Heddeland Instefjord, Hilde Elshaug, Rasmus Riis Kopperud, Hilde Synnøve Vollan, Karoline Bragstad, Olav Hungnes |
| EPI_ISL_549184 | Florida Bureau of Public Health Laboratories | Florida Bureau of Public Health Laboratories | Sarah Schmedes, Jason Blanton |

| | | | |
|---|---|---|---|
| EPI_ISL_550417, EPI_ISL_550418, EPI_ISL_550419, EPI_ISL_550420, EPI_ISL_550422, EPI_ISL_550424, EPI_ISL_550425, EPI_ISL_550426, EPI_ISL_550427, EPI_ISL_550428, EPI_ISL_550429, EPI_ISL_550430, EPI_ISL_550431, EPI_ISL_550432, EPI_ISL_550433, EPI_ISL_550434, EPI_ISL_550435, EPI_ISL_550436, EPI_ISL_550437, EPI_ISL_550440, EPI_ISL_550441, EPI_ISL_550443, EPI_ISL_550444, EPI_ISL_550445, EPI_ISL_550446, EPI_ISL_550447, EPI_ISL_550449, EPI_ISL_550450, EPI_ISL_550451, EPI_ISL_550452, EPI_ISL_550454 | | | |
| see above | Lighthouse Lab in Milton Keynes | Wellcome Sanger Institute for the COVID-19 Genomics UK (COG-UK) consortium | The Lighthouse Lab in Milton Keynes and Alex Alderton, Roberto Amato, Sonia Goncalves, Ewan Harrison, David K. Jackson, Ian Johnston, Dominic Kwiatkowski, Cordelia Langford, John Sillitoe on behalf of the Wellcome Sanger Institute COVID-19 Surveillance Team (http://www.sanger.ac.uk/covid-team) |
| EPI_ISL_550455 | Lighthouse Lab in Milton Keynes | Wellcome Sanger Institute for the COVID-19 Genomics UK (COG-UK) consortium | The Lighthouse Lab in Alderley Park and Alex Alderton, Roberto Amato, Sonia Goncalves, Ewan Harrison, David K. Jackson, Ian Johnston, Dominic Kwiatkowski, Cordelia Langford, John Sillitoe on behalf of the Wellcome Sanger Institute COVID-19 Surveillance Team |
| EPI_ISL_550456, EPI_ISL_550457 | Lighthouse Lab in Milton Keynes | Wellcome Sanger Institute for the COVID-19 Genomics UK (COG-UK) consortium | The Lighthouse Lab in Milton Keynes and Alex Alderton, Roberto Amato, Sonia Goncalves, Ewan Harrison, David K. Jackson, Ian Johnston, Dominic Kwiatkowski, Cordelia Langford, John Sillitoe on behalf of the Wellcome Sanger Institute COVID-19 Surveillance Team (http://www.sanger.ac.uk/covid-team) |
| EPI_ISL_550458 | Lighthouse Lab in Milton Keynes | Wellcome Sanger Institute for the COVID-19 Genomics UK (COG-UK) consortium | The Lighthouse Lab in Alderley Park and Alex Alderton, Roberto Amato, Sonia Goncalves, Ewan Harrison, David K. Jackson, Ian Johnston, Dominic Kwiatkowski, Cordelia Langford, John Sillitoe on behalf of the Wellcome Sanger Institute COVID-19 Surveillance Team |
| EPI_ISL_550459, EPI_ISL_550460, EPI_ISL_550461, EPI_ISL_550462, EPI_ISL_550463, EPI_ISL_550464, EPI_ISL_550465, EPI_ISL_550466, EPI_ISL_550467, EPI_ISL_550469, EPI_ISL_550471, EPI_ISL_550472, EPI_ISL_550473, EPI_ISL_550474, EPI_ISL_550475, EPI_ISL_550476, EPI_ISL_550479, EPI_ISL_550480, EPI_ISL_550481, EPI_ISL_550482, EPI_ISL_550483, EPI_ISL_550484, EPI_ISL_550486, EPI_ISL_550487, EPI_ISL_550488, EPI_ISL_550489, EPI_ISL_550490, EPI_ISL_550491, EPI_ISL_550492, EPI_ISL_550493, EPI_ISL_550494, EPI_ISL_550495, EPI_ISL_550496, EPI_ISL_550500, EPI_ISL_550501, EPI_ISL_550502, EPI_ISL_550504, EPI_ISL_550505, EPI_ISL_550506, EPI_ISL_550507, EPI_ISL_550508, EPI_ISL_550509, EPI_ISL_550510, EPI_ISL_550512, EPI_ISL_550513, EPI_ISL_550514, EPI_ISL_550515, EPI_ISL_550516, EPI_ISL_550517, EPI_ISL_552626 | | | |
| see above | Lighthouse Lab in Milton Keynes | Wellcome Sanger Institute for the COVID-19 Genomics UK (COG-UK) consortium | The Lighthouse Lab in Milton Keynes and Alex Alderton, Roberto Amato, Sonia Goncalves, Ewan Harrison, David K. Jackson, Ian Johnston, Dominic Kwiatkowski, Cordelia Langford, John Sillitoe on behalf of the Wellcome Sanger Institute COVID-19 Surveillance Team (http://www.sanger.ac.uk/covid-team) |
| EPI_ISL_555843, EPI_ISL_555864, EPI_ISL_555868, EPI_ISL_555875, EPI_ISL_555910, EPI_ISL_555939, EPI_ISL_555945, EPI_ISL_555948, EPI_ISL_555955, EPI_ISL_555973, EPI_ISL_555976, EPI_ISL_555993, EPI_ISL_556012, EPI_ISL_556021, EPI_ISL_556022, EPI_ISL_556026, EPI_ISL_556039, EPI_ISL_556041, EPI_ISL_556047, EPI_ISL_556058, EPI_ISL_556063, EPI_ISL_556066, EPI_ISL_556069, EPI_ISL_556305, EPI_ISL_556309, EPI_ISL_556317, EPI_ISL_556318, EPI_ISL_556321, EPI_ISL_556324, EPI_ISL_556338, EPI_ISL_556344, EPI_ISL_556349, EPI_ISL_556351, EPI_ISL_556357, EPI_ISL_556425, EPI_ISL_556434, EPI_ISL_556446, EPI_ISL_556459, EPI_ISL_556471, EPI_ISL_556482, EPI_ISL_556488, EPI_ISL_556494, EPI_ISL_556500, EPI_ISL_556508, EPI_ISL_556509, EPI_ISL_556528, EPI_ISL_556530, EPI_ISL_556531, EPI_ISL_556532, EPI_ISL_556535, EPI_ISL_556546, EPI_ISL_556557, EPI_ISL_556563, EPI_ISL_556565, EPI_ISL_556575, EPI_ISL_556650, EPI_ISL_556782, EPI_ISL_556825, EPI_ISL_556828, EPI_ISL_556829, EPI_ISL_556833, EPI_ISL_556835, EPI_ISL_556836, EPI_ISL_556839, EPI_ISL_556841, EPI_ISL_556843, EPI_ISL_556845, EPI_ISL_556849, EPI_ISL_556850, EPI_ISL_556851, EPI_ISL_556855, EPI_ISL_556856, EPI_ISL_556858, EPI_ISL_556859, EPI_ISL_556860, EPI_ISL_556861, EPI_ISL_556862, EPI_ISL_556865, EPI_ISL_556866, EPI_ISL_556867, EPI_ISL_556868, EPI_ISL_556869, EPI_ISL_556873, EPI_ISL_556874, EPI_ISL_556875, EPI_ISL_556878, EPI_ISL_556879, EPI_ISL_556882, EPI_ISL_556885 | | | |
| see above | Lighthouse Lab in Cambridge | Wellcome Sanger Institute for the COVID-19 Genomics UK (COG-UK) consortium | Rob Howes, The Lighthouse Lab in Cambridge and Alex Alderton, Roberto Amato, Sonia Goncalves, Ewan Harrison, David K. Jackson, Ian Johnston, Dominic Kwiatkowski, Cordelia Langford, John Sillitoe on behalf of the Wellcome Sanger Institute COVID-19 Surveillance Team |
| EPI_ISL_556887 | Lighthouse Lab in Cambridge | Wellcome Sanger Institute for the COVID-19 Genomics UK (COG-UK) Consortium | Rob Howes, The Lighthouse Lab in Cambridge and Alex Alderton, Roberto Amato, Sonia Goncalves, Ewan Harrison, David K. Jackson, Ian Johnston, Dominic Kwiatkowski, Cordelia Langford, John Sillitoe on behalf of the Wellcome Sanger Institute COVID-19 Surveillance Team |
| EPI_ISL_556890, EPI_ISL_556893, EPI_ISL_556896, EPI_ISL_556898, EPI_ISL_556899, EPI_ISL_556900 | Lighthouse Lab in Cambridge | Wellcome Sanger Institute for the COVID-19 Genomics UK (COG-UK) consortium | Rob Howes, The Lighthouse Lab in Cambridge and Alex Alderton, Roberto Amato, Sonia Goncalves, Ewan Harrison, David K. Jackson, Ian Johnston, Dominic Kwiatkowski, Cordelia Langford, John Sillitoe on behalf of the Wellcome Sanger Institute COVID-19 Surveillance Team |
| EPI_ISL_556904 | Lighthouse Lab in Cambridge | Wellcome Sanger Institute for the COVID-19 Genomics UK (COG-UK) consortium | Rob Howes, The Lighthouse Lab in Cambridge and Alex Alderton, Roberto Amato, Sonia Goncalves, Ewan Harrison, David K. Jackson, Ian Johnston, Dominic Kwiatkowski, Cordelia Langford, John Sillitoe on behalf of the Wellcome Sanger Institute COVID-19 Surveillance Team (http://www.sanger.ac.uk/covid-team) |
| EPI_ISL_556905, EPI_ISL_556907, EPI_ISL_556908, EPI_ISL_556910, EPI_ISL_556911, EPI_ISL_556913, EPI_ISL_556960 | Lighthouse Lab in Cambridge | Wellcome Sanger Institute for the COVID-19 Genomics UK (COG-UK) consortium | Rob Howes, The Lighthouse Lab in Cambridge and Alex Alderton, Roberto Amato, Sonia Goncalves, Ewan Harrison, David K. Jackson, Ian Johnston, Dominic Kwiatkowski, Cordelia Langford, John Sillitoe on behalf of the Wellcome Sanger Institute COVID-19 Surveillance Team |
| EPI_ISL_560408, EPI_ISL_560409, EPI_ISL_560410, EPI_ISL_560411, EPI_ISL_560412 | The National Institute of Public Health | State Veterinary Institute Prague | Nagy, A; Jirincova, H; Novakova, L; Trnka, D; Vecerova, J |
| EPI_ISL_560598, EPI_ISL_560599, EPI_ISL_560600, EPI_ISL_560601, EPI_ISL_560602, EPI_ISL_560603, EPI_ISL_560604, EPI_ISL_560605, EPI_ISL_560606 | Hospital | National Reference Center for Viruses of Respiratory Infections, Institut Pasteur, Paris | Sylvie Behillil, Fabiana Gambaro, Etienne Simon-Lorière, Vincent Enouf, Maud Vanpeene, Sylvie van der Werf |
| EPI_ISL_560992, EPI_ISL_560993, EPI_ISL_560994, EPI_ISL_560995, EPI_ISL_560996, EPI_ISL_560997, EPI_ISL_560998, EPI_ISL_560999, EPI_ISL_561000, EPI_ISL_561001, EPI_ISL_561002, EPI_ISL_561003, EPI_ISL_561004, EPI_ISL_561005, EPI_ISL_561006, EPI_ISL_561007, EPI_ISL_561008, EPI_ISL_561009, EPI_ISL_561010, EPI_ISL_561011, EPI_ISL_561012, EPI_ISL_561013, EPI_ISL_561025, EPI_ISL_561041, EPI_ISL_561042, EPI_ISL_561043, EPI_ISL_561044, EPI_ISL_561045, EPI_ISL_561046, EPI_ISL_561047, EPI_ISL_561048, EPI_ISL_561049, EPI_ISL_561050, EPI_ISL_561051, EPI_ISL_561052, EPI_ISL_561053, EPI_ISL_561054, EPI_ISL_561055, EPI_ISL_561056, EPI_ISL_561057, EPI_ISL_561058, EPI_ISL_561059, EPI_ISL_561060, EPI_ISL_561061, EPI_ISL_561062, EPI_ISL_561063, EPI_ISL_561064, EPI_ISL_561065, EPI_ISL_561066, EPI_ISL_561067, EPI_ISL_561068, EPI_ISL_561069, EPI_ISL_561070, EPI_ISL_561071, EPI_ISL_561072, EPI_ISL_561073, EPI_ISL_561074, EPI_ISL_561075, EPI_ISL_561076, EPI_ISL_561077, EPI_ISL_561078, EPI_ISL_561079, EPI_ISL_561080, EPI_ISL_561081, EPI_ISL_561082, EPI_ISL_561083, EPI_ISL_561084, EPI_ISL_561085, EPI_ISL_561086, EPI_ISL_561087, EPI_ISL_561088, EPI_ISL_561089, EPI_ISL_561090, EPI_ISL_561091, EPI_ISL_561092, EPI_ISL_561093, EPI_ISL_561094, EPI_ISL_561095, EPI_ISL_561096, EPI_ISL_561097, EPI_ISL_561098, EPI_ISL_561099, EPI_ISL_561100, EPI_ISL_561101, EPI_ISL_561102, EPI_ISL_561103, EPI_ISL_561104, EPI_ISL_561150, EPI_ISL_561151, EPI_ISL_561152, EPI_ISL_561153, EPI_ISL_561154, EPI_ISL_561155, EPI_ISL_561156, EPI_ISL_561157, EPI_ISL_561158, EPI_ISL_561159, EPI_ISL_561160, EPI_ISL_561161, EPI_ISL_561162, EPI_ISL_561163, EPI_ISL_561164, EPI_ISL_561165, EPI_ISL_561166, EPI_ISL_561167, EPI_ISL_561168, EPI_ISL_561169, EPI_ISL_561170, EPI_ISL_561171, EPI_ISL_561172, EPI_ISL_561173, EPI_ISL_561174, EPI_ISL_561175, EPI_ISL_561176, EPI_ISL_561177, EPI_ISL_561178, EPI_ISL_561179, EPI_ISL_561180, EPI_ISL_561181, EPI_ISL_561182, EPI_ISL_561183, EPI_ISL_561184, EPI_ISL_561185, EPI_ISL_561186, EPI_ISL_561187, EPI_ISL_561188, EPI_ISL_561189, EPI_ISL_561190, EPI_ISL_561191, EPI_ISL_561192, EPI_ISL_561193, EPI_ISL_561194, EPI_ISL_561195, EPI_ISL_561196, EPI_ISL_561197, EPI_ISL_561198, EPI_ISL_561199, EPI_ISL_561200, EPI_ISL_561201, EPI_ISL_561202, EPI_ISL_561203, EPI_ISL_561204, EPI_ISL_561235, EPI_ISL_561264, EPI_ISL_561266, EPI_ISL_561268, EPI_ISL_561281, EPI_ISL_561290, EPI_ISL_561291, EPI_ISL_561293, EPI_ISL_561294, EPI_ISL_561295, EPI_ISL_561297, EPI_ISL_561304, EPI_ISL_561309, EPI_ISL_561312, EPI_ISL_561315, EPI_ISL_561316 | | | |
| see above | MRCG at LSHTM Genomics lab | MRCG at LSHTM Genomics lab | Abdul Karim sesay, Abdoulie Kanteh, Jarra Manneh, Mariama Kujabi, Bakary Sanyang |
| EPI_ISL_561394 | Microbiological Diagnostic Unit - Public Health Laboratory (MDU-PHL) | MDU-PHL | Seemann, T., Schultz M. B., Sait, M., Sherry, N. |
| EPI_ISL_561397 | Victorian Infectious Diseases Reference Laboratory (VIDRL) | VIDRL and MDU-PHL | Caly, L., Seemann, T., Sait, M., Schultz, M. B., Druce J., Sherry, N. |
| EPI_ISL_561400 | Microbiological Diagnostic Unit - Public Health Laboratory (MDU-PHL) | MDU-PHL | Seemann, T., Schultz M. B., Sait, M., Sherry, N. |
| EPI_ISL_561403, EPI_ISL_561404, EPI_ISL_561409 | Victorian Infectious Diseases Reference Laboratory (VIDRL) | VIDRL and MDU-PHL | Caly, L., Seemann, T., Sait, M., Schultz, M. B., Druce J., Sherry, N. |
| EPI_ISL_561412 | Microbiological Diagnostic Unit - Public Health Laboratory (MDU-PHL) | MDU-PHL | Seemann, T., Schultz M. B., Sait, M., Sherry, N. |
| EPI_ISL_561414, EPI_ISL_561433 | Victorian Infectious Diseases Reference Laboratory (VIDRL) | VIDRL and MDU-PHL | Caly, L., Seemann, T., Sait, M., Schultz, M. B., Druce J., Sherry, N. |
| EPI_ISL_561438, EPI_ISL_561442, EPI_ISL_561464, EPI_ISL_561467, EPI_ISL_561481 | Microbiological Diagnostic Unit - Public Health Laboratory (MDU-PHL) | MDU-PHL | Seemann, T., Schultz M. B., Sait, M., Sherry, N. |
| EPI_ISL_561498 | Victorian Infectious Diseases Reference Laboratory (VIDRL) | VIDRL and MDU-PHL | Caly, L., Seemann, T., Sait, M., Schultz, M. B., Druce J., Sherry, N. |
| EPI_ISL_561501, EPI_ISL_561505, EPI_ISL_561506, EPI_ISL_561507 | Microbiological Diagnostic Unit - Public Health Laboratory (MDU-PHL) | MDU-PHL | Seemann, T., Schultz M. B., Sait, M., Sherry, N. |
| EPI_ISL_561508 | Victorian Infectious Diseases Reference Laboratory (VIDRL) | VIDRL and MDU-PHL | Caly, L., Seemann, T., Sait, M., Schultz, M. B., Druce J., Sherry, N. |
| EPI_ISL_561512, EPI_ISL_561513 | Microbiological Diagnostic Unit - Public Health Laboratory (MDU-PHL) | MDU-PHL | Seemann, T., Schultz M. B., Sait, M., Sherry, N. |
| EPI_ISL_561514, EPI_ISL_561515 | Victorian Infectious Diseases Reference Laboratory (VIDRL) | VIDRL and MDU-PHL | Caly, L., Seemann, T., Sait, M., Schultz, M. B., Druce J., Sherry, N. |
| EPI_ISL_561529 | Microbiological Diagnostic Unit - Public Health Laboratory (MDU-PHL) | MDU-PHL | Seemann, T., Schultz M. B., Sait, M., Sherry, N. |

| | | | |
|---|---|---|---|
| EPI_ISL_561533, EPI_ISL_561544 | Victorian Infectious Diseases Reference Laboratory (VIDRL) | VIDRL and MDU-PHL | Caly, L., Seemann, T., Sait, M., Schultz, M. B., Druce J., Sherry, N. |
| EPI_ISL_561545, EPI_ISL_561551, EPI_ISL_561554, EPI_ISL_561555, EPI_ISL_561558 | Microbiological Diagnostic Unit - Public Health Laboratory (MDU-PHL) | MDU-PHL | Seemann, T., Schultz M. B., Sait, M., Sherry, N. |
| EPI_ISL_561559, EPI_ISL_561565 | Victorian Infectious Diseases Reference Laboratory (VIDRL) | VIDRL and MDU-PHL | Caly, L., Seemann, T., Sait, M., Schultz, M. B., Druce J., Sherry, N. |
| EPI_ISL_561572, EPI_ISL_561576, EPI_ISL_561580 | Microbiological Diagnostic Unit - Public Health Laboratory (MDU-PHL) | MDU-PHL | Seemann, T., Schultz M. B., Sait, M., Sherry, N. |
| EPI_ISL_561582 | Victorian Infectious Diseases Reference Laboratory (VIDRL) | VIDRL and MDU-PHL | Caly, L., Seemann, T., Sait, M., Schultz, M. B., Druce J., Sherry, N. |
| EPI_ISL_561585 | Microbiological Diagnostic Unit - Public Health Laboratory (MDU-PHL) | MDU-PHL | Seemann, T., Schultz M. B., Sait, M., Sherry, N. |
| EPI_ISL_561586 | Victorian Infectious Diseases Reference Laboratory (VIDRL) | VIDRL and MDU-PHL | Caly, L., Seemann, T., Sait, M., Schultz, M. B., Druce J., Sherry, N. |
| EPI_ISL_561588, EPI_ISL_561597, EPI_ISL_561602 | Microbiological Diagnostic Unit - Public Health Laboratory (MDU-PHL) | MDU-PHL | Seemann, T., Schultz M. B., Sait, M., Sherry, N. |
| EPI_ISL_561612, EPI_ISL_561613 | Victorian Infectious Diseases Reference Laboratory (VIDRL) | VIDRL and MDU-PHL | Caly, L., Seemann, T., Sait, M., Schultz, M. B., Druce J., Sherry, N. |
| EPI_ISL_561619 | Microbiological Diagnostic Unit - Public Health Laboratory (MDU-PHL) | MDU-PHL | Seemann, T., Schultz M. B., Sait, M., Sherry, N. |
| EPI_ISL_561621, EPI_ISL_561622, EPI_ISL_561629 | Victorian Infectious Diseases Reference Laboratory (VIDRL) | VIDRL and MDU-PHL | Caly, L., Seemann, T., Sait, M., Schultz, M. B., Druce J., Sherry, N. |
| EPI_ISL_561633, EPI_ISL_561638, EPI_ISL_561641, EPI_ISL_561647 | Microbiological Diagnostic Unit - Public Health Laboratory (MDU-PHL) | MDU-PHL | Seemann, T., Schultz M. B., Sait, M., Sherry, N. |
| EPI_ISL_561657 | Victorian Infectious Diseases Reference Laboratory (VIDRL) | VIDRL and MDU-PHL | Caly, L., Seemann, T., Sait, M., Schultz, M. B., Druce J., Sherry, N. |
| EPI_ISL_561660 | Microbiological Diagnostic Unit - Public Health Laboratory (MDU-PHL) | MDU-PHL | Seemann, T., Schultz M. B., Sait, M., Sherry, N. |
| EPI_ISL_561661 | Victorian Infectious Diseases Reference Laboratory (VIDRL) | VIDRL and MDU-PHL | Caly, L., Seemann, T., Sait, M., Schultz, M. B., Druce J., Sherry, N. |
| EPI_ISL_561664, EPI_ISL_561670, EPI_ISL_561678, EPI_ISL_561680, EPI_ISL_561683, EPI_ISL_561685, EPI_ISL_561688, EPI_ISL_561694, EPI_ISL_561701, EPI_ISL_561703, EPI_ISL_561704 | | | |
| see above | Microbiological Diagnostic Unit - Public Health Laboratory (MDU-PHL) | MDU-PHL | Seemann, T., Schultz M. B., Sait, M., Sherry, N. |
| EPI_ISL_561707 | Victorian Infectious Diseases Reference Laboratory (VIDRL) | VIDRL and MDU-PHL | Caly, L., Seemann, T., Sait, M., Schultz, M. B., Druce J., Sherry, N. |
| EPI_ISL_561708 | Microbiological Diagnostic Unit - Public Health Laboratory (MDU-PHL) | MDU-PHL | Seemann, T., Schultz M. B., Sait, M., Sherry, N. |
| EPI_ISL_561715 | Victorian Infectious Diseases Reference Laboratory (VIDRL) | VIDRL and MDU-PHL | Caly, L., Seemann, T., Sait, M., Schultz, M. B., Druce J., Sherry, N. |
| EPI_ISL_561724, EPI_ISL_561725, EPI_ISL_561727, EPI_ISL_561730, EPI_ISL_561734, EPI_ISL_561739, EPI_ISL_561745, EPI_ISL_561748 | Microbiological Diagnostic Unit - Public Health Laboratory (MDU-PHL) | MDU-PHL | Seemann, T., Schultz M. B., Sait, M., Sherry, N. |
| EPI_ISL_561752 | Victorian Infectious Diseases Reference Laboratory (VIDRL) | VIDRL and MDU-PHL | Caly, L., Seemann, T., Sait, M., Schultz, M. B., Druce J., Sherry, N. |
| EPI_ISL_561761 | Microbiological Diagnostic Unit - Public Health Laboratory (MDU-PHL) | MDU-PHL | Seemann, T., Schultz M. B., Sait, M., Sherry, N. |
| EPI_ISL_561766 | Victorian Infectious Diseases Reference Laboratory (VIDRL) | VIDRL and MDU-PHL | Caly, L., Seemann, T., Sait, M., Schultz, M. B., Druce J., Sherry, N. |
| EPI_ISL_561769 | Microbiological Diagnostic Unit - Public Health Laboratory (MDU-PHL) | MDU-PHL | Seemann, T., Schultz M. B., Sait, M., Sherry, N. |
| EPI_ISL_561778 | Victorian Infectious Diseases Reference Laboratory (VIDRL) | VIDRL and MDU-PHL | Caly, L., Seemann, T., Sait, M., Schultz, M. B., Druce J., Sherry, N. |
| EPI_ISL_561779, EPI_ISL_561780, EPI_ISL_561783, EPI_ISL_561784 | Microbiological Diagnostic Unit - Public Health Laboratory (MDU-PHL) | MDU-PHL | Seemann, T., Schultz M. B., Sait, M., Sherry, N. |
| EPI_ISL_561787 | Victorian Infectious Diseases Reference Laboratory (VIDRL) | VIDRL and MDU-PHL | Caly, L., Seemann, T., Sait, M., Schultz, M. B., Druce J., Sherry, N. |
| EPI_ISL_561802 | Microbiological Diagnostic Unit - Public Health Laboratory (MDU-PHL) | MDU-PHL | Seemann, T., Schultz M. B., Sait, M., Sherry, N. |
| EPI_ISL_561803 | Victorian Infectious Diseases Reference Laboratory (VIDRL) | VIDRL and MDU-PHL | Caly, L., Seemann, T., Sait, M., Schultz, M. B., Druce J., Sherry, N. |
| EPI_ISL_561809, EPI_ISL_561815 | Microbiological Diagnostic Unit - Public Health Laboratory (MDU-PHL) | MDU-PHL | Seemann, T., Schultz M. B., Sait, M., Sherry, N. |
| EPI_ISL_561816 | Victorian Infectious Diseases Reference Laboratory (VIDRL) | VIDRL and MDU-PHL | Caly, L., Seemann, T., Sait, M., Schultz, M. B., Druce J., Sherry, N. |
| EPI_ISL_561827, EPI_ISL_561828, EPI_ISL_561830, EPI_ISL_561831, EPI_ISL_561835, EPI_ISL_561839, EPI_ISL_561842, EPI_ISL_561845, EPI_ISL_561846 | Microbiological Diagnostic Unit - Public Health Laboratory (MDU-PHL) | MDU-PHL | Seemann, T., Schultz M. B., Sait, M., Sherry, N. |
| EPI_ISL_561852 | Victorian Infectious Diseases Reference Laboratory (VIDRL) | VIDRL and MDU-PHL | Caly, L., Seemann, T., Sait, M., Schultz, M. B., Druce J., Sherry, N. |
| EPI_ISL_561854, EPI_ISL_561855, EPI_ISL_561857 | Microbiological Diagnostic Unit - Public Health Laboratory (MDU-PHL) | MDU-PHL | Seemann, T., Schultz M. B., Sait, M., Sherry, N. |
| EPI_ISL_561858 | Victorian Infectious Diseases Reference Laboratory (VIDRL) | VIDRL and MDU-PHL | Caly, L., Seemann, T., Sait, M., Schultz, M. B., Druce J., Sherry, N. |
| EPI_ISL_561872, EPI_ISL_561875, EPI_ISL_561876, EPI_ISL_561889, EPI_ISL_561891 | Microbiological Diagnostic Unit - Public Health Laboratory (MDU-PHL) | MDU-PHL | Seemann, T., Schultz M. B., Sait, M., Sherry, N. |
| EPI_ISL_561894 | Victorian Infectious Diseases Reference Laboratory (VIDRL) | VIDRL and MDU-PHL | Caly, L., Seemann, T., Sait, M., Schultz, M. B., Druce J., Sherry, N. |
| EPI_ISL_561896, EPI_ISL_561897, EPI_ISL_561898, EPI_ISL_561899, EPI_ISL_561909, EPI_ISL_561927, EPI_ISL_561948, EPI_ISL_561949, EPI_ISL_561950, EPI_ISL_561953, EPI_ISL_561960, EPI_ISL_561962, EPI_ISL_561969 | | | |
| see above | Microbiological Diagnostic Unit - Public Health Laboratory (MDU-PHL) | MDU-PHL | Seemann, T., Schultz M. B., Sait, M., Sherry, N. |

| | | | |
|---|---|---|---|
| EPI_ISL_561975 | Victorian Infectious Diseases Reference Laboratory (VIDRL) | VIDRL and MDU-PHL | Caly, L., Seemann, T., Sait, M., Schultz, M. B., Druce J., Sherry, N. |
| EPI_ISL_561979 | Microbiological Diagnostic Unit - Public Health Laboratory (MDU-PHL) | MDU-PHL | Seemann, T., Schultz M. B., Sait, M., Sherry, N. |
| EPI_ISL_561986, EPI_ISL_561987 | Victorian Infectious Diseases Reference Laboratory (VIDRL) | VIDRL and MDU-PHL | Caly, L., Seemann, T., Sait, M., Schultz, M. B., Druce J., Sherry, N. |
| EPI_ISL_561999, EPI_ISL_562000 | Microbiological Diagnostic Unit - Public Health Laboratory (MDU-PHL) | MDU-PHL | Seemann, T., Schultz M. B., Sait, M., Sherry, N. |
| EPI_ISL_562012, EPI_ISL_562016 | Victorian Infectious Diseases Reference Laboratory (VIDRL) | VIDRL and MDU-PHL | Caly, L., Seemann, T., Sait, M., Schultz, M. B., Druce J., Sherry, N. |
| EPI_ISL_562020, EPI_ISL_562022 | Microbiological Diagnostic Unit - Public Health Laboratory (MDU-PHL) | MDU-PHL | Seemann, T., Schultz M. B., Sait, M., Sherry, N. |
| EPI_ISL_562023 | Victorian Infectious Diseases Reference Laboratory (VIDRL) | VIDRL and MDU-PHL | Caly, L., Seemann, T., Sait, M., Schultz, M. B., Druce J., Sherry, N. |
| EPI_ISL_562028, EPI_ISL_562045, EPI_ISL_562046 | Microbiological Diagnostic Unit - Public Health Laboratory (MDU-PHL) | MDU-PHL | Seemann, T., Schultz M. B., Sait, M., Sherry, N. |
| EPI_ISL_562048, EPI_ISL_562057 | Victorian Infectious Diseases Reference Laboratory (VIDRL) | VIDRL and MDU-PHL | Caly, L., Seemann, T., Sait, M., Schultz, M. B., Druce J., Sherry, N. |
| EPI_ISL_562063, EPI_ISL_562073 | Microbiological Diagnostic Unit - Public Health Laboratory (MDU-PHL) | MDU-PHL | Seemann, T., Schultz M. B., Sait, M., Sherry, N. |
| EPI_ISL_562076 | Victorian Infectious Diseases Reference Laboratory (VIDRL) | VIDRL and MDU-PHL | Caly, L., Seemann, T., Sait, M., Schultz, M. B., Druce J., Sherry, N. |
| EPI_ISL_562077 | Microbiological Diagnostic Unit - Public Health Laboratory (MDU-PHL) | MDU-PHL | Seemann, T., Schultz M. B., Sait, M., Sherry, N. |
| EPI_ISL_562081, EPI_ISL_562085 | Victorian Infectious Diseases Reference Laboratory (VIDRL) | VIDRL and MDU-PHL | Caly, L., Seemann, T., Sait, M., Schultz, M. B., Druce J., Sherry, N. |
| EPI_ISL_562097, EPI_ISL_562100, EPI_ISL_562101, EPI_ISL_562107, EPI_ISL_562113 | Microbiological Diagnostic Unit - Public Health Laboratory (MDU-PHL) | MDU-PHL | Seemann, T., Schultz M. B., Sait, M., Sherry, N. |
| EPI_ISL_562115 | Victorian Infectious Diseases Reference Laboratory (VIDRL) | VIDRL and MDU-PHL | Caly, L., Seemann, T., Sait, M., Schultz, M. B., Druce J., Sherry, N. |
| EPI_ISL_562118, EPI_ISL_562120, EPI_ISL_562122 | Microbiological Diagnostic Unit - Public Health Laboratory (MDU-PHL) | MDU-PHL | Seemann, T., Schultz M. B., Sait, M., Sherry, N. |
| EPI_ISL_562123 | Victorian Infectious Diseases Reference Laboratory (VIDRL) | VIDRL and MDU-PHL | Caly, L., Seemann, T., Sait, M., Schultz, M. B., Druce J., Sherry, N. |
| EPI_ISL_562130, EPI_ISL_562132, EPI_ISL_562133, EPI_ISL_562149 | Microbiological Diagnostic Unit - Public Health Laboratory (MDU-PHL) | MDU-PHL | Seemann, T., Schultz M. B., Sait, M., Sherry, N. |
| EPI_ISL_562152 | Victorian Infectious Diseases Reference Laboratory (VIDRL) | VIDRL and MDU-PHL | Caly, L., Seemann, T., Sait, M., Schultz, M. B., Druce J., Sherry, N. |
| EPI_ISL_562153 | Microbiological Diagnostic Unit - Public Health Laboratory (MDU-PHL) | MDU-PHL | Seemann, T., Schultz M. B., Sait, M., Sherry, N. |
| EPI_ISL_562155 | Victorian Infectious Diseases Reference Laboratory (VIDRL) | VIDRL and MDU-PHL | Caly, L., Seemann, T., Sait, M., Schultz, M. B., Druce J., Sherry, N. |
| EPI_ISL_562156 | Microbiological Diagnostic Unit - Public Health Laboratory (MDU-PHL) | MDU-PHL | Seemann, T., Schultz M. B., Sait, M., Sherry, N. |
| EPI_ISL_562158, EPI_ISL_562164 | Victorian Infectious Diseases Reference Laboratory (VIDRL) | VIDRL and MDU-PHL | Caly, L., Seemann, T., Sait, M., Schultz, M. B., Druce J., Sherry, N. |
| EPI_ISL_562170 | Microbiological Diagnostic Unit - Public Health Laboratory (MDU-PHL) | MDU-PHL | Seemann, T., Schultz M. B., Sait, M., Sherry, N. |
| EPI_ISL_562175 | Victorian Infectious Diseases Reference Laboratory (VIDRL) | VIDRL and MDU-PHL | Caly, L., Seemann, T., Sait, M., Schultz, M. B., Druce J., Sherry, N. |
| EPI_ISL_562178, EPI_ISL_562181, EPI_ISL_562185, EPI_ISL_562188 | Microbiological Diagnostic Unit - Public Health Laboratory (MDU-PHL) | MDU-PHL | Seemann, T., Schultz M. B., Sait, M., Sherry, N. |
| EPI_ISL_562189, EPI_ISL_562197, EPI_ISL_562200 | Victorian Infectious Diseases Reference Laboratory (VIDRL) | VIDRL and MDU-PHL | Caly, L., Seemann, T., Sait, M., Schultz, M. B., Druce J., Sherry, N. |
| EPI_ISL_562206 | Microbiological Diagnostic Unit - Public Health Laboratory (MDU-PHL) | MDU-PHL | Seemann, T., Schultz M. B., Sait, M., Sherry, N. |
| EPI_ISL_562208, EPI_ISL_562209, EPI_ISL_562211 | Victorian Infectious Diseases Reference Laboratory (VIDRL) | VIDRL and MDU-PHL | Caly, L., Seemann, T., Sait, M., Schultz, M. B., Druce J., Sherry, N. |
| EPI_ISL_562212 | Microbiological Diagnostic Unit - Public Health Laboratory (MDU-PHL) | MDU-PHL | Seemann, T., Schultz M. B., Sait, M., Sherry, N. |
| EPI_ISL_562213, EPI_ISL_562217 | Victorian Infectious Diseases Reference Laboratory (VIDRL) | VIDRL and MDU-PHL | Caly, L., Seemann, T., Sait, M., Schultz, M. B., Druce J., Sherry, N. |
| EPI_ISL_562220 | Microbiological Diagnostic Unit - Public Health Laboratory (MDU-PHL) | MDU-PHL | Seemann, T., Schultz M. B., Sait, M., Sherry, N. |
| EPI_ISL_562231 | Victorian Infectious Diseases Reference Laboratory (VIDRL) | VIDRL and MDU-PHL | Caly, L., Seemann, T., Sait, M., Schultz, M. B., Druce J., Sherry, N. |
| EPI_ISL_562234, EPI_ISL_562236, EPI_ISL_562262 | Microbiological Diagnostic Unit - Public Health Laboratory (MDU-PHL) | MDU-PHL | Seemann, T., Schultz M. B., Sait, M., Sherry, N. |
| EPI_ISL_562267, EPI_ISL_562269, EPI_ISL_562275 | Victorian Infectious Diseases Reference Laboratory (VIDRL) | VIDRL and MDU-PHL | Caly, L., Seemann, T., Sait, M., Schultz, M. B., Druce J., Sherry, N. |
| EPI_ISL_562276, EPI_ISL_562284 | Microbiological Diagnostic Unit - Public Health Laboratory (MDU-PHL) | MDU-PHL | Seemann, T., Schultz M. B., Sait, M., Sherry, N. |
| EPI_ISL_562286 | Victorian Infectious Diseases Reference Laboratory (VIDRL) | VIDRL and MDU-PHL | Caly, L., Seemann, T., Sait, M., Schultz, M. B., Druce J., Sherry, N. |
| EPI_ISL_562291 | Microbiological Diagnostic Unit - Public Health Laboratory (MDU-PHL) | MDU-PHL | Seemann, T., Schultz M. B., Sait, M., Sherry, N. |
| EPI_ISL_562297 | Victorian Infectious Diseases Reference Laboratory (VIDRL) | VIDRL and MDU-PHL | Caly, L., Seemann, T., Sait, M., Schultz, M. B., Druce J., Sherry, N. |
| EPI_ISL_562306, EPI_ISL_562307 | Microbiological Diagnostic Unit - Public Health Laboratory (MDU-PHL) | MDU-PHL | Seemann, T., Schultz M. B., Sait, M., Sherry, N. |
| EPI_ISL_562317, EPI_ISL_562331, EPI_ISL_562333 | Victorian Infectious Diseases Reference Laboratory (VIDRL) | VIDRL and MDU-PHL | Caly, L., Seemann, T., Sait, M., Schultz, M. B., Druce J., Sherry, N. |
| EPI_ISL_562336, EPI_ISL_562338 | Microbiological Diagnostic Unit - Public Health Laboratory | MDU-PHL | Seemann, T., Schultz M. B., Sait, M., Sherry, N. |

| | | | |
|---|---|---|---|
| | (MDU-PHL) | | |
| EPI_ISL_562342 | Victorian Infectious Diseases Reference Laboratory (VIDRL) | VIDRL and MDU-PHL | Caly, L., Seemann, T., Sait, M., Schultz, M. B., Druce J., Sherry, N. |
| EPI_ISL_562343 | Microbiological Diagnostic Unit - Public Health Laboratory (MDU-PHL) | MDU-PHL | Seemann, T., Schultz M. B., Sait, M., Sherry, N. |
| EPI_ISL_562351, EPI_ISL_562357, EPI_ISL_562363 | Victorian Infectious Diseases Reference Laboratory (VIDRL) | VIDRL and MDU-PHL | Caly, L., Seemann, T., Sait, M., Schultz, M. B., Druce J., Sherry, N. |
| EPI_ISL_562370 | Microbiological Diagnostic Unit - Public Health Laboratory (MDU-PHL) | MDU-PHL | Seemann, T., Schultz M. B., Sait, M., Sherry, N. |
| EPI_ISL_562375 | Victorian Infectious Diseases Reference Laboratory (VIDRL) | VIDRL and MDU-PHL | Caly, L., Seemann, T., Sait, M., Schultz, M. B., Druce J., Sherry, N. |
| EPI_ISL_562378 | Microbiological Diagnostic Unit - Public Health Laboratory (MDU-PHL) | MDU-PHL | Seemann, T., Schultz M. B., Sait, M., Sherry, N. |
| EPI_ISL_562379 | Victorian Infectious Diseases Reference Laboratory (VIDRL) | VIDRL and MDU-PHL | Caly, L., Seemann, T., Sait, M., Schultz, M. B., Druce J., Sherry, N. |
| EPI_ISL_562386, EPI_ISL_562388 | Microbiological Diagnostic Unit - Public Health Laboratory (MDU-PHL) | MDU-PHL | Seemann, T., Schultz M. B., Sait, M., Sherry, N. |
| EPI_ISL_562389 | Victorian Infectious Diseases Reference Laboratory (VIDRL) | VIDRL and MDU-PHL | Caly, L., Seemann, T., Sait, M., Schultz, M. B., Druce J., Sherry, N. |
| EPI_ISL_562392 | Microbiological Diagnostic Unit - Public Health Laboratory (MDU-PHL) | MDU-PHL | Seemann, T., Schultz M. B., Sait, M., Sherry, N. |
| EPI_ISL_562393, EPI_ISL_562396 | Victorian Infectious Diseases Reference Laboratory (VIDRL) | VIDRL and MDU-PHL | Caly, L., Seemann, T., Sait, M., Schultz, M. B., Druce J., Sherry, N. |
| EPI_ISL_562404 | Microbiological Diagnostic Unit - Public Health Laboratory (MDU-PHL) | MDU-PHL | Seemann, T., Schultz M. B., Sait, M., Sherry, N. |
| EPI_ISL_562414, EPI_ISL_562415 | Victorian Infectious Diseases Reference Laboratory (VIDRL) | VIDRL and MDU-PHL | Caly, L., Seemann, T., Sait, M., Schultz, M. B., Druce J., Sherry, N. |
| EPI_ISL_562420, EPI_ISL_562429 | Microbiological Diagnostic Unit - Public Health Laboratory (MDU-PHL) | MDU-PHL | Seemann, T., Schultz M. B., Sait, M., Sherry, N. |
| EPI_ISL_562430, EPI_ISL_562437, EPI_ISL_562441 | Victorian Infectious Diseases Reference Laboratory (VIDRL) | VIDRL and MDU-PHL | Caly, L., Seemann, T., Sait, M., Schultz, M. B., Druce J., Sherry, N. |
| EPI_ISL_562443, EPI_ISL_562453, EPI_ISL_562457, EPI_ISL_562458, EPI_ISL_562459 | Microbiological Diagnostic Unit - Public Health Laboratory (MDU-PHL) | MDU-PHL | Seemann, T., Schultz M. B., Sait, M., Sherry, N. |
| EPI_ISL_562460, EPI_ISL_562463 | Victorian Infectious Diseases Reference Laboratory (VIDRL) | VIDRL and MDU-PHL | Caly, L., Seemann, T., Sait, M., Schultz, M. B., Druce J., Sherry, N. |
| EPI_ISL_562464 | Microbiological Diagnostic Unit - Public Health Laboratory (MDU-PHL) | MDU-PHL | Seemann, T., Schultz M. B., Sait, M., Sherry, N. |
| EPI_ISL_562466 | Victorian Infectious Diseases Reference Laboratory (VIDRL) | VIDRL and MDU-PHL | Caly, L., Seemann, T., Sait, M., Schultz, M. B., Druce J., Sherry, N. |
| EPI_ISL_562469, EPI_ISL_562470, EPI_ISL_562472, EPI_ISL_562477, EPI_ISL_562485, EPI_ISL_562498, EPI_ISL_562500 | Microbiological Diagnostic Unit - Public Health Laboratory (MDU-PHL) | MDU-PHL | Seemann, T., Schultz M. B., Sait, M., Sherry, N. |
| EPI_ISL_562503, EPI_ISL_562505, EPI_ISL_562509 | Victorian Infectious Diseases Reference Laboratory (VIDRL) | VIDRL and MDU-PHL | Caly, L., Seemann, T., Sait, M., Schultz, M. B., Druce J., Sherry, N. |
| EPI_ISL_562510, EPI_ISL_562512 | Microbiological Diagnostic Unit - Public Health Laboratory (MDU-PHL) | MDU-PHL | Seemann, T., Schultz M. B., Sait, M., Sherry, N. |
| EPI_ISL_562513, EPI_ISL_562514, EPI_ISL_562517, EPI_ISL_562524 | Victorian Infectious Diseases Reference Laboratory (VIDRL) | VIDRL and MDU-PHL | Caly, L., Seemann, T., Sait, M., Schultz, M. B., Druce J., Sherry, N. |
| EPI_ISL_562525 | Microbiological Diagnostic Unit - Public Health Laboratory (MDU-PHL) | MDU-PHL | Seemann, T., Schultz M. B., Sait, M., Sherry, N. |
| EPI_ISL_562526, EPI_ISL_562531, EPI_ISL_562548, EPI_ISL_562552 | Victorian Infectious Diseases Reference Laboratory (VIDRL) | VIDRL and MDU-PHL | Caly, L., Seemann, T., Sait, M., Schultz, M. B., Druce J., Sherry, N. |
| EPI_ISL_562554 | Microbiological Diagnostic Unit - Public Health Laboratory (MDU-PHL) | MDU-PHL | Seemann, T., Schultz M. B., Sait, M., Sherry, N. |
| EPI_ISL_562557, EPI_ISL_562559, EPI_ISL_562563 | Victorian Infectious Diseases Reference Laboratory (VIDRL) | VIDRL and MDU-PHL | Caly, L., Seemann, T., Sait, M., Schultz, M. B., Druce J., Sherry, N. |
| EPI_ISL_562566, EPI_ISL_562569 | Microbiological Diagnostic Unit - Public Health Laboratory (MDU-PHL) | MDU-PHL | Seemann, T., Schultz M. B., Sait, M., Sherry, N. |
| EPI_ISL_562583, EPI_ISL_562586, EPI_ISL_562587, EPI_ISL_562589 | Victorian Infectious Diseases Reference Laboratory (VIDRL) | VIDRL and MDU-PHL | Caly, L., Seemann, T., Sait, M., Schultz, M. B., Druce J., Sherry, N. |
| EPI_ISL_562606, EPI_ISL_562607, EPI_ISL_562608, EPI_ISL_562639, EPI_ISL_562647, EPI_ISL_562656 | Microbiological Diagnostic Unit - Public Health Laboratory (MDU-PHL) | MDU-PHL | Seemann, T., Schultz M. B., Sait, M., Sherry, N. |
| EPI_ISL_562658 | Victorian Infectious Diseases Reference Laboratory (VIDRL) | VIDRL and MDU-PHL | Caly, L., Seemann, T., Sait, M., Schultz, M. B., Druce J., Sherry, N. |
| EPI_ISL_562668, EPI_ISL_562703, EPI_ISL_562707, EPI_ISL_562755, EPI_ISL_562758, EPI_ISL_562785, EPI_ISL_562790, EPI_ISL_562848, EPI_ISL_562850, EPI_ISL_562852, EPI_ISL_562854, EPI_ISL_562855, EPI_ISL_562856, EPI_ISL_562857, EPI_ISL_562860, EPI_ISL_562862, EPI_ISL_562863, EPI_ISL_562867, EPI_ISL_562868, EPI_ISL_562869, EPI_ISL_562873, EPI_ISL_562874, EPI_ISL_562876, EPI_ISL_562877, EPI_ISL_562878, EPI_ISL_562879, EPI_ISL_562880, EPI_ISL_562881, EPI_ISL_562882, EPI_ISL_562883, EPI_ISL_562884, EPI_ISL_562885, EPI_ISL_562886, EPI_ISL_562888, EPI_ISL_562889, EPI_ISL_562890, EPI_ISL_562915, EPI_ISL_562954, EPI_ISL_563001, EPI_ISL_563005, EPI_ISL_563009, EPI_ISL_563016, EPI_ISL_563025, EPI_ISL_563026, EPI_ISL_563031, EPI_ISL_563036, EPI_ISL_563056, EPI_ISL_563057, EPI_ISL_563067, EPI_ISL_563072, EPI_ISL_563078, EPI_ISL_563089, EPI_ISL_563091, EPI_ISL_563092, EPI_ISL_563094, EPI_ISL_563096, EPI_ISL_563097, EPI_ISL_563104, EPI_ISL_563105, EPI_ISL_563106, EPI_ISL_563107, EPI_ISL_563108, EPI_ISL_563120, EPI_ISL_563121, EPI_ISL_563122, EPI_ISL_563123, EPI_ISL_563135, EPI_ISL_563163, EPI_ISL_563164, EPI_ISL_563165, EPI_ISL_563167, EPI_ISL_563170, EPI_ISL_563172, EPI_ISL_563181, EPI_ISL_563184, EPI_ISL_563190, EPI_ISL_563194, EPI_ISL_563199, EPI_ISL_563202, EPI_ISL_563205, EPI_ISL_563209, EPI_ISL_563216, EPI_ISL_563218, EPI_ISL_563222, EPI_ISL_563226, EPI_ISL_563229, EPI_ISL_563232, EPI_ISL_563234, EPI_ISL_563235, EPI_ISL_563236, EPI_ISL_563239, EPI_ISL_563240, EPI_ISL_563241, EPI_ISL_563255, EPI_ISL_563256, EPI_ISL_563263, EPI_ISL_563268, EPI_ISL_563271, EPI_ISL_563274, EPI_ISL_563275, EPI_ISL_563279, EPI_ISL_563281, EPI_ISL_563282, EPI_ISL_563284, EPI_ISL_563285, EPI_ISL_563286, EPI_ISL_563287, EPI_ISL_563290, EPI_ISL_563291, EPI_ISL_563292, EPI_ISL_563293, EPI_ISL_563294, EPI_ISL_563295, EPI_ISL_563329, EPI_ISL_563330, EPI_ISL_563424, EPI_ISL_563426, EPI_ISL_563427, EPI_ISL_563430, EPI_ISL_563431, EPI_ISL_563432, EPI_ISL_563433, EPI_ISL_563435, EPI_ISL_563436, EPI_ISL_563437, EPI_ISL_563441, EPI_ISL_563442, EPI_ISL_563443, EPI_ISL_563444, EPI_ISL_563445, EPI_ISL_563446, EPI_ISL_563447, EPI_ISL_563449, EPI_ISL_563450, EPI_ISL_563451, EPI_ISL_563453, EPI_ISL_563454, EPI_ISL_563455, EPI_ISL_563457, EPI_ISL_563459, EPI_ISL_563463, EPI_ISL_563465, EPI_ISL_563466, EPI_ISL_563467, EPI_ISL_563469, EPI_ISL_563471, EPI_ISL_563473, EPI_ISL_563474, EPI_ISL_563476, EPI_ISL_563479, EPI_ISL_563480, EPI_ISL_563486, EPI_ISL_563491, EPI_ISL_563496, EPI_ISL_563500, EPI_ISL_563503, EPI_ISL_563506, EPI_ISL_563511, EPI_ISL_563513, EPI_ISL_563514, EPI_ISL_563516, EPI_ISL_563522, EPI_ISL_563524, EPI_ISL_563530, EPI_ISL_563532, EPI_ISL_563536, EPI_ISL_563539, EPI_ISL_563541, EPI_ISL_563542, EPI_ISL_563544, EPI_ISL_563545, EPI_ISL_563546, EPI_ISL_563547, EPI_ISL_563549, EPI_ISL_563556, EPI_ISL_563559, EPI_ISL_563560, EPI_ISL_563561, EPI_ISL_563564, EPI_ISL_563565, EPI_ISL_563568, EPI_ISL_563570, EPI_ISL_563572, EPI_ISL_563574, EPI_ISL_563575, EPI_ISL_563577, EPI_ISL_563580, EPI_ISL_563581, EPI_ISL_563583, EPI_ISL_563586, EPI_ISL_563591, EPI_ISL_563598, EPI_ISL_563604, EPI_ISL_563606, EPI_ISL_563608, EPI_ISL_563610, EPI_ISL_563611, EPI_ISL_563612, EPI_ISL_563613, EPI_ISL_563614, EPI_ISL_563616, EPI_ISL_563617, EPI_ISL_563618, EPI_ISL_563619, EPI_ISL_563620, EPI_ISL_563621, EPI_ISL_563622, EPI_ISL_563623, EPI_ISL_563626, EPI_ISL_563629, EPI_ISL_563635, EPI_ISL_563637, EPI_ISL_563639, EPI_ISL_563642, EPI_ISL_563645, EPI_ISL_563646, EPI_ISL_563647, EPI_ISL_563648, EPI_ISL_563649, EPI_ISL_563650, EPI_ISL_563651, EPI_ISL_563652, EPI_ISL_563653, EPI_ISL_563654, EPI_ISL_563655, EPI_ISL_563656, EPI_ISL_563657, EPI_ISL_563658, EPI_ISL_563659, EPI_ISL_563660, EPI_ISL_563661, EPI_ISL_563662, EPI_ISL_563663, EPI_ISL_563664, EPI_ISL_563665, EPI_ISL_563666, EPI_ISL_563667, EPI_ISL_563668, EPI_ISL_563669, EPI_ISL_563670, EPI_ISL_563671, EPI_ISL_563672, EPI_ISL_563673, EPI_ISL_563677, EPI_ISL_563678, EPI_ISL_563679, EPI_ISL_563680, EPI_ISL_563681, EPI_ISL_563682, EPI_ISL_563683, EPI_ISL_563684, EPI_ISL_563685, | | | |

| | | | |
|---|---|---|---|
| EPI_ISL_563686, EPI_ISL_563687, EPI_ISL_563688, EPI_ISL_563689, EPI_ISL_563690, EPI_ISL_563691, EPI_ISL_563692, EPI_ISL_563693, EPI_ISL_563694, EPI_ISL_563695, EPI_ISL_563696, EPI_ISL_563697, EPI_ISL_563698, EPI_ISL_563700, EPI_ISL_563701, EPI_ISL_563702, EPI_ISL_563703, EPI_ISL_563704, EPI_ISL_563705, EPI_ISL_563706, EPI_ISL_563707, EPI_ISL_563708, EPI_ISL_563710, EPI_ISL_563711, EPI_ISL_563712, EPI_ISL_563713, EPI_ISL_563714, EPI_ISL_563715, EPI_ISL_563716, EPI_ISL_563717, EPI_ISL_563718, EPI_ISL_563719, EPI_ISL_563720, EPI_ISL_563743, EPI_ISL_563752, EPI_ISL_563768, EPI_ISL_563770, EPI_ISL_563789, EPI_ISL_563791, EPI_ISL_563794, EPI_ISL_563888, EPI_ISL_563892, EPI_ISL_563895, EPI_ISL_563898, EPI_ISL_563902, EPI_ISL_563905, EPI_ISL_563907, EPI_ISL_563910, EPI_ISL_563913, EPI_ISL_563916, EPI_ISL_563918, EPI_ISL_563921, EPI_ISL_563924, EPI_ISL_563928, EPI_ISL_563931, EPI_ISL_563932, EPI_ISL_563933, EPI_ISL_563935, EPI_ISL_563936, EPI_ISL_563937, EPI_ISL_563991, EPI_ISL_563992, EPI_ISL_563993, EPI_ISL_563995, EPI_ISL_563996 | | | |
| see above | Microbiological Diagnostic Unit - Public Health Laboratory (MDU-PHL) | MDU-PHL | Seemann, T., Schultz M. B., Sait, M., Sherry, N. |
| EPI_ISL_564003, EPI_ISL_564004, EPI_ISL_564005, EPI_ISL_564006, EPI_ISL_564007, EPI_ISL_564008, EPI_ISL_564009, EPI_ISL_564010, EPI_ISL_564011, EPI_ISL_564012, EPI_ISL_564013, EPI_ISL_564015, EPI_ISL_564016, EPI_ISL_564017, EPI_ISL_564018, EPI_ISL_564019, EPI_ISL_564020, EPI_ISL_564021, EPI_ISL_564022, EPI_ISL_564023, EPI_ISL_564024, EPI_ISL_564025, EPI_ISL_564026, EPI_ISL_564027, EPI_ISL_564028, EPI_ISL_564029, EPI_ISL_564030, EPI_ISL_564031, EPI_ISL_564032, EPI_ISL_564033, EPI_ISL_564034, EPI_ISL_564035, EPI_ISL_564036, EPI_ISL_564037, EPI_ISL_564038, EPI_ISL_564039, EPI_ISL_564040, EPI_ISL_564041, EPI_ISL_564042, EPI_ISL_564043, EPI_ISL_564044, EPI_ISL_564045, EPI_ISL_564046, EPI_ISL_564047, EPI_ISL_564048, EPI_ISL_564049 | | | |
| see above | Victorian Infectious Diseases Reference Laboratory (VIDRL) | VIDRL and MDU-PHL | Caly, L., Seemann, T., Sait, M., Schultz, M. B., Druce J., Sherry, N. |
| EPI_ISL_564050, EPI_ISL_564051, EPI_ISL_564052, EPI_ISL_564053, EPI_ISL_564054 | Microbiological Diagnostic Unit - Public Health Laboratory (MDU-PHL) | MDU-PHL | Seemann, T., Schultz M. B., Sait, M., Sherry, N. |
| EPI_ISL_564077, EPI_ISL_564078, EPI_ISL_564079, EPI_ISL_564080, EPI_ISL_564081, EPI_ISL_564082, EPI_ISL_564083, EPI_ISL_564084, EPI_ISL_564085, EPI_ISL_564086, EPI_ISL_564087, EPI_ISL_564088, EPI_ISL_564089, EPI_ISL_564090, EPI_ISL_564091, EPI_ISL_564092, EPI_ISL_564093, EPI_ISL_564094, EPI_ISL_564095, EPI_ISL_564096, EPI_ISL_564097, EPI_ISL_564098, EPI_ISL_564099, EPI_ISL_564100, EPI_ISL_564101, EPI_ISL_564102, EPI_ISL_564103, EPI_ISL_564104, EPI_ISL_564105, EPI_ISL_564106, EPI_ISL_564107, EPI_ISL_564108, EPI_ISL_564109, EPI_ISL_564110 | | | |
| see above | Victorian Infectious Diseases Reference Laboratory (VIDRL) | VIDRL and MDU-PHL | Caly, L., Seemann, T., Sait, M., Schultz, M. B., Druce J., Sherry, N. |
| EPI_ISL_564111, EPI_ISL_564112, EPI_ISL_564130, EPI_ISL_564136, EPI_ISL_564137, EPI_ISL_564138, EPI_ISL_564148, EPI_ISL_564153, EPI_ISL_564154, EPI_ISL_564157, EPI_ISL_564162, EPI_ISL_564164, EPI_ISL_564165, EPI_ISL_564166, EPI_ISL_564167, EPI_ISL_564168, EPI_ISL_564169, EPI_ISL_564170, EPI_ISL_564172, EPI_ISL_564175, EPI_ISL_564176, EPI_ISL_564177 | | | |
| see above | Microbiological Diagnostic Unit - Public Health Laboratory (MDU-PHL) | MDU-PHL | Seemann, T., Schultz M. B., Sait, M., Sherry, N. |
| EPI_ISL_564180, EPI_ISL_564182, EPI_ISL_564183, EPI_ISL_564184, EPI_ISL_564185, EPI_ISL_564186, EPI_ISL_564187, EPI_ISL_564188, EPI_ISL_564189, EPI_ISL_564190, EPI_ISL_564191, EPI_ISL_564192, EPI_ISL_564193, EPI_ISL_564194, EPI_ISL_564195, EPI_ISL_564196, EPI_ISL_564197, EPI_ISL_564198, EPI_ISL_564199, EPI_ISL_564200, EPI_ISL_564201, EPI_ISL_564202, EPI_ISL_564215, EPI_ISL_564216, EPI_ISL_564217, EPI_ISL_564221, EPI_ISL_564222, EPI_ISL_564244, EPI_ISL_564245, EPI_ISL_564246, EPI_ISL_564247, EPI_ISL_564248 | | | |
| see above | Victorian Infectious Diseases Reference Laboratory (VIDRL) | VIDRL and MDU-PHL | Caly, L., Seemann, T., Sait, M., Schultz, M. B., Druce J., Sherry, N. |
| EPI_ISL_564255, EPI_ISL_564260, EPI_ISL_564261, EPI_ISL_564262, EPI_ISL_564263, EPI_ISL_564264, EPI_ISL_564265, EPI_ISL_564266, EPI_ISL_564267, EPI_ISL_564272, EPI_ISL_564290, EPI_ISL_564293, EPI_ISL_564296, EPI_ISL_564297, EPI_ISL_564298, EPI_ISL_564299, EPI_ISL_564300, EPI_ISL_564301, EPI_ISL_564302, EPI_ISL_564303, EPI_ISL_564304, EPI_ISL_564305, EPI_ISL_564306, EPI_ISL_564307, EPI_ISL_564308, EPI_ISL_564309, EPI_ISL_564310, EPI_ISL_564311, EPI_ISL_564312, EPI_ISL_564313, EPI_ISL_564314, EPI_ISL_564315, EPI_ISL_564316, EPI_ISL_564317 | | | |
| see above | Microbiological Diagnostic Unit - Public Health Laboratory (MDU-PHL) | MDU-PHL | Seemann, T., Schultz M. B., Sait, M., Sherry, N. |
| EPI_ISL_564319, EPI_ISL_564320, EPI_ISL_564321, EPI_ISL_564322, EPI_ISL_564323, EPI_ISL_564324, EPI_ISL_564325, EPI_ISL_564326, EPI_ISL_564327, EPI_ISL_564328, EPI_ISL_564329, EPI_ISL_564335, EPI_ISL_564336, EPI_ISL_564337, EPI_ISL_564338, EPI_ISL_564339, EPI_ISL_564340, EPI_ISL_564342, EPI_ISL_564343 | | | |
| see above | Victorian Infectious Diseases Reference Laboratory (VIDRL) | VIDRL and MDU-PHL | Caly, L., Seemann, T., Sait, M., Schultz, M. B., Druce J., Sherry, N. |
| EPI_ISL_564351 | Microbiological Diagnostic Unit - Public Health Laboratory (MDU-PHL) | MDU-PHL | Seemann, T., Schultz M. B., Sait, M., Sherry, N. |
| EPI_ISL_564352, EPI_ISL_564353, EPI_ISL_564354, EPI_ISL_564355, EPI_ISL_564356, EPI_ISL_564357, EPI_ISL_564362, EPI_ISL_564363, EPI_ISL_564365, EPI_ISL_564366, EPI_ISL_564367, EPI_ISL_564368, EPI_ISL_564369, EPI_ISL_564370, EPI_ISL_564371, EPI_ISL_564372, EPI_ISL_564373, EPI_ISL_564374, EPI_ISL_564375, EPI_ISL_564376, EPI_ISL_564377, EPI_ISL_564378, EPI_ISL_564379, EPI_ISL_564380, EPI_ISL_564381, EPI_ISL_564382, EPI_ISL_564383, EPI_ISL_564384, EPI_ISL_564385, EPI_ISL_564386, EPI_ISL_564387, EPI_ISL_564388, EPI_ISL_564389, EPI_ISL_564390, EPI_ISL_564391, EPI_ISL_564392, EPI_ISL_564393, EPI_ISL_564397 | | | |
| see above | Victorian Infectious Diseases Reference Laboratory (VIDRL) | VIDRL and MDU-PHL | Caly, L., Seemann, T., Sait, M., Schultz, M. B., Druce J., Sherry, N. |
| EPI_ISL_564398, EPI_ISL_564399, EPI_ISL_564400, EPI_ISL_564401, EPI_ISL_564402, EPI_ISL_564403 | Microbiological Diagnostic Unit - Public Health Laboratory (MDU-PHL) | MDU-PHL | Seemann, T., Schultz M. B., Sait, M., Sherry, N. |
| EPI_ISL_564404 | Victorian Infectious Diseases Reference Laboratory (VIDRL) | VIDRL and MDU-PHL | Caly, L., Seemann, T., Sait, M., Schultz, M. B., Druce J., Sherry, N. |
| EPI_ISL_564406, EPI_ISL_564414, EPI_ISL_564416, EPI_ISL_564418, EPI_ISL_564419, EPI_ISL_564420, EPI_ISL_564422, EPI_ISL_564423, EPI_ISL_564424, EPI_ISL_564425, EPI_ISL_564426, EPI_ISL_564427, EPI_ISL_564429, EPI_ISL_564430, EPI_ISL_564432, EPI_ISL_564433, EPI_ISL_564434, EPI_ISL_564435, EPI_ISL_564437, EPI_ISL_564438, EPI_ISL_564441, EPI_ISL_564442, EPI_ISL_564443, EPI_ISL_564444, EPI_ISL_564445, EPI_ISL_564446, EPI_ISL_564447, EPI_ISL_564448, EPI_ISL_564449, EPI_ISL_564450, EPI_ISL_564452, EPI_ISL_564454, EPI_ISL_564472, EPI_ISL_564474, EPI_ISL_564475, EPI_ISL_564476, EPI_ISL_564477, EPI_ISL_564478, EPI_ISL_564479, EPI_ISL_564481, EPI_ISL_564482, EPI_ISL_564483, EPI_ISL_564484, EPI_ISL_564485, EPI_ISL_564486, EPI_ISL_564487, EPI_ISL_564488, EPI_ISL_564489, EPI_ISL_564490, EPI_ISL_564494, EPI_ISL_564495, EPI_ISL_564497, EPI_ISL_564508, EPI_ISL_564509, EPI_ISL_564510, EPI_ISL_564511, EPI_ISL_564512, EPI_ISL_564513, EPI_ISL_564515, EPI_ISL_564516, EPI_ISL_564517, EPI_ISL_564518, EPI_ISL_564520, EPI_ISL_564522, EPI_ISL_564523, EPI_ISL_564524, EPI_ISL_564525, EPI_ISL_564526, EPI_ISL_564527, EPI_ISL_564528, EPI_ISL_564535, EPI_ISL_564536, EPI_ISL_564537, EPI_ISL_564539, EPI_ISL_564540, EPI_ISL_564541, EPI_ISL_564542, EPI_ISL_564568, EPI_ISL_564569, EPI_ISL_564570, EPI_ISL_564571, EPI_ISL_564572, EPI_ISL_564573, EPI_ISL_564574, EPI_ISL_564575, EPI_ISL_564577, EPI_ISL_564578, EPI_ISL_564580, EPI_ISL_564581, EPI_ISL_564582, EPI_ISL_564583, EPI_ISL_564593, EPI_ISL_564594, EPI_ISL_564595, EPI_ISL_564602, EPI_ISL_564603, EPI_ISL_564625, EPI_ISL_564626, EPI_ISL_564627, EPI_ISL_564628, EPI_ISL_564629, EPI_ISL_564630, EPI_ISL_564631, EPI_ISL_564632, EPI_ISL_564676, EPI_ISL_564684, EPI_ISL_564685, EPI_ISL_564686, EPI_ISL_564687, EPI_ISL_564688, EPI_ISL_564689, EPI_ISL_564690, EPI_ISL_564691, EPI_ISL_564692, EPI_ISL_564693, EPI_ISL_564694, EPI_ISL_564695, EPI_ISL_564696, EPI_ISL_564697, EPI_ISL_564698, EPI_ISL_564699, EPI_ISL_564700, EPI_ISL_564701, EPI_ISL_564702, EPI_ISL_564703, EPI_ISL_564704, EPI_ISL_564705, EPI_ISL_564706, EPI_ISL_564707, EPI_ISL_564708, EPI_ISL_564709, EPI_ISL_564710, EPI_ISL_564711, EPI_ISL_564712, EPI_ISL_564713, EPI_ISL_564714, EPI_ISL_564715, EPI_ISL_564716, EPI_ISL_564717, EPI_ISL_564985, EPI_ISL_564986, EPI_ISL_564987, EPI_ISL_564988, EPI_ISL_564989, EPI_ISL_564990, EPI_ISL_564991, EPI_ISL_564992, EPI_ISL_564993, EPI_ISL_564994, EPI_ISL_564995, EPI_ISL_564996, EPI_ISL_564997, EPI_ISL_564998, EPI_ISL_564999, EPI_ISL_565000, EPI_ISL_565001, EPI_ISL_565002, EPI_ISL_565003, EPI_ISL_565009, EPI_ISL_565010, EPI_ISL_565011, EPI_ISL_565012, EPI_ISL_565013, EPI_ISL_565014, EPI_ISL_565015, EPI_ISL_565016, EPI_ISL_565017, EPI_ISL_565018, EPI_ISL_565019, EPI_ISL_565020, EPI_ISL_565021, EPI_ISL_565022, EPI_ISL_565023, EPI_ISL_565024, EPI_ISL_565025, EPI_ISL_565026, EPI_ISL_565027, EPI_ISL_565028, EPI_ISL_565029, EPI_ISL_565030, EPI_ISL_565031, EPI_ISL_565032, EPI_ISL_565033, EPI_ISL_565035, EPI_ISL_565036, EPI_ISL_565037, EPI_ISL_565038, EPI_ISL_565039, EPI_ISL_565042, EPI_ISL_565043, EPI_ISL_565044, EPI_ISL_565045, EPI_ISL_565046, EPI_ISL_565055, EPI_ISL_565056, EPI_ISL_565057, EPI_ISL_565058, EPI_ISL_565059, EPI_ISL_565060, EPI_ISL_565061, EPI_ISL_565062, EPI_ISL_565063, EPI_ISL_565064, EPI_ISL_565065, EPI_ISL_565066, EPI_ISL_565067, EPI_ISL_565068, EPI_ISL_565069, EPI_ISL_565070, EPI_ISL_565071, EPI_ISL_565111, EPI_ISL_565112, EPI_ISL_565113, EPI_ISL_565114, EPI_ISL_565115, EPI_ISL_565116 | | | |
| see above | Microbiological Diagnostic Unit - Public Health Laboratory (MDU-PHL) | MDU-PHL | Seemann, T., Schultz M. B., Sait, M., Sherry, N. |
| EPI_ISL_565136, EPI_ISL_565137, EPI_ISL_565138, EPI_ISL_565139, EPI_ISL_565140, EPI_ISL_565144, EPI_ISL_565145, EPI_ISL_565146, EPI_ISL_565147, EPI_ISL_565148, EPI_ISL_565149, EPI_ISL_565150, EPI_ISL_565151, EPI_ISL_565152, EPI_ISL_565153, EPI_ISL_565154, EPI_ISL_565155, EPI_ISL_565156, EPI_ISL_565157, EPI_ISL_565158, EPI_ISL_565159, EPI_ISL_565160, EPI_ISL_565161, EPI_ISL_565162, EPI_ISL_565163, EPI_ISL_565164, EPI_ISL_565165, EPI_ISL_565166, EPI_ISL_565167, EPI_ISL_565168, EPI_ISL_565169, EPI_ISL_565170, EPI_ISL_565171, EPI_ISL_565172, EPI_ISL_565173, EPI_ISL_565174, EPI_ISL_565175, EPI_ISL_565176, EPI_ISL_565177, EPI_ISL_565178, EPI_ISL_565179, EPI_ISL_565180, EPI_ISL_565181, EPI_ISL_565182, EPI_ISL_565183, EPI_ISL_565184, EPI_ISL_565185, EPI_ISL_565186, EPI_ISL_565187, EPI_ISL_565188, EPI_ISL_565189, EPI_ISL_565190, EPI_ISL_565191, EPI_ISL_565192, EPI_ISL_565193, EPI_ISL_565194, EPI_ISL_565195, EPI_ISL_565196, EPI_ISL_565197, EPI_ISL_565198, EPI_ISL_565199, EPI_ISL_565200, EPI_ISL_565201, EPI_ISL_565202, EPI_ISL_565203, EPI_ISL_565204, EPI_ISL_565205, EPI_ISL_565206, EPI_ISL_565207, EPI_ISL_565208, EPI_ISL_565212, EPI_ISL_565218, EPI_ISL_565219, EPI_ISL_565220, EPI_ISL_565221, EPI_ISL_565222 | | | |
| see above | Victorian Infectious Diseases Reference Laboratory (VIDRL) | VIDRL and MDU-PHL | Caly, L., Seemann, T., Sait, M., Schultz, M. B., Druce J., Sherry, N. |
| EPI_ISL_565223, EPI_ISL_565224, EPI_ISL_565225, EPI_ISL_565226, EPI_ISL_565228, EPI_ISL_565229, EPI_ISL_565231, EPI_ISL_565232, EPI_ISL_565233, EPI_ISL_565234, EPI_ISL_565235, EPI_ISL_565236, EPI_ISL_565237, EPI_ISL_565238, EPI_ISL_565239, EPI_ISL_565241, EPI_ISL_565244, EPI_ISL_565245, EPI_ISL_565246, EPI_ISL_565248, EPI_ISL_565249, EPI_ISL_565250, EPI_ISL_565251, EPI_ISL_565252, EPI_ISL_565253, EPI_ISL_565254, EPI_ISL_565255, EPI_ISL_565258, EPI_ISL_565259, EPI_ISL_565260, EPI_ISL_565261, EPI_ISL_565262, EPI_ISL_565263, EPI_ISL_565266, EPI_ISL_565267, EPI_ISL_565271, EPI_ISL_565281, EPI_ISL_565283, EPI_ISL_565284, EPI_ISL_565285, EPI_ISL_565286, EPI_ISL_565287, EPI_ISL_565288, EPI_ISL_565289, EPI_ISL_565290, EPI_ISL_565291, EPI_ISL_565292, EPI_ISL_565293, EPI_ISL_565294, EPI_ISL_565295, EPI_ISL_565296, EPI_ISL_565297, EPI_ISL_565298, EPI_ISL_565299, EPI_ISL_565300, EPI_ISL_565301, EPI_ISL_565302, EPI_ISL_565303, EPI_ISL_565304, EPI_ISL_565305, EPI_ISL_565306, EPI_ISL_565309, EPI_ISL_565315, EPI_ISL_565316, EPI_ISL_565317, EPI_ISL_565318, EPI_ISL_565319, EPI_ISL_565320, EPI_ISL_565321, EPI_ISL_565322, EPI_ISL_565323, EPI_ISL_565324, EPI_ISL_565325, EPI_ISL_565326, EPI_ISL_565327, EPI_ISL_565328, EPI_ISL_565329, EPI_ISL_565330, EPI_ISL_565331, EPI_ISL_565332, EPI_ISL_565333, EPI_ISL_565334, EPI_ISL_565335, EPI_ISL_565336, EPI_ISL_565337, EPI_ISL_565338, EPI_ISL_565339, EPI_ISL_565340, EPI_ISL_565341, EPI_ISL_565342, EPI_ISL_565343, EPI_ISL_565344, EPI_ISL_565345, EPI_ISL_565346, EPI_ISL_565347, EPI_ISL_565348, EPI_ISL_565349, EPI_ISL_565350, EPI_ISL_565353, EPI_ISL_565354, EPI_ISL_565355, EPI_ISL_565356, EPI_ISL_565357, EPI_ISL_565358, EPI_ISL_565359, EPI_ISL_565360, EPI_ISL_565361, EPI_ISL_565362, EPI_ISL_565363, EPI_ISL_565364, EPI_ISL_565365, EPI_ISL_565366, EPI_ISL_565367, EPI_ISL_565370, EPI_ISL_565514, EPI_ISL_565517, EPI_ISL_565518, EPI_ISL_565520, EPI_ISL_565621, EPI_ISL_565622, EPI_ISL_565626, EPI_ISL_565628, EPI_ISL_565630, EPI_ISL_565695, EPI_ISL_565696, EPI_ISL_565697, EPI_ISL_565699, EPI_ISL_565704, EPI_ISL_565707, EPI_ISL_565708, EPI_ISL_565709, EPI_ISL_565710, EPI_ISL_565711, EPI_ISL_565712, EPI_ISL_565713, EPI_ISL_565714, EPI_ISL_565718, EPI_ISL_565719, EPI_ISL_565720, EPI_ISL_565721, EPI_ISL_565723, EPI_ISL_565725, EPI_ISL_565743, EPI_ISL_565785, EPI_ISL_565786, EPI_ISL_565790, EPI_ISL_565791, EPI_ISL_565792, EPI_ISL_565793, EPI_ISL_565794, EPI_ISL_565795, EPI_ISL_565797, EPI_ISL_565798, EPI_ISL_565799, EPI_ISL_565801, EPI_ISL_565802, EPI_ISL_565803, EPI_ISL_565805, EPI_ISL_565806, EPI_ISL_565807, EPI_ISL_565808, EPI_ISL_565809, EPI_ISL_565810, EPI_ISL_565811, EPI_ISL_565814, EPI_ISL_565817, EPI_ISL_565819, EPI_ISL_565820, EPI_ISL_565821, EPI_ISL_565822 | | | |
| see above | Microbiological Diagnostic Unit - Public Health Laboratory (MDU-PHL) | MDU-PHL | Seemann, T., Schultz M. B., Sait, M., Sherry, N. |
| EPI_ISL_566035 | Michigan Department of Health and Human Services, Bureau of Laboratories | Michigan Department of Health and Human Services, Bureau of Laboratories | Blankenship HM, Riner D, Soehnlen MK |

| | | | |
|---|---|---|---|
| EPI_ISL_566058, EPI_ISL_566064 | Respiratory Virus Unit, Microbiology Services Colindale, Public Health England | Respiratory Virus Unit, Microbiology Services Colindale, Public Health England | PHE Covid Sequencing Team |
| EPI_ISL_568570, EPI_ISL_568571, EPI_ISL_568572 | Department of Infectious Diseases and Immunology, National Hospital Organization Nagoya Medical Center | Clinical Research Center, National Hospital Organization Nagoya Medical Center | Yoshihiro Nakata, Hirotaka Ode, Mai Kubota, Masakazu Matsuda, Kazuhiro Matsuoka, Nakasuji Miho, Mikiko Mori, Mayumi Imahashi, Yoshiyuki Yokomaku, Yasumasa Iwatani |
| EPI_ISL_568597, EPI_ISL_568598, EPI_ISL_568599, EPI_ISL_568600, EPI_ISL_568601, EPI_ISL_568602, EPI_ISL_568603, EPI_ISL_568604, EPI_ISL_568605, EPI_ISL_568606, EPI_ISL_568607, EPI_ISL_568608, EPI_ISL_568609, EPI_ISL_568610, EPI_ISL_568611, EPI_ISL_568613, EPI_ISL_568614, EPI_ISL_568615, EPI_ISL_568616, EPI_ISL_568617, EPI_ISL_568618, EPI_ISL_568619, EPI_ISL_568620, EPI_ISL_568621, EPI_ISL_568622, EPI_ISL_568623, EPI_ISL_568624, EPI_ISL_568625, EPI_ISL_568626, EPI_ISL_568627, EPI_ISL_568628 | | | |
| see above | Florida Bureau of Public Health Laboratories | Florida Bureau of Public Health Laboratories | Sarah Schmedes, Jason Blanton |
| EPI_ISL_569023, EPI_ISL_569024, EPI_ISL_569025, EPI_ISL_569026, EPI_ISL_569027, EPI_ISL_569028, EPI_ISL_569029, EPI_ISL_569033, EPI_ISL_569034, EPI_ISL_569035, EPI_ISL_569036, EPI_ISL_569037, EPI_ISL_569038, EPI_ISL_569039, EPI_ISL_569040 | | | |
| see above | MEPHI, Aix Marseille University | MEPHI, Aix Marseille University | Anthony LEVASSEUR |
| EPI_ISL_569617 | SD Urban Indian Health Pierre | South Dakota Public Health Laboratory | Matt Plumb, Jacob Garfin, Xiong Wang, and Chris Carlson |
| EPI_ISL_569618, EPI_ISL_569619 | Bethel Lutheran Home | South Dakota Public Health Laboratory | Matt Plumb, Jacob Garfin, Xiong Wang, and Chris Carlson |
| EPI_ISL_569620 | McCrossan Boys Ranch | South Dakota Public Health Laboratory | Matt Plumb, Jacob Garfin, Xiong Wang, and Chris Carlson |
| EPI_ISL_569791, EPI_ISL_569792, EPI_ISL_569793, EPI_ISL_569794, EPI_ISL_569795, EPI_ISL_569801, EPI_ISL_569808 | Omsk Research Institute of Natural Focal Infections | WHO National Influenza Centre Russian Federation | Artem Fadeev, Ekaterina Gradoboeva, Ekaterina Savkina, Daria Nashatyreva, Elena Poleshchuk, Aleksei Vasilenko, Valery Yakimenko, Andrey Komissarov |
| EPI_ISL_569980, EPI_ISL_569990, EPI_ISL_569991, EPI_ISL_569993, EPI_ISL_569994 | Unity Health Toronto | Ontario Institute for Cancer Research | Ramzi Fattouh, Larissa M. Matukas, Yan Chen,Mark Downing, Trina Otterman, Karel Boissinot, Wai Sum Siu, Zhi Cui, Le Luu, Samira Mubareka, TIBDN, Ilinca Lungu, Bernard Lam, Jeremy Johns, Paul Krzyzanowski, Richard de Borja, Felicia Vincelli, Philip Zuzarte, Jared T. Simpson |
| EPI_ISL_570803, EPI_ISL_570807, EPI_ISL_570835, EPI_ISL_570845, EPI_ISL_570981, EPI_ISL_570982 | UW Virology Lab | UW Virology Lab | Pavitra Roychoudhury, Hong Xie, Lasata Shrestha, Amin Addetia, Victoria M Rachleff, Meei-Li Huang, Keith R Jerome, Alexander Greninger |
| EPI_ISL_572246, EPI_ISL_572247, EPI_ISL_572250, EPI_ISL_572252, EPI_ISL_572253, EPI_ISL_572254, EPI_ISL_572255, EPI_ISL_572285, EPI_ISL_572286, EPI_ISL_572287, EPI_ISL_572288, EPI_ISL_572289, EPI_ISL_572290 | | | |
| see above | Virginia DCLS | Virginia DCLS | Virginia DCLS |
| EPI_ISL_572439, EPI_ISL_572664, EPI_ISL_572701, EPI_ISL_572744, EPI_ISL_573334, EPI_ISL_573335, EPI_ISL_573336, EPI_ISL_573337, EPI_ISL_573338, EPI_ISL_573339, EPI_ISL_573340, EPI_ISL_573341, EPI_ISL_573342, EPI_ISL_573343, EPI_ISL_573372 | | | |
| see above | Northumbria University / South Tees Hospitals NHS Foundation Trust / North Cumbria Integrated Care NHS Foundation Trust / North Tees and Hartlepool NHS Foundation Trust / Newcastle Hospitals NHS Foundation Trust | COVID-19 Genomics UK (COG-UK) Consortium | Darren L Smith,Andrew Nelson,Matthew Bashton,Greg R Young,Joshua Loh,John Allan,Mohammad A Tariq,Giles S Holt,Gary Black,Wen C Yew,Lynn Dover,Paul Baker,Steve Liggett,Sarah Essex,Jane Greenaway,Debra Padgett,Clive Graham,Garren Scott,Edward Barton,Emma Swindells,Brendan Payne,Jennifer Collins,Yusri Taha,Gary Eltringham |
| EPI_ISL_574326, EPI_ISL_574327 | LSUHS Emerging Viral Threat Laboratory | Microbial Genome Sequencing Center | Jeremy P. Kamil, Rona S. Scott, Maarten Van Diest, Malgorzata Bienkowska-Haba, Katarzyna Zwolinska, Andrew D. Yurochko, Christopher G. Kevil, Martin J. Sapp, Daniel J. Snyder, Vaughn S. Cooper, John A. Vanchiere |
| EPI_ISL_576145 | RSA Universitas Gadjah Mada | Genetics Working Group (Pokja Genetik) Faculty of Medicine, Public Health and Nursing Universitas Gadjah Mada (FK-KMK UGM); Disease Investigation Center Wates Ministry of Agriculture Indonesia; Department of Microbiology FK-KMK UGM; Laboratorium Diagnostik Yayasan Tahija World Mosquito Program (WMP) Yogyakarta Center for Tropical Medicine FK-KMK UGM; Integrated Research Center FK-KMK UGM; Department of Computer Science and Electronics FMIPA UGM | Gunadi, Hendra Wibawa, Marcellus, Mohamad S. Hakim, Edwin W. Daniwijaya, Ludhang P. Rizki, Endah Supriyati, Eggi Arguni, Titik Nuryastuti, Tri Wibawa, Dwi AA Nugrahaningsih, Afiahayati, Siswanto, Kristy Iskandar, Nungki Anggorowati, William Widitjiarso, Fadil Fahri |
| EPI_ISL_576146, EPI_ISL_576147 | Department of Respiratory & Other Viral Infections of L.V. Gromashevsky Institute of Epidemiology & Infectious Diseases NAMS of Ukraine | Department of Respiratory & Other Viral Infections of L.V. Gromashevsky Institute of Epidemiology & Infectious Diseases NAMS of Ukraine, JSC "Farmak" | Alla Mironenko, Ihor Kravchuk, Liudmyla Bolotova, Larysa Radchenko, Nataliia Teteriuk |
| EPI_ISL_576385 | Tirta Medical Center Angsana, Banjarmasin Kalsel | National Institute of Health Research and Development | Pawestri,HA;Subangkit;Puspa,KD;Nugraha,AA;Ikawati,HD;Pangesti,KNA;Soekarso,T;Paisal;Setiawaty,V |
| EPI_ISL_576386 | National Institute of Health Research and Development | National Institute of Health Research and Development | Pawestri,HA;Subangkit;Puspa,KD;Nugraha,AA;Ikawati,HD;Pangesti, KNA;Soekarso,T;Susilarini,NK;Hariastuti,NI;Nikmah,UA;Mursinah;Febriyani,A;Herman,R;Susanti,N;Herna;Febriyanti,T; Nurhadi,M; Paisal;Ramadhany,R;Agustiningsih;Kurniawati,J;Kipuw,NL;Muna,F;Indalau,IL;Adam,K;Wibowo,HA;Rizki,A;Puspandari,N;Setiawaty,V |
| EPI_ISL_577624, EPI_ISL_577625, EPI_ISL_577626 | The National Institute of Public Health | State Veterinary Institute Prague | Nagy,A;Jirincova,H;Novakova,L;Trnka,D;Vecerova,J |
| EPI_ISL_577729 | NIV Influenza | NIV Influenza | Potdar V |
| EPI_ISL_577843 | Dutch COVID-19 response team | Erasmus Medical Center | Bas Oude Munnink, Reina Sikkema, David Nieuwenhuijse, Irina Chestakova, Anne van der Linden, Marjan Boter, Emmanuelle Munger, Corine GeurtsvanKessel, Annemiek van der Eijk, Richard Molenkamp, Marion Koopmans, on behalf of the Dutch national COVID-19 response team. |
| EPI_ISL_578570, EPI_ISL_578571, EPI_ISL_578572, EPI_ISL_578573, EPI_ISL_578574, EPI_ISL_578575, EPI_ISL_578675, EPI_ISL_578676, EPI_ISL_578677 | Wisconsin State Laboratory of Hygiene Communicable Disease Division | Wisconsin State Laboratory of Hygiene Communicable Disease Division | Kelsey R. Florek, Abigail C. Shockey |
| EPI_ISL_578824, EPI_ISL_578825, EPI_ISL_578831, EPI_ISL_578847, EPI_ISL_578849, EPI_ISL_578850, EPI_ISL_578854, EPI_ISL_578861, EPI_ISL_578889, EPI_ISL_578891, EPI_ISL_578893, EPI_ISL_578895, EPI_ISL_578896, EPI_ISL_578897, EPI_ISL_578898, EPI_ISL_578900, EPI_ISL_578901, EPI_ISL_578902, EPI_ISL_578903, EPI_ISL_578904, EPI_ISL_578905, EPI_ISL_578906, EPI_ISL_578912, EPI_ISL_578913 | | | |
| see above | LSUHS Emerging Viral Threat Laboratory | Microbial Genome Sequencing Center | Jeremy P. Kamil, Rona S. Scott, Maarten Van Diest, Malgorzata Bienkowska-Haba, Katarzyna Zwolinska, Andrew D. Yurochko, Christopher G. Kevil, Martin J. Sapp, Daniel J. Snyder, Vaughn S. Cooper, John A. Vanchiere |
| EPI_ISL_581388, EPI_ISL_581389 | Lighthouse Lab in Milton Keynes | Wellcome Sanger Institute for the COVID-19 Genomics UK (COG-UK) consortium | The Lighthouse Lab in Milton Keynes and Alex Alderton, Roberto Amato, Sonia Goncalves, Ewan Harrison, David K. Jackson, Ian Johnston, Dominic Kwiatkowski, Cordelia Langford, John Sillitoe on behalf of the Wellcome Sanger Institute COVID-19 Surveillance Team |
| EPI_ISL_581508, EPI_ISL_581509, EPI_ISL_581510, EPI_ISL_581511, EPI_ISL_581512, EPI_ISL_581513, EPI_ISL_581514, EPI_ISL_581515, EPI_ISL_581516, EPI_ISL_581558, EPI_ISL_581559, EPI_ISL_581560, EPI_ISL_581561, EPI_ISL_581562, EPI_ISL_581563, EPI_ISL_581564, EPI_ISL_581565, EPI_ISL_581566, EPI_ISL_581567, EPI_ISL_581568, EPI_ISL_581569, EPI_ISL_581570, EPI_ISL_581571, EPI_ISL_581572, EPI_ISL_581573, EPI_ISL_581574 | | | |
| see above | Virginia DCLS | Virginia DCLS | Virginia DCLS |
| EPI_ISL_581930, EPI_ISL_581931, EPI_ISL_581932 | University Hospital Basel, Clinical Virology | University Hospital Basel, Clinical Bacteriology | Madlen Stange, Alfredo Mari, Tim Roloff, Helena MB Seth-Smith, Michael Schweitzer, Myrta Brunner, Karoline Leuzinger, Kirstine K. Soegaard, Alexander Gensch, Sarah Tschudin-Sutter, Simon Fuchs, Julia Bielicki, Hans Pargger, Martin Siegemund, Christian Nickel, Roland Bingisser, Michael Osthoff, Stefano Bassetti, Rita Schneider-Sliwa, Manuel Battegay, Hans Hirsch, Adrian Egli |
| EPI_ISL_582220, EPI_ISL_582222, EPI_ISL_582223, EPI_ISL_582228, | Wyoming Public Health Laboratory | Center for Global Health, University of New Mexico Health Sciences Center | Daryl Domman, Kurt Schwalm, Rob Christensen, Wanda Manley, Cari Sloma, Noah Hull, Darrell Dinwiddie |

| | | | |
|---|---|---|---|
| EPI_ISL_582229, EPI_ISL_582233, EPI_ISL_582238, EPI_ISL_582239 | | | |
| EPI_ISL_582359, EPI_ISL_582360, EPI_ISL_582361, EPI_ISL_582366, EPI_ISL_582369, EPI_ISL_582370, EPI_ISL_582371, EPI_ISL_582372, EPI_ISL_582377, EPI_ISL_582378, EPI_ISL_582380, EPI_ISL_582385, EPI_ISL_582386, EPI_ISL_582393, EPI_ISL_582399, EPI_ISL_582405, EPI_ISL_582406, EPI_ISL_582412, EPI_ISL_582414, EPI_ISL_582438, EPI_ISL_582479, EPI_ISL_582480, EPI_ISL_582484, EPI_ISL_582485, EPI_ISL_582493, EPI_ISL_582496, EPI_ISL_582497, EPI_ISL_582498, EPI_ISL_582499 | | | |
| see above | Cadham Provincial Laboratory | National Microbiology Laboratory (NML) | Anna Majer, Shari Tyson, Grace Seo, Philip Mabon, Elsie Grudeski, Rhiannon Huzarewich, Russell Mandes, Anneliese Landgraff, Jennifer Tanner, Natalie Knox, Morag Graham, Gary Van Domselaar, Paul Van Caeseele, Jared Bullard, David Alexander, Kerry Dust, Nathalie Bastien, Yan Li, Timothy Booth, Darian Hole, Madison Chapel, CanCOGeN's metadata curation team, Public Health Agency of Canada CanCOGeN team |
| EPI_ISL_582513 | Department of Respiratory and other Viral Infections of L.V.Gromashevsky Institute of Epidemiology & Infectious Diseases NAMS of Ukrain | Department of Respiratory and other Viral Infections of L.V.Gromashevsky Institute of Epidemiology & Infectious Diseases NAMS of Ukrain, JSC "Farmak" | Alla Mironenko, Andriy Goy, Ihor Kravchuk, Ludmyla Bolotova, Larysa Radchenko, Nataliia Teteriuk |
| EPI_ISL_582531, EPI_ISL_582532, EPI_ISL_582533, EPI_ISL_582534 | Veterinary Specialized Institute "Kraljevo", Serbia | Veterinary Specialized Institute "Kraljevo", Serbia | Vidanovic,D., Tesovic,B., Knezevic,A., Jovanovic,T., Jankovic,M., Sekler,M., Banovic Djeri,B., Petrovic,T., Volkening,J., Afonso,C. |
| EPI_ISL_582969, EPI_ISL_582970, EPI_ISL_582971, EPI_ISL_582972, EPI_ISL_582973 | San Luis Obispo Public Health Department | Chan-Zuckerberg Biohub | CZB Cliahub Consortium |
| EPI_ISL_583088, EPI_ISL_583089, EPI_ISL_583090, EPI_ISL_583091, EPI_ISL_583092, EPI_ISL_583093, EPI_ISL_583094, EPI_ISL_583095, EPI_ISL_583096, EPI_ISL_583097, EPI_ISL_583098, EPI_ISL_583099, EPI_ISL_583100, EPI_ISL_583101 | | | |
| see above | Humboldt County Public Health Laboratory | Chan-Zuckerberg Biohub | CZB Cliahub Consortium |
| EPI_ISL_583216 | UCSF Clinical Microbiology Laboratory | Chan-Zuckerberg Biohub | CZB Cliahub Consortium |
| EPI_ISL_584078 | The National Institute of Public Health | State Veterinary Institute Prague | Nagy,A;Jirincova,H;Novakova,L;Trnka,D;Vecerova,J |
| EPI_ISL_584621 | Liverpool Clinical Laboratories | COVID-19 Genomics UK (COG-UK) Consortium | Sam Haldenby, Anita Lucaci, Steve Paterson, Julian Hiscox, Alistair Darby, M Almsaud, A Alrezaihi, Muhannad Alruwaili, Stuart D Armstrong, Jones Benjamin, Eleanor G Bentley, Anu Chawla, Jordan J Clark, Angela Cowell, Richard Eccles, Isabel García-Dorival, Matthew Gemmell, Alessandro Gerada, PKF Gilmore, Richard Gregory, Ximeng Han, Catherine Hartley, Margaret Hughes, Miren Iturriza-Gomara, James Johnson, L Luu, Jenifer Manson, Charlotte Nelson, Elaine O'Toole, Cassie Olateju, Rebekah Penrice-Randal , Lucille Rainbow, N.P Randle, Trevor Ian Robinson, Parul Sharma, Ghada T Shawli, James P Stewart, Neil Swainston, Ecaterina Vamos, Joanne Watts, Mark Whitehead |
| EPI_ISL_590924, EPI_ISL_590925, EPI_ISL_590926, EPI_ISL_590939, EPI_ISL_590940, EPI_ISL_590947, EPI_ISL_590949, EPI_ISL_590950 | University Hospital of Northern Norway, Department for Microbiology and Infectious Disease Control | Norwegian Institute of Public Health, Department of Virology | Kathrine Stene-Johansen, Kamilla Heddeland Instefjord, Hilde Elshaug, Rasmus Riis Kopperud, Hilde Vollan, Karoline Bragstad, Olav Hungnes |
| EPI_ISL_591504 | South Eastern Area Laboratory Services (SEALS) | NSW Health Pathology - Institute of Clinical Pathology and Medical Research; Westmead Hospital; University of Sydney | CIDM-PH et al. |
| EPI_ISL_591518 | The Children's Hospital at Westmead | NSW Health Pathology - Institute of Clinical Pathology and Medical Research; Westmead Hospital; University of Sydney | CIDM-PH et al. |
| EPI_ISL_591527 | Medicina Norte U Chile - Servicio Medico Legal | Center for Mathematical Modeling and Center for Genome Regulation. Santiago, Chile | Gaggero A, Valiente F, Gaete A, Travisany D, Palma R, Urra C, Varas M, Allende ML, Maass A, González M, Ferres M. |
| EPI_ISL_591551, EPI_ISL_591552, EPI_ISL_591553, EPI_ISL_591554, EPI_ISL_591555 | Microbiological Diagnostic Unit - Public Health Laboratory (MDU-PHL) | MDU-PHL | Seemann T., Schultz, M. B., Sait, M., Sherry, N. |
| EPI_ISL_591558 | Victorian Infectious Diseases Reference Laboratory (VIDRL) | VIDRL and MDU-PHL | Caly L., Seemann T., Sait, M., Schultz, M. B., Druce J., Sherry, N. |
| EPI_ISL_591560 | Microbiological Diagnostic Unit - Public Health Laboratory (MDU-PHL) | MDU-PHL | Seemann T., Schultz, M. B., Sait, M., Sherry, N. |
| EPI_ISL_591562 | Victorian Infectious Diseases Reference Laboratory (VIDRL) | VIDRL and MDU-PHL | Caly L., Seemann T., Sait, M., Schultz, M. B., Druce J., Sherry, N. |
| EPI_ISL_591564, EPI_ISL_591566, EPI_ISL_591567, EPI_ISL_591570, EPI_ISL_591572, EPI_ISL_591575, EPI_ISL_591576, EPI_ISL_591579, EPI_ISL_591581, EPI_ISL_591583 | Microbiological Diagnostic Unit - Public Health Laboratory (MDU-PHL) | MDU-PHL | Seemann T., Schultz, M. B., Sait, M., Sherry, N. |
| EPI_ISL_591584 | Victorian Infectious Diseases Reference Laboratory (VIDRL) | VIDRL and MDU-PHL | Caly L., Seemann T., Sait, M., Schultz, M. B., Druce J., Sherry, N. |
| EPI_ISL_591586 | Microbiological Diagnostic Unit - Public Health Laboratory (MDU-PHL) | MDU-PHL | Seemann T., Schultz, M. B., Sait, M., Sherry, N. |
| EPI_ISL_591591 | Victorian Infectious Diseases Reference Laboratory (VIDRL) | VIDRL and MDU-PHL | Caly L., Seemann T., Sait, M., Schultz, M. B., Druce J., Sherry, N. |
| EPI_ISL_591594, EPI_ISL_591597, EPI_ISL_591598, EPI_ISL_591600, EPI_ISL_591601, EPI_ISL_591602, EPI_ISL_591605, EPI_ISL_591606, EPI_ISL_591608, EPI_ISL_591610, EPI_ISL_591611, EPI_ISL_591614, EPI_ISL_591616 | | | |
| see above | Microbiological Diagnostic Unit - Public Health Laboratory (MDU-PHL) | MDU-PHL | Seemann T., Schultz, M. B., Sait, M., Sherry, N. |
| EPI_ISL_591621 | Victorian Infectious Diseases Reference Laboratory (VIDRL) | VIDRL and MDU-PHL | Caly L., Seemann T., Sait, M., Schultz, M. B., Druce J., Sherry, N. |
| EPI_ISL_591622, EPI_ISL_591624, EPI_ISL_591625, EPI_ISL_591634, EPI_ISL_591635 | Microbiological Diagnostic Unit - Public Health Laboratory (MDU-PHL) | MDU-PHL | Seemann T., Schultz, M. B., Sait, M., Sherry, N. |
| EPI_ISL_591643 | Victorian Infectious Diseases Reference Laboratory (VIDRL) | VIDRL and MDU-PHL | Caly L., Seemann T., Sait, M., Schultz, M. B., Druce J., Sherry, N. |
| EPI_ISL_591647, EPI_ISL_591648, EPI_ISL_591649, EPI_ISL_591650 | Microbiological Diagnostic Unit - Public Health Laboratory (MDU-PHL) | MDU-PHL | Seemann T., Schultz, M. B., Sait, M., Sherry, N. |
| EPI_ISL_591654, EPI_ISL_591656 | Victorian Infectious Diseases Reference Laboratory (VIDRL) | VIDRL and MDU-PHL | Caly L., Seemann T., Sait, M., Schultz, M. B., Druce J., Sherry, N. |
| EPI_ISL_591661, EPI_ISL_591663 | Microbiological Diagnostic Unit - Public Health Laboratory (MDU-PHL) | MDU-PHL | Seemann T., Schultz, M. B., Sait, M., Sherry, N. |
| EPI_ISL_591666 | Victorian Infectious Diseases Reference Laboratory (VIDRL) | VIDRL and MDU-PHL | Caly L., Seemann T., Sait, M., Schultz, M. B., Druce J., Sherry, N. |
| EPI_ISL_591671, EPI_ISL_591673, EPI_ISL_591674, EPI_ISL_591682, EPI_ISL_591685, EPI_ISL_591687 | Microbiological Diagnostic Unit - Public Health Laboratory (MDU-PHL) | MDU-PHL | Seemann T., Schultz, M. B., Sait, M., Sherry, N. |
| EPI_ISL_591688 | Victorian Infectious Diseases Reference Laboratory (VIDRL) | VIDRL and MDU-PHL | Caly L., Seemann T., Sait, M., Schultz, M. B., Druce J., Sherry, N. |
| EPI_ISL_591690, EPI_ISL_591692, EPI_ISL_591693, EPI_ISL_591694, EPI_ISL_591695, EPI_ISL_591696, | Microbiological Diagnostic Unit - Public Health Laboratory (MDU-PHL) | MDU-PHL | Seemann T., Schultz, M. B., Sait, M., Sherry, N. |

| | | | |
|---|---|---|---|
| EPI_ISL_591699, EPI_ISL_591701, EPI_ISL_591705 | | | |
| EPI_ISL_591707 | Victorian Infectious Diseases Reference Laboratory (VIDRL) | VIDRL and MDU-PHL | Caly L., Seemann T., Sait, M., Schultz, M. B., Druce J., Sherry, N. |
| EPI_ISL_591712, EPI_ISL_591713, EPI_ISL_591716, EPI_ISL_591719, EPI_ISL_591720, EPI_ISL_591724, EPI_ISL_591731, EPI_ISL_591735, EPI_ISL_591741, EPI_ISL_591742, EPI_ISL_591745, EPI_ISL_591747, EPI_ISL_591749, EPI_ISL_591750, EPI_ISL_591753, EPI_ISL_591756, EPI_ISL_591762, EPI_ISL_591768 | | | |
| see above | Microbiological Diagnostic Unit - Public Health Laboratory (MDU-PHL) | MDU-PHL | Seemann T., Schultz, M. B., Sait, M., Sherry, N. |
| EPI_ISL_591778, EPI_ISL_591783 | Victorian Infectious Diseases Reference Laboratory (VIDRL) | VIDRL and MDU-PHL | Caly L., Seemann T., Sait, M., Schultz, M. B., Druce J., Sherry, N. |
| EPI_ISL_591787, EPI_ISL_591788, EPI_ISL_591793, EPI_ISL_591796, EPI_ISL_591797, EPI_ISL_591798, EPI_ISL_591804, EPI_ISL_591805, EPI_ISL_591806, EPI_ISL_591808, EPI_ISL_591810, EPI_ISL_591812, EPI_ISL_591826, EPI_ISL_591827, EPI_ISL_591828, EPI_ISL_591829, EPI_ISL_591830, EPI_ISL_591831, EPI_ISL_591832, EPI_ISL_591833, EPI_ISL_591834, EPI_ISL_591835, EPI_ISL_591836, EPI_ISL_591837, EPI_ISL_591838, EPI_ISL_591839, EPI_ISL_591840, EPI_ISL_591841, EPI_ISL_591842, EPI_ISL_591843, EPI_ISL_591844, EPI_ISL_591845, EPI_ISL_591846, EPI_ISL_591847, EPI_ISL_591848 | | | |
| see above | Microbiological Diagnostic Unit - Public Health Laboratory (MDU-PHL) | MDU-PHL | Seemann T., Schultz, M. B., Sait, M., Sherry, N. |
| EPI_ISL_591849, EPI_ISL_591860, EPI_ISL_591871, EPI_ISL_591896, EPI_ISL_591904 | Victorian Infectious Diseases Reference Laboratory (VIDRL) | VIDRL and MDU-PHL | Caly L., Seemann T., Sait, M., Schultz, M. B., Druce J., Sherry, N. |
| EPI_ISL_591906 | Microbiological Diagnostic Unit - Public Health Laboratory (MDU-PHL) | MDU-PHL | Seemann T., Schultz, M. B., Sait, M., Sherry, N. |
| EPI_ISL_591907, EPI_ISL_591908, EPI_ISL_591909 | Victorian Infectious Diseases Reference Laboratory (VIDRL) | VIDRL and MDU-PHL | Caly L., Seemann T., Sait, M., Schultz, M. B., Druce J., Sherry, N. |
| EPI_ISL_591910, EPI_ISL_591911 | Microbiological Diagnostic Unit - Public Health Laboratory (MDU-PHL) | MDU-PHL | Seemann T., Schultz, M. B., Sait, M., Sherry, N. |
| EPI_ISL_591912 | Victorian Infectious Diseases Reference Laboratory (VIDRL) | VIDRL and MDU-PHL | Caly L., Seemann T., Sait, M., Schultz, M. B., Druce J., Sherry, N. |
| EPI_ISL_591913, EPI_ISL_591915 | Microbiological Diagnostic Unit - Public Health Laboratory (MDU-PHL) | MDU-PHL | Seemann T., Schultz, M. B., Sait, M., Sherry, N. |
| EPI_ISL_591917 | Victorian Infectious Diseases Reference Laboratory (VIDRL) | VIDRL and MDU-PHL | Caly L., Seemann T., Sait, M., Schultz, M. B., Druce J., Sherry, N. |
| EPI_ISL_591918 | Microbiological Diagnostic Unit - Public Health Laboratory (MDU-PHL) | MDU-PHL | Seemann T., Schultz, M. B., Sait, M., Sherry, N. |
| EPI_ISL_591919 | Victorian Infectious Diseases Reference Laboratory (VIDRL) | VIDRL and MDU-PHL | Caly L., Seemann T., Sait, M., Schultz, M. B., Druce J., Sherry, N. |
| EPI_ISL_591920, EPI_ISL_591923, EPI_ISL_591924 | Microbiological Diagnostic Unit - Public Health Laboratory (MDU-PHL) | MDU-PHL | Seemann T., Schultz, M. B., Sait, M., Sherry, N. |
| EPI_ISL_591925 | Victorian Infectious Diseases Reference Laboratory (VIDRL) | VIDRL and MDU-PHL | Caly L., Seemann T., Sait, M., Schultz, M. B., Druce J., Sherry, N. |
| EPI_ISL_591926, EPI_ISL_591927 | Microbiological Diagnostic Unit - Public Health Laboratory (MDU-PHL) | MDU-PHL | Seemann T., Schultz, M. B., Sait, M., Sherry, N. |
| EPI_ISL_591928 | Victorian Infectious Diseases Reference Laboratory (VIDRL) | VIDRL and MDU-PHL | Caly L., Seemann T., Sait, M., Schultz, M. B., Druce J., Sherry, N. |
| EPI_ISL_591929, EPI_ISL_591930, EPI_ISL_591931 | Microbiological Diagnostic Unit - Public Health Laboratory (MDU-PHL) | MDU-PHL | Seemann T., Schultz, M. B., Sait, M., Sherry, N. |
| EPI_ISL_591932 | Victorian Infectious Diseases Reference Laboratory (VIDRL) | VIDRL and MDU-PHL | Caly L., Seemann T., Sait, M., Schultz, M. B., Druce J., Sherry, N. |
| EPI_ISL_591933, EPI_ISL_591936, EPI_ISL_591937, EPI_ISL_591938, EPI_ISL_591939 | Microbiological Diagnostic Unit - Public Health Laboratory (MDU-PHL) | MDU-PHL | Seemann T., Schultz, M. B., Sait, M., Sherry, N. |
| EPI_ISL_591956 | Victorian Infectious Diseases Reference Laboratory (VIDRL) | VIDRL and MDU-PHL | Caly L., Seemann T., Sait, M., Schultz, M. B., Druce J., Sherry, N. |
| EPI_ISL_591961, EPI_ISL_591962, EPI_ISL_591963, EPI_ISL_591964, EPI_ISL_591965, EPI_ISL_591966, EPI_ISL_591967, EPI_ISL_591969, EPI_ISL_591970, EPI_ISL_591971, EPI_ISL_591972, EPI_ISL_591973, EPI_ISL_591974, EPI_ISL_591975, EPI_ISL_591976, EPI_ISL_591977, EPI_ISL_591978, EPI_ISL_591979, EPI_ISL_591980, EPI_ISL_591981, EPI_ISL_591983, EPI_ISL_591984, EPI_ISL_591985, EPI_ISL_591986, EPI_ISL_591987, EPI_ISL_591988, EPI_ISL_591989, EPI_ISL_591990, EPI_ISL_591991, EPI_ISL_591992, EPI_ISL_591993, EPI_ISL_591994, EPI_ISL_591995, EPI_ISL_591996, EPI_ISL_591997, EPI_ISL_591998, EPI_ISL_592000, EPI_ISL_592001, EPI_ISL_592002, EPI_ISL_592003, EPI_ISL_592004, EPI_ISL_592005, EPI_ISL_592006, EPI_ISL_592007, EPI_ISL_592008, EPI_ISL_592009, EPI_ISL_592010, EPI_ISL_592011, EPI_ISL_592012, EPI_ISL_592014, EPI_ISL_592015, EPI_ISL_592016, EPI_ISL_592017, EPI_ISL_592018, EPI_ISL_592019, EPI_ISL_592020, EPI_ISL_592021, EPI_ISL_592022, EPI_ISL_592023, EPI_ISL_592024, EPI_ISL_592025, EPI_ISL_592026, EPI_ISL_592027, EPI_ISL_592029, EPI_ISL_592030, EPI_ISL_592031, EPI_ISL_592032, EPI_ISL_592033, EPI_ISL_592034, EPI_ISL_592035, EPI_ISL_592037, EPI_ISL_592038, EPI_ISL_592039, EPI_ISL_592040, EPI_ISL_592041, EPI_ISL_592042, EPI_ISL_592043, EPI_ISL_592044, EPI_ISL_592045, EPI_ISL_592046, EPI_ISL_592047, EPI_ISL_592048, EPI_ISL_592049, EPI_ISL_592051, EPI_ISL_592052, EPI_ISL_592053, EPI_ISL_592054, EPI_ISL_592055, EPI_ISL_592056, EPI_ISL_592057, EPI_ISL_592058, EPI_ISL_592059, EPI_ISL_592060, EPI_ISL_592061, EPI_ISL_592062, EPI_ISL_592063, EPI_ISL_592064, EPI_ISL_592065, EPI_ISL_592066, EPI_ISL_592067, EPI_ISL_592068, EPI_ISL_592070, EPI_ISL_592071, EPI_ISL_592072, EPI_ISL_592073, EPI_ISL_592074, EPI_ISL_592075, EPI_ISL_592076, EPI_ISL_592077, EPI_ISL_592078, EPI_ISL_592079, EPI_ISL_592080, EPI_ISL_592081, EPI_ISL_592082, EPI_ISL_592083, EPI_ISL_592084, EPI_ISL_592085, EPI_ISL_592086, EPI_ISL_592087, EPI_ISL_592088, EPI_ISL_592089, EPI_ISL_592090, EPI_ISL_592091, EPI_ISL_592092, EPI_ISL_592093, EPI_ISL_592094, EPI_ISL_592095, EPI_ISL_592096, EPI_ISL_592097, EPI_ISL_592098, EPI_ISL_592099, EPI_ISL_592100, EPI_ISL_592101, EPI_ISL_592102, EPI_ISL_592103, EPI_ISL_592106, EPI_ISL_592107, EPI_ISL_592108, EPI_ISL_592109, EPI_ISL_592110, EPI_ISL_592111, EPI_ISL_592112, EPI_ISL_592113, EPI_ISL_592114, EPI_ISL_592115, EPI_ISL_592116, EPI_ISL_592117, EPI_ISL_592118, EPI_ISL_592119, EPI_ISL_592120, EPI_ISL_592121, EPI_ISL_592122, EPI_ISL_592123, EPI_ISL_592125, EPI_ISL_592126, EPI_ISL_592127, EPI_ISL_592129, EPI_ISL_592130, EPI_ISL_592131, EPI_ISL_592132, EPI_ISL_592133, EPI_ISL_592134 | | | |
| see above | Microbiological Diagnostic Unit - Public Health Laboratory (MDU-PHL) | MDU-PHL | Seemann T., Schultz, M. B., Sait, M., Sherry, N. |
| EPI_ISL_592139, EPI_ISL_592146, EPI_ISL_592147, EPI_ISL_592150, EPI_ISL_592164, EPI_ISL_592165, EPI_ISL_592166, EPI_ISL_592186 | Victorian Infectious Diseases Reference Laboratory (VIDRL) | VIDRL and MDU-PHL | Caly L., Seemann T., Sait, M., Schultz, M. B., Druce J., Sherry, N. |
| EPI_ISL_592196, EPI_ISL_592197, EPI_ISL_592198, EPI_ISL_592199, EPI_ISL_592200, EPI_ISL_592201, EPI_ISL_592202, EPI_ISL_592203, EPI_ISL_592204, EPI_ISL_592205, EPI_ISL_592206, EPI_ISL_592207, EPI_ISL_592208, EPI_ISL_592209, EPI_ISL_592211, EPI_ISL_592212, EPI_ISL_592213, EPI_ISL_592214, EPI_ISL_592215, EPI_ISL_592216, EPI_ISL_592217, EPI_ISL_592218, EPI_ISL_592219, EPI_ISL_592220, EPI_ISL_592221, EPI_ISL_592222, EPI_ISL_592223, EPI_ISL_592224, EPI_ISL_592225, EPI_ISL_592226, EPI_ISL_592227, EPI_ISL_592228, EPI_ISL_592229, EPI_ISL_592230, EPI_ISL_592231, EPI_ISL_592232, EPI_ISL_592233, EPI_ISL_592234, EPI_ISL_592271, EPI_ISL_592273, EPI_ISL_592276, EPI_ISL_592277, EPI_ISL_592279, EPI_ISL_592281, EPI_ISL_592284, EPI_ISL_592286, EPI_ISL_592287, EPI_ISL_592288, EPI_ISL_592291, EPI_ISL_592353, EPI_ISL_592361, EPI_ISL_592363, EPI_ISL_592367, EPI_ISL_592368, EPI_ISL_592369, EPI_ISL_592371, EPI_ISL_592376, EPI_ISL_592393, EPI_ISL_592394, EPI_ISL_592398, EPI_ISL_592405, EPI_ISL_592406, EPI_ISL_592407, EPI_ISL_592411, EPI_ISL_592414, EPI_ISL_592423, EPI_ISL_592426, EPI_ISL_592449, EPI_ISL_592452, EPI_ISL_592455, EPI_ISL_592456, EPI_ISL_592461, EPI_ISL_592465, EPI_ISL_592466 | | | |
| see above | Microbiological Diagnostic Unit - Public Health Laboratory (MDU-PHL) | MDU-PHL | Seemann T., Schultz, M. B., Sait, M., Sherry, N. |
| EPI_ISL_592495 | Victorian Infectious Diseases Reference Laboratory (VIDRL) | VIDRL and MDU-PHL | Caly L., Seemann T., Sait, M., Schultz, M. B., Druce J., Sherry, N. |
| EPI_ISL_592538, EPI_ISL_592547, EPI_ISL_592553, EPI_ISL_592559, EPI_ISL_592561, EPI_ISL_592566, EPI_ISL_592573, EPI_ISL_592579, EPI_ISL_592583, EPI_ISL_592587, EPI_ISL_592611, EPI_ISL_592612, EPI_ISL_592622, EPI_ISL_592628, EPI_ISL_592632, EPI_ISL_592640, EPI_ISL_592648, EPI_ISL_592671, EPI_ISL_592715, EPI_ISL_592719, EPI_ISL_592721, EPI_ISL_592733, EPI_ISL_592765, EPI_ISL_592766, EPI_ISL_592773, EPI_ISL_592774, EPI_ISL_592775, EPI_ISL_592776, EPI_ISL_592794, EPI_ISL_592805, EPI_ISL_592811, EPI_ISL_592815, EPI_ISL_592852, EPI_ISL_592914, EPI_ISL_592917, EPI_ISL_592920, EPI_ISL_592921, EPI_ISL_592922, EPI_ISL_592924, EPI_ISL_592925, EPI_ISL_592929, EPI_ISL_592930, EPI_ISL_592931, EPI_ISL_592933, EPI_ISL_592941, EPI_ISL_592942, EPI_ISL_592948, EPI_ISL_592951, EPI_ISL_592952, EPI_ISL_592953, EPI_ISL_592954, EPI_ISL_592957, EPI_ISL_592963, EPI_ISL_592966, EPI_ISL_593022, EPI_ISL_593034, EPI_ISL_593038, EPI_ISL_593040, EPI_ISL_593044, EPI_ISL_593102, EPI_ISL_593103, EPI_ISL_593104, EPI_ISL_593106, EPI_ISL_593107, EPI_ISL_593109, EPI_ISL_593110, EPI_ISL_593111, EPI_ISL_593114, EPI_ISL_593115, EPI_ISL_593116, EPI_ISL_593117, EPI_ISL_593119, EPI_ISL_593120, EPI_ISL_593121, EPI_ISL_593122, EPI_ISL_593123, EPI_ISL_593258, EPI_ISL_593260, EPI_ISL_593265, EPI_ISL_593266, EPI_ISL_593276, EPI_ISL_593278, EPI_ISL_593285, EPI_ISL_593287, EPI_ISL_593289, EPI_ISL_593292, EPI_ISL_593293, EPI_ISL_593295, EPI_ISL_593297, EPI_ISL_593298, EPI_ISL_593299, EPI_ISL_593305, EPI_ISL_593309, EPI_ISL_593310, EPI_ISL_593313, EPI_ISL_593315, EPI_ISL_593317, EPI_ISL_593320, EPI_ISL_593321, EPI_ISL_593328, EPI_ISL_593332, EPI_ISL_593333, EPI_ISL_593344, EPI_ISL_593345, EPI_ISL_593346, EPI_ISL_593360, EPI_ISL_593361, EPI_ISL_593362, EPI_ISL_593363, EPI_ISL_593364, EPI_ISL_593365, EPI_ISL_593366, EPI_ISL_593367, EPI_ISL_593368, EPI_ISL_593370, EPI_ISL_593374, EPI_ISL_593378, EPI_ISL_593379, EPI_ISL_593380, EPI_ISL_593383, EPI_ISL_593384, EPI_ISL_593388, EPI_ISL_593393, EPI_ISL_593394, EPI_ISL_593401, EPI_ISL_593406, EPI_ISL_593408, EPI_ISL_593438 | | | |
| see above | Microbiological Diagnostic Unit - Public Health Laboratory | MDU-PHL | Seemann T., Schultz, M. B., Sait, M., Sherry, N. |

| | | | |
|---|---|---|---|
| | (MDU-PHL) | | |
| EPI_ISL_593671, EPI_ISL_593672 | Pathology West - NSW Health Pathology | NSW Health Pathology - Institute of Clinical Pathology and Medical Research; Westmead Hospital; University of Sydney | CIDM-PH et al. |
| EPI_ISL_593711, EPI_ISL_593748 | South Eastern Area Laboratory Services (SEALS) | NSW Health Pathology - Institute of Clinical Pathology and Medical Research; Westmead Hospital; University of Sydney | CIDM-PH et al. |
| EPI_ISL_593760 | Sydney South West Pathology Service (SSWPS) - Liverpool Hospital - NSW Health Pathology | NSW Health Pathology - Institute of Clinical Pathology and Medical Research; Westmead Hospital; University of Sydney | CIDM-PH et al. |
| EPI_ISL_593874, EPI_ISL_593875, EPI_ISL_593876, EPI_ISL_593877 | CHU Purpan - Laboratoire de Virologie - Institut Fédératif de Biologie | CHU Purpan - Laboratoire de Virologie - Institut Fédératif de Biologie | Latour J., Ranger N., Dubois M., Carcenac R., Harter A., Boyer P., Tremeaux P., Izopet J. |
| EPI_ISL_594139, EPI_ISL_594140, EPI_ISL_594141, EPI_ISL_594144 | MDU-PHL, The Peter Doherty Institute for Infection and Immunity | MDU-PHL, The Peter Doherty Institute for Infection and Immunity | Caly,L., Seemann,T., Sait,M.L., Schultz,M.B., Druce,J., Sherry,N.L. |
| EPI_ISL_594158, EPI_ISL_594162 | Israel Institute for Biological Research | Israel Institute for Biological Research | Galia Zaide, Inbar Cohen-Gihon, Ofir Israeli, Dana Stein, Shay Weiss, Orly Laskar, Yoav Gal, Libby Weiss, Emanuelle Mamroud, Adi Beth-Din and Anat Zvi |
| EPI_ISL_594297, EPI_ISL_594298, EPI_ISL_594299, EPI_ISL_594300 | Florida Bureau of Public Health Laboratories | Florida Bureau of Public Health Laboratories | Sarah Schmedes, Jason Blanton |
| EPI_ISL_596265 | WHO National Influenza Centre Russian Federation | WHO National Influenza Centre Russian Federation | Andrey Komissarov, Artem Fadeev, Anna Ivanova, Mariia Sergeeva, Kseniya Komissarova, Dmitry Bazhenov, Daria Danilenko |
| EPI_ISL_596266 | WHO National Influenza Centre Russian Federation | WHO National Influenza Centre Russian Federation | Andrey Komissarov, Artem Fadeev, Anna Ivanova, Kseniya Komissarova, Dmitry Bazhenov, Daria Danilenko |
| EPI_ISL_596449 | Institute for Medical Research, Infectious Disease Research Centre, National Institutes of Health, Ministry of Health Malaysia | Institute for Medical Research, Infectious Disease Research Centre, National Institutes of Health, Ministry of Health Malaysia | Suppiah J, Kamel K, Mohd-Zawawi Z, Thayan R |
| EPI_ISL_596520, EPI_ISL_596521 | Palestinian Ministry of Health | Molecular Genetics Lab | Nouar Qutob, Zaidoun Salah, Damien Richard, Hisham Darwish, Husam Sallam, Issa Shtayeh, Osama Najjar, Mahmoud Ruzayqat, Dana Najjar, Francois Balloux, Lucy van Dorp |
| EPI_ISL_596741, EPI_ISL_596795 | PathWest Laboratory Medicine WA | PathWest Laboratory Medicine WA Microbial Surveillance Unit | PathWest Laboratory Medicine WA Microbial Surveillance Unit |
| EPI_ISL_602318 | University of Miami Immunology and Histocompatibility Laboratory | University of Miami Immunology and Histocompatibility Laboratory | Emilio Margolles-Clark, PhD and Phillip Ruiz, MD, PhD |
| EPI_ISL_602596 | Pamela Youde Nethersole Eastern Hospital | Hong Kong Department of Health | Mak Gannon C.K., Lam Edman T.K., Chan Rickjason C.W., Tsang Dominic N.C. |
| EPI_ISL_602598 | Our Lady of Maryknoll Hospital | Hong Kong Department of Health | Mak Gannon C.K., Lam Edman T.K., Chan Rickjason C.W., Tsang Dominic N.C. |
| EPI_ISL_602599, EPI_ISL_602601 | Tseung Kwan O Hospital | Hong Kong Department of Health | Mak Gannon C.K., Lam Edman T.K., Chan Rickjason C.W., Tsang Dominic N.C. |
| EPI_ISL_602602 | Ruttonjee Hospital | Hong Kong Department of Health | Mak Gannon C.K., Lam Edman T.K., Chan Rickjason C.W., Tsang Dominic N.C. |
| EPI_ISL_602603, EPI_ISL_602604 | Queen Mary Hospital | Hong Kong Department of Health | Mak Gannon C.K., Lam Edman T.K., Chan Rickjason C.W., Tsang Dominic N.C. |
| EPI_ISL_602605 | Pamela Youde Nethersole Eastern Hospital | Hong Kong Department of Health | Mak Gannon C.K., Lam Edman T.K., Chan Rickjason C.W., Tsang Dominic N.C. |
| EPI_ISL_602606 | Queen Elizabeth Hospital | Hong Kong Department of Health | Mak Gannon C.K., Lam Edman T.K., Chan Rickjason C.W., Tsang Dominic N.C. |
| EPI_ISL_602607 | Tuen Mun Hospital | Hong Kong Department of Health | Mak Gannon C.K., Lam Edman T.K., Chan Rickjason C.W., Tsang Dominic N.C. |
| EPI_ISL_602608 | United Christian Hospital | Hong Kong Department of Health | Mak Gannon C.K., Lam Edman T.K., Chan Rickjason C.W., Tsang Dominic N.C. |
| EPI_ISL_602609 | Pamela Youde Nethersole Eastern Hospital | Hong Kong Department of Health | Mak Gannon C.K., Lam Edman T.K., Chan Rickjason C.W., Tsang Dominic N.C. |
| EPI_ISL_602610 | Queen Mary Hospital | Hong Kong Department of Health | Mak Gannon C.K., Lam Edman T.K., Chan Rickjason C.W., Tsang Dominic N.C. |
| EPI_ISL_602611 | Queen Elizabeth Hospital | Hong Kong Department of Health | Mak Gannon C.K., Lam Edman T.K., Chan Rickjason C.W., Tsang Dominic N.C. |
| EPI_ISL_602612 | Tseung Kwan O Hospital | Hong Kong Department of Health | Mak Gannon C.K., Lam Edman T.K., Chan Rickjason C.W., Tsang Dominic N.C. |
| EPI_ISL_602613, EPI_ISL_602614, EPI_ISL_602615 | Pamela Youde Nethersole Eastern Hospital | Hong Kong Department of Health | Mak Gannon C.K., Lam Edman T.K., Chan Rickjason C.W., Tsang Dominic N.C. |
| EPI_ISL_602616 | Kwong Wah Hospital | Hong Kong Department of Health | Mak Gannon C.K., Lam Edman T.K., Chan Rickjason C.W., Tsang Dominic N.C. |
| EPI_ISL_602630 | AHRI-Sigal | KRISP, KZN Research Innovation and Sequencing Platform | Gazy I, Sigl A, Karim F, Cele S, Giandhari J, Pillay S, Tegally H, Wilkinson E, de Oliveira T |
| EPI_ISL_603040, EPI_ISL_603042, EPI_ISL_603043 | MDU-PHL, The Peter Doherty Institute for Infection and Immunity | MDU-PHL, The Peter Doherty Institute for Infection and Immunity | Seemann,T., Caly,L., Sait,M., Schultz,M.B., Druce,J., Sherry,N. |
| EPI_ISL_603223, EPI_ISL_603224, EPI_ISL_603225 | National Institute of Laboratory Medicine and Referral Center | Genomic Research Lab, BCSIR | Md. Murshed Hasan Sarkar, Abu Sayeed Mohammad Mahmud, Mohammad Samir Uzzaman, Eshrar Osman, Md. Ahashan Habib, Shahina Akter, Tanjina Akhter Banu, Barna Goswami, Iffat Jahan, Md. Saddam Hossain, Tasnim Nafisa, Md. Maruf Ahmed Molla, Mahmuda Yeasmin, Asish Kumar Ghosh, A. K. M. Shamsuzzaman, Monira Parveen, Md. Masum Hossain Arif, Md. Salim Khan |
| EPI_ISL_605320, EPI_ISL_605321, EPI_ISL_605322, EPI_ISL_605323, EPI_ISL_605324, EPI_ISL_605325, EPI_ISL_605326, EPI_ISL_605327, EPI_ISL_605328, EPI_ISL_605329, EPI_ISL_605330, EPI_ISL_605331, EPI_ISL_605332, EPI_ISL_605333, EPI_ISL_605334, EPI_ISL_605335, EPI_ISL_605336, EPI_ISL_605337, EPI_ISL_605338, EPI_ISL_605339, EPI_ISL_605340, EPI_ISL_605341, EPI_ISL_605342, EPI_ISL_605343, EPI_ISL_605344, EPI_ISL_605345, EPI_ISL_605346, EPI_ISL_605347, EPI_ISL_605348, EPI_ISL_605349, EPI_ISL_605350, EPI_ISL_605351, EPI_ISL_605352, EPI_ISL_605353, EPI_ISL_605354, EPI_ISL_605355, EPI_ISL_605356, EPI_ISL_605357, EPI_ISL_605358, EPI_ISL_605359, EPI_ISL_605360, EPI_ISL_605361, EPI_ISL_605362, EPI_ISL_605363, EPI_ISL_605364, EPI_ISL_605365, EPI_ISL_605366, EPI_ISL_605367, EPI_ISL_605368, EPI_ISL_605369, EPI_ISL_605370, EPI_ISL_605371, EPI_ISL_605372, EPI_ISL_605373, EPI_ISL_605374, EPI_ISL_605375, EPI_ISL_605376, EPI_ISL_605377, EPI_ISL_605378, EPI_ISL_605379, EPI_ISL_605380, EPI_ISL_605381, EPI_ISL_605382, EPI_ISL_605383, EPI_ISL_605384, EPI_ISL_605385, EPI_ISL_605386, EPI_ISL_605387, EPI_ISL_605388, EPI_ISL_605389, EPI_ISL_605390, EPI_ISL_605391, EPI_ISL_605392, EPI_ISL_605393, EPI_ISL_605394, EPI_ISL_605395, EPI_ISL_605396, EPI_ISL_605397, EPI_ISL_605398, EPI_ISL_605399, EPI_ISL_605400, EPI_ISL_605401, EPI_ISL_605402, EPI_ISL_605403, EPI_ISL_605404 | | | |
| see above | Utah Public Health Laboratory | Utah Public Health Laboratory | Erin L. Young, Kelly Oakeson, Tara Gallagher, Michael T. Pyne, E. Susan Slechta, Melanie A. Mallory, Jeffrey B. Stevenson, Salika M. Shakir, David R. Hillyard |
| EPI_ISL_610126, EPI_ISL_610127 | Virginia DCLS | Virginia DCLS | Virginia DCLS |
| EPI_ISL_610208, EPI_ISL_610209, EPI_ISL_610212 | Department of Health Technology and Informatics, The Hong Kong Polytechnic University | Department of Health Technology and Informatics, The Hong Kong Polytechnic University | Siu,G.K.-H., Lee,L.-K., Leung,K.S.-S., Leung,J.S.-L., Ng,T.T.-L., Chan,C.T.-M., Tam,K.K.-G., Lao,H.-Y., Wu,A.K.-L., Yau,M.C.-Y., Lai,Y.W.-M., Fung,K.S.-C., Chau,S.K.-Y., Wong,B.K.-C., To,W.-K., Luk,K., Ho,A.Y.-M., Que,T.-L., Yip,K.-T., Yam,W.C., Shum,D.H.-K., Yip,S.P. |
| EPI_ISL_613463 | Public Health Laboratory - Infectious Disease Lab, Minnesota Department of Health Infectious Disease Laboratory Submission Group | Minnesota Department of Health, Public Health Laboratory | Plumb,M., Garfin,J., Lorentz,A., Wang,X. |
| EPI_ISL_613720, EPI_ISL_613723, EPI_ISL_613724, EPI_ISL_613747, EPI_ISL_613755, EPI_ISL_613780, EPI_ISL_613786, EPI_ISL_613789, EPI_ISL_613792, EPI_ISL_613794, EPI_ISL_613797, EPI_ISL_613820, EPI_ISL_613830, EPI_ISL_613840, EPI_ISL_613841, EPI_ISL_613842, EPI_ISL_613874, EPI_ISL_613875, EPI_ISL_613876, EPI_ISL_613877, EPI_ISL_613878, EPI_ISL_613879, EPI_ISL_613880, EPI_ISL_613881, EPI_ISL_613882, EPI_ISL_613883, EPI_ISL_613884, EPI_ISL_613885, EPI_ISL_613886, EPI_ISL_613887, EPI_ISL_613888, EPI_ISL_613889, EPI_ISL_613890, EPI_ISL_613891, EPI_ISL_613892, EPI_ISL_613893, EPI_ISL_613894, EPI_ISL_613895, EPI_ISL_613896, EPI_ISL_613897, EPI_ISL_613898, EPI_ISL_613950 | | | |
| see above | Florida Bureau of Public Health Laboratories | Florida Bureau of Public Health Laboratories | Sarah Schmedes, Jason Blanton |
| EPI_ISL_614307 | Faroese National Reference Laboratory for Fish and Animal Diseases | Faroese National Reference Laboratory for Fish and Animal Diseases | Maria Marjunardóttir Dahl, Petra Elisabeth Petersen, Debes Hammershaimb Christiansen |
| EPI_ISL_614381, EPI_ISL_614382, EPI_ISL_614383, EPI_ISL_614384 | Molecular diagnostic unit for viral haemorrhagic fevers and emerging viruses, Bouaké CHU Laboratory | Project group Epidemology of Highly Pathogenic Microorganisms, Robert Koch-Institute | Chantal Akoua-Koffi, Diané Bamourou, Etilé Anoh, Essia Belarbi, Safiatou Karidioula, Grit Schubert, Adjaratou Traoré, Soundélé Maïté, Monemo Pacome, Coulibaly Mbegnan, Bamba Fatoumata Touré, Kra Ouffoué, Fabian Leendertz |

| | | | |
|---|---|---|---|
| EPI_ISL_615908, EPI_ISL_615909, EPI_ISL_615910, EPI_ISL_617429, EPI_ISL_618231, EPI_ISL_618233, EPI_ISL_618234, EPI_ISL_618235, EPI_ISL_618236, EPI_ISL_618237, EPI_ISL_618238, EPI_ISL_618239, EPI_ISL_618242, EPI_ISL_618243, EPI_ISL_618244, EPI_ISL_618247, EPI_ISL_618248, EPI_ISL_618249, EPI_ISL_618252, EPI_ISL_618253, EPI_ISL_618254, EPI_ISL_618255, EPI_ISL_618256, EPI_ISL_618257, EPI_ISL_618263, EPI_ISL_618264, EPI_ISL_618265, EPI_ISL_618266, EPI_ISL_618267, EPI_ISL_618268, EPI_ISL_618269, EPI_ISL_618337, EPI_ISL_618338, EPI_ISL_618341, EPI_ISL_618348, EPI_ISL_618349, EPI_ISL_618350, EPI_ISL_618351, EPI_ISL_618352, EPI_ISL_618353, EPI_ISL_618354, EPI_ISL_618355, EPI_ISL_618356, EPI_ISL_618367, EPI_ISL_618368, EPI_ISL_618369, EPI_ISL_618370, EPI_ISL_618371, EPI_ISL_618376, EPI_ISL_618377, EPI_ISL_618378, EPI_ISL_618379, EPI_ISL_618381, EPI_ISL_618382, EPI_ISL_618383, EPI_ISL_618384, EPI_ISL_618385, EPI_ISL_618386, EPI_ISL_618387, EPI_ISL_618388, EPI_ISL_618389, EPI_ISL_618390, EPI_ISL_618391, EPI_ISL_618392, EPI_ISL_618393, EPI_ISL_618394, EPI_ISL_618395, EPI_ISL_618396, EPI_ISL_618397, EPI_ISL_618398, EPI_ISL_618399, EPI_ISL_618400, EPI_ISL_618401, EPI_ISL_618405, EPI_ISL_622667 | | | |
| see above | Department of Virus and Microbiological Special Diagnostics, Statens Serum Institut, Denmark | Albertsen lab, Department of Chemistry and Bioscience, Aalborg University, Denmark | Danish Covid-19 Genome Consortia |
| EPI_ISL_622943, EPI_ISL_622946, EPI_ISL_622947, EPI_ISL_622948, EPI_ISL_622949, EPI_ISL_622950, EPI_ISL_622951, EPI_ISL_622952, EPI_ISL_622953, EPI_ISL_622955, EPI_ISL_622957, EPI_ISL_622959, EPI_ISL_622960, EPI_ISL_622961, EPI_ISL_622962, EPI_ISL_622963, EPI_ISL_622964, EPI_ISL_622965, EPI_ISL_622966, EPI_ISL_622967, EPI_ISL_622968, EPI_ISL_622969, EPI_ISL_622978, EPI_ISL_622983, EPI_ISL_622985, EPI_ISL_622986, EPI_ISL_622988, EPI_ISL_622990, EPI_ISL_622994, EPI_ISL_623064, EPI_ISL_623068, EPI_ISL_623073 | | | |
| see above | Lancet Laboratories | National Institute for Communicable Diseases of the National Health Laboratory Service | Allam M, Ismail A, Khumalo Z, Kwenda S, Mtshali P, Mnyameni F, Mohale T, Subramoney K, Bhiman JN |
| EPI_ISL_625473, EPI_ISL_625475 | Child Health Research Foundation | Child Health Research Foundation | Senjuti Saha, Md Saiful Islam Sajib, Nikkon Sarkar, Syed Muktadir Al Sium, Afroza Akter Tanni, Roly Malaker, Arif Mohammad Tanmoy, Md Hafizur Rahman, Samir K Saha |
| EPI_ISL_625559, EPI_ISL_625560, EPI_ISL_625561, EPI_ISL_625562, EPI_ISL_625563, EPI_ISL_625564, EPI_ISL_625565, EPI_ISL_625566, EPI_ISL_625567, EPI_ISL_625568, EPI_ISL_625569, EPI_ISL_625570, EPI_ISL_625571, EPI_ISL_625572 | | | |
| see above | NaN | Chan-Zuckerberg Biohub | CZB Cliahub Consortium |
| EPI_ISL_625623 | County of San Luis Obispo Public Health Laboratory | Chan-Zuckerberg Biohub | CZB Cliahub Consortium |
| EPI_ISL_626540, EPI_ISL_626541, EPI_ISL_626542, EPI_ISL_626543, EPI_ISL_626544, EPI_ISL_626545, EPI_ISL_626546, EPI_ISL_626547, EPI_ISL_626548 | Northwestern Memorial Hospital | Ozer Lab | Ramon Lorenzo-Redondo, Hannah H. Nam, Scott C. Roberts, Lacy M. Simons, Chad J. Achenbach, Lawrence J. Jennings, Chao Qi, Alan R. Hauser, Michael G. Ison, Judd F. Hultquist, Egon A. Ozer |
| EPI_ISL_626549, EPI_ISL_626551, EPI_ISL_626553, EPI_ISL_626554, EPI_ISL_626555, EPI_ISL_626564, EPI_ISL_626565 | Laboratorio de Biología Molecular, Facultad de Medicina, Universidad de Atacama, Copiapo, Chile/ FONDAP CRG, Universidad Andrés Bello, Santiago, Chile | Center for Mathematical Modeling and Center for Genome Regulation. Santiago, Chile | Echeverría C, Manríquez R, Bastias M, Sanhueza D, Travisany D, Allende ML, Maass A, González M, Montecino, M, Orellana A, Castro E, Meneses C. |
| EPI_ISL_631399 | Wisconsin State Laboratory of Hygiene Communicable Disease Division | Wisconsin State Laboratory of Hygiene Communicable Disease Division | Kelsey R. Florek, Abigail C. Shockey |
| EPI_ISL_632267, EPI_ISL_632283, EPI_ISL_632284 | Communicable Disease Laboratory, Public Health Directorate | Communicable Disease Laboratory, Public Health Directorate | AlWasti,H., AlTaif,Z., AlHujairi,Z., AlAbbas,Z. |
| EPI_ISL_634877, EPI_ISL_634879 | Lab voor klinische biologie | Onderzoeksgroep Virologie | Laurens Lambrechts, Nick Vereecke, Marthe Pauwels, Bruno Verhasselt, Linos Vandekerckhove, Hans Nauwynck, Sebastiaan Theuns |
| EPI_ISL_635115, EPI_ISL_635117, EPI_ISL_635118, EPI_ISL_635147, EPI_ISL_635156, EPI_ISL_635157, EPI_ISL_635186 | University Hospital of Northern Norway, Department for Microbiology and Infectious Disease Control | Norwegian Institute of Public Health, Department of Virology | Kathrine Stene-Johansen, Kamilla Heddeland Instefjord, Hilde Elshaug, Marie Paulsen Madsen, Rasmus Riis Kopperud, Hilde Vollan, Karoline Bragstad, Olav Hungnes |
| EPI_ISL_635576 | Centro de Diagnstico COVID-19 UABC Tijuana | Andersen lab at Scripps Research | SEARCH Alliance San Diego with Idanya Rubí Serafín Higuera, Manuel Sánchez Alavez, Jorge Luis Jiménez Niebla, Germán Ibarra, Jonathan Vincent Baena, Oscar Efrén Zazueta Fierro |
| EPI_ISL_635680, EPI_ISL_635681, EPI_ISL_635682, EPI_ISL_635683, EPI_ISL_635684, EPI_ISL_635685, EPI_ISL_635687, EPI_ISL_635688, EPI_ISL_635692, EPI_ISL_635693, EPI_ISL_635696, EPI_ISL_635697, EPI_ISL_635698, EPI_ISL_635703, EPI_ISL_635704, EPI_ISL_635706, EPI_ISL_635710, EPI_ISL_635711, EPI_ISL_635712, EPI_ISL_635713, EPI_ISL_635715, EPI_ISL_635717, EPI_ISL_635723, EPI_ISL_635729, EPI_ISL_635791, EPI_ISL_635792, EPI_ISL_635804, EPI_ISL_635809, EPI_ISL_635813, EPI_ISL_635819, EPI_ISL_635822, EPI_ISL_635823, EPI_ISL_635910, EPI_ISL_635911, EPI_ISL_635913, EPI_ISL_635914, EPI_ISL_635915, EPI_ISL_635917, EPI_ISL_635918, EPI_ISL_635919, EPI_ISL_635922, EPI_ISL_635924, EPI_ISL_635926, EPI_ISL_635927, EPI_ISL_635928, EPI_ISL_635929, EPI_ISL_635930 | | | |
| see above | San Diego County Public Health Laboratory | Andersen lab at Scripps Research | SEARCH Alliance San Diego with Tracy Basler, Jovan Shephard, Brett Austin |
| EPI_ISL_636567, EPI_ISL_636587, EPI_ISL_636588 | Dutch COVID-19 response team | National Institute for Public Health and the Environment (RIVM) | Adam Meijer, Harry Vennema, Jeroen Cremer, Sharon van den Brink, Bas van der Veer, AnneMarie van den Brandt, Florian Zwagemaker, Dennis Schmitz, Chantal Reusken, on behalf of the national COVID-19 response team |
| EPI_ISL_636739, EPI_ISL_636748, EPI_ISL_636759, EPI_ISL_636770, EPI_ISL_636781, EPI_ISL_636792 | National Centre for Disease control (NCDC) | NCDC/CSIR-IGIB | Mahesh S. Dhar1*, Bharathram Uppili2*, Robin Marwal1*, Pooja Sharma2*, RadhaKrishnan VS, Vivekanand A, Nishu Tyagi, Shaista Khan, Simmi Tiwari, Manish Kumar, Ajit Shewale, Ishtaq Ahmed, Asangla Kamai, Aparna Swaminathan, Saruchi Wadhwa, Tushar Nale, Sandhya Kabra, Sujeet Singh, Mohammed Faruq#, Anurag Agrawal#, Partha Rakshit# |
| EPI_ISL_636841, EPI_ISL_636842, EPI_ISL_636843, EPI_ISL_636844 | Lithuanian University of Health Sciences Hospital, Department of Laboratory Medicine | Lithuanian University of Health Sciences, Molecular cardiology lab. | Lukas Zemaitis, Ingrida Olendrait, Arnoldas Pautienius, Kamile Tamusauskaite, Dovydas Gecys, Laura Pareckaite, Vaiva Lesauskaite, Astra Vitkauskiene |
| EPI_ISL_636997, EPI_ISL_636998, EPI_ISL_636999, EPI_ISL_637000, EPI_ISL_637001, EPI_ISL_637002, EPI_ISL_637003, EPI_ISL_637004 | Department of Infectious Diseases and Immunology, National Hospital Organization Nagoya Medical Center | Clinical Research Center, National Hospital Organization Nagoya Medical Center | Yoshihiro Nakata, Hirotaka Ode, Mai Kubota, Masakazu Matsuda, Kazuhiro Matsuoka, Miho Nakasuji, Mikiko Mori, Mayumi Imahashi, Yoshiyuki Yokomaku, Yasumasa Iwatani |
| EPI_ISL_640069, EPI_ISL_640070, EPI_ISL_640071 | Guguletu CHC wc GDH | NHLS/UCT | Arash Iranzadeh, Deelan Doolabh, Lynn Tyers, Bruna Galvao, Innocent Mudau, Marvin Hsiao, Kruger Marais, Diana Hardie, Stephen Korsman, Carolyn Williamson |
| EPI_ISL_640075 | Bothasig CDC wc BLD | NHLS/UCT | Arash Iranzadeh, Deelan Doolabh, Lynn Tyers, Bruna Galvao, Innocent Mudau, Marvin Hsiao, Kruger Marais, Diana Hardie, Stephen Korsman, Carolyn Williamson |
| EPI_ISL_640076 | Knysna Hospital wc KNY | NHLS/UCT | Arash Iranzadeh, Deelan Doolabh, Lynn Tyers, Bruna Galvao, Innocent Mudau, Marvin Hsiao, Kruger Marais, Diana Hardie, Stephen Korsman, Carolyn Williamson |
| EPI_ISL_640301, EPI_ISL_640303, EPI_ISL_640312, EPI_ISL_640315, EPI_ISL_640316, EPI_ISL_640322, EPI_ISL_640327, EPI_ISL_640328, EPI_ISL_640329, EPI_ISL_640331, EPI_ISL_640333, EPI_ISL_640335, EPI_ISL_640336, EPI_ISL_640339, EPI_ISL_640351, EPI_ISL_640353, EPI_ISL_640354, EPI_ISL_640360, EPI_ISL_640365, EPI_ISL_640367, EPI_ISL_640369, EPI_ISL_640377, EPI_ISL_640380, EPI_ISL_640381, EPI_ISL_640382, EPI_ISL_640384, EPI_ISL_640392, EPI_ISL_640395, EPI_ISL_640396, EPI_ISL_640397, EPI_ISL_640402, EPI_ISL_640403, EPI_ISL_640408, EPI_ISL_640411, EPI_ISL_640416, EPI_ISL_640417, EPI_ISL_640421, EPI_ISL_640423, EPI_ISL_640435, EPI_ISL_640439, EPI_ISL_640441, EPI_ISL_640444, EPI_ISL_640446, EPI_ISL_640449, EPI_ISL_640450, EPI_ISL_640451, EPI_ISL_640452, EPI_ISL_640454, EPI_ISL_640460, EPI_ISL_640462, EPI_ISL_640469, EPI_ISL_640479, EPI_ISL_640481, EPI_ISL_640485, EPI_ISL_640489, EPI_ISL_640490, EPI_ISL_640492, EPI_ISL_640494, EPI_ISL_640495, EPI_ISL_640496, EPI_ISL_640499, EPI_ISL_640500, EPI_ISL_640501, EPI_ISL_640503, EPI_ISL_640505, EPI_ISL_640508, EPI_ISL_640512, EPI_ISL_640514, EPI_ISL_640524, EPI_ISL_640527, EPI_ISL_640530, EPI_ISL_640532, EPI_ISL_640534, EPI_ISL_640536, EPI_ISL_640544, EPI_ISL_640546, EPI_ISL_640547, EPI_ISL_640549, EPI_ISL_640553, EPI_ISL_640554, EPI_ISL_640555, EPI_ISL_640557, EPI_ISL_640558, EPI_ISL_640560, EPI_ISL_640569, EPI_ISL_640570, EPI_ISL_640574, EPI_ISL_640575, EPI_ISL_640577, EPI_ISL_640581, EPI_ISL_640582, EPI_ISL_640590, EPI_ISL_640592, EPI_ISL_640594, EPI_ISL_640597, EPI_ISL_640598, EPI_ISL_640603, EPI_ISL_640605, EPI_ISL_640611, EPI_ISL_640615, EPI_ISL_640618, EPI_ISL_640620, EPI_ISL_640622, EPI_ISL_640629, EPI_ISL_640635, EPI_ISL_640640, EPI_ISL_640647, EPI_ISL_640651, EPI_ISL_640655, EPI_ISL_640659, EPI_ISL_640661, EPI_ISL_640663, EPI_ISL_640664, EPI_ISL_640666, EPI_ISL_640668, EPI_ISL_640669, EPI_ISL_640677, EPI_ISL_640678, EPI_ISL_640679, EPI_ISL_640681, EPI_ISL_640685, EPI_ISL_640687, EPI_ISL_640688, EPI_ISL_640691, EPI_ISL_640695, EPI_ISL_640697, EPI_ISL_640698, EPI_ISL_640700, EPI_ISL_640707, EPI_ISL_640708, EPI_ISL_640710, EPI_ISL_640712, EPI_ISL_640713, EPI_ISL_640717, EPI_ISL_640722, EPI_ISL_640723, EPI_ISL_640728, EPI_ISL_640730, EPI_ISL_640731, EPI_ISL_640732, EPI_ISL_640733, EPI_ISL_640743, EPI_ISL_640744, EPI_ISL_640746, EPI_ISL_640755, EPI_ISL_640759, EPI_ISL_640762, EPI_ISL_640765, EPI_ISL_640766, EPI_ISL_640767, EPI_ISL_640776, EPI_ISL_640778, EPI_ISL_640781, EPI_ISL_640782, EPI_ISL_640785, EPI_ISL_640787, EPI_ISL_640791, EPI_ISL_640793, EPI_ISL_640795, EPI_ISL_640796, EPI_ISL_640798, EPI_ISL_640801, EPI_ISL_640802, EPI_ISL_640805, EPI_ISL_640808, EPI_ISL_640816, EPI_ISL_640820, EPI_ISL_640821, EPI_ISL_640823, EPI_ISL_640826, EPI_ISL_640828, EPI_ISL_640829, EPI_ISL_640830, EPI_ISL_640834, EPI_ISL_640837, EPI_ISL_640844, EPI_ISL_640850, EPI_ISL_640855, EPI_ISL_640856, EPI_ISL_640859, EPI_ISL_640862, EPI_ISL_640863, EPI_ISL_640864, EPI_ISL_640865, EPI_ISL_640874, EPI_ISL_640875, EPI_ISL_640877, EPI_ISL_640878, EPI_ISL_640880, EPI_ISL_640884, EPI_ISL_640886, EPI_ISL_640888, EPI_ISL_640893, EPI_ISL_640897, EPI_ISL_640901, EPI_ISL_640904, EPI_ISL_640905, EPI_ISL_640910, EPI_ISL_640911, EPI_ISL_640912, EPI_ISL_640913, EPI_ISL_640916, EPI_ISL_640917, EPI_ISL_640918, EPI_ISL_640919, EPI_ISL_640922, EPI_ISL_640929, EPI_ISL_640934, EPI_ISL_640943, EPI_ISL_640946, EPI_ISL_640948, EPI_ISL_640949, EPI_ISL_640951, EPI_ISL_640953, EPI_ISL_640956, EPI_ISL_640957, EPI_ISL_640960, EPI_ISL_640974, EPI_ISL_640976, EPI_ISL_640981, EPI_ISL_640982, EPI_ISL_640985, EPI_ISL_641004, EPI_ISL_641005, EPI_ISL_641011, EPI_ISL_641018, EPI_ISL_641019, EPI_ISL_641020, EPI_ISL_641022, EPI_ISL_641023, EPI_ISL_641027, EPI_ISL_641030, EPI_ISL_641032, EPI_ISL_641038, EPI_ISL_641040, EPI_ISL_641043, EPI_ISL_641046, EPI_ISL_641062, EPI_ISL_641063, EPI_ISL_641064, EPI_ISL_641066, EPI_ISL_641068, EPI_ISL_641071, EPI_ISL_641075, EPI_ISL_641076, EPI_ISL_641081, EPI_ISL_641082, EPI_ISL_641084, EPI_ISL_641085, EPI_ISL_641087, EPI_ISL_641090, EPI_ISL_641093, EPI_ISL_641094, EPI_ISL_641095, EPI_ISL_641100, EPI_ISL_641101, EPI_ISL_641113, EPI_ISL_641114, EPI_ISL_641117, EPI_ISL_641120 | | | |
| see above | Microbiological Diagnostic Unit - Public Health Laboratory (MDU-PHL) | MDU-PHL | Seemann T., Schultz M.B., Sait, M.L., Sherry, N.L. |

| | | | |
|---|---|---|---|
| EPI_ISL_641125 | Victorian Infectious Diseases Reference Laboratory (VIDRL) | VIDRL and MDU-PHL | Caly L., Seemann T., Sait, M.L., Schultz M.B., Druce J., Sherry, N.L. |
| EPI_ISL_641129, EPI_ISL_641130, EPI_ISL_641132, EPI_ISL_641135, EPI_ISL_641136, EPI_ISL_641139, EPI_ISL_641145, EPI_ISL_641146, EPI_ISL_641151, EPI_ISL_641152, EPI_ISL_641154, EPI_ISL_641159, EPI_ISL_641160, EPI_ISL_641168, EPI_ISL_641173, EPI_ISL_641175, EPI_ISL_641183, EPI_ISL_641190, EPI_ISL_641192, EPI_ISL_641199, EPI_ISL_641201, EPI_ISL_641211, EPI_ISL_641219, EPI_ISL_641224, EPI_ISL_641228, EPI_ISL_641229, EPI_ISL_641235, EPI_ISL_641238, EPI_ISL_641242, EPI_ISL_641248, EPI_ISL_641249, EPI_ISL_641250, EPI_ISL_641251, EPI_ISL_641252, EPI_ISL_641259, EPI_ISL_641262, EPI_ISL_641263, EPI_ISL_641267, EPI_ISL_641271, EPI_ISL_641272, EPI_ISL_641274, EPI_ISL_641281, EPI_ISL_641282, EPI_ISL_641283, EPI_ISL_641284, EPI_ISL_641286, EPI_ISL_641290, EPI_ISL_641291, EPI_ISL_641293, EPI_ISL_641295, EPI_ISL_641296, EPI_ISL_641298, EPI_ISL_641299 | | | |
| see above | Microbiological Diagnostic Unit - Public Health Laboratory (MDU-PHL) | MDU-PHL | Seemann T., Schultz M.B., Sait, M.L., Sherry, N.L. |
| EPI_ISL_644574, EPI_ISL_644575, EPI_ISL_644576, EPI_ISL_644577, EPI_ISL_644578 | Veterinary Specialized Institute "Kraljevo", Serbia | Veterinary Specialized Institute "Kraljevo", Serbia | Vidanovic,D., Tesovic,B., Knezevic,A., Jovanovic,T., Jankovic,M., Sekler,M., Banovic Djeri,B., Petrovic,T., Volkening,J., Afonso,C. |
| EPI_ISL_644970, EPI_ISL_644971, EPI_ISL_644972, EPI_ISL_644973, EPI_ISL_644974, EPI_ISL_644975, EPI_ISL_644976, EPI_ISL_644977, EPI_ISL_644978, EPI_ISL_644979, EPI_ISL_644980 | | | |
| see above | Department of Infectious Diseases, Keio University School of Medicine, Tokyo, Japan | Center for Medical Genetics, Keio University School of Medicine, Tokyo, Japan | Kenjiro Kosaki, Yuka Iwasaki, Hirotsugu Ishizu, Haruhiko Siomi, Kodai Abe |
| EPI_ISL_648143 | The Public Health Agency of Sweden | The Public Health Agency of Sweden | Anna-Malin Linde, Maria Lind Karlberg, Mattias Haukland, Reza Advani, Olov Svartstrom, Oskar Karlsson Lindsjo, Sandra Broddesson, Petra Edquist, Mia Brytting, Anna Risberg, Karin Tegmark-Wisell |
| EPI_ISL_648378 | Laboratorio de Investigaciones de Baney | University Hospital Basel, Clinical Bacteriology | Carlos Cortes, Claudia Daubenberger, Adrian Egli, Guillermo Garcia, Salome Hosch, Bonifacio Manguire Nlavo, Alfredo Mari, Maximilian Mpina, Elizabeth Nyakarungu, Diosdado Odjama Nseng Ada, Mitoha Ondo O Ayekaba, Tim Roloff, Tobias Schindler, Helena Seth-Smith, Madlen Stange, Philip Wonder Phiri |
| EPI_ISL_648543, EPI_ISL_648544 | UCSF Clinical Microbiology Laboratory | Chan-Zuckerberg Biohub | CZB Cliahub Consortium |
| EPI_ISL_648741 | Department of Laboratory Medicine, Tan Tock Seng Hospital | Department of Laboratory Medicine, Tan Tock Seng Hospital | Chen YYC, Zair X, Lim JX, Li C, Tang WY, Maurer-Stroh S, Barkham TMS, Nagarajan N, Sessions OM |
| EPI_ISL_648981, EPI_ISL_648983, EPI_ISL_648985, EPI_ISL_648993, EPI_ISL_648994, EPI_ISL_649000, EPI_ISL_649001, EPI_ISL_649002, EPI_ISL_649003, EPI_ISL_649005 | San Diego County Public Health Laboratory | Andersen lab at Scripps Research | SEARCH Alliance San Diego with Tracy Basler, Jovan Shephard, Brett Austin |
| EPI_ISL_649150 | Microbiological Diagnostic Unit - Public Health Laboratory (MDU-PHL), The Peter Doherty Institute for Infection and Immunity | Microbiological Diagnostic Unit - Public Health Laboratory (MDU-PHL), The Peter Doherty Institute for Infection and Immunity | Seemann,T., Caly,L., Sait,M.L., Schultz,M.B., Druce,J., Sherry,N.L. |
| EPI_ISL_649170, EPI_ISL_649171 | Laboratorio de Investigaciones de Baney | University Hospital Basel, Clinical Bacteriology | Carlos Cortes, Claudia Daubenberger, Adrian Egli, Guillermo Garcia, Salome Hosch, Bonifacio Manguire Nlavo, Alfredo Mari, Maximilian Mpina, Elizabeth Nyakarungu, Diosdado Odjama Nseng Ada, Mitoha Ondo O Ayekaba, Tim Roloff, Tobias Schindler, Helena Seth-Smith, Madlen Stange, Philip Wonder Phiri |
| EPI_ISL_653234, EPI_ISL_653235, EPI_ISL_653236, EPI_ISL_653245, EPI_ISL_653246, EPI_ISL_653247, EPI_ISL_653318 | Florida Bureau of Public Health Laboratories | Florida Bureau of Public Health Laboratories | Sarah Schmedes, Jason Blanton |
| EPI_ISL_653475, EPI_ISL_653476, EPI_ISL_653477, EPI_ISL_653478, EPI_ISL_653479, EPI_ISL_653480, EPI_ISL_653481, EPI_ISL_653482, EPI_ISL_653483, EPI_ISL_653484, EPI_ISL_653485, EPI_ISL_653486, EPI_ISL_653487, EPI_ISL_653488, EPI_ISL_653489, EPI_ISL_653490, EPI_ISL_653491, EPI_ISL_653492, EPI_ISL_653493, EPI_ISL_653494, EPI_ISL_653495, EPI_ISL_653496, EPI_ISL_653497, EPI_ISL_653498, EPI_ISL_653499, EPI_ISL_653500, EPI_ISL_653501, EPI_ISL_653502, EPI_ISL_653503, EPI_ISL_653504, EPI_ISL_653505, EPI_ISL_653506, EPI_ISL_653507, EPI_ISL_653508, EPI_ISL_653509, EPI_ISL_653510, EPI_ISL_653511, EPI_ISL_653512, EPI_ISL_653513, EPI_ISL_653514, EPI_ISL_653515, EPI_ISL_653516, EPI_ISL_653517, EPI_ISL_653518, EPI_ISL_653519, EPI_ISL_653520, EPI_ISL_653521, EPI_ISL_653522, EPI_ISL_653523, EPI_ISL_653524, EPI_ISL_653525, EPI_ISL_653526, EPI_ISL_653527, EPI_ISL_653528, EPI_ISL_653529, EPI_ISL_653530, EPI_ISL_653531, EPI_ISL_653532, EPI_ISL_653533, EPI_ISL_653534, EPI_ISL_653535, EPI_ISL_653536, EPI_ISL_653537, EPI_ISL_653538, EPI_ISL_653539, EPI_ISL_653540, EPI_ISL_653541, EPI_ISL_653542, EPI_ISL_653543, EPI_ISL_653544, EPI_ISL_653545 | | | |
| see above | LSUHS Emerging Viral Threat Laboratory | Microbial Genome Sequencing Center | Jeremy P. Kamil, Rona S. Scott, Maarten Van Diest, Malgorzata Bienkowska-Haba, Katarzyna Zwolinska, Andrew D. Yurochko, Christopher G. Kevil, Martin J. Sapp, Daniel J. Snyder, Vaughn S. Cooper, John A. Vanchiere |
| EPI_ISL_654182, EPI_ISL_654278 | Hospital General Universitario Gregorio Marañón | SeqCOVID-SPAIN consortium/IBV(CSIC) | Darío García de Viedma, Laura Pérez-Lago, Marta Herranz, Jon Sicilia, Julia Suárez, Pilar Catalán, Patricia Muñoz and SeqCOVID-SPAIN consortium |
| EPI_ISL_654793 | Instituto Nacional de Salud, Bogotá, Colombia | Instituto Nacional de Salud, Bogotá, Colombia | Katherine Laiton-Donato, Diego A. Álvarez-Díaz, Carlos Franco-Muñoz, Mauricio Pacheco-Montealegre, Jonathan Reales, Diego Andrés Prada, Jose A. Usme-Ciro, Zulma M. Cucunubá, Christian Julian VillabonaArenas, Liz Villabona-Arenas, Sussy Echeverria, Astrid C. Flórez, Carolina Ferro, Diana Marcela Walteros-Acero, Franklin Prieto, Carlos Andrés Durán, Martha Lucia Ospina Martinez, Marcela Mercado-Reyes |
| EPI_ISL_660168, EPI_ISL_660171, EPI_ISL_660173, EPI_ISL_660174, EPI_ISL_660175, EPI_ISL_660176 | NHLS-IALCH | KRISP, KZN Research Innovation and Sequencing Platform | Gazy I, Sigal A, Karim F, Cele S, Giandhari J, Pillay S, Tegally H, Wilkinson E, de Oliveira T |
| EPI_ISL_660265, EPI_ISL_660266, EPI_ISL_660267, EPI_ISL_660268, EPI_ISL_660269, EPI_ISL_660270, EPI_ISL_660275, EPI_ISL_660276 | Servicio de Microbiología, Laboratori Clínic Metropolitana Nord. Hospital Universitari Germans Trias i Pujol. Institut d'Investigació en Ciències de la Salut Germans Trias i Pujol (IGTP) | SeqCOVID-SPAIN consortium/IBV(CSIC) | Elisa Martró, Antoni E. Bordoy, Anna Not, Adrián Antuori, Anabel Fernández, Nona Romaní and SeqCOVID-SPAIN consortium |
| EPI_ISL_660478, EPI_ISL_660479 | Laboratoire de Microbiologie CHU Souro Sanou | Centre Muraz | Abdoul-Salam Ouedraogo, Yacouba Sawadogo, Essia Belarbi, Grit Schubert, Fabian Leendertz, Arsène Zongo, Soumeya Ouangraoua, Zekiba Tarnagda, Lassana Sangaré, Halidou Tinto |
| EPI_ISL_660543 | Laboratory Medicine | Department of Laboratory Medicine, Lin-Kou Chang Gung Memorial Hospital, Taoyuan, Taiwan | Kuo-Chien Tsao, Yu-Nong Gong, Shu-Li Yang, Yi-Chun Liu, Chung-Guei Huang, Mei-Jen Hsiao, Po-Wei Huang, Cheng-Ta Yang, Cheng-Hsun Chiu, Peng-Nien Huang, Kuo-Ming Lee, Guang-Wu Chen, Shin-Ru Shih |
| EPI_ISL_661198 | Scientific Veterinary Institute Novi Sad | Veterinary Specialized Institute "Kraljevo", Serbia | Vidanovic,D., Tesovic,B., Knezevic,A., Jovanovic,T., Jankovic,M., Sekler,M., Banovic Djeri,B., Petrovic,T., Volkening,J., Afonso,C. |
| EPI_ISL_663291, EPI_ISL_663297, EPI_ISL_663299, EPI_ISL_663317, EPI_ISL_663328, EPI_ISL_663360, EPI_ISL_663384, EPI_ISL_663421, EPI_ISL_663472, EPI_ISL_663474, EPI_ISL_663494, EPI_ISL_663496, EPI_ISL_663547, EPI_ISL_663703, EPI_ISL_663704, EPI_ISL_663705, EPI_ISL_663785, EPI_ISL_663813, EPI_ISL_663814, EPI_ISL_663820, EPI_ISL_663822, EPI_ISL_663824, EPI_ISL_663825, EPI_ISL_663827, EPI_ISL_663829, EPI_ISL_663834, EPI_ISL_663835, EPI_ISL_663836, EPI_ISL_663837, EPI_ISL_663838, EPI_ISL_663839, EPI_ISL_663840, EPI_ISL_663843, EPI_ISL_663844, EPI_ISL_663846, EPI_ISL_663849, EPI_ISL_663918, EPI_ISL_663920, EPI_ISL_663922, EPI_ISL_663923, EPI_ISL_663924, EPI_ISL_663927, EPI_ISL_663928, EPI_ISL_663930, EPI_ISL_663931, EPI_ISL_663935, EPI_ISL_663941, EPI_ISL_663942, EPI_ISL_663944, EPI_ISL_663946, EPI_ISL_663947, EPI_ISL_663948, EPI_ISL_663950, EPI_ISL_663953, EPI_ISL_663954, EPI_ISL_663955, EPI_ISL_663964, EPI_ISL_663965, EPI_ISL_663969, EPI_ISL_663976, EPI_ISL_663977, EPI_ISL_663980, EPI_ISL_663983, EPI_ISL_663984 | | | |
| see above | Microbiological Diagnostic Unit - Public Health Laboratory (MDU-PHL) | MDU-PHL | Seemann T., Schultz M.B., Sait, M.L., Sherry, N.L. |
| EPI_ISL_666815, EPI_ISL_666816, EPI_ISL_666817, EPI_ISL_666818, EPI_ISL_666819, EPI_ISL_666830, EPI_ISL_666831, EPI_ISL_666865, EPI_ISL_666869 | Florida Bureau of Public Health Laboratories | Florida Bureau of Public Health Laboratories | Sarah Schmedes, Jason Blanton |
| EPI_ISL_667085, EPI_ISL_667086, EPI_ISL_667087, EPI_ISL_667088, EPI_ISL_667089, EPI_ISL_667090, EPI_ISL_667091, EPI_ISL_667092, EPI_ISL_667093, EPI_ISL_667094, EPI_ISL_667095, EPI_ISL_667096, EPI_ISL_667097 | | | |
| see above | OHSU Lab Services Molecular Microbiology Lab | Oregon SARS-CoV-2 Genome Sequencing Center | Brendan L. O'Connell, Ruth V. Nichols, Sally Grindstaff, Alec J. Hirsch, Donna Hansel, Guang Fan, Daniel N. Streblow, William B. Messer, Andrew C. Adey, Benjamin N. Bimber, Brian J. O'Roak |
| EPI_ISL_671245 | Department of Virus and Microbiological Special Diagnostics, Statens Serum Institut, Copenhagen, Denmark | Albertsen Lab, Department of Chemistry and Bioscience, Aalborg University, Denmark | Danish Covid-19 Genome Consortium |
| EPI_ISL_671794, EPI_ISL_671795, EPI_ISL_671796, EPI_ISL_671797, EPI_ISL_671798, EPI_ISL_671799, | Hospital Clínico Universitario Lozano Blesa de Zaragoza (España) | SeqCOVID-SPAIN consortium/IBV(CSIC) | Rafael Benito, Sonia Algarate, Jessica Bueno and SeqCOVID-SPAIN consortium |

| | | | |
|---|---|---|---|
| EPI_ISL_671800 | | | |
| EPI_ISL_671859, EPI_ISL_671860, EPI_ISL_671861, EPI_ISL_671862 | Servicio de Microbiología, Laboratori Clínic Metropolitana Nord. Hospital Universitari Germans Trias i Pujol. Institut d'Investigació en Ciències de la Salut Germans Trias i Pujol (IGTP) | SeqCOVID-SPAIN consortium/IBV(CSIC) | Elisa Martró, Antoni E. Bordoy, Anna Not, Adrián Antuori, Anabel Fernández, Nona Romaní, Verónica Saludes, Cristina Casañ and SeqCOVID-SPAIN consortium |
| EPI_ISL_672380, EPI_ISL_672381, EPI_ISL_672385 | San Francisco Public Health Laboratory | Chan-Zuckerberg Biohub | CZB Cliahub Consortium |
| EPI_ISL_672388 | UCSF Clinical Microbiology Laboratory | Chan-Zuckerberg Biohub | CZB Cliahub Consortium |
| EPI_ISL_672579 | Infectious Diseases and Tropical Medicine Research Center, Infectious Diseases and Tropical Medicine Research Center | Infectious Diseases and Tropical Medicine Research Center, Infectious Diseases and Tropical Medicine Research Center | Ahangarzadeh,S., Ataei,B., Shariati,L., Haghjooy Javanmard,S., Shoaei,P., Aboutalebian,S. |
| EPI_ISL_676514 | Gavleborg | The Public Health Agency of Sweden | Department of Microbiology, The Public Health Agency of Sweden |
| EPI_ISL_676528 | Uppsala klinisk mikrobiologi | The Public Health Agency of Sweden | Department of Microbiology, The Public Health Agency of Sweden |
| EPI_ISL_676615 | Texas Department of State Health Services | Texas Department of State Health Services | Rashmi Tuladhar, Bonnie Oh, Jenny Zhang, Maliha Rahman, Anita Pokharel, Myong Koag, Chung Wang, Rachel Lee, Grace Kubin, Mayela Pedrueza, James Daniel Bonser |
| EPI_ISL_676657 | Wadsworth Center, New York State Department.of Health | Wadsworth Center, New York State Department.of Health | Kirsten St. George, Daryl M. Lamson, Alexis Russel, Jonathan Plitnick, Navjot Singh, John Kelly, Sara Griesemer, Erasmus Schneider, Erica Lasek-Nesselquist |
| EPI_ISL_676665, EPI_ISL_676673 | Masonic Medical Research Institute | Wadsworth Center, New York State Department.of Health | Nathan Tucker, Kirsten St. George, Daryl M. Lamson, Alexis Russel, Jonathan Plitnick, Navjot Singh, John Kelly, Sara Griesemer, Erasmus Schneider, Erica Lasek-Nesselquist |
| EPI_ISL_676707, EPI_ISL_676994 | Wadsworth Center, New York State Department.of Health | Wadsworth Center, New York State Department.of Health | Kirsten St. George, Daryl M. Lamson, Alexis Russel, Jonathan Plitnick, Navjot Singh, John Kelly, Sara Griesemer, Erasmus Schneider, Erica Lasek-Nesselquist |
| EPI_ISL_677025, EPI_ISL_677115, EPI_ISL_677120, EPI_ISL_677121, EPI_ISL_677123 | Masonic Medical Research Institute | Wadsworth Center, New York State Department.of Health | Nathan Tucker, Kirsten St. George, Daryl M. Lamson, Alexis Russel, Jonathan Plitnick, Navjot Singh, John Kelly, Sara Griesemer, Erasmus Schneider, Erica Lasek-Nesselquist |
| EPI_ISL_677295, EPI_ISL_677296, EPI_ISL_677297 | Colorado Department of Public Health and Environment | Colorado Department of Puplic Health and Environment | Laura Bankers, Molly Hetherington-Rauth, Shannon Ely, Shannon R. Matzinger, Sarah Elizabeth Totten, Emily A. Travanty |
| EPI_ISL_677719 | General Hospital - Struga | Research Center for Genetic Engineering and Biotechnology "Georgi D. Efremov" , Macedonian Academy of Sciences and Arts | RCGEB - MASA |
| EPI_ISL_677720, EPI_ISL_677721 | Clinical Hospital - Shtip | Research Center for Genetic Engineering and Biotechnology "Georgi D. Efremov" , Macedonian Academy of Sciences and Arts | RCGEB - MASA |
| EPI_ISL_677826, EPI_ISL_677828, EPI_ISL_677829, EPI_ISL_677830, EPI_ISL_677832, EPI_ISL_677833, EPI_ISL_677834, EPI_ISL_677836, EPI_ISL_677840, EPI_ISL_677841, EPI_ISL_677843, EPI_ISL_677846, EPI_ISL_677847, EPI_ISL_677848, EPI_ISL_677849, EPI_ISL_677850, EPI_ISL_677851, EPI_ISL_677853, EPI_ISL_677854, EPI_ISL_677857, EPI_ISL_677858, EPI_ISL_677859, EPI_ISL_677860, EPI_ISL_677861, EPI_ISL_677863, EPI_ISL_677864, EPI_ISL_677866, EPI_ISL_677868, EPI_ISL_677869, EPI_ISL_677870, EPI_ISL_677871, EPI_ISL_677873, EPI_ISL_677876, EPI_ISL_677877, EPI_ISL_677878, EPI_ISL_677879, EPI_ISL_677881, EPI_ISL_677883, EPI_ISL_677884, EPI_ISL_677885, EPI_ISL_677886, EPI_ISL_677887, EPI_ISL_677906 | | | |
| see above | Innovative Genomics Institute, UC Berkeley | Innovative Genomics Institute, UC Berkeley | Stacia Wyman, Haridha Shivram, Phil Frankino, Liana Lareau, Shana McDevitt, Justin Choi |
| EPI_ISL_678164, EPI_ISL_678165 | Pathogen Genomics Lab King Abdullah University of Science and Technology(KAUST) | Pathogen Genomics Lab King Abdullah University of Science and Technology(KAUST) | Sara Mfarrej, Raushan Nugmanova, Olga Douvropoulou, Raeece Naeem, Sharif Hala, Luke Esau, Amanda Ooi, Awad Al-Omari, Samer Salih, Abbas Al Mutair, Arnab Pain |
| EPI_ISL_678170 | Pathogen Genomics Lab King Abdullah University of Science and Technology(KAUST) | Pathogen Genomics Lab King Abdullah University of Science and Technology(KAUST) | Muhammad Shuaib, Sara Mfarrej, Raushan Nugmanova, Olga Douvropoulou, Raeece Naeem, Sharif Hala, Luke Esau, Amanda Ooi, Awad Al-Omari, Samer Salih, Abbas Al Mutair, Arnab Pain |
| EPI_ISL_678174 | Pathogen Genomics Lab King Abdullah University of Science and Technology(KAUST) | Pathogen Genomics Lab King Abdullah University of Science and Technology(KAUST) | Muhammad Shuaib, Sara Mfarrej, Amanda Ooi, Luke Esau, Sharif Hala, Raeece Naeem, Awad Al-Omari, Samer Salih, Abbas Al Mutair, Arnab Pain |
| EPI_ISL_678178, EPI_ISL_678179 | Pathogen Genomics Lab King Abdullah University of Science and Technology(KAUST) | Pathogen Genomics Lab King Abdullah University of Science and Technology(KAUST) | Sara Mfarrej, Sharif Hala, Luke Esau, Amanda Ooi, Raeece Naeem, Awad Al-Omari, Samer Salih, Abbas Al Mutair, Arnab Pain |
| EPI_ISL_678182 | Pathogen Genomics Lab King Abdullah University of Science and Technology(KAUST) | Pathogen Genomics Lab King Abdullah University of Science and Technology(KAUST) | Sara Mfarrej, Olga Douvropoulou, Raushan Nugmanova, Raeece Naeem, Sharif Hala, Awad Al-Omari, Samer Salih, Abbas Al Mutair, Arnab Pain |
| EPI_ISL_678184 | Pathogen Genomics Lab King Abdullah University of Science and Technology(KAUST) | Pathogen Genomics Lab King Abdullah University of Science and Technology(KAUST) | Muhammad Shuaib, Raeece Naeem, Sara Mfarrej, Olga Douvropoulou, Raushan Nugmanova, Sharif Hala, Awad Al-Omari, Samer Salih, Abbas Al Mutair, Arnab Pain |
| EPI_ISL_678211 | Pathogen Genomics Lab King Abdullah University of Science and Technology(KAUST) | Pathogen Genomics Lab King Abdullah University of Science and Technology(KAUST) | Muhammad Shuaib, Sara Mfarrej, Amanda Ooi, Luke Esau, Sharif Hala, Raeece Naeem, Awad Al-Omari, Samer Salih, Abbas Al Mutair, Arnab Pain |
| EPI_ISL_678214 | Pathogen Genomics Lab King Abdullah University of Science and Technology(KAUST) | Pathogen Genomics Lab King Abdullah University of Science and Technology(KAUST) | Muhammad Shuaib, Raeece Naeem, Sara Mfarrej, Olga Douvropoulou, Raushan Nugmanova, Sharif Hala, Awad Al-Omari, Samer Salih, Abbas Al Mutair, Arnab Pain |
| EPI_ISL_678215 | Pathogen Genomics Lab King Abdullah University of Science and Technology(KAUST) | Pathogen Genomics Lab King Abdullah University of Science and Technology(KAUST) | Sara Mfarrej, Olga Douvropoulou, Raushan Nugmanova, Raeece Naeem, Sharif Hala, Awad Al-Omari, Samer Salih, Abbas Al Mutair, Arnab Pain |
| EPI_ISL_678218 | Pathogen Genomics Lab King Abdullah University of Science and Technology(KAUST) | Pathogen Genomics Lab King Abdullah University of Science and Technology(KAUST) | Sara Mfarrej, Sharif Hala, Luke Esau, Amanda Ooi, Raeece Naeem, Awad Al-Omari, Samer Salih, Abbas Al Mutair, Arnab Pain |
| EPI_ISL_678219 | Pathogen Genomics Lab King Abdullah University of Science and Technology(KAUST) | Pathogen Genomics Lab King Abdullah University of Science and Technology(KAUST) | Sara Mfarrej, Raeece Naeem, Luke Esau, Amanda Ooi, Sharif Hala, Awad Al-Omari, Samer Salih, Abbas Al Mutair, Arnab Pain |
| EPI_ISL_678239 | Pathogen Genomics Lab King Abdullah University of Science and Technology(KAUST) | Pathogen Genomics Lab King Abdullah University of Science and Technology(KAUST) | Muhammad Shuaib, Raeece Naeem, Sara Mfarrej, Olga Douvropoulou, Raushan Nugmanova, Sharif Hala, Awad Al-Omari, Samer Salih, Abbas Al Mutair, Arnab Pain |
| EPI_ISL_678240 | Pathogen Genomics Lab King Abdullah University of Science and Technology(KAUST) | Pathogen Genomics Lab King Abdullah University of Science and Technology(KAUST) | Sara Mfarrej, Raushan Nugmanova, Olga Douvropoulou, Raeece Naeem, Sharif Hala, Luke Esau, Amanda Ooi, Awad Al-Omari, Samer Salih, Abbas Al Mutair, Arnab Pain |
| EPI_ISL_678274, EPI_ISL_678281, EPI_ISL_678282, EPI_ISL_678284, EPI_ISL_678285 | Mikrobiologie, RARI | Mikrobiologie, RARI | Krasnov,Y.M., Naryshkina,E.A., Guseva,N.P., Sosedova,E.A., Fedorov,A.V., Badanin,D.V., Sharapova,N.A., Portenko,S.A., Shcherbakova,S.A., Kutyrev,V.V. |
| EPI_ISL_678326 | Area of Virology, Serology and Virology Division (SAViD), New South Wales Health Pathology Randwick | Virology Research Laboratory; Area of Virology, Serology and Virology Division (SAViD), New South Wales Health Pathology Randwick | Foster, C.; Au, J.; Ruiz Silva, M.; Deveson, I.; Bull, R.; Van Hal, S.; Rawlinson, W. |
| EPI_ISL_681695 | Molecular Medicine Laboratory, University of Magallanes | Centro Asistencial Docente y de Investigacion, Universidad de Magallanes | Jorge González, Jacqueline Aldridge, Diego Alvarez, Marco Montes de Oca, Hermy Alvarez, Roberto Uribe-Paredes, Marcelo Navarrete |
| EPI_ISL_681831, EPI_ISL_681839 | Molecular diagnostic unit for viral haemorrhagic fevers and emerging viruses, Bouaké CHU Laboratory | Project group Epidemology of Highly Pathogenic Microorganisms, Robert Koch-Institute | Chantal Akoua-Koffi, Diané Bamourou, Etilé Anoh, Essia Belarbi, Safiatou Karidioula, Grit Schubert, Adjaratou Traoré, Soundélé Maïté, Monemo Pacome, Coulibaly Mbegnan, Bamba Fatoumata Touré, Kra Ouffoué, Fabian Leendertz |

| | | | |
|---|---|---|---|
| EPI_ISL_681931, EPI_ISL_681932, EPI_ISL_681933, EPI_ISL_681934, EPI_ISL_681935 | UPMC Clinical Microbiology Laboratory | Microbial Genomic Epidemiology Laboratory, University of Pittsburgh | Mustapha M. Mustapha, Jane W. Marsh, Dan Snyder, Marissa P. Griffith, Stephanie L. Mitchell, Vatsala R. Srinivasa, Kady D. Waggle, Chinelo Ezeonwuku, Vaughn S. Cooper, Lee H. Harrison |
| EPI_ISL_682257 | HOSPITAL SAN JUAN DE DIOS | Inciensa, Instituto Costarricense de Investigación y Enseñanza en Nutrición y Salud | Francisco Duarte, Hebleen Porras, Claudio Soto-Garita, Estela Cordero, Adriana Godinez & Melany Calderon |
| EPI_ISL_682258 | AREA DE SALUD ALAJUELA NORTE - CLINICA DR. MARCIAL RODRIGUEZ | Inciensa, Instituto Costarricense de Investigación y Enseñanza en Nutrición y Salud | Francisco Duarte, Hebleen Porras, Claudio Soto-Garita, Estela Cordero, Adriana Godinez & Melany Calderon |
| EPI_ISL_682259 | AREA DE SALUD ALAJUELA NORTE - CLINICA DR. MARCIAL RODRIGUEZ | Inciensa, Instituto Costarricense de Investigación y Enseñanza en Nutrición y Salud | Francisco Duarte, Hebleen Porras, Claudio Soto-Garita, Estela Cordero, Adriana Godinez, Melany Calderon & Mariel López |
| EPI_ISL_682260 | AREA DE SALUD CATEDRAL NORESTE | Inciensa, Instituto Costarricense de Investigación y Enseñanza en Nutrición y Salud | Francisco Duarte, Hebleen Porras, Claudio Soto-Garita, Estela Cordero, Adriana Godinez, Melany Calderon & Mariel López |
| EPI_ISL_682261 | TAMIZAJE COMUNITARIO- PASO CANOAS | Inciensa, Instituto Costarricense de Investigación y Enseñanza en Nutrición y Salud | Francisco Duarte, Hebleen Porras, Claudio Soto-Garita, Estela Cordero, Adriana Godinez & Melany Calderon |
| EPI_ISL_683600, EPI_ISL_683633, EPI_ISL_683639, EPI_ISL_683640, EPI_ISL_683644 | Servicio de Microbiología, Laboratori Clínic Metropolitana Nord. Hospital Universitari Germans Trias i Pujol. Institut d'Investigació en Ciències de la Salut Germans Trias i Pujol (IGTP) | SeqCOVID-SPAIN consortium/IBV(CSIC) | Elisa Martró, Antoni E. Bordoy, Anna Not, Adrián Antuori, Anabel Fernández, Nona Romaní, Verónica Saludes, Cristina Casañ and SeqCOVID-SPAIN consortium |
| EPI_ISL_684002 | Utah Public Health Laboratory | Utah Public Health Laboratory | Erin Young, Kelly Oakeson |
| EPI_ISL_691628, EPI_ISL_691631, EPI_ISL_691632, EPI_ISL_691633, EPI_ISL_691636, EPI_ISL_691637 | Servicio de Microbiología, Hospital Universitario Son Espases | SeqCOVID-SPAIN consortium/IBV(CSIC) | Carla López-Causapé, Jordi Reina, Antonio Oliver and SeqCOVID-SPAIN consortium |
| EPI_ISL_693298 | Department of Microbiology, Yokohama City University School of Medicine | Department of Microbiology, Yokohama City University School of Medicine | Kei Miyakawa, Ryo Saji, Kazuya Sakai, Reo Matsumura, Mototsugu Nishii, Ichiro Takeuchi, Akihide Ryo |
| EPI_ISL_693697 | Delaware Public Health Laboratory | Delaware Public Health Laboratory | Gregory Hovan |
| EPI_ISL_695521, EPI_ISL_695522, EPI_ISL_695523, EPI_ISL_695524, EPI_ISL_695525, EPI_ISL_695526, EPI_ISL_695527, EPI_ISL_695528, EPI_ISL_695529, EPI_ISL_695530, EPI_ISL_695531, EPI_ISL_695532, EPI_ISL_695533, EPI_ISL_695534, EPI_ISL_695535, EPI_ISL_695536, EPI_ISL_695537, EPI_ISL_695538, EPI_ISL_695539, EPI_ISL_695540, EPI_ISL_695541, EPI_ISL_695542, EPI_ISL_695543, EPI_ISL_695544, EPI_ISL_695545, EPI_ISL_695546, EPI_ISL_695547, EPI_ISL_695548, EPI_ISL_695549, EPI_ISL_695550, EPI_ISL_695551, EPI_ISL_695552, EPI_ISL_695553, EPI_ISL_695554, EPI_ISL_695555, EPI_ISL_695556, EPI_ISL_695557, EPI_ISL_695558, EPI_ISL_695559, EPI_ISL_695560, EPI_ISL_695561 | | | |
| see above | TGen North | TGen North | Jolene Bowers, Megan Folkerts, Chris French, Hayley Yaglom, Ashlyn Pfeiffer, Darrin Lemmer, Dave Engelthaler, The Arizona COVID Genomics Union (ACGU) |
| EPI_ISL_695701, EPI_ISL_695712, EPI_ISL_695713, EPI_ISL_695714, EPI_ISL_695715, EPI_ISL_695716, EPI_ISL_695717, EPI_ISL_695718, EPI_ISL_695719, EPI_ISL_695720, EPI_ISL_695721, EPI_ISL_695722, EPI_ISL_695723, EPI_ISL_695724, EPI_ISL_695725, EPI_ISL_695726, EPI_ISL_695727, EPI_ISL_695728, EPI_ISL_695729, EPI_ISL_695730, EPI_ISL_695731, EPI_ISL_695732, EPI_ISL_695733, EPI_ISL_695734, EPI_ISL_695735, EPI_ISL_695736, EPI_ISL_695737, EPI_ISL_695738, EPI_ISL_695739, EPI_ISL_695740 | | | |
| see above | AZ SPHL, Arizona Department of Health Services | TGen North | Jolene Bowers, Megan Folkerts, Chris French, Hayley Yaglom, Ashlyn Pfeiffer, Darrin Lemmer, Dave Engelthaler, The Arizona COVID Genomics Union (ACGU) |
| EPI_ISL_699508, EPI_ISL_699509 | Diagnostic Virology Laboratory, USDA National Veterinary Services Laboratories | Diagnostic Virology Laboratory, USDA National Veterinary Services Laboratories | Hamer,S.A., Pauvolid-Correa,A., Zecca,I.B., Davila,E., Auckland,L.D., Roundy,C.M., Tang,W., Torchetti,M., Killian,M.L., Jenkins-Moore,M., Akpalu,Y., Ghai,R.R., Spengler,J., Barton Behravesh,C., Fischer,R.S., Hamer,G.L., Franzen,K.M., Love,E.R. |
| EPI_ISL_699650 | Douglas Hanly Moir | NSW Health Pathology - Institute of Clinical Pathology and Medical Research; Westmead Hospital; University of Sydney | CIDM-PH et al. |
| EPI_ISL_700005, EPI_ISL_700006, EPI_ISL_700007, EPI_ISL_700008, EPI_ISL_700009, EPI_ISL_700010, EPI_ISL_700011, EPI_ISL_700012, EPI_ISL_700013, EPI_ISL_700014, EPI_ISL_700015, EPI_ISL_700016, EPI_ISL_700017, EPI_ISL_700018, EPI_ISL_700019, EPI_ISL_700020, EPI_ISL_700021, EPI_ISL_700022, EPI_ISL_700023, EPI_ISL_700024, EPI_ISL_700025, EPI_ISL_700026, EPI_ISL_700027, EPI_ISL_700028, EPI_ISL_700029, EPI_ISL_700030, EPI_ISL_700031, EPI_ISL_700032, EPI_ISL_700033, EPI_ISL_700034, EPI_ISL_700035, EPI_ISL_700036, EPI_ISL_700037, EPI_ISL_700038, EPI_ISL_700039, EPI_ISL_700040, EPI_ISL_700041, EPI_ISL_700042, EPI_ISL_700043, EPI_ISL_700044, EPI_ISL_700045, EPI_ISL_700046, EPI_ISL_700047, EPI_ISL_700048, EPI_ISL_700049, EPI_ISL_700050, EPI_ISL_700051, EPI_ISL_700052, EPI_ISL_700053, EPI_ISL_700054, EPI_ISL_700055, EPI_ISL_700056, EPI_ISL_700057, EPI_ISL_700058, EPI_ISL_700059, EPI_ISL_700060, EPI_ISL_700061, EPI_ISL_700062, EPI_ISL_700063, EPI_ISL_700064, EPI_ISL_700065, EPI_ISL_700066, EPI_ISL_700067, EPI_ISL_700068, EPI_ISL_700069, EPI_ISL_700070, EPI_ISL_700071, EPI_ISL_700072, EPI_ISL_700073, EPI_ISL_700074, EPI_ISL_700075, EPI_ISL_700076, EPI_ISL_700077, EPI_ISL_700078 | | | |
| see above | Hematopathology Laboratory, ACTREC, TMC | Hematopathology Laboratory, ACTREC, TMC | Hematopathology Laboratory, ACTREC |
| EPI_ISL_707787 | Rwanda National Reference Laboratory | Rwanda National Reference Laboratory | Enatha Mukantwari,Jeanne d'Arc Umuringa |
| EPI_ISL_707937, EPI_ISL_707938, EPI_ISL_707939 | Pamukkale University Hospital | Pamukkale University Department of Medical Genetics | Onur TOKGUN et al. |
| EPI_ISL_708027 | University Hospital of Northern Norway, Department for Microbiology and Infectious Disease Control | Norwegian Institute of Public Health, Department of Virology | Kathrine Stene-Johansen, Kamilla Heddeland Instefjord, Hilde Elshaug, Marie Paulsen Madsen, Rasmus Riis Kopperud, Hilde Vollan, Karoline Bragstad, Olav Hungnes |
| EPI_ISL_708184, EPI_ISL_708185, EPI_ISL_708190, EPI_ISL_708191 | Pamukkale University Hospital | Pamukkale University Department of Medical Genetics | Onur TOKGUN et al. |
| EPI_ISL_708398 | Delaware Public Health Lab | Delaware Public Health Lab | Gregory Hovan |
| EPI_ISL_708737, EPI_ISL_708797, EPI_ISL_708800 | Regional medical sciences center 6 chonburi | National Institute of Health, Department of Medical Sciences, Ministry of Public Health, Thailand | Pilailuk Okada; Siripaporn Phuygun; Thanutsapa Thanadachakul; Sittiporn Parnmen; Warawan Wongboot; Sunthareeya Waicharoen; Malinee Chittaganpitch |
| EPI_ISL_708801, EPI_ISL_708802 | Regional medical sciences center 6 chonburi | National Institute of Health, Department of Medical Sciences, Ministry of Public Health, Thailand | Pilailuk Okada; Siripaporn Phuygun; Thanutsapa Thanadachakul; Sittiporn Parnmen; Pakorn Piromtong; Warawan Wongboot; Sunthareeya Waicharoen; Malinee Chittaganpitch |
| EPI_ISL_710099, EPI_ISL_710100, EPI_ISL_710101, EPI_ISL_710109 | Los Angeles County PHL | Los Angeles County PHL | P. Hemarajata et al. |
| EPI_ISL_710215, EPI_ISL_710319 | Colorado Department of Public Health and Environment | Colorado Department of Puplic Health and Environment | Laura Bankers, Molly C. Hetherington-Rauth, Shannon Ely, Shannon R. Matzinger, Sarah Elizabeth Totten, Emily A. Travanty |
| EPI_ISL_714931 | Department of Virus and Microbiological Special Diagnostics, Statens Serum Institut, Copenhagen, Denmark | Albertsen Lab, Department of Chemistry and Bioscience, Aalborg University, Denmark | Danish Covid-19 Genome Consortium |
| EPI_ISL_717785, EPI_ISL_717786, EPI_ISL_717788, EPI_ISL_717789, EPI_ISL_717790, EPI_ISL_717792, EPI_ISL_717794, EPI_ISL_717795, EPI_ISL_717796, EPI_ISL_717798, EPI_ISL_717799, EPI_ISL_717800, EPI_ISL_717803, EPI_ISL_717804 | | | |
| see above | LACEN RJ - Noel Nutels | Bioinformatics Laboratory / LNCC | Carolina M Voloch, Ronaldo da Silva F Jr, Luiz G P de Almeida, Cynthia C Cardoso, Otavio Bustrolini, Alexandra L Gerber, Ana Paula de C Guimarães, Diana Mariani, Andréa Cony Cavalcanti, Claudia dos Santos Rodrigues, Terezinha M P P Castiñeira, Amilcar Tanuri, Ana Tereza R de Vasconcelos |
| EPI_ISL_717841 | LACEN Dr. Francisco Rimolo Neto | Bioinformatics Laboratory / LNCC | Carolina M Voloch, Ronaldo da Silva F Jr, Luiz G P de Almeida, Cynthia C Cardoso, Otavio Bustrolini, Alexandra L Gerber, Ana Paula de C Guimarães, Diana Mariani, Andréa Cony Cavalcanti, Claudia dos Santos Rodrigues, Terezinha M P P Castiñeira, Amilcar Tanuri, Ana Tereza R de Vasconcelos |
| EPI_ISL_717899, EPI_ISL_717900, EPI_ISL_717901, EPI_ISL_717902, EPI_ISL_717903, EPI_ISL_717904, EPI_ISL_717905, EPI_ISL_717906, EPI_ISL_717909 | LACEN RJ - Noel Nutels | Bioinformatics Laboratory / LNCC | Carolina M Voloch, Ronaldo da Silva F Jr, Luiz G P de Almeida, Cynthia C Cardoso, Otavio Bustrolini, Alexandra L Gerber, Ana Paula de C Guimarães, Diana Mariani, Andréa Cony Cavalcanti, Claudia dos Santos Rodrigues, Terezinha M P P Castiñeira, Amilcar Tanuri, Ana Tereza R de Vasconcelos |
| EPI_ISL_717913, EPI_ISL_717914, | LACEN Dr. Francisco Rimolo Neto | Bioinformatics Laboratory / LNCC | Carolina M Voloch, Ronaldo da Silva F Jr, Luiz G P de Almeida, Cynthia C Cardoso, Otavio Bustrolini, Alexandra L Gerber, Ana Paula de C Guimarães, Diana |

| | | | |
|---|---|---|---|
| EPI_ISL_717915, EPI_ISL_717916, EPI_ISL_717917, EPI_ISL_717958 | | | Mariani, Andréa Cony Cavalcanti, Claudia dos Santos Rodrigues, Terezinha M P P Castiñeira, Amilcar Tanuri, Ana Tereza R de Vasconcelos |
| EPI_ISL_717962 | LACEN RJ - Noel Nutels | Bioinformatics Laboratory / LNCC | Carolina M Voloch, Ronaldo da Silva F Jr, Luiz G P de Almeida, Cynthia C Cardoso, Otavio Bustrolini, Alexandra L Gerber, Ana Paula de C Guimarães, Diana Mariani, Andréa Cony Cavalcanti, Claudia dos Santos Rodrigues, Terezinha M P P Castiñeira, Amilcar Tanuri, Ana Tereza R de Vasconcelos |
| EPI_ISL_717963, EPI_ISL_717964 | LACEN Dr. Francisco Rimolo Neto | Bioinformatics Laboratory / LNCC | Carolina M Voloch, Ronaldo da Silva F Jr, Luiz G P de Almeida, Cynthia C Cardoso, Otavio Bustrolini, Alexandra L Gerber, Ana Paula de C Guimarães, Diana Mariani, Andréa Cony Cavalcanti, Claudia dos Santos Rodrigues, Terezinha M P P Castiñeira, Amilcar Tanuri, Ana Tereza R de Vasconcelos |
| EPI_ISL_718267, EPI_ISL_718268, EPI_ISL_718269, EPI_ISL_718270, EPI_ISL_718283, EPI_ISL_718284 | Institute for Medical Research, Infectious Disease Research Centre, National Institutes of Health, Ministry of Health Malaysia | Institute for Medical Research, Infectious Disease Research Centre, National Institutes of Health, Ministry of Health Malaysia | Suppiah J, Kamel K, Mohd-Zawawi Z, Thayan R |
| EPI_ISL_721631, EPI_ISL_721632, EPI_ISL_721633, EPI_ISL_721634, EPI_ISL_721635, EPI_ISL_721636 | Armed Forces Medical College | National Centre For Cell Science | Dhiraj Paul, Kunal Jani, Radha Chauhan, Janesh Kumar, Vasudevan Seshadri, Girdhari Lal, Rajesh Karyakarte, Suvarna Joshi, Murlidhar Tambe, Sourav Sen, Santosh Karade, Kavita Bala Anand, Shelinder Pal Singh Shergill, Rajiv Mohan Gupta, Manoj Kumar Bhat, Arvind Sahu, Yogesh S Shouche |
| EPI_ISL_721654, EPI_ISL_721655, EPI_ISL_721656, EPI_ISL_722177, EPI_ISL_722179 | National Centre For Cell Science | National Centre For Cell Science | Dhiraj Paul, Kunal Jani, Radha Chauhan, Janesh Kumar, Vasudevan Seshadri, Girdhari Lal, Rajesh Karyakarte, Suvarna Joshi, Murlidhar Tambe, Sourav Sen, Santosh Karade, Kavita Bala Anand, Shelinder Pal Singh Shergill, Rajiv Mohan Gupta, Manoj Kumar Bhat, Arvind Sahu, Yogesh S Shouche |
| EPI_ISL_722198, EPI_ISL_722199, EPI_ISL_722200 | Armed Forces Medical College | National Centre For Cell Science | Dhiraj Paul, Kunal Jani, Radha Chauhan, Janesh Kumar, Vasudevan Seshadri, Girdhari Lal, Rajesh Karyakarte, Suvarna Joshi, Murlidhar Tambe, Sourav Sen, Santosh Karade, Kavita Bala Anand, Shelinder Pal Singh Shergill, Rajiv Mohan Gupta, Manoj Kumar Bhat, Arvind Sahu, Yogesh S Shouche |
| EPI_ISL_722332, EPI_ISL_722354, EPI_ISL_722810, EPI_ISL_722811, EPI_ISL_722812, EPI_ISL_722813, EPI_ISL_722814 | Dutch COVID-19 response team | Erasmus Medical Center | Bas Oude Munnink, Reina Sikkema, David Nieuwenhuijse, Irina Chestakova, Anne van der Linden, Marjan Boter, Emmanuelle Munger, Corine GeurtsvanKessel, Annemiek van der Eijk, Richard Molenkamp, Marion Koopmans, on behalf of the Dutch national COVID-19 response team. |
| EPI_ISL_722897 | Dipartimento di Scienze Biomediche e Oncologia Umana - Azienda Ospedaliero Universitaria Consorziale Policlinico | Istituto Zooprofilattico Sperimentale della Puglia e della Basilicata | Parisi A., Bianco A., Capozzi L., Del Sambro L., Chironna M., Loconsole D. |
| EPI_ISL_728326 | B.J. Govt. Medical College | National Centre For Cell Science | Dhiraj Paul, Kunal Jani, Radha Chauhan, Janesh Kumar, Vasudevan Seshadri, Girdhari Lal, Rajesh Karyakarte, Suvarna Joshi, Murlidhar Tambe, Sourav Sen, Santosh Karade, Kavita Bala Anand, Shelinder Pal Singh Shergill, Rajiv Mohan Gupta, Manoj Kumar Bhat, Arvind Sahu, Yogesh S Shouche |
| EPI_ISL_729572, EPI_ISL_729573, EPI_ISL_729574, EPI_ISL_729575, EPI_ISL_729576, EPI_ISL_729577, EPI_ISL_729578, EPI_ISL_729579, EPI_ISL_729580, EPI_ISL_729581, EPI_ISL_729582, EPI_ISL_729583 | | | |
| see above | A. Krumbholz, Labor Dr. Krause und Kollegen MVZ GmbH, Kiel | Charité Universitätsmedizin Berlin, Institut für Virologie | Victor M Corman, Barbara Mühlemann, Jörn Beheim-Schwarzbach, Talitha Veith, Julia Schneider, Terry Jones, Christian Drosten |
| EPI_ISL_729845, EPI_ISL_729848, EPI_ISL_729851 | Laboratorio Central de Saude Publica do Estado do Rio Grande do Sul (LACEN-RS) | Laboratory of Respiratory Viruses and Measles, Oswaldo Cruz Institute, FIOCRUZ | Paola Resende, Luciana Appolinario, Fernando Motta, Anna Carolina Paixão, Ana Carolina Mendonça, Tatiana Schaffer Gregianini, Marilda Tereza Mar da Rosa, Marilda Siqueira |
| EPI_ISL_729958, EPI_ISL_729994 | Nigeria Centre for Disease Control (NCDC) | African Centre of Excellence for Genomics of Infectious Diseases (ACEGID), Redeemer's University, Ede, Osun State, Nigeria | Oluniyi P.E. et al |
| EPI_ISL_732531 | Bundeswehr Institute of Microbiology | Bundeswehr Institute of Microbiology | Elham Khatamzas, Markus Antwerpen, Mathias Walter, Alexandra Rehn, Sabine Zange, Enrico Georgi, Michael von Bergwelt-Baildon, Roman Wölfel |
| EPI_ISL_732994, EPI_ISL_732995, EPI_ISL_732996, EPI_ISL_732997, EPI_ISL_732998, EPI_ISL_732999 | UMMC-Health | WHO National Influenza Centre Russian Federation | Andrey Komissarov, Artem Fadeev, Anna Ivanova, Kseniya Komissarova, Dmitry Bazhenov, Tatiana Platonova, Daria Danilenko, Ksenia Safina, Elena Nabieva, Georgii Bazykin, Dmitry Lioznov |
| EPI_ISL_733015 | WHO National Influenza Centre Russian Federation | WHO National Influenza Centre Russian Federation | Andrey Komissarov, Artem Fadeev, Anna Ivanova, Kseniya Komissarova, Dmitry Bazhenov, Daria Danilenko, Ksenia Safina, Elena Nabieva, Georgii Bazykin, Dmitry Lioznov |
| EPI_ISL_733170, EPI_ISL_733171 | Pathogenic Microorganisms Variability Laboratory | WHO National Influenza Centre Russian Federation | Andrey Komissarov, Artem Fadeev, Anna Ivanova, Kseniya Komissarova, Dmitry Bazhenov, Daria Danilenko, Ksenia Safina, Elena Nabieva, Georgii Bazykin, Nadezhda Kuznetsova, Elena Shidlovskaya, Sergey Alkhovsky, Tatyana Vishnevskaya, Elizaveta Divisenko, Alexey Shchetinin, Maria Nikiforova, Andrey Pochtovyy, Evgeny Usachev, Elena Vokalova, Maxim Rubalsky, Oleg Rubalsky, Artem Tkachuk, Vladimir Gushchin, Alexander Gintsburg, Dmitry Lioznov |
| EPI_ISL_734825, EPI_ISL_734826, EPI_ISL_734827, EPI_ISL_734828, EPI_ISL_734829, EPI_ISL_734830, EPI_ISL_734831, EPI_ISL_734832, EPI_ISL_734833, EPI_ISL_734834, EPI_ISL_734835, EPI_ISL_734836, EPI_ISL_734837, EPI_ISL_734838, EPI_ISL_734839 | | | |
| see above | UZ Leuven, National Reference Laboratory for Coronaviruses, Laboratory Medicine, Leuven, Belgium | KU Leuven, Rega Institute, Clinical and Epidemiological Virology | Tony Wawina-Bokalanga, Joan Marti-Carerras, Bert Vanmechelen, Piet Maes |
| EPI_ISL_737931, EPI_ISL_737965, EPI_ISL_737974, EPI_ISL_737975, EPI_ISL_737976 | Uganda Central Public Health Lab and Uganda Virus Research Institute | MRC/UVRI & LSHTM Uganda Research Unit | Matthew Cotten, Dan Lule Bugembe, My V.T. Phan, Pontiano Kaleebu et al. |
| EPI_ISL_738519, EPI_ISL_738547, EPI_ISL_738626, EPI_ISL_738813, EPI_ISL_738819, EPI_ISL_738904, EPI_ISL_739093, EPI_ISL_739338, EPI_ISL_739407 | Alameda County Public Health Lab | Chan-Zuckerberg Biohub | CZB Cliahub Consortium |
| EPI_ISL_739834, EPI_ISL_740048, EPI_ISL_740079, EPI_ISL_740098, EPI_ISL_740251, EPI_ISL_740360, EPI_ISL_740430, EPI_ISL_744198, EPI_ISL_744202, EPI_ISL_744235, EPI_ISL_744406, EPI_ISL_744434, EPI_ISL_744444, EPI_ISL_744598, EPI_ISL_744719, EPI_ISL_744751, EPI_ISL_744958, EPI_ISL_745013, EPI_ISL_745022 | | | |
| see above | Laboratoire national de santé, Microbiology, Virology | Laboratoire national de santé, Microbiology, Microbial Genomics Platform | Anke Wienecke-Baldacchino, Catherine Ragimbeau, Tamir Abdelrahman, Jessica Tapp, Fatu Djabi |
| EPI_ISL_745472, EPI_ISL_745483, EPI_ISL_745484, EPI_ISL_745486, EPI_ISL_745492, EPI_ISL_745501, EPI_ISL_745503, EPI_ISL_745510, EPI_ISL_745519, EPI_ISL_745527, EPI_ISL_745547, EPI_ISL_745558, EPI_ISL_745561, EPI_ISL_745568, EPI_ISL_745575, EPI_ISL_745591, EPI_ISL_745598, EPI_ISL_745603, EPI_ISL_745605, EPI_ISL_745609, EPI_ISL_745615, EPI_ISL_745620, EPI_ISL_745621, EPI_ISL_745633, EPI_ISL_745636, EPI_ISL_745644, EPI_ISL_745657, EPI_ISL_745659, EPI_ISL_745661, EPI_ISL_745662, EPI_ISL_745665, EPI_ISL_745671, EPI_ISL_745672, EPI_ISL_745673, EPI_ISL_745674, EPI_ISL_745675, EPI_ISL_745676, EPI_ISL_745677, EPI_ISL_745678, EPI_ISL_745679, EPI_ISL_745680, EPI_ISL_745681, EPI_ISL_745682, EPI_ISL_745683, EPI_ISL_745684, EPI_ISL_745685, EPI_ISL_745686, EPI_ISL_745687, EPI_ISL_745688, EPI_ISL_745689, EPI_ISL_745690, EPI_ISL_745691, EPI_ISL_745692, EPI_ISL_745693, EPI_ISL_745694, EPI_ISL_745695, EPI_ISL_745696, EPI_ISL_745697, EPI_ISL_745698, EPI_ISL_745699, EPI_ISL_745700, EPI_ISL_745701, EPI_ISL_745702, EPI_ISL_745703, EPI_ISL_745704, EPI_ISL_745705, EPI_ISL_745706, EPI_ISL_745707, EPI_ISL_745708, EPI_ISL_745709, EPI_ISL_745710, EPI_ISL_745711, EPI_ISL_745712, EPI_ISL_745713, EPI_ISL_745714, EPI_ISL_745715, EPI_ISL_745716, EPI_ISL_745717, EPI_ISL_745718, EPI_ISL_745719, EPI_ISL_745720, EPI_ISL_745730, EPI_ISL_745797, EPI_ISL_745821, EPI_ISL_746033, EPI_ISL_746050, EPI_ISL_746057, EPI_ISL_746058, EPI_ISL_746088 | | | |
| see above | Ginkgo Bioworks Clinical Laboratory | Utah Public Health Laboratory | Erin L. Young, Kelly Oakeson, Tara Gallagher, Michael T. Pyne, E. Susan Slechta, Melanie A. Mallory, Jeffrey B. Stevenson, Salika M. Shakir, David R. Hillyard, Malaika McKenzie-Bennett, James McGann, Jim Griffin, Keith Robison, Alex Plocik, Becky Schilling, Martha Pierson, Rebecca Littlefield, Michelle Spencer, Birgitte Simen |
| EPI_ISL_746625, EPI_ISL_746626, EPI_ISL_746628, EPI_ISL_746629, EPI_ISL_746630, EPI_ISL_746631, EPI_ISL_746632, EPI_ISL_746633, EPI_ISL_746634, EPI_ISL_746635, EPI_ISL_746636, EPI_ISL_746637, EPI_ISL_746638, EPI_ISL_746639, EPI_ISL_746640, EPI_ISL_746641, EPI_ISL_746642, EPI_ISL_746643 | | | |
| see above | Genetica Molecular and Subdepartamento de Virologia ISP Chile | Instituto de Salud Publica de Chile | Javier Tognarelli, Barbara Parra, Loredana Arata, Jaime Lagos, Gisselle Barra, Patricia Bustos, Rodrigo Fasce, Andres Castillo, Jorge Fernandez |
| EPI_ISL_751189, EPI_ISL_751190 | CENUR Litoral Norte - UdelaR, Salto, Uruguay | Institut Pasteur de Montevideo | Daiana Mir, Natalia Rego, Paola Cristina Resende, Fernando Lopez-Tort, Tamara Fernandez-Calero, Veronica Noya, Mariana Brandes, Tania Possi, Mailen Arleo, Natalia Reyes, Matias Victoria, Andres Lizasoain, Matias Castells, Leticia Maya, Matías Salvo, Tatiana Schäffer Gregianini, Marilda Tereza Mar da Rosa, Letícia Garay Martins, Cecilia Alonso, Yasser Vega, Cecilia Salazar, Ignacio Ferrés, Pablo Smircich, Jose Sotelo, Ighor Arantes, Luciana Appolinario, Ana Carolina Mendonça, Maria Jose Benitez-Galeano, Martín Graña, Camila Simoes, Fernando Motta, Marilda Mendonça Siqueira, Gonzalo Bello, Rodney Colina, Lucia Spangenberg |

| | | | |
|---|---|---|---|
| EPI_ISL_752719, EPI_ISL_752720, EPI_ISL_752721, EPI_ISL_752722, EPI_ISL_752723, EPI_ISL_752724, EPI_ISL_752725, EPI_ISL_752731, EPI_ISL_752732, EPI_ISL_752733, EPI_ISL_752734, EPI_ISL_752735, EPI_ISL_752736, EPI_ISL_752737, EPI_ISL_752738, EPI_ISL_752739, EPI_ISL_752740, EPI_ISL_752741, EPI_ISL_752742, EPI_ISL_752743, EPI_ISL_752744, EPI_ISL_752745, EPI_ISL_752746, EPI_ISL_752952 | | | |
| see above | State Laboratories Division, Hawaii State Department of Health | State Laboratories Division, Hawaii State Department of Health | Pamela O'Brien, Sabrina Diemert, Drew Kuwazaki, Razvan Sultana, Edward Desmond |
| EPI_ISL_753703, EPI_ISL_753708, EPI_ISL_753986, EPI_ISL_753989, EPI_ISL_753991, EPI_ISL_753993 | Charité Universitätsmedizin Berlin, Institut für Virologie/Labor Berlin | Charité Universitätsmedizin Berlin, Institut für Virologie | Victor M Corman, Jörn Beheim-Schwarzbach, Barbara Mühlemann, Julia Schneider, Talitha Veith, Terry Jones, Christian Drosten |
| EPI_ISL_754068, EPI_ISL_754069 | Nepal Korea Friendship Municipality Hospital | Nepal Health Research Council | Pradip Gyanwali, Meghnath Dhimal |
| EPI_ISL_754231 | The Republican Research and Practical Center for Epidemiology and Microbiology (RRPCEM) | WHO National Influenza Centre Russian Federation | Elena Gasich, Kirill Bulda, Anatoly Krasko, Andrey Komissarov, Artem Fadeev, Anna Ivanova, Kseniya Komissarova, Dmitry Bazhenov, Daria Danilenko, Ksenia Safina, Elena Nabieva, Georgii Bazykin, Dmitry Lioznov |
| EPI_ISL_754241 | Dinkes Tasikmalaya | "School of Life Sciences and Technology & School of Pharmacy-Institut Teknologi Bandung; Molecular Genetics Laboratory-Faculty of Medicine-Universitas Padjadjaran; Laboratorium Kesehatan Provinsi Jawa Barat" | Husna Nugrahapraja, Marselina Irasonia Tan, Yunia Sribudiani, Catur Riani, Azzania Fibriani, Tarwadi, Ema Rahmawati, Savira Ekawardhani, Hesti Lina Wiraswati, Ryan Bayusantika Ristandi, Rifky Waluyajati Rachman, Cut Nur Cinthia Alamanda, Lia Faridah, Miftahul Faridl, Karimatu Khoirunnisa, Hammam Riza, Soni Solistia Wirawan, Agung Eru Wibowo, Irvan Faizal |
| EPI_ISL_754936, EPI_ISL_755044, EPI_ISL_755045, EPI_ISL_755046 | California Department of Public Health | California Department of Public Health | CDPH IDLB COVIDNet |
| EPI_ISL_756330, EPI_ISL_756331, EPI_ISL_756332, EPI_ISL_756333, EPI_ISL_756334, EPI_ISL_756335, EPI_ISL_756336, EPI_ISL_756355 | Innovative Genomics Institute, UC Berkeley | Innovative Genomics Institute, UC Berkeley | Stacia Wyman, Haridha Shivram, Phil Frankino, Liana Lareau, Shana McDevitt, Justin Choi |
| EPI_ISL_760129, EPI_ISL_760130 | Division of Emerging Infectious Diseases, Bureau of Infectious Diseases Diagnosis Control, Korea Disease Control and Prevention Agency | Division of Emerging Infectious Diseases, Bureau of Infectious Diseases Diagnosis Control, Korea Disease Control and Prevention Agency | Ae Kyung Park, Il-Hwan Kim, Heui Man Kim, Jeong-Min Kim, Namjoo Lee, Chaeyoung Lee, Sang Hee Woo, Eun-Jin Kim |
| EPI_ISL_765680, EPI_ISL_765681, EPI_ISL_765683, EPI_ISL_765684, EPI_ISL_765685, EPI_ISL_765686 | Massachusetts General Hospital | Infectious Disease Program, Broad Institute of Harvard and MIT | Lemieux,J.E., Siddle,K.J., Shaw,B., Adams,G., Pierce,V., Turbett,S., Anahtar,M., Branda,J., Slater,D., Harris,J., Lin,A.E., Gladden-Young,A., Lagerborg,K., Rudy,M., DeRuff,K., Carter,A., Normandin,E., Bauer,M., Reilly,S., Tomkins-Tinch,C., Loreth,C., Chaluvadi,S., Neumann,A., Cusick,C., Chapman,S.B., Gnirke,A., Flowers,K., Cerrato,F., Birren,B.W., Gallagher,G., Smole,S., Park,D.J., MacInnis,B.L., Ryan,E., LaRocque,R., Rosenberg,E. and Sabeti,P.C. |
| EPI_ISL_766650, EPI_ISL_766676, EPI_ISL_766681, EPI_ISL_766687 | Texas Department of State Health Services | Texas Department of State Health Services | Rashmi Tuladhar, Bonnie Oh, Jenny Zhang, Maliha Rahman, Anita Pokharel, Myong Koag, Chung Wang, Rachel Lee, Grace Kubin, Mayela Pedrueza, James Daniel Bonser |
| EPI_ISL_768740 | Child Health Research Foundation | Child Health Research Foundation | Senjuti Saha, Afroza Akter Tanni, Roly Malaker, Sharmistha Goswami, Syed Muktadir Al Sium, Arif Mohammad Tanmoy, Md Hafizur Rahman, Samir K Saha |
| EPI_ISL_771190, EPI_ISL_771195 | Colorado Department of Public Health and Environment | Colorado Department of Puplic Health and Environment | Laura Bankers, Molly C. Hetherington-Rauth, Diana Ir, Shannon Ely, Shannon R. Matzinger, Sarah Elizabeth Totten, Emily A. Travanty |
| EPI_ISL_771361, EPI_ISL_771362, EPI_ISL_771363, EPI_ISL_771364, EPI_ISL_771365, EPI_ISL_771366 | Washington State Department of Health | Seattle Flu Study | Deborah A. Nickerson, Chris D. Frazar, Jover Lee, Benjamin Pelle, Matthew Richardson, Amanda Adler, Elisabeth Brandstetter, Peter D. Han, Kairsten Fay, Misja Ilcisin, Kirsten Lacombe, Thomas R. Sibley, Melissa Truong, Caitlin R. Wolf, Romesh Gautom, Geoff Melly, Brian Hiatt, Philip Dykema, Scott Lindquist, Michael Boeckh, Janet A. Englund, Michael Famulare, Barry R. Lutz, Mark J. Rieder, Lea M. Starita, Matthew Thompson, Helen Y. Chu, Jay Shendure, Trevor Bedford |
| EPI_ISL_776687 | UW Virology Lab | UW Virology Lab | Pavitra Roychoudhury, Hong Xie, Lasata Shrestha, Meei-Li Huang, Keith R Jerome, Alexander Greninger |
| EPI_ISL_779444, EPI_ISL_779445, EPI_ISL_779446, EPI_ISL_779448, EPI_ISL_779449, EPI_ISL_779450, EPI_ISL_779451, EPI_ISL_779452, EPI_ISL_779454, EPI_ISL_779456, EPI_ISL_779472, EPI_ISL_779473, EPI_ISL_779474, EPI_ISL_779475, EPI_ISL_779486, EPI_ISL_779487, EPI_ISL_779488, EPI_ISL_779489, EPI_ISL_779490, EPI_ISL_779491, EPI_ISL_779492, EPI_ISL_779493, EPI_ISL_779496, EPI_ISL_779497, EPI_ISL_779498, EPI_ISL_779499, EPI_ISL_779501, EPI_ISL_779502, EPI_ISL_779580, EPI_ISL_779589 | | | |
| see above | Microbiological Diagnostic Unit - Public Health Laboratory (MDU-PHL) | MDU-PHL | Seemann T., Sait, M.L., Sherry, N.L. |
| EPI_ISL_794609 | Virology, tehran university of medical sciences | Virology, tehran university of medical sciences | Soltani,S., Zandi,M. and Abbasi,S. |
| EPI_ISL_804469, EPI_ISL_804475, EPI_ISL_804478, EPI_ISL_804483, EPI_ISL_804484, EPI_ISL_804485, EPI_ISL_804486, EPI_ISL_804487, EPI_ISL_804585, EPI_ISL_804586, EPI_ISL_804587, EPI_ISL_804588, EPI_ISL_804589, EPI_ISL_804590, EPI_ISL_804591 | | | |
| see above | MEPHI, Aix Marseille University | MEPHI, Aix Marseille University | Anthony LEVASSEUR |
| EPI_ISL_804933 | DC Public Health Lab/ Dept. of Forensic Sciences | DC Public Health Lab/ Dept. of Forensic Sciences | Scott Nguyen, Elizabeth Zelaya, Connie Maza, Monica Mann, Brittany Hamilton, David Payne, Jocelyn Hauser |
| EPI_ISL_804952 | Hospital Comarcal de Melilla | Instituto de Salud Carlos III | Iglesias-Caballero, M. Molinero Calamita, M. González-Esguevillas, M. Camarero, S. Pozo, F. Casas, I. Jiménez, P. Jiménez, M. Zaballos, A. Monzón, S. Varona, S. Juliá, M. Cuesta, I, J. López |
| EPI_ISL_806592, EPI_ISL_806593, EPI_ISL_806698, EPI_ISL_806701 | KEMRI-Wellcome Trust Research Programme/KEMRI-CGMR-C Kilifi | KEMRI-Wellcome Trust Research Programme/KEMRI-CGMR-C Kilifi | Githinji et al |
| EPI_ISL_810987, EPI_ISL_810989, EPI_ISL_810991, EPI_ISL_811002, EPI_ISL_811003, EPI_ISL_811004, EPI_ISL_811005, EPI_ISL_811006, EPI_ISL_811007, EPI_ISL_811008 | MRCG at LSHTM Genomics lab | MRCG at LSHTM Genomics lab | Abdul Karim sesay, Abdoulie Kanteh, Jarra Manneh, Mariama Kujabi, Bakary Sanyang |
| EPI_ISL_811160, EPI_ISL_811170, EPI_ISL_811175, EPI_ISL_811191, EPI_ISL_811192, EPI_ISL_811193 | Dharwad | CSIR Institute of Genomics and Integrative Biology | Dr. Shivarudrapp B Bhairappanavar, Rahul Bhoyar, Mohammed Imran, Mohit Divakar, Disha Sharma, Dr. Vijay A Yenagi, Dr. Suresh B Arakera, Dr. Amit Ugargol, Dr. Rgavendra B Nayak, Bani Jolly, Abhinav Jain, Paras Sehgal, Gyan Ranjan, Vinod Scaria, Sridhar Sivasubbu |
| EPI_ISL_812161, EPI_ISL_812162 | GA Department of Public Health Laboratory | Pathogen Discovery, Respiratory Viruses Branch, Division of Viral Diseases, Centers for Disease Control and Prevention | Yan Li, Ying Tao, Anna Montmayeur, Jing Zhang, Brian Lynch, Krista Queen, Anna Uehara, Rachel Marine, Peter Cook, Clinton R. Paden, Haibin Wang, Suxiang Tong |
| EPI_ISL_812537, EPI_ISL_812538, EPI_ISL_812540, EPI_ISL_812552, EPI_ISL_812554, EPI_ISL_812555, EPI_ISL_812560, EPI_ISL_812561, EPI_ISL_812562, EPI_ISL_812563, EPI_ISL_812564, EPI_ISL_812565, EPI_ISL_812566, EPI_ISL_812567, EPI_ISL_812568, EPI_ISL_812569, EPI_ISL_812570, EPI_ISL_812571, EPI_ISL_812572, EPI_ISL_812573, EPI_ISL_812574, EPI_ISL_812575, EPI_ISL_812576, EPI_ISL_812577, EPI_ISL_812578, EPI_ISL_812579, EPI_ISL_812580 | | | |
| see above | United States Air Force School of Aerospace Medicine | United States Air Force School of Aerospace Medicine | Anthony Fries, Jennifer Meyer, Amanda Javorina, Sarah Purves, William Gruner, Clarise Starr, Elizabeth Macias |
| EPI_ISL_812805, EPI_ISL_812807, EPI_ISL_812815, EPI_ISL_812821, EPI_ISL_812823, EPI_ISL_812831, EPI_ISL_812835, EPI_ISL_812841, EPI_ISL_812843, EPI_ISL_812867, EPI_ISL_812869, EPI_ISL_812870 | | | |
| see above | Genomics Program, Children Cancer Hospital | Genomics Program, Children Cancer Hospital | Hatem,A., Hadad,A., Abouelnaga,S., Amer,K., Salah,H., Farawyla,H., Halafawy,A., Mansour,T., shalaby,L., Hassan,W., Soliman,M., Gomaa,C., Hassan,R., Soliman,S., Monuir,G., Hammad,M., Hussein,S., Abdo,I., Jalal,D., El-Zayat,M., El-Shaqnqery,H., Diab,A., Bakry,U., Samir,O., Magdeldin,S., Sayed,A. |
| EPI_ISL_812962, EPI_ISL_812963, EPI_ISL_812964, EPI_ISL_812965 | Division of Pathogen Resource Management, Korea National Institute of Health, Korea Disease Control and Prevention Agency | Division of Pathogen Resource Management, Korea National Institute of Health, Korea Disease Control and Prevention Agency | Kim,S.T., Kim,S.Y., Choi,Y.S., Kim,E.-J., Kim,J.-M., Yun,M.-r., Choi,C. |
| EPI_ISL_815318, EPI_ISL_815329, EPI_ISL_815391, EPI_ISL_815398, EPI_ISL_815399 | Centogene | Centogene | Peter Bauer, Krishna Kumar Kandaswamy, Vivi Hue-Trang Lieu |
| EPI_ISL_816717, EPI_ISL_816718, EPI_ISL_816719, EPI_ISL_816720, EPI_ISL_816721 | Bioinformatics and Biostatistics Lab, Advanced Sequencing Facility | COVID-19 Genomics UK (COG-UK) Consortium | Aengus Stewart,Jerome Nicod,Chelsea Sawyer,Laura Cubitt,Harshil Patel,Margaret Crawford |

| | | | |
|---|---|---|---|
| EPI_ISL_822343 | Lighthouse Lab in Glasgow | Wellcome Sanger Institute for the COVID-19 Genomics UK (COG-UK) Consortium | Harper VanSteenhouse, Yumi Kasai, David Gray, Carol Clugston, Anna Dominiczak and Alex Alderton, Roberto Amato, Sonia Goncalves, Ewan Harrison, David K. Jackson, Ian Johnston, Dominic Kwiatkowski, Cordelia Langford, John Sillitoe on behalf of the Wellcome Sanger Institute COVID-19 Surveillance Team |
| EPI_ISL_824414, EPI_ISL_824453, EPI_ISL_824454, EPI_ISL_824455, EPI_ISL_824456, EPI_ISL_824457, EPI_ISL_824458 | Hospital Universitari Vall d'Hebron - Vall d'Hebron Institut de Rercerca | Hospital Universitari Vall d'Hebron | Cristina Andrés, Maria Piñana, Josep F Abril, Damir Garcia-Cehic, Ariadna Rando, Juliana Esperalba, Maria Gema Codina, Carla Castillo, Maria Carmen Martin, Tomàs Pumarola, Josep Quer, Andrés Antón |
| EPI_ISL_825634, EPI_ISL_825687, EPI_ISL_825688, EPI_ISL_825689, EPI_ISL_825690, EPI_ISL_825691, EPI_ISL_825692, EPI_ISL_825693, EPI_ISL_825694, EPI_ISL_825695, EPI_ISL_825696, EPI_ISL_825697, EPI_ISL_825698, EPI_ISL_825699, EPI_ISL_825700, EPI_ISL_825701, EPI_ISL_825702, EPI_ISL_825703, EPI_ISL_825704, EPI_ISL_825705, EPI_ISL_825706, EPI_ISL_825707, EPI_ISL_826077, EPI_ISL_826123, EPI_ISL_826124, EPI_ISL_826125, EPI_ISL_826126, EPI_ISL_826127, EPI_ISL_826128, EPI_ISL_826129, EPI_ISL_826130, EPI_ISL_826131 | | | |
| see above | Laboratoire de santé publique du Québec | Laboratoire de santé publique du Québec | Sandrine Moreira, Ioannis Ragoussis, Guillaume Bourque, Jesse Shapiro, Mark Lathrop and Michel Roger on behalf of the CoVSeQ research group (http://covseq.ca/researchgroup) |
| EPI_ISL_826693 | The National University Hospital of Iceland | deCODE genetics | Daniel F Gudbjartsson; Agnar Helgason; Hakon Jonsson; Olafur T Magnusson; Pall Melsted; Gudmundur L Norddahl; Jona Saemundsdottir; Asgeir Sigurdsson; Patrick Sulem; Arna B Agustsdottir; Hannes Eggertsson; Berglind Eiriksdottir; Run Fridriksdottir; Elisabet E Gardarsdottir; Gudmundur Georgsson; Olafia S Gretarsdottir; Kjartan R Gudmundsson; Thora R Gunnarsdottir; Arnaldur Gylfason; Hilma Holm; Brynjar O Jensson; Aslaug Jonasdottir; Kamilla S Josefsdottir; Thordur Kristjansson; Droplaug N Magnusdottir; Solvi Rognvaldsson; Louise le Roux; Gudrun Sigmundsdottir; Gardar Sveinbjornsson; Kristin E Sveinsdottir; Maney Sveinsdottir; Emil A Thorarensen; Bjarni Thorbjornsson; Gisli Masson; Ingileif Jonsdottir; Alma Moller; Thorolfur Gudnason; Karl G Kristinsson; Unnur Thorsteinsdottir; Kari Stefansson |
| EPI_ISL_826825 | INSPI-CRN DE INFLUENZA Y OTROS VIRUS RESPIRATORIOS | Instituto de Salud Publica de Chile | Javier Tognarelli, Barbara Parra, Loredana Arata, Jaime Lagos, Gisselle Barra, Alfredo Bruno, Domenica de Mora, Solon Narvaez, Jimmy Garcez, Michelle Paez, Martiza Olmedo, Manuel Gonzalez, Patricia Bustos, Rodrigo Fasce, Andres Castillo, Jorge Fernandez |
| EPI_ISL_827011, EPI_ISL_827051, EPI_ISL_827288, EPI_ISL_827289, EPI_ISL_827290, EPI_ISL_827291, EPI_ISL_827293, EPI_ISL_827377, EPI_ISL_827379, EPI_ISL_827584, EPI_ISL_827588, EPI_ISL_827608, EPI_ISL_827609, EPI_ISL_827876, EPI_ISL_827878, EPI_ISL_827879, EPI_ISL_827880, EPI_ISL_827881, EPI_ISL_828009, EPI_ISL_828211, EPI_ISL_828212, EPI_ISL_828334, EPI_ISL_828503, EPI_ISL_828504, EPI_ISL_828505, EPI_ISL_828506, EPI_ISL_828507, EPI_ISL_828618, EPI_ISL_828993, EPI_ISL_829000 | | | |
| see above | The National University Hospital of Iceland | deCODE genetics | Daniel F Gudbjartsson; Agnar Helgason; Hakon Jonsson; Olafur T Magnusson; Pall Melsted; Gudmundur L Norddahl; Jona Saemundsdottir; Asgeir Sigurdsson; Patrick Sulem; Arna B Agustsdottir; Hannes Eggertsson; Berglind Eiriksdottir; Run Fridriksdottir; Elisabet E Gardarsdottir; Gudmundur Georgsson; Olafia S Gretarsdottir; Kjartan R Gudmundsson; Thora R Gunnarsdottir; Arnaldur Gylfason; Hilma Holm; Brynjar O Jensson; Aslaug Jonasdottir; Kamilla S Josefsdottir; Thordur Kristjansson; Droplaug N Magnusdottir; Solvi Rognvaldsson; Louise le Roux; Gudrun Sigmundsdottir; Gardar Sveinbjornsson; Kristin E Sveinsdottir; Maney Sveinsdottir; Emil A Thorarensen; Bjarni Thorbjornsson; Gisli Masson; Ingileif Jonsdottir; Alma Moller; Thorolfur Gudnason; Karl G Kristinsson; Unnur Thorsteinsdottir; Kari Stefansson |
| EPI_ISL_829073 | deCODE genetics | deCODE genetics | Daniel F Gudbjartsson; Agnar Helgason; Hakon Jonsson; Olafur T Magnusson; Pall Melsted; Gudmundur L Norddahl; Jona Saemundsdottir; Asgeir Sigurdsson; Patrick Sulem; Arna B Agustsdottir; Hannes Eggertsson; Berglind Eiriksdottir; Run Fridriksdottir; Elisabet E Gardarsdottir; Gudmundur Georgsson; Olafia S Gretarsdottir; Kjartan R Gudmundsson; Thora R Gunnarsdottir; Arnaldur Gylfason; Hilma Holm; Brynjar O Jensson; Aslaug Jonasdottir; Kamilla S Josefsdottir; Thordur Kristjansson; Droplaug N Magnusdottir; Solvi Rognvaldsson; Louise le Roux; Gudrun Sigmundsdottir; Gardar Sveinbjornsson; Kristin E Sveinsdottir; Maney Sveinsdottir; Emil A Thorarensen; Bjarni Thorbjornsson; Gisli Masson; Ingileif Jonsdottir; Alma Moller; Thorolfur Gudnason; Karl G Kristinsson; Unnur Thorsteinsdottir; Kari Stefansson |
| EPI_ISL_829241, EPI_ISL_829364, EPI_ISL_829378, EPI_ISL_829379, EPI_ISL_829477 | The National University Hospital of Iceland | deCODE genetics | Daniel F Gudbjartsson; Agnar Helgason; Hakon Jonsson; Olafur T Magnusson; Pall Melsted; Gudmundur L Norddahl; Jona Saemundsdottir; Asgeir Sigurdsson; Patrick Sulem; Arna B Agustsdottir; Hannes Eggertsson; Berglind Eiriksdottir; Run Fridriksdottir; Elisabet E Gardarsdottir; Gudmundur Georgsson; Olafia S Gretarsdottir; Kjartan R Gudmundsson; Thora R Gunnarsdottir; Arnaldur Gylfason; Hilma Holm; Brynjar O Jensson; Aslaug Jonasdottir; Kamilla S Josefsdottir; Thordur Kristjansson; Droplaug N Magnusdottir; Solvi Rognvaldsson; Louise le Roux; Gudrun Sigmundsdottir; Gardar Sveinbjornsson; Kristin E Sveinsdottir; Maney Sveinsdottir; Emil A Thorarensen; Bjarni Thorbjornsson; Gisli Masson; Ingileif Jonsdottir; Alma Moller; Thorolfur Gudnason; Karl G Kristinsson; Unnur Thorsteinsdottir; Kari Stefansson |
| EPI_ISL_829673 | deCODE genetics | deCODE genetics | Daniel F Gudbjartsson; Agnar Helgason; Hakon Jonsson; Olafur T Magnusson; Pall Melsted; Gudmundur L Norddahl; Jona Saemundsdottir; Asgeir Sigurdsson; Patrick Sulem; Arna B Agustsdottir; Hannes Eggertsson; Berglind Eiriksdottir; Run Fridriksdottir; Elisabet E Gardarsdottir; Gudmundur Georgsson; Olafia S Gretarsdottir; Kjartan R Gudmundsson; Thora R Gunnarsdottir; Arnaldur Gylfason; Hilma Holm; Brynjar O Jensson; Aslaug Jonasdottir; Kamilla S Josefsdottir; Thordur Kristjansson; Droplaug N Magnusdottir; Solvi Rognvaldsson; Louise le Roux; Gudrun Sigmundsdottir; Gardar Sveinbjornsson; Kristin E Sveinsdottir; Maney Sveinsdottir; Emil A Thorarensen; Bjarni Thorbjornsson; Gisli Masson; Ingileif Jonsdottir; Alma Moller; Thorolfur Gudnason; Karl G Kristinsson; Unnur Thorsteinsdottir; Kari Stefansson |
| EPI_ISL_829891, EPI_ISL_830191, EPI_ISL_830193, EPI_ISL_830387, EPI_ISL_830529, EPI_ISL_830530 | The National University Hospital of Iceland | deCODE genetics | Daniel F Gudbjartsson; Agnar Helgason; Hakon Jonsson; Olafur T Magnusson; Pall Melsted; Gudmundur L Norddahl; Jona Saemundsdottir; Asgeir Sigurdsson; Patrick Sulem; Arna B Agustsdottir; Hannes Eggertsson; Berglind Eiriksdottir; Run Fridriksdottir; Elisabet E Gardarsdottir; Gudmundur Georgsson; Olafia S Gretarsdottir; Kjartan R Gudmundsson; Thora R Gunnarsdottir; Arnaldur Gylfason; Hilma Holm; Brynjar O Jensson; Aslaug Jonasdottir; Kamilla S Josefsdottir; Thordur Kristjansson; Droplaug N Magnusdottir; Solvi Rognvaldsson; Louise le Roux; Gudrun Sigmundsdottir; Gardar Sveinbjornsson; Kristin E Sveinsdottir; Maney Sveinsdottir; Emil A Thorarensen; Bjarni Thorbjornsson; Gisli Masson; Ingileif Jonsdottir; Alma Moller; Thorolfur Gudnason; Karl G Kristinsson; Unnur Thorsteinsdottir; Kari Stefansson |
| EPI_ISL_833150 | National Institute of Laboratory Medicine and Referral Center | Genomic Research Lab, BCSIR | Md. Saddam Hossain, Mohammad Samir Uzzaman, Eshrar Osman, Md. Ahashan Habib, Shahina Akter, Tanjina Akhtar Banu, Abu Sayeed Mohammad Mahmud, Md. Murshed Hasan Sarkar, Barna Goswami, Iffat Jahan, Tasnim Nafisa, Md. Maruf Ahmed Molla, Mahmuda Yeasmin, Asish Kumar Ghosh, A. K. M. Shamsuzzaman, Monira Parveen, Md. Masum Hossain Arif, Md. Salim Khan |
| EPI_ISL_833191 | Hôpital Bichat Claude Bernard, Laboratoire de Virologie | IAME UMR1137 Inserm, Université de Paris, Hôpital Bichat | Antoine Bridier, Amélie Recoing, Quentin Le Hingrat, Lena Daniel, Siham Hamri, Gilles Collin, Alexandre Storto, Mélanie Bertine, Charlotte Charpentier, Nadhira Houhou-Fidouh, Diane Descamps, Benoit Visseaux |
| EPI_ISL_833512, EPI_ISL_833514, EPI_ISL_833515 | Veterinary Specialized Instute "Nis" | Veterinary Specialized Institute "Kraljevo", Serbia | Vidanovic,D., Tesovic,B., Manic,M., Petrovic,M.,Knezevic,A., Jovanovic,T., Jankovic,M., Sekler,M., Banovic Djeri,B., Petrovic,T., Volkening,J., Afonso,C. |
| EPI_ISL_833572 | Veterinary Specialized Instute "Nis" | Scientific Veterinary Institute "Novi Sad" | Vidanovic,D., Tesovic,B., Manic,M., Petrovic,M.,Knezevic,A., Jovanovic,T., Jankovic,M., Sekler,M., Banovic Djeri,B., Petrovic,T., Volkening,J., Afonso,C. |
| EPI_ISL_837590, EPI_ISL_837591, EPI_ISL_837592, EPI_ISL_837593, EPI_ISL_837594 | Laboratorio Nacional de Salud | Laboratory of Respiratory Viruses and Measles, Oswaldo Cruz Institute, FIOCRUZ | Paola Resende, Cesar Roberto Conde Pereira, Claudia Estrada, Luciana Appolinario, Fernando Motta, Anna Carolina Paixao, Ana Carolina Mendonca, Marilda Siqueira |
| EPI_ISL_837781, EPI_ISL_837782, EPI_ISL_837783, EPI_ISL_837784, EPI_ISL_837785, EPI_ISL_837786, EPI_ISL_837787, EPI_ISL_837788, EPI_ISL_837789, EPI_ISL_837790 | Instituto Nacional de Enfermedades Respiratorias (INER) | Instituto Nacional de Enfermedades Respiratorias (INER) | Celia Boukadida, Margarita Matías-Florentino, Alma Rincón-Rubio, Hector Esteban Paz-Juárez, Olivia Briceño, Edgar Sevilla-Reyes, Fidencio Mejía-Nepomuceno, Mario Mújica-Sánchez, Eduardo Becerril-Vargas, José Arturo Martínez-Orozco, Alejandra Hernández-Terán, Jorge Salas-Hernández, Santiago Ávila-Ríos, Joel Armando Vázquez-Pérez |
| EPI_ISL_845657, EPI_ISL_845658, EPI_ISL_845783 | Quest Diagnostics | Quest Diagnostics | Rosenthal,S.H., Gerasimova,A., Kagan,R.M., Anderson, B., Bernstein, L.E., Livingston, K.E., Hua, M., Liu Y., Shalhout, D.F., Shlyakhter, I.A., Owen, R., Lacbawan, F. |
| EPI_ISL_845808 | National Institute of Laboratory Medicine and Referral Center | Genomic Research Lab, BCSIR | Md. Saddam Hossain,Mohammad Samir Uzzaman, Eshrar Osman, Md. Ahashan Habib, Shahina Akter, Tanjina Akhtar Banu,Abu Sayeed Mohammad Mahmud, Md. Murshed Hasan Sarkar, Barna Goswami, Iffat Jahan, Tasnim Nafisa, Md. Maruf Ahmed Molla, Mahmuda Yeasmin, Asish Kumar Ghosh, A. K. M. Shamsuzzaman, Md. Salim Khan. |
| EPI_ISL_848300, EPI_ISL_848301, EPI_ISL_848302, EPI_ISL_848303, EPI_ISL_848304, EPI_ISL_848305, EPI_ISL_848306, EPI_ISL_848307, EPI_ISL_848308, EPI_ISL_848309, EPI_ISL_848310, EPI_ISL_848311, EPI_ISL_848312, EPI_ISL_848313, EPI_ISL_848314, EPI_ISL_848315, EPI_ISL_848316, EPI_ISL_848486, EPI_ISL_848487, EPI_ISL_848488, EPI_ISL_848489, EPI_ISL_848490, EPI_ISL_848491, EPI_ISL_848492, EPI_ISL_848493 | | | |
| see above | Illinois Department of Public Health | Gagnon Lab, Southern Illinois University | Keith Gagnon |
| EPI_ISL_849109, EPI_ISL_849110, EPI_ISL_849111, EPI_ISL_849112, EPI_ISL_849113, EPI_ISL_849114, EPI_ISL_849115, EPI_ISL_849116, EPI_ISL_849117, EPI_ISL_849118, EPI_ISL_849119 | | | |

| | | | |
|---|---|---|---|
| see above | Florida Bureau of Public Health Laboratories | Florida Bureau of Public Health Laboratories | Sarah Schmedes, Jason Blanton |
| EPI_ISL_849656, EPI_ISL_849657, EPI_ISL_849658, EPI_ISL_849659, EPI_ISL_849660, EPI_ISL_849661 | Servizio Igiene Epidemiologia e Sanità Pubblica (SIESP)-L'Aquila | Istituto Zooprofilattico Sperimentale dell'Abruzzo e Molise "G.Caporale" | Lorusso A, Marcacci M, Di Domenico M, Curini V, Ancora M, Cammà C, Rinaldi A, Mangone I, Di Pasquale A, Puglia I, Savini G. |
| EPI_ISL_849662 | Servizio di igiene epidemiologia e sanità pubblica (SIESP)-Chieti | Istituto Zooprofilattico Sperimentale dell'Abruzzo e Molise "G.Caporale" | Lorusso A, Marcacci M, Di Domenico M, Curini V, Ancora M, Cammà C, Rinaldi A, Mangone I, Di Pasquale A, Puglia I, Savini G. |
| EPI_ISL_849683 | unknown | PHV-FSS | Son Nguyen et al. |
| EPI_ISL_849928, EPI_ISL_849929, EPI_ISL_849930, EPI_ISL_849933, EPI_ISL_849934, EPI_ISL_849935, EPI_ISL_849936 | UC Davis- Department of Pathology and Laboratory Medicine | Chan-Zuckerberg Biohub | CZB Cliahub Consortium |
| EPI_ISL_850240, EPI_ISL_850241, EPI_ISL_850242, EPI_ISL_850243, EPI_ISL_850244, EPI_ISL_850245, EPI_ISL_850246, EPI_ISL_850247, EPI_ISL_850248, EPI_ISL_850249, EPI_ISL_850250, EPI_ISL_850251, EPI_ISL_850252, EPI_ISL_850253, EPI_ISL_850254, EPI_ISL_850255, EPI_ISL_850256 | | | |
| see above | Division of Emerging Infectious Diseases, Bureau of Infectious Diseases Diagnosis Control, Korea Disease Control and Prevention Agency | Division of Emerging Infectious Diseases, Bureau of Infectious Diseases Diagnosis Control, Korea Disease Control and Prevention Agency | Ae Kyung Park, Il-Hwan Kim, Heui Man Kim, Jeong-Min Kim, Namjoo Lee, Chaeyoung Lee, Sang Hee Woo, Eun-Jin Kim |
| EPI_ISL_850507 | Gandhi Medical College and Hospital | VRDL-Gandhi Medical College | Nagamani Kammili, Rajeshwar Rao, Madhavi Latha Manolla, Winnie Thomas, Shailaja VV, Sudhamadhuri Devara, Vanisree Rajoli, Archana GJ,Sushma Rajyalakshmi Gudiseva, Sunitha Pakalapati, Manisha Rani, Amrithesh Kumar, Raja Rao Mesepogu, Vinay Shekar Reddy, Thrilok Chander Bingi, Sofia Banu, Divya Tej Sowpati |
| EPI_ISL_853293 | UPMC Clinical Microbiology Laboratory | Microbial Genome Sequencing Center; Microbial Genomic Epidemiology Laboratory | Mustapha M. Mustapha, Jane W. Marsh, Dan Snyder, Marissa P. Griffith, Stephanie L. Mitchell, Vatsala R. Srinivasa, Kady D. Waggle, Chinelo Ezeonwuku, Vaughn S. Cooper, Lee H. Harrison |
| EPI_ISL_853842, EPI_ISL_853892, EPI_ISL_853893, EPI_ISL_853894 | Center for Virology, Medical University of Vienna | Bergthaler laboratory, CeMM Research Center for Molecular Medicine of the Austrian Academy of Sciences | Lukas Endler, Alexandra Popa, Benedikt Agerer, Jakob-Wendelin Genger, Alexander Lercher, Anna Schedl, Thomas Penz, Michael Schuster, Jan Laine, Martin Senekowitsch, Christoph Bock, Andreas Bergthaler |
| EPI_ISL_854237 | Institute of Legal Medicine, Medical University of Innsbruck | Bergthaler laboratory, CeMM Research Center for Molecular Medicine of the Austrian Academy of Sciences | Lukas Endler, Alexandra Popa, Benedikt Agerer, Jakob-Wendelin Genger, Alexander Lercher, Anna Schedl, Thomas Penz, Michael Schuster, Jan Laine, Martin Senekowitsch, Christoph Bock, Andreas Bergthaler |
| EPI_ISL_855349, EPI_ISL_855350, EPI_ISL_855351 | Department of Epidemiology, Infectious Disease Control and Prevention, Hiroshima University, Japan | Department of Epidemiology, Infectious Disease Control and Prevention, Hiroshima University, Japan | Junko Tanaka, Kazuaki Takahashi, Masao Kuwabara, Eisaku Kishita, Shintaro Nagashima, Ko Ko |
| EPI_ISL_861891, EPI_ISL_861895, EPI_ISL_861897, EPI_ISL_861901 | LATE - Laboratório de Técnicas Especiais - Hospital Israelita Albert Einstein | LATE - Laboratório de Técnicas Especiais - Hospital Israelita Albert Einstein | Deyvid Amgarten, Fernanda de Mello Malta, Raquel Riyuzo, Ana Paula Moreira Salles, Pedro Henrique Sebe Rodrigues, João Renato Rebello Pinho |
| EPI_ISL_862276, EPI_ISL_862284, EPI_ISL_862285, EPI_ISL_862286, EPI_ISL_862290, EPI_ISL_862300, EPI_ISL_862301, EPI_ISL_862344, EPI_ISL_862357, EPI_ISL_862370, EPI_ISL_862382, EPI_ISL_862385, EPI_ISL_862386, EPI_ISL_862387, EPI_ISL_862388, EPI_ISL_862389, EPI_ISL_862390, EPI_ISL_862391, EPI_ISL_862392, EPI_ISL_862393, EPI_ISL_862394, EPI_ISL_862395, EPI_ISL_862396, EPI_ISL_862397, EPI_ISL_862398, EPI_ISL_862399, EPI_ISL_862400, EPI_ISL_862401, EPI_ISL_862402, EPI_ISL_862403, EPI_ISL_862405, EPI_ISL_862416, EPI_ISL_862425, EPI_ISL_862468, EPI_ISL_862478, EPI_ISL_862481, EPI_ISL_862496, EPI_ISL_862497, EPI_ISL_862498, EPI_ISL_862499, EPI_ISL_862500, EPI_ISL_862501, EPI_ISL_862502, EPI_ISL_862503 | | | |
| see above | Kurnool Medical College (KMC) | CSIR Institute of Genomics and Integrative Biology | Pallavali Roja Rani, Mohamed Imran, J. Vijaya Lakshmi, Bani Jolly, S. Afsar, Abhinav Jain, Mohit Kumar Divakar, Panyam Suresh, Disha Sharma, Nambi Rajesh, Rahul C Bhoyar, Dasari Ankaiah, Sanaga Shanthi Kumari, Gyan Ranjan, Valluri Anitha Lavanya, Mercy Rophina, S. Umadevi, Paras Sehgal, Avula Renuka Devi, A. Surekha, Pulala Chandra, Rajamadugu Hymavathy, P R Vanaja, Vinod Scaria, Sridhar Sivasubbu |
| EPI_ISL_876038 | Montefiore Medical Center | Albert Einstein College of Medicine, Dept. of Microbiology & Immunology, Chandran lab | J. Maximilian Fels, Saad Khan, Ryan Forster, Karin A. Skalina, Surksha Sirichand, Amy S. Fox, Aviv Bergman, William B. Mitchell, Lucia R. Wolgast, Wendy Szymczak, Robert H. Bortz III, M. Eugenia Dieterle, Catalina Florez, Denise Haslwanter, Rohit K. Jangra, Ethan Laudermilch, Ariel S. Wirchnianski, Jason Barnhill, David L. Goldman, Hnin Khine, D. Yitzchak Goldstein, Johanna P. Daily, Kartik Chandran, Libusha Kelly |
| EPI_ISL_876953 | Quest Diagnostics | Quest Diagnostics | Rosenthal,S.H., Gerasimova,A., Kagan,R.M., Anderson, B., Hua, M., Liu Y., Bernstein, L.E., Livingston, K.E., Perez, A., Shalhout, D.F., Shlyakhter, I.A., Owen, R., Tanpaiboon, P., Lacbawan, F. |
| EPI_ISL_877654, EPI_ISL_877655, EPI_ISL_877656, EPI_ISL_877657, EPI_ISL_877658, EPI_ISL_877659, EPI_ISL_877660, EPI_ISL_877661, EPI_ISL_877662, EPI_ISL_877663 | Clinical Molecular Microbiology Laboratory, UNC Hospital | Dirk Dittmer | Razia Moorad , Justin T. Landis , Brent A. Eason, Melissa B. Miller, Linda Pluta, Dirk Dittmer, Angelica Juarez, Cecilia Thompson , Cameroon Grant, Evelyn Hoffman, Patricio Cano, Jason Wong, Carolina Caro-Vegas, Blossom Damania. |
| EPI_ISL_878552 | Robert Garry lab | Andersen lab at Scripps Research | Allison Smither, Gilberto Sabino-Santos, Patricia Snarski, Lilia Melnik, Antoinette Bell, Kaylynn Genemaras, Arnaud Drouin, Dahlene Fusco, Robert Garry with SEARCH Alliance San Diego |
| EPI_ISL_882657 | LACEN do Estado do Piaui, Dr. Costa Alvarenga | Instituto Adolfo Lutz, Interdiciplinary Procedures Center, Strategic Laboratory | Claudio Tavares Sacchi, Claudia Regina Gonçalves, Erica Valessa Ramos Gomes, Karoline Rodrigues Campos |
| EPI_ISL_884250 | LATE - Laboratório de Técnicas Especiais - Hospital Israelita Albert Einstein | LATE - Laboratório de Técnicas Especiais - Hospital Israelita Albert Einstein | Deyvid Amgarten, Fernanda de Mello Malta, Raquel Riyuzo, Ana Paula Moreira Salles, Pedro Henrique Sebe Rodrigues, João Renato Rebello Pinho |
| EPI_ISL_884304, EPI_ISL_884307, EPI_ISL_884328, EPI_ISL_884329, EPI_ISL_884330, EPI_ISL_884338, EPI_ISL_884346, EPI_ISL_884375, EPI_ISL_884405, EPI_ISL_884430 | Infectious Diseases, Quest Diagnostics | Infectious Diseases, Quest Diagnostics | Rosenthal,S.H., Gerasimova,A., Kagan,R.M., Anderson,B., Bernstein,L.E., Livingston,K.E., Hua,M., Liu,Y., Shalhout,D.F., Owen,R., Lacbawan,F. |
| EPI_ISL_884835, EPI_ISL_884840, EPI_ISL_884842, EPI_ISL_884843, EPI_ISL_884852, EPI_ISL_884853, EPI_ISL_884855 | Department of Biochemistry, Cell and Molecular Biology, West African Centre for Cell Biology of Infectious Pathogens (WACCBIP), University of Ghana | Department of Biochemistry, Cell and Molecular Biology, West African Centre for Cell Biology of Infectious Pathogens (WACCBIP), University of Ghana | Ngoi,J.M., Tei-Maya,F., Morang'a,C.M., Magnussen,V., Amuzu,D.S., Mohammed,A., Tapela,K., Kibinge,N., Diallo,A.B., Kumi-Ansah,F., Odoom,T., Boakye,O.D., Amoako,E., Abass,A.-K., Quashie,P., Amenga-Etego,L.N., Akoriyea,S.K., Awandare,G.A., Bediako,Y. |
| EPI_ISL_887147, EPI_ISL_887148, EPI_ISL_887149 | Massachusetts General Hospital | Infectious Disease Program, Broad Institute of Harvard and MIT | Lemieux,J.E., Siddle,K.J., Shaw,B., Adams,G., Pierce,V., Turbett,S., Anahtar,M., Branda,J., Slater,D., Harris,J., Lin,A.E., Gladden-Young,A., Lagerborg,K., Rudy,M., DeRuff,K., Carter,A., Normandin,E., Bauer,M., Reilly,S., Tomkins-Tinch,C., Loreth,C., Chaluvadi,S., Neumann,A., Cusick,C., Chapman,S.B., Gnirke,A., Flowers,K., Cerrato,F., Birren,B.W., Gallagher,G., Smole,S., Park,D.J., MacInnis,B.L., Ryan,E., LaRocque,R., Rosenberg,E. and Sabeti,P.C. |
| EPI_ISL_889336 | The University Hospital Brno | Institute of Applied Biotechnologies a.s. | Petr Klempt, Ondej Brzo, Martin Kašný, Kateina Kvapilová, Martina Lengerová, Petr Kvapil |
| EPI_ISL_889359, EPI_ISL_889360 | Motol University Hospital | Institute of Applied Biotechnologies a.s. | Petr Klempt, Ondej Brzo, Martin Kašný, Kateina Kvapilová, Pavel Devínek, Petr Kvapil |
| EPI_ISL_890111, EPI_ISL_890112, EPI_ISL_890113, EPI_ISL_890114, EPI_ISL_890115, EPI_ISL_890116 | Laboratoire de santé publique du Québec | Laboratoire de santé publique du Québec | Sandrine Moreira, Ioannis Ragoussis, Guillaume Bourque, Jesse Shapiro, Mark Lathrop and Michel Roger on behalf of the CoVSeQ research group |
| EPI_ISL_891224, EPI_ISL_891225, EPI_ISL_891226 | The Oncology Institute "Prof. Dr. Ion Chiricuta" Cluj Napoca | "Stefan cel Mare" University Metagenomics Lab | Lobiuc Andrei, Gheorghita Roxana |
| EPI_ISL_891258 | COVID lab, Mymensingh Medical College | Department of Pathology, Bangladesh Agricultural University and Department of Microbiology, Mymensingh Medical College | Afrin, S. Z. Paul, S. K. Parvin, R. |
| EPI_ISL_891269 | Institute of Biocides and Medical Ecology, Belgarde, Serbia | Virology department Institute of microbiology and | Banko Ana, Miljanovic Danijela, Milicevic Ognjen, Loncar Ana, Abazovic Dzihan, Despot Dragana |

| | | | |
|---|---|---|---|
| | | immunology Faculty of Medicine University of Belgrade | |
| EPI_ISL_896124, EPI_ISL_896129, EPI_ISL_896177, EPI_ISL_896180, EPI_ISL_900079, EPI_ISL_900102, EPI_ISL_900108, EPI_ISL_900120, EPI_ISL_900131, EPI_ISL_900149, EPI_ISL_900157, EPI_ISL_900189, EPI_ISL_900238, EPI_ISL_900293, EPI_ISL_900318, EPI_ISL_900325, EPI_ISL_900337, EPI_ISL_900347, EPI_ISL_900355, EPI_ISL_900378, EPI_ISL_900385, EPI_ISL_900440, EPI_ISL_900466 | | | |
| see above | MEPHI, Aix Marseille University | MEPHI, Aix Marseille University | Anthony LEVASSEUR |
| EPI_ISL_900734, EPI_ISL_900735, EPI_ISL_900736, EPI_ISL_900737, EPI_ISL_900738 | Bozeman Health Deaconess Hospital | Wiedenheft lab, Montana State University | Artem Nemudryi, Anna Nemudraia, Tanner Wiegand, Joseph Nichols, Deann T. Snyder, Jodi F. Hedges, Calvin Cicha, Helen Lee, Karl K. Vanderwood, Diane Bimczok, Mark A. Jutila and Blake Wiedenheft |
| EPI_ISL_903346 | Bozeman Health Deaconess Hospital | Wiedenheft lab, Montana State University | Artem Nemudryi, Anna Nemudraia, Tanner Wiegand, Joseph Nichols, Deann T. Snyder, Jodi F. Hedges, Calvin Cicha, Helen Lee, Karl K. Vanderwood, Diane Bimczok, Mark A. Jutila and Blake Wiedenheft |
| EPI_ISL_910294, EPI_ISL_910295, EPI_ISL_910296, EPI_ISL_910297, EPI_ISL_910298, EPI_ISL_910299, EPI_ISL_910300, EPI_ISL_910301, EPI_ISL_910302, EPI_ISL_910303, EPI_ISL_910304, EPI_ISL_910305, EPI_ISL_910314, EPI_ISL_910315, EPI_ISL_910316, EPI_ISL_910317, EPI_ISL_910318, EPI_ISL_910319, EPI_ISL_910320, EPI_ISL_910321, EPI_ISL_910322, EPI_ISL_910323 | | | |
| see above | CSIR-Centre for Cellular and Molecular Biology | CSIR-Centre for Cellular and Molecular Biology | Payel Mukherjee,Pratheusa Maccha,Namami Gaur,Lamuk Zaveri,Tulasi Nagabandi,Purushotham Vodnala,Blessy B John,Viswagithe S L,B Himasri,Sofia Banu,Priya Singh,Archana Bharadwaj Siva,Karthik Bharadwaj Tallapaka,Rakesh K Mishra,Divya Tej Sowpati |
| EPI_ISL_913913, EPI_ISL_913914, EPI_ISL_913956 | Instituto de Diagnostico y Referencia Epidemiologicos INDRE_RNLSP | Instituto de Diagnostico y Referencia Epidemiologicos (INDRE) | Claudia Wong-Arambula, Abril Rodriguez-Maldonado, Fabiola Garces-Ayala, Adnan Araiza-Rodriguez, David Fragoso-Fonseca, Sergio Rangel-Guerrero, Mayra Jimenez-Morales, Nancy Munoz-Hernandez, Natividad Cruz-Ortiz, Tatiana Nunez-Garcia, Gisela Barrera-Badillo, Lucia Hernandez-Rivas, Irma Lopez-Martinez, Ernesto Ramirez-Gonzalez. |
| EPI_ISL_914577, EPI_ISL_914578, EPI_ISL_914587 | TGen North | TGen North | "Jolene Bowers, Megan Folkerts, Chris French, Hayley Yaglom, Ashlyn Pfeiffer, Darrin Lemmer, Dave Engelthaler, The Arizona COVID Genomics Union (ACGU)" |
| EPI_ISL_914879 | Instituto de Diagnostico y Referencia Epidemiologicos INDRE_RNLSP | Instituto de Diagnostico y Referencia Epidemiologicos (INDRE) | Claudia Wong-Arambula, Abril Rodriguez-Maldonado, Fabiola Garces-Ayala, Adnan Araiza-Rodriguez, David Fragoso-Fonseca, Sergio Rangel-Guerrero, Mayra Jimenez-Morales, Nancy Munoz-Hernandez, Natividad Cruz-Ortiz, Tatiana Nunez-Garcia, Gisela Barrera-Badillo, Lucia Hernandez-Rivas, Irma Lopez-Martinez, Ernesto Ramirez-Gonzalez. |
| EPI_ISL_925406, EPI_ISL_925408, EPI_ISL_925413, EPI_ISL_925414 | Department of Clinical Microbiology | GIGA Medical Genomics | Keith Durkin, Maria Artesi, Sébastien Bontems, Raphaël Boreux, Bouchra Boujemla, Cécile Meex, Pierrette Melin, Marie-Pierre Hayette, Vincent Bours |
| EPI_ISL_930856, EPI_ISL_930857, EPI_ISL_930858 | Central Laboratory of Public Health of Rio Grande do Sul(Lacen_RS) | State Center for Health Surveillance of the Health Department of the State of Rio Grande do Sul(CEVS_SES-RS) | Barcellos R, Campos A, Dornelles C, Godinho F, Gonzalez A, Gregianini T, Molina C, Salvato R, Schaurich A, |
| EPI_ISL_933653, EPI_ISL_933654, EPI_ISL_933655, EPI_ISL_933656, EPI_ISL_933657 | Toronto Invasive Bacterial Diseases Network | McMaster University | Allison McGeer, Patryk Aftanas, Hooman Derakhshani, Angel Li, Kuganya Nirmalarajah, Emily Panousis, Ahmed Draia, Jalees Nasir, Michael Surette, Samira Mubareka, Andrew G. McArthur |
| EPI_ISL_933665 | Instituto de Diagnostico y Referencia Epidemiologicos INDRE_RNLSP | Instituto de Diagnostico y Referencia Epidemiologicos (INDRE) | Claudia Wong-Arambula, Abril Rodriguez-Maldonado, Fabiola Garces-Ayala, Adnan Araiza-Rodriguez, David Fragoso-Fonseca, Sergio Rangel-Guerrero, Mayra Jimenez-Morales, Nancy Munoz-Hernandez, Natividad Cruz-Ortiz, Tatiana Nunez-Garcia, Gisela Barrera-Badillo, Lucia Hernandez-Rivas, Irma Lopez-Martinez, Ernesto Ramirez-Gonzalez. |
| EPI_ISL_935754, EPI_ISL_935755, EPI_ISL_935756 | Cadham Provincial laboratory | National Microbiology Laboratory (NML) | Anna Majer, Shari Tyson, Grace Seo, Philip Mabon, Elsie Grudeski, Rhiannon Huzarewich, Russell Mandes, Anneliese Landgraff, Jennifer Tanner, Natalie Knox, Morag Graham, Gary Van Domselaar, Paul Van Caeseele, Jared Bullard, David Alexander, Kerry Dust, Nathalie Bastien, Yan Li, Timothy Booth, Darian Hole, Madison Chapel, Kirsten Biggar, CanCOGeN's metadata curation team, Public Health Agency of Canada CanCOGeN team |
| EPI_ISL_940547 | Hôpital Bichat Claude Bernard, Laboratoire de Virologie | IAME UMR1137 Inserm, Université de Paris, Hôpital Bichat | Antoine Bridier-Nahmias, Amélie Recoing, Quentin Le Hingrat, Lena Daniel, Siham Hamri, Gilles Collin, Alexandre Storto, Mélanie Bertine, Charlotte Charpentier, Nadhira Houhou-Fidouh, Diane Descamps, Benoit Visseaux |
| EPI_ISL_940894 | NCSLPH | NCSLPH | Chase K, Miller MC, Greene S, Glover W |
| EPI_ISL_940920, EPI_ISL_940921, EPI_ISL_940922, EPI_ISL_940923, EPI_ISL_940924, EPI_ISL_940925, EPI_ISL_940927, EPI_ISL_940933 | Centers for Disease Control and Prevention, Dengue Branch | Centers for Disease Control and Prevention, Dengue Branch | Gilberto A. Santiago, Glenda Gonzalez, Betzabel Flores, Keyla Charriez, Gabriela Paz-Bailey, Jorge L. Munoz-Jordan |
| EPI_ISL_941927, EPI_ISL_941928, EPI_ISL_941935 | Florida Bureau of Public Health Laboratories | Florida Bureau of Public Health Laboratories | Sarah Schmedes, Jason Blanton |
| EPI_ISL_942374, EPI_ISL_942375, EPI_ISL_942407, EPI_ISL_942896 | Lacen_RS | CEVS_SES_RS | Barcellos R, Campos A, Crescente L, Da Silva A, Dornelles C, Fonseca V, Garay L, Godinho F, Gonzalez A, Gregianini T, Molina C, Salvato R, Schaurich A |
| EPI_ISL_943601 | Lacen_RS | State Center for Health Surveillance. Rio Grande do Sul State Secretary of Health | Aline Campos, Amanda da Silva, Anelise Schaurich, Claudia Dornelles, Cynthia Molina, Fernanda Godinho, Lara Crescente, Leticia Garay, Regina Barcellos, Richard Salvato, Tatiana Gregianini, Vagner Fonseca |
| EPI_ISL_953404, EPI_ISL_953420 | Laboratorio de Investigaciones de Baney | "Swiss Tropical and Public Health Institute" | "Carlos Cortes, Claudia Daubenberger, Guillermo Garcia, Salome Hosch, Bonifacio Manguire Nlavo, Maximilian Mpina, Elizabeth Nyakarungu, Diosdado Odjama Nseng Ada, Mitoha Ondo O Ayekaba, Tobias Schindler, Philip Wonder Phiri" |
| EPI_ISL_954217 | 1.AO Universitaria 'S. Giovanni di Dio e Ruggi D'Aragona, Scuola Medica Salernitana' Hospital / 2.UOC di Virologia e Microbiologia, Università della Campania 'L. Vanvitelli' / 3.AO Universitaria 'Federico II' Napoli Hospital / 4.AORN 'San Giuseppe Moscati' Avellino Hospital / 5.AO 'San Pio - presidio G. Rummo' Benevento Hospital / 6.AO 'Sant'Anna e San Sebastiano' Caserta Hospital / 7.PO 'Maria Santissima Addolorata' Eboli Hospital / 8.Biogem Istituto di Ricerche Genetiche | 1. Genome Research Center for Health (CRGS) / 2. Laboratory of Molecular Medicine and Genomics(LMMGe) / 3. Center for Research in Pure and Applied Mathematics (CRMPA) | Giorgio Giurato, Francesca Rizzo, Alessandro Weisz, Gianluigi Franci, Giovanni Nassa, Pasquale Pagliano, Roberta Tarallo, Elena Alexandrova, Ylenia D'Agostino, Carlo Ferravante, Jessica Lamberti, Viola Melone, Domenico Memoli, Valeria Mirici Cappa, Domenico Palumbo, Giovanni Pecoraro, Assunta Sellitto, Oriana Strianese, Ilaria Terenzi, Giuseppe Fenza, Aniello Gentile, Antonello Saccomanno, Sonia Amabile, Teresa Rocco, Annamaria Salvati, Emilia Vaccaro, Massimiliano Galdiero, Michele Cennamo, Giuseppe Portella, Maria Grazia Foti, Mariarosaria Ingino, Maria Landi, Maurizio Fumi, Vincenzo Rocco, Rita Greco, Vittoria Letizia, Arnolfo Petruzziello, Maddalena Schioppa, Gregorio Goffredi, Francesca Marciano, Michele Caraglia, Alessia Cossu, Marianna Scrima, Edmondo Adorisio, Morena D'Avenia, Michela Iacobellis, Rosanna Piluscio, Giorgio Dirani, Vittorio Sambri, Simona Semprini, Silvia Zanoli, Francesco Curcio, Stefania Marzinotto, Andreina Baj, Fausto Sessa. |
| EPI_ISL_956278 | Siti Khodijah Hospital | Institute of Tropical Disease, Universitas Airlangga | Aldise M Nastri, Jezzy R Dewantari, Rima R Prasetya, Krisnoadi Rahardjo, Muhammad Hamdan, Gatot Soegiarto, Laksmi Wulandari, Resti Yudhawati, Soetjipto, Yasuko Mori, Maria I Lusida, Kazufumi Shimizu |
| EPI_ISL_959914, EPI_ISL_959919, EPI_ISL_960089, EPI_ISL_960090, EPI_ISL_960091, EPI_ISL_960092, EPI_ISL_960093 | University Medical Center Hamburg Eppendorf | Heinrich Pette Institute, Leibniz Institute for Experimental Virology | Alexis Robitaille, Thomas Günther, Johannes Knobloch, Martin Aepfelbacher, Nicole Fischer, Adam Grundhoff |
| EPI_ISL_960152 | Tshwaragano Hospital | National Health Laboratory Service/UCT | Arash Iranzadeh, Deelan Doolabh, Lynn Tyers, Bruna Galvao, Innocent Mudau, Marvin Hsiao, Kruger Marais, Diana Hardie, Stephen Korsman, Carolyn Williamson |
| EPI_ISL_960300, EPI_ISL_960301 | Nucleic Acid Testing, National Reference Laboratory | GIGA Medical Genomics | Yvan Butera, Keith Durkin, Maria Artesi, Bouchra Boujemla, Robert Rutayisire, Patrick Tuyisenge, Esperence Umumararungu, Sébastien Bontems, Marie-Pierre Hayette, Nathalie Renotte, Corinne Fasquelle, Swaibu Gatare, Jacob Souopgui, Sabin Nsanzimana, Vincent Bours, Léon Mutesa |
| EPI_ISL_961764, EPI_ISL_961765 | Laboratorio de Infectología, Servicio de Infectología, Hospital Universitario Dr. José Eleuterio González - Universidad Autónoma de Nuevo León | Laboratorio de Infectología Molecular, Departamento de Bioquímica y Medicina Molecular,Facultad de Medicina - Universidad Autónoma de Nuevo León | Kame A. Galán-Huerta, María F. Herrera-Saldivar, Natalia Martínez-Acuña, Sonia A. Lozano-Sepúlveda, Daniel Arellanos-Soto, Ana M. Rivas-Estilla, Paola Bocanegra-Ibarias, Samantha M. Flores-Treviño, Elvira Garza-González, Eduardo Perez-Alba, Laura Nuzzolo-Shihadeh, Adrian Camacho-Ortiz |
| EPI_ISL_962817 | Microbiological Diagnostic Unit - Public Health Laboratory | MDU-PHL | Seemann T., Sait, M.L., Sherry, N.L. |

| | | | |
|---|---|---|---|
| | (MDU-PHL) | | |
| EPI_ISL_964899 | Hospital Senillosa | Laboratorio Central Mg. Luis Alfredo Pianciola on behalf of 'Proyecto Argentino Interinstitucional de genomica de SARS-CoV-2' (PAIS Consortium) | L Pianciola, M Mazzeo, C Ziehm, C Pintos, M Fernandez, J Ousset, M Nabaes, M Viegas. |
| EPI_ISL_968164 | Clinical Molecular Microbiology Laboratory, UNC Hospital | Dirk Dittmer | Justin T. Landis , Razia Moorad , Brent A. Eason, Melissa B. Miller, Linda Pluta, Dirk Dittmer, Angelica Juarez, Cecilia Thompson, Shawn Hawken, Cameroon Grant, Evelyn Hoffman, Patricio Cano, Jason Wong, Carolina Caro-Vegas, Ryan McNamara, Blossom Damania. |
| EPI_ISL_968363, EPI_ISL_968364, EPI_ISL_968365, EPI_ISL_968366, EPI_ISL_968367, EPI_ISL_968368, EPI_ISL_968369, EPI_ISL_968370, EPI_ISL_968371, EPI_ISL_968372, EPI_ISL_968373, EPI_ISL_968374, EPI_ISL_968375, EPI_ISL_968376, EPI_ISL_968377, EPI_ISL_968378, EPI_ISL_968379, EPI_ISL_968380, EPI_ISL_968381, EPI_ISL_968382, EPI_ISL_968383, EPI_ISL_968384, EPI_ISL_968385, EPI_ISL_968386, EPI_ISL_968387, EPI_ISL_968388, EPI_ISL_968389, EPI_ISL_968390, EPI_ISL_968391, EPI_ISL_968392, EPI_ISL_968393, EPI_ISL_968394, EPI_ISL_968395, EPI_ISL_968396, EPI_ISL_968397, EPI_ISL_968398, EPI_ISL_968399, EPI_ISL_968400, EPI_ISL_968401, EPI_ISL_968402, EPI_ISL_968403, EPI_ISL_968404, EPI_ISL_968405, EPI_ISL_968406, EPI_ISL_968407, EPI_ISL_968408, EPI_ISL_968409, EPI_ISL_968410, EPI_ISL_968411, EPI_ISL_968412, EPI_ISL_968413, EPI_ISL_968414, EPI_ISL_968415, EPI_ISL_968416, EPI_ISL_968417, EPI_ISL_968418, EPI_ISL_968419, EPI_ISL_968420, EPI_ISL_968421, EPI_ISL_968422, EPI_ISL_968423, EPI_ISL_968424, EPI_ISL_968425, EPI_ISL_968426, EPI_ISL_968427, EPI_ISL_968428, EPI_ISL_968429, EPI_ISL_968430, EPI_ISL_968431, EPI_ISL_968432, EPI_ISL_968433, EPI_ISL_968434, EPI_ISL_968435, EPI_ISL_968436, EPI_ISL_968437, EPI_ISL_968438, EPI_ISL_968439, EPI_ISL_968440, EPI_ISL_968441, EPI_ISL_968442, EPI_ISL_968443, EPI_ISL_968444, EPI_ISL_968445, EPI_ISL_968446, EPI_ISL_968447, EPI_ISL_968448, EPI_ISL_968449, EPI_ISL_968450, EPI_ISL_968451, EPI_ISL_968452, EPI_ISL_968453, EPI_ISL_968454, EPI_ISL_968455, EPI_ISL_968456, EPI_ISL_968457, EPI_ISL_968458, EPI_ISL_968459, EPI_ISL_968460, EPI_ISL_968461, EPI_ISL_968462 | | | |
| see above | BCCDC Public Health Laboratory | BCCDC Public Health Laboratory | Prystajecky Natalie, Linda Hoang, Dan Fornika, John Tyson, Shannon Russell, Kim Macdonald, Kimia Kamelian, Ana Pacagnella, Corrinne Ng, Loretta Janz, Robert Azana Terry Snutch, Mel Krajden |
| EPI_ISL_977254, EPI_ISL_977260, EPI_ISL_977268, EPI_ISL_977269, EPI_ISL_977305, EPI_ISL_977309, EPI_ISL_977318, EPI_ISL_977319, EPI_ISL_977320, EPI_ISL_977431, EPI_ISL_977441, EPI_ISL_977442, EPI_ISL_977443, EPI_ISL_977444, EPI_ISL_977451, EPI_ISL_977458 | | | |
| see above | University of Zambia, School of Veterinary Medicine | UNZAVET and PATH | Mulenga Mwenda-Chimfwembe, Ngonda Saasa, Daniel Bridges |
| EPI_ISL_978497, EPI_ISL_978498 | Central Public Health Laboratory - LACEN -Bahia, Salvador, Brazil | Central Public Health Laboratory - LACEN -Bahia, Salvador, Brazil | Stephane Tosta, Luciana Oliveira, Vanessa Nardy,Patrícia Cajado,Marcela Gómez, Breno Dominguez, Jaqueline Gomes, Vagner Fonseca,Marta Giovanetti,Luiz Alcantara, Felicidade Pereira, Arabela Leal |
| EPI_ISL_979355 | Microbiological Diagnostic Unit - Public Health Laboratory (MDU-PHL) | MDU-PHL | Seemann T., Sait, M.L., Sherry, N.L. |
| EPI_ISL_981031 | Hospital Villa Regina | Laboratorio Central Mg. Luis Alfredo Pianciola on behalf of 'Proyecto Argentino Interinstitucional de genomica de SARS-CoV-2' (PAIS Consortium) | L Pianciola, M Mazzeo, C Ziehm, C Pintos, M Fernandez, J Ousset, M Nabaes, M Viegas. |
| EPI_ISL_981032 | hospital Cipolleti | Laboratorio Central Mg. Luis Alfredo Pianciola on behalf of 'Proyecto Argentino Interinstitucional de genomica de SARS-CoV-2' (PAIS Consortium) | L Pianciola, M Mazzeo, C Ziehm, C Pintos, M Fernandez, J Ousset, M Nabaes, M Viegas. |
| EPI_ISL_981035, EPI_ISL_981051 | Hospital Dr. Francisco López Lima | Laboratorio Central Mg. Luis Alfredo Pianciola on behalf of 'Proyecto Argentino Interinstitucional de genomica de SARS-CoV-2' (PAIS Consortium) | L Pianciola, M Mazzeo, C Ziehm, C Pintos, M Fernandez, J Ousset, M Nabaes, M Viegas. |
