## Supplementary material for "Detection and characterization of the SARS-CoV-2 lineage B.1.526 in New York": Supp. Table 3 GISAID Acknowledgment part 2: gisaid_hcov-19_acknowledgement_table_2021_02_12_23-3.pdf

We gratefully acknowledge the following Authors from the Originating laboratories responsible for obtaining the specimens, as well as the Submitting laboratories where the genome data were generated and shared via GISAID, on which this research is based.

All Submitters of data may be contacted directly via www.gisaid.org

Authors are sorted alphabetically.

| Accession ID | Originating Laboratory | Submitting Laboratory | Authors |
|---|---|---|---|
| EPI_ISL_510544 | SA Pathology | SA Pathology | Lex Leong, Chuan Kok Lim, Mark Turra, Ivan Bastian, Geoff Higgins |
| EPI_ISL_512842 | National Public Health Laboratory, National Centre for Infectious Diseases | National Public Health Laboratory, National Centre for Infectious Diseases | Mak TM, Octavia S, Zhou Z, Chavatte JM, Cui L, Lin RTP |
| EPI_ISL_512920, EPI_ISL_512921, EPI_ISL_512922, EPI_ISL_512923, EPI_ISL_512924, EPI_ISL_512925, EPI_ISL_512926, EPI_ISL_512927 | Pathogen Genomics Lab King Abdullah University of Science and Technology(KAUST) | Pathogen Genomics Lab King Abdullah University of Science and Technology(KAUST) | Fadwa Alofi, Sharif Hala, Rahul P Salunke, Sara Mfarrej, Amit Kumar Subudhi, Fathia Ben Rached, Amanda, Luke, Afrah Alsomali, Asim Khogeer, Jumana Taha, Abdulaziz Alahmadi, Kahled Algithami, Raeece Naeem, Anwar Hashem, Naif Almontashiri, Arnab Pain |
| EPI_ISL_513393 | Pathology West - NSW Health Pathology | NSW Health Pathology - Institute of Clinical Pathology and Medical Research; Westmead Hospital; University of Sydney | CIDM-PH et al. |
| EPI_ISL_513402 | 4Cyte Pathology | NSW Health Pathology - Institute of Clinical Pathology and Medical Research; Westmead Hospital; University of Sydney | CIDM-PH et al. |
| EPI_ISL_513411, EPI_ISL_513412 | Sydney South West Pathology Service (SSWPS) - Liverpool Hospital - NSW Health Pathology | NSW Health Pathology - Institute of Clinical Pathology and Medical Research; Westmead Hospital; University of Sydney | CIDM-PH et al. |
| EPI_ISL_513911, EPI_ISL_513912, EPI_ISL_513913 | SA Pathology | SA Pathology | Lex Leong, Chuan Kok Lim, Mark Turra, Ivan Bastian, Geoff Higgins |
| EPI_ISL_514345, EPI_ISL_514346, EPI_ISL_514347, EPI_ISL_514348, EPI_ISL_514349, EPI_ISL_514351, EPI_ISL_514352 | Respiratory Virus Unit, Microbiology Services Colindale, Public Health England | Respiratory Virus Unit, Microbiology Services Colindale, Public Health England | PHE Covid Sequencing Team |
| EPI_ISL_514443, EPI_ISL_514448, EPI_ISL_514449, EPI_ISL_514451 | Department of Pathology, University of Cambridge | COVID-19 Genomics UK (COG-UK) Consortium | Luke W Meredith, M. Estée Török, Myra Hosmillo, William L. Hamilton, Martin D. Curran, Theresa Feltwell, Grant Hall, Anna Yakovleva, Fahad A Khokhar, Charlotte J. Houldcroft, Laura G Caller, Aminu S. Jahun, Sarah L. Caddy, Yasmin Chaudhry, Malte Pinckert, Ian Goodfellow |
| EPI_ISL_514452, EPI_ISL_514453, EPI_ISL_514454 | Queens Medical Centre, Clinical Microbiology Department / DeepSeq Nottingham | COVID-19 Genomics UK (COG-UK) Consortium | Gemma Clark, Wendy Smith, Manjinder Khakh, Vicki M Fleming, Michelle M Lister, Hannah Howson-Wells, Jonathan Ball, Patrick McClure, Joseph Chappell, Theocharis Tsoleridis, Nadine Holmes, Matthew Carlisle, Christopher Moore, Fei Sang, Johnny Debebe, Victoria Wright, Matthew Loose |
| EPI_ISL_514514, EPI_ISL_514515 | West of Scotland Specialist Virology Centre, NHSGGC / MRC-University of Glasgow Centre for Virus Research | COVID-19 Genomics UK (COG-UK) Consortium | Ana da Silva Filipe, Natasha Johnson, Kathy Smollett, Daniel Mair, Stephen Carmichael, Lily Tong, Jenna Nichols, Elihu Aranday-Cortes, Kirstyn Brunker, Yasmin Parr, Alice Broos, Kyriaki Nomikou; Sarah McDonald, Marc Niebel, Patawee Asamaphan; Richard Orton, Joseph Hughes, Sreenu Vattipally, David L Robertson; Alasdair MacLean, Rory Gunson; Kathy Li, Natasha Jesudason, Rajiv Shah, James Shepherd, Antonia Ho, Emma Thomson |
| EPI_ISL_514516, EPI_ISL_514517, EPI_ISL_514518, EPI_ISL_514519, EPI_ISL_514520, EPI_ISL_514521, EPI_ISL_514522, EPI_ISL_514523, EPI_ISL_514524, EPI_ISL_514525, EPI_ISL_514526, EPI_ISL_514527, EPI_ISL_514528, EPI_ISL_514529, EPI_ISL_514530, EPI_ISL_514531, EPI_ISL_514532, EPI_ISL_514533, EPI_ISL_514534, EPI_ISL_514535, EPI_ISL_514536, EPI_ISL_514537, EPI_ISL_514538, EPI_ISL_514539, EPI_ISL_514540, EPI_ISL_514541, EPI_ISL_514542, EPI_ISL_514543, EPI_ISL_514544, EPI_ISL_514545, EPI_ISL_514546, EPI_ISL_514547, EPI_ISL_514548, EPI_ISL_514549, EPI_ISL_514550, EPI_ISL_514551, EPI_ISL_514552, EPI_ISL_514553, EPI_ISL_514554, EPI_ISL_514555, EPI_ISL_514556, EPI_ISL_514557, EPI_ISL_514558, EPI_ISL_514559, EPI_ISL_514560, EPI_ISL_514561, EPI_ISL_514562 | | | |
| see above | Virology Department, Royal Infirmary of Edinburgh, NHS Lothian / School of Biological Sciences, University of Edinburgh / Institute of Genetics and Molecular Medicine, University of Edinburgh | COVID-19 Genomics UK (COG-UK) Consortium | McHugh M, Dewar R, Rooke S, Gallagher M, Balcaza C, O'Toole Á, Scher E, Hill V, McCrone JT, Colquhoun R, Yu X, Jackson B, Rambaut A, Williams TC, Templeton K |
| EPI_ISL_514577, EPI_ISL_514579 | Wales Specialist Virology Centre Sequencing lab: Pathogen Genomics Unit | COVID-19 Genomics UK (COG-UK) Consortium | Catherine Moore, Johnathan Evans, Laura Gifford, Malorie Perry, Simon Cottrell, Angela Marchbank, Alec Birchley, Alexander Adams, Amy Gaskin, Bree Gatica-Wilcox, Jason Coombes, Joel Southgate, Lauren Gilbert, Lee Graham, Nicole Pacchiarini, Sara Kumziene-Summerhayes, Sarah Taylor, Sophie Jones, Sara Rey, Matthew Bull, Joanne Watkins, Sally Corden, Tom Connor |
| EPI_ISL_515187, EPI_ISL_515188, EPI_ISL_515189, EPI_ISL_515190, EPI_ISL_515191, EPI_ISL_515192, EPI_ISL_515193, EPI_ISL_515194, EPI_ISL_515195 | E. Gulbja Laboratorija | Latvian Biomedical Research and Study Centre | Ivars Silamielis, Kaspars Megnis, Monta Ustinova, ikita Zrelovs, Vita Rovte, Mikus Gavars, Dmitrijs Perminovs, Uga Dumpis, Jnis Kloviš |
| EPI_ISL_515196 | Centrl laboratorija | Latvian Biomedical Research and Study Centre | Ivars Silamielis, Kaspars Megnis, Monta Ustinova, ikita Zrelovs, Vita Rovte, Stella Lapia, Jana Oste, Marta Priedte, Uga Dumpis, Jnis Kloviš |
| EPI_ISL_516446, EPI_ISL_516449, EPI_ISL_516451, EPI_ISL_516452, EPI_ISL_516453, EPI_ISL_516454, EPI_ISL_516455, EPI_ISL_516456, EPI_ISL_516460, EPI_ISL_516461, EPI_ISL_516462, EPI_ISL_516463, EPI_ISL_516465, EPI_ISL_516467, EPI_ISL_516468, EPI_ISL_516469, EPI_ISL_516470, EPI_ISL_516473, EPI_ISL_516475, EPI_ISL_516477, EPI_ISL_516478, EPI_ISL_516496, EPI_ISL_516506, EPI_ISL_516520, EPI_ISL_516524 | | | |
| see above | University of Wisconsin-Madison AIDS Vaccine Research Laboratories | University of Wisconsin-Madison AIDS Vaccine Research Laboratories | Gage Moreno, Katarina Braun, et al. AIDS Vaccine Research Laboratories |
| EPI_ISL_516557, EPI_ISL_516558, EPI_ISL_516559, EPI_ISL_516560, EPI_ISL_516561, EPI_ISL_516562, EPI_ISL_516563, EPI_ISL_516564, EPI_ISL_516565, EPI_ISL_516566, EPI_ISL_516567, EPI_ISL_516568, EPI_ISL_516569, EPI_ISL_516570, EPI_ISL_516571, EPI_ISL_516572, EPI_ISL_516573, EPI_ISL_516574, EPI_ISL_516575, EPI_ISL_516576, EPI_ISL_516577, EPI_ISL_516578, EPI_ISL_516579, EPI_ISL_516580, EPI_ISL_516581, EPI_ISL_516582, EPI_ISL_516583, EPI_ISL_516584, EPI_ISL_516585, EPI_ISL_516586, EPI_ISL_516587, EPI_ISL_516588, EPI_ISL_516589, EPI_ISL_516590, EPI_ISL_516591, EPI_ISL_516592, EPI_ISL_516593, EPI_ISL_516594, EPI_ISL_516595, EPI_ISL_516596, EPI_ISL_516597, EPI_ISL_516598, EPI_ISL_516599, EPI_ISL_516600, EPI_ISL_516601, EPI_ISL_516602, EPI_ISL_516603, EPI_ISL_516604, EPI_ISL_516605, EPI_ISL_516606, EPI_ISL_516607 | | | |
| see above | Viollier AG | Department of Biosystems Science and Engineering, ETH Zürich | Christian Beisel, Sarah Nadeau, Ivan Topolsky, Pedro Ferreira, Philipp Jablonski, Susana Posada-Céspedes, Tobias Schär, Ina Nissen, Natascha Santacroce, Elodie Burcklen, Christiane Beckmann, Maurice Redondo, Olivier Kobel, Christoph Noppen, Sophie Seidel, Noemie Santamaria de Souza, Niko Beerenwinkel, Tanja Stadler |
| EPI_ISL_516813, EPI_ISL_516814, EPI_ISL_516828 | National Public Health Laboratory, National Centre for Infectious Diseases | National Public Health Laboratory, National Centre for Infectious Diseases | Mak TM, Octavia S, Zhou Z, Cui L, Lin RTP |
| EPI_ISL_516829 | RSUD Nyi Ageng Serang | Genetics Working Group (Pokja Genetik) Faculty of Medicine, Public Health and Nursing Universitas Gadjah Mada (FK-KMK UGM), Disease Investigation Center Wates Ministry of Agriculture Indonesia, Department of Microbiology FK-KMK UGM, Laboratorium Diagnostik Yayasan Tahija World Mosquito Program (WMP) Yogyakarta Center for Tropical Medicine FK-KMK UGM, Integrated Research Center FK-KMK UGM, Department of Computer Science and Electronics FMIPA UGM | Gunadi, Hendra Wibawa, . Marcellus, Mohamad S. Hakim, Edwin W. Daniwijaya, Ludhang P. Rizki, Endah Supriyati, Eggi Arguni, Titik Nuryastuti, Tri Wibawa, Dwi AA Nugrahaningsih, Afiahayati , . Siswanto, Beby Dewi Sartika, Dyah Ayu Puspitarani |
| EPI_ISL_516990 | Department of Pathology, University of Cambridge | COVID-19 Genomics UK (COG-UK) Consortium | Luke W Meredith, M. Estée Török, Myra Hosmillo, William L. Hamilton, Martin D. Curran, Theresa Feltwell, Grant Hall, Anna Yakovleva, Fahad A Khokhar, |

| | | | |
|---|---|---|---|
| | | | Charlotte J. Houldcroft, Laura G Caller, Aminu S. Jahun, Sarah L. Caddy, Yasmin Chaudhry, Malte Pinckert, Ian Goodfellow |
| EPI_ISL_517545, EPI_ISL_517555, EPI_ISL_517557, EPI_ISL_517558, EPI_ISL_517561, EPI_ISL_517567, EPI_ISL_517568, EPI_ISL_517571, EPI_ISL_517572, EPI_ISL_517576, EPI_ISL_517579 | | | |
| see above | Virology Department, Sheffield Teaching Hospitals NHS Foundation Trust/Department of Infection, Immunity and Cardiovascular Disease, The Medical School, University of Sheffield | COVID-19 Genomics UK (COG-UK) Consortium | Thushan de Silva, Matthew Parker, Nikki Smith, Adri Angyal, Rebecca Brown, Luke Green, Rachel Tucker, Paul Parsons, Danielle Groves, Katie Johnson, Laura Carrilero, Alex Keeley, Dave Partridge, Matthew Wyles, Benjamin Lindsey, Mehmet Yavuz, Mohammad Raza, Cariad Evans |
| EPI_ISL_517580, EPI_ISL_517581, EPI_ISL_517582 | West of Scotland Specialist Virology Centre, NHSGGC / MRC-University of Glasgow Centre for Virus Research | COVID-19 Genomics UK (COG-UK) Consortium | Ana da Silva Filipe, Natasha Johnson, Kathy Smollett, Daniel Mair, Stephen Carmichael, Lily Tong, Jenna Nichols, Elihu Aranday-Cortes, Kirstyn Brunker, Yasmin Parr, Alice Broos, Kyriaki Nomikou; Sarah McDonald, Marc Niebel, Patawee Asamaphan; Richard Orton, Joseph Hughes, Sreenu Vattipally, David L Robertson; Alasdair MacLean, Rory Gunson; Kathy Li, Natasha Jesudason, Rajiv Shah, James Shepherd, Antonia Ho, Emma Thomson |
| EPI_ISL_517588, EPI_ISL_517589 | Virology Department, Royal Infirmary of Edinburgh, NHS Lothian / School of Biological Sciences, University of Edinburgh / Institute of Genetics and Molecular Medicine, University of Edinburgh | COVID-19 Genomics UK (COG-UK) Consortium | McHugh M, Dewar R, Rooke S, Gallagher M, Balcaza C, O'Toole Á, Scher E, Hill V, McCrone JT, Colquhoun R, Yu X, Jackson B, Rambaut A, Williams TC, Templeton K |
| EPI_ISL_517592, EPI_ISL_517594, EPI_ISL_517596, EPI_ISL_517600, EPI_ISL_517601, EPI_ISL_517602, EPI_ISL_517603, EPI_ISL_517604, EPI_ISL_517610 | Wales Specialist Virology Centre Sequencing lab: Pathogen Genomics Unit | COVID-19 Genomics UK (COG-UK) Consortium | Catherine Moore, Johnathan Evans, Laura Gifford, Malorie Perry, Simon Cottrell, Angela Marchbank, Alec Birchley, Alexander Adams, Amy Gaskin, Bree Gatica-Wilcox, Jason Coombes, Joel Southgate, Lauren Gilbert, Lee Graham, Nicole Pacchiarini, Sara Kumziene-Summerhayes, Sarah Taylor, Sophie Jones, Sara Rey, Matthew Bull, Joanne Watkins, Sally Corden, Tom Connor |
| EPI_ISL_517661, EPI_ISL_517662 | Academic Hospital Paramaribo | Erasmus Medical Center | Bas Oude Munnink, Dion Gajadin, Ed Ijzerman, Emmanuelle Munger, Gary Gummels, Ingrid Krishnadath, Lycke Woittiez, Marion Koopmans, Mireille Van de Veer, Princes Wongsowidjojo, Radjesh Ori, Rohma Banwari, Stephen Vreden |
| EPI_ISL_517928, EPI_ISL_517929, EPI_ISL_517930, EPI_ISL_517931, EPI_ISL_517932 | Florida Bureau of Public Health Laboratories | Florida Bureau of Public Health Laboratories | Sarah Schmedes, Jason Blanton |
| EPI_ISL_522877 | Instituto Nacional de Medicina Genomica | Instituto Nacional de Medicina Genomica | Hidalgo-Miranda A, Mendoza-Vargas A, Reyes-Grajeda JP, Cisneros-Villanueva M, Cedro-Tanda A, Hurtado-Cordova E, Peñaloza-Figueroa F, Herrera-Montalvo LA |
| EPI_ISL_522878 | Instituto Nacional de Medicina Genomica | Instituto Nacional de Medicina Genomica | Hidalgo-Miranda A, Mendoza-Vargas A, Reyes-Grajeda JP, Cisneros-Villanueva M, Hurtado-Cordova E, Cedro-Tanda A, Peñaloza-Figueroa F, Herrera-Montalvo LA |
| EPI_ISL_522880 | Instituto Nacional de Medicina Genómica | Instituto Nacional de Medicina Genómica | Hidalgo-Miranda A, Mendoza-Vargas A, Reyes-Grajeda JP, Cisneros-Villanueva M, Cedro-Tanda A, Hurtado-Cordova E, Peñaloza-Figueroa F, Herrera-Montalvo LA |
| EPI_ISL_522881, EPI_ISL_522883, EPI_ISL_522901, EPI_ISL_522906, EPI_ISL_522915, EPI_ISL_522916, EPI_ISL_522917, EPI_ISL_522918, EPI_ISL_522919, EPI_ISL_522920, EPI_ISL_522921, EPI_ISL_522923, EPI_ISL_522924, EPI_ISL_522925, EPI_ISL_522926, EPI_ISL_522929, EPI_ISL_522930, EPI_ISL_522931, EPI_ISL_522932, EPI_ISL_522933, EPI_ISL_522934, EPI_ISL_522936, EPI_ISL_522937, EPI_ISL_522939 | | | |
| see above | University of Wisconsin-Madison AIDS Vaccine Research Laboratories | University of Wisconsin-Madison AIDS Vaccine Research Laboratories | Gage Moreno, Katarina Braun, et al. AIDS Vaccine Research Laboratories |
| EPI_ISL_522940, EPI_ISL_522942, EPI_ISL_522978, EPI_ISL_522979, EPI_ISL_522980, EPI_ISL_522981, EPI_ISL_522982, EPI_ISL_522983, EPI_ISL_522984, EPI_ISL_522985, EPI_ISL_522986 | | | |
| see above | Instituto Nacional de Medicina Genómica | Instituto Nacional de Medicina Genómica | Hidalgo-Miranda A, Mendoza-Vargas A, Reyes-Grajeda JP, Cisneros-Villanueva M, Cedro-Tanda A, Hurtado-Cordova E, Peñaloza-Figueroa F, Herrera-Montalvo LA |
| EPI_ISL_523275, EPI_ISL_523276, EPI_ISL_523277, EPI_ISL_523280 | Dutch COVID-19 response team | Erasmus Medical Center | Bas Oude Munnink, David Nieuwenhuijse, Reina Sikkema, Claudia Schapendonk, Irina Chestakova, Anne van der Linden, Theo Bestebroer, Stefan van Nieuwkoop, Mark Pronk, Pascal Lexmond, Corien Swaan, Manon Haverkate, Madelief Mollers, Mart Stein, Sandra Kengne Kamga Mobou, Jeroen van Kampen, Jolanda Voermans, Aura Timen, Corine GeurtsvanKessel, Annemiek van der Eijk, Richard Molenkamp, Marion Koopmans, on behalf of the Dutch national COVID-19 response team. |
| EPI_ISL_523500 | Instituto Nacional de Medicina Genómica | Instituto Nacional de Medicina Genómica | Hidalgo-Miranda A, Mendoza-Vargas A, Reyes-Grajeda JP, Cisneros-Villanueva M, Cedro-Tanda A, Hurtado-Cordova E, Peñaloza-Figueroa F, Herrera-Montalvo LA |
| EPI_ISL_523539, EPI_ISL_523540, EPI_ISL_523541, EPI_ISL_523542, EPI_ISL_523543, EPI_ISL_523544, EPI_ISL_523545, EPI_ISL_523546, EPI_ISL_523695, EPI_ISL_523696, EPI_ISL_523697, EPI_ISL_523698, EPI_ISL_523699, EPI_ISL_523700, EPI_ISL_523701, EPI_ISL_523702, EPI_ISL_523703, EPI_ISL_523704, EPI_ISL_523705, EPI_ISL_523706, EPI_ISL_523707, EPI_ISL_523708, EPI_ISL_523709, EPI_ISL_523710, EPI_ISL_523711, EPI_ISL_523712, EPI_ISL_523713, EPI_ISL_523714, EPI_ISL_523715, EPI_ISL_523716, EPI_ISL_523717, EPI_ISL_523718, EPI_ISL_523719, EPI_ISL_523720, EPI_ISL_523721, EPI_ISL_523722, EPI_ISL_523723, EPI_ISL_523724, EPI_ISL_523725, EPI_ISL_523726, EPI_ISL_523727, EPI_ISL_523728, EPI_ISL_523729, EPI_ISL_523730, EPI_ISL_523747, EPI_ISL_523749, EPI_ISL_523751, EPI_ISL_523752, EPI_ISL_523753, EPI_ISL_523755, EPI_ISL_523757, EPI_ISL_523758, EPI_ISL_523759, EPI_ISL_523762, EPI_ISL_523809 | | | |
| see above | Dutch COVID-19 response team | Erasmus Medical Center | Bas Oude Munnink, David Nieuwenhuijse, Reina Sikkema, Claudia Schapendonk, Irina Chestakova, Anne van der Linden, Theo Bestebroer, Stefan van Nieuwkoop, Mark Pronk, Pascal Lexmond, Corien Swaan, Manon Haverkate, Madelief Mollers, Mart Stein, Sandra Kengne Kamga Mobou, Jeroen van Kampen, Jolanda Voermans, Aura Timen, Corine GeurtsvanKessel, Annemiek van der Eijk, Richard Molenkamp, Marion Koopmans, on behalf of the Dutch national COVID-19 response team. |
| EPI_ISL_523856, EPI_ISL_523857, EPI_ISL_523858, EPI_ISL_523859, EPI_ISL_523860, EPI_ISL_523861, EPI_ISL_523862, EPI_ISL_523863, EPI_ISL_523864, EPI_ISL_523865, EPI_ISL_523866, EPI_ISL_523867, EPI_ISL_523868, EPI_ISL_523869, EPI_ISL_523870, EPI_ISL_523871, EPI_ISL_523872, EPI_ISL_523873, EPI_ISL_523874, EPI_ISL_523875, EPI_ISL_523876, EPI_ISL_523877, EPI_ISL_523878, EPI_ISL_523879, EPI_ISL_523907, EPI_ISL_523908, EPI_ISL_523909, EPI_ISL_523910, EPI_ISL_523911, EPI_ISL_523912, EPI_ISL_523913, EPI_ISL_523914, EPI_ISL_523915, EPI_ISL_523916, EPI_ISL_523917 | | | |
| see above | Viollier AG | Department of Biosystems Science and Engineering, ETH Zürich | Christian Beisel, Sarah Nadeau, Ivan Topolsky, Pedro Ferreira, Philipp Jablonski, Susana Posada-Céspedes, Tobias Schär, Ina Nissen, Natascha Santacroce, Elodie Burcklen, Christiane Beckmann, Maurice Redondo, Olivier Kobel, Christoph Noppen, Sophie Seidel, Noemie Santamaria de Souza, Niko Beerenwinkel, Tanja Stadler |
| EPI_ISL_524487, EPI_ISL_524488, EPI_ISL_524490, EPI_ISL_524491, EPI_ISL_524492 | PHE South West Regional Laboratory, National Infection Service | Wellcome Sanger Institute for the COVID-19 Genomics UK (COG-UK) consortium | Stephanie Hutchings, Hannah Pymont, Dr Peter Muir, Barry Vipond, Rich Hopes; and Alex Alderton, Roberto Amato, Sonia Goncalves, Ewan Harrison, David K. Jackson, Ian Johnston, Dominic Kwiatkowski, Cordelia Langford, John Sillitoe on behalf of the Wellcome Sanger Institute COVID-19 Surveillance Team (http://www.sanger.ac.uk/covid-team) |
| EPI_ISL_525371, EPI_ISL_525372, EPI_ISL_525373, EPI_ISL_525374, EPI_ISL_525375, EPI_ISL_525376, EPI_ISL_525377, EPI_ISL_525378, EPI_ISL_525379, EPI_ISL_525380, EPI_ISL_525381, EPI_ISL_525382, EPI_ISL_525383, EPI_ISL_525384, EPI_ISL_525385, EPI_ISL_525386, EPI_ISL_525387, EPI_ISL_525388, EPI_ISL_525389, EPI_ISL_525390, EPI_ISL_525391, EPI_ISL_525392, EPI_ISL_525393, EPI_ISL_525394, EPI_ISL_525395, EPI_ISL_525396, EPI_ISL_525397, EPI_ISL_525398, EPI_ISL_525399, EPI_ISL_525400, EPI_ISL_525401, EPI_ISL_525402, EPI_ISL_525403, EPI_ISL_525404, EPI_ISL_525405, EPI_ISL_525406, EPI_ISL_525407, EPI_ISL_525408, EPI_ISL_525409, EPI_ISL_525410, EPI_ISL_525411, EPI_ISL_525412, EPI_ISL_525413, EPI_ISL_525414, EPI_ISL_525415, EPI_ISL_525416, EPI_ISL_525417, EPI_ISL_525418 | | | |
| see above | National Virus Reference Laboratory | National Virus Reference Laboratory | Michael Carr, Gabriel Gonzalez, Jonathan Dean, Aditi Chaturvedi, Suzie Coughlan, Cillian F De Gascun |
| EPI_ISL_525755 | Seattle Flu Study | Seattle Flu Study | Deborah A. Nickerson, Chris D. Frazar, Jover Lee, Benjamin Pelle, Matthew Richardson, Amanda Adler, Elisabeth Brandstetter, Peter D. Han, Kairsten Fay, Misja Ilcisin, Kirsten Lacombe, Thomas R. Sibley, Melissa Truong, Caitlin R. Wolf, Karen Cowgill, Stephanie Schrag, Jeff Duchin, Michael Boeckh, Janet A. Englund, Michael Famulare, Barry R. Lutz, Mark J. Rieder, Lea M. Starita, Matthew Thompson, Helen Y. Chu, Trevor Bedford, Jay Shendure |
| EPI_ISL_526121, EPI_ISL_526122 | St Vincent's Pathology (SydPath) | NSW Health Pathology - Institute of Clinical Pathology and Medical Research; Westmead Hospital; University of Sydney | CIDM-PH et al. |
| EPI_ISL_526125 | Sydney South West Pathology Service (SSWPS) - Concord Repatriation General Hospital - NSW Health Pathology | NSW Health Pathology - Institute of Clinical Pathology and Medical Research; Westmead Hospital; University of Sydney | CIDM-PH et al. |
| EPI_ISL_526131 | Sydney South West Pathology Service (SSWPS) - Liverpool Hospital - NSW Health Pathology | NSW Health Pathology - Institute of Clinical Pathology and Medical Research; Westmead Hospital; University of Sydney | CIDM-PH et al. |
| EPI_ISL_526132, EPI_ISL_526133 | Pathology North - Royal North Shore Hospital - NSW Health Pathology | NSW Health Pathology - Institute of Clinical Pathology and Medical Research; Westmead Hospital; University of Sydney | CIDM-PH et al. |

| | | | |
|---|---|---|---|
| EPI_ISL_526138, EPI_ISL_526139, EPI_ISL_526140, EPI_ISL_526141 | Pathology West - NSW Health Pathology | NSW Health Pathology - Institute of Clinical Pathology and Medical Research; Westmead Hospital; University of Sydney | CIDM-PH et al. |
| EPI_ISL_526149 | South Eastern Area Laboratory Services (SEALS) | NSW Health Pathology - Institute of Clinical Pathology and Medical Research; Westmead Hospital; University of Sydney | CIDM-PH et al. |
| EPI_ISL_526150, EPI_ISL_526151 | Pathology West - NSW Health Pathology | NSW Health Pathology - Institute of Clinical Pathology and Medical Research; Westmead Hospital; University of Sydney | CIDM-PH et al. |
| EPI_ISL_526152 | Pathology North - Hunter - NSW Health Pathology | NSW Health Pathology - Institute of Clinical Pathology and Medical Research; Westmead Hospital; University of Sydney | CIDM-PH et al. |
| EPI_ISL_526153 | Douglass Hanly Moir Pathology | NSW Health Pathology - Institute of Clinical Pathology and Medical Research; Westmead Hospital; University of Sydney | CIDM-PH et al. |
| EPI_ISL_526154, EPI_ISL_526155 | Pathology North - Royal North Shore Hospital - NSW Health Pathology | NSW Health Pathology - Institute of Clinical Pathology and Medical Research; Westmead Hospital; University of Sydney | CIDM-PH et al. |
| EPI_ISL_526156 | Pathology West - NSW Health Pathology | NSW Health Pathology - Institute of Clinical Pathology and Medical Research; Westmead Hospital; University of Sydney | CIDM-PH et al. |
| EPI_ISL_526157, EPI_ISL_526158, EPI_ISL_526159, EPI_ISL_526160 | Sydney South West Pathology Service (SSWPS) - Liverpool Hospital - NSW Health Pathology | NSW Health Pathology - Institute of Clinical Pathology and Medical Research; Westmead Hospital; University of Sydney | CIDM-PH et al. |
| EPI_ISL_526161, EPI_ISL_526162, EPI_ISL_526163, EPI_ISL_526164, EPI_ISL_526165 | Pathology West - NSW Health Pathology | NSW Health Pathology - Institute of Clinical Pathology and Medical Research; Westmead Hospital; University of Sydney | CIDM-PH et al. |
| EPI_ISL_526166, EPI_ISL_526167, EPI_ISL_526168, EPI_ISL_526169 | Austech Medical Laboratories | NSW Health Pathology - Institute of Clinical Pathology and Medical Research; Westmead Hospital; University of Sydney | CIDM-PH et al. |
| EPI_ISL_526171 | St Vincent's Pathology (SydPath) | NSW Health Pathology - Institute of Clinical Pathology and Medical Research; Westmead Hospital; University of Sydney | CIDM-PH et al. |
| EPI_ISL_526172, EPI_ISL_526173, EPI_ISL_526174, EPI_ISL_526175, EPI_ISL_526176 | Sydney South West Pathology Service (SSWPS) - Liverpool Hospital - NSW Health Pathology | NSW Health Pathology - Institute of Clinical Pathology and Medical Research; Westmead Hospital; University of Sydney | CIDM-PH et al. |
| EPI_ISL_526177, EPI_ISL_526178, EPI_ISL_526179 | St Vincent's Pathology (SydPath) | NSW Health Pathology - Institute of Clinical Pathology and Medical Research; Westmead Hospital; University of Sydney | CIDM-PH et al. |
| EPI_ISL_526186 | 4Cyte Pathology | NSW Health Pathology - Institute of Clinical Pathology and Medical Research; Westmead Hospital; University of Sydney | CIDM-PH et al. |
| EPI_ISL_526187, EPI_ISL_526188, EPI_ISL_526189, EPI_ISL_526190 | St Vincent's Pathology (SydPath) | NSW Health Pathology - Institute of Clinical Pathology and Medical Research; Westmead Hospital; University of Sydney | CIDM-PH et al. |
| EPI_ISL_526192 | Medlab Pathology | NSW Health Pathology - Institute of Clinical Pathology and Medical Research; Westmead Hospital; University of Sydney | CIDM-PH et al. |
| EPI_ISL_526197, EPI_ISL_526198 | Sydney South West Pathology Service (SSWPS) - Liverpool Hospital - NSW Health Pathology | NSW Health Pathology - Institute of Clinical Pathology and Medical Research; Westmead Hospital; University of Sydney | CIDM-PH et al. |
| EPI_ISL_526202, EPI_ISL_526203 | Laverty Pathology | NSW Health Pathology - Institute of Clinical Pathology and Medical Research; Westmead Hospital; University of Sydney | CIDM-PH et al. |
| EPI_ISL_526205 | The Children's Hospital at Westmead | NSW Health Pathology - Institute of Clinical Pathology and Medical Research; Westmead Hospital; University of Sydney | CIDM-PH et al. |
| EPI_ISL_526337, EPI_ISL_526338, EPI_ISL_526339, EPI_ISL_526340, EPI_ISL_526341, EPI_ISL_526342, EPI_ISL_526343, EPI_ISL_526344, EPI_ISL_526345, EPI_ISL_526346, EPI_ISL_526347, EPI_ISL_526348, EPI_ISL_526349, EPI_ISL_526350, EPI_ISL_526351, EPI_ISL_526352, EPI_ISL_526353, EPI_ISL_526354, EPI_ISL_526355, EPI_ISL_526356, EPI_ISL_526357, EPI_ISL_526358, EPI_ISL_526359, EPI_ISL_526360, EPI_ISL_526361, EPI_ISL_526362, EPI_ISL_526363, EPI_ISL_526364, EPI_ISL_526365, EPI_ISL_526366, EPI_ISL_526423, EPI_ISL_526424 | | | |
| see above | Queens Medical Centre, Clinical Microbiology Department / DeepSeq Nottingham | COVID-19 Genomics UK (COG-UK) Consortium | Gemma Clark, Wendy Smith, Manjinder Khakh, Vicki M Fleming, Michelle M Lister, Hannah Howson-Wells, Jonathan Ball, Patrick McClure, Joseph Chappell, Theocharis Tsoleridis, Nadine Holmes, Matthew Carlisle, Christopher Moore, Fei Sang, Johnny Debebe, Victoria Wright, Matthew Loose |
| EPI_ISL_526442 | Virology Department, Sheffield Teaching Hospitals NHS Foundation Trust/Department of Infection, Immunity and Cardiovascular Disease, The Medical School, University of Sheffield | COVID-19 Genomics UK (COG-UK) Consortium | Thushan de Silva, Matthew Parker, Nikki Smith, Adri Angyal, Rebecca Brown, Luke Green, Rachel Tucker, Paul Parsons, Danielle Groves, Katie Johnson, Laura Carrilero, Alex Keeley, Dave Partridge, Matthew Wyles, Benjamin Lindsey, Mehmet Yavuz, Mohammad Raza, Cariad Evans |
| EPI_ISL_526518, EPI_ISL_526519, EPI_ISL_526520, EPI_ISL_526522, EPI_ISL_526528, EPI_ISL_526529 | Virology Department, Royal Infirmary of Edinburgh, NHS Lothian / School of Biological Sciences, University of Edinburgh / Institute of Genetics and Molecular Medicine, University of Edinburgh | COVID-19 Genomics UK (COG-UK) Consortium | McHugh M, Dewar R, Rooke S, Gallagher M, Balcaza C, O'Toole Á, Scher E, Hill V, McCrone JT, Colquhoun R, Yu X, Jackson B, Rambaut A, Williams TC, Templeton K |
| EPI_ISL_526587, EPI_ISL_526588, EPI_ISL_526589, EPI_ISL_526590, EPI_ISL_526591, EPI_ISL_526592, EPI_ISL_526593, EPI_ISL_526594, EPI_ISL_526595, EPI_ISL_526596, EPI_ISL_526597, EPI_ISL_526598, EPI_ISL_526599, EPI_ISL_526600, EPI_ISL_526601, EPI_ISL_526602, EPI_ISL_526603, EPI_ISL_526604, EPI_ISL_526605, EPI_ISL_526606, EPI_ISL_526607, EPI_ISL_526609, EPI_ISL_526611 | | | |
| see above | Florida Bureau of Public Health Laboratories | Florida Bureau of Public Health Laboratories | Sarah Schmedes, Jason Blanton |
| EPI_ISL_526745 | Center for Laboratory Control of Infectious Diseases, Korea Centers for Diseases Control and Prevention | Center for Laboratory Control of Infectious Diseases, Korea Centers for Diseases Control and Prevention | Junyoung Kim, Ae Kyung Park, Eunkyung Shin, Jin Sun No, Jeong-Min Kim, Yoon-Seok Chung, Heui Man Kim, Myung Guk Han |
| EPI_ISL_527382, EPI_ISL_527383, EPI_ISL_527384, EPI_ISL_527385, EPI_ISL_527386, EPI_ISL_527387, EPI_ISL_527388, EPI_ISL_527389 | University of Miami Immunology and Histocompatibility Laboratory | University of Miami Immunology and Histocompatibility Laboratory | Emilio Margolles-Clark, PhD and Phillip Ruiz, MD, PhD |
| EPI_ISL_527602, EPI_ISL_527603, EPI_ISL_527604, EPI_ISL_527605, EPI_ISL_527606, EPI_ISL_527607, EPI_ISL_527608, EPI_ISL_527609, EPI_ISL_527610, EPI_ISL_527611, EPI_ISL_527612, EPI_ISL_527613, EPI_ISL_527614, EPI_ISL_527615, EPI_ISL_527616, EPI_ISL_527617, EPI_ISL_527618, EPI_ISL_527619, EPI_ISL_527620, EPI_ISL_527621, EPI_ISL_527622, EPI_ISL_527623, EPI_ISL_527624, EPI_ISL_527625, EPI_ISL_527626, EPI_ISL_527627, EPI_ISL_527628, EPI_ISL_527629 | | | |
| see above | Minnesota Department of Health, Public Health Laboratory | Minnesota Department of Health, Public Health Laboratory | Matt Plumb, Jacob Garfin, and Xiong Wang |
| EPI_ISL_527794, EPI_ISL_527795, EPI_ISL_527796, EPI_ISL_527797, EPI_ISL_527798, EPI_ISL_527799, EPI_ISL_527801, EPI_ISL_527805, EPI_ISL_527806, EPI_ISL_527808 | University of Wisconsin-Madison AIDS Vaccine Research Laboratories | University of Wisconsin-Madison AIDS Vaccine Research Laboratories | Gage Moreno, Katarina Braun, et al. AIDS Vaccine Research Laboratories |
| EPI_ISL_527809, EPI_ISL_527810, EPI_ISL_527811 | Institute of Microbiology, Universidad San Francisco de Quito | Institute of Microbiology, Universidad San Francisco de Quito | Belén Prado-Vivar, Sully Márquez, Juan José Guadalupe, Monica Becerra-Wong, Bernardo Gutiérrez, Stephanie Arregui, Rene Bracho, Karina Barragan, Anita Garcia, Carlos Tobar, Verónica Barragán, Patricio Rojas-Silva, Gabriel Trueba, Michelle Grunauer, Paúl Cárdenas |
| EPI_ISL_528440, EPI_ISL_528441, EPI_ISL_528442, EPI_ISL_528443, EPI_ISL_528444, EPI_ISL_528445, EPI_ISL_528446, EPI_ISL_528447, EPI_ISL_528448, EPI_ISL_528449, EPI_ISL_528450, EPI_ISL_528451, EPI_ISL_528452, EPI_ISL_528454, EPI_ISL_528455 | | | |

| | | | |
|---|---|---|---|
| see above | National Virus Reference Laboratory | National Virus Reference Laboratory | Michael Carr, Gabriel Gonzalez, Jonathan Dean, Suzie Coughlan, Cillian F De Gascun |
| EPI_ISL_528651, EPI_ISL_528652, EPI_ISL_528653, EPI_ISL_528654 | Virginia DCLS | Virginia DCLS | Virginia DCLS |
| EPI_ISL_528753 | Dinkes Kota Bogor | School of Life Sciences and Technology & School of Pharmacy-Institut Teknologi Bandung; Molecular Genetics Laboratory-Faculty of Medicine-Universitas Padjadjaran; Laboratorium Kesehatan Provinsi Jawa Barat | Azzania Fibriani, Catur Riani, Marselina Irasonia Tan, Yunia Sribudiani, Husna Nugrahapraja, Tarwadi, Ema Rahmawati, Savira Ekawardhani, Hesti Lina Wiraswati, Ryan Bayusantika Ristandi, Rifky Waluyajati Rachman, Cut Nur Cinthia Alamanda, Lia Faridah, Davin H. E. Setiamarga, Rizki Mardian , Hammam Riza, Sony Solistia Wirawan, Agung Eru Wibowo, Irvan Faizal |
| EPI_ISL_529028 | Respiratory Virus Unit, Microbiology Services Colindale, Public Health England | Respiratory Virus Unit, Microbiology Services Colindale, Public Health England | PHE Covid Sequencing Team |
| EPI_ISL_529235 | University of Birmingham | COVID-19 Genomics UK (COG-UK) Consortium | Institute of Microbiology, University of Birmingham: Claire McMurray, Joanne Stockton, Samuel Nicholls, Radoslaw Poplawski, Will Rowe, Josh Quick, Nicholas Loman. University of Birmingham Testing Laboratory: Celina M Whalley, Andrew Bosworth, Charlotte Poxon, Kasun Wanigasooriya, Oliver Pickles, Mike Kidd, Alex Richter, Andrew D Beggs PHE Heartlands Lab: Husam Osman, Andrew Bosworth. Queen Elizabeth Hospital: Anna Casey |
| EPI_ISL_529258 | Liverpool Clinical Laboratories | COVID-19 Genomics UK (COG-UK) Consortium | Sam Haldenby, Anita Lucaci, Steve Paterson, Julian Hiscox, Alistair Darby, M Almsaud, A Alrezaihi, Muhannad Alruwaili, Stuart D Armstrong, Jones Benjamin, Eleanor G Bentley, Anu Chawla, Jordan J Clark, Angela Cowell, Richard Eccles, Isabel García-Dorival, Matthew Gemmell, Alessandro Gerada, PKF Gilmore, Richard Gregory, Ximeng Han, Catherine Hartley, Margaret Hughes, Miren Iturriza-Gomara, James Johnson, L Luu, Jenifer Manson, Charlotte Nelson, Elaine O'Toole, Cassie Olateju, Rebekah Penrice-Randal , Lucille Rainbow, N.P Randle, Trevor Ian Robinson, Parul Sharma, Ghada T Shawli, James P Stewart, Neil Swainston, Ecaterina Vamos, Joanne Watts, Mark Whitehead |
| EPI_ISL_529264, EPI_ISL_529266 | Queens Medical Centre, Clinical Microbiology Department / DeepSeq Nottingham | COVID-19 Genomics UK (COG-UK) Consortium | Gemma Clark, Wendy Smith, Manjinder Khakh, Vicki M Fleming, Michelle M Lister, Hannah Howson-Wells, Jonathan Ball, Patrick McClure, Joseph Chappell, Theocharis Tsoleridis, Nadine Holmes, Matthew Carlisle, Christopher Moore, Fei Sang, Johnny Debebe, Victoria Wright, Matthew Loose |
| EPI_ISL_529267 | Quadram Institute Bioscience | COVID-19 Genomics UK (COG-UK) Consortium | Dave J. Baker, Gemma L. Kay, Alp Aydin, Thanh Le-Viet, Steven Rudder, Ana P. Tedim, Anastasia Kolyva, Maria Diaz, Leonardo de Oliveira Martins, Nabil-Fareed Alikhan, Lizzie Meadows, Rachael Stanley, Ngozi Elumogo, Muhammed Yasir, Nicholas M. Thomson, Alexander J Trotter, Rachel Gilroy, Samuel Bloomfield, Claire Stuart, Andrew Bell, Reenesh Prakash, Samir Dervisevic, Alison E. Mather, John Wain, Mark Webber, Andrew J. Page, Justin O'Grady |
| EPI_ISL_529270, EPI_ISL_529271 | Liverpool Clinical Laboratories | COVID-19 Genomics UK (COG-UK) Consortium | Sam Haldenby, Anita Lucaci, Steve Paterson, Julian Hiscox, Alistair Darby, M Almsaud, A Alrezaihi, Muhannad Alruwaili, Stuart D Armstrong, Jones Benjamin, Eleanor G Bentley, Anu Chawla, Jordan J Clark, Angela Cowell, Richard Eccles, Isabel García-Dorival, Matthew Gemmell, Alessandro Gerada, PKF Gilmore, Richard Gregory, Ximeng Han, Catherine Hartley, Margaret Hughes, Miren Iturriza-Gomara, James Johnson, L Luu, Jenifer Manson, Charlotte Nelson, Elaine O'Toole, Cassie Olateju, Rebekah Penrice-Randal , Lucille Rainbow, N.P Randle, Trevor Ian Robinson, Parul Sharma, Ghada T Shawli, James P Stewart, Neil Swainston, Ecaterina Vamos, Joanne Watts, Mark Whitehead |
| EPI_ISL_529275 | University of Birmingham | COVID-19 Genomics UK (COG-UK) Consortium | Institute of Microbiology, University of Birmingham: Claire McMurray, Joanne Stockton, Samuel Nicholls, Radoslaw Poplawski, Will Rowe, Josh Quick, Nicholas Loman. University of Birmingham Testing Laboratory: Celina M Whalley, Andrew Bosworth, Charlotte Poxon, Kasun Wanigasooriya, Oliver Pickles, Mike Kidd, Alex Richter, Andrew D Beggs PHE Heartlands Lab: Husam Osman, Andrew Bosworth. Queen Elizabeth Hospital: Anna Casey |
| EPI_ISL_529279 | Queens Medical Centre, Clinical Microbiology Department / DeepSeq Nottingham | COVID-19 Genomics UK (COG-UK) Consortium | Gemma Clark, Wendy Smith, Manjinder Khakh, Vicki M Fleming, Michelle M Lister, Hannah Howson-Wells, Jonathan Ball, Patrick McClure, Joseph Chappell, Theocharis Tsoleridis, Nadine Holmes, Matthew Carlisle, Christopher Moore, Fei Sang, Johnny Debebe, Victoria Wright, Matthew Loose |
| EPI_ISL_529280 | Liverpool Clinical Laboratories | COVID-19 Genomics UK (COG-UK) Consortium | Sam Haldenby, Anita Lucaci, Steve Paterson, Julian Hiscox, Alistair Darby, M Almsaud, A Alrezaihi, Muhannad Alruwaili, Stuart D Armstrong, Jones Benjamin, Eleanor G Bentley, Anu Chawla, Jordan J Clark, Angela Cowell, Richard Eccles, Isabel García-Dorival, Matthew Gemmell, Alessandro Gerada, PKF Gilmore, Richard Gregory, Ximeng Han, Catherine Hartley, Margaret Hughes, Miren Iturriza-Gomara, James Johnson, L Luu, Jenifer Manson, Charlotte Nelson, Elaine O'Toole, Cassie Olateju, Rebekah Penrice-Randal , Lucille Rainbow, N.P Randle, Trevor Ian Robinson, Parul Sharma, Ghada T Shawli, James P Stewart, Neil Swainston, Ecaterina Vamos, Joanne Watts, Mark Whitehead |
| EPI_ISL_529288 | Queens Medical Centre, Clinical Microbiology Department / DeepSeq Nottingham | COVID-19 Genomics UK (COG-UK) Consortium | Gemma Clark, Wendy Smith, Manjinder Khakh, Vicki M Fleming, Michelle M Lister, Hannah Howson-Wells, Jonathan Ball, Patrick McClure, Joseph Chappell, Theocharis Tsoleridis, Nadine Holmes, Matthew Carlisle, Christopher Moore, Fei Sang, Johnny Debebe, Victoria Wright, Matthew Loose |
| EPI_ISL_529291 | University of Birmingham | COVID-19 Genomics UK (COG-UK) Consortium | Institute of Microbiology, University of Birmingham: Claire McMurray, Joanne Stockton, Samuel Nicholls, Radoslaw Poplawski, Will Rowe, Josh Quick, Nicholas Loman. University of Birmingham Testing Laboratory: Celina M Whalley, Andrew Bosworth, Charlotte Poxon, Kasun Wanigasooriya, Oliver Pickles, Mike Kidd, Alex Richter, Andrew D Beggs PHE Heartlands Lab: Husam Osman, Andrew Bosworth. Queen Elizabeth Hospital: Anna Casey |
| EPI_ISL_529293, EPI_ISL_529296, EPI_ISL_529302, EPI_ISL_529304 | Liverpool Clinical Laboratories | COVID-19 Genomics UK (COG-UK) Consortium | Sam Haldenby, Anita Lucaci, Steve Paterson, Julian Hiscox, Alistair Darby, M Almsaud, A Alrezaihi, Muhannad Alruwaili, Stuart D Armstrong, Jones Benjamin, Eleanor G Bentley, Anu Chawla, Jordan J Clark, Angela Cowell, Richard Eccles, Isabel García-Dorival, Matthew Gemmell, Alessandro Gerada, PKF Gilmore, Richard Gregory, Ximeng Han, Catherine Hartley, Margaret Hughes, Miren Iturriza-Gomara, James Johnson, L Luu, Jenifer Manson, Charlotte Nelson, Elaine O'Toole, Cassie Olateju, Rebekah Penrice-Randal , Lucille Rainbow, N.P Randle, Trevor Ian Robinson, Parul Sharma, Ghada T Shawli, James P Stewart, Neil Swainston, Ecaterina Vamos, Joanne Watts, Mark Whitehead |
| EPI_ISL_529308, EPI_ISL_529309, EPI_ISL_529311 | Queens Medical Centre, Clinical Microbiology Department / DeepSeq Nottingham | COVID-19 Genomics UK (COG-UK) Consortium | Gemma Clark, Wendy Smith, Manjinder Khakh, Vicki M Fleming, Michelle M Lister, Hannah Howson-Wells, Jonathan Ball, Patrick McClure, Joseph Chappell, Theocharis Tsoleridis, Nadine Holmes, Matthew Carlisle, Christopher Moore, Fei Sang, Johnny Debebe, Victoria Wright, Matthew Loose |
| EPI_ISL_529330, EPI_ISL_529359 | University of Birmingham | COVID-19 Genomics UK (COG-UK) Consortium | Institute of Microbiology, University of Birmingham: Claire McMurray, Joanne Stockton, Samuel Nicholls, Radoslaw Poplawski, Will Rowe, Josh Quick, Nicholas Loman. University of Birmingham Testing Laboratory: Celina M Whalley, Andrew Bosworth, Charlotte Poxon, Kasun Wanigasooriya, Oliver Pickles, Mike Kidd, Alex Richter, Andrew D Beggs PHE Heartlands Lab: Husam Osman, Andrew Bosworth. Queen Elizabeth Hospital: Anna Casey |
| EPI_ISL_529379, EPI_ISL_529380, EPI_ISL_529381 | Quadram Institute Bioscience | COVID-19 Genomics UK (COG-UK) Consortium | Dave J. Baker, Gemma L. Kay, Alp Aydin, Thanh Le-Viet, Steven Rudder, Ana P. Tedim, Anastasia Kolyva, Maria Diaz, Leonardo de Oliveira Martins, Nabil-Fareed Alikhan, Lizzie Meadows, Rachael Stanley, Ngozi Elumogo, Muhammed Yasir, Nicholas M. Thomson, Alexander J Trotter, Rachel Gilroy, Samuel Bloomfield, Claire Stuart, Andrew Bell, Reenesh Prakash, Samir Dervisevic, Alison E. Mather, John Wain, Mark Webber, Andrew J. Page, Justin O'Grady |
| EPI_ISL_529397, EPI_ISL_529406, EPI_ISL_529435 | University of Birmingham | COVID-19 Genomics UK (COG-UK) Consortium | Institute of Microbiology, University of Birmingham: Claire McMurray, Joanne Stockton, Samuel Nicholls, Radoslaw Poplawski, Will Rowe, Josh Quick, Nicholas Loman. University of Birmingham Testing Laboratory: Celina M Whalley, Andrew Bosworth, Charlotte Poxon, Kasun Wanigasooriya, Oliver Pickles, Mike Kidd, Alex Richter, Andrew D Beggs PHE Heartlands Lab: Husam Osman, Andrew Bosworth. Queen Elizabeth Hospital: Anna Casey |
| EPI_ISL_529437 | Quadram Institute Bioscience | COVID-19 Genomics UK (COG-UK) Consortium | Dave J. Baker, Gemma L. Kay, Alp Aydin, Thanh Le-Viet, Steven Rudder, Ana P. Tedim, Anastasia Kolyva, Maria Diaz, Leonardo de Oliveira Martins, Nabil-Fareed Alikhan, Lizzie Meadows, Rachael Stanley, Ngozi Elumogo, Muhammed Yasir, Nicholas M. Thomson, Alexander J Trotter, Rachel Gilroy, Samuel Bloomfield, Claire Stuart, Andrew Bell, Reenesh Prakash, Samir Dervisevic, Alison E. Mather, John Wain, Mark Webber, Andrew J. Page, Justin O'Grady |
| EPI_ISL_529450 | Queens Medical Centre, Clinical Microbiology Department / DeepSeq Nottingham | COVID-19 Genomics UK (COG-UK) Consortium | Gemma Clark, Wendy Smith, Manjinder Khakh, Vicki M Fleming, Michelle M Lister, Hannah Howson-Wells, Jonathan Ball, Patrick McClure, Joseph Chappell, Theocharis Tsoleridis, Nadine Holmes, Matthew Carlisle, Christopher Moore, Fei Sang, Johnny Debebe, Victoria Wright, Matthew Loose |
| EPI_ISL_529451, EPI_ISL_529458, EPI_ISL_529460 | Quadram Institute Bioscience | COVID-19 Genomics UK (COG-UK) Consortium | Dave J. Baker, Gemma L. Kay, Alp Aydin, Thanh Le-Viet, Steven Rudder, Ana P. Tedim, Anastasia Kolyva, Maria Diaz, Leonardo de Oliveira Martins, Nabil-Fareed Alikhan, Lizzie Meadows, Rachael Stanley, Ngozi Elumogo, Muhammed Yasir, Nicholas M. Thomson, Alexander J Trotter, Rachel Gilroy, Samuel Bloomfield, Claire Stuart, Andrew Bell, Reenesh Prakash, Samir Dervisevic, Alison E. Mather, John Wain, Mark Webber, Andrew J. Page, Justin O'Grady |
| EPI_ISL_529461, EPI_ISL_529462 | University of Birmingham | COVID-19 Genomics UK (COG-UK) Consortium | Institute of Microbiology, University of Birmingham: Claire McMurray, Joanne Stockton, Samuel Nicholls, Radoslaw Poplawski, Will Rowe, Josh Quick, Nicholas Loman. University of Birmingham Testing Laboratory: Celina M Whalley, Andrew Bosworth, Charlotte Poxon, Kasun Wanigasooriya, Oliver Pickles, Mike Kidd, Alex Richter, Andrew D Beggs PHE Heartlands Lab: Husam Osman, Andrew Bosworth. Queen Elizabeth Hospital: Anna Casey |
| EPI_ISL_529463, EPI_ISL_529467 | Queens Medical Centre, Clinical Microbiology Department / DeepSeq Nottingham | COVID-19 Genomics UK (COG-UK) Consortium | Gemma Clark, Wendy Smith, Manjinder Khakh, Vicki M Fleming, Michelle M Lister, Hannah Howson-Wells, Jonathan Ball, Patrick McClure, Joseph Chappell, Theocharis Tsoleridis, Nadine Holmes, Matthew Carlisle, Christopher Moore, Fei Sang, Johnny Debebe, Victoria Wright, Matthew Loose |

| | | | |
|---|---|---|---|
| EPI_ISL_529472, EPI_ISL_529474, EPI_ISL_529480 | University of Birmingham | COVID-19 Genomics UK (COG-UK) Consortium | Institute of Microbiology, University of Birmingham: Claire McMurray, Joanne Stockton, Samuel Nicholls, Radoslaw Poplawski, Will Rowe, Josh Quick, Nicholas Loman. University of Birmingham Testing Laboratory: Celina M Whalley, Andrew Bosworth, Charlotte Poxon, Kasun Wanigasooriya, Oliver Pickles, Mike Kidd, Alex Richter, Andrew D Beggs PHE Heartlands Lab: Husam Osman, Andrew Bosworth. Queen Elizabeth Hospital: Anna Casey |
| EPI_ISL_529484 | Liverpool Clinical Laboratories | COVID-19 Genomics UK (COG-UK) Consortium | Sam Haldenby, Anita Lucaci, Steve Paterson, Julian Hiscox, Alistair Darby, M Almsaud, A Alrezaihi, Muhannad Alruwaili, Stuart D Armstrong, Jones Benjamin, Eleanor G Bentley, Anu Chawla, Jordan J Clark, Angela Cowell, Richard Eccles, Isabel García-Dorival, Matthew Gemmell, Alessandro Gerada, PKF Gilmore, Richard Gregory, Ximeng Han, Catherine Hartley, Margaret Hughes, Miren Iturriza-Gomara, James Johnson, L Luu, Jenifer Manson, Charlotte Nelson, Elaine O'Toole, Cassie Olateju, Rebekah Penrice-Randal , Lucille Rainbow, N.P Randle, Trevor Ian Robinson, Parul Sharma, Ghada T Shawli, James P Stewart, Neil Swainston, Ecaterina Vamos, Joanne Watts, Mark Whitehead |
| EPI_ISL_529489 | Quadram Institute Bioscience | COVID-19 Genomics UK (COG-UK) Consortium | Dave J. Baker, Gemma L. Kay, Alp Aydin, Thanh Le-Viet, Steven Rudder, Ana P. Tedim, Anastasia Kolyva, Maria Diaz, Leonardo de Oliveira Martins, Nabil-Fareed Alikhan, Lizzie Meadows, Rachael Stanley, Ngozi Elumogo, Muhammed Yasir, Nicholas M. Thomson, Alexander J Trotter, Rachel Gilroy, Samuel Bloomfield, Claire Stuart, Andrew Bell, Reenesh Prakash, Samir Dervisevic, Alison E. Mather, John Wain, Mark Webber, Andrew J. Page, Justin O'Grady |
| EPI_ISL_529498 | Liverpool Clinical Laboratories | COVID-19 Genomics UK (COG-UK) Consortium | Sam Haldenby, Anita Lucaci, Steve Paterson, Julian Hiscox, Alistair Darby, M Almsaud, A Alrezaihi, Muhannad Alruwaili, Stuart D Armstrong, Jones Benjamin, Eleanor G Bentley, Anu Chawla, Jordan J Clark, Angela Cowell, Richard Eccles, Isabel García-Dorival, Matthew Gemmell, Alessandro Gerada, PKF Gilmore, Richard Gregory, Ximeng Han, Catherine Hartley, Margaret Hughes, Miren Iturriza-Gomara, James Johnson, L Luu, Jenifer Manson, Charlotte Nelson, Elaine O'Toole, Cassie Olateju, Rebekah Penrice-Randal , Lucille Rainbow, N.P Randle, Trevor Ian Robinson, Parul Sharma, Ghada T Shawli, James P Stewart, Neil Swainston, Ecaterina Vamos, Joanne Watts, Mark Whitehead |
| EPI_ISL_529499 | University of Birmingham | COVID-19 Genomics UK (COG-UK) Consortium | Institute of Microbiology, University of Birmingham: Claire McMurray, Joanne Stockton, Samuel Nicholls, Radoslaw Poplawski, Will Rowe, Josh Quick, Nicholas Loman. University of Birmingham Testing Laboratory: Celina M Whalley, Andrew Bosworth, Charlotte Poxon, Kasun Wanigasooriya, Oliver Pickles, Mike Kidd, Alex Richter, Andrew D Beggs PHE Heartlands Lab: Husam Osman, Andrew Bosworth. Queen Elizabeth Hospital: Anna Casey |
| EPI_ISL_529504 | Queens Medical Centre, Clinical Microbiology Department / DeepSeq Nottingham | COVID-19 Genomics UK (COG-UK) Consortium | Gemma Clark, Wendy Smith, Manjinder Khakh, Vicki M Fleming, Michelle M Lister, Hannah Howson-Wells, Jonathan Ball, Patrick McClure, Joseph Chappell, Theocharis Tsoleridis, Nadine Holmes, Matthew Carlisle, Christopher Moore, Fei Sang, Johnny Debebe, Victoria Wright, Matthew Loose |
| EPI_ISL_529508 | University of Birmingham | COVID-19 Genomics UK (COG-UK) Consortium | Institute of Microbiology, University of Birmingham: Claire McMurray, Joanne Stockton, Samuel Nicholls, Radoslaw Poplawski, Will Rowe, Josh Quick, Nicholas Loman. University of Birmingham Testing Laboratory: Celina M Whalley, Andrew Bosworth, Charlotte Poxon, Kasun Wanigasooriya, Oliver Pickles, Mike Kidd, Alex Richter, Andrew D Beggs PHE Heartlands Lab: Husam Osman, Andrew Bosworth. Queen Elizabeth Hospital: Anna Casey |
| EPI_ISL_529509 | Liverpool Clinical Laboratories | COVID-19 Genomics UK (COG-UK) Consortium | Sam Haldenby, Anita Lucaci, Steve Paterson, Julian Hiscox, Alistair Darby, M Almsaud, A Alrezaihi, Muhannad Alruwaili, Stuart D Armstrong, Jones Benjamin, Eleanor G Bentley, Anu Chawla, Jordan J Clark, Angela Cowell, Richard Eccles, Isabel García-Dorival, Matthew Gemmell, Alessandro Gerada, PKF Gilmore, Richard Gregory, Ximeng Han, Catherine Hartley, Margaret Hughes, Miren Iturriza-Gomara, James Johnson, L Luu, Jenifer Manson, Charlotte Nelson, Elaine O'Toole, Cassie Olateju, Rebekah Penrice-Randal , Lucille Rainbow, N.P Randle, Trevor Ian Robinson, Parul Sharma, Ghada T Shawli, James P Stewart, Neil Swainston, Ecaterina Vamos, Joanne Watts, Mark Whitehead |
| EPI_ISL_529512, EPI_ISL_529514 | University of Birmingham | COVID-19 Genomics UK (COG-UK) Consortium | Institute of Microbiology, University of Birmingham: Claire McMurray, Joanne Stockton, Samuel Nicholls, Radoslaw Poplawski, Will Rowe, Josh Quick, Nicholas Loman. University of Birmingham Testing Laboratory: Celina M Whalley, Andrew Bosworth, Charlotte Poxon, Kasun Wanigasooriya, Oliver Pickles, Mike Kidd, Alex Richter, Andrew D Beggs PHE Heartlands Lab: Husam Osman, Andrew Bosworth. Queen Elizabeth Hospital: Anna Casey |
| EPI_ISL_529516 | Liverpool Clinical Laboratories | COVID-19 Genomics UK (COG-UK) Consortium | Sam Haldenby, Anita Lucaci, Steve Paterson, Julian Hiscox, Alistair Darby, M Almsaud, A Alrezaihi, Muhannad Alruwaili, Stuart D Armstrong, Jones Benjamin, Eleanor G Bentley, Anu Chawla, Jordan J Clark, Angela Cowell, Richard Eccles, Isabel García-Dorival, Matthew Gemmell, Alessandro Gerada, PKF Gilmore, Richard Gregory, Ximeng Han, Catherine Hartley, Margaret Hughes, Miren Iturriza-Gomara, James Johnson, L Luu, Jenifer Manson, Charlotte Nelson, Elaine O'Toole, Cassie Olateju, Rebekah Penrice-Randal , Lucille Rainbow, N.P Randle, Trevor Ian Robinson, Parul Sharma, Ghada T Shawli, James P Stewart, Neil Swainston, Ecaterina Vamos, Joanne Watts, Mark Whitehead |
| EPI_ISL_529519 | Queens Medical Centre, Clinical Microbiology Department / DeepSeq Nottingham | COVID-19 Genomics UK (COG-UK) Consortium | Gemma Clark, Wendy Smith, Manjinder Khakh, Vicki M Fleming, Michelle M Lister, Hannah Howson-Wells, Jonathan Ball, Patrick McClure, Joseph Chappell, Theocharis Tsoleridis, Nadine Holmes, Matthew Carlisle, Christopher Moore, Fei Sang, Johnny Debebe, Victoria Wright, Matthew Loose |
| EPI_ISL_529523 | Quadram Institute Bioscience | COVID-19 Genomics UK (COG-UK) Consortium | Dave J. Baker, Gemma L. Kay, Alp Aydin, Thanh Le-Viet, Steven Rudder, Ana P. Tedim, Anastasia Kolyva, Maria Diaz, Leonardo de Oliveira Martins, Nabil-Fareed Alikhan, Lizzie Meadows, Rachael Stanley, Ngozi Elumogo, Muhammed Yasir, Nicholas M. Thomson, Alexander J Trotter, Rachel Gilroy, Samuel Bloomfield, Claire Stuart, Andrew Bell, Reenesh Prakash, Samir Dervisevic, Alison E. Mather, John Wain, Mark Webber, Andrew J. Page, Justin O'Grady |
| EPI_ISL_529533, EPI_ISL_529534, EPI_ISL_529535, EPI_ISL_529536, EPI_ISL_529537, EPI_ISL_529538, EPI_ISL_529539, EPI_ISL_529540, EPI_ISL_529541, EPI_ISL_529542, EPI_ISL_529543, EPI_ISL_529544, EPI_ISL_529545, EPI_ISL_529546, EPI_ISL_529547, EPI_ISL_529548, EPI_ISL_529549, EPI_ISL_529550, EPI_ISL_529551, EPI_ISL_529552 | | | |
| see above | Queens Medical Centre, Clinical Microbiology Department / DeepSeq Nottingham | COVID-19 Genomics UK (COG-UK) Consortium | Gemma Clark, Wendy Smith, Manjinder Khakh, Vicki M Fleming, Michelle M Lister, Hannah Howson-Wells, Jonathan Ball, Patrick McClure, Joseph Chappell, Theocharis Tsoleridis, Nadine Holmes, Matthew Carlisle, Christopher Moore, Fei Sang, Johnny Debebe, Victoria Wright, Matthew Loose |
| EPI_ISL_529567, EPI_ISL_529568, EPI_ISL_529572, EPI_ISL_529573, EPI_ISL_529574, EPI_ISL_529575, EPI_ISL_529576, EPI_ISL_529577, EPI_ISL_529578, EPI_ISL_529579, EPI_ISL_529580, EPI_ISL_529581, EPI_ISL_529587, EPI_ISL_529588, EPI_ISL_529589 | | | |
| see above | Quadram Institute Bioscience | COVID-19 Genomics UK (COG-UK) Consortium | Dave J. Baker, Gemma L. Kay, Alp Aydin, Thanh Le-Viet, Steven Rudder, Ana P. Tedim, Anastasia Kolyva, Maria Diaz, Leonardo de Oliveira Martins, Nabil-Fareed Alikhan, Lizzie Meadows, Rachael Stanley, Ngozi Elumogo, Muhammed Yasir, Nicholas M. Thomson, Alexander J Trotter, Rachel Gilroy, Samuel Bloomfield, Claire Stuart, Andrew Bell, Reenesh Prakash, Samir Dervisevic, Alison E. Mather, John Wain, Mark Webber, Andrew J. Page, Justin O'Grady |
| EPI_ISL_529644 | University of Birmingham | COVID-19 Genomics UK (COG-UK) Consortium | Institute of Microbiology, University of Birmingham: Claire McMurray, Joanne Stockton, Samuel Nicholls, Radoslaw Poplawski, Will Rowe, Josh Quick, Nicholas Loman. University of Birmingham Testing Laboratory: Celina M Whalley, Andrew Bosworth, Charlotte Poxon, Kasun Wanigasooriya, Oliver Pickles, Mike Kidd, Alex Richter, Andrew D Beggs PHE Heartlands Lab: Husam Osman, Andrew Bosworth. Queen Elizabeth Hospital: Anna Casey |
| EPI_ISL_529700, EPI_ISL_529704 | Wales Specialist Virology Centre Sequencing lab: Pathogen Genomics Unit | COVID-19 Genomics UK (COG-UK) Consortium | Catherine Moore, Johnathan Evans, Laura Gifford, Malorie Perry, Simon Cottrell, Angela Marchbank, Alec Birchley, Alexander Adams, Amy Gaskin, Bree Gatica-Wilcox, Jason Coombes, Joel Southgate, Lauren Gilbert, Lee Graham, Nicole Pacchiarini, Sara Kumziene-Summerhayes, Sarah Taylor, Sophie Jones, Sara Rey, Matthew Bull, Joanne Watkins, Sally Corden, Tom Connor |
| EPI_ISL_529844, EPI_ISL_529845, EPI_ISL_529846, EPI_ISL_529847, EPI_ISL_529848, EPI_ISL_529849, EPI_ISL_529850, EPI_ISL_529851, EPI_ISL_529855, EPI_ISL_529866, EPI_ISL_529867, EPI_ISL_529869, EPI_ISL_529870, EPI_ISL_529873, EPI_ISL_529874, EPI_ISL_529875, EPI_ISL_529876, EPI_ISL_529877, EPI_ISL_529880, EPI_ISL_529882, EPI_ISL_529883, EPI_ISL_529884, EPI_ISL_529907 | | | |
| see above | Michigan Department of Health and Human Services, Bureau of Laboratories | Michigan Department of Health and Human Services, Bureau of Laboratories | Blankenship HM, Riner D, Soehnlen MK |
| EPI_ISL_529943, EPI_ISL_529944, EPI_ISL_529945, EPI_ISL_529946, EPI_ISL_529947, EPI_ISL_529948, EPI_ISL_529949, EPI_ISL_529950, EPI_ISL_529951, EPI_ISL_529952, EPI_ISL_529953 | | | |
| see above | Virginia DCLS | Virginia DCLS | Virginia DCLS |
| EPI_ISL_530127, EPI_ISL_530128, EPI_ISL_530129, EPI_ISL_530130, EPI_ISL_530131 | Seattle Flu Study | Seattle Flu Study | Deborah A. Nickerson, Chris D. Frazar, Jover Lee, Benjamin Pelle, Matthew Richardson, Amanda Adler, Elisabeth Brandstetter, Peter D. Han, Kairsten Fay, Misja Ilcisin, Kirsten Lacombe, Thomas R. Sibley, Melissa Truong, Caitlin R. Wolf, Karen Cowgill, Stephanie Schrag, Jeff Duchin, Michael Boeckh, Janet A. Englund, Michael Famulare, Barry R. Lutz, Mark J. Rieder, Lea M. Starita, Matthew Thompson, Helen Y. Chu, Trevor Bedford, Jay Shendure |
| EPI_ISL_530132 | Seattle Flu Study | Seattle Flu Study | Deborah A. Nickerson, Chris D. Frazar, Jover Lee, Benjamin Pelle, Matthew Richardson, Amanda Adler, Elisabeth Brandstetter, Peter D. Han, Kairsten Fay, Misja Ilcisin, Kirsten Lacombe, Thomas R. Sibley, Melissa Truong, Caitlin R. Wolf, Michael Boeckh, Janet A. Englund, Michael Famulare, Barry R. Lutz, Mark J. Rieder, Lea M. Starita, Matthew Thompson, Jay Shendure, Trevor Bedford, Helen Y. Chu |
| EPI_ISL_530133, EPI_ISL_530134, EPI_ISL_530135, EPI_ISL_530136, EPI_ISL_530137, EPI_ISL_530138, EPI_ISL_530139, EPI_ISL_530140 | Seattle Flu Study | Seattle Flu Study | Deborah A. Nickerson, Chris D. Frazar, Jover Lee, Benjamin Pelle, Matthew Richardson, Amanda Adler, Elisabeth Brandstetter, Peter D. Han, Kairsten Fay, Misja Ilcisin, Kirsten Lacombe, Thomas R. Sibley, Melissa Truong, Caitlin R. Wolf, Karen Cowgill, Stephanie Schrag, Jeff Duchin, Michael Boeckh, Janet A. Englund, Michael Famulare, Barry R. Lutz, Mark J. Rieder, Lea M. Starita, Matthew Thompson, Helen Y. Chu, Trevor Bedford, Jay Shendure |

| | | | |
|---|---|---|---|
| EPI_ISL_530172 | Minnesota Department of Health, Public Health Laboratory | Minnesota Department of Health, Public Health Laboratory | Matt Plumb, Jacob Garfin, and Xiong Wang |
| EPI_ISL_530254, EPI_ISL_530280 | Queensland Health Forensic and Scientific Services, Public Health Virology | Public Health Virology Laboratory, Forensic and Scientific Services, Queensland Health | Son Nguyen et al |
| EPI_ISL_530805, EPI_ISL_530816, EPI_ISL_530818, EPI_ISL_530819, EPI_ISL_530837, EPI_ISL_530861 | Lighthouse Lab in Glasgow | Wellcome Sanger Institute for the COVID-19 Genomics UK (COG-UK) consortium | Harper VanSteenhouse, Yumi Kasai, David Gray, Carol Clugston, Anna Dominiczak and Alex Alderton, Roberto Amato, Sonia Goncalves, Ewan Harrison, David K. Jackson, Ian Johnston, Dominic Kwiatkowski, Cordelia Langford, John Sillitoe |
| EPI_ISL_530862 | NHSGGC West of Scotland Specialist Virology Centre / MRC-University of Glasgow Centre for Virus Research | Wellcome Sanger Institute for the COVID-19 Genomics UK (COG-UK) consortium | Ana da Silva Filipe, Natasha Johnson, Kathy Smollett, Daniel Mair, Stephen Carmichael, Lily Tong, Jenna Nichols, Elihu Aranday-Cortes, Kirstyn Brunker, Yasmin Parr, Kyriaki Nomikou; Sarah McDonald, Marc Niebel, Patawee Asamaphan; Richard Orton, Joseph Hughes, Sreenu Vattipally, David L Robertson; Alasdair MacLean, Rory Gunson; Kathy Li, Natasha Jesudason, Rajiv Shah, James Shepherd, Antonia Ho, Alice Broos, Emma Thomson and Alex Alderton, Roberto Amato, Sonia Goncalves, Ewan Harrison, David K. Jackson, Ian Johnston, Dominic Kwiatkowski, Cordelia Langford, John Sillitoe |
| EPI_ISL_530869, EPI_ISL_530875, EPI_ISL_530885, EPI_ISL_530902, EPI_ISL_530915 | Lighthouse Lab in Glasgow | Wellcome Sanger Institute for the COVID-19 Genomics UK (COG-UK) consortium | Harper VanSteenhouse, Yumi Kasai, David Gray, Carol Clugston, Anna Dominiczak and Alex Alderton, Roberto Amato, Sonia Goncalves, Ewan Harrison, David K. Jackson, Ian Johnston, Dominic Kwiatkowski, Cordelia Langford, John Sillitoe |
| EPI_ISL_530929 | NHSGGC West of Scotland Specialist Virology Centre / MRC-University of Glasgow Centre for Virus Research | Wellcome Sanger Institute for the COVID-19 Genomics UK (COG-UK) consortium | Ana da Silva Filipe, Natasha Johnson, Kathy Smollett, Daniel Mair, Stephen Carmichael, Lily Tong, Jenna Nichols, Elihu Aranday-Cortes, Kirstyn Brunker, Yasmin Parr, Kyriaki Nomikou; Sarah McDonald, Marc Niebel, Patawee Asamaphan; Richard Orton, Joseph Hughes, Sreenu Vattipally, David L Robertson; Alasdair MacLean, Rory Gunson; Kathy Li, Natasha Jesudason, Rajiv Shah, James Shepherd, Antonia Ho, Alice Broos, Emma Thomson and Alex Alderton, Roberto Amato, Sonia Goncalves, Ewan Harrison, David K. Jackson, Ian Johnston, Dominic Kwiatkowski, Cordelia Langford, John Sillitoe |
| EPI_ISL_530932, EPI_ISL_530938, EPI_ISL_530954, EPI_ISL_531050, EPI_ISL_531060 | Lighthouse Lab in Glasgow | Wellcome Sanger Institute for the COVID-19 Genomics UK (COG-UK) consortium | Harper VanSteenhouse, Yumi Kasai, David Gray, Carol Clugston, Anna Dominiczak and Alex Alderton, Roberto Amato, Sonia Goncalves, Ewan Harrison, David K. Jackson, Ian Johnston, Dominic Kwiatkowski, Cordelia Langford, John Sillitoe |
| EPI_ISL_531211 | Lighthouse Lab in Glasgow | Wellcome Sanger Institute for the COVID-19 Genomics UK (COG-UK) Consortium | Harper VanSteenhouse, Yumi Kasai, David Gray, Carol Clugston, Anna Dominiczak and Alex Alderton, Roberto Amato, Sonia Goncalves, Ewan Harrison, David K. Jackson, Ian Johnston, Dominic Kwiatkowski, Cordelia Langford, John Sillitoe on behalf of the Wellcome Sanger Institute COVID-19 Surveillance Team |
| EPI_ISL_531472, EPI_ISL_531479, EPI_ISL_531482, EPI_ISL_531488, EPI_ISL_531490, EPI_ISL_531491, EPI_ISL_531496, EPI_ISL_531499, EPI_ISL_531503, EPI_ISL_531512, EPI_ISL_531515, EPI_ISL_531516, EPI_ISL_531523, EPI_ISL_531527, EPI_ISL_531531, EPI_ISL_531535, EPI_ISL_531538, EPI_ISL_531541, EPI_ISL_531542, EPI_ISL_531549, EPI_ISL_531552 | | | |
| see above | Lighthouse Lab in Glasgow | Wellcome Sanger Institute for the COVID-19 Genomics UK (COG-UK) consortium | Harper VanSteenhouse, Yumi Kasai, David Gray, Carol Clugston, Anna Dominiczak and Alex Alderton, Roberto Amato, Sonia Goncalves, Ewan Harrison, David K. Jackson, Ian Johnston, Dominic Kwiatkowski, Cordelia Langford, John Sillitoe |
| EPI_ISL_531557 | NHSGGC West of Scotland Specialist Virology Centre / MRC-University of Glasgow Centre for Virus Research | Wellcome Sanger Institute for the COVID-19 Genomics UK (COG-UK) consortium | Ana da Silva Filipe, Natasha Johnson, Kathy Smollett, Daniel Mair, Stephen Carmichael, Lily Tong, Jenna Nichols, Elihu Aranday-Cortes, Kirstyn Brunker, Yasmin Parr, Kyriaki Nomikou; Sarah McDonald, Marc Niebel, Patawee Asamaphan; Richard Orton, Joseph Hughes, Sreenu Vattipally, David L Robertson; Alasdair MacLean, Rory Gunson; Kathy Li, Natasha Jesudason, Rajiv Shah, James Shepherd, Antonia Ho, Alice Broos, Emma Thomson and Alex Alderton, Roberto Amato, Sonia Goncalves, Ewan Harrison, David K. Jackson, Ian Johnston, Dominic Kwiatkowski, Cordelia Langford, John Sillitoe |
| EPI_ISL_531560, EPI_ISL_531567, EPI_ISL_531568, EPI_ISL_531576, EPI_ISL_531579, EPI_ISL_531580, EPI_ISL_531581, EPI_ISL_531582, EPI_ISL_531584 | Lighthouse Lab in Glasgow | Wellcome Sanger Institute for the COVID-19 Genomics UK (COG-UK) consortium | Harper VanSteenhouse, Yumi Kasai, David Gray, Carol Clugston, Anna Dominiczak and Alex Alderton, Roberto Amato, Sonia Goncalves, Ewan Harrison, David K. Jackson, Ian Johnston, Dominic Kwiatkowski, Cordelia Langford, John Sillitoe |
| EPI_ISL_531585 | Lighthouse Lab in Glasgow | Wellcome Sanger Institute for the COVID-19 Genomics UK (COG-UK) consortium | Harper VanSteenhouse, Yumi Kasai, David Gray, Carol Clugston, Anna Dominiczak and Alex Alderton, Roberto Amato, Sonia Goncalves, Ewan Harrison, David K. Jackson, Ian Johnston, Dominic Kwiatkowski, Cordelia Langford, John Sillitoe on behalf of the Wellcome Sanger Institute COVID-19 Surveillance Team (http://www.sanger.ac.uk/covid-team) |
| EPI_ISL_531586, EPI_ISL_531587, EPI_ISL_531588, EPI_ISL_531591, EPI_ISL_531592, EPI_ISL_531593, EPI_ISL_531594 | Lighthouse Lab in Glasgow | Wellcome Sanger Institute for the COVID-19 Genomics UK (COG-UK) consortium | Harper VanSteenhouse, Yumi Kasai, David Gray, Carol Clugston, Anna Dominiczak and Alex Alderton, Roberto Amato, Sonia Goncalves, Ewan Harrison, David K. Jackson, Ian Johnston, Dominic Kwiatkowski, Cordelia Langford, John Sillitoe |
| EPI_ISL_531595 | NHSGGC West of Scotland Specialist Virology Centre / MRC-University of Glasgow Centre for Virus Research | Wellcome Sanger Institute for the COVID-19 Genomics UK (COG-UK) consortium | Ana da Silva Filipe, Natasha Johnson, Kathy Smollett, Daniel Mair, Stephen Carmichael, Lily Tong, Jenna Nichols, Elihu Aranday-Cortes, Kirstyn Brunker, Yasmin Parr, Kyriaki Nomikou; Sarah McDonald, Marc Niebel, Patawee Asamaphan; Richard Orton, Joseph Hughes, Sreenu Vattipally, David L Robertson; Alasdair MacLean, Rory Gunson; Kathy Li, Natasha Jesudason, Rajiv Shah, James Shepherd, Antonia Ho, Alice Broos, Emma Thomson and Alex Alderton, Roberto Amato, Sonia Goncalves, Ewan Harrison, David K. Jackson, Ian Johnston, Dominic Kwiatkowski, Cordelia Langford, John Sillitoe |
| EPI_ISL_531596, EPI_ISL_531597 | Lighthouse Lab in Glasgow | Wellcome Sanger Institute for the COVID-19 Genomics UK (COG-UK) consortium | Harper VanSteenhouse, Yumi Kasai, David Gray, Carol Clugston, Anna Dominiczak and Alex Alderton, Roberto Amato, Sonia Goncalves, Ewan Harrison, David K. Jackson, Ian Johnston, Dominic Kwiatkowski, Cordelia Langford, John Sillitoe |
| EPI_ISL_531598, EPI_ISL_531599 | NHSGGC West of Scotland Specialist Virology Centre / MRC-University of Glasgow Centre for Virus Research | Wellcome Sanger Institute for the COVID-19 Genomics UK (COG-UK) consortium | Ana da Silva Filipe, Natasha Johnson, Kathy Smollett, Daniel Mair, Stephen Carmichael, Lily Tong, Jenna Nichols, Elihu Aranday-Cortes, Kirstyn Brunker, Yasmin Parr, Kyriaki Nomikou; Sarah McDonald, Marc Niebel, Patawee Asamaphan; Richard Orton, Joseph Hughes, Sreenu Vattipally, David L Robertson; Alasdair MacLean, Rory Gunson; Kathy Li, Natasha Jesudason, Rajiv Shah, James Shepherd, Antonia Ho, Alice Broos, Emma Thomson and Alex Alderton, Roberto Amato, Sonia Goncalves, Ewan Harrison, David K. Jackson, Ian Johnston, Dominic Kwiatkowski, Cordelia Langford, John Sillitoe |
| EPI_ISL_531600, EPI_ISL_531601, EPI_ISL_531602, EPI_ISL_531605, EPI_ISL_531606 | Lighthouse Lab in Glasgow | Wellcome Sanger Institute for the COVID-19 Genomics UK (COG-UK) consortium | Harper VanSteenhouse, Yumi Kasai, David Gray, Carol Clugston, Anna Dominiczak and Alex Alderton, Roberto Amato, Sonia Goncalves, Ewan Harrison, David K. Jackson, Ian Johnston, Dominic Kwiatkowski, Cordelia Langford, John Sillitoe |
| EPI_ISL_531607, EPI_ISL_531609 | NHSGGC West of Scotland Specialist Virology Centre / MRC-University of Glasgow Centre for Virus Research | Wellcome Sanger Institute for the COVID-19 Genomics UK (COG-UK) consortium | Ana da Silva Filipe, Natasha Johnson, Kathy Smollett, Daniel Mair, Stephen Carmichael, Lily Tong, Jenna Nichols, Elihu Aranday-Cortes, Kirstyn Brunker, Yasmin Parr, Kyriaki Nomikou; Sarah McDonald, Marc Niebel, Patawee Asamaphan; Richard Orton, Joseph Hughes, Sreenu Vattipally, David L Robertson; Alasdair MacLean, Rory Gunson; Kathy Li, Natasha Jesudason, Rajiv Shah, James Shepherd, Antonia Ho, Alice Broos, Emma Thomson and Alex Alderton, Roberto Amato, Sonia Goncalves, Ewan Harrison, David K. Jackson, Ian Johnston, Dominic Kwiatkowski, Cordelia Langford, John Sillitoe |
| EPI_ISL_531612, EPI_ISL_531613, EPI_ISL_531614, EPI_ISL_531615, EPI_ISL_531616, EPI_ISL_531619, EPI_ISL_531621, EPI_ISL_531623, EPI_ISL_531624, EPI_ISL_531626, EPI_ISL_531627, EPI_ISL_531630 | | | |
| see above | Lighthouse Lab in Glasgow | Wellcome Sanger Institute for the COVID-19 Genomics UK (COG-UK) consortium | Harper VanSteenhouse, Yumi Kasai, David Gray, Carol Clugston, Anna Dominiczak and Alex Alderton, Roberto Amato, Sonia Goncalves, Ewan Harrison, David K. Jackson, Ian Johnston, Dominic Kwiatkowski, Cordelia Langford, John Sillitoe |
| EPI_ISL_531631 | NHSGGC West of Scotland Specialist Virology Centre / MRC-University of Glasgow Centre for Virus Research | Wellcome Sanger Institute for the COVID-19 Genomics UK (COG-UK) consortium | Ana da Silva Filipe, Natasha Johnson, Kathy Smollett, Daniel Mair, Stephen Carmichael, Lily Tong, Jenna Nichols, Elihu Aranday-Cortes, Kirstyn Brunker, Yasmin Parr, Kyriaki Nomikou; Sarah McDonald, Marc Niebel, Patawee Asamaphan; Richard Orton, Joseph Hughes, Sreenu Vattipally, David L Robertson; Alasdair MacLean, Rory Gunson; Kathy Li, Natasha Jesudason, Rajiv Shah, James Shepherd, Antonia Ho, Alice Broos, Emma Thomson and Alex Alderton, Roberto Amato, Sonia Goncalves, Ewan Harrison, David K. Jackson, Ian Johnston, Dominic Kwiatkowski, Cordelia Langford, John Sillitoe |
| EPI_ISL_531632, EPI_ISL_531633, EPI_ISL_531635, EPI_ISL_531636, EPI_ISL_531637 | Lighthouse Lab in Glasgow | Wellcome Sanger Institute for the COVID-19 Genomics UK (COG-UK) consortium | Harper VanSteenhouse, Yumi Kasai, David Gray, Carol Clugston, Anna Dominiczak and Alex Alderton, Roberto Amato, Sonia Goncalves, Ewan Harrison, David K. Jackson, Ian Johnston, Dominic Kwiatkowski, Cordelia Langford, John Sillitoe |
| EPI_ISL_531638, EPI_ISL_531639 | NHSGGC West of Scotland Specialist Virology Centre / MRC-University of Glasgow Centre for Virus Research | Wellcome Sanger Institute for the COVID-19 Genomics UK (COG-UK) consortium | Ana da Silva Filipe, Natasha Johnson, Kathy Smollett, Daniel Mair, Stephen Carmichael, Lily Tong, Jenna Nichols, Elihu Aranday-Cortes, Kirstyn Brunker, Yasmin Parr, Kyriaki Nomikou; Sarah McDonald, Marc Niebel, Patawee Asamaphan; Richard Orton, Joseph Hughes, Sreenu Vattipally, David L Robertson; Alasdair MacLean, Rory Gunson; Kathy Li, Natasha Jesudason, Rajiv Shah, James Shepherd, Antonia Ho, Alice Broos, Emma Thomson and Alex Alderton, Roberto Amato, Sonia Goncalves, Ewan Harrison, David K. Jackson, Ian Johnston, Dominic Kwiatkowski, Cordelia Langford, John Sillitoe |
| EPI_ISL_531640, EPI_ISL_531642, EPI_ISL_531644, EPI_ISL_531645, EPI_ISL_531646, EPI_ISL_531647, EPI_ISL_531648, EPI_ISL_531650, EPI_ISL_531651, EPI_ISL_531652, EPI_ISL_531653, EPI_ISL_531654, EPI_ISL_531656, EPI_ISL_531657, EPI_ISL_531658, EPI_ISL_531659 | | | |

| | | | |
|---|---|---|---|
| see above | Lighthouse Lab in Glasgow | Wellcome Sanger Institute for the COVID-19 Genomics UK (COG-UK) consortium | Harper VanSteenhouse, Yumi Kasai, David Gray, Carol Clugston, Anna Dominiczak and Alex Alderton, Roberto Amato, Sonia Goncalves, Ewan Harrison, David K. Jackson, Ian Johnston, Dominic Kwiatkowski, Cordelia Langford, John Sillitoe |
| EPI_ISL_531660 | NHSGGC West of Scotland Specialist Virology Centre / MRC-University of Glasgow Centre for Virus Research | Wellcome Sanger Institute for the COVID-19 Genomics UK (COG-UK) consortium | Ana da Silva Filipe, Natasha Johnson, Kathy Smollett, Daniel Mair, Stephen Carmichael, Lily Tong, Jenna Nichols, Elihu Aranday-Cortes, Kirstyn Brunker, Yasmin Parr, Kyriaki Nomikou; Sarah McDonald, Marc Niebel, Patawee Asamaphan; Richard Orton, Joseph Hughes, Sreenu Vattipally, David L Robertson; Alasdair MacLean, Rory Gunson; Kathy Li, Natasha Jesudason, Rajiv Shah, James Shepherd, Antonia Ho, Alice Broos, Emma Thomson and Alex Alderton, Roberto Amato, Sonia Goncalves, Ewan Harrison, David K. Jackson, Ian Johnston, Dominic Kwiatkowski, Cordelia Langford, John Sillitoe |
| EPI_ISL_531661, EPI_ISL_531662, EPI_ISL_531663, EPI_ISL_531664, EPI_ISL_531665, EPI_ISL_531666, EPI_ISL_531667, EPI_ISL_531671, EPI_ISL_531673, EPI_ISL_531675, EPI_ISL_531676, EPI_ISL_531679, EPI_ISL_531680, EPI_ISL_531682, EPI_ISL_531683, EPI_ISL_531684, EPI_ISL_531686 | | | |
| see above | Lighthouse Lab in Glasgow | Wellcome Sanger Institute for the COVID-19 Genomics UK (COG-UK) consortium | Harper VanSteenhouse, Yumi Kasai, David Gray, Carol Clugston, Anna Dominiczak and Alex Alderton, Roberto Amato, Sonia Goncalves, Ewan Harrison, David K. Jackson, Ian Johnston, Dominic Kwiatkowski, Cordelia Langford, John Sillitoe |
| EPI_ISL_531688 | Lighthouse Lab in Glasgow | Wellcome Sanger Institute for the COVID-19 Genomics UK (COG-UK) Consortium | Harper VanSteenhouse, Yumi Kasai, David Gray, Carol Clugston, Anna Dominiczak and Alex Alderton, Roberto Amato, Sonia Goncalves, Ewan Harrison, David K. Jackson, Ian Johnston, Dominic Kwiatkowski, Cordelia Langford, John Sillitoe on behalf of the Wellcome Sanger Institute COVID-19 Surveillance Team |
| EPI_ISL_531689, EPI_ISL_531690 | NHSGGC West of Scotland Specialist Virology Centre / MRC-University of Glasgow Centre for Virus Research | Wellcome Sanger Institute for the COVID-19 Genomics UK (COG-UK) consortium | Ana da Silva Filipe, Natasha Johnson, Kathy Smollett, Daniel Mair, Stephen Carmichael, Lily Tong, Jenna Nichols, Elihu Aranday-Cortes, Kirstyn Brunker, Yasmin Parr, Kyriaki Nomikou; Sarah McDonald, Marc Niebel, Patawee Asamaphan; Richard Orton, Joseph Hughes, Sreenu Vattipally, David L Robertson; Alasdair MacLean, Rory Gunson; Kathy Li, Natasha Jesudason, Rajiv Shah, James Shepherd, Antonia Ho, Alice Broos, Emma Thomson and Alex Alderton, Roberto Amato, Sonia Goncalves, Ewan Harrison, David K. Jackson, Ian Johnston, Dominic Kwiatkowski, Cordelia Langford, John Sillitoe |
| EPI_ISL_531692, EPI_ISL_531694, EPI_ISL_531695, EPI_ISL_531696, EPI_ISL_531697, EPI_ISL_531698, EPI_ISL_531699, EPI_ISL_531700 | Lighthouse Lab in Glasgow | Wellcome Sanger Institute for the COVID-19 Genomics UK (COG-UK) consortium | Harper VanSteenhouse, Yumi Kasai, David Gray, Carol Clugston, Anna Dominiczak and Alex Alderton, Roberto Amato, Sonia Goncalves, Ewan Harrison, David K. Jackson, Ian Johnston, Dominic Kwiatkowski, Cordelia Langford, John Sillitoe |
| EPI_ISL_531702 | NHSGGC West of Scotland Specialist Virology Centre / MRC-University of Glasgow Centre for Virus Research | Wellcome Sanger Institute for the COVID-19 Genomics UK (COG-UK) consortium | Ana da Silva Filipe, Natasha Johnson, Kathy Smollett, Daniel Mair, Stephen Carmichael, Lily Tong, Jenna Nichols, Elihu Aranday-Cortes, Kirstyn Brunker, Yasmin Parr, Kyriaki Nomikou; Sarah McDonald, Marc Niebel, Patawee Asamaphan; Richard Orton, Joseph Hughes, Sreenu Vattipally, David L Robertson; Alasdair MacLean, Rory Gunson; Kathy Li, Natasha Jesudason, Rajiv Shah, James Shepherd, Antonia Ho, Alice Broos, Emma Thomson and Alex Alderton, Roberto Amato, Sonia Goncalves, Ewan Harrison, David K. Jackson, Ian Johnston, Dominic Kwiatkowski, Cordelia Langford, John Sillitoe |
| EPI_ISL_531704, EPI_ISL_531705, EPI_ISL_531708, EPI_ISL_531709, EPI_ISL_531710, EPI_ISL_531711, EPI_ISL_531712, EPI_ISL_531715, EPI_ISL_531716, EPI_ISL_531717, EPI_ISL_531718, EPI_ISL_531719, EPI_ISL_531720, EPI_ISL_531722, EPI_ISL_531723, EPI_ISL_531724 | | | |
| see above | Lighthouse Lab in Glasgow | Wellcome Sanger Institute for the COVID-19 Genomics UK (COG-UK) consortium | Harper VanSteenhouse, Yumi Kasai, David Gray, Carol Clugston, Anna Dominiczak and Alex Alderton, Roberto Amato, Sonia Goncalves, Ewan Harrison, David K. Jackson, Ian Johnston, Dominic Kwiatkowski, Cordelia Langford, John Sillitoe |
| EPI_ISL_531726 | Lighthouse Lab in Glasgow | Wellcome Sanger Institute for the COVID-19 Genomics UK (COG-UK) consortium | Harper VanSteenhouse, Yumi Kasai, David Gray, Carol Clugston, Anna Dominiczak and Alex Alderton, Roberto Amato, Sonia Goncalves, Ewan Harrison, David K. Jackson, Ian Johnston, Dominic Kwiatkowski, Cordelia Langford, John Sillitoe on behalf of the Wellcome Sanger Institute COVID-19 Surveillance Team (http://www.sanger.ac.uk/covid-team) |
| EPI_ISL_531729 | Lighthouse Lab in Glasgow | Wellcome Sanger Institute for the COVID-19 Genomics UK (COG-UK) consortium | Harper VanSteenhouse, Yumi Kasai, David Gray, Carol Clugston, Anna Dominiczak and Alex Alderton, Roberto Amato, Sonia Goncalves, Ewan Harrison, David K. Jackson, Ian Johnston, Dominic Kwiatkowski, Cordelia Langford, John Sillitoe |
| EPI_ISL_531732 | NHSGGC West of Scotland Specialist Virology Centre / MRC-University of Glasgow Centre for Virus Research | Wellcome Sanger Institute for the COVID-19 Genomics UK (COG-UK) consortium | Ana da Silva Filipe, Natasha Johnson, Kathy Smollett, Daniel Mair, Stephen Carmichael, Lily Tong, Jenna Nichols, Elihu Aranday-Cortes, Kirstyn Brunker, Yasmin Parr, Kyriaki Nomikou; Sarah McDonald, Marc Niebel, Patawee Asamaphan; Richard Orton, Joseph Hughes, Sreenu Vattipally, David L Robertson; Alasdair MacLean, Rory Gunson; Kathy Li, Natasha Jesudason, Rajiv Shah, James Shepherd, Antonia Ho, Alice Broos, Emma Thomson and Alex Alderton, Roberto Amato, Sonia Goncalves, Ewan Harrison, David K. Jackson, Ian Johnston, Dominic Kwiatkowski, Cordelia Langford, John Sillitoe |
| EPI_ISL_531733 | Lighthouse Lab in Glasgow | Wellcome Sanger Institute for the COVID-19 Genomics UK (COG-UK) consortium | Harper VanSteenhouse, Yumi Kasai, David Gray, Carol Clugston, Anna Dominiczak and Alex Alderton, Roberto Amato, Sonia Goncalves, Ewan Harrison, David K. Jackson, Ian Johnston, Dominic Kwiatkowski, Cordelia Langford, John Sillitoe |
| EPI_ISL_531734 | Lighthouse Lab in Glasgow | Wellcome Sanger Institute for the COVID-19 Genomics UK (COG-UK) consortium | Harper VanSteenhouse, Yumi Kasai, David Gray, Carol Clugston, Anna Dominiczak and Alex Alderton, Roberto Amato, Sonia Goncalves, Ewan Harrison, David K. Jackson, Ian Johnston, Dominic Kwiatkowski, Cordelia Langford, John Sillitoe on behalf of the Wellcome Sanger Institute COVID-19 Surveillance Team (http://www.sanger.ac.uk/covid-team) |
| EPI_ISL_531735, EPI_ISL_531737, EPI_ISL_531739, EPI_ISL_531740, EPI_ISL_531741 | Lighthouse Lab in Glasgow | Wellcome Sanger Institute for the COVID-19 Genomics UK (COG-UK) consortium | Harper VanSteenhouse, Yumi Kasai, David Gray, Carol Clugston, Anna Dominiczak and Alex Alderton, Roberto Amato, Sonia Goncalves, Ewan Harrison, David K. Jackson, Ian Johnston, Dominic Kwiatkowski, Cordelia Langford, John Sillitoe |
| EPI_ISL_531743 | NHSGGC West of Scotland Specialist Virology Centre / MRC-University of Glasgow Centre for Virus Research | Wellcome Sanger Institute for the COVID-19 Genomics UK (COG-UK) consortium | Ana da Silva Filipe, Natasha Johnson, Kathy Smollett, Daniel Mair, Stephen Carmichael, Lily Tong, Jenna Nichols, Elihu Aranday-Cortes, Kirstyn Brunker, Yasmin Parr, Kyriaki Nomikou; Sarah McDonald, Marc Niebel, Patawee Asamaphan; Richard Orton, Joseph Hughes, Sreenu Vattipally, David L Robertson; Alasdair MacLean, Rory Gunson; Kathy Li, Natasha Jesudason, Rajiv Shah, James Shepherd, Antonia Ho, Alice Broos, Emma Thomson and Alex Alderton, Roberto Amato, Sonia Goncalves, Ewan Harrison, David K. Jackson, Ian Johnston, Dominic Kwiatkowski, Cordelia Langford, John Sillitoe |
| EPI_ISL_531744, EPI_ISL_531746, EPI_ISL_531747, EPI_ISL_531748, EPI_ISL_531749, EPI_ISL_531750, EPI_ISL_531751 | Lighthouse Lab in Glasgow | Wellcome Sanger Institute for the COVID-19 Genomics UK (COG-UK) consortium | Harper VanSteenhouse, Yumi Kasai, David Gray, Carol Clugston, Anna Dominiczak and Alex Alderton, Roberto Amato, Sonia Goncalves, Ewan Harrison, David K. Jackson, Ian Johnston, Dominic Kwiatkowski, Cordelia Langford, John Sillitoe |
| EPI_ISL_531754, EPI_ISL_531755 | NHSGGC West of Scotland Specialist Virology Centre / MRC-University of Glasgow Centre for Virus Research | Wellcome Sanger Institute for the COVID-19 Genomics UK (COG-UK) consortium | Ana da Silva Filipe, Natasha Johnson, Kathy Smollett, Daniel Mair, Stephen Carmichael, Lily Tong, Jenna Nichols, Elihu Aranday-Cortes, Kirstyn Brunker, Yasmin Parr, Kyriaki Nomikou; Sarah McDonald, Marc Niebel, Patawee Asamaphan; Richard Orton, Joseph Hughes, Sreenu Vattipally, David L Robertson; Alasdair MacLean, Rory Gunson; Kathy Li, Natasha Jesudason, Rajiv Shah, James Shepherd, Antonia Ho, Alice Broos, Emma Thomson and Alex Alderton, Roberto Amato, Sonia Goncalves, Ewan Harrison, David K. Jackson, Ian Johnston, Dominic Kwiatkowski, Cordelia Langford, John Sillitoe |
| EPI_ISL_531756, EPI_ISL_531757 | Lighthouse Lab in Glasgow | Wellcome Sanger Institute for the COVID-19 Genomics UK (COG-UK) consortium | Harper VanSteenhouse, Yumi Kasai, David Gray, Carol Clugston, Anna Dominiczak and Alex Alderton, Roberto Amato, Sonia Goncalves, Ewan Harrison, David K. Jackson, Ian Johnston, Dominic Kwiatkowski, Cordelia Langford, John Sillitoe |
| EPI_ISL_531758 | NHSGGC West of Scotland Specialist Virology Centre / MRC-University of Glasgow Centre for Virus Research | Wellcome Sanger Institute for the COVID-19 Genomics UK (COG-UK) consortium | Ana da Silva Filipe, Natasha Johnson, Kathy Smollett, Daniel Mair, Stephen Carmichael, Lily Tong, Jenna Nichols, Elihu Aranday-Cortes, Kirstyn Brunker, Yasmin Parr, Kyriaki Nomikou; Sarah McDonald, Marc Niebel, Patawee Asamaphan; Richard Orton, Joseph Hughes, Sreenu Vattipally, David L Robertson; Alasdair MacLean, Rory Gunson; Kathy Li, Natasha Jesudason, Rajiv Shah, James Shepherd, Antonia Ho, Alice Broos, Emma Thomson and Alex Alderton, Roberto Amato, Sonia Goncalves, Ewan Harrison, David K. Jackson, Ian Johnston, Dominic Kwiatkowski, Cordelia Langford, John Sillitoe |
| EPI_ISL_531759, EPI_ISL_531760, EPI_ISL_531761, EPI_ISL_531762, EPI_ISL_531764, EPI_ISL_531765, EPI_ISL_531766, EPI_ISL_531767, EPI_ISL_531769, EPI_ISL_531770, EPI_ISL_531771, EPI_ISL_531772, EPI_ISL_531774, EPI_ISL_531775, EPI_ISL_531776, EPI_ISL_531777, EPI_ISL_531778 | | | |
| see above | Lighthouse Lab in Glasgow | Wellcome Sanger Institute for the COVID-19 Genomics UK (COG-UK) consortium | Harper VanSteenhouse, Yumi Kasai, David Gray, Carol Clugston, Anna Dominiczak and Alex Alderton, Roberto Amato, Sonia Goncalves, Ewan Harrison, David K. Jackson, Ian Johnston, Dominic Kwiatkowski, Cordelia Langford, John Sillitoe |
| EPI_ISL_531779 | NHSGGC West of Scotland Specialist Virology Centre / MRC-University of Glasgow Centre for Virus Research | Wellcome Sanger Institute for the COVID-19 Genomics UK (COG-UK) consortium | Ana da Silva Filipe, Natasha Johnson, Kathy Smollett, Daniel Mair, Stephen Carmichael, Lily Tong, Jenna Nichols, Elihu Aranday-Cortes, Kirstyn Brunker, Yasmin Parr, Kyriaki Nomikou; Sarah McDonald, Marc Niebel, Patawee Asamaphan; Richard Orton, Joseph Hughes, Sreenu Vattipally, David L Robertson; Alasdair MacLean, Rory Gunson; Kathy Li, Natasha Jesudason, Rajiv Shah, James Shepherd, Antonia Ho, Alice Broos, Emma Thomson and Alex Alderton, Roberto Amato, Sonia Goncalves, Ewan Harrison, David K. Jackson, Ian Johnston, Dominic Kwiatkowski, Cordelia Langford, John Sillitoe |
| EPI_ISL_531780, EPI_ISL_531781, EPI_ISL_531782, EPI_ISL_531783, EPI_ISL_531784, EPI_ISL_531786, EPI_ISL_531787 | Lighthouse Lab in Glasgow | Wellcome Sanger Institute for the COVID-19 Genomics UK (COG-UK) consortium | Harper VanSteenhouse, Yumi Kasai, David Gray, Carol Clugston, Anna Dominiczak and Alex Alderton, Roberto Amato, Sonia Goncalves, Ewan Harrison, David K. Jackson, Ian Johnston, Dominic Kwiatkowski, Cordelia Langford, John Sillitoe |
| EPI_ISL_531788 | NHSGGC West of Scotland Specialist Virology Centre / MRC-University of Glasgow Centre for Virus Research | Wellcome Sanger Institute for the COVID-19 Genomics UK (COG-UK) consortium | Ana da Silva Filipe, Natasha Johnson, Kathy Smollett, Daniel Mair, Stephen Carmichael, Lily Tong, Jenna Nichols, Elihu Aranday-Cortes, Kirstyn Brunker, Yasmin Parr, Kyriaki Nomikou; Sarah McDonald, Marc Niebel, Patawee Asamaphan; Richard Orton, Joseph Hughes, Sreenu Vattipally, David L Robertson; |

| | | | |
|---|---|---|---|
| | | | Alasdair MacLean, Rory Gunson; Kathy Li, Natasha Jesudason, Rajiv Shah, James Shepherd, Antonia Ho, Alice Broos, Emma Thomson and Alex Alderton, Roberto Amato, Sonia Goncalves, Ewan Harrison, David K. Jackson, Ian Johnston, Dominic Kwiatkowski, Cordelia Langford, John Sillitoe |
| EPI_ISL_531789, EPI_ISL_531791, EPI_ISL_531793, EPI_ISL_531794, EPI_ISL_531795, EPI_ISL_531796, EPI_ISL_531798, EPI_ISL_531799, EPI_ISL_531800, EPI_ISL_531803, EPI_ISL_531804, EPI_ISL_531806, EPI_ISL_531807, EPI_ISL_531808, EPI_ISL_531809, EPI_ISL_531810, EPI_ISL_531812, EPI_ISL_531813 | | | |
| see above | Lighthouse Lab in Glasgow | Wellcome Sanger Institute for the COVID-19 Genomics UK (COG-UK) consortium | Harper VanSteenhouse, Yumi Kasai, David Gray, Carol Clugston, Anna Dominiczak and Alex Alderton, Roberto Amato, Sonia Goncalves, Ewan Harrison, David K. Jackson, Ian Johnston, Dominic Kwiatkowski, Cordelia Langford, John Sillitoe |
| EPI_ISL_531814 | NHSGGC West of Scotland Specialist Virology Centre / MRC-University of Glasgow Centre for Virus Research | Wellcome Sanger Institute for the COVID-19 Genomics UK (COG-UK) consortium | Ana da Silva Filipe, Natasha Johnson, Kathy Smollett, Daniel Mair, Stephen Carmichael, Lily Tong, Jenna Nichols, Elihu Aranday-Cortes, Kirstyn Brunker, Yasmin Parr, Kyriaki Nomikou; Sarah McDonald, Marc Niebel, Patawee Asamaphan; Richard Orton, Joseph Hughes, Sreenu Vattipally, David L Robertson; Alasdair MacLean, Rory Gunson; Kathy Li, Natasha Jesudason, Rajiv Shah, James Shepherd, Antonia Ho, Alice Broos, Emma Thomson and Alex Alderton, Roberto Amato, Sonia Goncalves, Ewan Harrison, David K. Jackson, Ian Johnston, Dominic Kwiatkowski, Cordelia Langford, John Sillitoe |
| EPI_ISL_531816, EPI_ISL_531817, EPI_ISL_531818, EPI_ISL_531819, EPI_ISL_531820, EPI_ISL_531823 | Lighthouse Lab in Glasgow | Wellcome Sanger Institute for the COVID-19 Genomics UK (COG-UK) consortium | Harper VanSteenhouse, Yumi Kasai, David Gray, Carol Clugston, Anna Dominiczak and Alex Alderton, Roberto Amato, Sonia Goncalves, Ewan Harrison, David K. Jackson, Ian Johnston, Dominic Kwiatkowski, Cordelia Langford, John Sillitoe |
| EPI_ISL_531824 | NHSGGC West of Scotland Specialist Virology Centre / MRC-University of Glasgow Centre for Virus Research | Wellcome Sanger Institute for the COVID-19 Genomics UK (COG-UK) consortium | Ana da Silva Filipe, Natasha Johnson, Kathy Smollett, Daniel Mair, Stephen Carmichael, Lily Tong, Jenna Nichols, Elihu Aranday-Cortes, Kirstyn Brunker, Yasmin Parr, Kyriaki Nomikou; Sarah McDonald, Marc Niebel, Patawee Asamaphan; Richard Orton, Joseph Hughes, Sreenu Vattipally, David L Robertson; Alasdair MacLean, Rory Gunson; Kathy Li, Natasha Jesudason, Rajiv Shah, James Shepherd, Antonia Ho, Alice Broos, Emma Thomson and Alex Alderton, Roberto Amato, Sonia Goncalves, Ewan Harrison, David K. Jackson, Ian Johnston, Dominic Kwiatkowski, Cordelia Langford, John Sillitoe |
| EPI_ISL_531825 | Lighthouse Lab in Glasgow | Wellcome Sanger Institute for the COVID-19 Genomics UK (COG-UK) consortium | Harper VanSteenhouse, Yumi Kasai, David Gray, Carol Clugston, Anna Dominiczak and Alex Alderton, Roberto Amato, Sonia Goncalves, Ewan Harrison, David K. Jackson, Ian Johnston, Dominic Kwiatkowski, Cordelia Langford, John Sillitoe |
| EPI_ISL_531826 | Lighthouse Lab in Glasgow | Wellcome Sanger Institute for the COVID-19 Genomics UK (COG-UK) consortium | Harper VanSteenhouse, Yumi Kasai, David Gray, Carol Clugston, Anna Dominiczak and Alex Alderton, Roberto Amato, Sonia Goncalves, Ewan Harrison, David K. Jackson, Ian Johnston, Dominic Kwiatkowski, Cordelia Langford, John Sillitoe on behalf of the Wellcome Sanger Institute COVID-19 Surveillance Team (http://www.sanger.ac.uk/covid-team) |
| EPI_ISL_531827, EPI_ISL_531828, EPI_ISL_531830, EPI_ISL_531831, EPI_ISL_531833, EPI_ISL_531835, EPI_ISL_531836, EPI_ISL_531837, EPI_ISL_531839, EPI_ISL_531840, EPI_ISL_531841, EPI_ISL_531843, EPI_ISL_531844, EPI_ISL_531845, EPI_ISL_531846, EPI_ISL_531848, EPI_ISL_531849, EPI_ISL_531850 | | | |
| see above | Lighthouse Lab in Glasgow | Wellcome Sanger Institute for the COVID-19 Genomics UK (COG-UK) consortium | Harper VanSteenhouse, Yumi Kasai, David Gray, Carol Clugston, Anna Dominiczak and Alex Alderton, Roberto Amato, Sonia Goncalves, Ewan Harrison, David K. Jackson, Ian Johnston, Dominic Kwiatkowski, Cordelia Langford, John Sillitoe |
| EPI_ISL_531851 | NHSGGC West of Scotland Specialist Virology Centre / MRC-University of Glasgow Centre for Virus Research | Wellcome Sanger Institute for the COVID-19 Genomics UK (COG-UK) consortium | Ana da Silva Filipe, Natasha Johnson, Kathy Smollett, Daniel Mair, Stephen Carmichael, Lily Tong, Jenna Nichols, Elihu Aranday-Cortes, Kirstyn Brunker, Yasmin Parr, Kyriaki Nomikou; Sarah McDonald, Marc Niebel, Patawee Asamaphan; Richard Orton, Joseph Hughes, Sreenu Vattipally, David L Robertson; Alasdair MacLean, Rory Gunson; Kathy Li, Natasha Jesudason, Rajiv Shah, James Shepherd, Antonia Ho, Alice Broos, Emma Thomson and Alex Alderton, Roberto Amato, Sonia Goncalves, Ewan Harrison, David K. Jackson, Ian Johnston, Dominic Kwiatkowski, Cordelia Langford, John Sillitoe |
| EPI_ISL_531852, EPI_ISL_531853, EPI_ISL_531854 | Lighthouse Lab in Glasgow | Wellcome Sanger Institute for the COVID-19 Genomics UK (COG-UK) consortium | Harper VanSteenhouse, Yumi Kasai, David Gray, Carol Clugston, Anna Dominiczak and Alex Alderton, Roberto Amato, Sonia Goncalves, Ewan Harrison, David K. Jackson, Ian Johnston, Dominic Kwiatkowski, Cordelia Langford, John Sillitoe |
| EPI_ISL_531856, EPI_ISL_531857 | NHSGGC West of Scotland Specialist Virology Centre / MRC-University of Glasgow Centre for Virus Research | Wellcome Sanger Institute for the COVID-19 Genomics UK (COG-UK) consortium | Ana da Silva Filipe, Natasha Johnson, Kathy Smollett, Daniel Mair, Stephen Carmichael, Lily Tong, Jenna Nichols, Elihu Aranday-Cortes, Kirstyn Brunker, Yasmin Parr, Kyriaki Nomikou; Sarah McDonald, Marc Niebel, Patawee Asamaphan; Richard Orton, Joseph Hughes, Sreenu Vattipally, David L Robertson; Alasdair MacLean, Rory Gunson; Kathy Li, Natasha Jesudason, Rajiv Shah, James Shepherd, Antonia Ho, Alice Broos, Emma Thomson and Alex Alderton, Roberto Amato, Sonia Goncalves, Ewan Harrison, David K. Jackson, Ian Johnston, Dominic Kwiatkowski, Cordelia Langford, John Sillitoe |
| EPI_ISL_531859, EPI_ISL_531860, EPI_ISL_531861 | Lighthouse Lab in Glasgow | Wellcome Sanger Institute for the COVID-19 Genomics UK (COG-UK) consortium | Harper VanSteenhouse, Yumi Kasai, David Gray, Carol Clugston, Anna Dominiczak and Alex Alderton, Roberto Amato, Sonia Goncalves, Ewan Harrison, David K. Jackson, Ian Johnston, Dominic Kwiatkowski, Cordelia Langford, John Sillitoe |
| EPI_ISL_531862, EPI_ISL_531863 | NHSGGC West of Scotland Specialist Virology Centre / MRC-University of Glasgow Centre for Virus Research | Wellcome Sanger Institute for the COVID-19 Genomics UK (COG-UK) consortium | Ana da Silva Filipe, Natasha Johnson, Kathy Smollett, Daniel Mair, Stephen Carmichael, Lily Tong, Jenna Nichols, Elihu Aranday-Cortes, Kirstyn Brunker, Yasmin Parr, Kyriaki Nomikou; Sarah McDonald, Marc Niebel, Patawee Asamaphan; Richard Orton, Joseph Hughes, Sreenu Vattipally, David L Robertson; Alasdair MacLean, Rory Gunson; Kathy Li, Natasha Jesudason, Rajiv Shah, James Shepherd, Antonia Ho, Alice Broos, Emma Thomson and Alex Alderton, Roberto Amato, Sonia Goncalves, Ewan Harrison, David K. Jackson, Ian Johnston, Dominic Kwiatkowski, Cordelia Langford, John Sillitoe |
| EPI_ISL_531864 | Lighthouse Lab in Glasgow | Wellcome Sanger Institute for the COVID-19 Genomics UK (COG-UK) consortium | Harper VanSteenhouse, Yumi Kasai, David Gray, Carol Clugston, Anna Dominiczak and Alex Alderton, Roberto Amato, Sonia Goncalves, Ewan Harrison, David K. Jackson, Ian Johnston, Dominic Kwiatkowski, Cordelia Langford, John Sillitoe |
| EPI_ISL_531865 | NHSGGC West of Scotland Specialist Virology Centre / MRC-University of Glasgow Centre for Virus Research | Wellcome Sanger Institute for the COVID-19 Genomics UK (COG-UK) consortium | Ana da Silva Filipe, Natasha Johnson, Kathy Smollett, Daniel Mair, Stephen Carmichael, Lily Tong, Jenna Nichols, Elihu Aranday-Cortes, Kirstyn Brunker, Yasmin Parr, Kyriaki Nomikou; Sarah McDonald, Marc Niebel, Patawee Asamaphan; Richard Orton, Joseph Hughes, Sreenu Vattipally, David L Robertson; Alasdair MacLean, Rory Gunson; Kathy Li, Natasha Jesudason, Rajiv Shah, James Shepherd, Antonia Ho, Alice Broos, Emma Thomson and Alex Alderton, Roberto Amato, Sonia Goncalves, Ewan Harrison, David K. Jackson, Ian Johnston, Dominic Kwiatkowski, Cordelia Langford, John Sillitoe |
| EPI_ISL_531866, EPI_ISL_531867, EPI_ISL_531868, EPI_ISL_531869, EPI_ISL_531870, EPI_ISL_531871 | Lighthouse Lab in Glasgow | Wellcome Sanger Institute for the COVID-19 Genomics UK (COG-UK) consortium | Harper VanSteenhouse, Yumi Kasai, David Gray, Carol Clugston, Anna Dominiczak and Alex Alderton, Roberto Amato, Sonia Goncalves, Ewan Harrison, David K. Jackson, Ian Johnston, Dominic Kwiatkowski, Cordelia Langford, John Sillitoe |
| EPI_ISL_531872 | Lighthouse Lab in Glasgow | Wellcome Sanger Institute for the COVID-19 Genomics UK (COG-UK) Consortium | Harper VanSteenhouse, Yumi Kasai, David Gray, Carol Clugston, Anna Dominiczak and Alex Alderton, Roberto Amato, Sonia Goncalves, Ewan Harrison, David K. Jackson, Ian Johnston, Dominic Kwiatkowski, Cordelia Langford, John Sillitoe on behalf of the Wellcome Sanger Institute COVID-19 Surveillance Team |
| EPI_ISL_531873 | NHSGGC West of Scotland Specialist Virology Centre / MRC-University of Glasgow Centre for Virus Research | Wellcome Sanger Institute for the COVID-19 Genomics UK (COG-UK) consortium | Ana da Silva Filipe, Natasha Johnson, Kathy Smollett, Daniel Mair, Stephen Carmichael, Lily Tong, Jenna Nichols, Elihu Aranday-Cortes, Kirstyn Brunker, Yasmin Parr, Kyriaki Nomikou; Sarah McDonald, Marc Niebel, Patawee Asamaphan; Richard Orton, Joseph Hughes, Sreenu Vattipally, David L Robertson; Alasdair MacLean, Rory Gunson; Kathy Li, Natasha Jesudason, Rajiv Shah, James Shepherd, Antonia Ho, Alice Broos, Emma Thomson and Alex Alderton, Roberto Amato, Sonia Goncalves, Ewan Harrison, David K. Jackson, Ian Johnston, Dominic Kwiatkowski, Cordelia Langford, John Sillitoe |
| EPI_ISL_531874, EPI_ISL_531875, EPI_ISL_531876 | Lighthouse Lab in Glasgow | Wellcome Sanger Institute for the COVID-19 Genomics UK (COG-UK) consortium | Harper VanSteenhouse, Yumi Kasai, David Gray, Carol Clugston, Anna Dominiczak and Alex Alderton, Roberto Amato, Sonia Goncalves, Ewan Harrison, David K. Jackson, Ian Johnston, Dominic Kwiatkowski, Cordelia Langford, John Sillitoe |
| EPI_ISL_531877 | NHSGGC West of Scotland Specialist Virology Centre / MRC-University of Glasgow Centre for Virus Research | Wellcome Sanger Institute for the COVID-19 Genomics UK (COG-UK) consortium | Ana da Silva Filipe, Natasha Johnson, Kathy Smollett, Daniel Mair, Stephen Carmichael, Lily Tong, Jenna Nichols, Elihu Aranday-Cortes, Kirstyn Brunker, Yasmin Parr, Kyriaki Nomikou; Sarah McDonald, Marc Niebel, Patawee Asamaphan; Richard Orton, Joseph Hughes, Sreenu Vattipally, David L Robertson; Alasdair MacLean, Rory Gunson; Kathy Li, Natasha Jesudason, Rajiv Shah, James Shepherd, Antonia Ho, Alice Broos, Emma Thomson and Alex Alderton, Roberto Amato, Sonia Goncalves, Ewan Harrison, David K. Jackson, Ian Johnston, Dominic Kwiatkowski, Cordelia Langford, John Sillitoe |
| EPI_ISL_531878, EPI_ISL_531879 | Lighthouse Lab in Glasgow | Wellcome Sanger Institute for the COVID-19 Genomics UK (COG-UK) consortium | Harper VanSteenhouse, Yumi Kasai, David Gray, Carol Clugston, Anna Dominiczak and Alex Alderton, Roberto Amato, Sonia Goncalves, Ewan Harrison, David K. Jackson, Ian Johnston, Dominic Kwiatkowski, Cordelia Langford, John Sillitoe |
| EPI_ISL_531880, EPI_ISL_531881 | NHSGGC West of Scotland Specialist Virology Centre / MRC-University of Glasgow Centre for Virus Research | Wellcome Sanger Institute for the COVID-19 Genomics UK (COG-UK) consortium | Ana da Silva Filipe, Natasha Johnson, Kathy Smollett, Daniel Mair, Stephen Carmichael, Lily Tong, Jenna Nichols, Elihu Aranday-Cortes, Kirstyn Brunker, Yasmin Parr, Kyriaki Nomikou; Sarah McDonald, Marc Niebel, Patawee Asamaphan; Richard Orton, Joseph Hughes, Sreenu Vattipally, David L Robertson; Alasdair MacLean, Rory Gunson; Kathy Li, Natasha Jesudason, Rajiv Shah, James Shepherd, Antonia Ho, Alice Broos, Emma Thomson and Alex Alderton, Roberto Amato, Sonia Goncalves, Ewan Harrison, David K. Jackson, Ian Johnston, Dominic Kwiatkowski, Cordelia Langford, John Sillitoe |
| EPI_ISL_531882, EPI_ISL_531883 | Lighthouse Lab in Glasgow | Wellcome Sanger Institute for the COVID-19 Genomics UK (COG-UK) consortium | Harper VanSteenhouse, Yumi Kasai, David Gray, Carol Clugston, Anna Dominiczak and Alex Alderton, Roberto Amato, Sonia Goncalves, Ewan Harrison, David K. Jackson, Ian Johnston, Dominic Kwiatkowski, Cordelia Langford, John Sillitoe |
| EPI_ISL_531884 | NHSGGC West of Scotland Specialist Virology Centre / MRC-University of Glasgow Centre for Virus Research | Wellcome Sanger Institute for the COVID-19 Genomics UK (COG-UK) consortium | Ana da Silva Filipe, Natasha Johnson, Kathy Smollett, Daniel Mair, Stephen Carmichael, Lily Tong, Jenna Nichols, Elihu Aranday-Cortes, Kirstyn Brunker, Yasmin Parr, Kyriaki Nomikou; Sarah McDonald, Marc Niebel, Patawee Asamaphan; Richard Orton, Joseph Hughes, Sreenu Vattipally, David L Robertson; Alasdair MacLean, Rory Gunson; Kathy Li, Natasha Jesudason, Rajiv Shah, James Shepherd, Antonia Ho, Alice Broos, Emma Thomson and Alex Alderton, Roberto Amato, Sonia Goncalves, Ewan Harrison, David K. Jackson, Ian Johnston, Dominic Kwiatkowski, Cordelia Langford, John Sillitoe |

| | | | |
|---|---|---|---|
| EPI_ISL_531885, EPI_ISL_531886 | Lighthouse Lab in Glasgow | Wellcome Sanger Institute for the COVID-19 Genomics UK (COG-UK) consortium | Harper VanSteenhouse, Yumi Kasai, David Gray, Carol Clugston, Anna Dominiczak and Alex Alderton, Roberto Amato, Sonia Goncalves, Ewan Harrison, David K. Jackson, Ian Johnston, Dominic Kwiatkowski, Cordelia Langford, John Sillitoe |
| EPI_ISL_531887 | NHSGGC West of Scotland Specialist Virology Centre / MRC-University of Glasgow Centre for Virus Research | Wellcome Sanger Institute for the COVID-19 Genomics UK (COG-UK) consortium | Ana da Silva Filipe, Natasha Johnson, Kathy Smollett, Daniel Mair, Stephen Carmichael, Lily Tong, Jenna Nichols, Elihu Aranday-Cortes, Kirstyn Brunker, Yasmin Parr, Kyriaki Nomikou; Sarah McDonald, Marc Niebel, Patawee Asamaphan; Richard Orton, Joseph Hughes, Sreenu Vattipally, David L Robertson; Alasdair MacLean, Rory Gunson; Kathy Li, Natasha Jesudason, Rajiv Shah, James Shepherd, Antonia Ho, Alice Broos, Emma Thomson and Alex Alderton, Roberto Amato, Sonia Goncalves, Ewan Harrison, David K. Jackson, Ian Johnston, Dominic Kwiatkowski, Cordelia Langford, John Sillitoe |
| EPI_ISL_531888 | Lighthouse Lab in Glasgow | Wellcome Sanger Institute for the COVID-19 Genomics UK (COG-UK) consortium | Harper VanSteenhouse, Yumi Kasai, David Gray, Carol Clugston, Anna Dominiczak and Alex Alderton, Roberto Amato, Sonia Goncalves, Ewan Harrison, David K. Jackson, Ian Johnston, Dominic Kwiatkowski, Cordelia Langford, John Sillitoe |
| EPI_ISL_531889, EPI_ISL_531890 | NHSGGC West of Scotland Specialist Virology Centre / MRC-University of Glasgow Centre for Virus Research | Wellcome Sanger Institute for the COVID-19 Genomics UK (COG-UK) consortium | Ana da Silva Filipe, Natasha Johnson, Kathy Smollett, Daniel Mair, Stephen Carmichael, Lily Tong, Jenna Nichols, Elihu Aranday-Cortes, Kirstyn Brunker, Yasmin Parr, Kyriaki Nomikou; Sarah McDonald, Marc Niebel, Patawee Asamaphan; Richard Orton, Joseph Hughes, Sreenu Vattipally, David L Robertson; Alasdair MacLean, Rory Gunson; Kathy Li, Natasha Jesudason, Rajiv Shah, James Shepherd, Antonia Ho, Alice Broos, Emma Thomson and Alex Alderton, Roberto Amato, Sonia Goncalves, Ewan Harrison, David K. Jackson, Ian Johnston, Dominic Kwiatkowski, Cordelia Langford, John Sillitoe |
| EPI_ISL_531891 | Lighthouse Lab in Glasgow | Wellcome Sanger Institute for the COVID-19 Genomics UK (COG-UK) consortium | Harper VanSteenhouse, Yumi Kasai, David Gray, Carol Clugston, Anna Dominiczak and Alex Alderton, Roberto Amato, Sonia Goncalves, Ewan Harrison, David K. Jackson, Ian Johnston, Dominic Kwiatkowski, Cordelia Langford, John Sillitoe |
| EPI_ISL_531892 | NHSGGC West of Scotland Specialist Virology Centre / MRC-University of Glasgow Centre for Virus Research | Wellcome Sanger Institute for the COVID-19 Genomics UK (COG-UK) consortium | Ana da Silva Filipe, Natasha Johnson, Kathy Smollett, Daniel Mair, Stephen Carmichael, Lily Tong, Jenna Nichols, Elihu Aranday-Cortes, Kirstyn Brunker, Yasmin Parr, Kyriaki Nomikou; Sarah McDonald, Marc Niebel, Patawee Asamaphan; Richard Orton, Joseph Hughes, Sreenu Vattipally, David L Robertson; Alasdair MacLean, Rory Gunson; Kathy Li, Natasha Jesudason, Rajiv Shah, James Shepherd, Antonia Ho, Alice Broos, Emma Thomson and Alex Alderton, Roberto Amato, Sonia Goncalves, Ewan Harrison, David K. Jackson, Ian Johnston, Dominic Kwiatkowski, Cordelia Langford, John Sillitoe |
| EPI_ISL_531893 | Lighthouse Lab in Glasgow | Wellcome Sanger Institute for the COVID-19 Genomics UK (COG-UK) consortium | Harper VanSteenhouse, Yumi Kasai, David Gray, Carol Clugston, Anna Dominiczak and Alex Alderton, Roberto Amato, Sonia Goncalves, Ewan Harrison, David K. Jackson, Ian Johnston, Dominic Kwiatkowski, Cordelia Langford, John Sillitoe |
| EPI_ISL_531894 | NHSGGC West of Scotland Specialist Virology Centre / MRC-University of Glasgow Centre for Virus Research | Wellcome Sanger Institute for the COVID-19 Genomics UK (COG-UK) consortium | Ana da Silva Filipe, Natasha Johnson, Kathy Smollett, Daniel Mair, Stephen Carmichael, Lily Tong, Jenna Nichols, Elihu Aranday-Cortes, Kirstyn Brunker, Yasmin Parr, Kyriaki Nomikou; Sarah McDonald, Marc Niebel, Patawee Asamaphan; Richard Orton, Joseph Hughes, Sreenu Vattipally, David L Robertson; Alasdair MacLean, Rory Gunson; Kathy Li, Natasha Jesudason, Rajiv Shah, James Shepherd, Antonia Ho, Alice Broos, Emma Thomson and Alex Alderton, Roberto Amato, Sonia Goncalves, Ewan Harrison, David K. Jackson, Ian Johnston, Dominic Kwiatkowski, Cordelia Langford, John Sillitoe |
| EPI_ISL_531895, EPI_ISL_531896, EPI_ISL_531897, EPI_ISL_531898, EPI_ISL_531899, EPI_ISL_531900, EPI_ISL_531901, EPI_ISL_531902, EPI_ISL_531903, EPI_ISL_531904 | Lighthouse Lab in Glasgow | Wellcome Sanger Institute for the COVID-19 Genomics UK (COG-UK) consortium | Harper VanSteenhouse, Yumi Kasai, David Gray, Carol Clugston, Anna Dominiczak and Alex Alderton, Roberto Amato, Sonia Goncalves, Ewan Harrison, David K. Jackson, Ian Johnston, Dominic Kwiatkowski, Cordelia Langford, John Sillitoe |
| EPI_ISL_531905 | NHSGGC West of Scotland Specialist Virology Centre / MRC-University of Glasgow Centre for Virus Research | Wellcome Sanger Institute for the COVID-19 Genomics UK (COG-UK) consortium | Ana da Silva Filipe, Natasha Johnson, Kathy Smollett, Daniel Mair, Stephen Carmichael, Lily Tong, Jenna Nichols, Elihu Aranday-Cortes, Kirstyn Brunker, Yasmin Parr, Kyriaki Nomikou; Sarah McDonald, Marc Niebel, Patawee Asamaphan; Richard Orton, Joseph Hughes, Sreenu Vattipally, David L Robertson; Alasdair MacLean, Rory Gunson; Kathy Li, Natasha Jesudason, Rajiv Shah, James Shepherd, Antonia Ho, Alice Broos, Emma Thomson and Alex Alderton, Roberto Amato, Sonia Goncalves, Ewan Harrison, David K. Jackson, Ian Johnston, Dominic Kwiatkowski, Cordelia Langford, John Sillitoe |
| EPI_ISL_531906, EPI_ISL_531907, EPI_ISL_531908, EPI_ISL_531909, EPI_ISL_531910, EPI_ISL_531911 | Lighthouse Lab in Glasgow | Wellcome Sanger Institute for the COVID-19 Genomics UK (COG-UK) consortium | Harper VanSteenhouse, Yumi Kasai, David Gray, Carol Clugston, Anna Dominiczak and Alex Alderton, Roberto Amato, Sonia Goncalves, Ewan Harrison, David K. Jackson, Ian Johnston, Dominic Kwiatkowski, Cordelia Langford, John Sillitoe |
| EPI_ISL_531912, EPI_ISL_531913 | NHSGGC West of Scotland Specialist Virology Centre / MRC-University of Glasgow Centre for Virus Research | Wellcome Sanger Institute for the COVID-19 Genomics UK (COG-UK) consortium | Ana da Silva Filipe, Natasha Johnson, Kathy Smollett, Daniel Mair, Stephen Carmichael, Lily Tong, Jenna Nichols, Elihu Aranday-Cortes, Kirstyn Brunker, Yasmin Parr, Kyriaki Nomikou; Sarah McDonald, Marc Niebel, Patawee Asamaphan; Richard Orton, Joseph Hughes, Sreenu Vattipally, David L Robertson; Alasdair MacLean, Rory Gunson; Kathy Li, Natasha Jesudason, Rajiv Shah, James Shepherd, Antonia Ho, Alice Broos, Emma Thomson and Alex Alderton, Roberto Amato, Sonia Goncalves, Ewan Harrison, David K. Jackson, Ian Johnston, Dominic Kwiatkowski, Cordelia Langford, John Sillitoe |
| EPI_ISL_531914, EPI_ISL_531915, EPI_ISL_531916, EPI_ISL_531917 | Lighthouse Lab in Glasgow | Wellcome Sanger Institute for the COVID-19 Genomics UK (COG-UK) consortium | Harper VanSteenhouse, Yumi Kasai, David Gray, Carol Clugston, Anna Dominiczak and Alex Alderton, Roberto Amato, Sonia Goncalves, Ewan Harrison, David K. Jackson, Ian Johnston, Dominic Kwiatkowski, Cordelia Langford, John Sillitoe |
| EPI_ISL_531918, EPI_ISL_531919 | NHSGGC West of Scotland Specialist Virology Centre / MRC-University of Glasgow Centre for Virus Research | Wellcome Sanger Institute for the COVID-19 Genomics UK (COG-UK) consortium | Ana da Silva Filipe, Natasha Johnson, Kathy Smollett, Daniel Mair, Stephen Carmichael, Lily Tong, Jenna Nichols, Elihu Aranday-Cortes, Kirstyn Brunker, Yasmin Parr, Kyriaki Nomikou; Sarah McDonald, Marc Niebel, Patawee Asamaphan; Richard Orton, Joseph Hughes, Sreenu Vattipally, David L Robertson; Alasdair MacLean, Rory Gunson; Kathy Li, Natasha Jesudason, Rajiv Shah, James Shepherd, Antonia Ho, Alice Broos, Emma Thomson and Alex Alderton, Roberto Amato, Sonia Goncalves, Ewan Harrison, David K. Jackson, Ian Johnston, Dominic Kwiatkowski, Cordelia Langford, John Sillitoe |
| EPI_ISL_531920, EPI_ISL_531921, EPI_ISL_531922, EPI_ISL_531923, EPI_ISL_531924 | Lighthouse Lab in Glasgow | Wellcome Sanger Institute for the COVID-19 Genomics UK (COG-UK) consortium | Harper VanSteenhouse, Yumi Kasai, David Gray, Carol Clugston, Anna Dominiczak and Alex Alderton, Roberto Amato, Sonia Goncalves, Ewan Harrison, David K. Jackson, Ian Johnston, Dominic Kwiatkowski, Cordelia Langford, John Sillitoe |
| EPI_ISL_531925 | NHSGGC West of Scotland Specialist Virology Centre / MRC-University of Glasgow Centre for Virus Research | Wellcome Sanger Institute for the COVID-19 Genomics UK (COG-UK) consortium | Ana da Silva Filipe, Natasha Johnson, Kathy Smollett, Daniel Mair, Stephen Carmichael, Lily Tong, Jenna Nichols, Elihu Aranday-Cortes, Kirstyn Brunker, Yasmin Parr, Kyriaki Nomikou; Sarah McDonald, Marc Niebel, Patawee Asamaphan; Richard Orton, Joseph Hughes, Sreenu Vattipally, David L Robertson; Alasdair MacLean, Rory Gunson; Kathy Li, Natasha Jesudason, Rajiv Shah, James Shepherd, Antonia Ho, Alice Broos, Emma Thomson and Alex Alderton, Roberto Amato, Sonia Goncalves, Ewan Harrison, David K. Jackson, Ian Johnston, Dominic Kwiatkowski, Cordelia Langford, John Sillitoe |
| EPI_ISL_531926, EPI_ISL_531927, EPI_ISL_531928 | Lighthouse Lab in Glasgow | Wellcome Sanger Institute for the COVID-19 Genomics UK (COG-UK) consortium | Harper VanSteenhouse, Yumi Kasai, David Gray, Carol Clugston, Anna Dominiczak and Alex Alderton, Roberto Amato, Sonia Goncalves, Ewan Harrison, David K. Jackson, Ian Johnston, Dominic Kwiatkowski, Cordelia Langford, John Sillitoe |
| EPI_ISL_531929 | NHSGGC West of Scotland Specialist Virology Centre / MRC-University of Glasgow Centre for Virus Research | Wellcome Sanger Institute for the COVID-19 Genomics UK (COG-UK) consortium | Ana da Silva Filipe, Natasha Johnson, Kathy Smollett, Daniel Mair, Stephen Carmichael, Lily Tong, Jenna Nichols, Elihu Aranday-Cortes, Kirstyn Brunker, Yasmin Parr, Kyriaki Nomikou; Sarah McDonald, Marc Niebel, Patawee Asamaphan; Richard Orton, Joseph Hughes, Sreenu Vattipally, David L Robertson; Alasdair MacLean, Rory Gunson; Kathy Li, Natasha Jesudason, Rajiv Shah, James Shepherd, Antonia Ho, Alice Broos, Emma Thomson and Alex Alderton, Roberto Amato, Sonia Goncalves, Ewan Harrison, David K. Jackson, Ian Johnston, Dominic Kwiatkowski, Cordelia Langford, John Sillitoe |
| EPI_ISL_531930, EPI_ISL_531931 | Lighthouse Lab in Glasgow | Wellcome Sanger Institute for the COVID-19 Genomics UK (COG-UK) consortium | Harper VanSteenhouse, Yumi Kasai, David Gray, Carol Clugston, Anna Dominiczak and Alex Alderton, Roberto Amato, Sonia Goncalves, Ewan Harrison, David K. Jackson, Ian Johnston, Dominic Kwiatkowski, Cordelia Langford, John Sillitoe |
| EPI_ISL_531932, EPI_ISL_531933 | NHSGGC West of Scotland Specialist Virology Centre / MRC-University of Glasgow Centre for Virus Research | Wellcome Sanger Institute for the COVID-19 Genomics UK (COG-UK) consortium | Ana da Silva Filipe, Natasha Johnson, Kathy Smollett, Daniel Mair, Stephen Carmichael, Lily Tong, Jenna Nichols, Elihu Aranday-Cortes, Kirstyn Brunker, Yasmin Parr, Kyriaki Nomikou; Sarah McDonald, Marc Niebel, Patawee Asamaphan; Richard Orton, Joseph Hughes, Sreenu Vattipally, David L Robertson; Alasdair MacLean, Rory Gunson; Kathy Li, Natasha Jesudason, Rajiv Shah, James Shepherd, Antonia Ho, Alice Broos, Emma Thomson and Alex Alderton, Roberto Amato, Sonia Goncalves, Ewan Harrison, David K. Jackson, Ian Johnston, Dominic Kwiatkowski, Cordelia Langford, John Sillitoe |
| EPI_ISL_531934 | Lighthouse Lab in Glasgow | Wellcome Sanger Institute for the COVID-19 Genomics UK (COG-UK) consortium | Harper VanSteenhouse, Yumi Kasai, David Gray, Carol Clugston, Anna Dominiczak and Alex Alderton, Roberto Amato, Sonia Goncalves, Ewan Harrison, David K. Jackson, Ian Johnston, Dominic Kwiatkowski, Cordelia Langford, John Sillitoe |
| EPI_ISL_531935, EPI_ISL_531936 | NHSGGC West of Scotland Specialist Virology Centre / MRC-University of Glasgow Centre for Virus Research | Wellcome Sanger Institute for the COVID-19 Genomics UK (COG-UK) consortium | Ana da Silva Filipe, Natasha Johnson, Kathy Smollett, Daniel Mair, Stephen Carmichael, Lily Tong, Jenna Nichols, Elihu Aranday-Cortes, Kirstyn Brunker, Yasmin Parr, Kyriaki Nomikou; Sarah McDonald, Marc Niebel, Patawee Asamaphan; Richard Orton, Joseph Hughes, Sreenu Vattipally, David L Robertson; Alasdair MacLean, Rory Gunson; Kathy Li, Natasha Jesudason, Rajiv Shah, James Shepherd, Antonia Ho, Alice Broos, Emma Thomson and Alex Alderton, Roberto Amato, Sonia Goncalves, Ewan Harrison, David K. Jackson, Ian Johnston, Dominic Kwiatkowski, Cordelia Langford, John Sillitoe |
| EPI_ISL_531937, EPI_ISL_531938, EPI_ISL_531939, EPI_ISL_531940, EPI_ISL_531941, EPI_ISL_531942 | Lighthouse Lab in Glasgow | Wellcome Sanger Institute for the COVID-19 Genomics UK (COG-UK) consortium | Harper VanSteenhouse, Yumi Kasai, David Gray, Carol Clugston, Anna Dominiczak and Alex Alderton, Roberto Amato, Sonia Goncalves, Ewan Harrison, David K. Jackson, Ian Johnston, Dominic Kwiatkowski, Cordelia Langford, John Sillitoe |
| EPI_ISL_531943, EPI_ISL_531945 | NHSGGC West of Scotland Specialist Virology Centre / MRC-University of Glasgow Centre for Virus Research | Wellcome Sanger Institute for the COVID-19 Genomics UK (COG-UK) consortium | Ana da Silva Filipe, Natasha Johnson, Kathy Smollett, Daniel Mair, Stephen Carmichael, Lily Tong, Jenna Nichols, Elihu Aranday-Cortes, Kirstyn Brunker, Yasmin Parr, Kyriaki Nomikou; Sarah McDonald, Marc Niebel, Patawee Asamaphan; Richard Orton, Joseph Hughes, Sreenu Vattipally, David L Robertson; |

| | | | |
|---|---|---|---|
| | | | Alasdair MacLean, Rory Gunson; Kathy Li, Natasha Jesudason, Rajiv Shah, James Shepherd, Antonia Ho, Alice Broos, Emma Thomson and Alex Alderton, Roberto Amato, Sonia Goncalves, Ewan Harrison, David K. Jackson, Ian Johnston, Dominic Kwiatkowski, Cordelia Langford, John Sillitoe |
| EPI_ISL_531946, EPI_ISL_531947, EPI_ISL_531948, EPI_ISL_531949, EPI_ISL_531950 | Lighthouse Lab in Glasgow | Wellcome Sanger Institute for the COVID-19 Genomics UK (COG-UK) consortium | Harper VanSteenhouse, Yumi Kasai, David Gray, Carol Clugston, Anna Dominiczak and Alex Alderton, Roberto Amato, Sonia Goncalves, Ewan Harrison, David K. Jackson, Ian Johnston, Dominic Kwiatkowski, Cordelia Langford, John Sillitoe |
| EPI_ISL_531951 | NHSGGC West of Scotland Specialist Virology Centre / MRC-University of Glasgow Centre for Virus Research | Wellcome Sanger Institute for the COVID-19 Genomics UK (COG-UK) consortium | Ana da Silva Filipe, Natasha Johnson, Kathy Smollett, Daniel Mair, Stephen Carmichael, Lily Tong, Jenna Nichols, Elihu Aranday-Cortes, Kirstyn Brunker, Yasmin Parr, Kyriaki Nomikou; Sarah McDonald, Marc Niebel, Patawee Asamaphan; Richard Orton, Joseph Hughes, Sreenu Vattipally, David L Robertson; Alasdair MacLean, Rory Gunson; Kathy Li, Natasha Jesudason, Rajiv Shah, James Shepherd, Antonia Ho, Alice Broos, Emma Thomson and Alex Alderton, Roberto Amato, Sonia Goncalves, Ewan Harrison, David K. Jackson, Ian Johnston, Dominic Kwiatkowski, Cordelia Langford, John Sillitoe |
| EPI_ISL_531952, EPI_ISL_531953 | Lighthouse Lab in Glasgow | Wellcome Sanger Institute for the COVID-19 Genomics UK (COG-UK) consortium | Harper VanSteenhouse, Yumi Kasai, David Gray, Carol Clugston, Anna Dominiczak and Alex Alderton, Roberto Amato, Sonia Goncalves, Ewan Harrison, David K. Jackson, Ian Johnston, Dominic Kwiatkowski, Cordelia Langford, John Sillitoe |
| EPI_ISL_531954 | NHSGGC West of Scotland Specialist Virology Centre / MRC-University of Glasgow Centre for Virus Research | Wellcome Sanger Institute for the COVID-19 Genomics UK (COG-UK) consortium | Ana da Silva Filipe, Natasha Johnson, Kathy Smollett, Daniel Mair, Stephen Carmichael, Lily Tong, Jenna Nichols, Elihu Aranday-Cortes, Kirstyn Brunker, Yasmin Parr, Kyriaki Nomikou; Sarah McDonald, Marc Niebel, Patawee Asamaphan; Richard Orton, Joseph Hughes, Sreenu Vattipally, David L Robertson; Alasdair MacLean, Rory Gunson; Kathy Li, Natasha Jesudason, Rajiv Shah, James Shepherd, Antonia Ho, Alice Broos, Emma Thomson and Alex Alderton, Roberto Amato, Sonia Goncalves, Ewan Harrison, David K. Jackson, Ian Johnston, Dominic Kwiatkowski, Cordelia Langford, John Sillitoe |
| EPI_ISL_531955, EPI_ISL_531956, EPI_ISL_531957, EPI_ISL_531958, EPI_ISL_531959, EPI_ISL_531960, EPI_ISL_531961, EPI_ISL_531962 | Lighthouse Lab in Glasgow | Wellcome Sanger Institute for the COVID-19 Genomics UK (COG-UK) consortium | Harper VanSteenhouse, Yumi Kasai, David Gray, Carol Clugston, Anna Dominiczak and Alex Alderton, Roberto Amato, Sonia Goncalves, Ewan Harrison, David K. Jackson, Ian Johnston, Dominic Kwiatkowski, Cordelia Langford, John Sillitoe |
| EPI_ISL_531963 | NHSGGC West of Scotland Specialist Virology Centre / MRC-University of Glasgow Centre for Virus Research | Wellcome Sanger Institute for the COVID-19 Genomics UK (COG-UK) consortium | Ana da Silva Filipe, Natasha Johnson, Kathy Smollett, Daniel Mair, Stephen Carmichael, Lily Tong, Jenna Nichols, Elihu Aranday-Cortes, Kirstyn Brunker, Yasmin Parr, Kyriaki Nomikou; Sarah McDonald, Marc Niebel, Patawee Asamaphan; Richard Orton, Joseph Hughes, Sreenu Vattipally, David L Robertson; Alasdair MacLean, Rory Gunson; Kathy Li, Natasha Jesudason, Rajiv Shah, James Shepherd, Antonia Ho, Alice Broos, Emma Thomson and Alex Alderton, Roberto Amato, Sonia Goncalves, Ewan Harrison, David K. Jackson, Ian Johnston, Dominic Kwiatkowski, Cordelia Langford, John Sillitoe |
| EPI_ISL_531964, EPI_ISL_531965, EPI_ISL_531966, EPI_ISL_531967, EPI_ISL_531968, EPI_ISL_531969, EPI_ISL_531970, EPI_ISL_531971, EPI_ISL_531972 | Lighthouse Lab in Glasgow | Wellcome Sanger Institute for the COVID-19 Genomics UK (COG-UK) consortium | Harper VanSteenhouse, Yumi Kasai, David Gray, Carol Clugston, Anna Dominiczak and Alex Alderton, Roberto Amato, Sonia Goncalves, Ewan Harrison, David K. Jackson, Ian Johnston, Dominic Kwiatkowski, Cordelia Langford, John Sillitoe |
| EPI_ISL_531973 | NHSGGC West of Scotland Specialist Virology Centre / MRC-University of Glasgow Centre for Virus Research | Wellcome Sanger Institute for the COVID-19 Genomics UK (COG-UK) consortium | Ana da Silva Filipe, Natasha Johnson, Kathy Smollett, Daniel Mair, Stephen Carmichael, Lily Tong, Jenna Nichols, Elihu Aranday-Cortes, Kirstyn Brunker, Yasmin Parr, Kyriaki Nomikou; Sarah McDonald, Marc Niebel, Patawee Asamaphan; Richard Orton, Joseph Hughes, Sreenu Vattipally, David L Robertson; Alasdair MacLean, Rory Gunson; Kathy Li, Natasha Jesudason, Rajiv Shah, James Shepherd, Antonia Ho, Alice Broos, Emma Thomson and Alex Alderton, Roberto Amato, Sonia Goncalves, Ewan Harrison, David K. Jackson, Ian Johnston, Dominic Kwiatkowski, Cordelia Langford, John Sillitoe |
| EPI_ISL_531974, EPI_ISL_531975, EPI_ISL_531976 | Lighthouse Lab in Glasgow | Wellcome Sanger Institute for the COVID-19 Genomics UK (COG-UK) consortium | Harper VanSteenhouse, Yumi Kasai, David Gray, Carol Clugston, Anna Dominiczak and Alex Alderton, Roberto Amato, Sonia Goncalves, Ewan Harrison, David K. Jackson, Ian Johnston, Dominic Kwiatkowski, Cordelia Langford, John Sillitoe |
| EPI_ISL_531978, EPI_ISL_531979 | NHSGGC West of Scotland Specialist Virology Centre / MRC-University of Glasgow Centre for Virus Research | Wellcome Sanger Institute for the COVID-19 Genomics UK (COG-UK) consortium | Ana da Silva Filipe, Natasha Johnson, Kathy Smollett, Daniel Mair, Stephen Carmichael, Lily Tong, Jenna Nichols, Elihu Aranday-Cortes, Kirstyn Brunker, Yasmin Parr, Kyriaki Nomikou; Sarah McDonald, Marc Niebel, Patawee Asamaphan; Richard Orton, Joseph Hughes, Sreenu Vattipally, David L Robertson; Alasdair MacLean, Rory Gunson; Kathy Li, Natasha Jesudason, Rajiv Shah, James Shepherd, Antonia Ho, Alice Broos, Emma Thomson and Alex Alderton, Roberto Amato, Sonia Goncalves, Ewan Harrison, David K. Jackson, Ian Johnston, Dominic Kwiatkowski, Cordelia Langford, John Sillitoe |
| EPI_ISL_531980, EPI_ISL_531981 | Lighthouse Lab in Glasgow | Wellcome Sanger Institute for the COVID-19 Genomics UK (COG-UK) consortium | Harper VanSteenhouse, Yumi Kasai, David Gray, Carol Clugston, Anna Dominiczak and Alex Alderton, Roberto Amato, Sonia Goncalves, Ewan Harrison, David K. Jackson, Ian Johnston, Dominic Kwiatkowski, Cordelia Langford, John Sillitoe |
| EPI_ISL_531982 | NHSGGC West of Scotland Specialist Virology Centre / MRC-University of Glasgow Centre for Virus Research | Wellcome Sanger Institute for the COVID-19 Genomics UK (COG-UK) consortium | Ana da Silva Filipe, Natasha Johnson, Kathy Smollett, Daniel Mair, Stephen Carmichael, Lily Tong, Jenna Nichols, Elihu Aranday-Cortes, Kirstyn Brunker, Yasmin Parr, Kyriaki Nomikou; Sarah McDonald, Marc Niebel, Patawee Asamaphan; Richard Orton, Joseph Hughes, Sreenu Vattipally, David L Robertson; Alasdair MacLean, Rory Gunson; Kathy Li, Natasha Jesudason, Rajiv Shah, James Shepherd, Antonia Ho, Alice Broos, Emma Thomson and Alex Alderton, Roberto Amato, Sonia Goncalves, Ewan Harrison, David K. Jackson, Ian Johnston, Dominic Kwiatkowski, Cordelia Langford, John Sillitoe |
| EPI_ISL_531983 | Lighthouse Lab in Glasgow | Wellcome Sanger Institute for the COVID-19 Genomics UK (COG-UK) consortium | Harper VanSteenhouse, Yumi Kasai, David Gray, Carol Clugston, Anna Dominiczak and Alex Alderton, Roberto Amato, Sonia Goncalves, Ewan Harrison, David K. Jackson, Ian Johnston, Dominic Kwiatkowski, Cordelia Langford, John Sillitoe |
| EPI_ISL_531984 | NHSGGC West of Scotland Specialist Virology Centre / MRC-University of Glasgow Centre for Virus Research | Wellcome Sanger Institute for the COVID-19 Genomics UK (COG-UK) consortium | Ana da Silva Filipe, Natasha Johnson, Kathy Smollett, Daniel Mair, Stephen Carmichael, Lily Tong, Jenna Nichols, Elihu Aranday-Cortes, Kirstyn Brunker, Yasmin Parr, Kyriaki Nomikou; Sarah McDonald, Marc Niebel, Patawee Asamaphan; Richard Orton, Joseph Hughes, Sreenu Vattipally, David L Robertson; Alasdair MacLean, Rory Gunson; Kathy Li, Natasha Jesudason, Rajiv Shah, James Shepherd, Antonia Ho, Alice Broos, Emma Thomson and Alex Alderton, Roberto Amato, Sonia Goncalves, Ewan Harrison, David K. Jackson, Ian Johnston, Dominic Kwiatkowski, Cordelia Langford, John Sillitoe |
| EPI_ISL_531985, EPI_ISL_531986 | Lighthouse Lab in Glasgow | Wellcome Sanger Institute for the COVID-19 Genomics UK (COG-UK) consortium | Harper VanSteenhouse, Yumi Kasai, David Gray, Carol Clugston, Anna Dominiczak and Alex Alderton, Roberto Amato, Sonia Goncalves, Ewan Harrison, David K. Jackson, Ian Johnston, Dominic Kwiatkowski, Cordelia Langford, John Sillitoe |
| EPI_ISL_531987 | NHSGGC West of Scotland Specialist Virology Centre / MRC-University of Glasgow Centre for Virus Research | Wellcome Sanger Institute for the COVID-19 Genomics UK (COG-UK) consortium | Ana da Silva Filipe, Natasha Johnson, Kathy Smollett, Daniel Mair, Stephen Carmichael, Lily Tong, Jenna Nichols, Elihu Aranday-Cortes, Kirstyn Brunker, Yasmin Parr, Kyriaki Nomikou; Sarah McDonald, Marc Niebel, Patawee Asamaphan; Richard Orton, Joseph Hughes, Sreenu Vattipally, David L Robertson; Alasdair MacLean, Rory Gunson; Kathy Li, Natasha Jesudason, Rajiv Shah, James Shepherd, Antonia Ho, Alice Broos, Emma Thomson and Alex Alderton, Roberto Amato, Sonia Goncalves, Ewan Harrison, David K. Jackson, Ian Johnston, Dominic Kwiatkowski, Cordelia Langford, John Sillitoe |
| EPI_ISL_531988, EPI_ISL_531989 | Lighthouse Lab in Glasgow | Wellcome Sanger Institute for the COVID-19 Genomics UK (COG-UK) consortium | Harper VanSteenhouse, Yumi Kasai, David Gray, Carol Clugston, Anna Dominiczak and Alex Alderton, Roberto Amato, Sonia Goncalves, Ewan Harrison, David K. Jackson, Ian Johnston, Dominic Kwiatkowski, Cordelia Langford, John Sillitoe |
| EPI_ISL_531990 | NHSGGC West of Scotland Specialist Virology Centre / MRC-University of Glasgow Centre for Virus Research | Wellcome Sanger Institute for the COVID-19 Genomics UK (COG-UK) consortium | Ana da Silva Filipe, Natasha Johnson, Kathy Smollett, Daniel Mair, Stephen Carmichael, Lily Tong, Jenna Nichols, Elihu Aranday-Cortes, Kirstyn Brunker, Yasmin Parr, Kyriaki Nomikou; Sarah McDonald, Marc Niebel, Patawee Asamaphan; Richard Orton, Joseph Hughes, Sreenu Vattipally, David L Robertson; Alasdair MacLean, Rory Gunson; Kathy Li, Natasha Jesudason, Rajiv Shah, James Shepherd, Antonia Ho, Alice Broos, Emma Thomson and Alex Alderton, Roberto Amato, Sonia Goncalves, Ewan Harrison, David K. Jackson, Ian Johnston, Dominic Kwiatkowski, Cordelia Langford, John Sillitoe |
| EPI_ISL_531991, EPI_ISL_531992, EPI_ISL_531993, EPI_ISL_531994 | Lighthouse Lab in Glasgow | Wellcome Sanger Institute for the COVID-19 Genomics UK (COG-UK) consortium | Harper VanSteenhouse, Yumi Kasai, David Gray, Carol Clugston, Anna Dominiczak and Alex Alderton, Roberto Amato, Sonia Goncalves, Ewan Harrison, David K. Jackson, Ian Johnston, Dominic Kwiatkowski, Cordelia Langford, John Sillitoe |
| EPI_ISL_531995 | NHSGGC West of Scotland Specialist Virology Centre / MRC-University of Glasgow Centre for Virus Research | Wellcome Sanger Institute for the COVID-19 Genomics UK (COG-UK) consortium | Ana da Silva Filipe, Natasha Johnson, Kathy Smollett, Daniel Mair, Stephen Carmichael, Lily Tong, Jenna Nichols, Elihu Aranday-Cortes, Kirstyn Brunker, Yasmin Parr, Kyriaki Nomikou; Sarah McDonald, Marc Niebel, Patawee Asamaphan; Richard Orton, Joseph Hughes, Sreenu Vattipally, David L Robertson; Alasdair MacLean, Rory Gunson; Kathy Li, Natasha Jesudason, Rajiv Shah, James Shepherd, Antonia Ho, Alice Broos, Emma Thomson and Alex Alderton, Roberto Amato, Sonia Goncalves, Ewan Harrison, David K. Jackson, Ian Johnston, Dominic Kwiatkowski, Cordelia Langford, John Sillitoe |
| EPI_ISL_531996, EPI_ISL_531997, EPI_ISL_531998, EPI_ISL_531999, EPI_ISL_532000, EPI_ISL_532001, EPI_ISL_532002 | Lighthouse Lab in Glasgow | Wellcome Sanger Institute for the COVID-19 Genomics UK (COG-UK) consortium | Harper VanSteenhouse, Yumi Kasai, David Gray, Carol Clugston, Anna Dominiczak and Alex Alderton, Roberto Amato, Sonia Goncalves, Ewan Harrison, David K. Jackson, Ian Johnston, Dominic Kwiatkowski, Cordelia Langford, John Sillitoe |
| EPI_ISL_532003, EPI_ISL_532004, EPI_ISL_532005 | NHSGGC West of Scotland Specialist Virology Centre / MRC-University of Glasgow Centre for Virus Research | Wellcome Sanger Institute for the COVID-19 Genomics UK (COG-UK) consortium | Ana da Silva Filipe, Natasha Johnson, Kathy Smollett, Daniel Mair, Stephen Carmichael, Lily Tong, Jenna Nichols, Elihu Aranday-Cortes, Kirstyn Brunker, Yasmin Parr, Kyriaki Nomikou; Sarah McDonald, Marc Niebel, Patawee Asamaphan; Richard Orton, Joseph Hughes, Sreenu Vattipally, David L Robertson; Alasdair MacLean, Rory Gunson; Kathy Li, Natasha Jesudason, Rajiv Shah, James Shepherd, Antonia Ho, Alice Broos, Emma Thomson and Alex Alderton, Roberto Amato, Sonia Goncalves, Ewan Harrison, David K. Jackson, Ian Johnston, Dominic Kwiatkowski, Cordelia Langford, John Sillitoe |

| | | | |
|---|---|---|---|
| EPI_ISL_532006 | Lighthouse Lab in Glasgow | Wellcome Sanger Institute for the COVID-19 Genomics UK (COG-UK) consortium | Harper VanSteenhouse, Yumi Kasai, David Gray, Carol Clugston, Anna Dominiczak and Alex Alderton, Roberto Amato, Sonia Goncalves, Ewan Harrison, David K. Jackson, Ian Johnston, Dominic Kwiatkowski, Cordelia Langford, John Sillitoe |
| EPI_ISL_532007 | NHSGGC West of Scotland Specialist Virology Centre / MRC-University of Glasgow Centre for Virus Research | Wellcome Sanger Institute for the COVID-19 Genomics UK (COG-UK) consortium | Ana da Silva Filipe, Natasha Johnson, Kathy Smollett, Daniel Mair, Stephen Carmichael, Lily Tong, Jenna Nichols, Elihu Aranday-Cortes, Kirstyn Brunker, Yasmin Parr, Kyriaki Nomikou; Sarah McDonald, Marc Niebel, Patawee Asamaphan; Richard Orton, Joseph Hughes, Sreenu Vattipally, David L Robertson; Alasdair MacLean, Rory Gunson; Kathy Li, Natasha Jesudason, Rajiv Shah, James Shepherd, Antonia Ho, Alice Broos, Emma Thomson and Alex Alderton, Roberto Amato, Sonia Goncalves, Ewan Harrison, David K. Jackson, Ian Johnston, Dominic Kwiatkowski, Cordelia Langford, John Sillitoe |
| EPI_ISL_532008, EPI_ISL_532009, EPI_ISL_532010, EPI_ISL_532011 | Lighthouse Lab in Glasgow | Wellcome Sanger Institute for the COVID-19 Genomics UK (COG-UK) consortium | Harper VanSteenhouse, Yumi Kasai, David Gray, Carol Clugston, Anna Dominiczak and Alex Alderton, Roberto Amato, Sonia Goncalves, Ewan Harrison, David K. Jackson, Ian Johnston, Dominic Kwiatkowski, Cordelia Langford, John Sillitoe |
| EPI_ISL_532012 | Lighthouse Lab in Glasgow | Wellcome Sanger Institute for the COVID-19 Genomics UK (COG-UK) Consortium | Harper VanSteenhouse, Yumi Kasai, David Gray, Carol Clugston, Anna Dominiczak and Alex Alderton, Roberto Amato, Sonia Goncalves, Ewan Harrison, David K. Jackson, Ian Johnston, Dominic Kwiatkowski, Cordelia Langford, John Sillitoe on behalf of the Wellcome Sanger Institute COVID-19 Surveillance Team |
| EPI_ISL_532013, EPI_ISL_532014, EPI_ISL_532015, EPI_ISL_532016, EPI_ISL_532017 | Lighthouse Lab in Glasgow | Wellcome Sanger Institute for the COVID-19 Genomics UK (COG-UK) consortium | Harper VanSteenhouse, Yumi Kasai, David Gray, Carol Clugston, Anna Dominiczak and Alex Alderton, Roberto Amato, Sonia Goncalves, Ewan Harrison, David K. Jackson, Ian Johnston, Dominic Kwiatkowski, Cordelia Langford, John Sillitoe |
| EPI_ISL_532018 | NHSGGC West of Scotland Specialist Virology Centre / MRC-University of Glasgow Centre for Virus Research | Wellcome Sanger Institute for the COVID-19 Genomics UK (COG-UK) consortium | Ana da Silva Filipe, Natasha Johnson, Kathy Smollett, Daniel Mair, Stephen Carmichael, Lily Tong, Jenna Nichols, Elihu Aranday-Cortes, Kirstyn Brunker, Yasmin Parr, Kyriaki Nomikou; Sarah McDonald, Marc Niebel, Patawee Asamaphan; Richard Orton, Joseph Hughes, Sreenu Vattipally, David L Robertson; Alasdair MacLean, Rory Gunson; Kathy Li, Natasha Jesudason, Rajiv Shah, James Shepherd, Antonia Ho, Alice Broos, Emma Thomson and Alex Alderton, Roberto Amato, Sonia Goncalves, Ewan Harrison, David K. Jackson, Ian Johnston, Dominic Kwiatkowski, Cordelia Langford, John Sillitoe |
| EPI_ISL_532019, EPI_ISL_532020, EPI_ISL_532021, EPI_ISL_532022, EPI_ISL_532023 | Lighthouse Lab in Glasgow | Wellcome Sanger Institute for the COVID-19 Genomics UK (COG-UK) consortium | Harper VanSteenhouse, Yumi Kasai, David Gray, Carol Clugston, Anna Dominiczak and Alex Alderton, Roberto Amato, Sonia Goncalves, Ewan Harrison, David K. Jackson, Ian Johnston, Dominic Kwiatkowski, Cordelia Langford, John Sillitoe |
| EPI_ISL_532024 | NHSGGC West of Scotland Specialist Virology Centre / MRC-University of Glasgow Centre for Virus Research | Wellcome Sanger Institute for the COVID-19 Genomics UK (COG-UK) consortium | Ana da Silva Filipe, Natasha Johnson, Kathy Smollett, Daniel Mair, Stephen Carmichael, Lily Tong, Jenna Nichols, Elihu Aranday-Cortes, Kirstyn Brunker, Yasmin Parr, Kyriaki Nomikou; Sarah McDonald, Marc Niebel, Patawee Asamaphan; Richard Orton, Joseph Hughes, Sreenu Vattipally, David L Robertson; Alasdair MacLean, Rory Gunson; Kathy Li, Natasha Jesudason, Rajiv Shah, James Shepherd, Antonia Ho, Alice Broos, Emma Thomson and Alex Alderton, Roberto Amato, Sonia Goncalves, Ewan Harrison, David K. Jackson, Ian Johnston, Dominic Kwiatkowski, Cordelia Langford, John Sillitoe |
| EPI_ISL_532025, EPI_ISL_532026, EPI_ISL_532027, EPI_ISL_532028, EPI_ISL_532029, EPI_ISL_532030, EPI_ISL_532031 | Lighthouse Lab in Glasgow | Wellcome Sanger Institute for the COVID-19 Genomics UK (COG-UK) consortium | Harper VanSteenhouse, Yumi Kasai, David Gray, Carol Clugston, Anna Dominiczak and Alex Alderton, Roberto Amato, Sonia Goncalves, Ewan Harrison, David K. Jackson, Ian Johnston, Dominic Kwiatkowski, Cordelia Langford, John Sillitoe |
| EPI_ISL_532032 | NHSGGC West of Scotland Specialist Virology Centre / MRC-University of Glasgow Centre for Virus Research | Wellcome Sanger Institute for the COVID-19 Genomics UK (COG-UK) consortium | Ana da Silva Filipe, Natasha Johnson, Kathy Smollett, Daniel Mair, Stephen Carmichael, Lily Tong, Jenna Nichols, Elihu Aranday-Cortes, Kirstyn Brunker, Yasmin Parr, Kyriaki Nomikou; Sarah McDonald, Marc Niebel, Patawee Asamaphan; Richard Orton, Joseph Hughes, Sreenu Vattipally, David L Robertson; Alasdair MacLean, Rory Gunson; Kathy Li, Natasha Jesudason, Rajiv Shah, James Shepherd, Antonia Ho, Alice Broos, Emma Thomson and Alex Alderton, Roberto Amato, Sonia Goncalves, Ewan Harrison, David K. Jackson, Ian Johnston, Dominic Kwiatkowski, Cordelia Langford, John Sillitoe |
| EPI_ISL_532033, EPI_ISL_532034, EPI_ISL_532035, EPI_ISL_532036, EPI_ISL_532037 | Lighthouse Lab in Glasgow | Wellcome Sanger Institute for the COVID-19 Genomics UK (COG-UK) consortium | Harper VanSteenhouse, Yumi Kasai, David Gray, Carol Clugston, Anna Dominiczak and Alex Alderton, Roberto Amato, Sonia Goncalves, Ewan Harrison, David K. Jackson, Ian Johnston, Dominic Kwiatkowski, Cordelia Langford, John Sillitoe |
| EPI_ISL_532038 | NHSGGC West of Scotland Specialist Virology Centre / MRC-University of Glasgow Centre for Virus Research | Wellcome Sanger Institute for the COVID-19 Genomics UK (COG-UK) consortium | Ana da Silva Filipe, Natasha Johnson, Kathy Smollett, Daniel Mair, Stephen Carmichael, Lily Tong, Jenna Nichols, Elihu Aranday-Cortes, Kirstyn Brunker, Yasmin Parr, Kyriaki Nomikou; Sarah McDonald, Marc Niebel, Patawee Asamaphan; Richard Orton, Joseph Hughes, Sreenu Vattipally, David L Robertson; Alasdair MacLean, Rory Gunson; Kathy Li, Natasha Jesudason, Rajiv Shah, James Shepherd, Antonia Ho, Alice Broos, Emma Thomson and Alex Alderton, Roberto Amato, Sonia Goncalves, Ewan Harrison, David K. Jackson, Ian Johnston, Dominic Kwiatkowski, Cordelia Langford, John Sillitoe |
| EPI_ISL_532039, EPI_ISL_532040, EPI_ISL_532041, EPI_ISL_532042, EPI_ISL_532043, EPI_ISL_532044, EPI_ISL_532045, EPI_ISL_532046, EPI_ISL_532047, EPI_ISL_532048 | Lighthouse Lab in Glasgow | Wellcome Sanger Institute for the COVID-19 Genomics UK (COG-UK) consortium | Harper VanSteenhouse, Yumi Kasai, David Gray, Carol Clugston, Anna Dominiczak and Alex Alderton, Roberto Amato, Sonia Goncalves, Ewan Harrison, David K. Jackson, Ian Johnston, Dominic Kwiatkowski, Cordelia Langford, John Sillitoe |
| EPI_ISL_532049 | NHSGGC West of Scotland Specialist Virology Centre / MRC-University of Glasgow Centre for Virus Research | Wellcome Sanger Institute for the COVID-19 Genomics UK (COG-UK) consortium | Ana da Silva Filipe, Natasha Johnson, Kathy Smollett, Daniel Mair, Stephen Carmichael, Lily Tong, Jenna Nichols, Elihu Aranday-Cortes, Kirstyn Brunker, Yasmin Parr, Kyriaki Nomikou; Sarah McDonald, Marc Niebel, Patawee Asamaphan; Richard Orton, Joseph Hughes, Sreenu Vattipally, David L Robertson; Alasdair MacLean, Rory Gunson; Kathy Li, Natasha Jesudason, Rajiv Shah, James Shepherd, Antonia Ho, Alice Broos, Emma Thomson and Alex Alderton, Roberto Amato, Sonia Goncalves, Ewan Harrison, David K. Jackson, Ian Johnston, Dominic Kwiatkowski, Cordelia Langford, John Sillitoe |
| EPI_ISL_532050, EPI_ISL_532051, EPI_ISL_532052 | Lighthouse Lab in Glasgow | Wellcome Sanger Institute for the COVID-19 Genomics UK (COG-UK) consortium | Harper VanSteenhouse, Yumi Kasai, David Gray, Carol Clugston, Anna Dominiczak and Alex Alderton, Roberto Amato, Sonia Goncalves, Ewan Harrison, David K. Jackson, Ian Johnston, Dominic Kwiatkowski, Cordelia Langford, John Sillitoe |
| EPI_ISL_532053 | NHSGGC West of Scotland Specialist Virology Centre / MRC-University of Glasgow Centre for Virus Research | Wellcome Sanger Institute for the COVID-19 Genomics UK (COG-UK) consortium | Ana da Silva Filipe, Natasha Johnson, Kathy Smollett, Daniel Mair, Stephen Carmichael, Lily Tong, Jenna Nichols, Elihu Aranday-Cortes, Kirstyn Brunker, Yasmin Parr, Kyriaki Nomikou; Sarah McDonald, Marc Niebel, Patawee Asamaphan; Richard Orton, Joseph Hughes, Sreenu Vattipally, David L Robertson; Alasdair MacLean, Rory Gunson; Kathy Li, Natasha Jesudason, Rajiv Shah, James Shepherd, Antonia Ho, Alice Broos, Emma Thomson and Alex Alderton, Roberto Amato, Sonia Goncalves, Ewan Harrison, David K. Jackson, Ian Johnston, Dominic Kwiatkowski, Cordelia Langford, John Sillitoe |
| EPI_ISL_532054 | Lighthouse Lab in Glasgow | Wellcome Sanger Institute for the COVID-19 Genomics UK (COG-UK) consortium | Harper VanSteenhouse, Yumi Kasai, David Gray, Carol Clugston, Anna Dominiczak and Alex Alderton, Roberto Amato, Sonia Goncalves, Ewan Harrison, David K. Jackson, Ian Johnston, Dominic Kwiatkowski, Cordelia Langford, John Sillitoe |
| EPI_ISL_532055 | NHSGGC West of Scotland Specialist Virology Centre / MRC-University of Glasgow Centre for Virus Research | Wellcome Sanger Institute for the COVID-19 Genomics UK (COG-UK) consortium | Ana da Silva Filipe, Natasha Johnson, Kathy Smollett, Daniel Mair, Stephen Carmichael, Lily Tong, Jenna Nichols, Elihu Aranday-Cortes, Kirstyn Brunker, Yasmin Parr, Kyriaki Nomikou; Sarah McDonald, Marc Niebel, Patawee Asamaphan; Richard Orton, Joseph Hughes, Sreenu Vattipally, David L Robertson; Alasdair MacLean, Rory Gunson; Kathy Li, Natasha Jesudason, Rajiv Shah, James Shepherd, Antonia Ho, Alice Broos, Emma Thomson and Alex Alderton, Roberto Amato, Sonia Goncalves, Ewan Harrison, David K. Jackson, Ian Johnston, Dominic Kwiatkowski, Cordelia Langford, John Sillitoe |
| EPI_ISL_532056 | Lighthouse Lab in Glasgow | Wellcome Sanger Institute for the COVID-19 Genomics UK (COG-UK) consortium | Harper VanSteenhouse, Yumi Kasai, David Gray, Carol Clugston, Anna Dominiczak and Alex Alderton, Roberto Amato, Sonia Goncalves, Ewan Harrison, David K. Jackson, Ian Johnston, Dominic Kwiatkowski, Cordelia Langford, John Sillitoe |
| EPI_ISL_532057 | NHSGGC West of Scotland Specialist Virology Centre / MRC-University of Glasgow Centre for Virus Research | Wellcome Sanger Institute for the COVID-19 Genomics UK (COG-UK) consortium | Ana da Silva Filipe, Natasha Johnson, Kathy Smollett, Daniel Mair, Stephen Carmichael, Lily Tong, Jenna Nichols, Elihu Aranday-Cortes, Kirstyn Brunker, Yasmin Parr, Kyriaki Nomikou; Sarah McDonald, Marc Niebel, Patawee Asamaphan; Richard Orton, Joseph Hughes, Sreenu Vattipally, David L Robertson; Alasdair MacLean, Rory Gunson; Kathy Li, Natasha Jesudason, Rajiv Shah, James Shepherd, Antonia Ho, Alice Broos, Emma Thomson and Alex Alderton, Roberto Amato, Sonia Goncalves, Ewan Harrison, David K. Jackson, Ian Johnston, Dominic Kwiatkowski, Cordelia Langford, John Sillitoe |
| EPI_ISL_532058, EPI_ISL_532059, EPI_ISL_532060, EPI_ISL_532061 | Lighthouse Lab in Glasgow | Wellcome Sanger Institute for the COVID-19 Genomics UK (COG-UK) consortium | Harper VanSteenhouse, Yumi Kasai, David Gray, Carol Clugston, Anna Dominiczak and Alex Alderton, Roberto Amato, Sonia Goncalves, Ewan Harrison, David K. Jackson, Ian Johnston, Dominic Kwiatkowski, Cordelia Langford, John Sillitoe |
| EPI_ISL_532062 | NHSGGC West of Scotland Specialist Virology Centre / MRC-University of Glasgow Centre for Virus Research | Wellcome Sanger Institute for the COVID-19 Genomics UK (COG-UK) consortium | Ana da Silva Filipe, Natasha Johnson, Kathy Smollett, Daniel Mair, Stephen Carmichael, Lily Tong, Jenna Nichols, Elihu Aranday-Cortes, Kirstyn Brunker, Yasmin Parr, Kyriaki Nomikou; Sarah McDonald, Marc Niebel, Patawee Asamaphan; Richard Orton, Joseph Hughes, Sreenu Vattipally, David L Robertson; Alasdair MacLean, Rory Gunson; Kathy Li, Natasha Jesudason, Rajiv Shah, James Shepherd, Antonia Ho, Alice Broos, Emma Thomson and Alex Alderton, Roberto Amato, Sonia Goncalves, Ewan Harrison, David K. Jackson, Ian Johnston, Dominic Kwiatkowski, Cordelia Langford, John Sillitoe |
| EPI_ISL_532063, EPI_ISL_532064, EPI_ISL_532066, EPI_ISL_532067, EPI_ISL_532068, EPI_ISL_532069 | Lighthouse Lab in Glasgow | Wellcome Sanger Institute for the COVID-19 Genomics UK (COG-UK) consortium | Harper VanSteenhouse, Yumi Kasai, David Gray, Carol Clugston, Anna Dominiczak and Alex Alderton, Roberto Amato, Sonia Goncalves, Ewan Harrison, David K. Jackson, Ian Johnston, Dominic Kwiatkowski, Cordelia Langford, John Sillitoe |

| | | | |
|---|---|---|---|
| EPI_ISL_532070, EPI_ISL_532071 | NHSGGC West of Scotland Specialist Virology Centre / MRC-University of Glasgow Centre for Virus Research | Wellcome Sanger Institute for the COVID-19 Genomics UK (COG-UK) consortium | Ana da Silva Filipe, Natasha Johnson, Kathy Smollett, Daniel Mair, Stephen Carmichael, Lily Tong, Jenna Nichols, Elihu Aranday-Cortes, Kirstyn Brunker, Yasmin Parr, Kyriaki Nomikou; Sarah McDonald, Marc Niebel, Patawee Asamaphan; Richard Orton, Joseph Hughes, Sreenu Vattipally, David L Robertson; Alasdair MacLean, Rory Gunson; Kathy Li, Natasha Jesudason, Rajiv Shah, James Shepherd, Antonia Ho, Alice Broos, Emma Thomson and Alex Alderton, Roberto Amato, Sonia Goncalves, Ewan Harrison, David K. Jackson, Ian Johnston, Dominic Kwiatkowski, Cordelia Langford, John Sillitoe |
| EPI_ISL_532072, EPI_ISL_532073, EPI_ISL_532074, EPI_ISL_532075 | Lighthouse Lab in Glasgow | Wellcome Sanger Institute for the COVID-19 Genomics UK (COG-UK) consortium | Harper VanSteenhouse, Yumi Kasai, David Gray, Carol Clugston, Anna Dominiczak and Alex Alderton, Roberto Amato, Sonia Goncalves, Ewan Harrison, David K. Jackson, Ian Johnston, Dominic Kwiatkowski, Cordelia Langford, John Sillitoe |
| EPI_ISL_532076, EPI_ISL_532077, EPI_ISL_532078 | NHSGGC West of Scotland Specialist Virology Centre / MRC-University of Glasgow Centre for Virus Research | Wellcome Sanger Institute for the COVID-19 Genomics UK (COG-UK) consortium | Ana da Silva Filipe, Natasha Johnson, Kathy Smollett, Daniel Mair, Stephen Carmichael, Lily Tong, Jenna Nichols, Elihu Aranday-Cortes, Kirstyn Brunker, Yasmin Parr, Kyriaki Nomikou; Sarah McDonald, Marc Niebel, Patawee Asamaphan; Richard Orton, Joseph Hughes, Sreenu Vattipally, David L Robertson; Alasdair MacLean, Rory Gunson; Kathy Li, Natasha Jesudason, Rajiv Shah, James Shepherd, Antonia Ho, Alice Broos, Emma Thomson and Alex Alderton, Roberto Amato, Sonia Goncalves, Ewan Harrison, David K. Jackson, Ian Johnston, Dominic Kwiatkowski, Cordelia Langford, John Sillitoe |
| EPI_ISL_532079 | Lighthouse Lab in Glasgow | Wellcome Sanger Institute for the COVID-19 Genomics UK (COG-UK) consortium | Harper VanSteenhouse, Yumi Kasai, David Gray, Carol Clugston, Anna Dominiczak and Alex Alderton, Roberto Amato, Sonia Goncalves, Ewan Harrison, David K. Jackson, Ian Johnston, Dominic Kwiatkowski, Cordelia Langford, John Sillitoe |
| EPI_ISL_532080, EPI_ISL_532081 | NHSGGC West of Scotland Specialist Virology Centre / MRC-University of Glasgow Centre for Virus Research | Wellcome Sanger Institute for the COVID-19 Genomics UK (COG-UK) consortium | Ana da Silva Filipe, Natasha Johnson, Kathy Smollett, Daniel Mair, Stephen Carmichael, Lily Tong, Jenna Nichols, Elihu Aranday-Cortes, Kirstyn Brunker, Yasmin Parr, Kyriaki Nomikou; Sarah McDonald, Marc Niebel, Patawee Asamaphan; Richard Orton, Joseph Hughes, Sreenu Vattipally, David L Robertson; Alasdair MacLean, Rory Gunson; Kathy Li, Natasha Jesudason, Rajiv Shah, James Shepherd, Antonia Ho, Alice Broos, Emma Thomson and Alex Alderton, Roberto Amato, Sonia Goncalves, Ewan Harrison, David K. Jackson, Ian Johnston, Dominic Kwiatkowski, Cordelia Langford, John Sillitoe |
| EPI_ISL_532082, EPI_ISL_532083, EPI_ISL_532084 | Lighthouse Lab in Glasgow | Wellcome Sanger Institute for the COVID-19 Genomics UK (COG-UK) consortium | Harper VanSteenhouse, Yumi Kasai, David Gray, Carol Clugston, Anna Dominiczak and Alex Alderton, Roberto Amato, Sonia Goncalves, Ewan Harrison, David K. Jackson, Ian Johnston, Dominic Kwiatkowski, Cordelia Langford, John Sillitoe |
| EPI_ISL_532085 | NHSGGC West of Scotland Specialist Virology Centre / MRC-University of Glasgow Centre for Virus Research | Wellcome Sanger Institute for the COVID-19 Genomics UK (COG-UK) consortium | Ana da Silva Filipe, Natasha Johnson, Kathy Smollett, Daniel Mair, Stephen Carmichael, Lily Tong, Jenna Nichols, Elihu Aranday-Cortes, Kirstyn Brunker, Yasmin Parr, Kyriaki Nomikou; Sarah McDonald, Marc Niebel, Patawee Asamaphan; Richard Orton, Joseph Hughes, Sreenu Vattipally, David L Robertson; Alasdair MacLean, Rory Gunson; Kathy Li, Natasha Jesudason, Rajiv Shah, James Shepherd, Antonia Ho, Alice Broos, Emma Thomson and Alex Alderton, Roberto Amato, Sonia Goncalves, Ewan Harrison, David K. Jackson, Ian Johnston, Dominic Kwiatkowski, Cordelia Langford, John Sillitoe |
| EPI_ISL_532086, EPI_ISL_532087, EPI_ISL_532088, EPI_ISL_532089, EPI_ISL_532090, EPI_ISL_532091, EPI_ISL_532092 | Lighthouse Lab in Glasgow | Wellcome Sanger Institute for the COVID-19 Genomics UK (COG-UK) consortium | Harper VanSteenhouse, Yumi Kasai, David Gray, Carol Clugston, Anna Dominiczak and Alex Alderton, Roberto Amato, Sonia Goncalves, Ewan Harrison, David K. Jackson, Ian Johnston, Dominic Kwiatkowski, Cordelia Langford, John Sillitoe |
| EPI_ISL_532093, EPI_ISL_532094 | NHSGGC West of Scotland Specialist Virology Centre / MRC-University of Glasgow Centre for Virus Research | Wellcome Sanger Institute for the COVID-19 Genomics UK (COG-UK) consortium | Ana da Silva Filipe, Natasha Johnson, Kathy Smollett, Daniel Mair, Stephen Carmichael, Lily Tong, Jenna Nichols, Elihu Aranday-Cortes, Kirstyn Brunker, Yasmin Parr, Kyriaki Nomikou; Sarah McDonald, Marc Niebel, Patawee Asamaphan; Richard Orton, Joseph Hughes, Sreenu Vattipally, David L Robertson; Alasdair MacLean, Rory Gunson; Kathy Li, Natasha Jesudason, Rajiv Shah, James Shepherd, Antonia Ho, Alice Broos, Emma Thomson and Alex Alderton, Roberto Amato, Sonia Goncalves, Ewan Harrison, David K. Jackson, Ian Johnston, Dominic Kwiatkowski, Cordelia Langford, John Sillitoe |
| EPI_ISL_532095, EPI_ISL_532096, EPI_ISL_532097 | Lighthouse Lab in Glasgow | Wellcome Sanger Institute for the COVID-19 Genomics UK (COG-UK) consortium | Harper VanSteenhouse, Yumi Kasai, David Gray, Carol Clugston, Anna Dominiczak and Alex Alderton, Roberto Amato, Sonia Goncalves, Ewan Harrison, David K. Jackson, Ian Johnston, Dominic Kwiatkowski, Cordelia Langford, John Sillitoe |
| EPI_ISL_532098 | NHSGGC West of Scotland Specialist Virology Centre / MRC-University of Glasgow Centre for Virus Research | Wellcome Sanger Institute for the COVID-19 Genomics UK (COG-UK) consortium | Ana da Silva Filipe, Natasha Johnson, Kathy Smollett, Daniel Mair, Stephen Carmichael, Lily Tong, Jenna Nichols, Elihu Aranday-Cortes, Kirstyn Brunker, Yasmin Parr, Kyriaki Nomikou; Sarah McDonald, Marc Niebel, Patawee Asamaphan; Richard Orton, Joseph Hughes, Sreenu Vattipally, David L Robertson; Alasdair MacLean, Rory Gunson; Kathy Li, Natasha Jesudason, Rajiv Shah, James Shepherd, Antonia Ho, Alice Broos, Emma Thomson and Alex Alderton, Roberto Amato, Sonia Goncalves, Ewan Harrison, David K. Jackson, Ian Johnston, Dominic Kwiatkowski, Cordelia Langford, John Sillitoe |
| EPI_ISL_532099, EPI_ISL_532100, EPI_ISL_532101 | Lighthouse Lab in Glasgow | Wellcome Sanger Institute for the COVID-19 Genomics UK (COG-UK) consortium | Harper VanSteenhouse, Yumi Kasai, David Gray, Carol Clugston, Anna Dominiczak and Alex Alderton, Roberto Amato, Sonia Goncalves, Ewan Harrison, David K. Jackson, Ian Johnston, Dominic Kwiatkowski, Cordelia Langford, John Sillitoe |
| EPI_ISL_532102 | NHSGGC West of Scotland Specialist Virology Centre / MRC-University of Glasgow Centre for Virus Research | Wellcome Sanger Institute for the COVID-19 Genomics UK (COG-UK) consortium | Ana da Silva Filipe, Natasha Johnson, Kathy Smollett, Daniel Mair, Stephen Carmichael, Lily Tong, Jenna Nichols, Elihu Aranday-Cortes, Kirstyn Brunker, Yasmin Parr, Kyriaki Nomikou; Sarah McDonald, Marc Niebel, Patawee Asamaphan; Richard Orton, Joseph Hughes, Sreenu Vattipally, David L Robertson; Alasdair MacLean, Rory Gunson; Kathy Li, Natasha Jesudason, Rajiv Shah, James Shepherd, Antonia Ho, Alice Broos, Emma Thomson and Alex Alderton, Roberto Amato, Sonia Goncalves, Ewan Harrison, David K. Jackson, Ian Johnston, Dominic Kwiatkowski, Cordelia Langford, John Sillitoe |
| EPI_ISL_532103, EPI_ISL_532104 | Lighthouse Lab in Glasgow | Wellcome Sanger Institute for the COVID-19 Genomics UK (COG-UK) consortium | Harper VanSteenhouse, Yumi Kasai, David Gray, Carol Clugston, Anna Dominiczak and Alex Alderton, Roberto Amato, Sonia Goncalves, Ewan Harrison, David K. Jackson, Ian Johnston, Dominic Kwiatkowski, Cordelia Langford, John Sillitoe |
| EPI_ISL_532105 | NHSGGC West of Scotland Specialist Virology Centre / MRC-University of Glasgow Centre for Virus Research | Wellcome Sanger Institute for the COVID-19 Genomics UK (COG-UK) consortium | Ana da Silva Filipe, Natasha Johnson, Kathy Smollett, Daniel Mair, Stephen Carmichael, Lily Tong, Jenna Nichols, Elihu Aranday-Cortes, Kirstyn Brunker, Yasmin Parr, Kyriaki Nomikou; Sarah McDonald, Marc Niebel, Patawee Asamaphan; Richard Orton, Joseph Hughes, Sreenu Vattipally, David L Robertson; Alasdair MacLean, Rory Gunson; Kathy Li, Natasha Jesudason, Rajiv Shah, James Shepherd, Antonia Ho, Alice Broos, Emma Thomson and Alex Alderton, Roberto Amato, Sonia Goncalves, Ewan Harrison, David K. Jackson, Ian Johnston, Dominic Kwiatkowski, Cordelia Langford, John Sillitoe |
| EPI_ISL_532106, EPI_ISL_532107, EPI_ISL_532108, EPI_ISL_532109, EPI_ISL_532110, EPI_ISL_532111, EPI_ISL_532112, EPI_ISL_532113, EPI_ISL_532114, EPI_ISL_532115, EPI_ISL_532116, EPI_ISL_532117 | | | |
| see above | Lighthouse Lab in Glasgow | Wellcome Sanger Institute for the COVID-19 Genomics UK (COG-UK) consortium | Harper VanSteenhouse, Yumi Kasai, David Gray, Carol Clugston, Anna Dominiczak and Alex Alderton, Roberto Amato, Sonia Goncalves, Ewan Harrison, David K. Jackson, Ian Johnston, Dominic Kwiatkowski, Cordelia Langford, John Sillitoe |
| EPI_ISL_532118 | NHSGGC West of Scotland Specialist Virology Centre / MRC-University of Glasgow Centre for Virus Research | Wellcome Sanger Institute for the COVID-19 Genomics UK (COG-UK) consortium | Ana da Silva Filipe, Natasha Johnson, Kathy Smollett, Daniel Mair, Stephen Carmichael, Lily Tong, Jenna Nichols, Elihu Aranday-Cortes, Kirstyn Brunker, Yasmin Parr, Kyriaki Nomikou; Sarah McDonald, Marc Niebel, Patawee Asamaphan; Richard Orton, Joseph Hughes, Sreenu Vattipally, David L Robertson; Alasdair MacLean, Rory Gunson; Kathy Li, Natasha Jesudason, Rajiv Shah, James Shepherd, Antonia Ho, Alice Broos, Emma Thomson and Alex Alderton, Roberto Amato, Sonia Goncalves, Ewan Harrison, David K. Jackson, Ian Johnston, Dominic Kwiatkowski, Cordelia Langford, John Sillitoe |
| EPI_ISL_532119, EPI_ISL_532120, EPI_ISL_532121, EPI_ISL_532122 | Lighthouse Lab in Glasgow | Wellcome Sanger Institute for the COVID-19 Genomics UK (COG-UK) consortium | Harper VanSteenhouse, Yumi Kasai, David Gray, Carol Clugston, Anna Dominiczak and Alex Alderton, Roberto Amato, Sonia Goncalves, Ewan Harrison, David K. Jackson, Ian Johnston, Dominic Kwiatkowski, Cordelia Langford, John Sillitoe |
| EPI_ISL_532123, EPI_ISL_532125 | NHSGGC West of Scotland Specialist Virology Centre / MRC-University of Glasgow Centre for Virus Research | Wellcome Sanger Institute for the COVID-19 Genomics UK (COG-UK) consortium | Ana da Silva Filipe, Natasha Johnson, Kathy Smollett, Daniel Mair, Stephen Carmichael, Lily Tong, Jenna Nichols, Elihu Aranday-Cortes, Kirstyn Brunker, Yasmin Parr, Kyriaki Nomikou; Sarah McDonald, Marc Niebel, Patawee Asamaphan; Richard Orton, Joseph Hughes, Sreenu Vattipally, David L Robertson; Alasdair MacLean, Rory Gunson; Kathy Li, Natasha Jesudason, Rajiv Shah, James Shepherd, Antonia Ho, Alice Broos, Emma Thomson and Alex Alderton, Roberto Amato, Sonia Goncalves, Ewan Harrison, David K. Jackson, Ian Johnston, Dominic Kwiatkowski, Cordelia Langford, John Sillitoe |
| EPI_ISL_532126, EPI_ISL_532127, EPI_ISL_532128 | Lighthouse Lab in Glasgow | Wellcome Sanger Institute for the COVID-19 Genomics UK (COG-UK) consortium | Harper VanSteenhouse, Yumi Kasai, David Gray, Carol Clugston, Anna Dominiczak and Alex Alderton, Roberto Amato, Sonia Goncalves, Ewan Harrison, David K. Jackson, Ian Johnston, Dominic Kwiatkowski, Cordelia Langford, John Sillitoe |
| EPI_ISL_532129, EPI_ISL_532130, EPI_ISL_532131 | NHSGGC West of Scotland Specialist Virology Centre / MRC-University of Glasgow Centre for Virus Research | Wellcome Sanger Institute for the COVID-19 Genomics UK (COG-UK) consortium | Ana da Silva Filipe, Natasha Johnson, Kathy Smollett, Daniel Mair, Stephen Carmichael, Lily Tong, Jenna Nichols, Elihu Aranday-Cortes, Kirstyn Brunker, Yasmin Parr, Kyriaki Nomikou; Sarah McDonald, Marc Niebel, Patawee Asamaphan; Richard Orton, Joseph Hughes, Sreenu Vattipally, David L Robertson; Alasdair MacLean, Rory Gunson; Kathy Li, Natasha Jesudason, Rajiv Shah, James Shepherd, Antonia Ho, Alice Broos, Emma Thomson and Alex Alderton, Roberto Amato, Sonia Goncalves, Ewan Harrison, David K. Jackson, Ian Johnston, Dominic Kwiatkowski, Cordelia Langford, John Sillitoe |
| EPI_ISL_532132, EPI_ISL_532133, EPI_ISL_532134, EPI_ISL_532135, EPI_ISL_532136, EPI_ISL_532137 | Lighthouse Lab in Glasgow | Wellcome Sanger Institute for the COVID-19 Genomics UK (COG-UK) consortium | Harper VanSteenhouse, Yumi Kasai, David Gray, Carol Clugston, Anna Dominiczak and Alex Alderton, Roberto Amato, Sonia Goncalves, Ewan Harrison, David K. Jackson, Ian Johnston, Dominic Kwiatkowski, Cordelia Langford, John Sillitoe |
| EPI_ISL_532138 | NHSGGC West of Scotland Specialist Virology Centre / MRC-University of Glasgow Centre for Virus Research | Wellcome Sanger Institute for the COVID-19 Genomics UK (COG-UK) consortium | Ana da Silva Filipe, Natasha Johnson, Kathy Smollett, Daniel Mair, Stephen Carmichael, Lily Tong, Jenna Nichols, Elihu Aranday-Cortes, Kirstyn Brunker, Yasmin Parr, Kyriaki Nomikou; Sarah McDonald, Marc Niebel, Patawee Asamaphan; Richard Orton, Joseph Hughes, Sreenu Vattipally, David L Robertson; Alasdair MacLean, Rory Gunson; Kathy Li, Natasha Jesudason, Rajiv Shah, James Shepherd, Antonia Ho, Alice Broos, Emma Thomson and Alex Alderton, Roberto Amato, Sonia Goncalves, Ewan Harrison, David K. Jackson, Ian Johnston, Dominic Kwiatkowski, Cordelia Langford, John Sillitoe |
| EPI_ISL_532139 | Lighthouse Lab in Glasgow | Wellcome Sanger Institute for the COVID-19 Genomics UK | Harper VanSteenhouse, Yumi Kasai, David Gray, Carol Clugston, Anna Dominiczak and Alex Alderton, Roberto Amato, Sonia Goncalves, Ewan Harrison, |

| | | | |
|---|---|---|---|
| | | (COG-UK) consortium | David K. Jackson, Ian Johnston, Dominic Kwiatkowski, Cordelia Langford, John Sillitoe |
| EPI_ISL_532283 | NHSGGC West of Scotland Specialist Virology Centre / MRC-University of Glasgow Centre for Virus Research | Wellcome Sanger Institute for the COVID-19 Genomics UK (COG-UK) consortium | Ana da Silva Filipe, Natasha Johnson, Kathy Smollett, Daniel Mair, Stephen Carmichael, Lily Tong, Jenna Nichols, Elihu Aranday-Cortes, Kirstyn Brunker, Yasmin Parr, Kyriaki Nomikou; Sarah McDonald, Marc Niebel, Patawee Asamaphan; Richard Orton, Joseph Hughes, Sreenu Vattipally, David L Robertson; Alasdair MacLean, Rory Gunson; Kathy Li, Natasha Jesudason, Rajiv Shah, James Shepherd, Antonia Ho, Alice Broos, Emma Thomson and Alex Alderton, Roberto Amato, Sonia Goncalves, Ewan Harrison, David K. Jackson, Ian Johnston, Dominic Kwiatkowski, Cordelia Langford, John Sillitoe |
| EPI_ISL_532284, EPI_ISL_532285, EPI_ISL_532286, EPI_ISL_532287 | Lighthouse Lab in Glasgow | Wellcome Sanger Institute for the COVID-19 Genomics UK (COG-UK) consortium | Harper VanSteenhouse, Yumi Kasai, David Gray, Carol Clugston, Anna Dominiczak and Alex Alderton, Roberto Amato, Sonia Goncalves, Ewan Harrison, David K. Jackson, Ian Johnston, Dominic Kwiatkowski, Cordelia Langford, John Sillitoe |
| EPI_ISL_532288 | NHSGGC West of Scotland Specialist Virology Centre / MRC-University of Glasgow Centre for Virus Research | Wellcome Sanger Institute for the COVID-19 Genomics UK (COG-UK) consortium | Ana da Silva Filipe, Natasha Johnson, Kathy Smollett, Daniel Mair, Stephen Carmichael, Lily Tong, Jenna Nichols, Elihu Aranday-Cortes, Kirstyn Brunker, Yasmin Parr, Kyriaki Nomikou; Sarah McDonald, Marc Niebel, Patawee Asamaphan; Richard Orton, Joseph Hughes, Sreenu Vattipally, David L Robertson; Alasdair MacLean, Rory Gunson; Kathy Li, Natasha Jesudason, Rajiv Shah, James Shepherd, Antonia Ho, Alice Broos, Emma Thomson and Alex Alderton, Roberto Amato, Sonia Goncalves, Ewan Harrison, David K. Jackson, Ian Johnston, Dominic Kwiatkowski, Cordelia Langford, John Sillitoe |
| EPI_ISL_532289, EPI_ISL_532290, EPI_ISL_532291, EPI_ISL_532292 | Lighthouse Lab in Glasgow | Wellcome Sanger Institute for the COVID-19 Genomics UK (COG-UK) consortium | Harper VanSteenhouse, Yumi Kasai, David Gray, Carol Clugston, Anna Dominiczak and Alex Alderton, Roberto Amato, Sonia Goncalves, Ewan Harrison, David K. Jackson, Ian Johnston, Dominic Kwiatkowski, Cordelia Langford, John Sillitoe |
| EPI_ISL_532293 | NHSGGC West of Scotland Specialist Virology Centre / MRC-University of Glasgow Centre for Virus Research | Wellcome Sanger Institute for the COVID-19 Genomics UK (COG-UK) consortium | Ana da Silva Filipe, Natasha Johnson, Kathy Smollett, Daniel Mair, Stephen Carmichael, Lily Tong, Jenna Nichols, Elihu Aranday-Cortes, Kirstyn Brunker, Yasmin Parr, Kyriaki Nomikou; Sarah McDonald, Marc Niebel, Patawee Asamaphan; Richard Orton, Joseph Hughes, Sreenu Vattipally, David L Robertson; Alasdair MacLean, Rory Gunson; Kathy Li, Natasha Jesudason, Rajiv Shah, James Shepherd, Antonia Ho, Alice Broos, Emma Thomson and Alex Alderton, Roberto Amato, Sonia Goncalves, Ewan Harrison, David K. Jackson, Ian Johnston, Dominic Kwiatkowski, Cordelia Langford, John Sillitoe |
| EPI_ISL_532294 | Lighthouse Lab in Glasgow | Wellcome Sanger Institute for the COVID-19 Genomics UK (COG-UK) Consortium | Harper VanSteenhouse, Yumi Kasai, David Gray, Carol Clugston, Anna Dominiczak and Alex Alderton, Roberto Amato, Sonia Goncalves, Ewan Harrison, David K. Jackson, Ian Johnston, Dominic Kwiatkowski, Cordelia Langford, John Sillitoe on behalf of the Wellcome Sanger Institute COVID-19 Surveillance Team |
| EPI_ISL_532295, EPI_ISL_532296, EPI_ISL_532297, EPI_ISL_532298, EPI_ISL_532299, EPI_ISL_532300 | Lighthouse Lab in Glasgow | Wellcome Sanger Institute for the COVID-19 Genomics UK (COG-UK) consortium | Harper VanSteenhouse, Yumi Kasai, David Gray, Carol Clugston, Anna Dominiczak and Alex Alderton, Roberto Amato, Sonia Goncalves, Ewan Harrison, David K. Jackson, Ian Johnston, Dominic Kwiatkowski, Cordelia Langford, John Sillitoe |
| EPI_ISL_532301 | NHSGGC West of Scotland Specialist Virology Centre / MRC-University of Glasgow Centre for Virus Research | Wellcome Sanger Institute for the COVID-19 Genomics UK (COG-UK) consortium | Ana da Silva Filipe, Natasha Johnson, Kathy Smollett, Daniel Mair, Stephen Carmichael, Lily Tong, Jenna Nichols, Elihu Aranday-Cortes, Kirstyn Brunker, Yasmin Parr, Kyriaki Nomikou; Sarah McDonald, Marc Niebel, Patawee Asamaphan; Richard Orton, Joseph Hughes, Sreenu Vattipally, David L Robertson; Alasdair MacLean, Rory Gunson; Kathy Li, Natasha Jesudason, Rajiv Shah, James Shepherd, Antonia Ho, Alice Broos, Emma Thomson and Alex Alderton, Roberto Amato, Sonia Goncalves, Ewan Harrison, David K. Jackson, Ian Johnston, Dominic Kwiatkowski, Cordelia Langford, John Sillitoe |
| EPI_ISL_532302, EPI_ISL_532303, EPI_ISL_532304, EPI_ISL_532305, EPI_ISL_532306, EPI_ISL_532307, EPI_ISL_532308, EPI_ISL_532309, EPI_ISL_532310 | Lighthouse Lab in Glasgow | Wellcome Sanger Institute for the COVID-19 Genomics UK (COG-UK) consortium | Harper VanSteenhouse, Yumi Kasai, David Gray, Carol Clugston, Anna Dominiczak and Alex Alderton, Roberto Amato, Sonia Goncalves, Ewan Harrison, David K. Jackson, Ian Johnston, Dominic Kwiatkowski, Cordelia Langford, John Sillitoe |
| EPI_ISL_532311 | NHSGGC West of Scotland Specialist Virology Centre / MRC-University of Glasgow Centre for Virus Research | Wellcome Sanger Institute for the COVID-19 Genomics UK (COG-UK) consortium | Ana da Silva Filipe, Natasha Johnson, Kathy Smollett, Daniel Mair, Stephen Carmichael, Lily Tong, Jenna Nichols, Elihu Aranday-Cortes, Kirstyn Brunker, Yasmin Parr, Kyriaki Nomikou; Sarah McDonald, Marc Niebel, Patawee Asamaphan; Richard Orton, Joseph Hughes, Sreenu Vattipally, David L Robertson; Alasdair MacLean, Rory Gunson; Kathy Li, Natasha Jesudason, Rajiv Shah, James Shepherd, Antonia Ho, Alice Broos, Emma Thomson and Alex Alderton, Roberto Amato, Sonia Goncalves, Ewan Harrison, David K. Jackson, Ian Johnston, Dominic Kwiatkowski, Cordelia Langford, John Sillitoe |
| EPI_ISL_532312, EPI_ISL_532313, EPI_ISL_532314, EPI_ISL_532315 | Lighthouse Lab in Glasgow | Wellcome Sanger Institute for the COVID-19 Genomics UK (COG-UK) consortium | Harper VanSteenhouse, Yumi Kasai, David Gray, Carol Clugston, Anna Dominiczak and Alex Alderton, Roberto Amato, Sonia Goncalves, Ewan Harrison, David K. Jackson, Ian Johnston, Dominic Kwiatkowski, Cordelia Langford, John Sillitoe |
| EPI_ISL_532316, EPI_ISL_532317 | NHSGGC West of Scotland Specialist Virology Centre / MRC-University of Glasgow Centre for Virus Research | Wellcome Sanger Institute for the COVID-19 Genomics UK (COG-UK) consortium | Ana da Silva Filipe, Natasha Johnson, Kathy Smollett, Daniel Mair, Stephen Carmichael, Lily Tong, Jenna Nichols, Elihu Aranday-Cortes, Kirstyn Brunker, Yasmin Parr, Kyriaki Nomikou; Sarah McDonald, Marc Niebel, Patawee Asamaphan; Richard Orton, Joseph Hughes, Sreenu Vattipally, David L Robertson; Alasdair MacLean, Rory Gunson; Kathy Li, Natasha Jesudason, Rajiv Shah, James Shepherd, Antonia Ho, Alice Broos, Emma Thomson and Alex Alderton, Roberto Amato, Sonia Goncalves, Ewan Harrison, David K. Jackson, Ian Johnston, Dominic Kwiatkowski, Cordelia Langford, John Sillitoe |
| EPI_ISL_532318, EPI_ISL_532319, EPI_ISL_532320 | Lighthouse Lab in Glasgow | Wellcome Sanger Institute for the COVID-19 Genomics UK (COG-UK) consortium | Harper VanSteenhouse, Yumi Kasai, David Gray, Carol Clugston, Anna Dominiczak and Alex Alderton, Roberto Amato, Sonia Goncalves, Ewan Harrison, David K. Jackson, Ian Johnston, Dominic Kwiatkowski, Cordelia Langford, John Sillitoe |
| EPI_ISL_532321 | NHSGGC West of Scotland Specialist Virology Centre / MRC-University of Glasgow Centre for Virus Research | Wellcome Sanger Institute for the COVID-19 Genomics UK (COG-UK) consortium | Ana da Silva Filipe, Natasha Johnson, Kathy Smollett, Daniel Mair, Stephen Carmichael, Lily Tong, Jenna Nichols, Elihu Aranday-Cortes, Kirstyn Brunker, Yasmin Parr, Kyriaki Nomikou; Sarah McDonald, Marc Niebel, Patawee Asamaphan; Richard Orton, Joseph Hughes, Sreenu Vattipally, David L Robertson; Alasdair MacLean, Rory Gunson; Kathy Li, Natasha Jesudason, Rajiv Shah, James Shepherd, Antonia Ho, Alice Broos, Emma Thomson and Alex Alderton, Roberto Amato, Sonia Goncalves, Ewan Harrison, David K. Jackson, Ian Johnston, Dominic Kwiatkowski, Cordelia Langford, John Sillitoe |
| EPI_ISL_532322, EPI_ISL_532323, EPI_ISL_532324, EPI_ISL_532325, EPI_ISL_532326 | Lighthouse Lab in Glasgow | Wellcome Sanger Institute for the COVID-19 Genomics UK (COG-UK) consortium | Harper VanSteenhouse, Yumi Kasai, David Gray, Carol Clugston, Anna Dominiczak and Alex Alderton, Roberto Amato, Sonia Goncalves, Ewan Harrison, David K. Jackson, Ian Johnston, Dominic Kwiatkowski, Cordelia Langford, John Sillitoe |
| EPI_ISL_532327, EPI_ISL_532328 | NHSGGC West of Scotland Specialist Virology Centre / MRC-University of Glasgow Centre for Virus Research | Wellcome Sanger Institute for the COVID-19 Genomics UK (COG-UK) consortium | Ana da Silva Filipe, Natasha Johnson, Kathy Smollett, Daniel Mair, Stephen Carmichael, Lily Tong, Jenna Nichols, Elihu Aranday-Cortes, Kirstyn Brunker, Yasmin Parr, Kyriaki Nomikou; Sarah McDonald, Marc Niebel, Patawee Asamaphan; Richard Orton, Joseph Hughes, Sreenu Vattipally, David L Robertson; Alasdair MacLean, Rory Gunson; Kathy Li, Natasha Jesudason, Rajiv Shah, James Shepherd, Antonia Ho, Alice Broos, Emma Thomson and Alex Alderton, Roberto Amato, Sonia Goncalves, Ewan Harrison, David K. Jackson, Ian Johnston, Dominic Kwiatkowski, Cordelia Langford, John Sillitoe |
| EPI_ISL_532329, EPI_ISL_532330, EPI_ISL_532331, EPI_ISL_532332, EPI_ISL_532333 | Lighthouse Lab in Glasgow | Wellcome Sanger Institute for the COVID-19 Genomics UK (COG-UK) consortium | Harper VanSteenhouse, Yumi Kasai, David Gray, Carol Clugston, Anna Dominiczak and Alex Alderton, Roberto Amato, Sonia Goncalves, Ewan Harrison, David K. Jackson, Ian Johnston, Dominic Kwiatkowski, Cordelia Langford, John Sillitoe |
| EPI_ISL_532334, EPI_ISL_532335 | NHSGGC West of Scotland Specialist Virology Centre / MRC-University of Glasgow Centre for Virus Research | Wellcome Sanger Institute for the COVID-19 Genomics UK (COG-UK) consortium | Ana da Silva Filipe, Natasha Johnson, Kathy Smollett, Daniel Mair, Stephen Carmichael, Lily Tong, Jenna Nichols, Elihu Aranday-Cortes, Kirstyn Brunker, Yasmin Parr, Kyriaki Nomikou; Sarah McDonald, Marc Niebel, Patawee Asamaphan; Richard Orton, Joseph Hughes, Sreenu Vattipally, David L Robertson; Alasdair MacLean, Rory Gunson; Kathy Li, Natasha Jesudason, Rajiv Shah, James Shepherd, Antonia Ho, Alice Broos, Emma Thomson and Alex Alderton, Roberto Amato, Sonia Goncalves, Ewan Harrison, David K. Jackson, Ian Johnston, Dominic Kwiatkowski, Cordelia Langford, John Sillitoe |
| EPI_ISL_532336, EPI_ISL_532337 | Lighthouse Lab in Glasgow | Wellcome Sanger Institute for the COVID-19 Genomics UK (COG-UK) consortium | Harper VanSteenhouse, Yumi Kasai, David Gray, Carol Clugston, Anna Dominiczak and Alex Alderton, Roberto Amato, Sonia Goncalves, Ewan Harrison, David K. Jackson, Ian Johnston, Dominic Kwiatkowski, Cordelia Langford, John Sillitoe |
| EPI_ISL_532338 | NHSGGC West of Scotland Specialist Virology Centre / MRC-University of Glasgow Centre for Virus Research | Wellcome Sanger Institute for the COVID-19 Genomics UK (COG-UK) consortium | Ana da Silva Filipe, Natasha Johnson, Kathy Smollett, Daniel Mair, Stephen Carmichael, Lily Tong, Jenna Nichols, Elihu Aranday-Cortes, Kirstyn Brunker, Yasmin Parr, Kyriaki Nomikou; Sarah McDonald, Marc Niebel, Patawee Asamaphan; Richard Orton, Joseph Hughes, Sreenu Vattipally, David L Robertson; Alasdair MacLean, Rory Gunson; Kathy Li, Natasha Jesudason, Rajiv Shah, James Shepherd, Antonia Ho, Alice Broos, Emma Thomson and Alex Alderton, Roberto Amato, Sonia Goncalves, Ewan Harrison, David K. Jackson, Ian Johnston, Dominic Kwiatkowski, Cordelia Langford, John Sillitoe |
| EPI_ISL_532339 | Lighthouse Lab in Glasgow | Wellcome Sanger Institute for the COVID-19 Genomics UK (COG-UK) consortium | Harper VanSteenhouse, Yumi Kasai, David Gray, Carol Clugston, Anna Dominiczak and Alex Alderton, Roberto Amato, Sonia Goncalves, Ewan Harrison, David K. Jackson, Ian Johnston, Dominic Kwiatkowski, Cordelia Langford, John Sillitoe |
| EPI_ISL_532340 | NHSGGC West of Scotland Specialist Virology Centre / MRC-University of Glasgow Centre for Virus Research | Wellcome Sanger Institute for the COVID-19 Genomics UK (COG-UK) consortium | Ana da Silva Filipe, Natasha Johnson, Kathy Smollett, Daniel Mair, Stephen Carmichael, Lily Tong, Jenna Nichols, Elihu Aranday-Cortes, Kirstyn Brunker, Yasmin Parr, Kyriaki Nomikou; Sarah McDonald, Marc Niebel, Patawee Asamaphan; Richard Orton, Joseph Hughes, Sreenu Vattipally, David L Robertson; Alasdair MacLean, Rory Gunson; Kathy Li, Natasha Jesudason, Rajiv Shah, James Shepherd, Antonia Ho, Alice Broos, Emma Thomson and Alex Alderton, Roberto Amato, Sonia Goncalves, Ewan Harrison, David K. Jackson, Ian Johnston, Dominic Kwiatkowski, Cordelia Langford, John Sillitoe |
| EPI_ISL_532341 | Lighthouse Lab in Glasgow | Wellcome Sanger Institute for the COVID-19 Genomics UK (COG-UK) consortium | Harper VanSteenhouse, Yumi Kasai, David Gray, Carol Clugston, Anna Dominiczak and Alex Alderton, Roberto Amato, Sonia Goncalves, Ewan Harrison, David K. Jackson, Ian Johnston, Dominic Kwiatkowski, Cordelia Langford, John Sillitoe |

| | | | |
|---|---|---|---|
| EPI_ISL_532342 | NHSGGC West of Scotland Specialist Virology Centre / MRC-University of Glasgow Centre for Virus Research | Wellcome Sanger Institute for the COVID-19 Genomics UK (COG-UK) consortium | Ana da Silva Filipe, Natasha Johnson, Kathy Smollett, Daniel Mair, Stephen Carmichael, Lily Tong, Jenna Nichols, Elihu Aranday-Cortes, Kirstyn Brunker, Yasmin Parr, Kyriaki Nomikou; Sarah McDonald, Marc Niebel, Patawee Asamaphan; Richard Orton, Joseph Hughes, Sreenu Vattipally, David L Robertson; Alasdair MacLean, Rory Gunson; Kathy Li, Natasha Jesudason, Rajiv Shah, James Shepherd, Antonia Ho, Alice Broos, Emma Thomson and Alex Alderton, Roberto Amato, Sonia Goncalves, Ewan Harrison, David K. Jackson, Ian Johnston, Dominic Kwiatkowski, Cordelia Langford, John Sillitoe |
| EPI_ISL_532343, EPI_ISL_532344, EPI_ISL_532345, EPI_ISL_532346 | Lighthouse Lab in Glasgow | Wellcome Sanger Institute for the COVID-19 Genomics UK (COG-UK) consortium | Harper VanSteenhouse, Yumi Kasai, David Gray, Carol Clugston, Anna Dominiczak and Alex Alderton, Roberto Amato, Sonia Goncalves, Ewan Harrison, David K. Jackson, Ian Johnston, Dominic Kwiatkowski, Cordelia Langford, John Sillitoe |
| EPI_ISL_532347, EPI_ISL_532349 | NHSGGC West of Scotland Specialist Virology Centre / MRC-University of Glasgow Centre for Virus Research | Wellcome Sanger Institute for the COVID-19 Genomics UK (COG-UK) consortium | Ana da Silva Filipe, Natasha Johnson, Kathy Smollett, Daniel Mair, Stephen Carmichael, Lily Tong, Jenna Nichols, Elihu Aranday-Cortes, Kirstyn Brunker, Yasmin Parr, Kyriaki Nomikou; Sarah McDonald, Marc Niebel, Patawee Asamaphan; Richard Orton, Joseph Hughes, Sreenu Vattipally, David L Robertson; Alasdair MacLean, Rory Gunson; Kathy Li, Natasha Jesudason, Rajiv Shah, James Shepherd, Antonia Ho, Alice Broos, Emma Thomson and Alex Alderton, Roberto Amato, Sonia Goncalves, Ewan Harrison, David K. Jackson, Ian Johnston, Dominic Kwiatkowski, Cordelia Langford, John Sillitoe |
| EPI_ISL_532350 | Lighthouse Lab in Glasgow | Wellcome Sanger Institute for the COVID-19 Genomics UK (COG-UK) consortium | Harper VanSteenhouse, Yumi Kasai, David Gray, Carol Clugston, Anna Dominiczak and Alex Alderton, Roberto Amato, Sonia Goncalves, Ewan Harrison, David K. Jackson, Ian Johnston, Dominic Kwiatkowski, Cordelia Langford, John Sillitoe |
| EPI_ISL_532351 | NHSGGC West of Scotland Specialist Virology Centre / MRC-University of Glasgow Centre for Virus Research | Wellcome Sanger Institute for the COVID-19 Genomics UK (COG-UK) consortium | Ana da Silva Filipe, Natasha Johnson, Kathy Smollett, Daniel Mair, Stephen Carmichael, Lily Tong, Jenna Nichols, Elihu Aranday-Cortes, Kirstyn Brunker, Yasmin Parr, Kyriaki Nomikou; Sarah McDonald, Marc Niebel, Patawee Asamaphan; Richard Orton, Joseph Hughes, Sreenu Vattipally, David L Robertson; Alasdair MacLean, Rory Gunson; Kathy Li, Natasha Jesudason, Rajiv Shah, James Shepherd, Antonia Ho, Alice Broos, Emma Thomson and Alex Alderton, Roberto Amato, Sonia Goncalves, Ewan Harrison, David K. Jackson, Ian Johnston, Dominic Kwiatkowski, Cordelia Langford, John Sillitoe |
| EPI_ISL_532352 | Lighthouse Lab in Glasgow | Wellcome Sanger Institute for the COVID-19 Genomics UK (COG-UK) consortium | Harper VanSteenhouse, Yumi Kasai, David Gray, Carol Clugston, Anna Dominiczak and Alex Alderton, Roberto Amato, Sonia Goncalves, Ewan Harrison, David K. Jackson, Ian Johnston, Dominic Kwiatkowski, Cordelia Langford, John Sillitoe |
| EPI_ISL_532353 | NHSGGC West of Scotland Specialist Virology Centre / MRC-University of Glasgow Centre for Virus Research | Wellcome Sanger Institute for the COVID-19 Genomics UK (COG-UK) consortium | Ana da Silva Filipe, Natasha Johnson, Kathy Smollett, Daniel Mair, Stephen Carmichael, Lily Tong, Jenna Nichols, Elihu Aranday-Cortes, Kirstyn Brunker, Yasmin Parr, Kyriaki Nomikou; Sarah McDonald, Marc Niebel, Patawee Asamaphan; Richard Orton, Joseph Hughes, Sreenu Vattipally, David L Robertson; Alasdair MacLean, Rory Gunson; Kathy Li, Natasha Jesudason, Rajiv Shah, James Shepherd, Antonia Ho, Alice Broos, Emma Thomson and Alex Alderton, Roberto Amato, Sonia Goncalves, Ewan Harrison, David K. Jackson, Ian Johnston, Dominic Kwiatkowski, Cordelia Langford, John Sillitoe |
| EPI_ISL_532354 | Lighthouse Lab in Glasgow | Wellcome Sanger Institute for the COVID-19 Genomics UK (COG-UK) consortium | Harper VanSteenhouse, Yumi Kasai, David Gray, Carol Clugston, Anna Dominiczak and Alex Alderton, Roberto Amato, Sonia Goncalves, Ewan Harrison, David K. Jackson, Ian Johnston, Dominic Kwiatkowski, Cordelia Langford, John Sillitoe |
| EPI_ISL_532355, EPI_ISL_532356 | NHSGGC West of Scotland Specialist Virology Centre / MRC-University of Glasgow Centre for Virus Research | Wellcome Sanger Institute for the COVID-19 Genomics UK (COG-UK) consortium | Ana da Silva Filipe, Natasha Johnson, Kathy Smollett, Daniel Mair, Stephen Carmichael, Lily Tong, Jenna Nichols, Elihu Aranday-Cortes, Kirstyn Brunker, Yasmin Parr, Kyriaki Nomikou; Sarah McDonald, Marc Niebel, Patawee Asamaphan; Richard Orton, Joseph Hughes, Sreenu Vattipally, David L Robertson; Alasdair MacLean, Rory Gunson; Kathy Li, Natasha Jesudason, Rajiv Shah, James Shepherd, Antonia Ho, Alice Broos, Emma Thomson and Alex Alderton, Roberto Amato, Sonia Goncalves, Ewan Harrison, David K. Jackson, Ian Johnston, Dominic Kwiatkowski, Cordelia Langford, John Sillitoe |
| EPI_ISL_532357, EPI_ISL_532358, EPI_ISL_532359, EPI_ISL_532360, EPI_ISL_532361, EPI_ISL_532362 | Lighthouse Lab in Glasgow | Wellcome Sanger Institute for the COVID-19 Genomics UK (COG-UK) consortium | Harper VanSteenhouse, Yumi Kasai, David Gray, Carol Clugston, Anna Dominiczak and Alex Alderton, Roberto Amato, Sonia Goncalves, Ewan Harrison, David K. Jackson, Ian Johnston, Dominic Kwiatkowski, Cordelia Langford, John Sillitoe |
| EPI_ISL_532363, EPI_ISL_532364 | NHSGGC West of Scotland Specialist Virology Centre / MRC-University of Glasgow Centre for Virus Research | Wellcome Sanger Institute for the COVID-19 Genomics UK (COG-UK) consortium | Ana da Silva Filipe, Natasha Johnson, Kathy Smollett, Daniel Mair, Stephen Carmichael, Lily Tong, Jenna Nichols, Elihu Aranday-Cortes, Kirstyn Brunker, Yasmin Parr, Kyriaki Nomikou; Sarah McDonald, Marc Niebel, Patawee Asamaphan; Richard Orton, Joseph Hughes, Sreenu Vattipally, David L Robertson; Alasdair MacLean, Rory Gunson; Kathy Li, Natasha Jesudason, Rajiv Shah, James Shepherd, Antonia Ho, Alice Broos, Emma Thomson and Alex Alderton, Roberto Amato, Sonia Goncalves, Ewan Harrison, David K. Jackson, Ian Johnston, Dominic Kwiatkowski, Cordelia Langford, John Sillitoe |
| EPI_ISL_532365 | Lighthouse Lab in Glasgow | Wellcome Sanger Institute for the COVID-19 Genomics UK (COG-UK) consortium | Harper VanSteenhouse, Yumi Kasai, David Gray, Carol Clugston, Anna Dominiczak and Alex Alderton, Roberto Amato, Sonia Goncalves, Ewan Harrison, David K. Jackson, Ian Johnston, Dominic Kwiatkowski, Cordelia Langford, John Sillitoe on behalf of the Wellcome Sanger Institute COVID-19 Surveillance Team (http://www.sanger.ac.uk/covid-team) |
| EPI_ISL_532366, EPI_ISL_532367, EPI_ISL_532368, EPI_ISL_532369, EPI_ISL_532370, EPI_ISL_532371, EPI_ISL_532372, EPI_ISL_532373, EPI_ISL_532374, EPI_ISL_532375, EPI_ISL_532376, EPI_ISL_532377, EPI_ISL_532378 | | | |
| see above | Lighthouse Lab in Glasgow | Wellcome Sanger Institute for the COVID-19 Genomics UK (COG-UK) consortium | Harper VanSteenhouse, Yumi Kasai, David Gray, Carol Clugston, Anna Dominiczak and Alex Alderton, Roberto Amato, Sonia Goncalves, Ewan Harrison, David K. Jackson, Ian Johnston, Dominic Kwiatkowski, Cordelia Langford, John Sillitoe |
| EPI_ISL_532379 | NHSGGC West of Scotland Specialist Virology Centre / MRC-University of Glasgow Centre for Virus Research | Wellcome Sanger Institute for the COVID-19 Genomics UK (COG-UK) consortium | Ana da Silva Filipe, Natasha Johnson, Kathy Smollett, Daniel Mair, Stephen Carmichael, Lily Tong, Jenna Nichols, Elihu Aranday-Cortes, Kirstyn Brunker, Yasmin Parr, Kyriaki Nomikou; Sarah McDonald, Marc Niebel, Patawee Asamaphan; Richard Orton, Joseph Hughes, Sreenu Vattipally, David L Robertson; Alasdair MacLean, Rory Gunson; Kathy Li, Natasha Jesudason, Rajiv Shah, James Shepherd, Antonia Ho, Alice Broos, Emma Thomson and Alex Alderton, Roberto Amato, Sonia Goncalves, Ewan Harrison, David K. Jackson, Ian Johnston, Dominic Kwiatkowski, Cordelia Langford, John Sillitoe |
| EPI_ISL_532380, EPI_ISL_532381, EPI_ISL_532382, EPI_ISL_532383 | Lighthouse Lab in Glasgow | Wellcome Sanger Institute for the COVID-19 Genomics UK (COG-UK) consortium | Harper VanSteenhouse, Yumi Kasai, David Gray, Carol Clugston, Anna Dominiczak and Alex Alderton, Roberto Amato, Sonia Goncalves, Ewan Harrison, David K. Jackson, Ian Johnston, Dominic Kwiatkowski, Cordelia Langford, John Sillitoe |
| EPI_ISL_532384 | NHSGGC West of Scotland Specialist Virology Centre / MRC-University of Glasgow Centre for Virus Research | Wellcome Sanger Institute for the COVID-19 Genomics UK (COG-UK) consortium | Ana da Silva Filipe, Natasha Johnson, Kathy Smollett, Daniel Mair, Stephen Carmichael, Lily Tong, Jenna Nichols, Elihu Aranday-Cortes, Kirstyn Brunker, Yasmin Parr, Kyriaki Nomikou; Sarah McDonald, Marc Niebel, Patawee Asamaphan; Richard Orton, Joseph Hughes, Sreenu Vattipally, David L Robertson; Alasdair MacLean, Rory Gunson; Kathy Li, Natasha Jesudason, Rajiv Shah, James Shepherd, Antonia Ho, Alice Broos, Emma Thomson and Alex Alderton, Roberto Amato, Sonia Goncalves, Ewan Harrison, David K. Jackson, Ian Johnston, Dominic Kwiatkowski, Cordelia Langford, John Sillitoe |
| EPI_ISL_532385, EPI_ISL_532386, EPI_ISL_532387 | Lighthouse Lab in Glasgow | Wellcome Sanger Institute for the COVID-19 Genomics UK (COG-UK) consortium | Harper VanSteenhouse, Yumi Kasai, David Gray, Carol Clugston, Anna Dominiczak and Alex Alderton, Roberto Amato, Sonia Goncalves, Ewan Harrison, David K. Jackson, Ian Johnston, Dominic Kwiatkowski, Cordelia Langford, John Sillitoe |
| EPI_ISL_532388 | NHSGGC West of Scotland Specialist Virology Centre / MRC-University of Glasgow Centre for Virus Research | Wellcome Sanger Institute for the COVID-19 Genomics UK (COG-UK) consortium | Ana da Silva Filipe, Natasha Johnson, Kathy Smollett, Daniel Mair, Stephen Carmichael, Lily Tong, Jenna Nichols, Elihu Aranday-Cortes, Kirstyn Brunker, Yasmin Parr, Kyriaki Nomikou; Sarah McDonald, Marc Niebel, Patawee Asamaphan; Richard Orton, Joseph Hughes, Sreenu Vattipally, David L Robertson; Alasdair MacLean, Rory Gunson; Kathy Li, Natasha Jesudason, Rajiv Shah, James Shepherd, Antonia Ho, Alice Broos, Emma Thomson and Alex Alderton, Roberto Amato, Sonia Goncalves, Ewan Harrison, David K. Jackson, Ian Johnston, Dominic Kwiatkowski, Cordelia Langford, John Sillitoe |
| EPI_ISL_532389 | Lighthouse Lab in Glasgow | Wellcome Sanger Institute for the COVID-19 Genomics UK (COG-UK) consortium | Harper VanSteenhouse, Yumi Kasai, David Gray, Carol Clugston, Anna Dominiczak and Alex Alderton, Roberto Amato, Sonia Goncalves, Ewan Harrison, David K. Jackson, Ian Johnston, Dominic Kwiatkowski, Cordelia Langford, John Sillitoe |
| EPI_ISL_532390 | NHSGGC West of Scotland Specialist Virology Centre / MRC-University of Glasgow Centre for Virus Research | Wellcome Sanger Institute for the COVID-19 Genomics UK (COG-UK) consortium | Ana da Silva Filipe, Natasha Johnson, Kathy Smollett, Daniel Mair, Stephen Carmichael, Lily Tong, Jenna Nichols, Elihu Aranday-Cortes, Kirstyn Brunker, Yasmin Parr, Kyriaki Nomikou; Sarah McDonald, Marc Niebel, Patawee Asamaphan; Richard Orton, Joseph Hughes, Sreenu Vattipally, David L Robertson; Alasdair MacLean, Rory Gunson; Kathy Li, Natasha Jesudason, Rajiv Shah, James Shepherd, Antonia Ho, Alice Broos, Emma Thomson and Alex Alderton, Roberto Amato, Sonia Goncalves, Ewan Harrison, David K. Jackson, Ian Johnston, Dominic Kwiatkowski, Cordelia Langford, John Sillitoe |
| EPI_ISL_532391, EPI_ISL_532392, EPI_ISL_532393 | Lighthouse Lab in Glasgow | Wellcome Sanger Institute for the COVID-19 Genomics UK (COG-UK) consortium | Harper VanSteenhouse, Yumi Kasai, David Gray, Carol Clugston, Anna Dominiczak and Alex Alderton, Roberto Amato, Sonia Goncalves, Ewan Harrison, David K. Jackson, Ian Johnston, Dominic Kwiatkowski, Cordelia Langford, John Sillitoe |
| EPI_ISL_532394 | NHSGGC West of Scotland Specialist Virology Centre / MRC-University of Glasgow Centre for Virus Research | Wellcome Sanger Institute for the COVID-19 Genomics UK (COG-UK) consortium | Ana da Silva Filipe, Natasha Johnson, Kathy Smollett, Daniel Mair, Stephen Carmichael, Lily Tong, Jenna Nichols, Elihu Aranday-Cortes, Kirstyn Brunker, Yasmin Parr, Kyriaki Nomikou; Sarah McDonald, Marc Niebel, Patawee Asamaphan; Richard Orton, Joseph Hughes, Sreenu Vattipally, David L Robertson; Alasdair MacLean, Rory Gunson; Kathy Li, Natasha Jesudason, Rajiv Shah, James Shepherd, Antonia Ho, Alice Broos, Emma Thomson and Alex Alderton, Roberto Amato, Sonia Goncalves, Ewan Harrison, David K. Jackson, Ian Johnston, Dominic Kwiatkowski, Cordelia Langford, John Sillitoe |
| EPI_ISL_532395, EPI_ISL_532396, EPI_ISL_532397, EPI_ISL_532398 | Lighthouse Lab in Glasgow | Wellcome Sanger Institute for the COVID-19 Genomics UK (COG-UK) consortium | Harper VanSteenhouse, Yumi Kasai, David Gray, Carol Clugston, Anna Dominiczak and Alex Alderton, Roberto Amato, Sonia Goncalves, Ewan Harrison, David K. Jackson, Ian Johnston, Dominic Kwiatkowski, Cordelia Langford, John Sillitoe |
| EPI_ISL_532399 | NHSGGC West of Scotland Specialist Virology Centre / MRC-University of Glasgow Centre for Virus Research | Wellcome Sanger Institute for the COVID-19 Genomics UK (COG-UK) consortium | Ana da Silva Filipe, Natasha Johnson, Kathy Smollett, Daniel Mair, Stephen Carmichael, Lily Tong, Jenna Nichols, Elihu Aranday-Cortes, Kirstyn Brunker, Yasmin Parr, Kyriaki Nomikou; Sarah McDonald, Marc Niebel, Patawee Asamaphan; Richard Orton, Joseph Hughes, Sreenu Vattipally, David L Robertson; Alasdair MacLean, Rory Gunson; Kathy Li, Natasha Jesudason, Rajiv Shah, James Shepherd, Antonia Ho, Alice Broos, Emma Thomson and Alex Alderton, Roberto Amato, Sonia Goncalves, Ewan Harrison, David K. Jackson, Ian Johnston, Dominic Kwiatkowski, Cordelia Langford, John Sillitoe |

| | | | |
|---|---|---|---|
| EPI_ISL_532400, EPI_ISL_532401 | Lighthouse Lab in Glasgow | Wellcome Sanger Institute for the COVID-19 Genomics UK (COG-UK) consortium | Harper VanSteenhouse, Yumi Kasai, David Gray, Carol Clugston, Anna Dominiczak and Alex Alderton, Roberto Amato, Sonia Goncalves, Ewan Harrison, David K. Jackson, Ian Johnston, Dominic Kwiatkowski, Cordelia Langford, John Sillitoe |
| EPI_ISL_532402 | NHSGGC West of Scotland Specialist Virology Centre / MRC-University of Glasgow Centre for Virus Research | Wellcome Sanger Institute for the COVID-19 Genomics UK (COG-UK) consortium | Ana da Silva Filipe, Natasha Johnson, Kathy Smollett, Daniel Mair, Stephen Carmichael, Lily Tong, Jenna Nichols, Elihu Aranday-Cortes, Kirstyn Brunker, Yasmin Parr, Kyriaki Nomikou; Sarah McDonald, Marc Niebel, Patawee Asamaphan; Richard Orton, Joseph Hughes, Sreenu Vattipally, David L Robertson; Alasdair MacLean, Rory Gunson; Kathy Li, Natasha Jesudason, Rajiv Shah, James Shepherd, Antonia Ho, Alice Broos, Emma Thomson and Alex Alderton, Roberto Amato, Sonia Goncalves, Ewan Harrison, David K. Jackson, Ian Johnston, Dominic Kwiatkowski, Cordelia Langford, John Sillitoe |
| EPI_ISL_532403 | Lighthouse Lab in Glasgow | Wellcome Sanger Institute for the COVID-19 Genomics UK (COG-UK) consortium | Harper VanSteenhouse, Yumi Kasai, David Gray, Carol Clugston, Anna Dominiczak and Alex Alderton, Roberto Amato, Sonia Goncalves, Ewan Harrison, David K. Jackson, Ian Johnston, Dominic Kwiatkowski, Cordelia Langford, John Sillitoe |
| EPI_ISL_532404 | NHSGGC West of Scotland Specialist Virology Centre / MRC-University of Glasgow Centre for Virus Research | Wellcome Sanger Institute for the COVID-19 Genomics UK (COG-UK) consortium | Ana da Silva Filipe, Natasha Johnson, Kathy Smollett, Daniel Mair, Stephen Carmichael, Lily Tong, Jenna Nichols, Elihu Aranday-Cortes, Kirstyn Brunker, Yasmin Parr, Kyriaki Nomikou; Sarah McDonald, Marc Niebel, Patawee Asamaphan; Richard Orton, Joseph Hughes, Sreenu Vattipally, David L Robertson; Alasdair MacLean, Rory Gunson; Kathy Li, Natasha Jesudason, Rajiv Shah, James Shepherd, Antonia Ho, Alice Broos, Emma Thomson and Alex Alderton, Roberto Amato, Sonia Goncalves, Ewan Harrison, David K. Jackson, Ian Johnston, Dominic Kwiatkowski, Cordelia Langford, John Sillitoe |
| EPI_ISL_532405, EPI_ISL_532406, EPI_ISL_532407, EPI_ISL_532408, EPI_ISL_532409, EPI_ISL_532410, EPI_ISL_532411, EPI_ISL_532412, EPI_ISL_532413 | Lighthouse Lab in Glasgow | Wellcome Sanger Institute for the COVID-19 Genomics UK (COG-UK) consortium | Harper VanSteenhouse, Yumi Kasai, David Gray, Carol Clugston, Anna Dominiczak and Alex Alderton, Roberto Amato, Sonia Goncalves, Ewan Harrison, David K. Jackson, Ian Johnston, Dominic Kwiatkowski, Cordelia Langford, John Sillitoe |
| EPI_ISL_532414 | NHSGGC West of Scotland Specialist Virology Centre / MRC-University of Glasgow Centre for Virus Research | Wellcome Sanger Institute for the COVID-19 Genomics UK (COG-UK) consortium | Ana da Silva Filipe, Natasha Johnson, Kathy Smollett, Daniel Mair, Stephen Carmichael, Lily Tong, Jenna Nichols, Elihu Aranday-Cortes, Kirstyn Brunker, Yasmin Parr, Kyriaki Nomikou; Sarah McDonald, Marc Niebel, Patawee Asamaphan; Richard Orton, Joseph Hughes, Sreenu Vattipally, David L Robertson; Alasdair MacLean, Rory Gunson; Kathy Li, Natasha Jesudason, Rajiv Shah, James Shepherd, Antonia Ho, Alice Broos, Emma Thomson and Alex Alderton, Roberto Amato, Sonia Goncalves, Ewan Harrison, David K. Jackson, Ian Johnston, Dominic Kwiatkowski, Cordelia Langford, John Sillitoe |
| EPI_ISL_532415, EPI_ISL_532416, EPI_ISL_532417, EPI_ISL_532418, EPI_ISL_532419, EPI_ISL_532420, EPI_ISL_532421, EPI_ISL_532422, EPI_ISL_532423, EPI_ISL_532424, EPI_ISL_532425, EPI_ISL_532426, EPI_ISL_532427, EPI_ISL_532428, EPI_ISL_532429 | | | |
| see above | Lighthouse Lab in Glasgow | Wellcome Sanger Institute for the COVID-19 Genomics UK (COG-UK) consortium | Harper VanSteenhouse, Yumi Kasai, David Gray, Carol Clugston, Anna Dominiczak and Alex Alderton, Roberto Amato, Sonia Goncalves, Ewan Harrison, David K. Jackson, Ian Johnston, Dominic Kwiatkowski, Cordelia Langford, John Sillitoe |
| EPI_ISL_532430, EPI_ISL_532431, EPI_ISL_532432 | NHSGGC West of Scotland Specialist Virology Centre / MRC-University of Glasgow Centre for Virus Research | Wellcome Sanger Institute for the COVID-19 Genomics UK (COG-UK) consortium | Ana da Silva Filipe, Natasha Johnson, Kathy Smollett, Daniel Mair, Stephen Carmichael, Lily Tong, Jenna Nichols, Elihu Aranday-Cortes, Kirstyn Brunker, Yasmin Parr, Kyriaki Nomikou; Sarah McDonald, Marc Niebel, Patawee Asamaphan; Richard Orton, Joseph Hughes, Sreenu Vattipally, David L Robertson; Alasdair MacLean, Rory Gunson; Kathy Li, Natasha Jesudason, Rajiv Shah, James Shepherd, Antonia Ho, Alice Broos, Emma Thomson and Alex Alderton, Roberto Amato, Sonia Goncalves, Ewan Harrison, David K. Jackson, Ian Johnston, Dominic Kwiatkowski, Cordelia Langford, John Sillitoe |
| EPI_ISL_532433, EPI_ISL_532434, EPI_ISL_532435, EPI_ISL_532436, EPI_ISL_532437 | Lighthouse Lab in Glasgow | Wellcome Sanger Institute for the COVID-19 Genomics UK (COG-UK) consortium | Harper VanSteenhouse, Yumi Kasai, David Gray, Carol Clugston, Anna Dominiczak and Alex Alderton, Roberto Amato, Sonia Goncalves, Ewan Harrison, David K. Jackson, Ian Johnston, Dominic Kwiatkowski, Cordelia Langford, John Sillitoe |
| EPI_ISL_532438 | NHSGGC West of Scotland Specialist Virology Centre / MRC-University of Glasgow Centre for Virus Research | Wellcome Sanger Institute for the COVID-19 Genomics UK (COG-UK) consortium | Ana da Silva Filipe, Natasha Johnson, Kathy Smollett, Daniel Mair, Stephen Carmichael, Lily Tong, Jenna Nichols, Elihu Aranday-Cortes, Kirstyn Brunker, Yasmin Parr, Kyriaki Nomikou; Sarah McDonald, Marc Niebel, Patawee Asamaphan; Richard Orton, Joseph Hughes, Sreenu Vattipally, David L Robertson; Alasdair MacLean, Rory Gunson; Kathy Li, Natasha Jesudason, Rajiv Shah, James Shepherd, Antonia Ho, Alice Broos, Emma Thomson and Alex Alderton, Roberto Amato, Sonia Goncalves, Ewan Harrison, David K. Jackson, Ian Johnston, Dominic Kwiatkowski, Cordelia Langford, John Sillitoe |
| EPI_ISL_532439 | Lighthouse Lab in Glasgow | Wellcome Sanger Institute for the COVID-19 Genomics UK (COG-UK) consortium | Harper VanSteenhouse, Yumi Kasai, David Gray, Carol Clugston, Anna Dominiczak and Alex Alderton, Roberto Amato, Sonia Goncalves, Ewan Harrison, David K. Jackson, Ian Johnston, Dominic Kwiatkowski, Cordelia Langford, John Sillitoe |
| EPI_ISL_532440 | NHSGGC West of Scotland Specialist Virology Centre / MRC-University of Glasgow Centre for Virus Research | Wellcome Sanger Institute for the COVID-19 Genomics UK (COG-UK) consortium | Ana da Silva Filipe, Natasha Johnson, Kathy Smollett, Daniel Mair, Stephen Carmichael, Lily Tong, Jenna Nichols, Elihu Aranday-Cortes, Kirstyn Brunker, Yasmin Parr, Kyriaki Nomikou; Sarah McDonald, Marc Niebel, Patawee Asamaphan; Richard Orton, Joseph Hughes, Sreenu Vattipally, David L Robertson; Alasdair MacLean, Rory Gunson; Kathy Li, Natasha Jesudason, Rajiv Shah, James Shepherd, Antonia Ho, Alice Broos, Emma Thomson and Alex Alderton, Roberto Amato, Sonia Goncalves, Ewan Harrison, David K. Jackson, Ian Johnston, Dominic Kwiatkowski, Cordelia Langford, John Sillitoe |
| EPI_ISL_532441, EPI_ISL_532442, EPI_ISL_532443 | Lighthouse Lab in Glasgow | Wellcome Sanger Institute for the COVID-19 Genomics UK (COG-UK) consortium | Harper VanSteenhouse, Yumi Kasai, David Gray, Carol Clugston, Anna Dominiczak and Alex Alderton, Roberto Amato, Sonia Goncalves, Ewan Harrison, David K. Jackson, Ian Johnston, Dominic Kwiatkowski, Cordelia Langford, John Sillitoe |
| EPI_ISL_532444 | NHSGGC West of Scotland Specialist Virology Centre / MRC-University of Glasgow Centre for Virus Research | Wellcome Sanger Institute for the COVID-19 Genomics UK (COG-UK) consortium | Ana da Silva Filipe, Natasha Johnson, Kathy Smollett, Daniel Mair, Stephen Carmichael, Lily Tong, Jenna Nichols, Elihu Aranday-Cortes, Kirstyn Brunker, Yasmin Parr, Kyriaki Nomikou; Sarah McDonald, Marc Niebel, Patawee Asamaphan; Richard Orton, Joseph Hughes, Sreenu Vattipally, David L Robertson; Alasdair MacLean, Rory Gunson; Kathy Li, Natasha Jesudason, Rajiv Shah, James Shepherd, Antonia Ho, Alice Broos, Emma Thomson and Alex Alderton, Roberto Amato, Sonia Goncalves, Ewan Harrison, David K. Jackson, Ian Johnston, Dominic Kwiatkowski, Cordelia Langford, John Sillitoe |
| EPI_ISL_532445 | Lighthouse Lab in Glasgow | Wellcome Sanger Institute for the COVID-19 Genomics UK (COG-UK) consortium | Harper VanSteenhouse, Yumi Kasai, David Gray, Carol Clugston, Anna Dominiczak and Alex Alderton, Roberto Amato, Sonia Goncalves, Ewan Harrison, David K. Jackson, Ian Johnston, Dominic Kwiatkowski, Cordelia Langford, John Sillitoe |
| EPI_ISL_532446 | NHSGGC West of Scotland Specialist Virology Centre / MRC-University of Glasgow Centre for Virus Research | Wellcome Sanger Institute for the COVID-19 Genomics UK (COG-UK) consortium | Ana da Silva Filipe, Natasha Johnson, Kathy Smollett, Daniel Mair, Stephen Carmichael, Lily Tong, Jenna Nichols, Elihu Aranday-Cortes, Kirstyn Brunker, Yasmin Parr, Kyriaki Nomikou; Sarah McDonald, Marc Niebel, Patawee Asamaphan; Richard Orton, Joseph Hughes, Sreenu Vattipally, David L Robertson; Alasdair MacLean, Rory Gunson; Kathy Li, Natasha Jesudason, Rajiv Shah, James Shepherd, Antonia Ho, Alice Broos, Emma Thomson and Alex Alderton, Roberto Amato, Sonia Goncalves, Ewan Harrison, David K. Jackson, Ian Johnston, Dominic Kwiatkowski, Cordelia Langford, John Sillitoe |
| EPI_ISL_532447, EPI_ISL_532448, EPI_ISL_532449, EPI_ISL_532450, EPI_ISL_532451 | Lighthouse Lab in Glasgow | Wellcome Sanger Institute for the COVID-19 Genomics UK (COG-UK) consortium | Harper VanSteenhouse, Yumi Kasai, David Gray, Carol Clugston, Anna Dominiczak and Alex Alderton, Roberto Amato, Sonia Goncalves, Ewan Harrison, David K. Jackson, Ian Johnston, Dominic Kwiatkowski, Cordelia Langford, John Sillitoe |
| EPI_ISL_532452 | NHSGGC West of Scotland Specialist Virology Centre / MRC-University of Glasgow Centre for Virus Research | Wellcome Sanger Institute for the COVID-19 Genomics UK (COG-UK) consortium | Ana da Silva Filipe, Natasha Johnson, Kathy Smollett, Daniel Mair, Stephen Carmichael, Lily Tong, Jenna Nichols, Elihu Aranday-Cortes, Kirstyn Brunker, Yasmin Parr, Kyriaki Nomikou; Sarah McDonald, Marc Niebel, Patawee Asamaphan; Richard Orton, Joseph Hughes, Sreenu Vattipally, David L Robertson; Alasdair MacLean, Rory Gunson; Kathy Li, Natasha Jesudason, Rajiv Shah, James Shepherd, Antonia Ho, Alice Broos, Emma Thomson and Alex Alderton, Roberto Amato, Sonia Goncalves, Ewan Harrison, David K. Jackson, Ian Johnston, Dominic Kwiatkowski, Cordelia Langford, John Sillitoe |
| EPI_ISL_532453 | Lighthouse Lab in Glasgow | Wellcome Sanger Institute for the COVID-19 Genomics UK (COG-UK) consortium | Harper VanSteenhouse, Yumi Kasai, David Gray, Carol Clugston, Anna Dominiczak and Alex Alderton, Roberto Amato, Sonia Goncalves, Ewan Harrison, David K. Jackson, Ian Johnston, Dominic Kwiatkowski, Cordelia Langford, John Sillitoe |
| EPI_ISL_532454 | NHSGGC West of Scotland Specialist Virology Centre / MRC-University of Glasgow Centre for Virus Research | Wellcome Sanger Institute for the COVID-19 Genomics UK (COG-UK) consortium | Ana da Silva Filipe, Natasha Johnson, Kathy Smollett, Daniel Mair, Stephen Carmichael, Lily Tong, Jenna Nichols, Elihu Aranday-Cortes, Kirstyn Brunker, Yasmin Parr, Kyriaki Nomikou; Sarah McDonald, Marc Niebel, Patawee Asamaphan; Richard Orton, Joseph Hughes, Sreenu Vattipally, David L Robertson; Alasdair MacLean, Rory Gunson; Kathy Li, Natasha Jesudason, Rajiv Shah, James Shepherd, Antonia Ho, Alice Broos, Emma Thomson and Alex Alderton, Roberto Amato, Sonia Goncalves, Ewan Harrison, David K. Jackson, Ian Johnston, Dominic Kwiatkowski, Cordelia Langford, John Sillitoe |
| EPI_ISL_532455 | Lighthouse Lab in Glasgow | Wellcome Sanger Institute for the COVID-19 Genomics UK (COG-UK) consortium | Harper VanSteenhouse, Yumi Kasai, David Gray, Carol Clugston, Anna Dominiczak and Alex Alderton, Roberto Amato, Sonia Goncalves, Ewan Harrison, David K. Jackson, Ian Johnston, Dominic Kwiatkowski, Cordelia Langford, John Sillitoe |
| EPI_ISL_532456 | NHSGGC West of Scotland Specialist Virology Centre / MRC-University of Glasgow Centre for Virus Research | Wellcome Sanger Institute for the COVID-19 Genomics UK (COG-UK) consortium | Ana da Silva Filipe, Natasha Johnson, Kathy Smollett, Daniel Mair, Stephen Carmichael, Lily Tong, Jenna Nichols, Elihu Aranday-Cortes, Kirstyn Brunker, Yasmin Parr, Kyriaki Nomikou; Sarah McDonald, Marc Niebel, Patawee Asamaphan; Richard Orton, Joseph Hughes, Sreenu Vattipally, David L Robertson; Alasdair MacLean, Rory Gunson; Kathy Li, Natasha Jesudason, Rajiv Shah, James Shepherd, Antonia Ho, Alice Broos, Emma Thomson and Alex Alderton, Roberto Amato, Sonia Goncalves, Ewan Harrison, David K. Jackson, Ian Johnston, Dominic Kwiatkowski, Cordelia Langford, John Sillitoe |
| EPI_ISL_532457, EPI_ISL_532458, EPI_ISL_532459, EPI_ISL_532460, EPI_ISL_532461, EPI_ISL_532462 | Lighthouse Lab in Glasgow | Wellcome Sanger Institute for the COVID-19 Genomics UK (COG-UK) consortium | Harper VanSteenhouse, Yumi Kasai, David Gray, Carol Clugston, Anna Dominiczak and Alex Alderton, Roberto Amato, Sonia Goncalves, Ewan Harrison, David K. Jackson, Ian Johnston, Dominic Kwiatkowski, Cordelia Langford, John Sillitoe |

| | | | |
|---|---|---|---|
| EPI_ISL_532463, EPI_ISL_532464 | NHSGGC West of Scotland Specialist Virology Centre / MRC-University of Glasgow Centre for Virus Research | Wellcome Sanger Institute for the COVID-19 Genomics UK (COG-UK) consortium | Ana da Silva Filipe, Natasha Johnson, Kathy Smollett, Daniel Mair, Stephen Carmichael, Lily Tong, Jenna Nichols, Elihu Aranday-Cortes, Kirstyn Brunker, Yasmin Parr, Kyriaki Nomikou; Sarah McDonald, Marc Niebel, Patawee Asamaphan; Richard Orton, Joseph Hughes, Sreenu Vattipally, David L Robertson; Alasdair MacLean, Rory Gunson; Kathy Li, Natasha Jesudason, Rajiv Shah, James Shepherd, Antonia Ho, Alice Broos, Emma Thomson and Alex Alderton, Roberto Amato, Sonia Goncalves, Ewan Harrison, David K. Jackson, Ian Johnston, Dominic Kwiatkowski, Cordelia Langford, John Sillitoe |
| EPI_ISL_532465, EPI_ISL_532466, EPI_ISL_532467, EPI_ISL_532468 | Lighthouse Lab in Glasgow | Wellcome Sanger Institute for the COVID-19 Genomics UK (COG-UK) consortium | Harper VanSteenhouse, Yumi Kasai, David Gray, Carol Clugston, Anna Dominiczak and Alex Alderton, Roberto Amato, Sonia Goncalves, Ewan Harrison, David K. Jackson, Ian Johnston, Dominic Kwiatkowski, Cordelia Langford, John Sillitoe |
| EPI_ISL_532469 | NHSGGC West of Scotland Specialist Virology Centre / MRC-University of Glasgow Centre for Virus Research | Wellcome Sanger Institute for the COVID-19 Genomics UK (COG-UK) consortium | Ana da Silva Filipe, Natasha Johnson, Kathy Smollett, Daniel Mair, Stephen Carmichael, Lily Tong, Jenna Nichols, Elihu Aranday-Cortes, Kirstyn Brunker, Yasmin Parr, Kyriaki Nomikou; Sarah McDonald, Marc Niebel, Patawee Asamaphan; Richard Orton, Joseph Hughes, Sreenu Vattipally, David L Robertson; Alasdair MacLean, Rory Gunson; Kathy Li, Natasha Jesudason, Rajiv Shah, James Shepherd, Antonia Ho, Alice Broos, Emma Thomson and Alex Alderton, Roberto Amato, Sonia Goncalves, Ewan Harrison, David K. Jackson, Ian Johnston, Dominic Kwiatkowski, Cordelia Langford, John Sillitoe |
| EPI_ISL_532470, EPI_ISL_532471 | Lighthouse Lab in Glasgow | Wellcome Sanger Institute for the COVID-19 Genomics UK (COG-UK) consortium | Harper VanSteenhouse, Yumi Kasai, David Gray, Carol Clugston, Anna Dominiczak and Alex Alderton, Roberto Amato, Sonia Goncalves, Ewan Harrison, David K. Jackson, Ian Johnston, Dominic Kwiatkowski, Cordelia Langford, John Sillitoe |
| EPI_ISL_532472, EPI_ISL_532473 | NHSGGC West of Scotland Specialist Virology Centre / MRC-University of Glasgow Centre for Virus Research | Wellcome Sanger Institute for the COVID-19 Genomics UK (COG-UK) consortium | Ana da Silva Filipe, Natasha Johnson, Kathy Smollett, Daniel Mair, Stephen Carmichael, Lily Tong, Jenna Nichols, Elihu Aranday-Cortes, Kirstyn Brunker, Yasmin Parr, Kyriaki Nomikou; Sarah McDonald, Marc Niebel, Patawee Asamaphan; Richard Orton, Joseph Hughes, Sreenu Vattipally, David L Robertson; Alasdair MacLean, Rory Gunson; Kathy Li, Natasha Jesudason, Rajiv Shah, James Shepherd, Antonia Ho, Alice Broos, Emma Thomson and Alex Alderton, Roberto Amato, Sonia Goncalves, Ewan Harrison, David K. Jackson, Ian Johnston, Dominic Kwiatkowski, Cordelia Langford, John Sillitoe |
| EPI_ISL_532474, EPI_ISL_532475, EPI_ISL_532476, EPI_ISL_532477, EPI_ISL_532478, EPI_ISL_532479 | Lighthouse Lab in Glasgow | Wellcome Sanger Institute for the COVID-19 Genomics UK (COG-UK) consortium | Harper VanSteenhouse, Yumi Kasai, David Gray, Carol Clugston, Anna Dominiczak and Alex Alderton, Roberto Amato, Sonia Goncalves, Ewan Harrison, David K. Jackson, Ian Johnston, Dominic Kwiatkowski, Cordelia Langford, John Sillitoe |
| EPI_ISL_532480 | NHSGGC West of Scotland Specialist Virology Centre / MRC-University of Glasgow Centre for Virus Research | Wellcome Sanger Institute for the COVID-19 Genomics UK (COG-UK) consortium | Ana da Silva Filipe, Natasha Johnson, Kathy Smollett, Daniel Mair, Stephen Carmichael, Lily Tong, Jenna Nichols, Elihu Aranday-Cortes, Kirstyn Brunker, Yasmin Parr, Kyriaki Nomikou; Sarah McDonald, Marc Niebel, Patawee Asamaphan; Richard Orton, Joseph Hughes, Sreenu Vattipally, David L Robertson; Alasdair MacLean, Rory Gunson; Kathy Li, Natasha Jesudason, Rajiv Shah, James Shepherd, Antonia Ho, Alice Broos, Emma Thomson and Alex Alderton, Roberto Amato, Sonia Goncalves, Ewan Harrison, David K. Jackson, Ian Johnston, Dominic Kwiatkowski, Cordelia Langford, John Sillitoe |
| EPI_ISL_532481, EPI_ISL_532482, EPI_ISL_532483 | Lighthouse Lab in Glasgow | Wellcome Sanger Institute for the COVID-19 Genomics UK (COG-UK) consortium | Harper VanSteenhouse, Yumi Kasai, David Gray, Carol Clugston, Anna Dominiczak and Alex Alderton, Roberto Amato, Sonia Goncalves, Ewan Harrison, David K. Jackson, Ian Johnston, Dominic Kwiatkowski, Cordelia Langford, John Sillitoe |
| EPI_ISL_532484 | NHSGGC West of Scotland Specialist Virology Centre / MRC-University of Glasgow Centre for Virus Research | Wellcome Sanger Institute for the COVID-19 Genomics UK (COG-UK) consortium | Ana da Silva Filipe, Natasha Johnson, Kathy Smollett, Daniel Mair, Stephen Carmichael, Lily Tong, Jenna Nichols, Elihu Aranday-Cortes, Kirstyn Brunker, Yasmin Parr, Kyriaki Nomikou; Sarah McDonald, Marc Niebel, Patawee Asamaphan; Richard Orton, Joseph Hughes, Sreenu Vattipally, David L Robertson; Alasdair MacLean, Rory Gunson; Kathy Li, Natasha Jesudason, Rajiv Shah, James Shepherd, Antonia Ho, Alice Broos, Emma Thomson and Alex Alderton, Roberto Amato, Sonia Goncalves, Ewan Harrison, David K. Jackson, Ian Johnston, Dominic Kwiatkowski, Cordelia Langford, John Sillitoe |
| EPI_ISL_532485, EPI_ISL_532486, EPI_ISL_532487, EPI_ISL_532488, EPI_ISL_532489 | Lighthouse Lab in Glasgow | Wellcome Sanger Institute for the COVID-19 Genomics UK (COG-UK) consortium | Harper VanSteenhouse, Yumi Kasai, David Gray, Carol Clugston, Anna Dominiczak and Alex Alderton, Roberto Amato, Sonia Goncalves, Ewan Harrison, David K. Jackson, Ian Johnston, Dominic Kwiatkowski, Cordelia Langford, John Sillitoe |
| EPI_ISL_532490 | NHSGGC West of Scotland Specialist Virology Centre / MRC-University of Glasgow Centre for Virus Research | Wellcome Sanger Institute for the COVID-19 Genomics UK (COG-UK) consortium | Ana da Silva Filipe, Natasha Johnson, Kathy Smollett, Daniel Mair, Stephen Carmichael, Lily Tong, Jenna Nichols, Elihu Aranday-Cortes, Kirstyn Brunker, Yasmin Parr, Kyriaki Nomikou; Sarah McDonald, Marc Niebel, Patawee Asamaphan; Richard Orton, Joseph Hughes, Sreenu Vattipally, David L Robertson; Alasdair MacLean, Rory Gunson; Kathy Li, Natasha Jesudason, Rajiv Shah, James Shepherd, Antonia Ho, Alice Broos, Emma Thomson and Alex Alderton, Roberto Amato, Sonia Goncalves, Ewan Harrison, David K. Jackson, Ian Johnston, Dominic Kwiatkowski, Cordelia Langford, John Sillitoe |
| EPI_ISL_532491 | Lighthouse Lab in Glasgow | Wellcome Sanger Institute for the COVID-19 Genomics UK (COG-UK) consortium | Harper VanSteenhouse, Yumi Kasai, David Gray, Carol Clugston, Anna Dominiczak and Alex Alderton, Roberto Amato, Sonia Goncalves, Ewan Harrison, David K. Jackson, Ian Johnston, Dominic Kwiatkowski, Cordelia Langford, John Sillitoe |
| EPI_ISL_532492, EPI_ISL_532493 | NHSGGC West of Scotland Specialist Virology Centre / MRC-University of Glasgow Centre for Virus Research | Wellcome Sanger Institute for the COVID-19 Genomics UK (COG-UK) consortium | Ana da Silva Filipe, Natasha Johnson, Kathy Smollett, Daniel Mair, Stephen Carmichael, Lily Tong, Jenna Nichols, Elihu Aranday-Cortes, Kirstyn Brunker, Yasmin Parr, Kyriaki Nomikou; Sarah McDonald, Marc Niebel, Patawee Asamaphan; Richard Orton, Joseph Hughes, Sreenu Vattipally, David L Robertson; Alasdair MacLean, Rory Gunson; Kathy Li, Natasha Jesudason, Rajiv Shah, James Shepherd, Antonia Ho, Alice Broos, Emma Thomson and Alex Alderton, Roberto Amato, Sonia Goncalves, Ewan Harrison, David K. Jackson, Ian Johnston, Dominic Kwiatkowski, Cordelia Langford, John Sillitoe |
| EPI_ISL_532494, EPI_ISL_532495, EPI_ISL_532496, EPI_ISL_532497, EPI_ISL_532498, EPI_ISL_532499, EPI_ISL_532500, EPI_ISL_532501, EPI_ISL_532502, EPI_ISL_532503, EPI_ISL_532504, EPI_ISL_532505 | | | |
| see above | Lighthouse Lab in Glasgow | Wellcome Sanger Institute for the COVID-19 Genomics UK (COG-UK) consortium | Harper VanSteenhouse, Yumi Kasai, David Gray, Carol Clugston, Anna Dominiczak and Alex Alderton, Roberto Amato, Sonia Goncalves, Ewan Harrison, David K. Jackson, Ian Johnston, Dominic Kwiatkowski, Cordelia Langford, John Sillitoe |
| EPI_ISL_532506 | NHSGGC West of Scotland Specialist Virology Centre / MRC-University of Glasgow Centre for Virus Research | Wellcome Sanger Institute for the COVID-19 Genomics UK (COG-UK) consortium | Ana da Silva Filipe, Natasha Johnson, Kathy Smollett, Daniel Mair, Stephen Carmichael, Lily Tong, Jenna Nichols, Elihu Aranday-Cortes, Kirstyn Brunker, Yasmin Parr, Kyriaki Nomikou; Sarah McDonald, Marc Niebel, Patawee Asamaphan; Richard Orton, Joseph Hughes, Sreenu Vattipally, David L Robertson; Alasdair MacLean, Rory Gunson; Kathy Li, Natasha Jesudason, Rajiv Shah, James Shepherd, Antonia Ho, Alice Broos, Emma Thomson and Alex Alderton, Roberto Amato, Sonia Goncalves, Ewan Harrison, David K. Jackson, Ian Johnston, Dominic Kwiatkowski, Cordelia Langford, John Sillitoe |
| EPI_ISL_532507, EPI_ISL_532508, EPI_ISL_532509 | Lighthouse Lab in Glasgow | Wellcome Sanger Institute for the COVID-19 Genomics UK (COG-UK) consortium | Harper VanSteenhouse, Yumi Kasai, David Gray, Carol Clugston, Anna Dominiczak and Alex Alderton, Roberto Amato, Sonia Goncalves, Ewan Harrison, David K. Jackson, Ian Johnston, Dominic Kwiatkowski, Cordelia Langford, John Sillitoe |
| EPI_ISL_532510 | NHSGGC West of Scotland Specialist Virology Centre / MRC-University of Glasgow Centre for Virus Research | Wellcome Sanger Institute for the COVID-19 Genomics UK (COG-UK) consortium | Ana da Silva Filipe, Natasha Johnson, Kathy Smollett, Daniel Mair, Stephen Carmichael, Lily Tong, Jenna Nichols, Elihu Aranday-Cortes, Kirstyn Brunker, Yasmin Parr, Kyriaki Nomikou; Sarah McDonald, Marc Niebel, Patawee Asamaphan; Richard Orton, Joseph Hughes, Sreenu Vattipally, David L Robertson; Alasdair MacLean, Rory Gunson; Kathy Li, Natasha Jesudason, Rajiv Shah, James Shepherd, Antonia Ho, Alice Broos, Emma Thomson and Alex Alderton, Roberto Amato, Sonia Goncalves, Ewan Harrison, David K. Jackson, Ian Johnston, Dominic Kwiatkowski, Cordelia Langford, John Sillitoe |
| EPI_ISL_532511 | Lighthouse Lab in Glasgow | Wellcome Sanger Institute for the COVID-19 Genomics UK (COG-UK) Consortium | Harper VanSteenhouse, Yumi Kasai, David Gray, Carol Clugston, Anna Dominiczak and Alex Alderton, Roberto Amato, Sonia Goncalves, Ewan Harrison, David K. Jackson, Ian Johnston, Dominic Kwiatkowski, Cordelia Langford, John Sillitoe on behalf of the Wellcome Sanger Institute COVID-19 Surveillance Team |
| EPI_ISL_532512 | NHSGGC West of Scotland Specialist Virology Centre / MRC-University of Glasgow Centre for Virus Research | Wellcome Sanger Institute for the COVID-19 Genomics UK (COG-UK) consortium | Ana da Silva Filipe, Natasha Johnson, Kathy Smollett, Daniel Mair, Stephen Carmichael, Lily Tong, Jenna Nichols, Elihu Aranday-Cortes, Kirstyn Brunker, Yasmin Parr, Kyriaki Nomikou; Sarah McDonald, Marc Niebel, Patawee Asamaphan; Richard Orton, Joseph Hughes, Sreenu Vattipally, David L Robertson; Alasdair MacLean, Rory Gunson; Kathy Li, Natasha Jesudason, Rajiv Shah, James Shepherd, Antonia Ho, Alice Broos, Emma Thomson and Alex Alderton, Roberto Amato, Sonia Goncalves, Ewan Harrison, David K. Jackson, Ian Johnston, Dominic Kwiatkowski, Cordelia Langford, John Sillitoe |
| EPI_ISL_532513 | Lighthouse Lab in Glasgow | Wellcome Sanger Institute for the COVID-19 Genomics UK (COG-UK) consortium | Harper VanSteenhouse, Yumi Kasai, David Gray, Carol Clugston, Anna Dominiczak and Alex Alderton, Roberto Amato, Sonia Goncalves, Ewan Harrison, David K. Jackson, Ian Johnston, Dominic Kwiatkowski, Cordelia Langford, John Sillitoe |
| EPI_ISL_532514, EPI_ISL_532515 | NHSGGC West of Scotland Specialist Virology Centre / MRC-University of Glasgow Centre for Virus Research | Wellcome Sanger Institute for the COVID-19 Genomics UK (COG-UK) consortium | Ana da Silva Filipe, Natasha Johnson, Kathy Smollett, Daniel Mair, Stephen Carmichael, Lily Tong, Jenna Nichols, Elihu Aranday-Cortes, Kirstyn Brunker, Yasmin Parr, Kyriaki Nomikou; Sarah McDonald, Marc Niebel, Patawee Asamaphan; Richard Orton, Joseph Hughes, Sreenu Vattipally, David L Robertson; Alasdair MacLean, Rory Gunson; Kathy Li, Natasha Jesudason, Rajiv Shah, James Shepherd, Antonia Ho, Alice Broos, Emma Thomson and Alex Alderton, Roberto Amato, Sonia Goncalves, Ewan Harrison, David K. Jackson, Ian Johnston, Dominic Kwiatkowski, Cordelia Langford, John Sillitoe |
| EPI_ISL_532516, EPI_ISL_532517 | Lighthouse Lab in Glasgow | Wellcome Sanger Institute for the COVID-19 Genomics UK (COG-UK) consortium | Harper VanSteenhouse, Yumi Kasai, David Gray, Carol Clugston, Anna Dominiczak and Alex Alderton, Roberto Amato, Sonia Goncalves, Ewan Harrison, David K. Jackson, Ian Johnston, Dominic Kwiatkowski, Cordelia Langford, John Sillitoe |
| EPI_ISL_532518 | NHSGGC West of Scotland Specialist Virology Centre / MRC-University of Glasgow Centre for Virus Research | Wellcome Sanger Institute for the COVID-19 Genomics UK (COG-UK) consortium | Ana da Silva Filipe, Natasha Johnson, Kathy Smollett, Daniel Mair, Stephen Carmichael, Lily Tong, Jenna Nichols, Elihu Aranday-Cortes, Kirstyn Brunker, Yasmin Parr, Kyriaki Nomikou; Sarah McDonald, Marc Niebel, Patawee Asamaphan; Richard Orton, Joseph Hughes, Sreenu Vattipally, David L Robertson; Alasdair MacLean, Rory Gunson; Kathy Li, Natasha Jesudason, Rajiv Shah, James Shepherd, Antonia Ho, Alice Broos, Emma Thomson and Alex Alderton, Roberto Amato, Sonia Goncalves, Ewan Harrison, David K. Jackson, Ian Johnston, Dominic Kwiatkowski, Cordelia Langford, John Sillitoe |
| EPI_ISL_532519, EPI_ISL_532520, | Lighthouse Lab in Glasgow | Wellcome Sanger Institute for the COVID-19 Genomics UK | Harper VanSteenhouse, Yumi Kasai, David Gray, Carol Clugston, Anna Dominiczak and Alex Alderton, Roberto Amato, Sonia Goncalves, Ewan Harrison, |

| | | | |
|---|---|---|---|
| EPI_ISL_532521, EPI_ISL_532522 | | (COG-UK) consortium | David K. Jackson, Ian Johnston, Dominic Kwiatkowski, Cordelia Langford, John Sillitoe |
| EPI_ISL_532523 | NHSGGC West of Scotland Specialist Virology Centre / MRC-University of Glasgow Centre for Virus Research | Wellcome Sanger Institute for the COVID-19 Genomics UK (COG-UK) consortium | Ana da Silva Filipe, Natasha Johnson, Kathy Smollett, Daniel Mair, Stephen Carmichael, Lily Tong, Jenna Nichols, Elihu Aranday-Cortes, Kirstyn Brunker, Yasmin Parr, Kyriaki Nomikou; Sarah McDonald, Marc Niebel, Patawee Asamaphan; Richard Orton, Joseph Hughes, Sreenu Vattipally, David L Robertson; Alasdair MacLean, Rory Gunson; Kathy Li, Natasha Jesudason, Rajiv Shah, James Shepherd, Antonia Ho, Alice Broos, Emma Thomson and Alex Alderton, Roberto Amato, Sonia Goncalves, Ewan Harrison, David K. Jackson, Ian Johnston, Dominic Kwiatkowski, Cordelia Langford, John Sillitoe |
| EPI_ISL_532524, EPI_ISL_532525, EPI_ISL_532526, EPI_ISL_532527, EPI_ISL_532528, EPI_ISL_532529, EPI_ISL_532530 | Lighthouse Lab in Glasgow | Wellcome Sanger Institute for the COVID-19 Genomics UK (COG-UK) consortium | Harper VanSteenhouse, Yumi Kasai, David Gray, Carol Clugston, Anna Dominiczak and Alex Alderton, Roberto Amato, Sonia Goncalves, Ewan Harrison, David K. Jackson, Ian Johnston, Dominic Kwiatkowski, Cordelia Langford, John Sillitoe |
| EPI_ISL_532531 | NHSGGC West of Scotland Specialist Virology Centre / MRC-University of Glasgow Centre for Virus Research | Wellcome Sanger Institute for the COVID-19 Genomics UK (COG-UK) consortium | Ana da Silva Filipe, Natasha Johnson, Kathy Smollett, Daniel Mair, Stephen Carmichael, Lily Tong, Jenna Nichols, Elihu Aranday-Cortes, Kirstyn Brunker, Yasmin Parr, Kyriaki Nomikou; Sarah McDonald, Marc Niebel, Patawee Asamaphan; Richard Orton, Joseph Hughes, Sreenu Vattipally, David L Robertson; Alasdair MacLean, Rory Gunson; Kathy Li, Natasha Jesudason, Rajiv Shah, James Shepherd, Antonia Ho, Alice Broos, Emma Thomson and Alex Alderton, Roberto Amato, Sonia Goncalves, Ewan Harrison, David K. Jackson, Ian Johnston, Dominic Kwiatkowski, Cordelia Langford, John Sillitoe |
| EPI_ISL_532532, EPI_ISL_532533, EPI_ISL_532534, EPI_ISL_532535, EPI_ISL_532536, EPI_ISL_532537 | Lighthouse Lab in Glasgow | Wellcome Sanger Institute for the COVID-19 Genomics UK (COG-UK) consortium | Harper VanSteenhouse, Yumi Kasai, David Gray, Carol Clugston, Anna Dominiczak and Alex Alderton, Roberto Amato, Sonia Goncalves, Ewan Harrison, David K. Jackson, Ian Johnston, Dominic Kwiatkowski, Cordelia Langford, John Sillitoe |
| EPI_ISL_532538 | NHSGGC West of Scotland Specialist Virology Centre / MRC-University of Glasgow Centre for Virus Research | Wellcome Sanger Institute for the COVID-19 Genomics UK (COG-UK) consortium | Ana da Silva Filipe, Natasha Johnson, Kathy Smollett, Daniel Mair, Stephen Carmichael, Lily Tong, Jenna Nichols, Elihu Aranday-Cortes, Kirstyn Brunker, Yasmin Parr, Kyriaki Nomikou; Sarah McDonald, Marc Niebel, Patawee Asamaphan; Richard Orton, Joseph Hughes, Sreenu Vattipally, David L Robertson; Alasdair MacLean, Rory Gunson; Kathy Li, Natasha Jesudason, Rajiv Shah, James Shepherd, Antonia Ho, Alice Broos, Emma Thomson and Alex Alderton, Roberto Amato, Sonia Goncalves, Ewan Harrison, David K. Jackson, Ian Johnston, Dominic Kwiatkowski, Cordelia Langford, John Sillitoe |
| EPI_ISL_532539 | Lighthouse Lab in Glasgow | Wellcome Sanger Institute for the COVID-19 Genomics UK (COG-UK) consortium | Harper VanSteenhouse, Yumi Kasai, David Gray, Carol Clugston, Anna Dominiczak and Alex Alderton, Roberto Amato, Sonia Goncalves, Ewan Harrison, David K. Jackson, Ian Johnston, Dominic Kwiatkowski, Cordelia Langford, John Sillitoe |
| EPI_ISL_532540 | NHSGGC West of Scotland Specialist Virology Centre / MRC-University of Glasgow Centre for Virus Research | Wellcome Sanger Institute for the COVID-19 Genomics UK (COG-UK) consortium | Ana da Silva Filipe, Natasha Johnson, Kathy Smollett, Daniel Mair, Stephen Carmichael, Lily Tong, Jenna Nichols, Elihu Aranday-Cortes, Kirstyn Brunker, Yasmin Parr, Kyriaki Nomikou; Sarah McDonald, Marc Niebel, Patawee Asamaphan; Richard Orton, Joseph Hughes, Sreenu Vattipally, David L Robertson; Alasdair MacLean, Rory Gunson; Kathy Li, Natasha Jesudason, Rajiv Shah, James Shepherd, Antonia Ho, Alice Broos, Emma Thomson and Alex Alderton, Roberto Amato, Sonia Goncalves, Ewan Harrison, David K. Jackson, Ian Johnston, Dominic Kwiatkowski, Cordelia Langford, John Sillitoe |
| EPI_ISL_532546, EPI_ISL_532561 | Lighthouse Lab in Glasgow | Wellcome Sanger Institute for the COVID-19 Genomics UK (COG-UK) consortium | Harper VanSteenhouse, Yumi Kasai, David Gray, Carol Clugston, Anna Dominiczak and Alex Alderton, Roberto Amato, Sonia Goncalves, Ewan Harrison, David K. Jackson, Ian Johnston, Dominic Kwiatkowski, Cordelia Langford, John Sillitoe |
| EPI_ISL_532563, EPI_ISL_532564 | NHSGGC West of Scotland Specialist Virology Centre / MRC-University of Glasgow Centre for Virus Research | Wellcome Sanger Institute for the COVID-19 Genomics UK (COG-UK) consortium | Ana da Silva Filipe, Natasha Johnson, Kathy Smollett, Daniel Mair, Stephen Carmichael, Lily Tong, Jenna Nichols, Elihu Aranday-Cortes, Kirstyn Brunker, Yasmin Parr, Kyriaki Nomikou; Sarah McDonald, Marc Niebel, Patawee Asamaphan; Richard Orton, Joseph Hughes, Sreenu Vattipally, David L Robertson; Alasdair MacLean, Rory Gunson; Kathy Li, Natasha Jesudason, Rajiv Shah, James Shepherd, Antonia Ho, Alice Broos, Emma Thomson and Alex Alderton, Roberto Amato, Sonia Goncalves, Ewan Harrison, David K. Jackson, Ian Johnston, Dominic Kwiatkowski, Cordelia Langford, John Sillitoe |
| EPI_ISL_532569 | Lighthouse Lab in Glasgow | Wellcome Sanger Institute for the COVID-19 Genomics UK (COG-UK) consortium | Harper VanSteenhouse, Yumi Kasai, David Gray, Carol Clugston, Anna Dominiczak and Alex Alderton, Roberto Amato, Sonia Goncalves, Ewan Harrison, David K. Jackson, Ian Johnston, Dominic Kwiatkowski, Cordelia Langford, John Sillitoe |
| EPI_ISL_532572, EPI_ISL_532587 | NHSGGC West of Scotland Specialist Virology Centre / MRC-University of Glasgow Centre for Virus Research | Wellcome Sanger Institute for the COVID-19 Genomics UK (COG-UK) consortium | Ana da Silva Filipe, Natasha Johnson, Kathy Smollett, Daniel Mair, Stephen Carmichael, Lily Tong, Jenna Nichols, Elihu Aranday-Cortes, Kirstyn Brunker, Yasmin Parr, Kyriaki Nomikou; Sarah McDonald, Marc Niebel, Patawee Asamaphan; Richard Orton, Joseph Hughes, Sreenu Vattipally, David L Robertson; Alasdair MacLean, Rory Gunson; Kathy Li, Natasha Jesudason, Rajiv Shah, James Shepherd, Antonia Ho, Alice Broos, Emma Thomson and Alex Alderton, Roberto Amato, Sonia Goncalves, Ewan Harrison, David K. Jackson, Ian Johnston, Dominic Kwiatkowski, Cordelia Langford, John Sillitoe |
| EPI_ISL_532588 | Lighthouse Lab in Glasgow | Wellcome Sanger Institute for the COVID-19 Genomics UK (COG-UK) consortium | Harper VanSteenhouse, Yumi Kasai, David Gray, Carol Clugston, Anna Dominiczak and Alex Alderton, Roberto Amato, Sonia Goncalves, Ewan Harrison, David K. Jackson, Ian Johnston, Dominic Kwiatkowski, Cordelia Langford, John Sillitoe |
| EPI_ISL_532598 | NHSGGC West of Scotland Specialist Virology Centre / MRC-University of Glasgow Centre for Virus Research | Wellcome Sanger Institute for the COVID-19 Genomics UK (COG-UK) consortium | Ana da Silva Filipe, Natasha Johnson, Kathy Smollett, Daniel Mair, Stephen Carmichael, Lily Tong, Jenna Nichols, Elihu Aranday-Cortes, Kirstyn Brunker, Yasmin Parr, Kyriaki Nomikou; Sarah McDonald, Marc Niebel, Patawee Asamaphan; Richard Orton, Joseph Hughes, Sreenu Vattipally, David L Robertson; Alasdair MacLean, Rory Gunson; Kathy Li, Natasha Jesudason, Rajiv Shah, James Shepherd, Antonia Ho, Alice Broos, Emma Thomson and Alex Alderton, Roberto Amato, Sonia Goncalves, Ewan Harrison, David K. Jackson, Ian Johnston, Dominic Kwiatkowski, Cordelia Langford, John Sillitoe |
| EPI_ISL_532600, EPI_ISL_532615 | Lighthouse Lab in Glasgow | Wellcome Sanger Institute for the COVID-19 Genomics UK (COG-UK) consortium | Harper VanSteenhouse, Yumi Kasai, David Gray, Carol Clugston, Anna Dominiczak and Alex Alderton, Roberto Amato, Sonia Goncalves, Ewan Harrison, David K. Jackson, Ian Johnston, Dominic Kwiatkowski, Cordelia Langford, John Sillitoe |
| EPI_ISL_532620, EPI_ISL_532621, EPI_ISL_532626 | NHSGGC West of Scotland Specialist Virology Centre / MRC-University of Glasgow Centre for Virus Research | Wellcome Sanger Institute for the COVID-19 Genomics UK (COG-UK) consortium | Ana da Silva Filipe, Natasha Johnson, Kathy Smollett, Daniel Mair, Stephen Carmichael, Lily Tong, Jenna Nichols, Elihu Aranday-Cortes, Kirstyn Brunker, Yasmin Parr, Kyriaki Nomikou; Sarah McDonald, Marc Niebel, Patawee Asamaphan; Richard Orton, Joseph Hughes, Sreenu Vattipally, David L Robertson; Alasdair MacLean, Rory Gunson; Kathy Li, Natasha Jesudason, Rajiv Shah, James Shepherd, Antonia Ho, Alice Broos, Emma Thomson and Alex Alderton, Roberto Amato, Sonia Goncalves, Ewan Harrison, David K. Jackson, Ian Johnston, Dominic Kwiatkowski, Cordelia Langford, John Sillitoe |
| EPI_ISL_532650 | Lighthouse Lab in Glasgow | Wellcome Sanger Institute for the COVID-19 Genomics UK (COG-UK) consortium | Harper VanSteenhouse, Yumi Kasai, David Gray, Carol Clugston, Anna Dominiczak and Alex Alderton, Roberto Amato, Sonia Goncalves, Ewan Harrison, David K. Jackson, Ian Johnston, Dominic Kwiatkowski, Cordelia Langford, John Sillitoe |
| EPI_ISL_534199 | E. Gulbja Laboratorija | Latvian Biomedical Research and Study Centre | Ivars Silamielis, Jnis Pjalkovskis, Kaspars Megnis, Monta Ustinova, ikita Zrelovs, Vita Rovte, Mikus Gavars, Dmitrijs Perminovs, Uga Dumpis, Jnis Kloviš |
| EPI_ISL_534200, EPI_ISL_534201, EPI_ISL_534202 | Centrl laboratorija | Latvian Biomedical Research and Study Centre | Ivars Silamielis, Jnis Pjalkovskis, Kaspars Megnis, Monta Ustinova, ikita Zrelovs, Vita Rovte, Stella Lapia, Jana Oste, Marta Priedte, Uga Dumpis, Jnis Kloviš |
| EPI_ISL_534203 | E. Gulbja Laboratorija | Latvian Biomedical Research and Study Centre | Ivars Silamielis, Jnis Pjalkovskis, Kaspars Megnis, Monta Ustinova, ikita Zrelovs, Vita Rovte, Mikus Gavars, Dmitrijs Perminovs, Uga Dumpis, Jnis Kloviš |
| EPI_ISL_534204, EPI_ISL_534205 | Latvijas Infektoloijas centrs | Latvian Biomedical Research and Study Centre | Ivars Silamielis, Jnis Pjalkovskis, Kaspars Megnis, Monta Ustinova, ikita Zrelovs, Vita Rovte, Jeena Storoženko, Tatjana Kolupajeva, Oksana Savicka, Uga Dumpis, Jnis Kloviš |
| EPI_ISL_534206 | Centrl laboratorija | Latvian Biomedical Research and Study Centre | Ivars Silamielis, Jnis Pjalkovskis, Kaspars Megnis, Monta Ustinova, ikita Zrelovs, Vita Rovte, Stella Lapia, Jana Oste, Marta Priedte, Uga Dumpis, Jnis Kloviš |
| EPI_ISL_534230, EPI_ISL_534231, EPI_ISL_534232 | Capio S:t Gorans sjukhus | The Public Health Agency of Sweden | Anna-Malin Linde, Maria Lind Karlberg, Mattias Haukland, Reza Advani, Olov Svartstrom, Oskar Karlsson Lindsjo, Sandra Broddesson, Petra Edquist, Mia Brytting, Anna Risberg, Karin Tegmark-Wisell |
| EPI_ISL_534250 | Gavle Sjukhus | The Public Health Agency of Sweden | Anna-Malin Linde, Maria Lind Karlberg, Mattias Haukland, Reza Advani, Olov Svartstrom, Oskar Karlsson Lindsjo, Sandra Broddesson, Petra Edquist, Mia Brytting, Anna Risberg, Karin Tegmark-Wisell |
| EPI_ISL_534254, EPI_ISL_534255 | Laboratoriemedicin Vasternorrland | The Public Health Agency of Sweden | Anna-Malin Linde, Maria Lind Karlberg, Mattias Haukland, Reza Advani, Olov Svartstrom, Oskar Karlsson Lindsjo, Sandra Broddesson, Petra Edquist, Mia Brytting, Anna Risberg, Karin Tegmark-Wisell |
| EPI_ISL_534259 | Karolinska universitetslaboratoriet | The Public Health Agency of Sweden | Anna-Malin Linde, Maria Lind Karlberg, Mattias Haukland, Reza Advani, Olov Svartstrom, Oskar Karlsson Lindsjo, Sandra Broddesson, Petra Edquist, Mia Brytting, Anna Risberg, Karin Tegmark-Wisell |
| EPI_ISL_534328 | Hospital Universitario 12 de Octubre | Hospital Universitario 12 de Octubre | Raúl Recio, Sara González, Esther Viedma, Elias Dahdouh, Fernando Lázaro, Natalia Stella, Julio García, Juan Carlos Galán, Rafael Cantón, Ma Dolores Folgueira, Rafael Delgado, Jesús Mingorance |
| EPI_ISL_534406, EPI_ISL_534407, EPI_ISL_534408, EPI_ISL_534409, EPI_ISL_534410, EPI_ISL_534411, EPI_ISL_534412, EPI_ISL_534413, EPI_ISL_534414, EPI_ISL_534415, EPI_ISL_534416, EPI_ISL_534417, EPI_ISL_534418, EPI_ISL_534419, EPI_ISL_534420, EPI_ISL_534421, EPI_ISL_534422, EPI_ISL_534423, EPI_ISL_534424, EPI_ISL_534425, EPI_ISL_534426, EPI_ISL_534427, EPI_ISL_534428, EPI_ISL_534429, EPI_ISL_534430, EPI_ISL_534431, EPI_ISL_534432, EPI_ISL_534433, EPI_ISL_534434, EPI_ISL_534435, EPI_ISL_534436, EPI_ISL_534437, EPI_ISL_534438 | | | |
| see above | NHSGGC West of Scotland Specialist Virology Centre / MRC-University of Glasgow Centre for Virus Research | Wellcome Sanger Institute for the COVID-19 Genomics UK (COG-UK) consortium | Ana da Silva Filipe, Natasha Johnson, Kathy Smollett, Daniel Mair, Stephen Carmichael, Lily Tong, Jenna Nichols, Elihu Aranday-Cortes, Kirstyn Brunker, Yasmin Parr, Kyriaki Nomikou; Sarah McDonald, Marc Niebel, Patawee Asamaphan; Richard Orton, Joseph Hughes, Sreenu Vattipally, David L Robertson; |

| | | | |
|---|---|---|---|
| | | | Alasdair MacLean, Rory Gunson; Kathy Li, Natasha Jesudason, Rajiv Shah, James Shepherd, Antonia Ho, Alice Broos, Emma Thomson and Alex Alderton, Roberto Amato, Sonia Goncalves, Ewan Harrison, David K. Jackson, Ian Johnston, Dominic Kwiatkowski, Cordelia Langford, John Sillitoe on behalf of the Wellcome Sanger Institute COVID-19 Surveillance Team (http://www.sanger.ac.uk/covid-team) |
| EPI_ISL_534693, EPI_ISL_534694, EPI_ISL_534695 | University of Miami Immunology and Histocompatibility Laboratory | University of Miami Immunology and Histocompatibility Laboratory | Emilio Margolles-Clark, PhD and Phillip Ruiz, MD, PhD |
| EPI_ISL_534720, EPI_ISL_534721, EPI_ISL_534730 | Respiratory Virus Unit, Microbiology Services Colindale, Public Health England | Respiratory Virus Unit, Microbiology Services Colindale, Public Health England | PHE Covid Sequencing Team |
| EPI_ISL_535390, EPI_ISL_535391, EPI_ISL_535392, EPI_ISL_535393, EPI_ISL_535394, EPI_ISL_535395, EPI_ISL_535396, EPI_ISL_535397, EPI_ISL_535398, EPI_ISL_535399, EPI_ISL_535400, EPI_ISL_535401, EPI_ISL_535402, EPI_ISL_535403, EPI_ISL_535404, EPI_ISL_535448, EPI_ISL_535449, EPI_ISL_535451, EPI_ISL_535454, EPI_ISL_535458, EPI_ISL_535472, EPI_ISL_535480, EPI_ISL_535486, EPI_ISL_535487, EPI_ISL_535496, EPI_ISL_535501, EPI_ISL_535503, EPI_ISL_535504, EPI_ISL_535505, EPI_ISL_535506, EPI_ISL_535507, EPI_ISL_535508, EPI_ISL_535509, EPI_ISL_535510, EPI_ISL_535511, EPI_ISL_535512, EPI_ISL_535513, EPI_ISL_535514, EPI_ISL_535515, EPI_ISL_535516, EPI_ISL_535517, EPI_ISL_535518, EPI_ISL_535519, EPI_ISL_535520, EPI_ISL_535521, EPI_ISL_535522, EPI_ISL_535523, EPI_ISL_535524, EPI_ISL_535525, EPI_ISL_535526, EPI_ISL_535527, EPI_ISL_535528, EPI_ISL_535529, EPI_ISL_535530, EPI_ISL_535531, EPI_ISL_535532, EPI_ISL_535533, EPI_ISL_535534, EPI_ISL_535535, EPI_ISL_535536, EPI_ISL_535537, EPI_ISL_535538, EPI_ISL_535539, EPI_ISL_535540, EPI_ISL_535541, EPI_ISL_535542, EPI_ISL_535543, EPI_ISL_535544, EPI_ISL_535545, EPI_ISL_535546, EPI_ISL_535547, EPI_ISL_535548, EPI_ISL_535549, EPI_ISL_535550, EPI_ISL_535551 | | | |
| see above | NHLS-IALCH | KRISP, KZN Research Innovation and Sequencing Platform | Giandhari J, Pillay S, Lessells R, Mdlalose K, York D, Khan S, Tegally H, Wilkinson E, de Oliveira T |
| EPI_ISL_536411 | Medtimes Molecular Laboratory | Medtimes Molecular Laboratory | Eric Chan, Winsome Wong, Jacqueline Tam, Isaac Chow |
| EPI_ISL_536572, EPI_ISL_536573, EPI_ISL_536575, EPI_ISL_536576, EPI_ISL_536577, EPI_ISL_536579, EPI_ISL_536580, EPI_ISL_536581, EPI_ISL_536582, EPI_ISL_536583, EPI_ISL_536584 | | | |
| see above | University of Wisconsin-Madison AIDS Vaccine Research Laboratories | University of Wisconsin-Madison AIDS Vaccine Research Laboratories | Gage Moreno, Katarina Braun, et al. AIDS Vaccine Research Laboratories |
| EPI_ISL_536837, EPI_ISL_536862, EPI_ISL_536881, EPI_ISL_536913, EPI_ISL_536914, EPI_ISL_536916, EPI_ISL_536927, EPI_ISL_536943, EPI_ISL_536951, EPI_ISL_536963, EPI_ISL_536964, EPI_ISL_536965, EPI_ISL_536966, EPI_ISL_536968, EPI_ISL_536969, EPI_ISL_536970, EPI_ISL_536971, EPI_ISL_536973, EPI_ISL_536974, EPI_ISL_536975, EPI_ISL_536976, EPI_ISL_536977, EPI_ISL_536979, EPI_ISL_536981, EPI_ISL_536982, EPI_ISL_536983, EPI_ISL_536984, EPI_ISL_536985, EPI_ISL_536986, EPI_ISL_536987, EPI_ISL_536988, EPI_ISL_536990, EPI_ISL_536991, EPI_ISL_536993, EPI_ISL_536994, EPI_ISL_536995, EPI_ISL_536996, EPI_ISL_537000, EPI_ISL_537001, EPI_ISL_537003, EPI_ISL_537004, EPI_ISL_537005, EPI_ISL_537006, EPI_ISL_537008, EPI_ISL_537009, EPI_ISL_537010, EPI_ISL_537011, EPI_ISL_537012, EPI_ISL_537013, EPI_ISL_537014, EPI_ISL_537015, EPI_ISL_537016 | | | |
| see above | Lighthouse Lab in Glasgow | Wellcome Sanger Institute for the COVID-19 Genomics UK (COG-UK) consortium | Harper VanSteenhouse, Yumi Kasai, David Gray, Carol Clugston, Anna Dominiczak and Alex Alderton, Roberto Amato, Sonia Goncalves, Ewan Harrison, David K. Jackson, Ian Johnston, Dominic Kwiatkowski, Cordelia Langford, John Sillitoe on behalf of the Wellcome Sanger Institute COVID-19 Surveillance Team |
| EPI_ISL_537017 | Lighthouse Lab in Glasgow | Wellcome Sanger Institute for the COVID-19 Genomics UK (COG-UK) Consortium | Harper VanSteenhouse, Yumi Kasai, David Gray, Carol Clugston, Anna Dominiczak and Alex Alderton, Roberto Amato, Sonia Goncalves, Ewan Harrison, David K. Jackson, Ian Johnston, Dominic Kwiatkowski, Cordelia Langford, John Sillitoe on behalf of the Wellcome Sanger Institute COVID-19 Surveillance Team |
| EPI_ISL_537018, EPI_ISL_537019, EPI_ISL_537020, EPI_ISL_537021, EPI_ISL_537022, EPI_ISL_537023, EPI_ISL_537024, EPI_ISL_537025, EPI_ISL_537026, EPI_ISL_537028, EPI_ISL_537029, EPI_ISL_537030, EPI_ISL_537031, EPI_ISL_537032, EPI_ISL_537034, EPI_ISL_537035, EPI_ISL_537036, EPI_ISL_537037, EPI_ISL_537038, EPI_ISL_537039, EPI_ISL_537041, EPI_ISL_537042, EPI_ISL_537043, EPI_ISL_537044, EPI_ISL_537045, EPI_ISL_537046, EPI_ISL_537047, EPI_ISL_537048, EPI_ISL_537049, EPI_ISL_537050, EPI_ISL_537051, EPI_ISL_537052, EPI_ISL_537053, EPI_ISL_537054, EPI_ISL_537055, EPI_ISL_537056, EPI_ISL_537057, EPI_ISL_537058, EPI_ISL_537060, EPI_ISL_537061, EPI_ISL_537062, EPI_ISL_537064, EPI_ISL_537065, EPI_ISL_537066, EPI_ISL_537067, EPI_ISL_537068, EPI_ISL_537069, EPI_ISL_537071, EPI_ISL_537072, EPI_ISL_537073, EPI_ISL_537074, EPI_ISL_537075, EPI_ISL_537076, EPI_ISL_537077, EPI_ISL_537078, EPI_ISL_537079, EPI_ISL_537080, EPI_ISL_537082, EPI_ISL_537083, EPI_ISL_537084, EPI_ISL_537085, EPI_ISL_537086, EPI_ISL_537087, EPI_ISL_537088, EPI_ISL_537090, EPI_ISL_537091, EPI_ISL_537092, EPI_ISL_537093, EPI_ISL_537094, EPI_ISL_537095, EPI_ISL_537096, EPI_ISL_537097, EPI_ISL_537098, EPI_ISL_537099, EPI_ISL_537100, EPI_ISL_537101, EPI_ISL_537102, EPI_ISL_537105, EPI_ISL_537106, EPI_ISL_537110, EPI_ISL_537111, EPI_ISL_537112, EPI_ISL_537113, EPI_ISL_537114, EPI_ISL_537116, EPI_ISL_537117, EPI_ISL_537118, EPI_ISL_537119, EPI_ISL_537120, EPI_ISL_537121, EPI_ISL_537122, EPI_ISL_537123, EPI_ISL_537124, EPI_ISL_537125, EPI_ISL_537126, EPI_ISL_537127, EPI_ISL_537128, EPI_ISL_537129, EPI_ISL_537131, EPI_ISL_537133, EPI_ISL_537135, EPI_ISL_537136, EPI_ISL_537137, EPI_ISL_537138, EPI_ISL_537139, EPI_ISL_537140, EPI_ISL_537141, EPI_ISL_537143, EPI_ISL_537144, EPI_ISL_537145, EPI_ISL_537147 | | | |
| see above | Lighthouse Lab in Glasgow | Wellcome Sanger Institute for the COVID-19 Genomics UK (COG-UK) consortium | Harper VanSteenhouse, Yumi Kasai, David Gray, Carol Clugston, Anna Dominiczak and Alex Alderton, Roberto Amato, Sonia Goncalves, Ewan Harrison, David K. Jackson, Ian Johnston, Dominic Kwiatkowski, Cordelia Langford, John Sillitoe on behalf of the Wellcome Sanger Institute COVID-19 Surveillance Team |
| EPI_ISL_538237, EPI_ISL_538239, EPI_ISL_538240 | TriCore Reference Laboratories | Center for Global Health, University of New Mexico Health Sciences Center | Daryl Domman, Kurt Schwalm, Twila Kunde, Joseph Hicks, Michael Edwards, Darrell Dinwiddie |
| EPI_ISL_538500, EPI_ISL_538501 | RS Prima Husada Cipta Medan North Sumatra | National Institute of Health Research and Development | Pawestri, HA; Subangkit; Puspa, KD; Nugraha, AA; Ikawati, HD; Pangesti, KNA; Soekarso, T; Paisal; Setiawaty,V. |
| EPI_ISL_538510 | RSUD Ulin Banjarmasin South Kalimantan | National Institute of Health Research and Development | Pawestri, HA; Subangkit; Puspa, KD; Nugraha, AA; Ikawati, HD; Pangesti, KNA; Soekarso, T; Paisal; Pasaribu, M; Setiawaty,V. |
| EPI_ISL_538511 | RSUD Wahidin Sudirohusodo Mojokerto East Java | National Institute of Health Research and Development | Pawestri, HA; Subangkit; Puspa, KD; Nugraha, AA; Ikawati, HD; Pangesti, KNA; Soekarso, T; Paisal; Setiawaty,V. |
| EPI_ISL_538521 | Infectious Diseases, North Carolina State Laboratory of Public Health COVID-19 Response Team | Infectious Diseases, North Carolina State Laboratory of Public Health COVID-19 Response Team | Chase,K. |
| EPI_ISL_538522 | Infectious Diseases, North Carolina State Laboratory of Public Health COVID-19 Response Team | North Carolina State Laboratory of Public Health | Chase,K. |
| EPI_ISL_539567 | Hospital Comarcal de Melilla | Instituto de Salud Carlos III | Iglesias-Caballero, M. Molinero Calamita, M. González-Esguevillas, M. Camarero, S. Pozo, F. Casas, I. Jiménez, P. Jiménez, M. Zaballos, A. Monzón, S. Varona, S. Juliá, M. Cuesta, I, J. López |
| EPI_ISL_539568 | Hospital Comarcal de Melilla | Instituto de Salud Carlos III | Iglesias-Caballero, M. Molinero Calamita, M. González-Esguevillas, M. Camarero, S. Pozo, F. Casas, I. Jiménez, P. Jiménez, M. Zaballos, A. Monzón, S. Varona, S. Juliá, M. Cuesta, I, C. Ezpeleta |
| EPI_ISL_539616 | CSIR-Centre for Cellular and Molecular Biology | CSIR-Centre for Cellular and Molecular Biology | Lamuk Zaveri, Shagufta Khan, Namami Gaur, Sakshi Shambhavi, Nikhil Hajirnis, M Soujanya Reddy, Pratheusa Maccha, Tulasi Nagabandi, Purushotham Vodnala, Payel Mukherjee, Sofia Banu, Priya Singh, Onkar Kulkarni, Dhiviya Vedagiri, Divya Gupta, Vishal Sah, Santosh Kumar Kuncha, Krishnan Harinivas Harshan, Archana Bharadwaj Siva, Karthik Bharadwaj Tallapaka, Renu Sudhakar, Somesh Gorde, Gangumala Srinivas Reddy, Sujoy Deb, Swati Bayyana, Rakesh K Mishra, Divya Tej Sowpati |
| EPI_ISL_539620 | CSIR-Centre for Cellular and Molecular Biology | CSIR-Centre for Cellular and Molecular Biology | M Soujanya Reddy, Nikhil Hajirnis, Pratheusa Maccha, Payel Mukherjee, Sofia Banu, Priya Singh, Onkar Kulkarni,Tulasi Nagabandi, Namami Gaur, Sakshi Shambhavi, Lamuk Zaveri, Shagufta Khan, Purushotham Vodnala, Dhiviya Vedagiri, Divya Gupta, Vishal Sah, Santosh Kumar Kuncha, Krishnan Harinivas Harshan, Archana Bharadwaj Siva, Karthik Bharadwaj Tallapaka,Kezia J Ann, Radhika Khandelwal, Roshan Maku Venkata, Shemin Mansuri, Sonu Uday, Rakesh K Mishra, Divya Tej Sowpati |
| EPI_ISL_539621 | CSIR-Centre for Cellular and Molecular Biology | CSIR-Centre for Cellular and Molecular Biology | M Soujanya Reddy, Nikhil Hajirnis, Pratheusa Maccha, Sakshi Shambhavi, Lamuk Zaveri, Shagufta Khan, Namami Gaur, Tulasi Nagabandi, Purushotham Vodnala, Payel Mukherjee, Sofia Banu, Priya Singh,Onkar Kulkarni, Dhiviya Vedagiri, Divya Gupta, Vishal Sah, Santosh Kumar Kuncha, Krishnan Harinivas Harshan, Archana Bharadwaj Siva, Karthik Bharadwaj Tallapaka, G. Aditya Kumar, Koushick Sivakumar, Pooja Ramesh Gupta, Rajan Kumar Jha, Shraddha Vijay Lahoti, Rakesh K Mishra, Divya Tej Sowpati |
| EPI_ISL_539622 | CSIR-Centre for Cellular and Molecular Biology | CSIR-Centre for Cellular and Molecular Biology | Namami Gaur, Sakshi Shambhavi, Lamuk Zaveri, Shagufta Khan, Nikhil Hajirnis, M Soujanya Reddy, Pratheusa Maccha, Tulasi Nagabandi, Purushotham Vodnala, Payel Mukherjee, Sofia Banu, Priya Singh, Onkar Kulkarni, Dhiviya Vedagiri, Divya Gupta, Vishal Sah, Santosh Kumar Kuncha, Krishnan Harinivas Harshan, Archana Bharadwaj Siva, Karthik Bharadwaj Tallapaka, Zeba Rizvi, Zuberwasim Sayyad, Kakade Aishwarya Arun, Amrutha H C, Ananga Ghosh, Rakesh K Mishra, Divya Tej Sowpati |
| EPI_ISL_539623 | CSIR-Centre for Cellular and Molecular Biology | CSIR-Centre for Cellular and Molecular Biology | Namami Gaur, Sakshi Shambhavi, Lamuk Zaveri, Shagufta Khan, Nikhil Hajirnis, M Soujanya Reddy, Pratheusa Maccha, Tulasi Nagabandi, Purushotham Vodnala, Payel Mukherjee, Sofia Banu, Priya Singh, Onkar Kulkarni, Dhiviya Vedagiri, Divya Gupta, Vishal Sah, Santosh Kumar Kuncha, Krishnan Harinivas Harshan, Archana Bharadwaj Siva, Karthik Bharadwaj Tallapaka,G. Aditya Kumar, Koushick Sivakumar, Rakesh K Mishra, Divya Tej Sowpati |
| EPI_ISL_539624 | CSIR-Centre for Cellular and Molecular Biology | CSIR-Centre for Cellular and Molecular Biology | Namami Gaur, Sakshi Shambhavi, Lamuk Zaveri, Shagufta Khan, Nikhil Hajirnis, M Soujanya Reddy, Pratheusa Maccha, Tulasi Nagabandi, Purushotham Vodnala, Payel Mukherjee, Sofia Banu, Priya Singh,Onkar Kulkarni, Dhiviya Vedagiri, Divya Gupta, Vishal Sah, Santosh Kumar Kuncha, Krishnan Harinivas Harshan, Archana Bharadwaj Siva, Karthik Bharadwaj Tallapaka, Zeba Rizvi, Zuberwasim Sayyad, Kakade Aishwarya Arun, Amrutha H C, Ananga Ghosh, Rakesh K Mishra, Divya Tej Sowpati |
| EPI_ISL_539625 | CSIR-Centre for Cellular and Molecular Biology | CSIR-Centre for Cellular and Molecular Biology | Nikhil Hajirnis, M Soujanya Reddy, Pratheusa Maccha, Lamuk Zaveri, Shagufta Khan, Namami Gaur, Sakshi Shambhavi, Tulasi Nagabandi, Purushotham Vodnala, Payel Mukherjee, Sofia Banu, Priya Singh, Onkar Kulkarni, Dhiviya Vedagiri, Divya Gupta, Vishal Sah, Santosh Kumar Kuncha, Krishnan Harinivas |

| | | | |
|---|---|---|---|
| | | | Harshan, Archana Bharadwaj Siva, Karthik Bharadwaj Tallapaka,Zeba Rizvi, Zuberwasim Sayyad, Kakade Aishwarya Arun, Amrutha H C, Ananga Ghosh, Rakesh K Mishra, Divya Tej Sowpati |
| EPI_ISL_539626 | CSIR-Centre for Cellular and Molecular Biology | CSIR-Centre for Cellular and Molecular Biology | Nikhil Hajirnis, M Soujanya Reddy, Pratheusa Maccha, Namami Gaur, Sakshi Shambhavi, Lamuk Zaveri, Shagufta Khan, Tulasi Nagabandi, Purushotham Vodnala, Payel Mukherjee, Sofia Banu, Priya Singh,Onkar Kulkarni, Dhiviya Vedagiri, Divya Gupta, Vishal Sah, Santosh Kumar Kuncha, Krishnan Harinivas Harshan, Archana Bharadwaj Siva, Karthik Bharadwaj Tallapaka,Kezia J Ann, Radhika Khandelwal, Roshan Maku Venkata, Shemin Mansuri, Sonu Uday, Rakesh K Mishra, Divya Tej Sowpati |
| EPI_ISL_539630 | CSIR-Centre for Cellular and Molecular Biology | CSIR-Centre for Cellular and Molecular Biology | Payel Mukherjee, Sofia Banu, Priya Singh, Onkar Kulkarni, Dhiviya Vedagiri, Divya Gupta, Vishal Sah, Santosh Kumar Kuncha, Krishnan Harinivas Harshan, Archana Bharadwaj Siva, Karthik Bharadwaj Tallapaka, Shagufta Khan, Lamuk Zaveri, Nikhil Hajirnis, M Soujanya Reddy, Pratheusa Maccha, Namami Gaur, Sakshi Shambhavi, Tulasi Nagabandi, Purushotham Vodnala, Rakesh K Mishra, Sonu Uday, Sudipta Mondal, Annapoorna P Karthyayani, Debabrata Jana, Debrya Saha, Divya Tej Sowpati |
| EPI_ISL_539643 | CSIR-Centre for Cellular and Molecular Biology | CSIR-Centre for Cellular and Molecular Biology | Tulasi Nagabandi, Namami Gaur, Sakshi Shambhavi, Lamuk Zaveri, Shagufta Khan, Nikhil Hajirnis, M Soujanya Reddy, Pratheusa Maccha, Purushotham Vodnala, Payel Mukherjee, Sofia Banu, Priya Singh, Onkar Kulkarni, Dhiviya Vedagiri, Divya Gupta, Vishal Sah, Santosh Kumar Kuncha, Krishnan Harinivas Harshan, Archana Bharadwaj Siva, Karthik Bharadwaj Tallapaka,G. Aditya Kumar, Koushick Sivakumar, Pooja Ramesh Gupta, Rajan Kumar Jha, Shraddha Vijay Lahoti, Rakesh K Mishra, Divya Tej Sowpati |
| EPI_ISL_539644 | CSIR-Centre for Cellular and Molecular Biology | CSIR-Centre for Cellular and Molecular Biology | Tulasi Nagabandi, Namami Gaur, Sakshi Shambhavi, Lamuk Zaveri, Shagufta Khan, Nikhil Hajirnis, M Soujanya Reddy, Pratheusa Maccha, Purushotham Vodnala, Payel Mukherjee, Sofia Banu, Priya Singh,Onkar Kulkarni , Dhiviya Vedagiri, Divya Gupta, Vishal Sah, Santosh Kumar Kuncha, Krishnan Harinivas Harshan, Archana Bharadwaj Siva, Karthik Bharadwaj Tallapaka,G. Aditya Kumar, Koushick Sivakumar, Pooja Ramesh Gupta, Rajan Kumar Jha, Shraddha Vijay Lahoti, Rakesh K Mishra, Divya Tej Sowpati |
| EPI_ISL_539645 | CSIR-Centre for Cellular and Molecular Biology | CSIR-Centre for Cellular and Molecular Biology | Tulasi Nagabandi, Namami Gaur, Sakshi Shambhavi, Lamuk Zaveri, Shagufta Khan, Nikhil Hajirnis, M Soujanya Reddy, Pratheusa Maccha, Purushotham Vodnala, Payel Mukherjee, Sofia Banu, Priya Singh,Onkar Kulkarni, Dhiviya Vedagiri, Divya Gupta, Vishal Sah, Santosh Kumar Kuncha, Krishnan Harinivas Harshan, Archana Bharadwaj Siva, Karthik Bharadwaj Tallapaka,Kezia J Ann, Radhika Khandelwal, Roshan Maku Venkata, Shemin Mansuri, Sonu Uday, Rakesh K Mishra, Divya Tej Sowpati |
| EPI_ISL_539646 | CSIR-Centre for Cellular and Molecular Biology | CSIR-Centre for Cellular and Molecular Biology | Lamuk Zaveri, Shagufta Khan, Namami Gaur, Sakshi Shambhavi, Nikhil Hajirnis, M Soujanya Reddy, Pratheusa Maccha, Tulasi Nagabandi, Purushotham Vodnala, Payel Mukherjee, Sofia Banu, Priya Singh, Onkar Kulkarni, Dhiviya Vedagiri, Divya Gupta, Vishal Sah, Santosh Kumar Kuncha, Krishnan Harinivas Harshan, Archana Bharadwaj Siva, Karthik Bharadwaj Tallapaka, Renu Sudhakar, Somesh Gorde, Gangumala Srinivas Reddy, Sujoy Deb, Swati Bayyana, Rakesh K Mishra, Divya Tej Sowpati |
| EPI_ISL_539648 | CSIR-Centre for Cellular and Molecular Biology | CSIR-Centre for Cellular and Molecular Biology | Lamuk Zaveri, Shagufta Khan,Nikhil Hajirnis, M Soujanya Reddy, Pratheusa Maccha, Namami Gaur, Sakshi Shambhavi, Tulasi Nagabandi, Purushotham Vodnala, Payel Mukherjee, Sofia Banu, Priya Singh, Onkar Kulkarni, Dhiviya Vedagiri, Divya Gupta, Vishal Sah, Santosh Kumar Kuncha, Krishnan Harinivas Harshan, Archana Bharadwaj Siva, Karthik Bharadwaj Tallapaka,Zeba Rizvi, Zuberwasim Sayyad, Kakade Aishwarya Arun, Amrutha H C, Ananga Ghosh, Rakesh K Mishra, Divya Tej Sowpati |
| EPI_ISL_539651 | CSIR-Centre for Cellular and Molecular Biology | CSIR-Centre for Cellular and Molecular Biology | M Soujanya Reddy, Nikhil Hajirnis, Pratheusa Maccha, Sakshi Shambhavi, Lamuk Zaveri, Shagufta Khan, Namami Gaur, Tulasi Nagabandi, Purushotham Vodnala, Payel Mukherjee, Sofia Banu, Priya Singh,Onkar Kulkarni, Dhiviya Vedagiri, Divya Gupta, Vishal Sah, Santosh Kumar Kuncha, Krishnan Harinivas Harshan, Archana Bharadwaj Siva, Karthik Bharadwaj Tallapaka, G. Aditya Kumar, Koushick Sivakumar, Pooja Ramesh Gupta, Rajan Kumar Jha, Shraddha Vijay Lahoti, Rakesh K Mishra, Divya Tej Sowpati |
| EPI_ISL_539667 | CSIR-Centre for Cellular and Molecular Biology | CSIR-Centre for Cellular and Molecular Biology | Shagufta Khan, Lamuk Zaveri, Namami Gaur, Sakshi Shambhavi, Nikhil Hajirnis, M Soujanya Reddy, Pratheusa Maccha, Tulasi Nagabandi, Purushotham Vodnala, Payel Mukherjee, Sofia Banu, Priya Singh, Onkar Kulkarni, Dhiviya Vedagiri, Divya Gupta, Vishal Sah, Santosh Kumar Kuncha, Krishnan Harinivas Harshan, Archana Bharadwaj Siva, Karthik Bharadwaj Tallapaka, Renu Sudhakar, Somesh Gorde, Gangumala Srinivas Reddy, Sujoy Deb, Swati Bayyana, Rakesh K Mishra, Divya Tej Sowpati |
| EPI_ISL_539668 | CSIR-Centre for Cellular and Molecular Biology | CSIR-Centre for Cellular and Molecular Biology | Shagufta Khan, Lamuk Zaveri, Namami Gaur, Sakshi Shambhavi, Nikhil Hajirnis, M Soujanya Reddy, Pratheusa Maccha, Tulasi Nagabandi, Purushotham Vodnala, Payel Mukherjee, Sofia Banu, Priya Singh, Onkar Kulkarni, Dhiviya Vedagiri, Divya Gupta, Vishal Sah, Santosh Kumar Kuncha, Krishnan Harinivas Harshan, Archana Bharadwaj Siva, Karthik Bharadwaj Tallapaka,Preethi Jampala, Sharada Ravi Iyer, Sulagana Mukherjee, Swetha Sundar, Peddapuvala Sai Uday Kiran Rakesh K Mishra, Divya Tej Sowpati |
| EPI_ISL_539676 | CSIR-Centre for Cellular and Molecular Biology | CSIR-Centre for Cellular and Molecular Biology | Lamuk Zaveri, Shagufta Khan, Namami Gaur, Sakshi Shambhavi, Nikhil Hajirnis, M Soujanya Reddy, Pratheusa Maccha, Tulasi Nagabandi, Purushotham Vodnala, Payel Mukherjee, Sofia Banu, Priya Singh, Onkar Kulkarni, Dhiviya Vedagiri, Divya Gupta, Vishal Sah, Santosh Kumar Kuncha, Krishnan Harinivas Harshan, Archana Bharadwaj Siva, Karthik Bharadwaj Tallapaka, Renu Sudhakar, Somesh Gorde, Gangumala Srinivas Reddy, Sujoy Deb, Swati Bayyana, Rakesh K Mishra, Divya Tej Sowpati |
| EPI_ISL_539677 | CSIR-Centre for Cellular and Molecular Biology | CSIR-Centre for Cellular and Molecular Biology | Lamuk Zaveri, Shagufta Khan,Nikhil Hajirnis, M Soujanya Reddy, Pratheusa Maccha, Namami Gaur, Sakshi Shambhavi, Tulasi Nagabandi, Purushotham Vodnala, Payel Mukherjee, Sofia Banu, Priya Singh, Onkar Kulkarni, Dhiviya Vedagiri, Divya Gupta, Vishal Sah, Santosh Kumar Kuncha, Krishnan Harinivas Harshan, Archana Bharadwaj Siva, Karthik Bharadwaj Tallapaka,Umesh Kumar, Unis Ahmad Bhat, Ajay Sarawagi, Priyanka Pant, Rajkanwar Nathawat, Rakesh K Mishra, Divya Tej Sowpati |
| EPI_ISL_539679 | CSIR-Centre for Cellular and Molecular Biology | CSIR-Centre for Cellular and Molecular Biology | M Soujanya Reddy, Nikhil Hajirnis, Pratheusa Maccha, Namami Gaur, Sakshi Shambhavi, Lamuk Zaveri, Shagufta Khan, Tulasi Nagabandi, Purushotham Vodnala, Payel Mukherjee, Sofia Banu, Priya Singh, Onkar Kulkarni, Dhiviya Vedagiri, Divya Gupta, Vishal Sah, Santosh Kumar Kuncha, Krishnan Harinivas Harshan, Archana Bharadwaj Siva, Karthik Bharadwaj Tallapaka, Zeba Rizvi, Zuberwasim Sayyad, Kakade Aishwarya Arun, Amrutha H C, Ananga Ghosh, Rakesh K Mishra, Divya Tej Sowpati |
| EPI_ISL_539680 | CSIR-Centre for Cellular and Molecular Biology | CSIR-Centre for Cellular and Molecular Biology | M Soujanya Reddy, Nikhil Hajirnis, Pratheusa Maccha, Payel Mukherjee, Sofia Banu, Priya Singh, Onkar Kulkarni,Tulasi Nagabandi, Namami Gaur, Sakshi Shambhavi, Lamuk Zaveri, Shagufta Khan, Purushotham Vodnala, Dhiviya Vedagiri, Divya Gupta, Vishal Sah, Santosh Kumar Kuncha, Krishnan Harinivas Harshan, Archana Bharadwaj Siva, Karthik Bharadwaj Tallapaka,Kezia J Ann, Radhika Khandelwal, Roshan Maku Venkata, Shemin Mansuri, Sonu Uday, Rakesh K Mishra, Divya Tej Sowpati |
| EPI_ISL_539683 | CSIR-Centre for Cellular and Molecular Biology | CSIR-Centre for Cellular and Molecular Biology | Namami Gaur, Sakshi Shambhavi, Lamuk Zaveri, Shagufta Khan, Nikhil Hajirnis, M Soujanya Reddy, Pratheusa Maccha, Tulasi Nagabandi, Purushotham Vodnala, Payel Mukherjee, Sofia Banu, Priya Singh, Onkar Kulkarni, Dhiviya Vedagiri, Divya Gupta, Vishal Sah, Santosh Kumar Kuncha, Krishnan Harinivas Harshan, Archana Bharadwaj Siva, Karthik Bharadwaj Tallapaka,G. Aditya Kumar, Koushick Sivakumar, Rakesh K Mishra, Divya Tej Sowpati |
| EPI_ISL_539684 | CSIR-Centre for Cellular and Molecular Biology | CSIR-Centre for Cellular and Molecular Biology | Namami Gaur, Sakshi Shambhavi, Lamuk Zaveri, Shagufta Khan, Nikhil Hajirnis, M Soujanya Reddy, Pratheusa Maccha, Tulasi Nagabandi, Purushotham Vodnala, Payel Mukherjee, Sofia Banu, Priya Singh,Onkar Kulkarni, Dhiviya Vedagiri, Divya Gupta, Vishal Sah, Santosh Kumar Kuncha, Krishnan Harinivas Harshan, Archana Bharadwaj Siva, Karthik Bharadwaj Tallapaka, Zeba Rizvi, Zuberwasim Sayyad, Kakade Aishwarya Arun, Amrutha H C, Ananga Ghosh, Rakesh K Mishra, Divya Tej Sowpati |
| EPI_ISL_539711 | CSIR-Centre for Cellular and Molecular Biology | CSIR-Centre for Cellular and Molecular Biology | M Soujanya Reddy, Nikhil Hajirnis, Pratheusa Maccha, Sakshi Shambhavi, Lamuk Zaveri, Shagufta Khan, Namami Gaur, Tulasi Nagabandi, Purushotham Vodnala, Payel Mukherjee, Sofia Banu, Priya Singh,Onkar Kulkarni, Dhiviya Vedagiri, Divya Gupta, Vishal Sah, Santosh Kumar Kuncha, Krishnan Harinivas Harshan, Archana Bharadwaj Siva, Karthik Bharadwaj Tallapaka, G. Aditya Kumar, Koushick Sivakumar, Pooja Ramesh Gupta, Rajan Kumar Jha, Shraddha Vijay Lahoti, Rakesh K Mishra, Divya Tej Sowpati |
| EPI_ISL_539712 | CSIR-Centre for Cellular and Molecular Biology | CSIR-Centre for Cellular and Molecular Biology | Namami Gaur, Sakshi Shambhavi, Lamuk Zaveri, Shagufta Khan, Nikhil Hajirnis, M Soujanya Reddy, Pratheusa Maccha, Tulasi Nagabandi, Purushotham Vodnala, Payel Mukherjee, Sofia Banu, Priya Singh, Onkar Kulkarni, Dhiviya Vedagiri, Divya Gupta, Vishal Sah, Santosh Kumar Kuncha, Krishnan Harinivas Harshan, Archana Bharadwaj Siva, Karthik Bharadwaj Tallapaka, Zeba Rizvi, Zuberwasim Sayyad, Kakade Aishwarya Arun, Amrutha H C, Ananga Ghosh, Rakesh K Mishra, Divya Tej Sowpati |
| EPI_ISL_539720 | CSIR-Centre for Cellular and Molecular Biology | CSIR-Centre for Cellular and Molecular Biology | Payel Mukherjee, Sofia Banu, Priya Singh, Onkar Kulkarni, Dhiviya Vedagiri, Divya Gupta, Vishal Sah, Santosh Kumar Kuncha, Krishnan Harinivas Harshan, Archana Bharadwaj Siva, Karthik Bharadwaj Tallapaka, Shagufta Khan, Lamuk Zaveri, Nikhil Hajirnis, M Soujanya Reddy, Pratheusa Maccha, Namami Gaur, Sakshi Shambhavi, Tulasi Nagabandi, Purushotham Vodnala, Rakesh K Mishra, Sonu Uday, Sudipta Mondal, Annapoorna P Karthyayani, Debabrata Jana, Debrya Saha, Divya Tej Sowpati |

| | | | |
|---|---|---|---|
| EPI_ISL_539724 | CSIR-Centre for Cellular and Molecular Biology | CSIR-Centre for Cellular and Molecular Biology | Sakshi Shambhavi, Lamuk Zaveri, Shagufta Khan, Namami Gaur, Nikhil Hajirnis, M Soujanya Reddy, Pratheusa Maccha, Tulasi Nagabandi, Purushotham Vodnala, Payel Mukherjee, Sofia Banu, Priya Singh, Onkar Kulkarni, Dhiviya Vedagiri, Divya Gupta, Vishal Sah, Santosh Kumar Kuncha, Krishnan Harinivas Harshan, Archana Bharadwaj Siva, Karthik Bharadwaj Tallapaka, Deepak Kumar, Devi Prasad Vijayashankar, Disha Nanda, Divya Das, Jotin Gogoi, Manish Bhattacharjee, Rakesh K Mishra, Divya Tej Sowpati |
| EPI_ISL_539739 | CSIR-Centre for Cellular and Molecular Biology | CSIR-Centre for Cellular and Molecular Biology | M Soujanya Reddy, Nikhil Hajirnis, Pratheusa Maccha, Namami Gaur, Sakshi Shambhavi, Lamuk Zaveri, Shagufta Khan, Tulasi Nagabandi, Purushotham Vodnala, Payel Mukherjee, Sofia Banu, Priya Singh, Onkar Kulkarni, Dhiviya Vedagiri, Divya Gupta, Vishal Sah, Santosh Kumar Kuncha, Krishnan Harinivas Harshan, Archana Bharadwaj Siva, Karthik Bharadwaj Tallapaka, Zeba Rizvi, Zuberwasim Sayyad, Kakade Aishwarya Arun, Amrutha H C, Ananga Ghosh, Rakesh K Mishra, Divya Tej Sowpati |
| EPI_ISL_539745 | CSIR-Centre for Cellular and Molecular Biology | CSIR-Centre for Cellular and Molecular Biology | Nikhil Hajirnis, M Soujanya Reddy, Pratheusa Maccha, Lamuk Zaveri, Shagufta Khan, Namami Gaur, Sakshi Shambhavi, Tulasi Nagabandi, Purushotham Vodnala, Payel Mukherjee, Sofia Banu, Priya Singh, Onkar Kulkarni, Dhiviya Vedagiri, Divya Gupta, Vishal Sah, Santosh Kumar Kuncha, Krishnan Harinivas Harshan, Archana Bharadwaj Siva, Karthik Bharadwaj Tallapaka,Zeba Rizvi, Zuberwasim Sayyad, Kakade Aishwarya Arun, Amrutha H C, Ananga Ghosh, Rakesh K Mishra, Divya Tej Sowpati |
| EPI_ISL_539753 | CSIR-Centre for Cellular and Molecular Biology | CSIR-Centre for Cellular and Molecular Biology | Pratheusa Maccha, Sofia Banu, Payel Mukherjee, Priya Singh, Onkar Kulkarni, Dhiviya Vedagiri, Divya Gupta, Vishal Sah, Santosh Kumar Kuncha, Krishnan Harinivas Harshan, Archana Bharadwaj Siva, Karthik Bharadwaj Tallapaka, Shagufta Khan, Lamuk Zaveri, Namami Gaur, Sakshi Shambhavi, Nikhil Hajirnis, M Soujanya Reddy, Tulasi Nagabandi, Purushotham Vodnala,Preethi Jampala, Sharada Ravi Iyer, Sulagana Mukherjee, Swetha Sundar, Peddapuvala Sai Uday Kiran, Rakesh K Mishra, Divya Tej Sowpati |
| EPI_ISL_539755 | CSIR-Centre for Cellular and Molecular Biology | CSIR-Centre for Cellular and Molecular Biology | Sakshi Shambhavi, Lamuk Zaveri, Shagufta Khan, Namami Gaur, Nikhil Hajirnis, M Soujanya Reddy, Pratheusa Maccha,Tulasi Nagabandi, Purushotham Vodnala, Payel Mukherjee, Sofia Banu, Priya Singh,Onkar Kulkarni, Dhiviya Vedagiri, Divya Gupta, Vishal Sah, Santosh Kumar Kuncha, Krishnan Harinivas Harshan, Archana Bharadwaj Siva, Karthik Bharadwaj Tallapaka, G. Aditya Kumar, Koushick Sivakumar, Pooja Ramesh Gupta, Rajan Kumar Jha, Shraddha Vijay Lahoti, Rakesh K Mishra, Divya Tej Sowpati |
| EPI_ISL_539758 | CSIR-Centre for Cellular and Molecular Biology | CSIR-Centre for Cellular and Molecular Biology | Shagufta Khan, Lamuk Zaveri, Namami Gaur, Sakshi Shambhavi, Nikhil Hajirnis, M Soujanya Reddy, Pratheusa Maccha, Tulasi Nagabandi, Purushotham Vodnala, Payel Mukherjee, Sofia Banu, Priya Singh, Onkar Kulkarni, Dhiviya Vedagiri, Divya Gupta, Vishal Sah, Santosh Kumar Kuncha, Krishnan Harinivas Harshan, Archana Bharadwaj Siva, Karthik Bharadwaj Tallapaka,Preethi Jampala, Sharada Ravi Iyer, Sulagana Mukherjee, Swetha Sundar, Peddapuvala Sai Uday Kiran Rakesh K Mishra, Divya Tej Sowpati |
| EPI_ISL_539761 | CSIR-Centre for Cellular and Molecular Biology | CSIR-Centre for Cellular and Molecular Biology | Sofia Banu, Payel Mukherjee, Priya Singh,Onkar Kulkarni, Dhiviya Vedagiri, Divya Gupta, Vishal Sah, Santosh Kumar Kuncha, Krishnan Harinivas Harshan, Archana Bharadwaj Siva, Karthik Bharadwaj Tallapaka, Shagufta Khan, Lamuk Zaveri, Namami Gaur, Sakshi Shambhavi, Nikhil Hajirnis, M Soujanya Reddy, Pratheusa Maccha,Tulasi Nagabandi, Purushotham Vodnala, Deepak Kumar, Devi Prasad Vijayashankar, Disha Nanda, Divya Das, Jotin Gogoi, Manish Bhattacharjee, Rakesh K Mishra, Divya Tej Sowpati |
| EPI_ISL_539774 | CSIR-Centre for Cellular and Molecular Biology | CSIR-Centre for Cellular and Molecular Biology | Payel Mukherjee, Sofia Banu, Priya Singh, Onkar Kulkarni, Dhiviya Vedagiri, Divya Gupta, Vishal Sah, Santosh Kumar Kuncha, Krishnan Harinivas Harshan, Archana Bharadwaj Siva, Karthik Bharadwaj Tallapaka, Shagufta Khan, Lamuk Zaveri, Nikhil Hajirnis, M Soujanya Reddy, Pratheusa Maccha, Namami Gaur, Sakshi Shambhavi, Tulasi Nagabandi, Purushotham Vodnala, Rakesh K Mishra, Sonu Uday, Sudipta Mondal, Annapoorna P Karthyayani, Debabrata Jana, Debrya Saha, Divya Tej Sowpati |
| EPI_ISL_539775 | CSIR-Centre for Cellular and Molecular Biology | CSIR-Centre for Cellular and Molecular Biology | Payel Mukherjee, Sofia Banu, Priya Singh, Onkar Kulkarni, Dhiviya Vedagiri, Divya Gupta, Vishal Sah, Santosh Kumar Kuncha, Krishnan Harinivas Harshan, Archana Bharadwaj Siva, Karthik Bharadwaj Tallapaka, Shagufta Khan, Lamuk Zaveri, Nikhil Hajirnis, M Soujanya Reddy, Pratheusa Maccha, Namami Gaur, Sakshi Shambhavi, Tulasi Nagabandi, Purushotham Vodnala, Gokulan C G, Gunjan Purohit, Hanuman Tulashiram Kale, Pankaj Kumar, Prachand Issarapu, Rakesh K Mishra, Divya Tej Sowpati |
| EPI_ISL_539785, EPI_ISL_539786, EPI_ISL_539787, EPI_ISL_539788 | Institute of Microbiology, Universidad San Francisco de Quito | Institute of Microbiology, Universidad San Francisco de Quito | Andrea Macias, Belén Prado-Vivar, Sully Márquez, Juan José Guadalupe, Monica Becerra-Wong, Bernardo Gutiérrez, Verónica Barragán, Patricio Rojas-Silva, Gabriel Trueba, Michelle Grunauer, Paúl Cárdenas |
| EPI_ISL_539841 | Minnesota Department of Health, Public Health Laboratory | Minnesota Department of Health, Public Health Laboratory | Matt Plumb, Jacob Garfin, and Xiong Wang |
| EPI_ISL_539886 | Barnakuten | The Public Health Agency of Sweden | Anna-Malin Linde, Maria Lind Karlberg, Oskar Karlsson Lindsjo, Olov Svartstrom, Mattias Haukland, Reza Advani, Sandra Broddesson, Anna Risberg, Theresa Enkirch, Mia Brytting, Karin Tegmark-Wisell |
| EPI_ISL_539933 | Lighthouse Lab in Glasgow | Wellcome Sanger Institute for the COVID-19 Genomics UK (COG-UK) consortium | Harper VanSteenhouse, Yumi Kasai, David Gray, Carol Clugston, Anna Dominiczak and Alex Alderton, Roberto Amato, Sonia Goncalves, Ewan Harrison, David K. Jackson, Ian Johnston, Dominic Kwiatkowski, Cordelia Langford, John Sillitoe on behalf of the Wellcome Sanger Institute COVID-19 Surveillance Team |
| EPI_ISL_540507, EPI_ISL_540509, EPI_ISL_540510, EPI_ISL_540511, EPI_ISL_540512, EPI_ISL_540513, EPI_ISL_540514, EPI_ISL_540515, EPI_ISL_540516, EPI_ISL_540517, EPI_ISL_540518, EPI_ISL_540519, EPI_ISL_540520, EPI_ISL_540521, EPI_ISL_540522, EPI_ISL_540523, EPI_ISL_540524, EPI_ISL_540525, EPI_ISL_540526, EPI_ISL_540527, EPI_ISL_540528, EPI_ISL_540529, EPI_ISL_540530, EPI_ISL_540531, EPI_ISL_540532, EPI_ISL_540533, EPI_ISL_540534, EPI_ISL_540535, EPI_ISL_540536, EPI_ISL_540537, EPI_ISL_540538, EPI_ISL_540539, EPI_ISL_540540, EPI_ISL_540541, EPI_ISL_540542, EPI_ISL_540543, EPI_ISL_540544, EPI_ISL_540545, EPI_ISL_540546, EPI_ISL_540547, EPI_ISL_540548, EPI_ISL_540549, EPI_ISL_540550, EPI_ISL_540551, EPI_ISL_540552, EPI_ISL_540553, EPI_ISL_540554, EPI_ISL_540555, EPI_ISL_540556, EPI_ISL_540557, EPI_ISL_540558, EPI_ISL_540559, EPI_ISL_540560, EPI_ISL_540561, EPI_ISL_540562, EPI_ISL_540563, EPI_ISL_540564, EPI_ISL_540565, EPI_ISL_540566, EPI_ISL_540567 | | | |
| see above | Department of Clinical Microbiology | GIGA Medical Genomics | Keith Durkin, Maria Artesi, Sébastien Bontems, Raphaël Boreux, Bouchra Boujemla, Cécile Meex, Axelle Chaslain, Céline Fombellida-Lopez, Pierrette Melin, Marie-Pierre Hayette, Vincent Bours |
| EPI_ISL_540585, EPI_ISL_540586, EPI_ISL_540587, EPI_ISL_540588, EPI_ISL_540589, EPI_ISL_540590, EPI_ISL_540601, EPI_ISL_540604, EPI_ISL_540605, EPI_ISL_540608, EPI_ISL_540609, EPI_ISL_540610, EPI_ISL_540615, EPI_ISL_540628, EPI_ISL_540629, EPI_ISL_540630, EPI_ISL_540631, EPI_ISL_540632, EPI_ISL_540633, EPI_ISL_540634, EPI_ISL_540635, EPI_ISL_540636, EPI_ISL_540637 | | | |
| see above | Liverpool Clinical Laboratories | COVID-19 Genomics UK (COG-UK) Consortium | Sam Haldenby, Anita Lucaci, Steve Paterson, Julian Hiscox, Alistair Darby, M Almsaud, A Alrezaihi, Muhannad Alruwaili, Stuart D Armstrong, Jones Benjamin, Eleanor G Bentley, Anu Chawla, Jordan J Clark, Angela Cowell, Richard Eccles, Isabel García-Dorival, Matthew Gemmell, Alessandro Gerada, PKF Gilmore, Richard Gregory, Ximeng Han, Catherine Hartley, Margaret Hughes, Miren Iturriza-Gomara, James Johnson, L Luu, Jenifer Manson, Charlotte Nelson, Elaine O'Toole, Cassie Olateju, Rebekah Penrice-Randal , Lucille Rainbow, N.P Randle, Trevor Ian Robinson, Parul Sharma, Ghada T Shawli, James P Stewart, Neil Swainston, Ecaterina Vamos, Joanne Watts, Mark Whitehead |
| EPI_ISL_541130, EPI_ISL_541131, EPI_ISL_541132, EPI_ISL_541133, EPI_ISL_541134, EPI_ISL_541135, EPI_ISL_541136, EPI_ISL_541137 | Servicio de Microbiología. Hospital Universitario Donostia. OSI Donostialdea. Área de Enfermedades Infecciosas, Grupo de Infección Respiratoria y Resistencia Antimicrobiana. Instituto de Investigación Sanitaria Biodonostia | SeqCOVID-SPAIN consortium/Institute of Biomedicine of Valencia, IBV-CSIC | Gustavo Cilla, Milagrosa Montes, Luis Piñeiro, Jose Maria Marimón and SeqCOVID-SPAIN consortium |
| EPI_ISL_541139, EPI_ISL_541140, EPI_ISL_541141, EPI_ISL_541142, EPI_ISL_541199, EPI_ISL_541200, EPI_ISL_541201, EPI_ISL_541202, EPI_ISL_541203, EPI_ISL_541204, EPI_ISL_541205, EPI_ISL_541206, EPI_ISL_541207, EPI_ISL_541208, EPI_ISL_541209, EPI_ISL_541210 | | | |
| see above | Florida Bureau of Public Health Laboratories, Florida Department of Health | Florida Bureau of Public Health Laboratories, Florida Department of Health | Schmedes,S., Blanton,J. |
| EPI_ISL_541701, EPI_ISL_541702, EPI_ISL_541703, EPI_ISL_541704, EPI_ISL_541705, EPI_ISL_541706, EPI_ISL_541707, EPI_ISL_541708 | National Institute of Virology, NIV Influenza | National Institute of Virology, NIV Influenza | Potdar V |
| EPI_ISL_541753 | Microbiology Department, Barking Havering and Redbridge University Hospitals NHS trust | Wellcome Sanger Institute for the COVID-19 Genomics UK (COG-UK) consortium | Amy Ash, Fatima Ali, Cherian Koshy and Alex Alderton, Roberto Amato, Sonia Goncalves, Ewan Harrison, David K. Jackson, Ian Johnston, Dominic Kwiatkowski, Cordelia Langford, John Sillitoe on behalf of the Wellcome Sanger Institute COVID-19 Surveillance Team |
| EPI_ISL_541904, EPI_ISL_541905, EPI_ISL_541906, EPI_ISL_541907, EPI_ISL_541908, EPI_ISL_541909, EPI_ISL_541910, EPI_ISL_541911, EPI_ISL_541912, EPI_ISL_541913, EPI_ISL_541914, EPI_ISL_541915, EPI_ISL_541916, EPI_ISL_541917, EPI_ISL_541918, EPI_ISL_541919 | | | |
| see above | Hospital General Universitario Gregorio Marañón | SeqCOVID-SPAIN consortium/IBV(CSIC) | Laura Pérez-Lago, Marta Herranz, Jon Sicilia, Julia Suárez, Pilar Catalán, Patricia Muñoz, Darío García de Viedma and SeqCOVID-SPAIN consortium |
| EPI_ISL_542983, EPI_ISL_542984, EPI_ISL_542985, EPI_ISL_542986, EPI_ISL_542987, EPI_ISL_542988, | TriCore Reference Laboratories | Center for Global Health, University of New Mexico Health Sciences Center | Daryl Domman, Kurt Schwalm, Twila Kunde, Joseph Hicks, Michael Edwards, Darrell Dinwiddie |

| | | | |
|---|---|---|---|
| EPI_ISL_542989, EPI_ISL_542991 | | | |
| EPI_ISL_544958, EPI_ISL_544959 | Sydney South West Pathology Service (SSWPS) - Liverpool Hospital - NSW Health Pathology | NSW Health Pathology - Institute of Clinical Pathology and Medical Research; Westmead Hospital; University of Sydney | CIDM-PH et al. |
| EPI_ISL_544960 | Histopath | NSW Health Pathology - Institute of Clinical Pathology and Medical Research; Westmead Hospital; University of Sydney | CIDM-PH et al. |
| EPI_ISL_544963 | The Children's Hospital at Westmead | NSW Health Pathology - Institute of Clinical Pathology and Medical Research; Westmead Hospital; University of Sydney | CIDM-PH et al. |
| EPI_ISL_545016 | South Eastern Area Laboratory Services (SEALS) | NSW Health Pathology - Institute of Clinical Pathology and Medical Research; Westmead Hospital; University of Sydney | CIDM-PH et al. |
| EPI_ISL_545018, EPI_ISL_545019 | Sydney South West Pathology Service (SSWPS) - Royal Prince Alfred Hospital - NSW Health Pathology | NSW Health Pathology - Institute of Clinical Pathology and Medical Research; Westmead Hospital; University of Sydney | CIDM-PH et al. |
| EPI_ISL_546935, EPI_ISL_546936 | The National Institute of Public Health | State Veterinary Institute Prague | Nagy,A; Jirincova,H; Novakova,L; Trnka,D; Vecerova,J |
| EPI_ISL_547436, EPI_ISL_547438, EPI_ISL_547439, EPI_ISL_547440, EPI_ISL_547441, EPI_ISL_547442, EPI_ISL_547443 | Microbiology, Department of Pathology, St. Bernard's Hospital, Gibraltar Health Authority | Respiratory Virus Unit, Microbiology Services Colindale, Public Health England | PHE Covid Sequencing Team, Dr Nicholas Cortes (Gibraltar), Charlotte Gillborn-Jones (Gibraltar) |
| EPI_ISL_547655, EPI_ISL_547656, EPI_ISL_547657, EPI_ISL_547658, EPI_ISL_547659, EPI_ISL_547660, EPI_ISL_547661, EPI_ISL_547662, EPI_ISL_547663, EPI_ISL_547664, EPI_ISL_547665 | | | |
| see above | Gundersen Molecular Diagnostics Laboratory | Kabara Cancer Research Institute | Craig S. Richmond, Paraic A. Kenny |
| EPI_ISL_547676 | Gundersen Clinical Microbiology Laboratory | Kabara Cancer Research Institute | Craig S. Richmond, Paraic A. Kenny |
| EPI_ISL_547677, EPI_ISL_547678, EPI_ISL_547679, EPI_ISL_547680, EPI_ISL_547681, EPI_ISL_547682 | Gundersen Molecular Diagnostics Laboratory | Kabara Cancer Research Institute | Craig S. Richmond, Paraic A. Kenny |
| EPI_ISL_547683, EPI_ISL_547684 | Gundersen Clinical Microbiology Laboratory | Kabara Cancer Research Institute | Craig S. Richmond, Paraic A. Kenny |
| EPI_ISL_547685, EPI_ISL_547686, EPI_ISL_547687, EPI_ISL_547688 | Gundersen Molecular Diagnostics Laboratory | Kabara Cancer Research Institute | Craig S. Richmond, Paraic A. Kenny |
| EPI_ISL_547966, EPI_ISL_547967, EPI_ISL_547968 | The National Institute of Public Health | State Veterinary Institute Prague | Nagy,A;Jirincova,H;Novakova,L;Trnka,D;Vecerova,J |
| EPI_ISL_548144 | Middlemore Hospital | Institute of Environmental Science and Research (ESR) | Xiaoyun Ren, Matt Storey, Nikki Freed, Muhammad Faisal, Jing Wang, Hermes Perez, Anja Werno, Antje van der Linden, Arlo Upton, Chris Mansell, David Hammer, Dragana Drinkovic, Gary McAuliffe, Hana Sofia Andersson, James Ussher, Jill Sherwood, Josh Freeman, Julia Howard, Juliet Elvy, Mary DeAlmeida, Matt Blakiston, Matthew Rogers, Max Bloomfield, Michael Addidle, Michelle Balm, Sally Roberts, Sarah Jefferies, Sharmini Muttaiyah, Susan Morpeth, Susan Taylor, Timothy Blackmore, Vani Sathyendran, Veronica Playle, Virginia Hope, Erasmus Smit, Lauren Jelly, Olin Silander, Joep de Ligt |
| EPI_ISL_548258 | Karolinska universitetslaboratoriet | The Public Health Agency of Sweden | Anna-Malin Linde, Maria Lind Karlberg, Mattias Haukland, Reza Advani, Olov Svartstrom, Oskar Karlsson Lindsjo, Sandra Broddesson, Petra Edquist, Mia Brytting, Anna Risberg, Karin Tegmark-Wisell |
| EPI_ISL_548261 | Genome Center | Genome Center | Najuj Sakib, A. S. M. Rubayet- Ul- Alam, Tanay Chakrovarty, Md. Shazid Hasan, Hassan M. Al-Emran, Ovinu Kibria Islam, Md. Tanvir Islam, Pravas Chandra Roy, Md. Iqbal Kabir Jahid, Md. Anwar Hossain |
| EPI_ISL_548262 | Genome Center | Genome Center | Tanay Chakrovarty, Najuj Sakib, A. S. M. Rubayet- Ul- Alam, Md. Shazid Hasan, Hassan M. Al-Emran, Ovinu Kibria Islam, Md. Tanvir Islam, Pravas Chandra Roy, Md. Iqbal Kabir Jahid, Md. Anwar Hossain |
| EPI_ISL_548263 | Genome Center | Genome Center | A. S. M. Rubayet- Ul- Alam, Md. Shazid Hasan, Najuj Sakib, Ovinu Kibria Islam, Tanay Chakrovarty, Md. Tanvir Islam, Hassan M. Al-Emran, Pravas Chandra Roy, Md. Iqbal Kabir Jahid, Md. Anwar Hossain |
| EPI_ISL_548347 | County of San Luis Obispo Public Health Laboratory | Chan-Zuckerberg Biohub | CZB Cliahub Consortium |
| EPI_ISL_548420, EPI_ISL_548434, EPI_ISL_548470 | County of Santa Clara Public Health Department | Chan-Zuckerberg Biohub | CZB Cliahub Consortium |
| EPI_ISL_548520, EPI_ISL_548558, EPI_ISL_548577 | Contra Costa Public Health Lab | Chan-Zuckerberg Biohub | CZB Cliahub Consortium |
| EPI_ISL_548580, EPI_ISL_548581 | Orange County Public Health Laboratory | Chan-Zuckerberg Biohub | CZB Cliahub Consortium |
| EPI_ISL_548582, EPI_ISL_548584 | County of Santa Clara Public Health Department | Chan-Zuckerberg Biohub | CZB Cliahub Consortium |
| EPI_ISL_548585, EPI_ISL_548586 | Orange County Public Health Laboratory | Chan-Zuckerberg Biohub | CZB Cliahub Consortium |
| EPI_ISL_548588 | County of Santa Clara Public Health Department | Chan-Zuckerberg Biohub | CZB Cliahub Consortium |
| EPI_ISL_548592, EPI_ISL_548594 | Orange County Public Health Laboratory | Chan-Zuckerberg Biohub | CZB Cliahub Consortium |
| EPI_ISL_548595, EPI_ISL_548606 | County of Santa Clara Public Health Department | Chan-Zuckerberg Biohub | CZB Cliahub Consortium |
| EPI_ISL_548609, EPI_ISL_548611 | Orange County Public Health Laboratory | Chan-Zuckerberg Biohub | CZB Cliahub Consortium |
| EPI_ISL_548612, EPI_ISL_548616, EPI_ISL_548617, EPI_ISL_548618, EPI_ISL_548619, EPI_ISL_548621 | County of Santa Clara Public Health Department | Chan-Zuckerberg Biohub | CZB Cliahub Consortium |
| EPI_ISL_548623 | Orange County Public Health Laboratory | Chan-Zuckerberg Biohub | CZB Cliahub Consortium |
| EPI_ISL_548624 | County of Santa Clara Public Health Department | Chan-Zuckerberg Biohub | CZB Cliahub Consortium |
| EPI_ISL_548625, EPI_ISL_548627, EPI_ISL_548628 | Orange County Public Health Laboratory | Chan-Zuckerberg Biohub | CZB Cliahub Consortium |
| EPI_ISL_548629, EPI_ISL_548630, EPI_ISL_548633 | County of Santa Clara Public Health Department | Chan-Zuckerberg Biohub | CZB Cliahub Consortium |
| EPI_ISL_548635 | Orange County Public Health Laboratory | Chan-Zuckerberg Biohub | CZB Cliahub Consortium |
| EPI_ISL_548636 | County of Santa Clara Public Health Department | Chan-Zuckerberg Biohub | CZB Cliahub Consortium |
| EPI_ISL_548637 | Orange County Public Health Laboratory | Chan-Zuckerberg Biohub | CZB Cliahub Consortium |
| EPI_ISL_548639, EPI_ISL_548643, EPI_ISL_548645, EPI_ISL_548647 | County of Santa Clara Public Health Department | Chan-Zuckerberg Biohub | CZB Cliahub Consortium |
| EPI_ISL_548648 | Orange County Public Health Laboratory | Chan-Zuckerberg Biohub | CZB Cliahub Consortium |
| EPI_ISL_548650 | County of Santa Clara Public Health Department | Chan-Zuckerberg Biohub | CZB Cliahub Consortium |

| | | | |
|---|---|---|---|
| EPI_ISL_548651, EPI_ISL_548653, EPI_ISL_548654 | Orange County Public Health Laboratory | Chan-Zuckerberg Biohub | CZB Cliahub Consortium |
| EPI_ISL_548655, EPI_ISL_548656, EPI_ISL_548658 | County of Santa Clara Public Health Department | Chan-Zuckerberg Biohub | CZB Cliahub Consortium |
| EPI_ISL_548660, EPI_ISL_548662 | Orange County Public Health Laboratory | Chan-Zuckerberg Biohub | CZB Cliahub Consortium |
| EPI_ISL_548664 | County of Santa Clara Public Health Department | Chan-Zuckerberg Biohub | CZB Cliahub Consortium |
| EPI_ISL_548665, EPI_ISL_548667, EPI_ISL_548668 | Orange County Public Health Laboratory | Chan-Zuckerberg Biohub | CZB Cliahub Consortium |
| EPI_ISL_548669, EPI_ISL_548670, EPI_ISL_548672, EPI_ISL_548673 | County of Santa Clara Public Health Department | Chan-Zuckerberg Biohub | CZB Cliahub Consortium |
| EPI_ISL_548674, EPI_ISL_548675 | Orange County Public Health Laboratory | Chan-Zuckerberg Biohub | CZB Cliahub Consortium |
| EPI_ISL_548676 | County of Santa Clara Public Health Department | Chan-Zuckerberg Biohub | CZB Cliahub Consortium |
| EPI_ISL_548677 | Orange County Public Health Laboratory | Chan-Zuckerberg Biohub | CZB Cliahub Consortium |
| EPI_ISL_548678 | County of Santa Clara Public Health Department | Chan-Zuckerberg Biohub | CZB Cliahub Consortium |
| EPI_ISL_549032 | Oslo University Hospital, Department of Medical Microbiology | Norwegian Institute of Public Health, Department of Virology | Kathrine Stene-Johansen, Kamilla Heddeland Instefjord, Hilde Elshaug, Rasmus Riis Kopperud, Hilde Synnøve Vollan, Karoline Bragstad, Olav Hungnes |
| EPI_ISL_549042, EPI_ISL_549043, EPI_ISL_549044, EPI_ISL_549045, EPI_ISL_549046, EPI_ISL_549047 | Ostfold Hospital Trust - Kalnes, Centre for Laboratory Medicine, Section for gene technology and infection serology | Norwegian Institute of Public Health, Department of Virology | Kathrine Stene-Johansen, Kamilla Heddeland Instefjord, Hilde Elshaug, Rasmus Riis Kopperud, Hilde Synnøve Vollan, Karoline Bragstad, Olav Hungnes |
| EPI_ISL_549052 | Hospital of Southern Norway - Kristiansand, Department of Medical Microbiology | Norwegian Institute of Public Health, Department of Virology | Kathrine Stene-Johansen, Kamilla Heddeland Instefjord, Hilde Elshaug, Rasmus Riis Kopperud, Hilde Synnøve Vollan, Karoline Bragstad, Olav Hungnes |
| EPI_ISL_549053, EPI_ISL_549054, EPI_ISL_549055, EPI_ISL_549056, EPI_ISL_549057, EPI_ISL_549058 | Furst Medical Laboratory | Norwegian Institute of Public Health, Department of Virology | Kathrine Stene-Johansen, Kamilla Heddeland Instefjord, Hilde Elshaug, Rasmus Riis Kopperud, Hilde Synnøve Vollan, Karoline Bragstad, Olav Hungnes |
| EPI_ISL_549059 | Medical Microbiology Unit, Department for Laboratory Medicine, Drammen Hospital, Vestre Viken Health Trust, | Norwegian Institute of Public Health, Department of Virology | Kathrine Stene-Johansen, Kamilla Heddeland Instefjord, Hilde Elshaug, Rasmus Riis Kopperud, Hilde Synnøve Vollan, Karoline Bragstad, Olav Hungnes |
| EPI_ISL_549060, EPI_ISL_549061, EPI_ISL_549062, EPI_ISL_549063, EPI_ISL_549064, EPI_ISL_549065, EPI_ISL_549066, EPI_ISL_549067, EPI_ISL_549068, EPI_ISL_549072, EPI_ISL_549073, EPI_ISL_549074, EPI_ISL_549075, EPI_ISL_549076, EPI_ISL_549077, EPI_ISL_549078, EPI_ISL_549079, EPI_ISL_549080, EPI_ISL_549082 | | | |
| see above | Furst Medical Laboratory | Norwegian Institute of Public Health, Department of Virology | Kathrine Stene-Johansen, Kamilla Heddeland Instefjord, Hilde Elshaug, Rasmus Riis Kopperud, Hilde Synnøve Vollan, Karoline Bragstad, Olav Hungnes |
| EPI_ISL_549083, EPI_ISL_549084 | Akershus University Hospital, Department for Microbiology and Infectious Disease Control | Norwegian Institute of Public Health, Department of Virology | Kathrine Stene-Johansen, Kamilla Heddeland Instefjord, Hilde Elshaug, Rasmus Riis Kopperud, Hilde Synnøve Vollan, Karoline Bragstad, Olav Hungnes |
| EPI_ISL_549085, EPI_ISL_549086 | Medical Microbiology Unit, Department for Laboratory Medicine, Drammen Hospital, Vestre Viken Health Trust, | Norwegian Institute of Public Health, Department of Virology | Kathrine Stene-Johansen, Kamilla Heddeland Instefjord, Hilde Elshaug, Rasmus Riis Kopperud, Hilde Synnøve Vollan, Karoline Bragstad, Olav Hungnes |
| EPI_ISL_549087 | Vestfold Hospital, Toensberg Department of Microbiology | Norwegian Institute of Public Health, Department of Virology | Kathrine Stene-Johansen, Kamilla Heddeland Instefjord, Hilde Elshaug, Rasmus Riis Kopperud, Hilde Synnøve Vollan, Karoline Bragstad, Olav Hungnes |
| EPI_ISL_549088 | Furst Medical Laboratory | Norwegian Institute of Public Health, Department of Virology | Kathrine Stene-Johansen, Kamilla Heddeland Instefjord, Hilde Elshaug, Rasmus Riis Kopperud, Hilde Synnøve Vollan, Karoline Bragstad, Olav Hungnes |
| EPI_ISL_549090 | Akershus University Hospital, Department for Microbiology and Infectious Disease Control | Norwegian Institute of Public Health, Department of Virology | Kathrine Stene-Johansen, Kamilla Heddeland Instefjord, Hilde Elshaug, Rasmus Riis Kopperud, Hilde Synnøve Vollan, Karoline Bragstad, Olav Hungnes |
| EPI_ISL_549101 | Ostfold Hospital Trust - Kalnes, Centre for Laboratory Medicine, Section for gene technology and infection serology | Norwegian Institute of Public Health, Department of Virology | Kathrine Stene-Johansen, Kamilla Heddeland Instefjord, Hilde Elshaug, Rasmus Riis Kopperud, Hilde Synnøve Vollan, Karoline Bragstad, Olav Hungnes |
| EPI_ISL_549112, EPI_ISL_549119, EPI_ISL_549120, EPI_ISL_549121 | Furst Medical Laboratory | Norwegian Institute of Public Health, Department of Virology | Kathrine Stene-Johansen, Kamilla Heddeland Instefjord, Hilde Elshaug, Rasmus Riis Kopperud, Hilde Synnøve Vollan, Karoline Bragstad, Olav Hungnes |
| EPI_ISL_549123, EPI_ISL_549125 | Unilabs Laboratory Medicine | Norwegian Institute of Public Health, Department of Virology | Kathrine Stene-Johansen, Kamilla Heddeland Instefjord, Hilde Elshaug, Rasmus Riis Kopperud, Hilde Synnøve Vollan, Karoline Bragstad, Olav Hungnes |
| EPI_ISL_549126, EPI_ISL_549127, EPI_ISL_549128, EPI_ISL_549129, EPI_ISL_549130, EPI_ISL_549131, EPI_ISL_549133, EPI_ISL_549134, EPI_ISL_549135, EPI_ISL_549136, EPI_ISL_549137, EPI_ISL_549138, EPI_ISL_549139, EPI_ISL_549140, EPI_ISL_549141, EPI_ISL_549142, EPI_ISL_549144, EPI_ISL_549145, EPI_ISL_549146, EPI_ISL_549147, EPI_ISL_549148, EPI_ISL_549149, EPI_ISL_549150, EPI_ISL_549151, EPI_ISL_549152, EPI_ISL_549153, EPI_ISL_549155, EPI_ISL_549156, EPI_ISL_549157, EPI_ISL_549158, EPI_ISL_549159, EPI_ISL_549160, EPI_ISL_549161, EPI_ISL_549162, EPI_ISL_549163 | | | |
| see above | Ostfold Hospital Trust - Kalnes, Centre for Laboratory Medicine, Section for gene technology and infection serology | Norwegian Institute of Public Health, Department of Virology | Kathrine Stene-Johansen, Kamilla Heddeland Instefjord, Hilde Elshaug, Rasmus Riis Kopperud, Hilde Synnøve Vollan, Karoline Bragstad, Olav Hungnes |
| EPI_ISL_549192, EPI_ISL_549193, EPI_ISL_549194, EPI_ISL_549195, EPI_ISL_549196, EPI_ISL_549197, EPI_ISL_549198, EPI_ISL_549199 | Florida Bureau of Public Health Laboratories | Florida Bureau of Public Health Laboratories | Sarah Schmedes, Jason Blanton |
| EPI_ISL_549950 | Lighthouse Lab in Milton Keynes | Wellcome Sanger Institute for the COVID-19 Genomics UK (COG-UK) consortium | The Lighthouse Lab in Milton Keynes and Alex Alderton, Roberto Amato, Sonia Goncalves, Ewan Harrison, David K. Jackson, Ian Johnston, Dominic Kwiatkowski, Cordelia Langford, John Sillitoe on behalf of the Wellcome Sanger Institute COVID-19 Surveillance Team (http://www.sanger.ac.uk/covid-team) |
| EPI_ISL_550421 | Lighthouse Lab in Milton Keynes | Wellcome Sanger Institute for the COVID-19 Genomics UK (COG-UK) consortium | The Lighthouse Lab in Alderley Park and Alex Alderton, Roberto Amato, Sonia Goncalves, Ewan Harrison, David K. Jackson, Ian Johnston, Dominic Kwiatkowski, Cordelia Langford, John Sillitoe on behalf of the Wellcome Sanger Institute COVID-19 Surveillance Team |
| EPI_ISL_550423, EPI_ISL_550438, EPI_ISL_550439, EPI_ISL_550448, EPI_ISL_550453, EPI_ISL_550468, EPI_ISL_550470, EPI_ISL_550477, EPI_ISL_550478 | Lighthouse Lab in Milton Keynes | Wellcome Sanger Institute for the COVID-19 Genomics UK (COG-UK) consortium | The Lighthouse Lab in Milton Keynes and Alex Alderton, Roberto Amato, Sonia Goncalves, Ewan Harrison, David K. Jackson, Ian Johnston, Dominic Kwiatkowski, Cordelia Langford, John Sillitoe on behalf of the Wellcome Sanger Institute COVID-19 Surveillance Team (http://www.sanger.ac.uk/covid-team) |
| EPI_ISL_550485 | Lighthouse Lab in Milton Keynes | Wellcome Sanger Institute for the COVID-19 Genomics UK (COG-UK) Consortium | The Lighthouse Lab in Milton Keynes and Alex Alderton, Roberto Amato, Sonia Goncalves, Ewan Harrison, David K. Jackson, Ian Johnston, Dominic Kwiatkowski, Cordelia Langford, John Sillitoe on behalf of the Wellcome Sanger Institute COVID-19 Surveillance Team |
| EPI_ISL_550497, EPI_ISL_550498, EPI_ISL_550499, EPI_ISL_550503, EPI_ISL_550511 | Lighthouse Lab in Milton Keynes | Wellcome Sanger Institute for the COVID-19 Genomics UK (COG-UK) consortium | The Lighthouse Lab in Milton Keynes and Alex Alderton, Roberto Amato, Sonia Goncalves, Ewan Harrison, David K. Jackson, Ian Johnston, Dominic Kwiatkowski, Cordelia Langford, John Sillitoe on behalf of the Wellcome Sanger Institute COVID-19 Surveillance Team (http://www.sanger.ac.uk/covid-team) |
| EPI_ISL_552446, EPI_ISL_552449 | Lighthouse Lab in Alderley Park | Wellcome Sanger Institute for the COVID-19 Genomics UK (COG-UK) consortium | The Lighthouse Lab in Alderley Park and Alex Alderton, Roberto Amato, Sonia Goncalves, Ewan Harrison, David K. Jackson, Ian Johnston, Dominic Kwiatkowski, Cordelia Langford, John Sillitoe on behalf of the Wellcome Sanger Institute COVID-19 Surveillance Team |
| EPI_ISL_552463 | Lighthouse Lab in Alderley Park | Wellcome Sanger Institute for the COVID-19 Genomics UK (COG-UK) consortium | The Lighthouse Lab in Alderley Park and Alex Alderton, Roberto Amato, Sonia Goncalves, Ewan Harrison, David K. Jackson, Ian Johnston, Dominic Kwiatkowski, Cordelia Langford, John Sillitoe on behalf of the Wellcome Sanger Institute COVID-19 Surveillance Team (http://www.sanger.ac.uk/covid-team) |
| EPI_ISL_552464, EPI_ISL_552466, EPI_ISL_552468, EPI_ISL_552473, EPI_ISL_552476, EPI_ISL_552481, | Lighthouse Lab in Alderley Park | Wellcome Sanger Institute for the COVID-19 Genomics UK (COG-UK) consortium | The Lighthouse Lab in Alderley Park and Alex Alderton, Roberto Amato, Sonia Goncalves, Ewan Harrison, David K. Jackson, Ian Johnston, Dominic Kwiatkowski, Cordelia Langford, John Sillitoe on behalf of the Wellcome Sanger Institute COVID-19 Surveillance Team |

| | | | |
|---|---|---|---|
| EPI_ISL_552486 | | | |
| EPI_ISL_552488 | Lighthouse Lab in Alderley Park | Wellcome Sanger Institute for the COVID-19 Genomics UK (COG-UK) consortium | The Lighthouse Lab in Alderley Park and Alex Alderton, Roberto Amato, Sonia Goncalves, Ewan Harrison, David K. Jackson, Ian Johnston, Dominic Kwiatkowski, Cordelia Langford, John Sillitoe on behalf of the Wellcome Sanger Institute COVID-19 Surveillance Team (http://www.sanger.ac.uk/covid-team) |
| EPI_ISL_552492 | Lighthouse Lab in Alderley Park | Wellcome Sanger Institute for the COVID-19 Genomics UK (COG-UK) Consortium | The Lighthouse Lab in Alderley Park and Alex Alderton, Roberto Amato, Sonia Goncalves, Ewan Harrison, David K. Jackson, Ian Johnston, Dominic Kwiatkowski, Cordelia Langford, John Sillitoe on behalf of the Wellcome Sanger Institute COVID-19 Surveillance Team |
| EPI_ISL_552506, EPI_ISL_552507, EPI_ISL_552508, EPI_ISL_552512, EPI_ISL_552515, EPI_ISL_552516, EPI_ISL_552523, EPI_ISL_552526, EPI_ISL_552527, EPI_ISL_552530, EPI_ISL_552533 | | | |
| see above | Lighthouse Lab in Alderley Park | Wellcome Sanger Institute for the COVID-19 Genomics UK (COG-UK) consortium | The Lighthouse Lab in Alderley Park and Alex Alderton, Roberto Amato, Sonia Goncalves, Ewan Harrison, David K. Jackson, Ian Johnston, Dominic Kwiatkowski, Cordelia Langford, John Sillitoe on behalf of the Wellcome Sanger Institute COVID-19 Surveillance Team |
| EPI_ISL_552534 | Lighthouse Lab in Alderley Park | Wellcome Sanger Institute for the COVID-19 Genomics UK (COG-UK) consortium | The Lighthouse Lab in Alderley Park and Alex Alderton, Roberto Amato, Sonia Goncalves, Ewan Harrison, David K. Jackson, Ian Johnston, Dominic Kwiatkowski, Cordelia Langford, John Sillitoe on behalf of the Wellcome Sanger Institute COVID-19 Surveillance Team (http://www.sanger.ac.uk/covid-team) |
| EPI_ISL_552539, EPI_ISL_552540, EPI_ISL_552547, EPI_ISL_552556, EPI_ISL_552562, EPI_ISL_552565, EPI_ISL_552566, EPI_ISL_552572 | Lighthouse Lab in Alderley Park | Wellcome Sanger Institute for the COVID-19 Genomics UK (COG-UK) consortium | The Lighthouse Lab in Alderley Park and Alex Alderton, Roberto Amato, Sonia Goncalves, Ewan Harrison, David K. Jackson, Ian Johnston, Dominic Kwiatkowski, Cordelia Langford, John Sillitoe on behalf of the Wellcome Sanger Institute COVID-19 Surveillance Team |
| EPI_ISL_552574 | Lighthouse Lab in Alderley Park | Wellcome Sanger Institute for the COVID-19 Genomics UK (COG-UK) consortium | The Lighthouse Lab in Alderley Park and Alex Alderton, Roberto Amato, Sonia Goncalves, Ewan Harrison, David K. Jackson, Ian Johnston, Dominic Kwiatkowski, Cordelia Langford, John Sillitoe on behalf of the Wellcome Sanger Institute COVID-19 Surveillance Team (http://www.sanger.ac.uk/covid-team) |
| EPI_ISL_552576, EPI_ISL_552577, EPI_ISL_552579, EPI_ISL_552580, EPI_ISL_552583, EPI_ISL_552587, EPI_ISL_552590, EPI_ISL_552591, EPI_ISL_552595, EPI_ISL_552596, EPI_ISL_552598, EPI_ISL_552607, EPI_ISL_552608, EPI_ISL_552614, EPI_ISL_552617, EPI_ISL_552618, EPI_ISL_552620, EPI_ISL_552622, EPI_ISL_552629 | | | |
| see above | Lighthouse Lab in Alderley Park | Wellcome Sanger Institute for the COVID-19 Genomics UK (COG-UK) consortium | The Lighthouse Lab in Alderley Park and Alex Alderton, Roberto Amato, Sonia Goncalves, Ewan Harrison, David K. Jackson, Ian Johnston, Dominic Kwiatkowski, Cordelia Langford, John Sillitoe on behalf of the Wellcome Sanger Institute COVID-19 Surveillance Team |
| EPI_ISL_552639 | Lighthouse Lab in Alderley Park | Wellcome Sanger Institute for the COVID-19 Genomics UK (COG-UK) consortium | The Lighthouse Lab in Alderley Park and Alex Alderton, Roberto Amato, Sonia Goncalves, Ewan Harrison, David K. Jackson, Ian Johnston, Dominic Kwiatkowski, Cordelia Langford, John Sillitoe on behalf of the Wellcome Sanger Institute COVID-19 Surveillance Team (http://www.sanger.ac.uk/covid-team) |
| EPI_ISL_552642, EPI_ISL_552644, EPI_ISL_552646, EPI_ISL_552655, EPI_ISL_552659, EPI_ISL_552665, EPI_ISL_552666, EPI_ISL_552669, EPI_ISL_552671, EPI_ISL_552679, EPI_ISL_552681, EPI_ISL_552686 | | | |
| see above | Lighthouse Lab in Alderley Park | Wellcome Sanger Institute for the COVID-19 Genomics UK (COG-UK) consortium | The Lighthouse Lab in Alderley Park and Alex Alderton, Roberto Amato, Sonia Goncalves, Ewan Harrison, David K. Jackson, Ian Johnston, Dominic Kwiatkowski, Cordelia Langford, John Sillitoe on behalf of the Wellcome Sanger Institute COVID-19 Surveillance Team |
| EPI_ISL_552689 | Lighthouse Lab in Alderley Park | Wellcome Sanger Institute for the COVID-19 Genomics UK (COG-UK) consortium | The Lighthouse Lab in Alderley Park and Alex Alderton, Roberto Amato, Sonia Goncalves, Ewan Harrison, David K. Jackson, Ian Johnston, Dominic Kwiatkowski, Cordelia Langford, John Sillitoe on behalf of the Wellcome Sanger Institute COVID-19 Surveillance Team (http://www.sanger.ac.uk/covid-team) |
| EPI_ISL_552699, EPI_ISL_552703, EPI_ISL_552706, EPI_ISL_552707, EPI_ISL_552712, EPI_ISL_552713, EPI_ISL_552717, EPI_ISL_552719, EPI_ISL_552720, EPI_ISL_552721, EPI_ISL_552722, EPI_ISL_552723, EPI_ISL_552724, EPI_ISL_552726, EPI_ISL_552728, EPI_ISL_552729 | | | |
| see above | Lighthouse Lab in Alderley Park | Wellcome Sanger Institute for the COVID-19 Genomics UK (COG-UK) consortium | The Lighthouse Lab in Alderley Park and Alex Alderton, Roberto Amato, Sonia Goncalves, Ewan Harrison, David K. Jackson, Ian Johnston, Dominic Kwiatkowski, Cordelia Langford, John Sillitoe on behalf of the Wellcome Sanger Institute COVID-19 Surveillance Team |
| EPI_ISL_552730 | Lighthouse Lab in Alderley Park | Wellcome Sanger Institute for the COVID-19 Genomics UK (COG-UK) consortium | The Lighthouse Lab in Alderley Park and Alex Alderton, Roberto Amato, Sonia Goncalves, Ewan Harrison, David K. Jackson, Ian Johnston, Dominic Kwiatkowski, Cordelia Langford, John Sillitoe on behalf of the Wellcome Sanger Institute COVID-19 Surveillance Team (http://www.sanger.ac.uk/covid-team) |
| EPI_ISL_552731, EPI_ISL_552732, EPI_ISL_552735 | Lighthouse Lab in Alderley Park | Wellcome Sanger Institute for the COVID-19 Genomics UK (COG-UK) consortium | The Lighthouse Lab in Alderley Park and Alex Alderton, Roberto Amato, Sonia Goncalves, Ewan Harrison, David K. Jackson, Ian Johnston, Dominic Kwiatkowski, Cordelia Langford, John Sillitoe on behalf of the Wellcome Sanger Institute COVID-19 Surveillance Team |
| EPI_ISL_552741, EPI_ISL_552742, EPI_ISL_552745, EPI_ISL_552746, EPI_ISL_552748, EPI_ISL_552749 | Lighthouse Lab in Alderley Park | Wellcome Sanger Institute for the COVID-19 Genomics UK (COG-UK) consortium | The Lighthouse Lab in Alderley Park and Alex Alderton, Roberto Amato, Sonia Goncalves, Ewan Harrison, David K. Jackson, Ian Johnston, Dominic Kwiatkowski, Cordelia Langford, John Sillitoe on behalf of the Wellcome Sanger Institute COVID-19 Surveillance Team (http://www.sanger.ac.uk/covid-team) |
| EPI_ISL_552750, EPI_ISL_552760, EPI_ISL_552761, EPI_ISL_552768, EPI_ISL_552770, EPI_ISL_552771, EPI_ISL_552773 | Lighthouse Lab in Alderley Park | Wellcome Sanger Institute for the COVID-19 Genomics UK (COG-UK) consortium | The Lighthouse Lab in Alderley Park and Alex Alderton, Roberto Amato, Sonia Goncalves, Ewan Harrison, David K. Jackson, Ian Johnston, Dominic Kwiatkowski, Cordelia Langford, John Sillitoe on behalf of the Wellcome Sanger Institute COVID-19 Surveillance Team |
| EPI_ISL_552774 | Lighthouse Lab in Alderley Park | Wellcome Sanger Institute for the COVID-19 Genomics UK (COG-UK) consortium | The Lighthouse Lab in Alderley Park and Alex Alderton, Roberto Amato, Sonia Goncalves, Ewan Harrison, David K. Jackson, Ian Johnston, Dominic Kwiatkowski, Cordelia Langford, John Sillitoe on behalf of the Wellcome Sanger Institute COVID-19 Surveillance Team (http://www.sanger.ac.uk/covid-team) |
| EPI_ISL_552775, EPI_ISL_552776, EPI_ISL_552777, EPI_ISL_552779, EPI_ISL_552780, EPI_ISL_552782, EPI_ISL_552783, EPI_ISL_552785, EPI_ISL_552792, EPI_ISL_552793, EPI_ISL_552794, EPI_ISL_552796 | | | |
| see above | Lighthouse Lab in Alderley Park | Wellcome Sanger Institute for the COVID-19 Genomics UK (COG-UK) consortium | The Lighthouse Lab in Alderley Park and Alex Alderton, Roberto Amato, Sonia Goncalves, Ewan Harrison, David K. Jackson, Ian Johnston, Dominic Kwiatkowski, Cordelia Langford, John Sillitoe on behalf of the Wellcome Sanger Institute COVID-19 Surveillance Team |
| EPI_ISL_560317 | UMMC-Health | WHO National Influenza Centre Russian Federation | Andrey Komissarov, Artem Fadeev, Anna Ivanova, Tatiana Platonova, Daria Danilenko |
| EPI_ISL_560607, EPI_ISL_560608, EPI_ISL_560609, EPI_ISL_560610, EPI_ISL_560611, EPI_ISL_560612, EPI_ISL_560613, EPI_ISL_560614, EPI_ISL_560615, EPI_ISL_560616, EPI_ISL_560617, EPI_ISL_560618 | | | |
| see above | Hospital | National Reference Center for Viruses of Respiratory Infections, Institut Pasteur, Paris | Sylvie Behillil, Fabiana Gambaro, Etienne Simon-Lorière, Vincent Enouf, Maud Vanpeene, Sylvie van der Werf |
| EPI_ISL_560981 | Capio S:t Gorans sjukhus | The Public Health Agency of Sweden | Anna-Malin Linde, Maria Lind Karlberg, Mattias Haukland, Reza Advani, Olov Svartstrom, Oskar Karlsson Lindsjo, Sandra Broddesson, Petra Edquist, Mia Brytting, Anna Risberg, Karin Tegmark-Wisell |
| EPI_ISL_560989, EPI_ISL_560990 | Karolinska universitetslaboratoriet | The Public Health Agency of Sweden | Anna-Malin Linde, Maria Lind Karlberg, Mattias Haukland, Reza Advani, Olov Svartstrom, Oskar Karlsson Lindsjo, Sandra Broddesson, Petra Edquist, Mia Brytting, Anna Risberg, Karin Tegmark-Wisell |
| EPI_ISL_561236, EPI_ISL_561237, EPI_ISL_561288, EPI_ISL_561289, EPI_ISL_561292 | MRCG at LSHTM Genomics lab | MRCG at LSHTM Genomics lab | Abdul Karim sesay, Abdoulie Kanteh, Jarra Manneh, Mariama Kujabi, Bakary Sanyang |
| EPI_ISL_561402, EPI_ISL_561405, EPI_ISL_561413 | Microbiological Diagnostic Unit - Public Health Laboratory (MDU-PHL) | MDU-PHL | Seemann, T., Schultz M. B., Sait, M., Sherry, N. |
| EPI_ISL_561421 | Victorian Infectious Diseases Reference Laboratory (VIDRL) | VIDRL and MDU-PHL | Caly, L., Seemann, T., Sait, M., Schultz, M. B., Druce J., Sherry, N. |
| EPI_ISL_561424 | Microbiological Diagnostic Unit - Public Health Laboratory (MDU-PHL) | MDU-PHL | Seemann, T., Schultz M. B., Sait, M., Sherry, N. |
| EPI_ISL_561426 | Victorian Infectious Diseases Reference Laboratory (VIDRL) | VIDRL and MDU-PHL | Caly, L., Seemann, T., Sait, M., Schultz, M. B., Druce J., Sherry, N. |
| EPI_ISL_561432, EPI_ISL_561435, EPI_ISL_561443, EPI_ISL_561449, EPI_ISL_561460, EPI_ISL_561471, EPI_ISL_561478, EPI_ISL_561479, EPI_ISL_561487, EPI_ISL_561489, EPI_ISL_561490, EPI_ISL_561492, EPI_ISL_561493, EPI_ISL_561494, EPI_ISL_561503, EPI_ISL_561516, EPI_ISL_561526, EPI_ISL_561527, EPI_ISL_561534, EPI_ISL_561538, EPI_ISL_561539, EPI_ISL_561540, EPI_ISL_561560, EPI_ISL_561567, EPI_ISL_561574, EPI_ISL_561581, EPI_ISL_561583 | | | |
| see above | Microbiological Diagnostic Unit - Public Health Laboratory (MDU-PHL) | MDU-PHL | Seemann, T., Schultz M. B., Sait, M., Sherry, N. |
| EPI_ISL_561590, EPI_ISL_561591 | Victorian Infectious Diseases Reference Laboratory (VIDRL) | VIDRL and MDU-PHL | Caly, L., Seemann, T., Sait, M., Schultz, M. B., Druce J., Sherry, N. |
| EPI_ISL_561592, EPI_ISL_561595, EPI_ISL_561601, EPI_ISL_561623, | Microbiological Diagnostic Unit - Public Health Laboratory (MDU-PHL) | MDU-PHL | Seemann, T., Schultz M. B., Sait, M., Sherry, N. |

| | | | |
|---|---|---|---|
| EPI_ISL_561627, EPI_ISL_561634, EPI_ISL_561635, EPI_ISL_561642, EPI_ISL_561648 | | | |
| EPI_ISL_561653 | Victorian Infectious Diseases Reference Laboratory (VIDRL) | VIDRL and MDU-PHL | Caly, L., Seemann, T., Sait, M., Schultz, M. B., Druce J., Sherry, N. |
| EPI_ISL_561658, EPI_ISL_561665 | Microbiological Diagnostic Unit - Public Health Laboratory (MDU-PHL) | MDU-PHL | Seemann, T., Schultz M. B., Sait, M., Sherry, N. |
| EPI_ISL_561673, EPI_ISL_561674, EPI_ISL_561679 | Victorian Infectious Diseases Reference Laboratory (VIDRL) | VIDRL and MDU-PHL | Caly, L., Seemann, T., Sait, M., Schultz, M. B., Druce J., Sherry, N. |
| EPI_ISL_561696 | Microbiological Diagnostic Unit - Public Health Laboratory (MDU-PHL) | MDU-PHL | Seemann, T., Schultz M. B., Sait, M., Sherry, N. |
| EPI_ISL_561697, EPI_ISL_561711 | Victorian Infectious Diseases Reference Laboratory (VIDRL) | VIDRL and MDU-PHL | Caly, L., Seemann, T., Sait, M., Schultz, M. B., Druce J., Sherry, N. |
| EPI_ISL_561738, EPI_ISL_561740, EPI_ISL_561754, EPI_ISL_561770, EPI_ISL_561773, EPI_ISL_561785, EPI_ISL_561789, EPI_ISL_561792, EPI_ISL_561794, EPI_ISL_561813, EPI_ISL_561818, EPI_ISL_561822, EPI_ISL_561823, EPI_ISL_561826, EPI_ISL_561832, EPI_ISL_561859, EPI_ISL_561864, EPI_ISL_561870, EPI_ISL_561877 | | | |
| see above | Microbiological Diagnostic Unit - Public Health Laboratory (MDU-PHL) | MDU-PHL | Seemann, T., Schultz M. B., Sait, M., Sherry, N. |
| EPI_ISL_561879 | Victorian Infectious Diseases Reference Laboratory (VIDRL) | VIDRL and MDU-PHL | Caly, L., Seemann, T., Sait, M., Schultz, M. B., Druce J., Sherry, N. |
| EPI_ISL_561901, EPI_ISL_561903, EPI_ISL_561911, EPI_ISL_561934, EPI_ISL_561941, EPI_ISL_561943, EPI_ISL_561963, EPI_ISL_561970, EPI_ISL_561974, EPI_ISL_561982, EPI_ISL_561998, EPI_ISL_562001, EPI_ISL_562002 | | | |
| see above | Microbiological Diagnostic Unit - Public Health Laboratory (MDU-PHL) | MDU-PHL | Seemann, T., Schultz M. B., Sait, M., Sherry, N. |
| EPI_ISL_562011 | Victorian Infectious Diseases Reference Laboratory (VIDRL) | VIDRL and MDU-PHL | Caly, L., Seemann, T., Sait, M., Schultz, M. B., Druce J., Sherry, N. |
| EPI_ISL_562017, EPI_ISL_562030, EPI_ISL_562034, EPI_ISL_562036, EPI_ISL_562051, EPI_ISL_562061, EPI_ISL_562062 | Microbiological Diagnostic Unit - Public Health Laboratory (MDU-PHL) | MDU-PHL | Seemann, T., Schultz M. B., Sait, M., Sherry, N. |
| EPI_ISL_562069 | Victorian Infectious Diseases Reference Laboratory (VIDRL) | VIDRL and MDU-PHL | Caly, L., Seemann, T., Sait, M., Schultz, M. B., Druce J., Sherry, N. |
| EPI_ISL_562079, EPI_ISL_562086, EPI_ISL_562090, EPI_ISL_562102, EPI_ISL_562108 | Microbiological Diagnostic Unit - Public Health Laboratory (MDU-PHL) | MDU-PHL | Seemann, T., Schultz M. B., Sait, M., Sherry, N. |
| EPI_ISL_562110 | Victorian Infectious Diseases Reference Laboratory (VIDRL) | VIDRL and MDU-PHL | Caly, L., Seemann, T., Sait, M., Schultz, M. B., Druce J., Sherry, N. |
| EPI_ISL_562111, EPI_ISL_562119, EPI_ISL_562135 | Microbiological Diagnostic Unit - Public Health Laboratory (MDU-PHL) | MDU-PHL | Seemann, T., Schultz M. B., Sait, M., Sherry, N. |
| EPI_ISL_562138 | Victorian Infectious Diseases Reference Laboratory (VIDRL) | VIDRL and MDU-PHL | Caly, L., Seemann, T., Sait, M., Schultz, M. B., Druce J., Sherry, N. |
| EPI_ISL_562139, EPI_ISL_562140, EPI_ISL_562146, EPI_ISL_562150, EPI_ISL_562154, EPI_ISL_562165, EPI_ISL_562168 | Microbiological Diagnostic Unit - Public Health Laboratory (MDU-PHL) | MDU-PHL | Seemann, T., Schultz M. B., Sait, M., Sherry, N. |
| EPI_ISL_562194 | Victorian Infectious Diseases Reference Laboratory (VIDRL) | VIDRL and MDU-PHL | Caly, L., Seemann, T., Sait, M., Schultz, M. B., Druce J., Sherry, N. |
| EPI_ISL_562195, EPI_ISL_562210, EPI_ISL_562221 | Microbiological Diagnostic Unit - Public Health Laboratory (MDU-PHL) | MDU-PHL | Seemann, T., Schultz M. B., Sait, M., Sherry, N. |
| EPI_ISL_562225, EPI_ISL_562245 | Victorian Infectious Diseases Reference Laboratory (VIDRL) | VIDRL and MDU-PHL | Caly, L., Seemann, T., Sait, M., Schultz, M. B., Druce J., Sherry, N. |
| EPI_ISL_562249 | Microbiological Diagnostic Unit - Public Health Laboratory (MDU-PHL) | MDU-PHL | Seemann, T., Schultz M. B., Sait, M., Sherry, N. |
| EPI_ISL_562264 | Victorian Infectious Diseases Reference Laboratory (VIDRL) | VIDRL and MDU-PHL | Caly, L., Seemann, T., Sait, M., Schultz, M. B., Druce J., Sherry, N. |
| EPI_ISL_562277 | Microbiological Diagnostic Unit - Public Health Laboratory (MDU-PHL) | MDU-PHL | Seemann, T., Schultz M. B., Sait, M., Sherry, N. |
| EPI_ISL_562288, EPI_ISL_562289 | Victorian Infectious Diseases Reference Laboratory (VIDRL) | VIDRL and MDU-PHL | Caly, L., Seemann, T., Sait, M., Schultz, M. B., Druce J., Sherry, N. |
| EPI_ISL_562298, EPI_ISL_562300, EPI_ISL_562301 | Microbiological Diagnostic Unit - Public Health Laboratory (MDU-PHL) | MDU-PHL | Seemann, T., Schultz M. B., Sait, M., Sherry, N. |
| EPI_ISL_562308 | Victorian Infectious Diseases Reference Laboratory (VIDRL) | VIDRL and MDU-PHL | Caly, L., Seemann, T., Sait, M., Schultz, M. B., Druce J., Sherry, N. |
| EPI_ISL_562324, EPI_ISL_562341 | Microbiological Diagnostic Unit - Public Health Laboratory (MDU-PHL) | MDU-PHL | Seemann, T., Schultz M. B., Sait, M., Sherry, N. |
| EPI_ISL_562354 | Victorian Infectious Diseases Reference Laboratory (VIDRL) | VIDRL and MDU-PHL | Caly, L., Seemann, T., Sait, M., Schultz, M. B., Druce J., Sherry, N. |
| EPI_ISL_562364, EPI_ISL_562367, EPI_ISL_562371, EPI_ISL_562380 | Microbiological Diagnostic Unit - Public Health Laboratory (MDU-PHL) | MDU-PHL | Seemann, T., Schultz M. B., Sait, M., Sherry, N. |
| EPI_ISL_562390 | Victorian Infectious Diseases Reference Laboratory (VIDRL) | VIDRL and MDU-PHL | Caly, L., Seemann, T., Sait, M., Schultz, M. B., Druce J., Sherry, N. |
| EPI_ISL_562402, EPI_ISL_562405, EPI_ISL_562409, EPI_ISL_562416, EPI_ISL_562417, EPI_ISL_562427, EPI_ISL_562433 | Microbiological Diagnostic Unit - Public Health Laboratory (MDU-PHL) | MDU-PHL | Seemann, T., Schultz M. B., Sait, M., Sherry, N. |
| EPI_ISL_562442 | Victorian Infectious Diseases Reference Laboratory (VIDRL) | VIDRL and MDU-PHL | Caly, L., Seemann, T., Sait, M., Schultz, M. B., Druce J., Sherry, N. |
| EPI_ISL_562467, EPI_ISL_562468, EPI_ISL_562476 | Microbiological Diagnostic Unit - Public Health Laboratory (MDU-PHL) | MDU-PHL | Seemann, T., Schultz M. B., Sait, M., Sherry, N. |
| EPI_ISL_562484 | Victorian Infectious Diseases Reference Laboratory (VIDRL) | VIDRL and MDU-PHL | Caly, L., Seemann, T., Sait, M., Schultz, M. B., Druce J., Sherry, N. |
| EPI_ISL_562487, EPI_ISL_562507, EPI_ISL_562515, EPI_ISL_562529, EPI_ISL_562550 | Microbiological Diagnostic Unit - Public Health Laboratory (MDU-PHL) | MDU-PHL | Seemann, T., Schultz M. B., Sait, M., Sherry, N. |
| EPI_ISL_562558 | Victorian Infectious Diseases Reference Laboratory (VIDRL) | VIDRL and MDU-PHL | Caly, L., Seemann, T., Sait, M., Schultz, M. B., Druce J., Sherry, N. |
| EPI_ISL_562571, EPI_ISL_562601, EPI_ISL_562602, EPI_ISL_562611, | Microbiological Diagnostic Unit - Public Health Laboratory (MDU-PHL) | MDU-PHL | Seemann, T., Schultz M. B., Sait, M., Sherry, N. |

| | | | |
|---|---|---|---|
| EPI_ISL_562618, EPI_ISL_562619 | | | |
| EPI_ISL_562628 | Victorian Infectious Diseases Reference Laboratory (VIDRL) | VIDRL and MDU-PHL | Caly, L., Seemann, T., Sait, M., Schultz, M. B., Druce J., Sherry, N. |
| EPI_ISL_562638 | Microbiological Diagnostic Unit - Public Health Laboratory (MDU-PHL) | MDU-PHL | Seemann, T., Schultz M. B., Sait, M., Sherry, N. |
| EPI_ISL_562659 | Victorian Infectious Diseases Reference Laboratory (VIDRL) | VIDRL and MDU-PHL | Caly, L., Seemann, T., Sait, M., Schultz, M. B., Druce J., Sherry, N. |
| EPI_ISL_562660 | Microbiological Diagnostic Unit - Public Health Laboratory (MDU-PHL) | MDU-PHL | Seemann, T., Schultz M. B., Sait, M., Sherry, N. |
| EPI_ISL_562669, EPI_ISL_562705 | Victorian Infectious Diseases Reference Laboratory (VIDRL) | VIDRL and MDU-PHL | Caly, L., Seemann, T., Sait, M., Schultz, M. B., Druce J., Sherry, N. |
| EPI_ISL_562802, EPI_ISL_562903, EPI_ISL_562904, EPI_ISL_562907, EPI_ISL_562908, EPI_ISL_562909, EPI_ISL_562937, EPI_ISL_562945, EPI_ISL_562948, EPI_ISL_562953, EPI_ISL_562960, EPI_ISL_562962, EPI_ISL_562965, EPI_ISL_562966, EPI_ISL_562967, EPI_ISL_562968, EPI_ISL_562969, EPI_ISL_562970, EPI_ISL_562971, EPI_ISL_562972, EPI_ISL_562973, EPI_ISL_562975, EPI_ISL_562976, EPI_ISL_562977, EPI_ISL_562978, EPI_ISL_562979, EPI_ISL_562980, EPI_ISL_562981, EPI_ISL_562982, EPI_ISL_562983, EPI_ISL_562984, EPI_ISL_562985, EPI_ISL_562986, EPI_ISL_562987, EPI_ISL_562988, EPI_ISL_562989, EPI_ISL_562990, EPI_ISL_562991, EPI_ISL_562992, EPI_ISL_562993, EPI_ISL_562994, EPI_ISL_562995, EPI_ISL_562996, EPI_ISL_562997, EPI_ISL_562998, EPI_ISL_562999, EPI_ISL_563020, EPI_ISL_563029, EPI_ISL_563032, EPI_ISL_563035, EPI_ISL_563039, EPI_ISL_563051, EPI_ISL_563058, EPI_ISL_563069, EPI_ISL_563070, EPI_ISL_563081, EPI_ISL_563100, EPI_ISL_563101, EPI_ISL_563103, EPI_ISL_563109, EPI_ISL_563110, EPI_ISL_563117, EPI_ISL_563134, EPI_ISL_563145, EPI_ISL_563146, EPI_ISL_563147, EPI_ISL_563149, EPI_ISL_563151, EPI_ISL_563178, EPI_ISL_563185, EPI_ISL_563187, EPI_ISL_563188, EPI_ISL_563191, EPI_ISL_563192, EPI_ISL_563193, EPI_ISL_563206, EPI_ISL_563210, EPI_ISL_563211, EPI_ISL_563213, EPI_ISL_563215, EPI_ISL_563221, EPI_ISL_563227, EPI_ISL_563231, EPI_ISL_563238, EPI_ISL_563243, EPI_ISL_563254, EPI_ISL_563260, EPI_ISL_563261, EPI_ISL_563262, EPI_ISL_563264, EPI_ISL_563266, EPI_ISL_563267, EPI_ISL_563269, EPI_ISL_563272, EPI_ISL_563273, EPI_ISL_563276, EPI_ISL_563278, EPI_ISL_563280, EPI_ISL_563283, EPI_ISL_563298, EPI_ISL_563299, EPI_ISL_563300, EPI_ISL_563342, EPI_ISL_563343, EPI_ISL_563345, EPI_ISL_563396 | | | |
| see above | Microbiological Diagnostic Unit - Public Health Laboratory (MDU-PHL) | MDU-PHL | Seemann, T., Schultz M. B., Sait, M., Sherry, N. |
| EPI_ISL_563407, EPI_ISL_563408 | Victorian Infectious Diseases Reference Laboratory (VIDRL) | VIDRL and MDU-PHL | Caly, L., Seemann, T., Sait, M., Schultz, M. B., Druce J., Sherry, N. |
| EPI_ISL_563674, EPI_ISL_563675, EPI_ISL_563676, EPI_ISL_563709, EPI_ISL_563724, EPI_ISL_563726, EPI_ISL_563728, EPI_ISL_563729, EPI_ISL_563730, EPI_ISL_563731, EPI_ISL_563732, EPI_ISL_563734, EPI_ISL_563739, EPI_ISL_563740, EPI_ISL_563741, EPI_ISL_563742, EPI_ISL_563744, EPI_ISL_563745, EPI_ISL_563746, EPI_ISL_563748, EPI_ISL_563749, EPI_ISL_563750, EPI_ISL_563751, EPI_ISL_563753, EPI_ISL_563754, EPI_ISL_563755, EPI_ISL_563756, EPI_ISL_563757, EPI_ISL_563758, EPI_ISL_563759, EPI_ISL_563760, EPI_ISL_563761, EPI_ISL_563762, EPI_ISL_563763, EPI_ISL_563764, EPI_ISL_563766, EPI_ISL_563767, EPI_ISL_563771, EPI_ISL_563772, EPI_ISL_563773, EPI_ISL_563774, EPI_ISL_563775, EPI_ISL_563781, EPI_ISL_563782, EPI_ISL_563783, EPI_ISL_563784, EPI_ISL_563785, EPI_ISL_563786, EPI_ISL_563787, EPI_ISL_563788, EPI_ISL_563793, EPI_ISL_563795, EPI_ISL_563796, EPI_ISL_564119, EPI_ISL_564171, EPI_ISL_564173, EPI_ISL_564174, EPI_ISL_564179 | | | |
| see above | Microbiological Diagnostic Unit - Public Health Laboratory (MDU-PHL) | MDU-PHL | Seemann, T., Schultz M. B., Sait, M., Sherry, N. |
| EPI_ISL_564218, EPI_ISL_564219, EPI_ISL_564220, EPI_ISL_564223, EPI_ISL_564224, EPI_ISL_564225, EPI_ISL_564226, EPI_ISL_564227, EPI_ISL_564228, EPI_ISL_564229, EPI_ISL_564230, EPI_ISL_564231, EPI_ISL_564232, EPI_ISL_564233, EPI_ISL_564234, EPI_ISL_564235, EPI_ISL_564236, EPI_ISL_564237, EPI_ISL_564238, EPI_ISL_564239, EPI_ISL_564240, EPI_ISL_564241, EPI_ISL_564242, EPI_ISL_564243, EPI_ISL_564249, EPI_ISL_564250, EPI_ISL_564251, EPI_ISL_564252, EPI_ISL_564253, EPI_ISL_564254 | | | |
| see above | Victorian Infectious Diseases Reference Laboratory (VIDRL) | VIDRL and MDU-PHL | Caly, L., Seemann, T., Sait, M., Schultz, M. B., Druce J., Sherry, N. |
| EPI_ISL_564256, EPI_ISL_564257, EPI_ISL_564258, EPI_ISL_564259 | Microbiological Diagnostic Unit - Public Health Laboratory (MDU-PHL) | MDU-PHL | Seemann, T., Schultz M. B., Sait, M., Sherry, N. |
| EPI_ISL_564318, EPI_ISL_564341, EPI_ISL_564344, EPI_ISL_564345, EPI_ISL_564346, EPI_ISL_564347, EPI_ISL_564348, EPI_ISL_564349, EPI_ISL_564350, EPI_ISL_564394, EPI_ISL_564395, EPI_ISL_564396 | | | |
| see above | Victorian Infectious Diseases Reference Laboratory (VIDRL) | VIDRL and MDU-PHL | Caly, L., Seemann, T., Sait, M., Schultz, M. B., Druce J., Sherry, N. |
| EPI_ISL_564405, EPI_ISL_564407, EPI_ISL_564408, EPI_ISL_564409, EPI_ISL_564410, EPI_ISL_564411, EPI_ISL_564412, EPI_ISL_564413, EPI_ISL_564415 | Microbiological Diagnostic Unit - Public Health Laboratory (MDU-PHL) | MDU-PHL | Seemann, T., Schultz M. B., Sait, M., Sherry, N. |
| EPI_ISL_564544, EPI_ISL_564546, EPI_ISL_564547, EPI_ISL_564548, EPI_ISL_564549, EPI_ISL_564550, EPI_ISL_564551, EPI_ISL_564552, EPI_ISL_564553, EPI_ISL_564554, EPI_ISL_564557, EPI_ISL_564558, EPI_ISL_564559, EPI_ISL_564560, EPI_ISL_564561, EPI_ISL_564562, EPI_ISL_564563, EPI_ISL_564564 | | | |
| see above | Victorian Infectious Diseases Reference Laboratory (VIDRL) | VIDRL and MDU-PHL | Caly, L., Seemann, T., Sait, M., Schultz, M. B., Druce J., Sherry, N. |
| EPI_ISL_564576, EPI_ISL_564579, EPI_ISL_564584, EPI_ISL_564585, EPI_ISL_564586, EPI_ISL_564587, EPI_ISL_564588, EPI_ISL_564589, EPI_ISL_564590, EPI_ISL_564591, EPI_ISL_564592, EPI_ISL_564596, EPI_ISL_564597, EPI_ISL_564598, EPI_ISL_564599, EPI_ISL_564600, EPI_ISL_564601, EPI_ISL_564604, EPI_ISL_564605, EPI_ISL_564606, EPI_ISL_564607, EPI_ISL_564608, EPI_ISL_564609, EPI_ISL_564610, EPI_ISL_564611, EPI_ISL_564612, EPI_ISL_564613, EPI_ISL_564614, EPI_ISL_564615, EPI_ISL_564616, EPI_ISL_564617, EPI_ISL_564619, EPI_ISL_564620, EPI_ISL_564621, EPI_ISL_564622, EPI_ISL_564623, EPI_ISL_564624, EPI_ISL_564633, EPI_ISL_564634, EPI_ISL_564635, EPI_ISL_564636, EPI_ISL_564637, EPI_ISL_564638, EPI_ISL_564639, EPI_ISL_564640, EPI_ISL_564641, EPI_ISL_564642, EPI_ISL_564643, EPI_ISL_564644, EPI_ISL_564645, EPI_ISL_564646, EPI_ISL_564647, EPI_ISL_564648, EPI_ISL_564649, EPI_ISL_564650, EPI_ISL_564651, EPI_ISL_564652, EPI_ISL_564653, EPI_ISL_564654, EPI_ISL_564655, EPI_ISL_564656, EPI_ISL_564657, EPI_ISL_564658, EPI_ISL_564659, EPI_ISL_564660, EPI_ISL_564661, EPI_ISL_564662, EPI_ISL_564663, EPI_ISL_564664, EPI_ISL_564665, EPI_ISL_564666, EPI_ISL_564667, EPI_ISL_564668, EPI_ISL_564669, EPI_ISL_564670, EPI_ISL_564671, EPI_ISL_564672, EPI_ISL_564673, EPI_ISL_564674, EPI_ISL_564675, EPI_ISL_564677, EPI_ISL_564678, EPI_ISL_564679, EPI_ISL_564680, EPI_ISL_564681, EPI_ISL_564718, EPI_ISL_564719, EPI_ISL_564720, EPI_ISL_564722, EPI_ISL_564723, EPI_ISL_564724, EPI_ISL_564725, EPI_ISL_564726, EPI_ISL_564727, EPI_ISL_564728, EPI_ISL_564729, EPI_ISL_564733, EPI_ISL_564734, EPI_ISL_564735, EPI_ISL_564736, EPI_ISL_564739, EPI_ISL_564740, EPI_ISL_564741, EPI_ISL_564742, EPI_ISL_564743, EPI_ISL_564744, EPI_ISL_564745, EPI_ISL_564746, EPI_ISL_564747, EPI_ISL_564748, EPI_ISL_564749, EPI_ISL_564750, EPI_ISL_564751, EPI_ISL_564752, EPI_ISL_564753, EPI_ISL_564754, EPI_ISL_564762, EPI_ISL_564763, EPI_ISL_564765, EPI_ISL_564766, EPI_ISL_564768, EPI_ISL_564769, EPI_ISL_564770, EPI_ISL_564771, EPI_ISL_564772, EPI_ISL_564773, EPI_ISL_564774, EPI_ISL_564775, EPI_ISL_564776, EPI_ISL_564778, EPI_ISL_564779, EPI_ISL_564780, EPI_ISL_564789, EPI_ISL_564865, EPI_ISL_564866, EPI_ISL_564867, EPI_ISL_564868, EPI_ISL_564869, EPI_ISL_564870, EPI_ISL_564871, EPI_ISL_564872, EPI_ISL_564873, EPI_ISL_564874, EPI_ISL_564875, EPI_ISL_564876, EPI_ISL_564877, EPI_ISL_564896, EPI_ISL_564897, EPI_ISL_564898, EPI_ISL_564899, EPI_ISL_564900, EPI_ISL_564901, EPI_ISL_564902, EPI_ISL_564903, EPI_ISL_564906, EPI_ISL_564913, EPI_ISL_564914, EPI_ISL_564917, EPI_ISL_564930, EPI_ISL_564936, EPI_ISL_564937, EPI_ISL_564952, EPI_ISL_564953, EPI_ISL_564954, EPI_ISL_564957, EPI_ISL_564958, EPI_ISL_564959, EPI_ISL_564960, EPI_ISL_564962, EPI_ISL_564978, EPI_ISL_565072, EPI_ISL_565073, EPI_ISL_565074, EPI_ISL_565075, EPI_ISL_565076, EPI_ISL_565077, EPI_ISL_565078, EPI_ISL_565079, EPI_ISL_565080, EPI_ISL_565105, EPI_ISL_565107, EPI_ISL_565117, EPI_ISL_565118, EPI_ISL_565119, EPI_ISL_565120, EPI_ISL_565121, EPI_ISL_565122, EPI_ISL_565123, EPI_ISL_565124 | | | |
| see above | Microbiological Diagnostic Unit - Public Health Laboratory (MDU-PHL) | MDU-PHL | Seemann, T., Schultz M. B., Sait, M., Sherry, N. |
| EPI_ISL_565209, EPI_ISL_565210, EPI_ISL_565211, EPI_ISL_565213, EPI_ISL_565214, EPI_ISL_565215, EPI_ISL_565216, EPI_ISL_565217 | Victorian Infectious Diseases Reference Laboratory (VIDRL) | VIDRL and MDU-PHL | Caly, L., Seemann, T., Sait, M., Schultz, M. B., Druce J., Sherry, N. |
| EPI_ISL_565351, EPI_ISL_565352, EPI_ISL_565368, EPI_ISL_565369, EPI_ISL_565372 | Microbiological Diagnostic Unit - Public Health Laboratory (MDU-PHL) | MDU-PHL | Seemann, T., Schultz M. B., Sait, M., Sherry, N. |
| EPI_ISL_565382, EPI_ISL_565385, EPI_ISL_565394, EPI_ISL_565395, EPI_ISL_565396, EPI_ISL_565399, EPI_ISL_565401, EPI_ISL_565402, EPI_ISL_565403, EPI_ISL_565404, EPI_ISL_565409 | | | |
| see above | Victorian Infectious Diseases Reference Laboratory (VIDRL) | VIDRL and MDU-PHL | Caly, L., Seemann, T., Sait, M., Schultz, M. B., Druce J., Sherry, N. |
| EPI_ISL_565456, EPI_ISL_565510, EPI_ISL_565511, EPI_ISL_565512, EPI_ISL_565513, EPI_ISL_565515, EPI_ISL_565516, EPI_ISL_565519, EPI_ISL_565521, EPI_ISL_565603, EPI_ISL_565604, EPI_ISL_565605, EPI_ISL_565606, EPI_ISL_565625, EPI_ISL_565627, EPI_ISL_565629, EPI_ISL_565644, EPI_ISL_565645, EPI_ISL_565680, EPI_ISL_565681, EPI_ISL_565682, EPI_ISL_565683, EPI_ISL_565684, EPI_ISL_565685, EPI_ISL_565745, EPI_ISL_565746, EPI_ISL_565754, EPI_ISL_565755, EPI_ISL_565756 | | | |
| see above | Microbiological Diagnostic Unit - Public Health Laboratory (MDU-PHL) | MDU-PHL | Seemann, T., Schultz M. B., Sait, M., Sherry, N. |
| EPI_ISL_566031 | Michigan Department of Health and Human Services, Bureau of Laboratories | Michigan Department of Health and Human Services, Bureau of Laboratories | Blankenship HM, Riner D, Soehnlen MK |
| EPI_ISL_566065 | Respiratory Virus Unit, Microbiology Services Colindale, Public Health England | Respiratory Virus Unit, Microbiology Services Colindale, Public Health England | PHE Covid Sequencing Team |
| EPI_ISL_568573 | Department of Infectious Diseases and Immunology, National Hospital Organization Nagoya Medical Center | Clinical Research Center, National Hospital Organization Nagoya Medical Center | Yoshihiro Nakata, Hirotaka Ode, Mai Kubota, Masakazu Matsuda, Kazuhiro Matsuoka, Nakasuji Miho, Mikiko Mori, Mayumi Imahashi, Yoshiyuki Yokomaku, Yasumasa Iwatani |
| EPI_ISL_568587, EPI_ISL_568588, EPI_ISL_568589 | Florida Bureau of Public Health Laboratories | Florida Bureau of Public Health Laboratories | Sarah Schmedes, Jason Blanton |
| EPI_ISL_569030, EPI_ISL_569031, EPI_ISL_569032, EPI_ISL_569041, EPI_ISL_569042, EPI_ISL_569043, EPI_ISL_569044, EPI_ISL_569045, EPI_ISL_569046, EPI_ISL_569047, EPI_ISL_569048, EPI_ISL_569049, EPI_ISL_569050, EPI_ISL_569051, EPI_ISL_569052, EPI_ISL_569053, EPI_ISL_569054, EPI_ISL_569055, | | | |

| | | | |
|---|---|---|---|
| EPI_ISL_569056, EPI_ISL_569057, EPI_ISL_569058, EPI_ISL_569059, EPI_ISL_569060, EPI_ISL_569061, EPI_ISL_569062, EPI_ISL_569063, EPI_ISL_569064, EPI_ISL_569145, EPI_ISL_569146, EPI_ISL_569147, EPI_ISL_569148, EPI_ISL_569149, EPI_ISL_569150, EPI_ISL_569151, EPI_ISL_569152, EPI_ISL_569153, EPI_ISL_569154, EPI_ISL_569155, EPI_ISL_569156, EPI_ISL_569157, EPI_ISL_569158, EPI_ISL_569159, EPI_ISL_569160, EPI_ISL_569161, EPI_ISL_569162, EPI_ISL_569163, EPI_ISL_569164, EPI_ISL_569165, EPI_ISL_569166, EPI_ISL_569167, EPI_ISL_569168, EPI_ISL_569169, EPI_ISL_569170, EPI_ISL_569171, EPI_ISL_569180 | | | |
| see above | MEPHI, Aix Marseille University | MEPHI, Aix Marseille University | Anthony LEVASSEUR |
| EPI_ISL_569621 | Quick Care Watertown | South Dakota Public Health Laboratory | Matt Plumb, Jacob Garfin, Xiong Wang, and Chris Carlson |
| EPI_ISL_569622 | Woodrow Wilson Keeble Memorial Healthcare | South Dakota Public Health Laboratory | Matt Plumb, Jacob Garfin, Xiong Wang, and Chris Carlson |
| EPI_ISL_570000 | Unity Health Toronto | Ontario Institute for Cancer Research | Ramzi Fattouh, Larissa M. Matukas, Yan Chen,Mark Downing, Trina Otterman, Karel Boissinot, Wai Sum Siu, Zhi Cui, Le Luu, Samira Mubareka, TIBDN, Ilinca Lungu, Bernard Lam, Jeremy Johns, Paul Krzyzanowski, Richard de Borja, Felicia Vincelli, Philip Zuzarte, Jared T. Simpson |
| EPI_ISL_570548, EPI_ISL_570550, EPI_ISL_570551, EPI_ISL_570552, EPI_ISL_570558, EPI_ISL_570559, EPI_ISL_570560, EPI_ISL_570561, EPI_ISL_570562, EPI_ISL_570563, EPI_ISL_570564, EPI_ISL_570565, EPI_ISL_570566, EPI_ISL_570567, EPI_ISL_570568, EPI_ISL_570569, EPI_ISL_570570, EPI_ISL_570571, EPI_ISL_570572, EPI_ISL_570573, EPI_ISL_570576, EPI_ISL_570578, EPI_ISL_570582, EPI_ISL_570585, EPI_ISL_570586, EPI_ISL_570587, EPI_ISL_570588, EPI_ISL_570589, EPI_ISL_570590, EPI_ISL_570591, EPI_ISL_570592, EPI_ISL_570593, EPI_ISL_570594, EPI_ISL_570597, EPI_ISL_570598, EPI_ISL_570605, EPI_ISL_570606, EPI_ISL_570607, EPI_ISL_570608, EPI_ISL_570609, EPI_ISL_570610, EPI_ISL_570611, EPI_ISL_570613, EPI_ISL_570614, EPI_ISL_570615, EPI_ISL_570616, EPI_ISL_570617, EPI_ISL_570618, EPI_ISL_570619, EPI_ISL_570620, EPI_ISL_570621, EPI_ISL_570622, EPI_ISL_570623, EPI_ISL_570624, EPI_ISL_570625, EPI_ISL_570626, EPI_ISL_570627, EPI_ISL_570628, EPI_ISL_570629, EPI_ISL_570630, EPI_ISL_570631, EPI_ISL_570632, EPI_ISL_570633, EPI_ISL_570634, EPI_ISL_570635, EPI_ISL_570636, EPI_ISL_570637, EPI_ISL_570638, EPI_ISL_570639, EPI_ISL_570640, EPI_ISL_570641, EPI_ISL_570642, EPI_ISL_570643, EPI_ISL_570644, EPI_ISL_570645, EPI_ISL_570646, EPI_ISL_570647, EPI_ISL_570648, EPI_ISL_570649, EPI_ISL_570650, EPI_ISL_570651, EPI_ISL_570652, EPI_ISL_570653, EPI_ISL_570654, EPI_ISL_570655, EPI_ISL_570656, EPI_ISL_570657, EPI_ISL_570658, EPI_ISL_570659, EPI_ISL_570660, EPI_ISL_570661, EPI_ISL_570662, EPI_ISL_570663, EPI_ISL_570664, EPI_ISL_570665, EPI_ISL_570666, EPI_ISL_570667, EPI_ISL_570668, EPI_ISL_570669, EPI_ISL_570670, EPI_ISL_570671, EPI_ISL_570672, EPI_ISL_570673, EPI_ISL_570674, EPI_ISL_570675, EPI_ISL_570676, EPI_ISL_570677, EPI_ISL_570678, EPI_ISL_570679, EPI_ISL_570680, EPI_ISL_570681, EPI_ISL_570682, EPI_ISL_570683, EPI_ISL_570684, EPI_ISL_570685, EPI_ISL_570686, EPI_ISL_570687, EPI_ISL_570688, EPI_ISL_570689, EPI_ISL_570690, EPI_ISL_570691, EPI_ISL_570692, EPI_ISL_570693, EPI_ISL_570694, EPI_ISL_570695, EPI_ISL_570696, EPI_ISL_570697, EPI_ISL_570698, EPI_ISL_570699, EPI_ISL_570700, EPI_ISL_570701, EPI_ISL_570702, EPI_ISL_570703, EPI_ISL_570704, EPI_ISL_570705, EPI_ISL_570706, EPI_ISL_570707, EPI_ISL_570708, EPI_ISL_570709, EPI_ISL_570710, EPI_ISL_570711, EPI_ISL_570712, EPI_ISL_570713, EPI_ISL_570714, EPI_ISL_570715, EPI_ISL_570716, EPI_ISL_570717, EPI_ISL_570718, EPI_ISL_570719, EPI_ISL_570720, EPI_ISL_570721, EPI_ISL_570722, EPI_ISL_570723, EPI_ISL_570724, EPI_ISL_570725, EPI_ISL_570726, EPI_ISL_570727, EPI_ISL_570728, EPI_ISL_570729, EPI_ISL_570730, EPI_ISL_570731, EPI_ISL_570732, EPI_ISL_570733, EPI_ISL_570734, EPI_ISL_570736, EPI_ISL_570737, EPI_ISL_570738, EPI_ISL_570739, EPI_ISL_570740, EPI_ISL_570741, EPI_ISL_570742, EPI_ISL_570743, EPI_ISL_570744, EPI_ISL_570745, EPI_ISL_570746, EPI_ISL_570747, EPI_ISL_570748, EPI_ISL_570749, EPI_ISL_570750, EPI_ISL_570751, EPI_ISL_570752, EPI_ISL_570753, EPI_ISL_570754, EPI_ISL_570755, EPI_ISL_570756, EPI_ISL_570757, EPI_ISL_570758, EPI_ISL_570759, EPI_ISL_570760, EPI_ISL_570775, EPI_ISL_570804, EPI_ISL_570808, EPI_ISL_570810, EPI_ISL_570811, EPI_ISL_570812, EPI_ISL_570813, EPI_ISL_570815, EPI_ISL_570817, EPI_ISL_570820, EPI_ISL_570821, EPI_ISL_570822, EPI_ISL_570823, EPI_ISL_570826, EPI_ISL_570827, EPI_ISL_570828, EPI_ISL_570829, EPI_ISL_570830, EPI_ISL_570832, EPI_ISL_570838, EPI_ISL_570839, EPI_ISL_570840, EPI_ISL_570843, EPI_ISL_570846, EPI_ISL_570848, EPI_ISL_570852, EPI_ISL_570853, EPI_ISL_570854, EPI_ISL_570855, EPI_ISL_570856, EPI_ISL_570857, EPI_ISL_570861, EPI_ISL_570978, EPI_ISL_570980, EPI_ISL_570984, EPI_ISL_570985, EPI_ISL_570988, EPI_ISL_570989, EPI_ISL_570990, EPI_ISL_570991, EPI_ISL_570992, EPI_ISL_570993, EPI_ISL_570994, EPI_ISL_570995, EPI_ISL_570996, EPI_ISL_570997, EPI_ISL_570998, EPI_ISL_570999, EPI_ISL_571000, EPI_ISL_571001, EPI_ISL_571002, EPI_ISL_571003, EPI_ISL_571004, EPI_ISL_571005, EPI_ISL_571006, EPI_ISL_571007, EPI_ISL_571008, EPI_ISL_571009, EPI_ISL_571010, EPI_ISL_571011, EPI_ISL_571012, EPI_ISL_571013, EPI_ISL_571014, EPI_ISL_571015 | | | |
| see above | UW Virology Lab | UW Virology Lab | Pavitra Roychoudhury, Hong Xie, Lasata Shrestha, Amin Addetia, Victoria M Rachleff, Meei-Li Huang, Keith R Jerome, Alexander Greninger |
| EPI_ISL_572438, EPI_ISL_572488, EPI_ISL_572530, EPI_ISL_572532, EPI_ISL_572753, EPI_ISL_573314, EPI_ISL_573315, EPI_ISL_573344, EPI_ISL_573345, EPI_ISL_573346, EPI_ISL_573347, EPI_ISL_573348, EPI_ISL_573349, EPI_ISL_573350, EPI_ISL_573351, EPI_ISL_573352, EPI_ISL_573353, EPI_ISL_573354, EPI_ISL_573355, EPI_ISL_573356, EPI_ISL_573373, EPI_ISL_573374 | | | |
| see above | Northumbria University / South Tees Hospitals NHS Foundation Trust / North Cumbria Integrated Care NHS Foundation Trust / North Tees and Hartlepool NHS Foundation Trust / Newcastle Hospitals NHS Foundation Trust | COVID-19 Genomics UK (COG-UK) Consortium | Darren L Smith,Andrew Nelson,Matthew Bashton,Greg R Young,Joshua Loh,John Allan,Mohammad A Tariq,Giles S Holt,Gary Black,Wen C Yew,Lynn Dover,Paul Baker,Steve Liggett,Sarah Essex,Jane Greenaway,Debra Padgett,Clive Graham,Garren Scott,Edward Barton,Emma Swindells,Brendan Payne,Jennifer Collins,Yusri Taha,Gary Eltringham |
| EPI_ISL_574328 | LSUHS Emerging Viral Threat Laboratory | Microbial Genome Sequencing Center | Jeremy P. Kamil, Rona S. Scott, Maarten Van Diest, Malgorzata Bienkowska-Haba, Katarzyna Zwolinska, Andrew D. Yurochko, Christopher G. Kevil, Martin J. Sapp, Daniel J. Snyder, Vaughn S. Cooper, John A. Vanchiere |
| EPI_ISL_575238 | Utah Public Health Laboratory | Utah Public Health Laboratory | Erin Young, Kelly Oakeson |
| EPI_ISL_576148 | Department of Respiratory & Other Viral Infections of L.V. Gromashevsky Institute of Epidemiology & Infectious Diseases NAMS of Ukraine | Department of Respiratory & Other Viral Infections of L.V. Gromashevsky Institute of Epidemiology & Infectious Diseases NAMS of Ukraine, JSC "Farmak" | Alla Mironenko, Ihor Kravchuk, Liudmyla Bolotova, Larysa Radchenko, Nataliia Teteriuk |
| EPI_ISL_576149 | Department of Respiratory & Other Viral Infections of L.V. Gromashevsky Institute of Epidemiology & Infectious Diseases NAMS of Ukraine | Department of Respiratory & Other Viral Infections of L.V. Gromashevsky Institute of Epidemiology & Infectious Diseases NAMS of Ukraine, JSC "Farmak" | Alla Mironenko, Andriy Goy, Ihor Kravchuk, Liudmyla Bolotova, Larysa Radchenko, Nataliia Teteriuk |
| EPI_ISL_576261 | Instituto de Diagnostico y Referencia Epidemiologicos (INDRE) | Instituto de Diagnostico y Referencia Epidemiologicos (INDRE) | Gisela Barrera-Badillo , Abril Rodriguez-Maldonado, Claudia Wong-Arambula , Natividad Cruz-Ortiz, Tatiana Nunez-Garcia, Dayanira Arellano-Suarez, Adnan Araiza-Rodriguez, Edgar Mendieta-Condado, Lucia Hernandez-Rivas, Irma Lopez-Martinez, Ernesto Ramirez-Gonzalez. |
| EPI_ISL_576336, EPI_ISL_576337, EPI_ISL_576338, EPI_ISL_576339, EPI_ISL_576340, EPI_ISL_576341, EPI_ISL_576342, EPI_ISL_576343, EPI_ISL_576344, EPI_ISL_576345, EPI_ISL_576346 | | | |
| see above | Texas Department of State Health Services | Texas Department of State Health Services | Rashmi Tuladhar, Bonnie Oh, Mayela Pedrueza, Jenny Zhang, Maliha Rahman, Anita Pokharel, Myong Koag, Chun Wang, Rachel Lee, Grace Kubin |
| EPI_ISL_576407, EPI_ISL_576408, EPI_ISL_576409, EPI_ISL_576410, EPI_ISL_576411, EPI_ISL_576412, EPI_ISL_576413, EPI_ISL_576414, EPI_ISL_576415, EPI_ISL_576416, EPI_ISL_576417, EPI_ISL_576418, EPI_ISL_576419, EPI_ISL_576420, EPI_ISL_576421, EPI_ISL_576422, EPI_ISL_576423, EPI_ISL_576424, EPI_ISL_576425, EPI_ISL_576426, EPI_ISL_576427, EPI_ISL_576428, EPI_ISL_576429, EPI_ISL_576430, EPI_ISL_576431, EPI_ISL_576432, EPI_ISL_576433, EPI_ISL_576434, EPI_ISL_576435, EPI_ISL_576436, EPI_ISL_576437, EPI_ISL_576438, EPI_ISL_576439, EPI_ISL_576440, EPI_ISL_576444, EPI_ISL_576445, EPI_ISL_576446, EPI_ISL_576447, EPI_ISL_576448, EPI_ISL_576449, EPI_ISL_576450, EPI_ISL_576451, EPI_ISL_576452, EPI_ISL_576453, EPI_ISL_576455, EPI_ISL_576456, EPI_ISL_576457, EPI_ISL_576458, EPI_ISL_576461, EPI_ISL_576470, EPI_ISL_576471, EPI_ISL_576474, EPI_ISL_576477, EPI_ISL_576479, EPI_ISL_576482, EPI_ISL_576484 | | | |
| see above | UW Virology Lab | UW Virology Lab | Pavitra Roychoudhury, Hong Xie, Lasata Shrestha, Amin Addetia, Victoria M Rachleff, Meei-Li Huang, Keith R Jerome, Alexander Greninger |
| EPI_ISL_576568 | Texas Department of State Health Services | Texas Department of State Health Services | Rashmi Tuladhar, Bonnie Oh, Mayela Pedrueza, Jenny Zhang, Maliha Rahman, Anita Pokharel, Myong Koag, Chun Wang, Rachel Lee, Grace Kubin |
| EPI_ISL_577607 | Royal Hobart Hospital | Royal Hobart Hospital | Cooley L., van Haeften R. |
| EPI_ISL_577627, EPI_ISL_577628 | The National Institute of Public Health | State Veterinary Institute Prague | Nagy,A;Jirincova,H;Novakova,L;Trnka,D;Vecerova,J |
| EPI_ISL_577747 | Dutch COVID-19 response team | Erasmus Medical Center | Bas Oude Munnink, Reina Sikkema, David Nieuwenhuijse, Irina Chestakova, Anne van der Linden, Marjan Boter, Emmanuelle Munger, Corine GeurtsvanKessel, Annemiek van der Eijk, Richard Molenkamp, Marion Koopmans, on behalf of the Dutch national COVID-19 response team. |
| EPI_ISL_577749, EPI_ISL_577750, EPI_ISL_577751, EPI_ISL_577752, EPI_ISL_577753, EPI_ISL_577754 | Dutch COVID-19 response team | Erasmus Medical Center | OH consortium |
| EPI_ISL_577844, EPI_ISL_577910, EPI_ISL_577911, EPI_ISL_577912, EPI_ISL_577913 | Dutch COVID-19 response team | Erasmus Medical Center | Bas Oude Munnink, Reina Sikkema, David Nieuwenhuijse, Irina Chestakova, Anne van der Linden, Marjan Boter, Emmanuelle Munger, Corine GeurtsvanKessel, Annemiek van der Eijk, Richard Molenkamp, Marion Koopmans, on behalf of the Dutch national COVID-19 response team. |
| EPI_ISL_578919, EPI_ISL_578923, EPI_ISL_578927, EPI_ISL_578928, EPI_ISL_578929, EPI_ISL_578930, EPI_ISL_578933, EPI_ISL_578934, EPI_ISL_578935, EPI_ISL_578937, EPI_ISL_578938, EPI_ISL_578939, EPI_ISL_578949, EPI_ISL_578950, EPI_ISL_578951, EPI_ISL_578953, EPI_ISL_578955, EPI_ISL_578956, EPI_ISL_578957, EPI_ISL_578958, EPI_ISL_578959, EPI_ISL_578960, EPI_ISL_578961, EPI_ISL_578962, EPI_ISL_578963, EPI_ISL_578964, EPI_ISL_578965, EPI_ISL_578966, EPI_ISL_579008, EPI_ISL_579009, EPI_ISL_579012, EPI_ISL_579019, EPI_ISL_579021, EPI_ISL_579023, EPI_ISL_579026, EPI_ISL_579029, EPI_ISL_579032, EPI_ISL_579033, EPI_ISL_579035, EPI_ISL_579036, EPI_ISL_579037, EPI_ISL_579038, EPI_ISL_579039, EPI_ISL_579040, EPI_ISL_579042, EPI_ISL_579043, EPI_ISL_579044, EPI_ISL_579045, EPI_ISL_579046, EPI_ISL_579047, EPI_ISL_579048, EPI_ISL_579049, EPI_ISL_579050, EPI_ISL_579051, EPI_ISL_579052, EPI_ISL_579053, EPI_ISL_579054, EPI_ISL_579055, EPI_ISL_579056, EPI_ISL_579057 | | | |
| see above | LSUHS Emerging Viral Threat Laboratory | Microbial Genome Sequencing Center | Jeremy P. Kamil, Rona S. Scott, Maarten Van Diest, Malgorzata Bienkowska-Haba, Katarzyna Zwolinska, Andrew D. Yurochko, Christopher G. Kevil, Martin J. Sapp, Daniel J. Snyder, Vaughn S. Cooper, John A. Vanchiere |
| EPI_ISL_581410, EPI_ISL_581418, EPI_ISL_581419, EPI_ISL_581420 | Lighthouse Lab in Glasgow | Wellcome Sanger Institute for the COVID-19 Genomics UK (COG-UK) consortium | Harper VanSteenhouse, Yumi Kasai, David Gray, Carol Clugston, Anna Dominiczak and Alex Alderton, Roberto Amato, Sonia Goncalves, Ewan Harrison, David K. Jackson, Ian Johnston, Dominic Kwiatkowski, Cordelia Langford, John Sillitoe on behalf of the Wellcome Sanger Institute COVID-19 Surveillance Team |
| EPI_ISL_581449, EPI_ISL_581497 | CSIR-Indian Institute of Chemical Biology, MEDICA Supercpecialty Hospital Kolkata | CSIR-Indian Institute of Chemical Biology, MEDICA Supercpecialty Hospital Kolkata | Sujay Krishna Maity, Priyanka Mallick, Debaleena Bhowmik, Abhishake Lahiri, Dr. Aviral Roy, Dr. Soumen Saha, Dr. Arpita Ghosh Mitra, Dr. Rajesh Pandey, Dr. Sandip Paul, Dr. Partha Chakrabarti, Dr. Saikat Chakrabarti |
| EPI_ISL_581504 | CSIR-Indian Institute of Chemical Biology, MEDICA Supercpecialty Hospital Kolkata | CSIR-Indian Institute of Chemical Biology, MEDICA Supercpecialty Hospital Kolkata | Sujay Krishna Maity, Priyanka Mallick, Debaleena Bhowmik, Abhishake Lahiri, Dr. AviralRoy, Dr. Soumen Saha, Dr. Arpita Ghosh Mitra, Dr. Rajesh Pandey, Dr. Sandip Paul, Dr.Partha Chakrabarti, Dr. Saikat Chakrabarti |

| | | | |
|---|---|---|---|
| EPI_ISL_581517, EPI_ISL_581518, EPI_ISL_581520, EPI_ISL_581522, EPI_ISL_581523, EPI_ISL_581524, EPI_ISL_581525, EPI_ISL_581526, EPI_ISL_581527, EPI_ISL_581528, EPI_ISL_581529, EPI_ISL_581530, EPI_ISL_581531, EPI_ISL_581532, EPI_ISL_581533, EPI_ISL_581534, EPI_ISL_581535, EPI_ISL_581536, EPI_ISL_581537, EPI_ISL_581538, EPI_ISL_581539, EPI_ISL_581540, EPI_ISL_581541 | | | |
| see above | Virginia DCLS | Virginia DCLS | Virginia DCLS |
| EPI_ISL_581933, EPI_ISL_581934, EPI_ISL_581935, EPI_ISL_581936, EPI_ISL_581937, EPI_ISL_581938, EPI_ISL_581939, EPI_ISL_581940, EPI_ISL_581941, EPI_ISL_581942, EPI_ISL_581943, EPI_ISL_581944, EPI_ISL_581945, EPI_ISL_581961, EPI_ISL_581962, EPI_ISL_581963 | | | |
| see above | University Hospital Basel, Clinical Virology | University Hospital Basel, Clinical Bacteriology | Madlen Stange, Alfredo Mari, Tim Roloff, Helena MB Seth-Smith, Michael Schweitzer, Myrta Brunner, Karoline Leuzinger, Kirstine K. Soegaard, Alexander Gensch, Sarah Tschudin-Sutter, Simon Fuchs, Julia Bielicki, Hans Pargger, Martin Siegemund, Christian Nickel, Roland Bingisser, Michael Osthoff, Stefano Bassetti, Rita Schneider-Sliwa, Manuel Battegay, Hans Hirsch, Adrian Egli |
| EPI_ISL_582010 | Infectious Diseases and Tropical Medicine Research Center, Infectious Diseases and Tropical Medicine Research Center | Infectious Diseases and Tropical Medicine Research Center, Infectious Diseases and Tropical Medicine Research Center | Ahangarzadeh,S., Haghjooy Javanmard,S., Shariati,L., Ataei,B., Ranjbar,M.M., Shoaei,P. |
| EPI_ISL_582011 | Infectious Diseases and Tropical Medicine Research Center, Infectious Diseases and Tropical Medicine Research Center | Infectious Diseases and Tropical Medicine Research Center, Infectious Diseases and Tropical Medicine Research Center | Shariati,L., Haghjooy Javanmard,S., Ataei,B., Shoaei,P., Ranjbar,M.M., Ahangarzadeh,S. |
| EPI_ISL_582012 | Infectious Diseases and Tropical Medicine Research Center, Infectious Diseases and Tropical Medicine Research Center | Infectious Diseases and Tropical Medicine Research Center, Infectious Diseases and Tropical Medicine Research Center | Ataei,B., Haghjooy Javanmard,S., Ahangarzadeh,S., Shoaei,P., Shariati,L., Ranjbar,M.M. |
| EPI_ISL_582016 | Infectious Diseases and Tropical Medicine Research Center, Infectious Diseases and Tropical Medicine Research Center | Infectious Diseases and Tropical Medicine Research Center, Infectious Diseases and Tropical Medicine Research Center | Haghjooy Javanmard,S., Shariati,l., Ahangarzadeh,S., Ataei,B., Shoaei,P. |
| EPI_ISL_582121 | Centre Hospitalier de Bourg en Bresse | CNR Virus des Infections Respiratoires - France SUD | Antonin Bal, Gregory Destras, Gwendolyne Burfin, Hadrien Règue, Alexandre Gaymard, Maude Bouscambert-Duchamp, Florence Morfin-Sherpa, Martine Valette, Bruno Lina, Laurence Josset |
| EPI_ISL_582230, EPI_ISL_582234 | Wyoming Public Health Laboratory | Center for Global Health, University of New Mexico Health Sciences Center | Daryl Domman, Kurt Schwalm, Rob Christensen, Wanda Manley, Cari Sloma, Noah Hull, Darrell Dinwiddie |
| EPI_ISL_582416, EPI_ISL_582417, EPI_ISL_582418, EPI_ISL_582419, EPI_ISL_582420, EPI_ISL_582422, EPI_ISL_582424, EPI_ISL_582425, EPI_ISL_582426, EPI_ISL_582427, EPI_ISL_582428, EPI_ISL_582429, EPI_ISL_582430, EPI_ISL_582431, EPI_ISL_582432, EPI_ISL_582433, EPI_ISL_582434, EPI_ISL_582435, EPI_ISL_582436, EPI_ISL_582437, EPI_ISL_582439, EPI_ISL_582440, EPI_ISL_582441, EPI_ISL_582442, EPI_ISL_582443, EPI_ISL_582445, EPI_ISL_582446, EPI_ISL_582447, EPI_ISL_582448, EPI_ISL_582449, EPI_ISL_582450, EPI_ISL_582451, EPI_ISL_582452, EPI_ISL_582453, EPI_ISL_582454, EPI_ISL_582455, EPI_ISL_582456, EPI_ISL_582457, EPI_ISL_582458, EPI_ISL_582459, EPI_ISL_582460, EPI_ISL_582461, EPI_ISL_582462, EPI_ISL_582467, EPI_ISL_582469, EPI_ISL_582470, EPI_ISL_582471, EPI_ISL_582472, EPI_ISL_582473, EPI_ISL_582478, EPI_ISL_582481, EPI_ISL_582482, EPI_ISL_582483, EPI_ISL_582488, EPI_ISL_582489, EPI_ISL_582490, EPI_ISL_582491, EPI_ISL_582492, EPI_ISL_582495, EPI_ISL_582500, EPI_ISL_582501, EPI_ISL_582502, EPI_ISL_582503, EPI_ISL_582504 | | | |
| see above | Cadham Provincial Laboratory | National Microbiology Laboratory (NML) | Anna Majer, Shari Tyson, Grace Seo, Philip Mabon, Elsie Grudeski, Rhiannon Huzarewich, Russell Mandes, Anneliese Landgraff, Jennifer Tanner, Natalie Knox, Morag Graham, Gary Van Domselaar, Paul Van Caeseele, Jared Bullard, David Alexander, Kerry Dust, Nathalie Bastien, Yan Li, Timothy Booth, Darian Hole, Madison Chapel, CanCOGeN's metadata curation team, Public Health Agency of Canada CanCOGeN team |
| EPI_ISL_582509 | Department of Respiratory and other Viral Infections of L.V.Gromashevsky Institute of Epidemiology & Infectious Diseases NAMS of Ukraine | Department of Respiratory and other Viral Infections of L.V.Gromashevsky Institute of Epidemiology & Infectious Diseases NAMS of Ukraine, JSC "Farmak" | Alla Mironenko, Andriy Goy, Ihor Kravchuk, Ludmyla Bolotova, Larysa Radchenko, Nataliia Teteriuk |
| EPI_ISL_582510 | Department of Respiratory and other Viral Infections of L.V.Gromashevsky Institute of Epidemiology & Infectious Diseases NAMS of Ukraine | Department of Respiratory and other Viral Infections of L.V.Gromashevsky Institute of Epidemiology & Infectious Diseases NAMS of Ukrain, JSC "Farmak" | Alla Mironenko, Andriy Goy, Ihor Kravchuk, Ludmyla Bolotova, Larysa Radchenko, Nataliia Teteriuk |
| EPI_ISL_582511, EPI_ISL_582512 | Department of Respiratory and other Viral Infections of L.V.Gromashevsky Institute of Epidemiology & Infectious Diseases NAMS of Ukrain | Department of Respiratory and other Viral Infections of L.V.Gromashevsky Institute of Epidemiology & Infectious Diseases NAMS of Ukrain, JSC "Farmak" | Alla Mironenko, Andriy Goy, Ihor Kravchuk, Ludmyla Bolotova, Larysa Radchenko, Nataliia Teteriuk |
| EPI_ISL_582783, EPI_ISL_582784 | Uppsala klinisk mikrobiologi | The Public Health Agency of Sweden | Anna-Malin Linde, Maria Lind Karlberg, Mattias Haukland, Reza Advani, Olov Svartstrom, Oskar Karlsson Lindsjo, Sandra Broddesson, Petra Edquist, Mia Brytting, Anna Risberg, Karin Tegmark-Wisell |
| EPI_ISL_582801 | Gavle klinisk mikrobiologi | The Public Health Agency of Sweden | Anna-Malin Linde, Maria Lind Karlberg, Mattias Haukland, Reza Advani, Olov Svartstrom, Oskar Karlsson Lindsjo, Sandra Broddesson, Petra Edquist, Mia Brytting, Anna Risberg, Karin Tegmark-Wisell |
| EPI_ISL_582807 | Klinisk mikrobiologi SAS Boras | The Public Health Agency of Sweden | Anna-Malin Linde, Maria Lind Karlberg, Mattias Haukland, Reza Advani, Olov Svartstrom, Oskar Karlsson Lindsjo, Sandra Broddesson, Petra Edquist, Mia Brytting, Anna Risberg, Karin Tegmark-Wisell |
| EPI_ISL_582921, EPI_ISL_582924, EPI_ISL_582925, EPI_ISL_582926, EPI_ISL_582940, EPI_ISL_582941 | County of Santa Clara Public Health Department | Chan-Zuckerberg Biohub | CZB Cliahub Consortium |
| EPI_ISL_583102, EPI_ISL_583103, EPI_ISL_583104, EPI_ISL_583105, EPI_ISL_583106, EPI_ISL_583107, EPI_ISL_583108, EPI_ISL_583109, EPI_ISL_583110, EPI_ISL_583111, EPI_ISL_583112, EPI_ISL_583113, EPI_ISL_583114, EPI_ISL_583115, EPI_ISL_583116, EPI_ISL_583117, EPI_ISL_583118, EPI_ISL_583119 | | | |
| see above | Humboldt County Public Health Laboratory | Chan-Zuckerberg Biohub | CZB Cliahub Consortium |
| EPI_ISL_583197 | San Francisco Public Health Laboratory | Chan-Zuckerberg Biohub | CZB Cliahub Consortium |
| EPI_ISL_583217, EPI_ISL_583218 | UCSF Clinical Microbiology Laboratory | Chan-Zuckerberg Biohub | CZB Cliahub Consortium |
| EPI_ISL_583535 | Genome Centre | Genome Centre | Selina Akter, Pravas Chandra Roy, Amina Ferdaus manami, Habiba Ibnat, A. S. M. Rubayet Ul Alam, Shireen Nigar, Iqbal Kabir Jahid, M.Anwar Hossain |
| EPI_ISL_583887, EPI_ISL_583888, EPI_ISL_583889, EPI_ISL_583890, EPI_ISL_583891, EPI_ISL_583892 | Austrian Agency for Health and Food Safety (AGES) | Bergthaler laboratory, CeMM Research Center for Molecular Medicine of the Austrian Academy of Sciences | Alexandra Popa, Benedikt Agerer, Henrique Colaco, Lukas Endler, Jakob-Wendelin Genger, Alexander Lercher, Mark Smyth, Thomas Penz, Michael Schuster, Jan Laine, Martin Senekowitsch, Judith Aberle, Stephan Aberle, Peter Hufnagl, Daniela Schmid, Franz Allerberger, Elisabeth Puchhammer-Stoeckl, Manfred Nairz, Guenter Weiss, Gregor Hörmann, Kinga Rigler-Hohenwarter, Rainer Gattringer, Wegene Borena, Dorothee von Laer, Gernot Walder, Peter Obrist, Christian Paar, Sabine Sussitz-Rack, Gunther Vogl, Adi Steinrigl, Christoph Bock, Andreas Bergthaler |
| EPI_ISL_590323 | Lighthouse Lab in Glasgow | Wellcome Sanger Institute for the COVID-19 Genomics UK (COG-UK) consortium | Harper VanSteenhouse, Yumi Kasai, David Gray, Carol Clugston, Anna Dominiczak and Alex Alderton, Roberto Amato, Sonia Goncalves, Ewan Harrison, David K. Jackson, Ian Johnston, Dominic Kwiatkowski, Cordelia Langford, John Sillitoe on behalf of the Wellcome Sanger Institute COVID-19 Surveillance Team (http://www.sanger.ac.uk/covid-team) |
| EPI_ISL_590919, EPI_ISL_590920 | Vestfold Hospital, Toensberg Department of Microbiology | Norwegian Institute of Public Health, Department of Virology | Kathrine Stene-Johansen, Kamilla Heddeland Instefjord, Hilde Elshaug, Rasmus Riis Kopperud, Hilde Vollan, Karoline Bragstad, Olav Hungnes |
| EPI_ISL_590921, EPI_ISL_590922 | Ostfold Hospital Trust - Kalnes, Centre for Laboratory Medicine, Section for gene technology and infection serology | Norwegian Institute of Public Health, Department of Virology | Kathrine Stene-Johansen, Kamilla Heddeland Instefjord, Hilde Elshaug, Rasmus Riis Kopperud, Hilde Vollan, Karoline Bragstad, Olav Hungnes |
| EPI_ISL_590938, EPI_ISL_590941, EPI_ISL_590942, EPI_ISL_590943, EPI_ISL_590946 | University Hospital of Northern Norway, Department for Microbiology and Infectious Disease Control | Norwegian Institute of Public Health, Department of Virology | Kathrine Stene-Johansen, Kamilla Heddeland Instefjord, Hilde Elshaug, Rasmus Riis Kopperud, Hilde Vollan, Karoline Bragstad, Olav Hungnes |
| EPI_ISL_590986 | Ostfold Hospital Trust - Kalnes, Centre for Laboratory Medicine, Section for gene technology and infection serology | Norwegian Institute of Public Health, Department of Virology | Kathrine Stene-Johansen, Kamilla Heddeland Instefjord, Hilde Elshaug, Rasmus Riis Kopperud, Hilde Vollan, Karoline Bragstad, Olav Hungnes |
| EPI_ISL_591007 | Department of Medical Microbiology, St. Olavs hospital | Norwegian Institute of Public Health, Department of Virology | Kathrine Stene-Johansen, Kamilla Heddeland Instefjord, Hilde Elshaug, Rasmus Riis Kopperud, Hilde Vollan, Karoline Bragstad, Olav Hungnes |
| EPI_ISL_591021 | Dept. of Medical Microbiology, Stavanger University Hospital, Helse Stavanger HF | Norwegian Institute of Public Health, Department of Virology | Kathrine Stene-Johansen, Kamilla Heddeland Instefjord, Hilde Elshaug, Rasmus Riis Kopperud, Hilde Vollan, Karoline Bragstad, Olav Hungnes |
| EPI_ISL_591059 | Infectious Diseases and Tropical Medicine Research Center, Infectious Diseases and Tropical Medicine Research Center | Infectious Diseases and Tropical Medicine Research Center, Infectious Diseases and Tropical Medicine Research Center | Ahangarzadeh,S., Haghjooy Javanmard,S., Shariati,L. |
| EPI_ISL_591343 | Infectious Diseases and Tropical Medicine Research Center, Infectious Diseases and Tropical Medicine Research Center | Infectious Diseases and Tropical Medicine Research Center, Infectious Diseases and Tropical Medicine Research Center | Haghjooy Javanmard,S., Ahangarzadeh,S., Shariati,L., Ataei,B., Shoaei,P., Ranjbar,M.M., Abutalebian,S. |

| | | | |
|---|---|---|---|
| EPI_ISL_591346 | Infectious Diseases and Tropical Medicine Research Center, Infectious Diseases and Tropical Medicine Research Center | Infectious Diseases and Tropical Medicine Research Center, Infectious Diseases and Tropical Medicine Research Center | Haghjooy Javanmard,S., Abutalebian,S., Shoaei,P., Ataei,B., Shariati,L., Ahangarzadeh,S. |
| EPI_ISL_591509 | Sydney South West Pathology Service (SSWPS) - Liverpool Hospital - NSW Health Pathology | NSW Health Pathology - Institute of Clinical Pathology and Medical Research; Westmead Hospital; University of Sydney | CIDM-PH et al. |
| EPI_ISL_591556 | Microbiological Diagnostic Unit - Public Health Laboratory (MDU-PHL) | MDU-PHL | Seemann T., Schultz, M. B., Sait, M., Sherry, N. |
| EPI_ISL_591561, EPI_ISL_591565, EPI_ISL_591571, EPI_ISL_591573, EPI_ISL_591580, EPI_ISL_591585, EPI_ISL_591587 | Victorian Infectious Diseases Reference Laboratory (VIDRL) | VIDRL and MDU-PHL | Caly L., Seemann T., Sait, M., Schultz, M. B., Druce J., Sherry, N. |
| EPI_ISL_591588 | Microbiological Diagnostic Unit - Public Health Laboratory (MDU-PHL) | MDU-PHL | Seemann T., Schultz, M. B., Sait, M., Sherry, N. |
| EPI_ISL_591590, EPI_ISL_591592, EPI_ISL_591595 | Victorian Infectious Diseases Reference Laboratory (VIDRL) | VIDRL and MDU-PHL | Caly L., Seemann T., Sait, M., Schultz, M. B., Druce J., Sherry, N. |
| EPI_ISL_591596 | Microbiological Diagnostic Unit - Public Health Laboratory (MDU-PHL) | MDU-PHL | Seemann T., Schultz, M. B., Sait, M., Sherry, N. |
| EPI_ISL_591603, EPI_ISL_591609 | Victorian Infectious Diseases Reference Laboratory (VIDRL) | VIDRL and MDU-PHL | Caly L., Seemann T., Sait, M., Schultz, M. B., Druce J., Sherry, N. |
| EPI_ISL_591612 | Microbiological Diagnostic Unit - Public Health Laboratory (MDU-PHL) | MDU-PHL | Seemann T., Schultz, M. B., Sait, M., Sherry, N. |
| EPI_ISL_591613, EPI_ISL_591615, EPI_ISL_591619 | Victorian Infectious Diseases Reference Laboratory (VIDRL) | VIDRL and MDU-PHL | Caly L., Seemann T., Sait, M., Schultz, M. B., Druce J., Sherry, N. |
| EPI_ISL_591620 | Microbiological Diagnostic Unit - Public Health Laboratory (MDU-PHL) | MDU-PHL | Seemann T., Schultz, M. B., Sait, M., Sherry, N. |
| EPI_ISL_591623 | Victorian Infectious Diseases Reference Laboratory (VIDRL) | VIDRL and MDU-PHL | Caly L., Seemann T., Sait, M., Schultz, M. B., Druce J., Sherry, N. |
| EPI_ISL_591633, EPI_ISL_591639, EPI_ISL_591641, EPI_ISL_591644, EPI_ISL_591651 | Microbiological Diagnostic Unit - Public Health Laboratory (MDU-PHL) | MDU-PHL | Seemann T., Schultz, M. B., Sait, M., Sherry, N. |
| EPI_ISL_591652, EPI_ISL_591657 | Victorian Infectious Diseases Reference Laboratory (VIDRL) | VIDRL and MDU-PHL | Caly L., Seemann T., Sait, M., Schultz, M. B., Druce J., Sherry, N. |
| EPI_ISL_591665 | Microbiological Diagnostic Unit - Public Health Laboratory (MDU-PHL) | MDU-PHL | Seemann T., Schultz, M. B., Sait, M., Sherry, N. |
| EPI_ISL_591670 | Victorian Infectious Diseases Reference Laboratory (VIDRL) | VIDRL and MDU-PHL | Caly L., Seemann T., Sait, M., Schultz, M. B., Druce J., Sherry, N. |
| EPI_ISL_591676, EPI_ISL_591677, EPI_ISL_591678 | Microbiological Diagnostic Unit - Public Health Laboratory (MDU-PHL) | MDU-PHL | Seemann T., Schultz, M. B., Sait, M., Sherry, N. |
| EPI_ISL_591679, EPI_ISL_591681, EPI_ISL_591684, EPI_ISL_591698, EPI_ISL_591703, EPI_ISL_591706 | Victorian Infectious Diseases Reference Laboratory (VIDRL) | VIDRL and MDU-PHL | Caly L., Seemann T., Sait, M., Schultz, M. B., Druce J., Sherry, N. |
| EPI_ISL_591709, EPI_ISL_591717, EPI_ISL_591718 | Microbiological Diagnostic Unit - Public Health Laboratory (MDU-PHL) | MDU-PHL | Seemann T., Schultz, M. B., Sait, M., Sherry, N. |
| EPI_ISL_591723 | Victorian Infectious Diseases Reference Laboratory (VIDRL) | VIDRL and MDU-PHL | Caly L., Seemann T., Sait, M., Schultz, M. B., Druce J., Sherry, N. |
| EPI_ISL_591727, EPI_ISL_591728 | Microbiological Diagnostic Unit - Public Health Laboratory (MDU-PHL) | MDU-PHL | Seemann T., Schultz, M. B., Sait, M., Sherry, N. |
| EPI_ISL_591729, EPI_ISL_591734 | Victorian Infectious Diseases Reference Laboratory (VIDRL) | VIDRL and MDU-PHL | Caly L., Seemann T., Sait, M., Schultz, M. B., Druce J., Sherry, N. |
| EPI_ISL_591736 | Microbiological Diagnostic Unit - Public Health Laboratory (MDU-PHL) | MDU-PHL | Seemann T., Schultz, M. B., Sait, M., Sherry, N. |
| EPI_ISL_591737, EPI_ISL_591740, EPI_ISL_591743, EPI_ISL_591744, EPI_ISL_591746 | Victorian Infectious Diseases Reference Laboratory (VIDRL) | VIDRL and MDU-PHL | Caly L., Seemann T., Sait, M., Schultz, M. B., Druce J., Sherry, N. |
| EPI_ISL_591748, EPI_ISL_591752 | Microbiological Diagnostic Unit - Public Health Laboratory (MDU-PHL) | MDU-PHL | Seemann T., Schultz, M. B., Sait, M., Sherry, N. |
| EPI_ISL_591758 | Victorian Infectious Diseases Reference Laboratory (VIDRL) | VIDRL and MDU-PHL | Caly L., Seemann T., Sait, M., Schultz, M. B., Druce J., Sherry, N. |
| EPI_ISL_591759 | Microbiological Diagnostic Unit - Public Health Laboratory (MDU-PHL) | MDU-PHL | Seemann T., Schultz, M. B., Sait, M., Sherry, N. |
| EPI_ISL_591763 | Victorian Infectious Diseases Reference Laboratory (VIDRL) | VIDRL and MDU-PHL | Caly L., Seemann T., Sait, M., Schultz, M. B., Druce J., Sherry, N. |
| EPI_ISL_591765, EPI_ISL_591766 | Microbiological Diagnostic Unit - Public Health Laboratory (MDU-PHL) | MDU-PHL | Seemann T., Schultz, M. B., Sait, M., Sherry, N. |
| EPI_ISL_591767, EPI_ISL_591769, EPI_ISL_591771, EPI_ISL_591772, EPI_ISL_591773, EPI_ISL_591775 | Victorian Infectious Diseases Reference Laboratory (VIDRL) | VIDRL and MDU-PHL | Caly L., Seemann T., Sait, M., Schultz, M. B., Druce J., Sherry, N. |
| EPI_ISL_591776, EPI_ISL_591777 | Microbiological Diagnostic Unit - Public Health Laboratory (MDU-PHL) | MDU-PHL | Seemann T., Schultz, M. B., Sait, M., Sherry, N. |
| EPI_ISL_591779 | Victorian Infectious Diseases Reference Laboratory (VIDRL) | VIDRL and MDU-PHL | Caly L., Seemann T., Sait, M., Schultz, M. B., Druce J., Sherry, N. |
| EPI_ISL_591780, EPI_ISL_591784, EPI_ISL_591785 | Microbiological Diagnostic Unit - Public Health Laboratory (MDU-PHL) | MDU-PHL | Seemann T., Schultz, M. B., Sait, M., Sherry, N. |
| EPI_ISL_591786, EPI_ISL_591792, EPI_ISL_591799, EPI_ISL_591800, EPI_ISL_591822 | Victorian Infectious Diseases Reference Laboratory (VIDRL) | VIDRL and MDU-PHL | Caly L., Seemann T., Sait, M., Schultz, M. B., Druce J., Sherry, N. |
| EPI_ISL_591823 | Microbiological Diagnostic Unit - Public Health Laboratory (MDU-PHL) | MDU-PHL | Seemann T., Schultz, M. B., Sait, M., Sherry, N. |
| EPI_ISL_591824, EPI_ISL_591850, EPI_ISL_591851, EPI_ISL_591852 | Victorian Infectious Diseases Reference Laboratory (VIDRL) | VIDRL and MDU-PHL | Caly L., Seemann T., Sait, M., Schultz, M. B., Druce J., Sherry, N. |

| | | | |
|---|---|---|---|
| EPI_ISL_591853 | Microbiological Diagnostic Unit - Public Health Laboratory (MDU-PHL) | MDU-PHL | Seemann T., Schultz, M. B., Sait, M., Sherry, N. |
| EPI_ISL_591854, EPI_ISL_591856, EPI_ISL_591857, EPI_ISL_591858, EPI_ISL_591859, EPI_ISL_591861, EPI_ISL_591862, EPI_ISL_591863, EPI_ISL_591864, EPI_ISL_591865, EPI_ISL_591866, EPI_ISL_591867, EPI_ISL_591869, EPI_ISL_591870, EPI_ISL_591872, EPI_ISL_591873, EPI_ISL_591874, EPI_ISL_591875, EPI_ISL_591876, EPI_ISL_591878, EPI_ISL_591879, EPI_ISL_591880, EPI_ISL_591881, EPI_ISL_591882, EPI_ISL_591883, EPI_ISL_591884, EPI_ISL_591887, EPI_ISL_591888, EPI_ISL_591889, EPI_ISL_591890, EPI_ISL_591891, EPI_ISL_591893, EPI_ISL_591894, EPI_ISL_591895, EPI_ISL_591897, EPI_ISL_591898, EPI_ISL_591899, EPI_ISL_591900, EPI_ISL_591942, EPI_ISL_591943, EPI_ISL_591944, EPI_ISL_591945, EPI_ISL_591946, EPI_ISL_591947, EPI_ISL_591948, EPI_ISL_591949, EPI_ISL_591950, EPI_ISL_591951, EPI_ISL_591952, EPI_ISL_591954, EPI_ISL_591955, EPI_ISL_591959, EPI_ISL_591960 | | | |
| see above | Victorian Infectious Diseases Reference Laboratory (VIDRL) | VIDRL and MDU-PHL | Caly L., Seemann T., Sait, M., Schultz, M. B., Druce J., Sherry, N. |
| EPI_ISL_592124, EPI_ISL_592128, EPI_ISL_592135 | Microbiological Diagnostic Unit - Public Health Laboratory (MDU-PHL) | MDU-PHL | Seemann T., Schultz, M. B., Sait, M., Sherry, N. |
| EPI_ISL_592136, EPI_ISL_592137, EPI_ISL_592138, EPI_ISL_592140, EPI_ISL_592141, EPI_ISL_592142, EPI_ISL_592143, EPI_ISL_592144, EPI_ISL_592145, EPI_ISL_592148, EPI_ISL_592149, EPI_ISL_592151, EPI_ISL_592152, EPI_ISL_592153, EPI_ISL_592154, EPI_ISL_592155, EPI_ISL_592156, EPI_ISL_592157, EPI_ISL_592158, EPI_ISL_592159, EPI_ISL_592161, EPI_ISL_592162, EPI_ISL_592163, EPI_ISL_592167, EPI_ISL_592168, EPI_ISL_592169, EPI_ISL_592170, EPI_ISL_592171, EPI_ISL_592172, EPI_ISL_592173, EPI_ISL_592174, EPI_ISL_592175, EPI_ISL_592176, EPI_ISL_592177, EPI_ISL_592178, EPI_ISL_592179, EPI_ISL_592180, EPI_ISL_592181, EPI_ISL_592182, EPI_ISL_592183, EPI_ISL_592184, EPI_ISL_592185, EPI_ISL_592187, EPI_ISL_592189, EPI_ISL_592190, EPI_ISL_592191, EPI_ISL_592192, EPI_ISL_592193, EPI_ISL_592194, EPI_ISL_592195 | | | |
| see above | Victorian Infectious Diseases Reference Laboratory (VIDRL) | VIDRL and MDU-PHL | Caly L., Seemann T., Sait, M., Schultz, M. B., Druce J., Sherry, N. |
| EPI_ISL_592210, EPI_ISL_592244, EPI_ISL_592245, EPI_ISL_592246, EPI_ISL_592247, EPI_ISL_592248, EPI_ISL_592249, EPI_ISL_592250, EPI_ISL_592251, EPI_ISL_592252, EPI_ISL_592253, EPI_ISL_592254, EPI_ISL_592255, EPI_ISL_592256, EPI_ISL_592257, EPI_ISL_592258, EPI_ISL_592259, EPI_ISL_592260, EPI_ISL_592261, EPI_ISL_592262, EPI_ISL_592263, EPI_ISL_592264, EPI_ISL_592265, EPI_ISL_592266, EPI_ISL_592267, EPI_ISL_592268, EPI_ISL_592290, EPI_ISL_592292, EPI_ISL_592293, EPI_ISL_592294, EPI_ISL_592307, EPI_ISL_592315, EPI_ISL_592321, EPI_ISL_592323, EPI_ISL_592324, EPI_ISL_592352, EPI_ISL_592354, EPI_ISL_592356, EPI_ISL_592357, EPI_ISL_592358, EPI_ISL_592359, EPI_ISL_592360, EPI_ISL_592364, EPI_ISL_592365, EPI_ISL_592370, EPI_ISL_592372, EPI_ISL_592373, EPI_ISL_592374, EPI_ISL_592375, EPI_ISL_592377, EPI_ISL_592379, EPI_ISL_592381, EPI_ISL_592382, EPI_ISL_592384, EPI_ISL_592386, EPI_ISL_592387, EPI_ISL_592388, EPI_ISL_592389, EPI_ISL_592390, EPI_ISL_592391, EPI_ISL_592395, EPI_ISL_592397, EPI_ISL_592399, EPI_ISL_592400, EPI_ISL_592401, EPI_ISL_592402, EPI_ISL_592403, EPI_ISL_592408, EPI_ISL_592409, EPI_ISL_592410, EPI_ISL_592412, EPI_ISL_592413, EPI_ISL_592417, EPI_ISL_592418, EPI_ISL_592419, EPI_ISL_592420, EPI_ISL_592421, EPI_ISL_592422, EPI_ISL_592425, EPI_ISL_592427, EPI_ISL_592428, EPI_ISL_592429, EPI_ISL_592430, EPI_ISL_592431, EPI_ISL_592432, EPI_ISL_592433, EPI_ISL_592434, EPI_ISL_592436, EPI_ISL_592437, EPI_ISL_592438, EPI_ISL_592439, EPI_ISL_592440, EPI_ISL_592441, EPI_ISL_592442, EPI_ISL_592443, EPI_ISL_592444, EPI_ISL_592445, EPI_ISL_592447, EPI_ISL_592448, EPI_ISL_592450, EPI_ISL_592453, EPI_ISL_592454, EPI_ISL_592457, EPI_ISL_592458, EPI_ISL_592459, EPI_ISL_592460, EPI_ISL_592462, EPI_ISL_592463, EPI_ISL_592464, EPI_ISL_592467, EPI_ISL_592468, EPI_ISL_592518, EPI_ISL_592522, EPI_ISL_592524, EPI_ISL_592527, EPI_ISL_592541, EPI_ISL_592545, EPI_ISL_592546, EPI_ISL_592549, EPI_ISL_592550, EPI_ISL_592560, EPI_ISL_592563, EPI_ISL_592565, EPI_ISL_592569, EPI_ISL_592574, EPI_ISL_592575, EPI_ISL_592576, EPI_ISL_592593, EPI_ISL_592594, EPI_ISL_592595, EPI_ISL_592596, EPI_ISL_592599, EPI_ISL_592602, EPI_ISL_592603, EPI_ISL_592605, EPI_ISL_592606, EPI_ISL_592607, EPI_ISL_592613, EPI_ISL_592615, EPI_ISL_592619, EPI_ISL_592621, EPI_ISL_592623, EPI_ISL_592626, EPI_ISL_592627, EPI_ISL_592630, EPI_ISL_592634, EPI_ISL_592635, EPI_ISL_592636, EPI_ISL_592637, EPI_ISL_592639, EPI_ISL_592641, EPI_ISL_592642, EPI_ISL_592643, EPI_ISL_592645, EPI_ISL_592652, EPI_ISL_592653, EPI_ISL_592654, EPI_ISL_592657, EPI_ISL_592658, EPI_ISL_592659, EPI_ISL_592669, EPI_ISL_592670, EPI_ISL_592672, EPI_ISL_592674, EPI_ISL_592675, EPI_ISL_592680, EPI_ISL_592684, EPI_ISL_592685, EPI_ISL_592689, EPI_ISL_592690, EPI_ISL_592691, EPI_ISL_592693, EPI_ISL_592694, EPI_ISL_592695, EPI_ISL_592696, EPI_ISL_592697, EPI_ISL_592699, EPI_ISL_592700, EPI_ISL_592702, EPI_ISL_592704, EPI_ISL_592705, EPI_ISL_592708, EPI_ISL_592709, EPI_ISL_592712, EPI_ISL_592713, EPI_ISL_592717, EPI_ISL_592723, EPI_ISL_592727, EPI_ISL_592729, EPI_ISL_592735, EPI_ISL_592738, EPI_ISL_592741, EPI_ISL_592751, EPI_ISL_592752, EPI_ISL_592755, EPI_ISL_592756, EPI_ISL_592758, EPI_ISL_592759, EPI_ISL_592760, EPI_ISL_592761, EPI_ISL_592764, EPI_ISL_592768, EPI_ISL_592780, EPI_ISL_592786, EPI_ISL_592787, EPI_ISL_592791, EPI_ISL_592792, EPI_ISL_592793, EPI_ISL_592797, EPI_ISL_592808, EPI_ISL_592809, EPI_ISL_592812, EPI_ISL_592818, EPI_ISL_592832, EPI_ISL_592835, EPI_ISL_592838, EPI_ISL_592839, EPI_ISL_592840, EPI_ISL_592853, EPI_ISL_592854, EPI_ISL_592855, EPI_ISL_592857, EPI_ISL_592867, EPI_ISL_592869, EPI_ISL_592870, EPI_ISL_592874, EPI_ISL_592877, EPI_ISL_592882, EPI_ISL_592885, EPI_ISL_592886, EPI_ISL_592887, EPI_ISL_592889, EPI_ISL_592890, EPI_ISL_592891, EPI_ISL_592892, EPI_ISL_592894, EPI_ISL_592895, EPI_ISL_592896, EPI_ISL_592898, EPI_ISL_592899, EPI_ISL_592900, EPI_ISL_592902, EPI_ISL_592908, EPI_ISL_592909, EPI_ISL_592910, EPI_ISL_592911, EPI_ISL_592912, EPI_ISL_592913, EPI_ISL_592916, EPI_ISL_592918, EPI_ISL_592923, EPI_ISL_592927, EPI_ISL_592928, EPI_ISL_592934, EPI_ISL_592937, EPI_ISL_592938, EPI_ISL_592939, EPI_ISL_592940, EPI_ISL_592944, EPI_ISL_592945, EPI_ISL_592946, EPI_ISL_592949, EPI_ISL_592950, EPI_ISL_592955, EPI_ISL_592959, EPI_ISL_592961, EPI_ISL_592962, EPI_ISL_592964, EPI_ISL_592967, EPI_ISL_593015, EPI_ISL_593029, EPI_ISL_593031, EPI_ISL_593043, EPI_ISL_593046, EPI_ISL_593132, EPI_ISL_593136, EPI_ISL_593139, EPI_ISL_593140, EPI_ISL_593141, EPI_ISL_593144, EPI_ISL_593147, EPI_ISL_593152, EPI_ISL_593154, EPI_ISL_593156, EPI_ISL_593162, EPI_ISL_593167, EPI_ISL_593181, EPI_ISL_593187, EPI_ISL_593188, EPI_ISL_593189, EPI_ISL_593191, EPI_ISL_593192, EPI_ISL_593193, EPI_ISL_593194, EPI_ISL_593195, EPI_ISL_593196, EPI_ISL_593197, EPI_ISL_593198, EPI_ISL_593199, EPI_ISL_593200, EPI_ISL_593201, EPI_ISL_593202, EPI_ISL_593203, EPI_ISL_593204, EPI_ISL_593205, EPI_ISL_593206, EPI_ISL_593207, EPI_ISL_593208, EPI_ISL_593209, EPI_ISL_593210, EPI_ISL_593211, EPI_ISL_593212, EPI_ISL_593214, EPI_ISL_593225, EPI_ISL_593228, EPI_ISL_593232, EPI_ISL_593233, EPI_ISL_593235, EPI_ISL_593236, EPI_ISL_593240, EPI_ISL_593243, EPI_ISL_593248, EPI_ISL_593249, EPI_ISL_593250, EPI_ISL_593251, EPI_ISL_593252, EPI_ISL_593253, EPI_ISL_593254, EPI_ISL_593255, EPI_ISL_593259, EPI_ISL_593261, EPI_ISL_593262, EPI_ISL_593263, EPI_ISL_593264, EPI_ISL_593267, EPI_ISL_593268, EPI_ISL_593269, EPI_ISL_593270, EPI_ISL_593271, EPI_ISL_593272, EPI_ISL_593273, EPI_ISL_593274, EPI_ISL_593275, EPI_ISL_593277, EPI_ISL_593279, EPI_ISL_593280, EPI_ISL_593281, EPI_ISL_593282, EPI_ISL_593283, EPI_ISL_593286, EPI_ISL_593288, EPI_ISL_593290, EPI_ISL_593291, EPI_ISL_593296, EPI_ISL_593300, EPI_ISL_593301, EPI_ISL_593302, EPI_ISL_593303, EPI_ISL_593304, EPI_ISL_593306, EPI_ISL_593307, EPI_ISL_593308, EPI_ISL_593311, EPI_ISL_593312, EPI_ISL_593314, EPI_ISL_593316, EPI_ISL_593318, EPI_ISL_593319, EPI_ISL_593322, EPI_ISL_593323, EPI_ISL_593324, EPI_ISL_593325, EPI_ISL_593326, EPI_ISL_593327, EPI_ISL_593330, EPI_ISL_593331, EPI_ISL_593334, EPI_ISL_593335, EPI_ISL_593336, EPI_ISL_593337, EPI_ISL_593338, EPI_ISL_593339, EPI_ISL_593340, EPI_ISL_593341, EPI_ISL_593342, EPI_ISL_593343, EPI_ISL_593349, EPI_ISL_593350, EPI_ISL_593351, EPI_ISL_593352, EPI_ISL_593353, EPI_ISL_593354, EPI_ISL_593355, EPI_ISL_593356, EPI_ISL_593357, EPI_ISL_593358, EPI_ISL_593359, EPI_ISL_593371, EPI_ISL_593372, EPI_ISL_593373, EPI_ISL_593375, EPI_ISL_593376, EPI_ISL_593377, EPI_ISL_593381, EPI_ISL_593382, EPI_ISL_593386, EPI_ISL_593389, EPI_ISL_593390, EPI_ISL_593391, EPI_ISL_593392, EPI_ISL_593395, EPI_ISL_593396, EPI_ISL_593397, EPI_ISL_593398, EPI_ISL_593399, EPI_ISL_593400, EPI_ISL_593403, EPI_ISL_593404, EPI_ISL_593405, EPI_ISL_593407, EPI_ISL_593409, EPI_ISL_593414, EPI_ISL_593415, EPI_ISL_593416, EPI_ISL_593418, EPI_ISL_593419, EPI_ISL_593420, EPI_ISL_593421, EPI_ISL_593428, EPI_ISL_593429, EPI_ISL_593431, EPI_ISL_593432, EPI_ISL_593434, EPI_ISL_593435, EPI_ISL_593437, EPI_ISL_593439, EPI_ISL_593440, EPI_ISL_593441, EPI_ISL_593443, EPI_ISL_593444, EPI_ISL_593445, EPI_ISL_593446, EPI_ISL_593447, EPI_ISL_593448, EPI_ISL_593450, EPI_ISL_593454, EPI_ISL_593455, EPI_ISL_593459, EPI_ISL_593460, EPI_ISL_593461, EPI_ISL_593462, EPI_ISL_593465, EPI_ISL_593466, EPI_ISL_593467, EPI_ISL_593468, EPI_ISL_593469, EPI_ISL_593470, EPI_ISL_593471, EPI_ISL_593472, EPI_ISL_593473 | | | |
| see above | Microbiological Diagnostic Unit - Public Health Laboratory (MDU-PHL) | MDU-PHL | Seemann T., Schultz, M. B., Sait, M., Sherry, N. |
| EPI_ISL_593653 | Laverty Pathology | NSW Health Pathology - Institute of Clinical Pathology and Medical Research; Westmead Hospital; University of Sydney | CIDM-PH et al. |
| EPI_ISL_593656, EPI_ISL_593657 | Pathology North - Hunter - NSW Health Pathology | NSW Health Pathology - Institute of Clinical Pathology and Medical Research; Westmead Hospital; University of Sydney | CIDM-PH et al. |
| EPI_ISL_593659 | Pathology North - Royal North Shore Hospital - NSW Health Pathology | NSW Health Pathology - Institute of Clinical Pathology and Medical Research; Westmead Hospital; University of Sydney | CIDM-PH et al. |
| EPI_ISL_593673, EPI_ISL_593674, EPI_ISL_593675, EPI_ISL_593676, EPI_ISL_593677 | Pathology West - NSW Health Pathology | NSW Health Pathology - Institute of Clinical Pathology and Medical Research; Westmead Hospital; University of Sydney | CIDM-PH et al. |
| EPI_ISL_593708, EPI_ISL_593710, EPI_ISL_593712 | South Eastern Area Laboratory Services (SEALS) | NSW Health Pathology - Institute of Clinical Pathology and Medical Research; Westmead Hospital; University of Sydney | CIDM-PH et al. |
| EPI_ISL_593761 | Sydney South West Pathology Service (SSWPS) - Liverpool Hospital - NSW Health Pathology | NSW Health Pathology - Institute of Clinical Pathology and Medical Research; Westmead Hospital; University of Sydney | CIDM-PH et al. |
| EPI_ISL_593766 | Sydney South West Pathology Service (SSWPS) - Royal Prince Alfred Hospital - NSW Health Pathology | NSW Health Pathology - Institute of Clinical Pathology and Medical Research; Westmead Hospital; University of Sydney | CIDM-PH et al. |
| EPI_ISL_593769 | The Children's Hospital at Westmead | NSW Health Pathology - Institute of Clinical Pathology and Medical Research; Westmead Hospital; University of Sydney | CIDM-PH et al. |
| EPI_ISL_593878, EPI_ISL_593879, EPI_ISL_593880, EPI_ISL_593884, EPI_ISL_593885, EPI_ISL_593886 | CHU Purpan - Laboratoire de Virologie - Institut Fédératif de Biologie | CHU Purpan - Laboratoire de Virologie - Institut Fédératif de Biologie | Latour J., Ranger N., Dubois M., Carcenac R., Harter A., Boyer P., Tremeaux P., Izopet J. |
| EPI_ISL_594134, EPI_ISL_594137, EPI_ISL_594147 | MDU-PHL, The Peter Doherty Institute for Infection and Immunity | MDU-PHL, The Peter Doherty Institute for Infection and Immunity | Caly,L., Seemann,T., Sait,M.L., Schultz,M.B., Druce,J., Sherry,N.L. |
| EPI_ISL_594366, EPI_ISL_594367, EPI_ISL_594370 | Florida Bureau of Public Health Laboratories | Florida Bureau of Public Health Laboratories | Sarah Schmedes, Jason Blanton |
| EPI_ISL_596267, EPI_ISL_596353, EPI_ISL_596354, EPI_ISL_596355, EPI_ISL_596356 | WHO National Influenza Centre Russian Federation | WHO National Influenza Centre Russian Federation | Andrey Komissarov, Artem Fadeev, Anna Ivanova, Kseniya Komissarova, Dmitry Bazhenov, Daria Danilenko |
| EPI_ISL_596476 | National Public Health Laboratory, National Centre for Infectious Diseases | National Public Health Laboratory, National Centre for Infectious Diseases | Tze Minn Mak, Sophie Octavia, Zhenyang Zhou, Lin Cui, Raymond Tzer Pin Lin |
| EPI_ISL_602307, EPI_ISL_602308, EPI_ISL_602310, EPI_ISL_602312, EPI_ISL_602315, EPI_ISL_602316, EPI_ISL_602317, EPI_ISL_602319, EPI_ISL_602322, EPI_ISL_602325, EPI_ISL_602326, EPI_ISL_602327, EPI_ISL_602328 | | | |
| see above | University of Miami Immunology and Histocompatibility Laboratory | University of Miami Immunology and Histocompatibility Laboratory | Emilio Margolles-Clark, PhD and Phillip Ruiz, MD, PhD |
| EPI_ISL_602617, EPI_ISL_602618 | Queen Mary Hospital | Hong Kong Department of Health | Mak Gannon C.K., Lam Edman T.K., Chan Rickjason C.W., Tsang Dominic N.C. |
| EPI_ISL_602619 | United Christian Hospital | Hong Kong Department of Health | Mak Gannon C.K., Lam Edman T.K., Chan Rickjason C.W., Tsang Dominic N.C. |
| EPI_ISL_602620 | Queen Elizabeth Hospital | Hong Kong Department of Health | Mak Gannon C.K., Lam Edman T.K., Chan Rickjason C.W., Tsang Dominic N.C. |

| | | | |
|---|---|---|---|
| EPI_ISL_602621 | GLENEAGLES HONG KONG (CLINICAL LAB) | Hong Kong Department of Health | Mak Gannon C.K., Lam Edman T.K., Chan Rickjason C.W., Tsang Dominic N.C. |
| EPI_ISL_602956 | Minnesota Department of Health, Public Health Laboratory | Minnesota Department of Health, Public Health Laboratory | Matt Plumb, Jacob Garfin, Alexandra Lorentz, and Xiong Wang |
| EPI_ISL_603044, EPI_ISL_603045, EPI_ISL_603047 | MDU-PHL, The Peter Doherty Institute for Infection and Immunity | MDU-PHL, The Peter Doherty Institute for Infection and Immunity | Seemann,T., Caly,L., Sait,M., Schultz,M.B., Druce,J., Sherry,N. |
| EPI_ISL_603251, EPI_ISL_603253, EPI_ISL_603258, EPI_ISL_603261 | UW Virology Lab | UW Virology Lab | Pavitra Roychoudhury, Hong Xie, Lasata Shrestha, Meei-Li Huang, Keith R Jerome, Alexander Greninger |
| EPI_ISL_605147 | University of Miami Immunology and Histocompatibility Laboratory | University of Miami Immunology and Histocompatibility Laboratory | Emilio Margolles-Clark, PhD and Phillip Ruiz, MD, PhD |
| EPI_ISL_605783 | Genome Center | Genome Center | Md. Shazid Hasan, Hassan M. Al-Emran, Ovinu Kibria Islam, A. S. M. Rubayet- Ul- Alam, Selina Akter, Shireen Nigar, Md. Tanvir Islam, Pravas Chandra Roy, Shovon Lal Sarkar, Najmuj Sakib, S. M. Tanjil Shah, Md. Iqbal Kabir Jahid, Md. Anwar Hossain |
| EPI_ISL_610118, EPI_ISL_610119, EPI_ISL_610120, EPI_ISL_610121, EPI_ISL_610122, EPI_ISL_610123, EPI_ISL_610124, EPI_ISL_610125 | Virginia DCLS | Virginia DCLS | Virginia DCLS |
| EPI_ISL_612455 | University College London, Great Ormond Street Hospital for Children NHS Foundation Trust, Imperial College Healthcare NHS Trust | COVID-19 Genomics UK (COG-UK) Consortium | Sergi Castellano, Rachel Williams, Mark Kristiansen, Paola Resende Silva, Sunando Roy, Tony Brooks, Helena Tutill, Paola Niola, Patricia Dyal, Charlotte Williams, Leysa Forrest, Yasmin Panchbhaya, Jacqueline Findlay, Samuel Weeks, Julianne Brown, Kathryn Harris, Paul Randell, James Price, Alison Holmes, Judith Breuer |
| EPI_ISL_613728, EPI_ISL_613743, EPI_ISL_613753, EPI_ISL_613762, EPI_ISL_613812, EPI_ISL_613813, EPI_ISL_613827, EPI_ISL_613899, EPI_ISL_613900, EPI_ISL_613901, EPI_ISL_613902, EPI_ISL_613903 | | | |
| see above | Florida Bureau of Public Health Laboratories | Florida Bureau of Public Health Laboratories | Sarah Schmedes, Jason Blanton |
| EPI_ISL_614265 | Eurofins | National Reference Center for Viruses of Respiratory Infections, Institut Pasteur, Paris | Marion Barbet, Sylvie Behillil, Méline Bizard, Angela Brisebarre, Camille Capel, Etienne Simon-Lorière, Vincent Enouf, Maud Vanpeene, Sylvie van der Werf |
| EPI_ISL_614308, EPI_ISL_614309, EPI_ISL_614310, EPI_ISL_614311, EPI_ISL_614312, EPI_ISL_614313, EPI_ISL_614314, EPI_ISL_614315 | Faroese National Reference Laboratory for Fish and Animal Diseases | Faroese National Reference Laboratory for Fish and Animal Diseases | Maria Marjunardóttir Dahl, Petra Elisabeth Petersen, Debes Hammershaimb Christiansen |
| EPI_ISL_614385 | Molecular diagnostic unit for viral haemorrhagic fevers and emerging viruses, Bouaké CHU Laboratory | Project group Epidemology of Highly Pathogenic Microorganisms, Robert Koch-Institute | Chantal Akoua-Koffi, Diané Bamourou, Etilé Anoh, Essia Belarbi, Safiatou Karidioula, Grit Schubert, Adjaratou Traoré, Soundélé Maïté, Monemo Pacome, Coulibaly Mbegnan, Bamba Fatoumata Touré, Kra Ouffoué, Fabian Leendertz |
| EPI_ISL_615911, EPI_ISL_617406, EPI_ISL_617407, EPI_ISL_617408, EPI_ISL_617409, EPI_ISL_617410, EPI_ISL_617411, EPI_ISL_617412, EPI_ISL_617413, EPI_ISL_617414, EPI_ISL_617415, EPI_ISL_617416, EPI_ISL_617417, EPI_ISL_617418, EPI_ISL_617419, EPI_ISL_617420, EPI_ISL_617421, EPI_ISL_617422, EPI_ISL_617423, EPI_ISL_617424, EPI_ISL_617425, EPI_ISL_617426, EPI_ISL_617430, EPI_ISL_617431, EPI_ISL_617432, EPI_ISL_617433, EPI_ISL_617434, EPI_ISL_617435, EPI_ISL_617436, EPI_ISL_617437, EPI_ISL_617438, EPI_ISL_617439, EPI_ISL_617440, EPI_ISL_617441, EPI_ISL_617442, EPI_ISL_617443, EPI_ISL_617444, EPI_ISL_617445, EPI_ISL_617446, EPI_ISL_617447, EPI_ISL_617448, EPI_ISL_617449, EPI_ISL_617450, EPI_ISL_617451, EPI_ISL_617452, EPI_ISL_617453, EPI_ISL_617454, EPI_ISL_617455, EPI_ISL_617456, EPI_ISL_617457, EPI_ISL_617458, EPI_ISL_617459, EPI_ISL_617460, EPI_ISL_617461, EPI_ISL_617462, EPI_ISL_617463, EPI_ISL_617464, EPI_ISL_617465, EPI_ISL_617466, EPI_ISL_617467, EPI_ISL_617468, EPI_ISL_617469, EPI_ISL_617470, EPI_ISL_617471, EPI_ISL_617472, EPI_ISL_617473, EPI_ISL_617474, EPI_ISL_617475, EPI_ISL_617476, EPI_ISL_617477, EPI_ISL_617478, EPI_ISL_617479, EPI_ISL_617480, EPI_ISL_617481, EPI_ISL_617482, EPI_ISL_617483, EPI_ISL_617484, EPI_ISL_617485, EPI_ISL_617486, EPI_ISL_617487, EPI_ISL_617488, EPI_ISL_617489, EPI_ISL_617490, EPI_ISL_617491, EPI_ISL_617492, EPI_ISL_617493, EPI_ISL_617494, EPI_ISL_617495, EPI_ISL_617496, EPI_ISL_617497, EPI_ISL_617498, EPI_ISL_617499, EPI_ISL_617500, EPI_ISL_617501, EPI_ISL_617502, EPI_ISL_617503, EPI_ISL_617504, EPI_ISL_617505, EPI_ISL_617506, EPI_ISL_617507, EPI_ISL_617508, EPI_ISL_617509, EPI_ISL_617510, EPI_ISL_617511, EPI_ISL_617512, EPI_ISL_617513, EPI_ISL_617514, EPI_ISL_617515, EPI_ISL_617516, EPI_ISL_617517, EPI_ISL_617518, EPI_ISL_617519, EPI_ISL_617520, EPI_ISL_617521, EPI_ISL_617522, EPI_ISL_617523, EPI_ISL_617524, EPI_ISL_617525, EPI_ISL_617526, EPI_ISL_617527, EPI_ISL_617528, EPI_ISL_617529, EPI_ISL_617530, EPI_ISL_617531, EPI_ISL_617532, EPI_ISL_617533, EPI_ISL_617534, EPI_ISL_617535, EPI_ISL_617536, EPI_ISL_617537, EPI_ISL_617538, EPI_ISL_617539, EPI_ISL_617540, EPI_ISL_617541, EPI_ISL_617542, EPI_ISL_617543, EPI_ISL_617544, EPI_ISL_617545, EPI_ISL_617546, EPI_ISL_617547, EPI_ISL_617548, EPI_ISL_617549, EPI_ISL_617550, EPI_ISL_617551, EPI_ISL_617552, EPI_ISL_617553, EPI_ISL_617554, EPI_ISL_617555, EPI_ISL_617556, EPI_ISL_617557, EPI_ISL_617558, EPI_ISL_617559, EPI_ISL_617560, EPI_ISL_617561, EPI_ISL_617562, EPI_ISL_617563, EPI_ISL_617564, EPI_ISL_617565, EPI_ISL_617566, EPI_ISL_617567, EPI_ISL_617568, EPI_ISL_617569, EPI_ISL_617570, EPI_ISL_617571, EPI_ISL_617572, EPI_ISL_617573, EPI_ISL_617574, EPI_ISL_617575, EPI_ISL_617576, EPI_ISL_617577, EPI_ISL_617578, EPI_ISL_617579, EPI_ISL_617580, EPI_ISL_617581, EPI_ISL_617582, EPI_ISL_617583, EPI_ISL_617584, EPI_ISL_617585, EPI_ISL_617586, EPI_ISL_617587, EPI_ISL_617588, EPI_ISL_617589, EPI_ISL_617590, EPI_ISL_617591, EPI_ISL_617592, EPI_ISL_617593, EPI_ISL_617594, EPI_ISL_617595, EPI_ISL_617596, EPI_ISL_617597, EPI_ISL_617598, EPI_ISL_617599, EPI_ISL_617600, EPI_ISL_617601, EPI_ISL_617602, EPI_ISL_617603, EPI_ISL_617604, EPI_ISL_617605, EPI_ISL_617606, EPI_ISL_617607, EPI_ISL_617608, EPI_ISL_617609, EPI_ISL_617610, EPI_ISL_617611, EPI_ISL_617612, EPI_ISL_617613, EPI_ISL_617614, EPI_ISL_617615, EPI_ISL_617616, EPI_ISL_617617, EPI_ISL_617618, EPI_ISL_617619, EPI_ISL_617620, EPI_ISL_617621, EPI_ISL_617622, EPI_ISL_617623, EPI_ISL_617624, EPI_ISL_617625, EPI_ISL_617626, EPI_ISL_617627, EPI_ISL_617628, EPI_ISL_617629, EPI_ISL_617630, EPI_ISL_617631, EPI_ISL_617632, EPI_ISL_617633, EPI_ISL_617634, EPI_ISL_617635, EPI_ISL_617636, EPI_ISL_617637, EPI_ISL_617638, EPI_ISL_617639, EPI_ISL_617640, EPI_ISL_617641, EPI_ISL_617642, EPI_ISL_617643, EPI_ISL_617644, EPI_ISL_617645, EPI_ISL_617646, EPI_ISL_617647, EPI_ISL_617648, EPI_ISL_617649, EPI_ISL_617650, EPI_ISL_617651, EPI_ISL_617652, EPI_ISL_617653, EPI_ISL_617654, EPI_ISL_617655, EPI_ISL_617656, EPI_ISL_617657, EPI_ISL_617658, EPI_ISL_617659, EPI_ISL_617660, EPI_ISL_617661, EPI_ISL_617662, EPI_ISL_617663, EPI_ISL_617664, EPI_ISL_617665, EPI_ISL_617666, EPI_ISL_617667, EPI_ISL_617668, EPI_ISL_617669, EPI_ISL_617670, EPI_ISL_617671, EPI_ISL_617672, EPI_ISL_617673, EPI_ISL_617674, EPI_ISL_617675, EPI_ISL_617676, EPI_ISL_617677, EPI_ISL_617678, EPI_ISL_617679, EPI_ISL_617680, EPI_ISL_617681, EPI_ISL_617682, EPI_ISL_617683, EPI_ISL_617684, EPI_ISL_617685, EPI_ISL_617686, EPI_ISL_617687, EPI_ISL_617688, EPI_ISL_617689, EPI_ISL_617690, EPI_ISL_617691, EPI_ISL_617692, EPI_ISL_617693, EPI_ISL_617694, EPI_ISL_617695, EPI_ISL_617696, EPI_ISL_617697, EPI_ISL_617698, EPI_ISL_617699, EPI_ISL_617700, EPI_ISL_617701, EPI_ISL_617702, EPI_ISL_617703, EPI_ISL_617704, EPI_ISL_617705, EPI_ISL_617706, EPI_ISL_617707, EPI_ISL_617708, EPI_ISL_617709, EPI_ISL_617710, EPI_ISL_617711, EPI_ISL_617712, EPI_ISL_617713, EPI_ISL_617714, EPI_ISL_617715, EPI_ISL_617716, EPI_ISL_617717, EPI_ISL_617718, EPI_ISL_617719, EPI_ISL_617720, EPI_ISL_617721, EPI_ISL_617722, EPI_ISL_617723, EPI_ISL_617724, EPI_ISL_617725, EPI_ISL_617726, EPI_ISL_617727, EPI_ISL_617728, EPI_ISL_617729, EPI_ISL_617730, EPI_ISL_617731, EPI_ISL_617732, EPI_ISL_617733, EPI_ISL_617734, EPI_ISL_617735, EPI_ISL_617736, EPI_ISL_617737, EPI_ISL_617738, EPI_ISL_617739, EPI_ISL_617740, EPI_ISL_617741, EPI_ISL_617742, EPI_ISL_617743, EPI_ISL_617744, EPI_ISL_617745, EPI_ISL_617746, EPI_ISL_617747, EPI_ISL_617748, EPI_ISL_617749, EPI_ISL_617750, EPI_ISL_617751, EPI_ISL_617752, EPI_ISL_617753, EPI_ISL_617754, EPI_ISL_617756, EPI_ISL_617757, EPI_ISL_617758, EPI_ISL_617759, EPI_ISL_617760, EPI_ISL_617761, EPI_ISL_617762, EPI_ISL_617763, EPI_ISL_617764, EPI_ISL_617765, EPI_ISL_617766, EPI_ISL_617767, EPI_ISL_617768, EPI_ISL_617769, EPI_ISL_617770, EPI_ISL_617771, EPI_ISL_617772, EPI_ISL_617773, EPI_ISL_617774, EPI_ISL_617775, EPI_ISL_617776, EPI_ISL_617777, EPI_ISL_617778, EPI_ISL_617779, EPI_ISL_617780, EPI_ISL_617781, EPI_ISL_617782, EPI_ISL_617783, EPI_ISL_617784, EPI_ISL_617785, EPI_ISL_617786, EPI_ISL_617787, EPI_ISL_617788, EPI_ISL_617789, EPI_ISL_617792, EPI_ISL_617793, EPI_ISL_617797, EPI_ISL_617798, EPI_ISL_617810, EPI_ISL_617811, EPI_ISL_617812, EPI_ISL_617813, EPI_ISL_618339, EPI_ISL_618340, EPI_ISL_618343, EPI_ISL_618344, EPI_ISL_618346, EPI_ISL_618347, EPI_ISL_618363, EPI_ISL_618364, EPI_ISL_618365, EPI_ISL_618366, EPI_ISL_622666, EPI_ISL_622668, EPI_ISL_622669, EPI_ISL_622681, EPI_ISL_622729, EPI_ISL_622730, EPI_ISL_622731 | | | |
| see above | Department of Virus and Microbiological Special Diagnostics, Statens Serum Institut, Denmark | Albertsen lab, Department of Chemistry and Bioscience, Aalborg University, Denmark | Danish Covid-19 Genome Consortia |
| EPI_ISL_622998, EPI_ISL_623004, EPI_ISL_623011, EPI_ISL_623029, EPI_ISL_623039, EPI_ISL_623054 | National Health Laboratory Service | National Institute for Communicable Diseases of the National Health Laboratory Service | Allam M, Ismail A, Khumalo Z, Kwenda S, Mtshali P, Mnyameni F, Mohale T, Subramoney K, Bhiman JN |
| EPI_ISL_625472 | Child Health Research Foundation | Child Health Research Foundation | Senjuti Saha, Md Saiful Islam Sajib, Nikkon Sarkar, Syed Muktadir Al Sium, Afroza Akter Tanni, Roly Malaker, Arif Mohammad Tanmoy, Md Hafizur Rahman, Samir K Saha |
| EPI_ISL_625579, EPI_ISL_625580, EPI_ISL_625581, EPI_ISL_625582, EPI_ISL_625583, EPI_ISL_625584, EPI_ISL_625585, EPI_ISL_625586, EPI_ISL_625587, EPI_ISL_625588, EPI_ISL_625589, EPI_ISL_625590, EPI_ISL_625591, EPI_ISL_625592, EPI_ISL_625593 | | | |
| see above | NaN | Chan-Zuckerberg Biohub | CZB Cliahub Consortium |
| EPI_ISL_625681, EPI_ISL_625682 | Laboratory of Molecular Medicine, University of Magallanes | Centro Asistencial Docente y de Investigacion, Universidad de Magallanes | Jorge Gonzalez, Jacqueline Aldridge, Diego Alvarez, Marcelo Navarrete |
| EPI_ISL_626556, EPI_ISL_626557, EPI_ISL_626558, EPI_ISL_626559, EPI_ISL_626560, EPI_ISL_626561, EPI_ISL_626562, EPI_ISL_626563 | Laboratorio de Biología Molecular, Facultad de Medicina, Universidad de Atacama, Copiapo, Chile/ FONDAP CRG, Universidad Andrés Bello, Santiago, Chile | Center for Mathematical Modeling and Center for Genome Regulation. Santiago, Chile | Echeverría C, Manríquez R, Bastias M, Sanhueza D, Travisany D, Allende ML, Maass A, González M, Montecino, M, Orellana A, Castro E, Meneses C. |
| EPI_ISL_627465, EPI_ISL_627471 | University College London, Great Ormond Street Hospital for Children NHS Foundation Trust, Imperial College Healthcare NHS Trust | COVID-19 Genomics UK (COG-UK) Consortium | Sergi Castellano, Rachel Williams, Mark Kristiansen, Paola Resende Silva, Sunando Roy, Tony Brooks, Helena Tutill, Paola Niola, Patricia Dyal, Charlotte Williams, Leysa Forrest, Yasmin Panchbhaya, Jacqueline Findlay, Samuel Weeks, Julianne Brown, Kathryn Harris, Paul Randell, James Price, Alison Holmes, Judith Breuer |
| EPI_ISL_631490 | Wisconsin State Laboratory of Hygiene Communicable Disease Division | Wisconsin State Laboratory of Hygiene Communicable Disease Division | Kelsey R. Florek, Abigail C. Shockey |
| EPI_ISL_632543 | Dutch COVID-19 response team | Erasmus Medical Center | Bas Oude Munnink, David Nieuwenhuijse, Reina Sikkema, Claudia Schapendonk, Irina Chestakova, Anne van der Linden, Theo Bestebroer, Stefan van Nieuwkoop, Mark Pronk, Pascal Lexmond, Corien Swaan, Manon Haverkate, Madelief Mollers, Mart Stein, Sandra Kengne Kamga Mobou, Jeroen van Kampen, Jolanda Voermans, Aura Timen, Corine GeurtsvanKessel, Annemiek van der Eijk, Richard Molenkamp, Marion Koopmans, on behalf of the Dutch national COVID-19 response team. |
| EPI_ISL_635113, EPI_ISL_635114 | University Hospital of Northern Norway, Department for | Norwegian Institute of Public Health, Department of Virology | Kathrine Stene-Johansen, Kamilla Heddeland Instefjord, Hilde Elshaug, Marie Paulsen Madsen, Rasmus Riis Kopperud, Hilde Vollan, Karoline Bragstad, Olav |

| | | | |
|---|---|---|---|
| | Microbiology and Infectious Disease Control | | Hungnes |
| EPI_ISL_635146 | Dept. of Medical Microbiology, Stavanger University Hospital, Helse Stavanger HF | Norwegian Institute of Public Health, Department of Virology | Kathrine Stene-Johansen, Kamilla Heddeland Instefjord, Hilde Elshaug, Marie Paulsen Madsen, Rasmus Riis Kopperud, Hilde Vollan, Karoline Bragstad, Olav Hungnes |
| EPI_ISL_635148 | University Hospital of Northern Norway, Department for Microbiology and Infectious Disease Control | Norwegian Institute of Public Health, Department of Virology | Kathrine Stene-Johansen, Kamilla Heddeland Instefjord, Hilde Elshaug, Marie Paulsen Madsen, Rasmus Riis Kopperud, Hilde Vollan, Karoline Bragstad, Olav Hungnes |
| EPI_ISL_635701, EPI_ISL_635705, EPI_ISL_635707, EPI_ISL_635708, EPI_ISL_635709, EPI_ISL_635714, EPI_ISL_635716, EPI_ISL_635718, EPI_ISL_635719, EPI_ISL_635720, EPI_ISL_635721, EPI_ISL_635731, EPI_ISL_635733, EPI_ISL_635735, EPI_ISL_635789, EPI_ISL_635790, EPI_ISL_635820, EPI_ISL_635821, EPI_ISL_635824, EPI_ISL_635828, EPI_ISL_635831, EPI_ISL_635916, EPI_ISL_636002, EPI_ISL_636040, EPI_ISL_636049, EPI_ISL_636053, EPI_ISL_636055 | | | |
| see above | San Diego County Public Health Laboratory | Andersen lab at Scripps Research | SEARCH Alliance San Diego with Tracy Basler, Jovan Shephard, Brett Austin |
| EPI_ISL_636514, EPI_ISL_636515, EPI_ISL_636516, EPI_ISL_636517 | Dutch COVID-19 response team | National Institute for Public Health and the Environment (RIVM) | Adam Meijer, Harry Vennema, Jeroen Cremer, Sharon van den Brink, Bas van der Veer, AnneMarie van den Brandt, Florian Zwagemaker, Dennis Schmitz, Chantal Reusken, on behalf of the national COVID-19 response team |
| EPI_ISL_636738, EPI_ISL_636740, EPI_ISL_636742, EPI_ISL_636743, EPI_ISL_636744, EPI_ISL_636745, EPI_ISL_636746, EPI_ISL_636747, EPI_ISL_636750, EPI_ISL_636752, EPI_ISL_636753, EPI_ISL_636756, EPI_ISL_636758, EPI_ISL_636763, EPI_ISL_636764, EPI_ISL_636765, EPI_ISL_636766, EPI_ISL_636767, EPI_ISL_636769, EPI_ISL_636772, EPI_ISL_636773, EPI_ISL_636774, EPI_ISL_636775, EPI_ISL_636776, EPI_ISL_636778, EPI_ISL_636779, EPI_ISL_636780, EPI_ISL_636782, EPI_ISL_636783, EPI_ISL_636784, EPI_ISL_636785, EPI_ISL_636786, EPI_ISL_636787, EPI_ISL_636788, EPI_ISL_636789, EPI_ISL_636790, EPI_ISL_636791, EPI_ISL_636793, EPI_ISL_636794, EPI_ISL_636795, EPI_ISL_636796, EPI_ISL_636797, EPI_ISL_636798, EPI_ISL_636799, EPI_ISL_636800, EPI_ISL_636801, EPI_ISL_636802, EPI_ISL_636803, EPI_ISL_636804, EPI_ISL_636805, EPI_ISL_636806, EPI_ISL_636807, EPI_ISL_636808, EPI_ISL_636809, EPI_ISL_636810, EPI_ISL_636811, EPI_ISL_636812, EPI_ISL_636813, EPI_ISL_636814, EPI_ISL_636815, EPI_ISL_636816, EPI_ISL_636822, EPI_ISL_636823, EPI_ISL_636824, EPI_ISL_636825, EPI_ISL_636828, EPI_ISL_636829, EPI_ISL_636830, EPI_ISL_636831, EPI_ISL_636832, EPI_ISL_636833 | | | |
| see above | National Centre for Disease control (NCDC) | NCDC/CSIR-IGIB | Mahesh S. Dhar1*, Bharathram Uppili2*, Robin Marwal1*, Pooja Sharma2*, RadhaKrishnan VS, Vivekanand A, Nishu Tyagi, Shaista Khan, Simmi Tiwari, Manish Kumar, Ajit Shewale, Ishtaq Ahmed, Asangla Kamai, Aparna Swaminathan, Saruchi Wadhwa, Tushar Nale, Sandhya Kabra, Sujeet Singh, Mohammed Faruq#, Anurag Agrawal#, Partha Rakshit# |
| EPI_ISL_636845, EPI_ISL_636846, EPI_ISL_636847, EPI_ISL_636848, EPI_ISL_636849, EPI_ISL_636850, EPI_ISL_636851, EPI_ISL_636852, EPI_ISL_636853, EPI_ISL_636854, EPI_ISL_636855, EPI_ISL_636856, EPI_ISL_636857, EPI_ISL_636858 | | | |
| see above | Lithuanian University of Health Sciences Hospital, Department of Laboratory Medicine | Lithuanian University of Health Sciences, Molecular cardiology lab. | Lukas Zemaitis, Ingrida Olendrait, Arnoldas Pautienius, Kamile Tamusauskaite, Dovydas Gecys, Laura Pareckaite, Vaiva Lesauskaite, Astra Vitkauskiene |
| EPI_ISL_637005, EPI_ISL_637006, EPI_ISL_637007, EPI_ISL_637008, EPI_ISL_637009 | Department of Infectious Diseases and Immunology, National Hospital Organization Nagoya Medical Center | Clinical Research Center, National Hospital Organization Nagoya Medical Center | Yoshihiro Nakata, Hirotaka Ode, Mai Kubota, Masakazu Matsuda, Kazuhiro Matsuoka, Miho Nakasuji, Mikiko Mori, Mayumi Imahashi, Yoshiyuki Yokomaku, Yasumasa Iwatani |
| EPI_ISL_637022 | Tin Shui Wai Hospital | Hong Kong Department of Health | Mak Gannon C.K., Lam Edman T.K., Chan Rickjason C.W., Tsang Dominic N.C. |
| EPI_ISL_637023 | Queen Elizabeth Hospital | Hong Kong Department of Health | Mak Gannon C.K., Lam Edman T.K., Chan Rickjason C.W., Tsang Dominic N.C. |
| EPI_ISL_637024 | Pamela Youde Nethersole Eastern Hospital | Hong Kong Department of Health | Mak Gannon C.K., Lam Edman T.K., Chan Rickjason C.W., Tsang Dominic N.C. |
| EPI_ISL_637025 | Queen Elizabeth Hospital | Hong Kong Department of Health | Mak Gannon C.K., Lam Edman T.K., Chan Rickjason C.W., Tsang Dominic N.C. |
| EPI_ISL_637026 | Kwong Wah Hospital | Hong Kong Department of Health | Mak Gannon C.K., Lam Edman T.K., Chan Rickjason C.W., Tsang Dominic N.C. |
| EPI_ISL_637027 | Hong Kong Baptist Hospital | Hong Kong Department of Health | Mak Gannon C.K., Lam Edman T.K., Chan Rickjason C.W., Tsang Dominic N.C. |
| EPI_ISL_637029 | Caritas Medical Centre | Hong Kong Department of Health | Mak Gannon C.K., Lam Edman T.K., Chan Rickjason C.W., Tsang Dominic N.C. |
| EPI_ISL_639729, EPI_ISL_639730 | Sydney South West Pathology Service (SSWPS) - Liverpool Hospital - NSW Health Pathology | NSW Health Pathology - Institute of Clinical Pathology and Medical Research; Westmead Hospital; University of Sydney | CIDM-PH et al. |
| EPI_ISL_640016 | George Hospital wc GRH | NHLS/UCT | Arash Iranzadeh, Deelan Doolabh, Lynn Tyers, Bruna Galvao, Innocent Mudau, Marvin Hsiao, Kruger Marais, Diana Hardie, Stephen Korsman, Carolyn Williamson |
| EPI_ISL_640302, EPI_ISL_640304, EPI_ISL_640306, EPI_ISL_640307, EPI_ISL_640308, EPI_ISL_640309, EPI_ISL_640310, EPI_ISL_640311, EPI_ISL_640314, EPI_ISL_640320, EPI_ISL_640321, EPI_ISL_640323, EPI_ISL_640324, EPI_ISL_640325, EPI_ISL_640326, EPI_ISL_640332, EPI_ISL_640340, EPI_ISL_640341, EPI_ISL_640343, EPI_ISL_640345, EPI_ISL_640346, EPI_ISL_640348, EPI_ISL_640352, EPI_ISL_640355, EPI_ISL_640357, EPI_ISL_640359, EPI_ISL_640362, EPI_ISL_640363, EPI_ISL_640364, EPI_ISL_640368, EPI_ISL_640370, EPI_ISL_640371, EPI_ISL_640373, EPI_ISL_640374, EPI_ISL_640376, EPI_ISL_640378, EPI_ISL_640386, EPI_ISL_640387, EPI_ISL_640388, EPI_ISL_640389, EPI_ISL_640390, EPI_ISL_640393, EPI_ISL_640394, EPI_ISL_640399, EPI_ISL_640400, EPI_ISL_640401, EPI_ISL_640404, EPI_ISL_640405, EPI_ISL_640406, EPI_ISL_640409, EPI_ISL_640412, EPI_ISL_640413, EPI_ISL_640414, EPI_ISL_640415, EPI_ISL_640418, EPI_ISL_640420, EPI_ISL_640422, EPI_ISL_640424, EPI_ISL_640425, EPI_ISL_640426, EPI_ISL_640427, EPI_ISL_640428, EPI_ISL_640429, EPI_ISL_640430, EPI_ISL_640431, EPI_ISL_640432, EPI_ISL_640433, EPI_ISL_640434, EPI_ISL_640436, EPI_ISL_640438, EPI_ISL_640442, EPI_ISL_640443, EPI_ISL_640445, EPI_ISL_640447, EPI_ISL_640448, EPI_ISL_640455, EPI_ISL_640456, EPI_ISL_640457, EPI_ISL_640458, EPI_ISL_640459, EPI_ISL_640461, EPI_ISL_640464, EPI_ISL_640466, EPI_ISL_640467, EPI_ISL_640468, EPI_ISL_640470, EPI_ISL_640471, EPI_ISL_640472, EPI_ISL_640474, EPI_ISL_640475, EPI_ISL_640476, EPI_ISL_640477, EPI_ISL_640478, EPI_ISL_640482, EPI_ISL_640483, EPI_ISL_640484, EPI_ISL_640487, EPI_ISL_640488, EPI_ISL_640491, EPI_ISL_640493, EPI_ISL_640497, EPI_ISL_640498, EPI_ISL_640504, EPI_ISL_640506, EPI_ISL_640509, EPI_ISL_640510, EPI_ISL_640511, EPI_ISL_640513, EPI_ISL_640515, EPI_ISL_640516, EPI_ISL_640517, EPI_ISL_640518, EPI_ISL_640520, EPI_ISL_640521, EPI_ISL_640522, EPI_ISL_640526, EPI_ISL_640528, EPI_ISL_640531, EPI_ISL_640533, EPI_ISL_640535, EPI_ISL_640537, EPI_ISL_640539, EPI_ISL_640541, EPI_ISL_640542, EPI_ISL_640545, EPI_ISL_640551, EPI_ISL_640552, EPI_ISL_640556, EPI_ISL_640559, EPI_ISL_640563, EPI_ISL_640564, EPI_ISL_640565, EPI_ISL_640566, EPI_ISL_640568, EPI_ISL_640571, EPI_ISL_640572, EPI_ISL_640573, EPI_ISL_640576, EPI_ISL_640578, EPI_ISL_640584, EPI_ISL_640586, EPI_ISL_640587, EPI_ISL_640589, EPI_ISL_640591, EPI_ISL_640593, EPI_ISL_640599, EPI_ISL_640601, EPI_ISL_640602, EPI_ISL_640604, EPI_ISL_640606, EPI_ISL_640608, EPI_ISL_640612, EPI_ISL_640613, EPI_ISL_640616, EPI_ISL_640619, EPI_ISL_640621, EPI_ISL_640625, EPI_ISL_640626, EPI_ISL_640627, EPI_ISL_640628, EPI_ISL_640631, EPI_ISL_640632, EPI_ISL_640633, EPI_ISL_640634, EPI_ISL_640636, EPI_ISL_640637, EPI_ISL_640638, EPI_ISL_640642, EPI_ISL_640644, EPI_ISL_640645, EPI_ISL_640648, EPI_ISL_640649, EPI_ISL_640650, EPI_ISL_640652, EPI_ISL_640653, EPI_ISL_640654, EPI_ISL_640657, EPI_ISL_640658, EPI_ISL_640662, EPI_ISL_640670, EPI_ISL_640671, EPI_ISL_640672, EPI_ISL_640673, EPI_ISL_640674, EPI_ISL_640675, EPI_ISL_640676, EPI_ISL_640683, EPI_ISL_640684, EPI_ISL_640686, EPI_ISL_640690, EPI_ISL_640692, EPI_ISL_640693, EPI_ISL_640694, EPI_ISL_640696, EPI_ISL_640699, EPI_ISL_640701, EPI_ISL_640702, EPI_ISL_640703, EPI_ISL_640704, EPI_ISL_640705, EPI_ISL_640706, EPI_ISL_640709, EPI_ISL_640714, EPI_ISL_640715, EPI_ISL_640718, EPI_ISL_640719, EPI_ISL_640720, EPI_ISL_640721, EPI_ISL_640724, EPI_ISL_640727, EPI_ISL_640729, EPI_ISL_640734, EPI_ISL_640735, EPI_ISL_640736, EPI_ISL_640737, EPI_ISL_640738, EPI_ISL_640739, EPI_ISL_640740, EPI_ISL_640742, EPI_ISL_640745, EPI_ISL_640747, EPI_ISL_640748, EPI_ISL_640749, EPI_ISL_640752, EPI_ISL_640753, EPI_ISL_640756, EPI_ISL_640757, EPI_ISL_640758, EPI_ISL_640761, EPI_ISL_640764, EPI_ISL_640768, EPI_ISL_640769, EPI_ISL_640770, EPI_ISL_640771, EPI_ISL_640772, EPI_ISL_640773, EPI_ISL_640774, EPI_ISL_640777, EPI_ISL_640780, EPI_ISL_640783, EPI_ISL_640786, EPI_ISL_640788, EPI_ISL_640789, EPI_ISL_640790, EPI_ISL_640792, EPI_ISL_640794, EPI_ISL_640797, EPI_ISL_640799, EPI_ISL_640806, EPI_ISL_640807, EPI_ISL_640809, EPI_ISL_640810, EPI_ISL_640812, EPI_ISL_640813, EPI_ISL_640815, EPI_ISL_640817, EPI_ISL_640822, EPI_ISL_640824, EPI_ISL_640825, EPI_ISL_640827, EPI_ISL_640831, EPI_ISL_640832, EPI_ISL_640833, EPI_ISL_640836, EPI_ISL_640839, EPI_ISL_640840, EPI_ISL_640841, EPI_ISL_640842, EPI_ISL_640846, EPI_ISL_640848, EPI_ISL_640849, EPI_ISL_640851, EPI_ISL_640852, EPI_ISL_640853, EPI_ISL_640854, EPI_ISL_640857, EPI_ISL_640858, EPI_ISL_640860, EPI_ISL_640866, EPI_ISL_640868, EPI_ISL_640870, EPI_ISL_640871, EPI_ISL_640876, EPI_ISL_640879, EPI_ISL_640881, EPI_ISL_640882, EPI_ISL_640883, EPI_ISL_640885, EPI_ISL_640887, EPI_ISL_640889, EPI_ISL_640891, EPI_ISL_640895, EPI_ISL_640896, EPI_ISL_640899, EPI_ISL_640900, EPI_ISL_640902, EPI_ISL_640903, EPI_ISL_640906, EPI_ISL_640907, EPI_ISL_640914, EPI_ISL_640915, EPI_ISL_640920, EPI_ISL_640928, EPI_ISL_640930, EPI_ISL_640931, EPI_ISL_640932, EPI_ISL_640935, EPI_ISL_640936, EPI_ISL_640937, EPI_ISL_640938, EPI_ISL_640940, EPI_ISL_640941, EPI_ISL_640944, EPI_ISL_640945, EPI_ISL_640950, EPI_ISL_640952, EPI_ISL_640954, EPI_ISL_640955, EPI_ISL_640958, EPI_ISL_640962, EPI_ISL_640964, EPI_ISL_640966, EPI_ISL_640967, EPI_ISL_640968, EPI_ISL_640971, EPI_ISL_640972, EPI_ISL_640973, EPI_ISL_640977, EPI_ISL_640979, EPI_ISL_640980, EPI_ISL_640983, EPI_ISL_640986, EPI_ISL_640987, EPI_ISL_640989, EPI_ISL_640990, EPI_ISL_640993, EPI_ISL_640995, EPI_ISL_640996, EPI_ISL_640997, EPI_ISL_641001, EPI_ISL_641008, EPI_ISL_641009, EPI_ISL_641010, EPI_ISL_641013, EPI_ISL_641014, EPI_ISL_641015, EPI_ISL_641016, EPI_ISL_641021, EPI_ISL_641024, EPI_ISL_641025, EPI_ISL_641026, EPI_ISL_641028, EPI_ISL_641031, EPI_ISL_641037, EPI_ISL_641042, EPI_ISL_641045, EPI_ISL_641047, EPI_ISL_641049, EPI_ISL_641051, EPI_ISL_641055, EPI_ISL_641056, EPI_ISL_641058, EPI_ISL_641067, EPI_ISL_641069, EPI_ISL_641070, EPI_ISL_641072, EPI_ISL_641073, EPI_ISL_641074, EPI_ISL_641078, EPI_ISL_641079, EPI_ISL_641083, EPI_ISL_641089, EPI_ISL_641092, EPI_ISL_641096, EPI_ISL_641097, EPI_ISL_641098, EPI_ISL_641103, EPI_ISL_641106, EPI_ISL_641109, EPI_ISL_641111, EPI_ISL_641115, EPI_ISL_641118, EPI_ISL_641119, EPI_ISL_641122, EPI_ISL_641123, EPI_ISL_641124, EPI_ISL_641128, EPI_ISL_641131, EPI_ISL_641133, EPI_ISL_641134, EPI_ISL_641137, EPI_ISL_641138, EPI_ISL_641141, EPI_ISL_641142, EPI_ISL_641144, EPI_ISL_641147, EPI_ISL_641148, EPI_ISL_641149, EPI_ISL_641150, EPI_ISL_641153, EPI_ISL_641155, EPI_ISL_641158, EPI_ISL_641162, EPI_ISL_641163, EPI_ISL_641164, EPI_ISL_641165, EPI_ISL_641167, EPI_ISL_641169, EPI_ISL_641170, EPI_ISL_641171, EPI_ISL_641172, EPI_ISL_641176, EPI_ISL_641178, EPI_ISL_641179, EPI_ISL_641180, EPI_ISL_641184, EPI_ISL_641185, EPI_ISL_641186, EPI_ISL_641187, EPI_ISL_641188, EPI_ISL_641189, EPI_ISL_641191, EPI_ISL_641194, EPI_ISL_641195, EPI_ISL_641198, EPI_ISL_641202, EPI_ISL_641203, EPI_ISL_641204, EPI_ISL_641205, EPI_ISL_641206, EPI_ISL_641209, EPI_ISL_641212, EPI_ISL_641213, EPI_ISL_641215, EPI_ISL_641217, EPI_ISL_641221, EPI_ISL_641222, EPI_ISL_641223, EPI_ISL_641225, EPI_ISL_641226, EPI_ISL_641227, EPI_ISL_641230, EPI_ISL_641231, EPI_ISL_641234, EPI_ISL_641236, EPI_ISL_641237, EPI_ISL_641241, EPI_ISL_641243, EPI_ISL_641244, EPI_ISL_641245, EPI_ISL_641246, EPI_ISL_641247, EPI_ISL_641254, EPI_ISL_641255, EPI_ISL_641256, EPI_ISL_641257, EPI_ISL_641258, EPI_ISL_641260, EPI_ISL_641261, EPI_ISL_641265, EPI_ISL_641268, EPI_ISL_641269, EPI_ISL_641270, EPI_ISL_641275, EPI_ISL_641276, EPI_ISL_641278, EPI_ISL_641279, EPI_ISL_641280, EPI_ISL_641285, EPI_ISL_641288, EPI_ISL_641289, EPI_ISL_641292, EPI_ISL_641297, EPI_ISL_641300, EPI_ISL_641303, EPI_ISL_641306, EPI_ISL_641307, EPI_ISL_641308 | | | |
| see above | Microbiological Diagnostic Unit - Public Health Laboratory (MDU-PHL) | MDU-PHL | Seemann T., Schultz M.B., Sait, M.L., Sherry, N.L. |
| EPI_ISL_644532 | MEPHI, Aix Marseille University | MEPHI, Aix Marseille University | Anthony LEVASSEUR |
| EPI_ISL_644981, EPI_ISL_644982, EPI_ISL_644983, EPI_ISL_644984, EPI_ISL_644985, EPI_ISL_644986, EPI_ISL_644987, EPI_ISL_644988, EPI_ISL_644989, EPI_ISL_644990, EPI_ISL_644991 | | | |
| see above | Department of Infectious Diseases, Keio University School of Medicine, Tokyo, Japan | Center for Medical Genetics, Keio University School of Medicine, Tokyo, Japan | Kenjiro Kosaki, Yuka Iwasaki, Hirotsugu Ishizu, Haruhiko Siomi, Kodai Abe |
| EPI_ISL_648165 | The Public Health Agency of Sweden | The Public Health Agency of Sweden | Anna-Malin Linde, Maria Lind Karlberg, Mattias Haukland, Reza Advani, Olov Svartstrom, Oskar Karlsson Lindsjo, Sandra Broddesson, Petra Edquist, Mia Brytting, Anna Risberg, Karin Tegmark-Wisell |
| EPI_ISL_648169 | Klinisk mikrobiologi NAL Trollhattan | The Public Health Agency of Sweden | Anna-Malin Linde, Maria Lind Karlberg, Mattias Haukland, Reza Advani, Olov Svartstrom, Oskar Karlsson Lindsjo, Sandra Broddesson, Petra Edquist, Mia Brytting, Anna Risberg, Karin Tegmark-Wisell |
| EPI_ISL_648182, EPI_ISL_648183 | The Public Health Agency of Sweden | The Public Health Agency of Sweden | Anna-Malin Linde, Maria Lind Karlberg, Mattias Haukland, Reza Advani, Olov Svartstrom, Oskar Karlsson Lindsjo, Sandra Broddesson, Petra Edquist, Mia Brytting, Anna Risberg, Karin Tegmark-Wisell |

| | | | |
|---|---|---|---|
| EPI_ISL_648337, EPI_ISL_648379 | Laboratorio de Investigaciones de Baney | University Hospital Basel, Clinical Bacteriology | Carlos Cortes, Claudia Daubenberger, Adrian Egli, Guillermo Garcia, Salome Hosch, Bonifacio Manguire Nlavo, Alfredo Mari, Maximilian Mpina, Elizabeth Nyakarungu, Diosdado Odjama Nseng Ada, Mitoha Ondo O Ayekaba, Tim Roloff, Tobias Schindler, Helena Seth-Smith, Madlen Stange, Philip Wonder Phiri |
| EPI_ISL_648383 | Santa Clara County Public Health Laboratory | Chan-Zuckerberg Biohub | CZB Cliahub Consortium |
| EPI_ISL_648870, EPI_ISL_648877, EPI_ISL_648881, EPI_ISL_648882, EPI_ISL_648885, EPI_ISL_648886, EPI_ISL_648890, EPI_ISL_648902, EPI_ISL_648916, EPI_ISL_648988, EPI_ISL_648989, EPI_ISL_648990 | | | |
| see above | San Diego County Public Health Laboratory | Andersen lab at Scripps Research | SEARCH Alliance San Diego with Tracy Basler, Jovan Shephard, Brett Austin |
| EPI_ISL_649151 | Microbiological Diagnostic Unit - Public Health Laboratory (MDU-PHL), The Peter Doherty Institute for Infection and Immunity | Microbiological Diagnostic Unit - Public Health Laboratory (MDU-PHL), The Peter Doherty Institute for Infection and Immunity | Seemann,T., Caly,L., Sait,M.L., Schultz,M.B., Druce,J., Sherry,N.L. |
| EPI_ISL_649172 | Laboratorio de Investigaciones de Baney | University Hospital Basel, Clinical Bacteriology | Carlos Cortes, Claudia Daubenberger, Adrian Egli, Guillermo Garcia, Salome Hosch, Bonifacio Manguire Nlavo, Alfredo Mari, Maximilian Mpina, Elizabeth Nyakarungu, Diosdado Odjama Nseng Ada, Mitoha Ondo O Ayekaba, Tim Roloff, Tobias Schindler, Helena Seth-Smith, Madlen Stange, Philip Wonder Phiri |
| EPI_ISL_653319, EPI_ISL_653320, EPI_ISL_653321, EPI_ISL_653322 | Florida Bureau of Public Health Laboratories | Florida Bureau of Public Health Laboratories | Sarah Schmedes, Jason Blanton |
| EPI_ISL_653590, EPI_ISL_653594, EPI_ISL_653600, EPI_ISL_653607, EPI_ISL_653644, EPI_ISL_653645 | LSUHS Emerging Viral Threat Laboratory | Microbial Genome Sequencing Center | Jeremy P. Kamil, Rona S. Scott, Maarten Van Diest, Malgorzata Bienkowska-Haba, Katarzyna Zwolinska, Andrew D. Yurochko, Christopher G. Kevil, Martin J. Sapp, Daniel J. Snyder, Vaughn S. Cooper, John A. Vanchiere |
| EPI_ISL_653745, EPI_ISL_653746, EPI_ISL_653747 | Instituto Nacional de Salud, Bogotá, Colombia | Instituto Nacional de Salud, Bogotá, Colombia | Katherine Laiton-Donato, Diego A. Álvarez-Díaz, Carlos Franco-Muñoz, Mauricio Pacheco-Montealegre, Jonathan Reales, Diego Andrés Prada, Jose A. Usme-Ciro, Zulma M. Cucunubá, Christian Julian VillabonaArenas, Liz Villabona-Arenas, Sussy Echeverria, Astrid C. Flórez, Carolina Ferro, Diana Marcela Walteros-Acero, Franklin Prieto, Carlos Andrés Durán, Martha Lucia Ospina Martinez, Marcela Mercado-Reyes |
| EPI_ISL_654026, EPI_ISL_654032, EPI_ISL_654039, EPI_ISL_654043, EPI_ISL_654049, EPI_ISL_654053, EPI_ISL_654054, EPI_ISL_654055, EPI_ISL_654056, EPI_ISL_654057, EPI_ISL_654058, EPI_ISL_654059, EPI_ISL_654060, EPI_ISL_654061, EPI_ISL_654172, EPI_ISL_654330, EPI_ISL_654333, EPI_ISL_654335, EPI_ISL_654339, EPI_ISL_654362, EPI_ISL_654366, EPI_ISL_654367, EPI_ISL_654369 | | | |
| see above | Hospital General Universitario Gregorio Marañón | SeqCOVID-SPAIN consortium/IBV(CSIC) | Darío García de Viedma, Laura Pérez-Lago, Marta Herranz, Jon Sicilia, Julia Suárez, Pilar Catalán, Patricia Muñoz and SeqCOVID-SPAIN consortium |
| EPI_ISL_654506 | The Public Health Agency of Sweden | The Public Health Agency of Sweden | Anna-Malin Linde, Maria Lind Karlberg, Mattias Haukland, Reza Advani, Olov Svartstrom, Oskar Karlsson Lindsjo, Sandra Broddesson, Petra Edquist, Mia Brytting, Anna Risberg, Karin Tegmark-Wisell |
| EPI_ISL_654818 | Department of Respiratory and other Viral Infections of L.V.Gromashevsky Institute of Epidemiology & Infectious Diseases NAMS of Ukrain | Department of Respiratory and other Viral Infections of L.V.Gromashevsky Institute of Epidemiology & Infectious Diseases NAMS of Ukrain, JSC "Farmak" | Alla Mironenko, Andriy Goy, Ihor Kravchuk, Ludmyla Bolotova, Larysa Radchenko, Nataliia Teteriuk |
| EPI_ISL_654819, EPI_ISL_654820 | Department of Respiratory and other Viral Infections of L.V.Gromashevsky Institute of Epidemiology & Infectious Diseases NAMS of Ukrain | Department of Respiratory and other Viral Infections of L.V.Gromashevsky Institute of Epidemiology & Infectious Diseases NAMS of Ukrain, JSC "Farmak" | Alla Mironenko, Andriy Goy, Ihor Kravchuk, Ludmyla Bolotova, Larysa Radchenko, Nataliia Teteriuk |
| EPI_ISL_654892, EPI_ISL_654942, EPI_ISL_654946, EPI_ISL_654947, EPI_ISL_654948, EPI_ISL_654949, EPI_ISL_654950 | Klinisk mikrobiologi | The Public Health Agency of Sweden | Anna-Malin Linde, Maria Lind Karlberg, Mattias Haukland, Reza Advani, Olov Svartstrom, Oskar Karlsson Lindsjo, Sandra Broddesson, Petra Edquist, Mia Brytting, Anna Risberg, Karin Tegmark-Wisell |
| EPI_ISL_660177 | NHLS-IALCH | KRISP, KZN Research Innovation and Sequencing Platform | Gazy I, Sigal A, Karim F, Cele S, Giandhari J, Pillay S, Tegally H, Wilkinson E, de Oliveira T |
| EPI_ISL_660271, EPI_ISL_660272, EPI_ISL_660273, EPI_ISL_660274, EPI_ISL_660277, EPI_ISL_660278, EPI_ISL_660279, EPI_ISL_660280, EPI_ISL_660281, EPI_ISL_660282, EPI_ISL_660283, EPI_ISL_660284, EPI_ISL_660285, EPI_ISL_660286, EPI_ISL_660287, EPI_ISL_660288, EPI_ISL_660289, EPI_ISL_660290, EPI_ISL_660291, EPI_ISL_660292, EPI_ISL_660293, EPI_ISL_660294, EPI_ISL_660295, EPI_ISL_660296, EPI_ISL_660297, EPI_ISL_660298, EPI_ISL_660299, EPI_ISL_660300, EPI_ISL_660301, EPI_ISL_660302, EPI_ISL_660303, EPI_ISL_660305, EPI_ISL_660306, EPI_ISL_660307, EPI_ISL_660308, EPI_ISL_660309, EPI_ISL_660310, EPI_ISL_660311, EPI_ISL_660312 | | | |
| see above | Servicio de Microbiología, Laboratori Clínic Metropolitana Nord. Hospital Universitari Germans Trias i Pujol. Institut d'Investigació en Ciències de la Salut Germans Trias i Pujol (IGTP) | SeqCOVID-SPAIN consortium/IBV(CSIC) | Elisa Martró, Antoni E. Bordoy, Anna Not, Adrián Antuori, Anabel Fernández, Nona Romaní and SeqCOVID-SPAIN consortium |
| EPI_ISL_660447, EPI_ISL_660454 | Laboratoire de Microbiologie CHU Souro Sanou | Centre Muraz | Abdoul-Salam Ouedraogo, Yacouba Sawadogo, Essia Belarbi, Grit Schubert, Fabian Leendertz, Arsène Zongo, Soumeya Ouangraoua, Zekiba Tarnagda, Lassana Sangaré, Halidou Tinto |
| EPI_ISL_663292, EPI_ISL_663296, EPI_ISL_663304, EPI_ISL_663306, EPI_ISL_663308, EPI_ISL_663315, EPI_ISL_663320, EPI_ISL_663321, EPI_ISL_663322, EPI_ISL_663325, EPI_ISL_663326, EPI_ISL_663329, EPI_ISL_663333, EPI_ISL_663338, EPI_ISL_663339, EPI_ISL_663341, EPI_ISL_663343, EPI_ISL_663345, EPI_ISL_663347, EPI_ISL_663348, EPI_ISL_663356, EPI_ISL_663358, EPI_ISL_663361, EPI_ISL_663363, EPI_ISL_663368, EPI_ISL_663370, EPI_ISL_663371, EPI_ISL_663376, EPI_ISL_663380, EPI_ISL_663382, EPI_ISL_663387, EPI_ISL_663389, EPI_ISL_663390, EPI_ISL_663397, EPI_ISL_663404, EPI_ISL_663408, EPI_ISL_663414, EPI_ISL_663416, EPI_ISL_663418, EPI_ISL_663419, EPI_ISL_663420, EPI_ISL_663431, EPI_ISL_663436, EPI_ISL_663441, EPI_ISL_663443, EPI_ISL_663451, EPI_ISL_663456, EPI_ISL_663464, EPI_ISL_663465, EPI_ISL_663473, EPI_ISL_663475, EPI_ISL_663479, EPI_ISL_663488, EPI_ISL_663495, EPI_ISL_663499, EPI_ISL_663502, EPI_ISL_663506, EPI_ISL_663508, EPI_ISL_663510, EPI_ISL_663516, EPI_ISL_663518, EPI_ISL_663519, EPI_ISL_663522, EPI_ISL_663524, EPI_ISL_663526, EPI_ISL_663529, EPI_ISL_663530, EPI_ISL_663531, EPI_ISL_663532, EPI_ISL_663533, EPI_ISL_663534, EPI_ISL_663535, EPI_ISL_663536, EPI_ISL_663546, EPI_ISL_663548, EPI_ISL_663549, EPI_ISL_663550, EPI_ISL_663551, EPI_ISL_663552, EPI_ISL_663553, EPI_ISL_663554, EPI_ISL_663555, EPI_ISL_663556, EPI_ISL_663557, EPI_ISL_663558, EPI_ISL_663559, EPI_ISL_663560, EPI_ISL_663561, EPI_ISL_663562, EPI_ISL_663563, EPI_ISL_663564, EPI_ISL_663565, EPI_ISL_663566, EPI_ISL_663567, EPI_ISL_663568, EPI_ISL_663569, EPI_ISL_663570, EPI_ISL_663571, EPI_ISL_663572, EPI_ISL_663573, EPI_ISL_663574, EPI_ISL_663575, EPI_ISL_663576, EPI_ISL_663577, EPI_ISL_663578, EPI_ISL_663579, EPI_ISL_663580, EPI_ISL_663581, EPI_ISL_663596, EPI_ISL_663599, EPI_ISL_663600, EPI_ISL_663601, EPI_ISL_663602, EPI_ISL_663603, EPI_ISL_663611, EPI_ISL_663612, EPI_ISL_663617, EPI_ISL_663620, EPI_ISL_663621, EPI_ISL_663622, EPI_ISL_663623, EPI_ISL_663624, EPI_ISL_663625, EPI_ISL_663627, EPI_ISL_663628, EPI_ISL_663629, EPI_ISL_663630, EPI_ISL_663631, EPI_ISL_663633, EPI_ISL_663637, EPI_ISL_663639, EPI_ISL_663641, EPI_ISL_663642, EPI_ISL_663643, EPI_ISL_663644, EPI_ISL_663645, EPI_ISL_663647, EPI_ISL_663648, EPI_ISL_663649, EPI_ISL_663650, EPI_ISL_663651, EPI_ISL_663652, EPI_ISL_663653, EPI_ISL_663654, EPI_ISL_663655, EPI_ISL_663656, EPI_ISL_663657, EPI_ISL_663658, EPI_ISL_663676, EPI_ISL_663677, EPI_ISL_663679, EPI_ISL_663681, EPI_ISL_663682, EPI_ISL_663684, EPI_ISL_663686, EPI_ISL_663687, EPI_ISL_663688, EPI_ISL_663689, EPI_ISL_663691, EPI_ISL_663692, EPI_ISL_663694, EPI_ISL_663706, EPI_ISL_663707, EPI_ISL_663708, EPI_ISL_663709, EPI_ISL_663710, EPI_ISL_663711, EPI_ISL_663713, EPI_ISL_663714, EPI_ISL_663715, EPI_ISL_663717, EPI_ISL_663720, EPI_ISL_663723, EPI_ISL_663724, EPI_ISL_663725, EPI_ISL_663728, EPI_ISL_663729, EPI_ISL_663730, EPI_ISL_663731, EPI_ISL_663732, EPI_ISL_663733, EPI_ISL_663734, EPI_ISL_663735, EPI_ISL_663736, EPI_ISL_663737, EPI_ISL_663742, EPI_ISL_663743, EPI_ISL_663744, EPI_ISL_663745, EPI_ISL_663747, EPI_ISL_663751, EPI_ISL_663753, EPI_ISL_663756, EPI_ISL_663779, EPI_ISL_663780, EPI_ISL_663781, EPI_ISL_663783, EPI_ISL_663784, EPI_ISL_663786, EPI_ISL_663812, EPI_ISL_663815, EPI_ISL_663816, EPI_ISL_663817, EPI_ISL_663886, EPI_ISL_663959, EPI_ISL_663960, EPI_ISL_663961, EPI_ISL_663962, EPI_ISL_663963, EPI_ISL_663966, EPI_ISL_663967, EPI_ISL_663968, EPI_ISL_663970, EPI_ISL_663971, EPI_ISL_663972, EPI_ISL_663973, EPI_ISL_663974, EPI_ISL_663975, EPI_ISL_663978, EPI_ISL_663979, EPI_ISL_663981, EPI_ISL_663982, EPI_ISL_663985, EPI_ISL_664008, EPI_ISL_664009, EPI_ISL_664010, EPI_ISL_664011, EPI_ISL_664012, EPI_ISL_664013, EPI_ISL_664014 | | | |
| see above | Microbiological Diagnostic Unit - Public Health Laboratory (MDU-PHL) | MDU-PHL | Seemann T., Schultz M.B., Sait, M.L., Sherry, N.L. |
| EPI_ISL_666826, EPI_ISL_666827, EPI_ISL_666836, EPI_ISL_666837, EPI_ISL_666838, EPI_ISL_666839, EPI_ISL_666840, EPI_ISL_666841, EPI_ISL_666853, EPI_ISL_666854, EPI_ISL_666855 | | | |
| see above | Florida Bureau of Public Health Laboratories | Florida Bureau of Public Health Laboratories | Sarah Schmedes, Jason Blanton |
| EPI_ISL_667098, EPI_ISL_667099, EPI_ISL_667100, EPI_ISL_667101, EPI_ISL_667102, EPI_ISL_667103, EPI_ISL_667104, EPI_ISL_667105, EPI_ISL_667106, EPI_ISL_667107, EPI_ISL_667108, EPI_ISL_667109, EPI_ISL_667110, EPI_ISL_667111, EPI_ISL_667112, EPI_ISL_667113, EPI_ISL_667114, EPI_ISL_667115, EPI_ISL_667116, EPI_ISL_667117, EPI_ISL_667118, EPI_ISL_667119, EPI_ISL_667120, EPI_ISL_667121, EPI_ISL_667122, EPI_ISL_667123, EPI_ISL_667124, EPI_ISL_667125, EPI_ISL_667126, EPI_ISL_667127, EPI_ISL_667128, EPI_ISL_667129, EPI_ISL_667130, EPI_ISL_667131, EPI_ISL_667132, EPI_ISL_667133, EPI_ISL_667134, EPI_ISL_667135, EPI_ISL_667136, EPI_ISL_667137, EPI_ISL_667138, EPI_ISL_667139, EPI_ISL_667140, EPI_ISL_667141, EPI_ISL_667142, EPI_ISL_667143, EPI_ISL_667144, EPI_ISL_667145, EPI_ISL_667146, EPI_ISL_667147, EPI_ISL_667148, EPI_ISL_667149, EPI_ISL_667150, EPI_ISL_667151, EPI_ISL_667152, EPI_ISL_667153, EPI_ISL_667154, EPI_ISL_667155, EPI_ISL_667156, EPI_ISL_667157, EPI_ISL_667158, EPI_ISL_667159, EPI_ISL_667160, EPI_ISL_667161, EPI_ISL_667162, EPI_ISL_667163, EPI_ISL_667164, EPI_ISL_667165, EPI_ISL_667166, EPI_ISL_667167, EPI_ISL_667168, EPI_ISL_667169, EPI_ISL_667170, EPI_ISL_667171, EPI_ISL_667172, EPI_ISL_667173, EPI_ISL_667174, EPI_ISL_667175, EPI_ISL_667176, EPI_ISL_667177, EPI_ISL_667178, EPI_ISL_667179, EPI_ISL_667180, EPI_ISL_667181, EPI_ISL_667182, EPI_ISL_667183, EPI_ISL_667220, EPI_ISL_667221, EPI_ISL_667222, EPI_ISL_667223, EPI_ISL_667224, EPI_ISL_667551, EPI_ISL_667552, EPI_ISL_667553 | | | |
| see above | OHSU Lab Services Molecular Microbiology Lab | Oregon SARS-CoV-2 Genome Sequencing Center | Brendan L. O'Connell, Ruth V. Nichols, Sally Grindstaff, Alec J. Hirsch, Donna Hansel, Guang Fan, Daniel N. Streblow, William B. Messer, Andrew C. Adey, Benjamin N. Bimber, Brian J. O'Roak |
| EPI_ISL_671243, EPI_ISL_671244 | Department of Virus and Microbiological Special Diagnostics, Statens Serum Institut, Copenhagen, Denmark | Albertsen Lab, Department of Chemistry and Bioscience, Aalborg University, Denmark | Danish Covid-19 Genome Consortium |
| EPI_ISL_671975 | CHU Purpan - Laboratoire de Virologie - Institut Fédératif de Biologie | CHU Purpan - Laboratoire de Virologie - Institut Fédératif de Biologie | Latour J., Ranger N., Dubois M., Carcenac R., Harter A., Boyer P., Tremeaux P., Izopet J. |
| EPI_ISL_672378 | San Francisco Public Health Laboratory | Chan-Zuckerberg Biohub | CZB Cliahub Consortium |

| | | | |
|---|---|---|---|
| EPI_ISL_672584 | Infectious Diseases and Tropical Medicine Research Center, Infectious Diseases and Tropical Medicine Research Center | Infectious Diseases and Tropical Medicine Research Center, Infectious Diseases and Tropical Medicine Research Center | Ahangarzadeh,S., Haghjooy Javanmard,S., Shoaei,P., Ataei,B., Shariati,L. |
| EPI_ISL_672603, EPI_ISL_672604, EPI_ISL_672606, EPI_ISL_672607 | Infectious Diseases and Tropical Medicine Research Center, Infectious Diseases and Tropical Medicine Research Center | Infectious Diseases and Tropical Medicine Research Center, Infectious Diseases and Tropical Medicine Research Center | Ahangarzadeh,S., Haghjooy Javanmard,S., Shariati,L., Aboutalebian,S., Ataei,B., Shoaei,P. |
| EPI_ISL_676510 | Respiratory Virus Unit, Microbiology Services Colindale, Public Health England | COVID-19 Genomics UK (COG-UK) Consortium | PHE Covid Sequencing Team |
| EPI_ISL_676520 | Klinisk mikrobiologi | The Public Health Agency of Sweden | Department of Microbiology, The Public Health Agency of Sweden |
| EPI_ISL_676530 | Uppsala klinisk mikrobiologi | The Public Health Agency of Sweden | Department of Microbiology, The Public Health Agency of Sweden |
| EPI_ISL_676651 | Wadsworth Center, New York State Department.of Health | Wadsworth Center, New York State Department.of Health | Kirsten St. George, Daryl M. Lamson, Alexis Russel, Jonathan Plitnick, Navjot Singh, John Kelly, Sara Griesemer, Erasmus Schneider, Erica Lasek-Nesselquist |
| EPI_ISL_676652 | Masonic Medical Research Institute | Wadsworth Center, New York State Department.of Health | Nathan Tucker, Kirsten St. George, Daryl M. Lamson, Alexis Russel, Jonathan Plitnick, Navjot Singh, John Kelly, Sara Griesemer, Erasmus Schneider, Erica Lasek-Nesselquist |
| EPI_ISL_676653, EPI_ISL_676660, EPI_ISL_676662, EPI_ISL_676671, EPI_ISL_677026, EPI_ISL_677027, EPI_ISL_677028, EPI_ISL_677029, EPI_ISL_677030, EPI_ISL_677031, EPI_ISL_677032, EPI_ISL_677033, EPI_ISL_677034, EPI_ISL_677035, EPI_ISL_677036, EPI_ISL_677037, EPI_ISL_677038, EPI_ISL_677039, EPI_ISL_677040, EPI_ISL_677041 | | | |
| see above | Wadsworth Center, New York State Department.of Health | Wadsworth Center, New York State Department.of Health | Kirsten St. George, Daryl M. Lamson, Alexis Russel, Jonathan Plitnick, Navjot Singh, John Kelly, Sara Griesemer, Erasmus Schneider, Erica Lasek-Nesselquist |
| EPI_ISL_677113, EPI_ISL_677114, EPI_ISL_677116, EPI_ISL_677127, EPI_ISL_677128 | Masonic Medical Research Institute | Wadsworth Center, New York State Department.of Health | Nathan Tucker, Kirsten St. George, Daryl M. Lamson, Alexis Russel, Jonathan Plitnick, Navjot Singh, John Kelly, Sara Griesemer, Erasmus Schneider, Erica Lasek-Nesselquist |
| EPI_ISL_677256, EPI_ISL_677265, EPI_ISL_677298, EPI_ISL_677299 | Colorado Department of Public Health and Environment | Colorado Department of Puplic Health and Environment | Laura Bankers, Molly Hetherington-Rauth, Shannon Ely, Shannon R. Matzinger, Sarah Elizabeth Totten, Emily A. Travanty |
| EPI_ISL_677672 | Masonic Medical Research Institute | Wadsworth Center, New York State Department.of Health | Nathan Tucker, Kirsten St. George, Daryl M. Lamson, Alexis Russel, Jonathan Plitnick, Navjot Singh, John Kelly, Sara Griesemer, Erasmus Schneider, Erica Lasek-Nesselquist |
| EPI_ISL_677722 | General Hospital - Ohrid | Research Center for Genetic Engineering and Biotechnology "Georgi D. Efremov" , Macedonian Academy of Sciences and Arts | RCGEB - MASA |
| EPI_ISL_677723 | Institute for Lung Diseases in Children - Skopje | Research Center for Genetic Engineering and Biotechnology "Georgi D. Efremov" , Macedonian Academy of Sciences and Arts | RCGEB - MASA |
| EPI_ISL_677724 | Clinical Hospital - Shtip | Research Center for Genetic Engineering and Biotechnology "Georgi D. Efremov" , Macedonian Academy of Sciences and Arts | RCGEB - MASA |
| EPI_ISL_677725, EPI_ISL_677726 | General Hospital - Ohrid | Research Center for Genetic Engineering and Biotechnology "Georgi D. Efremov" , Macedonian Academy of Sciences and Arts | RCGEB - MASA |
| EPI_ISL_677827, EPI_ISL_677831, EPI_ISL_677835, EPI_ISL_677837, EPI_ISL_677838, EPI_ISL_677839, EPI_ISL_677842, EPI_ISL_677844, EPI_ISL_677845, EPI_ISL_677852, EPI_ISL_677855, EPI_ISL_677856, EPI_ISL_677862, EPI_ISL_677865, EPI_ISL_677867, EPI_ISL_677872, EPI_ISL_677874, EPI_ISL_677875, EPI_ISL_677880, EPI_ISL_677882, EPI_ISL_677888 | | | |
| see above | Innovative Genomics Institute, UC Berkeley | Innovative Genomics Institute, UC Berkeley | Stacia Wyman, Haridha Shivram, Phil Frankino, Liana Lareau, Shana McDevitt, Justin Choi |
| EPI_ISL_678248, EPI_ISL_678249 | Center for public health - Skopje | Research Center for Genetic Engineering and Biotechnology "Georgi D. Efremov" , Macedonian Academy of Sciences and Arts | RCGEB - MASA |
| EPI_ISL_678250 | Clinical Hospital - Shtip | Research Center for Genetic Engineering and Biotechnology "Georgi D. Efremov" , Macedonian Academy of Sciences and Arts | RCGEB - MASA |
| EPI_ISL_678336, EPI_ISL_678337 | Area of Virology, Serology and Virology Division (SAViD), New South Wales Health Pathology Randwick | Virology Research Laboratory; Area of Virology, Serology and Virology Division (SAViD), New South Wales Health Pathology Randwick | Foster, C.; Au, J.; Ruiz Silva, M.; Deveson, I.; Bull, R.; Van Hal, S.; Rawlinson, W. |
| EPI_ISL_678489 | Veterinary Specialized Institute "Sabac", Serbia | Veterinary Specialized Institute "Kraljevo", Serbia | Vidanovic,D., Tesovic,B., Knezevic,A., Jovanovic,T., Jankovic,M., Sekler,M., Banovic Djeri,B., Petrovic,T., Mrkovacki, S., Volkening,J., Afonso,C. |
| EPI_ISL_681840, EPI_ISL_681841 | Molecular diagnostic unit for viral haemorrhagic fevers and emerging viruses, Bouaké CHU Laboratory | Project group Epidemology of Highly Pathogenic Microorganisms, Robert Koch-Institute | Chantal Akoua-Koffi, Diané Bamourou, Etilé Anoh, Essia Belarbi, Safiatou Karidioula, Grit Schubert, Adjaratou Traoré, Soundélé Maïté, Monemo Pacome, Coulibaly Mbegnan, Bamba Fatoumata Touré, Kra Ouffoué, Fabian Leendertz |
| EPI_ISL_682262, EPI_ISL_682263, EPI_ISL_682264 | HOSPITAL SAN JUAN DE DIOS | Inciensa, Instituto Costarricense de Investigación y Enseñanza en Nutrición y Salud | Francisco Duarte, Hebleen Porras, Claudio Soto-Garita, Estela Cordero, Adriana Godinez & Melany Calderon |
| EPI_ISL_682265 | HOSPITAL UPALA | Inciensa, Instituto Costarricense de Investigación y Enseñanza en Nutrición y Salud | Francisco Duarte, Hebleen Porras, Claudio Soto-Garita, Estela Cordero, Adriana Godinez & Melany Calderon |
| EPI_ISL_682266, EPI_ISL_682267, EPI_ISL_682268, EPI_ISL_682270, EPI_ISL_682271 | HOSPITAL SAN JUAN DE DIOS | Inciensa, Instituto Costarricense de Investigación y Enseñanza en Nutrición y Salud | Francisco Duarte, Hebleen Porras, Claudio Soto-Garita, Estela Cordero, Adriana Godinez & Melany Calderon |
| EPI_ISL_683364, EPI_ISL_683365, EPI_ISL_683366, EPI_ISL_683367, EPI_ISL_683368 | CNR Virus des Infections Respiratoires - France SUD | CNR Virus des Infections Respiratoires - France SUD | Antonin Bal, Gregory Destras, Gwendolyne Burfin, Quentin Semanas, Martine Valette, Bruno Lina, Laurence Josset |
| EPI_ISL_684001 | Utah Public Health Laboratory | Utah Public Health Laboratory | Erin Young, Kelly Oakeson |
| EPI_ISL_691612, EPI_ISL_691618, EPI_ISL_691620, EPI_ISL_691622, EPI_ISL_691626, EPI_ISL_691629, EPI_ISL_691638, EPI_ISL_691641, EPI_ISL_691653, EPI_ISL_691658 | Servicio de Microbiología, Hospital Universitario Son Espases | SeqCOVID-SPAIN consortium/IBV(CSIC) | Carla López-Causapé, Jordi Reina, Antonio Oliver and SeqCOVID-SPAIN consortium |
| EPI_ISL_692733, EPI_ISL_692734, EPI_ISL_692735, EPI_ISL_692755 | CNR Virus des Infections Respiratoires - France SUD | CNR Virus des Infections Respiratoires - France SUD | Antonin Bal, Gregory Destras, Gwendolyne Burfin, Solenne Brun, Martine Valette, Bruno Lina, Laurence Josset |
| EPI_ISL_693756 | Delaware Public Health Laboratory | Delaware Public Health Laboratory | Gregory Hovan |
| EPI_ISL_695562, EPI_ISL_695614, EPI_ISL_695615, EPI_ISL_695616, EPI_ISL_695617, EPI_ISL_695618, EPI_ISL_695619, EPI_ISL_695620, EPI_ISL_695621, EPI_ISL_695622, EPI_ISL_695623, EPI_ISL_695624, EPI_ISL_695625, EPI_ISL_695626, EPI_ISL_695627, EPI_ISL_695628, EPI_ISL_695629, EPI_ISL_695630, EPI_ISL_695631, EPI_ISL_695632, EPI_ISL_695633, EPI_ISL_695634, EPI_ISL_695635, EPI_ISL_695636, EPI_ISL_695637, EPI_ISL_695638 | | | |
| see above | TGen North | TGen North | Jolene Bowers, Megan Folkerts, Chris French, Hayley Yaglom, Ashlyn Pfeiffer, Darrin Lemmer, Dave Engelthaler, The Arizona COVID Genomics Union |

| | | | |
|---|---|---|---|
| | | | (ACGU) |
| EPI_ISL_695741, EPI_ISL_695742, EPI_ISL_695743, EPI_ISL_695744, EPI_ISL_695745, EPI_ISL_695746, EPI_ISL_695747, EPI_ISL_695748, EPI_ISL_695749, EPI_ISL_695750, EPI_ISL_695751, EPI_ISL_695752, EPI_ISL_695830, EPI_ISL_695831, EPI_ISL_695832, EPI_ISL_695833, EPI_ISL_695834, EPI_ISL_695835, EPI_ISL_695836, EPI_ISL_695837, EPI_ISL_695838, EPI_ISL_695839 | | | |
| see above | AZ SPHL, Arizona Department of Health Services | TGen North | Jolene Bowers, Megan Folkerts, Chris French, Hayley Yaglom, Ashlyn Pfeiffer, Darrin Lemmer, Dave Engelthaler, The Arizona COVID Genomics Union (ACGU) |
| EPI_ISL_700079, EPI_ISL_700080, EPI_ISL_700081, EPI_ISL_700082, EPI_ISL_700083, EPI_ISL_700084, EPI_ISL_700085, EPI_ISL_700086, EPI_ISL_700087, EPI_ISL_700088, EPI_ISL_700089, EPI_ISL_700090, EPI_ISL_700091, EPI_ISL_700092, EPI_ISL_700093, EPI_ISL_700094, EPI_ISL_700095, EPI_ISL_700096, EPI_ISL_700097, EPI_ISL_700098, EPI_ISL_700099, EPI_ISL_700100, EPI_ISL_700101, EPI_ISL_700102, EPI_ISL_700103, EPI_ISL_700104, EPI_ISL_700105, EPI_ISL_700106, EPI_ISL_700107, EPI_ISL_700108, EPI_ISL_700109, EPI_ISL_700110, EPI_ISL_700111, EPI_ISL_700112, EPI_ISL_700113, EPI_ISL_700114, EPI_ISL_700115, EPI_ISL_700116 | | | |
| see above | Hematopathology Laboratory, ACTREC, TMC | Hematopathology Laboratory, ACTREC, TMC | Hematopathology Laboratory, ACTREC |
| EPI_ISL_700334, EPI_ISL_700338 | Child Health Research Foundation | Child Health Research Foundation | Senjuti Saha, Afroza Akter Tanni, Syed Muktadir Al Sium, Roly Malaker, Sharmistha Goswami, Arif Mohammad Tanmoy, Md Hafizur Rahman, Samir K Saha |
| EPI_ISL_700466 | Ladismith (Nissenville) Clinic wc LAF | NHLS/UCT | Arash Iranzadeh, Deelan Doolabh, Lynn Tyers, Bruna Galvao, Innocent Mudau, Marvin Hsiao, Kruger Marais, Diana Hardie, Stephen Korsman, Carolyn Williamson |
| EPI_ISL_700479 | Mowbray Maternity Hospital wc MMH | NHLS/UCT | Arash Iranzadeh, Deelan Doolabh, Lynn Tyers, Bruna Galvao, Innocent Mudau, Marvin Hsiao, Kruger Marais, Diana Hardie, Stephen Korsman, Carolyn Williamson |
| EPI_ISL_700495 | Kwanokuthula CDC wc KWA | NHLS/UCT | Arash Iranzadeh, Deelan Doolabh, Lynn Tyers, Bruna Galvao, Innocent Mudau, Marvin Hsiao, Kruger Marais, Diana Hardie, Stephen Korsman, Carolyn Williamson |
| EPI_ISL_700501 | Knysna Hospital wc KNY | NHLS/UCT | Arash Iranzadeh, Deelan Doolabh, Lynn Tyers, Bruna Galvao, Innocent Mudau, Marvin Hsiao, Kruger Marais, Diana Hardie, Stephen Korsman, Carolyn Williamson |
| EPI_ISL_700521 | Alma CDC wc AHC | NHLS/UCT | Arash Iranzadeh, Deelan Doolabh, Lynn Tyers, Bruna Galvao, Innocent Mudau, Marvin Hsiao, Kruger Marais, Diana Hardie, Stephen Korsman, Carolyn Williamson |
| EPI_ISL_700534 | Groote Schuur Hospital wc GSH | NHLS/UCT | Arash Iranzadeh, Deelan Doolabh, Lynn Tyers, Bruna Galvao, Innocent Mudau, Marvin Hsiao, Kruger Marais, Diana Hardie, Stephen Korsman, Carolyn Williamson |
| EPI_ISL_700546 | Victoria Hospital wc VHW | NHLS/UCT | Arash Iranzadeh, Deelan Doolabh, Lynn Tyers, Bruna Galvao, Innocent Mudau, Marvin Hsiao, Kruger Marais, Diana Hardie, Stephen Korsman, Carolyn Williamson |
| EPI_ISL_700577 | Groote Schuur Hospital wc GSH | NHLS/UCT | Arash Iranzadeh, Deelan Doolabh, Lynn Tyers, Bruna Galvao, Innocent Mudau, Marvin Hsiao, Kruger Marais, Diana Hardie, Stephen Korsman, Carolyn Williamson |
| EPI_ISL_700594 | Oudtshoorn Hospital wc OUD | NHLS/UCT | Arash Iranzadeh, Deelan Doolabh, Lynn Tyers, Bruna Galvao, Innocent Mudau, Marvin Hsiao, Kruger Marais, Diana Hardie, Stephen Korsman, Carolyn Williamson |
| EPI_ISL_707936 | Pamukkale University Hospital | Pamukkale University Department of Medical Genetics | Onur TOKGUN et al. |
| EPI_ISL_708026 | University Hospital of Northern Norway, Department for Microbiology and Infectious Disease Control | Norwegian Institute of Public Health, Department of Virology | Kathrine Stene-Johansen, Kamilla Heddeland Instefjord, Hilde Elshaug, Marie Paulsen Madsen, Rasmus Riis Kopperud, Hilde Vollan, Karoline Bragstad, Olav Hungnes |
| EPI_ISL_708394, EPI_ISL_708414 | Delaware Public Health Lab | Delaware Public Health Lab | Gregory Hovan |
| EPI_ISL_710102, EPI_ISL_710103, EPI_ISL_710104, EPI_ISL_710105, EPI_ISL_710106, EPI_ISL_710107, EPI_ISL_710108, EPI_ISL_710110, EPI_ISL_710111, EPI_ISL_710112, EPI_ISL_710113, EPI_ISL_710114, EPI_ISL_710115, EPI_ISL_710116, EPI_ISL_710117, EPI_ISL_710118 | | | |
| see above | Los Angeles County PHL | Los Angeles County PHL | P. Hemarajata et al. |
| EPI_ISL_710247, EPI_ISL_710320, EPI_ISL_710321, EPI_ISL_710322 | Colorado Department of Public Health and Environment | Colorado Department of Puplic Health and Environment | Laura Bankers, Molly C. Hetherington-Rauth, Shannon Ely, Shannon R. Matzinger, Sarah Elizabeth Totten, Emily A. Travanty |
| EPI_ISL_717769 | UW Virology Lab | UW Virology Lab | Pavitra Roychoudhury, Hong Xie, Lasata Shrestha, Michelle Lin, Meei-Li Huang, Keith R Jerome, Alexander Greninger |
| EPI_ISL_717863, EPI_ISL_717864, EPI_ISL_717865, EPI_ISL_717866, EPI_ISL_717867, EPI_ISL_717868 | Laboratorio de Virologia Molecular / UFRJ | Bioinformatics Laboratory / LNCC | Carolina M Voloch, Ronaldo da Silva F Jr, Luiz G P de Almeida, Cynthia C Cardoso, Otavio Bustrolini, Alexandra L Gerber, Ana Paula de C Guimarães, Diana Mariani, Andréa Cony Cavalcanti, Claudia dos Santos Rodrigues, Terezinha M P P Castiñeira, Amilcar Tanuri, Ana Tereza R de Vasconcelos |
| EPI_ISL_717908 | LACEN RJ - Noel Nutels | Bioinformatics Laboratory / LNCC | Carolina M Voloch, Ronaldo da Silva F Jr, Luiz G P de Almeida, Cynthia C Cardoso, Otavio Bustrolini, Alexandra L Gerber, Ana Paula de C Guimarães, Diana Mariani, Andréa Cony Cavalcanti, Claudia dos Santos Rodrigues, Terezinha M P P Castiñeira, Amilcar Tanuri, Ana Tereza R de Vasconcelos |
| EPI_ISL_717959 | Laboratorio de Virologia Molecular / UFRJ | Bioinformatics Laboratory / LNCC | Carolina M Voloch, Ronaldo da Silva F Jr, Luiz G P de Almeida, Cynthia C Cardoso, Otavio Bustrolini, Alexandra L Gerber, Ana Paula de C Guimarães, Diana Mariani, Andréa Cony Cavalcanti, Claudia dos Santos Rodrigues, Terezinha M P P Castiñeira, Amilcar Tanuri, Ana Tereza R de Vasconcelos |
| EPI_ISL_721678 | Viollier AG | Department of Biosystems Science and Engineering, ETH Zürich | Christian Beisel |
| EPI_ISL_723053 | Hematopathology Laboratory, ACTREC, TMC | Hematopathology Laboratory, ACTREC, TMC | Hematopathology Laboratory, ACTREC |
| EPI_ISL_729373, EPI_ISL_729584, EPI_ISL_729585, EPI_ISL_729586, EPI_ISL_729587 | A. Krumbholz, Labor Dr. Krause und Kollegen MVZ GmbH, Kiel | Charité Universitätsmedizin Berlin, Institut für Virologie | Victor M Corman, Barbara Mühlemann, Jörn Beheim-Schwarzbach, Talitha Veith, Julia Schneider, Terry Jones, Christian Drosten |
| EPI_ISL_729802, EPI_ISL_729852, EPI_ISL_729858 | Laboratorio Central de Saude Publica do Estado do Rio Grande do Sul (LACEN-RS) | Laboratory of Respiratory Viruses and Measles, Oswaldo Cruz Institute, FIOCRUZ | Paola Resende, Luciana Appolinario, Fernando Motta, Anna Carolina Paixão, Ana Carolina Mendonça, Tatiana Schaffer Gregianini, Marilda Tereza Mar da Rosa, Marilda Siqueira |
| EPI_ISL_729975, EPI_ISL_729976, EPI_ISL_729977 | Nigeria Centre for Disease Control (NCDC) | African Centre of Excellence for Genomics of Infectious Diseases (ACEGID), Redeemer's University, Ede, Osun State, Nigeria | Oluniyi P.E. et al |
| EPI_ISL_730209, EPI_ISL_730210, EPI_ISL_730211, EPI_ISL_730222, EPI_ISL_730223, EPI_ISL_730224, EPI_ISL_730225, EPI_ISL_730226, EPI_ISL_730227, EPI_ISL_730228 | Genomica Lab Molecular, M©xico | Andersen lab at Scripps Research | SEARCH Alliance San Diego with Jonathan Gonzalez Garcia, Jose Roman Chavez Mendez, Jose Horacio Reyna Verdugo, Martin Gonzalez Ibarra, Luis Alberto Rangel Gonzalez |
| EPI_ISL_732754, EPI_ISL_732811 | Centro de Investigación Biomédica de La Rioja - Hospital San Pedro Logroño | SeqCOVID-SPAIN consortium/IBV(CSIC) | María de Toro, José Manuel Azcona Gutiérrez, María Pilar Bea Escudero, Miriam Blasco Alberdi and SeqCOVID-SPAIN consortium |
| EPI_ISL_732972, EPI_ISL_732973 | WHO National Influenza Centre Russian Federation | WHO National Influenza Centre Russian Federation | Andrey Komissarov, Artem Fadeev, Anna Ivanova, Kseniya Komissarova, Dmitry Bazhenov, Daria Danilenko, Ksenia Safina, Elena Nabieva, Georgii Bazykin, Dmitry Lioznov |
| EPI_ISL_733000, EPI_ISL_733001, EPI_ISL_733002, EPI_ISL_733003, EPI_ISL_733004 | UMMC-Health | WHO National Influenza Centre Russian Federation | Andrey Komissarov, Artem Fadeev, Anna Ivanova, Kseniya Komissarova, Dmitry Bazhenov, Tatiana Platonova, Daria Danilenko, Ksenia Safina, Elena Nabieva, Georgii Bazykin, Dmitry Lioznov |
| EPI_ISL_733005, EPI_ISL_733006, EPI_ISL_733007, EPI_ISL_733008, | WHO National Influenza Centre Russian Federation | WHO National Influenza Centre Russian Federation | Andrey Komissarov, Artem Fadeev, Anna Ivanova, Kseniya Komissarova, Dmitry Bazhenov, Daria Danilenko, Ksenia Safina, Elena Nabieva, Georgii Bazykin, Dmitry Lioznov |

| | | | |
|---|---|---|---|
| EPI_ISL_733009, EPI_ISL_733016, EPI_ISL_733017, EPI_ISL_733018, EPI_ISL_733019, EPI_ISL_733021 | | | |
| EPI_ISL_733164, EPI_ISL_733165, EPI_ISL_733166, EPI_ISL_733167, EPI_ISL_733168, EPI_ISL_733169, EPI_ISL_733172, EPI_ISL_733173, EPI_ISL_733181, EPI_ISL_733182, EPI_ISL_733198, EPI_ISL_733210, EPI_ISL_733211 | | | |
| see above | Pathogenic Microorganisms Variability Laboratory | WHO National Influenza Centre Russian Federation | Andrey Komissarov, Artem Fadeev, Anna Ivanova, Kseniya Komissarova, Dmitry Bazhenov, Daria Danilenko, Ksenia Safina, Elena Nabieva, Georgii Bazykin, Nadezhda Kuznetsova, Elena Shidlovskaya, Sergey Alkhovsky, Tatyana Vishnevskaya, Elizaveta Divisenko, Alexey Shchetinin, Maria Nikiforova, Andrey Pochtovyy, Evgeny Usachev, Elena Vokalova, Maxim Rubalsky, Oleg Rubalsky, Artem Tkachuk, Vladimir Gushchin, Alexander Gintsburg, Dmitry Lioznov |
| EPI_ISL_733392 | WHO National Influenza Centre Russian Federation | WHO National Influenza Centre Russian Federation | Andrey Komissarov, Artem Fadeev, Anna Ivanova, Kseniya Komissarova, Dmitry Bazhenov, Daria Danilenko, Ksenia Safina, Elena Nabieva, Georgii Bazykin, Dmitry Lioznov |
| EPI_ISL_734840, EPI_ISL_734841, EPI_ISL_734842, EPI_ISL_734843, EPI_ISL_734844, EPI_ISL_734845, EPI_ISL_734846, EPI_ISL_734847, EPI_ISL_734848, EPI_ISL_734849, EPI_ISL_734850, EPI_ISL_734851, EPI_ISL_734852, EPI_ISL_734853, EPI_ISL_734854, EPI_ISL_734855, EPI_ISL_734856, EPI_ISL_734857, EPI_ISL_734858, EPI_ISL_734859, EPI_ISL_734860, EPI_ISL_734861, EPI_ISL_734862, EPI_ISL_734863, EPI_ISL_734864, EPI_ISL_734865, EPI_ISL_734866, EPI_ISL_734867, EPI_ISL_734868, EPI_ISL_734869, EPI_ISL_734871, EPI_ISL_734872, EPI_ISL_734873, EPI_ISL_734874, EPI_ISL_734875, EPI_ISL_734876, EPI_ISL_734877, EPI_ISL_734878, EPI_ISL_734879, EPI_ISL_734880, EPI_ISL_734881, EPI_ISL_734882, EPI_ISL_734883, EPI_ISL_734884, EPI_ISL_734885, EPI_ISL_734886, EPI_ISL_734887, EPI_ISL_734888, EPI_ISL_734889, EPI_ISL_734890, EPI_ISL_734891, EPI_ISL_734892, EPI_ISL_734893, EPI_ISL_734894, EPI_ISL_734895, EPI_ISL_734896, EPI_ISL_734897, EPI_ISL_734898, EPI_ISL_734899, EPI_ISL_734900, EPI_ISL_734901, EPI_ISL_734902, EPI_ISL_734903, EPI_ISL_734904, EPI_ISL_734905, EPI_ISL_734906, EPI_ISL_734907, EPI_ISL_734908, EPI_ISL_734909, EPI_ISL_734910, EPI_ISL_734911, EPI_ISL_734912, EPI_ISL_734913, EPI_ISL_734914, EPI_ISL_734915, EPI_ISL_734916 | | | |
| see above | UZ Leuven, National Reference Laboratory for Coronaviruses, Laboratory Medicine, Leuven, Belgium | KU Leuven, Rega Institute, Clinical and Epidemiological Virology | Tony Wawina-Bokalanga, Joan Marti-Carerras, Bert Vanmechelen, Piet Maes |
| EPI_ISL_737034, EPI_ISL_737035, EPI_ISL_737036 | Department of Virology and Immunology, University of Helsinki and Helsinki University Hospital, Huslab Finland | Department of Virology, Faculty of Medicine, University of Helsinki, Helsinki, Finland | Teemu Smura, Ravi Kant, Phuoc Truong, Hussein Alburkat, Hannimari Kallio-Kokko, Jenni Virtanen, Maija Suvanto, Sari Hannula, Harri Kangas, Pekka Ellonen, Olli Vapalahti |
| EPI_ISL_737971, EPI_ISL_738013, EPI_ISL_738014, EPI_ISL_738015 | Uganda Central Public Health Lab and Uganda Virus Research Institute | MRC/UVRI & LSHTM Uganda Research Unit | Matthew Cotten, Dan Lule Bugembe, My V.T. Phan, Pontiano Kaleebu et al. |
| EPI_ISL_738604, EPI_ISL_738748, EPI_ISL_738766, EPI_ISL_738800, EPI_ISL_738825, EPI_ISL_738919, EPI_ISL_738974, EPI_ISL_739049, EPI_ISL_739096, EPI_ISL_739150, EPI_ISL_739220, EPI_ISL_739226, EPI_ISL_739261, EPI_ISL_739262, EPI_ISL_739302, EPI_ISL_739308, EPI_ISL_739361, EPI_ISL_739449, EPI_ISL_739518, EPI_ISL_739519, EPI_ISL_739572 | | | |
| see above | Alameda County Public Health Lab | Chan-Zuckerberg Biohub | CZB Cliahub Consortium |
| EPI_ISL_739773, EPI_ISL_739833, EPI_ISL_740005 | Laboratoire national de santé, Microbiology, Virology | Laboratoire national de santé, Microbiology, Microbial Genomics Platform | Anke Wienecke-Baldacchino, Catherine Ragimbeau, Tamir Abdelrahman, Jessica Tapp, Fatu Djabi |
| EPI_ISL_740028 | Laboratoire national de santé, Microbiology, Virology | Laboratoire national de santé, Microbiology, Microbial Genomics Platform | Anke Wienecke-Baldacchino, Catherine Ragimbeau,Jessica Tapp, Fatu Djabi, Lise Pignon, Raoul Salmon, Tamir Abdelrahman |
| EPI_ISL_740051, EPI_ISL_740278, EPI_ISL_740471 | Laboratoire national de santé, Microbiology, Virology | Laboratoire national de santé, Microbiology, Microbial Genomics Platform | Anke Wienecke-Baldacchino, Catherine Ragimbeau, Tamir Abdelrahman, Jessica Tapp, Fatu Djabi |
| EPI_ISL_744221 | Laboratoire national de santé, Microbiology, Virology | Laboratoire national de santé, Microbiology, Microbial Genomics Platform | Anke Wienecke-Baldacchino, Catherine Ragimbeau,Jessica Tapp, Fatu Djabi, Lise Pignon, Raoul Salmon, Tamir Abdelrahman |
| EPI_ISL_744311, EPI_ISL_744490, EPI_ISL_744874, EPI_ISL_744886, EPI_ISL_744954 | Laboratoire national de santé, Microbiology, Virology | Laboratoire national de santé, Microbiology, Microbial Genomics Platform | Anke Wienecke-Baldacchino, Catherine Ragimbeau, Tamir Abdelrahman, Jessica Tapp, Fatu Djabi |
| EPI_ISL_745395 | CNR Virus des Infections Respiratoires - France SUD | CNR Virus des Infections Respiratoires - France SUD | Antonin Bal, Gregory Destras, Claudia Gonzalez, Gwendolyne Burfin, Quentin Semanas, Martine Valette, Bruno Lina, Laurence Josset |
| EPI_ISL_745471, EPI_ISL_745473, EPI_ISL_745475, EPI_ISL_745477, EPI_ISL_745482, EPI_ISL_745489, EPI_ISL_745490, EPI_ISL_745493, EPI_ISL_745496, EPI_ISL_745511, EPI_ISL_745514, EPI_ISL_745515, EPI_ISL_745518, EPI_ISL_745521, EPI_ISL_745524, EPI_ISL_745530, EPI_ISL_745531, EPI_ISL_745532, EPI_ISL_745534, EPI_ISL_745536, EPI_ISL_745537, EPI_ISL_745538, EPI_ISL_745543, EPI_ISL_745546, EPI_ISL_745548, EPI_ISL_745555, EPI_ISL_745556, EPI_ISL_745557, EPI_ISL_745559, EPI_ISL_745565, EPI_ISL_745566, EPI_ISL_745570, EPI_ISL_745571, EPI_ISL_745572, EPI_ISL_745573, EPI_ISL_745574, EPI_ISL_745577, EPI_ISL_745579, EPI_ISL_745584, EPI_ISL_745588, EPI_ISL_745589, EPI_ISL_745594, EPI_ISL_745595, EPI_ISL_745597, EPI_ISL_745599, EPI_ISL_745602, EPI_ISL_745606, EPI_ISL_745614, EPI_ISL_745616, EPI_ISL_745617, EPI_ISL_745618, EPI_ISL_745622, EPI_ISL_745624, EPI_ISL_745625, EPI_ISL_745628, EPI_ISL_745632, EPI_ISL_745637, EPI_ISL_745642, EPI_ISL_745643, EPI_ISL_745647, EPI_ISL_745651, EPI_ISL_745655, EPI_ISL_745656, EPI_ISL_745721, EPI_ISL_745722, EPI_ISL_745723, EPI_ISL_745724, EPI_ISL_745725, EPI_ISL_745726, EPI_ISL_745727, EPI_ISL_745728, EPI_ISL_745729, EPI_ISL_745731, EPI_ISL_745732, EPI_ISL_745733, EPI_ISL_745734, EPI_ISL_745735, EPI_ISL_745736, EPI_ISL_745737, EPI_ISL_745738, EPI_ISL_745739, EPI_ISL_745740, EPI_ISL_745741, EPI_ISL_745742, EPI_ISL_745743, EPI_ISL_745744, EPI_ISL_745745, EPI_ISL_745746, EPI_ISL_745747, EPI_ISL_745748, EPI_ISL_745749, EPI_ISL_745750, EPI_ISL_745751, EPI_ISL_745752, EPI_ISL_745753, EPI_ISL_745754, EPI_ISL_745755, EPI_ISL_745756, EPI_ISL_745757, EPI_ISL_745758, EPI_ISL_745759, EPI_ISL_745760, EPI_ISL_745761, EPI_ISL_745762, EPI_ISL_745763, EPI_ISL_745764, EPI_ISL_745765, EPI_ISL_745766, EPI_ISL_745767, EPI_ISL_745768, EPI_ISL_745769, EPI_ISL_745770, EPI_ISL_745771, EPI_ISL_745772, EPI_ISL_745773, EPI_ISL_745774, EPI_ISL_745775, EPI_ISL_745777, EPI_ISL_745778, EPI_ISL_745779, EPI_ISL_745780, EPI_ISL_745781, EPI_ISL_745782, EPI_ISL_745784, EPI_ISL_745785, EPI_ISL_745786, EPI_ISL_745787, EPI_ISL_745788, EPI_ISL_745789, EPI_ISL_745790, EPI_ISL_745791, EPI_ISL_745793, EPI_ISL_745794, EPI_ISL_745795, EPI_ISL_745796, EPI_ISL_745798, EPI_ISL_745800, EPI_ISL_745801, EPI_ISL_745802, EPI_ISL_745803, EPI_ISL_745805, EPI_ISL_745806, EPI_ISL_745807, EPI_ISL_745808, EPI_ISL_745809, EPI_ISL_745810, EPI_ISL_745811, EPI_ISL_745812, EPI_ISL_745813, EPI_ISL_745814, EPI_ISL_745815, EPI_ISL_745816, EPI_ISL_745817, EPI_ISL_745818, EPI_ISL_745819, EPI_ISL_745820, EPI_ISL_745822, EPI_ISL_745823, EPI_ISL_745824, EPI_ISL_745825, EPI_ISL_745826, EPI_ISL_745827, EPI_ISL_745828, EPI_ISL_745829, EPI_ISL_745830, EPI_ISL_745831, EPI_ISL_745832, EPI_ISL_745833, EPI_ISL_745834, EPI_ISL_745835, EPI_ISL_745836, EPI_ISL_745837, EPI_ISL_745838, EPI_ISL_745839, EPI_ISL_745840, EPI_ISL_745841, EPI_ISL_745842, EPI_ISL_745843, EPI_ISL_745844, EPI_ISL_745845, EPI_ISL_745846, EPI_ISL_745847, EPI_ISL_745848, EPI_ISL_745849, EPI_ISL_745850, EPI_ISL_745851, EPI_ISL_745852, EPI_ISL_745853, EPI_ISL_745854, EPI_ISL_745855, EPI_ISL_745856, EPI_ISL_745857, EPI_ISL_745858, EPI_ISL_745859, EPI_ISL_745860, EPI_ISL_745861, EPI_ISL_745862, EPI_ISL_745863, EPI_ISL_745864, EPI_ISL_745865, EPI_ISL_745866, EPI_ISL_745867, EPI_ISL_745868, EPI_ISL_745869, EPI_ISL_745870, EPI_ISL_745871, EPI_ISL_745872, EPI_ISL_745873, EPI_ISL_745874, EPI_ISL_745875, EPI_ISL_745876, EPI_ISL_745877, EPI_ISL_745882, EPI_ISL_745885, EPI_ISL_745892, EPI_ISL_745902, EPI_ISL_745907, EPI_ISL_745908, EPI_ISL_745912, EPI_ISL_745914, EPI_ISL_745919, EPI_ISL_745922, EPI_ISL_745958, EPI_ISL_745980, EPI_ISL_745981, EPI_ISL_745983, EPI_ISL_745984, EPI_ISL_745985, EPI_ISL_745986, EPI_ISL_745990, EPI_ISL_745991, EPI_ISL_745993, EPI_ISL_745996, EPI_ISL_745997, EPI_ISL_745999, EPI_ISL_746003, EPI_ISL_746005, EPI_ISL_746006, EPI_ISL_746011, EPI_ISL_746014, EPI_ISL_746018, EPI_ISL_746019, EPI_ISL_746021, EPI_ISL_746025, EPI_ISL_746028, EPI_ISL_746031, EPI_ISL_746032, EPI_ISL_746034, EPI_ISL_746040, EPI_ISL_746041, EPI_ISL_746042, EPI_ISL_746043, EPI_ISL_746044, EPI_ISL_746045, EPI_ISL_746047, EPI_ISL_746049, EPI_ISL_746055, EPI_ISL_746056, EPI_ISL_746060, EPI_ISL_746063, EPI_ISL_746065, EPI_ISL_746066, EPI_ISL_746069, EPI_ISL_746070, EPI_ISL_746071, EPI_ISL_746072, EPI_ISL_746075, EPI_ISL_746076, EPI_ISL_746077, EPI_ISL_746078, EPI_ISL_746079, EPI_ISL_746080, EPI_ISL_746083, EPI_ISL_746084, EPI_ISL_746085, EPI_ISL_746086, EPI_ISL_746087, EPI_ISL_746089, EPI_ISL_746093, EPI_ISL_746265, EPI_ISL_746273, EPI_ISL_746281, EPI_ISL_746289, EPI_ISL_746306, EPI_ISL_746313, EPI_ISL_746314 | | | |
| see above | Ginkgo Bioworks Clinical Laboratory | Utah Public Health Laboratory | Erin L. Young, Kelly Oakeson, Tara Gallagher, Michael T. Pyne, E. Susan Slechta, Melanie A. Mallory, Jeffrey B. Stevenson, Salika M. Shakir, David R. Hillyard, Malaika McKenzie-Bennett, James McGann, Jim Griffin, Keith Robison, Alex Plocik, Becky Schilling, Martha Pierson, Rebecca Littlefield, Michelle Spencer, Birgitte Simen |
| EPI_ISL_746644, EPI_ISL_746645, EPI_ISL_746646 | Genetica Molecular and Subdepartamento de Virologia ISP Chile | Instituto de Salud Publica de Chile | Javier Tognarelli, Barbara Parra, Loredana Arata, Jaime Lagos, Gisselle Barra, Patricia Bustos, Rodrigo Fasce, Andres Castillo, Jorge Fernandez |
| EPI_ISL_751324, EPI_ISL_751339, EPI_ISL_751342 | IRCCS Sacro Cuore Don Calabria Hospital, Department of Infectious, Tropical Diseases & Microbiology | University of Verona, Department of Biotechnology | Antonio Mori, Michela Deiana, Elena Pomari, Chiara Piubelli; Giulia Lopatriello, Luca Marcolungo, Cristina Beltrami, Chiara Degli Esposti, Emanuela Cosentino, Massimo Delledonne |
| EPI_ISL_752747, EPI_ISL_752748, EPI_ISL_752749, EPI_ISL_752750, EPI_ISL_752751, EPI_ISL_752752, EPI_ISL_752753, EPI_ISL_752754, EPI_ISL_752755, EPI_ISL_752756, EPI_ISL_752757, EPI_ISL_752758, EPI_ISL_752759, EPI_ISL_752760, EPI_ISL_752761, EPI_ISL_752772, EPI_ISL_752773, EPI_ISL_752774, EPI_ISL_752775, EPI_ISL_752776, EPI_ISL_752777, EPI_ISL_752778, EPI_ISL_752779, EPI_ISL_752846, EPI_ISL_752847, EPI_ISL_752848, EPI_ISL_752849, EPI_ISL_752850, EPI_ISL_752851, EPI_ISL_752953, EPI_ISL_752954, EPI_ISL_752955, EPI_ISL_752956, EPI_ISL_752957, EPI_ISL_752958, EPI_ISL_752959, EPI_ISL_752960, EPI_ISL_752961, EPI_ISL_753040, EPI_ISL_753041, EPI_ISL_753042, EPI_ISL_753043 | | | |
| see above | State Laboratories Division, Hawaii State Department of Health | State Laboratories Division, Hawaii State Department of Health | Pamela O'Brien, Sabrina Diemert, Drew Kuwazaki, Razvan Sultana, Edward Desmond |
| EPI_ISL_753699 | RSUD Ulin Banjarmasin Kalimantan Selatan | National Institute of Health Research and Development | Subangkit;Pawestri,HA;Puspa,KD;Nugraha,AA;Ikawati,HD;Pangesti,KNA;Soekarso,T;Puspandari,N;Pasaribu,M;Setiawaty,V |
| EPI_ISL_753734, EPI_ISL_753768, EPI_ISL_753858, EPI_ISL_753973, EPI_ISL_753974, EPI_ISL_753975, EPI_ISL_753976, EPI_ISL_753977, EPI_ISL_753978, EPI_ISL_753979, EPI_ISL_753980, EPI_ISL_753988, EPI_ISL_753994, EPI_ISL_753996, EPI_ISL_753998, EPI_ISL_753999, EPI_ISL_754000, EPI_ISL_754001, EPI_ISL_754002, EPI_ISL_754003, EPI_ISL_754004, EPI_ISL_754005, EPI_ISL_754016, EPI_ISL_754043, EPI_ISL_754044, EPI_ISL_754045, EPI_ISL_754054 | | | |
| see above | Charité Universitätsmedizin Berlin, Institut für Virologie/Labor Berlin | Charité Universitätsmedizin Berlin, Institut für Virologie | Victor M Corman, Jörn Beheim-Schwarzbach, Barbara Mühlemann, Julia Schneider, Talitha Veith, Terry Jones, Christian Drosten |
| EPI_ISL_754060, EPI_ISL_754061 | National Public Health Laboratory | Nepal Health Research Council | Pradip Gyanwali, Meghnath Dhimal |
| EPI_ISL_754062, EPI_ISL_754063, EPI_ISL_754064, EPI_ISL_754065, EPI_ISL_754066, EPI_ISL_754067 | Kathmandu University | Nepal Health Research Council | Pradip Gyanwali, Meghnath Dhimal |
| EPI_ISL_754070, EPI_ISL_754071 | Nepal Korea Friendship Municipality Hospital | Nepal Health Research Council | Pradip Gyanwali, Meghnath Dhimal |

| | | | |
|---|---|---|---|
| EPI_ISL_754072, EPI_ISL_754073 | National Public Health Laboratory | Nepal Health Research Council | Pradip Gyanwali, Meghnath Dhimal |
| EPI_ISL_754938, EPI_ISL_754963, EPI_ISL_755011, EPI_ISL_755047 | California Department of Public Health | California Department of Public Health | CDPH IDLB COVIDNet |
| EPI_ISL_755301, EPI_ISL_755303 | Laboratorio de Bioingeniería, Instituto de Ciencias de la Ingeniería, Universidad de O´Higgins | Center for Mathematical Modeling and Center for Genome Regulation. Santiago, Chile | M. Latorre, G. Galvez, J. Ortega, Bastias M, Sanhueza D, Travisany D, Allende ML, Maass A, González M, Bustos F, Arriagada G, Montecino, M, Orellana A, Castro E, Meneses C. |
| EPI_ISL_755969, EPI_ISL_755995, EPI_ISL_755996, EPI_ISL_755997, EPI_ISL_755998, EPI_ISL_755999, EPI_ISL_756000, EPI_ISL_756001, EPI_ISL_756002, EPI_ISL_756003, EPI_ISL_756004, EPI_ISL_756005, EPI_ISL_756006, EPI_ISL_756007, EPI_ISL_756008, EPI_ISL_756009, EPI_ISL_756010, EPI_ISL_756011, EPI_ISL_756012, EPI_ISL_756013 | | | |
| see above | Department of Virology and Immunology, University of Helsinki and Helsinki University Hospital, Huslab Finland | Department of Virology, Faculty of Medicine, University of Helsinki, Helsinki, Finland | Teemu Smura, Ravi Kant, Phuoc Truong, Hussein Alburkat, Hannimari Kallio-Kokko, Jenni Virtanen, Maija Suvanto, Sari Hannula, Harri Kangas, Pekka Ellonen, Olli Vapalahti |
| EPI_ISL_756337 | Innovative Genomics Institute, UC Berkeley | Innovative Genomics Institute, UC Berkeley | Stacia Wyman, Haridha Shivram, Phil Frankino, Liana Lareau, Shana McDevitt, Justin Choi |
| EPI_ISL_756372 | CUMC - CHI Bergan Mercy | Creighton University School of Medicine, Departments of Medical Microbiology and Pharmacology and Neuroscience | Michael Belshan, Morgan A. Raine, Anne V. Cheng, Christopher J. Destache, Richard V. Goering, Jacob A. Siedlik, Holly A. Stessman |
| EPI_ISL_760131, EPI_ISL_760132 | Division of Emerging Infectious Diseases, Bureau of Infectious Diseases Diagnosis Control, Korea Disease Control and Prevention Agency | Division of Emerging Infectious Diseases, Bureau of Infectious Diseases Diagnosis Control, Korea Disease Control and Prevention Agency | Ae Kyung Park, Il-Hwan Kim, Heui Man Kim, Jeong-Min Kim, Namjoo Lee, Chaeyoung Lee, Sang Hee Woo, Eun-Jin Kim |
| EPI_ISL_763137, EPI_ISL_763139, EPI_ISL_763140, EPI_ISL_763141, EPI_ISL_763142, EPI_ISL_763143, EPI_ISL_763144, EPI_ISL_763145 | Dutch COVID-19 response team | Erasmus Medical Center | Bas Oude Munnink, Reina Sikkema, David Nieuwenhuijse, Irina Chestakova, Anne van der Linden, Marjan Boter, Emmanuelle Munger, Corine GeurtsvanKessel, Annemiek van der Eijk, Richard Molenkamp, Marion Koopmans, on behalf of the Dutch national COVID-19 response team. |
| EPI_ISL_765689, EPI_ISL_765690, EPI_ISL_765691, EPI_ISL_765692, EPI_ISL_765693, EPI_ISL_765694, EPI_ISL_765695, EPI_ISL_765696, EPI_ISL_765697, EPI_ISL_765698 | Massachusetts General Hospital | Infectious Disease Program, Broad Institute of Harvard and MIT | Lemieux,J.E., Siddle,K.J., Shaw,B., Adams,G., Pierce,V., Turbett,S., Anahtar,M., Branda,J., Slater,D., Harris,J., Lin,A.E., Gladden-Young,A., Lagerborg,K., Rudy,M., DeRuff,K., Carter,A., Normandin,E., Bauer,M., Reilly,S., Tomkins-Tinch,C., Loreth,C., Chaluvadi,S., Neumann,A., Cusick,C., Chapman,S.B., Gnirke,A., Flowers,K., Cerrato,F., Birren,B.W., Gallagher,G., Smole,S., Park,D.J., MacInnis,B.L., Ryan,E., LaRocque,R., Rosenberg,E. and Sabeti,P.C. |
| EPI_ISL_766654, EPI_ISL_766662, EPI_ISL_766672 | Texas Department of State Health Services | Texas Department of State Health Services | Rashmi Tuladhar, Bonnie Oh, Jenny Zhang, Maliha Rahman, Anita Pokharel, Myong Koag, Chung Wang, Rachel Lee, Grace Kubin, Mayela Pedrueza, James Daniel Bonser |
| EPI_ISL_775596 | Lab. Biologi Molekular FKIK Unwar, Bali | National Institute of Health Research and Development | Ikawati,HD;Subangkit;Pawestri,HA;Nugraha,AA;Puspa,KD;Dewi,SR;Pangesti,KNA;Soekarso,T;Puspandari,N;Setiawaty,V |
| EPI_ISL_776666 | UW Virology Lab | UW Virology Lab | Pavitra Roychoudhury, Hong Xie, Lasata Shrestha, Meei-Li Huang, Keith R Jerome, Alexander Greninger |
| EPI_ISL_779273, EPI_ISL_779275, EPI_ISL_779277, EPI_ISL_779281, EPI_ISL_779282, EPI_ISL_779283, EPI_ISL_779284, EPI_ISL_779285, EPI_ISL_779286, EPI_ISL_779287 | Jamil-ur-Rahman Center for Genome Research, Dr. Panjwani Center for Molecular Medicine and Drug Research | Jamil-ur-Rahman Center for Genome Research, Dr. Panjwani Center for Molecular Medicine and Drug Research | Shakeel,M., Irfan,M., Nisa,Z., Rashid,M., Ansari,S., Khan,I. |
| EPI_ISL_779428, EPI_ISL_779429, EPI_ISL_779430, EPI_ISL_779431, EPI_ISL_779512, EPI_ISL_779565, EPI_ISL_779567, EPI_ISL_779568, EPI_ISL_779569, EPI_ISL_779570, EPI_ISL_779581 | | | |
| see above | Microbiological Diagnostic Unit - Public Health Laboratory (MDU-PHL) | MDU-PHL | Seemann T., Sait, M.L., Sherry, N.L. |
| EPI_ISL_792604, EPI_ISL_792605 | LACEN-PB | Laboratory of Respiratory Viruses and Measles, Oswaldo Cruz Institute, FIOCRUZ | Paola Resende, Luciana Appolinario, Fernando Motta, Anna Carolina Paixao, Ana Carolina Mendonca, João Felipe Bezerra, Romero Henrique Teixeira de Vasconcelos, Dalane Loudal Florentino Teixeira, Thiago Franco de Oliveira Carneiro, Marilda Siqueira |
| EPI_ISL_801872, EPI_ISL_801892, EPI_ISL_801937, EPI_ISL_801976, EPI_ISL_802140, EPI_ISL_802141, EPI_ISL_802142, EPI_ISL_802143, EPI_ISL_802144, EPI_ISL_802145, EPI_ISL_802146, EPI_ISL_802147, EPI_ISL_802148, EPI_ISL_802149, EPI_ISL_802150, EPI_ISL_802151, EPI_ISL_802152, EPI_ISL_802153, EPI_ISL_802154, EPI_ISL_802155, EPI_ISL_802156, EPI_ISL_802157, EPI_ISL_802158, EPI_ISL_802159, EPI_ISL_802160 | | | |
| see above | MSHS Clinical Microbiology Laboratories | MSHS Pathogen Surveillance Program | Ana S. Gonzalez-Reiche, Hala Alshammary, Mitchell J. Sullivan, Brianne Ciferri, Ajay Obla, Angela Amoako, Mahmoud Awawda, Elena Hirsch, Ashley S. Salimbangon, Levy Sominsky, Katherine Beach, Kayla Russo, Charles Gleason, Shelcie Fabre, Giulio Kleiner, Zenab Khan, Bremy Alburquerque, Adriana van de Guchte, Komal Srivastava, Matthew M. Hernandez, Jayeeta Dutta, Denise Jurczyszak, Emily Ferreri, Rachel Chernet, Nancy Francoeur, Betsaida Salom Melo, Irina Oussenko, Gintaras Deikus, Juan Soto, Shwetha Hara Sridhar, Ying-Chih Wang, Kathryn Twyman, Andrew Kasarskis, Deena R. Altman, Robert Sebra, Adolfo García-Sastre, Marta Luksza, Gopi Patel, Sarah Schaefer, Melissa Gitman, Michael D. Nowak, Alberto Paniz-Mondolfi, Emilia Mia Sordillo, Viviana Simon, Harm van Bakel |
| EPI_ISL_803741 | Wisconsin State Laboratory of Hygiene Communicable Disease Division | Wisconsin State Laboratory of Hygiene Communicable Disease Division | Kelsey R. Florek, Abigail C. Shockey |
| EPI_ISL_804949, EPI_ISL_804950, EPI_ISL_804951, EPI_ISL_804953, EPI_ISL_804954, EPI_ISL_804955, EPI_ISL_804956, EPI_ISL_804957, EPI_ISL_804958, EPI_ISL_804959, EPI_ISL_804960, EPI_ISL_804962, EPI_ISL_804965, EPI_ISL_804966, EPI_ISL_804967, EPI_ISL_804970, EPI_ISL_804977 | | | |
| see above | Hospital Comarcal de Melilla | Instituto de Salud Carlos III | Iglesias-Caballero, M. Molinero Calamita, M. González-Esguevillas, M. Camarero, S. Pozo, F. Casas, I. Jiménez, P. Jiménez, M. Zaballos, A. Monzón, S. Varona, S. Juliá, M. Cuesta, I, J. López |
| EPI_ISL_806594, EPI_ISL_806595, EPI_ISL_806596, EPI_ISL_806597, EPI_ISL_806699, EPI_ISL_806700, EPI_ISL_806702, EPI_ISL_806703, EPI_ISL_806704, EPI_ISL_806705, EPI_ISL_806706, EPI_ISL_806707, EPI_ISL_806709, EPI_ISL_806711, EPI_ISL_806712 | | | |
| see above | KEMRI-Wellcome Trust Research Programme/KEMRI-CGMR-C Kilifi | KEMRI-Wellcome Trust Research Programme/KEMRI-CGMR-C Kilifi | Githinji et al |
| EPI_ISL_810971, EPI_ISL_810972, EPI_ISL_810974, EPI_ISL_810975, EPI_ISL_810976, EPI_ISL_810983, EPI_ISL_810985, EPI_ISL_810988, EPI_ISL_811009, EPI_ISL_811014, EPI_ISL_811015, EPI_ISL_811036 | | | |
| see above | MRCG at LSHTM Genomics lab | MRCG at LSHTM Genomics lab | Abdul Karim sesay, Abdoulie Kanteh, Jarra Manneh, Mariama Kujabi, Bakary Sanyang |
| EPI_ISL_812581, EPI_ISL_812582, EPI_ISL_812583, EPI_ISL_812584, EPI_ISL_812585, EPI_ISL_812586, EPI_ISL_812587, EPI_ISL_812588, EPI_ISL_812589, EPI_ISL_812590, EPI_ISL_812591, EPI_ISL_812592, EPI_ISL_812593, EPI_ISL_812594, EPI_ISL_812595, EPI_ISL_812596, EPI_ISL_812597, EPI_ISL_812599, EPI_ISL_812604, EPI_ISL_812610, EPI_ISL_812611 | | | |
| see above | United States Air Force School of Aerospace Medicine | United States Air Force School of Aerospace Medicine | Anthony Fries, Jennifer Meyer, Amanda Javorina, Sarah Purves, William Gruner, Clarise Starr, Elizabeth Macias |
| EPI_ISL_812802, EPI_ISL_812810, EPI_ISL_812814, EPI_ISL_812819, EPI_ISL_812830, EPI_ISL_812834, EPI_ISL_812844, EPI_ISL_812846, EPI_ISL_812852, EPI_ISL_812865 | Genomics Program, Children Cancer Hospital | Genomics Program, Children Cancer Hospital | Hatem,A., Hadad,A., Abouelnaga,S., Amer,K., Salah,H., Farawyla,H., Halafawy,A., Mansour,T., shalaby,L., Hassan,W., Soliman,M., Gomaa,C., Hassan,R., Soliman,S., Monuir,G., Hammad,M., Hussein,S., Abdo,I., Jalal,D., El-Zayat,M., El-Shaqnqery,H., Diab,A., Bakry,U., Samir,O., Magdeldin,S., Sayed,A. |
| EPI_ISL_814078, EPI_ISL_814079, EPI_ISL_814080, EPI_ISL_814090, EPI_ISL_814091 | Ministry of Health Turkey | Ministry of Health Turkey | Fatma Bayrakdar, Yasemin Cogun, Süleyman Yalcin, Aye Baak Alta, Gülay Korukluolu |
| EPI_ISL_815269, EPI_ISL_815283, EPI_ISL_815299, EPI_ISL_815300, EPI_ISL_815301, EPI_ISL_815331, EPI_ISL_815350, EPI_ISL_815379 | Centogene | Centogene | Peter Bauer, Krishna Kumar Kandaswamy, Vivi Hue-Trang Lieu |
| EPI_ISL_816722 | Bioinformatics and Biostatistics Lab, Advanced Sequencing | COVID-19 Genomics UK (COG-UK) Consortium | Aengus Stewart,Jerome Nicod,Chelsea Sawyer,Laura Cubitt,Harshil Patel,Margaret Crawford |

| | | | |
|---|---|---|---|
| | Facility | | |
| EPI_ISL_822325, EPI_ISL_822342 | Lighthouse Lab in Glasgow | Wellcome Sanger Institute for the COVID-19 Genomics UK (COG-UK) Consortium | Harper VanSteenhouse, Yumi Kasai, David Gray, Carol Clugston, Anna Dominiczak and Alex Alderton, Roberto Amato, Sonia Goncalves, Ewan Harrison, David K. Jackson, Ian Johnston, Dominic Kwiatkowski, Cordelia Langford, John Sillitoe on behalf of the Wellcome Sanger Institute COVID-19 Surveillance Team |
| EPI_ISL_824459, EPI_ISL_824460, EPI_ISL_824461, EPI_ISL_824462 | Hospital Universitari Vall d'Hebron - Vall d'Hebron Institut de Rercerca | Hospital Universitari Vall d'Hebron | Cristina Andrés, Maria Piñana, Josep F Abril, Damir Garcia-Cehic, Ariadna Rando, Juliana Esperalba, Maria Gema Codina, Carla Castillo, Maria Carmen Martin, Tomàs Pumarola, Josep Quer, Andrés Antón |
| EPI_ISL_825635, EPI_ISL_825636, EPI_ISL_825648, EPI_ISL_825708, EPI_ISL_825709, EPI_ISL_825710, EPI_ISL_825711, EPI_ISL_825726, EPI_ISL_825727, EPI_ISL_825728, EPI_ISL_825729, EPI_ISL_825730, EPI_ISL_825731, EPI_ISL_825732, EPI_ISL_825733, EPI_ISL_825734, EPI_ISL_825735, EPI_ISL_825736, EPI_ISL_825737, EPI_ISL_825738, EPI_ISL_825739, EPI_ISL_825740, EPI_ISL_825741, EPI_ISL_825742, EPI_ISL_825743, EPI_ISL_825744, EPI_ISL_825745, EPI_ISL_825746, EPI_ISL_825747, EPI_ISL_825748, EPI_ISL_825749, EPI_ISL_825750, EPI_ISL_825751, EPI_ISL_825752, EPI_ISL_825753, EPI_ISL_825754, EPI_ISL_825755, EPI_ISL_825756 | | | |
| see above | Laboratoire de santé publique du Québec | Laboratoire de santé publique du Québec | Sandrine Moreira, Ioannis Ragoussis, Guillaume Bourque, Jesse Shapiro, Mark Lathrop and Michel Roger on behalf of the CoVSeQ research group (http://covseq.ca/researchgroup) |
| EPI_ISL_826589, EPI_ISL_826655, EPI_ISL_826656, EPI_ISL_826657 | Montefiore Medical Center | Albert Einstein College of Medicine, Dept. of Microbiology & Immunology, Chandran lab | J. Maximilian Fels, Saad Khan, Ryan Forster, Karin A. Skalina, Surksha Sirichand, Amy S. Fox, Aviv Bergman, William B. Mitchell, Lucia R. Wolgast, Wendy Szymczak, Robert H. Bortz III, M. Eugenia Dieterle, Catalina Florez, Denise Haslwanter, Rohit K. Jangra, Ethan Laudermilch, Ariel S. Wirchnianski, Jason Barnhill, David L. Goldman, Hnin Khine, D. Yitzchak Goldstein, Johanna P. Daily, Kartik Chandran, Libusha Kelly |
| EPI_ISL_826792, EPI_ISL_826854, EPI_ISL_826855, EPI_ISL_826856, EPI_ISL_826858 | The National University Hospital of Iceland | deCODE genetics | Daniel F Gudbjartsson; Agnar Helgason; Hakon Jonsson; Olafur T Magnusson; Pall Melsted; Gudmundur L Norddahl; Jona Saemundsdottir; Asgeir Sigurdsson; Patrick Sulem; Arna B Agustsdottir; Hannes Eggertsson; Berglind Eiriksdottir; Run Fridriksdottir; Elisabet E Gardarsdottir; Gudmundur Georgsson; Olafia S Gretarsdottir; Kjartan R Gudmundsson; Thora R Gunnarsdottir; Arnaldur Gylfason; Hilma Holm; Brynjar O Jensson; Aslaug Jonasdottir; Kamilla S Josefsdottir; Thordur Kristjansson; Droplaug N Magnusdottir; Solvi Rognvaldsson; Louise le Roux; Gudrun Sigmundsdottir; Gardar Sveinbjornsson; Kristin E Sveinsdottir; Maney Sveinsdottir; Emil A Thorarensen; Bjarni Thorbjornsson; Gisli Masson; Ingileif Jonsdottir; Alma Moller; Thorolfur Gudnason; Karl G Kristinsson; Unnur Thorsteinsdottir; Kari Stefansson |
| EPI_ISL_826871, EPI_ISL_826887, EPI_ISL_827107 | deCODE genetics | deCODE genetics | Daniel F Gudbjartsson; Agnar Helgason; Hakon Jonsson; Olafur T Magnusson; Pall Melsted; Gudmundur L Norddahl; Jona Saemundsdottir; Asgeir Sigurdsson; Patrick Sulem; Arna B Agustsdottir; Hannes Eggertsson; Berglind Eiriksdottir; Run Fridriksdottir; Elisabet E Gardarsdottir; Gudmundur Georgsson; Olafia S Gretarsdottir; Kjartan R Gudmundsson; Thora R Gunnarsdottir; Arnaldur Gylfason; Hilma Holm; Brynjar O Jensson; Aslaug Jonasdottir; Kamilla S Josefsdottir; Thordur Kristjansson; Droplaug N Magnusdottir; Solvi Rognvaldsson; Louise le Roux; Gudrun Sigmundsdottir; Gardar Sveinbjornsson; Kristin E Sveinsdottir; Maney Sveinsdottir; Emil A Thorarensen; Bjarni Thorbjornsson; Gisli Masson; Ingileif Jonsdottir; Alma Moller; Thorolfur Gudnason; Karl G Kristinsson; Unnur Thorsteinsdottir; Kari Stefansson |
| EPI_ISL_827270, EPI_ISL_827271 | The National University Hospital of Iceland | deCODE genetics | Daniel F Gudbjartsson; Agnar Helgason; Hakon Jonsson; Olafur T Magnusson; Pall Melsted; Gudmundur L Norddahl; Jona Saemundsdottir; Asgeir Sigurdsson; Patrick Sulem; Arna B Agustsdottir; Hannes Eggertsson; Berglind Eiriksdottir; Run Fridriksdottir; Elisabet E Gardarsdottir; Gudmundur Georgsson; Olafia S Gretarsdottir; Kjartan R Gudmundsson; Thora R Gunnarsdottir; Arnaldur Gylfason; Hilma Holm; Brynjar O Jensson; Aslaug Jonasdottir; Kamilla S Josefsdottir; Thordur Kristjansson; Droplaug N Magnusdottir; Solvi Rognvaldsson; Louise le Roux; Gudrun Sigmundsdottir; Gardar Sveinbjornsson; Kristin E Sveinsdottir; Maney Sveinsdottir; Emil A Thorarensen; Bjarni Thorbjornsson; Gisli Masson; Ingileif Jonsdottir; Alma Moller; Thorolfur Gudnason; Karl G Kristinsson; Unnur Thorsteinsdottir; Kari Stefansson |
| EPI_ISL_827382 | deCODE genetics | deCODE genetics | Daniel F Gudbjartsson; Agnar Helgason; Hakon Jonsson; Olafur T Magnusson; Pall Melsted; Gudmundur L Norddahl; Jona Saemundsdottir; Asgeir Sigurdsson; Patrick Sulem; Arna B Agustsdottir; Hannes Eggertsson; Berglind Eiriksdottir; Run Fridriksdottir; Elisabet E Gardarsdottir; Gudmundur Georgsson; Olafia S Gretarsdottir; Kjartan R Gudmundsson; Thora R Gunnarsdottir; Arnaldur Gylfason; Hilma Holm; Brynjar O Jensson; Aslaug Jonasdottir; Kamilla S Josefsdottir; Thordur Kristjansson; Droplaug N Magnusdottir; Solvi Rognvaldsson; Louise le Roux; Gudrun Sigmundsdottir; Gardar Sveinbjornsson; Kristin E Sveinsdottir; Maney Sveinsdottir; Emil A Thorarensen; Bjarni Thorbjornsson; Gisli Masson; Ingileif Jonsdottir; Alma Moller; Thorolfur Gudnason; Karl G Kristinsson; Unnur Thorsteinsdottir; Kari Stefansson |
| EPI_ISL_827437, EPI_ISL_827590, EPI_ISL_827737, EPI_ISL_827781, EPI_ISL_827783, EPI_ISL_827935, EPI_ISL_828191, EPI_ISL_828250, EPI_ISL_828251, EPI_ISL_828265, EPI_ISL_828266, EPI_ISL_828270, EPI_ISL_828539, EPI_ISL_828540, EPI_ISL_828541, EPI_ISL_828543, EPI_ISL_828555, EPI_ISL_828556, EPI_ISL_828557, EPI_ISL_828642, EPI_ISL_828663 | | | |
| see above | The National University Hospital of Iceland | deCODE genetics | Daniel F Gudbjartsson; Agnar Helgason; Hakon Jonsson; Olafur T Magnusson; Pall Melsted; Gudmundur L Norddahl; Jona Saemundsdottir; Asgeir Sigurdsson; Patrick Sulem; Arna B Agustsdottir; Hannes Eggertsson; Berglind Eiriksdottir; Run Fridriksdottir; Elisabet E Gardarsdottir; Gudmundur Georgsson; Olafia S Gretarsdottir; Kjartan R Gudmundsson; Thora R Gunnarsdottir; Arnaldur Gylfason; Hilma Holm; Brynjar O Jensson; Aslaug Jonasdottir; Kamilla S Josefsdottir; Thordur Kristjansson; Droplaug N Magnusdottir; Solvi Rognvaldsson; Louise le Roux; Gudrun Sigmundsdottir; Gardar Sveinbjornsson; Kristin E Sveinsdottir; Maney Sveinsdottir; Emil A Thorarensen; Bjarni Thorbjornsson; Gisli Masson; Ingileif Jonsdottir; Alma Moller; Thorolfur Gudnason; Karl G Kristinsson; Unnur Thorsteinsdottir; Kari Stefansson |
| EPI_ISL_828788 | deCODE genetics | deCODE genetics | Daniel F Gudbjartsson; Agnar Helgason; Hakon Jonsson; Olafur T Magnusson; Pall Melsted; Gudmundur L Norddahl; Jona Saemundsdottir; Asgeir Sigurdsson; Patrick Sulem; Arna B Agustsdottir; Hannes Eggertsson; Berglind Eiriksdottir; Run Fridriksdottir; Elisabet E Gardarsdottir; Gudmundur Georgsson; Olafia S Gretarsdottir; Kjartan R Gudmundsson; Thora R Gunnarsdottir; Arnaldur Gylfason; Hilma Holm; Brynjar O Jensson; Aslaug Jonasdottir; Kamilla S Josefsdottir; Thordur Kristjansson; Droplaug N Magnusdottir; Solvi Rognvaldsson; Louise le Roux; Gudrun Sigmundsdottir; Gardar Sveinbjornsson; Kristin E Sveinsdottir; Maney Sveinsdottir; Emil A Thorarensen; Bjarni Thorbjornsson; Gisli Masson; Ingileif Jonsdottir; Alma Moller; Thorolfur Gudnason; Karl G Kristinsson; Unnur Thorsteinsdottir; Kari Stefansson |
| EPI_ISL_829116, EPI_ISL_829236, EPI_ISL_829237 | The National University Hospital of Iceland | deCODE genetics | Daniel F Gudbjartsson; Agnar Helgason; Hakon Jonsson; Olafur T Magnusson; Pall Melsted; Gudmundur L Norddahl; Jona Saemundsdottir; Asgeir Sigurdsson; Patrick Sulem; Arna B Agustsdottir; Hannes Eggertsson; Berglind Eiriksdottir; Run Fridriksdottir; Elisabet E Gardarsdottir; Gudmundur Georgsson; Olafia S Gretarsdottir; Kjartan R Gudmundsson; Thora R Gunnarsdottir; Arnaldur Gylfason; Hilma Holm; Brynjar O Jensson; Aslaug Jonasdottir; Kamilla S Josefsdottir; Thordur Kristjansson; Droplaug N Magnusdottir; Solvi Rognvaldsson; Louise le Roux; Gudrun Sigmundsdottir; Gardar Sveinbjornsson; Kristin E Sveinsdottir; Maney Sveinsdottir; Emil A Thorarensen; Bjarni Thorbjornsson; Gisli Masson; Ingileif Jonsdottir; Alma Moller; Thorolfur Gudnason; Karl G Kristinsson; Unnur Thorsteinsdottir; Kari Stefansson |
| EPI_ISL_829362 | deCODE genetics | deCODE genetics | Daniel F Gudbjartsson; Agnar Helgason; Hakon Jonsson; Olafur T Magnusson; Pall Melsted; Gudmundur L Norddahl; Jona Saemundsdottir; Asgeir Sigurdsson; Patrick Sulem; Arna B Agustsdottir; Hannes Eggertsson; Berglind Eiriksdottir; Run Fridriksdottir; Elisabet E Gardarsdottir; Gudmundur Georgsson; Olafia S Gretarsdottir; Kjartan R Gudmundsson; Thora R Gunnarsdottir; Arnaldur Gylfason; Hilma Holm; Brynjar O Jensson; Aslaug Jonasdottir; Kamilla S Josefsdottir; Thordur Kristjansson; Droplaug N Magnusdottir; Solvi Rognvaldsson; Louise le Roux; Gudrun Sigmundsdottir; Gardar Sveinbjornsson; Kristin E Sveinsdottir; Maney Sveinsdottir; Emil A Thorarensen; Bjarni Thorbjornsson; Gisli Masson; Ingileif Jonsdottir; Alma Moller; Thorolfur Gudnason; Karl G Kristinsson; Unnur Thorsteinsdottir; Kari Stefansson |
| EPI_ISL_829524, EPI_ISL_829525, EPI_ISL_829526, EPI_ISL_829529 | The National University Hospital of Iceland | deCODE genetics | Daniel F Gudbjartsson; Agnar Helgason; Hakon Jonsson; Olafur T Magnusson; Pall Melsted; Gudmundur L Norddahl; Jona Saemundsdottir; Asgeir Sigurdsson; Patrick Sulem; Arna B Agustsdottir; Hannes Eggertsson; Berglind Eiriksdottir; Run Fridriksdottir; Elisabet E Gardarsdottir; Gudmundur Georgsson; Olafia S Gretarsdottir; Kjartan R Gudmundsson; Thora R Gunnarsdottir; Arnaldur Gylfason; Hilma Holm; Brynjar O Jensson; Aslaug Jonasdottir; Kamilla S Josefsdottir; Thordur Kristjansson; Droplaug N Magnusdottir; Solvi Rognvaldsson; Louise le Roux; Gudrun Sigmundsdottir; Gardar Sveinbjornsson; Kristin E Sveinsdottir; Maney Sveinsdottir; Emil A Thorarensen; Bjarni Thorbjornsson; Gisli Masson; Ingileif Jonsdottir; Alma Moller; Thorolfur Gudnason; Karl G Kristinsson; Unnur Thorsteinsdottir; Kari Stefansson |
| EPI_ISL_829549 | deCODE genetics | deCODE genetics | Daniel F Gudbjartsson; Agnar Helgason; Hakon Jonsson; Olafur T Magnusson; Pall Melsted; Gudmundur L Norddahl; Jona Saemundsdottir; Asgeir Sigurdsson; Patrick Sulem; Arna B Agustsdottir; Hannes Eggertsson; Berglind Eiriksdottir; Run Fridriksdottir; Elisabet E Gardarsdottir; Gudmundur Georgsson; Olafia S Gretarsdottir; Kjartan R Gudmundsson; Thora R Gunnarsdottir; Arnaldur Gylfason; Hilma Holm; Brynjar O Jensson; Aslaug Jonasdottir; Kamilla S Josefsdottir; Thordur Kristjansson; Droplaug N Magnusdottir; Solvi Rognvaldsson; Louise le Roux; Gudrun Sigmundsdottir; Gardar Sveinbjornsson; Kristin E Sveinsdottir; Maney Sveinsdottir; Emil A Thorarensen; Bjarni Thorbjornsson; Gisli Masson; Ingileif Jonsdottir; Alma Moller; Thorolfur Gudnason; Karl G Kristinsson; Unnur Thorsteinsdottir; Kari Stefansson |
| EPI_ISL_829601, EPI_ISL_829679, EPI_ISL_829849, EPI_ISL_829850, EPI_ISL_829970, EPI_ISL_830449 | The National University Hospital of Iceland | deCODE genetics | Daniel F Gudbjartsson; Agnar Helgason; Hakon Jonsson; Olafur T Magnusson; Pall Melsted; Gudmundur L Norddahl; Jona Saemundsdottir; Asgeir Sigurdsson; Patrick Sulem; Arna B Agustsdottir; Hannes Eggertsson; Berglind Eiriksdottir; Run Fridriksdottir; Elisabet E Gardarsdottir; Gudmundur Georgsson; Olafia S Gretarsdottir; Kjartan R Gudmundsson; Thora R Gunnarsdottir; Arnaldur Gylfason; Hilma Holm; Brynjar O Jensson; Aslaug Jonasdottir; Kamilla S |

| | | | |
|---|---|---|---|
| | | | Josefsdottir; Thordur Kristjansson; Droplaug N Magnusdottir; Solvi Rognvaldsson; Louise le Roux; Gudrun Sigmundsdottir; Gardar Sveinbjornsson; Kristin E Sveinsdottir; Maney Sveinsdottir; Emil A Thorarensen; Bjarni Thorbjornsson; Gisli Masson; Ingileif Jonsdottir; Alma Moller; Thorolfur Gudnason; Karl G Kristinsson; Unnur Thorsteinsdottir; Kari Stefansson |
| EPI_ISL_831085, EPI_ISL_831086, EPI_ISL_831093 | Hospital Universitario La Paz (Madrid) | SeqCOVID-SPAIN consortium/IBV(CSIC) | María Rodríguez-Tejedor, Elias Dahdouh, Fernando Lázaro-Perona, Jesús Mingorance and SeqCOVID-SPAIN consortium |
| EPI_ISL_831689 | Laboratório de Microbiologia Molecular - Universidade FEEVALE | Universidade Federal de Ciências da Saúde de Porto Alegre | Vinícius Bonetti Franceschi, Amanda de Menezes Mayer, Gabriel Dickin Caldana, Carla Andretta Moreira Neves, Patrícia Aline Gröhs Ferrareze, Gabriela Bettella Cybis, Ricardo Ariel Zimerman, Lívia Kmetzsch, Fernando Rosado Spilki, Claudia Elizabeth Thompson |
| EPI_ISL_831919 | New Mexico Department of Health Scientific Laboratory | New Mexico Department of Health Scientific Laboratory | Ellie Johnson, Anastacia Griego-Fisher, D'eldra Malone |
| EPI_ISL_831973 | Klinisk mikrobiologi | The Public Health Agency of Sweden | Department of Microbiology, The Public Health Agency of Sweden |
| EPI_ISL_831974 | Klinisk mikrobiologi NAL Trollhattan | The Public Health Agency of Sweden | Department of Microbiology, The Public Health Agency of Sweden |
| EPI_ISL_833511 | Veterinary Specialized Instute "Nis" | Veterinary Specialized Institute "Kraljevo", Serbia | Vidanovic,D., Tesovic,B., Manic,M., Petrovic,M.,Knezevic,A., Jovanovic,T., Jankovic,M., Sekler,M., Banovic Djeri,B., Petrovic,T., Volkening,J., Afonso,C. |
| EPI_ISL_837554, EPI_ISL_837555 | Centro Nacional de Enfermedades Tropicales (CENETROP) | Laboratory of Respiratory Viruses and Measles, Oswaldo Cruz Institute, FIOCRUZ | Paola Resende, Roxana Loayza, Cinthia Avila, Luciana Appolinario, Fernando Motta, Anna Carolina Paixao, Ana Carolina Mendonca, Marilda Siqueira |
| EPI_ISL_837595, EPI_ISL_837596 | Laboratorio Nacional de Salud | Laboratory of Respiratory Viruses and Measles, Oswaldo Cruz Institute, FIOCRUZ | Paola Resende, Cesar Roberto Conde Pereira, Claudia Estrada, Luciana Appolinario, Fernando Motta, Anna Carolina Paixao, Ana Carolina Mendonca, Marilda Siqueira |
| EPI_ISL_837601, EPI_ISL_837791, EPI_ISL_837792, EPI_ISL_837793, EPI_ISL_837794, EPI_ISL_837795, EPI_ISL_837796, EPI_ISL_837797 | Instituto Nacional de Enfermedades Respiratorias (INER) | Instituto Nacional de Enfermedades Respiratorias (INER) | Celia Boukadida, Margarita Matías-Florentino, Alma Rincón-Rubio, Hector Esteban Paz-Juárez, Olivia Briceño, Edgar Sevilla-Reyes, Fidencio Mejía-Nepomuceno, Mario Mújica-Sánchez, Eduardo Becerril-Vargas, José Arturo Martínez-Orozco, Alejandra Hernández-Terán, Jorge Salas-Hernández, Santiago Ávila-Ríos, Joel Armando Vázquez-Pérez |
| EPI_ISL_845662, EPI_ISL_845663, EPI_ISL_845668, EPI_ISL_845670, EPI_ISL_845672, EPI_ISL_845673, EPI_ISL_845674, EPI_ISL_845675 | Quest Diagnostics | Quest Diagnostics | Rosenthal,S.H., Gerasimova,A., Kagan,R.M., Anderson, B., Bernstein, L.E., Livingston, K.E., Hua, M., Liu Y., Shalhout, D.F., Shlyakhter, I.A., Owen, R., Lacbawan, F. |
| EPI_ISL_848207, EPI_ISL_848220, EPI_ISL_848234, EPI_ISL_848317, EPI_ISL_848318, EPI_ISL_848494 | Illinois Department of Public Health | Gagnon Lab, Southern Illinois University | Keith Gagnon |
| EPI_ISL_849085, EPI_ISL_849086, EPI_ISL_849087, EPI_ISL_849088, EPI_ISL_849089, EPI_ISL_849090, EPI_ISL_849091, EPI_ISL_849092, EPI_ISL_849093, EPI_ISL_849094, EPI_ISL_849095, EPI_ISL_849096, EPI_ISL_849097, EPI_ISL_849098, EPI_ISL_849099, EPI_ISL_849100, EPI_ISL_849101, EPI_ISL_849102, EPI_ISL_849103, EPI_ISL_849104, EPI_ISL_849105, EPI_ISL_849106, EPI_ISL_849107, EPI_ISL_849108 | | | |
| see above | Florida Bureau of Public Health Laboratories | Florida Bureau of Public Health Laboratories | Sarah Schmedes, Jason Blanton |
| EPI_ISL_849684 | unknown | PHV-FSS | Son Nguyen et al. |
| EPI_ISL_849939, EPI_ISL_849940, EPI_ISL_849942, EPI_ISL_849946, EPI_ISL_849950 | UC Davis- Department of Pathology and Laboratory Medicine | Chan-Zuckerberg Biohub | CZB Cliahub Consortium |
| EPI_ISL_850257, EPI_ISL_850258, EPI_ISL_850259, EPI_ISL_850260, EPI_ISL_850261, EPI_ISL_850262, EPI_ISL_850263, EPI_ISL_850264, EPI_ISL_850265, EPI_ISL_850266, EPI_ISL_850267, EPI_ISL_850268, EPI_ISL_850269, EPI_ISL_850270, EPI_ISL_850271, EPI_ISL_850272, EPI_ISL_850275, EPI_ISL_850276, EPI_ISL_850277, EPI_ISL_850278, EPI_ISL_850279, EPI_ISL_850280, EPI_ISL_850281, EPI_ISL_850282, EPI_ISL_850283, EPI_ISL_850284, EPI_ISL_850285, EPI_ISL_850286, EPI_ISL_850287, EPI_ISL_850288, EPI_ISL_850289, EPI_ISL_850290, EPI_ISL_850291, EPI_ISL_850292, EPI_ISL_850293, EPI_ISL_850294, EPI_ISL_850319, EPI_ISL_850331, EPI_ISL_850343 | | | |
| see above | Division of Emerging Infectious Diseases, Bureau of Infectious Diseases Diagnosis Control, Korea Disease Control and Prevention Agency | Division of Emerging Infectious Diseases, Bureau of Infectious Diseases Diagnosis Control, Korea Disease Control and Prevention Agency | Ae Kyung Park, Il-Hwan Kim, Heui Man Kim, Jeong-Min Kim, Namjoo Lee, Chaeyoung Lee, Sang Hee Woo, Eun-Jin Kim |
| EPI_ISL_850509 | Gandhi Medical College and Hospital | VRDL-Gandhi Medical College | Nagamani Kammili, Rajeshwar Rao, Winnie Thomas, Madhavi Latha Manolla, Shailaja VV, Sudhamadhuri Devara, Vanisree Rajoli, Archana GJ, Sushma Rajyalakshmi Gudiseva, Sunitha Pakalapati, Manisha Rani, Amrithesh Kumar, Raja Rao Mesepogu, Vinay Shekar Reddy, Thrilok Chander Bingi, Sofia Banu, Divya Tej Sowpati |
| EPI_ISL_850510 | Gandhi Medical College and Hospital | MDRU-DHR,Gandhi Medical College | Nagamani Kammili, Winnie Thomas, Madhavi Latha Manolla, Shailaja VV, Sudhamadhuri Devara, Vanisree Rajoli, Archana GJ, Sushma Rajyalakshmi Gudiseva, Sunitha Pakalapati, Manisha Rani, Amrithesh Kumar, Raja Rao Mesepogu, Vinay Shekar Reddy, Thrilok Chander Bingi, Sofia Banu, Divya Tej Sowpati |
| EPI_ISL_852806 | Institute of Virology, Medical Center, University of Freiburg, Freiburg, Germany | Institute of Virology, Clinial Virus Genomics, Medical Center, University of Freiburg, Freiburg, Germany | Jonas Fuchs, Lisa Kern, Sandra Reuter, Hajo Grundmann, Marcus Panning |
| EPI_ISL_853843, EPI_ISL_853844, EPI_ISL_853895, EPI_ISL_853896, EPI_ISL_853897, EPI_ISL_853898, EPI_ISL_853899, EPI_ISL_853900, EPI_ISL_853957 | Center for Virology, Medical University of Vienna | Bergthaler laboratory, CeMM Research Center for Molecular Medicine of the Austrian Academy of Sciences | Lukas Endler, Alexandra Popa, Benedikt Agerer, Jakob-Wendelin Genger, Alexander Lercher, Anna Schedl, Thomas Penz, Michael Schuster, Jan Laine, Martin Senekowitsch, Christoph Bock, Andreas Bergthaler |
| EPI_ISL_855352 | Department of Epidemiology, Infectious Disease Control and Prevention, Hiroshima University, Japan | Department of Epidemiology, Infectious Disease Control and Prevention, Hiroshima University, Japan | Junko Tanaka, Kazuaki Takahashi, Masao Kuwabara, Eisaku Kishita, Shintaro Nagashima, Ko Ko |
| EPI_ISL_859664, EPI_ISL_859681 | BTC, Khalifa University | BTC, Khalifa University | Al Safar et al |
| EPI_ISL_861871, EPI_ISL_861884 | LATE - Laboratório de Técnicas Especiais - Hospital Israelita Albert Einstein | LATE - Laboratório de Técnicas Especiais - Hospital Israelita Albert Einstein | Deyvid Amgarten, Fernanda de Mello Malta, Raquel Riyuzo, Ana Paula Moreira Salles, Pedro Henrique Sebe Rodrigues, João Renato Rebello Pinho |
| EPI_ISL_862270, EPI_ISL_862271, EPI_ISL_862272, EPI_ISL_862274, EPI_ISL_862275, EPI_ISL_862280, EPI_ISL_862288, EPI_ISL_862289, EPI_ISL_862295, EPI_ISL_862298, EPI_ISL_862299, EPI_ISL_862304, EPI_ISL_862306, EPI_ISL_862307, EPI_ISL_862314, EPI_ISL_862315, EPI_ISL_862316, EPI_ISL_862317, EPI_ISL_862318, EPI_ISL_862319, EPI_ISL_862320, EPI_ISL_862321, EPI_ISL_862322, EPI_ISL_862323, EPI_ISL_862324, EPI_ISL_862325, EPI_ISL_862326, EPI_ISL_862327, EPI_ISL_862328, EPI_ISL_862329, EPI_ISL_862330, EPI_ISL_862331, EPI_ISL_862332, EPI_ISL_862333, EPI_ISL_862334, EPI_ISL_862335, EPI_ISL_862336, EPI_ISL_862337, EPI_ISL_862338, EPI_ISL_862339, EPI_ISL_862340, EPI_ISL_862341, EPI_ISL_862342, EPI_ISL_862343, EPI_ISL_862345, EPI_ISL_862346, EPI_ISL_862347, EPI_ISL_862348, EPI_ISL_862349, EPI_ISL_862350, EPI_ISL_862351, EPI_ISL_862352, EPI_ISL_862353, EPI_ISL_862354, EPI_ISL_862355, EPI_ISL_862356, EPI_ISL_862358, EPI_ISL_862359, EPI_ISL_862360, EPI_ISL_862361, EPI_ISL_862362, EPI_ISL_862363, EPI_ISL_862364, EPI_ISL_862365, EPI_ISL_862366, EPI_ISL_862367, EPI_ISL_862368, EPI_ISL_862369, EPI_ISL_862371, EPI_ISL_862372, EPI_ISL_862373, EPI_ISL_862374, EPI_ISL_862375, EPI_ISL_862376, EPI_ISL_862377, EPI_ISL_862378, EPI_ISL_862379, EPI_ISL_862380, EPI_ISL_862381, EPI_ISL_862383, EPI_ISL_862384, EPI_ISL_862404, EPI_ISL_862406, EPI_ISL_862407, EPI_ISL_862408, EPI_ISL_862409, EPI_ISL_862410, EPI_ISL_862411, EPI_ISL_862412, EPI_ISL_862413, EPI_ISL_862414, EPI_ISL_862415, EPI_ISL_862417, EPI_ISL_862418, EPI_ISL_862419, EPI_ISL_862420, EPI_ISL_862421, EPI_ISL_862422, EPI_ISL_862423, EPI_ISL_862424, EPI_ISL_862426, EPI_ISL_862427, EPI_ISL_862428, EPI_ISL_862429, EPI_ISL_862430, EPI_ISL_862431, EPI_ISL_862432, EPI_ISL_862433, EPI_ISL_862434, EPI_ISL_862435, EPI_ISL_862436, EPI_ISL_862467, EPI_ISL_862469, EPI_ISL_862470, EPI_ISL_862471, EPI_ISL_862472, EPI_ISL_862473, EPI_ISL_862474, EPI_ISL_862475, EPI_ISL_862476, EPI_ISL_862477, EPI_ISL_862479, EPI_ISL_862480, EPI_ISL_862482, EPI_ISL_862483, EPI_ISL_862484, EPI_ISL_862485, EPI_ISL_862486, EPI_ISL_862487, EPI_ISL_862488, EPI_ISL_862489, EPI_ISL_862490, EPI_ISL_862491, EPI_ISL_862492, EPI_ISL_862493, EPI_ISL_862494, EPI_ISL_862495, EPI_ISL_862504, EPI_ISL_862505, EPI_ISL_862506, EPI_ISL_862507, EPI_ISL_862508 | | | |
| see above | Kurnool Medical College (KMC) | CSIR Institute of Genomics and Integrative Biology | Pallavali Roja Rani, Mohamed Imran, J. Vijaya Lakshmi, Bani Jolly, S. Afsar, Abhinav Jain, Mohit Kumar Divakar, Panyam Suresh, Disha Sharma, Nambi Rajesh, Rahul C Bhoyar, Dasari Ankaiah, Sanaga Shanthi Kumari, Gyan Ranjan, Valluri Anitha Lavanya, Mercy Rophina, S. Umadevi, Paras Sehgal, Avula Renuka Devi, A. Surekha, Pulala Chandra, Rajamadugu Hymavathy, P R Vanaja, Vinod Scaria, Sridhar Sivasubbu |
| EPI_ISL_865892, EPI_ISL_865898 | University College London, Great Ormond Street Hospital for Children NHS Foundation Trust, Imperial College Healthcare NHS Trust | COVID-19 Genomics UK (COG-UK) Consortium | Sergi Castellano, Rachel Williams, Mark Kristiansen, Paola Resende Silva, Sunando Roy, Tony Brooks, Helena Tutill, Paola Niola, Patricia Dyal, Charlotte Williams, Leysa Forrest, Yasmin Panchbhaya, Jacqueline Findlay, Samuel Weeks, Julianne Brown, Kathryn Harris, Paul Randell, James Price, Alison Holmes, Judith Breuer |
| EPI_ISL_872601, EPI_ISL_872602, EPI_ISL_872603 | Nigeria Centre for Disease Control (NCDC) | African Centre of Excellence for Genomics of Infectious Diseases (ACEGID), Redeemer's University | Oluniyi P.E. et al |

| | | | |
|---|---|---|---|
| EPI_ISL_875542, EPI_ISL_875546, EPI_ISL_875547 | Instituto de Biotecnologia - UNESP-Botucatu-SP | Instituto de Biotecnologia - UNESP-Botucatu-SP | Leila Sabrina Ullmann; Fábio Sossai Possebon, Camila Dantas Malossi, Paula Rahal, Paulo Inacio da Costa, João Pessoa Araújo Jr. |
| EPI_ISL_876620, EPI_ISL_876621, EPI_ISL_876622, EPI_ISL_876623, EPI_ISL_876624 | Toronto Invasive Bacterial Diseases Network | McMaster University | Allison McGeer, Patryk Aftanas, Hooman Derakhshani, Angel Li, Kuganya Nirmalarajah, Emily Panousis, Ahmed Draia, Jalees Nasir, Michael Surette, Samira Mubareka, Andrew G. McArthur |
| EPI_ISL_876828, EPI_ISL_876946, EPI_ISL_876947, EPI_ISL_876948, EPI_ISL_876949, EPI_ISL_876950, EPI_ISL_876951, EPI_ISL_876952 | Quest Diagnostics | Quest Diagnostics | Rosenthal,S.H., Gerasimova,A., Kagan,R.M., Anderson, B., Hua, M., Liu Y., Bernstein, L.E., Livingston, K.E., Perez, A., Shalhout, D.F., Shlyakhter, I.A., Owen, R., Tanpaiboon, P., Lacbawan, F. |
| EPI_ISL_877664, EPI_ISL_877665, EPI_ISL_877666, EPI_ISL_877667, EPI_ISL_877668, EPI_ISL_877669, EPI_ISL_877670, EPI_ISL_877671, EPI_ISL_877672, EPI_ISL_877673, EPI_ISL_877674, EPI_ISL_877675 | | | |
| see above | Clinical Molecular Microbiology Laboratory, UNC Hospital | Dirk Dittmer | Razia Moorad , Justin T. Landis , Brent A. Eason, Melissa B. Miller, Linda Pluta, Dirk Dittmer, Angelica Juarez, Cecilia Thompson , Cameroon Grant, Evelyn Hoffman, Patricio Cano, Jason Wong, Carolina Caro-Vegas, Blossom Damania. |
| EPI_ISL_878554 | Robert Garry lab | Andersen lab at Scripps Research | Allison Smither, Gilberto Sabino-Santos, Patricia Snarski, Lilia Melnik, Antoinette Bell, Kaylynn Genemaras, Arnaud Drouin, Dahlene Fusco, Robert Garry with SEARCH Alliance San Diego |
| EPI_ISL_884316, EPI_ISL_884339, EPI_ISL_884357, EPI_ISL_884386, EPI_ISL_884394, EPI_ISL_884395, EPI_ISL_884406 | Infectious Diseases, Quest Diagnostics | Infectious Diseases, Quest Diagnostics | Rosenthal,S.H., Gerasimova,A., Kagan,R.M., Anderson,B., Bernstein,L.E., Livingston,K.E., Hua,M., Liu,Y., Shalhout,D.F., Owen,R., Lacbawan,F. |
| EPI_ISL_884854 | Department of Biochemistry, Cell and Molecular Biology, West African Centre for Cell Biology of Infectious Pathogens (WACCBIP), University of Ghana | Department of Biochemistry, Cell and Molecular Biology, West African Centre for Cell Biology of Infectious Pathogens (WACCBIP), University of Ghana | Ngoi,J.M., Tei-Maya,F., Morang'a,C.M., Magnussen,V., Amuzu,D.S., Mohammed,A., Tapela,K., Kibinge,N., Diallo,A.B., Kumi-Ansah,F., Odoom,T., Boakye,O.D., Amoako,E., Abass,A.-K., Quashie,P., Amenga-Etego,L.N., Akoriyea,S.K., Awandare,G.A., Bediako,Y. |
| EPI_ISL_887150, EPI_ISL_887151 | Massachusetts General Hospital | Infectious Disease Program, Broad Institute of Harvard and MIT | Lemieux,J.E., Siddle,K.J., Shaw,B., Adams,G., Pierce,V., Turbett,S., Anahtar,M., Branda,J., Slater,D., Harris,J., Lin,A.E., Gladden-Young,A., Lagerborg,K., Rudy,M., DeRuff,K., Carter,A., Normandin,E., Bauer,M., Reilly,S., Tomkins-Tinch,C., Loreth,C., Chaluvadi,S., Neumann,A., Cusick,C., Chapman,S.B., Gnirke,A., Flowers,K., Cerrato,F., Birren,B.W., Gallagher,G., Smole,S., Park,D.J., MacInnis,B.L., Ryan,E., LaRocque,R., Rosenberg,E. and Sabeti,P.C. |
| EPI_ISL_889337, EPI_ISL_889338, EPI_ISL_889339, EPI_ISL_889345 | The University Hospital Brno | Institute of Applied Biotechnologies a.s. | Petr Klempt, Ondej Brzo, Martin Kašný, Kateina Kvapilová, Martina Lengerová, Petr Kvapil |
| EPI_ISL_889356, EPI_ISL_889357, EPI_ISL_889358, EPI_ISL_889364, EPI_ISL_889365, EPI_ISL_889366, EPI_ISL_889367, EPI_ISL_889368, EPI_ISL_889369 | Motol University Hospital | Institute of Applied Biotechnologies a.s. | Petr Klempt, Ondej Brzo, Martin Kašný, Kateina Kvapilová, Pavel Devínek, Petr Kvapil |
| EPI_ISL_890117, EPI_ISL_890118, EPI_ISL_890119, EPI_ISL_890120, EPI_ISL_890121, EPI_ISL_890122, EPI_ISL_890123, EPI_ISL_890124, EPI_ISL_890125, EPI_ISL_890126, EPI_ISL_890127, EPI_ISL_890128, EPI_ISL_890129, EPI_ISL_890130, EPI_ISL_890131, EPI_ISL_890132, EPI_ISL_890133, EPI_ISL_890134, EPI_ISL_890135, EPI_ISL_890136, EPI_ISL_890137, EPI_ISL_890138, EPI_ISL_890139, EPI_ISL_890140, EPI_ISL_890141, EPI_ISL_890142, EPI_ISL_890143, EPI_ISL_890144, EPI_ISL_890145, EPI_ISL_890146, EPI_ISL_890147, EPI_ISL_890148, EPI_ISL_890149, EPI_ISL_890150, EPI_ISL_890151, EPI_ISL_890152, EPI_ISL_890153, EPI_ISL_890154, EPI_ISL_890155, EPI_ISL_890156, EPI_ISL_890157, EPI_ISL_890158, EPI_ISL_890159 | | | |
| see above | Laboratoire de santé publique du Québec | Laboratoire de santé publique du Québec | Sandrine Moreira, Ioannis Ragoussis, Guillaume Bourque, Jesse Shapiro, Mark Lathrop and Michel Roger on behalf of the CoVSeQ research group |
| EPI_ISL_891272 | Institute of Biocides and Medical Ecology, Belgarde, Serbia | Virology department Institute of microbiology and immunology Faculty of Medicine University of Belgrade | Banko Ana, Miljanovic Danijela, Milicevic Ognjen, Loncar Ana, Abazovic Dzihan, Despot Dragana |
| EPI_ISL_892236 | Lighthouse Lab in Glasgow | Wellcome Sanger Institute for the COVID-19 Genomics UK (COG-UK) Consortium | Harper VanSteenhouse, Yumi Kasai, David Gray, Carol Clugston, Anna Dominiczak and Alex Alderton, Roberto Amato, Sonia Goncalves, Ewan Harrison, David K. Jackson, Ian Johnston, Dominic Kwiatkowski, Cordelia Langford, John Sillitoe on behalf of the Wellcome Sanger Institute COVID-19 Surveillance Team |
| EPI_ISL_896126, EPI_ISL_900090, EPI_ISL_900109, EPI_ISL_900150, EPI_ISL_900164, EPI_ISL_900184, EPI_ISL_900186, EPI_ISL_900192, EPI_ISL_900222, EPI_ISL_900232, EPI_ISL_900235, EPI_ISL_900239, EPI_ISL_900248, EPI_ISL_900250, EPI_ISL_900251, EPI_ISL_900255, EPI_ISL_900256, EPI_ISL_900272, EPI_ISL_900274, EPI_ISL_900281, EPI_ISL_900294, EPI_ISL_900322, EPI_ISL_900328, EPI_ISL_900333, EPI_ISL_900343, EPI_ISL_900345, EPI_ISL_900346, EPI_ISL_900352, EPI_ISL_900386, EPI_ISL_900387, EPI_ISL_900395, EPI_ISL_900404, EPI_ISL_900416, EPI_ISL_900428, EPI_ISL_900433, EPI_ISL_900437, EPI_ISL_900444, EPI_ISL_900459, EPI_ISL_900478, EPI_ISL_900481 | | | |
| see above | MEPHI, Aix Marseille University | MEPHI, Aix Marseille University | Anthony LEVASSEUR |
| EPI_ISL_900739, EPI_ISL_900740, EPI_ISL_900741 | Bozeman Health Deaconess Hospital | Wiedenheft lab, Montana State University | Artem Nemudryi, Anna Nemudraia, Tanner Wiegand, Joseph Nichols, Deann T. Snyder, Jodi F. Hedges, Calvin Cicha, Helen Lee, Karl K. Vanderwood, Diane Bimczok, Mark A. Jutila and Blake Wiedenheft |
| EPI_ISL_902741 | Hospital Universitari Germans Trias i Pujol (HUGTiP) / Fundació Lluita contra la SIDA (FLSida) | IrsiCaixa - Can Ruti CovidSeq | Fundació irsiCaixa. Hospital Universitari Germans Trias i Pujol(HUGTiP), 2a planta, maternal Ctra Canyet s/n, Badalona Marta Massanella, Ester Ballana, Lidia Ruiz, Nuria Izquierdo, Jorge Carrillo, Roger Paredes, Julia Blanco, Joaquim Segalés, Bonaventura Clotet |
| EPI_ISL_911633 | Tanjungpura University Hospital | Tanjungpura University Hospital | Mahyarudin; Andriani; Virhan Novianry; Delima Fajar Liana ; Sofi Siti Shofiyah; Puji Astuti ; Muhammad Ibnu Kahtan ; Ambar Rialita; Eka Ardiani Putri; Wiwik Windarti; Helmi Sastriawan |
| EPI_ISL_914588, EPI_ISL_914593, EPI_ISL_914594, EPI_ISL_914595, EPI_ISL_914596, EPI_ISL_914599, EPI_ISL_914600, EPI_ISL_914601, EPI_ISL_914602, EPI_ISL_914603, EPI_ISL_914604, EPI_ISL_914605, EPI_ISL_914606, EPI_ISL_914607, EPI_ISL_914609, EPI_ISL_914610, EPI_ISL_914611, EPI_ISL_914613, EPI_ISL_914614, EPI_ISL_914615, EPI_ISL_914616 | | | |
| see above | TGen North | TGen North | "Jolene Bowers, Megan Folkerts, Chris French, Hayley Yaglom, Ashlyn Pfeiffer, Darrin Lemmer, Dave Engelthaler, The Arizona COVID Genomics Union (ACGU)" |
| EPI_ISL_915392, EPI_ISL_915393, EPI_ISL_915394 | Keio University School of Medicine | Keio University School of Medicine | Kenjiro Kosaki, Yuka Iwasaki, Hirotsugu Ishizu, Haruhiko Siomi, Kodai Abe |
| EPI_ISL_918411 | Institute of Biocides and Medical Ecology | Virology Department Institute of Microbiology and Immunology Faculty of Medicine University of Belgrade | Banko Ana, Miljanovic Danijela, Milicevic Ognjen, Loncar Ana, Abazovic Dzihan, Despot Dragana |
| EPI_ISL_925407, EPI_ISL_925409, EPI_ISL_925410, EPI_ISL_925415, EPI_ISL_925416, EPI_ISL_925417 | Department of Clinical Microbiology | GIGA Medical Genomics | Keith Durkin, Maria Artesi, Sébastien Bontems, Raphaël Boreux, Bouchra Boujemla, Cécile Meex, Pierrette Melin, Marie-Pierre Hayette, Vincent Bours |
| EPI_ISL_933658, EPI_ISL_933659 | Toronto Invasive Bacterial Diseases Network | McMaster University | Allison McGeer, Patryk Aftanas, Hooman Derakhshani, Angel Li, Kuganya Nirmalarajah, Emily Panousis, Ahmed Draia, Jalees Nasir, Michael Surette, Samira Mubareka, Andrew G. McArthur |
| EPI_ISL_935790, EPI_ISL_935792, EPI_ISL_935795, EPI_ISL_935796, EPI_ISL_935801 | Cadham Provincial laboratory | National Microbiology Laboratory (NML) | Anna Majer, Shari Tyson, Grace Seo, Philip Mabon, Elsie Grudeski, Rhiannon Huzarewich, Russell Mandes, Anneliese Landgraff, Jennifer Tanner, Natalie Knox, Morag Graham, Gary Van Domselaar, Paul Van Caeseele, Jared Bullard, David Alexander, Kerry Dust, Nathalie Bastien, Yan Li, Timothy Booth, Darian Hole, Madison Chapel, Kirsten Biggar, CanCOGeN's metadata curation team, Public Health Agency of Canada CanCOGeN team |
| EPI_ISL_937097 | Quest Diagnostics | Quest Diagnostics | Rosenthal,S.H., Gerasimova,A., Kagan,R.M., Anderson, B., Livingston, K.E., Hua, M., Liu Y., Shalhout, D.F., Owen, R., Lacbawan, F. |
| EPI_ISL_937514 | Cadham Provincial laboratory | National Microbiology Laboratory (NML) | Anna Majer, Shari Tyson, Grace Seo, Philip Mabon, Elsie Grudeski, Rhiannon Huzarewich, Russell Mandes, Anneliese Landgraff, Jennifer Tanner, Natalie Knox, Morag Graham, Gary Van Domselaar, Paul Van Caeseele, Jared Bullard, David Alexander, Kerry Dust, Nathalie Bastien, Yan Li, Timothy Booth, Darian Hole, Madison Chapel, Kirsten Biggar, CanCOGeN's metadata curation team, Public Health Agency of Canada CanCOGeN team |
| EPI_ISL_940756, EPI_ISL_940895, EPI_ISL_940896, EPI_ISL_940897, EPI_ISL_940899, EPI_ISL_940926, EPI_ISL_940929, EPI_ISL_940930, EPI_ISL_940931, EPI_ISL_940932, EPI_ISL_940934, EPI_ISL_940935 | | | |
| see above | Centers for Disease Control and Prevention, Dengue Branch | Centers for Disease Control and Prevention, Dengue Branch | Gilberto A. Santiago, Glenda Gonzalez, Betzabel Flores, Keyla Charriez, Gabriela Paz-Bailey, Jorge L. Munoz-Jordan |

| | | | |
|---|---|---|---|
| EPI_ISL_942897, EPI_ISL_942898 | Lacen_RS | CEVS_SES_RS | Barcellos R, Campos A, Crescente L, Da Silva A, Dornelles C, Fonseca V, Garay L, Godinho F, Gonzalez A, Gregianini T, Molina C, Salvato R, Schaurich A |
| EPI_ISL_943581, EPI_ISL_943582, EPI_ISL_943583, EPI_ISL_943584, EPI_ISL_943585, EPI_ISL_943586, EPI_ISL_943588 | Lacen_RS | State Center for Health Surveillance. Rio Grande do Sul State Secretary of Health | Aline Campos, Amanda da Silva, Anelise Schaurich, Claudia Dornelles, Cynthia Molina, Fernanda Godinho, Lara Crescente, Leticia Garay, Regina Barcellos, Richard Salvato, Tatiana Gregianini, Vagner Fonseca |
| EPI_ISL_949202, EPI_ISL_949206, EPI_ISL_949213, EPI_ISL_949217, EPI_ISL_949219, EPI_ISL_949226, EPI_ISL_949227, EPI_ISL_949236, EPI_ISL_949238 | Departamento de Microbiología, CDB, Hospital Clínic, Barcelona | SeqCOVID-SPAIN consortium/IBV(CSIC) | Andrea Vergara, Mikel Martínez, Elisa Rubio, Jéssica Navero, Aida Peiró and SeqCOVID-SPAIN consortium |
| EPI_ISL_953320 | Bioinformatics and Biostatistics Lab, Advanced Sequencing Facility | COVID-19 Genomics UK (COG-UK) Consortium | Aengus Stewart,Jerome Nicod,Chelsea Sawyer,Laura Cubitt,Harshil Patel,Margaret Crawford |
| EPI_ISL_953411 | Laboratorio de Investigaciones de Baney | "Swiss Tropical and Public Health Institute" | "Carlos Cortes, Claudia Daubenberger, Guillermo Garcia, Salome Hosch, Bonifacio Manguire Nlavo, Maximilian Mpina, Elizabeth Nyakarungu, Diosdado Odjama Nseng Ada, Mitoha Ondo O Ayekaba, Tobias Schindler, Philip Wonder Phiri" |
| EPI_ISL_959901, EPI_ISL_959902, EPI_ISL_959906, EPI_ISL_960072, EPI_ISL_960073, EPI_ISL_960074, EPI_ISL_960075, EPI_ISL_960076, EPI_ISL_960077, EPI_ISL_960078, EPI_ISL_960079, EPI_ISL_960080, EPI_ISL_960081, EPI_ISL_960082, EPI_ISL_960083, EPI_ISL_960084, EPI_ISL_960085, EPI_ISL_960086, EPI_ISL_960087, EPI_ISL_960088 | | | |
| see above | University Medical Center Hamburg Eppendorf | Heinrich Pette Institute, Leibniz Institute for Experimental Virology | Alexis Robitaille, Thomas Günther, Johannes Knobloch, Martin Aepfelbacher, Nicole Fischer, Adam Grundhoff |
| EPI_ISL_961761, EPI_ISL_961762, EPI_ISL_961763 | Laboratorio de Infectología, Servicio de Infectología, Hospital Universitario Dr. José Eleuterio González - Universidad Autónoma de Nuevo León | Laboratorio de Infectología Molecular, Departamento de Bioquímica y Medicina Molecular,Facultad de Medicina - Universidad Autónoma de Nuevo León | Kame A. Galán-Huerta, María F. Herrera-Saldivar, Natalia Martínez-Acuña, Sonia A. Lozano-Sepúlveda, Daniel Arellanos-Soto, Ana M. Rivas-Estilla, Paola Bocanegra-Ibarias, Samantha M. Flores-Treviño, Elvira Garza-González, Eduardo Perez-Alba, Laura Nuzzolo-Shihadeh, Adrian Camacho-Ortiz |
| EPI_ISL_961886, EPI_ISL_961980 | Illinois Department of Public Health | Gagnon Lab, Southern Illinois University | Keith Gagnon |
| EPI_ISL_964906 | Hospital Castro Rendon | Laboratorio Central Mg. Luis Alfredo Pianciola on behalf of 'Proyecto Argentino Interinstitucional de genomica de SARS-CoV-2' (PAIS Consortium) | L Pianciola, M Mazzeo, C Ziehm, C Pintos, M Fernandez, J Ousset, M Nabaes, M Viegas. |
| EPI_ISL_964909 | Hospital Bouquet Roldan | Laboratorio Central Mg. Luis Alfredo Pianciola on behalf of 'Proyecto Argentino Interinstitucional de genomica de SARS-CoV-2' (PAIS Consortium) | L Pianciola, M Mazzeo, C Ziehm, C Pintos, M Fernandez, J Ousset, M Nabaes, M Viegas. |
| EPI_ISL_964913 | Sistema de Emergencias | Laboratorio Central Mg. Luis Alfredo Pianciola on behalf of 'Proyecto Argentino Interinstitucional de genomica de SARS-CoV-2' (PAIS Consortium) | L Pianciola, M Mazzeo, C Ziehm, C Pintos, M Fernandez, J Ousset, M Nabaes, M Viegas. |
| EPI_ISL_968165 | Clinical Molecular Microbiology Laboratory, UNC Hospital | Dirk Dittmer | Justin T. Landis , Razia Moorad , Brent A. Eason, Melissa B. Miller, Linda Pluta, Dirk Dittmer, Angelica Juarez, Cecilia Thompson, Shawn Hawken, Cameroon Grant, Evelyn Hoffman, Patricio Cano, Jason Wong, Carolina Caro-Vegas, Ryan McNamara, Blossom Damania. |
| EPI_ISL_968463, EPI_ISL_968464, EPI_ISL_968465, EPI_ISL_968466, EPI_ISL_968467, EPI_ISL_968468, EPI_ISL_968469, EPI_ISL_968470, EPI_ISL_968471, EPI_ISL_968472, EPI_ISL_968473, EPI_ISL_968474, EPI_ISL_968475, EPI_ISL_968476, EPI_ISL_968477, EPI_ISL_968478, EPI_ISL_968479, EPI_ISL_968480, EPI_ISL_968481, EPI_ISL_968482, EPI_ISL_968483, EPI_ISL_968484, EPI_ISL_968485, EPI_ISL_968486, EPI_ISL_968487, EPI_ISL_968488, EPI_ISL_968489, EPI_ISL_968490, EPI_ISL_968491, EPI_ISL_968492, EPI_ISL_968493, EPI_ISL_968494, EPI_ISL_968495, EPI_ISL_968496, EPI_ISL_968497, EPI_ISL_968498, EPI_ISL_968499, EPI_ISL_968500, EPI_ISL_968501, EPI_ISL_968502, EPI_ISL_968503, EPI_ISL_968504, EPI_ISL_968505, EPI_ISL_968506, EPI_ISL_968507, EPI_ISL_968508, EPI_ISL_968509, EPI_ISL_968510, EPI_ISL_968511, EPI_ISL_968512, EPI_ISL_968513, EPI_ISL_968514, EPI_ISL_968515, EPI_ISL_968516, EPI_ISL_968517, EPI_ISL_968518, EPI_ISL_968519, EPI_ISL_968520, EPI_ISL_968521, EPI_ISL_968522, EPI_ISL_968523, EPI_ISL_968524, EPI_ISL_968525, EPI_ISL_968526, EPI_ISL_968527, EPI_ISL_968528, EPI_ISL_968529, EPI_ISL_968530, EPI_ISL_968531, EPI_ISL_968532, EPI_ISL_968533, EPI_ISL_968534, EPI_ISL_968535, EPI_ISL_968536, EPI_ISL_968537, EPI_ISL_968538, EPI_ISL_968539, EPI_ISL_968540, EPI_ISL_968541, EPI_ISL_968542, EPI_ISL_968543, EPI_ISL_968544, EPI_ISL_968545, EPI_ISL_968546, EPI_ISL_968547, EPI_ISL_968548, EPI_ISL_968549, EPI_ISL_968550, EPI_ISL_968551, EPI_ISL_968552, EPI_ISL_968553, EPI_ISL_968554, EPI_ISL_968555, EPI_ISL_968556, EPI_ISL_968557, EPI_ISL_968558, EPI_ISL_968559, EPI_ISL_968560, EPI_ISL_968561, EPI_ISL_968562, EPI_ISL_968563, EPI_ISL_968564, EPI_ISL_968565, EPI_ISL_968566, EPI_ISL_968567, EPI_ISL_968568, EPI_ISL_968569, EPI_ISL_968570, EPI_ISL_968571, EPI_ISL_968572, EPI_ISL_968573, EPI_ISL_968574, EPI_ISL_968575, EPI_ISL_968576, EPI_ISL_968577, EPI_ISL_968578, EPI_ISL_968579, EPI_ISL_968580, EPI_ISL_968581, EPI_ISL_968582, EPI_ISL_968583, EPI_ISL_968584, EPI_ISL_968585, EPI_ISL_968586, EPI_ISL_968587 | | | |
| see above | BCCDC Public Health Laboratory | BCCDC Public Health Laboratory | Prystajecky Natalie, Linda Hoang, Dan Fornika, John Tyson, Shannon Russell, Kim Macdonald, Kimia Kamelian, Ana Pacagnella, Corrinne Ng, Loretta Janz, Robert Azana Terry Snutch, Mel Krajden |
| EPI_ISL_977255, EPI_ISL_977256, EPI_ISL_977277, EPI_ISL_977283, EPI_ISL_977284, EPI_ISL_977285, EPI_ISL_977300, EPI_ISL_977301, EPI_ISL_977302, EPI_ISL_977331, EPI_ISL_977332, EPI_ISL_977460, EPI_ISL_977461, EPI_ISL_977462, EPI_ISL_977463, EPI_ISL_977464, EPI_ISL_977465, EPI_ISL_977466 | | | |
| see above | University of Zambia, School of Veterinary Medicine | UNZAVET and PATH | Mulenga Mwenda-Chimfwembe, Ngonda Saasa, Daniel Bridges |
| EPI_ISL_977471 | Instituto Adolfo Lutz - Regional de Presidente Prudente | Instituto Adolfo Lutz, Interdiciplinary Procedures Center, Strategic Laboratory | Claudio Tavares Sacchi, Claudia Regina Gonçalves, Erica Valessa Ramos Gomes, Karoline Rodrigues Campos |
| EPI_ISL_977472 | Instituto Adolfo Lutz Central | Instituto Adolfo Lutz, Interdiciplinary Procedures Center, Strategic Laboratory | Claudio Tavares Sacchi, Claudia Regina Gonçalves, Erica Valessa Ramos Gomes, Karoline Rodrigues Campos |
| EPI_ISL_977478 | Instituto Adolfo Lutz - Regional de Presidente Prudente | Instituto Adolfo Lutz, Interdiciplinary Procedures Center, Strategic Laboratory | Claudio Tavares Sacchi, Claudia Regina Gonçalves, Erica Valessa Ramos Gomes, Karoline Rodrigues Campos |
| EPI_ISL_977479 | Lab Loc - Itapecerica da Serra | Instituto Adolfo Lutz, Interdiciplinary Procedures Center, Strategic Laboratory | Claudio Tavares Sacchi, Claudia Regina Gonçalves, Erica Valessa Ramos Gomes, Karoline Rodrigues Campos |
| EPI_ISL_977483 | Instituto Adolfo Lutz Central | Instituto Adolfo Lutz, Interdiciplinary Procedures Center, Strategic Laboratory | Claudio Tavares Sacchi, Claudia Regina Gonçalves, Erica Valessa Ramos Gomes, Karoline Rodrigues Campos |
| EPI_ISL_978300 | Texas Department of State Health Services | Texas Department of State Health Services | Bonnie Oh, Anita Pokharel, James Daniel Bonser, Myong Koag, Chung Wang, Rachel Lee, Grace Kubin, Rashmi Tuladhar, Mayela Pedrueza, Maliha Rahman, Jenny Zhang |
| EPI_ISL_978499 | Central Public Health Laboratory - LACEN -Bahia, Salvador, Brazil | Central Public Health Laboratory - LACEN -Bahia, Salvador, Brazil | Stephane Tosta, Luciana Oliveira, Vanessa Nardy,Patrícia Cajado,Marcela Gómez, Breno Dominguez, Jaqueline Gomes, Vagner Fonseca,Marta Giovanetti,Luiz Alcantara, Felicidade Pereira, Arabela Leal |
| EPI_ISL_981033 | hospital Cipolleti | Laboratorio Central Mg. Luis Alfredo Pianciola on behalf of 'Proyecto Argentino Interinstitucional de genomica de SARS-CoV-2' (PAIS Consortium) | L Pianciola, M Mazzeo, C Ziehm, C Pintos, M Fernandez, J Ousset, M Nabaes, M Viegas. |
| EPI_ISL_981036, EPI_ISL_981037 | Hospital Dr. Francisco López Lima | Laboratorio Central Mg. Luis Alfredo Pianciola on behalf of 'Proyecto Argentino Interinstitucional de genomica de SARS-CoV-2' (PAIS Consortium) | L Pianciola, M Mazzeo, C Ziehm, C Pintos, M Fernandez, J Ousset, M Nabaes, M Viegas. |
| EPI_ISL_981045 | Hospital Bariloche | Laboratorio Central Mg. Luis Alfredo Pianciola on behalf of 'Proyecto Argentino Interinstitucional de genomica de SARS-CoV-2' (PAIS Consortium) | L Pianciola, M Mazzeo, C Ziehm, C Pintos, M Fernandez, J Ousset, M Nabaes, M Viegas. |
| EPI_ISL_981047 | Hospital Jacobacci | Laboratorio Central Mg. Luis Alfredo Pianciola on behalf of 'Proyecto Argentino Interinstitucional de genomica de SARS-CoV-2' (PAIS Consortium) | L Pianciola, M Mazzeo, C Ziehm, C Pintos, M Fernandez, J Ousset, M Nabaes, M Viegas. |
