## Supplementary material for "Detection and characterization of the SARS-CoV-2 lineage B.1.526 in New York": Supp. Table 3 GISAID Acknowledgment part 2: gisaid_hcov-19_acknowledgement_table_2021_02_12_23-5.pdf

We gratefully acknowledge the following Authors from the Originating laboratories responsible for obtaining the specimens, as well as the Submitting laboratories where the genome data were generated and shared via GISAID, on which this research is based.

All Submitters of data may be contacted directly via www.gisaid.org

Authors are sorted alphabetically.

| Accession ID | Originating Laboratory | Submitting Laboratory | Authors |
|---|---|---|---|
| EPI_ISL_526678, EPI_ISL_526679, EPI_ISL_526680, EPI_ISL_526681, EPI_ISL_526682, EPI_ISL_526683, EPI_ISL_526684, EPI_ISL_526685 | Florida Bureau of Public Health Laboratories | Florida Bureau of Public Health Laboratories | Sarah Schmedes, Jason Blanton |
| EPI_ISL_526997, EPI_ISL_526998 | Biological prevention, army | Biological prevention, army | Seadawy, M.G., Harty,B.E., Gad,A.F., Elhoseiny,M.F., Shamel,M.D., Shabaan,A.E., Ageez,A.M. |
| EPI_ISL_526999, EPI_ISL_527000, EPI_ISL_527001 | Biological prevention, army | Biological prevention, army | Seadawy,M.G., Harty,B.E., Gad,A.F., Elhoseiny,M.F., Shamel,M.D., Shabaan,A.E., Ageez,A.M. |
| EPI_ISL_527002 | Biological prevention, army | Biological prevention, army | Seadawy, M.G., Harty,B.E., Gad,A.F., Elhoseiny,M.F., Shamel,M.D., Shabaan,A.E., Ageez,A.M. |
| EPI_ISL_527003, EPI_ISL_527004, EPI_ISL_527005, EPI_ISL_527006 | Biological prevention, army | Biological prevention, army | Seadawy,M.G., Harty,B.E., Gad,A.F., Elhoseiny,M.F., Shamel,M.D., Shabaan,A.E., Ageez,A.M. |
| EPI_ISL_527007 | Biological Prevention, Army | Biological Prevention, Army | Seadawy, M.G., Harty,B.E., Gad,A.F., Elhoseiny,M.F., Shamel,M.D., Shabaan,A.E., Ageez,A.M. |
| EPI_ISL_527179, EPI_ISL_527180 | Area of Virology, Serology and Virology Division (SAViD), New South Wales Health Pathology Randwick | Area of Virology, Serology and Virology Division (SAViD), New South Wales Health Pathology Randwick | Rawlinson, W. |
| EPI_ISL_527362, EPI_ISL_527363, EPI_ISL_527378, EPI_ISL_527379 | National Public Health Laboratory, National Centre for Infectious Diseases | National Public Health Laboratory, National Centre for Infectious Diseases | Mak TM, Octavia S, Zhou Z, Cui L, Lin RTP |
| EPI_ISL_527400 | Area of Virology, Serology and Virology Division (SAViD), New South Wales Health Pathology Randwick | Area of Virology, Serology and Virology Division (SAViD), New South Wales Health Pathology Randwick | Rawlinson, W., Deveson, I., Bull, R. |
| EPI_ISL_528386 | Viral vaccines, VSVRI- Veterinary serum and vaccine research institute | Viral vaccines, VSVRI- Veterinary serum and vaccine research institute | Saleh,A.A., Saad,M.A. |
| EPI_ISL_528395 | Respiratory Virus Unit, Microbiology Services Colindale, Public Health England | Respiratory Virus Unit, Microbiology Services Colindale, Public Health England | PHE Covid Sequencing Team |
| EPI_ISL_528683 | NCDC Institute of Genomics and Integrative Biology | NCDC Institute of Genomics and Integrative Biology | Vivekanand A, Mahesh S. Dhar, Bharathram Uppili, Akshay Kanakan, Simmi Tiwari, RadhaKrishnan VS, Robin Marwal, Azka Khan, Ajit Shewale, Pooja Sharma, Tushar Nale, Rajesh Pandey, Sandhya Kabra, Mohammed Faruq, Sujeet Singh, Anurag Agrawal, Partha Rakshit |
| EPI_ISL_529227, EPI_ISL_529228, EPI_ISL_529229, EPI_ISL_529230, EPI_ISL_529231, EPI_ISL_529232, EPI_ISL_529233 | Wales Specialist Virology Centre Sequencing lab: Pathogen Genomics Unit | COVID-19 Genomics UK (COG-UK) Consortium | Catherine Moore, Johnathan Evans, Laura Gifford, Malorie Perry, Simon Cottrell, Angela Marchbank, Alec Birchley, Alexander Adams, Amy Gaskin, Bree Gatica-Wilcox, Jason Coombes, Joel Southgate, Lauren Gilbert, Lee Graham, Nicole Pacchiarini, Sara Kumziene-Summerhayes, Sarah Taylor, Sophie Jones, Sara Rey, Matthew Bull, Joanne Watkins, Sally Corden, Tom Connor |
| EPI_ISL_529238, EPI_ISL_529240, EPI_ISL_529241, EPI_ISL_529242, EPI_ISL_529243, EPI_ISL_529244, EPI_ISL_529245, EPI_ISL_529246, EPI_ISL_529247, EPI_ISL_529248, EPI_ISL_529249, EPI_ISL_529250 | | | |
| see above | Quadram Institute Bioscience | COVID-19 Genomics UK (COG-UK) Consortium | Dave J. Baker, Gemma L. Kay, Alp Aydin, Thanh Le-Viet, Steven Rudder, Ana P. Tedim, Anastasia Kolyva, Maria Diaz, Leonardo de Oliveira Martins, Nabil-Fareed Alikhan, Lizzie Meadows, Rachael Stanley, Ngozi Elumogo, Muhammed Yasir, Nicholas M. Thomson, Alexander J Trotter, Rachel Gilroy, Samuel Bloomfield, Claire Stuart, Andrew Bell, Reenesh Prakash, Samir Dervisevic, Alison E. Mather, John Wain, Mark Webber, Andrew J. Page, Justin O'Grady |
| EPI_ISL_529253 | Wales Specialist Virology Centre Sequencing lab: Pathogen Genomics Unit | COVID-19 Genomics UK (COG-UK) Consortium | Catherine Moore, Johnathan Evans, Laura Gifford, Malorie Perry, Simon Cottrell, Angela Marchbank, Alec Birchley, Alexander Adams, Amy Gaskin, Bree Gatica-Wilcox, Jason Coombes, Joel Southgate, Lauren Gilbert, Lee Graham, Nicole Pacchiarini, Sara Kumziene-Summerhayes, Sarah Taylor, Sophie Jones, Sara Rey, Matthew Bull, Joanne Watkins, Sally Corden, Tom Connor |
| EPI_ISL_529268 | University of Birmingham | COVID-19 Genomics UK (COG-UK) Consortium | Institute of Microbiology, University of Birmingham: Claire McMurray, Joanne Stockton, Samuel Nicholls, Radoslaw Poplawski, Will Rowe, Josh Quick, Nicholas Loman. University of Birmingham Testing Laboratory: Celina M Whalley, Andrew Bosworth, Charlotte Poxon, Kasun Wanigasooriya, Oliver Pickles, Mike Kidd, Alex Richter, Andrew D Beggs PHE Heartlands Lab: Husam Osman, Andrew Bosworth. Queen Elizabeth Hospital: Anna Casey |
| EPI_ISL_529273 | Quadram Institute Bioscience | COVID-19 Genomics UK (COG-UK) Consortium | Dave J. Baker, Gemma L. Kay, Alp Aydin, Thanh Le-Viet, Steven Rudder, Ana P. Tedim, Anastasia Kolyva, Maria Diaz, Leonardo de Oliveira Martins, Nabil-Fareed Alikhan, Lizzie Meadows, Rachael Stanley, Ngozi Elumogo, Muhammed Yasir, Nicholas M. Thomson, Alexander J Trotter, Rachel Gilroy, Samuel Bloomfield, Claire Stuart, Andrew Bell, Reenesh Prakash, Samir Dervisevic, Alison E. Mather, John Wain, Mark Webber, Andrew J. Page, Justin O'Grady |
| EPI_ISL_529285 | Virology Department, Royal Infirmary of Edinburgh, NHS Lothian / School of Biological Sciences, University of Edinburgh / Institute of Genetics and Molecular Medicine, University of Edinburgh | COVID-19 Genomics UK (COG-UK) Consortium | McHugh M, Dewar R, Rooke S, Gallagher M, Balcaza C, O'Toole Á, Scher E, Hill V, McCrone JT, Colquhoun R, Yu X, Jackson B, Rambaut A, Williams TC, Templeton K |
| EPI_ISL_529286, EPI_ISL_529290, EPI_ISL_529292, EPI_ISL_529294 | Quadram Institute Bioscience | COVID-19 Genomics UK (COG-UK) Consortium | Dave J. Baker, Gemma L. Kay, Alp Aydin, Thanh Le-Viet, Steven Rudder, Ana P. Tedim, Anastasia Kolyva, Maria Diaz, Leonardo de Oliveira Martins, Nabil-Fareed Alikhan, Lizzie Meadows, Rachael Stanley, Ngozi Elumogo, Muhammed Yasir, Nicholas M. Thomson, Alexander J Trotter, Rachel Gilroy, Samuel Bloomfield, Claire Stuart, Andrew Bell, Reenesh Prakash, Samir Dervisevic, Alison E. Mather, John Wain, Mark Webber, Andrew J. Page, Justin O'Grady |
| EPI_ISL_529315, EPI_ISL_529316 | West of Scotland Specialist Virology Centre, NHSGGC / MRC-University of Glasgow Centre for Virus Research | COVID-19 Genomics UK (COG-UK) Consortium | Ana da Silva Filipe, Natasha Johnson, Kathy Smollett, Daniel Mair, Stephen Carmichael, Lily Tong, Jenna Nichols, Elihu Aranday-Cortes, Kirstyn Brunker, Yasmin Parr, Alice Broos, Kyriaki Nomikou; Sarah McDonald, Marc Niebel, Patawee Asamaphan; Richard Orton, Joseph Hughes, Sreenu Vattipally, David L Robertson; Alasdair MacLean, Rory Gunson; Kathy Li, Natasha Jesudason, Rajiv Shah, James Shepherd, Antonia Ho, Emma Thomson |
| EPI_ISL_529317, EPI_ISL_529318 | Virology Department, Royal Infirmary of Edinburgh, NHS Lothian / School of Biological Sciences, University of Edinburgh / Institute of Genetics and Molecular Medicine, University of Edinburgh | COVID-19 Genomics UK (COG-UK) Consortium | McHugh M, Dewar R, Rooke S, Gallagher M, Balcaza C, O'Toole Á, Scher E, Hill V, McCrone JT, Colquhoun R, Yu X, Jackson B, Rambaut A, Williams TC, Templeton K |
| EPI_ISL_529334, EPI_ISL_529335, EPI_ISL_529367 | University of Birmingham | COVID-19 Genomics UK (COG-UK) Consortium | Institute of Microbiology, University of Birmingham: Claire McMurray, Joanne Stockton, Samuel Nicholls, Radoslaw Poplawski, Will Rowe, Josh Quick, Nicholas Loman. University of Birmingham Testing Laboratory: Celina M Whalley, Andrew Bosworth, Charlotte Poxon, Kasun Wanigasooriya, Oliver Pickles, Mike Kidd, Alex Richter, Andrew D Beggs PHE Heartlands Lab: Husam Osman, Andrew Bosworth. Queen Elizabeth Hospital: Anna Casey |
| EPI_ISL_529388 | Quadram Institute Bioscience | COVID-19 Genomics UK (COG-UK) Consortium | Dave J. Baker, Gemma L. Kay, Alp Aydin, Thanh Le-Viet, Steven Rudder, Ana P. Tedim, Anastasia Kolyva, Maria Diaz, Leonardo de Oliveira Martins, Nabil-Fareed Alikhan, Lizzie Meadows, Rachael Stanley, Ngozi Elumogo, Muhammed Yasir, Nicholas M. Thomson, Alexander J Trotter, Rachel Gilroy, Samuel Bloomfield, Claire Stuart, Andrew Bell, Reenesh Prakash, Samir Dervisevic, Alison E. Mather, John Wain, Mark Webber, Andrew J. Page, Justin O'Grady |

| | | | |
|---|---|---|---|
| EPI_ISL_529409 | West of Scotland Specialist Virology Centre, NHSGGC / MRC-University of Glasgow Centre for Virus Research | COVID-19 Genomics UK (COG-UK) Consortium | Ana da Silva Filipe, Natasha Johnson, Kathy Smollett, Daniel Mair, Stephen Carmichael, Lily Tong, Jenna Nichols, Elihu Aranday-Cortes, Kirstyn Brunker, Yasmin Parr, Alice Broos, Kyriaki Nomikou; Sarah McDonald, Marc Niebel, Patawee Asamaphan; Richard Orton, Joseph Hughes, Sreenu Vattipally, David L Robertson; Alasdair MacLean, Rory Gunson; Kathy Li, Natasha Jesudason, Rajiv Shah, James Shepherd, Antonia Ho, Emma Thomson |
| EPI_ISL_529413, EPI_ISL_529415, EPI_ISL_529416 | Quadram Institute Bioscience | COVID-19 Genomics UK (COG-UK) Consortium | Dave J. Baker, Gemma L. Kay, Alp Aydin, Thanh Le-Viet, Steven Rudder, Ana P. Tedim, Anastasia Kolyva, Maria Diaz, Leonardo de Oliveira Martins, Nabil-Fareed Alikhan, Lizzie Meadows, Rachael Stanley, Ngozi Elumogo, Muhammed Yasir, Nicholas M. Thomson, Alexander J Trotter, Rachel Gilroy, Samuel Bloomfield, Claire Stuart, Andrew Bell, Reenesh Prakash, Samir Dervisevic, Alison E. Mather, John Wain, Mark Webber, Andrew J. Page, Justin O'Grady |
| EPI_ISL_529418 | Wales Specialist Virology Centre Sequencing lab: Pathogen Genomics Unit | COVID-19 Genomics UK (COG-UK) Consortium | Catherine Moore, Johnathan Evans, Laura Gifford, Malorie Perry, Simon Cottrell, Angela Marchbank, Alec Birchley, Alexander Adams, Amy Gaskin, Bree Gatica-Wilcox, Jason Coombes, Joel Southgate, Lauren Gilbert, Lee Graham, Nicole Pacchiarini, Sara Kumziene-Summerhayes, Sarah Taylor, Sophie Jones, Sara Rey, Matthew Bull, Joanne Watkins, Sally Corden, Tom Connor |
| EPI_ISL_529422 | Quadram Institute Bioscience | COVID-19 Genomics UK (COG-UK) Consortium | Dave J. Baker, Gemma L. Kay, Alp Aydin, Thanh Le-Viet, Steven Rudder, Ana P. Tedim, Anastasia Kolyva, Maria Diaz, Leonardo de Oliveira Martins, Nabil-Fareed Alikhan, Lizzie Meadows, Rachael Stanley, Ngozi Elumogo, Muhammed Yasir, Nicholas M. Thomson, Alexander J Trotter, Rachel Gilroy, Samuel Bloomfield, Claire Stuart, Andrew Bell, Reenesh Prakash, Samir Dervisevic, Alison E. Mather, John Wain, Mark Webber, Andrew J. Page, Justin O'Grady |
| EPI_ISL_529442 | University of Birmingham | COVID-19 Genomics UK (COG-UK) Consortium | Institute of Microbiology, University of Birmingham: Claire McMurray, Joanne Stockton, Samuel Nicholls, Radoslaw Poplawski, Will Rowe, Josh Quick, Nicholas Loman. University of Birmingham Testing Laboratory: Celina M Whalley, Andrew Bosworth, Charlotte Poxon, Kasun Wanigasooriya, Oliver Pickles, Mike Kidd, Alex Richter, Andrew D Beggs PHE Heartlands Lab: Husam Osman, Andrew Bosworth. Queen Elizabeth Hospital: Anna Casey |
| EPI_ISL_529445 | Department of Pathology, University of Cambridge | COVID-19 Genomics UK (COG-UK) Consortium | Luke W Meredith, M. Estée Török, Myra Hosmillo, William L. Hamilton, Martin D. Curran, Theresa Feltwell, Grant Hall, Anna Yakovleva, Fahad A Khokhar, Charlotte J. Houldcroft, Laura G Caller, Aminu S. Jahun, Sarah L. Caddy, Yasmin Chaudhry, Malte Pinckert, Ian Goodfellow |
| EPI_ISL_529457, EPI_ISL_529459, EPI_ISL_529469, EPI_ISL_529476, EPI_ISL_529477, EPI_ISL_529487, EPI_ISL_529488, EPI_ISL_529492 | Quadram Institute Bioscience | COVID-19 Genomics UK (COG-UK) Consortium | Dave J. Baker, Gemma L. Kay, Alp Aydin, Thanh Le-Viet, Steven Rudder, Ana P. Tedim, Anastasia Kolyva, Maria Diaz, Leonardo de Oliveira Martins, Nabil-Fareed Alikhan, Lizzie Meadows, Rachael Stanley, Ngozi Elumogo, Muhammed Yasir, Nicholas M. Thomson, Alexander J Trotter, Rachel Gilroy, Samuel Bloomfield, Claire Stuart, Andrew Bell, Reenesh Prakash, Samir Dervisevic, Alison E. Mather, John Wain, Mark Webber, Andrew J. Page, Justin O'Grady |
| EPI_ISL_529505 | Wales Specialist Virology Centre Sequencing lab: Pathogen Genomics Unit | COVID-19 Genomics UK (COG-UK) Consortium | Catherine Moore, Johnathan Evans, Laura Gifford, Malorie Perry, Simon Cottrell, Angela Marchbank, Alec Birchley, Alexander Adams, Amy Gaskin, Bree Gatica-Wilcox, Jason Coombes, Joel Southgate, Lauren Gilbert, Lee Graham, Nicole Pacchiarini, Sara Kumziene-Summerhayes, Sarah Taylor, Sophie Jones, Sara Rey, Matthew Bull, Joanne Watkins, Sally Corden, Tom Connor |
| EPI_ISL_529506, EPI_ISL_529507 | Quadram Institute Bioscience | COVID-19 Genomics UK (COG-UK) Consortium | Dave J. Baker, Gemma L. Kay, Alp Aydin, Thanh Le-Viet, Steven Rudder, Ana P. Tedim, Anastasia Kolyva, Maria Diaz, Leonardo de Oliveira Martins, Nabil-Fareed Alikhan, Lizzie Meadows, Rachael Stanley, Ngozi Elumogo, Muhammed Yasir, Nicholas M. Thomson, Alexander J Trotter, Rachel Gilroy, Samuel Bloomfield, Claire Stuart, Andrew Bell, Reenesh Prakash, Samir Dervisevic, Alison E. Mather, John Wain, Mark Webber, Andrew J. Page, Justin O'Grady |
| EPI_ISL_529517 | University of Birmingham | COVID-19 Genomics UK (COG-UK) Consortium | Institute of Microbiology, University of Birmingham: Claire McMurray, Joanne Stockton, Samuel Nicholls, Radoslaw Poplawski, Will Rowe, Josh Quick, Nicholas Loman. University of Birmingham Testing Laboratory: Celina M Whalley, Andrew Bosworth, Charlotte Poxon, Kasun Wanigasooriya, Oliver Pickles, Mike Kidd, Alex Richter, Andrew D Beggs PHE Heartlands Lab: Husam Osman, Andrew Bosworth. Queen Elizabeth Hospital: Anna Casey |
| EPI_ISL_529521 | Virology Department, Sheffield Teaching Hospitals NHS Foundation Trust/Department of Infection, Immunity and Cardiovascular Disease, The Medical School, University of Sheffield | COVID-19 Genomics UK (COG-UK) Consortium | Thushan de Silva, Matthew Parker, Nikki Smith, Adri Angyal, Rebecca Brown, Luke Green, Rachel Tucker, Paul Parsons, Danielle Groves, Katie Johnson, Laura Carrilero, Alex Keeley, Dave Partridge, Matthew Wyles, Benjamin Lindsey, Mehmet Yavuz, Mohammad Raza, Cariad Evans |
| EPI_ISL_529526 | Quadram Institute Bioscience | COVID-19 Genomics UK (COG-UK) Consortium | Dave J. Baker, Gemma L. Kay, Alp Aydin, Thanh Le-Viet, Steven Rudder, Ana P. Tedim, Anastasia Kolyva, Maria Diaz, Leonardo de Oliveira Martins, Nabil-Fareed Alikhan, Lizzie Meadows, Rachael Stanley, Ngozi Elumogo, Muhammed Yasir, Nicholas M. Thomson, Alexander J Trotter, Rachel Gilroy, Samuel Bloomfield, Claire Stuart, Andrew Bell, Reenesh Prakash, Samir Dervisevic, Alison E. Mather, John Wain, Mark Webber, Andrew J. Page, Justin O'Grady |
| EPI_ISL_529628, EPI_ISL_529629, EPI_ISL_529630, EPI_ISL_529631, EPI_ISL_529632, EPI_ISL_529633, EPI_ISL_529634, EPI_ISL_529635, EPI_ISL_529636, EPI_ISL_529637, EPI_ISL_529652, EPI_ISL_529653 | | | |
| see above | University of Birmingham | COVID-19 Genomics UK (COG-UK) Consortium | Institute of Microbiology, University of Birmingham: Claire McMurray, Joanne Stockton, Samuel Nicholls, Radoslaw Poplawski, Will Rowe, Josh Quick, Nicholas Loman. University of Birmingham Testing Laboratory: Celina M Whalley, Andrew Bosworth, Charlotte Poxon, Kasun Wanigasooriya, Oliver Pickles, Mike Kidd, Alex Richter, Andrew D Beggs PHE Heartlands Lab: Husam Osman, Andrew Bosworth. Queen Elizabeth Hospital: Anna Casey |
| EPI_ISL_529667 | Queens Medical Centre, Clinical Microbiology Department / DeepSeq Nottingham | COVID-19 Genomics UK (COG-UK) Consortium | Gemma Clark, Wendy Smith, Manjinder Khakh, Vicki M Fleming, Michelle M Lister, Hannah Howson-Wells, Jonathan Ball, Patrick McClure, Joseph Chappell, Theocharis Tsoleridis, Nadine Holmes, Matthew Carlisle, Christopher Moore, Fei Sang, Johnny Debebe, Victoria Wright, Matthew Loose |
| EPI_ISL_529684, EPI_ISL_529685, EPI_ISL_529686, EPI_ISL_529687, EPI_ISL_529688 | West of Scotland Specialist Virology Centre, NHSGGC / MRC-University of Glasgow Centre for Virus Research | COVID-19 Genomics UK (COG-UK) Consortium | Ana da Silva Filipe, Natasha Johnson, Kathy Smollett, Daniel Mair, Stephen Carmichael, Lily Tong, Jenna Nichols, Elihu Aranday-Cortes, Kirstyn Brunker, Yasmin Parr, Alice Broos, Kyriaki Nomikou; Sarah McDonald, Marc Niebel, Patawee Asamaphan; Richard Orton, Joseph Hughes, Sreenu Vattipally, David L Robertson; Alasdair MacLean, Rory Gunson; Kathy Li, Natasha Jesudason, Rajiv Shah, James Shepherd, Antonia Ho, Emma Thomson |
| EPI_ISL_529693, EPI_ISL_529695, EPI_ISL_529696 | Virology Department, Royal Infirmary of Edinburgh, NHS Lothian / School of Biological Sciences, University of Edinburgh / Institute of Genetics and Molecular Medicine, University of Edinburgh | COVID-19 Genomics UK (COG-UK) Consortium | McHugh M, Dewar R, Rooke S, Gallagher M, Balcaza C, O'Toole Á, Scher E, Hill V, McCrone JT, Colquhoun R, Yu X, Jackson B, Rambaut A, Williams TC, Templeton K |
| EPI_ISL_529706, EPI_ISL_529708, EPI_ISL_529709, EPI_ISL_529710, EPI_ISL_529711, EPI_ISL_529712, EPI_ISL_529713, EPI_ISL_529714, EPI_ISL_529715, EPI_ISL_529716 | Wales Specialist Virology Centre Sequencing lab: Pathogen Genomics Unit | COVID-19 Genomics UK (COG-UK) Consortium | Catherine Moore, Johnathan Evans, Laura Gifford, Malorie Perry, Simon Cottrell, Angela Marchbank, Alec Birchley, Alexander Adams, Amy Gaskin, Bree Gatica-Wilcox, Jason Coombes, Joel Southgate, Lauren Gilbert, Lee Graham, Nicole Pacchiarini, Sara Kumziene-Summerhayes, Sarah Taylor, Sophie Jones, Sara Rey, Matthew Bull, Joanne Watkins, Sally Corden, Tom Connor |
| EPI_ISL_529763, EPI_ISL_529764, EPI_ISL_529765, EPI_ISL_529766, EPI_ISL_529767, EPI_ISL_529768, EPI_ISL_529769, EPI_ISL_529770, EPI_ISL_529771, EPI_ISL_529772, EPI_ISL_529773, EPI_ISL_529775, EPI_ISL_529778, EPI_ISL_529779, EPI_ISL_529780, EPI_ISL_529781, EPI_ISL_529782, EPI_ISL_529783, EPI_ISL_529784, EPI_ISL_529785, EPI_ISL_529786, EPI_ISL_529787, EPI_ISL_529788, EPI_ISL_529789, EPI_ISL_529790, EPI_ISL_529791, EPI_ISL_529792, EPI_ISL_529793, EPI_ISL_529794, EPI_ISL_529795, EPI_ISL_529797, EPI_ISL_529798, EPI_ISL_529799, EPI_ISL_529800, EPI_ISL_529801, EPI_ISL_529802, EPI_ISL_529803 | | | |
| see above | NHLS-IALCH | KRISP, KZN Research Innovation and Sequencing Platform | Giandhari J, Pillay S, Lessells R, Mdlalose K, York D, Khan S, Tegally H, Wilkinson E, de Oliveira T |
| EPI_ISL_530155, EPI_ISL_530156, EPI_ISL_530157, EPI_ISL_530158, EPI_ISL_530159, EPI_ISL_530160, EPI_ISL_530161, EPI_ISL_530162 | Seattle Flu Study | Seattle Flu Study | Deborah A. Nickerson, Chris D. Frazar, Jover Lee, Benjamin Pelle, Matthew Richardson, Amanda Adler, Elisabeth Brandstetter, Peter D. Han, Kairsten Fay, Misja Ilcisin, Kirsten Lacombe, Thomas R. Sibley, Melissa Truong, Caitlin R. Wolf, Karen Cowgill, Stephanie Schrag, Jeff Duchin, Michael Boeckh, Janet A. Englund, Michael Famulare, Barry R. Lutz, Mark J. Rieder, Lea M. Starita, Matthew Thompson, Helen Y. Chu, Trevor Bedford, Jay Shendure |
| EPI_ISL_530163, EPI_ISL_530164, EPI_ISL_530165 | M Health Fairview | Minnesota Department of Health, Public Health Laboratory | Matt Plumb, Jacob Garfin, and Xiong Wang |
| EPI_ISL_530255, EPI_ISL_530258, EPI_ISL_530259, EPI_ISL_530278, EPI_ISL_530279, EPI_ISL_530281, EPI_ISL_530282, EPI_ISL_530283 | Queensland Health Forensic and Scientific Services, Public Health Virology | Public Health Virology Laboratory, Forensic and Scientific Services, Queensland Health | Son Nguyen et al |
| EPI_ISL_530334, EPI_ISL_530335, EPI_ISL_530337, EPI_ISL_530338 | Area of Virology, Serology and Virology Division (SAViD), New South Wales Health Pathology Randwick | Area of Virology, Serology and Virology Division (SAViD), New South Wales Health Pathology Randwick | Rawlinson, W., Deveson, I., Bull, R., Van Hal, S. |

| | | | |
|---|---|---|---|
| EPI_ISL_530351, EPI_ISL_530352, EPI_ISL_530353, EPI_ISL_530354, EPI_ISL_530355, EPI_ISL_530356, EPI_ISL_530357, EPI_ISL_530358, EPI_ISL_530359, EPI_ISL_530360 | Lighthouse Lab in Glasgow | Wellcome Sanger Institute for the COVID-19 Genomics UK (COG-UK) consortium | Harper VanSteenhouse, Yumi Kasai, David Gray, Carol Clugston, Anna Dominiczak and Alex Alderton, Roberto Amato, Sonia Goncalves, Ewan Harrison, David K. Jackson, Ian Johnston, Dominic Kwiatkowski, Cordelia Langford, John Sillitoe |
| EPI_ISL_530361 | NHSGGC West of Scotland Specialist Virology Centre / MRC-University of Glasgow Centre for Virus Research | Wellcome Sanger Institute for the COVID-19 Genomics UK (COG-UK) consortium | Ana da Silva Filipe, Natasha Johnson, Kathy Smollett, Daniel Mair, Stephen Carmichael, Lily Tong, Jenna Nichols, Elihu Aranday-Cortes, Kirstyn Brunker, Yasmin Parr, Kyriaki Nomikou; Sarah McDonald, Marc Niebel, Patawee Asamaphan; Richard Orton, Joseph Hughes, Sreenu Vattipally, David L Robertson; Alasdair MacLean, Rory Gunson; Kathy Li, Natasha Jesudason, Rajiv Shah, James Shepherd, Antonia Ho, Alice Broos, Emma Thomson and Alex Alderton, Roberto Amato, Sonia Goncalves, Ewan Harrison, David K. Jackson, Ian Johnston, Dominic Kwiatkowski, Cordelia Langford, John Sillitoe |
| EPI_ISL_530362, EPI_ISL_530363, EPI_ISL_530364, EPI_ISL_530365 | Lighthouse Lab in Glasgow | Wellcome Sanger Institute for the COVID-19 Genomics UK (COG-UK) consortium | Harper VanSteenhouse, Yumi Kasai, David Gray, Carol Clugston, Anna Dominiczak and Alex Alderton, Roberto Amato, Sonia Goncalves, Ewan Harrison, David K. Jackson, Ian Johnston, Dominic Kwiatkowski, Cordelia Langford, John Sillitoe |
| EPI_ISL_530366 | NHSGGC West of Scotland Specialist Virology Centre / MRC-University of Glasgow Centre for Virus Research | Wellcome Sanger Institute for the COVID-19 Genomics UK (COG-UK) consortium | Ana da Silva Filipe, Natasha Johnson, Kathy Smollett, Daniel Mair, Stephen Carmichael, Lily Tong, Jenna Nichols, Elihu Aranday-Cortes, Kirstyn Brunker, Yasmin Parr, Kyriaki Nomikou; Sarah McDonald, Marc Niebel, Patawee Asamaphan; Richard Orton, Joseph Hughes, Sreenu Vattipally, David L Robertson; Alasdair MacLean, Rory Gunson; Kathy Li, Natasha Jesudason, Rajiv Shah, James Shepherd, Antonia Ho, Alice Broos, Emma Thomson and Alex Alderton, Roberto Amato, Sonia Goncalves, Ewan Harrison, David K. Jackson, Ian Johnston, Dominic Kwiatkowski, Cordelia Langford, John Sillitoe |
| EPI_ISL_530367, EPI_ISL_530368, EPI_ISL_530369 | Lighthouse Lab in Glasgow | Wellcome Sanger Institute for the COVID-19 Genomics UK (COG-UK) consortium | Harper VanSteenhouse, Yumi Kasai, David Gray, Carol Clugston, Anna Dominiczak and Alex Alderton, Roberto Amato, Sonia Goncalves, Ewan Harrison, David K. Jackson, Ian Johnston, Dominic Kwiatkowski, Cordelia Langford, John Sillitoe |
| EPI_ISL_530370 | NHSGGC West of Scotland Specialist Virology Centre / MRC-University of Glasgow Centre for Virus Research | Wellcome Sanger Institute for the COVID-19 Genomics UK (COG-UK) consortium | Ana da Silva Filipe, Natasha Johnson, Kathy Smollett, Daniel Mair, Stephen Carmichael, Lily Tong, Jenna Nichols, Elihu Aranday-Cortes, Kirstyn Brunker, Yasmin Parr, Kyriaki Nomikou; Sarah McDonald, Marc Niebel, Patawee Asamaphan; Richard Orton, Joseph Hughes, Sreenu Vattipally, David L Robertson; Alasdair MacLean, Rory Gunson; Kathy Li, Natasha Jesudason, Rajiv Shah, James Shepherd, Antonia Ho, Alice Broos, Emma Thomson and Alex Alderton, Roberto Amato, Sonia Goncalves, Ewan Harrison, David K. Jackson, Ian Johnston, Dominic Kwiatkowski, Cordelia Langford, John Sillitoe |
| EPI_ISL_530371, EPI_ISL_530372, EPI_ISL_530373, EPI_ISL_530374, EPI_ISL_530375, EPI_ISL_530376 | Lighthouse Lab in Glasgow | Wellcome Sanger Institute for the COVID-19 Genomics UK (COG-UK) consortium | Harper VanSteenhouse, Yumi Kasai, David Gray, Carol Clugston, Anna Dominiczak and Alex Alderton, Roberto Amato, Sonia Goncalves, Ewan Harrison, David K. Jackson, Ian Johnston, Dominic Kwiatkowski, Cordelia Langford, John Sillitoe |
| EPI_ISL_530377 | NHSGGC West of Scotland Specialist Virology Centre / MRC-University of Glasgow Centre for Virus Research | Wellcome Sanger Institute for the COVID-19 Genomics UK (COG-UK) consortium | Ana da Silva Filipe, Natasha Johnson, Kathy Smollett, Daniel Mair, Stephen Carmichael, Lily Tong, Jenna Nichols, Elihu Aranday-Cortes, Kirstyn Brunker, Yasmin Parr, Kyriaki Nomikou; Sarah McDonald, Marc Niebel, Patawee Asamaphan; Richard Orton, Joseph Hughes, Sreenu Vattipally, David L Robertson; Alasdair MacLean, Rory Gunson; Kathy Li, Natasha Jesudason, Rajiv Shah, James Shepherd, Antonia Ho, Alice Broos, Emma Thomson and Alex Alderton, Roberto Amato, Sonia Goncalves, Ewan Harrison, David K. Jackson, Ian Johnston, Dominic Kwiatkowski, Cordelia Langford, John Sillitoe |
| EPI_ISL_530378, EPI_ISL_530379, EPI_ISL_530380, EPI_ISL_530381, EPI_ISL_530382, EPI_ISL_530383, EPI_ISL_530384, EPI_ISL_530385 | Lighthouse Lab in Glasgow | Wellcome Sanger Institute for the COVID-19 Genomics UK (COG-UK) consortium | Harper VanSteenhouse, Yumi Kasai, David Gray, Carol Clugston, Anna Dominiczak and Alex Alderton, Roberto Amato, Sonia Goncalves, Ewan Harrison, David K. Jackson, Ian Johnston, Dominic Kwiatkowski, Cordelia Langford, John Sillitoe |
| EPI_ISL_530386 | NHSGGC West of Scotland Specialist Virology Centre / MRC-University of Glasgow Centre for Virus Research | Wellcome Sanger Institute for the COVID-19 Genomics UK (COG-UK) consortium | Ana da Silva Filipe, Natasha Johnson, Kathy Smollett, Daniel Mair, Stephen Carmichael, Lily Tong, Jenna Nichols, Elihu Aranday-Cortes, Kirstyn Brunker, Yasmin Parr, Kyriaki Nomikou; Sarah McDonald, Marc Niebel, Patawee Asamaphan; Richard Orton, Joseph Hughes, Sreenu Vattipally, David L Robertson; Alasdair MacLean, Rory Gunson; Kathy Li, Natasha Jesudason, Rajiv Shah, James Shepherd, Antonia Ho, Alice Broos, Emma Thomson and Alex Alderton, Roberto Amato, Sonia Goncalves, Ewan Harrison, David K. Jackson, Ian Johnston, Dominic Kwiatkowski, Cordelia Langford, John Sillitoe |
| EPI_ISL_530387 | Lighthouse Lab in Glasgow | Wellcome Sanger Institute for the COVID-19 Genomics UK (COG-UK) consortium | Harper VanSteenhouse, Yumi Kasai, David Gray, Carol Clugston, Anna Dominiczak and Alex Alderton, Roberto Amato, Sonia Goncalves, Ewan Harrison, David K. Jackson, Ian Johnston, Dominic Kwiatkowski, Cordelia Langford, John Sillitoe |
| EPI_ISL_530388 | NHSGGC West of Scotland Specialist Virology Centre / MRC-University of Glasgow Centre for Virus Research | Wellcome Sanger Institute for the COVID-19 Genomics UK (COG-UK) consortium | Ana da Silva Filipe, Natasha Johnson, Kathy Smollett, Daniel Mair, Stephen Carmichael, Lily Tong, Jenna Nichols, Elihu Aranday-Cortes, Kirstyn Brunker, Yasmin Parr, Kyriaki Nomikou; Sarah McDonald, Marc Niebel, Patawee Asamaphan; Richard Orton, Joseph Hughes, Sreenu Vattipally, David L Robertson; Alasdair MacLean, Rory Gunson; Kathy Li, Natasha Jesudason, Rajiv Shah, James Shepherd, Antonia Ho, Alice Broos, Emma Thomson and Alex Alderton, Roberto Amato, Sonia Goncalves, Ewan Harrison, David K. Jackson, Ian Johnston, Dominic Kwiatkowski, Cordelia Langford, John Sillitoe |
| EPI_ISL_530389, EPI_ISL_530390 | Lighthouse Lab in Glasgow | Wellcome Sanger Institute for the COVID-19 Genomics UK (COG-UK) consortium | Harper VanSteenhouse, Yumi Kasai, David Gray, Carol Clugston, Anna Dominiczak and Alex Alderton, Roberto Amato, Sonia Goncalves, Ewan Harrison, David K. Jackson, Ian Johnston, Dominic Kwiatkowski, Cordelia Langford, John Sillitoe |
| EPI_ISL_530391 | NHSGGC West of Scotland Specialist Virology Centre / MRC-University of Glasgow Centre for Virus Research | Wellcome Sanger Institute for the COVID-19 Genomics UK (COG-UK) consortium | Ana da Silva Filipe, Natasha Johnson, Kathy Smollett, Daniel Mair, Stephen Carmichael, Lily Tong, Jenna Nichols, Elihu Aranday-Cortes, Kirstyn Brunker, Yasmin Parr, Kyriaki Nomikou; Sarah McDonald, Marc Niebel, Patawee Asamaphan; Richard Orton, Joseph Hughes, Sreenu Vattipally, David L Robertson; Alasdair MacLean, Rory Gunson; Kathy Li, Natasha Jesudason, Rajiv Shah, James Shepherd, Antonia Ho, Alice Broos, Emma Thomson and Alex Alderton, Roberto Amato, Sonia Goncalves, Ewan Harrison, David K. Jackson, Ian Johnston, Dominic Kwiatkowski, Cordelia Langford, John Sillitoe |
| EPI_ISL_530392 | Lighthouse Lab in Glasgow | Wellcome Sanger Institute for the COVID-19 Genomics UK (COG-UK) consortium | Harper VanSteenhouse, Yumi Kasai, David Gray, Carol Clugston, Anna Dominiczak and Alex Alderton, Roberto Amato, Sonia Goncalves, Ewan Harrison, David K. Jackson, Ian Johnston, Dominic Kwiatkowski, Cordelia Langford, John Sillitoe |
| EPI_ISL_530393 | Lighthouse Lab in Glasgow | Wellcome Sanger Institute for the COVID-19 Genomics UK (COG-UK) consortium | Harper VanSteenhouse, Yumi Kasai, David Gray, Carol Clugston, Anna Dominiczak and Alex Alderton, Roberto Amato, Sonia Goncalves, Ewan Harrison, David K. Jackson, Ian Johnston, Dominic Kwiatkowski, Cordelia Langford, John Sillitoe on behalf of the Wellcome Sanger Institute COVID-19 Surveillance Team (http://www.sanger.ac.uk/covid-team) |
| EPI_ISL_530623, EPI_ISL_530630, EPI_ISL_530632, EPI_ISL_530633, EPI_ISL_530636, EPI_ISL_530637, EPI_ISL_530639, EPI_ISL_530640, EPI_ISL_530641 | Lighthouse Lab in Glasgow | Wellcome Sanger Institute for the COVID-19 Genomics UK (COG-UK) consortium | Harper VanSteenhouse, Yumi Kasai, David Gray, Carol Clugston, Anna Dominiczak and Alex Alderton, Roberto Amato, Sonia Goncalves, Ewan Harrison, David K. Jackson, Ian Johnston, Dominic Kwiatkowski, Cordelia Langford, John Sillitoe |
| EPI_ISL_530642 | NHSGGC West of Scotland Specialist Virology Centre / MRC-University of Glasgow Centre for Virus Research | Wellcome Sanger Institute for the COVID-19 Genomics UK (COG-UK) consortium | Ana da Silva Filipe, Natasha Johnson, Kathy Smollett, Daniel Mair, Stephen Carmichael, Lily Tong, Jenna Nichols, Elihu Aranday-Cortes, Kirstyn Brunker, Yasmin Parr, Kyriaki Nomikou; Sarah McDonald, Marc Niebel, Patawee Asamaphan; Richard Orton, Joseph Hughes, Sreenu Vattipally, David L Robertson; Alasdair MacLean, Rory Gunson; Kathy Li, Natasha Jesudason, Rajiv Shah, James Shepherd, Antonia Ho, Alice Broos, Emma Thomson and Alex Alderton, Roberto Amato, Sonia Goncalves, Ewan Harrison, David K. Jackson, Ian Johnston, Dominic Kwiatkowski, Cordelia Langford, John Sillitoe |
| EPI_ISL_530644, EPI_ISL_530645, EPI_ISL_530646, EPI_ISL_530652, EPI_ISL_530654, EPI_ISL_530655, EPI_ISL_530656, EPI_ISL_530657, EPI_ISL_530658 | Lighthouse Lab in Glasgow | Wellcome Sanger Institute for the COVID-19 Genomics UK (COG-UK) consortium | Harper VanSteenhouse, Yumi Kasai, David Gray, Carol Clugston, Anna Dominiczak and Alex Alderton, Roberto Amato, Sonia Goncalves, Ewan Harrison, David K. Jackson, Ian Johnston, Dominic Kwiatkowski, Cordelia Langford, John Sillitoe |
| EPI_ISL_530659 | NHSGGC West of Scotland Specialist Virology Centre / MRC-University of Glasgow Centre for Virus Research | Wellcome Sanger Institute for the COVID-19 Genomics UK (COG-UK) consortium | Ana da Silva Filipe, Natasha Johnson, Kathy Smollett, Daniel Mair, Stephen Carmichael, Lily Tong, Jenna Nichols, Elihu Aranday-Cortes, Kirstyn Brunker, Yasmin Parr, Kyriaki Nomikou; Sarah McDonald, Marc Niebel, Patawee Asamaphan; Richard Orton, Joseph Hughes, Sreenu Vattipally, David L Robertson; Alasdair MacLean, Rory Gunson; Kathy Li, Natasha Jesudason, Rajiv Shah, James Shepherd, Antonia Ho, Alice Broos, Emma Thomson and Alex Alderton, Roberto Amato, Sonia Goncalves, Ewan Harrison, David K. Jackson, Ian Johnston, Dominic Kwiatkowski, Cordelia Langford, John Sillitoe |
| EPI_ISL_530660, EPI_ISL_530661, EPI_ISL_530668, EPI_ISL_530675, EPI_ISL_530676, EPI_ISL_530677, EPI_ISL_530678, EPI_ISL_530680, EPI_ISL_530681 | Lighthouse Lab in Glasgow | Wellcome Sanger Institute for the COVID-19 Genomics UK (COG-UK) consortium | Harper VanSteenhouse, Yumi Kasai, David Gray, Carol Clugston, Anna Dominiczak and Alex Alderton, Roberto Amato, Sonia Goncalves, Ewan Harrison, David K. Jackson, Ian Johnston, Dominic Kwiatkowski, Cordelia Langford, John Sillitoe |
| EPI_ISL_530682 | NHSGGC West of Scotland Specialist Virology Centre / | Wellcome Sanger Institute for the COVID-19 Genomics UK | Ana da Silva Filipe, Natasha Johnson, Kathy Smollett, Daniel Mair, Stephen Carmichael, Lily Tong, Jenna Nichols, Elihu Aranday-Cortes, Kirstyn Brunker, |

| | | | |
|---|---|---|---|
| | MRC-University of Glasgow Centre for Virus Research | (COG-UK) consortium | Yasmin Parr, Kyriaki Nomikou; Sarah McDonald, Marc Niebel, Patawee Asamaphan; Richard Orton, Joseph Hughes, Sreenu Vattipally, David L Robertson; Alasdair MacLean, Rory Gunson; Kathy Li, Natasha Jesudason, Rajiv Shah, James Shepherd, Antonia Ho, Alice Broos, Emma Thomson and Alex Alderton, Roberto Amato, Sonia Goncalves, Ewan Harrison, David K. Jackson, Ian Johnston, Dominic Kwiatkowski, Cordelia Langford, John Sillitoe |
| EPI_ISL_530683, EPI_ISL_530684, EPI_ISL_530688 | Lighthouse Lab in Glasgow | Wellcome Sanger Institute for the COVID-19 Genomics UK (COG-UK) consortium | Harper VanSteenhouse, Yumi Kasai, David Gray, Carol Clugston, Anna Dominiczak and Alex Alderton, Roberto Amato, Sonia Goncalves, Ewan Harrison, David K. Jackson, Ian Johnston, Dominic Kwiatkowski, Cordelia Langford, John Sillitoe |
| EPI_ISL_530689 | NHSGGC West of Scotland Specialist Virology Centre / MRC-University of Glasgow Centre for Virus Research | Wellcome Sanger Institute for the COVID-19 Genomics UK (COG-UK) consortium | Ana da Silva Filipe, Natasha Johnson, Kathy Smollett, Daniel Mair, Stephen Carmichael, Lily Tong, Jenna Nichols, Elihu Aranday-Cortes, Kirstyn Brunker, Yasmin Parr, Kyriaki Nomikou; Sarah McDonald, Marc Niebel, Patawee Asamaphan; Richard Orton, Joseph Hughes, Sreenu Vattipally, David L Robertson; Alasdair MacLean, Rory Gunson; Kathy Li, Natasha Jesudason, Rajiv Shah, James Shepherd, Antonia Ho, Alice Broos, Emma Thomson and Alex Alderton, Roberto Amato, Sonia Goncalves, Ewan Harrison, David K. Jackson, Ian Johnston, Dominic Kwiatkowski, Cordelia Langford, John Sillitoe on behalf of the Wellcome Sanger Institute COVID-19 Surveillance Team (http://www.sanger.ac.uk/covid-team) |
| EPI_ISL_530693, EPI_ISL_530694, EPI_ISL_530695, EPI_ISL_530698, EPI_ISL_530699, EPI_ISL_530701, EPI_ISL_530702, EPI_ISL_530704, EPI_ISL_530705, EPI_ISL_530707 | Lighthouse Lab in Glasgow | Wellcome Sanger Institute for the COVID-19 Genomics UK (COG-UK) consortium | Harper VanSteenhouse, Yumi Kasai, David Gray, Carol Clugston, Anna Dominiczak and Alex Alderton, Roberto Amato, Sonia Goncalves, Ewan Harrison, David K. Jackson, Ian Johnston, Dominic Kwiatkowski, Cordelia Langford, John Sillitoe |
| EPI_ISL_530708 | NHSGGC West of Scotland Specialist Virology Centre / MRC-University of Glasgow Centre for Virus Research | Wellcome Sanger Institute for the COVID-19 Genomics UK (COG-UK) consortium | Ana da Silva Filipe, Natasha Johnson, Kathy Smollett, Daniel Mair, Stephen Carmichael, Lily Tong, Jenna Nichols, Elihu Aranday-Cortes, Kirstyn Brunker, Yasmin Parr, Kyriaki Nomikou; Sarah McDonald, Marc Niebel, Patawee Asamaphan; Richard Orton, Joseph Hughes, Sreenu Vattipally, David L Robertson; Alasdair MacLean, Rory Gunson; Kathy Li, Natasha Jesudason, Rajiv Shah, James Shepherd, Antonia Ho, Alice Broos, Emma Thomson and Alex Alderton, Roberto Amato, Sonia Goncalves, Ewan Harrison, David K. Jackson, Ian Johnston, Dominic Kwiatkowski, Cordelia Langford, John Sillitoe |
| EPI_ISL_530711, EPI_ISL_530715, EPI_ISL_530717, EPI_ISL_530719, EPI_ISL_530721 | Lighthouse Lab in Glasgow | Wellcome Sanger Institute for the COVID-19 Genomics UK (COG-UK) consortium | Harper VanSteenhouse, Yumi Kasai, David Gray, Carol Clugston, Anna Dominiczak and Alex Alderton, Roberto Amato, Sonia Goncalves, Ewan Harrison, David K. Jackson, Ian Johnston, Dominic Kwiatkowski, Cordelia Langford, John Sillitoe |
| EPI_ISL_530722 | NHSGGC West of Scotland Specialist Virology Centre / MRC-University of Glasgow Centre for Virus Research | Wellcome Sanger Institute for the COVID-19 Genomics UK (COG-UK) consortium | Ana da Silva Filipe, Natasha Johnson, Kathy Smollett, Daniel Mair, Stephen Carmichael, Lily Tong, Jenna Nichols, Elihu Aranday-Cortes, Kirstyn Brunker, Yasmin Parr, Kyriaki Nomikou; Sarah McDonald, Marc Niebel, Patawee Asamaphan; Richard Orton, Joseph Hughes, Sreenu Vattipally, David L Robertson; Alasdair MacLean, Rory Gunson; Kathy Li, Natasha Jesudason, Rajiv Shah, James Shepherd, Antonia Ho, Alice Broos, Emma Thomson and Alex Alderton, Roberto Amato, Sonia Goncalves, Ewan Harrison, David K. Jackson, Ian Johnston, Dominic Kwiatkowski, Cordelia Langford, John Sillitoe |
| EPI_ISL_530725, EPI_ISL_530726, EPI_ISL_530728, EPI_ISL_530731, EPI_ISL_530732, EPI_ISL_530735, EPI_ISL_530736, EPI_ISL_530741, EPI_ISL_530742 | Lighthouse Lab in Glasgow | Wellcome Sanger Institute for the COVID-19 Genomics UK (COG-UK) consortium | Harper VanSteenhouse, Yumi Kasai, David Gray, Carol Clugston, Anna Dominiczak and Alex Alderton, Roberto Amato, Sonia Goncalves, Ewan Harrison, David K. Jackson, Ian Johnston, Dominic Kwiatkowski, Cordelia Langford, John Sillitoe |
| EPI_ISL_530743 | NHSGGC West of Scotland Specialist Virology Centre / MRC-University of Glasgow Centre for Virus Research | Wellcome Sanger Institute for the COVID-19 Genomics UK (COG-UK) consortium | Ana da Silva Filipe, Natasha Johnson, Kathy Smollett, Daniel Mair, Stephen Carmichael, Lily Tong, Jenna Nichols, Elihu Aranday-Cortes, Kirstyn Brunker, Yasmin Parr, Kyriaki Nomikou; Sarah McDonald, Marc Niebel, Patawee Asamaphan; Richard Orton, Joseph Hughes, Sreenu Vattipally, David L Robertson; Alasdair MacLean, Rory Gunson; Kathy Li, Natasha Jesudason, Rajiv Shah, James Shepherd, Antonia Ho, Alice Broos, Emma Thomson and Alex Alderton, Roberto Amato, Sonia Goncalves, Ewan Harrison, David K. Jackson, Ian Johnston, Dominic Kwiatkowski, Cordelia Langford, John Sillitoe |
| EPI_ISL_530745 | Lighthouse Lab in Glasgow | Wellcome Sanger Institute for the COVID-19 Genomics UK (COG-UK) consortium | Harper VanSteenhouse, Yumi Kasai, David Gray, Carol Clugston, Anna Dominiczak and Alex Alderton, Roberto Amato, Sonia Goncalves, Ewan Harrison, David K. Jackson, Ian Johnston, Dominic Kwiatkowski, Cordelia Langford, John Sillitoe |
| EPI_ISL_530746 | NHSGGC West of Scotland Specialist Virology Centre / MRC-University of Glasgow Centre for Virus Research | Wellcome Sanger Institute for the COVID-19 Genomics UK (COG-UK) consortium | Ana da Silva Filipe, Natasha Johnson, Kathy Smollett, Daniel Mair, Stephen Carmichael, Lily Tong, Jenna Nichols, Elihu Aranday-Cortes, Kirstyn Brunker, Yasmin Parr, Kyriaki Nomikou; Sarah McDonald, Marc Niebel, Patawee Asamaphan; Richard Orton, Joseph Hughes, Sreenu Vattipally, David L Robertson; Alasdair MacLean, Rory Gunson; Kathy Li, Natasha Jesudason, Rajiv Shah, James Shepherd, Antonia Ho, Alice Broos, Emma Thomson and Alex Alderton, Roberto Amato, Sonia Goncalves, Ewan Harrison, David K. Jackson, Ian Johnston, Dominic Kwiatkowski, Cordelia Langford, John Sillitoe on behalf of the Wellcome Sanger Institute COVID-19 Surveillance Team (http://www.sanger.ac.uk/covid-team) |
| EPI_ISL_530747, EPI_ISL_530749, EPI_ISL_530750, EPI_ISL_530752, EPI_ISL_530753 | Lighthouse Lab in Glasgow | Wellcome Sanger Institute for the COVID-19 Genomics UK (COG-UK) consortium | Harper VanSteenhouse, Yumi Kasai, David Gray, Carol Clugston, Anna Dominiczak and Alex Alderton, Roberto Amato, Sonia Goncalves, Ewan Harrison, David K. Jackson, Ian Johnston, Dominic Kwiatkowski, Cordelia Langford, John Sillitoe |
| EPI_ISL_530754 | NHSGGC West of Scotland Specialist Virology Centre / MRC-University of Glasgow Centre for Virus Research | Wellcome Sanger Institute for the COVID-19 Genomics UK (COG-UK) consortium | Ana da Silva Filipe, Natasha Johnson, Kathy Smollett, Daniel Mair, Stephen Carmichael, Lily Tong, Jenna Nichols, Elihu Aranday-Cortes, Kirstyn Brunker, Yasmin Parr, Kyriaki Nomikou; Sarah McDonald, Marc Niebel, Patawee Asamaphan; Richard Orton, Joseph Hughes, Sreenu Vattipally, David L Robertson; Alasdair MacLean, Rory Gunson; Kathy Li, Natasha Jesudason, Rajiv Shah, James Shepherd, Antonia Ho, Alice Broos, Emma Thomson and Alex Alderton, Roberto Amato, Sonia Goncalves, Ewan Harrison, David K. Jackson, Ian Johnston, Dominic Kwiatkowski, Cordelia Langford, John Sillitoe |
| EPI_ISL_530759, EPI_ISL_530760, EPI_ISL_530761, EPI_ISL_530763, EPI_ISL_530765 | Lighthouse Lab in Glasgow | Wellcome Sanger Institute for the COVID-19 Genomics UK (COG-UK) consortium | Harper VanSteenhouse, Yumi Kasai, David Gray, Carol Clugston, Anna Dominiczak and Alex Alderton, Roberto Amato, Sonia Goncalves, Ewan Harrison, David K. Jackson, Ian Johnston, Dominic Kwiatkowski, Cordelia Langford, John Sillitoe |
| EPI_ISL_530768 | NHSGGC West of Scotland Specialist Virology Centre / MRC-University of Glasgow Centre for Virus Research | Wellcome Sanger Institute for the COVID-19 Genomics UK (COG-UK) consortium | Ana da Silva Filipe, Natasha Johnson, Kathy Smollett, Daniel Mair, Stephen Carmichael, Lily Tong, Jenna Nichols, Elihu Aranday-Cortes, Kirstyn Brunker, Yasmin Parr, Kyriaki Nomikou; Sarah McDonald, Marc Niebel, Patawee Asamaphan; Richard Orton, Joseph Hughes, Sreenu Vattipally, David L Robertson; Alasdair MacLean, Rory Gunson; Kathy Li, Natasha Jesudason, Rajiv Shah, James Shepherd, Antonia Ho, Alice Broos, Emma Thomson and Alex Alderton, Roberto Amato, Sonia Goncalves, Ewan Harrison, David K. Jackson, Ian Johnston, Dominic Kwiatkowski, Cordelia Langford, John Sillitoe |
| EPI_ISL_530770, EPI_ISL_530771, EPI_ISL_530773, EPI_ISL_530775, EPI_ISL_530778, EPI_ISL_530779, EPI_ISL_530783, EPI_ISL_530784, EPI_ISL_530785, EPI_ISL_530786, EPI_ISL_530787, EPI_ISL_530788, EPI_ISL_530790, EPI_ISL_530797, EPI_ISL_530798, EPI_ISL_530799 | | | |
| see above | Lighthouse Lab in Glasgow | Wellcome Sanger Institute for the COVID-19 Genomics UK (COG-UK) consortium | Harper VanSteenhouse, Yumi Kasai, David Gray, Carol Clugston, Anna Dominiczak and Alex Alderton, Roberto Amato, Sonia Goncalves, Ewan Harrison, David K. Jackson, Ian Johnston, Dominic Kwiatkowski, Cordelia Langford, John Sillitoe |
| EPI_ISL_530800 | Lighthouse Lab in Glasgow | Wellcome Sanger Institute for the COVID-19 Genomics UK (COG-UK) consortium | Harper VanSteenhouse, Yumi Kasai, David Gray, Carol Clugston, Anna Dominiczak and Alex Alderton, Roberto Amato, Sonia Goncalves, Ewan Harrison, David K. Jackson, Ian Johnston, Dominic Kwiatkowski, Cordelia Langford, John Sillitoe on behalf of the Wellcome Sanger Institute COVID-19 Surveillance Team (http://www.sanger.ac.uk/covid-team) |
| EPI_ISL_534210, EPI_ISL_534211 | Centrl laboratorija | Latvian Biomedical Research and Study Centre | Ivars Silamielis, Jnis Pjalkovskis, Kaspars Megnis, Monta Ustinova, ikita Zrelovs, Vita Rovte, Stella Lapia, Jana Oste, Marta Priedte, Uga Dumpis, Jnis Kloviš |
| EPI_ISL_534212, EPI_ISL_534213, EPI_ISL_534214, EPI_ISL_534215, EPI_ISL_534216, EPI_ISL_534218, EPI_ISL_534219 | E. Gulbja Laboratorija | Latvian Biomedical Research and Study Centre | Ivars Silamielis, Jnis Pjalkovskis, Kaspars Megnis, Monta Ustinova, ikita Zrelovs, Vita Rovte, Mikus Gavars, Dmitrijs Perminovs, Uga Dumpis, Jnis Kloviš |
| EPI_ISL_534220 | Centrl laboratorija | Latvian Biomedical Research and Study Centre | Ivars Silamielis, Jnis Pjalkovskis, Kaspars Megnis, Monta Ustinova, ikita Zrelovs, Vita Rovte, Stella Lapia, Jana Oste, Marta Priedte, Uga Dumpis, Jnis Kloviš |
| EPI_ISL_534221 | E. Gulbja Laboratorija | Latvian Biomedical Research and Study Centre | Ivars Silamielis, Jnis Pjalkovskis, Kaspars Megnis, Monta Ustinova, ikita Zrelovs, Vita Rovte, Mikus Gavars, Dmitrijs Perminovs, Uga Dumpis, Jnis Kloviš |
| EPI_ISL_534222, EPI_ISL_534223 | Centrl laboratorija | Latvian Biomedical Research and Study Centre | Ivars Silamielis, Jnis Pjalkovskis, Kaspars Megnis, Monta Ustinova, ikita Zrelovs, Vita Rovte, Stella Lapia, Jana Oste, Marta Priedte, Uga Dumpis, Jnis Kloviš |
| EPI_ISL_534257 | Laboratoriemedicin Vasternorrland | The Public Health Agency of Sweden | Anna-Malin Linde, Maria Lind Karlberg, Mattias Haukland, Reza Advani, Olov Svartstrom, Oskar Karlsson Lindsjo, Sandra Broddesson, Petra Edquist, Mia Brytting, Anna Risberg, Karin Tegmark-Wisell |

| | | | |
|---|---|---|---|
| EPI_ISL_534439, EPI_ISL_534440, EPI_ISL_534441, EPI_ISL_534442, EPI_ISL_534443, EPI_ISL_534444, EPI_ISL_534445, EPI_ISL_534446, EPI_ISL_534447, EPI_ISL_534448, EPI_ISL_534449, EPI_ISL_534450, EPI_ISL_534451, EPI_ISL_534517, EPI_ISL_534524, EPI_ISL_534526, EPI_ISL_534527, EPI_ISL_534528, EPI_ISL_534529, EPI_ISL_534531, EPI_ISL_534532, EPI_ISL_534539, EPI_ISL_534551, EPI_ISL_534559, EPI_ISL_534561, EPI_ISL_534564, EPI_ISL_534568, EPI_ISL_534569, EPI_ISL_534570, EPI_ISL_534571, EPI_ISL_534573, EPI_ISL_534576, EPI_ISL_534577, EPI_ISL_534582, EPI_ISL_534584, EPI_ISL_534587, EPI_ISL_534588, EPI_ISL_534589, EPI_ISL_534590, EPI_ISL_534592, EPI_ISL_534594, EPI_ISL_534599, EPI_ISL_534601, EPI_ISL_534602, EPI_ISL_534608, EPI_ISL_534612, EPI_ISL_534616, EPI_ISL_534617, EPI_ISL_534618, EPI_ISL_534621, EPI_ISL_534623, EPI_ISL_534624, EPI_ISL_534629, EPI_ISL_534631, EPI_ISL_534632, EPI_ISL_534633, EPI_ISL_534636, EPI_ISL_534638, EPI_ISL_534639, EPI_ISL_534640 | | | |
| see above | NHSGGC West of Scotland Specialist Virology Centre / MRC-University of Glasgow Centre for Virus Research | Wellcome Sanger Institute for the COVID-19 Genomics UK (COG-UK) consortium | Ana da Silva Filipe, Natasha Johnson, Kathy Smollett, Daniel Mair, Stephen Carmichael, Lily Tong, Jenna Nichols, Elihu Aranday-Cortes, Kirstyn Brunker, Yasmin Parr, Kyriaki Nomikou; Sarah McDonald, Marc Niebel, Patawee Asamaphan; Richard Orton, Joseph Hughes, Sreenu Vattipally, David L Robertson; Alasdair MacLean, Rory Gunson; Kathy Li, Natasha Jesudason, Rajiv Shah, James Shepherd, Antonia Ho, Alice Broos, Emma Thomson and Alex Alderton, Roberto Amato, Sonia Goncalves, Ewan Harrison, David K. Jackson, Ian Johnston, Dominic Kwiatkowski, Cordelia Langford, John Sillitoe on behalf of the Wellcome Sanger Institute COVID-19 Surveillance Team (http://www.sanger.ac.uk/covid-team) |
| EPI_ISL_534733, EPI_ISL_534734, EPI_ISL_534736, EPI_ISL_534737, EPI_ISL_534738, EPI_ISL_534739, EPI_ISL_534743, EPI_ISL_534747, EPI_ISL_534748 | Liverpool Clinical Laboratories | COVID-19 Genomics UK (COG-UK) Consortium | Sam Haldenby, Anita Lucaci, Steve Paterson, Julian Hiscox, Alistair Darby, M Almsaud, A Alrezaihi, Muhannad Alruwaili, Stuart D Armstrong, Jones Benjamin, Eleanor G Bentley, Anu Chawla, Jordan J Clark, Angela Cowell, Richard Eccles, Isabel García-Dorival, Matthew Gemmell, Alessandro Gerada, PKF Gilmore, Richard Gregory, Ximeng Han, Catherine Hartley, Margaret Hughes, Miren Iturriza-Gomara, James Johnson, L Luu, Jenifer Manson, Charlotte Nelson, Elaine O'Toole, Cassie Olateju, Rebekah Penrice-Randal , Lucille Rainbow, N.P Randle, Trevor Ian Robinson, Parul Sharma, Ghada T Shawli, James P Stewart, Neil Swainston, Ecaterina Vamos, Joanne Watts, Mark Whitehead |
| EPI_ISL_535071, EPI_ISL_535089 | Virology Department, Sheffield Teaching Hospitals NHS Foundation Trust/Department of Infection, Immunity and Cardiovascular Disease, The Medical School, University of Sheffield | COVID-19 Genomics UK (COG-UK) Consortium | Thushan de Silva, Matthew Parker, Nikki Smith, Adri Angyal, Rebecca Brown, Luke Green, Rachel Tucker, Paul Parsons, Danielle Groves, Katie Johnson, Laura Carrilero, Alex Keeley, Dave Partridge, Matthew Wyles, Benjamin Lindsey, Mehmet Yavuz, Mohammad Raza, Cariad Evans |
| EPI_ISL_535102, EPI_ISL_535103, EPI_ISL_535104, EPI_ISL_535105, EPI_ISL_535106, EPI_ISL_535107, EPI_ISL_535108, EPI_ISL_535109, EPI_ISL_535110, EPI_ISL_535111, EPI_ISL_535112, EPI_ISL_535113, EPI_ISL_535114, EPI_ISL_535115, EPI_ISL_535116, EPI_ISL_535117, EPI_ISL_535118, EPI_ISL_535119, EPI_ISL_535120, EPI_ISL_535121, EPI_ISL_535122, EPI_ISL_535123, EPI_ISL_535124, EPI_ISL_535125, EPI_ISL_535126, EPI_ISL_535127, EPI_ISL_535128, EPI_ISL_535129, EPI_ISL_535130, EPI_ISL_535131, EPI_ISL_535132, EPI_ISL_535133, EPI_ISL_535134, EPI_ISL_535135, EPI_ISL_535136, EPI_ISL_535137, EPI_ISL_535138, EPI_ISL_535139, EPI_ISL_535140, EPI_ISL_535141, EPI_ISL_535142, EPI_ISL_535143, EPI_ISL_535144, EPI_ISL_535145, EPI_ISL_535146, EPI_ISL_535147, EPI_ISL_535148, EPI_ISL_535149, EPI_ISL_535150, EPI_ISL_535151, EPI_ISL_535152, EPI_ISL_535153, EPI_ISL_535154, EPI_ISL_535155, EPI_ISL_535156, EPI_ISL_535157, EPI_ISL_535158, EPI_ISL_535159, EPI_ISL_535160, EPI_ISL_535161, EPI_ISL_535162, EPI_ISL_535163, EPI_ISL_535164, EPI_ISL_535165, EPI_ISL_535166, EPI_ISL_535167, EPI_ISL_535168, EPI_ISL_535169, EPI_ISL_535170, EPI_ISL_535171, EPI_ISL_535172, EPI_ISL_535173, EPI_ISL_535174, EPI_ISL_535175, EPI_ISL_535176, EPI_ISL_535177, EPI_ISL_535178, EPI_ISL_535179, EPI_ISL_535180, EPI_ISL_535181, EPI_ISL_535182 | | | |
| see above | West of Scotland Specialist Virology Centre, NHSGGC / MRC-University of Glasgow Centre for Virus Research | COVID-19 Genomics UK (COG-UK) Consortium | Ana da Silva Filipe, Natasha Johnson, Kathy Smollett, Daniel Mair, Stephen Carmichael, Lily Tong, Jenna Nichols, Elihu Aranday-Cortes, Yasmin Parr, Alice Broos, Kyriaki Nomikou; Sarah McDonald, Marc Niebel, Patawee Asamaphan; Richard Orton, Joseph Hughes, Sreenu Vattipally, David L Robertson; Alasdair MacLean, Rory Gunson; Kathy Li, Natasha Jesudason, Rajiv Shah, James Shepherd, Antonia Ho, Emma Thomson |
| EPI_ISL_535183 | Queens Medical Centre, Clinical Microbiology Department / DeepSeq Nottingham | COVID-19 Genomics UK (COG-UK) Consortium | Gemma Clark, Wendy Smith, Manjinder Khakh, Vicki M Fleming, Michelle M Lister, Hannah Howson-Wells, Jonathan Ball, Patrick McClure, Joseph Chappell, Theocharis Tsoleridis, Nadine Holmes, Matthew Carlisle, Christopher Moore, Fei Sang, Johnny Debebe, Victoria Wright, Matthew Loose |
| EPI_ISL_535185, EPI_ISL_535191, EPI_ISL_535194, EPI_ISL_535200, EPI_ISL_535201, EPI_ISL_535202, EPI_ISL_535212 | Wales Specialist Virology Centre Sequencing lab: Pathogen Genomics Unit | COVID-19 Genomics UK (COG-UK) Consortium | Catherine Moore, Johnathan Evans, Laura Gifford, Malorie Perry, Simon Cottrell, Angela Marchbank, Alec Birchley, Alexander Adams, Amy Gaskin, Bree Gatica-Wilcox, Jason Coombes, Joel Southgate, Lauren Gilbert, Lee Graham, Nicole Pacchiarini, Sara Kumziene-Summerhayes, Sarah Taylor, Sophie Jones, Sara Rey, Matthew Bull, Joanne Watkins, Sally Corden, Tom Connor |
| EPI_ISL_535575 | Hospital Universitari Germans Trias i Pujol(HUGTiP)/Fundació Lluita contra la SIDA (FLSida) | IrsiCaixa AIDS Research Lab | Marc Noguera-Julian, Mariona Parera, Maria Pilar Armengol, Marta Massanella, Ester Ballana, Lidia Ruiz, Nuria Izquierdo, Jorge Carrillo, Roger Paredes, Julia Blanco, Joaquim Segalés, Bonaventura Clotet |
| EPI_ISL_536417, EPI_ISL_536418, EPI_ISL_536419, EPI_ISL_536420, EPI_ISL_536421, EPI_ISL_536423, EPI_ISL_536424, EPI_ISL_536425, EPI_ISL_536427, EPI_ISL_536428, EPI_ISL_536443, EPI_ISL_536453 | | | |
| see above | National Public Health Laboratory, National Centre for Infectious Diseases | National Public Health Laboratory, National Centre for Infectious Diseases | Mak TM, Octavia S, Zhou Z, Cui L, Lin RTP |
| EPI_ISL_536631, EPI_ISL_536632, EPI_ISL_536633, EPI_ISL_536634, EPI_ISL_536635, EPI_ISL_536636, EPI_ISL_536637, EPI_ISL_536638, EPI_ISL_536639, EPI_ISL_536640, EPI_ISL_536641, EPI_ISL_536642, EPI_ISL_536643, EPI_ISL_536644, EPI_ISL_536645, EPI_ISL_536646, EPI_ISL_536647, EPI_ISL_536648, EPI_ISL_536649, EPI_ISL_536650, EPI_ISL_536651, EPI_ISL_536652, EPI_ISL_536653, EPI_ISL_536654, EPI_ISL_536655, EPI_ISL_536658, EPI_ISL_536663, EPI_ISL_536669, EPI_ISL_536670, EPI_ISL_536671, EPI_ISL_536672, EPI_ISL_536673, EPI_ISL_536675, EPI_ISL_536677, EPI_ISL_536679, EPI_ISL_536681, EPI_ISL_536682, EPI_ISL_536683, EPI_ISL_536685, EPI_ISL_536686, EPI_ISL_536687, EPI_ISL_536688, EPI_ISL_536689, EPI_ISL_536690, EPI_ISL_536692, EPI_ISL_536693, EPI_ISL_536694, EPI_ISL_536695, EPI_ISL_536697, EPI_ISL_536698, EPI_ISL_536699, EPI_ISL_536700, EPI_ISL_536701, EPI_ISL_536702, EPI_ISL_536703, EPI_ISL_536704, EPI_ISL_536705, EPI_ISL_536706 | | | |
| see above | University of Wisconsin-Madison AIDS Vaccine Research Laboratories | University of Wisconsin-Madison AIDS Vaccine Research Laboratories | Gage Moreno, Katarina Braun, et al. AIDS Vaccine Research Laboratories |
| EPI_ISL_536785 | University of Wisconsin-Madison Campus AIDS Vaccine Research Laboratories | University of Wisconsin-Madison AIDS Vaccine Research Laboratories | Gage Moreno, Katarina Braun, et al. AIDS Vaccine Research Laboratories |
| EPI_ISL_537148, EPI_ISL_537149, EPI_ISL_537150, EPI_ISL_537151, EPI_ISL_537152, EPI_ISL_537153, EPI_ISL_537154, EPI_ISL_537155, EPI_ISL_537156, EPI_ISL_537157, EPI_ISL_537158, EPI_ISL_537159, EPI_ISL_537160, EPI_ISL_537161, EPI_ISL_537162, EPI_ISL_537163, EPI_ISL_537164, EPI_ISL_537165, EPI_ISL_537166, EPI_ISL_537167, EPI_ISL_537168, EPI_ISL_537169, EPI_ISL_537170, EPI_ISL_537171, EPI_ISL_537172, EPI_ISL_537173, EPI_ISL_537174, EPI_ISL_537175, EPI_ISL_537176, EPI_ISL_537177, EPI_ISL_537178, EPI_ISL_537179, EPI_ISL_537180, EPI_ISL_537181, EPI_ISL_537182, EPI_ISL_537183, EPI_ISL_537184, EPI_ISL_537185, EPI_ISL_537186, EPI_ISL_537187, EPI_ISL_537188, EPI_ISL_537189, EPI_ISL_537190, EPI_ISL_537191, EPI_ISL_537192, EPI_ISL_537193, EPI_ISL_537194, EPI_ISL_537195, EPI_ISL_537196, EPI_ISL_537197, EPI_ISL_537198, EPI_ISL_537199, EPI_ISL_537200, EPI_ISL_537201, EPI_ISL_537202, EPI_ISL_537203, EPI_ISL_537204, EPI_ISL_537205 | | | |
| see above | Lighthouse Lab in Glasgow | Wellcome Sanger Institute for the COVID-19 Genomics UK (COG-UK) consortium | Harper VanSteenhouse, Yumi Kasai, David Gray, Carol Clugston, Anna Dominiczak and Alex Alderton, Roberto Amato, Sonia Goncalves, Ewan Harrison, David K. Jackson, Ian Johnston, Dominic Kwiatkowski, Cordelia Langford, John Sillitoe on behalf of the Wellcome Sanger Institute COVID-19 Surveillance Team |
| EPI_ISL_538552 | Hospital Universitari Germans Trias i Pujol(HUGTiP)/Fundació Lluita contra la SIDA (FLSida) | IrsiCaixa AIDS Research Lab | Marc Noguera-Julian, Mariona Parera, Maria Pilar Armengol, Marta Massanella, Ester Ballana, Lidia Ruiz, Nuria Izquierdo, Jorge Carrillo, Roger Paredes, Julia Blanco, Joaquim Segalés, Bonaventura Clotet |
| EPI_ISL_539341, EPI_ISL_539342, EPI_ISL_539343, EPI_ISL_539344, EPI_ISL_539345, EPI_ISL_539346, EPI_ISL_539347, EPI_ISL_539348, EPI_ISL_539349, EPI_ISL_539350, EPI_ISL_539351, EPI_ISL_539352, EPI_ISL_539353, EPI_ISL_539354, EPI_ISL_539355, EPI_ISL_539356, EPI_ISL_539357, EPI_ISL_539358, EPI_ISL_539359, EPI_ISL_539360, EPI_ISL_539361, EPI_ISL_539362, EPI_ISL_539363, EPI_ISL_539364, EPI_ISL_539365, EPI_ISL_539366, EPI_ISL_539367, EPI_ISL_539368, EPI_ISL_539369, EPI_ISL_539370, EPI_ISL_539371, EPI_ISL_539372, EPI_ISL_539373, EPI_ISL_539374, EPI_ISL_539375, EPI_ISL_539376, EPI_ISL_539377, EPI_ISL_539378, EPI_ISL_539379, EPI_ISL_539380, EPI_ISL_539381, EPI_ISL_539382, EPI_ISL_539383, EPI_ISL_539384, EPI_ISL_539385, EPI_ISL_539386, EPI_ISL_539387, EPI_ISL_539388, EPI_ISL_539390, EPI_ISL_539391, EPI_ISL_539392, EPI_ISL_539393, EPI_ISL_539394, EPI_ISL_539395, EPI_ISL_539396, EPI_ISL_539397, EPI_ISL_539398, EPI_ISL_539399, EPI_ISL_539400, EPI_ISL_539401, EPI_ISL_539402, EPI_ISL_539403, EPI_ISL_539404, EPI_ISL_539405, EPI_ISL_539406, EPI_ISL_539407, EPI_ISL_539408, EPI_ISL_539409, EPI_ISL_539410, EPI_ISL_539411, EPI_ISL_539412, EPI_ISL_539413, EPI_ISL_539414, EPI_ISL_539415, EPI_ISL_539416, EPI_ISL_539417, EPI_ISL_539418, EPI_ISL_539419, EPI_ISL_539420, EPI_ISL_539421, EPI_ISL_539422, EPI_ISL_539423, EPI_ISL_539424, EPI_ISL_539425, EPI_ISL_539426, EPI_ISL_539427, EPI_ISL_539428, EPI_ISL_539429, EPI_ISL_539430, EPI_ISL_539431, EPI_ISL_539432, EPI_ISL_539433, EPI_ISL_539434, EPI_ISL_539435, EPI_ISL_539436, EPI_ISL_539437, EPI_ISL_539438, EPI_ISL_539439, EPI_ISL_539440, EPI_ISL_539441, EPI_ISL_539442, EPI_ISL_539443, EPI_ISL_539444, EPI_ISL_539445, EPI_ISL_539446, EPI_ISL_539447, EPI_ISL_539448, EPI_ISL_539449, EPI_ISL_539450, EPI_ISL_539451, EPI_ISL_539452, EPI_ISL_539453, EPI_ISL_539454, EPI_ISL_539455, EPI_ISL_539456, EPI_ISL_539457, EPI_ISL_539458, EPI_ISL_539459, EPI_ISL_539460, EPI_ISL_539461, EPI_ISL_539462, EPI_ISL_539463, EPI_ISL_539464, EPI_ISL_539465, EPI_ISL_539466, EPI_ISL_539467, EPI_ISL_539468, EPI_ISL_539469, EPI_ISL_539470, EPI_ISL_539471, EPI_ISL_539472, EPI_ISL_539473, EPI_ISL_539474, EPI_ISL_539475, EPI_ISL_539476, EPI_ISL_539477, EPI_ISL_539478, EPI_ISL_539479, EPI_ISL_539480, EPI_ISL_539481, EPI_ISL_539482 | | | |
| see above | Viollier AG | Department of Biosystems Science and Engineering, ETH Zürich | Christian Beisel, Sarah Nadeau, Ivan Topolsky, Pedro Ferreira, Philipp Jablonski, Susana Posada-Céspedes, Tobias Schär, Ina Nissen, Natascha Santacroce, Elodie Burcklen, Christiane Beckmann, Maurice Redondo, Olivier Kobel, Christoph Noppen, Sophie Seidel, Noemie Santamaria de Souza, Niko Beerenwinkel, Tanja Stadler |
| EPI_ISL_539577, EPI_ISL_539578, EPI_ISL_539579, EPI_ISL_539580, EPI_ISL_539581, EPI_ISL_539582, EPI_ISL_539583, EPI_ISL_539584, EPI_ISL_539585, EPI_ISL_539586, EPI_ISL_539587, EPI_ISL_539588, EPI_ISL_539589, EPI_ISL_539590, EPI_ISL_539591, EPI_ISL_539592, EPI_ISL_539593, EPI_ISL_539594, EPI_ISL_539595, EPI_ISL_539613, EPI_ISL_539614 | | | |
| see above | ZOTZ KLIMAS MVZ Düsseldorf-Centrum GbR ÜBAG für Labormedizin, Genetik, Zytologie, Pathologie | Center of Medical Microbiology, Virology, and Hospital Hygiene, University of Duesseldorf | Maximilian Damagnez, Alexander Dilthey, Ashley-Jane Duplessis, Patrick Finzer, Katrin Hoffmann, Torsten Houwaart, Malte Kohns Vasconcelos, Marek Korencak, Nadine Lübke, Jessica Nicolai, Klaus Pfeffer, Daniel Strelow, Jörg Timm, Andreas Walker, Tobias Wienemann, Rainer Zotz |
| EPI_ISL_539877 | Center for Microbiology and Cell Biology, Instituto Venezolano de Investigaciones Científicas (CMBC, IVIC) | Center for Microbiology and Cell Biology, Instituto Venezolano de Investigaciones Científicas (CMBC, IVIC) | Loureiro,C.L., Jaspe,R.C., D'Angelo,P., Garzaro,D., Rodriguez,L., Alarcon,V., Delgado,M., Aguilar,M., Rangel,H.R., Pujol,F.H. |
| EPI_ISL_539880 | Respiratory Virus Unit, Microbiology Services Colindale, Public Health England | Respiratory Virus Unit, Microbiology Services Colindale, Public Health England | PHE Covid Sequencing Team |
| EPI_ISL_539897, EPI_ISL_539898, EPI_ISL_539899, EPI_ISL_539900, EPI_ISL_539901, EPI_ISL_539902, | Lighthouse Lab in Glasgow | Wellcome Sanger Institute for the COVID-19 Genomics UK (COG-UK) consortium | Harper VanSteenhouse, Yumi Kasai, David Gray, Carol Clugston, Anna Dominiczak and Alex Alderton, Roberto Amato, Sonia Goncalves, Ewan Harrison, David K. Jackson, Ian Johnston, Dominic Kwiatkowski, Cordelia Langford, John Sillitoe on behalf of the Wellcome Sanger Institute COVID-19 Surveillance Team |

| | | | |
|---|---|---|---|
| EPI_ISL_539903, EPI_ISL_539904, EPI_ISL_539905 | | | |
| EPI_ISL_539906 | Lighthouse Lab in Glasgow | Wellcome Sanger Institute for the COVID-19 Genomics UK (COG-UK) Consortium | Harper VanSteenhouse, Yumi Kasai, David Gray, Carol Clugston, Anna Dominiczak and Alex Alderton, Roberto Amato, Sonia Goncalves, Ewan Harrison, David K. Jackson, Ian Johnston, Dominic Kwiatkowski, Cordelia Langford, John Sillitoe on behalf of the Wellcome Sanger Institute COVID-19 Surveillance Team |
| EPI_ISL_539907, EPI_ISL_539908, EPI_ISL_539909, EPI_ISL_539910, EPI_ISL_539911, EPI_ISL_539912, EPI_ISL_539913, EPI_ISL_539914, EPI_ISL_539915, EPI_ISL_539916, EPI_ISL_539917, EPI_ISL_539918, EPI_ISL_539919, EPI_ISL_539920, EPI_ISL_539921, EPI_ISL_539922, EPI_ISL_539923, EPI_ISL_539924, EPI_ISL_539925, EPI_ISL_539926, EPI_ISL_539927, EPI_ISL_539928, EPI_ISL_539929, EPI_ISL_539930, EPI_ISL_539931, EPI_ISL_539932, EPI_ISL_539934, EPI_ISL_539935, EPI_ISL_539936, EPI_ISL_539937, EPI_ISL_539938, EPI_ISL_539939, EPI_ISL_539940, EPI_ISL_539941, EPI_ISL_539942, EPI_ISL_539943, EPI_ISL_539944, EPI_ISL_539945, EPI_ISL_539946, EPI_ISL_539947, EPI_ISL_539948, EPI_ISL_539949, EPI_ISL_539950, EPI_ISL_539951, EPI_ISL_539952, EPI_ISL_539953, EPI_ISL_539954, EPI_ISL_539955, EPI_ISL_539956, EPI_ISL_539957, EPI_ISL_539958, EPI_ISL_539959, EPI_ISL_539960, EPI_ISL_539961, EPI_ISL_539962, EPI_ISL_539963, EPI_ISL_539964, EPI_ISL_539965, EPI_ISL_539966, EPI_ISL_539967, EPI_ISL_539968, EPI_ISL_539970, EPI_ISL_539971, EPI_ISL_539972, EPI_ISL_539973, EPI_ISL_539974, EPI_ISL_539975, EPI_ISL_539976, EPI_ISL_539977, EPI_ISL_539978, EPI_ISL_539979, EPI_ISL_539980, EPI_ISL_539981, EPI_ISL_539982, EPI_ISL_539983, EPI_ISL_539984, EPI_ISL_539985, EPI_ISL_539986, EPI_ISL_539987, EPI_ISL_539988, EPI_ISL_539989, EPI_ISL_539990, EPI_ISL_539991, EPI_ISL_539992, EPI_ISL_539993, EPI_ISL_539994, EPI_ISL_539995, EPI_ISL_539996, EPI_ISL_539997, EPI_ISL_539998, EPI_ISL_539999, EPI_ISL_540000, EPI_ISL_540001, EPI_ISL_540002, EPI_ISL_540003, EPI_ISL_540004, EPI_ISL_540005, EPI_ISL_540006, EPI_ISL_540007, EPI_ISL_540008, EPI_ISL_540009, EPI_ISL_540010, EPI_ISL_540011, EPI_ISL_540012, EPI_ISL_540013, EPI_ISL_540014, EPI_ISL_540015, EPI_ISL_540016, EPI_ISL_540017, EPI_ISL_540018, EPI_ISL_540019, EPI_ISL_540020, EPI_ISL_540021, EPI_ISL_540022, EPI_ISL_540023, EPI_ISL_540024, EPI_ISL_540025, EPI_ISL_540026, EPI_ISL_540027, EPI_ISL_540028, EPI_ISL_540029 | | | |
| see above | Lighthouse Lab in Glasgow | Wellcome Sanger Institute for the COVID-19 Genomics UK (COG-UK) consortium | Harper VanSteenhouse, Yumi Kasai, David Gray, Carol Clugston, Anna Dominiczak and Alex Alderton, Roberto Amato, Sonia Goncalves, Ewan Harrison, David K. Jackson, Ian Johnston, Dominic Kwiatkowski, Cordelia Langford, John Sillitoe on behalf of the Wellcome Sanger Institute COVID-19 Surveillance Team |
| EPI_ISL_540030 | Lighthouse Lab in Glasgow | Wellcome Sanger Institute for the COVID-19 Genomics UK (COG-UK) Consortium | Harper VanSteenhouse, Yumi Kasai, David Gray, Carol Clugston, Anna Dominiczak and Alex Alderton, Roberto Amato, Sonia Goncalves, Ewan Harrison, David K. Jackson, Ian Johnston, Dominic Kwiatkowski, Cordelia Langford, John Sillitoe on behalf of the Wellcome Sanger Institute COVID-19 Surveillance Team |
| EPI_ISL_540031, EPI_ISL_540032, EPI_ISL_540033, EPI_ISL_540034, EPI_ISL_540035, EPI_ISL_540036, EPI_ISL_540037, EPI_ISL_540038, EPI_ISL_540039, EPI_ISL_540040, EPI_ISL_540041, EPI_ISL_540042, EPI_ISL_540043, EPI_ISL_540044, EPI_ISL_540045, EPI_ISL_540046, EPI_ISL_540047, EPI_ISL_540048, EPI_ISL_540049, EPI_ISL_540050, EPI_ISL_540051, EPI_ISL_540052, EPI_ISL_540053, EPI_ISL_540054, EPI_ISL_540055, EPI_ISL_540056, EPI_ISL_540057, EPI_ISL_540058, EPI_ISL_540059, EPI_ISL_540060, EPI_ISL_540061, EPI_ISL_540062, EPI_ISL_540063, EPI_ISL_540064, EPI_ISL_540065, EPI_ISL_540066, EPI_ISL_540067, EPI_ISL_540068, EPI_ISL_540069, EPI_ISL_540070, EPI_ISL_540071, EPI_ISL_540072, EPI_ISL_540073, EPI_ISL_540074, EPI_ISL_540075, EPI_ISL_540076, EPI_ISL_540077, EPI_ISL_540078, EPI_ISL_540079, EPI_ISL_540080, EPI_ISL_540081, EPI_ISL_540082, EPI_ISL_540083, EPI_ISL_540084, EPI_ISL_540085, EPI_ISL_540086, EPI_ISL_540087, EPI_ISL_540088, EPI_ISL_540089, EPI_ISL_540090, EPI_ISL_540091, EPI_ISL_540092, EPI_ISL_540093, EPI_ISL_540094, EPI_ISL_540095, EPI_ISL_540096, EPI_ISL_540097, EPI_ISL_540098, EPI_ISL_540099, EPI_ISL_540100, EPI_ISL_540101, EPI_ISL_540102, EPI_ISL_540103, EPI_ISL_540104, EPI_ISL_540105, EPI_ISL_540106, EPI_ISL_540107, EPI_ISL_540108, EPI_ISL_540109, EPI_ISL_540110, EPI_ISL_540111, EPI_ISL_540112, EPI_ISL_540113, EPI_ISL_540114, EPI_ISL_540115, EPI_ISL_540116, EPI_ISL_540117, EPI_ISL_540118, EPI_ISL_540119, EPI_ISL_540120, EPI_ISL_540121, EPI_ISL_540122, EPI_ISL_540123, EPI_ISL_540124, EPI_ISL_540125, EPI_ISL_540126, EPI_ISL_540127, EPI_ISL_540128, EPI_ISL_540129, EPI_ISL_540130, EPI_ISL_540131, EPI_ISL_540132, EPI_ISL_540133, EPI_ISL_540134, EPI_ISL_540135, EPI_ISL_540136, EPI_ISL_540137, EPI_ISL_540138, EPI_ISL_540178, EPI_ISL_540342, EPI_ISL_540343, EPI_ISL_540344, EPI_ISL_540345, EPI_ISL_540346, EPI_ISL_540347, EPI_ISL_540348, EPI_ISL_540349, EPI_ISL_540350 | | | |
| see above | Lighthouse Lab in Glasgow | Wellcome Sanger Institute for the COVID-19 Genomics UK (COG-UK) consortium | Harper VanSteenhouse, Yumi Kasai, David Gray, Carol Clugston, Anna Dominiczak and Alex Alderton, Roberto Amato, Sonia Goncalves, Ewan Harrison, David K. Jackson, Ian Johnston, Dominic Kwiatkowski, Cordelia Langford, John Sillitoe on behalf of the Wellcome Sanger Institute COVID-19 Surveillance Team |
| EPI_ISL_540351 | Lighthouse Lab in Glasgow | Wellcome Sanger Institute for the COVID-19 Genomics UK (COG-UK) Consortium | Harper VanSteenhouse, Yumi Kasai, David Gray, Carol Clugston, Anna Dominiczak and Alex Alderton, Roberto Amato, Sonia Goncalves, Ewan Harrison, David K. Jackson, Ian Johnston, Dominic Kwiatkowski, Cordelia Langford, John Sillitoe on behalf of the Wellcome Sanger Institute COVID-19 Surveillance Team |
| EPI_ISL_540352, EPI_ISL_540353, EPI_ISL_540354, EPI_ISL_540355, EPI_ISL_540356, EPI_ISL_540357, EPI_ISL_540358, EPI_ISL_540359, EPI_ISL_540360, EPI_ISL_540361, EPI_ISL_540362, EPI_ISL_540363, EPI_ISL_540364, EPI_ISL_540365, EPI_ISL_540366, EPI_ISL_540367, EPI_ISL_540368, EPI_ISL_540369, EPI_ISL_540370, EPI_ISL_540371, EPI_ISL_540372, EPI_ISL_540373, EPI_ISL_540374, EPI_ISL_540375, EPI_ISL_540376, EPI_ISL_540377, EPI_ISL_540378, EPI_ISL_540379, EPI_ISL_540380, EPI_ISL_540381, EPI_ISL_540382, EPI_ISL_540383, EPI_ISL_540384, EPI_ISL_540385, EPI_ISL_540386, EPI_ISL_540387, EPI_ISL_540388, EPI_ISL_540389, EPI_ISL_540390, EPI_ISL_540391, EPI_ISL_540392, EPI_ISL_540393, EPI_ISL_540394, EPI_ISL_540395, EPI_ISL_540396, EPI_ISL_540397, EPI_ISL_540398, EPI_ISL_540399, EPI_ISL_540400, EPI_ISL_540401, EPI_ISL_540402, EPI_ISL_540403, EPI_ISL_540404, EPI_ISL_540405, EPI_ISL_540406, EPI_ISL_540407, EPI_ISL_540408, EPI_ISL_540409, EPI_ISL_540410, EPI_ISL_540411, EPI_ISL_540412, EPI_ISL_540413, EPI_ISL_540414, EPI_ISL_540415, EPI_ISL_540416, EPI_ISL_540417, EPI_ISL_540418 | | | |
| see above | Lighthouse Lab in Glasgow | Wellcome Sanger Institute for the COVID-19 Genomics UK (COG-UK) consortium | Harper VanSteenhouse, Yumi Kasai, David Gray, Carol Clugston, Anna Dominiczak and Alex Alderton, Roberto Amato, Sonia Goncalves, Ewan Harrison, David K. Jackson, Ian Johnston, Dominic Kwiatkowski, Cordelia Langford, John Sillitoe on behalf of the Wellcome Sanger Institute COVID-19 Surveillance Team |
| EPI_ISL_540573, EPI_ISL_540574, EPI_ISL_540575, EPI_ISL_540576, EPI_ISL_540577, EPI_ISL_540578 | Department of Clinical Microbiology | GIGA Medical Genomics | Keith Durkin, Maria Artesi, Sébastien Bontems, Raphaël Boreux, Bouchra Boujemla, Cécile Meex, Axelle Chaslain, Céline Fombellida-Lopez, Pierrette Melin, Marie-Pierre Hayette, Vincent Bours |
| EPI_ISL_540584 | Liverpool Clinical Laboratories | COVID-19 Genomics UK (COG-UK) Consortium | Sam Haldenby, Anita Lucaci, Steve Paterson, Julian Hiscox, Alistair Darby, M Almsaud, A Alrezaihi, Muhannad Alruwaili, Stuart D Armstrong, Jones Benjamin, Eleanor G Bentley, Anu Chawla, Jordan J Clark, Angela Cowell, Richard Eccles, Isabel García-Dorival, Matthew Gemmell, Alessandro Gerada, PKF Gilmore, Richard Gregory, Ximeng Han, Catherine Hartley, Margaret Hughes, Miren Iturriza-Gomara, James Johnson, L Luu, Jenifer Manson, Charlotte Nelson, Elaine O'Toole, Cassie Olateju, Rebekah Penrice-Randal , Lucille Rainbow, N.P Randle, Trevor Ian Robinson, Parul Sharma, Ghada T Shawli, James P Stewart, Neil Swainston, Ecaterina Vamos, Joanne Watts, Mark Whitehead |
| EPI_ISL_540640, EPI_ISL_540641, EPI_ISL_540642, EPI_ISL_540643, EPI_ISL_540644, EPI_ISL_540645, EPI_ISL_540646, EPI_ISL_540647, EPI_ISL_540648, EPI_ISL_540649, EPI_ISL_540650, EPI_ISL_540651, EPI_ISL_540652, EPI_ISL_540653, EPI_ISL_540654, EPI_ISL_540655, EPI_ISL_540656, EPI_ISL_540657, EPI_ISL_540658, EPI_ISL_540659, EPI_ISL_540660, EPI_ISL_540661, EPI_ISL_540662, EPI_ISL_540663, EPI_ISL_540664, EPI_ISL_540665, EPI_ISL_540666, EPI_ISL_540667, EPI_ISL_540668, EPI_ISL_540669, EPI_ISL_540670, EPI_ISL_540671, EPI_ISL_540672, EPI_ISL_540673, EPI_ISL_540674, EPI_ISL_540675, EPI_ISL_540676, EPI_ISL_540677, EPI_ISL_540678, EPI_ISL_540679, EPI_ISL_540680, EPI_ISL_540681, EPI_ISL_540682, EPI_ISL_540683, EPI_ISL_540684, EPI_ISL_540685, EPI_ISL_540686, EPI_ISL_540687, EPI_ISL_540688, EPI_ISL_540689, EPI_ISL_540690 | | | |
| see above | Queens Medical Centre, Clinical Microbiology Department / DeepSeq Nottingham | COVID-19 Genomics UK (COG-UK) Consortium | Gemma Clark, Wendy Smith, Manjinder Khakh, Vicki M Fleming, Michelle M Lister, Hannah Howson-Wells, Jonathan Ball, Patrick McClure, Joseph Chappell, Theocharis Tsoleridis, Nadine Holmes, Matthew Carlisle, Christopher Moore, Fei Sang, Johnny Debebe, Victoria Wright, Matthew Loose |
| EPI_ISL_540692, EPI_ISL_540693, EPI_ISL_540694, EPI_ISL_540695, EPI_ISL_540696 | Quadram Institute Bioscience | COVID-19 Genomics UK (COG-UK) Consortium | Dave J. Baker, Gemma L. Kay, Alp Aydin, Thanh Le-Viet, Steven Rudder, Ana P. Tedim, Anastasia Kolyva, Maria Diaz, Leonardo de Oliveira Martins, Nabil-Fareed Alikhan, Lizzie Meadows, Rachael Stanley, Ngozi Elumogo, Muhammed Yasir, Nicholas M. Thomson, Alexander J Trotter, Rachel Gilroy, Samuel Bloomfield, Claire Stuart, Andrew Bell, Reenesh Prakash, Samir Dervisevic, Alison E. Mather, John Wain, Mark Webber, Andrew J. Page, Justin O'Grady |
| EPI_ISL_540719, EPI_ISL_540720, EPI_ISL_540724, EPI_ISL_540725, EPI_ISL_540726, EPI_ISL_540727, EPI_ISL_540730, EPI_ISL_540733, EPI_ISL_540735, EPI_ISL_540736, EPI_ISL_540737, EPI_ISL_540738, EPI_ISL_540739, EPI_ISL_540742, EPI_ISL_540744, EPI_ISL_540745, EPI_ISL_540747, EPI_ISL_540749, EPI_ISL_540751, EPI_ISL_540754, EPI_ISL_540758, EPI_ISL_540759, EPI_ISL_540760, EPI_ISL_540761, EPI_ISL_540763, EPI_ISL_540764, EPI_ISL_540765, EPI_ISL_540766, EPI_ISL_540770, EPI_ISL_540771 | | | |
| see above | Virology Department, Sheffield Teaching Hospitals NHS Foundation Trust/Department of Infection, Immunity and Cardiovascular Disease, The Medical School, University of Sheffield | COVID-19 Genomics UK (COG-UK) Consortium | Thushan de Silva, Matthew Parker, Nikki Smith, Adri Angyal, Rebecca Brown, Luke Green, Rachel Tucker, Paul Parsons, Danielle Groves, Katie Johnson, Laura Carrilero, Alex Keeley, Dave Partridge, Matthew Wyles, Benjamin Lindsey, Mehmet Yavuz, Mohammad Raza, Cariad Evans |
| EPI_ISL_540775, EPI_ISL_540776, EPI_ISL_540777, EPI_ISL_540778, EPI_ISL_540779, EPI_ISL_540780, EPI_ISL_540781, EPI_ISL_540782, EPI_ISL_540783, EPI_ISL_540784, EPI_ISL_540785, EPI_ISL_540792, EPI_ISL_540816 | | | |
| see above | West of Scotland Specialist Virology Centre, NHSGGC / MRC-University of Glasgow Centre for Virus Research | COVID-19 Genomics UK (COG-UK) Consortium | Ana da Silva Filipe, Natasha Johnson, Kathy Smollett, Daniel Mair, Stephen Carmichael, Lily Tong, Jenna Nichols, Elihu Aranday-Cortes, Kyriaki Nomikou; Sarah McDonald, Marc Niebel, Patawee Asamaphan; Richard Orton, Joseph Hughes, Sreenu Vattipally, David L Robertson; Alasdair MacLean, Rory Gunson; Kathy Li, Igor Starinskij, Natasha Jesudason, Rajiv Shah, James Shepherd, Antonia Ho, Emma Thomson |
| EPI_ISL_540823, EPI_ISL_540824, EPI_ISL_540825, EPI_ISL_540826, EPI_ISL_540827, EPI_ISL_540828, EPI_ISL_540829, EPI_ISL_540830, EPI_ISL_540831, EPI_ISL_540832, EPI_ISL_540833, EPI_ISL_540834, EPI_ISL_540835, EPI_ISL_540836, EPI_ISL_540837, EPI_ISL_540838, EPI_ISL_540839, EPI_ISL_540840, EPI_ISL_540841, EPI_ISL_540842, EPI_ISL_540843, EPI_ISL_540844, EPI_ISL_540845, EPI_ISL_540846, EPI_ISL_540847, EPI_ISL_540848, EPI_ISL_540849, EPI_ISL_540850, EPI_ISL_540851, EPI_ISL_540852, EPI_ISL_540853, EPI_ISL_540854, EPI_ISL_540855, EPI_ISL_540856, EPI_ISL_540857, EPI_ISL_540858, EPI_ISL_540859, EPI_ISL_540860, EPI_ISL_540861, EPI_ISL_540862, EPI_ISL_540863, EPI_ISL_540864, EPI_ISL_540865, EPI_ISL_540866, EPI_ISL_540867, EPI_ISL_540868, EPI_ISL_540869, EPI_ISL_540870 | | | |
| see above | Lighthouse Lab in Glasgow / MRC-University of Glasgow Centre for Virus Research | COVID-19 Genomics UK (COG-UK) Consortium | Ana da Silva Filipe, Natasha Johnson, Kathy Smollett, Daniel Mair, Stephen Carmichael, Lily Tong, Jenna Nichols, Elihu Aranday-Cortes, Kyriaki Nomikou; Sarah McDonald, Marc Niebel, Patawee Asamaphan; Harper VanSteenhouse, Yumi Kasai, David Gray, Carol Clugston, Anna Dominiczak; Alasdair MacLean, Rory Gunson; Richard Orton, Joseph Hughes, Sreenu Vattipally, David L Robertson; Sharif Shaaban, Matthew Holden; Kathy Li, Natasha Jesudason, Rajiv Shah, James Shepherd, Antonia Ho, Emma Thomson |
| EPI_ISL_540872, EPI_ISL_540873 | Virology Department, Royal Infirmary of Edinburgh, NHS Lothian / School of Biological Sciences, University of Edinburgh / Institute of Genetics and Molecular Medicine, University of Edinburgh | COVID-19 Genomics UK (COG-UK) Consortium | McHugh M, Dewar R, Rooke S, Gallagher M, Balcaza C, O'Toole Á, Scher E, Hill V, McCrone JT, Colquhoun R, Yu X, Jackson B, Rambaut A, Williams TC, Templeton K |

| | | | |
|---|---|---|---|
| EPI_ISL_540894, EPI_ISL_540895, EPI_ISL_540896 | Queens Medical Centre, Clinical Microbiology Department / DeepSeq Nottingham | COVID-19 Genomics UK (COG-UK) Consortium | Gemma Clark, Wendy Smith, Manjinder Khakh, Vicki M Fleming, Michelle M Lister, Hannah Howson-Wells, Jonathan Ball, Patrick McClure, Joseph Chappell, Theocharis Tsoleridis, Nadine Holmes, Matthew Carlisle, Christopher Moore, Fei Sang, Johnny Debebe, Victoria Wright, Matthew Loose |
| EPI_ISL_541212, EPI_ISL_541213, EPI_ISL_541214, EPI_ISL_541215, EPI_ISL_541216, EPI_ISL_541217, EPI_ISL_541218, EPI_ISL_541219, EPI_ISL_541220, EPI_ISL_541221, EPI_ISL_541222, EPI_ISL_541223, EPI_ISL_541224, EPI_ISL_541225, EPI_ISL_541226, EPI_ISL_541227, EPI_ISL_541228, EPI_ISL_541229, EPI_ISL_541230, EPI_ISL_541231, EPI_ISL_541232, EPI_ISL_541233, EPI_ISL_541234, EPI_ISL_541235, EPI_ISL_541236 | | | |
| see above | Florida Bureau of Public Health Laboratories, Florida Department of Health | Florida Bureau of Public Health Laboratories, Florida Department of Health | Schmedes,S., Blanton,J. |
| EPI_ISL_541411, EPI_ISL_541412, EPI_ISL_541413, EPI_ISL_541414, EPI_ISL_541415, EPI_ISL_541436, EPI_ISL_541437, EPI_ISL_541438, EPI_ISL_541439, EPI_ISL_541440, EPI_ISL_541441, EPI_ISL_541442, EPI_ISL_541443, EPI_ISL_541444, EPI_ISL_541445, EPI_ISL_541446, EPI_ISL_541450, EPI_ISL_541451, EPI_ISL_541452, EPI_ISL_541453, EPI_ISL_541454, EPI_ISL_541455, EPI_ISL_541456, EPI_ISL_541457, EPI_ISL_541458, EPI_ISL_541459, EPI_ISL_541460, EPI_ISL_541461, EPI_ISL_541462, EPI_ISL_541463, EPI_ISL_541464, EPI_ISL_541469, EPI_ISL_541495, EPI_ISL_541496, EPI_ISL_541497, EPI_ISL_541498, EPI_ISL_541499, EPI_ISL_541500, EPI_ISL_541501, EPI_ISL_541502, EPI_ISL_541503, EPI_ISL_541504, EPI_ISL_541505, EPI_ISL_541506, EPI_ISL_541508, EPI_ISL_541509, EPI_ISL_541524, EPI_ISL_541525, EPI_ISL_541526, EPI_ISL_541527, EPI_ISL_541528, EPI_ISL_541529, EPI_ISL_541530, EPI_ISL_541531, EPI_ISL_541532, EPI_ISL_541533, EPI_ISL_541534, EPI_ISL_541535, EPI_ISL_541536, EPI_ISL_541537, EPI_ISL_541538, EPI_ISL_541539 | | | |
| see above | Viollier AG | Department of Biosystems Science and Engineering, ETH Zürich | Christian Beisel, Sarah Nadeau, Ivan Topolsky, Pedro Ferreira, Philipp Jablonski, Susana Posada-Céspedes, Tobias Schär, Ina Nissen, Natascha Santacroce, Elodie Burcklen, Christiane Beckmann, Maurice Redondo, Olivier Kobel, Christoph Noppen, Sophie Seidel, Noemie Santamaria de Souza, Niko Beerenwinkel, Tanja Stadler |
| EPI_ISL_541714, EPI_ISL_541715, EPI_ISL_541716, EPI_ISL_541717, EPI_ISL_541718, EPI_ISL_541719, EPI_ISL_541720, EPI_ISL_541721 | National Institute of Virology, NIV Influenza | National Institute of Virology, NIV Influenza | Potdar V |
| EPI_ISL_541756 | Microbiology Department, Barking Havering and Redbridge University Hospitals NHS trust | Wellcome Sanger Institute for the COVID-19 Genomics UK (COG-UK) consortium | Amy Ash, Fatima Ali, Cherian Koshy and Alex Alderton, Roberto Amato, Sonia Goncalves, Ewan Harrison, David K. Jackson, Ian Johnston, Dominic Kwiatkowski, Cordelia Langford, John Sillitoe on behalf of the Wellcome Sanger Institute COVID-19 Surveillance Team |
| EPI_ISL_541784, EPI_ISL_541785, EPI_ISL_541786, EPI_ISL_541787, EPI_ISL_541788, EPI_ISL_541789, EPI_ISL_541790, EPI_ISL_541791, EPI_ISL_541792, EPI_ISL_541793, EPI_ISL_541794, EPI_ISL_541795, EPI_ISL_541796, EPI_ISL_541797, EPI_ISL_541798, EPI_ISL_541799, EPI_ISL_541800, EPI_ISL_541801, EPI_ISL_541802, EPI_ISL_541803, EPI_ISL_541804, EPI_ISL_541805, EPI_ISL_541806, EPI_ISL_541807, EPI_ISL_541808, EPI_ISL_541809, EPI_ISL_541810, EPI_ISL_541811, EPI_ISL_541812, EPI_ISL_541813, EPI_ISL_541814, EPI_ISL_541815, EPI_ISL_541816, EPI_ISL_541817, EPI_ISL_541818, EPI_ISL_541819, EPI_ISL_541820, EPI_ISL_541821, EPI_ISL_541822, EPI_ISL_541823, EPI_ISL_541824, EPI_ISL_541825, EPI_ISL_541826, EPI_ISL_541827, EPI_ISL_541828, EPI_ISL_541829, EPI_ISL_541830, EPI_ISL_541831, EPI_ISL_541832, EPI_ISL_541833, EPI_ISL_541834, EPI_ISL_541835, EPI_ISL_541836, EPI_ISL_541837, EPI_ISL_541838, EPI_ISL_541839, EPI_ISL_541840, EPI_ISL_541841, EPI_ISL_541842, EPI_ISL_541843, EPI_ISL_541844, EPI_ISL_541845, EPI_ISL_541846 | | | |
| see above | Lighthouse Lab in Glasgow | Wellcome Sanger Institute for the COVID-19 Genomics UK (COG-UK) consortium | Harper VanSteenhouse, Yumi Kasai, David Gray, Carol Clugston, Anna Dominiczak and Alex Alderton, Roberto Amato, Sonia Goncalves, Ewan Harrison, David K. Jackson, Ian Johnston, Dominic Kwiatkowski, Cordelia Langford, John Sillitoe on behalf of the Wellcome Sanger Institute COVID-19 Surveillance Team |
| EPI_ISL_541936, EPI_ISL_541937, EPI_ISL_541938, EPI_ISL_541939, EPI_ISL_541940, EPI_ISL_541941 | Hospital General Universitario Gregorio Marañón | SeqCOVID-SPAIN consortium/IBV(CSIC) | Laura Pérez-Lago, Marta Herranz, Jon Sicilia, Julia Suárez, Pilar Catalán, Patricia Muñoz, Darío García de Viedma and SeqCOVID-SPAIN consortium |
| EPI_ISL_542041, EPI_ISL_542042, EPI_ISL_542043, EPI_ISL_542044, EPI_ISL_542045, EPI_ISL_542046, EPI_ISL_542047, EPI_ISL_542048, EPI_ISL_542049, EPI_ISL_542050, EPI_ISL_542051, EPI_ISL_542052, EPI_ISL_542053, EPI_ISL_542054, EPI_ISL_542055, EPI_ISL_542056, EPI_ISL_542057, EPI_ISL_542058, EPI_ISL_542059, EPI_ISL_542060, EPI_ISL_542061, EPI_ISL_542062, EPI_ISL_542087 | | | |
| see above | New Mexico Department of Health Scientific Laboratory | New Mexico Department of Health Scientific Laboratory | Ellie Johnson, Anastacia Griego-Fisher, D'Eldra Malone |
| EPI_ISL_544968 | St Vincent's Pathology (SydPath) | NSW Health Pathology - Institute of Clinical Pathology and Medical Research; Westmead Hospital; University of Sydney | CIDM-PH et al. |
| EPI_ISL_544969 | Pathology North - Royal North Shore Hospital - NSW Health Pathology | NSW Health Pathology - Institute of Clinical Pathology and Medical Research; Westmead Hospital; University of Sydney | CIDM-PH et al. |
| EPI_ISL_544970 | South Eastern Area Laboratory Services (SEALS) | NSW Health Pathology - Institute of Clinical Pathology and Medical Research; Westmead Hospital; University of Sydney | CIDM-PH et al. |
| EPI_ISL_544971, EPI_ISL_544972, EPI_ISL_544973 | St Vincent's Pathology (SydPath) | NSW Health Pathology - Institute of Clinical Pathology and Medical Research; Westmead Hospital; University of Sydney | CIDM-PH et al. |
| EPI_ISL_544974 | Sydney South West Pathology Service (SSWPS) - Royal Prince Alfred Hospital - NSW Health Pathology | NSW Health Pathology - Institute of Clinical Pathology and Medical Research; Westmead Hospital; University of Sydney | CIDM-PH et al. |
| EPI_ISL_544975 | Histopath | NSW Health Pathology - Institute of Clinical Pathology and Medical Research; Westmead Hospital; University of Sydney | CIDM-PH et al. |
| EPI_ISL_544976 | Pathology North - Royal North Shore Hospital - NSW Health Pathology | NSW Health Pathology - Institute of Clinical Pathology and Medical Research; Westmead Hospital; University of Sydney | CIDM-PH et al. |
| EPI_ISL_544977, EPI_ISL_544978, EPI_ISL_544979, EPI_ISL_544980, EPI_ISL_544981 | St Vincent's Pathology (SydPath) | NSW Health Pathology - Institute of Clinical Pathology and Medical Research; Westmead Hospital; University of Sydney | CIDM-PH et al. |
| EPI_ISL_544982 | Pathology North - Royal North Shore Hospital - NSW Health Pathology | NSW Health Pathology - Institute of Clinical Pathology and Medical Research; Westmead Hospital; University of Sydney | CIDM-PH et al. |
| EPI_ISL_544983, EPI_ISL_544984 | South Eastern Area Laboratory Services (SEALS) | NSW Health Pathology - Institute of Clinical Pathology and Medical Research; Westmead Hospital; University of Sydney | CIDM-PH et al. |
| EPI_ISL_544985 | Sydney South West Pathology Service (SSWPS) - Liverpool Hospital - NSW Health Pathology | NSW Health Pathology - Institute of Clinical Pathology and Medical Research; Westmead Hospital; University of Sydney | CIDM-PH et al. |
| EPI_ISL_544986 | Pathology West - NSW Health Pathology | NSW Health Pathology - Institute of Clinical Pathology and Medical Research; Westmead Hospital; University of Sydney | CIDM-PH et al. |
| EPI_ISL_544987 | Histopath | NSW Health Pathology - Institute of Clinical Pathology and Medical Research; Westmead Hospital; University of Sydney | CIDM-PH et al. |
| EPI_ISL_544988 | Australian Clinical Labs | NSW Health Pathology - Institute of Clinical Pathology and Medical Research; Westmead Hospital; University of Sydney | CIDM-PH et al. |
| EPI_ISL_544989, EPI_ISL_544990 | Pathology West - NSW Health Pathology | NSW Health Pathology - Institute of Clinical Pathology and Medical Research; Westmead Hospital; University of Sydney | CIDM-PH et al. |
| EPI_ISL_544991 | Pathology North - Royal North Shore Hospital - NSW Health Pathology | NSW Health Pathology - Institute of Clinical Pathology and Medical Research; Westmead Hospital; University of Sydney | CIDM-PH et al. |
| EPI_ISL_544992, EPI_ISL_544993, EPI_ISL_544994, EPI_ISL_544995, EPI_ISL_544996, EPI_ISL_544997, EPI_ISL_544998 | St Vincent's Pathology (SydPath) | NSW Health Pathology - Institute of Clinical Pathology and Medical Research; Westmead Hospital; University of Sydney | CIDM-PH et al. |
| EPI_ISL_545003 | Sydney South West Pathology Service (SSWPS) - Liverpool Hospital - NSW Health Pathology | NSW Health Pathology - Institute of Clinical Pathology and Medical Research; Westmead Hospital; University of Sydney | CIDM-PH et al. |
| EPI_ISL_545005 | Sydney South West Pathology Service (SSWPS) - Royal Prince Alfred Hospital - NSW Health Pathology | NSW Health Pathology - Institute of Clinical Pathology and Medical Research; Westmead Hospital; University of Sydney | CIDM-PH et al. |

| | | | |
|---|---|---|---|
| EPI_ISL_545009, EPI_ISL_545012, EPI_ISL_545013, EPI_ISL_545014 | South Eastern Area Laboratory Services (SEALS) | NSW Health Pathology - Institute of Clinical Pathology and Medical Research; Westmead Hospital; University of Sydney | CIDM-PH et al. |
| EPI_ISL_545017, EPI_ISL_545020, EPI_ISL_545021, EPI_ISL_545022 | Sydney South West Pathology Service (SSWPS) - Royal Prince Alfred Hospital - NSW Health Pathology | NSW Health Pathology - Institute of Clinical Pathology and Medical Research; Westmead Hospital; University of Sydney | CIDM-PH et al. |
| EPI_ISL_547581, EPI_ISL_547582 | University of Wisconsin-Madison AIDS Vaccine Research Laboratories | University of Wisconsin-Madison AIDS Vaccine Research Laboratories | Gage Moreno, Katarina Braun, et al. AIDS Vaccine Research Laboratories |
| EPI_ISL_547584 | CMS, Roorkee | CSIR-Institute of Microbial Technology | Kanika Bansal, Sanjeet Kumar, Anu Singh, Debarghya Ghose, Rajesh Kumar Mishra, Dipak Dutta, Sanjeev Khosla, Prabhu B. Patil |
| EPI_ISL_547585, EPI_ISL_547586, EPI_ISL_547587, EPI_ISL_547588, EPI_ISL_547589, EPI_ISL_547590, EPI_ISL_547591, EPI_ISL_547592, EPI_ISL_547593 | Civil Hospital, Panchkula | CSIR-Institute of Microbial Technology | Kanika Bansal, Sanjeet Kumar, Anu Singh, Debarghya Ghose, Rajesh Kumar Mishra, Dipak Dutta, Sanjeev Khosla, Prabhu B. Patil |
| EPI_ISL_547601 | South Eastern Area Laboratory Services (SEALS) | NSW Health Pathology - Institute of Clinical Pathology and Medical Research; Westmead Hospital; University of Sydney | CIDM-PH et al. |
| EPI_ISL_547725, EPI_ISL_547726, EPI_ISL_547727, EPI_ISL_547728, EPI_ISL_547729, EPI_ISL_547730, EPI_ISL_547731, EPI_ISL_547732, EPI_ISL_547733, EPI_ISL_547734, EPI_ISL_547735, EPI_ISL_547736, EPI_ISL_547737, EPI_ISL_547738, EPI_ISL_547739, EPI_ISL_547740, EPI_ISL_547741, EPI_ISL_547742, EPI_ISL_547743, EPI_ISL_547744, EPI_ISL_547745, EPI_ISL_547746, EPI_ISL_547747, EPI_ISL_547748, EPI_ISL_547749, EPI_ISL_547750, EPI_ISL_547751, EPI_ISL_547752, EPI_ISL_547753, EPI_ISL_547754 | | | |
| see above | Gundersen Molecular Diagnostics Laboratory | Kabara Cancer Research Institute | Craig S. Richmond, Paraic A. Kenny |
| EPI_ISL_547965 | Laboratorio Biologia Molecolare SarsCov2 UOC LAboratorio Analisi SErvizio Medicina di Laboratorio Ospedale San Francesco ATS-ASSL Nuoro | Laboratorio Specialistico UOC Ematologia Ospedale San Francesco - ATS ASSL NUORO | Piras Giovanna, Asproni Rosanna, Monne Maria Itria, Fancello Tatiana,Fiamma Maura,Toja Alessandro, Sanna Filomena, Floris Anna Rita, Sulis Vincenzo, Palmas Angelo Domenico, Casu Gavino, Lo Maglio Iana, Mameli Giuseppe. |
| EPI_ISL_548018, EPI_ISL_548020 | North Shore Hospital | Institute of Environmental Science and Research (ESR) | Xiaoyun Ren, Matt Storey, Nikki Freed, Muhammad Faisal, Jing Wang, Hermes Perez, Anja Werno, Antje van der Linden, Arlo Upton, Chris Mansell, David Hammer, Dragana Drinkovic, Gary McAuliffe, Hana Sofia Andersson, James Ussher, Jill Sherwood, Josh Freeman, Julia Howard, Juliet Elvy, Mary DeAlmeida, Matt Blakiston, Matthew Rogers, Max Bloomfield, Michael Addidle, Michelle Balm, Sally Roberts, Sarah Jefferies, Sharmini Muttaiyah, Susan Morpeth, Susan Taylor, Timothy Blackmore, Vani Sathyendran, Veronica Playle, Virginia Hope, Erasmus Smit, Lauren Jelly, Olin Silander, Joep de Ligt |
| EPI_ISL_548021, EPI_ISL_548022, EPI_ISL_548023, EPI_ISL_548024, EPI_ISL_548025, EPI_ISL_548026, EPI_ISL_548027 | LabPLUS | Institute of Environmental Science and Research (ESR) | Xiaoyun Ren, Matt Storey, Nikki Freed, Muhammad Faisal, Jing Wang, Hermes Perez, Anja Werno, Antje van der Linden, Arlo Upton, Chris Mansell, David Hammer, Dragana Drinkovic, Gary McAuliffe, Hana Sofia Andersson, James Ussher, Jill Sherwood, Josh Freeman, Julia Howard, Juliet Elvy, Mary DeAlmeida, Matt Blakiston, Matthew Rogers, Max Bloomfield, Michael Addidle, Michelle Balm, Sally Roberts, Sarah Jefferies, Sharmini Muttaiyah, Susan Morpeth, Susan Taylor, Timothy Blackmore, Vani Sathyendran, Veronica Playle, Virginia Hope, Erasmus Smit, Lauren Jelly, Olin Silander, Joep de Ligt |
| EPI_ISL_548028, EPI_ISL_548029, EPI_ISL_548030 | North Shore Hospital | Institute of Environmental Science and Research (ESR) | Xiaoyun Ren, Matt Storey, Nikki Freed, Muhammad Faisal, Jing Wang, Hermes Perez, Anja Werno, Antje van der Linden, Arlo Upton, Chris Mansell, David Hammer, Dragana Drinkovic, Gary McAuliffe, Hana Sofia Andersson, James Ussher, Jill Sherwood, Josh Freeman, Julia Howard, Juliet Elvy, Mary DeAlmeida, Matt Blakiston, Matthew Rogers, Max Bloomfield, Michael Addidle, Michelle Balm, Sally Roberts, Sarah Jefferies, Sharmini Muttaiyah, Susan Morpeth, Susan Taylor, Timothy Blackmore, Vani Sathyendran, Veronica Playle, Virginia Hope, Erasmus Smit, Lauren Jelly, Olin Silander, Joep de Ligt |
| EPI_ISL_548032, EPI_ISL_548033, EPI_ISL_548034, EPI_ISL_548035, EPI_ISL_548037, EPI_ISL_548038 | Middlemore Hospital | Institute of Environmental Science and Research (ESR) | Xiaoyun Ren, Matt Storey, Nikki Freed, Muhammad Faisal, Jing Wang, Hermes Perez, Anja Werno, Antje van der Linden, Arlo Upton, Chris Mansell, David Hammer, Dragana Drinkovic, Gary McAuliffe, Hana Sofia Andersson, James Ussher, Jill Sherwood, Josh Freeman, Julia Howard, Juliet Elvy, Mary DeAlmeida, Matt Blakiston, Matthew Rogers, Max Bloomfield, Michael Addidle, Michelle Balm, Sally Roberts, Sarah Jefferies, Sharmini Muttaiyah, Susan Morpeth, Susan Taylor, Timothy Blackmore, Vani Sathyendran, Veronica Playle, Virginia Hope, Erasmus Smit, Lauren Jelly, Olin Silander, Joep de Ligt |
| EPI_ISL_548040, EPI_ISL_548041 | LabPLUS | Institute of Environmental Science and Research (ESR) | Xiaoyun Ren, Matt Storey, Nikki Freed, Muhammad Faisal, Jing Wang, Hermes Perez, Anja Werno, Antje van der Linden, Arlo Upton, Chris Mansell, David Hammer, Dragana Drinkovic, Gary McAuliffe, Hana Sofia Andersson, James Ussher, Jill Sherwood, Josh Freeman, Julia Howard, Juliet Elvy, Mary DeAlmeida, Matt Blakiston, Matthew Rogers, Max Bloomfield, Michael Addidle, Michelle Balm, Sally Roberts, Sarah Jefferies, Sharmini Muttaiyah, Susan Morpeth, Susan Taylor, Timothy Blackmore, Vani Sathyendran, Veronica Playle, Virginia Hope, Erasmus Smit, Lauren Jelly, Olin Silander, Joep de Ligt |
| EPI_ISL_548042, EPI_ISL_548043, EPI_ISL_548044 | North Shore Hospital | Institute of Environmental Science and Research (ESR) | Xiaoyun Ren, Matt Storey, Nikki Freed, Muhammad Faisal, Jing Wang, Hermes Perez, Anja Werno, Antje van der Linden, Arlo Upton, Chris Mansell, David Hammer, Dragana Drinkovic, Gary McAuliffe, Hana Sofia Andersson, James Ussher, Jill Sherwood, Josh Freeman, Julia Howard, Juliet Elvy, Mary DeAlmeida, Matt Blakiston, Matthew Rogers, Max Bloomfield, Michael Addidle, Michelle Balm, Sally Roberts, Sarah Jefferies, Sharmini Muttaiyah, Susan Morpeth, Susan Taylor, Timothy Blackmore, Vani Sathyendran, Veronica Playle, Virginia Hope, Erasmus Smit, Lauren Jelly, Olin Silander, Joep de Ligt |
| EPI_ISL_548046 | LabTests | Institute of Environmental Science and Research (ESR) | Xiaoyun Ren, Matt Storey, Nikki Freed, Muhammad Faisal, Jing Wang, Hermes Perez, Anja Werno, Antje van der Linden, Arlo Upton, Chris Mansell, David Hammer, Dragana Drinkovic, Gary McAuliffe, Hana Sofia Andersson, James Ussher, Jill Sherwood, Josh Freeman, Julia Howard, Juliet Elvy, Mary DeAlmeida, Matt Blakiston, Matthew Rogers, Max Bloomfield, Michael Addidle, Michelle Balm, Sally Roberts, Sarah Jefferies, Sharmini Muttaiyah, Susan Morpeth, Susan Taylor, Timothy Blackmore, Vani Sathyendran, Veronica Playle, Virginia Hope, Erasmus Smit, Lauren Jelly, Olin Silander, Joep de Ligt |
| EPI_ISL_548047, EPI_ISL_548048 | North Shore Hospital | Institute of Environmental Science and Research (ESR) | Xiaoyun Ren, Matt Storey, Nikki Freed, Muhammad Faisal, Jing Wang, Hermes Perez, Anja Werno, Antje van der Linden, Arlo Upton, Chris Mansell, David Hammer, Dragana Drinkovic, Gary McAuliffe, Hana Sofia Andersson, James Ussher, Jill Sherwood, Josh Freeman, Julia Howard, Juliet Elvy, Mary DeAlmeida, Matt Blakiston, Matthew Rogers, Max Bloomfield, Michael Addidle, Michelle Balm, Sally Roberts, Sarah Jefferies, Sharmini Muttaiyah, Susan Morpeth, Susan Taylor, Timothy Blackmore, Vani Sathyendran, Veronica Playle, Virginia Hope, Erasmus Smit, Lauren Jelly, Olin Silander, Joep de Ligt |
| EPI_ISL_548049, EPI_ISL_548050, EPI_ISL_548051 | LabPLUS | Institute of Environmental Science and Research (ESR) | Xiaoyun Ren, Matt Storey, Nikki Freed, Muhammad Faisal, Jing Wang, Hermes Perez, Anja Werno, Antje van der Linden, Arlo Upton, Chris Mansell, David Hammer, Dragana Drinkovic, Gary McAuliffe, Hana Sofia Andersson, James Ussher, Jill Sherwood, Josh Freeman, Julia Howard, Juliet Elvy, Mary DeAlmeida, Matt Blakiston, Matthew Rogers, Max Bloomfield, Michael Addidle, Michelle Balm, Sally Roberts, Sarah Jefferies, Sharmini Muttaiyah, Susan Morpeth, Susan Taylor, Timothy Blackmore, Vani Sathyendran, Veronica Playle, Virginia Hope, Erasmus Smit, Lauren Jelly, Olin Silander, Joep de Ligt |
| EPI_ISL_548052, EPI_ISL_548053, EPI_ISL_548054, EPI_ISL_548055, EPI_ISL_548056 | Middlemore Hospital | Institute of Environmental Science and Research (ESR) | Xiaoyun Ren, Matt Storey, Nikki Freed, Muhammad Faisal, Jing Wang, Hermes Perez, Anja Werno, Antje van der Linden, Arlo Upton, Chris Mansell, David Hammer, Dragana Drinkovic, Gary McAuliffe, Hana Sofia Andersson, James Ussher, Jill Sherwood, Josh Freeman, Julia Howard, Juliet Elvy, Mary DeAlmeida, Matt Blakiston, Matthew Rogers, Max Bloomfield, Michael Addidle, Michelle Balm, Sally Roberts, Sarah Jefferies, Sharmini Muttaiyah, Susan Morpeth, Susan Taylor, Timothy Blackmore, Vani Sathyendran, Veronica Playle, Virginia Hope, Erasmus Smit, Lauren Jelly, Olin Silander, Joep de Ligt |
| EPI_ISL_548057, EPI_ISL_548058, EPI_ISL_548059, EPI_ISL_548060, EPI_ISL_548061, EPI_ISL_548062, EPI_ISL_548063 | North Shore Hospital | Institute of Environmental Science and Research (ESR) | Xiaoyun Ren, Matt Storey, Nikki Freed, Muhammad Faisal, Jing Wang, Hermes Perez, Anja Werno, Antje van der Linden, Arlo Upton, Chris Mansell, David Hammer, Dragana Drinkovic, Gary McAuliffe, Hana Sofia Andersson, James Ussher, Jill Sherwood, Josh Freeman, Julia Howard, Juliet Elvy, Mary DeAlmeida, Matt Blakiston, Matthew Rogers, Max Bloomfield, Michael Addidle, Michelle Balm, Sally Roberts, Sarah Jefferies, Sharmini Muttaiyah, Susan Morpeth, Susan Taylor, Timothy Blackmore, Vani Sathyendran, Veronica Playle, Virginia Hope, Erasmus Smit, Lauren Jelly, Olin Silander, Joep de Ligt |
| EPI_ISL_548064 | Waikato Hospital | Institute of Environmental Science and Research (ESR) | Xiaoyun Ren, Matt Storey, Nikki Freed, Muhammad Faisal, Jing Wang, Hermes Perez, Anja Werno, Antje van der Linden, Arlo Upton, Chris Mansell, David Hammer, Dragana Drinkovic, Gary McAuliffe, Hana Sofia Andersson, James Ussher, Jill Sherwood, Josh Freeman, Julia Howard, Juliet Elvy, Mary DeAlmeida, Matt Blakiston, Matthew Rogers, Max Bloomfield, Michael Addidle, Michelle Balm, Sally Roberts, Sarah Jefferies, Sharmini Muttaiyah, Susan Morpeth, Susan Taylor, Timothy Blackmore, Vani Sathyendran, Veronica Playle, Virginia Hope, Erasmus Smit, Lauren Jelly, Olin Silander, Joep de Ligt |
| EPI_ISL_548081 | Middlemore Hospital | Institute of Environmental Science and Research (ESR) | Xiaoyun Ren, Matt Storey, Nikki Freed, Muhammad Faisal, Jing Wang, Hermes Perez, Anja Werno, Antje van der Linden, Arlo Upton, Chris Mansell, David Hammer, Dragana Drinkovic, Gary McAuliffe, Hana Sofia Andersson, James Ussher, Jill Sherwood, Josh Freeman, Julia Howard, Juliet Elvy, Mary DeAlmeida, Matt Blakiston, Matthew Rogers, Max Bloomfield, Michael Addidle, Michelle Balm, Sally Roberts, Sarah Jefferies, Sharmini Muttaiyah, Susan Morpeth, Susan Taylor, Timothy Blackmore, Vani Sathyendran, Veronica Playle, Virginia Hope, Erasmus Smit, Lauren Jelly, Olin Silander, Joep de Ligt |
| EPI_ISL_548108, EPI_ISL_548109, EPI_ISL_548110, EPI_ISL_548111 | LabPLUS | Institute of Environmental Science and Research (ESR) | Xiaoyun Ren, Matt Storey, Nikki Freed, Muhammad Faisal, Jing Wang, Hermes Perez, Anja Werno, Antje van der Linden, Arlo Upton, Chris Mansell, David Hammer, Dragana Drinkovic, Gary McAuliffe, Hana Sofia Andersson, James Ussher, Jill Sherwood, Josh Freeman, Julia Howard, Juliet Elvy, Mary DeAlmeida, Matt Blakiston, Matthew Rogers, Max Bloomfield, Michael Addidle, Michelle Balm, Sally Roberts, Sarah Jefferies, Sharmini Muttaiyah, Susan Morpeth, Susan Taylor, Timothy Blackmore, Vani Sathyendran, Veronica Playle, Virginia Hope, Erasmus Smit, Lauren Jelly, Olin Silander, Joep de Ligt |

| | | | |
|---|---|---|---|
| EPI_ISL_548112, EPI_ISL_548113, EPI_ISL_548114, EPI_ISL_548115 | Middlemore Hospital | Institute of Environmental Science and Research (ESR) | Xiaoyun Ren, Matt Storey, Nikki Freed, Muhammad Faisal, Jing Wang, Hermes Perez, Anja Werno, Antje van der Linden, Arlo Upton, Chris Mansell, David Hammer, Dragana Drinkovic, Gary McAuliffe, Hana Sofia Andersson, James Ussher, Jill Sherwood, Josh Freeman, Julia Howard, Juliet Elvy, Mary DeAlmeida, Matt Blakiston, Matthew Rogers, Max Bloomfield, Michael Addidle, Michelle Balm, Sally Roberts, Sarah Jefferies, Sharmini Muttaiyah, Susan Morpeth, Susan Taylor, Timothy Blackmore, Vani Sathyendran, Veronica Playle, Virginia Hope, Erasmus Smit, Lauren Jelly, Olin Silander, Joep de Ligt |
| EPI_ISL_548116, EPI_ISL_548117, EPI_ISL_548118 | Canterbury Health Laboratories | Institute of Environmental Science and Research (ESR) | Xiaoyun Ren, Matt Storey, Nikki Freed, Muhammad Faisal, Jing Wang, Hermes Perez, Anja Werno, Antje van der Linden, Arlo Upton, Chris Mansell, David Hammer, Dragana Drinkovic, Gary McAuliffe, Hana Sofia Andersson, James Ussher, Jill Sherwood, Josh Freeman, Julia Howard, Juliet Elvy, Mary DeAlmeida, Matt Blakiston, Matthew Rogers, Max Bloomfield, Michael Addidle, Michelle Balm, Sally Roberts, Sarah Jefferies, Sharmini Muttaiyah, Susan Morpeth, Susan Taylor, Timothy Blackmore, Vani Sathyendran, Veronica Playle, Virginia Hope, Erasmus Smit, Lauren Jelly, Olin Silander, Joep de Ligt |
| EPI_ISL_548599 | Tuolumne County Public Health | Chan-Zuckerberg Biohub | CZB Cliahub Consortium |
| EPI_ISL_548602 | County of Santa Clara Public Health Department | Chan-Zuckerberg Biohub | CZB Cliahub Consortium |
| EPI_ISL_548626 | Tuolumne County Public Health | Chan-Zuckerberg Biohub | CZB Cliahub Consortium |
| EPI_ISL_549028 | Furst Medical Laboratory | Norwegian Institute of Public Health, Department of Virology | Kathrine Stene-Johansen, Kamilla Heddeland Instefjord, Hilde Elshaug, Rasmus Riis Kopperud, Hilde Synnøve Vollan, Karoline Bragstad, Olav Hungnes |
| EPI_ISL_549036, EPI_ISL_549037 | Hospital of Southern Norway - Kristiansand, Department of Medical Microbiology | Norwegian Institute of Public Health, Department of Virology | Kathrine Stene-Johansen, Kamilla Heddeland Instefjord, Hilde Elshaug, Rasmus Riis Kopperud, Hilde Synnøve Vollan, Karoline Bragstad, Olav Hungnes |
| EPI_ISL_549039, EPI_ISL_549040, EPI_ISL_549041 | Medical Microbiology Unit, Department for Laboratory Medicine, Drammen Hospital, Vestre Viken Health Trust, | Norwegian Institute of Public Health, Department of Virology | Kathrine Stene-Johansen, Kamilla Heddeland Instefjord, Hilde Elshaug, Rasmus Riis Kopperud, Hilde Synnøve Vollan, Karoline Bragstad, Olav Hungnes |
| EPI_ISL_549202, EPI_ISL_549203, EPI_ISL_549204, EPI_ISL_549205, EPI_ISL_549206, EPI_ISL_549207, EPI_ISL_549208, EPI_ISL_549213, EPI_ISL_549214 | Florida Bureau of Public Health Laboratories | Florida Bureau of Public Health Laboratories | Sarah Schmedes, Jason Blanton |
| EPI_ISL_549332, EPI_ISL_549338, EPI_ISL_549339, EPI_ISL_549340, EPI_ISL_549342, EPI_ISL_549345, EPI_ISL_549346, EPI_ISL_549348, EPI_ISL_549414, EPI_ISL_549415, EPI_ISL_549416, EPI_ISL_549417, EPI_ISL_549418, EPI_ISL_549419, EPI_ISL_549420, EPI_ISL_549421, EPI_ISL_549422, EPI_ISL_549423, EPI_ISL_549424, EPI_ISL_549425, EPI_ISL_549426, EPI_ISL_549427, EPI_ISL_549428, EPI_ISL_549429, EPI_ISL_549440, EPI_ISL_549441, EPI_ISL_549442, EPI_ISL_549443, EPI_ISL_549444, EPI_ISL_549445, EPI_ISL_549446, EPI_ISL_549447, EPI_ISL_549448, EPI_ISL_549449, EPI_ISL_549450, EPI_ISL_549451, EPI_ISL_549452, EPI_ISL_549453, EPI_ISL_549454, EPI_ISL_549455, EPI_ISL_549456, EPI_ISL_549457, EPI_ISL_549458, EPI_ISL_549459, EPI_ISL_549523, EPI_ISL_549524, EPI_ISL_549525, EPI_ISL_549526, EPI_ISL_549527, EPI_ISL_549528, EPI_ISL_549529, EPI_ISL_549530, EPI_ISL_549531, EPI_ISL_549532, EPI_ISL_549533, EPI_ISL_549534, EPI_ISL_549535, EPI_ISL_549536, EPI_ISL_549537, EPI_ISL_549538, EPI_ISL_549539, EPI_ISL_549540, EPI_ISL_549541, EPI_ISL_549542, EPI_ISL_549543, EPI_ISL_549544, EPI_ISL_549545, EPI_ISL_549546 | | | |
| see above | Oxford Viromics, NDM, University of Oxford; Oxford University Hospitals; Basingstoke and North Hampshire Hospital | COVID-19 Genomics UK (COG-UK) Consortium | Tanya Golubchik, David Bonsall, George Macintyre, Amy Trebes, Mariateresa de Cesare, Catrin Moore, Alex Mobbs, Anita Justice, Robert Shaw, Monique Andersson, Timothy Peto, Emma Wise, Nathan Moore, Jessica Lynch, Nick Cortes, Matilde Mori, Stephen Kidd, David Buck, John Todd, Christophe Fraser |
| EPI_ISL_549808, EPI_ISL_549809, EPI_ISL_549810, EPI_ISL_549811, EPI_ISL_549812, EPI_ISL_549813, EPI_ISL_549814, EPI_ISL_549815, EPI_ISL_549816, EPI_ISL_549817, EPI_ISL_549818, EPI_ISL_549819, EPI_ISL_549820 | | | |
| see above | Lighthouse Lab in Milton Keynes | Wellcome Sanger Institute for the COVID-19 Genomics UK (COG-UK) consortium | The Lighthouse Lab in Milton Keynes and Alex Alderton, Roberto Amato, Sonia Goncalves, Ewan Harrison, David K. Jackson, Ian Johnston, Dominic Kwiatkowski, Cordelia Langford, John Sillitoe on behalf of the Wellcome Sanger Institute COVID-19 Surveillance Team (http://www.sanger.ac.uk/covid-team) |
| EPI_ISL_549821 | Lighthouse Lab in Milton Keynes | Wellcome Sanger Institute for the COVID-19 Genomics UK (COG-UK) consortium | The Lighthouse Lab in Alderley Park and Alex Alderton, Roberto Amato, Sonia Goncalves, Ewan Harrison, David K. Jackson, Ian Johnston, Dominic Kwiatkowski, Cordelia Langford, John Sillitoe on behalf of the Wellcome Sanger Institute COVID-19 Surveillance Team |
| EPI_ISL_549822, EPI_ISL_549823, EPI_ISL_549824, EPI_ISL_549825, EPI_ISL_549826, EPI_ISL_549827, EPI_ISL_549828 | Lighthouse Lab in Milton Keynes | Wellcome Sanger Institute for the COVID-19 Genomics UK (COG-UK) consortium | The Lighthouse Lab in Milton Keynes and Alex Alderton, Roberto Amato, Sonia Goncalves, Ewan Harrison, David K. Jackson, Ian Johnston, Dominic Kwiatkowski, Cordelia Langford, John Sillitoe on behalf of the Wellcome Sanger Institute COVID-19 Surveillance Team (http://www.sanger.ac.uk/covid-team) |
| EPI_ISL_549829 | Lighthouse Lab in Milton Keynes | Wellcome Sanger Institute for the COVID-19 Genomics UK (COG-UK) Consortium | The Lighthouse Lab in Milton Keynes and Alex Alderton, Roberto Amato, Sonia Goncalves, Ewan Harrison, David K. Jackson, Ian Johnston, Dominic Kwiatkowski, Cordelia Langford, John Sillitoe on behalf of the Wellcome Sanger Institute COVID-19 Surveillance Team |
| EPI_ISL_549830, EPI_ISL_549831, EPI_ISL_549832, EPI_ISL_549833, EPI_ISL_549834, EPI_ISL_549835, EPI_ISL_549836, EPI_ISL_549837, EPI_ISL_549838, EPI_ISL_549839, EPI_ISL_549840, EPI_ISL_549841, EPI_ISL_549842, EPI_ISL_549843, EPI_ISL_549844, EPI_ISL_549845, EPI_ISL_549846, EPI_ISL_549847, EPI_ISL_549848, EPI_ISL_549849, EPI_ISL_549850, EPI_ISL_549851, EPI_ISL_549852, EPI_ISL_549853, EPI_ISL_549854, EPI_ISL_549855, EPI_ISL_549856, EPI_ISL_549857, EPI_ISL_549858, EPI_ISL_549859, EPI_ISL_549860 | | | |
| see above | Lighthouse Lab in Milton Keynes | Wellcome Sanger Institute for the COVID-19 Genomics UK (COG-UK) consortium | The Lighthouse Lab in Milton Keynes and Alex Alderton, Roberto Amato, Sonia Goncalves, Ewan Harrison, David K. Jackson, Ian Johnston, Dominic Kwiatkowski, Cordelia Langford, John Sillitoe on behalf of the Wellcome Sanger Institute COVID-19 Surveillance Team (http://www.sanger.ac.uk/covid-team) |
| EPI_ISL_549861 | Lighthouse Lab in Milton Keynes | Wellcome Sanger Institute for the COVID-19 Genomics UK (COG-UK) Consortium | The Lighthouse Lab in Milton Keynes and Alex Alderton, Roberto Amato, Sonia Goncalves, Ewan Harrison, David K. Jackson, Ian Johnston, Dominic Kwiatkowski, Cordelia Langford, John Sillitoe on behalf of the Wellcome Sanger Institute COVID-19 Surveillance Team |
| EPI_ISL_549862 | Lighthouse Lab in Milton Keynes | Wellcome Sanger Institute for the COVID-19 Genomics UK (COG-UK) consortium | The Lighthouse Lab in Milton Keynes and Alex Alderton, Roberto Amato, Sonia Goncalves, Ewan Harrison, David K. Jackson, Ian Johnston, Dominic Kwiatkowski, Cordelia Langford, John Sillitoe on behalf of the Wellcome Sanger Institute COVID-19 Surveillance Team (http://www.sanger.ac.uk/covid-team) |
| EPI_ISL_549863 | Lighthouse Lab in Milton Keynes | Wellcome Sanger Institute for the COVID-19 Genomics UK (COG-UK) consortium | The Lighthouse Lab in Alderley Park and Alex Alderton, Roberto Amato, Sonia Goncalves, Ewan Harrison, David K. Jackson, Ian Johnston, Dominic Kwiatkowski, Cordelia Langford, John Sillitoe on behalf of the Wellcome Sanger Institute COVID-19 Surveillance Team |
| EPI_ISL_549864, EPI_ISL_549865, EPI_ISL_549866, EPI_ISL_549867, EPI_ISL_549868, EPI_ISL_549869, EPI_ISL_549870, EPI_ISL_549871, EPI_ISL_549872, EPI_ISL_549873, EPI_ISL_549874, EPI_ISL_549875, EPI_ISL_549876, EPI_ISL_549877, EPI_ISL_549878, EPI_ISL_549879, EPI_ISL_549880, EPI_ISL_549881, EPI_ISL_549882, EPI_ISL_549883, EPI_ISL_549884, EPI_ISL_549885, EPI_ISL_549886, EPI_ISL_549887, EPI_ISL_549888, EPI_ISL_549889, EPI_ISL_549890, EPI_ISL_549891, EPI_ISL_549892, EPI_ISL_549893, EPI_ISL_549894, EPI_ISL_549895, EPI_ISL_549896, EPI_ISL_549897, EPI_ISL_549898, EPI_ISL_549899, EPI_ISL_549900, EPI_ISL_549901, EPI_ISL_549902, EPI_ISL_549903, EPI_ISL_549904, EPI_ISL_549905, EPI_ISL_549906, EPI_ISL_549907, EPI_ISL_549908, EPI_ISL_549909, EPI_ISL_549910, EPI_ISL_549911, EPI_ISL_549912, EPI_ISL_549913, EPI_ISL_549914, EPI_ISL_549915, EPI_ISL_549916, EPI_ISL_549917, EPI_ISL_549918, EPI_ISL_549919, EPI_ISL_549920, EPI_ISL_549921, EPI_ISL_549922, EPI_ISL_549923, EPI_ISL_549924, EPI_ISL_549925, EPI_ISL_549926, EPI_ISL_549927, EPI_ISL_549928, EPI_ISL_549929, EPI_ISL_549930, EPI_ISL_549931, EPI_ISL_549932, EPI_ISL_549933, EPI_ISL_549934, EPI_ISL_549935, EPI_ISL_549936, EPI_ISL_549937, EPI_ISL_549938, EPI_ISL_549939 | | | |
| see above | Lighthouse Lab in Milton Keynes | Wellcome Sanger Institute for the COVID-19 Genomics UK (COG-UK) consortium | The Lighthouse Lab in Milton Keynes and Alex Alderton, Roberto Amato, Sonia Goncalves, Ewan Harrison, David K. Jackson, Ian Johnston, Dominic Kwiatkowski, Cordelia Langford, John Sillitoe on behalf of the Wellcome Sanger Institute COVID-19 Surveillance Team (http://www.sanger.ac.uk/covid-team) |
| EPI_ISL_549940 | Lighthouse Lab in Milton Keynes | Wellcome Sanger Institute for the COVID-19 Genomics UK (COG-UK) consortium | The Lighthouse Lab in Alderley Park and Alex Alderton, Roberto Amato, Sonia Goncalves, Ewan Harrison, David K. Jackson, Ian Johnston, Dominic Kwiatkowski, Cordelia Langford, John Sillitoe on behalf of the Wellcome Sanger Institute COVID-19 Surveillance Team |
| EPI_ISL_549941, EPI_ISL_549942, EPI_ISL_549943, EPI_ISL_549944, EPI_ISL_549945, EPI_ISL_549946, EPI_ISL_549947, EPI_ISL_549968, EPI_ISL_549970, EPI_ISL_550058, EPI_ISL_550062, EPI_ISL_550066, EPI_ISL_550073, EPI_ISL_550075, EPI_ISL_550077, EPI_ISL_550104, EPI_ISL_550107, EPI_ISL_550108, EPI_ISL_550111, EPI_ISL_550120, EPI_ISL_550121, EPI_ISL_550122, EPI_ISL_550123, EPI_ISL_550124, EPI_ISL_550135, EPI_ISL_550169, EPI_ISL_550171, EPI_ISL_550209, EPI_ISL_550212, EPI_ISL_550223, EPI_ISL_550243, EPI_ISL_550262, EPI_ISL_550263, EPI_ISL_550264, EPI_ISL_550265, EPI_ISL_550266, EPI_ISL_550267, EPI_ISL_550268, EPI_ISL_550269, EPI_ISL_550270, EPI_ISL_550271, EPI_ISL_550272, EPI_ISL_550273, EPI_ISL_550274, EPI_ISL_550275, EPI_ISL_550276 | | | |
| see above | Lighthouse Lab in Milton Keynes | Wellcome Sanger Institute for the COVID-19 Genomics UK (COG-UK) consortium | The Lighthouse Lab in Milton Keynes and Alex Alderton, Roberto Amato, Sonia Goncalves, Ewan Harrison, David K. Jackson, Ian Johnston, Dominic Kwiatkowski, Cordelia Langford, John Sillitoe on behalf of the Wellcome Sanger Institute COVID-19 Surveillance Team (http://www.sanger.ac.uk/covid-team) |
| EPI_ISL_550277 | Lighthouse Lab in Milton Keynes | Wellcome Sanger Institute for the COVID-19 Genomics UK (COG-UK) consortium | The Lighthouse Lab in Alderley Park and Alex Alderton, Roberto Amato, Sonia Goncalves, Ewan Harrison, David K. Jackson, Ian Johnston, Dominic Kwiatkowski, Cordelia Langford, John Sillitoe on behalf of the Wellcome Sanger Institute COVID-19 Surveillance Team |
| EPI_ISL_550278, EPI_ISL_550279, EPI_ISL_550280, EPI_ISL_550281, EPI_ISL_550282, EPI_ISL_550283, EPI_ISL_550284, EPI_ISL_550285, EPI_ISL_550286, EPI_ISL_550287, EPI_ISL_550288, EPI_ISL_550289, EPI_ISL_550290, EPI_ISL_550291, EPI_ISL_550292, EPI_ISL_550293, EPI_ISL_550294, EPI_ISL_550295, EPI_ISL_550296, EPI_ISL_550297, EPI_ISL_550298 | | | |
| see above | Lighthouse Lab in Milton Keynes | Wellcome Sanger Institute for the COVID-19 Genomics UK (COG-UK) consortium | The Lighthouse Lab in Milton Keynes and Alex Alderton, Roberto Amato, Sonia Goncalves, Ewan Harrison, David K. Jackson, Ian Johnston, Dominic Kwiatkowski, Cordelia Langford, John Sillitoe on behalf of the Wellcome Sanger Institute COVID-19 Surveillance Team |

| | | | |
|---|---|---|---|
| | | | (http://www.sanger.ac.uk/covid-team) |
| EPI_ISL_550299 | Lighthouse Lab in Milton Keynes | Wellcome Sanger Institute for the COVID-19 Genomics UK (COG-UK) consortium | The Lighthouse Lab in Milton Keynes and Alex Alderton, Roberto Amato, Sonia Goncalves, Ewan Harrison, David K. Jackson, Ian Johnston, Dominic Kwiatkowski, Cordelia Langford, John Sillitoe on behalf of the Wellcome Sanger Institute COVID-19 Surveillance Team |
| EPI_ISL_550300, EPI_ISL_550301, EPI_ISL_550302, EPI_ISL_550303, EPI_ISL_550304, EPI_ISL_550305, EPI_ISL_550306, EPI_ISL_550307, EPI_ISL_550308, EPI_ISL_550309, EPI_ISL_550310, EPI_ISL_550311, EPI_ISL_550312, EPI_ISL_550313, EPI_ISL_550314, EPI_ISL_550315, EPI_ISL_550316, EPI_ISL_550317, EPI_ISL_550318 | | | |
| see above | Lighthouse Lab in Milton Keynes | Wellcome Sanger Institute for the COVID-19 Genomics UK (COG-UK) consortium | The Lighthouse Lab in Milton Keynes and Alex Alderton, Roberto Amato, Sonia Goncalves, Ewan Harrison, David K. Jackson, Ian Johnston, Dominic Kwiatkowski, Cordelia Langford, John Sillitoe on behalf of the Wellcome Sanger Institute COVID-19 Surveillance Team (http://www.sanger.ac.uk/covid-team) |
| EPI_ISL_550319 | Lighthouse Lab in Milton Keynes | Wellcome Sanger Institute for the COVID-19 Genomics UK (COG-UK) consortium | The Lighthouse Lab in Alderley Park and Alex Alderton, Roberto Amato, Sonia Goncalves, Ewan Harrison, David K. Jackson, Ian Johnston, Dominic Kwiatkowski, Cordelia Langford, John Sillitoe on behalf of the Wellcome Sanger Institute COVID-19 Surveillance Team |
| EPI_ISL_550320, EPI_ISL_550321, EPI_ISL_550322, EPI_ISL_550323, EPI_ISL_550324, EPI_ISL_550325, EPI_ISL_550326, EPI_ISL_550327 | Lighthouse Lab in Milton Keynes | Wellcome Sanger Institute for the COVID-19 Genomics UK (COG-UK) consortium | The Lighthouse Lab in Milton Keynes and Alex Alderton, Roberto Amato, Sonia Goncalves, Ewan Harrison, David K. Jackson, Ian Johnston, Dominic Kwiatkowski, Cordelia Langford, John Sillitoe on behalf of the Wellcome Sanger Institute COVID-19 Surveillance Team (http://www.sanger.ac.uk/covid-team) |
| EPI_ISL_550328 | Lighthouse Lab in Milton Keynes | Wellcome Sanger Institute for the COVID-19 Genomics UK (COG-UK) consortium | The Lighthouse Lab in Alderley Park and Alex Alderton, Roberto Amato, Sonia Goncalves, Ewan Harrison, David K. Jackson, Ian Johnston, Dominic Kwiatkowski, Cordelia Langford, John Sillitoe on behalf of the Wellcome Sanger Institute COVID-19 Surveillance Team |
| EPI_ISL_550329, EPI_ISL_550330, EPI_ISL_550331, EPI_ISL_550332, EPI_ISL_550334, EPI_ISL_550335, EPI_ISL_550336, EPI_ISL_550337, EPI_ISL_550338, EPI_ISL_550339, EPI_ISL_550340, EPI_ISL_550341, EPI_ISL_550342, EPI_ISL_550343, EPI_ISL_550344, EPI_ISL_550345, EPI_ISL_550346, EPI_ISL_550347, EPI_ISL_550348, EPI_ISL_550349, EPI_ISL_550350, EPI_ISL_550351, EPI_ISL_550352, EPI_ISL_550353, EPI_ISL_550354, EPI_ISL_550355, EPI_ISL_550356 | | | |
| see above | Lighthouse Lab in Milton Keynes | Wellcome Sanger Institute for the COVID-19 Genomics UK (COG-UK) consortium | The Lighthouse Lab in Milton Keynes and Alex Alderton, Roberto Amato, Sonia Goncalves, Ewan Harrison, David K. Jackson, Ian Johnston, Dominic Kwiatkowski, Cordelia Langford, John Sillitoe on behalf of the Wellcome Sanger Institute COVID-19 Surveillance Team (http://www.sanger.ac.uk/covid-team) |
| EPI_ISL_550357 | Lighthouse Lab in Milton Keynes | Wellcome Sanger Institute for the COVID-19 Genomics UK (COG-UK) consortium | The Lighthouse Lab in Alderley Park and Alex Alderton, Roberto Amato, Sonia Goncalves, Ewan Harrison, David K. Jackson, Ian Johnston, Dominic Kwiatkowski, Cordelia Langford, John Sillitoe on behalf of the Wellcome Sanger Institute COVID-19 Surveillance Team |
| EPI_ISL_550358, EPI_ISL_550359, EPI_ISL_550360, EPI_ISL_550361, EPI_ISL_550362, EPI_ISL_550363 | Lighthouse Lab in Milton Keynes | Wellcome Sanger Institute for the COVID-19 Genomics UK (COG-UK) consortium | The Lighthouse Lab in Milton Keynes and Alex Alderton, Roberto Amato, Sonia Goncalves, Ewan Harrison, David K. Jackson, Ian Johnston, Dominic Kwiatkowski, Cordelia Langford, John Sillitoe on behalf of the Wellcome Sanger Institute COVID-19 Surveillance Team (http://www.sanger.ac.uk/covid-team) |
| EPI_ISL_550364 | Lighthouse Lab in Milton Keynes | Wellcome Sanger Institute for the COVID-19 Genomics UK (COG-UK) consortium | The Lighthouse Lab in Alderley Park and Alex Alderton, Roberto Amato, Sonia Goncalves, Ewan Harrison, David K. Jackson, Ian Johnston, Dominic Kwiatkowski, Cordelia Langford, John Sillitoe on behalf of the Wellcome Sanger Institute COVID-19 Surveillance Team |
| EPI_ISL_550365, EPI_ISL_550366, EPI_ISL_550367, EPI_ISL_550368, EPI_ISL_550369, EPI_ISL_550370, EPI_ISL_550371, EPI_ISL_550372 | Lighthouse Lab in Milton Keynes | Wellcome Sanger Institute for the COVID-19 Genomics UK (COG-UK) consortium | The Lighthouse Lab in Milton Keynes and Alex Alderton, Roberto Amato, Sonia Goncalves, Ewan Harrison, David K. Jackson, Ian Johnston, Dominic Kwiatkowski, Cordelia Langford, John Sillitoe on behalf of the Wellcome Sanger Institute COVID-19 Surveillance Team (http://www.sanger.ac.uk/covid-team) |
| EPI_ISL_550373 | Lighthouse Lab in Milton Keynes | Wellcome Sanger Institute for the COVID-19 Genomics UK (COG-UK) consortium | The Lighthouse Lab in Alderley Park and Alex Alderton, Roberto Amato, Sonia Goncalves, Ewan Harrison, David K. Jackson, Ian Johnston, Dominic Kwiatkowski, Cordelia Langford, John Sillitoe on behalf of the Wellcome Sanger Institute COVID-19 Surveillance Team |
| EPI_ISL_550374, EPI_ISL_550375, EPI_ISL_550376, EPI_ISL_550377, EPI_ISL_550378, EPI_ISL_550379, EPI_ISL_550380, EPI_ISL_550381, EPI_ISL_550382, EPI_ISL_550383, EPI_ISL_550384, EPI_ISL_550385, EPI_ISL_550386, EPI_ISL_550387, EPI_ISL_550388, EPI_ISL_550389, EPI_ISL_550390, EPI_ISL_550391, EPI_ISL_550392, EPI_ISL_550393, EPI_ISL_550394, EPI_ISL_550395 | | | |
| see above | Lighthouse Lab in Milton Keynes | Wellcome Sanger Institute for the COVID-19 Genomics UK (COG-UK) consortium | The Lighthouse Lab in Milton Keynes and Alex Alderton, Roberto Amato, Sonia Goncalves, Ewan Harrison, David K. Jackson, Ian Johnston, Dominic Kwiatkowski, Cordelia Langford, John Sillitoe on behalf of the Wellcome Sanger Institute COVID-19 Surveillance Team (http://www.sanger.ac.uk/covid-team) |
| EPI_ISL_550396, EPI_ISL_550397 | Lighthouse Lab in Milton Keynes | Wellcome Sanger Institute for the COVID-19 Genomics UK (COG-UK) consortium | The Lighthouse Lab in Alderley Park and Alex Alderton, Roberto Amato, Sonia Goncalves, Ewan Harrison, David K. Jackson, Ian Johnston, Dominic Kwiatkowski, Cordelia Langford, John Sillitoe on behalf of the Wellcome Sanger Institute COVID-19 Surveillance Team |
| EPI_ISL_550398, EPI_ISL_550399 | Lighthouse Lab in Milton Keynes | Wellcome Sanger Institute for the COVID-19 Genomics UK (COG-UK) consortium | The Lighthouse Lab in Milton Keynes and Alex Alderton, Roberto Amato, Sonia Goncalves, Ewan Harrison, David K. Jackson, Ian Johnston, Dominic Kwiatkowski, Cordelia Langford, John Sillitoe on behalf of the Wellcome Sanger Institute COVID-19 Surveillance Team (http://www.sanger.ac.uk/covid-team) |
| EPI_ISL_550400 | Lighthouse Lab in Milton Keynes | Wellcome Sanger Institute for the COVID-19 Genomics UK (COG-UK) consortium | The Lighthouse Lab in Alderley Park and Alex Alderton, Roberto Amato, Sonia Goncalves, Ewan Harrison, David K. Jackson, Ian Johnston, Dominic Kwiatkowski, Cordelia Langford, John Sillitoe on behalf of the Wellcome Sanger Institute COVID-19 Surveillance Team |
| EPI_ISL_550401, EPI_ISL_550402, EPI_ISL_550403, EPI_ISL_550404, EPI_ISL_550405, EPI_ISL_550406, EPI_ISL_550407, EPI_ISL_550408, EPI_ISL_550409, EPI_ISL_550410 | Lighthouse Lab in Milton Keynes | Wellcome Sanger Institute for the COVID-19 Genomics UK (COG-UK) consortium | The Lighthouse Lab in Milton Keynes and Alex Alderton, Roberto Amato, Sonia Goncalves, Ewan Harrison, David K. Jackson, Ian Johnston, Dominic Kwiatkowski, Cordelia Langford, John Sillitoe on behalf of the Wellcome Sanger Institute COVID-19 Surveillance Team (http://www.sanger.ac.uk/covid-team) |
| EPI_ISL_550411 | Lighthouse Lab in Milton Keynes | Wellcome Sanger Institute for the COVID-19 Genomics UK (COG-UK) consortium | The Lighthouse Lab in Alderley Park and Alex Alderton, Roberto Amato, Sonia Goncalves, Ewan Harrison, David K. Jackson, Ian Johnston, Dominic Kwiatkowski, Cordelia Langford, John Sillitoe on behalf of the Wellcome Sanger Institute COVID-19 Surveillance Team |
| EPI_ISL_550412, EPI_ISL_550413, EPI_ISL_550414, EPI_ISL_550415, EPI_ISL_550416, EPI_ISL_550518, EPI_ISL_550519, EPI_ISL_550520, EPI_ISL_550521, EPI_ISL_550522, EPI_ISL_550523, EPI_ISL_550524, EPI_ISL_550525, EPI_ISL_550526, EPI_ISL_550527, EPI_ISL_550528, EPI_ISL_550529, EPI_ISL_550530, EPI_ISL_550531, EPI_ISL_550532, EPI_ISL_550533, EPI_ISL_550534 | | | |
| see above | Lighthouse Lab in Milton Keynes | Wellcome Sanger Institute for the COVID-19 Genomics UK (COG-UK) consortium | The Lighthouse Lab in Milton Keynes and Alex Alderton, Roberto Amato, Sonia Goncalves, Ewan Harrison, David K. Jackson, Ian Johnston, Dominic Kwiatkowski, Cordelia Langford, John Sillitoe on behalf of the Wellcome Sanger Institute COVID-19 Surveillance Team (http://www.sanger.ac.uk/covid-team) |
| EPI_ISL_550535, EPI_ISL_550536 | Lighthouse Lab in Milton Keynes | Wellcome Sanger Institute for the COVID-19 Genomics UK (COG-UK) consortium | The Lighthouse Lab in Alderley Park and Alex Alderton, Roberto Amato, Sonia Goncalves, Ewan Harrison, David K. Jackson, Ian Johnston, Dominic Kwiatkowski, Cordelia Langford, John Sillitoe on behalf of the Wellcome Sanger Institute COVID-19 Surveillance Team |
| EPI_ISL_550537, EPI_ISL_550538, EPI_ISL_550539, EPI_ISL_550540, EPI_ISL_550541, EPI_ISL_550542, EPI_ISL_550543, EPI_ISL_550544, EPI_ISL_550545, EPI_ISL_550546, EPI_ISL_550547, EPI_ISL_550548, EPI_ISL_550549, EPI_ISL_550550 | | | |
| see above | Lighthouse Lab in Milton Keynes | Wellcome Sanger Institute for the COVID-19 Genomics UK (COG-UK) consortium | The Lighthouse Lab in Milton Keynes and Alex Alderton, Roberto Amato, Sonia Goncalves, Ewan Harrison, David K. Jackson, Ian Johnston, Dominic Kwiatkowski, Cordelia Langford, John Sillitoe on behalf of the Wellcome Sanger Institute COVID-19 Surveillance Team (http://www.sanger.ac.uk/covid-team) |
| EPI_ISL_550551, EPI_ISL_550552 | Lighthouse Lab in Milton Keynes | Wellcome Sanger Institute for the COVID-19 Genomics UK (COG-UK) consortium | The Lighthouse Lab in Alderley Park and Alex Alderton, Roberto Amato, Sonia Goncalves, Ewan Harrison, David K. Jackson, Ian Johnston, Dominic Kwiatkowski, Cordelia Langford, John Sillitoe on behalf of the Wellcome Sanger Institute COVID-19 Surveillance Team |
| EPI_ISL_550553, EPI_ISL_550554, EPI_ISL_550555, EPI_ISL_550556, EPI_ISL_550557, EPI_ISL_550558, EPI_ISL_550559, EPI_ISL_550560, EPI_ISL_550561, EPI_ISL_550562, EPI_ISL_550563, EPI_ISL_550564, EPI_ISL_550565, EPI_ISL_550566, EPI_ISL_550567, EPI_ISL_550568, EPI_ISL_550569, EPI_ISL_550570, EPI_ISL_550571, EPI_ISL_550572, EPI_ISL_550573, EPI_ISL_550574 | | | |
| see above | Lighthouse Lab in Milton Keynes | Wellcome Sanger Institute for the COVID-19 Genomics UK (COG-UK) consortium | The Lighthouse Lab in Milton Keynes and Alex Alderton, Roberto Amato, Sonia Goncalves, Ewan Harrison, David K. Jackson, Ian Johnston, Dominic Kwiatkowski, Cordelia Langford, John Sillitoe on behalf of the Wellcome Sanger Institute COVID-19 Surveillance Team (http://www.sanger.ac.uk/covid-team) |
| EPI_ISL_550575 | Lighthouse Lab in Milton Keynes | Wellcome Sanger Institute for the COVID-19 Genomics UK (COG-UK) consortium | The Lighthouse Lab in Alderley Park and Alex Alderton, Roberto Amato, Sonia Goncalves, Ewan Harrison, David K. Jackson, Ian Johnston, Dominic Kwiatkowski, Cordelia Langford, John Sillitoe on behalf of the Wellcome Sanger Institute COVID-19 Surveillance Team |

| | | | |
|---|---|---|---|
| EPI_ISL_550576, EPI_ISL_550577, EPI_ISL_550578, EPI_ISL_550579, EPI_ISL_550580, EPI_ISL_550581, EPI_ISL_550582, EPI_ISL_550583, EPI_ISL_550584, EPI_ISL_550585, EPI_ISL_550586 | | | |
| see above | Lighthouse Lab in Milton Keynes | Wellcome Sanger Institute for the COVID-19 Genomics UK (COG-UK) consortium | The Lighthouse Lab in Milton Keynes and Alex Alderton, Roberto Amato, Sonia Goncalves, Ewan Harrison, David K. Jackson, Ian Johnston, Dominic Kwiatkowski, Cordelia Langford, John Sillitoe on behalf of the Wellcome Sanger Institute COVID-19 Surveillance Team (http://www.sanger.ac.uk/covid-team) |
| EPI_ISL_550587 | Lighthouse Lab in Milton Keynes | Wellcome Sanger Institute for the COVID-19 Genomics UK (COG-UK) consortium | The Lighthouse Lab in Alderley Park and Alex Alderton, Roberto Amato, Sonia Goncalves, Ewan Harrison, David K. Jackson, Ian Johnston, Dominic Kwiatkowski, Cordelia Langford, John Sillitoe on behalf of the Wellcome Sanger Institute COVID-19 Surveillance Team |
| EPI_ISL_550588, EPI_ISL_550589, EPI_ISL_550590, EPI_ISL_550591, EPI_ISL_550592, EPI_ISL_550593, EPI_ISL_550594, EPI_ISL_550595, EPI_ISL_550596, EPI_ISL_550597, EPI_ISL_550598, EPI_ISL_550599, EPI_ISL_550600, EPI_ISL_550601, EPI_ISL_550602, EPI_ISL_550603, EPI_ISL_550604, EPI_ISL_550605, EPI_ISL_550606, EPI_ISL_550607, EPI_ISL_550608, EPI_ISL_550609, EPI_ISL_550610, EPI_ISL_550611, EPI_ISL_550612, EPI_ISL_550613 | | | |
| see above | Lighthouse Lab in Milton Keynes | Wellcome Sanger Institute for the COVID-19 Genomics UK (COG-UK) consortium | The Lighthouse Lab in Milton Keynes and Alex Alderton, Roberto Amato, Sonia Goncalves, Ewan Harrison, David K. Jackson, Ian Johnston, Dominic Kwiatkowski, Cordelia Langford, John Sillitoe on behalf of the Wellcome Sanger Institute COVID-19 Surveillance Team (http://www.sanger.ac.uk/covid-team) |
| EPI_ISL_550614 | Lighthouse Lab in Milton Keynes | Wellcome Sanger Institute for the COVID-19 Genomics UK (COG-UK) consortium | The Lighthouse Lab in Alderley Park and Alex Alderton, Roberto Amato, Sonia Goncalves, Ewan Harrison, David K. Jackson, Ian Johnston, Dominic Kwiatkowski, Cordelia Langford, John Sillitoe on behalf of the Wellcome Sanger Institute COVID-19 Surveillance Team |
| EPI_ISL_550615, EPI_ISL_550616, EPI_ISL_550617, EPI_ISL_550618, EPI_ISL_550619, EPI_ISL_550620, EPI_ISL_550621, EPI_ISL_550622, EPI_ISL_550623, EPI_ISL_550624, EPI_ISL_550625, EPI_ISL_550626, EPI_ISL_550627, EPI_ISL_550628, EPI_ISL_550629, EPI_ISL_550630, EPI_ISL_550631, EPI_ISL_550632, EPI_ISL_550633, EPI_ISL_550634, EPI_ISL_550635, EPI_ISL_550636, EPI_ISL_550637 | | | |
| see above | Lighthouse Lab in Milton Keynes | Wellcome Sanger Institute for the COVID-19 Genomics UK (COG-UK) consortium | The Lighthouse Lab in Milton Keynes and Alex Alderton, Roberto Amato, Sonia Goncalves, Ewan Harrison, David K. Jackson, Ian Johnston, Dominic Kwiatkowski, Cordelia Langford, John Sillitoe on behalf of the Wellcome Sanger Institute COVID-19 Surveillance Team (http://www.sanger.ac.uk/covid-team) |
| EPI_ISL_550638 | Lighthouse Lab in Milton Keynes | Wellcome Sanger Institute for the COVID-19 Genomics UK (COG-UK) consortium | The Lighthouse Lab in Alderley Park and Alex Alderton, Roberto Amato, Sonia Goncalves, Ewan Harrison, David K. Jackson, Ian Johnston, Dominic Kwiatkowski, Cordelia Langford, John Sillitoe on behalf of the Wellcome Sanger Institute COVID-19 Surveillance Team |
| EPI_ISL_550639, EPI_ISL_550640, EPI_ISL_550641, EPI_ISL_550642, EPI_ISL_550643, EPI_ISL_550644, EPI_ISL_550645 | Lighthouse Lab in Milton Keynes | Wellcome Sanger Institute for the COVID-19 Genomics UK (COG-UK) consortium | The Lighthouse Lab in Milton Keynes and Alex Alderton, Roberto Amato, Sonia Goncalves, Ewan Harrison, David K. Jackson, Ian Johnston, Dominic Kwiatkowski, Cordelia Langford, John Sillitoe on behalf of the Wellcome Sanger Institute COVID-19 Surveillance Team (http://www.sanger.ac.uk/covid-team) |
| EPI_ISL_550646 | Lighthouse Lab in Milton Keynes | Wellcome Sanger Institute for the COVID-19 Genomics UK (COG-UK) consortium | The Lighthouse Lab in Alderley Park and Alex Alderton, Roberto Amato, Sonia Goncalves, Ewan Harrison, David K. Jackson, Ian Johnston, Dominic Kwiatkowski, Cordelia Langford, John Sillitoe on behalf of the Wellcome Sanger Institute COVID-19 Surveillance Team |
| EPI_ISL_550647, EPI_ISL_550648, EPI_ISL_550649, EPI_ISL_550650, EPI_ISL_550651, EPI_ISL_550652, EPI_ISL_550653, EPI_ISL_550654, EPI_ISL_550655, EPI_ISL_550656, EPI_ISL_550657, EPI_ISL_550658, EPI_ISL_550659, EPI_ISL_550660, EPI_ISL_550661, EPI_ISL_550662, EPI_ISL_550663, EPI_ISL_550664 | | | |
| see above | Lighthouse Lab in Milton Keynes | Wellcome Sanger Institute for the COVID-19 Genomics UK (COG-UK) consortium | The Lighthouse Lab in Milton Keynes and Alex Alderton, Roberto Amato, Sonia Goncalves, Ewan Harrison, David K. Jackson, Ian Johnston, Dominic Kwiatkowski, Cordelia Langford, John Sillitoe on behalf of the Wellcome Sanger Institute COVID-19 Surveillance Team (http://www.sanger.ac.uk/covid-team) |
| EPI_ISL_550665 | Lighthouse Lab in Milton Keynes | Wellcome Sanger Institute for the COVID-19 Genomics UK (COG-UK) consortium | The Lighthouse Lab in Alderley Park and Alex Alderton, Roberto Amato, Sonia Goncalves, Ewan Harrison, David K. Jackson, Ian Johnston, Dominic Kwiatkowski, Cordelia Langford, John Sillitoe on behalf of the Wellcome Sanger Institute COVID-19 Surveillance Team |
| EPI_ISL_550666, EPI_ISL_550667, EPI_ISL_550668, EPI_ISL_550669, EPI_ISL_550670, EPI_ISL_550671, EPI_ISL_550672 | Lighthouse Lab in Milton Keynes | Wellcome Sanger Institute for the COVID-19 Genomics UK (COG-UK) consortium | The Lighthouse Lab in Milton Keynes and Alex Alderton, Roberto Amato, Sonia Goncalves, Ewan Harrison, David K. Jackson, Ian Johnston, Dominic Kwiatkowski, Cordelia Langford, John Sillitoe on behalf of the Wellcome Sanger Institute COVID-19 Surveillance Team (http://www.sanger.ac.uk/covid-team) |
| EPI_ISL_550673 | Lighthouse Lab in Milton Keynes | Wellcome Sanger Institute for the COVID-19 Genomics UK (COG-UK) consortium | The Lighthouse Lab in Alderley Park and Alex Alderton, Roberto Amato, Sonia Goncalves, Ewan Harrison, David K. Jackson, Ian Johnston, Dominic Kwiatkowski, Cordelia Langford, John Sillitoe on behalf of the Wellcome Sanger Institute COVID-19 Surveillance Team |
| EPI_ISL_550674 | Lighthouse Lab in Milton Keynes | Wellcome Sanger Institute for the COVID-19 Genomics UK (COG-UK) consortium | The Lighthouse Lab in Milton Keynes and Alex Alderton, Roberto Amato, Sonia Goncalves, Ewan Harrison, David K. Jackson, Ian Johnston, Dominic Kwiatkowski, Cordelia Langford, John Sillitoe on behalf of the Wellcome Sanger Institute COVID-19 Surveillance Team (http://www.sanger.ac.uk/covid-team) |
| EPI_ISL_550675 | Lighthouse Lab in Milton Keynes | Wellcome Sanger Institute for the COVID-19 Genomics UK (COG-UK) consortium | The Lighthouse Lab in Alderley Park and Alex Alderton, Roberto Amato, Sonia Goncalves, Ewan Harrison, David K. Jackson, Ian Johnston, Dominic Kwiatkowski, Cordelia Langford, John Sillitoe on behalf of the Wellcome Sanger Institute COVID-19 Surveillance Team |
| EPI_ISL_550676, EPI_ISL_550677, EPI_ISL_550678, EPI_ISL_550679, EPI_ISL_550680, EPI_ISL_550681 | Lighthouse Lab in Milton Keynes | Wellcome Sanger Institute for the COVID-19 Genomics UK (COG-UK) consortium | The Lighthouse Lab in Milton Keynes and Alex Alderton, Roberto Amato, Sonia Goncalves, Ewan Harrison, David K. Jackson, Ian Johnston, Dominic Kwiatkowski, Cordelia Langford, John Sillitoe on behalf of the Wellcome Sanger Institute COVID-19 Surveillance Team (http://www.sanger.ac.uk/covid-team) |
| EPI_ISL_550682 | Lighthouse Lab in Milton Keynes | Wellcome Sanger Institute for the COVID-19 Genomics UK (COG-UK) consortium | The Lighthouse Lab in Alderley Park and Alex Alderton, Roberto Amato, Sonia Goncalves, Ewan Harrison, David K. Jackson, Ian Johnston, Dominic Kwiatkowski, Cordelia Langford, John Sillitoe on behalf of the Wellcome Sanger Institute COVID-19 Surveillance Team |
| EPI_ISL_550683, EPI_ISL_550684, EPI_ISL_550685, EPI_ISL_550686, EPI_ISL_550687, EPI_ISL_550688, EPI_ISL_550689, EPI_ISL_550690, EPI_ISL_550691, EPI_ISL_550692, EPI_ISL_550693, EPI_ISL_550694, EPI_ISL_550695, EPI_ISL_550696, EPI_ISL_550697, EPI_ISL_550698, EPI_ISL_550699, EPI_ISL_550700, EPI_ISL_550701, EPI_ISL_550702, EPI_ISL_550703, EPI_ISL_550704, EPI_ISL_550705, EPI_ISL_550706, EPI_ISL_550707, EPI_ISL_550708, EPI_ISL_550709, EPI_ISL_550710 | | | |
| see above | Lighthouse Lab in Milton Keynes | Wellcome Sanger Institute for the COVID-19 Genomics UK (COG-UK) consortium | The Lighthouse Lab in Milton Keynes and Alex Alderton, Roberto Amato, Sonia Goncalves, Ewan Harrison, David K. Jackson, Ian Johnston, Dominic Kwiatkowski, Cordelia Langford, John Sillitoe on behalf of the Wellcome Sanger Institute COVID-19 Surveillance Team (http://www.sanger.ac.uk/covid-team) |
| EPI_ISL_550711 | Lighthouse Lab in Milton Keynes | Wellcome Sanger Institute for the COVID-19 Genomics UK (COG-UK) consortium | The Lighthouse Lab in Alderley Park and Alex Alderton, Roberto Amato, Sonia Goncalves, Ewan Harrison, David K. Jackson, Ian Johnston, Dominic Kwiatkowski, Cordelia Langford, John Sillitoe on behalf of the Wellcome Sanger Institute COVID-19 Surveillance Team |
| EPI_ISL_550712, EPI_ISL_550713, EPI_ISL_550714, EPI_ISL_550715, EPI_ISL_550716, EPI_ISL_550717, EPI_ISL_550718, EPI_ISL_550719, EPI_ISL_550720, EPI_ISL_550721, EPI_ISL_550722, EPI_ISL_550723, EPI_ISL_550724, EPI_ISL_550725 | | | |
| see above | Lighthouse Lab in Milton Keynes | Wellcome Sanger Institute for the COVID-19 Genomics UK (COG-UK) consortium | The Lighthouse Lab in Milton Keynes and Alex Alderton, Roberto Amato, Sonia Goncalves, Ewan Harrison, David K. Jackson, Ian Johnston, Dominic Kwiatkowski, Cordelia Langford, John Sillitoe on behalf of the Wellcome Sanger Institute COVID-19 Surveillance Team (http://www.sanger.ac.uk/covid-team) |
| EPI_ISL_550726 | Lighthouse Lab in Milton Keynes | Wellcome Sanger Institute for the COVID-19 Genomics UK (COG-UK) consortium | The Lighthouse Lab in Alderley Park and Alex Alderton, Roberto Amato, Sonia Goncalves, Ewan Harrison, David K. Jackson, Ian Johnston, Dominic Kwiatkowski, Cordelia Langford, John Sillitoe on behalf of the Wellcome Sanger Institute COVID-19 Surveillance Team |
| EPI_ISL_550727, EPI_ISL_550728 | Lighthouse Lab in Milton Keynes | Wellcome Sanger Institute for the COVID-19 Genomics UK (COG-UK) consortium | The Lighthouse Lab in Milton Keynes and Alex Alderton, Roberto Amato, Sonia Goncalves, Ewan Harrison, David K. Jackson, Ian Johnston, Dominic Kwiatkowski, Cordelia Langford, John Sillitoe on behalf of the Wellcome Sanger Institute COVID-19 Surveillance Team (http://www.sanger.ac.uk/covid-team) |
| EPI_ISL_550729 | Lighthouse Lab in Milton Keynes | Wellcome Sanger Institute for the COVID-19 Genomics UK (COG-UK) consortium | The Lighthouse Lab in Alderley Park and Alex Alderton, Roberto Amato, Sonia Goncalves, Ewan Harrison, David K. Jackson, Ian Johnston, Dominic Kwiatkowski, Cordelia Langford, John Sillitoe on behalf of the Wellcome Sanger Institute COVID-19 Surveillance Team |
| EPI_ISL_550730, EPI_ISL_550731, EPI_ISL_550732, EPI_ISL_550733, EPI_ISL_550734, EPI_ISL_550735, EPI_ISL_550736, EPI_ISL_550737, EPI_ISL_550738, EPI_ISL_550739, EPI_ISL_550740, EPI_ISL_550741, EPI_ISL_550742, EPI_ISL_550743, EPI_ISL_550744, EPI_ISL_550745, EPI_ISL_550746, EPI_ISL_550747, EPI_ISL_550748, EPI_ISL_550749 | | | |
| see above | Lighthouse Lab in Milton Keynes | Wellcome Sanger Institute for the COVID-19 Genomics UK (COG-UK) consortium | The Lighthouse Lab in Milton Keynes and Alex Alderton, Roberto Amato, Sonia Goncalves, Ewan Harrison, David K. Jackson, Ian Johnston, Dominic Kwiatkowski, Cordelia Langford, John Sillitoe on behalf of the Wellcome Sanger Institute COVID-19 Surveillance Team (http://www.sanger.ac.uk/covid-team) |
| EPI_ISL_550750 | Lighthouse Lab in Milton Keynes | Wellcome Sanger Institute for the COVID-19 Genomics UK (COG-UK) consortium | The Lighthouse Lab in Alderley Park and Alex Alderton, Roberto Amato, Sonia Goncalves, Ewan Harrison, David K. Jackson, Ian Johnston, Dominic Kwiatkowski, Cordelia Langford, John Sillitoe on behalf of the Wellcome Sanger Institute COVID-19 Surveillance Team |

| | | | |
|---|---|---|---|
| EPI_ISL_550751, EPI_ISL_550752, EPI_ISL_550753, EPI_ISL_550754, EPI_ISL_550755, EPI_ISL_550756, EPI_ISL_550757, EPI_ISL_550758, EPI_ISL_550759, EPI_ISL_550760, EPI_ISL_550761, EPI_ISL_550762, EPI_ISL_550763, EPI_ISL_550764, EPI_ISL_550765, EPI_ISL_550766 | | | |
| see above | Lighthouse Lab in Milton Keynes | Wellcome Sanger Institute for the COVID-19 Genomics UK (COG-UK) consortium | The Lighthouse Lab in Milton Keynes and Alex Alderton, Roberto Amato, Sonia Goncalves, Ewan Harrison, David K. Jackson, Ian Johnston, Dominic Kwiatkowski, Cordelia Langford, John Sillitoe on behalf of the Wellcome Sanger Institute COVID-19 Surveillance Team (http://www.sanger.ac.uk/covid-team) |
| EPI_ISL_550767 | Lighthouse Lab in Milton Keynes | Wellcome Sanger Institute for the COVID-19 Genomics UK (COG-UK) consortium | The Lighthouse Lab in Alderley Park and Alex Alderton, Roberto Amato, Sonia Goncalves, Ewan Harrison, David K. Jackson, Ian Johnston, Dominic Kwiatkowski, Cordelia Langford, John Sillitoe on behalf of the Wellcome Sanger Institute COVID-19 Surveillance Team |
| EPI_ISL_550768, EPI_ISL_550769 | Lighthouse Lab in Milton Keynes | Wellcome Sanger Institute for the COVID-19 Genomics UK (COG-UK) consortium | The Lighthouse Lab in Milton Keynes and Alex Alderton, Roberto Amato, Sonia Goncalves, Ewan Harrison, David K. Jackson, Ian Johnston, Dominic Kwiatkowski, Cordelia Langford, John Sillitoe on behalf of the Wellcome Sanger Institute COVID-19 Surveillance Team (http://www.sanger.ac.uk/covid-team) |
| EPI_ISL_550770, EPI_ISL_550771 | Lighthouse Lab in Milton Keynes | Wellcome Sanger Institute for the COVID-19 Genomics UK (COG-UK) consortium | The Lighthouse Lab in Alderley Park and Alex Alderton, Roberto Amato, Sonia Goncalves, Ewan Harrison, David K. Jackson, Ian Johnston, Dominic Kwiatkowski, Cordelia Langford, John Sillitoe on behalf of the Wellcome Sanger Institute COVID-19 Surveillance Team |
| EPI_ISL_550772, EPI_ISL_550773, EPI_ISL_550774, EPI_ISL_550775, EPI_ISL_550776, EPI_ISL_550777, EPI_ISL_550778, EPI_ISL_550779, EPI_ISL_550780, EPI_ISL_550781, EPI_ISL_550782, EPI_ISL_550783, EPI_ISL_550784, EPI_ISL_550785, EPI_ISL_550786, EPI_ISL_550787, EPI_ISL_550788, EPI_ISL_550789, EPI_ISL_550790, EPI_ISL_550791, EPI_ISL_550792, EPI_ISL_550793, EPI_ISL_550794, EPI_ISL_550795, EPI_ISL_550799, EPI_ISL_550800, EPI_ISL_550801, EPI_ISL_550804, EPI_ISL_550809, EPI_ISL_550810 | | | |
| see above | Lighthouse Lab in Milton Keynes | Wellcome Sanger Institute for the COVID-19 Genomics UK (COG-UK) consortium | The Lighthouse Lab in Milton Keynes and Alex Alderton, Roberto Amato, Sonia Goncalves, Ewan Harrison, David K. Jackson, Ian Johnston, Dominic Kwiatkowski, Cordelia Langford, John Sillitoe on behalf of the Wellcome Sanger Institute COVID-19 Surveillance Team (http://www.sanger.ac.uk/covid-team) |
| EPI_ISL_550811 | Lighthouse Lab in Milton Keynes | Wellcome Sanger Institute for the COVID-19 Genomics UK (COG-UK) consortium | The Lighthouse Lab in Alderley Park and Alex Alderton, Roberto Amato, Sonia Goncalves, Ewan Harrison, David K. Jackson, Ian Johnston, Dominic Kwiatkowski, Cordelia Langford, John Sillitoe on behalf of the Wellcome Sanger Institute COVID-19 Surveillance Team |
| EPI_ISL_550813, EPI_ISL_550815, EPI_ISL_550816, EPI_ISL_550820, EPI_ISL_550821 | Lighthouse Lab in Milton Keynes | Wellcome Sanger Institute for the COVID-19 Genomics UK (COG-UK) consortium | The Lighthouse Lab in Milton Keynes and Alex Alderton, Roberto Amato, Sonia Goncalves, Ewan Harrison, David K. Jackson, Ian Johnston, Dominic Kwiatkowski, Cordelia Langford, John Sillitoe on behalf of the Wellcome Sanger Institute COVID-19 Surveillance Team (http://www.sanger.ac.uk/covid-team) |
| EPI_ISL_550826 | Lighthouse Lab in Milton Keynes | Wellcome Sanger Institute for the COVID-19 Genomics UK (COG-UK) consortium | The Lighthouse Lab in Alderley Park and Alex Alderton, Roberto Amato, Sonia Goncalves, Ewan Harrison, David K. Jackson, Ian Johnston, Dominic Kwiatkowski, Cordelia Langford, John Sillitoe on behalf of the Wellcome Sanger Institute COVID-19 Surveillance Team |
| EPI_ISL_550827, EPI_ISL_550829, EPI_ISL_550830, EPI_ISL_550831, EPI_ISL_550833 | Lighthouse Lab in Milton Keynes | Wellcome Sanger Institute for the COVID-19 Genomics UK (COG-UK) consortium | The Lighthouse Lab in Milton Keynes and Alex Alderton, Roberto Amato, Sonia Goncalves, Ewan Harrison, David K. Jackson, Ian Johnston, Dominic Kwiatkowski, Cordelia Langford, John Sillitoe on behalf of the Wellcome Sanger Institute COVID-19 Surveillance Team (http://www.sanger.ac.uk/covid-team) |
| EPI_ISL_550834 | Lighthouse Lab in Milton Keynes | Wellcome Sanger Institute for the COVID-19 Genomics UK (COG-UK) consortium | The Lighthouse Lab in Alderley Park and Alex Alderton, Roberto Amato, Sonia Goncalves, Ewan Harrison, David K. Jackson, Ian Johnston, Dominic Kwiatkowski, Cordelia Langford, John Sillitoe on behalf of the Wellcome Sanger Institute COVID-19 Surveillance Team |
| EPI_ISL_550838, EPI_ISL_550841, EPI_ISL_550843 | Lighthouse Lab in Milton Keynes | Wellcome Sanger Institute for the COVID-19 Genomics UK (COG-UK) consortium | The Lighthouse Lab in Milton Keynes and Alex Alderton, Roberto Amato, Sonia Goncalves, Ewan Harrison, David K. Jackson, Ian Johnston, Dominic Kwiatkowski, Cordelia Langford, John Sillitoe on behalf of the Wellcome Sanger Institute COVID-19 Surveillance Team (http://www.sanger.ac.uk/covid-team) |
| EPI_ISL_550846 | Lighthouse Lab in Milton Keynes | Wellcome Sanger Institute for the COVID-19 Genomics UK (COG-UK) consortium | The Lighthouse Lab in Alderley Park and Alex Alderton, Roberto Amato, Sonia Goncalves, Ewan Harrison, David K. Jackson, Ian Johnston, Dominic Kwiatkowski, Cordelia Langford, John Sillitoe on behalf of the Wellcome Sanger Institute COVID-19 Surveillance Team |
| EPI_ISL_550849 | Lighthouse Lab in Milton Keynes | Wellcome Sanger Institute for the COVID-19 Genomics UK (COG-UK) Consortium | The Lighthouse Lab in Milton Keynes and Alex Alderton, Roberto Amato, Sonia Goncalves, Ewan Harrison, David K. Jackson, Ian Johnston, Dominic Kwiatkowski, Cordelia Langford, John Sillitoe on behalf of the Wellcome Sanger Institute COVID-19 Surveillance Team |
| EPI_ISL_550851, EPI_ISL_550852, EPI_ISL_550854, EPI_ISL_550856, EPI_ISL_550857, EPI_ISL_550863, EPI_ISL_550864, EPI_ISL_550866, EPI_ISL_550867 | Lighthouse Lab in Milton Keynes | Wellcome Sanger Institute for the COVID-19 Genomics UK (COG-UK) consortium | The Lighthouse Lab in Milton Keynes and Alex Alderton, Roberto Amato, Sonia Goncalves, Ewan Harrison, David K. Jackson, Ian Johnston, Dominic Kwiatkowski, Cordelia Langford, John Sillitoe on behalf of the Wellcome Sanger Institute COVID-19 Surveillance Team (http://www.sanger.ac.uk/covid-team) |
| EPI_ISL_550870 | Lighthouse Lab in Milton Keynes | Wellcome Sanger Institute for the COVID-19 Genomics UK (COG-UK) consortium | The Lighthouse Lab in Alderley Park and Alex Alderton, Roberto Amato, Sonia Goncalves, Ewan Harrison, David K. Jackson, Ian Johnston, Dominic Kwiatkowski, Cordelia Langford, John Sillitoe on behalf of the Wellcome Sanger Institute COVID-19 Surveillance Team |
| EPI_ISL_550871, EPI_ISL_550874, EPI_ISL_550879, EPI_ISL_550880, EPI_ISL_550881, EPI_ISL_550883, EPI_ISL_550889, EPI_ISL_550890, EPI_ISL_550893, EPI_ISL_550895, EPI_ISL_550896, EPI_ISL_550897, EPI_ISL_550899, EPI_ISL_550900, EPI_ISL_550903, EPI_ISL_550905, EPI_ISL_550906, EPI_ISL_550908, EPI_ISL_550909, EPI_ISL_550910, EPI_ISL_550913 | | | |
| see above | Lighthouse Lab in Milton Keynes | Wellcome Sanger Institute for the COVID-19 Genomics UK (COG-UK) consortium | The Lighthouse Lab in Milton Keynes and Alex Alderton, Roberto Amato, Sonia Goncalves, Ewan Harrison, David K. Jackson, Ian Johnston, Dominic Kwiatkowski, Cordelia Langford, John Sillitoe on behalf of the Wellcome Sanger Institute COVID-19 Surveillance Team (http://www.sanger.ac.uk/covid-team) |
| EPI_ISL_550914 | Lighthouse Lab in Milton Keynes | Wellcome Sanger Institute for the COVID-19 Genomics UK (COG-UK) consortium | The Lighthouse Lab in Alderley Park and Alex Alderton, Roberto Amato, Sonia Goncalves, Ewan Harrison, David K. Jackson, Ian Johnston, Dominic Kwiatkowski, Cordelia Langford, John Sillitoe on behalf of the Wellcome Sanger Institute COVID-19 Surveillance Team |
| EPI_ISL_550918, EPI_ISL_550919, EPI_ISL_550920, EPI_ISL_550921, EPI_ISL_550923, EPI_ISL_550924, EPI_ISL_550925, EPI_ISL_550926, EPI_ISL_550934, EPI_ISL_550936, EPI_ISL_550939, EPI_ISL_550941, EPI_ISL_550943, EPI_ISL_550944 | | | |
| see above | Lighthouse Lab in Milton Keynes | Wellcome Sanger Institute for the COVID-19 Genomics UK (COG-UK) consortium | The Lighthouse Lab in Milton Keynes and Alex Alderton, Roberto Amato, Sonia Goncalves, Ewan Harrison, David K. Jackson, Ian Johnston, Dominic Kwiatkowski, Cordelia Langford, John Sillitoe on behalf of the Wellcome Sanger Institute COVID-19 Surveillance Team (http://www.sanger.ac.uk/covid-team) |
| EPI_ISL_550945 | Lighthouse Lab in Milton Keynes | Wellcome Sanger Institute for the COVID-19 Genomics UK (COG-UK) consortium | The Lighthouse Lab in Alderley Park and Alex Alderton, Roberto Amato, Sonia Goncalves, Ewan Harrison, David K. Jackson, Ian Johnston, Dominic Kwiatkowski, Cordelia Langford, John Sillitoe on behalf of the Wellcome Sanger Institute COVID-19 Surveillance Team |
| EPI_ISL_550947, EPI_ISL_550951, EPI_ISL_550952, EPI_ISL_550956, EPI_ISL_550957, EPI_ISL_550959, EPI_ISL_550960, EPI_ISL_550962, EPI_ISL_550963, EPI_ISL_550965, EPI_ISL_550968, EPI_ISL_550970, EPI_ISL_550972, EPI_ISL_550973, EPI_ISL_550974, EPI_ISL_550980, EPI_ISL_550982 | | | |
| see above | Lighthouse Lab in Milton Keynes | Wellcome Sanger Institute for the COVID-19 Genomics UK (COG-UK) consortium | The Lighthouse Lab in Milton Keynes and Alex Alderton, Roberto Amato, Sonia Goncalves, Ewan Harrison, David K. Jackson, Ian Johnston, Dominic Kwiatkowski, Cordelia Langford, John Sillitoe on behalf of the Wellcome Sanger Institute COVID-19 Surveillance Team (http://www.sanger.ac.uk/covid-team) |
| EPI_ISL_550987 | Lighthouse Lab in Milton Keynes | Wellcome Sanger Institute for the COVID-19 Genomics UK (COG-UK) consortium | The Lighthouse Lab in Alderley Park and Alex Alderton, Roberto Amato, Sonia Goncalves, Ewan Harrison, David K. Jackson, Ian Johnston, Dominic Kwiatkowski, Cordelia Langford, John Sillitoe on behalf of the Wellcome Sanger Institute COVID-19 Surveillance Team |
| EPI_ISL_550990, EPI_ISL_550991, EPI_ISL_550992, EPI_ISL_550998, EPI_ISL_551000, EPI_ISL_551004, EPI_ISL_551009, EPI_ISL_551010, EPI_ISL_551011, EPI_ISL_551012 | Lighthouse Lab in Milton Keynes | Wellcome Sanger Institute for the COVID-19 Genomics UK (COG-UK) consortium | The Lighthouse Lab in Milton Keynes and Alex Alderton, Roberto Amato, Sonia Goncalves, Ewan Harrison, David K. Jackson, Ian Johnston, Dominic Kwiatkowski, Cordelia Langford, John Sillitoe on behalf of the Wellcome Sanger Institute COVID-19 Surveillance Team (http://www.sanger.ac.uk/covid-team) |
| EPI_ISL_551013 | Lighthouse Lab in Milton Keynes | Wellcome Sanger Institute for the COVID-19 Genomics UK (COG-UK) Consortium | The Lighthouse Lab in Milton Keynes and Alex Alderton, Roberto Amato, Sonia Goncalves, Ewan Harrison, David K. Jackson, Ian Johnston, Dominic Kwiatkowski, Cordelia Langford, John Sillitoe on behalf of the Wellcome Sanger Institute COVID-19 Surveillance Team |
| EPI_ISL_551017, EPI_ISL_551018, EPI_ISL_551019, EPI_ISL_551024, EPI_ISL_551026, EPI_ISL_551027, EPI_ISL_551031, EPI_ISL_551032, EPI_ISL_551033, EPI_ISL_551037, EPI_ISL_551039, EPI_ISL_551040, EPI_ISL_551042, EPI_ISL_551043 | | | |
| see above | Lighthouse Lab in Milton Keynes | Wellcome Sanger Institute for the COVID-19 Genomics UK (COG-UK) consortium | The Lighthouse Lab in Milton Keynes and Alex Alderton, Roberto Amato, Sonia Goncalves, Ewan Harrison, David K. Jackson, Ian Johnston, Dominic Kwiatkowski, Cordelia Langford, John Sillitoe on behalf of the Wellcome Sanger Institute COVID-19 Surveillance Team (http://www.sanger.ac.uk/covid-team) |

| | | | |
|---|---|---|---|
| EPI_ISL_551044 | Lighthouse Lab in Milton Keynes | Wellcome Sanger Institute for the COVID-19 Genomics UK (COG-UK) consortium | The Lighthouse Lab in Alderley Park and Alex Alderton, Roberto Amato, Sonia Goncalves, Ewan Harrison, David K. Jackson, Ian Johnston, Dominic Kwiatkowski, Cordelia Langford, John Sillitoe on behalf of the Wellcome Sanger Institute COVID-19 Surveillance Team |
| EPI_ISL_551047, EPI_ISL_551049, EPI_ISL_551050, EPI_ISL_551053, EPI_ISL_551054, EPI_ISL_551055, EPI_ISL_551056, EPI_ISL_551059, EPI_ISL_551060, EPI_ISL_551061, EPI_ISL_551062, EPI_ISL_551066, EPI_ISL_551068, EPI_ISL_551069, EPI_ISL_551071, EPI_ISL_551072, EPI_ISL_551073, EPI_ISL_551075, EPI_ISL_551076, EPI_ISL_551080, EPI_ISL_551082, EPI_ISL_551083, EPI_ISL_551084, EPI_ISL_551085, EPI_ISL_551086, EPI_ISL_551090, EPI_ISL_551091, EPI_ISL_551093, EPI_ISL_551095, EPI_ISL_551097, EPI_ISL_551098, EPI_ISL_551099, EPI_ISL_551101, EPI_ISL_551102, EPI_ISL_551103, EPI_ISL_551105, EPI_ISL_551106, EPI_ISL_551107, EPI_ISL_551109, EPI_ISL_551110, EPI_ISL_551112, EPI_ISL_551113, EPI_ISL_551115, EPI_ISL_551116, EPI_ISL_551118, EPI_ISL_551120, EPI_ISL_551122, EPI_ISL_551123, EPI_ISL_551124, EPI_ISL_551126, EPI_ISL_551129, EPI_ISL_551130, EPI_ISL_551131, EPI_ISL_551136, EPI_ISL_551138, EPI_ISL_551142, EPI_ISL_551143, EPI_ISL_551149, EPI_ISL_551150, EPI_ISL_551151, EPI_ISL_551154, EPI_ISL_551155, EPI_ISL_551158, EPI_ISL_551162, EPI_ISL_551164, EPI_ISL_551167, EPI_ISL_551173, EPI_ISL_551175, EPI_ISL_551176, EPI_ISL_551177, EPI_ISL_551179, EPI_ISL_551180, EPI_ISL_551186, EPI_ISL_551190, EPI_ISL_551191, EPI_ISL_551194, EPI_ISL_551197, EPI_ISL_551198, EPI_ISL_551199, EPI_ISL_551200, EPI_ISL_551201, EPI_ISL_551202, EPI_ISL_551205, EPI_ISL_551206, EPI_ISL_551208, EPI_ISL_551210, EPI_ISL_551213 | | | |
| see above | Lighthouse Lab in Milton Keynes | Wellcome Sanger Institute for the COVID-19 Genomics UK (COG-UK) consortium | The Lighthouse Lab in Milton Keynes and Alex Alderton, Roberto Amato, Sonia Goncalves, Ewan Harrison, David K. Jackson, Ian Johnston, Dominic Kwiatkowski, Cordelia Langford, John Sillitoe on behalf of the Wellcome Sanger Institute COVID-19 Surveillance Team (http://www.sanger.ac.uk/covid-team) |
| EPI_ISL_551215 | Lighthouse Lab in Milton Keynes | Wellcome Sanger Institute for the COVID-19 Genomics UK (COG-UK) consortium | The Lighthouse Lab in Alderley Park and Alex Alderton, Roberto Amato, Sonia Goncalves, Ewan Harrison, David K. Jackson, Ian Johnston, Dominic Kwiatkowski, Cordelia Langford, John Sillitoe on behalf of the Wellcome Sanger Institute COVID-19 Surveillance Team |
| EPI_ISL_551216, EPI_ISL_551217 | Lighthouse Lab in Milton Keynes | Wellcome Sanger Institute for the COVID-19 Genomics UK (COG-UK) consortium | The Lighthouse Lab in Milton Keynes and Alex Alderton, Roberto Amato, Sonia Goncalves, Ewan Harrison, David K. Jackson, Ian Johnston, Dominic Kwiatkowski, Cordelia Langford, John Sillitoe on behalf of the Wellcome Sanger Institute COVID-19 Surveillance Team (http://www.sanger.ac.uk/covid-team) |
| EPI_ISL_551218 | Lighthouse Lab in Milton Keynes | Wellcome Sanger Institute for the COVID-19 Genomics UK (COG-UK) Consortium | The Lighthouse Lab in Milton Keynes and Alex Alderton, Roberto Amato, Sonia Goncalves, Ewan Harrison, David K. Jackson, Ian Johnston, Dominic Kwiatkowski, Cordelia Langford, John Sillitoe on behalf of the Wellcome Sanger Institute COVID-19 Surveillance Team |
| EPI_ISL_551220, EPI_ISL_551221, EPI_ISL_551222, EPI_ISL_551227, EPI_ISL_551229, EPI_ISL_551230, EPI_ISL_551233, EPI_ISL_551235, EPI_ISL_551237, EPI_ISL_551241, EPI_ISL_551245, EPI_ISL_551248, EPI_ISL_551249, EPI_ISL_551251, EPI_ISL_551252, EPI_ISL_551253, EPI_ISL_551254, EPI_ISL_551255, EPI_ISL_551256, EPI_ISL_551257, EPI_ISL_551259, EPI_ISL_551260 | | | |
| see above | Lighthouse Lab in Milton Keynes | Wellcome Sanger Institute for the COVID-19 Genomics UK (COG-UK) consortium | The Lighthouse Lab in Milton Keynes and Alex Alderton, Roberto Amato, Sonia Goncalves, Ewan Harrison, David K. Jackson, Ian Johnston, Dominic Kwiatkowski, Cordelia Langford, John Sillitoe on behalf of the Wellcome Sanger Institute COVID-19 Surveillance Team (http://www.sanger.ac.uk/covid-team) |
| EPI_ISL_551263 | Lighthouse Lab in Milton Keynes | Wellcome Sanger Institute for the COVID-19 Genomics UK (COG-UK) consortium | The Lighthouse Lab in Alderley Park and Alex Alderton, Roberto Amato, Sonia Goncalves, Ewan Harrison, David K. Jackson, Ian Johnston, Dominic Kwiatkowski, Cordelia Langford, John Sillitoe on behalf of the Wellcome Sanger Institute COVID-19 Surveillance Team |
| EPI_ISL_551267, EPI_ISL_551271, EPI_ISL_551272, EPI_ISL_551275, EPI_ISL_551276, EPI_ISL_551277, EPI_ISL_551278, EPI_ISL_551282, EPI_ISL_551577, EPI_ISL_551578, EPI_ISL_551579 | | | |
| see above | Lighthouse Lab in Milton Keynes | Wellcome Sanger Institute for the COVID-19 Genomics UK (COG-UK) consortium | The Lighthouse Lab in Milton Keynes and Alex Alderton, Roberto Amato, Sonia Goncalves, Ewan Harrison, David K. Jackson, Ian Johnston, Dominic Kwiatkowski, Cordelia Langford, John Sillitoe on behalf of the Wellcome Sanger Institute COVID-19 Surveillance Team (http://www.sanger.ac.uk/covid-team) |
| EPI_ISL_551580 | Lighthouse Lab in Milton Keynes | Wellcome Sanger Institute for the COVID-19 Genomics UK (COG-UK) consortium | The Lighthouse Lab in Alderley Park and Alex Alderton, Roberto Amato, Sonia Goncalves, Ewan Harrison, David K. Jackson, Ian Johnston, Dominic Kwiatkowski, Cordelia Langford, John Sillitoe on behalf of the Wellcome Sanger Institute COVID-19 Surveillance Team |
| EPI_ISL_551581, EPI_ISL_551583 | Lighthouse Lab in Milton Keynes | Wellcome Sanger Institute for the COVID-19 Genomics UK (COG-UK) consortium | The Lighthouse Lab in Milton Keynes and Alex Alderton, Roberto Amato, Sonia Goncalves, Ewan Harrison, David K. Jackson, Ian Johnston, Dominic Kwiatkowski, Cordelia Langford, John Sillitoe on behalf of the Wellcome Sanger Institute COVID-19 Surveillance Team (http://www.sanger.ac.uk/covid-team) |
| EPI_ISL_551584, EPI_ISL_551585 | Lighthouse Lab in Milton Keynes | Wellcome Sanger Institute for the COVID-19 Genomics UK (COG-UK) consortium | The Lighthouse Lab in Alderley Park and Alex Alderton, Roberto Amato, Sonia Goncalves, Ewan Harrison, David K. Jackson, Ian Johnston, Dominic Kwiatkowski, Cordelia Langford, John Sillitoe on behalf of the Wellcome Sanger Institute COVID-19 Surveillance Team |
| EPI_ISL_551586, EPI_ISL_551587, EPI_ISL_551588, EPI_ISL_551589, EPI_ISL_551590, EPI_ISL_551591, EPI_ISL_551592, EPI_ISL_551593, EPI_ISL_551594, EPI_ISL_551595, EPI_ISL_551596, EPI_ISL_551597, EPI_ISL_551598, EPI_ISL_551599, EPI_ISL_551600, EPI_ISL_551601, EPI_ISL_551602 | | | |
| see above | Lighthouse Lab in Milton Keynes | Wellcome Sanger Institute for the COVID-19 Genomics UK (COG-UK) consortium | The Lighthouse Lab in Milton Keynes and Alex Alderton, Roberto Amato, Sonia Goncalves, Ewan Harrison, David K. Jackson, Ian Johnston, Dominic Kwiatkowski, Cordelia Langford, John Sillitoe on behalf of the Wellcome Sanger Institute COVID-19 Surveillance Team (http://www.sanger.ac.uk/covid-team) |
| EPI_ISL_551603 | Lighthouse Lab in Milton Keynes | Wellcome Sanger Institute for the COVID-19 Genomics UK (COG-UK) consortium | The Lighthouse Lab in Alderley Park and Alex Alderton, Roberto Amato, Sonia Goncalves, Ewan Harrison, David K. Jackson, Ian Johnston, Dominic Kwiatkowski, Cordelia Langford, John Sillitoe on behalf of the Wellcome Sanger Institute COVID-19 Surveillance Team |
| EPI_ISL_551604, EPI_ISL_551605, EPI_ISL_551606, EPI_ISL_551607, EPI_ISL_551608, EPI_ISL_551609, EPI_ISL_551610, EPI_ISL_551611, EPI_ISL_551612, EPI_ISL_551613, EPI_ISL_551614, EPI_ISL_551615, EPI_ISL_551616, EPI_ISL_551617, EPI_ISL_551618, EPI_ISL_551619, EPI_ISL_551620, EPI_ISL_551621, EPI_ISL_551622, EPI_ISL_551623 | | | |
| see above | Lighthouse Lab in Milton Keynes | Wellcome Sanger Institute for the COVID-19 Genomics UK (COG-UK) consortium | The Lighthouse Lab in Milton Keynes and Alex Alderton, Roberto Amato, Sonia Goncalves, Ewan Harrison, David K. Jackson, Ian Johnston, Dominic Kwiatkowski, Cordelia Langford, John Sillitoe on behalf of the Wellcome Sanger Institute COVID-19 Surveillance Team (http://www.sanger.ac.uk/covid-team) |
| EPI_ISL_551624 | Lighthouse Lab in Milton Keynes | Wellcome Sanger Institute for the COVID-19 Genomics UK (COG-UK) consortium | The Lighthouse Lab in Alderley Park and Alex Alderton, Roberto Amato, Sonia Goncalves, Ewan Harrison, David K. Jackson, Ian Johnston, Dominic Kwiatkowski, Cordelia Langford, John Sillitoe on behalf of the Wellcome Sanger Institute COVID-19 Surveillance Team |
| EPI_ISL_551625, EPI_ISL_551626 | Lighthouse Lab in Milton Keynes | Wellcome Sanger Institute for the COVID-19 Genomics UK (COG-UK) consortium | The Lighthouse Lab in Milton Keynes and Alex Alderton, Roberto Amato, Sonia Goncalves, Ewan Harrison, David K. Jackson, Ian Johnston, Dominic Kwiatkowski, Cordelia Langford, John Sillitoe on behalf of the Wellcome Sanger Institute COVID-19 Surveillance Team (http://www.sanger.ac.uk/covid-team) |
| EPI_ISL_551627 | Lighthouse Lab in Milton Keynes | Wellcome Sanger Institute for the COVID-19 Genomics UK (COG-UK) consortium | The Lighthouse Lab in Alderley Park and Alex Alderton, Roberto Amato, Sonia Goncalves, Ewan Harrison, David K. Jackson, Ian Johnston, Dominic Kwiatkowski, Cordelia Langford, John Sillitoe on behalf of the Wellcome Sanger Institute COVID-19 Surveillance Team |
| EPI_ISL_551628, EPI_ISL_551629, EPI_ISL_551630, EPI_ISL_551631, EPI_ISL_551632, EPI_ISL_551633, EPI_ISL_551634, EPI_ISL_551635, EPI_ISL_551636 | Lighthouse Lab in Milton Keynes | Wellcome Sanger Institute for the COVID-19 Genomics UK (COG-UK) consortium | The Lighthouse Lab in Milton Keynes and Alex Alderton, Roberto Amato, Sonia Goncalves, Ewan Harrison, David K. Jackson, Ian Johnston, Dominic Kwiatkowski, Cordelia Langford, John Sillitoe on behalf of the Wellcome Sanger Institute COVID-19 Surveillance Team (http://www.sanger.ac.uk/covid-team) |
| EPI_ISL_551637 | Lighthouse Lab in Milton Keynes | Wellcome Sanger Institute for the COVID-19 Genomics UK (COG-UK) consortium | The Lighthouse Lab in Alderley Park and Alex Alderton, Roberto Amato, Sonia Goncalves, Ewan Harrison, David K. Jackson, Ian Johnston, Dominic Kwiatkowski, Cordelia Langford, John Sillitoe on behalf of the Wellcome Sanger Institute COVID-19 Surveillance Team |
| EPI_ISL_551638, EPI_ISL_551639, EPI_ISL_551640, EPI_ISL_551641 | Lighthouse Lab in Milton Keynes | Wellcome Sanger Institute for the COVID-19 Genomics UK (COG-UK) consortium | The Lighthouse Lab in Milton Keynes and Alex Alderton, Roberto Amato, Sonia Goncalves, Ewan Harrison, David K. Jackson, Ian Johnston, Dominic Kwiatkowski, Cordelia Langford, John Sillitoe on behalf of the Wellcome Sanger Institute COVID-19 Surveillance Team (http://www.sanger.ac.uk/covid-team) |
| EPI_ISL_551642 | Lighthouse Lab in Milton Keynes | Wellcome Sanger Institute for the COVID-19 Genomics UK (COG-UK) consortium | The Lighthouse Lab in Alderley Park and Alex Alderton, Roberto Amato, Sonia Goncalves, Ewan Harrison, David K. Jackson, Ian Johnston, Dominic Kwiatkowski, Cordelia Langford, John Sillitoe on behalf of the Wellcome Sanger Institute COVID-19 Surveillance Team |
| EPI_ISL_551643, EPI_ISL_551644, EPI_ISL_551645, EPI_ISL_551646, EPI_ISL_551648 | Lighthouse Lab in Milton Keynes | Wellcome Sanger Institute for the COVID-19 Genomics UK (COG-UK) consortium | The Lighthouse Lab in Milton Keynes and Alex Alderton, Roberto Amato, Sonia Goncalves, Ewan Harrison, David K. Jackson, Ian Johnston, Dominic Kwiatkowski, Cordelia Langford, John Sillitoe on behalf of the Wellcome Sanger Institute COVID-19 Surveillance Team (http://www.sanger.ac.uk/covid-team) |
| EPI_ISL_551649 | Lighthouse Lab in Milton Keynes | Wellcome Sanger Institute for the COVID-19 Genomics UK (COG-UK) consortium | The Lighthouse Lab in Alderley Park and Alex Alderton, Roberto Amato, Sonia Goncalves, Ewan Harrison, David K. Jackson, Ian Johnston, Dominic Kwiatkowski, Cordelia Langford, John Sillitoe on behalf of the Wellcome Sanger Institute COVID-19 Surveillance Team |
| EPI_ISL_551650, EPI_ISL_551651, EPI_ISL_551652, EPI_ISL_551653, EPI_ISL_551654, EPI_ISL_551655, EPI_ISL_551656, EPI_ISL_551657, EPI_ISL_551658, EPI_ISL_551659, EPI_ISL_551660, EPI_ISL_551661, EPI_ISL_551662, EPI_ISL_551663, EPI_ISL_551664, EPI_ISL_551665, EPI_ISL_551666, EPI_ISL_551667, EPI_ISL_551668, EPI_ISL_551669, EPI_ISL_551670, EPI_ISL_551671, EPI_ISL_551672, EPI_ISL_551673, EPI_ISL_551674, EPI_ISL_551675, EPI_ISL_551676, EPI_ISL_551677, EPI_ISL_551678, EPI_ISL_551679, EPI_ISL_551680 | | | |
| see above | Lighthouse Lab in Milton Keynes | Wellcome Sanger Institute for the COVID-19 Genomics UK | The Lighthouse Lab in Milton Keynes and Alex Alderton, Roberto Amato, Sonia Goncalves, Ewan Harrison, David K. Jackson, Ian Johnston, Dominic |

| | | | |
|---|---|---|---|
| | | (COG-UK) consortium | Kwiatkowski, Cordelia Langford, John Sillitoe on behalf of the Wellcome Sanger Institute COVID-19 Surveillance Team (http://www.sanger.ac.uk/covid-team) |
| EPI_ISL_551681 | Lighthouse Lab in Milton Keynes | Wellcome Sanger Institute for the COVID-19 Genomics UK (COG-UK) consortium | The Lighthouse Lab in Alderley Park and Alex Alderton, Roberto Amato, Sonia Goncalves, Ewan Harrison, David K. Jackson, Ian Johnston, Dominic Kwiatkowski, Cordelia Langford, John Sillitoe on behalf of the Wellcome Sanger Institute COVID-19 Surveillance Team |
| EPI_ISL_551682, EPI_ISL_551683, EPI_ISL_551684, EPI_ISL_551685, EPI_ISL_551686, EPI_ISL_551687, EPI_ISL_551688, EPI_ISL_551689, EPI_ISL_551690, EPI_ISL_551691 | Lighthouse Lab in Milton Keynes | Wellcome Sanger Institute for the COVID-19 Genomics UK (COG-UK) consortium | The Lighthouse Lab in Milton Keynes and Alex Alderton, Roberto Amato, Sonia Goncalves, Ewan Harrison, David K. Jackson, Ian Johnston, Dominic Kwiatkowski, Cordelia Langford, John Sillitoe on behalf of the Wellcome Sanger Institute COVID-19 Surveillance Team (http://www.sanger.ac.uk/covid-team) |
| EPI_ISL_551692 | Lighthouse Lab in Milton Keynes | Wellcome Sanger Institute for the COVID-19 Genomics UK (COG-UK) consortium | The Lighthouse Lab in Alderley Park and Alex Alderton, Roberto Amato, Sonia Goncalves, Ewan Harrison, David K. Jackson, Ian Johnston, Dominic Kwiatkowski, Cordelia Langford, John Sillitoe on behalf of the Wellcome Sanger Institute COVID-19 Surveillance Team |
| EPI_ISL_551693, EPI_ISL_551694 | Lighthouse Lab in Milton Keynes | Wellcome Sanger Institute for the COVID-19 Genomics UK (COG-UK) consortium | The Lighthouse Lab in Milton Keynes and Alex Alderton, Roberto Amato, Sonia Goncalves, Ewan Harrison, David K. Jackson, Ian Johnston, Dominic Kwiatkowski, Cordelia Langford, John Sillitoe on behalf of the Wellcome Sanger Institute COVID-19 Surveillance Team (http://www.sanger.ac.uk/covid-team) |
| EPI_ISL_551696 | Lighthouse Lab in Milton Keynes | Wellcome Sanger Institute for the COVID-19 Genomics UK (COG-UK) consortium | The Lighthouse Lab in Alderley Park and Alex Alderton, Roberto Amato, Sonia Goncalves, Ewan Harrison, David K. Jackson, Ian Johnston, Dominic Kwiatkowski, Cordelia Langford, John Sillitoe on behalf of the Wellcome Sanger Institute COVID-19 Surveillance Team |
| EPI_ISL_551697, EPI_ISL_551698, EPI_ISL_551699, EPI_ISL_551700, EPI_ISL_551701, EPI_ISL_551703, EPI_ISL_551704, EPI_ISL_551705, EPI_ISL_551706, EPI_ISL_551707, EPI_ISL_551708, EPI_ISL_551709, EPI_ISL_551710, EPI_ISL_551711, EPI_ISL_551712, EPI_ISL_551713, EPI_ISL_551714, EPI_ISL_551715 | | | |
| see above | Lighthouse Lab in Milton Keynes | Wellcome Sanger Institute for the COVID-19 Genomics UK (COG-UK) consortium | The Lighthouse Lab in Milton Keynes and Alex Alderton, Roberto Amato, Sonia Goncalves, Ewan Harrison, David K. Jackson, Ian Johnston, Dominic Kwiatkowski, Cordelia Langford, John Sillitoe on behalf of the Wellcome Sanger Institute COVID-19 Surveillance Team (http://www.sanger.ac.uk/covid-team) |
| EPI_ISL_551716 | Lighthouse Lab in Milton Keynes | Wellcome Sanger Institute for the COVID-19 Genomics UK (COG-UK) consortium | The Lighthouse Lab in Alderley Park and Alex Alderton, Roberto Amato, Sonia Goncalves, Ewan Harrison, David K. Jackson, Ian Johnston, Dominic Kwiatkowski, Cordelia Langford, John Sillitoe on behalf of the Wellcome Sanger Institute COVID-19 Surveillance Team |
| EPI_ISL_551717, EPI_ISL_551718, EPI_ISL_551719, EPI_ISL_551720, EPI_ISL_551722, EPI_ISL_551723, EPI_ISL_551724, EPI_ISL_551725, EPI_ISL_551726, EPI_ISL_551727, EPI_ISL_551728, EPI_ISL_551729, EPI_ISL_551730, EPI_ISL_551731, EPI_ISL_551732, EPI_ISL_551733, EPI_ISL_551734 | | | |
| see above | Lighthouse Lab in Milton Keynes | Wellcome Sanger Institute for the COVID-19 Genomics UK (COG-UK) consortium | The Lighthouse Lab in Milton Keynes and Alex Alderton, Roberto Amato, Sonia Goncalves, Ewan Harrison, David K. Jackson, Ian Johnston, Dominic Kwiatkowski, Cordelia Langford, John Sillitoe on behalf of the Wellcome Sanger Institute COVID-19 Surveillance Team (http://www.sanger.ac.uk/covid-team) |
| EPI_ISL_551835, EPI_ISL_551836, EPI_ISL_551837, EPI_ISL_551838, EPI_ISL_551839, EPI_ISL_551840, EPI_ISL_551841, EPI_ISL_551842, EPI_ISL_551843, EPI_ISL_551844, EPI_ISL_551845, EPI_ISL_551846, EPI_ISL_551847, EPI_ISL_551848, EPI_ISL_551849 | | | |
| see above | Lighthouse Lab in Cambridge | Wellcome Sanger Institute for the COVID-19 Genomics UK (COG-UK) consortium | Rob Howes, The Lighthouse Lab in Cambridge and Alex Alderton, Roberto Amato, Sonia Goncalves, Ewan Harrison, David K. Jackson, Ian Johnston, Dominic Kwiatkowski, Cordelia Langford, John Sillitoe on behalf of the Wellcome Sanger Institute COVID-19 Surveillance Team |
| EPI_ISL_551850 | Lighthouse Lab in Cambridge | Wellcome Sanger Institute for the COVID-19 Genomics UK (COG-UK) Consortium | Rob Howes, The Lighthouse Lab in Cambridge and Alex Alderton, Roberto Amato, Sonia Goncalves, Ewan Harrison, David K. Jackson, Ian Johnston, Dominic Kwiatkowski, Cordelia Langford, John Sillitoe on behalf of the Wellcome Sanger Institute COVID-19 Surveillance Team |
| EPI_ISL_551851, EPI_ISL_551852, EPI_ISL_551853, EPI_ISL_551854, EPI_ISL_551855, EPI_ISL_551856, EPI_ISL_551857, EPI_ISL_551858, EPI_ISL_551859, EPI_ISL_551860, EPI_ISL_551861, EPI_ISL_551862, EPI_ISL_551863, EPI_ISL_551864, EPI_ISL_551865, EPI_ISL_551866, EPI_ISL_551867, EPI_ISL_551868, EPI_ISL_551869, EPI_ISL_551870, EPI_ISL_551871, EPI_ISL_551872, EPI_ISL_551873, EPI_ISL_551874, EPI_ISL_551875, EPI_ISL_551876, EPI_ISL_551877, EPI_ISL_551878, EPI_ISL_551879, EPI_ISL_551880, EPI_ISL_551881, EPI_ISL_551882, EPI_ISL_551883, EPI_ISL_551884, EPI_ISL_551885, EPI_ISL_551886, EPI_ISL_551887, EPI_ISL_551888, EPI_ISL_551889, EPI_ISL_551890, EPI_ISL_551891, EPI_ISL_551892, EPI_ISL_551893, EPI_ISL_551894, EPI_ISL_551895, EPI_ISL_551896, EPI_ISL_551897, EPI_ISL_551898, EPI_ISL_551899, EPI_ISL_551900, EPI_ISL_551901, EPI_ISL_551902, EPI_ISL_551903, EPI_ISL_551904, EPI_ISL_551905, EPI_ISL_551906, EPI_ISL_551907, EPI_ISL_551908, EPI_ISL_551909, EPI_ISL_551910, EPI_ISL_551911, EPI_ISL_551912, EPI_ISL_551913, EPI_ISL_551914, EPI_ISL_551915, EPI_ISL_551916, EPI_ISL_551917, EPI_ISL_551918, EPI_ISL_551919, EPI_ISL_551920, EPI_ISL_551921 | | | |
| see above | Lighthouse Lab in Cambridge | Wellcome Sanger Institute for the COVID-19 Genomics UK (COG-UK) consortium | Rob Howes, The Lighthouse Lab in Cambridge and Alex Alderton, Roberto Amato, Sonia Goncalves, Ewan Harrison, David K. Jackson, Ian Johnston, Dominic Kwiatkowski, Cordelia Langford, John Sillitoe on behalf of the Wellcome Sanger Institute COVID-19 Surveillance Team |
| EPI_ISL_551922, EPI_ISL_551923, EPI_ISL_551924 | Lighthouse Lab in Milton Keynes | Wellcome Sanger Institute for the COVID-19 Genomics UK (COG-UK) consortium | The Lighthouse Lab in Milton Keynes and Alex Alderton, Roberto Amato, Sonia Goncalves, Ewan Harrison, David K. Jackson, Ian Johnston, Dominic Kwiatkowski, Cordelia Langford, John Sillitoe on behalf of the Wellcome Sanger Institute COVID-19 Surveillance Team (http://www.sanger.ac.uk/covid-team) |
| EPI_ISL_551925 | Lighthouse Lab in Cambridge | Wellcome Sanger Institute for the COVID-19 Genomics UK (COG-UK) consortium | Rob Howes, The Lighthouse Lab in Cambridge and Alex Alderton, Roberto Amato, Sonia Goncalves, Ewan Harrison, David K. Jackson, Ian Johnston, Dominic Kwiatkowski, Cordelia Langford, John Sillitoe on behalf of the Wellcome Sanger Institute COVID-19 Surveillance Team |
| EPI_ISL_551926 | Lighthouse Lab in Milton Keynes | Wellcome Sanger Institute for the COVID-19 Genomics UK (COG-UK) consortium | The Lighthouse Lab in Milton Keynes and Alex Alderton, Roberto Amato, Sonia Goncalves, Ewan Harrison, David K. Jackson, Ian Johnston, Dominic Kwiatkowski, Cordelia Langford, John Sillitoe on behalf of the Wellcome Sanger Institute COVID-19 Surveillance Team (http://www.sanger.ac.uk/covid-team) |
| EPI_ISL_551927 | Lighthouse Lab in Milton Keynes | Wellcome Sanger Institute for the COVID-19 Genomics UK (COG-UK) consortium | The Lighthouse Lab in Alderley Park and Alex Alderton, Roberto Amato, Sonia Goncalves, Ewan Harrison, David K. Jackson, Ian Johnston, Dominic Kwiatkowski, Cordelia Langford, John Sillitoe on behalf of the Wellcome Sanger Institute COVID-19 Surveillance Team |
| EPI_ISL_551928, EPI_ISL_551929, EPI_ISL_551930, EPI_ISL_551931 | Lighthouse Lab in Milton Keynes | Wellcome Sanger Institute for the COVID-19 Genomics UK (COG-UK) consortium | The Lighthouse Lab in Milton Keynes and Alex Alderton, Roberto Amato, Sonia Goncalves, Ewan Harrison, David K. Jackson, Ian Johnston, Dominic Kwiatkowski, Cordelia Langford, John Sillitoe on behalf of the Wellcome Sanger Institute COVID-19 Surveillance Team (http://www.sanger.ac.uk/covid-team) |
| EPI_ISL_551932 | Lighthouse Lab in Cambridge | Wellcome Sanger Institute for the COVID-19 Genomics UK (COG-UK) consortium | Rob Howes, The Lighthouse Lab in Cambridge and Alex Alderton, Roberto Amato, Sonia Goncalves, Ewan Harrison, David K. Jackson, Ian Johnston, Dominic Kwiatkowski, Cordelia Langford, John Sillitoe on behalf of the Wellcome Sanger Institute COVID-19 Surveillance Team |
| EPI_ISL_551933, EPI_ISL_551934, EPI_ISL_551935, EPI_ISL_551936, EPI_ISL_551937 | Lighthouse Lab in Milton Keynes | Wellcome Sanger Institute for the COVID-19 Genomics UK (COG-UK) consortium | The Lighthouse Lab in Milton Keynes and Alex Alderton, Roberto Amato, Sonia Goncalves, Ewan Harrison, David K. Jackson, Ian Johnston, Dominic Kwiatkowski, Cordelia Langford, John Sillitoe on behalf of the Wellcome Sanger Institute COVID-19 Surveillance Team (http://www.sanger.ac.uk/covid-team) |
| EPI_ISL_551938 | Lighthouse Lab in Milton Keynes | Wellcome Sanger Institute for the COVID-19 Genomics UK (COG-UK) consortium | The Lighthouse Lab in Alderley Park and Alex Alderton, Roberto Amato, Sonia Goncalves, Ewan Harrison, David K. Jackson, Ian Johnston, Dominic Kwiatkowski, Cordelia Langford, John Sillitoe on behalf of the Wellcome Sanger Institute COVID-19 Surveillance Team |
| EPI_ISL_551939, EPI_ISL_551940, EPI_ISL_551941, EPI_ISL_551942, EPI_ISL_551943, EPI_ISL_551944, EPI_ISL_551945, EPI_ISL_551946, EPI_ISL_551947, EPI_ISL_551948 | Lighthouse Lab in Milton Keynes | Wellcome Sanger Institute for the COVID-19 Genomics UK (COG-UK) consortium | The Lighthouse Lab in Milton Keynes and Alex Alderton, Roberto Amato, Sonia Goncalves, Ewan Harrison, David K. Jackson, Ian Johnston, Dominic Kwiatkowski, Cordelia Langford, John Sillitoe on behalf of the Wellcome Sanger Institute COVID-19 Surveillance Team (http://www.sanger.ac.uk/covid-team) |
| EPI_ISL_551949 | Lighthouse Lab in Milton Keynes | Wellcome Sanger Institute for the COVID-19 Genomics UK (COG-UK) consortium | The Lighthouse Lab in Alderley Park and Alex Alderton, Roberto Amato, Sonia Goncalves, Ewan Harrison, David K. Jackson, Ian Johnston, Dominic Kwiatkowski, Cordelia Langford, John Sillitoe on behalf of the Wellcome Sanger Institute COVID-19 Surveillance Team |
| EPI_ISL_551950 | Lighthouse Lab in Milton Keynes | Wellcome Sanger Institute for the COVID-19 Genomics UK (COG-UK) consortium | The Lighthouse Lab in Milton Keynes and Alex Alderton, Roberto Amato, Sonia Goncalves, Ewan Harrison, David K. Jackson, Ian Johnston, Dominic Kwiatkowski, Cordelia Langford, John Sillitoe on behalf of the Wellcome Sanger Institute COVID-19 Surveillance Team (http://www.sanger.ac.uk/covid-team) |
| EPI_ISL_551951 | Lighthouse Lab in Cambridge | Wellcome Sanger Institute for the COVID-19 Genomics UK (COG-UK) consortium | Rob Howes, The Lighthouse Lab in Cambridge and Alex Alderton, Roberto Amato, Sonia Goncalves, Ewan Harrison, David K. Jackson, Ian Johnston, Dominic Kwiatkowski, Cordelia Langford, John Sillitoe on behalf of the Wellcome Sanger Institute COVID-19 Surveillance Team |
| EPI_ISL_551952 | Lighthouse Lab in Milton Keynes | Wellcome Sanger Institute for the COVID-19 Genomics UK (COG-UK) consortium | The Lighthouse Lab in Milton Keynes and Alex Alderton, Roberto Amato, Sonia Goncalves, Ewan Harrison, David K. Jackson, Ian Johnston, Dominic Kwiatkowski, Cordelia Langford, John Sillitoe on behalf of the Wellcome Sanger Institute COVID-19 Surveillance Team (http://www.sanger.ac.uk/covid-team) |
| EPI_ISL_551953 | Lighthouse Lab in Cambridge | Wellcome Sanger Institute for the COVID-19 Genomics UK | Rob Howes, The Lighthouse Lab in Cambridge and Alex Alderton, Roberto Amato, Sonia Goncalves, Ewan Harrison, David K. Jackson, Ian Johnston, |

| | | | |
|---|---|---|---|
| | | (COG-UK) consortium | Dominic Kwiatkowski, Cordelia Langford, John Sillitoe on behalf of the Wellcome Sanger Institute COVID-19 Surveillance Team |
| EPI_ISL_551954, EPI_ISL_551955, EPI_ISL_551956, EPI_ISL_551957, EPI_ISL_551958, EPI_ISL_551959, EPI_ISL_551960, EPI_ISL_551961, EPI_ISL_551962, EPI_ISL_551963, EPI_ISL_551964, EPI_ISL_551965, EPI_ISL_551966 | | | |
| see above | Lighthouse Lab in Milton Keynes | Wellcome Sanger Institute for the COVID-19 Genomics UK (COG-UK) consortium | The Lighthouse Lab in Milton Keynes and Alex Alderton, Roberto Amato, Sonia Goncalves, Ewan Harrison, David K. Jackson, Ian Johnston, Dominic Kwiatkowski, Cordelia Langford, John Sillitoe on behalf of the Wellcome Sanger Institute COVID-19 Surveillance Team (http://www.sanger.ac.uk/covid-team) |
| EPI_ISL_551967 | Lighthouse Lab in Cambridge | Wellcome Sanger Institute for the COVID-19 Genomics UK (COG-UK) consortium | Rob Howes, The Lighthouse Lab in Cambridge and Alex Alderton, Roberto Amato, Sonia Goncalves, Ewan Harrison, David K. Jackson, Ian Johnston, Dominic Kwiatkowski, Cordelia Langford, John Sillitoe on behalf of the Wellcome Sanger Institute COVID-19 Surveillance Team |
| EPI_ISL_551968, EPI_ISL_551969, EPI_ISL_551970, EPI_ISL_551971, EPI_ISL_551972, EPI_ISL_551973 | Lighthouse Lab in Milton Keynes | Wellcome Sanger Institute for the COVID-19 Genomics UK (COG-UK) consortium | The Lighthouse Lab in Milton Keynes and Alex Alderton, Roberto Amato, Sonia Goncalves, Ewan Harrison, David K. Jackson, Ian Johnston, Dominic Kwiatkowski, Cordelia Langford, John Sillitoe on behalf of the Wellcome Sanger Institute COVID-19 Surveillance Team (http://www.sanger.ac.uk/covid-team) |
| EPI_ISL_551974 | Lighthouse Lab in Milton Keynes | Wellcome Sanger Institute for the COVID-19 Genomics UK (COG-UK) consortium | The Lighthouse Lab in Alderley Park and Alex Alderton, Roberto Amato, Sonia Goncalves, Ewan Harrison, David K. Jackson, Ian Johnston, Dominic Kwiatkowski, Cordelia Langford, John Sillitoe on behalf of the Wellcome Sanger Institute COVID-19 Surveillance Team |
| EPI_ISL_551975, EPI_ISL_551976, EPI_ISL_551977, EPI_ISL_551978, EPI_ISL_551979 | Lighthouse Lab in Milton Keynes | Wellcome Sanger Institute for the COVID-19 Genomics UK (COG-UK) consortium | The Lighthouse Lab in Milton Keynes and Alex Alderton, Roberto Amato, Sonia Goncalves, Ewan Harrison, David K. Jackson, Ian Johnston, Dominic Kwiatkowski, Cordelia Langford, John Sillitoe on behalf of the Wellcome Sanger Institute COVID-19 Surveillance Team (http://www.sanger.ac.uk/covid-team) |
| EPI_ISL_551980 | Lighthouse Lab in Cambridge | Wellcome Sanger Institute for the COVID-19 Genomics UK (COG-UK) consortium | Rob Howes, The Lighthouse Lab in Cambridge and Alex Alderton, Roberto Amato, Sonia Goncalves, Ewan Harrison, David K. Jackson, Ian Johnston, Dominic Kwiatkowski, Cordelia Langford, John Sillitoe on behalf of the Wellcome Sanger Institute COVID-19 Surveillance Team |
| EPI_ISL_551981, EPI_ISL_551982 | Lighthouse Lab in Milton Keynes | Wellcome Sanger Institute for the COVID-19 Genomics UK (COG-UK) consortium | The Lighthouse Lab in Milton Keynes and Alex Alderton, Roberto Amato, Sonia Goncalves, Ewan Harrison, David K. Jackson, Ian Johnston, Dominic Kwiatkowski, Cordelia Langford, John Sillitoe on behalf of the Wellcome Sanger Institute COVID-19 Surveillance Team (http://www.sanger.ac.uk/covid-team) |
| EPI_ISL_551983 | Lighthouse Lab in Cambridge | Wellcome Sanger Institute for the COVID-19 Genomics UK (COG-UK) consortium | Rob Howes, The Lighthouse Lab in Cambridge and Alex Alderton, Roberto Amato, Sonia Goncalves, Ewan Harrison, David K. Jackson, Ian Johnston, Dominic Kwiatkowski, Cordelia Langford, John Sillitoe on behalf of the Wellcome Sanger Institute COVID-19 Surveillance Team |
| EPI_ISL_551984, EPI_ISL_551985, EPI_ISL_551986, EPI_ISL_551987, EPI_ISL_551988, EPI_ISL_551989 | Lighthouse Lab in Milton Keynes | Wellcome Sanger Institute for the COVID-19 Genomics UK (COG-UK) consortium | The Lighthouse Lab in Milton Keynes and Alex Alderton, Roberto Amato, Sonia Goncalves, Ewan Harrison, David K. Jackson, Ian Johnston, Dominic Kwiatkowski, Cordelia Langford, John Sillitoe on behalf of the Wellcome Sanger Institute COVID-19 Surveillance Team (http://www.sanger.ac.uk/covid-team) |
| EPI_ISL_551990 | Lighthouse Lab in Cambridge | Wellcome Sanger Institute for the COVID-19 Genomics UK (COG-UK) consortium | Rob Howes, The Lighthouse Lab in Cambridge and Alex Alderton, Roberto Amato, Sonia Goncalves, Ewan Harrison, David K. Jackson, Ian Johnston, Dominic Kwiatkowski, Cordelia Langford, John Sillitoe on behalf of the Wellcome Sanger Institute COVID-19 Surveillance Team |
| EPI_ISL_551991, EPI_ISL_551992, EPI_ISL_551993 | Lighthouse Lab in Milton Keynes | Wellcome Sanger Institute for the COVID-19 Genomics UK (COG-UK) consortium | The Lighthouse Lab in Milton Keynes and Alex Alderton, Roberto Amato, Sonia Goncalves, Ewan Harrison, David K. Jackson, Ian Johnston, Dominic Kwiatkowski, Cordelia Langford, John Sillitoe on behalf of the Wellcome Sanger Institute COVID-19 Surveillance Team (http://www.sanger.ac.uk/covid-team) |
| EPI_ISL_551994 | Lighthouse Lab in Cambridge | Wellcome Sanger Institute for the COVID-19 Genomics UK (COG-UK) consortium | Rob Howes, The Lighthouse Lab in Cambridge and Alex Alderton, Roberto Amato, Sonia Goncalves, Ewan Harrison, David K. Jackson, Ian Johnston, Dominic Kwiatkowski, Cordelia Langford, John Sillitoe on behalf of the Wellcome Sanger Institute COVID-19 Surveillance Team |
| EPI_ISL_551995, EPI_ISL_551996, EPI_ISL_551997, EPI_ISL_551998, EPI_ISL_551999, EPI_ISL_552000 | Lighthouse Lab in Milton Keynes | Wellcome Sanger Institute for the COVID-19 Genomics UK (COG-UK) consortium | The Lighthouse Lab in Milton Keynes and Alex Alderton, Roberto Amato, Sonia Goncalves, Ewan Harrison, David K. Jackson, Ian Johnston, Dominic Kwiatkowski, Cordelia Langford, John Sillitoe on behalf of the Wellcome Sanger Institute COVID-19 Surveillance Team (http://www.sanger.ac.uk/covid-team) |
| EPI_ISL_552001 | Lighthouse Lab in Cambridge | Wellcome Sanger Institute for the COVID-19 Genomics UK (COG-UK) consortium | Rob Howes, The Lighthouse Lab in Cambridge and Alex Alderton, Roberto Amato, Sonia Goncalves, Ewan Harrison, David K. Jackson, Ian Johnston, Dominic Kwiatkowski, Cordelia Langford, John Sillitoe on behalf of the Wellcome Sanger Institute COVID-19 Surveillance Team |
| EPI_ISL_552002, EPI_ISL_552003 | Lighthouse Lab in Milton Keynes | Wellcome Sanger Institute for the COVID-19 Genomics UK (COG-UK) consortium | The Lighthouse Lab in Alderley Park and Alex Alderton, Roberto Amato, Sonia Goncalves, Ewan Harrison, David K. Jackson, Ian Johnston, Dominic Kwiatkowski, Cordelia Langford, John Sillitoe on behalf of the Wellcome Sanger Institute COVID-19 Surveillance Team |
| EPI_ISL_552004 | Lighthouse Lab in Milton Keynes | Wellcome Sanger Institute for the COVID-19 Genomics UK (COG-UK) consortium | The Lighthouse Lab in Milton Keynes and Alex Alderton, Roberto Amato, Sonia Goncalves, Ewan Harrison, David K. Jackson, Ian Johnston, Dominic Kwiatkowski, Cordelia Langford, John Sillitoe on behalf of the Wellcome Sanger Institute COVID-19 Surveillance Team (http://www.sanger.ac.uk/covid-team) |
| EPI_ISL_552005 | Lighthouse Lab in Cambridge | Wellcome Sanger Institute for the COVID-19 Genomics UK (COG-UK) consortium | Rob Howes, The Lighthouse Lab in Cambridge and Alex Alderton, Roberto Amato, Sonia Goncalves, Ewan Harrison, David K. Jackson, Ian Johnston, Dominic Kwiatkowski, Cordelia Langford, John Sillitoe on behalf of the Wellcome Sanger Institute COVID-19 Surveillance Team |
| EPI_ISL_552006, EPI_ISL_552007 | Lighthouse Lab in Milton Keynes | Wellcome Sanger Institute for the COVID-19 Genomics UK (COG-UK) consortium | The Lighthouse Lab in Milton Keynes and Alex Alderton, Roberto Amato, Sonia Goncalves, Ewan Harrison, David K. Jackson, Ian Johnston, Dominic Kwiatkowski, Cordelia Langford, John Sillitoe on behalf of the Wellcome Sanger Institute COVID-19 Surveillance Team (http://www.sanger.ac.uk/covid-team) |
| EPI_ISL_552008, EPI_ISL_552010 | Lighthouse Lab in Cambridge | Wellcome Sanger Institute for the COVID-19 Genomics UK (COG-UK) consortium | Rob Howes, The Lighthouse Lab in Cambridge and Alex Alderton, Roberto Amato, Sonia Goncalves, Ewan Harrison, David K. Jackson, Ian Johnston, Dominic Kwiatkowski, Cordelia Langford, John Sillitoe on behalf of the Wellcome Sanger Institute COVID-19 Surveillance Team |
| EPI_ISL_552011, EPI_ISL_552012 | Lighthouse Lab in Milton Keynes | Wellcome Sanger Institute for the COVID-19 Genomics UK (COG-UK) consortium | The Lighthouse Lab in Milton Keynes and Alex Alderton, Roberto Amato, Sonia Goncalves, Ewan Harrison, David K. Jackson, Ian Johnston, Dominic Kwiatkowski, Cordelia Langford, John Sillitoe on behalf of the Wellcome Sanger Institute COVID-19 Surveillance Team (http://www.sanger.ac.uk/covid-team) |
| EPI_ISL_552013 | Lighthouse Lab in Milton Keynes | Wellcome Sanger Institute for the COVID-19 Genomics UK (COG-UK) consortium | The Lighthouse Lab in Alderley Park and Alex Alderton, Roberto Amato, Sonia Goncalves, Ewan Harrison, David K. Jackson, Ian Johnston, Dominic Kwiatkowski, Cordelia Langford, John Sillitoe on behalf of the Wellcome Sanger Institute COVID-19 Surveillance Team |
| EPI_ISL_552014, EPI_ISL_552015, EPI_ISL_552016 | Lighthouse Lab in Milton Keynes | Wellcome Sanger Institute for the COVID-19 Genomics UK (COG-UK) consortium | The Lighthouse Lab in Milton Keynes and Alex Alderton, Roberto Amato, Sonia Goncalves, Ewan Harrison, David K. Jackson, Ian Johnston, Dominic Kwiatkowski, Cordelia Langford, John Sillitoe on behalf of the Wellcome Sanger Institute COVID-19 Surveillance Team (http://www.sanger.ac.uk/covid-team) |
| EPI_ISL_552017 | Lighthouse Lab in Cambridge | Wellcome Sanger Institute for the COVID-19 Genomics UK (COG-UK) consortium | Rob Howes, The Lighthouse Lab in Cambridge and Alex Alderton, Roberto Amato, Sonia Goncalves, Ewan Harrison, David K. Jackson, Ian Johnston, Dominic Kwiatkowski, Cordelia Langford, John Sillitoe on behalf of the Wellcome Sanger Institute COVID-19 Surveillance Team |
| EPI_ISL_552018, EPI_ISL_552019, EPI_ISL_552020, EPI_ISL_552021, EPI_ISL_552022, EPI_ISL_552023, EPI_ISL_552024, EPI_ISL_552025, EPI_ISL_552026, EPI_ISL_552027, EPI_ISL_552028, EPI_ISL_552029, EPI_ISL_552030, EPI_ISL_552031, EPI_ISL_552032, EPI_ISL_552033, EPI_ISL_552034, EPI_ISL_552035, EPI_ISL_552036, EPI_ISL_552037, EPI_ISL_552038, EPI_ISL_552039 | | | |
| see above | Lighthouse Lab in Milton Keynes | Wellcome Sanger Institute for the COVID-19 Genomics UK (COG-UK) consortium | The Lighthouse Lab in Milton Keynes and Alex Alderton, Roberto Amato, Sonia Goncalves, Ewan Harrison, David K. Jackson, Ian Johnston, Dominic Kwiatkowski, Cordelia Langford, John Sillitoe on behalf of the Wellcome Sanger Institute COVID-19 Surveillance Team (http://www.sanger.ac.uk/covid-team) |
| EPI_ISL_552040 | Lighthouse Lab in Milton Keynes | Wellcome Sanger Institute for the COVID-19 Genomics UK (COG-UK) consortium | The Lighthouse Lab in Alderley Park and Alex Alderton, Roberto Amato, Sonia Goncalves, Ewan Harrison, David K. Jackson, Ian Johnston, Dominic Kwiatkowski, Cordelia Langford, John Sillitoe on behalf of the Wellcome Sanger Institute COVID-19 Surveillance Team |
| EPI_ISL_552041, EPI_ISL_552042, EPI_ISL_552043, EPI_ISL_552044 | Lighthouse Lab in Milton Keynes | Wellcome Sanger Institute for the COVID-19 Genomics UK (COG-UK) consortium | The Lighthouse Lab in Milton Keynes and Alex Alderton, Roberto Amato, Sonia Goncalves, Ewan Harrison, David K. Jackson, Ian Johnston, Dominic Kwiatkowski, Cordelia Langford, John Sillitoe on behalf of the Wellcome Sanger Institute COVID-19 Surveillance Team (http://www.sanger.ac.uk/covid-team) |
| EPI_ISL_552045 | Lighthouse Lab in Milton Keynes | Wellcome Sanger Institute for the COVID-19 Genomics UK (COG-UK) consortium | The Lighthouse Lab in Alderley Park and Alex Alderton, Roberto Amato, Sonia Goncalves, Ewan Harrison, David K. Jackson, Ian Johnston, Dominic Kwiatkowski, Cordelia Langford, John Sillitoe on behalf of the Wellcome Sanger Institute COVID-19 Surveillance Team |
| EPI_ISL_552046, EPI_ISL_552047, EPI_ISL_552048, EPI_ISL_552049, | Lighthouse Lab in Milton Keynes | Wellcome Sanger Institute for the COVID-19 Genomics UK (COG-UK) consortium | The Lighthouse Lab in Milton Keynes and Alex Alderton, Roberto Amato, Sonia Goncalves, Ewan Harrison, David K. Jackson, Ian Johnston, Dominic Kwiatkowski, Cordelia Langford, John Sillitoe on behalf of the Wellcome Sanger Institute COVID-19 Surveillance Team |

| | | | |
|---|---|---|---|
| EPI_ISL_552050 | | | (http://www.sanger.ac.uk/covid-team) |
| EPI_ISL_552051 | Lighthouse Lab in Cambridge | Wellcome Sanger Institute for the COVID-19 Genomics UK (COG-UK) consortium | Rob Howes, The Lighthouse Lab in Cambridge and Alex Alderton, Roberto Amato, Sonia Goncalves, Ewan Harrison, David K. Jackson, Ian Johnston, Dominic Kwiatkowski, Cordelia Langford, John Sillitoe on behalf of the Wellcome Sanger Institute COVID-19 Surveillance Team |
| EPI_ISL_552052 | Lighthouse Lab in Milton Keynes | Wellcome Sanger Institute for the COVID-19 Genomics UK (COG-UK) consortium | The Lighthouse Lab in Milton Keynes and Alex Alderton, Roberto Amato, Sonia Goncalves, Ewan Harrison, David K. Jackson, Ian Johnston, Dominic Kwiatkowski, Cordelia Langford, John Sillitoe on behalf of the Wellcome Sanger Institute COVID-19 Surveillance Team (http://www.sanger.ac.uk/covid-team) |
| EPI_ISL_552053 | Lighthouse Lab in Milton Keynes | Wellcome Sanger Institute for the COVID-19 Genomics UK (COG-UK) consortium | The Lighthouse Lab in Alderley Park and Alex Alderton, Roberto Amato, Sonia Goncalves, Ewan Harrison, David K. Jackson, Ian Johnston, Dominic Kwiatkowski, Cordelia Langford, John Sillitoe on behalf of the Wellcome Sanger Institute COVID-19 Surveillance Team |
| EPI_ISL_552054, EPI_ISL_552055 | Lighthouse Lab in Milton Keynes | Wellcome Sanger Institute for the COVID-19 Genomics UK (COG-UK) consortium | The Lighthouse Lab in Milton Keynes and Alex Alderton, Roberto Amato, Sonia Goncalves, Ewan Harrison, David K. Jackson, Ian Johnston, Dominic Kwiatkowski, Cordelia Langford, John Sillitoe on behalf of the Wellcome Sanger Institute COVID-19 Surveillance Team (http://www.sanger.ac.uk/covid-team) |
| EPI_ISL_552056 | Lighthouse Lab in Milton Keynes | Wellcome Sanger Institute for the COVID-19 Genomics UK (COG-UK) consortium | The Lighthouse Lab in Alderley Park and Alex Alderton, Roberto Amato, Sonia Goncalves, Ewan Harrison, David K. Jackson, Ian Johnston, Dominic Kwiatkowski, Cordelia Langford, John Sillitoe on behalf of the Wellcome Sanger Institute COVID-19 Surveillance Team |
| EPI_ISL_552057 | Lighthouse Lab in Milton Keynes | Wellcome Sanger Institute for the COVID-19 Genomics UK (COG-UK) consortium | The Lighthouse Lab in Milton Keynes and Alex Alderton, Roberto Amato, Sonia Goncalves, Ewan Harrison, David K. Jackson, Ian Johnston, Dominic Kwiatkowski, Cordelia Langford, John Sillitoe on behalf of the Wellcome Sanger Institute COVID-19 Surveillance Team (http://www.sanger.ac.uk/covid-team) |
| EPI_ISL_552058 | Lighthouse Lab in Milton Keynes | Wellcome Sanger Institute for the COVID-19 Genomics UK (COG-UK) consortium | The Lighthouse Lab in Alderley Park and Alex Alderton, Roberto Amato, Sonia Goncalves, Ewan Harrison, David K. Jackson, Ian Johnston, Dominic Kwiatkowski, Cordelia Langford, John Sillitoe on behalf of the Wellcome Sanger Institute COVID-19 Surveillance Team |
| EPI_ISL_552059, EPI_ISL_552060, EPI_ISL_552061 | Lighthouse Lab in Milton Keynes | Wellcome Sanger Institute for the COVID-19 Genomics UK (COG-UK) consortium | The Lighthouse Lab in Milton Keynes and Alex Alderton, Roberto Amato, Sonia Goncalves, Ewan Harrison, David K. Jackson, Ian Johnston, Dominic Kwiatkowski, Cordelia Langford, John Sillitoe on behalf of the Wellcome Sanger Institute COVID-19 Surveillance Team (http://www.sanger.ac.uk/covid-team) |
| EPI_ISL_552062 | Lighthouse Lab in Milton Keynes | Wellcome Sanger Institute for the COVID-19 Genomics UK (COG-UK) consortium | The Lighthouse Lab in Alderley Park and Alex Alderton, Roberto Amato, Sonia Goncalves, Ewan Harrison, David K. Jackson, Ian Johnston, Dominic Kwiatkowski, Cordelia Langford, John Sillitoe on behalf of the Wellcome Sanger Institute COVID-19 Surveillance Team |
| EPI_ISL_552063, EPI_ISL_552064 | Lighthouse Lab in Milton Keynes | Wellcome Sanger Institute for the COVID-19 Genomics UK (COG-UK) consortium | The Lighthouse Lab in Milton Keynes and Alex Alderton, Roberto Amato, Sonia Goncalves, Ewan Harrison, David K. Jackson, Ian Johnston, Dominic Kwiatkowski, Cordelia Langford, John Sillitoe on behalf of the Wellcome Sanger Institute COVID-19 Surveillance Team (http://www.sanger.ac.uk/covid-team) |
| EPI_ISL_552065 | Lighthouse Lab in Cambridge | Wellcome Sanger Institute for the COVID-19 Genomics UK (COG-UK) consortium | Rob Howes, The Lighthouse Lab in Cambridge and Alex Alderton, Roberto Amato, Sonia Goncalves, Ewan Harrison, David K. Jackson, Ian Johnston, Dominic Kwiatkowski, Cordelia Langford, John Sillitoe on behalf of the Wellcome Sanger Institute COVID-19 Surveillance Team |
| EPI_ISL_552066 | Lighthouse Lab in Milton Keynes | Wellcome Sanger Institute for the COVID-19 Genomics UK (COG-UK) consortium | The Lighthouse Lab in Alderley Park and Alex Alderton, Roberto Amato, Sonia Goncalves, Ewan Harrison, David K. Jackson, Ian Johnston, Dominic Kwiatkowski, Cordelia Langford, John Sillitoe on behalf of the Wellcome Sanger Institute COVID-19 Surveillance Team |
| EPI_ISL_552067, EPI_ISL_552068, EPI_ISL_552069, EPI_ISL_552070, EPI_ISL_552071, EPI_ISL_552072 | Lighthouse Lab in Milton Keynes | Wellcome Sanger Institute for the COVID-19 Genomics UK (COG-UK) consortium | The Lighthouse Lab in Milton Keynes and Alex Alderton, Roberto Amato, Sonia Goncalves, Ewan Harrison, David K. Jackson, Ian Johnston, Dominic Kwiatkowski, Cordelia Langford, John Sillitoe on behalf of the Wellcome Sanger Institute COVID-19 Surveillance Team (http://www.sanger.ac.uk/covid-team) |
| EPI_ISL_552073 | Lighthouse Lab in Cambridge | Wellcome Sanger Institute for the COVID-19 Genomics UK (COG-UK) consortium | Rob Howes, The Lighthouse Lab in Cambridge and Alex Alderton, Roberto Amato, Sonia Goncalves, Ewan Harrison, David K. Jackson, Ian Johnston, Dominic Kwiatkowski, Cordelia Langford, John Sillitoe on behalf of the Wellcome Sanger Institute COVID-19 Surveillance Team |
| EPI_ISL_552074, EPI_ISL_552075, EPI_ISL_552076, EPI_ISL_552077, EPI_ISL_552078, EPI_ISL_552079, EPI_ISL_552080 | Lighthouse Lab in Milton Keynes | Wellcome Sanger Institute for the COVID-19 Genomics UK (COG-UK) consortium | The Lighthouse Lab in Milton Keynes and Alex Alderton, Roberto Amato, Sonia Goncalves, Ewan Harrison, David K. Jackson, Ian Johnston, Dominic Kwiatkowski, Cordelia Langford, John Sillitoe on behalf of the Wellcome Sanger Institute COVID-19 Surveillance Team (http://www.sanger.ac.uk/covid-team) |
| EPI_ISL_552081 | Lighthouse Lab in Milton Keynes | Wellcome Sanger Institute for the COVID-19 Genomics UK (COG-UK) consortium | The Lighthouse Lab in Alderley Park and Alex Alderton, Roberto Amato, Sonia Goncalves, Ewan Harrison, David K. Jackson, Ian Johnston, Dominic Kwiatkowski, Cordelia Langford, John Sillitoe on behalf of the Wellcome Sanger Institute COVID-19 Surveillance Team |
| EPI_ISL_552082, EPI_ISL_552083, EPI_ISL_552084, EPI_ISL_552085, EPI_ISL_552086, EPI_ISL_552087, EPI_ISL_552088, EPI_ISL_552089, EPI_ISL_552090 | Lighthouse Lab in Milton Keynes | Wellcome Sanger Institute for the COVID-19 Genomics UK (COG-UK) consortium | The Lighthouse Lab in Milton Keynes and Alex Alderton, Roberto Amato, Sonia Goncalves, Ewan Harrison, David K. Jackson, Ian Johnston, Dominic Kwiatkowski, Cordelia Langford, John Sillitoe on behalf of the Wellcome Sanger Institute COVID-19 Surveillance Team (http://www.sanger.ac.uk/covid-team) |
| EPI_ISL_552091 | Lighthouse Lab in Cambridge | Wellcome Sanger Institute for the COVID-19 Genomics UK (COG-UK) consortium | Rob Howes, The Lighthouse Lab in Cambridge and Alex Alderton, Roberto Amato, Sonia Goncalves, Ewan Harrison, David K. Jackson, Ian Johnston, Dominic Kwiatkowski, Cordelia Langford, John Sillitoe on behalf of the Wellcome Sanger Institute COVID-19 Surveillance Team |
| EPI_ISL_552092, EPI_ISL_552093 | Lighthouse Lab in Milton Keynes | Wellcome Sanger Institute for the COVID-19 Genomics UK (COG-UK) consortium | The Lighthouse Lab in Milton Keynes and Alex Alderton, Roberto Amato, Sonia Goncalves, Ewan Harrison, David K. Jackson, Ian Johnston, Dominic Kwiatkowski, Cordelia Langford, John Sillitoe on behalf of the Wellcome Sanger Institute COVID-19 Surveillance Team (http://www.sanger.ac.uk/covid-team) |
| EPI_ISL_552094 | Lighthouse Lab in Cambridge | Wellcome Sanger Institute for the COVID-19 Genomics UK (COG-UK) consortium | Rob Howes, The Lighthouse Lab in Cambridge and Alex Alderton, Roberto Amato, Sonia Goncalves, Ewan Harrison, David K. Jackson, Ian Johnston, Dominic Kwiatkowski, Cordelia Langford, John Sillitoe on behalf of the Wellcome Sanger Institute COVID-19 Surveillance Team |
| EPI_ISL_552095, EPI_ISL_552096, EPI_ISL_552097 | Lighthouse Lab in Milton Keynes | Wellcome Sanger Institute for the COVID-19 Genomics UK (COG-UK) consortium | The Lighthouse Lab in Milton Keynes and Alex Alderton, Roberto Amato, Sonia Goncalves, Ewan Harrison, David K. Jackson, Ian Johnston, Dominic Kwiatkowski, Cordelia Langford, John Sillitoe on behalf of the Wellcome Sanger Institute COVID-19 Surveillance Team (http://www.sanger.ac.uk/covid-team) |
| EPI_ISL_552098 | Lighthouse Lab in Cambridge | Wellcome Sanger Institute for the COVID-19 Genomics UK (COG-UK) consortium | Rob Howes, The Lighthouse Lab in Cambridge and Alex Alderton, Roberto Amato, Sonia Goncalves, Ewan Harrison, David K. Jackson, Ian Johnston, Dominic Kwiatkowski, Cordelia Langford, John Sillitoe on behalf of the Wellcome Sanger Institute COVID-19 Surveillance Team |
| EPI_ISL_552099, EPI_ISL_552100, EPI_ISL_552101, EPI_ISL_552102, EPI_ISL_552103, EPI_ISL_552104 | Lighthouse Lab in Milton Keynes | Wellcome Sanger Institute for the COVID-19 Genomics UK (COG-UK) consortium | The Lighthouse Lab in Milton Keynes and Alex Alderton, Roberto Amato, Sonia Goncalves, Ewan Harrison, David K. Jackson, Ian Johnston, Dominic Kwiatkowski, Cordelia Langford, John Sillitoe on behalf of the Wellcome Sanger Institute COVID-19 Surveillance Team (http://www.sanger.ac.uk/covid-team) |
| EPI_ISL_552105 | Lighthouse Lab in Cambridge | Wellcome Sanger Institute for the COVID-19 Genomics UK (COG-UK) consortium | Rob Howes, The Lighthouse Lab in Cambridge and Alex Alderton, Roberto Amato, Sonia Goncalves, Ewan Harrison, David K. Jackson, Ian Johnston, Dominic Kwiatkowski, Cordelia Langford, John Sillitoe on behalf of the Wellcome Sanger Institute COVID-19 Surveillance Team |
| EPI_ISL_552106 | Lighthouse Lab in Milton Keynes | Wellcome Sanger Institute for the COVID-19 Genomics UK (COG-UK) Consortium | The Lighthouse Lab in Milton Keynes and Alex Alderton, Roberto Amato, Sonia Goncalves, Ewan Harrison, David K. Jackson, Ian Johnston, Dominic Kwiatkowski, Cordelia Langford, John Sillitoe on behalf of the Wellcome Sanger Institute COVID-19 Surveillance Team |
| EPI_ISL_552107, EPI_ISL_552108, EPI_ISL_552109, EPI_ISL_552110, EPI_ISL_552111, EPI_ISL_552112, EPI_ISL_552113, EPI_ISL_552114, EPI_ISL_552115, EPI_ISL_552116 | Lighthouse Lab in Milton Keynes | Wellcome Sanger Institute for the COVID-19 Genomics UK (COG-UK) consortium | The Lighthouse Lab in Milton Keynes and Alex Alderton, Roberto Amato, Sonia Goncalves, Ewan Harrison, David K. Jackson, Ian Johnston, Dominic Kwiatkowski, Cordelia Langford, John Sillitoe on behalf of the Wellcome Sanger Institute COVID-19 Surveillance Team (http://www.sanger.ac.uk/covid-team) |
| EPI_ISL_552117 | Lighthouse Lab in Milton Keynes | Wellcome Sanger Institute for the COVID-19 Genomics UK (COG-UK) consortium | The Lighthouse Lab in Alderley Park and Alex Alderton, Roberto Amato, Sonia Goncalves, Ewan Harrison, David K. Jackson, Ian Johnston, Dominic Kwiatkowski, Cordelia Langford, John Sillitoe on behalf of the Wellcome Sanger Institute COVID-19 Surveillance Team |
| EPI_ISL_552118, EPI_ISL_552119, | Lighthouse Lab in Milton Keynes | Wellcome Sanger Institute for the COVID-19 Genomics UK | The Lighthouse Lab in Milton Keynes and Alex Alderton, Roberto Amato, Sonia Goncalves, Ewan Harrison, David K. Jackson, Ian Johnston, Dominic |

| | | | |
|---|---|---|---|
| EPI_ISL_552120, EPI_ISL_552121, EPI_ISL_552122, EPI_ISL_552123 | | (COG-UK) consortium | Kwiatkowski, Cordelia Langford, John Sillitoe on behalf of the Wellcome Sanger Institute COVID-19 Surveillance Team (http://www.sanger.ac.uk/covid-team) |
| EPI_ISL_552124 | Lighthouse Lab in Cambridge | Wellcome Sanger Institute for the COVID-19 Genomics UK (COG-UK) consortium | Rob Howes, The Lighthouse Lab in Cambridge and Alex Alderton, Roberto Amato, Sonia Goncalves, Ewan Harrison, David K. Jackson, Ian Johnston, Dominic Kwiatkowski, Cordelia Langford, John Sillitoe on behalf of the Wellcome Sanger Institute COVID-19 Surveillance Team |
| EPI_ISL_552125, EPI_ISL_552126, EPI_ISL_552127, EPI_ISL_552128, EPI_ISL_552129, EPI_ISL_552130, EPI_ISL_552131 | Lighthouse Lab in Milton Keynes | Wellcome Sanger Institute for the COVID-19 Genomics UK (COG-UK) consortium | The Lighthouse Lab in Milton Keynes and Alex Alderton, Roberto Amato, Sonia Goncalves, Ewan Harrison, David K. Jackson, Ian Johnston, Dominic Kwiatkowski, Cordelia Langford, John Sillitoe on behalf of the Wellcome Sanger Institute COVID-19 Surveillance Team (http://www.sanger.ac.uk/covid-team) |
| EPI_ISL_552132 | Lighthouse Lab in Milton Keynes | Wellcome Sanger Institute for the COVID-19 Genomics UK (COG-UK) consortium | The Lighthouse Lab in Alderley Park and Alex Alderton, Roberto Amato, Sonia Goncalves, Ewan Harrison, David K. Jackson, Ian Johnston, Dominic Kwiatkowski, Cordelia Langford, John Sillitoe on behalf of the Wellcome Sanger Institute COVID-19 Surveillance Team |
| EPI_ISL_552133, EPI_ISL_552134, EPI_ISL_552135, EPI_ISL_552136, EPI_ISL_552137, EPI_ISL_552138, EPI_ISL_552139, EPI_ISL_552140, EPI_ISL_552141, EPI_ISL_552142, EPI_ISL_552143 | | | |
| see above | Lighthouse Lab in Milton Keynes | Wellcome Sanger Institute for the COVID-19 Genomics UK (COG-UK) consortium | The Lighthouse Lab in Milton Keynes and Alex Alderton, Roberto Amato, Sonia Goncalves, Ewan Harrison, David K. Jackson, Ian Johnston, Dominic Kwiatkowski, Cordelia Langford, John Sillitoe on behalf of the Wellcome Sanger Institute COVID-19 Surveillance Team (http://www.sanger.ac.uk/covid-team) |
| EPI_ISL_552144, EPI_ISL_552145 | Lighthouse Lab in Milton Keynes | Wellcome Sanger Institute for the COVID-19 Genomics UK (COG-UK) consortium | The Lighthouse Lab in Alderley Park and Alex Alderton, Roberto Amato, Sonia Goncalves, Ewan Harrison, David K. Jackson, Ian Johnston, Dominic Kwiatkowski, Cordelia Langford, John Sillitoe on behalf of the Wellcome Sanger Institute COVID-19 Surveillance Team |
| EPI_ISL_552146, EPI_ISL_552147, EPI_ISL_552148, EPI_ISL_552149, EPI_ISL_552150 | Lighthouse Lab in Milton Keynes | Wellcome Sanger Institute for the COVID-19 Genomics UK (COG-UK) consortium | The Lighthouse Lab in Milton Keynes and Alex Alderton, Roberto Amato, Sonia Goncalves, Ewan Harrison, David K. Jackson, Ian Johnston, Dominic Kwiatkowski, Cordelia Langford, John Sillitoe on behalf of the Wellcome Sanger Institute COVID-19 Surveillance Team (http://www.sanger.ac.uk/covid-team) |
| EPI_ISL_552151 | Lighthouse Lab in Cambridge | Wellcome Sanger Institute for the COVID-19 Genomics UK (COG-UK) consortium | Rob Howes, The Lighthouse Lab in Cambridge and Alex Alderton, Roberto Amato, Sonia Goncalves, Ewan Harrison, David K. Jackson, Ian Johnston, Dominic Kwiatkowski, Cordelia Langford, John Sillitoe on behalf of the Wellcome Sanger Institute COVID-19 Surveillance Team |
| EPI_ISL_552152, EPI_ISL_552153, EPI_ISL_552154, EPI_ISL_552155, EPI_ISL_552156 | Lighthouse Lab in Milton Keynes | Wellcome Sanger Institute for the COVID-19 Genomics UK (COG-UK) consortium | The Lighthouse Lab in Milton Keynes and Alex Alderton, Roberto Amato, Sonia Goncalves, Ewan Harrison, David K. Jackson, Ian Johnston, Dominic Kwiatkowski, Cordelia Langford, John Sillitoe on behalf of the Wellcome Sanger Institute COVID-19 Surveillance Team (http://www.sanger.ac.uk/covid-team) |
| EPI_ISL_552157 | Lighthouse Lab in Milton Keynes | Wellcome Sanger Institute for the COVID-19 Genomics UK (COG-UK) Consortium | The Lighthouse Lab in Milton Keynes and Alex Alderton, Roberto Amato, Sonia Goncalves, Ewan Harrison, David K. Jackson, Ian Johnston, Dominic Kwiatkowski, Cordelia Langford, John Sillitoe on behalf of the Wellcome Sanger Institute COVID-19 Surveillance Team |
| EPI_ISL_552158, EPI_ISL_552159, EPI_ISL_552160, EPI_ISL_552161 | Lighthouse Lab in Milton Keynes | Wellcome Sanger Institute for the COVID-19 Genomics UK (COG-UK) consortium | The Lighthouse Lab in Milton Keynes and Alex Alderton, Roberto Amato, Sonia Goncalves, Ewan Harrison, David K. Jackson, Ian Johnston, Dominic Kwiatkowski, Cordelia Langford, John Sillitoe on behalf of the Wellcome Sanger Institute COVID-19 Surveillance Team (http://www.sanger.ac.uk/covid-team) |
| EPI_ISL_552162 | Lighthouse Lab in Milton Keynes | Wellcome Sanger Institute for the COVID-19 Genomics UK (COG-UK) consortium | The Lighthouse Lab in Alderley Park and Alex Alderton, Roberto Amato, Sonia Goncalves, Ewan Harrison, David K. Jackson, Ian Johnston, Dominic Kwiatkowski, Cordelia Langford, John Sillitoe on behalf of the Wellcome Sanger Institute COVID-19 Surveillance Team |
| EPI_ISL_552163 | Lighthouse Lab in Milton Keynes | Wellcome Sanger Institute for the COVID-19 Genomics UK (COG-UK) consortium | The Lighthouse Lab in Milton Keynes and Alex Alderton, Roberto Amato, Sonia Goncalves, Ewan Harrison, David K. Jackson, Ian Johnston, Dominic Kwiatkowski, Cordelia Langford, John Sillitoe on behalf of the Wellcome Sanger Institute COVID-19 Surveillance Team (http://www.sanger.ac.uk/covid-team) |
| EPI_ISL_552164 | Lighthouse Lab in Milton Keynes | Wellcome Sanger Institute for the COVID-19 Genomics UK (COG-UK) consortium | The Lighthouse Lab in Alderley Park and Alex Alderton, Roberto Amato, Sonia Goncalves, Ewan Harrison, David K. Jackson, Ian Johnston, Dominic Kwiatkowski, Cordelia Langford, John Sillitoe on behalf of the Wellcome Sanger Institute COVID-19 Surveillance Team |
| EPI_ISL_552165 | Lighthouse Lab in Milton Keynes | Wellcome Sanger Institute for the COVID-19 Genomics UK (COG-UK) consortium | The Lighthouse Lab in Milton Keynes and Alex Alderton, Roberto Amato, Sonia Goncalves, Ewan Harrison, David K. Jackson, Ian Johnston, Dominic Kwiatkowski, Cordelia Langford, John Sillitoe on behalf of the Wellcome Sanger Institute COVID-19 Surveillance Team (http://www.sanger.ac.uk/covid-team) |
| EPI_ISL_552166 | Lighthouse Lab in Cambridge | Wellcome Sanger Institute for the COVID-19 Genomics UK (COG-UK) consortium | Rob Howes, The Lighthouse Lab in Cambridge and Alex Alderton, Roberto Amato, Sonia Goncalves, Ewan Harrison, David K. Jackson, Ian Johnston, Dominic Kwiatkowski, Cordelia Langford, John Sillitoe on behalf of the Wellcome Sanger Institute COVID-19 Surveillance Team |
| EPI_ISL_552167, EPI_ISL_552168, EPI_ISL_552169 | Lighthouse Lab in Milton Keynes | Wellcome Sanger Institute for the COVID-19 Genomics UK (COG-UK) consortium | The Lighthouse Lab in Milton Keynes and Alex Alderton, Roberto Amato, Sonia Goncalves, Ewan Harrison, David K. Jackson, Ian Johnston, Dominic Kwiatkowski, Cordelia Langford, John Sillitoe on behalf of the Wellcome Sanger Institute COVID-19 Surveillance Team (http://www.sanger.ac.uk/covid-team) |
| EPI_ISL_552170 | Lighthouse Lab in Cambridge | Wellcome Sanger Institute for the COVID-19 Genomics UK (COG-UK) consortium | Rob Howes, The Lighthouse Lab in Cambridge and Alex Alderton, Roberto Amato, Sonia Goncalves, Ewan Harrison, David K. Jackson, Ian Johnston, Dominic Kwiatkowski, Cordelia Langford, John Sillitoe on behalf of the Wellcome Sanger Institute COVID-19 Surveillance Team |
| EPI_ISL_552171, EPI_ISL_552172 | Lighthouse Lab in Milton Keynes | Wellcome Sanger Institute for the COVID-19 Genomics UK (COG-UK) consortium | The Lighthouse Lab in Milton Keynes and Alex Alderton, Roberto Amato, Sonia Goncalves, Ewan Harrison, David K. Jackson, Ian Johnston, Dominic Kwiatkowski, Cordelia Langford, John Sillitoe on behalf of the Wellcome Sanger Institute COVID-19 Surveillance Team (http://www.sanger.ac.uk/covid-team) |
| EPI_ISL_552173 | Lighthouse Lab in Cambridge | Wellcome Sanger Institute for the COVID-19 Genomics UK (COG-UK) consortium | Rob Howes, The Lighthouse Lab in Cambridge and Alex Alderton, Roberto Amato, Sonia Goncalves, Ewan Harrison, David K. Jackson, Ian Johnston, Dominic Kwiatkowski, Cordelia Langford, John Sillitoe on behalf of the Wellcome Sanger Institute COVID-19 Surveillance Team |
| EPI_ISL_552174, EPI_ISL_552175, EPI_ISL_552176, EPI_ISL_552177, EPI_ISL_552178, EPI_ISL_552179 | Lighthouse Lab in Milton Keynes | Wellcome Sanger Institute for the COVID-19 Genomics UK (COG-UK) consortium | The Lighthouse Lab in Milton Keynes and Alex Alderton, Roberto Amato, Sonia Goncalves, Ewan Harrison, David K. Jackson, Ian Johnston, Dominic Kwiatkowski, Cordelia Langford, John Sillitoe on behalf of the Wellcome Sanger Institute COVID-19 Surveillance Team (http://www.sanger.ac.uk/covid-team) |
| EPI_ISL_552180 | Lighthouse Lab in Cambridge | Wellcome Sanger Institute for the COVID-19 Genomics UK (COG-UK) consortium | Rob Howes, The Lighthouse Lab in Cambridge and Alex Alderton, Roberto Amato, Sonia Goncalves, Ewan Harrison, David K. Jackson, Ian Johnston, Dominic Kwiatkowski, Cordelia Langford, John Sillitoe on behalf of the Wellcome Sanger Institute COVID-19 Surveillance Team |
| EPI_ISL_552181 | Lighthouse Lab in Milton Keynes | Wellcome Sanger Institute for the COVID-19 Genomics UK (COG-UK) consortium | The Lighthouse Lab in Milton Keynes and Alex Alderton, Roberto Amato, Sonia Goncalves, Ewan Harrison, David K. Jackson, Ian Johnston, Dominic Kwiatkowski, Cordelia Langford, John Sillitoe on behalf of the Wellcome Sanger Institute COVID-19 Surveillance Team (http://www.sanger.ac.uk/covid-team) |
| EPI_ISL_552182 | Lighthouse Lab in Milton Keynes | Wellcome Sanger Institute for the COVID-19 Genomics UK (COG-UK) consortium | The Lighthouse Lab in Alderley Park and Alex Alderton, Roberto Amato, Sonia Goncalves, Ewan Harrison, David K. Jackson, Ian Johnston, Dominic Kwiatkowski, Cordelia Langford, John Sillitoe on behalf of the Wellcome Sanger Institute COVID-19 Surveillance Team |
| EPI_ISL_552183, EPI_ISL_552184 | Lighthouse Lab in Milton Keynes | Wellcome Sanger Institute for the COVID-19 Genomics UK (COG-UK) consortium | The Lighthouse Lab in Milton Keynes and Alex Alderton, Roberto Amato, Sonia Goncalves, Ewan Harrison, David K. Jackson, Ian Johnston, Dominic Kwiatkowski, Cordelia Langford, John Sillitoe on behalf of the Wellcome Sanger Institute COVID-19 Surveillance Team (http://www.sanger.ac.uk/covid-team) |
| EPI_ISL_552185 | Lighthouse Lab in Cambridge | Wellcome Sanger Institute for the COVID-19 Genomics UK (COG-UK) consortium | Rob Howes, The Lighthouse Lab in Cambridge and Alex Alderton, Roberto Amato, Sonia Goncalves, Ewan Harrison, David K. Jackson, Ian Johnston, Dominic Kwiatkowski, Cordelia Langford, John Sillitoe on behalf of the Wellcome Sanger Institute COVID-19 Surveillance Team |
| EPI_ISL_552186, EPI_ISL_552187, EPI_ISL_552188, EPI_ISL_552189, EPI_ISL_552190 | Lighthouse Lab in Milton Keynes | Wellcome Sanger Institute for the COVID-19 Genomics UK (COG-UK) consortium | The Lighthouse Lab in Milton Keynes and Alex Alderton, Roberto Amato, Sonia Goncalves, Ewan Harrison, David K. Jackson, Ian Johnston, Dominic Kwiatkowski, Cordelia Langford, John Sillitoe on behalf of the Wellcome Sanger Institute COVID-19 Surveillance Team (http://www.sanger.ac.uk/covid-team) |
| EPI_ISL_552191 | Lighthouse Lab in Milton Keynes | Wellcome Sanger Institute for the COVID-19 Genomics UK (COG-UK) consortium | The Lighthouse Lab in Alderley Park and Alex Alderton, Roberto Amato, Sonia Goncalves, Ewan Harrison, David K. Jackson, Ian Johnston, Dominic Kwiatkowski, Cordelia Langford, John Sillitoe on behalf of the Wellcome Sanger Institute COVID-19 Surveillance Team |
| EPI_ISL_552192, EPI_ISL_552193, EPI_ISL_552194, EPI_ISL_552195, EPI_ISL_552196, EPI_ISL_552197, EPI_ISL_552198, EPI_ISL_552199, EPI_ISL_552200, EPI_ISL_552201, EPI_ISL_552202, EPI_ISL_552203, EPI_ISL_552204 | | | |
| see above | Lighthouse Lab in Milton Keynes | Wellcome Sanger Institute for the COVID-19 Genomics UK | The Lighthouse Lab in Milton Keynes and Alex Alderton, Roberto Amato, Sonia Goncalves, Ewan Harrison, David K. Jackson, Ian Johnston, Dominic |

| | | | |
|---|---|---|---|
| | | (COG-UK) consortium | Kwiatkowski, Cordelia Langford, John Sillitoe on behalf of the Wellcome Sanger Institute COVID-19 Surveillance Team (http://www.sanger.ac.uk/covid-team) |
| EPI_ISL_552205 | Lighthouse Lab in Milton Keynes | Wellcome Sanger Institute for the COVID-19 Genomics UK (COG-UK) consortium | The Lighthouse Lab in Alderley Park and Alex Alderton, Roberto Amato, Sonia Goncalves, Ewan Harrison, David K. Jackson, Ian Johnston, Dominic Kwiatkowski, Cordelia Langford, John Sillitoe on behalf of the Wellcome Sanger Institute COVID-19 Surveillance Team |
| EPI_ISL_552206, EPI_ISL_552207, EPI_ISL_552208, EPI_ISL_552209, EPI_ISL_552210, EPI_ISL_552211 | Lighthouse Lab in Milton Keynes | Wellcome Sanger Institute for the COVID-19 Genomics UK (COG-UK) consortium | The Lighthouse Lab in Milton Keynes and Alex Alderton, Roberto Amato, Sonia Goncalves, Ewan Harrison, David K. Jackson, Ian Johnston, Dominic Kwiatkowski, Cordelia Langford, John Sillitoe on behalf of the Wellcome Sanger Institute COVID-19 Surveillance Team (http://www.sanger.ac.uk/covid-team) |
| EPI_ISL_552212 | Lighthouse Lab in Milton Keynes | Wellcome Sanger Institute for the COVID-19 Genomics UK (COG-UK) consortium | The Lighthouse Lab in Alderley Park and Alex Alderton, Roberto Amato, Sonia Goncalves, Ewan Harrison, David K. Jackson, Ian Johnston, Dominic Kwiatkowski, Cordelia Langford, John Sillitoe on behalf of the Wellcome Sanger Institute COVID-19 Surveillance Team |
| EPI_ISL_552213, EPI_ISL_552214, EPI_ISL_552215, EPI_ISL_552216, EPI_ISL_552217, EPI_ISL_552218, EPI_ISL_552219, EPI_ISL_552220, EPI_ISL_552221, EPI_ISL_552222, EPI_ISL_552223, EPI_ISL_552224, EPI_ISL_552225, EPI_ISL_552226, EPI_ISL_552227, EPI_ISL_552228, EPI_ISL_552229, EPI_ISL_552230, EPI_ISL_552231, EPI_ISL_552232, EPI_ISL_552233, EPI_ISL_552234, EPI_ISL_552235, EPI_ISL_552236, EPI_ISL_552237, EPI_ISL_552238, EPI_ISL_552239 | | | |
| see above | Lighthouse Lab in Milton Keynes | Wellcome Sanger Institute for the COVID-19 Genomics UK (COG-UK) consortium | The Lighthouse Lab in Milton Keynes and Alex Alderton, Roberto Amato, Sonia Goncalves, Ewan Harrison, David K. Jackson, Ian Johnston, Dominic Kwiatkowski, Cordelia Langford, John Sillitoe on behalf of the Wellcome Sanger Institute COVID-19 Surveillance Team (http://www.sanger.ac.uk/covid-team) |
| EPI_ISL_552240 | Lighthouse Lab in Milton Keynes | Wellcome Sanger Institute for the COVID-19 Genomics UK (COG-UK) Consortium | The Lighthouse Lab in Milton Keynes and Alex Alderton, Roberto Amato, Sonia Goncalves, Ewan Harrison, David K. Jackson, Ian Johnston, Dominic Kwiatkowski, Cordelia Langford, John Sillitoe on behalf of the Wellcome Sanger Institute COVID-19 Surveillance Team |
| EPI_ISL_552241, EPI_ISL_552242, EPI_ISL_552243, EPI_ISL_552244, EPI_ISL_552245, EPI_ISL_552246, EPI_ISL_552247, EPI_ISL_552248, EPI_ISL_552249, EPI_ISL_552250, EPI_ISL_552251, EPI_ISL_552252, EPI_ISL_552253, EPI_ISL_552254, EPI_ISL_552255, EPI_ISL_552256, EPI_ISL_552257, EPI_ISL_552258, EPI_ISL_552259, EPI_ISL_552260, EPI_ISL_552261, EPI_ISL_552262, EPI_ISL_552263, EPI_ISL_552264, EPI_ISL_552265, EPI_ISL_552266, EPI_ISL_552267, EPI_ISL_552268, EPI_ISL_552269, EPI_ISL_552270, EPI_ISL_552271, EPI_ISL_552272, EPI_ISL_552273, EPI_ISL_552274, EPI_ISL_552275, EPI_ISL_552276, EPI_ISL_552277 | | | |
| see above | Lighthouse Lab in Milton Keynes | Wellcome Sanger Institute for the COVID-19 Genomics UK (COG-UK) consortium | The Lighthouse Lab in Milton Keynes and Alex Alderton, Roberto Amato, Sonia Goncalves, Ewan Harrison, David K. Jackson, Ian Johnston, Dominic Kwiatkowski, Cordelia Langford, John Sillitoe on behalf of the Wellcome Sanger Institute COVID-19 Surveillance Team (http://www.sanger.ac.uk/covid-team) |
| EPI_ISL_552278 | Lighthouse Lab in Milton Keynes | Wellcome Sanger Institute for the COVID-19 Genomics UK (COG-UK) consortium | The Lighthouse Lab in Alderley Park and Alex Alderton, Roberto Amato, Sonia Goncalves, Ewan Harrison, David K. Jackson, Ian Johnston, Dominic Kwiatkowski, Cordelia Langford, John Sillitoe on behalf of the Wellcome Sanger Institute COVID-19 Surveillance Team |
| EPI_ISL_552279, EPI_ISL_552280, EPI_ISL_552281, EPI_ISL_552282, EPI_ISL_552283, EPI_ISL_552284, EPI_ISL_552285, EPI_ISL_552286, EPI_ISL_552287, EPI_ISL_552288, EPI_ISL_552289, EPI_ISL_552290, EPI_ISL_552291, EPI_ISL_552292, EPI_ISL_552293, EPI_ISL_552294, EPI_ISL_552295, EPI_ISL_552296, EPI_ISL_552297, EPI_ISL_552298, EPI_ISL_552299, EPI_ISL_552300, EPI_ISL_552301, EPI_ISL_552302, EPI_ISL_552303, EPI_ISL_552304, EPI_ISL_552305 | | | |
| see above | Lighthouse Lab in Milton Keynes | Wellcome Sanger Institute for the COVID-19 Genomics UK (COG-UK) consortium | The Lighthouse Lab in Milton Keynes and Alex Alderton, Roberto Amato, Sonia Goncalves, Ewan Harrison, David K. Jackson, Ian Johnston, Dominic Kwiatkowski, Cordelia Langford, John Sillitoe on behalf of the Wellcome Sanger Institute COVID-19 Surveillance Team (http://www.sanger.ac.uk/covid-team) |
| EPI_ISL_552306 | Lighthouse Lab in Milton Keynes | Wellcome Sanger Institute for the COVID-19 Genomics UK (COG-UK) Consortium | The Lighthouse Lab in Milton Keynes and Alex Alderton, Roberto Amato, Sonia Goncalves, Ewan Harrison, David K. Jackson, Ian Johnston, Dominic Kwiatkowski, Cordelia Langford, John Sillitoe on behalf of the Wellcome Sanger Institute COVID-19 Surveillance Team |
| EPI_ISL_552307, EPI_ISL_552308, EPI_ISL_552309, EPI_ISL_552310, EPI_ISL_552311, EPI_ISL_552312, EPI_ISL_552313, EPI_ISL_552314, EPI_ISL_552315, EPI_ISL_552316, EPI_ISL_552317, EPI_ISL_552318, EPI_ISL_552319, EPI_ISL_552320, EPI_ISL_552321, EPI_ISL_552322, EPI_ISL_552323, EPI_ISL_552324, EPI_ISL_552325, EPI_ISL_552326, EPI_ISL_552327, EPI_ISL_552328, EPI_ISL_552329, EPI_ISL_552330, EPI_ISL_552331, EPI_ISL_552332, EPI_ISL_552333 | | | |
| see above | Lighthouse Lab in Milton Keynes | Wellcome Sanger Institute for the COVID-19 Genomics UK (COG-UK) consortium | The Lighthouse Lab in Milton Keynes and Alex Alderton, Roberto Amato, Sonia Goncalves, Ewan Harrison, David K. Jackson, Ian Johnston, Dominic Kwiatkowski, Cordelia Langford, John Sillitoe on behalf of the Wellcome Sanger Institute COVID-19 Surveillance Team (http://www.sanger.ac.uk/covid-team) |
| EPI_ISL_552334 | Lighthouse Lab in Milton Keynes | Wellcome Sanger Institute for the COVID-19 Genomics UK (COG-UK) consortium | The Lighthouse Lab in Alderley Park and Alex Alderton, Roberto Amato, Sonia Goncalves, Ewan Harrison, David K. Jackson, Ian Johnston, Dominic Kwiatkowski, Cordelia Langford, John Sillitoe on behalf of the Wellcome Sanger Institute COVID-19 Surveillance Team |
| EPI_ISL_552335, EPI_ISL_552336, EPI_ISL_552337, EPI_ISL_552338, EPI_ISL_552339, EPI_ISL_552340, EPI_ISL_552341, EPI_ISL_552342, EPI_ISL_552343, EPI_ISL_552344, EPI_ISL_552345, EPI_ISL_552346, EPI_ISL_552347, EPI_ISL_552348, EPI_ISL_552349, EPI_ISL_552350, EPI_ISL_552351, EPI_ISL_552352, EPI_ISL_552353, EPI_ISL_552354, EPI_ISL_552355, EPI_ISL_552356, EPI_ISL_552357 | | | |
| see above | Lighthouse Lab in Milton Keynes | Wellcome Sanger Institute for the COVID-19 Genomics UK (COG-UK) consortium | The Lighthouse Lab in Milton Keynes and Alex Alderton, Roberto Amato, Sonia Goncalves, Ewan Harrison, David K. Jackson, Ian Johnston, Dominic Kwiatkowski, Cordelia Langford, John Sillitoe on behalf of the Wellcome Sanger Institute COVID-19 Surveillance Team (http://www.sanger.ac.uk/covid-team) |
| EPI_ISL_552358, EPI_ISL_552359 | Lighthouse Lab in Milton Keynes | Wellcome Sanger Institute for the COVID-19 Genomics UK (COG-UK) consortium | The Lighthouse Lab in Alderley Park and Alex Alderton, Roberto Amato, Sonia Goncalves, Ewan Harrison, David K. Jackson, Ian Johnston, Dominic Kwiatkowski, Cordelia Langford, John Sillitoe on behalf of the Wellcome Sanger Institute COVID-19 Surveillance Team |
| EPI_ISL_552360, EPI_ISL_552361, EPI_ISL_552362, EPI_ISL_552363, EPI_ISL_552364, EPI_ISL_552365, EPI_ISL_552366, EPI_ISL_552367, EPI_ISL_552368, EPI_ISL_552369, EPI_ISL_552370, EPI_ISL_552371, EPI_ISL_552372, EPI_ISL_552373, EPI_ISL_552374, EPI_ISL_552375, EPI_ISL_552376, EPI_ISL_552377, EPI_ISL_552378, EPI_ISL_552379, EPI_ISL_552380, EPI_ISL_552381, EPI_ISL_552382, EPI_ISL_552383, EPI_ISL_552384 | | | |
| see above | Lighthouse Lab in Milton Keynes | Wellcome Sanger Institute for the COVID-19 Genomics UK (COG-UK) consortium | The Lighthouse Lab in Milton Keynes and Alex Alderton, Roberto Amato, Sonia Goncalves, Ewan Harrison, David K. Jackson, Ian Johnston, Dominic Kwiatkowski, Cordelia Langford, John Sillitoe on behalf of the Wellcome Sanger Institute COVID-19 Surveillance Team (http://www.sanger.ac.uk/covid-team) |
| EPI_ISL_552385 | Lighthouse Lab in Milton Keynes | Wellcome Sanger Institute for the COVID-19 Genomics UK (COG-UK) consortium | The Lighthouse Lab in Alderley Park and Alex Alderton, Roberto Amato, Sonia Goncalves, Ewan Harrison, David K. Jackson, Ian Johnston, Dominic Kwiatkowski, Cordelia Langford, John Sillitoe on behalf of the Wellcome Sanger Institute COVID-19 Surveillance Team |
| EPI_ISL_552386, EPI_ISL_552387, EPI_ISL_552388, EPI_ISL_552389, EPI_ISL_552390, EPI_ISL_552391, EPI_ISL_552392, EPI_ISL_552393, EPI_ISL_552394, EPI_ISL_552395, EPI_ISL_552396, EPI_ISL_552397, EPI_ISL_552398 | | | |
| see above | Lighthouse Lab in Milton Keynes | Wellcome Sanger Institute for the COVID-19 Genomics UK (COG-UK) consortium | The Lighthouse Lab in Milton Keynes and Alex Alderton, Roberto Amato, Sonia Goncalves, Ewan Harrison, David K. Jackson, Ian Johnston, Dominic Kwiatkowski, Cordelia Langford, John Sillitoe on behalf of the Wellcome Sanger Institute COVID-19 Surveillance Team (http://www.sanger.ac.uk/covid-team) |
| EPI_ISL_552399 | Lighthouse Lab in Milton Keynes | Wellcome Sanger Institute for the COVID-19 Genomics UK (COG-UK) consortium | The Lighthouse Lab in Alderley Park and Alex Alderton, Roberto Amato, Sonia Goncalves, Ewan Harrison, David K. Jackson, Ian Johnston, Dominic Kwiatkowski, Cordelia Langford, John Sillitoe on behalf of the Wellcome Sanger Institute COVID-19 Surveillance Team |
| EPI_ISL_552400, EPI_ISL_552401, EPI_ISL_552402, EPI_ISL_552403, EPI_ISL_552404, EPI_ISL_552405, EPI_ISL_552406, EPI_ISL_552407 | Lighthouse Lab in Milton Keynes | Wellcome Sanger Institute for the COVID-19 Genomics UK (COG-UK) consortium | The Lighthouse Lab in Milton Keynes and Alex Alderton, Roberto Amato, Sonia Goncalves, Ewan Harrison, David K. Jackson, Ian Johnston, Dominic Kwiatkowski, Cordelia Langford, John Sillitoe on behalf of the Wellcome Sanger Institute COVID-19 Surveillance Team (http://www.sanger.ac.uk/covid-team) |
| EPI_ISL_552408 | Lighthouse Lab in Milton Keynes | Wellcome Sanger Institute for the COVID-19 Genomics UK (COG-UK) consortium | The Lighthouse Lab in Alderley Park and Alex Alderton, Roberto Amato, Sonia Goncalves, Ewan Harrison, David K. Jackson, Ian Johnston, Dominic Kwiatkowski, Cordelia Langford, John Sillitoe on behalf of the Wellcome Sanger Institute COVID-19 Surveillance Team |
| EPI_ISL_552409, EPI_ISL_552410, EPI_ISL_552411, EPI_ISL_552412, EPI_ISL_552413 | Lighthouse Lab in Milton Keynes | Wellcome Sanger Institute for the COVID-19 Genomics UK (COG-UK) consortium | The Lighthouse Lab in Milton Keynes and Alex Alderton, Roberto Amato, Sonia Goncalves, Ewan Harrison, David K. Jackson, Ian Johnston, Dominic Kwiatkowski, Cordelia Langford, John Sillitoe on behalf of the Wellcome Sanger Institute COVID-19 Surveillance Team (http://www.sanger.ac.uk/covid-team) |
| EPI_ISL_552414 | Lighthouse Lab in Milton Keynes | Wellcome Sanger Institute for the COVID-19 Genomics UK (COG-UK) consortium | The Lighthouse Lab in Alderley Park and Alex Alderton, Roberto Amato, Sonia Goncalves, Ewan Harrison, David K. Jackson, Ian Johnston, Dominic Kwiatkowski, Cordelia Langford, John Sillitoe on behalf of the Wellcome Sanger Institute COVID-19 Surveillance Team |
| EPI_ISL_552415, EPI_ISL_552416 | Lighthouse Lab in Milton Keynes | Wellcome Sanger Institute for the COVID-19 Genomics UK (COG-UK) consortium | The Lighthouse Lab in Milton Keynes and Alex Alderton, Roberto Amato, Sonia Goncalves, Ewan Harrison, David K. Jackson, Ian Johnston, Dominic Kwiatkowski, Cordelia Langford, John Sillitoe on behalf of the Wellcome Sanger Institute COVID-19 Surveillance Team (http://www.sanger.ac.uk/covid-team) |
| EPI_ISL_552417 | Lighthouse Lab in Milton Keynes | Wellcome Sanger Institute for the COVID-19 Genomics UK | The Lighthouse Lab in Alderley Park and Alex Alderton, Roberto Amato, Sonia Goncalves, Ewan Harrison, David K. Jackson, Ian Johnston, Dominic |

| | | | |
|---|---|---|---|
| | | (COG-UK) consortium | Kwiatkowski, Cordelia Langford, John Sillitoe on behalf of the Wellcome Sanger Institute COVID-19 Surveillance Team |
| EPI_ISL_552418, EPI_ISL_552419, EPI_ISL_552420, EPI_ISL_552421, EPI_ISL_552422, EPI_ISL_552423, EPI_ISL_552424, EPI_ISL_552425, EPI_ISL_552426, EPI_ISL_552427, EPI_ISL_552428, EPI_ISL_552429, EPI_ISL_552430, EPI_ISL_552431, EPI_ISL_552432, EPI_ISL_552433, EPI_ISL_552434, EPI_ISL_552435, EPI_ISL_552436, EPI_ISL_552437, EPI_ISL_552438, EPI_ISL_552439, EPI_ISL_552440 | | | |
| see above | Lighthouse Lab in Milton Keynes | Wellcome Sanger Institute for the COVID-19 Genomics UK (COG-UK) consortium | The Lighthouse Lab in Milton Keynes and Alex Alderton, Roberto Amato, Sonia Goncalves, Ewan Harrison, David K. Jackson, Ian Johnston, Dominic Kwiatkowski, Cordelia Langford, John Sillitoe on behalf of the Wellcome Sanger Institute COVID-19 Surveillance Team (http://www.sanger.ac.uk/covid-team) |
| EPI_ISL_552441 | Lighthouse Lab in Milton Keynes | Wellcome Sanger Institute for the COVID-19 Genomics UK (COG-UK) consortium | The Lighthouse Lab in Alderley Park and Alex Alderton, Roberto Amato, Sonia Goncalves, Ewan Harrison, David K. Jackson, Ian Johnston, Dominic Kwiatkowski, Cordelia Langford, John Sillitoe on behalf of the Wellcome Sanger Institute COVID-19 Surveillance Team |
| EPI_ISL_552442, EPI_ISL_552443, EPI_ISL_552445, EPI_ISL_552447, EPI_ISL_552452 | Lighthouse Lab in Milton Keynes | Wellcome Sanger Institute for the COVID-19 Genomics UK (COG-UK) consortium | The Lighthouse Lab in Milton Keynes and Alex Alderton, Roberto Amato, Sonia Goncalves, Ewan Harrison, David K. Jackson, Ian Johnston, Dominic Kwiatkowski, Cordelia Langford, John Sillitoe on behalf of the Wellcome Sanger Institute COVID-19 Surveillance Team (http://www.sanger.ac.uk/covid-team) |
| EPI_ISL_552458 | Lighthouse Lab in Milton Keynes | Wellcome Sanger Institute for the COVID-19 Genomics UK (COG-UK) Consortium | The Lighthouse Lab in Milton Keynes and Alex Alderton, Roberto Amato, Sonia Goncalves, Ewan Harrison, David K. Jackson, Ian Johnston, Dominic Kwiatkowski, Cordelia Langford, John Sillitoe on behalf of the Wellcome Sanger Institute COVID-19 Surveillance Team |
| EPI_ISL_552460, EPI_ISL_552461, EPI_ISL_552462, EPI_ISL_552465, EPI_ISL_552467, EPI_ISL_552470, EPI_ISL_552471, EPI_ISL_552472, EPI_ISL_552480, EPI_ISL_552484, EPI_ISL_552485, EPI_ISL_552487, EPI_ISL_552489 | | | |
| see above | Lighthouse Lab in Milton Keynes | Wellcome Sanger Institute for the COVID-19 Genomics UK (COG-UK) consortium | The Lighthouse Lab in Milton Keynes and Alex Alderton, Roberto Amato, Sonia Goncalves, Ewan Harrison, David K. Jackson, Ian Johnston, Dominic Kwiatkowski, Cordelia Langford, John Sillitoe on behalf of the Wellcome Sanger Institute COVID-19 Surveillance Team (http://www.sanger.ac.uk/covid-team) |
| EPI_ISL_552496 | Lighthouse Lab in Milton Keynes | Wellcome Sanger Institute for the COVID-19 Genomics UK (COG-UK) consortium | The Lighthouse Lab in Alderley Park and Alex Alderton, Roberto Amato, Sonia Goncalves, Ewan Harrison, David K. Jackson, Ian Johnston, Dominic Kwiatkowski, Cordelia Langford, John Sillitoe on behalf of the Wellcome Sanger Institute COVID-19 Surveillance Team |
| EPI_ISL_552498, EPI_ISL_552499, EPI_ISL_552500, EPI_ISL_552502, EPI_ISL_552503, EPI_ISL_552504, EPI_ISL_552509, EPI_ISL_552510, EPI_ISL_552511, EPI_ISL_552514, EPI_ISL_552517, EPI_ISL_552522, EPI_ISL_552525, EPI_ISL_552528, EPI_ISL_552529, EPI_ISL_552535, EPI_ISL_552537, EPI_ISL_552538, EPI_ISL_552542, EPI_ISL_552543, EPI_ISL_552544, EPI_ISL_552545, EPI_ISL_552549, EPI_ISL_552551, EPI_ISL_552552, EPI_ISL_552557, EPI_ISL_552558, EPI_ISL_552560, EPI_ISL_552561, EPI_ISL_552564, EPI_ISL_552567, EPI_ISL_552570, EPI_ISL_552573, EPI_ISL_552575, EPI_ISL_552581, EPI_ISL_552582, EPI_ISL_552584, EPI_ISL_552585, EPI_ISL_552588, EPI_ISL_552589, EPI_ISL_552593, EPI_ISL_552594, EPI_ISL_552599, EPI_ISL_552601, EPI_ISL_552605, EPI_ISL_552606, EPI_ISL_552611, EPI_ISL_552612, EPI_ISL_552613, EPI_ISL_552615, EPI_ISL_552616, EPI_ISL_552621, EPI_ISL_552625, EPI_ISL_552631, EPI_ISL_552637, EPI_ISL_552638, EPI_ISL_552640, EPI_ISL_552643, EPI_ISL_552645, EPI_ISL_552651, EPI_ISL_552652, EPI_ISL_552654, EPI_ISL_552658, EPI_ISL_552662, EPI_ISL_552667, EPI_ISL_552670, EPI_ISL_552676, EPI_ISL_552677, EPI_ISL_552688, EPI_ISL_552694, EPI_ISL_552695, EPI_ISL_552708, EPI_ISL_552710, EPI_ISL_552715, EPI_ISL_552718, EPI_ISL_552733, EPI_ISL_552734 | | | |
| see above | Lighthouse Lab in Milton Keynes | Wellcome Sanger Institute for the COVID-19 Genomics UK (COG-UK) consortium | The Lighthouse Lab in Milton Keynes and Alex Alderton, Roberto Amato, Sonia Goncalves, Ewan Harrison, David K. Jackson, Ian Johnston, Dominic Kwiatkowski, Cordelia Langford, John Sillitoe on behalf of the Wellcome Sanger Institute COVID-19 Surveillance Team (http://www.sanger.ac.uk/covid-team) |
| EPI_ISL_552737 | Lighthouse Lab in Milton Keynes | Wellcome Sanger Institute for the COVID-19 Genomics UK (COG-UK) consortium | The Lighthouse Lab in Alderley Park and Alex Alderton, Roberto Amato, Sonia Goncalves, Ewan Harrison, David K. Jackson, Ian Johnston, Dominic Kwiatkowski, Cordelia Langford, John Sillitoe on behalf of the Wellcome Sanger Institute COVID-19 Surveillance Team |
| EPI_ISL_552753, EPI_ISL_552756, EPI_ISL_552758, EPI_ISL_552763 | Lighthouse Lab in Milton Keynes | Wellcome Sanger Institute for the COVID-19 Genomics UK (COG-UK) consortium | The Lighthouse Lab in Milton Keynes and Alex Alderton, Roberto Amato, Sonia Goncalves, Ewan Harrison, David K. Jackson, Ian Johnston, Dominic Kwiatkowski, Cordelia Langford, John Sillitoe on behalf of the Wellcome Sanger Institute COVID-19 Surveillance Team (http://www.sanger.ac.uk/covid-team) |
| EPI_ISL_552765, EPI_ISL_552766, EPI_ISL_552769 | Lighthouse Lab in Milton Keynes | Wellcome Sanger Institute for the COVID-19 Genomics UK (COG-UK) consortium | The Lighthouse Lab in Alderley Park and Alex Alderton, Roberto Amato, Sonia Goncalves, Ewan Harrison, David K. Jackson, Ian Johnston, Dominic Kwiatkowski, Cordelia Langford, John Sillitoe on behalf of the Wellcome Sanger Institute COVID-19 Surveillance Team |
| EPI_ISL_552778, EPI_ISL_552781, EPI_ISL_552788, EPI_ISL_552790 | Lighthouse Lab in Milton Keynes | Wellcome Sanger Institute for the COVID-19 Genomics UK (COG-UK) consortium | The Lighthouse Lab in Milton Keynes and Alex Alderton, Roberto Amato, Sonia Goncalves, Ewan Harrison, David K. Jackson, Ian Johnston, Dominic Kwiatkowski, Cordelia Langford, John Sillitoe on behalf of the Wellcome Sanger Institute COVID-19 Surveillance Team (http://www.sanger.ac.uk/covid-team) |
| EPI_ISL_559753, EPI_ISL_559754, EPI_ISL_559755, EPI_ISL_559756, EPI_ISL_559757, EPI_ISL_559758, EPI_ISL_559759, EPI_ISL_559760, EPI_ISL_559761, EPI_ISL_559762, EPI_ISL_559763, EPI_ISL_559764, EPI_ISL_559765, EPI_ISL_559766, EPI_ISL_559767, EPI_ISL_559768, EPI_ISL_559769, EPI_ISL_559770, EPI_ISL_559771, EPI_ISL_559772, EPI_ISL_559773, EPI_ISL_559774, EPI_ISL_559775, EPI_ISL_559776, EPI_ISL_559777, EPI_ISL_559778, EPI_ISL_559779, EPI_ISL_559780, EPI_ISL_559781, EPI_ISL_559782, EPI_ISL_559783, EPI_ISL_559784, EPI_ISL_559785, EPI_ISL_559786, EPI_ISL_559787, EPI_ISL_559788, EPI_ISL_559789, EPI_ISL_559790, EPI_ISL_559791, EPI_ISL_559792, EPI_ISL_559793, EPI_ISL_559794, EPI_ISL_559795, EPI_ISL_559796, EPI_ISL_559797, EPI_ISL_559798, EPI_ISL_559834, EPI_ISL_559835, EPI_ISL_559837, EPI_ISL_559840, EPI_ISL_559841, EPI_ISL_559843, EPI_ISL_559845, EPI_ISL_559848, EPI_ISL_559851, EPI_ISL_559984, EPI_ISL_559985, EPI_ISL_559986, EPI_ISL_559987, EPI_ISL_559988, EPI_ISL_559989, EPI_ISL_559990, EPI_ISL_559991, EPI_ISL_559992, EPI_ISL_559993, EPI_ISL_559994, EPI_ISL_559995, EPI_ISL_559996, EPI_ISL_559997, EPI_ISL_559998, EPI_ISL_559999, EPI_ISL_560000, EPI_ISL_560001, EPI_ISL_560002, EPI_ISL_560003, EPI_ISL_560305, EPI_ISL_560306 | | | |
| see above | Oxford Viromics, NDM, University of Oxford; Oxford University Hospitals; Basingstoke and North Hampshire Hospital | COVID-19 Genomics UK (COG-UK) Consortium | Tanya Golubchik, David Bonsall, George Macintyre, Amy Trebes, Mariateresa de Cesare, Catrin Moore, Alex Mobbs, Anita Justice, Robert Shaw, Monique Andersson, Timothy Peto, Emma Wise, Nathan Moore, Jessica Lynch, Nick Cortes, Matilde Mori, Stephen Kidd, David Buck, John Todd, Christophe Fraser |
| EPI_ISL_560322, EPI_ISL_560323, EPI_ISL_560324 | Civil Hospital, Panchkula | CSIR-Institute of Microbial Technology | Kanika Bansal, Sanjeet Kumar, Anu Singh, Debarghya Ghose, Rajesh Kumar Mishra, Dipak Dutta, Sanjeev Khosla, Prabhu B. Patil |
| EPI_ISL_560793, EPI_ISL_560794, EPI_ISL_560795, EPI_ISL_560796, EPI_ISL_560797 | Mayo Clinic & Mayo Clinic Laboratories | Minnesota Department of Health, Public Health Laboratory | Matt Plumb, Jacob Garfin, and Xiong Wang |
| EPI_ISL_560798, EPI_ISL_560800, EPI_ISL_560801, EPI_ISL_560802, EPI_ISL_560803, EPI_ISL_560804, EPI_ISL_560805 | M Health Fairview | Minnesota Department of Health, Public Health Laboratory | Matt Plumb, Jacob Garfin, and Xiong Wang |
| EPI_ISL_561337, EPI_ISL_561338, EPI_ISL_561339, EPI_ISL_561340, EPI_ISL_561341, EPI_ISL_561342, EPI_ISL_561343 | Civil Hospital, Panchkula | CSIR-Institute of Microbial Technology | Kanika Bansal, Sanjeet Kumar, Anu Singh, Debarghya Ghose, Rajesh Kumar Mishra, Dipak Dutta, Sanjeev Khosla, Prabhu B. Patil |
| EPI_ISL_561384, EPI_ISL_561386 | Microbiological Diagnostic Unit - Public Health Laboratory (MDU-PHL) | MDU-PHL | Seemann, T., Schultz M. B., Sait, M., Sherry, N. |
| EPI_ISL_561392 | Victorian Infectious Diseases Reference Laboratory (VIDRL) | VIDRL and MDU-PHL | Caly, L., Seemann, T., Sait, M., Schultz, M. B., Druce J., Sherry, N. |
| EPI_ISL_561395, EPI_ISL_561401, EPI_ISL_561408, EPI_ISL_561410, EPI_ISL_561417, EPI_ISL_561418, EPI_ISL_561429, EPI_ISL_561430, EPI_ISL_561436 | Microbiological Diagnostic Unit - Public Health Laboratory (MDU-PHL) | MDU-PHL | Seemann, T., Schultz M. B., Sait, M., Sherry, N. |
| EPI_ISL_561440 | Victorian Infectious Diseases Reference Laboratory (VIDRL) | VIDRL and MDU-PHL | Caly, L., Seemann, T., Sait, M., Schultz, M. B., Druce J., Sherry, N. |
| EPI_ISL_561454, EPI_ISL_561455, EPI_ISL_561457 | Microbiological Diagnostic Unit - Public Health Laboratory (MDU-PHL) | MDU-PHL | Seemann, T., Schultz M. B., Sait, M., Sherry, N. |
| EPI_ISL_561468 | Victorian Infectious Diseases Reference Laboratory (VIDRL) | VIDRL and MDU-PHL | Caly, L., Seemann, T., Sait, M., Schultz, M. B., Druce J., Sherry, N. |
| EPI_ISL_561476, EPI_ISL_561485, EPI_ISL_561486, EPI_ISL_561495, EPI_ISL_561497, EPI_ISL_561500, EPI_ISL_561511, EPI_ISL_561520, EPI_ISL_561521 | Microbiological Diagnostic Unit - Public Health Laboratory (MDU-PHL) | MDU-PHL | Seemann, T., Schultz M. B., Sait, M., Sherry, N. |

| | | | |
|---|---|---|---|
| EPI_ISL_561541 | Victorian Infectious Diseases Reference Laboratory (VIDRL) | VIDRL and MDU-PHL | Caly, L., Seemann, T., Sait, M., Schultz, M. B., Druce J., Sherry, N. |
| EPI_ISL_561550, EPI_ISL_561552, EPI_ISL_561553, EPI_ISL_561563, EPI_ISL_561564, EPI_ISL_561566, EPI_ISL_561571, EPI_ISL_561573, EPI_ISL_561575, EPI_ISL_561577 | Microbiological Diagnostic Unit - Public Health Laboratory (MDU-PHL) | MDU-PHL | Seemann, T., Schultz M. B., Sait, M., Sherry, N. |
| EPI_ISL_561578 | Victorian Infectious Diseases Reference Laboratory (VIDRL) | VIDRL and MDU-PHL | Caly, L., Seemann, T., Sait, M., Schultz, M. B., Druce J., Sherry, N. |
| EPI_ISL_561593, EPI_ISL_561594, EPI_ISL_561600, EPI_ISL_561607, EPI_ISL_561609, EPI_ISL_561610, EPI_ISL_561616, EPI_ISL_561620, EPI_ISL_561626, EPI_ISL_561632, EPI_ISL_561637, EPI_ISL_561639, EPI_ISL_561640, EPI_ISL_561645 | | | |
| see above | Microbiological Diagnostic Unit - Public Health Laboratory (MDU-PHL) | MDU-PHL | Seemann, T., Schultz M. B., Sait, M., Sherry, N. |
| EPI_ISL_561651, EPI_ISL_561656 | Victorian Infectious Diseases Reference Laboratory (VIDRL) | VIDRL and MDU-PHL | Caly, L., Seemann, T., Sait, M., Schultz, M. B., Druce J., Sherry, N. |
| EPI_ISL_561663, EPI_ISL_561668, EPI_ISL_561669, EPI_ISL_561675, EPI_ISL_561676, EPI_ISL_561681, EPI_ISL_561702, EPI_ISL_561712, EPI_ISL_561718, EPI_ISL_561723, EPI_ISL_561729, EPI_ISL_561733, EPI_ISL_561735, EPI_ISL_561736, EPI_ISL_561737, EPI_ISL_561743, EPI_ISL_561751, EPI_ISL_561756, EPI_ISL_561759, EPI_ISL_561762 | | | |
| see above | Microbiological Diagnostic Unit - Public Health Laboratory (MDU-PHL) | MDU-PHL | Seemann, T., Schultz M. B., Sait, M., Sherry, N. |
| EPI_ISL_561764 | Victorian Infectious Diseases Reference Laboratory (VIDRL) | VIDRL and MDU-PHL | Caly, L., Seemann, T., Sait, M., Schultz, M. B., Druce J., Sherry, N. |
| EPI_ISL_561767, EPI_ISL_561771, EPI_ISL_561772, EPI_ISL_561775 | Microbiological Diagnostic Unit - Public Health Laboratory (MDU-PHL) | MDU-PHL | Seemann, T., Schultz M. B., Sait, M., Sherry, N. |
| EPI_ISL_561793 | Victorian Infectious Diseases Reference Laboratory (VIDRL) | VIDRL and MDU-PHL | Caly, L., Seemann, T., Sait, M., Schultz, M. B., Druce J., Sherry, N. |
| EPI_ISL_561797, EPI_ISL_561806, EPI_ISL_561807, EPI_ISL_561814, EPI_ISL_561819, EPI_ISL_561820 | Microbiological Diagnostic Unit - Public Health Laboratory (MDU-PHL) | MDU-PHL | Seemann, T., Schultz M. B., Sait, M., Sherry, N. |
| EPI_ISL_561840 | Victorian Infectious Diseases Reference Laboratory (VIDRL) | VIDRL and MDU-PHL | Caly, L., Seemann, T., Sait, M., Schultz, M. B., Druce J., Sherry, N. |
| EPI_ISL_561848, EPI_ISL_561849 | Microbiological Diagnostic Unit - Public Health Laboratory (MDU-PHL) | MDU-PHL | Seemann, T., Schultz M. B., Sait, M., Sherry, N. |
| EPI_ISL_561856 | Victorian Infectious Diseases Reference Laboratory (VIDRL) | VIDRL and MDU-PHL | Caly, L., Seemann, T., Sait, M., Schultz, M. B., Druce J., Sherry, N. |
| EPI_ISL_561865, EPI_ISL_561881, EPI_ISL_561885, EPI_ISL_561886, EPI_ISL_561895, EPI_ISL_561900, EPI_ISL_561902, EPI_ISL_561906, EPI_ISL_561908, EPI_ISL_561922, EPI_ISL_561931, EPI_ISL_561932, EPI_ISL_561954, EPI_ISL_561955, EPI_ISL_561959, EPI_ISL_561961, EPI_ISL_561965, EPI_ISL_561967, EPI_ISL_561968, EPI_ISL_561971, EPI_ISL_561972, EPI_ISL_561977, EPI_ISL_561981 | | | |
| see above | Microbiological Diagnostic Unit - Public Health Laboratory (MDU-PHL) | MDU-PHL | Seemann, T., Schultz M. B., Sait, M., Sherry, N. |
| EPI_ISL_561988 | Victorian Infectious Diseases Reference Laboratory (VIDRL) | VIDRL and MDU-PHL | Caly, L., Seemann, T., Sait, M., Schultz, M. B., Druce J., Sherry, N. |
| EPI_ISL_561989, EPI_ISL_561992, EPI_ISL_561993 | Microbiological Diagnostic Unit - Public Health Laboratory (MDU-PHL) | MDU-PHL | Seemann, T., Schultz M. B., Sait, M., Sherry, N. |
| EPI_ISL_561994 | Victorian Infectious Diseases Reference Laboratory (VIDRL) | VIDRL and MDU-PHL | Caly, L., Seemann, T., Sait, M., Schultz, M. B., Druce J., Sherry, N. |
| EPI_ISL_562006, EPI_ISL_562025, EPI_ISL_562032, EPI_ISL_562039, EPI_ISL_562040, EPI_ISL_562041, EPI_ISL_562042, EPI_ISL_562043, EPI_ISL_562053, EPI_ISL_562065, EPI_ISL_562078, EPI_ISL_562083, EPI_ISL_562084, EPI_ISL_562089, EPI_ISL_562092, EPI_ISL_562098, EPI_ISL_562103, EPI_ISL_562116, EPI_ISL_562117, EPI_ISL_562121, EPI_ISL_562125, EPI_ISL_562142, EPI_ISL_562143, EPI_ISL_562148, EPI_ISL_562160, EPI_ISL_562161, EPI_ISL_562162 | | | |
| see above | Microbiological Diagnostic Unit - Public Health Laboratory (MDU-PHL) | MDU-PHL | Seemann, T., Schultz M. B., Sait, M., Sherry, N. |
| EPI_ISL_562163 | Victorian Infectious Diseases Reference Laboratory (VIDRL) | VIDRL and MDU-PHL | Caly, L., Seemann, T., Sait, M., Schultz, M. B., Druce J., Sherry, N. |
| EPI_ISL_562172, EPI_ISL_562174, EPI_ISL_562190, EPI_ISL_562191, EPI_ISL_562192, EPI_ISL_562196, EPI_ISL_562226 | Microbiological Diagnostic Unit - Public Health Laboratory (MDU-PHL) | MDU-PHL | Seemann, T., Schultz M. B., Sait, M., Sherry, N. |
| EPI_ISL_562229 | Victorian Infectious Diseases Reference Laboratory (VIDRL) | VIDRL and MDU-PHL | Caly, L., Seemann, T., Sait, M., Schultz, M. B., Druce J., Sherry, N. |
| EPI_ISL_562230, EPI_ISL_562232, EPI_ISL_562233 | Microbiological Diagnostic Unit - Public Health Laboratory (MDU-PHL) | MDU-PHL | Seemann, T., Schultz M. B., Sait, M., Sherry, N. |
| EPI_ISL_562239 | Victorian Infectious Diseases Reference Laboratory (VIDRL) | VIDRL and MDU-PHL | Caly, L., Seemann, T., Sait, M., Schultz, M. B., Druce J., Sherry, N. |
| EPI_ISL_562241 | Microbiological Diagnostic Unit - Public Health Laboratory (MDU-PHL) | MDU-PHL | Seemann, T., Schultz M. B., Sait, M., Sherry, N. |
| EPI_ISL_562242 | Victorian Infectious Diseases Reference Laboratory (VIDRL) | VIDRL and MDU-PHL | Caly, L., Seemann, T., Sait, M., Schultz, M. B., Druce J., Sherry, N. |
| EPI_ISL_562246, EPI_ISL_562247, EPI_ISL_562253, EPI_ISL_562254 | Microbiological Diagnostic Unit - Public Health Laboratory (MDU-PHL) | MDU-PHL | Seemann, T., Schultz M. B., Sait, M., Sherry, N. |
| EPI_ISL_562259 | Victorian Infectious Diseases Reference Laboratory (VIDRL) | VIDRL and MDU-PHL | Caly, L., Seemann, T., Sait, M., Schultz, M. B., Druce J., Sherry, N. |
| EPI_ISL_562265, EPI_ISL_562278, EPI_ISL_562285 | Microbiological Diagnostic Unit - Public Health Laboratory (MDU-PHL) | MDU-PHL | Seemann, T., Schultz M. B., Sait, M., Sherry, N. |
| EPI_ISL_562290 | Victorian Infectious Diseases Reference Laboratory (VIDRL) | VIDRL and MDU-PHL | Caly, L., Seemann, T., Sait, M., Schultz, M. B., Druce J., Sherry, N. |
| EPI_ISL_562299, EPI_ISL_562303 | Microbiological Diagnostic Unit - Public Health Laboratory (MDU-PHL) | MDU-PHL | Seemann, T., Schultz M. B., Sait, M., Sherry, N. |
| EPI_ISL_562304 | Victorian Infectious Diseases Reference Laboratory (VIDRL) | VIDRL and MDU-PHL | Caly, L., Seemann, T., Sait, M., Schultz, M. B., Druce J., Sherry, N. |
| EPI_ISL_562305, EPI_ISL_562309, EPI_ISL_562310, EPI_ISL_562311, EPI_ISL_562314, EPI_ISL_562316, EPI_ISL_562319, EPI_ISL_562321, EPI_ISL_562323, EPI_ISL_562325, EPI_ISL_562329, EPI_ISL_562330, EPI_ISL_562332, EPI_ISL_562339, EPI_ISL_562340, EPI_ISL_562346, EPI_ISL_562358, EPI_ISL_562359, EPI_ISL_562365 | | | |
| see above | Microbiological Diagnostic Unit - Public Health Laboratory (MDU-PHL) | MDU-PHL | Seemann, T., Schultz M. B., Sait, M., Sherry, N. |
| EPI_ISL_562366 | Victorian Infectious Diseases Reference Laboratory (VIDRL) | VIDRL and MDU-PHL | Caly, L., Seemann, T., Sait, M., Schultz, M. B., Druce J., Sherry, N. |
| EPI_ISL_562368, EPI_ISL_562372, EPI_ISL_562374, EPI_ISL_562382, EPI_ISL_562394, EPI_ISL_562399, EPI_ISL_562408, EPI_ISL_562419, EPI_ISL_562422, EPI_ISL_562424, EPI_ISL_562426 | | | |
| see above | Microbiological Diagnostic Unit - Public Health Laboratory (MDU-PHL) | MDU-PHL | Seemann, T., Schultz M. B., Sait, M., Sherry, N. |
| EPI_ISL_562440 | Victorian Infectious Diseases Reference Laboratory (VIDRL) | VIDRL and MDU-PHL | Caly, L., Seemann, T., Sait, M., Schultz, M. B., Druce J., Sherry, N. |

| | | | |
|---|---|---|---|
| EPI_ISL_562444 | Microbiological Diagnostic Unit - Public Health Laboratory (MDU-PHL) | MDU-PHL | Seemann, T., Schultz M. B., Sait, M., Sherry, N. |
| EPI_ISL_562447 | Victorian Infectious Diseases Reference Laboratory (VIDRL) | VIDRL and MDU-PHL | Caly, L., Seemann, T., Sait, M., Schultz, M. B., Druce J., Sherry, N. |
| EPI_ISL_562452, EPI_ISL_562454, EPI_ISL_562455, EPI_ISL_562461, EPI_ISL_562465 | Microbiological Diagnostic Unit - Public Health Laboratory (MDU-PHL) | MDU-PHL | Seemann, T., Schultz M. B., Sait, M., Sherry, N. |
| EPI_ISL_562474 | Victorian Infectious Diseases Reference Laboratory (VIDRL) | VIDRL and MDU-PHL | Caly, L., Seemann, T., Sait, M., Schultz, M. B., Druce J., Sherry, N. |
| EPI_ISL_562478, EPI_ISL_562480, EPI_ISL_562486, EPI_ISL_562497, EPI_ISL_562501 | Microbiological Diagnostic Unit - Public Health Laboratory (MDU-PHL) | MDU-PHL | Seemann, T., Schultz M. B., Sait, M., Sherry, N. |
| EPI_ISL_562502 | Victorian Infectious Diseases Reference Laboratory (VIDRL) | VIDRL and MDU-PHL | Caly, L., Seemann, T., Sait, M., Schultz, M. B., Druce J., Sherry, N. |
| EPI_ISL_562504, EPI_ISL_562506, EPI_ISL_562511, EPI_ISL_562520, EPI_ISL_562522, EPI_ISL_562523, EPI_ISL_562530, EPI_ISL_562532, EPI_ISL_562533 | Microbiological Diagnostic Unit - Public Health Laboratory (MDU-PHL) | MDU-PHL | Seemann, T., Schultz M. B., Sait, M., Sherry, N. |
| EPI_ISL_562545 | Victorian Infectious Diseases Reference Laboratory (VIDRL) | VIDRL and MDU-PHL | Caly, L., Seemann, T., Sait, M., Schultz, M. B., Druce J., Sherry, N. |
| EPI_ISL_562551, EPI_ISL_562555, EPI_ISL_562560, EPI_ISL_562564, EPI_ISL_562567, EPI_ISL_562570, EPI_ISL_562573, EPI_ISL_562596, EPI_ISL_562597, EPI_ISL_562612, EPI_ISL_562613, EPI_ISL_562614, EPI_ISL_562633, EPI_ISL_562635, EPI_ISL_562636, EPI_ISL_562637, EPI_ISL_562642, EPI_ISL_562643, EPI_ISL_562644 | | | |
| see above | Microbiological Diagnostic Unit - Public Health Laboratory (MDU-PHL) | MDU-PHL | Seemann, T., Schultz M. B., Sait, M., Sherry, N. |
| EPI_ISL_562645, EPI_ISL_562646 | Victorian Infectious Diseases Reference Laboratory (VIDRL) | VIDRL and MDU-PHL | Caly, L., Seemann, T., Sait, M., Schultz, M. B., Druce J., Sherry, N. |
| EPI_ISL_562648, EPI_ISL_562649, EPI_ISL_562651, EPI_ISL_562654, EPI_ISL_562655, EPI_ISL_562657, EPI_ISL_562670, EPI_ISL_562681, EPI_ISL_562682, EPI_ISL_562683, EPI_ISL_562684, EPI_ISL_562685, EPI_ISL_562687, EPI_ISL_562688, EPI_ISL_562690, EPI_ISL_562702, EPI_ISL_562708, EPI_ISL_562709, EPI_ISL_562710, EPI_ISL_562711, EPI_ISL_562713, EPI_ISL_562714, EPI_ISL_562715, EPI_ISL_562722, EPI_ISL_562723, EPI_ISL_562727, EPI_ISL_562728, EPI_ISL_562729, EPI_ISL_562730, EPI_ISL_562731, EPI_ISL_562732, EPI_ISL_562733, EPI_ISL_562734, EPI_ISL_562735, EPI_ISL_562737, EPI_ISL_562738, EPI_ISL_562739, EPI_ISL_562740, EPI_ISL_562741, EPI_ISL_562743, EPI_ISL_562744, EPI_ISL_562745, EPI_ISL_562746, EPI_ISL_562747, EPI_ISL_562748, EPI_ISL_562749, EPI_ISL_562750, EPI_ISL_562751, EPI_ISL_562752, EPI_ISL_562753, EPI_ISL_562761, EPI_ISL_562762, EPI_ISL_562763, EPI_ISL_562764, EPI_ISL_562765, EPI_ISL_562766, EPI_ISL_562768, EPI_ISL_562770, EPI_ISL_562771, EPI_ISL_562772, EPI_ISL_562773, EPI_ISL_562774, EPI_ISL_562775, EPI_ISL_562776, EPI_ISL_562777, EPI_ISL_562778, EPI_ISL_562779, EPI_ISL_562780, EPI_ISL_562781, EPI_ISL_562782, EPI_ISL_562783, EPI_ISL_562786, EPI_ISL_562787, EPI_ISL_562788, EPI_ISL_562789, EPI_ISL_562793, EPI_ISL_562794, EPI_ISL_562796, EPI_ISL_562797, EPI_ISL_562798, EPI_ISL_562799, EPI_ISL_562800, EPI_ISL_562810, EPI_ISL_562811, EPI_ISL_562813, EPI_ISL_562814, EPI_ISL_562815, EPI_ISL_562816, EPI_ISL_562817, EPI_ISL_562818, EPI_ISL_562819, EPI_ISL_562820, EPI_ISL_562821, EPI_ISL_562822, EPI_ISL_562823, EPI_ISL_562824, EPI_ISL_562825, EPI_ISL_562826, EPI_ISL_562827, EPI_ISL_562828, EPI_ISL_562829, EPI_ISL_562830, EPI_ISL_562831, EPI_ISL_562832, EPI_ISL_562834, EPI_ISL_562835, EPI_ISL_562836, EPI_ISL_562837, EPI_ISL_562846, EPI_ISL_562847, EPI_ISL_562853, EPI_ISL_562864, EPI_ISL_562866, EPI_ISL_562870, EPI_ISL_562871, EPI_ISL_562887, EPI_ISL_562892, EPI_ISL_562899, EPI_ISL_562900, EPI_ISL_562917 | | | |
| see above | Microbiological Diagnostic Unit - Public Health Laboratory (MDU-PHL) | MDU-PHL | Seemann, T., Schultz M. B., Sait, M., Sherry, N. |
| EPI_ISL_562924 | Victorian Infectious Diseases Reference Laboratory (VIDRL) | VIDRL and MDU-PHL | Caly, L., Seemann, T., Sait, M., Schultz, M. B., Druce J., Sherry, N. |
| EPI_ISL_562925, EPI_ISL_562926 | Microbiological Diagnostic Unit - Public Health Laboratory (MDU-PHL) | MDU-PHL | Seemann, T., Schultz M. B., Sait, M., Sherry, N. |
| EPI_ISL_562929 | Victorian Infectious Diseases Reference Laboratory (VIDRL) | VIDRL and MDU-PHL | Caly, L., Seemann, T., Sait, M., Schultz, M. B., Druce J., Sherry, N. |
| EPI_ISL_562930 | Microbiological Diagnostic Unit - Public Health Laboratory (MDU-PHL) | MDU-PHL | Seemann, T., Schultz M. B., Sait, M., Sherry, N. |
| EPI_ISL_562936 | Victorian Infectious Diseases Reference Laboratory (VIDRL) | VIDRL and MDU-PHL | Caly, L., Seemann, T., Sait, M., Schultz, M. B., Druce J., Sherry, N. |
| EPI_ISL_562939 | Microbiological Diagnostic Unit - Public Health Laboratory (MDU-PHL) | MDU-PHL | Seemann, T., Schultz M. B., Sait, M., Sherry, N. |
| EPI_ISL_562946 | Victorian Infectious Diseases Reference Laboratory (VIDRL) | VIDRL and MDU-PHL | Caly, L., Seemann, T., Sait, M., Schultz, M. B., Druce J., Sherry, N. |
| EPI_ISL_562950 | Microbiological Diagnostic Unit - Public Health Laboratory (MDU-PHL) | MDU-PHL | Seemann, T., Schultz M. B., Sait, M., Sherry, N. |
| EPI_ISL_562951 | Victorian Infectious Diseases Reference Laboratory (VIDRL) | VIDRL and MDU-PHL | Caly, L., Seemann, T., Sait, M., Schultz, M. B., Druce J., Sherry, N. |
| EPI_ISL_562958 | Microbiological Diagnostic Unit - Public Health Laboratory (MDU-PHL) | MDU-PHL | Seemann, T., Schultz M. B., Sait, M., Sherry, N. |
| EPI_ISL_562961 | Victorian Infectious Diseases Reference Laboratory (VIDRL) | VIDRL and MDU-PHL | Caly, L., Seemann, T., Sait, M., Schultz, M. B., Druce J., Sherry, N. |
| EPI_ISL_562974, EPI_ISL_563002, EPI_ISL_563064, EPI_ISL_563065, EPI_ISL_563082, EPI_ISL_563087, EPI_ISL_563113 | Microbiological Diagnostic Unit - Public Health Laboratory (MDU-PHL) | MDU-PHL | Seemann, T., Schultz M. B., Sait, M., Sherry, N. |
| EPI_ISL_563139, EPI_ISL_563143 | Victorian Infectious Diseases Reference Laboratory (VIDRL) | VIDRL and MDU-PHL | Caly, L., Seemann, T., Sait, M., Schultz, M. B., Druce J., Sherry, N. |
| EPI_ISL_563253, EPI_ISL_563257, EPI_ISL_563259, EPI_ISL_563289, EPI_ISL_563301, EPI_ISL_563302, EPI_ISL_563303, EPI_ISL_563304, EPI_ISL_563305, EPI_ISL_563306, EPI_ISL_563307, EPI_ISL_563311, EPI_ISL_563323, EPI_ISL_563324, EPI_ISL_563326, EPI_ISL_563327, EPI_ISL_563332, EPI_ISL_563333, EPI_ISL_563334, EPI_ISL_563335, EPI_ISL_563336, EPI_ISL_563337, EPI_ISL_563338, EPI_ISL_563339, EPI_ISL_563340, EPI_ISL_563341, EPI_ISL_563346, EPI_ISL_563348, EPI_ISL_563349, EPI_ISL_563350, EPI_ISL_563351, EPI_ISL_563352, EPI_ISL_563353, EPI_ISL_563354, EPI_ISL_563355, EPI_ISL_563356, EPI_ISL_563357, EPI_ISL_563358, EPI_ISL_563359, EPI_ISL_563360, EPI_ISL_563361, EPI_ISL_563362, EPI_ISL_563363, EPI_ISL_563364, EPI_ISL_563365, EPI_ISL_563367, EPI_ISL_563368, EPI_ISL_563370, EPI_ISL_563371, EPI_ISL_563372, EPI_ISL_563373, EPI_ISL_563374, EPI_ISL_563375, EPI_ISL_563376, EPI_ISL_563377, EPI_ISL_563378, EPI_ISL_563379, EPI_ISL_563381, EPI_ISL_563382, EPI_ISL_563383, EPI_ISL_563384, EPI_ISL_563385, EPI_ISL_563386, EPI_ISL_563387, EPI_ISL_563388, EPI_ISL_563389, EPI_ISL_563390, EPI_ISL_563391, EPI_ISL_563392, EPI_ISL_563393, EPI_ISL_563394, EPI_ISL_563395, EPI_ISL_563398, EPI_ISL_563399, EPI_ISL_563404, EPI_ISL_563406 | | | |
| see above | Microbiological Diagnostic Unit - Public Health Laboratory (MDU-PHL) | MDU-PHL | Seemann, T., Schultz M. B., Sait, M., Sherry, N. |
| EPI_ISL_563409, EPI_ISL_563410, EPI_ISL_563411, EPI_ISL_563412, EPI_ISL_563413, EPI_ISL_563414, EPI_ISL_563415, EPI_ISL_563416, EPI_ISL_563417, EPI_ISL_563418, EPI_ISL_563419, EPI_ISL_563420, EPI_ISL_563421, EPI_ISL_563422, EPI_ISL_563721, EPI_ISL_563722 | | | |
| see above | Victorian Infectious Diseases Reference Laboratory (VIDRL) | VIDRL and MDU-PHL | Caly, L., Seemann, T., Sait, M., Schultz, M. B., Druce J., Sherry, N. |
| EPI_ISL_563798, EPI_ISL_563800, EPI_ISL_563801, EPI_ISL_563803, EPI_ISL_563819, EPI_ISL_563822, EPI_ISL_563824, EPI_ISL_563826, EPI_ISL_563833, EPI_ISL_563836, EPI_ISL_563838, EPI_ISL_563839, EPI_ISL_563845, EPI_ISL_563848, EPI_ISL_563853, EPI_ISL_563854, EPI_ISL_563855, EPI_ISL_563858, EPI_ISL_563859, EPI_ISL_563860, EPI_ISL_563862, EPI_ISL_563876, EPI_ISL_563879, EPI_ISL_563891, EPI_ISL_563940, EPI_ISL_563943, EPI_ISL_563952, EPI_ISL_563953, EPI_ISL_563958, EPI_ISL_563962, EPI_ISL_563963, EPI_ISL_563971, EPI_ISL_563972, EPI_ISL_565377, EPI_ISL_565383, EPI_ISL_565384, EPI_ISL_565405, EPI_ISL_565406, EPI_ISL_565407, EPI_ISL_565408, EPI_ISL_565410, EPI_ISL_565412, EPI_ISL_565413, EPI_ISL_565414, EPI_ISL_565416, EPI_ISL_565417, EPI_ISL_565419, EPI_ISL_565424, EPI_ISL_565425, EPI_ISL_565436, EPI_ISL_565437, EPI_ISL_565438, EPI_ISL_565440, EPI_ISL_565441, EPI_ISL_565442, EPI_ISL_565443, EPI_ISL_565445, EPI_ISL_565446, EPI_ISL_565447, EPI_ISL_565448, EPI_ISL_565450, EPI_ISL_565452, EPI_ISL_565468, EPI_ISL_565470, EPI_ISL_565471, EPI_ISL_565472, EPI_ISL_565474, EPI_ISL_565475, EPI_ISL_565477, EPI_ISL_565500, EPI_ISL_565501, EPI_ISL_565502, EPI_ISL_565504, EPI_ISL_565528, EPI_ISL_565530, EPI_ISL_565535, EPI_ISL_565589, EPI_ISL_565610, EPI_ISL_565611, EPI_ISL_565612, EPI_ISL_565613, EPI_ISL_565614, EPI_ISL_565615, EPI_ISL_565616, EPI_ISL_565617, EPI_ISL_565633, EPI_ISL_565634, EPI_ISL_565635, EPI_ISL_565638, EPI_ISL_565639, EPI_ISL_565643, EPI_ISL_565652, EPI_ISL_565653, EPI_ISL_565654, EPI_ISL_565655, EPI_ISL_565656, EPI_ISL_565657, EPI_ISL_565658, EPI_ISL_565659, EPI_ISL_565660, EPI_ISL_565661, EPI_ISL_565662, EPI_ISL_565663, EPI_ISL_565667, EPI_ISL_565668, EPI_ISL_565669, EPI_ISL_565670, EPI_ISL_565671, EPI_ISL_565672, EPI_ISL_565673, EPI_ISL_565674, EPI_ISL_565675, EPI_ISL_565676, EPI_ISL_565677, EPI_ISL_565678, EPI_ISL_565679, EPI_ISL_565686, EPI_ISL_565687, EPI_ISL_565688, EPI_ISL_565689, EPI_ISL_565690, EPI_ISL_565691, EPI_ISL_565692, EPI_ISL_565693, EPI_ISL_565694, EPI_ISL_565715 | | | |
| see above | Microbiological Diagnostic Unit - Public Health Laboratory (MDU-PHL) | MDU-PHL | Seemann, T., Schultz M. B., Sait, M., Sherry, N. |
| EPI_ISL_565742 | Victorian Infectious Diseases Reference Laboratory (VIDRL) | VIDRL and MDU-PHL | Caly, L., Seemann, T., Sait, M., Schultz, M. B., Druce J., Sherry, N. |

| | | | |
|---|---|---|---|
| EPI_ISL_565762, EPI_ISL_565763, EPI_ISL_565764, EPI_ISL_565765, EPI_ISL_565766, EPI_ISL_565767, EPI_ISL_565768, EPI_ISL_565769, EPI_ISL_565770, EPI_ISL_565771, EPI_ISL_565772, EPI_ISL_565773, EPI_ISL_565774, EPI_ISL_565775, EPI_ISL_565776, EPI_ISL_565777, EPI_ISL_565778, EPI_ISL_565779, EPI_ISL_565780, EPI_ISL_565781, EPI_ISL_565782, EPI_ISL_565783, EPI_ISL_565784, EPI_ISL_565787, EPI_ISL_565813, EPI_ISL_565815, EPI_ISL_565816, EPI_ISL_565823, EPI_ISL_565824, EPI_ISL_565825, EPI_ISL_565826, EPI_ISL_565827, EPI_ISL_565828, EPI_ISL_565829 | | | |
| see above | Microbiological Diagnostic Unit - Public Health Laboratory (MDU-PHL) | MDU-PHL | Seemann, T., Schultz M. B., Sait, M., Sherry, N. |
| EPI_ISL_565938, EPI_ISL_566005, EPI_ISL_566009, EPI_ISL_566020, EPI_ISL_566026, EPI_ISL_566033, EPI_ISL_566034 | Michigan Department of Health and Human Services, Bureau of Laboratories | Michigan Department of Health and Human Services, Bureau of Laboratories | Blankenship HM, Riner D, Soehnlen MK |
| EPI_ISL_566045, EPI_ISL_566069, EPI_ISL_566070, EPI_ISL_566071, EPI_ISL_566072, EPI_ISL_566073 | Respiratory Virus Unit, Microbiology Services Colindale, Public Health England | Respiratory Virus Unit, Microbiology Services Colindale, Public Health England | PHE Covid Sequencing Team |
| EPI_ISL_566576 | Lighthouse Lab in Cambridge | Wellcome Sanger Institute for the COVID-19 Genomics UK (COG-UK) consortium | Rob Howes, The Lighthouse Lab in Cambridge and Alex Alderton, Roberto Amato, Sonia Goncalves, Ewan Harrison, David K. Jackson, Ian Johnston, Dominic Kwiatkowski, Cordelia Langford, John Sillitoe on behalf of the Wellcome Sanger Institute COVID-19 Surveillance Team |
| EPI_ISL_568291, EPI_ISL_568292, EPI_ISL_568293, EPI_ISL_568294, EPI_ISL_568295, EPI_ISL_568296, EPI_ISL_568297, EPI_ISL_568298, EPI_ISL_568299, EPI_ISL_568300 | Lighthouse Lab in Glasgow | Wellcome Sanger Institute for the COVID-19 Genomics UK (COG-UK) consortium | Harper VanSteenhouse, Yumi Kasai, David Gray, Carol Clugston, Anna Dominiczak and Alex Alderton, Roberto Amato, Sonia Goncalves, Ewan Harrison, David K. Jackson, Ian Johnston, Dominic Kwiatkowski, Cordelia Langford, John Sillitoe on behalf of the Wellcome Sanger Institute COVID-19 Surveillance Team |
| EPI_ISL_568301 | Lighthouse Lab in Glasgow | Wellcome Sanger Institute for the COVID-19 Genomics UK (COG-UK) Consortium | Harper VanSteenhouse, Yumi Kasai, David Gray, Carol Clugston, Anna Dominiczak and Alex Alderton, Roberto Amato, Sonia Goncalves, Ewan Harrison, David K. Jackson, Ian Johnston, Dominic Kwiatkowski, Cordelia Langford, John Sillitoe on behalf of the Wellcome Sanger Institute COVID-19 Surveillance Team |
| EPI_ISL_568302, EPI_ISL_568303, EPI_ISL_568304, EPI_ISL_568305, EPI_ISL_568306, EPI_ISL_568307, EPI_ISL_568308, EPI_ISL_568309, EPI_ISL_568310, EPI_ISL_568311, EPI_ISL_568312, EPI_ISL_568313, EPI_ISL_568314 | | | |
| see above | Lighthouse Lab in Glasgow | Wellcome Sanger Institute for the COVID-19 Genomics UK (COG-UK) consortium | Harper VanSteenhouse, Yumi Kasai, David Gray, Carol Clugston, Anna Dominiczak and Alex Alderton, Roberto Amato, Sonia Goncalves, Ewan Harrison, David K. Jackson, Ian Johnston, Dominic Kwiatkowski, Cordelia Langford, John Sillitoe on behalf of the Wellcome Sanger Institute COVID-19 Surveillance Team |
| EPI_ISL_568315 | Lighthouse Lab in Glasgow | Wellcome Sanger Institute for the COVID-19 Genomics UK (COG-UK) Consortium | Harper VanSteenhouse, Yumi Kasai, David Gray, Carol Clugston, Anna Dominiczak and Alex Alderton, Roberto Amato, Sonia Goncalves, Ewan Harrison, David K. Jackson, Ian Johnston, Dominic Kwiatkowski, Cordelia Langford, John Sillitoe on behalf of the Wellcome Sanger Institute COVID-19 Surveillance Team |
| EPI_ISL_568316 | Lighthouse Lab in Glasgow | Wellcome Sanger Institute for the COVID-19 Genomics UK (COG-UK) consortium | Harper VanSteenhouse, Yumi Kasai, David Gray, Carol Clugston, Anna Dominiczak and Alex Alderton, Roberto Amato, Sonia Goncalves, Ewan Harrison, David K. Jackson, Ian Johnston, Dominic Kwiatkowski, Cordelia Langford, John Sillitoe on behalf of the Wellcome Sanger Institute COVID-19 Surveillance Team |
| EPI_ISL_568317 | Lighthouse Lab in Glasgow | Wellcome Sanger Institute for the COVID-19 Genomics UK (COG-UK) Consortium | Harper VanSteenhouse, Yumi Kasai, David Gray, Carol Clugston, Anna Dominiczak and Alex Alderton, Roberto Amato, Sonia Goncalves, Ewan Harrison, David K. Jackson, Ian Johnston, Dominic Kwiatkowski, Cordelia Langford, John Sillitoe on behalf of the Wellcome Sanger Institute COVID-19 Surveillance Team |
| EPI_ISL_568318, EPI_ISL_568319, EPI_ISL_568320, EPI_ISL_568321, EPI_ISL_568322, EPI_ISL_568323, EPI_ISL_568324, EPI_ISL_568325, EPI_ISL_568326, EPI_ISL_568327, EPI_ISL_568328, EPI_ISL_568329, EPI_ISL_568330, EPI_ISL_568331, EPI_ISL_568332, EPI_ISL_568333, EPI_ISL_568334, EPI_ISL_568335, EPI_ISL_568336, EPI_ISL_568337, EPI_ISL_568338, EPI_ISL_568339, EPI_ISL_568340, EPI_ISL_568341, EPI_ISL_568342, EPI_ISL_568343 | | | |
| see above | Lighthouse Lab in Glasgow | Wellcome Sanger Institute for the COVID-19 Genomics UK (COG-UK) consortium | Harper VanSteenhouse, Yumi Kasai, David Gray, Carol Clugston, Anna Dominiczak and Alex Alderton, Roberto Amato, Sonia Goncalves, Ewan Harrison, David K. Jackson, Ian Johnston, Dominic Kwiatkowski, Cordelia Langford, John Sillitoe on behalf of the Wellcome Sanger Institute COVID-19 Surveillance Team |
| EPI_ISL_568344 | Lighthouse Lab in Glasgow | Wellcome Sanger Institute for the COVID-19 Genomics UK (COG-UK) Consortium | Harper VanSteenhouse, Yumi Kasai, David Gray, Carol Clugston, Anna Dominiczak and Alex Alderton, Roberto Amato, Sonia Goncalves, Ewan Harrison, David K. Jackson, Ian Johnston, Dominic Kwiatkowski, Cordelia Langford, John Sillitoe on behalf of the Wellcome Sanger Institute COVID-19 Surveillance Team |
| EPI_ISL_568345, EPI_ISL_568346, EPI_ISL_568347, EPI_ISL_568348, EPI_ISL_568349, EPI_ISL_568350, EPI_ISL_568351, EPI_ISL_568352, EPI_ISL_568353, EPI_ISL_568354, EPI_ISL_568355, EPI_ISL_568356, EPI_ISL_568357, EPI_ISL_568358, EPI_ISL_568359, EPI_ISL_568360, EPI_ISL_568361, EPI_ISL_568362, EPI_ISL_568363 | | | |
| see above | Lighthouse Lab in Glasgow | Wellcome Sanger Institute for the COVID-19 Genomics UK (COG-UK) consortium | Harper VanSteenhouse, Yumi Kasai, David Gray, Carol Clugston, Anna Dominiczak and Alex Alderton, Roberto Amato, Sonia Goncalves, Ewan Harrison, David K. Jackson, Ian Johnston, Dominic Kwiatkowski, Cordelia Langford, John Sillitoe on behalf of the Wellcome Sanger Institute COVID-19 Surveillance Team |
| EPI_ISL_568364, EPI_ISL_568365 | Lighthouse Lab in Milton Keynes | Wellcome Sanger Institute for the COVID-19 Genomics UK (COG-UK) consortium | The Lighthouse Lab in Milton Keynes and Alex Alderton, Roberto Amato, Sonia Goncalves, Ewan Harrison, David K. Jackson, Ian Johnston, Dominic Kwiatkowski, Cordelia Langford, John Sillitoe on behalf of the Wellcome Sanger Institute COVID-19 Surveillance Team |
| EPI_ISL_568366, EPI_ISL_568367 | Lighthouse Lab in Glasgow | Wellcome Sanger Institute for the COVID-19 Genomics UK (COG-UK) consortium | Harper VanSteenhouse, Yumi Kasai, David Gray, Carol Clugston, Anna Dominiczak and Alex Alderton, Roberto Amato, Sonia Goncalves, Ewan Harrison, David K. Jackson, Ian Johnston, Dominic Kwiatkowski, Cordelia Langford, John Sillitoe on behalf of the Wellcome Sanger Institute COVID-19 Surveillance Team |
| EPI_ISL_568385 | Lighthouse Lab in Cambridge | Wellcome Sanger Institute for the COVID-19 Genomics UK (COG-UK) consortium | Rob Howes, The Lighthouse Lab in Cambridge and Alex Alderton, Roberto Amato, Sonia Goncalves, Ewan Harrison, David K. Jackson, Ian Johnston, Dominic Kwiatkowski, Cordelia Langford, John Sillitoe on behalf of the Wellcome Sanger Institute COVID-19 Surveillance Team |
| EPI_ISL_568386, EPI_ISL_568387 | Lighthouse Lab in Milton Keynes | Wellcome Sanger Institute for the COVID-19 Genomics UK (COG-UK) consortium | The Lighthouse Lab in Milton Keynes and Alex Alderton, Roberto Amato, Sonia Goncalves, Ewan Harrison, David K. Jackson, Ian Johnston, Dominic Kwiatkowski, Cordelia Langford, John Sillitoe on behalf of the Wellcome Sanger Institute COVID-19 Surveillance Team |
| EPI_ISL_568388, EPI_ISL_568389, EPI_ISL_568390, EPI_ISL_568391 | Lighthouse Lab in Glasgow | Wellcome Sanger Institute for the COVID-19 Genomics UK (COG-UK) consortium | Harper VanSteenhouse, Yumi Kasai, David Gray, Carol Clugston, Anna Dominiczak and Alex Alderton, Roberto Amato, Sonia Goncalves, Ewan Harrison, David K. Jackson, Ian Johnston, Dominic Kwiatkowski, Cordelia Langford, John Sillitoe on behalf of the Wellcome Sanger Institute COVID-19 Surveillance Team |
| EPI_ISL_568392 | Lighthouse Lab in Glasgow | Wellcome Sanger Institute for the COVID-19 Genomics UK (COG-UK) Consortium | Harper VanSteenhouse, Yumi Kasai, David Gray, Carol Clugston, Anna Dominiczak and Alex Alderton, Roberto Amato, Sonia Goncalves, Ewan Harrison, David K. Jackson, Ian Johnston, Dominic Kwiatkowski, Cordelia Langford, John Sillitoe on behalf of the Wellcome Sanger Institute COVID-19 Surveillance Team |
| EPI_ISL_568393, EPI_ISL_568394, EPI_ISL_568396, EPI_ISL_568397 | Lighthouse Lab in Glasgow | Wellcome Sanger Institute for the COVID-19 Genomics UK (COG-UK) consortium | Harper VanSteenhouse, Yumi Kasai, David Gray, Carol Clugston, Anna Dominiczak and Alex Alderton, Roberto Amato, Sonia Goncalves, Ewan Harrison, David K. Jackson, Ian Johnston, Dominic Kwiatkowski, Cordelia Langford, John Sillitoe on behalf of the Wellcome Sanger Institute COVID-19 Surveillance Team |
| EPI_ISL_568398 | Lighthouse Lab in Glasgow | Wellcome Sanger Institute for the COVID-19 Genomics UK (COG-UK) Consortium | Harper VanSteenhouse, Yumi Kasai, David Gray, Carol Clugston, Anna Dominiczak and Alex Alderton, Roberto Amato, Sonia Goncalves, Ewan Harrison, David K. Jackson, Ian Johnston, Dominic Kwiatkowski, Cordelia Langford, John Sillitoe on behalf of the Wellcome Sanger Institute COVID-19 Surveillance Team |
| EPI_ISL_568431, EPI_ISL_568432, EPI_ISL_568433, EPI_ISL_568434, EPI_ISL_568435 | Lighthouse Lab in Glasgow | Wellcome Sanger Institute for the COVID-19 Genomics UK (COG-UK) consortium | Harper VanSteenhouse, Yumi Kasai, David Gray, Carol Clugston, Anna Dominiczak and Alex Alderton, Roberto Amato, Sonia Goncalves, Ewan Harrison, David K. Jackson, Ian Johnston, Dominic Kwiatkowski, Cordelia Langford, John Sillitoe on behalf of the Wellcome Sanger Institute COVID-19 Surveillance Team |
| EPI_ISL_568436 | Lighthouse Lab in Cambridge | Wellcome Sanger Institute for the COVID-19 Genomics UK (COG-UK) consortium | Rob Howes, The Lighthouse Lab in Cambridge and Alex Alderton, Roberto Amato, Sonia Goncalves, Ewan Harrison, David K. Jackson, Ian Johnston, Dominic Kwiatkowski, Cordelia Langford, John Sillitoe on behalf of the Wellcome Sanger Institute COVID-19 Surveillance Team |
| EPI_ISL_568437, EPI_ISL_568438, EPI_ISL_568439, EPI_ISL_568440, EPI_ISL_568441, EPI_ISL_568442, EPI_ISL_568443, EPI_ISL_568444, EPI_ISL_568445, EPI_ISL_568446, EPI_ISL_568447, EPI_ISL_568448, EPI_ISL_568449, EPI_ISL_568450, EPI_ISL_568451, EPI_ISL_568452 | | | |

| | | | |
|---|---|---|---|
| see above | Lighthouse Lab in Glasgow | Wellcome Sanger Institute for the COVID-19 Genomics UK (COG-UK) consortium | Harper VanSteenhouse, Yumi Kasai, David Gray, Carol Clugston, Anna Dominiczak and Alex Alderton, Roberto Amato, Sonia Goncalves, Ewan Harrison, David K. Jackson, Ian Johnston, Dominic Kwiatkowski, Cordelia Langford, John Sillitoe on behalf of the Wellcome Sanger Institute COVID-19 Surveillance Team |
| EPI_ISL_568453, EPI_ISL_568454 | Lighthouse Lab in Cambridge | Wellcome Sanger Institute for the COVID-19 Genomics UK (COG-UK) consortium | Rob Howes, The Lighthouse Lab in Cambridge and Alex Alderton, Roberto Amato, Sonia Goncalves, Ewan Harrison, David K. Jackson, Ian Johnston, Dominic Kwiatkowski, Cordelia Langford, John Sillitoe on behalf of the Wellcome Sanger Institute COVID-19 Surveillance Team |
| EPI_ISL_568455, EPI_ISL_568456, EPI_ISL_568457 | Lighthouse Lab in Glasgow | Wellcome Sanger Institute for the COVID-19 Genomics UK (COG-UK) consortium | Harper VanSteenhouse, Yumi Kasai, David Gray, Carol Clugston, Anna Dominiczak and Alex Alderton, Roberto Amato, Sonia Goncalves, Ewan Harrison, David K. Jackson, Ian Johnston, Dominic Kwiatkowski, Cordelia Langford, John Sillitoe on behalf of the Wellcome Sanger Institute COVID-19 Surveillance Team |
| EPI_ISL_568458 | Lighthouse Lab in Cambridge | Wellcome Sanger Institute for the COVID-19 Genomics UK (COG-UK) consortium | Rob Howes, The Lighthouse Lab in Cambridge and Alex Alderton, Roberto Amato, Sonia Goncalves, Ewan Harrison, David K. Jackson, Ian Johnston, Dominic Kwiatkowski, Cordelia Langford, John Sillitoe on behalf of the Wellcome Sanger Institute COVID-19 Surveillance Team |
| EPI_ISL_568459, EPI_ISL_568460 | Lighthouse Lab in Glasgow | Wellcome Sanger Institute for the COVID-19 Genomics UK (COG-UK) consortium | Harper VanSteenhouse, Yumi Kasai, David Gray, Carol Clugston, Anna Dominiczak and Alex Alderton, Roberto Amato, Sonia Goncalves, Ewan Harrison, David K. Jackson, Ian Johnston, Dominic Kwiatkowski, Cordelia Langford, John Sillitoe on behalf of the Wellcome Sanger Institute COVID-19 Surveillance Team |
| EPI_ISL_568461 | Lighthouse Lab in Cambridge | Wellcome Sanger Institute for the COVID-19 Genomics UK (COG-UK) consortium | Rob Howes, The Lighthouse Lab in Cambridge and Alex Alderton, Roberto Amato, Sonia Goncalves, Ewan Harrison, David K. Jackson, Ian Johnston, Dominic Kwiatkowski, Cordelia Langford, John Sillitoe on behalf of the Wellcome Sanger Institute COVID-19 Surveillance Team |
| EPI_ISL_568462, EPI_ISL_568463, EPI_ISL_568464, EPI_ISL_568465 | Lighthouse Lab in Glasgow | Wellcome Sanger Institute for the COVID-19 Genomics UK (COG-UK) consortium | Harper VanSteenhouse, Yumi Kasai, David Gray, Carol Clugston, Anna Dominiczak and Alex Alderton, Roberto Amato, Sonia Goncalves, Ewan Harrison, David K. Jackson, Ian Johnston, Dominic Kwiatkowski, Cordelia Langford, John Sillitoe on behalf of the Wellcome Sanger Institute COVID-19 Surveillance Team |
| EPI_ISL_568518, EPI_ISL_568519, EPI_ISL_568520, EPI_ISL_568521, EPI_ISL_568522, EPI_ISL_568523, EPI_ISL_568524, EPI_ISL_568525, EPI_ISL_568526, EPI_ISL_568527, EPI_ISL_568528, EPI_ISL_568529, EPI_ISL_568530, EPI_ISL_568531, EPI_ISL_568532, EPI_ISL_568533, EPI_ISL_568534, EPI_ISL_568535, EPI_ISL_568536, EPI_ISL_568537, EPI_ISL_568538, EPI_ISL_568539, EPI_ISL_568540, EPI_ISL_568541, EPI_ISL_568542, EPI_ISL_568543, EPI_ISL_568544, EPI_ISL_568545, EPI_ISL_568546, EPI_ISL_568547, EPI_ISL_568548, EPI_ISL_568549, EPI_ISL_568550, EPI_ISL_568551, EPI_ISL_568552, EPI_ISL_568553, EPI_ISL_568554, EPI_ISL_568555 | | | |
| see above | Laboratorio de Referencia Nacional de Virus Respiratorios, Instituto Nacional de Salud Peru | Laboratorio de Genómica Microbiana, Universidad Peruana Cayetano Heredia | Pablo Tsukayama, Alejandra Dávila-Barclay, Luis González, Pedro E. Romero, Brenda Ayzanoa, Janet Huancachoque, Pool Marcos, Maribel Huaringa, Camila Castillo-Vilcahuaman, Guillermo Salvatierra |
| EPI_ISL_568638, EPI_ISL_568639, EPI_ISL_568640, EPI_ISL_568641, EPI_ISL_568642, EPI_ISL_568643 | Florida Bureau of Public Health Laboratories | Florida Bureau of Public Health Laboratories | Sarah Schmedes, Jason Blanton |
| EPI_ISL_570778 | UW Virology Lab | UW Virology Lab | Pavitra Roychoudhury, Hong Xie, Lasata Shrestha, Amin Addetia, Victoria M Rachleff, Meei-Li Huang, Keith R Jerome, Alexander Greninger |
| EPI_ISL_572201, EPI_ISL_572202, EPI_ISL_572203, EPI_ISL_572204, EPI_ISL_572205, EPI_ISL_572206, EPI_ISL_572207, EPI_ISL_572208, EPI_ISL_572209, EPI_ISL_572210, EPI_ISL_572211, EPI_ISL_572212, EPI_ISL_572213, EPI_ISL_572214, EPI_ISL_572215, EPI_ISL_572216, EPI_ISL_572217 | | | |
| see above | Virginia DCLS | Virginia DCLS | Virginia DCLS |
| EPI_ISL_572424, EPI_ISL_572452 | Northumbria University / South Tees Hospitals NHS Foundation Trust / North Cumbria Integrated Care NHS Foundation Trust / North Tees and Hartlepool NHS Foundation Trust / Newcastle Hospitals NHS Foundation Trust | COVID-19 Genomics UK (COG-UK) Consortium | Darren L Smith,Andrew Nelson,Matthew Bashton,Greg R Young,Joshua Loh,John Allan,Mohammad A Tariq,Giles S Holt,Gary Black,Wen C Yew,Lynn Dover,Paul Baker,Steve Liggett,Sarah Essex,Jane Greenaway,Debra Padgett,Clive Graham,Garren Scott,Edward Barton,Emma Swindells,Brendan Payne,Jennifer Collins,Yusri Taha,Gary Eltringham |
| EPI_ISL_572563 | Quadram Institute Bioscience | COVID-19 Genomics UK (COG-UK) Consortium | Dave J. Baker, Gemma L. Kay, Alp Aydin, Thanh Le-Viet, Steven Rudder, Ana P. Tedim, Anastasia Kolyva, Maria Diaz, Leonardo de Oliveira Martins, Nabil-Fareed Alikhan, Lizzie Meadows, Rachael Stanley, Ngozi Elumogo, Muhammed Yasir, Nicholas M. Thomson, Alexander J Trotter, Rachel Gilroy, Samuel Bloomfield, Claire Stuart, Andrew Bell, Reenesh Prakash, Samir Dervisevic, Alison E. Mather, John Wain, Mark Webber, Andrew J. Page, Justin O'Grady |
| EPI_ISL_572700, EPI_ISL_572729 | Northumbria University / South Tees Hospitals NHS Foundation Trust / North Cumbria Integrated Care NHS Foundation Trust / North Tees and Hartlepool NHS Foundation Trust / Newcastle Hospitals NHS Foundation Trust | COVID-19 Genomics UK (COG-UK) Consortium | Darren L Smith,Andrew Nelson,Matthew Bashton,Greg R Young,Joshua Loh,John Allan,Mohammad A Tariq,Giles S Holt,Gary Black,Wen C Yew,Lynn Dover,Paul Baker,Steve Liggett,Sarah Essex,Jane Greenaway,Debra Padgett,Clive Graham,Garren Scott,Edward Barton,Emma Swindells,Brendan Payne,Jennifer Collins,Yusri Taha,Gary Eltringham |
| EPI_ISL_572849, EPI_ISL_572857, EPI_ISL_572858, EPI_ISL_572859, EPI_ISL_572893, EPI_ISL_572894, EPI_ISL_572895, EPI_ISL_572896, EPI_ISL_572897, EPI_ISL_572933, EPI_ISL_572953 | | | |
| see above | Oxford Viromics, NDM, University of Oxford; Oxford University Hospitals; Basingstoke and North Hampshire Hospital | COVID-19 Genomics UK (COG-UK) Consortium | Tanya Golubchik, David Bonsall, George Macintyre, Amy Trebes, Mariateresa de Cesare, Catrin Moore, Alex Mobbs, Anita Justice, Robert Shaw, Monique Andersson, Timothy Peto, Emma Wise, Nathan Moore, Jessica Lynch, Nick Cortes, Matilde Mori, Stephen Kidd, David Buck, John Todd, Christophe Fraser |
| EPI_ISL_572957 | University College London, Great Ormond Street Hospital for Children NHS Foundation Trust, Imperial College Healthcare NHS Trust | COVID-19 Genomics UK (COG-UK) Consortium | Sergi Castellano, Rachel Williams, Mark Kristiansen, Paola Resende Silva, Sunando Roy, Tony Brooks, Helena Tutill, Paola Niola, Patricia Dyal, Charlotte Williams, Leysa Forrest, Yasmin Panchbhaya, Jacqueline Findlay, Samuel Weeks, Julianne Brown, Kathryn Harris, Paul Randell, James Price, Alison Holmes, Judith Breuer |
| EPI_ISL_572958, EPI_ISL_572962, EPI_ISL_572963, EPI_ISL_572964, EPI_ISL_572965, EPI_ISL_572966, EPI_ISL_572967, EPI_ISL_572968, EPI_ISL_572969, EPI_ISL_572970, EPI_ISL_572971, EPI_ISL_572972, EPI_ISL_572973, EPI_ISL_572974, EPI_ISL_572975, EPI_ISL_572976, EPI_ISL_572977, EPI_ISL_572978, EPI_ISL_572979, EPI_ISL_572980, EPI_ISL_572981, EPI_ISL_572982, EPI_ISL_572983, EPI_ISL_572984, EPI_ISL_572985, EPI_ISL_572986, EPI_ISL_572987, EPI_ISL_572988, EPI_ISL_572989, EPI_ISL_572990, EPI_ISL_572991, EPI_ISL_572992, EPI_ISL_572993, EPI_ISL_572994, EPI_ISL_572995, EPI_ISL_572996, EPI_ISL_572997, EPI_ISL_572998, EPI_ISL_572999, EPI_ISL_573000, EPI_ISL_573001, EPI_ISL_573002, EPI_ISL_573003, EPI_ISL_573004, EPI_ISL_573005, EPI_ISL_573006, EPI_ISL_573007, EPI_ISL_573008, EPI_ISL_573009, EPI_ISL_573010, EPI_ISL_573011, EPI_ISL_573012, EPI_ISL_573013, EPI_ISL_573014, EPI_ISL_573015, EPI_ISL_573016, EPI_ISL_573017, EPI_ISL_573018, EPI_ISL_573019, EPI_ISL_573020, EPI_ISL_573021, EPI_ISL_573022, EPI_ISL_573023, EPI_ISL_573024, EPI_ISL_573025, EPI_ISL_573026, EPI_ISL_573027, EPI_ISL_573028, EPI_ISL_573029, EPI_ISL_573030, EPI_ISL_573031, EPI_ISL_573032, EPI_ISL_573033, EPI_ISL_573034, EPI_ISL_573035, EPI_ISL_573036, EPI_ISL_573037 | | | |
| see above | Oxford Viromics, NDM, University of Oxford; Oxford University Hospitals; Basingstoke and North Hampshire Hospital | COVID-19 Genomics UK (COG-UK) Consortium | Tanya Golubchik, David Bonsall, George Macintyre, Amy Trebes, Mariateresa de Cesare, Catrin Moore, Alex Mobbs, Anita Justice, Robert Shaw, Monique Andersson, Timothy Peto, Emma Wise, Nathan Moore, Jessica Lynch, Nick Cortes, Matilde Mori, Stephen Kidd, David Buck, John Todd, Christophe Fraser |
| EPI_ISL_573316, EPI_ISL_573317, EPI_ISL_573318, EPI_ISL_573319, EPI_ISL_573375 | Northumbria University / South Tees Hospitals NHS Foundation Trust / North Cumbria Integrated Care NHS Foundation Trust / North Tees and Hartlepool NHS Foundation Trust / Newcastle Hospitals NHS Foundation Trust | COVID-19 Genomics UK (COG-UK) Consortium | Darren L Smith,Andrew Nelson,Matthew Bashton,Greg R Young,Joshua Loh,John Allan,Mohammad A Tariq,Giles S Holt,Gary Black,Wen C Yew,Lynn Dover,Paul Baker,Steve Liggett,Sarah Essex,Jane Greenaway,Debra Padgett,Clive Graham,Garren Scott,Edward Barton,Emma Swindells,Brendan Payne,Jennifer Collins,Yusri Taha,Gary Eltringham |
| EPI_ISL_573512 | University College London, Great Ormond Street Hospital for Children NHS Foundation Trust, Imperial College Healthcare NHS Trust | COVID-19 Genomics UK (COG-UK) Consortium | Sergi Castellano, Rachel Williams, Mark Kristiansen, Paola Resende Silva, Sunando Roy, Tony Brooks, Helena Tutill, Paola Niola, Patricia Dyal, Charlotte Williams, Leysa Forrest, Yasmin Panchbhaya, Jacqueline Findlay, Samuel Weeks, Julianne Brown, Kathryn Harris, Paul Randell, James Price, Alison Holmes, Judith Breuer |
| EPI_ISL_573773, EPI_ISL_573774 | West of Scotland Specialist Virology Centre, NHSGGC / MRC-University of Glasgow Centre for Virus Research | COVID-19 Genomics UK (COG-UK) Consortium | Ana da Silva Filipe, Natasha Johnson, Kathy Smollett, Daniel Mair, Stephen Carmichael, Lily Tong, Jenna Nichols, Elihu Aranday-Cortes, Kyriaki Nomikou; Sarah McDonald, Marc Niebel, Patawee Asamaphan; Richard Orton, Joseph Hughes, Sreenu Vattipally, David L Robertson; Alasdair MacLean, Rory Gunson; Kathy Li, Igor Starinskij, Natasha Jesudason, Rajiv Shah, James Shepherd, Antonia Ho, Emma Thomson |
| EPI_ISL_574260, EPI_ISL_574261, EPI_ISL_574262, EPI_ISL_574263, EPI_ISL_574264, EPI_ISL_574265, EPI_ISL_574266, EPI_ISL_574267, EPI_ISL_574268, EPI_ISL_574269, EPI_ISL_574270, EPI_ISL_574271, EPI_ISL_574272, EPI_ISL_574273, EPI_ISL_574274, EPI_ISL_574275, EPI_ISL_574276, EPI_ISL_574277, EPI_ISL_574278, EPI_ISL_574280, EPI_ISL_574281, EPI_ISL_574282, EPI_ISL_574283, EPI_ISL_574284, EPI_ISL_574285, EPI_ISL_574286, EPI_ISL_574287, EPI_ISL_574288, EPI_ISL_574289, EPI_ISL_574290, EPI_ISL_574291 | | | |
| see above | New Mexico Department of Health Scientific Laboratory | New Mexico Department of Health Scientific Laboratory | Ellie Johnson, Anastacia Griego-Fisher, D'Eldra Malone |
| EPI_ISL_574354, EPI_ISL_574355, EPI_ISL_574356, EPI_ISL_574357, EPI_ISL_574358, EPI_ISL_574359, EPI_ISL_574360, EPI_ISL_574361, EPI_ISL_574362, EPI_ISL_574363, EPI_ISL_574364, EPI_ISL_574365, EPI_ISL_574366, EPI_ISL_574367, EPI_ISL_574368, EPI_ISL_574369, EPI_ISL_574370, EPI_ISL_574371, EPI_ISL_574372, EPI_ISL_574373, EPI_ISL_574374, EPI_ISL_574375, EPI_ISL_574376, EPI_ISL_574377, EPI_ISL_574378, EPI_ISL_574379, EPI_ISL_574380, EPI_ISL_574381, EPI_ISL_574382, EPI_ISL_574383, EPI_ISL_574384, EPI_ISL_574385, EPI_ISL_574386, EPI_ISL_574387, EPI_ISL_574388, EPI_ISL_574389, EPI_ISL_574390, EPI_ISL_574391, EPI_ISL_574392, EPI_ISL_574393, EPI_ISL_574394, EPI_ISL_574395, EPI_ISL_574396, EPI_ISL_574397, EPI_ISL_574398, EPI_ISL_574399, EPI_ISL_574400, EPI_ISL_574401, EPI_ISL_574402, EPI_ISL_574403, EPI_ISL_574404, EPI_ISL_574405, EPI_ISL_574406, EPI_ISL_574407, EPI_ISL_574408, EPI_ISL_574409, EPI_ISL_574410, EPI_ISL_574411, EPI_ISL_574412, EPI_ISL_574413, EPI_ISL_574414, EPI_ISL_574415, EPI_ISL_574416, EPI_ISL_574417, EPI_ISL_574418, EPI_ISL_574419, EPI_ISL_574420, EPI_ISL_574421, EPI_ISL_574422, EPI_ISL_574423, EPI_ISL_574424, EPI_ISL_574425, EPI_ISL_574426, EPI_ISL_574427, EPI_ISL_574428, EPI_ISL_574429, EPI_ISL_574430 | | | |
| see above | LSUHS Emerging Viral Threat Laboratory | Microbial Genome Sequencing Center | Jeremy P. Kamil, Rona S. Scott, Maarten Van Diest, Malgorzata Bienkowska-Haba, Katarzyna Zwolinska, Andrew D. Yurochko, Christopher G. Kevil, Martin J. Sapp, Daniel J. Snyder, Vaughn S. Cooper, John A. Vanchiere |

| | | | |
|---|---|---|---|
| EPI_ISL_574649, EPI_ISL_574650, EPI_ISL_574651 | Seattle Flu Study | Seattle Flu Study | Deborah A. Nickerson, Chris D. Frazar, Jover Lee, Benjamin Pelle, Matthew Richardson, Amanda Adler, Elisabeth Brandstetter, Peter D. Han, Kairsten Fay, Misja Ilcisin, Kirsten Lacombe, Thomas R. Sibley, Melissa Truong, Caitlin R. Wolf, Karen Cowgill, Stephanie Schrag, Jeff Duchin, Michael Boeckh, Janet A. Englund, Michael Famulare, Barry R. Lutz, Mark J. Rieder, Lea M. Starita, Matthew Thompson, Helen Y. Chu, Trevor Bedford, Jay Shendure |
| EPI_ISL_574786, EPI_ISL_574787, EPI_ISL_574788, EPI_ISL_574789 | Dutch COVID-19 response team | Erasmus Medical Center | Bas Oude Munnink, Reina Sikkema, David Nieuwenhuijse, Irina Chestakova, Anne van der Linden, Marjan Boter, Emmanuelle Munger, Corine GeurtsvanKessel, Annemiek van der Eijk, Richard Molenkamp, Marion Koopmans, on behalf of the Dutch national COVID-19 response team. |
| EPI_ISL_575029, EPI_ISL_575031 | Seattle Flu Study | Seattle Flu Study | Deborah A. Nickerson, Chris D. Frazar, Jover Lee, Benjamin Pelle, Matthew Richardson, Amanda Adler, Elisabeth Brandstetter, Peter D. Han, Kairsten Fay, Misja Ilcisin, Kirsten Lacombe, Thomas R. Sibley, Melissa Truong, Caitlin R. Wolf, Karen Cowgill, Stephanie Schrag, Jeff Duchin, Michael Boeckh, Janet A. Englund, Michael Famulare, Barry R. Lutz, Mark J. Rieder, Lea M. Starita, Matthew Thompson, Helen Y. Chu, Trevor Bedford, Jay Shendure |
| EPI_ISL_575761, EPI_ISL_575765, EPI_ISL_575772, EPI_ISL_575773, EPI_ISL_575776, EPI_ISL_575781, EPI_ISL_575785, EPI_ISL_575788, EPI_ISL_575791, EPI_ISL_575794, EPI_ISL_575795, EPI_ISL_575796, EPI_ISL_575797, EPI_ISL_575810, EPI_ISL_575814, EPI_ISL_575817, EPI_ISL_575822, EPI_ISL_575824, EPI_ISL_575825, EPI_ISL_575828, EPI_ISL_575862, EPI_ISL_575878, EPI_ISL_575890, EPI_ISL_575895, EPI_ISL_575902, EPI_ISL_575908, EPI_ISL_575913, EPI_ISL_575914, EPI_ISL_575917, EPI_ISL_575918, EPI_ISL_575920, EPI_ISL_575928, EPI_ISL_575929, EPI_ISL_575938, EPI_ISL_575941, EPI_ISL_575943, EPI_ISL_575945, EPI_ISL_575946, EPI_ISL_575948, EPI_ISL_575952, EPI_ISL_575953, EPI_ISL_575954, EPI_ISL_575955, EPI_ISL_575960, EPI_ISL_575961, EPI_ISL_575963, EPI_ISL_575966, EPI_ISL_575968, EPI_ISL_575970, EPI_ISL_575971 | | | |
| see above | Lighthouse Lab in Milton Keynes | Wellcome Sanger Institute for the COVID-19 Genomics UK (COG-UK) consortium | The Lighthouse Lab in Milton Keynes and Alex Alderton, Roberto Amato, Sonia Goncalves, Ewan Harrison, David K. Jackson, Ian Johnston, Dominic Kwiatkowski, Cordelia Langford, John Sillitoe on behalf of the Wellcome Sanger Institute COVID-19 Surveillance Team |
| EPI_ISL_575979, EPI_ISL_576094 | Lighthouse Lab in Milton Keynes | Wellcome Sanger Institute for the COVID-19 Genomics UK (COG-UK) consortium | The Lighthouse Lab in Milton Keynes and Alex Alderton, Roberto Amato, Sonia Goncalves, Ewan Harrison, David K. Jackson, Ian Johnston, Dominic Kwiatkowski, Cordelia Langford, John Sillitoe on behalf of the Wellcome Sanger Institute COVID-19 Surveillance Team (http://www.sanger.ac.uk/covid-team) |
| EPI_ISL_576128 | RSUP Dr. Sardjito | Genetics Working Group (Pokja Genetik) Faculty of Medicine, Public Health and Nursing Universitas Gadjah Mada (FK-KMK UGM); Disease Investigation Center Wates Ministry of Agriculture Indonesia; Department of Microbiology FK-KMK UGM; Laboratorium Diagnostik Yayasan Tahija World Mosquito Program (WMP) Yogyakarta Center for Tropical Medicine FK-KMK UGM; Integrated Research Center FK-KMK UGM; Department of Computer Science and Electronics FMIPA UGM | Gunadi, Hendra Wibawa, Marcellus, Mohamad S. Hakim, Edwin W. Daniwijaya, Ludhang P. Rizki, Endah Supriyati, Eggi Arguni, Titik Nuryastuti, Tri Wibawa, Dwi AA Nugrahaningsih, Afiahayati, Siswanto, Kristy Iskandar, Nungki Anggorowati, Ika Trisnawati, Sumardi, Eko Budiono, Bambang Sigit Riyanto, Heni Retnowulan, Munawar Gani, Satria Maulana, Nur Rahmi Ananda, Riat El Khair, Yunika Puspadewi, Osman Sianipar, Umi Solekhah Intansari, Elizabeth Henny Herningtiyas, Ira Puspitawati, Nur Imma Fatimah Harahap, Dyah Ayu Puspitarani, Kemala Athollah |
| EPI_ISL_576131 | unknown | Public Health Virology Laboratory, Forensic and Scientific Services (PHV-FSS) | Son Nguyen et al. |
| EPI_ISL_576265 | Instituto de Diagnostico y Referencia Epidemiologicos (INDRE) | Instituto de Diagnostico y Referencia Epidemiologicos (INDRE) | Ernesto Ramirez-Gonzalez, Abril Rodriguez-Maldonado, Claudia Wong-Arambula , Natividad Cruz-Ortiz, Tatiana Nunez-Garcia, Dayanira Arellano-Suarez, Adnan Araiza-Rodriguez, Edgar Mendieta-Condado, Lucia Hernandez-Rivas, Irma Lopez-Martinez, Gisela Barrera-Badillo. |
| EPI_ISL_576266, EPI_ISL_576267, EPI_ISL_576268 | Instituto de Diagnostico y Referencia Epidemiologicos (INDRE) | Instituto de Diagnostico y Referencia Epidemiologicos (INDRE) | Ernesto Ramirez-Gonzalez, Abril Rodriguez-Maldonado, Claudia Wong-Arambula , Natividad Cruz-Ortiz, Tatiana Nunez-Garcia, Dayanira Arellano-Suarez, Adnan Araiza-Rodriguez, Fabiola Garces-Ayala, Lucia Hernandez-Rivas, Irma Lopez-Martinez, Gisela Barrera-Badillo. |
| EPI_ISL_576272 | Instituto de Diagnostico y Referencia Epidemiologicos (INDRE) | Instituto de Diagnostico y Referencia Epidemiologicos (INDRE) | Gisela Barrera-Badillo , Abril Rodriguez-Maldonado, Claudia Wong-Arambula , Natividad Cruz-Ortiz, Tatiana Nunez-Garcia, Dayanira Arellano-Suarez, Fabiola Garces-Ayala, Edgar Mendieta-Condado, Lucia Hernandez-Rivas, Irma Lopez-Martinez, Ernesto Ramirez-Gonzalez. |
| EPI_ISL_576273 | Instituto de Diagnostico y Referencia Epidemiologicos (INDRE) | Instituto de Diagnostico y Referencia Epidemiologicos (INDRE) | Ernesto Ramirez-Gonzalez, Abril Rodriguez-Maldonado, Claudia Wong-Arambula , Natividad Cruz-Ortiz, Tatiana Nunez-Garcia, Dayanira Arellano-Suarez, Adnan Araiza-Rodriguez, Fabiola Garces-Ayala, Lucia Hernandez-Rivas, Irma Lopez-Martinez, Gisela Barrera-Badillo. |
| EPI_ISL_576274 | Instituto de Diagnostico y Referencia Epidemiologicos (INDRE) | Instituto de Diagnostico y Referencia Epidemiologicos (INDRE) | Gisela Barrera-Badillo , Abril Rodriguez-Maldonado, Claudia Wong-Arambula , Natividad Cruz-Ortiz, Tatiana Nunez-Garcia, Dayanira Arellano-Suarez, Adnan Araiza-Rodriguez, Edgar Mendieta-Condado, Lucia Hernandez-Rivas, Irma Lopez-Martinez, Ernesto Ramirez-Gonzalez. |
| EPI_ISL_576278 | Instituto de Diagnostico y Referencia Epidemiologicos (INDRE) | Instituto de Diagnostico y Referencia Epidemiologicos (INDRE) | Ernesto Ramirez-Gonzalez, Abril Rodriguez-Maldonado, Claudia Wong-Arambula , Natividad Cruz-Ortiz, Tatiana Nunez-Garcia, Dayanira Arellano-Suarez, Adnan Araiza-Rodriguez, Edgar Mendieta-Condado, Lucia Hernandez-Rivas, Irma Lopez-Martinez, Gisela Barrera-Badillo. |
| EPI_ISL_576279 | Instituto de Diagnostico y Referencia Epidemiologicos (INDRE) | Instituto de Diagnostico y Referencia Epidemiologicos (INDRE) | Ernesto Ramirez-Gonzalez, Abril Rodriguez-Maldonado, Claudia Wong-Arambula , Natividad Cruz-Ortiz, Tatiana Nunez-Garcia, Dayanira Arellano-Suarez, Adnan Araiza-Rodriguez, Fabiola Garces-Ayala, Lucia Hernandez-Rivas, Irma Lopez-Martinez, Gisela Barrera-Badillo. |
| EPI_ISL_576877, EPI_ISL_576878, EPI_ISL_576879, EPI_ISL_576880, EPI_ISL_576881 | University of Birmingham | COVID-19 Genomics UK (COG-UK) Consortium | Institute of Microbiology, University of Birmingham: Claire McMurray, Joanne Stockton, Samuel Nicholls, Radoslaw Poplawski, Will Rowe, Josh Quick, Nicholas Loman. University of Birmingham Testing Laboratory: Celina M Whalley, Andrew Bosworth, Charlotte Poxon, Kasun Wanigasooriya, Oliver Pickles, Mike Kidd, Alex Richter, Andrew D Beggs PHE Heartlands Lab: Husam Osman, Andrew Bosworth. Queen Elizabeth Hospital: Anna Casey |
| EPI_ISL_576986, EPI_ISL_576988, EPI_ISL_576989, EPI_ISL_576990, EPI_ISL_576991, EPI_ISL_576992, EPI_ISL_576993, EPI_ISL_576994, EPI_ISL_576995, EPI_ISL_576996, EPI_ISL_576997, EPI_ISL_576998, EPI_ISL_576999, EPI_ISL_577000, EPI_ISL_577001, EPI_ISL_577002, EPI_ISL_577003, EPI_ISL_577004, EPI_ISL_577005, EPI_ISL_577006, EPI_ISL_577007, EPI_ISL_577008, EPI_ISL_577009, EPI_ISL_577010, EPI_ISL_577011, EPI_ISL_577012, EPI_ISL_577013, EPI_ISL_577014, EPI_ISL_577015, EPI_ISL_577016, EPI_ISL_577017, EPI_ISL_577018, EPI_ISL_577019, EPI_ISL_577020, EPI_ISL_577021, EPI_ISL_577022, EPI_ISL_577023, EPI_ISL_577024, EPI_ISL_577025, EPI_ISL_577026, EPI_ISL_577027, EPI_ISL_577028, EPI_ISL_577029, EPI_ISL_577030, EPI_ISL_577031, EPI_ISL_577032, EPI_ISL_577033, EPI_ISL_577034, EPI_ISL_577035, EPI_ISL_577036, EPI_ISL_577037, EPI_ISL_577038, EPI_ISL_577039 | | | |
| see above | Northumbria University / South Tees Hospitals NHS Foundation Trust / North Cumbria Integrated Care NHS Foundation Trust / North Tees and Hartlepool NHS Foundation Trust / Newcastle Hospitals NHS Foundation Trust | COVID-19 Genomics UK (COG-UK) Consortium | Darren L Smith,Andrew Nelson,Matthew Bashton,Greg R Young,Joshua Loh,John Allan,Mohammad A Tariq,Giles S Holt,Gary Black,Wen C Yew,Lynn Dover,Paul Baker,Steve Liggett,Sarah Essex,Jane Greenaway,Debra Padgett,Clive Graham,Garren Scott,Edward Barton,Emma Swindells,Brendan Payne,Jennifer Collins,Yusri Taha,Gary Eltringham |
| EPI_ISL_577547, EPI_ISL_577551, EPI_ISL_577557, EPI_ISL_577560, EPI_ISL_577568, EPI_ISL_577569, EPI_ISL_577570, EPI_ISL_577571, EPI_ISL_577573 | Michigan Department of Health and Human Services, Bureau of Laboratories | Michigan Department of Health and Human Services, Bureau of Laboratories | Blankenship HM, Riner D, Soehnlen MK |
| EPI_ISL_577634, EPI_ISL_577635, EPI_ISL_577636, EPI_ISL_577637, EPI_ISL_577638 | The National Institute of Public Health | State Veterinary Institute Prague | Nagy,A;Jirincova,H;Novakova,L;Trnka,D;Vecerova,J |
| EPI_ISL_577671 | NIV Influenza | NIV Influenza | Potdar V |
| EPI_ISL_577782, EPI_ISL_577783, EPI_ISL_577784, EPI_ISL_577785, EPI_ISL_577786, EPI_ISL_577787, EPI_ISL_577788, EPI_ISL_577789, EPI_ISL_577790, EPI_ISL_577791, EPI_ISL_577792, EPI_ISL_577793, EPI_ISL_577794, EPI_ISL_577795 | | | |
| see above | Dutch COVID-19 response team | Erasmus Medical Center | OH consortium |
| EPI_ISL_577837, EPI_ISL_577838, EPI_ISL_577839, EPI_ISL_577840, EPI_ISL_577841, EPI_ISL_577842, EPI_ISL_577847, EPI_ISL_577848, EPI_ISL_577849, EPI_ISL_577850, EPI_ISL_577853, EPI_ISL_577854, EPI_ISL_577855, EPI_ISL_577856, EPI_ISL_577860, EPI_ISL_577861, EPI_ISL_577862, EPI_ISL_577886, EPI_ISL_577887, EPI_ISL_577888, EPI_ISL_577889, EPI_ISL_577890, EPI_ISL_577891, EPI_ISL_577892, EPI_ISL_577893, EPI_ISL_577894, EPI_ISL_577895, EPI_ISL_577896, EPI_ISL_577897, EPI_ISL_577898, EPI_ISL_577899, EPI_ISL_577900, EPI_ISL_577901, EPI_ISL_577902, EPI_ISL_577903, EPI_ISL_577904, EPI_ISL_577905, EPI_ISL_577915, EPI_ISL_577916, EPI_ISL_577926, EPI_ISL_577927, EPI_ISL_577928, EPI_ISL_577929, EPI_ISL_577930, EPI_ISL_577931, EPI_ISL_577932, EPI_ISL_577933, EPI_ISL_577934, EPI_ISL_577935, EPI_ISL_577936, EPI_ISL_577937, EPI_ISL_577938, EPI_ISL_577939, EPI_ISL_577940, EPI_ISL_577941, EPI_ISL_577942, EPI_ISL_577943, EPI_ISL_577948, EPI_ISL_577949, EPI_ISL_577952 | | | |
| see above | Dutch COVID-19 response team | Erasmus Medical Center | Bas Oude Munnink, Reina Sikkema, David Nieuwenhuijse, Irina Chestakova, Anne van der Linden, Marjan Boter, Emmanuelle Munger, Corine GeurtsvanKessel, Annemiek van der Eijk, Richard Molenkamp, Marion Koopmans, on behalf of the Dutch national COVID-19 response team. |
| EPI_ISL_578205, EPI_ISL_578243 | National Virus Reference Laboratory | National Virus Reference Laboratory | Michael Carr, Gabriel Gonzalez, Jonathan Dean, Suzie Coughlan, Cillian F De Gascun |
| EPI_ISL_581369, EPI_ISL_581382 | Lighthouse Lab in Milton Keynes | Wellcome Sanger Institute for the COVID-19 Genomics UK (COG-UK) consortium | The Lighthouse Lab in Milton Keynes and Alex Alderton, Roberto Amato, Sonia Goncalves, Ewan Harrison, David K. Jackson, Ian Johnston, Dominic Kwiatkowski, Cordelia Langford, John Sillitoe on behalf of the Wellcome Sanger Institute COVID-19 Surveillance Team |
| EPI_ISL_581384 | Lighthouse Lab in Glasgow | Wellcome Sanger Institute for the COVID-19 Genomics UK (COG-UK) consortium | Harper VanSteenhouse, Yumi Kasai, David Gray, Carol Clugston, Anna Dominiczak and Alex Alderton, Roberto Amato, Sonia Goncalves, Ewan Harrison, David K. Jackson, Ian Johnston, Dominic Kwiatkowski, Cordelia Langford, John Sillitoe on behalf of the Wellcome Sanger Institute COVID-19 Surveillance Team |

| | | | |
|---|---|---|---|
| EPI_ISL_581386 | Lighthouse Lab in Milton Keynes | Wellcome Sanger Institute for the COVID-19 Genomics UK (COG-UK) consortium | The Lighthouse Lab in Milton Keynes and Alex Alderton, Roberto Amato, Sonia Goncalves, Ewan Harrison, David K. Jackson, Ian Johnston, Dominic Kwiatkowski, Cordelia Langford, John Sillitoe on behalf of the Wellcome Sanger Institute COVID-19 Surveillance Team |
| EPI_ISL_581387 | Lighthouse Lab in Glasgow | Wellcome Sanger Institute for the COVID-19 Genomics UK (COG-UK) consortium | Harper VanSteenhouse, Yumi Kasai, David Gray, Carol Clugston, Anna Dominiczak and Alex Alderton, Roberto Amato, Sonia Goncalves, Ewan Harrison, David K. Jackson, Ian Johnston, Dominic Kwiatkowski, Cordelia Langford, John Sillitoe on behalf of the Wellcome Sanger Institute COVID-19 Surveillance Team |
| EPI_ISL_581390 | Lighthouse Lab in Milton Keynes | Wellcome Sanger Institute for the COVID-19 Genomics UK (COG-UK) consortium | The Lighthouse Lab in Milton Keynes and Alex Alderton, Roberto Amato, Sonia Goncalves, Ewan Harrison, David K. Jackson, Ian Johnston, Dominic Kwiatkowski, Cordelia Langford, John Sillitoe on behalf of the Wellcome Sanger Institute COVID-19 Surveillance Team |
| EPI_ISL_581392 | Lighthouse Lab in Glasgow | Wellcome Sanger Institute for the COVID-19 Genomics UK (COG-UK) consortium | Harper VanSteenhouse, Yumi Kasai, David Gray, Carol Clugston, Anna Dominiczak and Alex Alderton, Roberto Amato, Sonia Goncalves, Ewan Harrison, David K. Jackson, Ian Johnston, Dominic Kwiatkowski, Cordelia Langford, John Sillitoe on behalf of the Wellcome Sanger Institute COVID-19 Surveillance Team |
| EPI_ISL_581394 | Lighthouse Lab in Cambridge | Wellcome Sanger Institute for the COVID-19 Genomics UK (COG-UK) consortium | Rob Howes, The Lighthouse Lab in Cambridge and Alex Alderton, Roberto Amato, Sonia Goncalves, Ewan Harrison, David K. Jackson, Ian Johnston, Dominic Kwiatkowski, Cordelia Langford, John Sillitoe on behalf of the Wellcome Sanger Institute COVID-19 Surveillance Team |
| EPI_ISL_581395 | Lighthouse Lab in Glasgow | Wellcome Sanger Institute for the COVID-19 Genomics UK (COG-UK) consortium | Harper VanSteenhouse, Yumi Kasai, David Gray, Carol Clugston, Anna Dominiczak and Alex Alderton, Roberto Amato, Sonia Goncalves, Ewan Harrison, David K. Jackson, Ian Johnston, Dominic Kwiatkowski, Cordelia Langford, John Sillitoe on behalf of the Wellcome Sanger Institute COVID-19 Surveillance Team |
| EPI_ISL_581396 | Lighthouse Lab in Cambridge | Wellcome Sanger Institute for the COVID-19 Genomics UK (COG-UK) consortium | Rob Howes, The Lighthouse Lab in Cambridge and Alex Alderton, Roberto Amato, Sonia Goncalves, Ewan Harrison, David K. Jackson, Ian Johnston, Dominic Kwiatkowski, Cordelia Langford, John Sillitoe on behalf of the Wellcome Sanger Institute COVID-19 Surveillance Team |
| EPI_ISL_581506, EPI_ISL_581507 | NCDC/IGIB | NCDC/IGIB | Vivekanand A, Mahesh S. Dhar, Bharathram Uppili, Nishu Tyagi, Pooja Sharma, Akshay Kanakan, Simmi Tiwari, RadhaKrishnan VS, Robin Marwal, Azka Khan, Ajit Shewale, Tushar Nale, Rajesh Pandey, Sandhya Kabra, Mohammed Faruq, Sujeet Singh, Anurag Agrawal, Partha Rakshit |
| EPI_ISL_581576, EPI_ISL_581577, EPI_ISL_581580, EPI_ISL_581581, EPI_ISL_581588, EPI_ISL_581589, EPI_ISL_581622, EPI_ISL_581623, EPI_ISL_581624, EPI_ISL_581625, EPI_ISL_581626, EPI_ISL_581627, EPI_ISL_581628, EPI_ISL_581629, EPI_ISL_581630, EPI_ISL_581631, EPI_ISL_581632, EPI_ISL_581633, EPI_ISL_581634, EPI_ISL_581635, EPI_ISL_581636 | | | |
| see above | Department of Clinical Microbiology | GIGA Medical Genomics | Keith Durkin, Maria Artesi, Sébastien Bontems, Raphaël Boreux, Bouchra Boujemla, Cécile Meex, Pierrette Melin, Marie-Pierre Hayette, Vincent Bours |
| EPI_ISL_581979, EPI_ISL_581980, EPI_ISL_581981, EPI_ISL_581982, EPI_ISL_581983, EPI_ISL_581984, EPI_ISL_581985, EPI_ISL_581986, EPI_ISL_581987, EPI_ISL_581988, EPI_ISL_581989, EPI_ISL_581990, EPI_ISL_581991, EPI_ISL_581992, EPI_ISL_581993, EPI_ISL_581994, EPI_ISL_581995, EPI_ISL_581996, EPI_ISL_581997, EPI_ISL_581998, EPI_ISL_582002 | | | |
| see above | University Hospital Basel, Clinical Virology | University Hospital Basel, Clinical Bacteriology | Madlen Stange, Alfredo Mari, Tim Roloff, Helena MB Seth-Smith, Michael Schweitzer, Myrta Brunner, Karoline Leuzinger, Kirstine K. Soegaard, Alexander Gensch, Sarah Tschudin-Sutter, Simon Fuchs, Julia Bielicki, Hans Pargger, Martin Siegemund, Christian Nickel, Roland Bingisser, Michael Osthoff, Stefano Bassetti, Rita Schneider-Sliwa, Manuel Battegay, Hans Hirsch, Adrian Egli |
| EPI_ISL_582027 | Department of Clinical Microbiology | GIGA Medical Genomics | Keith Durkin, Maria Artesi, Sébastien Bontems, Raphaël Boreux, Bouchra Boujemla, Cécile Meex, Pierrette Melin, Marie-Pierre Hayette, Vincent Bours |
| EPI_ISL_582114, EPI_ISL_582115 | CNR Virus des Infections Respiratoires - France SUD | CNR Virus des Infections Respiratoires - France SUD | Antonin Bal, Gregory Destras, Gwendolyne Burfin, Hadrien Règue, Alexandre Gaymard, Maude Bouscambert-Duchamp, Florence Morfin-Sherpa, Martine Valette, Bruno Lina, Laurence Josset |
| EPI_ISL_582224, EPI_ISL_582225, EPI_ISL_582231, EPI_ISL_582236, EPI_ISL_582240 | Wyoming Public Health Laboratory | Center for Global Health, University of New Mexico Health Sciences Center | Daryl Domman, Kurt Schwalm, Rob Christensen, Wanda Manley, Cari Sloma, Noah Hull, Darrell Dinwiddie |
| EPI_ISL_582642, EPI_ISL_582643, EPI_ISL_582644, EPI_ISL_582645, EPI_ISL_582646, EPI_ISL_582647, EPI_ISL_582648, EPI_ISL_582649, EPI_ISL_582650, EPI_ISL_582651, EPI_ISL_582652, EPI_ISL_582653, EPI_ISL_582654, EPI_ISL_582655, EPI_ISL_582656, EPI_ISL_582657 | | | |
| see above | Sheikh Khalifa Medical City | Molecular/Surveillance lab Sheikh Khalifa Medical City | Amirtharaj Francis, Sajeed Abdul, Hala Imambaccus, Sahar Almarzooqi, Hiba Saud, Stefan Weber |
| EPI_ISL_582808 | Klinisk Mikrobiologi | The Public Health Agency of Sweden | Anna-Malin Linde, Maria Lind Karlberg, Mattias Haukland, Reza Advani, Olov Svartstrom, Oskar Karlsson Lindsjo, Sandra Broddesson, Petra Edquist, Mia Brytting, Anna Risberg, Karin Tegmark-Wisell |
| EPI_ISL_582821, EPI_ISL_582822, EPI_ISL_582823, EPI_ISL_582824, EPI_ISL_582825, EPI_ISL_582826, EPI_ISL_582827, EPI_ISL_582828, EPI_ISL_582829, EPI_ISL_582830, EPI_ISL_582831 | | | |
| see above | Hospital General Universitario Gregorio Marañón | SeqCOVID-SPAIN consortium/IBV(CSIC) | Darío García de Viedma, Laura Pérez-Lago, Marta Herranz, Jon Sicilia, Julia Suárez, Pilar Catalán, Patricia Muñoz and SeqCOVID-SPAIN consortium |
| EPI_ISL_582841 | Orebro klinisk mikrobiologi | The Public Health Agency of Sweden | Anna-Malin Linde, Maria Lind Karlberg, Mattias Haukland, Reza Advani, Olov Svartstrom, Oskar Karlsson Lindsjo, Sandra Broddesson, Petra Edquist, Mia Brytting, Anna Risberg, Karin Tegmark-Wisell |
| EPI_ISL_582844 | Hospital General Universitario Gregorio Marañón | SeqCOVID-SPAIN consortium/IBV(CSIC) | Darío García de Viedma, Laura Pérez-Lago, Marta Herranz, Jon Sicilia, Julia Suárez, Pilar Catalán, Patricia Muñoz and SeqCOVID-SPAIN consortium |
| EPI_ISL_582890, EPI_ISL_582891, EPI_ISL_582892, EPI_ISL_582893, EPI_ISL_582894, EPI_ISL_582895, EPI_ISL_582896, EPI_ISL_582920, EPI_ISL_582929, EPI_ISL_582939, EPI_ISL_582942, EPI_ISL_582957, EPI_ISL_582958 | | | |
| see above | County of Santa Clara Public Health Department | Chan-Zuckerberg Biohub | CZB Cliahub Consortium |
| EPI_ISL_582999, EPI_ISL_583000, EPI_ISL_583001, EPI_ISL_583002, EPI_ISL_583003, EPI_ISL_583004, EPI_ISL_583005, EPI_ISL_583006, EPI_ISL_583007, EPI_ISL_583008, EPI_ISL_583009, EPI_ISL_583010 | | | |
| see above | Orange County Public Health Lab | Chan-Zuckerberg Biohub | CZB Cliahub Consortium |
| EPI_ISL_583133, EPI_ISL_583134, EPI_ISL_583135, EPI_ISL_583136, EPI_ISL_583137, EPI_ISL_583138, EPI_ISL_583139, EPI_ISL_583140, EPI_ISL_583141, EPI_ISL_583142, EPI_ISL_583143, EPI_ISL_583144, EPI_ISL_583145, EPI_ISL_583146, EPI_ISL_583147, EPI_ISL_583148, EPI_ISL_583149, EPI_ISL_583150, EPI_ISL_583151, EPI_ISL_583152, EPI_ISL_583153, EPI_ISL_583154, EPI_ISL_583155, EPI_ISL_583156, EPI_ISL_583157, EPI_ISL_583158, EPI_ISL_583159, EPI_ISL_583160 | | | |
| see above | Humboldt County Public Health Laboratory | Chan-Zuckerberg Biohub | CZB Cliahub Consortium |
| EPI_ISL_583199, EPI_ISL_583200, EPI_ISL_583201, EPI_ISL_583202, EPI_ISL_583203, EPI_ISL_583208, EPI_ISL_583209, EPI_ISL_583225 | UCSF Clinical Microbiology Laboratory | Chan-Zuckerberg Biohub | CZB Cliahub Consortium |
| EPI_ISL_583249, EPI_ISL_583250, EPI_ISL_583251, EPI_ISL_583252, EPI_ISL_583253, EPI_ISL_583254, EPI_ISL_583255, EPI_ISL_583256, EPI_ISL_583257, EPI_ISL_583259, EPI_ISL_583260, EPI_ISL_583261, EPI_ISL_583262, EPI_ISL_583263, EPI_ISL_583265, EPI_ISL_583266, EPI_ISL_583267, EPI_ISL_583268, EPI_ISL_583269, EPI_ISL_583270, EPI_ISL_583271, EPI_ISL_583272, EPI_ISL_583273, EPI_ISL_583274, EPI_ISL_583276, EPI_ISL_583277, EPI_ISL_583278, EPI_ISL_583279, EPI_ISL_583280, EPI_ISL_583281, EPI_ISL_583282, EPI_ISL_583283, EPI_ISL_583284, EPI_ISL_583285, EPI_ISL_583286, EPI_ISL_583287, EPI_ISL_583288, EPI_ISL_583289, EPI_ISL_583291, EPI_ISL_583294, EPI_ISL_583295, EPI_ISL_583296, EPI_ISL_583298, EPI_ISL_583299, EPI_ISL_583302, EPI_ISL_583305, EPI_ISL_583308, EPI_ISL_583309, EPI_ISL_583311, EPI_ISL_583314, EPI_ISL_583315 | | | |
| see above | University of Michigan Clinical Microbiology Laboratory | Lauring Lab, University of Michigan, Department of Microbiology and Immunology | Valesano |
| EPI_ISL_583506 | Michigan Department of Health and Human Services, Bureau of Laboratories | Michigan Department of Health and Human Services, Bureau of Laboratories | Blankenship HM, Riner D, Soehnlen MK |
| EPI_ISL_583537, EPI_ISL_583538, EPI_ISL_583539, EPI_ISL_583540, EPI_ISL_583541, EPI_ISL_583542 | Genome Centre | Genome Centre | Selina Akter, Pravas Chandra Roy, Amina Ferdaus manami, Habiba Ibnat, A. S. M. Rubayet Ul Alam, Shireen Nigar, Iqbal Kabir Jahid, M.Anwar Hossain |
| EPI_ISL_583894, EPI_ISL_583895 | Singapore General Hospital | Department of Microbiology | Nurdyana Abdul Rahman, Kun Lee Lim, Chenhao Li, Sui Sin Goh, Kenneth Xin Long Chan, Kian Sing Chan, Lynette Oon, Kern Rei Chng, Niranjan Nagarajan, Karrie Ko |
| EPI_ISL_584095 | University of Michigan Clinical Microbiology Laboratory | Lauring Lab, University of Michigan, Department of Microbiology and Immunology | Valesano |
| EPI_ISL_584673 | University College London, Great Ormond Street Hospital for Children NHS Foundation Trust, Imperial College Healthcare NHS Trust | COVID-19 Genomics UK (COG-UK) Consortium | Sergi Castellano, Rachel Williams, Mark Kristiansen, Paola Resende Silva, Sunando Roy, Tony Brooks, Helena Tutill, Paola Niola, Patricia Dyal, Charlotte Williams, Leysa Forrest, Yasmin Panchbhaya, Jacqueline Findlay, Samuel Weeks, Julianne Brown, Kathryn Harris, Paul Randell, James Price, Alison Holmes, Judith Breuer |

| | | | |
|---|---|---|---|
| EPI_ISL_585223, EPI_ISL_585224, EPI_ISL_585225, EPI_ISL_585227, EPI_ISL_585228, EPI_ISL_585229, EPI_ISL_585230, EPI_ISL_585232, EPI_ISL_585234, EPI_ISL_585236, EPI_ISL_585237, EPI_ISL_585238, EPI_ISL_585239, EPI_ISL_585240, EPI_ISL_585241, EPI_ISL_585242, EPI_ISL_585244, EPI_ISL_585245 | | | |
| see above | Regional Virus Laboratory, Belfast Health and Social Care Trust | COVID-19 Genomics UK (COG-UK) Consortium | Conall McCaughey, James McKenna, Tanya Curran, Susan Feeney, Alison Watt, Ciara Cox, Mairead Connor, Zoltan Molnar, David Simpson, Derek Fairley |
| EPI_ISL_586362, EPI_ISL_586365 | Toronto Invasive Bacterial Diseases Network | McMaster University | Allison McGeer, Patryk Aftanas, Hooman Derakhshani, Angel Li, Kuganya Nirmalarajah, Emily Panousis, Ahmed Draia, Jalees Nasir, Michael Surette, Samira Mubareka, Andrew G. McArthur |
| EPI_ISL_588539 | Lighthouse Lab in Glasgow | Wellcome Sanger Institute for the COVID-19 Genomics UK (COG-UK) consortium | Harper VanSteenhouse, Yumi Kasai, David Gray, Carol Clugston, Anna Dominiczak and Alex Alderton, Roberto Amato, Sonia Goncalves, Ewan Harrison, David K. Jackson, Ian Johnston, Dominic Kwiatkowski, Cordelia Langford, John Sillitoe on behalf of the Wellcome Sanger Institute COVID-19 Surveillance Team |
| EPI_ISL_590181, EPI_ISL_590186, EPI_ISL_590193, EPI_ISL_590194, EPI_ISL_590198, EPI_ISL_590200, EPI_ISL_590206, EPI_ISL_590210, EPI_ISL_590213, EPI_ISL_590214, EPI_ISL_590215, EPI_ISL_590220, EPI_ISL_590224, EPI_ISL_590225, EPI_ISL_590226, EPI_ISL_590228, EPI_ISL_590234, EPI_ISL_590235, EPI_ISL_590239, EPI_ISL_590246, EPI_ISL_590250, EPI_ISL_590255, EPI_ISL_590257, EPI_ISL_590259, EPI_ISL_590260, EPI_ISL_590261, EPI_ISL_590263, EPI_ISL_590265, EPI_ISL_590272, EPI_ISL_590274, EPI_ISL_590275, EPI_ISL_590276, EPI_ISL_590277, EPI_ISL_590282, EPI_ISL_590283, EPI_ISL_590284, EPI_ISL_590286, EPI_ISL_590288, EPI_ISL_590295, EPI_ISL_590296, EPI_ISL_590300, EPI_ISL_590306, EPI_ISL_590307, EPI_ISL_590309, EPI_ISL_590314, EPI_ISL_590319, EPI_ISL_590321, EPI_ISL_590327, EPI_ISL_590328, EPI_ISL_590329, EPI_ISL_590335, EPI_ISL_590337, EPI_ISL_590346, EPI_ISL_590347, EPI_ISL_590351, EPI_ISL_590352, EPI_ISL_590355, EPI_ISL_590357, EPI_ISL_590359, EPI_ISL_590361, EPI_ISL_590364, EPI_ISL_590366, EPI_ISL_590367, EPI_ISL_590369, EPI_ISL_590373, EPI_ISL_590379, EPI_ISL_590380, EPI_ISL_590383, EPI_ISL_590386, EPI_ISL_590388, EPI_ISL_590390, EPI_ISL_590392, EPI_ISL_590397, EPI_ISL_590398, EPI_ISL_590399, EPI_ISL_590403, EPI_ISL_590405, EPI_ISL_590419, EPI_ISL_590422, EPI_ISL_590423, EPI_ISL_590438, EPI_ISL_590441, EPI_ISL_590442, EPI_ISL_590446, EPI_ISL_590448, EPI_ISL_590452, EPI_ISL_590453, EPI_ISL_590462 | | | |
| see above | Lighthouse Lab in Alderley Park | Wellcome Sanger Institute for the COVID-19 Genomics UK (COG-UK) consortium | Jacquelyn Wynn, Mairead Hyland, The Lighthouse Lab in Alderley Park and Alex Alderton, Roberto Amato, Sonia Goncalves, Ewan Harrison, David K. Jackson, Ian Johnston, Dominic Kwiatkowski, Cordelia Langford, John Sillitoe on behalf of the Wellcome Sanger Institute COVID-19 Surveillance Team (http://www.sanger.ac.uk/covid-team) |
| EPI_ISL_590885, EPI_ISL_590886 | Vestfold Hospital, Toensberg Department of Microbiology | Norwegian Institute of Public Health, Department of Virology | Kathrine Stene-Johansen, Kamilla Heddeland Instefjord, Hilde Elshaug, Rasmus Riis Kopperud, Hilde Vollan, Karoline Bragstad, Olav Hungnes |
| EPI_ISL_590904, EPI_ISL_590905, EPI_ISL_590906 | Hospital of Southern Norway - Kristiansand, Department of Medical Microbiology | Norwegian Institute of Public Health, Department of Virology | Kathrine Stene-Johansen, Kamilla Heddeland Instefjord, Hilde Elshaug, Rasmus Riis Kopperud, Hilde Vollan, Karoline Bragstad, Olav Hungnes |
| EPI_ISL_590908, EPI_ISL_590909, EPI_ISL_590910, EPI_ISL_590913 | Oslo University Hospital, Department of Medical Microbiology | Norwegian Institute of Public Health, Department of Virology | Kathrine Stene-Johansen, Kamilla Heddeland Instefjord, Hilde Elshaug, Rasmus Riis Kopperud, Hilde Vollan, Karoline Bragstad, Olav Hungnes |
| EPI_ISL_590988, EPI_ISL_590992, EPI_ISL_590993, EPI_ISL_590994, EPI_ISL_590997, EPI_ISL_591016, EPI_ISL_591017 | Ostfold Hospital Trust - Kalnes, Centre for Laboratory Medicine, Section for gene technology and infection serology | Norwegian Institute of Public Health, Department of Virology | Kathrine Stene-Johansen, Kamilla Heddeland Instefjord, Hilde Elshaug, Rasmus Riis Kopperud, Hilde Vollan, Karoline Bragstad, Olav Hungnes |
| EPI_ISL_591100, EPI_ISL_591101, EPI_ISL_591102, EPI_ISL_591103, EPI_ISL_591104, EPI_ISL_591105, EPI_ISL_591106, EPI_ISL_591107, EPI_ISL_591108, EPI_ISL_591109, EPI_ISL_591110, EPI_ISL_591111, EPI_ISL_591112, EPI_ISL_591113, EPI_ISL_591114, EPI_ISL_591115, EPI_ISL_591116, EPI_ISL_591117, EPI_ISL_591185, EPI_ISL_591186, EPI_ISL_591187, EPI_ISL_591188, EPI_ISL_591189, EPI_ISL_591190, EPI_ISL_591191, EPI_ISL_591192, EPI_ISL_591193, EPI_ISL_591194, EPI_ISL_591195, EPI_ISL_591196, EPI_ISL_591197, EPI_ISL_591198, EPI_ISL_591199, EPI_ISL_591200, EPI_ISL_591201, EPI_ISL_591202, EPI_ISL_591203, EPI_ISL_591204, EPI_ISL_591205, EPI_ISL_591206, EPI_ISL_591207 | | | |
| see above | Toronto Invasive Bacterial Diseases Network | McMaster University | Allison McGeer, Patryk Aftanas, Hooman Derakhshani, Angel Li, Kuganya Nirmalarajah, Emily Panousis, Ahmed Draia, Jalees Nasir, Michael Surette, Samira Mubareka, Andrew G. McArthur |
| EPI_ISL_591531, EPI_ISL_591532, EPI_ISL_591533, EPI_ISL_591534 | Laboratorio de Infectologia y virologia molecular | Center for Mathematical Modeling and Center for Genome Regulation. Santiago, Chile | Valiente F, Gaete A, Travisany D, Palma R, Urra C, Varas M, Allende ML, Maass A, González M, Ferres M. |
| EPI_ISL_591557, EPI_ISL_591577, EPI_ISL_591607, EPI_ISL_591632, EPI_ISL_591637, EPI_ISL_591680, EPI_ISL_591702, EPI_ISL_591711, EPI_ISL_591730 | Microbiological Diagnostic Unit - Public Health Laboratory (MDU-PHL) | MDU-PHL | Seemann T., Schultz, M. B., Sait, M., Sherry, N. |
| EPI_ISL_591782 | Victorian Infectious Diseases Reference Laboratory (VIDRL) | VIDRL and MDU-PHL | Caly L., Seemann T., Sait, M., Schultz, M. B., Druce J., Sherry, N. |
| EPI_ISL_591794, EPI_ISL_591802 | Microbiological Diagnostic Unit - Public Health Laboratory (MDU-PHL) | MDU-PHL | Seemann T., Schultz, M. B., Sait, M., Sherry, N. |
| EPI_ISL_591807 | Victorian Infectious Diseases Reference Laboratory (VIDRL) | VIDRL and MDU-PHL | Caly L., Seemann T., Sait, M., Schultz, M. B., Druce J., Sherry, N. |
| EPI_ISL_591816, EPI_ISL_591817, EPI_ISL_592278, EPI_ISL_592289, EPI_ISL_592308, EPI_ISL_592325, EPI_ISL_592326, EPI_ISL_592327, EPI_ISL_592328, EPI_ISL_592330, EPI_ISL_592331, EPI_ISL_592333, EPI_ISL_592335, EPI_ISL_592336, EPI_ISL_592338, EPI_ISL_592339, EPI_ISL_592340, EPI_ISL_592378, EPI_ISL_592446, EPI_ISL_592483, EPI_ISL_592484 | | | |
| see above | Microbiological Diagnostic Unit - Public Health Laboratory (MDU-PHL) | MDU-PHL | Seemann T., Schultz, M. B., Sait, M., Sherry, N. |
| EPI_ISL_592489, EPI_ISL_592498, EPI_ISL_592515 | Victorian Infectious Diseases Reference Laboratory (VIDRL) | VIDRL and MDU-PHL | Caly L., Seemann T., Sait, M., Schultz, M. B., Druce J., Sherry, N. |
| EPI_ISL_592536 | Microbiological Diagnostic Unit - Public Health Laboratory (MDU-PHL) | MDU-PHL | Seemann T., Schultz, M. B., Sait, M., Sherry, N. |
| EPI_ISL_592537 | Victorian Infectious Diseases Reference Laboratory (VIDRL) | VIDRL and MDU-PHL | Caly L., Seemann T., Sait, M., Schultz, M. B., Druce J., Sherry, N. |
| EPI_ISL_592551, EPI_ISL_592558, EPI_ISL_592562 | Microbiological Diagnostic Unit - Public Health Laboratory (MDU-PHL) | MDU-PHL | Seemann T., Schultz, M. B., Sait, M., Sherry, N. |
| EPI_ISL_592564, EPI_ISL_592568 | Victorian Infectious Diseases Reference Laboratory (VIDRL) | VIDRL and MDU-PHL | Caly L., Seemann T., Sait, M., Schultz, M. B., Druce J., Sherry, N. |
| EPI_ISL_592572 | Microbiological Diagnostic Unit - Public Health Laboratory (MDU-PHL) | MDU-PHL | Seemann T., Schultz, M. B., Sait, M., Sherry, N. |
| EPI_ISL_592577 | Victorian Infectious Diseases Reference Laboratory (VIDRL) | VIDRL and MDU-PHL | Caly L., Seemann T., Sait, M., Schultz, M. B., Druce J., Sherry, N. |
| EPI_ISL_592582, EPI_ISL_592586, EPI_ISL_592589 | Microbiological Diagnostic Unit - Public Health Laboratory (MDU-PHL) | MDU-PHL | Seemann T., Schultz, M. B., Sait, M., Sherry, N. |
| EPI_ISL_592597 | Victorian Infectious Diseases Reference Laboratory (VIDRL) | VIDRL and MDU-PHL | Caly L., Seemann T., Sait, M., Schultz, M. B., Druce J., Sherry, N. |
| EPI_ISL_592620, EPI_ISL_592625 | Microbiological Diagnostic Unit - Public Health Laboratory (MDU-PHL) | MDU-PHL | Seemann T., Schultz, M. B., Sait, M., Sherry, N. |
| EPI_ISL_592660, EPI_ISL_592661 | Victorian Infectious Diseases Reference Laboratory (VIDRL) | VIDRL and MDU-PHL | Caly L., Seemann T., Sait, M., Schultz, M. B., Druce J., Sherry, N. |
| EPI_ISL_592665, EPI_ISL_592677, EPI_ISL_592679, EPI_ISL_592682, EPI_ISL_592688, EPI_ISL_592707, EPI_ISL_592714, EPI_ISL_592718 | Microbiological Diagnostic Unit - Public Health Laboratory (MDU-PHL) | MDU-PHL | Seemann T., Schultz, M. B., Sait, M., Sherry, N. |
| EPI_ISL_592720 | Victorian Infectious Diseases Reference Laboratory (VIDRL) | VIDRL and MDU-PHL | Caly L., Seemann T., Sait, M., Schultz, M. B., Druce J., Sherry, N. |
| EPI_ISL_592736, EPI_ISL_592746, EPI_ISL_592748, EPI_ISL_592777, EPI_ISL_592779 | Microbiological Diagnostic Unit - Public Health Laboratory (MDU-PHL) | MDU-PHL | Seemann T., Schultz, M. B., Sait, M., Sherry, N. |
| EPI_ISL_592781 | Victorian Infectious Diseases Reference Laboratory (VIDRL) | VIDRL and MDU-PHL | Caly L., Seemann T., Sait, M., Schultz, M. B., Druce J., Sherry, N. |

| | | | |
|---|---|---|---|
| EPI_ISL_592784, EPI_ISL_592785 | Microbiological Diagnostic Unit - Public Health Laboratory (MDU-PHL) | MDU-PHL | Seemann T., Schultz, M. B., Sait, M., Sherry, N. |
| EPI_ISL_592790 | Victorian Infectious Diseases Reference Laboratory (VIDRL) | VIDRL and MDU-PHL | Caly L., Seemann T., Sait, M., Schultz, M. B., Druce J., Sherry, N. |
| EPI_ISL_592799, EPI_ISL_592803, EPI_ISL_592807, EPI_ISL_592813 | Microbiological Diagnostic Unit - Public Health Laboratory (MDU-PHL) | MDU-PHL | Seemann T., Schultz, M. B., Sait, M., Sherry, N. |
| EPI_ISL_592814 | Victorian Infectious Diseases Reference Laboratory (VIDRL) | VIDRL and MDU-PHL | Caly L., Seemann T., Sait, M., Schultz, M. B., Druce J., Sherry, N. |
| EPI_ISL_592822, EPI_ISL_592823 | Microbiological Diagnostic Unit - Public Health Laboratory (MDU-PHL) | MDU-PHL | Seemann T., Schultz, M. B., Sait, M., Sherry, N. |
| EPI_ISL_592824 | Victorian Infectious Diseases Reference Laboratory (VIDRL) | VIDRL and MDU-PHL | Caly L., Seemann T., Sait, M., Schultz, M. B., Druce J., Sherry, N. |
| EPI_ISL_592827 | Microbiological Diagnostic Unit - Public Health Laboratory (MDU-PHL) | MDU-PHL | Seemann T., Schultz, M. B., Sait, M., Sherry, N. |
| EPI_ISL_592828 | Victorian Infectious Diseases Reference Laboratory (VIDRL) | VIDRL and MDU-PHL | Caly L., Seemann T., Sait, M., Schultz, M. B., Druce J., Sherry, N. |
| EPI_ISL_592829, EPI_ISL_592830, EPI_ISL_592833, EPI_ISL_592834, EPI_ISL_592841, EPI_ISL_592843, EPI_ISL_592845, EPI_ISL_592846, EPI_ISL_592847, EPI_ISL_592848, EPI_ISL_592849, EPI_ISL_592850, EPI_ISL_592851, EPI_ISL_592866, EPI_ISL_592883, EPI_ISL_592884, EPI_ISL_592893, EPI_ISL_592897, EPI_ISL_592901, EPI_ISL_592904, EPI_ISL_592905, EPI_ISL_592906 | | | |
| see above | Microbiological Diagnostic Unit - Public Health Laboratory (MDU-PHL) | MDU-PHL | Seemann T., Schultz, M. B., Sait, M., Sherry, N. |
| EPI_ISL_592970, EPI_ISL_592973, EPI_ISL_592976, EPI_ISL_592985, EPI_ISL_593007 | Victorian Infectious Diseases Reference Laboratory (VIDRL) | VIDRL and MDU-PHL | Caly L., Seemann T., Sait, M., Schultz, M. B., Druce J., Sherry, N. |
| EPI_ISL_593021, EPI_ISL_593027 | Microbiological Diagnostic Unit - Public Health Laboratory (MDU-PHL) | MDU-PHL | Seemann T., Schultz, M. B., Sait, M., Sherry, N. |
| EPI_ISL_593033, EPI_ISL_593035 | Victorian Infectious Diseases Reference Laboratory (VIDRL) | VIDRL and MDU-PHL | Caly L., Seemann T., Sait, M., Schultz, M. B., Druce J., Sherry, N. |
| EPI_ISL_593036, EPI_ISL_593045, EPI_ISL_593047, EPI_ISL_593049, EPI_ISL_593051, EPI_ISL_593068, EPI_ISL_593072, EPI_ISL_593080, EPI_ISL_593083, EPI_ISL_593089, EPI_ISL_593093, EPI_ISL_593105, EPI_ISL_593108, EPI_ISL_593112, EPI_ISL_593113, EPI_ISL_593118 | | | |
| see above | Microbiological Diagnostic Unit - Public Health Laboratory (MDU-PHL) | MDU-PHL | Seemann T., Schultz, M. B., Sait, M., Sherry, N. |
| EPI_ISL_593126 | Victorian Infectious Diseases Reference Laboratory (VIDRL) | VIDRL and MDU-PHL | Caly L., Seemann T., Sait, M., Schultz, M. B., Druce J., Sherry, N. |
| EPI_ISL_593131, EPI_ISL_593134, EPI_ISL_593135, EPI_ISL_593137, EPI_ISL_593148, EPI_ISL_593149, EPI_ISL_593151, EPI_ISL_593153, EPI_ISL_593157, EPI_ISL_593158, EPI_ISL_593159, EPI_ISL_593160, EPI_ISL_593165, EPI_ISL_593166, EPI_ISL_593173, EPI_ISL_593174, EPI_ISL_593178, EPI_ISL_593182, EPI_ISL_593186 | | | |
| see above | Microbiological Diagnostic Unit - Public Health Laboratory (MDU-PHL) | MDU-PHL | Seemann T., Schultz, M. B., Sait, M., Sherry, N. |
| EPI_ISL_593215, EPI_ISL_593216 | Victorian Infectious Diseases Reference Laboratory (VIDRL) | VIDRL and MDU-PHL | Caly L., Seemann T., Sait, M., Schultz, M. B., Druce J., Sherry, N. |
| EPI_ISL_593217, EPI_ISL_593219, EPI_ISL_593221, EPI_ISL_593223, EPI_ISL_593224, EPI_ISL_593226, EPI_ISL_593229, EPI_ISL_593234, EPI_ISL_593237, EPI_ISL_593242, EPI_ISL_593244, EPI_ISL_593257 | | | |
| see above | Microbiological Diagnostic Unit - Public Health Laboratory (MDU-PHL) | MDU-PHL | Seemann T., Schultz, M. B., Sait, M., Sherry, N. |
| EPI_ISL_593556 | Brigham and Women's Hospital | Jonathan Li Laboratory | Manish C. Choudhary, James Regan, Jonathan Z. Li |
| EPI_ISL_593640, EPI_ISL_593641, EPI_ISL_593642, EPI_ISL_593643, EPI_ISL_593645 | unknown | Public Health Virology Laboratory, Forensic and Scientific Services (PHV-FSS) | Son Nguyen et al. |
| EPI_ISL_593647, EPI_ISL_593648 | 4Cyte Pathology | NSW Health Pathology - Institute of Clinical Pathology and Medical Research; Westmead Hospital; University of Sydney | CIDM-PH et al. |
| EPI_ISL_593652 | Histopath | NSW Health Pathology - Institute of Clinical Pathology and Medical Research; Westmead Hospital; University of Sydney | CIDM-PH et al. |
| EPI_ISL_593658 | Pathology North - Hunter - NSW Health Pathology | NSW Health Pathology - Institute of Clinical Pathology and Medical Research; Westmead Hospital; University of Sydney | CIDM-PH et al. |
| EPI_ISL_593682 | Pathology West - NSW Health Pathology | NSW Health Pathology - Institute of Clinical Pathology and Medical Research; Westmead Hospital; University of Sydney | CIDM-PH et al. |
| EPI_ISL_593713, EPI_ISL_593714, EPI_ISL_593715, EPI_ISL_593716, EPI_ISL_593717, EPI_ISL_593718, EPI_ISL_593719, EPI_ISL_593725 | South Eastern Area Laboratory Services (SEALS) | NSW Health Pathology - Institute of Clinical Pathology and Medical Research; Westmead Hospital; University of Sydney | CIDM-PH et al. |
| EPI_ISL_593763 | Sydney South West Pathology Service (SSWPS) - Liverpool Hospital - NSW Health Pathology | NSW Health Pathology - Institute of Clinical Pathology and Medical Research; Westmead Hospital; University of Sydney | CIDM-PH et al. |
| EPI_ISL_593882, EPI_ISL_593887, EPI_ISL_593889 | CHU Purpan - Laboratoire de Virologie - Institut Fédératif de Biologie | CHU Purpan - Laboratoire de Virologie - Institut Fédératif de Biologie | Latour J., Ranger N., Dubois M., Carcenac R., Harter A., Boyer P., Tremeaux P., Izopet J. |
| EPI_ISL_593912 | Hospital, Marange Silvange | National Reference Center for Viruses of Respiratory Infections, Institut Pasteur, Paris | Sylvie Behillil, Fabiana Gambaro, Etienne Simon-Lorière, Vincent Enouf, Maud Vanpeene, Sylvie van der Werf |
| EPI_ISL_593917 | Hospital, Montigny les Metz | National Reference Center for Viruses of Respiratory Infections, Institut Pasteur, Paris | Sylvie Behillil, Fabiana Gambaro, Etienne Simon-Lorière, Vincent Enouf, Maud Vanpeene, Sylvie van der Werf |
| EPI_ISL_593931, EPI_ISL_593932 | Labo Analyses Med, Puteaux | National Reference Center for Viruses of Respiratory Infections, Institut Pasteur, Paris | Sylvie Behillil, Fabiana Gambaro, Etienne Simon-Lorière, Vincent Enouf, Maud Vanpeene, Sylvie van der Werf |
| EPI_ISL_594145 | MDU-PHL, The Peter Doherty Institute for Infection and Immunity | MDU-PHL, The Peter Doherty Institute for Infection and Immunity | Caly,L., Seemann,T., Sait,M.L., Schultz,M.B., Druce,J., Sherry,N.L. |
| EPI_ISL_594163, EPI_ISL_594164 | hopital | National Reference Center for Viruses of Respiratory Infections, Institut Pasteur, Paris | Sylvie Behillil, Fabiana Gambaro, Etienne Simon-Lorière, Vincent Enouf, Maud Vanpeene, Sylvie van der Werf |
| EPI_ISL_594167, EPI_ISL_594168, EPI_ISL_594169, EPI_ISL_594170, EPI_ISL_594171, EPI_ISL_594172, EPI_ISL_594173, EPI_ISL_594174 | PathWest Laboratory Medicine WA | PathWest Laboratory Medicine WA Microbial Surveillance Unit | PathWest Laboratory Medicine WA Microbial Surveillance Unit |
| EPI_ISL_594387, EPI_ISL_594388, EPI_ISL_594390, EPI_ISL_594391, | Florida Bureau of Public Health Laboratories | Florida Bureau of Public Health Laboratories | Sarah Schmedes, Jason Blanton |

| | | | |
|---|---|---|---|
| EPI_ISL_594392, EPI_ISL_594393 | | | |
| EPI_ISL_596229, EPI_ISL_596230 | WHO National Influenza Centre Russian Federation | WHO National Influenza Centre Russian Federation | Andrey Komissarov, Artem Fadeev, Anna Ivanova, Kseniya Komissarova, Dmitry Bazhenov, Daria Danilenko |
| EPI_ISL_596569 | University of Michigan Clinical Microbiology Laboratory | Lauring Lab, University of Michigan, Department of Microbiology and Immunology | Valesano |
| EPI_ISL_596691, EPI_ISL_596694, EPI_ISL_596703 | PathWest Laboratory Medicine WA | PathWest Laboratory Medicine WA Microbial Surveillance Unit | PathWest Laboratory Medicine WA Microbial Surveillance Unit |
| EPI_ISL_602146, EPI_ISL_602147, EPI_ISL_602148, EPI_ISL_602149, EPI_ISL_602150, EPI_ISL_602151, EPI_ISL_602152, EPI_ISL_602153 | Lighthouse Lab in Milton Keynes | Wellcome Sanger Institute for the COVID-19 Genomics UK (COG-UK) consortium | The Lighthouse Lab in Milton Keynes and Alex Alderton, Roberto Amato, Sonia Goncalves, Ewan Harrison, David K. Jackson, Ian Johnston, Dominic Kwiatkowski, Cordelia Langford, John Sillitoe on behalf of the Wellcome Sanger Institute COVID-19 Surveillance Team (http://www.sanger.ac.uk/covid-team) |
| EPI_ISL_602154 | Lighthouse Lab in Cambridge | Wellcome Sanger Institute for the COVID-19 Genomics UK (COG-UK) consortium | Rob Howes, The Lighthouse Lab in Cambridge and Alex Alderton, Roberto Amato, Sonia Goncalves, Ewan Harrison, David K. Jackson, Ian Johnston, Dominic Kwiatkowski, Cordelia Langford, John Sillitoe on behalf of the Wellcome Sanger Institute COVID-19 Surveillance Team (http://www.sanger.ac.uk/covid-team) |
| EPI_ISL_602162, EPI_ISL_602163, EPI_ISL_602164, EPI_ISL_602165, EPI_ISL_602166, EPI_ISL_602167, EPI_ISL_602168, EPI_ISL_602169, EPI_ISL_602170, EPI_ISL_602171, EPI_ISL_602172, EPI_ISL_602173, EPI_ISL_602174, EPI_ISL_602175, EPI_ISL_602176, EPI_ISL_602177, EPI_ISL_602178, EPI_ISL_602179, EPI_ISL_602180, EPI_ISL_602181, EPI_ISL_602182, EPI_ISL_602183, EPI_ISL_602184, EPI_ISL_602185, EPI_ISL_602186, EPI_ISL_602187, EPI_ISL_602188, EPI_ISL_602189, EPI_ISL_602190, EPI_ISL_602191, EPI_ISL_602192, EPI_ISL_602193, EPI_ISL_602194, EPI_ISL_602195, EPI_ISL_602196, EPI_ISL_602197, EPI_ISL_602198, EPI_ISL_602199, EPI_ISL_602200, EPI_ISL_602201, EPI_ISL_602202, EPI_ISL_602203, EPI_ISL_602204, EPI_ISL_602205, EPI_ISL_602206, EPI_ISL_602207 | | | |
| see above | New Mexico Department of Health Scientific Laboratory | New Mexico Department of Health Scientific Laboratory | Ellie Johnson, Anastacia Griego-Fisher, D'Eldra Malone |
| EPI_ISL_602306, EPI_ISL_602313, EPI_ISL_602320 | University of Miami Immunology and Histocompatibility Laboratory | University of Miami Immunology and Histocompatibility Laboratory | Emilio Margolles-Clark, PhD and Phillip Ruiz, MD, PhD |
| EPI_ISL_602632, EPI_ISL_602633, EPI_ISL_602634, EPI_ISL_602637, EPI_ISL_602640, EPI_ISL_602641, EPI_ISL_602644, EPI_ISL_602645, EPI_ISL_602650, EPI_ISL_602654, EPI_ISL_602659, EPI_ISL_602660, EPI_ISL_602661, EPI_ISL_602666, EPI_ISL_602667, EPI_ISL_602668, EPI_ISL_602669, EPI_ISL_602670, EPI_ISL_602671, EPI_ISL_602672, EPI_ISL_602673, EPI_ISL_602674, EPI_ISL_602675, EPI_ISL_602676, EPI_ISL_602677, EPI_ISL_602678, EPI_ISL_602679, EPI_ISL_602680, EPI_ISL_602681 | | | |
| see above | NHLS-IALCH | KRISP, KZN Research Innovation and Sequencing Platform | Giandhari J, Pillay S, Lessells R, Mdlalose K, York D, Khan S, Tegally H, Wilkinson E, de Oliveira T |
| EPI_ISL_609993 | INMI Lazzaro Spallanzani IRCCS | INMI Lazzaro Spallanzani IRCCS | E Giombini, M Rueca, B Bartolini, C.E.M Gruber, F Messina, A Di Caro, MR Capobianchi |
| EPI_ISL_610158 | RS Panti Rini | Genetics Working Group (Pokja Genetik) Faculty of Medicine, Public Health and Nursing Universitas Gadjah Mada (FK-KMK UGM); Disease Investigation Center Wates Ministry of Agriculture Indonesia; Department of Microbiology FK-KMK UGM; Laboratorium Diagnostik Yayasan Tahija World Mosquito Program (WMP) Yogyakarta Center for Tropical Medicine FK-KMK UGM; Integrated Research Center FK-KMK UGM; Department of Computer Science and Electronics FMIPA UGM | Gunadi, Hendra Wibawa, Marcellus, Mohamad S. Hakim, Edwin W. Daniwijaya, Ludhang P. Rizki, Endah Supriyati, Eggi Arguni, Titik Nuryastuti, Tri Wibawa, Dwi AA Nugrahaningsih, Afiahayati, Siswanto, Kristy Iskandar, Nungki Anggorowati, V. Noegroho Isti Donodjati, Wawan Triharjanto, Dwiki Afandy, Dyah Ayu Puspitarani |
| EPI_ISL_610162 | RSUD Dr. Tjitrowardojo | Genetics Working Group (Pokja Genetik) Faculty of Medicine, Public Health and Nursing Universitas Gadjah Mada (FK-KMK UGM); Disease Investigation Center Wates Ministry of Agriculture Indonesia; Department of Microbiology FK-KMK UGM; Laboratorium Diagnostik Yayasan Tahija World Mosquito Program (WMP) Yogyakarta Center for Tropical Medicine FK-KMK UGM; Integrated Research Center FK-KMK UGM; Department of Computer Science and Electronics FMIPA UGM; Balai Besar Teknik Kesehatan Lingkungan dan Pengendalian Penyakit (BBTKLPP) Yogyakarta | Gunadi, Hendra Wibawa, Marcellus, Mohamad S. Hakim, Edwin W. Daniwijaya, Ludhang P. Rizki, Endah Supriyati, Eggi Arguni, Titik Nuryastuti, Tri Wibawa, Dwi AA Nugrahaningsih, Afiahayati, Siswanto, Kristy Iskandar, Nungki Anggorowati, Irene, Indaryati, Havid Setyawan, Wuryanto, Susan Simanjaya, Alvin Santoso Kalim |
| EPI_ISL_610205 | Department of Health Technology and Informatics, The Hong Kong Polytechnic University | Department of Health Technology and Informatics, The Hong Kong Polytechnic University | Siu,G.K.-H., Lee,L.-K., Leung,K.S.-S., Leung,J.S.-L., Ng,T.T.-L., Chan,C.T.-M., Tam,K.K.-G., Lao,H.-Y., Wu,A.K.-L., Yau,M.C.-Y., Lai,Y.W.-M., Fung,K.S.-C., Chau,S.K.-Y., Wong,B.K.-C., To,W.-K., Luk,K., Ho,A.Y.-M., Que,T.-L., Yip,K.-T., Yam,W.C., Shum,D.H.-K., Yip,S.P. |
| EPI_ISL_610239, EPI_ISL_610240, EPI_ISL_610241, EPI_ISL_610242, EPI_ISL_610243, EPI_ISL_610244 | Molecular diagnostic laboratory of Federal Budget Institution of Science "Central Research Institute of Epidemiology" of The Federal Service on Customers' Rights Protection and Human Well-being Surveillance | Group of Genomics and Postgenomic Technologies of Central Research Institute of Epidemilology | Samoilov AE, Kaptelova VV, Valdokhina AV, Bulanenko VP, Speranskaya AS, Tivanova EV, Shipulina OY, Akimkin VG |
| EPI_ISL_611579 | Liverpool Clinical Laboratories | COVID-19 Genomics UK (COG-UK) Consortium | Sam Haldenby, Anita Lucaci, Steve Paterson, Julian Hiscox, Alistair Darby, M Almsaud, A Alrezaihi, Muhannad Alruwaili, Stuart D Armstrong, Jones Benjamin, Eleanor G Bentley, Anu Chawla, Jordan J Clark, Angela Cowell, Richard Eccles, Isabel García-Dorival, Matthew Gemmell, Alessandro Gerada, PKF Gilmore, Richard Gregory, Ximeng Han, Catherine Hartley, Margaret Hughes, Miren Iturriza-Gomara, James Johnson, L Luu, Jenifer Manson, Charlotte Nelson, Elaine O'Toole, Cassie Olateju, Rebekah Penrice-Randal , Lucille Rainbow, N.P Randle, Trevor Ian Robinson, Parul Sharma, Ghada T Shawli, James P Stewart, Neil Swainston, Ecaterina Vamos, Joanne Watts, Mark Whitehead |
| EPI_ISL_612131, EPI_ISL_612456 | University College London, Great Ormond Street Hospital for Children NHS Foundation Trust, Imperial College Healthcare NHS Trust | COVID-19 Genomics UK (COG-UK) Consortium | Sergi Castellano, Rachel Williams, Mark Kristiansen, Paola Resende Silva, Sunando Roy, Tony Brooks, Helena Tutill, Paola Niola, Patricia Dyal, Charlotte Williams, Leysa Forrest, Yasmin Panchbhaya, Jacqueline Findlay, Samuel Weeks, Julianne Brown, Kathryn Harris, Paul Randell, James Price, Alison Holmes, Judith Breuer |
| EPI_ISL_613467, EPI_ISL_613468, EPI_ISL_613469, EPI_ISL_613470, EPI_ISL_613471, EPI_ISL_613472, EPI_ISL_613473, EPI_ISL_613474, EPI_ISL_613475, EPI_ISL_613476 | Public Health Laboratory - Infectious Disease Lab, Minnesota Department of Health Infectious Disease Laboratory Submission Group | Minnesota Department of Health, Public Health Laboratory | Plumb,M., Garfin,J., Lorentz,A., Wang,X. |
| EPI_ISL_613547, EPI_ISL_613548, EPI_ISL_613550 | CHRU Pontchaillou - Laboratoire de Virologie 2, rue Henri Le Guilloux | National Reference Center for Viruses of Respiratory Infections, Institut Pasteur, Paris | Marion Barbet, Sylvie Behillil, Méline Bizard, Angela Brisebarre, Camille Capel, Etienne Simon-Lorière, Vincent Enouf, Maud Vanpeene, Sylvie van der Werf, Gisèle Lagathu |
| EPI_ISL_614046, EPI_ISL_614047 | Virginia DCLS | Virginia DCLS | Virginia DCLS |
| EPI_ISL_614390, EPI_ISL_614391, EPI_ISL_614392 | Molecular diagnostic unit for viral haemorrhagic fevers and emerging viruses, Bouaké CHU Laboratory | Project group Epidemology of Highly Pathogenic Microorganisms, Robert Koch-Institute | Chantal Akoua-Koffi, Diané Bamourou, Etilé Anoh, Essia Belarbi, Safiatou Karidioula, Grit Schubert, Adjaratou Traoré, Soundélé Maïté, Monemo Pacome, Coulibaly Mbegnan, Bamba Fatoumata Touré, Kra Ouffoué, Fabian Leendertz |
| EPI_ISL_615101, EPI_ISL_615102 | Gavle klinisk mikrobiologi | The Public Health Agency of Sweden | Anna-Malin Linde, Maria Lind Karlberg, Mattias Haukland, Reza Advani, Olov Svartstrom, Oskar Karlsson Lindsjo, Sandra Broddesson, Petra Edquist, Mia Brytting, Anna Risberg, Karin Tegmark-Wisell |
| EPI_ISL_615108 | Klinisk Mikrobiologi | The Public Health Agency of Sweden | Anna-Malin Linde, Maria Lind Karlberg, Mattias Haukland, Reza Advani, Olov Svartstrom, Oskar Karlsson Lindsjo, Sandra Broddesson, Petra Edquist, Mia Brytting, Anna Risberg, Karin Tegmark-Wisell |
| EPI_ISL_615109 | Klinisk mikrobiologi NAL Trollhattan | The Public Health Agency of Sweden | Anna-Malin Linde, Maria Lind Karlberg, Mattias Haukland, Reza Advani, Olov Svartstrom, Oskar Karlsson Lindsjo, Sandra Broddesson, Petra Edquist, Mia Brytting, Anna Risberg, Karin Tegmark-Wisell |
| EPI_ISL_615110 | Klinsisk mikrobiologi Linkoping | The Public Health Agency of Sweden | Anna-Malin Linde, Maria Lind Karlberg, Mattias Haukland, Reza Advani, Olov Svartstrom, Oskar Karlsson Lindsjo, Sandra Broddesson, Petra Edquist, Mia Brytting, Anna Risberg, Karin Tegmark-Wisell |

| | | | |
|---|---|---|---|
| EPI_ISL_615111, EPI_ISL_615112, EPI_ISL_615113, EPI_ISL_615114 | Orebro klinisk mikrobiologi | The Public Health Agency of Sweden | Anna-Malin Linde, Maria Lind Karlberg, Mattias Haukland, Reza Advani, Olov Svartstrom, Oskar Karlsson Lindsjo, Sandra Broddesson, Petra Edquist, Mia Brytting, Anna Risberg, Karin Tegmark-Wisell |
| EPI_ISL_615322, EPI_ISL_615323, EPI_ISL_615324, EPI_ISL_615325, EPI_ISL_615332, EPI_ISL_615406, EPI_ISL_615407, EPI_ISL_615408, EPI_ISL_615409, EPI_ISL_615410, EPI_ISL_615411, EPI_ISL_615412, EPI_ISL_615413, EPI_ISL_615414, EPI_ISL_615415, EPI_ISL_615416, EPI_ISL_615417, EPI_ISL_615418, EPI_ISL_615419, EPI_ISL_615420, EPI_ISL_615421, EPI_ISL_615422, EPI_ISL_615423, EPI_ISL_615424, EPI_ISL_615425, EPI_ISL_615426, EPI_ISL_615427, EPI_ISL_615428, EPI_ISL_615429, EPI_ISL_615430, EPI_ISL_615431, EPI_ISL_615432, EPI_ISL_615433, EPI_ISL_615434, EPI_ISL_615435, EPI_ISL_615436, EPI_ISL_615437, EPI_ISL_615438, EPI_ISL_615439, EPI_ISL_615440, EPI_ISL_615441, EPI_ISL_615442, EPI_ISL_615443, EPI_ISL_615444, EPI_ISL_615445, EPI_ISL_615446, EPI_ISL_615447, EPI_ISL_615448, EPI_ISL_615449, EPI_ISL_615450, EPI_ISL_615451, EPI_ISL_615452, EPI_ISL_615473, EPI_ISL_615474, EPI_ISL_615475, EPI_ISL_615476, EPI_ISL_615477, EPI_ISL_615478, EPI_ISL_615479, EPI_ISL_615480, EPI_ISL_615481, EPI_ISL_615482, EPI_ISL_615483, EPI_ISL_615484, EPI_ISL_615485, EPI_ISL_615486, EPI_ISL_615487, EPI_ISL_615488, EPI_ISL_615489, EPI_ISL_615490, EPI_ISL_615491, EPI_ISL_615492, EPI_ISL_615493, EPI_ISL_615494, EPI_ISL_615495, EPI_ISL_615496, EPI_ISL_615497, EPI_ISL_615498, EPI_ISL_615499, EPI_ISL_615500, EPI_ISL_615501, EPI_ISL_615502, EPI_ISL_615503, EPI_ISL_615504, EPI_ISL_615505, EPI_ISL_615506, EPI_ISL_615507, EPI_ISL_615508, EPI_ISL_615509, EPI_ISL_615510, EPI_ISL_615511, EPI_ISL_615512, EPI_ISL_615513, EPI_ISL_615514, EPI_ISL_615515, EPI_ISL_615523, EPI_ISL_615524, EPI_ISL_615525, EPI_ISL_615526, EPI_ISL_615527, EPI_ISL_615528, EPI_ISL_615552, EPI_ISL_615563, EPI_ISL_615567, EPI_ISL_615568, EPI_ISL_615569, EPI_ISL_615570, EPI_ISL_615571, EPI_ISL_615573, EPI_ISL_615581, EPI_ISL_615638, EPI_ISL_615639, EPI_ISL_615640, EPI_ISL_615641, EPI_ISL_615642, EPI_ISL_615643, EPI_ISL_615644, EPI_ISL_615906, EPI_ISL_615907, EPI_ISL_615912, EPI_ISL_615913, EPI_ISL_615914, EPI_ISL_615915, EPI_ISL_615916, EPI_ISL_615917, EPI_ISL_615918, EPI_ISL_615919, EPI_ISL_615920, EPI_ISL_615921, EPI_ISL_615922, EPI_ISL_615923, EPI_ISL_615924, EPI_ISL_615925, EPI_ISL_615926, EPI_ISL_615927, EPI_ISL_615928, EPI_ISL_615929, EPI_ISL_615930, EPI_ISL_615931, EPI_ISL_615932, EPI_ISL_615933, EPI_ISL_615934, EPI_ISL_615935, EPI_ISL_615936, EPI_ISL_615937, EPI_ISL_615938, EPI_ISL_615939, EPI_ISL_615940, EPI_ISL_615941, EPI_ISL_615942, EPI_ISL_615943, EPI_ISL_615944, EPI_ISL_615945, EPI_ISL_615946, EPI_ISL_615947, EPI_ISL_615948, EPI_ISL_615949, EPI_ISL_615950, EPI_ISL_615951, EPI_ISL_615952, EPI_ISL_615953, EPI_ISL_615954, EPI_ISL_615955, EPI_ISL_615956, EPI_ISL_615957, EPI_ISL_615958, EPI_ISL_615959, EPI_ISL_615960, EPI_ISL_615961, EPI_ISL_615962, EPI_ISL_615963, EPI_ISL_615964, EPI_ISL_615965, EPI_ISL_615966, EPI_ISL_615967, EPI_ISL_615968, EPI_ISL_615969, EPI_ISL_615970, EPI_ISL_615971, EPI_ISL_615972, EPI_ISL_615973, EPI_ISL_615974, EPI_ISL_615975, EPI_ISL_615976, EPI_ISL_615977, EPI_ISL_615978, EPI_ISL_615979, EPI_ISL_615980, EPI_ISL_615981, EPI_ISL_615982, EPI_ISL_615983, EPI_ISL_615984, EPI_ISL_615985, EPI_ISL_615986, EPI_ISL_615987, EPI_ISL_615988, EPI_ISL_615989, EPI_ISL_615990, EPI_ISL_615991, EPI_ISL_615992, EPI_ISL_615993, EPI_ISL_615994, EPI_ISL_615995, EPI_ISL_615996, EPI_ISL_615997, EPI_ISL_615998, EPI_ISL_615999, EPI_ISL_616000, EPI_ISL_616001, EPI_ISL_616002, EPI_ISL_616003, EPI_ISL_616004, EPI_ISL_616005, EPI_ISL_616006, EPI_ISL_616007, EPI_ISL_616008, EPI_ISL_616009, EPI_ISL_616010, EPI_ISL_616011, EPI_ISL_616012, EPI_ISL_616013, EPI_ISL_616014, EPI_ISL_616015, EPI_ISL_616016, EPI_ISL_616017, EPI_ISL_616018, EPI_ISL_616019, EPI_ISL_616020, EPI_ISL_616021, EPI_ISL_616022, EPI_ISL_616023, EPI_ISL_616024, EPI_ISL_616025, EPI_ISL_616026, EPI_ISL_616027, EPI_ISL_616028, EPI_ISL_616029, EPI_ISL_616030, EPI_ISL_616031, EPI_ISL_616032, EPI_ISL_616033, EPI_ISL_616034, EPI_ISL_616035, EPI_ISL_616036, EPI_ISL_616037, EPI_ISL_616038, EPI_ISL_616039, EPI_ISL_616040, EPI_ISL_616041, EPI_ISL_616042, EPI_ISL_616043, EPI_ISL_616044, EPI_ISL_616045, EPI_ISL_616046, EPI_ISL_616047, EPI_ISL_616048, EPI_ISL_616049, EPI_ISL_616050, EPI_ISL_616051, EPI_ISL_616052, EPI_ISL_616053, EPI_ISL_616054, EPI_ISL_616055, EPI_ISL_616056, EPI_ISL_616057, EPI_ISL_616058, EPI_ISL_616059, EPI_ISL_616060, EPI_ISL_616061, EPI_ISL_616062, EPI_ISL_616063, EPI_ISL_616064, EPI_ISL_616065, EPI_ISL_616066, EPI_ISL_616067, EPI_ISL_616068, EPI_ISL_616069, EPI_ISL_616070, EPI_ISL_616071, EPI_ISL_616072, EPI_ISL_616073, EPI_ISL_616074, EPI_ISL_616075, EPI_ISL_616076, EPI_ISL_616077, EPI_ISL_616078, EPI_ISL_616079, EPI_ISL_616080, EPI_ISL_616081, EPI_ISL_616082, EPI_ISL_616083, EPI_ISL_616084, EPI_ISL_616085, EPI_ISL_616086, EPI_ISL_616087, EPI_ISL_616088, EPI_ISL_616089, EPI_ISL_616090, EPI_ISL_616091, EPI_ISL_616092, EPI_ISL_616093, EPI_ISL_616094, EPI_ISL_616095, EPI_ISL_616096, EPI_ISL_616097, EPI_ISL_616098, EPI_ISL_616099, EPI_ISL_616100, EPI_ISL_616101, EPI_ISL_616102, EPI_ISL_616103, EPI_ISL_616104, EPI_ISL_616105, EPI_ISL_616106, EPI_ISL_616107, EPI_ISL_616108, EPI_ISL_616109, EPI_ISL_616110, EPI_ISL_616111, EPI_ISL_616112, EPI_ISL_616113, EPI_ISL_616114, EPI_ISL_616115, EPI_ISL_616116, EPI_ISL_616117, EPI_ISL_616118, EPI_ISL_616119, EPI_ISL_616120, EPI_ISL_616121, EPI_ISL_616122, EPI_ISL_616123, EPI_ISL_616124, EPI_ISL_616125, EPI_ISL_616126, EPI_ISL_616127, EPI_ISL_616128, EPI_ISL_616129, EPI_ISL_616130, EPI_ISL_616131, EPI_ISL_616132, EPI_ISL_616133, EPI_ISL_616134, EPI_ISL_616135, EPI_ISL_616136, EPI_ISL_616137, EPI_ISL_616138, EPI_ISL_616139, EPI_ISL_616140, EPI_ISL_616141, EPI_ISL_616142, EPI_ISL_616143, EPI_ISL_616144, EPI_ISL_616145, EPI_ISL_616146, EPI_ISL_616147, EPI_ISL_616148, EPI_ISL_616149, EPI_ISL_616150, EPI_ISL_616151, EPI_ISL_616152, EPI_ISL_616153, EPI_ISL_616154, EPI_ISL_616155, EPI_ISL_616156, EPI_ISL_616157, EPI_ISL_616158, EPI_ISL_616159, EPI_ISL_616160, EPI_ISL_616161, EPI_ISL_616162, EPI_ISL_616163, EPI_ISL_616164, EPI_ISL_616165, EPI_ISL_616166, EPI_ISL_616167, EPI_ISL_616168, EPI_ISL_616169, EPI_ISL_616170, EPI_ISL_616171, EPI_ISL_616172, EPI_ISL_616173, EPI_ISL_616174, EPI_ISL_616175, EPI_ISL_616176, EPI_ISL_616177, EPI_ISL_616178, EPI_ISL_616179, EPI_ISL_616180, EPI_ISL_616181, EPI_ISL_616182, EPI_ISL_616183, EPI_ISL_616184, EPI_ISL_616185, EPI_ISL_616186, EPI_ISL_616187, EPI_ISL_616188, EPI_ISL_616189, EPI_ISL_616190, EPI_ISL_616191, EPI_ISL_616192, EPI_ISL_616193, EPI_ISL_616194, EPI_ISL_616195, EPI_ISL_616196, EPI_ISL_616197, EPI_ISL_616198, EPI_ISL_616199, EPI_ISL_616200, EPI_ISL_616201, EPI_ISL_616202, EPI_ISL_616203, EPI_ISL_616204, EPI_ISL_616205, EPI_ISL_616206, EPI_ISL_616207, EPI_ISL_616208, EPI_ISL_616209, EPI_ISL_616210, EPI_ISL_616211, EPI_ISL_616212, EPI_ISL_616213, EPI_ISL_616214, EPI_ISL_616215, EPI_ISL_616216, EPI_ISL_616217, EPI_ISL_616218, EPI_ISL_616219, EPI_ISL_616220, EPI_ISL_616221, EPI_ISL_616222, EPI_ISL_616223, EPI_ISL_616224, EPI_ISL_616225, EPI_ISL_616226, EPI_ISL_616227, EPI_ISL_616228, EPI_ISL_616229, EPI_ISL_616230, EPI_ISL_616231, EPI_ISL_616232, EPI_ISL_616233, EPI_ISL_616234, EPI_ISL_616235, EPI_ISL_616236, EPI_ISL_616237, EPI_ISL_616238, EPI_ISL_616239, EPI_ISL_616240, EPI_ISL_616241, EPI_ISL_616242, EPI_ISL_616243, EPI_ISL_616244, EPI_ISL_616245, EPI_ISL_616246, EPI_ISL_616247, EPI_ISL_616248, EPI_ISL_616249, EPI_ISL_616250, EPI_ISL_616251, EPI_ISL_616252, EPI_ISL_616253, EPI_ISL_616254, EPI_ISL_616255, EPI_ISL_616256, EPI_ISL_616257, EPI_ISL_616258, EPI_ISL_616259, EPI_ISL_616260, EPI_ISL_616261, EPI_ISL_616262, EPI_ISL_616263, EPI_ISL_616264, EPI_ISL_616265, EPI_ISL_616266, EPI_ISL_616267, EPI_ISL_616268, EPI_ISL_616269, EPI_ISL_616270, EPI_ISL_616271, EPI_ISL_616272, EPI_ISL_616273, EPI_ISL_616274, EPI_ISL_616275, EPI_ISL_616276, EPI_ISL_616277, EPI_ISL_616278, EPI_ISL_616279, EPI_ISL_616280, EPI_ISL_616281, EPI_ISL_616282, EPI_ISL_616283, EPI_ISL_616284, EPI_ISL_616285, EPI_ISL_616286, EPI_ISL_616287, EPI_ISL_616288, EPI_ISL_616289, EPI_ISL_616290, EPI_ISL_616291, EPI_ISL_616292, EPI_ISL_616293, EPI_ISL_616294, EPI_ISL_616295, EPI_ISL_616296, EPI_ISL_616297, EPI_ISL_616298, EPI_ISL_616299, EPI_ISL_616300, EPI_ISL_616301, EPI_ISL_616302, EPI_ISL_616303, EPI_ISL_616304, EPI_ISL_616305, EPI_ISL_616306, EPI_ISL_616307, EPI_ISL_616308, EPI_ISL_616309, EPI_ISL_616310, EPI_ISL_616311, EPI_ISL_616312, EPI_ISL_616313, EPI_ISL_616314, EPI_ISL_616315, EPI_ISL_616316, EPI_ISL_616317, EPI_ISL_616318, EPI_ISL_616319, EPI_ISL_616320, EPI_ISL_616321, EPI_ISL_616322, EPI_ISL_616323, EPI_ISL_616324, EPI_ISL_616325, EPI_ISL_616326, EPI_ISL_616327, EPI_ISL_616328, EPI_ISL_616329, EPI_ISL_616330, EPI_ISL_616331, EPI_ISL_616332, EPI_ISL_616333, EPI_ISL_616334, EPI_ISL_616335, EPI_ISL_616336, EPI_ISL_616337, EPI_ISL_616338, EPI_ISL_616339, EPI_ISL_616340, EPI_ISL_616341, EPI_ISL_616342, EPI_ISL_616343, EPI_ISL_616344, EPI_ISL_616345, EPI_ISL_616346, EPI_ISL_616347, EPI_ISL_616348, EPI_ISL_616349, EPI_ISL_616350, EPI_ISL_616351, EPI_ISL_616352, EPI_ISL_616353, EPI_ISL_616354, EPI_ISL_616355, EPI_ISL_616356, EPI_ISL_616357, EPI_ISL_616358, EPI_ISL_616359, EPI_ISL_616360, EPI_ISL_616361, EPI_ISL_616362, EPI_ISL_616363, EPI_ISL_616364, EPI_ISL_616365, EPI_ISL_616366, EPI_ISL_616367, EPI_ISL_616368, EPI_ISL_616369, EPI_ISL_616370, EPI_ISL_616373, EPI_ISL_616380, EPI_ISL_616382, EPI_ISL_616384, EPI_ISL_616385, EPI_ISL_616386, EPI_ISL_616387, EPI_ISL_616388, EPI_ISL_616389, EPI_ISL_616390, EPI_ISL_616391, EPI_ISL_616392, EPI_ISL_616393, EPI_ISL_616394, EPI_ISL_616395, EPI_ISL_616396, EPI_ISL_616397, EPI_ISL_616398, EPI_ISL_616399, EPI_ISL_616400, EPI_ISL_616401, EPI_ISL_616402, EPI_ISL_616403, EPI_ISL_616404, EPI_ISL_616405, EPI_ISL_616536, EPI_ISL_616537, EPI_ISL_616540, EPI_ISL_616548, EPI_ISL_616556, EPI_ISL_616562, EPI_ISL_616563, EPI_ISL_616564, EPI_ISL_616565, EPI_ISL_616566, EPI_ISL_616567, EPI_ISL_616568, EPI_ISL_616570, EPI_ISL_616571, EPI_ISL_616572, EPI_ISL_616573, EPI_ISL_616575, EPI_ISL_616576, EPI_ISL_616577, EPI_ISL_616578, EPI_ISL_616579, EPI_ISL_616580, EPI_ISL_616581, EPI_ISL_616582, EPI_ISL_616583, EPI_ISL_616584, EPI_ISL_616585, EPI_ISL_616586, EPI_ISL_616587, EPI_ISL_616588, EPI_ISL_616589, EPI_ISL_616590, EPI_ISL_616591, EPI_ISL_616592, EPI_ISL_616593, EPI_ISL_616594, EPI_ISL_616595, EPI_ISL_616596, EPI_ISL_616597, EPI_ISL_616598, EPI_ISL_616599, EPI_ISL_616600, EPI_ISL_616601, EPI_ISL_616602, EPI_ISL_616603, EPI_ISL_616604, EPI_ISL_616605, EPI_ISL_616606, EPI_ISL_616607, EPI_ISL_616608, EPI_ISL_616609, EPI_ISL_616610, EPI_ISL_616611, EPI_ISL_616612, EPI_ISL_616613, EPI_ISL_616614, EPI_ISL_616615, EPI_ISL_616616, EPI_ISL_616617, EPI_ISL_616618, EPI_ISL_616619, EPI_ISL_616620, EPI_ISL_616621, EPI_ISL_616622, EPI_ISL_616623, EPI_ISL_616624, EPI_ISL_616625, EPI_ISL_616626, EPI_ISL_616627, EPI_ISL_616628, EPI_ISL_616629, EPI_ISL_616630, EPI_ISL_616631, EPI_ISL_616632, EPI_ISL_616639, EPI_ISL_616640, EPI_ISL_616652, EPI_ISL_616653, EPI_ISL_616654, EPI_ISL_616655, EPI_ISL_616656, EPI_ISL_616657, EPI_ISL_616658, EPI_ISL_616659, EPI_ISL_616660, EPI_ISL_616661, EPI_ISL_616662, EPI_ISL_616663, EPI_ISL_616664, EPI_ISL_616665, EPI_ISL_616666, EPI_ISL_616667, EPI_ISL_616668, EPI_ISL_616669, EPI_ISL_616670, EPI_ISL_616671, EPI_ISL_616672, EPI_ISL_616673, EPI_ISL_616674, EPI_ISL_616675, EPI_ISL_616676, EPI_ISL_616694, EPI_ISL_616695, EPI_ISL_616696, EPI_ISL_616697, EPI_ISL_616698, EPI_ISL_616699, EPI_ISL_616700, EPI_ISL_616701, EPI_ISL_616702, EPI_ISL_616703, EPI_ISL_616704, EPI_ISL_616705, EPI_ISL_616706, EPI_ISL_616707, EPI_ISL_616708, EPI_ISL_616709, EPI_ISL_616710, EPI_ISL_616711, EPI_ISL_616712, EPI_ISL_616713, EPI_ISL_616714, EPI_ISL_616715, EPI_ISL_616716, EPI_ISL_616717, EPI_ISL_616718, EPI_ISL_616719, EPI_ISL_616720, EPI_ISL_616721, EPI_ISL_616722, EPI_ISL_616723, EPI_ISL_616724, EPI_ISL_616725, EPI_ISL_616726, EPI_ISL_616727, EPI_ISL_616728, EPI_ISL_616729, EPI_ISL_616730, EPI_ISL_616731, EPI_ISL_616732, EPI_ISL_616733, EPI_ISL_616734, EPI_ISL_616735, EPI_ISL_616736, EPI_ISL_616737, EPI_ISL_616738, EPI_ISL_616739, EPI_ISL_616740, EPI_ISL_616741, EPI_ISL_616742, EPI_ISL_616743, EPI_ISL_616744, EPI_ISL_616745, EPI_ISL_616746, EPI_ISL_616747, EPI_ISL_616748, EPI_ISL_616749, EPI_ISL_616750, EPI_ISL_616751, EPI_ISL_616752, EPI_ISL_616753, EPI_ISL_616754, EPI_ISL_616755, EPI_ISL_616756, EPI_ISL_616757, EPI_ISL_616758, EPI_ISL_616759, EPI_ISL_616760, EPI_ISL_616761, EPI_ISL_616762, EPI_ISL_616763, EPI_ISL_616764, EPI_ISL_616765, EPI_ISL_616766, EPI_ISL_616767, EPI_ISL_616768, EPI_ISL_616769, EPI_ISL_616770, EPI_ISL_616771, EPI_ISL_616772, EPI_ISL_616773, EPI_ISL_616774, EPI_ISL_616775, EPI_ISL_616776, EPI_ISL_616777, EPI_ISL_616778, EPI_ISL_616779, EPI_ISL_616780, EPI_ISL_616781, EPI_ISL_616782, EPI_ISL_616783, EPI_ISL_616784, EPI_ISL_616785, EPI_ISL_616786, EPI_ISL_616787, EPI_ISL_616788, EPI_ISL_616789, EPI_ISL_616790, EPI_ISL_616791, EPI_ISL_616792, EPI_ISL_616793, EPI_ISL_616794, EPI_ISL_616795, EPI_ISL_616796, EPI_ISL_616797, EPI_ISL_616798, EPI_ISL_616799, EPI_ISL_616800, EPI_ISL_616801, EPI_ISL_616802, EPI_ISL_616803, EPI_ISL_616804, EPI_ISL_616805, EPI_ISL_616806, EPI_ISL_616807, EPI_ISL_616808, EPI_ISL_616809, EPI_ISL_616810, EPI_ISL_616811, EPI_ISL_616812, EPI_ISL_616813, EPI_ISL_616814, EPI_ISL_616815, EPI_ISL_616816, EPI_ISL_616817, EPI_ISL_616818, EPI_ISL_616819, EPI_ISL_616820, EPI_ISL_616821, EPI_ISL_616822, EPI_ISL_616823, EPI_ISL_616824, EPI_ISL_616825, EPI_ISL_616826, EPI_ISL_616827, EPI_ISL_616828, EPI_ISL_616829, EPI_ISL_616830, EPI_ISL_616831, EPI_ISL_616832, EPI_ISL_616833, EPI_ISL_616834, EPI_ISL_616835, EPI_ISL_616836, EPI_ISL_616837, EPI_ISL_616838, EPI_ISL_616839, EPI_ISL_616840, EPI_ISL_616841, EPI_ISL_616842, EPI_ISL_616843, EPI_ISL_616844, EPI_ISL_616845, EPI_ISL_616846, EPI_ISL_616847, EPI_ISL_616848, EPI_ISL_616849, EPI_ISL_616850, EPI_ISL_616851, EPI_ISL_616852, EPI_ISL_616853, EPI_ISL_616854, EPI_ISL_616855, EPI_ISL_616856, EPI_ISL_616857, EPI_ISL_616858, EPI_ISL_616859, EPI_ISL_616860, EPI_ISL_616861, EPI_ISL_616862, EPI_ISL_616863, EPI_ISL_616864, EPI_ISL_616865, EPI_ISL_616866, EPI_ISL_616867, EPI_ISL_616868, EPI_ISL_616869, EPI_ISL_616870, EPI_ISL_616871, EPI_ISL_616872, EPI_ISL_616873, EPI_ISL_616874, EPI_ISL_616875, EPI_ISL_616876, EPI_ISL_616877, EPI_ISL_616878, EPI_ISL_616879, EPI_ISL_616880, EPI_ISL_616881, EPI_ISL_616882, EPI_ISL_616883, EPI_ISL_616884, EPI_ISL_616885, EPI_ISL_616886, EPI_ISL_616887, EPI_ISL_616888, EPI_ISL_616889, EPI_ISL_616890, EPI_ISL_616891, EPI_ISL_616892, EPI_ISL_616893, EPI_ISL_616894, EPI_ISL_616895, EPI_ISL_616896, EPI_ISL_616897, EPI_ISL_616898, EPI_ISL_616899, EPI_ISL_616900, EPI_ISL_616901, EPI_ISL_616902, EPI_ISL_616903, EPI_ISL_616904, EPI_ISL_616905, EPI_ISL_621139, EPI_ISL_621140, EPI_ISL_621143, EPI_ISL_621144, EPI_ISL_621146, EPI_ISL_621152, EPI_ISL_621153, EPI_ISL_621191, EPI_ISL_621192, EPI_ISL_622676, EPI_ISL_622682, EPI_ISL_622683, EPI_ISL_622684, EPI_ISL_622685, EPI_ISL_622686, EPI_ISL_622687, EPI_ISL_622688, EPI_ISL_622689, EPI_ISL_622690, EPI_ISL_622691, EPI_ISL_622692, EPI_ISL_622693, EPI_ISL_622694, EPI_ISL_622695, EPI_ISL_622696, EPI_ISL_622697, EPI_ISL_622698, EPI_ISL_622699, EPI_ISL_622700, EPI_ISL_622701, EPI_ISL_622702, EPI_ISL_622703, EPI_ISL_622704, EPI_ISL_622705, EPI_ISL_622706, EPI_ISL_622707, EPI_ISL_622708, EPI_ISL_622709, EPI_ISL_622710, EPI_ISL_622711, EPI_ISL_622712, EPI_ISL_622713, EPI_ISL_622714, EPI_ISL_622715, EPI_ISL_622716, EPI_ISL_622717, EPI_ISL_622718, EPI_ISL_622719, EPI_ISL_622720, EPI_ISL_622721, EPI_ISL_622722, EPI_ISL_622723, EPI_ISL_622725, EPI_ISL_622732, EPI_ISL_622733, EPI_ISL_622735, EPI_ISL_622737, EPI_ISL_622738 | | | |
| see above | Department of Virus and Microbiological Special Diagnostics, Statens Serum Institut, Denmark | Albertsen lab, Department of Chemistry and Bioscience, Aalborg University, Denmark | Danish Covid-19 Genome Consortia |
| EPI_ISL_625680 | Laboratory of Molecular Medicine, University of Magallanes | Centro Asistencial Docente y de Investigacion, Universidad de Magallanes | Jorge Gonzalez, Jacqueline Aldridge, Diego Alvarez, Marcelo Navarrete |
| EPI_ISL_626237, EPI_ISL_626238, EPI_ISL_626239, EPI_ISL_626240, EPI_ISL_626241 | Department of Clinical Microbiology | GIGA Medical Genomics | Keith Durkin, Maria Artesi, Sébastien Bontems, Raphaël Boreux, Bouchra Boujemla, Cécile Meex, Pierrette Melin, Marie-Pierre Hayette, Vincent Bours |
| EPI_ISL_627467, EPI_ISL_627468, EPI_ISL_627469, EPI_ISL_627470 | University College London, Great Ormond Street Hospital for Children NHS Foundation Trust, Imperial College Healthcare NHS Trust | COVID-19 Genomics UK (COG-UK) Consortium | Sergi Castellano, Rachel Williams, Mark Kristiansen, Paola Resende Silva, Sunando Roy, Tony Brooks, Helena Tutill, Paola Niola, Patricia Dyal, Charlotte Williams, Leysa Forrest, Yasmin Panchbhaya, Jacqueline Findlay, Samuel Weeks, Julianne Brown, Kathryn Harris, Paul Randell, James Price, Alison Holmes, Judith Breuer |
| EPI_ISL_631427, EPI_ISL_631438, EPI_ISL_631439, EPI_ISL_631440, EPI_ISL_631491, EPI_ISL_631492 | Wisconsin State Laboratory of Hygiene Communicable Disease Division | Wisconsin State Laboratory of Hygiene Communicable Disease Division | Kelsey R. Florek, Abigail C. Shockey |
| EPI_ISL_632263 | Communicable Disease Laboratory, Public Health Directorate | Communicable Disease Laboratory, Public Health Directorate | AlWasti,H., AlTaif,Z., AlHujairi,Z., AlAbbas,Z. |
| EPI_ISL_632312 | NU-sjukvården | Clinical microbiology, Sahlgrenska University Hospital | Johan Ringlander, Josefin Olausson, Hedvig Engström Jakobsson, Magnus Lindh |
| EPI_ISL_632664 | Dutch COVID-19 response team | Erasmus Medical Center | Bas Oude Munnink, David Nieuwenhuijse, Reina Sikkema, Claudia Schapendonk, Irina Chestakova, Anne van der Linden, Theo Bestebroer, Stefan van Nieuwkoop, Mark Pronk, Pascal Lexmond, Corien Swaan, Manon Haverkate, Madelief Mollers, Mart Stein, Sandra Kengne Kamga Mobou, Jeroen van Kampen, Jolanda Voermans, Aura Timen, Corine GeurtsvanKessel, Annemiek van der Eijk, Richard Molenkamp, Marion Koopmans, on behalf of the Dutch national COVID-19 response team. |
| EPI_ISL_632936 | Puskesmas Mlati 1 Sleman | Genetics Working Group (Pokja Genetik) Faculty of Medicine, Public Health and Nursing Universitas Gadjah Mada (FK-KMK UGM); Disease Investigation Center Wates Ministry of Agriculture Indonesia; Department of Microbiology FK-KMK UGM; Laboratorium Diagnostik Yayasan Tahija World Mosquito Program (WMP) Yogyakarta Center for Tropical | Gunadi, Hendra Wibawa, Marcellus, Mohamad S. Hakim, Edwin W. Daniwijaya, Ludhang P. Rizki, Endah Supriyati, Eggi Arguni, Titik Nuryastuti, Tri Wibawa, Dwi AA Nugrahaningsih, Afiahayati, Siswanto, Kristy Iskandar, Nungki Anggorowati, Ika Trisnawati, Sumardi, Eko Budiono, Bambang Sigit Riyanto, Heni Retnowulan, Munawar Gani, Satria Maulana, Nur Rahmi Ananda, Riat El Khair, Yunika Puspadewi, Osman Sianipar, Umi Solekhah Intansari, Elizabeth Henny Herningtiyas, Ira Puspitawati, Nur Imma Fatimah Harahap, Ernawati, Untung Wirawan , William Widitjiarso |

| | | | |
|---|---|---|---|
| | | Medicine FK-KMK UGM; Integrated Research Center FK-KMK UGM; Department of Computer Science and Electronics FMIPA UGM; RSUP Dr. Sardjito | |
| EPI_ISL_632999, EPI_ISL_633000, EPI_ISL_633001, EPI_ISL_633002 | DOHMH Jamaica | New York City Public Health Laboratory | Jade Wang, et al. |
| EPI_ISL_633012 | DOHMH Riverside | New York City Public Health Laboratory | Jade Wang, et al. |
| EPI_ISL_633013 | DOHMH PHL | New York City Public Health Laboratory | Jade Wang, et al. |
| EPI_ISL_633014 | DOHMH Chelsea | New York City Public Health Laboratory | Jade Wang, et al. |
| EPI_ISL_633015 | DOHMH Morrisania | New York City Public Health Laboratory | Jade Wang, et al. |
| EPI_ISL_633016, EPI_ISL_633017 | DOHMH Chelsea | New York City Public Health Laboratory | Jade Wang, et al. |
| EPI_ISL_635107, EPI_ISL_635149 | Ostfold Hospital Trust - Kalnes, Centre for Laboratory Medicine, Section for gene technology and infection serology | Norwegian Institute of Public Health, Department of Virology | Kathrine Stene-Johansen, Kamilla Heddeland Instefjord, Hilde Elshaug, Marie Paulsen Madsen, Rasmus Riis Kopperud, Hilde Vollan, Karoline Bragstad, Olav Hungnes |
| EPI_ISL_635772, EPI_ISL_635773, EPI_ISL_635774, EPI_ISL_635775, EPI_ISL_635776, EPI_ISL_635800, EPI_ISL_635801, EPI_ISL_635802, EPI_ISL_635805, EPI_ISL_635808, EPI_ISL_635810, EPI_ISL_635811, EPI_ISL_635812, EPI_ISL_635815, EPI_ISL_635816, EPI_ISL_635826, EPI_ISL_635832, EPI_ISL_635835, EPI_ISL_635836, EPI_ISL_635854, EPI_ISL_635859, EPI_ISL_635860, EPI_ISL_635861, EPI_ISL_635862, EPI_ISL_635863, EPI_ISL_635865, EPI_ISL_635867, EPI_ISL_635868, EPI_ISL_635873, EPI_ISL_635876, EPI_ISL_635879, EPI_ISL_635882, EPI_ISL_635884, EPI_ISL_635885, EPI_ISL_635891, EPI_ISL_635892, EPI_ISL_635986, EPI_ISL_635987 | | | |
| see above | San Diego County Public Health Laboratory | Andersen lab at Scripps Research | SEARCH Alliance San Diego with Tracy Basler, Jovan Shephard, Brett Austin |
| EPI_ISL_636518 | Dutch COVID-19 response team | National Institute for Public Health and the Environment (RIVM) | Adam Meijer, Harry Vennema, Jeroen Cremer, Sharon van den Brink, Bas van der Veer, AnneMarie van den Brandt, Florian Zwagemaker, Dennis Schmitz, Chantal Reusken, on behalf of the national COVID-19 response team |
| EPI_ISL_636872, EPI_ISL_636873, EPI_ISL_636874, EPI_ISL_636875, EPI_ISL_636876, EPI_ISL_636877, EPI_ISL_636878, EPI_ISL_636879, EPI_ISL_636880, EPI_ISL_636881 | Lithuanian University of Health Sciences Hospital, Department of Laboratory Medicine | Lithuanian University of Health Sciences, Molecular cardiology lab. | Lukas Zemaitis, Ingrida Olendrait, Arnoldas Pautienius, Kamile Tamusauskaite, Dovydas Gecys, Laura Pareckaite, Vaiva Lesauskaite, Astra Vitkauskiene |
| EPI_ISL_636973 | Public Health Lab | Public Health Lab | Alwasti, H |
| EPI_ISL_637013, EPI_ISL_637014 | Department of Infectious Diseases and Immunology, National Hospital Organization Nagoya Medical Center | Clinical Research Center, National Hospital Organization Nagoya Medical Center | Yoshihiro Nakata, Hirotaka Ode, Mai Kubota, Masakazu Matsuda, Kazuhiro Matsuoka, Miho Nakasuji, Mikiko Mori, Mayumi Imahashi, Yoshiyuki Yokomaku, Yasumasa Iwatani |
| EPI_ISL_637041 | Private medical practitioner | Hong Kong Department of Health | Mak Gannon C.K., Lam Edman T.K., Chan Rickjason C.W., Tsang Dominic N.C. |
| EPI_ISL_637042 | Prince of Wales Hospital | Hong Kong Department of Health | Mak Gannon C.K., Lam Edman T.K., Chan Rickjason C.W., Tsang Dominic N.C. |
| EPI_ISL_637043 | Temporary Specimen Collection Centre at the AsiaWorld-Expo | Hong Kong Department of Health | Mak Gannon C.K., Lam Edman T.K., Chan Rickjason C.W., Tsang Dominic N.C. |
| EPI_ISL_638020 | Department of Pathology, University of Cambridge | COVID-19 Genomics UK (COG-UK) Consortium | Aminu S. Jahun, Yasmin Chaudhry, Grant Hall, Iliana Georgana, Myra Hosmillo, Martin D. Curran, Malte Pinckert, Surendra Parmar, Ian Goodfellow |
| EPI_ISL_639631 | E. Gulbja Laboratorija | Latvian Biomedical Research and Study Centre | Ivars Silamielis, Kaspars Megnis, Monta Ustinova, ikita Zrelovs, Vita Rovte, Mikus Gavars, Dmitrijs Perminovs, Uga Dumpis, Jnis Kloviš |
| EPI_ISL_639632, EPI_ISL_639633 | Centrl Laboratorija | Latvian Biomedical Research and Study Centre | Ivars Silamielis, Kaspars Megnis, Monta Ustinova, ikita Zrelovs, Vita Rovte, Stella Lapia, Jana Oste, Marta Priedte, Uga Dumpis, Jnis Kloviš |
| EPI_ISL_639638 | E. Gulbja Laboratorija | Latvian Biomedical Research and Study Centre | Ivars Silamielis, Kaspars Megnis, Monta Ustinova, ikita Zrelovs, Vita Rovte, Mikus Gavars, Dmitrijs Perminovs, Uga Dumpis, Jnis Kloviš |
| EPI_ISL_639639 | Latvijas Infektoloijas centrs | Latvian Biomedical Research and Study Centre | Ivars Silamielis, Kaspars Megnis, Monta Ustinova, ikita Zrelovs, Vita Rovte, Jeena Storoženko, Tatjana Kolupajeva, Oksana Savicka, Uga Dumpis, Jnis Kloviš |
| EPI_ISL_639649 | E. Gulbja Laboratorija | Latvian Biomedical Research and Study Centre | Ivars Silamielis, Kaspars Megnis, Monta Ustinova, ikita Zrelovs, Vita Rovte, Mikus Gavars, Dmitrijs Perminovs, Uga Dumpis, Jnis Kloviš |
| EPI_ISL_639657 | Centrl Laboratorija | Latvian Biomedical Research and Study Centre | Ivars Silamielis, Kaspars Megnis, Monta Ustinova, ikita Zrelovs, Vita Rovte, Stella Lapia, Jana Oste, Marta Priedte, Uga Dumpis, Jnis Kloviš |
| EPI_ISL_639658, EPI_ISL_639659, EPI_ISL_639660 | E. Gulbja Laboratorija | Latvian Biomedical Research and Study Centre | Ivars Silamielis, Kaspars Megnis, Monta Ustinova, ikita Zrelovs, Vita Rovte, Mikus Gavars, Dmitrijs Perminovs, Uga Dumpis, Jnis Kloviš |
| EPI_ISL_640036 | Bridgeton CDC wc BTC | NHLS/UCT | Arash Iranzadeh, Deelan Doolabh, Lynn Tyers, Bruna Galvao, Innocent Mudau, Marvin Hsiao, Kruger Marais, Diana Hardie, Stephen Korsman, Carolyn Williamson |
| EPI_ISL_640037, EPI_ISL_640046 | Valkenberg Hospital wc VBH | NHLS/UCT | Arash Iranzadeh, Deelan Doolabh, Lynn Tyers, Bruna Galvao, Innocent Mudau, Marvin Hsiao, Kruger Marais, Diana Hardie, Stephen Korsman, Carolyn Williamson |
| EPI_ISL_640047 | Beaufort West Hospital wc BWH | NHLS/UCT | Arash Iranzadeh, Deelan Doolabh, Lynn Tyers, Bruna Galvao, Innocent Mudau, Marvin Hsiao, Kruger Marais, Diana Hardie, Stephen Korsman, Carolyn Williamson |
| EPI_ISL_640048 | D'Almeida Clinic wc DAL | NHLS/UCT | Arash Iranzadeh, Deelan Doolabh, Lynn Tyers, Bruna Galvao, Innocent Mudau, Marvin Hsiao, Kruger Marais, Diana Hardie, Stephen Korsman, Carolyn Williamson |
| EPI_ISL_640049 | George Road Sat Clinic wc GWM | NHLS/UCT | Arash Iranzadeh, Deelan Doolabh, Lynn Tyers, Bruna Galvao, Innocent Mudau, Marvin Hsiao, Kruger Marais, Diana Hardie, Stephen Korsman, Carolyn Williamson |
| EPI_ISL_640077 | Oudtshoorn Hospital wc OUD | NHLS/UCT | Arash Iranzadeh, Deelan Doolabh, Lynn Tyers, Bruna Galvao, Innocent Mudau, Marvin Hsiao, Kruger Marais, Diana Hardie, Stephen Korsman, Carolyn Williamson |
| EPI_ISL_640751, EPI_ISL_641266 | Microbiological Diagnostic Unit - Public Health Laboratory (MDU-PHL) | MDU-PHL | Seemann T., Schultz M.B., Sait, M.L., Sherry, N.L. |
| EPI_ISL_641496, EPI_ISL_641497, EPI_ISL_641498 | Department of Virus and Microbiological Special Diagnostics, Statens Serum Institut, Copenhagen, Denmark | Albertsen lab, Department of Chemistry and Bioscience, Aalborg University, Denmark | Thomas Bruun Rasmussen, Jannik Fonager, Morten Rasmussen |
| EPI_ISL_644375, EPI_ISL_644379, EPI_ISL_644384, EPI_ISL_644387, EPI_ISL_644388, EPI_ISL_644447, EPI_ISL_644452, EPI_ISL_644453, EPI_ISL_644454, EPI_ISL_644455, EPI_ISL_644456, EPI_ISL_644457, EPI_ISL_644458, EPI_ISL_644459, EPI_ISL_644460, EPI_ISL_644461, EPI_ISL_644462, EPI_ISL_644463, EPI_ISL_644464, EPI_ISL_644465, EPI_ISL_644466, EPI_ISL_644467, EPI_ISL_644468, EPI_ISL_644469, EPI_ISL_644470, EPI_ISL_644471, EPI_ISL_644472, EPI_ISL_644473, EPI_ISL_644474, EPI_ISL_644475, EPI_ISL_644476, EPI_ISL_644477, EPI_ISL_644478, EPI_ISL_644479, EPI_ISL_644480, EPI_ISL_644481, EPI_ISL_644482, EPI_ISL_644483, EPI_ISL_644484, EPI_ISL_644491, EPI_ISL_644492, EPI_ISL_644493 | | | |
| see above | MEPHI, Aix Marseille University | MEPHI, Aix Marseille University | Anthony LEVASSEUR |
| EPI_ISL_644825, EPI_ISL_644826, EPI_ISL_644827, EPI_ISL_644828, EPI_ISL_644829, EPI_ISL_644830, EPI_ISL_644831, EPI_ISL_644832, EPI_ISL_644833, EPI_ISL_644834, EPI_ISL_644835, EPI_ISL_644836, EPI_ISL_644837, EPI_ISL_644838, EPI_ISL_644839, EPI_ISL_644840, EPI_ISL_644841 | | | |
| see above | Virginia DCLS | Virginia DCLS | Virginia DCLS |
| EPI_ISL_644998, EPI_ISL_644999 | Department of Infectious Diseases, Keio University School of Medicine, Tokyo, Japan | Center for Medical Genetics, Keio University School of Medicine, Tokyo, Japan | Kenjiro Kosaki, Yuka Iwasaki, Hirotsugu Ishizu, Haruhiko Siomi, Kodai Abe |
| EPI_ISL_648004, EPI_ISL_648005, EPI_ISL_648006, EPI_ISL_648034, EPI_ISL_648037, EPI_ISL_648038, EPI_ISL_648039 | MS Public Health Laboratory | Pathogen Discovery, Respiratory Viruses Branch, Division of Viral Diseases, Centers for Disease Control and Prevention | Yan Li, Jing Zhang, Ying Tao, Brian Lynch, Krista Queen, Anna Montmayeur, Anna Uehara, Clinton R. Paden, Rachel Marine, Haibin Wang, Suxiang Tong |
| EPI_ISL_648142 | Gavle klinisk mikrobiologi | The Public Health Agency of Sweden | Anna-Malin Linde, Maria Lind Karlberg, Mattias Haukland, Reza Advani, Olov Svartstrom, Oskar Karlsson Lindsjo, Sandra Broddesson, Petra Edquist, Mia |

| | | | |
|---|---|---|---|
| | | | Brytting, Anna Risberg, Karin Tegmark-Wisell |
| EPI_ISL_648204 | Orebro klinisk mikrobiologi | The Public Health Agency of Sweden | Anna-Malin Linde, Maria Lind Karlberg, Mattias Haukland, Reza Advani, Olov Svartstrom, Oskar Karlsson Lindsjo, Sandra Broddesson, Petra Edquist, Mia Brytting, Anna Risberg, Karin Tegmark-Wisell |
| EPI_ISL_648496 | Orange County Public Health Lab | Chan-Zuckerberg Biohub | CZB Cliahub Consortium |
| EPI_ISL_648822 | Virus Ecology Section, RML | Virus Ecology Section, RML | van Doremalen,N., Holbrook,M.G., Barbian,K.D., Bushmaker,C., Bushmaker,T., Martens,C.A. and Munster,V.J |
| EPI_ISL_648873, EPI_ISL_648921 | San Diego County Public Health Laboratory | Andersen lab at Scripps Research | SEARCH Alliance San Diego with Tracy Basler, Jovan Shephard, Brett Austin |
| EPI_ISL_649145, EPI_ISL_649146, EPI_ISL_649147, EPI_ISL_649148 | Animal health and puplic health, Escola de Medicina Veterinaria | Animal health and puplic health, Escola de Medicina Veterinaria | Ribeiro Junior,J.C., da Silva,M.O., Carvalho,R.F., dos Santos,I.G.C., Ribeiro,J. |
| EPI_ISL_649785 | I.R.C.C.S. "S. De Bellis" - Ente Ospedaliero | Istituto Zooprofilattico Sperimentale della Puglia e della Basilicata | Parisi A., Bianco A., Capozzi L., Del Sambro L., Lippolis A., Notarnicola M., Manzulli V, Rondinone V., Pace L. |
| EPI_ISL_649938 | I.R.C.C.S. "S. De Bellis" - Ente Ospedaliero | Istituto Zooprofilattico Sperimentale della Puglia e della Basilicata | Parisi A., Bianco A., Capozzi L., Del Sambro L., Lippolis A., Notarnicola M., Cipolletta D., Galante D. |
| EPI_ISL_649939 | I.R.C.C.S. "S. De Bellis" - Ente Ospedaliero | Istituto Zooprofilattico Sperimentale della Puglia e della Basilicata | Parisi A., Bianco A., Capozzi L., Del Sambro L., Lippolis A., Notarnicola M., Manzulli V, Rondinone V., Pace L. |
| EPI_ISL_649940 | Istituto Zooprofilattico Sperimentale della Puglia e della Basilicata | Istituto Zooprofilattico Sperimentale della Puglia e della Basilicata | Parisi A., Bianco A., Capozzi L., Del Sambro L., Manzulli V, Rondinone V., Pace L., Cipolletta D., Galante D. |
| EPI_ISL_653309, EPI_ISL_653339, EPI_ISL_653340, EPI_ISL_653341 | Florida Bureau of Public Health Laboratories | Florida Bureau of Public Health Laboratories | Sarah Schmedes, Jason Blanton |
| EPI_ISL_653696, EPI_ISL_653697, EPI_ISL_653698, EPI_ISL_653699, EPI_ISL_653700, EPI_ISL_653701, EPI_ISL_653702, EPI_ISL_653703, EPI_ISL_653704, EPI_ISL_653705, EPI_ISL_653706, EPI_ISL_653707, EPI_ISL_653708, EPI_ISL_653709, EPI_ISL_653710, EPI_ISL_653711, EPI_ISL_653712, EPI_ISL_653713, EPI_ISL_653714, EPI_ISL_653715, EPI_ISL_653716, EPI_ISL_653717, EPI_ISL_653718, EPI_ISL_653719, EPI_ISL_653720, EPI_ISL_653721, EPI_ISL_653722, EPI_ISL_653723, EPI_ISL_653724, EPI_ISL_653725, EPI_ISL_653726, EPI_ISL_653727, EPI_ISL_653728, EPI_ISL_653729, EPI_ISL_653730, EPI_ISL_653731, EPI_ISL_653732, EPI_ISL_653733, EPI_ISL_653734, EPI_ISL_653735, EPI_ISL_653736, EPI_ISL_653737, EPI_ISL_653738, EPI_ISL_653739, EPI_ISL_653740, EPI_ISL_653741, EPI_ISL_653742, EPI_ISL_653743, EPI_ISL_653744 | | | |
| see above | LSUHS Emerging Viral Threat Laboratory | Microbial Genome Sequencing Center | Jeremy P. Kamil, Rona S. Scott, Maarten Van Diest, Malgorzata Bienkowska-Haba, Katarzyna Zwolinska, Andrew D. Yurochko, Christopher G. Kevil, Martin J. Sapp, Daniel J. Snyder, Vaughn S. Cooper, John A. Vanchiere |
| EPI_ISL_653758 | Instituto Nacional de Salud, Bogotá, Colombia | Instituto Nacional de Salud, Bogotá, Colombia | Katherine Laiton-Donato, Diego A. Álvarez-Díaz, Carlos Franco-Muñoz, Mauricio Pacheco-Montealegre, Jonathan Reales, Diego Andrés Prada, Jose A. Usme-Ciro, Zulma M. Cucunubá, Christian Julian VillabonaArenas, Liz Villabona-Arenas, Sussy Echeverria, Astrid C. Flórez, Carolina Ferro, Diana Marcela Walteros-Acero, Franklin Prieto, Carlos Andrés Durán, Martha Lucia Ospina Martinez, Marcela Mercado-Reyes |
| EPI_ISL_653763, EPI_ISL_653764 | I.R.C.C.S. "S. De Bellis" - Ente Ospedaliero | Istituto Zooprofilattico Sperimentale della Puglia e della Basilicata | Parisi A., Bianco A., Capozzi L., Del Sambro L., Lippolis A., Notarnicola M., Manzulli V, Rondinone V., Pace L. |
| EPI_ISL_653922, EPI_ISL_653923, EPI_ISL_653924, EPI_ISL_653925, EPI_ISL_653926, EPI_ISL_653927, EPI_ISL_653928, EPI_ISL_653929, EPI_ISL_653930, EPI_ISL_653931, EPI_ISL_653932 | | | |
| see above | Molecular diagnostic laboratory of Federal Budget Institution of Science "Central Research Institute of Epidemiology" of The Federal Service on Customers' Rights Protection and Human Well-being Surveillance | Group of Genomics and Postgenomic Technologies of Central Research Institute of Epidemilology | Samoilov AE, Kaptelova VV, Dudorova A.V., Speranskaya AS, Tivanova EV, Shipulina OY, Akimkin VG |
| EPI_ISL_654281, EPI_ISL_654346, EPI_ISL_654364, EPI_ISL_654368, EPI_ISL_654370 | Hospital General Universitario Gregorio Marañón | SeqCOVID-SPAIN consortium/IBV(CSIC) | Darío García de Viedma, Laura Pérez-Lago, Marta Herranz, Jon Sicilia, Julia Suárez, Pilar Catalán, Patricia Muñoz and SeqCOVID-SPAIN consortium |
| EPI_ISL_654519 | Servicio de Microbiología, Laboratori Clínic Metropolitana Nord. Hospital Universitari Germans Trias i Pujol. Institut d'Investigació en Ciències de la Salut Germans Trias i Pujol (IGTP) | SeqCOVID-SPAIN consortium/IBV(CSIC) | Elisa Martró, Antoni E. Bordoy, Anna Not, Adrián Antuori, Anabel Fernández, Nona Romaní and SeqCOVID-SPAIN consortium |
| EPI_ISL_654698 | Mayo Clinic & Mayo Clinic Laboratories | Minnesota Department of Health, Public Health Laboratory | Matt Plumb, Jacob Garfin, Alexandra Lorentz, and Xiong Wang |
| EPI_ISL_654952 | Klinsisk mikrobiologi Linkoping | The Public Health Agency of Sweden | Anna-Malin Linde, Maria Lind Karlberg, Mattias Haukland, Reza Advani, Olov Svartstrom, Oskar Karlsson Lindsjo, Sandra Broddesson, Petra Edquist, Mia Brytting, Anna Risberg, Karin Tegmark-Wisell |
| EPI_ISL_654954 | Klinisk Mikrobiologi | The Public Health Agency of Sweden | Anna-Malin Linde, Maria Lind Karlberg, Mattias Haukland, Reza Advani, Olov Svartstrom, Oskar Karlsson Lindsjo, Sandra Broddesson, Petra Edquist, Mia Brytting, Anna Risberg, Karin Tegmark-Wisell |
| EPI_ISL_660183 | NHLS-IALCH | KRISP, KZN Research Innovation and Sequencing Platform | Gazy I, Sigal A, Karim F, Cele S, Giandhari J, Pillay S, Tegally H, Wilkinson E, de Oliveira T |
| EPI_ISL_660379, EPI_ISL_660426 | Orebro klinisk mikrobiologi | The Public Health Agency of Sweden | Anna-Malin Linde, Maria Lind Karlberg, Mattias Haukland, Reza Advani, Olov Svartstrom, Oskar Karlsson Lindsjo, Sandra Broddesson, Petra Edquist, Mia Brytting, Anna Risberg, Karin Tegmark-Wisell |
| EPI_ISL_660437, EPI_ISL_660438 | Molecular diagnostic laboratory of Federal Budget Institution of Science "Central Research Institute of Epidemiology" of The Federal Service on Customers' Rights Protection and Human Well-being Surveillance | Group of Genomics and Postgenomic Technologies of Central Research Institute of Epidemilology | Samoilov AE, Kaptelova VV, Valdokhina AV, Bulanenko VP, Speranskaya AS, Tivanova EV, Shipulina OY, Akimkin VG |
| EPI_ISL_660500, EPI_ISL_660505, EPI_ISL_660526 | Laboratoire de Microbiologie CHU Souro Sanou | Centre Muraz | Abdoul-Salam Ouedraogo, Yacouba Sawadogo, Essia Belarbi, Grit Schubert, Fabian Leendertz, Arsène Zongo, Soumeya Ouangraoua, Zekiba Tarnagda, Lassana Sangaré, Halidou Tinto |
| EPI_ISL_661266 | Russian Academy of Sciences, Federal Research Center for Virology and Microbiology | Russian Academy of Sciences, Federal Research Center for Virology and Microbiology | Titov,I., Nefedeva,M., Egorova,I., Malogolovkin,A. |
| EPI_ISL_661282 | Orebro klinisk mikrobiologi | The Public Health Agency of Sweden | Department of Microbiology, The Public Health Agency of Sweden |
| EPI_ISL_663293, EPI_ISL_663298, EPI_ISL_663309, EPI_ISL_663323, EPI_ISL_663324, EPI_ISL_663327, EPI_ISL_663332, EPI_ISL_663336, EPI_ISL_663337, EPI_ISL_663342, EPI_ISL_663344, EPI_ISL_663351, EPI_ISL_663354, EPI_ISL_663355, EPI_ISL_663395, EPI_ISL_663399, EPI_ISL_663400, EPI_ISL_663402, EPI_ISL_663410, EPI_ISL_663411, EPI_ISL_663417, EPI_ISL_663423, EPI_ISL_663430, EPI_ISL_663442, EPI_ISL_663445, EPI_ISL_663446, EPI_ISL_663452, EPI_ISL_663458, EPI_ISL_663460, EPI_ISL_663466, EPI_ISL_663480, EPI_ISL_663493, EPI_ISL_663498, EPI_ISL_663503, EPI_ISL_663504, EPI_ISL_663509, EPI_ISL_663511, EPI_ISL_663513, EPI_ISL_663527, EPI_ISL_663543, EPI_ISL_663854, EPI_ISL_663856, EPI_ISL_663857, EPI_ISL_663862, EPI_ISL_663894, EPI_ISL_663895, EPI_ISL_663896, EPI_ISL_663897, EPI_ISL_663900, EPI_ISL_663906, EPI_ISL_663909, EPI_ISL_663910, EPI_ISL_663916, EPI_ISL_663917, EPI_ISL_663919, EPI_ISL_663925, EPI_ISL_663926, EPI_ISL_663929, EPI_ISL_663932, EPI_ISL_663937, EPI_ISL_663938, EPI_ISL_663940, EPI_ISL_663943, EPI_ISL_663949, EPI_ISL_663952, EPI_ISL_663987, EPI_ISL_663997, EPI_ISL_663998, EPI_ISL_663999, EPI_ISL_664000, EPI_ISL_664001, EPI_ISL_664002, EPI_ISL_664003, EPI_ISL_664004, EPI_ISL_664005, EPI_ISL_664006, EPI_ISL_664007, EPI_ISL_664015, EPI_ISL_664016, EPI_ISL_664017, EPI_ISL_664018, EPI_ISL_664019 | | | |
| see above | Microbiological Diagnostic Unit - Public Health Laboratory (MDU-PHL) | MDU-PHL | Seemann T., Schultz M.B., Sait, M.L., Sherry, N.L. |
| EPI_ISL_666613 | LSUHS Emerging Viral Threat Laboratory | Microbial Genome Sequencing Center | Jeremy P. Kamil, Rona S. Scott, Maarten Van Diest, Malgorzata Bienkowska-Haba, Katarzyna Zwolinska, Andrew D. Yurochko, Christopher G. Kevil, Martin J. Sapp, Daniel J. Snyder, Vaughn S. Cooper, John A. Vanchiere |
| EPI_ISL_666630, EPI_ISL_666631 | ZOTZ KLIMAS MVZ Düsseldorf-Centrum GbR ÜBAG für Labormedizin, Genetik, Zytologie, Pathologie | Center of Medical Microbiology, Virology, and Hospital Hygiene, University of Duesseldorf | Maximilian Damagnez, Alexander Dilthey, Ashley-Jane Duplessis, Patrick Finzer, Katrin Hoffmann, Torsten Houwaart, Lisanna Hülse, Malte Kohns Vasconcelos, Marek Korencak, Nadine Lübke, Jessica Nicolai, Klaus Pfeffer, Daniel Strelow, Jörg Timm, Andreas Walker, Tobias Wienemann, Rainer Zotz |
| EPI_ISL_666828, EPI_ISL_666829, EPI_ISL_666843, EPI_ISL_666844, EPI_ISL_666845, EPI_ISL_666846, EPI_ISL_666847 | Florida Bureau of Public Health Laboratories | Florida Bureau of Public Health Laboratories | Sarah Schmedes, Jason Blanton |

| | | | |
|---|---|---|---|
| EPI_ISL_666993 | San Diego County Public Health Laboratory | Andersen lab at Scripps Research | SEARCH Alliance San Diego with Tracy Basler, Jovan Shephard, Brett Austin |

EPI_ISL_667260, EPI_ISL_667261, EPI_ISL_667262, EPI_ISL_667263, EPI_ISL_667264, EPI_ISL_667265, EPI_ISL_667266, EPI_ISL_667267, EPI_ISL_667269, EPI_ISL_667270, EPI_ISL_667271, EPI_ISL_667272, EPI_ISL_667273, EPI_ISL_667274, EPI_ISL_667275, EPI_ISL_667276, EPI_ISL_667277, EPI_ISL_667278, EPI_ISL_667279, EPI_ISL_667280, EPI_ISL_667281, EPI_ISL_667282, EPI_ISL_667283, EPI_ISL_667284, EPI_ISL_667285, EPI_ISL_667286, EPI_ISL_667287, EPI_ISL_667288, EPI_ISL_667289, EPI_ISL_667290, EPI_ISL_667291, EPI_ISL_667292, EPI_ISL_667293, EPI_ISL_667294, EPI_ISL_667295, EPI_ISL_667296, EPI_ISL_667297, EPI_ISL_667298, EPI_ISL_667299, EPI_ISL_667300, EPI_ISL_667301, EPI_ISL_667302, EPI_ISL_667303, EPI_ISL_667304, EPI_ISL_667305, EPI_ISL_667306, EPI_ISL_667307, EPI_ISL_667308, EPI_ISL_667309, EPI_ISL_667310, EPI_ISL_667311, EPI_ISL_667312, EPI_ISL_667313, EPI_ISL_667314, EPI_ISL_667315, EPI_ISL_667316, EPI_ISL_667317, EPI_ISL_667428, EPI_ISL_667429, EPI_ISL_667430, EPI_ISL_667432, EPI_ISL_667433, EPI_ISL_667434, EPI_ISL_667554

| | | | |
|---|---|---|---|
| see above | OHSU Lab Services Molecular Microbiology Lab | Oregon SARS-CoV-2 Genome Sequencing Center | Brendan L. O'Connell, Ruth V. Nichols, Sally Grindstaff, Alec J. Hirsch, Donna Hansel, Guang Fan, Daniel N. Streblow, William B. Messer, Andrew C. Adey, Benjamin N. Bimber, Brian J. O'Roak |
| EPI_ISL_667777 | Pathology West - NSW Health Pathology | NSW Health Pathology - Institute of Clinical Pathology and Medical Research; Westmead Hospital; University of Sydney | CIDM-PH et al. |
| EPI_ISL_668438 | Ostfold Hospital Trust - Kalnes, Centre for Laboratory Medicine, Section for gene technology and infection serology | Norwegian Institute of Public Health, Department of Virology | Kathrine Stene-Johansen, Kamilla Heddeland Instefjord, Hilde Elshaug, Marie Paulsen Madsen, Rasmus Riis Kopperud, Hilde Vollan, Karoline Bragstad, Olav Hungnes |
| EPI_ISL_670477, EPI_ISL_670729, EPI_ISL_671239, EPI_ISL_671240, EPI_ISL_671241, EPI_ISL_671242 | Department of Virus and Microbiological Special Diagnostics, Statens Serum Institut, Copenhagen, Denmark | Albertsen Lab, Department of Chemistry and Bioscience, Aalborg University, Denmark | Danish Covid-19 Genome Consortium |

EPI_ISL_671419, EPI_ISL_671420, EPI_ISL_671421, EPI_ISL_671422, EPI_ISL_671423, EPI_ISL_671424, EPI_ISL_671425, EPI_ISL_671426, EPI_ISL_671427, EPI_ISL_671429, EPI_ISL_671437

| | | | |
|---|---|---|---|
| see above | University of Debrecen, Department of Medical Microbiology | National Laboratory of Virology, Szentágothai Research Centre | Endre Gábor Tóth, Balázs Somogyi, Brigitta Zana, Eszter Csoma, Ferenc Jakab, Gábor Kemenesi |
| EPI_ISL_671801, EPI_ISL_671802 | Hospital Clínico Universitario Lozano Blesa de Zaragoza (España) | SeqCOVID-SPAIN consortium/IBV(CSIC) | Rafael Benito, Sonia Algarate, Jessica Bueno and SeqCOVID-SPAIN consortium |
| EPI_ISL_671805, EPI_ISL_671806, EPI_ISL_671807, EPI_ISL_671808, EPI_ISL_671809, EPI_ISL_671810, EPI_ISL_671811, EPI_ISL_671812, EPI_ISL_671813 | Hospital de la Santa Creu i Sant Pau. Servicio de Microbiología | SeqCOVID-SPAIN consortium/IBV(CSIC) | Ferran Navarro, Núria Rabella, Elisenda Miró and SeqCOVID-SPAIN consortium |
| EPI_ISL_671968, EPI_ISL_671972 | CHU Purpan - Laboratoire de Virologie - Institut Fédératif de Biologie | CHU Purpan - Laboratoire de Virologie - Institut Fédératif de Biologie | Latour J., Ranger N., Dubois M., Carcenac R., Harter A., Boyer P., Tremeaux P., Izopet J. |
| EPI_ISL_672028 | County of San Luis Obispo Public Health Laboratory | Chan-Zuckerberg Biohub | CZB Cliahub Consortium |
| EPI_ISL_672062, EPI_ISL_672076, EPI_ISL_672086, EPI_ISL_672454, EPI_ISL_672455 | Alameda County Public Health Lab | Chan-Zuckerberg Biohub | CZB Cliahub Consortium |
| EPI_ISL_676492 | Klinisk Mikrobiologi | The Public Health Agency of Sweden | Department of Microbiology, The Public Health Agency of Sweden |
| EPI_ISL_676513 | Orebro klinisk mikrobiologi | The Public Health Agency of Sweden | Department of Microbiology, The Public Health Agency of Sweden |
| EPI_ISL_676519 | Klinsisk mikrobiologi Linkoping | The Public Health Agency of Sweden | Department of Microbiology, The Public Health Agency of Sweden |
| EPI_ISL_676625 | Texas Department of State Health Services | Texas Department of State Health Services | Rashmi Tuladhar, Bonnie Oh, Jenny Zhang, Maliha Rahman, Anita Pokharel, Myong Koag, Chung Wang, Rachel Lee, Grace Kubin, Mayela Pedrueza, James Daniel Bonser |
| EPI_ISL_677079, EPI_ISL_677080, EPI_ISL_677081, EPI_ISL_677082, EPI_ISL_677083, EPI_ISL_677084, EPI_ISL_677085, EPI_ISL_677086, EPI_ISL_677087, EPI_ISL_677088 | Wadsworth Center, New York State Department.of Health | Wadsworth Center, New York State Department.of Health | Kirsten St. George, Daryl M. Lamson, Alexis Russel, Jonathan Plitnick, Navjot Singh, John Kelly, Sara Griesemer, Erasmus Schneider, Erica Lasek-Nesselquist |
| EPI_ISL_677111, EPI_ISL_677119, EPI_ISL_677124 | Masonic Medical Research Institute | Wadsworth Center, New York State Department.of Health | Nathan Tucker, Kirsten St. George, Daryl M. Lamson, Alexis Russel, Jonathan Plitnick, Navjot Singh, John Kelly, Sara Griesemer, Erasmus Schneider, Erica Lasek-Nesselquist |

EPI_ISL_677728, EPI_ISL_677729, EPI_ISL_677730, EPI_ISL_677731, EPI_ISL_677732, EPI_ISL_677733, EPI_ISL_677734, EPI_ISL_677735, EPI_ISL_677736, EPI_ISL_677737, EPI_ISL_677738

| | | | |
|---|---|---|---|
| see above | University of Szeged, Institute of Clinical Microbiology | National Laboratory of Virology, Szentágothai Research Centre | Endre Gábor Tóth, Balázs Somogyi, Brigitta, Gabriella Terhes, Ferenc Jakab, Gábor Kemenesi |
| EPI_ISL_678340 | Area of Virology, Serology and Virology Division (SAViD), New South Wales Health Pathology Randwick | Virology Research Laboratory; Area of Virology, Serology and Virology Division (SAViD), New South Wales Health Pathology Randwick | Foster, C.; Au, J.; Ruiz Silva, M.; Deveson, I.; Bull, R.; Van Hal, S.; Rawlinson, W. |
| EPI_ISL_682026, EPI_ISL_682027 | UPMC Clinical Microbiology Laboratory | Microbial Genomic Epidemiology Laboratory, University of Pittsburgh | Mustapha M. Mustapha, Jane W. Marsh, Dan Snyder, Marissa P. Griffith, Stephanie L. Mitchell, Vatsala R. Srinivasa, Kady D. Waggle, Chinelo Ezeonwuku, Vaughn S. Cooper, Lee H. Harrison |
| EPI_ISL_683092, EPI_ISL_683093, EPI_ISL_683094 | Department of Virus and Microbiological Special Diagnostics, Statens Serum Institut, Copenhagen, Denmark | Albertsen Lab, Department of Chemistry and Bioscience, Aalborg University, Denmark | Danish Covid-19 Genome Consortium |
| EPI_ISL_683376, EPI_ISL_683377, EPI_ISL_683378, EPI_ISL_683379, EPI_ISL_683380, EPI_ISL_683381, EPI_ISL_683382, EPI_ISL_683383 | CNR Virus des Infections Respiratoires - France SUD | CNR Virus des Infections Respiratoires - France SUD | Antonin Bal, Gregory Destras, Gwendolyne Burfin, Quentin Semanas, Martine Valette, Bruno Lina, Laurence Josset |
| EPI_ISL_683629, EPI_ISL_683634, EPI_ISL_683643, EPI_ISL_683645, EPI_ISL_683653 | Servicio de Microbiología, Laboratori Clínic Metropolitana Nord. Hospital Universitari Germans Trias i Pujol. Institut d'Investigació en Ciències de la Salut Germans Trias i Pujol (IGTP) | SeqCOVID-SPAIN consortium/IBV(CSIC) | Elisa Martró, Antoni E. Bordoy, Anna Not, Adrián Antuori, Anabel Fernández, Nona Romaní, Verónica Saludes, Cristina Casañ and SeqCOVID-SPAIN consortium |
| EPI_ISL_691645 | Servicio de Microbiología, Hospital Universitario Son Espases | SeqCOVID-SPAIN consortium/IBV(CSIC) | Carla López-Causapé, Jordi Reina, Antonio Oliver and SeqCOVID-SPAIN consortium |
| EPI_ISL_691723, EPI_ISL_691724, EPI_ISL_691725, EPI_ISL_691726, EPI_ISL_691727, EPI_ISL_691728 | Hospital Clínico San Carlos | Instituto de Salud Carlos III | Iglesias-Caballero, M. Camarero, S. Molinero Calamita, M. González-Esguevillas, M. Pozo, F. Casas, I. Jiménez, P. Jiménez, M. Zaballos, A. Monzón, S. Varona, S. Juliá, M. Cuesta, I. Rodríguez, I. |
| EPI_ISL_692756, EPI_ISL_692757, EPI_ISL_692758, EPI_ISL_692759, EPI_ISL_692760, EPI_ISL_692761 | CNR Virus des Infections Respiratoires - France SUD | CNR Virus des Infections Respiratoires - France SUD | Antonin Bal, Gregory Destras, Gwendolyne Burfin, Solenne Brun, Martine Valette, Bruno Lina, Laurence Josset |
| EPI_ISL_693473, EPI_ISL_693477 | Central Public Health Laboratory | National Public Health Laboratory, National Centre for Infectious Diseases | Tze Minn Mak, Sophie Octavia, Zhenyang Zhou, Esorom Daoni, Theresa Palou, Lin Cui, Raymond Tzer Pin Lin |

EPI_ISL_693516, EPI_ISL_693518, EPI_ISL_693519, EPI_ISL_693520, EPI_ISL_693523, EPI_ISL_693524, EPI_ISL_693534, EPI_ISL_693536, EPI_ISL_693537, EPI_ISL_693539, EPI_ISL_693630, EPI_ISL_693631, EPI_ISL_693632, EPI_ISL_693633, EPI_ISL_693634, EPI_ISL_693635, EPI_ISL_693636, EPI_ISL_693637, EPI_ISL_693638, EPI_ISL_693639, EPI_ISL_693640, EPI_ISL_693641, EPI_ISL_693642, EPI_ISL_693643, EPI_ISL_693644, EPI_ISL_693645, EPI_ISL_693646, EPI_ISL_693647, EPI_ISL_693648, EPI_ISL_693649, EPI_ISL_693650, EPI_ISL_693651, EPI_ISL_693652, EPI_ISL_693653, EPI_ISL_693654, EPI_ISL_693655, EPI_ISL_693656, EPI_ISL_693657

| | | | |
|---|---|---|---|
| see above | Hospital Vila Franca de Xira | Instituto Nacional de Saude (INSA) | Borges et al |
| EPI_ISL_693725, EPI_ISL_693743 | Delaware Public Health Laboratory | Delaware Public Health Laboratory | Gregory Hovan |
| EPI_ISL_695710, EPI_ISL_695753, EPI_ISL_695754 | TGen North | TGen North | Jolene Bowers, Megan Folkerts, Chris French, Hayley Yaglom, Ashlyn Pfeiffer, Darrin Lemmer, Dave Engelthaler, The Arizona COVID Genomics Union (ACGU) |
| EPI_ISL_700164, EPI_ISL_700165, EPI_ISL_700166, EPI_ISL_700167, EPI_ISL_700168, EPI_ISL_700169, EPI_ISL_700170, EPI_ISL_700171, EPI_ISL_700172, EPI_ISL_700173, EPI_ISL_700174, EPI_ISL_700175, EPI_ISL_700176, EPI_ISL_700177, EPI_ISL_700178, EPI_ISL_700179, EPI_ISL_700180, EPI_ISL_700181, EPI_ISL_700182, EPI_ISL_700183, EPI_ISL_700184, EPI_ISL_700185, EPI_ISL_700186, EPI_ISL_700187, EPI_ISL_700188, EPI_ISL_700189, EPI_ISL_700190, EPI_ISL_700191, EPI_ISL_700192, EPI_ISL_700193, EPI_ISL_700194, EPI_ISL_700195, EPI_ISL_700196, EPI_ISL_700197, EPI_ISL_700198, EPI_ISL_700199, EPI_ISL_700200, EPI_ISL_700201, EPI_ISL_700202, EPI_ISL_700203, EPI_ISL_700204, EPI_ISL_700205, EPI_ISL_700206, EPI_ISL_700207, EPI_ISL_700208, EPI_ISL_700209, EPI_ISL_700210, EPI_ISL_700211, EPI_ISL_700212, EPI_ISL_700213, EPI_ISL_700214, EPI_ISL_700215, EPI_ISL_700216, EPI_ISL_700217 | | | |
| see above | Hematopathology Laboratory, ACTREC, TMC | Hematopathology Laboratory, ACTREC, TMC | Hematopathology Laboratory, ACTREC |
| EPI_ISL_700329, EPI_ISL_700330, EPI_ISL_700333, EPI_ISL_700335 | Child Health Research Foundation | Child Health Research Foundation | Senjuti Saha, Afroza Akter Tanni, Syed Muktadir Al Sium, Roly Malaker, Sharmistha Goswami, Arif Mohammad Tanmoy, Md Hafizur Rahman, Samir K Saha |
| EPI_ISL_700528 | Plettenberg Bay Clinic wc PLC | NHLS/UCT | Arash Iranzadeh, Deelan Doolabh, Lynn Tyers, Bruna Galvao, Innocent Mudau, Marvin Hsiao, Kruger Marais, Diana Hardie, Stephen Korsman, Carolyn Williamson |
| EPI_ISL_700572 | Mitchells Plain Hospital wc MPH | NHLS/UCT | Arash Iranzadeh, Deelan Doolabh, Lynn Tyers, Bruna Galvao, Innocent Mudau, Marvin Hsiao, Kruger Marais, Diana Hardie, Stephen Korsman, Carolyn Williamson |
| EPI_ISL_700596 | Victoria Hospital wc VHW | NHLS/UCT | Arash Iranzadeh, Deelan Doolabh, Lynn Tyers, Bruna Galvao, Innocent Mudau, Marvin Hsiao, Kruger Marais, Diana Hardie, Stephen Korsman, Carolyn Williamson |
| EPI_ISL_700597 | Mitchells Plain Hospital wc MPH | NHLS/UCT | Arash Iranzadeh, Deelan Doolabh, Lynn Tyers, Bruna Galvao, Innocent Mudau, Marvin Hsiao, Kruger Marais, Diana Hardie, Stephen Korsman, Carolyn Williamson |
| EPI_ISL_700704, EPI_ISL_700716, EPI_ISL_700722, EPI_ISL_700735 | Texas Department of State Health Services | Texas Department of State Health Services | Rashmi Tuladhar, Bonnie Oh, Jenny Zhang, Maliha Rahman, Anita Pokharel, Myong Koag, Chung Wang, Rachel Lee, Grace Kubin, Mayela Pedrueza, James Daniel Bonser |
| EPI_ISL_708186, EPI_ISL_708188, EPI_ISL_708189, EPI_ISL_708192, EPI_ISL_708193, EPI_ISL_708194, EPI_ISL_708195 | Pamukkale University Hospital | Pamukkale University Department of Medical Genetics | Onur TOKGUN et al. |
| EPI_ISL_708418, EPI_ISL_708422, EPI_ISL_708423, EPI_ISL_708424, EPI_ISL_708426, EPI_ISL_708448 | Delaware Public Health Lab | Delaware Public Health Lab | Gregory Hovan |
| EPI_ISL_710250, EPI_ISL_710255, EPI_ISL_710258, EPI_ISL_710281, EPI_ISL_710299, EPI_ISL_710326, EPI_ISL_710327, EPI_ISL_710328, EPI_ISL_710329 | Colorado Department of Public Health and Environment | Colorado Department of Puplic Health and Environment | Laura Bankers, Molly C. Hetherington-Rauth, Shannon Ely, Shannon R. Matzinger, Sarah Elizabeth Totten, Emily A. Travanty |
| EPI_ISL_710548, EPI_ISL_710549, EPI_ISL_710556 | University Hospital Dubrava | Ruer Boškovic Institute; Forensic Science Centre Ivan Vueti; University of Zagreb Faculty of Science | Robert Beluži, Marina Korolija, Ana Livun, Vjekoslav Tomai, Dunja Glavaš, Maja Kuzman, Paula Štancl, Lucija Markulin, Lucija Basi, Antonela Blažekovi, Fran Boroveki, Lidija Cvetko-Krajinovi, Ivana elap, Fuad osovi, Mirjana Domazet-Lošo, Tomislav Domazet-Lošo, Valentina umlijan-Combaj, Kristina Gotovac Jerei, Jasna Kašman, Vladimir Krajinovi, Danilo Licastro, Boris Maek, Željka Maak Šafranko, Gordana Maravi Vlahoviek, Senica Pejša, Josipa Skelin, Ivan Šamija, Mario Štefanovi, Sanja Tadinac, Katarina Marija Tupek, Petra Vrabec, Rosa Karli, Kristian Vlahoviek |
| EPI_ISL_717694, EPI_ISL_717695, EPI_ISL_717699 | Trinidad Public Health Laboratory | Carrington Lab, Department of PreClinical Sciences, Faculty of Medical Sciences, The University of the West Indies | Nikita S. D. Sahadeo, Arianne Brown-Jordan, Sarah Hill, Vernie Ramkissoon, Naresh Nandram, Avery Hinds, Jerome Foster, Stanley Giddings, Karla Georges, Marsha Ivey, Rahul Naidu, Risha Singh, SueMin Nathaniel, Rajini Haraksingh, Jaya Jayaraman, Chinna Chinnadurai, Adesh Ramsubhag, Nuno Faria, Oliver Pybus, Christopher Oura, Gabriel Escobar, Christine V. F. Carrington |
| EPI_ISL_722815 | Dutch COVID-19 response team | Erasmus Medical Center | Bas Oude Munnink, Reina Sikkema, David Nieuwenhuijse, Irina Chestakova, Anne van der Linden, Marjan Boter, Emmanuelle Munger, Corine GeurtsvanKessel, Annemiek van der Eijk, Richard Molenkamp, Marion Koopmans, on behalf of the Dutch national COVID-19 response team. |
| EPI_ISL_722851 | I.R.C.C.S. "S. De Bellis" - Ente Ospedaliero | Istituto Zooprofilattico Sperimentale della Puglia e della Basilicata | Parisi A., Bianco A., Capozzi L., Del Sambro L., Lippolis A., Notarnicola M., Manzulli V, Rondinone V., Pace L. |
| EPI_ISL_722852 | I.R.C.C.S. "S. De Bellis" - Ente Ospedaliero | Istituto Zooprofilattico Sperimentale della Puglia e della Basilicata | Parisi A., Bianco A., Capozzi L., Del Sambro L., Lippolis A., Notarnicola M., Cipolletta D., Galante D. |
| EPI_ISL_722853 | I.R.C.C.S. "S. De Bellis" - Ente Ospedaliero | Istituto Zooprofilattico Sperimentale della Puglia e della Basilicata | Parisi A., Bianco A., Capozzi L., Del Sambro L., Lippolis A., Notarnicola M., Manzulli V, Rondinone V., Pace L. |
| EPI_ISL_722854 | I.R.C.C.S. "S. De Bellis" - Ente Ospedaliero | Istituto Zooprofilattico Sperimentale della Puglia e della Basilicata | Parisi A., Bianco A., Capozzi L., Del Sambro L., Lippolis A., Notarnicola M., Cipolletta D., Galante D. |
| EPI_ISL_722859, EPI_ISL_722860 | I.R.C.C.S. "S. De Bellis" - Ente Ospedaliero | Istituto Zooprofilattico Sperimentale della Puglia e della Basilicata | Parisi A., Bianco A., Capozzi L., Del Sambro L., Lippolis A., Notarnicola M., Manzulli V, Rondinone V., Pace L. |
| EPI_ISL_722861 | I.R.C.C.S. "S. De Bellis" - Ente Ospedaliero | Istituto Zooprofilattico Sperimentale della Puglia e della Basilicata | Parisi A., Bianco A., Capozzi L., Del Sambro L., Lippolis A., Notarnicola M., Cipolletta D., Galante D. |
| EPI_ISL_722862 | I.R.C.C.S. "S. De Bellis" - Ente Ospedaliero | Istituto Zooprofilattico Sperimentale della Puglia e della Basilicata | Parisi A., Bianco A., Capozzi L., Del Sambro L., Lippolis A., Notarnicola M., Manzulli V, Rondinone V., Pace L. |
| EPI_ISL_722863 | I.R.C.C.S. "S. De Bellis" - Ente Ospedaliero | Istituto Zooprofilattico Sperimentale della Puglia e della Basilicata | Parisi A., Bianco A., Capozzi L., Del Sambro L., Lippolis A., Notarnicola M., Cipolletta D., Galante D. |
| EPI_ISL_722864, EPI_ISL_722865 | I.R.C.C.S. "S. De Bellis" - Ente Ospedaliero | Istituto Zooprofilattico Sperimentale della Puglia e della Basilicata | Parisi A., Bianco A., Capozzi L., Del Sambro L., Lippolis A., Notarnicola M., Manzulli V, Rondinone V., Pace L. |
| EPI_ISL_722866 | I.R.C.C.S. "S. De Bellis" - Ente Ospedaliero | Istituto Zooprofilattico Sperimentale della Puglia e della Basilicata | Parisi A., Bianco A., Capozzi L., Del Sambro L., Lippolis A., Notarnicola M., Cipolletta D., Galante D. |
| EPI_ISL_722867, EPI_ISL_722868 | I.R.C.C.S. "S. De Bellis" - Ente Ospedaliero | Istituto Zooprofilattico Sperimentale della Puglia e della Basilicata | Parisi A., Bianco A., Capozzi L., Del Sambro L., Lippolis A., Notarnicola M., Manzulli V, Rondinone V., Pace L. |
| EPI_ISL_722869 | I.R.C.C.S. "S. De Bellis" - Ente Ospedaliero | Istituto Zooprofilattico Sperimentale della Puglia e della Basilicata | Parisi A., Bianco A., Capozzi L., Del Sambro L., Lippolis A., Notarnicola M., Cipolletta D., Galante D. |
| EPI_ISL_722870 | I.R.C.C.S. "S. De Bellis" - Ente Ospedaliero | Istituto Zooprofilattico Sperimentale della Puglia e della Basilicata | Parisi A., Bianco A., Capozzi L., Del Sambro L., Lippolis A., Notarnicola M., Manzulli V, Rondinone V., Pace L. |
| EPI_ISL_722871 | I.R.C.C.S. "S. De Bellis" - Ente Ospedaliero | Istituto Zooprofilattico Sperimentale della Puglia e della Basilicata | Parisi A., Bianco A., Capozzi L., Del Sambro L., Lippolis A., Notarnicola M., Cipolletta D., Galante D. |
| EPI_ISL_722892 | Istituto Zooprofilattico Sperimentale della Puglia e della Basilicata | Istituto Zooprofilattico Sperimentale della Puglia e della Basilicata | Parisi A., Bianco A., Capozzi L., Del Sambro L., Manzulli V, Rondinone V., Pace L., Cipolletta D., Galante D. |
| EPI_ISL_722895, EPI_ISL_722898, | I.R.C.C.S. "S. De Bellis" - Ente Ospedaliero | Istituto Zooprofilattico Sperimentale della Puglia e della | Parisi A., Bianco A., Capozzi L., Del Sambro L., Lippolis A., Notarnicola M., Cipolletta D., Galante D. |

| | | | |
|---|---|---|---|
| EPI_ISL_722900 | | Basilicata | |
| EPI_ISL_723056, EPI_ISL_723057 | Hematopathology Laboratory, ACTREC, TMC | Hematopathology Laboratory, ACTREC, TMC | Hematopathology Laboratory, ACTREC |
| EPI_ISL_729592, EPI_ISL_729593, EPI_ISL_729594, EPI_ISL_729595 | A. Krumbholz, Labor Dr. Krause und Kollegen MVZ GmbH, Kiel | Charité Universitätsmedizin Berlin, Institut für Virologie | Victor M Corman, Barbara Mühlemann, Jörn Beheim-Schwarzbach, Talitha Veith, Julia Schneider, Terry Jones, Christian Drosten |
| EPI_ISL_729902, EPI_ISL_729912, EPI_ISL_729919 | Instituto de Medicina Tropical, Universidad Nacional Toribio Rodríguez de Mendoza de Amazonas | Laboratorio de Genómica Microbiana, Universidad Peruana Cayetano Heredia | Pablo Tsukayama, Alejandra Dávila-Barclay, Luis González, Pedro E. Romero, Brenda Ayzanoa, Janet Huancachoque, Pool Marcos, Stella Chenet, Rafael Tapia, Cecilia Pajuelo, Carla Montenegro |
| EPI_ISL_729978 | Nigeria Centre for Disease Control (NCDC) | African Centre of Excellence for Genomics of Infectious Diseases (ACEGID), Redeemer's University, Ede, Osun State, Nigeria | Oluniyi P.E. et al |
| EPI_ISL_730127, EPI_ISL_730128, EPI_ISL_730129, EPI_ISL_730130, EPI_ISL_730131, EPI_ISL_730132, EPI_ISL_730133, EPI_ISL_730134, EPI_ISL_730135, EPI_ISL_730136 | Sharp HealthCare Laboratory | Andersen lab at Scripps Research | SEARCH Alliance San Diego with Aaron Harding, Jacquelyn Berumen, Cathy Woerle, Liam McGinnis, Art Mendoza, Omid Bakhtar |
| EPI_ISL_730215, EPI_ISL_730216, EPI_ISL_730217, EPI_ISL_730218, EPI_ISL_730219, EPI_ISL_730220, EPI_ISL_730221 | Genomica Lab Molecular, M©xico | Andersen lab at Scripps Research | SEARCH Alliance San Diego with Jonathan Gonzalez Garcia, Jose Roman Chavez Mendez, Jose Horacio Reyna Verdugo, Martin Gonzalez Ibarra, Luis Alberto Rangel Gonzalez |
| EPI_ISL_730571, EPI_ISL_730572 | Gazi University Faculty of Medicine, Medical Virology Laboratory | Gazi University Faculty of Medicine, Medical Virology Laboratory | Erdem ahin, Gülendam Bozday, Hager Muftah, Selin Yiit, Shaknoza Sarzhanova, Özlem Güzel Tunçcan, Murat Dizbay, Il Fidan, Kayhan Çalar |
| EPI_ISL_730578 | Home Quarantine Taskforce | Hong Kong Department of Health | Mak Gannon C.K., Lam Edman T.K., Chan Rickjason C.W., Tsang Dominic N.C. |
| EPI_ISL_730601 | Temporary Specimen Collection Centre at the AsiaWorld-Expo | Hong Kong Department of Health | Mak Gannon C.K., Lam Edman T.K., Chan Rickjason C.W., Tsang Dominic N.C. |
| EPI_ISL_732704, EPI_ISL_732705, EPI_ISL_732706, EPI_ISL_732707, EPI_ISL_732708, EPI_ISL_732709, EPI_ISL_732710, EPI_ISL_732711, EPI_ISL_732712, EPI_ISL_732713, EPI_ISL_732714, EPI_ISL_732715, EPI_ISL_732716, EPI_ISL_732717, EPI_ISL_732718, EPI_ISL_732719, EPI_ISL_732720, EPI_ISL_732721, EPI_ISL_732722, EPI_ISL_732723, EPI_ISL_732724, EPI_ISL_732725, EPI_ISL_732726, EPI_ISL_732727, EPI_ISL_732728, EPI_ISL_732729, EPI_ISL_732730, EPI_ISL_732731, EPI_ISL_732732, EPI_ISL_732733, EPI_ISL_732734, EPI_ISL_732735, EPI_ISL_732736, EPI_ISL_732737, EPI_ISL_732738, EPI_ISL_732739, EPI_ISL_732740, EPI_ISL_732741, EPI_ISL_732742, EPI_ISL_732743, EPI_ISL_732744, EPI_ISL_732745, EPI_ISL_732746, EPI_ISL_732747 | | | |
| see above | New Mexico Department of Health Scientific Laboratory | New Mexico Department of Health Scientific Laboratory | Ellie Johnson, Anastacia Griego-Fisher, D'Eldra Malone |
| EPI_ISL_732767, EPI_ISL_732782, EPI_ISL_732810 | Centro de Investigación Biomédica de La Rioja - Hospital San Pedro Logroño | SeqCOVID-SPAIN consortium/IBV(CSIC) | María de Toro, José Manuel Azcona Gutiérrez, María Pilar Bea Escudero, Miriam Blasco Alberdi and SeqCOVID-SPAIN consortium |
| EPI_ISL_733209, EPI_ISL_733212, EPI_ISL_733222 | Pathogenic Microorganisms Variability Laboratory | WHO National Influenza Centre Russian Federation | Andrey Komissarov, Artem Fadeev, Anna Ivanova, Kseniya Komissarova, Dmitry Bazhenov, Daria Danilenko, Ksenia Safina, Elena Nabieva, Georgii Bazykin, Nadezhda Kuznetsova, Elena Shidlovskaya, Sergey Alkhovsky, Tatyana Vishnevskaya, Elizaveta Divisenko, Alexey Shchetinin, Maria Nikiforova, Andrey Pochtovyy, Evgeny Usachev, Elena Vokalova, Maxim Rubalsky, Oleg Rubalsky, Artem Tkachuk, Vladimir Gushchin, Alexander Gintsburg, Dmitry Lioznov |
| EPI_ISL_733223, EPI_ISL_733224, EPI_ISL_733225, EPI_ISL_733226, EPI_ISL_733227 | UMMC-Health | WHO National Influenza Centre Russian Federation | Andrey Komissarov, Artem Fadeev, Anna Ivanova, Kseniya Komissarova, Dmitry Bazhenov, Tatiana Platonova, Daria Danilenko, Ksenia Safina, Elena Nabieva, Georgii Bazykin, Dmitry Lioznov |
| EPI_ISL_735436 | Nucleic Acid Testing - Rwanda National Reference Laboratory | GIGA Medical Genomics | Yvan Butera,Keith Durkin, Maria Artesi, Bouchra Boujemla, Robert Rutayisire, Patrick Tuyisenge, Esperence Umumararungu, Sébastien Bontems, Marie-Pierre Hayette, Swaibu Gatare, Jacob Souopgui, Sabin Nsanzimana, Vincent Bours, Léon Mutesa |
| EPI_ISL_735437 | Nucleic Acid Testing - Rwanda National Reference Laboratory | GIGA Medical Genomics | Yvan Butera, Keith Durkin, Maria Artesi, Bouchra Boujemla, Robert Rutayisire, Patrick Tuyisenge, Esperence Umumararungu, Sébastien Bontems, Marie-Pierre Hayette, Swaibu Gatare, Jacob Souopgui, Sabin Nsanzimana, Vincent Bours, Léon Mutesa |
| EPI_ISL_735438 | Nuclei Acid Testing - Rwanda National Reference Laboratory | GIGA Medical Genomics | Yvan Butera, Keith Durkin, Maria Artesi, Bouchra Boujemla, Robert Rutayisire, Patrick Tuyisenge, Esperence Umumararungu, Sébastien Bontems, Marie-Pierre Hayette, Swaibu Gatare, Jacob Souopgui, Sabin Nsanzimana, Vincent Bours, Léon Mutesa |
| EPI_ISL_735501, EPI_ISL_735502 | Abdul Malek Ukil Medical College, Noakhali | Central Biological Research Laboratory and Department of Biochemistry and Molecular Biology | H. M. Abdullah Al Masud, Mohammad Omar Faruque, Sajib Rudra, Md. Khondakar Raziur Rahman, Imam Hossen, Md. Arif Hossain, Shanta Paul, Md. Omer Faruq, Md. Imranul Hoq, Robiul Hasan Bhuiyan |
| EPI_ISL_737200 | National Reference Laboratory, Nigeria Centre for Disease Control. | National Reference Laboratory, Nigeria Centre for Disease Control, Gaduwa,Abuja, Nigeria | Dr Ndodo Nnaemeka, Olusola Akanbi, Chimaobi Chukwu, Dr Adesuyi Omoare, Nwando Mba, Shirlee Wohl, Dr Chikwe Ihekweazu |
| EPI_ISL_737207 | National Reference Laboratory, Nigeria Centre for Disease Control. | National Reference Laboratory, Nigeria Centre for Disease Control, Gaduwa,Abuja, Nigeria | Dr Ndodo Nnaemeka, Olusola Akanbi, Chimaobi Chukwu, Dr Adesuyi Omoare, Shirlee Wohl, Anthony Ahumibe, Abdulmajid Musa, Nneamaka Uba, Bamidele Olorunfemi, Kingsley Madubuike, Dr Sikiru Badaru, Adama Ahmad, Michael Popoola, Dr Chikwe Ihekweazu |
| EPI_ISL_737940, EPI_ISL_737941, EPI_ISL_737942, EPI_ISL_737943, EPI_ISL_737944, EPI_ISL_737980, EPI_ISL_737981, EPI_ISL_737982, EPI_ISL_737983, EPI_ISL_737985, EPI_ISL_737986, EPI_ISL_737987, EPI_ISL_737988, EPI_ISL_737989, EPI_ISL_737997, EPI_ISL_737998, EPI_ISL_737999, EPI_ISL_738000, EPI_ISL_738003, EPI_ISL_738004, EPI_ISL_738006, EPI_ISL_738022, EPI_ISL_738023 | | | |
| see above | Uganda Central Public Health Lab and Uganda Virus Research Institute | MRC/UVRI & LSHTM Uganda Research Unit | Matthew Cotten, Dan Lule Bugembe, My V.T. Phan, Pontiano Kaleebu et al. |
| EPI_ISL_738122, EPI_ISL_738123, EPI_ISL_738124, EPI_ISL_738125, EPI_ISL_738126, EPI_ISL_738127, EPI_ISL_738128, EPI_ISL_738130, EPI_ISL_738131, EPI_ISL_738132 | IZSM-U.O.C. Virologia | Istituto Zooprofilattico Sperimentale del Mezzogiorno | Maurizio Viscardi, Lorena Cardillo, Giovanna Fusco |
| EPI_ISL_738513, EPI_ISL_738523, EPI_ISL_738527, EPI_ISL_738531, EPI_ISL_738537, EPI_ISL_738562, EPI_ISL_738573, EPI_ISL_738592, EPI_ISL_738609, EPI_ISL_738615, EPI_ISL_738711, EPI_ISL_738774, EPI_ISL_738846, EPI_ISL_739059, EPI_ISL_739082, EPI_ISL_739088, EPI_ISL_739179, EPI_ISL_739218, EPI_ISL_739231, EPI_ISL_739312, EPI_ISL_739328, EPI_ISL_739368, EPI_ISL_739426, EPI_ISL_739427, EPI_ISL_739473, EPI_ISL_739483, EPI_ISL_739541, EPI_ISL_739565, EPI_ISL_739575, EPI_ISL_739610 | | | |
| see above | Alameda County Public Health Lab | Chan-Zuckerberg Biohub | CZB Cliahub Consortium |
| EPI_ISL_739733, EPI_ISL_739934, EPI_ISL_740092, EPI_ISL_744187, EPI_ISL_744846 | Laboratoire national de santé, Microbiology, Virology | Laboratoire national de santé, Microbiology, Microbial Genomics Platform | Anke Wienecke-Baldacchino, Catherine Ragimbeau,Jessica Tapp, Fatu Djabi, Lise Pignon, Raoul Salmon, Tamir Abdelrahman |
| EPI_ISL_745223, EPI_ISL_745259 | Molecular diagnostic laboratory of Federal Budget Institution of Science "Central Research Institute of Epidemiology" of The Federal Service on Customers' Rights Protection and Human Well-being Surveillance | Group of Genomics and Postgenomic Technologies of Central Research Institute of Epidemilology | Samoilov AE, Kaptelova VV, Korneenko EV, Dudorova AV, Saenko SS, Speranskaya AS, Tivanova EV, Shipulina OY, Akimkin VG |
| EPI_ISL_746494, EPI_ISL_746495, EPI_ISL_746520, EPI_ISL_746669, EPI_ISL_746671, EPI_ISL_746672, EPI_ISL_746673, EPI_ISL_746675, EPI_ISL_746676, EPI_ISL_746677, EPI_ISL_746678, EPI_ISL_746679, EPI_ISL_746680, EPI_ISL_746681, EPI_ISL_746682, EPI_ISL_746683, EPI_ISL_746684, EPI_ISL_746685, EPI_ISL_746686, EPI_ISL_746687, EPI_ISL_746688, EPI_ISL_746689, EPI_ISL_746690, EPI_ISL_746691, EPI_ISL_746692, EPI_ISL_746693, EPI_ISL_746694, EPI_ISL_746695, EPI_ISL_746696, EPI_ISL_746697 | | | |
| see above | Genetica Molecular and Subdepartamento de Virologia ISP Chile | Instituto de Salud Publica de Chile | Javier Tognarelli, Barbara Parra, Loredana Arata, Jaime Lagos, Gisselle Barra, Patricia Bustos, Rodrigo Fasce, Andres Castillo, Jorge Fernandez |
| EPI_ISL_747242 | Pathogen Lab (BSL3), Biomedical Innovation Department, Applied and Experimental Biology Division, Scientific Research Center and High Education from Ensenada | Pathogen Laboratory (BSL3), Biomedical Innovation Department, Experimental and Applied Biology Division, Scientific Research Center and High Education from | Cervantes-Luevano K, Martinez M, Saavedra A, Galindo C and Licea-Navarro A |

| | | | |
|---|---|---|---|
| | (CICESE) | Ensenada (CICESE) | |
| EPI_ISL_747256, EPI_ISL_747257, EPI_ISL_747262, EPI_ISL_747264, EPI_ISL_747271, EPI_ISL_747272, EPI_ISL_747273, EPI_ISL_747279 | Division of Emerging Infectious Diseases, Bureau of Infectious Diseases Diagnosis Control, Korea Disease Control and Prevention Agency | Division of Emerging Infectious Diseases, Bureau of Infectious Diseases Diagnosis Control, Korea Disease Control and Prevention Agency | Ae Kyung Park, Il-Hwan Kim, Heui Man Kim, Jeong-Min Kim, Namjoo Lee, Chaeyoung Lee, Sang Hee Woo, Eun-Jin Kim |
| EPI_ISL_751209 | Pathogen Genomics Lab King Abdullah University of Science and Technology(KAUST) | Pathogen Genomics Lab King Abdullah University of Science and Technology(KAUST) | Sara Mfarrej, Olga Douvropoulou, Raushan Nugmanova, Raeece Naeem, Sharif Hala, Fadwa Alofi, Asim Khogeer, Afrah Alsomali, Jumana Taha, Abdulaziz Alahmadi, Kahled Algithami, Anwar Hashem, Naif Almontashiri, Arnab Pain |
| EPI_ISL_751215 | Pathogen Genomics Lab King Abdullah University of Science and Technology(KAUST) | Pathogen Genomics Lab King Abdullah University of Science and Technology(KAUST) | Sara Mfarrej, Raushan Nugmanova, Olga Douvropoulou, Sharif Hala, Raeece Naeem, Fadwa Alofi, Asim Khogeer, Afrah Alsomali, Jumana Taha, Abdulaziz Alahmadi, Kahled Algithami, Anwar Hashem, Naif Almontashiri, Arnab Pain |
| EPI_ISL_751216 | Pathogen Genomics Lab King Abdullah University of Science and Technology(KAUST) | Pathogen Genomics Lab King Abdullah University of Science and Technology(KAUST) | Sara Mfarrej, Olga Douvropoulou, Raushan Nugmanova, Raeece Naeem, Sharif Hala, Fadwa Alofi, Asim Khogeer, Afrah Alsomali, Jumana Taha, Abdulaziz Alahmadi, Kahled Algithami, Anwar Hashem, Naif Almontashiri, Arnab Pain |
| EPI_ISL_751217 | Pathogen Genomics Lab King Abdullah University of Science and Technology(KAUST) | Pathogen Genomics Lab King Abdullah University of Science and Technology(KAUST) | Sara Mfarrej, Sharif Hala, Raushan Nugmanova, Olga Douvropoulou, Raeece Naeem, Fadwa Alofi, Asim Khogeer, Afrah Alsomali, Jumana Taha, Abdulaziz Alahmadi, Kahled Algithami, Anwar Hashem, Naif Almontashiri, Arnab Pain |
| EPI_ISL_751225 | Pathogen Genomics Lab King Abdullah University of Science and Technology(KAUST) | Pathogen Genomics Lab King Abdullah University of Science and Technology(KAUST) | Raushan Nugmanova, Olga Douvropoulou, Sara Mfarrej, Raeece Naeem, Sharif Hala, Fadwa Alofi, Asim Khogeer, Afrah Alsomali, Jumana Taha, Abdulaziz Alahmadi, Kahled Algithami, Anwar Hashem, Naif Almontashiri, Arnab Pain |
| EPI_ISL_751227, EPI_ISL_751229 | Pathogen Genomics Lab King Abdullah University of Science and Technology(KAUST) | Pathogen Genomics Lab King Abdullah University of Science and Technology(KAUST) | Sara Mfarrej, Sharif Hala, Raushan Nugmanova, Olga Douvropoulou, Raeece Naeem, Fadwa Alofi, Asim Khogeer, Afrah Alsomali, Jumana Taha, Abdulaziz Alahmadi, Kahled Algithami, Anwar Hashem, Naif Almontashiri, Arnab Pain |
| EPI_ISL_751230 | Pathogen Genomics Lab King Abdullah University of Science and Technology(KAUST) | Pathogen Genomics Lab King Abdullah University of Science and Technology(KAUST) | Sara Mfarrej, Olga Douvropoulou, Raushan Nugmanova, Raeece Naeem, Sharif Hala, Fadwa Alofi, Asim Khogeer, Afrah Alsomali, Jumana Taha, Abdulaziz Alahmadi, Kahled Algithami, Anwar Hashem, Naif Almontashiri, Arnab Pain |
| EPI_ISL_751231 | Pathogen Genomics Lab King Abdullah University of Science and Technology(KAUST) | Pathogen Genomics Lab King Abdullah University of Science and Technology(KAUST) | Sara Mfarrej, Raushan Nugmanova, Olga Douvropoulou, Sharif Hala, Raeece Naeem, Fadwa Alofi, Asim Khogeer, Afrah Alsomali, Jumana Taha, Abdulaziz Alahmadi, Kahled Algithami, Anwar Hashem, Naif Almontashiri, Arnab Pain |
| EPI_ISL_751232 | Pathogen Genomics Lab King Abdullah University of Science and Technology(KAUST) | Pathogen Genomics Lab King Abdullah University of Science and Technology(KAUST) | Sara Mfarrej, Sharif Hala, Raushan Nugmanova, Olga Douvropoulou, Raeece Naeem, Fadwa Alofi, Asim Khogeer, Afrah Alsomali, Jumana Taha, Abdulaziz Alahmadi, Kahled Algithami, Anwar Hashem, Naif Almontashiri, Arnab Pain |
| EPI_ISL_751233 | Pathogen Genomics Lab King Abdullah University of Science and Technology(KAUST) | Pathogen Genomics Lab King Abdullah University of Science and Technology(KAUST) | Olga Douvropoulou, Raushan Nugmanova, Sara Mfarrej, Raeece Naeem, Sharif Hala, Fadwa Alofi, Asim Khogeer, Afrah Alsomali, Jumana Taha, Abdulaziz Alahmadi, Kahled Algithami, Anwar Hashem, Naif Almontashiri, Arnab Pain |
| EPI_ISL_751235, EPI_ISL_751236 | Pathogen Genomics Lab King Abdullah University of Science and Technology(KAUST) | Pathogen Genomics Lab King Abdullah University of Science and Technology(KAUST) | Sara Mfarrej, Olga Douvropoulou, Raushan Nugmanova, Raeece Naeem, Sharif Hala, Fadwa Alofi, Asim Khogeer, Afrah Alsomali, Jumana Taha, Abdulaziz Alahmadi, Kahled Algithami, Anwar Hashem, Naif Almontashiri, Arnab Pain |
| EPI_ISL_751325, EPI_ISL_751335, EPI_ISL_751340, EPI_ISL_751341, EPI_ISL_751344, EPI_ISL_751345, EPI_ISL_751346 | IRCCS Sacro Cuore Don Calabria Hospital, Department of Infectious, Tropical Diseases & Microbiology | University of Verona, Department of Biotechnology | Antonio Mori, Michela Deiana, Elena Pomari, Chiara Piubelli; Giulia Lopatriello, Luca Marcolungo, Cristina Beltrami, Chiara Degli Esposti, Emanuela Cosentino, Massimo Delledonne |
| EPI_ISL_751501 | Pathogen Genomics Lab King Abdullah University of Science and Technology(KAUST) | Pathogen Genomics Lab King Abdullah University of Science and Technology(KAUST) | Sara Mfarrej, Raushan Nugmanova, Olga Douvropoulou, Sharif Hala, Raeece Naeem, Fadwa Alofi, Asim Khogeer, Afrah Alsomali, Jumana Taha, Abdulaziz Alahmadi, Kahled Algithami, Anwar Hashem, Naif Almontashiri, Arnab Pain |
| EPI_ISL_752795, EPI_ISL_752815, EPI_ISL_752816, EPI_ISL_752817, EPI_ISL_752818, EPI_ISL_752819, EPI_ISL_752820, EPI_ISL_752821, EPI_ISL_752822, EPI_ISL_752823, EPI_ISL_752824, EPI_ISL_752825, EPI_ISL_752888, EPI_ISL_752889, EPI_ISL_752890, EPI_ISL_752897, EPI_ISL_752898, EPI_ISL_752899, EPI_ISL_752900, EPI_ISL_752901 | | | |
| see above | State Laboratories Division, Hawaii State Department of Health | State Laboratories Division, Hawaii State Department of Health | Pamela O'Brien, Sabrina Diemert, Drew Kuwazaki, Razvan Sultana, Edward Desmond |
| EPI_ISL_754904, EPI_ISL_754906 | Laboratory Diagnostics and Clinical Immunology of Developmental Age, Medical University of Warsaw | genXone SA, Research & Development Laboratory; The Faculty of Mathematics, Informatics and Mechanics of the University of Warsaw | Maciej Sykulski, Grzegorz Nowicki, Monika Makowska-Woniak, Jakub Grabowski, Natalia Drwska-Matelska, ukasz Krych, Micha Kaszuba, Anna Gambin, Urszula Demkow |
| EPI_ISL_755008 | California Department of Public Health | California Department of Public Health | CDPH IDLB COVIDNet |
| EPI_ISL_755124, EPI_ISL_755126, EPI_ISL_755145, EPI_ISL_755233, EPI_ISL_755234, EPI_ISL_755237, EPI_ISL_755238, EPI_ISL_755240, EPI_ISL_755253, EPI_ISL_755254, EPI_ISL_755255 | | | |
| see above | Biolab Diagnostic Laboratories | Andersen lab at Scripps Research | Issa Abu-Dayyeh, Ahmad Tibi, Lama Hussein, Lina Mohammad, Zein Naber, Amid Abdelnour with SEARCH Alliance San Diego |
| EPI_ISL_755306, EPI_ISL_755307, EPI_ISL_755308, EPI_ISL_755309, EPI_ISL_755310, EPI_ISL_755311, EPI_ISL_755312, EPI_ISL_755313, EPI_ISL_755314, EPI_ISL_755315 | Maine Health and Environmental Testing Laboratory | Tewhey Lab, The Jackson Laboratory | Matluk,N., Dewey,H., Iosue,F., Barter,M., Lynch,R., Munger,H. and Tewhey,R. |
| EPI_ISL_755882, EPI_ISL_755883, EPI_ISL_755884, EPI_ISL_755885, EPI_ISL_755886, EPI_ISL_755887, EPI_ISL_755888, EPI_ISL_755889, EPI_ISL_755890, EPI_ISL_755891, EPI_ISL_755892, EPI_ISL_755893, EPI_ISL_755894, EPI_ISL_755895, EPI_ISL_755896, EPI_ISL_755897, EPI_ISL_755898, EPI_ISL_755899, EPI_ISL_755900, EPI_ISL_755901, EPI_ISL_755902, EPI_ISL_755903, EPI_ISL_755904, EPI_ISL_755905 | | | |
| see above | Toronto Invasive Bacterial Diseases Network | McMaster University | Allison McGeer, Patryk Aftanas, Hooman Derakhshani, Angel Li, Kuganya Nirmalarajah, Emily Panousis, Ahmed Draia, Jalees Nasir, Michael Surette, Samira Mubareka, Andrew G. McArthur |
| EPI_ISL_756362 | Trinidad Public Health Laboratory | Carrington Lab, Department of PreClinical Sciences, Faculty of Medical Sciences, The University of the West Indies | Nikita S. D. Sahadeo, Arianne Brown-Jordan, Sarah Hill, Vernie Ramkissoon, Roshan Parasram, Naresh Nandram, Avery Hinds, Jerome Foster, Stanley Giddings, Karla Georges, Marsha Ivey, Rahul Naidu, Risha Singh, SueMin Nathaniel, Rajini Haraksingh, Jaya Jayaraman, Chinna Chinnadurai, Adesh Ramsubhag, Nuno Faria, Oliver Pybus, Christopher Oura, Gabriel Escobar, Christine V. F. Carrington |
| EPI_ISL_756369, EPI_ISL_756370, EPI_ISL_756371, EPI_ISL_756373 | CUMC - CHI Bergan Mercy | Creighton University School of Medicine, Departments of Medical Microbiology and Pharmacology and Neuroscience | Michael Belshan, Morgan A. Raine, Anne V. Cheng, Christopher J. Destache, Richard V. Goering, Jacob A. Siedlik, Holly A. Stessman |
| EPI_ISL_756401 | National Institute of Health Research and Development | National Institute of Health Research and Development | Adam,K; Wibowo,HA; Ramadhany,R; Rukminiati,Y; Agustiningsih; Pawestri,HA; Subangkit; Puspa,KD; Nugraha,AA; Ikawati,HD; Pangesti,KNA; Soekarso,T; Susilarini,NK; Hariastuti,NI; Nikmah,UA; Mursinah; Febriyani,A; Herman,R; Susanti,N; Herna; Febriyanti,T; Nurhadi,M; Kurniawati,J; Kipuw,NL; Muna,F; Indalao,IL; Rizki,A; Puspandari,N; Setiawaty,Vivi |
| EPI_ISL_757377, EPI_ISL_757383, EPI_ISL_757384, EPI_ISL_759763, EPI_ISL_759764, EPI_ISL_759809, EPI_ISL_759810, EPI_ISL_759811, EPI_ISL_759899 | Department of Virology and Immunology, University of Helsinki and Helsinki University Hospital, Huslab Finland | Department of Virology, Faculty of Medicine, University of Helsinki, Helsinki, Finland | Teemu Smura, Ravi Kant, Phuoc Truong, Hussein Alburkat, Hannimari Kallio-Kokko, Jenni Virtanen, Maija Suvanto, Sari Hannula, Harri Kangas, Pekka Ellonen, Olli Vapalahti |
| EPI_ISL_760003, EPI_ISL_760004, EPI_ISL_760005, EPI_ISL_760006, EPI_ISL_760008, EPI_ISL_760009, EPI_ISL_760010, EPI_ISL_760114, EPI_ISL_760133, EPI_ISL_760134 | Division of Emerging Infectious Diseases, Bureau of Infectious Diseases Diagnosis Control, Korea Disease Control and Prevention Agency | Division of Emerging Infectious Diseases, Bureau of Infectious Diseases Diagnosis Control, Korea Disease Control and Prevention Agency | Ae Kyung Park, Il-Hwan Kim, Heui Man Kim, Jeong-Min Kim, Namjoo Lee, Chaeyoung Lee, Sang Hee Woo, Eun-Jin Kim |
| EPI_ISL_765704, EPI_ISL_765724, EPI_ISL_765725, EPI_ISL_765727 | Massachusetts General Hospital | Infectious Disease Program, Broad Institute of Harvard and MIT | Lemieux,J.E., Siddle,K.J., Shaw,B., Adams,G., Pierce,V., Turbett,S., Anahtar,M., Branda,J., Slater,D., Harris,J., Lin,A.E., Gladden-Young,A., Lagerborg,K., Rudy,M., DeRuff,K., Carter,A., Normandin,E., Bauer,M., Reilly,S., Tomkins-Tinch,C., Loreth,C., Chaluvadi,S., Neumann,A., Cusick,C., Chapman,S.B., Gnirke,A., Flowers,K., Cerrato,F., Birren,B.W., Gallagher,G., Smole,S., Park,D.J., MacInnis,B.L., Ryan,E., LaRocque,R., Rosenberg,E. and Sabeti,P.C. |

| | | | |
|---|---|---|---|
| EPI_ISL_768742 | Child Health Research Foundation | Child Health Research Foundation | Senjuti Saha, Afroza Akter Tanni, Roly Malaker, Sharmistha Goswami, Syed Muktadir Al Sium, Arif Mohammad Tanmoy, Md Hafizur Rahman, Samir K Saha |
| EPI_ISL_770797 | Hennepin County Medical Center | Minnesota Department of Health, Public Health Laboratory | Alexandra Lorentz, Jacob Garfin, Matt Plumb, and Xiong Wang |
| EPI_ISL_771204, EPI_ISL_771221 | Colorado Department of Public Health and Environment | Colorado Department of Puplic Health and Environment | Laura Bankers, Molly C. Hetherington-Rauth, Diana Ir, Shannon Ely, Shannon R. Matzinger, Sarah Elizabeth Totten, Emily A. Travanty |
| EPI_ISL_776689, EPI_ISL_776690, EPI_ISL_776691, EPI_ISL_776692 | UW Virology Lab | UW Virology Lab | Pavitra Roychoudhury, Hong Xie, Lasata Shrestha, Meei-Li Huang, Keith R Jerome, Alexander Greninger |
| EPI_ISL_779414, EPI_ISL_779415, EPI_ISL_779416, EPI_ISL_779417, EPI_ISL_779423, EPI_ISL_779424, EPI_ISL_779425, EPI_ISL_779426, EPI_ISL_779427, EPI_ISL_779437, EPI_ISL_779438, EPI_ISL_779439, EPI_ISL_779440, EPI_ISL_779441, EPI_ISL_779463, EPI_ISL_779465, EPI_ISL_779485, EPI_ISL_779494, EPI_ISL_779495, EPI_ISL_779500, EPI_ISL_779503, EPI_ISL_779504, EPI_ISL_779505, EPI_ISL_779506, EPI_ISL_779507, EPI_ISL_779508, EPI_ISL_779509, EPI_ISL_779510, EPI_ISL_779511, EPI_ISL_779513, EPI_ISL_779514, EPI_ISL_779515, EPI_ISL_779516, EPI_ISL_779517, EPI_ISL_779518, EPI_ISL_779519, EPI_ISL_779520, EPI_ISL_779521, EPI_ISL_779522, EPI_ISL_779523, EPI_ISL_779524, EPI_ISL_779525, EPI_ISL_779526, EPI_ISL_779527, EPI_ISL_779528, EPI_ISL_779529, EPI_ISL_779530, EPI_ISL_779531, EPI_ISL_779532, EPI_ISL_779533, EPI_ISL_779534, EPI_ISL_779535, EPI_ISL_779536, EPI_ISL_779537, EPI_ISL_779538, EPI_ISL_779539, EPI_ISL_779540, EPI_ISL_779541, EPI_ISL_779542, EPI_ISL_779543, EPI_ISL_779545, EPI_ISL_779546, EPI_ISL_779547, EPI_ISL_779548, EPI_ISL_779549, EPI_ISL_779550, EPI_ISL_779551, EPI_ISL_779552, EPI_ISL_779553, EPI_ISL_779554, EPI_ISL_779555, EPI_ISL_779556, EPI_ISL_779557, EPI_ISL_779558, EPI_ISL_779559, EPI_ISL_779560, EPI_ISL_779564, EPI_ISL_779582, EPI_ISL_779583, EPI_ISL_779584, EPI_ISL_779585, EPI_ISL_779586, EPI_ISL_779587, EPI_ISL_779588, EPI_ISL_779590, EPI_ISL_779591, EPI_ISL_779592, EPI_ISL_779593, EPI_ISL_779594, EPI_ISL_779595, EPI_ISL_779596, EPI_ISL_779597, EPI_ISL_779598, EPI_ISL_779599, EPI_ISL_779600, EPI_ISL_779601, EPI_ISL_779602, EPI_ISL_779603, EPI_ISL_779604, EPI_ISL_779634, EPI_ISL_779636, EPI_ISL_779637 | | | |
| see above | Microbiological Diagnostic Unit - Public Health Laboratory (MDU-PHL) | MDU-PHL | Seemann T., Sait, M.L., Sherry, N.L. |
| EPI_ISL_779703 | The Foundation for Medical Research | The Foundation for Medical Research | Ayan Mandal, Kayzad Nilgiriwala, Kalpana Sriraman, Ambreen Shaikh, Grishma Patel, Tejal Mestry, Smriti Vaswani, Swapneil Parikh, Shreevatsa Udupa, Nirjhar Chatterjee, Jayanthi Shastri, Nerges Mistry |
| EPI_ISL_788103, EPI_ISL_788104, EPI_ISL_788105, EPI_ISL_788106, EPI_ISL_788107, EPI_ISL_788108, EPI_ISL_788109, EPI_ISL_788110, EPI_ISL_788111, EPI_ISL_788112, EPI_ISL_788113, EPI_ISL_788114, EPI_ISL_788115, EPI_ISL_788116, EPI_ISL_788117, EPI_ISL_788118, EPI_ISL_788119, EPI_ISL_788120, EPI_ISL_788121, EPI_ISL_788122, EPI_ISL_788123, EPI_ISL_788124, EPI_ISL_788125, EPI_ISL_788126, EPI_ISL_788127, EPI_ISL_788128, EPI_ISL_788129, EPI_ISL_788130, EPI_ISL_788131, EPI_ISL_788132, EPI_ISL_788133, EPI_ISL_788134, EPI_ISL_788135, EPI_ISL_788136, EPI_ISL_788137, EPI_ISL_788138, EPI_ISL_788139, EPI_ISL_788140, EPI_ISL_788141, EPI_ISL_788142, EPI_ISL_788143, EPI_ISL_788144, EPI_ISL_788145, EPI_ISL_788146, EPI_ISL_788147, EPI_ISL_788148, EPI_ISL_788149, EPI_ISL_788150, EPI_ISL_788151, EPI_ISL_788152, EPI_ISL_788153, EPI_ISL_788154, EPI_ISL_788155, EPI_ISL_788156, EPI_ISL_788157, EPI_ISL_788158, EPI_ISL_788159, EPI_ISL_788160, EPI_ISL_788161, EPI_ISL_788162, EPI_ISL_788163, EPI_ISL_788164, EPI_ISL_788165, EPI_ISL_788166, EPI_ISL_788167, EPI_ISL_788168, EPI_ISL_788169, EPI_ISL_788170, EPI_ISL_788171, EPI_ISL_788172, EPI_ISL_788173, EPI_ISL_788174, EPI_ISL_788175, EPI_ISL_788176, EPI_ISL_788177, EPI_ISL_788178, EPI_ISL_788179, EPI_ISL_788180, EPI_ISL_788181, EPI_ISL_788182, EPI_ISL_788183, EPI_ISL_788184, EPI_ISL_788186, EPI_ISL_788187, EPI_ISL_788188, EPI_ISL_788189, EPI_ISL_788190, EPI_ISL_788191, EPI_ISL_788192, EPI_ISL_788193, EPI_ISL_788194, EPI_ISL_788195, EPI_ISL_788196, EPI_ISL_788197, EPI_ISL_788200, EPI_ISL_788201, EPI_ISL_788202, EPI_ISL_788203, EPI_ISL_788204, EPI_ISL_788205, EPI_ISL_788206, EPI_ISL_788207, EPI_ISL_788208, EPI_ISL_788209, EPI_ISL_788210, EPI_ISL_788211, EPI_ISL_788212, EPI_ISL_788213, EPI_ISL_788214, EPI_ISL_788215, EPI_ISL_788216, EPI_ISL_788217, EPI_ISL_788218, EPI_ISL_788219, EPI_ISL_788220, EPI_ISL_788221, EPI_ISL_788222, EPI_ISL_788223, EPI_ISL_788224, EPI_ISL_788225, EPI_ISL_788226, EPI_ISL_788227, EPI_ISL_788228, EPI_ISL_788229, EPI_ISL_788230, EPI_ISL_788231, EPI_ISL_788232, EPI_ISL_788233, EPI_ISL_788234, EPI_ISL_788235, EPI_ISL_788236, EPI_ISL_788237, EPI_ISL_788238, EPI_ISL_788239, EPI_ISL_788240, EPI_ISL_788241, EPI_ISL_788242, EPI_ISL_788243, EPI_ISL_788244, EPI_ISL_788245, EPI_ISL_788246, EPI_ISL_788247, EPI_ISL_788248, EPI_ISL_788249, EPI_ISL_788250, EPI_ISL_788251, EPI_ISL_788252, EPI_ISL_788253, EPI_ISL_788254, EPI_ISL_788255, EPI_ISL_788256, EPI_ISL_788257, EPI_ISL_788258, EPI_ISL_788259, EPI_ISL_788260, EPI_ISL_788261, EPI_ISL_788262, EPI_ISL_788263, EPI_ISL_788264, EPI_ISL_788265, EPI_ISL_788266, EPI_ISL_788267, EPI_ISL_788268, EPI_ISL_788269, EPI_ISL_788270, EPI_ISL_788271, EPI_ISL_788272, EPI_ISL_788273, EPI_ISL_788274, EPI_ISL_788275, EPI_ISL_788276, EPI_ISL_788277, EPI_ISL_788278, EPI_ISL_788279, EPI_ISL_788280, EPI_ISL_788281, EPI_ISL_788282, EPI_ISL_788283, EPI_ISL_788284, EPI_ISL_788285, EPI_ISL_788286, EPI_ISL_788287, EPI_ISL_788288, EPI_ISL_788289, EPI_ISL_788290, EPI_ISL_788291, EPI_ISL_788292, EPI_ISL_788293, EPI_ISL_788294, EPI_ISL_788295, EPI_ISL_788296, EPI_ISL_788297, EPI_ISL_788298, EPI_ISL_788299, EPI_ISL_788300, EPI_ISL_788301, EPI_ISL_788302, EPI_ISL_788303, EPI_ISL_788304, EPI_ISL_788305, EPI_ISL_788306, EPI_ISL_788307, EPI_ISL_788308, EPI_ISL_788309, EPI_ISL_788310, EPI_ISL_788311, EPI_ISL_788312, EPI_ISL_788313, EPI_ISL_788314, EPI_ISL_788315, EPI_ISL_788316, EPI_ISL_788317, EPI_ISL_788318, EPI_ISL_788319, EPI_ISL_788320, EPI_ISL_788321, EPI_ISL_788322, EPI_ISL_788323, EPI_ISL_788324, EPI_ISL_788325, EPI_ISL_788326, EPI_ISL_788327, EPI_ISL_788328, EPI_ISL_788329, EPI_ISL_788330, EPI_ISL_788331, EPI_ISL_788332, EPI_ISL_788333, EPI_ISL_788334, EPI_ISL_788335, EPI_ISL_788336, EPI_ISL_788337, EPI_ISL_788338, EPI_ISL_788339, EPI_ISL_788340, EPI_ISL_788341, EPI_ISL_788342, EPI_ISL_788343, EPI_ISL_788344, EPI_ISL_788345, EPI_ISL_788346, EPI_ISL_788347, EPI_ISL_788348, EPI_ISL_788349, EPI_ISL_788350, EPI_ISL_788351, EPI_ISL_788352, EPI_ISL_788353, EPI_ISL_788354, EPI_ISL_788355, EPI_ISL_788356, EPI_ISL_788357, EPI_ISL_788358, EPI_ISL_788359, EPI_ISL_788360, EPI_ISL_788361, EPI_ISL_788362, EPI_ISL_788363, EPI_ISL_788364, EPI_ISL_788365, EPI_ISL_788366, EPI_ISL_788367, EPI_ISL_788368, EPI_ISL_788369, EPI_ISL_788370, EPI_ISL_788371, EPI_ISL_788372, EPI_ISL_788373, EPI_ISL_788374, EPI_ISL_788375, EPI_ISL_788376, EPI_ISL_788377, EPI_ISL_788378, EPI_ISL_788379, EPI_ISL_788380, EPI_ISL_788381, EPI_ISL_788382, EPI_ISL_788383, EPI_ISL_788384, EPI_ISL_788385, EPI_ISL_788386, EPI_ISL_788387, EPI_ISL_788388, EPI_ISL_788389, EPI_ISL_788390, EPI_ISL_788391, EPI_ISL_788392, EPI_ISL_788393, EPI_ISL_788394, EPI_ISL_788395, EPI_ISL_788396, EPI_ISL_788397, EPI_ISL_788398, EPI_ISL_788399, EPI_ISL_788400, EPI_ISL_788401, EPI_ISL_788402, EPI_ISL_788403, EPI_ISL_788404, EPI_ISL_788405, EPI_ISL_788406, EPI_ISL_788407, EPI_ISL_788408, EPI_ISL_788409, EPI_ISL_788410, EPI_ISL_788411, EPI_ISL_788412, EPI_ISL_788413, EPI_ISL_788414, EPI_ISL_788415, EPI_ISL_788416, EPI_ISL_788417, EPI_ISL_788418, EPI_ISL_788419, EPI_ISL_788420, EPI_ISL_788421, EPI_ISL_788422, EPI_ISL_788423, EPI_ISL_788424, EPI_ISL_788425, EPI_ISL_788426, EPI_ISL_788427, EPI_ISL_788428, EPI_ISL_788429, EPI_ISL_788430, EPI_ISL_788431, EPI_ISL_788432, EPI_ISL_788433, EPI_ISL_788434, EPI_ISL_788435, EPI_ISL_788436, EPI_ISL_788437, EPI_ISL_788438, EPI_ISL_788439, EPI_ISL_788440, EPI_ISL_788441, EPI_ISL_788442, EPI_ISL_788443, EPI_ISL_788444, EPI_ISL_788445, EPI_ISL_788446, EPI_ISL_788447, EPI_ISL_788448, EPI_ISL_788449, EPI_ISL_788450, EPI_ISL_788451, EPI_ISL_788452, EPI_ISL_788453, EPI_ISL_788454, EPI_ISL_788455, EPI_ISL_788456, EPI_ISL_788457, EPI_ISL_788458, EPI_ISL_788459, EPI_ISL_788460, EPI_ISL_788461, EPI_ISL_788462, EPI_ISL_788463, EPI_ISL_788464, EPI_ISL_788465, EPI_ISL_788466, EPI_ISL_788467, EPI_ISL_788468, EPI_ISL_788469, EPI_ISL_788470, EPI_ISL_788471, EPI_ISL_788472, EPI_ISL_788473, EPI_ISL_788474, EPI_ISL_788475, EPI_ISL_788476, EPI_ISL_788477, EPI_ISL_788478, EPI_ISL_788479, EPI_ISL_788480, EPI_ISL_788481, EPI_ISL_788482, EPI_ISL_788483, EPI_ISL_788484, EPI_ISL_788485, EPI_ISL_788486, EPI_ISL_788487, EPI_ISL_788488, EPI_ISL_788489, EPI_ISL_788490, EPI_ISL_788491, EPI_ISL_788492, EPI_ISL_788493, EPI_ISL_788494, EPI_ISL_788495, EPI_ISL_788496, EPI_ISL_788497, EPI_ISL_788498, EPI_ISL_788499, EPI_ISL_788500, EPI_ISL_788501, EPI_ISL_788502, EPI_ISL_788503, EPI_ISL_788504, EPI_ISL_788505, EPI_ISL_788506, EPI_ISL_788507, EPI_ISL_788508, EPI_ISL_788509, EPI_ISL_788510, EPI_ISL_788511, EPI_ISL_788512, EPI_ISL_788513, EPI_ISL_788514, EPI_ISL_788515, EPI_ISL_788516, EPI_ISL_788517, EPI_ISL_788518, EPI_ISL_788519, EPI_ISL_788520, EPI_ISL_788521, EPI_ISL_788522, EPI_ISL_788523, EPI_ISL_788524, EPI_ISL_788525, EPI_ISL_788526, EPI_ISL_788527, EPI_ISL_788528, EPI_ISL_788529, EPI_ISL_788530, EPI_ISL_788531, EPI_ISL_788532, EPI_ISL_788533, EPI_ISL_788534, EPI_ISL_788535, EPI_ISL_788536, EPI_ISL_788537, EPI_ISL_788538, EPI_ISL_788539, EPI_ISL_788540, EPI_ISL_788541, EPI_ISL_788542, EPI_ISL_788543, EPI_ISL_788544, EPI_ISL_788545, EPI_ISL_788546, EPI_ISL_788547, EPI_ISL_788548, EPI_ISL_788549, EPI_ISL_788550, EPI_ISL_788551, EPI_ISL_788552, EPI_ISL_788553, EPI_ISL_788554, EPI_ISL_788555, EPI_ISL_788556, EPI_ISL_788557, EPI_ISL_788558, EPI_ISL_788559, EPI_ISL_788560, EPI_ISL_788561, EPI_ISL_788562, EPI_ISL_788563, EPI_ISL_788564, EPI_ISL_788565, EPI_ISL_788566, EPI_ISL_788567, EPI_ISL_788568, EPI_ISL_788569, EPI_ISL_788570, EPI_ISL_788571, EPI_ISL_788572, EPI_ISL_788573, EPI_ISL_788574, EPI_ISL_788575, EPI_ISL_788576, EPI_ISL_788577, EPI_ISL_788578, EPI_ISL_788579, EPI_ISL_788580, EPI_ISL_788581, EPI_ISL_788582, EPI_ISL_788583, EPI_ISL_788584, EPI_ISL_788585, EPI_ISL_788586, EPI_ISL_788587, EPI_ISL_788588, EPI_ISL_788589, EPI_ISL_788590, EPI_ISL_788591, EPI_ISL_788592, EPI_ISL_788593, EPI_ISL_788594, EPI_ISL_788595, EPI_ISL_788596, EPI_ISL_788597, EPI_ISL_788598, EPI_ISL_788599, EPI_ISL_788600, EPI_ISL_788601, EPI_ISL_788602, EPI_ISL_788603, EPI_ISL_788604, EPI_ISL_788605, EPI_ISL_788606, EPI_ISL_788607, EPI_ISL_788608, EPI_ISL_788609, EPI_ISL_788610, EPI_ISL_788611, EPI_ISL_788612, EPI_ISL_788613, EPI_ISL_788614, EPI_ISL_788615, EPI_ISL_788616, EPI_ISL_788617, EPI_ISL_788618, EPI_ISL_788619, EPI_ISL_788620, EPI_ISL_788621, EPI_ISL_788622, EPI_ISL_788623, EPI_ISL_788624, EPI_ISL_788625, EPI_ISL_788626, EPI_ISL_788627, EPI_ISL_788628, EPI_ISL_788629, EPI_ISL_788630, EPI_ISL_788631, EPI_ISL_788632, EPI_ISL_788633, EPI_ISL_788634, EPI_ISL_788635, EPI_ISL_788636, EPI_ISL_788637, EPI_ISL_788638, EPI_ISL_788639, EPI_ISL_788640, EPI_ISL_788641, EPI_ISL_788642, EPI_ISL_788643, EPI_ISL_788644, EPI_ISL_788645, EPI_ISL_788646, EPI_ISL_788647, EPI_ISL_788648, EPI_ISL_788649, EPI_ISL_788650, EPI_ISL_788651, EPI_ISL_788652, EPI_ISL_788653, EPI_ISL_788654, EPI_ISL_788655, EPI_ISL_788656, EPI_ISL_788657, EPI_ISL_788658, EPI_ISL_788659, EPI_ISL_788660, EPI_ISL_788662, EPI_ISL_788663, EPI_ISL_788664, EPI_ISL_788665, EPI_ISL_788666, EPI_ISL_788667, EPI_ISL_788668, EPI_ISL_788669, EPI_ISL_788670, EPI_ISL_788671, EPI_ISL_788672, EPI_ISL_788673, EPI_ISL_788674, EPI_ISL_788675, EPI_ISL_788676, EPI_ISL_788677, EPI_ISL_788680, EPI_ISL_788681, EPI_ISL_788682, EPI_ISL_788683, EPI_ISL_788685, EPI_ISL_788686, EPI_ISL_788687, EPI_ISL_788688, EPI_ISL_788689, EPI_ISL_788690, EPI_ISL_788691, EPI_ISL_788692, EPI_ISL_788693, EPI_ISL_788694, EPI_ISL_788697, EPI_ISL_788698, EPI_ISL_788699, EPI_ISL_788700, EPI_ISL_788701, EPI_ISL_788702, EPI_ISL_788703, EPI_ISL_788704, EPI_ISL_788705, EPI_ISL_788709, EPI_ISL_788710, EPI_ISL_788711, EPI_ISL_788712, EPI_ISL_788713, EPI_ISL_788714, EPI_ISL_788715, EPI_ISL_788716, EPI_ISL_788717, EPI_ISL_788721, EPI_ISL_788722, EPI_ISL_788723, EPI_ISL_788724, EPI_ISL_788725, EPI_ISL_788726, EPI_ISL_788727, EPI_ISL_788745, EPI_ISL_788786 | | | |
| see above | Houston Methodist Hospital | Houston Methodist Hospital | S. Wesley Long, Randall J. Olsen, Paul A. Christensen, David W. Bernard, James J. Davis, Maulik Shukla, Marcus Nguyen, Matthew Ojeda Saavedra, Prasanti Yerramilli, Layne Pruitt, Sishir Subedi, Heather Hendrickson, and James M. Musser |
| EPI_ISL_788953, EPI_ISL_788955, EPI_ISL_788956, EPI_ISL_788957, EPI_ISL_788958, EPI_ISL_788959, EPI_ISL_788960, EPI_ISL_788961, EPI_ISL_788973, EPI_ISL_788974, EPI_ISL_788975, EPI_ISL_788976, EPI_ISL_788977, EPI_ISL_788978 | | | |
| see above | Ospedale "Di Venere" | Beaconlab (Bioinformatics, Evolution and Comparative Genomics lab), Dept of Biosciences, University on Milan | Iacobellis M, d'Avenia M, Piluscio R, Parisi A, Chiara M, Manzari C, Pesole G |
| EPI_ISL_789780, EPI_ISL_789782, EPI_ISL_789784, EPI_ISL_789786, EPI_ISL_789787, EPI_ISL_789788, EPI_ISL_789790, EPI_ISL_789792, EPI_ISL_789793, EPI_ISL_789795, EPI_ISL_789796, EPI_ISL_789797, EPI_ISL_789798, EPI_ISL_789800, EPI_ISL_789801, EPI_ISL_789802, EPI_ISL_789803, EPI_ISL_789804, EPI_ISL_789807, EPI_ISL_789810, EPI_ISL_789811, EPI_ISL_789812, EPI_ISL_789813, EPI_ISL_789816, EPI_ISL_789817, EPI_ISL_789819, EPI_ISL_789826, EPI_ISL_789827, EPI_ISL_789828, EPI_ISL_789831, EPI_ISL_789832, EPI_ISL_789833, EPI_ISL_789835, EPI_ISL_789836, EPI_ISL_789839, EPI_ISL_789840, EPI_ISL_789841, EPI_ISL_789843, EPI_ISL_789844, EPI_ISL_789845, EPI_ISL_789846, EPI_ISL_789848, EPI_ISL_789849, EPI_ISL_789850, EPI_ISL_789851, EPI_ISL_789854, EPI_ISL_789858, EPI_ISL_789860, EPI_ISL_789861, EPI_ISL_789863, EPI_ISL_789869, EPI_ISL_789870, EPI_ISL_789871, EPI_ISL_789875, EPI_ISL_789876, EPI_ISL_789896, EPI_ISL_789904, EPI_ISL_789905, EPI_ISL_789906, EPI_ISL_789907, EPI_ISL_789908, EPI_ISL_789909, EPI_ISL_789911, EPI_ISL_789912, EPI_ISL_789913, EPI_ISL_789914, EPI_ISL_789915, EPI_ISL_789917, EPI_ISL_789922, EPI_ISL_789923, EPI_ISL_789925, EPI_ISL_789926, EPI_ISL_789928, EPI_ISL_789930, EPI_ISL_789931, EPI_ISL_789938, EPI_ISL_790048, EPI_ISL_790091, EPI_ISL_790396, EPI_ISL_790397, EPI_ISL_790398, EPI_ISL_790399, EPI_ISL_790401, EPI_ISL_790402, EPI_ISL_790403, EPI_ISL_790404, EPI_ISL_790405, EPI_ISL_790406, EPI_ISL_790407, EPI_ISL_790408, EPI_ISL_790409, EPI_ISL_790410, EPI_ISL_790411, EPI_ISL_790412, EPI_ISL_790413, EPI_ISL_790414, EPI_ISL_790415, EPI_ISL_790416, EPI_ISL_790417, EPI_ISL_790418, EPI_ISL_790419, EPI_ISL_790420, EPI_ISL_790421, EPI_ISL_790422, EPI_ISL_790423, EPI_ISL_790424, EPI_ISL_790425, EPI_ISL_790426, EPI_ISL_790427, EPI_ISL_790428, EPI_ISL_790429, EPI_ISL_790430, EPI_ISL_790431, EPI_ISL_790432, EPI_ISL_790433, EPI_ISL_790434, EPI_ISL_790435, EPI_ISL_790436, EPI_ISL_790437, EPI_ISL_790438, EPI_ISL_790439, EPI_ISL_790440, EPI_ISL_790441, EPI_ISL_790442, EPI_ISL_790443, EPI_ISL_790444, EPI_ISL_790445, EPI_ISL_790446, EPI_ISL_790447, EPI_ISL_790448, EPI_ISL_790449, EPI_ISL_790450, EPI_ISL_790451, EPI_ISL_790452, EPI_ISL_790453, EPI_ISL_790454, EPI_ISL_790455, EPI_ISL_790456, EPI_ISL_790457, EPI_ISL_790458, EPI_ISL_790459, EPI_ISL_790460, EPI_ISL_790461, EPI_ISL_790462, EPI_ISL_790463, EPI_ISL_790464, EPI_ISL_790465, EPI_ISL_790466, EPI_ISL_790467, EPI_ISL_790468, EPI_ISL_790469, EPI_ISL_790470, EPI_ISL_790471, EPI_ISL_790472, EPI_ISL_790473, EPI_ISL_790474, EPI_ISL_790475, EPI_ISL_790476, EPI_ISL_790477, EPI_ISL_790478, EPI_ISL_790479, EPI_ISL_790480, EPI_ISL_790481, EPI_ISL_790482, EPI_ISL_790483, EPI_ISL_790484, EPI_ISL_790485, EPI_ISL_790486, EPI_ISL_790488, EPI_ISL_790489, EPI_ISL_790490, EPI_ISL_790491, EPI_ISL_790492, EPI_ISL_790493, EPI_ISL_790494, EPI_ISL_790495, EPI_ISL_790496, EPI_ISL_790497, EPI_ISL_790498, EPI_ISL_790499, EPI_ISL_790500, EPI_ISL_790501, EPI_ISL_790502, EPI_ISL_790504, EPI_ISL_790505, EPI_ISL_790506, EPI_ISL_790507 | | | |
| see above | Houston Methodist Hospital | Houston Methodist Hospital | S. Wesley Long, Randall J. Olsen, Paul A. Christensen, David W. Bernard, James J. Davis, Maulik Shukla, Marcus Nguyen, Matthew Ojeda Saavedra, Prasanti Yerramilli, Layne Pruitt, Sishir Subedi, Heather Hendrickson, and James M. Musser |
| EPI_ISL_792611 | LACEN-PB | Laboratory of Respiratory Viruses and Measles, Oswaldo Cruz Institute, FIOCRUZ | Paola Resende, Luciana Appolinario, Fernando Motta, Anna Carolina Paixao, Ana Carolina Mendonca, João Felipe Bezerra, Romero Henrique Teixeira de Vasconcelos, Dalane Loudal Florentino Teixeira, Thiago Franco de Oliveira Carneiro, Marilda Siqueira |
| EPI_ISL_801397, EPI_ISL_801398 | Laboratório Central de Saúde Pública do Amazonas - LACEN-AM | Laboratorio de Ecologia de Doencas Transmissiveis na Amazonia, Instituto Leonidas e Maria Deane - Fiocruz Amazonia | Valdinete Nascimento, Victor Souza, André Corado, Fernanda Nascimento, George Silva, Ágatha Costa, Debora Duarte, Luciana Gonçalves, Maria Júlia Brandão, Michele Jesus, Felipe Naveca |
| EPI_ISL_801878, EPI_ISL_801886, EPI_ISL_801904, EPI_ISL_801925, EPI_ISL_801927, EPI_ISL_801930, EPI_ISL_801936, EPI_ISL_801968, EPI_ISL_802189, EPI_ISL_802190, EPI_ISL_802191, EPI_ISL_802192, EPI_ISL_802193, EPI_ISL_802194, EPI_ISL_802195, EPI_ISL_802196, EPI_ISL_802197, EPI_ISL_802198, EPI_ISL_802199, EPI_ISL_802200, EPI_ISL_802201, EPI_ISL_802202, EPI_ISL_802203, EPI_ISL_802204, EPI_ISL_802205, EPI_ISL_802206, EPI_ISL_802207, EPI_ISL_802208, EPI_ISL_802209, EPI_ISL_802210, EPI_ISL_802211, EPI_ISL_802212, EPI_ISL_802213, EPI_ISL_802214, EPI_ISL_802215, EPI_ISL_802216, EPI_ISL_802217, EPI_ISL_802218 | | | |
| see above | MSHS Clinical Microbiology Laboratories | MSHS Pathogen Surveillance Program | Ana S. Gonzalez-Reiche, Hala Alshammary, Mitchell J. Sullivan, Brianne Ciferri, Ajay Obla, Angela Amoako, Mahmoud Awawda, Elena Hirsch, Ashley S. Salimbangon, Levy Sominsky, Katherine Beach, Kayla Russo, Charles Gleason, Shelcie Fabre, Giulio Kleiner, Zenab Khan, Bremy Alburquerque, Adriana van de Guchte, Komal Srivastava, Matthew M. Hernandez, Jayeeta Dutta, Denise Jurczyszak, Emily Ferreri, Rachel Chernet, Nancy Francoeur, Betsaida |

| | | | |
|---|---|---|---|
| | | | Salom Melo, Irina Oussenko, Gintaras Deikus, Juan Soto, Shwetha Hara Sridhar, Ying-Chih Wang, Kathryn Twyman, Andrew Kasarskis, Deena R. Altman, Robert Sebra, Adolfo García-Sastre, Marta Luksza, Gopi Patel, Sarah Schaefer, Melissa Gitman, Michael D. Nowak, Alberto Paniz-Mondolfi, Emilia Mia Sordillo, Viviana Simon, Harm van Bakel |
| EPI_ISL_803123, EPI_ISL_803124, EPI_ISL_803125, EPI_ISL_803126, EPI_ISL_803127, EPI_ISL_803128, EPI_ISL_803129, EPI_ISL_803130, EPI_ISL_803131, EPI_ISL_803132, EPI_ISL_803133, EPI_ISL_803141, EPI_ISL_803216, EPI_ISL_803217, EPI_ISL_803218, EPI_ISL_803219, EPI_ISL_803220, EPI_ISL_803221, EPI_ISL_803222, EPI_ISL_803223, EPI_ISL_803224, EPI_ISL_803225, EPI_ISL_803226, EPI_ISL_803232, EPI_ISL_803233, EPI_ISL_803234, EPI_ISL_803235, EPI_ISL_803236, EPI_ISL_803251, EPI_ISL_803252, EPI_ISL_803253 | | | |
| see above | Wisconsin State Laboratory of Hygiene Communicable Disease Division | Wisconsin State Laboratory of Hygiene Communicable Disease Division | Kelsey R. Florek, Abigail C. Shockey |
| EPI_ISL_804011, EPI_ISL_804012, EPI_ISL_804013, EPI_ISL_804014, EPI_ISL_804015 | Grupo de Resistencia Antimicrobiana en bacterias patógenas y ambientales GRABPA | Facultad de Ciencias de la Vida, UNAB | Dayán Sanhueza, Macarena Bastías, César Echeverría, Sebastián Wolter, Waldo Díaz, Jorge Olivares, Eduardo Castro, Claudio Meneses |
| EPI_ISL_804961 | Hospital Comarcal de Melilla | Instituto de Salud Carlos III | Iglesias-Caballero, M. Molinero Calamita, M. González-Esguevillas, M. Camarero, S. Pozo, F. Casas, I. Jiménez, P. Jiménez, M. Zaballos, A. Monzón, S. Varona, S. Juliá, M. Cuesta, I, J. López |
| EPI_ISL_806543 | Santa Clara County Public Health Laboratory | Santa Clara County Public Health Laboratory | Santa Clara Co Public Health Department |
| EPI_ISL_806545 | Santa Clara County Public Health Laboratory | Santa Clara County Public Health Laboratory | Santa CLara County Public Health Department |
| EPI_ISL_806583, EPI_ISL_806588, EPI_ISL_806589, EPI_ISL_806590, EPI_ISL_806599, EPI_ISL_806601, EPI_ISL_806602, EPI_ISL_806603, EPI_ISL_806604, EPI_ISL_806605, EPI_ISL_806606, EPI_ISL_806607, EPI_ISL_806608, EPI_ISL_806609, EPI_ISL_806610, EPI_ISL_806613, EPI_ISL_806615, EPI_ISL_806616, EPI_ISL_806618 | | | |
| see above | KEMRI-Wellcome Trust Research Programme/KEMRI-CGMR-C Kilifi | KEMRI-Wellcome Trust Research Programme/KEMRI-CGMR-C Kilifi | Githinji et al |
| EPI_ISL_806853, EPI_ISL_806854, EPI_ISL_806855, EPI_ISL_806856, EPI_ISL_806857, EPI_ISL_806858, EPI_ISL_806859, EPI_ISL_806860, EPI_ISL_806861, EPI_ISL_806862, EPI_ISL_806863, EPI_ISL_806864, EPI_ISL_806865, EPI_ISL_806866, EPI_ISL_806965, EPI_ISL_806975, EPI_ISL_806976, EPI_ISL_806977, EPI_ISL_806978, EPI_ISL_806979, EPI_ISL_806980, EPI_ISL_806981, EPI_ISL_806982, EPI_ISL_806983, EPI_ISL_806984, EPI_ISL_806985, EPI_ISL_806986, EPI_ISL_806987, EPI_ISL_806988, EPI_ISL_806989 | | | |
| see above | Washington State Department of Health | Seattle Flu Study | Deborah A. Nickerson, Chris D. Frazar, Jover Lee, Benjamin Pelle, Matthew Richardson, Amanda Adler, Elisabeth Brandstetter, Peter D. Han, Kairsten Fay, Misja Ilcisin, Kirsten Lacombe, Thomas R. Sibley, Melissa Truong, Caitlin R. Wolf, Romesh Gautom, Geoff Melly, Brian Hiatt, Philip Dykema, Scott Lindquist, Michael Boeckh, Janet A. Englund, Michael Famulare, Barry R. Lutz, Mark J. Rieder, Lea M. Starita, Matthew Thompson, Helen Y. Chu, Jay Shendure, Trevor Bedford |
| EPI_ISL_810977, EPI_ISL_810978, EPI_ISL_810979, EPI_ISL_811028, EPI_ISL_811029, EPI_ISL_811030, EPI_ISL_811031 | MRCG at LSHTM Genomics lab | MRCG at LSHTM Genomics lab | Abdul Karim sesay, Abdoulie Kanteh, Jarra Manneh, Mariama Kujabi, Bakary Sanyang |
| EPI_ISL_812115, EPI_ISL_812116, EPI_ISL_812119, EPI_ISL_812120 | Santa Clara County Public Health Laboratory | Santa Clara County Public Health Laboratory | Santa Clara County Public Health Department |
| EPI_ISL_812135 | UT-Unified State Labs: Public Health Utah DOH | Pathogen Discovery, Respiratory Viruses Branch, Division of Viral Diseases, Centers for Disease Control and Prevention | Yan Li, Ying Tao, Anna Montmayeur, Jing Zhang, Brian Lynch, Krista Queen, Anna Uehara, Rachel Marine, Peter Cook, Clinton R. Paden, Haibin Wang, Suxiang Tong |
| EPI_ISL_812615, EPI_ISL_812617, EPI_ISL_812625, EPI_ISL_812636, EPI_ISL_812637, EPI_ISL_812638, EPI_ISL_812639, EPI_ISL_812640, EPI_ISL_812641, EPI_ISL_812642, EPI_ISL_812643, EPI_ISL_812644, EPI_ISL_812645, EPI_ISL_812646, EPI_ISL_812647, EPI_ISL_812648, EPI_ISL_812649, EPI_ISL_812650, EPI_ISL_812651, EPI_ISL_812652, EPI_ISL_812653, EPI_ISL_812654, EPI_ISL_812655, EPI_ISL_812656, EPI_ISL_812657, EPI_ISL_812658, EPI_ISL_812659, EPI_ISL_812660, EPI_ISL_812661, EPI_ISL_812662, EPI_ISL_812663, EPI_ISL_812664, EPI_ISL_812666, EPI_ISL_812667, EPI_ISL_812669, EPI_ISL_812670, EPI_ISL_812671, EPI_ISL_812672, EPI_ISL_812677, EPI_ISL_812680, EPI_ISL_812681, EPI_ISL_812682, EPI_ISL_812684 | | | |
| see above | United States Air Force School of Aerospace Medicine | United States Air Force School of Aerospace Medicine | Anthony Fries, Jennifer Meyer, Amanda Javorina, Sarah Purves, William Gruner, Clarise Starr, Elizabeth Macias |
| EPI_ISL_812857 | Genomics Program, Children Cancer Hospital | Genomics Program, Children Cancer Hospital | Hatem,A., Hadad,A., Abouelnaga,S., Amer,K., Salah,H., Farawyla,H., Halafawy,A., Mansour,T., shalaby,L., Hassan,W., Soliman,M., Gomaa,C., Hassan,R., Soliman,S., Monuir,G., Hammad,M., Hussein,S., Abdo,I., Jalal,D., El-Zayat,M., El-Shaqnqery,H., Diab,A., Bakry,U., Samir,O., Magdeldin,S., Sayed,A. |
| EPI_ISL_812970 | Oxford Viromics, NDM, University of Oxford; Oxford University Hospitals; Basingstoke and North Hampshire Hospital | COVID-19 Genomics UK (COG-UK) Consortium | Tanya Golubchik, David Bonsall, George Macintyre, Amy Trebes, Mariateresa de Cesare, Catrin Moore, Alex Mobbs, Anita Justice, Robert Shaw, Monique Andersson, Timothy Peto, Emma Wise, Nathan Moore, Jessica Lynch, Nick Cortes, Matilde Mori, Stephen Kidd, David Buck, John Todd, Christophe Fraser |
| EPI_ISL_815280, EPI_ISL_815305, EPI_ISL_815306, EPI_ISL_815309, EPI_ISL_815335, EPI_ISL_815345, EPI_ISL_815370 | Centogene | Centogene | Peter Bauer, Krishna Kumar Kandaswamy, Vivi Hue-Trang Lieu |
| EPI_ISL_822328, EPI_ISL_822329 | Lighthouse Lab in Glasgow | Wellcome Sanger Institute for the COVID-19 Genomics UK (COG-UK) Consortium | Harper VanSteenhouse, Yumi Kasai, David Gray, Carol Clugston, Anna Dominiczak and Alex Alderton, Roberto Amato, Sonia Goncalves, Ewan Harrison, David K. Jackson, Ian Johnston, Dominic Kwiatkowski, Cordelia Langford, John Sillitoe on behalf of the Wellcome Sanger Institute COVID-19 Surveillance Team |
| EPI_ISL_824470, EPI_ISL_824471, EPI_ISL_824472, EPI_ISL_824473, EPI_ISL_824474, EPI_ISL_824475, EPI_ISL_824476, EPI_ISL_824477, EPI_ISL_824478, EPI_ISL_824479, EPI_ISL_824480, EPI_ISL_824481 | | | |
| see above | Hospital Universitari Vall d'Hebron - Vall d'Hebron Institut de Rercerca | Hospital Universitari Vall d'Hebron | Cristina Andrés, Maria Piñana, Josep F Abril, Damir Garcia-Cehic, Ariadna Rando, Juliana Esperalba, Maria Gema Codina, Carla Castillo, Maria Carmen Martin, Tomàs Pumarola, Josep Quer, Andrés Antón |
| EPI_ISL_825716, EPI_ISL_825717, EPI_ISL_825720, EPI_ISL_825762, EPI_ISL_825763, EPI_ISL_825765, EPI_ISL_825766, EPI_ISL_825767, EPI_ISL_825769, EPI_ISL_825770, EPI_ISL_825771, EPI_ISL_825772, EPI_ISL_825773, EPI_ISL_825774, EPI_ISL_825775, EPI_ISL_825776, EPI_ISL_825777, EPI_ISL_825778, EPI_ISL_825779, EPI_ISL_825783 | | | |
| see above | Laboratoire de santé publique du Québec | Laboratoire de santé publique du Québec | Sandrine Moreira, Ioannis Ragoussis, Guillaume Bourque, Jesse Shapiro, Mark Lathrop and Michel Roger on behalf of the CoVSeQ research group (http://covseq.ca/researchgroup) |
| EPI_ISL_826520 | University of Bari Biomedical Sciences and Human Oncology | University of Bari Biomedical Sciences and Human Oncology | Chironna M, Sallustio A., Loconsole D., Accogli M. |
| EPI_ISL_826590, EPI_ISL_826593, EPI_ISL_826658, EPI_ISL_826665 | Montefiore Medical Center | Albert Einstein College of Medicine, Dept. of Microbiology & Immunology, Chandran lab | J. Maximilian Fels, Saad Khan, Ryan Forster, Karin A. Skalina, Surksha Sirichand, Amy S. Fox, Aviv Bergman, William B. Mitchell, Lucia R. Wolgast, Wendy Szymczak, Robert H. Bortz III, M. Eugenia Dieterle, Catalina Florez, Denise Haslwanter, Rohit K. Jangra, Ethan Laudermilch, Ariel S. Wirchnianski, Jason Barnhill, David L. Goldman, Hnin Khine, D. Yitzchak Goldstein, Johanna P. Daily, Kartik Chandran, Libusha Kelly |
| EPI_ISL_826827 | INSPI-CRN DE INFLUENZA Y OTROS VIRUS RESPIRATORIOS | Instituto de Salud Publica de Chile | Javier Tognarelli, Barbara Parra, Loredana Arata, Jaime Lagos, Gisselle Barra, Alfredo Bruno, Domenica de Mora, Solon Narvaez, Jimmy Garcez, Michelle Paez, Martiza Olmedo, Manuel Gonzalez, Patricia Bustos, Rodrigo Fasce, Andres Castillo, Jorge Fernandez |
| EPI_ISL_826959, EPI_ISL_827000 | deCODE genetics | deCODE genetics | Daniel F Gudbjartsson; Agnar Helgason; Hakon Jonsson; Olafur T Magnusson; Pall Melsted; Gudmundur L Norddahl; Jona Saemundsdottir; Asgeir Sigurdsson; Patrick Sulem; Arna B Agustsdottir; Hannes Eggertsson; Berglind Eiriksdottir; Run Fridriksdottir; Elisabet E Gardarsdottir; Gudmundur Georgsson; Olafia S Gretarsdottir; Kjartan R Gudmundsson; Thora R Gunnarsdottir; Arnaldur Gylfason; Hilma Holm; Brynjar O Jensson; Aslaug Jonasdottir; Kamilla S Josefsdottir; Thordur Kristjansson; Droplaug N Magnusdottir; Solvi Rognvaldsson; Louise le Roux; Gudrun Sigmundsdottir; Gardar Sveinbjornsson; Kristin E Sveinsdottir; Maney Sveinsdottir; Emil A Thorarensen; Bjarni Thorbjornsson; Gisli Masson; Ingileif Jonsdottir; Alma Moller; Thorolfur Gudnason; Karl G Kristinsson; Unnur Thorsteinsdottir; Kari Stefansson |
| EPI_ISL_827383, EPI_ISL_827549, EPI_ISL_827551 | The National University Hospital of Iceland | deCODE genetics | Daniel F Gudbjartsson; Agnar Helgason; Hakon Jonsson; Olafur T Magnusson; Pall Melsted; Gudmundur L Norddahl; Jona Saemundsdottir; Asgeir Sigurdsson; Patrick Sulem; Arna B Agustsdottir; Hannes Eggertsson; Berglind Eiriksdottir; Run Fridriksdottir; Elisabet E Gardarsdottir; Gudmundur Georgsson; Olafia S Gretarsdottir; Kjartan R Gudmundsson; Thora R Gunnarsdottir; Arnaldur Gylfason; Hilma Holm; Brynjar O Jensson; Aslaug Jonasdottir; Kamilla S Josefsdottir; Thordur Kristjansson; Droplaug N Magnusdottir; Solvi Rognvaldsson; Louise le Roux; Gudrun Sigmundsdottir; Gardar Sveinbjornsson; Kristin E Sveinsdottir; Maney Sveinsdottir; Emil A Thorarensen; Bjarni Thorbjornsson; Gisli Masson; Ingileif Jonsdottir; Alma Moller; Thorolfur Gudnason; Karl G Kristinsson; Unnur Thorsteinsdottir; Kari Stefansson |
| EPI_ISL_827552 | deCODE genetics | deCODE genetics | Daniel F Gudbjartsson; Agnar Helgason; Hakon Jonsson; Olafur T Magnusson; Pall Melsted; Gudmundur L Norddahl; Jona Saemundsdottir; Asgeir |

| | | | |
|---|---|---|---|
| | | | Sigurdsson; Patrick Sulem; Arna B Agustsdottir; Hannes Eggertsson; Berglind Eiriksdottir; Run Fridriksdottir; Elisabet E Gardarsdottir; Gudmundur Georgsson; Olafia S Gretarsdottir; Kjartan R Gudmundsson; Thora R Gunnarsdottir; Arnaldur Gylfason; Hilma Holm; Brynjar O Jensson; Aslaug Jonasdottir; Kamilla S Josefsdottir; Thordur Kristjansson; Droplaug N Magnusdottir; Solvi Rognvaldsson; Louise le Roux; Gudrun Sigmundsdottir; Gardar Sveinbjornsson; Kristin E Sveinsdottir; Maney Sveinsdottir; Emil A Thorarensen; Bjarni Thorbjornsson; Gisli Masson; Ingileif Jonsdottir; Alma Moller; Thorolfur Gudnason; Karl G Kristinsson; Unnur Thorsteinsdottir; Kari Stefansson |
| EPI_ISL_827822, EPI_ISL_827825 | The National University Hospital of Iceland | deCODE genetics | Daniel F Gudbjartsson; Agnar Helgason; Hakon Jonsson; Olafur T Magnusson; Pall Melsted; Gudmundur L Norddahl; Jona Saemundsdottir; Asgeir Sigurdsson; Patrick Sulem; Arna B Agustsdottir; Hannes Eggertsson; Berglind Eiriksdottir; Run Fridriksdottir; Elisabet E Gardarsdottir; Gudmundur Georgsson; Olafia S Gretarsdottir; Kjartan R Gudmundsson; Thora R Gunnarsdottir; Arnaldur Gylfason; Hilma Holm; Brynjar O Jensson; Aslaug Jonasdottir; Kamilla S Josefsdottir; Thordur Kristjansson; Droplaug N Magnusdottir; Solvi Rognvaldsson; Louise le Roux; Gudrun Sigmundsdottir; Gardar Sveinbjornsson; Kristin E Sveinsdottir; Maney Sveinsdottir; Emil A Thorarensen; Bjarni Thorbjornsson; Gisli Masson; Ingileif Jonsdottir; Alma Moller; Thorolfur Gudnason; Karl G Kristinsson; Unnur Thorsteinsdottir; Kari Stefansson |
| EPI_ISL_827853, EPI_ISL_827854, EPI_ISL_827855, EPI_ISL_827856, EPI_ISL_827857, EPI_ISL_827873, EPI_ISL_827874, EPI_ISL_827924 | deCODE genetics | deCODE genetics | Daniel F Gudbjartsson; Agnar Helgason; Hakon Jonsson; Olafur T Magnusson; Pall Melsted; Gudmundur L Norddahl; Jona Saemundsdottir; Asgeir Sigurdsson; Patrick Sulem; Arna B Agustsdottir; Hannes Eggertsson; Berglind Eiriksdottir; Run Fridriksdottir; Elisabet E Gardarsdottir; Gudmundur Georgsson; Olafia S Gretarsdottir; Kjartan R Gudmundsson; Thora R Gunnarsdottir; Arnaldur Gylfason; Hilma Holm; Brynjar O Jensson; Aslaug Jonasdottir; Kamilla S Josefsdottir; Thordur Kristjansson; Droplaug N Magnusdottir; Solvi Rognvaldsson; Louise le Roux; Gudrun Sigmundsdottir; Gardar Sveinbjornsson; Kristin E Sveinsdottir; Maney Sveinsdottir; Emil A Thorarensen; Bjarni Thorbjornsson; Gisli Masson; Ingileif Jonsdottir; Alma Moller; Thorolfur Gudnason; Karl G Kristinsson; Unnur Thorsteinsdottir; Kari Stefansson |
| EPI_ISL_828215 | The National University Hospital of Iceland | deCODE genetics | Daniel F Gudbjartsson; Agnar Helgason; Hakon Jonsson; Olafur T Magnusson; Pall Melsted; Gudmundur L Norddahl; Jona Saemundsdottir; Asgeir Sigurdsson; Patrick Sulem; Arna B Agustsdottir; Hannes Eggertsson; Berglind Eiriksdottir; Run Fridriksdottir; Elisabet E Gardarsdottir; Gudmundur Georgsson; Olafia S Gretarsdottir; Kjartan R Gudmundsson; Thora R Gunnarsdottir; Arnaldur Gylfason; Hilma Holm; Brynjar O Jensson; Aslaug Jonasdottir; Kamilla S Josefsdottir; Thordur Kristjansson; Droplaug N Magnusdottir; Solvi Rognvaldsson; Louise le Roux; Gudrun Sigmundsdottir; Gardar Sveinbjornsson; Kristin E Sveinsdottir; Maney Sveinsdottir; Emil A Thorarensen; Bjarni Thorbjornsson; Gisli Masson; Ingileif Jonsdottir; Alma Moller; Thorolfur Gudnason; Karl G Kristinsson; Unnur Thorsteinsdottir; Kari Stefansson |
| EPI_ISL_828217, EPI_ISL_828218 | deCODE genetics | deCODE genetics | Daniel F Gudbjartsson; Agnar Helgason; Hakon Jonsson; Olafur T Magnusson; Pall Melsted; Gudmundur L Norddahl; Jona Saemundsdottir; Asgeir Sigurdsson; Patrick Sulem; Arna B Agustsdottir; Hannes Eggertsson; Berglind Eiriksdottir; Run Fridriksdottir; Elisabet E Gardarsdottir; Gudmundur Georgsson; Olafia S Gretarsdottir; Kjartan R Gudmundsson; Thora R Gunnarsdottir; Arnaldur Gylfason; Hilma Holm; Brynjar O Jensson; Aslaug Jonasdottir; Kamilla S Josefsdottir; Thordur Kristjansson; Droplaug N Magnusdottir; Solvi Rognvaldsson; Louise le Roux; Gudrun Sigmundsdottir; Gardar Sveinbjornsson; Kristin E Sveinsdottir; Maney Sveinsdottir; Emil A Thorarensen; Bjarni Thorbjornsson; Gisli Masson; Ingileif Jonsdottir; Alma Moller; Thorolfur Gudnason; Karl G Kristinsson; Unnur Thorsteinsdottir; Kari Stefansson |
| EPI_ISL_828320, EPI_ISL_828322, EPI_ISL_828595, EPI_ISL_828596 | The National University Hospital of Iceland | deCODE genetics | Daniel F Gudbjartsson; Agnar Helgason; Hakon Jonsson; Olafur T Magnusson; Pall Melsted; Gudmundur L Norddahl; Jona Saemundsdottir; Asgeir Sigurdsson; Patrick Sulem; Arna B Agustsdottir; Hannes Eggertsson; Berglind Eiriksdottir; Run Fridriksdottir; Elisabet E Gardarsdottir; Gudmundur Georgsson; Olafia S Gretarsdottir; Kjartan R Gudmundsson; Thora R Gunnarsdottir; Arnaldur Gylfason; Hilma Holm; Brynjar O Jensson; Aslaug Jonasdottir; Kamilla S Josefsdottir; Thordur Kristjansson; Droplaug N Magnusdottir; Solvi Rognvaldsson; Louise le Roux; Gudrun Sigmundsdottir; Gardar Sveinbjornsson; Kristin E Sveinsdottir; Maney Sveinsdottir; Emil A Thorarensen; Bjarni Thorbjornsson; Gisli Masson; Ingileif Jonsdottir; Alma Moller; Thorolfur Gudnason; Karl G Kristinsson; Unnur Thorsteinsdottir; Kari Stefansson |
| EPI_ISL_828597, EPI_ISL_828598 | deCODE genetics | deCODE genetics | Daniel F Gudbjartsson; Agnar Helgason; Hakon Jonsson; Olafur T Magnusson; Pall Melsted; Gudmundur L Norddahl; Jona Saemundsdottir; Asgeir Sigurdsson; Patrick Sulem; Arna B Agustsdottir; Hannes Eggertsson; Berglind Eiriksdottir; Run Fridriksdottir; Elisabet E Gardarsdottir; Gudmundur Georgsson; Olafia S Gretarsdottir; Kjartan R Gudmundsson; Thora R Gunnarsdottir; Arnaldur Gylfason; Hilma Holm; Brynjar O Jensson; Aslaug Jonasdottir; Kamilla S Josefsdottir; Thordur Kristjansson; Droplaug N Magnusdottir; Solvi Rognvaldsson; Louise le Roux; Gudrun Sigmundsdottir; Gardar Sveinbjornsson; Kristin E Sveinsdottir; Maney Sveinsdottir; Emil A Thorarensen; Bjarni Thorbjornsson; Gisli Masson; Ingileif Jonsdottir; Alma Moller; Thorolfur Gudnason; Karl G Kristinsson; Unnur Thorsteinsdottir; Kari Stefansson |
| EPI_ISL_828630, EPI_ISL_828634 | The National University Hospital of Iceland | deCODE genetics | Daniel F Gudbjartsson; Agnar Helgason; Hakon Jonsson; Olafur T Magnusson; Pall Melsted; Gudmundur L Norddahl; Jona Saemundsdottir; Asgeir Sigurdsson; Patrick Sulem; Arna B Agustsdottir; Hannes Eggertsson; Berglind Eiriksdottir; Run Fridriksdottir; Elisabet E Gardarsdottir; Gudmundur Georgsson; Olafia S Gretarsdottir; Kjartan R Gudmundsson; Thora R Gunnarsdottir; Arnaldur Gylfason; Hilma Holm; Brynjar O Jensson; Aslaug Jonasdottir; Kamilla S Josefsdottir; Thordur Kristjansson; Droplaug N Magnusdottir; Solvi Rognvaldsson; Louise le Roux; Gudrun Sigmundsdottir; Gardar Sveinbjornsson; Kristin E Sveinsdottir; Maney Sveinsdottir; Emil A Thorarensen; Bjarni Thorbjornsson; Gisli Masson; Ingileif Jonsdottir; Alma Moller; Thorolfur Gudnason; Karl G Kristinsson; Unnur Thorsteinsdottir; Kari Stefansson |
| EPI_ISL_828636, EPI_ISL_828637, EPI_ISL_828638, EPI_ISL_828640, EPI_ISL_829348 | deCODE genetics | deCODE genetics | Daniel F Gudbjartsson; Agnar Helgason; Hakon Jonsson; Olafur T Magnusson; Pall Melsted; Gudmundur L Norddahl; Jona Saemundsdottir; Asgeir Sigurdsson; Patrick Sulem; Arna B Agustsdottir; Hannes Eggertsson; Berglind Eiriksdottir; Run Fridriksdottir; Elisabet E Gardarsdottir; Gudmundur Georgsson; Olafia S Gretarsdottir; Kjartan R Gudmundsson; Thora R Gunnarsdottir; Arnaldur Gylfason; Hilma Holm; Brynjar O Jensson; Aslaug Jonasdottir; Kamilla S Josefsdottir; Thordur Kristjansson; Droplaug N Magnusdottir; Solvi Rognvaldsson; Louise le Roux; Gudrun Sigmundsdottir; Gardar Sveinbjornsson; Kristin E Sveinsdottir; Maney Sveinsdottir; Emil A Thorarensen; Bjarni Thorbjornsson; Gisli Masson; Ingileif Jonsdottir; Alma Moller; Thorolfur Gudnason; Karl G Kristinsson; Unnur Thorsteinsdottir; Kari Stefansson |
| EPI_ISL_829376 | The National University Hospital of Iceland | deCODE genetics | Daniel F Gudbjartsson; Agnar Helgason; Hakon Jonsson; Olafur T Magnusson; Pall Melsted; Gudmundur L Norddahl; Jona Saemundsdottir; Asgeir Sigurdsson; Patrick Sulem; Arna B Agustsdottir; Hannes Eggertsson; Berglind Eiriksdottir; Run Fridriksdottir; Elisabet E Gardarsdottir; Gudmundur Georgsson; Olafia S Gretarsdottir; Kjartan R Gudmundsson; Thora R Gunnarsdottir; Arnaldur Gylfason; Hilma Holm; Brynjar O Jensson; Aslaug Jonasdottir; Kamilla S Josefsdottir; Thordur Kristjansson; Droplaug N Magnusdottir; Solvi Rognvaldsson; Louise le Roux; Gudrun Sigmundsdottir; Gardar Sveinbjornsson; Kristin E Sveinsdottir; Maney Sveinsdottir; Emil A Thorarensen; Bjarni Thorbjornsson; Gisli Masson; Ingileif Jonsdottir; Alma Moller; Thorolfur Gudnason; Karl G Kristinsson; Unnur Thorsteinsdottir; Kari Stefansson |
| EPI_ISL_829535, EPI_ISL_829567, EPI_ISL_829720, EPI_ISL_829739, EPI_ISL_829740, EPI_ISL_829821, EPI_ISL_829975, EPI_ISL_830032 | deCODE genetics | deCODE genetics | Daniel F Gudbjartsson; Agnar Helgason; Hakon Jonsson; Olafur T Magnusson; Pall Melsted; Gudmundur L Norddahl; Jona Saemundsdottir; Asgeir Sigurdsson; Patrick Sulem; Arna B Agustsdottir; Hannes Eggertsson; Berglind Eiriksdottir; Run Fridriksdottir; Elisabet E Gardarsdottir; Gudmundur Georgsson; Olafia S Gretarsdottir; Kjartan R Gudmundsson; Thora R Gunnarsdottir; Arnaldur Gylfason; Hilma Holm; Brynjar O Jensson; Aslaug Jonasdottir; Kamilla S Josefsdottir; Thordur Kristjansson; Droplaug N Magnusdottir; Solvi Rognvaldsson; Louise le Roux; Gudrun Sigmundsdottir; Gardar Sveinbjornsson; Kristin E Sveinsdottir; Maney Sveinsdottir; Emil A Thorarensen; Bjarni Thorbjornsson; Gisli Masson; Ingileif Jonsdottir; Alma Moller; Thorolfur Gudnason; Karl G Kristinsson; Unnur Thorsteinsdottir; Kari Stefansson |
| EPI_ISL_830037, EPI_ISL_830038 | The National University Hospital of Iceland | deCODE genetics | Daniel F Gudbjartsson; Agnar Helgason; Hakon Jonsson; Olafur T Magnusson; Pall Melsted; Gudmundur L Norddahl; Jona Saemundsdottir; Asgeir Sigurdsson; Patrick Sulem; Arna B Agustsdottir; Hannes Eggertsson; Berglind Eiriksdottir; Run Fridriksdottir; Elisabet E Gardarsdottir; Gudmundur Georgsson; Olafia S Gretarsdottir; Kjartan R Gudmundsson; Thora R Gunnarsdottir; Arnaldur Gylfason; Hilma Holm; Brynjar O Jensson; Aslaug Jonasdottir; Kamilla S Josefsdottir; Thordur Kristjansson; Droplaug N Magnusdottir; Solvi Rognvaldsson; Louise le Roux; Gudrun Sigmundsdottir; Gardar Sveinbjornsson; Kristin E Sveinsdottir; Maney Sveinsdottir; Emil A Thorarensen; Bjarni Thorbjornsson; Gisli Masson; Ingileif Jonsdottir; Alma Moller; Thorolfur Gudnason; Karl G Kristinsson; Unnur Thorsteinsdottir; Kari Stefansson |
| EPI_ISL_830183, EPI_ISL_830273, EPI_ISL_830275, EPI_ISL_830277, EPI_ISL_830279 | deCODE genetics | deCODE genetics | Daniel F Gudbjartsson; Agnar Helgason; Hakon Jonsson; Olafur T Magnusson; Pall Melsted; Gudmundur L Norddahl; Jona Saemundsdottir; Asgeir Sigurdsson; Patrick Sulem; Arna B Agustsdottir; Hannes Eggertsson; Berglind Eiriksdottir; Run Fridriksdottir; Elisabet E Gardarsdottir; Gudmundur Georgsson; Olafia S Gretarsdottir; Kjartan R Gudmundsson; Thora R Gunnarsdottir; Arnaldur Gylfason; Hilma Holm; Brynjar O Jensson; Aslaug Jonasdottir; Kamilla S Josefsdottir; Thordur Kristjansson; Droplaug N Magnusdottir; Solvi Rognvaldsson; Louise le Roux; Gudrun Sigmundsdottir; Gardar Sveinbjornsson; Kristin E Sveinsdottir; Maney Sveinsdottir; Emil A Thorarensen; Bjarni Thorbjornsson; Gisli Masson; Ingileif Jonsdottir; Alma Moller; Thorolfur Gudnason; Karl G Kristinsson; Unnur Thorsteinsdottir; Kari Stefansson |
| EPI_ISL_830377, EPI_ISL_830378, EPI_ISL_830379, EPI_ISL_830380, EPI_ISL_830381, EPI_ISL_830382 | The National University Hospital of Iceland | deCODE genetics | Daniel F Gudbjartsson; Agnar Helgason; Hakon Jonsson; Olafur T Magnusson; Pall Melsted; Gudmundur L Norddahl; Jona Saemundsdottir; Asgeir Sigurdsson; Patrick Sulem; Arna B Agustsdottir; Hannes Eggertsson; Berglind Eiriksdottir; Run Fridriksdottir; Elisabet E Gardarsdottir; Gudmundur Georgsson; Olafia S Gretarsdottir; Kjartan R Gudmundsson; Thora R Gunnarsdottir; Arnaldur Gylfason; Hilma Holm; Brynjar O Jensson; Aslaug Jonasdottir; Kamilla S Josefsdottir; Thordur Kristjansson; Droplaug N Magnusdottir; Solvi Rognvaldsson; Louise le Roux; Gudrun Sigmundsdottir; Gardar |

| | | | |
|---|---|---|---|
| | | | Sveinbjornsson; Kristin E Sveinsdottir; Maney Sveinsdottir; Emil A Thorarensen; Bjarni Thorbjornsson; Gisli Masson; Ingileif Jonsdottir; Alma Moller; Thorolfur Gudnason; Karl G Kristinsson; Unnur Thorsteinsdottir; Kari Stefansson |
| EPI_ISL_831090 | Hospital Universitario La Paz (Madrid) | SeqCOVID-SPAIN consortium/IBV(CSIC) | María Rodríguez-Tejedor, Elias Dahdouh, Fernando Lázaro-Perona, Jesús Mingorance and SeqCOVID-SPAIN consortium |
| EPI_ISL_831898, EPI_ISL_831913, EPI_ISL_831938 | Laboratório de Microbiologia Molecular - Universidade FEEVALE | Universidade Federal de Ciências da Saúde de Porto Alegre | Vinícius Bonetti Franceschi, Amanda de Menezes Mayer, Gabriel Dickin Caldana, Carla Andretta Moreira Neves, Patrícia Aline Gröhs Ferrareze, Gabriela Bettella Cybis, Ricardo Ariel Zimerman, Lívia Kmetzsch, Fernando Rosado Spilki, Claudia Elizabeth Thompson |
| EPI_ISL_831969 | unknown | The Public Health Agency of Sweden | Department of Microbiology, The Public Health Agency of Sweden |
| EPI_ISL_831971 | Orebro klinisk mikrobiologi | The Public Health Agency of Sweden | Department of Microbiology, The Public Health Agency of Sweden |
| EPI_ISL_832101 | Hospital Universitari Germans Trias i Pujol(HUGTiP)/Fundació Lluita contra la SIDA (FLSida) | IrsiCaixa AIDS Research Lab | Marc Noguera-Julian, Mariona Parera, Maria Pilar Armengol, Marta Massanella, Ester Ballana, Lidia Ruiz, Nuria Izquierdo, Jorge Carrillo, Roger Paredes, Julia Blanco, Joaquim Segalés, Bonaventura Clotet |
| EPI_ISL_832412, EPI_ISL_832413 | OHSU Lab Services Molecular Microbiology Lab | Oregon SARS-CoV-2 Genome Sequencing Center | Brendan L. O'Connell, Ruth V. Nichols, Sally Grindstaff, Alec J. Hirsch, Donna Hansel, Guang Fan, Daniel N. Streblow, William B. Messer, Andrew C. Adey, Benjamin N. Bimber, Brian J. O'Roak |
| EPI_ISL_833497 | Klinik Apotek Dein, Jakarta, Indonesia | Biosafety Level-3 Laboratory, Indonesian Institute of Sciences (LIPI) | Syam Budi Iryanto, Andri Wardiana, Anggia Prasetyoputri, Anik Budhi Dharmayanthi, Isa Nuryana, Ade Andriani, Ahmad Fathoni, Saiful anwar, Pamella Apriliana, Ratih Asmana Ningrum |
| EPI_ISL_833499 | RS Qadr, Tangerang, Indonesia | Biosafety Level-3 Laboratory, Indonesian Institute of Sciences (LIPI) | Isa Nuryana, Ade Andriani, Ahmad Fathoni, Anik Budhi Dharmayanthi, Syam Budi Iryanto, Andri Wardiana, Anggia Prasetyoputri, Ratna Dwi Ramadani, Sugiyono Saputra, Ratih Asmana Ningrum |
| EPI_ISL_833519 | Veterinary Specialized Instute "Nis" | Veterinary Specialized Institute "Kraljevo", Serbia | Vidanovic,D., Tesovic,B., Manic,M., Petrovic,M.,Knezevic,A., Jovanovic,T., Jankovic,M., Sekler,M., Banovic Djeri,B., Petrovic,T., Volkening,J., Afonso,C. |
| EPI_ISL_837269, EPI_ISL_837270, EPI_ISL_837271 | Istituto Zooprofilattico Sperimentale del Mezzogiorno | TIGEM | Patrizia Annunziata, Andrea Ballabio, Valentina Bouche, Davide Cacchiarelli (CorrespAuthor), Pellegrino Cerino, Chiara Colantuono, Lucio Di Filippo, Antonio Grimaldi, Antonio Limone, Gabriella Loconte, Anna Manfredi, Francesco Panariello, Biancamaria Pierri, Marcello Salvi, Lucia Vassallo |
| EPI_ISL_837602, EPI_ISL_837603, EPI_ISL_837805, EPI_ISL_837806, EPI_ISL_837807, EPI_ISL_837808, EPI_ISL_837809 | Instituto Nacional de Enfermedades Respiratorias (INER) | Instituto Nacional de Enfermedades Respiratorias (INER) | Celia Boukadida, Margarita Matías-Florentino, Alma Rincón-Rubio, Hector Esteban Paz-Juárez, Olivia Briceño, Edgar Sevilla-Reyes, Fidencio Mejía-Nepomuceno, Mario Mújica-Sánchez, Eduardo Becerril-Vargas, José Arturo Martínez-Orozco, Alejandra Hernández-Terán, Jorge Salas-Hernández, Santiago Ávila-Ríos, Joel Armando Vázquez-Pérez |
| EPI_ISL_845660, EPI_ISL_845667 | Quest Diagnostics | Quest Diagnostics | Rosenthal,S.H., Gerasimova,A., Kagan,R.M., Anderson, B., Bernstein, L.E., Livingston, K.E., Hua, M., Liu Y., Shalhout, D.F., Shlyakhter, I.A., Owen, R., Lacbawan, F. |
| EPI_ISL_849941, EPI_ISL_849943 | UC Davis- Department of Pathology and Laboratory Medicine | Chan-Zuckerberg Biohub | CZB Cliahub Consortium |
| EPI_ISL_850396, EPI_ISL_850401, EPI_ISL_850409, EPI_ISL_850410, EPI_ISL_850411, EPI_ISL_850413, EPI_ISL_850414, EPI_ISL_850415, EPI_ISL_850416, EPI_ISL_850417, EPI_ISL_850418, EPI_ISL_850419, EPI_ISL_850420, EPI_ISL_850421, EPI_ISL_850422, EPI_ISL_850423, EPI_ISL_850424, EPI_ISL_850425, EPI_ISL_850426, EPI_ISL_850427, EPI_ISL_850428, EPI_ISL_850429, EPI_ISL_850430, EPI_ISL_850431, EPI_ISL_850432, EPI_ISL_850433, EPI_ISL_850434, EPI_ISL_850435, EPI_ISL_850436, EPI_ISL_850437, EPI_ISL_850438, EPI_ISL_850439, EPI_ISL_850440, EPI_ISL_850441, EPI_ISL_850442, EPI_ISL_850443, EPI_ISL_850444, EPI_ISL_850445, EPI_ISL_850446, EPI_ISL_850447, EPI_ISL_850448, EPI_ISL_850449, EPI_ISL_850450, EPI_ISL_850451, EPI_ISL_850452, EPI_ISL_850453, EPI_ISL_850454, EPI_ISL_850455, EPI_ISL_850457, EPI_ISL_850458, EPI_ISL_850459, EPI_ISL_850460, EPI_ISL_850461, EPI_ISL_850462, EPI_ISL_850463, EPI_ISL_850464, EPI_ISL_850465, EPI_ISL_850466, EPI_ISL_850467, EPI_ISL_850468, EPI_ISL_850469, EPI_ISL_850470, EPI_ISL_850471, EPI_ISL_850472, EPI_ISL_850473, EPI_ISL_850474, EPI_ISL_850475, EPI_ISL_850476, EPI_ISL_850477, EPI_ISL_850478, EPI_ISL_850479, EPI_ISL_850480, EPI_ISL_850481, EPI_ISL_850482, EPI_ISL_850483, EPI_ISL_850484, EPI_ISL_850485, EPI_ISL_850486, EPI_ISL_850487, EPI_ISL_850488, EPI_ISL_850489, EPI_ISL_850490, EPI_ISL_850491, EPI_ISL_850492, EPI_ISL_850493, EPI_ISL_850495 | | | |
| see above | Division of Emerging Infectious Diseases, Bureau of Infectious Diseases Diagnosis Control, Korea Disease Control and Prevention Agency | Division of Emerging Infectious Diseases, Bureau of Infectious Diseases Diagnosis Control, Korea Disease Control and Prevention Agency | Ae Kyung Park, Il-Hwan Kim, Heui Man Kim, Jeong-Min Kim, Namjoo Lee, Chaeyoung Lee, Sang Hee Woo, Eun-Jin Kim |
| EPI_ISL_852586, EPI_ISL_852588, EPI_ISL_852591, EPI_ISL_852592 | Max von Pettenkofer Institute, Virology, National Reference Center for Retroviruses, LMU München | Laboratory for Functional Genome Analysis, Dept. Genomics, Gene Center of the LMU Munich | Max Muenchhoff, Stefan Krebs, Alexander Graf, Oliver Keppler, Helmut Blum |
| EPI_ISL_852655, EPI_ISL_852656 | Institute of Virology, Medical Center, University of Freiburg, Freiburg, Germany | Institute of Virology, Clinial Virus Genomics, Medical Center, University of Freiburg, Freiburg, Germany | Jonas Fuchs, Lisa Kern, Sandra Reuter, Hajo Grundmann, Marcus Panning |
| EPI_ISL_853785 | Department of Microbiology, University Innsbruck | Bergthaler laboratory, CeMM Research Center for Molecular Medicine of the Austrian Academy of Sciences | Lukas Endler, Alexandra Popa, Benedikt Agerer, Jakob-Wendelin Genger, Alexander Lercher, Anna Schedl, Thomas Penz, Michael Schuster, Jan Laine, Martin Senekowitsch, Christoph Bock, Andreas Bergthaler |
| EPI_ISL_853846, EPI_ISL_853847, EPI_ISL_853848, EPI_ISL_853849, EPI_ISL_853850, EPI_ISL_853851, EPI_ISL_853852, EPI_ISL_853853, EPI_ISL_853904, EPI_ISL_853905, EPI_ISL_853906, EPI_ISL_853907, EPI_ISL_853908, EPI_ISL_853909, EPI_ISL_853910, EPI_ISL_853911, EPI_ISL_853912, EPI_ISL_853913, EPI_ISL_853958, EPI_ISL_853961, EPI_ISL_854218, EPI_ISL_854219 | | | |
| see above | Center for Virology, Medical University of Vienna | Bergthaler laboratory, CeMM Research Center for Molecular Medicine of the Austrian Academy of Sciences | Lukas Endler, Alexandra Popa, Benedikt Agerer, Jakob-Wendelin Genger, Alexander Lercher, Anna Schedl, Thomas Penz, Michael Schuster, Jan Laine, Martin Senekowitsch, Christoph Bock, Andreas Bergthaler |
| EPI_ISL_854229 | Institute of Legal Medicine, Medical University of Innsbruck | Bergthaler laboratory, CeMM Research Center for Molecular Medicine of the Austrian Academy of Sciences | Lukas Endler, Alexandra Popa, Benedikt Agerer, Jakob-Wendelin Genger, Alexander Lercher, Anna Schedl, Thomas Penz, Michael Schuster, Jan Laine, Martin Senekowitsch, Christoph Bock, Andreas Bergthaler |
| EPI_ISL_854241, EPI_ISL_854253, EPI_ISL_854258, EPI_ISL_854295 | Center for Virology, Medical University of Vienna | Bergthaler laboratory, CeMM Research Center for Molecular Medicine of the Austrian Academy of Sciences | Lukas Endler, Alexandra Popa, Benedikt Agerer, Jakob-Wendelin Genger, Alexander Lercher, Anna Schedl, Thomas Penz, Michael Schuster, Jan Laine, Martin Senekowitsch, Christoph Bock, Andreas Bergthaler |
| EPI_ISL_854760 | Microbiological Diagnostic Unit - Public Health Laboratory (MDU-PHL) | MDU-PHL | Seemann T., Sait, M.L., Sherry, N.L. |
| EPI_ISL_859561, EPI_ISL_859562, EPI_ISL_859563, EPI_ISL_859564, EPI_ISL_859565, EPI_ISL_859566, EPI_ISL_859567, EPI_ISL_859568, EPI_ISL_859569, EPI_ISL_859570, EPI_ISL_859571, EPI_ISL_859572, EPI_ISL_859573, EPI_ISL_859574, EPI_ISL_859575, EPI_ISL_859576, EPI_ISL_859577, EPI_ISL_859601, EPI_ISL_859602, EPI_ISL_859603, EPI_ISL_859604, EPI_ISL_859605, EPI_ISL_859607, EPI_ISL_859609, EPI_ISL_859610, EPI_ISL_859619, EPI_ISL_859622, EPI_ISL_859637, EPI_ISL_859656, EPI_ISL_859657, EPI_ISL_859658, EPI_ISL_859659, EPI_ISL_859660, EPI_ISL_859662, EPI_ISL_859666, EPI_ISL_859667, EPI_ISL_859668, EPI_ISL_859669, EPI_ISL_859670, EPI_ISL_859671, EPI_ISL_859672, EPI_ISL_859673, EPI_ISL_859674 | | | |
| see above | BTC, Khalifa University | BTC, Khalifa University | Al Safar et al |
| EPI_ISL_861910 | LATE - Laboratório de Técnicas Especiais - Hospital Israelita Albert Einstein | LATE - Laboratório de Técnicas Especiais - Hospital Israelita Albert Einstein | Deyvid Amgarten, Fernanda de Mello Malta, Raquel Riyuzo, Ana Paula Moreira Salles, Pedro Henrique Sebe Rodrigues, João Renato Rebello Pinho |
| EPI_ISL_862052, EPI_ISL_862053, EPI_ISL_862054, EPI_ISL_862055 | Department of Virology and Immunology, University of Helsinki and Helsinki University Hospital, Huslab Finland | Department of Virology, Faculty of Medicine, University of Helsinki, Helsinki, Finland | Teemu Smura, Ravi Kant, Phuoc Truong, Hussein Alburkat, Hanna Liimatainen, Hannimari Kallio-Kokko, Jenni Virtanen, Maija Suvanto, Sari Hannula, Harri Kangas, Pekka Ellonen, Olli Vapalahti |
| EPI_ISL_862594 | Hospital Comarcal de Melilla | Instituto de Salud Carlos III | Iglesias-Caballero, M. Camarero, S. Molinero Calamita, M. González-Esguevillas, M. Pozo, F. Casas, I. Jiménez, P. Jiménez, M. Zaballos, A. Monzón, S. Varona, S. Juliá, M. Cuesta, I. López, J. |
| EPI_ISL_871790 | Multidisciplinary Research Unit, DHR-ICMR, Institute of Medical Sciences, Banaras Hindu University | Multidisciplinary Research Unit, DHR-ICMR, Institute of Medical Sciences, Banaras Hindu University | Royana Singh, Priyoneel Basu, Ashish, Nitish Kumar Singh, Abhay Kumar Yadav, Manpreet Kaur, Arup Acharjee, Deepa Devadas, Chetan Sahni, Gulshan Kumar, Tribhuwan Mohan Mohapatra, Richa Arya, Prashant Singh, Jay Prakash Maurya, Surendra Pratap Mishra |
| EPI_ISL_871990 | Hospital Clínico San Carlos | Instituto de Salud Carlos III | Iglesias-Caballero, M. Camarero, S. Molinero Calamita, M. González-Esguevillas, M. Pozo, F. Casas, I. Jiménez, P. Jiménez, M. Zaballos, A. Monzón, S. Varona, S. Juliá, M. Cuesta, I. Rodríguez, I. |
| EPI_ISL_871991 | Hospital Universitario Severo Ochoa | Instituto de Salud Carlos III | Iglesias-Caballero, M. Camarero, S. Molinero Calamita, M. González-Esguevillas, M. Pozo, F. Casas, I. Jiménez, P. Jiménez, M. Zaballos, A. Monzón, S. Varona, S. Juliá, M. Cuesta, I. García, M.L. |
| EPI_ISL_871992, EPI_ISL_871993 | Hospital Clínico San Carlos | Instituto de Salud Carlos III | Iglesias-Caballero, M. Camarero, S. Molinero Calamita, M. González-Esguevillas, M. Pozo, F. Casas, I. Jiménez, P. Jiménez, M. Zaballos, A. Monzón, S. Varona, S. Juliá, M. Cuesta, I. Rodríguez, I. |
| EPI_ISL_872009, EPI_ISL_872010, EPI_ISL_872011, EPI_ISL_872012, EPI_ISL_872013, EPI_ISL_872014, EPI_ISL_872015, EPI_ISL_872016, EPI_ISL_872017, EPI_ISL_872018, EPI_ISL_872019, EPI_ISL_872020, EPI_ISL_872021, EPI_ISL_872022, EPI_ISL_872023, EPI_ISL_872024, EPI_ISL_872025, EPI_ISL_872026 | | | |
| see above | Ospedale "Di Venere" | Beaconlab (Bioinformatics, Evolution and Comparative Genomics lab), Dept of Biosciences, University on Milan | Iacobellis M, d'Avenia M, Piluscio R, Parisi A, Chiara M, Manzari C, Pesole G |

| | | | |
|---|---|---|---|
| EPI_ISL_875515 | Influenza etiology and epidemiology laboratory | Pathogenic Microorganisms Variability Laboratory | Alexey Shchetinin, Olesya Venchakova, Maria Nikiforova, Andrei Siniavin, Nadezhda Kuznetsova, Elena Shidlovskaya, Elizaveta Divisenko, Kirill Krasnoslobotsev, Evgeniya Mukasheva, Anna Ignatieva, Svetlana Trushakova, Andrey Pochtovyy, Valeria Bacalin, Evgeny Usachev, Olga Burgasova, Ludmila Kolobukhina, Svetlana Smetanina, Elena Burtseva, Denis Logunov, Vladimir Gushchin, Alexander Gintsburg |
| EPI_ISL_876573, EPI_ISL_876574, EPI_ISL_876575 | Florida Bureau of Public Health Laboratories | Florida Bureau of Public Health Laboratories | Sarah Schmedes, Jason Blanton |
| EPI_ISL_876598, EPI_ISL_876599, EPI_ISL_876600, EPI_ISL_876601, EPI_ISL_876602, EPI_ISL_876603, EPI_ISL_876604, EPI_ISL_876605, EPI_ISL_876606, EPI_ISL_876607, EPI_ISL_876608 | | | |
| see above | Toronto Invasive Bacterial Diseases Network | McMaster University | Allison McGeer, Patryk Aftanas, Hooman Derakhshani, Angel Li, Kuganya Nirmalarajah, Emily Panousis, Ahmed Draia, Jalees Nasir, Michael Surette, Samira Mubareka, Andrew G. McArthur |
| EPI_ISL_876817, EPI_ISL_876818, EPI_ISL_876822, EPI_ISL_876830, EPI_ISL_876913, EPI_ISL_876914, EPI_ISL_876915, EPI_ISL_876916, EPI_ISL_876917, EPI_ISL_876918, EPI_ISL_876919, EPI_ISL_876920, EPI_ISL_876921, EPI_ISL_876922, EPI_ISL_876923, EPI_ISL_876924, EPI_ISL_876925, EPI_ISL_876926, EPI_ISL_877121 | | | |
| see above | Quest Diagnostics | Quest Diagnostics | Rosenthal,S.H., Gerasimova,A., Kagan,R.M., Anderson, B., Hua, M., Liu Y., Bernstein, L.E., Livingston, K.E., Perez, A., Shalhout, D.F., Shlyakhter, I.A., Owen, R., Tanpaiboon, P., Lacbawan, F. |
| EPI_ISL_877676, EPI_ISL_877677, EPI_ISL_877678, EPI_ISL_877679, EPI_ISL_877680, EPI_ISL_877681, EPI_ISL_877682, EPI_ISL_877683, EPI_ISL_877684, EPI_ISL_877685, EPI_ISL_877686, EPI_ISL_877687, EPI_ISL_877688, EPI_ISL_877689, EPI_ISL_877690, EPI_ISL_877691, EPI_ISL_877692, EPI_ISL_877693, EPI_ISL_877694, EPI_ISL_877695, EPI_ISL_877696, EPI_ISL_877697, EPI_ISL_877698, EPI_ISL_877699, EPI_ISL_877700, EPI_ISL_877701, EPI_ISL_877702, EPI_ISL_877703, EPI_ISL_877704, EPI_ISL_877705, EPI_ISL_877706, EPI_ISL_877707, EPI_ISL_877708, EPI_ISL_877709, EPI_ISL_877710, EPI_ISL_877711, EPI_ISL_877712, EPI_ISL_877713, EPI_ISL_877714, EPI_ISL_877715, EPI_ISL_877716, EPI_ISL_877717, EPI_ISL_877718, EPI_ISL_877719, EPI_ISL_877720, EPI_ISL_877721, EPI_ISL_877722, EPI_ISL_877723, EPI_ISL_877724, EPI_ISL_877725, EPI_ISL_877726, EPI_ISL_877727, EPI_ISL_877728, EPI_ISL_877729, EPI_ISL_877730, EPI_ISL_877731, EPI_ISL_877732, EPI_ISL_877733, EPI_ISL_877734, EPI_ISL_877735, EPI_ISL_877736, EPI_ISL_877737, EPI_ISL_877738, EPI_ISL_877739, EPI_ISL_877740, EPI_ISL_877741, EPI_ISL_877742, EPI_ISL_877743, EPI_ISL_877744, EPI_ISL_877745, EPI_ISL_877746, EPI_ISL_877747, EPI_ISL_877748, EPI_ISL_877749, EPI_ISL_877750, EPI_ISL_877751, EPI_ISL_877752, EPI_ISL_877753, EPI_ISL_877754, EPI_ISL_877755, EPI_ISL_877756, EPI_ISL_877757, EPI_ISL_877758, EPI_ISL_877759, EPI_ISL_877760 | | | |
| see above | Clinical Molecular Microbiology Laboratory, UNC Hospital | Dirk Dittmer | Razia Moorad , Justin T. Landis , Brent A. Eason, Melissa B. Miller, Linda Pluta, Dirk Dittmer, Angelica Juarez, Cecilia Thompson , Cameroon Grant, Evelyn Hoffman, Patricio Cano, Jason Wong, Carolina Caro-Vegas, Blossom Damania. |
| EPI_ISL_882619 | COVID lab, Mymensingh Medical College | Department of Pathology, Bangladesh Agricultural University & Department of Microbiology, Mymensingh Medical College | Afrin, S. Z. Paul, S. K. Parvin, R. |
| EPI_ISL_884293 | Clinical Molecular Microbiology Laboratory, UNC Hospital | Dirk Dittmer | Razia Moorad , Justin T. Landis , Brent A. Eason, Melissa B. Miller, Linda Pluta, Dirk Dittmer, Angelica Juarez, Cecilia Thompson , Cameroon Grant, Evelyn Hoffman, Patricio Cano, Jason Wong, Carolina Caro-Vegas, Blossom Damania. |
| EPI_ISL_884331, EPI_ISL_884341, EPI_ISL_884358, EPI_ISL_884407, EPI_ISL_884417 | Infectious Diseases, Quest Diagnostics | Infectious Diseases, Quest Diagnostics | Rosenthal,S.H., Gerasimova,A., Kagan,R.M., Anderson,B., Bernstein,L.E., Livingston,K.E., Hua,M., Liu,Y., Shalhout,D.F., Owen,R., Lacbawan,F. |
| EPI_ISL_884839, EPI_ISL_884841 | Department of Biochemistry, Cell and Molecular Biology, West African Centre for Cell Biology of Infectious Pathogens (WACCBIP), University of Ghana | Department of Biochemistry, Cell and Molecular Biology, West African Centre for Cell Biology of Infectious Pathogens (WACCBIP), University of Ghana | Ngoi,J.M., Tei-Maya,F., Morang'a,C.M., Magnussen,V., Amuzu,D.S., Mohammed,A., Tapela,K., Kibinge,N., Diallo,A.B., Kumi-Ansah,F., Odoom,T., Boakye,O.D., Amoako,E., Abass,A.-K., Quashie,P., Amenga-Etego,L.N., Akoriyea,S.K., Awandare,G.A., Bediako,Y. |
| EPI_ISL_884868, EPI_ISL_884869 | CA DPH Viral and Rickettsial Disease Laboratory | Chan-Zuckerberg Biohub | CZB Cliahub Consortium |
| EPI_ISL_887429, EPI_ISL_887500, EPI_ISL_887503 | Instituto Nacional de Saude (INS), Mozambique | KRISP, KZN Research Innovation and Sequencing Platform | Nalia Ismael, Nadia Sitoe, Paulo Arnaldo, Nedio Mabunda, Giandhari J, Pillay S, Tegally H, Wilkinson E, de Oliveira T |
| EPI_ISL_889371 | Olomouc University Hospital | Institute of Applied Biotechnologies a.s. | Petr Klempt, Ondej Brzo, Martin Kašný, Kateina Kvapilová, Petr Kvapil |
| EPI_ISL_890164, EPI_ISL_890165, EPI_ISL_890166, EPI_ISL_890167, EPI_ISL_890168, EPI_ISL_890169, EPI_ISL_890170, EPI_ISL_890171, EPI_ISL_890172, EPI_ISL_890173 | Laboratoire de santé publique du Québec | Laboratoire de santé publique du Québec | Sandrine Moreira, Ioannis Ragoussis, Guillaume Bourque, Jesse Shapiro, Mark Lathrop and Michel Roger on behalf of the CoVSeQ research group |
| EPI_ISL_891260 | COVID lab, Mymensingh Medical College | Department of Pathology, Bangladesh Agricultural University and Department of Microbiology, Mymensingh Medical College | Afrin, S. Z. Paul, S. k. Parvin, R. |
| EPI_ISL_892231 | Lighthouse Lab in Milton Keynes | Wellcome Sanger Institute for the COVID-19 Genomics UK (COG-UK) Consortium | The Lighthouse Lab in Milton Keynes and Alex Alderton, Roberto Amato, Sonia Goncalves, Ewan Harrison, David K. Jackson, Ian Johnston, Dominic Kwiatkowski, Cordelia Langford, John Sillitoe on behalf of the Wellcome Sanger Institute COVID-19 Surveillance Team |
| EPI_ISL_896127, EPI_ISL_896133, EPI_ISL_896136, EPI_ISL_896174, EPI_ISL_896181, EPI_ISL_900060, EPI_ISL_900066, EPI_ISL_900077, EPI_ISL_900085, EPI_ISL_900113, EPI_ISL_900121, EPI_ISL_900133, EPI_ISL_900155, EPI_ISL_900160, EPI_ISL_900166, EPI_ISL_900168, EPI_ISL_900193, EPI_ISL_900209, EPI_ISL_900210, EPI_ISL_900214, EPI_ISL_900221, EPI_ISL_900233, EPI_ISL_900246, EPI_ISL_900254, EPI_ISL_900258, EPI_ISL_900261, EPI_ISL_900265, EPI_ISL_900268, EPI_ISL_900275, EPI_ISL_900278, EPI_ISL_900285, EPI_ISL_900288, EPI_ISL_900300, EPI_ISL_900309, EPI_ISL_900315, EPI_ISL_900324, EPI_ISL_900340, EPI_ISL_900341, EPI_ISL_900349, EPI_ISL_900351, EPI_ISL_900361, EPI_ISL_900367, EPI_ISL_900380, EPI_ISL_900384, EPI_ISL_900398, EPI_ISL_900415, EPI_ISL_900421, EPI_ISL_900439, EPI_ISL_900454, EPI_ISL_900460 | | | |
| see above | MEPHI, Aix Marseille University | MEPHI, Aix Marseille University | Anthony LEVASSEUR |
| EPI_ISL_902740, EPI_ISL_902743, EPI_ISL_902746, EPI_ISL_902748 | Hospital Universitari Germans Trias i Pujol (HUGTiP) / Fundació Lluita contra la SIDA (FLSida) | IrsiCaixa - Can Ruti CovidSeq | Fundació irsiCaixa. Hospital Universitari Germans Trias i Pujol(HUGTiP), 2a planta, maternal Ctra Canyet s/n, Badalona Marta Massanella, Ester Ballana, Lidia Ruiz, Nuria Izquierdo, Jorge Carrillo, Roger Paredes, Julia Blanco, Joaquim Segalés, Bonaventura Clotet |
| EPI_ISL_902917 | Tanjungpura University Hospital | Tanjungpura University Hospital | Andriani; Mahyarudin; Novianry V; Liana DF; Shofiyah SS Astuti P; Kahtan MI; Rialita A; Putri EA; Windarti W; Sastriawan W |
| EPI_ISL_906060 | Tilia Laboratories s.r.o. | Tilia Laboratories s.r.o. | Sona Pekova, MD, PhD. |
| EPI_ISL_906752 | Hematology Laboratory, Section of Molecular Diagnostics, University Clinical Centre, Medical University of Gdansk | Laboratory of Recombinant Vaccines | Lukasz Rabalski, Maciej Kosinski, Maciej Grzybek, Adam Sodol, Aneta Szulc, Krzysztof Lewandowski, Ewa Milosz, Marlena Robakowska, Boguslaw Szewczyk, Krystyna Bienkowska-Szewczyk |
| EPI_ISL_909955, EPI_ISL_909959 | Apollo Hospitals | CSIR-Centre for Cellular and Molecular Biology | Onkar Kulkarni, Suneetha Narreddy, Lamuk Zaveri, Irawathy Goud, Sofia Banu, Payel Mukherjee, Karthik Bharadwaj Tallapaka, Divya Tej Sowpati |
| EPI_ISL_910031, EPI_ISL_910032, EPI_ISL_910033, EPI_ISL_910034, EPI_ISL_910035, EPI_ISL_910036, EPI_ISL_910037, EPI_ISL_910038, EPI_ISL_910039, EPI_ISL_910040, EPI_ISL_910041, EPI_ISL_910042, EPI_ISL_910043, EPI_ISL_910044, EPI_ISL_910045, EPI_ISL_910046, EPI_ISL_910047, EPI_ISL_910048, EPI_ISL_910049, EPI_ISL_910050, EPI_ISL_910051, EPI_ISL_910052, EPI_ISL_910053, EPI_ISL_910054, EPI_ISL_910055, EPI_ISL_910056, EPI_ISL_910057, EPI_ISL_910058, EPI_ISL_910059, EPI_ISL_910060, EPI_ISL_910061, EPI_ISL_910062, EPI_ISL_910063, EPI_ISL_910064, EPI_ISL_910065, EPI_ISL_910066, EPI_ISL_910067, EPI_ISL_910068, EPI_ISL_910069, EPI_ISL_910070, EPI_ISL_910071, EPI_ISL_910072, EPI_ISL_910073, EPI_ISL_910074, EPI_ISL_910075, EPI_ISL_910076, EPI_ISL_910077, EPI_ISL_910078, EPI_ISL_910079, EPI_ISL_910080, EPI_ISL_910081, EPI_ISL_910082, EPI_ISL_910083, EPI_ISL_910084, EPI_ISL_910085, EPI_ISL_910086, EPI_ISL_910087, EPI_ISL_910088, EPI_ISL_910089, EPI_ISL_910090, EPI_ISL_910091, EPI_ISL_910092, EPI_ISL_910093, EPI_ISL_910094, EPI_ISL_910095, EPI_ISL_910096, EPI_ISL_910097, EPI_ISL_910098, EPI_ISL_910099, EPI_ISL_910100, EPI_ISL_910101, EPI_ISL_910102, EPI_ISL_910103, EPI_ISL_910104, EPI_ISL_910105, EPI_ISL_910106, EPI_ISL_910107, EPI_ISL_910108, EPI_ISL_910109, EPI_ISL_910110, EPI_ISL_910111, EPI_ISL_910113, EPI_ISL_910114, EPI_ISL_910115, EPI_ISL_910116, EPI_ISL_910117, EPI_ISL_910118, EPI_ISL_910119, EPI_ISL_910120, EPI_ISL_910121, EPI_ISL_910122 | | | |
| see above | CSIR-Centre for Cellular and Molecular Biology | CSIR-Centre for Cellular and Molecular Biology | Payel Mukherjee,Pratheusa Maccha,Namami Gaur,Lamuk Zaveri,Tulasi Nagabandi,Purushotham Vodnala,Blessy B John,Viswagithe S L,B Himasri,Sofia Banu,Priya Singh,Archana Bharadwaj Siva,Karthik Bharadwaj Tallapaka,Rakesh K Mishra,Divya Tej Sowpati |
| EPI_ISL_913476 | Klinisk mikrobiologi | The Public Health Agency of Sweden | Anna-Malin Linde, Maria Lind Karlberg, Carlo Berg, Oskar Karlsson Lindsjo, Sofia Stamouli, Reza Advani, Mattias Haukland, Petra Holmstrom, Noura Walai, Petra Edquist, Mia Brytting, Anna Risberg, Karin Tegmark-Wisell |
| EPI_ISL_925411, EPI_ISL_925421, EPI_ISL_925422, EPI_ISL_925423, EPI_ISL_925428 | Department of Clinical Microbiology | GIGA Medical Genomics | Keith Durkin, Maria Artesi, Sébastien Bontems, Raphaël Boreux, Bouchra Boujemla, Cécile Meex, Pierrette Melin, Marie-Pierre Hayette, Vincent Bours |
| EPI_ISL_934330, EPI_ISL_934331, EPI_ISL_934332, EPI_ISL_934333, EPI_ISL_934334 | Klinisk mikrobiologi | The Public Health Agency of Sweden | Anna-Malin Linde, Maria Lind Karlberg, Carlo Berg, Oskar Karlsson Lindsjo, Sofia Stamouli, Reza Advani, Mattias Haukland, Petra Holmstrom, Noura Walai, Petra Edquist, Mia Brytting, Anna Risberg, Karin Tegmark-Wisell |
| EPI_ISL_935523, EPI_ISL_935524 | University of Massachusetts Medical School | Infectious Disease Program, Broad Institute of Harvard and MIT | Tomkins-Tinch, Christopher H.; Daly, Jennifer S.; Gladden-Young, Adrianne; Theodoropoulos, Nicole M.; Madaio, Michael; Yu, Neng; Vanguri, Vijay K; Siddle, Katherine J.; Adams, Gordon; Krasilnikova, Lydia A.; Movahedi, Babak; Bozorgzadeh, Adel; Simin, Karl; Lemieux, Jacob E.; Luban, Jeremy; Park, Daniel J.; MacInnis, Bronwyn L.; Sabeti, Pardis C.; Levitz, Stuart M. |

| | | | |
|---|---|---|---|
| EPI_ISL_935762, EPI_ISL_935763, EPI_ISL_935764, EPI_ISL_935765, EPI_ISL_935766, EPI_ISL_935767, EPI_ISL_935768, EPI_ISL_935769, EPI_ISL_935770, EPI_ISL_935771, EPI_ISL_935772, EPI_ISL_935773, EPI_ISL_935774, EPI_ISL_935775, EPI_ISL_935776, EPI_ISL_935777, EPI_ISL_935778, EPI_ISL_935779, EPI_ISL_935780, EPI_ISL_935781, EPI_ISL_935782, EPI_ISL_935783, EPI_ISL_935784, EPI_ISL_935785, EPI_ISL_935786, EPI_ISL_935787, EPI_ISL_935809, EPI_ISL_935810, EPI_ISL_935811, EPI_ISL_935812, EPI_ISL_935813, EPI_ISL_935814, EPI_ISL_935815, EPI_ISL_935816, EPI_ISL_935823, EPI_ISL_935831, EPI_ISL_935856, EPI_ISL_935860, EPI_ISL_935861 | | | |
| see above | Cadham Provincial laboratory | National Microbiology Laboratory (NML) | Anna Majer, Shari Tyson, Grace Seo, Philip Mabon, Elsie Grudeski, Rhiannon Huzarewich, Russell Mandes, Anneliese Landgraff, Jennifer Tanner, Natalie Knox, Morag Graham, Gary Van Domselaar, Paul Van Caeseele, Jared Bullard, David Alexander, Kerry Dust, Nathalie Bastien, Yan Li, Timothy Booth, Darian Hole, Madison Chapel, Kirsten Biggar, CanCOGeN's metadata curation team, Public Health Agency of Canada CanCOGeN team |
| EPI_ISL_936828 | Northwestern Memorial Hospital | Ozer Lab | Ramon Lorenzo-Redondo, Lacy M. Simons, Chad J. Achenbach, Lawrence J. Jennings, Michael G. Ison, Judd F. Hultquist, Egon A. Ozer |
| EPI_ISL_937026, EPI_ISL_937035, EPI_ISL_937041, EPI_ISL_937042, EPI_ISL_937067, EPI_ISL_937079, EPI_ISL_937090, EPI_ISL_937091, EPI_ISL_937096, EPI_ISL_937110, EPI_ISL_937111, EPI_ISL_937112, EPI_ISL_937120 | | | |
| see above | Quest Diagnostics | Quest Diagnostics | Rosenthal,S.H., Gerasimova,A., Kagan,R.M., Anderson, B., Livingston, K.E., Hua, M., Liu Y., Shalhout, D.F., Owen, R., Lacbawan, F. |
| EPI_ISL_939625 | USC Clinical Lab | Los Angeles County PHL | P. Hemarajata et al. |
| EPI_ISL_940191, EPI_ISL_940193, EPI_ISL_940195, EPI_ISL_940196, EPI_ISL_940499, EPI_ISL_940500 | Hôpital Bichat Claude Bernard, Laboratoire de Virologie | IAME UMR1137 Inserm, Université de Paris, Hôpital Bichat | Antoine Bridier-Nahmias, Amélie Recoing, Quentin Le Hingrat, Lena Daniel, Siham Hamri, Gilles Collin, Alexandre Storto, Mélanie Bertine, Charlotte Charpentier, Nadhira Houhou-Fidouh, Diane Descamps, Benoit Visseaux |
| EPI_ISL_940950, EPI_ISL_940955, EPI_ISL_940956, EPI_ISL_940957, EPI_ISL_940958, EPI_ISL_940959 | Centers for Disease Control and Prevention, Dengue Branch | Centers for Disease Control and Prevention, Dengue Branch | Gilberto A. Santiago, Glenda Gonzalez, Betzabel Flores, Keyla Charriez, Gabriela Paz-Bailey, Jorge L. Munoz-Jordan |
| EPI_ISL_949204, EPI_ISL_949212, EPI_ISL_949215, EPI_ISL_949228, EPI_ISL_949241 | Departamento de Microbiología, CDB, Hospital Clínic, Barcelona | SeqCOVID-SPAIN consortium/IBV(CSIC) | Andrea Vergara, Mikel Martínez, Elisa Rubio, Jéssica Navero, Aida Peiró and SeqCOVID-SPAIN consortium |
| EPI_ISL_959281 | National Influenza Center, Virology Department | National Influenza Center | NZ Shafiei Jandaghi, V Salimi, A Nejati, K Sadeghi, J Yavarian, F Ajaminejad,N Ghavvami and T Mokhtari Azad |
| EPI_ISL_959903, EPI_ISL_960042, EPI_ISL_960043, EPI_ISL_960044, EPI_ISL_960045, EPI_ISL_960046, EPI_ISL_960047, EPI_ISL_960048, EPI_ISL_960049, EPI_ISL_960050, EPI_ISL_960051, EPI_ISL_960052, EPI_ISL_960053, EPI_ISL_960054, EPI_ISL_960055, EPI_ISL_960056, EPI_ISL_960057, EPI_ISL_960058, EPI_ISL_960059, EPI_ISL_960060 | | | |
| see above | University Medical Center Hamburg Eppendorf | Heinrich Pette Institute, Leibniz Institute for Experimental Virology | Alexis Robitaille, Thomas Günther, Johannes Knobloch, Martin Aepfelbacher, Nicole Fischer, Adam Grundhoff |
| EPI_ISL_960227, EPI_ISL_960229, EPI_ISL_960230, EPI_ISL_960231, EPI_ISL_960232, EPI_ISL_960233, EPI_ISL_960238, EPI_ISL_960239, EPI_ISL_960240, EPI_ISL_960241, EPI_ISL_960242, EPI_ISL_960243, EPI_ISL_960244, EPI_ISL_960245, EPI_ISL_960246, EPI_ISL_960247, EPI_ISL_960248, EPI_ISL_960249, EPI_ISL_960250, EPI_ISL_960251, EPI_ISL_960252, EPI_ISL_960253, EPI_ISL_960254, EPI_ISL_960255, EPI_ISL_960256, EPI_ISL_960257, EPI_ISL_960258, EPI_ISL_960259, EPI_ISL_960260, EPI_ISL_960261, EPI_ISL_960262, EPI_ISL_960263, EPI_ISL_960264, EPI_ISL_960265, EPI_ISL_960266, EPI_ISL_960267, EPI_ISL_960268, EPI_ISL_960269, EPI_ISL_960270, EPI_ISL_960271, EPI_ISL_960272, EPI_ISL_960273, EPI_ISL_960274, EPI_ISL_960275, EPI_ISL_960276, EPI_ISL_960277, EPI_ISL_960278, EPI_ISL_960279, EPI_ISL_960280, EPI_ISL_960281 | | | |
| see above | Nucleic Acid Testing, National Reference Laboratory | GIGA Medical Genomics | Yvan Butera, Keith Durkin, Maria Artesi, Bouchra Boujemla, Robert Rutayisire, Patrick Tuyisenge, Esperence Umumararungu, Sébastien Bontems, Marie-Pierre Hayette, Nathalie Renotte, Corinne Fasquelle, Swaibu Gatare, Jacob Souopgui, Sabin Nsanzimana, Vincent Bours, Léon Mutesa |
| EPI_ISL_962816 | Microbiological Diagnostic Unit - Public Health Laboratory (MDU-PHL) | MDU-PHL | Seemann T., Sait, M.L., Sherry, N.L. |
| EPI_ISL_964907 | Hospital Zapala | Laboratorio Central Mg. Luis Alfredo Pianciola on behalf of 'Proyecto Argentino Interinstitucional de genomica de SARS-CoV-2' (PAIS Consortium) | L Pianciola, M Mazzeo, C Ziehm, C Pintos, M Fernandez, J Ousset, M Nabaes, M Viegas. |
| EPI_ISL_964908 | Hospital Las Ovejas | Laboratorio Central Mg. Luis Alfredo Pianciola on behalf of 'Proyecto Argentino Interinstitucional de genomica de SARS-CoV-2' (PAIS Consortium) | L Pianciola, M Mazzeo, C Ziehm, C Pintos, M Fernandez, J Ousset, M Nabaes, M Viegas. |
| EPI_ISL_965629, EPI_ISL_965654, EPI_ISL_965783 | Dutch COVID-19 response team | Medical Microbiology, Maastricht University Medical Centre | Jozef Dingemans*, Brian van der Veer*, Erik Beuken, Carmen Reumkens, Lieke van Alphen, Christian Hoebe, Paul Savelkoul |
| EPI_ISL_968169, EPI_ISL_968170, EPI_ISL_968171, EPI_ISL_968172, EPI_ISL_968185 | Clinical Molecular Microbiology Laboratory, UNC Hospital | Dirk Dittmer | Justin T. Landis , Razia Moorad , Brent A. Eason, Melissa B. Miller, Linda Pluta, Dirk Dittmer, Angelica Juarez, Cecilia Thompson, Shawn Hawken, Cameroon Grant, Evelyn Hoffman, Patricio Cano, Jason Wong, Carolina Caro-Vegas, Ryan McNamara, Blossom Damania. |
| EPI_ISL_969099, EPI_ISL_969100, EPI_ISL_969101, EPI_ISL_969102, EPI_ISL_969103, EPI_ISL_969104, EPI_ISL_969105, EPI_ISL_969106, EPI_ISL_969107, EPI_ISL_969108, EPI_ISL_969109, EPI_ISL_969110, EPI_ISL_969111, EPI_ISL_969112, EPI_ISL_969113, EPI_ISL_969114, EPI_ISL_969115, EPI_ISL_969116, EPI_ISL_969117, EPI_ISL_969118, EPI_ISL_969119, EPI_ISL_969120, EPI_ISL_969121, EPI_ISL_969122, EPI_ISL_969123, EPI_ISL_969124, EPI_ISL_969125, EPI_ISL_969126, EPI_ISL_969127, EPI_ISL_969128, EPI_ISL_969129, EPI_ISL_969130, EPI_ISL_969132, EPI_ISL_969133, EPI_ISL_969134, EPI_ISL_969135, EPI_ISL_969136, EPI_ISL_969137, EPI_ISL_969138, EPI_ISL_969139, EPI_ISL_969140, EPI_ISL_969141, EPI_ISL_969142, EPI_ISL_969143, EPI_ISL_969144, EPI_ISL_969145, EPI_ISL_969146, EPI_ISL_969147, EPI_ISL_969148, EPI_ISL_969149, EPI_ISL_969150, EPI_ISL_969151, EPI_ISL_969152, EPI_ISL_969153, EPI_ISL_969154, EPI_ISL_969155, EPI_ISL_969156, EPI_ISL_969157, EPI_ISL_969158, EPI_ISL_969159, EPI_ISL_969160, EPI_ISL_969161, EPI_ISL_969162, EPI_ISL_969163, EPI_ISL_969164, EPI_ISL_969165, EPI_ISL_969166, EPI_ISL_969167, EPI_ISL_969168, EPI_ISL_969169, EPI_ISL_969170, EPI_ISL_969171, EPI_ISL_969172, EPI_ISL_969173, EPI_ISL_969174, EPI_ISL_969175, EPI_ISL_969176, EPI_ISL_969177, EPI_ISL_969178, EPI_ISL_969179, EPI_ISL_969180, EPI_ISL_969181, EPI_ISL_969182, EPI_ISL_969183, EPI_ISL_969184, EPI_ISL_969185, EPI_ISL_969186, EPI_ISL_969187, EPI_ISL_969188, EPI_ISL_969189, EPI_ISL_969190, EPI_ISL_969191, EPI_ISL_969192, EPI_ISL_969193, EPI_ISL_969194, EPI_ISL_969195, EPI_ISL_969196, EPI_ISL_969197, EPI_ISL_969198, EPI_ISL_969199, EPI_ISL_969200, EPI_ISL_969201, EPI_ISL_969202, EPI_ISL_969203, EPI_ISL_969204, EPI_ISL_969205, EPI_ISL_969206, EPI_ISL_969207, EPI_ISL_969208, EPI_ISL_969209, EPI_ISL_969210, EPI_ISL_969211, EPI_ISL_969212, EPI_ISL_969213, EPI_ISL_969214, EPI_ISL_969215, EPI_ISL_969216, EPI_ISL_969217, EPI_ISL_969219, EPI_ISL_969220, EPI_ISL_969221, EPI_ISL_969222, EPI_ISL_969223, EPI_ISL_969224, EPI_ISL_969225, EPI_ISL_969226, EPI_ISL_969228, EPI_ISL_969229, EPI_ISL_969230, EPI_ISL_969231, EPI_ISL_969232, EPI_ISL_969233, EPI_ISL_969234, EPI_ISL_969235, EPI_ISL_969236, EPI_ISL_969237, EPI_ISL_969238, EPI_ISL_969239, EPI_ISL_969240, EPI_ISL_969241, EPI_ISL_969242, EPI_ISL_969243, EPI_ISL_969244, EPI_ISL_969246, EPI_ISL_969247, EPI_ISL_969248, EPI_ISL_969249, EPI_ISL_969250, EPI_ISL_969251, EPI_ISL_969252, EPI_ISL_969253, EPI_ISL_969254, EPI_ISL_969255, EPI_ISL_969256, EPI_ISL_969257, EPI_ISL_969259, EPI_ISL_969260, EPI_ISL_969261, EPI_ISL_969262, EPI_ISL_969263, EPI_ISL_969264, EPI_ISL_969265, EPI_ISL_969266, EPI_ISL_969267, EPI_ISL_969268, EPI_ISL_969269, EPI_ISL_969270, EPI_ISL_969271, EPI_ISL_969272, EPI_ISL_969273, EPI_ISL_969274, EPI_ISL_969275, EPI_ISL_969276, EPI_ISL_969277, EPI_ISL_969278, EPI_ISL_969279, EPI_ISL_969280, EPI_ISL_969281, EPI_ISL_969282, EPI_ISL_969283, EPI_ISL_969284, EPI_ISL_969285, EPI_ISL_969286, EPI_ISL_969287, EPI_ISL_969288, EPI_ISL_969289, EPI_ISL_969291, EPI_ISL_969292, EPI_ISL_969293, EPI_ISL_969294, EPI_ISL_969295, EPI_ISL_969296, EPI_ISL_969298, EPI_ISL_969299, EPI_ISL_969300, EPI_ISL_969301, EPI_ISL_969302, EPI_ISL_969303, EPI_ISL_969304, EPI_ISL_969305, EPI_ISL_969306, EPI_ISL_969307, EPI_ISL_969308, EPI_ISL_969309, EPI_ISL_969310, EPI_ISL_969311, EPI_ISL_969312, EPI_ISL_969313, EPI_ISL_969314, EPI_ISL_969315, EPI_ISL_969316, EPI_ISL_969317, EPI_ISL_969318, EPI_ISL_969319, EPI_ISL_969320, EPI_ISL_969321, EPI_ISL_969322, EPI_ISL_969323, EPI_ISL_969324, EPI_ISL_969325, EPI_ISL_969326, EPI_ISL_969327, EPI_ISL_969328, EPI_ISL_969329, EPI_ISL_969330, EPI_ISL_969331, EPI_ISL_969333, EPI_ISL_969334, EPI_ISL_969335, EPI_ISL_969336, EPI_ISL_969337, EPI_ISL_969338, EPI_ISL_969339, EPI_ISL_969340, EPI_ISL_969341, EPI_ISL_969342, EPI_ISL_969343, EPI_ISL_969344, EPI_ISL_969345, EPI_ISL_969346, EPI_ISL_969347, EPI_ISL_969348, EPI_ISL_969349, EPI_ISL_969350, EPI_ISL_969351, EPI_ISL_969352, EPI_ISL_969353, EPI_ISL_969354, EPI_ISL_969355, EPI_ISL_969356, EPI_ISL_969357, EPI_ISL_969358, EPI_ISL_969359, EPI_ISL_969360, EPI_ISL_969361, EPI_ISL_969362, EPI_ISL_969363, EPI_ISL_969364, EPI_ISL_969366, EPI_ISL_969367, EPI_ISL_969368, EPI_ISL_969369, EPI_ISL_969370, EPI_ISL_969371, EPI_ISL_969372, EPI_ISL_969373, EPI_ISL_969374, EPI_ISL_969375, EPI_ISL_969376, EPI_ISL_969377, EPI_ISL_969378, EPI_ISL_969379, EPI_ISL_969380, EPI_ISL_969381, EPI_ISL_969382, EPI_ISL_969383, EPI_ISL_969384, EPI_ISL_969385, EPI_ISL_969386, EPI_ISL_969387, EPI_ISL_969388, EPI_ISL_969389, EPI_ISL_969390, EPI_ISL_969391, EPI_ISL_969392, EPI_ISL_969393, EPI_ISL_969394, EPI_ISL_969395, EPI_ISL_969396, EPI_ISL_969398, EPI_ISL_969399, EPI_ISL_969400, EPI_ISL_969401, EPI_ISL_969402, EPI_ISL_969403, EPI_ISL_969404, EPI_ISL_969405, EPI_ISL_969406, EPI_ISL_969407, EPI_ISL_969408, EPI_ISL_969409, EPI_ISL_969410, EPI_ISL_969411, EPI_ISL_969412, EPI_ISL_969413, EPI_ISL_969414, EPI_ISL_969415, EPI_ISL_969416, EPI_ISL_969417, EPI_ISL_969418, EPI_ISL_969419, EPI_ISL_969420, EPI_ISL_969421, EPI_ISL_969422, EPI_ISL_969423, EPI_ISL_969424, EPI_ISL_969425, EPI_ISL_969426, EPI_ISL_969427, EPI_ISL_969428, EPI_ISL_969429, EPI_ISL_969430, EPI_ISL_969431, EPI_ISL_969432, EPI_ISL_969433, EPI_ISL_969434, EPI_ISL_969435, EPI_ISL_969436, EPI_ISL_969437, EPI_ISL_969438, EPI_ISL_969439, EPI_ISL_969440, EPI_ISL_969441, EPI_ISL_969442, EPI_ISL_969443, EPI_ISL_969444, EPI_ISL_969445, EPI_ISL_969446, EPI_ISL_969447, EPI_ISL_969448, EPI_ISL_969449, EPI_ISL_969450, EPI_ISL_969451, EPI_ISL_969452, EPI_ISL_969453, EPI_ISL_969470, EPI_ISL_969499, EPI_ISL_969547, EPI_ISL_969590, EPI_ISL_969601, EPI_ISL_969648, EPI_ISL_969676, EPI_ISL_969678, EPI_ISL_969680, EPI_ISL_969681, EPI_ISL_969684, EPI_ISL_969686, EPI_ISL_969687, EPI_ISL_969689, EPI_ISL_969692, EPI_ISL_969694, EPI_ISL_969696, EPI_ISL_969698, EPI_ISL_969700, EPI_ISL_969703, EPI_ISL_969705, EPI_ISL_969707, EPI_ISL_969709, EPI_ISL_969711, EPI_ISL_969713, EPI_ISL_969715, EPI_ISL_969717, EPI_ISL_969719, EPI_ISL_969721, EPI_ISL_969723, EPI_ISL_969725, EPI_ISL_969727, EPI_ISL_969729, EPI_ISL_969731, EPI_ISL_969733, EPI_ISL_969735, EPI_ISL_969737, EPI_ISL_969738, EPI_ISL_969741, EPI_ISL_969743, EPI_ISL_969745, EPI_ISL_969747, EPI_ISL_969749, EPI_ISL_969751, EPI_ISL_969753, EPI_ISL_969755, EPI_ISL_969757 | | | |
| see above | BCCDC Public Health Laboratory | BCCDC Public Health Laboratory | Prystajecky Natalie, Linda Hoang, Dan Fornika, John Tyson, Shannon Russell, Kim Macdonald, Kimia Kamelian, Ana Pacagnella, Corinne Ng, Loretta Janz, Robert Azana Terry Snutch, Mel Krajden |
| EPI_ISL_976910, EPI_ISL_976920, EPI_ISL_976923, EPI_ISL_976927, EPI_ISL_976932, EPI_ISL_976934, EPI_ISL_976935, EPI_ISL_976936, EPI_ISL_976937 | University of Massachusetts Medical School | Infectious Disease Program, Broad Institute of Harvard and MIT | Lemieux,J.E., Siddle,K.J., Ward,D., Ellison,R., Adams,G., Gladden-Young,A., Lagerborg,K., Rudy,M., DeRuff,K., Carter,A., Normandin,E., Bauer,M., Reilly,S., Tomkins-Tinch,C., Loreth,C., Chaluvadi,S., Birren,B.W., Gallagher,G., Smole,S., Park,D.J., MacInnis,B.L., and Sabeti,P.C. |
| EPI_ISL_977173, EPI_ISL_977174 | Microbiologia e Virologia | Istituto Zooprofilattico Sperimentale delle Venezie | Adelaide Milani, Alessia Schivo, Annalisa Salviato, Erika Giorgia Quaranta, Ambra Pastori, Bianca Zecchin, Alice Fusaro, Isabella Monne, Calogero Terregino, Antonia Ricci |
| EPI_ISL_978501 | Central Public Health Laboratory - LACEN -Bahia, Salvador, Brazil | Central Public Health Laboratory - LACEN -Bahia, Salvador, Brazil | Stephane Tosta, Luciana Oliveira, Vanessa Nardy,Patrícia Cajado,Marcela Gómez, Breno Dominguez, Jaqueline Gomes, Vagner Fonseca,Marta Giovanetti,Luiz Alcantara, Felicidade Pereira, Arabela Leal |

| | | | |
|---|---|---|---|
| EPI_ISL_981048, EPI_ISL_981049, EPI_ISL_981056 | Hospital Viedma | Laboratorio Central Mg. Luis Alfredo Pianciola on behalf of 'Proyecto Argentino Interinstitucional de genomica de SARS-CoV-2' (PAIS Consortium) | L Pianciola, M Mazzeo, C Ziehm, C Pintos, M Fernandez, J Ousset, M Nabaes, M Viegas. |
