## Supplementary material for "Detection and characterization of the SARS-CoV-2 lineage B.1.526 in New York": Supp. Table 3 GISAID Acknowledgment part 2: gisaid_hcov-19_acknowledgement_table_2021_02_12_23-6.pdf

We gratefully acknowledge the following Authors from the Originating laboratories responsible for obtaining the specimens, as well as the Submitting laboratories where the genome data were generated and shared via GISAID, on which this research is based.

All Submitters of data may be contacted directly via www.gisaid.org

Authors are sorted alphabetically.

| Accession ID | Originating Laboratory | Submitting Laboratory | Authors |
| --- | --- | --- | --- |
| EPI_ISL_529638, EPI_ISL_529639, EPI_ISL_529640, EPI_ISL_529641, EPI_ISL_529642, EPI_ISL_529643 | University of Birmingham | COVID-19 Genomics UK (COG-UK) Consortium | Institute of Microbiology, University of Birmingham: Claire McMurray, Joanne Stockton, Samuel Nicholls, Radoslaw Poplawski, Will Rowe, Josh Quick, Nicholas Loman. University of Birmingham Testing Laboratory: Celina M Whalley, Andrew Bosworth, Charlotte Poxon, Kasun Wanigasooriya, Oliver Pickles, Mike Kidd, Alex Richter, Andrew D Beggs PHE Heartlands Lab: Husam Osman, Andrew Bosworth. Queen Elizabeth Hospital: Anna Casey |
| EPI_ISL_530336, EPI_ISL_530339, EPI_ISL_530340 | Area of Virology, Serology and Virology Division (SAViD), New South Wales Health Pathology Randwick | Area of Virology, Serology and Virology Division (SAViD), New South Wales Health Pathology Randwick | Rawlinson, W., Deveson, I., Bull, R., Van Hal, S. |
| EPI_ISL_534723, EPI_ISL_534724, EPI_ISL_534725, EPI_ISL_534726, EPI_ISL_534727, EPI_ISL_534728, EPI_ISL_534729 | Respiratory Virus Unit, Microbiology Services Colindale, Public Health England | Respiratory Virus Unit, Microbiology Services Colindale, Public Health England | PHE Covid Sequencing Team |
| EPI_ISL_534756, EPI_ISL_534757, EPI_ISL_534758 | Queens Medical Centre, Clinical Microbiology Department / DeepSeq Nottingham | COVID-19 Genomics UK (COG-UK) Consortium | Gemma Clark, Wendy Smith, Manjinder Khakh, Vicki M Fleming, Michelle M Lister, Hannah Howson-Wells, Jonathan Ball, Patrick McClure, Joseph Chappell, Theocharis Tsoleridis, Nadine Holmes, Matthew Carlisle, Christopher Moore, Fei Sang, Johnny Debebe, Victoria Wright, Matthew Loose |
| EPI_ISL_535184, EPI_ISL_535186, EPI_ISL_535187, EPI_ISL_535188, EPI_ISL_535189, EPI_ISL_535190, EPI_ISL_535192, EPI_ISL_535193, EPI_ISL_535195, EPI_ISL_535196, EPI_ISL_535197, EPI_ISL_535198, EPI_ISL_535199, EPI_ISL_535203, EPI_ISL_535204, EPI_ISL_535205, EPI_ISL_535206, EPI_ISL_535207, EPI_ISL_535209, EPI_ISL_535210, EPI_ISL_535211, EPI_ISL_535213, EPI_ISL_535214, EPI_ISL_535215, EPI_ISL_535216, EPI_ISL_535217, EPI_ISL_535218, EPI_ISL_535219, EPI_ISL_535220, EPI_ISL_535221, EPI_ISL_535222, EPI_ISL_535223, EPI_ISL_535224, EPI_ISL_535225, EPI_ISL_535226, EPI_ISL_535227, EPI_ISL_535228, EPI_ISL_535229, EPI_ISL_535230, EPI_ISL_535231, EPI_ISL_535232, EPI_ISL_535233, EPI_ISL_535234, EPI_ISL_535235, EPI_ISL_535236, EPI_ISL_535237, EPI_ISL_535238, EPI_ISL_535239, EPI_ISL_535240, EPI_ISL_535241, EPI_ISL_535242, EPI_ISL_535243, EPI_ISL_535244, EPI_ISL_535245, EPI_ISL_535246, EPI_ISL_535247, EPI_ISL_535248, EPI_ISL_535249, EPI_ISL_535250, EPI_ISL_535251, EPI_ISL_535252, EPI_ISL_535253, EPI_ISL_535254, EPI_ISL_535255, EPI_ISL_535256, EPI_ISL_535257, EPI_ISL_535258, EPI_ISL_535259, EPI_ISL_535260, EPI_ISL_535261, EPI_ISL_535262 | | | |
| see above | Wales Specialist Virology Centre Sequencing lab: Pathogen Genomics Unit | COVID-19 Genomics UK (COG-UK) Consortium | Catherine Moore, Johnathan Evans, Laura Gifford, Malorie Perry, Simon Cottrell, Angela Marchbank, Alec Birchley, Alexander Adams, Amy Gaskin, Bree Gatica-Wilcox, Jason Coombes, Joel Southgate, Lauren Gilbert, Lee Graham, Nicole Pacchiarini, Sara Kumziene-Summerhayes, Sarah Taylor, Sophie Jones, Sara Rey, Matthew Bull, Joanne Watkins, Sally Corden, Tom Connor |
| EPI_ISL_536429, EPI_ISL_536430, EPI_ISL_536431, EPI_ISL_536432, EPI_ISL_536433, EPI_ISL_536434, EPI_ISL_536438, EPI_ISL_536442, EPI_ISL_536444, EPI_ISL_536445, EPI_ISL_536446, EPI_ISL_536449, EPI_ISL_536451, EPI_ISL_536454 | | | |
| see above | National Public Health Laboratory, National Centre for Infectious Diseases | National Public Health Laboratory, National Centre for Infectious Diseases | Mak TM, Octavia S, Zhou Z, Cui L, Lin RTP |
| EPI_ISL_536659, EPI_ISL_536660, EPI_ISL_536661, EPI_ISL_536662, EPI_ISL_536664, EPI_ISL_536665, EPI_ISL_536666, EPI_ISL_536667, EPI_ISL_536668, EPI_ISL_536674, EPI_ISL_536676, EPI_ISL_536678, EPI_ISL_536680, EPI_ISL_536684, EPI_ISL_536691, EPI_ISL_536696, EPI_ISL_536707, EPI_ISL_536708, EPI_ISL_536709, EPI_ISL_536710, EPI_ISL_536711, EPI_ISL_536712, EPI_ISL_536713, EPI_ISL_536714, EPI_ISL_536715, EPI_ISL_536716, EPI_ISL_536717, EPI_ISL_536718, EPI_ISL_536719, EPI_ISL_536720, EPI_ISL_536721, EPI_ISL_536722, EPI_ISL_536723, EPI_ISL_536724, EPI_ISL_536725, EPI_ISL_536726, EPI_ISL_536727, EPI_ISL_536728, EPI_ISL_536729, EPI_ISL_536730, EPI_ISL_536731, EPI_ISL_536732, EPI_ISL_536733, EPI_ISL_536734, EPI_ISL_536735, EPI_ISL_536736, EPI_ISL_536737, EPI_ISL_536738, EPI_ISL_536739, EPI_ISL_536740, EPI_ISL_536741, EPI_ISL_536742, EPI_ISL_536743, EPI_ISL_536744, EPI_ISL_536745, EPI_ISL_536746, EPI_ISL_536747, EPI_ISL_536748, EPI_ISL_536749, EPI_ISL_536750, EPI_ISL_536751, EPI_ISL_536752, EPI_ISL_536753, EPI_ISL_536754, EPI_ISL_536755, EPI_ISL_536756, EPI_ISL_536757, EPI_ISL_536758, EPI_ISL_536759, EPI_ISL_536760, EPI_ISL_536761, EPI_ISL_536762, EPI_ISL_536763, EPI_ISL_536764, EPI_ISL_536765, EPI_ISL_536766, EPI_ISL_536767, EPI_ISL_536768, EPI_ISL_536769, EPI_ISL_536770, EPI_ISL_536771, EPI_ISL_536772, EPI_ISL_536773, EPI_ISL_536774, EPI_ISL_536775, EPI_ISL_536776, EPI_ISL_536777, EPI_ISL_536778, EPI_ISL_536779, EPI_ISL_536780, EPI_ISL_536781, EPI_ISL_536782, EPI_ISL_536783, EPI_ISL_536784, EPI_ISL_536786 | | | |
| see above | University of Wisconsin-Madison AIDS Vaccine Research Laboratories | University of Wisconsin-Madison AIDS Vaccine Research Laboratories | Gage Moreno, Katarina Braun, et al. AIDS Vaccine Research Laboratories |
| EPI_ISL_537286, EPI_ISL_537287 | Area of Virology, Serology and Virology Division (SAViD), New South Wales Health Pathology Randwick | Area of Virology, Serology and Virology Division (SAViD), New South Wales Health Pathology Randwick | Rawlinson, W., Deveson, I., Bull, R., Van Hal, S. |
| EPI_ISL_540139, EPI_ISL_540140, EPI_ISL_540141, EPI_ISL_540142, EPI_ISL_540143, EPI_ISL_540144, EPI_ISL_540145, EPI_ISL_540146, EPI_ISL_540147, EPI_ISL_540148, EPI_ISL_540149, EPI_ISL_540150, EPI_ISL_540151, EPI_ISL_540152, EPI_ISL_540153, EPI_ISL_540154, EPI_ISL_540155, EPI_ISL_540156, EPI_ISL_540157, EPI_ISL_540158, EPI_ISL_540159, EPI_ISL_540160, EPI_ISL_540161, EPI_ISL_540162, EPI_ISL_540163, EPI_ISL_540164, EPI_ISL_540165, EPI_ISL_540166, EPI_ISL_540167, EPI_ISL_540168, EPI_ISL_540169, EPI_ISL_540170, EPI_ISL_540171, EPI_ISL_540172, EPI_ISL_540173, EPI_ISL_540174, EPI_ISL_540175, EPI_ISL_540176, EPI_ISL_540177, EPI_ISL_540179, EPI_ISL_540180, EPI_ISL_540181, EPI_ISL_540182, EPI_ISL_540183, EPI_ISL_540184, EPI_ISL_540185, EPI_ISL_540186, EPI_ISL_540187, EPI_ISL_540188, EPI_ISL_540189, EPI_ISL_540190, EPI_ISL_540191, EPI_ISL_540192, EPI_ISL_540193, EPI_ISL_540194, EPI_ISL_540195, EPI_ISL_540196, EPI_ISL_540197, EPI_ISL_540198, EPI_ISL_540199, EPI_ISL_540200, EPI_ISL_540201, EPI_ISL_540202, EPI_ISL_540203, EPI_ISL_540204, EPI_ISL_540205, EPI_ISL_540206, EPI_ISL_540207, EPI_ISL_540208, EPI_ISL_540209, EPI_ISL_540210, EPI_ISL_540211, EPI_ISL_540212, EPI_ISL_540213, EPI_ISL_540214, EPI_ISL_540215, EPI_ISL_540216, EPI_ISL_540218, EPI_ISL_540219, EPI_ISL_540220, EPI_ISL_540221, EPI_ISL_540222, EPI_ISL_540223, EPI_ISL_540224, EPI_ISL_540225, EPI_ISL_540226, EPI_ISL_540227, EPI_ISL_540228, EPI_ISL_540229, EPI_ISL_540230, EPI_ISL_540231, EPI_ISL_540232 | | | |
| see above | Lighthouse Lab in Glasgow | Wellcome Sanger Institute for the COVID-19 Genomics UK (COG-UK) consortium | Harper VanSteenhouse, Yumi Kasai, David Gray, Carol Clugston, Anna Dominiczak and Alex Alderton, Roberto Amato, Sonia Goncalves, Ewan Harrison, David K. Jackson, Ian Johnston, Dominic Kwiatkowski, Cordelia Langford, John Sillitoe on behalf of the Wellcome Sanger Institute COVID-19 Surveillance Team |
| EPI_ISL_540691, EPI_ISL_540697, EPI_ISL_540698 | Quadram Institute Bioscience | COVID-19 Genomics UK (COG-UK) Consortium | Dave J. Baker, Gemma L. Kay, Alp Aydin, Thanh Le-Viet, Steven Rudder, Ana P. Tedim, Anastasia Kolyva, Maria Diaz, Leonardo de Oliveira Martins, Nabil-Fareed Alikhan, Lizzie Meadows, Rachael Stanley, Ngozi Elumogo, Muhammed Yasir, Nicholas M. Thomson, Alexander J Trotter, Rachel Gilroy, Samuel Bloomfield, Claire Stuart, Andrew Bell, Reenesh Prakash, Samir Dervisevic, Alison E. Mather, John Wain, Mark Webber, Andrew J. Page, Justin O'Grady |
| EPI_ISL_540699, EPI_ISL_540700 | Queens Medical Centre, Clinical Microbiology Department / DeepSeq Nottingham | COVID-19 Genomics UK (COG-UK) Consortium | Gemma Clark, Wendy Smith, Manjinder Khakh, Vicki M Fleming, Michelle M Lister, Hannah Howson-Wells, Jonathan Ball, Patrick McClure, Joseph Chappell, Theocharis Tsoleridis, Nadine Holmes, Matthew Carlisle, Christopher Moore, Fei Sang, Johnny Debebe, Victoria Wright, Matthew Loose |
| EPI_ISL_540721, EPI_ISL_540722, EPI_ISL_540729, EPI_ISL_540732, EPI_ISL_540740, EPI_ISL_540746, EPI_ISL_540748, EPI_ISL_540752, EPI_ISL_540753, EPI_ISL_540755, EPI_ISL_540756, EPI_ISL_540757, EPI_ISL_540762, EPI_ISL_540767, EPI_ISL_540768, EPI_ISL_540769, EPI_ISL_540772, EPI_ISL_540774 | | | |
| see above | Virology Department, Sheffield Teaching Hospitals NHS Foundation Trust/Department of Infection, Immunity and Cardiovascular Disease, The Medical School, University of Sheffield | COVID-19 Genomics UK (COG-UK) Consortium | Thushan de Silva, Matthew Parker, Nikki Smith, Adri Angyal, Rebecca Brown, Luke Green, Rachel Tucker, Paul Parsons, Danielle Groves, Katie Johnson, Laura Carrilero, Alex Keeley, Dave Partridge, Matthew Wyles, Benjamin Lindsey, Mehmet Yavuz, Mohammad Raza, Cariad Evans |
| EPI_ISL_540786, EPI_ISL_540787, EPI_ISL_540788, EPI_ISL_540789, EPI_ISL_540790, EPI_ISL_540791, EPI_ISL_540794, EPI_ISL_540795, EPI_ISL_540796, EPI_ISL_540797, EPI_ISL_540798, EPI_ISL_540799, EPI_ISL_540800, EPI_ISL_540801, EPI_ISL_540802, EPI_ISL_540803, EPI_ISL_540804, EPI_ISL_540805, EPI_ISL_540806, EPI_ISL_540807, EPI_ISL_540808, EPI_ISL_540809, EPI_ISL_540810, EPI_ISL_540811, EPI_ISL_540812, EPI_ISL_540813, EPI_ISL_540814, EPI_ISL_540815, EPI_ISL_540817, EPI_ISL_540818, EPI_ISL_540819, EPI_ISL_540820, EPI_ISL_540821, EPI_ISL_540822 | | | |
| see above | West of Scotland Specialist Virology Centre, NHSGGC / MRC-University of Glasgow Centre for Virus Research | COVID-19 Genomics UK (COG-UK) Consortium | Ana da Silva Filipe, Natasha Johnson, Kathy Smollett, Daniel Mair, Stephen Carmichael, Lily Tong, Jenna Nichols, Elihu Aranday-Cortes, Kyriaki Nomikou; Sarah McDonald, Marc Niebel, Patawee Asamaphan; Richard Orton, Joseph Hughes, Sreenu Vattipally, David L Robertson; Alasdair MacLean, Rory Gunson; Kathy Li, Igor Starinskij, Natasha Jesudason, Rajiv Shah, James Shepherd, Antonia Ho, Emma Thomson |
| EPI_ISL_540878, EPI_ISL_540879, EPI_ISL_540880, EPI_ISL_540881, EPI_ISL_540882, EPI_ISL_540883, EPI_ISL_540884, EPI_ISL_540885, EPI_ISL_540890 | Virology Department, Royal Infirmary of Edinburgh, NHS Lothian / School of Biological Sciences, University of Edinburgh / Institute of Genetics and Molecular Medicine, University of Edinburgh | COVID-19 Genomics UK (COG-UK) Consortium | McHugh M, Dewar R, Rooke S, Gallagher M, Balcaza C, O'Toole Á, Scher E, Hill V, McCrone JT, Colquhoun R, Yu X, Jackson B, Rambaut A, Williams TC, Templeton K |
| EPI_ISL_540899, EPI_ISL_540900, EPI_ISL_540901, EPI_ISL_540902, EPI_ISL_540903, EPI_ISL_540904, EPI_ISL_540905, EPI_ISL_540906, EPI_ISL_540907, EPI_ISL_540908, EPI_ISL_540909, EPI_ISL_540910, EPI_ISL_540911, EPI_ISL_540912, EPI_ISL_540913, EPI_ISL_540914, EPI_ISL_540915, EPI_ISL_540916, EPI_ISL_540917, EPI_ISL_540918, EPI_ISL_540919, EPI_ISL_540920 | | | |
| see above | Wales Specialist Virology Centre Sequencing lab: Pathogen Genomics Unit | COVID-19 Genomics UK (COG-UK) Consortium | Catherine Moore, Johnathan Evans, Laura Gifford, Malorie Perry, Simon Cottrell, Angela Marchbank, Alec Birchley, Alexander Adams, Amy Gaskin, Bree Gatica-Wilcox, Jason Coombes, Joel Southgate, Lauren Gilbert, Lee Graham, Nicole Pacchiarini, Sara Kumziene-Summerhayes, Sarah Taylor, Sophie |

| | | | |
|---|---|---|---|
| | | | Jones, Sara Rey, Matthew Bull, Joanne Watkins, Sally Corden, Tom Connor |
| EPI_ISL_541401, EPI_ISL_541402, EPI_ISL_541403, EPI_ISL_541404, EPI_ISL_541405, EPI_ISL_541406, EPI_ISL_541407, EPI_ISL_541408, EPI_ISL_541409, EPI_ISL_541410, EPI_ISL_541416, EPI_ISL_541417, EPI_ISL_541418, EPI_ISL_541419, EPI_ISL_541420, EPI_ISL_541421, EPI_ISL_541422, EPI_ISL_541423, EPI_ISL_541424, EPI_ISL_541425, EPI_ISL_541426, EPI_ISL_541427, EPI_ISL_541428, EPI_ISL_541429, EPI_ISL_541430, EPI_ISL_541431, EPI_ISL_541432, EPI_ISL_541433, EPI_ISL_541434, EPI_ISL_541435, EPI_ISL_541447, EPI_ISL_541448, EPI_ISL_541449, EPI_ISL_541465, EPI_ISL_541466, EPI_ISL_541467, EPI_ISL_541468, EPI_ISL_541470, EPI_ISL_541471, EPI_ISL_541472, EPI_ISL_541473, EPI_ISL_541474, EPI_ISL_541475, EPI_ISL_541476, EPI_ISL_541477, EPI_ISL_541478, EPI_ISL_541479, EPI_ISL_541480, EPI_ISL_541481, EPI_ISL_541482, EPI_ISL_541483, EPI_ISL_541484, EPI_ISL_541485, EPI_ISL_541486, EPI_ISL_541487, EPI_ISL_541488, EPI_ISL_541489, EPI_ISL_541490, EPI_ISL_541491, EPI_ISL_541492, EPI_ISL_541493, EPI_ISL_541494, EPI_ISL_541507, EPI_ISL_541510, EPI_ISL_541511, EPI_ISL_541512, EPI_ISL_541513, EPI_ISL_541514, EPI_ISL_541515, EPI_ISL_541516, EPI_ISL_541517, EPI_ISL_541518, EPI_ISL_541519, EPI_ISL_541520, EPI_ISL_541521, EPI_ISL_541522, EPI_ISL_541523 | | | |
| see above | Viollier AG | Department of Biosystems Science and Engineering, ETH Zürich | Christian Beisel, Sarah Nadeau, Ivan Topolsky, Pedro Ferreira, Philipp Jablonski, Susana Posada-Céspedes, Tobias Schär, Ina Nissen, Natascha Santacroce, Elodie Burcklen, Christiane Beckmann, Maurice Redondo, Olivier Kobel, Christoph Noppen, Sophie Seidel, Noemie Santamaria de Souza, Niko Beerenwinkel, Tanja Stadler |
| EPI_ISL_541568 | University of Wisconsin-Madison Campus AIDS Vaccine Research Laboratories | University of Wisconsin-Madison AIDS Vaccine Research Laboratories | Gage Moreno, Katarina Braun, et al. AIDS Vaccine Research Laboratories |
| EPI_ISL_541723, EPI_ISL_541724 | National Institute of Virology, NIV Influenza | National Institute of Virology, NIV Influenza | Potdar V |
| EPI_ISL_542063, EPI_ISL_542064, EPI_ISL_542065, EPI_ISL_542066, EPI_ISL_542067, EPI_ISL_542068, EPI_ISL_542069, EPI_ISL_542070, EPI_ISL_542071, EPI_ISL_542072, EPI_ISL_542073, EPI_ISL_542074, EPI_ISL_542075, EPI_ISL_542076, EPI_ISL_542077, EPI_ISL_542078, EPI_ISL_542079, EPI_ISL_542080, EPI_ISL_542081, EPI_ISL_542082, EPI_ISL_542083, EPI_ISL_542084, EPI_ISL_542085, EPI_ISL_542086, EPI_ISL_542088, EPI_ISL_542089, EPI_ISL_542090, EPI_ISL_542091, EPI_ISL_542092, EPI_ISL_542093, EPI_ISL_542094, EPI_ISL_542095, EPI_ISL_542096, EPI_ISL_542097 | | | |
| see above | New Mexico Department of Health Scientific Laboratory | New Mexico Department of Health Scientific Laboratory | Ellie Johnson, Anastacia Griego-Fisher, D'Eldra Malone |
| EPI_ISL_544999, EPI_ISL_545000, EPI_ISL_545001 | St Vincent's Pathology (SydPath) | NSW Health Pathology - Institute of Clinical Pathology and Medical Research; Westmead Hospital; University of Sydney | CIDM-PH et al. |
| EPI_ISL_545002 | The Children's Hospital at Westmead | NSW Health Pathology - Institute of Clinical Pathology and Medical Research; Westmead Hospital; University of Sydney | CIDM-PH et al. |
| EPI_ISL_545004 | Sydney South West Pathology Service (SSWPS) - Royal Prince Alfred Hospital - NSW Health Pathology | NSW Health Pathology - Institute of Clinical Pathology and Medical Research; Westmead Hospital; University of Sydney | CIDM-PH et al. |
| EPI_ISL_545006 | Pathology West - NSW Health Pathology | NSW Health Pathology - Institute of Clinical Pathology and Medical Research; Westmead Hospital; University of Sydney | CIDM-PH et al. |
| EPI_ISL_545007 | Histopath | NSW Health Pathology - Institute of Clinical Pathology and Medical Research; Westmead Hospital; University of Sydney | CIDM-PH et al. |
| EPI_ISL_545008 | Pathology North - Royal North Shore Hospital - NSW Health Pathology | NSW Health Pathology - Institute of Clinical Pathology and Medical Research; Westmead Hospital; University of Sydney | CIDM-PH et al. |
| EPI_ISL_545011 | South Eastern Area Laboratory Services (SEALS) | NSW Health Pathology - Institute of Clinical Pathology and Medical Research; Westmead Hospital; University of Sydney | CIDM-PH et al. |
| EPI_ISL_545024 | Australian Clinical Labs | NSW Health Pathology - Institute of Clinical Pathology and Medical Research; Westmead Hospital; University of Sydney | CIDM-PH et al. |
| EPI_ISL_545025 | Sydney South West Pathology Service (SSWPS) - Liverpool Hospital - NSW Health Pathology | NSW Health Pathology - Institute of Clinical Pathology and Medical Research; Westmead Hospital; University of Sydney | CIDM-PH et al. |
| EPI_ISL_545026 | St Vincent's Pathology (SydPath) | NSW Health Pathology - Institute of Clinical Pathology and Medical Research; Westmead Hospital; University of Sydney | CIDM-PH et al. |
| EPI_ISL_545027 | Douglass Hanly Moir Pathology | NSW Health Pathology - Institute of Clinical Pathology and Medical Research; Westmead Hospital; University of Sydney | CIDM-PH et al. |
| EPI_ISL_545028 | Sydney South West Pathology Service (SSWPS) - Royal Prince Alfred Hospital - NSW Health Pathology | NSW Health Pathology - Institute of Clinical Pathology and Medical Research; Westmead Hospital; University of Sydney | CIDM-PH et al. |
| EPI_ISL_545029 | Pathology West - NSW Health Pathology | NSW Health Pathology - Institute of Clinical Pathology and Medical Research; Westmead Hospital; University of Sydney | CIDM-PH et al. |
| EPI_ISL_545030 | St Vincent's Pathology (SydPath) | NSW Health Pathology - Institute of Clinical Pathology and Medical Research; Westmead Hospital; University of Sydney | CIDM-PH et al. |
| EPI_ISL_545031 | South Eastern Area Laboratory Services (SEALS) | NSW Health Pathology - Institute of Clinical Pathology and Medical Research; Westmead Hospital; University of Sydney | CIDM-PH et al. |
| EPI_ISL_545040 | Laverty Pathology | NSW Health Pathology - Institute of Clinical Pathology and Medical Research; Westmead Hospital; University of Sydney | CIDM-PH et al. |
| EPI_ISL_545041, EPI_ISL_545043 | Sydney South West Pathology Service (SSWPS) - Royal Prince Alfred Hospital - NSW Health Pathology | NSW Health Pathology - Institute of Clinical Pathology and Medical Research; Westmead Hospital; University of Sydney | CIDM-PH et al. |
| EPI_ISL_545044 | Sydney South West Pathology Service (SSWPS) - Concord Repatriation General Hospital - NSW Health Pathology | NSW Health Pathology - Institute of Clinical Pathology and Medical Research; Westmead Hospital; University of Sydney | CIDM-PH et al. |
| EPI_ISL_545045 | Sydney South West Pathology Service (SSWPS) - Royal Prince Alfred Hospital - NSW Health Pathology | NSW Health Pathology - Institute of Clinical Pathology and Medical Research; Westmead Hospital; University of Sydney | CIDM-PH et al. |
| EPI_ISL_547583 | University of Wisconsin-Madison AIDS Vaccine Research Laboratories | University of Wisconsin-Madison AIDS Vaccine Research Laboratories | Gage Moreno, Katarina Braun, et al. AIDS Vaccine Research Laboratories |
| EPI_ISL_547594 | Sydney South West Pathology Service (SSWPS) - Liverpool Hospital - NSW Health Pathology | NSW Health Pathology - Institute of Clinical Pathology and Medical Research; Westmead Hospital; University of Sydney | CIDM-PH et al. |
| EPI_ISL_547595, EPI_ISL_547596 | St Vincent's Pathology (SydPath) | NSW Health Pathology - Institute of Clinical Pathology and Medical Research; Westmead Hospital; University of Sydney | CIDM-PH et al. |
| EPI_ISL_547597 | Australian Clinical Labs | NSW Health Pathology - Institute of Clinical Pathology and Medical Research; Westmead Hospital; University of Sydney | CIDM-PH et al. |
| EPI_ISL_547598 | Sydney South West Pathology Service (SSWPS) - Concord Repatriation General Hospital - NSW Health Pathology | NSW Health Pathology - Institute of Clinical Pathology and Medical Research; Westmead Hospital; University of Sydney | CIDM-PH et al. |
| EPI_ISL_547599 | Sydney South West Pathology Service (SSWPS) - Royal Prince Alfred Hospital - NSW Health Pathology | NSW Health Pathology - Institute of Clinical Pathology and Medical Research; Westmead Hospital; University of Sydney | CIDM-PH et al. |
| EPI_ISL_547600 | Histopath | NSW Health Pathology - Institute of Clinical Pathology and Medical Research; Westmead Hospital; University of Sydney | CIDM-PH et al. |
| EPI_ISL_547755, EPI_ISL_547756, EPI_ISL_547757, EPI_ISL_547758, EPI_ISL_547759, EPI_ISL_547760, EPI_ISL_547761, EPI_ISL_547762, EPI_ISL_547763, EPI_ISL_547764, EPI_ISL_547765, EPI_ISL_547766, EPI_ISL_547767, EPI_ISL_547768, EPI_ISL_547769, EPI_ISL_547770, EPI_ISL_547771, EPI_ISL_547772, EPI_ISL_547773, EPI_ISL_547774, EPI_ISL_547775, EPI_ISL_547776, EPI_ISL_547777, EPI_ISL_547778, EPI_ISL_547779, EPI_ISL_547780, EPI_ISL_547781, EPI_ISL_547782, EPI_ISL_547783, EPI_ISL_547784, EPI_ISL_547785, EPI_ISL_547786, EPI_ISL_547787, EPI_ISL_547788, EPI_ISL_547789, EPI_ISL_547790, EPI_ISL_547791, EPI_ISL_547792, EPI_ISL_547793, EPI_ISL_547794, EPI_ISL_547795, EPI_ISL_547796, EPI_ISL_547797, EPI_ISL_547798, EPI_ISL_547799, EPI_ISL_547800, EPI_ISL_547801, EPI_ISL_547802, EPI_ISL_547803, EPI_ISL_547804, EPI_ISL_547805, EPI_ISL_547806, EPI_ISL_547807, EPI_ISL_547808, EPI_ISL_547809, EPI_ISL_547810, EPI_ISL_547811, EPI_ISL_547812, EPI_ISL_547813, EPI_ISL_547814, EPI_ISL_547815, EPI_ISL_547816, EPI_ISL_547817, EPI_ISL_547818 | | | |

| | | | |
|---|---|---|---|
| see above | Gundersen Molecular Diagnostics Laboratory | Kabara Cancer Research Institute | Craig S. Richmond, Paraic A. Kenny |
| EPI_ISL_548065, EPI_ISL_548066, EPI_ISL_548067 | North Shore Hospital | Institute of Environmental Science and Research (ESR) | Xiaoyun Ren, Matt Storey, Nikki Freed, Muhammad Faisal, Jing Wang, Hermes Perez, Anja Werno, Antje van der Linden, Arlo Upton, Chris Mansell, David Hammer, Dragana Drinkovic, Gary McAuliffe, Hana Sofia Andersson, James Ussher, Jill Sherwood, Josh Freeman, Julia Howard, Juliet Elvy, Mary DeAlmeida, Matt Blakiston, Matthew Rogers, Max Bloomfield, Michael Addidle, Michelle Balm, Sally Roberts, Sarah Jefferies, Sharmini Muttaiyah, Susan Morpeth, Susan Taylor, Timothy Blackmore, Vani Sathyendran, Veronica Playle, Virginia Hope, Erasmus Smit, Lauren Jelly, Olin Silander, Joep de Ligt |
| EPI_ISL_548068, EPI_ISL_548069, EPI_ISL_548070, EPI_ISL_548071, EPI_ISL_548072, EPI_ISL_548073 | LabPLUS | Institute of Environmental Science and Research (ESR) | Xiaoyun Ren, Matt Storey, Nikki Freed, Muhammad Faisal, Jing Wang, Hermes Perez, Anja Werno, Antje van der Linden, Arlo Upton, Chris Mansell, David Hammer, Dragana Drinkovic, Gary McAuliffe, Hana Sofia Andersson, James Ussher, Jill Sherwood, Josh Freeman, Julia Howard, Juliet Elvy, Mary DeAlmeida, Matt Blakiston, Matthew Rogers, Max Bloomfield, Michael Addidle, Michelle Balm, Sally Roberts, Sarah Jefferies, Sharmini Muttaiyah, Susan Morpeth, Susan Taylor, Timothy Blackmore, Vani Sathyendran, Veronica Playle, Virginia Hope, Erasmus Smit, Lauren Jelly, Olin Silander, Joep de Ligt |
| EPI_ISL_548074 | Middlemore Hospital | Institute of Environmental Science and Research (ESR) | Xiaoyun Ren, Matt Storey, Nikki Freed, Muhammad Faisal, Jing Wang, Hermes Perez, Anja Werno, Antje van der Linden, Arlo Upton, Chris Mansell, David Hammer, Dragana Drinkovic, Gary McAuliffe, Hana Sofia Andersson, James Ussher, Jill Sherwood, Josh Freeman, Julia Howard, Juliet Elvy, Mary DeAlmeida, Matt Blakiston, Matthew Rogers, Max Bloomfield, Michael Addidle, Michelle Balm, Sally Roberts, Sarah Jefferies, Sharmini Muttaiyah, Susan Morpeth, Susan Taylor, Timothy Blackmore, Vani Sathyendran, Veronica Playle, Virginia Hope, Erasmus Smit, Lauren Jelly, Olin Silander, Joep de Ligt |
| EPI_ISL_548075, EPI_ISL_548076 | North Shore Hospital | Institute of Environmental Science and Research (ESR) | Xiaoyun Ren, Matt Storey, Nikki Freed, Muhammad Faisal, Jing Wang, Hermes Perez, Anja Werno, Antje van der Linden, Arlo Upton, Chris Mansell, David Hammer, Dragana Drinkovic, Gary McAuliffe, Hana Sofia Andersson, James Ussher, Jill Sherwood, Josh Freeman, Julia Howard, Juliet Elvy, Mary DeAlmeida, Matt Blakiston, Matthew Rogers, Max Bloomfield, Michael Addidle, Michelle Balm, Sally Roberts, Sarah Jefferies, Sharmini Muttaiyah, Susan Morpeth, Susan Taylor, Timothy Blackmore, Vani Sathyendran, Veronica Playle, Virginia Hope, Erasmus Smit, Lauren Jelly, Olin Silander, Joep de Ligt |
| EPI_ISL_548077, EPI_ISL_548078, EPI_ISL_548079, EPI_ISL_548080, EPI_ISL_548083, EPI_ISL_548119, EPI_ISL_548120 | LabPLUS | Institute of Environmental Science and Research (ESR) | Xiaoyun Ren, Matt Storey, Nikki Freed, Muhammad Faisal, Jing Wang, Hermes Perez, Anja Werno, Antje van der Linden, Arlo Upton, Chris Mansell, David Hammer, Dragana Drinkovic, Gary McAuliffe, Hana Sofia Andersson, James Ussher, Jill Sherwood, Josh Freeman, Julia Howard, Juliet Elvy, Mary DeAlmeida, Matt Blakiston, Matthew Rogers, Max Bloomfield, Michael Addidle, Michelle Balm, Sally Roberts, Sarah Jefferies, Sharmini Muttaiyah, Susan Morpeth, Susan Taylor, Timothy Blackmore, Vani Sathyendran, Veronica Playle, Virginia Hope, Erasmus Smit, Lauren Jelly, Olin Silander, Joep de Ligt |
| EPI_ISL_548121 | Middlemore Hospital | Institute of Environmental Science and Research (ESR) | Xiaoyun Ren, Matt Storey, Nikki Freed, Muhammad Faisal, Jing Wang, Hermes Perez, Anja Werno, Antje van der Linden, Arlo Upton, Chris Mansell, David Hammer, Dragana Drinkovic, Gary McAuliffe, Hana Sofia Andersson, James Ussher, Jill Sherwood, Josh Freeman, Julia Howard, Juliet Elvy, Mary DeAlmeida, Matt Blakiston, Matthew Rogers, Max Bloomfield, Michael Addidle, Michelle Balm, Sally Roberts, Sarah Jefferies, Sharmini Muttaiyah, Susan Morpeth, Susan Taylor, Timothy Blackmore, Vani Sathyendran, Veronica Playle, Virginia Hope, Erasmus Smit, Lauren Jelly, Olin Silander, Joep de Ligt |
| EPI_ISL_548122 | LabPLUS | Institute of Environmental Science and Research (ESR) | Xiaoyun Ren, Matt Storey, Nikki Freed, Muhammad Faisal, Jing Wang, Hermes Perez, Anja Werno, Antje van der Linden, Arlo Upton, Chris Mansell, David Hammer, Dragana Drinkovic, Gary McAuliffe, Hana Sofia Andersson, James Ussher, Jill Sherwood, Josh Freeman, Julia Howard, Juliet Elvy, Mary DeAlmeida, Matt Blakiston, Matthew Rogers, Max Bloomfield, Michael Addidle, Michelle Balm, Sally Roberts, Sarah Jefferies, Sharmini Muttaiyah, Susan Morpeth, Susan Taylor, Timothy Blackmore, Vani Sathyendran, Veronica Playle, Virginia Hope, Erasmus Smit, Lauren Jelly, Olin Silander, Joep de Ligt |
| EPI_ISL_548973 | National Public Health Laboratory, National Centre for Infectious Diseases | National Public Health Laboratory, National Centre for Infectious Diseases | Mak TM, Octavia S, Zhou Z, Cui L, Lin RTP |
| EPI_ISL_549201, EPI_ISL_549209, EPI_ISL_549210, EPI_ISL_549211, EPI_ISL_549212, EPI_ISL_549215, EPI_ISL_549216, EPI_ISL_549217, EPI_ISL_549218, EPI_ISL_549219, EPI_ISL_549220, EPI_ISL_549221, EPI_ISL_549222, EPI_ISL_549223, EPI_ISL_549224, EPI_ISL_549225, EPI_ISL_549226, EPI_ISL_549227 | | | |
| see above | Florida Bureau of Public Health Laboratories | Florida Bureau of Public Health Laboratories | Sarah Schmedes, Jason Blanton |
| EPI_ISL_549329, EPI_ISL_549330, EPI_ISL_549331, EPI_ISL_549333, EPI_ISL_549335, EPI_ISL_549337 | Oxford Viromics, NDM, University of Oxford; Oxford University Hospitals; Basingstoke and North Hampshire Hospital | COVID-19 Genomics UK (COG-UK) Consortium | Tanya Golubchik, David Bonsall, George Macintyre, Amy Trebes, Mariateresa de Cesare, Catrin Moore, Alex Mobbs, Anita Justice, Robert Shaw, Monique Andersson, Timothy Peto, Emma Wise, Nathan Moore, Jessica Lynch, Nick Cortes, Matilde Mori, Stephen Kidd, David Buck, John Todd, Christophe Fraser |
| EPI_ISL_549343 | Centre for Enzyme Innovation, University of Portsmouth / Translational Research Laboratory, Portsmouth Hospitals NHS Trust | COVID-19 Genomics UK (COG-UK) Consortium | Angela Beckett,Yann Bourgeois,Garry Scarlett,Sharon Glaysher,Scott Elliott,Kelly Bicknell,Robert Impey,Allyson Lloyd,Sarah Wyllie,Ethan Butcher,Anoop Chauhan,Samuel Robson |
| EPI_ISL_549344 | Quadram Institute Bioscience | COVID-19 Genomics UK (COG-UK) Consortium | Dave J. Baker, Gemma L. Kay, Alp Aydin, Thanh Le-Viet, Steven Rudder, Ana P. Tedim, Anastasia Kolyva, Maria Diaz, Leonardo de Oliveira Martins, Nabil-Fareed Alikhan, Lizzie Meadows, Rachael Stanley, Ngozi Elumogo, Muhammed Yasir, Nicholas M. Thomson, Alexander J Trotter, Rachel Gilroy, Samuel Bloomfield, Claire Stuart, Andrew Bell, Reenesh Prakash, Samir Dervisevic, Alison E. Mather, John Wain, Mark Webber, Andrew J. Page, Justin O'Grady |
| EPI_ISL_549347 | Oxford Viromics, NDM, University of Oxford; Oxford University Hospitals; Basingstoke and North Hampshire Hospital | COVID-19 Genomics UK (COG-UK) Consortium | Tanya Golubchik, David Bonsall, George Macintyre, Amy Trebes, Mariateresa de Cesare, Catrin Moore, Alex Mobbs, Anita Justice, Robert Shaw, Monique Andersson, Timothy Peto, Emma Wise, Nathan Moore, Jessica Lynch, Nick Cortes, Matilde Mori, Stephen Kidd, David Buck, John Todd, Christophe Fraser |
| EPI_ISL_549350, EPI_ISL_549351 | Quadram Institute Bioscience | COVID-19 Genomics UK (COG-UK) Consortium | Dave J. Baker, Gemma L. Kay, Alp Aydin, Thanh Le-Viet, Steven Rudder, Ana P. Tedim, Anastasia Kolyva, Maria Diaz, Leonardo de Oliveira Martins, Nabil-Fareed Alikhan, Lizzie Meadows, Rachael Stanley, Ngozi Elumogo, Muhammed Yasir, Nicholas M. Thomson, Alexander J Trotter, Rachel Gilroy, Samuel Bloomfield, Claire Stuart, Andrew Bell, Reenesh Prakash, Samir Dervisevic, Alison E. Mather, John Wain, Mark Webber, Andrew J. Page, Justin O'Grady |
| EPI_ISL_549352 | Lincolnshire Hospitals and DeepSeq Nottingham | COVID-19 Genomics UK (COG-UK) Consortium | Nichola Duckworth, Tim Sloan, Sarah Walsh, Jonathan Ball, Patrick McClure, Joeseph Chappell, Nadine Holmes, Matthew Carlisle, Christopher Moore, Fei Sang, Johnny Debebe, Victoria Wright, Matthew Loose |
| EPI_ISL_549358 | Liverpool Clinical Laboratories | COVID-19 Genomics UK (COG-UK) Consortium | Sam Haldenby, Anita Lucaci, Steve Paterson, Julian Hiscox, Alistair Darby, M Almsaud, A Alrezaihi, Muhannad Alruwaili, Stuart D Armstrong, Jones Benjamin, Eleanor G Bentley, Anu Chawla, Jordan J Clark, Angela Cowell, Richard Eccles, Isabel García-Dorival, Matthew Gemmell, Alessandro Gerada, PKF Gilmore, Richard Gregory, Ximeng Han, Catherine Hartley, Margaret Hughes, Miren Iturriza-Gomara, James Johnson, L Luu, Jenifer Manson, Charlotte Nelson, Elaine O'Toole, Cassie Olateju, Rebekah Penrice-Randal , Lucille Rainbow, N.P Randle, Trevor Ian Robinson, Parul Sharma, Ghada T Shawli, James P Stewart, Neil Swainston, Ecaterina Vamos, Joanne Watts, Mark Whitehead |
| EPI_ISL_549362 | Lincolnshire Hospitals and DeepSeq Nottingham | COVID-19 Genomics UK (COG-UK) Consortium | Nichola Duckworth, Tim Sloan, Sarah Walsh, Jonathan Ball, Patrick McClure, Joeseph Chappell, Nadine Holmes, Matthew Carlisle, Christopher Moore, Fei Sang, Johnny Debebe, Victoria Wright, Matthew Loose |
| EPI_ISL_549363 | Centre for Enzyme Innovation, University of Portsmouth / Translational Research Laboratory, Portsmouth Hospitals NHS Trust | COVID-19 Genomics UK (COG-UK) Consortium | Angela Beckett,Yann Bourgeois,Garry Scarlett,Sharon Glaysher,Scott Elliott,Kelly Bicknell,Robert Impey,Allyson Lloyd,Sarah Wyllie,Ethan Butcher,Anoop Chauhan,Samuel Robson |
| EPI_ISL_549365 | Lincolnshire Hospitals and DeepSeq Nottingham | COVID-19 Genomics UK (COG-UK) Consortium | Nichola Duckworth, Tim Sloan, Sarah Walsh, Jonathan Ball, Patrick McClure, Joeseph Chappell, Nadine Holmes, Matthew Carlisle, Christopher Moore, Fei Sang, Johnny Debebe, Victoria Wright, Matthew Loose |
| EPI_ISL_549367 | Quadram Institute Bioscience | COVID-19 Genomics UK (COG-UK) Consortium | Dave J. Baker, Gemma L. Kay, Alp Aydin, Thanh Le-Viet, Steven Rudder, Ana P. Tedim, Anastasia Kolyva, Maria Diaz, Leonardo de Oliveira Martins, Nabil-Fareed Alikhan, Lizzie Meadows, Rachael Stanley, Ngozi Elumogo, Muhammed Yasir, Nicholas M. Thomson, Alexander J Trotter, Rachel Gilroy, Samuel Bloomfield, Claire Stuart, Andrew Bell, Reenesh Prakash, Samir Dervisevic, Alison E. Mather, John Wain, Mark Webber, Andrew J. Page, Justin O'Grady |
| EPI_ISL_549374 | Department of Pathology, University of Cambridge | COVID-19 Genomics UK (COG-UK) Consortium | Aminu S. Jahun, Yasmin Chaudhry, Grant Hall, Iliana Georgana, Myra Hosmillo, Martin D. Curran, Malte Pinckert, Surendra Parmar, Ian Goodfellow |
| EPI_ISL_549375, EPI_ISL_549381 | Lincolnshire Hospitals and DeepSeq Nottingham | COVID-19 Genomics UK (COG-UK) Consortium | Nichola Duckworth, Tim Sloan, Sarah Walsh, Jonathan Ball, Patrick McClure, Joeseph Chappell, Nadine Holmes, Matthew Carlisle, Christopher Moore, Fei Sang, Johnny Debebe, Victoria Wright, Matthew Loose |
| EPI_ISL_549382 | Quadram Institute Bioscience | COVID-19 Genomics UK (COG-UK) Consortium | Dave J. Baker, Gemma L. Kay, Alp Aydin, Thanh Le-Viet, Steven Rudder, Ana P. Tedim, Anastasia Kolyva, Maria Diaz, Leonardo de Oliveira Martins, Nabil-Fareed Alikhan, Lizzie Meadows, Rachael Stanley, Ngozi Elumogo, Muhammed Yasir, Nicholas M. Thomson, Alexander J Trotter, Rachel Gilroy, Samuel Bloomfield, Claire Stuart, Andrew Bell, Reenesh Prakash, Samir Dervisevic, Alison E. Mather, John Wain, Mark Webber, Andrew J. Page, Justin O'Grady |
| EPI_ISL_549386, EPI_ISL_549388, | Department of Pathology, University of Cambridge | COVID-19 Genomics UK (COG-UK) Consortium | Aminu S. Jahun, Yasmin Chaudhry, Grant Hall, Iliana Georgana, Myra Hosmillo, Martin D. Curran, Malte Pinckert, Surendra Parmar, Ian Goodfellow |

| | | | |
|---|---|---|---|
| EPI_ISL_549389, EPI_ISL_549390, EPI_ISL_549391, EPI_ISL_549392, EPI_ISL_549393, EPI_ISL_549394 | | | |
| EPI_ISL_549396, EPI_ISL_549397, EPI_ISL_549398, EPI_ISL_549399, EPI_ISL_549400, EPI_ISL_549401, EPI_ISL_549402, EPI_ISL_549403, EPI_ISL_549404, EPI_ISL_549405, EPI_ISL_549406, EPI_ISL_549407, EPI_ISL_549408, EPI_ISL_549409, EPI_ISL_549410, EPI_ISL_549411, EPI_ISL_549412, EPI_ISL_549413, EPI_ISL_549430, EPI_ISL_549431, EPI_ISL_549432, EPI_ISL_549433, EPI_ISL_549434, EPI_ISL_549435, EPI_ISL_549436, EPI_ISL_549437, EPI_ISL_549438, EPI_ISL_549439 | | | |
| see above | Oxford Viromics, NDM, University of Oxford; Oxford University Hospitals; Basingstoke and North Hampshire Hospital | COVID-19 Genomics UK (COG-UK) Consortium | Tanya Golubchik, David Bonsall, George Macintyre, Amy Trebes, Mariateresa de Cesare, Catrin Moore, Alex Mobbs, Anita Justice, Robert Shaw, Monique Andersson, Timothy Peto, Emma Wise, Nathan Moore, Jessica Lynch, Nick Cortes, Matilde Mori, Stephen Kidd, David Buck, John Todd, Christophe Fraser |
| EPI_ISL_549460, EPI_ISL_549462, EPI_ISL_549463, EPI_ISL_549464, EPI_ISL_549465, EPI_ISL_549467, EPI_ISL_549468 | Centre for Enzyme Innovation, University of Portsmouth / Translational Research Laboratory, Portsmouth Hospitals NHS Trust | COVID-19 Genomics UK (COG-UK) Consortium | Angela Beckett,Yann Bourgeois,Garry Scarlett,Sharon Glaysher,Scott Elliott,Kelly Bicknell,Robert Impey,Allyson Lloyd,Sarah Wyllie,Ethan Butcher,Anoop Chauhan,Samuel Robson |
| EPI_ISL_549488 | Quadram Institute Bioscience | COVID-19 Genomics UK (COG-UK) Consortium | Dave J. Baker, Gemma L. Kay, Alp Aydin, Thanh Le-Viet, Steven Rudder, Ana P. Tedim, Anastasia Kolyva, Maria Diaz, Leonardo de Oliveira Martins, Nabil-Fareed Alikhan, Lizzie Meadows, Rachael Stanley, Ngozi Elumogo, Muhammed Yasir, Nicholas M. Thomson, Alexander J Trotter, Rachel Gilroy, Samuel Bloomfield, Claire Stuart, Andrew Bell, Reenesh Prakash, Samir Dervisevic, Alison E. Mather, John Wain, Mark Webber, Andrew J. Page, Justin O'Grady |
| EPI_ISL_549517, EPI_ISL_549518 | Lincolnshire Hospitals and DeepSeq Nottingham | COVID-19 Genomics UK (COG-UK) Consortium | Nichola Duckworth, Tim Sloan, Sarah Walsh, Jonathan Ball, Patrick McClure, Joeseph Chappell, Nadine Holmes, Matthew Carlisle, Christopher Moore, Fei Sang, Johnny Debebe, Victoria Wright, Matthew Loose |
| EPI_ISL_549547, EPI_ISL_549548, EPI_ISL_549549, EPI_ISL_549550, EPI_ISL_549551, EPI_ISL_549552, EPI_ISL_549553, EPI_ISL_549554, EPI_ISL_549556, EPI_ISL_549557, EPI_ISL_549558, EPI_ISL_549559, EPI_ISL_549560, EPI_ISL_549561, EPI_ISL_549562, EPI_ISL_549563, EPI_ISL_549564, EPI_ISL_549565, EPI_ISL_549567, EPI_ISL_549568, EPI_ISL_549569, EPI_ISL_549570, EPI_ISL_549571, EPI_ISL_549572, EPI_ISL_549573, EPI_ISL_549574, EPI_ISL_549575, EPI_ISL_549577, EPI_ISL_549578, EPI_ISL_549579, EPI_ISL_549580, EPI_ISL_549581, EPI_ISL_549582, EPI_ISL_549583, EPI_ISL_549584, EPI_ISL_549585, EPI_ISL_549587, EPI_ISL_549588, EPI_ISL_549589, EPI_ISL_549590, EPI_ISL_549591, EPI_ISL_549592, EPI_ISL_549593, EPI_ISL_549594, EPI_ISL_549596, EPI_ISL_549597, EPI_ISL_549598, EPI_ISL_549599, EPI_ISL_549600, EPI_ISL_549601, EPI_ISL_549602, EPI_ISL_549603, EPI_ISL_549604, EPI_ISL_549605, EPI_ISL_549606, EPI_ISL_549607, EPI_ISL_549608, EPI_ISL_549609, EPI_ISL_549611, EPI_ISL_549612, EPI_ISL_549613, EPI_ISL_549614, EPI_ISL_549615, EPI_ISL_549616, EPI_ISL_549618, EPI_ISL_549619, EPI_ISL_549620, EPI_ISL_549621, EPI_ISL_549622, EPI_ISL_549623, EPI_ISL_549624, EPI_ISL_549625, EPI_ISL_549626, EPI_ISL_549627, EPI_ISL_549628, EPI_ISL_549629, EPI_ISL_549630, EPI_ISL_549631, EPI_ISL_549632, EPI_ISL_549633, EPI_ISL_549634, EPI_ISL_549635, EPI_ISL_549636, EPI_ISL_549637, EPI_ISL_549638, EPI_ISL_549639, EPI_ISL_549640, EPI_ISL_549641, EPI_ISL_549642, EPI_ISL_549643, EPI_ISL_549644, EPI_ISL_549645, EPI_ISL_549646, EPI_ISL_549647, EPI_ISL_549648, EPI_ISL_549649, EPI_ISL_549650, EPI_ISL_549651, EPI_ISL_549652, EPI_ISL_549653, EPI_ISL_549654, EPI_ISL_549655, EPI_ISL_549656, EPI_ISL_549658, EPI_ISL_549659, EPI_ISL_549660, EPI_ISL_549661, EPI_ISL_549662, EPI_ISL_549663, EPI_ISL_549664, EPI_ISL_549665, EPI_ISL_549666, EPI_ISL_549667, EPI_ISL_549668, EPI_ISL_549669, EPI_ISL_549670, EPI_ISL_549671, EPI_ISL_549672, EPI_ISL_549673, EPI_ISL_549674, EPI_ISL_549675, EPI_ISL_549676, EPI_ISL_549677, EPI_ISL_549678, EPI_ISL_549679, EPI_ISL_549680, EPI_ISL_549681, EPI_ISL_549682, EPI_ISL_549683, EPI_ISL_549684, EPI_ISL_549685, EPI_ISL_549686, EPI_ISL_549687, EPI_ISL_549688, EPI_ISL_549689, EPI_ISL_549690, EPI_ISL_549691, EPI_ISL_549692, EPI_ISL_549693, EPI_ISL_549694, EPI_ISL_549695, EPI_ISL_549696, EPI_ISL_549697, EPI_ISL_549698, EPI_ISL_549699, EPI_ISL_549700, EPI_ISL_549701, EPI_ISL_549702, EPI_ISL_549703, EPI_ISL_549704, EPI_ISL_549705, EPI_ISL_549706, EPI_ISL_549707, EPI_ISL_549709, EPI_ISL_549710, EPI_ISL_549711, EPI_ISL_549712, EPI_ISL_549713, EPI_ISL_549714, EPI_ISL_549715, EPI_ISL_549716, EPI_ISL_549717, EPI_ISL_549718, EPI_ISL_549719, EPI_ISL_549720, EPI_ISL_549722, EPI_ISL_549723, EPI_ISL_549725, EPI_ISL_549726, EPI_ISL_549727, EPI_ISL_549728, EPI_ISL_549729, EPI_ISL_549730, EPI_ISL_549731, EPI_ISL_549732, EPI_ISL_549733, EPI_ISL_549734, EPI_ISL_549735, EPI_ISL_549736, EPI_ISL_549739, EPI_ISL_549740, EPI_ISL_549741, EPI_ISL_549742, EPI_ISL_549743, EPI_ISL_549744, EPI_ISL_549745, EPI_ISL_549746, EPI_ISL_549747, EPI_ISL_549748, EPI_ISL_549749, EPI_ISL_549750, EPI_ISL_549751, EPI_ISL_549752, EPI_ISL_549753, EPI_ISL_549754, EPI_ISL_549755, EPI_ISL_549756, EPI_ISL_549757, EPI_ISL_549758, EPI_ISL_549759, EPI_ISL_549760, EPI_ISL_549761, EPI_ISL_549762, EPI_ISL_549763, EPI_ISL_549764, EPI_ISL_549765, EPI_ISL_549766, EPI_ISL_549767, EPI_ISL_549768, EPI_ISL_549769, EPI_ISL_549770, EPI_ISL_549771 | | | |
| see above | Lighthouse Lab in Glasgow | Wellcome Sanger Institute for the COVID-19 Genomics UK (COG-UK) consortium | Harper VanSteenhouse, Yumi Kasai, David Gray, Carol Clugston, Anna Dominiczak and Alex Alderton, Roberto Amato, Sonia Goncalves, Ewan Harrison, David K. Jackson, Ian Johnston, Dominic Kwiatkowski, Cordelia Langford, John Sillitoe on behalf of the Wellcome Sanger Institute COVID-19 Surveillance Team |
| EPI_ISL_549772 | Lighthouse Lab in Glasgow | Wellcome Sanger Institute for the COVID-19 Genomics UK (COG-UK) Consortium | Harper VanSteenhouse, Yumi Kasai, David Gray, Carol Clugston, Anna Dominiczak and Alex Alderton, Roberto Amato, Sonia Goncalves, Ewan Harrison, David K. Jackson, Ian Johnston, Dominic Kwiatkowski, Cordelia Langford, John Sillitoe on behalf of the Wellcome Sanger Institute COVID-19 Surveillance Team |
| EPI_ISL_549773, EPI_ISL_549774, EPI_ISL_549775, EPI_ISL_549776, EPI_ISL_549777, EPI_ISL_549778, EPI_ISL_549779, EPI_ISL_549780, EPI_ISL_549781, EPI_ISL_549782, EPI_ISL_549783 | | | |
| see above | Lighthouse Lab in Glasgow | Wellcome Sanger Institute for the COVID-19 Genomics UK (COG-UK) consortium | Harper VanSteenhouse, Yumi Kasai, David Gray, Carol Clugston, Anna Dominiczak and Alex Alderton, Roberto Amato, Sonia Goncalves, Ewan Harrison, David K. Jackson, Ian Johnston, Dominic Kwiatkowski, Cordelia Langford, John Sillitoe on behalf of the Wellcome Sanger Institute COVID-19 Surveillance Team |
| EPI_ISL_549784 | Lighthouse Lab in Glasgow | Wellcome Sanger Institute for the COVID-19 Genomics UK (COG-UK) Consortium | Harper VanSteenhouse, Yumi Kasai, David Gray, Carol Clugston, Anna Dominiczak and Alex Alderton, Roberto Amato, Sonia Goncalves, Ewan Harrison, David K. Jackson, Ian Johnston, Dominic Kwiatkowski, Cordelia Langford, John Sillitoe on behalf of the Wellcome Sanger Institute COVID-19 Surveillance Team |
| EPI_ISL_549785, EPI_ISL_549786, EPI_ISL_549787, EPI_ISL_549788, EPI_ISL_549789, EPI_ISL_549790, EPI_ISL_549792, EPI_ISL_549793, EPI_ISL_549794, EPI_ISL_549795, EPI_ISL_549796, EPI_ISL_549797, EPI_ISL_549798, EPI_ISL_549799, EPI_ISL_549800, EPI_ISL_549801, EPI_ISL_549802, EPI_ISL_549803, EPI_ISL_549804, EPI_ISL_549805, EPI_ISL_549806, EPI_ISL_549807 | | | |
| see above | Lighthouse Lab in Glasgow | Wellcome Sanger Institute for the COVID-19 Genomics UK (COG-UK) consortium | Harper VanSteenhouse, Yumi Kasai, David Gray, Carol Clugston, Anna Dominiczak and Alex Alderton, Roberto Amato, Sonia Goncalves, Ewan Harrison, David K. Jackson, Ian Johnston, Dominic Kwiatkowski, Cordelia Langford, John Sillitoe on behalf of the Wellcome Sanger Institute COVID-19 Surveillance Team |
| EPI_ISL_549948, EPI_ISL_549949, EPI_ISL_549951, EPI_ISL_549952, EPI_ISL_549953, EPI_ISL_549954, EPI_ISL_549955 | Lighthouse Lab in Milton Keynes | Wellcome Sanger Institute for the COVID-19 Genomics UK (COG-UK) consortium | The Lighthouse Lab in Milton Keynes and Alex Alderton, Roberto Amato, Sonia Goncalves, Ewan Harrison, David K. Jackson, Ian Johnston, Dominic Kwiatkowski, Cordelia Langford, John Sillitoe on behalf of the Wellcome Sanger Institute COVID-19 Surveillance Team (http://www.sanger.ac.uk/covid-team) |
| EPI_ISL_549956 | Lighthouse Lab in Milton Keynes | Wellcome Sanger Institute for the COVID-19 Genomics UK (COG-UK) consortium | The Lighthouse Lab in Alderley Park and Alex Alderton, Roberto Amato, Sonia Goncalves, Ewan Harrison, David K. Jackson, Ian Johnston, Dominic Kwiatkowski, Cordelia Langford, John Sillitoe on behalf of the Wellcome Sanger Institute COVID-19 Surveillance Team |
| EPI_ISL_549957, EPI_ISL_549958, EPI_ISL_549959, EPI_ISL_549960, EPI_ISL_549961, EPI_ISL_549962, EPI_ISL_549963, EPI_ISL_549964, EPI_ISL_549965, EPI_ISL_549966, EPI_ISL_549967, EPI_ISL_549969, EPI_ISL_549971, EPI_ISL_549972, EPI_ISL_549973, EPI_ISL_549974, EPI_ISL_549975, EPI_ISL_549976, EPI_ISL_549977, EPI_ISL_549978 | | | |
| see above | Lighthouse Lab in Milton Keynes | Wellcome Sanger Institute for the COVID-19 Genomics UK (COG-UK) consortium | The Lighthouse Lab in Milton Keynes and Alex Alderton, Roberto Amato, Sonia Goncalves, Ewan Harrison, David K. Jackson, Ian Johnston, Dominic Kwiatkowski, Cordelia Langford, John Sillitoe on behalf of the Wellcome Sanger Institute COVID-19 Surveillance Team (http://www.sanger.ac.uk/covid-team) |
| EPI_ISL_549979 | Lighthouse Lab in Milton Keynes | Wellcome Sanger Institute for the COVID-19 Genomics UK (COG-UK) consortium | The Lighthouse Lab in Alderley Park and Alex Alderton, Roberto Amato, Sonia Goncalves, Ewan Harrison, David K. Jackson, Ian Johnston, Dominic Kwiatkowski, Cordelia Langford, John Sillitoe on behalf of the Wellcome Sanger Institute COVID-19 Surveillance Team |
| EPI_ISL_549980, EPI_ISL_549981, EPI_ISL_549982, EPI_ISL_549983, EPI_ISL_549984, EPI_ISL_549985, EPI_ISL_549986, EPI_ISL_549987, EPI_ISL_549988, EPI_ISL_549989, EPI_ISL_549990, EPI_ISL_549991, EPI_ISL_549992, EPI_ISL_549993, EPI_ISL_549994, EPI_ISL_549995, EPI_ISL_549996, EPI_ISL_549997, EPI_ISL_549998, EPI_ISL_549999 | | | |
| see above | Lighthouse Lab in Milton Keynes | Wellcome Sanger Institute for the COVID-19 Genomics UK (COG-UK) consortium | The Lighthouse Lab in Milton Keynes and Alex Alderton, Roberto Amato, Sonia Goncalves, Ewan Harrison, David K. Jackson, Ian Johnston, Dominic Kwiatkowski, Cordelia Langford, John Sillitoe on behalf of the Wellcome Sanger Institute COVID-19 Surveillance Team (http://www.sanger.ac.uk/covid-team) |
| EPI_ISL_550000 | Lighthouse Lab in Milton Keynes | Wellcome Sanger Institute for the COVID-19 Genomics UK (COG-UK) consortium | The Lighthouse Lab in Alderley Park and Alex Alderton, Roberto Amato, Sonia Goncalves, Ewan Harrison, David K. Jackson, Ian Johnston, Dominic Kwiatkowski, Cordelia Langford, John Sillitoe on behalf of the Wellcome Sanger Institute COVID-19 Surveillance Team |
| EPI_ISL_550001, EPI_ISL_550002 | Lighthouse Lab in Milton Keynes | Wellcome Sanger Institute for the COVID-19 Genomics UK (COG-UK) consortium | The Lighthouse Lab in Milton Keynes and Alex Alderton, Roberto Amato, Sonia Goncalves, Ewan Harrison, David K. Jackson, Ian Johnston, Dominic Kwiatkowski, Cordelia Langford, John Sillitoe on behalf of the Wellcome Sanger Institute COVID-19 Surveillance Team (http://www.sanger.ac.uk/covid-team) |
| EPI_ISL_550003 | Lighthouse Lab in Milton Keynes | Wellcome Sanger Institute for the COVID-19 Genomics UK (COG-UK) consortium | The Lighthouse Lab in Alderley Park and Alex Alderton, Roberto Amato, Sonia Goncalves, Ewan Harrison, David K. Jackson, Ian Johnston, Dominic Kwiatkowski, Cordelia Langford, John Sillitoe on behalf of the Wellcome Sanger Institute COVID-19 Surveillance Team |
| EPI_ISL_550004, EPI_ISL_550005, EPI_ISL_550006, EPI_ISL_550007, EPI_ISL_550008, EPI_ISL_550009, EPI_ISL_550010, EPI_ISL_550011, EPI_ISL_550012, EPI_ISL_550013, EPI_ISL_550014, EPI_ISL_550015, EPI_ISL_550016 | | | |
| see above | Lighthouse Lab in Milton Keynes | Wellcome Sanger Institute for the COVID-19 Genomics UK (COG-UK) consortium | The Lighthouse Lab in Milton Keynes and Alex Alderton, Roberto Amato, Sonia Goncalves, Ewan Harrison, David K. Jackson, Ian Johnston, Dominic Kwiatkowski, Cordelia Langford, John Sillitoe on behalf of the Wellcome Sanger Institute COVID-19 Surveillance Team |

| | | | |
|---|---|---|---|
| | | | (http://www.sanger.ac.uk/covid-team) |
| EPI_ISL_550017 | Lighthouse Lab in Milton Keynes | Wellcome Sanger Institute for the COVID-19 Genomics UK (COG-UK) consortium | The Lighthouse Lab in Alderley Park and Alex Alderton, Roberto Amato, Sonia Goncalves, Ewan Harrison, David K. Jackson, Ian Johnston, Dominic Kwiatkowski, Cordelia Langford, John Sillitoe on behalf of the Wellcome Sanger Institute COVID-19 Surveillance Team |
| EPI_ISL_550018 | Lighthouse Lab in Milton Keynes | Wellcome Sanger Institute for the COVID-19 Genomics UK (COG-UK) consortium | The Lighthouse Lab in Milton Keynes and Alex Alderton, Roberto Amato, Sonia Goncalves, Ewan Harrison, David K. Jackson, Ian Johnston, Dominic Kwiatkowski, Cordelia Langford, John Sillitoe on behalf of the Wellcome Sanger Institute COVID-19 Surveillance Team (http://www.sanger.ac.uk/covid-team) |
| EPI_ISL_550019 | Lighthouse Lab in Milton Keynes | Wellcome Sanger Institute for the COVID-19 Genomics UK (COG-UK) consortium | The Lighthouse Lab in Alderley Park and Alex Alderton, Roberto Amato, Sonia Goncalves, Ewan Harrison, David K. Jackson, Ian Johnston, Dominic Kwiatkowski, Cordelia Langford, John Sillitoe on behalf of the Wellcome Sanger Institute COVID-19 Surveillance Team |
| EPI_ISL_550020, EPI_ISL_550021, EPI_ISL_550022 | Lighthouse Lab in Milton Keynes | Wellcome Sanger Institute for the COVID-19 Genomics UK (COG-UK) consortium | The Lighthouse Lab in Milton Keynes and Alex Alderton, Roberto Amato, Sonia Goncalves, Ewan Harrison, David K. Jackson, Ian Johnston, Dominic Kwiatkowski, Cordelia Langford, John Sillitoe on behalf of the Wellcome Sanger Institute COVID-19 Surveillance Team (http://www.sanger.ac.uk/covid-team) |
| EPI_ISL_550023 | Lighthouse Lab in Milton Keynes | Wellcome Sanger Institute for the COVID-19 Genomics UK (COG-UK) consortium | The Lighthouse Lab in Alderley Park and Alex Alderton, Roberto Amato, Sonia Goncalves, Ewan Harrison, David K. Jackson, Ian Johnston, Dominic Kwiatkowski, Cordelia Langford, John Sillitoe on behalf of the Wellcome Sanger Institute COVID-19 Surveillance Team |
| EPI_ISL_550024, EPI_ISL_550025, EPI_ISL_550026, EPI_ISL_550027, EPI_ISL_550028, EPI_ISL_550029 | Lighthouse Lab in Milton Keynes | Wellcome Sanger Institute for the COVID-19 Genomics UK (COG-UK) consortium | The Lighthouse Lab in Milton Keynes and Alex Alderton, Roberto Amato, Sonia Goncalves, Ewan Harrison, David K. Jackson, Ian Johnston, Dominic Kwiatkowski, Cordelia Langford, John Sillitoe on behalf of the Wellcome Sanger Institute COVID-19 Surveillance Team (http://www.sanger.ac.uk/covid-team) |
| EPI_ISL_550030 | Lighthouse Lab in Milton Keynes | Wellcome Sanger Institute for the COVID-19 Genomics UK (COG-UK) consortium | The Lighthouse Lab in Alderley Park and Alex Alderton, Roberto Amato, Sonia Goncalves, Ewan Harrison, David K. Jackson, Ian Johnston, Dominic Kwiatkowski, Cordelia Langford, John Sillitoe on behalf of the Wellcome Sanger Institute COVID-19 Surveillance Team |
| EPI_ISL_550031, EPI_ISL_550032, EPI_ISL_550033, EPI_ISL_550034, EPI_ISL_550035 | Lighthouse Lab in Milton Keynes | Wellcome Sanger Institute for the COVID-19 Genomics UK (COG-UK) consortium | The Lighthouse Lab in Milton Keynes and Alex Alderton, Roberto Amato, Sonia Goncalves, Ewan Harrison, David K. Jackson, Ian Johnston, Dominic Kwiatkowski, Cordelia Langford, John Sillitoe on behalf of the Wellcome Sanger Institute COVID-19 Surveillance Team (http://www.sanger.ac.uk/covid-team) |
| EPI_ISL_550036, EPI_ISL_550037 | Lighthouse Lab in Milton Keynes | Wellcome Sanger Institute for the COVID-19 Genomics UK (COG-UK) consortium | The Lighthouse Lab in Alderley Park and Alex Alderton, Roberto Amato, Sonia Goncalves, Ewan Harrison, David K. Jackson, Ian Johnston, Dominic Kwiatkowski, Cordelia Langford, John Sillitoe on behalf of the Wellcome Sanger Institute COVID-19 Surveillance Team |
| EPI_ISL_550038, EPI_ISL_550039, EPI_ISL_550040 | Lighthouse Lab in Milton Keynes | Wellcome Sanger Institute for the COVID-19 Genomics UK (COG-UK) consortium | The Lighthouse Lab in Milton Keynes and Alex Alderton, Roberto Amato, Sonia Goncalves, Ewan Harrison, David K. Jackson, Ian Johnston, Dominic Kwiatkowski, Cordelia Langford, John Sillitoe on behalf of the Wellcome Sanger Institute COVID-19 Surveillance Team (http://www.sanger.ac.uk/covid-team) |
| EPI_ISL_550041 | Lighthouse Lab in Milton Keynes | Wellcome Sanger Institute for the COVID-19 Genomics UK (COG-UK) consortium | The Lighthouse Lab in Alderley Park and Alex Alderton, Roberto Amato, Sonia Goncalves, Ewan Harrison, David K. Jackson, Ian Johnston, Dominic Kwiatkowski, Cordelia Langford, John Sillitoe on behalf of the Wellcome Sanger Institute COVID-19 Surveillance Team |
| EPI_ISL_550042, EPI_ISL_550043, EPI_ISL_550044, EPI_ISL_550045 | Lighthouse Lab in Milton Keynes | Wellcome Sanger Institute for the COVID-19 Genomics UK (COG-UK) consortium | The Lighthouse Lab in Milton Keynes and Alex Alderton, Roberto Amato, Sonia Goncalves, Ewan Harrison, David K. Jackson, Ian Johnston, Dominic Kwiatkowski, Cordelia Langford, John Sillitoe on behalf of the Wellcome Sanger Institute COVID-19 Surveillance Team (http://www.sanger.ac.uk/covid-team) |
| EPI_ISL_550046 | Lighthouse Lab in Milton Keynes | Wellcome Sanger Institute for the COVID-19 Genomics UK (COG-UK) consortium | The Lighthouse Lab in Alderley Park and Alex Alderton, Roberto Amato, Sonia Goncalves, Ewan Harrison, David K. Jackson, Ian Johnston, Dominic Kwiatkowski, Cordelia Langford, John Sillitoe on behalf of the Wellcome Sanger Institute COVID-19 Surveillance Team |
| EPI_ISL_550047, EPI_ISL_550048, EPI_ISL_550049, EPI_ISL_550050, EPI_ISL_550051, EPI_ISL_550052, EPI_ISL_550053, EPI_ISL_550054, EPI_ISL_550055, EPI_ISL_550056, EPI_ISL_550057, EPI_ISL_550059, EPI_ISL_550060, EPI_ISL_550061, EPI_ISL_550063, EPI_ISL_550064, EPI_ISL_550065, EPI_ISL_550067, EPI_ISL_550068 | | | |
| see above | Lighthouse Lab in Milton Keynes | Wellcome Sanger Institute for the COVID-19 Genomics UK (COG-UK) consortium | The Lighthouse Lab in Milton Keynes and Alex Alderton, Roberto Amato, Sonia Goncalves, Ewan Harrison, David K. Jackson, Ian Johnston, Dominic Kwiatkowski, Cordelia Langford, John Sillitoe on behalf of the Wellcome Sanger Institute COVID-19 Surveillance Team (http://www.sanger.ac.uk/covid-team) |
| EPI_ISL_550069 | Lighthouse Lab in Milton Keynes | Wellcome Sanger Institute for the COVID-19 Genomics UK (COG-UK) Consortium | The Lighthouse Lab in Milton Keynes and Alex Alderton, Roberto Amato, Sonia Goncalves, Ewan Harrison, David K. Jackson, Ian Johnston, Dominic Kwiatkowski, Cordelia Langford, John Sillitoe on behalf of the Wellcome Sanger Institute COVID-19 Surveillance Team |
| EPI_ISL_550070 | Lighthouse Lab in Milton Keynes | Wellcome Sanger Institute for the COVID-19 Genomics UK (COG-UK) consortium | The Lighthouse Lab in Milton Keynes and Alex Alderton, Roberto Amato, Sonia Goncalves, Ewan Harrison, David K. Jackson, Ian Johnston, Dominic Kwiatkowski, Cordelia Langford, John Sillitoe on behalf of the Wellcome Sanger Institute COVID-19 Surveillance Team (http://www.sanger.ac.uk/covid-team) |
| EPI_ISL_550071 | Lighthouse Lab in Milton Keynes | Wellcome Sanger Institute for the COVID-19 Genomics UK (COG-UK) consortium | The Lighthouse Lab in Alderley Park and Alex Alderton, Roberto Amato, Sonia Goncalves, Ewan Harrison, David K. Jackson, Ian Johnston, Dominic Kwiatkowski, Cordelia Langford, John Sillitoe on behalf of the Wellcome Sanger Institute COVID-19 Surveillance Team |
| EPI_ISL_550072, EPI_ISL_550074, EPI_ISL_550076, EPI_ISL_550078, EPI_ISL_550079, EPI_ISL_550080, EPI_ISL_550081, EPI_ISL_550082, EPI_ISL_550083, EPI_ISL_550084, EPI_ISL_550085, EPI_ISL_550086, EPI_ISL_550087, EPI_ISL_550088, EPI_ISL_550089, EPI_ISL_550090, EPI_ISL_550091, EPI_ISL_550092, EPI_ISL_550093, EPI_ISL_550094, EPI_ISL_550095, EPI_ISL_550096, EPI_ISL_550097, EPI_ISL_550098, EPI_ISL_550099, EPI_ISL_550100, EPI_ISL_550101, EPI_ISL_550102, EPI_ISL_550103, EPI_ISL_550105, EPI_ISL_550106, EPI_ISL_550109, EPI_ISL_550110, EPI_ISL_550112, EPI_ISL_550113, EPI_ISL_550114, EPI_ISL_550115, EPI_ISL_550116, EPI_ISL_550117, EPI_ISL_550118, EPI_ISL_550119, EPI_ISL_550125, EPI_ISL_550126, EPI_ISL_550127, EPI_ISL_550129, EPI_ISL_550130, EPI_ISL_550131, EPI_ISL_550132, EPI_ISL_550133, EPI_ISL_550134, EPI_ISL_550136, EPI_ISL_550137 | | | |
| see above | Lighthouse Lab in Milton Keynes | Wellcome Sanger Institute for the COVID-19 Genomics UK (COG-UK) consortium | The Lighthouse Lab in Milton Keynes and Alex Alderton, Roberto Amato, Sonia Goncalves, Ewan Harrison, David K. Jackson, Ian Johnston, Dominic Kwiatkowski, Cordelia Langford, John Sillitoe on behalf of the Wellcome Sanger Institute COVID-19 Surveillance Team (http://www.sanger.ac.uk/covid-team) |
| EPI_ISL_550138 | Lighthouse Lab in Milton Keynes | Wellcome Sanger Institute for the COVID-19 Genomics UK (COG-UK) consortium | The Lighthouse Lab in Alderley Park and Alex Alderton, Roberto Amato, Sonia Goncalves, Ewan Harrison, David K. Jackson, Ian Johnston, Dominic Kwiatkowski, Cordelia Langford, John Sillitoe on behalf of the Wellcome Sanger Institute COVID-19 Surveillance Team |
| EPI_ISL_550139, EPI_ISL_550140, EPI_ISL_550141, EPI_ISL_550142, EPI_ISL_550143, EPI_ISL_550144 | Lighthouse Lab in Milton Keynes | Wellcome Sanger Institute for the COVID-19 Genomics UK (COG-UK) consortium | The Lighthouse Lab in Milton Keynes and Alex Alderton, Roberto Amato, Sonia Goncalves, Ewan Harrison, David K. Jackson, Ian Johnston, Dominic Kwiatkowski, Cordelia Langford, John Sillitoe on behalf of the Wellcome Sanger Institute COVID-19 Surveillance Team (http://www.sanger.ac.uk/covid-team) |
| EPI_ISL_550145, EPI_ISL_550146 | Lighthouse Lab in Milton Keynes | Wellcome Sanger Institute for the COVID-19 Genomics UK (COG-UK) consortium | The Lighthouse Lab in Alderley Park and Alex Alderton, Roberto Amato, Sonia Goncalves, Ewan Harrison, David K. Jackson, Ian Johnston, Dominic Kwiatkowski, Cordelia Langford, John Sillitoe on behalf of the Wellcome Sanger Institute COVID-19 Surveillance Team |
| EPI_ISL_550147, EPI_ISL_550148, EPI_ISL_550149, EPI_ISL_550150, EPI_ISL_550151, EPI_ISL_550152, EPI_ISL_550153, EPI_ISL_550154, EPI_ISL_550155, EPI_ISL_550156, EPI_ISL_550157, EPI_ISL_550158 | | | |
| see above | Lighthouse Lab in Milton Keynes | Wellcome Sanger Institute for the COVID-19 Genomics UK (COG-UK) consortium | The Lighthouse Lab in Milton Keynes and Alex Alderton, Roberto Amato, Sonia Goncalves, Ewan Harrison, David K. Jackson, Ian Johnston, Dominic Kwiatkowski, Cordelia Langford, John Sillitoe on behalf of the Wellcome Sanger Institute COVID-19 Surveillance Team (http://www.sanger.ac.uk/covid-team) |
| EPI_ISL_550159 | Lighthouse Lab in Milton Keynes | Wellcome Sanger Institute for the COVID-19 Genomics UK (COG-UK) consortium | The Lighthouse Lab in Alderley Park and Alex Alderton, Roberto Amato, Sonia Goncalves, Ewan Harrison, David K. Jackson, Ian Johnston, Dominic Kwiatkowski, Cordelia Langford, John Sillitoe on behalf of the Wellcome Sanger Institute COVID-19 Surveillance Team |
| EPI_ISL_550160, EPI_ISL_550161, EPI_ISL_550162 | Lighthouse Lab in Milton Keynes | Wellcome Sanger Institute for the COVID-19 Genomics UK (COG-UK) consortium | The Lighthouse Lab in Milton Keynes and Alex Alderton, Roberto Amato, Sonia Goncalves, Ewan Harrison, David K. Jackson, Ian Johnston, Dominic Kwiatkowski, Cordelia Langford, John Sillitoe on behalf of the Wellcome Sanger Institute COVID-19 Surveillance Team (http://www.sanger.ac.uk/covid-team) |
| EPI_ISL_550163 | Lighthouse Lab in Milton Keynes | Wellcome Sanger Institute for the COVID-19 Genomics UK (COG-UK) consortium | The Lighthouse Lab in Alderley Park and Alex Alderton, Roberto Amato, Sonia Goncalves, Ewan Harrison, David K. Jackson, Ian Johnston, Dominic Kwiatkowski, Cordelia Langford, John Sillitoe on behalf of the Wellcome Sanger Institute COVID-19 Surveillance Team |
| EPI_ISL_550164, EPI_ISL_550165, EPI_ISL_550166, EPI_ISL_550167, | Lighthouse Lab in Milton Keynes | Wellcome Sanger Institute for the COVID-19 Genomics UK (COG-UK) consortium | The Lighthouse Lab in Milton Keynes and Alex Alderton, Roberto Amato, Sonia Goncalves, Ewan Harrison, David K. Jackson, Ian Johnston, Dominic Kwiatkowski, Cordelia Langford, John Sillitoe on behalf of the Wellcome Sanger Institute COVID-19 Surveillance Team (http://www.sanger.ac.uk/covid-team) |

| | | | |
|---|---|---|---|
| EPI_ISL_550168, EPI_ISL_550170, EPI_ISL_550172, EPI_ISL_550173, EPI_ISL_550174 | | | |
| EPI_ISL_550175 | Lighthouse Lab in Milton Keynes | Wellcome Sanger Institute for the COVID-19 Genomics UK (COG-UK) consortium | The Lighthouse Lab in Alderley Park and Alex Alderton, Roberto Amato, Sonia Goncalves, Ewan Harrison, David K. Jackson, Ian Johnston, Dominic Kwiatkowski, Cordelia Langford, John Sillitoe on behalf of the Wellcome Sanger Institute COVID-19 Surveillance Team |
| EPI_ISL_550176, EPI_ISL_550177, EPI_ISL_550178, EPI_ISL_550179, EPI_ISL_550180, EPI_ISL_550181, EPI_ISL_550182, EPI_ISL_550183, EPI_ISL_550184 | Lighthouse Lab in Milton Keynes | Wellcome Sanger Institute for the COVID-19 Genomics UK (COG-UK) consortium | The Lighthouse Lab in Milton Keynes and Alex Alderton, Roberto Amato, Sonia Goncalves, Ewan Harrison, David K. Jackson, Ian Johnston, Dominic Kwiatkowski, Cordelia Langford, John Sillitoe on behalf of the Wellcome Sanger Institute COVID-19 Surveillance Team (http://www.sanger.ac.uk/covid-team) |
| EPI_ISL_550185, EPI_ISL_550186, EPI_ISL_550187 | Lighthouse Lab in Milton Keynes | Wellcome Sanger Institute for the COVID-19 Genomics UK (COG-UK) consortium | The Lighthouse Lab in Alderley Park and Alex Alderton, Roberto Amato, Sonia Goncalves, Ewan Harrison, David K. Jackson, Ian Johnston, Dominic Kwiatkowski, Cordelia Langford, John Sillitoe on behalf of the Wellcome Sanger Institute COVID-19 Surveillance Team |
| EPI_ISL_550188, EPI_ISL_550189, EPI_ISL_550190, EPI_ISL_550191, EPI_ISL_550192, EPI_ISL_550193, EPI_ISL_550194, EPI_ISL_550195, EPI_ISL_550196, EPI_ISL_550197, EPI_ISL_550198, EPI_ISL_550199, EPI_ISL_550201, EPI_ISL_550202, EPI_ISL_550203, EPI_ISL_550204, EPI_ISL_550205, EPI_ISL_550207, EPI_ISL_550208, EPI_ISL_550210, EPI_ISL_550211, EPI_ISL_550213, EPI_ISL_550214, EPI_ISL_550215, EPI_ISL_550216, EPI_ISL_550217, EPI_ISL_550218, EPI_ISL_550219, EPI_ISL_550220, EPI_ISL_550221, EPI_ISL_550222, EPI_ISL_550224, EPI_ISL_550225, EPI_ISL_550226, EPI_ISL_550227, EPI_ISL_550228, EPI_ISL_550229, EPI_ISL_550230, EPI_ISL_550231, EPI_ISL_550232, EPI_ISL_550233, EPI_ISL_550234, EPI_ISL_550235, EPI_ISL_550237, EPI_ISL_550238, EPI_ISL_550239, EPI_ISL_550240, EPI_ISL_550241, EPI_ISL_550242 | | | |
| see above | Lighthouse Lab in Milton Keynes | Wellcome Sanger Institute for the COVID-19 Genomics UK (COG-UK) consortium | The Lighthouse Lab in Milton Keynes and Alex Alderton, Roberto Amato, Sonia Goncalves, Ewan Harrison, David K. Jackson, Ian Johnston, Dominic Kwiatkowski, Cordelia Langford, John Sillitoe on behalf of the Wellcome Sanger Institute COVID-19 Surveillance Team (http://www.sanger.ac.uk/covid-team) |
| EPI_ISL_550244 | Lighthouse Lab in Milton Keynes | Wellcome Sanger Institute for the COVID-19 Genomics UK (COG-UK) consortium | The Lighthouse Lab in Alderley Park and Alex Alderton, Roberto Amato, Sonia Goncalves, Ewan Harrison, David K. Jackson, Ian Johnston, Dominic Kwiatkowski, Cordelia Langford, John Sillitoe on behalf of the Wellcome Sanger Institute COVID-19 Surveillance Team |
| EPI_ISL_550245, EPI_ISL_550246, EPI_ISL_550248, EPI_ISL_550249, EPI_ISL_550250 | Lighthouse Lab in Milton Keynes | Wellcome Sanger Institute for the COVID-19 Genomics UK (COG-UK) consortium | The Lighthouse Lab in Milton Keynes and Alex Alderton, Roberto Amato, Sonia Goncalves, Ewan Harrison, David K. Jackson, Ian Johnston, Dominic Kwiatkowski, Cordelia Langford, John Sillitoe on behalf of the Wellcome Sanger Institute COVID-19 Surveillance Team (http://www.sanger.ac.uk/covid-team) |
| EPI_ISL_550251 | Lighthouse Lab in Milton Keynes | Wellcome Sanger Institute for the COVID-19 Genomics UK (COG-UK) consortium | The Lighthouse Lab in Alderley Park and Alex Alderton, Roberto Amato, Sonia Goncalves, Ewan Harrison, David K. Jackson, Ian Johnston, Dominic Kwiatkowski, Cordelia Langford, John Sillitoe on behalf of the Wellcome Sanger Institute COVID-19 Surveillance Team |
| EPI_ISL_550252, EPI_ISL_550253, EPI_ISL_550255, EPI_ISL_550256 | Lighthouse Lab in Milton Keynes | Wellcome Sanger Institute for the COVID-19 Genomics UK (COG-UK) consortium | The Lighthouse Lab in Milton Keynes and Alex Alderton, Roberto Amato, Sonia Goncalves, Ewan Harrison, David K. Jackson, Ian Johnston, Dominic Kwiatkowski, Cordelia Langford, John Sillitoe on behalf of the Wellcome Sanger Institute COVID-19 Surveillance Team (http://www.sanger.ac.uk/covid-team) |
| EPI_ISL_550257 | Lighthouse Lab in Milton Keynes | Wellcome Sanger Institute for the COVID-19 Genomics UK (COG-UK) consortium | The Lighthouse Lab in Alderley Park and Alex Alderton, Roberto Amato, Sonia Goncalves, Ewan Harrison, David K. Jackson, Ian Johnston, Dominic Kwiatkowski, Cordelia Langford, John Sillitoe on behalf of the Wellcome Sanger Institute COVID-19 Surveillance Team |
| EPI_ISL_550258, EPI_ISL_550259, EPI_ISL_550260, EPI_ISL_550261, EPI_ISL_550796, EPI_ISL_550797, EPI_ISL_550798, EPI_ISL_550802, EPI_ISL_550803, EPI_ISL_550805, EPI_ISL_550806, EPI_ISL_550807, EPI_ISL_550808, EPI_ISL_550812, EPI_ISL_550814, EPI_ISL_550817, EPI_ISL_550818, EPI_ISL_550819, EPI_ISL_550822, EPI_ISL_550823, EPI_ISL_550824 | | | |
| see above | Lighthouse Lab in Milton Keynes | Wellcome Sanger Institute for the COVID-19 Genomics UK (COG-UK) consortium | The Lighthouse Lab in Milton Keynes and Alex Alderton, Roberto Amato, Sonia Goncalves, Ewan Harrison, David K. Jackson, Ian Johnston, Dominic Kwiatkowski, Cordelia Langford, John Sillitoe on behalf of the Wellcome Sanger Institute COVID-19 Surveillance Team (http://www.sanger.ac.uk/covid-team) |
| EPI_ISL_550825 | Lighthouse Lab in Milton Keynes | Wellcome Sanger Institute for the COVID-19 Genomics UK (COG-UK) consortium | The Lighthouse Lab in Alderley Park and Alex Alderton, Roberto Amato, Sonia Goncalves, Ewan Harrison, David K. Jackson, Ian Johnston, Dominic Kwiatkowski, Cordelia Langford, John Sillitoe on behalf of the Wellcome Sanger Institute COVID-19 Surveillance Team |
| EPI_ISL_550828, EPI_ISL_550832, EPI_ISL_550835, EPI_ISL_550836, EPI_ISL_550837, EPI_ISL_550839, EPI_ISL_550840, EPI_ISL_550842, EPI_ISL_550844, EPI_ISL_550845, EPI_ISL_550847, EPI_ISL_550848 | | | |
| see above | Lighthouse Lab in Milton Keynes | Wellcome Sanger Institute for the COVID-19 Genomics UK (COG-UK) consortium | The Lighthouse Lab in Milton Keynes and Alex Alderton, Roberto Amato, Sonia Goncalves, Ewan Harrison, David K. Jackson, Ian Johnston, Dominic Kwiatkowski, Cordelia Langford, John Sillitoe on behalf of the Wellcome Sanger Institute COVID-19 Surveillance Team (http://www.sanger.ac.uk/covid-team) |
| EPI_ISL_550850 | Lighthouse Lab in Milton Keynes | Wellcome Sanger Institute for the COVID-19 Genomics UK (COG-UK) consortium | The Lighthouse Lab in Alderley Park and Alex Alderton, Roberto Amato, Sonia Goncalves, Ewan Harrison, David K. Jackson, Ian Johnston, Dominic Kwiatkowski, Cordelia Langford, John Sillitoe on behalf of the Wellcome Sanger Institute COVID-19 Surveillance Team |
| EPI_ISL_550853, EPI_ISL_550855, EPI_ISL_550858, EPI_ISL_550859, EPI_ISL_550860, EPI_ISL_550861, EPI_ISL_550862, EPI_ISL_550865, EPI_ISL_550868, EPI_ISL_550869, EPI_ISL_550872, EPI_ISL_550873, EPI_ISL_550875, EPI_ISL_550876, EPI_ISL_550877, EPI_ISL_550878, EPI_ISL_550882, EPI_ISL_550884, EPI_ISL_550885, EPI_ISL_550886, EPI_ISL_550887, EPI_ISL_550888, EPI_ISL_550891, EPI_ISL_550892, EPI_ISL_550894, EPI_ISL_550898 | | | |
| see above | Lighthouse Lab in Milton Keynes | Wellcome Sanger Institute for the COVID-19 Genomics UK (COG-UK) consortium | The Lighthouse Lab in Milton Keynes and Alex Alderton, Roberto Amato, Sonia Goncalves, Ewan Harrison, David K. Jackson, Ian Johnston, Dominic Kwiatkowski, Cordelia Langford, John Sillitoe on behalf of the Wellcome Sanger Institute COVID-19 Surveillance Team (http://www.sanger.ac.uk/covid-team) |
| EPI_ISL_550901 | Lighthouse Lab in Milton Keynes | Wellcome Sanger Institute for the COVID-19 Genomics UK (COG-UK) consortium | The Lighthouse Lab in Alderley Park and Alex Alderton, Roberto Amato, Sonia Goncalves, Ewan Harrison, David K. Jackson, Ian Johnston, Dominic Kwiatkowski, Cordelia Langford, John Sillitoe on behalf of the Wellcome Sanger Institute COVID-19 Surveillance Team |
| EPI_ISL_550902, EPI_ISL_550904, EPI_ISL_550907, EPI_ISL_550911, EPI_ISL_550912, EPI_ISL_550915, EPI_ISL_550916, EPI_ISL_550917, EPI_ISL_550922, EPI_ISL_550927, EPI_ISL_550928, EPI_ISL_550929, EPI_ISL_550930, EPI_ISL_550931, EPI_ISL_550932, EPI_ISL_550933, EPI_ISL_550935, EPI_ISL_550937, EPI_ISL_550938, EPI_ISL_550940, EPI_ISL_550942, EPI_ISL_550946, EPI_ISL_550948, EPI_ISL_550949, EPI_ISL_550950, EPI_ISL_550953, EPI_ISL_550954, EPI_ISL_550955, EPI_ISL_550958, EPI_ISL_550961, EPI_ISL_550964, EPI_ISL_550966, EPI_ISL_550967, EPI_ISL_550969, EPI_ISL_550971, EPI_ISL_550975, EPI_ISL_550976, EPI_ISL_550977 | | | |
| see above | Lighthouse Lab in Milton Keynes | Wellcome Sanger Institute for the COVID-19 Genomics UK (COG-UK) consortium | The Lighthouse Lab in Milton Keynes and Alex Alderton, Roberto Amato, Sonia Goncalves, Ewan Harrison, David K. Jackson, Ian Johnston, Dominic Kwiatkowski, Cordelia Langford, John Sillitoe on behalf of the Wellcome Sanger Institute COVID-19 Surveillance Team (http://www.sanger.ac.uk/covid-team) |
| EPI_ISL_550978 | Lighthouse Lab in Milton Keynes | Wellcome Sanger Institute for the COVID-19 Genomics UK (COG-UK) consortium | The Lighthouse Lab in Alderley Park and Alex Alderton, Roberto Amato, Sonia Goncalves, Ewan Harrison, David K. Jackson, Ian Johnston, Dominic Kwiatkowski, Cordelia Langford, John Sillitoe on behalf of the Wellcome Sanger Institute COVID-19 Surveillance Team |
| EPI_ISL_550979 | Lighthouse Lab in Milton Keynes | Wellcome Sanger Institute for the COVID-19 Genomics UK (COG-UK) consortium | The Lighthouse Lab in Milton Keynes and Alex Alderton, Roberto Amato, Sonia Goncalves, Ewan Harrison, David K. Jackson, Ian Johnston, Dominic Kwiatkowski, Cordelia Langford, John Sillitoe on behalf of the Wellcome Sanger Institute COVID-19 Surveillance Team (http://www.sanger.ac.uk/covid-team) |
| EPI_ISL_550981 | Lighthouse Lab in Milton Keynes | Wellcome Sanger Institute for the COVID-19 Genomics UK (COG-UK) consortium | The Lighthouse Lab in Alderley Park and Alex Alderton, Roberto Amato, Sonia Goncalves, Ewan Harrison, David K. Jackson, Ian Johnston, Dominic Kwiatkowski, Cordelia Langford, John Sillitoe on behalf of the Wellcome Sanger Institute COVID-19 Surveillance Team |
| EPI_ISL_550983 | Lighthouse Lab in Milton Keynes | Wellcome Sanger Institute for the COVID-19 Genomics UK (COG-UK) Consortium | The Lighthouse Lab in Milton Keynes and Alex Alderton, Roberto Amato, Sonia Goncalves, Ewan Harrison, David K. Jackson, Ian Johnston, Dominic Kwiatkowski, Cordelia Langford, John Sillitoe on behalf of the Wellcome Sanger Institute COVID-19 Surveillance Team |
| EPI_ISL_550984, EPI_ISL_550985, EPI_ISL_550986, EPI_ISL_550988, EPI_ISL_550989, EPI_ISL_550993, EPI_ISL_550994, EPI_ISL_550995, EPI_ISL_550996, EPI_ISL_550997, EPI_ISL_550999 | | | |
| see above | Lighthouse Lab in Milton Keynes | Wellcome Sanger Institute for the COVID-19 Genomics UK (COG-UK) consortium | The Lighthouse Lab in Milton Keynes and Alex Alderton, Roberto Amato, Sonia Goncalves, Ewan Harrison, David K. Jackson, Ian Johnston, Dominic Kwiatkowski, Cordelia Langford, John Sillitoe on behalf of the Wellcome Sanger Institute COVID-19 Surveillance Team (http://www.sanger.ac.uk/covid-team) |
| EPI_ISL_551001 | Lighthouse Lab in Milton Keynes | Wellcome Sanger Institute for the COVID-19 Genomics UK (COG-UK) Consortium | The Lighthouse Lab in Milton Keynes and Alex Alderton, Roberto Amato, Sonia Goncalves, Ewan Harrison, David K. Jackson, Ian Johnston, Dominic Kwiatkowski, Cordelia Langford, John Sillitoe on behalf of the Wellcome Sanger Institute COVID-19 Surveillance Team |
| EPI_ISL_551002, EPI_ISL_551003 | Lighthouse Lab in Milton Keynes | Wellcome Sanger Institute for the COVID-19 Genomics UK (COG-UK) consortium | The Lighthouse Lab in Milton Keynes and Alex Alderton, Roberto Amato, Sonia Goncalves, Ewan Harrison, David K. Jackson, Ian Johnston, Dominic Kwiatkowski, Cordelia Langford, John Sillitoe on behalf of the Wellcome Sanger Institute COVID-19 Surveillance Team |

| | | | |
|---|---|---|---|
| | | | (http://www.sanger.ac.uk/covid-team) |
| EPI_ISL_551005 | Lighthouse Lab in Milton Keynes | Wellcome Sanger Institute for the COVID-19 Genomics UK (COG-UK) consortium | The Lighthouse Lab in Alderley Park and Alex Alderton, Roberto Amato, Sonia Goncalves, Ewan Harrison, David K. Jackson, Ian Johnston, Dominic Kwiatkowski, Cordelia Langford, John Sillitoe on behalf of the Wellcome Sanger Institute COVID-19 Surveillance Team |
| EPI_ISL_551006 | Lighthouse Lab in Milton Keynes | Wellcome Sanger Institute for the COVID-19 Genomics UK (COG-UK) consortium | The Lighthouse Lab in Milton Keynes and Alex Alderton, Roberto Amato, Sonia Goncalves, Ewan Harrison, David K. Jackson, Ian Johnston, Dominic Kwiatkowski, Cordelia Langford, John Sillitoe on behalf of the Wellcome Sanger Institute COVID-19 Surveillance Team (http://www.sanger.ac.uk/covid-team) |
| EPI_ISL_551007 | Lighthouse Lab in Milton Keynes | Wellcome Sanger Institute for the COVID-19 Genomics UK (COG-UK) Consortium | The Lighthouse Lab in Milton Keynes and Alex Alderton, Roberto Amato, Sonia Goncalves, Ewan Harrison, David K. Jackson, Ian Johnston, Dominic Kwiatkowski, Cordelia Langford, John Sillitoe on behalf of the Wellcome Sanger Institute COVID-19 Surveillance Team |
| EPI_ISL_551008, EPI_ISL_551014, EPI_ISL_551015, EPI_ISL_551016, EPI_ISL_551020 | Lighthouse Lab in Milton Keynes | Wellcome Sanger Institute for the COVID-19 Genomics UK (COG-UK) consortium | The Lighthouse Lab in Milton Keynes and Alex Alderton, Roberto Amato, Sonia Goncalves, Ewan Harrison, David K. Jackson, Ian Johnston, Dominic Kwiatkowski, Cordelia Langford, John Sillitoe on behalf of the Wellcome Sanger Institute COVID-19 Surveillance Team (http://www.sanger.ac.uk/covid-team) |
| EPI_ISL_551021 | Lighthouse Lab in Milton Keynes | Wellcome Sanger Institute for the COVID-19 Genomics UK (COG-UK) consortium | The Lighthouse Lab in Alderley Park and Alex Alderton, Roberto Amato, Sonia Goncalves, Ewan Harrison, David K. Jackson, Ian Johnston, Dominic Kwiatkowski, Cordelia Langford, John Sillitoe on behalf of the Wellcome Sanger Institute COVID-19 Surveillance Team |
| EPI_ISL_551022, EPI_ISL_551023, EPI_ISL_551025, EPI_ISL_551028, EPI_ISL_551029, EPI_ISL_551030, EPI_ISL_551034, EPI_ISL_551035, EPI_ISL_551036, EPI_ISL_551038, EPI_ISL_551041, EPI_ISL_551045, EPI_ISL_551046, EPI_ISL_551048, EPI_ISL_551051, EPI_ISL_551052 | | | |
| see above | Lighthouse Lab in Milton Keynes | Wellcome Sanger Institute for the COVID-19 Genomics UK (COG-UK) consortium | The Lighthouse Lab in Milton Keynes and Alex Alderton, Roberto Amato, Sonia Goncalves, Ewan Harrison, David K. Jackson, Ian Johnston, Dominic Kwiatkowski, Cordelia Langford, John Sillitoe on behalf of the Wellcome Sanger Institute COVID-19 Surveillance Team (http://www.sanger.ac.uk/covid-team) |
| EPI_ISL_551057, EPI_ISL_551058 | Lighthouse Lab in Milton Keynes | Wellcome Sanger Institute for the COVID-19 Genomics UK (COG-UK) consortium | The Lighthouse Lab in Alderley Park and Alex Alderton, Roberto Amato, Sonia Goncalves, Ewan Harrison, David K. Jackson, Ian Johnston, Dominic Kwiatkowski, Cordelia Langford, John Sillitoe on behalf of the Wellcome Sanger Institute COVID-19 Surveillance Team |
| EPI_ISL_551063, EPI_ISL_551064, EPI_ISL_551065, EPI_ISL_551067, EPI_ISL_551070, EPI_ISL_551074, EPI_ISL_551077, EPI_ISL_551078, EPI_ISL_551079, EPI_ISL_551081, EPI_ISL_551087, EPI_ISL_551088, EPI_ISL_551089 | | | |
| see above | Lighthouse Lab in Milton Keynes | Wellcome Sanger Institute for the COVID-19 Genomics UK (COG-UK) consortium | The Lighthouse Lab in Milton Keynes and Alex Alderton, Roberto Amato, Sonia Goncalves, Ewan Harrison, David K. Jackson, Ian Johnston, Dominic Kwiatkowski, Cordelia Langford, John Sillitoe on behalf of the Wellcome Sanger Institute COVID-19 Surveillance Team (http://www.sanger.ac.uk/covid-team) |
| EPI_ISL_551092 | Lighthouse Lab in Milton Keynes | Wellcome Sanger Institute for the COVID-19 Genomics UK (COG-UK) Consortium | The Lighthouse Lab in Milton Keynes and Alex Alderton, Roberto Amato, Sonia Goncalves, Ewan Harrison, David K. Jackson, Ian Johnston, Dominic Kwiatkowski, Cordelia Langford, John Sillitoe on behalf of the Wellcome Sanger Institute COVID-19 Surveillance Team |
| EPI_ISL_551094, EPI_ISL_551096, EPI_ISL_551100, EPI_ISL_551104, EPI_ISL_551108, EPI_ISL_551111 | Lighthouse Lab in Milton Keynes | Wellcome Sanger Institute for the COVID-19 Genomics UK (COG-UK) consortium | The Lighthouse Lab in Milton Keynes and Alex Alderton, Roberto Amato, Sonia Goncalves, Ewan Harrison, David K. Jackson, Ian Johnston, Dominic Kwiatkowski, Cordelia Langford, John Sillitoe on behalf of the Wellcome Sanger Institute COVID-19 Surveillance Team (http://www.sanger.ac.uk/covid-team) |
| EPI_ISL_551114 | Lighthouse Lab in Milton Keynes | Wellcome Sanger Institute for the COVID-19 Genomics UK (COG-UK) consortium | The Lighthouse Lab in Alderley Park and Alex Alderton, Roberto Amato, Sonia Goncalves, Ewan Harrison, David K. Jackson, Ian Johnston, Dominic Kwiatkowski, Cordelia Langford, John Sillitoe on behalf of the Wellcome Sanger Institute COVID-19 Surveillance Team |
| EPI_ISL_551117, EPI_ISL_551119, EPI_ISL_551121, EPI_ISL_551125, EPI_ISL_551127, EPI_ISL_551128, EPI_ISL_551132, EPI_ISL_551133, EPI_ISL_551134, EPI_ISL_551135, EPI_ISL_551137, EPI_ISL_551139, EPI_ISL_551140, EPI_ISL_551141, EPI_ISL_551144, EPI_ISL_551145, EPI_ISL_551146, EPI_ISL_551147, EPI_ISL_551148, EPI_ISL_551152, EPI_ISL_551153, EPI_ISL_551156, EPI_ISL_551157, EPI_ISL_551159, EPI_ISL_551160, EPI_ISL_551161, EPI_ISL_551163, EPI_ISL_551165 | | | |
| see above | Lighthouse Lab in Milton Keynes | Wellcome Sanger Institute for the COVID-19 Genomics UK (COG-UK) consortium | The Lighthouse Lab in Milton Keynes and Alex Alderton, Roberto Amato, Sonia Goncalves, Ewan Harrison, David K. Jackson, Ian Johnston, Dominic Kwiatkowski, Cordelia Langford, John Sillitoe on behalf of the Wellcome Sanger Institute COVID-19 Surveillance Team (http://www.sanger.ac.uk/covid-team) |
| EPI_ISL_551166 | Lighthouse Lab in Milton Keynes | Wellcome Sanger Institute for the COVID-19 Genomics UK (COG-UK) consortium | The Lighthouse Lab in Alderley Park and Alex Alderton, Roberto Amato, Sonia Goncalves, Ewan Harrison, David K. Jackson, Ian Johnston, Dominic Kwiatkowski, Cordelia Langford, John Sillitoe on behalf of the Wellcome Sanger Institute COVID-19 Surveillance Team |
| EPI_ISL_551168, EPI_ISL_551169, EPI_ISL_551170, EPI_ISL_551171, EPI_ISL_551172, EPI_ISL_551174, EPI_ISL_551178, EPI_ISL_551181, EPI_ISL_551182, EPI_ISL_551183, EPI_ISL_551184, EPI_ISL_551185, EPI_ISL_551187, EPI_ISL_551188, EPI_ISL_551189, EPI_ISL_551192, EPI_ISL_551193, EPI_ISL_551195, EPI_ISL_551196, EPI_ISL_551203, EPI_ISL_551204, EPI_ISL_551207, EPI_ISL_551209, EPI_ISL_551211, EPI_ISL_551212, EPI_ISL_551214, EPI_ISL_551219, EPI_ISL_551223, EPI_ISL_551224, EPI_ISL_551225, EPI_ISL_551226, EPI_ISL_551228, EPI_ISL_551231, EPI_ISL_551232, EPI_ISL_551234, EPI_ISL_551236, EPI_ISL_551238, EPI_ISL_551239, EPI_ISL_551240, EPI_ISL_551242, EPI_ISL_551243 | | | |
| see above | Lighthouse Lab in Milton Keynes | Wellcome Sanger Institute for the COVID-19 Genomics UK (COG-UK) consortium | The Lighthouse Lab in Milton Keynes and Alex Alderton, Roberto Amato, Sonia Goncalves, Ewan Harrison, David K. Jackson, Ian Johnston, Dominic Kwiatkowski, Cordelia Langford, John Sillitoe on behalf of the Wellcome Sanger Institute COVID-19 Surveillance Team (http://www.sanger.ac.uk/covid-team) |
| EPI_ISL_551244 | Lighthouse Lab in Milton Keynes | Wellcome Sanger Institute for the COVID-19 Genomics UK (COG-UK) consortium | The Lighthouse Lab in Alderley Park and Alex Alderton, Roberto Amato, Sonia Goncalves, Ewan Harrison, David K. Jackson, Ian Johnston, Dominic Kwiatkowski, Cordelia Langford, John Sillitoe on behalf of the Wellcome Sanger Institute COVID-19 Surveillance Team |
| EPI_ISL_551246, EPI_ISL_551247, EPI_ISL_551250, EPI_ISL_551258, EPI_ISL_551261, EPI_ISL_551262, EPI_ISL_551264, EPI_ISL_551265, EPI_ISL_551266, EPI_ISL_551268, EPI_ISL_551269, EPI_ISL_551270, EPI_ISL_551273, EPI_ISL_551274, EPI_ISL_551279, EPI_ISL_551280, EPI_ISL_551281, EPI_ISL_551283, EPI_ISL_551284 | | | |
| see above | Lighthouse Lab in Milton Keynes | Wellcome Sanger Institute for the COVID-19 Genomics UK (COG-UK) consortium | The Lighthouse Lab in Milton Keynes and Alex Alderton, Roberto Amato, Sonia Goncalves, Ewan Harrison, David K. Jackson, Ian Johnston, Dominic Kwiatkowski, Cordelia Langford, John Sillitoe on behalf of the Wellcome Sanger Institute COVID-19 Surveillance Team (http://www.sanger.ac.uk/covid-team) |
| EPI_ISL_552009 | Lighthouse Lab in Cambridge | Wellcome Sanger Institute for the COVID-19 Genomics UK (COG-UK) consortium | Rob Howes, The Lighthouse Lab in Cambridge and Alex Alderton, Roberto Amato, Sonia Goncalves, Ewan Harrison, David K. Jackson, Ian Johnston, Dominic Kwiatkowski, Cordelia Langford, John Sillitoe on behalf of the Wellcome Sanger Institute COVID-19 Surveillance Team |
| EPI_ISL_559799, EPI_ISL_559800, EPI_ISL_559801, EPI_ISL_559802, EPI_ISL_559803, EPI_ISL_559804, EPI_ISL_559805, EPI_ISL_559806, EPI_ISL_559807, EPI_ISL_559808, EPI_ISL_559809, EPI_ISL_559810, EPI_ISL_559811, EPI_ISL_559812, EPI_ISL_559813, EPI_ISL_559814, EPI_ISL_559815, EPI_ISL_559816, EPI_ISL_559817, EPI_ISL_559818, EPI_ISL_559819, EPI_ISL_559820, EPI_ISL_559821, EPI_ISL_559822, EPI_ISL_559823, EPI_ISL_559824, EPI_ISL_559825, EPI_ISL_559826, EPI_ISL_559827, EPI_ISL_559828, EPI_ISL_559829, EPI_ISL_559830, EPI_ISL_559831, EPI_ISL_559832, EPI_ISL_559833, EPI_ISL_559836, EPI_ISL_559838, EPI_ISL_559839, EPI_ISL_559842, EPI_ISL_559844, EPI_ISL_559846, EPI_ISL_559847, EPI_ISL_559849, EPI_ISL_559850, EPI_ISL_559852, EPI_ISL_559853, EPI_ISL_559854, EPI_ISL_559855, EPI_ISL_559856, EPI_ISL_559857, EPI_ISL_559858, EPI_ISL_559859, EPI_ISL_559860, EPI_ISL_559861, EPI_ISL_559862, EPI_ISL_559863, EPI_ISL_559864, EPI_ISL_559865, EPI_ISL_559866, EPI_ISL_559867, EPI_ISL_559868, EPI_ISL_559869, EPI_ISL_559870, EPI_ISL_559871, EPI_ISL_559872, EPI_ISL_559873, EPI_ISL_559874, EPI_ISL_559875, EPI_ISL_559876, EPI_ISL_559877, EPI_ISL_559878, EPI_ISL_559879, EPI_ISL_559880, EPI_ISL_559881, EPI_ISL_559882, EPI_ISL_559883, EPI_ISL_559884, EPI_ISL_559885, EPI_ISL_559886, EPI_ISL_559887, EPI_ISL_559888, EPI_ISL_559889, EPI_ISL_559890, EPI_ISL_559891, EPI_ISL_559892, EPI_ISL_559893, EPI_ISL_559894, EPI_ISL_559895, EPI_ISL_559896, EPI_ISL_559897 | | | |
| see above | Oxford Viromics, NDM, University of Oxford; Oxford University Hospitals; Basingstoke and North Hampshire Hospital | COVID-19 Genomics UK (COG-UK) Consortium | Tanya Golubchik, David Bonsall, George Macintyre, Amy Trebes, Mariateresa de Cesare, Catrin Moore, Alex Mobbs, Anita Justice, Robert Shaw, Monique Andersson, Timothy Peto, Emma Wise, Nathan Moore, Jessica Lynch, Nick Cortes, Matilde Mori, Stephen Kidd, David Buck, John Todd, Christophe Fraser |
| EPI_ISL_559898, EPI_ISL_559900, EPI_ISL_559901, EPI_ISL_559902, EPI_ISL_559904, EPI_ISL_559910, EPI_ISL_559912, EPI_ISL_559914, EPI_ISL_559920, EPI_ISL_559921, EPI_ISL_559922, EPI_ISL_559923, EPI_ISL_559927, EPI_ISL_559928, EPI_ISL_559930, EPI_ISL_559931, EPI_ISL_559932, EPI_ISL_559935, EPI_ISL_559937, EPI_ISL_559938, EPI_ISL_559939 | | | |
| see above | Virology Department, Sheffield Teaching Hospitals NHS Foundation Trust/Department of Infection, Immunity and Cardiovascular Disease, The Medical School, University of Sheffield | COVID-19 Genomics UK (COG-UK) Consortium | Thushan de Silva, Matthew Parker, Nikki Smith, Adri Angyal, Rebecca Brown, Luke Green, Rachel Tucker, Paul Parsons, Danielle Groves, Katie Johnson, Laura Carrilero, Alex Keeley, Dave Partridge, Matthew Wyles, Benjamin Lindsey, Mehmet Yavuz, Mohammad Raza, Cariad Evans |
| EPI_ISL_559947, EPI_ISL_559948, EPI_ISL_559949, EPI_ISL_559950, EPI_ISL_559951, EPI_ISL_559952, EPI_ISL_559953, EPI_ISL_559954, EPI_ISL_559955, EPI_ISL_559956, EPI_ISL_559957, EPI_ISL_559958, EPI_ISL_559959, EPI_ISL_559960, EPI_ISL_559961, EPI_ISL_559962, EPI_ISL_559963, EPI_ISL_559964, EPI_ISL_559965, EPI_ISL_559966, EPI_ISL_559967, EPI_ISL_559968, EPI_ISL_559969, EPI_ISL_559970 | | | |
| see above | Lighthouse Lab in Glasgow / MRC-University of Glasgow Centre for Virus Research | COVID-19 Genomics UK (COG-UK) Consortium | Ana da Silva Filipe, Natasha Johnson, Kathy Smollett, Daniel Mair, Stephen Carmichael, Lily Tong, Jenna Nichols, Elihu Aranday-Cortes, Kyriaki Nomikou; Sarah McDonald, Marc Niebel, Patawee Asamaphan; Harper VanSteenhouse, Yumi Kasai, David Gray, Carol Clugston, Anna Dominiczak; Alasdair MacLean, Rory Gunson; Richard Orton, Joseph Hughes, Sreenu Vattipally, David L Robertson; Sharif Shaaban, Matthew Holden; Kathy Li, Natasha Jesudason, Rajiv Shah, James Shepherd, Antonia Ho, Emma Thomson |
| EPI_ISL_560004, EPI_ISL_560005, EPI_ISL_560006, EPI_ISL_560007, EPI_ISL_560008, EPI_ISL_560009, EPI_ISL_560010, EPI_ISL_560011, EPI_ISL_560012, EPI_ISL_560013, EPI_ISL_560014, EPI_ISL_560015, EPI_ISL_560016, EPI_ISL_560017, EPI_ISL_560018, EPI_ISL_560019, EPI_ISL_560020, EPI_ISL_560021, EPI_ISL_560022, EPI_ISL_560023, EPI_ISL_560024, EPI_ISL_560025, EPI_ISL_560026, EPI_ISL_560027, EPI_ISL_560028, EPI_ISL_560029, EPI_ISL_560030, EPI_ISL_560031, EPI_ISL_560032, EPI_ISL_560033, EPI_ISL_560034, EPI_ISL_560035, EPI_ISL_560036, EPI_ISL_560037, EPI_ISL_560038, EPI_ISL_560039, | | | |

| | | | |
|---|---|---|---|
| EPI_ISL_560040, EPI_ISL_560041, EPI_ISL_560042, EPI_ISL_560043, EPI_ISL_560044, EPI_ISL_560045, EPI_ISL_560046, EPI_ISL_560047, EPI_ISL_560048, EPI_ISL_560049, EPI_ISL_560050, EPI_ISL_560051, EPI_ISL_560052, EPI_ISL_560053 | | | |
| see above | Oxford Viromics, NDM, University of Oxford; Oxford University Hospitals; Basingstoke and North Hampshire Hospital | COVID-19 Genomics UK (COG-UK) Consortium | Tanya Golubchik, David Bonsall, George Macintyre, Amy Trebes, Mariateresa de Cesare, Catrin Moore, Alex Mobbs, Anita Justice, Robert Shaw, Monique Andersson, Timothy Peto, Emma Wise, Nathan Moore, Jessica Lynch, Nick Cortes, Matilde Mori, Stephen Kidd, David Buck, John Todd, Christophe Fraser |
| EPI_ISL_560054, EPI_ISL_560055, EPI_ISL_560056, EPI_ISL_560057, EPI_ISL_560058, EPI_ISL_560059, EPI_ISL_560060, EPI_ISL_560061, EPI_ISL_560062, EPI_ISL_560063, EPI_ISL_560064, EPI_ISL_560065, EPI_ISL_560066, EPI_ISL_560067, EPI_ISL_560068 | | | |
| see above | Virology Department, Sheffield Teaching Hospitals NHS Foundation Trust/Department of Infection, Immunity and Cardiovascular Disease, The Medical School, University of Sheffield | COVID-19 Genomics UK (COG-UK) Consortium | Thushan de Silva, Matthew Parker, Nikki Smith, Adri Angyal, Rebecca Brown, Luke Green, Rachel Tucker, Paul Parsons, Danielle Groves, Katie Johnson, Laura Carrilero, Alex Keeley, Dave Partridge, Matthew Wyles, Benjamin Lindsey, Mehmet Yavuz, Mohammad Raza, Cariad Evans |
| EPI_ISL_560069, EPI_ISL_560070, EPI_ISL_560071, EPI_ISL_560072, EPI_ISL_560073, EPI_ISL_560074, EPI_ISL_560075, EPI_ISL_560076, EPI_ISL_560077, EPI_ISL_560078, EPI_ISL_560079, EPI_ISL_560080, EPI_ISL_560081, EPI_ISL_560082 | | | |
| see above | Oxford Viromics, NDM, University of Oxford; Oxford University Hospitals; Basingstoke and North Hampshire Hospital | COVID-19 Genomics UK (COG-UK) Consortium | Tanya Golubchik, David Bonsall, George Macintyre, Amy Trebes, Mariateresa de Cesare, Catrin Moore, Alex Mobbs, Anita Justice, Robert Shaw, Monique Andersson, Timothy Peto, Emma Wise, Nathan Moore, Jessica Lynch, Nick Cortes, Matilde Mori, Stephen Kidd, David Buck, John Todd, Christophe Fraser |
| EPI_ISL_560084, EPI_ISL_560085, EPI_ISL_560088, EPI_ISL_560091, EPI_ISL_560092, EPI_ISL_560093, EPI_ISL_560094, EPI_ISL_560096, EPI_ISL_560103, EPI_ISL_560104, EPI_ISL_560109, EPI_ISL_560110, EPI_ISL_560111, EPI_ISL_560112, EPI_ISL_560113, EPI_ISL_560114, EPI_ISL_560117, EPI_ISL_560122, EPI_ISL_560125, EPI_ISL_560126, EPI_ISL_560128, EPI_ISL_560129, EPI_ISL_560131, EPI_ISL_560137, EPI_ISL_560139, EPI_ISL_560140, EPI_ISL_560143, EPI_ISL_560147, EPI_ISL_560150, EPI_ISL_560154, EPI_ISL_560155, EPI_ISL_560156, EPI_ISL_560159, EPI_ISL_560163, EPI_ISL_560164, EPI_ISL_560176, EPI_ISL_560185, EPI_ISL_560190, EPI_ISL_560195, EPI_ISL_560197, EPI_ISL_560199, EPI_ISL_560201, EPI_ISL_560202, EPI_ISL_560205, EPI_ISL_560206, EPI_ISL_560207, EPI_ISL_560219, EPI_ISL_560278 | | | |
| see above | Wales Specialist Virology Centre Sequencing lab: Pathogen Genomics Unit | COVID-19 Genomics UK (COG-UK) Consortium | Catherine Moore, Johnathan Evans, Laura Gifford, Malorie Perry, Simon Cottrell, Angela Marchbank, Alec Birchley, Alexander Adams, Amy Gaskin, Bree Gatica-Wilcox, Jason Coombes, Joel Southgate, Lauren Gilbert, Lee Graham, Nicole Pacchiarini, Sara Kumziene-Summerhayes, Sarah Taylor, Sophie Jones, Sara Rey, Matthew Bull, Joanne Watkins, Sally Corden, Tom Connor |
| EPI_ISL_560416, EPI_ISL_560417, EPI_ISL_560418, EPI_ISL_560419, EPI_ISL_560420, EPI_ISL_560421, EPI_ISL_560422, EPI_ISL_560423, EPI_ISL_560424, EPI_ISL_560425, EPI_ISL_560427, EPI_ISL_560429, EPI_ISL_560430, EPI_ISL_560431, EPI_ISL_560432, EPI_ISL_560433, EPI_ISL_560434, EPI_ISL_560435, EPI_ISL_560436, EPI_ISL_560437, EPI_ISL_560438, EPI_ISL_560439, EPI_ISL_560440, EPI_ISL_560441, EPI_ISL_560442, EPI_ISL_560443, EPI_ISL_560444, EPI_ISL_560445, EPI_ISL_560455, EPI_ISL_560456, EPI_ISL_560457, EPI_ISL_560458, EPI_ISL_560459, EPI_ISL_560460, EPI_ISL_560461, EPI_ISL_560462, EPI_ISL_560463, EPI_ISL_560468, EPI_ISL_560470, EPI_ISL_560471, EPI_ISL_560472, EPI_ISL_560476, EPI_ISL_560477, EPI_ISL_560478, EPI_ISL_560479, EPI_ISL_560481, EPI_ISL_560483, EPI_ISL_560484, EPI_ISL_560485, EPI_ISL_560486, EPI_ISL_560487, EPI_ISL_560488, EPI_ISL_560489, EPI_ISL_560490, EPI_ISL_560491, EPI_ISL_560492, EPI_ISL_560493, EPI_ISL_560494, EPI_ISL_560495, EPI_ISL_560496, EPI_ISL_560497, EPI_ISL_560498, EPI_ISL_560499, EPI_ISL_560500, EPI_ISL_560501, EPI_ISL_560502, EPI_ISL_560503, EPI_ISL_560504, EPI_ISL_560505, EPI_ISL_560506, EPI_ISL_560507, EPI_ISL_560508, EPI_ISL_560509, EPI_ISL_560510, EPI_ISL_560511, EPI_ISL_560512, EPI_ISL_560513, EPI_ISL_560514, EPI_ISL_560515, EPI_ISL_560516, EPI_ISL_560517, EPI_ISL_560518, EPI_ISL_560519, EPI_ISL_560520, EPI_ISL_560521, EPI_ISL_560522, EPI_ISL_560524, EPI_ISL_560525, EPI_ISL_560527, EPI_ISL_560534, EPI_ISL_560535, EPI_ISL_560541, EPI_ISL_560542, EPI_ISL_560543, EPI_ISL_560544, EPI_ISL_560545, EPI_ISL_560546, EPI_ISL_560547, EPI_ISL_560548, EPI_ISL_560549, EPI_ISL_560550, EPI_ISL_560551, EPI_ISL_560552 | | | |
| see above | Viollier AG | Department of Biosystems Science and Engineering, ETH Zürich | Christian Beisel, Sarah Nadeau, Ivan Topolsky, Pedro Ferreira, Philipp Jablonski, Susana Posada-Céspedes, Tobias Schär, Ina Nissen, Natascha Santacroce, Elodie Burcklen, Christiane Beckmann, Maurice Redondo, Olivier Kobel, Christoph Noppen, Sophie Seidel, Noemie Santamaria de Souza, Niko Beerenwinkel, Tanja Stadler |
| EPI_ISL_561302 | MRCG at LSHTM Genomics lab | MRCG at LSHTM Genomics lab | Abdul Karim sesay, Abdoulie Kanteh, Jarra Manneh, Mariama Kujabi, Bakary Sanyang |
| EPI_ISL_561378, EPI_ISL_561381, EPI_ISL_561390, EPI_ISL_561411, EPI_ISL_561431, EPI_ISL_561448, EPI_ISL_561488, EPI_ISL_561546, EPI_ISL_561548, EPI_ISL_561549, EPI_ISL_561589, EPI_ISL_561604, EPI_ISL_561636, EPI_ISL_561650, EPI_ISL_561667, EPI_ISL_561693, EPI_ISL_561709, EPI_ISL_561713, EPI_ISL_561747 | | | |
| see above | Microbiological Diagnostic Unit - Public Health Laboratory (MDU-PHL) | MDU-PHL | Seemann, T., Schultz M. B., Sait, M., Sherry, N. |
| EPI_ISL_561750 | Victorian Infectious Diseases Reference Laboratory (VIDRL) | VIDRL and MDU-PHL | Caly, L., Seemann, T., Sait, M., Schultz, M. B., Druce J., Sherry, N. |
| EPI_ISL_561760, EPI_ISL_561768, EPI_ISL_561776, EPI_ISL_561810, EPI_ISL_561890, EPI_ISL_561915, EPI_ISL_561916, EPI_ISL_561942, EPI_ISL_561951, EPI_ISL_561983, EPI_ISL_562004, EPI_ISL_562005, EPI_ISL_562014 | | | |
| see above | Microbiological Diagnostic Unit - Public Health Laboratory (MDU-PHL) | MDU-PHL | Seemann, T., Schultz M. B., Sait, M., Sherry, N. |
| EPI_ISL_562021 | Victorian Infectious Diseases Reference Laboratory (VIDRL) | VIDRL and MDU-PHL | Caly, L., Seemann, T., Sait, M., Schultz, M. B., Druce J., Sherry, N. |
| EPI_ISL_562026, EPI_ISL_562059, EPI_ISL_562087, EPI_ISL_562127, EPI_ISL_562173, EPI_ISL_562177, EPI_ISL_562187, EPI_ISL_562204, EPI_ISL_562214, EPI_ISL_562227, EPI_ISL_562248, EPI_ISL_562251, EPI_ISL_562256, EPI_ISL_562257, EPI_ISL_562263 | | | |
| see above | Microbiological Diagnostic Unit - Public Health Laboratory (MDU-PHL) | MDU-PHL | Seemann, T., Schultz M. B., Sait, M., Sherry, N. |
| EPI_ISL_562274 | Victorian Infectious Diseases Reference Laboratory (VIDRL) | VIDRL and MDU-PHL | Caly, L., Seemann, T., Sait, M., Schultz, M. B., Druce J., Sherry, N. |
| EPI_ISL_562293, EPI_ISL_562312, EPI_ISL_562318 | Microbiological Diagnostic Unit - Public Health Laboratory (MDU-PHL) | MDU-PHL | Seemann, T., Schultz M. B., Sait, M., Sherry, N. |
| EPI_ISL_562326 | Victorian Infectious Diseases Reference Laboratory (VIDRL) | VIDRL and MDU-PHL | Caly, L., Seemann, T., Sait, M., Schultz, M. B., Druce J., Sherry, N. |
| EPI_ISL_562344, EPI_ISL_562349, EPI_ISL_562373, EPI_ISL_562383, EPI_ISL_562384, EPI_ISL_562395, EPI_ISL_562403, EPI_ISL_562432, EPI_ISL_562434 | Microbiological Diagnostic Unit - Public Health Laboratory (MDU-PHL) | MDU-PHL | Seemann, T., Schultz M. B., Sait, M., Sherry, N. |
| EPI_ISL_562435 | Victorian Infectious Diseases Reference Laboratory (VIDRL) | VIDRL and MDU-PHL | Caly, L., Seemann, T., Sait, M., Schultz, M. B., Druce J., Sherry, N. |
| EPI_ISL_562448, EPI_ISL_562456, EPI_ISL_562489, EPI_ISL_562495, EPI_ISL_562537, EPI_ISL_562543, EPI_ISL_562561, EPI_ISL_562591, EPI_ISL_562592 | Microbiological Diagnostic Unit - Public Health Laboratory (MDU-PHL) | MDU-PHL | Seemann, T., Schultz M. B., Sait, M., Sherry, N. |
| EPI_ISL_562593, EPI_ISL_562595 | Victorian Infectious Diseases Reference Laboratory (VIDRL) | VIDRL and MDU-PHL | Caly, L., Seemann, T., Sait, M., Schultz, M. B., Druce J., Sherry, N. |
| EPI_ISL_562598, EPI_ISL_562599, EPI_ISL_562603, EPI_ISL_562604, EPI_ISL_562605, EPI_ISL_562609, EPI_ISL_562616, EPI_ISL_562691, EPI_ISL_562692, EPI_ISL_562693, EPI_ISL_562694, EPI_ISL_562695 | | | |
| see above | Microbiological Diagnostic Unit - Public Health Laboratory (MDU-PHL) | MDU-PHL | Seemann, T., Schultz M. B., Sait, M., Sherry, N. |
| EPI_ISL_562700 | Victorian Infectious Diseases Reference Laboratory (VIDRL) | VIDRL and MDU-PHL | Caly, L., Seemann, T., Sait, M., Schultz, M. B., Druce J., Sherry, N. |
| EPI_ISL_562701, EPI_ISL_562712, EPI_ISL_562716, EPI_ISL_562717, EPI_ISL_562718, EPI_ISL_562719, EPI_ISL_562720, EPI_ISL_562721, EPI_ISL_562736, EPI_ISL_562849, EPI_ISL_562851, EPI_ISL_562858, EPI_ISL_562859, EPI_ISL_562861, EPI_ISL_562865, EPI_ISL_562872, EPI_ISL_562875 | | | |
| see above | Microbiological Diagnostic Unit - Public Health Laboratory (MDU-PHL) | MDU-PHL | Seemann, T., Schultz M. B., Sait, M., Sherry, N. |
| EPI_ISL_562910, EPI_ISL_562911, EPI_ISL_562912, EPI_ISL_562913 | Victorian Infectious Diseases Reference Laboratory (VIDRL) | VIDRL and MDU-PHL | Caly, L., Seemann, T., Sait, M., Schultz, M. B., Druce J., Sherry, N. |
| EPI_ISL_562914, EPI_ISL_562916, EPI_ISL_562918, EPI_ISL_562920, EPI_ISL_562932, EPI_ISL_562933, EPI_ISL_562934, EPI_ISL_562935, EPI_ISL_562938, EPI_ISL_562940, EPI_ISL_562941, EPI_ISL_562942, EPI_ISL_562943, EPI_ISL_562944, EPI_ISL_562952, EPI_ISL_562955, EPI_ISL_562957, EPI_ISL_562963 | | | |
| see above | Microbiological Diagnostic Unit - Public Health Laboratory (MDU-PHL) | MDU-PHL | Seemann, T., Schultz M. B., Sait, M., Sherry, N. |
| EPI_ISL_563000 | Victorian Infectious Diseases Reference Laboratory (VIDRL) | VIDRL and MDU-PHL | Caly, L., Seemann, T., Sait, M., Schultz, M. B., Druce J., Sherry, N. |
| EPI_ISL_563003, EPI_ISL_563004, EPI_ISL_563007, EPI_ISL_563008, EPI_ISL_563011, EPI_ISL_563013, EPI_ISL_563014, EPI_ISL_563015, EPI_ISL_563018, EPI_ISL_563019, EPI_ISL_563021, EPI_ISL_563022, EPI_ISL_563024, EPI_ISL_563027, EPI_ISL_563028, EPI_ISL_563030, EPI_ISL_563033, EPI_ISL_563034, EPI_ISL_563037, EPI_ISL_563038, EPI_ISL_563040, EPI_ISL_563042, EPI_ISL_563043, EPI_ISL_563044, EPI_ISL_563045, EPI_ISL_563046, EPI_ISL_563047, EPI_ISL_563048, EPI_ISL_563049, EPI_ISL_563050, EPI_ISL_563053, EPI_ISL_563054, EPI_ISL_563055, EPI_ISL_563059, EPI_ISL_563060, EPI_ISL_563061, | | | |

| | | | |
|---|---|---|---|
| EPI_ISL_563062, EPI_ISL_563063, EPI_ISL_563068, EPI_ISL_563071, EPI_ISL_563074, EPI_ISL_563075, EPI_ISL_563076, EPI_ISL_563077, EPI_ISL_563079, EPI_ISL_563080, EPI_ISL_563083, EPI_ISL_563084, EPI_ISL_563085, EPI_ISL_563086, EPI_ISL_563088, EPI_ISL_563090, EPI_ISL_563093, EPI_ISL_563095, EPI_ISL_563098, EPI_ISL_563099, EPI_ISL_563102, EPI_ISL_563111, EPI_ISL_563118, EPI_ISL_563119, EPI_ISL_563124, EPI_ISL_563125, EPI_ISL_563126, EPI_ISL_563127, EPI_ISL_563128, EPI_ISL_563129, EPI_ISL_563130 | | | |
| see above | Microbiological Diagnostic Unit - Public Health Laboratory (MDU-PHL) | MDU-PHL | Seemann, T., Schultz M. B., Sait, M., Sherry, N. |
| EPI_ISL_563131, EPI_ISL_563132, EPI_ISL_563144 | Victorian Infectious Diseases Reference Laboratory (VIDRL) | VIDRL and MDU-PHL | Caly, L., Seemann, T., Sait, M., Schultz, M. B., Druce J., Sherry, N. |
| EPI_ISL_563148, EPI_ISL_563150, EPI_ISL_563152, EPI_ISL_563153, EPI_ISL_563154, EPI_ISL_563155, EPI_ISL_563156, EPI_ISL_563157, EPI_ISL_563158, EPI_ISL_563159, EPI_ISL_563160, EPI_ISL_563162, EPI_ISL_563166, EPI_ISL_563168, EPI_ISL_563171, EPI_ISL_563174, EPI_ISL_563175, EPI_ISL_563176, EPI_ISL_563177, EPI_ISL_563179, EPI_ISL_563180, EPI_ISL_563182, EPI_ISL_563183, EPI_ISL_563195, EPI_ISL_563198, EPI_ISL_563200, EPI_ISL_563204, EPI_ISL_563208, EPI_ISL_563212, EPI_ISL_563214, EPI_ISL_563224, EPI_ISL_563228, EPI_ISL_563230, EPI_ISL_563233, EPI_ISL_563237, EPI_ISL_563258, EPI_ISL_563288, EPI_ISL_563296, EPI_ISL_563297, EPI_ISL_563400, EPI_ISL_563401, EPI_ISL_563405 | | | |
| see above | Microbiological Diagnostic Unit - Public Health Laboratory (MDU-PHL) | MDU-PHL | Seemann, T., Schultz M. B., Sait, M., Sherry, N. |
| EPI_ISL_563423, EPI_ISL_563497, EPI_ISL_563498 | Victorian Infectious Diseases Reference Laboratory (VIDRL) | VIDRL and MDU-PHL | Caly, L., Seemann, T., Sait, M., Schultz, M. B., Druce J., Sherry, N. |
| EPI_ISL_563508, EPI_ISL_563510, EPI_ISL_563550, EPI_ISL_563555, EPI_ISL_563567, EPI_ISL_563569, EPI_ISL_563578, EPI_ISL_563579, EPI_ISL_563588 | Microbiological Diagnostic Unit - Public Health Laboratory (MDU-PHL) | MDU-PHL | Seemann, T., Schultz M. B., Sait, M., Sherry, N. |
| EPI_ISL_563790 | Victorian Infectious Diseases Reference Laboratory (VIDRL) | VIDRL and MDU-PHL | Caly, L., Seemann, T., Sait, M., Schultz, M. B., Druce J., Sherry, N. |
| EPI_ISL_563810, EPI_ISL_563813, EPI_ISL_563827, EPI_ISL_563828, EPI_ISL_563881, EPI_ISL_563883 | Microbiological Diagnostic Unit - Public Health Laboratory (MDU-PHL) | MDU-PHL | Seemann, T., Schultz M. B., Sait, M., Sherry, N. |
| EPI_ISL_566074, EPI_ISL_566076 | Respiratory Virus Unit, Microbiology Services Colindale, Public Health England | Respiratory Virus Unit, Microbiology Services Colindale, Public Health England | PHE Covid Sequencing Team |
| EPI_ISL_566422 | Lighthouse Lab in Milton Keynes | Wellcome Sanger Institute for the COVID-19 Genomics UK (COG-UK) consortium | The Lighthouse Lab in Milton Keynes and Alex Alderton, Roberto Amato, Sonia Goncalves, Ewan Harrison, David K. Jackson, Ian Johnston, Dominic Kwiatkowski, Cordelia Langford, John Sillitoe on behalf of the Wellcome Sanger Institute COVID-19 Surveillance Team |
| EPI_ISL_566423 | Lighthouse Lab in Cambridge | Wellcome Sanger Institute for the COVID-19 Genomics UK (COG-UK) consortium | Rob Howes, The Lighthouse Lab in Cambridge and Alex Alderton, Roberto Amato, Sonia Goncalves, Ewan Harrison, David K. Jackson, Ian Johnston, Dominic Kwiatkowski, Cordelia Langford, John Sillitoe on behalf of the Wellcome Sanger Institute COVID-19 Surveillance Team |
| EPI_ISL_566427 | Lighthouse Lab in Milton Keynes | Wellcome Sanger Institute for the COVID-19 Genomics UK (COG-UK) consortium | The Lighthouse Lab in Milton Keynes and Alex Alderton, Roberto Amato, Sonia Goncalves, Ewan Harrison, David K. Jackson, Ian Johnston, Dominic Kwiatkowski, Cordelia Langford, John Sillitoe on behalf of the Wellcome Sanger Institute COVID-19 Surveillance Team |
| EPI_ISL_566428, EPI_ISL_566429, EPI_ISL_566444, EPI_ISL_566445 | Lighthouse Lab in Cambridge | Wellcome Sanger Institute for the COVID-19 Genomics UK (COG-UK) consortium | Rob Howes, The Lighthouse Lab in Cambridge and Alex Alderton, Roberto Amato, Sonia Goncalves, Ewan Harrison, David K. Jackson, Ian Johnston, Dominic Kwiatkowski, Cordelia Langford, John Sillitoe on behalf of the Wellcome Sanger Institute COVID-19 Surveillance Team |
| EPI_ISL_566447, EPI_ISL_566451 | Lighthouse Lab in Milton Keynes | Wellcome Sanger Institute for the COVID-19 Genomics UK (COG-UK) consortium | The Lighthouse Lab in Milton Keynes and Alex Alderton, Roberto Amato, Sonia Goncalves, Ewan Harrison, David K. Jackson, Ian Johnston, Dominic Kwiatkowski, Cordelia Langford, John Sillitoe on behalf of the Wellcome Sanger Institute COVID-19 Surveillance Team |
| EPI_ISL_566453 | Lighthouse Lab in Cambridge | Wellcome Sanger Institute for the COVID-19 Genomics UK (COG-UK) consortium | Rob Howes, The Lighthouse Lab in Cambridge and Alex Alderton, Roberto Amato, Sonia Goncalves, Ewan Harrison, David K. Jackson, Ian Johnston, Dominic Kwiatkowski, Cordelia Langford, John Sillitoe on behalf of the Wellcome Sanger Institute COVID-19 Surveillance Team |
| EPI_ISL_566454 | Lighthouse Lab in Milton Keynes | Wellcome Sanger Institute for the COVID-19 Genomics UK (COG-UK) consortium | The Lighthouse Lab in Milton Keynes and Alex Alderton, Roberto Amato, Sonia Goncalves, Ewan Harrison, David K. Jackson, Ian Johnston, Dominic Kwiatkowski, Cordelia Langford, John Sillitoe on behalf of the Wellcome Sanger Institute COVID-19 Surveillance Team |
| EPI_ISL_566455 | Lighthouse Lab in Cambridge | Wellcome Sanger Institute for the COVID-19 Genomics UK (COG-UK) Consortium | Rob Howes, The Lighthouse Lab in Cambridge and Alex Alderton, Roberto Amato, Sonia Goncalves, Ewan Harrison, David K. Jackson, Ian Johnston, Dominic Kwiatkowski, Cordelia Langford, John Sillitoe on behalf of the Wellcome Sanger Institute COVID-19 Surveillance Team |
| EPI_ISL_566456, EPI_ISL_566457 | Lighthouse Lab in Cambridge | Wellcome Sanger Institute for the COVID-19 Genomics UK (COG-UK) consortium | Rob Howes, The Lighthouse Lab in Cambridge and Alex Alderton, Roberto Amato, Sonia Goncalves, Ewan Harrison, David K. Jackson, Ian Johnston, Dominic Kwiatkowski, Cordelia Langford, John Sillitoe on behalf of the Wellcome Sanger Institute COVID-19 Surveillance Team |
| EPI_ISL_566460, EPI_ISL_566465 | Lighthouse Lab in Milton Keynes | Wellcome Sanger Institute for the COVID-19 Genomics UK (COG-UK) consortium | The Lighthouse Lab in Milton Keynes and Alex Alderton, Roberto Amato, Sonia Goncalves, Ewan Harrison, David K. Jackson, Ian Johnston, Dominic Kwiatkowski, Cordelia Langford, John Sillitoe on behalf of the Wellcome Sanger Institute COVID-19 Surveillance Team |
| EPI_ISL_566467 | Lighthouse Lab in Cambridge | Wellcome Sanger Institute for the COVID-19 Genomics UK (COG-UK) consortium | Rob Howes, The Lighthouse Lab in Cambridge and Alex Alderton, Roberto Amato, Sonia Goncalves, Ewan Harrison, David K. Jackson, Ian Johnston, Dominic Kwiatkowski, Cordelia Langford, John Sillitoe on behalf of the Wellcome Sanger Institute COVID-19 Surveillance Team |
| EPI_ISL_566469 | Lighthouse Lab in Milton Keynes | Wellcome Sanger Institute for the COVID-19 Genomics UK (COG-UK) consortium | The Lighthouse Lab in Milton Keynes and Alex Alderton, Roberto Amato, Sonia Goncalves, Ewan Harrison, David K. Jackson, Ian Johnston, Dominic Kwiatkowski, Cordelia Langford, John Sillitoe on behalf of the Wellcome Sanger Institute COVID-19 Surveillance Team |
| EPI_ISL_566473 | Lighthouse Lab in Cambridge | Wellcome Sanger Institute for the COVID-19 Genomics UK (COG-UK) consortium | Rob Howes, The Lighthouse Lab in Cambridge and Alex Alderton, Roberto Amato, Sonia Goncalves, Ewan Harrison, David K. Jackson, Ian Johnston, Dominic Kwiatkowski, Cordelia Langford, John Sillitoe on behalf of the Wellcome Sanger Institute COVID-19 Surveillance Team |
| EPI_ISL_566474 | Lighthouse Lab in Cambridge | Wellcome Sanger Institute for the COVID-19 Genomics UK (COG-UK) Consortium | Rob Howes, The Lighthouse Lab in Cambridge and Alex Alderton, Roberto Amato, Sonia Goncalves, Ewan Harrison, David K. Jackson, Ian Johnston, Dominic Kwiatkowski, Cordelia Langford, John Sillitoe on behalf of the Wellcome Sanger Institute COVID-19 Surveillance Team |
| EPI_ISL_566479, EPI_ISL_566485, EPI_ISL_566490 | Lighthouse Lab in Milton Keynes | Wellcome Sanger Institute for the COVID-19 Genomics UK (COG-UK) consortium | The Lighthouse Lab in Milton Keynes and Alex Alderton, Roberto Amato, Sonia Goncalves, Ewan Harrison, David K. Jackson, Ian Johnston, Dominic Kwiatkowski, Cordelia Langford, John Sillitoe on behalf of the Wellcome Sanger Institute COVID-19 Surveillance Team |
| EPI_ISL_566492, EPI_ISL_566493 | Lighthouse Lab in Cambridge | Wellcome Sanger Institute for the COVID-19 Genomics UK (COG-UK) consortium | Rob Howes, The Lighthouse Lab in Cambridge and Alex Alderton, Roberto Amato, Sonia Goncalves, Ewan Harrison, David K. Jackson, Ian Johnston, Dominic Kwiatkowski, Cordelia Langford, John Sillitoe on behalf of the Wellcome Sanger Institute COVID-19 Surveillance Team |
| EPI_ISL_566494 | Lighthouse Lab in Milton Keynes | Wellcome Sanger Institute for the COVID-19 Genomics UK (COG-UK) consortium | The Lighthouse Lab in Milton Keynes and Alex Alderton, Roberto Amato, Sonia Goncalves, Ewan Harrison, David K. Jackson, Ian Johnston, Dominic Kwiatkowski, Cordelia Langford, John Sillitoe on behalf of the Wellcome Sanger Institute COVID-19 Surveillance Team |
| EPI_ISL_566495, EPI_ISL_566500 | Lighthouse Lab in Cambridge | Wellcome Sanger Institute for the COVID-19 Genomics UK (COG-UK) consortium | Rob Howes, The Lighthouse Lab in Cambridge and Alex Alderton, Roberto Amato, Sonia Goncalves, Ewan Harrison, David K. Jackson, Ian Johnston, Dominic Kwiatkowski, Cordelia Langford, John Sillitoe on behalf of the Wellcome Sanger Institute COVID-19 Surveillance Team |
| EPI_ISL_566502, EPI_ISL_566504 | Lighthouse Lab in Milton Keynes | Wellcome Sanger Institute for the COVID-19 Genomics UK (COG-UK) consortium | The Lighthouse Lab in Milton Keynes and Alex Alderton, Roberto Amato, Sonia Goncalves, Ewan Harrison, David K. Jackson, Ian Johnston, Dominic Kwiatkowski, Cordelia Langford, John Sillitoe on behalf of the Wellcome Sanger Institute COVID-19 Surveillance Team |
| EPI_ISL_566508 | Lighthouse Lab in Cambridge | Wellcome Sanger Institute for the COVID-19 Genomics UK (COG-UK) consortium | Rob Howes, The Lighthouse Lab in Cambridge and Alex Alderton, Roberto Amato, Sonia Goncalves, Ewan Harrison, David K. Jackson, Ian Johnston, Dominic Kwiatkowski, Cordelia Langford, John Sillitoe on behalf of the Wellcome Sanger Institute COVID-19 Surveillance Team |
| EPI_ISL_566510, EPI_ISL_566515 | Lighthouse Lab in Milton Keynes | Wellcome Sanger Institute for the COVID-19 Genomics UK (COG-UK) consortium | The Lighthouse Lab in Milton Keynes and Alex Alderton, Roberto Amato, Sonia Goncalves, Ewan Harrison, David K. Jackson, Ian Johnston, Dominic Kwiatkowski, Cordelia Langford, John Sillitoe on behalf of the Wellcome Sanger Institute COVID-19 Surveillance Team |
| EPI_ISL_566518, EPI_ISL_566519, EPI_ISL_566524, EPI_ISL_566528 | Lighthouse Lab in Cambridge | Wellcome Sanger Institute for the COVID-19 Genomics UK (COG-UK) consortium | Rob Howes, The Lighthouse Lab in Cambridge and Alex Alderton, Roberto Amato, Sonia Goncalves, Ewan Harrison, David K. Jackson, Ian Johnston, Dominic Kwiatkowski, Cordelia Langford, John Sillitoe on behalf of the Wellcome Sanger Institute COVID-19 Surveillance Team |
| EPI_ISL_566530, EPI_ISL_566531 | Lighthouse Lab in Milton Keynes | Wellcome Sanger Institute for the COVID-19 Genomics UK (COG-UK) consortium | The Lighthouse Lab in Milton Keynes and Alex Alderton, Roberto Amato, Sonia Goncalves, Ewan Harrison, David K. Jackson, Ian Johnston, Dominic Kwiatkowski, Cordelia Langford, John Sillitoe on behalf of the Wellcome Sanger Institute COVID-19 Surveillance Team |
| EPI_ISL_566534, EPI_ISL_566537, | Lighthouse Lab in Cambridge | Wellcome Sanger Institute for the COVID-19 Genomics UK | Rob Howes, The Lighthouse Lab in Cambridge and Alex Alderton, Roberto Amato, Sonia Goncalves, Ewan Harrison, David K. Jackson, Ian Johnston, |

| | | | |
|---|---|---|---|
| EPI_ISL_566543, EPI_ISL_566544 | | (COG-UK) consortium | Dominic Kwiatkowski, Cordelia Langford, John Sillitoe on behalf of the Wellcome Sanger Institute COVID-19 Surveillance Team |
| EPI_ISL_566547, EPI_ISL_566548, EPI_ISL_566559, EPI_ISL_566560 | Lighthouse Lab in Milton Keynes | Wellcome Sanger Institute for the COVID-19 Genomics UK (COG-UK) consortium | The Lighthouse Lab in Milton Keynes and Alex Alderton, Roberto Amato, Sonia Goncalves, Ewan Harrison, David K. Jackson, Ian Johnston, Dominic Kwiatkowski, Cordelia Langford, John Sillitoe on behalf of the Wellcome Sanger Institute COVID-19 Surveillance Team |
| EPI_ISL_566567 | Lighthouse Lab in Cambridge | Wellcome Sanger Institute for the COVID-19 Genomics UK (COG-UK) consortium | Rob Howes, The Lighthouse Lab in Cambridge and Alex Alderton, Roberto Amato, Sonia Goncalves, Ewan Harrison, David K. Jackson, Ian Johnston, Dominic Kwiatkowski, Cordelia Langford, John Sillitoe on behalf of the Wellcome Sanger Institute COVID-19 Surveillance Team |
| EPI_ISL_566573, EPI_ISL_566575, EPI_ISL_566579, EPI_ISL_566580, EPI_ISL_566582, EPI_ISL_566583, EPI_ISL_566585 | Lighthouse Lab in Milton Keynes | Wellcome Sanger Institute for the COVID-19 Genomics UK (COG-UK) consortium | The Lighthouse Lab in Milton Keynes and Alex Alderton, Roberto Amato, Sonia Goncalves, Ewan Harrison, David K. Jackson, Ian Johnston, Dominic Kwiatkowski, Cordelia Langford, John Sillitoe on behalf of the Wellcome Sanger Institute COVID-19 Surveillance Team |
| EPI_ISL_566586 | Lighthouse Lab in Cambridge | Wellcome Sanger Institute for the COVID-19 Genomics UK (COG-UK) consortium | Rob Howes, The Lighthouse Lab in Cambridge and Alex Alderton, Roberto Amato, Sonia Goncalves, Ewan Harrison, David K. Jackson, Ian Johnston, Dominic Kwiatkowski, Cordelia Langford, John Sillitoe on behalf of the Wellcome Sanger Institute COVID-19 Surveillance Team |
| EPI_ISL_566587 | Lighthouse Lab in Milton Keynes | Wellcome Sanger Institute for the COVID-19 Genomics UK (COG-UK) consortium | The Lighthouse Lab in Milton Keynes and Alex Alderton, Roberto Amato, Sonia Goncalves, Ewan Harrison, David K. Jackson, Ian Johnston, Dominic Kwiatkowski, Cordelia Langford, John Sillitoe on behalf of the Wellcome Sanger Institute COVID-19 Surveillance Team |
| EPI_ISL_566589 | Lighthouse Lab in Cambridge | Wellcome Sanger Institute for the COVID-19 Genomics UK (COG-UK) consortium | Rob Howes, The Lighthouse Lab in Cambridge and Alex Alderton, Roberto Amato, Sonia Goncalves, Ewan Harrison, David K. Jackson, Ian Johnston, Dominic Kwiatkowski, Cordelia Langford, John Sillitoe on behalf of the Wellcome Sanger Institute COVID-19 Surveillance Team |
| EPI_ISL_566591, EPI_ISL_566603, EPI_ISL_566605, EPI_ISL_566608, EPI_ISL_566609 | Lighthouse Lab in Milton Keynes | Wellcome Sanger Institute for the COVID-19 Genomics UK (COG-UK) consortium | The Lighthouse Lab in Milton Keynes and Alex Alderton, Roberto Amato, Sonia Goncalves, Ewan Harrison, David K. Jackson, Ian Johnston, Dominic Kwiatkowski, Cordelia Langford, John Sillitoe on behalf of the Wellcome Sanger Institute COVID-19 Surveillance Team |
| EPI_ISL_566610, EPI_ISL_566611 | Lighthouse Lab in Cambridge | Wellcome Sanger Institute for the COVID-19 Genomics UK (COG-UK) consortium | Rob Howes, The Lighthouse Lab in Cambridge and Alex Alderton, Roberto Amato, Sonia Goncalves, Ewan Harrison, David K. Jackson, Ian Johnston, Dominic Kwiatkowski, Cordelia Langford, John Sillitoe on behalf of the Wellcome Sanger Institute COVID-19 Surveillance Team |
| EPI_ISL_566612, EPI_ISL_566622 | Lighthouse Lab in Milton Keynes | Wellcome Sanger Institute for the COVID-19 Genomics UK (COG-UK) consortium | The Lighthouse Lab in Milton Keynes and Alex Alderton, Roberto Amato, Sonia Goncalves, Ewan Harrison, David K. Jackson, Ian Johnston, Dominic Kwiatkowski, Cordelia Langford, John Sillitoe on behalf of the Wellcome Sanger Institute COVID-19 Surveillance Team |
| EPI_ISL_566625 | Lighthouse Lab in Cambridge | Wellcome Sanger Institute for the COVID-19 Genomics UK (COG-UK) consortium | Rob Howes, The Lighthouse Lab in Cambridge and Alex Alderton, Roberto Amato, Sonia Goncalves, Ewan Harrison, David K. Jackson, Ian Johnston, Dominic Kwiatkowski, Cordelia Langford, John Sillitoe on behalf of the Wellcome Sanger Institute COVID-19 Surveillance Team |
| EPI_ISL_566626, EPI_ISL_566628 | Lighthouse Lab in Milton Keynes | Wellcome Sanger Institute for the COVID-19 Genomics UK (COG-UK) consortium | The Lighthouse Lab in Milton Keynes and Alex Alderton, Roberto Amato, Sonia Goncalves, Ewan Harrison, David K. Jackson, Ian Johnston, Dominic Kwiatkowski, Cordelia Langford, John Sillitoe on behalf of the Wellcome Sanger Institute COVID-19 Surveillance Team |
| EPI_ISL_566631 | Lighthouse Lab in Cambridge | Wellcome Sanger Institute for the COVID-19 Genomics UK (COG-UK) consortium | Rob Howes, The Lighthouse Lab in Cambridge and Alex Alderton, Roberto Amato, Sonia Goncalves, Ewan Harrison, David K. Jackson, Ian Johnston, Dominic Kwiatkowski, Cordelia Langford, John Sillitoe on behalf of the Wellcome Sanger Institute COVID-19 Surveillance Team |
| EPI_ISL_566637 | Lighthouse Lab in Milton Keynes | Wellcome Sanger Institute for the COVID-19 Genomics UK (COG-UK) consortium | The Lighthouse Lab in Milton Keynes and Alex Alderton, Roberto Amato, Sonia Goncalves, Ewan Harrison, David K. Jackson, Ian Johnston, Dominic Kwiatkowski, Cordelia Langford, John Sillitoe on behalf of the Wellcome Sanger Institute COVID-19 Surveillance Team |
| EPI_ISL_566638 | Lighthouse Lab in Cambridge | Wellcome Sanger Institute for the COVID-19 Genomics UK (COG-UK) consortium | Rob Howes, The Lighthouse Lab in Cambridge and Alex Alderton, Roberto Amato, Sonia Goncalves, Ewan Harrison, David K. Jackson, Ian Johnston, Dominic Kwiatkowski, Cordelia Langford, John Sillitoe on behalf of the Wellcome Sanger Institute COVID-19 Surveillance Team |
| EPI_ISL_566642, EPI_ISL_566643, EPI_ISL_566645, EPI_ISL_566657, EPI_ISL_566658 | Lighthouse Lab in Milton Keynes | Wellcome Sanger Institute for the COVID-19 Genomics UK (COG-UK) consortium | The Lighthouse Lab in Milton Keynes and Alex Alderton, Roberto Amato, Sonia Goncalves, Ewan Harrison, David K. Jackson, Ian Johnston, Dominic Kwiatkowski, Cordelia Langford, John Sillitoe on behalf of the Wellcome Sanger Institute COVID-19 Surveillance Team |
| EPI_ISL_566659, EPI_ISL_566661, EPI_ISL_566662 | Lighthouse Lab in Cambridge | Wellcome Sanger Institute for the COVID-19 Genomics UK (COG-UK) consortium | Rob Howes, The Lighthouse Lab in Cambridge and Alex Alderton, Roberto Amato, Sonia Goncalves, Ewan Harrison, David K. Jackson, Ian Johnston, Dominic Kwiatkowski, Cordelia Langford, John Sillitoe on behalf of the Wellcome Sanger Institute COVID-19 Surveillance Team |
| EPI_ISL_566665 | Lighthouse Lab in Milton Keynes | Wellcome Sanger Institute for the COVID-19 Genomics UK (COG-UK) consortium | The Lighthouse Lab in Milton Keynes and Alex Alderton, Roberto Amato, Sonia Goncalves, Ewan Harrison, David K. Jackson, Ian Johnston, Dominic Kwiatkowski, Cordelia Langford, John Sillitoe on behalf of the Wellcome Sanger Institute COVID-19 Surveillance Team |
| EPI_ISL_566666 | Lighthouse Lab in Cambridge | Wellcome Sanger Institute for the COVID-19 Genomics UK (COG-UK) consortium | Rob Howes, The Lighthouse Lab in Cambridge and Alex Alderton, Roberto Amato, Sonia Goncalves, Ewan Harrison, David K. Jackson, Ian Johnston, Dominic Kwiatkowski, Cordelia Langford, John Sillitoe on behalf of the Wellcome Sanger Institute COVID-19 Surveillance Team |
| EPI_ISL_566667 | Lighthouse Lab in Milton Keynes | Wellcome Sanger Institute for the COVID-19 Genomics UK (COG-UK) consortium | The Lighthouse Lab in Milton Keynes and Alex Alderton, Roberto Amato, Sonia Goncalves, Ewan Harrison, David K. Jackson, Ian Johnston, Dominic Kwiatkowski, Cordelia Langford, John Sillitoe on behalf of the Wellcome Sanger Institute COVID-19 Surveillance Team |
| EPI_ISL_566672 | Lighthouse Lab in Cambridge | Wellcome Sanger Institute for the COVID-19 Genomics UK (COG-UK) consortium | Rob Howes, The Lighthouse Lab in Cambridge and Alex Alderton, Roberto Amato, Sonia Goncalves, Ewan Harrison, David K. Jackson, Ian Johnston, Dominic Kwiatkowski, Cordelia Langford, John Sillitoe on behalf of the Wellcome Sanger Institute COVID-19 Surveillance Team |
| EPI_ISL_566673 | Lighthouse Lab in Milton Keynes | Wellcome Sanger Institute for the COVID-19 Genomics UK (COG-UK) consortium | The Lighthouse Lab in Milton Keynes and Alex Alderton, Roberto Amato, Sonia Goncalves, Ewan Harrison, David K. Jackson, Ian Johnston, Dominic Kwiatkowski, Cordelia Langford, John Sillitoe on behalf of the Wellcome Sanger Institute COVID-19 Surveillance Team |
| EPI_ISL_566674 | Lighthouse Lab in Cambridge | Wellcome Sanger Institute for the COVID-19 Genomics UK (COG-UK) consortium | Rob Howes, The Lighthouse Lab in Cambridge and Alex Alderton, Roberto Amato, Sonia Goncalves, Ewan Harrison, David K. Jackson, Ian Johnston, Dominic Kwiatkowski, Cordelia Langford, John Sillitoe on behalf of the Wellcome Sanger Institute COVID-19 Surveillance Team |
| EPI_ISL_566675, EPI_ISL_566676, EPI_ISL_566677, EPI_ISL_566680, EPI_ISL_566681, EPI_ISL_566682, EPI_ISL_566683, EPI_ISL_566685, EPI_ISL_566687, EPI_ISL_566688, EPI_ISL_566690, EPI_ISL_566692, EPI_ISL_566693, EPI_ISL_566694, EPI_ISL_566697, EPI_ISL_566698, EPI_ISL_566699, EPI_ISL_566700, EPI_ISL_566701, EPI_ISL_566703, EPI_ISL_566705, EPI_ISL_566707, EPI_ISL_566708, EPI_ISL_566710, EPI_ISL_566711, EPI_ISL_566713, EPI_ISL_566721, EPI_ISL_566725, EPI_ISL_566726, EPI_ISL_566727, EPI_ISL_566728, EPI_ISL_566729, EPI_ISL_566730, EPI_ISL_566731, EPI_ISL_566732, EPI_ISL_566734, EPI_ISL_566735, EPI_ISL_566740, EPI_ISL_566741, EPI_ISL_566742, EPI_ISL_566744, EPI_ISL_566746, EPI_ISL_566748, EPI_ISL_566749, EPI_ISL_566751, EPI_ISL_566752, EPI_ISL_566755, EPI_ISL_566757, EPI_ISL_566759, EPI_ISL_566763, EPI_ISL_566764, EPI_ISL_566765, EPI_ISL_566768, EPI_ISL_566771, EPI_ISL_566772, EPI_ISL_566773, EPI_ISL_566774, EPI_ISL_566775, EPI_ISL_566776, EPI_ISL_566777, EPI_ISL_566778, EPI_ISL_566780, EPI_ISL_566781, EPI_ISL_566782, EPI_ISL_566784, EPI_ISL_566788, EPI_ISL_566791, EPI_ISL_566793, EPI_ISL_566794, EPI_ISL_566795, EPI_ISL_566796, EPI_ISL_566797, EPI_ISL_566798, EPI_ISL_566799, EPI_ISL_566800, EPI_ISL_566801, EPI_ISL_566803, EPI_ISL_566805, EPI_ISL_566807, EPI_ISL_566809, EPI_ISL_566811, EPI_ISL_566813, EPI_ISL_566814, EPI_ISL_566818, EPI_ISL_566820, EPI_ISL_566825, EPI_ISL_566827, EPI_ISL_566828, EPI_ISL_566829, EPI_ISL_566830, EPI_ISL_566831, EPI_ISL_566835, EPI_ISL_566836, EPI_ISL_566837, EPI_ISL_566838, EPI_ISL_566839, EPI_ISL_566843, EPI_ISL_566846, EPI_ISL_566847, EPI_ISL_566850, EPI_ISL_566852, EPI_ISL_566853, EPI_ISL_566854, EPI_ISL_566855, EPI_ISL_566856, EPI_ISL_566857, EPI_ISL_566864, EPI_ISL_566865, EPI_ISL_566866, EPI_ISL_566869, EPI_ISL_566870, EPI_ISL_566871, EPI_ISL_566874, EPI_ISL_566875, EPI_ISL_566877, EPI_ISL_566878, EPI_ISL_566879, EPI_ISL_566880, EPI_ISL_566882, EPI_ISL_566883, EPI_ISL_566884, EPI_ISL_566885, EPI_ISL_566887, EPI_ISL_566889, EPI_ISL_566890, EPI_ISL_566891, EPI_ISL_566892, EPI_ISL_566893, EPI_ISL_566894, EPI_ISL_566896, EPI_ISL_566900, EPI_ISL_566901, EPI_ISL_566902, EPI_ISL_566906, EPI_ISL_566907, EPI_ISL_566908, EPI_ISL_566909, EPI_ISL_566912, EPI_ISL_566913, EPI_ISL_566917, EPI_ISL_566918, EPI_ISL_566919, EPI_ISL_566920, EPI_ISL_566921, EPI_ISL_566923, EPI_ISL_566924, EPI_ISL_566926, EPI_ISL_566928, EPI_ISL_566929, EPI_ISL_566930, EPI_ISL_566931, EPI_ISL_566934, EPI_ISL_566935, EPI_ISL_566936, EPI_ISL_566938, EPI_ISL_566939, EPI_ISL_566941, EPI_ISL_566942, EPI_ISL_566943, EPI_ISL_566944, EPI_ISL_566946, EPI_ISL_566948, EPI_ISL_566949 | | | |
| see above | Lighthouse Lab in Milton Keynes | Wellcome Sanger Institute for the COVID-19 Genomics UK (COG-UK) consortium | The Lighthouse Lab in Milton Keynes and Alex Alderton, Roberto Amato, Sonia Goncalves, Ewan Harrison, David K. Jackson, Ian Johnston, Dominic Kwiatkowski, Cordelia Langford, John Sillitoe on behalf of the Wellcome Sanger Institute COVID-19 Surveillance Team |
| EPI_ISL_566951 | Lighthouse Lab in Milton Keynes | Wellcome Sanger Institute for the COVID-19 Genomics UK (COG-UK) Consortium | The Lighthouse Lab in Milton Keynes and Alex Alderton, Roberto Amato, Sonia Goncalves, Ewan Harrison, David K. Jackson, Ian Johnston, Dominic Kwiatkowski, Cordelia Langford, John Sillitoe on behalf of the Wellcome Sanger Institute COVID-19 Surveillance Team |
| EPI_ISL_566952, EPI_ISL_566954, EPI_ISL_566955, EPI_ISL_566956, EPI_ISL_566958, EPI_ISL_566960, EPI_ISL_566961, EPI_ISL_566962, EPI_ISL_566963, EPI_ISL_566966, EPI_ISL_566967, EPI_ISL_566968, EPI_ISL_566970, EPI_ISL_566973, EPI_ISL_566974, EPI_ISL_566975, EPI_ISL_566976, EPI_ISL_566978, EPI_ISL_566979, EPI_ISL_566980, EPI_ISL_566981, EPI_ISL_566982, EPI_ISL_566983, EPI_ISL_566984, EPI_ISL_566985 | | | |
| see above | Lighthouse Lab in Milton Keynes | Wellcome Sanger Institute for the COVID-19 Genomics UK (COG-UK) consortium | The Lighthouse Lab in Milton Keynes and Alex Alderton, Roberto Amato, Sonia Goncalves, Ewan Harrison, David K. Jackson, Ian Johnston, Dominic Kwiatkowski, Cordelia Langford, John Sillitoe on behalf of the Wellcome Sanger Institute COVID-19 Surveillance Team |
| EPI_ISL_566986 | Lighthouse Lab in Milton Keynes | Wellcome Sanger Institute for the COVID-19 Genomics UK (COG-UK) Consortium | The Lighthouse Lab in Milton Keynes and Alex Alderton, Roberto Amato, Sonia Goncalves, Ewan Harrison, David K. Jackson, Ian Johnston, Dominic Kwiatkowski, Cordelia Langford, John Sillitoe on behalf of the Wellcome Sanger Institute COVID-19 Surveillance Team |
| EPI_ISL_566987, EPI_ISL_566988, EPI_ISL_566989, EPI_ISL_566990, EPI_ISL_566991, EPI_ISL_566993, EPI_ISL_566995, EPI_ISL_566997 | Lighthouse Lab in Milton Keynes | Wellcome Sanger Institute for the COVID-19 Genomics UK (COG-UK) consortium | The Lighthouse Lab in Milton Keynes and Alex Alderton, Roberto Amato, Sonia Goncalves, Ewan Harrison, David K. Jackson, Ian Johnston, Dominic Kwiatkowski, Cordelia Langford, John Sillitoe on behalf of the Wellcome Sanger Institute COVID-19 Surveillance Team |

| | | | |
|---|---|---|---|
| EPI_ISL_566998, EPI_ISL_567065, EPI_ISL_567080, EPI_ISL_567115, EPI_ISL_567150, EPI_ISL_567165, EPI_ISL_567243, EPI_ISL_567248, EPI_ISL_567250, EPI_ISL_567254, EPI_ISL_567260, EPI_ISL_567274, EPI_ISL_567289, EPI_ISL_567294, EPI_ISL_567338, EPI_ISL_567355, EPI_ISL_567358, EPI_ISL_567363, EPI_ISL_567375, EPI_ISL_567382, EPI_ISL_567390, EPI_ISL_567396, EPI_ISL_567398, EPI_ISL_567427, EPI_ISL_567429, EPI_ISL_567435, EPI_ISL_567446, EPI_ISL_567453, EPI_ISL_567464, EPI_ISL_567473, EPI_ISL_567482, EPI_ISL_567489, EPI_ISL_567501, EPI_ISL_567522, EPI_ISL_567524 | | | |
| see above | Lighthouse Lab in Glasgow | Wellcome Sanger Institute for the COVID-19 Genomics UK (COG-UK) consortium | Harper VanSteenhouse, Yumi Kasai, David Gray, Carol Clugston, Anna Dominiczak and Alex Alderton, Roberto Amato, Sonia Goncalves, Ewan Harrison, David K. Jackson, Ian Johnston, Dominic Kwiatkowski, Cordelia Langford, John Sillitoe on behalf of the Wellcome Sanger Institute COVID-19 Surveillance Team |
| EPI_ISL_567563, EPI_ISL_567886, EPI_ISL_567887, EPI_ISL_567888, EPI_ISL_567889, EPI_ISL_567890, EPI_ISL_567891, EPI_ISL_567892, EPI_ISL_567894, EPI_ISL_567895, EPI_ISL_567896, EPI_ISL_567897, EPI_ISL_567901, EPI_ISL_567902, EPI_ISL_567903, EPI_ISL_567906, EPI_ISL_567907, EPI_ISL_567909, EPI_ISL_567910, EPI_ISL_567911, EPI_ISL_567913, EPI_ISL_567915, EPI_ISL_567917, EPI_ISL_567918, EPI_ISL_567919, EPI_ISL_567920, EPI_ISL_567921, EPI_ISL_567924, EPI_ISL_567927, EPI_ISL_567928, EPI_ISL_567931, EPI_ISL_567932, EPI_ISL_567933, EPI_ISL_567936, EPI_ISL_567937, EPI_ISL_567938, EPI_ISL_567939, EPI_ISL_567940, EPI_ISL_567942, EPI_ISL_567943, EPI_ISL_567944, EPI_ISL_567945, EPI_ISL_567946, EPI_ISL_567949, EPI_ISL_567950, EPI_ISL_567951, EPI_ISL_567953, EPI_ISL_567957, EPI_ISL_567960, EPI_ISL_567961, EPI_ISL_567963, EPI_ISL_567964, EPI_ISL_567965, EPI_ISL_567967, EPI_ISL_567968, EPI_ISL_567969, EPI_ISL_567970, EPI_ISL_567974, EPI_ISL_567975, EPI_ISL_567977, EPI_ISL_567981, EPI_ISL_567982, EPI_ISL_567986, EPI_ISL_567987, EPI_ISL_567988, EPI_ISL_567989, EPI_ISL_567990, EPI_ISL_567991, EPI_ISL_567993, EPI_ISL_567994, EPI_ISL_567995, EPI_ISL_567996, EPI_ISL_567997, EPI_ISL_567998, EPI_ISL_568001, EPI_ISL_568002, EPI_ISL_568003, EPI_ISL_568005, EPI_ISL_568006, EPI_ISL_568007, EPI_ISL_568008, EPI_ISL_568009, EPI_ISL_568010, EPI_ISL_568012, EPI_ISL_568013, EPI_ISL_568014, EPI_ISL_568015, EPI_ISL_568016, EPI_ISL_568017, EPI_ISL_568019, EPI_ISL_568021, EPI_ISL_568022, EPI_ISL_568025, EPI_ISL_568026, EPI_ISL_568028, EPI_ISL_568030, EPI_ISL_568031, EPI_ISL_568032, EPI_ISL_568033, EPI_ISL_568034, EPI_ISL_568035, EPI_ISL_568036, EPI_ISL_568038, EPI_ISL_568039, EPI_ISL_568041, EPI_ISL_568042, EPI_ISL_568044, EPI_ISL_568045, EPI_ISL_568046, EPI_ISL_568047, EPI_ISL_568049, EPI_ISL_568050, EPI_ISL_568051, EPI_ISL_568053, EPI_ISL_568056, EPI_ISL_568058, EPI_ISL_568061, EPI_ISL_568062, EPI_ISL_568063, EPI_ISL_568065, EPI_ISL_568066, EPI_ISL_568067, EPI_ISL_568068, EPI_ISL_568069, EPI_ISL_568070, EPI_ISL_568071, EPI_ISL_568076, EPI_ISL_568078, EPI_ISL_568079, EPI_ISL_568081, EPI_ISL_568082, EPI_ISL_568084, EPI_ISL_568085, EPI_ISL_568086, EPI_ISL_568087, EPI_ISL_568090, EPI_ISL_568091, EPI_ISL_568092, EPI_ISL_568096, EPI_ISL_568099, EPI_ISL_568100, EPI_ISL_568101, EPI_ISL_568103, EPI_ISL_568104, EPI_ISL_568105, EPI_ISL_568106, EPI_ISL_568107, EPI_ISL_568108, EPI_ISL_568110, EPI_ISL_568111, EPI_ISL_568112, EPI_ISL_568113, EPI_ISL_568114, EPI_ISL_568115, EPI_ISL_568116, EPI_ISL_568117, EPI_ISL_568118, EPI_ISL_568119, EPI_ISL_568120, EPI_ISL_568121, EPI_ISL_568122, EPI_ISL_568123, EPI_ISL_568125, EPI_ISL_568126, EPI_ISL_568128, EPI_ISL_568129, EPI_ISL_568130, EPI_ISL_568132, EPI_ISL_568133, EPI_ISL_568134, EPI_ISL_568135, EPI_ISL_568137, EPI_ISL_568138, EPI_ISL_568140, EPI_ISL_568141, EPI_ISL_568146, EPI_ISL_568149, EPI_ISL_568150, EPI_ISL_568152, EPI_ISL_568154, EPI_ISL_568156, EPI_ISL_568157, EPI_ISL_568158, EPI_ISL_568160, EPI_ISL_568161, EPI_ISL_568162, EPI_ISL_568163, EPI_ISL_568165, EPI_ISL_568166, EPI_ISL_568167, EPI_ISL_568168, EPI_ISL_568169, EPI_ISL_568170, EPI_ISL_568171, EPI_ISL_568172, EPI_ISL_568173, EPI_ISL_568174, EPI_ISL_568175, EPI_ISL_568177, EPI_ISL_568178, EPI_ISL_568179, EPI_ISL_568180, EPI_ISL_568181, EPI_ISL_568182, EPI_ISL_568183, EPI_ISL_568184, EPI_ISL_568185, EPI_ISL_568186, EPI_ISL_568188, EPI_ISL_568189, EPI_ISL_568190, EPI_ISL_568192, EPI_ISL_568193, EPI_ISL_568196, EPI_ISL_568198, EPI_ISL_568199, EPI_ISL_568200, EPI_ISL_568201, EPI_ISL_568203, EPI_ISL_568205, EPI_ISL_568206, EPI_ISL_568207, EPI_ISL_568208, EPI_ISL_568209, EPI_ISL_568211, EPI_ISL_568218, EPI_ISL_568221, EPI_ISL_568222 | | | |
| see above | Lighthouse Lab in Alderley Park | Wellcome Sanger Institute for the COVID-19 Genomics UK (COG-UK) consortium | Jacquelyn Wynn, Mairead Hyland, The Lighthouse Lab in Alderley Park and Alex Alderton, Roberto Amato, Sonia Goncalves, Ewan Harrison, David K. Jackson, Ian Johnston, Dominic Kwiatkowski, Cordelia Langford, John Sillitoe on behalf of the Wellcome Sanger Institute COVID-19 Surveillance Team |
| EPI_ISL_568223, EPI_ISL_568225, EPI_ISL_568228, EPI_ISL_568229, EPI_ISL_568230, EPI_ISL_568232, EPI_ISL_568233, EPI_ISL_568234, EPI_ISL_568236, EPI_ISL_568239, EPI_ISL_568240, EPI_ISL_568241, EPI_ISL_568243, EPI_ISL_568244, EPI_ISL_568245, EPI_ISL_568248, EPI_ISL_568251, EPI_ISL_568252, EPI_ISL_568253, EPI_ISL_568254, EPI_ISL_568256, EPI_ISL_568258, EPI_ISL_568259, EPI_ISL_568260, EPI_ISL_568261, EPI_ISL_568264, EPI_ISL_568267, EPI_ISL_568272, EPI_ISL_568274, EPI_ISL_568276, EPI_ISL_568280, EPI_ISL_568281, EPI_ISL_568282, EPI_ISL_568283, EPI_ISL_568284, EPI_ISL_568285, EPI_ISL_568288 | | | |
| see above | Lighthouse Lab in Glasgow | Wellcome Sanger Institute for the COVID-19 Genomics UK (COG-UK) consortium | Harper VanSteenhouse, Yumi Kasai, David Gray, Carol Clugston, Anna Dominiczak and Alex Alderton, Roberto Amato, Sonia Goncalves, Ewan Harrison, David K. Jackson, Ian Johnston, Dominic Kwiatkowski, Cordelia Langford, John Sillitoe on behalf of the Wellcome Sanger Institute COVID-19 Surveillance Team |
| EPI_ISL_568289, EPI_ISL_568290 | Lighthouse Lab in Milton Keynes | Wellcome Sanger Institute for the COVID-19 Genomics UK (COG-UK) consortium | The Lighthouse Lab in Milton Keynes and Alex Alderton, Roberto Amato, Sonia Goncalves, Ewan Harrison, David K. Jackson, Ian Johnston, Dominic Kwiatkowski, Cordelia Langford, John Sillitoe on behalf of the Wellcome Sanger Institute COVID-19 Surveillance Team |
| EPI_ISL_568368, EPI_ISL_568369, EPI_ISL_568370, EPI_ISL_568371, EPI_ISL_568372, EPI_ISL_568373, EPI_ISL_568374, EPI_ISL_568375, EPI_ISL_568376, EPI_ISL_568377, EPI_ISL_568378, EPI_ISL_568379, EPI_ISL_568380, EPI_ISL_568381, EPI_ISL_568382, EPI_ISL_568383 | | | |
| see above | Lighthouse Lab in Glasgow | Wellcome Sanger Institute for the COVID-19 Genomics UK (COG-UK) consortium | Harper VanSteenhouse, Yumi Kasai, David Gray, Carol Clugston, Anna Dominiczak and Alex Alderton, Roberto Amato, Sonia Goncalves, Ewan Harrison, David K. Jackson, Ian Johnston, Dominic Kwiatkowski, Cordelia Langford, John Sillitoe on behalf of the Wellcome Sanger Institute COVID-19 Surveillance Team |
| EPI_ISL_568577 | The National Institute of Public Health | State Veterinary Institute Prague | Nagy, A; Jirincova, H; Novakova, L; Trnka,D; Vecerova, J |
| EPI_ISL_568644, EPI_ISL_568645, EPI_ISL_568646, EPI_ISL_568647, EPI_ISL_568648, EPI_ISL_568649, EPI_ISL_568650, EPI_ISL_568651, EPI_ISL_568652, EPI_ISL_568653, EPI_ISL_568654, EPI_ISL_568655, EPI_ISL_568656, EPI_ISL_568657 | | | |
| see above | Florida Bureau of Public Health Laboratories | Florida Bureau of Public Health Laboratories | Sarah Schmedes, Jason Blanton |
| EPI_ISL_569197, EPI_ISL_569198, EPI_ISL_569199, EPI_ISL_569200, EPI_ISL_569201, EPI_ISL_569202, EPI_ISL_569203, EPI_ISL_569204, EPI_ISL_569205, EPI_ISL_569206, EPI_ISL_569207, EPI_ISL_569208, EPI_ISL_569209, EPI_ISL_569210, EPI_ISL_569211, EPI_ISL_569212, EPI_ISL_569213, EPI_ISL_569214, EPI_ISL_569215, EPI_ISL_569216, EPI_ISL_569217, EPI_ISL_569218, EPI_ISL_569219, EPI_ISL_569221, EPI_ISL_569222, EPI_ISL_569223, EPI_ISL_569224, EPI_ISL_569225, EPI_ISL_569226, EPI_ISL_569227, EPI_ISL_569228, EPI_ISL_569229 | | | |
| see above | MEPHI, Aix Marseille University | MEPHI, Aix Marseille University | Anthony LEVASSEUR |
| EPI_ISL_569625, EPI_ISL_569626, EPI_ISL_569627, EPI_ISL_569628, EPI_ISL_569629, EPI_ISL_569630, EPI_ISL_569631, EPI_ISL_569632, EPI_ISL_569633, EPI_ISL_569634, EPI_ISL_569635, EPI_ISL_569636, EPI_ISL_569637, EPI_ISL_569638, EPI_ISL_569639, EPI_ISL_569640, EPI_ISL_569641, EPI_ISL_569642, EPI_ISL_569643, EPI_ISL_569644, EPI_ISL_569645, EPI_ISL_569646, EPI_ISL_569647, EPI_ISL_569648, EPI_ISL_569649, EPI_ISL_569650, EPI_ISL_569651, EPI_ISL_569652, EPI_ISL_569653, EPI_ISL_569654, EPI_ISL_569655, EPI_ISL_569656, EPI_ISL_569657, EPI_ISL_569658, EPI_ISL_569659, EPI_ISL_569660, EPI_ISL_569661, EPI_ISL_569662, EPI_ISL_569663 | | | |
| see above | New Mexico Department of Health Scientific Laboratory | New Mexico Department of Health Scientific Laboratory | Ellie Johnson, Anastacia Griego-Fisher, D'Eldra Malone |
| EPI_ISL_569799 | Omsk Research Institute of Natural Focal Infections | WHO National Influenza Centre Russian Federation | Artem Fadeev, Ekaterina Gradoboeva, Ekaterina Savkina, Daria Nashatyreva, Elena Poleshchuk, Aleksei Vasilenko, Valery Yakimenko, Andrey Komissarov |
| EPI_ISL_570553, EPI_ISL_570554, EPI_ISL_570555, EPI_ISL_570556, EPI_ISL_570557, EPI_ISL_570574, EPI_ISL_570577, EPI_ISL_570579, EPI_ISL_570604, EPI_ISL_570779, EPI_ISL_570780, EPI_ISL_570781, EPI_ISL_570782, EPI_ISL_570783, EPI_ISL_570784, EPI_ISL_570785, EPI_ISL_570786, EPI_ISL_570787, EPI_ISL_570788, EPI_ISL_570789, EPI_ISL_570790, EPI_ISL_570791, EPI_ISL_570792, EPI_ISL_570793, EPI_ISL_570794, EPI_ISL_570795, EPI_ISL_570796, EPI_ISL_570797 | | | |
| see above | UW Virology Lab | UW Virology Lab | Pavitra Roychoudhury, Hong Xie, Lasata Shrestha, Amin Addetia, Victoria M Rachleff, Meei-Li Huang, Keith R Jerome, Alexander Greninger |
| EPI_ISL_572218, EPI_ISL_572219, EPI_ISL_572269 | Virginia DCLS | Virginia DCLS | Virginia DCLS |
| EPI_ISL_572412 | Northumbria University / South Tees Hospitals NHS Foundation Trust / North Cumbria Integrated Care NHS Foundation Trust / North Tees and Hartlepool NHS Foundation Trust / Newcastle Hospitals NHS Foundation Trust | COVID-19 Genomics UK (COG-UK) Consortium | Darren L Smith,Andrew Nelson,Matthew Bashton,Greg R Young,Joshua Loh,John Allan,Mohammad A Tariq,Giles S Holt,Gary Black,Wen C Yew,Lynn Dover,Paul Baker,Steve Liggett,Sarah Essex,Jane Greenaway,Debra Padgett,Clive Graham,Garren Scott,Edward Barton,Emma Swindells,Brendan Payne,Jennifer Collins,Yusri Taha,Gary Eltringham |
| EPI_ISL_572417 | Virology Department, Sheffield Teaching Hospitals NHS Foundation Trust/Department of Infection, Immunity and Cardiovascular Disease, The Medical School, University of Sheffield | COVID-19 Genomics UK (COG-UK) Consortium | Thushan de Silva, Matthew Parker, Nikki Smith, Adri Angyal, Rebecca Brown, Luke Green, Rachel Tucker, Paul Parsons, Danielle Groves, Katie Johnson, Laura Carrilero, Alex Keeley, Dave Partridge, Matthew Wyles, Benjamin Lindsey, Mehmet Yavuz, Mohammad Raza, Cariad Evans |
| EPI_ISL_572421 | Virology Department, Royal Infirmary of Edinburgh, NHS Lothian / School of Biological Sciences, University of Edinburgh / Institute of Genetics and Molecular Medicine, University of Edinburgh | COVID-19 Genomics UK (COG-UK) Consortium | McHugh M, Dewar R, Rooke S, Gallagher M, Balcaza C, O'Toole Á, Scher E, Hill V, McCrone JT, Colquhoun R, Yu X, Jackson B, Rambaut A, Williams TC, Templeton K |
| EPI_ISL_572426 | Liverpool Clinical Laboratories | COVID-19 Genomics UK (COG-UK) Consortium | Sam Haldenby, Anita Lucaci, Steve Paterson, Julian Hiscox, Alistair Darby, M Almsaud, A Alrezaihi, Muhannad Alruwaili, Stuart D Armstrong, Jones Benjamin, Eleanor G Bentley, Anu Chawla, Jordan J Clark, Angela Cowell, Richard Eccles, Isabel García-Dorival, Matthew Gemmell, Alessandro Gerada, PKF Gilmore, Richard Gregory, Ximeng Han, Catherine Hartley, Margaret Hughes, Miren Iturriza-Gomara, James Johnson, L Luu, Jenifer Manson, Charlotte Nelson, Elaine O'Toole, Cassie Olateju, Rebekah Penrice-Randal , Lucille Rainbow, N.P Randle, Trevor Ian Robinson, Parul Sharma, Ghada T Shawli, James P Stewart, Neil Swainston, Ecaterina Vamos, Joanne Watts, Mark Whitehead |
| EPI_ISL_572435 | Virology Department, Royal Infirmary of Edinburgh, NHS Lothian / School of Biological Sciences, University of Edinburgh / Institute of Genetics and Molecular Medicine, University of Edinburgh | COVID-19 Genomics UK (COG-UK) Consortium | McHugh M, Dewar R, Rooke S, Gallagher M, Balcaza C, O'Toole Á, Scher E, Hill V, McCrone JT, Colquhoun R, Yu X, Jackson B, Rambaut A, Williams TC, Templeton K |
| EPI_ISL_572450 | Northumbria University / South Tees Hospitals NHS Foundation Trust / North Cumbria Integrated Care NHS Foundation Trust / North Tees and Hartlepool NHS Foundation Trust / Newcastle Hospitals NHS Foundation Trust | COVID-19 Genomics UK (COG-UK) Consortium | Darren L Smith,Andrew Nelson,Matthew Bashton,Greg R Young,Joshua Loh,John Allan,Mohammad A Tariq,Giles S Holt,Gary Black,Wen C Yew,Lynn Dover,Paul Baker,Steve Liggett,Sarah Essex,Jane Greenaway,Debra Padgett,Clive Graham,Garren Scott,Edward Barton,Emma Swindells,Brendan Payne,Jennifer Collins,Yusri Taha,Gary Eltringham |

| | | | |
|---|---|---|---|
| EPI_ISL_572457, EPI_ISL_572458 | Virology Department, Sheffield Teaching Hospitals NHS Foundation Trust/Department of Infection, Immunity and Cardiovascular Disease, The Medical School, University of Sheffield | COVID-19 Genomics UK (COG-UK) Consortium | Thushan de Silva, Matthew Parker, Nikki Smith, Adri Angyal, Rebecca Brown, Luke Green, Rachel Tucker, Paul Parsons, Danielle Groves, Katie Johnson, Laura Carrilero, Alex Keeley, Dave Partridge, Matthew Wyles, Benjamin Lindsey, Mehmet Yavuz, Mohammad Raza, Cariad Evans |
| EPI_ISL_572478 | Liverpool Clinical Laboratories | COVID-19 Genomics UK (COG-UK) Consortium | Sam Haldenby, Anita Lucaci, Steve Paterson, Julian Hiscox, Alistair Darby, M Almsaud, A Alrezaihi, Muhannad Alruwaili, Stuart D Armstrong, Jones Benjamin, Eleanor G Bentley, Anu Chawla, Jordan J Clark, Angela Cowell, Richard Eccles, Isabel García-Dorival, Matthew Gemmell, Alessandro Gerada, PKF Gilmore, Richard Gregory, Ximeng Han, Catherine Hartley, Margaret Hughes, Miren Iturriza-Gomara, James Johnson, L Luu, Jenifer Manson, Charlotte Nelson, Elaine O'Toole, Cassie Olateju, Rebekah Penrice-Randal , Lucille Rainbow, N.P Randle, Trevor Ian Robinson, Parul Sharma, Ghada T Shawli, James P Stewart, Neil Swainston, Ecaterina Vamos, Joanne Watts, Mark Whitehead |
| EPI_ISL_572513 | Northumbria University / South Tees Hospitals NHS Foundation Trust / North Cumbria Integrated Care NHS Foundation Trust / North Tees and Hartlepool NHS Foundation Trust / Newcastle Hospitals NHS Foundation Trust | COVID-19 Genomics UK (COG-UK) Consortium | Darren L Smith,Andrew Nelson,Matthew Bashton,Greg R Young,Joshua Loh,John Allan,Mohammad A Tariq,Giles S Holt,Gary Black,Wen C Yew,Lynn Dover,Paul Baker,Steve Liggett,Sarah Essex,Jane Greenaway,Debra Padgett,Clive Graham,Garren Scott,Edward Barton,Emma Swindells,Brendan Payne,Jennifer Collins,Yusri Taha,Gary Eltringham |
| EPI_ISL_572514 | Virology Department, Royal Infirmary of Edinburgh, NHS Lothian / School of Biological Sciences, University of Edinburgh / Institute of Genetics and Molecular Medicine, University of Edinburgh | COVID-19 Genomics UK (COG-UK) Consortium | McHugh M, Dewar R, Rooke S, Gallagher M, Balcaza C, O'Toole Á, Scher E, Hill V, McCrone JT, Colquhoun R, Yu X, Jackson B, Rambaut A, Williams TC, Templeton K |
| EPI_ISL_572515, EPI_ISL_572517, EPI_ISL_572525 | Northumbria University / South Tees Hospitals NHS Foundation Trust / North Cumbria Integrated Care NHS Foundation Trust / North Tees and Hartlepool NHS Foundation Trust / Newcastle Hospitals NHS Foundation Trust | COVID-19 Genomics UK (COG-UK) Consortium | Darren L Smith,Andrew Nelson,Matthew Bashton,Greg R Young,Joshua Loh,John Allan,Mohammad A Tariq,Giles S Holt,Gary Black,Wen C Yew,Lynn Dover,Paul Baker,Steve Liggett,Sarah Essex,Jane Greenaway,Debra Padgett,Clive Graham,Garren Scott,Edward Barton,Emma Swindells,Brendan Payne,Jennifer Collins,Yusri Taha,Gary Eltringham |
| EPI_ISL_572529 | Virology Department, Sheffield Teaching Hospitals NHS Foundation Trust/Department of Infection, Immunity and Cardiovascular Disease, The Medical School, University of Sheffield | COVID-19 Genomics UK (COG-UK) Consortium | Thushan de Silva, Matthew Parker, Nikki Smith, Adri Angyal, Rebecca Brown, Luke Green, Rachel Tucker, Paul Parsons, Danielle Groves, Katie Johnson, Laura Carrilero, Alex Keeley, Dave Partridge, Matthew Wyles, Benjamin Lindsey, Mehmet Yavuz, Mohammad Raza, Cariad Evans |
| EPI_ISL_572573 | Virology Department, Royal Infirmary of Edinburgh, NHS Lothian / School of Biological Sciences, University of Edinburgh / Institute of Genetics and Molecular Medicine, University of Edinburgh | COVID-19 Genomics UK (COG-UK) Consortium | McHugh M, Dewar R, Rooke S, Gallagher M, Balcaza C, O'Toole Á, Scher E, Hill V, McCrone JT, Colquhoun R, Yu X, Jackson B, Rambaut A, Williams TC, Templeton K |
| EPI_ISL_572574 | Wales Specialist Virology Centre Sequencing lab: Pathogen Genomics Unit | COVID-19 Genomics UK (COG-UK) Consortium | Catherine Moore, Johnathan Evans, Laura Gifford, Malorie Perry, Simon Cottrell, Angela Marchbank, Alec Birchley, Alexander Adams, Amy Gaskin, Bree Gatica-Wilcox, Jason Coombes, Joel Southgate, Lauren Gilbert, Lee Graham, Nicole Pacchiarini, Sara Kumziene-Summerhayes, Sarah Taylor, Sophie Jones, Sara Rey, Matthew Bull, Joanne Watkins, Sally Corden, Tom Connor |
| EPI_ISL_572590, EPI_ISL_572593 | Virology Department, Royal Infirmary of Edinburgh, NHS Lothian / School of Biological Sciences, University of Edinburgh / Institute of Genetics and Molecular Medicine, University of Edinburgh | COVID-19 Genomics UK (COG-UK) Consortium | McHugh M, Dewar R, Rooke S, Gallagher M, Balcaza C, O'Toole Á, Scher E, Hill V, McCrone JT, Colquhoun R, Yu X, Jackson B, Rambaut A, Williams TC, Templeton K |
| EPI_ISL_572594, EPI_ISL_572596 | Northumbria University / South Tees Hospitals NHS Foundation Trust / North Cumbria Integrated Care NHS Foundation Trust / North Tees and Hartlepool NHS Foundation Trust / Newcastle Hospitals NHS Foundation Trust | COVID-19 Genomics UK (COG-UK) Consortium | Darren L Smith,Andrew Nelson,Matthew Bashton,Greg R Young,Joshua Loh,John Allan,Mohammad A Tariq,Giles S Holt,Gary Black,Wen C Yew,Lynn Dover,Paul Baker,Steve Liggett,Sarah Essex,Jane Greenaway,Debra Padgett,Clive Graham,Garren Scott,Edward Barton,Emma Swindells,Brendan Payne,Jennifer Collins,Yusri Taha,Gary Eltringham |
| EPI_ISL_572608, EPI_ISL_572617 | Virology Department, Royal Infirmary of Edinburgh, NHS Lothian / School of Biological Sciences, University of Edinburgh / Institute of Genetics and Molecular Medicine, University of Edinburgh | COVID-19 Genomics UK (COG-UK) Consortium | McHugh M, Dewar R, Rooke S, Gallagher M, Balcaza C, O'Toole Á, Scher E, Hill V, McCrone JT, Colquhoun R, Yu X, Jackson B, Rambaut A, Williams TC, Templeton K |
| EPI_ISL_572619 | Virology Department, Sheffield Teaching Hospitals NHS Foundation Trust/Department of Infection, Immunity and Cardiovascular Disease, The Medical School, University of Sheffield | COVID-19 Genomics UK (COG-UK) Consortium | Thushan de Silva, Matthew Parker, Nikki Smith, Adri Angyal, Rebecca Brown, Luke Green, Rachel Tucker, Paul Parsons, Danielle Groves, Katie Johnson, Laura Carrilero, Alex Keeley, Dave Partridge, Matthew Wyles, Benjamin Lindsey, Mehmet Yavuz, Mohammad Raza, Cariad Evans |
| EPI_ISL_572621, EPI_ISL_572622, EPI_ISL_572627, EPI_ISL_572628, EPI_ISL_572629, EPI_ISL_572630, EPI_ISL_572631, EPI_ISL_572632 | West of Scotland Specialist Virology Centre, NHSGGC / MRC-University of Glasgow Centre for Virus Research | COVID-19 Genomics UK (COG-UK) Consortium | Ana da Silva Filipe, Natasha Johnson, Kathy Smollett, Daniel Mair, Stephen Carmichael, Lily Tong, Jenna Nichols, Elihu Aranday-Cortes, Kyriaki Nomikou; Sarah McDonald, Marc Niebel, Patawee Asamaphan; Richard Orton, Joseph Hughes, Sreenu Vattipally, David L Robertson; Alasdair MacLean, Rory Gunson; Kathy Li, Igor Starinskij, Natasha Jesudason, Rajiv Shah, James Shepherd, Antonia Ho, Emma Thomson |
| EPI_ISL_572644 | Northumbria University / South Tees Hospitals NHS Foundation Trust / North Cumbria Integrated Care NHS Foundation Trust / North Tees and Hartlepool NHS Foundation Trust / Newcastle Hospitals NHS Foundation Trust | COVID-19 Genomics UK (COG-UK) Consortium | Darren L Smith,Andrew Nelson,Matthew Bashton,Greg R Young,Joshua Loh,John Allan,Mohammad A Tariq,Giles S Holt,Gary Black,Wen C Yew,Lynn Dover,Paul Baker,Steve Liggett,Sarah Essex,Jane Greenaway,Debra Padgett,Clive Graham,Garren Scott,Edward Barton,Emma Swindells,Brendan Payne,Jennifer Collins,Yusri Taha,Gary Eltringham |
| EPI_ISL_572678 | Virology Department, Sheffield Teaching Hospitals NHS Foundation Trust/Department of Infection, Immunity and Cardiovascular Disease, The Medical School, University of Sheffield | COVID-19 Genomics UK (COG-UK) Consortium | Thushan de Silva, Matthew Parker, Nikki Smith, Adri Angyal, Rebecca Brown, Luke Green, Rachel Tucker, Paul Parsons, Danielle Groves, Katie Johnson, Laura Carrilero, Alex Keeley, Dave Partridge, Matthew Wyles, Benjamin Lindsey, Mehmet Yavuz, Mohammad Raza, Cariad Evans |
| EPI_ISL_572695 | Liverpool Clinical Laboratories | COVID-19 Genomics UK (COG-UK) Consortium | Sam Haldenby, Anita Lucaci, Steve Paterson, Julian Hiscox, Alistair Darby, M Almsaud, A Alrezaihi, Muhannad Alruwaili, Stuart D Armstrong, Jones Benjamin, Eleanor G Bentley, Anu Chawla, Jordan J Clark, Angela Cowell, Richard Eccles, Isabel García-Dorival, Matthew Gemmell, Alessandro Gerada, PKF Gilmore, Richard Gregory, Ximeng Han, Catherine Hartley, Margaret Hughes, Miren Iturriza-Gomara, James Johnson, L Luu, Jenifer Manson, Charlotte Nelson, Elaine O'Toole, Cassie Olateju, Rebekah Penrice-Randal , Lucille Rainbow, N.P Randle, Trevor Ian Robinson, Parul Sharma, Ghada T Shawli, James P Stewart, Neil Swainston, Ecaterina Vamos, Joanne Watts, Mark Whitehead |
| EPI_ISL_572702 | Northumbria University / South Tees Hospitals NHS Foundation Trust / North Cumbria Integrated Care NHS Foundation Trust / North Tees and Hartlepool NHS Foundation Trust / Newcastle Hospitals NHS Foundation Trust | COVID-19 Genomics UK (COG-UK) Consortium | Darren L Smith,Andrew Nelson,Matthew Bashton,Greg R Young,Joshua Loh,John Allan,Mohammad A Tariq,Giles S Holt,Gary Black,Wen C Yew,Lynn Dover,Paul Baker,Steve Liggett,Sarah Essex,Jane Greenaway,Debra Padgett,Clive Graham,Garren Scott,Edward Barton,Emma Swindells,Brendan Payne,Jennifer Collins,Yusri Taha,Gary Eltringham |
| EPI_ISL_572709 | University College London, Great Ormond Street Hospital for Children NHS Foundation Trust, Imperial College Healthcare NHS Trust | COVID-19 Genomics UK (COG-UK) Consortium | Sergi Castellano, Rachel Williams, Mark Kristiansen, Paola Resende Silva, Sunando Roy, Tony Brooks, Helena Tutill, Paola Niola, Patricia Dyal, Charlotte Williams, Leysa Forrest, Yasmin Panchbhaya, Jacqueline Findlay, Samuel Weeks, Julianne Brown, Kathryn Harris, Paul Randell, James Price, Alison Holmes, Judith Breuer |
| EPI_ISL_572712 | Northumbria University / South Tees Hospitals NHS Foundation Trust / North Cumbria Integrated Care NHS Foundation Trust / North Tees and Hartlepool NHS Foundation Trust / Newcastle Hospitals NHS Foundation Trust | COVID-19 Genomics UK (COG-UK) Consortium | Darren L Smith,Andrew Nelson,Matthew Bashton,Greg R Young,Joshua Loh,John Allan,Mohammad A Tariq,Giles S Holt,Gary Black,Wen C Yew,Lynn Dover,Paul Baker,Steve Liggett,Sarah Essex,Jane Greenaway,Debra Padgett,Clive Graham,Garren Scott,Edward Barton,Emma Swindells,Brendan Payne,Jennifer Collins,Yusri Taha,Gary Eltringham |
| EPI_ISL_572715 | Centre for Enzyme Innovation, University of Portsmouth / Translational Research Laboratory, Portsmouth Hospitals | COVID-19 Genomics UK (COG-UK) Consortium | Angela Beckett,Yann Bourgeois,Garry Scarlett,Sharon Glaysher,Scott Elliott,Kelly Bicknell,Robert Impey,Allyson Lloyd,Sarah Wyllie,Ethan Butcher,Anoop Chauhan,Samuel Robson |

| | | | |
|---|---|---|---|
| | NHS Trust | | |
| EPI_ISL_572722 | Virology Department, Sheffield Teaching Hospitals NHS Foundation Trust/Department of Infection, Immunity and Cardiovascular Disease, The Medical School, University of Sheffield | COVID-19 Genomics UK (COG-UK) Consortium | Thushan de Silva, Matthew Parker, Nikki Smith, Adri Angyal, Rebecca Brown, Luke Green, Rachel Tucker, Paul Parsons, Danielle Groves, Katie Johnson, Laura Carrilero, Alex Keeley, Dave Partridge, Matthew Wyles, Benjamin Lindsey, Mehmet Yavuz, Mohammad Raza, Cariad Evans |
| EPI_ISL_572723, EPI_ISL_572724, EPI_ISL_572726, EPI_ISL_572727, EPI_ISL_572728 | Oxford Viromics, NDM, University of Oxford; Oxford University Hospitals; Basingstoke and North Hampshire Hospital | COVID-19 Genomics UK (COG-UK) Consortium | Tanya Golubchik, David Bonsall, George Macintyre, Amy Trebes, Mariateresa de Cesare, Catrin Moore, Alex Mobbs, Anita Justice, Robert Shaw, Monique Andersson, Timothy Peto, Emma Wise, Nathan Moore, Jessica Lynch, Nick Cortes, Matilde Mori, Stephen Kidd, David Buck, John Todd, Christophe Fraser |
| EPI_ISL_572741 | West of Scotland Specialist Virology Centre, NHSGGC / MRC-University of Glasgow Centre for Virus Research | COVID-19 Genomics UK (COG-UK) Consortium | Ana da Silva Filipe, Natasha Johnson, Kathy Smollett, Daniel Mair, Stephen Carmichael, Lily Tong, Jenna Nichols, Elihu Aranday-Cortes, Kyriaki Nomikou; Sarah McDonald, Marc Niebel, Patawee Asamaphan; Richard Orton, Joseph Hughes, Sreenu Vattipally, David L Robertson; Alasdair MacLean, Rory Gunson; Kathy Li, Igor Starinskij, Natasha Jesudason, Rajiv Shah, James Shepherd, Antonia Ho, Emma Thomson |
| EPI_ISL_572747, EPI_ISL_572751 | Northumbria University / South Tees Hospitals NHS Foundation Trust / North Cumbria Integrated Care NHS Foundation Trust / North Tees and Hartlepool NHS Foundation Trust / Newcastle Hospitals NHS Foundation Trust | COVID-19 Genomics UK (COG-UK) Consortium | Darren L Smith,Andrew Nelson,Matthew Bashton,Greg R Young,Joshua Loh,John Allan,Mohammad A Tariq,Giles S Holt,Gary Black,Wen C Yew,Lynn Dover,Paul Baker,Steve Liggett,Sarah Essex,Jane Greenaway,Debra Padgett,Clive Graham,Garren Scott,Edward Barton,Emma Swindells,Brendan Payne,Jennifer Collins,Yusri Taha,Gary Eltringham |
| EPI_ISL_572752 | Virology Department, Sheffield Teaching Hospitals NHS Foundation Trust/Department of Infection, Immunity and Cardiovascular Disease, The Medical School, University of Sheffield | COVID-19 Genomics UK (COG-UK) Consortium | Thushan de Silva, Matthew Parker, Nikki Smith, Adri Angyal, Rebecca Brown, Luke Green, Rachel Tucker, Paul Parsons, Danielle Groves, Katie Johnson, Laura Carrilero, Alex Keeley, Dave Partridge, Matthew Wyles, Benjamin Lindsey, Mehmet Yavuz, Mohammad Raza, Cariad Evans |
| EPI_ISL_572770 | University College London, Great Ormond Street Hospital for Children NHS Foundation Trust, Imperial College Healthcare NHS Trust | COVID-19 Genomics UK (COG-UK) Consortium | Sergi Castellano, Rachel Williams, Mark Kristiansen, Paola Resende Silva, Sunando Roy, Tony Brooks, Helena Tutill, Paola Niola, Patricia Dyal, Charlotte Williams, Leysa Forrest, Yasmin Panchbhaya, Jacqueline Findlay, Samuel Weeks, Julianne Brown, Kathryn Harris, Paul Randell, James Price, Alison Holmes, Judith Breuer |
| EPI_ISL_572795 | Northumbria University / South Tees Hospitals NHS Foundation Trust / North Cumbria Integrated Care NHS Foundation Trust / North Tees and Hartlepool NHS Foundation Trust / Newcastle Hospitals NHS Foundation Trust | COVID-19 Genomics UK (COG-UK) Consortium | Darren L Smith,Andrew Nelson,Matthew Bashton,Greg R Young,Joshua Loh,John Allan,Mohammad A Tariq,Giles S Holt,Gary Black,Wen C Yew,Lynn Dover,Paul Baker,Steve Liggett,Sarah Essex,Jane Greenaway,Debra Padgett,Clive Graham,Garren Scott,Edward Barton,Emma Swindells,Brendan Payne,Jennifer Collins,Yusri Taha,Gary Eltringham |
| EPI_ISL_572796 | Virology Department, Sheffield Teaching Hospitals NHS Foundation Trust/Department of Infection, Immunity and Cardiovascular Disease, The Medical School, University of Sheffield | COVID-19 Genomics UK (COG-UK) Consortium | Thushan de Silva, Matthew Parker, Nikki Smith, Adri Angyal, Rebecca Brown, Luke Green, Rachel Tucker, Paul Parsons, Danielle Groves, Katie Johnson, Laura Carrilero, Alex Keeley, Dave Partridge, Matthew Wyles, Benjamin Lindsey, Mehmet Yavuz, Mohammad Raza, Cariad Evans |
| EPI_ISL_572807 | University College London, Great Ormond Street Hospital for Children NHS Foundation Trust, Imperial College Healthcare NHS Trust | COVID-19 Genomics UK (COG-UK) Consortium | Sergi Castellano, Rachel Williams, Mark Kristiansen, Paola Resende Silva, Sunando Roy, Tony Brooks, Helena Tutill, Paola Niola, Patricia Dyal, Charlotte Williams, Leysa Forrest, Yasmin Panchbhaya, Jacqueline Findlay, Samuel Weeks, Julianne Brown, Kathryn Harris, Paul Randell, James Price, Alison Holmes, Judith Breuer |
| EPI_ISL_572821 | Virology Department, Royal Infirmary of Edinburgh, NHS Lothian / School of Biological Sciences, University of Edinburgh / Institute of Genetics and Molecular Medicine, University of Edinburgh | COVID-19 Genomics UK (COG-UK) Consortium | McHugh M, Dewar R, Rooke S, Gallagher M, Balcaza C, O'Toole Á, Scher E, Hill V, McCrone JT, Colquhoun R, Yu X, Jackson B, Rambaut A, Williams TC, Templeton K |
| EPI_ISL_572828, EPI_ISL_572838 | University College London, Great Ormond Street Hospital for Children NHS Foundation Trust, Imperial College Healthcare NHS Trust | COVID-19 Genomics UK (COG-UK) Consortium | Sergi Castellano, Rachel Williams, Mark Kristiansen, Paola Resende Silva, Sunando Roy, Tony Brooks, Helena Tutill, Paola Niola, Patricia Dyal, Charlotte Williams, Leysa Forrest, Yasmin Panchbhaya, Jacqueline Findlay, Samuel Weeks, Julianne Brown, Kathryn Harris, Paul Randell, James Price, Alison Holmes, Judith Breuer |
| EPI_ISL_572840, EPI_ISL_572845 | Northumbria University / South Tees Hospitals NHS Foundation Trust / North Cumbria Integrated Care NHS Foundation Trust / North Tees and Hartlepool NHS Foundation Trust / Newcastle Hospitals NHS Foundation Trust | COVID-19 Genomics UK (COG-UK) Consortium | Darren L Smith,Andrew Nelson,Matthew Bashton,Greg R Young,Joshua Loh,John Allan,Mohammad A Tariq,Giles S Holt,Gary Black,Wen C Yew,Lynn Dover,Paul Baker,Steve Liggett,Sarah Essex,Jane Greenaway,Debra Padgett,Clive Graham,Garren Scott,Edward Barton,Emma Swindells,Brendan Payne,Jennifer Collins,Yusri Taha,Gary Eltringham |
| EPI_ISL_572846, EPI_ISL_572850, EPI_ISL_572869, EPI_ISL_572870 | Oxford Viromics, NDM, University of Oxford; Oxford University Hospitals; Basingstoke and North Hampshire Hospital | COVID-19 Genomics UK (COG-UK) Consortium | Tanya Golubchik, David Bonsall, George Macintyre, Amy Trebes, Mariateresa de Cesare, Catrin Moore, Alex Mobbs, Anita Justice, Robert Shaw, Monique Andersson, Timothy Peto, Emma Wise, Nathan Moore, Jessica Lynch, Nick Cortes, Matilde Mori, Stephen Kidd, David Buck, John Todd, Christophe Fraser |
| EPI_ISL_572871 | Virology Department, Royal Infirmary of Edinburgh, NHS Lothian / School of Biological Sciences, University of Edinburgh / Institute of Genetics and Molecular Medicine, University of Edinburgh | COVID-19 Genomics UK (COG-UK) Consortium | McHugh M, Dewar R, Rooke S, Gallagher M, Balcaza C, O'Toole Á, Scher E, Hill V, McCrone JT, Colquhoun R, Yu X, Jackson B, Rambaut A, Williams TC, Templeton K |
| EPI_ISL_572873 | Oxford Viromics, NDM, University of Oxford; Oxford University Hospitals; Basingstoke and North Hampshire Hospital | COVID-19 Genomics UK (COG-UK) Consortium | Tanya Golubchik, David Bonsall, George Macintyre, Amy Trebes, Mariateresa de Cesare, Catrin Moore, Alex Mobbs, Anita Justice, Robert Shaw, Monique Andersson, Timothy Peto, Emma Wise, Nathan Moore, Jessica Lynch, Nick Cortes, Matilde Mori, Stephen Kidd, David Buck, John Todd, Christophe Fraser |
| EPI_ISL_572874 | Virology Department, Royal Infirmary of Edinburgh, NHS Lothian / School of Biological Sciences, University of Edinburgh / Institute of Genetics and Molecular Medicine, University of Edinburgh | COVID-19 Genomics UK (COG-UK) Consortium | McHugh M, Dewar R, Rooke S, Gallagher M, Balcaza C, O'Toole Á, Scher E, Hill V, McCrone JT, Colquhoun R, Yu X, Jackson B, Rambaut A, Williams TC, Templeton K |
| EPI_ISL_572892, EPI_ISL_572902 | Oxford Viromics, NDM, University of Oxford; Oxford University Hospitals; Basingstoke and North Hampshire Hospital | COVID-19 Genomics UK (COG-UK) Consortium | Tanya Golubchik, David Bonsall, George Macintyre, Amy Trebes, Mariateresa de Cesare, Catrin Moore, Alex Mobbs, Anita Justice, Robert Shaw, Monique Andersson, Timothy Peto, Emma Wise, Nathan Moore, Jessica Lynch, Nick Cortes, Matilde Mori, Stephen Kidd, David Buck, John Todd, Christophe Fraser |
| EPI_ISL_572909, EPI_ISL_572910 | Northumbria University / South Tees Hospitals NHS Foundation Trust / North Cumbria Integrated Care NHS Foundation Trust / North Tees and Hartlepool NHS Foundation Trust / Newcastle Hospitals NHS Foundation Trust | COVID-19 Genomics UK (COG-UK) Consortium | Darren L Smith,Andrew Nelson,Matthew Bashton,Greg R Young,Joshua Loh,John Allan,Mohammad A Tariq,Giles S Holt,Gary Black,Wen C Yew,Lynn Dover,Paul Baker,Steve Liggett,Sarah Essex,Jane Greenaway,Debra Padgett,Clive Graham,Garren Scott,Edward Barton,Emma Swindells,Brendan Payne,Jennifer Collins,Yusri Taha,Gary Eltringham |
| EPI_ISL_572912, EPI_ISL_572913, EPI_ISL_572914, EPI_ISL_572915, EPI_ISL_572916 | Virology Department, Sheffield Teaching Hospitals NHS Foundation Trust/Department of Infection, Immunity and Cardiovascular Disease, The Medical School, University of Sheffield | COVID-19 Genomics UK (COG-UK) Consortium | Thushan de Silva, Matthew Parker, Nikki Smith, Adri Angyal, Rebecca Brown, Luke Green, Rachel Tucker, Paul Parsons, Danielle Groves, Katie Johnson, Laura Carrilero, Alex Keeley, Dave Partridge, Matthew Wyles, Benjamin Lindsey, Mehmet Yavuz, Mohammad Raza, Cariad Evans |
| EPI_ISL_572917, EPI_ISL_572918, EPI_ISL_572919, EPI_ISL_572920, EPI_ISL_572921, EPI_ISL_572922, EPI_ISL_572923, EPI_ISL_572924, EPI_ISL_572925, EPI_ISL_572926, EPI_ISL_572934, EPI_ISL_572935, EPI_ISL_572936 | | | |
| see above | Oxford Viromics, NDM, University of Oxford; Oxford University Hospitals; Basingstoke and North Hampshire Hospital | COVID-19 Genomics UK (COG-UK) Consortium | Tanya Golubchik, David Bonsall, George Macintyre, Amy Trebes, Mariateresa de Cesare, Catrin Moore, Alex Mobbs, Anita Justice, Robert Shaw, Monique Andersson, Timothy Peto, Emma Wise, Nathan Moore, Jessica Lynch, Nick Cortes, Matilde Mori, Stephen Kidd, David Buck, John Todd, Christophe Fraser |
| EPI_ISL_572945, EPI_ISL_572946 | Virology Department, Sheffield Teaching Hospitals NHS Foundation Trust/Department of Infection, Immunity and Cardiovascular Disease, The Medical School, University of Sheffield | COVID-19 Genomics UK (COG-UK) Consortium | Thushan de Silva, Matthew Parker, Nikki Smith, Adri Angyal, Rebecca Brown, Luke Green, Rachel Tucker, Paul Parsons, Danielle Groves, Katie Johnson, Laura Carrilero, Alex Keeley, Dave Partridge, Matthew Wyles, Benjamin Lindsey, Mehmet Yavuz, Mohammad Raza, Cariad Evans |
| EPI_ISL_572947 | Oxford Viromics, NDM, University of Oxford; Oxford University Hospitals; Basingstoke and North Hampshire Hospital | COVID-19 Genomics UK (COG-UK) Consortium | Tanya Golubchik, David Bonsall, George Macintyre, Amy Trebes, Mariateresa de Cesare, Catrin Moore, Alex Mobbs, Anita Justice, Robert Shaw, Monique Andersson, Timothy Peto, Emma Wise, Nathan Moore, Jessica Lynch, Nick Cortes, Matilde Mori, Stephen Kidd, David Buck, John Todd, Christophe Fraser |

| | | | |
|---|---|---|---|
| EPI_ISL_572950 | Northumbria University / South Tees Hospitals NHS Foundation Trust / North Cumbria Integrated Care NHS Foundation Trust / North Tees and Hartlepool NHS Foundation Trust / Newcastle Hospitals NHS Foundation Trust | COVID-19 Genomics UK (COG-UK) Consortium | Darren L Smith,Andrew Nelson,Matthew Bashton,Greg R Young,Joshua Loh,John Allan,Mohammad A Tariq,Giles S Holt,Gary Black,Wen C Yew,Lynn Dover,Paul Baker,Steve Liggett,Sarah Essex,Jane Greenaway,Debra Padgett,Clive Graham,Garren Scott,Edward Barton,Emma Swindells,Brendan Payne,Jennifer Collins,Yusri Taha,Gary Eltringham |
| EPI_ISL_572952 | Oxford Viromics, NDM, University of Oxford; Oxford University Hospitals; Basingstoke and North Hampshire Hospital | COVID-19 Genomics UK (COG-UK) Consortium | Tanya Golubchik, David Bonsall, George Macintyre, Amy Trebes, Mariateresa de Cesare, Catrin Moore, Alex Mobbs, Anita Justice, Robert Shaw, Monique Andersson, Timothy Peto, Emma Wise, Nathan Moore, Jessica Lynch, Nick Cortes, Matilde Mori, Stephen Kidd, David Buck, John Todd, Christophe Fraser |
| EPI_ISL_572959 | Virology Department, Sheffield Teaching Hospitals NHS Foundation Trust/Department of Infection, Immunity and Cardiovascular Disease, The Medical School, University of Sheffield | COVID-19 Genomics UK (COG-UK) Consortium | Thushan de Silva, Matthew Parker, Nikki Smith, Adri Angyal, Rebecca Brown, Luke Green, Rachel Tucker, Paul Parsons, Danielle Groves, Katie Johnson, Laura Carrilero, Alex Keeley, Dave Partridge, Matthew Wyles, Benjamin Lindsey, Mehmet Yavuz, Mohammad Raza, Cariad Evans |
| EPI_ISL_572960, EPI_ISL_573038, EPI_ISL_573039, EPI_ISL_573040, EPI_ISL_573041, EPI_ISL_573042, EPI_ISL_573043, EPI_ISL_573044, EPI_ISL_573045, EPI_ISL_573046, EPI_ISL_573047, EPI_ISL_573048, EPI_ISL_573049, EPI_ISL_573050, EPI_ISL_573051, EPI_ISL_573052, EPI_ISL_573053, EPI_ISL_573054, EPI_ISL_573055, EPI_ISL_573056, EPI_ISL_573057, EPI_ISL_573058, EPI_ISL_573059, EPI_ISL_573060, EPI_ISL_573061, EPI_ISL_573062, EPI_ISL_573063, EPI_ISL_573064, EPI_ISL_573065, EPI_ISL_573066, EPI_ISL_573067, EPI_ISL_573068, EPI_ISL_573069, EPI_ISL_573070, EPI_ISL_573071, EPI_ISL_573072, EPI_ISL_573073, EPI_ISL_573074, EPI_ISL_573075, EPI_ISL_573076, EPI_ISL_573077, EPI_ISL_573078, EPI_ISL_573079, EPI_ISL_573080, EPI_ISL_573084, EPI_ISL_573154, EPI_ISL_573155, EPI_ISL_573156, EPI_ISL_573157, EPI_ISL_573158, EPI_ISL_573159, EPI_ISL_573160, EPI_ISL_573161, EPI_ISL_573162, EPI_ISL_573163, EPI_ISL_573164, EPI_ISL_573165, EPI_ISL_573166, EPI_ISL_573167, EPI_ISL_573168, EPI_ISL_573169, EPI_ISL_573170, EPI_ISL_573171, EPI_ISL_573172, EPI_ISL_573173, EPI_ISL_573174, EPI_ISL_573175, EPI_ISL_573176, EPI_ISL_573177, EPI_ISL_573178, EPI_ISL_573179, EPI_ISL_573180, EPI_ISL_573181, EPI_ISL_573182, EPI_ISL_573183, EPI_ISL_573184, EPI_ISL_573185, EPI_ISL_573186, EPI_ISL_573187, EPI_ISL_573188, EPI_ISL_573189, EPI_ISL_573190, EPI_ISL_573191, EPI_ISL_573192, EPI_ISL_573193, EPI_ISL_573194, EPI_ISL_573196, EPI_ISL_573197, EPI_ISL_573202, EPI_ISL_573204, EPI_ISL_573205, EPI_ISL_573209, EPI_ISL_573220, EPI_ISL_573226, EPI_ISL_573228, EPI_ISL_573229, EPI_ISL_573230, EPI_ISL_573233, EPI_ISL_573234, EPI_ISL_573235 | | | |
| see above | Oxford Viromics, NDM, University of Oxford; Oxford University Hospitals; Basingstoke and North Hampshire Hospital | COVID-19 Genomics UK (COG-UK) Consortium | Tanya Golubchik, David Bonsall, George Macintyre, Amy Trebes, Mariateresa de Cesare, Catrin Moore, Alex Mobbs, Anita Justice, Robert Shaw, Monique Andersson, Timothy Peto, Emma Wise, Nathan Moore, Jessica Lynch, Nick Cortes, Matilde Mori, Stephen Kidd, David Buck, John Todd, Christophe Fraser |
| EPI_ISL_573246 | Liverpool Clinical Laboratories | COVID-19 Genomics UK (COG-UK) Consortium | Sam Haldenby, Anita Lucaci, Steve Paterson, Julian Hiscox, Alistair Darby, M Almsaud, A Alrezaihi, Muhannad Alruwaili, Stuart D Armstrong, Jones Benjamin, Eleanor G Bentley, Anu Chawla, Jordan J Clark, Angela Cowell, Richard Eccles, Isabel García-Dorival, Matthew Gemmell, Alessandro Gerada, PKF Gilmore, Richard Gregory, Ximeng Han, Catherine Hartley, Margaret Hughes, Miren Iturriza-Gomara, James Johnson, L Luu, Jenifer Manson, Charlotte Nelson, Elaine O'Toole, Cassie Olateju, Rebekah Penrice-Randal , Lucille Rainbow, N.P Randle, Trevor Ian Robinson, Parul Sharma, Ghada T Shawli, James P Stewart, Neil Swainston, Ecaterina Vamos, Joanne Watts, Mark Whitehead |
| EPI_ISL_573247, EPI_ISL_573248, EPI_ISL_573249, EPI_ISL_573250, EPI_ISL_573251, EPI_ISL_573252 | Virology Department, Royal Infirmary of Edinburgh, NHS Lothian / School of Biological Sciences, University of Edinburgh / Institute of Genetics and Molecular Medicine, University of Edinburgh | COVID-19 Genomics UK (COG-UK) Consortium | McHugh M, Dewar R, Rooke S, Gallagher M, Balcaza C, O'Toole Á, Scher E, Hill V, McCrone JT, Colquhoun R, Yu X, Jackson B, Rambaut A, Williams TC, Templeton K |
| EPI_ISL_573253, EPI_ISL_573254, EPI_ISL_573256, EPI_ISL_573258, EPI_ISL_573259, EPI_ISL_573260, EPI_ISL_573265 | Centre for Enzyme Innovation, University of Portsmouth / Translational Research Laboratory, Portsmouth Hospitals NHS Trust | COVID-19 Genomics UK (COG-UK) Consortium | Angela Beckett,Yann Bourgeois,Garry Scarlett,Sharon Glaysher,Scott Elliott,Kelly Bicknell,Robert Impey,Allyson Lloyd,Sarah Wyllie,Ethan Butcher,Anoop Chauhan,Samuel Robson |
| EPI_ISL_573266, EPI_ISL_573267, EPI_ISL_573268, EPI_ISL_573269, EPI_ISL_573270, EPI_ISL_573271, EPI_ISL_573272, EPI_ISL_573273, EPI_ISL_573274, EPI_ISL_573275, EPI_ISL_573276, EPI_ISL_573279, EPI_ISL_573280, EPI_ISL_573281, EPI_ISL_573282, EPI_ISL_573283, EPI_ISL_573284, EPI_ISL_573285, EPI_ISL_573286, EPI_ISL_573287, EPI_ISL_573288, EPI_ISL_573289, EPI_ISL_573290, EPI_ISL_573291, EPI_ISL_573292, EPI_ISL_573294, EPI_ISL_573295, EPI_ISL_573296, EPI_ISL_573297, EPI_ISL_573299, EPI_ISL_573300, EPI_ISL_573301, EPI_ISL_573302, EPI_ISL_573303, EPI_ISL_573304, EPI_ISL_573306, EPI_ISL_573307, EPI_ISL_573308, EPI_ISL_573309, EPI_ISL_573310, EPI_ISL_573320, EPI_ISL_573321, EPI_ISL_573322, EPI_ISL_573376, EPI_ISL_573377, EPI_ISL_573378, EPI_ISL_573379, EPI_ISL_573380, EPI_ISL_573381 | | | |
| see above | Northumbria University / South Tees Hospitals NHS Foundation Trust / North Cumbria Integrated Care NHS Foundation Trust / North Tees and Hartlepool NHS Foundation Trust / Newcastle Hospitals NHS Foundation Trust | COVID-19 Genomics UK (COG-UK) Consortium | Darren L Smith,Andrew Nelson,Matthew Bashton,Greg R Young,Joshua Loh,John Allan,Mohammad A Tariq,Giles S Holt,Gary Black,Wen C Yew,Lynn Dover,Paul Baker,Steve Liggett,Sarah Essex,Jane Greenaway,Debra Padgett,Clive Graham,Garren Scott,Edward Barton,Emma Swindells,Brendan Payne,Jennifer Collins,Yusri Taha,Gary Eltringham |
| EPI_ISL_573478, EPI_ISL_573479, EPI_ISL_573480, EPI_ISL_573481, EPI_ISL_573482, EPI_ISL_573483, EPI_ISL_573484, EPI_ISL_573485, EPI_ISL_573486, EPI_ISL_573487, EPI_ISL_573488, EPI_ISL_573489, EPI_ISL_573490, EPI_ISL_573491, EPI_ISL_573492, EPI_ISL_573493, EPI_ISL_573494, EPI_ISL_573495, EPI_ISL_573496, EPI_ISL_573497, EPI_ISL_573498, EPI_ISL_573499, EPI_ISL_573500, EPI_ISL_573501, EPI_ISL_573502, EPI_ISL_573503, EPI_ISL_573504, EPI_ISL_573505, EPI_ISL_573506, EPI_ISL_573507, EPI_ISL_573508, EPI_ISL_573509, EPI_ISL_573510, EPI_ISL_573511, EPI_ISL_573513, EPI_ISL_573514, EPI_ISL_573515, EPI_ISL_573516, EPI_ISL_573517, EPI_ISL_573518, EPI_ISL_573519, EPI_ISL_573520, EPI_ISL_573521, EPI_ISL_573522, EPI_ISL_573523, EPI_ISL_573524, EPI_ISL_573525 | | | |
| see above | University College London, Great Ormond Street Hospital for Children NHS Foundation Trust, Imperial College Healthcare NHS Trust | COVID-19 Genomics UK (COG-UK) Consortium | Sergi Castellano, Rachel Williams, Mark Kristiansen, Paola Resende Silva, Sunando Roy, Tony Brooks, Helena Tutill, Paola Niola, Patricia Dyal, Charlotte Williams, Leysa Forrest, Yasmin Panchbhaya, Jacqueline Findlay, Samuel Weeks, Julianne Brown, Kathryn Harris, Paul Randell, James Price, Alison Holmes, Judith Breuer |
| EPI_ISL_573526, EPI_ISL_573527, EPI_ISL_573528, EPI_ISL_573529, EPI_ISL_573530, EPI_ISL_573531, EPI_ISL_573532, EPI_ISL_573533, EPI_ISL_573534, EPI_ISL_573535, EPI_ISL_573536, EPI_ISL_573537, EPI_ISL_573538, EPI_ISL_573539, EPI_ISL_573540, EPI_ISL_573541, EPI_ISL_573542, EPI_ISL_573543, EPI_ISL_573544, EPI_ISL_573545, EPI_ISL_573546, EPI_ISL_573547, EPI_ISL_573548, EPI_ISL_573549, EPI_ISL_573550, EPI_ISL_573551, EPI_ISL_573552, EPI_ISL_573553, EPI_ISL_573554, EPI_ISL_573555, EPI_ISL_573556, EPI_ISL_573557, EPI_ISL_573558, EPI_ISL_573559, EPI_ISL_573560, EPI_ISL_573561, EPI_ISL_573562, EPI_ISL_573563, EPI_ISL_573564, EPI_ISL_573565 | | | |
| see above | Virology Department, Sheffield Teaching Hospitals NHS Foundation Trust/Department of Infection, Immunity and Cardiovascular Disease, The Medical School, University of Sheffield | COVID-19 Genomics UK (COG-UK) Consortium | Thushan de Silva, Matthew Parker, Nikki Smith, Adri Angyal, Rebecca Brown, Luke Green, Rachel Tucker, Paul Parsons, Danielle Groves, Katie Johnson, Laura Carrilero, Alex Keeley, Dave Partridge, Matthew Wyles, Benjamin Lindsey, Mehmet Yavuz, Mohammad Raza, Cariad Evans |
| EPI_ISL_573566, EPI_ISL_573567, EPI_ISL_573568, EPI_ISL_573569, EPI_ISL_573570, EPI_ISL_573571, EPI_ISL_573572, EPI_ISL_573573, EPI_ISL_573574, EPI_ISL_573575, EPI_ISL_573576, EPI_ISL_573577, EPI_ISL_573578, EPI_ISL_573579, EPI_ISL_573580, EPI_ISL_573581, EPI_ISL_573582, EPI_ISL_573583, EPI_ISL_573584, EPI_ISL_573585, EPI_ISL_573586, EPI_ISL_573587, EPI_ISL_573588, EPI_ISL_573589, EPI_ISL_573590, EPI_ISL_573591, EPI_ISL_573592, EPI_ISL_573593, EPI_ISL_573594, EPI_ISL_573595, EPI_ISL_573596, EPI_ISL_573600, EPI_ISL_573597, EPI_ISL_573598, EPI_ISL_573599, EPI_ISL_573600, EPI_ISL_573601, EPI_ISL_573602, EPI_ISL_573603, EPI_ISL_573604, EPI_ISL_573605, EPI_ISL_573606, EPI_ISL_573607, EPI_ISL_573608, EPI_ISL_573609, EPI_ISL_573610, EPI_ISL_573611, EPI_ISL_573612, EPI_ISL_573613, EPI_ISL_573614, EPI_ISL_573615, EPI_ISL_573616, EPI_ISL_573617, EPI_ISL_573618, EPI_ISL_573619, EPI_ISL_573620, EPI_ISL_573621, EPI_ISL_573622, EPI_ISL_573623, EPI_ISL_573624, EPI_ISL_573625, EPI_ISL_573626, EPI_ISL_573627, EPI_ISL_573628, EPI_ISL_573629, EPI_ISL_573630, EPI_ISL_573631, EPI_ISL_573632, EPI_ISL_573633, EPI_ISL_573634, EPI_ISL_573635, EPI_ISL_573636, EPI_ISL_573637, EPI_ISL_573638, EPI_ISL_573639, EPI_ISL_573640, EPI_ISL_573641, EPI_ISL_573642, EPI_ISL_573643, EPI_ISL_573644, EPI_ISL_573645, EPI_ISL_573646, EPI_ISL_573647, EPI_ISL_573648, EPI_ISL_573649, EPI_ISL_573650, EPI_ISL_573651, EPI_ISL_573759 | | | |
| see above | Oxford Viromics, NDM, University of Oxford; Oxford University Hospitals; Basingstoke and North Hampshire Hospital | COVID-19 Genomics UK (COG-UK) Consortium | Tanya Golubchik, David Bonsall, George Macintyre, Amy Trebes, Mariateresa de Cesare, Catrin Moore, Alex Mobbs, Anita Justice, Robert Shaw, Monique Andersson, Timothy Peto, Emma Wise, Nathan Moore, Jessica Lynch, Nick Cortes, Matilde Mori, Stephen Kidd, David Buck, John Todd, Christophe Fraser |
| EPI_ISL_573760, EPI_ISL_573761 | Northumbria University / South Tees Hospitals NHS Foundation Trust / North Cumbria Integrated Care NHS Foundation Trust / North Tees and Hartlepool NHS Foundation Trust / Newcastle Hospitals NHS Foundation Trust | COVID-19 Genomics UK (COG-UK) Consortium | Darren L Smith,Andrew Nelson,Matthew Bashton,Greg R Young,Joshua Loh,John Allan,Mohammad A Tariq,Giles S Holt,Gary Black,Wen C Yew,Lynn Dover,Paul Baker,Steve Liggett,Sarah Essex,Jane Greenaway,Debra Padgett,Clive Graham,Garren Scott,Edward Barton,Emma Swindells,Brendan Payne,Jennifer Collins,Yusri Taha,Gary Eltringham |
| EPI_ISL_573762, EPI_ISL_573763 | Virology Department, Sheffield Teaching Hospitals NHS Foundation Trust/Department of Infection, Immunity and Cardiovascular Disease, The Medical School, University of Sheffield | COVID-19 Genomics UK (COG-UK) Consortium | Thushan de Silva, Matthew Parker, Nikki Smith, Adri Angyal, Rebecca Brown, Luke Green, Rachel Tucker, Paul Parsons, Danielle Groves, Katie Johnson, Laura Carrilero, Alex Keeley, Dave Partridge, Matthew Wyles, Benjamin Lindsey, Mehmet Yavuz, Mohammad Raza, Cariad Evans |
| EPI_ISL_573765, EPI_ISL_573766, EPI_ISL_573767, EPI_ISL_573768 | West of Scotland Specialist Virology Centre, NHSGGC / MRC-University of Glasgow Centre for Virus Research | COVID-19 Genomics UK (COG-UK) Consortium | Ana da Silva Filipe, Natasha Johnson, Kathy Smollett, Daniel Mair, Stephen Carmichael, Lily Tong, Jenna Nichols, Elihu Aranday-Cortes, Kyriaki Nomikou; Sarah McDonald, Marc Niebel, Patawee Asamaphan; Richard Orton, Joseph Hughes, Sreenu Vattipally, David L Robertson; Alasdair MacLean, Rory Gunson; Kathy Li, Igor Starinskij, Natasha Jesudason, Rajiv Shah, James Shepherd, Antonia Ho, Emma Thomson |
| EPI_ISL_573769 | Lighthouse Lab in Glasgow / MRC-University of Glasgow Centre for Virus Research | COVID-19 Genomics UK (COG-UK) Consortium | Ana da Silva Filipe, Natasha Johnson, Kathy Smollett, Daniel Mair, Stephen Carmichael, Lily Tong, Jenna Nichols, Elihu Aranday-Cortes, Kyriaki Nomikou; Sarah McDonald, Marc Niebel, Patawee Asamaphan; Harper VanSteenhouse, Yumi Kasai, David Gray, Carol Clugston, Anna Dominiczak; Alasdair MacLean, Rory Gunson; Richard Orton, Joseph Hughes, Sreenu Vattipally, David L Robertson; Sharif Shaaban, Matthew Holden; Kathy Li, Natasha Jesudason, Rajiv Shah, James Shepherd, Antonia Ho, Emma Thomson |
| EPI_ISL_573772, EPI_ISL_573775, EPI_ISL_573776, EPI_ISL_573777, EPI_ISL_573778, EPI_ISL_573779 | West of Scotland Specialist Virology Centre, NHSGGC / MRC-University of Glasgow Centre for Virus Research | COVID-19 Genomics UK (COG-UK) Consortium | Ana da Silva Filipe, Natasha Johnson, Kathy Smollett, Daniel Mair, Stephen Carmichael, Lily Tong, Jenna Nichols, Elihu Aranday-Cortes, Kyriaki Nomikou; Sarah McDonald, Marc Niebel, Patawee Asamaphan; Richard Orton, Joseph Hughes, Sreenu Vattipally, David L Robertson; Alasdair MacLean, Rory Gunson; Kathy Li, Igor Starinskij, Natasha Jesudason, Rajiv Shah, James Shepherd, Antonia Ho, Emma Thomson |
| EPI_ISL_573799 | Northumbria University / South Tees Hospitals NHS | COVID-19 Genomics UK (COG-UK) Consortium | Darren L Smith,Andrew Nelson,Matthew Bashton,Greg R Young,Joshua Loh,John Allan,Mohammad A Tariq,Giles S Holt,Gary Black,Wen C Yew,Lynn |

| | | | |
|---|---|---|---|
| | Foundation Trust / North Cumbria Integrated Care NHS Foundation Trust / North Tees and Hartlepool NHS Foundation Trust / Newcastle Hospitals NHS Foundation Trust | | Dover,Paul Baker,Steve Liggett,Sarah Essex,Jane Greenaway,Debra Padgett,Clive Graham,Garren Scott,Edward Barton,Emma Swindells,Brendan Payne,Jennifer Collins,Yusri Taha,Gary Eltringham |
| EPI_ISL_573800, EPI_ISL_573801, EPI_ISL_573802, EPI_ISL_573803, EPI_ISL_573804, EPI_ISL_573805 | University College London, Great Ormond Street Hospital for Children NHS Foundation Trust, Imperial College Healthcare NHS Trust | COVID-19 Genomics UK (COG-UK) Consortium | Sergi Castellano, Rachel Williams, Mark Kristiansen, Paola Resende Silva, Sunando Roy, Tony Brooks, Helena Tutill, Paola Niola, Patricia Dyal, Charlotte Williams, Leysa Forrest, Yasmin Panchbhaya, Jacqueline Findlay, Samuel Weeks, Julianne Brown, Kathryn Harris, Paul Randell, James Price, Alison Holmes, Judith Breuer |
| EPI_ISL_573806, EPI_ISL_573807, EPI_ISL_573808, EPI_ISL_573809, EPI_ISL_573810 | Virology Department, Sheffield Teaching Hospitals NHS Foundation Trust/Department of Infection, Immunity and Cardiovascular Disease, The Medical School, University of Sheffield | COVID-19 Genomics UK (COG-UK) Consortium | Thushan de Silva, Matthew Parker, Nikki Smith, Adri Angyal, Rebecca Brown, Luke Green, Rachel Tucker, Paul Parsons, Danielle Groves, Katie Johnson, Laura Carrilero, Alex Keeley, Dave Partridge, Matthew Wyles, Benjamin Lindsey, Mehmet Yavuz, Mohammad Raza, Cariad Evans |
| EPI_ISL_573811, EPI_ISL_573812, EPI_ISL_573813, EPI_ISL_573814, EPI_ISL_573815, EPI_ISL_573816, EPI_ISL_573817, EPI_ISL_573818, EPI_ISL_573819, EPI_ISL_573820, EPI_ISL_573821, EPI_ISL_573822, EPI_ISL_573823, EPI_ISL_573824, EPI_ISL_573825, EPI_ISL_573826, EPI_ISL_573827, EPI_ISL_573828, EPI_ISL_573829, EPI_ISL_573830, EPI_ISL_573831, EPI_ISL_573832, EPI_ISL_573833, EPI_ISL_573834, EPI_ISL_573835, EPI_ISL_573836, EPI_ISL_573837, EPI_ISL_573838, EPI_ISL_573839, EPI_ISL_573840, EPI_ISL_573841, EPI_ISL_573842, EPI_ISL_573843, EPI_ISL_573844, EPI_ISL_573845, EPI_ISL_573846, EPI_ISL_573847, EPI_ISL_573848, EPI_ISL_573849, EPI_ISL_573850, EPI_ISL_573851, EPI_ISL_573852, EPI_ISL_573853, EPI_ISL_573854, EPI_ISL_573855, EPI_ISL_573866, EPI_ISL_573867, EPI_ISL_573868 | | | |
| see above | Oxford Viromics, NDM, University of Oxford; Oxford University Hospitals; Basingstoke and North Hampshire Hospital | COVID-19 Genomics UK (COG-UK) Consortium | Tanya Golubchik, David Bonsall, George Macintyre, Amy Trebes, Mariateresa de Cesare, Catrin Moore, Alex Mobbs, Anita Justice, Robert Shaw, Monique Andersson, Timothy Peto, Emma Wise, Nathan Moore, Jessica Lynch, Nick Cortes, Matilde Mori, Stephen Kidd, David Buck, John Todd, Christophe Fraser |
| EPI_ISL_573872 | Northumbria University / South Tees Hospitals NHS Foundation Trust / North Cumbria Integrated Care NHS Foundation Trust / North Tees and Hartlepool NHS Foundation Trust / Newcastle Hospitals NHS Foundation Trust | COVID-19 Genomics UK (COG-UK) Consortium | Darren L Smith,Andrew Nelson,Matthew Bashton,Greg R Young,Joshua Loh,John Allan,Mohammad A Tariq,Giles S Holt,Gary Black,Wen C Yew,Lynn Dover,Paul Baker,Steve Liggett,Sarah Essex,Jane Greenaway,Debra Padgett,Clive Graham,Garren Scott,Edward Barton,Emma Swindells,Brendan Payne,Jennifer Collins,Yusri Taha,Gary Eltringham |
| EPI_ISL_573947, EPI_ISL_573977, EPI_ISL_573993, EPI_ISL_574015, EPI_ISL_574038, EPI_ISL_574053, EPI_ISL_574059, EPI_ISL_574132, EPI_ISL_574175 | Wales Specialist Virology Centre Sequencing lab: Pathogen Genomics Unit | COVID-19 Genomics UK (COG-UK) Consortium | Catherine Moore, Johnathan Evans, Laura Gifford, Malorie Perry, Simon Cottrell, Angela Marchbank, Alec Birchley, Alexander Adams, Amy Gaskin, Bree Gatica-Wilcox, Jason Coombes, Joel Southgate, Lauren Gilbert, Lee Graham, Nicole Pacchiarini, Sara Kumziene-Summerhayes, Sarah Taylor, Sophie Jones, Sara Rey, Matthew Bull, Joanne Watkins, Sally Corden, Tom Connor |
| EPI_ISL_574257 | Oxford Viromics, NDM, University of Oxford; Oxford University Hospitals; Basingstoke and North Hampshire Hospital | COVID-19 Genomics UK (COG-UK) Consortium | Tanya Golubchik, David Bonsall, George Macintyre, Amy Trebes, Mariateresa de Cesare, Catrin Moore, Alex Mobbs, Anita Justice, Robert Shaw, Monique Andersson, Timothy Peto, Emma Wise, Nathan Moore, Jessica Lynch, Nick Cortes, Matilde Mori, Stephen Kidd, David Buck, John Todd, Christophe Fraser |
| EPI_ISL_574652, EPI_ISL_574653, EPI_ISL_574654, EPI_ISL_574655, EPI_ISL_574656, EPI_ISL_574657, EPI_ISL_574658, EPI_ISL_574659, EPI_ISL_574660, EPI_ISL_574661, EPI_ISL_574662 | | | |
| see above | Seattle Flu Study | Seattle Flu Study | Deborah A. Nickerson, Chris D. Frazar, Jover Lee, Benjamin Pelle, Matthew Richardson, Amanda Adler, Elisabeth Brandstetter, Peter D. Han, Kairsten Fay, Misja Ilcisin, Kirsten Lacombe, Thomas R. Sibley, Melissa Truong, Caitlin R. Wolf, Karen Cowgill, Stephanie Schrag, Jeff Duchin, Michael Boeckh, Janet A. Englund, Michael Famulare, Barry R. Lutz, Mark J. Rieder, Lea M. Starita, Matthew Thompson, Helen Y. Chu, Trevor Bedford, Jay Shendure |
| EPI_ISL_574663, EPI_ISL_574664, EPI_ISL_574665, EPI_ISL_574666, EPI_ISL_574667 | Seattle Flu Study | Seattle Flu Study | Deborah A. Nickerson, Chris D. Frazar, Jover Lee, Benjamin Pelle, Matthew Richardson, Amanda Adler, Elisabeth Brandstetter, Peter D. Han, Kairsten Fay, Misja Ilcisin, Kirsten Lacombe, Thomas R. Sibley, Melissa Truong, Caitlin R. Wolf, Michael Boeckh, Janet A. Englund, Michael Famulare, Barry R. Lutz, Mark J. Rieder, Lea M. Starita, Matthew Thompson, Jay Shendure, Trevor Bedford, Helen Y. Chu |
| EPI_ISL_574668, EPI_ISL_574669, EPI_ISL_574670, EPI_ISL_574671, EPI_ISL_574672, EPI_ISL_574673, EPI_ISL_574674, EPI_ISL_574675, EPI_ISL_575028, EPI_ISL_575030, EPI_ISL_575032 | | | |
| see above | Seattle Flu Study | Seattle Flu Study | Deborah A. Nickerson, Chris D. Frazar, Jover Lee, Benjamin Pelle, Matthew Richardson, Amanda Adler, Elisabeth Brandstetter, Peter D. Han, Kairsten Fay, Misja Ilcisin, Kirsten Lacombe, Thomas R. Sibley, Melissa Truong, Caitlin R. Wolf, Karen Cowgill, Stephanie Schrag, Jeff Duchin, Michael Boeckh, Janet A. Englund, Michael Famulare, Barry R. Lutz, Mark J. Rieder, Lea M. Starita, Matthew Thompson, Helen Y. Chu, Trevor Bedford, Jay Shendure |
| EPI_ISL_575289, EPI_ISL_575290, EPI_ISL_575291, EPI_ISL_575292, EPI_ISL_575293, EPI_ISL_575294, EPI_ISL_575295, EPI_ISL_575296, EPI_ISL_575297, EPI_ISL_575298, EPI_ISL_575299, EPI_ISL_575300, EPI_ISL_575301, EPI_ISL_575302, EPI_ISL_575303, EPI_ISL_575304, EPI_ISL_575305, EPI_ISL_575306, EPI_ISL_575307, EPI_ISL_575308, EPI_ISL_575309, EPI_ISL_575310, EPI_ISL_575311, EPI_ISL_575314, EPI_ISL_575322 | | | |
| see above | Molecular Biology, New Mexico Department of Health Scientific Laboratory | Molecular Biology, New Mexico Department of Health Scientific Laboratory | Johnson,E.J., Griego-Fisher,A.M., Malone,D. |
| EPI_ISL_575332, EPI_ISL_575333, EPI_ISL_575334 | Israel Central Virology laboratory | Israel Central Virology laboratory | Neta Zuckerman, Efrat Dahan Bucris, Oran Erster, Ella Mendelson, Michal Mandelboim |
| EPI_ISL_575986, EPI_ISL_575987, EPI_ISL_575988, EPI_ISL_575989, EPI_ISL_575990, EPI_ISL_575991, EPI_ISL_575992, EPI_ISL_575993, EPI_ISL_575994 | Lighthouse Lab in Milton Keynes | Wellcome Sanger Institute for the COVID-19 Genomics UK (COG-UK) consortium | The Lighthouse Lab in Milton Keynes and Alex Alderton, Roberto Amato, Sonia Goncalves, Ewan Harrison, David K. Jackson, Ian Johnston, Dominic Kwiatkowski, Cordelia Langford, John Sillitoe on behalf of the Wellcome Sanger Institute COVID-19 Surveillance Team (http://www.sanger.ac.uk/covid-team) |
| EPI_ISL_576053, EPI_ISL_576076, EPI_ISL_576082 | Lighthouse Lab in Glasgow | Wellcome Sanger Institute for the COVID-19 Genomics UK (COG-UK) consortium | Harper VanSteenhouse, Yumi Kasai, David Gray, Carol Clugston, Anna Dominiczak and Alex Alderton, Roberto Amato, Sonia Goncalves, Ewan Harrison, David K. Jackson, Ian Johnston, Dominic Kwiatkowski, Cordelia Langford, John Sillitoe on behalf of the Wellcome Sanger Institute COVID-19 Surveillance Team (http://www.sanger.ac.uk/covid-team) |
| EPI_ISL_576118, EPI_ISL_576120, EPI_ISL_576122, EPI_ISL_576123, EPI_ISL_576124, EPI_ISL_576125, EPI_ISL_576126 | Laboratory, The Bio Arte Limited | Laboratory, The Bio Arte Limited | Biazzo,M., Madeddu,S., Santoro,F., Pinzauti,D. |
| EPI_ISL_576132, EPI_ISL_576133, EPI_ISL_576134, EPI_ISL_576135, EPI_ISL_576136, EPI_ISL_576137, EPI_ISL_576138, EPI_ISL_576139, EPI_ISL_576140, EPI_ISL_576141 | unknown | Public Health Virology Laboratory, Forensic and Scientific Services (PHV-FSS) | Son Nguyen et al. |
| EPI_ISL_576173 | FL Department of Health-Bureau of Epidemiology | Pathogen Discovery, Respiratory Viruses Branch, Division of Viral Diseases, Centers for Disease Control and Prevention | Ying Tao, Jing Zhang, Brian Lynch, Yan Li, Krista Queen, Anna Uehara, Clinton R. Paden, Peter Cook, Haibin Wang, Suxiang Tong |
| EPI_ISL_576347, EPI_ISL_576348, EPI_ISL_576349, EPI_ISL_576350, EPI_ISL_576351, EPI_ISL_576352, EPI_ISL_576353, EPI_ISL_576354, EPI_ISL_576355, EPI_ISL_576356, EPI_ISL_576357 | | | |
| see above | Texas Department of State Health Services | Texas Department of State Health Services | Rashmi Tuladhar, Bonnie Oh, Mayela Pedrueza, Jenny Zhang, Maliha Rahman, Anita Pokharel, Myong Koag, Chun Wang, Rachel Lee, Grace Kubin |
| EPI_ISL_576383 | RSUD Budi Rahayu Kota Magelang | Genetics Working Group (Pokja Genetik) Faculty of Medicine, Public Health and Nursing Universitas Gadjah Mada (FK-KMK UGM); Disease Investigation Center Wates Ministry of Agriculture Indonesia; Department of Microbiology FK-KMK UGM; Laboratorium Diagnostik Yayasan Tahija World Mosquito Program (WMP) Yogyakarta Center for Tropical Medicine FK-KMK UGM; Integrated Research Center FK-KMK UGM; Department of Computer Science and Electronics FMIPA UGM; Balai Besar Teknik Kesehatan Lingkungan dan Pengendalian Penyakit (BBTKLPP) Yogyakarta | Gunadi, Hendra Wibawa, Marcellus, Mohamad S. Hakim, Edwin W. Daniwijaya, Ludhang P. Rizki, Endah Supriyati, Eggi Arguni, Titik Nuryastuti, Tri Wibawa, Dwi AA Nugrahaningsih, Afiahayati, Siswanto, Kristy Iskandar, Nungki Anggorowati, Irene, Indaryati, Havid Setyawan, Ari Meliyanti, Merliana Sari Situmeang, Audric Kenny Tedja, Aditya Rifqi Fauzi |
| EPI_ISL_576882, EPI_ISL_576883, EPI_ISL_576884, EPI_ISL_576885, EPI_ISL_576886, EPI_ISL_576887, EPI_ISL_576888, EPI_ISL_576889, EPI_ISL_576890, EPI_ISL_576891, EPI_ISL_576892, EPI_ISL_576893, EPI_ISL_576894, EPI_ISL_576895, EPI_ISL_576896 | | | |

| | | | |
|---|---|---|---|
| see above | University of Birmingham | COVID-19 Genomics UK (COG-UK) Consortium | Institute of Microbiology, University of Birmingham: Claire McMurray, Joanne Stockton, Samuel Nicholls, Radoslaw Poplawski, Will Rowe, Josh Quick, Nicholas Loman. University of Birmingham Testing Laboratory: Celina M Whalley, Andrew Bosworth, Charlotte Poxon, Kasun Wanigasooriya, Oliver Pickles, Mike Kidd, Alex Richter, Andrew D Beggs PHE Heartlands Lab: Husam Osman, Andrew Bosworth. Queen Elizabeth Hospital: Anna Casey |
| EPI_ISL_577212, EPI_ISL_577213, EPI_ISL_577214, EPI_ISL_577215, EPI_ISL_577217, EPI_ISL_577218, EPI_ISL_577219 | Centre for Enzyme Innovation, University of Portsmouth / Translational Research Laboratory, Portsmouth Hospitals NHS Trust | COVID-19 Genomics UK (COG-UK) Consortium | Angela Beckett,Yann Bourgeois,Garry Scarlett,Sharon Glaysher,Scott Elliott,Kelly Bicknell,Robert Impey,Allyson Lloyd,Sarah Wyllie,Ethan Butcher,Anoop Chauhan,Samuel Robson |
| EPI_ISL_577220, EPI_ISL_577221, EPI_ISL_577222, EPI_ISL_577223, EPI_ISL_577224, EPI_ISL_577225, EPI_ISL_577226, EPI_ISL_577227, EPI_ISL_577228, EPI_ISL_577229, EPI_ISL_577230, EPI_ISL_577231, EPI_ISL_577232, EPI_ISL_577233, EPI_ISL_577234, EPI_ISL_577235, EPI_ISL_577236, EPI_ISL_577237, EPI_ISL_577238, EPI_ISL_577239, EPI_ISL_577240, EPI_ISL_577241, EPI_ISL_577242, EPI_ISL_577243, EPI_ISL_577244, EPI_ISL_577245, EPI_ISL_577246, EPI_ISL_577247, EPI_ISL_577248, EPI_ISL_577249, EPI_ISL_577250, EPI_ISL_577251, EPI_ISL_577252, EPI_ISL_577253, EPI_ISL_577254, EPI_ISL_577255, EPI_ISL_577256, EPI_ISL_577257, EPI_ISL_577258, EPI_ISL_577259, EPI_ISL_577260, EPI_ISL_577261, EPI_ISL_577262, EPI_ISL_577263, EPI_ISL_577264, EPI_ISL_577265, EPI_ISL_577266, EPI_ISL_577267, EPI_ISL_577268, EPI_ISL_577269, EPI_ISL_577270, EPI_ISL_577271, EPI_ISL_577272, EPI_ISL_577298, EPI_ISL_577346 | | | |
| see above | University of Exeter | COVID-19 Genomics UK (COG-UK) Consortium | Ben Temperton,Aaron Jeffries,Michelle Michelsen,Joanna Warwick-Dugdale,Audrey Farbos,Robyn Manley,Stephen Michell,Jane Masoli |
| EPI_ISL_577349 | Virology Department, Royal Infirmary of Edinburgh, NHS Lothian / School of Biological Sciences, University of Edinburgh / Institute of Genetics and Molecular Medicine, University of Edinburgh | COVID-19 Genomics UK (COG-UK) Consortium | McHugh M, Dewar R, Rooke S, Gallagher M, Balcaza C, O'Toole Á, Scher E, Hill V, McCrone JT, Colquhoun R, Yu X, Jackson B, Rambaut A, Williams TC, Templeton K |
| EPI_ISL_577359, EPI_ISL_577360, EPI_ISL_577361, EPI_ISL_577362, EPI_ISL_577363, EPI_ISL_577364, EPI_ISL_577365, EPI_ISL_577366, EPI_ISL_577367, EPI_ISL_577368, EPI_ISL_577369, EPI_ISL_577370, EPI_ISL_577371, EPI_ISL_577372, EPI_ISL_577373, EPI_ISL_577544 | | | |
| see above | University of Exeter | COVID-19 Genomics UK (COG-UK) Consortium | Ben Temperton,Aaron Jeffries,Michelle Michelsen,Joanna Warwick-Dugdale,Audrey Farbos,Robyn Manley,Stephen Michell,Jane Masoli |
| EPI_ISL_577549, EPI_ISL_577552, EPI_ISL_577553, EPI_ISL_577555, EPI_ISL_577561, EPI_ISL_577562, EPI_ISL_577564, EPI_ISL_577572, EPI_ISL_577574, EPI_ISL_577575, EPI_ISL_577576, EPI_ISL_577577, EPI_ISL_577578, EPI_ISL_577579, EPI_ISL_577580, EPI_ISL_577581, EPI_ISL_577582, EPI_ISL_577583, EPI_ISL_577584, EPI_ISL_577588, EPI_ISL_577589, EPI_ISL_577590, EPI_ISL_577594 | | | |
| see above | Michigan Department of Health and Human Services, Bureau of Laboratories | Michigan Department of Health and Human Services, Bureau of Laboratories | Blankenship HM, Riner D, Soehnlen MK |
| EPI_ISL_577639, EPI_ISL_577672, EPI_ISL_577673, EPI_ISL_577727, EPI_ISL_577728 | NIV Influenza | NIV Influenza | Potdar V |
| EPI_ISL_577796, EPI_ISL_577797, EPI_ISL_577798, EPI_ISL_577799, EPI_ISL_577800, EPI_ISL_577801, EPI_ISL_577802, EPI_ISL_577803, EPI_ISL_577805, EPI_ISL_577807, EPI_ISL_577808, EPI_ISL_577809, EPI_ISL_577810, EPI_ISL_577811, EPI_ISL_577812, EPI_ISL_577813, EPI_ISL_577814, EPI_ISL_577815 | | | |
| see above | Dutch COVID-19 response team | Erasmus Medical Center | OH consortium |
| EPI_ISL_577944, EPI_ISL_577950, EPI_ISL_577951, EPI_ISL_577953, EPI_ISL_577954, EPI_ISL_577980, EPI_ISL_577981, EPI_ISL_577982, EPI_ISL_578042 | Dutch COVID-19 response team | Erasmus Medical Center | Bas Oude Munnink, Reina Sikkema, David Nieuwenhuijse, Irina Chestakova, Anne van der Linden, Marjan Boter, Emmanuelle Munger, Corine GeurtsvanKessel, Annemiek van der Eijk, Richard Molenkamp, Marion Koopmans, on behalf of the Dutch national COVID-19 response team. |
| EPI_ISL_578203, EPI_ISL_578204, EPI_ISL_578206, EPI_ISL_578207, EPI_ISL_578208, EPI_ISL_578209, EPI_ISL_578210, EPI_ISL_578211, EPI_ISL_578212, EPI_ISL_578213, EPI_ISL_578217, EPI_ISL_578220, EPI_ISL_578226, EPI_ISL_578228, EPI_ISL_578233, EPI_ISL_578234, EPI_ISL_578236, EPI_ISL_578241, EPI_ISL_578245, EPI_ISL_578248, EPI_ISL_578251, EPI_ISL_578252, EPI_ISL_578253, EPI_ISL_578260, EPI_ISL_578262, EPI_ISL_578264, EPI_ISL_578265, EPI_ISL_578272, EPI_ISL_578275, EPI_ISL_578278, EPI_ISL_578279, EPI_ISL_578295, EPI_ISL_578297, EPI_ISL_578298, EPI_ISL_578308, EPI_ISL_578309, EPI_ISL_578312, EPI_ISL_578313, EPI_ISL_578315, EPI_ISL_578316, EPI_ISL_578318, EPI_ISL_578328, EPI_ISL_578329, EPI_ISL_578330, EPI_ISL_578331, EPI_ISL_578332 | | | |
| see above | National Virus Reference Laboratory | National Virus Reference Laboratory | Michael Carr, Gabriel Gonzalez, Jonathan Dean, Suzie Coughlan, Cillian F De Gascun |
| EPI_ISL_579092 | Canterbury Health Laboratories | Institute of Environmental Science and Research (ESR) | Xiaoyun Ren, Matt Storey, Nikki Freed, Muhammad Faisal, Jing Wang, Hermes Perez, Anja Werno, Antje van der Linden, Arlo Upton, Chris Mansell, David Hammer, Dragana Drinkovic, Gary McAuliffe, Hana Sofia Andersson, James Ussher, Jill Sherwood, Josh Freeman, Julia Howard, Juliet Elvy, Mary DeAlmeida, Matt Blakiston, Matthew Rogers, Max Bloomfield, Michael Addidle, Michelle Balm, Sally Roberts, Sarah Jefferies, Sharmini Muttaiyah, Susan Morpeth, Susan Taylor, Timothy Blackmore, Vani Sathyendran, Veronica Playle, Virginia Hope, Erasmus Smit, Lauren Jelly, Olin Silander, Joep de Ligt |
| EPI_ISL_579096 | North Shore Hospital | Institute of Environmental Science and Research (ESR) | Xiaoyun Ren, Matt Storey, Nikki Freed, Muhammad Faisal, Jing Wang, Hermes Perez, Anja Werno, Antje van der Linden, Arlo Upton, Chris Mansell, David Hammer, Dragana Drinkovic, Gary McAuliffe, Hana Sofia Andersson, James Ussher, Jill Sherwood, Josh Freeman, Julia Howard, Juliet Elvy, Mary DeAlmeida, Matt Blakiston, Matthew Rogers, Max Bloomfield, Michael Addidle, Michelle Balm, Sally Roberts, Sarah Jefferies, Sharmini Muttaiyah, Susan Morpeth, Susan Taylor, Timothy Blackmore, Vani Sathyendran, Veronica Playle, Virginia Hope, Erasmus Smit, Lauren Jelly, Olin Silander, Joep de Ligt |
| EPI_ISL_579097 | LabPLUS | Institute of Environmental Science and Research (ESR) | Xiaoyun Ren, Matt Storey, Nikki Freed, Muhammad Faisal, Jing Wang, Hermes Perez, Anja Werno, Antje van der Linden, Arlo Upton, Chris Mansell, David Hammer, Dragana Drinkovic, Gary McAuliffe, Hana Sofia Andersson, James Ussher, Jill Sherwood, Josh Freeman, Julia Howard, Juliet Elvy, Mary DeAlmeida, Matt Blakiston, Matthew Rogers, Max Bloomfield, Michael Addidle, Michelle Balm, Sally Roberts, Sarah Jefferies, Sharmini Muttaiyah, Susan Morpeth, Susan Taylor, Timothy Blackmore, Vani Sathyendran, Veronica Playle, Virginia Hope, Erasmus Smit, Lauren Jelly, Olin Silander, Joep de Ligt |
| EPI_ISL_581381 | Lighthouse Lab in Glasgow | Wellcome Sanger Institute for the COVID-19 Genomics UK (COG-UK) consortium | Harper VanSteenhouse, Yumi Kasai, David Gray, Carol Clugston, Anna Dominiczak and Alex Alderton, Roberto Amato, Sonia Goncalves, Ewan Harrison, David K. Jackson, Ian Johnston, Dominic Kwiatkowski, Cordelia Langford, John Sillitoe on behalf of the Wellcome Sanger Institute COVID-19 Surveillance Team |
| EPI_ISL_581383 | Lighthouse Lab in Milton Keynes | Wellcome Sanger Institute for the COVID-19 Genomics UK (COG-UK) consortium | The Lighthouse Lab in Milton Keynes and Alex Alderton, Roberto Amato, Sonia Goncalves, Ewan Harrison, David K. Jackson, Ian Johnston, Dominic Kwiatkowski, Cordelia Langford, John Sillitoe on behalf of the Wellcome Sanger Institute COVID-19 Surveillance Team |
| EPI_ISL_581999 | University Hospital Basel, Clinical Virology | University Hospital Basel, Clinical Bacteriology | Madlen Stange, Alfredo Mari, Tim Roloff, Helena MB Seth-Smith, Michael Schweitzer, Myrta Brunner, Karoline Leuzinger, Kirstine K. Soegaard, Alexander Gensch, Sarah Tschudin-Sutter, Simon Fuchs, Julia Bielicki, Hans Pargger, Martin Siegemund, Christian Nickel, Roland Bingisser, Michael Osthoff, Stefano Bassetti, Rita Schneider-Sliwa, Manuel Battegay, Hans Hirsch, Adrian Egli |
| EPI_ISL_582031, EPI_ISL_582032 | Institute of Human Genetics, Polish Academy of Sciences | Institute of Human Genetics, Polish Academy of Sciences | Szymon Hryhorowicz, Adam Ustaszewski, Marta Kaczmarek-Ry, Emilia Lis, Ewa Zitkiewicz, Micha Witt, Andrzej Pawski |
| EPI_ISL_582036, EPI_ISL_582037, EPI_ISL_582039, EPI_ISL_582043, EPI_ISL_582046, EPI_ISL_582047, EPI_ISL_582048, EPI_ISL_582049, EPI_ISL_582054, EPI_ISL_582055, EPI_ISL_582056, EPI_ISL_582057, EPI_ISL_582058, EPI_ISL_582059, EPI_ISL_582060, EPI_ISL_582061, EPI_ISL_582062, EPI_ISL_582063, EPI_ISL_582064, EPI_ISL_582065, EPI_ISL_582066, EPI_ISL_582067, EPI_ISL_582068, EPI_ISL_582069, EPI_ISL_582070, EPI_ISL_582071 | | | |
| see above | Servicio de Microbiología. Hospital Universitario Donostia. OSI Donostialdea. Área de Enfermedades Infecciosas, Grupo de Infección Respiratoria y Resistencia Antimicrobiana. Instituto de Investigación Sanitaria Biodonostia | SeqCOVID-SPAIN consortium/IBV(CSIC) | Gustavo Cilla, Milagrosa Montes, Luis Piñeiro, Jose Maria Marimón and SeqCOVID-SPAIN consortium |
| EPI_ISL_582091, EPI_ISL_582092, EPI_ISL_582093, EPI_ISL_582094, EPI_ISL_582095, EPI_ISL_582096, EPI_ISL_582100, EPI_ISL_582101, EPI_ISL_582102, EPI_ISL_582103, EPI_ISL_582104, EPI_ISL_582105, EPI_ISL_582106, EPI_ISL_582107, EPI_ISL_582108 | | | |
| see above | Hospital Universitario Marqués de Valdecilla - IDIVAL (Santander, Cantabria) | SeqCOVID-SPAIN consortium/IBV(CSIC) | María Eliecer Cano García, Mónica Gozalo Margüello, Jose Manuel Méndez Legaza, Daniel Pablo Marcos, Jesús Rodríguez Rodríguez, María Siller Ruiz and SeqCOVID-SPAIN consortium |
| EPI_ISL_582110, EPI_ISL_582112, EPI_ISL_582117, EPI_ISL_582118, EPI_ISL_582119, EPI_ISL_582120 | CNR Virus des Infections Respiratoires - France SUD | CNR Virus des Infections Respiratoires - France SUD | Antonin Bal, Gregory Destras, Gwendolyne Burfin, Hadrien Règue, Alexandre Gaymard, Maude Bouscambert-Duchamp, Florence Morfin-Sherpa, Martine Valette, Bruno Lina, Laurence Josset |
| EPI_ISL_582658, EPI_ISL_582659, EPI_ISL_582660, EPI_ISL_582661, EPI_ISL_582662, EPI_ISL_582663 | Sheikh Khalifa Medical City | Molecular/Surveillance lab Sheikh Khalifa Medical City | Amirtharaj Francis, Sajeed Abdul, Hala Imambaccus, Sahar Almarzooqi, Hiba Saud, Stefan Weber |
| EPI_ISL_582832 | Hospital General Universitario Gregorio Marañón | SeqCOVID-SPAIN consortium/IBV(CSIC) | Darío García de Viedma, Laura Pérez-Lago, Marta Herranz, Jon Sicilia, Julia Suárez, Pilar Catalán, Patricia Muñoz and SeqCOVID-SPAIN consortium |

| | | | |
|---|---|---|---|
| EPI_ISL_582840 | Narhalsan Fjallbacka VC | The Public Health Agency of Sweden | Anna-Malin Linde, Maria Lind Karlberg, Mattias Haukland, Reza Advani, Olov Svartstrom, Oskar Karlsson Lindsjo, Sandra Broddesson, Petra Edquist, Mia Brytting, Anna Risberg, Karin Tegmark-Wisell |
| EPI_ISL_582897, EPI_ISL_582898, EPI_ISL_582899, EPI_ISL_582900, EPI_ISL_582901, EPI_ISL_582902, EPI_ISL_582903, EPI_ISL_582904, EPI_ISL_582905, EPI_ISL_582906, EPI_ISL_582907, EPI_ISL_582908, EPI_ISL_582909, EPI_ISL_582910, EPI_ISL_582911, EPI_ISL_582912, EPI_ISL_582913, EPI_ISL_582914, EPI_ISL_582915, EPI_ISL_582916, EPI_ISL_582917, EPI_ISL_582918, EPI_ISL_582919, EPI_ISL_582922, EPI_ISL_582923, EPI_ISL_582927, EPI_ISL_582928, EPI_ISL_582930, EPI_ISL_582931, EPI_ISL_582932, EPI_ISL_582933, EPI_ISL_582934, EPI_ISL_582935, EPI_ISL_582936, EPI_ISL_582937, EPI_ISL_582952, EPI_ISL_582953 | | | |
| see above | County of Santa Clara Public Health Department | Chan-Zuckerberg Biohub | CZB Cliahub Consortium |
| EPI_ISL_583011, EPI_ISL_583012, EPI_ISL_583013, EPI_ISL_583014, EPI_ISL_583015, EPI_ISL_583016, EPI_ISL_583017, EPI_ISL_583018, EPI_ISL_583019, EPI_ISL_583020, EPI_ISL_583021, EPI_ISL_583022, EPI_ISL_583023, EPI_ISL_583024, EPI_ISL_583025, EPI_ISL_583026 | | | |
| see above | Orange County Public Health Lab | Chan-Zuckerberg Biohub | CZB Cliahub Consortium |
| EPI_ISL_583161, EPI_ISL_583162, EPI_ISL_583163, EPI_ISL_583164, EPI_ISL_583165, EPI_ISL_583166, EPI_ISL_583167, EPI_ISL_583168, EPI_ISL_583169, EPI_ISL_583170 | Humboldt County Public Health Laboratory | Chan-Zuckerberg Biohub | CZB Cliahub Consortium |
| EPI_ISL_583190, EPI_ISL_583191, EPI_ISL_583192, EPI_ISL_583193, EPI_ISL_583194, EPI_ISL_583195, EPI_ISL_583196 | San Francisco Public Health Laboratory | Chan-Zuckerberg Biohub | CZB Cliahub Consortium |
| EPI_ISL_583198 | Marin County Public Health Department | Chan-Zuckerberg Biohub | CZB Cliahub Consortium |
| EPI_ISL_583204, EPI_ISL_583205, EPI_ISL_583206, EPI_ISL_583207, EPI_ISL_583229, EPI_ISL_583230, EPI_ISL_583231 | UCSF Clinical Microbiology Laboratory | Chan-Zuckerberg Biohub | CZB Cliahub Consortium |
| EPI_ISL_583290, EPI_ISL_583292, EPI_ISL_583293, EPI_ISL_583297, EPI_ISL_583300, EPI_ISL_583301, EPI_ISL_583303, EPI_ISL_583304, EPI_ISL_583306, EPI_ISL_583307, EPI_ISL_583310, EPI_ISL_583312, EPI_ISL_583313, EPI_ISL_583316, EPI_ISL_583317, EPI_ISL_583318, EPI_ISL_583319, EPI_ISL_583320, EPI_ISL_583321, EPI_ISL_583322, EPI_ISL_583323, EPI_ISL_583324, EPI_ISL_583325, EPI_ISL_583326, EPI_ISL_583327, EPI_ISL_583328, EPI_ISL_583330, EPI_ISL_583332, EPI_ISL_583333, EPI_ISL_583334, EPI_ISL_583335, EPI_ISL_583336, EPI_ISL_583338, EPI_ISL_583339, EPI_ISL_583340, EPI_ISL_583341, EPI_ISL_583342, EPI_ISL_583343, EPI_ISL_583345, EPI_ISL_583346 | | | |
| see above | University of Michigan Clinical Microbiology Laboratory | Lauring Lab, University of Michigan, Department of Microbiology and Immunology | Valesano |
| EPI_ISL_584081, EPI_ISL_584082 | The National Institute of Public Health | State Veterinary Institute Prague | Nagy,A;Jirincova,H;Novakova,L;Trnka,D;Vecerova,J |
| EPI_ISL_584096 | University of Michigan Clinical Microbiology Laboratory | Lauring Lab, University of Michigan, Department of Microbiology and Immunology | Valesano |
| EPI_ISL_584674 | University College London, Great Ormond Street Hospital for Children NHS Foundation Trust, Imperial College Healthcare NHS Trust | COVID-19 Genomics UK (COG-UK) Consortium | Sergi Castellano, Rachel Williams, Mark Kristiansen, Paola Resende Silva, Sunando Roy, Tony Brooks, Helena Tutill, Paola Niola, Patricia Dyal, Charlotte Williams, Leysa Forrest, Yasmin Panchbhaya, Jacqueline Findlay, Samuel Weeks, Julianne Brown, Kathryn Harris, Paul Randell, James Price, Alison Holmes, Judith Breuer |
| EPI_ISL_585185, EPI_ISL_585186, EPI_ISL_585188, EPI_ISL_585189, EPI_ISL_585190, EPI_ISL_585195, EPI_ISL_585201, EPI_ISL_585202, EPI_ISL_585203, EPI_ISL_585210, EPI_ISL_585212, EPI_ISL_585226, EPI_ISL_585243, EPI_ISL_585249, EPI_ISL_585251, EPI_ISL_585253 | | | |
| see above | Regional Virus Laboratory, Belfast Health and Social Care Trust | COVID-19 Genomics UK (COG-UK) Consortium | Conall McCaughey, James McKenna, Tanya Curran, Susan Feeney, Alison Watt, Ciara Cox, Mairead Connor, Zoltan Molnar, David Simpson, Derek Fairley |
| EPI_ISL_585422, EPI_ISL_585423 | West of Scotland Specialist Virology Centre, NHSGGC / MRC-University of Glasgow Centre for Virus Research | COVID-19 Genomics UK (COG-UK) Consortium | Ana da Silva Filipe, Natasha Johnson, Kathy Smollett, Daniel Mair, Stephen Carmichael, Lily Tong, Jenna Nichols, Elihu Aranday-Cortes, Kyriaki Nomikou; Sarah McDonald, Marc Niebel, Patawee Asamaphan; Richard Orton, Joseph Hughes, Sreenu Vattipally, David L Robertson; Alasdair MacLean, Rory Gunson; Kathy Li, Igor Starinskij, Natasha Jesudason, Rajiv Shah, James Shepherd, Antonia Ho, Emma Thomson |
| EPI_ISL_585653 | Wales Specialist Virology Centre Sequencing lab: Pathogen Genomics Unit | COVID-19 Genomics UK (COG-UK) Consortium | Catherine Moore, Johnathan Evans, Laura Gifford, Malorie Perry, Simon Cottrell, Angela Marchbank, Alec Birchley, Alexander Adams, Amy Gaskin, Bree Gatica-Wilcox, Jason Coombes, Joel Southgate, Lauren Gilbert, Lee Graham, Nicole Pacchiarini, Sara Kumziene-Summerhayes, Sarah Taylor, Sophie Jones, Sara Rey, Matthew Bull, Joanne Watkins, Sally Corden, Tom Connor |
| EPI_ISL_586569 | GCRI,Ahmedabad | Gujarat Biotechnology Research Centre | Shashank Pandya, Harsha Panchal, Apurvasinh Puvar, Janvi Raval, Zarna Patel, Monika Gandhi, Pinal Trivedi, Maharshi Pandya, Nidhi Patel, Nitin Savaliya, Raghawendra Kumar, Dinesh Kumar, Zuber Saiyed, Komal Patel, Labdhi Pandya, Afzal Ansari, Nikha Trivedi, R D Dixit, A M Kadri, Harsh Bakshi, Chaitanya Joshi, Madhvi Joshi |
| EPI_ISL_586570 | GCRI,Ahmedabad | Gujarat Biotechnology Research Centre | Harsha Panchal, Apurvasinh Puvar, Janvi Raval, Zarna Patel, Monika Gandhi, Pinal Trivedi, Maharshi Pandya, Nidhi Patel, Nitin Savaliya, Raghawendra Kumar, Dinesh Kumar, Zuber Saiyed, Komal Patel, Labdhi Pandya, Afzal Ansari, Nikha Trivedi, Shashank Pandya, R D Dixit, A M Kadri, Harsh Bakshi, Chaitanya Joshi, Madhvi Joshi |
| EPI_ISL_590173 | Lighthouse Lab in Cambridge | Wellcome Sanger Institute for the COVID-19 Genomics UK (COG-UK) consortium | Rob Howes, The Lighthouse Lab in Cambridge and Alex Alderton, Roberto Amato, Sonia Goncalves, Ewan Harrison, David K. Jackson, Ian Johnston, Dominic Kwiatkowski, Cordelia Langford, John Sillitoe on behalf of the Wellcome Sanger Institute COVID-19 Surveillance Team (http://www.sanger.ac.uk/covid-team) |
| EPI_ISL_590174, EPI_ISL_590175 | Lighthouse Lab in Milton Keynes | Wellcome Sanger Institute for the COVID-19 Genomics UK (COG-UK) consortium | The Lighthouse Lab in Milton Keynes and Alex Alderton, Roberto Amato, Sonia Goncalves, Ewan Harrison, David K. Jackson, Ian Johnston, Dominic Kwiatkowski, Cordelia Langford, John Sillitoe on behalf of the Wellcome Sanger Institute COVID-19 Surveillance Team (http://www.sanger.ac.uk/covid-team) |
| EPI_ISL_590178, EPI_ISL_590179, EPI_ISL_590180, EPI_ISL_590182, EPI_ISL_590183, EPI_ISL_590184, EPI_ISL_590185, EPI_ISL_590187, EPI_ISL_590188, EPI_ISL_590189, EPI_ISL_590190, EPI_ISL_590191, EPI_ISL_590192, EPI_ISL_590195, EPI_ISL_590196, EPI_ISL_590197, EPI_ISL_590199, EPI_ISL_590201, EPI_ISL_590202, EPI_ISL_590203, EPI_ISL_590204, EPI_ISL_590205, EPI_ISL_590207, EPI_ISL_590208, EPI_ISL_590209, EPI_ISL_590211, EPI_ISL_590212, EPI_ISL_590216, EPI_ISL_590217, EPI_ISL_590218, EPI_ISL_590219, EPI_ISL_590221, EPI_ISL_590222, EPI_ISL_590223, EPI_ISL_590227, EPI_ISL_590229, EPI_ISL_590230, EPI_ISL_590231, EPI_ISL_590232, EPI_ISL_590233, EPI_ISL_590236, EPI_ISL_590237, EPI_ISL_590238, EPI_ISL_590240, EPI_ISL_590241, EPI_ISL_590242, EPI_ISL_590243, EPI_ISL_590244, EPI_ISL_590245, EPI_ISL_590247, EPI_ISL_590248, EPI_ISL_590249, EPI_ISL_590251, EPI_ISL_590252, EPI_ISL_590253, EPI_ISL_590254, EPI_ISL_590256, EPI_ISL_590258, EPI_ISL_590262, EPI_ISL_590264, EPI_ISL_590266, EPI_ISL_590267, EPI_ISL_590268, EPI_ISL_590269, EPI_ISL_590270, EPI_ISL_590271, EPI_ISL_590273, EPI_ISL_590278, EPI_ISL_590279, EPI_ISL_590280, EPI_ISL_590281, EPI_ISL_590285, EPI_ISL_590287, EPI_ISL_590289, EPI_ISL_590290, EPI_ISL_590291, EPI_ISL_590292, EPI_ISL_590293, EPI_ISL_590294, EPI_ISL_590297, EPI_ISL_590298, EPI_ISL_590299, EPI_ISL_590301, EPI_ISL_590302, EPI_ISL_590303, EPI_ISL_590304, EPI_ISL_590305, EPI_ISL_590308, EPI_ISL_590310, EPI_ISL_590311, EPI_ISL_590312, EPI_ISL_590313, EPI_ISL_590315, EPI_ISL_590316, EPI_ISL_590317, EPI_ISL_590318, EPI_ISL_590320, EPI_ISL_590322, EPI_ISL_590324, EPI_ISL_590325, EPI_ISL_590326, EPI_ISL_590330, EPI_ISL_590331, EPI_ISL_590332, EPI_ISL_590333, EPI_ISL_590334, EPI_ISL_590336, EPI_ISL_590338, EPI_ISL_590339, EPI_ISL_590340, EPI_ISL_590341, EPI_ISL_590342, EPI_ISL_590343, EPI_ISL_590344, EPI_ISL_590345, EPI_ISL_590348, EPI_ISL_590349, EPI_ISL_590350, EPI_ISL_590353, EPI_ISL_590354, EPI_ISL_590356, EPI_ISL_590358, EPI_ISL_590360, EPI_ISL_590362, EPI_ISL_590363, EPI_ISL_590365, EPI_ISL_590368, EPI_ISL_590370, EPI_ISL_590371, EPI_ISL_590372, EPI_ISL_590374, EPI_ISL_590375, EPI_ISL_590376, EPI_ISL_590377, EPI_ISL_590378, EPI_ISL_590381, EPI_ISL_590382, EPI_ISL_590384, EPI_ISL_590385, EPI_ISL_590387, EPI_ISL_590389, EPI_ISL_590391, EPI_ISL_590393, EPI_ISL_590394, EPI_ISL_590395, EPI_ISL_590396, EPI_ISL_590400, EPI_ISL_590401, EPI_ISL_590402, EPI_ISL_590404, EPI_ISL_590406, EPI_ISL_590407, EPI_ISL_590408, EPI_ISL_590409, EPI_ISL_590410, EPI_ISL_590411, EPI_ISL_590412, EPI_ISL_590413, EPI_ISL_590414, EPI_ISL_590415, EPI_ISL_590416, EPI_ISL_590417, EPI_ISL_590418, EPI_ISL_590420, EPI_ISL_590421, EPI_ISL_590424, EPI_ISL_590425, EPI_ISL_590426, EPI_ISL_590427, EPI_ISL_590428, EPI_ISL_590429, EPI_ISL_590430, EPI_ISL_590431, EPI_ISL_590432, EPI_ISL_590433, EPI_ISL_590434, EPI_ISL_590435, EPI_ISL_590436, EPI_ISL_590437, EPI_ISL_590439, EPI_ISL_590440, EPI_ISL_590443, EPI_ISL_590444, EPI_ISL_590445, EPI_ISL_590447, EPI_ISL_590449, EPI_ISL_590450, EPI_ISL_590451, EPI_ISL_590454, EPI_ISL_590455, EPI_ISL_590456, EPI_ISL_590457, EPI_ISL_590458, EPI_ISL_590459, EPI_ISL_590460, EPI_ISL_590461, EPI_ISL_590463, EPI_ISL_590464, EPI_ISL_590465, EPI_ISL_590466, EPI_ISL_590467, EPI_ISL_590468, EPI_ISL_590469, EPI_ISL_590470, EPI_ISL_590471, EPI_ISL_590472, EPI_ISL_590473, EPI_ISL_590474, EPI_ISL_590475, EPI_ISL_590476, EPI_ISL_590477, EPI_ISL_590478, EPI_ISL_590479, EPI_ISL_590480, EPI_ISL_590481, EPI_ISL_590482, EPI_ISL_590483, EPI_ISL_590484, EPI_ISL_590485, EPI_ISL_590486, EPI_ISL_590487, EPI_ISL_590488, EPI_ISL_590489, EPI_ISL_590490, EPI_ISL_590491, EPI_ISL_590492, EPI_ISL_590493, EPI_ISL_590494, EPI_ISL_590495, EPI_ISL_590496 | | | |
| see above | Lighthouse Lab in Glasgow | Wellcome Sanger Institute for the COVID-19 Genomics UK (COG-UK) consortium | Harper VanSteenhouse, Yumi Kasai, David Gray, Carol Clugston, Anna Dominiczak and Alex Alderton, Roberto Amato, Sonia Goncalves, Ewan Harrison, David K. Jackson, Ian Johnston, Dominic Kwiatkowski, Cordelia Langford, John Sillitoe on behalf of the Wellcome Sanger Institute COVID-19 Surveillance Team (http://www.sanger.ac.uk/covid-team) |
| EPI_ISL_590497 | Lighthouse Lab in Glasgow | Wellcome Sanger Institute for the COVID-19 Genomics UK (COG-UK) Consortium | Harper VanSteenhouse, Yumi Kasai, David Gray, Carol Clugston, Anna Dominiczak and Alex Alderton, Roberto Amato, Sonia Goncalves, Ewan Harrison, David K. Jackson, Ian Johnston, Dominic Kwiatkowski, Cordelia Langford, John Sillitoe on behalf of the Wellcome Sanger Institute COVID-19 Surveillance Team |
| EPI_ISL_590498, EPI_ISL_590499, EPI_ISL_590500, EPI_ISL_590501, EPI_ISL_590502, EPI_ISL_590503, EPI_ISL_590504, EPI_ISL_590505, EPI_ISL_590506, EPI_ISL_590507, EPI_ISL_590508, EPI_ISL_590509, EPI_ISL_590510, EPI_ISL_590511, EPI_ISL_590512, EPI_ISL_590513, EPI_ISL_590514, EPI_ISL_590515, EPI_ISL_590516, EPI_ISL_590517, EPI_ISL_590518, EPI_ISL_590519, EPI_ISL_590520, EPI_ISL_590521, EPI_ISL_590522, EPI_ISL_590523, EPI_ISL_590524, EPI_ISL_590525, EPI_ISL_590526, EPI_ISL_590527, EPI_ISL_590528, EPI_ISL_590529, EPI_ISL_590530, EPI_ISL_590531, EPI_ISL_590532, EPI_ISL_590533, | | | |

| | | | |
|---|---|---|---|
| EPI_ISL_590534, EPI_ISL_590535, EPI_ISL_590536, EPI_ISL_590537, EPI_ISL_590538, EPI_ISL_590539, EPI_ISL_590540, EPI_ISL_590541, EPI_ISL_590542, EPI_ISL_590543, EPI_ISL_590544, EPI_ISL_590545, EPI_ISL_590546, EPI_ISL_590547, EPI_ISL_590548, EPI_ISL_590549, EPI_ISL_590550, EPI_ISL_590551, EPI_ISL_590552, EPI_ISL_590553, EPI_ISL_590554, EPI_ISL_590555, EPI_ISL_590556, EPI_ISL_590557, EPI_ISL_590558, EPI_ISL_590559, EPI_ISL_590560, EPI_ISL_590561, EPI_ISL_590562, EPI_ISL_590563, EPI_ISL_590564, EPI_ISL_590565, EPI_ISL_590566, EPI_ISL_590567, EPI_ISL_590568, EPI_ISL_590569, EPI_ISL_590570, EPI_ISL_590571, EPI_ISL_590572, EPI_ISL_590573, EPI_ISL_590574, EPI_ISL_590575, EPI_ISL_590576, EPI_ISL_590577, EPI_ISL_590578, EPI_ISL_590579, EPI_ISL_590580, EPI_ISL_590581, EPI_ISL_590582, EPI_ISL_590583, EPI_ISL_590584, EPI_ISL_590585, EPI_ISL_590586, EPI_ISL_590587, EPI_ISL_590588, EPI_ISL_590589, EPI_ISL_590590, EPI_ISL_590591, EPI_ISL_590592, EPI_ISL_590593, EPI_ISL_590594, EPI_ISL_590595, EPI_ISL_590596, EPI_ISL_590597, EPI_ISL_590598, EPI_ISL_590599, EPI_ISL_590600, EPI_ISL_590601, EPI_ISL_590602, EPI_ISL_590603, EPI_ISL_590604, EPI_ISL_590605, EPI_ISL_590606, EPI_ISL_590607, EPI_ISL_590608, EPI_ISL_590609, EPI_ISL_590610, EPI_ISL_590611, EPI_ISL_590612, EPI_ISL_590613, EPI_ISL_590614, EPI_ISL_590615, EPI_ISL_590616, EPI_ISL_590617, EPI_ISL_590618, EPI_ISL_590619, EPI_ISL_590620, EPI_ISL_590621, EPI_ISL_590622, EPI_ISL_590623, EPI_ISL_590624, EPI_ISL_590625, EPI_ISL_590626, EPI_ISL_590627, EPI_ISL_590628, EPI_ISL_590629, EPI_ISL_590630, EPI_ISL_590631, EPI_ISL_590632, EPI_ISL_590633, EPI_ISL_590634, EPI_ISL_590635, EPI_ISL_590636, EPI_ISL_590637, EPI_ISL_590638, EPI_ISL_590639, EPI_ISL_590640, EPI_ISL_590641, EPI_ISL_590642, EPI_ISL_590643, EPI_ISL_590644, EPI_ISL_590645, EPI_ISL_590646, EPI_ISL_590647, EPI_ISL_590648, EPI_ISL_590649, EPI_ISL_590650, EPI_ISL_590651, EPI_ISL_590652, EPI_ISL_590653, EPI_ISL_590654, EPI_ISL_590655, EPI_ISL_590656, EPI_ISL_590657, EPI_ISL_590658, EPI_ISL_590659, EPI_ISL_590660, EPI_ISL_590661, EPI_ISL_590662, EPI_ISL_590663, EPI_ISL_590664, EPI_ISL_590665, EPI_ISL_590666, EPI_ISL_590667, EPI_ISL_590668, EPI_ISL_590669, EPI_ISL_590670, EPI_ISL_590671, EPI_ISL_590672, EPI_ISL_590673, EPI_ISL_590674, EPI_ISL_590675, EPI_ISL_590676, EPI_ISL_590677, EPI_ISL_590678, EPI_ISL_590679, EPI_ISL_590680, EPI_ISL_590681, EPI_ISL_590682, EPI_ISL_590683, EPI_ISL_590684, EPI_ISL_590685, EPI_ISL_590686, EPI_ISL_590687, EPI_ISL_590688 | | | |
| see above | Lighthouse Lab in Glasgow | Wellcome Sanger Institute for the COVID-19 Genomics UK (COG-UK) consortium | Harper VanSteenhouse, Yumi Kasai, David Gray, Carol Clugston, Anna Dominiczak and Alex Alderton, Roberto Amato, Sonia Goncalves, Ewan Harrison, David K. Jackson, Ian Johnston, Dominic Kwiatkowski, Cordelia Langford, John Sillitoe on behalf of the Wellcome Sanger Institute COVID-19 Surveillance Team (http://www.sanger.ac.uk/covid-team) |
| EPI_ISL_590878 | Oslo University Hospital, Department of Medical Microbiology | Norwegian Institute of Public Health, Department of Virology | Kathrine Stene-Johansen, Kamilla Heddeland Instefjord, Hilde Elshaug, Rasmus Riis Kopperud, Hilde Vollan, Karoline Bragstad, Olav Hungnes |
| EPI_ISL_590879, EPI_ISL_590880 | Medical Microbiology Unit, Department for Laboratory Medicine, Drammen Hospital, Vestre Viken Health Trust, | Norwegian Institute of Public Health, Department of Virology | Kathrine Stene-Johansen, Kamilla Heddeland Instefjord, Hilde Elshaug, Rasmus Riis Kopperud, Hilde Vollan, Karoline Bragstad, Olav Hungnes |
| EPI_ISL_590881 | Furst Medical Laboratory | Norwegian Institute of Public Health, Department of Virology | Kathrine Stene-Johansen, Kamilla Heddeland Instefjord, Hilde Elshaug, Rasmus Riis Kopperud, Hilde Vollan, Karoline Bragstad, Olav Hungnes |
| EPI_ISL_590882 | Unilabs Laboratory Medicine | Norwegian Institute of Public Health, Department of Virology | Kathrine Stene-Johansen, Kamilla Heddeland Instefjord, Hilde Elshaug, Rasmus Riis Kopperud, Hilde Vollan, Karoline Bragstad, Olav Hungnes |
| EPI_ISL_590883, EPI_ISL_590884 | Foerde Hospital, Department of Microbiology | Norwegian Institute of Public Health, Department of Virology | Kathrine Stene-Johansen, Kamilla Heddeland Instefjord, Hilde Elshaug, Rasmus Riis Kopperud, Hilde Vollan, Karoline Bragstad, Olav Hungnes |
| EPI_ISL_590894, EPI_ISL_590895 | Ostfold Hospital Trust - Kalnes, Centre for Laboratory Medicine, Section for gene technology and infection serology | Norwegian Institute of Public Health, Department of Virology | Kathrine Stene-Johansen, Kamilla Heddeland Instefjord, Hilde Elshaug, Rasmus Riis Kopperud, Hilde Vollan, Karoline Bragstad, Olav Hungnes |
| EPI_ISL_590898 | Furst Medical Laboratory | Norwegian Institute of Public Health, Department of Virology | Kathrine Stene-Johansen, Kamilla Heddeland Instefjord, Hilde Elshaug, Rasmus Riis Kopperud, Hilde Vollan, Karoline Bragstad, Olav Hungnes |
| EPI_ISL_590899 | Foerde Hospital, Department of Microbiology | Norwegian Institute of Public Health, Department of Virology | Kathrine Stene-Johansen, Kamilla Heddeland Instefjord, Hilde Elshaug, Rasmus Riis Kopperud, Hilde Vollan, Karoline Bragstad, Olav Hungnes |
| EPI_ISL_590900 | Oslo University Hospital, Department of Medical Microbiology | Norwegian Institute of Public Health, Department of Virology | Kathrine Stene-Johansen, Kamilla Heddeland Instefjord, Hilde Elshaug, Rasmus Riis Kopperud, Hilde Vollan, Karoline Bragstad, Olav Hungnes |
| EPI_ISL_590901, EPI_ISL_590902, EPI_ISL_590903 | Furst Medical Laboratory | Norwegian Institute of Public Health, Department of Virology | Kathrine Stene-Johansen, Kamilla Heddeland Instefjord, Hilde Elshaug, Rasmus Riis Kopperud, Hilde Vollan, Karoline Bragstad, Olav Hungnes |
| EPI_ISL_590907 | Hospital of Southern Norway - Kristiansand, Department of Medical Microbiology | Norwegian Institute of Public Health, Department of Virology | Kathrine Stene-Johansen, Kamilla Heddeland Instefjord, Hilde Elshaug, Rasmus Riis Kopperud, Hilde Vollan, Karoline Bragstad, Olav Hungnes |
| EPI_ISL_590911 | Unilabs Laboratory Medicine | Norwegian Institute of Public Health, Department of Virology | Kathrine Stene-Johansen, Kamilla Heddeland Instefjord, Hilde Elshaug, Rasmus Riis Kopperud, Hilde Vollan, Karoline Bragstad, Olav Hungnes |
| EPI_ISL_590912 | Furst Medical Laboratory | Norwegian Institute of Public Health, Department of Virology | Kathrine Stene-Johansen, Kamilla Heddeland Instefjord, Hilde Elshaug, Rasmus Riis Kopperud, Hilde Vollan, Karoline Bragstad, Olav Hungnes |
| EPI_ISL_590934 | Oslo University Hospital, Department of Medical Microbiology | Norwegian Institute of Public Health, Department of Virology | Kathrine Stene-Johansen, Kamilla Heddeland Instefjord, Hilde Elshaug, Rasmus Riis Kopperud, Hilde Vollan, Karoline Bragstad, Olav Hungnes |
| EPI_ISL_590987, EPI_ISL_590989, EPI_ISL_590990, EPI_ISL_590991, EPI_ISL_590995, EPI_ISL_590996, EPI_ISL_590998, EPI_ISL_590999, EPI_ISL_591000, EPI_ISL_591001, EPI_ISL_591002, EPI_ISL_591003, EPI_ISL_591004, EPI_ISL_591005, EPI_ISL_591015 | | | |
| see above | Ostfold Hospital Trust - Kalnes, Centre for Laboratory Medicine, Section for gene technology and infection serology | Norwegian Institute of Public Health, Department of Virology | Kathrine Stene-Johansen, Kamilla Heddeland Instefjord, Hilde Elshaug, Rasmus Riis Kopperud, Hilde Vollan, Karoline Bragstad, Olav Hungnes |
| EPI_ISL_591020 | Oslo University Hospital, Department of Medical Microbiology | Norwegian Institute of Public Health, Department of Virology | Kathrine Stene-Johansen, Kamilla Heddeland Instefjord, Hilde Elshaug, Rasmus Riis Kopperud, Hilde Vollan, Karoline Bragstad, Olav Hungnes |
| EPI_ISL_591487 | Laverty Pathology | NSW Health Pathology - Institute of Clinical Pathology and Medical Research; Westmead Hospital; University of Sydney | CIDM-PH et al. |
| EPI_ISL_591507 | Sydney South West Pathology Service (SSWPS) - Concord Repatriation General Hospital - NSW Health Pathology | NSW Health Pathology - Institute of Clinical Pathology and Medical Research; Westmead Hospital; University of Sydney | CIDM-PH et al. |
| EPI_ISL_591568, EPI_ISL_591593, EPI_ISL_591631, EPI_ISL_591660, EPI_ISL_591675, EPI_ISL_591689, EPI_ISL_591697, EPI_ISL_591710, EPI_ISL_591714, EPI_ISL_591733, EPI_ISL_591738, EPI_ISL_591751, EPI_ISL_591781, EPI_ISL_591795, EPI_ISL_591801, EPI_ISL_591803, EPI_ISL_591813, EPI_ISL_591818, EPI_ISL_591819, EPI_ISL_591820, EPI_ISL_592235, EPI_ISL_592236, EPI_ISL_592238, EPI_ISL_592239, EPI_ISL_592240, EPI_ISL_592241, EPI_ISL_592242, EPI_ISL_592243, EPI_ISL_592269, EPI_ISL_592270, EPI_ISL_592272, EPI_ISL_592274, EPI_ISL_592275, EPI_ISL_592280, EPI_ISL_592282, EPI_ISL_592285, EPI_ISL_592311, EPI_ISL_592312, EPI_ISL_592313, EPI_ISL_592329, EPI_ISL_592332, EPI_ISL_592334, EPI_ISL_592337, EPI_ISL_592350, EPI_ISL_592351, EPI_ISL_592355, EPI_ISL_592366, EPI_ISL_592385, EPI_ISL_592396, EPI_ISL_592404, EPI_ISL_592416, EPI_ISL_592424, EPI_ISL_592435, EPI_ISL_592451, EPI_ISL_592480, EPI_ISL_592481, EPI_ISL_592482, EPI_ISL_592540, EPI_ISL_592543, EPI_ISL_592618, EPI_ISL_592631, EPI_ISL_592638, EPI_ISL_592647, EPI_ISL_592650, EPI_ISL_592651, EPI_ISL_592656, EPI_ISL_592664, EPI_ISL_592706, EPI_ISL_592728 | | | |
| see above | Microbiological Diagnostic Unit - Public Health Laboratory (MDU-PHL) | MDU-PHL | Seemann T., Schultz, M. B., Sait, M., Sherry, N. |
| EPI_ISL_592730 | Victorian Infectious Diseases Reference Laboratory (VIDRL) | VIDRL and MDU-PHL | Caly L., Seemann T., Sait, M., Schultz, M. B., Druce J., Sherry, N. |
| EPI_ISL_592731, EPI_ISL_592732, EPI_ISL_592739, EPI_ISL_592750, EPI_ISL_592819 | Microbiological Diagnostic Unit - Public Health Laboratory (MDU-PHL) | MDU-PHL | Seemann T., Schultz, M. B., Sait, M., Sherry, N. |
| EPI_ISL_592820, EPI_ISL_592821 | Victorian Infectious Diseases Reference Laboratory (VIDRL) | VIDRL and MDU-PHL | Caly L., Seemann T., Sait, M., Schultz, M. B., Druce J., Sherry, N. |
| EPI_ISL_592861 | Microbiological Diagnostic Unit - Public Health Laboratory (MDU-PHL) | MDU-PHL | Seemann T., Schultz, M. B., Sait, M., Sherry, N. |
| EPI_ISL_592862, EPI_ISL_592863, EPI_ISL_592864, EPI_ISL_592865 | Victorian Infectious Diseases Reference Laboratory (VIDRL) | VIDRL and MDU-PHL | Caly L., Seemann T., Sait, M., Schultz, M. B., Druce J., Sherry, N. |
| EPI_ISL_592996, EPI_ISL_593001, EPI_ISL_593002 | Microbiological Diagnostic Unit - Public Health Laboratory (MDU-PHL) | MDU-PHL | Seemann T., Schultz, M. B., Sait, M., Sherry, N. |
| EPI_ISL_593004, EPI_ISL_593006 | Victorian Infectious Diseases Reference Laboratory (VIDRL) | VIDRL and MDU-PHL | Caly L., Seemann T., Sait, M., Schultz, M. B., Druce J., Sherry, N. |
| EPI_ISL_593039, EPI_ISL_593041, EPI_ISL_593052, EPI_ISL_593053, EPI_ISL_593055, EPI_ISL_593061, EPI_ISL_593063, EPI_ISL_593069, EPI_ISL_593070, EPI_ISL_593071, EPI_ISL_593091, EPI_ISL_593095, EPI_ISL_593097, EPI_ISL_593098, EPI_ISL_593100, EPI_ISL_593101 | | | |
| see above | Microbiological Diagnostic Unit - Public Health Laboratory (MDU-PHL) | MDU-PHL | Seemann T., Schultz, M. B., Sait, M., Sherry, N. |
| EPI_ISL_593256 | Victorian Infectious Diseases Reference Laboratory (VIDRL) | VIDRL and MDU-PHL | Caly L., Seemann T., Sait, M., Schultz, M. B., Druce J., Sherry, N. |
| EPI_ISL_593557, EPI_ISL_593558 | Brigham and Women's Hospital | Jonathan Li Laboratory | Manish C. Choudhary, James Regan, Jonathan Z. Li |
| EPI_ISL_593616, EPI_ISL_593635, EPI_ISL_593639 | unknown | Public Health Virology Laboratory, Forensic and Scientific Services (PHV-FSS) | Son Nguyen et al. |
| EPI_ISL_593727, EPI_ISL_593728, EPI_ISL_593738, EPI_ISL_593741, EPI_ISL_593742, EPI_ISL_593743 | South Eastern Area Laboratory Services (SEALS) | NSW Health Pathology - Institute of Clinical Pathology and Medical Research; Westmead Hospital; University of Sydney | CIDM-PH et al. |
| EPI_ISL_593764 | Sydney South West Pathology Service (SSWPS) - Liverpool Hospital - NSW Health Pathology | NSW Health Pathology - Institute of Clinical Pathology and Medical Research; Westmead Hospital; University of Sydney | CIDM-PH et al. |
| EPI_ISL_593805, EPI_ISL_593837 | Respiratory Virus Unit, Microbiology Services Colindale, Public | Respiratory Virus Unit, Microbiology Services Colindale, Public | PHE Covid Sequencing Team |

| | | | |
|---|---|---|---|
| | Health England | Health England | |
| EPI_ISL_593888 | CHU Purpan - Laboratoire de Virologie - Institut Fédératif de Biologie | CHU Purpan - Laboratoire de Virologie - Institut Fédératif de Biologie | Latour J., Ranger N., Dubois M., Carcenac R., Harter A., Boyer P., Tremeaux P., Izopet J. |
| EPI_ISL_593903 | Hospital, Argenteuil | National Reference Center for Viruses of Respiratory Infections, Institut Pasteur, Paris | Sylvie Behillil, Fabiana Gambaro, Etienne Simon-Lorière, Vincent Enouf, Maud Vanpeene, Sylvie van der Werf |
| EPI_ISL_593908 | Labo Analyses Med, Sarcelles | National Reference Center for Viruses of Respiratory Infections, Institut Pasteur, Paris | Sylvie Behillil, Fabiana Gambaro, Etienne Simon-Lorière, Vincent Enouf, Maud Vanpeene, Sylvie van der Werf |
| EPI_ISL_593911 | Hospital, Le Chesnay Cédex | National Reference Center for Viruses of Respiratory Infections, Institut Pasteur, Paris | Sylvie Behillil, Fabiana Gambaro, Etienne Simon-Lorière, Vincent Enouf, Maud Vanpeene, Sylvie van der Werf |
| EPI_ISL_593913 | Hospital, Talange | National Reference Center for Viruses of Respiratory Infections, Institut Pasteur, Paris | Sylvie Behillil, Fabiana Gambaro, Etienne Simon-Lorière, Vincent Enouf, Maud Vanpeene, Sylvie van der Werf |
| EPI_ISL_593914 | Hospital, Metz | National Reference Center for Viruses of Respiratory Infections, Institut Pasteur, Paris | Sylvie Behillil, Fabiana Gambaro, Etienne Simon-Lorière, Vincent Enouf, Maud Vanpeene, Sylvie van der Werf |
| EPI_ISL_593915 | Hospital, Amneville | National Reference Center for Viruses of Respiratory Infections, Institut Pasteur, Paris | Sylvie Behillil, Fabiana Gambaro, Etienne Simon-Lorière, Vincent Enouf, Maud Vanpeene, Sylvie van der Werf |
| EPI_ISL_593916 | Hospital, Sarreguemines | National Reference Center for Viruses of Respiratory Infections, Institut Pasteur, Paris | Sylvie Behillil, Fabiana Gambaro, Etienne Simon-Lorière, Vincent Enouf, Maud Vanpeene, Sylvie van der Werf |
| EPI_ISL_593918 | Hospital, Joeuf | National Reference Center for Viruses of Respiratory Infections, Institut Pasteur, Paris | Sylvie Behillil, Fabiana Gambaro, Etienne Simon-Lorière, Vincent Enouf, Maud Vanpeene, Sylvie van der Werf |
| EPI_ISL_593919 | Hospital, Ottonville | National Reference Center for Viruses of Respiratory Infections, Institut Pasteur, Paris | Sylvie Behillil, Fabiana Gambaro, Etienne Simon-Lorière, Vincent Enouf, Maud Vanpeene, Sylvie van der Werf |
| EPI_ISL_593920 | Hospital, Metz | National Reference Center for Viruses of Respiratory Infections, Institut Pasteur, Paris | Sylvie Behillil, Fabiana Gambaro, Etienne Simon-Lorière, Vincent Enouf, Maud Vanpeene, Sylvie van der Werf |
| EPI_ISL_593921 | Hospital, Fameck | National Reference Center for Viruses of Respiratory Infections, Institut Pasteur, Paris | Sylvie Behillil, Fabiana Gambaro, Etienne Simon-Lorière, Vincent Enouf, Maud Vanpeene, Sylvie van der Werf |
| EPI_ISL_593922, EPI_ISL_593923 | Hospital, Maizières les Metz | National Reference Center for Viruses of Respiratory Infections, Institut Pasteur, Paris | Sylvie Behillil, Fabiana Gambaro, Etienne Simon-Lorière, Vincent Enouf, Maud Vanpeene, Sylvie van der Werf |
| EPI_ISL_593925, EPI_ISL_593929, EPI_ISL_593930, EPI_ISL_593933 | Labo Analyses Med, Puteaux | National Reference Center for Viruses of Respiratory Infections, Institut Pasteur, Paris | Sylvie Behillil, Fabiana Gambaro, Etienne Simon-Lorière, Vincent Enouf, Maud Vanpeene, Sylvie van der Werf |
| EPI_ISL_594142 | MDU-PHL, The Peter Doherty Institute for Infection and Immunity | MDU-PHL, The Peter Doherty Institute for Infection and Immunity | Caly,L., Seemann,T., Sait,M.L., Schultz,M.B., Druce,J., Sherry,N.L. |
| EPI_ISL_594165 | hopital | National Reference Center for Viruses of Respiratory Infections, Institut Pasteur, Paris | Sylvie Behillil, Fabiana Gambaro, Etienne Simon-Lorière, Vincent Enouf, Maud Vanpeene, Sylvie van der Werf |
| EPI_ISL_594301, EPI_ISL_594302, EPI_ISL_594303, EPI_ISL_594304, EPI_ISL_594305, EPI_ISL_594306, EPI_ISL_594307, EPI_ISL_594308, EPI_ISL_594309, EPI_ISL_594310, EPI_ISL_594311 | | | |
| see above | Florida Bureau of Public Health Laboratories | Florida Bureau of Public Health Laboratories | Sarah Schmedes, Jason Blanton |
| EPI_ISL_596231 | WHO National Influenza Centre Russian Federation | WHO National Influenza Centre Russian Federation | Andrey Komissarov, Artem Fadeev, Anna Ivanova, Kseniya Komissarova, Dmitry Bazhenov, Daria Danilenko |
| EPI_ISL_596250, EPI_ISL_596251, EPI_ISL_596252, EPI_ISL_596253, EPI_ISL_596254, EPI_ISL_596255, EPI_ISL_596256, EPI_ISL_596257, EPI_ISL_596258, EPI_ISL_596259, EPI_ISL_596260, EPI_ISL_596261, EPI_ISL_596262, EPI_ISL_596263, EPI_ISL_596270, EPI_ISL_596271, EPI_ISL_596275, EPI_ISL_596276, EPI_ISL_596277, EPI_ISL_596278, EPI_ISL_596279, EPI_ISL_596283, EPI_ISL_596295, EPI_ISL_596296, EPI_ISL_596297, EPI_ISL_596298, EPI_ISL_596299, EPI_ISL_596300, EPI_ISL_596301, EPI_ISL_596302, EPI_ISL_596305, EPI_ISL_596306, EPI_ISL_596307, EPI_ISL_596308, EPI_ISL_596310, EPI_ISL_596311, EPI_ISL_596312, EPI_ISL_596313, EPI_ISL_596314, EPI_ISL_596315, EPI_ISL_596316, EPI_ISL_596317, EPI_ISL_596357, EPI_ISL_596358, EPI_ISL_596359, EPI_ISL_596360, EPI_ISL_596361, EPI_ISL_596362, EPI_ISL_596363, EPI_ISL_596364, EPI_ISL_596365, EPI_ISL_596366, EPI_ISL_596367, EPI_ISL_596368, EPI_ISL_596369, EPI_ISL_596370, EPI_ISL_596371, EPI_ISL_596372, EPI_ISL_596373, EPI_ISL_596374, EPI_ISL_596375, EPI_ISL_596376, EPI_ISL_596377 | | | |
| see above | HELIX LCC | WHO National Influenza Centre Russian Federation | Andrey Komissarov, Artem Fadeev, Anna Ivanova, Kseniya Komissarova, Dmitry Bazhenov, Daria Danilenko |
| EPI_ISL_596702, EPI_ISL_596764, EPI_ISL_596766 | PathWest Laboratory Medicine WA | PathWest Laboratory Medicine WA Microbial Surveillance Unit | PathWest Laboratory Medicine WA Microbial Surveillance Unit |
| EPI_ISL_596889, EPI_ISL_596891, EPI_ISL_596894, EPI_ISL_596896, EPI_ISL_596903, EPI_ISL_596905, EPI_ISL_596906, EPI_ISL_596907, EPI_ISL_596908 | National Virus Reference Laboratory | National Virus Reference Laboratory | Michael Carr, Gabriel Gonzalez, Jonathan Dean, Daniel Hare, Cillian F De Gascun |
| EPI_ISL_602027, EPI_ISL_602028, EPI_ISL_602029 | Lighthouse Lab in Milton Keynes | Wellcome Sanger Institute for the COVID-19 Genomics UK (COG-UK) consortium | The Lighthouse Lab in Milton Keynes and Alex Alderton, Roberto Amato, Sonia Goncalves, Ewan Harrison, David K. Jackson, Ian Johnston, Dominic Kwiatkowski, Cordelia Langford, John Sillitoe on behalf of the Wellcome Sanger Institute COVID-19 Surveillance Team (http://www.sanger.ac.uk/covid-team) |
| EPI_ISL_602043, EPI_ISL_602044, EPI_ISL_602045, EPI_ISL_602046, EPI_ISL_602047, EPI_ISL_602048, EPI_ISL_602049, EPI_ISL_602050, EPI_ISL_602051, EPI_ISL_602052, EPI_ISL_602053, EPI_ISL_602054, EPI_ISL_602055, EPI_ISL_602056, EPI_ISL_602057, EPI_ISL_602058, EPI_ISL_602059, EPI_ISL_602060, EPI_ISL_602061, EPI_ISL_602062, EPI_ISL_602063, EPI_ISL_602064, EPI_ISL_602065, EPI_ISL_602066, EPI_ISL_602067, EPI_ISL_602068, EPI_ISL_602069, EPI_ISL_602070, EPI_ISL_602071, EPI_ISL_602072, EPI_ISL_602073, EPI_ISL_602074, EPI_ISL_602075, EPI_ISL_602076 | | | |
| see above | Lighthouse Lab in Glasgow | Wellcome Sanger Institute for the COVID-19 Genomics UK (COG-UK) consortium | Harper VanSteenhouse, Yumi Kasai, David Gray, Carol Clugston, Anna Dominiczak and Alex Alderton, Roberto Amato, Sonia Goncalves, Ewan Harrison, David K. Jackson, Ian Johnston, Dominic Kwiatkowski, Cordelia Langford, John Sillitoe on behalf of the Wellcome Sanger Institute COVID-19 Surveillance Team (http://www.sanger.ac.uk/covid-team) |
| EPI_ISL_602077 | Lighthouse Lab in Glasgow | Wellcome Sanger Institute for the COVID-19 Genomics UK (COG-UK) Consortium | Harper VanSteenhouse, Yumi Kasai, David Gray, Carol Clugston, Anna Dominiczak and Alex Alderton, Roberto Amato, Sonia Goncalves, Ewan Harrison, David K. Jackson, Ian Johnston, Dominic Kwiatkowski, Cordelia Langford, John Sillitoe on behalf of the Wellcome Sanger Institute COVID-19 Surveillance Team |
| EPI_ISL_602078, EPI_ISL_602079, EPI_ISL_602080, EPI_ISL_602081, EPI_ISL_602082 | Lighthouse Lab in Glasgow | Wellcome Sanger Institute for the COVID-19 Genomics UK (COG-UK) consortium | Harper VanSteenhouse, Yumi Kasai, David Gray, Carol Clugston, Anna Dominiczak and Alex Alderton, Roberto Amato, Sonia Goncalves, Ewan Harrison, David K. Jackson, Ian Johnston, Dominic Kwiatkowski, Cordelia Langford, John Sillitoe on behalf of the Wellcome Sanger Institute COVID-19 Surveillance Team (http://www.sanger.ac.uk/covid-team) |
| EPI_ISL_602083 | Lighthouse Lab in Glasgow | Wellcome Sanger Institute for the COVID-19 Genomics UK (COG-UK) Consortium | Harper VanSteenhouse, Yumi Kasai, David Gray, Carol Clugston, Anna Dominiczak and Alex Alderton, Roberto Amato, Sonia Goncalves, Ewan Harrison, David K. Jackson, Ian Johnston, Dominic Kwiatkowski, Cordelia Langford, John Sillitoe on behalf of the Wellcome Sanger Institute COVID-19 Surveillance Team |
| EPI_ISL_602084, EPI_ISL_602085, EPI_ISL_602086, EPI_ISL_602087, EPI_ISL_602088, EPI_ISL_602089 | Lighthouse Lab in Glasgow | Wellcome Sanger Institute for the COVID-19 Genomics UK (COG-UK) consortium | Harper VanSteenhouse, Yumi Kasai, David Gray, Carol Clugston, Anna Dominiczak and Alex Alderton, Roberto Amato, Sonia Goncalves, Ewan Harrison, David K. Jackson, Ian Johnston, Dominic Kwiatkowski, Cordelia Langford, John Sillitoe on behalf of the Wellcome Sanger Institute COVID-19 Surveillance Team (http://www.sanger.ac.uk/covid-team) |
| EPI_ISL_602090, EPI_ISL_602091 | Lighthouse Lab in Glasgow | Wellcome Sanger Institute for the COVID-19 Genomics UK (COG-UK) Consortium | Harper VanSteenhouse, Yumi Kasai, David Gray, Carol Clugston, Anna Dominiczak and Alex Alderton, Roberto Amato, Sonia Goncalves, Ewan Harrison, David K. Jackson, Ian Johnston, Dominic Kwiatkowski, Cordelia Langford, John Sillitoe on behalf of the Wellcome Sanger Institute COVID-19 Surveillance Team |
| EPI_ISL_602092, EPI_ISL_602093, | Lighthouse Lab in Glasgow | Wellcome Sanger Institute for the COVID-19 Genomics UK | Harper VanSteenhouse, Yumi Kasai, David Gray, Carol Clugston, Anna Dominiczak and Alex Alderton, Roberto Amato, Sonia Goncalves, Ewan Harrison, |

| | | | |
|---|---|---|---|
| EPI_ISL_602094, EPI_ISL_602095, EPI_ISL_602096, EPI_ISL_602097, EPI_ISL_602098, EPI_ISL_602099, EPI_ISL_602100, EPI_ISL_602101 | | (COG-UK) consortium | David K. Jackson, Ian Johnston, Dominic Kwiatkowski, Cordelia Langford, John Sillitoe on behalf of the Wellcome Sanger Institute COVID-19 Surveillance Team (http://www.sanger.ac.uk/covid-team) |
| EPI_ISL_602102 | Lighthouse Lab in Glasgow | Wellcome Sanger Institute for the COVID-19 Genomics UK (COG-UK) Consortium | Harper VanSteenhouse, Yumi Kasai, David Gray, Carol Clugston, Anna Dominiczak and Alex Alderton, Roberto Amato, Sonia Goncalves, Ewan Harrison, David K. Jackson, Ian Johnston, Dominic Kwiatkowski, Cordelia Langford, John Sillitoe on behalf of the Wellcome Sanger Institute COVID-19 Surveillance Team |
| EPI_ISL_602103, EPI_ISL_602104, EPI_ISL_602105 | Lighthouse Lab in Glasgow | Wellcome Sanger Institute for the COVID-19 Genomics UK (COG-UK) consortium | Harper VanSteenhouse, Yumi Kasai, David Gray, Carol Clugston, Anna Dominiczak and Alex Alderton, Roberto Amato, Sonia Goncalves, Ewan Harrison, David K. Jackson, Ian Johnston, Dominic Kwiatkowski, Cordelia Langford, John Sillitoe on behalf of the Wellcome Sanger Institute COVID-19 Surveillance Team (http://www.sanger.ac.uk/covid-team) |
| EPI_ISL_602106 | Lighthouse Lab in Glasgow | Wellcome Sanger Institute for the COVID-19 Genomics UK (COG-UK) Consortium | Harper VanSteenhouse, Yumi Kasai, David Gray, Carol Clugston, Anna Dominiczak and Alex Alderton, Roberto Amato, Sonia Goncalves, Ewan Harrison, David K. Jackson, Ian Johnston, Dominic Kwiatkowski, Cordelia Langford, John Sillitoe on behalf of the Wellcome Sanger Institute COVID-19 Surveillance Team |
| EPI_ISL_602107, EPI_ISL_602108 | Lighthouse Lab in Glasgow | Wellcome Sanger Institute for the COVID-19 Genomics UK (COG-UK) consortium | Harper VanSteenhouse, Yumi Kasai, David Gray, Carol Clugston, Anna Dominiczak and Alex Alderton, Roberto Amato, Sonia Goncalves, Ewan Harrison, David K. Jackson, Ian Johnston, Dominic Kwiatkowski, Cordelia Langford, John Sillitoe on behalf of the Wellcome Sanger Institute COVID-19 Surveillance Team (http://www.sanger.ac.uk/covid-team) |
| EPI_ISL_602109 | Lighthouse Lab in Glasgow | Wellcome Sanger Institute for the COVID-19 Genomics UK (COG-UK) Consortium | Harper VanSteenhouse, Yumi Kasai, David Gray, Carol Clugston, Anna Dominiczak and Alex Alderton, Roberto Amato, Sonia Goncalves, Ewan Harrison, David K. Jackson, Ian Johnston, Dominic Kwiatkowski, Cordelia Langford, John Sillitoe on behalf of the Wellcome Sanger Institute COVID-19 Surveillance Team |
| EPI_ISL_602110, EPI_ISL_602111, EPI_ISL_602112, EPI_ISL_602113, EPI_ISL_602114, EPI_ISL_602115, EPI_ISL_602116, EPI_ISL_602117, EPI_ISL_602118, EPI_ISL_602119, EPI_ISL_602120, EPI_ISL_602121 | | | |
| see above | Lighthouse Lab in Glasgow | Wellcome Sanger Institute for the COVID-19 Genomics UK (COG-UK) consortium | Harper VanSteenhouse, Yumi Kasai, David Gray, Carol Clugston, Anna Dominiczak and Alex Alderton, Roberto Amato, Sonia Goncalves, Ewan Harrison, David K. Jackson, Ian Johnston, Dominic Kwiatkowski, Cordelia Langford, John Sillitoe on behalf of the Wellcome Sanger Institute COVID-19 Surveillance Team (http://www.sanger.ac.uk/covid-team) |
| EPI_ISL_602122 | Lighthouse Lab in Glasgow | Wellcome Sanger Institute for the COVID-19 Genomics UK (COG-UK) Consortium | Harper VanSteenhouse, Yumi Kasai, David Gray, Carol Clugston, Anna Dominiczak and Alex Alderton, Roberto Amato, Sonia Goncalves, Ewan Harrison, David K. Jackson, Ian Johnston, Dominic Kwiatkowski, Cordelia Langford, John Sillitoe on behalf of the Wellcome Sanger Institute COVID-19 Surveillance Team |
| EPI_ISL_602123, EPI_ISL_602124, EPI_ISL_602125, EPI_ISL_602126, EPI_ISL_602127, EPI_ISL_602128, EPI_ISL_602129, EPI_ISL_602130, EPI_ISL_602131, EPI_ISL_602132, EPI_ISL_602133, EPI_ISL_602134, EPI_ISL_602135, EPI_ISL_602136, EPI_ISL_602137, EPI_ISL_602138, EPI_ISL_602139, EPI_ISL_602140, EPI_ISL_602141, EPI_ISL_602142, EPI_ISL_602143, EPI_ISL_602144 | | | |
| see above | Lighthouse Lab in Glasgow | Wellcome Sanger Institute for the COVID-19 Genomics UK (COG-UK) consortium | Harper VanSteenhouse, Yumi Kasai, David Gray, Carol Clugston, Anna Dominiczak and Alex Alderton, Roberto Amato, Sonia Goncalves, Ewan Harrison, David K. Jackson, Ian Johnston, Dominic Kwiatkowski, Cordelia Langford, John Sillitoe on behalf of the Wellcome Sanger Institute COVID-19 Surveillance Team (http://www.sanger.ac.uk/covid-team) |
| EPI_ISL_602145 | Lighthouse Lab in Milton Keynes | Wellcome Sanger Institute for the COVID-19 Genomics UK (COG-UK) consortium | The Lighthouse Lab in Milton Keynes and Alex Alderton, Roberto Amato, Sonia Goncalves, Ewan Harrison, David K. Jackson, Ian Johnston, Dominic Kwiatkowski, Cordelia Langford, John Sillitoe on behalf of the Wellcome Sanger Institute COVID-19 Surveillance Team (http://www.sanger.ac.uk/covid-team) |
| EPI_ISL_602329, EPI_ISL_602330, EPI_ISL_602331, EPI_ISL_602332, EPI_ISL_602333, EPI_ISL_602334, EPI_ISL_602335, EPI_ISL_602341 | HELIX LLC | WHO National Influenza Centre Russian Federation | Andrey Komissarov, Artem Fadeev, Kseniya Komissarova, Anna Ivanova, Dmitry Bazhenov, Daria Danilenko |
| EPI_ISL_602639, EPI_ISL_602642, EPI_ISL_602643, EPI_ISL_602646, EPI_ISL_602647, EPI_ISL_602648, EPI_ISL_602649, EPI_ISL_602651, EPI_ISL_602652, EPI_ISL_602653, EPI_ISL_602655, EPI_ISL_602656, EPI_ISL_602657, EPI_ISL_602658, EPI_ISL_602662, EPI_ISL_602663, EPI_ISL_602664, EPI_ISL_602665, EPI_ISL_602682, EPI_ISL_602683, EPI_ISL_602684, EPI_ISL_602685, EPI_ISL_602686, EPI_ISL_602687, EPI_ISL_602688, EPI_ISL_602689, EPI_ISL_602690, EPI_ISL_602691, EPI_ISL_602692, EPI_ISL_602693, EPI_ISL_602694, EPI_ISL_602695, EPI_ISL_602696, EPI_ISL_602697, EPI_ISL_602698, EPI_ISL_602699, EPI_ISL_602700, EPI_ISL_602701, EPI_ISL_602702, EPI_ISL_602703, EPI_ISL_602704, EPI_ISL_602705, EPI_ISL_602706, EPI_ISL_602707, EPI_ISL_602708, EPI_ISL_602709, EPI_ISL_602710, EPI_ISL_602711, EPI_ISL_602712, EPI_ISL_602713, EPI_ISL_602714, EPI_ISL_602715, EPI_ISL_602716, EPI_ISL_602717, EPI_ISL_602718, EPI_ISL_602719, EPI_ISL_602720, EPI_ISL_602721, EPI_ISL_602722, EPI_ISL_602723, EPI_ISL_602724, EPI_ISL_602725, EPI_ISL_602726, EPI_ISL_602727, EPI_ISL_602728, EPI_ISL_602729, EPI_ISL_602730, EPI_ISL_602731, EPI_ISL_602732, EPI_ISL_602733, EPI_ISL_602734, EPI_ISL_602735, EPI_ISL_602736, EPI_ISL_602737, EPI_ISL_602738, EPI_ISL_602739, EPI_ISL_602740, EPI_ISL_602741, EPI_ISL_602742, EPI_ISL_602743, EPI_ISL_602744, EPI_ISL_602745, EPI_ISL_602746, EPI_ISL_602747, EPI_ISL_602748, EPI_ISL_602749, EPI_ISL_602750, EPI_ISL_602751, EPI_ISL_602752, EPI_ISL_602753, EPI_ISL_602754, EPI_ISL_602755, EPI_ISL_602756, EPI_ISL_602757, EPI_ISL_602758, EPI_ISL_602759, EPI_ISL_602760, EPI_ISL_602766, EPI_ISL_602768, EPI_ISL_602779, EPI_ISL_602780, EPI_ISL_602781 | | | |
| see above | NHLS-IALCH | KRISP, KZN Research Innovation and Sequencing Platform | Giandhari J, Pillay S, Lessells R, Mdlalose K, York D, Khan S, Tegally H, Wilkinson E, de Oliveira T |
| EPI_ISL_603216, EPI_ISL_603217 | CHU Purpan - Laboratoire de Virologie - Institut Fédératif de Biologie | CHU Purpan - Laboratoire de Virologie - Institut Fédératif de Biologie | Latour J., Ranger N., Dubois M., Carcenac R., Harter A., Boyer P., Tremeaux P., Izopet J. |
| EPI_ISL_605405, EPI_ISL_605406, EPI_ISL_605407, EPI_ISL_605504, EPI_ISL_605505, EPI_ISL_605506, EPI_ISL_605507, EPI_ISL_605508, EPI_ISL_605509, EPI_ISL_605510, EPI_ISL_605520, EPI_ISL_605521, EPI_ISL_605522, EPI_ISL_605523, EPI_ISL_605528, EPI_ISL_605565, EPI_ISL_605567, EPI_ISL_605570, EPI_ISL_605573, EPI_ISL_605574, EPI_ISL_605578, EPI_ISL_605583, EPI_ISL_605585, EPI_ISL_605587, EPI_ISL_605594, EPI_ISL_605596, EPI_ISL_605603, EPI_ISL_605604, EPI_ISL_605605, EPI_ISL_605609, EPI_ISL_605610, EPI_ISL_605671, EPI_ISL_605676, EPI_ISL_605677, EPI_ISL_605682, EPI_ISL_605684, EPI_ISL_605685, EPI_ISL_605686, EPI_ISL_605687, EPI_ISL_605688, EPI_ISL_605689, EPI_ISL_605690, EPI_ISL_605691, EPI_ISL_605693, EPI_ISL_605694, EPI_ISL_605695, EPI_ISL_605696, EPI_ISL_605697, EPI_ISL_605699, EPI_ISL_605701, EPI_ISL_605705, EPI_ISL_605706, EPI_ISL_605714, EPI_ISL_605718, EPI_ISL_605721, EPI_ISL_605737, EPI_ISL_605738, EPI_ISL_605739, EPI_ISL_605740, EPI_ISL_605741, EPI_ISL_605742 | | | |
| see above | University of Wisconsin-Madison AIDS Vaccine Research Laboratories | University of Wisconsin-Madison AIDS Vaccine Research Laboratories | Gage Moreno, Katarina Braun, et al. AIDS Vaccine Research Laboratories |
| EPI_ISL_609801, EPI_ISL_609802, EPI_ISL_609805, EPI_ISL_609806, EPI_ISL_609807, EPI_ISL_609809, EPI_ISL_609813, EPI_ISL_609816, EPI_ISL_609817, EPI_ISL_609818, EPI_ISL_609820, EPI_ISL_609821, EPI_ISL_609824, EPI_ISL_609825, EPI_ISL_609826 | | | |
| see above | Unity Health Toronto | Ontario Institute for Cancer Research | Ramzi Fattouh, Larissa M. Matukas, Yan Chen,Mark Downing, Trina Otterman, Karel Boissinot, Wai Sum Siu, Zhi Cui, Le Luu, Samira Mubareka, TIBDN, Ilinca Lungu, Bernard Lam, Jeremy Johns, Paul Krzyzanowski, Richard de Borja, Felicia Vincelli, Philip Zuzarte, Jared T. Simpson |
| EPI_ISL_609971, EPI_ISL_609972, EPI_ISL_609973, EPI_ISL_609974, EPI_ISL_609975, EPI_ISL_609976, EPI_ISL_609977, EPI_ISL_609978, EPI_ISL_609979, EPI_ISL_609980, EPI_ISL_609981, EPI_ISL_609982, EPI_ISL_609983 | | | |
| see above | Virginia DCLS | Virginia DCLS | Virginia DCLS |
| EPI_ISL_610155 | Puskesmas Cangkrep | Genetics Working Group (Pokja Genetik) Faculty of Medicine, Public Health and Nursing Universitas Gadjah Mada (FK-KMK UGM); Disease Investigation Center Wates Ministry of Agriculture Indonesia; Department of Microbiology FK-KMK UGM; Laboratorium Diagnostik Yayasan Tahija World Mosquito Program (WMP) Yogyakarta Center for Tropical Medicine FK-KMK UGM; Integrated Research Center FK-KMK UGM; Department of Computer Science and Electronics FMIPA UGM; Balai Besar Teknik Kesehatan Lingkungan dan Pengendalian Penyakit (BBTKLPP) Yogyakarta | Gunadi, Hendra Wibawa, Marcellus, Mohamad S. Hakim, Edwin W. Daniwijaya, Ludhang P. Rizki, Endah Supriyati, Eggi Arguni, Titik Nuryastuti, Tri Wibawa, Dwi AA Nugrahaningsih, Afiahayati, Siswanto, Kristy Iskandar, Nungki Anggorowati, Irene, Indaryati, Havid Setyawan, Nawang Sukestiningsih, Fadil Fahri, Ririn Enggy Yuliyanti |
| EPI_ISL_610161 | RSUD Dr. Tjitrowardojo | Genetics Working Group (Pokja Genetik) Faculty of Medicine, Public Health and Nursing Universitas Gadjah Mada (FK-KMK UGM); Disease Investigation Center Wates Ministry of Agriculture Indonesia; Department of Microbiology FK-KMK UGM; Laboratorium Diagnostik Yayasan Tahija World Mosquito Program (WMP) Yogyakarta Center for Tropical Medicine FK-KMK UGM; Integrated Research Center FK-KMK UGM; Department of Computer Science and Electronics | Gunadi, Hendra Wibawa, Marcellus, Mohamad S. Hakim, Edwin W. Daniwijaya, Ludhang P. Rizki, Endah Supriyati, Eggi Arguni, Titik Nuryastuti, Tri Wibawa, Dwi AA Nugrahaningsih, Afiahayati, Siswanto, Kristy Iskandar, Nungki Anggorowati, Irene, Indaryati, Havid Setyawan, Wuryanto, Kemala Athollah, Desyifa Mursalin |

| | | | |
|---|---|---|---|
| | | FMIPA UGM; Balai Besar Teknik Kesehatan Lingkungan dan Pengendalian Penyakit (BBTKLPP) Yogyakarta | |
| EPI_ISL_610216, EPI_ISL_610217 | Department of Health Technology and Informatics, The Hong Kong Polytechnic University | Department of Health Technology and Informatics, The Hong Kong Polytechnic University | Siu,G.K.-H., Lee,L.-K., Leung,K.S.-S., Leung,J.S.-L., Ng,T.T.-L., Chan,C.T.-M., Tam,K.K.-G., Lao,H.-Y., Wu,A.K.-L., Yau,M.C.-Y., Lai,Y.W.-M., Fung,K.S.-C., Chau,S.K.-Y., Wong,B.K.-C., To,W.-K., Luk,K., Ho,A.Y.-M., Que,T.-L., Yip,K.-T., Yam,W.C., Shum,D.H.-K., Yip,S.P. |
| EPI_ISL_611521, EPI_ISL_611629, EPI_ISL_611723, EPI_ISL_612065 | Liverpool Clinical Laboratories | COVID-19 Genomics UK (COG-UK) Consortium | Sam Haldenby, Anita Lucaci, Steve Paterson, Julian Hiscox, Alistair Darby, M Almsaud, A Alrezaihi, Muhannad Alruwaili, Stuart D Armstrong, Jones Benjamin, Eleanor G Bentley, Anu Chawla, Jordan J Clark, Angela Cowell, Richard Eccles, Isabel García-Dorival, Matthew Gemmell, Alessandro Gerada, PKF Gilmore, Richard Gregory, Ximeng Han, Catherine Hartley, Margaret Hughes, Miren Iturriza-Gomara, James Johnson, L Luu, Jenifer Manson, Charlotte Nelson, Elaine O'Toole, Cassie Olateju, Rebekah Penrice-Randal , Lucille Rainbow, N.P Randle, Trevor Ian Robinson, Parul Sharma, Ghada T Shawli, James P Stewart, Neil Swainston, Ecaterina Vamos, Joanne Watts, Mark Whitehead |
| EPI_ISL_612284 | West of Scotland Specialist Virology Centre, NHSGGC / MRC-University of Glasgow Centre for Virus Research | COVID-19 Genomics UK (COG-UK) Consortium | Ana da Silva Filipe, Natasha Johnson, Kathy Smollett, Daniel Mair, Stephen Carmichael, Lily Tong, Jenna Nichols, Elihu Aranday-Cortes, Kyriaki Nomikou; Sarah McDonald, Marc Niebel, Patawee Asamaphan; Richard Orton, Joseph Hughes, Sreenu Vattipally, David L Robertson; Alasdair MacLean, Rory Gunson; Kathy Li, Igor Starinskij, Natasha Jesudason, Rajiv Shah, James Shepherd, Antonia Ho, Emma Thomson |
| EPI_ISL_613549, EPI_ISL_613551, EPI_ISL_613552, EPI_ISL_613553, EPI_ISL_613554, EPI_ISL_613555, EPI_ISL_613556, EPI_ISL_613557, EPI_ISL_613558, EPI_ISL_613559 | CHRU Pontchaillou - Laboratoire de Virologie 2, rue Henri Le Guilloux | National Reference Center for Viruses of Respiratory Infections, Institut Pasteur, Paris | Marion Barbet, Sylvie Behillil, Méline Bizard, Angela Brisebarre, Camille Capel, Etienne Simon-Lorière, Vincent Enouf, Maud Vanpeene, Sylvie van der Werf, Gisèle Lagathu |
| EPI_ISL_614050, EPI_ISL_614051, EPI_ISL_614052, EPI_ISL_614053, EPI_ISL_614054, EPI_ISL_614075, EPI_ISL_614123, EPI_ISL_614124, EPI_ISL_614125, EPI_ISL_614126, EPI_ISL_614127 | | | |
| see above | Virginia DCLS | Virginia DCLS | Virginia DCLS |
| EPI_ISL_614393, EPI_ISL_614394, EPI_ISL_614395 | Molecular diagnostic unit for viral haemorrhagic fevers and emerging viruses, Bouaké CHU Laboratory | Project group Epidemology of Highly Pathogenic Microorganisms, Robert Koch-Institute | Chantal Akoua-Koffi, Diané Bamourou, Etilé Anoh, Essia Belarbi, Safiatou Karidioula, Grit Schubert, Adjaratou Traoré, Soundélé Maïté, Monemo Pacome, Coulibaly Mbegnan, Bamba Fatoumata Touré, Kra Ouffoué, Fabian Leendertz |
| EPI_ISL_615104 | Halmstad klinisk mikrobiologi | The Public Health Agency of Sweden | Anna-Malin Linde, Maria Lind Karlberg, Mattias Haukland, Reza Advani, Olov Svartstrom, Oskar Karlsson Lindsjo, Sandra Broddesson, Petra Edquist, Mia Brytting, Anna Risberg, Karin Tegmark-Wisell |
| EPI_ISL_615115, EPI_ISL_615116 | Skovde/Unilabs | The Public Health Agency of Sweden | Anna-Malin Linde, Maria Lind Karlberg, Mattias Haukland, Reza Advani, Olov Svartstrom, Oskar Karlsson Lindsjo, Sandra Broddesson, Petra Edquist, Mia Brytting, Anna Risberg, Karin Tegmark-Wisell |
| EPI_ISL_615178, EPI_ISL_615181, EPI_ISL_615187, EPI_ISL_615188, EPI_ISL_615189, EPI_ISL_615190, EPI_ISL_615191, EPI_ISL_615192, EPI_ISL_615193, EPI_ISL_615326, EPI_ISL_615327, EPI_ISL_615397, EPI_ISL_615398, EPI_ISL_615399, EPI_ISL_615400, EPI_ISL_615401, EPI_ISL_615402, EPI_ISL_615403, EPI_ISL_615453, EPI_ISL_615454, EPI_ISL_615455, EPI_ISL_615456, EPI_ISL_615457, EPI_ISL_615458, EPI_ISL_615459, EPI_ISL_615460, EPI_ISL_615461, EPI_ISL_615462, EPI_ISL_615463, EPI_ISL_615464, EPI_ISL_615465, EPI_ISL_615466, EPI_ISL_615468, EPI_ISL_615469, EPI_ISL_615470, EPI_ISL_615471, EPI_ISL_615472, EPI_ISL_615516, EPI_ISL_615517, EPI_ISL_615518, EPI_ISL_615519, EPI_ISL_615520, EPI_ISL_615521, EPI_ISL_615522, EPI_ISL_615529, EPI_ISL_615530, EPI_ISL_615531, EPI_ISL_615532, EPI_ISL_615533, EPI_ISL_615534, EPI_ISL_615535, EPI_ISL_615536, EPI_ISL_615537, EPI_ISL_615538, EPI_ISL_615539, EPI_ISL_615540, EPI_ISL_615541, EPI_ISL_615542, EPI_ISL_615543, EPI_ISL_615544, EPI_ISL_615545, EPI_ISL_615546, EPI_ISL_615547, EPI_ISL_615548, EPI_ISL_615549, EPI_ISL_615550, EPI_ISL_615551, EPI_ISL_615553, EPI_ISL_615554, EPI_ISL_615555, EPI_ISL_615556, EPI_ISL_615557, EPI_ISL_615558, EPI_ISL_615559, EPI_ISL_615560, EPI_ISL_615561, EPI_ISL_615562, EPI_ISL_615564, EPI_ISL_615565, EPI_ISL_615566, EPI_ISL_615572, EPI_ISL_615574, EPI_ISL_615575, EPI_ISL_615576, EPI_ISL_615578, EPI_ISL_615579, EPI_ISL_615580, EPI_ISL_615582, EPI_ISL_615583, EPI_ISL_615584, EPI_ISL_615585, EPI_ISL_615586, EPI_ISL_615587, EPI_ISL_615588, EPI_ISL_615589, EPI_ISL_615590, EPI_ISL_615591, EPI_ISL_615592, EPI_ISL_615593, EPI_ISL_615594, EPI_ISL_615595, EPI_ISL_615596, EPI_ISL_615597, EPI_ISL_615598, EPI_ISL_615599, EPI_ISL_615607, EPI_ISL_615609, EPI_ISL_615610, EPI_ISL_615611, EPI_ISL_615612, EPI_ISL_615613, EPI_ISL_615614, EPI_ISL_615615, EPI_ISL_615616, EPI_ISL_615618, EPI_ISL_615619, EPI_ISL_615620, EPI_ISL_615621, EPI_ISL_615622, EPI_ISL_615623, EPI_ISL_615624, EPI_ISL_615625, EPI_ISL_615627, EPI_ISL_615633, EPI_ISL_615634, EPI_ISL_615645, EPI_ISL_615646, EPI_ISL_615647, EPI_ISL_615648, EPI_ISL_615649, EPI_ISL_615650, EPI_ISL_615651, EPI_ISL_615670, EPI_ISL_615671, EPI_ISL_615672, EPI_ISL_615673, EPI_ISL_615675, EPI_ISL_615676, EPI_ISL_615677, EPI_ISL_615678, EPI_ISL_615679, EPI_ISL_615680, EPI_ISL_615681, EPI_ISL_615682, EPI_ISL_615683, EPI_ISL_615687, EPI_ISL_615688, EPI_ISL_615689, EPI_ISL_615690, EPI_ISL_615691, EPI_ISL_615692, EPI_ISL_615693, EPI_ISL_615694, EPI_ISL_615695, EPI_ISL_615696, EPI_ISL_615697, EPI_ISL_615698, EPI_ISL_615699, EPI_ISL_615700, EPI_ISL_615701, EPI_ISL_615702, EPI_ISL_615703, EPI_ISL_615704, EPI_ISL_615705, EPI_ISL_615706, EPI_ISL_615707, EPI_ISL_615708, EPI_ISL_615709, EPI_ISL_615710, EPI_ISL_615711, EPI_ISL_615712, EPI_ISL_615713, EPI_ISL_615714, EPI_ISL_615715, EPI_ISL_615716, EPI_ISL_615717, EPI_ISL_615718, EPI_ISL_615719, EPI_ISL_615720, EPI_ISL_615721, EPI_ISL_615722, EPI_ISL_615723, EPI_ISL_615724, EPI_ISL_615725, EPI_ISL_615726, EPI_ISL_615727, EPI_ISL_615728, EPI_ISL_615729, EPI_ISL_615730, EPI_ISL_615731, EPI_ISL_615732, EPI_ISL_615733, EPI_ISL_615734, EPI_ISL_615735, EPI_ISL_615736, EPI_ISL_615737, EPI_ISL_615738, EPI_ISL_615739, EPI_ISL_615740, EPI_ISL_615741, EPI_ISL_615742, EPI_ISL_615743, EPI_ISL_615744, EPI_ISL_615745, EPI_ISL_615746, EPI_ISL_615747, EPI_ISL_615748, EPI_ISL_615749, EPI_ISL_615750, EPI_ISL_615751, EPI_ISL_615752, EPI_ISL_616372, EPI_ISL_616374, EPI_ISL_616375, EPI_ISL_616376, EPI_ISL_616377, EPI_ISL_616378, EPI_ISL_616379, EPI_ISL_616381, EPI_ISL_616383, EPI_ISL_621186, EPI_ISL_621187, EPI_ISL_621188, EPI_ISL_621189, EPI_ISL_621190, EPI_ISL_621826, EPI_ISL_622726, EPI_ISL_622734, EPI_ISL_622736, EPI_ISL_622739, EPI_ISL_622740, EPI_ISL_622741, EPI_ISL_622742, EPI_ISL_622743, EPI_ISL_622744, EPI_ISL_622745, EPI_ISL_622746, EPI_ISL_622747 | | | |
| see above | Department of Virus and Microbiological Special Diagnostics, Statens Serum Institut, Denmark | Albertsen lab, Department of Chemistry and Bioscience, Aalborg University, Denmark | Danish Covid-19 Genome Consortia |
| EPI_ISL_623098, EPI_ISL_623100, EPI_ISL_623101, EPI_ISL_623102 | CNR Virus des Infections Respiratoires - France SUD | CNR Virus des Infections Respiratoires - France SUD | Antonin Bal, Gregory Destras, Gwendolyne Burfin, Hadrien Règue, Alexandre Gaymard, Maude Bouscambert-Duchamp, Florence Morfin-Sherpa, Martine Valette, Bruno Lina, Laurence Josset |
| EPI_ISL_625622 | Alameda County Public Health Lab | Chan-Zuckerberg Biohub | CZB Cliahub Consortium |
| EPI_ISL_625675, EPI_ISL_625676 | Laboratory of Molecular Medicine, University of Magallanes | Centro Asistencial Docente y de Investigacion, Universidad de Magallanes | Jorge Gonzalez, Jacqueline Aldridge, Diego Alvarez, Marcelo Navarrete |
| EPI_ISL_626242, EPI_ISL_626243, EPI_ISL_626244, EPI_ISL_626245, EPI_ISL_626246, EPI_ISL_626247, EPI_ISL_626248, EPI_ISL_626249, EPI_ISL_626250, EPI_ISL_626251, EPI_ISL_626252, EPI_ISL_626253, EPI_ISL_626254 | | | |
| see above | Department of Clinical Microbiology | GIGA Medical Genomics | Keith Durkin, Maria Artesi, Sébastien Bontems, Raphaël Boreux, Bouchra Boujemla, Cécile Meex, Pierrette Melin, Marie-Pierre Hayette, Vincent Bours |
| EPI_ISL_626575 | The National Institute of Public Health | State Veterinary Institute Prague | Nagy,A;Jirincova,H;Novakova,L;Trnka,D;Vecerova,J |
| EPI_ISL_626937, EPI_ISL_627052 | University of Exeter | COVID-19 Genomics UK (COG-UK) Consortium | Ben Temperton,Aaron Jeffries,Michelle Michelsen,Joanna Warwick-Dugdale,Audrey Farbos,Robyn Manley,Stephen Michell,Jane Masoli |
| EPI_ISL_627811 | Wales Specialist Virology Centre Sequencing lab: Pathogen Genomics Unit | COVID-19 Genomics UK (COG-UK) Consortium | Catherine Moore, Johnathan Evans, Laura Gifford, Malorie Perry, Simon Cottrell, Angela Marchbank, Alec Birchley, Alexander Adams, Amy Gaskin, Bree Gatica-Wilcox, Jason Coombes, Joel Southgate, Lauren Gilbert, Lee Graham, Nicole Pacchiarini, Sara Kumziene-Summerhayes, Sarah Taylor, Sophie Jones, Sara Rey, Matthew Bull, Joanne Watkins, Sally Corden, Tom Connor |
| EPI_ISL_628543 | Oxford Viromics, NDM, University of Oxford; Oxford University Hospitals; Basingstoke and North Hampshire Hospital | COVID-19 Genomics UK (COG-UK) Consortium | Tanya Golubchik, David Bonsall, George Macintyre, Amy Trebes, Mariateresa de Cesare, Catrin Moore, Alex Mobbs, Anita Justice, Robert Shaw, Monique Andersson, Timothy Peto, Emma Wise, Nathan Moore, Jessica Lynch, Nick Cortes, Matilde Mori, Stephen Kidd, David Buck, John Todd, Christophe Fraser |
| EPI_ISL_631493 | Wisconsin State Laboratory of Hygiene Communicable Disease Division | Wisconsin State Laboratory of Hygiene Communicable Disease Division | Kelsey R. Florek, Abigail C. Shockey |
| EPI_ISL_632377, EPI_ISL_632386, EPI_ISL_632415, EPI_ISL_632422, EPI_ISL_632520, EPI_ISL_632521, EPI_ISL_632522, EPI_ISL_632523, EPI_ISL_632524, EPI_ISL_632525, EPI_ISL_632526, EPI_ISL_632527, EPI_ISL_632528, EPI_ISL_632529, EPI_ISL_632532, EPI_ISL_632533, EPI_ISL_632534, EPI_ISL_632577, EPI_ISL_632588 | | | |
| see above | Dutch COVID-19 response team | Erasmus Medical Center | Bas Oude Munnink, David Nieuwenhuijse, Reina Sikkema, Claudia Schapendonk, Irina Chestakova, Anne van der Linden, Theo Bestebroer, Stefan van Nieuwkoop, Mark Pronk, Pascal Lexmond, Corien Swaan, Manon Haverkate, Madelief Mollers, Mart Stein, Sandra Kengne Kamga Mobou, Jeroen van Kampen, Jolanda Voermans, Aura Timen, Corine GeurtsvanKessel, Annemiek van der Eijk, Richard Molenkamp, Marion Koopmans, on behalf of the Dutch national COVID-19 response team. |
| EPI_ISL_632823, EPI_ISL_632824, EPI_ISL_632825, EPI_ISL_632826, EPI_ISL_632827, EPI_ISL_632828, EPI_ISL_632829, EPI_ISL_632830, EPI_ISL_632831, EPI_ISL_632832, EPI_ISL_632833, EPI_ISL_632834, EPI_ISL_632835, EPI_ISL_632836, EPI_ISL_632837, EPI_ISL_632838, EPI_ISL_632839, EPI_ISL_632840, EPI_ISL_632841, EPI_ISL_632842, EPI_ISL_632843, EPI_ISL_632844, EPI_ISL_632845 | | | |
| see above | New Mexico Department of Health Scientific Laboratory | New Mexico Department of Health Scientific Laboratory | Ellie Johnson, Anastacia Griego-Fisher, D'Eldra Malone |
| EPI_ISL_632937 | RSUD Saptosari Gunung Kidul | Genetics Working Group (Pokja Genetik) Faculty of Medicine, Public Health and Nursing Universitas Gadjah Mada (FK-KMK UGM); Disease Investigation Center Wates Ministry of Agriculture Indonesia; Department of Microbiology FK-KMK UGM; Laboratorium Diagnostik Yayasan Tahija World Mosquito Program (WMP) Yogyakarta Center for Tropical Medicine FK-KMK UGM; Integrated Research Center FK-KMK UGM; Department of Computer Science and Electronics | Gunadi, Hendra Wibawa, Marcellus, Mohamad S. Hakim, Edwin W. Daniwijaya, Ludhang P. Rizki, Endah Supriyati, Eggi Arguni, Titik Nuryastuti, Tri Wibawa, Dwi AA Nugrahaningsih, Afiahayati, Siswanto, Kristy Iskandar, Nungki Anggorowati, Irene, Indaryati, Havid Setyawan, Eko Darmawan, Maria Patricia Inggriani, Audric Kenny Tedja |

| | | | |
|---|---|---|---|
| | | FMIPA UGM; Balai Besar Teknik Kesehatan Lingkungan dan Pengendalian Penyakit (BBTKLPP) Yogyakarta | |
| EPI_ISL_633053 | DOHMH Morrisania | New York City Public Health Laboratory | Jade Wang, et al. |
| EPI_ISL_633054, EPI_ISL_633055 | DOHMH PHL | New York City Public Health Laboratory | Jade Wang, et al. |
| EPI_ISL_634880 | Lab voor klinische biologie | Onderzoeksgroep Virologie | Laurens Lambrechts, Nick Vereecke, Marthe Pauwels, Bruno Verhasselt, Linos Vandekerckhove, Hans Nauwynck, Sebastiaan Theuns |
| EPI_ISL_635104 | Unilabs Laboratory Medicine | Norwegian Institute of Public Health, Department of Virology | Kathrine Stene-Johansen, Kamilla Heddeland Instefjord, Hilde Elshaug, Marie Paulsen Madsen, Rasmus Riis Kopperud, Hilde Vollan, Karoline Bragstad, Olav Hungnes |
| EPI_ISL_635108, EPI_ISL_635109, EPI_ISL_635150, EPI_ISL_635151, EPI_ISL_635152 | Ostfold Hospital Trust - Kalnes, Centre for Laboratory Medicine, Section for gene technology and infection serology | Norwegian Institute of Public Health, Department of Virology | Kathrine Stene-Johansen, Kamilla Heddeland Instefjord, Hilde Elshaug, Marie Paulsen Madsen, Rasmus Riis Kopperud, Hilde Vollan, Karoline Bragstad, Olav Hungnes |
| EPI_ISL_635749, EPI_ISL_635750, EPI_ISL_635751, EPI_ISL_635752, EPI_ISL_635753, EPI_ISL_635754, EPI_ISL_635755, EPI_ISL_635756, EPI_ISL_635757, EPI_ISL_635758, EPI_ISL_635759, EPI_ISL_635760, EPI_ISL_635761, EPI_ISL_635762, EPI_ISL_635763, EPI_ISL_635764, EPI_ISL_635765, EPI_ISL_635766, EPI_ISL_635767, EPI_ISL_635768, EPI_ISL_635769, EPI_ISL_635770, EPI_ISL_635771, EPI_ISL_635793, EPI_ISL_635794, EPI_ISL_635795, EPI_ISL_635796, EPI_ISL_635797, EPI_ISL_635798, EPI_ISL_635799, EPI_ISL_635803, EPI_ISL_635806, EPI_ISL_635807, EPI_ISL_635814, EPI_ISL_635817, EPI_ISL_635818, EPI_ISL_635825, EPI_ISL_635827, EPI_ISL_635829, EPI_ISL_635830, EPI_ISL_635833, EPI_ISL_635834, EPI_ISL_635837, EPI_ISL_635838, EPI_ISL_635839, EPI_ISL_635840, EPI_ISL_635841, EPI_ISL_635842, EPI_ISL_635843, EPI_ISL_635932, EPI_ISL_635933, EPI_ISL_635934, EPI_ISL_635935, EPI_ISL_635936, EPI_ISL_635937, EPI_ISL_635938, EPI_ISL_635939, EPI_ISL_635940, EPI_ISL_635941, EPI_ISL_635942, EPI_ISL_635943, EPI_ISL_635944, EPI_ISL_635945, EPI_ISL_635994, EPI_ISL_636050, EPI_ISL_636054, EPI_ISL_636056, EPI_ISL_636060, EPI_ISL_636061, EPI_ISL_636062, EPI_ISL_636069, EPI_ISL_636070, EPI_ISL_636071 | | | |
| see above | San Diego County Public Health Laboratory | Andersen lab at Scripps Research | SEARCH Alliance San Diego with Tracy Basler, Jovan Shephard, Brett Austin |
| EPI_ISL_636548, EPI_ISL_636578, EPI_ISL_636590, EPI_ISL_636600, EPI_ISL_636601, EPI_ISL_636602, EPI_ISL_636603 | Dutch COVID-19 response team | National Institute for Public Health and the Environment (RIVM) | Adam Meijer, Harry Vennema, Jeroen Cremer, Sharon van den Brink, Bas van der Veer, AnneMarie van den Brandt, Florian Zwagemaker, Dennis Schmitz, Chantal Reusken, on behalf of the national COVID-19 response team |
| EPI_ISL_636882, EPI_ISL_636883, EPI_ISL_636884, EPI_ISL_636885, EPI_ISL_636886, EPI_ISL_636887, EPI_ISL_636888, EPI_ISL_636889, EPI_ISL_636890, EPI_ISL_636891 | Lithuanian University of Health Sciences Hospital, Department of Laboratory Medicine | Lithuanian University of Health Sciences, Molecular cardiology lab. | Lukas Zemaitis, Ingrida Olendrait, Arnoldas Pautienius, Kamile Tamusauskaite, Dovydas Gecys, Laura Pareckaite, Vaiva Lesauskaite, Astra Vitkauskiene |
| EPI_ISL_637261, EPI_ISL_637282, EPI_ISL_637292, EPI_ISL_637324, EPI_ISL_637334, EPI_ISL_637335, EPI_ISL_637341, EPI_ISL_637345 | Department of Pathology, University of Cambridge | COVID-19 Genomics UK (COG-UK) Consortium | Aminu S. Jahun, Yasmin Chaudhry, Grant Hall, Iliana Georgana, Myra Hosmillo, Martin D. Curran, Malte Pinckert, Surendra Parmar, Ian Goodfellow |
| EPI_ISL_637351 | Wales Specialist Virology Centre Sequencing lab: Pathogen Genomics Unit | COVID-19 Genomics UK (COG-UK) Consortium | Catherine Moore, Johnathan Evans, Laura Gifford, Malorie Perry, Simon Cottrell, Angela Marchbank, Alec Birchley, Alexander Adams, Amy Gaskin, Bree Gatica-Wilcox, Jason Coombes, Joel Southgate, Lauren Gilbert, Lee Graham, Nicole Pacchiarini, Sara Kumziene-Summerhayes, Sarah Taylor, Sophie Jones, Sara Rey, Matthew Bull, Joanne Watkins, Sally Corden, Tom Connor |
| EPI_ISL_637429, EPI_ISL_637430, EPI_ISL_637431, EPI_ISL_637440, EPI_ISL_637545, EPI_ISL_637547, EPI_ISL_637570, EPI_ISL_637590, EPI_ISL_637591, EPI_ISL_637592, EPI_ISL_637595, EPI_ISL_637632, EPI_ISL_637633, EPI_ISL_637634, EPI_ISL_637635, EPI_ISL_637662, EPI_ISL_637663, EPI_ISL_637664, EPI_ISL_637665, EPI_ISL_637669, EPI_ISL_637670, EPI_ISL_637671, EPI_ISL_637801, EPI_ISL_637802, EPI_ISL_637913, EPI_ISL_637955, EPI_ISL_638079, EPI_ISL_638614, EPI_ISL_638615, EPI_ISL_638616, EPI_ISL_638617, EPI_ISL_638618, EPI_ISL_638619, EPI_ISL_638620, EPI_ISL_638621, EPI_ISL_638622, EPI_ISL_638623, EPI_ISL_638624, EPI_ISL_638625, EPI_ISL_638626, EPI_ISL_638627, EPI_ISL_638628, EPI_ISL_638629, EPI_ISL_638630, EPI_ISL_638631, EPI_ISL_638632, EPI_ISL_638633, EPI_ISL_638634, EPI_ISL_638635, EPI_ISL_638636, EPI_ISL_638637, EPI_ISL_638638, EPI_ISL_638639, EPI_ISL_638640, EPI_ISL_638895, EPI_ISL_638896, EPI_ISL_638897, EPI_ISL_638898, EPI_ISL_638899, EPI_ISL_638900, EPI_ISL_638901, EPI_ISL_638902, EPI_ISL_638903, EPI_ISL_638904, EPI_ISL_638905, EPI_ISL_638906, EPI_ISL_638907, EPI_ISL_638908, EPI_ISL_638909, EPI_ISL_638910, EPI_ISL_639624, EPI_ISL_639625 | | | |
| see above | Department of Pathology, University of Cambridge | COVID-19 Genomics UK (COG-UK) Consortium | Aminu S. Jahun, Yasmin Chaudhry, Grant Hall, Iliana Georgana, Myra Hosmillo, Martin D. Curran, Malte Pinckert, Surendra Parmar, Ian Goodfellow |
| EPI_ISL_639634 | Latvijas Infektoloijas centrs | Latvian Biomedical Research and Study Centre | Ivars Silamielis, Kaspars Megnis, Monta Ustinova, ikita Zrelovs, Vita Rovte, Jeena Storoženko, Tatjana Kolupajeva, Oksana Savicka, Uga Dumpis, Jnis Kloviš |
| EPI_ISL_639635 | E. Gulbja Laboratorija | Latvian Biomedical Research and Study Centre | Ivars Silamielis, Kaspars Megnis, Monta Ustinova, ikita Zrelovs, Vita Rovte, Mikus Gavars, Dmitrijs Perminovs, Uga Dumpis, Jnis Kloviš |
| EPI_ISL_639636 | Centrl Laboratorija | Latvian Biomedical Research and Study Centre | Ivars Silamielis, Kaspars Megnis, Monta Ustinova, ikita Zrelovs, Vita Rovte, Stella Lapia, Jana Oste, Marta Priedte, Uga Dumpis, Jnis Kloviš |
| EPI_ISL_639637 | E. Gulbja Laboratorija | Latvian Biomedical Research and Study Centre | Ivars Silamielis, Kaspars Megnis, Monta Ustinova, ikita Zrelovs, Vita Rovte, Mikus Gavars, Dmitrijs Perminovs, Uga Dumpis, Jnis Kloviš |
| EPI_ISL_639640 | Centrl Laboratorija | Latvian Biomedical Research and Study Centre | Ivars Silamielis, Kaspars Megnis, Monta Ustinova, ikita Zrelovs, Vita Rovte, Stella Lapia, Jana Oste, Marta Priedte, Uga Dumpis, Jnis Kloviš |
| EPI_ISL_639641 | E. Gulbja Laboratorija | Latvian Biomedical Research and Study Centre | Ivars Silamielis, Kaspars Megnis, Monta Ustinova, ikita Zrelovs, Vita Rovte, Mikus Gavars, Dmitrijs Perminovs, Uga Dumpis, Jnis Kloviš |
| EPI_ISL_639642, EPI_ISL_639650 | Centrl Laboratorija | Latvian Biomedical Research and Study Centre | Ivars Silamielis, Kaspars Megnis, Monta Ustinova, ikita Zrelovs, Vita Rovte, Stella Lapia, Jana Oste, Marta Priedte, Uga Dumpis, Jnis Kloviš |
| EPI_ISL_639661 | E. Gulbja Laboratorija | Latvian Biomedical Research and Study Centre | Ivars Silamielis, Kaspars Megnis, Monta Ustinova, ikita Zrelovs, Vita Rovte, Mikus Gavars, Dmitrijs Perminovs, Uga Dumpis, Jnis Kloviš |
| EPI_ISL_639826, EPI_ISL_639827, EPI_ISL_639833, EPI_ISL_639836, EPI_ISL_639838, EPI_ISL_639839, EPI_ISL_639844, EPI_ISL_639848, EPI_ISL_639850, EPI_ISL_639855, EPI_ISL_639856, EPI_ISL_639857, EPI_ISL_639858, EPI_ISL_639859, EPI_ISL_639861, EPI_ISL_639862 | | | |
| see above | National Virus Reference Laboratory | National Virus Reference Laboratory | Michael Carr, Gabriel Gonzalez, Jonathan Dean, Daniel Hare, Cillian F De Gascun |
| EPI_ISL_639950 | HELIX LLC | WHO National Influenza Centre Russian Federation | Andrey Komissarov, Artem Fadeev, Kseniya Komissarova, Anna Ivanova, Dmitry Bazhenov, Daria Danilenko |
| EPI_ISL_639994, EPI_ISL_639999 | CNR Virus des Infections Respiratoires - France SUD | CNR Virus des Infections Respiratoires - France SUD | Antonin Bal, Gregory Destras, Gwendolyne Burfin, Hadrien Règue, Alexandre Gaymard, Maude Bouscambert-Duchamp, Florence Morfin-Sherpa, Martine Valette, Bruno Lina, Laurence Josset |
| EPI_ISL_640107 | 2 Military Hospital wc MAA | NHLS/UCT | Arash Iranzadeh, Deelan Doolabh, Lynn Tyers, Bruna Galvao, Innocent Mudau, Marvin Hsiao, Kruger Marais, Diana Hardie, Stephen Korsman, Carolyn Williamson |
| EPI_ISL_640108, EPI_ISL_640109, EPI_ISL_640110 | Victoria Hospital wc VHW | NHLS/UCT | Arash Iranzadeh, Deelan Doolabh, Lynn Tyers, Bruna Galvao, Innocent Mudau, Marvin Hsiao, Kruger Marais, Diana Hardie, Stephen Korsman, Carolyn Williamson |
| EPI_ISL_640111, EPI_ISL_640112 | Groote Schuur Hospital wc GSH | NHLS/UCT | Arash Iranzadeh, Deelan Doolabh, Lynn Tyers, Bruna Galvao, Innocent Mudau, Marvin Hsiao, Kruger Marais, Diana Hardie, Stephen Korsman, Carolyn Williamson |
| EPI_ISL_640113 | Victoria Hospital wc VHW | NHLS/UCT | Arash Iranzadeh, Deelan Doolabh, Lynn Tyers, Bruna Galvao, Innocent Mudau, Marvin Hsiao, Kruger Marais, Diana Hardie, Stephen Korsman, Carolyn Williamson |
| EPI_ISL_640114 | Valkenberg Hospital wc VBH | NHLS/UCT | Arash Iranzadeh, Deelan Doolabh, Lynn Tyers, Bruna Galvao, Innocent Mudau, Marvin Hsiao, Kruger Marais, Diana Hardie, Stephen Korsman, Carolyn Williamson |
| EPI_ISL_640115 | 2 Military Hospital wc MAA | NHLS/UCT | Arash Iranzadeh, Deelan Doolabh, Lynn Tyers, Bruna Galvao, Innocent Mudau, Marvin Hsiao, Kruger Marais, Diana Hardie, Stephen Korsman, Carolyn Williamson |
| EPI_ISL_640118 | Conville CDC wc CVC | NHLS/UCT | Arash Iranzadeh, Deelan Doolabh, Lynn Tyers, Bruna Galvao, Innocent Mudau, Marvin Hsiao, Kruger Marais, Diana Hardie, Stephen Korsman, Carolyn Williamson |
| EPI_ISL_640305, EPI_ISL_640317, EPI_ISL_640334, EPI_ISL_640453, EPI_ISL_640523, EPI_ISL_640561, EPI_ISL_640610, EPI_ISL_640665, EPI_ISL_640711, EPI_ISL_640775, EPI_ISL_640818, EPI_ISL_640963, EPI_ISL_641048, EPI_ISL_641053, EPI_ISL_641054 | | | |
| see above | Microbiological Diagnostic Unit - Public Health Laboratory (MDU-PHL) | MDU-PHL | Seemann T., Schultz M.B., Sait, M.L., Sherry, N.L. |
| EPI_ISL_641121 | Victorian Infectious Diseases Reference Laboratory (VIDRL) | VIDRL and MDU-PHL | Caly L., Seemann T., Sait, M.L., Schultz M.B., Druce J., Sherry, N.L. |
| EPI_ISL_641126, EPI_ISL_641193, | Microbiological Diagnostic Unit - Public Health Laboratory | MDU-PHL | Seemann T., Schultz M.B., Sait, M.L., Sherry, N.L. |

| | | | |
|---|---|---|---|
| EPI_ISL_641200, EPI_ISL_641305 | (MDU-PHL) | | |
| EPI_ISL_641470, EPI_ISL_641471, EPI_ISL_641472, EPI_ISL_641473, EPI_ISL_641474, EPI_ISL_641475 | Department of Virus and Microbiological Special Diagnostics, Statens Serum Institut, Copenhagen, Denmark | Albertsen lab, Department of Chemistry and Bioscience, Aalborg University, Denmark | Thomas Bruun Rasmussen, Jannik Fonager, Morten Rasmussen |
| EPI_ISL_644411, EPI_ISL_644412, EPI_ISL_644485, EPI_ISL_644486 | MEPHI, Aix Marseille University | MEPHI, Aix Marseille University | Anthony LEVASSEUR |
| EPI_ISL_644584, EPI_ISL_644585, EPI_ISL_644586, EPI_ISL_644587, EPI_ISL_644588, EPI_ISL_644589, EPI_ISL_644590, EPI_ISL_644591, EPI_ISL_644592, EPI_ISL_644593, EPI_ISL_644594, EPI_ISL_644595, EPI_ISL_644596, EPI_ISL_644597, EPI_ISL_644598, EPI_ISL_644599, EPI_ISL_644600, EPI_ISL_644601, EPI_ISL_644602, EPI_ISL_644603, EPI_ISL_644604, EPI_ISL_644605, EPI_ISL_644606, EPI_ISL_644607, EPI_ISL_644608, EPI_ISL_644609 | | | |
| see above | New Mexico Department of Health Scientific Laboratory | New Mexico Department of Health Scientific Laboratory | Ellie Johnson, Anastacia Griego-Fisher, D'Eldra Malone |
| EPI_ISL_644671, EPI_ISL_644672 | Institute for Medical Research, Infectious Disease Research Centre, National Institutes of Health, Ministry of Health Malaysia | Institute for Medical Research, Infectious Disease Research Centre, National Institutes of Health, Ministry of Health Malaysia | Suppiah J, Kamel K, Mohd-Zawawi Z, Thayan R |
| EPI_ISL_644856, EPI_ISL_644857, EPI_ISL_644859, EPI_ISL_644860, EPI_ISL_644861, EPI_ISL_644862, EPI_ISL_644865 | Virginia DCLS | Virginia DCLS | Virginia DCLS |
| EPI_ISL_645303 | Lighthouse Lab in Alderley Park | Wellcome Sanger Institute for the COVID-19 Genomics UK (COG-UK) Consortium | Jacquelyn Wynn, Mairead Hyland, The Lighthouse Lab in Alderley Park and Alex Alderton, Roberto Amato, Sonia Goncalves, Ewan Harrison, David K. Jackson, Ian Johnston, Dominic Kwiatkowski, Cordelia Langford, John Sillitoe on behalf of the Wellcome Sanger Institute COVID-19 Surveillance Team |
| EPI_ISL_645517 | Lighthouse Lab in Milton Keynes | Wellcome Sanger Institute for the COVID-19 Genomics UK (COG-UK) Consortium | The Lighthouse Lab in Milton Keynes and Alex Alderton, Roberto Amato, Sonia Goncalves, Ewan Harrison, David K. Jackson, Ian Johnston, Dominic Kwiatkowski, Cordelia Langford, John Sillitoe on behalf of the Wellcome Sanger Institute COVID-19 Surveillance Team |
| EPI_ISL_647987, EPI_ISL_647988, EPI_ISL_647989 | GA Department of Public Health Laboratory | Pathogen Discovery, Respiratory Viruses Branch, Division of Viral Diseases, Centers for Disease Control and Prevention | Yan Li, Jing Zhang, Ying Tao, Brian Lynch, Krista Queen, Anna Montmayeur, Anna Uehara, Clinton R. Paden, Rachel Marine, Haibin Wang, Suxiang Tong |
| EPI_ISL_648007, EPI_ISL_648008, EPI_ISL_648009, EPI_ISL_648010, EPI_ISL_648011, EPI_ISL_648012, EPI_ISL_648013, EPI_ISL_648014, EPI_ISL_648015, EPI_ISL_648040 | MS Public Health Laboratory | Pathogen Discovery, Respiratory Viruses Branch, Division of Viral Diseases, Centers for Disease Control and Prevention | Yan Li, Jing Zhang, Ying Tao, Brian Lynch, Krista Queen, Anna Montmayeur, Anna Uehara, Clinton R. Paden, Rachel Marine, Haibin Wang, Suxiang Tong |
| EPI_ISL_648123 | GA Department of Public Health Laboratory | Pathogen Discovery, Respiratory Viruses Branch, Division of Viral Diseases, Centers for Disease Control and Prevention | Yan Li, Jing Zhang, Ying Tao, Brian Lynch, Krista Queen, Anna Montmayeur, Anna Uehara, Clinton R. Paden, Rachel Marine, Haibin Wang, Suxiang Tong |
| EPI_ISL_648127 | UHAS COVID-19 Lab | UHAS COVID-19 Lab | Kwabena O. Duedu, Jones Gyamfi, Reuben Ayivor-Djanie, John O. Gyapong and the UHAS COVID-19 Lab Team |
| EPI_ISL_648136 | The Public Health Agency of Sweden | The Public Health Agency of Sweden | Anna-Malin Linde, Maria Lind Karlberg, Mattias Haukland, Reza Advani, Olov Svartstrom, Oskar Karlsson Lindsjo, Sandra Broddesson, Petra Edquist, Mia Brytting, Anna Risberg, Karin Tegmark-Wisell |
| EPI_ISL_648140 | Gavle klinisk mikrobiologi | The Public Health Agency of Sweden | Anna-Malin Linde, Maria Lind Karlberg, Mattias Haukland, Reza Advani, Olov Svartstrom, Oskar Karlsson Lindsjo, Sandra Broddesson, Petra Edquist, Mia Brytting, Anna Risberg, Karin Tegmark-Wisell |
| EPI_ISL_648148 | Halmstad klinisk mikrobiologi | The Public Health Agency of Sweden | Anna-Malin Linde, Maria Lind Karlberg, Mattias Haukland, Reza Advani, Olov Svartstrom, Oskar Karlsson Lindsjo, Sandra Broddesson, Petra Edquist, Mia Brytting, Anna Risberg, Karin Tegmark-Wisell |
| EPI_ISL_648171 | The Public Health Agency of Sweden | The Public Health Agency of Sweden | Anna-Malin Linde, Maria Lind Karlberg, Mattias Haukland, Reza Advani, Olov Svartstrom, Oskar Karlsson Lindsjo, Sandra Broddesson, Petra Edquist, Mia Brytting, Anna Risberg, Karin Tegmark-Wisell |
| EPI_ISL_648198, EPI_ISL_648199 | Skovde/Unilabs | The Public Health Agency of Sweden | Anna-Malin Linde, Maria Lind Karlberg, Mattias Haukland, Reza Advani, Olov Svartstrom, Oskar Karlsson Lindsjo, Sandra Broddesson, Petra Edquist, Mia Brytting, Anna Risberg, Karin Tegmark-Wisell |
| EPI_ISL_648460, EPI_ISL_648461, EPI_ISL_648462, EPI_ISL_648464 | Santa Clara County Public Health Laboratory | Chan-Zuckerberg Biohub | CZB Cliahub Consortium |
| EPI_ISL_648497, EPI_ISL_648498, EPI_ISL_648499, EPI_ISL_648500 | Orange County Public Health Lab | Chan-Zuckerberg Biohub | CZB Cliahub Consortium |
| EPI_ISL_648550, EPI_ISL_648551, EPI_ISL_648552, EPI_ISL_648553, EPI_ISL_648554 | Madera County Department of Public Health | Chan-Zuckerberg Biohub | CZB Cliahub Consortium |
| EPI_ISL_648871 | San Diego County Public Health Laboratory | Andersen lab at Scripps Research | SEARCH Alliance San Diego with Tracy Basler, Jovan Shephard, Brett Austin |
| EPI_ISL_651590 | University of Birmingham | COVID-19 Genomics UK (COG-UK) Consortium | Institute of Microbiology, University of Birmingham: Claire McMurray, Joanne Stockton, Samuel Nicholls, Radoslaw Poplawski, Will Rowe, Josh Quick, Nicholas Loman. University of Birmingham Testing Laboratory: Celina M Whalley, Andrew Bosworth, Charlotte Poxon, Kasun Wanigasooriya, Oliver Pickles, Mike Kidd, Alex Richter, Andrew D Beggs PHE Heartlands Lab: Husam Osman, Andrew Bosworth. Queen Elizabeth Hospital: Anna Casey |
| EPI_ISL_653754 | Instituto Nacional de Salud, Bogotá, Colombia | Instituto Nacional de Salud, Bogotá, Colombia | Katherine Laiton-Donato, Diego A. Álvarez-Díaz, Carlos Franco-Muñoz, Mauricio Pacheco-Montealegre, Jonathan Reales, Diego Andrés Prada, Jose A. Usme-Ciro, Zulma M. Cucunubá, Christian Julian VillabonaArenas, Liz Villabona-Arenas, Sussy Echeverria, Astrid C. Flórez, Carolina Ferro, Diana Marcela Walteros-Acero, Franklin Prieto, Carlos Andrés Durán, Martha Lucia Ospina Martinez, Marcela Mercado-Reyes |
| EPI_ISL_654086, EPI_ISL_654087, EPI_ISL_654089, EPI_ISL_654090, EPI_ISL_654170, EPI_ISL_654282, EPI_ISL_654365 | Hospital General Universitario Gregorio Marañón | SeqCOVID-SPAIN consortium/IBV(CSIC) | Darío García de Viedma, Laura Pérez-Lago, Marta Herranz, Jon Sicilia, Julia Suárez, Pilar Catalán, Patricia Muñoz and SeqCOVID-SPAIN consortium |
| EPI_ISL_654515 | Servicio de Microbiología, Laboratori Clínic Metropolitana Nord. Hospital Universitari Germans Trias i Pujol. Institut d'Investigació en Ciències de la Salut Germans Trias i Pujol (IGTP) | SeqCOVID-SPAIN consortium/IBV(CSIC) | Elisa Martró, Antoni E. Bordoy, Anna Not, Adrián Antuori, Anabel Fernández, Nona Romaní and SeqCOVID-SPAIN consortium |
| EPI_ISL_654956 | Skovde/Unilabs | The Public Health Agency of Sweden | Anna-Malin Linde, Maria Lind Karlberg, Mattias Haukland, Reza Advani, Olov Svartstrom, Oskar Karlsson Lindsjo, Sandra Broddesson, Petra Edquist, Mia Brytting, Anna Risberg, Karin Tegmark-Wisell |
| EPI_ISL_658901 | Instituto de Diagnostico y Referencia Epidemiologicos (INDRE) | Instituto de Diagnostico y Referencia Epidemiologicos (INDRE) | Ernesto Ramirez-Gonzalez, Abril Rodriguez-Maldonado, Claudia Wong-Arambula , Natividad Cruz-Ortiz, Tatiana Nunez-Garcia, Dayanira Arellano-Suarez, Fabiola Garces-Ayala, Lucia Hernandez-Rivas, Irma Lopez-Martinez, Gisela Barrera-Badillo. |
| EPI_ISL_660304, EPI_ISL_660322 | Servicio de Microbiología, Laboratori Clínic Metropolitana Nord. Hospital Universitari Germans Trias i Pujol. Institut d'Investigació en Ciències de la Salut Germans Trias i Pujol (IGTP) | SeqCOVID-SPAIN consortium/IBV(CSIC) | Elisa Martró, Antoni E. Bordoy, Anna Not, Adrián Antuori, Anabel Fernández, Nona Romaní and SeqCOVID-SPAIN consortium |
| EPI_ISL_660429, EPI_ISL_660430 | Klinisk Mikrobiologi | The Public Health Agency of Sweden | Anna-Malin Linde, Maria Lind Karlberg, Mattias Haukland, Reza Advani, Olov Svartstrom, Oskar Karlsson Lindsjo, Sandra Broddesson, Petra Edquist, Mia Brytting, Anna Risberg, Karin Tegmark-Wisell |
| EPI_ISL_660457, EPI_ISL_660480, | Laboratoire de Microbiologie CHU Souro Sanou | Centre Muraz | Abdoul-Salam Ouedraogo, Yacouba Sawadogo, Essia Belarbi, Grit Schubert, Fabian Leendertz, Arsène Zongo, Soumeya Ouangraoua, Zekiba Tarnagda, |

| | | | |
|---|---|---|---|
| EPI_ISL_660491, EPI_ISL_660508, EPI_ISL_660512, EPI_ISL_660523 | | | Lassana Sangaré, Halidou Tinto |
| EPI_ISL_660544 | Laboratory Medicine | Department of Laboratory Medicine, Lin-Kou Chang Gung Memorial Hospital, Taoyuan, Taiwan | Kuo-Chien Tsao, Yu-Nong Gong, Shu-Li Yang, Yi-Chun Liu, Chung-Guei Huang, Mei-Jen Hsiao, Po-Wei Huang, Cheng-Ta Yang, Cheng-Hsun Chiu, Peng-Nien Huang, Kuo-Ming Lee, Guang-Wu Chen, Shin-Ru Shih |
| EPI_ISL_661269 | Russian Academy of Sciences, Federal Research Center for Virology and Microbiology | Russian Academy of Sciences, Federal Research Center for Virology and Microbiology | Titov,I., Nefedeva,M., Egorova,I., Malogolovkin,A. |
| EPI_ISL_663285 | Microbiological Diagnostic Unit - Public Health Laboratory (MDU-PHL) | MDU-PHL | Seemann T., Schultz M.B., Sait, M.L., Sherry, N.L. |
| EPI_ISL_664197 | University College London Hospital | COVID-19 Genomics UK (COG-UK) Consortium | Judith Heaney, Matthew Byott, Catherine Houlihan, Dan Frampton, Stuart Kirk, Moira Spyer and Eleni Nastouli |
| EPI_ISL_665285, EPI_ISL_665470, EPI_ISL_665956 | University of Exeter | COVID-19 Genomics UK (COG-UK) Consortium | Ben Temperton,Aaron Jeffries,Michelle Michelsen,Joanna Warwick-Dugdale,Audrey Farbos,Robyn Manley,Stephen Michell,Jane Masoli |
| EPI_ISL_666632, EPI_ISL_666633, EPI_ISL_666634, EPI_ISL_666635, EPI_ISL_666636 | ZOTZ KLIMAS MVZ Düsseldorf-Centrum GbR ÜBAG für Labormedizin, Genetik, Zytologie, Pathologie | Center of Medical Microbiology, Virology, and Hospital Hygiene, University of Duesseldorf | Maximilian Damagnez, Alexander Dilthey, Ashley-Jane Duplessis, Patrick Finzer, Katrin Hoffmann, Torsten Houwaart, Lisanna Hülse, Malte Kohns Vasconcelos, Marek Korencak, Nadine Lübke, Jessica Nicolai, Klaus Pfeffer, Daniel Strelow, Jörg Timm, Andreas Walker, Tobias Wienemann, Rainer Zotz |
| EPI_ISL_666848, EPI_ISL_666849, EPI_ISL_666850, EPI_ISL_666851, EPI_ISL_666852, EPI_ISL_666866, EPI_ISL_666867, EPI_ISL_666868 | Florida Bureau of Public Health Laboratories | Florida Bureau of Public Health Laboratories | Sarah Schmedes, Jason Blanton |
| EPI_ISL_666994, EPI_ISL_667056 | San Diego County Public Health Laboratory | Andersen lab at Scripps Research | SEARCH Alliance San Diego with Tracy Basler, Jovan Shephard, Brett Austin |
| EPI_ISL_667318, EPI_ISL_667319, EPI_ISL_667320, EPI_ISL_667321, EPI_ISL_667322, EPI_ISL_667323, EPI_ISL_667324, EPI_ISL_667325, EPI_ISL_667326, EPI_ISL_667327, EPI_ISL_667328, EPI_ISL_667329, EPI_ISL_667330, EPI_ISL_667331, EPI_ISL_667332, EPI_ISL_667333, EPI_ISL_667334, EPI_ISL_667335, EPI_ISL_667336, EPI_ISL_667337, EPI_ISL_667338, EPI_ISL_667339, EPI_ISL_667340, EPI_ISL_667341, EPI_ISL_667342, EPI_ISL_667343, EPI_ISL_667344, EPI_ISL_667345, EPI_ISL_667346, EPI_ISL_667347, EPI_ISL_667348, EPI_ISL_667349, EPI_ISL_667350, EPI_ISL_667351, EPI_ISL_667352, EPI_ISL_667353, EPI_ISL_667354, EPI_ISL_667355, EPI_ISL_667356, EPI_ISL_667357, EPI_ISL_667358, EPI_ISL_667359, EPI_ISL_667360, EPI_ISL_667361, EPI_ISL_667362, EPI_ISL_667363, EPI_ISL_667364, EPI_ISL_667365, EPI_ISL_667366, EPI_ISL_667367, EPI_ISL_667368, EPI_ISL_667369, EPI_ISL_667370, EPI_ISL_667371, EPI_ISL_667372, EPI_ISL_667373, EPI_ISL_667374, EPI_ISL_667375, EPI_ISL_667376, EPI_ISL_667377, EPI_ISL_667378, EPI_ISL_667379, EPI_ISL_667380, EPI_ISL_667381, EPI_ISL_667382, EPI_ISL_667383, EPI_ISL_667384, EPI_ISL_667385, EPI_ISL_667386, EPI_ISL_667387, EPI_ISL_667388, EPI_ISL_667389, EPI_ISL_667390, EPI_ISL_667391, EPI_ISL_667392, EPI_ISL_667393, EPI_ISL_667394, EPI_ISL_667395, EPI_ISL_667396, EPI_ISL_667397, EPI_ISL_667398, EPI_ISL_667399, EPI_ISL_667400, EPI_ISL_667401, EPI_ISL_667402, EPI_ISL_667403, EPI_ISL_667404, EPI_ISL_667405, EPI_ISL_667406, EPI_ISL_667407, EPI_ISL_667408, EPI_ISL_667409, EPI_ISL_667410, EPI_ISL_667411, EPI_ISL_667412, EPI_ISL_667413, EPI_ISL_667414, EPI_ISL_667415, EPI_ISL_667416, EPI_ISL_667417, EPI_ISL_667418, EPI_ISL_667419, EPI_ISL_667420, EPI_ISL_667421, EPI_ISL_667422, EPI_ISL_667423, EPI_ISL_667424, EPI_ISL_667425, EPI_ISL_667426, EPI_ISL_667427, EPI_ISL_667435, EPI_ISL_667436, EPI_ISL_667437, EPI_ISL_667438, EPI_ISL_667439, EPI_ISL_667440, EPI_ISL_667441, EPI_ISL_667442, EPI_ISL_667443, EPI_ISL_667444, EPI_ISL_667445, EPI_ISL_667446, EPI_ISL_667447, EPI_ISL_667448, EPI_ISL_667555, EPI_ISL_667556, EPI_ISL_667557 | | | |
| see above | OHSU Lab Services Molecular Microbiology Lab | Oregon SARS-CoV-2 Genome Sequencing Center | Brendan L. O'Connell, Ruth V. Nichols, Sally Grindstaff, Alec J. Hirsch, Donna Hansel, Guang Fan, Daniel N. Streblow, William B. Messer, Andrew C. Adey, Benjamin N. Bimber, Brian J. O'Roak |
| EPI_ISL_667758, EPI_ISL_667759, EPI_ISL_667760, EPI_ISL_667761, EPI_ISL_667762, EPI_ISL_667763, EPI_ISL_667764, EPI_ISL_667765, EPI_ISL_667766, EPI_ISL_667767, EPI_ISL_667768, EPI_ISL_667769, EPI_ISL_667770, EPI_ISL_667771, EPI_ISL_667772, EPI_ISL_667773, EPI_ISL_667774, EPI_ISL_667775 | | | |
| see above | Microbiology, Infectious Diseases and Immunology, Centre de Recherche du Centre Hospitalier de l'Universite de Montreal | Microbiology, Infectious Diseases and Immunology, Centre de Recherche du Centre Hospitalier de l'Universite de Montreal | Benoit,P., Point,F., Gagnon,S., Hardy,I., Kaufmann,D., Tremblay,C., Coutlee,F., Grandjean-Lapierre,S. |
| EPI_ISL_668440 | Ostfold Hospital Trust - Kalnes, Centre for Laboratory Medicine, Section for gene technology and infection serology | Norwegian Institute of Public Health, Department of Virology | Kathrine Stene-Johansen, Kamilla Heddeland Instefjord, Hilde Elshaug, Marie Paulsen Madsen, Rasmus Riis Kopperud, Hilde Vollan, Karoline Bragstad, Olav Hungnes |
| EPI_ISL_670517, EPI_ISL_670659, EPI_ISL_670660, EPI_ISL_670661, EPI_ISL_670662, EPI_ISL_670663, EPI_ISL_670664, EPI_ISL_670665, EPI_ISL_670701, EPI_ISL_670728, EPI_ISL_670745, EPI_ISL_671234, EPI_ISL_671235, EPI_ISL_671236, EPI_ISL_671237, EPI_ISL_671238 | | | |
| see above | Department of Virus and Microbiological Special Diagnostics, Statens Serum Institut, Copenhagen, Denmark | Albertsen Lab, Department of Chemistry and Bioscience, Aalborg University, Denmark | Danish Covid-19 Genome Consortium |
| EPI_ISL_671414 | National Laboratory of Virology, Szentágothai Research Centre | National Laboratory of Virology, Szentágothai Research Centre | Endre Gábor Tóth, Balázs Somogyi, Brigitta, Ferenc Jakab, Gábor Kemenesi |
| EPI_ISL_671428, EPI_ISL_671430, EPI_ISL_671431, EPI_ISL_671432, EPI_ISL_671433, EPI_ISL_671434, EPI_ISL_671435, EPI_ISL_671436, EPI_ISL_671438, EPI_ISL_671439, EPI_ISL_671440, EPI_ISL_671441, EPI_ISL_671442, EPI_ISL_671443, EPI_ISL_671444, EPI_ISL_671445, EPI_ISL_671446, EPI_ISL_671447, EPI_ISL_671448, EPI_ISL_671449, EPI_ISL_671450, EPI_ISL_671451, EPI_ISL_671452, EPI_ISL_671453, EPI_ISL_671454 | | | |
| see above | University of Debrecen, Department of Medical Microbiology | National Laboratory of Virology, Szentágothai Research Centre | Endre Gábor Tóth, Balázs Somogyi, Brigitta Zana, Eszter Csoma, Ferenc Jakab, Gábor Kemenesi |
| EPI_ISL_671814, EPI_ISL_671815, EPI_ISL_671816, EPI_ISL_671817, EPI_ISL_671818, EPI_ISL_671819, EPI_ISL_671820, EPI_ISL_671821, EPI_ISL_671822, EPI_ISL_671823, EPI_ISL_671824 | | | |
| see above | Hospital de la Santa Creu i Sant Pau. Servicio de Microbiología | SeqCOVID-SPAIN consortium/IBV(CSIC) | Ferran Navarro, Núria Rabella, Elisenda Miró and SeqCOVID-SPAIN consortium |
| EPI_ISL_671941, EPI_ISL_671942 | CHU Purpan - Laboratoire de Virologie - Institut Fédératif de Biologie | CHU Purpan - Laboratoire de Virologie - Institut Fédératif de Biologie | Latour J., Ranger N., Dubois M., Carcenac R., Harter A., Boyer P., Tremeaux P., Izopet J. |
| EPI_ISL_672024, EPI_ISL_672029, EPI_ISL_672036, EPI_ISL_672045, EPI_ISL_672091, EPI_ISL_672113 | Alameda County Public Health Lab | Chan-Zuckerberg Biohub | CZB Cliahub Consortium |
| EPI_ISL_672423, EPI_ISL_672425 | Fresno County Public Health Laboratory | Chan-Zuckerberg Biohub | CZB Cliahub Consortium |
| EPI_ISL_672456, EPI_ISL_672457, EPI_ISL_672458, EPI_ISL_672459, EPI_ISL_672460, EPI_ISL_672461, EPI_ISL_672462, EPI_ISL_672463, EPI_ISL_672464, EPI_ISL_672465, EPI_ISL_672466, EPI_ISL_672467, EPI_ISL_672468, EPI_ISL_672469, EPI_ISL_672470, EPI_ISL_672471, EPI_ISL_672472, EPI_ISL_672473, EPI_ISL_672474, EPI_ISL_672475, EPI_ISL_672476 | | | |
| see above | Alameda County Public Health Lab | Chan-Zuckerberg Biohub | CZB Cliahub Consortium |
| EPI_ISL_676509 | TNMC & Nair ch. Hospital | CSIR-Institute of Genomics and Integrative Biology | Rajesh Pandey, Jayanthi Shastri, Akshay Kanakan, Janani Srinivasa Vasudevan, Ranjeet Maurya, Sachee Agrawal, Nirjhar Chatterjee, Swapneil Parikh, Manish Pathak, Subrat Thanapati, Jasmina Savak, Suresh Poojari, Mahesh Sangar, Amol Borse, Shweta Kawankar, Vasil Nachan, Mayuresh Vishwanathan, Shruthi Sachidanandan, Shrutika Pophale, Utkarsha Yelve |
| EPI_ISL_676512 | Klinsisk mikrobiologi Linkoping | The Public Health Agency of Sweden | Department of Microbiology, The Public Health Agency of Sweden |
| EPI_ISL_676619 | Texas Department of State Health Services | Texas Department of State Health Services | Rashmi Tuladhar, Bonnie Oh, Jenny Zhang, Maliha Rahman, Anita Pokharel, Myong Koag, Chung Wang, Rachel Lee, Grace Kubin, Mayela Pedrueza, James Daniel Bonser |
| EPI_ISL_677089, EPI_ISL_677090, EPI_ISL_677091, EPI_ISL_677092, EPI_ISL_677093, EPI_ISL_677094, EPI_ISL_677095, EPI_ISL_677096, EPI_ISL_677097, EPI_ISL_677098 | Wadsworth Center, New York State Department.of Health | Wadsworth Center, New York State Department.of Health | Kirsten St. George, Daryl M. Lamson, Alexis Russel, Jonathan Plitnick, Navjot Singh, John Kelly, Sara Griesemer, Erasmus Schneider, Erica Lasek-Nesselquist |
| EPI_ISL_677244, EPI_ISL_677245, EPI_ISL_677247, EPI_ISL_677248, EPI_ISL_677257, EPI_ISL_677270, EPI_ISL_677271, EPI_ISL_677280, EPI_ISL_677281, EPI_ISL_677282, EPI_ISL_677283, EPI_ISL_677284, EPI_ISL_677285, EPI_ISL_677311, EPI_ISL_677312, EPI_ISL_677313, EPI_ISL_677314 | | | |
| see above | Colorado Department of Public Health and Environment | Colorado Department of Puplic Health and Environment | Laura Bankers, Molly Hetherington-Rauth, Shannon Ely, Shannon R. Matzinger, Sarah Elizabeth Totten, Emily A. Travanty |
| EPI_ISL_677409, EPI_ISL_677410, EPI_ISL_677411, EPI_ISL_677412 | University of Wisconsin-Madison AIDS Vaccine Research Laboratories | University of Wisconsin-Madison AIDS Vaccine Research Laboratories | Gage Moreno, Katarina Braun, et al. AIDS Vaccine Research Laboratories |
| EPI_ISL_677638, EPI_ISL_677639, | Colorado Department of Public Health and Environment | Colorado Department of Puplic Health and Environment | Laura Bankers, Molly C. Hetherington-Rauth, Shannon Ely, Shannon R. Matzinger, Sarah Elizabeth Totten, Emily A. Travanty |

| | | | |
|---|---|---|---|
| EPI_ISL_677640, EPI_ISL_677641, EPI_ISL_677642, EPI_ISL_677643, EPI_ISL_677644, EPI_ISL_677645, EPI_ISL_677646, EPI_ISL_677647 | | | |
| EPI_ISL_677670 | Wadsworth Center, New York State Department.of Health | Wadsworth Center, New York State Department.of Health | Kirsten St. George, Daryl M. Lamson, Alexis Russel, Jonathan Plitnick, Navjot Singh, John Kelly, Sara Griesemer, Erasmus Schneider, Erica Lasek-Nesselquist |
| EPI_ISL_677695 | Vanda Pharmaceuticals Clinical Site | Vanda Pharmaceuticals | Vanda Pharmaceuticals |
| EPI_ISL_677739, EPI_ISL_677740, EPI_ISL_677741, EPI_ISL_677742, EPI_ISL_677743, EPI_ISL_677744, EPI_ISL_677745, EPI_ISL_677746, EPI_ISL_677747, EPI_ISL_677748, EPI_ISL_677749, EPI_ISL_677750, EPI_ISL_677751, EPI_ISL_677752, EPI_ISL_677753, EPI_ISL_677754, EPI_ISL_677755, EPI_ISL_677756, EPI_ISL_677757, EPI_ISL_677758, EPI_ISL_677759, EPI_ISL_677760, EPI_ISL_677761, EPI_ISL_677762, EPI_ISL_677763, EPI_ISL_677764, EPI_ISL_677765, EPI_ISL_677766, EPI_ISL_677767, EPI_ISL_677768 | | | |
| see above | University of Szeged, Institute of Clinical Microbiology | National Laboratory of Virology, Szentágothai Research Centre | Endre Gábor Tóth, Balázs Somogyi, Brigitta, Gabriella Terhes, Ferenc Jakab, Gábor Kemenesi |
| EPI_ISL_678253 | Institute for Lung Diseases in Children - Skopje | Research Center for Genetic Engineering and Biotechnology "Georgi D. Efremov" , Macedonian Academy of Sciences and Arts | RCGEB - MASA |
| EPI_ISL_678254 | General Hospital - Strumica | Research Center for Genetic Engineering and Biotechnology "Georgi D. Efremov" , Macedonian Academy of Sciences and Arts | RCGEB - MASA |
| EPI_ISL_678255, EPI_ISL_678256 | General Hospital - Ohrid | Research Center for Genetic Engineering and Biotechnology "Georgi D. Efremov" , Macedonian Academy of Sciences and Arts | RCGEB - MASA |
| EPI_ISL_678257 | PHI Institute of Nephrology - Struga | Research Center for Genetic Engineering and Biotechnology "Georgi D. Efremov" , Macedonian Academy of Sciences and Arts | RCGEB - MASA |
| EPI_ISL_678258 | General Hospital - Prilep | Research Center for Genetic Engineering and Biotechnology "Georgi D. Efremov" , Macedonian Academy of Sciences and Arts | RCGEB - MASA |
| EPI_ISL_678259 | Clinical Hospital - Shtip | Research Center for Genetic Engineering and Biotechnology "Georgi D. Efremov" , Macedonian Academy of Sciences and Arts | RCGEB - MASA |
| EPI_ISL_678345, EPI_ISL_678347 | Area of Virology, Serology and Virology Division (SAViD), New South Wales Health Pathology Randwick | Virology Research Laboratory; Area of Virology, Serology and Virology Division (SAViD), New South Wales Health Pathology Randwick | Foster, C.; Au, J.; Ruiz Silva, M.; Deveson, I.; Bull, R.; Van Hal, S.; Rawlinson, W. |
| EPI_ISL_679408 | University College London Hospital | COVID-19 Genomics UK (COG-UK) Consortium | Judith Heaney, Matthew Byott, Catherine Houlihan, Dan Frampton, Stuart Kirk, Moira Spyer and Eleni Nastouli |
| EPI_ISL_680300 | Regional Virus Laboratory, Belfast Health and Social Care Trust | COVID-19 Genomics UK (COG-UK) Consortium | Conall McCaughey, James McKenna, Tanya Curran, Susan Feeney, Alison Watt, Ciara Cox, Mairead Connor, Zoltan Molnar, David Simpson, Derek Fairley |
| EPI_ISL_681938, EPI_ISL_682035, EPI_ISL_682037 | UPMC Clinical Microbiology Laboratory | Microbial Genomic Epidemiology Laboratory, University of Pittsburgh | Mustapha M. Mustapha, Jane W. Marsh, Dan Snyder, Marissa P. Griffith, Stephanie L. Mitchell, Vatsala R. Srinivasa, Kady D. Waggle, Chinelo Ezeonwuku, Vaughn S. Cooper, Lee H. Harrison |
| EPI_ISL_683004, EPI_ISL_683095 | Department of Virus and Microbiological Special Diagnostics, Statens Serum Institut, Copenhagen, Denmark | Albertsen Lab, Department of Chemistry and Bioscience, Aalborg University, Denmark | Danish Covid-19 Genome Consortium |
| EPI_ISL_683384, EPI_ISL_683385, EPI_ISL_683386, EPI_ISL_683387, EPI_ISL_683388, EPI_ISL_683389, EPI_ISL_683390, EPI_ISL_683391, EPI_ISL_683392, EPI_ISL_683393, EPI_ISL_683394, EPI_ISL_683395, EPI_ISL_683396, EPI_ISL_683397 | | | |
| see above | CNR Virus des Infections Respiratoires - France SUD | CNR Virus des Infections Respiratoires - France SUD | Antonin Bal, Gregory Destras, Gwendolyne Burfin, Quentin Semanas, Martine Valette, Bruno Lina, Laurence Josset |
| EPI_ISL_683628, EPI_ISL_683630 | Servicio de Microbiología, Laboratori Clínic Metropolitana Nord. Hospital Universitari Germans Trias i Pujol. Institut d'Investigació en Ciències de la Salut Germans Trias i Pujol (IGTP) | SeqCOVID-SPAIN consortium/IBV(CSIC) | Elisa Martró, Antoni E. Bordoy, Anna Not, Adrián Antuori, Anabel Fernández, Nona Romaní, Verónica Saludes, Cristina Casañ and SeqCOVID-SPAIN consortium |
| EPI_ISL_691614, EPI_ISL_691642, EPI_ISL_691646, EPI_ISL_691648, EPI_ISL_691652, EPI_ISL_691655 | Servicio de Microbiología, Hospital Universitario Son Espases | SeqCOVID-SPAIN consortium/IBV(CSIC) | Carla López-Causapé, Jordi Reina, Antonio Oliver and SeqCOVID-SPAIN consortium |
| EPI_ISL_691729, EPI_ISL_691730 | Hospital Universitario Severo Ochoa | Instituto de Salud Carlos III | Iglesias-Caballero, M. Camarero, S. Molinero Calamita, M. González-Esguevillas, M. Pozo, F. Casas, I. Jiménez, P. Jiménez, M. Zaballos, A. Monzón, S. Varona, S. Juliá, M. Cuesta, I. García, M. |
| EPI_ISL_692737, EPI_ISL_692738, EPI_ISL_692739, EPI_ISL_692740, EPI_ISL_692741, EPI_ISL_692742, EPI_ISL_692743, EPI_ISL_692744, EPI_ISL_692745, EPI_ISL_692762, EPI_ISL_692763, EPI_ISL_692764, EPI_ISL_692765, EPI_ISL_692766, EPI_ISL_692767, EPI_ISL_692768 | | | |
| see above | CNR Virus des Infections Respiratoires - France SUD | CNR Virus des Infections Respiratoires - France SUD | Antonin Bal, Gregory Destras, Gwendolyne Burfin, Solenne Brun, Martine Valette, Bruno Lina, Laurence Josset |
| EPI_ISL_692948 | Massachusetts State Public Health Laboratory | Massachusetts State Public Health Laboratory | Andrew Lang, Timelia Fink, Glen Gallagher, Sandra Smole |
| EPI_ISL_693478 | Central Public Health Laboratory | National Public Health Laboratory, National Centre for Infectious Diseases | Tze Minn Mak, Sophie Octavia, Zhenyang Zhou, Esorom Daoni, Theresa Palou, Lin Cui, Raymond Tzer Pin Lin |
| EPI_ISL_693490 | CNR Virus des Infections Respiratoires - France SUD | CNR Virus des Infections Respiratoires - France SUD | Antonin Bal, Gregory Destras, Gwendolyne Burfin, Quentin Semanas, Martine Valette, Bruno Lina, Laurence Josset |
| EPI_ISL_693535 | Hospital Vila Franca de Xira | Instituto Nacional de Saude (INSA) | Borges et al |
| EPI_ISL_693723, EPI_ISL_693757 | Delaware Public Health Laboratory | Delaware Public Health Laboratory | Gregory Hovan |
| EPI_ISL_693758, EPI_ISL_693759, EPI_ISL_693760, EPI_ISL_693761 | Hospital | National Reference Center for Viruses of Respiratory Infections, Institut Pasteur, Paris | Marion Barbet, Sylvie Behillil, Méline Bizard, Angela Brisebarre, Camille Capel, Etienne Simon-Lorière, Vincent Enouf, Maud Vanpeene, Sylvie van der Werf, Gisèle Lagathu |
| EPI_ISL_697789, EPI_ISL_697790, EPI_ISL_697792, EPI_ISL_697793 | Institute of Microbiology, Universidad San Francisco de Quito | Institute of Microbiology, Universidad San Francisco de Quito | Belén Prado-Vivar, Sully Márquez, Juan José Guadalupe, Monica Becerra-Wong, Bernardo Gutiérrez, Jonathan Araujo, Verónica Barragán, Patricio Rojas-Silva, Gabriel Trueba, Michelle Grunauer, Paúl Cárdenas |
| EPI_ISL_700218, EPI_ISL_700219, EPI_ISL_700220, EPI_ISL_700221, EPI_ISL_700222, EPI_ISL_700223, EPI_ISL_700224, EPI_ISL_700225, EPI_ISL_700226, EPI_ISL_700227, EPI_ISL_700228, EPI_ISL_700229, EPI_ISL_700230, EPI_ISL_700231, EPI_ISL_700232, EPI_ISL_700233, EPI_ISL_700234, EPI_ISL_700235, EPI_ISL_700236, EPI_ISL_700237, EPI_ISL_700238, EPI_ISL_700239, EPI_ISL_700240, EPI_ISL_700241, EPI_ISL_700242, EPI_ISL_700243, EPI_ISL_700244, EPI_ISL_700245, EPI_ISL_700246, EPI_ISL_700247, EPI_ISL_700248, EPI_ISL_700249, EPI_ISL_700250 | | | |
| see above | Hematopathology Laboratory, ACTREC, TMC | Hematopathology Laboratory, ACTREC, TMC | Hematopathology Laboratory, ACTREC |
| EPI_ISL_700327 | CNR Virus des Infections Respiratoires - France SUD | CNR Virus des Infections Respiratoires - France SUD | Antonin Bal, Gregory Destras, Gwendolyne Burfin, Solenne Brun, Martine Valette, Bruno Lina, Laurence Josset |
| EPI_ISL_700331, EPI_ISL_700342, EPI_ISL_700343 | Child Health Research Foundation | Child Health Research Foundation | Senjuti Saha, Afroza Akter Tanni, Syed Muktadir Al Sium, Roly Malaker, Sharmistha Goswami, Arif Mohammad Tanmoy, Md Hafizur Rahman, Samir K Saha |
| EPI_ISL_700414 | Conville CDC wc CVC | NHLS/UCT | Arash Iranzadeh, Deelan Doolabh, Lynn Tyers, Bruna Galvao, Innocent Mudau, Marvin Hsiao, Kruger Marais, Diana Hardie, Stephen Korsman, Carolyn Williamson |

| | | | |
|---|---|---|---|
| EPI_ISL_700453 | Beaufort West Hospital wc BWH | NHLS/UCT | Arash Iranzadeh, Deelan Doolabh, Lynn Tyers, Bruna Galvao, Innocent Mudau, Marvin Hsiao, Kruger Marais, Diana Hardie, Stephen Korsman, Carolyn Williamson |
| EPI_ISL_700558 | Hornlee Clinic wc HLC | NHLS/UCT | Arash Iranzadeh, Deelan Doolabh, Lynn Tyers, Bruna Galvao, Innocent Mudau, Marvin Hsiao, Kruger Marais, Diana Hardie, Stephen Korsman, Carolyn Williamson |
| EPI_ISL_700700, EPI_ISL_700709, EPI_ISL_700723, EPI_ISL_700741 | Texas Department of State Health Services | Texas Department of State Health Services | Rashmi Tuladhar, Bonnie Oh, Jenny Zhang, Maliha Rahman, Anita Pokharel, Myong Koag, Chung Wang, Rachel Lee, Grace Kubin, Mayela Pedrueza, James Daniel Bonser |
| EPI_ISL_707715, EPI_ISL_707716, EPI_ISL_707722, EPI_ISL_707723, EPI_ISL_707724, EPI_ISL_707725, EPI_ISL_707727, EPI_ISL_707728, EPI_ISL_707729, EPI_ISL_707730, EPI_ISL_707731, EPI_ISL_707732, EPI_ISL_707733, EPI_ISL_707734, EPI_ISL_707735, EPI_ISL_707736, EPI_ISL_707737, EPI_ISL_707738, EPI_ISL_707739, EPI_ISL_707740, EPI_ISL_707741, EPI_ISL_707742, EPI_ISL_707743, EPI_ISL_707744, EPI_ISL_707745, EPI_ISL_707746, EPI_ISL_707747, EPI_ISL_707748, EPI_ISL_707749, EPI_ISL_707750, EPI_ISL_707751, EPI_ISL_707752, EPI_ISL_707753, EPI_ISL_707754, EPI_ISL_707755, EPI_ISL_707756 | | | |
| see above | Department of Clinical Microbiology | GIGA Medical Genomics | Keith Durkin, Maria Artesi, Sébastien Bontems, Raphaël Boreux, Bouchra Boujemla, Cécile Meex, Pierrette Melin, Marie-Pierre Hayette, Vincent Bours |
| EPI_ISL_707788, EPI_ISL_707790 | Rwanda National Reference Laboratory | Rwanda National Reference Laboratory | Enatha Mukantwari,Jeanne d'Arc Umuringa |
| EPI_ISL_708182, EPI_ISL_708183 | Los Angeles County PHL | Los Angeles County PHL | P. Hemarajata et al. |
| EPI_ISL_708313, EPI_ISL_708328 | Michigan Department of Health and Human Services, Bureau of Laboratories | Michigan Department of Health and Human Services, Bureau of Laboratories | Blankenship HM, Riner D, Soehnlen MK |
| EPI_ISL_708389, EPI_ISL_708403, EPI_ISL_708406, EPI_ISL_708407, EPI_ISL_708408, EPI_ISL_708412, EPI_ISL_708436, EPI_ISL_708446 | Delaware Public Health Lab | Delaware Public Health Lab | Gregory Hovan |
| EPI_ISL_708453, EPI_ISL_708454 | Department of Clinical Microbiology | GIGA Medical Genomics | Keith Durkin, Maria Artesi, Sébastien Bontems, Raphaël Boreux, Bouchra Boujemla, Cécile Meex, Pierrette Melin, Marie-Pierre Hayette, Vincent Bours |
| EPI_ISL_708727, EPI_ISL_708728 | Los Angeles County Public Health Laboratory | Los Angeles County Public Health Laboratory | P. Hemarajata et al. |
| EPI_ISL_710130, EPI_ISL_710149, EPI_ISL_710158, EPI_ISL_710222, EPI_ISL_710223, EPI_ISL_710245, EPI_ISL_710262, EPI_ISL_710263, EPI_ISL_710283, EPI_ISL_710284, EPI_ISL_710330, EPI_ISL_710331, EPI_ISL_710332 | | | |
| see above | Colorado Department of Public Health and Environment | Colorado Department of Puplic Health and Environment | Laura Bankers, Molly C. Hetherington-Rauth, Shannon Ely, Shannon R. Matzinger, Sarah Elizabeth Totten, Emily A. Travanty |
| EPI_ISL_710547 | Sestre Milosrdnice University Hospital Center | Ruer Boškovic Institute; Forensic Science Centre Ivan Vueti; University of Zagreb Faculty of Science | Robert Beluži, Marina Korolija, Ivana elap, Vjekoslav Tomai, Maja Kuzman, Dunja Glavaš, Maja Kuzman, Paula Štancl, Lucija Markulin, Antonela Blažekovi, Fran Boroveki, Lidija Cvetko-Krajinovi, Mirjana Domazet-Lošo, Tomislav Domazet-Lošo, Kristina Gotovac Jerei, Vladimir Krajinovi, Danilo Licastro, Ana Livun, Boris Maek, Željka Maak Šafranko, Gordana Maravi Vlahoviek, Josipa Skelin, Ivan Šamija, Mario Štefanovi, Sanja Tadinac, Rosa Karli, Kristian Vlahoviek |
| EPI_ISL_710550, EPI_ISL_710551, EPI_ISL_710557, EPI_ISL_710558, EPI_ISL_710559, EPI_ISL_710560, EPI_ISL_710561, EPI_ISL_710562 | University Hospital Dubrava | Ruer Boškovic Institute; Forensic Science Centre Ivan Vueti; University of Zagreb Faculty of Science | Robert Beluži, Marina Korolija, Ana Livun, Vjekoslav Tomai, Dunja Glavaš, Maja Kuzman, Paula Štancl, Lucija Markulin, Lucija Basi, Antonela Blažekovi, Fran Boroveki, Lidija Cvetko-Krajinovi, Ivana elap, Fuad osovi, Mirjana Domazet-Lošo, Tomislav Domazet-Lošo, Valentina umlijan-Combaj, Kristina Gotovac Jerei, Jasna Kašman, Vladimir Krajinovi, Danilo Licastro, Boris Maek, Željka Maak Šafranko, Gordana Maravi Vlahoviek, Senica Pejša, Josipa Skelin, Ivan Šamija, Mario Štefanovi, Sanja Tadinac, Katarina Marija Tupek, Petra Vrabec, Rosa Karli, Kristian Vlahoviek |
| EPI_ISL_710563 | Sestre Milosrdnice University Hospital Center | Ruer Boškovic Institute; Forensic Science Centre Ivan Vueti; University of Zagreb Faculty of Science | Robert Beluži, Marina Korolija, Ivana elap, Vjekoslav Tomai, Maja Kuzman, Dunja Glavaš, Maja Kuzman, Paula Štancl, Lucija Markulin, Antonela Blažekovi, Fran Boroveki, Lidija Cvetko-Krajinovi, Mirjana Domazet-Lošo, Tomislav Domazet-Lošo, Kristina Gotovac Jerei, Vladimir Krajinovi, Danilo Licastro, Ana Livun, Boris Maek, Željka Maak Šafranko, Gordana Maravi Vlahoviek, Josipa Skelin, Ivan Šamija, Mario Štefanovi, Sanja Tadinac, Rosa Karli, Kristian Vlahoviek |
| EPI_ISL_710609 | Gavle klinisk mikrobiologi | The Public Health Agency of Sweden | Department of Microbiology, The Public Health Agency of Sweden |
| EPI_ISL_714930 | Department of Virus and Microbiological Special Diagnostics, Statens Serum Institut, Copenhagen, Denmark | Albertsen Lab, Department of Chemistry and Bioscience, Aalborg University, Denmark | Danish Covid-19 Genome Consortium |
| EPI_ISL_717692, EPI_ISL_717697 | Trinidad Public Health Laboratory | Carrington Lab, Department of PreClinical Sciences, Faculty of Medical Sciences, The University of the West Indies | Nikita S. D. Sahadeo, Arianne Brown-Jordan, Sarah Hill, Vernie Ramkissoon, Naresh Nandram, Avery Hinds, Jerome Foster, Stanley Giddings, Karla Georges, Marsha Ivey, Rahul Naidu, Risha Singh, SueMin Nathaniel, Rajini Haraksingh, Jaya Jayaraman, Chinna Chinnadurai, Adesh Ramsubhag, Nuno Faria, Oliver Pybus, Christopher Oura, Gabriel Escobar, Christine V. F. Carrington |
| EPI_ISL_717770, EPI_ISL_717771, EPI_ISL_717772 | UW Virology Lab | UW Virology Lab | Pavitra Roychoudhury, Hong Xie, Lasata Shrestha, Michelle Lin, Meei-Li Huang, Keith R Jerome, Alexander Greninger |
| EPI_ISL_717806, EPI_ISL_717814, EPI_ISL_717815, EPI_ISL_717816, EPI_ISL_717817, EPI_ISL_717869, EPI_ISL_717870, EPI_ISL_717871, EPI_ISL_717872, EPI_ISL_717873 | Laboratorio de Virologia Molecular / UFRJ | Bioinformatics Laboratory / LNCC | Carolina M Voloch, Ronaldo da Silva F Jr, Luiz G P de Almeida, Cynthia C Cardoso, Otavio Bustrolini, Alexandra L Gerber, Ana Paula de C Guimarães, Diana Mariani, Andréa Cony Cavalcanti, Claudia dos Santos Rodrigues, Terezinha M P P Castiñeira, Amilcar Tanuri, Ana Tereza R de Vasconcelos |
| EPI_ISL_718287, EPI_ISL_718288, EPI_ISL_718289, EPI_ISL_718290, EPI_ISL_718291, EPI_ISL_718292, EPI_ISL_718293, EPI_ISL_718294, EPI_ISL_718295, EPI_ISL_718296, EPI_ISL_718297, EPI_ISL_718298, EPI_ISL_718299, EPI_ISL_718300, EPI_ISL_718301, EPI_ISL_718302, EPI_ISL_718303, EPI_ISL_718304, EPI_ISL_718305, EPI_ISL_718306 | | | |
| see above | Institute for Medical Research, Infectious Disease Research Centre, National Institutes of Health, Ministry of Health Malaysia | Institute for Medical Research, Infectious Disease Research Centre, National Institutes of Health, Ministry of Health Malaysia | Suppiah J, Kamel K, Mohd-Zawawi Z, Thayan R |
| EPI_ISL_722300, EPI_ISL_722436, EPI_ISL_722463, EPI_ISL_722842, EPI_ISL_722843, EPI_ISL_722844, EPI_ISL_722845, EPI_ISL_722846, EPI_ISL_722847, EPI_ISL_722848, EPI_ISL_722849 | | | |
| see above | Dutch COVID-19 response team | Erasmus Medical Center | Bas Oude Munnink, Reina Sikkema, David Nieuwenhuijse, Irina Chestakova, Anne van der Linden, Marjan Boter, Emmanuelle Munger, Corine GeurtsvanKessel, Annemiek van der Eijk, Richard Molenkamp, Marion Koopmans, on behalf of the Dutch national COVID-19 response team. |
| EPI_ISL_723058, EPI_ISL_723059, EPI_ISL_723060, EPI_ISL_723061, EPI_ISL_723062, EPI_ISL_723063, EPI_ISL_723064, EPI_ISL_723065, EPI_ISL_723066 | Hematopathology Laboratory, ACTREC, TMC | Hematopathology Laboratory, ACTREC, TMC | Hematopathology Laboratory, ACTREC |
| EPI_ISL_729904, EPI_ISL_729905 | Instituto de Medicina Tropical, Universidad Nacional Toribio Rodríguez de Mendoza de Amazonas | Laboratorio de Genómica Microbiana, Universidad Peruana Cayetano Heredia | Pablo Tsukayama, Alejandra Dávila-Barclay, Luis González, Pedro E. Romero, Brenda Ayzanoa, Janet Huancachoque, Pool Marcos, Stella Chenet, Rafael Tapia, Cecilia Pajuelo, Carla Montenegro |
| EPI_ISL_729979, EPI_ISL_729980 | Nigeria Centre for Disease Control (NCDC) | African Centre of Excellence for Genomics of Infectious Diseases (ACEGID), Redeemer's University, Ede, Osun State, Nigeria | Oluniyi P.E. et al |
| EPI_ISL_730091, EPI_ISL_730181 | San Diego County Public Health Laboratory | Andersen lab at Scripps Research | SEARCH Alliance San Diego with Tracy Basler, Jovan Shephard, Brett Austin |
| EPI_ISL_730568 | Gazi University Faculty of Medicine, Medical Virology Laboratory | Gazi University Faculty of Medicine, Medical Virology Laboratory | Erdem ahin, Hager Muftah, Selin Yiit, Shaknoza Sarzhanova, Özlem Güzel Tunçcan, Murat Dizbay, Il Fidan, Kayhan Çalar, Gülendam Bozday |
| EPI_ISL_730573 | Gazi University Faculty of Medicine, Medical Virology Laboratory | Gazi University Faculty of Medicine, Medical Virology Laboratory | Erdem ahin, Gülendam Bozday, Hager Muftah, Selin Yiit, Shaknoza Sarzhanova, Özlem Güzel Tunçcan, Murat Dizbay, Il Fidan, Kayhan Çalar |
| EPI_ISL_730582 | Hong Kong Adventist Hospital - Tsuen Wan | Hong Kong Department of Health | Mak Gannon C.K., Lam Edman T.K., Chan Rickjason C.W., Tsang Dominic N.C. |
| EPI_ISL_730589 | Prince of Wales Hospital | Hong Kong Department of Health | Mak Gannon C.K., Lam Edman T.K., Chan Rickjason C.W., Tsang Dominic N.C. |

| | | | |
|---|---|---|---|
| EPI_ISL_730596 | Queen Mary Hospital | Hong Kong Department of Health | Mak Gannon C.K., Lam Edman T.K., Chan Rickjason C.W., Tsang Dominic N.C. |
| EPI_ISL_730617 | Universal Community Testing Programme | Hong Kong Department of Health | Mak Gannon C.K., Lam Edman T.K., Chan Rickjason C.W., Tsang Dominic N.C. |
| EPI_ISL_730618 | Yan Chai Hospital | Hong Kong Department of Health | Mak Gannon C.K., Lam Edman T.K., Chan Rickjason C.W., Tsang Dominic N.C. |
| EPI_ISL_732760, EPI_ISL_732768, EPI_ISL_732819 | Centro de Investigación Biomédica de La Rioja - Hospital San Pedro Logroño | SeqCOVID-SPAIN consortium/IBV(CSIC) | María de Toro, José Manuel Azcona Gutiérrez, María Pilar Bea Escudero, Miriam Blasco Alberdi and SeqCOVID-SPAIN consortium |
| EPI_ISL_733183 | WHO National Influenza Centre Russian Federation | WHO National Influenza Centre Russian Federation | Andrey Komissarov, Artem Fadeev, Anna Ivanova, Kseniya Komissarova, Dmitry Bazhenov, Daria Danilenko, Ksenia Safina, Elena Nabieva, Georgii Bazykin, Dmitry Lioznov |
| EPI_ISL_733191, EPI_ISL_733193, EPI_ISL_733213 | Pathogenic Microorganisms Variability Laboratory | WHO National Influenza Centre Russian Federation | Andrey Komissarov, Artem Fadeev, Anna Ivanova, Kseniya Komissarova, Dmitry Bazhenov, Daria Danilenko, Ksenia Safina, Elena Nabieva, Georgii Bazykin, Nadezhda Kuznetsova, Elena Shidlovskaya, Sergey Alkhovsky, Tatyana Vishnevskaya, Elizaveta Divisenko, Alexey Shchetinin, Maria Nikiforova, Andrey Pochtovyy, Evgeny Usachev, Elena Vokalova, Maxim Rubalsky, Oleg Rubalsky, Artem Tkachuk, Vladimir Gushchin, Alexander Gintsburg, Dmitry Lioznov |
| EPI_ISL_733228, EPI_ISL_733229 | UMMC-Health | WHO National Influenza Centre Russian Federation | Andrey Komissarov, Artem Fadeev, Anna Ivanova, Kseniya Komissarova, Dmitry Bazhenov, Tatiana Platonova, Daria Danilenko, Ksenia Safina, Elena Nabieva, Georgii Bazykin, Dmitry Lioznov |
| EPI_ISL_733393 | HELIX LLC | WHO National Influenza Centre Russian Federation | Andrey Komissarov, Artem Fadeev, Anna Ivanova, Kseniya Komissarova, Dmitry Bazhenov, Daria Danilenko, Ksenia Safina, Elena Nabieva, Georgii Bazykin, Dmitry Lioznov |
| EPI_ISL_733402, EPI_ISL_733404 | WHO National Influenza Centre Russian Federation | WHO National Influenza Centre Russian Federation | Andrey Komissarov, Artem Fadeev, Anna Ivanova, Kseniya Komissarova, Dmitry Bazhenov, Daria Danilenko, Ksenia Safina, Elena Nabieva, Georgii Bazykin, Dmitry Lioznov |
| EPI_ISL_734287, EPI_ISL_734307 | Wadsworth Center, New York State Department.of Health | Wadsworth Center, New York State Department.of Health | Kirsten St. George, Daryl M. Lamson, Alexis Russel, Jonathan Plitnick, Navjot Singh, John Kelly, Sara Griesemer, Erasmus Schneider, Erica Lasek-Nesselquist |
| EPI_ISL_734870, EPI_ISL_734917, EPI_ISL_734918, EPI_ISL_734919, EPI_ISL_734920, EPI_ISL_734921, EPI_ISL_734922 | UZ Leuven, National Reference Laboratory for Coronaviruses, Laboratory Medicine, Leuven, Belgium | KU Leuven, Rega Institute, Clinical and Epidemiological Virology | Tony Wawina-Bokalanga, Joan Marti-Carerras, Bert Vanmechelen, Piet Maes |
| EPI_ISL_737990, EPI_ISL_738025, EPI_ISL_738026, EPI_ISL_738027, EPI_ISL_738028, EPI_ISL_738029, EPI_ISL_738031, EPI_ISL_738032, EPI_ISL_738033, EPI_ISL_738034, EPI_ISL_738035, EPI_ISL_738036, EPI_ISL_738037 | | | |
| see above | Uganda Central Public Health Lab and Uganda Virus Research Institute | MRC/UVRI & LSHTM Uganda Research Unit | Matthew Cotten, Dan Lule Bugembe, My V.T. Phan, Pontiano Kaleebu et al. |
| EPI_ISL_738692, EPI_ISL_738745, EPI_ISL_738868 | Alameda County Public Health Lab | Chan-Zuckerberg Biohub | CZB Cliahub Consortium |
| EPI_ISL_738870 | Napa-Solano-Yolo- Marin County (NSYM) Public Health Laboratories | Chan-Zuckerberg Biohub | CZB Cliahub Consortium |
| EPI_ISL_738890, EPI_ISL_739095 | County of San Luis Obispo Public Health Laboratory | Chan-Zuckerberg Biohub | CZB Cliahub Consortium |
| EPI_ISL_739127, EPI_ISL_739288 | Alameda County Public Health Lab | Chan-Zuckerberg Biohub | CZB Cliahub Consortium |
| EPI_ISL_739363 | County of San Luis Obispo Public Health Laboratory | Chan-Zuckerberg Biohub | CZB Cliahub Consortium |
| EPI_ISL_739372 | Monterey County Public Health Lab | Chan-Zuckerberg Biohub | CZB Cliahub Consortium |
| EPI_ISL_739645 | Napa-Solano-Yolo- Marin County (NSYM) Public Health Laboratories | Chan-Zuckerberg Biohub | CZB Cliahub Consortium |
| EPI_ISL_740488, EPI_ISL_744149, EPI_ISL_744418 | Laboratoire national de santé, Microbiology, Virology | Laboratoire national de santé, Microbiology, Microbial Genomics Platform | Anke Wienecke-Baldacchino, Catherine Ragimbeau,Jessica Tapp, Fatu Djabi, Lise Pignon, Raoul Salmon, Tamir Abdelrahman |
| EPI_ISL_745315 | CHU Clermont-Ferrand | CNR Virus des Infections Respiratoires - France SUD | Antonin Bal, Gregory Destras, Gwendolyne Burfin, Hadrien Règue, Quentin Semanas, Martine Valette, Bruno Lina, Christine Archimbaud, Amélie Brebion, Hélène Chabrolles, Martine Chambon, Audrey Mirand, Christel Regagnon, Maxime Bisseux, Patricia Combes, Cécile Henquell, Laurence Josset |
| EPI_ISL_745498, EPI_ISL_745601, EPI_ISL_746096, EPI_ISL_746104, EPI_ISL_746110, EPI_ISL_746113, EPI_ISL_746127, EPI_ISL_746128, EPI_ISL_746129, EPI_ISL_746135, EPI_ISL_746137, EPI_ISL_746143, EPI_ISL_746144 | | | |
| see above | Ginkgo Bioworks Clinical Laboratory | Utah Public Health Laboratory | Erin L. Young, Kelly Oakeson, Tara Gallagher, Michael T. Pyne, E. Susan Slechta, Melanie A. Mallory, Jeffrey B. Stevenson, Salika M. Shakir, David R. Hillyard, Malaika McKenzie-Bennett, James McGann, Jim Griffin, Keith Robison, Alex Plocik, Becky Schilling, Martha Pierson, Rebecca Littlefield, Michelle Spencer, Birgitte Simen |
| EPI_ISL_746478, EPI_ISL_746496, EPI_ISL_746499, EPI_ISL_746507, EPI_ISL_746508, EPI_ISL_746698, EPI_ISL_746699, EPI_ISL_746700, EPI_ISL_746701, EPI_ISL_746702, EPI_ISL_746703, EPI_ISL_746704, EPI_ISL_746705, EPI_ISL_746706, EPI_ISL_746707, EPI_ISL_746708, EPI_ISL_746709, EPI_ISL_746710, EPI_ISL_746711, EPI_ISL_746712, EPI_ISL_746713, EPI_ISL_746714, EPI_ISL_746715, EPI_ISL_746716, EPI_ISL_746717, EPI_ISL_746718, EPI_ISL_746719, EPI_ISL_746720, EPI_ISL_746721, EPI_ISL_746722, EPI_ISL_746723, EPI_ISL_746724, EPI_ISL_746725, EPI_ISL_746726, EPI_ISL_746727, EPI_ISL_746728, EPI_ISL_746729, EPI_ISL_746730 | | | |
| see above | Genetica Molecular and Subdepartamento de Virologia ISP Chile | Instituto de Salud Publica de Chile | Javier Tognarelli, Barbara Parra, Loredana Arata, Jaime Lagos, Gisselle Barra, Patricia Bustos, Rodrigo Fasce, Andres Castillo, Jorge Fernandez |
| EPI_ISL_746997 | Utah Public Health Laboratory | Utah Public Health Laboratory | Erin Young, Kelly Oakeson, Tara Gallagher |
| EPI_ISL_747245, EPI_ISL_747246, EPI_ISL_747247, EPI_ISL_747248, EPI_ISL_747249, EPI_ISL_747250, EPI_ISL_747251, EPI_ISL_747252, EPI_ISL_747253, EPI_ISL_747254, EPI_ISL_747255, EPI_ISL_747258, EPI_ISL_747259, EPI_ISL_747260, EPI_ISL_747261, EPI_ISL_747263, EPI_ISL_747267, EPI_ISL_747269, EPI_ISL_747270, EPI_ISL_747277, EPI_ISL_747278, EPI_ISL_747280, EPI_ISL_747289, EPI_ISL_747300, EPI_ISL_747301, EPI_ISL_747326, EPI_ISL_747334, EPI_ISL_747335 | | | |
| see above | Division of Emerging Infectious Diseases, Bureau of Infectious Diseases Diagnosis Control, Korea Disease Control and Prevention Agency | Division of Emerging Infectious Diseases, Bureau of Infectious Diseases Diagnosis Control, Korea Disease Control and Prevention Agency | Ae Kyung Park, Il-Hwan Kim, Heui Man Kim, Jeong-Min Kim, Namjoo Lee, Chaeyoung Lee, Sang Hee Woo, Eun-Jin Kim |
| EPI_ISL_751574, EPI_ISL_751652, EPI_ISL_751697, EPI_ISL_751698, EPI_ISL_751699, EPI_ISL_751723, EPI_ISL_751731, EPI_ISL_751760, EPI_ISL_751764, EPI_ISL_751765 | AR Dept. of Health-Public Health Lab | Genomics and Discovery, Respiratory Viruses Branch, Division of Viral Diseases, Centers for Disease Control and Prevention | Krista Queen, Yan Li, Ying Tao, Jing Zhang, Anna Uehara, Anna Montmayeur, Clinton R. Paden, Peter W. Cook,Rachel Marine, Mili Sheth, Haibin Wang, Justin Lee, Suxiang Tong |
| EPI_ISL_752826, EPI_ISL_752827, EPI_ISL_752828, EPI_ISL_752829, EPI_ISL_752830, EPI_ISL_752831, EPI_ISL_752832, EPI_ISL_752833, EPI_ISL_752834, EPI_ISL_752835, EPI_ISL_752836, EPI_ISL_752837, EPI_ISL_752838, EPI_ISL_752839, EPI_ISL_752840, EPI_ISL_752841, EPI_ISL_752902, EPI_ISL_752903, EPI_ISL_752923, EPI_ISL_752924, EPI_ISL_752941, EPI_ISL_752942, EPI_ISL_752943, EPI_ISL_752944, EPI_ISL_752945, EPI_ISL_752948, EPI_ISL_752996, EPI_ISL_752997, EPI_ISL_753100 | | | |
| see above | State Laboratories Division, Hawaii State Department of Health | State Laboratories Division, Hawaii State Department of Health | Pamela O'Brien, Sabrina Diemert, Drew Kuwazaki, Razvan Sultana, Edward Desmond |
| EPI_ISL_754136 | CHU Purpan - Laboratoire de Virologie - Institut Fédératif de Biologie | CHU Purpan - Laboratoire de Virologie - Institut Fédératif de Biologie | Latour J., Ranger N., Dubois M., Carcenac R., Harter A., Boyer P., Tremeaux P., Izopet J. |
| EPI_ISL_754195 | Department for Virology, Molecular Biology and Genome Research, R. G. Lugar Center for Public Health Research, National Center for Disease Control and Public Health (NCDC) of Georgia. | Department for Virology, Molecular Biology and Genome Research, R. G. Lugar Center for Public Health Research, National Center for Disease Control and Public Health (NCDC) of Georgia. | Gvantsa Brachveli, Meri Pantsulaia, Nino Berishvili, Tata Imnadze, Giorgi Tomashvili, Ana Papkiauri, Gvantsa Chanturia, Ann Machablishvili, Nato Kotaria, Marine Murtskhvaladze, Lela Sabadze, Mari Gavashelidze, Tamar Jashiashvili, Tea Tevdoradze, Ketevan Sidamonidze, Ekaterine Khmaladze, Ekaterine Zhghenti, Roena Sukhiashvili, Mariam Zakalashvili, Lela Urushadze, Magda Dgebuadze, Davit Tsaguria, Ekaterine Zangaladze, Adam Kotorashvili, Maia Alkhazashvili, Irma Burjanadze, Anna Kasradze, Khatuna Zakhashvili, Paata Imnadze, Amiran Gamkrelidze. |
| EPI_ISL_754392 | Genetica Molecular and Subdepartamento de Virologia ISP | Insituto de Salud Publica de Chile | Javier Tognarelli, Barbara Parra, Loredana Arata, Jaime Lagos, Gisselle Barra, Patricia Bustos, Rodrigo Fasce, Andres Castillo, Jorge Fernandez |

| | | | |
|---|---|---|---|
| | Chile | | |
| EPI_ISL_754905, EPI_ISL_754907, EPI_ISL_754913 | Laboratory Diagnostics and Clinical Immunology of Developmental Age, Medical University of Warsaw | genXone SA, Research & Development Laboratory; The Faculty of Mathematics, Informatics and Mechanics of the University of Warsaw | Maciej Sykulski, Grzegorz Nowicki, Monika Makowska-Woniak, Jakub Grabowski, Natalia Drwska-Matelska, ukasz Krych, Micha Kaszuba, Anna Gambin, Urszula Demkow |
| EPI_ISL_755121, EPI_ISL_755127, EPI_ISL_755131, EPI_ISL_755141, EPI_ISL_755143, EPI_ISL_755235, EPI_ISL_755239, EPI_ISL_755241, EPI_ISL_755243, EPI_ISL_755245, EPI_ISL_755249, EPI_ISL_755250, EPI_ISL_755256, EPI_ISL_755261, EPI_ISL_755263, EPI_ISL_755264, EPI_ISL_755265 | | | |
| see above | Biolab Diagnostic Laboratories | Andersen lab at Scripps Research | Issa Abu-Dayyeh, Ahmad Tibi, Lama Hussein, Lina Mohammad, Zein Naber, Amid Abdelnour with SEARCH Alliance San Diego |
| EPI_ISL_755316, EPI_ISL_755317, EPI_ISL_755318, EPI_ISL_755319, EPI_ISL_755320, EPI_ISL_755321, EPI_ISL_755322, EPI_ISL_755323, EPI_ISL_755324, EPI_ISL_755325, EPI_ISL_755326, EPI_ISL_755327, EPI_ISL_755328, EPI_ISL_755329, EPI_ISL_755330, EPI_ISL_755331, EPI_ISL_755332, EPI_ISL_755333, EPI_ISL_755335 | | | |
| see above | Maine Health and Environmental Testing Laboratory | Tewhey Lab, The Jackson Laboratory | Matluk,N., Dewey,H., Iosue,F., Barter,M., Lynch,R., Munger,H. and Tewhey,R. |
| EPI_ISL_755808, EPI_ISL_755906, EPI_ISL_755907, EPI_ISL_755908, EPI_ISL_755909, EPI_ISL_755910, EPI_ISL_755911, EPI_ISL_755912, EPI_ISL_755913, EPI_ISL_755914, EPI_ISL_755915, EPI_ISL_755916, EPI_ISL_755917, EPI_ISL_755918, EPI_ISL_755919, EPI_ISL_755920, EPI_ISL_755921, EPI_ISL_755922, EPI_ISL_755923, EPI_ISL_755924, EPI_ISL_755925, EPI_ISL_755926, EPI_ISL_755927, EPI_ISL_755928, EPI_ISL_755929, EPI_ISL_755930, EPI_ISL_755931, EPI_ISL_755932, EPI_ISL_755933, EPI_ISL_755934, EPI_ISL_755935, EPI_ISL_755936, EPI_ISL_755937, EPI_ISL_755938, EPI_ISL_755939 | | | |
| see above | Toronto Invasive Bacterial Diseases Network | McMaster University | Allison McGeer, Patryk Aftanas, Hooman Derakhshani, Angel Li, Kuganya Nirmalarajah, Emily Panousis, Ahmed Draia, Jalees Nasir, Michael Surette, Samira Mubareka, Andrew G. McArthur |
| EPI_ISL_755993, EPI_ISL_755994, EPI_ISL_756088, EPI_ISL_756089 | Department of Virology and Immunology, University of Helsinki and Helsinki University Hospital, Huslab Finland | Department of Virology, Faculty of Medicine, University of Helsinki, Helsinki, Finland | Teemu Smura, Ravi Kant, Phuoc Truong, Hussein Alburkat, Hannimari Kallio-Kokko, Jenni Virtanen, Maija Suvanto, Sari Hannula, Harri Kangas, Pekka Ellonen, Olli Vapalahti |
| EPI_ISL_756275 | State Laboratories Division, Hawaii State Department of Health | State Laboratories Division, Hawaii State Department of Health | Pamela O'Brien, Sabrina Diemert, Drew Kuwazaki, Razvan Sultana, Edward Desmond |
| EPI_ISL_756363 | Trinidad Public Health Laboratory | Carrington Lab, Department of PreClinical Sciences, Faculty of Medical Sciences, The University of the West Indies | Nikita S. D. Sahadeo, Arianne Brown-Jordan, Sarah Hill, Vernie Ramkissoon, Roshan Parasram, Naresh Nandram, Avery Hinds, Jerome Foster, Stanley Giddings, Karla Georges, Marsha Ivey, Rahul Naidu, Risha Singh, SueMin Nathaniel, Rajini Haraksingh, Jaya Jayaraman, Chinna Chinnadurai, Adesh Ramsubhag, Nuno Faria, Oliver Pybus, Christopher Oura, Gabriel Escobar, Christine V. F. Carrington |
| EPI_ISL_757365, EPI_ISL_757366, EPI_ISL_757367 | Department of Virology and Immunology, University of Helsinki and Helsinki University Hospital, Huslab Finland | Department of Virology, Faculty of Medicine, University of Helsinki, Helsinki, Finland | Teemu Smura, Ravi Kant, Phuoc Truong, Hussein Alburkat, Hannimari Kallio-Kokko, Jenni Virtanen, Maija Suvanto, Sari Hannula, Harri Kangas, Pekka Ellonen, Olli Vapalahti |
| EPI_ISL_765480, EPI_ISL_765484, EPI_ISL_765487, EPI_ISL_765488, EPI_ISL_765496, EPI_ISL_765497, EPI_ISL_765498, EPI_ISL_765499, EPI_ISL_765500, EPI_ISL_765501 | Wadsworth Center, New York State Department.of Health | Wadsworth Center, New York State Department.of Health | Kirsten St. George, Daryl M. Lamson, Alexis Russel, Matthew Shudt, Melissa A Leisner, Jonathan Plitnick, Navjot Singh, John Kelly, Sara Griesemer, Erasmus Schneider, Erica Lasek-Nesselquist |
| EPI_ISL_765713, EPI_ISL_765714, EPI_ISL_765715, EPI_ISL_765716, EPI_ISL_765717, EPI_ISL_765718, EPI_ISL_765719, EPI_ISL_765720, EPI_ISL_765721, EPI_ISL_765723 | Massachusetts General Hospital | Infectious Disease Program, Broad Institute of Harvard and MIT | Lemieux,J.E., Siddle,K.J., Shaw,B., Adams,G., Pierce,V., Turbett,S., Anahtar,M., Branda,J., Slater,D., Harris,J., Lin,A.E., Gladden-Young,A., Lagerborg,K., Rudy,M., DeRuff,K., Carter,A., Normandin,E., Bauer,M., Reilly,S., Tomkins-Tinch,C., Loreth,C., Chaluvadi,S., Neumann,A., Cusick,C., Chapman,S.B., Gnirke,A., Flowers,K., Cerrato,F., Birren,B.W., Gallagher,G., Smole,S., Park,D.J., MacInnis,B.L., Ryan,E., LaRocque,R., Rosenberg,E. and Sabeti,P.C. |
| EPI_ISL_769990 | Hospital Dr. Rafael A. Calderon Guardia | Inciensa, Instituto Costarricense de Investigación y Enseñanza en Nutrición y Salud | Francisco Duarte, Hebleen Porras, Claudio Soto-Garita, Estela Cordero, Adriana Godínez, Melany Calderón & Mariel López |
| EPI_ISL_769993 | Area De Salud El Guarco | Inciensa, Instituto Costarricense de Investigación y Enseñanza en Nutrición y Salud | Francisco Duarte, Hebleen Porras, Claudio Soto-Garita, Estela Cordero, Adriana Godínez, Melany Calderón & Mariel López |
| EPI_ISL_770006 | Hle - Asociacion Hogar De Ancianos De Palmar Sur De Osa | Inciensa, Instituto Costarricense de Investigación y Enseñanza en Nutrición y Salud | Francisco Duarte, Hebleen Porras, Claudio Soto-Garita, Estela Cordero, Adriana Godínez, Melany Calderón & Mariel López |
| EPI_ISL_770010 | Area De Salud Escazu (Coopesana) | Inciensa, Instituto Costarricense de Investigación y Enseñanza en Nutrición y Salud | Francisco Duarte, Hebleen Porras, Claudio Soto-Garita, Estela Cordero, Adriana Godínez, Melany Calderón & Mariel López |
| EPI_ISL_770011 | Area De Salud Coronado | Inciensa, Instituto Costarricense de Investigación y Enseñanza en Nutrición y Salud | Francisco Duarte, Hebleen Porras, Claudio Soto-Garita, Estela Cordero, Adriana Godínez, Melany Calderón & Mariel López |
| EPI_ISL_770047 | Latvijas Infektoloijas centrs | Latvian Biomedical Research and Study Centre | Ivars Silamielis, Kaspars Megnis, Monta Ustinova, Jnis Pjalkovskis, ikita Zrelovs, Vita Rovte, Jeena Storoženko, Tatjana Kolupajeva, Oksana Savicka, Uga Dumpis, Jnis Kloviš |
| EPI_ISL_770050 | E. Gulbja Laboratorija | Latvian Biomedical Research and Study Centre | Ivars Silamielis, Kaspars Megnis, Monta Ustinova, Jnis Pjalkovskis, ikita Zrelovs, Vita Rovte, Mikus Gavars, Dmitrijs Perminovs, Uga Dumpis, Jnis Kloviš |
| EPI_ISL_775217 | Gonoshasthya-RNA Molecular Research Center | Gonoshasthya-RNA Molecular Research Center | Nihad Adnan, Mohd. Raeed Jamiruddin, Md. Ahsanul Haq, Mohib Ullah Khondoker, Nafisa Azmuda, Firoz Ahmed, Shahana Sharmin, Salma Akter, Taslin Jahan Mou, Mahfuza Marzan, Sayeda Moriam Liza, Nowshin Jahan, Tamanna Ali, Maha Jamiruddin, Mousumi Chaity, Shahad Saif Khandker, Mumtarin Jannat Oishee |
| EPI_ISL_776968, EPI_ISL_776969, EPI_ISL_776970, EPI_ISL_776971, EPI_ISL_776972, EPI_ISL_776973, EPI_ISL_776974, EPI_ISL_776975, EPI_ISL_776976, EPI_ISL_776977, EPI_ISL_776978, EPI_ISL_776979, EPI_ISL_776980, EPI_ISL_776981 | | | |
| see above | Istituto Zooprofilattico Sperimentale del Mezzogiorno | TIGEM | Patrizia Annunziata, Andrea Ballabio, Valentina Bouche, Davide Cacchiarelli (CorrespAuthor), Pellegrino Cerino, Chiara Colantuono, Lucio Di Filippo, Antonio Grimaldi, Antonio Limone, Gabriella Loconte, Anna Manfredi, Francesco Panariello, Biancamaria Pierri, Marcello Salvi, Lucia Vassallo |
| EPI_ISL_779316 | Utah Public Health Laboratory, Utah Public Health Laboratory Infectious Disease submission group | Utah Public Health Laboratory, Utah Public Health Laboratory Infectious Disease submission group | Gallagher,T., Young,E.L., Oakeson,K.F. |
| EPI_ISL_779561, EPI_ISL_779562, EPI_ISL_779563 | Microbiological Diagnostic Unit - Public Health Laboratory (MDU-PHL) | MDU-PHL | Seemann T., Sait, M.L., Sherry, N.L. |
| EPI_ISL_779705, EPI_ISL_779706, EPI_ISL_779708, EPI_ISL_779710 | The Foundation for Medical Research | The Foundation for Medical Research | Ayan Mandal, Kayzad Nilgiriwala, Kalpana Sriraman, Ambreen Shaikh, Grishma Patel, Tejal Mestry, Smriti Vaswani, Swapneil Parikh, Shreevatsa Udupa, Nirjhar Chatterjee, Jayanthi Shastri, Nerges Mistry |
| EPI_ISL_780275, EPI_ISL_780683, EPI_ISL_783775, EPI_ISL_784585, EPI_ISL_785485, EPI_ISL_786385, EPI_ISL_787285, EPI_ISL_788185 | Houston Methodist Hospital | Houston Methodist Hospital | S. Wesley Long, Randall J. Olsen, Paul A. Christensen, David W. Bernard, James J. Davis, Maulik Shukla, Marcus Nguyen, Matthew Ojeda Saavedra, Prasanti Yerramilli, Layne Pruitt, Sishir Subedi, Heather Hendrickson, and James M. Musser |
| EPI_ISL_788943, EPI_ISL_788948, EPI_ISL_788949, EPI_ISL_788950, EPI_ISL_788951, EPI_ISL_788952, EPI_ISL_788968, EPI_ISL_788969 | Ospedale "Di Venere" | Beaconlab (Bioinformatics, Evolution and Comparative Genomics lab), Dept of Biosciences, University on Milan | Iacobellis M, d'Avenia M, Piluscio R, Parisi A, Chiara M, Manzari C, Pesole G |
| EPI_ISL_789105, EPI_ISL_789108, EPI_ISL_789115, EPI_ISL_789117, EPI_ISL_789121, EPI_ISL_789123, EPI_ISL_789124, EPI_ISL_789125, EPI_ISL_789129, EPI_ISL_789134, EPI_ISL_789135, EPI_ISL_789140, EPI_ISL_789141, EPI_ISL_789144, EPI_ISL_789145, EPI_ISL_789148, EPI_ISL_789149, EPI_ISL_789155, EPI_ISL_789158, EPI_ISL_789159, EPI_ISL_789162, EPI_ISL_789164, EPI_ISL_789170, EPI_ISL_789171, EPI_ISL_789172, EPI_ISL_789179, EPI_ISL_789180, EPI_ISL_789193, EPI_ISL_789200, EPI_ISL_789217, EPI_ISL_789218, EPI_ISL_789221, EPI_ISL_789231, EPI_ISL_789232, EPI_ISL_789233, EPI_ISL_789260, EPI_ISL_789268, EPI_ISL_789272, EPI_ISL_789273, EPI_ISL_789697, EPI_ISL_789698, EPI_ISL_789699, EPI_ISL_789700, EPI_ISL_789701, EPI_ISL_789702, EPI_ISL_789703, EPI_ISL_789704, EPI_ISL_789705, EPI_ISL_789706, EPI_ISL_789707, EPI_ISL_789708, EPI_ISL_789709, EPI_ISL_789710, EPI_ISL_789711, EPI_ISL_789712, EPI_ISL_789713, EPI_ISL_789714, EPI_ISL_789715, EPI_ISL_789716, EPI_ISL_789717, EPI_ISL_789718, EPI_ISL_789719, EPI_ISL_789720, EPI_ISL_789721, EPI_ISL_789722, EPI_ISL_789723, EPI_ISL_789724, EPI_ISL_789725, EPI_ISL_789726, EPI_ISL_789727, EPI_ISL_789728, EPI_ISL_789729, EPI_ISL_789730, EPI_ISL_789731, EPI_ISL_789732, EPI_ISL_789733, EPI_ISL_789734, EPI_ISL_789735, EPI_ISL_789736, EPI_ISL_789737, EPI_ISL_789738, EPI_ISL_789739, EPI_ISL_789740, EPI_ISL_789741, EPI_ISL_789742, EPI_ISL_789743, EPI_ISL_789744, EPI_ISL_789745, EPI_ISL_789746, EPI_ISL_789747, EPI_ISL_789748, EPI_ISL_789749, EPI_ISL_789750, EPI_ISL_789751, EPI_ISL_789752, EPI_ISL_789753, EPI_ISL_789754, EPI_ISL_789755, EPI_ISL_789756, EPI_ISL_789757, EPI_ISL_789758, EPI_ISL_789759, EPI_ISL_789760, EPI_ISL_789761, EPI_ISL_789762, EPI_ISL_789763, EPI_ISL_789764, EPI_ISL_789765, EPI_ISL_789766, EPI_ISL_789767, EPI_ISL_789768, EPI_ISL_789769, EPI_ISL_789770, EPI_ISL_789771, EPI_ISL_789772, EPI_ISL_789773, EPI_ISL_789774, EPI_ISL_789775, EPI_ISL_789776, EPI_ISL_789777, EPI_ISL_789778, EPI_ISL_789779, EPI_ISL_789822, EPI_ISL_789825, EPI_ISL_789852, EPI_ISL_789886, EPI_ISL_789897, EPI_ISL_789929, EPI_ISL_789933, EPI_ISL_789943, EPI_ISL_789944, EPI_ISL_789956, EPI_ISL_789959, EPI_ISL_789964, EPI_ISL_789978, EPI_ISL_789981, EPI_ISL_789983, EPI_ISL_789987, EPI_ISL_789990, EPI_ISL_789991, EPI_ISL_789994, EPI_ISL_789995, EPI_ISL_789996, EPI_ISL_789997, | | | |

| | | | |
|---|---|---|---|
| EPI_ISL_789998, EPI_ISL_789999, EPI_ISL_790000, EPI_ISL_790001, EPI_ISL_790002, EPI_ISL_790003, EPI_ISL_790004, EPI_ISL_790005, EPI_ISL_790006, EPI_ISL_790007, EPI_ISL_790008, EPI_ISL_790009, EPI_ISL_790010, EPI_ISL_790011, EPI_ISL_790012, EPI_ISL_790013, EPI_ISL_790014, EPI_ISL_790015, EPI_ISL_790016, EPI_ISL_790017, EPI_ISL_790018, EPI_ISL_790019, EPI_ISL_790020, EPI_ISL_790021, EPI_ISL_790022, EPI_ISL_790023, EPI_ISL_790024, EPI_ISL_790025, EPI_ISL_790026, EPI_ISL_790027, EPI_ISL_790028, EPI_ISL_790030, EPI_ISL_790032, EPI_ISL_790033, EPI_ISL_790034, EPI_ISL_790035, EPI_ISL_790036, EPI_ISL_790037, EPI_ISL_790038, EPI_ISL_790039, EPI_ISL_790040, EPI_ISL_790041, EPI_ISL_790042, EPI_ISL_790043, EPI_ISL_790044, EPI_ISL_790045, EPI_ISL_790046, EPI_ISL_790071, EPI_ISL_790072, EPI_ISL_790073, EPI_ISL_790074, EPI_ISL_790075, EPI_ISL_790077, EPI_ISL_790078, EPI_ISL_790082, EPI_ISL_790083, EPI_ISL_790087, EPI_ISL_790088, EPI_ISL_790089, EPI_ISL_790090, EPI_ISL_790093, EPI_ISL_790094, EPI_ISL_790096, EPI_ISL_790099, EPI_ISL_790100, EPI_ISL_790104, EPI_ISL_790105, EPI_ISL_790106, EPI_ISL_790107, EPI_ISL_790108, EPI_ISL_790109, EPI_ISL_790110, EPI_ISL_790111, EPI_ISL_790114, EPI_ISL_790133, EPI_ISL_790400, EPI_ISL_790487, EPI_ISL_790503 | | | |
| see above | Houston Methodist Hospital | Houston Methodist Hospital | S. Wesley Long, Randall J. Olsen, Paul A. Christensen, David W. Bernard, James J. Davis, Maulik Shukla, Marcus Nguyen, Matthew Ojeda Saavedra, Prasanti Yerramilli, Layne Pruitt, Sishir Subedi, Heather Hendrickson, and James M. Musser |
| EPI_ISL_794014, EPI_ISL_794015, EPI_ISL_794016, EPI_ISL_794017, EPI_ISL_794018, EPI_ISL_794019, EPI_ISL_794020, EPI_ISL_794021, EPI_ISL_794022, EPI_ISL_794023, EPI_ISL_794024, EPI_ISL_794025, EPI_ISL_794026, EPI_ISL_794027, EPI_ISL_794028, EPI_ISL_794029, EPI_ISL_794030, EPI_ISL_794031, EPI_ISL_794032, EPI_ISL_794033, EPI_ISL_794034 | | | |
| see above | Wadsworth Center, New York State Department.of Health | Wadsworth Center, New York State Department.of Health | Kirsten St. George, Daryl M. Lamson, Alexis Russel, Matthew Shudt, Melissa A Leisner, Jonathan Plitnick, Navjot Singh, John Kelly, Sara Griesemer, Erasmus Schneider, Erica Lasek-Nesselquist |
| EPI_ISL_801862, EPI_ISL_801866, EPI_ISL_801870, EPI_ISL_801874, EPI_ISL_801880, EPI_ISL_801881, EPI_ISL_801893, EPI_ISL_801897, EPI_ISL_801905, EPI_ISL_801914, EPI_ISL_801915, EPI_ISL_801920, EPI_ISL_801922, EPI_ISL_801923, EPI_ISL_801926, EPI_ISL_801931, EPI_ISL_801940, EPI_ISL_801941, EPI_ISL_801969, EPI_ISL_801970, EPI_ISL_802219, EPI_ISL_802220, EPI_ISL_802221, EPI_ISL_802222, EPI_ISL_802223, EPI_ISL_802224, EPI_ISL_802225, EPI_ISL_802226, EPI_ISL_802227, EPI_ISL_802228, EPI_ISL_802229, EPI_ISL_802230, EPI_ISL_802231, EPI_ISL_802232, EPI_ISL_802233, EPI_ISL_802234, EPI_ISL_802235, EPI_ISL_802236, EPI_ISL_802237 | | | |
| see above | MSHS Clinical Microbiology Laboratories | MSHS Pathogen Surveillance Program | Ana S. Gonzalez-Reiche, Hala Alshammary, Mitchell J. Sullivan, Brianne Ciferri, Ajay Obla, Angela Amoako, Mahmoud Awawda, Elena Hirsch, Ashley S. Salimbangon, Levy Sominsky, Katherine Beach, Kayla Russo, Charles Gleason, Shelcie Fabre, Giulio Kleiner, Zenab Khan, Bremy Alburquerque, Adriana van de Guchte, Komal Srivastava, Matthew M. Hernandez, Jayeeta Dutta, Denise Jurczyszak, Emily Ferreri, Rachel Chernet, Nancy Francoeur, Betsaida Salom Melo, Irina Oussenko, Gintaras Deikus, Juan Soto, Shwetha Hara Sridhar, Ying-Chih Wang, Kathryn Twyman, Andrew Kasarskis, Deena R. Altman, Robert Sebra, Adolfo García-Sastre, Marta Luksza, Gopi Patel, Sarah Schaefer, Melissa Gitman, Michael D. Nowak, Alberto Paniz-Mondolfi, Emilia Mia Sordillo, Viviana Simon, Harm van Bakel |
| EPI_ISL_803134, EPI_ISL_803135, EPI_ISL_803136, EPI_ISL_803137, EPI_ISL_803227, EPI_ISL_803228, EPI_ISL_803237, EPI_ISL_803333, EPI_ISL_803334, EPI_ISL_803335, EPI_ISL_803336, EPI_ISL_803337, EPI_ISL_803338, EPI_ISL_803339, EPI_ISL_803340, EPI_ISL_803486, EPI_ISL_803487, EPI_ISL_803488, EPI_ISL_803489, EPI_ISL_803490, EPI_ISL_803491, EPI_ISL_803492, EPI_ISL_803493, EPI_ISL_803508, EPI_ISL_803518, EPI_ISL_803519, EPI_ISL_803520, EPI_ISL_803521, EPI_ISL_803522, EPI_ISL_803535, EPI_ISL_803536, EPI_ISL_803537 | | | |
| see above | Wisconsin State Laboratory of Hygiene Communicable Disease Division | Wisconsin State Laboratory of Hygiene Communicable Disease Division | Kelsey R. Florek, Abigail C. Shockey |
| EPI_ISL_804378 | CHU Purpan - Laboratoire de Virologie - Institut Fédératif de Biologie | CHU Purpan - Laboratoire de Virologie - Institut Fédératif de Biologie | Latour J., Ranger N., Dubois M., Carcenac R., Harter A., Boyer P., Tremeaux P., Izopet J. |
| EPI_ISL_806611, EPI_ISL_806612, EPI_ISL_806614, EPI_ISL_806619, EPI_ISL_806620, EPI_ISL_806621 | KEMRI-Wellcome Trust Research Programme/KEMRI-CGMR-C Kilifi | KEMRI-Wellcome Trust Research Programme/KEMRI-CGMR-C Kilifi | Githinji et al |
| EPI_ISL_806867, EPI_ISL_806868, EPI_ISL_806869, EPI_ISL_806870, EPI_ISL_806871, EPI_ISL_806872, EPI_ISL_806873, EPI_ISL_806874, EPI_ISL_806875, EPI_ISL_806876, EPI_ISL_806877, EPI_ISL_806878, EPI_ISL_806879, EPI_ISL_806880, EPI_ISL_806881, EPI_ISL_806882, EPI_ISL_806883, EPI_ISL_806884, EPI_ISL_806885, EPI_ISL_806886, EPI_ISL_806887, EPI_ISL_806888, EPI_ISL_806889, EPI_ISL_806890, EPI_ISL_806891, EPI_ISL_806892, EPI_ISL_806893, EPI_ISL_806894, EPI_ISL_806895, EPI_ISL_806896, EPI_ISL_806897, EPI_ISL_806898 | | | |
| see above | Washington State Department of Health | Seattle Flu Study | Deborah A. Nickerson, Chris D. Frazar, Jover Lee, Benjamin Pelle, Matthew Richardson, Amanda Adler, Elisabeth Brandstetter, Peter D. Han, Kairsten Fay, Misja Ilcisin, Kirsten Lacombe, Thomas R. Sibley, Melissa Truong, Caitlin R. Wolf, Romesh Gautom, Geoff Melly, Brian Hiatt, Philip Dykema, Scott Lindquist, Michael Boeckh, Janet A. Englund, Michael Famulare, Barry R. Lutz, Mark J. Rieder, Lea M. Starita, Matthew Thompson, Helen Y. Chu, Jay Shendure, Trevor Bedford |
| EPI_ISL_811032, EPI_ISL_811033 | MRCG at LSHTM Genomics lab | MRCG at LSHTM Genomics lab | Abdul Karim sesay, Abdoulie Kanteh, Jarra Manneh, Mariama Kujabi, Bakary Sanyang |
| EPI_ISL_812665, EPI_ISL_812668, EPI_ISL_812673, EPI_ISL_812674, EPI_ISL_812675, EPI_ISL_812676, EPI_ISL_812678, EPI_ISL_812679, EPI_ISL_812683, EPI_ISL_812685, EPI_ISL_812686, EPI_ISL_812687, EPI_ISL_812691, EPI_ISL_812693, EPI_ISL_812695, EPI_ISL_812696, EPI_ISL_812697, EPI_ISL_812698, EPI_ISL_812701, EPI_ISL_812703 | | | |
| see above | United States Air Force School of Aerospace Medicine | United States Air Force School of Aerospace Medicine | Anthony Fries, Jennifer Meyer, Amanda Javorina, Sarah Purves, William Gruner, Clarise Starr, Elizabeth Macias |
| EPI_ISL_813356 | Oxford Viromics, NDM, University of Oxford; Oxford University Hospitals; Basingstoke and North Hampshire Hospital | COVID-19 Genomics UK (COG-UK) Consortium | Tanya Golubchik, David Bonsall, George Macintyre, Amy Trebes, Mariateresa de Cesare, Catrin Moore, Alex Mobbs, Anita Justice, Robert Shaw, Monique Andersson, Timothy Peto, Emma Wise, Nathan Moore, Jessica Lynch, Nick Cortes, Matilde Mori, Stephen Kidd, David Buck, John Todd, Christophe Fraser |
| EPI_ISL_813983, EPI_ISL_813999, EPI_ISL_814022 | Hospital General Universitario Gregorio Marañón | SeqCOVID-SPAIN consortium/IBV(CSIC) | Darío García de Viedma, Laura Pérez-Lago, Marta Herranz, Jon Sicilia, Julia Suárez, Pilar Catalán, Patricia Muñoz and SeqCOVID-SPAIN consortium |
| EPI_ISL_815257 | Centogene | Centogene | Peter Bauer, Krishna Kumar Kandaswamy, Vivi Hue-Trang Lieu |
| EPI_ISL_824482, EPI_ISL_824483, EPI_ISL_824484, EPI_ISL_824485, EPI_ISL_824486 | Hospital Universitari Vall d'Hebron - Vall d'Hebron Institut de Rercerca | Hospital Universitari Vall d'Hebron | Cristina Andrés, Maria Piñana, Josep F Abril, Damir Garcia-Cehic, Ariadna Rando, Juliana Esperalba, Maria Gema Codina, Carla Castillo, Maria Carmen Martin, Tomàs Pumarola, Josep Quer, Andrés Antón |
| EPI_ISL_825633 | Nigeria Centre For Disease Control | National reference Laboratory, NCDC, Gaduwa, Abuja | Dr Ndodo Nnaemeka, Olusola Anuoluwapo Akanbi, Chimaobi Chukwu, Dr Omoare Adesuyi, Esebanmen Grace, Anthony Ahumibe, Naidoo Dhamari, Nwando Mba, Dr Chikwe Ihekweazu |
| EPI_ISL_825715, EPI_ISL_825718, EPI_ISL_825719, EPI_ISL_825721, EPI_ISL_825722, EPI_ISL_825723, EPI_ISL_825724, EPI_ISL_825725, EPI_ISL_825780, EPI_ISL_825781, EPI_ISL_825782, EPI_ISL_825784, EPI_ISL_825785, EPI_ISL_825786, EPI_ISL_825787, EPI_ISL_825788, EPI_ISL_825789, EPI_ISL_825790, EPI_ISL_825791, EPI_ISL_825792, EPI_ISL_825793, EPI_ISL_825794, EPI_ISL_825795, EPI_ISL_825796, EPI_ISL_825797, EPI_ISL_825798, EPI_ISL_825799, EPI_ISL_825800, EPI_ISL_825801, EPI_ISL_825802, EPI_ISL_825803, EPI_ISL_825804, EPI_ISL_825805, EPI_ISL_825806 | | | |
| see above | Laboratoire de santé publique du Québec | Laboratoire de santé publique du Québec | Sandrine Moreira, Ioannis Ragoussis, Guillaume Bourque, Jesse Shapiro, Mark Lathrop and Michel Roger on behalf of the CoVSeQ research group (http://covseq.ca/researchgroup) |
| EPI_ISL_826588, EPI_ISL_826659, EPI_ISL_826660, EPI_ISL_826661, EPI_ISL_826662, EPI_ISL_826663, EPI_ISL_826664 | Montefiore Medical Center | Albert Einstein College of Medicine, Dept. of Microbiology & Immunology, Chandran lab | J. Maximilian Fels, Saad Khan, Ryan Forster, Karin A. Skalina, Surksha Sirichand, Amy S. Fox, Aviv Bergman, William B. Mitchell, Lucia R. Wolgast, Wendy Szymczak, Robert H. Bortz III, M. Eugenia Dieterle, Catalina Florez, Denise Haslwanter, Rohit K. Jangra, Ethan Laudermilch, Ariel S. Wirchnianski, Jason Barnhill, David L. Goldman, Hnin Khine, D. Yitzchak Goldstein, Johanna P. Daily, Kartik Chandran, Libusha Kelly |
| EPI_ISL_826817 | The National University Hospital of Iceland | deCODE genetics | Daniel F Gudbjartsson; Agnar Helgason; Hakon Jonsson; Olafur T Magnusson; Pall Melsted; Gudmundur L Norddahl; Jona Saemundsdottir; Asgeir Sigurdsson; Patrick Sulem; Arna B Agustsdottir; Hannes Eggertsson; Berglind Eiriksdottir; Run Fridriksdottir; Elisabet E Gardarsdottir; Gudmundur Georgsson; Olafia S Gretarsdottir; Kjartan R Gudmundsson; Thora R Gunnarsdottir; Arnaldur Gylfason; Hilma Holm; Brynjar O Jensson; Aslaug Jonasdottir; Kamilla S Josefsdottir; Thordur Kristjansson; Droplaug N Magnusdottir; Solvi Rognvaldsson; Louise le Roux; Gudrun Sigmundsdottir; Gardar Sveinbjornsson; Kristin E Sveinsdottir; Maney Sveinsdottir; Emil A Thorarensen; Bjarni Thorbjornsson; Gisli Masson; Ingileif Jonsdottir; Alma Moller; Thorolfur Gudnason; Karl G Kristinsson; Unnur Thorsteinsdottir; Kari Stefansson |
| EPI_ISL_826829, EPI_ISL_826830, EPI_ISL_826832 | INSPI-CRN DE INFLUENZA Y OTROS VIRUS RESPIRATORIOS | Instituto de Salud Publica de Chile | Javier Tognarelli, Barbara Parra, Loredana Arata, Jaime Lagos, Gisselle Barra, Alfredo Bruno, Domenica de Mora, Solon Narvaez, Jimmy Garcez, Michelle Paez, Martiza Olmedo, Manuel Gonzalez, Patricia Bustos, Rodrigo Fasce, Andres Castillo, Jorge Fernandez |
| EPI_ISL_826885 | The National University Hospital of Iceland | deCODE genetics | Daniel F Gudbjartsson; Agnar Helgason; Hakon Jonsson; Olafur T Magnusson; Pall Melsted; Gudmundur L Norddahl; Jona Saemundsdottir; Asgeir Sigurdsson; Patrick Sulem; Arna B Agustsdottir; Hannes Eggertsson; Berglind Eiriksdottir; Run Fridriksdottir; Elisabet E Gardarsdottir; Gudmundur Georgsson; Olafia S Gretarsdottir; Kjartan R Gudmundsson; Thora R Gunnarsdottir; Arnaldur Gylfason; Hilma Holm; Brynjar O Jensson; Aslaug Jonasdottir; Kamilla S Josefsdottir; Thordur Kristjansson; Droplaug N Magnusdottir; Solvi Rognvaldsson; Louise le Roux; Gudrun Sigmundsdottir; Gardar Sveinbjornsson; Kristin E Sveinsdottir; Maney Sveinsdottir; Emil A Thorarensen; Bjarni Thorbjornsson; Gisli Masson; Ingileif Jonsdottir; Alma Moller; Thorolfur Gudnason; Karl G Kristinsson; Unnur Thorsteinsdottir; Kari Stefansson |
| EPI_ISL_827006, EPI_ISL_827393, EPI_ISL_827394, EPI_ISL_827504, EPI_ISL_827505 | deCODE genetics | deCODE genetics | Daniel F Gudbjartsson; Agnar Helgason; Hakon Jonsson; Olafur T Magnusson; Pall Melsted; Gudmundur L Norddahl; Jona Saemundsdottir; Asgeir Sigurdsson; Patrick Sulem; Arna B Agustsdottir; Hannes Eggertsson; Berglind Eiriksdottir; Run Fridriksdottir; Elisabet E Gardarsdottir; Gudmundur Georgsson; Olafia S Gretarsdottir; Kjartan R Gudmundsson; Thora R Gunnarsdottir; Arnaldur Gylfason; Hilma Holm; Brynjar O Jensson; Aslaug |

| | | | |
|---|---|---|---|
| | | | Jonasdottir; Kamilla S Josefsdottir; Thordur Kristjansson; Droplaug N Magnusdottir; Solvi Rognvaldsson; Louise le Roux; Gudrun Sigmundsdottir; Gardar Sveinbjornsson; Kristin E Sveinsdottir; Maney Sveinsdottir; Emil A Thorarensen; Bjarni Thorbjornsson; Gisli Masson; Ingileif Jonsdottir; Alma Moller; Thorolfur Gudnason; Karl G Kristinsson; Unnur Thorsteinsdottir; Kari Stefansson |
| EPI_ISL_827513, EPI_ISL_827514 | The National University Hospital of Iceland | deCODE genetics | Daniel F Gudbjartsson; Agnar Helgason; Hakon Jonsson; Olafur T Magnusson; Pall Melsted; Gudmundur L Norddahl; Jona Saemundsdottir; Asgeir Sigurdsson; Patrick Sulem; Arna B Agustsdottir; Hannes Eggertsson; Berglind Eiriksdottir; Run Fridriksdottir; Elisabet E Gardarsdottir; Gudmundur Georgsson; Olafia S Gretarsdottir; Kjartan R Gudmundsson; Thora R Gunnarsdottir; Arnaldur Gylfason; Hilma Holm; Brynjar O Jensson; Aslaug Jonasdottir; Kamilla S Josefsdottir; Thordur Kristjansson; Droplaug N Magnusdottir; Solvi Rognvaldsson; Louise le Roux; Gudrun Sigmundsdottir; Gardar Sveinbjornsson; Kristin E Sveinsdottir; Maney Sveinsdottir; Emil A Thorarensen; Bjarni Thorbjornsson; Gisli Masson; Ingileif Jonsdottir; Alma Moller; Thorolfur Gudnason; Karl G Kristinsson; Unnur Thorsteinsdottir; Kari Stefansson |
| EPI_ISL_827685, EPI_ISL_827686 | deCODE genetics | deCODE genetics | Daniel F Gudbjartsson; Agnar Helgason; Hakon Jonsson; Olafur T Magnusson; Pall Melsted; Gudmundur L Norddahl; Jona Saemundsdottir; Asgeir Sigurdsson; Patrick Sulem; Arna B Agustsdottir; Hannes Eggertsson; Berglind Eiriksdottir; Run Fridriksdottir; Elisabet E Gardarsdottir; Gudmundur Georgsson; Olafia S Gretarsdottir; Kjartan R Gudmundsson; Thora R Gunnarsdottir; Arnaldur Gylfason; Hilma Holm; Brynjar O Jensson; Aslaug Jonasdottir; Kamilla S Josefsdottir; Thordur Kristjansson; Droplaug N Magnusdottir; Solvi Rognvaldsson; Louise le Roux; Gudrun Sigmundsdottir; Gardar Sveinbjornsson; Kristin E Sveinsdottir; Maney Sveinsdottir; Emil A Thorarensen; Bjarni Thorbjornsson; Gisli Masson; Ingileif Jonsdottir; Alma Moller; Thorolfur Gudnason; Karl G Kristinsson; Unnur Thorsteinsdottir; Kari Stefansson |
| EPI_ISL_827687, EPI_ISL_827688, EPI_ISL_827831 | The National University Hospital of Iceland | deCODE genetics | Daniel F Gudbjartsson; Agnar Helgason; Hakon Jonsson; Olafur T Magnusson; Pall Melsted; Gudmundur L Norddahl; Jona Saemundsdottir; Asgeir Sigurdsson; Patrick Sulem; Arna B Agustsdottir; Hannes Eggertsson; Berglind Eiriksdottir; Run Fridriksdottir; Elisabet E Gardarsdottir; Gudmundur Georgsson; Olafia S Gretarsdottir; Kjartan R Gudmundsson; Thora R Gunnarsdottir; Arnaldur Gylfason; Hilma Holm; Brynjar O Jensson; Aslaug Jonasdottir; Kamilla S Josefsdottir; Thordur Kristjansson; Droplaug N Magnusdottir; Solvi Rognvaldsson; Louise le Roux; Gudrun Sigmundsdottir; Gardar Sveinbjornsson; Kristin E Sveinsdottir; Maney Sveinsdottir; Emil A Thorarensen; Bjarni Thorbjornsson; Gisli Masson; Ingileif Jonsdottir; Alma Moller; Thorolfur Gudnason; Karl G Kristinsson; Unnur Thorsteinsdottir; Kari Stefansson |
| EPI_ISL_828477, EPI_ISL_828902 | deCODE genetics | deCODE genetics | Daniel F Gudbjartsson; Agnar Helgason; Hakon Jonsson; Olafur T Magnusson; Pall Melsted; Gudmundur L Norddahl; Jona Saemundsdottir; Asgeir Sigurdsson; Patrick Sulem; Arna B Agustsdottir; Hannes Eggertsson; Berglind Eiriksdottir; Run Fridriksdottir; Elisabet E Gardarsdottir; Gudmundur Georgsson; Olafia S Gretarsdottir; Kjartan R Gudmundsson; Thora R Gunnarsdottir; Arnaldur Gylfason; Hilma Holm; Brynjar O Jensson; Aslaug Jonasdottir; Kamilla S Josefsdottir; Thordur Kristjansson; Droplaug N Magnusdottir; Solvi Rognvaldsson; Louise le Roux; Gudrun Sigmundsdottir; Gardar Sveinbjornsson; Kristin E Sveinsdottir; Maney Sveinsdottir; Emil A Thorarensen; Bjarni Thorbjornsson; Gisli Masson; Ingileif Jonsdottir; Alma Moller; Thorolfur Gudnason; Karl G Kristinsson; Unnur Thorsteinsdottir; Kari Stefansson |
| EPI_ISL_829165 | The National University Hospital of Iceland | deCODE genetics | Daniel F Gudbjartsson; Agnar Helgason; Hakon Jonsson; Olafur T Magnusson; Pall Melsted; Gudmundur L Norddahl; Jona Saemundsdottir; Asgeir Sigurdsson; Patrick Sulem; Arna B Agustsdottir; Hannes Eggertsson; Berglind Eiriksdottir; Run Fridriksdottir; Elisabet E Gardarsdottir; Gudmundur Georgsson; Olafia S Gretarsdottir; Kjartan R Gudmundsson; Thora R Gunnarsdottir; Arnaldur Gylfason; Hilma Holm; Brynjar O Jensson; Aslaug Jonasdottir; Kamilla S Josefsdottir; Thordur Kristjansson; Droplaug N Magnusdottir; Solvi Rognvaldsson; Louise le Roux; Gudrun Sigmundsdottir; Gardar Sveinbjornsson; Kristin E Sveinsdottir; Maney Sveinsdottir; Emil A Thorarensen; Bjarni Thorbjornsson; Gisli Masson; Ingileif Jonsdottir; Alma Moller; Thorolfur Gudnason; Karl G Kristinsson; Unnur Thorsteinsdottir; Kari Stefansson |
| EPI_ISL_829167, EPI_ISL_829388, EPI_ISL_829627, EPI_ISL_829682, EPI_ISL_829683 | deCODE genetics | deCODE genetics | Daniel F Gudbjartsson; Agnar Helgason; Hakon Jonsson; Olafur T Magnusson; Pall Melsted; Gudmundur L Norddahl; Jona Saemundsdottir; Asgeir Sigurdsson; Patrick Sulem; Arna B Agustsdottir; Hannes Eggertsson; Berglind Eiriksdottir; Run Fridriksdottir; Elisabet E Gardarsdottir; Gudmundur Georgsson; Olafia S Gretarsdottir; Kjartan R Gudmundsson; Thora R Gunnarsdottir; Arnaldur Gylfason; Hilma Holm; Brynjar O Jensson; Aslaug Jonasdottir; Kamilla S Josefsdottir; Thordur Kristjansson; Droplaug N Magnusdottir; Solvi Rognvaldsson; Louise le Roux; Gudrun Sigmundsdottir; Gardar Sveinbjornsson; Kristin E Sveinsdottir; Maney Sveinsdottir; Emil A Thorarensen; Bjarni Thorbjornsson; Gisli Masson; Ingileif Jonsdottir; Alma Moller; Thorolfur Gudnason; Karl G Kristinsson; Unnur Thorsteinsdottir; Kari Stefansson |
| EPI_ISL_829697 | The National University Hospital of Iceland | deCODE genetics | Daniel F Gudbjartsson; Agnar Helgason; Hakon Jonsson; Olafur T Magnusson; Pall Melsted; Gudmundur L Norddahl; Jona Saemundsdottir; Asgeir Sigurdsson; Patrick Sulem; Arna B Agustsdottir; Hannes Eggertsson; Berglind Eiriksdottir; Run Fridriksdottir; Elisabet E Gardarsdottir; Gudmundur Georgsson; Olafia S Gretarsdottir; Kjartan R Gudmundsson; Thora R Gunnarsdottir; Arnaldur Gylfason; Hilma Holm; Brynjar O Jensson; Aslaug Jonasdottir; Kamilla S Josefsdottir; Thordur Kristjansson; Droplaug N Magnusdottir; Solvi Rognvaldsson; Louise le Roux; Gudrun Sigmundsdottir; Gardar Sveinbjornsson; Kristin E Sveinsdottir; Maney Sveinsdottir; Emil A Thorarensen; Bjarni Thorbjornsson; Gisli Masson; Ingileif Jonsdottir; Alma Moller; Thorolfur Gudnason; Karl G Kristinsson; Unnur Thorsteinsdottir; Kari Stefansson |
| EPI_ISL_830043, EPI_ISL_830044, EPI_ISL_830045, EPI_ISL_830046, EPI_ISL_830047, EPI_ISL_830048, EPI_ISL_830049 | deCODE genetics | deCODE genetics | Daniel F Gudbjartsson; Agnar Helgason; Hakon Jonsson; Olafur T Magnusson; Pall Melsted; Gudmundur L Norddahl; Jona Saemundsdottir; Asgeir Sigurdsson; Patrick Sulem; Arna B Agustsdottir; Hannes Eggertsson; Berglind Eiriksdottir; Run Fridriksdottir; Elisabet E Gardarsdottir; Gudmundur Georgsson; Olafia S Gretarsdottir; Kjartan R Gudmundsson; Thora R Gunnarsdottir; Arnaldur Gylfason; Hilma Holm; Brynjar O Jensson; Aslaug Jonasdottir; Kamilla S Josefsdottir; Thordur Kristjansson; Droplaug N Magnusdottir; Solvi Rognvaldsson; Louise le Roux; Gudrun Sigmundsdottir; Gardar Sveinbjornsson; Kristin E Sveinsdottir; Maney Sveinsdottir; Emil A Thorarensen; Bjarni Thorbjornsson; Gisli Masson; Ingileif Jonsdottir; Alma Moller; Thorolfur Gudnason; Karl G Kristinsson; Unnur Thorsteinsdottir; Kari Stefansson |
| EPI_ISL_830393 | The National University Hospital of Iceland | deCODE genetics | Daniel F Gudbjartsson; Agnar Helgason; Hakon Jonsson; Olafur T Magnusson; Pall Melsted; Gudmundur L Norddahl; Jona Saemundsdottir; Asgeir Sigurdsson; Patrick Sulem; Arna B Agustsdottir; Hannes Eggertsson; Berglind Eiriksdottir; Run Fridriksdottir; Elisabet E Gardarsdottir; Gudmundur Georgsson; Olafia S Gretarsdottir; Kjartan R Gudmundsson; Thora R Gunnarsdottir; Arnaldur Gylfason; Hilma Holm; Brynjar O Jensson; Aslaug Jonasdottir; Kamilla S Josefsdottir; Thordur Kristjansson; Droplaug N Magnusdottir; Solvi Rognvaldsson; Louise le Roux; Gudrun Sigmundsdottir; Gardar Sveinbjornsson; Kristin E Sveinsdottir; Maney Sveinsdottir; Emil A Thorarensen; Bjarni Thorbjornsson; Gisli Masson; Ingileif Jonsdottir; Alma Moller; Thorolfur Gudnason; Karl G Kristinsson; Unnur Thorsteinsdottir; Kari Stefansson |
| EPI_ISL_830414 | deCODE genetics | deCODE genetics | Daniel F Gudbjartsson; Agnar Helgason; Hakon Jonsson; Olafur T Magnusson; Pall Melsted; Gudmundur L Norddahl; Jona Saemundsdottir; Asgeir Sigurdsson; Patrick Sulem; Arna B Agustsdottir; Hannes Eggertsson; Berglind Eiriksdottir; Run Fridriksdottir; Elisabet E Gardarsdottir; Gudmundur Georgsson; Olafia S Gretarsdottir; Kjartan R Gudmundsson; Thora R Gunnarsdottir; Arnaldur Gylfason; Hilma Holm; Brynjar O Jensson; Aslaug Jonasdottir; Kamilla S Josefsdottir; Thordur Kristjansson; Droplaug N Magnusdottir; Solvi Rognvaldsson; Louise le Roux; Gudrun Sigmundsdottir; Gardar Sveinbjornsson; Kristin E Sveinsdottir; Maney Sveinsdottir; Emil A Thorarensen; Bjarni Thorbjornsson; Gisli Masson; Ingileif Jonsdottir; Alma Moller; Thorolfur Gudnason; Karl G Kristinsson; Unnur Thorsteinsdottir; Kari Stefansson |
| EPI_ISL_830417, EPI_ISL_830501, EPI_ISL_830502 | The National University Hospital of Iceland | deCODE genetics | Daniel F Gudbjartsson; Agnar Helgason; Hakon Jonsson; Olafur T Magnusson; Pall Melsted; Gudmundur L Norddahl; Jona Saemundsdottir; Asgeir Sigurdsson; Patrick Sulem; Arna B Agustsdottir; Hannes Eggertsson; Berglind Eiriksdottir; Run Fridriksdottir; Elisabet E Gardarsdottir; Gudmundur Georgsson; Olafia S Gretarsdottir; Kjartan R Gudmundsson; Thora R Gunnarsdottir; Arnaldur Gylfason; Hilma Holm; Brynjar O Jensson; Aslaug Jonasdottir; Kamilla S Josefsdottir; Thordur Kristjansson; Droplaug N Magnusdottir; Solvi Rognvaldsson; Louise le Roux; Gudrun Sigmundsdottir; Gardar Sveinbjornsson; Kristin E Sveinsdottir; Maney Sveinsdottir; Emil A Thorarensen; Bjarni Thorbjornsson; Gisli Masson; Ingileif Jonsdottir; Alma Moller; Thorolfur Gudnason; Karl G Kristinsson; Unnur Thorsteinsdottir; Kari Stefansson |
| EPI_ISL_830728, EPI_ISL_830740, EPI_ISL_830743, EPI_ISL_830744, EPI_ISL_830772, EPI_ISL_830775, EPI_ISL_830779, EPI_ISL_830780, EPI_ISL_830781, EPI_ISL_830782, EPI_ISL_830783, EPI_ISL_830784, EPI_ISL_830785, EPI_ISL_830786, EPI_ISL_830787, EPI_ISL_830788, EPI_ISL_830789, EPI_ISL_830790, EPI_ISL_830791, EPI_ISL_830792, EPI_ISL_830793, EPI_ISL_830794, EPI_ISL_830795, EPI_ISL_830796, EPI_ISL_830797, EPI_ISL_830798, EPI_ISL_830985, EPI_ISL_830986, EPI_ISL_831012 | | | |
| see above | University Hospital Basel, Clinical Virology | University Hospital Basel, Clinical Bacteriology | Tim Roloff, Madlen Stange, Helena MB Seth-Smith, Alfredo Mari, Karoline Leuzinger, Julia Bielicki, Manuel Battegay, Hans Hirsch, Adrian Egli |
| EPI_ISL_831091, EPI_ISL_831092, EPI_ISL_831107, EPI_ISL_831108, EPI_ISL_831109, EPI_ISL_831115, EPI_ISL_831116, EPI_ISL_831117, EPI_ISL_831118, EPI_ISL_831119, EPI_ISL_831120, EPI_ISL_831121, EPI_ISL_831122, EPI_ISL_831123, EPI_ISL_831124, EPI_ISL_831125, EPI_ISL_831126, EPI_ISL_831127, EPI_ISL_831128, EPI_ISL_831129, EPI_ISL_831130, EPI_ISL_831131, EPI_ISL_831132, EPI_ISL_831133, EPI_ISL_831134, EPI_ISL_831135 | | | |
| see above | Hospital Universitario La Paz (Madrid) | SeqCOVID-SPAIN consortium/IBV(CSIC) | María Rodríguez-Tejedor, Elias Dahdouh, Fernando Lázaro-Perona, Jesús Mingorance and SeqCOVID-SPAIN consortium |
| EPI_ISL_831939 | Laboratório de Microbiologia Molecular - Universidade FEEVALE | Universidade Federal de Ciências da Saúde de Porto Alegre | Vinícius Bonetti Franceschi, Amanda de Menezes Mayer, Gabriel Dickin Caldana, Carla Andretta Moreira Neves, Patrícia Aline Gröhs Ferrareze, Gabriela Bettela Cybis, Ricardo Ariel Zimerman, Lívia Kmetzsch, Fernando Rosado Spilki, Claudia Elizabeth Thompson |
| EPI_ISL_831963, EPI_ISL_831964, EPI_ISL_831965, EPI_ISL_831967, | Orebro klinisk mikrobiologi | The Public Health Agency of Sweden | Department of Microbiology, The Public Health Agency of Sweden |

| | | | |
|---|---|---|---|
| EPI_ISL_831972 | | | |
| EPI_ISL_833337 | Christ the Saviour General Hospital - Batangas | Research Institute for Tropical Medicine | Hannah Leah Morito, Othoniel Jan Onza, John Leonard Chan, Ma Angelica Tujan, Francisco Gerardo Polotan, Inez Andrea Medado, Kirstyn Brunker, Edelwisa Mercado, Daria Manalo, Catalino Demetria |
| EPI_ISL_833509 | RS Melania, Bogor, Indonesia | Biosafety Level-3 Laboratory, Indonesian Institute of Sciences (LIPI) | Syam Budi Iryanto, Andri Wardiana, Anik Budhi Dharmayanthi, Anggia Prasetyoputri, Isa Nuryana, Ade Andriani, Ahmad Fathoni, Ratih Asmana Ningrum |
| EPI_ISL_833510 | RS Melania, Bogor, Indonesia | Biosafety Level-3 Laboratory, Indonesian Institute of Sciences (LIPI) | Anik Budhi Dharmayanthi, Syam Budi Iryanto, Andri Wardiana, Anggia Prasetyoputri, Isa Nuryana, Ade Andriani, Ahmad Fathoni, Ratih Asmana Ningrum |
| EPI_ISL_837272, EPI_ISL_837273, EPI_ISL_837274, EPI_ISL_837275, EPI_ISL_837276, EPI_ISL_837277, EPI_ISL_837278, EPI_ISL_837279, EPI_ISL_837280, EPI_ISL_837281, EPI_ISL_837282, EPI_ISL_837283, EPI_ISL_837284 | | | |
| see above | Istituto Zooprofilattico Sperimentale del Mezzogiorno | TIGEM | Patrizia Annunziata, Andrea Ballabio, Valentina Bouche, Davide Cacchiarelli (CorrespAuthor), Pellegrino Cerino, Chiara Colantuono, Lucio Di Filippo, Antonio Grimaldi, Antonio Limone, Gabriella Loconte, Anna Manfredi, Francesco Panariello, Biancamaria Pierri, Marcello Salvi, Lucia Vassallo |
| EPI_ISL_837810, EPI_ISL_837811, EPI_ISL_837812, EPI_ISL_837813, EPI_ISL_837814 | Instituto Nacional de Enfermedades Respiratorias (INER) | Instituto Nacional de Enfermedades Respiratorias (INER) | Celia Boukadida, Margarita Matías-Florentino, Alma Rincón-Rubio, Hector Esteban Paz-Juárez, Olivia Briceño, Edgar Sevilla-Reyes, Fidencio Mejía-Nepomuceno, Mario Mújica-Sánchez, Eduardo Becerril-Vargas, José Arturo Martínez-Orozco, Alejandra Hernández-Terán, Jorge Salas-Hernández, Santiago Ávila-Ríos, Joel Armando Vázquez-Pérez |
| EPI_ISL_845659, EPI_ISL_845671 | Quest Diagnostics | Quest Diagnostics | Rosenthal,S.H., Gerasimova,A., Kagan,R.M., Anderson, B., Bernstein, L.E., Livingston, K.E., Hua, M., Liu Y., Shalhout, D.F., Shlyakhter, I.A., Owen, R., Lacbawan, F. |
| EPI_ISL_845754 | Emory Molecular Diagnostics Laboratory, Emory Healthcare | Piantadosi Lab, Emory Department of Pathology | Ahmed Babiker, Anne Piantadosi |
| EPI_ISL_850494, EPI_ISL_850496, EPI_ISL_850497, EPI_ISL_850498, EPI_ISL_850499, EPI_ISL_850500, EPI_ISL_850501, EPI_ISL_850502, EPI_ISL_850503, EPI_ISL_850504, EPI_ISL_850505 | | | |
| see above | Division of Emerging Infectious Diseases, Bureau of Infectious Diseases Diagnosis Control, Korea Disease Control and Prevention Agency | Division of Emerging Infectious Diseases, Bureau of Infectious Diseases Diagnosis Control, Korea Disease Control and Prevention Agency | Ae Kyung Park, Il-Hwan Kim, Heui Man Kim, Jeong-Min Kim, Namjoo Lee, Chaeyoung Lee, Sang Hee Woo, Eun-Jin Kim |
| EPI_ISL_852587, EPI_ISL_852589, EPI_ISL_852590, EPI_ISL_852593 | Max von Pettenkofer Institute, Virology, National Reference Center for Retroviruses, LMU München | Laboratory for Functional Genome Analysis, Dept. Genomics, Gene Center of the LMU Munich | Max Muenchhoff, Stefan Krebs, Alexander Graf, Oliver Keppler, Helmut Blum |
| EPI_ISL_853299, EPI_ISL_853300 | UPMC Clinical Microbiology Laboratory | Microbial Genome Sequencing Center; Microbial Genomic Epidemiology Laboratory | Mustapha M. Mustapha, Jane W. Marsh, Dan Snyder, Marissa P. Griffith, Stephanie L. Mitchell, Vatsala R. Srinivasa, Kady D. Waggle, Chinelo Ezeonwuku, Vaughn S. Cooper, Lee H. Harrison |
| EPI_ISL_854299 | Center for Virology, Medical University of Vienna | Bergthaler laboratory, CeMM Research Center for Molecular Medicine of the Austrian Academy of Sciences | Lukas Endler, Alexandra Popa, Benedikt Agerer, Jakob-Wendelin Genger, Alexander Lercher, Anna Schedl, Thomas Penz, Michael Schuster, Jan Laine, Martin Senekowitsch, Christoph Bock, Andreas Bergthaler |
| EPI_ISL_859684, EPI_ISL_859692, EPI_ISL_859693, EPI_ISL_859694, EPI_ISL_859763, EPI_ISL_859764, EPI_ISL_859765, EPI_ISL_859766, EPI_ISL_859767 | BTC, Khalifa University | BTC, Khalifa University | Al Safar et al |
| EPI_ISL_860138, EPI_ISL_860139 | Keio University School of Medicine | Keio University School of Medicine | Kenjiro Kosaki, Yuka Iwasaki, Hirotsugu Ishizu, Haruhiko Siomi, Kodai Abe |
| EPI_ISL_861242 | Instituto de Biotecnologia - UNESP-Botucatu-SP | Instituto de Biotecnologia - UNESP-Botucatu-SP | Leila Sabrina Ullmann; Fábio Sossai Possebon, Camila Dantas Malossi, Paula Rahal, Paulo Inacio da Costa, João Pessoa Araújo Jr. |
| EPI_ISL_861885 | LATE - Laboratório de Técnicas Especiais - Hospital Israelita Albert Einstein | LATE - Laboratório de Técnicas Especiais - Hospital Israelita Albert Einstein | Deyvid Amgarten, Fernanda de Mello Malta, Raquel Riyuzo, Ana Paula Moreira Salles, Pedro Henrique Sebe Rodrigues, João Renato Rebello Pinho |
| EPI_ISL_872003, EPI_ISL_872004, EPI_ISL_872005, EPI_ISL_872006, EPI_ISL_872007, EPI_ISL_872008 | Ospedale "Di Venere" | Beaconlab (Bioinformatics, Evolution and Comparative Genomics lab), Dept of Biosciences, University on Milan | Iacobellis M, d'Avenia M, Piluscio R, Parisi A, Chiara M, Manzari C, Pesole G |
| EPI_ISL_872604 | Nigeria Centre for Disease Control (NCDC) | African Centre of Excellence for Genomics of Infectious Diseases (ACEGID), Redeemer's University | Oluniyi P.E. et al |
| EPI_ISL_876900, EPI_ISL_876901, EPI_ISL_876902, EPI_ISL_876903, EPI_ISL_876904, EPI_ISL_876905, EPI_ISL_876906, EPI_ISL_876907, EPI_ISL_876908, EPI_ISL_876909, EPI_ISL_876910, EPI_ISL_876911, EPI_ISL_876912 | | | |
| see above | Quest Diagnostics | Quest Diagnostics | Rosenthal,S.H., Gerasimova,A., Kagan,R.M., Anderson, B., Hua, M., Liu Y., Bernstein, L.E., Livingston, K.E., Perez, A., Shalhout, D.F., Shlyakhter, I.A., Owen, R., Tanpaiboon, P., Lacbawan, F. |
| EPI_ISL_877126 | Academic Hospital of Gajdah Mada University (RSA UGM) | Genetics Working Group (Pokja Genetik) Faculty of Medicine, Public Health and Nursing Universitas Gadjah Mada (FK-KMK UGM); Disease Investigation Center Wates Ministry of Agriculture Indonesia; Department of Microbiology FK-KMK UGM; Laboratorium Diagnostik Yayasan Tahija World Mosquito Program (WMP) Yogyakarta Center for Tropical Medicine FK-KMK UGM; Integrated Research center FK-KMK UGM; Department of Computer Science and Electronics FMIPA UGM | Gunadi, Hendra Wibawa, Marcellus, Mohamad S. Hakim, Edwin W. Daniwijaya, Ludhang P. Rizki, Endah Supriyati, Eggi Arguni, Titik Nuryastuti, Tri Wibawa, Dwi AA Nugrahaningsih, Afiahayati, Siswanto,Untung Riawan, William Widitjiarso |
| EPI_ISL_877127 | Academic Hospital of Gajdah Mada University (RSA UGM) | Genetics Working Group (Pokja Genetik) Faculty of Medicine, Public Health and Nursing Universitas Gadjah Mada (FK-KMK UGM); Disease Investigation Center Wates Ministry of Agriculture Indonesia; Department of Microbiology FK-KMK UGM; Laboratorium Diagnostik Yayasan Tahija World Mosquito Program (WMP) Yogyakarta Center for Tropical Medicine FK-KMK UGM; Integrated Research center FK-KMK UGM; Department of Computer Science and Electronics FMIPA UGM | Gunadi, Hendra Wibawa, Marcellus, Mohamad S. Hakim, Edwin W. Daniwijaya, Ludhang P. Rizki, Endah Supriyati, Eggi Arguni, Titik Nuryastuti, Tri Wibawa, Dwi AA Nugrahaningsih, Afiahayati, Siswanto,Audric Kenny Tedja, Maria Patricia Inggriani |
| EPI_ISL_877128 | Academic Hospital of Gajdah Mada University (RSA UGM) | Genetics Working Group (Pokja Genetik) Faculty of Medicine, Public Health and Nursing Universitas Gadjah Mada (FK-KMK UGM); Disease Investigation Center Wates Ministry of Agriculture Indonesia; Department of Microbiology FK-KMK UGM; Laboratorium Diagnostik Yayasan Tahija World Mosquito Program (WMP) Yogyakarta Center for Tropical Medicine FK-KMK UGM; Integrated Research center FK-KMK UGM; Department of Computer Science and Electronics FMIPA UGM | Gunadi, Hendra Wibawa, Marcellus, Mohamad S. Hakim, Edwin W. Daniwijaya, Ludhang P. Rizki, Endah Supriyati, Eggi Arguni, Titik Nuryastuti, Tri Wibawa, Dwi AA Nugrahaningsih, Afiahayati, Siswanto, Kristy Iskandar, Nungki Anggorowati, Aditya Rifqi Fauzi, Fadil Fahri |
| EPI_ISL_877129 | Academic Hospital of Gajdah Mada University (RSA UGM) | Genetics Working Group (Pokja Genetik) Faculty of Medicine, Public Health and Nursing Universitas Gadjah Mada (FK-KMK UGM); Disease Investigation Center Wates Ministry of Agriculture Indonesia; Department of Microbiology FK-KMK UGM; Laboratorium Diagnostik Yayasan Tahija World | Gunadi, Hendra Wibawa, Marcellus, Mohamad S. Hakim, Edwin W. Daniwijaya, Ludhang P. Rizki, Endah Supriyati, Eggi Arguni, Titik Nuryastuti, Tri Wibawa, Dwi AA Nugrahaningsih, Afiahayati, Siswanto, Kristy Iskandar, Nungki Anggorowati, Dwiki afandy, Dyah Ayu Puspitarani |

| | | | |
|---|---|---|---|
| | | Mosquito Program (WMP) Yogyakarta Center for Tropical Medicine FK-KMK UGM; Integrated Research Center FK-KMK UGM; Department of Computer Science and Electronics FMIPA UGM | |
| EPI_ISL_877130 | Academic Hospital of Gajdah Mada University (RSA UGM) | Genetics Working Group (Pokja Genetik) Faculty of Medicine, Public Health and Nursing Universitas Gadjah Mada (FK-KMK UGM); Disease Investigation Center Wates Ministry of Agriculture Indonesia; Department of Microbiology FK-KMK UGM; Laboratorium Diagnostik Yayasan Tahija World Mosquito Program (WMP) Yogyakarta Center for Tropical Medicine FK-KMK UGM; Integrated Research center FK-KMK UGM; Department of Computer Science and Electronics FMIPA UGM | Gunadi, Hendra Wibawa, Marcellus, Mohamad S. Hakim, Edwin W. Daniwijaya, Ludhang P. Rizki, Endah Supriyati, Eggi Arguni, Titik Nuryastuti, Tri Wibawa, Dwi AA Nugrahaningsih, Afiahayati, Siswanto, Kristy Iskandar, Nungki Anggorowati, Kemala Athollah, Desyifa Mursalin |
| EPI_ISL_877131 | Academic Hospital of Gajdah Mada University (RSA UGM) | Genetics Working Group (Pokja Genetik) Faculty of Medicine, Public Health and Nursing Universitas Gadjah Mada (FK-KMK UGM); Disease Investigation Center Wates Ministry of Agriculture Indonesia; Department of Microbiology FK-KMK UGM; Laboratorium Diagnostik Yayasan Tahija World Mosquito Program (WMP) Yogyakarta Center for Tropical Medicine FK-KMK UGM; Integrated Research center FK-KMK UGM; Department of Computer Science and Electronics FMIPA UGM | Gunadi, Hendra Wibawa, Marcellus, Mohamad S. Hakim, Edwin W. Daniwijaya, Ludhang P. Rizki, Endah Supriyati, Eggi Arguni, Titik Nuryastuti, Tri Wibawa, Dwi AA Nugrahaningsih, Afiahayati, Siswanto, Kristy Iskandar, Nungki Anggorowati, Susan Simanjaya, Alvin Santoso Kalim |
| EPI_ISL_882614 | COVID lab, Mymensingh Medical College | Department of Pathology, Bangladesh Agricultural University & Department of Microbiology, Mymensingh Medical College | Afrin, S. Z. Paul, S. K. Parvin, R. |
| EPI_ISL_884340, EPI_ISL_884433, EPI_ISL_884434 | Infectious Diseases, Quest Diagnostics | Infectious Diseases, Quest Diagnostics | Rosenthal,S.H., Gerasimova,A., Kagan,R.M., Anderson,B., Bernstein,L.E., Livingston,K.E., Hua,M., Liu,Y., Shalhout,D.F., Owen,R., Lacbawan,F. |
| EPI_ISL_887152 | Massachusetts General Hospital | Infectious Disease Program, Broad Institute of Harvard and MIT | Lemieux,J.E., Siddle,K.J., Shaw,B., Adams,G., Pierce,V., Turbett,S., Anahtar,M., Branda,J., Slater,D., Harris,J., Lin,A.E., Gladden-Young,A., Lagerborg,K., Rudy,M., DeRuff,K., Carter,A., Normandin,E., Bauer,M., Reilly,S., Tomkins-Tinch,C., Loreth,C., Chaluvadi,S., Neumann,A., Cusick,C., Chapman,S.B., Gnirke,A., Flowers,K., Cerrato,F., Birren,B.W., Gallagher,G., Smole,S., Park,D.J., MacInnis,B.L., Ryan,E., LaRocque,R., Rosenberg,E. and Sabeti,P.C. |
| EPI_ISL_887420, EPI_ISL_887426, EPI_ISL_887502 | Instituto Nacional de Saude (INS), Mozambique | KRISP, KZN Research Innovation and Sequencing Platform | Nalia Ismael, Nadia Sitoe, Paulo Arnaldo, Nedio Mabunda, Giandhari J, Pillay S, Tegally H, Wilkinson E, de Oliveira T |
| EPI_ISL_890186 | Academic Hospital of Gadjah Mada University (RSA UGM) | Genetics Working Group (Pokja Genetik) Faculty of Medicine, Public Health and Nursing Universitas Gadjah Mada (FK-KMK UGM); Disease Investigation Center Wates Ministry of Agriculture Indonesia; Department of Microbiology FK-KMK UGM; Laboratorium Diagnostik Yayasan Tahija World Mosquito Program (WMP) Yogyakarta Center for Tropical Medicine FK-KMK UGM; Integrated Research center FK-KMK UGM; Department of Computer Science and Electronics FMIPA UGM | Gunadi, Hendra Wibawa, Marcellus, Mohamad S. Hakim, Edwin W. Daniwijaya, Ludhang P. Rizki, Endah Supriyati, Eggi Arguni, Titik Nuryastuti, Tri Wibawa, Dwi AA Nugrahaningsih, Afiahayati, Siswanto, Kristy Iskandar, Nungki Anggorowati, Audric Kenny Tedja, Maria Patricia Inggriani |
| EPI_ISL_890187 | Academic Hospital of Gadjah Mada University (RSA UGM) | Genetics Working Group (Pokja Genetik) Faculty of Medicine, Public Health and Nursing Universitas Gadjah Mada (FK-KMK UGM); Disease Investigation Center Wates Ministry of Agriculture Indonesia; Department of Microbiology FK-KMK UGM; Laboratorium Diagnostik Yayasan Tahija World Mosquito Program (WMP) Yogyakarta Center for Tropical Medicine FK-KMK UGM; Integrated Research center FK-KMK UGM; Department of Computer Science and Electronics FMIPA UGM | Gunadi, Hendra Wibawa, Marcellus, Mohamad S. Hakim, Edwin W. Daniwijaya, Ludhang P. Rizki, Endah Supriyati, Eggi Arguni, Titik Nuryastuti, Tri Wibawa, Dwi AA Nugrahaningsih, Afiahayati, Siswanto, Kristy Iskandar, Nungki Anggorowati, Untung Riawan, William Widitjiarso |
| EPI_ISL_890193 | Gonoshasthaya-RNA Research Center, Gonoshasthaya-RNA Molecular Diagnostics and Research Center | Gonoshasthaya-RNA Research Center, Gonoshasthaya-RNA Molecular Diagnostics and Research Center | Jamiruddin,M.R., Khondoker,M.U., Sharif,N., Azmuda,N., Ahmed,M.F., Sharmin,S., Akter,S., Mou,T.J., Marzan,M., Liza,S.M., Nahar,S., Jahan,N., Ali,T., Khandker,S.S., Jamiruddin,M., Haq,M.A., Adnan,N., Chaity,M., Oishee,M. |
| EPI_ISL_891259 | COVID lab, Department of Microbiology | Department of Pathology, Bangladesh Agricultural University and Department of Microbiology, Mymensingh Medical College | Afrin, S.Z. Paul, S. K. Parvin, R. |
| EPI_ISL_892230 | Lighthouse Lab in Alderley Park | Wellcome Sanger Institute for the COVID-19 Genomics UK (COG-UK) Consortium | Jacquelyn Wynn, Mairead Hyland, The Lighthouse Lab in Alderley Park and Alex Alderton, Roberto Amato, Sonia Goncalves, Ewan Harrison, David K. Jackson, Ian Johnston, Dominic Kwiatkowski, Cordelia Langford, John Sillitoe on behalf of the Wellcome Sanger Institute COVID-19 Surveillance Team |
| EPI_ISL_900082, EPI_ISL_900123, EPI_ISL_900170, EPI_ISL_900219, EPI_ISL_900303, EPI_ISL_900312, EPI_ISL_900448, EPI_ISL_900462 | MEPHI, Aix Marseille University | MEPHI, Aix Marseille University | Anthony LEVASSEUR |
| EPI_ISL_902918 | Tanjungpura University Hospital | Tanjungpura University Hospital | Andriani; Mahyarudin; Virhan Novianry ; Delima Fajar Liana; Sofi Siti Shofiyah; Puji Astuti P; Muhammad Ibnu Kahtan; Ambar Rialita; Eka Ardiani Putri EA; Wiwik Windarti ; Helmi Sastriawan; Willy Handoko |
| EPI_ISL_903328, EPI_ISL_903329, EPI_ISL_903330, EPI_ISL_903331, EPI_ISL_903332, EPI_ISL_903333, EPI_ISL_903334, EPI_ISL_903335 | University of Iowa Hospitals & Clinics, Microbiology Laboratory | University or Iowa, Lung Biology and Cystic Fibrosis Research Center, Pezzulo Lab | Miguel E Ortiz, Alejandro A Pezzulo |
| EPI_ISL_906056 | Tilia Laboratories s.r.o. | Tilia Laboratories s.r.o. | Sona Pekova, MD, PhD. |
| EPI_ISL_906103 | Child Health Research Foundation | Child Health Research Foundation | Senjuti Saha, Arif Mohammad Tanmoy, Sharmistha Goswami, Afroza Akter Tanni, Syed Muktadir Al Sium, Roly Malaker, Md Hafizur Rahman, Samir K Saha |
| EPI_ISL_910112, EPI_ISL_910123, EPI_ISL_910124, EPI_ISL_910125, EPI_ISL_910126, EPI_ISL_910127, EPI_ISL_910128, EPI_ISL_910129, EPI_ISL_910130, EPI_ISL_910131, EPI_ISL_910132, EPI_ISL_910133, EPI_ISL_910134, EPI_ISL_910135, EPI_ISL_910136, EPI_ISL_910137, EPI_ISL_910138, EPI_ISL_910139, EPI_ISL_910140, EPI_ISL_910141, EPI_ISL_910142, EPI_ISL_910143, EPI_ISL_910144, EPI_ISL_910145, EPI_ISL_910146, EPI_ISL_910147, EPI_ISL_910148, EPI_ISL_910149, EPI_ISL_910150, EPI_ISL_910151, EPI_ISL_910152, EPI_ISL_910153, EPI_ISL_910154, EPI_ISL_910155, EPI_ISL_910156, EPI_ISL_910157, EPI_ISL_910158, EPI_ISL_910159, EPI_ISL_910160 | | | |
| see above | CSIR-Centre for Cellular and Molecular Biology | CSIR-Centre for Cellular and Molecular Biology | Payel Mukherjee,Pratheusa Maccha,Namami Gaur,Lamuk Zaveri,Tulasi Nagabandi,Purushotham Vodnala,Blessy B John,Viswagithe S L,B Himasri,Sofia Banu,Priya Singh,Archana Bharadwaj Siva,Karthik Bharadwaj Tallapaka,Rakesh K Mishra,Divya Tej Sowpati |
| EPI_ISL_913082, EPI_ISL_913083, EPI_ISL_913084, EPI_ISL_913093 | Center for Virology | Center for Virology | Jeremy V. Camp, Irene Goerzer, Monika Redlberger-Fritz, Stephan W. Aberle |
| EPI_ISL_920975 | Northumbria University / South Tees Hospitals NHS | COVID-19 Genomics UK (COG-UK) Consortium | Darren L Smith,Andrew Nelson,Matthew Bashton,Greg R Young,Joshua Loh,John Allan,Mohammad A Tariq,Giles S Holt,Gary Black,Wen C Yew,Lynn |

| | | | |
|---|---|---|---|
| | Foundation Trust / North Cumbria Integrated Care NHS Foundation Trust / North Tees and Hartlepool NHS Foundation Trust / Newcastle Hospitals NHS Foundation Trust | | Dover,Paul Baker,Steve Liggett,Sarah Essex,Jane Greenaway,Debra Padgett,Clive Graham,Garren Scott,Edward Barton,Emma Swindells,Brendan Payne,Jennifer Collins,Yusri Taha,Gary Eltringham |
| EPI_ISL_925098, EPI_ISL_925109, EPI_ISL_925110, EPI_ISL_925120, EPI_ISL_925131, EPI_ISL_925132 | 1.AO Universitaria 'S. Giovanni di Dio e Ruggi D'Aragona, Scuola Medica Salernitana' Hospital / 2.UOC di Virologia e Microbiologia, Università della Campania 'L. Vanvitelli' / 3.AO Universitaria 'Federico II' Napoli Hospital / 4.AORN 'San Giuseppe Moscati' Avellino Hospital / 5.AO 'San Pio - presidio G. Rummo' Benevento Hospital / 6.AO 'Sant'Anna e San Sebastiano' Caserta Hospital / 7.PO 'Maria Santissima Addolorata' Eboli Hospital / 8.Biogem Istituto di Ricerche Genetiche | 1. Genome Research Center for Health (CRGS) / 2. Laboratory of Molecular Medicine and Genomics(LMMGe) / 3. Center for Research in Pure and Applied Mathematics (CRMPA) | Giorgio Giurato, Francesca Rizzo, Alessandro Weisz, Gianluigi Franci, Giovanni Nassa, Pasquale Pagliano, Roberta Tarallo, Elena Alexandrova, Ylenia D'Agostino, Carlo Ferravante, Jessica Lamberti, Viola Melone, Domenico Memoli, Valeria Mirici Cappa, Domenico Palumbo, Giovanni Pecoraro, Assunta Sellitto, Oriana Strianese, Ilaria Terenzi, Giuseppe Fenza, Aniello Gentile, Antonello Saccomanno, Sonia Amabile, Teresa Rocco, Annamaria Salvati, Emilia Vaccaro, Massimiliano Galdiero, Michele Cennamo, Giuseppe Portella, Maria Grazia Foti, Mariarosaria Ingino, Maria Landi, Maurizio Fumi, Vincenzo Rocco, Rita Greco, Vittoria Letizia, Arnolfo Petruzziello, Maddalena Schioppa, Gregorio Goffredi, Francesca Marciano, Michele Caraglia, Alessia Cossu, Marianna Scrima |
| EPI_ISL_925369, EPI_ISL_925370, EPI_ISL_925381 | Department of Clinical Microbiology | GIGA Medical Genomics | Keith Durkin, Maria Artesi, Sébastien Bontems, Raphaël Boreux, Bouchra Boujemla, Cécile Meex, Pierrette Melin, Marie-Pierre Hayette, Vincent Bours |
| EPI_ISL_935526 | University of Massachusetts Medical School | Infectious Disease Program, Broad Institute of Harvard and MIT | Tomkins-Tinch, Christopher H.; Daly, Jennifer S.; Gladden-Young, Adrianne; Theodoropoulos, Nicole M.; Madaio, Michael; Yu, Neng; Vanguri, Vijay K; Siddle, Katherine J.; Adams, Gordon; Krasilnikova, Lydia A.; Movahedi, Babak; Bozorgzadeh, Adel; Simin, Karl; Lemieux, Jacob E.; Luban, Jeremy; Park, Daniel J.; MacInnis, Bronwyn L.; Sabeti, Pardis C.; Levitz, Stuart M. |
| EPI_ISL_935822, EPI_ISL_935824, EPI_ISL_935825, EPI_ISL_935826, EPI_ISL_935827, EPI_ISL_935840, EPI_ISL_935841, EPI_ISL_935852 | Cadham Provincial laboratory | National Microbiology Laboratory (NML) | Anna Majer, Shari Tyson, Grace Seo, Philip Mabon, Elsie Grudeski, Rhiannon Huzarewich, Russell Mandes, Anneliese Landgraff, Jennifer Tanner, Natalie Knox, Morag Graham, Gary Van Domselaar, Paul Van Caeseele, Jared Bullard, David Alexander, Kerry Dust, Nathalie Bastien, Yan Li, Timothy Booth, Darian Hole, Madison Chapel, Kirsten Biggar, CanCOGeN's metadata curation team, Public Health Agency of Canada CanCOGeN team |
| EPI_ISL_936829, EPI_ISL_936830, EPI_ISL_936831, EPI_ISL_936832, EPI_ISL_936833, EPI_ISL_936834, EPI_ISL_936835, EPI_ISL_936836 | Northwestern Memorial Hospital | Ozer Lab | Ramon Lorenzo-Redondo, Lacy M. Simons, Chad J. Achenbach, Lawrence J. Jennings, Michael G. Ison, Judd F. Hultquist, Egon A. Ozer |
| EPI_ISL_937028, EPI_ISL_937029, EPI_ISL_937032, EPI_ISL_937045, EPI_ISL_937053 | Quest Diagnostics | Quest Diagnostics | Rosenthal,S.H., Gerasimova,A., Kagan,R.M., Anderson, B., Livingston, K.E., Hua, M., Liu Y., Shalhout, D.F., Owen, R., Lacbawan, F. |
| EPI_ISL_940198, EPI_ISL_940199, EPI_ISL_940200, EPI_ISL_940201, EPI_ISL_940202, EPI_ISL_940203, EPI_ISL_940204, EPI_ISL_940503, EPI_ISL_940504, EPI_ISL_940506 | Hôpital Bichat Claude Bernard, Laboratoire de Virologie | IAME UMR1137 Inserm, Université de Paris, Hôpital Bichat | Antoine Bridier-Nahmias, Amélie Recoing, Quentin Le Hingrat, Lena Daniel, Siham Hamri, Gilles Collin, Alexandre Storto, Mélanie Bertine, Charlotte Charpentier, Nadhira Houhou-Fidouh, Diane Descamps, Benoit Visseaux |
| EPI_ISL_940851 | Vaccines and Infectious Diseases Analytics Research Unit (VIDA) | KRISP, KZN Research Innovation and Sequencing Platform | Baillie Vicky, du Plessis Jeanine, Giandhari Jennifer, Pillay Sureshnee, Naidoo Yeshnee, Tegally Houriiyah, de Oliveira Tulio, Madhi Shabir |
| EPI_ISL_940954, EPI_ISL_940963, EPI_ISL_940964, EPI_ISL_940965, EPI_ISL_940966, EPI_ISL_940967, EPI_ISL_940968, EPI_ISL_940969, EPI_ISL_940970, EPI_ISL_940971, EPI_ISL_940972, EPI_ISL_942011, EPI_ISL_942012 | | | |
| see above | Centers for Disease Control and Prevention, Dengue Branch | Centers for Disease Control and Prevention, Dengue Branch | Gilberto A. Santiago, Glenda Gonzalez, Betzabel Flores, Keyla Charriez, Gabriela Paz-Bailey, Jorge L. Munoz-Jordan |
| EPI_ISL_949195, EPI_ISL_949196, EPI_ISL_949209, EPI_ISL_949222, EPI_ISL_949237 | Departamento de Microbiología, CDB, Hospital Clínic, Barcelona | SeqCOVID-SPAIN consortium/IBV(CSIC) | Andrea Vergara, Mikel Martínez, Elisa Rubio, Jéssica Navero, Aida Peiró and SeqCOVID-SPAIN consortium |
| EPI_ISL_954242, EPI_ISL_954243, EPI_ISL_954244, EPI_ISL_954245, EPI_ISL_954246, EPI_ISL_954247, EPI_ISL_954248, EPI_ISL_954249, EPI_ISL_954250, EPI_ISL_954251, EPI_ISL_954252, EPI_ISL_954253, EPI_ISL_954254 | | | |
| see above | MRC/UVRI & LSHTM Uganda Research Unit | Where sequence data have been generated and submitted to GISAID | Matthew Cotten, Dan Lule Bugembe, My V.T. Phan, Isaac Sseeewanyana, Patrick Semanda, Susan Nabadda, Pontiano Kaleebu |
| EPI_ISL_955146 | University of Sarajevo, Veterinary Faculty, Laboratory for Molecular Diagnostic and Research Laboratory | University of Sarajevo, Veterinary Faculty, Laboratory for Molecular Diagnostic and Research Laboratory | Goleti T., Goleti Š., Softi A., Ali-Šeho A., Nicevi M., Jaži A., Šabi E., Terzi I., Hodži A. |
| EPI_ISL_959503 | Division of Emerging Infectious Diseases, Bureau of Infectious Diseases Diagnosis Control, Korea Disease Control and Prevention Agency | Division of Emerging Infectious Diseases, Bureau of Infectious Diseases Diagnosis Control, Korea Disease Control and Prevention Agency | Ae Kyung Park, Il-Hwan Kim, Heui Man Kim, Jeong-Min Kim, Namjoo Lee, Chae Young Lee, Sang Hee Woo, Eun-Jin Kim |
| EPI_ISL_959911, EPI_ISL_959917, EPI_ISL_959918, EPI_ISL_960022, EPI_ISL_960023, EPI_ISL_960024, EPI_ISL_960025, EPI_ISL_960026, EPI_ISL_960027, EPI_ISL_960028, EPI_ISL_960029, EPI_ISL_960030, EPI_ISL_960031, EPI_ISL_960032, EPI_ISL_960033, EPI_ISL_960034, EPI_ISL_960035, EPI_ISL_960036, EPI_ISL_960037, EPI_ISL_960038, EPI_ISL_960039, EPI_ISL_960040, EPI_ISL_960041 | | | |
| see above | University Medical Center Hamburg Eppendorf | Heinrich Pette Institute, Leibniz Institute for Experimental Virology | Alexis Robitaille, Thomas Günther, Johannes Knobloch, Martin Aepfelbacher, Nicole Fischer, Adam Grundhoff |
| EPI_ISL_960150 | Victoria Hospital wc VHW | National Health Laboratory Service/UCT | Arash Iranzadeh, Deelan Doolabh, Lynn Tyers, Bruna Galvao, Innocent Mudau, Marvin Hsiao, Kruger Marais, Diana Hardie, Stephen Korsman, Carolyn Williamson |
| EPI_ISL_968173 | Clinical Molecular Microbiology Laboratory, UNC Hospital | Dirk Dittmer | Justin T. Landis , Razia Moorad , Brent A. Eason, Melissa B. Miller, Linda Pluta, Dirk Dittmer, Angelica Juarez, Cecilia Thompson, Shawn Hawken, Cameroon Grant, Evelyn Hoffman, Patricio Cano, Jason Wong, Carolina Caro-Vegas, Ryan McNamara, Blossom Damania. |
| EPI_ISL_969758, EPI_ISL_969761, EPI_ISL_969763, EPI_ISL_969765, EPI_ISL_969767, EPI_ISL_969770, EPI_ISL_969772, EPI_ISL_969773, EPI_ISL_969775, EPI_ISL_969777, EPI_ISL_969778, EPI_ISL_969781, EPI_ISL_969783, EPI_ISL_969784, EPI_ISL_969787, EPI_ISL_969788, EPI_ISL_969790, EPI_ISL_969792, EPI_ISL_969794, EPI_ISL_969796, EPI_ISL_969798, EPI_ISL_969800, EPI_ISL_969803, EPI_ISL_969805, EPI_ISL_969807, EPI_ISL_969810, EPI_ISL_969811, EPI_ISL_969814, EPI_ISL_969816, EPI_ISL_969818, EPI_ISL_969820, EPI_ISL_969822, EPI_ISL_969825, EPI_ISL_969827, EPI_ISL_969829, EPI_ISL_969830, EPI_ISL_969832, EPI_ISL_969835, EPI_ISL_969837, EPI_ISL_969839, EPI_ISL_969842, EPI_ISL_969844, EPI_ISL_969845, EPI_ISL_969847, EPI_ISL_969850, EPI_ISL_969851, EPI_ISL_969854, EPI_ISL_969855, EPI_ISL_969857, EPI_ISL_969860, EPI_ISL_969862, EPI_ISL_969864, EPI_ISL_969866, EPI_ISL_969868, EPI_ISL_969870, EPI_ISL_969872, EPI_ISL_969874, EPI_ISL_969876, EPI_ISL_969878, EPI_ISL_969880, EPI_ISL_969882, EPI_ISL_969885, EPI_ISL_969888, EPI_ISL_969890, EPI_ISL_969892, EPI_ISL_969894, EPI_ISL_969896, EPI_ISL_969898, EPI_ISL_969900, EPI_ISL_969902, EPI_ISL_969904, EPI_ISL_969905, EPI_ISL_969908, EPI_ISL_969910, EPI_ISL_969912, EPI_ISL_969913, EPI_ISL_969915, EPI_ISL_969918, EPI_ISL_969920, EPI_ISL_969922, EPI_ISL_969924, EPI_ISL_969927, EPI_ISL_969929, EPI_ISL_969931, EPI_ISL_969933, EPI_ISL_969935, EPI_ISL_969938, EPI_ISL_969941, EPI_ISL_969942, EPI_ISL_969944, EPI_ISL_969946, EPI_ISL_969949, EPI_ISL_969950, EPI_ISL_969953, EPI_ISL_969954, EPI_ISL_969957, EPI_ISL_969959, EPI_ISL_969960, EPI_ISL_969962, EPI_ISL_969964, EPI_ISL_969966, EPI_ISL_969968, EPI_ISL_969970, EPI_ISL_969972, EPI_ISL_969973, EPI_ISL_969976, EPI_ISL_969978, EPI_ISL_969980, EPI_ISL_969981, EPI_ISL_969984, EPI_ISL_969985, EPI_ISL_969987, EPI_ISL_969989, EPI_ISL_969991, EPI_ISL_969993, EPI_ISL_969995, EPI_ISL_969997, EPI_ISL_969999, EPI_ISL_970001, EPI_ISL_970003, EPI_ISL_970006, EPI_ISL_970008, EPI_ISL_970010, EPI_ISL_970012, EPI_ISL_970014, EPI_ISL_970016, EPI_ISL_970018, EPI_ISL_970020, EPI_ISL_970021, EPI_ISL_970023, EPI_ISL_970025, EPI_ISL_970027, EPI_ISL_970029, EPI_ISL_970031, EPI_ISL_970034, EPI_ISL_970036, EPI_ISL_970038, EPI_ISL_970040, EPI_ISL_970042, EPI_ISL_970044, EPI_ISL_970046, EPI_ISL_970048, EPI_ISL_970051, EPI_ISL_970053, EPI_ISL_970054, EPI_ISL_970056, EPI_ISL_970058, EPI_ISL_970060, EPI_ISL_970062, EPI_ISL_970065, EPI_ISL_970067, EPI_ISL_970069, EPI_ISL_970071, EPI_ISL_970072, EPI_ISL_970074, EPI_ISL_970077, EPI_ISL_970078, EPI_ISL_970081, EPI_ISL_970083, EPI_ISL_970085, EPI_ISL_970088, EPI_ISL_970090, EPI_ISL_970093, EPI_ISL_970095, EPI_ISL_970097, EPI_ISL_970100, EPI_ISL_970102, EPI_ISL_970104, EPI_ISL_970106, EPI_ISL_970108, EPI_ISL_970110, EPI_ISL_970112, EPI_ISL_970114, EPI_ISL_970116, EPI_ISL_970118, EPI_ISL_970121, EPI_ISL_970123, EPI_ISL_970125, EPI_ISL_970127, EPI_ISL_970129, EPI_ISL_970131, EPI_ISL_970133, EPI_ISL_970135, EPI_ISL_970137, EPI_ISL_970139, EPI_ISL_970141, EPI_ISL_970143, EPI_ISL_970144, EPI_ISL_970146, EPI_ISL_970149, EPI_ISL_970151, EPI_ISL_970154, EPI_ISL_970156, EPI_ISL_970158, EPI_ISL_970159, EPI_ISL_970162, EPI_ISL_970164, EPI_ISL_970167, EPI_ISL_970169, EPI_ISL_970171, EPI_ISL_970174, EPI_ISL_970176, EPI_ISL_970178, EPI_ISL_970180, EPI_ISL_970182, EPI_ISL_970185, EPI_ISL_970187, EPI_ISL_970189, EPI_ISL_970191, EPI_ISL_970193, EPI_ISL_970194, EPI_ISL_970196, EPI_ISL_970199, EPI_ISL_970201, EPI_ISL_970203, EPI_ISL_970204, EPI_ISL_970207, EPI_ISL_970208, EPI_ISL_970210, EPI_ISL_970212, EPI_ISL_970214, EPI_ISL_970216, EPI_ISL_970218, EPI_ISL_970220, EPI_ISL_970222, EPI_ISL_970223, EPI_ISL_970226, EPI_ISL_970228, EPI_ISL_970230, EPI_ISL_970232, EPI_ISL_970234, EPI_ISL_970236, EPI_ISL_970238, EPI_ISL_970240, EPI_ISL_970241, EPI_ISL_970243, EPI_ISL_970244, EPI_ISL_970245, EPI_ISL_970246, EPI_ISL_970248, EPI_ISL_970250, EPI_ISL_970253, EPI_ISL_970255, EPI_ISL_970257, EPI_ISL_970259, EPI_ISL_970261, EPI_ISL_970263, EPI_ISL_970265, EPI_ISL_970267, EPI_ISL_970269, EPI_ISL_970271, EPI_ISL_970273, EPI_ISL_970276, EPI_ISL_970277, EPI_ISL_970280, EPI_ISL_970282, EPI_ISL_970284, EPI_ISL_970286, EPI_ISL_970289, EPI_ISL_970290, EPI_ISL_970293, EPI_ISL_970294, EPI_ISL_970296, EPI_ISL_970298, EPI_ISL_970300, EPI_ISL_970302, EPI_ISL_970304, EPI_ISL_970305, EPI_ISL_970308, EPI_ISL_970310, EPI_ISL_970311, EPI_ISL_970313, EPI_ISL_970315, EPI_ISL_970317, EPI_ISL_970318, EPI_ISL_970320, EPI_ISL_970322, EPI_ISL_970325, EPI_ISL_970327, EPI_ISL_970329, EPI_ISL_970331, EPI_ISL_970333, EPI_ISL_970335, EPI_ISL_970336, EPI_ISL_970339, EPI_ISL_970341, EPI_ISL_970343, EPI_ISL_970345, EPI_ISL_970347, EPI_ISL_970349, EPI_ISL_970351, EPI_ISL_970353 | | | |
| see above | BCCDC Public Health Laboratory | BCCDC Public Health Laboratory | Prystajecky Natalie, Linda Hoang, Dan Fornika, John Tyson, Shannon Russell, Kim Macdonald, Kimia Kamelian, Ana Pacagnella, Corrinne Ng, Loretta Janz, Robert Azana Terry Snutch, Mel Krajden |
| EPI_ISL_982228 | MEPHI, Aix Marseille University | MEPHI, Aix Marseille University | Anthony LEVASSEUR |
