## Supplementary material for "Detection and characterization of the SARS-CoV-2 lineage B.1.526 in New York": Supp. Table 3 GISAID Acknowledgment part 2: gisaid_hcov-19_acknowledgement_table_2021_02_12_23-7.pdf

We gratefully acknowledge the following Authors from the Originating laboratories responsible for obtaining the specimens, as well as the Submitting laboratories where the genome data were generated and shared via GISAID, on which this research is based.

All Submitters of data may be contacted directly via www.gisaid.org

Authors are sorted alphabetically.

| Accession ID | Originating Laboratory | Submitting Laboratory | Authors |
|---|---|---|---|
| EPI_ISL_534759, EPI_ISL_534760, EPI_ISL_534761, EPI_ISL_534762 | Queens Medical Centre, Clinical Microbiology Department / DeepSeq Nottingham | COVID-19 Genomics UK (COG-UK) Consortium | Gemma Clark, Wendy Smith, Manjinder Khakh, Vicki M Fleming, Michelle M Lister, Hannah Howson-Wells, Jonathan Ball, Patrick McClure, Joseph Chappell, Theocharis Tsoleridis, Nadine Holmes, Matthew Carlisle, Christopher Moore, Fei Sang, Johnny Debebe, Victoria Wright, Matthew Loose |
| EPI_ISL_536656, EPI_ISL_536657 | University of Wisconsin-Madison Campus AIDS Vaccine Research Laboratories | University of Wisconsin-Madison AIDS Vaccine Research Laboratories | Gage Moreno, Katarina Braun, et al. AIDS Vaccine Research Laboratories |
| EPI_ISL_539327 | Area of Virology, Serology and Virology Division (SAViD), New South Wales Health Pathology Randwick | Area of Virology, Serology and Virology Division (SAViD), New South Wales Health Pathology Randwick | Rawlinson, W., Bull, R., Deveson, I., Van Hal, S. |
| EPI_ISL_540701, EPI_ISL_540702, EPI_ISL_540703, EPI_ISL_540704, EPI_ISL_540705, EPI_ISL_540706, EPI_ISL_540707, EPI_ISL_540708, EPI_ISL_540709, EPI_ISL_540710, EPI_ISL_540711, EPI_ISL_540712, EPI_ISL_540713, EPI_ISL_540714, EPI_ISL_540715, EPI_ISL_540716, EPI_ISL_540717, EPI_ISL_540718 | | | |
| see above | Queens Medical Centre, Clinical Microbiology Department / DeepSeq Nottingham | COVID-19 Genomics UK (COG-UK) Consortium | Gemma Clark, Wendy Smith, Manjinder Khakh, Vicki M Fleming, Michelle M Lister, Hannah Howson-Wells, Jonathan Ball, Patrick McClure, Joseph Chappell, Theocharis Tsoleridis, Nadine Holmes, Matthew Carlisle, Christopher Moore, Fei Sang, Johnny Debebe, Victoria Wright, Matthew Loose |
| EPI_ISL_540731, EPI_ISL_540743, EPI_ISL_540750, EPI_ISL_540773 | Virology Department, Sheffield Teaching Hospitals NHS Foundation Trust/Department of Infection, Immunity and Cardiovascular Disease, The Medical School, University of Sheffield | COVID-19 Genomics UK (COG-UK) Consortium | Thushan de Silva, Matthew Parker, Nikki Smith, Adri Angyal, Rebecca Brown, Luke Green, Rachel Tucker, Paul Parsons, Danielle Groves, Katie Johnson, Laura Carrilero, Alex Keeley, Dave Partridge, Matthew Wyles, Benjamin Lindsey, Mehmet Yavuz, Mohammad Raza, Cariad Evans |
| EPI_ISL_540886, EPI_ISL_540887, EPI_ISL_540888, EPI_ISL_540889, EPI_ISL_540891, EPI_ISL_540892, EPI_ISL_540893 | Virology Department, Royal Infirmary of Edinburgh, NHS Lothian / School of Biological Sciences, University of Edinburgh / Institute of Genetics and Molecular Medicine, University of Edinburgh | COVID-19 Genomics UK (COG-UK) Consortium | McHugh M, Dewar R, Rooke S, Gallagher M, Balcaza C, O'Toole Á, Scher E, Hill V, McCrone JT, Colquhoun R, Yu X, Jackson B, Rambaut A, Williams TC, Templeton K |
| EPI_ISL_541540, EPI_ISL_541541, EPI_ISL_541542, EPI_ISL_541543, EPI_ISL_541544, EPI_ISL_541545, EPI_ISL_541546, EPI_ISL_541547, EPI_ISL_541548, EPI_ISL_541549, EPI_ISL_541550 | | | |
| see above | University of Wisconsin-Madison AIDS Vaccine Research Laboratories | University of Wisconsin-Madison AIDS Vaccine Research Laboratories | Gage Moreno, Katarina Braun, et al. AIDS Vaccine Research Laboratories |
| EPI_ISL_541551, EPI_ISL_541552, EPI_ISL_541553, EPI_ISL_541554, EPI_ISL_541555, EPI_ISL_541556, EPI_ISL_541557, EPI_ISL_541558, EPI_ISL_541559, EPI_ISL_541560, EPI_ISL_541561, EPI_ISL_541562, EPI_ISL_541563, EPI_ISL_541564, EPI_ISL_541565 | | | |
| see above | University of Wisconsin-Madison Campus AIDS Vaccine Research Laboratories | University of Wisconsin-Madison AIDS Vaccine Research Laboratories | Gage Moreno, Katarina Braun, et al. AIDS Vaccine Research Laboratories |
| EPI_ISL_541566, EPI_ISL_541567, EPI_ISL_541569, EPI_ISL_541570, EPI_ISL_541571, EPI_ISL_541572, EPI_ISL_541573, EPI_ISL_541574, EPI_ISL_541575, EPI_ISL_541576, EPI_ISL_541577, EPI_ISL_541578, EPI_ISL_541579, EPI_ISL_541580, EPI_ISL_541581, EPI_ISL_541582, EPI_ISL_541583, EPI_ISL_541584, EPI_ISL_541585, EPI_ISL_541586, EPI_ISL_541587, EPI_ISL_541588, EPI_ISL_541589, EPI_ISL_541590, EPI_ISL_541591, EPI_ISL_541592, EPI_ISL_541593, EPI_ISL_541594, EPI_ISL_541595, EPI_ISL_541596, EPI_ISL_541597, EPI_ISL_541598, EPI_ISL_541599, EPI_ISL_541600, EPI_ISL_541601, EPI_ISL_541602, EPI_ISL_541603, EPI_ISL_541604, EPI_ISL_541605, EPI_ISL_541606, EPI_ISL_541607, EPI_ISL_541608, EPI_ISL_541609, EPI_ISL_541610, EPI_ISL_541611, EPI_ISL_541612, EPI_ISL_541613, EPI_ISL_541614, EPI_ISL_541615, EPI_ISL_541616, EPI_ISL_541617, EPI_ISL_541618, EPI_ISL_541619, EPI_ISL_541620, EPI_ISL_541621, EPI_ISL_541622, EPI_ISL_541623, EPI_ISL_541624, EPI_ISL_541625, EPI_ISL_541626, EPI_ISL_541627, EPI_ISL_541628, EPI_ISL_541629, EPI_ISL_541630, EPI_ISL_541631, EPI_ISL_541632, EPI_ISL_541633, EPI_ISL_541634, EPI_ISL_541635, EPI_ISL_541636, EPI_ISL_541637, EPI_ISL_541638, EPI_ISL_541639, EPI_ISL_541640, EPI_ISL_541641, EPI_ISL_541642, EPI_ISL_541643, EPI_ISL_541644, EPI_ISL_541645, EPI_ISL_541646, EPI_ISL_541647, EPI_ISL_541648 | | | |
| see above | University of Wisconsin-Madison AIDS Vaccine Research Laboratories | University of Wisconsin-Madison AIDS Vaccine Research Laboratories | Gage Moreno, Katarina Braun, et al. AIDS Vaccine Research Laboratories |
| EPI_ISL_545032, EPI_ISL_545033 | Pathology West - NSW Health Pathology | NSW Health Pathology - Institute of Clinical Pathology and Medical Research; Westmead Hospital; University of Sydney | CIDM-PH et al. |
| EPI_ISL_545034, EPI_ISL_545035, EPI_ISL_545036 | Histopath | NSW Health Pathology - Institute of Clinical Pathology and Medical Research; Westmead Hospital; University of Sydney | CIDM-PH et al. |
| EPI_ISL_545037 | Sydney South West Pathology Service (SSWPS) - Concord Repatriation General Hospital - NSW Health Pathology | NSW Health Pathology - Institute of Clinical Pathology and Medical Research; Westmead Hospital; University of Sydney | CIDM-PH et al. |

| | | | |
|---|---|---|---|
| EPI_ISL_545038 | Pathology West - NSW Health Pathology | NSW Health Pathology - Institute of Clinical Pathology and Medical Research; Westmead Hospital; University of Sydney | CIDM-PH et al. |
| EPI_ISL_545039 | Sydney South West Pathology Service (SSWPS) - Royal Prince Alfred Hospital - NSW Health Pathology | NSW Health Pathology - Institute of Clinical Pathology and Medical Research; Westmead Hospital; University of Sydney | CIDM-PH et al. |
| EPI_ISL_545042 | Sydney South West Pathology Service (SSWPS) - Liverpool Hospital - NSW Health Pathology | NSW Health Pathology - Institute of Clinical Pathology and Medical Research; Westmead Hospital; University of Sydney | CIDM-PH et al. |
| EPI_ISL_545046, EPI_ISL_545047 | Pathology West - NSW Health Pathology | NSW Health Pathology - Institute of Clinical Pathology and Medical Research; Westmead Hospital; University of Sydney | CIDM-PH et al. |
| EPI_ISL_545048 | South Eastern Area Laboratory Services (SEALS) | NSW Health Pathology - Institute of Clinical Pathology and Medical Research; Westmead Hospital; University of Sydney | CIDM-PH et al. |
| EPI_ISL_545049 | Sydney South West Pathology Service (SSWPS) - Concord Repatriation General Hospital - NSW Health Pathology | NSW Health Pathology - Institute of Clinical Pathology and Medical Research; Westmead Hospital; University of Sydney | CIDM-PH et al. |
| EPI_ISL_547819, EPI_ISL_547820, EPI_ISL_547821, EPI_ISL_547822, EPI_ISL_547823, EPI_ISL_547824, EPI_ISL_547825, EPI_ISL_547826, EPI_ISL_547827, EPI_ISL_547828, EPI_ISL_547829, EPI_ISL_547830, EPI_ISL_547831, EPI_ISL_547832, EPI_ISL_547833, EPI_ISL_547834, EPI_ISL_547835, EPI_ISL_547836, EPI_ISL_547837, EPI_ISL_547838, EPI_ISL_547839, EPI_ISL_547840, EPI_ISL_547841, EPI_ISL_547842, EPI_ISL_547843, EPI_ISL_547844, EPI_ISL_547845, EPI_ISL_547846, EPI_ISL_547847, EPI_ISL_547848, EPI_ISL_547849, EPI_ISL_547850, EPI_ISL_547851, EPI_ISL_547852, EPI_ISL_547853, EPI_ISL_547854, EPI_ISL_547855, EPI_ISL_547856, EPI_ISL_547857, EPI_ISL_547858, EPI_ISL_547859, EPI_ISL_547860, EPI_ISL_547861, EPI_ISL_547862, EPI_ISL_547863, EPI_ISL_547864, EPI_ISL_547865, EPI_ISL_547866, EPI_ISL_547867, EPI_ISL_547868, EPI_ISL_547869 | | | |
| see above | Gundersen Molecular Diagnostics Laboratory | Kabara Cancer Research Institute | Craig S. Richmond, Paraic A. Kenny |
| EPI_ISL_548082 | Middlemore Hospital | Institute of Environmental Science and Research (ESR) | Xiaoyun Ren, Matt Storey, Nikki Freed, Muhammad Faisal, Jing Wang, Hermes Perez, Anja Werno, Antje van der Linden, Arlo Upton, Chris Mansell, David Hammer, Dragana Drinkovic, Gary McAuliffe, Hana Sofia Andersson, James Ussher, Jill Sherwood, Josh Freeman, Julia Howard, Juliet Elvy, Mary DeAlmeida, Matt Blakiston, Matthew Rogers, Max Bloomfield, Michael Addidle, Michelle Balm, Sally Roberts, Sarah Jefferies, Sharmini Muttaiyah, Susan Morpeth, Susan Taylor, Timothy Blackmore, Vani Sathyendran, Veronica Playle, Virginia Hope, Erasmus Smit, Lauren Jelly, Olin Silander, Joep de Ligt |
| EPI_ISL_548084 | LabPLUS | Institute of Environmental Science and Research (ESR) | Xiaoyun Ren, Matt Storey, Nikki Freed, Muhammad Faisal, Jing Wang, Hermes Perez, Anja Werno, Antje van der Linden, Arlo Upton, Chris Mansell, David Hammer, Dragana Drinkovic, Gary McAuliffe, Hana Sofia Andersson, James Ussher, Jill Sherwood, Josh Freeman, Julia Howard, Juliet Elvy, Mary DeAlmeida, Matt Blakiston, Matthew Rogers, Max Bloomfield, Michael Addidle, Michelle Balm, Sally Roberts, Sarah Jefferies, Sharmini Muttaiyah, Susan Morpeth, Susan Taylor, Timothy Blackmore, Vani Sathyendran, Veronica Playle, Virginia Hope, Erasmus Smit, Lauren Jelly, Olin Silander, Joep de Ligt |
| EPI_ISL_548972, EPI_ISL_548974, EPI_ISL_548975, EPI_ISL_548976 | National Public Health Laboratory, National Centre for Infectious Diseases | National Public Health Laboratory, National Centre for Infectious Diseases | Mak TM, Octavia S, Zhou Z, Cui L, Lin RTP |
| EPI_ISL_549334 | Queens Medical Centre, Clinical Microbiology Department / DeepSeq Nottingham | COVID-19 Genomics UK (COG-UK) Consortium | Gemma Clark, Wendy Smith, Manjinder Khakh, Vicki M Fleming, Michelle M Lister, Hannah Howson-Wells, Jonathan Ball, Patrick McClure, Joseph Chappell, Theocharis Tsoleridis, Nadine Holmes, Matthew Carlisle, Christopher Moore, Fei Sang, Johnny Debebe, Victoria Wright, Matthew Loose |
| EPI_ISL_549341 | Centre for Enzyme Innovation, University of Portsmouth / Translational Research Laboratory, Portsmouth Hospitals NHS Trust | COVID-19 Genomics UK (COG-UK) Consortium | Angela Beckett,Yann Bourgeois,Garry Scarlett,Sharon Glaysher,Scott Elliott,Kelly Bicknell,Robert Impey,Allyson Lloyd,Sarah Wyllie,Ethan Butcher,Anoop Chauhan,Samuel Robson |
| EPI_ISL_549354 | Liverpool Clinical Laboratories | COVID-19 Genomics UK (COG-UK) Consortium | Sam Haldenby, Anita Lucaci, Steve Paterson, Julian Hiscox, Alistair Darby, M Almsaud, A Alrezaihi, Muhannad Alruwaili, Stuart D Armstrong, Jones Benjamin, Eleanor G Bentley, Anu Chawla, Jordan J Clark, Angela Cowell, Richard Eccles, Isabel García-Dorival, Matthew Gemmell, Alessandro Gerada, PKF Gilmore, Richard Gregory, Ximeng Han, Catherine Hartley, Margaret Hughes, Miren Iturriza-Gomara, James Johnson, L Luu, Jenifer Manson, Charlotte Nelson, Elaine O'Toole, Cassie Olateju, Rebekah Penrice-Randal , Lucille Rainbow, N.P Randle, Trevor Ian Robinson, Parul Sharma, Ghada T Shawli, James P Stewart, Neil Swainston, Ecaterina Vamos, Joanne Watts, Mark Whitehead |
| EPI_ISL_549355 | Quadram Institute Bioscience | COVID-19 Genomics UK (COG-UK) Consortium | Dave J. Baker, Gemma L. Kay, Alp Aydin, Thanh Le-Viet, Steven Rudder, Ana P. Tedim, Anastasia Kolyva, Maria Diaz, Leonardo de Oliveira Martins, Nabil-Fareed Alikhan, Lizzie Meadows, Rachael Stanley, Ngozi Elumogo, Muhammed Yasir, Nicholas M. Thomson, Alexander J Trotter, Rachel Gilroy, Samuel Bloomfield, Claire Stuart, Andrew Bell, Reenesh Prakash, Samir Dervisevic, Alison E. Mather, John Wain, Mark Webber, Andrew J. Page, Justin O'Grady |
| EPI_ISL_549356 | Lincolnshire Hospitals and DeepSeq Nottingham | COVID-19 Genomics UK (COG-UK) Consortium | Nichola Duckworth, Tim Sloan, Sarah Walsh, Jonathan Ball, Patrick McClure, Joeseph Chappell, Nadine Holmes, Matthew Carlisle, Christopher Moore, Fei Sang, Johnny Debebe, Victoria Wright, Matthew Loose |
| EPI_ISL_549360, EPI_ISL_549364, EPI_ISL_549366 | Centre for Enzyme Innovation, University of Portsmouth / Translational Research Laboratory, Portsmouth Hospitals NHS Trust | COVID-19 Genomics UK (COG-UK) Consortium | Angela Beckett,Yann Bourgeois,Garry Scarlett,Sharon Glaysher,Scott Elliott,Kelly Bicknell,Robert Impey,Allyson Lloyd,Sarah Wyllie,Ethan Butcher,Anoop Chauhan,Samuel Robson |
| EPI_ISL_549371, EPI_ISL_549372 | Lincolnshire Hospitals and DeepSeq Nottingham | COVID-19 Genomics UK (COG-UK) Consortium | Nichola Duckworth, Tim Sloan, Sarah Walsh, Jonathan Ball, Patrick McClure, Joeseph Chappell, Nadine Holmes, Matthew Carlisle, Christopher Moore, Fei Sang, Johnny Debebe, Victoria Wright, Matthew Loose |
| EPI_ISL_549380 | Queens Medical Centre, Clinical Microbiology Department / DeepSeq Nottingham | COVID-19 Genomics UK (COG-UK) Consortium | Gemma Clark, Wendy Smith, Manjinder Khakh, Vicki M Fleming, Michelle M Lister, Hannah Howson-Wells, Jonathan Ball, Patrick McClure, Joseph Chappell, Theocharis Tsoleridis, Nadine Holmes, Matthew Carlisle, Christopher Moore, Fei Sang, Johnny Debebe, Victoria Wright, Matthew Loose |
| EPI_ISL_549387 | Lincolnshire Hospitals and DeepSeq Nottingham | COVID-19 Genomics UK (COG-UK) Consortium | Nichola Duckworth, Tim Sloan, Sarah Walsh, Jonathan Ball, Patrick McClure, Joeseph Chappell, Nadine Holmes, Matthew Carlisle, Christopher Moore, Fei Sang, Johnny Debebe, Victoria Wright, Matthew Loose |
| EPI_ISL_549461, | Centre for Enzyme | COVID-19 Genomics UK | Angela Beckett,Yann Bourgeois,Garry Scarlett,Sharon Glaysher,Scott Elliott,Kelly Bicknell,Robert Impey,Allyson Lloyd,Sarah Wyllie,Ethan Butcher,Anoop Chauhan,Samuel Robson |

| | | | |
|---|---|---|---|
| EPI_ISL_549466, EPI_ISL_549469, EPI_ISL_549470, EPI_ISL_549471, EPI_ISL_549472, EPI_ISL_549473, EPI_ISL_549474, EPI_ISL_549475, EPI_ISL_549476 | Innovation, University of Portsmouth / Translational Research Laboratory, Portsmouth Hospitals NHS Trust | (COG-UK) Consortium | |
| EPI_ISL_549479, EPI_ISL_549480, EPI_ISL_549481 | Quadram Institute Bioscience | COVID-19 Genomics UK (COG-UK) Consortium | Dave J. Baker, Gemma L. Kay, Alp Aydin, Thanh Le-Viet, Steven Rudder, Ana P. Tedim, Anastasia Kolyva, Maria Diaz, Leonardo de Oliveira Martins, Nabil-Fareed Alikhan, Lizzie Meadows, Rachael Stanley, Ngozi Elumogo, Muhammed Yasir, Nicholas M. Thomson, Alexander J Trotter, Rachel Gilroy, Samuel Bloomfield, Claire Stuart, Andrew Bell, Reenesh Prakash, Samir Dervisevic, Alison E. Mather, John Wain, Mark Webber, Andrew J. Page, Justin O'Grady |
| EPI_ISL_549491, EPI_ISL_549492, EPI_ISL_549493, EPI_ISL_549494, EPI_ISL_549495, EPI_ISL_549496, EPI_ISL_549497, EPI_ISL_549498, EPI_ISL_549499, EPI_ISL_549500, EPI_ISL_549501, EPI_ISL_549502, EPI_ISL_549503, EPI_ISL_549504 | | | |
| see above | Queens Medical Centre, Clinical Microbiology Department / DeepSeq Nottingham | COVID-19 Genomics UK (COG-UK) Consortium | Gemma Clark, Wendy Smith, Manjinder Khakh, Vicki M Fleming, Michelle M Lister, Hannah Howson-Wells, Jonathan Ball, Patrick McClure, Joseph Chappell, Theocharis Tsoleridis, Nadine Holmes, Matthew Carlisle, Christopher Moore, Fei Sang, Johnny Debebe, Victoria Wright, Matthew Loose |
| EPI_ISL_549519, EPI_ISL_549520, EPI_ISL_549521, EPI_ISL_549522 | Lincolnshire Hospitals and DeepSeq Nottingham | COVID-19 Genomics UK (COG-UK) Consortium | Nichola Duckworth, Tim Sloan, Sarah Walsh, Jonathan Ball, Patrick McClure, Joeseph Chappell, Nadine Holmes, Matthew Carlisle, Christopher Moore, Fei Sang, Johnny Debebe, Victoria Wright, Matthew Loose |
| EPI_ISL_549555, EPI_ISL_549566, EPI_ISL_549576, EPI_ISL_549586, EPI_ISL_549595, EPI_ISL_549610, EPI_ISL_549617, EPI_ISL_549657, EPI_ISL_549708, EPI_ISL_549721, EPI_ISL_549724, EPI_ISL_549737, EPI_ISL_549738, EPI_ISL_549791 | | | |
| see above | Lighthouse Lab in Glasgow | Wellcome Sanger Institute for the COVID-19 Genomics UK (COG-UK) consortium | Harper VanSteenhouse, Yumi Kasai, David Gray, Carol Clugston, Anna Dominiczak and Alex Alderton, Roberto Amato, Sonia Goncalves, Ewan Harrison, David K. Jackson, Ian Johnston, Dominic Kwiatkowski, Cordelia Langford, John Sillitoe on behalf of the Wellcome Sanger Institute COVID-19 Surveillance Team |
| EPI_ISL_559899, EPI_ISL_559903, EPI_ISL_559905, EPI_ISL_559906, EPI_ISL_559907, EPI_ISL_559908, EPI_ISL_559909, EPI_ISL_559913, EPI_ISL_559915, EPI_ISL_559916, EPI_ISL_559917, EPI_ISL_559918, EPI_ISL_559919, EPI_ISL_559924, EPI_ISL_559925, EPI_ISL_559926, EPI_ISL_559929, EPI_ISL_559933, EPI_ISL_559934, EPI_ISL_559936, EPI_ISL_559940, EPI_ISL_559941, EPI_ISL_559942, EPI_ISL_559943, EPI_ISL_559944, EPI_ISL_559945, EPI_ISL_559946 | | | |
| see above | Virology Department, Sheffield Teaching Hospitals NHS Foundation Trust/Department of Infection, Immunity and Cardiovascular Disease, The Medical School, University of Sheffield | COVID-19 Genomics UK (COG-UK) Consortium | Thushan de Silva, Matthew Parker, Nikki Smith, Adri Angyal, Rebecca Brown, Luke Green, Rachel Tucker, Paul Parsons, Danielle Groves, Katie Johnson, Laura Carrilero, Alex Keeley, Dave Partridge, Matthew Wyles, Benjamin Lindsey, Mehmet Yavuz, Mohammad Raza, Cariad Evans |
| EPI_ISL_559979, EPI_ISL_559980, EPI_ISL_559981, EPI_ISL_559982 | Virology Department, Royal Infirmary of Edinburgh, NHS Lothian / School of Biological Sciences, University of Edinburgh / Institute of Genetics and Molecular Medicine, University of Edinburgh | COVID-19 Genomics UK (COG-UK) Consortium | McHugh M, Dewar R, Rooke S, Gallagher M, Balcaza C, O'Toole Á, Scher E, Hill V, McCrone JT, Colquhoun R, Yu X, Jackson B, Rambaut A, Williams TC, Templeton K |
| EPI_ISL_559983 | Centre for Enzyme Innovation, University of Portsmouth / Translational Research Laboratory, Portsmouth Hospitals NHS Trust | COVID-19 Genomics UK (COG-UK) Consortium | Angela Beckett,Yann Bourgeois,Garry Scarlett,Sharon Glaysher,Scott Elliott,Kelly Bicknell,Robert Impey,Allyson Lloyd,Sarah Wyllie,Ethan Butcher,Anoop Chauhan,Samuel Robson |
| EPI_ISL_560083, EPI_ISL_560086, EPI_ISL_560087, EPI_ISL_560089, EPI_ISL_560090, EPI_ISL_560095, EPI_ISL_560097, EPI_ISL_560098, EPI_ISL_560099, EPI_ISL_560100, EPI_ISL_560101, EPI_ISL_560102, EPI_ISL_560105, EPI_ISL_560106, EPI_ISL_560107, EPI_ISL_560108, EPI_ISL_560115, EPI_ISL_560116, EPI_ISL_560118, EPI_ISL_560119, EPI_ISL_560120, EPI_ISL_560121, EPI_ISL_560123, EPI_ISL_560130, EPI_ISL_560132, EPI_ISL_560133, EPI_ISL_560134, EPI_ISL_560135, EPI_ISL_560136, EPI_ISL_560138, EPI_ISL_560141, EPI_ISL_560142, EPI_ISL_560144, EPI_ISL_560145, EPI_ISL_560146, EPI_ISL_560148, EPI_ISL_560149, EPI_ISL_560151, EPI_ISL_560152, EPI_ISL_560153, EPI_ISL_560157, EPI_ISL_560158, EPI_ISL_560160, EPI_ISL_560161, EPI_ISL_560162, EPI_ISL_560165, EPI_ISL_560166, EPI_ISL_560167, EPI_ISL_560168, EPI_ISL_560169, EPI_ISL_560170, EPI_ISL_560171, EPI_ISL_560172, EPI_ISL_560173, EPI_ISL_560174, EPI_ISL_560175, EPI_ISL_560177, EPI_ISL_560178, EPI_ISL_560179, EPI_ISL_560180, EPI_ISL_560181, EPI_ISL_560182, EPI_ISL_560183, EPI_ISL_560184, EPI_ISL_560186, EPI_ISL_560187, EPI_ISL_560188, EPI_ISL_560189, EPI_ISL_560191, EPI_ISL_560192, EPI_ISL_560193, EPI_ISL_560194, EPI_ISL_560196, EPI_ISL_560198, EPI_ISL_560200, EPI_ISL_560203, EPI_ISL_560204, EPI_ISL_560208, EPI_ISL_560209, EPI_ISL_560210, EPI_ISL_560211, EPI_ISL_560212, EPI_ISL_560213, EPI_ISL_560214, EPI_ISL_560215, EPI_ISL_560216, EPI_ISL_560217, EPI_ISL_560218, EPI_ISL_560220, EPI_ISL_560221, EPI_ISL_560222, EPI_ISL_560223, EPI_ISL_560224, EPI_ISL_560225, EPI_ISL_560226, EPI_ISL_560227, EPI_ISL_560228, EPI_ISL_560229, EPI_ISL_560230, EPI_ISL_560231, EPI_ISL_560232, EPI_ISL_560233, EPI_ISL_560234, EPI_ISL_560235, EPI_ISL_560236, EPI_ISL_560237, EPI_ISL_560238, EPI_ISL_560239, EPI_ISL_560240, EPI_ISL_560241, EPI_ISL_560242, EPI_ISL_560243, EPI_ISL_560244, EPI_ISL_560245, EPI_ISL_560246, EPI_ISL_560247, EPI_ISL_560248, EPI_ISL_560249, EPI_ISL_560250, EPI_ISL_560251, EPI_ISL_560252, EPI_ISL_560253, EPI_ISL_560254, EPI_ISL_560255, EPI_ISL_560256, EPI_ISL_560257, EPI_ISL_560258, EPI_ISL_560259, EPI_ISL_560260, EPI_ISL_560261, EPI_ISL_560262, EPI_ISL_560263, EPI_ISL_560264, EPI_ISL_560265, EPI_ISL_560266, EPI_ISL_560267, EPI_ISL_560268, EPI_ISL_560269, EPI_ISL_560270, EPI_ISL_560271, EPI_ISL_560272, EPI_ISL_560273, EPI_ISL_560274, EPI_ISL_560275, EPI_ISL_560276, EPI_ISL_560277, EPI_ISL_560279, EPI_ISL_560280, EPI_ISL_560281, EPI_ISL_560282, EPI_ISL_560283, EPI_ISL_560284, EPI_ISL_560285, EPI_ISL_560286, EPI_ISL_560287, EPI_ISL_560288, EPI_ISL_560289, EPI_ISL_560290, EPI_ISL_560291, EPI_ISL_560292, EPI_ISL_560293, EPI_ISL_560294, EPI_ISL_560295, EPI_ISL_560296, EPI_ISL_560297, EPI_ISL_560298, EPI_ISL_560299, EPI_ISL_560300, EPI_ISL_560301, EPI_ISL_560302, EPI_ISL_560303, EPI_ISL_560304 | | | |
| see above | Wales Specialist Virology Centre Sequencing lab: Pathogen Genomics Unit | COVID-19 Genomics UK (COG-UK) Consortium | Catherine Moore, Johnathan Evans, Laura Gifford, Malorie Perry, Simon Cottrell, Angela Marchbank, Alec Birchley, Alexander Adams, Amy Gaskin, Bree Gatica-Wilcox, Jason Coombes, Joel Southgate, Lauren Gilbert, Lee Graham, Nicole Pacchiarini, Sara Kumziene-Summerhayes, Sarah Taylor, Sophie Jones, Sara Rey, Matthew Bull, Joanne Watkins, Sally Corden, Tom Connor |
| EPI_ISL_560426, EPI_ISL_560428, EPI_ISL_560446, EPI_ISL_560447, EPI_ISL_560448, EPI_ISL_560449, EPI_ISL_560450, EPI_ISL_560451, EPI_ISL_560452, EPI_ISL_560453, EPI_ISL_560454, EPI_ISL_560464, EPI_ISL_560465, EPI_ISL_560466, EPI_ISL_560467, EPI_ISL_560469, EPI_ISL_560473, EPI_ISL_560474, EPI_ISL_560475, EPI_ISL_560480, EPI_ISL_560482, EPI_ISL_560523, EPI_ISL_560526, EPI_ISL_560528, EPI_ISL_560529, EPI_ISL_560530, EPI_ISL_560531, EPI_ISL_560532, EPI_ISL_560533, EPI_ISL_560536, EPI_ISL_560537, EPI_ISL_560538, EPI_ISL_560539, EPI_ISL_560540, EPI_ISL_560553 | | | |
| see above | Viollier AG | Department of Biosystems Science and Engineering, ETH Zürich | Christian Beisel, Sarah Nadeau, Ivan Topolsky, Pedro Ferreira, Philipp Jablonski, Susana Posada-Céspedes, Tobias Schär, Ina Nissen, Natascha Santacroce, Elodie Burcklen, Christiane Beckmann, Maurice Redondo, Olivier Kobel, Christoph Noppen, Sophie Seidel, Noemie Santamaria de Souza, Niko Beerenwinkel, Tanja Stadler |
| EPI_ISL_560817, EPI_ISL_560818, EPI_ISL_560819, EPI_ISL_560820, EPI_ISL_560821, EPI_ISL_560822, EPI_ISL_560823, EPI_ISL_560824, EPI_ISL_560825, EPI_ISL_560826, EPI_ISL_560827, EPI_ISL_560828, EPI_ISL_560829 | | | |
| see above | Maryland Public Health Laboratory | Maryland Public Health Laboratory | Maryland Department of Health Laboratories Administration |
| EPI_ISL_561398, EPI_ISL_561427, EPI_ISL_561439, EPI_ISL_561441, EPI_ISL_561447, EPI_ISL_561451, EPI_ISL_561452, EPI_ISL_561461, EPI_ISL_561463, EPI_ISL_561473, EPI_ISL_561484, EPI_ISL_561518, EPI_ISL_561537, EPI_ISL_561587, EPI_ISL_561598, EPI_ISL_561608, EPI_ISL_561659 | | | |
| see above | Microbiological Diagnostic Unit - Public Health Laboratory (MDU-PHL) | MDU-PHL | Seemann, T., Schultz M. B., Sait, M., Sherry, N. |
| EPI_ISL_561682 | Victorian Infectious Diseases Reference Laboratory (VIDRL) | VIDRL and MDU-PHL | Caly, L., Seemann, T., Sait, M., Schultz, M. B., Druce J., Sherry, N. |

| | | | |
|---|---|---|---|
| EPI_ISL_561691, EPI_ISL_561744, EPI_ISL_561795, EPI_ISL_561812, EPI_ISL_561834, EPI_ISL_561874, EPI_ISL_561892, EPI_ISL_561926, EPI_ISL_561929, EPI_ISL_561958, EPI_ISL_561984, EPI_ISL_561985, EPI_ISL_562064, EPI_ISL_562075 | | | |
| see above | Microbiological Diagnostic Unit - Public Health Laboratory (MDU-PHL) | MDU-PHL | Seemann, T., Schultz M. B., Sait, M., Sherry, N. |
| EPI_ISL_562104 | Victorian Infectious Diseases Reference Laboratory (VIDRL) | VIDRL and MDU-PHL | Caly, L., Seemann, T., Sait, M., Schultz, M. B., Druce J., Sherry, N. |
| EPI_ISL_562128, EPI_ISL_562151, EPI_ISL_562201, EPI_ISL_562218, EPI_ISL_562243, EPI_ISL_562252 | Microbiological Diagnostic Unit - Public Health Laboratory (MDU-PHL) | MDU-PHL | Seemann, T., Schultz M. B., Sait, M., Sherry, N. |
| EPI_ISL_562261, EPI_ISL_562272, EPI_ISL_562280 | Victorian Infectious Diseases Reference Laboratory (VIDRL) | VIDRL and MDU-PHL | Caly, L., Seemann, T., Sait, M., Schultz, M. B., Druce J., Sherry, N. |
| EPI_ISL_562295 | Microbiological Diagnostic Unit - Public Health Laboratory (MDU-PHL) | MDU-PHL | Seemann, T., Schultz M. B., Sait, M., Sherry, N. |
| EPI_ISL_562335 | Victorian Infectious Diseases Reference Laboratory (VIDRL) | VIDRL and MDU-PHL | Caly, L., Seemann, T., Sait, M., Schultz, M. B., Druce J., Sherry, N. |
| EPI_ISL_562418, EPI_ISL_562421, EPI_ISL_562423, EPI_ISL_562450, EPI_ISL_562481, EPI_ISL_562493, EPI_ISL_562494, EPI_ISL_562496, EPI_ISL_562508, EPI_ISL_562516, EPI_ISL_562519, EPI_ISL_562556, EPI_ISL_562575, EPI_ISL_562594, EPI_ISL_562615, EPI_ISL_562620, EPI_ISL_562622, EPI_ISL_562724, EPI_ISL_562725, EPI_ISL_562726, EPI_ISL_562756 | | | |
| see above | Microbiological Diagnostic Unit - Public Health Laboratory (MDU-PHL) | MDU-PHL | Seemann, T., Schultz M. B., Sait, M., Sherry, N. |
| EPI_ISL_563499 | Victorian Infectious Diseases Reference Laboratory (VIDRL) | VIDRL and MDU-PHL | Caly, L., Seemann, T., Sait, M., Schultz, M. B., Druce J., Sherry, N. |
| EPI_ISL_563505, EPI_ISL_563507, EPI_ISL_563509, EPI_ISL_563517, EPI_ISL_563518, EPI_ISL_563519, EPI_ISL_563521, EPI_ISL_563526, EPI_ISL_563534, EPI_ISL_563535, EPI_ISL_563538, EPI_ISL_563540, EPI_ISL_563543, EPI_ISL_563548, EPI_ISL_563552, EPI_ISL_563553, EPI_ISL_563554, EPI_ISL_563557, EPI_ISL_563558, EPI_ISL_563562, EPI_ISL_563563, EPI_ISL_563566, EPI_ISL_563571, EPI_ISL_563573, EPI_ISL_563576, EPI_ISL_563582, EPI_ISL_563584, EPI_ISL_563585, EPI_ISL_563587, EPI_ISL_563589, EPI_ISL_563590, EPI_ISL_563592, EPI_ISL_563593, EPI_ISL_563595, EPI_ISL_563596, EPI_ISL_563597, EPI_ISL_563599, EPI_ISL_563600, EPI_ISL_563601, EPI_ISL_563602, EPI_ISL_563603, EPI_ISL_563605, EPI_ISL_563607, EPI_ISL_563615 | | | |
| see above | Microbiological Diagnostic Unit - Public Health Laboratory (MDU-PHL) | MDU-PHL | Seemann, T., Schultz M. B., Sait, M., Sherry, N. |
| EPI_ISL_563723 | Victorian Infectious Diseases Reference Laboratory (VIDRL) | VIDRL and MDU-PHL | Caly, L., Seemann, T., Sait, M., Schultz, M. B., Druce J., Sherry, N. |
| EPI_ISL_563725, EPI_ISL_563727, EPI_ISL_563733, EPI_ISL_563735, EPI_ISL_563736, EPI_ISL_563737, EPI_ISL_563738, EPI_ISL_563747, EPI_ISL_563765, EPI_ISL_563769 | Microbiological Diagnostic Unit - Public Health Laboratory (MDU-PHL) | MDU-PHL | Seemann, T., Schultz M. B., Sait, M., Sherry, N. |
| EPI_ISL_563792 | Victorian Infectious Diseases Reference Laboratory (VIDRL) | VIDRL and MDU-PHL | Caly, L., Seemann, T., Sait, M., Schultz, M. B., Druce J., Sherry, N. |
| EPI_ISL_563809 | Microbiological Diagnostic Unit - Public Health Laboratory (MDU-PHL) | MDU-PHL | Seemann, T., Schultz M. B., Sait, M., Sherry, N. |
| EPI_ISL_563811 | Victorian Infectious Diseases Reference Laboratory (VIDRL) | VIDRL and MDU-PHL | Caly, L., Seemann, T., Sait, M., Schultz, M. B., Druce J., Sherry, N. |
| EPI_ISL_563821, EPI_ISL_563823, EPI_ISL_563825, EPI_ISL_563829, EPI_ISL_563834, EPI_ISL_563835 | Microbiological Diagnostic Unit - Public Health Laboratory (MDU-PHL) | MDU-PHL | Seemann, T., Schultz M. B., Sait, M., Sherry, N. |
| EPI_ISL_563840 | Victorian Infectious Diseases Reference Laboratory (VIDRL) | VIDRL and MDU-PHL | Caly, L., Seemann, T., Sait, M., Schultz, M. B., Druce J., Sherry, N. |
| EPI_ISL_563841, EPI_ISL_563842, EPI_ISL_563843, EPI_ISL_563844, EPI_ISL_563846, EPI_ISL_563849, EPI_ISL_563850, EPI_ISL_563851, EPI_ISL_563852, EPI_ISL_563856, EPI_ISL_563857, EPI_ISL_563861, EPI_ISL_563863, EPI_ISL_563864, EPI_ISL_563865, EPI_ISL_563867, EPI_ISL_563868, EPI_ISL_563869, EPI_ISL_563870, EPI_ISL_563871, EPI_ISL_563872, EPI_ISL_563873, EPI_ISL_563874, EPI_ISL_563875, EPI_ISL_563877, EPI_ISL_563878, EPI_ISL_563882, EPI_ISL_563884, EPI_ISL_563901, EPI_ISL_563927, EPI_ISL_563929 | | | |
| see above | Microbiological Diagnostic Unit - Public Health Laboratory (MDU-PHL) | MDU-PHL | Seemann, T., Schultz M. B., Sait, M., Sherry, N. |
| EPI_ISL_563938 | Victorian Infectious Diseases Reference Laboratory (VIDRL) | VIDRL and MDU-PHL | Caly, L., Seemann, T., Sait, M., Schultz, M. B., Druce J., Sherry, N. |
| EPI_ISL_563939, EPI_ISL_563941, EPI_ISL_563942, EPI_ISL_563945, EPI_ISL_563948, EPI_ISL_563949, EPI_ISL_563950, EPI_ISL_563951, EPI_ISL_563954, EPI_ISL_563955, EPI_ISL_563956, EPI_ISL_563957, EPI_ISL_563959, EPI_ISL_563960, EPI_ISL_563961, EPI_ISL_563964 | | | |

| | | | |
|---|---|---|---|
| see above | Microbiological Diagnostic Unit - Public Health Laboratory (MDU-PHL) | MDU-PHL | Seemann, T., Schultz M. B., Sait, M., Sherry, N. |
| EPI_ISL_563973, EPI_ISL_563974, EPI_ISL_563975 | Victorian Infectious Diseases Reference Laboratory (VIDRL) | VIDRL and MDU-PHL | Caly, L., Seemann, T., Sait, M., Schultz, M. B., Druce J., Sherry, N. |
| EPI_ISL_566110, EPI_ISL_566111, EPI_ISL_566112, EPI_ISL_566113, EPI_ISL_566114, EPI_ISL_566115, EPI_ISL_566116, EPI_ISL_566117, EPI_ISL_566118, EPI_ISL_566119, EPI_ISL_566120, EPI_ISL_566121, EPI_ISL_566122, EPI_ISL_566123, EPI_ISL_566124, EPI_ISL_566125, EPI_ISL_566126, EPI_ISL_566127, EPI_ISL_566128, EPI_ISL_566129, EPI_ISL_566130, EPI_ISL_566131, EPI_ISL_566132, EPI_ISL_566133, EPI_ISL_566134, EPI_ISL_566135, EPI_ISL_566136, EPI_ISL_566137, EPI_ISL_566138, EPI_ISL_566139, EPI_ISL_566140, EPI_ISL_566141, EPI_ISL_566142, EPI_ISL_566143, EPI_ISL_566144, EPI_ISL_566145, EPI_ISL_566146, EPI_ISL_566147, EPI_ISL_566148, EPI_ISL_566149, EPI_ISL_566150, EPI_ISL_566151, EPI_ISL_566152, EPI_ISL_566153, EPI_ISL_566154, EPI_ISL_566155, EPI_ISL_566156, EPI_ISL_566157, EPI_ISL_566158, EPI_ISL_566159, EPI_ISL_566160, EPI_ISL_566161, EPI_ISL_566162, EPI_ISL_566163, EPI_ISL_566164, EPI_ISL_566165, EPI_ISL_566166, EPI_ISL_566167, EPI_ISL_566168, EPI_ISL_566169, EPI_ISL_566170, EPI_ISL_566171, EPI_ISL_566172, EPI_ISL_566173, EPI_ISL_566174, EPI_ISL_566175, EPI_ISL_566176, EPI_ISL_566177, EPI_ISL_566178, EPI_ISL_566179, EPI_ISL_566180, EPI_ISL_566181, EPI_ISL_566182, EPI_ISL_566183, EPI_ISL_566184, EPI_ISL_566185, EPI_ISL_566186, EPI_ISL_566187, EPI_ISL_566188, EPI_ISL_566189, EPI_ISL_566190, EPI_ISL_566191, EPI_ISL_566192, EPI_ISL_566193, EPI_ISL_566194, EPI_ISL_566195, EPI_ISL_566196, EPI_ISL_566197, EPI_ISL_566198, EPI_ISL_566199, EPI_ISL_566200, EPI_ISL_566201, EPI_ISL_566202, EPI_ISL_566203, EPI_ISL_566204, EPI_ISL_566205, EPI_ISL_566206, EPI_ISL_566207, EPI_ISL_566208, EPI_ISL_566209, EPI_ISL_566210, EPI_ISL_566211, EPI_ISL_566212, EPI_ISL_566213, EPI_ISL_566214, EPI_ISL_566215, EPI_ISL_566216, EPI_ISL_566217, EPI_ISL_566218, EPI_ISL_566219, EPI_ISL_566220, EPI_ISL_566221, EPI_ISL_566222, EPI_ISL_566223, EPI_ISL_566224, EPI_ISL_566225, EPI_ISL_566226, EPI_ISL_566227, EPI_ISL_566228, EPI_ISL_566229, EPI_ISL_566230, EPI_ISL_566231, EPI_ISL_566232, EPI_ISL_566233, EPI_ISL_566234, EPI_ISL_566235, EPI_ISL_566236, EPI_ISL_566237, EPI_ISL_566238, EPI_ISL_566239, EPI_ISL_566240, EPI_ISL_566241, EPI_ISL_566242, EPI_ISL_566243, EPI_ISL_566244, EPI_ISL_566245, EPI_ISL_566246, EPI_ISL_566247, EPI_ISL_566248, EPI_ISL_566249, EPI_ISL_566250, EPI_ISL_566251, EPI_ISL_566252, EPI_ISL_566253, EPI_ISL_566254, EPI_ISL_566255, EPI_ISL_566256, EPI_ISL_566257, EPI_ISL_566258, EPI_ISL_566259, EPI_ISL_566260, EPI_ISL_566261, EPI_ISL_566262, EPI_ISL_566263, EPI_ISL_566264, EPI_ISL_566265, EPI_ISL_566266, EPI_ISL_566267, EPI_ISL_566268, EPI_ISL_566269, EPI_ISL_566270, EPI_ISL_566271, EPI_ISL_566272, EPI_ISL_566273, EPI_ISL_566274, EPI_ISL_566275, EPI_ISL_566276, EPI_ISL_566277, EPI_ISL_566278, EPI_ISL_566279, EPI_ISL_566280, EPI_ISL_566281, EPI_ISL_566282, EPI_ISL_566283, EPI_ISL_566284, EPI_ISL_566285, EPI_ISL_566286, EPI_ISL_566287, EPI_ISL_566288, EPI_ISL_566289, EPI_ISL_566290, EPI_ISL_566291, EPI_ISL_566292, EPI_ISL_566293, EPI_ISL_566294, EPI_ISL_566295, EPI_ISL_566296, EPI_ISL_566297, EPI_ISL_566298, EPI_ISL_566299, EPI_ISL_566300, EPI_ISL_566301, EPI_ISL_566302, EPI_ISL_566303, EPI_ISL_566304, EPI_ISL_566305, EPI_ISL_566306, EPI_ISL_566307, EPI_ISL_566308, EPI_ISL_566309, EPI_ISL_566310, EPI_ISL_566311, EPI_ISL_566312, EPI_ISL_566313, EPI_ISL_566314, EPI_ISL_566315, EPI_ISL_566316, EPI_ISL_566317, EPI_ISL_566318, EPI_ISL_566319, EPI_ISL_566320, EPI_ISL_566321, EPI_ISL_566322, EPI_ISL_566323, EPI_ISL_566324, EPI_ISL_566325, EPI_ISL_566326, EPI_ISL_566327, EPI_ISL_566328, EPI_ISL_566329, EPI_ISL_566330, EPI_ISL_566331, EPI_ISL_566332, EPI_ISL_566333, EPI_ISL_566334, EPI_ISL_566335, EPI_ISL_566336, EPI_ISL_566337, EPI_ISL_566338, EPI_ISL_566339, EPI_ISL_566340, EPI_ISL_566341, EPI_ISL_566342, EPI_ISL_566343, EPI_ISL_566344, EPI_ISL_566345, EPI_ISL_566346, EPI_ISL_566347, EPI_ISL_566348, EPI_ISL_566349, EPI_ISL_566350, EPI_ISL_566351, EPI_ISL_566352, EPI_ISL_566353, EPI_ISL_566354, EPI_ISL_566355, EPI_ISL_566356, EPI_ISL_566357, EPI_ISL_566358, EPI_ISL_566359, EPI_ISL_566360, EPI_ISL_566361, EPI_ISL_566362, EPI_ISL_566363, EPI_ISL_566364, EPI_ISL_566365, EPI_ISL_566366, EPI_ISL_566367, EPI_ISL_566368, EPI_ISL_566369, EPI_ISL_566370, EPI_ISL_566371, EPI_ISL_566372, EPI_ISL_566373, EPI_ISL_566374, EPI_ISL_566375, EPI_ISL_566376, EPI_ISL_566377, EPI_ISL_566378, EPI_ISL_566379, EPI_ISL_566380, EPI_ISL_566381, EPI_ISL_566382, EPI_ISL_566383, EPI_ISL_566384, EPI_ISL_566385, EPI_ISL_566386, EPI_ISL_566387, EPI_ISL_566388, EPI_ISL_566389, EPI_ISL_566390, EPI_ISL_566391, EPI_ISL_566392, EPI_ISL_566393, EPI_ISL_566394, EPI_ISL_566395, EPI_ISL_566396, EPI_ISL_566397, EPI_ISL_566398, EPI_ISL_566399, EPI_ISL_566400, EPI_ISL_566401, EPI_ISL_566402, EPI_ISL_566403, EPI_ISL_566404, EPI_ISL_566405, EPI_ISL_566406, EPI_ISL_566407, EPI_ISL_566408, EPI_ISL_566409, EPI_ISL_566410, EPI_ISL_566411, EPI_ISL_566412, EPI_ISL_566413, EPI_ISL_566414, EPI_ISL_566415, EPI_ISL_566416, EPI_ISL_566417, EPI_ISL_566418, EPI_ISL_566419, EPI_ISL_566420 | | | |
| see above | Lighthouse Lab in Alderley Park | Wellcome Sanger Institute for the COVID-19 Genomics UK (COG-UK) consortium | Jacquelyn Wynn, Mairead Hyland, The Lighthouse Lab in Alderley Park and Alex Alderton, Roberto Amato, Sonia Goncalves, Ewan Harrison, David K. Jackson, Ian Johnston, Dominic Kwiatkowski, Cordelia Langford, John Sillitoe on behalf of the Wellcome Sanger Institute COVID-19 Surveillance Team |
| EPI_ISL_566421, EPI_ISL_566424, EPI_ISL_566425, EPI_ISL_566426, EPI_ISL_566430, EPI_ISL_566431, EPI_ISL_566432, EPI_ISL_566433, EPI_ISL_566434, EPI_ISL_566435, EPI_ISL_566436, EPI_ISL_566437, EPI_ISL_566438, EPI_ISL_566439, EPI_ISL_566440, EPI_ISL_566441, EPI_ISL_566442, EPI_ISL_566443, EPI_ISL_566446, EPI_ISL_566448, EPI_ISL_566449, EPI_ISL_566450, EPI_ISL_566452, EPI_ISL_566458, EPI_ISL_566459, EPI_ISL_566461, EPI_ISL_566462, EPI_ISL_566463, EPI_ISL_566464, EPI_ISL_566466, EPI_ISL_566468, EPI_ISL_566470, EPI_ISL_566471, EPI_ISL_566472, EPI_ISL_566475, EPI_ISL_566476, EPI_ISL_566477, EPI_ISL_566478, EPI_ISL_566480, EPI_ISL_566481, EPI_ISL_566482, EPI_ISL_566483, EPI_ISL_566484, EPI_ISL_566486, EPI_ISL_566487, EPI_ISL_566488, EPI_ISL_566489, EPI_ISL_566491, EPI_ISL_566496, EPI_ISL_566497, EPI_ISL_566498, EPI_ISL_566499, EPI_ISL_566501, EPI_ISL_566503, EPI_ISL_566505, EPI_ISL_566506, EPI_ISL_566507, EPI_ISL_566509, EPI_ISL_566511, EPI_ISL_566512, EPI_ISL_566513, EPI_ISL_566514, EPI_ISL_566516, EPI_ISL_566517, EPI_ISL_566520, EPI_ISL_566521, EPI_ISL_566522, EPI_ISL_566523, EPI_ISL_566525, EPI_ISL_566526, EPI_ISL_566527, EPI_ISL_566529, EPI_ISL_566532, EPI_ISL_566533, EPI_ISL_566535, EPI_ISL_566536, EPI_ISL_566538, EPI_ISL_566539, EPI_ISL_566540, EPI_ISL_566541, EPI_ISL_566542, EPI_ISL_566545, EPI_ISL_566546, EPI_ISL_566549, EPI_ISL_566550, EPI_ISL_566551, EPI_ISL_566552, EPI_ISL_566553, EPI_ISL_566554, EPI_ISL_566555, EPI_ISL_566556, EPI_ISL_566557, EPI_ISL_566558, EPI_ISL_566561, EPI_ISL_566562, EPI_ISL_566563, EPI_ISL_566564, EPI_ISL_566565, EPI_ISL_566566, EPI_ISL_566568, EPI_ISL_566569, EPI_ISL_566570, EPI_ISL_566571, EPI_ISL_566572, EPI_ISL_566574, EPI_ISL_566577, EPI_ISL_566578, EPI_ISL_566581, EPI_ISL_566584, EPI_ISL_566588, EPI_ISL_566590, EPI_ISL_566592, EPI_ISL_566593, EPI_ISL_566594, EPI_ISL_566595, EPI_ISL_566596, EPI_ISL_566597, EPI_ISL_566598, EPI_ISL_566599, EPI_ISL_566600, EPI_ISL_566601, EPI_ISL_566602, EPI_ISL_566604, EPI_ISL_566606, EPI_ISL_566607, EPI_ISL_566613, EPI_ISL_566614, EPI_ISL_566615, EPI_ISL_566616, EPI_ISL_566617, EPI_ISL_566618, EPI_ISL_566619, EPI_ISL_566620, EPI_ISL_566621, EPI_ISL_566623, EPI_ISL_566624, EPI_ISL_566627, EPI_ISL_566629, EPI_ISL_566630, EPI_ISL_566632, EPI_ISL_566633, EPI_ISL_566634, EPI_ISL_566635, EPI_ISL_566636, EPI_ISL_566639, EPI_ISL_566640, EPI_ISL_566641, EPI_ISL_566644, EPI_ISL_566646, EPI_ISL_566647, EPI_ISL_566648, EPI_ISL_566649, EPI_ISL_566650, EPI_ISL_566651, EPI_ISL_566652, EPI_ISL_566653, EPI_ISL_566654, EPI_ISL_566655, EPI_ISL_566656, EPI_ISL_566660, EPI_ISL_566663, EPI_ISL_566664, EPI_ISL_566668, EPI_ISL_566669, EPI_ISL_566670, EPI_ISL_566671, EPI_ISL_566678, EPI_ISL_566679, EPI_ISL_566684, EPI_ISL_566686, EPI_ISL_566689, EPI_ISL_566691, EPI_ISL_566695, EPI_ISL_566696, EPI_ISL_566702, EPI_ISL_566704, EPI_ISL_566706, EPI_ISL_566709, EPI_ISL_566712, EPI_ISL_566714, EPI_ISL_566715, EPI_ISL_566716, EPI_ISL_566717, EPI_ISL_566718, EPI_ISL_566719, EPI_ISL_566720, EPI_ISL_566722, EPI_ISL_566723, EPI_ISL_566724, EPI_ISL_566733, EPI_ISL_566736, EPI_ISL_566737, EPI_ISL_566738, EPI_ISL_566739, EPI_ISL_566743, EPI_ISL_566745, EPI_ISL_566747, EPI_ISL_566750, EPI_ISL_566753, EPI_ISL_566754, EPI_ISL_566756, EPI_ISL_566758, EPI_ISL_566760, EPI_ISL_566761, EPI_ISL_566762, EPI_ISL_566766, EPI_ISL_566767, EPI_ISL_566769, EPI_ISL_566770, EPI_ISL_566779, EPI_ISL_566783, EPI_ISL_566785, EPI_ISL_566786, EPI_ISL_566787, EPI_ISL_566789, EPI_ISL_566790, EPI_ISL_566792, EPI_ISL_566802, EPI_ISL_566804, EPI_ISL_566806, EPI_ISL_566808, EPI_ISL_566810, EPI_ISL_566812, EPI_ISL_566815, EPI_ISL_566816, EPI_ISL_566817, EPI_ISL_566819, EPI_ISL_566821, EPI_ISL_566822, EPI_ISL_566823, EPI_ISL_566824, EPI_ISL_566826, EPI_ISL_566832, EPI_ISL_566833, EPI_ISL_566834, EPI_ISL_566840, EPI_ISL_566841, EPI_ISL_566842, EPI_ISL_566844, EPI_ISL_566845, EPI_ISL_566848, EPI_ISL_566849, EPI_ISL_566851, EPI_ISL_566858, EPI_ISL_566859, EPI_ISL_566860, EPI_ISL_566861, EPI_ISL_566862, EPI_ISL_566863, EPI_ISL_566867, EPI_ISL_566868, EPI_ISL_566872, EPI_ISL_566873, EPI_ISL_566876, EPI_ISL_566881, EPI_ISL_566886, EPI_ISL_566888, EPI_ISL_566895, EPI_ISL_566897, EPI_ISL_566898, EPI_ISL_566899, EPI_ISL_566903, EPI_ISL_566904, EPI_ISL_566905, EPI_ISL_566910, EPI_ISL_566911, EPI_ISL_566914, EPI_ISL_566915, EPI_ISL_566916, EPI_ISL_566922, EPI_ISL_566925, EPI_ISL_566927, EPI_ISL_566932, EPI_ISL_566933, EPI_ISL_566937, EPI_ISL_566940, EPI_ISL_566945, EPI_ISL_566947, EPI_ISL_566950, EPI_ISL_566953, EPI_ISL_566957, EPI_ISL_566959, EPI_ISL_566964, EPI_ISL_566965, EPI_ISL_566969, EPI_ISL_566971, EPI_ISL_566972, EPI_ISL_566977, EPI_ISL_566992, EPI_ISL_566994, EPI_ISL_566996 | | | |
| see above | Lighthouse Lab in Milton Keynes | Wellcome Sanger Institute for the COVID-19 Genomics UK (COG-UK) consortium | The Lighthouse Lab in Milton Keynes and Alex Alderton, Roberto Amato, Sonia Goncalves, Ewan Harrison, David K. Jackson, Ian Johnston, Dominic Kwiatkowski, Cordelia Langford, John Sillitoe on behalf of the Wellcome Sanger Institute COVID-19 Surveillance Team |
| EPI_ISL_566999, EPI_ISL_567000, EPI_ISL_567001, EPI_ISL_567002, EPI_ISL_567003, EPI_ISL_567004, EPI_ISL_567005, EPI_ISL_567006, EPI_ISL_567007, EPI_ISL_567008, EPI_ISL_567009, EPI_ISL_567010, EPI_ISL_567011, EPI_ISL_567012, EPI_ISL_567013, EPI_ISL_567014, EPI_ISL_567015, EPI_ISL_567016, EPI_ISL_567017, EPI_ISL_567018, EPI_ISL_567019, EPI_ISL_567020, EPI_ISL_567021, EPI_ISL_567022, EPI_ISL_567023 | | | |
| see above | Lighthouse Lab in Glasgow | Wellcome Sanger Institute for the COVID-19 Genomics UK (COG-UK) consortium | Harper VanSteenhouse, Yumi Kasai, David Gray, Carol Clugston, Anna Dominiczak and Alex Alderton, Roberto Amato, Sonia Goncalves, Ewan Harrison, David K. Jackson, Ian Johnston, Dominic Kwiatkowski, Cordelia Langford, John Sillitoe on behalf of the Wellcome Sanger Institute COVID-19 Surveillance Team |
| EPI_ISL_567024 | Lighthouse Lab in Glasgow | Wellcome Sanger Institute for the COVID-19 Genomics UK (COG-UK) Consortium | Harper VanSteenhouse, Yumi Kasai, David Gray, Carol Clugston, Anna Dominiczak and Alex Alderton, Roberto Amato, Sonia Goncalves, Ewan Harrison, David K. Jackson, Ian Johnston, Dominic Kwiatkowski, Cordelia Langford, John Sillitoe on behalf of the Wellcome Sanger Institute COVID-19 Surveillance Team |
| EPI_ISL_567025, EPI_ISL_567026, EPI_ISL_567027, EPI_ISL_567028, EPI_ISL_567029, EPI_ISL_567030, EPI_ISL_567031, EPI_ISL_567032, EPI_ISL_567033, EPI_ISL_567034, EPI_ISL_567035, EPI_ISL_567036, EPI_ISL_567037, EPI_ISL_567038, EPI_ISL_567039, EPI_ISL_567040, EPI_ISL_567041, EPI_ISL_567042, EPI_ISL_567043, EPI_ISL_567044, EPI_ISL_567045, EPI_ISL_567046, EPI_ISL_567047, EPI_ISL_567048, EPI_ISL_567049, EPI_ISL_567050, EPI_ISL_567051, EPI_ISL_567052, EPI_ISL_567053, EPI_ISL_567054, EPI_ISL_567055, EPI_ISL_567056, EPI_ISL_567057, EPI_ISL_567058 | | | |
| see above | Lighthouse Lab in Glasgow | Wellcome Sanger Institute for the COVID-19 Genomics UK (COG-UK) consortium | Harper VanSteenhouse, Yumi Kasai, David Gray, Carol Clugston, Anna Dominiczak and Alex Alderton, Roberto Amato, Sonia Goncalves, Ewan Harrison, David K. Jackson, Ian Johnston, Dominic Kwiatkowski, Cordelia Langford, John Sillitoe on behalf of the Wellcome Sanger Institute COVID-19 Surveillance Team |
| EPI_ISL_567059 | Lighthouse Lab in Glasgow | Wellcome Sanger Institute for the COVID-19 Genomics UK (COG-UK) Consortium | Harper VanSteenhouse, Yumi Kasai, David Gray, Carol Clugston, Anna Dominiczak and Alex Alderton, Roberto Amato, Sonia Goncalves, Ewan Harrison, David K. Jackson, Ian Johnston, Dominic Kwiatkowski, Cordelia Langford, John Sillitoe on behalf of the Wellcome Sanger Institute COVID-19 Surveillance Team |
| EPI_ISL_567060, EPI_ISL_567061, EPI_ISL_567062, EPI_ISL_567063, EPI_ISL_567064, EPI_ISL_567066, EPI_ISL_567067, EPI_ISL_567068, EPI_ISL_567069, EPI_ISL_567070, EPI_ISL_567071, EPI_ISL_567072, EPI_ISL_567073, EPI_ISL_567074, EPI_ISL_567075, EPI_ISL_567076, EPI_ISL_567077, EPI_ISL_567078, EPI_ISL_567079, EPI_ISL_567081, EPI_ISL_567082, EPI_ISL_567083, EPI_ISL_567084, EPI_ISL_567085, EPI_ISL_567086, EPI_ISL_567087, EPI_ISL_567088, EPI_ISL_567089, EPI_ISL_567090, EPI_ISL_567091, EPI_ISL_567092, EPI_ISL_567093, EPI_ISL_567094, EPI_ISL_567095, EPI_ISL_567096, EPI_ISL_567097, EPI_ISL_567098, EPI_ISL_567099, EPI_ISL_567100, EPI_ISL_567101, EPI_ISL_567102, EPI_ISL_567103, EPI_ISL_567104, EPI_ISL_567105, EPI_ISL_567106, EPI_ISL_567107, EPI_ISL_567108, EPI_ISL_567109, EPI_ISL_567110, EPI_ISL_567111, EPI_ISL_567112, EPI_ISL_567113, EPI_ISL_567114, EPI_ISL_567116, EPI_ISL_567117, EPI_ISL_567118, EPI_ISL_567119, EPI_ISL_567120, EPI_ISL_567121, EPI_ISL_567122, EPI_ISL_567123, EPI_ISL_567124, EPI_ISL_567125, EPI_ISL_567126, EPI_ISL_567127, EPI_ISL_567128, EPI_ISL_567129, EPI_ISL_567130, EPI_ISL_567131, EPI_ISL_567132, EPI_ISL_567133, EPI_ISL_567134, EPI_ISL_567135, EPI_ISL_567136, EPI_ISL_567137, EPI_ISL_567138, EPI_ISL_567139, EPI_ISL_567140, EPI_ISL_567141, EPI_ISL_567142, EPI_ISL_567143, EPI_ISL_567144, EPI_ISL_567145, EPI_ISL_567146, EPI_ISL_567147, EPI_ISL_567148, EPI_ISL_567149, EPI_ISL_567151, EPI_ISL_567152, EPI_ISL_567153, EPI_ISL_567154, EPI_ISL_567155, EPI_ISL_567156, EPI_ISL_567157, EPI_ISL_567158, EPI_ISL_567159, EPI_ISL_567160, EPI_ISL_567161, EPI_ISL_567162, EPI_ISL_567163, EPI_ISL_567164, EPI_ISL_567166, EPI_ISL_567167, EPI_ISL_567168, EPI_ISL_567169, EPI_ISL_567170, EPI_ISL_567171, EPI_ISL_567172, EPI_ISL_567173, EPI_ISL_567174, EPI_ISL_567175, EPI_ISL_567176, EPI_ISL_567177, EPI_ISL_567178, EPI_ISL_567179, EPI_ISL_567180, EPI_ISL_567181, EPI_ISL_567182, EPI_ISL_567183 | | | |
| see above | Lighthouse Lab in Glasgow | Wellcome Sanger Institute for the COVID-19 Genomics UK (COG-UK) consortium | Harper VanSteenhouse, Yumi Kasai, David Gray, Carol Clugston, Anna Dominiczak and Alex Alderton, Roberto Amato, Sonia Goncalves, Ewan Harrison, David K. Jackson, Ian Johnston, Dominic Kwiatkowski, Cordelia Langford, John Sillitoe on behalf of the Wellcome Sanger Institute COVID-19 Surveillance Team |
| EPI_ISL_567184 | Lighthouse Lab in Glasgow | Wellcome Sanger Institute for the COVID-19 Genomics UK (COG-UK) Consortium | Harper VanSteenhouse, Yumi Kasai, David Gray, Carol Clugston, Anna Dominiczak and Alex Alderton, Roberto Amato, Sonia Goncalves, Ewan Harrison, David K. Jackson, Ian Johnston, Dominic Kwiatkowski, Cordelia Langford, John Sillitoe on behalf of the Wellcome Sanger Institute COVID-19 Surveillance Team |
| EPI_ISL_567185, EPI_ISL_567186, EPI_ISL_567187, EPI_ISL_567188, EPI_ISL_567189, EPI_ISL_567190, EPI_ISL_567191, EPI_ISL_567192, EPI_ISL_567193, EPI_ISL_567194, EPI_ISL_567195, EPI_ISL_567196, EPI_ISL_567197, EPI_ISL_567198, EPI_ISL_567199, EPI_ISL_567200, EPI_ISL_567201 | | | |
| see above | Lighthouse Lab in Glasgow | Wellcome Sanger Institute for the COVID-19 Genomics | Harper VanSteenhouse, Yumi Kasai, David Gray, Carol Clugston, Anna Dominiczak and Alex Alderton, Roberto Amato, Sonia Goncalves, Ewan Harrison, David K. Jackson, Ian Johnston, Dominic Kwiatkowski, Cordelia Langford, John Sillitoe on behalf of the Wellcome Sanger Institute COVID-19 Surveillance Team |

| | | | |
|---|---|---|---|
| | | UK (COG-UK) consortium | |
| EPI_ISL_567202 | Lighthouse Lab in Glasgow | Wellcome Sanger Institute for the COVID-19 Genomics UK (COG-UK) Consortium | Harper VanSteenhouse, Yumi Kasai, David Gray, Carol Clugston, Anna Dominiczak and Alex Alderton, Roberto Amato, Sonia Goncalves, Ewan Harrison, David K. Jackson, Ian Johnston, Dominic Kwiatkowski, Cordelia Langford, John Sillitoe on behalf of the Wellcome Sanger Institute COVID-19 Surveillance Team |
| EPI_ISL_567203, EPI_ISL_567204, EPI_ISL_567205, EPI_ISL_567206, EPI_ISL_567207, EPI_ISL_567208, EPI_ISL_567209, EPI_ISL_567210, EPI_ISL_567211, EPI_ISL_567212, EPI_ISL_567213, EPI_ISL_567214, EPI_ISL_567215, EPI_ISL_567216, EPI_ISL_567217, EPI_ISL_567218, EPI_ISL_567219, EPI_ISL_567220, EPI_ISL_567221, EPI_ISL_567222, EPI_ISL_567223, EPI_ISL_567224, EPI_ISL_567225, EPI_ISL_567226, EPI_ISL_567227, EPI_ISL_567228, EPI_ISL_567229, EPI_ISL_567230 | | | |
| see above | Lighthouse Lab in Glasgow | Wellcome Sanger Institute for the COVID-19 Genomics UK (COG-UK) consortium | Harper VanSteenhouse, Yumi Kasai, David Gray, Carol Clugston, Anna Dominiczak and Alex Alderton, Roberto Amato, Sonia Goncalves, Ewan Harrison, David K. Jackson, Ian Johnston, Dominic Kwiatkowski, Cordelia Langford, John Sillitoe on behalf of the Wellcome Sanger Institute COVID-19 Surveillance Team |
| EPI_ISL_567231 | Lighthouse Lab in Glasgow | Wellcome Sanger Institute for the COVID-19 Genomics UK (COG-UK) Consortium | Harper VanSteenhouse, Yumi Kasai, David Gray, Carol Clugston, Anna Dominiczak and Alex Alderton, Roberto Amato, Sonia Goncalves, Ewan Harrison, David K. Jackson, Ian Johnston, Dominic Kwiatkowski, Cordelia Langford, John Sillitoe on behalf of the Wellcome Sanger Institute COVID-19 Surveillance Team |
| EPI_ISL_567232, EPI_ISL_567233, EPI_ISL_567234, EPI_ISL_567235, EPI_ISL_567236, EPI_ISL_567237, EPI_ISL_567238, EPI_ISL_567239, EPI_ISL_567240, EPI_ISL_567241, EPI_ISL_567242, EPI_ISL_567244, EPI_ISL_567245, EPI_ISL_567246, EPI_ISL_567247, EPI_ISL_567249, EPI_ISL_567251, EPI_ISL_567252, EPI_ISL_567253, EPI_ISL_567255, EPI_ISL_567256, EPI_ISL_567257, EPI_ISL_567258, EPI_ISL_567259, EPI_ISL_567261, EPI_ISL_567262, EPI_ISL_567263, EPI_ISL_567264, EPI_ISL_567265, EPI_ISL_567266, EPI_ISL_567267, EPI_ISL_567268, EPI_ISL_567269, EPI_ISL_567270, EPI_ISL_567271, EPI_ISL_567272, EPI_ISL_567273, EPI_ISL_567275, EPI_ISL_567276, EPI_ISL_567277, EPI_ISL_567278, EPI_ISL_567279, EPI_ISL_567280, EPI_ISL_567281, EPI_ISL_567282, EPI_ISL_567283, EPI_ISL_567284, EPI_ISL_567285, EPI_ISL_567286, EPI_ISL_567287, EPI_ISL_567288, EPI_ISL_567290, EPI_ISL_567291, EPI_ISL_567292, EPI_ISL_567293, EPI_ISL_567295, EPI_ISL_567296, EPI_ISL_567297, EPI_ISL_567298, EPI_ISL_567299, EPI_ISL_567300, EPI_ISL_567301, EPI_ISL_567302, EPI_ISL_567303, EPI_ISL_567304, EPI_ISL_567305, EPI_ISL_567306, EPI_ISL_567307, EPI_ISL_567308, EPI_ISL_567309, EPI_ISL_567310, EPI_ISL_567311, EPI_ISL_567312, EPI_ISL_567313, EPI_ISL_567314, EPI_ISL_567315, EPI_ISL_567316, EPI_ISL_567317, EPI_ISL_567318, EPI_ISL_567319, EPI_ISL_567320, EPI_ISL_567321, EPI_ISL_567322, EPI_ISL_567323, EPI_ISL_567324, EPI_ISL_567325, EPI_ISL_567326, EPI_ISL_567327, EPI_ISL_567328, EPI_ISL_567329, EPI_ISL_567330, EPI_ISL_567331, EPI_ISL_567332, EPI_ISL_567333, EPI_ISL_567334, EPI_ISL_567335, EPI_ISL_567336, EPI_ISL_567337, EPI_ISL_567339, EPI_ISL_567340, EPI_ISL_567341, EPI_ISL_567342, EPI_ISL_567343, EPI_ISL_567344, EPI_ISL_567345, EPI_ISL_567346, EPI_ISL_567347, EPI_ISL_567348, EPI_ISL_567349, EPI_ISL_567350, EPI_ISL_567351, EPI_ISL_567352, EPI_ISL_567353, EPI_ISL_567354, EPI_ISL_567356, EPI_ISL_567357, EPI_ISL_567359, EPI_ISL_567360, EPI_ISL_567361, EPI_ISL_567362, EPI_ISL_567364, EPI_ISL_567365, EPI_ISL_567366, EPI_ISL_567367, EPI_ISL_567368, EPI_ISL_567369, EPI_ISL_567370, EPI_ISL_567371, EPI_ISL_567372, EPI_ISL_567373, EPI_ISL_567374, EPI_ISL_567376, EPI_ISL_567377, EPI_ISL_567378, EPI_ISL_567379, EPI_ISL_567380, EPI_ISL_567381, EPI_ISL_567383, EPI_ISL_567384, EPI_ISL_567385, EPI_ISL_567386, EPI_ISL_567387, EPI_ISL_567388, EPI_ISL_567389, EPI_ISL_567391, EPI_ISL_567392, EPI_ISL_567393, EPI_ISL_567394, EPI_ISL_567395, EPI_ISL_567397, EPI_ISL_567399, EPI_ISL_567400, EPI_ISL_567401, EPI_ISL_567402, EPI_ISL_567404, EPI_ISL_567405, EPI_ISL_567406, EPI_ISL_567407, EPI_ISL_567408, EPI_ISL_567409, EPI_ISL_567410, EPI_ISL_567411, EPI_ISL_567412, EPI_ISL_567413, EPI_ISL_567414, EPI_ISL_567415, EPI_ISL_567416, EPI_ISL_567417, EPI_ISL_567418, EPI_ISL_567419, EPI_ISL_567420, EPI_ISL_567421, EPI_ISL_567422, EPI_ISL_567423, EPI_ISL_567424, EPI_ISL_567425, EPI_ISL_567426, EPI_ISL_567428, EPI_ISL_567430, EPI_ISL_567431, EPI_ISL_567432, EPI_ISL_567433, EPI_ISL_567434, EPI_ISL_567436, EPI_ISL_567437, EPI_ISL_567438, EPI_ISL_567439, EPI_ISL_567440, EPI_ISL_567441, EPI_ISL_567442, EPI_ISL_567443, EPI_ISL_567444, EPI_ISL_567445, EPI_ISL_567447, EPI_ISL_567448, EPI_ISL_567449, EPI_ISL_567450, EPI_ISL_567451, EPI_ISL_567452, EPI_ISL_567454, EPI_ISL_567455, EPI_ISL_567456, EPI_ISL_567457, EPI_ISL_567458, EPI_ISL_567459, EPI_ISL_567460, EPI_ISL_567461, EPI_ISL_567462, EPI_ISL_567463, EPI_ISL_567465, EPI_ISL_567466, EPI_ISL_567467, EPI_ISL_567468, EPI_ISL_567469, EPI_ISL_567470, EPI_ISL_567471, EPI_ISL_567472, EPI_ISL_567474, EPI_ISL_567475, EPI_ISL_567476, EPI_ISL_567477, EPI_ISL_567478, EPI_ISL_567479, EPI_ISL_567480, EPI_ISL_567481, EPI_ISL_567483, EPI_ISL_567484, EPI_ISL_567485, EPI_ISL_567486, EPI_ISL_567487, EPI_ISL_567488, EPI_ISL_567490, EPI_ISL_567491, EPI_ISL_567492, EPI_ISL_567493, EPI_ISL_567494, EPI_ISL_567495, EPI_ISL_567496, EPI_ISL_567497, EPI_ISL_567498, EPI_ISL_567499, EPI_ISL_567500, EPI_ISL_567502, EPI_ISL_567503, EPI_ISL_567504, EPI_ISL_567505, EPI_ISL_567506, EPI_ISL_567507, EPI_ISL_567508, EPI_ISL_567509, EPI_ISL_567510, EPI_ISL_567511, EPI_ISL_567512, EPI_ISL_567513, EPI_ISL_567514, EPI_ISL_567515, EPI_ISL_567516, EPI_ISL_567517, EPI_ISL_567518, EPI_ISL_567519, EPI_ISL_567520, EPI_ISL_567521, EPI_ISL_567523, EPI_ISL_567525, EPI_ISL_567526, EPI_ISL_567527, EPI_ISL_567528, EPI_ISL_567529, EPI_ISL_567530, EPI_ISL_567531, EPI_ISL_567532 | | | |
| see above | Lighthouse Lab in Glasgow | Wellcome Sanger Institute for the COVID-19 Genomics UK (COG-UK) consortium | Harper VanSteenhouse, Yumi Kasai, David Gray, Carol Clugston, Anna Dominiczak and Alex Alderton, Roberto Amato, Sonia Goncalves, Ewan Harrison, David K. Jackson, Ian Johnston, Dominic Kwiatkowski, Cordelia Langford, John Sillitoe on behalf of the Wellcome Sanger Institute COVID-19 Surveillance Team |
| EPI_ISL_567533, EPI_ISL_567534, EPI_ISL_567535, EPI_ISL_567536, EPI_ISL_567537, EPI_ISL_567538, EPI_ISL_567539, EPI_ISL_567540, EPI_ISL_567541, EPI_ISL_567542, EPI_ISL_567543, EPI_ISL_567544, EPI_ISL_567545, EPI_ISL_567546, EPI_ISL_567547, EPI_ISL_567548, EPI_ISL_567549, EPI_ISL_567550, EPI_ISL_567551, EPI_ISL_567552, EPI_ISL_567553, EPI_ISL_567554, EPI_ISL_567555, EPI_ISL_567556, EPI_ISL_567557, EPI_ISL_567558, EPI_ISL_567559, EPI_ISL_567560, EPI_ISL_567561, EPI_ISL_567562, EPI_ISL_567564, EPI_ISL_567565, EPI_ISL_567566, EPI_ISL_567567, EPI_ISL_567568, EPI_ISL_567569, EPI_ISL_567570, EPI_ISL_567571, EPI_ISL_567572, EPI_ISL_567573, EPI_ISL_567574, EPI_ISL_567575, EPI_ISL_567576, EPI_ISL_567577, EPI_ISL_567578, EPI_ISL_567579, EPI_ISL_567580, EPI_ISL_567581, EPI_ISL_567582, EPI_ISL_567583, EPI_ISL_567584, EPI_ISL_567585, EPI_ISL_567586, EPI_ISL_567587, EPI_ISL_567588, EPI_ISL_567589, EPI_ISL_567590, EPI_ISL_567591, EPI_ISL_567592, EPI_ISL_567593, EPI_ISL_567594, EPI_ISL_567595, EPI_ISL_567596, EPI_ISL_567597, EPI_ISL_567598, EPI_ISL_567599, EPI_ISL_567600, EPI_ISL_567601, EPI_ISL_567602, EPI_ISL_567603, EPI_ISL_567604, EPI_ISL_567605, EPI_ISL_567606, EPI_ISL_567607, EPI_ISL_567608, EPI_ISL_567609, EPI_ISL_567610, EPI_ISL_567611, EPI_ISL_567612, EPI_ISL_567613, EPI_ISL_567614, EPI_ISL_567615, EPI_ISL_567616, EPI_ISL_567617, EPI_ISL_567618, EPI_ISL_567619, EPI_ISL_567620, EPI_ISL_567621, EPI_ISL_567622, EPI_ISL_567623, EPI_ISL_567624, EPI_ISL_567625, EPI_ISL_567626, EPI_ISL_567627, EPI_ISL_567628, EPI_ISL_567629, EPI_ISL_567630, EPI_ISL_567631, EPI_ISL_567632, EPI_ISL_567633, EPI_ISL_567634, EPI_ISL_567635, EPI_ISL_567636, EPI_ISL_567637, EPI_ISL_567638, EPI_ISL_567639, EPI_ISL_567640, EPI_ISL_567641, EPI_ISL_567642, EPI_ISL_567643, EPI_ISL_567644, EPI_ISL_567645, EPI_ISL_567646, EPI_ISL_567647, EPI_ISL_567648, EPI_ISL_567649, EPI_ISL_567650, EPI_ISL_567651, EPI_ISL_567652, EPI_ISL_567653, EPI_ISL_567654, EPI_ISL_567655, EPI_ISL_567656, EPI_ISL_567657, EPI_ISL_567658, EPI_ISL_567659, EPI_ISL_567660, EPI_ISL_567661, EPI_ISL_567662, EPI_ISL_567663, EPI_ISL_567664, EPI_ISL_567665, EPI_ISL_567666, EPI_ISL_567667, EPI_ISL_567668, EPI_ISL_567669, EPI_ISL_567670, EPI_ISL_567671, EPI_ISL_567672, EPI_ISL_567673, EPI_ISL_567674, EPI_ISL_567675, EPI_ISL_567676, EPI_ISL_567677, EPI_ISL_567678, EPI_ISL_567679, EPI_ISL_567680, EPI_ISL_567681, EPI_ISL_567682, EPI_ISL_567683, EPI_ISL_567684, EPI_ISL_567685, EPI_ISL_567686, EPI_ISL_567687, EPI_ISL_567688, EPI_ISL_567689, EPI_ISL_567690, EPI_ISL_567691, EPI_ISL_567692, EPI_ISL_567693, EPI_ISL_567694, EPI_ISL_567695, EPI_ISL_567696, EPI_ISL_567697, EPI_ISL_567698, EPI_ISL_567699, EPI_ISL_567700, EPI_ISL_567701, EPI_ISL_567702, EPI_ISL_567703, EPI_ISL_567704, EPI_ISL_567705, EPI_ISL_567706, EPI_ISL_567707, EPI_ISL_567708, EPI_ISL_567709, EPI_ISL_567710, EPI_ISL_567711, EPI_ISL_567712, EPI_ISL_567713, EPI_ISL_567714, EPI_ISL_567715, EPI_ISL_567716, EPI_ISL_567717, EPI_ISL_567718, EPI_ISL_567719, EPI_ISL_567720, EPI_ISL_567721, EPI_ISL_567722, EPI_ISL_567723, EPI_ISL_567724, EPI_ISL_567725, EPI_ISL_567726, EPI_ISL_567727, EPI_ISL_567728, EPI_ISL_567729, EPI_ISL_567730, EPI_ISL_567731, EPI_ISL_567732, EPI_ISL_567733, EPI_ISL_567734, EPI_ISL_567735, EPI_ISL_567736, EPI_ISL_567737, EPI_ISL_567738, EPI_ISL_567739, EPI_ISL_567740, EPI_ISL_567741, EPI_ISL_567742, EPI_ISL_567743, EPI_ISL_567744, EPI_ISL_567745, EPI_ISL_567746, EPI_ISL_567747, EPI_ISL_567748, EPI_ISL_567749, EPI_ISL_567750, EPI_ISL_567751, EPI_ISL_567752, EPI_ISL_567753, EPI_ISL_567754, EPI_ISL_567755, EPI_ISL_567756, EPI_ISL_567757, EPI_ISL_567758, EPI_ISL_567759, EPI_ISL_567760, EPI_ISL_567761, EPI_ISL_567762, EPI_ISL_567763, EPI_ISL_567764, EPI_ISL_567765, EPI_ISL_567766, EPI_ISL_567767, EPI_ISL_567768, EPI_ISL_567769, EPI_ISL_567770, EPI_ISL_567771, EPI_ISL_567772, EPI_ISL_567773, EPI_ISL_567774, EPI_ISL_567775, EPI_ISL_567776, EPI_ISL_567777, EPI_ISL_567778, EPI_ISL_567779, EPI_ISL_567780, EPI_ISL_567781, EPI_ISL_567782, EPI_ISL_567783, EPI_ISL_567784, EPI_ISL_567785, EPI_ISL_567786, EPI_ISL_567787, EPI_ISL_567788, EPI_ISL_567789, EPI_ISL_567790, EPI_ISL_567791, EPI_ISL_567792, EPI_ISL_567793, EPI_ISL_567794, EPI_ISL_567795, EPI_ISL_567796, EPI_ISL_567797, EPI_ISL_567798, EPI_ISL_567799, EPI_ISL_567800, EPI_ISL_567801, EPI_ISL_567802 | | | |
| see above | Lighthouse Lab in Alderley Park | Wellcome Sanger Institute for the COVID-19 Genomics UK (COG-UK) consortium | Jacquelyn Wynn, Mairead Hyland, The Lighthouse Lab in Alderley Park and Alex Alderton, Roberto Amato, Sonia Goncalves, Ewan Harrison, David K. Jackson, Ian Johnston, Dominic Kwiatkowski, Cordelia Langford, John Sillitoe on behalf of the Wellcome Sanger Institute COVID-19 Surveillance Team |
| EPI_ISL_567803 | Lighthouse Lab in Alderley Park | Wellcome Sanger Institute for the COVID-19 Genomics UK (COG-UK) Consortium | Jacquelyn Wynn, Mairead Hyland, The Lighthouse Lab in Alderley Park and Alex Alderton, Roberto Amato, Sonia Goncalves, Ewan Harrison, David K. Jackson, Ian Johnston, Dominic Kwiatkowski, Cordelia Langford, John Sillitoe on behalf of the Wellcome Sanger Institute COVID-19 Surveillance Team |
| EPI_ISL_567804, EPI_ISL_567805, EPI_ISL_567806, EPI_ISL_567807, EPI_ISL_567808, EPI_ISL_567809, EPI_ISL_567810 | Lighthouse Lab in Alderley Park | Wellcome Sanger Institute for the COVID-19 Genomics UK (COG-UK) consortium | Jacquelyn Wynn, Mairead Hyland, The Lighthouse Lab in Alderley Park and Alex Alderton, Roberto Amato, Sonia Goncalves, Ewan Harrison, David K. Jackson, Ian Johnston, Dominic Kwiatkowski, Cordelia Langford, John Sillitoe on behalf of the Wellcome Sanger Institute COVID-19 Surveillance Team |
| EPI_ISL_567811 | Lighthouse Lab in Alderley Park | Wellcome Sanger Institute for the COVID-19 Genomics UK (COG-UK) Consortium | Jacquelyn Wynn, Mairead Hyland, The Lighthouse Lab in Alderley Park and Alex Alderton, Roberto Amato, Sonia Goncalves, Ewan Harrison, David K. Jackson, Ian Johnston, Dominic Kwiatkowski, Cordelia Langford, John Sillitoe on behalf of the Wellcome Sanger Institute COVID-19 Surveillance Team |
| EPI_ISL_567812, EPI_ISL_567813, EPI_ISL_567814, EPI_ISL_567815, EPI_ISL_567816, EPI_ISL_567817, EPI_ISL_567818, EPI_ISL_567819, EPI_ISL_567820, EPI_ISL_567821, EPI_ISL_567822, EPI_ISL_567823, EPI_ISL_567824, EPI_ISL_567825, EPI_ISL_567826, EPI_ISL_567827, EPI_ISL_567828, EPI_ISL_567829, EPI_ISL_567830, EPI_ISL_567831, EPI_ISL_567832, EPI_ISL_567833, EPI_ISL_567834, EPI_ISL_567835, EPI_ISL_567836, EPI_ISL_567837, EPI_ISL_567838, EPI_ISL_567839, EPI_ISL_567840, EPI_ISL_567841, EPI_ISL_567842, EPI_ISL_567843, EPI_ISL_567844, EPI_ISL_567845, EPI_ISL_567846, EPI_ISL_567847, EPI_ISL_567848, EPI_ISL_567849, EPI_ISL_567850, EPI_ISL_567851, EPI_ISL_567852, EPI_ISL_567853, EPI_ISL_567854, EPI_ISL_567855, EPI_ISL_567856, EPI_ISL_567857, EPI_ISL_567858, EPI_ISL_567859, EPI_ISL_567860, EPI_ISL_567861, EPI_ISL_567862, EPI_ISL_567863, EPI_ISL_567864, EPI_ISL_567865, EPI_ISL_567866, EPI_ISL_567867, EPI_ISL_567868, EPI_ISL_567869, EPI_ISL_567870, EPI_ISL_567871, EPI_ISL_567872, EPI_ISL_567873, EPI_ISL_567874, EPI_ISL_567875, EPI_ISL_567876, EPI_ISL_567877, EPI_ISL_567878, EPI_ISL_567879, EPI_ISL_567880, EPI_ISL_567881, EPI_ISL_567882, EPI_ISL_567883, EPI_ISL_567884, EPI_ISL_567885, EPI_ISL_567893, EPI_ISL_567898, EPI_ISL_567899, EPI_ISL_567900, EPI_ISL_567904, EPI_ISL_567905, EPI_ISL_567908, EPI_ISL_567912, EPI_ISL_567914, EPI_ISL_567916, EPI_ISL_567922, EPI_ISL_567923, EPI_ISL_567925, EPI_ISL_567926, EPI_ISL_567929, EPI_ISL_567930, EPI_ISL_567934, EPI_ISL_567935, EPI_ISL_567941, EPI_ISL_567947, EPI_ISL_567948, EPI_ISL_567952, EPI_ISL_567954, EPI_ISL_567955, EPI_ISL_567956, EPI_ISL_567958, EPI_ISL_567959, EPI_ISL_567962, EPI_ISL_567966, EPI_ISL_567971, EPI_ISL_567972, EPI_ISL_567973, EPI_ISL_567976, EPI_ISL_567978, EPI_ISL_567979, EPI_ISL_567980, EPI_ISL_567983, EPI_ISL_567984, EPI_ISL_567985, EPI_ISL_567992, EPI_ISL_567999, EPI_ISL_568000, EPI_ISL_568004, EPI_ISL_568011, EPI_ISL_568018, EPI_ISL_568020, EPI_ISL_568023, EPI_ISL_568024, EPI_ISL_568027, EPI_ISL_568029, EPI_ISL_568037, EPI_ISL_568040, EPI_ISL_568043, EPI_ISL_568048, EPI_ISL_568052, EPI_ISL_568054, EPI_ISL_568055, EPI_ISL_568057, EPI_ISL_568059, EPI_ISL_568060, EPI_ISL_568064, EPI_ISL_568072, EPI_ISL_568073, EPI_ISL_568074, EPI_ISL_568075, EPI_ISL_568077, EPI_ISL_568080, EPI_ISL_568083, EPI_ISL_568088, EPI_ISL_568089, EPI_ISL_568093, EPI_ISL_568094, EPI_ISL_568095, EPI_ISL_568097, EPI_ISL_568098, EPI_ISL_568102, EPI_ISL_568109, EPI_ISL_568124, EPI_ISL_568127, EPI_ISL_568131, EPI_ISL_568136, EPI_ISL_568139, EPI_ISL_568142, EPI_ISL_568143, EPI_ISL_568144, EPI_ISL_568145, EPI_ISL_568147, EPI_ISL_568148, EPI_ISL_568151, EPI_ISL_568153, EPI_ISL_568155, EPI_ISL_568159, EPI_ISL_568164, EPI_ISL_568176, EPI_ISL_568187, EPI_ISL_568191, EPI_ISL_568194, EPI_ISL_568195, EPI_ISL_568197, EPI_ISL_568202, EPI_ISL_568204, EPI_ISL_568210, EPI_ISL_568212, EPI_ISL_568213, EPI_ISL_568214, EPI_ISL_568215, EPI_ISL_568216, EPI_ISL_568217, EPI_ISL_568219, EPI_ISL_568220 | | | |
| see above | Lighthouse Lab in Alderley Park | Wellcome Sanger Institute for the COVID-19 Genomics UK (COG-UK) consortium | Jacquelyn Wynn, Mairead Hyland, The Lighthouse Lab in Alderley Park and Alex Alderton, Roberto Amato, Sonia Goncalves, Ewan Harrison, David K. Jackson, Ian Johnston, Dominic Kwiatkowski, Cordelia Langford, John Sillitoe on behalf of the Wellcome Sanger Institute COVID-19 Surveillance Team |
| EPI_ISL_568224, EPI_ISL_568226, EPI_ISL_568227, EPI_ISL_568231, EPI_ISL_568235, EPI_ISL_568237, EPI_ISL_568238, EPI_ISL_568242, EPI_ISL_568246, EPI_ISL_568247, EPI_ISL_568249, EPI_ISL_568250, EPI_ISL_568255, EPI_ISL_568257, EPI_ISL_568262, EPI_ISL_568263, EPI_ISL_568265, EPI_ISL_568266, EPI_ISL_568268, EPI_ISL_568269, EPI_ISL_568270, EPI_ISL_568271, EPI_ISL_568273, EPI_ISL_568275, EPI_ISL_568277, EPI_ISL_568278, EPI_ISL_568279, EPI_ISL_568286, EPI_ISL_568287 | | | |
| see above | Lighthouse Lab in Glasgow | Wellcome Sanger Institute for the COVID-19 Genomics | Harper VanSteenhouse, Yumi Kasai, David Gray, Carol Clugston, Anna Dominiczak and Alex Alderton, Roberto Amato, Sonia Goncalves, Ewan Harrison, David K. Jackson, Ian Johnston, Dominic Kwiatkowski, Cordelia Langford, John Sillitoe on behalf of the Wellcome Sanger Institute COVID-19 Surveillance Team |

| | | | |
|---|---|---|---|
| | | UK (COG-UK) consortium | |
| EPI_ISL_568590, EPI_ISL_568591, EPI_ISL_568592, EPI_ISL_568593, EPI_ISL_568594, EPI_ISL_568595, EPI_ISL_568596, EPI_ISL_568658, EPI_ISL_568659, EPI_ISL_568660, EPI_ISL_568661, EPI_ISL_568662, EPI_ISL_568663, EPI_ISL_568664, EPI_ISL_568665, EPI_ISL_568666, EPI_ISL_568667, EPI_ISL_568668, EPI_ISL_568669, EPI_ISL_568670, EPI_ISL_568671, EPI_ISL_568672, EPI_ISL_568673, EPI_ISL_568674, EPI_ISL_568675, EPI_ISL_568676, EPI_ISL_568677, EPI_ISL_568678, EPI_ISL_568679 | | | |
| see above | Florida Bureau of Public Health Laboratories | Florida Bureau of Public Health Laboratories | Sarah Schmedes, Jason Blanton |
| EPI_ISL_569220, EPI_ISL_569230, EPI_ISL_569231, EPI_ISL_569232, EPI_ISL_569233, EPI_ISL_569234, EPI_ISL_569235, EPI_ISL_569236, EPI_ISL_569237, EPI_ISL_569238, EPI_ISL_569239, EPI_ISL_569240, EPI_ISL_569241 | | | |
| see above | MEPHI, Aix Marseille University | MEPHI, Aix Marseille University | Anthony LEVASSEUR |
| EPI_ISL_569858 | Area of Virology, Serology and Virology Division (SAViD), New South Wales Health Pathology Randwick | Area of Virology, Serology and Virology Division (SAViD), New South Wales Health Pathology Randwick | Rawlinson, W., Deveson, I., Bull, R., Van Hal, S. |
| EPI_ISL_569859, EPI_ISL_569860, EPI_ISL_569861, EPI_ISL_569862 | CSIR-Indian Institute of Chemical Biology, MEDICA Supercpecialty Hospital Kolkata | CSIR-Indian Institute of Chemical Biology, MEDICA Supercpecialty Hospital Kolkata | Sujay Krishna Maity, Priyanka Mallick, Debaleena Bhowmik, Abhishake Lahiri, Dr. Aviral Roy, Dr. Soumen Saha, Dr. Arpita Ghosh Mitra, Dr. Rajesh Pandey, Dr. Sandip Paul, Dr. Partha Chakrabarti, Dr. Saikat Chakrabarti |
| EPI_ISL_572320, EPI_ISL_572321, EPI_ISL_572322, EPI_ISL_572323 | IZSM | IZSM | Maurizio Viscardi, Lorena Cardillo, Giovanna Fusco |
| EPI_ISL_572399, EPI_ISL_572401, EPI_ISL_572403, EPI_ISL_572406 | Quadram Institute Bioscience | COVID-19 Genomics UK (COG-UK) Consortium | Dave J. Baker, Gemma L. Kay, Alp Aydin, Thanh Le-Viet, Steven Rudder, Ana P. Tedim, Anastasia Kolyva, Maria Diaz, Leonardo de Oliveira Martins, Nabil-Fareed Alikhan, Lizzie Meadows, Rachael Stanley, Ngozi Elumogo, Muhammed Yasir, Nicholas M. Thomson, Alexander J Trotter, Rachel Gilroy, Samuel Bloomfield, Claire Stuart, Andrew Bell, Reenesh Prakash, Samir Dervisevic, Alison E. Mather, John Wain, Mark Webber, Andrew J. Page, Justin O'Grady |
| EPI_ISL_572409, EPI_ISL_572410 | Northumbria University / South Tees Hospitals NHS Foundation Trust / North Cumbria Integrated Care NHS Foundation Trust / North Tees and Hartlepool NHS Foundation Trust / Newcastle Hospitals NHS Foundation Trust | COVID-19 Genomics UK (COG-UK) Consortium | Darren L Smith,Andrew Nelson,Matthew Bashton,Greg R Young,Joshua Loh,John Allan,Mohammad A Tariq,Giles S Holt,Gary Black,Wen C Yew,Lynn Dover,Paul Baker,Steve Liggett,Sarah Essex,Jane Greenaway,Debra Padgett,Clive Graham,Garren Scott,Edward Barton,Emma Swindells,Brendan Payne,Jennifer Collins,Yusri Taha,Gary Eltringham |
| EPI_ISL_572415 | West of Scotland Specialist Virology Centre, NHSGGC / MRC-University of Glasgow Centre for Virus Research | COVID-19 Genomics UK (COG-UK) Consortium | Ana da Silva Filipe, Natasha Johnson, Kathy Smollett, Daniel Mair, Stephen Carmichael, Lily Tong, Jenna Nichols, Elihu Aranday-Cortes, Kyriaki Nomikou; Sarah McDonald, Marc Niebel, Patawee Asamaphan; Richard Orton, Joseph Hughes, Sreenu Vattipally, David L Robertson; Alasdair MacLean, Rory Gunson; Kathy Li, Igor Starinskij, Natasha Jesudason, Rajiv Shah, James Shepherd, Antonia Ho, Emma Thomson |
| EPI_ISL_572423, EPI_ISL_572430 | Quadram Institute Bioscience | COVID-19 Genomics UK (COG-UK) Consortium | Dave J. Baker, Gemma L. Kay, Alp Aydin, Thanh Le-Viet, Steven Rudder, Ana P. Tedim, Anastasia Kolyva, Maria Diaz, Leonardo de Oliveira Martins, Nabil-Fareed Alikhan, Lizzie Meadows, Rachael Stanley, Ngozi Elumogo, Muhammed Yasir, Nicholas M. Thomson, Alexander J Trotter, Rachel Gilroy, Samuel Bloomfield, Claire Stuart, Andrew Bell, Reenesh Prakash, Samir Dervisevic, Alison E. Mather, John Wain, Mark Webber, Andrew J. Page, Justin O'Grady |
| EPI_ISL_572432, EPI_ISL_572433 | Virology Department, Sheffield Teaching Hospitals NHS Foundation Trust/Department of Infection, Immunity and Cardiovascular Disease, The Medical School, University of Sheffield | COVID-19 Genomics UK (COG-UK) Consortium | Thushan de Silva, Matthew Parker, Nikki Smith, Adri Angyal, Rebecca Brown, Luke Green, Rachel Tucker, Paul Parsons, Danielle Groves, Katie Johnson, Laura Carrilero, Alex Keeley, Dave Partridge, Matthew Wyles, Benjamin Lindsey, Mehmet Yavuz, Mohammad Raza, Cariad Evans |
| EPI_ISL_572436 | Liverpool Clinical Laboratories | COVID-19 Genomics UK (COG-UK) Consortium | Sam Haldenby, Anita Lucaci, Steve Paterson, Julian Hiscox, Alistair Darby, M Almsaud, A Alrezaihi, Muhannad Alruwaili, Stuart D Armstrong, Jones Benjamin, Eleanor G Bentley, Anu Chawla, Jordan J Clark, Angela Cowell, Richard Eccles, Isabel García-Dorival, Matthew Gemmell, Alessandro Gerada, PKF Gilmore, Richard Gregory, Ximeng Han, Catherine Hartley, Margaret Hughes, Miren Iturriza-Gomara, James Johnson, L Luu, Jenifer Manson, Charlotte Nelson, Elaine O'Toole, Cassie Olateju, Rebekah Penrice-Randal , Lucille Rainbow, N.P Randle, Trevor Ian Robinson, Parul Sharma, Ghada T Shawli, James P Stewart, Neil Swainston, Ecaterina Vamos, Joanne Watts, Mark Whitehead |
| EPI_ISL_572437, EPI_ISL_572440, EPI_ISL_572441, EPI_ISL_572447, EPI_ISL_572448, EPI_ISL_572449, EPI_ISL_572453, EPI_ISL_572454 | Quadram Institute Bioscience | COVID-19 Genomics UK (COG-UK) Consortium | Dave J. Baker, Gemma L. Kay, Alp Aydin, Thanh Le-Viet, Steven Rudder, Ana P. Tedim, Anastasia Kolyva, Maria Diaz, Leonardo de Oliveira Martins, Nabil-Fareed Alikhan, Lizzie Meadows, Rachael Stanley, Ngozi Elumogo, Muhammed Yasir, Nicholas M. Thomson, Alexander J Trotter, Rachel Gilroy, Samuel Bloomfield, Claire Stuart, Andrew Bell, Reenesh Prakash, Samir Dervisevic, Alison E. Mather, John Wain, Mark Webber, Andrew J. Page, Justin O'Grady |
| EPI_ISL_572455 | Wales Specialist Virology Centre Sequencing lab: Pathogen Genomics Unit | COVID-19 Genomics UK (COG-UK) Consortium | Catherine Moore, Johnathan Evans, Laura Gifford, Malorie Perry, Simon Cottrell, Angela Marchbank, Alec Birchley, Alexander Adams, Amy Gaskin, Bree Gatica-Wilcox, Jason Coombes, Joel Southgate, Lauren Gilbert, Lee Graham, Nicole Pacchiarini, Sara Kumziene-Summerhayes, Sarah Taylor, Sophie Jones, Sara Rey, Matthew Bull, Joanne Watkins, Sally Corden, Tom Connor |
| EPI_ISL_572456, EPI_ISL_572459, EPI_ISL_572463 | Quadram Institute Bioscience | COVID-19 Genomics UK (COG-UK) Consortium | Dave J. Baker, Gemma L. Kay, Alp Aydin, Thanh Le-Viet, Steven Rudder, Ana P. Tedim, Anastasia Kolyva, Maria Diaz, Leonardo de Oliveira Martins, Nabil-Fareed Alikhan, Lizzie Meadows, Rachael Stanley, Ngozi Elumogo, Muhammed Yasir, Nicholas M. Thomson, Alexander J Trotter, Rachel Gilroy, Samuel Bloomfield, Claire Stuart, Andrew Bell, Reenesh Prakash, Samir Dervisevic, Alison E. Mather, John Wain, Mark Webber, Andrew J. Page, Justin O'Grady |
| EPI_ISL_572464 | Northumbria University / South Tees Hospitals NHS Foundation Trust / North Cumbria Integrated Care NHS Foundation Trust / North Tees and Hartlepool NHS Foundation Trust / Newcastle Hospitals NHS Foundation Trust | COVID-19 Genomics UK (COG-UK) Consortium | Darren L Smith,Andrew Nelson,Matthew Bashton,Greg R Young,Joshua Loh,John Allan,Mohammad A Tariq,Giles S Holt,Gary Black,Wen C Yew,Lynn Dover,Paul Baker,Steve Liggett,Sarah Essex,Jane Greenaway,Debra Padgett,Clive Graham,Garren Scott,Edward Barton,Emma Swindells,Brendan Payne,Jennifer Collins,Yusri Taha,Gary Eltringham |
| EPI_ISL_572466 | Liverpool Clinical Laboratories | COVID-19 Genomics UK (COG-UK) Consortium | Sam Haldenby, Anita Lucaci, Steve Paterson, Julian Hiscox, Alistair Darby, M Almsaud, A Alrezaihi, Muhannad Alruwaili, Stuart D Armstrong, Jones Benjamin, Eleanor G Bentley, Anu Chawla, Jordan J Clark, Angela Cowell, Richard Eccles, Isabel García-Dorival, Matthew Gemmell, Alessandro Gerada, PKF Gilmore, Richard Gregory, Ximeng Han, Catherine Hartley, Margaret Hughes, Miren Iturriza-Gomara, James Johnson, L Luu, Jenifer Manson, Charlotte Nelson, Elaine O'Toole, Cassie Olateju, Rebekah Penrice-Randal , Lucille Rainbow, N.P Randle, Trevor Ian Robinson, Parul Sharma, Ghada T Shawli, James P Stewart, Neil Swainston, Ecaterina Vamos, Joanne Watts, Mark Whitehead |
| EPI_ISL_572468 | Wales Specialist Virology Centre Sequencing lab: | COVID-19 Genomics UK (COG-UK) Consortium | Catherine Moore, Johnathan Evans, Laura Gifford, Malorie Perry, Simon Cottrell, Angela Marchbank, Alec Birchley, Alexander Adams, Amy Gaskin, Bree Gatica-Wilcox, Jason Coombes, Joel Southgate, Lauren Gilbert, Lee Graham, Nicole Pacchiarini, Sara Kumziene-Summerhayes, Sarah Taylor, Sophie Jones, Sara Rey, Matthew Bull, Joanne Watkins, Sally Corden, Tom Connor |

| | | | |
|---|---|---|---|
| | Pathogen Genomics Unit | | |
| EPI_ISL_572470 | West of Scotland Specialist Virology Centre, NHSGGC / MRC-University of Glasgow Centre for Virus Research | COVID-19 Genomics UK (COG-UK) Consortium | Ana da Silva Filipe, Natasha Johnson, Kathy Smollett, Daniel Mair, Stephen Carmichael, Lily Tong, Jenna Nichols, Elihu Aranday-Cortes, Kyriaki Nomikou; Sarah McDonald, Marc Niebel, Patawee Asamaphan; Richard Orton, Joseph Hughes, Sreenu Vattipally, David L Robertson; Alasdair MacLean, Rory Gunson; Kathy Li, Igor Starinskij, Natasha Jesudason, Rajiv Shah, James Shepherd, Antonia Ho, Emma Thomson |
| EPI_ISL_572471, EPI_ISL_572472 | Department of Pathology, University of Cambridge | COVID-19 Genomics UK (COG-UK) Consortium | Aminu S. Jahun, Yasmin Chaudhry, Grant Hall, Iliana Georgana, Myra Hosmillo, Martin D. Curran, Malte Pinckert, Surendra Parmar, Ian Goodfellow |
| EPI_ISL_572477 | Wales Specialist Virology Centre Sequencing lab: Pathogen Genomics Unit | COVID-19 Genomics UK (COG-UK) Consortium | Catherine Moore, Johnathan Evans, Laura Gifford, Malorie Perry, Simon Cottrell, Angela Marchbank, Alec Birchley, Alexander Adams, Amy Gaskin, Bree Gatica-Wilcox, Jason Coombes, Joel Southgate, Lauren Gilbert, Lee Graham, Nicole Pacchiarini, Sara Kumziene-Summerhayes, Sarah Taylor, Sophie Jones, Sara Rey, Matthew Bull, Joanne Watkins, Sally Corden, Tom Connor |
| EPI_ISL_572479 | Liverpool Clinical Laboratories | COVID-19 Genomics UK (COG-UK) Consortium | Sam Haldenby, Anita Lucaci, Steve Paterson, Julian Hiscox, Alistair Darby, M Almsaud, A Alrezaihi, Muhannad Alruwaili, Stuart D Armstrong, Jones Benjamin, Eleanor G Bentley, Anu Chawla, Jordan J Clark, Angela Cowell, Richard Eccles, Isabel García-Dorival, Matthew Gemmell, Alessandro Gerada, PKF Gilmore, Richard Gregory, Ximeng Han, Catherine Hartley, Margaret Hughes, Miren Iturriza-Gomara, James Johnson, L Luu, Jenifer Manson, Charlotte Nelson, Elaine O'Toole, Cassie Olateju, Rebekah Penrice-Randal , Lucille Rainbow, N.P Randle, Trevor Ian Robinson, Parul Sharma, Ghada T Shawli, James P Stewart, Neil Swainston, Ecaterina Vamos, Joanne Watts, Mark Whitehead |
| EPI_ISL_572484 | Department of Pathology, University of Cambridge | COVID-19 Genomics UK (COG-UK) Consortium | Aminu S. Jahun, Yasmin Chaudhry, Grant Hall, Iliana Georgana, Myra Hosmillo, Martin D. Curran, Malte Pinckert, Surendra Parmar, Ian Goodfellow |
| EPI_ISL_572485 | Quadram Institute Bioscience | COVID-19 Genomics UK (COG-UK) Consortium | Dave J. Baker, Gemma L. Kay, Alp Aydin, Thanh Le-Viet, Steven Rudder, Ana P. Tedim, Anastasia Kolyva, Maria Diaz, Leonardo de Oliveira Martins, Nabil-Fareed Alikhan, Lizzie Meadows, Rachael Stanley, Ngozi Elumogo, Muhammed Yasir, Nicholas M. Thomson, Alexander J Trotter, Rachel Gilroy, Samuel Bloomfield, Claire Stuart, Andrew Bell, Reenesh Prakash, Samir Dervisevic, Alison E. Mather, John Wain, Mark Webber, Andrew J. Page, Justin O'Grady |
| EPI_ISL_572489 | Northumbria University / South Tees Hospitals NHS Foundation Trust / North Cumbria Integrated Care NHS Foundation Trust / North Tees and Hartlepool NHS Foundation Trust / Newcastle Hospitals NHS Foundation Trust | COVID-19 Genomics UK (COG-UK) Consortium | Darren L Smith,Andrew Nelson,Matthew Bashton,Greg R Young,Joshua Loh,John Allan,Mohammad A Tariq,Giles S Holt,Gary Black,Wen C Yew,Lynn Dover,Paul Baker,Steve Liggett,Sarah Essex,Jane Greenaway,Debra Padgett,Clive Graham,Garren Scott,Edward Barton,Emma Swindells,Brendan Payne,Jennifer Collins,Yusri Taha,Gary Eltringham |
| EPI_ISL_572492, EPI_ISL_572493, EPI_ISL_572495, EPI_ISL_572497, EPI_ISL_572499, EPI_ISL_572503, EPI_ISL_572504, EPI_ISL_572505 | Quadram Institute Bioscience | COVID-19 Genomics UK (COG-UK) Consortium | Dave J. Baker, Gemma L. Kay, Alp Aydin, Thanh Le-Viet, Steven Rudder, Ana P. Tedim, Anastasia Kolyva, Maria Diaz, Leonardo de Oliveira Martins, Nabil-Fareed Alikhan, Lizzie Meadows, Rachael Stanley, Ngozi Elumogo, Muhammed Yasir, Nicholas M. Thomson, Alexander J Trotter, Rachel Gilroy, Samuel Bloomfield, Claire Stuart, Andrew Bell, Reenesh Prakash, Samir Dervisevic, Alison E. Mather, John Wain, Mark Webber, Andrew J. Page, Justin O'Grady |
| EPI_ISL_572510 | Lincolnshire Hospitals and DeepSeq Nottingham | COVID-19 Genomics UK (COG-UK) Consortium | Nichola Duckworth, Tim Sloan, Sarah Walsh, Jonathan Ball, Patrick McClure, Joeseph Chappell, Nadine Holmes, Matthew Carlisle, Christopher Moore, Fei Sang, Johnny Debebe, Victoria Wright, Matthew Loose |
| EPI_ISL_572516 | Virology Department, Sheffield Teaching Hospitals NHS Foundation Trust/Department of Infection, Immunity and Cardiovascular Disease, The Medical School, University of Sheffield | COVID-19 Genomics UK (COG-UK) Consortium | Thushan de Silva, Matthew Parker, Nikki Smith, Adri Angyal, Rebecca Brown, Luke Green, Rachel Tucker, Paul Parsons, Danielle Groves, Katie Johnson, Laura Carrilero, Alex Keeley, Dave Partridge, Matthew Wyles, Benjamin Lindsey, Mehmet Yavuz, Mohammad Raza, Cariad Evans |
| EPI_ISL_572518 | West of Scotland Specialist Virology Centre, NHSGGC / MRC-University of Glasgow Centre for Virus Research | COVID-19 Genomics UK (COG-UK) Consortium | Ana da Silva Filipe, Natasha Johnson, Kathy Smollett, Daniel Mair, Stephen Carmichael, Lily Tong, Jenna Nichols, Elihu Aranday-Cortes, Kyriaki Nomikou; Sarah McDonald, Marc Niebel, Patawee Asamaphan; Richard Orton, Joseph Hughes, Sreenu Vattipally, David L Robertson; Alasdair MacLean, Rory Gunson; Kathy Li, Igor Starinskij, Natasha Jesudason, Rajiv Shah, James Shepherd, Antonia Ho, Emma Thomson |
| EPI_ISL_572519 | Department of Pathology, University of Cambridge | COVID-19 Genomics UK (COG-UK) Consortium | Aminu S. Jahun, Yasmin Chaudhry, Grant Hall, Iliana Georgana, Myra Hosmillo, Martin D. Curran, Malte Pinckert, Surendra Parmar, Ian Goodfellow |
| EPI_ISL_572520 | Quadram Institute Bioscience | COVID-19 Genomics UK (COG-UK) Consortium | Dave J. Baker, Gemma L. Kay, Alp Aydin, Thanh Le-Viet, Steven Rudder, Ana P. Tedim, Anastasia Kolyva, Maria Diaz, Leonardo de Oliveira Martins, Nabil-Fareed Alikhan, Lizzie Meadows, Rachael Stanley, Ngozi Elumogo, Muhammed Yasir, Nicholas M. Thomson, Alexander J Trotter, Rachel Gilroy, Samuel Bloomfield, Claire Stuart, Andrew Bell, Reenesh Prakash, Samir Dervisevic, Alison E. Mather, John Wain, Mark Webber, Andrew J. Page, Justin O'Grady |
| EPI_ISL_572523 | West of Scotland Specialist Virology Centre, NHSGGC / MRC-University of Glasgow Centre for Virus Research | COVID-19 Genomics UK (COG-UK) Consortium | Ana da Silva Filipe, Natasha Johnson, Kathy Smollett, Daniel Mair, Stephen Carmichael, Lily Tong, Jenna Nichols, Elihu Aranday-Cortes, Kyriaki Nomikou; Sarah McDonald, Marc Niebel, Patawee Asamaphan; Richard Orton, Joseph Hughes, Sreenu Vattipally, David L Robertson; Alasdair MacLean, Rory Gunson; Kathy Li, Igor Starinskij, Natasha Jesudason, Rajiv Shah, James Shepherd, Antonia Ho, Emma Thomson |
| EPI_ISL_572524 | Department of Pathology, University of Cambridge | COVID-19 Genomics UK (COG-UK) Consortium | Aminu S. Jahun, Yasmin Chaudhry, Grant Hall, Iliana Georgana, Myra Hosmillo, Martin D. Curran, Malte Pinckert, Surendra Parmar, Ian Goodfellow |
| EPI_ISL_572526 | Quadram Institute Bioscience | COVID-19 Genomics UK (COG-UK) Consortium | Dave J. Baker, Gemma L. Kay, Alp Aydin, Thanh Le-Viet, Steven Rudder, Ana P. Tedim, Anastasia Kolyva, Maria Diaz, Leonardo de Oliveira Martins, Nabil-Fareed Alikhan, Lizzie Meadows, Rachael Stanley, Ngozi Elumogo, Muhammed Yasir, Nicholas M. Thomson, Alexander J Trotter, Rachel Gilroy, Samuel Bloomfield, Claire Stuart, Andrew Bell, Reenesh Prakash, Samir Dervisevic, Alison E. Mather, John Wain, Mark Webber, Andrew J. Page, Justin O'Grady |
| EPI_ISL_572527 | Lincolnshire Hospitals and DeepSeq Nottingham | COVID-19 Genomics UK (COG-UK) Consortium | Nichola Duckworth, Tim Sloan, Sarah Walsh, Jonathan Ball, Patrick McClure, Joeseph Chappell, Nadine Holmes, Matthew Carlisle, Christopher Moore, Fei Sang, Johnny Debebe, Victoria Wright, Matthew Loose |
| EPI_ISL_572528 | West of Scotland Specialist Virology Centre, NHSGGC / MRC-University of Glasgow Centre for Virus Research | COVID-19 Genomics UK (COG-UK) Consortium | Ana da Silva Filipe, Natasha Johnson, Kathy Smollett, Daniel Mair, Stephen Carmichael, Lily Tong, Jenna Nichols, Elihu Aranday-Cortes, Kyriaki Nomikou; Sarah McDonald, Marc Niebel, Patawee Asamaphan; Richard Orton, Joseph Hughes, Sreenu Vattipally, David L Robertson; Alasdair MacLean, Rory Gunson; Kathy Li, Igor Starinskij, Natasha Jesudason, Rajiv Shah, James Shepherd, Antonia Ho, Emma Thomson |
| EPI_ISL_572533 | Quadram Institute Bioscience | COVID-19 Genomics UK (COG-UK) Consortium | Dave J. Baker, Gemma L. Kay, Alp Aydin, Thanh Le-Viet, Steven Rudder, Ana P. Tedim, Anastasia Kolyva, Maria Diaz, Leonardo de Oliveira Martins, Nabil-Fareed Alikhan, Lizzie Meadows, Rachael Stanley, Ngozi Elumogo, Muhammed Yasir, Nicholas M. Thomson, Alexander J Trotter, Rachel Gilroy, Samuel Bloomfield, Claire Stuart, Andrew Bell, Reenesh Prakash, Samir Dervisevic, Alison E. Mather, John Wain, Mark Webber, Andrew J. Page, Justin O'Grady |
| EPI_ISL_572534 | West of Scotland Specialist Virology Centre, NHSGGC / MRC-University of Glasgow Centre for Virus Research | COVID-19 Genomics UK (COG-UK) Consortium | Ana da Silva Filipe, Natasha Johnson, Kathy Smollett, Daniel Mair, Stephen Carmichael, Lily Tong, Jenna Nichols, Elihu Aranday-Cortes, Kyriaki Nomikou; Sarah McDonald, Marc Niebel, Patawee Asamaphan; Richard Orton, Joseph Hughes, Sreenu Vattipally, David L Robertson; Alasdair MacLean, Rory Gunson; Kathy Li, Igor Starinskij, Natasha Jesudason, Rajiv Shah, James Shepherd, Antonia Ho, Emma Thomson |
| EPI_ISL_572541 | Department of Pathology, University of Cambridge | COVID-19 Genomics UK (COG-UK) Consortium | Aminu S. Jahun, Yasmin Chaudhry, Grant Hall, Iliana Georgana, Myra Hosmillo, Martin D. Curran, Malte Pinckert, Surendra Parmar, Ian Goodfellow |
| EPI_ISL_572546, EPI_ISL_572548, | Quadram Institute Bioscience | COVID-19 Genomics UK (COG-UK) Consortium | Dave J. Baker, Gemma L. Kay, Alp Aydin, Thanh Le-Viet, Steven Rudder, Ana P. Tedim, Anastasia Kolyva, Maria Diaz, Leonardo de Oliveira Martins, Nabil-Fareed Alikhan, Lizzie Meadows, Rachael Stanley, Ngozi Elumogo, Muhammed Yasir, Nicholas M. Thomson, Alexander J Trotter, Rachel Gilroy, Samuel Bloomfield, Claire Stuart, Andrew Bell, Reenesh Prakash, Samir Dervisevic, Alison E. Mather, John Wain, Mark Webber, Andrew J. Page, Justin O'Grady |

| | | | |
|---|---|---|---|
| EPI_ISL_572549, EPI_ISL_572550, EPI_ISL_572551, EPI_ISL_572553, EPI_ISL_572556 | | | |
| EPI_ISL_572558, EPI_ISL_572564 | Wales Specialist Virology Centre Sequencing lab: Pathogen Genomics Unit | COVID-19 Genomics UK (COG-UK) Consortium | Catherine Moore, Johnathan Evans, Laura Gifford, Malorie Perry, Simon Cottrell, Angela Marchbank, Alec Birchley, Alexander Adams, Amy Gaskin, Bree Gatica-Wilcox, Jason Coombes, Joel Southgate, Lauren Gilbert, Lee Graham, Nicole Pacchiarini, Sara Kumziene-Summerhayes, Sarah Taylor, Sophie Jones, Sara Rey, Matthew Bull, Joanne Watkins, Sally Corden, Tom Connor |
| EPI_ISL_572568 | Northumbria University / South Tees Hospitals NHS Foundation Trust / North Cumbria Integrated Care NHS Foundation Trust / North Tees and Hartlepool NHS Foundation Trust / Newcastle Hospitals NHS Foundation Trust | COVID-19 Genomics UK (COG-UK) Consortium | Darren L Smith,Andrew Nelson,Matthew Bashton,Greg R Young,Joshua Loh,John Allan,Mohammad A Tariq,Giles S Holt,Gary Black,Wen C Yew,Lynn Dover,Paul Baker,Steve Liggett,Sarah Essex,Jane Greenaway,Debra Padgett,Clive Graham,Garren Scott,Edward Barton,Emma Swindells,Brendan Payne,Jennifer Collins,Yusri Taha,Gary Eltringham |
| EPI_ISL_572569 | Quadram Institute Bioscience | COVID-19 Genomics UK (COG-UK) Consortium | Dave J. Baker, Gemma L. Kay, Alp Aydin, Thanh Le-Viet, Steven Rudder, Ana P. Tedim, Anastasia Kolyva, Maria Diaz, Leonardo de Oliveira Martins, Nabil-Fareed Alikhan, Lizzie Meadows, Rachael Stanley, Ngozi Elumogo, Muhammed Yasir, Nicholas M. Thomson, Alexander J Trotter, Rachel Gilroy, Samuel Bloomfield, Claire Stuart, Andrew Bell, Reenesh Prakash, Samir Dervisevic, Alison E. Mather, John Wain, Mark Webber, Andrew J. Page, Justin O'Grady |
| EPI_ISL_572570, EPI_ISL_572571 | Northumbria University / South Tees Hospitals NHS Foundation Trust / North Cumbria Integrated Care NHS Foundation Trust / North Tees and Hartlepool NHS Foundation Trust / Newcastle Hospitals NHS Foundation Trust | COVID-19 Genomics UK (COG-UK) Consortium | Darren L Smith,Andrew Nelson,Matthew Bashton,Greg R Young,Joshua Loh,John Allan,Mohammad A Tariq,Giles S Holt,Gary Black,Wen C Yew,Lynn Dover,Paul Baker,Steve Liggett,Sarah Essex,Jane Greenaway,Debra Padgett,Clive Graham,Garren Scott,Edward Barton,Emma Swindells,Brendan Payne,Jennifer Collins,Yusri Taha,Gary Eltringham |
| EPI_ISL_572580 | Wales Specialist Virology Centre Sequencing lab: Pathogen Genomics Unit | COVID-19 Genomics UK (COG-UK) Consortium | Catherine Moore, Johnathan Evans, Laura Gifford, Malorie Perry, Simon Cottrell, Angela Marchbank, Alec Birchley, Alexander Adams, Amy Gaskin, Bree Gatica-Wilcox, Jason Coombes, Joel Southgate, Lauren Gilbert, Lee Graham, Nicole Pacchiarini, Sara Kumziene-Summerhayes, Sarah Taylor, Sophie Jones, Sara Rey, Matthew Bull, Joanne Watkins, Sally Corden, Tom Connor |
| EPI_ISL_572581 | Northumbria University / South Tees Hospitals NHS Foundation Trust / North Cumbria Integrated Care NHS Foundation Trust / North Tees and Hartlepool NHS Foundation Trust / Newcastle Hospitals NHS Foundation Trust | COVID-19 Genomics UK (COG-UK) Consortium | Darren L Smith,Andrew Nelson,Matthew Bashton,Greg R Young,Joshua Loh,John Allan,Mohammad A Tariq,Giles S Holt,Gary Black,Wen C Yew,Lynn Dover,Paul Baker,Steve Liggett,Sarah Essex,Jane Greenaway,Debra Padgett,Clive Graham,Garren Scott,Edward Barton,Emma Swindells,Brendan Payne,Jennifer Collins,Yusri Taha,Gary Eltringham |
| EPI_ISL_572583 | Quadram Institute Bioscience | COVID-19 Genomics UK (COG-UK) Consortium | Dave J. Baker, Gemma L. Kay, Alp Aydin, Thanh Le-Viet, Steven Rudder, Ana P. Tedim, Anastasia Kolyva, Maria Diaz, Leonardo de Oliveira Martins, Nabil-Fareed Alikhan, Lizzie Meadows, Rachael Stanley, Ngozi Elumogo, Muhammed Yasir, Nicholas M. Thomson, Alexander J Trotter, Rachel Gilroy, Samuel Bloomfield, Claire Stuart, Andrew Bell, Reenesh Prakash, Samir Dervisevic, Alison E. Mather, John Wain, Mark Webber, Andrew J. Page, Justin O'Grady |
| EPI_ISL_572585 | West of Scotland Specialist Virology Centre, NHSGGC / MRC-University of Glasgow Centre for Virus Research | COVID-19 Genomics UK (COG-UK) Consortium | Ana da Silva Filipe, Natasha Johnson, Kathy Smollett, Daniel Mair, Stephen Carmichael, Lily Tong, Jenna Nichols, Elihu Aranday-Cortes, Kyriaki Nomikou; Sarah McDonald, Marc Niebel, Patawee Asamaphan; Richard Orton, Joseph Hughes, Sreenu Vattipally, David L Robertson; Alasdair MacLean, Rory Gunson; Kathy Li, Igor Starinskij, Natasha Jesudason, Rajiv Shah, James Shepherd, Antonia Ho, Emma Thomson |
| EPI_ISL_572591 | Quadram Institute Bioscience | COVID-19 Genomics UK (COG-UK) Consortium | Dave J. Baker, Gemma L. Kay, Alp Aydin, Thanh Le-Viet, Steven Rudder, Ana P. Tedim, Anastasia Kolyva, Maria Diaz, Leonardo de Oliveira Martins, Nabil-Fareed Alikhan, Lizzie Meadows, Rachael Stanley, Ngozi Elumogo, Muhammed Yasir, Nicholas M. Thomson, Alexander J Trotter, Rachel Gilroy, Samuel Bloomfield, Claire Stuart, Andrew Bell, Reenesh Prakash, Samir Dervisevic, Alison E. Mather, John Wain, Mark Webber, Andrew J. Page, Justin O'Grady |
| EPI_ISL_572592 | Department of Pathology, University of Cambridge | COVID-19 Genomics UK (COG-UK) Consortium | Aminu S. Jahun, Yasmin Chaudhry, Grant Hall, Iliana Georgana, Myra Hosmillo, Martin D. Curran, Malte Pinckert, Surendra Parmar, Ian Goodfellow |
| EPI_ISL_572597 | Northumbria University / South Tees Hospitals NHS Foundation Trust / North Cumbria Integrated Care NHS Foundation Trust / North Tees and Hartlepool NHS Foundation Trust / Newcastle Hospitals NHS Foundation Trust | COVID-19 Genomics UK (COG-UK) Consortium | Darren L Smith,Andrew Nelson,Matthew Bashton,Greg R Young,Joshua Loh,John Allan,Mohammad A Tariq,Giles S Holt,Gary Black,Wen C Yew,Lynn Dover,Paul Baker,Steve Liggett,Sarah Essex,Jane Greenaway,Debra Padgett,Clive Graham,Garren Scott,Edward Barton,Emma Swindells,Brendan Payne,Jennifer Collins,Yusri Taha,Gary Eltringham |
| EPI_ISL_572600, EPI_ISL_572602 | Quadram Institute Bioscience | COVID-19 Genomics UK (COG-UK) Consortium | Dave J. Baker, Gemma L. Kay, Alp Aydin, Thanh Le-Viet, Steven Rudder, Ana P. Tedim, Anastasia Kolyva, Maria Diaz, Leonardo de Oliveira Martins, Nabil-Fareed Alikhan, Lizzie Meadows, Rachael Stanley, Ngozi Elumogo, Muhammed Yasir, Nicholas M. Thomson, Alexander J Trotter, Rachel Gilroy, Samuel Bloomfield, Claire Stuart, Andrew Bell, Reenesh Prakash, Samir Dervisevic, Alison E. Mather, John Wain, Mark Webber, Andrew J. Page, Justin O'Grady |
| EPI_ISL_572604 | Liverpool Clinical Laboratories | COVID-19 Genomics UK (COG-UK) Consortium | Sam Haldenby, Anita Lucaci, Steve Paterson, Julian Hiscox, Alistair Darby, M Almsaud, A Alrezaihi, Muhannad Alruwaili, Stuart D Armstrong, Jones Benjamin, Eleanor G Bentley, Anu Chawla, Jordan J Clark, Angela Cowell, Richard Eccles, Isabel García-Dorival, Matthew Gemmell, Alessandro Gerada, PKF Gilmore, Richard Gregory, Ximeng Han, Catherine Hartley, Margaret Hughes, Miren Iturriza-Gomara, James Johnson, L Luu, Jenifer Manson, Charlotte Nelson, Elaine O'Toole, Cassie Olateju, Rebekah Penrice-Randal , Lucille Rainbow, N.P Randle, Trevor Ian Robinson, Parul Sharma, Ghada T Shawli, James P Stewart, Neil Swainston, Ecaterina Vamos, Joanne Watts, Mark Whitehead |
| EPI_ISL_572615 | Quadram Institute Bioscience | COVID-19 Genomics UK (COG-UK) Consortium | Dave J. Baker, Gemma L. Kay, Alp Aydin, Thanh Le-Viet, Steven Rudder, Ana P. Tedim, Anastasia Kolyva, Maria Diaz, Leonardo de Oliveira Martins, Nabil-Fareed Alikhan, Lizzie Meadows, Rachael Stanley, Ngozi Elumogo, Muhammed Yasir, Nicholas M. Thomson, Alexander J Trotter, Rachel Gilroy, Samuel Bloomfield, Claire Stuart, Andrew Bell, Reenesh Prakash, Samir Dervisevic, Alison E. Mather, John Wain, Mark Webber, Andrew J. Page, Justin O'Grady |
| EPI_ISL_572616 | Department of Pathology, University of Cambridge | COVID-19 Genomics UK (COG-UK) Consortium | Aminu S. Jahun, Yasmin Chaudhry, Grant Hall, Iliana Georgana, Myra Hosmillo, Martin D. Curran, Malte Pinckert, Surendra Parmar, Ian Goodfellow |
| EPI_ISL_572620, EPI_ISL_572623, EPI_ISL_572624, EPI_ISL_572625, EPI_ISL_572626, EPI_ISL_572633, EPI_ISL_572634, EPI_ISL_572635, EPI_ISL_572636, EPI_ISL_572637, EPI_ISL_572638, EPI_ISL_572639, EPI_ISL_572640, EPI_ISL_572641 | | | |
| see above | West of Scotland Specialist Virology Centre, NHSGGC / MRC-University of Glasgow Centre for Virus Research | COVID-19 Genomics UK (COG-UK) Consortium | Ana da Silva Filipe, Natasha Johnson, Kathy Smollett, Daniel Mair, Stephen Carmichael, Lily Tong, Jenna Nichols, Elihu Aranday-Cortes, Kyriaki Nomikou; Sarah McDonald, Marc Niebel, Patawee Asamaphan; Richard Orton, Joseph Hughes, Sreenu Vattipally, David L Robertson; Alasdair MacLean, Rory Gunson; Kathy Li, Igor Starinskij, Natasha Jesudason, Rajiv Shah, James Shepherd, Antonia Ho, Emma Thomson |
| EPI_ISL_572642 | Department of Pathology, University of Cambridge | COVID-19 Genomics UK (COG-UK) Consortium | Aminu S. Jahun, Yasmin Chaudhry, Grant Hall, Iliana Georgana, Myra Hosmillo, Martin D. Curran, Malte Pinckert, Surendra Parmar, Ian Goodfellow |
| EPI_ISL_572643 | Wales Specialist Virology Centre Sequencing lab: Pathogen Genomics Unit | COVID-19 Genomics UK (COG-UK) Consortium | Catherine Moore, Johnathan Evans, Laura Gifford, Malorie Perry, Simon Cottrell, Angela Marchbank, Alec Birchley, Alexander Adams, Amy Gaskin, Bree Gatica-Wilcox, Jason Coombes, Joel Southgate, Lauren Gilbert, Lee Graham, Nicole Pacchiarini, Sara Kumziene-Summerhayes, Sarah Taylor, Sophie Jones, Sara Rey, Matthew Bull, Joanne Watkins, Sally Corden, Tom Connor |

| | | | |
|---|---|---|---|
| EPI_ISL_572646 | Quadram Institute Bioscience | COVID-19 Genomics UK (COG-UK) Consortium | Dave J. Baker, Gemma L. Kay, Alp Aydin, Thanh Le-Viet, Steven Rudder, Ana P. Tedim, Anastasia Kolyva, Maria Diaz, Leonardo de Oliveira Martins, Nabil-Fareed Alikhan, Lizzie Meadows, Rachael Stanley, Ngozi Elumogo, Muhammed Yasir, Nicholas M. Thomson, Alexander J Trotter, Rachel Gilroy, Samuel Bloomfield, Claire Stuart, Andrew Bell, Reenesh Prakash, Samir Dervisevic, Alison E. Mather, John Wain, Mark Webber, Andrew J. Page, Justin O'Grady |
| EPI_ISL_572647 | Wales Specialist Virology Centre Sequencing lab: Pathogen Genomics Unit | COVID-19 Genomics UK (COG-UK) Consortium | Catherine Moore, Johnathan Evans, Laura Gifford, Malorie Perry, Simon Cottrell, Angela Marchbank, Alec Birchley, Alexander Adams, Amy Gaskin, Bree Gatica-Wilcox, Jason Coombes, Joel Southgate, Lauren Gilbert, Lee Graham, Nicole Pacchiarini, Sara Kumziene-Summerhayes, Sarah Taylor, Sophie Jones, Sara Rey, Matthew Bull, Joanne Watkins, Sally Corden, Tom Connor |
| EPI_ISL_572649, EPI_ISL_572653, EPI_ISL_572655, EPI_ISL_572656, EPI_ISL_572658, EPI_ISL_572660, EPI_ISL_572665, EPI_ISL_572666, EPI_ISL_572670, EPI_ISL_572671, EPI_ISL_572673 | | | |
| see above | Quadram Institute Bioscience | COVID-19 Genomics UK (COG-UK) Consortium | Dave J. Baker, Gemma L. Kay, Alp Aydin, Thanh Le-Viet, Steven Rudder, Ana P. Tedim, Anastasia Kolyva, Maria Diaz, Leonardo de Oliveira Martins, Nabil-Fareed Alikhan, Lizzie Meadows, Rachael Stanley, Ngozi Elumogo, Muhammed Yasir, Nicholas M. Thomson, Alexander J Trotter, Rachel Gilroy, Samuel Bloomfield, Claire Stuart, Andrew Bell, Reenesh Prakash, Samir Dervisevic, Alison E. Mather, John Wain, Mark Webber, Andrew J. Page, Justin O'Grady |
| EPI_ISL_572684, EPI_ISL_572685 | University College London, Great Ormond Street Hospital for Children NHS Foundation Trust, Imperial College Healthcare NHS Trust | COVID-19 Genomics UK (COG-UK) Consortium | Sergi Castellano, Rachel Williams, Mark Kristiansen, Paola Resende Silva, Sunando Roy, Tony Brooks, Helena Tutill, Paola Niola, Patricia Dyal, Charlotte Williams, Leysa Forrest, Yasmin Panchbhaya, Jacqueline Findlay, Samuel Weeks, Julianne Brown, Kathryn Harris, Paul Randell, James Price, Alison Holmes, Judith Breuer |
| EPI_ISL_572686 | West of Scotland Specialist Virology Centre, NHSGGC / MRC-University of Glasgow Centre for Virus Research | COVID-19 Genomics UK (COG-UK) Consortium | Ana da Silva Filipe, Natasha Johnson, Kathy Smollett, Daniel Mair, Stephen Carmichael, Lily Tong, Jenna Nichols, Elihu Aranday-Cortes, Kyriaki Nomikou; Sarah McDonald, Marc Niebel, Patawee Asamaphan; Richard Orton, Joseph Hughes, Sreenu Vattipally, David L Robertson; Alasdair MacLean, Rory Gunson; Kathy Li, Igor Starinskij, Natasha Jesudason, Rajiv Shah, James Shepherd, Antonia Ho, Emma Thomson |
| EPI_ISL_572703, EPI_ISL_572704 | Northumbria University / South Tees Hospitals NHS Foundation Trust / North Cumbria Integrated Care NHS Foundation Trust / North Tees and Hartlepool NHS Foundation Trust / Newcastle Hospitals NHS Foundation Trust | COVID-19 Genomics UK (COG-UK) Consortium | Darren L Smith,Andrew Nelson,Matthew Bashton,Greg R Young,Joshua Loh,John Allan,Mohammad A Tariq,Giles S Holt,Gary Black,Wen C Yew,Lynn Dover,Paul Baker,Steve Liggett,Sarah Essex,Jane Greenaway,Debra Padgett,Clive Graham,Garren Scott,Edward Barton,Emma Swindells,Brendan Payne,Jennifer Collins,Yusri Taha,Gary Eltringham |
| EPI_ISL_572711 | Quadram Institute Bioscience | COVID-19 Genomics UK (COG-UK) Consortium | Dave J. Baker, Gemma L. Kay, Alp Aydin, Thanh Le-Viet, Steven Rudder, Ana P. Tedim, Anastasia Kolyva, Maria Diaz, Leonardo de Oliveira Martins, Nabil-Fareed Alikhan, Lizzie Meadows, Rachael Stanley, Ngozi Elumogo, Muhammed Yasir, Nicholas M. Thomson, Alexander J Trotter, Rachel Gilroy, Samuel Bloomfield, Claire Stuart, Andrew Bell, Reenesh Prakash, Samir Dervisevic, Alison E. Mather, John Wain, Mark Webber, Andrew J. Page, Justin O'Grady |
| EPI_ISL_572713 | Northumbria University / South Tees Hospitals NHS Foundation Trust / North Cumbria Integrated Care NHS Foundation Trust / North Tees and Hartlepool NHS Foundation Trust / Newcastle Hospitals NHS Foundation Trust | COVID-19 Genomics UK (COG-UK) Consortium | Darren L Smith,Andrew Nelson,Matthew Bashton,Greg R Young,Joshua Loh,John Allan,Mohammad A Tariq,Giles S Holt,Gary Black,Wen C Yew,Lynn Dover,Paul Baker,Steve Liggett,Sarah Essex,Jane Greenaway,Debra Padgett,Clive Graham,Garren Scott,Edward Barton,Emma Swindells,Brendan Payne,Jennifer Collins,Yusri Taha,Gary Eltringham |
| EPI_ISL_572714 | Oxford Viromics, NDM, University of Oxford; Oxford University Hospitals; Basingstoke and North Hampshire Hospital | COVID-19 Genomics UK (COG-UK) Consortium | Tanya Golubchik, David Bonsall, George Macintyre, Amy Trebes, Mariateresa de Cesare, Catrin Moore, Alex Mobbs, Anita Justice, Robert Shaw, Monique Andersson, Timothy Peto, Emma Wise, Nathan Moore, Jessica Lynch, Nick Cortes, Matilde Mori, Stephen Kidd, David Buck, John Todd, Christophe Fraser |
| EPI_ISL_572716 | Centre for Enzyme Innovation, University of Portsmouth / Translational Research Laboratory, Portsmouth Hospitals NHS Trust | COVID-19 Genomics UK (COG-UK) Consortium | Angela Beckett,Yann Bourgeois,Garry Scarlett,Sharon Glaysher,Scott Elliott,Kelly Bicknell,Robert Impey,Allyson Lloyd,Sarah Wyllie,Ethan Butcher,Anoop Chauhan,Samuel Robson |
| EPI_ISL_572717, EPI_ISL_572718 | Quadram Institute Bioscience | COVID-19 Genomics UK (COG-UK) Consortium | Dave J. Baker, Gemma L. Kay, Alp Aydin, Thanh Le-Viet, Steven Rudder, Ana P. Tedim, Anastasia Kolyva, Maria Diaz, Leonardo de Oliveira Martins, Nabil-Fareed Alikhan, Lizzie Meadows, Rachael Stanley, Ngozi Elumogo, Muhammed Yasir, Nicholas M. Thomson, Alexander J Trotter, Rachel Gilroy, Samuel Bloomfield, Claire Stuart, Andrew Bell, Reenesh Prakash, Samir Dervisevic, Alison E. Mather, John Wain, Mark Webber, Andrew J. Page, Justin O'Grady |
| EPI_ISL_572721 | Centre for Enzyme Innovation, University of Portsmouth / Translational Research Laboratory, Portsmouth Hospitals NHS Trust | COVID-19 Genomics UK (COG-UK) Consortium | Angela Beckett,Yann Bourgeois,Garry Scarlett,Sharon Glaysher,Scott Elliott,Kelly Bicknell,Robert Impey,Allyson Lloyd,Sarah Wyllie,Ethan Butcher,Anoop Chauhan,Samuel Robson |
| EPI_ISL_572725 | University College London, Great Ormond Street Hospital for Children NHS Foundation Trust, Imperial College Healthcare NHS Trust | COVID-19 Genomics UK (COG-UK) Consortium | Sergi Castellano, Rachel Williams, Mark Kristiansen, Paola Resende Silva, Sunando Roy, Tony Brooks, Helena Tutill, Paola Niola, Patricia Dyal, Charlotte Williams, Leysa Forrest, Yasmin Panchbhaya, Jacqueline Findlay, Samuel Weeks, Julianne Brown, Kathryn Harris, Paul Randell, James Price, Alison Holmes, Judith Breuer |
| EPI_ISL_572730, EPI_ISL_572735 | Quadram Institute Bioscience | COVID-19 Genomics UK (COG-UK) Consortium | Dave J. Baker, Gemma L. Kay, Alp Aydin, Thanh Le-Viet, Steven Rudder, Ana P. Tedim, Anastasia Kolyva, Maria Diaz, Leonardo de Oliveira Martins, Nabil-Fareed Alikhan, Lizzie Meadows, Rachael Stanley, Ngozi Elumogo, Muhammed Yasir, Nicholas M. Thomson, Alexander J Trotter, Rachel Gilroy, Samuel Bloomfield, Claire Stuart, Andrew Bell, Reenesh Prakash, Samir Dervisevic, Alison E. Mather, John Wain, Mark Webber, Andrew J. Page, Justin O'Grady |
| EPI_ISL_572737, EPI_ISL_572738 | Liverpool Clinical Laboratories | COVID-19 Genomics UK (COG-UK) Consortium | Sam Haldenby, Anita Lucaci, Steve Paterson, Julian Hiscox, Alistair Darby, M Almsaud, A Alrezaihi, Muhannad Alruwaili, Stuart D Armstrong, Jones Benjamin, Eleanor G Bentley, Anu Chawla, Jordan J Clark, Angela Cowell, Richard Eccles, Isabel García-Dorival, Matthew Gemmell, Alessandro Gerada, PKF Gilmore, Richard Gregory, Ximeng Han, Catherine Hartley, Margaret Hughes, Miren Iturriza-Gomara, James Johnson, L Luu, Jenifer Manson, Charlotte Nelson, Elaine O'Toole, Cassie Olateju, Rebekah Penrice-Randal , Lucille Rainbow, N.P Randle, Trevor Ian Robinson, Parul Sharma, Ghada T Shawli, James P Stewart, Neil Swainston, Ecaterina Vamos, Joanne Watts, Mark Whitehead |
| EPI_ISL_572742, EPI_ISL_572745 | Quadram Institute Bioscience | COVID-19 Genomics UK (COG-UK) Consortium | Dave J. Baker, Gemma L. Kay, Alp Aydin, Thanh Le-Viet, Steven Rudder, Ana P. Tedim, Anastasia Kolyva, Maria Diaz, Leonardo de Oliveira Martins, Nabil-Fareed Alikhan, Lizzie Meadows, Rachael Stanley, Ngozi Elumogo, Muhammed Yasir, Nicholas M. Thomson, Alexander J Trotter, Rachel Gilroy, Samuel Bloomfield, Claire Stuart, Andrew Bell, Reenesh Prakash, Samir Dervisevic, Alison E. Mather, John Wain, Mark Webber, Andrew J. Page, Justin O'Grady |
| EPI_ISL_572746 | West of Scotland Specialist Virology Centre, NHSGGC / MRC-University of Glasgow Centre for Virus Research | COVID-19 Genomics UK (COG-UK) Consortium | Ana da Silva Filipe, Natasha Johnson, Kathy Smollett, Daniel Mair, Stephen Carmichael, Lily Tong, Jenna Nichols, Elihu Aranday-Cortes, Kyriaki Nomikou; Sarah McDonald, Marc Niebel, Patawee Asamaphan; Richard Orton, Joseph Hughes, Sreenu Vattipally, David L Robertson; Alasdair MacLean, Rory Gunson; Kathy Li, Igor Starinskij, Natasha Jesudason, Rajiv Shah, James Shepherd, Antonia Ho, Emma Thomson |
| EPI_ISL_572749, EPI_ISL_572755, EPI_ISL_572758, EPI_ISL_572759 | Quadram Institute Bioscience | COVID-19 Genomics UK (COG-UK) Consortium | Dave J. Baker, Gemma L. Kay, Alp Aydin, Thanh Le-Viet, Steven Rudder, Ana P. Tedim, Anastasia Kolyva, Maria Diaz, Leonardo de Oliveira Martins, Nabil-Fareed Alikhan, Lizzie Meadows, Rachael Stanley, Ngozi Elumogo, Muhammed Yasir, Nicholas M. Thomson, Alexander J Trotter, Rachel Gilroy, Samuel Bloomfield, Claire Stuart, Andrew Bell, Reenesh Prakash, Samir Dervisevic, Alison E. Mather, John Wain, Mark Webber, Andrew J. Page, Justin O'Grady |

| | | | |
|---|---|---|---|
| EPI_ISL_572763, EPI_ISL_572765 | West of Scotland Specialist Virology Centre, NHSGGC / MRC-University of Glasgow Centre for Virus Research | COVID-19 Genomics UK (COG-UK) Consortium | Ana da Silva Filipe, Natasha Johnson, Kathy Smollett, Daniel Mair, Stephen Carmichael, Lily Tong, Jenna Nichols, Elihu Aranday-Cortes, Kyriaki Nomikou; Sarah McDonald, Marc Niebel, Patawee Asamaphan; Richard Orton, Joseph Hughes, Sreenu Vattipally, David L Robertson; Alasdair MacLean, Rory Gunson; Kathy Li, Igor Starinskij, Natasha Jesudason, Rajiv Shah, James Shepherd, Antonia Ho, Emma Thomson |
| EPI_ISL_572767 | Virology Department, Sheffield Teaching Hospitals NHS Foundation Trust/Department of Infection, Immunity and Cardiovascular Disease, The Medical School, University of Sheffield | COVID-19 Genomics UK (COG-UK) Consortium | Thushan de Silva, Matthew Parker, Nikki Smith, Adri Angyal, Rebecca Brown, Luke Green, Rachel Tucker, Paul Parsons, Danielle Groves, Katie Johnson, Laura Carrilero, Alex Keeley, Dave Partridge, Matthew Wyles, Benjamin Lindsey, Mehmet Yavuz, Mohammad Raza, Cariad Evans |
| EPI_ISL_572768, EPI_ISL_572775, EPI_ISL_572780 | Quadram Institute Bioscience | COVID-19 Genomics UK (COG-UK) Consortium | Dave J. Baker, Gemma L. Kay, Alp Aydin, Thanh Le-Viet, Steven Rudder, Ana P. Tedim, Anastasia Kolyva, Maria Diaz, Leonardo de Oliveira Martins, Nabil-Fareed Alikhan, Lizzie Meadows, Rachael Stanley, Ngozi Elumogo, Muhammed Yasir, Nicholas M. Thomson, Alexander J Trotter, Rachel Gilroy, Samuel Bloomfield, Claire Stuart, Andrew Bell, Reenesh Prakash, Samir Dervisevic, Alison E. Mather, John Wain, Mark Webber, Andrew J. Page, Justin O'Grady |
| EPI_ISL_572781, EPI_ISL_572784 | Wales Specialist Virology Centre Sequencing lab: Pathogen Genomics Unit | COVID-19 Genomics UK (COG-UK) Consortium | Catherine Moore, Johnathan Evans, Laura Gifford, Malorie Perry, Simon Cottrell, Angela Marchbank, Alec Birchley, Alexander Adams, Amy Gaskin, Bree Gatica-Wilcox, Jason Coombes, Joel Southgate, Lauren Gilbert, Lee Graham, Nicole Pacchiarini, Sara Kumziene-Summerhayes, Sarah Taylor, Sophie Jones, Sara Rey, Matthew Bull, Joanne Watkins, Sally Corden, Tom Connor |
| EPI_ISL_572786, EPI_ISL_572787 | Quadram Institute Bioscience | COVID-19 Genomics UK (COG-UK) Consortium | Dave J. Baker, Gemma L. Kay, Alp Aydin, Thanh Le-Viet, Steven Rudder, Ana P. Tedim, Anastasia Kolyva, Maria Diaz, Leonardo de Oliveira Martins, Nabil-Fareed Alikhan, Lizzie Meadows, Rachael Stanley, Ngozi Elumogo, Muhammed Yasir, Nicholas M. Thomson, Alexander J Trotter, Rachel Gilroy, Samuel Bloomfield, Claire Stuart, Andrew Bell, Reenesh Prakash, Samir Dervisevic, Alison E. Mather, John Wain, Mark Webber, Andrew J. Page, Justin O'Grady |
| EPI_ISL_572788 | West of Scotland Specialist Virology Centre, NHSGGC / MRC-University of Glasgow Centre for Virus Research | COVID-19 Genomics UK (COG-UK) Consortium | Ana da Silva Filipe, Natasha Johnson, Kathy Smollett, Daniel Mair, Stephen Carmichael, Lily Tong, Jenna Nichols, Elihu Aranday-Cortes, Kyriaki Nomikou; Sarah McDonald, Marc Niebel, Patawee Asamaphan; Richard Orton, Joseph Hughes, Sreenu Vattipally, David L Robertson; Alasdair MacLean, Rory Gunson; Kathy Li, Igor Starinskij, Natasha Jesudason, Rajiv Shah, James Shepherd, Antonia Ho, Emma Thomson |
| EPI_ISL_572789, EPI_ISL_572791 | Quadram Institute Bioscience | COVID-19 Genomics UK (COG-UK) Consortium | Dave J. Baker, Gemma L. Kay, Alp Aydin, Thanh Le-Viet, Steven Rudder, Ana P. Tedim, Anastasia Kolyva, Maria Diaz, Leonardo de Oliveira Martins, Nabil-Fareed Alikhan, Lizzie Meadows, Rachael Stanley, Ngozi Elumogo, Muhammed Yasir, Nicholas M. Thomson, Alexander J Trotter, Rachel Gilroy, Samuel Bloomfield, Claire Stuart, Andrew Bell, Reenesh Prakash, Samir Dervisevic, Alison E. Mather, John Wain, Mark Webber, Andrew J. Page, Justin O'Grady |
| EPI_ISL_572792 | Department of Pathology, University of Cambridge | COVID-19 Genomics UK (COG-UK) Consortium | Aminu S. Jahun, Yasmin Chaudhry, Grant Hall, Iliana Georgana, Myra Hosmillo, Martin D. Curran, Malte Pinckert, Surendra Parmar, Ian Goodfellow |
| EPI_ISL_572794 | West of Scotland Specialist Virology Centre, NHSGGC / MRC-University of Glasgow Centre for Virus Research | COVID-19 Genomics UK (COG-UK) Consortium | Ana da Silva Filipe, Natasha Johnson, Kathy Smollett, Daniel Mair, Stephen Carmichael, Lily Tong, Jenna Nichols, Elihu Aranday-Cortes, Kyriaki Nomikou; Sarah McDonald, Marc Niebel, Patawee Asamaphan; Richard Orton, Joseph Hughes, Sreenu Vattipally, David L Robertson; Alasdair MacLean, Rory Gunson; Kathy Li, Igor Starinskij, Natasha Jesudason, Rajiv Shah, James Shepherd, Antonia Ho, Emma Thomson |
| EPI_ISL_572797 | Quadram Institute Bioscience | COVID-19 Genomics UK (COG-UK) Consortium | Dave J. Baker, Gemma L. Kay, Alp Aydin, Thanh Le-Viet, Steven Rudder, Ana P. Tedim, Anastasia Kolyva, Maria Diaz, Leonardo de Oliveira Martins, Nabil-Fareed Alikhan, Lizzie Meadows, Rachael Stanley, Ngozi Elumogo, Muhammed Yasir, Nicholas M. Thomson, Alexander J Trotter, Rachel Gilroy, Samuel Bloomfield, Claire Stuart, Andrew Bell, Reenesh Prakash, Samir Dervisevic, Alison E. Mather, John Wain, Mark Webber, Andrew J. Page, Justin O'Grady |
| EPI_ISL_572798 | Northumbria University / South Tees Hospitals NHS Foundation Trust / North Cumbria Integrated Care NHS Foundation Trust / North Tees and Hartlepool NHS Foundation Trust / Newcastle Hospitals NHS Foundation Trust | COVID-19 Genomics UK (COG-UK) Consortium | Darren L Smith,Andrew Nelson,Matthew Bashton,Greg R Young,Joshua Loh,John Allan,Mohammad A Tariq,Giles S Holt,Gary Black,Wen C Yew,Lynn Dover,Paul Baker,Steve Liggett,Sarah Essex,Jane Greenaway,Debra Padgett,Clive Graham,Garren Scott,Edward Barton,Emma Swindells,Brendan Payne,Jennifer Collins,Yusri Taha,Gary Eltringham |
| EPI_ISL_572800 | University College London, Great Ormond Street Hospital for Children NHS Foundation Trust, Imperial College Healthcare NHS Trust | COVID-19 Genomics UK (COG-UK) Consortium | Sergi Castellano, Rachel Williams, Mark Kristiansen, Paola Resende Silva, Sunando Roy, Tony Brooks, Helena Tutill, Paola Niola, Patricia Dyal, Charlotte Williams, Leysa Forrest, Yasmin Panchbhaya, Jacqueline Findlay, Samuel Weeks, Julianne Brown, Kathryn Harris, Paul Randell, James Price, Alison Holmes, Judith Breuer |
| EPI_ISL_572804, EPI_ISL_572806 | Quadram Institute Bioscience | COVID-19 Genomics UK (COG-UK) Consortium | Dave J. Baker, Gemma L. Kay, Alp Aydin, Thanh Le-Viet, Steven Rudder, Ana P. Tedim, Anastasia Kolyva, Maria Diaz, Leonardo de Oliveira Martins, Nabil-Fareed Alikhan, Lizzie Meadows, Rachael Stanley, Ngozi Elumogo, Muhammed Yasir, Nicholas M. Thomson, Alexander J Trotter, Rachel Gilroy, Samuel Bloomfield, Claire Stuart, Andrew Bell, Reenesh Prakash, Samir Dervisevic, Alison E. Mather, John Wain, Mark Webber, Andrew J. Page, Justin O'Grady |
| EPI_ISL_572808, EPI_ISL_572809 | University College London, Great Ormond Street Hospital for Children NHS Foundation Trust, Imperial College Healthcare NHS Trust | COVID-19 Genomics UK (COG-UK) Consortium | Sergi Castellano, Rachel Williams, Mark Kristiansen, Paola Resende Silva, Sunando Roy, Tony Brooks, Helena Tutill, Paola Niola, Patricia Dyal, Charlotte Williams, Leysa Forrest, Yasmin Panchbhaya, Jacqueline Findlay, Samuel Weeks, Julianne Brown, Kathryn Harris, Paul Randell, James Price, Alison Holmes, Judith Breuer |
| EPI_ISL_572811, EPI_ISL_572812, EPI_ISL_572819 | Quadram Institute Bioscience | COVID-19 Genomics UK (COG-UK) Consortium | Dave J. Baker, Gemma L. Kay, Alp Aydin, Thanh Le-Viet, Steven Rudder, Ana P. Tedim, Anastasia Kolyva, Maria Diaz, Leonardo de Oliveira Martins, Nabil-Fareed Alikhan, Lizzie Meadows, Rachael Stanley, Ngozi Elumogo, Muhammed Yasir, Nicholas M. Thomson, Alexander J Trotter, Rachel Gilroy, Samuel Bloomfield, Claire Stuart, Andrew Bell, Reenesh Prakash, Samir Dervisevic, Alison E. Mather, John Wain, Mark Webber, Andrew J. Page, Justin O'Grady |
| EPI_ISL_572822 | Liverpool Clinical Laboratories | COVID-19 Genomics UK (COG-UK) Consortium | Sam Haldenby, Anita Lucaci, Steve Paterson, Julian Hiscox, Alistair Darby, M Almsaud, A Alrezaihi, Muhannad Alruwaili, Stuart D Armstrong, Jones Benjamin, Eleanor G Bentley, Anu Chawla, Jordan J Clark, Angela Cowell, Richard Eccles, Isabel García-Dorival, Matthew Gemmell, Alessandro Gerada, PKF Gilmore, Richard Gregory, Ximeng Han, Catherine Hartley, Margaret Hughes, Miren Iturriza-Gomara, James Johnson, L Luu, Jenifer Manson, Charlotte Nelson, Elaine O'Toole, Cassie Olateju, Rebekah Penrice-Randal , Lucille Rainbow, N.P Randle, Trevor Ian Robinson, Parul Sharma, Ghada T Shawli, James P Stewart, Neil Swainston, Ecaterina Vamos, Joanne Watts, Mark Whitehead |
| EPI_ISL_572823, EPI_ISL_572825, EPI_ISL_572826 | Quadram Institute Bioscience | COVID-19 Genomics UK (COG-UK) Consortium | Dave J. Baker, Gemma L. Kay, Alp Aydin, Thanh Le-Viet, Steven Rudder, Ana P. Tedim, Anastasia Kolyva, Maria Diaz, Leonardo de Oliveira Martins, Nabil-Fareed Alikhan, Lizzie Meadows, Rachael Stanley, Ngozi Elumogo, Muhammed Yasir, Nicholas M. Thomson, Alexander J Trotter, Rachel Gilroy, Samuel Bloomfield, Claire Stuart, Andrew Bell, Reenesh Prakash, Samir Dervisevic, Alison E. Mather, John Wain, Mark Webber, Andrew J. Page, Justin O'Grady |
| EPI_ISL_572829 | University College London, Great Ormond Street Hospital for Children NHS Foundation Trust, Imperial College Healthcare NHS Trust | COVID-19 Genomics UK (COG-UK) Consortium | Sergi Castellano, Rachel Williams, Mark Kristiansen, Paola Resende Silva, Sunando Roy, Tony Brooks, Helena Tutill, Paola Niola, Patricia Dyal, Charlotte Williams, Leysa Forrest, Yasmin Panchbhaya, Jacqueline Findlay, Samuel Weeks, Julianne Brown, Kathryn Harris, Paul Randell, James Price, Alison Holmes, Judith Breuer |
| EPI_ISL_572831 | Quadram Institute Bioscience | COVID-19 Genomics UK (COG-UK) Consortium | Dave J. Baker, Gemma L. Kay, Alp Aydin, Thanh Le-Viet, Steven Rudder, Ana P. Tedim, Anastasia Kolyva, Maria Diaz, Leonardo de Oliveira Martins, Nabil-Fareed Alikhan, Lizzie Meadows, Rachael Stanley, Ngozi Elumogo, Muhammed Yasir, Nicholas M. Thomson, Alexander J Trotter, Rachel Gilroy, Samuel Bloomfield, Claire Stuart, Andrew Bell, Reenesh Prakash, Samir Dervisevic, Alison E. Mather, John Wain, Mark Webber, Andrew J. Page, Justin O'Grady |
| EPI_ISL_572832 | West of Scotland Specialist Virology Centre, NHSGGC / MRC-University of Glasgow Centre for Virus Research | COVID-19 Genomics UK (COG-UK) Consortium | Ana da Silva Filipe, Natasha Johnson, Kathy Smollett, Daniel Mair, Stephen Carmichael, Lily Tong, Jenna Nichols, Elihu Aranday-Cortes, Kyriaki Nomikou; Sarah McDonald, Marc Niebel, Patawee Asamaphan; Richard Orton, Joseph Hughes, Sreenu Vattipally, David L Robertson; Alasdair MacLean, Rory Gunson; Kathy Li, Igor Starinskij, Natasha Jesudason, Rajiv Shah, James Shepherd, Antonia Ho, Emma Thomson |

| | | | |
|---|---|---|---|
| EPI_ISL_572835 | Quadram Institute Bioscience | COVID-19 Genomics UK (COG-UK) Consortium | Dave J. Baker, Gemma L. Kay, Alp Aydin, Thanh Le-Viet, Steven Rudder, Ana P. Tedim, Anastasia Kolyva, Maria Diaz, Leonardo de Oliveira Martins, Nabil-Fareed Alikhan, Lizzie Meadows, Rachael Stanley, Ngozi Elumogo, Muhammed Yasir, Nicholas M. Thomson, Alexander J Trotter, Rachel Gilroy, Samuel Bloomfield, Claire Stuart, Andrew Bell, Reenesh Prakash, Samir Dervisevic, Alison E. Mather, John Wain, Mark Webber, Andrew J. Page, Justin O'Grady |
| EPI_ISL_572837 | University College London, Great Ormond Street Hospital for Children NHS Foundation Trust, Imperial College Healthcare NHS Trust | COVID-19 Genomics UK (COG-UK) Consortium | Sergi Castellano, Rachel Williams, Mark Kristiansen, Paola Resende Silva, Sunando Roy, Tony Brooks, Helena Tutill, Paola Niola, Patricia Dyal, Charlotte Williams, Leysa Forrest, Yasmin Panchbhaya, Jacqueline Findlay, Samuel Weeks, Julianne Brown, Kathryn Harris, Paul Randell, James Price, Alison Holmes, Judith Breuer |
| EPI_ISL_572844, EPI_ISL_572847 | Quadram Institute Bioscience | COVID-19 Genomics UK (COG-UK) Consortium | Dave J. Baker, Gemma L. Kay, Alp Aydin, Thanh Le-Viet, Steven Rudder, Ana P. Tedim, Anastasia Kolyva, Maria Diaz, Leonardo de Oliveira Martins, Nabil-Fareed Alikhan, Lizzie Meadows, Rachael Stanley, Ngozi Elumogo, Muhammed Yasir, Nicholas M. Thomson, Alexander J Trotter, Rachel Gilroy, Samuel Bloomfield, Claire Stuart, Andrew Bell, Reenesh Prakash, Samir Dervisevic, Alison E. Mather, John Wain, Mark Webber, Andrew J. Page, Justin O'Grady |
| EPI_ISL_572851, EPI_ISL_572852 | Department of Pathology, University of Cambridge | COVID-19 Genomics UK (COG-UK) Consortium | Aminu S. Jahun, Yasmin Chaudhry, Grant Hall, Iliana Georgana, Myra Hosmillo, Martin D. Curran, Malte Pinckert, Surendra Parmar, Ian Goodfellow |
| EPI_ISL_572860, EPI_ISL_572861 | Oxford Viromics, NDM, University of Oxford; Oxford University Hospitals; Basingstoke and North Hampshire Hospital | COVID-19 Genomics UK (COG-UK) Consortium | Tanya Golubchik, David Bonsall, George Macintyre, Amy Trebes, Mariateresa de Cesare, Catrin Moore, Alex Mobbs, Anita Justice, Robert Shaw, Monique Andersson, Timothy Peto, Emma Wise, Nathan Moore, Jessica Lynch, Nick Cortes, Matilde Mori, Stephen Kidd, David Buck, John Todd, Christophe Fraser |
| EPI_ISL_572862, EPI_ISL_572863, EPI_ISL_572864, EPI_ISL_572865, EPI_ISL_572866, EPI_ISL_572867, EPI_ISL_572868 | Department of Pathology, University of Cambridge | COVID-19 Genomics UK (COG-UK) Consortium | Aminu S. Jahun, Yasmin Chaudhry, Grant Hall, Iliana Georgana, Myra Hosmillo, Martin D. Curran, Malte Pinckert, Surendra Parmar, Ian Goodfellow |
| EPI_ISL_572877, EPI_ISL_572878 | Wales Specialist Virology Centre Sequencing lab: Pathogen Genomics Unit | COVID-19 Genomics UK (COG-UK) Consortium | Catherine Moore, Johnathan Evans, Laura Gifford, Malorie Perry, Simon Cottrell, Angela Marchbank, Alec Birchley, Alexander Adams, Amy Gaskin, Bree Gatica-Wilcox, Jason Coombes, Joel Southgate, Lauren Gilbert, Lee Graham, Nicole Pacchiarini, Sara Kumziene-Summerhayes, Sarah Taylor, Sophie Jones, Sara Rey, Matthew Bull, Joanne Watkins, Sally Corden, Tom Connor |
| EPI_ISL_572891 | Oxford Viromics, NDM, University of Oxford; Oxford University Hospitals; Basingstoke and North Hampshire Hospital | COVID-19 Genomics UK (COG-UK) Consortium | Tanya Golubchik, David Bonsall, George Macintyre, Amy Trebes, Mariateresa de Cesare, Catrin Moore, Alex Mobbs, Anita Justice, Robert Shaw, Monique Andersson, Timothy Peto, Emma Wise, Nathan Moore, Jessica Lynch, Nick Cortes, Matilde Mori, Stephen Kidd, David Buck, John Todd, Christophe Fraser |
| EPI_ISL_572898, EPI_ISL_572899, EPI_ISL_572900, EPI_ISL_572901 | Quadram Institute Bioscience | COVID-19 Genomics UK (COG-UK) Consortium | Dave J. Baker, Gemma L. Kay, Alp Aydin, Thanh Le-Viet, Steven Rudder, Ana P. Tedim, Anastasia Kolyva, Maria Diaz, Leonardo de Oliveira Martins, Nabil-Fareed Alikhan, Lizzie Meadows, Rachael Stanley, Ngozi Elumogo, Muhammed Yasir, Nicholas M. Thomson, Alexander J Trotter, Rachel Gilroy, Samuel Bloomfield, Claire Stuart, Andrew Bell, Reenesh Prakash, Samir Dervisevic, Alison E. Mather, John Wain, Mark Webber, Andrew J. Page, Justin O'Grady |
| EPI_ISL_572903 | Oxford Viromics, NDM, University of Oxford; Oxford University Hospitals; Basingstoke and North Hampshire Hospital | COVID-19 Genomics UK (COG-UK) Consortium | Tanya Golubchik, David Bonsall, George Macintyre, Amy Trebes, Mariateresa de Cesare, Catrin Moore, Alex Mobbs, Anita Justice, Robert Shaw, Monique Andersson, Timothy Peto, Emma Wise, Nathan Moore, Jessica Lynch, Nick Cortes, Matilde Mori, Stephen Kidd, David Buck, John Todd, Christophe Fraser |
| EPI_ISL_572905, EPI_ISL_572906, EPI_ISL_572907, EPI_ISL_572908 | Liverpool Clinical Laboratories | COVID-19 Genomics UK (COG-UK) Consortium | Sam Haldenby, Anita Lucaci, Steve Paterson, Julian Hiscox, Alistair Darby, M Almsaud, A Alrezaihi, Muhannad Alruwaili, Stuart D Armstrong, Jones Benjamin, Eleanor G Bentley, Anu Chawla, Jordan J Clark, Angela Cowell, Richard Eccles, Isabel García-Dorival, Matthew Gemmell, Alessandro Gerada, PKF Gilmore, Richard Gregory, Ximeng Han, Catherine Hartley, Margaret Hughes, Miren Iturriza-Gomara, James Johnson, L Luu, Jenifer Manson, Charlotte Nelson, Elaine O'Toole, Cassie Olateju, Rebekah Penrice-Randal , Lucille Rainbow, N.P Randle, Trevor Ian Robinson, Parul Sharma, Ghada T Shawli, James P Stewart, Neil Swainston, Ecaterina Vamos, Joanne Watts, Mark Whitehead |
| EPI_ISL_572928 | Virology Department, Sheffield Teaching Hospitals NHS Foundation Trust/Department of Infection, Immunity and Cardiovascular Disease, The Medical School, University of Sheffield | COVID-19 Genomics UK (COG-UK) Consortium | Thushan de Silva, Matthew Parker, Nikki Smith, Adri Angyal, Rebecca Brown, Luke Green, Rachel Tucker, Paul Parsons, Danielle Groves, Katie Johnson, Laura Carrilero, Alex Keeley, Dave Partridge, Matthew Wyles, Benjamin Lindsey, Mehmet Yavuz, Mohammad Raza, Cariad Evans |
| EPI_ISL_572931 | Wales Specialist Virology Centre Sequencing lab: Pathogen Genomics Unit | COVID-19 Genomics UK (COG-UK) Consortium | Catherine Moore, Johnathan Evans, Laura Gifford, Malorie Perry, Simon Cottrell, Angela Marchbank, Alec Birchley, Alexander Adams, Amy Gaskin, Bree Gatica-Wilcox, Jason Coombes, Joel Southgate, Lauren Gilbert, Lee Graham, Nicole Pacchiarini, Sara Kumziene-Summerhayes, Sarah Taylor, Sophie Jones, Sara Rey, Matthew Bull, Joanne Watkins, Sally Corden, Tom Connor |
| EPI_ISL_572937 | Oxford Viromics, NDM, University of Oxford; Oxford University Hospitals; Basingstoke and North Hampshire Hospital | COVID-19 Genomics UK (COG-UK) Consortium | Tanya Golubchik, David Bonsall, George Macintyre, Amy Trebes, Mariateresa de Cesare, Catrin Moore, Alex Mobbs, Anita Justice, Robert Shaw, Monique Andersson, Timothy Peto, Emma Wise, Nathan Moore, Jessica Lynch, Nick Cortes, Matilde Mori, Stephen Kidd, David Buck, John Todd, Christophe Fraser |
| EPI_ISL_572939, EPI_ISL_572940, EPI_ISL_572941 | Quadram Institute Bioscience | COVID-19 Genomics UK (COG-UK) Consortium | Dave J. Baker, Gemma L. Kay, Alp Aydin, Thanh Le-Viet, Steven Rudder, Ana P. Tedim, Anastasia Kolyva, Maria Diaz, Leonardo de Oliveira Martins, Nabil-Fareed Alikhan, Lizzie Meadows, Rachael Stanley, Ngozi Elumogo, Muhammed Yasir, Nicholas M. Thomson, Alexander J Trotter, Rachel Gilroy, Samuel Bloomfield, Claire Stuart, Andrew Bell, Reenesh Prakash, Samir Dervisevic, Alison E. Mather, John Wain, Mark Webber, Andrew J. Page, Justin O'Grady |
| EPI_ISL_572942 | Northumbria University / South Tees Hospitals NHS Foundation Trust / North Cumbria Integrated Care NHS Foundation Trust / North Tees and Hartlepool NHS Foundation Trust / Newcastle Hospitals NHS Foundation Trust | COVID-19 Genomics UK (COG-UK) Consortium | Darren L Smith,Andrew Nelson,Matthew Bashton,Greg R Young,Joshua Loh,John Allan,Mohammad A Tariq,Giles S Holt,Gary Black,Wen C Yew,Lynn Dover,Paul Baker,Steve Liggett,Sarah Essex,Jane Greenaway,Debra Padgett,Clive Graham,Garren Scott,Edward Barton,Emma Swindells,Brendan Payne,Jennifer Collins,Yusri Taha,Gary Eltringham |
| EPI_ISL_572943, EPI_ISL_572944 | Lincolnshire Hospitals and DeepSeq Nottingham | COVID-19 Genomics UK (COG-UK) Consortium | Nichola Duckworth, Tim Sloan, Sarah Walsh, Jonathan Ball, Patrick McClure, Joeseph Chappell, Nadine Holmes, Matthew Carlisle, Christopher Moore, Fei Sang, Johnny Debebe, Victoria Wright, Matthew Loose |
| EPI_ISL_572951 | Wales Specialist Virology Centre Sequencing lab: Pathogen Genomics Unit | COVID-19 Genomics UK (COG-UK) Consortium | Catherine Moore, Johnathan Evans, Laura Gifford, Malorie Perry, Simon Cottrell, Angela Marchbank, Alec Birchley, Alexander Adams, Amy Gaskin, Bree Gatica-Wilcox, Jason Coombes, Joel Southgate, Lauren Gilbert, Lee Graham, Nicole Pacchiarini, Sara Kumziene-Summerhayes, Sarah Taylor, Sophie Jones, Sara Rey, Matthew Bull, Joanne Watkins, Sally Corden, Tom Connor |

| | | | |
|---|---|---|---|
| EPI_ISL_572954 | Quadram Institute Bioscience | COVID-19 Genomics UK (COG-UK) Consortium | Dave J. Baker, Gemma L. Kay, Alp Aydin, Thanh Le-Viet, Steven Rudder, Ana P. Tedim, Anastasia Kolyva, Maria Diaz, Leonardo de Oliveira Martins, Nabil-Fareed Alikhan, Lizzie Meadows, Rachael Stanley, Ngozi Elumogo, Muhammed Yasir, Nicholas M. Thomson, Alexander J Trotter, Rachel Gilroy, Samuel Bloomfield, Claire Stuart, Andrew Bell, Reenesh Prakash, Samir Dervisevic, Alison E. Mather, John Wain, Mark Webber, Andrew J. Page, Justin O'Grady |
| EPI_ISL_572956 | Virology Department, Sheffield Teaching Hospitals NHS Foundation Trust/Department of Infection, Immunity and Cardiovascular Disease, The Medical School, University of Sheffield | COVID-19 Genomics UK (COG-UK) Consortium | Thushan de Silva, Matthew Parker, Nikki Smith, Adri Angyal, Rebecca Brown, Luke Green, Rachel Tucker, Paul Parsons, Danielle Groves, Katie Johnson, Laura Carrilero, Alex Keeley, Dave Partridge, Matthew Wyles, Benjamin Lindsey, Mehmet Yavuz, Mohammad Raza, Cariad Evans |
| EPI_ISL_573081, EPI_ISL_573082, EPI_ISL_573083 | Oxford Viromics, NDM, University of Oxford; Oxford University Hospitals; Basingstoke and North Hampshire Hospital | COVID-19 Genomics UK (COG-UK) Consortium | Tanya Golubchik, David Bonsall, George Macintyre, Amy Trebes, Mariateresa de Cesare, Catrin Moore, Alex Mobbs, Anita Justice, Robert Shaw, Monique Andersson, Timothy Peto, Emma Wise, Nathan Moore, Jessica Lynch, Nick Cortes, Matilde Mori, Stephen Kidd, David Buck, John Todd, Christophe Fraser |
| EPI_ISL_573085, EPI_ISL_573086, EPI_ISL_573087, EPI_ISL_573088, EPI_ISL_573089, EPI_ISL_573090, EPI_ISL_573091, EPI_ISL_573092, EPI_ISL_573093, EPI_ISL_573094, EPI_ISL_573095, EPI_ISL_573096, EPI_ISL_573097, EPI_ISL_573098, EPI_ISL_573099, EPI_ISL_573100, EPI_ISL_573101, EPI_ISL_573102, EPI_ISL_573103, EPI_ISL_573104, EPI_ISL_573105, EPI_ISL_573106, EPI_ISL_573107, EPI_ISL_573108, EPI_ISL_573109, EPI_ISL_573110, EPI_ISL_573111, EPI_ISL_573112, EPI_ISL_573113, EPI_ISL_573114, EPI_ISL_573115, EPI_ISL_573116, EPI_ISL_573117, EPI_ISL_573118, EPI_ISL_573119, EPI_ISL_573120, EPI_ISL_573121, EPI_ISL_573122, EPI_ISL_573123, EPI_ISL_573124, EPI_ISL_573125, EPI_ISL_573126, EPI_ISL_573127, EPI_ISL_573128, EPI_ISL_573129, EPI_ISL_573130, EPI_ISL_573131 | | | |
| see above | Quadram Institute Bioscience | COVID-19 Genomics UK (COG-UK) Consortium | Dave J. Baker, Gemma L. Kay, Alp Aydin, Thanh Le-Viet, Steven Rudder, Ana P. Tedim, Anastasia Kolyva, Maria Diaz, Leonardo de Oliveira Martins, Nabil-Fareed Alikhan, Lizzie Meadows, Rachael Stanley, Ngozi Elumogo, Muhammed Yasir, Nicholas M. Thomson, Alexander J Trotter, Rachel Gilroy, Samuel Bloomfield, Claire Stuart, Andrew Bell, Reenesh Prakash, Samir Dervisevic, Alison E. Mather, John Wain, Mark Webber, Andrew J. Page, Justin O'Grady |
| EPI_ISL_573138, EPI_ISL_573139, EPI_ISL_573140, EPI_ISL_573141, EPI_ISL_573142, EPI_ISL_573143, EPI_ISL_573144, EPI_ISL_573146, EPI_ISL_573148, EPI_ISL_573149, EPI_ISL_573153 | | | |
| see above | Department of Pathology, University of Cambridge | COVID-19 Genomics UK (COG-UK) Consortium | Aminu S. Jahun, Yasmin Chaudhry, Grant Hall, Iliana Georgana, Myra Hosmillo, Martin D. Curran, Malte Pinckert, Surendra Parmar, Ian Goodfellow |
| EPI_ISL_573195, EPI_ISL_573198, EPI_ISL_573199, EPI_ISL_573200, EPI_ISL_573201, EPI_ISL_573203, EPI_ISL_573206, EPI_ISL_573207, EPI_ISL_573208, EPI_ISL_573210, EPI_ISL_573211, EPI_ISL_573212, EPI_ISL_573213, EPI_ISL_573214, EPI_ISL_573215, EPI_ISL_573216, EPI_ISL_573217, EPI_ISL_573218, EPI_ISL_573219, EPI_ISL_573221, EPI_ISL_573222, EPI_ISL_573223, EPI_ISL_573224, EPI_ISL_573225, EPI_ISL_573227, EPI_ISL_573231, EPI_ISL_573232 | | | |
| see above | Oxford Viromics, NDM, University of Oxford; Oxford University Hospitals; Basingstoke and North Hampshire Hospital | COVID-19 Genomics UK (COG-UK) Consortium | Tanya Golubchik, David Bonsall, George Macintyre, Amy Trebes, Mariateresa de Cesare, Catrin Moore, Alex Mobbs, Anita Justice, Robert Shaw, Monique Andersson, Timothy Peto, Emma Wise, Nathan Moore, Jessica Lynch, Nick Cortes, Matilde Mori, Stephen Kidd, David Buck, John Todd, Christophe Fraser |
| EPI_ISL_573255, EPI_ISL_573257, EPI_ISL_573261, EPI_ISL_573262, EPI_ISL_573263, EPI_ISL_573264 | Centre for Enzyme Innovation, University of Portsmouth / Translational Research Laboratory, Portsmouth Hospitals NHS Trust | COVID-19 Genomics UK (COG-UK) Consortium | Angela Beckett,Yann Bourgeois,Garry Scarlett,Sharon Glaysher,Scott Elliott,Kelly Bicknell,Robert Impey,Allyson Lloyd,Sarah Wyllie,Ethan Butcher,Anoop Chauhan,Samuel Robson |
| EPI_ISL_573277, EPI_ISL_573278, EPI_ISL_573293, EPI_ISL_573298, EPI_ISL_573305, EPI_ISL_573382, EPI_ISL_573383, EPI_ISL_573384, EPI_ISL_573385, EPI_ISL_573386, EPI_ISL_573387, EPI_ISL_573388, EPI_ISL_573389, EPI_ISL_573390, EPI_ISL_573391, EPI_ISL_573392, EPI_ISL_573393, EPI_ISL_573394, EPI_ISL_573395, EPI_ISL_573396, EPI_ISL_573397, EPI_ISL_573398, EPI_ISL_573399, EPI_ISL_573400, EPI_ISL_573401, EPI_ISL_573402, EPI_ISL_573403, EPI_ISL_573404, EPI_ISL_573405, EPI_ISL_573406, EPI_ISL_573407 | | | |
| see above | Northumbria University / South Tees Hospitals NHS Foundation Trust / North Cumbria Integrated Care NHS Foundation Trust / North Tees and Hartlepool NHS Foundation Trust / Newcastle Hospitals NHS Foundation Trust | COVID-19 Genomics UK (COG-UK) Consortium | Darren L Smith,Andrew Nelson,Matthew Bashton,Greg R Young,Joshua Loh,John Allan,Mohammad A Tariq,Giles S Holt,Gary Black,Wen C Yew,Lynn Dover,Paul Baker,Steve Liggett,Sarah Essex,Jane Greenaway,Debra Padgett,Clive Graham,Garren Scott,Edward Barton,Emma Swindells,Brendan Payne,Jennifer Collins,Yusri Taha,Gary Eltringham |
| EPI_ISL_573416 | Quadram Institute Bioscience | COVID-19 Genomics UK (COG-UK) Consortium | Dave J. Baker, Gemma L. Kay, Alp Aydin, Thanh Le-Viet, Steven Rudder, Ana P. Tedim, Anastasia Kolyva, Maria Diaz, Leonardo de Oliveira Martins, Nabil-Fareed Alikhan, Lizzie Meadows, Rachael Stanley, Ngozi Elumogo, Muhammed Yasir, Nicholas M. Thomson, Alexander J Trotter, Rachel Gilroy, Samuel Bloomfield, Claire Stuart, Andrew Bell, Reenesh Prakash, Samir Dervisevic, Alison E. Mather, John Wain, Mark Webber, Andrew J. Page, Justin O'Grady |
| EPI_ISL_573437, EPI_ISL_573438, EPI_ISL_573439, EPI_ISL_573440, EPI_ISL_573441, EPI_ISL_573442, EPI_ISL_573443, EPI_ISL_573444, EPI_ISL_573445, EPI_ISL_573446 | Lincolnshire Hospitals and DeepSeq Nottingham | COVID-19 Genomics UK (COG-UK) Consortium | Nichola Duckworth, Tim Sloan, Sarah Walsh, Jonathan Ball, Patrick McClure, Joeseph Chappell, Nadine Holmes, Matthew Carlisle, Christopher Moore, Fei Sang, Johnny Debebe, Victoria Wright, Matthew Loose |
| EPI_ISL_573652, EPI_ISL_573653, EPI_ISL_573654, EPI_ISL_573655, EPI_ISL_573656, EPI_ISL_573657, EPI_ISL_573658, EPI_ISL_573659, EPI_ISL_573660, EPI_ISL_573661, EPI_ISL_573662, EPI_ISL_573663, EPI_ISL_573664, EPI_ISL_573665, EPI_ISL_573666, EPI_ISL_573667, EPI_ISL_573668, EPI_ISL_573669, EPI_ISL_573670, EPI_ISL_573671, EPI_ISL_573672, EPI_ISL_573673, EPI_ISL_573674, EPI_ISL_573675, EPI_ISL_573676, EPI_ISL_573677, EPI_ISL_573678, EPI_ISL_573679, EPI_ISL_573680, EPI_ISL_573681, EPI_ISL_573682, EPI_ISL_573683, EPI_ISL_573684, EPI_ISL_573685 | | | |
| see above | University College London, Great Ormond Street Hospital for Children NHS Foundation Trust, Imperial College Healthcare NHS Trust | COVID-19 Genomics UK (COG-UK) Consortium | Sergi Castellano, Rachel Williams, Mark Kristiansen, Paola Resende Silva, Sunando Roy, Tony Brooks, Helena Tutill, Paola Niola, Patricia Dyal, Charlotte Williams, Leysa Forrest, Yasmin Panchbhaya, Jacqueline Findlay, Samuel Weeks, Julianne Brown, Kathryn Harris, Paul Randell, James Price, Alison Holmes, Judith Breuer |
| EPI_ISL_573692, EPI_ISL_573694, EPI_ISL_573695, EPI_ISL_573699, EPI_ISL_573700, EPI_ISL_573709, EPI_ISL_573713, EPI_ISL_573719, EPI_ISL_573726, EPI_ISL_573727, EPI_ISL_573735, EPI_ISL_573741, EPI_ISL_573750, EPI_ISL_573753, EPI_ISL_573754 | | | |
| see above | Virology Department, Sheffield Teaching Hospitals NHS Foundation Trust/Department of Infection, Immunity and Cardiovascular Disease, The Medical School, University of Sheffield | COVID-19 Genomics UK (COG-UK) Consortium | Thushan de Silva, Matthew Parker, Nikki Smith, Adri Angyal, Rebecca Brown, Luke Green, Rachel Tucker, Paul Parsons, Danielle Groves, Katie Johnson, Laura Carrilero, Alex Keeley, Dave Partridge, Matthew Wyles, Benjamin Lindsey, Mehmet Yavuz, Mohammad Raza, Cariad Evans |
| EPI_ISL_573764 | University College London, Great Ormond Street | COVID-19 Genomics UK (COG-UK) Consortium | Sergi Castellano, Rachel Williams, Mark Kristiansen, Paola Resende Silva, Sunando Roy, Tony Brooks, Helena Tutill, Paola Niola, Patricia Dyal, Charlotte Williams, Leysa Forrest, Yasmin Panchbhaya, Jacqueline Findlay, Samuel Weeks, Julianne Brown, Kathryn Harris, Paul Randell, James Price, Alison Holmes, Judith Breuer |

| | | | |
|---|---|---|---|
| | Hospital for Children NHS Foundation Trust, Imperial College Healthcare NHS Trust | | |
| EPI_ISL_573770, EPI_ISL_573771, EPI_ISL_573780, EPI_ISL_573781, EPI_ISL_573782, EPI_ISL_573783, EPI_ISL_573784, EPI_ISL_573785, EPI_ISL_573786, EPI_ISL_573787, EPI_ISL_573788, EPI_ISL_573789, EPI_ISL_573790, EPI_ISL_573791, EPI_ISL_573792, EPI_ISL_573793 | | | |
| see above | West of Scotland Specialist Virology Centre, NHSGGC / MRC-University of Glasgow Centre for Virus Research | COVID-19 Genomics UK (COG-UK) Consortium | Ana da Silva Filipe, Natasha Johnson, Kathy Smollett, Daniel Mair, Stephen Carmichael, Lily Tong, Jenna Nichols, Elihu Aranday-Cortes, Kyriaki Nomikou; Sarah McDonald, Marc Niebel, Patawee Asamaphan; Richard Orton, Joseph Hughes, Sreenu Vattipally, David L Robertson; Alasdair MacLean, Rory Gunson; Kathy Li, Igor Starinskij, Natasha Jesudason, Rajiv Shah, James Shepherd, Antonia Ho, Emma Thomson |
| EPI_ISL_573794 | Virology Department, Royal Infirmary of Edinburgh, NHS Lothian / School of Biological Sciences, University of Edinburgh / Institute of Genetics and Molecular Medicine, University of Edinburgh | COVID-19 Genomics UK (COG-UK) Consortium | McHugh M, Dewar R, Rooke S, Gallagher M, Balcaza C, O'Toole Á, Scher E, Hill V, McCrone JT, Colquhoun R, Yu X, Jackson B, Rambaut A, Williams TC, Templeton K |
| EPI_ISL_573856, EPI_ISL_573857, EPI_ISL_573858, EPI_ISL_573859, EPI_ISL_573860, EPI_ISL_573861, EPI_ISL_573862, EPI_ISL_573863, EPI_ISL_573864, EPI_ISL_573865 | University College London, Great Ormond Street Hospital for Children NHS Foundation Trust, Imperial College Healthcare NHS Trust | COVID-19 Genomics UK (COG-UK) Consortium | Sergi Castellano, Rachel Williams, Mark Kristiansen, Paola Resende Silva, Sunando Roy, Tony Brooks, Helena Tutill, Paola Niola, Patricia Dyal, Charlotte Williams, Leysa Forrest, Yasmin Panchbhaya, Jacqueline Findlay, Samuel Weeks, Julianne Brown, Kathryn Harris, Paul Randell, James Price, Alison Holmes, Judith Breuer |
| EPI_ISL_573869 | Oxford Viromics, NDM, University of Oxford; Oxford University Hospitals; Basingstoke and North Hampshire Hospital | COVID-19 Genomics UK (COG-UK) Consortium | Tanya Golubchik, David Bonsall, George Macintyre, Amy Trebes, Mariateresa de Cesare, Catrin Moore, Alex Mobbs, Anita Justice, Robert Shaw, Monique Andersson, Timothy Peto, Emma Wise, Nathan Moore, Jessica Lynch, Nick Cortes, Matilde Mori, Stephen Kidd, David Buck, John Todd, Christophe Fraser |
| EPI_ISL_573870 | Centre for Enzyme Innovation, University of Portsmouth / Translational Research Laboratory, Portsmouth Hospitals NHS Trust | COVID-19 Genomics UK (COG-UK) Consortium | Angela Beckett,Yann Bourgeois,Garry Scarlett,Sharon Glaysher,Scott Elliott,Kelly Bicknell,Robert Impey,Allyson Lloyd,Sarah Wyllie,Ethan Butcher,Anoop Chauhan,Samuel Robson |
| EPI_ISL_573871 | Northumbria University / South Tees Hospitals NHS Foundation Trust / North Cumbria Integrated Care NHS Foundation Trust / North Tees and Hartlepool NHS Foundation Trust / Newcastle Hospitals NHS Foundation Trust | COVID-19 Genomics UK (COG-UK) Consortium | Darren L Smith,Andrew Nelson,Matthew Bashton,Greg R Young,Joshua Loh,John Allan,Mohammad A Tariq,Giles S Holt,Gary Black,Wen C Yew,Lynn Dover,Paul Baker,Steve Liggett,Sarah Essex,Jane Greenaway,Debra Padgett,Clive Graham,Garren Scott,Edward Barton,Emma Swindells,Brendan Payne,Jennifer Collins,Yusri Taha,Gary Eltringham |
| EPI_ISL_573875, EPI_ISL_573877, EPI_ISL_573880, EPI_ISL_573883, EPI_ISL_573884, EPI_ISL_573885, EPI_ISL_573887, EPI_ISL_573888, EPI_ISL_573889, EPI_ISL_573891, EPI_ISL_573893, EPI_ISL_573894, EPI_ISL_573895, EPI_ISL_573896, EPI_ISL_573898, EPI_ISL_573900, EPI_ISL_573901, EPI_ISL_573903, EPI_ISL_573904, EPI_ISL_573905, EPI_ISL_573908, EPI_ISL_573909, EPI_ISL_573910, EPI_ISL_573911, EPI_ISL_573912, EPI_ISL_573914, EPI_ISL_573915, EPI_ISL_573918, EPI_ISL_573919, EPI_ISL_573920, EPI_ISL_573921, EPI_ISL_573923, EPI_ISL_573924, EPI_ISL_573925, EPI_ISL_573926, EPI_ISL_573927, EPI_ISL_573928, EPI_ISL_573929, EPI_ISL_573930, EPI_ISL_573931, EPI_ISL_573932, EPI_ISL_573934, EPI_ISL_573938, EPI_ISL_573939, EPI_ISL_573940, EPI_ISL_573941, EPI_ISL_573944, EPI_ISL_573948, EPI_ISL_573950, EPI_ISL_573951, EPI_ISL_573954, EPI_ISL_573956, EPI_ISL_573958, EPI_ISL_573960, EPI_ISL_573961, EPI_ISL_573962, EPI_ISL_573963, EPI_ISL_573964, EPI_ISL_573967, EPI_ISL_573969, EPI_ISL_573970, EPI_ISL_573971, EPI_ISL_573972, EPI_ISL_573973, EPI_ISL_573974, EPI_ISL_573975, EPI_ISL_573976, EPI_ISL_573978, EPI_ISL_573982, EPI_ISL_573983, EPI_ISL_573988, EPI_ISL_573989, EPI_ISL_573990, EPI_ISL_573991, EPI_ISL_573992, EPI_ISL_573994, EPI_ISL_573995, EPI_ISL_573996, EPI_ISL_573998, EPI_ISL_573999, EPI_ISL_574001, EPI_ISL_574006, EPI_ISL_574007, EPI_ISL_574008, EPI_ISL_574009, EPI_ISL_574011, EPI_ISL_574013, EPI_ISL_574016, EPI_ISL_574031, EPI_ISL_574037, EPI_ISL_574041, EPI_ISL_574058, EPI_ISL_574065, EPI_ISL_574066, EPI_ISL_574068, EPI_ISL_574072, EPI_ISL_574074, EPI_ISL_574080, EPI_ISL_574091, EPI_ISL_574092, EPI_ISL_574099, EPI_ISL_574100, EPI_ISL_574116, EPI_ISL_574138, EPI_ISL_574166, EPI_ISL_574184, EPI_ISL_574191, EPI_ISL_574193, EPI_ISL_574217, EPI_ISL_574224, EPI_ISL_574228, EPI_ISL_574230, EPI_ISL_574237, EPI_ISL_574244 | | | |
| see above | Wales Specialist Virology Centre Sequencing lab: Pathogen Genomics Unit | COVID-19 Genomics UK (COG-UK) Consortium | Catherine Moore, Johnathan Evans, Laura Gifford, Malorie Perry, Simon Cottrell, Angela Marchbank, Alec Birchley, Alexander Adams, Amy Gaskin, Bree Gatica-Wilcox, Jason Coombes, Joel Southgate, Lauren Gilbert, Lee Graham, Nicole Pacchiarini, Sara Kumziene-Summerhayes, Sarah Taylor, Sophie Jones, Sara Rey, Matthew Bull, Joanne Watkins, Sally Corden, Tom Connor |
| EPI_ISL_574258 | University College London, Great Ormond Street Hospital for Children NHS Foundation Trust, Imperial College Healthcare NHS Trust | COVID-19 Genomics UK (COG-UK) Consortium | Sergi Castellano, Rachel Williams, Mark Kristiansen, Paola Resende Silva, Sunando Roy, Tony Brooks, Helena Tutill, Paola Niola, Patricia Dyal, Charlotte Williams, Leysa Forrest, Yasmin Panchbhaya, Jacqueline Findlay, Samuel Weeks, Julianne Brown, Kathryn Harris, Paul Randell, James Price, Alison Holmes, Judith Breuer |
| EPI_ISL_574431 | Hospital IESS Babahoyo | Institute of Microbiology, Universidad San Francisco de Quito | Belén Prado-Vivar, Sully Márquez, Juan José Guadalupe, Monica Becerra-Wong, Fernanda Zurita, Bernardo Gutiérrez, Francisco Cordova, Ninfa Henriquez, Killen Briones-Zamora, Killen Briones-Claudette, Verónica Barragán, Patricio Rojas-Silva, Gabriel Trueba, Michelle Grunauer, Paúl Cárdenas |
| EPI_ISL_574676, EPI_ISL_574677, EPI_ISL_574678 | Seattle Flu Study | Seattle Flu Study | Deborah A. Nickerson, Chris D. Frazar, Jover Lee, Benjamin Pelle, Matthew Richardson, Amanda Adler, Elisabeth Brandstetter, Peter D. Han, Kairsten Fay, Misja Ilcisin, Kirsten Lacombe, Thomas R. Sibley, Melissa Truong, Caitlin R. Wolf, Karen Cowgill, Stephanie Schrag, Jeff Duchin, Michael Boeckh, Janet A. Englund, Michael Famulare, Barry R. Lutz, Mark J. Rieder, Lea M. Starita, Matthew Thompson, Helen Y. Chu, Trevor Bedford, Jay Shendure |
| EPI_ISL_574679 | Seattle Flu Study | Seattle Flu Study | Deborah A. Nickerson, Chris D. Frazar, Jover Lee, Benjamin Pelle, Matthew Richardson, Amanda Adler, Elisabeth Brandstetter, Peter D. Han, Kairsten Fay, Misja Ilcisin, Kirsten Lacombe, Thomas R. Sibley, Melissa Truong, Caitlin R. Wolf, Michael Boeckh, Janet A. Englund, Michael Famulare, Barry R. Lutz, Mark J. Rieder, Lea M. Starita, Matthew Thompson, Jay Shendure, Trevor Bedford, Helen Y. Chu |
| EPI_ISL_574680, EPI_ISL_574681 | Seattle Flu Study | Seattle Flu Study | Deborah A. Nickerson, Chris D. Frazar, Jover Lee, Benjamin Pelle, Matthew Richardson, Amanda Adler, Elisabeth Brandstetter, Peter D. Han, Kairsten Fay, Misja Ilcisin, Kirsten Lacombe, Thomas R. Sibley, Melissa Truong, Caitlin R. Wolf, Karen Cowgill, Stephanie Schrag, Jeff Duchin, Michael Boeckh, Janet A. Englund, Michael Famulare, Barry R. Lutz, Mark J. Rieder, Lea M. Starita, Matthew Thompson, Helen Y. Chu, Trevor Bedford, Jay Shendure |
| EPI_ISL_574682 | Seattle Flu Study | Seattle Flu Study | Deborah A. Nickerson, Chris D. Frazar, Jover Lee, Benjamin Pelle, Matthew Richardson, Amanda Adler, Elisabeth Brandstetter, Peter D. Han, Kairsten Fay, Misja Ilcisin, Kirsten Lacombe, Thomas R. Sibley, Melissa Truong, Caitlin R. Wolf, Michael Boeckh, Janet A. Englund, Michael Famulare, Barry R. Lutz, Mark J. Rieder, Lea M. Starita, Matthew Thompson, Jay Shendure, Trevor Bedford, Helen Y. Chu |
| EPI_ISL_574683 | Seattle Flu Study | Seattle Flu Study | Deborah A. Nickerson, Chris D. Frazar, Jover Lee, Benjamin Pelle, Matthew Richardson, Amanda Adler, Elisabeth Brandstetter, Peter D. Han, Kairsten Fay, Misja Ilcisin, Kirsten Lacombe, Thomas R. Sibley, Melissa Truong, Caitlin R. Wolf, Karen Cowgill, Stephanie Schrag, Jeff Duchin, Michael Boeckh, Janet A. Englund, Michael Famulare, Barry R. Lutz, Mark J. Rieder, Lea M. Starita, Matthew Thompson, Helen Y. Chu, Trevor Bedford, Jay Shendure |
| EPI_ISL_574684 | Seattle Flu Study | Seattle Flu Study | Deborah A. Nickerson, Chris D. Frazar, Jover Lee, Benjamin Pelle, Matthew Richardson, Amanda Adler, Elisabeth Brandstetter, Peter D. Han, Kairsten Fay, Misja Ilcisin, Kirsten Lacombe, Thomas R. Sibley, Melissa Truong, Caitlin R. Wolf, Michael Boeckh, Janet A. Englund, Michael Famulare, Barry R. Lutz, Mark J. Rieder, Lea M. Starita, Matthew Thompson, Jay Shendure, Trevor Bedford, Helen Y. Chu |

| | | | |
|---|---|---|---|
| EPI_ISL_574685, EPI_ISL_574686, EPI_ISL_574687, EPI_ISL_574688 | Seattle Flu Study | Seattle Flu Study | Deborah A. Nickerson, Chris D. Frazar, Jover Lee, Benjamin Pelle, Matthew Richardson, Amanda Adler, Elisabeth Brandstetter, Peter D. Han, Kairsten Fay, Misja Ilcisin, Kirsten Lacombe, Thomas R. Sibley, Melissa Truong, Caitlin R. Wolf, Karen Cowgill, Stephanie Schrag, Jeff Duchin, Michael Boeckh, Janet A. Englund, Michael Famulare, Barry R. Lutz, Mark J. Rieder, Lea M. Starita, Matthew Thompson, Helen Y. Chu, Trevor Bedford, Jay Shendure |
| EPI_ISL_574697 | Respiratory Virus Unit, Microbiology Services Colindale, Public Health England | Respiratory Virus Unit, Microbiology Services Colindale, Public Health England | PHE Covid Sequencing Team |
| EPI_ISL_575399, EPI_ISL_575400, EPI_ISL_575401, EPI_ISL_575402, EPI_ISL_575403, EPI_ISL_575404, EPI_ISL_575405, EPI_ISL_575406, EPI_ISL_575407, EPI_ISL_575408, EPI_ISL_575409, EPI_ISL_575410, EPI_ISL_575411, EPI_ISL_575412, EPI_ISL_575413, EPI_ISL_575414, EPI_ISL_575415, EPI_ISL_575416, EPI_ISL_575417, EPI_ISL_575418, EPI_ISL_575419, EPI_ISL_575420, EPI_ISL_575421, EPI_ISL_575422, EPI_ISL_575423, EPI_ISL_575424, EPI_ISL_575425, EPI_ISL_575426, EPI_ISL_575427, EPI_ISL_575428, EPI_ISL_575429, EPI_ISL_575430, EPI_ISL_575431, EPI_ISL_575432, EPI_ISL_575433, EPI_ISL_575434, EPI_ISL_575435, EPI_ISL_575436, EPI_ISL_575437, EPI_ISL_575438, EPI_ISL_575439, EPI_ISL_575440, EPI_ISL_575441, EPI_ISL_575442, EPI_ISL_575443, EPI_ISL_575444, EPI_ISL_575445, EPI_ISL_575446, EPI_ISL_575447, EPI_ISL_575448, EPI_ISL_575449, EPI_ISL_575450, EPI_ISL_575451, EPI_ISL_575452, EPI_ISL_575453, EPI_ISL_575454, EPI_ISL_575455, EPI_ISL_575456, EPI_ISL_575457, EPI_ISL_575458, EPI_ISL_575459, EPI_ISL_575460, EPI_ISL_575461, EPI_ISL_575462, EPI_ISL_575463, EPI_ISL_575464, EPI_ISL_575465, EPI_ISL_575466, EPI_ISL_575467, EPI_ISL_575468, EPI_ISL_575469, EPI_ISL_575470, EPI_ISL_575471, EPI_ISL_575472, EPI_ISL_575473, EPI_ISL_575474, EPI_ISL_575475, EPI_ISL_575476, EPI_ISL_575477, EPI_ISL_575478, EPI_ISL_575479, EPI_ISL_575480, EPI_ISL_575481, EPI_ISL_575482, EPI_ISL_575483, EPI_ISL_575484, EPI_ISL_575485, EPI_ISL_575486, EPI_ISL_575487, EPI_ISL_575488, EPI_ISL_575489, EPI_ISL_575490, EPI_ISL_575491, EPI_ISL_575492, EPI_ISL_575493, EPI_ISL_575494, EPI_ISL_575495, EPI_ISL_575496, EPI_ISL_575497, EPI_ISL_575498, EPI_ISL_575499, EPI_ISL_575500, EPI_ISL_575501, EPI_ISL_575502, EPI_ISL_575503, EPI_ISL_575504, EPI_ISL_575505, EPI_ISL_575506, EPI_ISL_575507, EPI_ISL_575508, EPI_ISL_575509, EPI_ISL_575510, EPI_ISL_575511, EPI_ISL_575512, EPI_ISL_575513, EPI_ISL_575514, EPI_ISL_575515, EPI_ISL_575516, EPI_ISL_575517, EPI_ISL_575518, EPI_ISL_575519, EPI_ISL_575520, EPI_ISL_575521, EPI_ISL_575522, EPI_ISL_575523, EPI_ISL_575524, EPI_ISL_575525, EPI_ISL_575526, EPI_ISL_575527, EPI_ISL_575528, EPI_ISL_575529, EPI_ISL_575530, EPI_ISL_575531, EPI_ISL_575532, EPI_ISL_575533, EPI_ISL_575534, EPI_ISL_575535 | | | |
| see above | Lighthouse Lab in Alderley Park | Wellcome Sanger Institute for the COVID-19 Genomics UK (COG-UK) consortium | Jacquelyn Wynn, Mairead Hyland, The Lighthouse Lab in Alderley Park and Alex Alderton, Roberto Amato, Sonia Goncalves, Ewan Harrison, David K. Jackson, Ian Johnston, Dominic Kwiatkowski, Cordelia Langford, John Sillitoe on behalf of the Wellcome Sanger Institute COVID-19 Surveillance Team |
| EPI_ISL_575536 | Lighthouse Lab in Alderley Park | Wellcome Sanger Institute for the COVID-19 Genomics UK (COG-UK) Consortium | Jacquelyn Wynn, Mairead Hyland, The Lighthouse Lab in Alderley Park and Alex Alderton, Roberto Amato, Sonia Goncalves, Ewan Harrison, David K. Jackson, Ian Johnston, Dominic Kwiatkowski, Cordelia Langford, John Sillitoe on behalf of the Wellcome Sanger Institute COVID-19 Surveillance Team |
| EPI_ISL_575537, EPI_ISL_575538, EPI_ISL_575539, EPI_ISL_575540, EPI_ISL_575541, EPI_ISL_575542, EPI_ISL_575543, EPI_ISL_575544, EPI_ISL_575545, EPI_ISL_575546, EPI_ISL_575547, EPI_ISL_575548, EPI_ISL_575549, EPI_ISL_575550, EPI_ISL_575551, EPI_ISL_575552, EPI_ISL_575553, EPI_ISL_575554, EPI_ISL_575555, EPI_ISL_575556, EPI_ISL_575557, EPI_ISL_575558, EPI_ISL_575559, EPI_ISL_575560, EPI_ISL_575561, EPI_ISL_575562, EPI_ISL_575563, EPI_ISL_575564, EPI_ISL_575565, EPI_ISL_575566, EPI_ISL_575567, EPI_ISL_575568, EPI_ISL_575569, EPI_ISL_575570, EPI_ISL_575571, EPI_ISL_575572, EPI_ISL_575573, EPI_ISL_575574, EPI_ISL_575575, EPI_ISL_575576, EPI_ISL_575577, EPI_ISL_575578, EPI_ISL_575579, EPI_ISL_575580, EPI_ISL_575581, EPI_ISL_575582, EPI_ISL_575583, EPI_ISL_575584, EPI_ISL_575585, EPI_ISL_575586, EPI_ISL_575587, EPI_ISL_575588, EPI_ISL_575589, EPI_ISL_575590, EPI_ISL_575591, EPI_ISL_575592, EPI_ISL_575593, EPI_ISL_575594, EPI_ISL_575595, EPI_ISL_575596, EPI_ISL_575597, EPI_ISL_575598, EPI_ISL_575599, EPI_ISL_575600, EPI_ISL_575601, EPI_ISL_575602, EPI_ISL_575603, EPI_ISL_575604, EPI_ISL_575605, EPI_ISL_575606, EPI_ISL_575607, EPI_ISL_575608, EPI_ISL_575609, EPI_ISL_575610, EPI_ISL_575611, EPI_ISL_575612, EPI_ISL_575613, EPI_ISL_575614, EPI_ISL_575615, EPI_ISL_575616, EPI_ISL_575617, EPI_ISL_575618, EPI_ISL_575619, EPI_ISL_575620, EPI_ISL_575621, EPI_ISL_575622, EPI_ISL_575623, EPI_ISL_575624, EPI_ISL_575625, EPI_ISL_575626, EPI_ISL_575627, EPI_ISL_575628, EPI_ISL_575629, EPI_ISL_575630, EPI_ISL_575631, EPI_ISL_575632, EPI_ISL_575633, EPI_ISL_575634, EPI_ISL_575635, EPI_ISL_575636, EPI_ISL_575637, EPI_ISL_575638, EPI_ISL_575639, EPI_ISL_575640, EPI_ISL_575641, EPI_ISL_575642, EPI_ISL_575643, EPI_ISL_575644, EPI_ISL_575645, EPI_ISL_575646, EPI_ISL_575647, EPI_ISL_575648, EPI_ISL_575649, EPI_ISL_575650, EPI_ISL_575651, EPI_ISL_575652, EPI_ISL_575653, EPI_ISL_575654, EPI_ISL_575655, EPI_ISL_575656, EPI_ISL_575657, EPI_ISL_575658, EPI_ISL_575659, EPI_ISL_575660, EPI_ISL_575661, EPI_ISL_575662, EPI_ISL_575663, EPI_ISL_575664, EPI_ISL_575665, EPI_ISL_575666, EPI_ISL_575667, EPI_ISL_575668, EPI_ISL_575669, EPI_ISL_575670, EPI_ISL_575671, EPI_ISL_575672, EPI_ISL_575673, EPI_ISL_575674, EPI_ISL_575675, EPI_ISL_575676, EPI_ISL_575677, EPI_ISL_575678, EPI_ISL_575679, EPI_ISL_575680, EPI_ISL_575681, EPI_ISL_575682, EPI_ISL_575683, EPI_ISL_575684, EPI_ISL_575685, EPI_ISL_575686, EPI_ISL_575687, EPI_ISL_575688, EPI_ISL_575689, EPI_ISL_575690, EPI_ISL_575691, EPI_ISL_575692, EPI_ISL_575693, EPI_ISL_575694, EPI_ISL_575695, EPI_ISL_575696, EPI_ISL_575697, EPI_ISL_575698, EPI_ISL_575699, EPI_ISL_575700, EPI_ISL_575701, EPI_ISL_575702, EPI_ISL_575703, EPI_ISL_575704, EPI_ISL_575705, EPI_ISL_575706, EPI_ISL_575707, EPI_ISL_575708, EPI_ISL_575709, EPI_ISL_575710, EPI_ISL_575711, EPI_ISL_575712, EPI_ISL_575713, EPI_ISL_575714, EPI_ISL_575715, EPI_ISL_575716, EPI_ISL_575717, EPI_ISL_575718, EPI_ISL_575719, EPI_ISL_575720, EPI_ISL_575721, EPI_ISL_575722, EPI_ISL_575723, EPI_ISL_575724, EPI_ISL_575725, EPI_ISL_575726, EPI_ISL_575727, EPI_ISL_575728, EPI_ISL_575729, EPI_ISL_575730, EPI_ISL_575731, EPI_ISL_575732, EPI_ISL_575733, EPI_ISL_575734, EPI_ISL_575735, EPI_ISL_575736, EPI_ISL_575737, EPI_ISL_575738, EPI_ISL_575739, EPI_ISL_575740, EPI_ISL_575741, EPI_ISL_575742, EPI_ISL_575743, EPI_ISL_575744, EPI_ISL_575745, EPI_ISL_575746, EPI_ISL_575747, EPI_ISL_575748, EPI_ISL_575749, EPI_ISL_575750, EPI_ISL_575751, EPI_ISL_575752, EPI_ISL_575753, EPI_ISL_575754, EPI_ISL_575755, EPI_ISL_575756, EPI_ISL_575757, EPI_ISL_575758, EPI_ISL_575759, EPI_ISL_575760 | | | |
| see above | Lighthouse Lab in Alderley Park | Wellcome Sanger Institute for the COVID-19 Genomics UK (COG-UK) consortium | Jacquelyn Wynn, Mairead Hyland, The Lighthouse Lab in Alderley Park and Alex Alderton, Roberto Amato, Sonia Goncalves, Ewan Harrison, David K. Jackson, Ian Johnston, Dominic Kwiatkowski, Cordelia Langford, John Sillitoe on behalf of the Wellcome Sanger Institute COVID-19 Surveillance Team |
| EPI_ISL_575982, EPI_ISL_575983 | Lighthouse Lab in Alderley Park | Wellcome Sanger Institute for the COVID-19 Genomics UK (COG-UK) consortium | Jacquelyn Wynn, Mairead Hyland, The Lighthouse Lab in Alderley Park and Alex Alderton, Roberto Amato, Sonia Goncalves, Ewan Harrison, David K. Jackson, Ian Johnston, Dominic Kwiatkowski, Cordelia Langford, John Sillitoe on behalf of the Wellcome Sanger Institute COVID-19 Surveillance Team (http://www.sanger.ac.uk/covid-team) |
| EPI_ISL_575984, EPI_ISL_575985 | Lighthouse Lab in Milton Keynes | Wellcome Sanger Institute for the COVID-19 Genomics UK (COG-UK) consortium | The Lighthouse Lab in Milton Keynes and Alex Alderton, Roberto Amato, Sonia Goncalves, Ewan Harrison, David K. Jackson, Ian Johnston, Dominic Kwiatkowski, Cordelia Langford, John Sillitoe on behalf of the Wellcome Sanger Institute COVID-19 Surveillance Team (http://www.sanger.ac.uk/covid-team) |
| EPI_ISL_575995, EPI_ISL_575996, EPI_ISL_575997, EPI_ISL_575998, EPI_ISL_575999, EPI_ISL_576000, EPI_ISL_576001, EPI_ISL_576002, EPI_ISL_576003, EPI_ISL_576004, EPI_ISL_576005, EPI_ISL_576006, EPI_ISL_576007, EPI_ISL_576008, EPI_ISL_576009, EPI_ISL_576010, EPI_ISL_576011, EPI_ISL_576012, EPI_ISL_576013, EPI_ISL_576014, EPI_ISL_576015, EPI_ISL_576016, EPI_ISL_576017, EPI_ISL_576018, EPI_ISL_576019, EPI_ISL_576020, EPI_ISL_576021, EPI_ISL_576022, EPI_ISL_576023, EPI_ISL_576024, EPI_ISL_576025, EPI_ISL_576026, EPI_ISL_576027, EPI_ISL_576028, EPI_ISL_576029, EPI_ISL_576030, EPI_ISL_576031, EPI_ISL_576032, EPI_ISL_576033, EPI_ISL_576034, EPI_ISL_576035, EPI_ISL_576036, EPI_ISL_576037, EPI_ISL_576038, EPI_ISL_576039, EPI_ISL_576040, EPI_ISL_576041, EPI_ISL_576042, EPI_ISL_576043, EPI_ISL_576044, EPI_ISL_576045, EPI_ISL_576046, EPI_ISL_576047, EPI_ISL_576048, EPI_ISL_576049 | | | |
| see above | Lighthouse Lab in Glasgow | Wellcome Sanger Institute for the COVID-19 Genomics UK (COG-UK) consortium | Harper VanSteenhouse, Yumi Kasai, David Gray, Carol Clugston, Anna Dominiczak and Alex Alderton, Roberto Amato, Sonia Goncalves, Ewan Harrison, David K. Jackson, Ian Johnston, Dominic Kwiatkowski, Cordelia Langford, John Sillitoe on behalf of the Wellcome Sanger Institute COVID-19 Surveillance Team (http://www.sanger.ac.uk/covid-team) |
| EPI_ISL_576050 | Lighthouse Lab in Glasgow | Wellcome Sanger Institute for the COVID-19 Genomics UK (COG-UK) Consortium | Harper VanSteenhouse, Yumi Kasai, David Gray, Carol Clugston, Anna Dominiczak and Alex Alderton, Roberto Amato, Sonia Goncalves, Ewan Harrison, David K. Jackson, Ian Johnston, Dominic Kwiatkowski, Cordelia Langford, John Sillitoe on behalf of the Wellcome Sanger Institute COVID-19 Surveillance Team |
| EPI_ISL_576051, EPI_ISL_576052, EPI_ISL_576054, EPI_ISL_576055, EPI_ISL_576056, EPI_ISL_576057, EPI_ISL_576058, EPI_ISL_576059, EPI_ISL_576060, EPI_ISL_576061, EPI_ISL_576062, EPI_ISL_576063 | | | |
| see above | Lighthouse Lab in Glasgow | Wellcome Sanger Institute for the COVID-19 Genomics UK (COG-UK) consortium | Harper VanSteenhouse, Yumi Kasai, David Gray, Carol Clugston, Anna Dominiczak and Alex Alderton, Roberto Amato, Sonia Goncalves, Ewan Harrison, David K. Jackson, Ian Johnston, Dominic Kwiatkowski, Cordelia Langford, John Sillitoe on behalf of the Wellcome Sanger Institute COVID-19 Surveillance Team (http://www.sanger.ac.uk/covid-team) |
| EPI_ISL_576064 | Lighthouse Lab in Glasgow | Wellcome Sanger Institute for the COVID-19 Genomics UK (COG-UK) Consortium | Harper VanSteenhouse, Yumi Kasai, David Gray, Carol Clugston, Anna Dominiczak and Alex Alderton, Roberto Amato, Sonia Goncalves, Ewan Harrison, David K. Jackson, Ian Johnston, Dominic Kwiatkowski, Cordelia Langford, John Sillitoe on behalf of the Wellcome Sanger Institute COVID-19 Surveillance Team |
| EPI_ISL_576065, EPI_ISL_576066, EPI_ISL_576067, EPI_ISL_576068, EPI_ISL_576069, EPI_ISL_576071, EPI_ISL_576072, EPI_ISL_576073, EPI_ISL_576074, EPI_ISL_576075, EPI_ISL_576077, EPI_ISL_576078, EPI_ISL_576079, EPI_ISL_576080, EPI_ISL_576081, EPI_ISL_576083, EPI_ISL_576084, EPI_ISL_576085, EPI_ISL_576086, EPI_ISL_576087 | | | |
| see above | Lighthouse Lab in Glasgow | Wellcome Sanger Institute for the COVID-19 Genomics UK (COG-UK) consortium | Harper VanSteenhouse, Yumi Kasai, David Gray, Carol Clugston, Anna Dominiczak and Alex Alderton, Roberto Amato, Sonia Goncalves, Ewan Harrison, David K. Jackson, Ian Johnston, Dominic Kwiatkowski, Cordelia Langford, John Sillitoe on behalf of the Wellcome Sanger Institute COVID-19 Surveillance Team (http://www.sanger.ac.uk/covid-team) |
| EPI_ISL_576088 | Lighthouse Lab in Glasgow | Wellcome Sanger Institute for the COVID-19 Genomics UK (COG-UK) Consortium | Harper VanSteenhouse, Yumi Kasai, David Gray, Carol Clugston, Anna Dominiczak and Alex Alderton, Roberto Amato, Sonia Goncalves, Ewan Harrison, David K. Jackson, Ian Johnston, Dominic Kwiatkowski, Cordelia Langford, John Sillitoe on behalf of the Wellcome Sanger Institute COVID-19 Surveillance Team |
| EPI_ISL_576089, EPI_ISL_576090, EPI_ISL_576091, EPI_ISL_576092, EPI_ISL_576093 | Lighthouse Lab in Glasgow | Wellcome Sanger Institute for the COVID-19 Genomics UK (COG-UK) consortium | Harper VanSteenhouse, Yumi Kasai, David Gray, Carol Clugston, Anna Dominiczak and Alex Alderton, Roberto Amato, Sonia Goncalves, Ewan Harrison, David K. Jackson, Ian Johnston, Dominic Kwiatkowski, Cordelia Langford, John Sillitoe on behalf of the Wellcome Sanger Institute COVID-19 Surveillance Team (http://www.sanger.ac.uk/covid-team) |
| EPI_ISL_576142, EPI_ISL_576143, EPI_ISL_576144 | unknown | Public Health Virology Laboratory, Forensic and Scientific Services (PHV-FSS) | Son Nguyen et al. |

| | | | |
|---|---|---|---|
| EPI_ISL_576189, EPI_ISL_576190, EPI_ISL_576191 | GA Department of Public Health Laboratory | Pathogen Discovery, Respiratory Viruses Branch, Division of Viral Diseases, Centers for Disease Control and Prevention | Ying Tao, Jing Zhang, Brian Lynch, Yan Li, Krista Queen, Anna Uehara, Clinton R. Paden, Peter Cook, Haibin Wang, Suxiang Tong |
| EPI_ISL_576228, EPI_ISL_576229, EPI_ISL_576230, EPI_ISL_576231, EPI_ISL_576232, EPI_ISL_576233, EPI_ISL_576234, EPI_ISL_576235, EPI_ISL_576236, EPI_ISL_576238, EPI_ISL_576240, EPI_ISL_576241, EPI_ISL_576242, EPI_ISL_576244, EPI_ISL_576245, EPI_ISL_576246, EPI_ISL_576247, EPI_ISL_576248, EPI_ISL_576249, EPI_ISL_576250, EPI_ISL_576251, EPI_ISL_576256 | | | |
| see above | Minnesota Department of Health, Public Health Laboratory | Minnesota Department of Health, Public Health Laboratory | Matt Plumb, Jacob Garfin, Alexandra Lorentz, and Xiong Wang |
| EPI_ISL_576358, EPI_ISL_576359, EPI_ISL_576360, EPI_ISL_576361, EPI_ISL_576362 | Texas Department of State Health Services | Texas Department of State Health Services | Rashmi Tuladhar, Bonnie Oh, Mayela Pedrueza, Jenny Zhang, Maliha Rahman, Anita Pokharel, Myong Koag, Chun Wang, Rachel Lee, Grace Kubin |
| EPI_ISL_576609, EPI_ISL_576610, EPI_ISL_576612, EPI_ISL_576617, EPI_ISL_576618, EPI_ISL_576629, EPI_ISL_576630, EPI_ISL_576631, EPI_ISL_576632, EPI_ISL_576633, EPI_ISL_576634, EPI_ISL_576635, EPI_ISL_576646, EPI_ISL_576659, EPI_ISL_576831, EPI_ISL_576832, EPI_ISL_576833 | | | |
| see above | Oxford Viromics, NDM, University of Oxford; Oxford University Hospitals; Basingstoke and North Hampshire Hospital | COVID-19 Genomics UK (COG-UK) Consortium | Tanya Golubchik, David Bonsall, George Macintyre, Amy Trebes, Mariateresa de Cesare, Catrin Moore, Alex Mobbs, Anita Justice, Robert Shaw, Monique Andersson, Timothy Peto, Emma Wise, Nathan Moore, Jessica Lynch, Nick Cortes, Matilde Mori, Stephen Kidd, David Buck, John Todd, Christophe Fraser |
| EPI_ISL_576965, EPI_ISL_576966, EPI_ISL_576967, EPI_ISL_576968, EPI_ISL_576969, EPI_ISL_576970, EPI_ISL_576971, EPI_ISL_576972, EPI_ISL_576973, EPI_ISL_576974, EPI_ISL_576975, EPI_ISL_576976, EPI_ISL_576977, EPI_ISL_576980 | | | |
| see above | Liverpool Clinical Laboratories | COVID-19 Genomics UK (COG-UK) Consortium | Sam Haldenby, Anita Lucaci, Steve Paterson, Julian Hiscox, Alistair Darby, M Almsaud, A Alrezaihi, Muhannad Alruwaili, Stuart D Armstrong, Jones Benjamin, Eleanor G Bentley, Anu Chawla, Jordan J Clark, Angela Cowell, Richard Eccles, Isabel García-Dorival, Matthew Gemmell, Alessandro Gerada, PKF Gilmore, Richard Gregory, Ximeng Han, Catherine Hartley, Margaret Hughes, Miren Iturriza-Gomara, James Johnson, L Luu, Jenifer Manson, Charlotte Nelson, Elaine O'Toole, Cassie Olateju, Rebekah Penrice-Randal , Lucille Rainbow, N.P Randle, Trevor Ian Robinson, Parul Sharma, Ghada T Shawli, James P Stewart, Neil Swainston, Ecaterina Vamos, Joanne Watts, Mark Whitehead |
| EPI_ISL_577040, EPI_ISL_577044, EPI_ISL_577046, EPI_ISL_577049, EPI_ISL_577051, EPI_ISL_577066, EPI_ISL_577196, EPI_ISL_577197, EPI_ISL_577199, EPI_ISL_577200, EPI_ISL_577204, EPI_ISL_577207, EPI_ISL_577209, EPI_ISL_577216 | | | |
| see above | Centre for Enzyme Innovation, University of Portsmouth / Translational Research Laboratory, Portsmouth Hospitals NHS Trust | COVID-19 Genomics UK (COG-UK) Consortium | Angela Beckett,Yann Bourgeois,Garry Scarlett,Sharon Glaysher,Scott Elliott,Kelly Bicknell,Robert Impey,Allyson Lloyd,Sarah Wyllie,Ethan Butcher,Anoop Chauhan,Samuel Robson |
| EPI_ISL_577273, EPI_ISL_577274, EPI_ISL_577275, EPI_ISL_577276, EPI_ISL_577277, EPI_ISL_577278, EPI_ISL_577279, EPI_ISL_577280, EPI_ISL_577281, EPI_ISL_577282, EPI_ISL_577283, EPI_ISL_577284, EPI_ISL_577285, EPI_ISL_577286, EPI_ISL_577287, EPI_ISL_577288, EPI_ISL_577289, EPI_ISL_577290, EPI_ISL_577291, EPI_ISL_577292, EPI_ISL_577293, EPI_ISL_577294, EPI_ISL_577295, EPI_ISL_577296, EPI_ISL_577297 | | | |
| see above | University of Exeter | COVID-19 Genomics UK (COG-UK) Consortium | Ben Temperton,Aaron Jeffries,Michelle Michelsen,Joanna Warwick-Dugdale,Audrey Farbos,Robyn Manley,Stephen Michell,Jane Masoli |
| EPI_ISL_577303, EPI_ISL_577312, EPI_ISL_577322, EPI_ISL_577331 | Centre for Enzyme Innovation, University of Portsmouth / Translational Research Laboratory, Portsmouth Hospitals NHS Trust | COVID-19 Genomics UK (COG-UK) Consortium | Angela Beckett,Yann Bourgeois,Garry Scarlett,Sharon Glaysher,Scott Elliott,Kelly Bicknell,Robert Impey,Allyson Lloyd,Sarah Wyllie,Ethan Butcher,Anoop Chauhan,Samuel Robson |
| EPI_ISL_577347, EPI_ISL_577348, EPI_ISL_577350, EPI_ISL_577351, EPI_ISL_577352, EPI_ISL_577353, EPI_ISL_577354 | Virology Department, Royal Infirmary of Edinburgh, NHS Lothian / School of Biological Sciences, University of Edinburgh / Institute of Genetics and Molecular Medicine, University of Edinburgh | COVID-19 Genomics UK (COG-UK) Consortium | McHugh M, Dewar R, Rooke S, Gallagher M, Balcaza C, O'Toole Á, Scher E, Hill V, McCrone JT, Colquhoun R, Yu X, Jackson B, Rambaut A, Williams TC, Templeton K |
| EPI_ISL_577374, EPI_ISL_577375, EPI_ISL_577376, EPI_ISL_577377, EPI_ISL_577378, EPI_ISL_577379, EPI_ISL_577380 | University of Exeter | COVID-19 Genomics UK (COG-UK) Consortium | Ben Temperton,Aaron Jeffries,Michelle Michelsen,Joanna Warwick-Dugdale,Audrey Farbos,Robyn Manley,Stephen Michell,Jane Masoli |
| EPI_ISL_577419, EPI_ISL_577460, EPI_ISL_577482, EPI_ISL_577485 | Wales Specialist Virology Centre Sequencing lab: Pathogen Genomics Unit | COVID-19 Genomics UK (COG-UK) Consortium | Catherine Moore, Johnathan Evans, Laura Gifford, Malorie Perry, Simon Cottrell, Angela Marchbank, Alec Birchley, Alexander Adams, Amy Gaskin, Bree Gatica-Wilcox, Jason Coombes, Joel Southgate, Lauren Gilbert, Lee Graham, Nicole Pacchiarini, Sara Kumziene-Summerhayes, Sarah Taylor, Sophie Jones, Sara Rey, Matthew Bull, Joanne Watkins, Sally Corden, Tom Connor |
| EPI_ISL_577548, EPI_ISL_577558, EPI_ISL_577559, EPI_ISL_577563, EPI_ISL_577585, EPI_ISL_577586, EPI_ISL_577587, EPI_ISL_577591, EPI_ISL_577592, EPI_ISL_577593, EPI_ISL_577595, EPI_ISL_577596 | | | |
| see above | Michigan Department of Health and Human Services, Bureau of Laboratories | Michigan Department of Health and Human Services, Bureau of Laboratories | Blankenship HM, Riner D, Soehnlen MK |
| EPI_ISL_577630 | The National Institute of Public Health | State Veterinary Institute Prague | Nagy,A;Jirincova,H;Novakova,L;Trnka,D;Vecerova,J |
| EPI_ISL_577641, EPI_ISL_577642, EPI_ISL_577643, EPI_ISL_577644, EPI_ISL_577645, EPI_ISL_577646, EPI_ISL_577648, EPI_ISL_577661, EPI_ISL_577663, EPI_ISL_577664, EPI_ISL_577666, EPI_ISL_577667, EPI_ISL_577668, EPI_ISL_577669, EPI_ISL_577670, EPI_ISL_577674, EPI_ISL_577675, EPI_ISL_577676, EPI_ISL_577678, EPI_ISL_577703, EPI_ISL_577704, EPI_ISL_577705, EPI_ISL_577708, EPI_ISL_577716, EPI_ISL_577730, EPI_ISL_577731 | | | |
| see above | NIV Influenza | NIV Influenza | Potdar V |
| EPI_ISL_577816, EPI_ISL_577817, EPI_ISL_577818, EPI_ISL_577819, EPI_ISL_577820, EPI_ISL_577821, EPI_ISL_577822, EPI_ISL_577823, EPI_ISL_577824, EPI_ISL_577825, EPI_ISL_577826, EPI_ISL_577827, EPI_ISL_577828 | | | |
| see above | Dutch COVID-19 response team | Erasmus Medical Center | OH consortium |
| EPI_ISL_577863, EPI_ISL_577864, EPI_ISL_577865, EPI_ISL_577866, EPI_ISL_577964, EPI_ISL_577965, EPI_ISL_577966, EPI_ISL_577967, EPI_ISL_577968, EPI_ISL_577969, EPI_ISL_577970, EPI_ISL_577971, EPI_ISL_577972, EPI_ISL_577973, EPI_ISL_577974, EPI_ISL_577975, EPI_ISL_577976, EPI_ISL_577977, EPI_ISL_577978, EPI_ISL_577979, EPI_ISL_578035, EPI_ISL_578043, EPI_ISL_578044, EPI_ISL_578047, EPI_ISL_578048 | | | |

| | | | |
|---|---|---|---|
| see above | Dutch COVID-19 response team | Erasmus Medical Center | Bas Oude Munnink, Reina Sikkema, David Nieuwenhuijse, Irina Chestakova, Anne van der Linden, Marjan Boter, Emmanuelle Munger, Corine GeurtsvanKessel, Annemiek van der Eijk, Richard Molenkamp, Marion Koopmans, on behalf of the Dutch national COVID-19 response team. |
| EPI_ISL_578080 | CSIR-Indian Institute of Chemical Biology, MEDICA Supercpecialty Hospital Kolkata | CSIR-Indian Institute of Chemical Biology, MEDICA Supercpecialty Hospital Kolkata | Sujay Krishna Maity, Priyanka Mallick, Debaleena Bhowmik, Abhishake Lahiri, Dr. AviralRoy, Dr. Soumen Saha, Dr. Arpita Ghosh Mitra, Dr. Rajesh Pandey, Dr. Sandip Paul, Dr.Partha Chakrabarti, Dr. Saikat Chakrabarti |
| EPI_ISL_578081, EPI_ISL_578168, EPI_ISL_578175, EPI_ISL_578178, EPI_ISL_578179, EPI_ISL_578180, EPI_ISL_578181, EPI_ISL_578182, EPI_ISL_578183, EPI_ISL_578184 | CSIR-Indian Institute of Chemical Biology, MEDICA Supercpecialty Hospital Kolkata | CSIR-Indian Institute of Chemical Biology, MEDICA Supercpecialty Hospital Kolkata | Sujay Krishna Maity, Priyanka Mallick, Debaleena Bhowmik, Abhishake Lahiri, Dr. Aviral Roy, Dr. Soumen Saha, Dr. Arpita Ghosh Mitra, Dr. Rajesh Pandey, Dr. Sandip Paul, Dr. Partha Chakrabarti, Dr. Saikat Chakrabarti |
| EPI_ISL_578202, EPI_ISL_578214, EPI_ISL_578215, EPI_ISL_578216, EPI_ISL_578218, EPI_ISL_578221, EPI_ISL_578222, EPI_ISL_578223, EPI_ISL_578224, EPI_ISL_578225, EPI_ISL_578231, EPI_ISL_578239, EPI_ISL_578244, EPI_ISL_578247, EPI_ISL_578259, EPI_ISL_578261, EPI_ISL_578263, EPI_ISL_578266, EPI_ISL_578267, EPI_ISL_578268, EPI_ISL_578269, EPI_ISL_578270, EPI_ISL_578271, EPI_ISL_578273, EPI_ISL_578274, EPI_ISL_578280, EPI_ISL_578281, EPI_ISL_578284, EPI_ISL_578285, EPI_ISL_578286, EPI_ISL_578287, EPI_ISL_578288, EPI_ISL_578289, EPI_ISL_578290, EPI_ISL_578291, EPI_ISL_578292, EPI_ISL_578293, EPI_ISL_578294, EPI_ISL_578296, EPI_ISL_578299, EPI_ISL_578300, EPI_ISL_578301, EPI_ISL_578311, EPI_ISL_578314, EPI_ISL_578319, EPI_ISL_578320 | | | |
| see above | National Virus Reference Laboratory | National Virus Reference Laboratory | Michael Carr, Gabriel Gonzalez, Jonathan Dean, Suzie Coughlan, Cillian F De Gascun |
| EPI_ISL_579093, EPI_ISL_579094, EPI_ISL_579095 | North Shore Hospital | Institute of Environmental Science and Research (ESR) | Xiaoyun Ren, Matt Storey, Nikki Freed, Muhammad Faisal, Jing Wang, Hermes Perez, Anja Werno, Antje van der Linden, Arlo Upton, Chris Mansell, David Hammer, Dragana Drinkovic, Gary McAuliffe, Hana Sofia Andersson, James Ussher, Jill Sherwood, Josh Freeman, Julia Howard, Juliet Elvy, Mary DeAlmeida, Matt Blakiston, Matthew Rogers, Max Bloomfield, Michael Addidle, Michelle Balm, Sally Roberts, Sarah Jefferies, Sharmini Muttaiyah, Susan Morpeth, Susan Taylor, Timothy Blackmore, Vani Sathyendran, Veronica Playle, Virginia Hope, Erasmus Smit, Lauren Jelly, Olin Silander, Joep de Ligt |
| EPI_ISL_579098 | LabPLUS | Institute of Environmental Science and Research (ESR) | Xiaoyun Ren, Matt Storey, Nikki Freed, Muhammad Faisal, Jing Wang, Hermes Perez, Anja Werno, Antje van der Linden, Arlo Upton, Chris Mansell, David Hammer, Dragana Drinkovic, Gary McAuliffe, Hana Sofia Andersson, James Ussher, Jill Sherwood, Josh Freeman, Julia Howard, Juliet Elvy, Mary DeAlmeida, Matt Blakiston, Matthew Rogers, Max Bloomfield, Michael Addidle, Michelle Balm, Sally Roberts, Sarah Jefferies, Sharmini Muttaiyah, Susan Morpeth, Susan Taylor, Timothy Blackmore, Vani Sathyendran, Veronica Playle, Virginia Hope, Erasmus Smit, Lauren Jelly, Olin Silander, Joep de Ligt |
| EPI_ISL_579099 | LabTests | Institute of Environmental Science and Research (ESR) | Xiaoyun Ren, Matt Storey, Nikki Freed, Muhammad Faisal, Jing Wang, Hermes Perez, Anja Werno, Antje van der Linden, Arlo Upton, Chris Mansell, David Hammer, Dragana Drinkovic, Gary McAuliffe, Hana Sofia Andersson, James Ussher, Jill Sherwood, Josh Freeman, Julia Howard, Juliet Elvy, Mary DeAlmeida, Matt Blakiston, Matthew Rogers, Max Bloomfield, Michael Addidle, Michelle Balm, Sally Roberts, Sarah Jefferies, Sharmini Muttaiyah, Susan Morpeth, Susan Taylor, Timothy Blackmore, Vani Sathyendran, Veronica Playle, Virginia Hope, Erasmus Smit, Lauren Jelly, Olin Silander, Joep de Ligt |
| EPI_ISL_579101, EPI_ISL_579102 | Middlemore Hospital | Institute of Environmental Science and Research (ESR) | Xiaoyun Ren, Matt Storey, Nikki Freed, Muhammad Faisal, Jing Wang, Hermes Perez, Anja Werno, Antje van der Linden, Arlo Upton, Chris Mansell, David Hammer, Dragana Drinkovic, Gary McAuliffe, Hana Sofia Andersson, James Ussher, Jill Sherwood, Josh Freeman, Julia Howard, Juliet Elvy, Mary DeAlmeida, Matt Blakiston, Matthew Rogers, Max Bloomfield, Michael Addidle, Michelle Balm, Sally Roberts, Sarah Jefferies, Sharmini Muttaiyah, Susan Morpeth, Susan Taylor, Timothy Blackmore, Vani Sathyendran, Veronica Playle, Virginia Hope, Erasmus Smit, Lauren Jelly, Olin Silander, Joep de Ligt |
| EPI_ISL_581318 | Lighthouse Lab in Milton Keynes | Wellcome Sanger Institute for the COVID-19 Genomics UK (COG-UK) consortium | The Lighthouse Lab in Milton Keynes and Alex Alderton, Roberto Amato, Sonia Goncalves, Ewan Harrison, David K. Jackson, Ian Johnston, Dominic Kwiatkowski, Cordelia Langford, John Sillitoe on behalf of the Wellcome Sanger Institute COVID-19 Surveillance Team |
| EPI_ISL_581367 | Lighthouse Lab in Alderley Park | Wellcome Sanger Institute for the COVID-19 Genomics UK (COG-UK) consortium | Jacquelyn Wynn, Mairead Hyland, The Lighthouse Lab in Alderley Park and Alex Alderton, Roberto Amato, Sonia Goncalves, Ewan Harrison, David K. Jackson, Ian Johnston, Dominic Kwiatkowski, Cordelia Langford, John Sillitoe on behalf of the Wellcome Sanger Institute COVID-19 Surveillance Team |
| EPI_ISL_581371, EPI_ISL_581374, EPI_ISL_581375, EPI_ISL_581376 | Lighthouse Lab in Milton Keynes | Wellcome Sanger Institute for the COVID-19 Genomics UK (COG-UK) consortium | The Lighthouse Lab in Milton Keynes and Alex Alderton, Roberto Amato, Sonia Goncalves, Ewan Harrison, David K. Jackson, Ian Johnston, Dominic Kwiatkowski, Cordelia Langford, John Sillitoe on behalf of the Wellcome Sanger Institute COVID-19 Surveillance Team |
| EPI_ISL_581377 | Lighthouse Lab in Glasgow | Wellcome Sanger Institute for the COVID-19 Genomics UK (COG-UK) consortium | Harper VanSteenhouse, Yumi Kasai, David Gray, Carol Clugston, Anna Dominiczak and Alex Alderton, Roberto Amato, Sonia Goncalves, Ewan Harrison, David K. Jackson, Ian Johnston, Dominic Kwiatkowski, Cordelia Langford, John Sillitoe on behalf of the Wellcome Sanger Institute COVID-19 Surveillance Team |
| EPI_ISL_581378, EPI_ISL_581379, EPI_ISL_581380 | Lighthouse Lab in Alderley Park | Wellcome Sanger Institute for the COVID-19 Genomics UK (COG-UK) consortium | Jacquelyn Wynn, Mairead Hyland, The Lighthouse Lab in Alderley Park and Alex Alderton, Roberto Amato, Sonia Goncalves, Ewan Harrison, David K. Jackson, Ian Johnston, Dominic Kwiatkowski, Cordelia Langford, John Sillitoe on behalf of the Wellcome Sanger Institute COVID-19 Surveillance Team |
| EPI_ISL_582034, EPI_ISL_582035, EPI_ISL_582038, EPI_ISL_582041, EPI_ISL_582042, EPI_ISL_582044, EPI_ISL_582045, EPI_ISL_582050, EPI_ISL_582051, EPI_ISL_582072, EPI_ISL_582073, EPI_ISL_582074, EPI_ISL_582075, EPI_ISL_582076, EPI_ISL_582077, EPI_ISL_582078, EPI_ISL_582079, EPI_ISL_582080, EPI_ISL_582081, EPI_ISL_582082, EPI_ISL_582083, EPI_ISL_582084, EPI_ISL_582085, EPI_ISL_582086, EPI_ISL_582087 | | | |
| see above | Servicio de Microbiología. Hospital Universitario Donostia. OSI Donostialdea. Área de Enfermedades Infecciosas, Grupo de Infección Respiratoria y Resistencia Antimicrobiana. Instituto de Investigación Sanitaria Biodonostia | SeqCOVID-SPAIN consortium/IBV(CSIC) | Gustavo Cilla, Milagrosa Montes, Luis Piñeiro, Jose Maria Marimón and SeqCOVID-SPAIN consortium |
| EPI_ISL_582111, EPI_ISL_582113, EPI_ISL_582116, EPI_ISL_582122 | CNR Virus des Infections Respiratoires - France SUD | CNR Virus des Infections Respiratoires - France SUD | Antonin Bal, Gregory Destras, Gwendolyne Burfin, Hadrien Règue, Alexandre Gaymard, Maude Bouscambert-Duchamp, Florence Morfin-Sherpa, Martine Valette, Bruno Lina, Laurence Josset |
| EPI_ISL_582128, EPI_ISL_582129, EPI_ISL_582130, EPI_ISL_582131, EPI_ISL_582132 | Antwerp University Hospital | Institute of Tropical Medicine | Philippe Selhorst, Colin Anthony |
| EPI_ISL_582508 | CNR Virus des Infections Respiratoires - France SUD | CNR Virus des Infections Respiratoires - France SUD | Antonin Bal, Gregory Destras, Gwendolyne Burfin, Hadrien Règue, Alexandre Gaymard, Maude Bouscambert-Duchamp, Florence Morfin-Sherpa, Martine Valette, Bruno Lina, Laurence Josset |
| EPI_ISL_582664, EPI_ISL_582665, EPI_ISL_582666, EPI_ISL_582667, EPI_ISL_582668, EPI_ISL_582669, EPI_ISL_582670, EPI_ISL_582671, EPI_ISL_582672, EPI_ISL_582673, EPI_ISL_582674, EPI_ISL_582675, EPI_ISL_582676 | | | |
| see above | Sheikh Khalifa Medical City | Molecular/Surveillance lab Sheikh Khalifa Medical City | Amirtharaj Francis, Sajeed Abdul, Hala Imambaccus, Sahar Almarzooqi, Hiba Saud, Stefan Weber |
| EPI_ISL_582771, EPI_ISL_582772, EPI_ISL_582773, | Klinisk mikrobiologi Linkoping | The Public Health Agency of Sweden | Anna-Malin Linde, Maria Lind Karlberg, Mattias Haukland, Reza Advani, Olov Svartstrom, Oskar Karlsson Lindsjo, Sandra Broddesson, Petra Edquist, Mia Brytting, Anna Risberg, Karin Tegmark-Wisell |

| | | | |
|---|---|---|---|
| EPI_ISL_582774, EPI_ISL_582775 | | | |
| EPI_ISL_582938, EPI_ISL_582943, EPI_ISL_582944, EPI_ISL_582945, EPI_ISL_582946, EPI_ISL_582947, EPI_ISL_582948, EPI_ISL_582951, EPI_ISL_582954 | County of Santa Clara Public Health Department | Chan-Zuckerberg Biohub | CZB Cliahub Consortium |
| EPI_ISL_583027, EPI_ISL_583028, EPI_ISL_583029, EPI_ISL_583030, EPI_ISL_583031, EPI_ISL_583032, EPI_ISL_583033, EPI_ISL_583034, EPI_ISL_583035, EPI_ISL_583036, EPI_ISL_583037, EPI_ISL_583038, EPI_ISL_583039, EPI_ISL_583040, EPI_ISL_583041, EPI_ISL_583042, EPI_ISL_583043, EPI_ISL_583044 | | | |
| see above | Orange County Public Health Lab | Chan-Zuckerberg Biohub | CZB Cliahub Consortium |
| EPI_ISL_583329, EPI_ISL_583331, EPI_ISL_583337, EPI_ISL_583344, EPI_ISL_583347, EPI_ISL_583348, EPI_ISL_583349, EPI_ISL_583350, EPI_ISL_583351, EPI_ISL_583352, EPI_ISL_583353, EPI_ISL_583354, EPI_ISL_583355, EPI_ISL_583356, EPI_ISL_583357, EPI_ISL_583358, EPI_ISL_583359, EPI_ISL_583360, EPI_ISL_583361, EPI_ISL_583363, EPI_ISL_583364, EPI_ISL_583365, EPI_ISL_583366, EPI_ISL_583367, EPI_ISL_583368, EPI_ISL_583369, EPI_ISL_583370, EPI_ISL_583371, EPI_ISL_583372, EPI_ISL_583373, EPI_ISL_583374, EPI_ISL_583378, EPI_ISL_583382, EPI_ISL_583384, EPI_ISL_583385, EPI_ISL_583386, EPI_ISL_583387, EPI_ISL_583388, EPI_ISL_583389, EPI_ISL_583390, EPI_ISL_583391, EPI_ISL_583392, EPI_ISL_583393, EPI_ISL_583394, EPI_ISL_583395, EPI_ISL_583396, EPI_ISL_583397, EPI_ISL_583398, EPI_ISL_583399, EPI_ISL_583400, EPI_ISL_583401, EPI_ISL_583402, EPI_ISL_583403 | | | |
| see above | University of Michigan Clinical Microbiology Laboratory | Lauring Lab, University of Michigan, Department of Microbiology and Immunology | Valesano |
| EPI_ISL_583481 | Institute of Virology, Biomedical Research Center of the Slovak Academy of Sciences, Bratislava | Faculty of Natural Sciences, Comenius University, Bratislava | Viktória Hodorová, Kristína Boršová, Broa Brejová, Viktória abanová, Dominika Friová, Sabina Fumaová Havlíková, Juraj Kopáek, Martina Liková, ubomíra Lukáiková, Martina Neboháová, Monika Sláviková, Edita Staroová, Elena Tichá, Tomáš Vina, Jozef Nosek, Boris Klempa |
| EPI_ISL_583482 | Institute of Virology, Biomedical Research Center of the Slovak Academy of Sciences, Bratislava | Faculty of Natural Sciences, Comenius University, Bratislava | Kristína Boršová, Viktória Hodorová, Broa Brejová, Viktória abanová, Dominika Friová, Sabina Fumaová Havlíková, Juraj Kopáek, Martina Liková, ubomíra Lukáiková, Martina Neboháová, Monika Sláviková, Edita Staroová, Elena Tichá, Tomáš Vina, Boris Klempa, Jozef Nosek |
| EPI_ISL_583483 | Institute of Virology, Biomedical Research Center of the Slovak Academy of Sciences, Bratislava | Faculty of Natural Sciences, Comenius University, Bratislava | Viktória Hodorová, Kristína Boršová, Broa Brejová, Viktória abanová, Dominika Friová, Sabina Fumaová Havlíková, Juraj Kopáek, Martina Liková, ubomíra Lukáiková, Martina Neboháová, Monika Sláviková, Edita Staroová, Elena Tichá, Tomáš Vina, Jozef Nosek, Boris Klempa |
| EPI_ISL_583484 | Institute of Virology, Biomedical Research Center of the Slovak Academy of Sciences, Bratislava | Faculty of Natural Sciences, Comenius University, Bratislava | Kristína Boršová, Viktória Hodorová, Broa Brejová, Viktória abanová, Dominika Friová, Sabina Fumaová Havlíková, Juraj Kopáek, Martina Liková, ubomíra Lukáiková, Martina Neboháová, Monika Sláviková, Edita Staroová, Elena Tichá, Tomáš Vina, Boris Klempa, Jozef Nosek |
| EPI_ISL_583485 | Institute of Virology, Biomedical Research Center of the Slovak Academy of Sciences, Bratislava | Faculty of Natural Sciences, Comenius University, Bratislava | Viktória Hodorová, Kristína Boršová, Broa Brejová, Viktória abanová, Dominika Friová, Sabina Fumaová Havlíková, Juraj Kopáek, Martina Liková, ubomíra Lukáiková, Martina Neboháová, Monika Sláviková, Edita Staroová, Elena Tichá, Tomáš Vina, Jozef Nosek, Boris Klempa |
| EPI_ISL_583486 | Institute of Virology, Biomedical Research Center of the Slovak Academy of Sciences, Bratislava | Faculty of Natural Sciences, Comenius University, Bratislava | Kristína Boršová, Viktória Hodorová, Broa Brejová, Viktória abanová, Dominika Friová, Sabina Fumaová Havlíková, Juraj Kopáek, Martina Liková, ubomíra Lukáiková, Martina Neboháová, Monika Sláviková, Edita Staroová, Elena Tichá, Tomáš Vina, Boris Klempa, Jozef Nosek |
| EPI_ISL_583487, EPI_ISL_583488 | Institute of Virology, Biomedical Research Center of the Slovak Academy of Sciences, Bratislava | Faculty of Natural Sciences, Comenius University, Bratislava | Broa Brejová, Viktória Hodorová, Kristína Boršová, Viktória abanová, Dominika Friová, Sabina Fumaová Havlíková, Juraj Kopáek, Martina Liková, ubomíra Lukáiková, Martina Neboháová, Monika Sláviková, Edita Staroová, Elena Tichá, Tomáš Vina, Jozef Nosek, Boris Klempa |
| EPI_ISL_583489 | Institute of Virology, Biomedical Research Center of the Slovak Academy of Sciences, Bratislava | Faculty of Natural Sciences, Comenius University, Bratislava | Viktória Hodorová, Kristína Boršová, Broa Brejová, Viktória abanová, Dominika Friová, Sabina Fumaová Havlíková, Juraj Kopáek, Martina Liková, ubomíra Lukáiková, Martina Neboháová, Monika Sláviková, Edita Staroová, Elena Tichá, Tomáš Vina, Jozef Nosek, Boris Klempa |
| EPI_ISL_583524 | Michigan Department of Health and Human Services, Bureau of Laboratories | Michigan Department of Health and Human Services, Bureau of Laboratories | Blankenship HM, Riner D, Soehnlen MK |
| EPI_ISL_583993 | Respiratory Virus Unit, Microbiology Services Colindale, Public Health England | Respiratory Virus Unit, Microbiology Services Colindale, Public Health England | PHE Covid Sequencing Team |
| EPI_ISL_584072 | IZSM | IZSM | Maurizio Viscardi, Lorena Cardillo, Giovanna Fusco |
| EPI_ISL_584097 | University of Michigan Clinical Microbiology Laboratory | Lauring Lab, University of Michigan, Department of Microbiology and Immunology | Valesano |
| EPI_ISL_584165, EPI_ISL_584166 | Quadram Institute Bioscience | COVID-19 Genomics UK (COG-UK) Consortium | Dave J. Baker, Gemma L. Kay, Alp Aydin, Thanh Le-Viet, Steven Rudder, Ana P. Tedim, Anastasia Kolyva, Maria Diaz, Leonardo de Oliveira Martins, Nabil-Fareed Alikhan, Lizzie Meadows, Rachael Stanley, Ngozi Elumogo, Muhammed Yasir, Nicholas M. Thomson, Alexander J Trotter, Rachel Gilroy, Samuel Bloomfield, Claire Stuart, Andrew Bell, Reenesh Prakash, Samir Dervisevic, Alison E. Mather, John Wain, Mark Webber, Andrew J. Page, Justin O'Grady |
| EPI_ISL_584370, EPI_ISL_584371, EPI_ISL_584372, EPI_ISL_584377 | Virology Department, Sheffield Teaching Hospitals NHS Foundation Trust/Department of Infection, Immunity and Cardiovascular Disease, The Medical School, University of Sheffield | COVID-19 Genomics UK (COG-UK) Consortium | Thushan de Silva, Matthew Parker, Nikki Smith, Adri Angyal, Rebecca Brown, Luke Green, Rachel Tucker, Paul Parsons, Danielle Groves, Katie Johnson, Laura Carrilero, Alex Keeley, Dave Partridge, Matthew Wyles, Benjamin Lindsey, Mehmet Yavuz, Mohammad Raza, Cariad Evans |
| EPI_ISL_584675, EPI_ISL_584676, | University College London, Great Ormond Street | COVID-19 Genomics UK (COG-UK) Consortium | Sergi Castellano, Rachel Williams, Mark Kristiansen, Paola Resende Silva, Sunando Roy, Tony Brooks, Helena Tutill, Paola Niola, Patricia Dyal, Charlotte Williams, Leysa Forrest, Yasmin Panchbhaya, Jacqueline Findlay, Samuel Weeks, Julianne Brown, Kathryn Harris, Paul Randell, James Price, Alison Holmes, Judith Breuer |

| | | | |
|---|---|---|---|
| EPI_ISL_584677 | Hospital for Children NHS Foundation Trust, Imperial College Healthcare NHS Trust | | |
| EPI_ISL_585024, EPI_ISL_585026, EPI_ISL_585035, EPI_ISL_585036, EPI_ISL_585037, EPI_ISL_585038, EPI_ISL_585041, EPI_ISL_585071, EPI_ISL_585090 | Virology Department, Sheffield Teaching Hospitals NHS Foundation Trust/Department of Infection, Immunity and Cardiovascular Disease, The Medical School, University of Sheffield | COVID-19 Genomics UK (COG-UK) Consortium | Thushan de Silva, Matthew Parker, Nikki Smith, Adri Angyal, Rebecca Brown, Luke Green, Rachel Tucker, Paul Parsons, Danielle Groves, Katie Johnson, Laura Carrilero, Alex Keeley, Dave Partridge, Matthew Wyles, Benjamin Lindsey, Mehmet Yavuz, Mohammad Raza, Cariad Evans |
| EPI_ISL_585187, EPI_ISL_585191, EPI_ISL_585192, EPI_ISL_585193, EPI_ISL_585194, EPI_ISL_585196, EPI_ISL_585197, EPI_ISL_585198, EPI_ISL_585199, EPI_ISL_585200, EPI_ISL_585204, EPI_ISL_585205, EPI_ISL_585206, EPI_ISL_585207, EPI_ISL_585208, EPI_ISL_585209, EPI_ISL_585211, EPI_ISL_585213, EPI_ISL_585214, EPI_ISL_585215, EPI_ISL_585216, EPI_ISL_585217, EPI_ISL_585218, EPI_ISL_585219, EPI_ISL_585220, EPI_ISL_585221, EPI_ISL_585222 | | | |
| see above | Regional Virus Laboratory, Belfast Health and Social Care Trust | COVID-19 Genomics UK (COG-UK) Consortium | Conall McCaughey, James McKenna, Tanya Curran, Susan Feeney, Alison Watt, Ciara Cox, Mairead Connor, Zoltan Molnar, David Simpson, Derek Fairley |
| EPI_ISL_585261, EPI_ISL_585262, EPI_ISL_585263, EPI_ISL_585264, EPI_ISL_585265, EPI_ISL_585266, EPI_ISL_585272 | West of Scotland Specialist Virology Centre, NHSGGC / MRC-University of Glasgow Centre for Virus Research | COVID-19 Genomics UK (COG-UK) Consortium | Ana da Silva Filipe, Natasha Johnson, Kathy Smollett, Daniel Mair, Stephen Carmichael, Lily Tong, Jenna Nichols, Elihu Aranday-Cortes, Kyriaki Nomikou; Sarah McDonald, Marc Niebel, Patawee Asamaphan; Richard Orton, Joseph Hughes, Sreenu Vattipally, David L Robertson; Alasdair MacLean, Rory Gunson; Kathy Li, Igor Starinskij, Natasha Jesudason, Rajiv Shah, James Shepherd, Antonia Ho, Emma Thomson |
| EPI_ISL_585297, EPI_ISL_585298, EPI_ISL_585299, EPI_ISL_585300, EPI_ISL_585301, EPI_ISL_585302, EPI_ISL_585303, EPI_ISL_585304, EPI_ISL_585421 | Lighthouse Lab in Glasgow / MRC-University of Glasgow Centre for Virus Research | COVID-19 Genomics UK (COG-UK) Consortium | Ana da Silva Filipe, Natasha Johnson, Kathy Smollett, Daniel Mair, Stephen Carmichael, Lily Tong, Jenna Nichols, Elihu Aranday-Cortes, Kyriaki Nomikou; Sarah McDonald, Marc Niebel, Patawee Asamaphan; Harper VanSteenhouse, Yumi Kasai, David Gray, Carol Clugston, Anna Dominiczak; Alasdair MacLean, Rory Gunson; Richard Orton, Joseph Hughes, Sreenu Vattipally, David L Robertson; Sharif Shaaban, Matthew Holden; Kathy Li, Natasha Jesudason, Rajiv Shah, James Shepherd, Antonia Ho, Emma Thomson |
| EPI_ISL_585424 | West of Scotland Specialist Virology Centre, NHSGGC / MRC-University of Glasgow Centre for Virus Research | COVID-19 Genomics UK (COG-UK) Consortium | Ana da Silva Filipe, Natasha Johnson, Kathy Smollett, Daniel Mair, Stephen Carmichael, Lily Tong, Jenna Nichols, Elihu Aranday-Cortes, Kyriaki Nomikou; Sarah McDonald, Marc Niebel, Patawee Asamaphan; Richard Orton, Joseph Hughes, Sreenu Vattipally, David L Robertson; Alasdair MacLean, Rory Gunson; Kathy Li, Igor Starinskij, Natasha Jesudason, Rajiv Shah, James Shepherd, Antonia Ho, Emma Thomson |
| EPI_ISL_585426 | Virology Department, Royal Infirmary of Edinburgh, NHS Lothian / School of Biological Sciences, University of Edinburgh / Institute of Genetics and Molecular Medicine, University of Edinburgh | COVID-19 Genomics UK (COG-UK) Consortium | McHugh M, Dewar R, Rooke S, Gallagher M, Balcaza C, O'Toole Á, Scher E, Hill V, McCrone JT, Colquhoun R, Yu X, Jackson B, Rambaut A, Williams TC, Templeton K |
| EPI_ISL_585626 | Virology Department, Sheffield Teaching Hospitals NHS Foundation Trust/Department of Infection, Immunity and Cardiovascular Disease, The Medical School, University of Sheffield | COVID-19 Genomics UK (COG-UK) Consortium | Thushan de Silva, Matthew Parker, Nikki Smith, Adri Angyal, Rebecca Brown, Luke Green, Rachel Tucker, Paul Parsons, Danielle Groves, Katie Johnson, Laura Carrilero, Alex Keeley, Dave Partridge, Matthew Wyles, Benjamin Lindsey, Mehmet Yavuz, Mohammad Raza, Cariad Evans |
| EPI_ISL_585797 | Wales Specialist Virology Centre Sequencing lab: Pathogen Genomics Unit | COVID-19 Genomics UK (COG-UK) Consortium | Catherine Moore, Johnathan Evans, Laura Gifford, Malorie Perry, Simon Cottrell, Angela Marchbank, Alec Birchley, Alexander Adams, Amy Gaskin, Bree Gatica-Wilcox, Jason Coombes, Joel Southgate, Lauren Gilbert, Lee Graham, Nicole Pacchiarini, Sara Kumziene-Summerhayes, Sarah Taylor, Sophie Jones, Sara Rey, Matthew Bull, Joanne Watkins, Sally Corden, Tom Connor |
| EPI_ISL_589801 | Lighthouse Lab in Milton Keynes | Wellcome Sanger Institute for the COVID-19 Genomics UK (COG-UK) consortium | The Lighthouse Lab in Milton Keynes and Alex Alderton, Roberto Amato, Sonia Goncalves, Ewan Harrison, David K. Jackson, Ian Johnston, Dominic Kwiatkowski, Cordelia Langford, John Sillitoe on behalf of the Wellcome Sanger Institute COVID-19 Surveillance Team (http://www.sanger.ac.uk/covid-team) |
| EPI_ISL_589802 | Lighthouse Lab in Alderley Park | Wellcome Sanger Institute for the COVID-19 Genomics UK (COG-UK) consortium | Jacquelyn Wynn, Mairead Hyland, The Lighthouse Lab in Alderley Park and Alex Alderton, Roberto Amato, Sonia Goncalves, Ewan Harrison, David K. Jackson, Ian Johnston, Dominic Kwiatkowski, Cordelia Langford, John Sillitoe on behalf of the Wellcome Sanger Institute COVID-19 Surveillance Team (http://www.sanger.ac.uk/covid-team) |
| EPI_ISL_589803, EPI_ISL_589804, EPI_ISL_589805, EPI_ISL_589806, EPI_ISL_589807, EPI_ISL_589808, EPI_ISL_589809, EPI_ISL_589810, EPI_ISL_589811, EPI_ISL_589812, EPI_ISL_589813, EPI_ISL_589814, EPI_ISL_589815, EPI_ISL_589816, EPI_ISL_589817, EPI_ISL_589818, EPI_ISL_589819, EPI_ISL_589820, EPI_ISL_589821, EPI_ISL_589822, EPI_ISL_589823, EPI_ISL_589824, EPI_ISL_589825, EPI_ISL_589826, EPI_ISL_589827, EPI_ISL_589828, EPI_ISL_589829, EPI_ISL_589830, EPI_ISL_589831, EPI_ISL_589832, EPI_ISL_589833, EPI_ISL_589834, EPI_ISL_589835, EPI_ISL_589836, EPI_ISL_589837, EPI_ISL_589838, EPI_ISL_589839, EPI_ISL_589840, EPI_ISL_589841, EPI_ISL_589842, EPI_ISL_589843, EPI_ISL_589844, EPI_ISL_589845, EPI_ISL_589846, EPI_ISL_589847, EPI_ISL_589848, EPI_ISL_589849, EPI_ISL_589850, EPI_ISL_589851, EPI_ISL_589852, EPI_ISL_589853, EPI_ISL_589854, EPI_ISL_589855, EPI_ISL_589856, EPI_ISL_589857, EPI_ISL_589858, EPI_ISL_589859, EPI_ISL_589860, EPI_ISL_589861, EPI_ISL_589862, EPI_ISL_589863, EPI_ISL_589864, EPI_ISL_589865, EPI_ISL_589866, EPI_ISL_589867, EPI_ISL_589868, EPI_ISL_589869, EPI_ISL_589870, EPI_ISL_589871, EPI_ISL_589872, EPI_ISL_589873, EPI_ISL_589874, EPI_ISL_589875, EPI_ISL_589876, EPI_ISL_589877, EPI_ISL_589878, EPI_ISL_589879, EPI_ISL_589880, EPI_ISL_589881, EPI_ISL_589882, EPI_ISL_589883, EPI_ISL_589884, EPI_ISL_589885, EPI_ISL_589886, EPI_ISL_589887, EPI_ISL_589888, EPI_ISL_589889, EPI_ISL_589890, EPI_ISL_589891, EPI_ISL_589892, EPI_ISL_589893, EPI_ISL_589894, EPI_ISL_589895, EPI_ISL_589896, EPI_ISL_589897, EPI_ISL_589898, EPI_ISL_589899, EPI_ISL_589900, EPI_ISL_589901, EPI_ISL_589902, EPI_ISL_589903, EPI_ISL_589904, EPI_ISL_589905, EPI_ISL_589906, EPI_ISL_589907, EPI_ISL_589908, EPI_ISL_589909, EPI_ISL_589910, EPI_ISL_589911, EPI_ISL_589912, EPI_ISL_589913, EPI_ISL_589914, EPI_ISL_589915, EPI_ISL_589916, EPI_ISL_589917, EPI_ISL_589918, EPI_ISL_589919, EPI_ISL_589920, EPI_ISL_589921, EPI_ISL_589922, EPI_ISL_589923, EPI_ISL_589924, EPI_ISL_589925, EPI_ISL_589926, EPI_ISL_589927, EPI_ISL_589928, EPI_ISL_589929, EPI_ISL_589930, EPI_ISL_589931, EPI_ISL_589932, EPI_ISL_589933, EPI_ISL_589934, EPI_ISL_589935 | | | |
| see above | Lighthouse Lab in Glasgow | Wellcome Sanger Institute for the COVID-19 Genomics UK (COG-UK) consortium | Harper VanSteenhouse, Yumi Kasai, David Gray, Carol Clugston, Anna Dominiczak and Alex Alderton, Roberto Amato, Sonia Goncalves, Ewan Harrison, David K. Jackson, Ian Johnston, Dominic Kwiatkowski, Cordelia Langford, John Sillitoe on behalf of the Wellcome Sanger Institute COVID-19 Surveillance Team (http://www.sanger.ac.uk/covid-team) |
| EPI_ISL_589936 | Lighthouse Lab in Glasgow | Wellcome Sanger Institute for the COVID-19 Genomics UK (COG-UK) Consortium | Harper VanSteenhouse, Yumi Kasai, David Gray, Carol Clugston, Anna Dominiczak and Alex Alderton, Roberto Amato, Sonia Goncalves, Ewan Harrison, David K. Jackson, Ian Johnston, Dominic Kwiatkowski, Cordelia Langford, John Sillitoe on behalf of the Wellcome Sanger Institute COVID-19 Surveillance Team |
| EPI_ISL_589937, EPI_ISL_589938, EPI_ISL_589939, EPI_ISL_589940, EPI_ISL_589941, EPI_ISL_589942, EPI_ISL_589943, EPI_ISL_589944, EPI_ISL_589945, EPI_ISL_589946, EPI_ISL_589947, EPI_ISL_589948, EPI_ISL_589949, EPI_ISL_589950, EPI_ISL_589951, EPI_ISL_589952, EPI_ISL_589953, EPI_ISL_589954, EPI_ISL_589955, EPI_ISL_589956, EPI_ISL_589957, EPI_ISL_589958, EPI_ISL_589959, EPI_ISL_589960, EPI_ISL_589961, EPI_ISL_589962, EPI_ISL_589963, EPI_ISL_589964, EPI_ISL_589965, EPI_ISL_589966, EPI_ISL_589967, EPI_ISL_589968, EPI_ISL_589969, EPI_ISL_589970, EPI_ISL_589971, EPI_ISL_589972, EPI_ISL_589973, EPI_ISL_589974, EPI_ISL_589975, EPI_ISL_589976, EPI_ISL_589977, EPI_ISL_589978, EPI_ISL_589979, EPI_ISL_589980, EPI_ISL_589981, EPI_ISL_589982, EPI_ISL_589983, EPI_ISL_589984, EPI_ISL_589985, EPI_ISL_589986, EPI_ISL_589987, EPI_ISL_589988, EPI_ISL_589990, EPI_ISL_589991, | | | |

| | | | |
|---|---|---|---|
| EPI_ISL_589992, EPI_ISL_589993, EPI_ISL_589994, EPI_ISL_589995, EPI_ISL_589996, EPI_ISL_589997, EPI_ISL_589998, EPI_ISL_589999, EPI_ISL_590000, EPI_ISL_590001, EPI_ISL_590002, EPI_ISL_590003, EPI_ISL_590004, EPI_ISL_590005, EPI_ISL_590006, EPI_ISL_590007, EPI_ISL_590008, EPI_ISL_590009, EPI_ISL_590010, EPI_ISL_590011, EPI_ISL_590012, EPI_ISL_590013, EPI_ISL_590014, EPI_ISL_590015, EPI_ISL_590016, EPI_ISL_590017, EPI_ISL_590018, EPI_ISL_590019, EPI_ISL_590020, EPI_ISL_590021, EPI_ISL_590022, EPI_ISL_590023, EPI_ISL_590024, EPI_ISL_590025, EPI_ISL_590026, EPI_ISL_590027, EPI_ISL_590028, EPI_ISL_590029, EPI_ISL_590030, EPI_ISL_590031, EPI_ISL_590032, EPI_ISL_590033, EPI_ISL_590034, EPI_ISL_590035, EPI_ISL_590036, EPI_ISL_590037, EPI_ISL_590038, EPI_ISL_590039, EPI_ISL_590040, EPI_ISL_590041, EPI_ISL_590042, EPI_ISL_590043, EPI_ISL_590044, EPI_ISL_590045 | | | |
| see above | Lighthouse Lab in Glasgow | Wellcome Sanger Institute for the COVID-19 Genomics UK (COG-UK) consortium | Harper VanSteenhouse, Yumi Kasai, David Gray, Carol Clugston, Anna Dominiczak and Alex Alderton, Roberto Amato, Sonia Goncalves, Ewan Harrison, David K. Jackson, Ian Johnston, Dominic Kwiatkowski, Cordelia Langford, John Sillitoe on behalf of the Wellcome Sanger Institute COVID-19 Surveillance Team (http://www.sanger.ac.uk/covid-team) |
| EPI_ISL_590046 | Lighthouse Lab in Glasgow | Wellcome Sanger Institute for the COVID-19 Genomics UK (COG-UK) Consortium | Harper VanSteenhouse, Yumi Kasai, David Gray, Carol Clugston, Anna Dominiczak and Alex Alderton, Roberto Amato, Sonia Goncalves, Ewan Harrison, David K. Jackson, Ian Johnston, Dominic Kwiatkowski, Cordelia Langford, John Sillitoe on behalf of the Wellcome Sanger Institute COVID-19 Surveillance Team |
| EPI_ISL_590047, EPI_ISL_590048, EPI_ISL_590049, EPI_ISL_590050, EPI_ISL_590051, EPI_ISL_590052, EPI_ISL_590053, EPI_ISL_590054, EPI_ISL_590055, EPI_ISL_590056, EPI_ISL_590057, EPI_ISL_590058, EPI_ISL_590059, EPI_ISL_590060, EPI_ISL_590061, EPI_ISL_590062, EPI_ISL_590063, EPI_ISL_590064, EPI_ISL_590065, EPI_ISL_590066, EPI_ISL_590067, EPI_ISL_590068, EPI_ISL_590069, EPI_ISL_590070, EPI_ISL_590071, EPI_ISL_590072, EPI_ISL_590073, EPI_ISL_590074, EPI_ISL_590075, EPI_ISL_590076, EPI_ISL_590077, EPI_ISL_590078, EPI_ISL_590079, EPI_ISL_590080, EPI_ISL_590081, EPI_ISL_590082, EPI_ISL_590083, EPI_ISL_590084, EPI_ISL_590085, EPI_ISL_590086, EPI_ISL_590087, EPI_ISL_590088, EPI_ISL_590089, EPI_ISL_590090, EPI_ISL_590091, EPI_ISL_590092, EPI_ISL_590093, EPI_ISL_590094, EPI_ISL_590095, EPI_ISL_590096, EPI_ISL_590097, EPI_ISL_590098, EPI_ISL_590099, EPI_ISL_590100, EPI_ISL_590101, EPI_ISL_590102, EPI_ISL_590103, EPI_ISL_590104, EPI_ISL_590105, EPI_ISL_590106, EPI_ISL_590107, EPI_ISL_590108, EPI_ISL_590109, EPI_ISL_590110, EPI_ISL_590111, EPI_ISL_590112, EPI_ISL_590113, EPI_ISL_590114, EPI_ISL_590115, EPI_ISL_590116, EPI_ISL_590117, EPI_ISL_590118, EPI_ISL_590119, EPI_ISL_590120, EPI_ISL_590121, EPI_ISL_590122, EPI_ISL_590123, EPI_ISL_590124, EPI_ISL_590125, EPI_ISL_590126, EPI_ISL_590127, EPI_ISL_590128, EPI_ISL_590129, EPI_ISL_590130, EPI_ISL_590131, EPI_ISL_590132, EPI_ISL_590133, EPI_ISL_590135, EPI_ISL_590136, EPI_ISL_590137, EPI_ISL_590138, EPI_ISL_590139, EPI_ISL_590140, EPI_ISL_590141, EPI_ISL_590142, EPI_ISL_590143, EPI_ISL_590144, EPI_ISL_590145, EPI_ISL_590146, EPI_ISL_590147, EPI_ISL_590148, EPI_ISL_590149, EPI_ISL_590150, EPI_ISL_590151, EPI_ISL_590152, EPI_ISL_590153, EPI_ISL_590154, EPI_ISL_590155, EPI_ISL_590156, EPI_ISL_590157, EPI_ISL_590158, EPI_ISL_590159, EPI_ISL_590160, EPI_ISL_590161, EPI_ISL_590162, EPI_ISL_590163, EPI_ISL_590164, EPI_ISL_590165, EPI_ISL_590166, EPI_ISL_590167, EPI_ISL_590168, EPI_ISL_590169, EPI_ISL_590170, EPI_ISL_590171 | | | |
| see above | Lighthouse Lab in Glasgow | Wellcome Sanger Institute for the COVID-19 Genomics UK (COG-UK) consortium | Harper VanSteenhouse, Yumi Kasai, David Gray, Carol Clugston, Anna Dominiczak and Alex Alderton, Roberto Amato, Sonia Goncalves, Ewan Harrison, David K. Jackson, Ian Johnston, Dominic Kwiatkowski, Cordelia Langford, John Sillitoe on behalf of the Wellcome Sanger Institute COVID-19 Surveillance Team (http://www.sanger.ac.uk/covid-team) |
| EPI_ISL_590172 | Lighthouse Lab in Milton Keynes | Wellcome Sanger Institute for the COVID-19 Genomics UK (COG-UK) consortium | The Lighthouse Lab in Milton Keynes and Alex Alderton, Roberto Amato, Sonia Goncalves, Ewan Harrison, David K. Jackson, Ian Johnston, Dominic Kwiatkowski, Cordelia Langford, John Sillitoe on behalf of the Wellcome Sanger Institute COVID-19 Surveillance Team (http://www.sanger.ac.uk/covid-team) |
| EPI_ISL_590176, EPI_ISL_590177 | Lighthouse Lab in Glasgow | Wellcome Sanger Institute for the COVID-19 Genomics UK (COG-UK) consortium | Harper VanSteenhouse, Yumi Kasai, David Gray, Carol Clugston, Anna Dominiczak and Alex Alderton, Roberto Amato, Sonia Goncalves, Ewan Harrison, David K. Jackson, Ian Johnston, Dominic Kwiatkowski, Cordelia Langford, John Sillitoe on behalf of the Wellcome Sanger Institute COVID-19 Surveillance Team (http://www.sanger.ac.uk/covid-team) |
| EPI_ISL_590887, EPI_ISL_590888 | Vestfold Hospital, Toensberg Department of Microbiology | Norwegian Institute of Public Health, Department of Virology | Kathrine Stene-Johansen, Kamilla Heddeland Instefjord, Hilde Elshaug, Rasmus Riis Kopperud, Hilde Vollan, Karoline Bragstad, Olav Hungnes |
| EPI_ISL_590889, EPI_ISL_590890, EPI_ISL_590891, EPI_ISL_590892 | Akershus University Hospital, Department for Microbiology and Infectious Disease Control | Norwegian Institute of Public Health, Department of Virology | Kathrine Stene-Johansen, Kamilla Heddeland Instefjord, Hilde Elshaug, Rasmus Riis Kopperud, Hilde Vollan, Karoline Bragstad, Olav Hungnes |
| EPI_ISL_590893 | Medical Microbiology Unit, Department for Laboratory Medicine, Drammen Hospital, Vestre Viken Health Trust, | Norwegian Institute of Public Health, Department of Virology | Kathrine Stene-Johansen, Kamilla Heddeland Instefjord, Hilde Elshaug, Rasmus Riis Kopperud, Hilde Vollan, Karoline Bragstad, Olav Hungnes |
| EPI_ISL_590896, EPI_ISL_590897 | Ostfold Hospital Trust - Kalnes, Centre for Laboratory Medicine, Section for gene technology and infection serology | Norwegian Institute of Public Health, Department of Virology | Kathrine Stene-Johansen, Kamilla Heddeland Instefjord, Hilde Elshaug, Rasmus Riis Kopperud, Hilde Vollan, Karoline Bragstad, Olav Hungnes |
| EPI_ISL_590927, EPI_ISL_590928, EPI_ISL_590930 | Oslo University Hospital, Department of Medical Microbiology | Norwegian Institute of Public Health, Department of Virology | Kathrine Stene-Johansen, Kamilla Heddeland Instefjord, Hilde Elshaug, Rasmus Riis Kopperud, Hilde Vollan, Karoline Bragstad, Olav Hungnes |
| EPI_ISL_590932 | Medical Microbiology Unit, Department for Laboratory Medicine, Drammen Hospital, Vestre Viken Health Trust, | Norwegian Institute of Public Health, Department of Virology | Kathrine Stene-Johansen, Kamilla Heddeland Instefjord, Hilde Elshaug, Rasmus Riis Kopperud, Hilde Vollan, Karoline Bragstad, Olav Hungnes |
| EPI_ISL_590935 | Oslo University Hospital, Department of Medical Microbiology | Norwegian Institute of Public Health, Department of Virology | Kathrine Stene-Johansen, Kamilla Heddeland Instefjord, Hilde Elshaug, Rasmus Riis Kopperud, Hilde Vollan, Karoline Bragstad, Olav Hungnes |
| EPI_ISL_590936 | Hospital of Southern Norway - Kristiansand, Department of Medical Microbiology | Norwegian Institute of Public Health, Department of Virology | Kathrine Stene-Johansen, Kamilla Heddeland Instefjord, Hilde Elshaug, Rasmus Riis Kopperud, Hilde Vollan, Karoline Bragstad, Olav Hungnes |
| EPI_ISL_590982 | Vestfold Hospital, Toensberg Department of Microbiology | Norwegian Institute of Public Health, Department of Virology | Kathrine Stene-Johansen, Kamilla Heddeland Instefjord, Hilde Elshaug, Rasmus Riis Kopperud, Hilde Vollan, Karoline Bragstad, Olav Hungnes |
| EPI_ISL_591493 | Pathology North - Royal North Shore Hospital - NSW Health Pathology | NSW Health Pathology - Institute of Clinical Pathology and Medical Research; Westmead Hospital; University of Sydney | CIDM-PH et al. |
| EPI_ISL_591496 | Pathology West - NSW Health Pathology | NSW Health Pathology - Institute of Clinical Pathology and Medical Research; Westmead Hospital; University of Sydney | CIDM-PH et al. |
| EPI_ISL_591502 | South Eastern Area Laboratory Services (SEALS) | NSW Health Pathology - Institute of Clinical Pathology and Medical Research; Westmead Hospital; University of Sydney | CIDM-PH et al. |
| EPI_ISL_591582, EPI_ISL_591589, EPI_ISL_591604, EPI_ISL_591629 | Microbiological Diagnostic Unit - Public Health Laboratory (MDU-PHL) | MDU-PHL | Seemann T., Schultz, M. B., Sait, M., Sherry, N. |

| | | | |
|---|---|---|---|
| EPI_ISL_591645 | Victorian Infectious Diseases Reference Laboratory (VIDRL) | VIDRL and MDU-PHL | Caly L., Seemann T., Sait, M., Schultz, M. B., Druce J., Sherry, N. |
| EPI_ISL_591646 | Microbiological Diagnostic Unit - Public Health Laboratory (MDU-PHL) | MDU-PHL | Seemann T., Schultz, M. B., Sait, M., Sherry, N. |
| EPI_ISL_591653, EPI_ISL_591667 | Victorian Infectious Diseases Reference Laboratory (VIDRL) | VIDRL and MDU-PHL | Caly L., Seemann T., Sait, M., Schultz, M. B., Druce J., Sherry, N. |
| EPI_ISL_591757 | Microbiological Diagnostic Unit - Public Health Laboratory (MDU-PHL) | MDU-PHL | Seemann T., Schultz, M. B., Sait, M., Sherry, N. |
| EPI_ISL_591760 | Victorian Infectious Diseases Reference Laboratory (VIDRL) | VIDRL and MDU-PHL | Caly L., Seemann T., Sait, M., Schultz, M. B., Druce J., Sherry, N. |
| EPI_ISL_591814, EPI_ISL_592310, EPI_ISL_592342, EPI_ISL_592344, EPI_ISL_592345, EPI_ISL_592347, EPI_ISL_592348, EPI_ISL_592349 | Microbiological Diagnostic Unit - Public Health Laboratory (MDU-PHL) | MDU-PHL | Seemann T., Schultz, M. B., Sait, M., Sherry, N. |
| EPI_ISL_592509, EPI_ISL_592600 | Victorian Infectious Diseases Reference Laboratory (VIDRL) | VIDRL and MDU-PHL | Caly L., Seemann T., Sait, M., Schultz, M. B., Druce J., Sherry, N. |
| EPI_ISL_592646, EPI_ISL_592681, EPI_ISL_592734, EPI_ISL_592762, EPI_ISL_592788, EPI_ISL_592935, EPI_ISL_592965 | Microbiological Diagnostic Unit - Public Health Laboratory (MDU-PHL) | MDU-PHL | Seemann T., Schultz, M. B., Sait, M., Sherry, N. |
| EPI_ISL_592979 | Victorian Infectious Diseases Reference Laboratory (VIDRL) | VIDRL and MDU-PHL | Caly L., Seemann T., Sait, M., Schultz, M. B., Druce J., Sherry, N. |
| EPI_ISL_592988, EPI_ISL_592989, EPI_ISL_593227, EPI_ISL_593230, EPI_ISL_593231, EPI_ISL_593241 | Microbiological Diagnostic Unit - Public Health Laboratory (MDU-PHL) | MDU-PHL | Seemann T., Schultz, M. B., Sait, M., Sherry, N. |
| EPI_ISL_593617, EPI_ISL_593644 | unknown | Public Health Virology Laboratory, Forensic and Scientific Services (PHV-FSS) | Son Nguyen et al. |
| EPI_ISL_593729, EPI_ISL_593730, EPI_ISL_593731, EPI_ISL_593732, EPI_ISL_593733, EPI_ISL_593734, EPI_ISL_593735, EPI_ISL_593736, EPI_ISL_593737, EPI_ISL_593739, EPI_ISL_593740, EPI_ISL_593750 | | | |
| see above | South Eastern Area Laboratory Services (SEALS) | NSW Health Pathology - Institute of Clinical Pathology and Medical Research; Westmead Hospital; University of Sydney | CIDM-PH et al. |
| EPI_ISL_593770 | Pathology West - NSW Health Pathology | NSW Health Pathology - Institute of Clinical Pathology and Medical Research; Westmead Hospital; University of Sydney | CIDM-PH et al. |
| EPI_ISL_593829, EPI_ISL_593831, EPI_ISL_593834, EPI_ISL_593843, EPI_ISL_593846, EPI_ISL_593848, EPI_ISL_593850 | Respiratory Virus Unit, Microbiology Services Colindale, Public Health England | Respiratory Virus Unit, Microbiology Services Colindale, Public Health England | PHE Covid Sequencing Team |
| EPI_ISL_593902 | Sentinelles, Limay | National Reference Center for Viruses of Respiratory Infections, Institut Pasteur, Paris | Sylvie Behillil, Fabiana Gambaro, Etienne Simon-Lorière, Vincent Enouf, Maud Vanpeene, Sylvie van der Werf |
| EPI_ISL_593904, EPI_ISL_593905, EPI_ISL_593906, EPI_ISL_593907, EPI_ISL_593909 | Labo Analyses Med, Sarcelles | National Reference Center for Viruses of Respiratory Infections, Institut Pasteur, Paris | Sylvie Behillil, Fabiana Gambaro, Etienne Simon-Lorière, Vincent Enouf, Maud Vanpeene, Sylvie van der Werf |
| EPI_ISL_593910 | Hospital, Boulogne-Billancourt | National Reference Center for Viruses of Respiratory Infections, Institut Pasteur, Paris | Sylvie Behillil, Fabiana Gambaro, Etienne Simon-Lorière, Vincent Enouf, Maud Vanpeene, Sylvie van der Werf |
| EPI_ISL_593924, | Labo Analyses Med, | National Reference Center | Sylvie Behillil, Fabiana Gambaro, Etienne Simon-Lorière, Vincent Enouf, Maud Vanpeene, Sylvie van der Werf |

| | | | |
|---|---|---|---|
| EPI_ISL_593926, EPI_ISL_593927, EPI_ISL_593928 | Puteaux | for Viruses of Respiratory Infections, Institut Pasteur, Paris | |
| EPI_ISL_594149, EPI_ISL_594150, EPI_ISL_594151, EPI_ISL_594152, EPI_ISL_594153 | Klinisk mikrobiologi Linkoping | The Public Health Agency of Sweden | Anna-Malin Linde, Maria Lind Karlberg, Mattias Haukland, Reza Advani, Olov Svartstrom, Oskar Karlsson Lindsjo, Sandra Broddesson, Petra Edquist, Mia Brytting, Anna Risberg, Karin Tegmark-Wisell |
| EPI_ISL_594312, EPI_ISL_594321, EPI_ISL_594322, EPI_ISL_594323, EPI_ISL_594324, EPI_ISL_594325, EPI_ISL_594326 | Florida Bureau of Public Health Laboratories | Florida Bureau of Public Health Laboratories | Sarah Schmedes, Jason Blanton |
| EPI_ISL_594516, EPI_ISL_594517, EPI_ISL_594518, EPI_ISL_594519, EPI_ISL_594520, EPI_ISL_594521 | University of Birmingham | COVID-19 Genomics UK (COG-UK) Consortium | Institute of Microbiology, University of Birmingham: Claire McMurray, Joanne Stockton, Samuel Nicholls, Radoslaw Poplawski, Will Rowe, Josh Quick, Nicholas Loman. University of Birmingham Testing Laboratory: Celina M Whalley, Andrew Bosworth, Charlotte Poxon, Kasun Wanigasooriya, Oliver Pickles, Mike Kidd, Alex Richter, Andrew D Beggs PHE Heartlands Lab: Husam Osman, Andrew Bosworth. Queen Elizabeth Hospital: Anna Casey |
| EPI_ISL_594849 | Liverpool Clinical Laboratories | COVID-19 Genomics UK (COG-UK) Consortium | Sam Haldenby, Anita Lucaci, Steve Paterson, Julian Hiscox, Alistair Darby, M Almsaud, A Alrezaihi, Muhannad Alruwaili, Stuart D Armstrong, Jones Benjamin, Eleanor G Bentley, Anu Chawla, Jordan J Clark, Angela Cowell, Richard Eccles, Isabel García-Dorival, Matthew Gemmell, Alessandro Gerada, PKF Gilmore, Richard Gregory, Ximeng Han, Catherine Hartley, Margaret Hughes, Miren Iturriza-Gomara, James Johnson, L Luu, Jenifer Manson, Charlotte Nelson, Elaine O'Toole, Cassie Olateju, Rebekah Penrice-Randal , Lucille Rainbow, N.P Randle, Trevor Ian Robinson, Parul Sharma, Ghada T Shawli, James P Stewart, Neil Swainston, Ecaterina Vamos, Joanne Watts, Mark Whitehead |
| EPI_ISL_596232 | WHO National Influenza Centre Russian Federation | WHO National Influenza Centre Russian Federation | Andrey Komissarov, Artem Fadeev, Anna Ivanova, Mariia Sergeeva, Kseniya Komissarova, Dmitry Bazhenov, Daria Danilenko |
| EPI_ISL_596235, EPI_ISL_596236 | WHO National Influenza Centre Russian Federation | WHO National Influenza Centre Russian Federation | Andrey Komissarov, Artem Fadeev, Anna Ivanova, Kseniya Komissarova, Dmitry Bazhenov, Daria Danilenko |
| EPI_ISL_596268, EPI_ISL_596269, EPI_ISL_596272, EPI_ISL_596273, EPI_ISL_596274, EPI_ISL_596280, EPI_ISL_596281, EPI_ISL_596282, EPI_ISL_596284, EPI_ISL_596285, EPI_ISL_596286, EPI_ISL_596287, EPI_ISL_596288, EPI_ISL_596289, EPI_ISL_596290, EPI_ISL_596291, EPI_ISL_596292, EPI_ISL_596293, EPI_ISL_596294, EPI_ISL_596303, EPI_ISL_596304, EPI_ISL_596309 | | | |
| see above | HELIX LCC | WHO National Influenza Centre Russian Federation | Andrey Komissarov, Artem Fadeev, Anna Ivanova, Kseniya Komissarova, Dmitry Bazhenov, Daria Danilenko |
| EPI_ISL_596387 | Seattle Flu Study | Seattle Flu Study | Deborah A. Nickerson, Chris D. Frazar, Jover Lee, Benjamin Pelle, Matthew Richardson, Amanda Adler, Elisabeth Brandstetter, Peter D. Han, Kairsten Fay, Misja Ilcisin, Kirsten Lacombe, Thomas R. Sibley, Melissa Truong, Caitlin R. Wolf, Michael Boeckh, Janet A. Englund, Michael Famulare, Barry R. Lutz, Mark J. Rieder, Lea M. Starita, Matthew Thompson, Jay Shendure, Trevor Bedford, Helen Y. Chu |
| EPI_ISL_596388, EPI_ISL_596389, EPI_ISL_596390, EPI_ISL_596391 | Seattle Flu Study | Seattle Flu Study | Deborah A. Nickerson, Chris D. Frazar, Jover Lee, Benjamin Pelle, Matthew Richardson, Amanda Adler, Elisabeth Brandstetter, Peter D. Han, Kairsten Fay, Misja Ilcisin, Kirsten Lacombe, Thomas R. Sibley, Melissa Truong, Caitlin R. Wolf, Karen Cowgill, Stephanie Schrag, Jeff Duchin, Michael Boeckh, Janet A. Englund, Michael Famulare, Barry R. Lutz, Mark J. Rieder, Lea M. Starita, Matthew Thompson, Helen Y. Chu, Trevor Bedford, Jay Shendure |
| EPI_ISL_596734 | PathWest Laboratory Medicine WA | PathWest Laboratory Medicine WA Microbial Surveillance Unit | PathWest Laboratory Medicine WA Microbial Surveillance Unit |
| EPI_ISL_596890, EPI_ISL_596892, EPI_ISL_596893, EPI_ISL_596895, EPI_ISL_596897, EPI_ISL_596898, EPI_ISL_596899, EPI_ISL_596900, EPI_ISL_596901, EPI_ISL_596902, EPI_ISL_596904, EPI_ISL_596909, EPI_ISL_596910, EPI_ISL_596911, EPI_ISL_596912, EPI_ISL_596913, EPI_ISL_596914, EPI_ISL_596915, EPI_ISL_596916, EPI_ISL_596917, EPI_ISL_596918, EPI_ISL_596919, EPI_ISL_596920, EPI_ISL_596921, EPI_ISL_596922, EPI_ISL_596923, EPI_ISL_596924, EPI_ISL_596925, EPI_ISL_596926, EPI_ISL_596927 | | | |
| see above | National Virus Reference Laboratory | National Virus Reference Laboratory | Michael Carr, Gabriel Gonzalez, Jonathan Dean, Daniel Hare, Cillian F De Gascun |
| EPI_ISL_601478, EPI_ISL_601490, EPI_ISL_601491, EPI_ISL_601507, EPI_ISL_601513, EPI_ISL_601520, EPI_ISL_601529, EPI_ISL_601536, EPI_ISL_601539, EPI_ISL_601546, EPI_ISL_601564, EPI_ISL_601597, EPI_ISL_601599, EPI_ISL_601612, EPI_ISL_601630, EPI_ISL_601635, EPI_ISL_601637, EPI_ISL_601640, EPI_ISL_601645, EPI_ISL_601647, EPI_ISL_601649 | | | |
| see above | Lighthouse Lab in Milton Keynes | Wellcome Sanger Institute for the COVID-19 Genomics UK (COG-UK) consortium | The Lighthouse Lab in Milton Keynes and Alex Alderton, Roberto Amato, Sonia Goncalves, Ewan Harrison, David K. Jackson, Ian Johnston, Dominic Kwiatkowski, Cordelia Langford, John Sillitoe on behalf of the Wellcome Sanger Institute COVID-19 Surveillance Team (http://www.sanger.ac.uk/covid-team) |
| EPI_ISL_601667 | Lighthouse Lab in Milton Keynes | Wellcome Sanger Institute for the COVID-19 Genomics UK (COG-UK) Consortium | The Lighthouse Lab in Milton Keynes and Alex Alderton, Roberto Amato, Sonia Goncalves, Ewan Harrison, David K. Jackson, Ian Johnston, Dominic Kwiatkowski, Cordelia Langford, John Sillitoe on behalf of the Wellcome Sanger Institute COVID-19 Surveillance Team |
| EPI_ISL_601669, EPI_ISL_601671, EPI_ISL_601686, EPI_ISL_601694, EPI_ISL_601706, EPI_ISL_601711, EPI_ISL_601737, EPI_ISL_601745, EPI_ISL_601746, EPI_ISL_601751, EPI_ISL_601752, EPI_ISL_601761, EPI_ISL_601774, EPI_ISL_601784 | | | |
| see above | Lighthouse Lab in Milton Keynes | Wellcome Sanger Institute for the COVID-19 Genomics UK (COG-UK) consortium | The Lighthouse Lab in Milton Keynes and Alex Alderton, Roberto Amato, Sonia Goncalves, Ewan Harrison, David K. Jackson, Ian Johnston, Dominic Kwiatkowski, Cordelia Langford, John Sillitoe on behalf of the Wellcome Sanger Institute COVID-19 Surveillance Team (http://www.sanger.ac.uk/covid-team) |
| EPI_ISL_601785 | Lighthouse Lab in Cambridge | Wellcome Sanger Institute for the COVID-19 Genomics UK (COG-UK) consortium | Rob Howes, The Lighthouse Lab in Cambridge and Alex Alderton, Roberto Amato, Sonia Goncalves, Ewan Harrison, David K. Jackson, Ian Johnston, Dominic Kwiatkowski, Cordelia Langford, John Sillitoe on behalf of the Wellcome Sanger Institute COVID-19 Surveillance Team (http://www.sanger.ac.uk/covid-team) |
| EPI_ISL_601787, EPI_ISL_601788, EPI_ISL_601789, EPI_ISL_601790, EPI_ISL_601791, EPI_ISL_601792, EPI_ISL_601793, EPI_ISL_601794, EPI_ISL_601795, EPI_ISL_601796 | Lighthouse Lab in Glasgow | Wellcome Sanger Institute for the COVID-19 Genomics UK (COG-UK) consortium | Harper VanSteenhouse, Yumi Kasai, David Gray, Carol Clugston, Anna Dominiczak and Alex Alderton, Roberto Amato, Sonia Goncalves, Ewan Harrison, David K. Jackson, Ian Johnston, Dominic Kwiatkowski, Cordelia Langford, John Sillitoe on behalf of the Wellcome Sanger Institute COVID-19 Surveillance Team (http://www.sanger.ac.uk/covid-team) |
| EPI_ISL_601797 | Lighthouse Lab in Glasgow | Wellcome Sanger Institute for the COVID-19 Genomics UK (COG-UK) Consortium | Harper VanSteenhouse, Yumi Kasai, David Gray, Carol Clugston, Anna Dominiczak and Alex Alderton, Roberto Amato, Sonia Goncalves, Ewan Harrison, David K. Jackson, Ian Johnston, Dominic Kwiatkowski, Cordelia Langford, John Sillitoe on behalf of the Wellcome Sanger Institute COVID-19 Surveillance Team |
| EPI_ISL_601798, EPI_ISL_601799, EPI_ISL_601800, EPI_ISL_601801, EPI_ISL_601802, EPI_ISL_601803, EPI_ISL_601804, EPI_ISL_601805, EPI_ISL_601806, EPI_ISL_601807, EPI_ISL_601808, EPI_ISL_601809, EPI_ISL_601810, EPI_ISL_601811, EPI_ISL_601812, EPI_ISL_601813, EPI_ISL_601814, EPI_ISL_601815, EPI_ISL_601816, EPI_ISL_601817, EPI_ISL_601818, EPI_ISL_601819, EPI_ISL_601820, EPI_ISL_601821, EPI_ISL_601822, EPI_ISL_601823, EPI_ISL_601824, EPI_ISL_601825, EPI_ISL_601826, EPI_ISL_601827, EPI_ISL_601828, EPI_ISL_601829, EPI_ISL_601830, EPI_ISL_601831, EPI_ISL_601832, EPI_ISL_601833 | | | |
| see above | Lighthouse Lab in Glasgow | Wellcome Sanger Institute | Harper VanSteenhouse, Yumi Kasai, David Gray, Carol Clugston, Anna Dominiczak and Alex Alderton, Roberto Amato, Sonia Goncalves, Ewan Harrison, David K. Jackson, Ian Johnston, Dominic Kwiatkowski, Cordelia Langford, John Sillitoe on |

| | | | |
|---|---|---|---|
| | | for the COVID-19 Genomics UK (COG-UK) consortium | behalf of the Wellcome Sanger Institute COVID-19 Surveillance Team (http://www.sanger.ac.uk/covid-team) |
| EPI_ISL_601834 | Lighthouse Lab in Glasgow | Wellcome Sanger Institute for the COVID-19 Genomics UK (COG-UK) Consortium | Harper VanSteenhouse, Yumi Kasai, David Gray, Carol Clugston, Anna Dominiczak and Alex Alderton, Roberto Amato, Sonia Goncalves, Ewan Harrison, David K. Jackson, Ian Johnston, Dominic Kwiatkowski, Cordelia Langford, John Sillitoe on behalf of the Wellcome Sanger Institute COVID-19 Surveillance Team |
| EPI_ISL_601835, EPI_ISL_601836, EPI_ISL_601837, EPI_ISL_601838, EPI_ISL_601839, EPI_ISL_601840, EPI_ISL_601841, EPI_ISL_601842, EPI_ISL_601843, EPI_ISL_601844, EPI_ISL_601845, EPI_ISL_601846, EPI_ISL_601847, EPI_ISL_601848, EPI_ISL_601849, EPI_ISL_601850, EPI_ISL_601851, EPI_ISL_601852, EPI_ISL_601853, EPI_ISL_601854, EPI_ISL_601855, EPI_ISL_601856, EPI_ISL_601857, EPI_ISL_601858, EPI_ISL_601859, EPI_ISL_601860, EPI_ISL_601861, EPI_ISL_601862, EPI_ISL_601863, EPI_ISL_601864, EPI_ISL_601865, EPI_ISL_601866, EPI_ISL_601867, EPI_ISL_601868, EPI_ISL_601869, EPI_ISL_601870, EPI_ISL_601871, EPI_ISL_601872, EPI_ISL_601873, EPI_ISL_601874, EPI_ISL_601875, EPI_ISL_601876, EPI_ISL_601877, EPI_ISL_601878, EPI_ISL_601879, EPI_ISL_601880, EPI_ISL_601881, EPI_ISL_601882, EPI_ISL_601883, EPI_ISL_601884, EPI_ISL_601885, EPI_ISL_601886, EPI_ISL_601887, EPI_ISL_601888, EPI_ISL_601889, EPI_ISL_601890, EPI_ISL_601891, EPI_ISL_601892, EPI_ISL_601893, EPI_ISL_601894, EPI_ISL_601895, EPI_ISL_601896, EPI_ISL_601897, EPI_ISL_601898, EPI_ISL_601899, EPI_ISL_601900, EPI_ISL_601901, EPI_ISL_601902, EPI_ISL_601903, EPI_ISL_601904, EPI_ISL_601905, EPI_ISL_601906, EPI_ISL_601907, EPI_ISL_601908, EPI_ISL_601909, EPI_ISL_601910, EPI_ISL_601911, EPI_ISL_601912, EPI_ISL_601913, EPI_ISL_601914, EPI_ISL_601915, EPI_ISL_601916, EPI_ISL_601917, EPI_ISL_601918, EPI_ISL_601919, EPI_ISL_601920, EPI_ISL_601921, EPI_ISL_601923, EPI_ISL_601924, EPI_ISL_601925, EPI_ISL_601926, EPI_ISL_601927, EPI_ISL_601928, EPI_ISL_601929, EPI_ISL_601930, EPI_ISL_601931, EPI_ISL_601932, EPI_ISL_601933, EPI_ISL_601934, EPI_ISL_601935, EPI_ISL_601936, EPI_ISL_601937, EPI_ISL_601938, EPI_ISL_601939, EPI_ISL_601940, EPI_ISL_601941, EPI_ISL_601942, EPI_ISL_601943, EPI_ISL_601944, EPI_ISL_601945, EPI_ISL_601946, EPI_ISL_601947, EPI_ISL_601948, EPI_ISL_601949 | | | |
| see above | Lighthouse Lab in Glasgow | Wellcome Sanger Institute for the COVID-19 Genomics UK (COG-UK) consortium | Harper VanSteenhouse, Yumi Kasai, David Gray, Carol Clugston, Anna Dominiczak and Alex Alderton, Roberto Amato, Sonia Goncalves, Ewan Harrison, David K. Jackson, Ian Johnston, Dominic Kwiatkowski, Cordelia Langford, John Sillitoe on behalf of the Wellcome Sanger Institute COVID-19 Surveillance Team (http://www.sanger.ac.uk/covid-team) |
| EPI_ISL_601950 | Lighthouse Lab in Glasgow | Wellcome Sanger Institute for the COVID-19 Genomics UK (COG-UK) Consortium | Harper VanSteenhouse, Yumi Kasai, David Gray, Carol Clugston, Anna Dominiczak and Alex Alderton, Roberto Amato, Sonia Goncalves, Ewan Harrison, David K. Jackson, Ian Johnston, Dominic Kwiatkowski, Cordelia Langford, John Sillitoe on behalf of the Wellcome Sanger Institute COVID-19 Surveillance Team |
| EPI_ISL_601951, EPI_ISL_601952, EPI_ISL_601953, EPI_ISL_601954, EPI_ISL_601955, EPI_ISL_601956, EPI_ISL_601957, EPI_ISL_601958, EPI_ISL_601959, EPI_ISL_601960, EPI_ISL_601961, EPI_ISL_601962, EPI_ISL_601963, EPI_ISL_601964, EPI_ISL_601965, EPI_ISL_601966, EPI_ISL_601967, EPI_ISL_601968, EPI_ISL_601969, EPI_ISL_601970, EPI_ISL_601971, EPI_ISL_601972, EPI_ISL_601973, EPI_ISL_601974, EPI_ISL_601975, EPI_ISL_601976, EPI_ISL_601977, EPI_ISL_601978, EPI_ISL_601979, EPI_ISL_601980, EPI_ISL_601981, EPI_ISL_601982, EPI_ISL_601983, EPI_ISL_601984, EPI_ISL_601985, EPI_ISL_601986, EPI_ISL_601987, EPI_ISL_601988, EPI_ISL_601989, EPI_ISL_601990, EPI_ISL_601991, EPI_ISL_601992, EPI_ISL_601993, EPI_ISL_601994, EPI_ISL_601995, EPI_ISL_601996, EPI_ISL_601997, EPI_ISL_601998, EPI_ISL_601999, EPI_ISL_602000, EPI_ISL_602001, EPI_ISL_602002, EPI_ISL_602003, EPI_ISL_602004, EPI_ISL_602005 | | | |
| see above | Lighthouse Lab in Glasgow | Wellcome Sanger Institute for the COVID-19 Genomics UK (COG-UK) consortium | Harper VanSteenhouse, Yumi Kasai, David Gray, Carol Clugston, Anna Dominiczak and Alex Alderton, Roberto Amato, Sonia Goncalves, Ewan Harrison, David K. Jackson, Ian Johnston, Dominic Kwiatkowski, Cordelia Langford, John Sillitoe on behalf of the Wellcome Sanger Institute COVID-19 Surveillance Team (http://www.sanger.ac.uk/covid-team) |
| EPI_ISL_602006 | Lighthouse Lab in Glasgow | Wellcome Sanger Institute for the COVID-19 Genomics UK (COG-UK) Consortium | Harper VanSteenhouse, Yumi Kasai, David Gray, Carol Clugston, Anna Dominiczak and Alex Alderton, Roberto Amato, Sonia Goncalves, Ewan Harrison, David K. Jackson, Ian Johnston, Dominic Kwiatkowski, Cordelia Langford, John Sillitoe on behalf of the Wellcome Sanger Institute COVID-19 Surveillance Team |
| EPI_ISL_602007, EPI_ISL_602008, EPI_ISL_602009, EPI_ISL_602010, EPI_ISL_602011, EPI_ISL_602012, EPI_ISL_602013, EPI_ISL_602014 | Lighthouse Lab in Glasgow | Wellcome Sanger Institute for the COVID-19 Genomics UK (COG-UK) consortium | Harper VanSteenhouse, Yumi Kasai, David Gray, Carol Clugston, Anna Dominiczak and Alex Alderton, Roberto Amato, Sonia Goncalves, Ewan Harrison, David K. Jackson, Ian Johnston, Dominic Kwiatkowski, Cordelia Langford, John Sillitoe on behalf of the Wellcome Sanger Institute COVID-19 Surveillance Team (http://www.sanger.ac.uk/covid-team) |
| EPI_ISL_602015 | Lighthouse Lab in Glasgow | Wellcome Sanger Institute for the COVID-19 Genomics UK (COG-UK) Consortium | Harper VanSteenhouse, Yumi Kasai, David Gray, Carol Clugston, Anna Dominiczak and Alex Alderton, Roberto Amato, Sonia Goncalves, Ewan Harrison, David K. Jackson, Ian Johnston, Dominic Kwiatkowski, Cordelia Langford, John Sillitoe on behalf of the Wellcome Sanger Institute COVID-19 Surveillance Team |
| EPI_ISL_602016, EPI_ISL_602017, EPI_ISL_602018, EPI_ISL_602019, EPI_ISL_602020, EPI_ISL_602021, EPI_ISL_602022, EPI_ISL_602023, EPI_ISL_602024, EPI_ISL_602025, EPI_ISL_602026, EPI_ISL_602030, EPI_ISL_602031, EPI_ISL_602032, EPI_ISL_602033, EPI_ISL_602034, EPI_ISL_602035, EPI_ISL_602036, EPI_ISL_602037, EPI_ISL_602038, EPI_ISL_602039, EPI_ISL_602040, EPI_ISL_602041, EPI_ISL_602042 | | | |
| see above | Lighthouse Lab in Glasgow | Wellcome Sanger Institute for the COVID-19 Genomics UK (COG-UK) consortium | Harper VanSteenhouse, Yumi Kasai, David Gray, Carol Clugston, Anna Dominiczak and Alex Alderton, Roberto Amato, Sonia Goncalves, Ewan Harrison, David K. Jackson, Ian Johnston, Dominic Kwiatkowski, Cordelia Langford, John Sillitoe on behalf of the Wellcome Sanger Institute COVID-19 Surveillance Team (http://www.sanger.ac.uk/covid-team) |
| EPI_ISL_602336, EPI_ISL_602337, EPI_ISL_602338, EPI_ISL_602339, EPI_ISL_602340, EPI_ISL_602342, EPI_ISL_602344, EPI_ISL_602345, EPI_ISL_602348, EPI_ISL_602349, EPI_ISL_602350, EPI_ISL_602351, EPI_ISL_602352, EPI_ISL_602353, EPI_ISL_602360, EPI_ISL_602361, EPI_ISL_602362, EPI_ISL_602363, EPI_ISL_602364, EPI_ISL_602365, EPI_ISL_602366, EPI_ISL_602367, EPI_ISL_602368, EPI_ISL_602369, EPI_ISL_602370, EPI_ISL_602371, EPI_ISL_602372, EPI_ISL_602374, EPI_ISL_602375, EPI_ISL_602376, EPI_ISL_602404 | | | |
| see above | HELIX LLC | WHO National Influenza Centre Russian Federation | Andrey Komissarov, Artem Fadeev, Kseniya Komissarova, Anna Ivanova, Dmitry Bazhenov, Daria Danilenko |
| EPI_ISL_602413, EPI_ISL_602414, EPI_ISL_602415, EPI_ISL_602416 | WHO National Influenza Centre Russian Federation | WHO National Influenza Centre Russian Federation | Andrey Komissarov, Artem Fadeev, Kseniya Komissarova, Anna Ivanova, Dmitry Bazhenov, Daria Danilenko |
| EPI_ISL_602417, EPI_ISL_602418, EPI_ISL_602424, EPI_ISL_602427, EPI_ISL_602428, EPI_ISL_602429, EPI_ISL_602442, EPI_ISL_602443, EPI_ISL_602453, EPI_ISL_602457, EPI_ISL_602460 | | | |
| see above | HELIX LLC | WHO National Influenza Centre Russian Federation | Andrey Komissarov, Artem Fadeev, Kseniya Komissarova, Anna Ivanova, Dmitry Bazhenov, Daria Danilenko |
| EPI_ISL_602761, EPI_ISL_602762, EPI_ISL_602763, EPI_ISL_602764, EPI_ISL_602765, EPI_ISL_602767, EPI_ISL_602769, EPI_ISL_602770, EPI_ISL_602771, EPI_ISL_602772, EPI_ISL_602773, EPI_ISL_602774, EPI_ISL_602775, EPI_ISL_602776, EPI_ISL_602777, EPI_ISL_602778, EPI_ISL_602782, EPI_ISL_602783, EPI_ISL_602784, EPI_ISL_602785, EPI_ISL_602786, EPI_ISL_602787, EPI_ISL_602788, EPI_ISL_602789, EPI_ISL_602790, EPI_ISL_602791, EPI_ISL_602792, EPI_ISL_602793, EPI_ISL_602794, EPI_ISL_602795, EPI_ISL_602796, EPI_ISL_602797, EPI_ISL_602798, EPI_ISL_602799, EPI_ISL_602800, EPI_ISL_602801, EPI_ISL_602802, EPI_ISL_602803, EPI_ISL_602804, EPI_ISL_602805, EPI_ISL_602806, EPI_ISL_602807, EPI_ISL_602808, EPI_ISL_602809, EPI_ISL_602810, EPI_ISL_602811, EPI_ISL_602812, EPI_ISL_602813, EPI_ISL_602814, EPI_ISL_602815, EPI_ISL_602816, EPI_ISL_602817, EPI_ISL_602818, EPI_ISL_602819, EPI_ISL_602820, EPI_ISL_602821, EPI_ISL_602822, EPI_ISL_602823, EPI_ISL_602824, EPI_ISL_602825, EPI_ISL_602826, EPI_ISL_602827, EPI_ISL_602828, EPI_ISL_602829 | | | |
| see above | NHLS-IALCH | KRISP, KZN Research Innovation and Sequencing Platform | Giandhari J, Pillay S, Lessells R, Mdlalose K, York D, Khan S, Tegally H, Wilkinson E, de Oliveira T |
| EPI_ISL_602958, EPI_ISL_602959, EPI_ISL_602960, EPI_ISL_602961, EPI_ISL_602966, EPI_ISL_602967, EPI_ISL_602968, EPI_ISL_602969, EPI_ISL_602970, EPI_ISL_602971 | Minnesota Department of Health, Public Health Laboratory | Minnesota Department of Health, Public Health Laboratory | Matt Plumb, Jacob Garfin, Alexandra Lorentz, and Xiong Wang |
| EPI_ISL_603046 | MDU-PHL, The Peter Doherty Institute for Infection and Immunity | MDU-PHL, The Peter Doherty Institute for Infection and Immunity | Seemann,T., Caly,L., Sait,M., Schultz,M.B., Druce,J., Sherry,N. |
| EPI_ISL_605408, EPI_ISL_605409, EPI_ISL_605410, EPI_ISL_605411, EPI_ISL_605412, EPI_ISL_605413, EPI_ISL_605414, EPI_ISL_605415, EPI_ISL_605416, EPI_ISL_605417, EPI_ISL_605418, EPI_ISL_605419, EPI_ISL_605420, EPI_ISL_605421, EPI_ISL_605422, EPI_ISL_605423, EPI_ISL_605424, EPI_ISL_605425, EPI_ISL_605426, EPI_ISL_605427, EPI_ISL_605428, EPI_ISL_605429, EPI_ISL_605430, EPI_ISL_605431, EPI_ISL_605432, EPI_ISL_605433, EPI_ISL_605434, EPI_ISL_605435, EPI_ISL_605436, EPI_ISL_605437, EPI_ISL_605438, EPI_ISL_605439, EPI_ISL_605440, EPI_ISL_605441, EPI_ISL_605442, EPI_ISL_605443, | | | |

EPI_ISL_605444, EPI_ISL_605445, EPI_ISL_605446, EPI_ISL_605447, EPI_ISL_605448, EPI_ISL_605449, EPI_ISL_605450, EPI_ISL_605451, EPI_ISL_605452, EPI_ISL_605453, EPI_ISL_605454, EPI_ISL_605455, EPI_ISL_605456, EPI_ISL_605457, EPI_ISL_605458, EPI_ISL_605459, EPI_ISL_605460, EPI_ISL_605461, EPI_ISL_605462, EPI_ISL_605485, EPI_ISL_605486, EPI_ISL_605487, EPI_ISL_605488, EPI_ISL_605489, EPI_ISL_605490, EPI_ISL_605511, EPI_ISL_605512, EPI_ISL_605513, EPI_ISL_605514, EPI_ISL_605515, EPI_ISL_605516, EPI_ISL_605517, EPI_ISL_605518, EPI_ISL_605519, EPI_ISL_605524, EPI_ISL_605525, EPI_ISL_605526, EPI_ISL_605527, EPI_ISL_605529, EPI_ISL_605530, EPI_ISL_605531, EPI_ISL_605532, EPI_ISL_605533, EPI_ISL_605535, EPI_ISL_605536, EPI_ISL_605537, EPI_ISL_605538, EPI_ISL_605539, EPI_ISL_605540, EPI_ISL_605541, EPI_ISL_605542, EPI_ISL_605543, EPI_ISL_605544, EPI_ISL_605545, EPI_ISL_605546, EPI_ISL_605547, EPI_ISL_605548, EPI_ISL_605549, EPI_ISL_605550, EPI_ISL_605551, EPI_ISL_605552, EPI_ISL_605553, EPI_ISL_605554, EPI_ISL_605555, EPI_ISL_605556, EPI_ISL_605557, EPI_ISL_605558, EPI_ISL_605559, EPI_ISL_605560, EPI_ISL_605561, EPI_ISL_605562, EPI_ISL_605563, EPI_ISL_605564, EPI_ISL_605566, EPI_ISL_605568, EPI_ISL_605569, EPI_ISL_605571, EPI_ISL_605572, EPI_ISL_605575, EPI_ISL_605576, EPI_ISL_605577, EPI_ISL_605579, EPI_ISL_605580, EPI_ISL_605581, EPI_ISL_605582, EPI_ISL_605584, EPI_ISL_605586, EPI_ISL_605588, EPI_ISL_605589, EPI_ISL_605590, EPI_ISL_605591, EPI_ISL_605592, EPI_ISL_605593, EPI_ISL_605595, EPI_ISL_605597, EPI_ISL_605598, EPI_ISL_605599, EPI_ISL_605600, EPI_ISL_605601, EPI_ISL_605602, EPI_ISL_605606, EPI_ISL_605607, EPI_ISL_605608, EPI_ISL_605611, EPI_ISL_605612, EPI_ISL_605613, EPI_ISL_605617, EPI_ISL_605618, EPI_ISL_605619, EPI_ISL_605620, EPI_ISL_605621, EPI_ISL_605622, EPI_ISL_605623, EPI_ISL_605624, EPI_ISL_605625, EPI_ISL_605626, EPI_ISL_605627, EPI_ISL_605628, EPI_ISL_605629, EPI_ISL_605630, EPI_ISL_605631, EPI_ISL_605632, EPI_ISL_605633, EPI_ISL_605635, EPI_ISL_605636, EPI_ISL_605637, EPI_ISL_605638, EPI_ISL_605639, EPI_ISL_605640, EPI_ISL_605641, EPI_ISL_605642, EPI_ISL_605643, EPI_ISL_605644, EPI_ISL_605645, EPI_ISL_605646, EPI_ISL_605647, EPI_ISL_605648, EPI_ISL_605649, EPI_ISL_605650, EPI_ISL_605651, EPI_ISL_605652, EPI_ISL_605653, EPI_ISL_605654, EPI_ISL_605655, EPI_ISL_605656, EPI_ISL_605657, EPI_ISL_605658, EPI_ISL_605659, EPI_ISL_605660, EPI_ISL_605661, EPI_ISL_605662, EPI_ISL_605663, EPI_ISL_605664, EPI_ISL_605665, EPI_ISL_605666, EPI_ISL_605667, EPI_ISL_605668, EPI_ISL_605669, EPI_ISL_605670, EPI_ISL_605672, EPI_ISL_605673, EPI_ISL_605674, EPI_ISL_605675, EPI_ISL_605678, EPI_ISL_605679, EPI_ISL_605680, EPI_ISL_605681, EPI_ISL_605683, EPI_ISL_605692, EPI_ISL_605698, EPI_ISL_605700, EPI_ISL_605702, EPI_ISL_605703, EPI_ISL_605704, EPI_ISL_605707, EPI_ISL_605708, EPI_ISL_605709, EPI_ISL_605710, EPI_ISL_605711, EPI_ISL_605712, EPI_ISL_605713, EPI_ISL_605715, EPI_ISL_605716, EPI_ISL_605717, EPI_ISL_605719, EPI_ISL_605720, EPI_ISL_605722, EPI_ISL_605723, EPI_ISL_605724, EPI_ISL_605725, EPI_ISL_605726, EPI_ISL_605727, EPI_ISL_605728, EPI_ISL_605729, EPI_ISL_605730, EPI_ISL_605731, EPI_ISL_605732, EPI_ISL_605733, EPI_ISL_605734, EPI_ISL_605735, EPI_ISL_605736, EPI_ISL_605743, EPI_ISL_605744, EPI_ISL_605745, EPI_ISL_605746, EPI_ISL_605747, EPI_ISL_605748, EPI_ISL_605749, EPI_ISL_605750, EPI_ISL_605751, EPI_ISL_605752, EPI_ISL_605753, EPI_ISL_605754, EPI_ISL_605755, EPI_ISL_605756, EPI_ISL_605757, EPI_ISL_605764, EPI_ISL_605769, EPI_ISL_605771, EPI_ISL_605772, EPI_ISL_605773, EPI_ISL_605774, EPI_ISL_605775, EPI_ISL_605776, EPI_ISL_605777, EPI_ISL_605778, EPI_ISL_605779

| | | | |
|---|---|---|---|
| see above | University of Wisconsin-Madison AIDS Vaccine Research Laboratories | University of Wisconsin-Madison AIDS Vaccine Research Laboratories | Gage Moreno, Katarina Braun, et al. AIDS Vaccine Research Laboratories |
| EPI_ISL_609803, EPI_ISL_609804, EPI_ISL_609808, EPI_ISL_609810, EPI_ISL_609812, EPI_ISL_609814, EPI_ISL_609815, EPI_ISL_609819, EPI_ISL_609823 | Unity Health Toronto | Ontario Institute for Cancer Research | Ramzi Fattouh, Larissa M. Matukas, Yan Chen,Mark Downing, Trina Otterman, Karel Boissinot, Wai Sum Siu, Zhi Cui, Le Luu, Samira Mubareka, TIBDN, Ilinca Lungu, Bernard Lam, Jeremy Johns, Paul Krzyzanowski, Richard de Borja, Felicia Vincelli, Philip Zuzarte, Jared T. Simpson |
| EPI_ISL_610218, EPI_ISL_610219 | Department of Health Technology and Informatics, The Hong Kong Polytechnic University | Department of Health Technology and Informatics, The Hong Kong Polytechnic University | Siu,G.K.-H., Lee,L.-K., Leung,K.S.-S., Leung,J.S.-L., Ng,T.T.-L., Chan,C.T.-M., Tam,K.K.-G., Lao,H.-Y., Wu,A.K.-L., Yau,M.C.-Y., Lai,Y.W.-M., Fung,K.S.-C., Chau,S.K.-Y., Wong,B.K.-C., To,W.-K., Luk,K., Ho,A.Y.-M., Que,T.-L., Yip,K.-T., Yam,W.C., Shum,D.H.-K., Yip,S.P. |

EPI_ISL_611539, EPI_ISL_611540, EPI_ISL_611551, EPI_ISL_611552, EPI_ISL_611607, EPI_ISL_611615, EPI_ISL_611616, EPI_ISL_611617, EPI_ISL_611618, EPI_ISL_611619, EPI_ISL_611620, EPI_ISL_611641, EPI_ISL_611701, EPI_ISL_611702, EPI_ISL_611703

| | | | |
|---|---|---|---|
| see above | University of Exeter | COVID-19 Genomics UK (COG-UK) Consortium | Ben Temperton,Aaron Jeffries,Michelle Michelsen,Joanna Warwick-Dugdale,Audrey Farbos,Robyn Manley,Stephen Michell,Jane Masoli |
| EPI_ISL_611766 | Northumbria University / South Tees Hospitals NHS Foundation Trust / North Cumbria Integrated Care NHS Foundation Trust / North Tees and Hartlepool NHS Foundation Trust / Newcastle Hospitals NHS Foundation Trust | COVID-19 Genomics UK (COG-UK) Consortium | Darren L Smith,Andrew Nelson,Matthew Bashton,Greg R Young,Joshua Loh,John Allan,Mohammad A Tariq,Giles S Holt,Gary Black,Wen C Yew,Lynn Dover,Paul Baker,Steve Liggett,Sarah Essex,Jane Greenaway,Debra Padgett,Clive Graham,Garren Scott,Edward Barton,Emma Swindells,Brendan Payne,Jennifer Collins,Yusri Taha,Gary Eltringham |
| EPI_ISL_612056, EPI_ISL_612057, EPI_ISL_612095, EPI_ISL_612096, EPI_ISL_612395, EPI_ISL_612396, EPI_ISL_612397, EPI_ISL_612398, EPI_ISL_612399 | University of Exeter | COVID-19 Genomics UK (COG-UK) Consortium | Ben Temperton,Aaron Jeffries,Michelle Michelsen,Joanna Warwick-Dugdale,Audrey Farbos,Robyn Manley,Stephen Michell,Jane Masoli |
| EPI_ISL_612548 | Regional Virus Laboratory, Belfast Health and Social Care Trust | COVID-19 Genomics UK (COG-UK) Consortium | Conall McCaughey, James McKenna, Tanya Curran, Susan Feeney, Alison Watt, Ciara Cox, Mairead Connor, Zoltan Molnar, David Simpson, Derek Fairley |
| EPI_ISL_613477 | Public Health Laboratory - Infectious Disease Lab, Minnesota Department of Health Infectious Disease Laboratory Submission Group | Minnesota Department of Health, Public Health Laboratory | Plumb,M., Garfin,J., Lorentz,A., Wang,X. |

EPI_ISL_614048, EPI_ISL_614055, EPI_ISL_614056, EPI_ISL_614057, EPI_ISL_614058, EPI_ISL_614059, EPI_ISL_614060, EPI_ISL_614061, EPI_ISL_614062, EPI_ISL_614128, EPI_ISL_614130, EPI_ISL_614131, EPI_ISL_614132, EPI_ISL_614133, EPI_ISL_614134

| | | | |
|---|---|---|---|
| see above | Virginia DCLS | Virginia DCLS | Virginia DCLS |
| EPI_ISL_614301 | Faroese National Reference Laboratory for Fish and Animal Diseases | Faroese National Reference Laboratory for Fish and Animal Diseases | Maria Marjunardóttir Dahl, Petra Elisabeth Petersen, Debes Hammershaimb Christiansen |
| EPI_ISL_615105, EPI_ISL_615106, EPI_ISL_615107 | Halmstad klinisk mikrobiologi | The Public Health Agency of Sweden | Anna-Malin Linde, Maria Lind Karlberg, Mattias Haukland, Reza Advani, Olov Svartstrom, Oskar Karlsson Lindsjo, Sandra Broddesson, Petra Edquist, Mia Brytting, Anna Risberg, Karin Tegmark-Wisell |

EPI_ISL_615166, EPI_ISL_615167, EPI_ISL_615168, EPI_ISL_615169, EPI_ISL_615170, EPI_ISL_615171, EPI_ISL_615172, EPI_ISL_615173, EPI_ISL_615174, EPI_ISL_615175, EPI_ISL_615176, EPI_ISL_615177, EPI_ISL_615179, EPI_ISL_615180, EPI_ISL_615182, EPI_ISL_615183, EPI_ISL_615184, EPI_ISL_615185, EPI_ISL_615186, EPI_ISL_615194, EPI_ISL_615195, EPI_ISL_615196, EPI_ISL_615197, EPI_ISL_615198, EPI_ISL_615199, EPI_ISL_615200, EPI_ISL_615201, EPI_ISL_615202, EPI_ISL_615203, EPI_ISL_615204, EPI_ISL_615205, EPI_ISL_615206, EPI_ISL_615207, EPI_ISL_615208, EPI_ISL_615209, EPI_ISL_615210, EPI_ISL_615211, EPI_ISL_615212, EPI_ISL_615214, EPI_ISL_615215, EPI_ISL_615216, EPI_ISL_615217, EPI_ISL_615218, EPI_ISL_615219, EPI_ISL_615220, EPI_ISL_615221, EPI_ISL_615222, EPI_ISL_615223, EPI_ISL_615224, EPI_ISL_615225, EPI_ISL_615226, EPI_ISL_615227, EPI_ISL_615228, EPI_ISL_615229, EPI_ISL_615230, EPI_ISL_615231, EPI_ISL_615232, EPI_ISL_615233, EPI_ISL_615234, EPI_ISL_615235, EPI_ISL_615236, EPI_ISL_615237, EPI_ISL_615238, EPI_ISL_615239, EPI_ISL_615240, EPI_ISL_615241, EPI_ISL_615242, EPI_ISL_615243, EPI_ISL_615244, EPI_ISL_615245, EPI_ISL_615246, EPI_ISL_615247, EPI_ISL_615248, EPI_ISL_615249, EPI_ISL_615250, EPI_ISL_615251, EPI_ISL_615252, EPI_ISL_615253, EPI_ISL_615254, EPI_ISL_615255, EPI_ISL_615256, EPI_ISL_615257, EPI_ISL_615258, EPI_ISL_615259, EPI_ISL_615260, EPI_ISL_615261, EPI_ISL_615262, EPI_ISL_615263, EPI_ISL_615264, EPI_ISL_615265, EPI_ISL_615266, EPI_ISL_615267, EPI_ISL_615268, EPI_ISL_615269, EPI_ISL_615270, EPI_ISL_615271, EPI_ISL_615272, EPI_ISL_615273, EPI_ISL_615274, EPI_ISL_615275, EPI_ISL_615276, EPI_ISL_615277, EPI_ISL_615278, EPI_ISL_615279, EPI_ISL_615280, EPI_ISL_615281, EPI_ISL_615282, EPI_ISL_615283, EPI_ISL_615284, EPI_ISL_615285, EPI_ISL_615286, EPI_ISL_615287, EPI_ISL_615288, EPI_ISL_615289, EPI_ISL_615290, EPI_ISL_615291, EPI_ISL_615292, EPI_ISL_615293, EPI_ISL_615294, EPI_ISL_615295, EPI_ISL_615296, EPI_ISL_615297, EPI_ISL_615298, EPI_ISL_615299, EPI_ISL_615300, EPI_ISL_615302, EPI_ISL_615303, EPI_ISL_615304, EPI_ISL_615305, EPI_ISL_615306, EPI_ISL_615307, EPI_ISL_615308, EPI_ISL_615309, EPI_ISL_615310, EPI_ISL_615311, EPI_ISL_615312, EPI_ISL_615313, EPI_ISL_615314, EPI_ISL_615315, EPI_ISL_615316, EPI_ISL_615317, EPI_ISL_615318, EPI_ISL_615319, EPI_ISL_615320, EPI_ISL_615321, EPI_ISL_615328, EPI_ISL_615329, EPI_ISL_615330, EPI_ISL_615331, EPI_ISL_615333, EPI_ISL_615334, EPI_ISL_615335, EPI_ISL_615336, EPI_ISL_615337, EPI_ISL_615338, EPI_ISL_615339, EPI_ISL_615340, EPI_ISL_615341, EPI_ISL_615342, EPI_ISL_615343, EPI_ISL_615344, EPI_ISL_615345, EPI_ISL_615346, EPI_ISL_615347, EPI_ISL_615348, EPI_ISL_615349, EPI_ISL_615350, EPI_ISL_615351, EPI_ISL_615352, EPI_ISL_615353, EPI_ISL_615354, EPI_ISL_615355, EPI_ISL_615356, EPI_ISL_615357, EPI_ISL_615358, EPI_ISL_615359, EPI_ISL_615360, EPI_ISL_615361, EPI_ISL_615362, EPI_ISL_615363, EPI_ISL_615364, EPI_ISL_615365, EPI_ISL_615366, EPI_ISL_615367, EPI_ISL_615368, EPI_ISL_615369, EPI_ISL_615370, EPI_ISL_615371, EPI_ISL_615372, EPI_ISL_615373, EPI_ISL_615374, EPI_ISL_615375, EPI_ISL_615376, EPI_ISL_615377, EPI_ISL_615378, EPI_ISL_615379, EPI_ISL_615380, EPI_ISL_615381, EPI_ISL_615382, EPI_ISL_615383, EPI_ISL_615384, EPI_ISL_615385, EPI_ISL_615386, EPI_ISL_615387, EPI_ISL_615388, EPI_ISL_615389, EPI_ISL_615390, EPI_ISL_615391, EPI_ISL_615392, EPI_ISL_615393, EPI_ISL_615394, EPI_ISL_615395, EPI_ISL_615396, EPI_ISL_615600, EPI_ISL_615601, EPI_ISL_615602,

| | | | |
|---|---|---|---|
| EPI_ISL_615603, EPI_ISL_615604, EPI_ISL_615605, EPI_ISL_615606, EPI_ISL_615608, EPI_ISL_615626, EPI_ISL_615628, EPI_ISL_615629, EPI_ISL_615630, EPI_ISL_615631, EPI_ISL_615632, EPI_ISL_615635, EPI_ISL_615652, EPI_ISL_615653, EPI_ISL_615654, EPI_ISL_615655, EPI_ISL_615656, EPI_ISL_615657, EPI_ISL_615658, EPI_ISL_615659, EPI_ISL_615660, EPI_ISL_615661, EPI_ISL_615662, EPI_ISL_615663, EPI_ISL_615664, EPI_ISL_615665, EPI_ISL_615666, EPI_ISL_615667, EPI_ISL_615668, EPI_ISL_615669, EPI_ISL_615674, EPI_ISL_615684, EPI_ISL_615685, EPI_ISL_615686, EPI_ISL_615753, EPI_ISL_615754, EPI_ISL_615755, EPI_ISL_615756, EPI_ISL_615757, EPI_ISL_615758, EPI_ISL_615759, EPI_ISL_615760, EPI_ISL_615761, EPI_ISL_615762, EPI_ISL_615763, EPI_ISL_615764, EPI_ISL_615765, EPI_ISL_615766, EPI_ISL_615767, EPI_ISL_615768, EPI_ISL_615769, EPI_ISL_615770, EPI_ISL_615771, EPI_ISL_615772, EPI_ISL_615773, EPI_ISL_615774, EPI_ISL_615775, EPI_ISL_615776, EPI_ISL_615777, EPI_ISL_615778, EPI_ISL_615779, EPI_ISL_615780, EPI_ISL_615781, EPI_ISL_615782, EPI_ISL_615783, EPI_ISL_615784, EPI_ISL_615785, EPI_ISL_615786, EPI_ISL_615787, EPI_ISL_615788, EPI_ISL_615789, EPI_ISL_615790, EPI_ISL_615791, EPI_ISL_615792, EPI_ISL_615793, EPI_ISL_615794, EPI_ISL_615795, EPI_ISL_615796, EPI_ISL_615797, EPI_ISL_615798, EPI_ISL_615799, EPI_ISL_615800, EPI_ISL_615801, EPI_ISL_615802, EPI_ISL_615803, EPI_ISL_615804, EPI_ISL_615805, EPI_ISL_615806, EPI_ISL_615807, EPI_ISL_615808, EPI_ISL_615809, EPI_ISL_615810, EPI_ISL_615811, EPI_ISL_615812, EPI_ISL_615813, EPI_ISL_615814, EPI_ISL_615815, EPI_ISL_615816, EPI_ISL_615817, EPI_ISL_615818, EPI_ISL_615819, EPI_ISL_615820, EPI_ISL_615821, EPI_ISL_615822, EPI_ISL_615823, EPI_ISL_615824, EPI_ISL_615825, EPI_ISL_615826, EPI_ISL_615827, EPI_ISL_615828, EPI_ISL_615829, EPI_ISL_615830, EPI_ISL_615831, EPI_ISL_615832, EPI_ISL_615833, EPI_ISL_615834, EPI_ISL_615835, EPI_ISL_615836, EPI_ISL_615837, EPI_ISL_615838, EPI_ISL_615839, EPI_ISL_615840, EPI_ISL_615841, EPI_ISL_615842, EPI_ISL_615843, EPI_ISL_615844, EPI_ISL_615845, EPI_ISL_615846, EPI_ISL_615847, EPI_ISL_615848, EPI_ISL_615849, EPI_ISL_615850, EPI_ISL_615851, EPI_ISL_615852, EPI_ISL_615853, EPI_ISL_615854, EPI_ISL_615855, EPI_ISL_615856, EPI_ISL_615857, EPI_ISL_615858, EPI_ISL_615859, EPI_ISL_615860, EPI_ISL_615861, EPI_ISL_615862, EPI_ISL_615863, EPI_ISL_615864, EPI_ISL_615865, EPI_ISL_615866, EPI_ISL_615867, EPI_ISL_615868, EPI_ISL_615869, EPI_ISL_615870, EPI_ISL_615871, EPI_ISL_615872, EPI_ISL_615873, EPI_ISL_615874, EPI_ISL_615875, EPI_ISL_615876, EPI_ISL_615877, EPI_ISL_615878, EPI_ISL_615879, EPI_ISL_615880, EPI_ISL_615881, EPI_ISL_615882, EPI_ISL_615883, EPI_ISL_615884, EPI_ISL_615885, EPI_ISL_615886, EPI_ISL_615887, EPI_ISL_615888, EPI_ISL_615889, EPI_ISL_615890, EPI_ISL_615891, EPI_ISL_615892, EPI_ISL_615893, EPI_ISL_615894, EPI_ISL_615895, EPI_ISL_615896, EPI_ISL_615897, EPI_ISL_615898, EPI_ISL_615899, EPI_ISL_615900, EPI_ISL_615901, EPI_ISL_615902, EPI_ISL_615903, EPI_ISL_615904, EPI_ISL_615905, EPI_ISL_620853, EPI_ISL_621177, EPI_ISL_621178, EPI_ISL_621179, EPI_ISL_621180, EPI_ISL_621181, EPI_ISL_621182, EPI_ISL_621184, EPI_ISL_621185, EPI_ISL_621536, EPI_ISL_621537, EPI_ISL_621541, EPI_ISL_621548, EPI_ISL_621568, EPI_ISL_621569, EPI_ISL_621570, EPI_ISL_621736, EPI_ISL_621737, EPI_ISL_621738, EPI_ISL_621745, EPI_ISL_621752, EPI_ISL_621753, EPI_ISL_621754, EPI_ISL_621755, EPI_ISL_621756, EPI_ISL_621757, EPI_ISL_621758, EPI_ISL_621761, EPI_ISL_621762, EPI_ISL_621763, EPI_ISL_621764, EPI_ISL_621765, EPI_ISL_621766, EPI_ISL_621767, EPI_ISL_621768, EPI_ISL_621769, EPI_ISL_621770, EPI_ISL_621771, EPI_ISL_621772, EPI_ISL_621773, EPI_ISL_621774, EPI_ISL_621775, EPI_ISL_621776, EPI_ISL_621777, EPI_ISL_621779, EPI_ISL_621780, EPI_ISL_621781, EPI_ISL_621782, EPI_ISL_621783, EPI_ISL_621784, EPI_ISL_621785, EPI_ISL_621786, EPI_ISL_621787, EPI_ISL_621788, EPI_ISL_621789, EPI_ISL_621790, EPI_ISL_621791, EPI_ISL_621792, EPI_ISL_621793, EPI_ISL_621794, EPI_ISL_621795, EPI_ISL_621798, EPI_ISL_621803, EPI_ISL_621814, EPI_ISL_621818, EPI_ISL_621819, EPI_ISL_621827, EPI_ISL_621833, EPI_ISL_621835, EPI_ISL_621836, EPI_ISL_621837, EPI_ISL_621838, EPI_ISL_621839, EPI_ISL_621846, EPI_ISL_621847, EPI_ISL_621848, EPI_ISL_621849, EPI_ISL_621850, EPI_ISL_621851, EPI_ISL_621866, EPI_ISL_621870, EPI_ISL_621871, EPI_ISL_621872, EPI_ISL_621873, EPI_ISL_621874, EPI_ISL_621875, EPI_ISL_621876, EPI_ISL_621877, EPI_ISL_621878, EPI_ISL_621879, EPI_ISL_621880, EPI_ISL_621881, EPI_ISL_621882, EPI_ISL_621883, EPI_ISL_621884, EPI_ISL_621886, EPI_ISL_621887, EPI_ISL_621889, EPI_ISL_621890, EPI_ISL_621891, EPI_ISL_621893, EPI_ISL_621894, EPI_ISL_621895, EPI_ISL_621896, EPI_ISL_621897, EPI_ISL_621898, EPI_ISL_621899, EPI_ISL_621900, EPI_ISL_621901, EPI_ISL_621902, EPI_ISL_621903, EPI_ISL_621904, EPI_ISL_621905, EPI_ISL_621906, EPI_ISL_621907, EPI_ISL_621908, EPI_ISL_621909, EPI_ISL_621910, EPI_ISL_621911, EPI_ISL_621912, EPI_ISL_621913, EPI_ISL_621914, EPI_ISL_621915, EPI_ISL_621917, EPI_ISL_621918, EPI_ISL_621919, EPI_ISL_621920, EPI_ISL_621921, EPI_ISL_621927, EPI_ISL_621930, EPI_ISL_621948, EPI_ISL_621986, EPI_ISL_621987, EPI_ISL_621988, EPI_ISL_621990, EPI_ISL_621991, EPI_ISL_621992, EPI_ISL_621993, EPI_ISL_621994, EPI_ISL_621995, EPI_ISL_621997, EPI_ISL_621998, EPI_ISL_621999, EPI_ISL_622000, EPI_ISL_622001, EPI_ISL_622002, EPI_ISL_622003, EPI_ISL_622004, EPI_ISL_622005, EPI_ISL_622006, EPI_ISL_622007, EPI_ISL_622008, EPI_ISL_622009, EPI_ISL_622010, EPI_ISL_622011, EPI_ISL_622012, EPI_ISL_622013, EPI_ISL_622014, EPI_ISL_622015, EPI_ISL_622016, EPI_ISL_622017, EPI_ISL_622018, EPI_ISL_622019, EPI_ISL_622020, EPI_ISL_622021, EPI_ISL_622022, EPI_ISL_622023, EPI_ISL_622024, EPI_ISL_622025, EPI_ISL_622026, EPI_ISL_622027, EPI_ISL_622028, EPI_ISL_622029, EPI_ISL_622030, EPI_ISL_622031, EPI_ISL_622032, EPI_ISL_622033, EPI_ISL_622034, EPI_ISL_622035, EPI_ISL_622036, EPI_ISL_622037, EPI_ISL_622038, EPI_ISL_622039, EPI_ISL_622040, EPI_ISL_622041, EPI_ISL_622042, EPI_ISL_622043, EPI_ISL_622044, EPI_ISL_622045, EPI_ISL_622046, EPI_ISL_622047, EPI_ISL_622048, EPI_ISL_622049, EPI_ISL_622050, EPI_ISL_622051, EPI_ISL_622052, EPI_ISL_622053, EPI_ISL_622054, EPI_ISL_622055, EPI_ISL_622056, EPI_ISL_622057, EPI_ISL_622058, EPI_ISL_622059, EPI_ISL_622060, EPI_ISL_622061, EPI_ISL_622062, EPI_ISL_622063, EPI_ISL_622064, EPI_ISL_622065, EPI_ISL_622066, EPI_ISL_622067, EPI_ISL_622068, EPI_ISL_622069, EPI_ISL_622070, EPI_ISL_622071, EPI_ISL_622072, EPI_ISL_622073, EPI_ISL_622074, EPI_ISL_622075, EPI_ISL_622076, EPI_ISL_622077, EPI_ISL_622090, EPI_ISL_622091, EPI_ISL_622092, EPI_ISL_622093, EPI_ISL_622094, EPI_ISL_622095, EPI_ISL_622096, EPI_ISL_622097, EPI_ISL_622098, EPI_ISL_622099, EPI_ISL_622100, EPI_ISL_622101, EPI_ISL_622102, EPI_ISL_622103, EPI_ISL_622104, EPI_ISL_622105, EPI_ISL_622106, EPI_ISL_622107, EPI_ISL_622108, EPI_ISL_622109, EPI_ISL_622110, EPI_ISL_622111, EPI_ISL_622112, EPI_ISL_622113, EPI_ISL_622114, EPI_ISL_622115, EPI_ISL_622116, EPI_ISL_622117, EPI_ISL_622118, EPI_ISL_622119, EPI_ISL_622120, EPI_ISL_622121, EPI_ISL_622122, EPI_ISL_622123, EPI_ISL_622124, EPI_ISL_622125, EPI_ISL_622126, EPI_ISL_622127, EPI_ISL_622128, EPI_ISL_622129, EPI_ISL_622130, EPI_ISL_622131, EPI_ISL_622132, EPI_ISL_622133, EPI_ISL_622134, EPI_ISL_622135, EPI_ISL_622136, EPI_ISL_622137, EPI_ISL_622138, EPI_ISL_622139, EPI_ISL_622140, EPI_ISL_622141, EPI_ISL_622142, EPI_ISL_622143, EPI_ISL_622144, EPI_ISL_622145, EPI_ISL_622146, EPI_ISL_622147, EPI_ISL_622148, EPI_ISL_622149, EPI_ISL_622165, EPI_ISL_622166, EPI_ISL_622167, EPI_ISL_622168, EPI_ISL_622186, EPI_ISL_622187, EPI_ISL_622188, EPI_ISL_622189, EPI_ISL_622190, EPI_ISL_622191, EPI_ISL_622192, EPI_ISL_622193, EPI_ISL_622194, EPI_ISL_622195, EPI_ISL_622196, EPI_ISL_622197, EPI_ISL_622198, EPI_ISL_622199, EPI_ISL_622200, EPI_ISL_622243, EPI_ISL_622244, EPI_ISL_622245, EPI_ISL_622246, EPI_ISL_622247, EPI_ISL_622248, EPI_ISL_622249, EPI_ISL_622250, EPI_ISL_622251, EPI_ISL_622252, EPI_ISL_622253, EPI_ISL_622254, EPI_ISL_622255, EPI_ISL_622256, EPI_ISL_622257, EPI_ISL_622258, EPI_ISL_622259, EPI_ISL_622748, EPI_ISL_622749, EPI_ISL_622750, EPI_ISL_622751, EPI_ISL_622752, EPI_ISL_622753, EPI_ISL_622754, EPI_ISL_622755, EPI_ISL_622756, EPI_ISL_622757, EPI_ISL_622758, EPI_ISL_622759, EPI_ISL_622760 | | | |
| see above | Department of Virus and Microbiological Special Diagnostics, Statens Serum Institut, Denmark | Albertsen lab, Department of Chemistry and Bioscience, Aalborg University, Denmark | Danish Covid-19 Genome Consortia |
| EPI_ISL_622805, EPI_ISL_622806 | PathLab Bay of Plenty | Institute of Environmental Science and Research (ESR) | Xiaoyun Ren, Matt Storey, Nikki Freed, Muhammad Faisal, Jing Wang, Hermes Perez, Anja Werno, Antje van der Linden, Arlo Upton, Chris Mansell, David Hammer, Dragana Drinkovic, Gary McAuliffe, Hana Sofia Andersson, James Ussher, Jill Sherwood, Josh Freeman, Julia Howard, Juliet Elvy, Mary DeAlmeida, Matt Blakiston, Matthew Rogers, Max Bloomfield, Michael Addidle, Michelle Balm, Sally Roberts, Sarah Jefferies, Sharmini Muttaiyah, Susan Morpeth, Susan Taylor, Timothy Blackmore, Vani Sathyendran, Veronica Playle, Virginia Hope, Erasmus Smit, Lauren Jelly, Olin Silander, Joep de Ligt |
| EPI_ISL_625522, EPI_ISL_625523, EPI_ISL_625524, EPI_ISL_625525, EPI_ISL_625526, EPI_ISL_625527, EPI_ISL_625528, EPI_ISL_625529, EPI_ISL_625530, EPI_ISL_625531, EPI_ISL_625532, EPI_ISL_625533, EPI_ISL_625534, EPI_ISL_625535, EPI_ISL_625536, EPI_ISL_625537, EPI_ISL_625538, EPI_ISL_625539, EPI_ISL_625540, EPI_ISL_625541, EPI_ISL_625542, EPI_ISL_625544, EPI_ISL_625545 | | | |
| see above | Alameda County Public Health Lab | Chan-Zuckerberg Biohub | CZB Cliahub Consortium |
| EPI_ISL_625660 | San Joaquin County Public Health Lab | Chan-Zuckerberg Biohub | CZB Cliahub Consortium |
| EPI_ISL_625669 | UCSF Clinical Microbiology Laboratory | Chan-Zuckerberg Biohub | CZB Cliahub Consortium |
| EPI_ISL_625673 | Laboratory of Molecular Medicine, University of Magallanes | Centro Asistencial Docente y de Investigacion, Universidad de Magallanes | Jorge Gonzalez, Jacqueline Aldridge, Diego Alvarez, Marcelo Navarrete |
| EPI_ISL_626574, EPI_ISL_626576, EPI_ISL_626577, EPI_ISL_626578, EPI_ISL_626579, EPI_ISL_626580, EPI_ISL_626581, EPI_ISL_626582, EPI_ISL_626583, EPI_ISL_626584 | The National Institute of Public Health | State Veterinary Institute Prague | Nagy,A;Jirincova,H;Novakova,L;Trnka,D;Vecerova,J |
| EPI_ISL_626806, EPI_ISL_627064, EPI_ISL_627312 | University of Exeter | COVID-19 Genomics UK (COG-UK) Consortium | Ben Temperton,Aaron Jeffries,Michelle Michelsen,Joanna Warwick-Dugdale,Audrey Farbos,Robyn Manley,Stephen Michell,Jane Masoli |
| EPI_ISL_627350, EPI_ISL_627355, EPI_ISL_627359, EPI_ISL_627360 | West of Scotland Specialist Virology Centre, NHSGGC / MRC-University of Glasgow Centre for Virus Research | COVID-19 Genomics UK (COG-UK) Consortium | Ana da Silva Filipe, Natasha Johnson, Kathy Smollett, Daniel Mair, Stephen Carmichael, Lily Tong, Jenna Nichols, Elihu Aranday-Cortes, Kyriaki Nomikou; Sarah McDonald, Marc Niebel, Patawee Asamaphan; Richard Orton, Joseph Hughes, Sreenu Vattipally, David L Robertson; Alasdair MacLean, Rory Gunson; Kathy Li, Igor Starinskij, Natasha Jesudason, Rajiv Shah, James Shepherd, Antonia Ho, Emma Thomson |
| EPI_ISL_627441, EPI_ISL_627444 | University of Exeter | COVID-19 Genomics UK (COG-UK) Consortium | Ben Temperton,Aaron Jeffries,Michelle Michelsen,Joanna Warwick-Dugdale,Audrey Farbos,Robyn Manley,Stephen Michell,Jane Masoli |
| EPI_ISL_629001 | South Eastern Area Laboratory Services (SEALS) | NSW Health Pathology - Institute of Clinical Pathology and Medical Research; Westmead Hospital; University of Sydney | CIDM-PH et al. |
| EPI_ISL_631365, EPI_ISL_631366, EPI_ISL_631367, EPI_ISL_631368, EPI_ISL_631369, EPI_ISL_631370, EPI_ISL_631371, EPI_ISL_631372, EPI_ISL_631373, EPI_ISL_631374, EPI_ISL_631375, EPI_ISL_631376 | | | |
| see above | ZOTZ KLIMAS MVZ Düsseldorf-Centrum GbR ÜBAG für Labormedizin, Genetik, Zytologie, Pathologie | Center of Medical Microbiology, Virology, and Hospital Hygiene, University of Duesseldorf | Maximilian Damagnez, Alexander Dilthey, Ashley-Jane Duplessis, Patrick Finzer, Katrin Hoffmann, Torsten Houwaart, Lisanna Hülse, Malte Kohns Vasconcelos, Marek Korencak, Nadine Lübke, Jessica Nicolai, Klaus Pfeffer, Daniel Strelow, Jörg Timm, Andreas Walker, Tobias Wienemann, Rainer Zotz |
| EPI_ISL_631378, EPI_ISL_631379, EPI_ISL_631383 | University Hospital Cologne | Center of Medical Microbiology, Virology, and Hospital Hygiene, University | Maximilian Damagnez, Alexander Dilthey, Ashley-Jane Duplessis, Eva Heger, Torsten Houwaart, Rolf Kaiser, Florian Klein, Elena Knops, Malte Kohns Vasconcelos, Jessica Nicolai, Klaus Pfeffer, Gibran Rubio Quintanares, Saleta Sierra-Aragón, Daniel Strelow, Jörg Timm, Andreas Walker, Tobias Wienemann |

| | | | |
|---|---|---|---|
| | | of Duesseldorf | |
| EPI_ISL_631387 | ZOTZ KLIMAS MVZ Düsseldorf-Centrum GbR ÜBAG für Labormedizin, Genetik, Zytologie, Pathologie | Center of Medical Microbiology, Virology, and Hospital Hygiene, University of Duesseldorf | Maximilian Damagnez, Alexander Dilthey, Ashley-Jane Duplessis, Patrick Finzer, Katrin Hoffmann, Torsten Houwaart, Lisanna Hülse, Malte Kohns Vasconcelos, Marek Korencak, Nadine Lübke, Jessica Nicolai, Klaus Pfeffer, Daniel Strelow, Jörg Timm, Andreas Walker, Tobias Wienemann, Rainer Zotz |
| EPI_ISL_631441, EPI_ISL_631442, EPI_ISL_631443, EPI_ISL_631444, EPI_ISL_631445, EPI_ISL_631446, EPI_ISL_631447, EPI_ISL_631448, EPI_ISL_631449, EPI_ISL_631450, EPI_ISL_631451, EPI_ISL_631452, EPI_ISL_631453, EPI_ISL_631454, EPI_ISL_631494, EPI_ISL_631495, EPI_ISL_631496 | | | |
| see above | Wisconsin State Laboratory of Hygiene Communicable Disease Division | Wisconsin State Laboratory of Hygiene Communicable Disease Division | Kelsey R. Florek, Abigail C. Shockey |
| EPI_ISL_632327, EPI_ISL_632346, EPI_ISL_632387, EPI_ISL_632388, EPI_ISL_632391, EPI_ISL_632405, EPI_ISL_632409, EPI_ISL_632417, EPI_ISL_632460, EPI_ISL_632495, EPI_ISL_632496, EPI_ISL_632497, EPI_ISL_632498, EPI_ISL_632499, EPI_ISL_632500, EPI_ISL_632501, EPI_ISL_632503, EPI_ISL_632504, EPI_ISL_632547, EPI_ISL_632548, EPI_ISL_632578, EPI_ISL_632579, EPI_ISL_632580, EPI_ISL_632581, EPI_ISL_632582, EPI_ISL_632583, EPI_ISL_632589, EPI_ISL_632762, EPI_ISL_632763 | | | |
| see above | Dutch COVID-19 response team | Erasmus Medical Center | Bas Oude Munnink, David Nieuwenhuijse, Reina Sikkema, Claudia Schapendonk, Irina Chestakova, Anne van der Linden, Theo Bestebroer, Stefan van Nieuwkoop, Mark Pronk, Pascal Lexmond, Corien Swaan, Manon Haverkate, Madelief Mollers, Mart Stein, Sandra Kengne Kamga Mobou, Jeroen van Kampen, Jolanda Voermans, Aura Timen, Corine GeurtsvanKessel, Annemiek van der Eijk, Richard Molenkamp, Marion Koopmans, on behalf of the Dutch national COVID-19 response team. |
| EPI_ISL_632890, EPI_ISL_632891, EPI_ISL_632892, EPI_ISL_632893, EPI_ISL_632894, EPI_ISL_632895, EPI_ISL_632896, EPI_ISL_632897, EPI_ISL_632898 | University of Wisconsin-Madison AIDS Vaccine Research Laboratories | University of Wisconsin-Madison AIDS Vaccine Research Laboratories | Gage Moreno, Katarina Braun, et al. AIDS Vaccine Research Laboratories |
| EPI_ISL_632934 | Department of Acute Infectious Diseases Control and Prevention,Yunnan Provincial Center for Disease Control and Prevention | Department of Acute Infectious Diseases Control and Prevention,Yunnan Provincial Center for Disease Control and Prevention | Meiling Zhang,Jienan Zhou,Senquan Jia,Xiaonan Zhao,Xiaoqing Fu |
| EPI_ISL_632935 | Department of Acute Infectious Diseases Control and Prevention, Yunnan Provincial Center for Disease Control and Prevention | Department of Acute Infectious Diseases Control and Prevention, Yunnan Provincial Center for Disease Control and Prevention | Meiling Zhang, Jienan Zhou, Senquan Jia, Xiaonan Zhao, Xiaoqing Fu |
| EPI_ISL_632976, EPI_ISL_632977, EPI_ISL_632978, EPI_ISL_632979 | Empire City Laboratories | New York City Public Health Laboratory | Jade Wang, et al. |
| EPI_ISL_632981 | DOHMH Corona | New York City Public Health Laboratory | Jade Wang, et al. |
| EPI_ISL_632994 | DOHMH Crown Heights | New York City Public Health Laboratory | Jade Wang, et al. |
| EPI_ISL_633035 | DOHMH PHL | New York City Public Health Laboratory | Jade Wang, et al. |
| EPI_ISL_633036 | DOHMH Morrisania | New York City Public Health Laboratory | Jade Wang, et al. |
| EPI_ISL_633037 | DOHMH Central Harlem | New York City Public Health Laboratory | Jade Wang, et al. |
| EPI_ISL_633038 | DOHMH PHL | New York City Public Health Laboratory | Jade Wang, et al. |
| EPI_ISL_635946, EPI_ISL_635947, EPI_ISL_635948, EPI_ISL_635949, EPI_ISL_635950, EPI_ISL_635951, EPI_ISL_635952, EPI_ISL_635967, EPI_ISL_635968, EPI_ISL_635969, EPI_ISL_635970, EPI_ISL_635971, EPI_ISL_635972, EPI_ISL_635976, EPI_ISL_635977, EPI_ISL_635982, EPI_ISL_636031, EPI_ISL_636032, EPI_ISL_636033, EPI_ISL_636034, EPI_ISL_636036, EPI_ISL_636037, EPI_ISL_636038, EPI_ISL_636039, EPI_ISL_636041, EPI_ISL_636046, EPI_ISL_636047, EPI_ISL_636051, EPI_ISL_636052, EPI_ISL_636057, EPI_ISL_636063, EPI_ISL_636066, EPI_ISL_636067, EPI_ISL_636068, EPI_ISL_636125 | | | |
| see above | San Diego County Public Health Laboratory | Andersen lab at Scripps Research | SEARCH Alliance San Diego with Tracy Basler, Jovan Shephard, Brett Austin |
| EPI_ISL_636589 | Dutch COVID-19 response team | National Institute for Public Health and the Environment (RIVM) | Adam Meijer, Harry Vennema, Jeroen Cremer, Sharon van den Brink, Bas van der Veer, AnneMarie van den Brandt, Florian Zwagemaker, Dennis Schmitz, Chantal Reusken, on behalf of the national COVID-19 response team |
| EPI_ISL_636892, EPI_ISL_636893, EPI_ISL_636894, EPI_ISL_636895, EPI_ISL_636896, EPI_ISL_636897, EPI_ISL_636898, EPI_ISL_636899, EPI_ISL_636900, EPI_ISL_636901, EPI_ISL_636902, EPI_ISL_636903, EPI_ISL_636904, EPI_ISL_636905, EPI_ISL_636906 | | | |
| see above | Lithuanian University of Health Sciences Hospital, Department of Laboratory Medicine | Lithuanian University of Health Sciences, Molecular cardiology lab. | Lukas Zemaitis, Ingrida Olendrait, Arnoldas Pautienius, Kamile Tamusauskaite, Dovydas Gecys, Laura Pareckaite, Vaiva Lesauskaite, Astra Vitkauskiene |
| EPI_ISL_637015 | Department of Infectious Diseases and Immunology, National Hospital Organization Nagoya Medical Center | Clinical Research Center, National Hospital Organization Nagoya Medical Center | Yoshihiro Nakata, Hirotaka Ode, Mai Kubota, Masakazu Matsuda, Kazuhiro Matsuoka, Miho Nakasuji, Mikiko Mori, Mayumi Imahashi, Yoshiyuki Yokomaku, Yasumasa Iwatani |
| EPI_ISL_637495 | Centre for Enzyme Innovation, University of Portsmouth / Translational Research Laboratory, Portsmouth Hospitals NHS Trust | COVID-19 Genomics UK (COG-UK) Consortium | Angela Beckett,Yann Bourgeois,Garry Scarlett,Sharon Glaysher,Scott Elliott,Kelly Bicknell,Robert Impey,Allyson Lloyd,Sarah Wyllie,Ethan Butcher,Anoop Chauhan,Samuel Robson |
| EPI_ISL_639645, EPI_ISL_639647 | E. Gulbja Laboratorija | Latvian Biomedical Research and Study Centre | Ivars Silamielis, Kaspars Megnis, Monta Ustinova, ikita Zrelovs, Vita Rovte, Mikus Gavars, Dmitrijs Perminovs, Uga Dumpis, Jnis Kloviš |

| | | | |
|---|---|---|---|
| EPI_ISL_639651 | Centrl Laboratorija | Latvian Biomedical Research and Study Centre | Ivars Silamielis, Kaspars Megnis, Monta Ustinova, ikita Zrelovs, Vita Rovte, Stella Lapia, Jana Oste, Marta Priedte, Uga Dumpis, Jnis Kloviš |
| EPI_ISL_639664, EPI_ISL_639665 | E. Gulbja Laboratorija | Latvian Biomedical Research and Study Centre | Ivars Silamielis, Kaspars Megnis, Monta Ustinova, ikita Zrelovs, Vita Rovte, Mikus Gavars, Dmitrijs Perminovs, Uga Dumpis, Jnis Kloviš |
| EPI_ISL_639825, EPI_ISL_639828, EPI_ISL_639829, EPI_ISL_639830, EPI_ISL_639831, EPI_ISL_639834, EPI_ISL_639835, EPI_ISL_639837, EPI_ISL_639841, EPI_ISL_639843, EPI_ISL_639845, EPI_ISL_639846, EPI_ISL_639847, EPI_ISL_639849, EPI_ISL_639851, EPI_ISL_639852, EPI_ISL_639854, EPI_ISL_639860, EPI_ISL_639863, EPI_ISL_639864 | | | |
| see above | National Virus Reference Laboratory | National Virus Reference Laboratory | Michael Carr, Gabriel Gonzalez, Jonathan Dean, Daniel Hare, Cillian F De Gascun |
| EPI_ISL_639948, EPI_ISL_639949 | HELIX LLC | WHO National Influenza Centre Russian Federation | Andrey Komissarov, Artem Fadeev, Kseniya Komissarova, Anna Ivanova, Dmitry Bazhenov, Daria Danilenko |
| EPI_ISL_639978 | Centre Hospitalier Saint Joseph Saint Luc | CNR Virus des Infections Respiratoires - France SUD | Antonin Bal, Gregory Destras, Gwendolyne Burfin, Hadrien Règue, Alexandre Gaymard, Maude Bouscambert-Duchamp, Florence Morfin-Sherpa, Martine Valette, Bruno Lina, Laurence Josset |
| EPI_ISL_640116 | False Bay Hospital wc FBH | NHLS/UCT | Arash Iranzadeh, Deelan Doolabh, Lynn Tyers, Bruna Galvao, Innocent Mudau, Marvin Hsiao, Kruger Marais, Diana Hardie, Stephen Korsman, Carolyn Williamson |
| EPI_ISL_640117 | 2 Military Hospital wc MAA | NHLS/UCT | Arash Iranzadeh, Deelan Doolabh, Lynn Tyers, Bruna Galvao, Innocent Mudau, Marvin Hsiao, Kruger Marais, Diana Hardie, Stephen Korsman, Carolyn Williamson |
| EPI_ISL_640119, EPI_ISL_640120 | Groote Schuur Hospital wc GSH | NHLS/UCT | Arash Iranzadeh, Deelan Doolabh, Lynn Tyers, Bruna Galvao, Innocent Mudau, Marvin Hsiao, Kruger Marais, Diana Hardie, Stephen Korsman, Carolyn Williamson |
| EPI_ISL_640121 | False Bay Hospital wc FBH | NHLS/UCT | Arash Iranzadeh, Deelan Doolabh, Lynn Tyers, Bruna Galvao, Innocent Mudau, Marvin Hsiao, Kruger Marais, Diana Hardie, Stephen Korsman, Carolyn Williamson |
| EPI_ISL_640583 | Victorian Infectious Diseases Reference Laboratory (VIDRL) | VIDRL and MDU-PHL | Caly L., Seemann T., Sait, M.L., Schultz M.B., Druce J., Sherry, N.L. |
| EPI_ISL_640969, EPI_ISL_641143, EPI_ISL_641166, EPI_ISL_641197 | Microbiological Diagnostic Unit - Public Health Laboratory (MDU-PHL) | MDU-PHL | Seemann T., Schultz M.B., Sait, M.L., Sherry, N.L. |
| EPI_ISL_641447, EPI_ISL_641448, EPI_ISL_641476, EPI_ISL_641477, EPI_ISL_641478, EPI_ISL_641479, EPI_ISL_641480, EPI_ISL_641481, EPI_ISL_641482, EPI_ISL_641483, EPI_ISL_641484, EPI_ISL_641485, EPI_ISL_641486, EPI_ISL_641487, EPI_ISL_641488, EPI_ISL_641489, EPI_ISL_641490, EPI_ISL_641491, EPI_ISL_641492, EPI_ISL_641493, EPI_ISL_641494, EPI_ISL_641495 | | | |
| see above | Department of Virus and Microbiological Special Diagnostics, Statens Serum Institut, Copenhagen, Denmark | Albertsen lab, Department of Chemistry and Bioscience, Aalborg University, Denmark | Thomas Bruun Rasmussen, Jannik Fonager, Morten Rasmussen |
| EPI_ISL_641557, EPI_ISL_641558, EPI_ISL_641559, EPI_ISL_641560, EPI_ISL_641561, EPI_ISL_641562, EPI_ISL_641563, EPI_ISL_641564, EPI_ISL_641565, EPI_ISL_641566, EPI_ISL_641567, EPI_ISL_641568, EPI_ISL_641569, EPI_ISL_641570, EPI_ISL_641571, EPI_ISL_641572 | | | |
| see above | Department of Clinical Microbiology | GIGA Medical Genomics | Keith Durkin, Maria Artesi, Sébastien Bontems, Raphaël Boreux, Bouchra Boujemla, Cécile Meex, Pierrette Melin, Marie-Pierre Hayette, Vincent Bours |
| EPI_ISL_644382, EPI_ISL_644393, EPI_ISL_644394, EPI_ISL_644395, EPI_ISL_644400, EPI_ISL_644401, EPI_ISL_644413, EPI_ISL_644414, EPI_ISL_644415, EPI_ISL_644416, EPI_ISL_644417, EPI_ISL_644418, EPI_ISL_644419, EPI_ISL_644420, EPI_ISL_644421, EPI_ISL_644422, EPI_ISL_644423, EPI_ISL_644424, EPI_ISL_644426, EPI_ISL_644427, EPI_ISL_644428, EPI_ISL_644429, EPI_ISL_644430, EPI_ISL_644431, EPI_ISL_644432, EPI_ISL_644433, EPI_ISL_644434, EPI_ISL_644435, EPI_ISL_644436, EPI_ISL_644437, EPI_ISL_644438 | | | |
| see above | MEPHI, Aix Marseille University | MEPHI, Aix Marseille University | Anthony LEVASSEUR |
| EPI_ISL_644610 | National Virus Reference Laboratory | National Virus Reference Laboratory | Michael Carr, Gabriel Gonzalez, Jonathan Dean, Daniel Hare, Cillian F De Gascun |
| EPI_ISL_644842, EPI_ISL_644843, EPI_ISL_644844, EPI_ISL_644845, EPI_ISL_644846, EPI_ISL_644847, EPI_ISL_644848, EPI_ISL_644849, EPI_ISL_644850, EPI_ISL_644851, EPI_ISL_644852, EPI_ISL_644853, EPI_ISL_644854, EPI_ISL_644855, EPI_ISL_644858, EPI_ISL_644863, EPI_ISL_644864, EPI_ISL_644866, EPI_ISL_644867, EPI_ISL_644868, EPI_ISL_644869, EPI_ISL_644870, EPI_ISL_644871, EPI_ISL_644872, EPI_ISL_644873, EPI_ISL_644874, EPI_ISL_644875, EPI_ISL_644876, EPI_ISL_644877, EPI_ISL_644878, EPI_ISL_644879, EPI_ISL_644880, EPI_ISL_644881 | | | |
| see above | Virginia DCLS | Virginia DCLS | Virginia DCLS |
| EPI_ISL_645116 | National Public Health Laboratory, National Centre for Infectious Diseases | National Public Health Laboratory, National Centre for Infectious Diseases | Tze Minn Mak, Sophie Octavia, Zhenyang Zhou, Lin Cui, Raymond Tzer Pin Lin |
| EPI_ISL_648016, EPI_ISL_648017 | MS Public Health Laboratory | Pathogen Discovery, Respiratory Viruses Branch, Division of Viral Diseases, Centers for Disease Control and Prevention | Yan Li, Jing Zhang, Ying Tao, Brian Lynch, Krista Queen, Anna Montmayeur, Anna Uehara, Clinton R. Paden, Rachel Marine, Haibin Wang, Suxiang Tong |
| EPI_ISL_648128 | UHAS COVID-19 Lab | UHAS COVID-19 Lab | Kwabena O. Duedu, Jones Gyamfi, Reuben Ayivor-Djanie, John O. Gyapong and the UHAS COVID-19 Lab Team |
| EPI_ISL_648170 | The Public Health Agency of Sweden | The Public Health Agency of Sweden | Anna-Malin Linde, Maria Lind Karlberg, Mattias Haukland, Reza Advani, Olov Svartstrom, Oskar Karlsson Lindsjo, Sandra Broddesson, Petra Edquist, Mia Brytting, Anna Risberg, Karin Tegmark-Wisell |
| EPI_ISL_648381, EPI_ISL_648382, EPI_ISL_648463, EPI_ISL_648465, EPI_ISL_648466 | Santa Clara County Public Health Laboratory | Chan-Zuckerberg Biohub | CZB Cliahub Consortium |
| EPI_ISL_648501, EPI_ISL_648502, EPI_ISL_648503 | Orange County Public Health Lab | Chan-Zuckerberg Biohub | CZB Cliahub Consortium |
| EPI_ISL_648545, EPI_ISL_648546, EPI_ISL_648547, EPI_ISL_648548, EPI_ISL_648549 | Madera County Department of Public Health | Chan-Zuckerberg Biohub | CZB Cliahub Consortium |
| EPI_ISL_649048 | San Diego County Public Health Laboratory | Andersen lab at Scripps Research | SEARCH Alliance San Diego with Tracy Basler, Jovan Shephard, Brett Austin |
| EPI_ISL_650850 | West of Scotland Specialist Virology Centre, NHSGGC / MRC-University of Glasgow Centre for Virus Research | COVID-19 Genomics UK (COG-UK) Consortium | Ana da Silva Filipe, Natasha Johnson, Kathy Smollett, Daniel Mair, Stephen Carmichael, Alice Broos, Lily Tong, Jenna Nichols, Kyriaki Nomikou; Sarah McDonald; Richard Orton, Joseph Hughes, Sreenu Vattipally, David L Robertson; Alasdair MacLean, Rory Gunson; Sharif Shaaban, Matthew Holden; Rachel Blacow, Guy Mollett, Kathy Li, James Shepherd, Antonia Ho, Emma Thomson |

| | | | |
|---|---|---|---|
| EPI_ISL_653916 | Diagnostic- and Research Institute of Pathology, Medical University of Graz | Diagnostic- and Research Institute of Pathology, Medical University of Graz | Karl Kashofer, Peter Regitnig, Martin Zacharias, Gregor Gorkiewicz |
| EPI_ISL_654021, EPI_ISL_654022, EPI_ISL_654023, EPI_ISL_654024, EPI_ISL_654025, EPI_ISL_654027, EPI_ISL_654028, EPI_ISL_654029, EPI_ISL_654030, EPI_ISL_654031, EPI_ISL_654033, EPI_ISL_654034, EPI_ISL_654036, EPI_ISL_654037, EPI_ISL_654038, EPI_ISL_654040, EPI_ISL_654041, EPI_ISL_654042, EPI_ISL_654044, EPI_ISL_654045, EPI_ISL_654046, EPI_ISL_654047, EPI_ISL_654050, EPI_ISL_654051, EPI_ISL_654052, EPI_ISL_654063, EPI_ISL_654064, EPI_ISL_654065, EPI_ISL_654066, EPI_ISL_654067, EPI_ISL_654068, EPI_ISL_654069, EPI_ISL_654070, EPI_ISL_654071, EPI_ISL_654072, EPI_ISL_654073, EPI_ISL_654074, EPI_ISL_654075, EPI_ISL_654076, EPI_ISL_654077, EPI_ISL_654078, EPI_ISL_654079, EPI_ISL_654080, EPI_ISL_654091, EPI_ISL_654092, EPI_ISL_654093, EPI_ISL_654094, EPI_ISL_654095, EPI_ISL_654096, EPI_ISL_654181, EPI_ISL_654196, EPI_ISL_654197, EPI_ISL_654198, EPI_ISL_654199, EPI_ISL_654200, EPI_ISL_654348 | | | |
| see above | Hospital General Universitario Gregorio Marañón | SeqCOVID-SPAIN consortium/IBV(CSIC) | Darío García de Viedma, Laura Pérez-Lago, Marta Herranz, Jon Sicilia, Julia Suárez, Pilar Catalán, Patricia Muñoz and SeqCOVID-SPAIN consortium |
| EPI_ISL_654483, EPI_ISL_654484, EPI_ISL_654485, EPI_ISL_654486, EPI_ISL_654487, EPI_ISL_654488 | Servicio de Microbiología. Hospital Clínico Universitario de Valencia | SeqCOVID-SPAIN consortium/IBV(CSIC) | David Navarro Ortega, Eliseo Albert Vicent, Ignacio Torres and SeqCOVID-SPAIN consortium |
| EPI_ISL_654513, EPI_ISL_654524 | Servicio de Microbiología, Laboratori Clínic Metropolitana Nord. Hospital Universitari Germans Trias i Pujol. Institut d'Investigació en Ciències de la Salut Germans Trias i Pujol (IGTP) | SeqCOVID-SPAIN consortium/IBV(CSIC) | Elisa Martró, Antoni E. Bordoy, Anna Not, Adrián Antuori, Anabel Fernández, Nona Romaní and SeqCOVID-SPAIN consortium |
| EPI_ISL_654893, EPI_ISL_654894, EPI_ISL_654895, EPI_ISL_654896, EPI_ISL_654897, EPI_ISL_654898 | Klinisk mikrobiologi | The Public Health Agency of Sweden | Anna-Malin Linde, Maria Lind Karlberg, Mattias Haukland, Reza Advani, Olov Svartstrom, Oskar Karlsson Lindsjo, Sandra Broddesson, Petra Edquist, Mia Brytting, Anna Risberg, Karin Tegmark-Wisell |
| EPI_ISL_660314 | Servicio de Microbiología, Laboratori Clínic Metropolitana Nord. Hospital Universitari Germans Trias i Pujol. Institut d'Investigació en Ciències de la Salut Germans Trias i Pujol (IGTP) | SeqCOVID-SPAIN consortium/IBV(CSIC) | Elisa Martró, Antoni E. Bordoy, Anna Not, Adrián Antuori, Anabel Fernández, Nona Romaní and SeqCOVID-SPAIN consortium |
| EPI_ISL_660427, EPI_ISL_660428, EPI_ISL_660431 | Klinisk Mikrobiologi | The Public Health Agency of Sweden | Anna-Malin Linde, Maria Lind Karlberg, Mattias Haukland, Reza Advani, Olov Svartstrom, Oskar Karlsson Lindsjo, Sandra Broddesson, Petra Edquist, Mia Brytting, Anna Risberg, Karin Tegmark-Wisell |
| EPI_ISL_660456, EPI_ISL_660458, EPI_ISL_660459, EPI_ISL_660460, EPI_ISL_660461, EPI_ISL_660462, EPI_ISL_660463, EPI_ISL_660481, EPI_ISL_660482, EPI_ISL_660483, EPI_ISL_660484, EPI_ISL_660485, EPI_ISL_660486, EPI_ISL_660487, EPI_ISL_660494, EPI_ISL_660496, EPI_ISL_660498, EPI_ISL_660501, EPI_ISL_660502, EPI_ISL_660509, EPI_ISL_660510, EPI_ISL_660517, EPI_ISL_660524, EPI_ISL_660525, EPI_ISL_660528 | | | |
| see above | Laboratoire de Microbiologie CHU Souro Sanou | Centre Muraz | Abdoul-Salam Ouedraogo, Yacouba Sawadogo, Essia Belarbi, Grit Schubert, Fabian Leendertz, Arsène Zongo, Soumeya Ouangraoua, Zekiba Tarnagda, Lassana Sangaré, Halidou Tinto |
| EPI_ISL_660795, EPI_ISL_660796, EPI_ISL_660797, EPI_ISL_660798, EPI_ISL_660799, EPI_ISL_660800, EPI_ISL_660801 | Gundersen Molecular Diagnostics Laboratory | Kabara Cancer Research Institute | Craig S. Richmond, Paraic A. Kenny |
| EPI_ISL_660802, EPI_ISL_660803, EPI_ISL_660804, EPI_ISL_660805 | Gundersen Clinical Microbiology Laboratory | Kabara Cancer Research Institute | Craig S. Richmond, Paraic A. Kenny |
| EPI_ISL_660806, EPI_ISL_660807, EPI_ISL_660808, EPI_ISL_660809, EPI_ISL_660810, EPI_ISL_660811, EPI_ISL_660812, EPI_ISL_660813, EPI_ISL_660814, EPI_ISL_660815, EPI_ISL_660816, EPI_ISL_660817, EPI_ISL_660818, EPI_ISL_660819 | | | |
| see above | Gundersen Molecular Diagnostics Laboratory | Kabara Cancer Research Institute | Craig S. Richmond, Paraic A. Kenny |
| EPI_ISL_661278 | Klinisk mikrobiologi | The Public Health Agency of Sweden | Department of Microbiology, The Public Health Agency of Sweden |
| EPI_ISL_661286 | Gavle klinisk mikrobiologi | The Public Health Agency of Sweden | Department of Microbiology, The Public Health Agency of Sweden |
| EPI_ISL_664233, EPI_ISL_664449, EPI_ISL_665153, EPI_ISL_665240 | University College London Hospital | COVID-19 Genomics UK (COG-UK) Consortium | Judith Heaney, Matthew Byott, Catherine Houlihan, Dan Frampton, Stuart Kirk, Moira Spyer and Eleni Nastouli |
| EPI_ISL_666637, EPI_ISL_666638, EPI_ISL_666639, EPI_ISL_666640, EPI_ISL_666641, EPI_ISL_666642, EPI_ISL_666643, EPI_ISL_666644, EPI_ISL_666645, EPI_ISL_666646, EPI_ISL_666647, EPI_ISL_666648, EPI_ISL_666649, EPI_ISL_666650, EPI_ISL_666651, EPI_ISL_666652 | | | |
| see above | ZOTZ KLIMAS MVZ Düsseldorf-Centrum GbR ÜBAG für Labormedizin, Genetik, Zytologie, Pathologie | Center of Medical Microbiology, Virology, and Hospital Hygiene, University of Duesseldorf | Maximilian Damagnez, Alexander Dilthey, Ashley-Jane Duplessis, Patrick Finzer, Katrin Hoffmann, Torsten Houwaart, Lisanna Hülse, Malte Kohns Vasconcelos, Marek Korencak, Nadine Lübke, Jessica Nicolai, Klaus Pfeffer, Daniel Strelow, Jörg Timm, Andreas Walker, Tobias Wienemann, Rainer Zotz |
| EPI_ISL_666856 | Florida Bureau of Public Health Laboratories | Florida Bureau of Public Health Laboratories | Sarah Schmedes, Jason Blanton |
| EPI_ISL_666995, EPI_ISL_666996, EPI_ISL_666997, EPI_ISL_666998 | San Diego County Public Health Laboratory | Andersen lab at Scripps Research | SEARCH Alliance San Diego with Tracy Basler, Jovan Shephard, Brett Austin |

| | | | |
|---|---|---|---|
| EPI_ISL_667449, EPI_ISL_667450, EPI_ISL_667451, EPI_ISL_667452, EPI_ISL_667453, EPI_ISL_667558 | OHSU Lab Services Molecular Microbiology Lab | Oregon SARS-CoV-2 Genome Sequencing Center | Brendan L. O'Connell, Ruth V. Nichols, Sally Grindstaff, Alec J. Hirsch, Donna Hansel, Guang Fan, Daniel N. Streblow, William B. Messer, Andrew C. Adey, Benjamin N. Bimber, Brian J. O'Roak |
| EPI_ISL_667809 | Laboratory Medicine | Department of Laboratory Medicine, Lin-Kou Chang Gung Memorial Hospital, Taoyuan, Taiwan | Kuo-Chien Tsao, Yu-Nong Gong, Shu-Li Yang, Yi-Chun Liu, Chung-Guei Huang, Mei-Jen Hsiao, Po-Wei Huang, Cheng-Ta Yang, Cheng-Hsun Chiu, Peng-Nien Huang, Kuo-Ming Lee, Guang-Wu Chen, Shin-Ru Shih |
| EPI_ISL_670651, EPI_ISL_670652, EPI_ISL_670653, EPI_ISL_670654, EPI_ISL_670655, EPI_ISL_670656, EPI_ISL_670657, EPI_ISL_670658, EPI_ISL_670672, EPI_ISL_670673, EPI_ISL_670674, EPI_ISL_670700, EPI_ISL_670767, EPI_ISL_671232, EPI_ISL_671233 | | | |
| see above | Department of Virus and Microbiological Special Diagnostics, Statens Serum Institut, Copenhagen, Denmark | Albertsen Lab, Department of Chemistry and Bioscience, Aalborg University, Denmark | Danish Covid-19 Genome Consortium |
| EPI_ISL_671674 | DOHMH Corona | New York City Public Health Laboratory | Jade Wang, et al. |
| EPI_ISL_671675 | DOHMH Jamaica | New York City Public Health Laboratory | Jade Wang, et al. |
| EPI_ISL_671825, EPI_ISL_671826, EPI_ISL_671827, EPI_ISL_671828, EPI_ISL_671829, EPI_ISL_671830, EPI_ISL_671831, EPI_ISL_671832, EPI_ISL_671833 | Hospital de la Santa Creu i Sant Pau. Servicio de Microbiología | SeqCOVID-SPAIN consortium/IBV(CSIC) | Ferran Navarro, Núria Rabella, Elisenda Miró and SeqCOVID-SPAIN consortium |
| EPI_ISL_671964 | CHU Purpan - Laboratoire de Virologie - Institut Fédératif de Biologie | CHU Purpan - Laboratoire de Virologie - Institut Fédératif de Biologie | Latour J., Ranger N., Dubois M., Carcenac R., Harter A., Boyer P., Tremeaux P., Izopet J. |
| EPI_ISL_671974 | Laboratorio de Virología y Microbiología Molecular, Depto. de Microbiología, Facultad de Medicina, Universidad de El Salvador/INS-laboratorio de Ref. Ministerio de Salud | Laboratorio de Virología y Microbiología Molecular, Depto. de Microbiología, Facultad de Medicina, Universidad de El Salvador/INS-laboratorio de Ref. Ministerio de Salud | Rivera NR, Ortega-Pérez C A, Xochitl Sandoval López, Carlos Hernández Ávila |
| EPI_ISL_672489 | Contra Costa County Public Health Lab | Chan-Zuckerberg Biohub | CZB Cliahub Consortium |
| EPI_ISL_672637 | PathWest Laboratory Medicine WA | PathWest Laboratory Medicine WA Microbial Surveillance Unit | PathWest Laboratory Medicine WA Microbial Surveillance Unit |
| EPI_ISL_676531, EPI_ISL_676532 | Klinisk mikrobiologi | The Public Health Agency of Sweden | Department of Microbiology, The Public Health Agency of Sweden |
| EPI_ISL_676610 | Texas Department of State Health Services | Texas Department of State Health Services | Rashmi Tuladhar, Bonnie Oh, Jenny Zhang, Maliha Rahman, Anita Pokharel, Myong Koag, Chung Wang, Rachel Lee, Grace Kubin, Mayela Pedrueza, James Daniel Bonser |
| EPI_ISL_677099, EPI_ISL_677100, EPI_ISL_677101, EPI_ISL_677102, EPI_ISL_677103 | Wadsworth Center, New York State Department.of Health | Wadsworth Center, New York State Department.of Health | Kirsten St. George, Daryl M. Lamson, Alexis Russel, Jonathan Plitnick, Navjot Singh, John Kelly, Sara Griesemer, Erasmus Schneider, Erica Lasek-Nesselquist |
| EPI_ISL_677243, EPI_ISL_677262, EPI_ISL_677277, EPI_ISL_677279, EPI_ISL_677315, EPI_ISL_677316, EPI_ISL_677317 | Colorado Department of Public Health and Environment | Colorado Department of Puplic Health and Environment | Laura Bankers, Molly Hetherington-Rauth, Shannon Ely, Shannon R. Matzinger, Sarah Elizabeth Totten, Emily A. Travanty |
| EPI_ISL_677343, EPI_ISL_677350, EPI_ISL_677413, EPI_ISL_677414, EPI_ISL_677415, EPI_ISL_677416, EPI_ISL_677417, EPI_ISL_677418, EPI_ISL_677419, EPI_ISL_677420, EPI_ISL_677421, EPI_ISL_677450 | | | |
| see above | University of Wisconsin-Madison AIDS Vaccine Research Laboratories | University of Wisconsin-Madison AIDS Vaccine Research Laboratories | Gage Moreno, Katarina Braun, et al. AIDS Vaccine Research Laboratories |
| EPI_ISL_677634 | Virology Unit, Institut Pasteur de Madagascar | Virology Unit, Institut Pasteur de Madagascar | Christian Ranaivoson, Cara E. Brook, Vida Ahyong, Soa Fy Andriamandimby, Vololoniaina Raharinosy, Tsiry Randriambolamanantsoa, Helisoa Razafimanjato, Norosoa Razanajatovo, Michelle Tan, Cristina M. Tato, Joseph L. DeRisi, Jean-Michel Heraud, Philippe Dussart |
| EPI_ISL_677648, EPI_ISL_677649, EPI_ISL_677650, EPI_ISL_677651, EPI_ISL_677652, EPI_ISL_677653, EPI_ISL_677654, EPI_ISL_677655, EPI_ISL_677656, EPI_ISL_677657 | Colorado Department of Public Health and Environment | Colorado Department of Puplic Health and Environment | Laura Bankers, Molly C. Hetherington-Rauth, Shannon Ely, Shannon R. Matzinger, Sarah Elizabeth Totten, Emily A. Travanty |
| EPI_ISL_677659 | University of | University of | Gage Moreno, Katarina Braun, et al. AIDS Vaccine Research Laboratories |

| | | | |
|---|---|---|---|
| | Wisconsin-Madison AIDS Vaccine Research Laboratories | Wisconsin-Madison AIDS Vaccine Research Laboratories | |
| EPI_ISL_677671 | Wadsworth Center, New York State Department.of Health | Wadsworth Center, New York State Department.of Health | Kirsten St. George, Daryl M. Lamson, Alexis Russel, Jonathan Plitnick, Navjot Singh, John Kelly, Sara Griesemer, Erasmus Schneider, Erica Lasek-Nesselquist |
| EPI_ISL_677696, EPI_ISL_677699 | Vanda Pharmaceuticals Clinical Site | Vanda Pharmaceuticals | Vanda Pharmaceuticals |
| EPI_ISL_678260 | General Hospital - Struga | Research Center for Genetic Engineering and Biotechnology "Georgi D. Efremov" , Macedonian Academy of Sciences and Arts | RCGEB - MASA |
| EPI_ISL_678343, EPI_ISL_678344, EPI_ISL_678346, EPI_ISL_678358 | Area of Virology, Serology and Virology Division (SAViD), New South Wales Health Pathology Randwick | Virology Research Laboratory; Area of Virology, Serology and Virology Division (SAViD), New South Wales Health Pathology Randwick | Foster, C.; Au, J.; Ruiz Silva, M.; Deveson, I.; Bull, R.; Van Hal, S.; Rawlinson, W. |
| EPI_ISL_678829 | Respiratory Virus Unit, Microbiology Services Colindale, Public Health England | COVID-19 Genomics UK (COG-UK) Consortium | PHE Covid Sequencing Team |
| EPI_ISL_679409, EPI_ISL_679410, EPI_ISL_679411 | University College London Hospital | COVID-19 Genomics UK (COG-UK) Consortium | Judith Heaney, Matthew Byott, Catherine Houlihan, Dan Frampton, Stuart Kirk, Moira Spyer and Eleni Nastouli |
| EPI_ISL_680258, EPI_ISL_680265, EPI_ISL_680301, EPI_ISL_680303 | Regional Virus Laboratory, Belfast Health and Social Care Trust | COVID-19 Genomics UK (COG-UK) Consortium | Conall McCaughey, James McKenna, Tanya Curran, Susan Feeney, Alison Watt, Ciara Cox, Mairead Connor, Zoltan Molnar, David Simpson, Derek Fairley |
| EPI_ISL_680497 | Virology Department, Royal Infirmary of Edinburgh, NHS Lothian / School of Biological Sciences, University of Edinburgh / Institute of Genetics and Molecular Medicine, University of Edinburgh | COVID-19 Genomics UK (COG-UK) Consortium | McHugh M, Dewar R, Rooke S, Gallagher M, Balcaza C, O'Toole Á, Scher E, Hill V, McCrone JT, Colquhoun R, Yu X, Jackson B, Rambaut A, Williams TC, Templeton K |
| EPI_ISL_681301, EPI_ISL_681309 | Communicable Disease Laboratory, Public Health Directorate | Communicable Disease Laboratory, Public Health Directorate | Alwasti,H., Altaif,Z., AlHujairi,Z., AlAbbas,Z. |
| EPI_ISL_681936, EPI_ISL_682034, EPI_ISL_682036, EPI_ISL_682038, EPI_ISL_682044, EPI_ISL_682045, EPI_ISL_682046, EPI_ISL_682047, EPI_ISL_682048, EPI_ISL_682049, EPI_ISL_682055, EPI_ISL_682056 | | | |
| see above | UPMC Clinical Microbiology Laboratory | Microbial Genomic Epidemiology Laboratory, University of Pittsburgh | Mustapha M. Mustapha, Jane W. Marsh, Dan Snyder, Marissa P. Griffith, Stephanie L. Mitchell, Vatsala R. Srinivasa, Kady D. Waggle, Chinelo Ezeonwuku, Vaughn S. Cooper, Lee H. Harrison |
| EPI_ISL_682300, EPI_ISL_682303 | Communicable Disease Laboratory, Public Health Directorate | Communicable Disease Laboratory, Public Health Directorate | Alwasti,H., Altaif,Z., AlHujairi,Z., AlAbbas,Z. |
| EPI_ISL_683601, EPI_ISL_683602, EPI_ISL_683603 | Servicio de Microbiología, Laboratori Clínic Metropolitana Nord. Hospital Universitari Germans Trias i Pujol. Institut d'Investigació en Ciències de la Salut Germans Trias i Pujol (IGTP) | SeqCOVID-SPAIN consortium/IBV(CSIC) | Elisa Martró, Antoni E. Bordoy, Anna Not, Adrián Antuori, Anabel Fernández, Nona Romaní, Verónica Saludes, Cristina Casañ and SeqCOVID-SPAIN consortium |
| EPI_ISL_684004 | Utah Public Health Laboratory | Utah Public Health Laboratory | Erin Young, Kelly Oakeson |
| EPI_ISL_692746, EPI_ISL_692747, EPI_ISL_692748, EPI_ISL_692749, EPI_ISL_692750, EPI_ISL_692751, EPI_ISL_692752, EPI_ISL_692753, EPI_ISL_692754 | CNR Virus des Infections Respiratoires - France SUD | CNR Virus des Infections Respiratoires - France SUD | Antonin Bal, Gregory Destras, Gwendolyne Burfin, Solenne Brun, Martine Valette, Bruno Lina, Laurence Josset |
| EPI_ISL_693299 | Department of Microbiology, Yokohama City University School of Medicine | Department of Microbiology, Yokohama City University School of Medicine | Kei Miyakawa, Ryo Saji, Kazuya Sakai, Reo Matsumura, Mototsugu Nishii, Ichiro Takeuchi, Akihide Ryo |
| EPI_ISL_693479, EPI_ISL_693481 | Central Public Health Laboratory | National Public Health Laboratory, National Centre for Infectious Diseases | Tze Minn Mak, Sophie Octavia, Zhenyang Zhou, Esorom Daoni, Theresa Palou, Lin Cui, Raymond Tzer Pin Lin |
| EPI_ISL_693488, EPI_ISL_693489, EPI_ISL_693491, EPI_ISL_693492, EPI_ISL_693493, EPI_ISL_693494, EPI_ISL_693495, EPI_ISL_693496, EPI_ISL_693497, EPI_ISL_693498, EPI_ISL_693499, EPI_ISL_693500 | | | |
| see above | CNR Virus des Infections Respiratoires - France SUD | CNR Virus des Infections Respiratoires - France SUD | Antonin Bal, Gregory Destras, Gwendolyne Burfin, Quentin Semanas, Martine Valette, Bruno Lina, Laurence Josset |
| EPI_ISL_693718, | Delaware Public Health | Delaware Public Health | Gregory Hovan |

| | | | |
|---|---|---|---|
| EPI_ISL_693720, EPI_ISL_693745 | Laboratory | Laboratory | |
| EPI_ISL_693763 | Hospital | National Reference Center for Viruses of Respiratory Infections, Institut Pasteur, Paris | Marion Barbet, Sylvie Behillil, Méline Bizard, Angela Brisebarre, Camille Capel, Etienne Simon-Lorière, Vincent Enouf, Maud Vanpeene, Sylvie van der Werf, Gisèle Lagathu |
| EPI_ISL_699642 | South Eastern Area Laboratory Services (SEALS) | NSW Health Pathology - Institute of Clinical Pathology and Medical Research; Westmead Hospital; University of Sydney | CIDM-PH et al. |
| EPI_ISL_700251, EPI_ISL_700252, EPI_ISL_700253, EPI_ISL_700254, EPI_ISL_700255, EPI_ISL_700256, EPI_ISL_700257, EPI_ISL_700258, EPI_ISL_700259, EPI_ISL_700260, EPI_ISL_700261, EPI_ISL_700262, EPI_ISL_700263, EPI_ISL_700264, EPI_ISL_700265, EPI_ISL_700266, EPI_ISL_700267, EPI_ISL_700268, EPI_ISL_700269, EPI_ISL_700270, EPI_ISL_700271, EPI_ISL_700272, EPI_ISL_700273, EPI_ISL_700274, EPI_ISL_700275, EPI_ISL_700276, EPI_ISL_700277, EPI_ISL_700278, EPI_ISL_700279, EPI_ISL_700280, EPI_ISL_700281, EPI_ISL_700282, EPI_ISL_700283, EPI_ISL_700284, EPI_ISL_700285, EPI_ISL_700286, EPI_ISL_700287, EPI_ISL_700288, EPI_ISL_700289, EPI_ISL_700290, EPI_ISL_700291, EPI_ISL_700292, EPI_ISL_700293, EPI_ISL_700294, EPI_ISL_700295 | | | |
| see above | Hematopathology Laboratory, ACTREC, TMC | Hematopathology Laboratory, ACTREC, TMC | Hematopathology Laboratory, ACTREC |
| EPI_ISL_700328, EPI_ISL_700332, EPI_ISL_700344, EPI_ISL_700345, EPI_ISL_700346 | Child Health Research Foundation | Child Health Research Foundation | Senjuti Saha, Afroza Akter Tanni, Syed Muktadir Al Sium, Roly Malaker, Sharmistha Goswami, Arif Mohammad Tanmoy, Md Hafizur Rahman, Samir K Saha |
| EPI_ISL_700532 | Kwamandlenkosi Clinic wc KMN | NHLS/UCT | Arash Iranzadeh, Deelan Doolabh, Lynn Tyers, Bruna Galvao, Innocent Mudau, Marvin Hsiao, Kruger Marais, Diana Hardie, Stephen Korsman, Carolyn Williamson |
| EPI_ISL_700699, EPI_ISL_700701, EPI_ISL_700724, EPI_ISL_700732, EPI_ISL_700734 | Texas Department of State Health Services | Texas Department of State Health Services | Rashmi Tuladhar, Bonnie Oh, Jenny Zhang, Maliha Rahman, Anita Pokharel, Myong Koag, Chung Wang, Rachel Lee, Grace Kubin, Mayela Pedrueza, James Daniel Bonser |
| EPI_ISL_706464 | Wales Specialist Virology Centre Sequencing lab: Pathogen Genomics Unit | COVID-19 Genomics UK (COG-UK) Consortium | Catherine Moore, Johnathan Evans, Laura Gifford, Malorie Perry, Simon Cottrell, Angela Marchbank, Alec Birchley, Alexander Adams, Amy Gaskin, Bree Gatica-Wilcox, Jason Coombes, Joel Southgate, Lauren Gilbert, Lee Graham, Nicole Pacchiarini, Sara Kumziene-Summerhayes, Sarah Taylor, Sophie Jones, Sara Rey, Matthew Bull, Joanne Watkins, Sally Corden, Tom Connor |
| EPI_ISL_707887 | Los Angeles County Public Health Laboratory | Los Angeles County Public Health Laboratory | P. Hemarajata et al. |
| EPI_ISL_708342 | Michigan Department of Health and Human Services, Bureau of Laboratories | Michigan Department of Health and Human Services, Bureau of Laboratories | Blankenship HM, Riner D, Soehnlen MK |
| EPI_ISL_708396, EPI_ISL_708397, EPI_ISL_708433, EPI_ISL_708434, EPI_ISL_708435 | Delaware Public Health Lab | Delaware Public Health Lab | Gregory Hovan |
| EPI_ISL_708812 | World Medical Hospital | National Institute of Health, Department of Medical Sciences, Ministry of Public Health, Thailand | Pilailuk Okada; Siripaporn Phuygun; Thanutsapa Thanadachakul; Sittiporn Parnmen; Pakorn Piromtong; Warawan Wongboot; Sunthareeya Waicharoen; Malinee Chittaganpitch |
| EPI_ISL_710132, EPI_ISL_710142, EPI_ISL_710152, EPI_ISL_710154, EPI_ISL_710167, EPI_ISL_710169, EPI_ISL_710170, EPI_ISL_710224, EPI_ISL_710225, EPI_ISL_710226, EPI_ISL_710227, EPI_ISL_710228, EPI_ISL_710229, EPI_ISL_710230, EPI_ISL_710231, EPI_ISL_710232, EPI_ISL_710233, EPI_ISL_710234, EPI_ISL_710235, EPI_ISL_710236, EPI_ISL_710252, EPI_ISL_710254, EPI_ISL_710256, EPI_ISL_710257, EPI_ISL_710264, EPI_ISL_710272, EPI_ISL_710273, EPI_ISL_710275, EPI_ISL_710276, EPI_ISL_710280, EPI_ISL_710282, EPI_ISL_710294, EPI_ISL_710298, EPI_ISL_710333, EPI_ISL_710334, EPI_ISL_710335, EPI_ISL_710336, EPI_ISL_710337, EPI_ISL_710338, EPI_ISL_710339, EPI_ISL_710340, EPI_ISL_710341, EPI_ISL_710342, EPI_ISL_710343, EPI_ISL_710344, EPI_ISL_710345, EPI_ISL_710346, EPI_ISL_710347 | | | |
| see above | Colorado Department of Public Health and Environment | Colorado Department of Puplic Health and Environment | Laura Bankers, Molly C. Hetherington-Rauth, Shannon Ely, Shannon R. Matzinger, Sarah Elizabeth Totten, Emily A. Travanty |
| EPI_ISL_714929 | Department of Virus and Microbiological Special Diagnostics, Statens Serum Institut, Copenhagen, Denmark | Albertsen Lab, Department of Chemistry and Bioscience, Aalborg University, Denmark | Danish Covid-19 Genome Consortium |
| EPI_ISL_717700 | Trinidad Public Health Laboratory | Carrington Lab, Department of PreClinical Sciences, Faculty of Medical Sciences, The University of the West Indies | Nikita S. D. Sahadeo, Arianne Brown-Jordan, Sarah Hill, Vernie Ramkissoon, Naresh Nandram, Avery Hinds, Jerome Foster, Stanley Giddings, Karla Georges, Marsha Ivey, Rahul Naidu, Risha Singh, SueMin Nathaniel, Rajini Haraksingh, Jaya Jayaraman, Chinna Chinnadurai, Adesh Ramsubhag, Nuno Faria, Oliver Pybus, Christopher Oura, Gabriel Escobar, Christine V. F. Carrington |
| EPI_ISL_717773, EPI_ISL_717774, EPI_ISL_717775 | UW Virology Lab | UW Virology Lab | Pavitra Roychoudhury, Hong Xie, Lasata Shrestha, Michelle Lin, Meei-Li Huang, Keith R Jerome, Alexander Greninger |
| EPI_ISL_717975 | Institute of Virology, Biomedical Research Center of the Slovak Academy of Sciences, Bratislava | Faculty of Natural Sciences, Comenius University, Bratislava | Broa Brejová, Viktória Hodorová, Kristína Boršová, Viktória abanová, Dominika Friová, Sabina Fumaová Havlíková, Juraj Kopáek, Martina Liková, ubomíra Lukáiková, Martina Neboháová, Monika Sláviková, Edita Staroová, Elena Tichá, Tomáš Vina, Jozef Nosek, Boris Klempa |
| EPI_ISL_718229, EPI_ISL_718233 | Hospital | National Reference Center for Viruses of Respiratory Infections, Institut Pasteur, Paris | Marion Barbet, Sylvie Behillil, Méline Bizard, Angela Brisebarre, Camille Capel, Etienne Simon-Lorière, Vincent Enouf, Maud Vanpeene, Sylvie van der Werf, Gisèle Lagathu |
| EPI_ISL_718282 | Institute for Medical Research, Infectious Disease Research Centre, National Institutes of Health, Ministry of Health Malaysia | Institute for Medical Research, Infectious Disease Research Centre, National Institutes of Health, Ministry of Health Malaysia | Suppiah J, Kamel K, Mohd-Zawawi Z, Thayan R |

| | | | |
|---|---|---|---|
| EPI_ISL_722375, EPI_ISL_722465, EPI_ISL_722816, EPI_ISL_722817, EPI_ISL_722818, EPI_ISL_722819, EPI_ISL_722820, EPI_ISL_722821, EPI_ISL_722822, EPI_ISL_722823 | Dutch COVID-19 response team | Erasmus Medical Center | Bas Oude Munnink, Reina Sikkema, David Nieuwenhuijse, Irina Chestakova, Anne van der Linden, Marjan Boter, Emmanuelle Munger, Corine GeurtsvanKessel, Annemiek van der Eijk, Richard Molenkamp, Marion Koopmans, on behalf of the Dutch national COVID-19 response team. |
| EPI_ISL_723067, EPI_ISL_723068, EPI_ISL_723069, EPI_ISL_723070, EPI_ISL_723071, EPI_ISL_723072, EPI_ISL_723073, EPI_ISL_723074, EPI_ISL_723075 | Hematopathology Laboratory, ACTREC, TMC | Hematopathology Laboratory, ACTREC, TMC | Hematopathology Laboratory, ACTREC |
| EPI_ISL_729343, EPI_ISL_729351, EPI_ISL_729352, EPI_ISL_729359, EPI_ISL_729375, EPI_ISL_729608 | A. Krumbholz, Labor Dr. Krause und Kollegen MVZ GmbH, Kiel | Charité Universitätsmedizin Berlin, Institut für Virologie | Victor M Corman, Barbara Mühlemann, Jörn Beheim-Schwarzbach, Talitha Veith, Julia Schneider, Terry Jones, Christian Drosten |
| EPI_ISL_729735, EPI_ISL_729736, EPI_ISL_729741, EPI_ISL_729743, EPI_ISL_729744, EPI_ISL_729745, EPI_ISL_729748, EPI_ISL_729749, EPI_ISL_729750, EPI_ISL_729752, EPI_ISL_729754, EPI_ISL_729760, EPI_ISL_729773, EPI_ISL_729779, EPI_ISL_729780 | | | |
| see above | Connecticut Department of Health | Grubaugh Lab - Yale School of Public Health | Joseph Fauver, Tara Alpert, Anderson Brito, Annie Watkins, Anne Wyllie, Chantal Vogels, Mary Petrone, Chaney Kalinich, Isabel Ott, Arnau Casanovas, Catherine Muenker, Adam Moore, Alice Lu, Maria Tokuyama, Patrick Wong, Peiwen Lu, Saad Omer, Richard Martinello, Allison Nelson, Shelli Farhadian, Akiko Iwasaki, Charlese Dela Cruz, Albert Ko, Nathan Grubaugh |
| EPI_ISL_729871, EPI_ISL_729906 | Instituto de Medicina Tropical, Universidad Nacional Toribio Rodríguez de Mendoza de Amazonas | Laboratorio de Genómica Microbiana, Universidad Peruana Cayetano Heredia | Pablo Tsukayama, Alejandra Dávila-Barclay, Luis González, Pedro E. Romero, Brenda Ayzanoa, Janet Huancachoque, Pool Marcos, Stella Chenet, Rafael Tapia, Cecilia Pajuelo, Carla Montenegro |
| EPI_ISL_730115, EPI_ISL_730116 | San Diego County Public Health Laboratory | Andersen lab at Scripps Research | SEARCH Alliance San Diego with Tracy Basler, Jovan Shephard, Brett Austin |
| EPI_ISL_730229 | Biolab Diagnostic Laboratories | Andersen lab at Scripps Research | Issa Abu-Dayyeh, Ahmad Tibi, Lama Hussein, Lina Mohammad, Zein Naber, Amid Abdelnour with SEARCH Alliance San Diego |
| EPI_ISL_730341 | San Diego County Public Health Laboratory | Andersen lab at Scripps Research | SEARCH Alliance San Diego with Tracy Basler, Jovan Shephard, Brett Austin |
| EPI_ISL_730569 | Gazi University Faculty of Medicine, Medical Virology Laboratory | Gazi University Faculty of Medicine, Medical Virology Laboratory | Erdem ahin, Hager Muftah, Selin Yiit, Shaknoza Sarzhanova, Özlem Güzel Tunçcan, Murat Dizbay, Il Fidan, Kayhan Çalar, Gülendam Bozday |
| EPI_ISL_730591 | Princess Margaret Hospital | Hong Kong Department of Health | Mak Gannon C.K., Lam Edman T.K., Chan Rickjason C.W., Tsang Dominic N.C. |
| EPI_ISL_730602 | Tseung Kwan O Hospital | Hong Kong Department of Health | Mak Gannon C.K., Lam Edman T.K., Chan Rickjason C.W., Tsang Dominic N.C. |
| EPI_ISL_730604, EPI_ISL_730605, EPI_ISL_730609 | Tuen Mun Hospital | Hong Kong Department of Health | Mak Gannon C.K., Lam Edman T.K., Chan Rickjason C.W., Tsang Dominic N.C. |
| EPI_ISL_730612, EPI_ISL_730613 | United Christian Hospital | Hong Kong Department of Health | Mak Gannon C.K., Lam Edman T.K., Chan Rickjason C.W., Tsang Dominic N.C. |
| EPI_ISL_732750, EPI_ISL_732769, EPI_ISL_732770, EPI_ISL_732771, EPI_ISL_732783, EPI_ISL_732815 | Centro de Investigación Biomédica de La Rioja - Hospital San Pedro Logroño | SeqCOVID-SPAIN consortium/IBV(CSIC) | María de Toro, José Manuel Azcona Gutiérrez, María Pilar Bea Escudero, Miriam Blasco Alberdi and SeqCOVID-SPAIN consortium |
| EPI_ISL_733046, EPI_ISL_733071, EPI_ISL_733075 | HELIX LLC | WHO National Influenza Centre Russian Federation | Andrey Komissarov, Artem Fadeev, Anna Ivanova, Kseniya Komissarova, Dmitry Bazhenov, Daria Danilenko, Ksenia Safina, Elena Nabieva, Georgii Bazykin, Dmitry Lioznov |
| EPI_ISL_733186, EPI_ISL_733194, EPI_ISL_733203, EPI_ISL_733204, EPI_ISL_733214 | Pathogenic Microorganisms Variability Laboratory | WHO National Influenza Centre Russian Federation | Andrey Komissarov, Artem Fadeev, Anna Ivanova, Kseniya Komissarova, Dmitry Bazhenov, Daria Danilenko, Ksenia Safina, Elena Nabieva, Georgii Bazykin, Nadezhda Kuznetsova, Elena Shidlovskaya, Sergey Alkhovsky, Tatyana Vishnevskaya, Elizaveta Divisenko, Alexey Shchetinin, Maria Nikiforova, Andrey Pochtovyy, Evgeny Usachev, Elena Vokalova, Maxim Rubalsky, Oleg Rubalsky, Artem Tkachuk, Vladimir Gushchin, Alexander Gintsburg, Dmitry Lioznov |
| EPI_ISL_733230 | UMMC-Health | WHO National Influenza Centre Russian Federation | Andrey Komissarov, Artem Fadeev, Anna Ivanova, Kseniya Komissarova, Dmitry Bazhenov, Tatiana Platonova, Daria Danilenko, Ksenia Safina, Elena Nabieva, Georgii Bazykin, Dmitry Lioznov |
| EPI_ISL_733298, EPI_ISL_733299, EPI_ISL_733395 | WHO National Influenza Centre Russian Federation | WHO National Influenza Centre Russian Federation | Andrey Komissarov, Artem Fadeev, Anna Ivanova, Kseniya Komissarova, Dmitry Bazhenov, Daria Danilenko, Ksenia Safina, Elena Nabieva, Georgii Bazykin, Dmitry Lioznov |
| EPI_ISL_734923, EPI_ISL_734924, EPI_ISL_734925, EPI_ISL_734926, EPI_ISL_734927, EPI_ISL_734928, EPI_ISL_734929, EPI_ISL_734930, EPI_ISL_734931, EPI_ISL_734932, EPI_ISL_734933, EPI_ISL_734934, EPI_ISL_734935, EPI_ISL_734936, EPI_ISL_734937, EPI_ISL_734938, EPI_ISL_734939, EPI_ISL_734940, EPI_ISL_734941, EPI_ISL_734942, EPI_ISL_734943, EPI_ISL_734944, EPI_ISL_734945, EPI_ISL_734946, EPI_ISL_734947, EPI_ISL_734948, EPI_ISL_734949, EPI_ISL_734950, EPI_ISL_734951, EPI_ISL_734952, EPI_ISL_734953, EPI_ISL_734954, EPI_ISL_734955, EPI_ISL_734956, EPI_ISL_734957, EPI_ISL_734958, EPI_ISL_734959, EPI_ISL_734960, EPI_ISL_734961, EPI_ISL_734962, EPI_ISL_734963, EPI_ISL_734964, EPI_ISL_734965, EPI_ISL_734966, EPI_ISL_734967, EPI_ISL_734968, EPI_ISL_734969, EPI_ISL_734970, EPI_ISL_734971, EPI_ISL_734972, EPI_ISL_734973, EPI_ISL_734974, EPI_ISL_734975, EPI_ISL_734976, EPI_ISL_734977, EPI_ISL_734978, EPI_ISL_734979, EPI_ISL_734980, EPI_ISL_734981, EPI_ISL_734982, EPI_ISL_734983, EPI_ISL_734984, EPI_ISL_734985, EPI_ISL_734986, EPI_ISL_734987, EPI_ISL_734988, EPI_ISL_734989, EPI_ISL_734990 | | | |
| see above | UZ Leuven, National Reference Laboratory for Coronaviruses, Laboratory Medicine, Leuven, Belgium | KU Leuven, Rega Institute, Clinical and Epidemiological Virology | Tony Wawina-Bokalanga, Joan Marti-Carerras, Bert Vanmechelen, Piet Maes |

| | | | |
|---|---|---|---|
| EPI_ISL_737061, EPI_ISL_737062, EPI_ISL_737063, EPI_ISL_737064, EPI_ISL_737065, EPI_ISL_737066, EPI_ISL_737067, EPI_ISL_737068, EPI_ISL_737234, EPI_ISL_737235 | Department of Virology and Immunology, University of Helsinki and Helsinki University Hospital, Huslab Finland | Department of Virology, Faculty of Medicine, University of Helsinki, Helsinki, Finland | Teemu Smura, Ravi Kant, Phuoc Truong, Hussein Alburkat, Hannimari Kallio-Kokko, Jenni Virtanen, Maija Suvanto, Sari Hannula, Harri Kangas, Pekka Ellonen, Olli Vapalahti |
| EPI_ISL_738223, EPI_ISL_738224, EPI_ISL_738225, EPI_ISL_738226 | UZ Leuven, National Reference Laboratory for Coronaviruses, Laboratory Medicine, Leuven, Belgium | KU Leuven, Rega Institute, Clinical and Epidemiological Virology | Tony Wawina-Bokalanga, Joan Marti-Carerras, Bert Vanmechelen, Piet Maes |
| EPI_ISL_738541 | Napa-Solano-Yolo- Marin County (NSYM) Public Health Laboratories | Chan-Zuckerberg Biohub | CZB Cliahub Consortium |
| EPI_ISL_738554, EPI_ISL_738797, EPI_ISL_738932 | Alameda County Public Health Lab | Chan-Zuckerberg Biohub | CZB Cliahub Consortium |
| EPI_ISL_738952 | Napa-Solano-Yolo- Marin County (NSYM) Public Health Laboratories | Chan-Zuckerberg Biohub | CZB Cliahub Consortium |
| EPI_ISL_738990 | County of San Luis Obispo Public Health Laboratory | Chan-Zuckerberg Biohub | CZB Cliahub Consortium |
| EPI_ISL_739062, EPI_ISL_739110 | Alameda County Public Health Lab | Chan-Zuckerberg Biohub | CZB Cliahub Consortium |
| EPI_ISL_739169 | Napa-Solano-Yolo- Marin County (NSYM) Public Health Laboratories | Chan-Zuckerberg Biohub | CZB Cliahub Consortium |
| EPI_ISL_739173 | Alameda County Public Health Lab | Chan-Zuckerberg Biohub | CZB Cliahub Consortium |
| EPI_ISL_739181, EPI_ISL_739223, EPI_ISL_739339 | Napa-Solano-Yolo- Marin County (NSYM) Public Health Laboratories | Chan-Zuckerberg Biohub | CZB Cliahub Consortium |
| EPI_ISL_739444 | Alameda County Public Health Lab | Chan-Zuckerberg Biohub | CZB Cliahub Consortium |
| EPI_ISL_739469 | Napa-Solano-Yolo- Marin County (NSYM) Public Health Laboratories | Chan-Zuckerberg Biohub | CZB Cliahub Consortium |
| EPI_ISL_739492 | County of San Luis Obispo Public Health Laboratory | Chan-Zuckerberg Biohub | CZB Cliahub Consortium |
| EPI_ISL_739507, EPI_ISL_739535 | Alameda County Public Health Lab | Chan-Zuckerberg Biohub | CZB Cliahub Consortium |
| EPI_ISL_739546 | Monterey County Public Health Lab | Chan-Zuckerberg Biohub | CZB Cliahub Consortium |
| EPI_ISL_739557 | Napa-Solano-Yolo- Marin County (NSYM) Public Health Laboratories | Chan-Zuckerberg Biohub | CZB Cliahub Consortium |
| EPI_ISL_739583 | Alameda County Public Health Lab | Chan-Zuckerberg Biohub | CZB Cliahub Consortium |
| EPI_ISL_739588 | County of San Luis Obispo Public Health Laboratory | Chan-Zuckerberg Biohub | CZB Cliahub Consortium |
| EPI_ISL_739623, EPI_ISL_739634 | Alameda County Public Health Lab | Chan-Zuckerberg Biohub | CZB Cliahub Consortium |
| EPI_ISL_739947, EPI_ISL_744291, EPI_ISL_744528, EPI_ISL_744531, EPI_ISL_744822, EPI_ISL_744936, EPI_ISL_744941 | Laboratoire national de santé, Microbiology, Virology | Laboratoire national de santé, Microbiology, Microbial Genomics Platform | Anke Wienecke-Baldacchino, Catherine Ragimbeau,Jessica Tapp, Fatu Djabi, Lise Pignon, Raoul Salmon, Tamir Abdelrahman |
| EPI_ISL_745554, EPI_ISL_745664 | Ginkgo Bioworks Clinical Laboratory | Utah Public Health Laboratory | Erin L. Young, Kelly Oakeson, Tara Gallagher, Michael T. Pyne, E. Susan Slechta, Melanie A. Mallory, Jeffrey B. Stevenson, Salika M. Shakir, David R. Hillyard, Malaika McKenzie-Bennett, James McGann, Jim Griffin, Keith Robison, Alex Plocik, Becky Schilling, Martha Pierson, Rebecca Littlefield, Michelle Spencer, Birgitte Simen |
| EPI_ISL_746481, EPI_ISL_746500, EPI_ISL_746509, EPI_ISL_746536, EPI_ISL_746537 | Genetica Molecular and Subdepartamento de Virologia ISP Chile | Instituto de Salud Publica de Chile | Javier Tognarelli, Barbara Parra, Loredana Arata, Jaime Lagos, Gisselle Barra, Patricia Bustos, Rodrigo Fasce, Andres Castillo, Jorge Fernandez |
| EPI_ISL_747265, EPI_ISL_747266, EPI_ISL_747268, EPI_ISL_747274, EPI_ISL_747275, EPI_ISL_747276, EPI_ISL_747281, EPI_ISL_747285, EPI_ISL_747286, EPI_ISL_747288, EPI_ISL_747291, EPI_ISL_747292, EPI_ISL_747293, EPI_ISL_747294, EPI_ISL_747298, EPI_ISL_747299, EPI_ISL_747303, EPI_ISL_747304, EPI_ISL_747305, EPI_ISL_747306, EPI_ISL_747319, EPI_ISL_747336, EPI_ISL_747337, EPI_ISL_747338, EPI_ISL_747339, EPI_ISL_747374 | | | |
| see above | Division of Emerging Infectious Diseases, Bureau of Infectious Diseases | Division of Emerging Infectious Diseases, Bureau of Infectious Diseases | Ae Kyung Park, Il-Hwan Kim, Heui Man Kim, Jeong-Min Kim, Namjoo Lee, Chaeyoung Lee, Sang Hee Woo, Eun-Jin Kim |

| | | | |
|---|---|---|---|
| | Diagnosis Control, Korea Disease Control and Prevention Agency | Diagnosis Control, Korea Disease Control and Prevention Agency | |
| EPI_ISL_748136, EPI_ISL_748137 | Department of Virus and Microbiological Special Diagnostics, Statens Serum Institut, Copenhagen, Denmark | Albertsen Lab, Department of Chemistry and Bioscience, Aalborg University, Denmark | Danish Covid-19 Genome Consortium |
| EPI_ISL_752842, EPI_ISL_752843, EPI_ISL_752904, EPI_ISL_752905, EPI_ISL_752906, EPI_ISL_752907, EPI_ISL_752908, EPI_ISL_752909, EPI_ISL_752910, EPI_ISL_752927, EPI_ISL_752946, EPI_ISL_752998, EPI_ISL_752999, EPI_ISL_753000, EPI_ISL_753001, EPI_ISL_753022, EPI_ISL_753023 | | | |
| see above | State Laboratories Division, Hawaii State Department of Health | State Laboratories Division, Hawaii State Department of Health | Pamela O'Brien, Sabrina Diemert, Drew Kuwazaki, Razvan Sultana, Edward Desmond |
| EPI_ISL_754180 | Department for Virology, Molecular Biology and Genome Research, R. G. Lugar Center for Public Health Research, National Center for Disease Control and Public Health (NCDC) of Georgia. | Department for Virology, Molecular Biology and Genome Research, R. G. Lugar Center for Public Health Research, National Center for Disease Control and Public Health (NCDC) of Georgia. | Salome Javashvili, Nino Berishvili, Tata Imnadze, Giorgi Tomashvili, Ana Papkiauri, Meri Pantsulaia, Gvantsa Brachveli, Gvantsa Chanturia, Ann Machablishvili, Nato Kotaria, Marine Murtskhvaladze, Lela Sabadze, Mari Gavashelidze, Tamar Jashiashvili, Tea Tevdoradze, Ketevan Sidamonidze, Ekaterine Khmaladze, Ekaterine Zhghenti, Roena Sukhiashvili, Mariam Zakalashvili, Lela Urushadze, Magda Dgebuadze, Davit Tsaguria, Ekaterine Zangaladze, Adam Kotorashvili, Maia Alkhazashvili, Irma Burjanadze, Anna Kasradze, Khatuna Zakhashvili, Paata Imnadze, Amiran Gamkrelidze. |
| EPI_ISL_754856 | CHU - Hôpital Cavale Blanche - Labo. de Virologie | National Reference Center for Viruses of Respiratory Infections, Institut Pasteur, Paris | Marion Barbet, Sylvie Behillil, Méline Bizard, Angela Brisebarre, Camille Capel, Etienne Simon-Lorière, Vincent Enouf, Maud Vanpeene, Sylvie van der Werf, Pilorge léa |
| EPI_ISL_754912 | Laboratory Diagnostics and Clinical Immunology of Developmental Age, Medical University of Warsaw | genXone SA, Research & Development Laboratory; The Faculty of Mathematics, Informatics and Mechanics of the University of Warsaw | Maciej Sykulski, Grzegorz Nowicki, Monika Makowska-Woniak, Jakub Grabowski, Natalia Drwska-Matelska, ukasz Krych, Micha Kaszuba, Anna Gambin, Urszula Demkow |
| EPI_ISL_754925, EPI_ISL_754958, EPI_ISL_754987, EPI_ISL_755049, EPI_ISL_755050 | California Department of Public Health | California Department of Public Health | CDPH IDLB COVIDNet |
| EPI_ISL_755118, EPI_ISL_755120, EPI_ISL_755122, EPI_ISL_755123, EPI_ISL_755125, EPI_ISL_755128, EPI_ISL_755142, EPI_ISL_755144, EPI_ISL_755214, EPI_ISL_755242, EPI_ISL_755244, EPI_ISL_755246, EPI_ISL_755247, EPI_ISL_755248, EPI_ISL_755251, EPI_ISL_755257, EPI_ISL_755258, EPI_ISL_755259, EPI_ISL_755260, EPI_ISL_755262, EPI_ISL_755266, EPI_ISL_755267, EPI_ISL_755268 | | | |
| see above | Biolab Diagnostic Laboratories | Andersen lab at Scripps Research | Issa Abu-Dayyeh, Ahmad Tibi, Lama Hussein, Lina Mohammad, Zein Naber, Amid Abdelnour with SEARCH Alliance San Diego |
| EPI_ISL_755334, EPI_ISL_755336, EPI_ISL_755337, EPI_ISL_755338, EPI_ISL_755339, EPI_ISL_755340, EPI_ISL_755341, EPI_ISL_755342, EPI_ISL_755343, EPI_ISL_755344, EPI_ISL_755345, EPI_ISL_755346, EPI_ISL_755347, EPI_ISL_755348, EPI_ISL_755349, EPI_ISL_755350, EPI_ISL_755351, EPI_ISL_755352, EPI_ISL_755353, EPI_ISL_755354, EPI_ISL_755355, EPI_ISL_755356, EPI_ISL_755357, EPI_ISL_755358, EPI_ISL_755359, EPI_ISL_755360, EPI_ISL_755361, EPI_ISL_755362, EPI_ISL_755363, EPI_ISL_755364, EPI_ISL_755365, EPI_ISL_755366, EPI_ISL_755367, EPI_ISL_755368, EPI_ISL_755369 | | | |
| see above | Maine Health and Environmental Testing Laboratory | Tewhey Lab, The Jackson Laboratory | Matluk,N., Dewey,H., Iosue,F., Barter,M., Lynch,R., Munger,H. and Tewhey,R. |
| EPI_ISL_755755, EPI_ISL_755756, EPI_ISL_755757, EPI_ISL_755758, EPI_ISL_755759, EPI_ISL_755809 | Toronto Invasive Bacterial Diseases Network | McMaster University | Allison McGeer, Patryk Aftanas, Hooman Derakhshani, Angel Li, Kuganya Nirmalarajah, Emily Panousis, Ahmed Draia, Jalees Nasir, Michael Surette, Samira Mubareka, Andrew G. McArthur |
| EPI_ISL_755966, EPI_ISL_755967, EPI_ISL_755968, EPI_ISL_756181, EPI_ISL_756182, EPI_ISL_756183, EPI_ISL_756249, EPI_ISL_756250, EPI_ISL_756251, EPI_ISL_756252 | Department of Virology and Immunology, University of Helsinki and Helsinki University Hospital, Huslab Finland | Department of Virology, Faculty of Medicine, University of Helsinki, Helsinki, Finland | Teemu Smura, Ravi Kant, Phuoc Truong, Hussein Alburkat, Hannimari Kallio-Kokko, Jenni Virtanen, Maija Suvanto, Sari Hannula, Harri Kangas, Pekka Ellonen, Olli Vapalahti |
| EPI_ISL_756374 | CUMC - CHI Bergan Mercy | Creighton University School of Medicine, Departments of Medical Microbiology and Pharmacology and Neuroscience | Michael Belshan, Morgan A. Raine, Anne V. Cheng, Christopher J. Destache, Richard V. Goering, Jacob A. Siedlik, Holly A. Stessman |
| EPI_ISL_756376 | National Institute of Health Research and Development | National Institute of Health Research and Development | Ramadhany,R; Rukminiati,Y; Agustiningsih; Adam,K; Wibowo,HA; Pawestri,HA; Subangkit;Puspa,KD; Nugraha,AA; Ikawati,HD; Pangesti,KNA; Soekarso,T; Susilarini,NK; Hariastuti,NI; Nikmah,UA;Mursinah; Febriyani,A; Herman,R; Susanti,N;Herna; Febriyanti,T; Nurhadi,M; Kurniawati,J; Kipuw,NL; Muna,F; Indalao,IL; Rizki,A; Puspandari,N; Setiawaty,Vivi |
| EPI_ISL_757287 | National Institute of Health Research and Development | National Institute of Health Research and Development | Rukminiati,Y;Agustiningsih;Adam,K;Wibowo,HA;Ramadhany,R;Pawestri,HA;Subangkit;Puspa,KD;Nugraha,AA;Ikawati,HD;Pangesti,KNA;Soekarso,T;Susilarini,NK;Hariastuti,NI;Nikmah,UA;Mursinah;Febriyani,A;Herman,R;Susanti,N;Herna;Febriyanti,T; Nurhadi,M;Kurniawati,J;Kipuw,NL;Muna,F;Indalao,IL;Rizki,A;Puspandari,N;Setiawaty,V |
| EPI_ISL_759702 | RS UNS Surakarta | Universitas Sebelas Maret (UNS), Rumah sakit Universitas Sebelas Maret (RS-UNS) Surakarta, National Institute of Health Research and Development, Indonesian Ministry of Health | Betty Suryawati, Yulia sari, hartono, maryani, Afif Avicena Gufron, Hana Apsari Pawestri, Kartika, Dewi Puspa, Hartanti Dian Ika, Arie Ardiansyah Nugraha, Vivi Setiawaty |
| EPI_ISL_760135, EPI_ISL_760157 | Division of Emerging Infectious Diseases, Bureau of Infectious Diseases | Division of Emerging Infectious Diseases, Bureau of Infectious Diseases | Ae Kyung Park, Il-Hwan Kim, Heui Man Kim, Jeong-Min Kim, Namjoo Lee, Chaeyoung Lee, Sang Hee Woo, Eun-Jin Kim |

| | | | |
|---|---|---|---|
| | Diagnosis Control, Korea Disease Control and Prevention Agency | Diagnosis Control, Korea Disease Control and Prevention Agency | |
| EPI_ISL_763064 | RS Kasih Ibu, Surakarta | Universitas Sebelas Maret (UNS); Rumah Sakit UNS (RS-UNS); National Institute of Health Research and Development, Indonesian Ministry of Health. | Betty Suryawati, Yulia Sari, Hartono, Maryani, Afif A Ghufron, Revi G H Novika, Hana A Pawestri, Kartika, Dewi Puspa, Hartanti Dian Ika, Arie A Nugraha, Vivi Setiawaty. |
| EPI_ISL_763306 | Dutch COVID-19 response team | Erasmus Medical Center | Bas Oude Munnink, Reina Sikkema, David Nieuwenhuijse, Irina Chestakova, Anne van der Linden, Marjan Boter, Emmanuelle Munger, Corine GeurtsvanKessel, Annemiek van der Eijk, Richard Molenkamp, Marion Koopmans, on behalf of the Dutch national COVID-19 response team. |
| EPI_ISL_765482, EPI_ISL_765486, EPI_ISL_765502, EPI_ISL_765503, EPI_ISL_765504, EPI_ISL_765505, EPI_ISL_765506, EPI_ISL_765507, EPI_ISL_765508 | Wadsworth Center, New York State Department.of Health | Wadsworth Center, New York State Department.of Health | Kirsten St. George, Daryl M. Lamson, Alexis Russel, Matthew Shudt, Melissa A Leisner, Jonathan Plitnick, Navjot Singh, John Kelly, Sara Griesemer, Erasmus Schneider, Erica Lasek-Nesselquist |
| EPI_ISL_765584, EPI_ISL_765585, EPI_ISL_765587, EPI_ISL_765588, EPI_ISL_765590 | Brigham and Womens Hospital | Infectious Disease Program, Broad Institute of Harvard and MIT | Lemieux,J.E., Siddle,K.J., Shaw,B., Adams,G., Pierce,V., Turbett,S., Anahtar,M., Branda,J., Slater,D., Harris,J., Lin,A.E., Gladden-Young,A., Lagerborg,K., Rudy,M., DeRuff,K., Carter,A., Normandin,E., Bauer,M., Reilly,S., Tomkins-Tinch,C., Loreth,C., Chaluvadi,S., Neumann,A., Cusick,C., Chapman,S.B., Gnirke,A., Flowers,K., Cerrato,F., Birren,B.W., Gallagher,G., Smole,S., Park,D.J., MacInnis,B.L., Ryan,E., LaRocque,R., Rosenberg,E. and Sabeti,P.C. |
| EPI_ISL_765722, EPI_ISL_765726, EPI_ISL_765728 | Massachusetts General Hospital | Infectious Disease Program, Broad Institute of Harvard and MIT | Lemieux,J.E., Siddle,K.J., Shaw,B., Adams,G., Pierce,V., Turbett,S., Anahtar,M., Branda,J., Slater,D., Harris,J., Lin,A.E., Gladden-Young,A., Lagerborg,K., Rudy,M., DeRuff,K., Carter,A., Normandin,E., Bauer,M., Reilly,S., Tomkins-Tinch,C., Loreth,C., Chaluvadi,S., Neumann,A., Cusick,C., Chapman,S.B., Gnirke,A., Flowers,K., Cerrato,F., Birren,B.W., Gallagher,G., Smole,S., Park,D.J., MacInnis,B.L., Ryan,E., LaRocque,R., Rosenberg,E. and Sabeti,P.C. |
| EPI_ISL_766592, EPI_ISL_766593, EPI_ISL_766625, EPI_ISL_766626, EPI_ISL_766627, EPI_ISL_766628, EPI_ISL_766629 | Klinisk mikrobiologi | The Public Health Agency of Sweden | Department of Microbiology, The Public Health Agency of Sweden |
| EPI_ISL_768765, EPI_ISL_768767, EPI_ISL_768768 | AIID | Irish Coronavirus Sequencing Consortium - National Virus Reference Laboratory | Michael Carr, Gabriel Gonzalez, Alejandro Abner Garcia Leon, Patrick Mallon |
| EPI_ISL_769991 | Laboratorio Clinico Labin | Inciensa, Instituto Costarricense de Investigación y Enseñanza en Nutrición y Salud | Francisco Duarte, Hebleen Porras, Claudio Soto-Garita, Estela Cordero, Adriana Godínez, Melany Calderón & Mariel López |
| EPI_ISL_769994 | Area De Salud Catedral Noreste | Inciensa, Instituto Costarricense de Investigación y Enseñanza en Nutrición y Salud | Francisco Duarte, Hebleen Porras, Claudio Soto-Garita, Estela Cordero, Adriana Godínez, Melany Calderón & Mariel López |
| EPI_ISL_770028 | Hospital Dr. Raul Blanco Cervantes | Inciensa, Instituto Costarricense de Investigación y Enseñanza en Nutrición y Salud | Francisco Duarte, Hebleen Porras, Claudio Soto-Garita, Estela Cordero, Adriana Godínez, Melany Calderón & Mariel López |
| EPI_ISL_770035 | E. Gulbja Laboratorija | Latvian Biomedical Research and Study Centre | Ivars Silamielis, Kaspars Megnis, Monta Ustinova, Jnis Pjalkovskis, ikita Zrelovs, Vita Rovte, Mikus Gavars, Dmitrijs Perminovs, Uga Dumpis, Jnis Kloviš |
| EPI_ISL_770039, EPI_ISL_770042 | Centrl Laboratorija | Latvian Biomedical Research and Study Centre | Ivars Silamielis, Kaspars Megnis, Monta Ustinova, Jnis Pjalkovskis, ikita Zrelovs, Vita Rovte, Stella Lapia, Jana Oste, Marta Priedte, Uga Dumpis, Jnis Kloviš |
| EPI_ISL_770043, EPI_ISL_770044, EPI_ISL_770046 | E. Gulbja Laboratorija | Latvian Biomedical Research and Study Centre | Ivars Silamielis, Kaspars Megnis, Monta Ustinova, Jnis Pjalkovskis, ikita Zrelovs, Vita Rovte, Mikus Gavars, Dmitrijs Perminovs, Uga Dumpis, Jnis Kloviš |
| EPI_ISL_770048, EPI_ISL_770049, EPI_ISL_770052 | Centrl Laboratorija | Latvian Biomedical Research and Study Centre | Ivars Silamielis, Kaspars Megnis, Monta Ustinova, Jnis Pjalkovskis, ikita Zrelovs, Vita Rovte, Stella Lapia, Jana Oste, Marta Priedte, Uga Dumpis, Jnis Kloviš |
| EPI_ISL_770053 | E. Gulbja Laboratorija | Latvian Biomedical Research and Study Centre | Ivars Silamielis, Kaspars Megnis, Monta Ustinova, Jnis Pjalkovskis, ikita Zrelovs, Vita Rovte, Mikus Gavars, Dmitrijs Perminovs, Uga Dumpis, Jnis Kloviš |
| EPI_ISL_770054 | Centrl Laboratorija | Latvian Biomedical Research and Study Centre | Ivars Silamielis, Kaspars Megnis, Monta Ustinova, Jnis Pjalkovskis, ikita Zrelovs, Vita Rovte, Stella Lapia, Jana Oste, Marta Priedte, Uga Dumpis, Jnis Kloviš |
| EPI_ISL_770055 | E. Gulbja Laboratorija | Latvian Biomedical Research and Study Centre | Ivars Silamielis, Kaspars Megnis, Monta Ustinova, Jnis Pjalkovskis, ikita Zrelovs, Vita Rovte, Mikus Gavars, Dmitrijs Perminovs, Uga Dumpis, Jnis Kloviš |
| EPI_ISL_770056 | Centrl Laboratorija | Latvian Biomedical Research and Study Centre | Ivars Silamielis, Kaspars Megnis, Monta Ustinova, Jnis Pjalkovskis, ikita Zrelovs, Vita Rovte, Stella Lapia, Jana Oste, Marta Priedte, Uga Dumpis, Jnis Kloviš |
| EPI_ISL_770057 | E. Gulbja Laboratorija | Latvian Biomedical Research and Study Centre | Ivars Silamielis, Kaspars Megnis, Monta Ustinova, Jnis Pjalkovskis, ikita Zrelovs, Vita Rovte, Mikus Gavars, Dmitrijs Perminovs, Uga Dumpis, Jnis Kloviš |
| EPI_ISL_770058 | Centrl Laboratorija | Latvian Biomedical Research and Study Centre | Ivars Silamielis, Kaspars Megnis, Monta Ustinova, Jnis Pjalkovskis, ikita Zrelovs, Vita Rovte, Stella Lapia, Jana Oste, Marta Priedte, Uga Dumpis, Jnis Kloviš |
| EPI_ISL_770060 | Latvijas Infektoloijas centrs | Latvian Biomedical Research and Study Centre | Ivars Silamielis, Kaspars Megnis, Monta Ustinova, Jnis Pjalkovskis, ikita Zrelovs, Vita Rovte, Jeena Storoženko, Tatjana Kolupajeva, Oksana Savicka, Uga Dumpis, Jnis Kloviš |
| EPI_ISL_770720, | ZOTZ KLIMAS MVZ | Center of Medical | Maximilian Damagnez, Alexander Dilthey, Ashley-Jane Duplessis, Patrick Finzer, Katrin Hoffmann, Torsten Houwaart, Lisanna Hülse, Malte Kohns Vasconcelos, Marek Korencak, Nadine Lübke, Jessica Nicolai, Klaus Pfeffer, Daniel Strelow, Jörg |

| | | | |
|---|---|---|---|
| EPI_ISL_770721, EPI_ISL_770722, EPI_ISL_770723, EPI_ISL_770724 | Düsseldorf-Centrum GbR ÜBAG für Labormedizin, Genetik, Zytologie, Pathologie | Microbiology, Virology, and Hospital Hygiene, University of Duesseldorf | Timm, Andreas Walker, Tobias Wienemann, Rainer Zotz |
| EPI_ISL_776982, EPI_ISL_776983, EPI_ISL_776984, EPI_ISL_776985, EPI_ISL_776986, EPI_ISL_776987, EPI_ISL_776988, EPI_ISL_776989, EPI_ISL_776990, EPI_ISL_776991, EPI_ISL_776992, EPI_ISL_776993, EPI_ISL_776995, EPI_ISL_776996, EPI_ISL_778704, EPI_ISL_778705, EPI_ISL_778706, EPI_ISL_778707, EPI_ISL_778708, EPI_ISL_778709, EPI_ISL_778710, EPI_ISL_778711, EPI_ISL_778712, EPI_ISL_778713, EPI_ISL_778714, EPI_ISL_778715, EPI_ISL_778716, EPI_ISL_778717, EPI_ISL_778718, EPI_ISL_778719, EPI_ISL_778720, EPI_ISL_778721 | | | |
| see above | Istituto Zooprofilattico Sperimentale del Mezzogiorno | TIGEM | Patrizia Annunziata, Andrea Ballabio, Valentina Bouche, Davide Cacchiarelli (CorrespAuthor), Pellegrino Cerino, Chiara Colantuono, Lucio Di Filippo, Antonio Grimaldi, Antonio Limone, Gabriella Loconte, Anna Manfredi, Francesco Panariello, Biancamaria Pierri, Marcello Salvi, Lucia Vassallo |
| EPI_ISL_779566 | Microbiological Diagnostic Unit - Public Health Laboratory (MDU-PHL) | MDU-PHL | Seemann T., Sait, M.L., Sherry, N.L. |
| EPI_ISL_789781, EPI_ISL_789791, EPI_ISL_789830, EPI_ISL_789842, EPI_ISL_789847, EPI_ISL_789853, EPI_ISL_789856, EPI_ISL_789866, EPI_ISL_789872, EPI_ISL_789879, EPI_ISL_789884, EPI_ISL_789891, EPI_ISL_789892, EPI_ISL_789898, EPI_ISL_789910, EPI_ISL_789918, EPI_ISL_789932, EPI_ISL_789939, EPI_ISL_789940, EPI_ISL_789947, EPI_ISL_789948, EPI_ISL_789957, EPI_ISL_789958, EPI_ISL_789960, EPI_ISL_789967, EPI_ISL_789971, EPI_ISL_789973, EPI_ISL_789974, EPI_ISL_789979, EPI_ISL_789980, EPI_ISL_789984, EPI_ISL_789985, EPI_ISL_789986, EPI_ISL_789989, EPI_ISL_790029, EPI_ISL_790031, EPI_ISL_790047, EPI_ISL_790049, EPI_ISL_790050, EPI_ISL_790051, EPI_ISL_790052, EPI_ISL_790053, EPI_ISL_790054, EPI_ISL_790055, EPI_ISL_790056, EPI_ISL_790057, EPI_ISL_790058, EPI_ISL_790059, EPI_ISL_790060, EPI_ISL_790061, EPI_ISL_790062, EPI_ISL_790063, EPI_ISL_790064, EPI_ISL_790065, EPI_ISL_790066, EPI_ISL_790067, EPI_ISL_790068, EPI_ISL_790069, EPI_ISL_790070, EPI_ISL_790076, EPI_ISL_790079, EPI_ISL_790080, EPI_ISL_790081, EPI_ISL_790084, EPI_ISL_790085, EPI_ISL_790086, EPI_ISL_790092, EPI_ISL_790095, EPI_ISL_790097, EPI_ISL_790098, EPI_ISL_790101, EPI_ISL_790102, EPI_ISL_790103, EPI_ISL_790112, EPI_ISL_790113, EPI_ISL_790115, EPI_ISL_790116, EPI_ISL_790117, EPI_ISL_790118, EPI_ISL_790119, EPI_ISL_790120, EPI_ISL_790121, EPI_ISL_790122, EPI_ISL_790123, EPI_ISL_790124, EPI_ISL_790125, EPI_ISL_790126, EPI_ISL_790127, EPI_ISL_790128, EPI_ISL_790129, EPI_ISL_790130, EPI_ISL_790131, EPI_ISL_790132, EPI_ISL_790134, EPI_ISL_790137, EPI_ISL_790138, EPI_ISL_790139, EPI_ISL_790141, EPI_ISL_790142, EPI_ISL_790143, EPI_ISL_790144, EPI_ISL_790145, EPI_ISL_790146, EPI_ISL_790148, EPI_ISL_790150, EPI_ISL_790151, EPI_ISL_790154, EPI_ISL_790155, EPI_ISL_790156, EPI_ISL_790157, EPI_ISL_790159, EPI_ISL_790160, EPI_ISL_790161, EPI_ISL_790162, EPI_ISL_790163, EPI_ISL_790164, EPI_ISL_790165, EPI_ISL_790166, EPI_ISL_790168, EPI_ISL_790171, EPI_ISL_790187, EPI_ISL_790195, EPI_ISL_790200, EPI_ISL_790201, EPI_ISL_790247, EPI_ISL_790252, EPI_ISL_790263 | | | |
| see above | Houston Methodist Hospital | Houston Methodist Hospital | S. Wesley Long, Randall J. Olsen, Paul A. Christensen, David W. Bernard, James J. Davis, Maulik Shukla, Marcus Nguyen, Matthew Ojeda Saavedra, Prasanti Yerramilli, Layne Pruitt, Sishir Subedi, Heather Hendrickson, and James M. Musser |
| EPI_ISL_792023 | Plateforme COVID IDF | National Reference Center for Viruses of Respiratory Infections, Institut Pasteur, Paris | Marion Barbet, Sylvie Behillil, Méline Bizard, Angela Brisebarre, Camille Capel, Etienne Simon-Lorière, Vincent Enouf, Maud Vanpeene, Sylvie van der Werf,Jacques Fourgeaud |
| EPI_ISL_792612, EPI_ISL_792613 | LACEN-PB | Laboratory of Respiratory Viruses and Measles, Oswaldo Cruz Institute, FIOCRUZ | Paola Resende, Luciana Appolinario, Fernando Motta, Anna Carolina Paixao, Ana Carolina Mendonca, João Felipe Bezerra, Romero Henrique Teixeira de Vasconcelos, Dalane Loudal Florentino Teixeira, Thiago Franco de Oliveira Carneiro, Marilda Siqueira |
| EPI_ISL_792687, EPI_ISL_792700 | The National Institute of Public Health | State Veterinary Institute Prague | Nagy,A;Jirincova,H;Trnka,D;Vecerova,J |
| EPI_ISL_796028, EPI_ISL_796029, EPI_ISL_796033, EPI_ISL_796039, EPI_ISL_796040, EPI_ISL_796041, EPI_ISL_796042, EPI_ISL_796054 | Institute of Virology, University of Cologne | Institute of Virology, University of Cologne | Saleta Sierra, Gibran Rubio, Zevanya Tessalonica, Dominik Aschenmeier, Eva Heger, Elena Knops, Rolf Kaiser, Maximilian Damagnez, Andreas Walker, Jörg Timm, Alexander Dilthey, Martin Däumer, Alex Thielen |
| EPI_ISL_801387, EPI_ISL_801388 | Laboratorio de Ecologia de Doencas Transmissiveis na Amazonia, Instituto Leonidas e Maria Deane - Fiocruz Amazonia | Laboratorio de Ecologia de Doencas Transmissiveis na Amazonia, Instituto Leonidas e Maria Deane - Fiocruz Amazonia | Valdinete Nascimento, Victor Souza, André Corado, Fernanda Nascimento, George Silva, Ágatha Costa, Debora Duarte, Luciana Gonçalves, Maria Júlia Brandão, Michele Jesus, Felipe Naveca |
| EPI_ISL_801400, EPI_ISL_801401 | Laboratório Central de Saúde Pública do Amazonas - LACEN-AM | Laboratorio de Ecologia de Doencas Transmissiveis na Amazonia, Instituto Leonidas e Maria Deane - Fiocruz Amazonia | Valdinete Nascimento, Victor Souza, André Corado, Fernanda Nascimento, George Silva, Ágatha Costa, Debora Duarte, Luciana Gonçalves, Maria Júlia Brandão, Michele Jesus, Felipe Naveca |
| EPI_ISL_801871, EPI_ISL_801873, EPI_ISL_801883, EPI_ISL_801899, EPI_ISL_801900, EPI_ISL_801934, EPI_ISL_802238, EPI_ISL_802239, EPI_ISL_802240, EPI_ISL_802241, EPI_ISL_802242, EPI_ISL_802243, EPI_ISL_802244, EPI_ISL_802245, EPI_ISL_802246, EPI_ISL_802247, EPI_ISL_802248, EPI_ISL_802249, EPI_ISL_802250, EPI_ISL_802251, EPI_ISL_802252, EPI_ISL_802253, EPI_ISL_802254, EPI_ISL_802255 | | | |
| see above | MSHS Clinical Microbiology Laboratories | MSHS Pathogen Surveillance Program | Ana S. Gonzalez-Reiche, Hala Alshammary, Mitchell J. Sullivan, Brianne Ciferri, Ajay Obla, Angela Amoako, Mahmoud Awawda, Elena Hirsch, Ashley S. Salimbangon, Levy Sominsky, Katherine Beach, Kayla Russo, Charles Gleason, Shelcie Fabre, Giulio Kleiner, Zenab Khan, Bremy Alburquerque, Adriana van de Guchte, Komal Srivastava, Matthew M. Hernandez, Jayeeta Dutta, Denise Jurczyszak, Emily Ferreri, Rachel Chernet, Nancy Francoeur, Betsaida Salom Melo, Irina Oussenko, Gintaras Deikus, Juan Soto, Shwetha Hara Sridhar, Ying-Chih Wang, Kathryn Twyman, Andrew Kasarskis, Deena R. Altman, Robert Sebra, Adolfo García-Sastre, Marta Luksza, Gopi Patel, Sarah Schaefer, Melissa Gitman, Michael D. Nowak, Alberto Paniz-Mondolfi, Emilia Mia Sordillo, Viviana Simon, Harm van Bakel |
| EPI_ISL_803142, EPI_ISL_803143, EPI_ISL_803144, EPI_ISL_803145, EPI_ISL_803146, EPI_ISL_803147, EPI_ISL_803148, EPI_ISL_803149, EPI_ISL_803150, EPI_ISL_803151, EPI_ISL_803152, EPI_ISL_803153, EPI_ISL_803154, EPI_ISL_803155, EPI_ISL_803156, EPI_ISL_803157, EPI_ISL_803158, EPI_ISL_803159, EPI_ISL_803160, EPI_ISL_803161, EPI_ISL_803162, EPI_ISL_803163, EPI_ISL_803164, EPI_ISL_803165, EPI_ISL_803166, EPI_ISL_803167, EPI_ISL_803168, EPI_ISL_803169, EPI_ISL_803170, EPI_ISL_803171, EPI_ISL_803172, EPI_ISL_803173, EPI_ISL_803174, EPI_ISL_803175, EPI_ISL_803176, EPI_ISL_803177, EPI_ISL_803178, EPI_ISL_803179, EPI_ISL_803180, EPI_ISL_803181, EPI_ISL_803182, EPI_ISL_803183, EPI_ISL_803184, EPI_ISL_803185, EPI_ISL_803186, EPI_ISL_803187, EPI_ISL_803188, EPI_ISL_803189, EPI_ISL_803190, EPI_ISL_803191, EPI_ISL_803192, EPI_ISL_803193, EPI_ISL_803194, EPI_ISL_803195, EPI_ISL_803229, EPI_ISL_803230, EPI_ISL_803231, EPI_ISL_803238, EPI_ISL_803239, EPI_ISL_803240, EPI_ISL_803241, EPI_ISL_803242, EPI_ISL_803243, EPI_ISL_803244, EPI_ISL_803245, EPI_ISL_803246, EPI_ISL_803247, EPI_ISL_803341, EPI_ISL_803342, EPI_ISL_803343, EPI_ISL_803344, EPI_ISL_803345, EPI_ISL_803346, EPI_ISL_803347, EPI_ISL_803348, EPI_ISL_803349, EPI_ISL_803350, EPI_ISL_803351, EPI_ISL_803494, EPI_ISL_803495, EPI_ISL_803496, EPI_ISL_803497, EPI_ISL_803498, EPI_ISL_803499, EPI_ISL_803500, EPI_ISL_803501, EPI_ISL_803502, EPI_ISL_803503, EPI_ISL_803504, EPI_ISL_803505, EPI_ISL_803506, EPI_ISL_803509, EPI_ISL_803510, EPI_ISL_803511, EPI_ISL_803523, EPI_ISL_803524, EPI_ISL_803525, EPI_ISL_803526, EPI_ISL_803527, EPI_ISL_803528, EPI_ISL_803529, EPI_ISL_803538, EPI_ISL_803539, EPI_ISL_803540, EPI_ISL_803742, EPI_ISL_803743, EPI_ISL_803744, EPI_ISL_803745, EPI_ISL_803746, EPI_ISL_803747 | | | |
| see above | Wisconsin State Laboratory of Hygiene Communicable Disease Division | Wisconsin State Laboratory of Hygiene Communicable Disease Division | Kelsey R. Florek, Abigail C. Shockey |
| EPI_ISL_804964, EPI_ISL_804968, EPI_ISL_804969, EPI_ISL_804971, EPI_ISL_804972, EPI_ISL_804973, EPI_ISL_804974, EPI_ISL_804975, EPI_ISL_804976, EPI_ISL_804978, EPI_ISL_804979 | | | |
| see above | Hospital Comarcal de Melilla | Instituto de Salud Carlos III | Iglesias-Caballero, M. Molinero Calamita, M. González-Esguevillas, M. Camarero, S. Pozo, F. Casas, I. Jiménez, P. Jiménez, M. Zaballos, A. Monzón, S. Varona, S. Juliá, M. Cuesta, I, J. López |
| EPI_ISL_806899, EPI_ISL_806900, EPI_ISL_806901, EPI_ISL_806902, EPI_ISL_806903, EPI_ISL_806904, EPI_ISL_806905, EPI_ISL_806906, EPI_ISL_806907, EPI_ISL_806908, EPI_ISL_806909, EPI_ISL_806910, EPI_ISL_806911, EPI_ISL_806912, EPI_ISL_806913, EPI_ISL_806914, EPI_ISL_806915, EPI_ISL_806916 | | | |
| see above | Washington State Department of Health | Seattle Flu Study | Deborah A. Nickerson, Chris D. Frazar, Jover Lee, Benjamin Pelle, Matthew Richardson, Amanda Adler, Elisabeth Brandstetter, Peter D. Han, Kairsten Fay, Misja Ilcisin, Kirsten Lacombe, Thomas R. Sibley, Melissa Truong, Caitlin R. Wolf, Romesh Gautom, Geoff Melly, Brian Hiatt, Philip Dykema, Scott Lindquist, Michael Boeckh, Janet A. Englund, Michael Famulare, Barry R. Lutz, Mark J. Rieder, Lea M. Starita, Matthew Thompson, Helen Y. Chu, Jay Shendure, Trevor Bedford |
| EPI_ISL_810980, EPI_ISL_811013 | MRCG at LSHTM Genomics lab | MRCG at LSHTM Genomics lab | Abdul Karim sesay, Abdoulie Kanteh, Jarra Manneh, Mariama Kujabi, Bakary Sanyang |
| EPI_ISL_812688, EPI_ISL_812689, EPI_ISL_812690, EPI_ISL_812692, EPI_ISL_812694, EPI_ISL_812699, EPI_ISL_812700, EPI_ISL_812702, EPI_ISL_812704, EPI_ISL_812705, EPI_ISL_812706, EPI_ISL_812707, EPI_ISL_812708, EPI_ISL_812709, EPI_ISL_812710, EPI_ISL_812711 | | | |
| see above | United States Air Force School of Aerospace Medicine | United States Air Force School of Aerospace Medicine | Anthony Fries, Jennifer Meyer, Amanda Javorina, Sarah Purves, William Gruner, Clarise Starr, Elizabeth Macias |

| | | | |
|---|---|---|---|
| EPI_ISL_813052 | University of Birmingham | COVID-19 Genomics UK (COG-UK) Consortium | Institute of Microbiology, University of Birmingham: Claire McMurray, Joanne Stockton, Samuel Nicholls, Radoslaw Poplawski, Will Rowe, Josh Quick, Nicholas Loman. University of Birmingham Testing Laboratory: Celina M Whalley, Andrew Bosworth, Charlotte Poxon, Kasun Wanigasooriya, Oliver Pickles, Mike Kidd, Alex Richter, Andrew D Beggs PHE Heartlands Lab: Husam Osman, Andrew Bosworth. Queen Elizabeth Hospital: Anna Casey |
| EPI_ISL_813154, EPI_ISL_813155, EPI_ISL_813156, EPI_ISL_813157, EPI_ISL_813158, EPI_ISL_813159, EPI_ISL_813162, EPI_ISL_813163, EPI_ISL_813164, EPI_ISL_813165 | Department of Pathology, University of Cambridge | COVID-19 Genomics UK (COG-UK) Consortium | Aminu S. Jahun, Yasmin Chaudhry, Grant Hall, Iliana Georgana, Myra Hosmillo, Martin D. Curran, Malte Pinckert, Surendra Parmar, Ian Goodfellow |
| EPI_ISL_815356 | Centogene | Centogene | Peter Bauer, Krishna Kumar Kandaswamy, Vivi Hue-Trang Lieu |
| EPI_ISL_824487, EPI_ISL_824488, EPI_ISL_824489, EPI_ISL_824490, EPI_ISL_824491 | Hospital Universitari Vall d'Hebron - Vall d'Hebron Institut de Rercerca | Hospital Universitari Vall d'Hebron | Cristina Andrés, Maria Piñana, Josep F Abril, Damir Garcia-Cehic, Ariadna Rando, Juliana Esperalba, Maria Gema Codina, Carla Castillo, Maria Carmen Martin, Tomàs Pumarola, Josep Quer, Andrés Antón |
| EPI_ISL_825028, EPI_ISL_825029, EPI_ISL_825030, EPI_ISL_825031, EPI_ISL_825032 | Infectious Diseases, NC SLPH COVID-19 Response Team | Infectious Diseases, NC SLPH COVID-19 Response Team | Miller,M.C., Chase,K. |
| EPI_ISL_825553 | Respiratory Virus Unit, National Infection Service, Public Health England | COVID-19 Genomics UK (COG-UK) Consortium | PHE Covid Sequencing Team |
| EPI_ISL_826834, EPI_ISL_826836 | INSPI-CRN DE INFLUENZA Y OTROS VIRUS RESPIRATORIOS | Instituto de Salud Publica de Chile | Javier Tognarelli, Barbara Parra, Loredana Arata, Jaime Lagos, Gisselle Barra, Alfredo Bruno, Domenica de Mora, Solon Narvaez, Jimmy Garcez, Michelle Paez, Martiza Olmedo, Manuel Gonzalez, Patricia Bustos, Rodrigo Fasce, Andres Castillo, Jorge Fernandez |
| EPI_ISL_826898, EPI_ISL_826899, EPI_ISL_826900, EPI_ISL_826901, EPI_ISL_826902 | deCODE genetics | deCODE genetics | Daniel F Gudbjartsson; Agnar Helgason; Hakon Jonsson; Olafur T Magnusson; Pall Melsted; Gudmundur L Norddahl; Jona Saemundsdottir; Asgeir Sigurdsson; Patrick Sulem; Arna B Agustsdottir; Hannes Eggertsson; Berglind Eiriksdottir; Run Fridriksdottir; Elisabet E Gardarsdottir; Gudmundur Georgsson; Olafia S Gretarsdottir; Kjartan R Gudmundsson; Thora R Gunnarsdottir; Arnaldur Gylfason; Hilma Holm; Brynjar O Jensson; Aslaug Jonasdottir; Kamilla S Josefsdottir; Thordur Kristjansson; Droplaug N Magnusdottir; Solvi Rognvaldsson; Louise le Roux; Gudrun Sigmundsdottir; Gardar Sveinbjornsson; Kristin E Sveinsdottir; Maney Sveinsdottir; Emil A Thorarensen; Bjarni Thorbjornsson; Gisli Masson; Ingileif Jonsdottir; Alma Moller; Thorolfur Gudnason; Karl G Kristinsson; Unnur Thorsteinsdottir; Kari Stefansson |
| EPI_ISL_826916 | The National University Hospital of Iceland | deCODE genetics | Daniel F Gudbjartsson; Agnar Helgason; Hakon Jonsson; Olafur T Magnusson; Pall Melsted; Gudmundur L Norddahl; Jona Saemundsdottir; Asgeir Sigurdsson; Patrick Sulem; Arna B Agustsdottir; Hannes Eggertsson; Berglind Eiriksdottir; Run Fridriksdottir; Elisabet E Gardarsdottir; Gudmundur Georgsson; Olafia S Gretarsdottir; Kjartan R Gudmundsson; Thora R Gunnarsdottir; Arnaldur Gylfason; Hilma Holm; Brynjar O Jensson; Aslaug Jonasdottir; Kamilla S Josefsdottir; Thordur Kristjansson; Droplaug N Magnusdottir; Solvi Rognvaldsson; Louise le Roux; Gudrun Sigmundsdottir; Gardar Sveinbjornsson; Kristin E Sveinsdottir; Maney Sveinsdottir; Emil A Thorarensen; Bjarni Thorbjornsson; Gisli Masson; Ingileif Jonsdottir; Alma Moller; Thorolfur Gudnason; Karl G Kristinsson; Unnur Thorsteinsdottir; Kari Stefansson |
| EPI_ISL_826922, EPI_ISL_826923, EPI_ISL_826924, EPI_ISL_827060, EPI_ISL_827061, EPI_ISL_827062, EPI_ISL_827072, EPI_ISL_827073, EPI_ISL_827338, EPI_ISL_827341, EPI_ISL_827639, EPI_ISL_827643, EPI_ISL_827790, EPI_ISL_827791, EPI_ISL_828210, EPI_ISL_828254, EPI_ISL_828304, EPI_ISL_828308 | | | |
| see above | deCODE genetics | deCODE genetics | Daniel F Gudbjartsson; Agnar Helgason; Hakon Jonsson; Olafur T Magnusson; Pall Melsted; Gudmundur L Norddahl; Jona Saemundsdottir; Asgeir Sigurdsson; Patrick Sulem; Arna B Agustsdottir; Hannes Eggertsson; Berglind Eiriksdottir; Run Fridriksdottir; Elisabet E Gardarsdottir; Gudmundur Georgsson; Olafia S Gretarsdottir; Kjartan R Gudmundsson; Thora R Gunnarsdottir; Arnaldur Gylfason; Hilma Holm; Brynjar O Jensson; Aslaug Jonasdottir; Kamilla S Josefsdottir; Thordur Kristjansson; Droplaug N Magnusdottir; Solvi Rognvaldsson; Louise le Roux; Gudrun Sigmundsdottir; Gardar Sveinbjornsson; Kristin E Sveinsdottir; Maney Sveinsdottir; Emil A Thorarensen; Bjarni Thorbjornsson; Gisli Masson; Ingileif Jonsdottir; Alma Moller; Thorolfur Gudnason; Karl G Kristinsson; Unnur Thorsteinsdottir; Kari Stefansson |
| EPI_ISL_828326, EPI_ISL_828327 | The National University Hospital of Iceland | deCODE genetics | Daniel F Gudbjartsson; Agnar Helgason; Hakon Jonsson; Olafur T Magnusson; Pall Melsted; Gudmundur L Norddahl; Jona Saemundsdottir; Asgeir Sigurdsson; Patrick Sulem; Arna B Agustsdottir; Hannes Eggertsson; Berglind Eiriksdottir; Run Fridriksdottir; Elisabet E Gardarsdottir; Gudmundur Georgsson; Olafia S Gretarsdottir; Kjartan R Gudmundsson; Thora R Gunnarsdottir; Arnaldur Gylfason; Hilma Holm; Brynjar O Jensson; Aslaug Jonasdottir; Kamilla S Josefsdottir; Thordur Kristjansson; Droplaug N Magnusdottir; Solvi Rognvaldsson; Louise le Roux; Gudrun Sigmundsdottir; Gardar Sveinbjornsson; Kristin E Sveinsdottir; Maney Sveinsdottir; Emil A Thorarensen; Bjarni Thorbjornsson; Gisli Masson; Ingileif Jonsdottir; Alma Moller; Thorolfur Gudnason; Karl G Kristinsson; Unnur Thorsteinsdottir; Kari Stefansson |
| EPI_ISL_828534 | deCODE genetics | deCODE genetics | Daniel F Gudbjartsson; Agnar Helgason; Hakon Jonsson; Olafur T Magnusson; Pall Melsted; Gudmundur L Norddahl; Jona Saemundsdottir; Asgeir Sigurdsson; Patrick Sulem; Arna B Agustsdottir; Hannes Eggertsson; Berglind Eiriksdottir; Run Fridriksdottir; Elisabet E Gardarsdottir; Gudmundur Georgsson; Olafia S Gretarsdottir; Kjartan R Gudmundsson; Thora R Gunnarsdottir; Arnaldur Gylfason; Hilma Holm; Brynjar O Jensson; Aslaug Jonasdottir; Kamilla S Josefsdottir; Thordur Kristjansson; Droplaug N Magnusdottir; Solvi Rognvaldsson; Louise le Roux; Gudrun Sigmundsdottir; Gardar Sveinbjornsson; Kristin E Sveinsdottir; Maney Sveinsdottir; Emil A Thorarensen; Bjarni Thorbjornsson; Gisli Masson; Ingileif Jonsdottir; Alma Moller; Thorolfur Gudnason; Karl G Kristinsson; Unnur Thorsteinsdottir; Kari Stefansson |
| EPI_ISL_828641, EPI_ISL_828775, EPI_ISL_828776 | The National University Hospital of Iceland | deCODE genetics | Daniel F Gudbjartsson; Agnar Helgason; Hakon Jonsson; Olafur T Magnusson; Pall Melsted; Gudmundur L Norddahl; Jona Saemundsdottir; Asgeir Sigurdsson; Patrick Sulem; Arna B Agustsdottir; Hannes Eggertsson; Berglind Eiriksdottir; Run Fridriksdottir; Elisabet E Gardarsdottir; Gudmundur Georgsson; Olafia S Gretarsdottir; Kjartan R Gudmundsson; Thora R Gunnarsdottir; Arnaldur Gylfason; Hilma Holm; Brynjar O Jensson; Aslaug Jonasdottir; Kamilla S Josefsdottir; Thordur Kristjansson; Droplaug N Magnusdottir; Solvi Rognvaldsson; Louise le Roux; Gudrun Sigmundsdottir; Gardar Sveinbjornsson; Kristin E Sveinsdottir; Maney Sveinsdottir; Emil A Thorarensen; Bjarni Thorbjornsson; Gisli Masson; Ingileif Jonsdottir; Alma Moller; Thorolfur Gudnason; Karl G Kristinsson; Unnur Thorsteinsdottir; Kari Stefansson |
| EPI_ISL_828777, EPI_ISL_828778 | deCODE genetics | deCODE genetics | Daniel F Gudbjartsson; Agnar Helgason; Hakon Jonsson; Olafur T Magnusson; Pall Melsted; Gudmundur L Norddahl; Jona Saemundsdottir; Asgeir Sigurdsson; Patrick Sulem; Arna B Agustsdottir; Hannes Eggertsson; Berglind Eiriksdottir; Run Fridriksdottir; Elisabet E Gardarsdottir; Gudmundur Georgsson; Olafia S Gretarsdottir; Kjartan R Gudmundsson; Thora R Gunnarsdottir; Arnaldur Gylfason; Hilma Holm; Brynjar O Jensson; Aslaug Jonasdottir; Kamilla S Josefsdottir; Thordur Kristjansson; Droplaug N Magnusdottir; Solvi Rognvaldsson; Louise le Roux; Gudrun Sigmundsdottir; Gardar Sveinbjornsson; Kristin E Sveinsdottir; Maney Sveinsdottir; Emil A Thorarensen; Bjarni Thorbjornsson; Gisli Masson; Ingileif Jonsdottir; Alma Moller; Thorolfur Gudnason; Karl G Kristinsson; Unnur Thorsteinsdottir; Kari Stefansson |
| EPI_ISL_828779 | The National University Hospital of Iceland | deCODE genetics | Daniel F Gudbjartsson; Agnar Helgason; Hakon Jonsson; Olafur T Magnusson; Pall Melsted; Gudmundur L Norddahl; Jona Saemundsdottir; Asgeir Sigurdsson; Patrick Sulem; Arna B Agustsdottir; Hannes Eggertsson; Berglind Eiriksdottir; Run Fridriksdottir; Elisabet E Gardarsdottir; Gudmundur Georgsson; Olafia S Gretarsdottir; Kjartan R Gudmundsson; Thora R Gunnarsdottir; Arnaldur Gylfason; Hilma Holm; Brynjar O Jensson; Aslaug Jonasdottir; Kamilla S Josefsdottir; Thordur Kristjansson; Droplaug N Magnusdottir; Solvi Rognvaldsson; Louise le Roux; Gudrun Sigmundsdottir; Gardar Sveinbjornsson; Kristin E Sveinsdottir; Maney Sveinsdottir; Emil A Thorarensen; Bjarni Thorbjornsson; Gisli Masson; Ingileif Jonsdottir; Alma Moller; Thorolfur Gudnason; Karl G Kristinsson; Unnur Thorsteinsdottir; Kari Stefansson |
| EPI_ISL_828781, EPI_ISL_828782, EPI_ISL_828783, EPI_ISL_828784, EPI_ISL_829025, EPI_ISL_829026, EPI_ISL_829031, EPI_ISL_829032, EPI_ISL_829033, EPI_ISL_829034, EPI_ISL_829035, EPI_ISL_829430 | | | |
| see above | deCODE genetics | deCODE genetics | Daniel F Gudbjartsson; Agnar Helgason; Hakon Jonsson; Olafur T Magnusson; Pall Melsted; Gudmundur L Norddahl; Jona Saemundsdottir; Asgeir Sigurdsson; Patrick Sulem; Arna B Agustsdottir; Hannes Eggertsson; Berglind Eiriksdottir; Run Fridriksdottir; Elisabet E Gardarsdottir; Gudmundur Georgsson; Olafia S Gretarsdottir; Kjartan R Gudmundsson; Thora R Gunnarsdottir; Arnaldur Gylfason; Hilma Holm; Brynjar O Jensson; Aslaug Jonasdottir; Kamilla S Josefsdottir; Thordur Kristjansson; Droplaug N Magnusdottir; Solvi Rognvaldsson; Louise le Roux; Gudrun Sigmundsdottir; Gardar Sveinbjornsson; Kristin E Sveinsdottir; Maney Sveinsdottir; Emil A Thorarensen; Bjarni Thorbjornsson; Gisli Masson; Ingileif Jonsdottir; Alma Moller; Thorolfur Gudnason; Karl G Kristinsson; Unnur Thorsteinsdottir; Kari Stefansson |
| EPI_ISL_829471, EPI_ISL_829707, EPI_ISL_829708, EPI_ISL_829709 | The National University Hospital of Iceland | deCODE genetics | Daniel F Gudbjartsson; Agnar Helgason; Hakon Jonsson; Olafur T Magnusson; Pall Melsted; Gudmundur L Norddahl; Jona Saemundsdottir; Asgeir Sigurdsson; Patrick Sulem; Arna B Agustsdottir; Hannes Eggertsson; Berglind Eiriksdottir; Run Fridriksdottir; Elisabet E Gardarsdottir; Gudmundur Georgsson; Olafia S Gretarsdottir; Kjartan R Gudmundsson; Thora R Gunnarsdottir; Arnaldur Gylfason; Hilma Holm; Brynjar O Jensson; Aslaug Jonasdottir; Kamilla S Josefsdottir; Thordur Kristjansson; Droplaug N Magnusdottir; Solvi Rognvaldsson; Louise le Roux; Gudrun Sigmundsdottir; Gardar Sveinbjornsson; Kristin E Sveinsdottir; Maney Sveinsdottir; Emil A Thorarensen; Bjarni Thorbjornsson; Gisli Masson; Ingileif Jonsdottir; Alma Moller; Thorolfur Gudnason; Karl G Kristinsson; Unnur Thorsteinsdottir; Kari Stefansson |
| EPI_ISL_829919, EPI_ISL_829920, EPI_ISL_829921, EPI_ISL_829922, EPI_ISL_829924, EPI_ISL_829926, EPI_ISL_829927, EPI_ISL_829928, EPI_ISL_830189, EPI_ISL_830190, EPI_ISL_830281, EPI_ISL_830283, EPI_ISL_830287, EPI_ISL_830290, EPI_ISL_830292, EPI_ISL_830295, EPI_ISL_830297, EPI_ISL_830485, EPI_ISL_830491, EPI_ISL_830495 | | | |
| see above | deCODE genetics | deCODE genetics | Daniel F Gudbjartsson; Agnar Helgason; Hakon Jonsson; Olafur T Magnusson; Pall Melsted; Gudmundur L Norddahl; Jona Saemundsdottir; Asgeir Sigurdsson; Patrick Sulem; Arna B Agustsdottir; Hannes Eggertsson; Berglind Eiriksdottir; Run Fridriksdottir; Elisabet E Gardarsdottir; Gudmundur Georgsson; Olafia S Gretarsdottir; Kjartan R Gudmundsson; Thora R Gunnarsdottir; Arnaldur Gylfason; Hilma Holm; Brynjar O Jensson; Aslaug Jonasdottir; Kamilla S Josefsdottir; Thordur Kristjansson; Droplaug N Magnusdottir; Solvi Rognvaldsson; Louise le Roux; Gudrun Sigmundsdottir; Gardar Sveinbjornsson; Kristin E Sveinsdottir; Maney Sveinsdottir; Emil A Thorarensen; Bjarni Thorbjornsson; Gisli Masson; Ingileif Jonsdottir; Alma |

| | | | |
|---|---|---|---|
| | | | Moller; Thorolfur Gudnason; Karl G Kristinsson; Unnur Thorsteinsdottir; Kari Stefansson |
| EPI_ISL_830737, EPI_ISL_830799, EPI_ISL_830800, EPI_ISL_830801, EPI_ISL_830802, EPI_ISL_830803, EPI_ISL_830804, EPI_ISL_830805, EPI_ISL_830806, EPI_ISL_830807, EPI_ISL_830808, EPI_ISL_830809, EPI_ISL_830810, EPI_ISL_830811, EPI_ISL_830987, EPI_ISL_830988, EPI_ISL_831005 | | | |
| see above | University Hospital Basel, Clinical Virology | University Hospital Basel, Clinical Bacteriology | Tim Roloff, Madlen Stange, Helena MB Seth-Smith, Alfredo Mari, Karoline Leuzinger, Julia Bielicki, Manuel Battegay, Hans Hirsch, Adrian Egli |
| EPI_ISL_831033, EPI_ISL_831136, EPI_ISL_831137, EPI_ISL_831138, EPI_ISL_831139, EPI_ISL_831140, EPI_ISL_831141, EPI_ISL_831142, EPI_ISL_831143, EPI_ISL_831144, EPI_ISL_831145, EPI_ISL_831146, EPI_ISL_831147, EPI_ISL_831148, EPI_ISL_831149, EPI_ISL_831150, EPI_ISL_831151, EPI_ISL_831152, EPI_ISL_831153, EPI_ISL_831154, EPI_ISL_831155, EPI_ISL_831156, EPI_ISL_831157, EPI_ISL_831158, EPI_ISL_831159, EPI_ISL_831160, EPI_ISL_831161, EPI_ISL_831162, EPI_ISL_831163, EPI_ISL_831164, EPI_ISL_831165, EPI_ISL_831166, EPI_ISL_831167, EPI_ISL_831168, EPI_ISL_831169 | | | |
| see above | Hospital Universitario La Paz (Madrid) | SeqCOVID-SPAIN consortium/IBV(CSIC) | María Rodríguez-Tejedor, Elias Dahdouh, Fernando Lázaro-Perona, Jesús Mingorance and SeqCOVID-SPAIN consortium |
| EPI_ISL_831940, EPI_ISL_832009 | Laboratório de Microbiologia Molecular - Universidade FEEVALE | Universidade Federal de Ciências da Saúde de Porto Alegre | Vinícius Bonetti Franceschi, Amanda de Menezes Mayer, Gabriel Dickin Caldana, Carla Andretta Moreira Neves, Patrícia Aline Gröhs Ferrareze, Gabriela Bettella Cybis, Ricardo Ariel Zimerman, Lívia Kmetzsch, Fernando Rosado Spilki, Claudia Elizabeth Thompson |
| EPI_ISL_832103, EPI_ISL_832104 | Hospital Universitari Germans Trias i Pujol(HUGTiP)/Fundació Lluita contra la SIDA (FLSida) | IrsiCaixa AIDS Research Lab | Marc Noguera-Julian, Mariona Parera, Maria Pilar Armengol, Marta Massanella, Ester Ballana, Lidia Ruiz, Nuria Izquierdo, Jorge Carrillo, Roger Paredes, Julia Blanco, Joaquim Segalés, Bonaventura Clotet |
| EPI_ISL_833040 | RSUD Dr. Abdul Aziz Singkawang | National Institute of Health Research and Development | Subangkit;Pawestri,HA;Ikawati,HD;Nugraha,AA;Puspa,KD;Pangesti,KNA;Soekarso,T;Junus,HN;Puspandari,N;Setiawaty,V |
| EPI_ISL_833332 | National Jail Management and Penology Training Institute | Research Institute for Tropical Medicine | Hannah Leah Morito, Othoniel Jan Onza, John Leonard Chan, Ma Angelica Tujan, Francisco Gerardo Polotan, Inez Andrea Medado, Kirstyn Brunker, Edelwisa Mercado, Daria Manalo, Catalino Demetria |
| EPI_ISL_837247, EPI_ISL_837248, EPI_ISL_837250, EPI_ISL_837252, EPI_ISL_837255, EPI_ISL_837285, EPI_ISL_837286, EPI_ISL_837287, EPI_ISL_837288, EPI_ISL_837289, EPI_ISL_837290, EPI_ISL_837291, EPI_ISL_837292, EPI_ISL_837293, EPI_ISL_837294, EPI_ISL_837295, EPI_ISL_837296, EPI_ISL_837297, EPI_ISL_837298, EPI_ISL_837299, EPI_ISL_837300, EPI_ISL_837301, EPI_ISL_837302, EPI_ISL_837303, EPI_ISL_837304, EPI_ISL_837305, EPI_ISL_837306, EPI_ISL_837307, EPI_ISL_837308, EPI_ISL_837309, EPI_ISL_837310, EPI_ISL_837311, EPI_ISL_837312, EPI_ISL_837313, EPI_ISL_837314, EPI_ISL_837315, EPI_ISL_837316, EPI_ISL_837317 | | | |
| see above | Istituto Zooprofilattico Sperimentale del Mezzogiorno | TIGEM | Patrizia Annunziata, Andrea Ballabio, Valentina Bouche, Davide Cacchiarelli (CorrespAuthor), Pellegrino Cerino, Chiara Colantuono, Lucio Di Filippo, Antonio Grimaldi, Antonio Limone, Gabriella Loconte, Anna Manfredi, Francesco Panariello, Biancamaria Pierri, Marcello Salvi, Lucia Vassallo |
| EPI_ISL_842838, EPI_ISL_842839, EPI_ISL_842840, EPI_ISL_842841, EPI_ISL_842842, EPI_ISL_842843, EPI_ISL_842858, EPI_ISL_842859 | Barts Health NHS Trust | COVID-19 Genomics UK (COG-UK) Consortium | CUTINO-MOGUEL, Maria-Teresa; HARRINGTON, David; OWOYEMI, Dola; SHYLINI, Raghavendran; BROAD, Claire; KELE, Beatrix |
| EPI_ISL_849033, EPI_ISL_849034, EPI_ISL_849035, EPI_ISL_849036, EPI_ISL_849037, EPI_ISL_849038, EPI_ISL_849039, EPI_ISL_849040, EPI_ISL_849041, EPI_ISL_849042, EPI_ISL_849043, EPI_ISL_849044, EPI_ISL_849045, EPI_ISL_849046, EPI_ISL_849047, EPI_ISL_849048, EPI_ISL_849049, EPI_ISL_849050, EPI_ISL_849051, EPI_ISL_849052, EPI_ISL_849053, EPI_ISL_849054, EPI_ISL_849055, EPI_ISL_849056, EPI_ISL_849057, EPI_ISL_849058, EPI_ISL_849059, EPI_ISL_849060, EPI_ISL_849061, EPI_ISL_849062, EPI_ISL_849063, EPI_ISL_849064, EPI_ISL_849065, EPI_ISL_849066, EPI_ISL_849067, EPI_ISL_849068, EPI_ISL_849069, EPI_ISL_849070, EPI_ISL_849071, EPI_ISL_849072, EPI_ISL_849073, EPI_ISL_849074, EPI_ISL_849075, EPI_ISL_849076, EPI_ISL_849077, EPI_ISL_849078, EPI_ISL_849079, EPI_ISL_849080, EPI_ISL_849081, EPI_ISL_849082, EPI_ISL_849083, EPI_ISL_849084 | | | |
| see above | Florida Bureau of Public Health Laboratories | Florida Bureau of Public Health Laboratories | Sarah Schmedes, Jason Blanton |
| EPI_ISL_849693 | unknown | PHV-FSS | Son Nguyen et al. |
| EPI_ISL_852584, EPI_ISL_852594 | Max von Pettenkofer Institute, Virology, National Reference Center for Retroviruses, LMU München | Laboratory for Functional Genome Analysis, Dept. Genomics, Gene Center of the LMU Munich | Max Muenchhoff, Stefan Krebs, Alexander Graf, Oliver Keppler, Helmut Blum |
| EPI_ISL_853298 | UPMC Clinical Microbiology Laboratory | Microbial Genome Sequencing Center; Microbial Genomic Epidemiology Laboratory | Mustapha M. Mustapha, Jane W. Marsh, Dan Snyder, Marissa P. Griffith, Stephanie L. Mitchell, Vatsala R. Srinivasa, Kady D. Waggle, Chinelo Ezeonwuku, Vaughn S. Cooper, Lee H. Harrison |
| EPI_ISL_855479, EPI_ISL_855480, EPI_ISL_855481, EPI_ISL_855482, EPI_ISL_855483, EPI_ISL_855484, EPI_ISL_855485, EPI_ISL_855487 | Servicio de Microbiología, Laboratori Clínic Metropolitana Nord. Hospital Universitari Germans Trias i Pujol. Institut d'Investigació en Ciències de la Salut Germans Trias i Pujol (IGTP) | SeqCOVID-SPAIN consortium/IBV(CSIC) | Elisa Martró, Antoni E. Bordoy, Anna Not, Adrián Antuori, Anabel Fernández, Nona Romaní, Verónica Saludes, Cristina Casañ and SeqCOVID-SPAIN consortium |
| EPI_ISL_855557, EPI_ISL_855558, EPI_ISL_855559, EPI_ISL_855560, EPI_ISL_855561, EPI_ISL_855562, EPI_ISL_855563, EPI_ISL_855564, EPI_ISL_855565, EPI_ISL_855566, EPI_ISL_855567, EPI_ISL_855568, EPI_ISL_855569, EPI_ISL_855570, EPI_ISL_855571, EPI_ISL_855572 | | | |
| see above | Department of Virology, Principal Military Hospital of Instruction of Tunis | Bundeswehr Institute of Microbiology | Susann Handrick, Malena Bestehorn-Willmann, Simone Eckstein, Mathias C. Walter, Markus H. Antwerpen, Habiba Naija, Kilian Stoecker, Roman Wölfel & Mohamed Ben Moussa |
| EPI_ISL_856682, EPI_ISL_856694, EPI_ISL_856700 | Lab. Microbiologia e Virologia, Cotugno, A.O. dei Colli | Lab. Microbiologia e Virologia, Cotugno, A.O. dei Colli | Luigi Atripaldi, Claudia Tiberio, Anna Perfetti |
| EPI_ISL_859682, EPI_ISL_859683, EPI_ISL_859695, EPI_ISL_859696, EPI_ISL_859697, EPI_ISL_859755, EPI_ISL_859756, EPI_ISL_859757 | BTC, Khalifa University | BTC, Khalifa University | Al Safar et al |
| EPI_ISL_860095 | Lab. Microbiologia e Virologia, Cotugno, A.O. dei Colli | Lab. Microbiologia e Virologia, Cotugno, A.O. dei Colli | Luigi Atripaldi, Claudia Tiberio, Anna Perfetti |
| EPI_ISL_860140, | Keio University School of | Keio University School of | Kenjiro Kosaki, Yuka Iwasaki, Hirotsugu Ishizu, Haruhiko Siomi, Kodai Abe |

| | | | |
|---|---|---|---|
| EPI_ISL_860142 | Medicine | Medicine | |
| EPI_ISL_860550 | St.Olavs hospital/NTNU | IKOM, NTNU | Aleksandr Ianevski, Svein Arne Nordbo, Denis Kainov |
| EPI_ISL_860794 | Lab. Microbiologia e Virologia, Cotugno, A.O. dei Colli | Lab. Microbiologia e Virologia, Cotugno, A.O. dei Colli | Luigi Atripaldi, Claudia Tiberio, Anna Perfetti |
| EPI_ISL_862589, EPI_ISL_862590, EPI_ISL_862591, EPI_ISL_862592, EPI_ISL_862593, EPI_ISL_862595, EPI_ISL_862596, EPI_ISL_862597, EPI_ISL_862598, EPI_ISL_862599, EPI_ISL_862600, EPI_ISL_862601, EPI_ISL_862602, EPI_ISL_862603, EPI_ISL_862604, EPI_ISL_862605, EPI_ISL_862606, EPI_ISL_862607, EPI_ISL_862608, EPI_ISL_862609, EPI_ISL_862610, EPI_ISL_862611, EPI_ISL_862612, EPI_ISL_862613, EPI_ISL_862614, EPI_ISL_862615, EPI_ISL_862616, EPI_ISL_862617, EPI_ISL_862618, EPI_ISL_862619, EPI_ISL_862620, EPI_ISL_862621, EPI_ISL_862622, EPI_ISL_862623, EPI_ISL_862624, EPI_ISL_862625, EPI_ISL_862626 | | | |
| see above | Hospital Comarcal de Melilla | Instituto de Salud Carlos III | Iglesias-Caballero, M. Camarero, S. Molinero Calamita, M. González-Esguevillas, M. Pozo, F. Casas, I. Jiménez, P. Jiménez, M. Zaballos, A. Monzón, S. Varona, S. Juliá, M. Cuesta, I. López, J. |
| EPI_ISL_862815 | Multidisciplinary Research Unit, DHR-ICMR, Institute of Medical Sciences, Banaras Hindu University, Varanasi | Multidisciplinary Research Unit, DHR-ICMR, Institute of Medical Sciences, Banaras Hindu University, Varanasi | Royana Singh, Priyoneel Basu, Ashish, Nitish Kumar Singh, Abhay Kumar Yadav, Manpreet Kaur, Deepa Devadas, Chetan Sahni, Gulshan Kumar, Tribhuwan Mohan Mohapatra, Richa Arya, Prashant Singh, Jay Prakash Maurya, Surendra Pratap Mishra |
| EPI_ISL_876820, EPI_ISL_876821, EPI_ISL_876886, EPI_ISL_876887, EPI_ISL_876888, EPI_ISL_876889, EPI_ISL_876890, EPI_ISL_876891, EPI_ISL_876892, EPI_ISL_876893, EPI_ISL_876894, EPI_ISL_876895, EPI_ISL_876896, EPI_ISL_876897, EPI_ISL_876898, EPI_ISL_876899, EPI_ISL_877120, EPI_ISL_877124, EPI_ISL_877125 | | | |
| see above | Quest Diagnostics | Quest Diagnostics | Rosenthal,S.H., Gerasimova,A., Kagan,R.M., Anderson, B., Hua, M., Liu Y., Bernstein, L.E., Livingston, K.E., Perez, A., Shalhout, D.F., Shlyakhter, I.A., Owen, R., Tanpaiboon, P., Lacbawan, F. |
| EPI_ISL_882684, EPI_ISL_882686, EPI_ISL_882748, EPI_ISL_882754 | 1.AO Universitaria 'S. Giovanni di Dio e Ruggi D'Aragona, Scuola Medica Salernitana' Hospital / 2.UOC di Virologia e Microbiologia, Università della Campania 'L. Vanvitelli' / 3.AO Universitaria 'Federico II' Napoli Hospital / 4.AORN 'San Giuseppe Moscati' Avellino Hospital / 5.AO 'San Pio - presidio G. Rummo' Benevento Hospital / 6.AO 'Sant'Anna e San Sebastiano' Caserta Hospital / 7.PO 'Maria Santissima Addolorata' Eboli Hospital / 8.Biogem Istituto di Ricerche Genetiche | 1. Genome Research Center for Health (CRGS) / 2. Laboratory of Molecular Medicine and Genomics(LMMGe) / 3. Center for Research in Pure and Applied Mathematics (CRMPA) | Giorgio Giurato, Francesca Rizzo, Alessandro Weisz, Gianluigi Franci, Giovanni Nassa, Pasquale Pagliano, Roberta Tarallo, Elena Alexandrova, Ylenia D'Agostino, Carlo Ferravante, Jessica Lamberti, Viola Melone, Domenico Memoli, Valeria Mirici Cappa, Domenico Palumbo, Giovanni Pecoraro, Assunta Sellitto, Oriana Strianese, Ilaria Terenzi, Giuseppe Fenza, Aniello Gentile, Antonello Saccomanno, Sonia Amabile, Teresa Rocco, Annamaria Salvati, Emilia Vaccaro, Massimiliano Galdiero, Michele Cennamo, Giuseppe Portella, Maria Grazia Foti, Mariarosaria Ingino, Maria Landi, Maurizio Fumi, Vincenzo Rocco, Rita Greco, Vittoria Letizia, Arnolfo Petruzziello, Maddalena Schioppa, Gregorio Goffredi, Francesca Marciano, Michele Caraglia, Alessia Cossu, Marianna Scrima |
| EPI_ISL_884308, EPI_ISL_884309, EPI_ISL_884310, EPI_ISL_884311, EPI_ISL_884408, EPI_ISL_884409, EPI_ISL_884410, EPI_ISL_884411, EPI_ISL_884412 | Infectious Diseases, Quest Diagnostics | Infectious Diseases, Quest Diagnostics | Rosenthal,S.H., Gerasimova,A., Kagan,R.M., Anderson,B., Bernstein,L.E., Livingston,K.E., Hua,M., Liu,Y., Shalhout,D.F., Owen,R., Lacbawan,F. |
| EPI_ISL_885150 | Hospital General Universitario Gregorio Marañón | Hospital General Universitario Gregorio Marañón | Sergio Buenestado Serrano, Pedro Sola Campoy, Laura Pérez-Lago, Pilar Catalán, Patricia Muñoz, Darío García de Viedma. |
| EPI_ISL_887109 | Institute of Medical Microbiology and Virology, University Hospital Carl Gustav Carus, TU Dresden | DRESDEN-concept Genome Center, CMCB, TU Dresden | Beil J., Brown T., Büttner L., Gscheidel N., Hochauf-Stange K., Klemroth S., Lindemann D., Mehnert G., Petzold A., Reinhardt S., Rost F., Sameith K., Winkler S. |
| EPI_ISL_889372 | Olomouc University Hospital | Institute of Applied Biotechnologies a.s. | Petr Klempt, Ondej Brzo, Martin Kašný, Kateina Kvapilová, Petr Kvapil |
| EPI_ISL_891267 | Institute of Biocides and Medical Ecology, Belgarde, Serbia | Virology department Institute of microbiology and immunology Faculty of Medicine University of Belgrade | Banko Ana, Miljanovic Danijela, Milicevic Ognjen, Loncar Ana, Abazovic Dzihan, Despot Dragana |
| EPI_ISL_892228, EPI_ISL_892229 | Lighthouse Lab in Glasgow | Wellcome Sanger Institute for the COVID-19 Genomics UK (COG-UK) Consortium | Harper VanSteenhouse, Yumi Kasai, David Gray, Carol Clugston, Anna Dominiczak and Alex Alderton, Roberto Amato, Sonia Goncalves, Ewan Harrison, David K. Jackson, Ian Johnston, Dominic Kwiatkowski, Cordelia Langford, John Sillitoe on behalf of the Wellcome Sanger Institute COVID-19 Surveillance Team |
| EPI_ISL_896182, EPI_ISL_900058, EPI_ISL_900069, EPI_ISL_900096, EPI_ISL_900100, EPI_ISL_900475, EPI_ISL_900489 | MEPHI, Aix Marseille University | MEPHI, Aix Marseille University | Anthony LEVASSEUR |
| EPI_ISL_902742, EPI_ISL_902745 | Hospital Universitari Germans Trias i Pujol (HUGTiP) / Fundació Lluita contra la SIDA (FLSida) | IrsiCaixa - Can Ruti CovidSeq | Fundació irsiCaixa. Hospital Universitari Germans Trias i Pujol(HUGTiP), 2a planta, maternal Ctra Canyet s/n, Badalona Marta Massanella, Ester Ballana, Lidia Ruiz, Nuria Izquierdo, Jorge Carrillo, Roger Paredes, Julia Blanco, Joaquim Segalés, Bonaventura Clotet |
| EPI_ISL_903336, EPI_ISL_903337, EPI_ISL_903338 | University of Iowa Hospitals & Clinics, Microbiology Laboratory | University or Iowa, Lung Biology and Cystic Fibrosis Research Center, Pezzulo Lab | Miguel E Ortiz, Alejandro A Pezzulo |
| EPI_ISL_906063, EPI_ISL_906064 | Tilia Laboratories s.r.o. | Tilia Laboratories s.r.o. | Sona Pekova, MD, PhD. |

| | | | |
|---|---|---|---|
| EPI_ISL_909772 | National Institute of Health Research and Development | National Institute of Health Research and Development | Holy Arif Wibowo; Ririn Ramadhany; Yuni Rukminiati; Agustiningsih; Kindi Adam; Hana Apsari Pawestri; Subangkit; Kartika Dewi Puspa; Arie Ardiansyah Nugraha; Hartanti Dian Ikawati; Krisna Nur Andriana Pangesti; Triyani Soekarso; Ni Ketut Susilarini; Nur Ika Hariastuti; Uli Alfi Nikmah; Mursinah; Asri Febriyani; Reni Herman; Nike Susanti; Herna; Tati Febriyanti; Juwita Kurniawati; Natalie Laurencia Kipuw; Fauzul Muna; Irene Lorinda Indalao; Aulia Rizki; Nelly Puspandari; vivi Setiawaty |
| EPI_ISL_910161, EPI_ISL_910162, EPI_ISL_910163, EPI_ISL_910164, EPI_ISL_910165, EPI_ISL_910166, EPI_ISL_910167, EPI_ISL_910168, EPI_ISL_910169, EPI_ISL_910170, EPI_ISL_910171, EPI_ISL_910172, EPI_ISL_910173, EPI_ISL_910174, EPI_ISL_910175, EPI_ISL_910176, EPI_ISL_910177, EPI_ISL_910178, EPI_ISL_910179, EPI_ISL_910180, EPI_ISL_910181, EPI_ISL_910182, EPI_ISL_910183, EPI_ISL_910184, EPI_ISL_910185, EPI_ISL_910186, EPI_ISL_910187, EPI_ISL_910188, EPI_ISL_910189, EPI_ISL_910190, EPI_ISL_910191, EPI_ISL_910192, EPI_ISL_910193, EPI_ISL_910194, EPI_ISL_910195, EPI_ISL_910196, EPI_ISL_910197, EPI_ISL_910198, EPI_ISL_910199, EPI_ISL_910200, EPI_ISL_910201, EPI_ISL_910202, EPI_ISL_910203, EPI_ISL_910204, EPI_ISL_910205, EPI_ISL_910206, EPI_ISL_910207, EPI_ISL_910208, EPI_ISL_910209, EPI_ISL_910210 | | | |
| see above | CSIR-Centre for Cellular and Molecular Biology | CSIR-Centre for Cellular and Molecular Biology | Payel Mukherjee,Pratheusa Maccha,Namami Gaur,Lamuk Zaveri,Tulasi Nagabandi,Purushotham Vodnala,Blessy B John,Viswagithe S L,B Himasri,Sofia Banu,Priya Singh,Archana Bharadwaj Siva,Karthik Bharadwaj Tallapaka,Rakesh K Mishra,Divya Tej Sowpati |
| EPI_ISL_912187, EPI_ISL_912188, EPI_ISL_912189 | California Institute of Technology | Chan-Zuckerberg Biohub | CZB Cliahub Consortium |
| EPI_ISL_915395, EPI_ISL_915396, EPI_ISL_915397 | Keio University School of Medicine | Keio University School of Medicine | Kenjiro Kosaki, Yuka Iwasaki, Hirotsugu Ishizu, Haruhiko Siomi, Kodai Abe |
| EPI_ISL_918126 | Lighthouse Lab in Glasgow | Wellcome Sanger Institute for the COVID-19 Genomics UK (COG-UK) Consortium | Harper VanSteenhouse, Yumi Kasai, David Gray, Carol Clugston, Anna Dominiczak and Alex Alderton, Roberto Amato, Sonia Goncalves, Ewan Harrison, David K. Jackson, Ian Johnston, Dominic Kwiatkowski, Cordelia Langford, John Sillitoe on behalf of the Wellcome Sanger Institute COVID-19 Surveillance Team |
| EPI_ISL_924428 | Virology Department, Sheffield Teaching Hospitals NHS Foundation Trust/Department of Infection, Immunity and Cardiovascular Disease, The Medical School, University of Sheffield | COVID-19 Genomics UK (COG-UK) Consortium | Thushan de Silva, Matthew Parker, Nikki Smith, Adri Angyal, Rebecca Brown, Luke Green, Rachel Tucker, Paul Parsons, Danielle Groves, Katie Johnson, Laura Carrilero, Alex Keeley, Dave Partridge, Matthew Wyles, Benjamin Lindsey, Mehmet Yavuz, Mohammad Raza, Cariad Evans |
| EPI_ISL_925429, EPI_ISL_925430, EPI_ISL_925431, EPI_ISL_925432, EPI_ISL_925433, EPI_ISL_925434 | Department of Clinical Microbiology | GIGA Medical Genomics | Keith Durkin, Maria Artesi, Sébastien Bontems, Raphaël Boreux, Bouchra Boujemla, Cécile Meex, Pierrette Melin, Marie-Pierre Hayette, Vincent Bours |
| EPI_ISL_935367, EPI_ISL_935368, EPI_ISL_935369, EPI_ISL_935370, EPI_ISL_935371, EPI_ISL_935372, EPI_ISL_935373, EPI_ISL_935374, EPI_ISL_935375, EPI_ISL_935376, EPI_ISL_935377, EPI_ISL_935378, EPI_ISL_935379, EPI_ISL_935380, EPI_ISL_935381, EPI_ISL_935382, EPI_ISL_935385, EPI_ISL_935386, EPI_ISL_935387, EPI_ISL_935388, EPI_ISL_935399, EPI_ISL_935400, EPI_ISL_935402, EPI_ISL_935403, EPI_ISL_935404 | | | |
| see above | Florida Bureau of Public Health Laboratories | Florida Bureau of Public Health Laboratories | Sarah Schmedes, Jason Blanton |
| EPI_ISL_935791, EPI_ISL_935817, EPI_ISL_935818, EPI_ISL_935819, EPI_ISL_935820, EPI_ISL_935821, EPI_ISL_935829, EPI_ISL_935830, EPI_ISL_935834, EPI_ISL_935837, EPI_ISL_935838, EPI_ISL_935839, EPI_ISL_935842, EPI_ISL_935843, EPI_ISL_935844, EPI_ISL_935845, EPI_ISL_935847, EPI_ISL_935848, EPI_ISL_935851 | | | |
| see above | Cadham Provincial laboratory | National Microbiology Laboratory (NML) | Anna Majer, Shari Tyson, Grace Seo, Philip Mabon, Elsie Grudeski, Rhiannon Huzarewich, Russell Mandes, Anneliese Landgraff, Jennifer Tanner, Natalie Knox, Morag Graham, Gary Van Domselaar, Paul Van Caeseele, Jared Bullard, David Alexander, Kerry Dust, Nathalie Bastien, Yan Li, Timothy Booth, Darian Hole, Madison Chapel, Kirsten Biggar, CanCOGeN's metadata curation team, Public Health Agency of Canada CanCOGeN team |
| EPI_ISL_936837, EPI_ISL_936838, EPI_ISL_936839, EPI_ISL_936840, EPI_ISL_936841, EPI_ISL_936842, EPI_ISL_936843 | Northwestern Memorial Hospital | Ozer Lab | Ramon Lorenzo-Redondo, Lacy M. Simons, Chad J. Achenbach, Lawrence J. Jennings, Michael G. Ison, Judd F. Hultquist, Egon A. Ozer |
| EPI_ISL_937030, EPI_ISL_937031, EPI_ISL_937036, EPI_ISL_937043, EPI_ISL_937044, EPI_ISL_937051, EPI_ISL_937052, EPI_ISL_937059, EPI_ISL_937064, EPI_ISL_937069, EPI_ISL_937080, EPI_ISL_937092, EPI_ISL_937094, EPI_ISL_937113 | | | |
| see above | Quest Diagnostics | Quest Diagnostics | Rosenthal,S.H., Gerasimova,A., Kagan,R.M., Anderson, B., Livingston, K.E., Hua, M., Liu Y., Shalhout, D.F., Owen, R., Lacbawan, F. |
| EPI_ISL_940205, EPI_ISL_940206, EPI_ISL_940207, EPI_ISL_940394 | Hôpital Bichat Claude Bernard, Laboratoire de Virologie | IAME UMR1137 Inserm, Université de Paris, Hôpital Bichat | Antoine Bridier-Nahmias, Amélie Recoing, Quentin Le Hingrat, Lena Daniel, Siham Hamri, Gilles Collin, Alexandre Storto, Mélanie Bertine, Charlotte Charpentier, Nadhira Houhou-Fidouh, Diane Descamps, Benoit Visseaux |
| EPI_ISL_940973, EPI_ISL_940974, EPI_ISL_940975 | Centers for Disease Control and Prevention, Dengue Branch | Centers for Disease Control and Prevention, Dengue Branch | Gilberto A. Santiago, Glenda Gonzalez, Betzabel Flores, Keyla Charriez, Gabriela Paz-Bailey, Jorge L. Munoz-Jordan |
| EPI_ISL_941273 | NCSLPH | NCSLPH | Chase K, Miller MC, Greene S, Glover W |
| EPI_ISL_941929, EPI_ISL_941940 | Florida Bureau of Public Health Laboratories | Florida Bureau of Public Health Laboratories | Sarah Schmedes, Jason Blanton |
| EPI_ISL_942010 | Centers for Disease Control and Prevention, Dengue Branch | Centers for Disease Control and Prevention, Dengue Branch | Gilberto A. Santiago, Glenda Gonzalez, Betzabel Flores, Keyla Charriez, Gabriela Paz-Bailey, Jorge L. Munoz-Jordan |
| EPI_ISL_949197, EPI_ISL_949218, EPI_ISL_949225, EPI_ISL_949242, EPI_ISL_949243 | Departamento de Microbiología, CDB, Hospital Clínic, Barcelona | SeqCOVID-SPAIN consortium/IBV(CSIC) | Andrea Vergara, Mikel Martínez, Elisa Rubio, Jéssica Navero, Aida Peiró and SeqCOVID-SPAIN consortium |
| EPI_ISL_955148 | University of Sarajevo, Veterinary Faculty, Laboratory for Molecular Diagnostic and Research Laboratory | University of Sarajevo, Veterinary Faculty, Laboratory for Molecular Diagnostic and Research Laboratory | Goleti T., Goleti Š., Softi A., Ali-Šeho A., Jaži A., Šabi E., Terzi I., Nicevi M., Hodži A. |
| EPI_ISL_956404 | General Hospital - Strumica | Research Center for Genetic Engineering and Biotechnology "Georgi D. Efremov" , Macedonian Academy of Sciences and Arts | Aleksandar J. Dimovski, Dijana Plasheska-Karanfilska, Predrag Noveski, Gjorgji Bozinovski, Milena Jakimovska |

| | | | |
|---|---|---|---|
| EPI_ISL_959907, EPI_ISL_959910, EPI_ISL_959912, EPI_ISL_959998, EPI_ISL_959999, EPI_ISL_960000, EPI_ISL_960001, EPI_ISL_960002, EPI_ISL_960003, EPI_ISL_960004, EPI_ISL_960005, EPI_ISL_960006, EPI_ISL_960007, EPI_ISL_960008, EPI_ISL_960009, EPI_ISL_960010, EPI_ISL_960011, EPI_ISL_960012, EPI_ISL_960013, EPI_ISL_960014, EPI_ISL_960015, EPI_ISL_960016, EPI_ISL_960017, EPI_ISL_960018, EPI_ISL_960019, EPI_ISL_960020, EPI_ISL_960021 | | | |
| see above | University Medical Center Hamburg Eppendorf | Heinrich Pette Institute, Leibniz Institute for Experimental Virology | Alexis Robitaille, Thomas Günther, Johannes Knobloch, Martin Aepfelbacher, Nicole Fischer, Adam Grundhoff |
| EPI_ISL_970355, EPI_ISL_970357, EPI_ISL_970359, EPI_ISL_970361, EPI_ISL_970363, EPI_ISL_970366, EPI_ISL_970367, EPI_ISL_970369, EPI_ISL_970372, EPI_ISL_970374, EPI_ISL_970375, EPI_ISL_970378, EPI_ISL_970379, EPI_ISL_970381, EPI_ISL_970384, EPI_ISL_970385, EPI_ISL_970388, EPI_ISL_970389, EPI_ISL_970391, EPI_ISL_970393, EPI_ISL_970395, EPI_ISL_970397, EPI_ISL_970399, EPI_ISL_970401, EPI_ISL_970403, EPI_ISL_970405, EPI_ISL_970407, EPI_ISL_970408, EPI_ISL_970410, EPI_ISL_970413, EPI_ISL_970414, EPI_ISL_970417, EPI_ISL_970418, EPI_ISL_970420, EPI_ISL_970423, EPI_ISL_970425, EPI_ISL_970427, EPI_ISL_970429, EPI_ISL_970431, EPI_ISL_970433, EPI_ISL_970434, EPI_ISL_970437, EPI_ISL_970438, EPI_ISL_970440, EPI_ISL_970442, EPI_ISL_970445, EPI_ISL_970447, EPI_ISL_970449, EPI_ISL_970451, EPI_ISL_970453, EPI_ISL_970455, EPI_ISL_970456, EPI_ISL_970458, EPI_ISL_970459, EPI_ISL_970461, EPI_ISL_970463, EPI_ISL_970465, EPI_ISL_970467, EPI_ISL_970471, EPI_ISL_970472, EPI_ISL_970474, EPI_ISL_970476, EPI_ISL_970479, EPI_ISL_970480, EPI_ISL_970482, EPI_ISL_970484, EPI_ISL_970485, EPI_ISL_970487, EPI_ISL_970489, EPI_ISL_970491, EPI_ISL_970494, EPI_ISL_970496, EPI_ISL_970498, EPI_ISL_970501, EPI_ISL_970503, EPI_ISL_970505, EPI_ISL_970507, EPI_ISL_970509, EPI_ISL_970511, EPI_ISL_970514, EPI_ISL_970516, EPI_ISL_970518, EPI_ISL_970520, EPI_ISL_970522, EPI_ISL_970524, EPI_ISL_970526, EPI_ISL_970529, EPI_ISL_970531, EPI_ISL_970532, EPI_ISL_970534, EPI_ISL_970536, EPI_ISL_970538, EPI_ISL_970539, EPI_ISL_970542, EPI_ISL_970544, EPI_ISL_970545, EPI_ISL_970547, EPI_ISL_970549, EPI_ISL_970550, EPI_ISL_970552, EPI_ISL_970554, EPI_ISL_970556, EPI_ISL_970557, EPI_ISL_970559, EPI_ISL_970561, EPI_ISL_970563, EPI_ISL_970565, EPI_ISL_970567, EPI_ISL_970570, EPI_ISL_970572, EPI_ISL_970574, EPI_ISL_970576, EPI_ISL_970578, EPI_ISL_970580, EPI_ISL_970582, EPI_ISL_970584, EPI_ISL_970586, EPI_ISL_970588, EPI_ISL_970590, EPI_ISL_970592, EPI_ISL_970594, EPI_ISL_970596, EPI_ISL_970598, EPI_ISL_970600, EPI_ISL_970603, EPI_ISL_970604, EPI_ISL_970606, EPI_ISL_970608, EPI_ISL_970610, EPI_ISL_970611, EPI_ISL_970613, EPI_ISL_970615, EPI_ISL_970617, EPI_ISL_970620, EPI_ISL_970622, EPI_ISL_970624, EPI_ISL_970626, EPI_ISL_970627, EPI_ISL_970629, EPI_ISL_970631, EPI_ISL_970634, EPI_ISL_970636, EPI_ISL_970638, EPI_ISL_970640, EPI_ISL_970642, EPI_ISL_970644, EPI_ISL_970646, EPI_ISL_970649, EPI_ISL_970651, EPI_ISL_970653, EPI_ISL_970655, EPI_ISL_970657, EPI_ISL_970659, EPI_ISL_970662, EPI_ISL_970663, EPI_ISL_970665, EPI_ISL_970667, EPI_ISL_970669, EPI_ISL_970671, EPI_ISL_970672, EPI_ISL_970674, EPI_ISL_970676, EPI_ISL_970678, EPI_ISL_970679, EPI_ISL_970682, EPI_ISL_970684, EPI_ISL_970686, EPI_ISL_970688, EPI_ISL_970691, EPI_ISL_970693, EPI_ISL_970695, EPI_ISL_970696, EPI_ISL_970698, EPI_ISL_970700, EPI_ISL_970702, EPI_ISL_970705, EPI_ISL_970707, EPI_ISL_970710, EPI_ISL_970712, EPI_ISL_970714, EPI_ISL_970716, EPI_ISL_970718, EPI_ISL_970719, EPI_ISL_970721, EPI_ISL_970723, EPI_ISL_970725, EPI_ISL_970728, EPI_ISL_970730, EPI_ISL_970731, EPI_ISL_970733, EPI_ISL_970735, EPI_ISL_970737, EPI_ISL_970740, EPI_ISL_970742, EPI_ISL_970743, EPI_ISL_970745, EPI_ISL_970748, EPI_ISL_970750, EPI_ISL_970752, EPI_ISL_970754, EPI_ISL_970756, EPI_ISL_970759, EPI_ISL_970761, EPI_ISL_970763, EPI_ISL_970765, EPI_ISL_970767, EPI_ISL_970769, EPI_ISL_970771, EPI_ISL_970773, EPI_ISL_970776, EPI_ISL_970778, EPI_ISL_970780, EPI_ISL_970783, EPI_ISL_970785, EPI_ISL_970787, EPI_ISL_970789, EPI_ISL_970791, EPI_ISL_970793, EPI_ISL_970795, EPI_ISL_970798, EPI_ISL_970799, EPI_ISL_970801, EPI_ISL_970804, EPI_ISL_970806, EPI_ISL_970809, EPI_ISL_970818, EPI_ISL_970840, EPI_ISL_970856, EPI_ISL_970878, EPI_ISL_970902, EPI_ISL_970931, EPI_ISL_970947, EPI_ISL_970965, EPI_ISL_970995, EPI_ISL_971012, EPI_ISL_971036, EPI_ISL_971040, EPI_ISL_971043, EPI_ISL_971046, EPI_ISL_971049, EPI_ISL_971052, EPI_ISL_971054, EPI_ISL_971057, EPI_ISL_971061, EPI_ISL_971064, EPI_ISL_971068, EPI_ISL_971072, EPI_ISL_971075, EPI_ISL_971077, EPI_ISL_971080, EPI_ISL_971084, EPI_ISL_971086, EPI_ISL_971089, EPI_ISL_971091, EPI_ISL_971094, EPI_ISL_971097, EPI_ISL_971101, EPI_ISL_971104, EPI_ISL_971107, EPI_ISL_971111, EPI_ISL_971114, EPI_ISL_971116, EPI_ISL_971120, EPI_ISL_971122, EPI_ISL_971126, EPI_ISL_971129, EPI_ISL_971132, EPI_ISL_971137, EPI_ISL_971139, EPI_ISL_971141, EPI_ISL_971144, EPI_ISL_971147, EPI_ISL_971150, EPI_ISL_971154, EPI_ISL_971157, EPI_ISL_971160, EPI_ISL_971162, EPI_ISL_971167, EPI_ISL_971170, EPI_ISL_971173, EPI_ISL_971177, EPI_ISL_971179, EPI_ISL_971182, EPI_ISL_971187, EPI_ISL_971189, EPI_ISL_971192, EPI_ISL_971196, EPI_ISL_971199, EPI_ISL_971202, EPI_ISL_971205, EPI_ISL_971208, EPI_ISL_971212, EPI_ISL_971216, EPI_ISL_971219, EPI_ISL_971222, EPI_ISL_971225, EPI_ISL_971227, EPI_ISL_971230, EPI_ISL_971233 | | | |
| see above | BCCDC Public Health Laboratory | BCCDC Public Health Laboratory | Prystajecky Natalie, Linda Hoang, Dan Fornika, John Tyson, Shannon Russell, Kim Macdonald, Kimia Kamelian, Ana Pacagnella, Corrinne Ng, Loretta Janz, Robert Azana Terry Snutch, Mel Krajden |
| EPI_ISL_976912, EPI_ISL_976913, EPI_ISL_976914, EPI_ISL_976915, EPI_ISL_976917, EPI_ISL_976918, EPI_ISL_976921, EPI_ISL_976922, EPI_ISL_976924, EPI_ISL_976926, EPI_ISL_976928, EPI_ISL_976929, EPI_ISL_976930, EPI_ISL_976931 | | | |
| see above | University of Massachusetts Medical School | Infectious Disease Program, Broad Institute of Harvard and MIT | Lemieux,J.E., Siddle,K.J., Ward,D., Ellison,R., Adams,G., Gladden-Young,A., Lagerborg,K., Rudy,M., DeRuff,K., Carter,A., Normandin,E., Bauer,M., Reilly,S., Tomkins-Tinch,C., Loreth,C., Chaluvadi,S., Birren,B.W., Gallagher,G., Smole,S., Park,D.J., MacInnis,B.L., and Sabeti,P.C. |
| EPI_ISL_979292, EPI_ISL_979293 | Cadham Provincial laboratory | National Microbiology Laboratory (NML) | Anna Majer, Shari Tyson, Grace Seo, Philip Mabon, Elsie Grudeski, Rhiannon Huzarewich, Russell Mandes, Anneliese Landgraff, Jennifer Tanner, Natalie Knox, Morag Graham, Gary Van Domselaar, Paul Van Caeseele, Jared Bullard, David Alexander, Kerry Dust, Nathalie Bastien, Yan Li, Timothy Booth, Darian Hole, Madison Chapel, Kirsten Biggar, CanCOGeN's metadata curation team, Public Health Agency of Canada CanCOGeN team |
