## Supplementary material for "Detection and characterization of the SARS-CoV-2 lineage B.1.526 in New York": Supp. Table 3 GISAID Acknowledgment part 2: gisaid_hcov-19_acknowledgement_table_2021_02_12_23-9.pdf

We gratefully acknowledge the following Authors from the Originating laboratories responsible for obtaining the specimens, as well as the Submitting laboratories where the genome data were generated and shared via GISAID, on which this research is based.

All Submitters of data may be contacted directly via www.gisaid.org

Authors are sorted alphabetically.

| Accession ID | Originating Laboratory | Submitting Laboratory | Authors |
|---|---|---|---|
| EPI_ISL_572405 | Wales Specialist Virology Centre Sequencing lab: Pathogen Genomics Unit | COVID-19 Genomics UK (COG-UK) Consortium | Catherine Moore, Johnathan Evans, Laura Gifford, Malorie Perry, Simon Cottrell, Angela Marchbank, Alec Birchley, Alexander Adams, Amy Gaskin, Bree Gatica-Wilcox, Jason Coombes, Joel Southgate, Lauren Gilbert, Lee Graham, Nicole Pacchiarini, Sara Kumziene-Summerhayes, Sarah Taylor, Sophie Jones, Sara Rey, Matthew Bull, Joanne Watkins, Sally Corden, Tom Connor |
| EPI_ISL_572419 | Quadram Institute Bioscience | COVID-19 Genomics UK (COG-UK) Consortium | Dave J. Baker, Gemma L. Kay, Alp Aydin, Thanh Le-Viet, Steven Rudder, Ana P. Tedim, Anastasia Kolyva, Maria Diaz, Leonardo de Oliveira Martins, Nabil-Fareed Alikhan, Lizzie Meadows, Rachael Stanley, Ngozi Elumogo, Muhammed Yasir, Nicholas M. Thomson, Alexander J Trotter, Rachel Gilroy, Samuel Bloomfield, Claire Stuart, Andrew Bell, Reenesh Prakash, Samir Dervisevic, Alison E. Mather, John Wain, Mark Webber, Andrew J. Page, Justin O'Grady |
| EPI_ISL_572444, EPI_ISL_572460 | Queens Medical Centre, Clinical Microbiology Department / DeepSeq Nottingham | COVID-19 Genomics UK (COG-UK) Consortium | Gemma Clark, Wendy Smith, Manjinder Khakh, Vicki M Fleming, Michelle M Lister, Hannah Howson-Wells, Jonathan Ball, Patrick McClure, Joseph Chappell, Theocharis Tsoleridis, Nadine Holmes, Matthew Carlisle, Christopher Moore, Fei Sang, Johnny Debebe, Victoria Wright, Matthew Loose |
| EPI_ISL_572462 | Quadram Institute Bioscience | COVID-19 Genomics UK (COG-UK) Consortium | Dave J. Baker, Gemma L. Kay, Alp Aydin, Thanh Le-Viet, Steven Rudder, Ana P. Tedim, Anastasia Kolyva, Maria Diaz, Leonardo de Oliveira Martins, Nabil-Fareed Alikhan, Lizzie Meadows, Rachael Stanley, Ngozi Elumogo, Muhammed Yasir, Nicholas M. Thomson, Alexander J Trotter, Rachel Gilroy, Samuel Bloomfield, Claire Stuart, Andrew Bell, Reenesh Prakash, Samir Dervisevic, Alison E. Mather, John Wain, Mark Webber, Andrew J. Page, Justin O'Grady |
| EPI_ISL_572474 | Wales Specialist Virology Centre Sequencing lab: Pathogen Genomics Unit | COVID-19 Genomics UK (COG-UK) Consortium | Catherine Moore, Johnathan Evans, Laura Gifford, Malorie Perry, Simon Cottrell, Angela Marchbank, Alec Birchley, Alexander Adams, Amy Gaskin, Bree Gatica-Wilcox, Jason Coombes, Joel Southgate, Lauren Gilbert, Lee Graham, Nicole Pacchiarini, Sara Kumziene-Summerhayes, Sarah Taylor, Sophie Jones, Sara Rey, Matthew Bull, Joanne Watkins, Sally Corden, Tom Connor |
| EPI_ISL_572475, EPI_ISL_572476 | Queens Medical Centre, Clinical Microbiology Department / DeepSeq Nottingham | COVID-19 Genomics UK (COG-UK) Consortium | Gemma Clark, Wendy Smith, Manjinder Khakh, Vicki M Fleming, Michelle M Lister, Hannah Howson-Wells, Jonathan Ball, Patrick McClure, Joseph Chappell, Theocharis Tsoleridis, Nadine Holmes, Matthew Carlisle, Christopher Moore, Fei Sang, Johnny Debebe, Victoria Wright, Matthew Loose |
| EPI_ISL_572481 | Virology Department, Sheffield Teaching Hospitals NHS Foundation Trust/Department of Infection, Immunity and Cardiovascular Disease, The Medical School, University of Sheffield | COVID-19 Genomics UK (COG-UK) Consortium | Thushan de Silva, Matthew Parker, Nikki Smith, Adri Angyal, Rebecca Brown, Luke Green, Rachel Tucker, Paul Parsons, Danielle Groves, Katie Johnson, Laura Carrilero, Alex Keeley, Dave Partridge, Matthew Wyles, Benjamin Lindsey, Mehmet Yavuz, Mohammad Raza, Cariad Evans |
| EPI_ISL_572501 | Wales Specialist Virology Centre Sequencing lab: Pathogen Genomics Unit | COVID-19 Genomics UK (COG-UK) Consortium | Catherine Moore, Johnathan Evans, Laura Gifford, Malorie Perry, Simon Cottrell, Angela Marchbank, Alec Birchley, Alexander Adams, Amy Gaskin, Bree Gatica-Wilcox, Jason Coombes, Joel Southgate, Lauren Gilbert, Lee Graham, Nicole Pacchiarini, Sara Kumziene-Summerhayes, Sarah Taylor, Sophie Jones, Sara Rey, Matthew Bull, Joanne Watkins, Sally Corden, Tom Connor |
| EPI_ISL_572522 | Virology Department, Sheffield Teaching Hospitals NHS Foundation Trust/Department of Infection, Immunity and Cardiovascular Disease, The Medical School, University of Sheffield | COVID-19 Genomics UK (COG-UK) Consortium | Thushan de Silva, Matthew Parker, Nikki Smith, Adri Angyal, Rebecca Brown, Luke Green, Rachel Tucker, Paul Parsons, Danielle Groves, Katie Johnson, Laura Carrilero, Alex Keeley, Dave Partridge, Matthew Wyles, Benjamin Lindsey, Mehmet Yavuz, Mohammad Raza, Cariad Evans |
| EPI_ISL_572555, EPI_ISL_572560, EPI_ISL_572565, EPI_ISL_572575 | Wales Specialist Virology Centre Sequencing lab: Pathogen Genomics Unit | COVID-19 Genomics UK (COG-UK) Consortium | Catherine Moore, Johnathan Evans, Laura Gifford, Malorie Perry, Simon Cottrell, Angela Marchbank, Alec Birchley, Alexander Adams, Amy Gaskin, Bree Gatica-Wilcox, Jason Coombes, Joel Southgate, Lauren Gilbert, Lee Graham, Nicole Pacchiarini, Sara Kumziene-Summerhayes, Sarah Taylor, Sophie Jones, Sara Rey, Matthew Bull, Joanne Watkins, Sally Corden, Tom Connor |
| EPI_ISL_572582 | Queens Medical Centre, Clinical Microbiology Department / DeepSeq Nottingham | COVID-19 Genomics UK (COG-UK) Consortium | Gemma Clark, Wendy Smith, Manjinder Khakh, Vicki M Fleming, Michelle M Lister, Hannah Howson-Wells, Jonathan Ball, Patrick McClure, Joseph Chappell, Theocharis Tsoleridis, Nadine Holmes, Matthew Carlisle, Christopher Moore, Fei Sang, Johnny Debebe, Victoria Wright, Matthew Loose |
| EPI_ISL_572589 | Virology Department, Sheffield Teaching Hospitals NHS Foundation Trust/Department of Infection, Immunity and Cardiovascular Disease, The Medical School, University of Sheffield | COVID-19 Genomics UK (COG-UK) Consortium | Thushan de Silva, Matthew Parker, Nikki Smith, Adri Angyal, Rebecca Brown, Luke Green, Rachel Tucker, Paul Parsons, Danielle Groves, Katie Johnson, Laura Carrilero, Alex Keeley, Dave Partridge, Matthew Wyles, Benjamin Lindsey, Mehmet Yavuz, Mohammad Raza, Cariad Evans |
| EPI_ISL_572598, EPI_ISL_572603 | Queens Medical Centre, Clinical Microbiology Department / DeepSeq Nottingham | COVID-19 Genomics UK (COG-UK) Consortium | Gemma Clark, Wendy Smith, Manjinder Khakh, Vicki M Fleming, Michelle M Lister, Hannah Howson-Wells, Jonathan Ball, Patrick McClure, Joseph Chappell, Theocharis Tsoleridis, Nadine Holmes, Matthew Carlisle, Christopher Moore, Fei Sang, Johnny Debebe, Victoria Wright, Matthew Loose |
| EPI_ISL_572605 | Wales Specialist Virology Centre Sequencing lab: Pathogen Genomics Unit | COVID-19 Genomics UK (COG-UK) Consortium | Catherine Moore, Johnathan Evans, Laura Gifford, Malorie Perry, Simon Cottrell, Angela Marchbank, Alec Birchley, Alexander Adams, Amy Gaskin, Bree Gatica-Wilcox, Jason Coombes, Joel Southgate, Lauren Gilbert, Lee Graham, Nicole Pacchiarini, Sara Kumziene-Summerhayes, Sarah Taylor, Sophie Jones, Sara Rey, Matthew Bull, Joanne Watkins, Sally Corden, Tom Connor |
| EPI_ISL_572613, EPI_ISL_572618 | Virology Department, Sheffield Teaching Hospitals NHS Foundation Trust/Department of Infection, Immunity and Cardiovascular Disease, The Medical School, University of Sheffield | COVID-19 Genomics UK (COG-UK) Consortium | Thushan de Silva, Matthew Parker, Nikki Smith, Adri Angyal, Rebecca Brown, Luke Green, Rachel Tucker, Paul Parsons, Danielle Groves, Katie Johnson, Laura Carrilero, Alex Keeley, Dave Partridge, Matthew Wyles, Benjamin Lindsey, Mehmet Yavuz, Mohammad Raza, Cariad Evans |
| EPI_ISL_572651 | Quadram Institute Bioscience | COVID-19 Genomics UK (COG-UK) Consortium | Dave J. Baker, Gemma L. Kay, Alp Aydin, Thanh Le-Viet, Steven Rudder, Ana P. Tedim, Anastasia Kolyva, Maria Diaz, Leonardo de Oliveira Martins, Nabil-Fareed Alikhan, Lizzie Meadows, Rachael Stanley, Ngozi Elumogo, Muhammed Yasir, Nicholas M. Thomson, Alexander J Trotter, Rachel Gilroy, Samuel Bloomfield, Claire Stuart, Andrew Bell, Reenesh Prakash, Samir Dervisevic, Alison E. Mather, John Wain, Mark Webber, Andrew J. Page, Justin O'Grady |
| EPI_ISL_572652, EPI_ISL_572654, EPI_ISL_572668 | Wales Specialist Virology Centre Sequencing lab: Pathogen Genomics Unit | COVID-19 Genomics UK (COG-UK) Consortium | Catherine Moore, Johnathan Evans, Laura Gifford, Malorie Perry, Simon Cottrell, Angela Marchbank, Alec Birchley, Alexander Adams, Amy Gaskin, Bree Gatica-Wilcox, Jason Coombes, Joel Southgate, Lauren Gilbert, Lee Graham, Nicole Pacchiarini, Sara Kumziene-Summerhayes, Sarah Taylor, Sophie Jones, Sara Rey, Matthew Bull, Joanne Watkins, Sally Corden, Tom Connor |
| EPI_ISL_572706 | Queens Medical Centre, Clinical Microbiology Department / DeepSeq Nottingham | COVID-19 Genomics UK (COG-UK) Consortium | Gemma Clark, Wendy Smith, Manjinder Khakh, Vicki M Fleming, Michelle M Lister, Hannah Howson-Wells, Jonathan Ball, Patrick McClure, Joseph Chappell, Theocharis Tsoleridis, Nadine Holmes, Matthew Carlisle, Christopher Moore, Fei Sang, Johnny Debebe, Victoria Wright, Matthew Loose |
| EPI_ISL_572732 | Quadram Institute Bioscience | COVID-19 Genomics UK (COG-UK) Consortium | Dave J. Baker, Gemma L. Kay, Alp Aydin, Thanh Le-Viet, Steven Rudder, Ana P. Tedim, Anastasia Kolyva, Maria Diaz, Leonardo de Oliveira Martins, Nabil-Fareed Alikhan, Lizzie Meadows, Rachael Stanley, Ngozi Elumogo, Muhammed Yasir, Nicholas M. Thomson, Alexander J Trotter, Rachel Gilroy, Samuel Bloomfield, Claire Stuart, Andrew Bell, Reenesh Prakash, Samir Dervisevic, Alison E. Mather, John Wain, Mark Webber, Andrew J. Page, Justin O'Grady |
| EPI_ISL_572754 | Wales Specialist Virology Centre Sequencing lab: Pathogen Genomics Unit | COVID-19 Genomics UK (COG-UK) Consortium | Catherine Moore, Johnathan Evans, Laura Gifford, Malorie Perry, Simon Cottrell, Angela Marchbank, Alec Birchley, Alexander Adams, Amy Gaskin, Bree Gatica-Wilcox, Jason Coombes, Joel Southgate, Lauren Gilbert, Lee Graham, Nicole Pacchiarini, Sara Kumziene-Summerhayes, Sarah Taylor, Sophie Jones, Sara Rey, Matthew Bull, Joanne Watkins, Sally Corden, Tom Connor |
| EPI_ISL_572764 | Quadram Institute Bioscience | COVID-19 Genomics UK (COG-UK) Consortium | Dave J. Baker, Gemma L. Kay, Alp Aydin, Thanh Le-Viet, Steven Rudder, Ana P. Tedim, Anastasia Kolyva, Maria Diaz, Leonardo de Oliveira Martins, Nabil-Fareed Alikhan, Lizzie Meadows, Rachael Stanley, Ngozi Elumogo, Muhammed Yasir, Nicholas M. Thomson, Alexander J Trotter, Rachel Gilroy, Samuel Bloomfield, Claire Stuart, Andrew Bell, Reenesh Prakash, Samir Dervisevic, Alison E. Mather, John Wain, Mark Webber, Andrew J. Page, Justin O'Grady |
| EPI_ISL_572778 | Wales Specialist Virology Centre Sequencing lab: Pathogen Genomics Unit | COVID-19 Genomics UK (COG-UK) Consortium | Catherine Moore, Johnathan Evans, Laura Gifford, Malorie Perry, Simon Cottrell, Angela Marchbank, Alec Birchley, Alexander Adams, Amy Gaskin, Bree Gatica-Wilcox, Jason Coombes, Joel Southgate, Lauren Gilbert, Lee Graham, Nicole Pacchiarini, Sara Kumziene-Summerhayes, Sarah Taylor, Sophie Jones, Sara Rey, Matthew Bull, Joanne Watkins, Sally Corden, Tom Connor |

| | | | |
|---|---|---|---|
| EPI_ISL_572799 | Queens Medical Centre, Clinical Microbiology Department / DeepSeq Nottingham | COVID-19 Genomics UK (COG-UK) Consortium | Gemma Clark, Wendy Smith, Manjinder Khakh, Vicki M Fleming, Michelle M Lister, Hannah Howson-Wells, Jonathan Ball, Patrick McClure, Joseph Chappell, Theocharis Tsoleridis, Nadine Holmes, Matthew Carlisle, Christopher Moore, Fei Sang, Johnny Debebe, Victoria Wright, Matthew Loose |
| EPI_ISL_572805, EPI_ISL_572810 | Wales Specialist Virology Centre Sequencing lab: Pathogen Genomics Unit | COVID-19 Genomics UK (COG-UK) Consortium | Catherine Moore, Johnathan Evans, Laura Gifford, Malorie Perry, Simon Cottrell, Angela Marchbank, Alec Birchley, Alexander Adams, Amy Gaskin, Bree Gatica-Wilcox, Jason Coombes, Joel Southgate, Lauren Gilbert, Lee Graham, Nicole Pacchiarini, Sara Kumziene-Summerhayes, Sarah Taylor, Sophie Jones, Sara Rey, Matthew Bull, Joanne Watkins, Sally Corden, Tom Connor |
| EPI_ISL_572814 | Quadram Institute Bioscience | COVID-19 Genomics UK (COG-UK) Consortium | Dave J. Baker, Gemma L. Kay, Alp Aydin, Thanh Le-Viet, Steven Rudder, Ana P. Tedim, Anastasia Kolyva, Maria Diaz, Leonardo de Oliveira Martins, Nabil-Fareed Alikhan, Lizzie Meadows, Rachael Stanley, Ngozi Elumogo, Muhammed Yasir, Nicholas M. Thomson, Alexander J Trotter, Rachel Gilroy, Samuel Bloomfield, Claire Stuart, Andrew Bell, Reenesh Prakash, Samir Dervisevic, Alison E. Mather, John Wain, Mark Webber, Andrew J. Page, Justin O'Grady |
| EPI_ISL_572833 | Queens Medical Centre, Clinical Microbiology Department / DeepSeq Nottingham | COVID-19 Genomics UK (COG-UK) Consortium | Gemma Clark, Wendy Smith, Manjinder Khakh, Vicki M Fleming, Michelle M Lister, Hannah Howson-Wells, Jonathan Ball, Patrick McClure, Joseph Chappell, Theocharis Tsoleridis, Nadine Holmes, Matthew Carlisle, Christopher Moore, Fei Sang, Johnny Debebe, Victoria Wright, Matthew Loose |
| EPI_ISL_572842 | Quadram Institute Bioscience | COVID-19 Genomics UK (COG-UK) Consortium | Dave J. Baker, Gemma L. Kay, Alp Aydin, Thanh Le-Viet, Steven Rudder, Ana P. Tedim, Anastasia Kolyva, Maria Diaz, Leonardo de Oliveira Martins, Nabil-Fareed Alikhan, Lizzie Meadows, Rachael Stanley, Ngozi Elumogo, Muhammed Yasir, Nicholas M. Thomson, Alexander J Trotter, Rachel Gilroy, Samuel Bloomfield, Claire Stuart, Andrew Bell, Reenesh Prakash, Samir Dervisevic, Alison E. Mather, John Wain, Mark Webber, Andrew J. Page, Justin O'Grady |
| EPI_ISL_572881, EPI_ISL_572882, EPI_ISL_572886, EPI_ISL_572888 | Wales Specialist Virology Centre Sequencing lab: Pathogen Genomics Unit | COVID-19 Genomics UK (COG-UK) Consortium | Catherine Moore, Johnathan Evans, Laura Gifford, Malorie Perry, Simon Cottrell, Angela Marchbank, Alec Birchley, Alexander Adams, Amy Gaskin, Bree Gatica-Wilcox, Jason Coombes, Joel Southgate, Lauren Gilbert, Lee Graham, Nicole Pacchiarini, Sara Kumziene-Summerhayes, Sarah Taylor, Sophie Jones, Sara Rey, Matthew Bull, Joanne Watkins, Sally Corden, Tom Connor |
| EPI_ISL_572929, EPI_ISL_572948 | Virology Department, Sheffield Teaching Hospitals NHS Foundation Trust/Department of Infection, Immunity and Cardiovascular Disease, The Medical School, University of Sheffield | COVID-19 Genomics UK (COG-UK) Consortium | Thushan de Silva, Matthew Parker, Nikki Smith, Adri Angyal, Rebecca Brown, Luke Green, Rachel Tucker, Paul Parsons, Danielle Groves, Katie Johnson, Laura Carrilero, Alex Keeley, Dave Partridge, Matthew Wyles, Benjamin Lindsey, Mehmet Yavuz, Mohammad Raza, Cariad Evans |
| EPI_ISL_573450, EPI_ISL_573451, EPI_ISL_573452, EPI_ISL_573453, EPI_ISL_573454, EPI_ISL_573455, EPI_ISL_573456, EPI_ISL_573457, EPI_ISL_573458, EPI_ISL_573459, EPI_ISL_573460, EPI_ISL_573461, EPI_ISL_573462, EPI_ISL_573463, EPI_ISL_573464, EPI_ISL_573465, EPI_ISL_573466, EPI_ISL_573467, EPI_ISL_573468, EPI_ISL_573469, EPI_ISL_573470, EPI_ISL_573471, EPI_ISL_573472, EPI_ISL_573473, EPI_ISL_573474, EPI_ISL_573475, EPI_ISL_573476, EPI_ISL_573477 | | | |
| see above | Queens Medical Centre, Clinical Microbiology Department / DeepSeq Nottingham | COVID-19 Genomics UK (COG-UK) Consortium | Gemma Clark, Wendy Smith, Manjinder Khakh, Vicki M Fleming, Michelle M Lister, Hannah Howson-Wells, Jonathan Ball, Patrick McClure, Joseph Chappell, Theocharis Tsoleridis, Nadine Holmes, Matthew Carlisle, Christopher Moore, Fei Sang, Johnny Debebe, Victoria Wright, Matthew Loose |
| EPI_ISL_573687, EPI_ISL_573691, EPI_ISL_573697, EPI_ISL_573703, EPI_ISL_573707, EPI_ISL_573710, EPI_ISL_573712, EPI_ISL_573724, EPI_ISL_573730, EPI_ISL_573731, EPI_ISL_573738, EPI_ISL_573744, EPI_ISL_573751, EPI_ISL_573752 | | | |
| see above | Virology Department, Sheffield Teaching Hospitals NHS Foundation Trust/Department of Infection, Immunity and Cardiovascular Disease, The Medical School, University of Sheffield | COVID-19 Genomics UK (COG-UK) Consortium | Thushan de Silva, Matthew Parker, Nikki Smith, Adri Angyal, Rebecca Brown, Luke Green, Rachel Tucker, Paul Parsons, Danielle Groves, Katie Johnson, Laura Carrilero, Alex Keeley, Dave Partridge, Matthew Wyles, Benjamin Lindsey, Mehmet Yavuz, Mohammad Raza, Cariad Evans |
| EPI_ISL_574018, EPI_ISL_574019, EPI_ISL_574030, EPI_ISL_574032, EPI_ISL_574034, EPI_ISL_574036, EPI_ISL_574039, EPI_ISL_574050, EPI_ISL_574054, EPI_ISL_574055, EPI_ISL_574061, EPI_ISL_574062, EPI_ISL_574064, EPI_ISL_574067, EPI_ISL_574069, EPI_ISL_574070, EPI_ISL_574087, EPI_ISL_574090, EPI_ISL_574094, EPI_ISL_574102, EPI_ISL_574108, EPI_ISL_574112, EPI_ISL_574113, EPI_ISL_574118, EPI_ISL_574121, EPI_ISL_574125, EPI_ISL_574129, EPI_ISL_574143, EPI_ISL_574146, EPI_ISL_574150, EPI_ISL_574151, EPI_ISL_574155, EPI_ISL_574171, EPI_ISL_574179, EPI_ISL_574183, EPI_ISL_574185, EPI_ISL_574187, EPI_ISL_574190, EPI_ISL_574192, EPI_ISL_574199, EPI_ISL_574201, EPI_ISL_574206, EPI_ISL_574208, EPI_ISL_574211, EPI_ISL_574213, EPI_ISL_574222, EPI_ISL_574229, EPI_ISL_574234, EPI_ISL_574241, EPI_ISL_574246, EPI_ISL_574249, EPI_ISL_574253, EPI_ISL_574256 | | | |
| see above | Wales Specialist Virology Centre Sequencing lab: Pathogen Genomics Unit | COVID-19 Genomics UK (COG-UK) Consortium | Catherine Moore, Johnathan Evans, Laura Gifford, Malorie Perry, Simon Cottrell, Angela Marchbank, Alec Birchley, Alexander Adams, Amy Gaskin, Bree Gatica-Wilcox, Jason Coombes, Joel Southgate, Lauren Gilbert, Lee Graham, Nicole Pacchiarini, Sara Kumziene-Summerhayes, Sarah Taylor, Sophie Jones, Sara Rey, Matthew Bull, Joanne Watkins, Sally Corden, Tom Connor |
| EPI_ISL_574494, EPI_ISL_574519 | National Public Health Laboratory, National Centre for Infectious Diseases | National Public Health Laboratory, National Centre for Infectious Diseases | Tze Minn Mak, Sophie Octavia, Zhenyang Zhou, Lin Cui, Raymond Tzer Pin Lin |
| EPI_ISL_576198, EPI_ISL_576199, EPI_ISL_576200, EPI_ISL_576201, EPI_ISL_576202, EPI_ISL_576203, EPI_ISL_576204, EPI_ISL_576205, EPI_ISL_576206, EPI_ISL_576207, EPI_ISL_576208, EPI_ISL_576209, EPI_ISL_576210, EPI_ISL_576211, EPI_ISL_576212, EPI_ISL_576213, EPI_ISL_576214, EPI_ISL_576215, EPI_ISL_576216, EPI_ISL_576217, EPI_ISL_576218, EPI_ISL_576219, EPI_ISL_576220, EPI_ISL_576221 | | | |
| see above | Michigan Department of Health and Human Services, Bureau of Laboratories | Michigan Department of Health and Human Services, Bureau of Laboratories | Blankenship HM, Riner D, Soehnlen MK |
| EPI_ISL_576396 | Genome Centre | Genome Centre | Selina Akter Pravas Chandra Roy Amina Ferdaus manami Habiba Ibnat A. S. M. Rubayet Ul Alam Shireen Nigar Iqbal Kabir Jahid and M. Anwar Hossain |
| EPI_ISL_576487, EPI_ISL_576488, EPI_ISL_576557 | UW Virology Lab | UW Virology Lab | Pavitra Roychoudhury, Hong Xie, Lasata Shrestha, Amin Addetia, Victoria M Rachleff, Meei-Li Huang, Keith R Jerome, Alexander Greninger |
| EPI_ISL_576902 | Department of Pathology, University of Cambridge | COVID-19 Genomics UK (COG-UK) Consortium | Aminu S. Jahun, Yasmin Chaudhry, Grant Hall, Iliana Georgana, Myra Hosmillo, Martin D. Curran, Malte Pinckert, Surendra Parmar, Ian Goodfellow |
| EPI_ISL_576981, EPI_ISL_576982, EPI_ISL_576983 | Liverpool Clinical Laboratories | COVID-19 Genomics UK (COG-UK) Consortium | Sam Haldenby, Anita Lucaci, Steve Paterson, Julian Hiscox, Alistair Darby, M Almsaud, A Alrezaihi, Muhannad Alruwaili, Stuart D Armstrong, Jones Benjamin, Eleanor G Bentley, Anu Chawla, Jordan J Clark, Angela Cowell, Richard Eccles, Isabel García-Dorival, Matthew Gemmell, Alessandro Gerada, PKF Gilmore, Richard Gregory, Ximeng Han, Catherine Hartley, Margaret Hughes, Miren Iturriza-Gomara, James Johnson, L Luu, Jenifer Manson, Charlotte Nelson, Elaine O'Toole, Cassie Olateju, Rebekah Penrice-Randal , Lucille Rainbow, N.P Randle, Trevor Ian Robinson, Parul Sharma, Ghada T Shawli, James P Stewart, Neil Swainston, Ecaterina Vamos, Joanne Watts, Mark Whitehead |
| EPI_ISL_577096, EPI_ISL_577097, EPI_ISL_577100, EPI_ISL_577101, EPI_ISL_577102, EPI_ISL_577105, EPI_ISL_577106, EPI_ISL_577107, EPI_ISL_577108, EPI_ISL_577109 | Quadram Institute Bioscience | COVID-19 Genomics UK (COG-UK) Consortium | Dave J. Baker, Gemma L. Kay, Alp Aydin, Thanh Le-Viet, Steven Rudder, Ana P. Tedim, Anastasia Kolyva, Maria Diaz, Leonardo de Oliveira Martins, Nabil-Fareed Alikhan, Lizzie Meadows, Rachael Stanley, Ngozi Elumogo, Muhammed Yasir, Nicholas M. Thomson, Alexander J Trotter, Rachel Gilroy, Samuel Bloomfield, Claire Stuart, Andrew Bell, Reenesh Prakash, Samir Dervisevic, Alison E. Mather, John Wain, Mark Webber, Andrew J. Page, Justin O'Grady |
| EPI_ISL_577110, EPI_ISL_577111, EPI_ISL_577112, EPI_ISL_577113, EPI_ISL_577114, EPI_ISL_577115, EPI_ISL_577116, EPI_ISL_577117, EPI_ISL_577118, EPI_ISL_577119, EPI_ISL_577120, EPI_ISL_577121, EPI_ISL_577122, EPI_ISL_577123 | | | |
| see above | Queens Medical Centre, Clinical Microbiology Department / DeepSeq Nottingham | COVID-19 Genomics UK (COG-UK) Consortium | Gemma Clark, Wendy Smith, Manjinder Khakh, Vicki M Fleming, Michelle M Lister, Hannah Howson-Wells, Jonathan Ball, Patrick McClure, Joseph Chappell, Theocharis Tsoleridis, Nadine Holmes, Matthew Carlisle, Christopher Moore, Fei Sang, Johnny Debebe, Victoria Wright, Matthew Loose |
| EPI_ISL_577418, EPI_ISL_577421, EPI_ISL_577423, EPI_ISL_577424, EPI_ISL_577425, EPI_ISL_577426, EPI_ISL_577428, EPI_ISL_577429, EPI_ISL_577431, EPI_ISL_577434, EPI_ISL_577435, EPI_ISL_577436, EPI_ISL_577438, EPI_ISL_577442, EPI_ISL_577443, EPI_ISL_577444, EPI_ISL_577445, EPI_ISL_577446, EPI_ISL_577447, EPI_ISL_577448, EPI_ISL_577450, EPI_ISL_577451, EPI_ISL_577452, EPI_ISL_577453, EPI_ISL_577454, EPI_ISL_577458, EPI_ISL_577459, EPI_ISL_577462, EPI_ISL_577463, EPI_ISL_577465, EPI_ISL_577466, EPI_ISL_577469, EPI_ISL_577472, EPI_ISL_577475, EPI_ISL_577476, EPI_ISL_577479, EPI_ISL_577481, EPI_ISL_577492, EPI_ISL_577493, EPI_ISL_577494, EPI_ISL_577495, EPI_ISL_577497, EPI_ISL_577499, EPI_ISL_577500, EPI_ISL_577501, EPI_ISL_577503, EPI_ISL_577504, EPI_ISL_577505, EPI_ISL_577507, EPI_ISL_577508, EPI_ISL_577509, EPI_ISL_577511, EPI_ISL_577512, EPI_ISL_577513, EPI_ISL_577515, EPI_ISL_577517, EPI_ISL_577521, EPI_ISL_577522, EPI_ISL_577523, EPI_ISL_577526, EPI_ISL_577527, EPI_ISL_577528, EPI_ISL_577529, EPI_ISL_577530, EPI_ISL_577531, EPI_ISL_577532, EPI_ISL_577534, EPI_ISL_577535, EPI_ISL_577536, EPI_ISL_577537, EPI_ISL_577539, EPI_ISL_577540, EPI_ISL_577541, EPI_ISL_577542, EPI_ISL_577543 | | | |
| see above | Wales Specialist Virology Centre Sequencing lab: Pathogen Genomics Unit | COVID-19 Genomics UK (COG-UK) Consortium | Catherine Moore, Johnathan Evans, Laura Gifford, Malorie Perry, Simon Cottrell, Angela Marchbank, Alec Birchley, Alexander Adams, Amy Gaskin, Bree Gatica-Wilcox, Jason Coombes, Joel Southgate, Lauren Gilbert, Lee Graham, Nicole Pacchiarini, Sara Kumziene-Summerhayes, Sarah Taylor, Sophie Jones, Sara Rey, Matthew Bull, Joanne Watkins, Sally Corden, Tom Connor |
| EPI_ISL_577640, EPI_ISL_577677, EPI_ISL_577679, EPI_ISL_577680, EPI_ISL_577681, EPI_ISL_577682, EPI_ISL_577683, EPI_ISL_577684, EPI_ISL_577685, EPI_ISL_577686, EPI_ISL_577687, EPI_ISL_577688, EPI_ISL_577689, EPI_ISL_577690, EPI_ISL_577691, EPI_ISL_577692, EPI_ISL_577693, EPI_ISL_577694, EPI_ISL_577695, EPI_ISL_577696, EPI_ISL_577697, EPI_ISL_577698, EPI_ISL_577699, EPI_ISL_577700, EPI_ISL_577701, EPI_ISL_577702, EPI_ISL_577706, EPI_ISL_577709, EPI_ISL_577710, EPI_ISL_577711, EPI_ISL_577712, EPI_ISL_577713, EPI_ISL_577717, EPI_ISL_577720, EPI_ISL_577721, EPI_ISL_577722, EPI_ISL_577723 | | | |
| see above | NIV Influenza | NIV Influenza | Potdar V |
| EPI_ISL_577834, EPI_ISL_577835, EPI_ISL_577836, EPI_ISL_578052, EPI_ISL_578073, EPI_ISL_578074, EPI_ISL_578075, EPI_ISL_578076, EPI_ISL_578077, EPI_ISL_578078, EPI_ISL_578079 | | | |
| see above | Dutch COVID-19 response team | Erasmus Medical Center | Bas Oude Munnink, Reina Sikkema, David Nieuwenhuijse, Irina Chestakova, Anne van der Linden, Marjan Boter, Emmanuelle Munger, Corine GeurtsvanKessel, Annemiek van der Eijk, Richard Molenkamp, Marion Koopmans, on behalf of the Dutch national COVID-19 response team. |

| | | | |
|---|---|---|---|
| EPI_ISL_579113, EPI_ISL_579114, EPI_ISL_579115 | LabPLUS | Institute of Environmental Science and Research (ESR) | Xiaoyun Ren, Matt Storey, Nikki Freed, Muhammad Faisal, Jing Wang, Hermes Perez, Anja Werno, Antje van der Linden, Arlo Upton, Chris Mansell, David Hammer, Dragana Drinkovic, Gary McAuliffe, Hana Sofia Andersson, James Ussher, Jill Sherwood, Josh Freeman, Julia Howard, Juliet Elvy, Mary DeAlmeida, Matt Blakiston, Matthew Rogers, Max Bloomfield, Michael Addidle, Michelle Balm, Sally Roberts, Sarah Jefferies, Sharmini Muttaiyah, Susan Morpeth, Susan Taylor, Timothy Blackmore, Vani Sathyendran, Veronica Playle, Virginia Hope, Erasmus Smit, Lauren Jelly, Olin Silander, Joep de Ligt |
| EPI_ISL_580302, EPI_ISL_580337, EPI_ISL_580405, EPI_ISL_580411, EPI_ISL_580424, EPI_ISL_580443, EPI_ISL_580501, EPI_ISL_580514, EPI_ISL_580526, EPI_ISL_580602 | Lighthouse Lab in Alderley Park | Wellcome Sanger Institute for the COVID-19 Genomics UK (COG-UK) consortium | Jacquelyn Wynn, Mairead Hyland, The Lighthouse Lab in Alderley Park and Alex Alderton, Roberto Amato, Sonia Goncalves, Ewan Harrison, David K. Jackson, Ian Johnston, Dominic Kwiatkowski, Cordelia Langford, John Sillitoe on behalf of the Wellcome Sanger Institute COVID-19 Surveillance Team |
| EPI_ISL_580654, EPI_ISL_580661, EPI_ISL_580665, EPI_ISL_580707, EPI_ISL_580711, EPI_ISL_580717, EPI_ISL_580725, EPI_ISL_580729, EPI_ISL_580739, EPI_ISL_580748, EPI_ISL_580760, EPI_ISL_580773, EPI_ISL_580777, EPI_ISL_580786, EPI_ISL_580812, EPI_ISL_580818, EPI_ISL_580826, EPI_ISL_580830, EPI_ISL_580831, EPI_ISL_580839 | | | |
| see above | Lighthouse Lab in Milton Keynes | Wellcome Sanger Institute for the COVID-19 Genomics UK (COG-UK) consortium | The Lighthouse Lab in Milton Keynes and Alex Alderton, Roberto Amato, Sonia Goncalves, Ewan Harrison, David K. Jackson, Ian Johnston, Dominic Kwiatkowski, Cordelia Langford, John Sillitoe on behalf of the Wellcome Sanger Institute COVID-19 Surveillance Team |
| EPI_ISL_581664, EPI_ISL_581665, EPI_ISL_581666 | Department of Clinical Microbiology | GIGA Medical Genomics | Keith Durkin, Maria Artesi, Sébastien Bontems, Raphaël Boreux, Bouchra Boujemla, Cécile Meex, Pierrette Melin, Marie-Pierre Hayette, Vincent Bours |
| EPI_ISL_582241 | NIV Influenza | NIV Influenza | Potdar V |
| EPI_ISL_582689, EPI_ISL_582690, EPI_ISL_582691 | Sheikh Khalifa Medical City | Molecular/Surveillance lab Sheikh Khalifa Medical City | Amirtharaj Francis, Sajeed Abdul, Hala Imambaccus, Sahar Almarzooqi, Hiba Saud, Stefan Weber |
| EPI_ISL_583421, EPI_ISL_583422, EPI_ISL_583423, EPI_ISL_583425 | University of Michigan Clinical Microbiology Laboratory | Lauring Lab, University of Michigan, Department of Microbiology and Immunology | Valesano |
| EPI_ISL_583514, EPI_ISL_583515, EPI_ISL_583516, EPI_ISL_583517, EPI_ISL_583518, EPI_ISL_583519, EPI_ISL_583520, EPI_ISL_583521, EPI_ISL_583525, EPI_ISL_583526, EPI_ISL_583527, EPI_ISL_583529, EPI_ISL_583532 | | | |
| see above | Michigan Department of Health and Human Services, Bureau of Laboratories | Michigan Department of Health and Human Services, Bureau of Laboratories | Blankenship HM, Riner D, Soehnlen MK |
| EPI_ISL_583554, EPI_ISL_583555, EPI_ISL_583556 | Genome Centre | Genome Centre | Selina Akter, Pravas Chandra Roy, Amina Ferdaus manami, Habiba Ibnat, A. S. M. Rubayet Ul Alam, Shireen Nigar, Iqbal Kabir Jahid, M.Anwar Hossain |
| EPI_ISL_583893 | Singapore General Hospital | Department of Microbiology | Nurdyana Abdul Rahman, Kun Lee Lim, Chenhao Li, Sui Sin Goh, Kenneth Xin Long Chan, Kian Sing Chan, Lynette Oon, Kern Rei Chng, Niranjan Nagarajan, Karrie Ko |
| EPI_ISL_583968, EPI_ISL_583970, EPI_ISL_583971, EPI_ISL_583972, EPI_ISL_583973, EPI_ISL_583974, EPI_ISL_583982, EPI_ISL_583983, EPI_ISL_583984, EPI_ISL_583985, EPI_ISL_583986, EPI_ISL_583987, EPI_ISL_583990, EPI_ISL_583991, EPI_ISL_584066 | | | |
| see above | Respiratory Virus Unit, Microbiology Services Colindale, Public Health England | Respiratory Virus Unit, Microbiology Services Colindale, Public Health England | PHE Covid Sequencing Team |
| EPI_ISL_584069, EPI_ISL_584071 | IZSM | IZSM | Maurizio Viscardi, Lorena Cardillo, Giovanna Fusco |
| EPI_ISL_584286, EPI_ISL_584287, EPI_ISL_584288, EPI_ISL_584289, EPI_ISL_584290, EPI_ISL_584294, EPI_ISL_584295, EPI_ISL_584296, EPI_ISL_584297, EPI_ISL_584298 | Department of Pathology, University of Cambridge | COVID-19 Genomics UK (COG-UK) Consortium | Aminu S. Jahun, Yasmin Chaudhry, Grant Hall, Iliana Georgana, Myra Hosmillo, Martin D. Curran, Malte Pinckert, Surendra Parmar, Ian Goodfellow |
| EPI_ISL_584592, EPI_ISL_584595, EPI_ISL_584596, EPI_ISL_584598, EPI_ISL_584599, EPI_ISL_584600, EPI_ISL_584601, EPI_ISL_584602, EPI_ISL_584603, EPI_ISL_584604, EPI_ISL_584605 | | | |
| see above | Liverpool Clinical Laboratories | COVID-19 Genomics UK (COG-UK) Consortium | Sam Haldenby, Anita Lucaci, Steve Paterson, Julian Hiscox, Alistair Darby, M Almsaud, A Alrezaihi, Muhannad Alruwaili, Stuart D Armstrong, Jones Benjamin, Eleanor G Bentley, Anu Chawla, Jordan J Clark, Angela Cowell, Richard Eccles, Isabel García-Dorival, Matthew Gemmell, Alessandro Gerada, PKF Gilmore, Richard Gregory, Ximeng Han, Catherine Hartley, Margaret Hughes, Miren Iturriza-Gomara, James Johnson, L Luu, Jenifer Manson, Charlotte Nelson, Elaine O'Toole, Cassie Olateju, Rebekah Penrice-Randal , Lucille Rainbow, N.P Randle, Trevor Ian Robinson, Parul Sharma, Ghada T Shawli, James P Stewart, Neil Swainston, Ecaterina Vamos, Joanne Watts, Mark Whitehead |
| EPI_ISL_584672 | University College London, Great Ormond Street Hospital for Children NHS Foundation Trust, Imperial College Healthcare NHS Trust | COVID-19 Genomics UK (COG-UK) Consortium | Sergi Castellano, Rachel Williams, Mark Kristiansen, Paola Resende Silva, Sunando Roy, Tony Brooks, Helena Tutill, Paola Niola, Patricia Dyal, Charlotte Williams, Leysa Forrest, Yasmin Panchbhaya, Jacqueline Findlay, Samuel Weeks, Julianne Brown, Kathryn Harris, Paul Randell, James Price, Alison Holmes, Judith Breuer |
| EPI_ISL_584690, EPI_ISL_584691, EPI_ISL_584692, EPI_ISL_584693, EPI_ISL_584694, EPI_ISL_584695, EPI_ISL_584696, EPI_ISL_584697, EPI_ISL_584698, EPI_ISL_584699, EPI_ISL_584700, EPI_ISL_584701, EPI_ISL_584702, EPI_ISL_584703, EPI_ISL_584704, EPI_ISL_584705 | | | |
| see above | Northumbria University / South Tees Hospitals NHS Foundation Trust / North Cumbria Integrated Care NHS Foundation Trust / North Tees and Hartlepool NHS Foundation Trust / Newcastle Hospitals NHS Foundation Trust | COVID-19 Genomics UK (COG-UK) Consortium | Darren L Smith,Andrew Nelson,Matthew Bashton,Greg R Young,Joshua Loh,John Allan,Mohammad A Tariq,Giles S Holt,Gary Black,Wen C Yew,Lynn Dover,Paul Baker,Steve Liggett,Sarah Essex,Jane Greenaway,Debra Padgett,Clive Graham,Garren Scott,Edward Barton,Emma Swindells,Brendan Payne,Jennifer Collins,Yusri Taha,Gary Eltringham |
| EPI_ISL_584716, EPI_ISL_584719, EPI_ISL_584722, EPI_ISL_584723 | Quadram Institute Bioscience | COVID-19 Genomics UK (COG-UK) Consortium | Dave J. Baker, Gemma L. Kay, Alp Aydin, Thanh Le-Viet, Steven Rudder, Ana P. Tedim, Anastasia Kolyva, Maria Diaz, Leonardo de Oliveira Martins, Nabil-Fareed Alikhan, Lizzie Meadows, Rachael Stanley, Ngozi Elumogo, Muhammed Yasir, Nicholas M. Thomson, Alexander J Trotter, Rachel Gilroy, Samuel Bloomfield, Claire Stuart, Andrew Bell, Reenesh Prakash, Samir Dervisevic, Alison E. Mather, John Wain, Mark Webber, Andrew J. Page, Justin O'Grady |
| EPI_ISL_584898, EPI_ISL_584899, EPI_ISL_584901, EPI_ISL_584902 | University of Exeter | COVID-19 Genomics UK (COG-UK) Consortium | Ben Temperton,Aaron Jeffries,Michelle Michelsen,Joanna Warwick-Dugdale,Audrey Farbos,Robyn Manley,Stephen Michell,Jane Masoli |
| EPI_ISL_584906, EPI_ISL_584907, EPI_ISL_584908, EPI_ISL_584909, EPI_ISL_584910, EPI_ISL_584911, EPI_ISL_584912, EPI_ISL_584913, EPI_ISL_584914, EPI_ISL_584915, EPI_ISL_584916, EPI_ISL_584917, EPI_ISL_584918, EPI_ISL_584919, EPI_ISL_584920, EPI_ISL_584921, EPI_ISL_584922, EPI_ISL_584923, EPI_ISL_584924, EPI_ISL_584925, EPI_ISL_584926, EPI_ISL_584927, EPI_ISL_584928, EPI_ISL_584929, EPI_ISL_584930, EPI_ISL_584931, EPI_ISL_584932, EPI_ISL_584933, EPI_ISL_584934, EPI_ISL_584935, EPI_ISL_584936, EPI_ISL_584937, EPI_ISL_584938, EPI_ISL_584939, EPI_ISL_584940, EPI_ISL_584941, EPI_ISL_584942, EPI_ISL_584943, EPI_ISL_584944, EPI_ISL_584945, EPI_ISL_584946, EPI_ISL_584947, EPI_ISL_584948, EPI_ISL_584949, EPI_ISL_584950, EPI_ISL_584951, EPI_ISL_584952, EPI_ISL_584956, EPI_ISL_584961, EPI_ISL_584962, EPI_ISL_584963, EPI_ISL_584964, EPI_ISL_584965, EPI_ISL_584966, EPI_ISL_584967, EPI_ISL_584968, EPI_ISL_584969, EPI_ISL_584970, EPI_ISL_584971, EPI_ISL_584972, EPI_ISL_584973, EPI_ISL_584981, EPI_ISL_584982, EPI_ISL_584983, EPI_ISL_584984, EPI_ISL_584985, EPI_ISL_584986, EPI_ISL_584987, EPI_ISL_584988, EPI_ISL_584989, EPI_ISL_584990, EPI_ISL_584991, EPI_ISL_584992, EPI_ISL_584993, EPI_ISL_584994, EPI_ISL_584995, EPI_ISL_584996, EPI_ISL_584997, EPI_ISL_584998, EPI_ISL_584999, EPI_ISL_585001, EPI_ISL_585006, EPI_ISL_585007, EPI_ISL_585008 | | | |
| see above | Quadram Institute Bioscience | COVID-19 Genomics UK (COG-UK) Consortium | Dave J. Baker, Gemma L. Kay, Alp Aydin, Thanh Le-Viet, Steven Rudder, Ana P. Tedim, Anastasia Kolyva, Maria Diaz, Leonardo de Oliveira Martins, Nabil-Fareed Alikhan, Lizzie Meadows, Rachael Stanley, Ngozi Elumogo, Muhammed Yasir, Nicholas M. Thomson, Alexander J Trotter, Rachel Gilroy, Samuel Bloomfield, Claire Stuart, Andrew Bell, Reenesh Prakash, Samir Dervisevic, Alison E. Mather, John Wain, Mark Webber, Andrew J. Page, Justin O'Grady |
| EPI_ISL_585011, EPI_ISL_585013, EPI_ISL_585014, EPI_ISL_585017, EPI_ISL_585019, EPI_ISL_585020, EPI_ISL_585021, EPI_ISL_585022, EPI_ISL_585030, EPI_ISL_585031, EPI_ISL_585032, EPI_ISL_585034, EPI_ISL_585039, EPI_ISL_585040, EPI_ISL_585043, EPI_ISL_585049, EPI_ISL_585051, EPI_ISL_585052, EPI_ISL_585053, EPI_ISL_585055, EPI_ISL_585057, EPI_ISL_585058, EPI_ISL_585059, EPI_ISL_585060, EPI_ISL_585061, EPI_ISL_585062, EPI_ISL_585064, EPI_ISL_585066, EPI_ISL_585068, EPI_ISL_585070, EPI_ISL_585076, EPI_ISL_585078, EPI_ISL_585080, EPI_ISL_585082, EPI_ISL_585083, EPI_ISL_585087, EPI_ISL_585089 | | | |
| see above | Virology Department, Sheffield Teaching Hospitals NHS Foundation Trust/Department of Infection, Immunity and Cardiovascular Disease, The Medical School, University of Sheffield | COVID-19 Genomics UK (COG-UK) Consortium | Thushan de Silva, Matthew Parker, Nikki Smith, Adri Angyal, Rebecca Brown, Luke Green, Rachel Tucker, Paul Parsons, Danielle Groves, Katie Johnson, Laura Carrilero, Alex Keeley, Dave Partridge, Matthew Wyles, Benjamin Lindsey, Mehmet Yavuz, Mohammad Raza, Cariad Evans |
| EPI_ISL_585260 | Quadram Institute Bioscience | COVID-19 Genomics UK (COG-UK) Consortium | Dave J. Baker, Gemma L. Kay, Alp Aydin, Thanh Le-Viet, Steven Rudder, Ana P. Tedim, Anastasia Kolyva, Maria Diaz, Leonardo de Oliveira Martins, Nabil-Fareed Alikhan, Lizzie Meadows, Rachael Stanley, Ngozi Elumogo, Muhammed Yasir, Nicholas M. Thomson, Alexander J Trotter, Rachel Gilroy, Samuel Bloomfield, Claire Stuart, Andrew Bell, Reenesh Prakash, Samir Dervisevic, Alison E. Mather, John Wain, Mark Webber, Andrew J. Page, Justin O'Grady |

| | | | |
|---|---|---|---|
| EPI_ISL_585425 | West of Scotland Specialist Virology Centre, NHSGGC / MRC-University of Glasgow Centre for Virus Research | COVID-19 Genomics UK (COG-UK) Consortium | Ana da Silva Filipe, Natasha Johnson, Kathy Smollett, Daniel Mair, Stephen Carmichael, Lily Tong, Jenna Nichols, Elihu Aranday-Cortes, Kyriaki Nomikou; Sarah McDonald, Marc Niebel, Patawee Asamaphan; Richard Orton, Joseph Hughes, Sreenu Vattipally, David L Robertson; Alasdair MacLean, Rory Gunson; Kathy Li, Igor Starinskij, Natasha Jesudason, Rajiv Shah, James Shepherd, Antonia Ho, Emma Thomson |
| EPI_ISL_585451, EPI_ISL_585452, EPI_ISL_585453, EPI_ISL_585454, EPI_ISL_585455, EPI_ISL_585456, EPI_ISL_585458, EPI_ISL_585459, EPI_ISL_585460, EPI_ISL_585461, EPI_ISL_585462, EPI_ISL_585463, EPI_ISL_585464, EPI_ISL_585465, EPI_ISL_585466, EPI_ISL_585467, EPI_ISL_585468, EPI_ISL_585469, EPI_ISL_585470, EPI_ISL_585471, EPI_ISL_585472, EPI_ISL_585473, EPI_ISL_585488, EPI_ISL_585492, EPI_ISL_585493, EPI_ISL_585496, EPI_ISL_585497 | | | |
| see above | Virology Department, Royal Infirmary of Edinburgh, NHS Lothian / School of Biological Sciences, University of Edinburgh / Institute of Genetics and Molecular Medicine, University of Edinburgh | COVID-19 Genomics UK (COG-UK) Consortium | McHugh M, Dewar R, Rooke S, Gallagher M, Balcaza C, O'Toole Á, Scher E, Hill V, McCrone JT, Colquhoun R, Yu X, Jackson B, Rambaut A, Williams TC, Templeton K |
| EPI_ISL_585583, EPI_ISL_585584, EPI_ISL_585585, EPI_ISL_585586, EPI_ISL_585587, EPI_ISL_585588, EPI_ISL_585589, EPI_ISL_585590, EPI_ISL_585591, EPI_ISL_585592, EPI_ISL_585593, EPI_ISL_585594, EPI_ISL_585595, EPI_ISL_585596, EPI_ISL_585597, EPI_ISL_585606, EPI_ISL_585607, EPI_ISL_585608, EPI_ISL_585619 | | | |
| see above | Quadram Institute Bioscience | COVID-19 Genomics UK (COG-UK) Consortium | Dave J. Baker, Gemma L. Kay, Alp Aydin, Thanh Le-Viet, Steven Rudder, Ana P. Tedim, Anastasia Kolyva, Maria Diaz, Leonardo de Oliveira Martins, Nabil-Fareed Alikhan, Lizzie Meadows, Rachael Stanley, Ngozi Elumogo, Muhammed Yasir, Nicholas M. Thomson, Alexander J Trotter, Rachel Gilroy, Samuel Bloomfield, Claire Stuart, Andrew Bell, Reenesh Prakash, Samir Dervisevic, Alison E. Mather, John Wain, Mark Webber, Andrew J. Page, Justin O'Grady |
| EPI_ISL_585631, EPI_ISL_585633, EPI_ISL_585634, EPI_ISL_585635, EPI_ISL_585637, EPI_ISL_585638, EPI_ISL_585639, EPI_ISL_585640, EPI_ISL_585641, EPI_ISL_585642, EPI_ISL_585643, EPI_ISL_585645, EPI_ISL_585646, EPI_ISL_585647, EPI_ISL_585649, EPI_ISL_585650, EPI_ISL_585651, EPI_ISL_585654, EPI_ISL_585655, EPI_ISL_585657, EPI_ISL_585658, EPI_ISL_585660, EPI_ISL_585661, EPI_ISL_585663, EPI_ISL_585664, EPI_ISL_585665, EPI_ISL_585666, EPI_ISL_585667, EPI_ISL_585668, EPI_ISL_585669, EPI_ISL_585672, EPI_ISL_585674, EPI_ISL_585675, EPI_ISL_585676, EPI_ISL_585677, EPI_ISL_585678, EPI_ISL_585679, EPI_ISL_585681, EPI_ISL_585683, EPI_ISL_585684, EPI_ISL_585685, EPI_ISL_585686, EPI_ISL_585687, EPI_ISL_585689, EPI_ISL_585690, EPI_ISL_585691, EPI_ISL_585692, EPI_ISL_585694, EPI_ISL_585698, EPI_ISL_585699, EPI_ISL_585700, EPI_ISL_585702, EPI_ISL_585703, EPI_ISL_585705, EPI_ISL_585707, EPI_ISL_585708, EPI_ISL_585709, EPI_ISL_585710, EPI_ISL_585711, EPI_ISL_585712, EPI_ISL_585713, EPI_ISL_585714, EPI_ISL_585715, EPI_ISL_585716, EPI_ISL_585718, EPI_ISL_585719, EPI_ISL_585721, EPI_ISL_585722, EPI_ISL_585723, EPI_ISL_585725, EPI_ISL_585726, EPI_ISL_585728, EPI_ISL_585729, EPI_ISL_585730, EPI_ISL_585732, EPI_ISL_585733, EPI_ISL_585735, EPI_ISL_585736, EPI_ISL_585738, EPI_ISL_585739, EPI_ISL_585741, EPI_ISL_585746, EPI_ISL_585747, EPI_ISL_585749, EPI_ISL_585750, EPI_ISL_585751, EPI_ISL_585752, EPI_ISL_585754, EPI_ISL_585755, EPI_ISL_585756, EPI_ISL_585757, EPI_ISL_585758, EPI_ISL_585759, EPI_ISL_585760, EPI_ISL_585762, EPI_ISL_585763, EPI_ISL_585765, EPI_ISL_585779, EPI_ISL_585781, EPI_ISL_585782, EPI_ISL_585783, EPI_ISL_585791, EPI_ISL_585794, EPI_ISL_585796, EPI_ISL_585798, EPI_ISL_585800, EPI_ISL_585801, EPI_ISL_585804, EPI_ISL_585806, EPI_ISL_585807, EPI_ISL_585825, EPI_ISL_585830, EPI_ISL_585833, EPI_ISL_585834, EPI_ISL_585835, EPI_ISL_585857, EPI_ISL_585862, EPI_ISL_585864, EPI_ISL_585867, EPI_ISL_585876, EPI_ISL_585878, EPI_ISL_585879, EPI_ISL_585886, EPI_ISL_585887, EPI_ISL_585888, EPI_ISL_585892, EPI_ISL_585893, EPI_ISL_585895, EPI_ISL_585899, EPI_ISL_585903, EPI_ISL_585904, EPI_ISL_585908, EPI_ISL_585918, EPI_ISL_585919, EPI_ISL_585920, EPI_ISL_585926, EPI_ISL_585929, EPI_ISL_585930, EPI_ISL_585931, EPI_ISL_585937, EPI_ISL_585938, EPI_ISL_585940, EPI_ISL_585942, EPI_ISL_585955, EPI_ISL_585962, EPI_ISL_585964, EPI_ISL_585975, EPI_ISL_585977, EPI_ISL_585978, EPI_ISL_585980, EPI_ISL_585985, EPI_ISL_585986, EPI_ISL_585988, EPI_ISL_585990, EPI_ISL_585998, EPI_ISL_586005, EPI_ISL_586011, EPI_ISL_586012, EPI_ISL_586013, EPI_ISL_586017, EPI_ISL_586020, EPI_ISL_586023, EPI_ISL_586024, EPI_ISL_586057, EPI_ISL_586074, EPI_ISL_586078, EPI_ISL_586080, EPI_ISL_586092, EPI_ISL_586128, EPI_ISL_586135, EPI_ISL_586162, EPI_ISL_586164, EPI_ISL_586167, EPI_ISL_586213, EPI_ISL_586225 | | | |
| see above | Wales Specialist Virology Centre Sequencing lab: Pathogen Genomics Unit | COVID-19 Genomics UK (COG-UK) Consortium | Catherine Moore, Johnathan Evans, Laura Gifford, Malorie Perry, Simon Cottrell, Angela Marchbank, Alec Birchley, Alexander Adams, Amy Gaskin, Bree Gatica-Wilcox, Jason Coombes, Joel Southgate, Lauren Gilbert, Lee Graham, Nicole Pacchiarini, Sara Kumziene-Summerhayes, Sarah Taylor, Sophie Jones, Sara Rey, Matthew Bull, Joanne Watkins, Sally Corden, Tom Connor |
| EPI_ISL_586240, EPI_ISL_586241, EPI_ISL_586242 | Quadram Institute Bioscience | COVID-19 Genomics UK (COG-UK) Consortium | Dave J. Baker, Gemma L. Kay, Alp Aydin, Thanh Le-Viet, Steven Rudder, Ana P. Tedim, Anastasia Kolyva, Maria Diaz, Leonardo de Oliveira Martins, Nabil-Fareed Alikhan, Lizzie Meadows, Rachael Stanley, Ngozi Elumogo, Muhammed Yasir, Nicholas M. Thomson, Alexander J Trotter, Rachel Gilroy, Samuel Bloomfield, Claire Stuart, Andrew Bell, Reenesh Prakash, Samir Dervisevic, Alison E. Mather, John Wain, Mark Webber, Andrew J. Page, Justin O'Grady |
| EPI_ISL_587303 | Lighthouse Lab in Glasgow | Wellcome Sanger Institute for the COVID-19 Genomics UK (COG-UK) Consortium | Harper VanSteenhouse, Yumi Kasai, David Gray, Carol Clugston, Anna Dominiczak and Alex Alderton, Roberto Amato, Sonia Goncalves, Ewan Harrison, David K. Jackson, Ian Johnston, Dominic Kwiatkowski, Cordelia Langford, John Sillitoe on behalf of the Wellcome Sanger Institute COVID-19 Surveillance Team |
| EPI_ISL_587304, EPI_ISL_587305, EPI_ISL_587306, EPI_ISL_587307, EPI_ISL_587308, EPI_ISL_587309, EPI_ISL_587310, EPI_ISL_587311, EPI_ISL_587312, EPI_ISL_587313, EPI_ISL_587314, EPI_ISL_587315, EPI_ISL_587316, EPI_ISL_587317, EPI_ISL_587318, EPI_ISL_587319, EPI_ISL_587320, EPI_ISL_587321, EPI_ISL_587322, EPI_ISL_587323, EPI_ISL_587324, EPI_ISL_587325, EPI_ISL_587326, EPI_ISL_587327, EPI_ISL_587328, EPI_ISL_587329, EPI_ISL_587330, EPI_ISL_587331, EPI_ISL_587332, EPI_ISL_587333, EPI_ISL_587334, EPI_ISL_587335, EPI_ISL_587336, EPI_ISL_587337, EPI_ISL_587338, EPI_ISL_587339, EPI_ISL_587340, EPI_ISL_587341, EPI_ISL_587342, EPI_ISL_587343, EPI_ISL_587344, EPI_ISL_587345, EPI_ISL_587346, EPI_ISL_587347, EPI_ISL_587348, EPI_ISL_587349, EPI_ISL_587350, EPI_ISL_587351, EPI_ISL_587352, EPI_ISL_587353, EPI_ISL_587354, EPI_ISL_587355, EPI_ISL_587356, EPI_ISL_587357, EPI_ISL_587358, EPI_ISL_587359, EPI_ISL_587360, EPI_ISL_587361, EPI_ISL_587362, EPI_ISL_587363, EPI_ISL_587364, EPI_ISL_587365, EPI_ISL_587366, EPI_ISL_587367, EPI_ISL_587368, EPI_ISL_587369, EPI_ISL_587370, EPI_ISL_587371, EPI_ISL_587372, EPI_ISL_587373, EPI_ISL_587374, EPI_ISL_587375, EPI_ISL_587376, EPI_ISL_587377, EPI_ISL_587378, EPI_ISL_587379, EPI_ISL_587380, EPI_ISL_587381, EPI_ISL_587382, EPI_ISL_587383, EPI_ISL_587384, EPI_ISL_587385 | | | |
| see above | Lighthouse Lab in Glasgow | Wellcome Sanger Institute for the COVID-19 Genomics UK (COG-UK) consortium | Harper VanSteenhouse, Yumi Kasai, David Gray, Carol Clugston, Anna Dominiczak and Alex Alderton, Roberto Amato, Sonia Goncalves, Ewan Harrison, David K. Jackson, Ian Johnston, Dominic Kwiatkowski, Cordelia Langford, John Sillitoe on behalf of the Wellcome Sanger Institute COVID-19 Surveillance Team |
| EPI_ISL_587386 | Lighthouse Lab in Glasgow | Wellcome Sanger Institute for the COVID-19 Genomics UK (COG-UK) Consortium | Harper VanSteenhouse, Yumi Kasai, David Gray, Carol Clugston, Anna Dominiczak and Alex Alderton, Roberto Amato, Sonia Goncalves, Ewan Harrison, David K. Jackson, Ian Johnston, Dominic Kwiatkowski, Cordelia Langford, John Sillitoe on behalf of the Wellcome Sanger Institute COVID-19 Surveillance Team |
| EPI_ISL_587387, EPI_ISL_587388, EPI_ISL_587389, EPI_ISL_587390, EPI_ISL_587391, EPI_ISL_587392, EPI_ISL_587393, EPI_ISL_587394, EPI_ISL_587395, EPI_ISL_587396, EPI_ISL_587397, EPI_ISL_587398, EPI_ISL_587399, EPI_ISL_587400, EPI_ISL_587401, EPI_ISL_587402, EPI_ISL_587403, EPI_ISL_587404, EPI_ISL_587405, EPI_ISL_587406, EPI_ISL_587407, EPI_ISL_587408, EPI_ISL_587409, EPI_ISL_587410, EPI_ISL_587411, EPI_ISL_587412, EPI_ISL_587413, EPI_ISL_587414, EPI_ISL_587415, EPI_ISL_587416, EPI_ISL_587417, EPI_ISL_587418, EPI_ISL_587419, EPI_ISL_587420, EPI_ISL_587421, EPI_ISL_587422, EPI_ISL_587423, EPI_ISL_587424, EPI_ISL_587425, EPI_ISL_587426, EPI_ISL_587427, EPI_ISL_587428, EPI_ISL_587429, EPI_ISL_587430, EPI_ISL_587431, EPI_ISL_587432, EPI_ISL_587433, EPI_ISL_587434, EPI_ISL_587435, EPI_ISL_587436, EPI_ISL_587437, EPI_ISL_587438, EPI_ISL_587439, EPI_ISL_587440, EPI_ISL_587441, EPI_ISL_587442, EPI_ISL_587443, EPI_ISL_587444, EPI_ISL_587445, EPI_ISL_587446, EPI_ISL_587447, EPI_ISL_587448, EPI_ISL_587449, EPI_ISL_587450, EPI_ISL_587451, EPI_ISL_587452, EPI_ISL_587453, EPI_ISL_587454, EPI_ISL_587455, EPI_ISL_587456, EPI_ISL_587457, EPI_ISL_587458, EPI_ISL_587459, EPI_ISL_587460, EPI_ISL_587461, EPI_ISL_587462, EPI_ISL_587463, EPI_ISL_587464, EPI_ISL_587465, EPI_ISL_587466, EPI_ISL_587467, EPI_ISL_587468, EPI_ISL_587469, EPI_ISL_587470, EPI_ISL_587471, EPI_ISL_587472, EPI_ISL_587473, EPI_ISL_587474, EPI_ISL_587475, EPI_ISL_587476 | | | |
| see above | Lighthouse Lab in Glasgow | Wellcome Sanger Institute for the COVID-19 Genomics UK (COG-UK) consortium | Harper VanSteenhouse, Yumi Kasai, David Gray, Carol Clugston, Anna Dominiczak and Alex Alderton, Roberto Amato, Sonia Goncalves, Ewan Harrison, David K. Jackson, Ian Johnston, Dominic Kwiatkowski, Cordelia Langford, John Sillitoe on behalf of the Wellcome Sanger Institute COVID-19 Surveillance Team |
| EPI_ISL_587477 | Lighthouse Lab in Glasgow | Wellcome Sanger Institute for the COVID-19 Genomics UK (COG-UK) Consortium | Harper VanSteenhouse, Yumi Kasai, David Gray, Carol Clugston, Anna Dominiczak and Alex Alderton, Roberto Amato, Sonia Goncalves, Ewan Harrison, David K. Jackson, Ian Johnston, Dominic Kwiatkowski, Cordelia Langford, John Sillitoe on behalf of the Wellcome Sanger Institute COVID-19 Surveillance Team |
| EPI_ISL_587478, EPI_ISL_587479, EPI_ISL_587480, EPI_ISL_587481, EPI_ISL_587482, EPI_ISL_587483, EPI_ISL_587484, EPI_ISL_587485, EPI_ISL_587486, EPI_ISL_587487, EPI_ISL_587488, EPI_ISL_587489, EPI_ISL_587490, EPI_ISL_587491, EPI_ISL_587492, EPI_ISL_587493, EPI_ISL_587494, EPI_ISL_587495, EPI_ISL_587496, EPI_ISL_587497, EPI_ISL_587498, EPI_ISL_587499, EPI_ISL_587500, EPI_ISL_587501, EPI_ISL_587502, EPI_ISL_587503, EPI_ISL_587504, EPI_ISL_587505, EPI_ISL_587506, EPI_ISL_587507, EPI_ISL_587508, EPI_ISL_587509, EPI_ISL_587510, EPI_ISL_587511, EPI_ISL_587512, EPI_ISL_587513, EPI_ISL_587514, EPI_ISL_587515, EPI_ISL_587516, EPI_ISL_587517, EPI_ISL_587518, EPI_ISL_587519, EPI_ISL_587520, EPI_ISL_587521, EPI_ISL_587522, EPI_ISL_587523, EPI_ISL_587524, EPI_ISL_587525, EPI_ISL_587526, EPI_ISL_587527, EPI_ISL_587528, EPI_ISL_587529, EPI_ISL_587530, EPI_ISL_587531, EPI_ISL_587532, EPI_ISL_587533, EPI_ISL_587534, EPI_ISL_587535, EPI_ISL_587536, EPI_ISL_587537, EPI_ISL_587538, EPI_ISL_587539, EPI_ISL_587540, EPI_ISL_587541, EPI_ISL_587542, EPI_ISL_587543, EPI_ISL_587544, EPI_ISL_587545, EPI_ISL_587546, EPI_ISL_587547, EPI_ISL_587548, EPI_ISL_587549, EPI_ISL_587550, EPI_ISL_587551, EPI_ISL_587552, EPI_ISL_587553, EPI_ISL_587554, EPI_ISL_587555, EPI_ISL_587556, EPI_ISL_587557, EPI_ISL_587558, EPI_ISL_587559, EPI_ISL_587560, EPI_ISL_587561, EPI_ISL_587562, EPI_ISL_587563, EPI_ISL_587564, EPI_ISL_587565, EPI_ISL_587566, EPI_ISL_587567, EPI_ISL_587568, EPI_ISL_587569, EPI_ISL_587570, EPI_ISL_587571, EPI_ISL_587572, EPI_ISL_587573, EPI_ISL_587574, EPI_ISL_587575, EPI_ISL_587576, EPI_ISL_587577, EPI_ISL_587578, EPI_ISL_587579, EPI_ISL_587580, EPI_ISL_587581, EPI_ISL_587582, EPI_ISL_587583, EPI_ISL_587584, EPI_ISL_587585, EPI_ISL_587586, EPI_ISL_587587, EPI_ISL_587588, EPI_ISL_587589, EPI_ISL_587590, EPI_ISL_587591, EPI_ISL_587592, EPI_ISL_587593, EPI_ISL_587594, EPI_ISL_587595, EPI_ISL_587596, EPI_ISL_587597, EPI_ISL_587598, EPI_ISL_587599, EPI_ISL_587600, EPI_ISL_587601, EPI_ISL_587602, EPI_ISL_587603, EPI_ISL_587604, EPI_ISL_587605, EPI_ISL_587606, EPI_ISL_587607, EPI_ISL_587608, EPI_ISL_587609, EPI_ISL_587610, EPI_ISL_587611, EPI_ISL_587612, EPI_ISL_587613, EPI_ISL_587614, EPI_ISL_587615, EPI_ISL_587616, EPI_ISL_587617, EPI_ISL_587618, EPI_ISL_587619, EPI_ISL_587620, EPI_ISL_587621, EPI_ISL_587622, EPI_ISL_587623, EPI_ISL_587624, EPI_ISL_587625, EPI_ISL_587626, EPI_ISL_587627, EPI_ISL_587628, EPI_ISL_587629, EPI_ISL_587630, EPI_ISL_587631, EPI_ISL_587632, EPI_ISL_587633, EPI_ISL_587634, EPI_ISL_587635, EPI_ISL_587636, EPI_ISL_587637, EPI_ISL_587638, EPI_ISL_587639, EPI_ISL_587640, EPI_ISL_587641, EPI_ISL_587642, EPI_ISL_587643, EPI_ISL_587644, EPI_ISL_587645, EPI_ISL_587646, EPI_ISL_587647, EPI_ISL_587648, EPI_ISL_587649, EPI_ISL_587650, EPI_ISL_587651, EPI_ISL_587652, EPI_ISL_587653, EPI_ISL_587654, EPI_ISL_587655, EPI_ISL_587656, EPI_ISL_587657, EPI_ISL_587658, EPI_ISL_587659, EPI_ISL_587660, EPI_ISL_587661, EPI_ISL_587662, EPI_ISL_587663, EPI_ISL_587664, EPI_ISL_587665, EPI_ISL_587666, EPI_ISL_587667, EPI_ISL_587668, EPI_ISL_587669, EPI_ISL_587670, EPI_ISL_587671, EPI_ISL_587672, EPI_ISL_587673, EPI_ISL_587674, EPI_ISL_587675, EPI_ISL_587676, EPI_ISL_587677, EPI_ISL_587678, EPI_ISL_587679, EPI_ISL_587680, EPI_ISL_587681, EPI_ISL_587682, EPI_ISL_587683, EPI_ISL_587684, EPI_ISL_587685, EPI_ISL_587686, EPI_ISL_587687, EPI_ISL_587688, EPI_ISL_587689, EPI_ISL_587690, EPI_ISL_587691, EPI_ISL_587692, EPI_ISL_587693, EPI_ISL_587694, EPI_ISL_587695, EPI_ISL_587696, EPI_ISL_587697, EPI_ISL_587698, EPI_ISL_587699, EPI_ISL_587700, EPI_ISL_587701, EPI_ISL_587702, EPI_ISL_587703, EPI_ISL_587704, EPI_ISL_587705, EPI_ISL_587706, EPI_ISL_587707, EPI_ISL_587708, EPI_ISL_587709, EPI_ISL_587710 | | | |
| see above | Lighthouse Lab in Glasgow | Wellcome Sanger Institute for the COVID-19 Genomics UK (COG-UK) consortium | Harper VanSteenhouse, Yumi Kasai, David Gray, Carol Clugston, Anna Dominiczak and Alex Alderton, Roberto Amato, Sonia Goncalves, Ewan Harrison, David K. Jackson, Ian Johnston, Dominic Kwiatkowski, Cordelia Langford, John Sillitoe on behalf of the Wellcome Sanger Institute COVID-19 Surveillance Team |
| EPI_ISL_587711, EPI_ISL_587712, EPI_ISL_587713, EPI_ISL_587714, EPI_ISL_587718, EPI_ISL_587720, EPI_ISL_587721, EPI_ISL_587722, EPI_ISL_587724, EPI_ISL_587726, EPI_ISL_587727, EPI_ISL_587729, EPI_ISL_587730, EPI_ISL_587731, EPI_ISL_587734, EPI_ISL_587738, EPI_ISL_587741, EPI_ISL_587744, EPI_ISL_587746, EPI_ISL_587748, EPI_ISL_587750, EPI_ISL_587751, EPI_ISL_587753, EPI_ISL_587754, EPI_ISL_587755, EPI_ISL_587758, EPI_ISL_587760, EPI_ISL_587761, EPI_ISL_587762, EPI_ISL_587763, EPI_ISL_587764, EPI_ISL_587765, EPI_ISL_587766, EPI_ISL_587768, EPI_ISL_587769, EPI_ISL_587770, EPI_ISL_587772, EPI_ISL_587773, EPI_ISL_587774, EPI_ISL_587775, EPI_ISL_587776, EPI_ISL_587777, EPI_ISL_587778, EPI_ISL_587779, EPI_ISL_587781, EPI_ISL_587782, EPI_ISL_587784, EPI_ISL_587785, EPI_ISL_587786, EPI_ISL_587787, EPI_ISL_587788, EPI_ISL_587789, EPI_ISL_587791, EPI_ISL_587792, EPI_ISL_587793, EPI_ISL_587794, EPI_ISL_587796, EPI_ISL_587797, EPI_ISL_587799, EPI_ISL_587800, EPI_ISL_587801, EPI_ISL_587803, EPI_ISL_587806, EPI_ISL_587807, EPI_ISL_587808, EPI_ISL_587809, EPI_ISL_587814, EPI_ISL_587816, EPI_ISL_587818, EPI_ISL_587819, EPI_ISL_587820, EPI_ISL_587822, EPI_ISL_587824, EPI_ISL_587825, EPI_ISL_587828, EPI_ISL_587829, EPI_ISL_587830, EPI_ISL_587833, EPI_ISL_587834, EPI_ISL_587835, EPI_ISL_587837, EPI_ISL_587838, EPI_ISL_587840, EPI_ISL_587843, EPI_ISL_587844, EPI_ISL_587846, EPI_ISL_587847, EPI_ISL_587849, EPI_ISL_587852, EPI_ISL_587853, EPI_ISL_587856, EPI_ISL_587859, EPI_ISL_587860, EPI_ISL_587861, EPI_ISL_587862, EPI_ISL_587864, EPI_ISL_587865, EPI_ISL_587866, EPI_ISL_587867, EPI_ISL_587869, EPI_ISL_587870, EPI_ISL_587873, EPI_ISL_587874, EPI_ISL_587875, EPI_ISL_587876, EPI_ISL_587883, EPI_ISL_587884, EPI_ISL_587885, EPI_ISL_587886, EPI_ISL_587887, EPI_ISL_587888, EPI_ISL_587890, EPI_ISL_587891, EPI_ISL_587897, EPI_ISL_587898, EPI_ISL_587899, EPI_ISL_587900, EPI_ISL_587901, EPI_ISL_587902, EPI_ISL_587904, EPI_ISL_587905, EPI_ISL_587906, EPI_ISL_587907, EPI_ISL_587908, EPI_ISL_587910, EPI_ISL_587911, EPI_ISL_587912, EPI_ISL_587913, EPI_ISL_587916, EPI_ISL_587917, EPI_ISL_587920, EPI_ISL_587922, EPI_ISL_587923, EPI_ISL_587924, EPI_ISL_587927, EPI_ISL_587928, EPI_ISL_587929, EPI_ISL_587930, EPI_ISL_587931, EPI_ISL_587932, EPI_ISL_587933, EPI_ISL_587934, EPI_ISL_587935, EPI_ISL_587937, | | | |

| | | | |
|---|---|---|---|
| EPI_ISL_587938, EPI_ISL_587941, EPI_ISL_587944, EPI_ISL_587946, EPI_ISL_587947, EPI_ISL_587950, EPI_ISL_587951, EPI_ISL_587952, EPI_ISL_587953, EPI_ISL_587957, EPI_ISL_587958, EPI_ISL_587960, EPI_ISL_587961, EPI_ISL_587965, EPI_ISL_587966, EPI_ISL_587967, EPI_ISL_587971, EPI_ISL_587972, EPI_ISL_587973, EPI_ISL_587976 | | | |
| see above | Lighthouse Lab in Alderley Park | Wellcome Sanger Institute for the COVID-19 Genomics UK (COG-UK) consortium | Jacquelyn Wynn, Mairead Hyland, The Lighthouse Lab in Alderley Park and Alex Alderton, Roberto Amato, Sonia Goncalves, Ewan Harrison, David K. Jackson, Ian Johnston, Dominic Kwiatkowski, Cordelia Langford, John Sillitoe on behalf of the Wellcome Sanger Institute COVID-19 Surveillance Team |
| EPI_ISL_587977 | Lighthouse Lab in Alderley Park | Wellcome Sanger Institute for the COVID-19 Genomics UK (COG-UK) Consortium | Jacquelyn Wynn, Mairead Hyland, The Lighthouse Lab in Alderley Park and Alex Alderton, Roberto Amato, Sonia Goncalves, Ewan Harrison, David K. Jackson, Ian Johnston, Dominic Kwiatkowski, Cordelia Langford, John Sillitoe on behalf of the Wellcome Sanger Institute COVID-19 Surveillance Team |
| EPI_ISL_587978, EPI_ISL_587981, EPI_ISL_587984, EPI_ISL_587986, EPI_ISL_587987, EPI_ISL_587990, EPI_ISL_587992, EPI_ISL_587993, EPI_ISL_587996, EPI_ISL_587997, EPI_ISL_587998, EPI_ISL_587999, EPI_ISL_588000, EPI_ISL_588001, EPI_ISL_588002, EPI_ISL_588004 | | | |
| see above | Lighthouse Lab in Alderley Park | Wellcome Sanger Institute for the COVID-19 Genomics UK (COG-UK) consortium | Jacquelyn Wynn, Mairead Hyland, The Lighthouse Lab in Alderley Park and Alex Alderton, Roberto Amato, Sonia Goncalves, Ewan Harrison, David K. Jackson, Ian Johnston, Dominic Kwiatkowski, Cordelia Langford, John Sillitoe on behalf of the Wellcome Sanger Institute COVID-19 Surveillance Team |
| EPI_ISL_588007 | Lighthouse Lab in Alderley Park | Wellcome Sanger Institute for the COVID-19 Genomics UK (COG-UK) Consortium | Jacquelyn Wynn, Mairead Hyland, The Lighthouse Lab in Alderley Park and Alex Alderton, Roberto Amato, Sonia Goncalves, Ewan Harrison, David K. Jackson, Ian Johnston, Dominic Kwiatkowski, Cordelia Langford, John Sillitoe on behalf of the Wellcome Sanger Institute COVID-19 Surveillance Team |
| EPI_ISL_588009, EPI_ISL_588010, EPI_ISL_588011, EPI_ISL_588012, EPI_ISL_588013, EPI_ISL_588014, EPI_ISL_588015, EPI_ISL_588017, EPI_ISL_588018, EPI_ISL_588019, EPI_ISL_588021, EPI_ISL_588022, EPI_ISL_588024, EPI_ISL_588025 | | | |
| see above | Lighthouse Lab in Alderley Park | Wellcome Sanger Institute for the COVID-19 Genomics UK (COG-UK) consortium | Jacquelyn Wynn, Mairead Hyland, The Lighthouse Lab in Alderley Park and Alex Alderton, Roberto Amato, Sonia Goncalves, Ewan Harrison, David K. Jackson, Ian Johnston, Dominic Kwiatkowski, Cordelia Langford, John Sillitoe on behalf of the Wellcome Sanger Institute COVID-19 Surveillance Team |
| EPI_ISL_588026 | Lighthouse Lab in Alderley Park | Wellcome Sanger Institute for the COVID-19 Genomics UK (COG-UK) Consortium | Jacquelyn Wynn, Mairead Hyland, The Lighthouse Lab in Alderley Park and Alex Alderton, Roberto Amato, Sonia Goncalves, Ewan Harrison, David K. Jackson, Ian Johnston, Dominic Kwiatkowski, Cordelia Langford, John Sillitoe on behalf of the Wellcome Sanger Institute COVID-19 Surveillance Team |
| EPI_ISL_588028, EPI_ISL_588029, EPI_ISL_588031, EPI_ISL_588032, EPI_ISL_588034, EPI_ISL_588036, EPI_ISL_588037, EPI_ISL_588038, EPI_ISL_588040, EPI_ISL_588042, EPI_ISL_588044, EPI_ISL_588045, EPI_ISL_588046, EPI_ISL_588048, EPI_ISL_588049, EPI_ISL_588051, EPI_ISL_588054, EPI_ISL_588055, EPI_ISL_588057, EPI_ISL_588059 | | | |
| see above | Lighthouse Lab in Alderley Park | Wellcome Sanger Institute for the COVID-19 Genomics UK (COG-UK) consortium | Jacquelyn Wynn, Mairead Hyland, The Lighthouse Lab in Alderley Park and Alex Alderton, Roberto Amato, Sonia Goncalves, Ewan Harrison, David K. Jackson, Ian Johnston, Dominic Kwiatkowski, Cordelia Langford, John Sillitoe on behalf of the Wellcome Sanger Institute COVID-19 Surveillance Team |
| EPI_ISL_588060, EPI_ISL_588061, EPI_ISL_588062, EPI_ISL_588063, EPI_ISL_588064, EPI_ISL_588065, EPI_ISL_588066, EPI_ISL_588067, EPI_ISL_588068, EPI_ISL_588069, EPI_ISL_588070, EPI_ISL_588071, EPI_ISL_588072, EPI_ISL_588073, EPI_ISL_588074 | | | |
| see above | Lighthouse Lab in Glasgow | Wellcome Sanger Institute for the COVID-19 Genomics UK (COG-UK) consortium | Harper VanSteenhouse, Yumi Kasai, David Gray, Carol Clugston, Anna Dominiczak and Alex Alderton, Roberto Amato, Sonia Goncalves, Ewan Harrison, David K. Jackson, Ian Johnston, Dominic Kwiatkowski, Cordelia Langford, John Sillitoe on behalf of the Wellcome Sanger Institute COVID-19 Surveillance Team |
| EPI_ISL_588075 | Lighthouse Lab in Glasgow | Wellcome Sanger Institute for the COVID-19 Genomics UK (COG-UK) Consortium | Harper VanSteenhouse, Yumi Kasai, David Gray, Carol Clugston, Anna Dominiczak and Alex Alderton, Roberto Amato, Sonia Goncalves, Ewan Harrison, David K. Jackson, Ian Johnston, Dominic Kwiatkowski, Cordelia Langford, John Sillitoe on behalf of the Wellcome Sanger Institute COVID-19 Surveillance Team |
| EPI_ISL_588076, EPI_ISL_588077, EPI_ISL_588078, EPI_ISL_588079, EPI_ISL_588080, EPI_ISL_588081, EPI_ISL_588082, EPI_ISL_588083, EPI_ISL_588084, EPI_ISL_588085, EPI_ISL_588086, EPI_ISL_588087, EPI_ISL_588088, EPI_ISL_588089, EPI_ISL_588090, EPI_ISL_588091, EPI_ISL_588092, EPI_ISL_588093, EPI_ISL_588094, EPI_ISL_588095, EPI_ISL_588096, EPI_ISL_588097, EPI_ISL_588098, EPI_ISL_588099, EPI_ISL_588100, EPI_ISL_588101, EPI_ISL_588102, EPI_ISL_588103, EPI_ISL_588104, EPI_ISL_588105, EPI_ISL_588106, EPI_ISL_588107, EPI_ISL_588108, EPI_ISL_588109, EPI_ISL_588110, EPI_ISL_588111, EPI_ISL_588112, EPI_ISL_588113, EPI_ISL_588114, EPI_ISL_588115, EPI_ISL_588116, EPI_ISL_588117, EPI_ISL_588118, EPI_ISL_588119, EPI_ISL_588120, EPI_ISL_588121, EPI_ISL_588122, EPI_ISL_588123, EPI_ISL_588124 | | | |
| see above | Lighthouse Lab in Glasgow | Wellcome Sanger Institute for the COVID-19 Genomics UK (COG-UK) consortium | Harper VanSteenhouse, Yumi Kasai, David Gray, Carol Clugston, Anna Dominiczak and Alex Alderton, Roberto Amato, Sonia Goncalves, Ewan Harrison, David K. Jackson, Ian Johnston, Dominic Kwiatkowski, Cordelia Langford, John Sillitoe on behalf of the Wellcome Sanger Institute COVID-19 Surveillance Team |
| EPI_ISL_588125 | Lighthouse Lab in Glasgow | Wellcome Sanger Institute for the COVID-19 Genomics UK (COG-UK) Consortium | Harper VanSteenhouse, Yumi Kasai, David Gray, Carol Clugston, Anna Dominiczak and Alex Alderton, Roberto Amato, Sonia Goncalves, Ewan Harrison, David K. Jackson, Ian Johnston, Dominic Kwiatkowski, Cordelia Langford, John Sillitoe on behalf of the Wellcome Sanger Institute COVID-19 Surveillance Team |
| EPI_ISL_588126, EPI_ISL_588127 | Lighthouse Lab in Glasgow | Wellcome Sanger Institute for the COVID-19 Genomics UK (COG-UK) consortium | Harper VanSteenhouse, Yumi Kasai, David Gray, Carol Clugston, Anna Dominiczak and Alex Alderton, Roberto Amato, Sonia Goncalves, Ewan Harrison, David K. Jackson, Ian Johnston, Dominic Kwiatkowski, Cordelia Langford, John Sillitoe on behalf of the Wellcome Sanger Institute COVID-19 Surveillance Team |
| EPI_ISL_588128 | Lighthouse Lab in Glasgow | Wellcome Sanger Institute for the COVID-19 Genomics UK (COG-UK) Consortium | Harper VanSteenhouse, Yumi Kasai, David Gray, Carol Clugston, Anna Dominiczak and Alex Alderton, Roberto Amato, Sonia Goncalves, Ewan Harrison, David K. Jackson, Ian Johnston, Dominic Kwiatkowski, Cordelia Langford, John Sillitoe on behalf of the Wellcome Sanger Institute COVID-19 Surveillance Team |
| EPI_ISL_588129, EPI_ISL_588130, EPI_ISL_588131, EPI_ISL_588132, EPI_ISL_588133, EPI_ISL_588134, EPI_ISL_588135, EPI_ISL_588136, EPI_ISL_588137, EPI_ISL_588138, EPI_ISL_588139, EPI_ISL_588140, EPI_ISL_588141, EPI_ISL_588142, EPI_ISL_588143, EPI_ISL_588144, EPI_ISL_588145, EPI_ISL_588146, EPI_ISL_588147, EPI_ISL_588148, EPI_ISL_588149, EPI_ISL_588150, EPI_ISL_588151, EPI_ISL_588152, EPI_ISL_588153, EPI_ISL_588154, EPI_ISL_588155, EPI_ISL_588156, EPI_ISL_588157, EPI_ISL_588158, EPI_ISL_588159, EPI_ISL_588160, EPI_ISL_588161, EPI_ISL_588162, EPI_ISL_588163, EPI_ISL_588164, EPI_ISL_588165, EPI_ISL_588166, EPI_ISL_588167, EPI_ISL_588168, EPI_ISL_588169, EPI_ISL_588170, EPI_ISL_588171, EPI_ISL_588172, EPI_ISL_588173, EPI_ISL_588174 | | | |
| see above | Lighthouse Lab in Glasgow | Wellcome Sanger Institute for the COVID-19 Genomics UK (COG-UK) consortium | Harper VanSteenhouse, Yumi Kasai, David Gray, Carol Clugston, Anna Dominiczak and Alex Alderton, Roberto Amato, Sonia Goncalves, Ewan Harrison, David K. Jackson, Ian Johnston, Dominic Kwiatkowski, Cordelia Langford, John Sillitoe on behalf of the Wellcome Sanger Institute COVID-19 Surveillance Team |
| EPI_ISL_588175 | Lighthouse Lab in Glasgow | Wellcome Sanger Institute for the COVID-19 Genomics UK (COG-UK) Consortium | Harper VanSteenhouse, Yumi Kasai, David Gray, Carol Clugston, Anna Dominiczak and Alex Alderton, Roberto Amato, Sonia Goncalves, Ewan Harrison, David K. Jackson, Ian Johnston, Dominic Kwiatkowski, Cordelia Langford, John Sillitoe on behalf of the Wellcome Sanger Institute COVID-19 Surveillance Team |
| EPI_ISL_588176, EPI_ISL_588177, EPI_ISL_588178, EPI_ISL_588179, EPI_ISL_588180, EPI_ISL_588181, EPI_ISL_588182, EPI_ISL_588183, EPI_ISL_588184, EPI_ISL_588185, EPI_ISL_588186, EPI_ISL_588187, EPI_ISL_588188, EPI_ISL_588189, EPI_ISL_588190, EPI_ISL_588191, EPI_ISL_588192, EPI_ISL_588193, EPI_ISL_588194, EPI_ISL_588195, EPI_ISL_588196, EPI_ISL_588197, EPI_ISL_588198, EPI_ISL_588199, EPI_ISL_588200, EPI_ISL_588201, EPI_ISL_588202, EPI_ISL_588203, EPI_ISL_588204, EPI_ISL_588205, EPI_ISL_588207, EPI_ISL_588208, EPI_ISL_588209, EPI_ISL_588210, EPI_ISL_588211, EPI_ISL_588212, EPI_ISL_588213, EPI_ISL_588214, EPI_ISL_588215, EPI_ISL_588216, EPI_ISL_588217, EPI_ISL_588218, EPI_ISL_588219, EPI_ISL_588220, EPI_ISL_588221, EPI_ISL_588222, EPI_ISL_588223, EPI_ISL_588224, EPI_ISL_588225, EPI_ISL_588226, EPI_ISL_588227, EPI_ISL_588228, EPI_ISL_588229, EPI_ISL_588230, EPI_ISL_588231, EPI_ISL_588232, EPI_ISL_588233, EPI_ISL_588234, EPI_ISL_588235, EPI_ISL_588236, EPI_ISL_588237, EPI_ISL_588238, EPI_ISL_588239, EPI_ISL_588240, EPI_ISL_588241, EPI_ISL_588242, EPI_ISL_588243, EPI_ISL_588244, EPI_ISL_588245, EPI_ISL_588246, EPI_ISL_588247, EPI_ISL_588248, EPI_ISL_588249, EPI_ISL_588250, EPI_ISL_588251, EPI_ISL_588252, EPI_ISL_588253, EPI_ISL_588254, EPI_ISL_588255, EPI_ISL_588256, EPI_ISL_588257, EPI_ISL_588258, EPI_ISL_588259, EPI_ISL_588260, EPI_ISL_588261, EPI_ISL_588262, EPI_ISL_588263, EPI_ISL_588264, EPI_ISL_588265, EPI_ISL_588266, EPI_ISL_588267, EPI_ISL_588268, EPI_ISL_588269, EPI_ISL_588270, EPI_ISL_588271, EPI_ISL_588272, EPI_ISL_588273, EPI_ISL_588274, EPI_ISL_588275, EPI_ISL_588276, EPI_ISL_588278, EPI_ISL_588279, EPI_ISL_588280, EPI_ISL_588281, EPI_ISL_588282, EPI_ISL_588283, EPI_ISL_588285, EPI_ISL_588286, EPI_ISL_588287, EPI_ISL_588288, EPI_ISL_588289, EPI_ISL_588290, EPI_ISL_588291, EPI_ISL_588292, EPI_ISL_588293, EPI_ISL_588294, EPI_ISL_588295, EPI_ISL_588296, EPI_ISL_588297, EPI_ISL_588298, EPI_ISL_588299, EPI_ISL_588300, EPI_ISL_588301, EPI_ISL_588302, EPI_ISL_588303, EPI_ISL_588304, EPI_ISL_588305, EPI_ISL_588306, EPI_ISL_588307, EPI_ISL_588309, EPI_ISL_588310, EPI_ISL_588311, EPI_ISL_588312, EPI_ISL_588315, EPI_ISL_588316, EPI_ISL_588317, EPI_ISL_588318, EPI_ISL_588319, EPI_ISL_588320, EPI_ISL_588321, EPI_ISL_588322, EPI_ISL_588323, EPI_ISL_588324, EPI_ISL_588325, EPI_ISL_588326, EPI_ISL_588327, EPI_ISL_588328, EPI_ISL_588329, EPI_ISL_588331, EPI_ISL_588332, EPI_ISL_588333, EPI_ISL_588334, EPI_ISL_588335, EPI_ISL_588336, EPI_ISL_588337, EPI_ISL_588338, EPI_ISL_588339, EPI_ISL_588340, EPI_ISL_588341, EPI_ISL_588342, EPI_ISL_588343, EPI_ISL_588344, EPI_ISL_588345, EPI_ISL_588348, EPI_ISL_588349, EPI_ISL_588350, EPI_ISL_588351, EPI_ISL_588353, EPI_ISL_588354, EPI_ISL_588355, EPI_ISL_588356, EPI_ISL_588357, EPI_ISL_588358, EPI_ISL_588359, EPI_ISL_588360, EPI_ISL_588361, EPI_ISL_588362, EPI_ISL_588363, EPI_ISL_588364, EPI_ISL_588365, EPI_ISL_588366, EPI_ISL_588367, EPI_ISL_588368, EPI_ISL_588369, EPI_ISL_588370, EPI_ISL_588371, EPI_ISL_588372, EPI_ISL_588373, EPI_ISL_588374, EPI_ISL_588375, EPI_ISL_588376, EPI_ISL_588377, EPI_ISL_588379, EPI_ISL_588380, EPI_ISL_588382, EPI_ISL_588383, EPI_ISL_588384, EPI_ISL_588385, EPI_ISL_588386, EPI_ISL_588388, EPI_ISL_588390, EPI_ISL_588391, EPI_ISL_588392, EPI_ISL_588395, EPI_ISL_588396, EPI_ISL_588397, EPI_ISL_588398, EPI_ISL_588399, EPI_ISL_588400, EPI_ISL_588401, EPI_ISL_588402, EPI_ISL_588403, EPI_ISL_588404, EPI_ISL_588405, EPI_ISL_588406, EPI_ISL_588407, EPI_ISL_588408, EPI_ISL_588409, EPI_ISL_588410, EPI_ISL_588411, EPI_ISL_588412, EPI_ISL_588413, EPI_ISL_588414, EPI_ISL_588415, EPI_ISL_588417, EPI_ISL_588418, EPI_ISL_588419, EPI_ISL_588420, EPI_ISL_588421, EPI_ISL_588423, EPI_ISL_588425, EPI_ISL_588426, EPI_ISL_588427, EPI_ISL_588429, EPI_ISL_588431, EPI_ISL_588432, EPI_ISL_588433, EPI_ISL_588434, EPI_ISL_588435, EPI_ISL_588436, EPI_ISL_588440, EPI_ISL_588441, EPI_ISL_588442, EPI_ISL_588444, EPI_ISL_588445, EPI_ISL_588446, EPI_ISL_588447, EPI_ISL_588448, EPI_ISL_588449, EPI_ISL_588450, EPI_ISL_588451, EPI_ISL_588452, EPI_ISL_588453, EPI_ISL_588454, EPI_ISL_588455, EPI_ISL_588457, EPI_ISL_588458, EPI_ISL_588459, EPI_ISL_588460, EPI_ISL_588461, EPI_ISL_588462, EPI_ISL_588464, EPI_ISL_588465, EPI_ISL_588466, EPI_ISL_588467, EPI_ISL_588468, EPI_ISL_588469, EPI_ISL_588470, EPI_ISL_588471, EPI_ISL_588472, EPI_ISL_588473, EPI_ISL_588474, EPI_ISL_588475, EPI_ISL_588476, EPI_ISL_588479, EPI_ISL_588488, EPI_ISL_588490 | | | |
| see above | Lighthouse Lab in Glasgow | Wellcome Sanger Institute for the COVID-19 Genomics UK (COG-UK) consortium | Harper VanSteenhouse, Yumi Kasai, David Gray, Carol Clugston, Anna Dominiczak and Alex Alderton, Roberto Amato, Sonia Goncalves, Ewan Harrison, David K. Jackson, Ian Johnston, Dominic Kwiatkowski, Cordelia Langford, John Sillitoe on behalf of the Wellcome Sanger Institute COVID-19 Surveillance Team |
| EPI_ISL_588496 | Lighthouse Lab in Glasgow | Wellcome Sanger Institute for the COVID-19 Genomics UK (COG-UK) Consortium | Harper VanSteenhouse, Yumi Kasai, David Gray, Carol Clugston, Anna Dominiczak and Alex Alderton, Roberto Amato, Sonia Goncalves, Ewan Harrison, David K. Jackson, Ian Johnston, Dominic Kwiatkowski, Cordelia Langford, John Sillitoe on behalf of the Wellcome Sanger Institute COVID-19 Surveillance Team |
| EPI_ISL_588523, EPI_ISL_588533, EPI_ISL_588537, EPI_ISL_588542, EPI_ISL_588546, EPI_ISL_588548, EPI_ISL_588552, EPI_ISL_588553, EPI_ISL_588556, EPI_ISL_588558, EPI_ISL_588563, EPI_ISL_588585, EPI_ISL_588589, EPI_ISL_588592, EPI_ISL_588598, EPI_ISL_588601, EPI_ISL_588608, EPI_ISL_588609, EPI_ISL_588614, EPI_ISL_588619, EPI_ISL_588626, EPI_ISL_588628, EPI_ISL_588633, EPI_ISL_588643, EPI_ISL_588656, EPI_ISL_588659, EPI_ISL_588661, EPI_ISL_588668, EPI_ISL_588672, EPI_ISL_588675, EPI_ISL_588682, EPI_ISL_588684, EPI_ISL_588691, EPI_ISL_588698, EPI_ISL_588699, EPI_ISL_588702, | | | |

| | | | |
|---|---|---|---|
| EPI_ISL_588706, EPI_ISL_588712 | | | |
| see above | Lighthouse Lab in Glasgow | Wellcome Sanger Institute for the COVID-19 Genomics UK (COG-UK) consortium | Harper VanSteenhouse, Yumi Kasai, David Gray, Carol Clugston, Anna Dominiczak and Alex Alderton, Roberto Amato, Sonia Goncalves, Ewan Harrison, David K. Jackson, Ian Johnston, Dominic Kwiatkowski, Cordelia Langford, John Sillitoe on behalf of the Wellcome Sanger Institute COVID-19 Surveillance Team |
| EPI_ISL_588717 | Lighthouse Lab in Glasgow | Wellcome Sanger Institute for the COVID-19 Genomics UK (COG-UK) Consortium | Harper VanSteenhouse, Yumi Kasai, David Gray, Carol Clugston, Anna Dominiczak and Alex Alderton, Roberto Amato, Sonia Goncalves, Ewan Harrison, David K. Jackson, Ian Johnston, Dominic Kwiatkowski, Cordelia Langford, John Sillitoe on behalf of the Wellcome Sanger Institute COVID-19 Surveillance Team |
| EPI_ISL_588725, EPI_ISL_588726, EPI_ISL_588727, EPI_ISL_588728, EPI_ISL_588729, EPI_ISL_588732, EPI_ISL_588733, EPI_ISL_588734, EPI_ISL_588735, EPI_ISL_588736, EPI_ISL_588737, EPI_ISL_588738, EPI_ISL_588739, EPI_ISL_588740, EPI_ISL_588741, EPI_ISL_588742, EPI_ISL_588743, EPI_ISL_588744, EPI_ISL_588745, EPI_ISL_588746, EPI_ISL_588747, EPI_ISL_588748, EPI_ISL_588749, EPI_ISL_588750, EPI_ISL_588751, EPI_ISL_588753, EPI_ISL_588754, EPI_ISL_588755, EPI_ISL_588756, EPI_ISL_588757, EPI_ISL_588758, EPI_ISL_588759, EPI_ISL_588760, EPI_ISL_588761, EPI_ISL_588762, EPI_ISL_588764, EPI_ISL_588765, EPI_ISL_588766, EPI_ISL_588767, EPI_ISL_588768, EPI_ISL_588769, EPI_ISL_588770, EPI_ISL_588771, EPI_ISL_588772, EPI_ISL_588773, EPI_ISL_588774, EPI_ISL_588775, EPI_ISL_588776, EPI_ISL_588777, EPI_ISL_588778, EPI_ISL_588779, EPI_ISL_588780, EPI_ISL_588781, EPI_ISL_588782, EPI_ISL_588783, EPI_ISL_588784, EPI_ISL_588785, EPI_ISL_588786, EPI_ISL_588787, EPI_ISL_588788, EPI_ISL_588789, EPI_ISL_588790, EPI_ISL_588791, EPI_ISL_588792, EPI_ISL_588793, EPI_ISL_588794, EPI_ISL_588795, EPI_ISL_588796, EPI_ISL_588797, EPI_ISL_588798, EPI_ISL_588799, EPI_ISL_588800, EPI_ISL_588801, EPI_ISL_588802, EPI_ISL_588804, EPI_ISL_588805, EPI_ISL_588807, EPI_ISL_588808, EPI_ISL_588809, EPI_ISL_588810, EPI_ISL_588811, EPI_ISL_588812, EPI_ISL_588813, EPI_ISL_588814, EPI_ISL_588815, EPI_ISL_588816, EPI_ISL_588817, EPI_ISL_588818, EPI_ISL_588819, EPI_ISL_588820, EPI_ISL_588821, EPI_ISL_588822, EPI_ISL_588823, EPI_ISL_588824, EPI_ISL_588825, EPI_ISL_588826, EPI_ISL_588827, EPI_ISL_588828, EPI_ISL_588829, EPI_ISL_588830, EPI_ISL_588831, EPI_ISL_588832, EPI_ISL_588833, EPI_ISL_588834, EPI_ISL_588835, EPI_ISL_588836, EPI_ISL_588837, EPI_ISL_588838, EPI_ISL_588839, EPI_ISL_588840, EPI_ISL_588841, EPI_ISL_588842, EPI_ISL_588843 | | | |
| see above | Lighthouse Lab in Glasgow | Wellcome Sanger Institute for the COVID-19 Genomics UK (COG-UK) consortium | Harper VanSteenhouse, Yumi Kasai, David Gray, Carol Clugston, Anna Dominiczak and Alex Alderton, Roberto Amato, Sonia Goncalves, Ewan Harrison, David K. Jackson, Ian Johnston, Dominic Kwiatkowski, Cordelia Langford, John Sillitoe on behalf of the Wellcome Sanger Institute COVID-19 Surveillance Team |
| EPI_ISL_588844, EPI_ISL_588845 | Lighthouse Lab in Glasgow | Wellcome Sanger Institute for the COVID-19 Genomics UK (COG-UK) Consortium | Harper VanSteenhouse, Yumi Kasai, David Gray, Carol Clugston, Anna Dominiczak and Alex Alderton, Roberto Amato, Sonia Goncalves, Ewan Harrison, David K. Jackson, Ian Johnston, Dominic Kwiatkowski, Cordelia Langford, John Sillitoe on behalf of the Wellcome Sanger Institute COVID-19 Surveillance Team |
| EPI_ISL_588846, EPI_ISL_588847, EPI_ISL_588848, EPI_ISL_588849, EPI_ISL_588850, EPI_ISL_588851, EPI_ISL_588852, EPI_ISL_588853, EPI_ISL_588854, EPI_ISL_588855, EPI_ISL_588856, EPI_ISL_588857, EPI_ISL_588858, EPI_ISL_588859, EPI_ISL_588860, EPI_ISL_588861, EPI_ISL_588862, EPI_ISL_588863, EPI_ISL_588864, EPI_ISL_588865, EPI_ISL_588866, EPI_ISL_588867, EPI_ISL_588868, EPI_ISL_588869, EPI_ISL_588870, EPI_ISL_588871, EPI_ISL_588872, EPI_ISL_588873, EPI_ISL_588874, EPI_ISL_588875, EPI_ISL_588876, EPI_ISL_588877, EPI_ISL_588878, EPI_ISL_588879, EPI_ISL_588880, EPI_ISL_588881, EPI_ISL_588882, EPI_ISL_588883, EPI_ISL_588884, EPI_ISL_588885, EPI_ISL_588886, EPI_ISL_588887, EPI_ISL_588888, EPI_ISL_588890, EPI_ISL_588891, EPI_ISL_588892, EPI_ISL_588893, EPI_ISL_588894, EPI_ISL_588895, EPI_ISL_588896, EPI_ISL_588897, EPI_ISL_588898, EPI_ISL_588899, EPI_ISL_588900, EPI_ISL_588901, EPI_ISL_588902, EPI_ISL_588903, EPI_ISL_588904, EPI_ISL_588905, EPI_ISL_588906, EPI_ISL_588907, EPI_ISL_588908, EPI_ISL_588909, EPI_ISL_588910, EPI_ISL_588911, EPI_ISL_588913, EPI_ISL_588914, EPI_ISL_588915, EPI_ISL_588916, EPI_ISL_588917, EPI_ISL_588918, EPI_ISL_588919, EPI_ISL_588920, EPI_ISL_588921, EPI_ISL_588922, EPI_ISL_588923, EPI_ISL_588924, EPI_ISL_588925, EPI_ISL_588926 | | | |
| see above | Lighthouse Lab in Glasgow | Wellcome Sanger Institute for the COVID-19 Genomics UK (COG-UK) consortium | Harper VanSteenhouse, Yumi Kasai, David Gray, Carol Clugston, Anna Dominiczak and Alex Alderton, Roberto Amato, Sonia Goncalves, Ewan Harrison, David K. Jackson, Ian Johnston, Dominic Kwiatkowski, Cordelia Langford, John Sillitoe on behalf of the Wellcome Sanger Institute COVID-19 Surveillance Team |
| EPI_ISL_588927 | Lighthouse Lab in Glasgow | Wellcome Sanger Institute for the COVID-19 Genomics UK (COG-UK) Consortium | Harper VanSteenhouse, Yumi Kasai, David Gray, Carol Clugston, Anna Dominiczak and Alex Alderton, Roberto Amato, Sonia Goncalves, Ewan Harrison, David K. Jackson, Ian Johnston, Dominic Kwiatkowski, Cordelia Langford, John Sillitoe on behalf of the Wellcome Sanger Institute COVID-19 Surveillance Team |
| EPI_ISL_588928, EPI_ISL_588929, EPI_ISL_588930, EPI_ISL_588931, EPI_ISL_588932, EPI_ISL_588933, EPI_ISL_588934, EPI_ISL_588935, EPI_ISL_588936, EPI_ISL_588937, EPI_ISL_588938, EPI_ISL_588939 | | | |
| see above | Lighthouse Lab in Glasgow | Wellcome Sanger Institute for the COVID-19 Genomics UK (COG-UK) consortium | Harper VanSteenhouse, Yumi Kasai, David Gray, Carol Clugston, Anna Dominiczak and Alex Alderton, Roberto Amato, Sonia Goncalves, Ewan Harrison, David K. Jackson, Ian Johnston, Dominic Kwiatkowski, Cordelia Langford, John Sillitoe on behalf of the Wellcome Sanger Institute COVID-19 Surveillance Team |
| EPI_ISL_588940 | Lighthouse Lab in Glasgow | Wellcome Sanger Institute for the COVID-19 Genomics UK (COG-UK) Consortium | Harper VanSteenhouse, Yumi Kasai, David Gray, Carol Clugston, Anna Dominiczak and Alex Alderton, Roberto Amato, Sonia Goncalves, Ewan Harrison, David K. Jackson, Ian Johnston, Dominic Kwiatkowski, Cordelia Langford, John Sillitoe on behalf of the Wellcome Sanger Institute COVID-19 Surveillance Team |
| EPI_ISL_588941, EPI_ISL_588942, EPI_ISL_588943, EPI_ISL_588944, EPI_ISL_588945, EPI_ISL_588946, EPI_ISL_588947, EPI_ISL_588948, EPI_ISL_588949, EPI_ISL_588950, EPI_ISL_588951, EPI_ISL_588952, EPI_ISL_588953, EPI_ISL_588954, EPI_ISL_588955, EPI_ISL_588956, EPI_ISL_588957, EPI_ISL_588958, EPI_ISL_588959, EPI_ISL_588960, EPI_ISL_588961 | | | |
| see above | Lighthouse Lab in Glasgow | Wellcome Sanger Institute for the COVID-19 Genomics UK (COG-UK) consortium | Harper VanSteenhouse, Yumi Kasai, David Gray, Carol Clugston, Anna Dominiczak and Alex Alderton, Roberto Amato, Sonia Goncalves, Ewan Harrison, David K. Jackson, Ian Johnston, Dominic Kwiatkowski, Cordelia Langford, John Sillitoe on behalf of the Wellcome Sanger Institute COVID-19 Surveillance Team |
| EPI_ISL_588962 | Lighthouse Lab in Glasgow | Wellcome Sanger Institute for the COVID-19 Genomics UK (COG-UK) Consortium | Harper VanSteenhouse, Yumi Kasai, David Gray, Carol Clugston, Anna Dominiczak and Alex Alderton, Roberto Amato, Sonia Goncalves, Ewan Harrison, David K. Jackson, Ian Johnston, Dominic Kwiatkowski, Cordelia Langford, John Sillitoe on behalf of the Wellcome Sanger Institute COVID-19 Surveillance Team |
| EPI_ISL_588963, EPI_ISL_588964, EPI_ISL_588965, EPI_ISL_588966, EPI_ISL_588967, EPI_ISL_588968, EPI_ISL_588969, EPI_ISL_588970, EPI_ISL_588971, EPI_ISL_588972, EPI_ISL_588974, EPI_ISL_588975, EPI_ISL_588976, EPI_ISL_588977, EPI_ISL_588978, EPI_ISL_588979, EPI_ISL_588980, EPI_ISL_588981, EPI_ISL_588982, EPI_ISL_588983, EPI_ISL_588984, EPI_ISL_588985, EPI_ISL_588986, EPI_ISL_588987, EPI_ISL_588988, EPI_ISL_588989, EPI_ISL_588990, EPI_ISL_588991, EPI_ISL_588992, EPI_ISL_588993, EPI_ISL_588994, EPI_ISL_588996, EPI_ISL_588997, EPI_ISL_588998, EPI_ISL_588999, EPI_ISL_589000, EPI_ISL_589001, EPI_ISL_589003, EPI_ISL_589004, EPI_ISL_589005, EPI_ISL_589006, EPI_ISL_589007, EPI_ISL_589008, EPI_ISL_589009, EPI_ISL_589010, EPI_ISL_589011, EPI_ISL_589012, EPI_ISL_589013, EPI_ISL_589014, EPI_ISL_589015, EPI_ISL_589016 | | | |
| see above | Lighthouse Lab in Glasgow | Wellcome Sanger Institute for the COVID-19 Genomics UK (COG-UK) consortium | Harper VanSteenhouse, Yumi Kasai, David Gray, Carol Clugston, Anna Dominiczak and Alex Alderton, Roberto Amato, Sonia Goncalves, Ewan Harrison, David K. Jackson, Ian Johnston, Dominic Kwiatkowski, Cordelia Langford, John Sillitoe on behalf of the Wellcome Sanger Institute COVID-19 Surveillance Team |
| EPI_ISL_589017, EPI_ISL_589018 | Lighthouse Lab in Milton Keynes | Wellcome Sanger Institute for the COVID-19 Genomics UK (COG-UK) consortium | The Lighthouse Lab in Milton Keynes and Alex Alderton, Roberto Amato, Sonia Goncalves, Ewan Harrison, David K. Jackson, Ian Johnston, Dominic Kwiatkowski, Cordelia Langford, John Sillitoe on behalf of the Wellcome Sanger Institute COVID-19 Surveillance Team |
| EPI_ISL_589020, EPI_ISL_589022, EPI_ISL_589023, EPI_ISL_589025 | Lighthouse Lab in Glasgow | Wellcome Sanger Institute for the COVID-19 Genomics UK (COG-UK) consortium | Harper VanSteenhouse, Yumi Kasai, David Gray, Carol Clugston, Anna Dominiczak and Alex Alderton, Roberto Amato, Sonia Goncalves, Ewan Harrison, David K. Jackson, Ian Johnston, Dominic Kwiatkowski, Cordelia Langford, John Sillitoe on behalf of the Wellcome Sanger Institute COVID-19 Surveillance Team |
| EPI_ISL_589026 | Lighthouse Lab in Milton Keynes | Wellcome Sanger Institute for the COVID-19 Genomics UK (COG-UK) consortium | The Lighthouse Lab in Milton Keynes and Alex Alderton, Roberto Amato, Sonia Goncalves, Ewan Harrison, David K. Jackson, Ian Johnston, Dominic Kwiatkowski, Cordelia Langford, John Sillitoe on behalf of the Wellcome Sanger Institute COVID-19 Surveillance Team |
| EPI_ISL_589027, EPI_ISL_589031, EPI_ISL_589032, EPI_ISL_589033, EPI_ISL_589035, EPI_ISL_589036, EPI_ISL_589037, EPI_ISL_589038 | Lighthouse Lab in Glasgow | Wellcome Sanger Institute for the COVID-19 Genomics UK (COG-UK) consortium | Harper VanSteenhouse, Yumi Kasai, David Gray, Carol Clugston, Anna Dominiczak and Alex Alderton, Roberto Amato, Sonia Goncalves, Ewan Harrison, David K. Jackson, Ian Johnston, Dominic Kwiatkowski, Cordelia Langford, John Sillitoe on behalf of the Wellcome Sanger Institute COVID-19 Surveillance Team |
| EPI_ISL_589041 | Lighthouse Lab in Milton Keynes | Wellcome Sanger Institute for the COVID-19 Genomics UK (COG-UK) consortium | The Lighthouse Lab in Milton Keynes and Alex Alderton, Roberto Amato, Sonia Goncalves, Ewan Harrison, David K. Jackson, Ian Johnston, Dominic Kwiatkowski, Cordelia Langford, John Sillitoe on behalf of the Wellcome Sanger Institute COVID-19 Surveillance Team |
| EPI_ISL_589042 | Lighthouse Lab in Glasgow | Wellcome Sanger Institute for the COVID-19 Genomics UK (COG-UK) consortium | Harper VanSteenhouse, Yumi Kasai, David Gray, Carol Clugston, Anna Dominiczak and Alex Alderton, Roberto Amato, Sonia Goncalves, Ewan Harrison, David K. Jackson, Ian Johnston, Dominic Kwiatkowski, Cordelia Langford, John Sillitoe on behalf of the Wellcome Sanger Institute COVID-19 Surveillance Team |
| EPI_ISL_589043 | Lighthouse Lab in Milton Keynes | Wellcome Sanger Institute for the COVID-19 Genomics UK (COG-UK) consortium | The Lighthouse Lab in Milton Keynes and Alex Alderton, Roberto Amato, Sonia Goncalves, Ewan Harrison, David K. Jackson, Ian Johnston, Dominic Kwiatkowski, Cordelia Langford, John Sillitoe on behalf of the Wellcome Sanger Institute COVID-19 Surveillance Team |
| EPI_ISL_589044, EPI_ISL_589045, EPI_ISL_589046, EPI_ISL_589047, EPI_ISL_589048, EPI_ISL_589049 | Lighthouse Lab in Glasgow | Wellcome Sanger Institute for the COVID-19 Genomics UK (COG-UK) consortium | Harper VanSteenhouse, Yumi Kasai, David Gray, Carol Clugston, Anna Dominiczak and Alex Alderton, Roberto Amato, Sonia Goncalves, Ewan Harrison, David K. Jackson, Ian Johnston, Dominic Kwiatkowski, Cordelia Langford, John Sillitoe on behalf of the Wellcome Sanger Institute COVID-19 Surveillance Team |
| EPI_ISL_589050 | Lighthouse Lab in Milton Keynes | Wellcome Sanger Institute for the COVID-19 Genomics UK (COG-UK) Consortium | The Lighthouse Lab in Milton Keynes and Alex Alderton, Roberto Amato, Sonia Goncalves, Ewan Harrison, David K. Jackson, Ian Johnston, Dominic Kwiatkowski, Cordelia Langford, John Sillitoe on behalf of the Wellcome Sanger Institute COVID-19 Surveillance Team |

| | | | |
|---|---|---|---|
| EPI_ISL_589053, EPI_ISL_589054, EPI_ISL_589055, EPI_ISL_589056 | Lighthouse Lab in Glasgow | Wellcome Sanger Institute for the COVID-19 Genomics UK (COG-UK) consortium | Harper VanSteenhouse, Yumi Kasai, David Gray, Carol Clugston, Anna Dominiczak and Alex Alderton, Roberto Amato, Sonia Goncalves, Ewan Harrison, David K. Jackson, Ian Johnston, Dominic Kwiatkowski, Cordelia Langford, John Sillitoe on behalf of the Wellcome Sanger Institute COVID-19 Surveillance Team |
| EPI_ISL_589057 | Lighthouse Lab in Milton Keynes | Wellcome Sanger Institute for the COVID-19 Genomics UK (COG-UK) consortium | The Lighthouse Lab in Milton Keynes and Alex Alderton, Roberto Amato, Sonia Goncalves, Ewan Harrison, David K. Jackson, Ian Johnston, Dominic Kwiatkowski, Cordelia Langford, John Sillitoe on behalf of the Wellcome Sanger Institute COVID-19 Surveillance Team |
| EPI_ISL_589058, EPI_ISL_589059, EPI_ISL_589060, EPI_ISL_589061, EPI_ISL_589062, EPI_ISL_589064, EPI_ISL_589065, EPI_ISL_589066, EPI_ISL_589068, EPI_ISL_589069 | Lighthouse Lab in Glasgow | Wellcome Sanger Institute for the COVID-19 Genomics UK (COG-UK) consortium | Harper VanSteenhouse, Yumi Kasai, David Gray, Carol Clugston, Anna Dominiczak and Alex Alderton, Roberto Amato, Sonia Goncalves, Ewan Harrison, David K. Jackson, Ian Johnston, Dominic Kwiatkowski, Cordelia Langford, John Sillitoe on behalf of the Wellcome Sanger Institute COVID-19 Surveillance Team |
| EPI_ISL_589070 | Lighthouse Lab in Milton Keynes | Wellcome Sanger Institute for the COVID-19 Genomics UK (COG-UK) consortium | The Lighthouse Lab in Milton Keynes and Alex Alderton, Roberto Amato, Sonia Goncalves, Ewan Harrison, David K. Jackson, Ian Johnston, Dominic Kwiatkowski, Cordelia Langford, John Sillitoe on behalf of the Wellcome Sanger Institute COVID-19 Surveillance Team |
| EPI_ISL_589071, EPI_ISL_589072, EPI_ISL_589073 | Lighthouse Lab in Glasgow | Wellcome Sanger Institute for the COVID-19 Genomics UK (COG-UK) consortium | Harper VanSteenhouse, Yumi Kasai, David Gray, Carol Clugston, Anna Dominiczak and Alex Alderton, Roberto Amato, Sonia Goncalves, Ewan Harrison, David K. Jackson, Ian Johnston, Dominic Kwiatkowski, Cordelia Langford, John Sillitoe on behalf of the Wellcome Sanger Institute COVID-19 Surveillance Team |
| EPI_ISL_589075 | Lighthouse Lab in Milton Keynes | Wellcome Sanger Institute for the COVID-19 Genomics UK (COG-UK) consortium | The Lighthouse Lab in Milton Keynes and Alex Alderton, Roberto Amato, Sonia Goncalves, Ewan Harrison, David K. Jackson, Ian Johnston, Dominic Kwiatkowski, Cordelia Langford, John Sillitoe on behalf of the Wellcome Sanger Institute COVID-19 Surveillance Team |
| EPI_ISL_589076, EPI_ISL_589078, EPI_ISL_589082 | Lighthouse Lab in Glasgow | Wellcome Sanger Institute for the COVID-19 Genomics UK (COG-UK) consortium | Harper VanSteenhouse, Yumi Kasai, David Gray, Carol Clugston, Anna Dominiczak and Alex Alderton, Roberto Amato, Sonia Goncalves, Ewan Harrison, David K. Jackson, Ian Johnston, Dominic Kwiatkowski, Cordelia Langford, John Sillitoe on behalf of the Wellcome Sanger Institute COVID-19 Surveillance Team |
| EPI_ISL_589083 | Lighthouse Lab in Milton Keynes | Wellcome Sanger Institute for the COVID-19 Genomics UK (COG-UK) consortium | The Lighthouse Lab in Milton Keynes and Alex Alderton, Roberto Amato, Sonia Goncalves, Ewan Harrison, David K. Jackson, Ian Johnston, Dominic Kwiatkowski, Cordelia Langford, John Sillitoe on behalf of the Wellcome Sanger Institute COVID-19 Surveillance Team |
| EPI_ISL_589084, EPI_ISL_589085, EPI_ISL_589087, EPI_ISL_589088, EPI_ISL_589089, EPI_ISL_589092, EPI_ISL_589093, EPI_ISL_589094 | Lighthouse Lab in Glasgow | Wellcome Sanger Institute for the COVID-19 Genomics UK (COG-UK) consortium | Harper VanSteenhouse, Yumi Kasai, David Gray, Carol Clugston, Anna Dominiczak and Alex Alderton, Roberto Amato, Sonia Goncalves, Ewan Harrison, David K. Jackson, Ian Johnston, Dominic Kwiatkowski, Cordelia Langford, John Sillitoe on behalf of the Wellcome Sanger Institute COVID-19 Surveillance Team |
| EPI_ISL_589095 | Lighthouse Lab in Milton Keynes | Wellcome Sanger Institute for the COVID-19 Genomics UK (COG-UK) consortium | The Lighthouse Lab in Milton Keynes and Alex Alderton, Roberto Amato, Sonia Goncalves, Ewan Harrison, David K. Jackson, Ian Johnston, Dominic Kwiatkowski, Cordelia Langford, John Sillitoe on behalf of the Wellcome Sanger Institute COVID-19 Surveillance Team |
| EPI_ISL_589096, EPI_ISL_589100, EPI_ISL_589102, EPI_ISL_589105, EPI_ISL_589106, EPI_ISL_589107, EPI_ISL_589112, EPI_ISL_589113, EPI_ISL_589116 | Lighthouse Lab in Glasgow | Wellcome Sanger Institute for the COVID-19 Genomics UK (COG-UK) consortium | Harper VanSteenhouse, Yumi Kasai, David Gray, Carol Clugston, Anna Dominiczak and Alex Alderton, Roberto Amato, Sonia Goncalves, Ewan Harrison, David K. Jackson, Ian Johnston, Dominic Kwiatkowski, Cordelia Langford, John Sillitoe on behalf of the Wellcome Sanger Institute COVID-19 Surveillance Team |
| EPI_ISL_589118 | Lighthouse Lab in Milton Keynes | Wellcome Sanger Institute for the COVID-19 Genomics UK (COG-UK) consortium | The Lighthouse Lab in Milton Keynes and Alex Alderton, Roberto Amato, Sonia Goncalves, Ewan Harrison, David K. Jackson, Ian Johnston, Dominic Kwiatkowski, Cordelia Langford, John Sillitoe on behalf of the Wellcome Sanger Institute COVID-19 Surveillance Team |
| EPI_ISL_589119, EPI_ISL_589120 | Lighthouse Lab in Glasgow | Wellcome Sanger Institute for the COVID-19 Genomics UK (COG-UK) consortium | Harper VanSteenhouse, Yumi Kasai, David Gray, Carol Clugston, Anna Dominiczak and Alex Alderton, Roberto Amato, Sonia Goncalves, Ewan Harrison, David K. Jackson, Ian Johnston, Dominic Kwiatkowski, Cordelia Langford, John Sillitoe on behalf of the Wellcome Sanger Institute COVID-19 Surveillance Team |
| EPI_ISL_589123 | Lighthouse Lab in Milton Keynes | Wellcome Sanger Institute for the COVID-19 Genomics UK (COG-UK) consortium | The Lighthouse Lab in Milton Keynes and Alex Alderton, Roberto Amato, Sonia Goncalves, Ewan Harrison, David K. Jackson, Ian Johnston, Dominic Kwiatkowski, Cordelia Langford, John Sillitoe on behalf of the Wellcome Sanger Institute COVID-19 Surveillance Team |
| EPI_ISL_589124, EPI_ISL_589125, EPI_ISL_589126 | Lighthouse Lab in Glasgow | Wellcome Sanger Institute for the COVID-19 Genomics UK (COG-UK) consortium | Harper VanSteenhouse, Yumi Kasai, David Gray, Carol Clugston, Anna Dominiczak and Alex Alderton, Roberto Amato, Sonia Goncalves, Ewan Harrison, David K. Jackson, Ian Johnston, Dominic Kwiatkowski, Cordelia Langford, John Sillitoe on behalf of the Wellcome Sanger Institute COVID-19 Surveillance Team |
| EPI_ISL_589127 | Lighthouse Lab in Milton Keynes | Wellcome Sanger Institute for the COVID-19 Genomics UK (COG-UK) consortium | The Lighthouse Lab in Milton Keynes and Alex Alderton, Roberto Amato, Sonia Goncalves, Ewan Harrison, David K. Jackson, Ian Johnston, Dominic Kwiatkowski, Cordelia Langford, John Sillitoe on behalf of the Wellcome Sanger Institute COVID-19 Surveillance Team |
| EPI_ISL_589128, EPI_ISL_589129 | Lighthouse Lab in Glasgow | Wellcome Sanger Institute for the COVID-19 Genomics UK (COG-UK) consortium | Harper VanSteenhouse, Yumi Kasai, David Gray, Carol Clugston, Anna Dominiczak and Alex Alderton, Roberto Amato, Sonia Goncalves, Ewan Harrison, David K. Jackson, Ian Johnston, Dominic Kwiatkowski, Cordelia Langford, John Sillitoe on behalf of the Wellcome Sanger Institute COVID-19 Surveillance Team |
| EPI_ISL_589130 | Lighthouse Lab in Milton Keynes | Wellcome Sanger Institute for the COVID-19 Genomics UK (COG-UK) Consortium | The Lighthouse Lab in Milton Keynes and Alex Alderton, Roberto Amato, Sonia Goncalves, Ewan Harrison, David K. Jackson, Ian Johnston, Dominic Kwiatkowski, Cordelia Langford, John Sillitoe on behalf of the Wellcome Sanger Institute COVID-19 Surveillance Team |
| EPI_ISL_589131, EPI_ISL_589132 | Lighthouse Lab in Glasgow | Wellcome Sanger Institute for the COVID-19 Genomics UK (COG-UK) consortium | Harper VanSteenhouse, Yumi Kasai, David Gray, Carol Clugston, Anna Dominiczak and Alex Alderton, Roberto Amato, Sonia Goncalves, Ewan Harrison, David K. Jackson, Ian Johnston, Dominic Kwiatkowski, Cordelia Langford, John Sillitoe on behalf of the Wellcome Sanger Institute COVID-19 Surveillance Team |
| EPI_ISL_589133 | Lighthouse Lab in Milton Keynes | Wellcome Sanger Institute for the COVID-19 Genomics UK (COG-UK) consortium | The Lighthouse Lab in Milton Keynes and Alex Alderton, Roberto Amato, Sonia Goncalves, Ewan Harrison, David K. Jackson, Ian Johnston, Dominic Kwiatkowski, Cordelia Langford, John Sillitoe on behalf of the Wellcome Sanger Institute COVID-19 Surveillance Team |
| EPI_ISL_589134 | Lighthouse Lab in Glasgow | Wellcome Sanger Institute for the COVID-19 Genomics UK (COG-UK) consortium | Harper VanSteenhouse, Yumi Kasai, David Gray, Carol Clugston, Anna Dominiczak and Alex Alderton, Roberto Amato, Sonia Goncalves, Ewan Harrison, David K. Jackson, Ian Johnston, Dominic Kwiatkowski, Cordelia Langford, John Sillitoe on behalf of the Wellcome Sanger Institute COVID-19 Surveillance Team |
| EPI_ISL_589135 | Lighthouse Lab in Milton Keynes | Wellcome Sanger Institute for the COVID-19 Genomics UK (COG-UK) consortium | The Lighthouse Lab in Milton Keynes and Alex Alderton, Roberto Amato, Sonia Goncalves, Ewan Harrison, David K. Jackson, Ian Johnston, Dominic Kwiatkowski, Cordelia Langford, John Sillitoe on behalf of the Wellcome Sanger Institute COVID-19 Surveillance Team |
| EPI_ISL_589136, EPI_ISL_589138, EPI_ISL_589139, EPI_ISL_589140, EPI_ISL_589141, EPI_ISL_589142, EPI_ISL_589143, EPI_ISL_589144, EPI_ISL_589145, EPI_ISL_589146 | Lighthouse Lab in Glasgow | Wellcome Sanger Institute for the COVID-19 Genomics UK (COG-UK) consortium | Harper VanSteenhouse, Yumi Kasai, David Gray, Carol Clugston, Anna Dominiczak and Alex Alderton, Roberto Amato, Sonia Goncalves, Ewan Harrison, David K. Jackson, Ian Johnston, Dominic Kwiatkowski, Cordelia Langford, John Sillitoe on behalf of the Wellcome Sanger Institute COVID-19 Surveillance Team |
| EPI_ISL_589147 | Lighthouse Lab in Milton Keynes | Wellcome Sanger Institute for the COVID-19 Genomics UK (COG-UK) consortium | The Lighthouse Lab in Milton Keynes and Alex Alderton, Roberto Amato, Sonia Goncalves, Ewan Harrison, David K. Jackson, Ian Johnston, Dominic Kwiatkowski, Cordelia Langford, John Sillitoe on behalf of the Wellcome Sanger Institute COVID-19 Surveillance Team |
| EPI_ISL_589148, EPI_ISL_589149, EPI_ISL_589150, EPI_ISL_589151, EPI_ISL_589152, EPI_ISL_589154, EPI_ISL_589155, EPI_ISL_589156 | Lighthouse Lab in Glasgow | Wellcome Sanger Institute for the COVID-19 Genomics UK (COG-UK) consortium | Harper VanSteenhouse, Yumi Kasai, David Gray, Carol Clugston, Anna Dominiczak and Alex Alderton, Roberto Amato, Sonia Goncalves, Ewan Harrison, David K. Jackson, Ian Johnston, Dominic Kwiatkowski, Cordelia Langford, John Sillitoe on behalf of the Wellcome Sanger Institute COVID-19 Surveillance Team |
| EPI_ISL_589157 | Lighthouse Lab in Milton Keynes | Wellcome Sanger Institute for the COVID-19 Genomics UK (COG-UK) consortium | The Lighthouse Lab in Milton Keynes and Alex Alderton, Roberto Amato, Sonia Goncalves, Ewan Harrison, David K. Jackson, Ian Johnston, Dominic Kwiatkowski, Cordelia Langford, John Sillitoe on behalf of the Wellcome Sanger Institute COVID-19 Surveillance Team |
| EPI_ISL_589158, EPI_ISL_589159 | Lighthouse Lab in Glasgow | Wellcome Sanger Institute for the COVID-19 Genomics UK | Harper VanSteenhouse, Yumi Kasai, David Gray, Carol Clugston, Anna Dominiczak and Alex Alderton, Roberto Amato, Sonia Goncalves, Ewan Harrison, |

| | | | |
|---|---|---|---|
| | | (COG-UK) consortium | David K. Jackson, Ian Johnston, Dominic Kwiatkowski, Cordelia Langford, John Sillitoe on behalf of the Wellcome Sanger Institute COVID-19 Surveillance Team |
| EPI_ISL_589160, EPI_ISL_589161 | Lighthouse Lab in Milton Keynes | Wellcome Sanger Institute for the COVID-19 Genomics UK (COG-UK) consortium | The Lighthouse Lab in Milton Keynes and Alex Alderton, Roberto Amato, Sonia Goncalves, Ewan Harrison, David K. Jackson, Ian Johnston, Dominic Kwiatkowski, Cordelia Langford, John Sillitoe on behalf of the Wellcome Sanger Institute COVID-19 Surveillance Team |
| EPI_ISL_589162, EPI_ISL_589163, EPI_ISL_589164, EPI_ISL_589165 | Lighthouse Lab in Glasgow | Wellcome Sanger Institute for the COVID-19 Genomics UK (COG-UK) consortium | Harper VanSteenhouse, Yumi Kasai, David Gray, Carol Clugston, Anna Dominiczak and Alex Alderton, Roberto Amato, Sonia Goncalves, Ewan Harrison, David K. Jackson, Ian Johnston, Dominic Kwiatkowski, Cordelia Langford, John Sillitoe on behalf of the Wellcome Sanger Institute COVID-19 Surveillance Team |
| EPI_ISL_589168 | Lighthouse Lab in Milton Keynes | Wellcome Sanger Institute for the COVID-19 Genomics UK (COG-UK) consortium | The Lighthouse Lab in Milton Keynes and Alex Alderton, Roberto Amato, Sonia Goncalves, Ewan Harrison, David K. Jackson, Ian Johnston, Dominic Kwiatkowski, Cordelia Langford, John Sillitoe on behalf of the Wellcome Sanger Institute COVID-19 Surveillance Team |
| EPI_ISL_589175, EPI_ISL_589177, EPI_ISL_589178 | Lighthouse Lab in Glasgow | Wellcome Sanger Institute for the COVID-19 Genomics UK (COG-UK) consortium | Harper VanSteenhouse, Yumi Kasai, David Gray, Carol Clugston, Anna Dominiczak and Alex Alderton, Roberto Amato, Sonia Goncalves, Ewan Harrison, David K. Jackson, Ian Johnston, Dominic Kwiatkowski, Cordelia Langford, John Sillitoe on behalf of the Wellcome Sanger Institute COVID-19 Surveillance Team |
| EPI_ISL_589179 | Lighthouse Lab in Milton Keynes | Wellcome Sanger Institute for the COVID-19 Genomics UK (COG-UK) consortium | The Lighthouse Lab in Milton Keynes and Alex Alderton, Roberto Amato, Sonia Goncalves, Ewan Harrison, David K. Jackson, Ian Johnston, Dominic Kwiatkowski, Cordelia Langford, John Sillitoe on behalf of the Wellcome Sanger Institute COVID-19 Surveillance Team |
| EPI_ISL_589180, EPI_ISL_589181 | Lighthouse Lab in Glasgow | Wellcome Sanger Institute for the COVID-19 Genomics UK (COG-UK) consortium | Harper VanSteenhouse, Yumi Kasai, David Gray, Carol Clugston, Anna Dominiczak and Alex Alderton, Roberto Amato, Sonia Goncalves, Ewan Harrison, David K. Jackson, Ian Johnston, Dominic Kwiatkowski, Cordelia Langford, John Sillitoe on behalf of the Wellcome Sanger Institute COVID-19 Surveillance Team |
| EPI_ISL_589182 | Lighthouse Lab in Milton Keynes | Wellcome Sanger Institute for the COVID-19 Genomics UK (COG-UK) consortium | The Lighthouse Lab in Milton Keynes and Alex Alderton, Roberto Amato, Sonia Goncalves, Ewan Harrison, David K. Jackson, Ian Johnston, Dominic Kwiatkowski, Cordelia Langford, John Sillitoe on behalf of the Wellcome Sanger Institute COVID-19 Surveillance Team |
| EPI_ISL_589184 | Lighthouse Lab in Milton Keynes | Wellcome Sanger Institute for the COVID-19 Genomics UK (COG-UK) Consortium | The Lighthouse Lab in Milton Keynes and Alex Alderton, Roberto Amato, Sonia Goncalves, Ewan Harrison, David K. Jackson, Ian Johnston, Dominic Kwiatkowski, Cordelia Langford, John Sillitoe on behalf of the Wellcome Sanger Institute COVID-19 Surveillance Team |
| EPI_ISL_589185 | Lighthouse Lab in Milton Keynes | Wellcome Sanger Institute for the COVID-19 Genomics UK (COG-UK) consortium | The Lighthouse Lab in Milton Keynes and Alex Alderton, Roberto Amato, Sonia Goncalves, Ewan Harrison, David K. Jackson, Ian Johnston, Dominic Kwiatkowski, Cordelia Langford, John Sillitoe on behalf of the Wellcome Sanger Institute COVID-19 Surveillance Team |
| EPI_ISL_589186 | Lighthouse Lab in Glasgow | Wellcome Sanger Institute for the COVID-19 Genomics UK (COG-UK) consortium | Harper VanSteenhouse, Yumi Kasai, David Gray, Carol Clugston, Anna Dominiczak and Alex Alderton, Roberto Amato, Sonia Goncalves, Ewan Harrison, David K. Jackson, Ian Johnston, Dominic Kwiatkowski, Cordelia Langford, John Sillitoe on behalf of the Wellcome Sanger Institute COVID-19 Surveillance Team |
| EPI_ISL_589187 | Lighthouse Lab in Milton Keynes | Wellcome Sanger Institute for the COVID-19 Genomics UK (COG-UK) consortium | The Lighthouse Lab in Milton Keynes and Alex Alderton, Roberto Amato, Sonia Goncalves, Ewan Harrison, David K. Jackson, Ian Johnston, Dominic Kwiatkowski, Cordelia Langford, John Sillitoe on behalf of the Wellcome Sanger Institute COVID-19 Surveillance Team |
| EPI_ISL_589188, EPI_ISL_589189, EPI_ISL_589190, EPI_ISL_589191, EPI_ISL_589192, EPI_ISL_589193, EPI_ISL_589194 | Lighthouse Lab in Glasgow | Wellcome Sanger Institute for the COVID-19 Genomics UK (COG-UK) consortium | Harper VanSteenhouse, Yumi Kasai, David Gray, Carol Clugston, Anna Dominiczak and Alex Alderton, Roberto Amato, Sonia Goncalves, Ewan Harrison, David K. Jackson, Ian Johnston, Dominic Kwiatkowski, Cordelia Langford, John Sillitoe on behalf of the Wellcome Sanger Institute COVID-19 Surveillance Team |
| EPI_ISL_589195, EPI_ISL_589196 | Lighthouse Lab in Milton Keynes | Wellcome Sanger Institute for the COVID-19 Genomics UK (COG-UK) consortium | The Lighthouse Lab in Milton Keynes and Alex Alderton, Roberto Amato, Sonia Goncalves, Ewan Harrison, David K. Jackson, Ian Johnston, Dominic Kwiatkowski, Cordelia Langford, John Sillitoe on behalf of the Wellcome Sanger Institute COVID-19 Surveillance Team |
| EPI_ISL_589197, EPI_ISL_589201, EPI_ISL_589202, EPI_ISL_589203, EPI_ISL_589204, EPI_ISL_589206, EPI_ISL_589208, EPI_ISL_589209, EPI_ISL_589210 | Lighthouse Lab in Glasgow | Wellcome Sanger Institute for the COVID-19 Genomics UK (COG-UK) consortium | Harper VanSteenhouse, Yumi Kasai, David Gray, Carol Clugston, Anna Dominiczak and Alex Alderton, Roberto Amato, Sonia Goncalves, Ewan Harrison, David K. Jackson, Ian Johnston, Dominic Kwiatkowski, Cordelia Langford, John Sillitoe on behalf of the Wellcome Sanger Institute COVID-19 Surveillance Team |
| EPI_ISL_589211 | Lighthouse Lab in Milton Keynes | Wellcome Sanger Institute for the COVID-19 Genomics UK (COG-UK) consortium | The Lighthouse Lab in Milton Keynes and Alex Alderton, Roberto Amato, Sonia Goncalves, Ewan Harrison, David K. Jackson, Ian Johnston, Dominic Kwiatkowski, Cordelia Langford, John Sillitoe on behalf of the Wellcome Sanger Institute COVID-19 Surveillance Team |
| EPI_ISL_589213, EPI_ISL_589214, EPI_ISL_589215, EPI_ISL_589216, EPI_ISL_589217, EPI_ISL_589218, EPI_ISL_589219, EPI_ISL_589220 | Lighthouse Lab in Glasgow | Wellcome Sanger Institute for the COVID-19 Genomics UK (COG-UK) consortium | Harper VanSteenhouse, Yumi Kasai, David Gray, Carol Clugston, Anna Dominiczak and Alex Alderton, Roberto Amato, Sonia Goncalves, Ewan Harrison, David K. Jackson, Ian Johnston, Dominic Kwiatkowski, Cordelia Langford, John Sillitoe on behalf of the Wellcome Sanger Institute COVID-19 Surveillance Team |
| EPI_ISL_589221 | Lighthouse Lab in Milton Keynes | Wellcome Sanger Institute for the COVID-19 Genomics UK (COG-UK) consortium | The Lighthouse Lab in Milton Keynes and Alex Alderton, Roberto Amato, Sonia Goncalves, Ewan Harrison, David K. Jackson, Ian Johnston, Dominic Kwiatkowski, Cordelia Langford, John Sillitoe on behalf of the Wellcome Sanger Institute COVID-19 Surveillance Team |
| EPI_ISL_589222, EPI_ISL_589223 | Lighthouse Lab in Glasgow | Wellcome Sanger Institute for the COVID-19 Genomics UK (COG-UK) consortium | Harper VanSteenhouse, Yumi Kasai, David Gray, Carol Clugston, Anna Dominiczak and Alex Alderton, Roberto Amato, Sonia Goncalves, Ewan Harrison, David K. Jackson, Ian Johnston, Dominic Kwiatkowski, Cordelia Langford, John Sillitoe on behalf of the Wellcome Sanger Institute COVID-19 Surveillance Team |
| EPI_ISL_589226 | Lighthouse Lab in Milton Keynes | Wellcome Sanger Institute for the COVID-19 Genomics UK (COG-UK) consortium | The Lighthouse Lab in Milton Keynes and Alex Alderton, Roberto Amato, Sonia Goncalves, Ewan Harrison, David K. Jackson, Ian Johnston, Dominic Kwiatkowski, Cordelia Langford, John Sillitoe on behalf of the Wellcome Sanger Institute COVID-19 Surveillance Team |
| EPI_ISL_589228 | Lighthouse Lab in Glasgow | Wellcome Sanger Institute for the COVID-19 Genomics UK (COG-UK) consortium | Harper VanSteenhouse, Yumi Kasai, David Gray, Carol Clugston, Anna Dominiczak and Alex Alderton, Roberto Amato, Sonia Goncalves, Ewan Harrison, David K. Jackson, Ian Johnston, Dominic Kwiatkowski, Cordelia Langford, John Sillitoe on behalf of the Wellcome Sanger Institute COVID-19 Surveillance Team |
| EPI_ISL_589229 | Lighthouse Lab in Milton Keynes | Wellcome Sanger Institute for the COVID-19 Genomics UK (COG-UK) consortium | The Lighthouse Lab in Milton Keynes and Alex Alderton, Roberto Amato, Sonia Goncalves, Ewan Harrison, David K. Jackson, Ian Johnston, Dominic Kwiatkowski, Cordelia Langford, John Sillitoe on behalf of the Wellcome Sanger Institute COVID-19 Surveillance Team |
| EPI_ISL_589230, EPI_ISL_589232 | Lighthouse Lab in Glasgow | Wellcome Sanger Institute for the COVID-19 Genomics UK (COG-UK) consortium | Harper VanSteenhouse, Yumi Kasai, David Gray, Carol Clugston, Anna Dominiczak and Alex Alderton, Roberto Amato, Sonia Goncalves, Ewan Harrison, David K. Jackson, Ian Johnston, Dominic Kwiatkowski, Cordelia Langford, John Sillitoe on behalf of the Wellcome Sanger Institute COVID-19 Surveillance Team |
| EPI_ISL_589233 | Lighthouse Lab in Milton Keynes | Wellcome Sanger Institute for the COVID-19 Genomics UK (COG-UK) consortium | The Lighthouse Lab in Milton Keynes and Alex Alderton, Roberto Amato, Sonia Goncalves, Ewan Harrison, David K. Jackson, Ian Johnston, Dominic Kwiatkowski, Cordelia Langford, John Sillitoe on behalf of the Wellcome Sanger Institute COVID-19 Surveillance Team |
| EPI_ISL_589234, EPI_ISL_589236, EPI_ISL_589238, EPI_ISL_589239, EPI_ISL_589240, EPI_ISL_589241 | Lighthouse Lab in Glasgow | Wellcome Sanger Institute for the COVID-19 Genomics UK (COG-UK) consortium | Harper VanSteenhouse, Yumi Kasai, David Gray, Carol Clugston, Anna Dominiczak and Alex Alderton, Roberto Amato, Sonia Goncalves, Ewan Harrison, David K. Jackson, Ian Johnston, Dominic Kwiatkowski, Cordelia Langford, John Sillitoe on behalf of the Wellcome Sanger Institute COVID-19 Surveillance Team |
| EPI_ISL_589242, EPI_ISL_589243 | Lighthouse Lab in Milton Keynes | Wellcome Sanger Institute for the COVID-19 Genomics UK (COG-UK) consortium | The Lighthouse Lab in Milton Keynes and Alex Alderton, Roberto Amato, Sonia Goncalves, Ewan Harrison, David K. Jackson, Ian Johnston, Dominic Kwiatkowski, Cordelia Langford, John Sillitoe on behalf of the Wellcome Sanger Institute COVID-19 Surveillance Team |
| EPI_ISL_589244, EPI_ISL_589245, EPI_ISL_589246 | Lighthouse Lab in Glasgow | Wellcome Sanger Institute for the COVID-19 Genomics UK (COG-UK) consortium | Harper VanSteenhouse, Yumi Kasai, David Gray, Carol Clugston, Anna Dominiczak and Alex Alderton, Roberto Amato, Sonia Goncalves, Ewan Harrison, David K. Jackson, Ian Johnston, Dominic Kwiatkowski, Cordelia Langford, John Sillitoe on behalf of the Wellcome Sanger Institute COVID-19 Surveillance Team |
| EPI_ISL_589248 | Lighthouse Lab in Glasgow | Wellcome Sanger Institute for the COVID-19 Genomics UK (COG-UK) Consortium | Harper VanSteenhouse, Yumi Kasai, David Gray, Carol Clugston, Anna Dominiczak and Alex Alderton, Roberto Amato, Sonia Goncalves, Ewan Harrison, David K. Jackson, Ian Johnston, Dominic Kwiatkowski, Cordelia Langford, John Sillitoe on behalf of the Wellcome Sanger Institute COVID-19 Surveillance Team |

| | | | |
|---|---|---|---|
| EPI_ISL_589249, EPI_ISL_589250 | Lighthouse Lab in Glasgow | Wellcome Sanger Institute for the COVID-19 Genomics UK (COG-UK) consortium | Harper VanSteenhouse, Yumi Kasai, David Gray, Carol Clugston, Anna Dominiczak and Alex Alderton, Roberto Amato, Sonia Goncalves, Ewan Harrison, David K. Jackson, Ian Johnston, Dominic Kwiatkowski, Cordelia Langford, John Sillitoe on behalf of the Wellcome Sanger Institute COVID-19 Surveillance Team |
| EPI_ISL_589251 | Lighthouse Lab in Milton Keynes | Wellcome Sanger Institute for the COVID-19 Genomics UK (COG-UK) consortium | The Lighthouse Lab in Milton Keynes and Alex Alderton, Roberto Amato, Sonia Goncalves, Ewan Harrison, David K. Jackson, Ian Johnston, Dominic Kwiatkowski, Cordelia Langford, John Sillitoe on behalf of the Wellcome Sanger Institute COVID-19 Surveillance Team |
| EPI_ISL_589252 | Lighthouse Lab in Glasgow | Wellcome Sanger Institute for the COVID-19 Genomics UK (COG-UK) consortium | Harper VanSteenhouse, Yumi Kasai, David Gray, Carol Clugston, Anna Dominiczak and Alex Alderton, Roberto Amato, Sonia Goncalves, Ewan Harrison, David K. Jackson, Ian Johnston, Dominic Kwiatkowski, Cordelia Langford, John Sillitoe on behalf of the Wellcome Sanger Institute COVID-19 Surveillance Team |
| EPI_ISL_589253 | Lighthouse Lab in Milton Keynes | Wellcome Sanger Institute for the COVID-19 Genomics UK (COG-UK) consortium | The Lighthouse Lab in Milton Keynes and Alex Alderton, Roberto Amato, Sonia Goncalves, Ewan Harrison, David K. Jackson, Ian Johnston, Dominic Kwiatkowski, Cordelia Langford, John Sillitoe on behalf of the Wellcome Sanger Institute COVID-19 Surveillance Team |
| EPI_ISL_589254, EPI_ISL_589255, EPI_ISL_589258 | Lighthouse Lab in Glasgow | Wellcome Sanger Institute for the COVID-19 Genomics UK (COG-UK) consortium | Harper VanSteenhouse, Yumi Kasai, David Gray, Carol Clugston, Anna Dominiczak and Alex Alderton, Roberto Amato, Sonia Goncalves, Ewan Harrison, David K. Jackson, Ian Johnston, Dominic Kwiatkowski, Cordelia Langford, John Sillitoe on behalf of the Wellcome Sanger Institute COVID-19 Surveillance Team |
| EPI_ISL_589259, EPI_ISL_589261 | Lighthouse Lab in Milton Keynes | Wellcome Sanger Institute for the COVID-19 Genomics UK (COG-UK) consortium | The Lighthouse Lab in Milton Keynes and Alex Alderton, Roberto Amato, Sonia Goncalves, Ewan Harrison, David K. Jackson, Ian Johnston, Dominic Kwiatkowski, Cordelia Langford, John Sillitoe on behalf of the Wellcome Sanger Institute COVID-19 Surveillance Team |
| EPI_ISL_589263 | Lighthouse Lab in Glasgow | Wellcome Sanger Institute for the COVID-19 Genomics UK (COG-UK) consortium | Harper VanSteenhouse, Yumi Kasai, David Gray, Carol Clugston, Anna Dominiczak and Alex Alderton, Roberto Amato, Sonia Goncalves, Ewan Harrison, David K. Jackson, Ian Johnston, Dominic Kwiatkowski, Cordelia Langford, John Sillitoe on behalf of the Wellcome Sanger Institute COVID-19 Surveillance Team |
| EPI_ISL_589266 | Lighthouse Lab in Milton Keynes | Wellcome Sanger Institute for the COVID-19 Genomics UK (COG-UK) consortium | The Lighthouse Lab in Milton Keynes and Alex Alderton, Roberto Amato, Sonia Goncalves, Ewan Harrison, David K. Jackson, Ian Johnston, Dominic Kwiatkowski, Cordelia Langford, John Sillitoe on behalf of the Wellcome Sanger Institute COVID-19 Surveillance Team |
| EPI_ISL_589577, EPI_ISL_589585, EPI_ISL_589586, EPI_ISL_589588, EPI_ISL_589591, EPI_ISL_589592, EPI_ISL_589594, EPI_ISL_589600, EPI_ISL_589602, EPI_ISL_589603, EPI_ISL_589608, EPI_ISL_589613, EPI_ISL_589614, EPI_ISL_589618, EPI_ISL_589621, EPI_ISL_589625, EPI_ISL_589626, EPI_ISL_589627, EPI_ISL_589628, EPI_ISL_589629, EPI_ISL_589636, EPI_ISL_589639, EPI_ISL_589642, EPI_ISL_589647, EPI_ISL_589649, EPI_ISL_589654, EPI_ISL_589655, EPI_ISL_589656, EPI_ISL_589657, EPI_ISL_589658, EPI_ISL_589664, EPI_ISL_589668, EPI_ISL_589669, EPI_ISL_589670, EPI_ISL_589671, EPI_ISL_589673, EPI_ISL_589674, EPI_ISL_589675, EPI_ISL_589678, EPI_ISL_589679, EPI_ISL_589681, EPI_ISL_589685, EPI_ISL_589688, EPI_ISL_589695, EPI_ISL_589698, EPI_ISL_589700, EPI_ISL_589709, EPI_ISL_589710, EPI_ISL_589713, EPI_ISL_589716, EPI_ISL_589717, EPI_ISL_589726, EPI_ISL_589730, EPI_ISL_589733, EPI_ISL_589734, EPI_ISL_589735, EPI_ISL_589739, EPI_ISL_589742, EPI_ISL_589743, EPI_ISL_589745, EPI_ISL_589746, EPI_ISL_589748, EPI_ISL_589754, EPI_ISL_589755, EPI_ISL_589757, EPI_ISL_589758, EPI_ISL_589759, EPI_ISL_589766 | | | |
| see above | Lighthouse Lab in Milton Keynes | Wellcome Sanger Institute for the COVID-19 Genomics UK (COG-UK) consortium | The Lighthouse Lab in Milton Keynes and Alex Alderton, Roberto Amato, Sonia Goncalves, Ewan Harrison, David K. Jackson, Ian Johnston, Dominic Kwiatkowski, Cordelia Langford, John Sillitoe on behalf of the Wellcome Sanger Institute COVID-19 Surveillance Team (http://www.sanger.ac.uk/covid-team) |
| EPI_ISL_590699, EPI_ISL_590702, EPI_ISL_590704, EPI_ISL_590705, EPI_ISL_590707, EPI_ISL_590709, EPI_ISL_590710, EPI_ISL_590711, EPI_ISL_590715, EPI_ISL_590716, EPI_ISL_590719, EPI_ISL_590720, EPI_ISL_590722, EPI_ISL_590723, EPI_ISL_590725, EPI_ISL_590726, EPI_ISL_590727, EPI_ISL_590728, EPI_ISL_590729, EPI_ISL_590730, EPI_ISL_590731, EPI_ISL_590732, EPI_ISL_590733, EPI_ISL_590734, EPI_ISL_590735, EPI_ISL_590736, EPI_ISL_590737, EPI_ISL_590738, EPI_ISL_590739, EPI_ISL_590740, EPI_ISL_590742, EPI_ISL_590743, EPI_ISL_590744, EPI_ISL_590745, EPI_ISL_590746 | | | |
| see above | University of Michigan Clinical Microbiology Laboratory | Lauring Lab, University of Michigan, Department of Microbiology and Immunology | Valesano |
| EPI_ISL_590877 | Dept. of Medical Microbiology, Stavanger University Hospital, Helse Stavanger HF | Norwegian Institute of Public Health, Department of Virology | Kathrine Stene-Johansen, Kamilla Heddeland Instefjord, Hilde Elshaug, Marie Paulsen Madsen, Rasmus Riis Kopperud, Hilde Vollan, Karoline Bragstad, Olav Hungnes |
| EPI_ISL_590955 | Dept. of Medical Microbiology, Stavanger University Hospital, Helse Stavanger HF | Norwegian Institute of Public Health, Department of Virology | Kathrine Stene-Johansen, Iren Löhr, Kamilla Heddeland Instefjord, Hilde Elshaug, Rasmus Riis Kopperud, Hilde Vollan, Karoline Bragstad, Olav Hungnes |
| EPI_ISL_590979, EPI_ISL_590980 | Oslo University Hospital, Department of Medical Microbiology | Norwegian Institute of Public Health, Department of Virology | Kathrine Stene-Johansen, Kamilla Heddeland Instefjord, Hilde Elshaug, Rasmus Riis Kopperud, Hilde Vollan, Karoline Bragstad, Olav Hungnes |
| EPI_ISL_590981, EPI_ISL_591008 | Foerde Hospital, Department of Microbiology | Norwegian Institute of Public Health, Department of Virology | Kathrine Stene-Johansen, Kamilla Heddeland Instefjord, Hilde Elshaug, Rasmus Riis Kopperud, Hilde Vollan, Karoline Bragstad, Olav Hungnes |
| EPI_ISL_591009 | Ostfold Hospital Trust - Kalnes, Centre for Laboratory Medicine, Section for gene technology and infection serology | Norwegian Institute of Public Health, Department of Virology | Kathrine Stene-Johansen, Kamilla Heddeland Instefjord, Hilde Elshaug, Rasmus Riis Kopperud, Hilde Vollan, Karoline Bragstad, Olav Hungnes |
| EPI_ISL_591011 | Oslo University Hospital, Department of Medical Microbiology | Norwegian Institute of Public Health, Department of Virology | Kathrine Stene-Johansen, Kamilla Heddeland Instefjord, Hilde Elshaug, Rasmus Riis Kopperud, Hilde Vollan, Karoline Bragstad, Olav Hungnes |
| EPI_ISL_591012 | Department of Medical Microbiology - section Molde, Molde Hospital | Norwegian Institute of Public Health, Department of Virology | Kathrine Stene-Johansen, Kamilla Heddeland Instefjord, Hilde Elshaug, Rasmus Riis Kopperud, Hilde Vollan, Karoline Bragstad, Olav Hungnes |
| EPI_ISL_591014 | Department of Medical Microbiology, St. Olavs hospital | Norwegian Institute of Public Health, Department of Virology | Kathrine Stene-Johansen, Kamilla Heddeland Instefjord, Hilde Elshaug, Rasmus Riis Kopperud, Hilde Vollan, Karoline Bragstad, Olav Hungnes |
| EPI_ISL_591018, EPI_ISL_591019 | Department of Medical Microbiology - section Molde, Molde Hospital | Norwegian Institute of Public Health, Department of Virology | Kathrine Stene-Johansen, Kamilla Heddeland Instefjord, Hilde Elshaug, Rasmus Riis Kopperud, Hilde Vollan, Karoline Bragstad, Olav Hungnes |
| EPI_ISL_591033, EPI_ISL_591034, EPI_ISL_591035, EPI_ISL_591036 | MD PHL | MD PHL | Maryland Department of Health Laboratories Administration |
| EPI_ISL_591279 | National Institute for Viral Disease Control and Prevention, China CDC | National Institute for Viral Disease Control and Prevention, China CDC | Huilai Ma, Zhaoguo Wang, Xiang Zhao, Jun Han, Yong Zhang, Hong Wang, Cao Chen, Ji Wang, Jingdong Song, Yao Meng, Yuchao Wu, Zhixiao Chen, Dayan Wang, Ruqin Gao, George F.Gao, Wenbo Xu |
| EPI_ISL_591517 | Sydney South West Pathology Service (SSWPS) - Royal Prince Alfred Hospital - NSW Health Pathology | NSW Health Pathology - Institute of Clinical Pathology and Medical Research; Westmead Hospital; University of Sydney | CIDM-PH et al. |
| EPI_ISL_592584, EPI_ISL_592588, EPI_ISL_592609, EPI_ISL_592749 | Microbiological Diagnostic Unit - Public Health Laboratory (MDU-PHL) | MDU-PHL | Seemann T., Schultz, M. B., Sait, M., Sherry, N. |
| EPI_ISL_592795 | Victorian Infectious Diseases Reference Laboratory (VIDRL) | VIDRL and MDU-PHL | Caly L., Seemann T., Sait, M., Schultz, M. B., Druce J., Sherry, N. |
| EPI_ISL_592801, EPI_ISL_592802, EPI_ISL_592817, EPI_ISL_593013, EPI_ISL_593014, EPI_ISL_593016, EPI_ISL_593099, EPI_ISL_593218, EPI_ISL_593220, EPI_ISL_593222, EPI_ISL_593238, EPI_ISL_593239, EPI_ISL_593284, EPI_ISL_593347, EPI_ISL_593348 | | | |
| see above | Microbiological Diagnostic Unit - Public Health Laboratory (MDU-PHL) | MDU-PHL | Seemann T., Schultz, M. B., Sait, M., Sherry, N. |
| EPI_ISL_593744, EPI_ISL_593753 | South Eastern Area Laboratory Services (SEALS) | NSW Health Pathology - Institute of Clinical Pathology and Medical Research; Westmead Hospital; University of Sydney | CIDM-PH et al. |
| EPI_ISL_593804 | Respiratory Virus Unit, Microbiology Services Colindale, Public Health England | Respiratory Virus Unit, Microbiology Services Colindale, Public Health England | PHE Covid Sequencing Team |
| EPI_ISL_593893, EPI_ISL_593894, EPI_ISL_593901 | CHU Purpan - Laboratoire de Virologie - Institut Fédératif de Biologie | CHU Purpan - Laboratoire de Virologie - Institut Fédératif de Biologie | Latour J., Ranger N., Dubois M., Carcenac R., Harter A., Boyer P., Tremeaux P., Izopet J. |
| EPI_ISL_593935 | Sentinelles, Plessis-Trevise | National Reference Center for Viruses of Respiratory Infections, Institut Pasteur, Paris | Sylvie Behillil, Fabiana Gambaro, Etienne Simon-Lorière, Vincent Enouf, Maud Vanpeene, Sylvie van der Werf |
| EPI_ISL_593936 | Sentinelles, Fondettes | National Reference Center for Viruses of Respiratory Infections, Institut Pasteur, Paris | Sylvie Behillil, Fabiana Gambaro, Etienne Simon-Lorière, Vincent Enouf, Maud Vanpeene, Sylvie van der Werf |
| EPI_ISL_593980 | Delaware Public Health Lab | Delaware Public Health Lab | Gregory Hovan |
| EPI_ISL_594062 | Utah Public Health Laboratory | Utah Public Health Laboratory | Erin Young, Kelly Oakeson |
| EPI_ISL_594175, EPI_ISL_594176, EPI_ISL_594177, EPI_ISL_594178, | PathWest Laboratory Medicine WA | PathWest Laboratory Medicine WA Microbial Surveillance Unit | PathWest Laboratory Medicine WA Microbial Surveillance Unit |

| | | | |
|---|---|---|---|
| EPI_ISL_594179 | | | |
| EPI_ISL_594478, EPI_ISL_594479, EPI_ISL_594480, EPI_ISL_594481, EPI_ISL_594482, EPI_ISL_594483 | Oxford Viromics, NDM, University of Oxford; Oxford University Hospitals; Basingstoke and North Hampshire Hospital | COVID-19 Genomics UK (COG-UK) Consortium | Tanya Golubchik, David Bonsall, George Macintyre, Amy Trebes, Mariateresa de Cesare, Catrin Moore, Alex Mobbs, Anita Justice, Robert Shaw, Monique Andersson, Timothy Peto, Emma Wise, Nathan Moore, Jessica Lynch, Nick Cortes, Matilde Mori, Stephen Kidd, David Buck, John Todd, Christophe Fraser |
| EPI_ISL_594484, EPI_ISL_594532, EPI_ISL_594533, EPI_ISL_594534, EPI_ISL_594535, EPI_ISL_594536, EPI_ISL_594537, EPI_ISL_594538, EPI_ISL_594539, EPI_ISL_594540, EPI_ISL_594541, EPI_ISL_594542, EPI_ISL_594543, EPI_ISL_594544, EPI_ISL_594545, EPI_ISL_594546, EPI_ISL_594547, EPI_ISL_594548, EPI_ISL_594549, EPI_ISL_594550, EPI_ISL_594551, EPI_ISL_594552, EPI_ISL_594553, EPI_ISL_594554, EPI_ISL_594555, EPI_ISL_594556, EPI_ISL_594557, EPI_ISL_594558, EPI_ISL_594597, EPI_ISL_594598, EPI_ISL_594601, EPI_ISL_594602, EPI_ISL_594603, EPI_ISL_594604, EPI_ISL_594605, EPI_ISL_594606, EPI_ISL_594608 | | | |
| see above | University of Birmingham | COVID-19 Genomics UK (COG-UK) Consortium | Institute of Microbiology, University of Birmingham: Claire McMurray, Joanne Stockton, Samuel Nicholls, Radoslaw Poplawski, Will Rowe, Josh Quick, Nicholas Loman. University of Birmingham Testing Laboratory: Celina M Whalley, Andrew Bosworth, Charlotte Poxon, Kasun Wanigasooriya, Oliver Pickles, Mike Kidd, Alex Richter, Andrew D Beggs PHE Heartlands Lab: Husam Osman, Andrew Bosworth. Queen Elizabeth Hospital: Anna Casey |
| EPI_ISL_594655, EPI_ISL_594659, EPI_ISL_594667, EPI_ISL_594671, EPI_ISL_594681, EPI_ISL_594692, EPI_ISL_594700, EPI_ISL_594710, EPI_ISL_594713, EPI_ISL_594714, EPI_ISL_594721, EPI_ISL_594733 | | | |
| see above | West of Scotland Specialist Virology Centre, NHSGGC / MRC-University of Glasgow Centre for Virus Research | COVID-19 Genomics UK (COG-UK) Consortium | Ana da Silva Filipe, Natasha Johnson, Kathy Smollett, Daniel Mair, Stephen Carmichael, Lily Tong, Jenna Nichols, Elihu Aranday-Cortes, Kyriaki Nomikou; Sarah McDonald, Marc Niebel, Patawee Asamaphan; Richard Orton, Joseph Hughes, Sreenu Vattipally, David L Robertson; Alasdair MacLean, Rory Gunson; Kathy Li, Igor Starinskij, Natasha Jesudason, Rajiv Shah, James Shepherd, Antonia Ho, Emma Thomson |
| EPI_ISL_594816, EPI_ISL_594817, EPI_ISL_594818, EPI_ISL_594819, EPI_ISL_594820, EPI_ISL_594821, EPI_ISL_594822, EPI_ISL_594823, EPI_ISL_594824, EPI_ISL_594825, EPI_ISL_594826, EPI_ISL_594827 | | | |
| see above | Virology Department, Royal Infirmary of Edinburgh, NHS Lothian / School of Biological Sciences, University of Edinburgh / Institute of Genetics and Molecular Medicine, University of Edinburgh | COVID-19 Genomics UK (COG-UK) Consortium | McHugh M, Dewar R, Rooke S, Gallagher M, Balcaza C, O'Toole Á, Scher E, Hill V, McCrone JT, Colquhoun R, Yu X, Jackson B, Rambaut A, Williams TC, Templeton K |
| EPI_ISL_594853, EPI_ISL_594854, EPI_ISL_594855, EPI_ISL_594856, EPI_ISL_594859, EPI_ISL_594860 | Liverpool Clinical Laboratories | COVID-19 Genomics UK (COG-UK) Consortium | Sam Haldenby, Anita Lucaci, Steve Paterson, Julian Hiscox, Alistair Darby, M Almsaud, A Alrezaihi, Muhannad Alruwaili, Stuart D Armstrong, Jones Benjamin, Eleanor G Bentley, Anu Chawla, Jordan J Clark, Angela Cowell, Richard Eccles, Isabel García-Dorival, Matthew Gemmell, Alessandro Gerada, PKF Gilmore, Richard Gregory, Ximeng Han, Catherine Hartley, Margaret Hughes, Miren Iturriza-Gomara, James Johnson, L Luu, Jenifer Manson, Charlotte Nelson, Elaine O'Toole, Cassie Olateju, Rebekah Penrice-Randal , Lucille Rainbow, N.P Randle, Trevor Ian Robinson, Parul Sharma, Ghada T Shawli, James P Stewart, Neil Swainston, Ecaterina Vamos, Joanne Watts, Mark Whitehead |
| EPI_ISL_594861, EPI_ISL_594862, EPI_ISL_594863, EPI_ISL_594864, EPI_ISL_594867, EPI_ISL_594868, EPI_ISL_594871 | University College London, Great Ormond Street Hospital for Children NHS Foundation Trust, Imperial College Healthcare NHS Trust | COVID-19 Genomics UK (COG-UK) Consortium | Sergi Castellano, Rachel Williams, Mark Kristiansen, Paola Resende Silva, Sunando Roy, Tony Brooks, Helena Tutill, Paola Niola, Patricia Dyal, Charlotte Williams, Leysa Forrest, Yasmin Panchbhaya, Jacqueline Findlay, Samuel Weeks, Julianne Brown, Kathryn Harris, Paul Randell, James Price, Alison Holmes, Judith Breuer |
| EPI_ISL_594900, EPI_ISL_594901, EPI_ISL_594913, EPI_ISL_594914, EPI_ISL_594915, EPI_ISL_594918, EPI_ISL_594919, EPI_ISL_594920, EPI_ISL_594921, EPI_ISL_594922 | Oxford Viromics, NDM, University of Oxford; Oxford University Hospitals; Basingstoke and North Hampshire Hospital | COVID-19 Genomics UK (COG-UK) Consortium | Tanya Golubchik, David Bonsall, George Macintyre, Amy Trebes, Mariateresa de Cesare, Catrin Moore, Alex Mobbs, Anita Justice, Robert Shaw, Monique Andersson, Timothy Peto, Emma Wise, Nathan Moore, Jessica Lynch, Nick Cortes, Matilde Mori, Stephen Kidd, David Buck, John Todd, Christophe Fraser |
| EPI_ISL_594987, EPI_ISL_594988, EPI_ISL_594989, EPI_ISL_594990, EPI_ISL_594991, EPI_ISL_594992, EPI_ISL_594993, EPI_ISL_594994, EPI_ISL_594995, EPI_ISL_594996, EPI_ISL_594997, EPI_ISL_594998, EPI_ISL_594999, EPI_ISL_595000, EPI_ISL_595001, EPI_ISL_595002, EPI_ISL_595003, EPI_ISL_595004, EPI_ISL_595005, EPI_ISL_595006, EPI_ISL_595007, EPI_ISL_595008, EPI_ISL_595009, EPI_ISL_595010, EPI_ISL_595011, EPI_ISL_595012, EPI_ISL_595013, EPI_ISL_595014, EPI_ISL_595015, EPI_ISL_595016, EPI_ISL_595017, EPI_ISL_595018, EPI_ISL_595019, EPI_ISL_595020, EPI_ISL_595021, EPI_ISL_595022, EPI_ISL_595023, EPI_ISL_595024, EPI_ISL_595025, EPI_ISL_595026, EPI_ISL_595027, EPI_ISL_595028, EPI_ISL_595029, EPI_ISL_595030, EPI_ISL_595031, EPI_ISL_595032, EPI_ISL_595033, EPI_ISL_595034, EPI_ISL_595035, EPI_ISL_595036, EPI_ISL_595037, EPI_ISL_595038, EPI_ISL_595039, EPI_ISL_595040, EPI_ISL_595041, EPI_ISL_595042, EPI_ISL_595043, EPI_ISL_595044, EPI_ISL_595045 | | | |
| see above | Northumbria University / South Tees Hospitals NHS Foundation Trust / North Cumbria Integrated Care NHS Foundation Trust / North Tees and Hartlepool NHS Foundation Trust / Newcastle Hospitals NHS Foundation Trust | COVID-19 Genomics UK (COG-UK) Consortium | Darren L Smith,Andrew Nelson,Matthew Bashton,Greg R Young,Joshua Loh,John Allan,Mohammad A Tariq,Giles S Holt,Gary Black,Wen C Yew,Lynn Dover,Paul Baker,Steve Liggett,Sarah Essex,Jane Greenaway,Debra Padgett,Clive Graham,Garren Scott,Edward Barton,Emma Swindells,Brendan Payne,Jennifer Collins,Yusri Taha,Gary Eltringham |
| EPI_ISL_595095, EPI_ISL_595096, EPI_ISL_595097, EPI_ISL_595098 | Queens Medical Centre, Clinical Microbiology Department / DeepSeq Nottingham | COVID-19 Genomics UK (COG-UK) Consortium | Gemma Clark, Wendy Smith, Manjinder Khakh, Vicki M Fleming, Michelle M Lister, Hannah Howson-Wells, Jonathan Ball, Patrick McClure, Joseph Chappell, Theocharis Tsoleridis, Nadine Holmes, Matthew Carlisle, Christopher Moore, Fei Sang, Johnny Debebe, Victoria Wright, Matthew Loose |
| EPI_ISL_595326, EPI_ISL_595327, EPI_ISL_595328 | Oxford Viromics, NDM, University of Oxford; Oxford University Hospitals; Basingstoke and North Hampshire Hospital | COVID-19 Genomics UK (COG-UK) Consortium | Tanya Golubchik, David Bonsall, George Macintyre, Amy Trebes, Mariateresa de Cesare, Catrin Moore, Alex Mobbs, Anita Justice, Robert Shaw, Monique Andersson, Timothy Peto, Emma Wise, Nathan Moore, Jessica Lynch, Nick Cortes, Matilde Mori, Stephen Kidd, David Buck, John Todd, Christophe Fraser |
| EPI_ISL_595369, EPI_ISL_595377, EPI_ISL_595380, EPI_ISL_595381, EPI_ISL_595382, EPI_ISL_595388, EPI_ISL_595391, EPI_ISL_595397, EPI_ISL_595402, EPI_ISL_595403, EPI_ISL_595404, EPI_ISL_595405, EPI_ISL_595407, EPI_ISL_595408, EPI_ISL_595412, EPI_ISL_595413, EPI_ISL_595414, EPI_ISL_595417, EPI_ISL_595424, EPI_ISL_595426, EPI_ISL_595427, EPI_ISL_595430, EPI_ISL_595435, EPI_ISL_595441, EPI_ISL_595444, EPI_ISL_595448, EPI_ISL_595452, EPI_ISL_595455, EPI_ISL_595457, EPI_ISL_595459, EPI_ISL_595468, EPI_ISL_595471, EPI_ISL_595473, EPI_ISL_595476, EPI_ISL_595478, EPI_ISL_595483, EPI_ISL_595491, EPI_ISL_595493, EPI_ISL_595494, EPI_ISL_595495, EPI_ISL_595496, EPI_ISL_595497, EPI_ISL_595498, EPI_ISL_595500, EPI_ISL_595501, EPI_ISL_595504, EPI_ISL_595505, EPI_ISL_595514, EPI_ISL_595520, EPI_ISL_595525, EPI_ISL_595527, EPI_ISL_595530, EPI_ISL_595532, EPI_ISL_595534, EPI_ISL_595538, EPI_ISL_595550, EPI_ISL_595554, EPI_ISL_595555, EPI_ISL_595556, EPI_ISL_595559, EPI_ISL_595562, EPI_ISL_595564, EPI_ISL_595565, EPI_ISL_595571, EPI_ISL_595572, EPI_ISL_595573, EPI_ISL_595579, EPI_ISL_595580, EPI_ISL_595586, EPI_ISL_595591, EPI_ISL_595592, EPI_ISL_595607, EPI_ISL_595610, EPI_ISL_595612 | | | |
| see above | Wales Specialist Virology Centre Sequencing lab: Pathogen Genomics Unit | COVID-19 Genomics UK (COG-UK) Consortium | Catherine Moore, Johnathan Evans, Laura Gifford, Malorie Perry, Simon Cottrell, Angela Marchbank, Alec Birchley, Alexander Adams, Amy Gaskin, Bree Gatica-Wilcox, Jason Coombes, Joel Southgate, Lauren Gilbert, Lee Graham, Nicole Pacchiarini, Sara Kumziene-Summerhayes, Sarah Taylor, Sophie Jones, Sara Rey, Matthew Bull, Joanne Watkins, Sally Corden, Tom Connor |
| EPI_ISL_595617, EPI_ISL_595618, EPI_ISL_595619, EPI_ISL_595620, EPI_ISL_595621, EPI_ISL_595622, EPI_ISL_595623, EPI_ISL_595624, EPI_ISL_595625, EPI_ISL_595626, EPI_ISL_595627, EPI_ISL_595628, EPI_ISL_595629, EPI_ISL_595630, EPI_ISL_595631, EPI_ISL_595632, EPI_ISL_595633, EPI_ISL_595634, EPI_ISL_595635, EPI_ISL_595636, EPI_ISL_595637, EPI_ISL_595638, EPI_ISL_595639, EPI_ISL_595640, EPI_ISL_595641, EPI_ISL_595642, EPI_ISL_595643, EPI_ISL_595644, EPI_ISL_595645, EPI_ISL_595646, EPI_ISL_595647, EPI_ISL_595648, EPI_ISL_595649, EPI_ISL_595650, EPI_ISL_595651, EPI_ISL_595652, EPI_ISL_595653, EPI_ISL_595654, EPI_ISL_595655, EPI_ISL_595656, EPI_ISL_595657, EPI_ISL_595658, EPI_ISL_595659, EPI_ISL_595660, EPI_ISL_595661, EPI_ISL_595662, EPI_ISL_595663, EPI_ISL_595664, EPI_ISL_595665, EPI_ISL_595666, EPI_ISL_595667, EPI_ISL_595668, EPI_ISL_595669, EPI_ISL_595670, EPI_ISL_595671, EPI_ISL_595672, EPI_ISL_595673, EPI_ISL_595674, EPI_ISL_595675, EPI_ISL_595676, EPI_ISL_595677, EPI_ISL_595678, EPI_ISL_595679, EPI_ISL_595680, EPI_ISL_595681, EPI_ISL_595682, EPI_ISL_595683, EPI_ISL_595684, EPI_ISL_595685, EPI_ISL_595686, EPI_ISL_595687, EPI_ISL_595688, EPI_ISL_595689, EPI_ISL_595690, EPI_ISL_595691, EPI_ISL_595692, EPI_ISL_595693, EPI_ISL_595694, EPI_ISL_595695, EPI_ISL_595696, EPI_ISL_595697, EPI_ISL_595698, EPI_ISL_595699, EPI_ISL_595700, EPI_ISL_595701, EPI_ISL_595702, EPI_ISL_595703, EPI_ISL_595704, EPI_ISL_595705, EPI_ISL_595706, EPI_ISL_595707, EPI_ISL_595708, EPI_ISL_595709, EPI_ISL_595710, EPI_ISL_595711, EPI_ISL_595712, EPI_ISL_595713, EPI_ISL_595714, EPI_ISL_595715, EPI_ISL_595716, EPI_ISL_595717, EPI_ISL_595718, EPI_ISL_595719, EPI_ISL_595720, EPI_ISL_595721, EPI_ISL_595722, EPI_ISL_595723, EPI_ISL_595724, EPI_ISL_595725, EPI_ISL_595726, EPI_ISL_595727, EPI_ISL_595728, EPI_ISL_595729, EPI_ISL_595730, EPI_ISL_595731, EPI_ISL_595732, EPI_ISL_595733, EPI_ISL_595734, EPI_ISL_595735, EPI_ISL_595736, EPI_ISL_595737, EPI_ISL_595738, EPI_ISL_595740, EPI_ISL_595741, EPI_ISL_595742, EPI_ISL_595743, EPI_ISL_595744, EPI_ISL_595745, EPI_ISL_595746, EPI_ISL_595747, EPI_ISL_595748, EPI_ISL_595749, EPI_ISL_595750, EPI_ISL_595751, EPI_ISL_595752, EPI_ISL_595753, EPI_ISL_595754, EPI_ISL_595755, EPI_ISL_595756, EPI_ISL_595757, EPI_ISL_595758, EPI_ISL_595759, EPI_ISL_595760, EPI_ISL_595761, EPI_ISL_595762, EPI_ISL_595763, EPI_ISL_595764, EPI_ISL_595765, EPI_ISL_595766, EPI_ISL_595767, EPI_ISL_595768, EPI_ISL_595769, EPI_ISL_595770, EPI_ISL_595771, EPI_ISL_595772, EPI_ISL_595773, EPI_ISL_595774, EPI_ISL_595775, EPI_ISL_595776, EPI_ISL_595777, EPI_ISL_595778, EPI_ISL_595779, EPI_ISL_595780, EPI_ISL_595781, EPI_ISL_595782, EPI_ISL_595783, EPI_ISL_595784, EPI_ISL_595785, EPI_ISL_595786, EPI_ISL_595787, EPI_ISL_595788, EPI_ISL_595789, EPI_ISL_595790, EPI_ISL_595791, EPI_ISL_595792, EPI_ISL_595793, EPI_ISL_595794, EPI_ISL_595795, EPI_ISL_595796, EPI_ISL_595797, EPI_ISL_595798 | | | |
| see above | Oxford Viromics, NDM, University of Oxford; Oxford University Hospitals; Basingstoke and North Hampshire Hospital | COVID-19 Genomics UK (COG-UK) Consortium | Tanya Golubchik, David Bonsall, George Macintyre, Amy Trebes, Mariateresa de Cesare, Catrin Moore, Alex Mobbs, Anita Justice, Robert Shaw, Monique Andersson, Timothy Peto, Emma Wise, Nathan Moore, Jessica Lynch, Nick Cortes, Matilde Mori, Stephen Kidd, David Buck, John Todd, Christophe Fraser |
| EPI_ISL_595799 | Liverpool Clinical Laboratories | COVID-19 Genomics UK (COG-UK) Consortium | Sam Haldenby, Anita Lucaci, Steve Paterson, Julian Hiscox, Alistair Darby, M Almsaud, A Alrezaihi, Muhannad Alruwaili, Stuart D Armstrong, Jones Benjamin, Eleanor G Bentley, Anu Chawla, Jordan J Clark, Angela Cowell, Richard Eccles, Isabel García-Dorival, Matthew Gemmell, Alessandro Gerada, PKF Gilmore, Richard Gregory, Ximeng Han, Catherine Hartley, Margaret Hughes, Miren Iturriza-Gomara, James Johnson, L Luu, Jenifer Manson, Charlotte Nelson, Elaine O'Toole, Cassie Olateju, Rebekah Penrice-Randal , Lucille Rainbow, N.P Randle, Trevor Ian Robinson, Parul Sharma, Ghada T Shawli, James P Stewart, Neil Swainston, Ecaterina Vamos, Joanne Watts, Mark Whitehead |
| EPI_ISL_595803, EPI_ISL_595813, EPI_ISL_595814, EPI_ISL_595815, EPI_ISL_595816, EPI_ISL_595817, EPI_ISL_595818, EPI_ISL_595823, EPI_ISL_595824, EPI_ISL_595825, EPI_ISL_595826, EPI_ISL_595827, EPI_ISL_595831 | | | |
| see above | Virology Department, Sheffield Teaching Hospitals NHS Foundation Trust/Department of Infection, Immunity and | COVID-19 Genomics UK (COG-UK) Consortium | Thushan de Silva, Matthew Parker, Nikki Smith, Adri Angyal, Rebecca Brown, Luke Green, Rachel Tucker, Paul Parsons, Danielle Groves, Katie Johnson, Laura Carrilero, Alex Keeley, Dave Partridge, Matthew Wyles, Benjamin Lindsey, Mehmet Yavuz, Mohammad Raza, Cariad Evans |

| | | | |
|---|---|---|---|
| | Cardiovascular Disease, The Medical School, University of Sheffield | | |
| EPI_ISL_595837, EPI_ISL_595838, EPI_ISL_595839, EPI_ISL_595840, EPI_ISL_595841, EPI_ISL_595842, EPI_ISL_595843, EPI_ISL_595844, EPI_ISL_595845, EPI_ISL_595846, EPI_ISL_595847 | | | |
| see above | Quadram Institute Bioscience | COVID-19 Genomics UK (COG-UK) Consortium | Dave J. Baker, Gemma L. Kay, Alp Aydin, Thanh Le-Viet, Steven Rudder, Ana P. Tedim, Anastasia Kolyva, Maria Diaz, Leonardo de Oliveira Martins, Nabil-Fareed Alikhan, Lizzie Meadows, Rachel Stanley, Ngozi Elumogo, Muhammed Yasir, Nicholas M. Thomson, Alexander J Trotter, Rachel Gilroy, Samuel Bloomfield, Claire Stuart, Andrew Bell, Reenesh Prakash, Samir Dervisevic, Alison E. Mather, John Wain, Mark Webber, Andrew J. Page, Justin O'Grady |
| EPI_ISL_596201, EPI_ISL_596202, EPI_ISL_596205, EPI_ISL_596209, EPI_ISL_596213, EPI_ISL_596215, EPI_ISL_596218, EPI_ISL_596220 | Virology Department, Sheffield Teaching Hospitals NHS Foundation Trust/Department of Infection, Immunity and Cardiovascular Disease, The Medical School, University of Sheffield | COVID-19 Genomics UK (COG-UK) Consortium | Thushan de Silva, Matthew Parker, Nikki Smith, Adri Angyal, Rebecca Brown, Luke Green, Rachel Tucker, Paul Parsons, Danielle Groves, Katie Johnson, Laura Carrilero, Alex Keeley, Dave Partridge, Matthew Wyles, Benjamin Lindsey, Mehmet Yavuz, Mohammad Raza, Cariad Evans |
| EPI_ISL_596246, EPI_ISL_596247, EPI_ISL_596248, EPI_ISL_596249 | WHO National Influenza Centre Russian Federation | WHO National Influenza Centre Russian Federation | Andrey Komissarov, Artem Fadeev, Anna Ivanova, Kseniya Komissarova, Dmitry Bazhenov, Daria Danilenko |
| EPI_ISL_596409 | Seattle Flu Study | Seattle Flu Study | Deborah A. Nickerson, Chris D. Frazar, Jover Lee, Benjamin Pelle, Matthew Richardson, Amanda Adler, Elisabeth Brandstetter, Peter D. Han, Kairsten Fay, Misja Ilcisin, Kirsten Lacombe, Thomas R. Sibley, Melissa Truong, Caitlin R. Wolf, Karen Cowgill, Stephanie Schrag, Jeff Duchin, Michael Boeckh, Janet A. Englund, Michael Famulare, Barry R. Lutz, Mark J. Rieder, Lea M. Starita, Matthew Thompson, Helen Y. Chu, Trevor Bedford, Jay Shendure |
| EPI_ISL_596410 | Seattle Flu Study | Seattle Flu Study | Deborah A. Nickerson, Chris D. Frazar, Jover Lee, Benjamin Pelle, Matthew Richardson, Amanda Adler, Elisabeth Brandstetter, Peter D. Han, Kairsten Fay, Misja Ilcisin, Kirsten Lacombe, Thomas R. Sibley, Melissa Truong, Caitlin R. Wolf, Michael Boeckh, Janet A. Englund, Michael Famulare, Barry R. Lutz, Mark J. Rieder, Lea M. Starita, Matthew Thompson, Jay Shendure, Trevor Bedford, Helen Y. Chu |
| EPI_ISL_596411 | Seattle Flu Study | Seattle Flu Study | Deborah A. Nickerson, Chris D. Frazar, Jover Lee, Benjamin Pelle, Matthew Richardson, Amanda Adler, Elisabeth Brandstetter, Peter D. Han, Kairsten Fay, Misja Ilcisin, Kirsten Lacombe, Thomas R. Sibley, Melissa Truong, Caitlin R. Wolf, Karen Cowgill, Stephanie Schrag, Jeff Duchin, Michael Boeckh, Janet A. Englund, Michael Famulare, Barry R. Lutz, Mark J. Rieder, Lea M. Starita, Matthew Thompson, Helen Y. Chu, Trevor Bedford, Jay Shendure |
| EPI_ISL_596412, EPI_ISL_596413, EPI_ISL_596414 | Seattle Flu Study | Seattle Flu Study | Deborah A. Nickerson, Chris D. Frazar, Jover Lee, Benjamin Pelle, Matthew Richardson, Amanda Adler, Elisabeth Brandstetter, Peter D. Han, Kairsten Fay, Misja Ilcisin, Kirsten Lacombe, Thomas R. Sibley, Melissa Truong, Caitlin R. Wolf, Michael Boeckh, Janet A. Englund, Michael Famulare, Barry R. Lutz, Mark J. Rieder, Lea M. Starita, Matthew Thompson, Jay Shendure, Trevor Bedford, Helen Y. Chu |
| EPI_ISL_596416 | Seattle Flu Study | Seattle Flu Study | Deborah A. Nickerson, Chris D. Frazar, Jover Lee, Benjamin Pelle, Matthew Richardson, Amanda Adler, Elisabeth Brandstetter, Peter D. Han, Kairsten Fay, Misja Ilcisin, Kirsten Lacombe, Thomas R. Sibley, Melissa Truong, Caitlin R. Wolf, Karen Cowgill, Stephanie Schrag, Jeff Duchin, Michael Boeckh, Janet A. Englund, Michael Famulare, Barry R. Lutz, Mark J. Rieder, Lea M. Starita, Matthew Thompson, Helen Y. Chu, Trevor Bedford, Jay Shendure |
| EPI_ISL_596417, EPI_ISL_596418, EPI_ISL_596419, EPI_ISL_596420 | Seattle Flu Study | Seattle Flu Study | Deborah A. Nickerson, Chris D. Frazar, Jover Lee, Benjamin Pelle, Matthew Richardson, Amanda Adler, Elisabeth Brandstetter, Peter D. Han, Kairsten Fay, Misja Ilcisin, Kirsten Lacombe, Thomas R. Sibley, Melissa Truong, Caitlin R. Wolf, Michael Boeckh, Janet A. Englund, Michael Famulare, Barry R. Lutz, Mark J. Rieder, Lea M. Starita, Matthew Thompson, Jay Shendure, Trevor Bedford, Helen Y. Chu |
| EPI_ISL_596421, EPI_ISL_596422, EPI_ISL_596423, EPI_ISL_596424, EPI_ISL_596425 | Seattle Flu Study | Seattle Flu Study | Deborah A. Nickerson, Chris D. Frazar, Jover Lee, Benjamin Pelle, Matthew Richardson, Amanda Adler, Elisabeth Brandstetter, Peter D. Han, Kairsten Fay, Misja Ilcisin, Kirsten Lacombe, Thomas R. Sibley, Melissa Truong, Caitlin R. Wolf, Karen Cowgill, Stephanie Schrag, Jeff Duchin, Michael Boeckh, Janet A. Englund, Michael Famulare, Barry R. Lutz, Mark J. Rieder, Lea M. Starita, Matthew Thompson, Helen Y. Chu, Trevor Bedford, Jay Shendure |
| EPI_ISL_596426, EPI_ISL_596427 | Seattle Flu Study | Seattle Flu Study | Deborah A. Nickerson, Chris D. Frazar, Jover Lee, Benjamin Pelle, Matthew Richardson, Amanda Adler, Elisabeth Brandstetter, Peter D. Han, Kairsten Fay, Misja Ilcisin, Kirsten Lacombe, Thomas R. Sibley, Melissa Truong, Caitlin R. Wolf, Michael Boeckh, Janet A. Englund, Michael Famulare, Barry R. Lutz, Mark J. Rieder, Lea M. Starita, Matthew Thompson, Jay Shendure, Trevor Bedford, Helen Y. Chu |
| EPI_ISL_596463, EPI_ISL_596468, EPI_ISL_596477, EPI_ISL_596481 | National Public Health Laboratory, National Centre for Infectious Diseases | National Public Health Laboratory, National Centre for Infectious Diseases | Tze Minn Mak, Sophie Octavia, Zhenyang Zhou, Lin Cui, Raymond Tzer Pin Lin |
| EPI_ISL_596572, EPI_ISL_596581 | University of Michigan Clinical Microbiology Laboratory | Lauring Lab, University of Michigan, Department of Microbiology and Immunology | Valesano |
| EPI_ISL_596883 | PathWest Laboratory Medicine WA | PathWest Laboratory Medicine WA Microbial Surveillance Unit | PathWest Laboratory Medicine WA Microbial Surveillance Unit |
| EPI_ISL_598831, EPI_ISL_598832, EPI_ISL_598833, EPI_ISL_598834, EPI_ISL_598835, EPI_ISL_598836, EPI_ISL_598837, EPI_ISL_598838, EPI_ISL_598839, EPI_ISL_598840, EPI_ISL_598841, EPI_ISL_598842, EPI_ISL_598843, EPI_ISL_598844, EPI_ISL_598845, EPI_ISL_598846, EPI_ISL_598847, EPI_ISL_598848, EPI_ISL_598849, EPI_ISL_598850, EPI_ISL_598851, EPI_ISL_598852, EPI_ISL_598853, EPI_ISL_598854, EPI_ISL_598855, EPI_ISL_598856, EPI_ISL_598857, EPI_ISL_598858, EPI_ISL_598859, EPI_ISL_598860, EPI_ISL_598861, EPI_ISL_598862, EPI_ISL_598863, EPI_ISL_598864, EPI_ISL_598865, EPI_ISL_598866, EPI_ISL_598867, EPI_ISL_598868, EPI_ISL_598869, EPI_ISL_598870, EPI_ISL_598871, EPI_ISL_598872, EPI_ISL_598873, EPI_ISL_598874, EPI_ISL_598875, EPI_ISL_598876, EPI_ISL_598877, EPI_ISL_598878, EPI_ISL_598879, EPI_ISL_598880, EPI_ISL_598881, EPI_ISL_598882, EPI_ISL_598883, EPI_ISL_598884, EPI_ISL_598885, EPI_ISL_598886, EPI_ISL_598887, EPI_ISL_598888, EPI_ISL_598889, EPI_ISL_598890, EPI_ISL_598891, EPI_ISL_598892, EPI_ISL_598893, EPI_ISL_598894, EPI_ISL_598895, EPI_ISL_598896, EPI_ISL_598897, EPI_ISL_598898, EPI_ISL_598899, EPI_ISL_598900, EPI_ISL_598901, EPI_ISL_598902, EPI_ISL_598903, EPI_ISL_598904, EPI_ISL_598905, EPI_ISL_598906, EPI_ISL_598907, EPI_ISL_598908, EPI_ISL_598909, EPI_ISL_598910, EPI_ISL_598911, EPI_ISL_598912, EPI_ISL_598913, EPI_ISL_598914, EPI_ISL_598915, EPI_ISL_598916, EPI_ISL_598917, EPI_ISL_598918, EPI_ISL_598919, EPI_ISL_598920, EPI_ISL_598921, EPI_ISL_598922, EPI_ISL_598923, EPI_ISL_598924, EPI_ISL_598925, EPI_ISL_598926, EPI_ISL_598927, EPI_ISL_598928, EPI_ISL_598929, EPI_ISL_598930, EPI_ISL_598931, EPI_ISL_598932, EPI_ISL_598933, EPI_ISL_598934, EPI_ISL_598935, EPI_ISL_598936, EPI_ISL_598937, EPI_ISL_598938, EPI_ISL_598939, EPI_ISL_598940, EPI_ISL_598941, EPI_ISL_598942, EPI_ISL_598943, EPI_ISL_598944, EPI_ISL_598945, EPI_ISL_598946, EPI_ISL_598947, EPI_ISL_598948, EPI_ISL_598949, EPI_ISL_598950, EPI_ISL_598951, EPI_ISL_598952, EPI_ISL_598953, EPI_ISL_598954, EPI_ISL_598955, EPI_ISL_598956, EPI_ISL_598957, EPI_ISL_598958, EPI_ISL_598959, EPI_ISL_598960, EPI_ISL_598961, EPI_ISL_598962, EPI_ISL_598963, EPI_ISL_598964, EPI_ISL_598965, EPI_ISL_598966, EPI_ISL_598967, EPI_ISL_598968, EPI_ISL_598969 | | | |
| see above | Lighthouse Lab in Glasgow | Wellcome Sanger Institute for the COVID-19 Genomics UK (COG-UK) consortium | Harper VanSteenhouse, Yumi Kasai, David Gray, Carol Clugston, Anna Dominiczak and Alex Alderton, Roberto Amato, Sonia Goncalves, Ewan Harrison, David K. Jackson, Ian Johnston, Dominic Kwiatkowski, Cordelia Langford, John Sillitoe on behalf of the Wellcome Sanger Institute COVID-19 Surveillance Team (http://www.sanger.ac.uk/covid-team) |
| EPI_ISL_598970 | Lighthouse Lab in Glasgow | Wellcome Sanger Institute for the COVID-19 Genomics UK (COG-UK) Consortium | Harper VanSteenhouse, Yumi Kasai, David Gray, Carol Clugston, Anna Dominiczak and Alex Alderton, Roberto Amato, Sonia Goncalves, Ewan Harrison, David K. Jackson, Ian Johnston, Dominic Kwiatkowski, Cordelia Langford, John Sillitoe on behalf of the Wellcome Sanger Institute COVID-19 Surveillance Team |
| EPI_ISL_598971, EPI_ISL_598972, EPI_ISL_598973, EPI_ISL_598974, EPI_ISL_598975, EPI_ISL_598976, EPI_ISL_598977, EPI_ISL_598978, EPI_ISL_598979, EPI_ISL_598980, EPI_ISL_598981, EPI_ISL_598982, EPI_ISL_598983, EPI_ISL_598984, EPI_ISL_598985, EPI_ISL_598986, EPI_ISL_598987, EPI_ISL_598988, EPI_ISL_598989, EPI_ISL_598990, EPI_ISL_598991, EPI_ISL_598992, EPI_ISL_598993, EPI_ISL_598994, EPI_ISL_598995, EPI_ISL_598996, EPI_ISL_598997, EPI_ISL_598998, EPI_ISL_598999, EPI_ISL_599000, EPI_ISL_599001, EPI_ISL_599002, EPI_ISL_599003, EPI_ISL_599004, EPI_ISL_599005, EPI_ISL_599006, EPI_ISL_599007, EPI_ISL_599008, EPI_ISL_599009 | | | |
| see above | Lighthouse Lab in Glasgow | Wellcome Sanger Institute for the COVID-19 Genomics UK (COG-UK) consortium | Harper VanSteenhouse, Yumi Kasai, David Gray, Carol Clugston, Anna Dominiczak and Alex Alderton, Roberto Amato, Sonia Goncalves, Ewan Harrison, David K. Jackson, Ian Johnston, Dominic Kwiatkowski, Cordelia Langford, John Sillitoe on behalf of the Wellcome Sanger Institute COVID-19 Surveillance Team (http://www.sanger.ac.uk/covid-team) |
| EPI_ISL_599010 | Lighthouse Lab in Glasgow | Wellcome Sanger Institute for the COVID-19 Genomics UK (COG-UK) Consortium | Harper VanSteenhouse, Yumi Kasai, David Gray, Carol Clugston, Anna Dominiczak and Alex Alderton, Roberto Amato, Sonia Goncalves, Ewan Harrison, David K. Jackson, Ian Johnston, Dominic Kwiatkowski, Cordelia Langford, John Sillitoe on behalf of the Wellcome Sanger Institute COVID-19 Surveillance Team |
| EPI_ISL_599011, EPI_ISL_599012, EPI_ISL_599013, EPI_ISL_599014, EPI_ISL_599015, EPI_ISL_599016, EPI_ISL_599017, EPI_ISL_599018, EPI_ISL_599019, EPI_ISL_599020, EPI_ISL_599021, EPI_ISL_599022, EPI_ISL_599023, EPI_ISL_599024, EPI_ISL_599025, EPI_ISL_599026, EPI_ISL_599027, EPI_ISL_599028, EPI_ISL_599029, EPI_ISL_599030, EPI_ISL_599031, EPI_ISL_599032, EPI_ISL_599033, EPI_ISL_599034, EPI_ISL_599035, EPI_ISL_599036, EPI_ISL_599037, EPI_ISL_599038, EPI_ISL_599039, EPI_ISL_599040, EPI_ISL_599041, EPI_ISL_599042, EPI_ISL_599043, EPI_ISL_599044, EPI_ISL_599045, EPI_ISL_599046, EPI_ISL_599047, EPI_ISL_599048, EPI_ISL_599049, EPI_ISL_599050, EPI_ISL_599051, EPI_ISL_599052, EPI_ISL_599053, EPI_ISL_599054, EPI_ISL_599055, EPI_ISL_599056, EPI_ISL_599057, EPI_ISL_599058, EPI_ISL_599059, EPI_ISL_599060, EPI_ISL_599061, EPI_ISL_599062, EPI_ISL_599063, EPI_ISL_599064, EPI_ISL_599065, EPI_ISL_599066, EPI_ISL_599067, EPI_ISL_599068, EPI_ISL_599069, EPI_ISL_599070, EPI_ISL_599071, EPI_ISL_599072, EPI_ISL_599073, EPI_ISL_599074, EPI_ISL_599075, EPI_ISL_599076 | | | |
| see above | Lighthouse Lab in Glasgow | Wellcome Sanger Institute for the COVID-19 Genomics UK (COG-UK) consortium | Harper VanSteenhouse, Yumi Kasai, David Gray, Carol Clugston, Anna Dominiczak and Alex Alderton, Roberto Amato, Sonia Goncalves, Ewan Harrison, David K. Jackson, Ian Johnston, Dominic Kwiatkowski, Cordelia Langford, John Sillitoe on behalf of the Wellcome Sanger Institute COVID-19 Surveillance Team (http://www.sanger.ac.uk/covid-team) |
| EPI_ISL_599077 | Lighthouse Lab in Glasgow | Wellcome Sanger Institute for the COVID-19 Genomics UK (COG-UK) Consortium | Harper VanSteenhouse, Yumi Kasai, David Gray, Carol Clugston, Anna Dominiczak and Alex Alderton, Roberto Amato, Sonia Goncalves, Ewan Harrison, David K. Jackson, Ian Johnston, Dominic Kwiatkowski, Cordelia Langford, John Sillitoe on behalf of the Wellcome Sanger Institute COVID-19 Surveillance Team |

| | | | |
|---|---|---|---|
| EPI_ISL_599078, EPI_ISL_599079, EPI_ISL_599080, EPI_ISL_599081, EPI_ISL_599082, EPI_ISL_599083, EPI_ISL_599084, EPI_ISL_599085, EPI_ISL_599086, EPI_ISL_599087, EPI_ISL_599088, EPI_ISL_599089, EPI_ISL_599090, EPI_ISL_599091, EPI_ISL_599092, EPI_ISL_599093, EPI_ISL_599094, EPI_ISL_599095, EPI_ISL_599096, EPI_ISL_599097, EPI_ISL_599098, EPI_ISL_599099, EPI_ISL_599100, EPI_ISL_599101, EPI_ISL_599102, EPI_ISL_599103, EPI_ISL_599104, EPI_ISL_599105, EPI_ISL_599106, EPI_ISL_599107, EPI_ISL_599108, EPI_ISL_599109, EPI_ISL_599110, EPI_ISL_599111, EPI_ISL_599112, EPI_ISL_599113, EPI_ISL_599114, EPI_ISL_599115, EPI_ISL_599116, EPI_ISL_599117, EPI_ISL_599118, EPI_ISL_599119, EPI_ISL_599120, EPI_ISL_599121, EPI_ISL_599122, EPI_ISL_599123, EPI_ISL_599124, EPI_ISL_599125, EPI_ISL_599126, EPI_ISL_599127, EPI_ISL_599128, EPI_ISL_599129, EPI_ISL_599130, EPI_ISL_599131, EPI_ISL_599132, EPI_ISL_599133, EPI_ISL_599134, EPI_ISL_599135, EPI_ISL_599136, EPI_ISL_599137, EPI_ISL_599139, EPI_ISL_599141, EPI_ISL_599142, EPI_ISL_599143, EPI_ISL_599144, EPI_ISL_599147, EPI_ISL_599150, EPI_ISL_599151, EPI_ISL_599152, EPI_ISL_599153, EPI_ISL_599155, EPI_ISL_599156, EPI_ISL_599157, EPI_ISL_599158, EPI_ISL_599159, EPI_ISL_599163, EPI_ISL_599164, EPI_ISL_599167, EPI_ISL_599168, EPI_ISL_599172, EPI_ISL_599174, EPI_ISL_599180, EPI_ISL_599182 | | | |
| see above | Lighthouse Lab in Glasgow | Wellcome Sanger Institute for the COVID-19 Genomics UK (COG-UK) consortium | Harper VanSteenhouse, Yumi Kasai, David Gray, Carol Clugston, Anna Dominiczak and Alex Alderton, Roberto Amato, Sonia Goncalves, Ewan Harrison, David K. Jackson, Ian Johnston, Dominic Kwiatkowski, Cordelia Langford, John Sillitoe on behalf of the Wellcome Sanger Institute COVID-19 Surveillance Team (http://www.sanger.ac.uk/covid-team) |
| EPI_ISL_599183 | Lighthouse Lab in Glasgow | Wellcome Sanger Institute for the COVID-19 Genomics UK (COG-UK) Consortium | Harper VanSteenhouse, Yumi Kasai, David Gray, Carol Clugston, Anna Dominiczak and Alex Alderton, Roberto Amato, Sonia Goncalves, Ewan Harrison, David K. Jackson, Ian Johnston, Dominic Kwiatkowski, Cordelia Langford, John Sillitoe on behalf of the Wellcome Sanger Institute COVID-19 Surveillance Team |
| EPI_ISL_599184, EPI_ISL_599185, EPI_ISL_599186, EPI_ISL_599190, EPI_ISL_599192, EPI_ISL_599193, EPI_ISL_599195, EPI_ISL_599198, EPI_ISL_599200, EPI_ISL_599205, EPI_ISL_599208, EPI_ISL_599213, EPI_ISL_599217, EPI_ISL_599218, EPI_ISL_599221, EPI_ISL_599224, EPI_ISL_599226, EPI_ISL_599229, EPI_ISL_599230, EPI_ISL_599232, EPI_ISL_599233, EPI_ISL_599240, EPI_ISL_599242, EPI_ISL_599243, EPI_ISL_599244, EPI_ISL_599245, EPI_ISL_599247, EPI_ISL_599251, EPI_ISL_599252, EPI_ISL_599254, EPI_ISL_599258, EPI_ISL_599259, EPI_ISL_599260, EPI_ISL_599261, EPI_ISL_599263, EPI_ISL_599264, EPI_ISL_599265 | | | |
| see above | Lighthouse Lab in Glasgow | Wellcome Sanger Institute for the COVID-19 Genomics UK (COG-UK) consortium | Harper VanSteenhouse, Yumi Kasai, David Gray, Carol Clugston, Anna Dominiczak and Alex Alderton, Roberto Amato, Sonia Goncalves, Ewan Harrison, David K. Jackson, Ian Johnston, Dominic Kwiatkowski, Cordelia Langford, John Sillitoe on behalf of the Wellcome Sanger Institute COVID-19 Surveillance Team (http://www.sanger.ac.uk/covid-team) |
| EPI_ISL_599270 | Lighthouse Lab in Glasgow | Wellcome Sanger Institute for the COVID-19 Genomics UK (COG-UK) Consortium | Harper VanSteenhouse, Yumi Kasai, David Gray, Carol Clugston, Anna Dominiczak and Alex Alderton, Roberto Amato, Sonia Goncalves, Ewan Harrison, David K. Jackson, Ian Johnston, Dominic Kwiatkowski, Cordelia Langford, John Sillitoe on behalf of the Wellcome Sanger Institute COVID-19 Surveillance Team |
| EPI_ISL_599272, EPI_ISL_599273, EPI_ISL_599274, EPI_ISL_599275, EPI_ISL_599278, EPI_ISL_599281, EPI_ISL_599282, EPI_ISL_599283, EPI_ISL_599284, EPI_ISL_599285, EPI_ISL_599287, EPI_ISL_599289, EPI_ISL_599290, EPI_ISL_599292, EPI_ISL_599296, EPI_ISL_599297, EPI_ISL_599298, EPI_ISL_599300, EPI_ISL_599301, EPI_ISL_599302, EPI_ISL_599303, EPI_ISL_599304, EPI_ISL_599305, EPI_ISL_599308, EPI_ISL_599309, EPI_ISL_599312, EPI_ISL_599313 | | | |
| see above | Lighthouse Lab in Glasgow | Wellcome Sanger Institute for the COVID-19 Genomics UK (COG-UK) consortium | Harper VanSteenhouse, Yumi Kasai, David Gray, Carol Clugston, Anna Dominiczak and Alex Alderton, Roberto Amato, Sonia Goncalves, Ewan Harrison, David K. Jackson, Ian Johnston, Dominic Kwiatkowski, Cordelia Langford, John Sillitoe on behalf of the Wellcome Sanger Institute COVID-19 Surveillance Team (http://www.sanger.ac.uk/covid-team) |
| EPI_ISL_599314 | Lighthouse Lab in Glasgow | Wellcome Sanger Institute for the COVID-19 Genomics UK (COG-UK) Consortium | Harper VanSteenhouse, Yumi Kasai, David Gray, Carol Clugston, Anna Dominiczak and Alex Alderton, Roberto Amato, Sonia Goncalves, Ewan Harrison, David K. Jackson, Ian Johnston, Dominic Kwiatkowski, Cordelia Langford, John Sillitoe on behalf of the Wellcome Sanger Institute COVID-19 Surveillance Team |
| EPI_ISL_599315, EPI_ISL_599317, EPI_ISL_599319, EPI_ISL_599322, EPI_ISL_599324, EPI_ISL_599327, EPI_ISL_599329, EPI_ISL_599334, EPI_ISL_599335, EPI_ISL_599336, EPI_ISL_599337, EPI_ISL_599338, EPI_ISL_599346, EPI_ISL_599348, EPI_ISL_599349, EPI_ISL_599350, EPI_ISL_599351, EPI_ISL_599355, EPI_ISL_599356, EPI_ISL_599357, EPI_ISL_599358, EPI_ISL_599359, EPI_ISL_599360, EPI_ISL_599363, EPI_ISL_599364, EPI_ISL_599365, EPI_ISL_599367, EPI_ISL_599368, EPI_ISL_599370, EPI_ISL_599371, EPI_ISL_599373, EPI_ISL_599375, EPI_ISL_599380, EPI_ISL_599381, EPI_ISL_599382, EPI_ISL_599383, EPI_ISL_599386, EPI_ISL_599387, EPI_ISL_599388, EPI_ISL_599389, EPI_ISL_599391, EPI_ISL_599392, EPI_ISL_599395, EPI_ISL_599396, EPI_ISL_599400, EPI_ISL_599402, EPI_ISL_599405, EPI_ISL_599406, EPI_ISL_599407, EPI_ISL_599415, EPI_ISL_599416, EPI_ISL_599417, EPI_ISL_599421, EPI_ISL_599422, EPI_ISL_599423, EPI_ISL_599424, EPI_ISL_599425, EPI_ISL_599427, EPI_ISL_599428, EPI_ISL_599430, EPI_ISL_599432, EPI_ISL_599433, EPI_ISL_599434, EPI_ISL_599439, EPI_ISL_599441, EPI_ISL_599442, EPI_ISL_599443, EPI_ISL_599446, EPI_ISL_599448, EPI_ISL_599451, EPI_ISL_599453, EPI_ISL_599454, EPI_ISL_599455, EPI_ISL_599456, EPI_ISL_599458, EPI_ISL_599460, EPI_ISL_599461, EPI_ISL_599462, EPI_ISL_599464, EPI_ISL_599467, EPI_ISL_599469, EPI_ISL_599473, EPI_ISL_599476, EPI_ISL_599478, EPI_ISL_599479, EPI_ISL_599483 | | | |
| see above | Lighthouse Lab in Glasgow | Wellcome Sanger Institute for the COVID-19 Genomics UK (COG-UK) consortium | Harper VanSteenhouse, Yumi Kasai, David Gray, Carol Clugston, Anna Dominiczak and Alex Alderton, Roberto Amato, Sonia Goncalves, Ewan Harrison, David K. Jackson, Ian Johnston, Dominic Kwiatkowski, Cordelia Langford, John Sillitoe on behalf of the Wellcome Sanger Institute COVID-19 Surveillance Team (http://www.sanger.ac.uk/covid-team) |
| EPI_ISL_599818, EPI_ISL_599819, EPI_ISL_599820, EPI_ISL_599821, EPI_ISL_599822, EPI_ISL_599823, EPI_ISL_599824, EPI_ISL_599825, EPI_ISL_599826, EPI_ISL_599827, EPI_ISL_599828, EPI_ISL_599829, EPI_ISL_599830, EPI_ISL_599831, EPI_ISL_599832, EPI_ISL_599833, EPI_ISL_599834, EPI_ISL_599835, EPI_ISL_599836, EPI_ISL_599837, EPI_ISL_599838, EPI_ISL_599839, EPI_ISL_599840, EPI_ISL_599841, EPI_ISL_599842, EPI_ISL_599843, EPI_ISL_599844 | | | |
| see above | Lighthouse Lab in Milton Keynes | Wellcome Sanger Institute for the COVID-19 Genomics UK (COG-UK) consortium | The Lighthouse Lab in Milton Keynes and Alex Alderton, Roberto Amato, Sonia Goncalves, Ewan Harrison, David K. Jackson, Ian Johnston, Dominic Kwiatkowski, Cordelia Langford, John Sillitoe on behalf of the Wellcome Sanger Institute COVID-19 Surveillance Team (http://www.sanger.ac.uk/covid-team) |
| EPI_ISL_599845 | Lighthouse Lab in Milton Keynes | Wellcome Sanger Institute for the COVID-19 Genomics UK (COG-UK) Consortium | The Lighthouse Lab in Milton Keynes and Alex Alderton, Roberto Amato, Sonia Goncalves, Ewan Harrison, David K. Jackson, Ian Johnston, Dominic Kwiatkowski, Cordelia Langford, John Sillitoe on behalf of the Wellcome Sanger Institute COVID-19 Surveillance Team |
| EPI_ISL_599846, EPI_ISL_599847, EPI_ISL_599848, EPI_ISL_599849, EPI_ISL_599850, EPI_ISL_599851, EPI_ISL_599852, EPI_ISL_599853, EPI_ISL_599854, EPI_ISL_599855, EPI_ISL_599856, EPI_ISL_599857, EPI_ISL_599858, EPI_ISL_599859, EPI_ISL_599860, EPI_ISL_599861, EPI_ISL_599862, EPI_ISL_599863, EPI_ISL_599864, EPI_ISL_599865, EPI_ISL_599866, EPI_ISL_599867, EPI_ISL_599868, EPI_ISL_599869, EPI_ISL_599870, EPI_ISL_599871, EPI_ISL_599872, EPI_ISL_599873, EPI_ISL_599874, EPI_ISL_599875, EPI_ISL_599876, EPI_ISL_599877, EPI_ISL_599878, EPI_ISL_599879, EPI_ISL_599880, EPI_ISL_599881, EPI_ISL_599882, EPI_ISL_599883, EPI_ISL_599884, EPI_ISL_599885, EPI_ISL_599886, EPI_ISL_599887 | | | |
| see above | Lighthouse Lab in Milton Keynes | Wellcome Sanger Institute for the COVID-19 Genomics UK (COG-UK) consortium | The Lighthouse Lab in Milton Keynes and Alex Alderton, Roberto Amato, Sonia Goncalves, Ewan Harrison, David K. Jackson, Ian Johnston, Dominic Kwiatkowski, Cordelia Langford, John Sillitoe on behalf of the Wellcome Sanger Institute COVID-19 Surveillance Team (http://www.sanger.ac.uk/covid-team) |
| EPI_ISL_599888 | Lighthouse Lab in Milton Keynes | Wellcome Sanger Institute for the COVID-19 Genomics UK (COG-UK) Consortium | The Lighthouse Lab in Milton Keynes and Alex Alderton, Roberto Amato, Sonia Goncalves, Ewan Harrison, David K. Jackson, Ian Johnston, Dominic Kwiatkowski, Cordelia Langford, John Sillitoe on behalf of the Wellcome Sanger Institute COVID-19 Surveillance Team |
| EPI_ISL_599889, EPI_ISL_599890, EPI_ISL_599891, EPI_ISL_599892, EPI_ISL_599893, EPI_ISL_599894, EPI_ISL_599895, EPI_ISL_599896, EPI_ISL_599897, EPI_ISL_599898, EPI_ISL_599899, EPI_ISL_599900, EPI_ISL_599901, EPI_ISL_599902, EPI_ISL_599903, EPI_ISL_599904, EPI_ISL_599905, EPI_ISL_599906, EPI_ISL_599907, EPI_ISL_599908, EPI_ISL_599909, EPI_ISL_599910, EPI_ISL_599911, EPI_ISL_599912, EPI_ISL_599913, EPI_ISL_599914, EPI_ISL_599915, EPI_ISL_599916, EPI_ISL_599917, EPI_ISL_599918, EPI_ISL_599919, EPI_ISL_599920, EPI_ISL_599921, EPI_ISL_599922, EPI_ISL_599923, EPI_ISL_599924, EPI_ISL_599925, EPI_ISL_599926, EPI_ISL_599927, EPI_ISL_599928, EPI_ISL_599929, EPI_ISL_599930, EPI_ISL_599931, EPI_ISL_599932, EPI_ISL_599933, EPI_ISL_599934, EPI_ISL_599935, EPI_ISL_599936, EPI_ISL_599937, EPI_ISL_599938, EPI_ISL_599939, EPI_ISL_599940, EPI_ISL_599941, EPI_ISL_599942, EPI_ISL_599943, EPI_ISL_599944, EPI_ISL_599945, EPI_ISL_599946, EPI_ISL_599947 | | | |
| see above | Lighthouse Lab in Milton Keynes | Wellcome Sanger Institute for the COVID-19 Genomics UK (COG-UK) consortium | The Lighthouse Lab in Milton Keynes and Alex Alderton, Roberto Amato, Sonia Goncalves, Ewan Harrison, David K. Jackson, Ian Johnston, Dominic Kwiatkowski, Cordelia Langford, John Sillitoe on behalf of the Wellcome Sanger Institute COVID-19 Surveillance Team (http://www.sanger.ac.uk/covid-team) |
| EPI_ISL_599948 | Lighthouse Lab in Milton Keynes | Wellcome Sanger Institute for the COVID-19 Genomics UK (COG-UK) Consortium | The Lighthouse Lab in Milton Keynes and Alex Alderton, Roberto Amato, Sonia Goncalves, Ewan Harrison, David K. Jackson, Ian Johnston, Dominic Kwiatkowski, Cordelia Langford, John Sillitoe on behalf of the Wellcome Sanger Institute COVID-19 Surveillance Team |
| EPI_ISL_599949, EPI_ISL_599950, EPI_ISL_599951, EPI_ISL_599952, EPI_ISL_599953, EPI_ISL_599954, EPI_ISL_599955, EPI_ISL_599956, EPI_ISL_599957, EPI_ISL_599958, EPI_ISL_599959 | | | |
| see above | Lighthouse Lab in Milton Keynes | Wellcome Sanger Institute for the COVID-19 Genomics UK (COG-UK) consortium | The Lighthouse Lab in Milton Keynes and Alex Alderton, Roberto Amato, Sonia Goncalves, Ewan Harrison, David K. Jackson, Ian Johnston, Dominic Kwiatkowski, Cordelia Langford, John Sillitoe on behalf of the Wellcome Sanger Institute COVID-19 Surveillance Team (http://www.sanger.ac.uk/covid-team) |
| EPI_ISL_599960 | Lighthouse Lab in Milton Keynes | Wellcome Sanger Institute for the COVID-19 Genomics UK (COG-UK) Consortium | The Lighthouse Lab in Milton Keynes and Alex Alderton, Roberto Amato, Sonia Goncalves, Ewan Harrison, David K. Jackson, Ian Johnston, Dominic Kwiatkowski, Cordelia Langford, John Sillitoe on behalf of the Wellcome Sanger Institute COVID-19 Surveillance Team |
| EPI_ISL_599961, EPI_ISL_599962, EPI_ISL_599963, EPI_ISL_599964, EPI_ISL_599965, EPI_ISL_599966, EPI_ISL_599967, EPI_ISL_599968, EPI_ISL_599969, EPI_ISL_599970, EPI_ISL_599971, EPI_ISL_599972, EPI_ISL_599973, EPI_ISL_599974, EPI_ISL_599975, EPI_ISL_599976, EPI_ISL_599977, EPI_ISL_599978, EPI_ISL_599979, EPI_ISL_599980, EPI_ISL_599981, EPI_ISL_599982, EPI_ISL_599983, EPI_ISL_599984, EPI_ISL_599985, EPI_ISL_599986, EPI_ISL_599987, EPI_ISL_599988, EPI_ISL_599989, EPI_ISL_599990, EPI_ISL_599991, EPI_ISL_599992, EPI_ISL_599993, EPI_ISL_599994, EPI_ISL_599995, EPI_ISL_599996, EPI_ISL_599997, EPI_ISL_599998, EPI_ISL_599999, EPI_ISL_600000, EPI_ISL_600001, EPI_ISL_600002, EPI_ISL_600003, EPI_ISL_600004, EPI_ISL_600005, EPI_ISL_600006, EPI_ISL_600007, EPI_ISL_600008, EPI_ISL_600009, EPI_ISL_600010, EPI_ISL_600011, EPI_ISL_600012, EPI_ISL_600013, EPI_ISL_600014, EPI_ISL_600015, EPI_ISL_600016, EPI_ISL_600017, EPI_ISL_600018, EPI_ISL_600019, EPI_ISL_600020, EPI_ISL_600021, EPI_ISL_600022, EPI_ISL_600023, EPI_ISL_600024, EPI_ISL_600025, EPI_ISL_600026, EPI_ISL_600027, EPI_ISL_600028, EPI_ISL_600029, EPI_ISL_600030, EPI_ISL_600031, EPI_ISL_600032, EPI_ISL_600033, EPI_ISL_600034, EPI_ISL_600035, EPI_ISL_600036, EPI_ISL_600037, EPI_ISL_600038, EPI_ISL_600039, EPI_ISL_600040, EPI_ISL_600041, EPI_ISL_600042, EPI_ISL_600043, EPI_ISL_600044, EPI_ISL_600045, EPI_ISL_600046, EPI_ISL_600047, EPI_ISL_600048, EPI_ISL_600049, EPI_ISL_600050, EPI_ISL_600051, EPI_ISL_600052, EPI_ISL_600053, EPI_ISL_600054, EPI_ISL_600055 | | | |
| see above | Lighthouse Lab in Milton Keynes | Wellcome Sanger Institute for the COVID-19 Genomics UK (COG-UK) consortium | The Lighthouse Lab in Milton Keynes and Alex Alderton, Roberto Amato, Sonia Goncalves, Ewan Harrison, David K. Jackson, Ian Johnston, Dominic Kwiatkowski, Cordelia Langford, John Sillitoe on behalf of the Wellcome Sanger Institute COVID-19 Surveillance Team (http://www.sanger.ac.uk/covid-team) |
| EPI_ISL_600056 | Lighthouse Lab in Milton Keynes | Wellcome Sanger Institute for the COVID-19 Genomics UK (COG-UK) Consortium | The Lighthouse Lab in Milton Keynes and Alex Alderton, Roberto Amato, Sonia Goncalves, Ewan Harrison, David K. Jackson, Ian Johnston, Dominic Kwiatkowski, Cordelia Langford, John Sillitoe on behalf of the Wellcome Sanger Institute COVID-19 Surveillance Team |
| EPI_ISL_600057, EPI_ISL_600058, EPI_ISL_600059, EPI_ISL_600060, EPI_ISL_600061, EPI_ISL_600062, EPI_ISL_600063, EPI_ISL_600064, EPI_ISL_600065, EPI_ISL_600066, EPI_ISL_600067, EPI_ISL_600068, EPI_ISL_600069, EPI_ISL_600070, EPI_ISL_600071, EPI_ISL_600072, EPI_ISL_600073, EPI_ISL_600074, EPI_ISL_600075, EPI_ISL_600076, EPI_ISL_600077, EPI_ISL_600078, EPI_ISL_600079, EPI_ISL_600080, EPI_ISL_600081, EPI_ISL_600082, EPI_ISL_600083, EPI_ISL_600084, EPI_ISL_600085, EPI_ISL_600086, EPI_ISL_600087, EPI_ISL_600088, EPI_ISL_600089, EPI_ISL_600090, EPI_ISL_600091, EPI_ISL_600092, | | | |

| | | | |
|---|---|---|---|
| EPI_ISL_600093, EPI_ISL_600094, EPI_ISL_600095, EPI_ISL_600096, EPI_ISL_600097, EPI_ISL_600098, EPI_ISL_600099, EPI_ISL_600100, EPI_ISL_600101, EPI_ISL_600102, EPI_ISL_600103, EPI_ISL_600104, EPI_ISL_600105, EPI_ISL_600106, EPI_ISL_600107, EPI_ISL_600108, EPI_ISL_600109, EPI_ISL_600110, EPI_ISL_600111, EPI_ISL_600112, EPI_ISL_600113, EPI_ISL_600114, EPI_ISL_600115, EPI_ISL_600116, EPI_ISL_600117, EPI_ISL_600118, EPI_ISL_600119, EPI_ISL_600120, EPI_ISL_600121, EPI_ISL_600122, EPI_ISL_600123, EPI_ISL_600124, EPI_ISL_600125, EPI_ISL_600126, EPI_ISL_600127 | | | |
| see above | Lighthouse Lab in Milton Keynes | Wellcome Sanger Institute for the COVID-19 Genomics UK (COG-UK) consortium | The Lighthouse Lab in Milton Keynes and Alex Alderton, Roberto Amato, Sonia Goncalves, Ewan Harrison, David K. Jackson, Ian Johnston, Dominic Kwiatkowski, Cordelia Langford, John Sillitoe on behalf of the Wellcome Sanger Institute COVID-19 Surveillance Team (http://www.sanger.ac.uk/covid-team) |
| EPI_ISL_600128, EPI_ISL_600130, EPI_ISL_600133, EPI_ISL_600134, EPI_ISL_600135, EPI_ISL_600136 | Lighthouse Lab in Glasgow | Wellcome Sanger Institute for the COVID-19 Genomics UK (COG-UK) consortium | Harper VanSteenhouse, Yumi Kasai, David Gray, Carol Clugston, Anna Dominiczak and Alex Alderton, Roberto Amato, Sonia Goncalves, Ewan Harrison, David K. Jackson, Ian Johnston, Dominic Kwiatkowski, Cordelia Langford, John Sillitoe on behalf of the Wellcome Sanger Institute COVID-19 Surveillance Team (http://www.sanger.ac.uk/covid-team) |
| EPI_ISL_600139 | Lighthouse Lab in Glasgow | Wellcome Sanger Institute for the COVID-19 Genomics UK (COG-UK) Consortium | Harper VanSteenhouse, Yumi Kasai, David Gray, Carol Clugston, Anna Dominiczak and Alex Alderton, Roberto Amato, Sonia Goncalves, Ewan Harrison, David K. Jackson, Ian Johnston, Dominic Kwiatkowski, Cordelia Langford, John Sillitoe on behalf of the Wellcome Sanger Institute COVID-19 Surveillance Team |
| EPI_ISL_600144, EPI_ISL_600147, EPI_ISL_600148, EPI_ISL_600150, EPI_ISL_600151, EPI_ISL_600152, EPI_ISL_600157, EPI_ISL_600158, EPI_ISL_600159, EPI_ISL_600160, EPI_ISL_600161, EPI_ISL_600163, EPI_ISL_600166, EPI_ISL_600167, EPI_ISL_600172, EPI_ISL_600177, EPI_ISL_600179, EPI_ISL_600180, EPI_ISL_600182, EPI_ISL_600185, EPI_ISL_600192, EPI_ISL_600193, EPI_ISL_600194, EPI_ISL_600196, EPI_ISL_600197, EPI_ISL_600200, EPI_ISL_600201, EPI_ISL_600204, EPI_ISL_600205, EPI_ISL_600209, EPI_ISL_600210, EPI_ISL_600212, EPI_ISL_600215, EPI_ISL_600217, EPI_ISL_600223, EPI_ISL_600224, EPI_ISL_600225, EPI_ISL_600231, EPI_ISL_600232, EPI_ISL_600233, EPI_ISL_600238, EPI_ISL_600239, EPI_ISL_600240, EPI_ISL_600241, EPI_ISL_600243, EPI_ISL_600249, EPI_ISL_600250, EPI_ISL_600252, EPI_ISL_600254, EPI_ISL_600257, EPI_ISL_600258, EPI_ISL_600262, EPI_ISL_600263, EPI_ISL_600266, EPI_ISL_600267, EPI_ISL_600269, EPI_ISL_600271, EPI_ISL_600272, EPI_ISL_600273, EPI_ISL_600279, EPI_ISL_600281, EPI_ISL_600283, EPI_ISL_600285, EPI_ISL_600286, EPI_ISL_600287, EPI_ISL_600290, EPI_ISL_600292, EPI_ISL_600293, EPI_ISL_600298, EPI_ISL_600299, EPI_ISL_600303, EPI_ISL_600304, EPI_ISL_600306, EPI_ISL_600309, EPI_ISL_600311, EPI_ISL_600313, EPI_ISL_600314, EPI_ISL_600315, EPI_ISL_600317, EPI_ISL_600325, EPI_ISL_600332, EPI_ISL_600334, EPI_ISL_600338, EPI_ISL_600342, EPI_ISL_600344, EPI_ISL_600345, EPI_ISL_600352, EPI_ISL_600355, EPI_ISL_600356, EPI_ISL_600357, EPI_ISL_600359, EPI_ISL_600361, EPI_ISL_600366, EPI_ISL_600369, EPI_ISL_600375, EPI_ISL_600378, EPI_ISL_600380, EPI_ISL_600381, EPI_ISL_600389, EPI_ISL_600391, EPI_ISL_600393, EPI_ISL_600396, EPI_ISL_600402, EPI_ISL_600404, EPI_ISL_600406, EPI_ISL_600407, EPI_ISL_600413, EPI_ISL_600414, EPI_ISL_600415, EPI_ISL_600416, EPI_ISL_600417, EPI_ISL_600419, EPI_ISL_600421, EPI_ISL_600422, EPI_ISL_600424, EPI_ISL_600427, EPI_ISL_600447, EPI_ISL_600448, EPI_ISL_600452, EPI_ISL_600455, EPI_ISL_600472, EPI_ISL_600475, EPI_ISL_600477, EPI_ISL_600479, EPI_ISL_600485, EPI_ISL_600496, EPI_ISL_600501, EPI_ISL_600503, EPI_ISL_600504, EPI_ISL_600509, EPI_ISL_600520 | | | |
| see above | Lighthouse Lab in Glasgow | Wellcome Sanger Institute for the COVID-19 Genomics UK (COG-UK) consortium | Harper VanSteenhouse, Yumi Kasai, David Gray, Carol Clugston, Anna Dominiczak and Alex Alderton, Roberto Amato, Sonia Goncalves, Ewan Harrison, David K. Jackson, Ian Johnston, Dominic Kwiatkowski, Cordelia Langford, John Sillitoe on behalf of the Wellcome Sanger Institute COVID-19 Surveillance Team (http://www.sanger.ac.uk/covid-team) |
| EPI_ISL_600521, EPI_ISL_600523, EPI_ISL_600525, EPI_ISL_600528, EPI_ISL_600529, EPI_ISL_600532, EPI_ISL_600536, EPI_ISL_600537 | Lighthouse Lab in Alderley Park | Wellcome Sanger Institute for the COVID-19 Genomics UK (COG-UK) consortium | Jacquelyn Wynn, Mairead Hyland, The Lighthouse Lab in Alderley Park and Alex Alderton, Roberto Amato, Sonia Goncalves, Ewan Harrison, David K. Jackson, Ian Johnston, Dominic Kwiatkowski, Cordelia Langford, John Sillitoe on behalf of the Wellcome Sanger Institute COVID-19 Surveillance Team (http://www.sanger.ac.uk/covid-team) |
| EPI_ISL_600539, EPI_ISL_600540, EPI_ISL_600542, EPI_ISL_600544, EPI_ISL_600545, EPI_ISL_600547, EPI_ISL_600548, EPI_ISL_600550, EPI_ISL_600551, EPI_ISL_600553, EPI_ISL_600555, EPI_ISL_600556, EPI_ISL_600558, EPI_ISL_600560, EPI_ISL_600561, EPI_ISL_600563, EPI_ISL_600565, EPI_ISL_600566, EPI_ISL_600568, EPI_ISL_600569, EPI_ISL_600571, EPI_ISL_600572, EPI_ISL_600573, EPI_ISL_600574, EPI_ISL_600575, EPI_ISL_600576, EPI_ISL_600577, EPI_ISL_600578, EPI_ISL_600579, EPI_ISL_600580, EPI_ISL_600581, EPI_ISL_600582, EPI_ISL_600583, EPI_ISL_600584, EPI_ISL_600585, EPI_ISL_600586, EPI_ISL_600587, EPI_ISL_600588, EPI_ISL_600589, EPI_ISL_600590, EPI_ISL_600591, EPI_ISL_600592, EPI_ISL_600593, EPI_ISL_600594, EPI_ISL_600595, EPI_ISL_600596, EPI_ISL_600597, EPI_ISL_600598, EPI_ISL_600599, EPI_ISL_600600, EPI_ISL_600601, EPI_ISL_600602, EPI_ISL_600603, EPI_ISL_600604, EPI_ISL_600605, EPI_ISL_600606, EPI_ISL_600607, EPI_ISL_600608, EPI_ISL_600609, EPI_ISL_600610, EPI_ISL_600611, EPI_ISL_600612, EPI_ISL_600613, EPI_ISL_600614, EPI_ISL_600615, EPI_ISL_600616, EPI_ISL_600617, EPI_ISL_600618, EPI_ISL_600619, EPI_ISL_600620, EPI_ISL_600621, EPI_ISL_600622, EPI_ISL_600623, EPI_ISL_600624, EPI_ISL_600625, EPI_ISL_600626, EPI_ISL_600627, EPI_ISL_600628, EPI_ISL_600629, EPI_ISL_600630, EPI_ISL_600631, EPI_ISL_600632, EPI_ISL_600633, EPI_ISL_600634, EPI_ISL_600635, EPI_ISL_600636, EPI_ISL_600637, EPI_ISL_600638, EPI_ISL_600639, EPI_ISL_600640, EPI_ISL_600641, EPI_ISL_600642 | | | |
| see above | Lighthouse Lab in Glasgow | Wellcome Sanger Institute for the COVID-19 Genomics UK (COG-UK) consortium | Harper VanSteenhouse, Yumi Kasai, David Gray, Carol Clugston, Anna Dominiczak and Alex Alderton, Roberto Amato, Sonia Goncalves, Ewan Harrison, David K. Jackson, Ian Johnston, Dominic Kwiatkowski, Cordelia Langford, John Sillitoe on behalf of the Wellcome Sanger Institute COVID-19 Surveillance Team (http://www.sanger.ac.uk/covid-team) |
| EPI_ISL_600643 | Lighthouse Lab in Glasgow | Wellcome Sanger Institute for the COVID-19 Genomics UK (COG-UK) Consortium | Harper VanSteenhouse, Yumi Kasai, David Gray, Carol Clugston, Anna Dominiczak and Alex Alderton, Roberto Amato, Sonia Goncalves, Ewan Harrison, David K. Jackson, Ian Johnston, Dominic Kwiatkowski, Cordelia Langford, John Sillitoe on behalf of the Wellcome Sanger Institute COVID-19 Surveillance Team |
| EPI_ISL_600644, EPI_ISL_600645, EPI_ISL_600646, EPI_ISL_600647, EPI_ISL_600648, EPI_ISL_600649, EPI_ISL_600650, EPI_ISL_600651, EPI_ISL_600652, EPI_ISL_600653, EPI_ISL_600654, EPI_ISL_600655, EPI_ISL_600656, EPI_ISL_600657, EPI_ISL_600658, EPI_ISL_600659, EPI_ISL_600660, EPI_ISL_600661, EPI_ISL_600662, EPI_ISL_600663, EPI_ISL_600664, EPI_ISL_600665, EPI_ISL_600666, EPI_ISL_600667, EPI_ISL_600668, EPI_ISL_600669, EPI_ISL_600670, EPI_ISL_600671, EPI_ISL_600672, EPI_ISL_600673, EPI_ISL_600674, EPI_ISL_600675, EPI_ISL_600676, EPI_ISL_600677, EPI_ISL_600678, EPI_ISL_600679, EPI_ISL_600680, EPI_ISL_600681, EPI_ISL_600682, EPI_ISL_600683, EPI_ISL_600684, EPI_ISL_600685, EPI_ISL_600686, EPI_ISL_600687, EPI_ISL_600688, EPI_ISL_600689, EPI_ISL_600690, EPI_ISL_600691, EPI_ISL_600692, EPI_ISL_600693, EPI_ISL_600694, EPI_ISL_600695, EPI_ISL_600696, EPI_ISL_600697, EPI_ISL_600698, EPI_ISL_600699, EPI_ISL_600700, EPI_ISL_600701, EPI_ISL_600702, EPI_ISL_600703, EPI_ISL_600704, EPI_ISL_600705, EPI_ISL_600706, EPI_ISL_600707, EPI_ISL_600708, EPI_ISL_600709, EPI_ISL_600710, EPI_ISL_600711, EPI_ISL_600712, EPI_ISL_600713, EPI_ISL_600714, EPI_ISL_600715, EPI_ISL_600716, EPI_ISL_600717, EPI_ISL_600718, EPI_ISL_600719, EPI_ISL_600720, EPI_ISL_600721, EPI_ISL_600722, EPI_ISL_600723, EPI_ISL_600724, EPI_ISL_600725, EPI_ISL_600726, EPI_ISL_600727, EPI_ISL_600728, EPI_ISL_600729, EPI_ISL_600730, EPI_ISL_600731, EPI_ISL_600732, EPI_ISL_600733, EPI_ISL_600734, EPI_ISL_600735, EPI_ISL_600736, EPI_ISL_600737, EPI_ISL_600738, EPI_ISL_600739, EPI_ISL_600740, EPI_ISL_600741, EPI_ISL_600742, EPI_ISL_600743, EPI_ISL_600744, EPI_ISL_600745, EPI_ISL_600746, EPI_ISL_600747, EPI_ISL_600748, EPI_ISL_600749, EPI_ISL_600750, EPI_ISL_600751, EPI_ISL_600752, EPI_ISL_600753, EPI_ISL_600754, EPI_ISL_600755, EPI_ISL_600756, EPI_ISL_600757, EPI_ISL_600758, EPI_ISL_600759, EPI_ISL_600761, EPI_ISL_600766, EPI_ISL_600768, EPI_ISL_600770, EPI_ISL_600773, EPI_ISL_600778, EPI_ISL_600780, EPI_ISL_600782, EPI_ISL_600783, EPI_ISL_600784, EPI_ISL_600785, EPI_ISL_600787, EPI_ISL_600788, EPI_ISL_600789, EPI_ISL_600790, EPI_ISL_600791, EPI_ISL_600792, EPI_ISL_600793, EPI_ISL_600794, EPI_ISL_600796, EPI_ISL_600797, EPI_ISL_600798, EPI_ISL_600803, EPI_ISL_600805, EPI_ISL_600807, EPI_ISL_600808, EPI_ISL_600809, EPI_ISL_600810, EPI_ISL_600812, EPI_ISL_600817, EPI_ISL_600818, EPI_ISL_600819, EPI_ISL_600820, EPI_ISL_600821, EPI_ISL_600823 | | | |
| see above | Lighthouse Lab in Glasgow | Wellcome Sanger Institute for the COVID-19 Genomics UK (COG-UK) consortium | Harper VanSteenhouse, Yumi Kasai, David Gray, Carol Clugston, Anna Dominiczak and Alex Alderton, Roberto Amato, Sonia Goncalves, Ewan Harrison, David K. Jackson, Ian Johnston, Dominic Kwiatkowski, Cordelia Langford, John Sillitoe on behalf of the Wellcome Sanger Institute COVID-19 Surveillance Team (http://www.sanger.ac.uk/covid-team) |
| EPI_ISL_600824 | Lighthouse Lab in Glasgow | Wellcome Sanger Institute for the COVID-19 Genomics UK (COG-UK) Consortium | Harper VanSteenhouse, Yumi Kasai, David Gray, Carol Clugston, Anna Dominiczak and Alex Alderton, Roberto Amato, Sonia Goncalves, Ewan Harrison, David K. Jackson, Ian Johnston, Dominic Kwiatkowski, Cordelia Langford, John Sillitoe on behalf of the Wellcome Sanger Institute COVID-19 Surveillance Team |
| EPI_ISL_600825, EPI_ISL_600826, EPI_ISL_600829, EPI_ISL_600830, EPI_ISL_600831, EPI_ISL_600832, EPI_ISL_600833, EPI_ISL_600834, EPI_ISL_600835, EPI_ISL_600836, EPI_ISL_600837, EPI_ISL_600838, EPI_ISL_600839, EPI_ISL_600840, EPI_ISL_600841, EPI_ISL_600842, EPI_ISL_600843, EPI_ISL_600844, EPI_ISL_600845, EPI_ISL_600846, EPI_ISL_600847, EPI_ISL_600848, EPI_ISL_600849, EPI_ISL_600850, EPI_ISL_600851, EPI_ISL_600852, EPI_ISL_600853, EPI_ISL_600854, EPI_ISL_600855, EPI_ISL_600856, EPI_ISL_600857, EPI_ISL_600858, EPI_ISL_600859, EPI_ISL_600860, EPI_ISL_600861, EPI_ISL_600862, EPI_ISL_600863, EPI_ISL_600864, EPI_ISL_600865, EPI_ISL_600866, EPI_ISL_600867, EPI_ISL_600868, EPI_ISL_600869, EPI_ISL_600870, EPI_ISL_600871, EPI_ISL_600872, EPI_ISL_600873, EPI_ISL_600874, EPI_ISL_600875, EPI_ISL_600876, EPI_ISL_600877, EPI_ISL_600878, EPI_ISL_600879, EPI_ISL_600880, EPI_ISL_600881, EPI_ISL_600882, EPI_ISL_600883, EPI_ISL_600884, EPI_ISL_600885, EPI_ISL_600886, EPI_ISL_600887, EPI_ISL_600888, EPI_ISL_600889, EPI_ISL_600890, EPI_ISL_600891, EPI_ISL_600893, EPI_ISL_600894, EPI_ISL_600895, EPI_ISL_600896, EPI_ISL_600897, EPI_ISL_600898, EPI_ISL_600899, EPI_ISL_600900, EPI_ISL_600902, EPI_ISL_600903 | | | |
| see above | Lighthouse Lab in Glasgow | Wellcome Sanger Institute for the COVID-19 Genomics UK (COG-UK) consortium | Harper VanSteenhouse, Yumi Kasai, David Gray, Carol Clugston, Anna Dominiczak and Alex Alderton, Roberto Amato, Sonia Goncalves, Ewan Harrison, David K. Jackson, Ian Johnston, Dominic Kwiatkowski, Cordelia Langford, John Sillitoe on behalf of the Wellcome Sanger Institute COVID-19 Surveillance Team (http://www.sanger.ac.uk/covid-team) |
| EPI_ISL_600904 | Lighthouse Lab in Glasgow | Wellcome Sanger Institute for the COVID-19 Genomics UK (COG-UK) Consortium | Harper VanSteenhouse, Yumi Kasai, David Gray, Carol Clugston, Anna Dominiczak and Alex Alderton, Roberto Amato, Sonia Goncalves, Ewan Harrison, David K. Jackson, Ian Johnston, Dominic Kwiatkowski, Cordelia Langford, John Sillitoe on behalf of the Wellcome Sanger Institute COVID-19 Surveillance Team |
| EPI_ISL_600905, EPI_ISL_600906, EPI_ISL_600907, EPI_ISL_600908, EPI_ISL_600909, EPI_ISL_600910, EPI_ISL_600911, EPI_ISL_600914, EPI_ISL_600916, EPI_ISL_600917, EPI_ISL_600918, EPI_ISL_600919, EPI_ISL_600920, EPI_ISL_600921, EPI_ISL_600922, EPI_ISL_600924 | | | |
| see above | Lighthouse Lab in Glasgow | Wellcome Sanger Institute for the COVID-19 Genomics UK (COG-UK) consortium | Harper VanSteenhouse, Yumi Kasai, David Gray, Carol Clugston, Anna Dominiczak and Alex Alderton, Roberto Amato, Sonia Goncalves, Ewan Harrison, David K. Jackson, Ian Johnston, Dominic Kwiatkowski, Cordelia Langford, John Sillitoe on behalf of the Wellcome Sanger Institute COVID-19 Surveillance Team (http://www.sanger.ac.uk/covid-team) |
| EPI_ISL_600925, EPI_ISL_600928, EPI_ISL_600929 | Lighthouse Lab in Milton Keynes | Wellcome Sanger Institute for the COVID-19 Genomics UK (COG-UK) consortium | The Lighthouse Lab in Milton Keynes and Alex Alderton, Roberto Amato, Sonia Goncalves, Ewan Harrison, David K. Jackson, Ian Johnston, Dominic Kwiatkowski, Cordelia Langford, John Sillitoe on behalf of the Wellcome Sanger Institute COVID-19 Surveillance Team (http://www.sanger.ac.uk/covid-team) |
| EPI_ISL_602258, EPI_ISL_602259, EPI_ISL_602260, EPI_ISL_602261, EPI_ISL_602262, EPI_ISL_602263 | MD PHL | MD PHL | Maryland Department of Health Laboratories Administration |
| EPI_ISL_602304 | Istituto Zooprofilattico Sperimentale del Mezzogiorno | U.O. Diagnostica Virologica Dip. Sanità Animale IZSM | Maurizio Viscardi, Lorena Cardillo , e Giovanna Fusco |
| EPI_ISL_602379, EPI_ISL_602390, EPI_ISL_602394, EPI_ISL_602395, EPI_ISL_602396, EPI_ISL_602405, EPI_ISL_602407, EPI_ISL_602408, EPI_ISL_602419, EPI_ISL_602421, EPI_ISL_602422, EPI_ISL_602431, EPI_ISL_602432, EPI_ISL_602435, EPI_ISL_602436, EPI_ISL_602437, EPI_ISL_602448, EPI_ISL_602454, EPI_ISL_602455, EPI_ISL_602458 | | | |
| see above | HELIX LLC | WHO National Influenza Centre Russian Federation | Andrey Komissarov, Artem Fadeev, Kseniya Komissarova, Anna Ivanova, Dmitry Bazhenov, Daria Danilenko |

| | | | |
|---|---|---|---|
| EPI_ISL_602830, EPI_ISL_602831, EPI_ISL_602832, EPI_ISL_602833, EPI_ISL_602834, EPI_ISL_602835, EPI_ISL_602836, EPI_ISL_602837, EPI_ISL_602838, EPI_ISL_602839, EPI_ISL_602840, EPI_ISL_602841, EPI_ISL_602842, EPI_ISL_602843, EPI_ISL_602844, EPI_ISL_602852, EPI_ISL_602853, EPI_ISL_602854, EPI_ISL_602855, EPI_ISL_602856, EPI_ISL_602857, EPI_ISL_602858, EPI_ISL_602859, EPI_ISL_602860, EPI_ISL_602861, EPI_ISL_602862, EPI_ISL_602863, EPI_ISL_602864, EPI_ISL_602866, EPI_ISL_602912, EPI_ISL_602913, EPI_ISL_602914, EPI_ISL_602915, EPI_ISL_602916, EPI_ISL_602917, EPI_ISL_602918, EPI_ISL_602919, EPI_ISL_602920, EPI_ISL_602921, EPI_ISL_602922, EPI_ISL_602923, EPI_ISL_602924, EPI_ISL_602925, EPI_ISL_602926, EPI_ISL_602927, EPI_ISL_602928, EPI_ISL_602929, EPI_ISL_602930 | | | |
| see above | NHLS-IALCH | KRISP, KZN Research Innovation and Sequencing Platform | Giandhari J, Pillay S, Lessells R, Mdlalose K, York D, Khan S, Tegally H, Wilkinson E, de Oliveira T |
| EPI_ISL_602977, EPI_ISL_602978, EPI_ISL_602979, EPI_ISL_602980, EPI_ISL_602981, EPI_ISL_602982, EPI_ISL_602983, EPI_ISL_602984 | Minnesota Department of Health, Public Health Laboratory | Minnesota Department of Health, Public Health Laboratory | Matt Plumb, Jacob Garfin, Alexandra Lorentz, and Xiong Wang |
| EPI_ISL_603002, EPI_ISL_603003 | Sanford South University Medical Center | Minnesota Department of Health, Public Health Laboratory | Matt Plumb, Jacob Garfin, Alexandra Lorentz, and Xiong Wang |
| EPI_ISL_603004 | Essentia Health-St. Mary's Medical Center | Minnesota Department of Health, Public Health Laboratory | Matt Plumb, Jacob Garfin, Alexandra Lorentz, and Xiong Wang |
| EPI_ISL_603011 | Mayo Clinic & Mayo Clinic Laboratories | Minnesota Department of Health, Public Health Laboratory | Matt Plumb, Jacob Garfin, Alexandra Lorentz, and Xiong Wang |
| EPI_ISL_603218, EPI_ISL_603219, EPI_ISL_603220 | CHU Purpan - Laboratoire de Virologie - Institut Fédératif de Biologie | CHU Purpan - Laboratoire de Virologie - Institut Fédératif de Biologie | Latour J., Ranger N., Dubois M., Carcenac R., Harter A., Boyer P., Tremeaux P., Izopet J. |
| EPI_ISL_603252, EPI_ISL_603254, EPI_ISL_603255 | UW Virology Lab | UW Virology Lab | Pavitra Roychoudhury, Hong Xie, Lasata Shrestha, Meei-Li Huang, Keith R Jerome, Alexander Greninger |
| EPI_ISL_603267, EPI_ISL_603268, EPI_ISL_603269, EPI_ISL_603270, EPI_ISL_603271, EPI_ISL_603272, EPI_ISL_603273, EPI_ISL_603274, EPI_ISL_603275, EPI_ISL_603276, EPI_ISL_603277, EPI_ISL_603278, EPI_ISL_603279, EPI_ISL_603280, EPI_ISL_603281, EPI_ISL_603282, EPI_ISL_603283, EPI_ISL_603284, EPI_ISL_603285, EPI_ISL_603286, EPI_ISL_603287, EPI_ISL_603288, EPI_ISL_603289, EPI_ISL_603290, EPI_ISL_603291, EPI_ISL_603292, EPI_ISL_603293, EPI_ISL_603294, EPI_ISL_603295, EPI_ISL_603296, EPI_ISL_603297, EPI_ISL_603298, EPI_ISL_603299, EPI_ISL_603300, EPI_ISL_603301, EPI_ISL_603302, EPI_ISL_603303, EPI_ISL_603304, EPI_ISL_603305, EPI_ISL_603306, EPI_ISL_603307, EPI_ISL_603308, EPI_ISL_603309, EPI_ISL_603310, EPI_ISL_603311, EPI_ISL_603312, EPI_ISL_603313, EPI_ISL_603314, EPI_ISL_603315, EPI_ISL_603316, EPI_ISL_603317, EPI_ISL_603318, EPI_ISL_603319, EPI_ISL_603320, EPI_ISL_603321, EPI_ISL_603322, EPI_ISL_603323, EPI_ISL_603324, EPI_ISL_603325, EPI_ISL_603326, EPI_ISL_603327, EPI_ISL_603328, EPI_ISL_603329, EPI_ISL_603330, EPI_ISL_603331, EPI_ISL_603332, EPI_ISL_603333, EPI_ISL_603334, EPI_ISL_603335, EPI_ISL_603336, EPI_ISL_603337, EPI_ISL_603338, EPI_ISL_603339, EPI_ISL_603340, EPI_ISL_603341, EPI_ISL_603342, EPI_ISL_603343, EPI_ISL_603344, EPI_ISL_603345, EPI_ISL_603346, EPI_ISL_603347, EPI_ISL_603348, EPI_ISL_603349, EPI_ISL_603350, EPI_ISL_603351, EPI_ISL_603352, EPI_ISL_603353, EPI_ISL_603354, EPI_ISL_603355, EPI_ISL_603356, EPI_ISL_603357, EPI_ISL_603358, EPI_ISL_603359, EPI_ISL_603360, EPI_ISL_603361, EPI_ISL_603362, EPI_ISL_603363, EPI_ISL_603364, EPI_ISL_603365, EPI_ISL_603366, EPI_ISL_603367, EPI_ISL_603368, EPI_ISL_603369, EPI_ISL_603370, EPI_ISL_603371, EPI_ISL_603372, EPI_ISL_603373, EPI_ISL_603374, EPI_ISL_603375, EPI_ISL_603376, EPI_ISL_603377, EPI_ISL_603378, EPI_ISL_603379, EPI_ISL_603380, EPI_ISL_603381, EPI_ISL_603382, EPI_ISL_603383, EPI_ISL_603384, EPI_ISL_603385, EPI_ISL_603386, EPI_ISL_603387, EPI_ISL_603388, EPI_ISL_603389, EPI_ISL_603390, EPI_ISL_603391, EPI_ISL_603392, EPI_ISL_603393, EPI_ISL_603394, EPI_ISL_603395, EPI_ISL_603396, EPI_ISL_603397, EPI_ISL_603398, EPI_ISL_603399, EPI_ISL_603400, EPI_ISL_603401, EPI_ISL_603402, EPI_ISL_603403, EPI_ISL_603404, EPI_ISL_603405, EPI_ISL_603406, EPI_ISL_603407, EPI_ISL_603408, EPI_ISL_603409, EPI_ISL_603410, EPI_ISL_603411, EPI_ISL_603412, EPI_ISL_603413, EPI_ISL_603414, EPI_ISL_603415, EPI_ISL_603416, EPI_ISL_603417, EPI_ISL_603418, EPI_ISL_603419, EPI_ISL_603420, EPI_ISL_603421, EPI_ISL_603422 | | | |
| see above | Viollier AG | Department of Biosystems Science and Engineering, ETH Zürich | Christian Beisel, Sarah Nadeau, Ivan Topolsky, Pedro Ferreira, Philipp Jablonski, Susana Posada-Céspedes, Tobias Schär, Ina Nissen, Natascha Santacroce, Elodie Burcklen, Christiane Beckmann, Maurice Redondo, Olivier Kobel, Christoph Noppen, Sophie Seidel, Noemie Santamaria de Souza, Niko Beerenwinkel, Tanja Stadler |
| EPI_ISL_605483, EPI_ISL_605484, EPI_ISL_605495, EPI_ISL_605496, EPI_ISL_605498, EPI_ISL_605499, EPI_ISL_605500, EPI_ISL_605501, EPI_ISL_605763, EPI_ISL_605767 | University of Wisconsin-Madison AIDS Vaccine Research Laboratories | University of Wisconsin-Madison AIDS Vaccine Research Laboratories | Gage Moreno, Katarina Braun, et al. AIDS Vaccine Research Laboratories |
| EPI_ISL_605786, EPI_ISL_605789, EPI_ISL_605790 | NHLS-IALCH | KRISP, KZN Research Innovation and Sequencing Platform | Giandhari J, Pillay S, Lessells R, Mdlalose K, York D, Khan S, Tegally H, Wilkinson E, de Oliveira T |
| EPI_ISL_609539 | Lighthouse Lab in Cambridge | Wellcome Sanger Institute for the COVID-19 Genomics UK (COG-UK) consortium | Rob Howes, The Lighthouse Lab in Cambridge and Alex Alderton, Roberto Amato, Sonia Goncalves, Ewan Harrison, David K. Jackson, Ian Johnston, Dominic Kwiatkowski, Cordelia Langford, John Sillitoe on behalf of the Wellcome Sanger Institute COVID-19 Surveillance Team |
| EPI_ISL_609827, EPI_ISL_609828, EPI_ISL_609829, EPI_ISL_609839, EPI_ISL_609857 | Respiratory Virus Unit, Microbiology Services Colindale, Public Health England | Respiratory Virus Unit, Microbiology Services Colindale, Public Health England | PHE Covid Sequencing Team |
| EPI_ISL_610153 | Singapore General Hospital | Department of Microbiology | Nurdyana Abdul Rahman, Kun Lee Lim, Chenhao Li, Sui Sin Goh, Kenneth Xin Long Chan, Kian Sing Chan, Lynette Oon, Kern Rei Chng, Niranjan Nagarajan, Karrie Ko |
| EPI_ISL_610196, EPI_ISL_610197, EPI_ISL_610198, EPI_ISL_610204 | Department of Health Technology and Informatics, The Hong Kong Polytechnic University | Department of Health Technology and Informatics, The Hong Kong Polytechnic University | Siu,G.K.-H., Lee,L.-K., Leung,K.S.-S., Leung,J.S.-L., Ng,T.T.-L., Chan,C.T.-M., Tam,K.K.-G., Lao,H.-Y., Wu,A.K.-L., Yau,M.C.-Y., Lai,Y.W.-M., Fung,K.S.-C., Chau,S.K.-Y., Wong,B.K.-C., To,W.-K., Luk,K., Ho,A.Y.-M., Que,T.-L., Yip,K.-T., Yam,W.C., Shum,D.H.-K., Yip,S.P. |
| EPI_ISL_611517 | Liverpool Clinical Laboratories | COVID-19 Genomics UK (COG-UK) Consortium | Sam Haldenby, Anita Lucaci, Steve Paterson, Julian Hiscox, Alistair Darby, M Almsaud, A Alrezaihi, Muhannad Alruwaili, Stuart D Armstrong, Jones Benjamin, Eleanor G Bentley, Anu Chawla, Jordan J Clark, Angela Cowell, Richard Eccles, Isabel García-Dorival, Matthew Gemmell, Alessandro Gerada, PKF Gilmore, Richard Gregory, Ximeng Han, Catherine Hartley, Margaret Hughes, Miren Iturriza-Gomara, James Johnson, L Luu, Jenifer Manson, Charlotte Nelson, Elaine O'Toole, Cassie Olateju, Rebekah Penrice-Randal , Lucille Rainbow, N.P Randle, Trevor Ian Robinson, Parul Sharma, Ghada T Shawli, James P Stewart, Neil Swainston, Ecaterina Vamos, Joanne Watts, Mark Whitehead |
| EPI_ISL_611533, EPI_ISL_611547, EPI_ISL_611548 | West of Scotland Specialist Virology Centre, NHSGGC / MRC-University of Glasgow Centre for Virus Research | COVID-19 Genomics UK (COG-UK) Consortium | Ana da Silva Filipe, Natasha Johnson, Kathy Smollett, Daniel Mair, Stephen Carmichael, Lily Tong, Jenna Nichols, Elihu Aranday-Cortes, Kyriaki Nomikou; Sarah McDonald, Marc Niebel, Patawee Asamaphan; Richard Orton, Joseph Hughes, Sreenu Vattipally, David L Robertson; Alasdair MacLean, Rory Gunson; Kathy Li, Igor Starinskij, Natasha Jesudason, Rajiv Shah, James Shepherd, Antonia Ho, Emma Thomson |
| EPI_ISL_611572 | Northumbria University / South Tees Hospitals NHS Foundation Trust / North Cumbria Integrated Care NHS Foundation Trust / North Tees and Hartlepool NHS Foundation Trust / Newcastle Hospitals NHS Foundation Trust | COVID-19 Genomics UK (COG-UK) Consortium | Darren L Smith,Andrew Nelson,Matthew Bashton,Greg R Young,Joshua Loh,John Allan,Mohammad A Tariq,Giles S Holt,Gary Black,Wen C Yew,Lynn Dover,Paul Baker,Steve Liggett,Sarah Essex,Jane Greenaway,Debra Padgett,Clive Graham,Garren Scott,Edward Barton,Emma Swindells,Brendan Payne,Jennifer Collins,Yusri Taha,Gary Eltringham |
| EPI_ISL_611603 | Liverpool Clinical Laboratories | COVID-19 Genomics UK (COG-UK) Consortium | Sam Haldenby, Anita Lucaci, Steve Paterson, Julian Hiscox, Alistair Darby, M Almsaud, A Alrezaihi, Muhannad Alruwaili, Stuart D Armstrong, Jones Benjamin, Eleanor G Bentley, Anu Chawla, Jordan J Clark, Angela Cowell, Richard Eccles, Isabel García-Dorival, Matthew Gemmell, Alessandro Gerada, PKF Gilmore, Richard Gregory, Ximeng Han, Catherine Hartley, Margaret Hughes, Miren Iturriza-Gomara, James Johnson, L Luu, Jenifer Manson, Charlotte Nelson, Elaine O'Toole, Cassie Olateju, Rebekah Penrice-Randal , Lucille Rainbow, N.P Randle, Trevor Ian Robinson, Parul Sharma, Ghada T Shawli, James P Stewart, Neil Swainston, Ecaterina Vamos, Joanne Watts, Mark Whitehead |
| EPI_ISL_611610, EPI_ISL_611611, EPI_ISL_611614 | West of Scotland Specialist Virology Centre, NHSGGC / MRC-University of Glasgow Centre for Virus Research | COVID-19 Genomics UK (COG-UK) Consortium | Ana da Silva Filipe, Natasha Johnson, Kathy Smollett, Daniel Mair, Stephen Carmichael, Lily Tong, Jenna Nichols, Elihu Aranday-Cortes, Kyriaki Nomikou; Sarah McDonald, Marc Niebel, Patawee Asamaphan; Richard Orton, Joseph Hughes, Sreenu Vattipally, David L Robertson; Alasdair MacLean, Rory Gunson; Kathy Li, Igor Starinskij, Natasha Jesudason, Rajiv Shah, James Shepherd, Antonia Ho, Emma Thomson |
| EPI_ISL_611631 | Wales Specialist Virology Centre Sequencing lab: Pathogen Genomics Unit | COVID-19 Genomics UK (COG-UK) Consortium | Catherine Moore, Johnathan Evans, Laura Gifford, Malorie Perry, Simon Cottrell, Angela Marchbank, Alec Birchley, Alexander Adams, Amy Gaskin, Bree Gatica-Wilcox, Jason Coombes, Joel Southgate, Lauren Gilbert, Lee Graham, Nicole Pacchiarini, Sara Kumziene-Summerhayes, Sarah Taylor, Sophie Jones, Sara Rey, Matthew Bull, Joanne Watkins, Sally Corden, Tom Connor |
| EPI_ISL_611632 | Liverpool Clinical Laboratories | COVID-19 Genomics UK (COG-UK) Consortium | Sam Haldenby, Anita Lucaci, Steve Paterson, Julian Hiscox, Alistair Darby, M Almsaud, A Alrezaihi, Muhannad Alruwaili, Stuart D Armstrong, Jones Benjamin, Eleanor G Bentley, Anu Chawla, Jordan J Clark, Angela Cowell, Richard Eccles, Isabel García-Dorival, Matthew Gemmell, Alessandro Gerada, PKF Gilmore, Richard Gregory, Ximeng Han, Catherine Hartley, Margaret Hughes, Miren Iturriza-Gomara, James Johnson, L Luu, Jenifer Manson, Charlotte Nelson, Elaine O'Toole, Cassie Olateju, Rebekah Penrice-Randal , Lucille Rainbow, N.P Randle, Trevor Ian Robinson, Parul Sharma, Ghada T Shawli, James P Stewart, Neil Swainston, Ecaterina Vamos, Joanne Watts, Mark Whitehead |
| EPI_ISL_611636 | Wales Specialist Virology Centre Sequencing lab: Pathogen Genomics Unit | COVID-19 Genomics UK (COG-UK) Consortium | Catherine Moore, Johnathan Evans, Laura Gifford, Malorie Perry, Simon Cottrell, Angela Marchbank, Alec Birchley, Alexander Adams, Amy Gaskin, Bree Gatica-Wilcox, Jason Coombes, Joel Southgate, Lauren Gilbert, Lee Graham, Nicole Pacchiarini, Sara Kumziene-Summerhayes, Sarah Taylor, Sophie Jones, Sara Rey, Matthew Bull, Joanne Watkins, Sally Corden, Tom Connor |

| | | | |
|---|---|---|---|
| EPI_ISL_611642 | Liverpool Clinical Laboratories | COVID-19 Genomics UK (COG-UK) Consortium | Sam Haldenby, Anita Lucaci, Steve Paterson, Julian Hiscox, Alistair Darby, M Almsaud, A Alrezaihi, Muhannad Alruwaili, Stuart D Armstrong, Jones Benjamin, Eleanor G Bentley, Anu Chawla, Jordan J Clark, Angela Cowell, Richard Eccles, Isabel García-Dorival, Matthew Gemmell, Alessandro Gerada, PKF Gilmore, Richard Gregory, Ximeng Han, Catherine Hartley, Margaret Hughes, Miren Iturriza-Gomara, James Johnson, L Luu, Jenifer Manson, Charlotte Nelson, Elaine O'Toole, Cassie Olateju, Rebekah Penrice-Randal , Lucille Rainbow, N.P Randle, Trevor Ian Robinson, Parul Sharma, Ghada T Shawli, James P Stewart, Neil Swainston, Ecaterina Vamos, Joanne Watts, Mark Whitehead |
| EPI_ISL_611644 | Northumbria University / South Tees Hospitals NHS Foundation Trust / North Cumbria Integrated Care NHS Foundation Trust / North Tees and Hartlepool NHS Foundation Trust / Newcastle Hospitals NHS Foundation Trust | COVID-19 Genomics UK (COG-UK) Consortium | Darren L Smith,Andrew Nelson,Matthew Bashton,Greg R Young,Joshua Loh,John Allan,Mohammad A Tariq,Giles S Holt,Gary Black,Wen C Yew,Lynn Dover,Paul Baker,Steve Liggett,Sarah Essex,Jane Greenaway,Debra Padgett,Clive Graham,Garren Scott,Edward Barton,Emma Swindells,Brendan Payne,Jennifer Collins,Yusri Taha,Gary Eltringham |
| EPI_ISL_611660 | West of Scotland Specialist Virology Centre, NHSGGC / MRC-University of Glasgow Centre for Virus Research | COVID-19 Genomics UK (COG-UK) Consortium | Ana da Silva Filipe, Natasha Johnson, Kathy Smollett, Daniel Mair, Stephen Carmichael, Lily Tong, Jenna Nichols, Elihu Aranday-Cortes, Kyriaki Nomikou; Sarah McDonald, Marc Niebel, Patawee Asamaphan; Richard Orton, Joseph Hughes, Sreenu Vattipally, David L Robertson; Alasdair MacLean, Rory Gunson; Kathy Li, Igor Starinskij, Natasha Jesudason, Rajiv Shah, James Shepherd, Antonia Ho, Emma Thomson |
| EPI_ISL_611664 | Liverpool Clinical Laboratories | COVID-19 Genomics UK (COG-UK) Consortium | Sam Haldenby, Anita Lucaci, Steve Paterson, Julian Hiscox, Alistair Darby, M Almsaud, A Alrezaihi, Muhannad Alruwaili, Stuart D Armstrong, Jones Benjamin, Eleanor G Bentley, Anu Chawla, Jordan J Clark, Angela Cowell, Richard Eccles, Isabel García-Dorival, Matthew Gemmell, Alessandro Gerada, PKF Gilmore, Richard Gregory, Ximeng Han, Catherine Hartley, Margaret Hughes, Miren Iturriza-Gomara, James Johnson, L Luu, Jenifer Manson, Charlotte Nelson, Elaine O'Toole, Cassie Olateju, Rebekah Penrice-Randal , Lucille Rainbow, N.P Randle, Trevor Ian Robinson, Parul Sharma, Ghada T Shawli, James P Stewart, Neil Swainston, Ecaterina Vamos, Joanne Watts, Mark Whitehead |
| EPI_ISL_611679 | Virology Department, Sheffield Teaching Hospitals NHS Foundation Trust/Department of Infection, Immunity and Cardiovascular Disease, The Medical School, University of Sheffield | COVID-19 Genomics UK (COG-UK) Consortium | Thushan de Silva, Matthew Parker, Nikki Smith, Adri Angyal, Rebecca Brown, Luke Green, Rachel Tucker, Paul Parsons, Danielle Groves, Katie Johnson, Laura Carrilero, Alex Keeley, Dave Partridge, Matthew Wyles, Benjamin Lindsey, Mehmet Yavuz, Mohammad Raza, Cariad Evans |
| EPI_ISL_611695 | Liverpool Clinical Laboratories | COVID-19 Genomics UK (COG-UK) Consortium | Sam Haldenby, Anita Lucaci, Steve Paterson, Julian Hiscox, Alistair Darby, M Almsaud, A Alrezaihi, Muhannad Alruwaili, Stuart D Armstrong, Jones Benjamin, Eleanor G Bentley, Anu Chawla, Jordan J Clark, Angela Cowell, Richard Eccles, Isabel García-Dorival, Matthew Gemmell, Alessandro Gerada, PKF Gilmore, Richard Gregory, Ximeng Han, Catherine Hartley, Margaret Hughes, Miren Iturriza-Gomara, James Johnson, L Luu, Jenifer Manson, Charlotte Nelson, Elaine O'Toole, Cassie Olateju, Rebekah Penrice-Randal , Lucille Rainbow, N.P Randle, Trevor Ian Robinson, Parul Sharma, Ghada T Shawli, James P Stewart, Neil Swainston, Ecaterina Vamos, Joanne Watts, Mark Whitehead |
| EPI_ISL_611709, EPI_ISL_611712, EPI_ISL_611713 | West of Scotland Specialist Virology Centre, NHSGGC / MRC-University of Glasgow Centre for Virus Research | COVID-19 Genomics UK (COG-UK) Consortium | Ana da Silva Filipe, Natasha Johnson, Kathy Smollett, Daniel Mair, Stephen Carmichael, Lily Tong, Jenna Nichols, Elihu Aranday-Cortes, Kyriaki Nomikou; Sarah McDonald, Marc Niebel, Patawee Asamaphan; Richard Orton, Joseph Hughes, Sreenu Vattipally, David L Robertson; Alasdair MacLean, Rory Gunson; Kathy Li, Igor Starinskij, Natasha Jesudason, Rajiv Shah, James Shepherd, Antonia Ho, Emma Thomson |
| EPI_ISL_611721, EPI_ISL_611728, EPI_ISL_611730, EPI_ISL_611736, EPI_ISL_611743 | Liverpool Clinical Laboratories | COVID-19 Genomics UK (COG-UK) Consortium | Sam Haldenby, Anita Lucaci, Steve Paterson, Julian Hiscox, Alistair Darby, M Almsaud, A Alrezaihi, Muhannad Alruwaili, Stuart D Armstrong, Jones Benjamin, Eleanor G Bentley, Anu Chawla, Jordan J Clark, Angela Cowell, Richard Eccles, Isabel García-Dorival, Matthew Gemmell, Alessandro Gerada, PKF Gilmore, Richard Gregory, Ximeng Han, Catherine Hartley, Margaret Hughes, Miren Iturriza-Gomara, James Johnson, L Luu, Jenifer Manson, Charlotte Nelson, Elaine O'Toole, Cassie Olateju, Rebekah Penrice-Randal , Lucille Rainbow, N.P Randle, Trevor Ian Robinson, Parul Sharma, Ghada T Shawli, James P Stewart, Neil Swainston, Ecaterina Vamos, Joanne Watts, Mark Whitehead |
| EPI_ISL_611746, EPI_ISL_611747 | Northumbria University / South Tees Hospitals NHS Foundation Trust / North Cumbria Integrated Care NHS Foundation Trust / North Tees and Hartlepool NHS Foundation Trust / Newcastle Hospitals NHS Foundation Trust | COVID-19 Genomics UK (COG-UK) Consortium | Darren L Smith,Andrew Nelson,Matthew Bashton,Greg R Young,Joshua Loh,John Allan,Mohammad A Tariq,Giles S Holt,Gary Black,Wen C Yew,Lynn Dover,Paul Baker,Steve Liggett,Sarah Essex,Jane Greenaway,Debra Padgett,Clive Graham,Garren Scott,Edward Barton,Emma Swindells,Brendan Payne,Jennifer Collins,Yusri Taha,Gary Eltringham |
| EPI_ISL_611784, EPI_ISL_611785, EPI_ISL_611790, EPI_ISL_611815 | Liverpool Clinical Laboratories | COVID-19 Genomics UK (COG-UK) Consortium | Sam Haldenby, Anita Lucaci, Steve Paterson, Julian Hiscox, Alistair Darby, M Almsaud, A Alrezaihi, Muhannad Alruwaili, Stuart D Armstrong, Jones Benjamin, Eleanor G Bentley, Anu Chawla, Jordan J Clark, Angela Cowell, Richard Eccles, Isabel García-Dorival, Matthew Gemmell, Alessandro Gerada, PKF Gilmore, Richard Gregory, Ximeng Han, Catherine Hartley, Margaret Hughes, Miren Iturriza-Gomara, James Johnson, L Luu, Jenifer Manson, Charlotte Nelson, Elaine O'Toole, Cassie Olateju, Rebekah Penrice-Randal , Lucille Rainbow, N.P Randle, Trevor Ian Robinson, Parul Sharma, Ghada T Shawli, James P Stewart, Neil Swainston, Ecaterina Vamos, Joanne Watts, Mark Whitehead |
| EPI_ISL_611818, EPI_ISL_611820, EPI_ISL_611826 | West of Scotland Specialist Virology Centre, NHSGGC / MRC-University of Glasgow Centre for Virus Research | COVID-19 Genomics UK (COG-UK) Consortium | Ana da Silva Filipe, Natasha Johnson, Kathy Smollett, Daniel Mair, Stephen Carmichael, Lily Tong, Jenna Nichols, Elihu Aranday-Cortes, Kyriaki Nomikou; Sarah McDonald, Marc Niebel, Patawee Asamaphan; Richard Orton, Joseph Hughes, Sreenu Vattipally, David L Robertson; Alasdair MacLean, Rory Gunson; Kathy Li, Igor Starinskij, Natasha Jesudason, Rajiv Shah, James Shepherd, Antonia Ho, Emma Thomson |
| EPI_ISL_611827 | Liverpool Clinical Laboratories | COVID-19 Genomics UK (COG-UK) Consortium | Sam Haldenby, Anita Lucaci, Steve Paterson, Julian Hiscox, Alistair Darby, M Almsaud, A Alrezaihi, Muhannad Alruwaili, Stuart D Armstrong, Jones Benjamin, Eleanor G Bentley, Anu Chawla, Jordan J Clark, Angela Cowell, Richard Eccles, Isabel García-Dorival, Matthew Gemmell, Alessandro Gerada, PKF Gilmore, Richard Gregory, Ximeng Han, Catherine Hartley, Margaret Hughes, Miren Iturriza-Gomara, James Johnson, L Luu, Jenifer Manson, Charlotte Nelson, Elaine O'Toole, Cassie Olateju, Rebekah Penrice-Randal , Lucille Rainbow, N.P Randle, Trevor Ian Robinson, Parul Sharma, Ghada T Shawli, James P Stewart, Neil Swainston, Ecaterina Vamos, Joanne Watts, Mark Whitehead |
| EPI_ISL_611844 | University of Exeter | COVID-19 Genomics UK (COG-UK) Consortium | Ben Temperton,Aaron Jeffries,Michelle Michelsen,Joanna Warwick-Dugdale,Audrey Farbos,Robyn Manley,Stephen Michell,Jane Masoli |
| EPI_ISL_611848 | West of Scotland Specialist Virology Centre, NHSGGC / MRC-University of Glasgow Centre for Virus Research | COVID-19 Genomics UK (COG-UK) Consortium | Ana da Silva Filipe, Natasha Johnson, Kathy Smollett, Daniel Mair, Stephen Carmichael, Lily Tong, Jenna Nichols, Elihu Aranday-Cortes, Kyriaki Nomikou; Sarah McDonald, Marc Niebel, Patawee Asamaphan; Richard Orton, Joseph Hughes, Sreenu Vattipally, David L Robertson; Alasdair MacLean, Rory Gunson; Kathy Li, Igor Starinskij, Natasha Jesudason, Rajiv Shah, James Shepherd, Antonia Ho, Emma Thomson |
| EPI_ISL_611849 | Wales Specialist Virology Centre Sequencing lab: Pathogen Genomics Unit | COVID-19 Genomics UK (COG-UK) Consortium | Catherine Moore, Johnathan Evans, Laura Gifford, Malorie Perry, Simon Cottrell, Angela Marchbank, Alec Birchley, Alexander Adams, Amy Gaskin, Bree Gatica-Wilcox, Jason Coombes, Joel Southgate, Lauren Gilbert, Lee Graham, Nicole Pacchiarini, Sara Kumziene-Summerhayes, Sarah Taylor, Sophie Jones, Sara Rey, Matthew Bull, Joanne Watkins, Sally Corden, Tom Connor |
| EPI_ISL_611851 | Liverpool Clinical Laboratories | COVID-19 Genomics UK (COG-UK) Consortium | Sam Haldenby, Anita Lucaci, Steve Paterson, Julian Hiscox, Alistair Darby, M Almsaud, A Alrezaihi, Muhannad Alruwaili, Stuart D Armstrong, Jones Benjamin, Eleanor G Bentley, Anu Chawla, Jordan J Clark, Angela Cowell, Richard Eccles, Isabel García-Dorival, Matthew Gemmell, Alessandro Gerada, PKF Gilmore, Richard Gregory, Ximeng Han, Catherine Hartley, Margaret Hughes, Miren Iturriza-Gomara, James Johnson, L Luu, Jenifer Manson, Charlotte Nelson, Elaine O'Toole, Cassie Olateju, Rebekah Penrice-Randal , Lucille Rainbow, N.P Randle, Trevor Ian Robinson, Parul Sharma, Ghada T Shawli, James P Stewart, Neil Swainston, Ecaterina Vamos, Joanne Watts, Mark Whitehead |
| EPI_ISL_611852 | West of Scotland Specialist Virology Centre, NHSGGC / MRC-University of Glasgow Centre for Virus Research | COVID-19 Genomics UK (COG-UK) Consortium | Ana da Silva Filipe, Natasha Johnson, Kathy Smollett, Daniel Mair, Stephen Carmichael, Lily Tong, Jenna Nichols, Elihu Aranday-Cortes, Kyriaki Nomikou; Sarah McDonald, Marc Niebel, Patawee Asamaphan; Richard Orton, Joseph Hughes, Sreenu Vattipally, David L Robertson; Alasdair MacLean, Rory Gunson; Kathy Li, Igor Starinskij, Natasha Jesudason, Rajiv Shah, James Shepherd, Antonia Ho, Emma Thomson |
| EPI_ISL_611857 | Liverpool Clinical Laboratories | COVID-19 Genomics UK (COG-UK) Consortium | Sam Haldenby, Anita Lucaci, Steve Paterson, Julian Hiscox, Alistair Darby, M Almsaud, A Alrezaihi, Muhannad Alruwaili, Stuart D Armstrong, Jones Benjamin, Eleanor G Bentley, Anu Chawla, Jordan J Clark, Angela Cowell, Richard Eccles, Isabel García-Dorival, Matthew Gemmell, Alessandro Gerada, PKF Gilmore, Richard Gregory, Ximeng Han, Catherine Hartley, Margaret Hughes, Miren Iturriza-Gomara, James Johnson, L Luu, Jenifer Manson, Charlotte Nelson, Elaine O'Toole, Cassie Olateju, Rebekah Penrice-Randal , Lucille Rainbow, N.P Randle, Trevor Ian Robinson, Parul Sharma, Ghada T Shawli, James P Stewart, Neil Swainston, Ecaterina Vamos, Joanne Watts, Mark Whitehead |
| EPI_ISL_611867, EPI_ISL_611872, EPI_ISL_611873, EPI_ISL_611877, EPI_ISL_611878, EPI_ISL_611879, EPI_ISL_611880, EPI_ISL_611881, EPI_ISL_611882, EPI_ISL_611883, EPI_ISL_611884, EPI_ISL_611890 | | | |
| see above | West of Scotland Specialist Virology Centre, NHSGGC / MRC-University of Glasgow Centre for Virus Research | COVID-19 Genomics UK (COG-UK) Consortium | Ana da Silva Filipe, Natasha Johnson, Kathy Smollett, Daniel Mair, Stephen Carmichael, Lily Tong, Jenna Nichols, Elihu Aranday-Cortes, Kyriaki Nomikou; Sarah McDonald, Marc Niebel, Patawee Asamaphan; Richard Orton, Joseph Hughes, Sreenu Vattipally, David L Robertson; Alasdair MacLean, Rory Gunson; Kathy Li, Igor Starinskij, Natasha Jesudason, Rajiv Shah, James Shepherd, Antonia Ho, Emma Thomson |

| | | | |
|---|---|---|---|
| EPI_ISL_611892 | Liverpool Clinical Laboratories | COVID-19 Genomics UK (COG-UK) Consortium | Sam Haldenby, Anita Lucaci, Steve Paterson, Julian Hiscox, Alistair Darby, M Almsaud, A Alrezaihi, Muhannad Alruwaili, Stuart D Armstrong, Jones Benjamin, Eleanor G Bentley, Anu Chawla, Jordan J Clark, Angela Cowell, Richard Eccles, Isabel García-Dorival, Matthew Gemmell, Alessandro Gerada, PKF Gilmore, Richard Gregory, Ximeng Han, Catherine Hartley, Margaret Hughes, Miren Iturriza-Gomara, James Johnson, L Luu, Jenifer Manson, Charlotte Nelson, Elaine O'Toole, Cassie Olateju, Rebekah Penrice-Randal , Lucille Rainbow, N.P Randle, Trevor Ian Robinson, Parul Sharma, Ghada T Shawli, James P Stewart, Neil Swainston, Ecaterina Vamos, Joanne Watts, Mark Whitehead |
| EPI_ISL_611917, EPI_ISL_611952 | Northumbria University / South Tees Hospitals NHS Foundation Trust / North Cumbria Integrated Care NHS Foundation Trust / North Tees and Hartlepool NHS Foundation Trust / Newcastle Hospitals NHS Foundation Trust | COVID-19 Genomics UK (COG-UK) Consortium | Darren L Smith,Andrew Nelson,Matthew Bashton,Greg R Young,Joshua Loh,John Allan,Mohammad A Tariq,Giles S Holt,Gary Black,Wen C Yew,Lynn Dover,Paul Baker,Steve Liggett,Sarah Essex,Jane Greenaway,Debra Padgett,Clive Graham,Garren Scott,Edward Barton,Emma Swindells,Brendan Payne,Jennifer Collins,Yusri Taha,Gary Eltringham |
| EPI_ISL_611969, EPI_ISL_611970 | Wales Specialist Virology Centre Sequencing lab: Pathogen Genomics Unit | COVID-19 Genomics UK (COG-UK) Consortium | Catherine Moore, Johnathan Evans, Laura Gifford, Malorie Perry, Simon Cottrell, Angela Marchbank, Alec Birchley, Alexander Adams, Amy Gaskin, Bree Gatica-Wilcox, Jason Coombes, Joel Southgate, Lauren Gilbert, Lee Graham, Nicole Pacchiarini, Sara Kumziene-Summerhayes, Sarah Taylor, Sophie Jones, Sara Rey, Matthew Bull, Joanne Watkins, Sally Corden, Tom Connor |
| EPI_ISL_611978, EPI_ISL_611984, EPI_ISL_612001, EPI_ISL_612009, EPI_ISL_612010, EPI_ISL_612039, EPI_ISL_612071, EPI_ISL_612101 | Liverpool Clinical Laboratories | COVID-19 Genomics UK (COG-UK) Consortium | Sam Haldenby, Anita Lucaci, Steve Paterson, Julian Hiscox, Alistair Darby, M Almsaud, A Alrezaihi, Muhannad Alruwaili, Stuart D Armstrong, Jones Benjamin, Eleanor G Bentley, Anu Chawla, Jordan J Clark, Angela Cowell, Richard Eccles, Isabel García-Dorival, Matthew Gemmell, Alessandro Gerada, PKF Gilmore, Richard Gregory, Ximeng Han, Catherine Hartley, Margaret Hughes, Miren Iturriza-Gomara, James Johnson, L Luu, Jenifer Manson, Charlotte Nelson, Elaine O'Toole, Cassie Olateju, Rebekah Penrice-Randal , Lucille Rainbow, N.P Randle, Trevor Ian Robinson, Parul Sharma, Ghada T Shawli, James P Stewart, Neil Swainston, Ecaterina Vamos, Joanne Watts, Mark Whitehead |
| EPI_ISL_612121, EPI_ISL_612132, EPI_ISL_612133 | Wales Specialist Virology Centre Sequencing lab: Pathogen Genomics Unit | COVID-19 Genomics UK (COG-UK) Consortium | Catherine Moore, Johnathan Evans, Laura Gifford, Malorie Perry, Simon Cottrell, Angela Marchbank, Alec Birchley, Alexander Adams, Amy Gaskin, Bree Gatica-Wilcox, Jason Coombes, Joel Southgate, Lauren Gilbert, Lee Graham, Nicole Pacchiarini, Sara Kumziene-Summerhayes, Sarah Taylor, Sophie Jones, Sara Rey, Matthew Bull, Joanne Watkins, Sally Corden, Tom Connor |
| EPI_ISL_612262, EPI_ISL_612263, EPI_ISL_612264 | Department of Pathology, University of Cambridge | COVID-19 Genomics UK (COG-UK) Consortium | Aminu S. Jahun, Yasmin Chaudhry, Grant Hall, Iliana Georgana, Myra Hosmillo, Martin D. Curran, Malte Pinckert, Surendra Parmar, Ian Goodfellow |
| EPI_ISL_612288, EPI_ISL_612289, EPI_ISL_612290, EPI_ISL_612291, EPI_ISL_612292, EPI_ISL_612293, EPI_ISL_612294, EPI_ISL_612297, EPI_ISL_612298, EPI_ISL_612299, EPI_ISL_612300, EPI_ISL_612301, EPI_ISL_612302, EPI_ISL_612303, EPI_ISL_612304, EPI_ISL_612305, EPI_ISL_612306, EPI_ISL_612307, EPI_ISL_612308, EPI_ISL_612309, EPI_ISL_612310, EPI_ISL_612311, EPI_ISL_612312, EPI_ISL_612313 | | | |
| see above | West of Scotland Specialist Virology Centre, NHSGGC / MRC-University of Glasgow Centre for Virus Research | COVID-19 Genomics UK (COG-UK) Consortium | Ana da Silva Filipe, Natasha Johnson, Kathy Smollett, Daniel Mair, Stephen Carmichael, Lily Tong, Jenna Nichols, Elihu Aranday-Cortes, Kyriaki Nomikou; Sarah McDonald, Marc Niebel, Patawee Asamaphan; Richard Orton, Joseph Hughes, Sreenu Vattipally, David L Robertson; Alasdair MacLean, Rory Gunson; Kathy Li, Igor Starinskij, Natasha Jesudason, Rajiv Shah, James Shepherd, Antonia Ho, Emma Thomson |
| EPI_ISL_612320, EPI_ISL_612321, EPI_ISL_612322, EPI_ISL_612323, EPI_ISL_612324 | Virology Department, Royal Infirmary of Edinburgh, NHS Lothian / School of Biological Sciences, University of Edinburgh / Institute of Genetics and Molecular Medicine, University of Edinburgh | COVID-19 Genomics UK (COG-UK) Consortium | McHugh M, Dewar R, Rooke S, Gallagher M, Balcaza C, O'Toole Á, Scher E, Hill V, McCrone JT, Colquhoun R, Yu X, Jackson B, Rambaut A, Williams TC, Templeton K |
| EPI_ISL_612375, EPI_ISL_612376, EPI_ISL_612377, EPI_ISL_612378, EPI_ISL_612379 | University of Exeter | COVID-19 Genomics UK (COG-UK) Consortium | Ben Temperton,Aaron Jeffries,Michelle Michelsen,Joanna Warwick-Dugdale,Audrey Farbos,Robyn Manley,Stephen Michell,Jane Masoli |
| EPI_ISL_612433, EPI_ISL_612435, EPI_ISL_612436 | Liverpool Clinical Laboratories | COVID-19 Genomics UK (COG-UK) Consortium | Sam Haldenby, Anita Lucaci, Steve Paterson, Julian Hiscox, Alistair Darby, M Almsaud, A Alrezaihi, Muhannad Alruwaili, Stuart D Armstrong, Jones Benjamin, Eleanor G Bentley, Anu Chawla, Jordan J Clark, Angela Cowell, Richard Eccles, Isabel García-Dorival, Matthew Gemmell, Alessandro Gerada, PKF Gilmore, Richard Gregory, Ximeng Han, Catherine Hartley, Margaret Hughes, Miren Iturriza-Gomara, James Johnson, L Luu, Jenifer Manson, Charlotte Nelson, Elaine O'Toole, Cassie Olateju, Rebekah Penrice-Randal , Lucille Rainbow, N.P Randle, Trevor Ian Robinson, Parul Sharma, Ghada T Shawli, James P Stewart, Neil Swainston, Ecaterina Vamos, Joanne Watts, Mark Whitehead |
| EPI_ISL_612458, EPI_ISL_612459, EPI_ISL_612460, EPI_ISL_612461, EPI_ISL_612462, EPI_ISL_612463, EPI_ISL_612464, EPI_ISL_612465, EPI_ISL_612466, EPI_ISL_612467, EPI_ISL_612468, EPI_ISL_612469, EPI_ISL_612470, EPI_ISL_612471, EPI_ISL_612472, EPI_ISL_612473, EPI_ISL_612474, EPI_ISL_612475, EPI_ISL_612476, EPI_ISL_612477, EPI_ISL_612478, EPI_ISL_612479, EPI_ISL_612480, EPI_ISL_612481, EPI_ISL_612482, EPI_ISL_612483, EPI_ISL_612484, EPI_ISL_612485, EPI_ISL_612486, EPI_ISL_612487, EPI_ISL_612493, EPI_ISL_612494, EPI_ISL_612495, EPI_ISL_612496, EPI_ISL_612497, EPI_ISL_612498, EPI_ISL_612499, EPI_ISL_612500 | | | |
| see above | Wales Specialist Virology Centre Sequencing lab: Pathogen Genomics Unit | COVID-19 Genomics UK (COG-UK) Consortium | Catherine Moore, Johnathan Evans, Laura Gifford, Malorie Perry, Simon Cottrell, Angela Marchbank, Alec Birchley, Alexander Adams, Amy Gaskin, Bree Gatica-Wilcox, Jason Coombes, Joel Southgate, Lauren Gilbert, Lee Graham, Nicole Pacchiarini, Sara Kumziene-Summerhayes, Sarah Taylor, Sophie Jones, Sara Rey, Matthew Bull, Joanne Watkins, Sally Corden, Tom Connor |
| EPI_ISL_612554, EPI_ISL_612555, EPI_ISL_612556, EPI_ISL_612558, EPI_ISL_612559, EPI_ISL_612560, EPI_ISL_612561, EPI_ISL_612562, EPI_ISL_612563 | Northumbria University / South Tees Hospitals NHS Foundation Trust / North Cumbria Integrated Care NHS Foundation Trust / North Tees and Hartlepool NHS Foundation Trust / Newcastle Hospitals NHS Foundation Trust | COVID-19 Genomics UK (COG-UK) Consortium | Darren L Smith,Andrew Nelson,Matthew Bashton,Greg R Young,Joshua Loh,John Allan,Mohammad A Tariq,Giles S Holt,Gary Black,Wen C Yew,Lynn Dover,Paul Baker,Steve Liggett,Sarah Essex,Jane Greenaway,Debra Padgett,Clive Graham,Garren Scott,Edward Barton,Emma Swindells,Brendan Payne,Jennifer Collins,Yusri Taha,Gary Eltringham |
| EPI_ISL_612880, EPI_ISL_612978, EPI_ISL_613003, EPI_ISL_613010, EPI_ISL_613014, EPI_ISL_613020, EPI_ISL_613045, EPI_ISL_613061, EPI_ISL_613078, EPI_ISL_613098, EPI_ISL_613105, EPI_ISL_613149, EPI_ISL_613151 | | | |
| see above | Wales Specialist Virology Centre Sequencing lab: Pathogen Genomics Unit | COVID-19 Genomics UK (COG-UK) Consortium | Catherine Moore, Johnathan Evans, Laura Gifford, Malorie Perry, Simon Cottrell, Angela Marchbank, Alec Birchley, Alexander Adams, Amy Gaskin, Bree Gatica-Wilcox, Jason Coombes, Joel Southgate, Lauren Gilbert, Lee Graham, Nicole Pacchiarini, Sara Kumziene-Summerhayes, Sarah Taylor, Sophie Jones, Sara Rey, Matthew Bull, Joanne Watkins, Sally Corden, Tom Connor |
| EPI_ISL_613379 | Virology Department, Sheffield Teaching Hospitals NHS Foundation Trust/Department of Infection, Immunity and Cardiovascular Disease, The Medical School, University of Sheffield | COVID-19 Genomics UK (COG-UK) Consortium | Thushan de Silva, Matthew Parker, Nikki Smith, Adri Angyal, Rebecca Brown, Luke Green, Rachel Tucker, Paul Parsons, Danielle Groves, Katie Johnson, Laura Carrilero, Alex Keeley, Dave Partridge, Matthew Wyles, Benjamin Lindsey, Mehmet Yavuz, Mohammad Raza, Cariad Evans |
| EPI_ISL_613489, EPI_ISL_613490, EPI_ISL_613491, EPI_ISL_613492 | Public Health Laboratory - Infectious Disease Lab, Minnesota Department of Health Infectious Disease Laboratory Submission Group | Minnesota Department of Health, Public Health Laboratory | Plumb,M., Garfin,J., Lorentz,A., Wang,X. |
| EPI_ISL_613718, EPI_ISL_613721, EPI_ISL_613722, EPI_ISL_613733, EPI_ISL_613737, EPI_ISL_613741, EPI_ISL_613752, EPI_ISL_613754, EPI_ISL_613756, EPI_ISL_613760, EPI_ISL_613778, EPI_ISL_613790, EPI_ISL_613796, EPI_ISL_613822, EPI_ISL_613843, EPI_ISL_613844, EPI_ISL_613845, EPI_ISL_613846, EPI_ISL_613847, EPI_ISL_613848, EPI_ISL_613849, EPI_ISL_613850, EPI_ISL_613851, EPI_ISL_613852, EPI_ISL_613853, EPI_ISL_613854, EPI_ISL_613855 | | | |
| see above | Florida Bureau of Public Health Laboratories | Florida Bureau of Public Health Laboratories | Sarah Schmedes, Jason Blanton |
| EPI_ISL_614049, EPI_ISL_614065, EPI_ISL_614071, EPI_ISL_614072, EPI_ISL_614145, EPI_ISL_614146, EPI_ISL_614147, EPI_ISL_614148 | Virginia DCLS | Virginia DCLS | Virginia DCLS |
| EPI_ISL_614160, EPI_ISL_614161, EPI_ISL_614165, EPI_ISL_614166, EPI_ISL_614168, EPI_ISL_614171, EPI_ISL_614175, EPI_ISL_614178 | Michigan Department of Health and Human Services, Bureau of Laboratories | Michigan Department of Health and Human Services, Bureau of Laboratories | Blankenship HM, Riner D, Soehnlen MK |
| EPI_ISL_614295, EPI_ISL_614304, EPI_ISL_614316, EPI_ISL_614317 | Faroese National Reference Laboratory for Fish and Animal Diseases | Faroese National Reference Laboratory for Fish and Animal Diseases | Maria Marjunardóttir Dahl, Petra Elisabeth Petersen, Debes Hammershaimb Christiansen |
| EPI_ISL_615117, EPI_ISL_615118, | Klinisk mikrobiologi Linkoping | The Public Health Agency of Sweden | Anna-Malin Linde, Maria Lind Karlberg, Mattias Haukland, Reza Advani, Olov Svartstrom, Oskar Karlsson Lindsjo, Sandra Broddesson, Petra Edquist, Mia |

| | | | |
|---|---|---|---|
| EPI_ISL_615119 | | | Brytting, Anna Risberg, Karin Tegmark-Wisell |
| EPI_ISL_615120 | Orebro klinisk mikrobiologi | The Public Health Agency of Sweden | Anna-Malin Linde, Maria Lind Karlberg, Mattias Haukland, Reza Advani, Olov Svartstrom, Oskar Karlsson Lindsjo, Sandra Broddesson, Petra Edquist, Mia Brytting, Anna Risberg, Karin Tegmark-Wisell |
| EPI_ISL_619906, EPI_ISL_619907, EPI_ISL_619908, EPI_ISL_619909, EPI_ISL_619910, EPI_ISL_619911, EPI_ISL_619912, EPI_ISL_619913, EPI_ISL_619914, EPI_ISL_619915, EPI_ISL_619916, EPI_ISL_619917, EPI_ISL_619918, EPI_ISL_619919, EPI_ISL_619920, EPI_ISL_619921, EPI_ISL_619922, EPI_ISL_619923, EPI_ISL_619924, EPI_ISL_619925, EPI_ISL_619926, EPI_ISL_619927, EPI_ISL_619928, EPI_ISL_619929, EPI_ISL_619930, EPI_ISL_619931, EPI_ISL_619932, EPI_ISL_619933, EPI_ISL_619934, EPI_ISL_619935, EPI_ISL_619936, EPI_ISL_619937, EPI_ISL_619938, EPI_ISL_619939, EPI_ISL_619940, EPI_ISL_619941, EPI_ISL_619942, EPI_ISL_619943, EPI_ISL_619944, EPI_ISL_619945, EPI_ISL_619946, EPI_ISL_619947, EPI_ISL_619948, EPI_ISL_619949, EPI_ISL_619950, EPI_ISL_619951, EPI_ISL_619952, EPI_ISL_619953, EPI_ISL_619954, EPI_ISL_619955, EPI_ISL_619956, EPI_ISL_619957, EPI_ISL_619958, EPI_ISL_619959, EPI_ISL_619960, EPI_ISL_619961, EPI_ISL_619962, EPI_ISL_619963, EPI_ISL_619964, EPI_ISL_619965, EPI_ISL_619966, EPI_ISL_619967, EPI_ISL_619968, EPI_ISL_619969, EPI_ISL_619970, EPI_ISL_619971, EPI_ISL_619972, EPI_ISL_619973, EPI_ISL_619974, EPI_ISL_619975, EPI_ISL_619976, EPI_ISL_620783, EPI_ISL_620800, EPI_ISL_620804, EPI_ISL_620805, EPI_ISL_620806, EPI_ISL_620807, EPI_ISL_620808, EPI_ISL_620809, EPI_ISL_620810, EPI_ISL_620811, EPI_ISL_620812, EPI_ISL_620813, EPI_ISL_620829, EPI_ISL_620850, EPI_ISL_620851, EPI_ISL_620852, EPI_ISL_620854, EPI_ISL_620855, EPI_ISL_620856, EPI_ISL_620857, EPI_ISL_620858, EPI_ISL_620859, EPI_ISL_620860, EPI_ISL_620861, EPI_ISL_620862, EPI_ISL_620863, EPI_ISL_620864, EPI_ISL_620865, EPI_ISL_620866, EPI_ISL_620867, EPI_ISL_620868, EPI_ISL_620869, EPI_ISL_620870, EPI_ISL_620871, EPI_ISL_620872, EPI_ISL_620873, EPI_ISL_620874, EPI_ISL_620875, EPI_ISL_620876, EPI_ISL_620877, EPI_ISL_620878, EPI_ISL_620879, EPI_ISL_620880, EPI_ISL_620881, EPI_ISL_620882, EPI_ISL_620883, EPI_ISL_620884, EPI_ISL_620885, EPI_ISL_620886, EPI_ISL_620887, EPI_ISL_620888, EPI_ISL_620889, EPI_ISL_620890, EPI_ISL_620891, EPI_ISL_620892, EPI_ISL_620893, EPI_ISL_620894, EPI_ISL_620895, EPI_ISL_620896, EPI_ISL_620897, EPI_ISL_620898, EPI_ISL_620899, EPI_ISL_620900, EPI_ISL_620901, EPI_ISL_620902, EPI_ISL_620903, EPI_ISL_620904, EPI_ISL_620905, EPI_ISL_620906, EPI_ISL_620907, EPI_ISL_620908, EPI_ISL_620909, EPI_ISL_620910, EPI_ISL_620911, EPI_ISL_620912, EPI_ISL_620913, EPI_ISL_620914, EPI_ISL_620915, EPI_ISL_620916, EPI_ISL_620917, EPI_ISL_620918, EPI_ISL_620919, EPI_ISL_620920, EPI_ISL_620922, EPI_ISL_620923, EPI_ISL_620924, EPI_ISL_620925, EPI_ISL_620927, EPI_ISL_620928, EPI_ISL_620929, EPI_ISL_620930, EPI_ISL_620931, EPI_ISL_620932, EPI_ISL_620933, EPI_ISL_620934, EPI_ISL_620935, EPI_ISL_620936, EPI_ISL_620937, EPI_ISL_620938, EPI_ISL_620939, EPI_ISL_620940, EPI_ISL_620941, EPI_ISL_620942, EPI_ISL_620943, EPI_ISL_620944, EPI_ISL_620945, EPI_ISL_620946, EPI_ISL_620947, EPI_ISL_620948, EPI_ISL_620949, EPI_ISL_620950, EPI_ISL_620951, EPI_ISL_620952, EPI_ISL_620953, EPI_ISL_620954, EPI_ISL_620955, EPI_ISL_620956, EPI_ISL_620957, EPI_ISL_620958, EPI_ISL_620959, EPI_ISL_620960, EPI_ISL_620961, EPI_ISL_620962, EPI_ISL_620963, EPI_ISL_620964, EPI_ISL_620965, EPI_ISL_620966, EPI_ISL_620967, EPI_ISL_620968, EPI_ISL_620969, EPI_ISL_620970, EPI_ISL_620971, EPI_ISL_620972, EPI_ISL_620973, EPI_ISL_620974, EPI_ISL_620975, EPI_ISL_620976, EPI_ISL_620977, EPI_ISL_620978, EPI_ISL_620979, EPI_ISL_620980, EPI_ISL_620981, EPI_ISL_620982, EPI_ISL_620983, EPI_ISL_620984, EPI_ISL_620985, EPI_ISL_620986, EPI_ISL_620987, EPI_ISL_620988, EPI_ISL_620989, EPI_ISL_620990, EPI_ISL_620991, EPI_ISL_620992, EPI_ISL_620993, EPI_ISL_620994, EPI_ISL_620995, EPI_ISL_620996, EPI_ISL_620997, EPI_ISL_620998, EPI_ISL_620999, EPI_ISL_621000, EPI_ISL_621001, EPI_ISL_621002, EPI_ISL_621003, EPI_ISL_621004, EPI_ISL_621005, EPI_ISL_621006, EPI_ISL_621007, EPI_ISL_621008, EPI_ISL_621009, EPI_ISL_621010, EPI_ISL_621011, EPI_ISL_621012, EPI_ISL_621013, EPI_ISL_621014, EPI_ISL_621015, EPI_ISL_621016, EPI_ISL_621017, EPI_ISL_621018, EPI_ISL_621019, EPI_ISL_621020, EPI_ISL_621021, EPI_ISL_621022, EPI_ISL_621023, EPI_ISL_621024, EPI_ISL_621025, EPI_ISL_621026, EPI_ISL_621027, EPI_ISL_621028, EPI_ISL_621029, EPI_ISL_621030, EPI_ISL_621031, EPI_ISL_621032, EPI_ISL_621033, EPI_ISL_621034, EPI_ISL_621035, EPI_ISL_621036, EPI_ISL_621037, EPI_ISL_621038, EPI_ISL_621039, EPI_ISL_621040, EPI_ISL_621041, EPI_ISL_621042, EPI_ISL_621043, EPI_ISL_621044, EPI_ISL_621045, EPI_ISL_621046, EPI_ISL_621047, EPI_ISL_621048, EPI_ISL_621049, EPI_ISL_621050, EPI_ISL_621051, EPI_ISL_621052, EPI_ISL_621053, EPI_ISL_621054, EPI_ISL_621055, EPI_ISL_621056, EPI_ISL_621057, EPI_ISL_621058, EPI_ISL_621059, EPI_ISL_621060, EPI_ISL_621061, EPI_ISL_621062, EPI_ISL_621063, EPI_ISL_621064, EPI_ISL_621065, EPI_ISL_621066, EPI_ISL_621067, EPI_ISL_621068, EPI_ISL_621069, EPI_ISL_621070, EPI_ISL_621071, EPI_ISL_621072, EPI_ISL_621073, EPI_ISL_621074, EPI_ISL_621075, EPI_ISL_621076, EPI_ISL_621077, EPI_ISL_621078, EPI_ISL_621079, EPI_ISL_621080, EPI_ISL_621081, EPI_ISL_621082, EPI_ISL_621090, EPI_ISL_621095, EPI_ISL_621096, EPI_ISL_621097, EPI_ISL_621098, EPI_ISL_621099, EPI_ISL_621100, EPI_ISL_621101, EPI_ISL_621102, EPI_ISL_621103, EPI_ISL_621104, EPI_ISL_621105, EPI_ISL_621106, EPI_ISL_621107, EPI_ISL_621108, EPI_ISL_621109, EPI_ISL_621110, EPI_ISL_621111, EPI_ISL_621112, EPI_ISL_621113, EPI_ISL_621118, EPI_ISL_621119, EPI_ISL_621120, EPI_ISL_621121, EPI_ISL_621122, EPI_ISL_621123, EPI_ISL_621124, EPI_ISL_621125, EPI_ISL_621150, EPI_ISL_621211, EPI_ISL_621212, EPI_ISL_621213, EPI_ISL_621214, EPI_ISL_621215, EPI_ISL_621216, EPI_ISL_621217, EPI_ISL_621218, EPI_ISL_621219, EPI_ISL_621220, EPI_ISL_621233, EPI_ISL_621234, EPI_ISL_621235, EPI_ISL_621236, EPI_ISL_621237, EPI_ISL_621238, EPI_ISL_621239, EPI_ISL_621240, EPI_ISL_621241, EPI_ISL_621242, EPI_ISL_621243, EPI_ISL_621244, EPI_ISL_621245, EPI_ISL_621246, EPI_ISL_621247, EPI_ISL_621248, EPI_ISL_621249, EPI_ISL_621250, EPI_ISL_621251, EPI_ISL_621252, EPI_ISL_621253, EPI_ISL_621254, EPI_ISL_621255, EPI_ISL_621256, EPI_ISL_621257, EPI_ISL_621258, EPI_ISL_621259, EPI_ISL_621260, EPI_ISL_621261, EPI_ISL_622302, EPI_ISL_622306, EPI_ISL_622307, EPI_ISL_622308, EPI_ISL_622309, EPI_ISL_622310, EPI_ISL_622311, EPI_ISL_622312, EPI_ISL_622313, EPI_ISL_622314, EPI_ISL_622315, EPI_ISL_622316, EPI_ISL_622317, EPI_ISL_622318, EPI_ISL_622319, EPI_ISL_622320, EPI_ISL_622321, EPI_ISL_622322, EPI_ISL_622323, EPI_ISL_622324, EPI_ISL_622325, EPI_ISL_622326, EPI_ISL_622327, EPI_ISL_622328, EPI_ISL_622329, EPI_ISL_622330, EPI_ISL_622331, EPI_ISL_622333, EPI_ISL_622339, EPI_ISL_622340 | | | |
| see above | Department of Virus and Microbiological Special Diagnostics, Statens Serum Institut, Denmark | Albertsen lab, Department of Chemistry and Bioscience, Aalborg University, Denmark | Danish Covid-19 Genome Consortia |
| EPI_ISL_622833, EPI_ISL_622834, EPI_ISL_622835, EPI_ISL_622836, EPI_ISL_622837, EPI_ISL_622838, EPI_ISL_622839, EPI_ISL_622840, EPI_ISL_622841, EPI_ISL_622842, EPI_ISL_622892, EPI_ISL_622893 | | | |
| see above | Respiratory Virus Unit, Microbiology Services Colindale, Public Health England | Respiratory Virus Unit, Microbiology Services Colindale, Public Health England | PHE Covid Sequencing Team |
| EPI_ISL_623179 | Utah Public Health Laboratory | Utah Public Health Laboratory | Erin Young, Kelly Oakeson |
| EPI_ISL_625485, EPI_ISL_625492, EPI_ISL_625495, EPI_ISL_625496, EPI_ISL_625497, EPI_ISL_625498, EPI_ISL_625499 | Santa Clara County Public Health Laboratory | Chan-Zuckerberg Biohub | CZB Cliahub Consortium |
| EPI_ISL_625546 | Alameda County Public Health Lab | Chan-Zuckerberg Biohub | CZB Cliahub Consortium |
| EPI_ISL_625601 | Santa Clara County Public Health Laboratory | Chan-Zuckerberg Biohub | CZB Cliahub Consortium |
| EPI_ISL_625611, EPI_ISL_625612, EPI_ISL_625613 | Alameda County Public Health Lab | Chan-Zuckerberg Biohub | CZB Cliahub Consortium |
| EPI_ISL_625647, EPI_ISL_625648, EPI_ISL_625649, EPI_ISL_625650, EPI_ISL_625651, EPI_ISL_625652, EPI_ISL_625653, EPI_ISL_625654, EPI_ISL_625655, EPI_ISL_625656, EPI_ISL_625657 | | | |
| see above | Orange County Public Health Lab | Chan-Zuckerberg Biohub | CZB Cliahub Consortium |
| EPI_ISL_625668 | UCSF Clinical Microbiology Laboratory | Chan-Zuckerberg Biohub | CZB Cliahub Consortium |
| EPI_ISL_625674 | Laboratory of Molecular Medicine, University of Magallanes | Centro Asistencial Docente y de Investigacion, Universidad de Magallanes | Jorge Gonzalez, Jacqueline Aldridge, Diego Alvarez, Marcelo Navarrete |
| EPI_ISL_626586, EPI_ISL_626587, EPI_ISL_626588, EPI_ISL_626589, EPI_ISL_626590, EPI_ISL_626591, EPI_ISL_626599, EPI_ISL_626600, EPI_ISL_626601 | The National Institute of Public Health | State Veterinary Institute Prague | Nagy,A;Jirincova,H;Novakova,L;Trnka,D;Vecerova,J |
| EPI_ISL_626794 | University of Exeter | COVID-19 Genomics UK (COG-UK) Consortium | Ben Temperton,Aaron Jeffries,Michelle Michelsen,Joanna Warwick-Dugdale,Audrey Farbos,Robyn Manley,Stephen Michell,Jane Masoli |
| EPI_ISL_626833, EPI_ISL_627026, EPI_ISL_627146, EPI_ISL_627203, EPI_ISL_627212, EPI_ISL_627220 | Wales Specialist Virology Centre Sequencing lab: Pathogen Genomics Unit | COVID-19 Genomics UK (COG-UK) Consortium | Catherine Moore, Johnathan Evans, Laura Gifford, Malorie Perry, Simon Cottrell, Angela Marchbank, Alec Birchley, Alexander Adams, Amy Gaskin, Bree Gatica-Wilcox, Jason Coombes, Joel Southgate, Lauren Gilbert, Lee Graham, Nicole Pacchiarini, Sara Kumziene-Summerhayes, Sarah Taylor, Sophie Jones, Sara Rey, Matthew Bull, Joanne Watkins, Sally Corden, Tom Connor |
| EPI_ISL_627436, EPI_ISL_627438, EPI_ISL_627439 | University of Exeter | COVID-19 Genomics UK (COG-UK) Consortium | Ben Temperton,Aaron Jeffries,Michelle Michelsen,Joanna Warwick-Dugdale,Audrey Farbos,Robyn Manley,Stephen Michell,Jane Masoli |
| EPI_ISL_627482, EPI_ISL_627483, EPI_ISL_627484, EPI_ISL_627485, EPI_ISL_627486, EPI_ISL_627487 | Wales Specialist Virology Centre Sequencing lab: Pathogen Genomics Unit | COVID-19 Genomics UK (COG-UK) Consortium | Catherine Moore, Johnathan Evans, Laura Gifford, Malorie Perry, Simon Cottrell, Angela Marchbank, Alec Birchley, Alexander Adams, Amy Gaskin, Bree Gatica-Wilcox, Jason Coombes, Joel Southgate, Lauren Gilbert, Lee Graham, Nicole Pacchiarini, Sara Kumziene-Summerhayes, Sarah Taylor, Sophie Jones, Sara Rey, Matthew Bull, Joanne Watkins, Sally Corden, Tom Connor |
| EPI_ISL_627491 | University College London, Great Ormond Street Hospital for Children NHS Foundation Trust, Imperial College Healthcare NHS Trust | COVID-19 Genomics UK (COG-UK) Consortium | Sergi Castellano, Rachel Williams, Mark Kristiansen, Paola Resende Silva, Sunando Roy, Tony Brooks, Helena Tutill, Paola Niola, Patricia Dyal, Charlotte Williams, Leysa Forrest, Yasmin Panchbhaya, Jacqueline Findlay, Samuel Weeks, Julianne Brown, Kathryn Harris, Paul Randell, James Price, Alison Holmes, Judith Breuer |
| EPI_ISL_627730, EPI_ISL_627731, EPI_ISL_627871, EPI_ISL_627899 | Wales Specialist Virology Centre Sequencing lab: Pathogen Genomics Unit | COVID-19 Genomics UK (COG-UK) Consortium | Catherine Moore, Johnathan Evans, Laura Gifford, Malorie Perry, Simon Cottrell, Angela Marchbank, Alec Birchley, Alexander Adams, Amy Gaskin, Bree Gatica-Wilcox, Jason Coombes, Joel Southgate, Lauren Gilbert, Lee Graham, Nicole Pacchiarini, Sara Kumziene-Summerhayes, Sarah Taylor, Sophie Jones, Sara Rey, Matthew Bull, Joanne Watkins, Sally Corden, Tom Connor |
| EPI_ISL_628374 | Virology Department, Sheffield Teaching Hospitals NHS Foundation Trust/Department of Infection, Immunity and Cardiovascular Disease, The Medical School, University of Sheffield | COVID-19 Genomics UK (COG-UK) Consortium | Thushan de Silva, Matthew Parker, Nikki Smith, Adri Angyal, Rebecca Brown, Luke Green, Rachel Tucker, Paul Parsons, Danielle Groves, Katie Johnson, Laura Carrilero, Alex Keeley, Dave Partridge, Matthew Wyles, Benjamin Lindsey, Mehmet Yavuz, Mohammad Raza, Cariad Evans |
| EPI_ISL_628939, EPI_ISL_628940, EPI_ISL_628941, EPI_ISL_628946, EPI_ISL_628947, EPI_ISL_628948, EPI_ISL_628949, EPI_ISL_628956, EPI_ISL_628960 | Utah Public Health Laboratory | Utah Public Health Laboratory | Erin Young, Kelly Oakeson |
| EPI_ISL_629030 | South Eastern Area Laboratory Services (SEALS) | NSW Health Pathology - Institute of Clinical Pathology and | CIDM-PH et al. |

| | | | |
|---|---|---|---|
| | | Medical Research; Westmead Hospital; University of Sydney | |
| EPI_ISL_631362, EPI_ISL_631363, EPI_ISL_631364 | ZOTZ KLIMAS MVZ Düsseldorf-Centrum GbR ÜBAG für Labormedizin, Genetik, Zytologie, Pathologie | Center of Medical Microbiology, Virology, and Hospital Hygiene, University of Duesseldorf | Maximilian Damagnez, Alexander Dilthey, Ashley-Jane Duplessis, Patrick Finzer, Katrin Hoffmann, Torsten Houwaart, Lisanna Hülse, Malte Kohns Vasconcelos, Marek Korencak, Nadine Lübke, Jessica Nicolai, Klaus Pfeffer, Daniel Strelow, Jörg Timm, Andreas Walker, Tobias Wienemann, Rainer Zotz |
| EPI_ISL_631377, EPI_ISL_631382, EPI_ISL_631386, EPI_ISL_631388 | University Hospital Cologne | Center of Medical Microbiology, Virology, and Hospital Hygiene, University of Duesseldorf | Maximilian Damagnez, Alexander Dilthey, Ashley-Jane Duplessis, Eva Heger, Torsten Houwaart, Rolf Kaiser, Florian Klein, Elena Knops, Malte Kohns Vasconcelos, Jessica Nicolai, Klaus Pfeffer, Gibran Rubio Quintanares, Saleta Sierra-Aragón, Daniel Strelow, Jörg Timm, Andreas Walker, Tobias Wienemann |
| EPI_ISL_632323, EPI_ISL_632345, EPI_ISL_632352, EPI_ISL_632355, EPI_ISL_632356, EPI_ISL_632360, EPI_ISL_632363, EPI_ISL_632367, EPI_ISL_632378 | Dutch COVID-19 response team | Erasmus Medical Center | Bas Oude Munnink, David Nieuwenhuijse, Reina Sikkema, Claudia Schapendonk, Irina Chestakova, Anne van der Linden, Theo Bestebroer, Stefan van Nieuwkoop, Mark Pronk, Pascal Lexmond, Corien Swaan, Manon Haverkate, Madelief Mollers, Mart Stein, Sandra Kengne Kamga Mobou, Jeroen van Kampen, Jolanda Voermans, Aura Timen, Corine GeurtsvanKessel, Annemiek van der Eijk, Richard Molenkamp, Marion Koopmans, on behalf of the Dutch national COVID-19 response team. |
| EPI_ISL_632385 | Dutch COVID-19 response team | Erasmus Medical Center | OH consortium |
| EPI_ISL_632389, EPI_ISL_632390, EPI_ISL_632399, EPI_ISL_632403, EPI_ISL_632410, EPI_ISL_632421 | Dutch COVID-19 response team | Erasmus Medical Center | Bas Oude Munnink, David Nieuwenhuijse, Reina Sikkema, Claudia Schapendonk, Irina Chestakova, Anne van der Linden, Theo Bestebroer, Stefan van Nieuwkoop, Mark Pronk, Pascal Lexmond, Corien Swaan, Manon Haverkate, Madelief Mollers, Mart Stein, Sandra Kengne Kamga Mobou, Jeroen van Kampen, Jolanda Voermans, Aura Timen, Corine GeurtsvanKessel, Annemiek van der Eijk, Richard Molenkamp, Marion Koopmans, on behalf of the Dutch national COVID-19 response team. |
| EPI_ISL_632456, EPI_ISL_632457, EPI_ISL_632458, EPI_ISL_632459 | Dutch COVID-19 response team | Erasmus Medical Center | OH consortium |
| EPI_ISL_632461, EPI_ISL_632493, EPI_ISL_632494, EPI_ISL_632502, EPI_ISL_632506, EPI_ISL_632510, EPI_ISL_632511, EPI_ISL_632513, EPI_ISL_632530, EPI_ISL_632531, EPI_ISL_632535, EPI_ISL_632544, EPI_ISL_632545, EPI_ISL_632546, EPI_ISL_632551, EPI_ISL_632552, EPI_ISL_632553, EPI_ISL_632554, EPI_ISL_632555, EPI_ISL_632556, EPI_ISL_632557, EPI_ISL_632558, EPI_ISL_632568, EPI_ISL_632569, EPI_ISL_632570, EPI_ISL_632571, EPI_ISL_632572, EPI_ISL_632592, EPI_ISL_632593, EPI_ISL_632596, EPI_ISL_632598, EPI_ISL_632599, EPI_ISL_632612, EPI_ISL_632613, EPI_ISL_632614, EPI_ISL_632615, EPI_ISL_632616, EPI_ISL_632617, EPI_ISL_632618, EPI_ISL_632619, EPI_ISL_632620, EPI_ISL_632621, EPI_ISL_632622, EPI_ISL_632665, EPI_ISL_632670, EPI_ISL_632671, EPI_ISL_632697, EPI_ISL_632716, EPI_ISL_632717, EPI_ISL_632718, EPI_ISL_632733, EPI_ISL_632754, EPI_ISL_632785 | | | |
| see above | Dutch COVID-19 response team | Erasmus Medical Center | Bas Oude Munnink, David Nieuwenhuijse, Reina Sikkema, Claudia Schapendonk, Irina Chestakova, Anne van der Linden, Theo Bestebroer, Stefan van Nieuwkoop, Mark Pronk, Pascal Lexmond, Corien Swaan, Manon Haverkate, Madelief Mollers, Mart Stein, Sandra Kengne Kamga Mobou, Jeroen van Kampen, Jolanda Voermans, Aura Timen, Corine GeurtsvanKessel, Annemiek van der Eijk, Richard Molenkamp, Marion Koopmans, on behalf of the Dutch national COVID-19 response team. |
| EPI_ISL_632983, EPI_ISL_632984, EPI_ISL_632985, EPI_ISL_632986, EPI_ISL_632987, EPI_ISL_632988, EPI_ISL_632989, EPI_ISL_632990, EPI_ISL_632991 | DOHMH Corona | New York City Public Health Laboratory | Jade Wang, et al. |
| EPI_ISL_632995 | DOHMH Crown Heights | New York City Public Health Laboratory | Jade Wang, et al. |
| EPI_ISL_633004, EPI_ISL_633005, EPI_ISL_633006, EPI_ISL_633007 | DOHMH Jamaica | New York City Public Health Laboratory | Jade Wang, et al. |
| EPI_ISL_633009, EPI_ISL_633010, EPI_ISL_633018 | DOHMH Central Harlem | New York City Public Health Laboratory | Jade Wang, et al. |
| EPI_ISL_633019, EPI_ISL_633020, EPI_ISL_633021, EPI_ISL_633022 | DOHMH Morrisania | New York City Public Health Laboratory | Jade Wang, et al. |
| EPI_ISL_633023 | DOHMH Chelsea | New York City Public Health Laboratory | Jade Wang, et al. |
| EPI_ISL_633024 | DOHMH Morrisania | New York City Public Health Laboratory | Jade Wang, et al. |
| EPI_ISL_633043 | DOHMH Riverside | New York City Public Health Laboratory | Jade Wang, et al. |
| EPI_ISL_633044 | DOHMH Morrisania | New York City Public Health Laboratory | Jade Wang, et al. |
| EPI_ISL_633045 | DOHMH PHL | New York City Public Health Laboratory | Jade Wang, et al. |
| EPI_ISL_633046, EPI_ISL_633047 | DOHMH Chelsea | New York City Public Health Laboratory | Jade Wang, et al. |
| EPI_ISL_633048, EPI_ISL_633049 | DOHMH Morrisania | New York City Public Health Laboratory | Jade Wang, et al. |
| EPI_ISL_633050, EPI_ISL_633051 | DOHMH PHL | New York City Public Health Laboratory | Jade Wang, et al. |
| EPI_ISL_633052 | DOHMH Chelsea | New York City Public Health Laboratory | Jade Wang, et al. |
| EPI_ISL_634884 | Lab voor klinische biologie | Onderzoeksgroep Virologie | Laurens Lambrechts, Nick Vereecke, Marthe Pauwels, Bruno Verhasselt, Linos Vandekerckhove, Hans Nauwynck, Sebastiaan Theuns |
| EPI_ISL_634923 | Utah Public Health Laboratory | Utah Public Health Laboratory | Erin L. Young, Kelly F. Oakeson |
| EPI_ISL_634978, EPI_ISL_634979, EPI_ISL_634980, EPI_ISL_634981, EPI_ISL_634982, EPI_ISL_634983, EPI_ISL_634984, EPI_ISL_634985, EPI_ISL_634986, EPI_ISL_634988, EPI_ISL_634989, EPI_ISL_634990, EPI_ISL_634991, EPI_ISL_634992 | | | |
| see above | National Health Laboratory Service - Inkosi Albert Luthuli Central Hospital (NHLS-IALCH) | KRISP, KZN Research Innovation and Sequencing Platform | Giandhari J, Pillay S, Lessells R, Mdlalose K, York D, Khan S, Tegally H, Wilkinson E, de Oliveira T |
| EPI_ISL_635064, EPI_ISL_635065 | Foerde Hospital, Department of Microbiology | Norwegian Institute of Public Health, Department of Virology | Kathrine Stene-Johansen, Kamilla Heddeland Instefjord, Hilde Elshaug, Marie Paulsen Madsen, Rasmus Riis Kopperud, Hilde Vollan, Karoline Bragstad, Olav Hungnes |
| EPI_ISL_635092 | Medical Microbiology Unit, Department for Laboratory Medicine, Drammen Hospital, Vestre Viken Health Trust, | Norwegian Institute of Public Health, Department of Virology | Kathrine Stene-Johansen, Kamilla Heddeland Instefjord, Hilde Elshaug, Marie Paulsen Madsen, Rasmus Riis Kopperud, Hilde Vollan, Karoline Bragstad, Olav Hungnes |
| EPI_ISL_635094 | Oslo University Hospital, Department of Medical Microbiology | Norwegian Institute of Public Health, Department of Virology | Kathrine Stene-Johansen, Kamilla Heddeland Instefjord, Hilde Elshaug, Marie Paulsen Madsen, Rasmus Riis Kopperud, Hilde Vollan, Karoline Bragstad, Olav Hungnes |
| EPI_ISL_635098 | Foerde Hospital, Department of Microbiology | Norwegian Institute of Public Health, Department of Virology | Kathrine Stene-Johansen, Kamilla Heddeland Instefjord, Hilde Elshaug, Marie Paulsen Madsen, Rasmus Riis Kopperud, Hilde Vollan, Karoline Bragstad, Olav Hungnes |
| EPI_ISL_635110, EPI_ISL_635111 | Department of Medical Microbiology, St. Olavs hospital | Norwegian Institute of Public Health, Department of Virology | Kathrine Stene-Johansen, Kamilla Heddeland Instefjord, Hilde Elshaug, Marie Paulsen Madsen, Rasmus Riis Kopperud, Hilde Vollan, Karoline Bragstad, Olav Hungnes |
| EPI_ISL_635120 | Dept. of Medical Microbiology, Stavanger University Hospital, Helse Stavanger HF | Norwegian Institute of Public Health, Department of Virology | Kathrine Stene-Johansen, Kamilla Heddeland Instefjord, Hilde Elshaug, Marie Paulsen Madsen, Rasmus Riis Kopperud, Hilde Vollan, Karoline Bragstad, Olav Hungnes |
| EPI_ISL_635121, EPI_ISL_635122, EPI_ISL_635123 | Hospital of Southern Norway - Kristiansand, Department of Medical Microbiology | Norwegian Institute of Public Health, Department of Virology | Kathrine Stene-Johansen, Kamilla Heddeland Instefjord, Hilde Elshaug, Marie Paulsen Madsen, Rasmus Riis Kopperud, Hilde Vollan, Karoline Bragstad, Olav Hungnes |
| EPI_ISL_635126 | Department of Medical Microbiology, St. Olavs hospital | Norwegian Institute of Public Health, Department of Virology | Kathrine Stene-Johansen, Kamilla Heddeland Instefjord, Hilde Elshaug, Marie Paulsen Madsen, Rasmus Riis Kopperud, Hilde Vollan, Karoline Bragstad, Olav Hungnes |
| EPI_ISL_635127 | Foerde Hospital, Department of Microbiology | Norwegian Institute of Public Health, Department of Virology | Kathrine Stene-Johansen, Kamilla Heddeland Instefjord, Hilde Elshaug, Marie Paulsen Madsen, Rasmus Riis Kopperud, Hilde Vollan, Karoline Bragstad, Olav Hungnes |
| EPI_ISL_635132 | Unilabs Laboratory Medicine | Norwegian Institute of Public Health, Department of Virology | Kathrine Stene-Johansen, Kamilla Heddeland Instefjord, Hilde Elshaug, Marie Paulsen Madsen, Rasmus Riis Kopperud, Hilde Vollan, Karoline Bragstad, Olav Hungnes |

| | | | |
|---|---|---|---|
| EPI_ISL_635135, EPI_ISL_635136 | Department of Medical Microbiology, St. Olavs hospital | Norwegian Institute of Public Health, Department of Virology | Kathrine Stene-Johansen, Kamilla Heddeland Instefjord, Hilde Elshaug, Marie Paulsen Madsen, Rasmus Riis Kopperud, Hilde Vollan, Karoline Bragstad, Olav Hungnes |
| EPI_ISL_635137 | University Hospital of Northern Norway, Department for Microbiology and Infectious Disease Control | Norwegian Institute of Public Health, Department of Virology | Kathrine Stene-Johansen, Kamilla Heddeland Instefjord, Hilde Elshaug, Marie Paulsen Madsen, Rasmus Riis Kopperud, Hilde Vollan, Karoline Bragstad, Olav Hungnes |
| EPI_ISL_635155 | Oslo University Hospital, Department of Medical Microbiology | Norwegian Institute of Public Health, Department of Virology | Kathrine Stene-Johansen, Kamilla Heddeland Instefjord, Hilde Elshaug, Marie Paulsen Madsen, Rasmus Riis Kopperud, Hilde Vollan, Karoline Bragstad, Olav Hungnes |
| EPI_ISL_635158 | Unilabs Laboratory Medicine | Norwegian Institute of Public Health, Department of Virology | Kathrine Stene-Johansen, Kamilla Heddeland Instefjord, Hilde Elshaug, Marie Paulsen Madsen, Rasmus Riis Kopperud, Hilde Vollan, Karoline Bragstad, Olav Hungnes |
| EPI_ISL_635188 | Norwegian Institute of Public Health, Department of Virology | Norwegian Institute of Public Health, Department of Virology | Kathrine Stene-Johansen, Kamilla Heddeland Instefjord, Hilde Elshaug, Marie Paulsen Madsen, Rasmus Riis Kopperud, Hilde Vollan, Karoline Bragstad, Olav Hungnes |
| EPI_ISL_636121, EPI_ISL_636122, EPI_ISL_636123, EPI_ISL_636126, EPI_ISL_636127, EPI_ISL_636130, EPI_ISL_636134, EPI_ISL_636140, EPI_ISL_636141, EPI_ISL_636144, EPI_ISL_636146, EPI_ISL_636147, EPI_ISL_636150, EPI_ISL_636153, EPI_ISL_636156, EPI_ISL_636157, EPI_ISL_636158, EPI_ISL_636160, EPI_ISL_636161 | | | |
| see above | San Diego County Public Health Laboratory | Andersen lab at Scripps Research | SEARCH Alliance San Diego with Tracy Basler, Jovan Shephard, Brett Austin |
| EPI_ISL_636267, EPI_ISL_636268, EPI_ISL_636269, EPI_ISL_636270, EPI_ISL_636271, EPI_ISL_636272, EPI_ISL_636273, EPI_ISL_636274, EPI_ISL_636275, EPI_ISL_636276, EPI_ISL_636277, EPI_ISL_636278, EPI_ISL_636279, EPI_ISL_636280, EPI_ISL_636281, EPI_ISL_636282, EPI_ISL_636283, EPI_ISL_636284, EPI_ISL_636285, EPI_ISL_636286, EPI_ISL_636287, EPI_ISL_636288, EPI_ISL_636289, EPI_ISL_636290, EPI_ISL_636291, EPI_ISL_636292, EPI_ISL_636293, EPI_ISL_636294, EPI_ISL_636295, EPI_ISL_636296, EPI_ISL_636297, EPI_ISL_636298, EPI_ISL_636299, EPI_ISL_636300, EPI_ISL_636301, EPI_ISL_636302, EPI_ISL_636303, EPI_ISL_636304, EPI_ISL_636305, EPI_ISL_636306, EPI_ISL_636307, EPI_ISL_636308, EPI_ISL_636309, EPI_ISL_636310, EPI_ISL_636311, EPI_ISL_636312, EPI_ISL_636313, EPI_ISL_636314, EPI_ISL_636315, EPI_ISL_636316, EPI_ISL_636317, EPI_ISL_636318, EPI_ISL_636319, EPI_ISL_636320, EPI_ISL_636321, EPI_ISL_636322, EPI_ISL_636323, EPI_ISL_636324, EPI_ISL_636325, EPI_ISL_636326, EPI_ISL_636327, EPI_ISL_636328 | | | |
| see above | Biolab Diagnostic Laboratories | Andersen lab at Scripps Research | Issa Abu-Dayyeh, Ahmad Tibi, Lama Hussein, Lina Mohammad, Zein Naber, Amid Abdelnour with SEARCH Alliance San Diego |
| EPI_ISL_636534, EPI_ISL_636535, EPI_ISL_636579 | Dutch COVID-19 response team | National Institute for Public Health and the Environment (RIVM) | Adam Meijer, Harry Vennema, Jeroen Cremer, Sharon van den Brink, Bas van der Veer, AnneMarie van den Brandt, Florian Zwagemaker, Dennis Schmitz, Chantal Reusken, on behalf of the national COVID-19 response team |
| EPI_ISL_636688, EPI_ISL_636689 | Respiratory Virus Unit, Microbiology Services Colindale, Public Health England | Respiratory Virus Unit, Microbiology Services Colindale, Public Health England | PHE Covid Sequencing Team |
| EPI_ISL_637017 | Department of Infectious Diseases and Immunology, National Hospital Organization Nagoya Medical Center | Clinical Research Center, National Hospital Organization Nagoya Medical Center | Yoshihiro Nakata, Hirotaka Ode, Mai Kubota, Masakazu Matsuda, Kazuhiro Matsuoka, Miho Nakasuji, Mikiko Mori, Mayumi Imahashi, Yoshiyuki Yokomaku, Yasumasa Iwatani |
| EPI_ISL_637307, EPI_ISL_637354, EPI_ISL_637554, EPI_ISL_637598, EPI_ISL_637666 | Department of Pathology, University of Cambridge | COVID-19 Genomics UK (COG-UK) Consortium | Aminu S. Jahun, Yasmin Chaudhry, Grant Hall, Iliana Georgana, Myra Hosmillo, Martin D. Curran, Malte Pinckert, Surendra Parmar, Ian Goodfellow |
| EPI_ISL_637793 | Northumbria University / South Tees Hospitals NHS Foundation Trust / North Cumbria Integrated Care NHS Foundation Trust / North Tees and Hartlepool NHS Foundation Trust / Newcastle Hospitals NHS Foundation Trust | COVID-19 Genomics UK (COG-UK) Consortium | Darren L Smith,Andrew Nelson,Matthew Bashton,Greg R Young,Joshua Loh,John Allan,Mohammad A Tariq,Giles S Holt,Gary Black,Wen C Yew,Lynn Dover,Paul Baker,Steve Liggett,Sarah Essex,Jane Greenaway,Debra Padgett,Clive Graham,Garren Scott,Edward Barton,Emma Swindells,Brendan Payne,Jennifer Collins,Yusri Taha,Gary Eltringham |
| EPI_ISL_637803, EPI_ISL_637804, EPI_ISL_637805, EPI_ISL_637806, EPI_ISL_637807, EPI_ISL_637808, EPI_ISL_637809, EPI_ISL_637856, EPI_ISL_637880, EPI_ISL_637888, EPI_ISL_637916 | | | |
| see above | Department of Pathology, University of Cambridge | COVID-19 Genomics UK (COG-UK) Consortium | Aminu S. Jahun, Yasmin Chaudhry, Grant Hall, Iliana Georgana, Myra Hosmillo, Martin D. Curran, Malte Pinckert, Surendra Parmar, Ian Goodfellow |
| EPI_ISL_637995 | University College London, Great Ormond Street Hospital for Children NHS Foundation Trust, Imperial College Healthcare NHS Trust | COVID-19 Genomics UK (COG-UK) Consortium | Sergi Castellano, Rachel Williams, Mark Kristiansen, Paola Resende Silva, Sunando Roy, Tony Brooks, Helena Tutill, Paola Niola, Patricia Dyal, Charlotte Williams, Leysa Forrest, Yasmin Panchbhaya, Jacqueline Findlay, Samuel Weeks, Julianne Brown, Kathryn Harris, Paul Randell, James Price, Alison Holmes, Judith Breuer |
| EPI_ISL_638080, EPI_ISL_638081, EPI_ISL_638082, EPI_ISL_638083, EPI_ISL_638174 | Department of Pathology, University of Cambridge | COVID-19 Genomics UK (COG-UK) Consortium | Aminu S. Jahun, Yasmin Chaudhry, Grant Hall, Iliana Georgana, Myra Hosmillo, Martin D. Curran, Malte Pinckert, Surendra Parmar, Ian Goodfellow |
| EPI_ISL_638562 | Northumbria University / South Tees Hospitals NHS Foundation Trust / North Cumbria Integrated Care NHS Foundation Trust / North Tees and Hartlepool NHS Foundation Trust / Newcastle Hospitals NHS Foundation Trust | COVID-19 Genomics UK (COG-UK) Consortium | Darren L Smith,Andrew Nelson,Matthew Bashton,Greg R Young,Joshua Loh,John Allan,Mohammad A Tariq,Giles S Holt,Gary Black,Wen C Yew,Lynn Dover,Paul Baker,Steve Liggett,Sarah Essex,Jane Greenaway,Debra Padgett,Clive Graham,Garren Scott,Edward Barton,Emma Swindells,Brendan Payne,Jennifer Collins,Yusri Taha,Gary Eltringham |
| EPI_ISL_638642, EPI_ISL_638643, EPI_ISL_638645, EPI_ISL_638646, EPI_ISL_638647, EPI_ISL_638648, EPI_ISL_638649, EPI_ISL_638650, EPI_ISL_638651, EPI_ISL_638652, EPI_ISL_638653, EPI_ISL_638654, EPI_ISL_638655, EPI_ISL_638656, EPI_ISL_638657, EPI_ISL_638658, EPI_ISL_638659, EPI_ISL_638660, EPI_ISL_638661, EPI_ISL_638662, EPI_ISL_638663, EPI_ISL_638664, EPI_ISL_638665, EPI_ISL_638666, EPI_ISL_638667, EPI_ISL_638668, EPI_ISL_638669, EPI_ISL_638670, EPI_ISL_638671, EPI_ISL_638672, EPI_ISL_638673, EPI_ISL_638674, EPI_ISL_638675, EPI_ISL_638676, EPI_ISL_638677, EPI_ISL_638678, EPI_ISL_638679, EPI_ISL_638681, EPI_ISL_638914, EPI_ISL_638915, EPI_ISL_638916, EPI_ISL_638917, EPI_ISL_638918, EPI_ISL_638919, EPI_ISL_638920, EPI_ISL_638921, EPI_ISL_638922, EPI_ISL_638923, EPI_ISL_638924, EPI_ISL_638925, EPI_ISL_638926, EPI_ISL_638927 | | | |
| see above | Department of Pathology, University of Cambridge | COVID-19 Genomics UK (COG-UK) Consortium | Aminu S. Jahun, Yasmin Chaudhry, Grant Hall, Iliana Georgana, Myra Hosmillo, Martin D. Curran, Malte Pinckert, Surendra Parmar, Ian Goodfellow |
| EPI_ISL_639668, EPI_ISL_639670, EPI_ISL_639673, EPI_ISL_639675 | E. Gulbja Laboratorija | Latvian Biomedical Research and Study Centre | Ivars Silamielis, Kaspars Megnis, Monta Ustinova, ikita Zrelovs, Vita Rovte, Mikus Gavars, Dmitrijs Perminovs, Uga Dumpis, Jnis Kloviš |
| EPI_ISL_639676, EPI_ISL_639677, EPI_ISL_639678, EPI_ISL_639679, EPI_ISL_639680 | Centrl Laboratorija | Latvian Biomedical Research and Study Centre | Ivars Silamielis, Kaspars Megnis, Monta Ustinova, ikita Zrelovs, Vita Rovte, Stella Lapia, Jana Oste, Marta Priedte, Uga Dumpis, Jnis Kloviš |
| EPI_ISL_639682 | E. Gulbja Laboratorija | Latvian Biomedical Research and Study Centre | Ivars Silamielis, Kaspars Megnis, Monta Ustinova, ikita Zrelovs, Vita Rovte, Mikus Gavars, Dmitrijs Perminovs, Uga Dumpis, Jnis Kloviš |
| EPI_ISL_639892, EPI_ISL_639893, EPI_ISL_639894, EPI_ISL_639895, EPI_ISL_639896, EPI_ISL_639897, EPI_ISL_639898, EPI_ISL_639899, EPI_ISL_639900, EPI_ISL_639901, EPI_ISL_639902, EPI_ISL_639903, EPI_ISL_639905, EPI_ISL_639906, EPI_ISL_639907, EPI_ISL_639908 | | | |
| see above | National Virus Reference Laboratory | National Virus Reference Laboratory | Michael Carr, Gabriel Gonzalez, Jonathan Dean, Daniel Hare, Cillian F De Gascun |
| EPI_ISL_639953, EPI_ISL_639954, EPI_ISL_639955, EPI_ISL_639956, EPI_ISL_639957, EPI_ISL_639958, EPI_ISL_639959, EPI_ISL_639962, EPI_ISL_639968, EPI_ISL_639972 | HELIX LLC | WHO National Influenza Centre Russian Federation | Andrey Komissarov, Artem Fadeev, Kseniya Komissarova, Anna Ivanova, Dmitry Bazhenov, Daria Danilenko |
| EPI_ISL_639991, EPI_ISL_639993, EPI_ISL_639995 | CNR Virus des Infections Respiratoires - France SUD | CNR Virus des Infections Respiratoires - France SUD | Antonin Bal, Gregory Destras, Gwendolyne Burfin, Hadrien Règue, Alexandre Gaymard, Maude Bouscambert-Duchamp, Florence Morfin-Sherpa, Martine Valette, Bruno Lina, Laurence Josset |
| EPI_ISL_640084 | Groote Schuur Hospital wc GSH | NHLS/UCT | Arash Iranzadeh, Deelan Doolabh, Lynn Tyers, Bruna Galvao, Innocent Mudau, Marvin Hsiao, Kruger Marais, Diana Hardie, Stephen Korsman, Carolyn Williamson |
| EPI_ISL_640085 | Victoria Hospital wc VHW | NHLS/UCT | Arash Iranzadeh, Deelan Doolabh, Lynn Tyers, Bruna Galvao, Innocent Mudau, Marvin Hsiao, Kruger Marais, Diana Hardie, Stephen Korsman, Carolyn Williamson |
| EPI_ISL_640086 | Red Cross Children's Hospital wc RXH | NHLS/UCT | Arash Iranzadeh, Deelan Doolabh, Lynn Tyers, Bruna Galvao, Innocent Mudau, Marvin Hsiao, Kruger Marais, Diana Hardie, Stephen Korsman, Carolyn Williamson |
| EPI_ISL_640131, EPI_ISL_640132, EPI_ISL_640133, EPI_ISL_640134, | Groote Schuur Hospital wc GSH | NHLS/UCT | Arash Iranzadeh, Deelan Doolabh, Lynn Tyers, Bruna Galvao, Innocent Mudau, Marvin Hsiao, Kruger Marais, Diana Hardie, Stephen Korsman, Carolyn Williamson |

| | | | |
|---|---|---|---|
| EPI_ISL_640135 | | | |
| EPI_ISL_640136 | Red Cross Children's Hospital wc RXH | NHLS/UCT | Arash Iranzadeh, Deelan Doolabh, Lynn Tyers, Bruna Galvao, Innocent Mudau, Marvin Hsiao, Kruger Marais, Diana Hardie, Stephen Korsman, Carolyn Williamson |
| EPI_ISL_641424, EPI_ISL_641425, EPI_ISL_641426, EPI_ISL_641427, EPI_ISL_641428, EPI_ISL_641429, EPI_ISL_641430, EPI_ISL_641431, EPI_ISL_641432, EPI_ISL_641433, EPI_ISL_641434, EPI_ISL_641435, EPI_ISL_641436, EPI_ISL_641437, EPI_ISL_641438, EPI_ISL_641439, EPI_ISL_641440, EPI_ISL_641441, EPI_ISL_641442, EPI_ISL_641443 | | | |
| see above | Department of Virus and Microbiological Special Diagnostics, Statens Serum Institut, Copenhagen, Denmark | Albertsen lab, Department of Chemistry and Bioscience, Aalborg University, Denmark | Thomas Bruun Rasmussen, Jannik Fonager, Morten Rasmussen |
| EPI_ISL_644383, EPI_ISL_644386, EPI_ISL_644389, EPI_ISL_644495, EPI_ISL_644497, EPI_ISL_644498, EPI_ISL_644499, EPI_ISL_644500, EPI_ISL_644501, EPI_ISL_644503, EPI_ISL_644504, EPI_ISL_644505, EPI_ISL_644507, EPI_ISL_644509, EPI_ISL_644510, EPI_ISL_644512, EPI_ISL_644514, EPI_ISL_644515, EPI_ISL_644516, EPI_ISL_644517, EPI_ISL_644518, EPI_ISL_644519, EPI_ISL_644520, EPI_ISL_644521, EPI_ISL_644522, EPI_ISL_644523, EPI_ISL_644524, EPI_ISL_644525, EPI_ISL_644526, EPI_ISL_644527 | | | |
| see above | MEPHI, Aix Marseille University | MEPHI, Aix Marseille University | Anthony LEVASSEUR |
| EPI_ISL_644900, EPI_ISL_644901, EPI_ISL_644921, EPI_ISL_644922, EPI_ISL_644923, EPI_ISL_644924, EPI_ISL_644925, EPI_ISL_644926, EPI_ISL_644927, EPI_ISL_644928, EPI_ISL_644929, EPI_ISL_644930, EPI_ISL_644931 | | | |
| see above | Virginia DCLS | Virginia DCLS | Virginia DCLS |
| EPI_ISL_644936, EPI_ISL_644938, EPI_ISL_644939 | Mayo Clinic & Mayo Clinic Laboratories | Minnesota Department of Health, Public Health Laboratory | Matt Plumb, Jacob Garfin, Alexandra Lorentz, and Xiong Wang |
| EPI_ISL_645240, EPI_ISL_645473, EPI_ISL_645499, EPI_ISL_645508 | Lighthouse Lab in Alderley Park | Wellcome Sanger Institute for the COVID-19 Genomics UK (COG-UK) Consortium | Jacquelyn Wynn, Mairead Hyland, The Lighthouse Lab in Alderley Park and Alex Alderton, Roberto Amato, Sonia Goncalves, Ewan Harrison, David K. Jackson, Ian Johnston, Dominic Kwiatkowski, Cordelia Langford, John Sillitoe on behalf of the Wellcome Sanger Institute COVID-19 Surveillance Team |
| EPI_ISL_648141 | Gavle klinisk mikrobiologi | The Public Health Agency of Sweden | Anna-Malin Linde, Maria Lind Karlberg, Mattias Haukland, Reza Advani, Olov Svartstrom, Oskar Karlsson Lindsjo, Sandra Broddesson, Petra Edquist, Mia Brytting, Anna Risberg, Karin Tegmark-Wisell |
| EPI_ISL_648146 | The Public Health Agency of Sweden | The Public Health Agency of Sweden | Anna-Malin Linde, Maria Lind Karlberg, Mattias Haukland, Reza Advani, Olov Svartstrom, Oskar Karlsson Lindsjo, Sandra Broddesson, Petra Edquist, Mia Brytting, Anna Risberg, Karin Tegmark-Wisell |
| EPI_ISL_648162 | Klinisk mikrobiologi Vasternorrland | The Public Health Agency of Sweden | Anna-Malin Linde, Maria Lind Karlberg, Mattias Haukland, Reza Advani, Olov Svartstrom, Oskar Karlsson Lindsjo, Sandra Broddesson, Petra Edquist, Mia Brytting, Anna Risberg, Karin Tegmark-Wisell |
| EPI_ISL_648164 | Kalmar klinisk mikrobiologi | The Public Health Agency of Sweden | Anna-Malin Linde, Maria Lind Karlberg, Mattias Haukland, Reza Advani, Olov Svartstrom, Oskar Karlsson Lindsjo, Sandra Broddesson, Petra Edquist, Mia Brytting, Anna Risberg, Karin Tegmark-Wisell |
| EPI_ISL_648174, EPI_ISL_648175, EPI_ISL_648197 | The Public Health Agency of Sweden | The Public Health Agency of Sweden | Anna-Malin Linde, Maria Lind Karlberg, Mattias Haukland, Reza Advani, Olov Svartstrom, Oskar Karlsson Lindsjo, Sandra Broddesson, Petra Edquist, Mia Brytting, Anna Risberg, Karin Tegmark-Wisell |
| EPI_ISL_648205, EPI_ISL_648206 | Orebro klinisk mikrobiologi | The Public Health Agency of Sweden | Anna-Malin Linde, Maria Lind Karlberg, Mattias Haukland, Reza Advani, Olov Svartstrom, Oskar Karlsson Lindsjo, Sandra Broddesson, Petra Edquist, Mia Brytting, Anna Risberg, Karin Tegmark-Wisell |
| EPI_ISL_648468, EPI_ISL_648469, EPI_ISL_648470, EPI_ISL_648471, EPI_ISL_648472, EPI_ISL_648473 | Santa Clara County Public Health Laboratory | Chan-Zuckerberg Biohub | CZB Cliahub Consortium |
| EPI_ISL_648537, EPI_ISL_648538 | Tulare County Public Health Lab | Chan-Zuckerberg Biohub | CZB Cliahub Consortium |
| EPI_ISL_648922, EPI_ISL_648931, EPI_ISL_648933, EPI_ISL_648938, EPI_ISL_648945, EPI_ISL_648946, EPI_ISL_648952, EPI_ISL_648958, EPI_ISL_648959, EPI_ISL_648963, EPI_ISL_649026, EPI_ISL_649028, EPI_ISL_649032, EPI_ISL_649034, EPI_ISL_649035, EPI_ISL_649039, EPI_ISL_649045, EPI_ISL_649054 | | | |
| see above | San Diego County Public Health Laboratory | Andersen lab at Scripps Research | SEARCH Alliance San Diego with Tracy Basler, Jovan Shephard, Brett Austin |
| EPI_ISL_650503 | West of Scotland Specialist Virology Centre, NHSGGC / MRC-University of Glasgow Centre for Virus Research | COVID-19 Genomics UK (COG-UK) Consortium | Ana da Silva Filipe, Natasha Johnson, Kathy Smollett, Daniel Mair, Stephen Carmichael, Alice Broos, Lily Tong, Jenna Nichols, Kyriaki Nomikou; Sarah McDonald; Richard Orton, Joseph Hughes, Sreenu Vattipally, David L Robertson; Alasdair MacLean, Rory Gunson; Sharif Shaaban, Matthew Holden; Rachel Blacow, Guy Mollett, Kathy Li, James Shepherd, Antonia Ho, Emma Thomson |
| EPI_ISL_650684 | Northumbria University / South Tees Hospitals NHS Foundation Trust / North Cumbria Integrated Care NHS Foundation Trust / North Tees and Hartlepool NHS Foundation Trust / Newcastle Hospitals NHS Foundation Trust | COVID-19 Genomics UK (COG-UK) Consortium | Darren L Smith,Andrew Nelson,Matthew Bashton,Greg R Young,Joshua Loh,John Allan,Mohammad A Tariq,Giles S Holt,Gary Black,Wen C Yew,Lynn Dover,Paul Baker,Steve Liggett,Sarah Essex,Jane Greenaway,Debra Padgett,Clive Graham,Garren Scott,Edward Barton,Emma Swindells,Brendan Payne,Jennifer Collins,Yusri Taha,Gary Eltringham |
| EPI_ISL_650762 | West of Scotland Specialist Virology Centre, NHSGGC / MRC-University of Glasgow Centre for Virus Research | COVID-19 Genomics UK (COG-UK) Consortium | Ana da Silva Filipe, Natasha Johnson, Kathy Smollett, Daniel Mair, Stephen Carmichael, Alice Broos, Lily Tong, Jenna Nichols, Kyriaki Nomikou; Sarah McDonald; Richard Orton, Joseph Hughes, Sreenu Vattipally, David L Robertson; Alasdair MacLean, Rory Gunson; Sharif Shaaban, Matthew Holden; Rachel Blacow, Guy Mollett, Kathy Li, James Shepherd, Antonia Ho, Emma Thomson |
| EPI_ISL_651126, EPI_ISL_651356 | Northumbria University / South Tees Hospitals NHS Foundation Trust / North Cumbria Integrated Care NHS Foundation Trust / North Tees and Hartlepool NHS Foundation Trust / Newcastle Hospitals NHS Foundation Trust | COVID-19 Genomics UK (COG-UK) Consortium | Darren L Smith,Andrew Nelson,Matthew Bashton,Greg R Young,Joshua Loh,John Allan,Mohammad A Tariq,Giles S Holt,Gary Black,Wen C Yew,Lynn Dover,Paul Baker,Steve Liggett,Sarah Essex,Jane Greenaway,Debra Padgett,Clive Graham,Garren Scott,Edward Barton,Emma Swindells,Brendan Payne,Jennifer Collins,Yusri Taha,Gary Eltringham |
| EPI_ISL_651498, EPI_ISL_651583 | West of Scotland Specialist Virology Centre, NHSGGC / MRC-University of Glasgow Centre for Virus Research | COVID-19 Genomics UK (COG-UK) Consortium | Ana da Silva Filipe, Natasha Johnson, Kathy Smollett, Daniel Mair, Stephen Carmichael, Alice Broos, Lily Tong, Jenna Nichols, Kyriaki Nomikou; Sarah McDonald; Richard Orton, Joseph Hughes, Sreenu Vattipally, David L Robertson; Alasdair MacLean, Rory Gunson; Sharif Shaaban, Matthew Holden; Rachel Blacow, Guy Mollett, Kathy Li, James Shepherd, Antonia Ho, Emma Thomson |
| EPI_ISL_652344 | Northumbria University / South Tees Hospitals NHS Foundation Trust / North Cumbria Integrated Care NHS Foundation Trust / North Tees and Hartlepool NHS Foundation Trust / Newcastle Hospitals NHS Foundation Trust | COVID-19 Genomics UK (COG-UK) Consortium | Darren L Smith,Andrew Nelson,Matthew Bashton,Greg R Young,Joshua Loh,John Allan,Mohammad A Tariq,Giles S Holt,Gary Black,Wen C Yew,Lynn Dover,Paul Baker,Steve Liggett,Sarah Essex,Jane Greenaway,Debra Padgett,Clive Graham,Garren Scott,Edward Barton,Emma Swindells,Brendan Payne,Jennifer Collins,Yusri Taha,Gary Eltringham |
| EPI_ISL_653785, EPI_ISL_653786 | Istituto Zooprofilattico Sperimentale della Puglia e della Basilicata | Istituto Zooprofilattico Sperimentale della Puglia e della Basilicata | Parisi A., Bianco A., Capozzi L., Del Sambro L., Manzulli V, Rondinone V., Pace L., Cipolletta D., Galante D. |
| EPI_ISL_654162, EPI_ISL_654163, EPI_ISL_654164, EPI_ISL_654165, EPI_ISL_654169, EPI_ISL_654171, EPI_ISL_654174, EPI_ISL_654175, EPI_ISL_654176, EPI_ISL_654177, EPI_ISL_654178, EPI_ISL_654180, EPI_ISL_654183, EPI_ISL_654185, EPI_ISL_654194, EPI_ISL_654205, EPI_ISL_654206, EPI_ISL_654207, EPI_ISL_654208, EPI_ISL_654209, EPI_ISL_654210, EPI_ISL_654211, EPI_ISL_654212, EPI_ISL_654213, EPI_ISL_654214, EPI_ISL_654215, EPI_ISL_654216, EPI_ISL_654217, EPI_ISL_654218, EPI_ISL_654219, EPI_ISL_654220, EPI_ISL_654221, EPI_ISL_654222, EPI_ISL_654223, EPI_ISL_654224, EPI_ISL_654225, EPI_ISL_654226, EPI_ISL_654227, EPI_ISL_654228, EPI_ISL_654229, EPI_ISL_654230, EPI_ISL_654283 | | | |
| see above | Hospital General Universitario Gregorio Marañón | SeqCOVID-SPAIN consortium/IBV(CSIC) | Darío García de Viedma, Laura Pérez-Lago, Marta Herranz, Jon Sicilia, Julia Suárez, Pilar Catalán, Patricia Muñoz and SeqCOVID-SPAIN consortium |
| EPI_ISL_654401, EPI_ISL_654404, EPI_ISL_654406, EPI_ISL_654412, EPI_ISL_654414, EPI_ISL_654416, EPI_ISL_654421, EPI_ISL_654422, EPI_ISL_654423, EPI_ISL_654424, EPI_ISL_654425, EPI_ISL_654433, EPI_ISL_654435, EPI_ISL_654437, EPI_ISL_654438, EPI_ISL_654439, EPI_ISL_654441, EPI_ISL_654442, EPI_ISL_654443, EPI_ISL_654444, EPI_ISL_654445, EPI_ISL_654446, EPI_ISL_654447, EPI_ISL_654448, EPI_ISL_654449, EPI_ISL_654450, EPI_ISL_654451, EPI_ISL_654452, EPI_ISL_654453, EPI_ISL_654454 | | | |
| see above | Servicio de Microbiología, Hospital Miguel Servet, Zaragoza | SeqCOVID-SPAIN consortium/IBV(CSIC) | Antonio Rezusta López, Alexander Tristancho Baró, Ana Milagro, Yolanda Gracia Grataloup, Nieves Martínez Cameo and SeqCOVID-SPAIN consortium |
| EPI_ISL_654510, EPI_ISL_654521 | Servicio de Microbiología, Laboratori Clínic Metropolitana Nord. Hospital Universitari Germans Trias i Pujol. Institut d'Investigació en Ciències de la Salut Germans Trias i Pujol (IGTP) | SeqCOVID-SPAIN consortium/IBV(CSIC) | Elisa Martró, Antoni E. Bordoy, Anna Not, Adrián Antuori, Anabel Fernández, Nona Romaní and SeqCOVID-SPAIN consortium |
| EPI_ISL_654553, EPI_ISL_654554 | Servicio de Microbiología. Hospital Universitario Donostia. | SeqCOVID-SPAIN consortium/IBV(CSIC) | Gustavo Cilla Eguiluz, Milagrosa Montes Ros, Luis Piñeiro Vázquez, Ane Sorrarain, Jose Maria Marimón and SeqCOVID-SPAIN consortium |

| | | | |
|---|---|---|---|
| | OSI Donostialdea. Área de Enfermedades Infecciosas, Grupo de Infección Respiratoria y Resistencia Antimicrobiana. Instituto de Investigación Sanitaria Biodonostia | | |
| EPI_ISL_654696, EPI_ISL_654697 | Essentia Health-St. Mary's Medical Center | Minnesota Department of Health, Public Health Laboratory | Matt Plumb, Jacob Garfin, Alexandra Lorentz, and Xiong Wang |
| EPI_ISL_660466, EPI_ISL_660467, EPI_ISL_660504 | Laboratoire de Microbiologie CHU Souro Sanou | Centre Muraz | Abdoul-Salam Ouedraogo, Yacouba Sawadogo, Essia Belarbi, Grit Schubert, Fabian Leendertz, Arsène Zongo, Soumeya Ouangraoua, Zekiba Tarnagda, Lassana Sangaré, Halidou Tinto |
| EPI_ISL_660529, EPI_ISL_660530, EPI_ISL_660531 | Institute of Microbiology, Universidad San Francisco de Quito | Institute of Microbiology, Universidad San Francisco de Quito | Sully Márquez, Belén Prado-Vivar, Juan José Guadalupe, Monica Becerra-Wong, Bernardo Gutiérrez, Manuel Jaramillo, Verónica Barragán, Patricio Rojas-Silva, Gabriel Trueba, Michelle Grunauer, Paúl Cárdenas |
| EPI_ISL_660558 | The National Institute of Public Health | State Veterinary Institute Prague | Nagy,A;Jirincova,H;Novakova,L;Trnka,D;Vecerova,J |
| EPI_ISL_660854, EPI_ISL_660855, EPI_ISL_660856, EPI_ISL_660857, EPI_ISL_660858, EPI_ISL_660859, EPI_ISL_660860, EPI_ISL_660861, EPI_ISL_660862, EPI_ISL_660863, EPI_ISL_660864, EPI_ISL_660865, EPI_ISL_660866, EPI_ISL_660867, EPI_ISL_660868 | | | |
| see above | Gundersen Molecular Diagnostics Laboratory | Kabara Cancer Research Institute | Craig S. Richmond, Paraic A. Kenny |
| EPI_ISL_660869, EPI_ISL_660870, EPI_ISL_660871, EPI_ISL_660872, EPI_ISL_660873, EPI_ISL_660874 | Gundersen Clinical Microbiology Laboratory | Kabara Cancer Research Institute | Craig S. Richmond, Paraic A. Kenny |
| EPI_ISL_661284, EPI_ISL_661285 | Klinisk mikrobiologi Linkoping | The Public Health Agency of Sweden | Department of Microbiology, The Public Health Agency of Sweden |
| EPI_ISL_664281, EPI_ISL_665158, EPI_ISL_665159, EPI_ISL_665160, EPI_ISL_665183, EPI_ISL_665184, EPI_ISL_665185, EPI_ISL_665223 | University College London Hospital | COVID-19 Genomics UK (COG-UK) Consortium | Judith Heaney, Matthew Byott, Catherine Houlihan, Dan Frampton, Stuart Kirk, Moira Spyer and Eleni Nastouli |
| EPI_ISL_665278 | Northumbria University / South Tees Hospitals NHS Foundation Trust / North Cumbria Integrated Care NHS Foundation Trust / North Tees and Hartlepool NHS Foundation Trust / Newcastle Hospitals NHS Foundation Trust | COVID-19 Genomics UK (COG-UK) Consortium | Darren L Smith,Andrew Nelson,Matthew Bashton,Greg R Young,Joshua Loh,John Allan,Mohammad A Tariq,Giles S Holt,Gary Black,Wen C Yew,Lynn Dover,Paul Baker,Steve Liggett,Sarah Essex,Jane Greenaway,Debra Padgett,Clive Graham,Garren Scott,Edward Barton,Emma Swindells,Brendan Payne,Jennifer Collins,Yusri Taha,Gary Eltringham |
| EPI_ISL_667454, EPI_ISL_667455, EPI_ISL_667456, EPI_ISL_667457, EPI_ISL_667458, EPI_ISL_667459, EPI_ISL_667460, EPI_ISL_667461, EPI_ISL_667462, EPI_ISL_667463, EPI_ISL_667464, EPI_ISL_667465, EPI_ISL_667466, EPI_ISL_667467, EPI_ISL_667468, EPI_ISL_667469, EPI_ISL_667470, EPI_ISL_667471, EPI_ISL_667472, EPI_ISL_667473, EPI_ISL_667474, EPI_ISL_667475, EPI_ISL_667476, EPI_ISL_667477, EPI_ISL_667478, EPI_ISL_667479, EPI_ISL_667480, EPI_ISL_667481, EPI_ISL_667482, EPI_ISL_667484 | | | |
| see above | OHSU Lab Services Molecular Microbiology Lab | Oregon SARS-CoV-2 Genome Sequencing Center | Brendan L. O'Connell, Ruth V. Nichols, Sally Grindstaff, Alec J. Hirsch, Donna Hansel, Guang Fan, Daniel N. Streblow, William B. Messer, Andrew C. Adey, Benjamin N. Bimber, Brian J. O'Roak |
| EPI_ISL_667611, EPI_ISL_667612, EPI_ISL_667613, EPI_ISL_667614, EPI_ISL_667615, EPI_ISL_667616, EPI_ISL_667617, EPI_ISL_667618, EPI_ISL_667619, EPI_ISL_667620, EPI_ISL_667637, EPI_ISL_667638, EPI_ISL_667667, EPI_ISL_667668, EPI_ISL_667669, EPI_ISL_667670, EPI_ISL_667671, EPI_ISL_667672, EPI_ISL_667673, EPI_ISL_667674, EPI_ISL_667675, EPI_ISL_667676, EPI_ISL_667677, EPI_ISL_667678, EPI_ISL_667679, EPI_ISL_667680, EPI_ISL_667681, EPI_ISL_667682, EPI_ISL_667683, EPI_ISL_667684, EPI_ISL_667685, EPI_ISL_667686, EPI_ISL_667687, EPI_ISL_667688, EPI_ISL_667689, EPI_ISL_667690, EPI_ISL_667691, EPI_ISL_667692, EPI_ISL_667693, EPI_ISL_667737, EPI_ISL_667738, EPI_ISL_667739 | | | |
| see above | Pathogen Genomics Center, National Institute of Infectious Diseases | Pathogen Genomics Center, National Institute of Infectious Diseases | Tsuyoshi Sekizuka, Kentaro Itokawa, Rina Tanaka, Masanori Hashino, Makoto Kuroda |
| EPI_ISL_668390 | Vestfold Hospital, Toensberg Department of Microbiology | Norwegian Institute of Public Health, Department of Virology | Kathrine Stene-Johansen, Kamilla Heddeland Instefjord, Hilde Elshaug, Marie Paulsen Madsen, Rasmus Riis Kopperud, Hilde Vollan, Karoline Bragstad, Olav Hungnes |
| EPI_ISL_668393 | Akershus University Hospital, Department for Microbiology and Infectious Disease Control | Norwegian Institute of Public Health, Department of Virology | Kathrine Stene-Johansen, Kamilla Heddeland Instefjord, Hilde Elshaug, Marie Paulsen Madsen, Rasmus Riis Kopperud, Hilde Vollan, Karoline Bragstad, Olav Hungnes |
| EPI_ISL_668439 | Ostfold Hospital Trust - Kalnes, Centre for Laboratory Medicine, Section for gene technology and infection serology | Norwegian Institute of Public Health, Department of Virology | Kathrine Stene-Johansen, Kamilla Heddeland Instefjord, Hilde Elshaug, Marie Paulsen Madsen, Rasmus Riis Kopperud, Hilde Vollan, Karoline Bragstad, Olav Hungnes |
| EPI_ISL_671652, EPI_ISL_671655, EPI_ISL_671660, EPI_ISL_671661, EPI_ISL_671662, EPI_ISL_671663, EPI_ISL_671664, EPI_ISL_671668, EPI_ISL_671670, EPI_ISL_671671 | Unity Health Toronto | Ontario Institute for Cancer Research | Ramzi Fattouh, Larissa M. Matukas, Yan Chen,Mark Downing, Trina Otterman, Karel Boissinot, Wai Sum Siu, Zhi Cui, Le Luu, Samira Mubareka, TIBDN, Ilinca Lungu, Bernard Lam, Jeremy Johns, Paul Krzyzanowski, Richard de Borja, Felicia Vincelli, Philip Zuzarte, Jared T. Simpson |
| EPI_ISL_671978 | Laboratorio de Virología y Microbiología Molecular, Depto. de Microbiología, Facultad de Medicina, Universidad de El Salvador/INS-laboratorio de Ref. Ministerio de Salud | Laboratorio de Virología y Microbiología Molecular, Depto. de Microbiología, Facultad de Medicina, Universidad de El Salvador/INS-laboratorio de Ref. Ministerio de Salud | Rivera NR et al |
| EPI_ISL_672012 | Laboratorio de Virología y Microbiología Molecular, Depto. de Microbiología, Facultad de Medicina, Universidad de El Salvador/INS-laboratorio de Ref. Ministerio de Salud. 1*: Dr. Noé Rigoberto Rivera, profesor del Departamento de Bioquímica e Investigador adjunto de la Sección de Virología y Microbiología Molecular; director de la Unidad de Investigaciones Científicas (UNICA), Facultad de Medicina Universidad de El Salvador, El Salvador C.A 1**: Dr. Carlos Alexander Ortega Pérez, Profesor del Departamento de Microbiología; investigador y Jefe de la Sección de Virología, Director Metodológico de la Unidad de Investigaciones Científicas (UNICA), Facultad de Medicina, Universidad de El Salvador, El Salvador C.A 2*: Dra. Xochitl Sandoval López: Directora e investigadora del Instituto Nacional de Salud (INS) El Salvador C.A 2** Dr. Hernandez Avila Carlos E Director de Gobernanza e investigador del Instituto Nacional de Salud (INS) El Salvador C.A | Laboratorio de Virología y Microbiología Molecular, Depto. de Microbiología, Facultad de Medicina, Universidad de El Salvador/INS-laboratorio de Ref. Ministerio de Salud. | Ortega Pérez CA et al |
| EPI_ISL_672033, EPI_ISL_672089 | Santa Clara County Public Health Laboratory | Chan-Zuckerberg Biohub | CZB Cliahub Consortium |
| EPI_ISL_672109 | Orange County Public Health Lab | Chan-Zuckerberg Biohub | CZB Cliahub Consortium |
| EPI_ISL_672111, EPI_ISL_672277, EPI_ISL_672278, EPI_ISL_672279, EPI_ISL_672280, EPI_ISL_672281, EPI_ISL_672282, EPI_ISL_672283 | Santa Clara County Public Health Laboratory | Chan-Zuckerberg Biohub | CZB Cliahub Consortium |
| EPI_ISL_672570 | Laboratorio de Virología y Microbiología Molecular, Depto. de Microbiología, Facultad de Medicina, Universidad de El Salvador/INS-laboratorio de Ref. Ministerio de Salud | Laboratorio de Virología y Microbiología Molecular, Depto. de Microbiología, Facultad de Medicina, Universidad de El Salvador/INS-laboratorio de Ref. Ministerio de Salud | Ortega Pérez CA et al |
| EPI_ISL_672572 | Laboratorio de Virología y Microbiología Molecular, Depto. de | Laboratorio de Virología y Microbiología Molecular, Depto. de | Rivera NR et al |

| | | | |
|---|---|---|---|
| | Microbiología, Facultad de Medicina, Universidad de El Salvador/INS-laboratorio de Ref. Ministerio de Salud. 1*: Dr. Noé Rigoberto Rivera, profesor del Departamento de Bioquímica e Investigador adjunto de la Sección de Virología y Microbiología Molecular; director de la Unidad de Investigaciones Científicas (UNICA), Facultad de Medicina Universidad de El Salvador, El Salvador C.A 1**: Dr. Carlos Alexander Ortega Pérez, Profesor del Departamento de Microbiología; investigador y Jefe de la Sección de Virología, Director Metodológico de la Unidad de Investigaciones Científicas (UNICA), Facultad de Medicina, Universidad de El Salvador, El Salvador C.A 2*: Dra. Xochitl Sandoval López: Directora e investigadora del Instituto Nacional de Salud (INS) El Salvador C.A 2** Dr. Hernandez Avila Carlos E Director de Gobernanza e investigador del Instituto Nacional de Salud (INS) El Salvador C.A | Microbiología, Facultad de Medicina, Universidad de El Salvador/INS-laboratorio de Ref. Ministerio de Salud. | |
| EPI_ISL_672644, EPI_ISL_672645, EPI_ISL_672646, EPI_ISL_672647, EPI_ISL_672648, EPI_ISL_672649, EPI_ISL_672650, EPI_ISL_672651, EPI_ISL_672652, EPI_ISL_672653 | PathWest Laboratory Medicine WA | PathWest Laboratory Medicine WA Microbial Surveillance Unit | PathWest Laboratory Medicine WA Microbial Surveillance Unit |
| EPI_ISL_676493 | Orebro klinisk mikrobiologi | The Public Health Agency of Sweden | Department of Microbiology, The Public Health Agency of Sweden |
| EPI_ISL_676506 | Klinisk mikrobiologi | The Public Health Agency of Sweden | Department of Microbiology, The Public Health Agency of Sweden |
| EPI_ISL_677131, EPI_ISL_677132, EPI_ISL_677133 | Wadsworth Center, New York State Department.of Health | Wadsworth Center, New York State Department.of Health | Kirsten St. George, Daryl M. Lamson, Alexis Russel, Jonathan Plitnick, Navjot Singh, John Kelly, Sara Griesemer, Erasmus Schneider, Erica Lasek-Nesselquist |
| EPI_ISL_677344, EPI_ISL_677346, EPI_ISL_677353, EPI_ISL_677354, EPI_ISL_677355, EPI_ISL_677357, EPI_ISL_677358, EPI_ISL_677359, EPI_ISL_677362, EPI_ISL_677366, EPI_ISL_677368, EPI_ISL_677370, EPI_ISL_677373, EPI_ISL_677375, EPI_ISL_677380, EPI_ISL_677384, EPI_ISL_677385, EPI_ISL_677388, EPI_ISL_677390, EPI_ISL_677394, EPI_ISL_677395, EPI_ISL_677396, EPI_ISL_677397, EPI_ISL_677399, EPI_ISL_677400, EPI_ISL_677401, EPI_ISL_677402, EPI_ISL_677403, EPI_ISL_677404, EPI_ISL_677405, EPI_ISL_677406, EPI_ISL_677407, EPI_ISL_677408, EPI_ISL_677427, EPI_ISL_677429, EPI_ISL_677432, EPI_ISL_677437, EPI_ISL_677438, EPI_ISL_677443, EPI_ISL_677444, EPI_ISL_677446, EPI_ISL_677451, EPI_ISL_677453, EPI_ISL_677465, EPI_ISL_677474, EPI_ISL_677493, EPI_ISL_677494, EPI_ISL_677495, EPI_ISL_677497, EPI_ISL_677509, EPI_ISL_677510, EPI_ISL_677511, EPI_ISL_677512, EPI_ISL_677514, EPI_ISL_677516, EPI_ISL_677517, EPI_ISL_677518, EPI_ISL_677520, EPI_ISL_677525, EPI_ISL_677529, EPI_ISL_677530, EPI_ISL_677534, EPI_ISL_677535, EPI_ISL_677536, EPI_ISL_677620, EPI_ISL_677621, EPI_ISL_677622, EPI_ISL_677623, EPI_ISL_677625, EPI_ISL_677660 | | | |
| see above | University of Wisconsin-Madison AIDS Vaccine Research Laboratories | University of Wisconsin-Madison AIDS Vaccine Research Laboratories | Gage Moreno, Katarina Braun, et al. AIDS Vaccine Research Laboratories |
| EPI_ISL_677727, EPI_ISL_677769, EPI_ISL_677772, EPI_ISL_677773, EPI_ISL_677774, EPI_ISL_677777, EPI_ISL_677782, EPI_ISL_677789, EPI_ISL_677790, EPI_ISL_677795, EPI_ISL_677797, EPI_ISL_677798, EPI_ISL_677799, EPI_ISL_677801, EPI_ISL_677802, EPI_ISL_677803, EPI_ISL_677804, EPI_ISL_677805, EPI_ISL_677815 | | | |
| see above | University of Szeged, Institute of Clinical Microbiology | National Laboratory of Virology, Szentágothai Research Centre | Endre Gábor Tóth, Balázs Somogyi, Brigitta, Gabriella Terhes, Ferenc Jakab, Gábor Kemenesi |
| EPI_ISL_678351 | Area of Virology, Serology and Virology Division (SAViD), New South Wales Health Pathology Randwick | Virology Research Laboratory; Area of Virology, Serology and Virology Division (SAViD), New South Wales Health Pathology Randwick | Foster, C.; Au, J.; Ruiz Silva, M.; Deveson, I.; Bull, R.; Van Hal, S.; Rawlinson, W. |
| EPI_ISL_679627, EPI_ISL_679628 | University College London, Great Ormond Street Hospital for Children NHS Foundation Trust, Imperial College Healthcare NHS Trust | COVID-19 Genomics UK (COG-UK) Consortium | Sergi Castellano, Rachel Williams, Mark Kristiansen, Paola Resende Silva, Sunando Roy, Tony Brooks, Helena Tutill, Paola Niola, Patricia Dyal, Charlotte Williams, Leysa Forrest, Yasmin Panchbhaya, Jacqueline Findlay, Samuel Weeks, Julianne Brown, Kathryn Harris, Paul Randell, James Price, Alison Holmes, Judith Breuer |
| EPI_ISL_679986, EPI_ISL_679987, EPI_ISL_679988, EPI_ISL_679989, EPI_ISL_679990, EPI_ISL_679991, EPI_ISL_679992, EPI_ISL_679993, EPI_ISL_679994, EPI_ISL_679995, EPI_ISL_679996, EPI_ISL_679997, EPI_ISL_679998, EPI_ISL_679999, EPI_ISL_680000, EPI_ISL_680001, EPI_ISL_680002, EPI_ISL_680003, EPI_ISL_680004, EPI_ISL_680005, EPI_ISL_680006, EPI_ISL_680007, EPI_ISL_680008, EPI_ISL_680009, EPI_ISL_680010, EPI_ISL_680011, EPI_ISL_680012, EPI_ISL_680013, EPI_ISL_680014, EPI_ISL_680015 | | | |
| see above | Virology Department, Sheffield Teaching Hospitals NHS Foundation Trust/Department of Infection, Immunity and Cardiovascular Disease, The Medical School, University of Sheffield | COVID-19 Genomics UK (COG-UK) Consortium | Thushan de Silva, Matthew Parker, Nikki Smith, Adri Angyal, Rebecca Brown, Luke Green, Rachel Tucker, Paul Parsons, Danielle Groves, Katie Johnson, Laura Carrilero, Alex Keeley, Dave Partridge, Matthew Wyles, Benjamin Lindsey, Mehmet Yavuz, Mohammad Raza, Cariad Evans |
| EPI_ISL_680231, EPI_ISL_680232, EPI_ISL_680233, EPI_ISL_680234, EPI_ISL_680235, EPI_ISL_680236, EPI_ISL_680237, EPI_ISL_680238, EPI_ISL_680239, EPI_ISL_680240, EPI_ISL_680241, EPI_ISL_680242, EPI_ISL_680243, EPI_ISL_680244, EPI_ISL_680245, EPI_ISL_680246, EPI_ISL_680247, EPI_ISL_680248, EPI_ISL_680249, EPI_ISL_680250, EPI_ISL_680251, EPI_ISL_680252, EPI_ISL_680253, EPI_ISL_680254, EPI_ISL_680255, EPI_ISL_680256, EPI_ISL_680257, EPI_ISL_680259, EPI_ISL_680260, EPI_ISL_680261, EPI_ISL_680262, EPI_ISL_680263, EPI_ISL_680264, EPI_ISL_680274, EPI_ISL_680275, EPI_ISL_680276, EPI_ISL_680277, EPI_ISL_680278, EPI_ISL_680282, EPI_ISL_680292, EPI_ISL_680293, EPI_ISL_680294, EPI_ISL_680295, EPI_ISL_680298, EPI_ISL_680299, EPI_ISL_680305, EPI_ISL_680306, EPI_ISL_680307, EPI_ISL_680308, EPI_ISL_680309, EPI_ISL_680310, EPI_ISL_680311, EPI_ISL_680312, EPI_ISL_680313, EPI_ISL_680314, EPI_ISL_680315, EPI_ISL_680441 | | | |
| see above | Regional Virus Laboratory, Belfast Health and Social Care Trust | COVID-19 Genomics UK (COG-UK) Consortium | Conall McCaughey, James McKenna, Tanya Curran, Susan Feeney, Alison Watt, Ciara Cox, Mairead Connor, Zoltan Molnar, David Simpson, Derek Fairley |
| EPI_ISL_681321 | Pathology and Laboratory Medicine, UW-Madison | Pathology and Laboratory Medicine, UW-Madison | Moreno,G., Braun,K., Baczenas,J.J., Baker,D. |
| EPI_ISL_682330, EPI_ISL_682332, EPI_ISL_682334, EPI_ISL_682335, EPI_ISL_682337, EPI_ISL_682342, EPI_ISL_682344 | NHLS Universitas Academic | UFS Virology | PA Bester, MM Nyaga, P Nthiga, MT Mogotsi, D Goedhals, T de Oliveira |
| EPI_ISL_684014 | Utah Public Health Laboratory | Utah Public Health Laboratory | Erin Young, Kelly Oakeson |
| EPI_ISL_686535, EPI_ISL_686536, EPI_ISL_686537, EPI_ISL_686538, EPI_ISL_686539, EPI_ISL_686540, EPI_ISL_686597 | Respiratory Virus Unit, Microbiology Services Colindale, Public Health England | COVID-19 Genomics UK (COG-UK) Consortium | PHE Covid Sequencing Team |
| EPI_ISL_691616, EPI_ISL_691624, EPI_ISL_691644, EPI_ISL_691651, EPI_ISL_691667, EPI_ISL_691668, EPI_ISL_691678 | Servicio de Microbiología, Hospital Universitario Son Espases | SeqCOVID-SPAIN consortium/IBV(CSIC) | Carla López-Causapé, Jordi Reina, Antonio Oliver and SeqCOVID-SPAIN consortium |
| EPI_ISL_692866, EPI_ISL_692867, EPI_ISL_692901, EPI_ISL_692903, EPI_ISL_692904, EPI_ISL_692906, EPI_ISL_692907, EPI_ISL_692908, EPI_ISL_692909, EPI_ISL_692911, EPI_ISL_692912, EPI_ISL_692913, EPI_ISL_692914, EPI_ISL_692915, EPI_ISL_692916, EPI_ISL_692917, EPI_ISL_692918, EPI_ISL_692919, EPI_ISL_692920, EPI_ISL_692921, EPI_ISL_692922, EPI_ISL_692923, EPI_ISL_692924, EPI_ISL_692925 | | | |
| see above | Massachusetts State Public Health Laboratory | Massachusetts State Public Health Laboratory | Andrew Lang, Timelia Fink, Glen Gallagher, Sandra Smole |
| EPI_ISL_693480 | Central Public Health Laboratory | National Public Health Laboratory, National Centre for Infectious Diseases | Tze Minn Mak, Sophie Octavia, Zhenyang Zhou, Esorom Daoni, Theresa Palou, Lin Cui, Raymond Tzer Pin Lin |
| EPI_ISL_693512 | CNR Virus des Infections Respiratoires - France SUD | CNR Virus des Infections Respiratoires - France SUD | Antonin Bal, Gregory Destras, Gwendolyne Burfin, Quentin Semanas, Martine Valette, Bruno Lina, Laurence Josset |
| EPI_ISL_693692, EPI_ISL_693721, EPI_ISL_693746 | Delaware Public Health Laboratory | Delaware Public Health Laboratory | Gregory Hovan |

| | | | |
|---|---|---|---|
| EPI_ISL_694046 | Los Angeles County Public Health Laboratory | Los Angeles County Public Health Laboratory | Peera Hemarajata et al. |
| EPI_ISL_697794, EPI_ISL_697795 | Institute of Microbiology, Universidad San Francisco de Quito | Institute of Microbiology, Universidad San Francisco de Quito | Andrea Macias, Belén Prado-Vivar, Sully Márquez, Juan José Guadalupe, Monica Becerra-Wong, Bernardo Gutiérrez, Verónica Barragán, Patricio Rojas-Silva, Gabriel Trueba, Michelle Grunauer, Paúl Cárdenas |
| EPI_ISL_700319, EPI_ISL_700320, EPI_ISL_700321, EPI_ISL_700322, EPI_ISL_700323, EPI_ISL_700324, EPI_ISL_700325 | Hematopathology Laboratory, ACTREC, TMC | Hematopathology Laboratory, ACTREC, TMC | Hematopathology Laboratory, ACTREC |
| EPI_ISL_700347 | Child Health Research Foundation | Child Health Research Foundation | Senjuti Saha, Afroza Akter Tanni, Syed Muktadir Al Sium, Roly Malaker, Sharmistha Goswami, Arif Mohammad Tanmoy, Md Hafizur Rahman, Samir K Saha |
| EPI_ISL_700727, EPI_ISL_700731, EPI_ISL_700739 | Texas Department of State Health Services | Texas Department of State Health Services | Rashmi Tuladhar, Bonnie Oh, Jenny Zhang, Maliha Rahman, Anita Pokharel, Myong Koag, Chung Wang, Rachel Lee, Grace Kubin, Mayela Pedrueza, James Daniel Bonser |
| EPI_ISL_705476 | Oxford Viromics, NDM, University of Oxford; Oxford University Hospitals; Basingstoke and North Hampshire Hospital | COVID-19 Genomics UK (COG-UK) Consortium | Tanya Golubchik, David Bonsall, George Macintyre, Amy Trebes, Mariateresa de Cesare, Catrin Moore, Alex Mobbs, Anita Justice, Robert Shaw, Monique Andersson, Timothy Peto, Emma Wise, Nathan Moore, Jessica Lynch, Nick Cortes, Matilde Mori, Stephen Kidd, David Buck, John Todd, Christophe Fraser |
| EPI_ISL_707694 | Los Angeles County Public Health Laboratory | Los Angeles County Public Health Laboratory | P. Hemarajata et al. |
| EPI_ISL_707778 | Medical Research Center, Faculty of Medicine, Syarif Hidayatullah State Islamic University Jakarta | Medical Research Center, Faculty of Medicine, Syarif Hidayatullah State Islamic University Jakarta | Erike A Suwarsono, Chris Adhiyanto, Laifa Hendarmin, Zeti Harriyati, Rini Puspitaningrum, Ferania Mela, Dennis Nurjadi |
| EPI_ISL_708395, EPI_ISL_708429, EPI_ISL_708430 | Delaware Public Health Lab | Delaware Public Health Lab | Gregory Hovan |
| EPI_ISL_709947 | Lighthouse Lab in Milton Keynes | Wellcome Sanger Institute for the COVID-19 Genomics UK (COG-UK) Consortium | The Lighthouse Lab in Milton Keynes and Alex Alderton, Roberto Amato, Sonia Goncalves, Ewan Harrison, David K. Jackson, Ian Johnston, Dominic Kwiatkowski, Cordelia Langford, John Sillitoe on behalf of the Wellcome Sanger Institute COVID-19 Surveillance Team |
| EPI_ISL_710265, EPI_ISL_710278, EPI_ISL_710348 | Colorado Department of Public Health and Environment | Colorado Department of Puplic Health and Environment | Laura Bankers, Molly C. Hetherington-Rauth, Shannon Ely, Shannon R. Matzinger, Sarah Elizabeth Totten, Emily A. Travanty |
| EPI_ISL_710552, EPI_ISL_710553, EPI_ISL_710565, EPI_ISL_710566, EPI_ISL_710567, EPI_ISL_710568, EPI_ISL_710569, EPI_ISL_710570 | University Hospital Dubrava | Ruer Boškovic Institute; Forensic Science Centre Ivan Vueti; University of Zagreb Faculty of Science | Robert Beluži, Marina Korolija, Ana Livun, Vjekoslav Tomai, Dunja Glavaš, Maja Kuzman, Paula Štancl, Lucija Markulin, Lucija Basi, Antonela Blažekovi, Fran Boroveki, Lidija Cvetko-Krajinovi, Ivana elap, Fuad osovi, Mirjana Domazet-Lošo, Tomislav Domazet-Lošo, Valentina umlijan-Combaj, Kristina Gotovac Jerei, Jasna Kašman, Vladimir Krajinovi, Danilo Licastro, Boris Maek, Željka Maak Šafranko, Gordana Maravi Vlahoviek, Senica Pejša, Josipa Skelin, Ivan Šamija, Mario Štefanovi, Sanja Tadinac, Katarina Marija Tupek, Petra Vrabec, Rosa Karli, Kristian Vlahoviek |
| EPI_ISL_712077, EPI_ISL_712078, EPI_ISL_712092 | East London NHLS Laboratory, Eastern Cape, South Africa | National Institute for Communicable Diseases of the National Health Laboratory Service | Mohale T, Ntuli N, Mahlangu B, Allam M, Ismail A, Bhiman JN |
| EPI_ISL_712568 | Laboratoire de Microbiologie- CHU Habib Bourguiba - Sfax adresse | Laboratoire des Procédés de Criblage Moléculaire et Cellulaire-Centre de Biotechnologie de Sfax | Souissi,A., Abid,N., Ben Ayed,I., Gargouri,S., Abdelmoulah,F.,Elargoubi,A., Smeti,I., Bensaid,M., Stambouli,N., Kharat,N., Ajili,F., Fki-berrajah,L., Mhalla,S., Chtourou,A., Gaaloul,I., Nabli,A., Turki,M., Aouni,M., Hammami,A., Mastouri,M., Karray Hakim,H., Kamoun,S., Rebai,A. and Masmoudi,S. |
| EPI_ISL_717777, EPI_ISL_717778, EPI_ISL_717779, EPI_ISL_717780, EPI_ISL_717781 | UW Virology Lab | UW Virology Lab | Pavitra Roychoudhury, Hong Xie, Lasata Shrestha, Michelle Lin, Meei-Li Huang, Keith R Jerome, Alexander Greninger |
| EPI_ISL_717809 | LACEN Dr. Francisco Rimolo Neto | Bioinformatics Laboratory / LNCC | Carolina M Voloch, Ronaldo da Silva F Jr, Luiz G P de Almeida, Cynthia C Cardoso, Otavio Bustrolini, Alexandra L Gerber, Ana Paula de C Guimarães, Diana Mariani, Andréa Cony Cavalcanti, Claudia dos Santos Rodrigues, Terezinha M P P Castiñeira, Amilcar Tanuri, Ana Tereza R de Vasconcelos |
| EPI_ISL_718214, EPI_ISL_718215 | Ministry of Health Hospitals | Institute of Health and Community Medicine | David Perera, Ooi Mong How, Chua Hock Hin, Tonnii Sia Loong Loong, Wong Jyn Shan, Wong Kiing Aik, Chan Chia Jui |
| EPI_ISL_718230 | Hospital | National Reference Center for Viruses of Respiratory Infections, Institut Pasteur, Paris | Marion Barbet, Sylvie Behillil, Méline Bizard, Angela Brisebarre, Camille Capel, Etienne Simon-Lorière, Vincent Enouf, Maud Vanpeene, Sylvie van der Werf, Gisèle Lagathu |
| EPI_ISL_718285, EPI_ISL_718286, EPI_ISL_718310, EPI_ISL_718312, EPI_ISL_718313 | Institute for Medical Research, Infectious Disease Research Centre, National Institutes of Health, Ministry of Health Malaysia | Institute for Medical Research, Infectious Disease Research Centre, National Institutes of Health, Ministry of Health Malaysia | Suppiah J, Kamel K, Mohd-Zawawi Z, Thayan R |
| EPI_ISL_722224, EPI_ISL_722225, EPI_ISL_722226, EPI_ISL_722227, EPI_ISL_722255, EPI_ISL_722256, EPI_ISL_722257, EPI_ISL_722260, EPI_ISL_722261 | Servicio de Microbiología, Hospital Miguel Servet, Zaragoza | SeqCOVID-SPAIN consortium/IBV(CSIC) | Antonio Rezusta López, Alexander Tristancho Baró, Ana Milagro, Yolanda Gracia Grataloup, Nieves Martínez Cameo and SeqCOVID-SPAIN consortium |
| EPI_ISL_722274, EPI_ISL_722278, EPI_ISL_722294, EPI_ISL_722297, EPI_ISL_722310, EPI_ISL_722353, EPI_ISL_722403, EPI_ISL_722413, EPI_ISL_722492, EPI_ISL_722493, EPI_ISL_722494, EPI_ISL_722495, EPI_ISL_722829, EPI_ISL_722830, EPI_ISL_722831, EPI_ISL_722832, EPI_ISL_722833, EPI_ISL_722834, EPI_ISL_722835, EPI_ISL_722836, EPI_ISL_722837, EPI_ISL_722838, EPI_ISL_722839, EPI_ISL_722840, EPI_ISL_722841 | | | |
| see above | Dutch COVID-19 response team | Erasmus Medical Center | Bas Oude Munnink, Reina Sikkema, David Nieuwenhuijse, Irina Chestakova, Anne van der Linden, Marjan Boter, Emmanuelle Munger, Corine GeurtsvanKessel, Annemiek van der Eijk, Richard Molenkamp, Marion Koopmans, on behalf of the Dutch national COVID-19 response team. |
| EPI_ISL_724967, EPI_ISL_724968, EPI_ISL_724969, EPI_ISL_724978 | Northumbria University / South Tees Hospitals NHS Foundation Trust / North Cumbria Integrated Care NHS Foundation Trust / North Tees and Hartlepool NHS Foundation Trust / Newcastle Hospitals NHS Foundation Trust | COVID-19 Genomics UK (COG-UK) Consortium | Darren L Smith,Andrew Nelson,Matthew Bashton,Greg R Young,Joshua Loh,John Allan,Mohammad A Tariq,Giles S Holt,Gary Black,Wen C Yew,Lynn Dover,Paul Baker,Steve Liggett,Sarah Essex,Jane Greenaway,Debra Padgett,Clive Graham,Garren Scott,Edward Barton,Emma Swindells,Brendan Payne,Jennifer Collins,Yusri Taha,Gary Eltringham |
| EPI_ISL_727705, EPI_ISL_727707, EPI_ISL_727712 | Centre for Enzyme Innovation, University of Portsmouth / Translational Research Laboratory, Portsmouth Hospitals NHS Trust | COVID-19 Genomics UK (COG-UK) Consortium | Angela Beckett,Yann Bourgeois,Garry Scarlett,Sharon Glaysher,Scott Elliott,Kelly Bicknell,Robert Impey,Allyson Lloyd,Sarah Wyllie,Ethan Butcher,Anoop Chauhan,Samuel Robson |
| EPI_ISL_728013, EPI_ISL_728018, EPI_ISL_728019, EPI_ISL_728024, EPI_ISL_728025, EPI_ISL_728026, EPI_ISL_728093 | University of Wisconsin-Madison AIDS Vaccine Research Laboratories | University of Wisconsin-Madison AIDS Vaccine Research Laboratories | Gage Moreno, Katarina Braun, et al. AIDS Vaccine Research Laboratories |
| EPI_ISL_728157, EPI_ISL_728158, EPI_ISL_728161, EPI_ISL_728162, EPI_ISL_728163 | Institute for Medical Research, Infectious Disease Research Centre, National Institutes of Health, Ministry of Health Malaysia | Institute for Medical Research, Infectious Disease Research Centre, National Institutes of Health, Ministry of Health Malaysia | Suppiah J, Kamel K, Mohd-Zawawi Z, Thayan R |
| EPI_ISL_729374, EPI_ISL_729403, EPI_ISL_729419, EPI_ISL_729610, EPI_ISL_729611, EPI_ISL_729615, EPI_ISL_729616 | A. Krumbholz, Labor Dr. Krause und Kollegen MVZ GmbH, Kiel | Charité Universitätsmedizin Berlin, Institut für Virologie | Victor M Corman, Barbara Mühlemann, Jörn Beheim-Schwarzbach, Talitha Veith, Julia Schneider, Terry Jones, Christian Drosten |
| EPI_ISL_729870, EPI_ISL_729903, EPI_ISL_729907 | Instituto de Medicina Tropical, Universidad Nacional Toribio Rodríguez de Mendoza de Amazonas | Laboratorio de Genómica Microbiana, Universidad Peruana Cayetano Heredia | Pablo Tsukayama, Alejandra Dávila-Barclay, Luis González, Pedro E. Romero, Brenda Ayzanoa, Janet Huancachoque, Pool Marcos, Stella Chenet, Rafael Tapia, Cecilia Pajuelo, Carla Montenegro |
| EPI_ISL_730085, EPI_ISL_730092, EPI_ISL_730097, EPI_ISL_730098, | San Diego County Public Health Laboratory | Andersen lab at Scripps Research | SEARCH Alliance San Diego with Tracy Basler, Jovan Shephard, Brett Austin |

| | | | |
|---|---|---|---|
| EPI_ISL_730105, EPI_ISL_730106, EPI_ISL_730107, EPI_ISL_730120, EPI_ISL_730345 | | | |
| EPI_ISL_730567 | Gazi University Faculty of Medicine, Medical Virology Laboratory | Gazi University Faculty of Medicine, Medical Virology Laboratory | Erdem ahin, Hager Muftah, Selin Yiit, Shaknoza Sarzhanova, Özlem Güzel Tunçcan, Murat Dizbay, Il Fidan, Kayhan Çalar, Gülendam Bozday |
| EPI_ISL_730580 | Home Quarantine Taskforce | Hong Kong Department of Health | Mak Gannon C.K., Lam Edman T.K., Chan Rickjason C.W., Tsang Dominic N.C. |
| EPI_ISL_730587 | Pamela Youde Nethersole Eastern Hospital | Hong Kong Department of Health | Mak Gannon C.K., Lam Edman T.K., Chan Rickjason C.W., Tsang Dominic N.C. |
| EPI_ISL_730588, EPI_ISL_730590 | Prince of Wales Hospital | Hong Kong Department of Health | Mak Gannon C.K., Lam Edman T.K., Chan Rickjason C.W., Tsang Dominic N.C. |
| EPI_ISL_730592 | Princess Margaret Hospital | Hong Kong Department of Health | Mak Gannon C.K., Lam Edman T.K., Chan Rickjason C.W., Tsang Dominic N.C. |
| EPI_ISL_730597 | Queen Mary Hospital | Hong Kong Department of Health | Mak Gannon C.K., Lam Edman T.K., Chan Rickjason C.W., Tsang Dominic N.C. |
| EPI_ISL_730614 | United Christian Hospital | Hong Kong Department of Health | Mak Gannon C.K., Lam Edman T.K., Chan Rickjason C.W., Tsang Dominic N.C. |
| EPI_ISL_732517 | National Virus Reference Laboratory | National Virus Reference Laboratory | Michael Carr, Gabriel Gonzalez, Jonathan Dean, Daniel Hare, Cillian F De Gascun |
| EPI_ISL_732694, EPI_ISL_732695 | CNR Virus des Infections Respiratoires - France SUD | CNR Virus des Infections Respiratoires - France SUD | Antonin Bal, Gregory Destras, Claudia Gonzalez, Gwendolyne Burfin, Quentin Semanas, Martine Valette, Bruno Lina, Laurence Josset |
| EPI_ISL_732784, EPI_ISL_732794, EPI_ISL_732804 | Centro de Investigación Biomédica de La Rioja - Hospital San Pedro Logroño | SeqCOVID-SPAIN consortium/IBV(CSIC) | María de Toro, José Manuel Azcona Gutiérrez, María Pilar Bea Escudero, Miriam Blasco Alberdi and SeqCOVID-SPAIN consortium |
| EPI_ISL_732820, EPI_ISL_732821, EPI_ISL_732822, EPI_ISL_732823, EPI_ISL_732824, EPI_ISL_732825, EPI_ISL_732826 | Nebraska Public Health Laboratory | UNMC COVID-19 Response Team | UNMC COVID-19 Response Team |
| EPI_ISL_733026, EPI_ISL_733028, EPI_ISL_733029, EPI_ISL_733030, EPI_ISL_733031, EPI_ISL_733033, EPI_ISL_733038, EPI_ISL_733040, EPI_ISL_733042, EPI_ISL_733047, EPI_ISL_733051, EPI_ISL_733052, EPI_ISL_733056, EPI_ISL_733059, EPI_ISL_733060, EPI_ISL_733064, EPI_ISL_733065, EPI_ISL_733066, EPI_ISL_733068, EPI_ISL_733074, EPI_ISL_733079, EPI_ISL_733080, EPI_ISL_733081, EPI_ISL_733082, EPI_ISL_733093, EPI_ISL_733094, EPI_ISL_733095, EPI_ISL_733096, EPI_ISL_733097 | | | |
| see above | HELIX LLC | WHO National Influenza Centre Russian Federation | Andrey Komissarov, Artem Fadeev, Anna Ivanova, Kseniya Komissarova, Dmitry Bazhenov, Daria Danilenko, Ksenia Safina, Elena Nabieva, Georgii Bazykin, Dmitry Lioznov |
| EPI_ISL_733190, EPI_ISL_733192, EPI_ISL_733208, EPI_ISL_733220 | Pathogenic Microorganisms Variability Laboratory | WHO National Influenza Centre Russian Federation | Andrey Komissarov, Artem Fadeev, Anna Ivanova, Kseniya Komissarova, Dmitry Bazhenov, Daria Danilenko, Ksenia Safina, Elena Nabieva, Georgii Bazykin, Nadezhda Kuznetsova, Elena Shidlovskaya, Sergey Alkhovsky, Tatyana Vishnevskaya, Elizaveta Divisenko, Alexey Shchetinin, Maria Nikiforova, Andrey Pochtovyy, Evgeny Usachev, Elena Vokalova, Maxim Rubalsky, Oleg Rubalsky, Artem Tkachuk, Vladimir Gushchin, Alexander Gintsburg, Dmitry Lioznov |
| EPI_ISL_733233, EPI_ISL_733235, EPI_ISL_733236 | UMMC-Health | WHO National Influenza Centre Russian Federation | Andrey Komissarov, Artem Fadeev, Anna Ivanova, Kseniya Komissarova, Dmitry Bazhenov, Tatiana Platonova, Daria Danilenko, Ksenia Safina, Elena Nabieva, Georgii Bazykin, Dmitry Lioznov |
| EPI_ISL_733242, EPI_ISL_733261, EPI_ISL_733262, EPI_ISL_733263, EPI_ISL_733296, EPI_ISL_733297, EPI_ISL_733394, EPI_ISL_733403 | WHO National Influenza Centre Russian Federation | WHO National Influenza Centre Russian Federation | Andrey Komissarov, Artem Fadeev, Anna Ivanova, Kseniya Komissarova, Dmitry Bazhenov, Daria Danilenko, Ksenia Safina, Elena Nabieva, Georgii Bazykin, Dmitry Lioznov |
| EPI_ISL_733405, EPI_ISL_733406, EPI_ISL_733407, EPI_ISL_733408, EPI_ISL_733409, EPI_ISL_733410, EPI_ISL_733411, EPI_ISL_733412 | UMMC-Health | WHO National Influenza Centre Russian Federation | Andrey Komissarov, Artem Fadeev, Anna Ivanova, Kseniya Komissarova, Dmitry Bazhenov, Tatiana Platonova, Daria Danilenko, Ksenia Safina, Elena Nabieva, Georgii Bazykin, Dmitry Lioznov |
| EPI_ISL_734309, EPI_ISL_734310, EPI_ISL_734311, EPI_ISL_734312, EPI_ISL_734313 | Wadsworth Center, New York State Department.of Health | Wadsworth Center, New York State Department.of Health | Kirsten St. George, Daryl M. Lamson, Alexis Russel, Jonathan Plitnick, Navjot Singh, John Kelly, Sara Griesemer, Erasmus Schneider, Erica Lasek-Nesselquist |
| EPI_ISL_735451 | UW Virology Lab | UW Virology Lab | Pavitra Roychoudhury, Hong Xie, Lasata Shrestha, Meei-Li Huang, Keith R Jerome, Alexander Greninger |
| EPI_ISL_738038, EPI_ISL_738039, EPI_ISL_738040, EPI_ISL_738041, EPI_ISL_738042, EPI_ISL_738043 | Uganda Central Public Health Lab and Uganda Virus Research Institute | MRC/UVRI & LSHTM Uganda Research Unit | Matthew Cotten, Dan Lule Bugembe, My V.T. Phan, Pontiano Kaleebu et al. |
| EPI_ISL_739086, EPI_ISL_739264, EPI_ISL_739297, EPI_ISL_739489 | Humboldt County Public Health Laboratory | Chan-Zuckerberg Biohub | CZB Cliahub Consortium |
| EPI_ISL_739795, EPI_ISL_740027, EPI_ISL_740222, EPI_ISL_740226, EPI_ISL_740291 | Laboratoire national de santé, Microbiology, Virology | Laboratoire national de santé, Microbiology, Microbial Genomics Platform | Anke Wienecke-Baldacchino, Catherine Ragimbeau,Jessica Tapp, Fatu Djabi, Lise Pignon, Raoul Salmon, Tamir Abdelrahman |
| EPI_ISL_740548, EPI_ISL_740549, EPI_ISL_740550, EPI_ISL_740551, EPI_ISL_740552, EPI_ISL_740553, EPI_ISL_740554, EPI_ISL_740555, EPI_ISL_740556, EPI_ISL_740557, EPI_ISL_740558, EPI_ISL_740559, EPI_ISL_740560, EPI_ISL_740561, EPI_ISL_740562, EPI_ISL_740563, EPI_ISL_740564, EPI_ISL_740565, EPI_ISL_740566, EPI_ISL_740567, EPI_ISL_740568, EPI_ISL_740569, EPI_ISL_740570, EPI_ISL_740571, EPI_ISL_740572, EPI_ISL_740573, EPI_ISL_740574, EPI_ISL_740575, EPI_ISL_740576, EPI_ISL_740577, EPI_ISL_740578, EPI_ISL_740579, EPI_ISL_740580, EPI_ISL_740581, EPI_ISL_740582, EPI_ISL_740583, EPI_ISL_740584, EPI_ISL_740585, EPI_ISL_740586, EPI_ISL_740587, EPI_ISL_740588, EPI_ISL_740589, EPI_ISL_740590, EPI_ISL_740591, EPI_ISL_740592, EPI_ISL_740593, EPI_ISL_740594, EPI_ISL_740595, EPI_ISL_740596, EPI_ISL_740597, EPI_ISL_740598, EPI_ISL_740599, EPI_ISL_740600, EPI_ISL_740601, EPI_ISL_740602, EPI_ISL_740603, EPI_ISL_740604, EPI_ISL_740605, EPI_ISL_740606, EPI_ISL_740607, EPI_ISL_740608, EPI_ISL_740609, EPI_ISL_740610, EPI_ISL_740611, EPI_ISL_740612, EPI_ISL_740613, EPI_ISL_740614, EPI_ISL_740615, EPI_ISL_740616, EPI_ISL_740617, EPI_ISL_740618, EPI_ISL_740619, EPI_ISL_740620, EPI_ISL_740621, EPI_ISL_740622, EPI_ISL_740655, EPI_ISL_740677, EPI_ISL_740681, EPI_ISL_740682, EPI_ISL_740684, EPI_ISL_740767, EPI_ISL_740777, EPI_ISL_740778, EPI_ISL_740779, EPI_ISL_740780, EPI_ISL_740781, EPI_ISL_740784, EPI_ISL_740785, EPI_ISL_740786, EPI_ISL_740787, EPI_ISL_740788, EPI_ISL_740789 | | | |
| see above | BCCDC Public Health Laboratory | BCCDC Public Health Laboratory | Prystajecky Natalie, Linda Hoang, Dan Fornika, Shannon Russell, Kim Macdonald, Kimia Kamelian, John Tyson, Inna Sekirov, Mel Krajden |
| EPI_ISL_741347 | Wales Specialist Virology Centre Sequencing lab: Pathogen Genomics Unit | COVID-19 Genomics UK (COG-UK) Consortium | Catherine Moore, Johnathan Evans, Laura Gifford, Malorie Perry, Simon Cottrell, Angela Marchbank, Alec Birchley, Alexander Adams, Amy Gaskin, Bree Gatica-Wilcox, Jason Coombes, Joel Southgate, Lauren Gilbert, Lee Graham, Nicole Pacchiarini, Sara Kumziene-Summerhayes, Sarah Taylor, Sophie Jones, Sara Rey, Matthew Bull, Joanne Watkins, Sally Corden, Tom Connor |
| EPI_ISL_744484, EPI_ISL_744521, EPI_ISL_744761, EPI_ISL_744861, EPI_ISL_744994, EPI_ISL_745023 | Laboratoire national de santé, Microbiology, Virology | Laboratoire national de santé, Microbiology, Microbial Genomics Platform | Anke Wienecke-Baldacchino, Catherine Ragimbeau,Jessica Tapp, Fatu Djabi, Lise Pignon, Raoul Salmon, Tamir Abdelrahman |
| EPI_ISL_745639, EPI_ISL_746121, EPI_ISL_746124 | Ginkgo Bioworks Clinical Laboratory | Utah Public Health Laboratory | Erin L. Young, Kelly Oakeson, Tara Gallagher, Michael T. Pyne, E. Susan Slechta, Melanie A. Mallory, Jeffrey B. Stevenson, Salika M. Shakir, David R. Hillyard, Malaika McKenzie-Bennett, James McGann, Jim Griffin, Keith Robison, Alex Plocik, Becky Schilling, Martha Pierson, Rebecca Littlefield, Michelle Spencer, Birgitte Simen |
| EPI_ISL_746480, EPI_ISL_746483, EPI_ISL_746491, EPI_ISL_746492, EPI_ISL_746498, EPI_ISL_746501, EPI_ISL_746502, EPI_ISL_746511, EPI_ISL_746512, EPI_ISL_746521, EPI_ISL_746522, EPI_ISL_746523, EPI_ISL_746524, EPI_ISL_746525, EPI_ISL_746526, EPI_ISL_746527, EPI_ISL_746528, EPI_ISL_746529, EPI_ISL_746742, EPI_ISL_746743, EPI_ISL_746744, EPI_ISL_746745, EPI_ISL_746746, EPI_ISL_746747, EPI_ISL_746748, EPI_ISL_746749, EPI_ISL_746750, EPI_ISL_746751, EPI_ISL_746752, EPI_ISL_746753, EPI_ISL_746754, EPI_ISL_746755, EPI_ISL_746756 | | | |
| see above | Genetica Molecular and Subdepartamento de Virologia ISP Chile | Instituto de Salud Publica de Chile | Javier Tognarelli, Barbara Parra, Loredana Arata, Jaime Lagos, Gisselle Barra, Patricia Bustos, Rodrigo Fasce, Andres Castillo, Jorge Fernandez |
| EPI_ISL_747039, EPI_ISL_747040 | Respiratory Viruses Branch, Centers for Disease Control and Prevention | Respiratory Viruses Branch, Centers for Disease Control and Prevention | Queen,K., Li,Y., Tao,Y., Uehara,A., Montmayeur,A., Paden,C.R., Cook,P.W., Marine,R., Sheth,M., Wang,H., Lee,J., Tong,S. |
| EPI_ISL_747237 | National Institute of Health Research and Development | National Institute of Health Research and Development | Subangkit,Nafiandi,Estaria,V;Pawestri,HA;Puspa,KD;Nugraha,AA;Ikawati,HD;Pangesti,KNA;Soekarso,T;Paisal;Puspandari,N;Setiawaty,V |
| EPI_ISL_747307, EPI_ISL_747315, EPI_ISL_747316, EPI_ISL_747328, EPI_ISL_747331, EPI_ISL_747343, EPI_ISL_747344, EPI_ISL_747345, EPI_ISL_747346, EPI_ISL_747347, EPI_ISL_747348, EPI_ISL_747349, EPI_ISL_747350, EPI_ISL_747351, EPI_ISL_747352, EPI_ISL_747353, EPI_ISL_747354, EPI_ISL_747355 | | | |

| | | | |
|---|---|---|---|
| see above | Division of Emerging Infectious Diseases, Bureau of Infectious Diseases Diagnosis Control, Korea Disease Control and Prevention Agency | Division of Emerging Infectious Diseases, Bureau of Infectious Diseases Diagnosis Control, Korea Disease Control and Prevention Agency | Ae Kyung Park, Il-Hwan Kim, Heui Man Kim, Jeong-Min Kim, Namjoo Lee, Chaeyoung Lee, Sang Hee Woo, Eun-Jin Kim |
| EPI_ISL_747597, EPI_ISL_748132, EPI_ISL_748133, EPI_ISL_748134, EPI_ISL_748135 | Department of Virus and Microbiological Special Diagnostics, Statens Serum Institut, Copenhagen, Denmark | Albertsen Lab, Department of Chemistry and Bioscience, Aalborg University, Denmark | Danish Covid-19 Genome Consortium |
| EPI_ISL_751499 | CHU Purpan - Laboratoire de Virologie - Institut Fédératif de Biologie | CHU Purpan - Laboratoire de Virologie - Institut Fédératif de Biologie | Latour J., Ranger N., Dubois M., Carcenac R., Harter A., Boyer P., Tremeaux P., Izopet J. |
| EPI_ISL_752912, EPI_ISL_752913, EPI_ISL_752914, EPI_ISL_752915, EPI_ISL_752916, EPI_ISL_752917, EPI_ISL_752918, EPI_ISL_752919, EPI_ISL_752920, EPI_ISL_752921, EPI_ISL_752922, EPI_ISL_752933, EPI_ISL_752934, EPI_ISL_752935, EPI_ISL_752936, EPI_ISL_752937, EPI_ISL_752938, EPI_ISL_752939, EPI_ISL_752940, EPI_ISL_752972, EPI_ISL_752973, EPI_ISL_752974, EPI_ISL_752975, EPI_ISL_752976, EPI_ISL_752977, EPI_ISL_753029, EPI_ISL_753030, EPI_ISL_753123, EPI_ISL_753149 | | | |
| see above | State Laboratories Division, Hawaii State Department of Health | State Laboratories Division, Hawaii State Department of Health | Pamela O'Brien, Sabrina Diemert, Drew Kuwazaki, Razvan Sultana, Edward Desmond |
| EPI_ISL_753731, EPI_ISL_753743, EPI_ISL_753751, EPI_ISL_753777, EPI_ISL_753786, EPI_ISL_753787, EPI_ISL_753788, EPI_ISL_753819, EPI_ISL_753820, EPI_ISL_753821, EPI_ISL_753837, EPI_ISL_753860, EPI_ISL_753876, EPI_ISL_753879, EPI_ISL_753884, EPI_ISL_753885, EPI_ISL_753886, EPI_ISL_753907, EPI_ISL_754021 | | | |
| see above | Charité Universitätsmedizin Berlin, Institut für Virologie/Labor Berlin | Charité Universitätsmedizin Berlin, Institut für Virologie | Victor M Corman, Jörn Beheim-Schwarzbach, Barbara Mühlemann, Julia Schneider, Talitha Veith, Terry Jones, Christian Drosten |
| EPI_ISL_754182 | Wyoming Public Health Laboratory | Wyoming Public Health Laboratory | Noah Hull, Taylor Fearing, Channing Weber, Ashley Norberg, Bailey Bowcutt, and Wanda Manley |
| EPI_ISL_754185 | Charité Universitätsmedizin Berlin, Institut für Virologie/Labor Berlin | Charité Universitätsmedizin Berlin, Institut für Virologie | Victor M Corman, Jörn Beheim-Schwarzbach, Barbara Mühlemann, Julia Schneider, Talitha Veith, Terry Jones, Christian Drosten |
| EPI_ISL_754393 | Genetica Molecular and Subdepartamento de Virologia ISP Chile | Insituto de Salud Publica de Chile | Javier Tognarelli, Barbara Parra, Loredana Arata, Jaime Lagos, Gisselle Barra, Patricia Bustos, Rodrigo Fasce, Andres Castillo, Jorge Fernandez |
| EPI_ISL_754526, EPI_ISL_754527, EPI_ISL_754528, EPI_ISL_754529, EPI_ISL_754530, EPI_ISL_754531, EPI_ISL_754532 | Wadsworth Center, New York State Department.of Health | Wadsworth Center, New York State Department.of Health | Kirsten St. George, Daryl M. Lamson, Alexis Russel, Matthew Shudt, Melissa A Leisner, Jonathan Plitnick, Navjot Singh, John Kelly, Sara Griesemer, Erasmus Schneider, Erica Lasek-Nesselquist |
| EPI_ISL_754902 | Laboratory Diagnostics and Clinical Immunology of Developmental Age, Medical University of Warsaw | genXone SA, Research & Development Laboratory; The Faculty of Mathematics, Informatics and Mechanics of the University of Warsaw | Maciej Sykulski, Grzegorz Nowicki, Monika Makowska-Woniak, Jakub Grabowski, Natalia Drwska-Matelska, ukasz Krych, Micha Kaszuba, Anna Gambin, Urszula Demkow |
| EPI_ISL_754915, EPI_ISL_754917, EPI_ISL_754921, EPI_ISL_754923, EPI_ISL_754931, EPI_ISL_754934, EPI_ISL_754952, EPI_ISL_754974, EPI_ISL_754975, EPI_ISL_754979, EPI_ISL_754981, EPI_ISL_754995, EPI_ISL_755016, EPI_ISL_755027, EPI_ISL_755031, EPI_ISL_755032, EPI_ISL_755057, EPI_ISL_755058, EPI_ISL_755059 | | | |
| see above | California Department of Public Health | California Department of Public Health | CDPH IDLB COVIDNet |
| EPI_ISL_755386, EPI_ISL_755387, EPI_ISL_755388, EPI_ISL_755389, EPI_ISL_755390, EPI_ISL_755391, EPI_ISL_755392, EPI_ISL_755393, EPI_ISL_755394, EPI_ISL_755395, EPI_ISL_755396, EPI_ISL_755397, EPI_ISL_755398, EPI_ISL_755399, EPI_ISL_755400, EPI_ISL_755401, EPI_ISL_755402, EPI_ISL_755403, EPI_ISL_755404, EPI_ISL_755405, EPI_ISL_755406, EPI_ISL_755407, EPI_ISL_755408, EPI_ISL_755409, EPI_ISL_755410, EPI_ISL_755411, EPI_ISL_755412, EPI_ISL_755413, EPI_ISL_755414, EPI_ISL_755415, EPI_ISL_755416, EPI_ISL_755417, EPI_ISL_755418, EPI_ISL_755419 | | | |
| see above | Maine Health and Environmental Testing Laboratory | Tewhey Lab, The Jackson Laboratory | Matluk,N., Dewey,H., Iosue,F., Barter,M., Lynch,R., Munger,H. and Tewhey,R. |
| EPI_ISL_755746 | Toronto Invasive Bacterial Diseases Network | McMaster University | Allison McGeer, Patryk Aftanas, Hooman Derakhshani, Angel Li, Kuganya Nirmalarajah, Emily Panousis, Ahmed Draia, Jalees Nasir, Michael Surette, Samira Mubareka, Andrew G. McArthur |
| EPI_ISL_756278 | State Laboratories Division, Hawaii State Department of Health | State Laboratories Division, Hawaii State Department of Health | Pamela O'Brien, Sabrina Diemert, Drew Kuwazaki, Razvan Sultana, Edward Desmond |
| EPI_ISL_757343, EPI_ISL_757344, EPI_ISL_757345, EPI_ISL_757346, EPI_ISL_757347, EPI_ISL_757348, EPI_ISL_757349, EPI_ISL_757350, EPI_ISL_757351, EPI_ISL_757352, EPI_ISL_757353, EPI_ISL_757354, EPI_ISL_757355, EPI_ISL_757356, EPI_ISL_757357, EPI_ISL_757358, EPI_ISL_757359, EPI_ISL_757360, EPI_ISL_757361, EPI_ISL_757362, EPI_ISL_757363, EPI_ISL_757364, EPI_ISL_759916, EPI_ISL_759917, EPI_ISL_759918, EPI_ISL_759919, EPI_ISL_759920, EPI_ISL_759921, EPI_ISL_759922, EPI_ISL_759923, EPI_ISL_759924, EPI_ISL_759925, EPI_ISL_759926, EPI_ISL_759927, EPI_ISL_759928, EPI_ISL_759945, EPI_ISL_759946, EPI_ISL_759947, EPI_ISL_759948, EPI_ISL_759949, EPI_ISL_759950, EPI_ISL_759951, EPI_ISL_759952, EPI_ISL_759953, EPI_ISL_759954 | | | |
| see above | Department of Virology and Immunology, University of Helsinki and Helsinki University Hospital, Huslab Finland | Department of Virology, Faculty of Medicine, University of Helsinki, Helsinki, Finland | Teemu Smura, Ravi Kant, Phuoc Truong, Hussein Alburkat, Hannimari Kallio-Kokko, Jenni Virtanen, Maija Suvanto, Sari Hannula, Harri Kangas, Pekka Ellonen, Olli Vapalahti |
| EPI_ISL_760037, EPI_ISL_760038, EPI_ISL_760043, EPI_ISL_760044, EPI_ISL_760048 | Hong Kong Department of Health | School of Public Health, The University of Hong Kong | Daniel Chu, Haogao Gu, Pavithra Krishnan, Daisy Ng, Gigi Liu, Carrie Wan, Malik Peiris, Leo Poon |
| EPI_ISL_760117 | Division of Emerging Infectious Diseases, Bureau of Infectious Diseases Diagnosis Control, Korea Disease Control and Prevention Agency | Division of Emerging Infectious Diseases, Bureau of Infectious Diseases Diagnosis Control, Korea Disease Control and Prevention Agency | Ae Kyung Park, Il-Hwan Kim, Heui Man Kim, Jeong-Min Kim, Namjoo Lee, Chaeyoung Lee, Sang Hee Woo, Eun-Jin Kim |
| EPI_ISL_765586, EPI_ISL_765589, EPI_ISL_765592, EPI_ISL_765595, EPI_ISL_765596, EPI_ISL_765597, EPI_ISL_765598, EPI_ISL_765599, EPI_ISL_765600, EPI_ISL_765601, EPI_ISL_765602, EPI_ISL_765603, EPI_ISL_765604, EPI_ISL_765605, EPI_ISL_765606, EPI_ISL_765607, EPI_ISL_765608, EPI_ISL_765609, EPI_ISL_765610, EPI_ISL_765612 | | | |
| see above | Brigham and Womens Hospital | Infectious Disease Program, Broad Institute of Harvard and MIT | Lemieux,J.E., Siddle,K.J., Shaw,B., Adams,G., Pierce,V., Turbett,S., Anahtar,M., Branda,J., Slater,D., Harris,J., Lin,A.E., Gladden-Young,A., Lagerborg,K., Rudy,M., DeRuff,K., Carter,A., Normandin,E., Bauer,M., Reilly,S., Tomkins-Tinch,C., Loreth,C., Chaluvadi,S., Neumann,A., Cusick,C., Chapman,S.B., Gnirke,A., Flowers,K., Cerrato,F., Birren,B.W., Gallagher,G., Smole,S., Park,D.J., MacInnis,B.L., Ryan,E., LaRocque,R., Rosenberg,E. and Sabeti,P.C. |
| EPI_ISL_765613, EPI_ISL_765615, EPI_ISL_765620, EPI_ISL_765624, EPI_ISL_765641, EPI_ISL_765642, EPI_ISL_765735, EPI_ISL_765737, EPI_ISL_765738, EPI_ISL_765739, EPI_ISL_765740, EPI_ISL_765741, EPI_ISL_765742, EPI_ISL_765743, EPI_ISL_765744, EPI_ISL_765745, EPI_ISL_765746, EPI_ISL_765747 | | | |
| see above | Massachusetts General Hospital | Infectious Disease Program, Broad Institute of Harvard and MIT | Lemieux,J.E., Siddle,K.J., Shaw,B., Adams,G., Pierce,V., Turbett,S., Anahtar,M., Branda,J., Slater,D., Harris,J., Lin,A.E., Gladden-Young,A., Lagerborg,K., Rudy,M., DeRuff,K., Carter,A., Normandin,E., Bauer,M., Reilly,S., Tomkins-Tinch,C., Loreth,C., Chaluvadi,S., Neumann,A., Cusick,C., Chapman,S.B., Gnirke,A., Flowers,K., Cerrato,F., Birren,B.W., Gallagher,G., Smole,S., Park,D.J., MacInnis,B.L., Ryan,E., LaRocque,R., Rosenberg,E. and Sabeti,P.C. |
| EPI_ISL_765945, EPI_ISL_765946, EPI_ISL_765947, EPI_ISL_765948, EPI_ISL_765949, EPI_ISL_765950, EPI_ISL_765951 | Worobey Lab, Department of Ecology and Evolutionary Biology, University of Arizona | Worobey Lab, Department of Ecology and Evolutionary Biology, University of Arizona | Brendan Larsen, Grace Quirk, Thomas Watts, David Baltrus, Michael Worobey |
| EPI_ISL_767439, EPI_ISL_767440, EPI_ISL_767441, EPI_ISL_767442, EPI_ISL_767443, EPI_ISL_767444, EPI_ISL_767445, EPI_ISL_767446, EPI_ISL_767447, EPI_ISL_767448, EPI_ISL_767449, EPI_ISL_767578 | | | |
| see above | Wadsworth Center, New York State Department.of Health | Wadsworth Center, New York State Department.of Health | Kirsten St. George, Daryl M. Lamson, Alexis Russel, Matthew Shudt, Melissa A Leisner, Jonathan Plitnick, Navjot Singh, John Kelly, Sara Griesemer, Erasmus Schneider, Erica Lasek-Nesselquist |
| EPI_ISL_768646, EPI_ISL_768647, EPI_ISL_768648, EPI_ISL_768649, EPI_ISL_768650, EPI_ISL_768659, EPI_ISL_768660 | Pathogen Genomics Center, National Institute of Infectious Diseases | Pathogen Genomics Center, National Institute of Infectious Diseases | Tsuyoshi Sekizuka, Kentaro Itokawa, Rina Tanaka, Masanori Hashino, Makoto Kuroda |
| EPI_ISL_768773 | AIID | Irish Coronavirus Sequencing Consortium - National Virus Reference Laboratory | Michael Carr, Gabriel Gonzalez, Alejandro Abner Garcia Leon, Patrick Mallon |
| EPI_ISL_769992, EPI_ISL_769995 | Hospital Metropolitano | Inciensa, Instituto Costarricense de Investigación y | Francisco Duarte, Hebleen Porras, Claudio Soto-Garita, Estela Cordero, Adriana Godínez, Melany Calderón & Margarita Lee Lui |

| | | | |
|---|---|---|---|
| | | Enseñanza en Nutrición y Salud | |
| EPI_ISL_770007 | Area De Salud San Rafael | Inciensa, Instituto Costarricense de Investigación y Enseñanza en Nutrición y Salud | Francisco Duarte, Hebleen Porras, Claudio Soto-Garita, Estela Cordero, Adriana Godínez, Melany Calderón & Mariel López |
| EPI_ISL_770027 | Area De Salud La Cruz | Inciensa, Instituto Costarricense de Investigación y Enseñanza en Nutrición y Salud | Francisco Duarte, Hebleen Porras, Claudio Soto-Garita, Estela Cordero, Adriana Godínez, Melany Calderón & Mariel López |
| EPI_ISL_770031, EPI_ISL_770032 | Hle-Asociacion De Atencion Integral Del Anciano San Cayetano | Inciensa, Instituto Costarricense de Investigación y Enseñanza en Nutrición y Salud | Francisco Duarte, Hebleen Porras, Claudio Soto-Garita, Estela Cordero, Adriana Godínez, Melany Calderón & Mariel López |
| EPI_ISL_770638, EPI_ISL_770642, EPI_ISL_770647, EPI_ISL_770648, EPI_ISL_770649, EPI_ISL_770650, EPI_ISL_770651, EPI_ISL_770652, EPI_ISL_770654, EPI_ISL_770655, EPI_ISL_770657, EPI_ISL_770662, EPI_ISL_770663, EPI_ISL_770665, EPI_ISL_770667, EPI_ISL_770668, EPI_ISL_770671, EPI_ISL_770672 | | | |
| see above | UMMC-Health | WHO National Influenza Centre Russian Federation | Andrey Komissarov, Artem Fadeev, Anna Ivanova, Kseniya Komissarova, Dmitry Bazhenov, Tatiana Platonova, Daria Danilenko, Ksenia Safina, Elena Nabieva, Georgii Bazykin, Dmitry Lioznov |
| EPI_ISL_770752, EPI_ISL_770753, EPI_ISL_770754, EPI_ISL_770755, EPI_ISL_770756, EPI_ISL_770757, EPI_ISL_770758, EPI_ISL_770759, EPI_ISL_770760, EPI_ISL_770761, EPI_ISL_770762, EPI_ISL_770763, EPI_ISL_770764, EPI_ISL_770765, EPI_ISL_770766, EPI_ISL_770767, EPI_ISL_770768, EPI_ISL_770769, EPI_ISL_770770, EPI_ISL_770771 | | | |
| see above | ZOTZ KLIMAS MVZ Düsseldorf-Centrum GbR ÜBAG für Labormedizin, Genetik, Zytologie, Pathologie | Center of Medical Microbiology, Virology, and Hospital Hygiene, University of Duesseldorf | Maximilian Damagnez, Alexander Dilthey, Ashley-Jane Duplessis, Patrick Finzer, Katrin Hoffmann, Torsten Houwaart, Lisanna Hülse, Malte Kohns Vasconcelos, Marek Korencak, Nadine Lübke, Jessica Nicolai, Klaus Pfeffer, Daniel Strelow, Jörg Timm, Andreas Walker, Tobias Wienemann, Rainer Zotz |
| EPI_ISL_770876 | Laboratoire national de santé, Microbiology, Virology | Laboratoire national de santé, Microbiology, Microbial Genomics Platform | Anke Wienecke-Baldacchino, Catherine Ragimbeau,Jessica Tapp, Fatu Djabi, Lise Pignon, Raoul Salmon, Tamir Abdelrahman |
| EPI_ISL_775307, EPI_ISL_775308 | Oslo University Hospital, Department of Medical Microbiology | Norwegian Institute of Public Health, Department of Virology | Kathrine Stene-Johansen, Kamilla Heddeland Instefjord, Hilde Elshaug, Atiya R Ali,Marie Paulsen Madsen, Rasmus Riis Kopperud, Hilde Vollan, Karoline Bragstad, Olav Hungnes |
| EPI_ISL_775310 | Medical Microbiology Unit, Department for Laboratory Medicine, Drammen Hospital, Vestre Viken Health Trust, | Norwegian Institute of Public Health, Department of Virology | Kathrine Stene-Johansen, Kamilla Heddeland Instefjord, Hilde Elshaug, Atiya R Ali,Marie Paulsen Madsen, Rasmus Riis Kopperud, Hilde Vollan, Karoline Bragstad, Olav Hungnes |
| EPI_ISL_775464, EPI_ISL_775465, EPI_ISL_775466 | Department of Medical Microbiology, St. Olavs hospital | Norwegian Institute of Public Health, Department of Virology | Kathrine Stene-Johansen, Kamilla Heddeland Instefjord, Hilde Elshaug, Atiya R Ali,Marie Paulsen Madsen, Rasmus Riis Kopperud, Hilde Vollan, Karoline Bragstad, Olav Hungnes |
| EPI_ISL_775467 | Department of Medical Microbiology - section Molde, Molde Hospital | Norwegian Institute of Public Health, Department of Virology | Kathrine Stene-Johansen, Kamilla Heddeland Instefjord, Hilde Elshaug, Atiya R Ali,Marie Paulsen Madsen, Rasmus Riis Kopperud, Hilde Vollan, Karoline Bragstad, Olav Hungnes |
| EPI_ISL_775468 | Unilabs Laboratory Medicine | Norwegian Institute of Public Health, Department of Virology | Kathrine Stene-Johansen, Kamilla Heddeland Instefjord, Hilde Elshaug, Atiya R Ali,Marie Paulsen Madsen, Rasmus Riis Kopperud, Hilde Vollan, Karoline Bragstad, Olav Hungnes |
| EPI_ISL_775479 | Akershus University Hospital, Department for Microbiology and Infectious Disease Control | Norwegian Institute of Public Health, Department of Virology | Kathrine Stene-Johansen, Kamilla Heddeland Instefjord, Hilde Elshaug, Atiya R Ali,Marie Paulsen Madsen, Rasmus Riis Kopperud, Hilde Vollan, Karoline Bragstad, Olav Hungnes |
| EPI_ISL_775481 | Department of Medical Microbiology, St. Olavs hospital | Norwegian Institute of Public Health, Department of Virology | Kathrine Stene-Johansen, Kamilla Heddeland Instefjord, Hilde Elshaug, Atiya R Ali,Marie Paulsen Madsen, Rasmus Riis Kopperud, Hilde Vollan, Karoline Bragstad, Olav Hungnes |
| EPI_ISL_775523 | Department of Medical Microbiology - section Molde, Molde Hospital | Norwegian Institute of Public Health, Department of Virology | Kathrine Stene-Johansen, Kamilla Heddeland Instefjord, Hilde Elshaug, Atiya R Ali,Marie Paulsen Madsen, Rasmus Riis Kopperud, Hilde Vollan, Karoline Bragstad, Olav Hungnes |
| EPI_ISL_775597 | RS Pertamina Sorong | National Institute of Health Research and Development | Puspa,KD;Subangkit;Pawestri,HA;Ikawati,HD;Nugraha,AA;Yosepin;Pangesti,KNA;Soekarso,T;Puspandari,N;Setiawaty,V |
| EPI_ISL_776960, EPI_ISL_776961, EPI_ISL_776962, EPI_ISL_776963, EPI_ISL_776964, EPI_ISL_776965, EPI_ISL_776966, EPI_ISL_776967, EPI_ISL_778691, EPI_ISL_778692, EPI_ISL_778693, EPI_ISL_778694, EPI_ISL_778695 | | | |
| see above | Istituto Zooprofilattico Sperimentale del Mezzogiorno | TIGEM | Patrizia Annunziata, Andrea Ballabio, Valentina Bouche, Davide Cacchiarelli (CorrespAuthor), Pellegrino Cerino, Chiara Colantuono, Lucio Di Filippo, Antonio Grimaldi, Antonio Limone, Gabriella Loconte, Anna Manfredi, Francesco Panariello, Biancamaria Pierri, Marcello Salvi, Lucia Vassallo |
| EPI_ISL_779221, EPI_ISL_779222, EPI_ISL_779223, EPI_ISL_779224, EPI_ISL_779225, EPI_ISL_779226, EPI_ISL_779227, EPI_ISL_779251, EPI_ISL_779252, EPI_ISL_779253, EPI_ISL_779666, EPI_ISL_779667, EPI_ISL_779668, EPI_ISL_779669 | | | |
| see above | Pathogen Genomics Center, National Institute of Infectious Diseases | Pathogen Genomics Center, National Institute of Infectious Diseases | Tsuyoshi Sekizuka, Kentaro Itokawa, Rina Tanaka, Masanori Hashino, Makoto Kuroda |
| EPI_ISL_786200, EPI_ISL_786201, EPI_ISL_786202, EPI_ISL_786203, EPI_ISL_786212, EPI_ISL_786213, EPI_ISL_786214, EPI_ISL_786215, EPI_ISL_786224, EPI_ISL_786225, EPI_ISL_786226, EPI_ISL_786227, EPI_ISL_786236, EPI_ISL_786237, EPI_ISL_786238, EPI_ISL_786239, EPI_ISL_786248, EPI_ISL_786249, EPI_ISL_786250, EPI_ISL_787389, EPI_ISL_787390, EPI_ISL_787391, EPI_ISL_787401, EPI_ISL_787402, EPI_ISL_787403, EPI_ISL_787412, EPI_ISL_787413, EPI_ISL_787414, EPI_ISL_787424, EPI_ISL_787425, EPI_ISL_787426, EPI_ISL_787436, EPI_ISL_787437, EPI_ISL_787438, EPI_ISL_787448, EPI_ISL_787449, EPI_ISL_787450, EPI_ISL_787460, EPI_ISL_787461, EPI_ISL_787462, EPI_ISL_787472, EPI_ISL_787473, EPI_ISL_787474, EPI_ISL_787964, EPI_ISL_787965, EPI_ISL_787966, EPI_ISL_787967, EPI_ISL_787976, EPI_ISL_787977, EPI_ISL_787978, EPI_ISL_787979, EPI_ISL_787988, EPI_ISL_787989, EPI_ISL_787990, EPI_ISL_787991, EPI_ISL_789789, EPI_ISL_789799, EPI_ISL_789806, EPI_ISL_789808, EPI_ISL_789814, EPI_ISL_789823, EPI_ISL_789834, EPI_ISL_789838, EPI_ISL_789874, EPI_ISL_789883, EPI_ISL_789888, EPI_ISL_789890, EPI_ISL_789894, EPI_ISL_789895, EPI_ISL_789903, EPI_ISL_789916, EPI_ISL_789934, EPI_ISL_789935, EPI_ISL_789936, EPI_ISL_789937, EPI_ISL_789941, EPI_ISL_789954, EPI_ISL_789961, EPI_ISL_789962, EPI_ISL_789963, EPI_ISL_789965, EPI_ISL_789966, EPI_ISL_789969, EPI_ISL_789970, EPI_ISL_789977, EPI_ISL_789988, EPI_ISL_790208, EPI_ISL_790220, EPI_ISL_790222, EPI_ISL_790229, EPI_ISL_790231, EPI_ISL_790268, EPI_ISL_790290, EPI_ISL_790291, EPI_ISL_790292, EPI_ISL_790293, EPI_ISL_790294, EPI_ISL_790295, EPI_ISL_790296, EPI_ISL_790297, EPI_ISL_790298, EPI_ISL_790299, EPI_ISL_790300, EPI_ISL_790301, EPI_ISL_790302, EPI_ISL_790303, EPI_ISL_790304, EPI_ISL_790305, EPI_ISL_790307, EPI_ISL_790308, EPI_ISL_790309, EPI_ISL_790310, EPI_ISL_790311, EPI_ISL_790312, EPI_ISL_790313, EPI_ISL_790314, EPI_ISL_790315, EPI_ISL_790316, EPI_ISL_790317, EPI_ISL_790318, EPI_ISL_790319, EPI_ISL_790320, EPI_ISL_790321, EPI_ISL_790322, EPI_ISL_790323, EPI_ISL_790324, EPI_ISL_790325, EPI_ISL_790326, EPI_ISL_790327, EPI_ISL_790328, EPI_ISL_790329, EPI_ISL_790330, EPI_ISL_790331, EPI_ISL_790332, EPI_ISL_790333, EPI_ISL_790334, EPI_ISL_790335, EPI_ISL_790336, EPI_ISL_790337, EPI_ISL_790338, EPI_ISL_790339, EPI_ISL_790340, EPI_ISL_790341, EPI_ISL_790342, EPI_ISL_790343, EPI_ISL_790344, EPI_ISL_790345, EPI_ISL_790346, EPI_ISL_790347, EPI_ISL_790348, EPI_ISL_790349, EPI_ISL_790350, EPI_ISL_790351, EPI_ISL_790352, EPI_ISL_790353, EPI_ISL_790354, EPI_ISL_790355, EPI_ISL_790356, EPI_ISL_790357, EPI_ISL_790358, EPI_ISL_790359, EPI_ISL_790360, EPI_ISL_790361, EPI_ISL_790362, EPI_ISL_790363, EPI_ISL_790364, EPI_ISL_790365, EPI_ISL_790366, EPI_ISL_790367, EPI_ISL_790368, EPI_ISL_790369, EPI_ISL_790370, EPI_ISL_790371, EPI_ISL_790372, EPI_ISL_790373, EPI_ISL_790374, EPI_ISL_790375, EPI_ISL_790376, EPI_ISL_790377, EPI_ISL_790378, EPI_ISL_790379, EPI_ISL_790380, EPI_ISL_790381, EPI_ISL_790382, EPI_ISL_790383, EPI_ISL_790384, EPI_ISL_790385, EPI_ISL_790386, EPI_ISL_790387, EPI_ISL_790388, EPI_ISL_790389, EPI_ISL_790390, EPI_ISL_790391, EPI_ISL_790392, EPI_ISL_790393, EPI_ISL_790394, EPI_ISL_790395 | | | |
| see above | Houston Methodist Hospital | Houston Methodist Hospital | S. Wesley Long, Randall J. Olsen, Paul A. Christensen, David W. Bernard, James J. Davis, Maulik Shukla, Marcus Nguyen, Matthew Ojeda Saavedra, Prasanti Yerramilli, Layne Pruitt, Sishir Subedi, Heather Hendrickson, and James M. Musser |
| EPI_ISL_790927, EPI_ISL_790942, EPI_ISL_790953 | Dutch COVID-19 response team | National Institute for Public Health and the Environment (RIVM) | Adam Meijer, Harry Vennema, Jeroen Cremer, Sharon van den Brink, Bas van der Veer, AnneMarie van den Brandt, Florian Zwagemaker, Dennis Schmitz, Chantal Reusken, on behalf of the national COVID-19 response team |
| EPI_ISL_791316 | National Virus Reference Laboratory | Irish Coronavirus Sequencing Consortium - Teagasc Moorepark | Alejandro Abner Garcia Leon, Paul Cotter, Fiona Crispie, John Kenny, Paddy Mallon, Calum Walsh |
| EPI_ISL_792602, EPI_ISL_792620, EPI_ISL_792621 | LACEN-PB | Laboratory of Respiratory Viruses and Measles, Oswaldo Cruz Institute, FIOCRUZ | Paola Resende, Luciana Appolinario, Fernando Motta, Anna Carolina Paixao, Ana Carolina Mendonca, João Felipe Bezerra, Romero Henrique Teixeira de Vasconcelos, Dalane Loudal Florentino Teixeira, Thiago Franco de Oliveira Carneiro, Marilda Siqueira |
| EPI_ISL_794035, EPI_ISL_794038, EPI_ISL_794039, EPI_ISL_794040, EPI_ISL_794041 | Wadsworth Center, New York State Department.of Health | Wadsworth Center, New York State Department.of Health | Kirsten St. George, Daryl M. Lamson, Alexis Russel, Matthew Shudt, Melissa A Leisner, Jonathan Plitnick, Navjot Singh, John Kelly, Sara Griesemer, Erasmus Schneider, Erica Lasek-Nesselquist |
| EPI_ISL_796032, EPI_ISL_796036, EPI_ISL_796037, EPI_ISL_796038, EPI_ISL_796048, EPI_ISL_796049, EPI_ISL_796050, EPI_ISL_796056, EPI_ISL_796057, EPI_ISL_796058 | Institute of Virology, University of Cologne | Institute of Virology, University of Cologne | Saleta Sierra, Gibran Rubio, Zevanya Tessalonica, Dominik Aschenmeier, Eva Heger, Elena Knops, Rolf Kaiser, Maximilian Damagnez, Andreas Walker, Jörg Timm, Alexander Dilthey, Martin Däumer, Alex Thielen |
| EPI_ISL_801885, EPI_ISL_801916, EPI_ISL_801918, EPI_ISL_801933, EPI_ISL_801942, EPI_ISL_801943, EPI_ISL_801947, EPI_ISL_801954, EPI_ISL_801959, EPI_ISL_801971, EPI_ISL_801972, EPI_ISL_802270, EPI_ISL_802271, EPI_ISL_802272, EPI_ISL_802273, EPI_ISL_802274, EPI_ISL_802275, EPI_ISL_802276, EPI_ISL_802277, EPI_ISL_802278, EPI_ISL_802279, EPI_ISL_802280, EPI_ISL_802281, EPI_ISL_802282, EPI_ISL_802283, EPI_ISL_802284, EPI_ISL_802285, EPI_ISL_802286, EPI_ISL_802287, EPI_ISL_802288, EPI_ISL_802289, EPI_ISL_802290, EPI_ISL_802291, EPI_ISL_802292, EPI_ISL_802293, EPI_ISL_802294, EPI_ISL_802295, EPI_ISL_802296, EPI_ISL_802297, EPI_ISL_802298, EPI_ISL_802299, EPI_ISL_802300, EPI_ISL_802301, EPI_ISL_802302, EPI_ISL_802303, EPI_ISL_802304, EPI_ISL_802305, EPI_ISL_802306, EPI_ISL_802307, EPI_ISL_802308, EPI_ISL_802309, EPI_ISL_802310, EPI_ISL_802311, EPI_ISL_802312, EPI_ISL_802313, EPI_ISL_802314, EPI_ISL_802315, EPI_ISL_802316, EPI_ISL_802317, EPI_ISL_802318, EPI_ISL_802319, EPI_ISL_802320, EPI_ISL_802321, EPI_ISL_802322, EPI_ISL_802323, EPI_ISL_802324, EPI_ISL_802325 | | | |
| see above | MSHS Clinical Microbiology Laboratories | MSHS Pathogen Surveillance Program | Ana S. Gonzalez-Reiche, Hala Alshammary, Mitchell J. Sullivan, Brianne Ciferri, Ajay Obla, Angela Amoako, Mahmoud Awawda, Elena Hirsch, Ashley S. Salimbangon, Levy Sominsky, Katherine Beach, Kayla Russo, Charles Gleason, Shelcie Fabre, Giulio Kleiner, Zenab Khan, Bremy Alburquerque, Adriana van de Guchte, Komal Srivastava, Matthew M. Hernandez, Jayeeta Dutta, Denise Jurczyszak, Emily Ferreri, Rachel Chernet, Nancy Francoeur, Betsaida Salom |

| | | | |
|---|---|---|---|
| | | | Melo, Irina Oussenko, Gintaras Deikus, Juan Soto, Shwetha Hara Sridhar, Ying-Chih Wang, Kathryn Twyman, Andrew Kasarskis, Deena R. Altman, Robert Sebra, Adolfo García-Sastre, Marta Luksza, Gopi Patel, Sarah Schaefer, Melissa Gitman, Michael D. Nowak, Alberto Paniz-Mondolfi, Emilia Mia Sordillo, Viviana Simon, Harm van Bakel |
| EPI_ISL_802860 | Vilnius University Hospital Santaros Klinikos, Vilnius University | Institute of Biotechnology, Life Sciences Center, Vilnius University | Emilija Vasiliunaite, Milda Norkiene, Albertas Timinskas, Alma Gedvilaite, Aurelija Zvirbliene, Daniel Naumovas, Laimonas Griskevicius |
| EPI_ISL_803308, EPI_ISL_803319, EPI_ISL_803320, EPI_ISL_803321, EPI_ISL_803323, EPI_ISL_803324, EPI_ISL_803325, EPI_ISL_803327, EPI_ISL_803328, EPI_ISL_803329, EPI_ISL_803330, EPI_ISL_803331, EPI_ISL_803332, EPI_ISL_803353, EPI_ISL_803559, EPI_ISL_803560, EPI_ISL_803561, EPI_ISL_803562, EPI_ISL_803563, EPI_ISL_803564, EPI_ISL_803565, EPI_ISL_803566, EPI_ISL_803567, EPI_ISL_803568, EPI_ISL_803569, EPI_ISL_803570, EPI_ISL_803571, EPI_ISL_803572, EPI_ISL_803573, EPI_ISL_803574, EPI_ISL_803575, EPI_ISL_803576, EPI_ISL_803587, EPI_ISL_803588, EPI_ISL_803589, EPI_ISL_803590, EPI_ISL_803591, EPI_ISL_803592, EPI_ISL_803593, EPI_ISL_803594, EPI_ISL_803595, EPI_ISL_803596, EPI_ISL_803597, EPI_ISL_803601, EPI_ISL_803605, EPI_ISL_803606, EPI_ISL_803607, EPI_ISL_803608, EPI_ISL_803609, EPI_ISL_803610, EPI_ISL_803611, EPI_ISL_803612, EPI_ISL_803613, EPI_ISL_803614, EPI_ISL_803615, EPI_ISL_803616, EPI_ISL_803617, EPI_ISL_803618, EPI_ISL_803619, EPI_ISL_803620, EPI_ISL_803621, EPI_ISL_803622, EPI_ISL_803623, EPI_ISL_803624, EPI_ISL_803625, EPI_ISL_803626, EPI_ISL_803628, EPI_ISL_803629, EPI_ISL_803725, EPI_ISL_803726, EPI_ISL_803750, EPI_ISL_803751, EPI_ISL_803752, EPI_ISL_803753, EPI_ISL_803754, EPI_ISL_803755 | | | |
| see above | Wisconsin State Laboratory of Hygiene Communicable Disease Division | Wisconsin State Laboratory of Hygiene Communicable Disease Division | Kelsey R. Florek, Abigail C. Shockey |
| EPI_ISL_804261, EPI_ISL_804262, EPI_ISL_804263 | Respiratory Virus Unit, National Infection Service, Public Health England | COVID-19 Genomics UK (COG-UK) Consortium | PHE Covid Sequencing Team |
| EPI_ISL_804963 | Hospital Comarcal de Melilla | Instituto de Salud Carlos III | Iglesias-Caballero, M. Molinero Calamita, M. González-Esguevillas, M. Camarero, S. Pozo, F. Casas, I. Jiménez, P. Jiménez, M. Zaballos, A. Monzón, S. Varona, S. Juliá, M. Cuesta, I, J. López |
| EPI_ISL_810981, EPI_ISL_810982, EPI_ISL_810986, EPI_ISL_811011, EPI_ISL_811037 | MRCG at LSHTM Genomics lab | MRCG at LSHTM Genomics lab | Abdul Karim sesay, Abdoulie Kanteh, Jarra Manneh, Mariama Kujabi, Bakary Sanyang |
| EPI_ISL_812121 | Santa Clara County Public Health Laboratory | Santa Clara County Public Health Laboratory | Santa Clara County Public Health Department |
| EPI_ISL_812134, EPI_ISL_812232 | UT-Unified State Labs: Public Health Utah DOH | Pathogen Discovery, Respiratory Viruses Branch, Division of Viral Diseases, Centers for Disease Control and Prevention | Yan Li, Ying Tao, Anna Montmayeur, Jing Zhang, Brian Lynch, Krista Queen, Anna Uehara, Rachel Marine, Peter Cook, Clinton R. Paden, Haibin Wang, Suxiang Tong |
| EPI_ISL_813661 | Liverpool Clinical Laboratories | COVID-19 Genomics UK (COG-UK) Consortium | Sam Haldenby, Anita Lucaci, Steve Paterson, Julian Hiscox, Alistair Darby, M Almsaud, A Alrezaihi, Muhannad Alruwaili, Stuart D Armstrong, Jones Benjamin, Eleanor G Bentley, Anu Chawla, Jordan J Clark, Angela Cowell, Richard Eccles, Isabel García-Dorival, Matthew Gemmell, Alessandro Gerada, PKF Gilmore, Richard Gregory, Ximeng Han, Catherine Hartley, Margaret Hughes, Miren Iturriza-Gomara, James Johnson, L Luu, Jenifer Manson, Charlotte Nelson, Elaine O'Toole, Cassie Olateju, Rebekah Penrice-Randal , Lucille Rainbow, N.P Randle, Trevor Ian Robinson, Parul Sharma, Ghada T Shawli, James P Stewart, Neil Swainston, Ecaterina Vamos, Joanne Watts, Mark Whitehead |
| EPI_ISL_813993 | Hospital General Universitario Gregorio Marañón | SeqCOVID-SPAIN consortium/IBV(CSIC) | Darío García de Viedma, Laura Pérez-Lago, Marta Herranz, Jon Sicilia, Julia Suárez, Pilar Catalán, Patricia Muñoz and SeqCOVID-SPAIN consortium |
| EPI_ISL_815258, EPI_ISL_815290, EPI_ISL_815294, EPI_ISL_815295, EPI_ISL_815298, EPI_ISL_815312, EPI_ISL_815332, EPI_ISL_815340, EPI_ISL_815344, EPI_ISL_815348, EPI_ISL_815349, EPI_ISL_815354, EPI_ISL_815359, EPI_ISL_815361, EPI_ISL_815362, EPI_ISL_815363, EPI_ISL_815368, EPI_ISL_815369, EPI_ISL_815373, EPI_ISL_815376, EPI_ISL_815382, EPI_ISL_815383, EPI_ISL_815389 | | | |
| see above | Centogene | Centogene | Peter Bauer, Krishna Kumar Kandaswamy, Vivi Hue-Trang Lieu |
| EPI_ISL_822318, EPI_ISL_822319, EPI_ISL_822320, EPI_ISL_822321 | Lighthouse Lab in Glasgow | Wellcome Sanger Institute for the COVID-19 Genomics UK (COG-UK) Consortium | Harper VanSteenhouse, Yumi Kasai, David Gray, Carol Clugston, Anna Dominiczak and Alex Alderton, Roberto Amato, Sonia Goncalves, Ewan Harrison, David K. Jackson, Ian Johnston, Dominic Kwiatkowski, Cordelia Langford, John Sillitoe on behalf of the Wellcome Sanger Institute COVID-19 Surveillance Team |
| EPI_ISL_824003, EPI_ISL_824157 | Dutch COVID-19 response team | National Institute for Public Health and the Environment (RIVM) | Adam Meijer, Harry Vennema, Jeroen Cremer, Sharon van den Brink, Bas van der Veer, AnneMarie van den Brandt, Florian Zwagemaker, Dennis Schmitz, Chantal Reusken, on behalf of the national COVID-19 response team |
| EPI_ISL_824495, EPI_ISL_824496, EPI_ISL_824497, EPI_ISL_824498, EPI_ISL_824499, EPI_ISL_824500, EPI_ISL_824501, EPI_ISL_824502 | Hospital Universitari Vall d'Hebron - Vall d'Hebron Institut de Rercerca | Hospital Universitari Vall d'Hebron | Cristina Andrés, Maria Piñana, Josep F Abril, Damir Garcia-Cehic, Ariadna Rando, Juliana Esperalba, Maria Gema Codina, Carla Castillo, Maria Carmen Martin, Tomàs Pumarola, Josep Quer, Andrés Antón |
| EPI_ISL_824513 | New Mexico Department of Health Scientific Laboratory | New Mexico Department of Health Scientific Laboratory | Ellie Johnson, Anastacia Griego-Fisher, D'eldra Malone |
| EPI_ISL_825493, EPI_ISL_825575 | Respiratory Virus Unit, National Infection Service, Public Health England | COVID-19 Genomics UK (COG-UK) Consortium | PHE Covid Sequencing Team |
| EPI_ISL_826934, EPI_ISL_826935, EPI_ISL_826936, EPI_ISL_826938, EPI_ISL_826939, EPI_ISL_826940, EPI_ISL_826941, EPI_ISL_826942, EPI_ISL_826943 | deCODE genetics | deCODE genetics | Daniel F Gudbjartsson; Agnar Helgason; Hakon Jonsson; Olafur T Magnusson; Pall Melsted; Gudmundur L Norddahl; Jona Saemundsdottir; Asgeir Sigurdsson; Patrick Sulem; Arna B Agustsdottir; Hannes Eggertsson; Berglind Eiriksdottir; Run Fridriksdottir; Elisabet E Gardarsdottir; Gudmundur Georgsson; Olafia S Gretarsdottir; Kjartan R Gudmundsson; Thora R Gunnarsdottir; Arnaldur Gylfason; Hilma Holm; Brynjar O Jensson; Aslaug Jonasdottir; Kamilla S Josefsdottir; Thordur Kristjansson; Droplaug N Magnusdottir; Solvi Rognvaldsson; Louise le Roux; Gudrun Sigmundsdottir; Gardar Sveinbjornsson; Kristin E Sveinsdottir; Maney Sveinsdottir; Emil A Thorarensen; Bjarni Thorbjornsson; Gisli Masson; Ingileif Jonsdottir; Alma Moller; Thorolfur Gudnason; Karl G Kristinsson; Unnur Thorsteinsdottir; Kari Stefansson |
| EPI_ISL_826955, EPI_ISL_827045, EPI_ISL_827069, EPI_ISL_827076, EPI_ISL_827082, EPI_ISL_827110, EPI_ISL_827126, EPI_ISL_827142 | The National University Hospital of Iceland | deCODE genetics | Daniel F Gudbjartsson; Agnar Helgason; Hakon Jonsson; Olafur T Magnusson; Pall Melsted; Gudmundur L Norddahl; Jona Saemundsdottir; Asgeir Sigurdsson; Patrick Sulem; Arna B Agustsdottir; Hannes Eggertsson; Berglind Eiriksdottir; Run Fridriksdottir; Elisabet E Gardarsdottir; Gudmundur Georgsson; Olafia S Gretarsdottir; Kjartan R Gudmundsson; Thora R Gunnarsdottir; Arnaldur Gylfason; Hilma Holm; Brynjar O Jensson; Aslaug Jonasdottir; Kamilla S Josefsdottir; Thordur Kristjansson; Droplaug N Magnusdottir; Solvi Rognvaldsson; Louise le Roux; Gudrun Sigmundsdottir; Gardar Sveinbjornsson; Kristin E Sveinsdottir; Maney Sveinsdottir; Emil A Thorarensen; Bjarni Thorbjornsson; Gisli Masson; Ingileif Jonsdottir; Alma Moller; Thorolfur Gudnason; Karl G Kristinsson; Unnur Thorsteinsdottir; Kari Stefansson |
| EPI_ISL_827322, EPI_ISL_827323, EPI_ISL_827325, EPI_ISL_827326, EPI_ISL_827327, EPI_ISL_827328, EPI_ISL_827329, EPI_ISL_827332 | deCODE genetics | deCODE genetics | Daniel F Gudbjartsson; Agnar Helgason; Hakon Jonsson; Olafur T Magnusson; Pall Melsted; Gudmundur L Norddahl; Jona Saemundsdottir; Asgeir Sigurdsson; Patrick Sulem; Arna B Agustsdottir; Hannes Eggertsson; Berglind Eiriksdottir; Run Fridriksdottir; Elisabet E Gardarsdottir; Gudmundur Georgsson; Olafia S Gretarsdottir; Kjartan R Gudmundsson; Thora R Gunnarsdottir; Arnaldur Gylfason; Hilma Holm; Brynjar O Jensson; Aslaug Jonasdottir; Kamilla S Josefsdottir; Thordur Kristjansson; Droplaug N Magnusdottir; Solvi Rognvaldsson; Louise le Roux; Gudrun Sigmundsdottir; Gardar Sveinbjornsson; Kristin E Sveinsdottir; Maney Sveinsdottir; Emil A Thorarensen; Bjarni Thorbjornsson; Gisli Masson; Ingileif Jonsdottir; Alma Moller; Thorolfur Gudnason; Karl G Kristinsson; Unnur Thorsteinsdottir; Kari Stefansson |
| EPI_ISL_827354 | The National University Hospital of Iceland | deCODE genetics | Daniel F Gudbjartsson; Agnar Helgason; Hakon Jonsson; Olafur T Magnusson; Pall Melsted; Gudmundur L Norddahl; Jona Saemundsdottir; Asgeir Sigurdsson; Patrick Sulem; Arna B Agustsdottir; Hannes Eggertsson; Berglind Eiriksdottir; Run Fridriksdottir; Elisabet E Gardarsdottir; Gudmundur Georgsson; Olafia S Gretarsdottir; Kjartan R Gudmundsson; Thora R Gunnarsdottir; Arnaldur Gylfason; Hilma Holm; Brynjar O Jensson; Aslaug Jonasdottir; Kamilla S Josefsdottir; Thordur Kristjansson; Droplaug N Magnusdottir; Solvi Rognvaldsson; Louise le Roux; Gudrun Sigmundsdottir; Gardar Sveinbjornsson; Kristin E Sveinsdottir; Maney Sveinsdottir; Emil A Thorarensen; Bjarni Thorbjornsson; Gisli Masson; Ingileif Jonsdottir; Alma Moller; Thorolfur Gudnason; Karl G Kristinsson; Unnur Thorsteinsdottir; Kari Stefansson |
| EPI_ISL_827610, EPI_ISL_827611, EPI_ISL_827612, EPI_ISL_827613, EPI_ISL_827614, EPI_ISL_827615, EPI_ISL_827616, EPI_ISL_827617, EPI_ISL_827618, EPI_ISL_827619, EPI_ISL_827630, EPI_ISL_827631, EPI_ISL_827632, EPI_ISL_827634, EPI_ISL_827635, EPI_ISL_827636, EPI_ISL_827738, EPI_ISL_827740, EPI_ISL_827741, EPI_ISL_827742, EPI_ISL_827743 | | | |
| see above | deCODE genetics | deCODE genetics | Daniel F Gudbjartsson; Agnar Helgason; Hakon Jonsson; Olafur T Magnusson; Pall Melsted; Gudmundur L Norddahl; Jona Saemundsdottir; Asgeir Sigurdsson; Patrick Sulem; Arna B Agustsdottir; Hannes Eggertsson; Berglind Eiriksdottir; Run Fridriksdottir; Elisabet E Gardarsdottir; Gudmundur Georgsson; Olafia S Gretarsdottir; Kjartan R Gudmundsson; Thora R Gunnarsdottir; Arnaldur Gylfason; Hilma Holm; Brynjar O Jensson; Aslaug Jonasdottir; Kamilla S Josefsdottir; Thordur Kristjansson; Droplaug N Magnusdottir; Solvi Rognvaldsson; Louise le Roux; Gudrun Sigmundsdottir; Gardar Sveinbjornsson; Kristin E Sveinsdottir; Maney Sveinsdottir; Emil A Thorarensen; Bjarni Thorbjornsson; Gisli Masson; Ingileif Jonsdottir; Alma Moller; Thorolfur Gudnason; Karl G Kristinsson; Unnur Thorsteinsdottir; Kari Stefansson |

| | | | |
|---|---|---|---|
| EPI_ISL_827746 | The National University Hospital of Iceland | deCODE genetics | Daniel F Gudbjartsson; Agnar Helgason; Hakon Jonsson; Olafur T Magnusson; Pall Melsted; Gudmundur L Norddahl; Jona Saemundsdottir; Asgeir Sigurdsson; Patrick Sulem; Arna B Agustsdottir; Hannes Eggertsson; Berglind Eiriksdottir; Run Fridriksdottir; Elisabet E Gardarsdottir; Gudmundur Georgsson; Olafia S Gretarsdottir; Kjartan R Gudmundsson; Thora R Gunnarsdottir; Arnaldur Gylfason; Hilma Holm; Brynjar O Jensson; Aslaug Jonasdottir; Kamilla S Josefsdottir; Thordur Kristjansson; Droplaug N Magnusdottir; Solvi Rognvaldsson; Louise le Roux; Gudrun Sigmundsdottir; Gardar Sveinbjornsson; Kristin E Sveinsdottir; Maney Sveinsdottir; Emil A Thorarensen; Bjarni Thorbjornsson; Gisli Masson; Ingileif Jonsdottir; Alma Moller; Thorolfur Gudnason; Karl G Kristinsson; Unnur Thorsteinsdottir; Kari Stefansson |
| EPI_ISL_827755, EPI_ISL_827756 | deCODE genetics | deCODE genetics | Daniel F Gudbjartsson; Agnar Helgason; Hakon Jonsson; Olafur T Magnusson; Pall Melsted; Gudmundur L Norddahl; Jona Saemundsdottir; Asgeir Sigurdsson; Patrick Sulem; Arna B Agustsdottir; Hannes Eggertsson; Berglind Eiriksdottir; Run Fridriksdottir; Elisabet E Gardarsdottir; Gudmundur Georgsson; Olafia S Gretarsdottir; Kjartan R Gudmundsson; Thora R Gunnarsdottir; Arnaldur Gylfason; Hilma Holm; Brynjar O Jensson; Aslaug Jonasdottir; Kamilla S Josefsdottir; Thordur Kristjansson; Droplaug N Magnusdottir; Solvi Rognvaldsson; Louise le Roux; Gudrun Sigmundsdottir; Gardar Sveinbjornsson; Kristin E Sveinsdottir; Maney Sveinsdottir; Emil A Thorarensen; Bjarni Thorbjornsson; Gisli Masson; Ingileif Jonsdottir; Alma Moller; Thorolfur Gudnason; Karl G Kristinsson; Unnur Thorsteinsdottir; Kari Stefansson |
| EPI_ISL_827800, EPI_ISL_827802, EPI_ISL_827803, EPI_ISL_827804, EPI_ISL_827805 | The National University Hospital of Iceland | deCODE genetics | Daniel F Gudbjartsson; Agnar Helgason; Hakon Jonsson; Olafur T Magnusson; Pall Melsted; Gudmundur L Norddahl; Jona Saemundsdottir; Asgeir Sigurdsson; Patrick Sulem; Arna B Agustsdottir; Hannes Eggertsson; Berglind Eiriksdottir; Run Fridriksdottir; Elisabet E Gardarsdottir; Gudmundur Georgsson; Olafia S Gretarsdottir; Kjartan R Gudmundsson; Thora R Gunnarsdottir; Arnaldur Gylfason; Hilma Holm; Brynjar O Jensson; Aslaug Jonasdottir; Kamilla S Josefsdottir; Thordur Kristjansson; Droplaug N Magnusdottir; Solvi Rognvaldsson; Louise le Roux; Gudrun Sigmundsdottir; Gardar Sveinbjornsson; Kristin E Sveinsdottir; Maney Sveinsdottir; Emil A Thorarensen; Bjarni Thorbjornsson; Gisli Masson; Ingileif Jonsdottir; Alma Moller; Thorolfur Gudnason; Karl G Kristinsson; Unnur Thorsteinsdottir; Kari Stefansson |
| EPI_ISL_827883, EPI_ISL_827914, EPI_ISL_827915, EPI_ISL_827916, EPI_ISL_827917, EPI_ISL_827918, EPI_ISL_827919 | deCODE genetics | deCODE genetics | Daniel F Gudbjartsson; Agnar Helgason; Hakon Jonsson; Olafur T Magnusson; Pall Melsted; Gudmundur L Norddahl; Jona Saemundsdottir; Asgeir Sigurdsson; Patrick Sulem; Arna B Agustsdottir; Hannes Eggertsson; Berglind Eiriksdottir; Run Fridriksdottir; Elisabet E Gardarsdottir; Gudmundur Georgsson; Olafia S Gretarsdottir; Kjartan R Gudmundsson; Thora R Gunnarsdottir; Arnaldur Gylfason; Hilma Holm; Brynjar O Jensson; Aslaug Jonasdottir; Kamilla S Josefsdottir; Thordur Kristjansson; Droplaug N Magnusdottir; Solvi Rognvaldsson; Louise le Roux; Gudrun Sigmundsdottir; Gardar Sveinbjornsson; Kristin E Sveinsdottir; Maney Sveinsdottir; Emil A Thorarensen; Bjarni Thorbjornsson; Gisli Masson; Ingileif Jonsdottir; Alma Moller; Thorolfur Gudnason; Karl G Kristinsson; Unnur Thorsteinsdottir; Kari Stefansson |
| EPI_ISL_827942, EPI_ISL_827943, EPI_ISL_828031, EPI_ISL_828032, EPI_ISL_828035, EPI_ISL_828036 | The National University Hospital of Iceland | deCODE genetics | Daniel F Gudbjartsson; Agnar Helgason; Hakon Jonsson; Olafur T Magnusson; Pall Melsted; Gudmundur L Norddahl; Jona Saemundsdottir; Asgeir Sigurdsson; Patrick Sulem; Arna B Agustsdottir; Hannes Eggertsson; Berglind Eiriksdottir; Run Fridriksdottir; Elisabet E Gardarsdottir; Gudmundur Georgsson; Olafia S Gretarsdottir; Kjartan R Gudmundsson; Thora R Gunnarsdottir; Arnaldur Gylfason; Hilma Holm; Brynjar O Jensson; Aslaug Jonasdottir; Kamilla S Josefsdottir; Thordur Kristjansson; Droplaug N Magnusdottir; Solvi Rognvaldsson; Louise le Roux; Gudrun Sigmundsdottir; Gardar Sveinbjornsson; Kristin E Sveinsdottir; Maney Sveinsdottir; Emil A Thorarensen; Bjarni Thorbjornsson; Gisli Masson; Ingileif Jonsdottir; Alma Moller; Thorolfur Gudnason; Karl G Kristinsson; Unnur Thorsteinsdottir; Kari Stefansson |
| EPI_ISL_828040, EPI_ISL_828041, EPI_ISL_828042, EPI_ISL_828043, EPI_ISL_828044, EPI_ISL_828045, EPI_ISL_828046, EPI_ISL_828048, EPI_ISL_828049 | deCODE genetics | deCODE genetics | Daniel F Gudbjartsson; Agnar Helgason; Hakon Jonsson; Olafur T Magnusson; Pall Melsted; Gudmundur L Norddahl; Jona Saemundsdottir; Asgeir Sigurdsson; Patrick Sulem; Arna B Agustsdottir; Hannes Eggertsson; Berglind Eiriksdottir; Run Fridriksdottir; Elisabet E Gardarsdottir; Gudmundur Georgsson; Olafia S Gretarsdottir; Kjartan R Gudmundsson; Thora R Gunnarsdottir; Arnaldur Gylfason; Hilma Holm; Brynjar O Jensson; Aslaug Jonasdottir; Kamilla S Josefsdottir; Thordur Kristjansson; Droplaug N Magnusdottir; Solvi Rognvaldsson; Louise le Roux; Gudrun Sigmundsdottir; Gardar Sveinbjornsson; Kristin E Sveinsdottir; Maney Sveinsdottir; Emil A Thorarensen; Bjarni Thorbjornsson; Gisli Masson; Ingileif Jonsdottir; Alma Moller; Thorolfur Gudnason; Karl G Kristinsson; Unnur Thorsteinsdottir; Kari Stefansson |
| EPI_ISL_828053, EPI_ISL_828064, EPI_ISL_828070, EPI_ISL_828099 | The National University Hospital of Iceland | deCODE genetics | Daniel F Gudbjartsson; Agnar Helgason; Hakon Jonsson; Olafur T Magnusson; Pall Melsted; Gudmundur L Norddahl; Jona Saemundsdottir; Asgeir Sigurdsson; Patrick Sulem; Arna B Agustsdottir; Hannes Eggertsson; Berglind Eiriksdottir; Run Fridriksdottir; Elisabet E Gardarsdottir; Gudmundur Georgsson; Olafia S Gretarsdottir; Kjartan R Gudmundsson; Thora R Gunnarsdottir; Arnaldur Gylfason; Hilma Holm; Brynjar O Jensson; Aslaug Jonasdottir; Kamilla S Josefsdottir; Thordur Kristjansson; Droplaug N Magnusdottir; Solvi Rognvaldsson; Louise le Roux; Gudrun Sigmundsdottir; Gardar Sveinbjornsson; Kristin E Sveinsdottir; Maney Sveinsdottir; Emil A Thorarensen; Bjarni Thorbjornsson; Gisli Masson; Ingileif Jonsdottir; Alma Moller; Thorolfur Gudnason; Karl G Kristinsson; Unnur Thorsteinsdottir; Kari Stefansson |
| EPI_ISL_828104 | deCODE genetics | deCODE genetics | Daniel F Gudbjartsson; Agnar Helgason; Hakon Jonsson; Olafur T Magnusson; Pall Melsted; Gudmundur L Norddahl; Jona Saemundsdottir; Asgeir Sigurdsson; Patrick Sulem; Arna B Agustsdottir; Hannes Eggertsson; Berglind Eiriksdottir; Run Fridriksdottir; Elisabet E Gardarsdottir; Gudmundur Georgsson; Olafia S Gretarsdottir; Kjartan R Gudmundsson; Thora R Gunnarsdottir; Arnaldur Gylfason; Hilma Holm; Brynjar O Jensson; Aslaug Jonasdottir; Kamilla S Josefsdottir; Thordur Kristjansson; Droplaug N Magnusdottir; Solvi Rognvaldsson; Louise le Roux; Gudrun Sigmundsdottir; Gardar Sveinbjornsson; Kristin E Sveinsdottir; Maney Sveinsdottir; Emil A Thorarensen; Bjarni Thorbjornsson; Gisli Masson; Ingileif Jonsdottir; Alma Moller; Thorolfur Gudnason; Karl G Kristinsson; Unnur Thorsteinsdottir; Kari Stefansson |
| EPI_ISL_828142, EPI_ISL_828143, EPI_ISL_828144 | The National University Hospital of Iceland | deCODE genetics | Daniel F Gudbjartsson; Agnar Helgason; Hakon Jonsson; Olafur T Magnusson; Pall Melsted; Gudmundur L Norddahl; Jona Saemundsdottir; Asgeir Sigurdsson; Patrick Sulem; Arna B Agustsdottir; Hannes Eggertsson; Berglind Eiriksdottir; Run Fridriksdottir; Elisabet E Gardarsdottir; Gudmundur Georgsson; Olafia S Gretarsdottir; Kjartan R Gudmundsson; Thora R Gunnarsdottir; Arnaldur Gylfason; Hilma Holm; Brynjar O Jensson; Aslaug Jonasdottir; Kamilla S Josefsdottir; Thordur Kristjansson; Droplaug N Magnusdottir; Solvi Rognvaldsson; Louise le Roux; Gudrun Sigmundsdottir; Gardar Sveinbjornsson; Kristin E Sveinsdottir; Maney Sveinsdottir; Emil A Thorarensen; Bjarni Thorbjornsson; Gisli Masson; Ingileif Jonsdottir; Alma Moller; Thorolfur Gudnason; Karl G Kristinsson; Unnur Thorsteinsdottir; Kari Stefansson |
| EPI_ISL_828153, EPI_ISL_828155, EPI_ISL_828156 | deCODE genetics | deCODE genetics | Daniel F Gudbjartsson; Agnar Helgason; Hakon Jonsson; Olafur T Magnusson; Pall Melsted; Gudmundur L Norddahl; Jona Saemundsdottir; Asgeir Sigurdsson; Patrick Sulem; Arna B Agustsdottir; Hannes Eggertsson; Berglind Eiriksdottir; Run Fridriksdottir; Elisabet E Gardarsdottir; Gudmundur Georgsson; Olafia S Gretarsdottir; Kjartan R Gudmundsson; Thora R Gunnarsdottir; Arnaldur Gylfason; Hilma Holm; Brynjar O Jensson; Aslaug Jonasdottir; Kamilla S Josefsdottir; Thordur Kristjansson; Droplaug N Magnusdottir; Solvi Rognvaldsson; Louise le Roux; Gudrun Sigmundsdottir; Gardar Sveinbjornsson; Kristin E Sveinsdottir; Maney Sveinsdottir; Emil A Thorarensen; Bjarni Thorbjornsson; Gisli Masson; Ingileif Jonsdottir; Alma Moller; Thorolfur Gudnason; Karl G Kristinsson; Unnur Thorsteinsdottir; Kari Stefansson |
| EPI_ISL_828157, EPI_ISL_828158, EPI_ISL_828159, EPI_ISL_828160, EPI_ISL_828162 | The National University Hospital of Iceland | deCODE genetics | Daniel F Gudbjartsson; Agnar Helgason; Hakon Jonsson; Olafur T Magnusson; Pall Melsted; Gudmundur L Norddahl; Jona Saemundsdottir; Asgeir Sigurdsson; Patrick Sulem; Arna B Agustsdottir; Hannes Eggertsson; Berglind Eiriksdottir; Run Fridriksdottir; Elisabet E Gardarsdottir; Gudmundur Georgsson; Olafia S Gretarsdottir; Kjartan R Gudmundsson; Thora R Gunnarsdottir; Arnaldur Gylfason; Hilma Holm; Brynjar O Jensson; Aslaug Jonasdottir; Kamilla S Josefsdottir; Thordur Kristjansson; Droplaug N Magnusdottir; Solvi Rognvaldsson; Louise le Roux; Gudrun Sigmundsdottir; Gardar Sveinbjornsson; Kristin E Sveinsdottir; Maney Sveinsdottir; Emil A Thorarensen; Bjarni Thorbjornsson; Gisli Masson; Ingileif Jonsdottir; Alma Moller; Thorolfur Gudnason; Karl G Kristinsson; Unnur Thorsteinsdottir; Kari Stefansson |
| EPI_ISL_828176, EPI_ISL_828180, EPI_ISL_828181, EPI_ISL_828189, EPI_ISL_828193, EPI_ISL_828207, EPI_ISL_828237, EPI_ISL_828238, EPI_ISL_828239, EPI_ISL_828240, EPI_ISL_828280, EPI_ISL_828284 | | | |
| see above | deCODE genetics | deCODE genetics | Daniel F Gudbjartsson; Agnar Helgason; Hakon Jonsson; Olafur T Magnusson; Pall Melsted; Gudmundur L Norddahl; Jona Saemundsdottir; Asgeir Sigurdsson; Patrick Sulem; Arna B Agustsdottir; Hannes Eggertsson; Berglind Eiriksdottir; Run Fridriksdottir; Elisabet E Gardarsdottir; Gudmundur Georgsson; Olafia S Gretarsdottir; Kjartan R Gudmundsson; Thora R Gunnarsdottir; Arnaldur Gylfason; Hilma Holm; Brynjar O Jensson; Aslaug Jonasdottir; Kamilla S Josefsdottir; Thordur Kristjansson; Droplaug N Magnusdottir; Solvi Rognvaldsson; Louise le Roux; Gudrun Sigmundsdottir; Gardar Sveinbjornsson; Kristin E Sveinsdottir; Maney Sveinsdottir; Emil A Thorarensen; Bjarni Thorbjornsson; Gisli Masson; Ingileif Jonsdottir; Alma Moller; Thorolfur Gudnason; Karl G Kristinsson; Unnur Thorsteinsdottir; Kari Stefansson |
| EPI_ISL_828287 | The National University Hospital of Iceland | deCODE genetics | Daniel F Gudbjartsson; Agnar Helgason; Hakon Jonsson; Olafur T Magnusson; Pall Melsted; Gudmundur L Norddahl; Jona Saemundsdottir; Asgeir Sigurdsson; Patrick Sulem; Arna B Agustsdottir; Hannes Eggertsson; Berglind Eiriksdottir; Run Fridriksdottir; Elisabet E Gardarsdottir; Gudmundur Georgsson; Olafia S Gretarsdottir; Kjartan R Gudmundsson; Thora R Gunnarsdottir; Arnaldur Gylfason; Hilma Holm; Brynjar O Jensson; Aslaug Jonasdottir; Kamilla S Josefsdottir; Thordur Kristjansson; Droplaug N Magnusdottir; Solvi Rognvaldsson; Louise le Roux; Gudrun Sigmundsdottir; Gardar Sveinbjornsson; Kristin E Sveinsdottir; Maney Sveinsdottir; Emil A Thorarensen; Bjarni Thorbjornsson; Gisli Masson; Ingileif Jonsdottir; Alma Moller; Thorolfur Gudnason; Karl G Kristinsson; Unnur Thorsteinsdottir; Kari Stefansson |
| EPI_ISL_828289, EPI_ISL_828293, | deCODE genetics | deCODE genetics | Daniel F Gudbjartsson; Agnar Helgason; Hakon Jonsson; Olafur T Magnusson; Pall Melsted; Gudmundur L Norddahl; Jona Saemundsdottir; Asgeir |

| | | | |
|---|---|---|---|
| EPI_ISL_828297, EPI_ISL_828302, EPI_ISL_828307, EPI_ISL_828311, EPI_ISL_828316, EPI_ISL_828318, EPI_ISL_828319 | | | Sigurdsson; Patrick Sulem; Arna B Agustsdottir; Hannes Eggertsson; Berglind Eiriksdottir; Run Fridriksdottir; Elisabet E Gardarsdottir; Gudmundur Georgsson; Olafia S Gretarsdottir; Kjartan R Gudmundsson; Thora R Gunnarsdottir; Arnaldur Gylfason; Hilma Holm; Brynjar O Jensson; Aslaug Jonasdottir; Kamilla S Josefsdottir; Thordur Kristjansson; Droplaug N Magnusdottir; Solvi Rognvaldsson; Louise le Roux; Gudrun Sigmundsdottir; Gardar Sveinbjornsson; Kristin E Sveinsdottir; Maney Sveinsdottir; Emil A Thorarensen; Bjarni Thorbjornsson; Gisli Masson; Ingileif Jonsdottir; Alma Moller; Thorolfur Gudnason; Karl G Kristinsson; Unnur Thorsteinsdottir; Kari Stefansson |
| EPI_ISL_828323 | The National University Hospital of Iceland | deCODE genetics | Daniel F Gudbjartsson; Agnar Helgason; Hakon Jonsson; Olafur T Magnusson; Pall Melsted; Gudmundur L Norddahl; Jona Saemundsdottir; Asgeir Sigurdsson; Patrick Sulem; Arna B Agustsdottir; Hannes Eggertsson; Berglind Eiriksdottir; Run Fridriksdottir; Elisabet E Gardarsdottir; Gudmundur Georgsson; Olafia S Gretarsdottir; Kjartan R Gudmundsson; Thora R Gunnarsdottir; Arnaldur Gylfason; Hilma Holm; Brynjar O Jensson; Aslaug Jonasdottir; Kamilla S Josefsdottir; Thordur Kristjansson; Droplaug N Magnusdottir; Solvi Rognvaldsson; Louise le Roux; Gudrun Sigmundsdottir; Gardar Sveinbjornsson; Kristin E Sveinsdottir; Maney Sveinsdottir; Emil A Thorarensen; Bjarni Thorbjornsson; Gisli Masson; Ingileif Jonsdottir; Alma Moller; Thorolfur Gudnason; Karl G Kristinsson; Unnur Thorsteinsdottir; Kari Stefansson |
| EPI_ISL_828417, EPI_ISL_828419, EPI_ISL_828420, EPI_ISL_828421, EPI_ISL_828422, EPI_ISL_828423, EPI_ISL_828424, EPI_ISL_828425, EPI_ISL_828426, EPI_ISL_828559, EPI_ISL_828560, EPI_ISL_828561, EPI_ISL_828563, EPI_ISL_828564, EPI_ISL_828579, EPI_ISL_828580, EPI_ISL_828581, EPI_ISL_828582, EPI_ISL_828583, EPI_ISL_828584, EPI_ISL_828585, EPI_ISL_828586 | | | |
| see above | deCODE genetics | deCODE genetics | Daniel F Gudbjartsson; Agnar Helgason; Hakon Jonsson; Olafur T Magnusson; Pall Melsted; Gudmundur L Norddahl; Jona Saemundsdottir; Asgeir Sigurdsson; Patrick Sulem; Arna B Agustsdottir; Hannes Eggertsson; Berglind Eiriksdottir; Run Fridriksdottir; Elisabet E Gardarsdottir; Gudmundur Georgsson; Olafia S Gretarsdottir; Kjartan R Gudmundsson; Thora R Gunnarsdottir; Arnaldur Gylfason; Hilma Holm; Brynjar O Jensson; Aslaug Jonasdottir; Kamilla S Josefsdottir; Thordur Kristjansson; Droplaug N Magnusdottir; Solvi Rognvaldsson; Louise le Roux; Gudrun Sigmundsdottir; Gardar Sveinbjornsson; Kristin E Sveinsdottir; Maney Sveinsdottir; Emil A Thorarensen; Bjarni Thorbjornsson; Gisli Masson; Ingileif Jonsdottir; Alma Moller; Thorolfur Gudnason; Karl G Kristinsson; Unnur Thorsteinsdottir; Kari Stefansson |
| EPI_ISL_828589, EPI_ISL_828591, EPI_ISL_828593, EPI_ISL_828594 | The National University Hospital of Iceland | deCODE genetics | Daniel F Gudbjartsson; Agnar Helgason; Hakon Jonsson; Olafur T Magnusson; Pall Melsted; Gudmundur L Norddahl; Jona Saemundsdottir; Asgeir Sigurdsson; Patrick Sulem; Arna B Agustsdottir; Hannes Eggertsson; Berglind Eiriksdottir; Run Fridriksdottir; Elisabet E Gardarsdottir; Gudmundur Georgsson; Olafia S Gretarsdottir; Kjartan R Gudmundsson; Thora R Gunnarsdottir; Arnaldur Gylfason; Hilma Holm; Brynjar O Jensson; Aslaug Jonasdottir; Kamilla S Josefsdottir; Thordur Kristjansson; Droplaug N Magnusdottir; Solvi Rognvaldsson; Louise le Roux; Gudrun Sigmundsdottir; Gardar Sveinbjornsson; Kristin E Sveinsdottir; Maney Sveinsdottir; Emil A Thorarensen; Bjarni Thorbjornsson; Gisli Masson; Ingileif Jonsdottir; Alma Moller; Thorolfur Gudnason; Karl G Kristinsson; Unnur Thorsteinsdottir; Kari Stefansson |
| EPI_ISL_828695, EPI_ISL_828703, EPI_ISL_828716, EPI_ISL_828799, EPI_ISL_828803, EPI_ISL_828805, EPI_ISL_828827, EPI_ISL_828830, EPI_ISL_828831, EPI_ISL_828832, EPI_ISL_828833, EPI_ISL_828939, EPI_ISL_828941, EPI_ISL_828942, EPI_ISL_828943 | | | |
| see above | deCODE genetics | deCODE genetics | Daniel F Gudbjartsson; Agnar Helgason; Hakon Jonsson; Olafur T Magnusson; Pall Melsted; Gudmundur L Norddahl; Jona Saemundsdottir; Asgeir Sigurdsson; Patrick Sulem; Arna B Agustsdottir; Hannes Eggertsson; Berglind Eiriksdottir; Run Fridriksdottir; Elisabet E Gardarsdottir; Gudmundur Georgsson; Olafia S Gretarsdottir; Kjartan R Gudmundsson; Thora R Gunnarsdottir; Arnaldur Gylfason; Hilma Holm; Brynjar O Jensson; Aslaug Jonasdottir; Kamilla S Josefsdottir; Thordur Kristjansson; Droplaug N Magnusdottir; Solvi Rognvaldsson; Louise le Roux; Gudrun Sigmundsdottir; Gardar Sveinbjornsson; Kristin E Sveinsdottir; Maney Sveinsdottir; Emil A Thorarensen; Bjarni Thorbjornsson; Gisli Masson; Ingileif Jonsdottir; Alma Moller; Thorolfur Gudnason; Karl G Kristinsson; Unnur Thorsteinsdottir; Kari Stefansson |
| EPI_ISL_828970, EPI_ISL_828971, EPI_ISL_828976, EPI_ISL_828977, EPI_ISL_829010, EPI_ISL_829011 | The National University Hospital of Iceland | deCODE genetics | Daniel F Gudbjartsson; Agnar Helgason; Hakon Jonsson; Olafur T Magnusson; Pall Melsted; Gudmundur L Norddahl; Jona Saemundsdottir; Asgeir Sigurdsson; Patrick Sulem; Arna B Agustsdottir; Hannes Eggertsson; Berglind Eiriksdottir; Run Fridriksdottir; Elisabet E Gardarsdottir; Gudmundur Georgsson; Olafia S Gretarsdottir; Kjartan R Gudmundsson; Thora R Gunnarsdottir; Arnaldur Gylfason; Hilma Holm; Brynjar O Jensson; Aslaug Jonasdottir; Kamilla S Josefsdottir; Thordur Kristjansson; Droplaug N Magnusdottir; Solvi Rognvaldsson; Louise le Roux; Gudrun Sigmundsdottir; Gardar Sveinbjornsson; Kristin E Sveinsdottir; Maney Sveinsdottir; Emil A Thorarensen; Bjarni Thorbjornsson; Gisli Masson; Ingileif Jonsdottir; Alma Moller; Thorolfur Gudnason; Karl G Kristinsson; Unnur Thorsteinsdottir; Kari Stefansson |
| EPI_ISL_829096, EPI_ISL_829097, EPI_ISL_829098, EPI_ISL_829099, EPI_ISL_829101, EPI_ISL_829102, EPI_ISL_829200, EPI_ISL_829201, EPI_ISL_829202, EPI_ISL_829203, EPI_ISL_829204, EPI_ISL_829205, EPI_ISL_829206, EPI_ISL_829207, EPI_ISL_829208 | | | |
| see above | deCODE genetics | deCODE genetics | Daniel F Gudbjartsson; Agnar Helgason; Hakon Jonsson; Olafur T Magnusson; Pall Melsted; Gudmundur L Norddahl; Jona Saemundsdottir; Asgeir Sigurdsson; Patrick Sulem; Arna B Agustsdottir; Hannes Eggertsson; Berglind Eiriksdottir; Run Fridriksdottir; Elisabet E Gardarsdottir; Gudmundur Georgsson; Olafia S Gretarsdottir; Kjartan R Gudmundsson; Thora R Gunnarsdottir; Arnaldur Gylfason; Hilma Holm; Brynjar O Jensson; Aslaug Jonasdottir; Kamilla S Josefsdottir; Thordur Kristjansson; Droplaug N Magnusdottir; Solvi Rognvaldsson; Louise le Roux; Gudrun Sigmundsdottir; Gardar Sveinbjornsson; Kristin E Sveinsdottir; Maney Sveinsdottir; Emil A Thorarensen; Bjarni Thorbjornsson; Gisli Masson; Ingileif Jonsdottir; Alma Moller; Thorolfur Gudnason; Karl G Kristinsson; Unnur Thorsteinsdottir; Kari Stefansson |
| EPI_ISL_829239, EPI_ISL_829290, EPI_ISL_829291, EPI_ISL_829292, EPI_ISL_829293 | The National University Hospital of Iceland | deCODE genetics | Daniel F Gudbjartsson; Agnar Helgason; Hakon Jonsson; Olafur T Magnusson; Pall Melsted; Gudmundur L Norddahl; Jona Saemundsdottir; Asgeir Sigurdsson; Patrick Sulem; Arna B Agustsdottir; Hannes Eggertsson; Berglind Eiriksdottir; Run Fridriksdottir; Elisabet E Gardarsdottir; Gudmundur Georgsson; Olafia S Gretarsdottir; Kjartan R Gudmundsson; Thora R Gunnarsdottir; Arnaldur Gylfason; Hilma Holm; Brynjar O Jensson; Aslaug Jonasdottir; Kamilla S Josefsdottir; Thordur Kristjansson; Droplaug N Magnusdottir; Solvi Rognvaldsson; Louise le Roux; Gudrun Sigmundsdottir; Gardar Sveinbjornsson; Kristin E Sveinsdottir; Maney Sveinsdottir; Emil A Thorarensen; Bjarni Thorbjornsson; Gisli Masson; Ingileif Jonsdottir; Alma Moller; Thorolfur Gudnason; Karl G Kristinsson; Unnur Thorsteinsdottir; Kari Stefansson |
| EPI_ISL_829466, EPI_ISL_829467, EPI_ISL_829468, EPI_ISL_829469, EPI_ISL_829472, EPI_ISL_829473, EPI_ISL_829474, EPI_ISL_829475 | deCODE genetics | deCODE genetics | Daniel F Gudbjartsson; Agnar Helgason; Hakon Jonsson; Olafur T Magnusson; Pall Melsted; Gudmundur L Norddahl; Jona Saemundsdottir; Asgeir Sigurdsson; Patrick Sulem; Arna B Agustsdottir; Hannes Eggertsson; Berglind Eiriksdottir; Run Fridriksdottir; Elisabet E Gardarsdottir; Gudmundur Georgsson; Olafia S Gretarsdottir; Kjartan R Gudmundsson; Thora R Gunnarsdottir; Arnaldur Gylfason; Hilma Holm; Brynjar O Jensson; Aslaug Jonasdottir; Kamilla S Josefsdottir; Thordur Kristjansson; Droplaug N Magnusdottir; Solvi Rognvaldsson; Louise le Roux; Gudrun Sigmundsdottir; Gardar Sveinbjornsson; Kristin E Sveinsdottir; Maney Sveinsdottir; Emil A Thorarensen; Bjarni Thorbjornsson; Gisli Masson; Ingileif Jonsdottir; Alma Moller; Thorolfur Gudnason; Karl G Kristinsson; Unnur Thorsteinsdottir; Kari Stefansson |
| EPI_ISL_829725, EPI_ISL_829735 | The National University Hospital of Iceland | deCODE genetics | Daniel F Gudbjartsson; Agnar Helgason; Hakon Jonsson; Olafur T Magnusson; Pall Melsted; Gudmundur L Norddahl; Jona Saemundsdottir; Asgeir Sigurdsson; Patrick Sulem; Arna B Agustsdottir; Hannes Eggertsson; Berglind Eiriksdottir; Run Fridriksdottir; Elisabet E Gardarsdottir; Gudmundur Georgsson; Olafia S Gretarsdottir; Kjartan R Gudmundsson; Thora R Gunnarsdottir; Arnaldur Gylfason; Hilma Holm; Brynjar O Jensson; Aslaug Jonasdottir; Kamilla S Josefsdottir; Thordur Kristjansson; Droplaug N Magnusdottir; Solvi Rognvaldsson; Louise le Roux; Gudrun Sigmundsdottir; Gardar Sveinbjornsson; Kristin E Sveinsdottir; Maney Sveinsdottir; Emil A Thorarensen; Bjarni Thorbjornsson; Gisli Masson; Ingileif Jonsdottir; Alma Moller; Thorolfur Gudnason; Karl G Kristinsson; Unnur Thorsteinsdottir; Kari Stefansson |
| EPI_ISL_829831, EPI_ISL_829832, EPI_ISL_829833, EPI_ISL_829834, EPI_ISL_829836, EPI_ISL_829837 | deCODE genetics | deCODE genetics | Daniel F Gudbjartsson; Agnar Helgason; Hakon Jonsson; Olafur T Magnusson; Pall Melsted; Gudmundur L Norddahl; Jona Saemundsdottir; Asgeir Sigurdsson; Patrick Sulem; Arna B Agustsdottir; Hannes Eggertsson; Berglind Eiriksdottir; Run Fridriksdottir; Elisabet E Gardarsdottir; Gudmundur Georgsson; Olafia S Gretarsdottir; Kjartan R Gudmundsson; Thora R Gunnarsdottir; Arnaldur Gylfason; Hilma Holm; Brynjar O Jensson; Aslaug Jonasdottir; Kamilla S Josefsdottir; Thordur Kristjansson; Droplaug N Magnusdottir; Solvi Rognvaldsson; Louise le Roux; Gudrun Sigmundsdottir; Gardar Sveinbjornsson; Kristin E Sveinsdottir; Maney Sveinsdottir; Emil A Thorarensen; Bjarni Thorbjornsson; Gisli Masson; Ingileif Jonsdottir; Alma Moller; Thorolfur Gudnason; Karl G Kristinsson; Unnur Thorsteinsdottir; Kari Stefansson |
| EPI_ISL_829916 | The National University Hospital of Iceland | deCODE genetics | Daniel F Gudbjartsson; Agnar Helgason; Hakon Jonsson; Olafur T Magnusson; Pall Melsted; Gudmundur L Norddahl; Jona Saemundsdottir; Asgeir Sigurdsson; Patrick Sulem; Arna B Agustsdottir; Hannes Eggertsson; Berglind Eiriksdottir; Run Fridriksdottir; Elisabet E Gardarsdottir; Gudmundur Georgsson; Olafia S Gretarsdottir; Kjartan R Gudmundsson; Thora R Gunnarsdottir; Arnaldur Gylfason; Hilma Holm; Brynjar O Jensson; Aslaug Jonasdottir; Kamilla S Josefsdottir; Thordur Kristjansson; Droplaug N Magnusdottir; Solvi Rognvaldsson; Louise le Roux; Gudrun Sigmundsdottir; Gardar Sveinbjornsson; Kristin E Sveinsdottir; Maney Sveinsdottir; Emil A Thorarensen; Bjarni Thorbjornsson; Gisli Masson; Ingileif Jonsdottir; Alma Moller; Thorolfur Gudnason; Karl G Kristinsson; Unnur Thorsteinsdottir; Kari Stefansson |
| EPI_ISL_830087, EPI_ISL_830089, EPI_ISL_830090, EPI_ISL_830091, EPI_ISL_830092 | deCODE genetics | deCODE genetics | Daniel F Gudbjartsson; Agnar Helgason; Hakon Jonsson; Olafur T Magnusson; Pall Melsted; Gudmundur L Norddahl; Jona Saemundsdottir; Asgeir Sigurdsson; Patrick Sulem; Arna B Agustsdottir; Hannes Eggertsson; Berglind Eiriksdottir; Run Fridriksdottir; Elisabet E Gardarsdottir; Gudmundur Georgsson; Olafia S Gretarsdottir; Kjartan R Gudmundsson; Thora R Gunnarsdottir; Arnaldur Gylfason; Hilma Holm; Brynjar O Jensson; Aslaug Jonasdottir; Kamilla S |

| | | | |
|---|---|---|---|
| | | | Josefsdottir; Thordur Kristjansson; Droplaug N Magnusdottir; Solvi Rognvaldsson; Louise le Roux; Gudrun Sigmundsdottir; Gardar Sveinbjornsson; Kristin E Sveinsdottir; Maney Sveinsdottir; Emil A Thorarensen; Bjarni Thorbjornsson; Gisli Masson; Ingileif Jonsdottir; Alma Moller; Thorolfur Gudnason; Karl G Kristinsson; Unnur Thorsteinsdottir; Kari Stefansson |
| EPI_ISL_830097, EPI_ISL_830098, EPI_ISL_830100 | The National University Hospital of Iceland | deCODE genetics | Daniel F Gudbjartsson; Agnar Helgason; Hakon Jonsson; Olafur T Magnusson; Pall Melsted; Gudmundur L Norddahl; Jona Saemundsdottir; Asgeir Sigurdsson; Patrick Sulem; Arna B Agustsdottir; Hannes Eggertsson; Berglind Eiriksdottir; Run Fridriksdottir; Elisabet E Gardarsdottir; Gudmundur Georgsson; Olafia S Gretarsdottir; Kjartan R Gudmundsson; Thora R Gunnarsdottir; Arnaldur Gylfason; Hilma Holm; Brynjar O Jensson; Aslaug Jonasdottir; Kamilla S Josefsdottir; Thordur Kristjansson; Droplaug N Magnusdottir; Solvi Rognvaldsson; Louise le Roux; Gudrun Sigmundsdottir; Gardar Sveinbjornsson; Kristin E Sveinsdottir; Maney Sveinsdottir; Emil A Thorarensen; Bjarni Thorbjornsson; Gisli Masson; Ingileif Jonsdottir; Alma Moller; Thorolfur Gudnason; Karl G Kristinsson; Unnur Thorsteinsdottir; Kari Stefansson |
| EPI_ISL_830106, EPI_ISL_830107, EPI_ISL_830108 | deCODE genetics | deCODE genetics | Daniel F Gudbjartsson; Agnar Helgason; Hakon Jonsson; Olafur T Magnusson; Pall Melsted; Gudmundur L Norddahl; Jona Saemundsdottir; Asgeir Sigurdsson; Patrick Sulem; Arna B Agustsdottir; Hannes Eggertsson; Berglind Eiriksdottir; Run Fridriksdottir; Elisabet E Gardarsdottir; Gudmundur Georgsson; Olafia S Gretarsdottir; Kjartan R Gudmundsson; Thora R Gunnarsdottir; Arnaldur Gylfason; Hilma Holm; Brynjar O Jensson; Aslaug Jonasdottir; Kamilla S Josefsdottir; Thordur Kristjansson; Droplaug N Magnusdottir; Solvi Rognvaldsson; Louise le Roux; Gudrun Sigmundsdottir; Gardar Sveinbjornsson; Kristin E Sveinsdottir; Maney Sveinsdottir; Emil A Thorarensen; Bjarni Thorbjornsson; Gisli Masson; Ingileif Jonsdottir; Alma Moller; Thorolfur Gudnason; Karl G Kristinsson; Unnur Thorsteinsdottir; Kari Stefansson |
| EPI_ISL_830184, EPI_ISL_830266, EPI_ISL_830309, EPI_ISL_830314, EPI_ISL_830368, EPI_ISL_830445 | The National University Hospital of Iceland | deCODE genetics | Daniel F Gudbjartsson; Agnar Helgason; Hakon Jonsson; Olafur T Magnusson; Pall Melsted; Gudmundur L Norddahl; Jona Saemundsdottir; Asgeir Sigurdsson; Patrick Sulem; Arna B Agustsdottir; Hannes Eggertsson; Berglind Eiriksdottir; Run Fridriksdottir; Elisabet E Gardarsdottir; Gudmundur Georgsson; Olafia S Gretarsdottir; Kjartan R Gudmundsson; Thora R Gunnarsdottir; Arnaldur Gylfason; Hilma Holm; Brynjar O Jensson; Aslaug Jonasdottir; Kamilla S Josefsdottir; Thordur Kristjansson; Droplaug N Magnusdottir; Solvi Rognvaldsson; Louise le Roux; Gudrun Sigmundsdottir; Gardar Sveinbjornsson; Kristin E Sveinsdottir; Maney Sveinsdottir; Emil A Thorarensen; Bjarni Thorbjornsson; Gisli Masson; Ingileif Jonsdottir; Alma Moller; Thorolfur Gudnason; Karl G Kristinsson; Unnur Thorsteinsdottir; Kari Stefansson |
| EPI_ISL_830448, EPI_ISL_830450, EPI_ISL_830496, EPI_ISL_830550 | deCODE genetics | deCODE genetics | Daniel F Gudbjartsson; Agnar Helgason; Hakon Jonsson; Olafur T Magnusson; Pall Melsted; Gudmundur L Norddahl; Jona Saemundsdottir; Asgeir Sigurdsson; Patrick Sulem; Arna B Agustsdottir; Hannes Eggertsson; Berglind Eiriksdottir; Run Fridriksdottir; Elisabet E Gardarsdottir; Gudmundur Georgsson; Olafia S Gretarsdottir; Kjartan R Gudmundsson; Thora R Gunnarsdottir; Arnaldur Gylfason; Hilma Holm; Brynjar O Jensson; Aslaug Jonasdottir; Kamilla S Josefsdottir; Thordur Kristjansson; Droplaug N Magnusdottir; Solvi Rognvaldsson; Louise le Roux; Gudrun Sigmundsdottir; Gardar Sveinbjornsson; Kristin E Sveinsdottir; Maney Sveinsdottir; Emil A Thorarensen; Bjarni Thorbjornsson; Gisli Masson; Ingileif Jonsdottir; Alma Moller; Thorolfur Gudnason; Karl G Kristinsson; Unnur Thorsteinsdottir; Kari Stefansson |
| EPI_ISL_830739, EPI_ISL_830764, EPI_ISL_830765, EPI_ISL_830773, EPI_ISL_830815, EPI_ISL_830816, EPI_ISL_830817, EPI_ISL_830818, EPI_ISL_830819, EPI_ISL_830820, EPI_ISL_830821, EPI_ISL_830822, EPI_ISL_830823, EPI_ISL_830824, EPI_ISL_830825, EPI_ISL_830826, EPI_ISL_830827, EPI_ISL_831003, EPI_ISL_831004 | | | |
| see above | University Hospital Basel, Clinical Virology | University Hospital Basel, Clinical Bacteriology | Tim Roloff, Madlen Stange, Helena MB Seth-Smith, Alfredo Mari, Karoline Leuzinger, Julia Bielicki, Manuel Battegay, Hans Hirsch, Adrian Egli |
| EPI_ISL_831180, EPI_ISL_831181, EPI_ISL_831182, EPI_ISL_831183, EPI_ISL_831184, EPI_ISL_831185, EPI_ISL_831186, EPI_ISL_831187, EPI_ISL_831188, EPI_ISL_831189 | Hospital Universitario La Paz (Madrid) | SeqCOVID-SPAIN consortium/IBV(CSIC) | María Rodríguez-Tejedor, Elias Dahdouh, Fernando Lázaro-Perona, Jesús Mingorance and SeqCOVID-SPAIN consortium |
| EPI_ISL_831976 | Orebro klinisk mikrobiologi | The Public Health Agency of Sweden | Department of Microbiology, The Public Health Agency of Sweden |
| EPI_ISL_831977, EPI_ISL_831978 | Gavle klinisk mikrobiologi | The Public Health Agency of Sweden | Department of Microbiology, The Public Health Agency of Sweden |
| EPI_ISL_833333 | Batangas City Health Office | Research Institute for Tropical Medicine | Hannah Leah Morito, Othoniel Jan Onza, John Leonard Chan, Ma Angelica Tujan, Francisco Gerardo Polotan, Inez Andrea Medado, Kirstyn Brunker, Edelwisa Mercado, Daria Manalo, Catalino Demetria |
| EPI_ISL_833339 | St. Pascual De Baylon Hospital | Research Institute for Tropical Medicine | Hannah Leah Morito, Othoniel Jan Onza, John Leonard Chan, Ma Angelica Tujan, Francisco Gerardo Polotan, Inez Andrea Medado, Kirstyn Brunker, Edelwisa Mercado, Daria Manalo, Catalino Demetria |
| EPI_ISL_833341 | Batangas City Health Office | Research Institute for Tropical Medicine | Hannah Leah Morito, Othoniel Jan Onza, John Leonard Chan, Ma Angelica Tujan, Francisco Gerardo Polotan, Inez Andrea Medado, Kirstyn Brunker, Edelwisa Mercado, Daria Manalo, Catalino Demetria |
| EPI_ISL_839732 | Northumbria University / South Tees Hospitals NHS Foundation Trust / North Cumbria Integrated Care NHS Foundation Trust / North Tees and Hartlepool NHS Foundation Trust / Newcastle Hospitals NHS Foundation Trust | COVID-19 Genomics UK (COG-UK) Consortium | Darren L Smith,Andrew Nelson,Matthew Bashton,Greg R Young,Joshua Loh,John Allan,Mohammad A Tariq,Giles S Holt,Gary Black,Wen C Yew,Lynn Dover,Paul Baker,Steve Liggett,Sarah Essex,Jane Greenaway,Debra Padgett,Clive Graham,Garren Scott,Edward Barton,Emma Swindells,Brendan Payne,Jennifer Collins,Yusri Taha,Gary Eltringham |
| EPI_ISL_842100 | Quadram Institute Bioscience | COVID-19 Genomics UK (COG-UK) Consortium | Dave J. Baker, Gemma L. Kay, Alp Aydin, Thanh Le-Viet, Steven Rudder, Ana P. Tedim, Anastasia Kolyva, Maria Diaz, Leonardo de Oliveira Martins, Nabil-Fareed Alikhan, Lizzie Meadows, Rachael Stanley, Ngozi Elumogo, Muhammed Yasir, Nicholas M. Thomson, Alexander J Trotter, Rachel Gilroy, Samuel Bloomfield, Claire Stuart, Andrew Bell, Reenesh Prakash, Samir Dervisevic, Alison E. Mather, John Wain, Mark Webber, Andrew J. Page, Justin O'Grady |
| EPI_ISL_842651 | University Hospital Basel, Clinical Virology | University Hospital Basel, Clinical Bacteriology | Tim Roloff, Madlen Stange, Helena MB Seth-Smith, Alfredo Mari, Karoline Leuzinger, Julia Bielicki, Manuel Battegay, Hans Hirsch, Adrian Egli |
| EPI_ISL_842847, EPI_ISL_842848, EPI_ISL_842849, EPI_ISL_842850, EPI_ISL_842851, EPI_ISL_842852, EPI_ISL_842853, EPI_ISL_842854, EPI_ISL_842855, EPI_ISL_842856, EPI_ISL_842857 | | | |
| see above | Barts Health NHS Trust | COVID-19 Genomics UK (COG-UK) Consortium | CUTINO-MOGUEL, Maria-Teresa; HARRINGTON, David; OWOYEMI, Dola; SHYLINI, Raghavendran; BROAD, Claire; KELE, Beatrix |
| EPI_ISL_848078, EPI_ISL_848081 | Laboratory of Clinical Research on Dermatozoonoses in Domestic Animals, Evandro Chagas National Institute of Infectious Diseases, Oswaldo Cruz Foundation (Fiocruz), Rio de Janeiro, RJ, Brazil. | Laboratory of Respiratory Viruses and Measles, Oswaldo Cruz Institute, FIOCRUZ | Guilherme Amaral Calvet, Michelle Fernanda Borges da Silva, Anielle de Pina Costa, Ezequias Batista Martins, Isabella Campos Vargas de Moraes, Lusiele Guaraldo, Patrícia Brasil, Sandro Antônio Pereira, Rodrigo Caldas Menezes, Isabella Dib Ferreira Gremião, Lucas Oliveira Keidel, Shanna Araujo dos Santos, Artur Augusto Velho Mendes Junior, Renato Orsini Ornellas, Maria Ogrzewalska, PaolaCristina Resende, Alex Pauvolid-Corrêa, Fernando do Couto Motta, Alice Sampaio Barreto da Rocha, Thiago C. Souza, Marilda Mendonça Siqueira |
| EPI_ISL_848319 | Illinois Department of Public Health | Gagnon Lab, Southern Illinois University | Keith Gagnon |
| EPI_ISL_849011, EPI_ISL_849012, EPI_ISL_849013, EPI_ISL_849014, EPI_ISL_849015, EPI_ISL_849016, EPI_ISL_849017, EPI_ISL_849018, EPI_ISL_849019, EPI_ISL_849020, EPI_ISL_849021, EPI_ISL_849022, EPI_ISL_849023, EPI_ISL_849024, EPI_ISL_849025, EPI_ISL_849026 | | | |
| see above | Florida Bureau of Public Health Laboratories | Florida Bureau of Public Health Laboratories | Sarah Schmedes, Jason Blanton |
| EPI_ISL_849483 | Seattle Flu Study | Seattle Flu Study | Deborah A. Nickerson, Chris D. Frazar, Jover Lee, Benjamin Pelle, Matthew Richardson, Amanda Adler, Elisabeth Brandstetter, Peter D. Han, Kairsten Fay, Misja Ilcisin, Kirsten Lacombe, Thomas R. Sibley, Melissa Truong, Caitlin R. Wolf, Michael Boeckh, Janet A. Englund, Michael Famulare, Barry R. Lutz, Mark J. Rieder, Lea M. Starita, Matthew Thompson, Jay Shendure, Trevor Bedford, Helen Y. Chu |
| EPI_ISL_853294, EPI_ISL_853306, EPI_ISL_853347, EPI_ISL_853348, EPI_ISL_853349, EPI_ISL_853350, EPI_ISL_853351, EPI_ISL_853390 | UPMC Clinical Microbiology Laboratory | Microbial Genome Sequencing Center; Microbial Genomic Epidemiology Laboratory | Mustapha M. Mustapha, Jane W. Marsh, Dan Snyder, Marissa P. Griffith, Stephanie L. Mitchell, Vatsala R. Srinivasa, Kady D. Waggle, Chinelo Ezeonwuku, Vaughn S. Cooper, Lee H. Harrison |
| EPI_ISL_854234 | Department of Microbiology, University Innsbruck | Bergthaler laboratory, CeMM Research Center for Molecular Medicine of the Austrian Academy of Sciences | Lukas Endler, Alexandra Popa, Benedikt Agerer, Jakob-Wendelin Genger, Alexander Lercher, Anna Schedl, Thomas Penz, Michael Schuster, Jan Laine, Martin Senekowitsch, Christoph Bock, Andreas Bergthaler |
| EPI_ISL_854714 | Public Health Ontario Laboratory | Public Health Ontario Laboratory | Vanessa G Allen, Philip Banh, Yao Chen, Richard de Borja, Alireza Eshaghi, Nahuel Fittipaldi, Christine Frantz, Jonathan B Gubbay, Jennifer L Guthrie, Lawrence Heisler, Esha Joshi, Michael Laszloffy, Aimin Li, Michael CY Li, Dean Maxwell, Sandeep Nagra, Samir N Patel, Jared Simpson, Karthikeyan Sivaraman, Ashleigh Sullivan, Yogi Sundaravadanam, Sarah Teatero, Matthew Watson, Andre Villegas, Sandra Zittermann |
| EPI_ISL_855356 | Hospital | National Reference Center for Viruses of Respiratory | Marion Barbet, Sylvie Behillil, Méline Bizard, Angela Brisebarre, Camille Capel, Etienne Simon-Lorière, Vincent Enouf, Maud Vanpeene, Sylvie van der Werf |

| | | | |
|---|---|---|---|
| | | Infections, Institut Pasteur, Paris | |
| EPI_ISL_855527 | KEMRI-Wellcome Trust Research Programme/KEMRI-CGMR-C Kilifi | KEMRI-Wellcome Trust Research Programme/KEMRI-CGMR-C Kilifi | Githinji et al |
| EPI_ISL_859685, EPI_ISL_859686, EPI_ISL_859688, EPI_ISL_859690, EPI_ISL_859691, EPI_ISL_859698, EPI_ISL_859699, EPI_ISL_859700, EPI_ISL_859701, EPI_ISL_859702, EPI_ISL_859703, EPI_ISL_859704, EPI_ISL_859705, EPI_ISL_859706, EPI_ISL_859707, EPI_ISL_859708, EPI_ISL_859709, EPI_ISL_859710, EPI_ISL_859711, EPI_ISL_859712, EPI_ISL_859713, EPI_ISL_859714, EPI_ISL_859715, EPI_ISL_859716, EPI_ISL_859717, EPI_ISL_859718, EPI_ISL_859719, EPI_ISL_859720, EPI_ISL_859721, EPI_ISL_859722, EPI_ISL_859723, EPI_ISL_859724, EPI_ISL_859725, EPI_ISL_859726, EPI_ISL_859727, EPI_ISL_859728, EPI_ISL_859729, EPI_ISL_859730, EPI_ISL_859731, EPI_ISL_859732, EPI_ISL_859733, EPI_ISL_859734, EPI_ISL_859735, EPI_ISL_859736, EPI_ISL_859737, EPI_ISL_859738, EPI_ISL_859739, EPI_ISL_859740, EPI_ISL_859741, EPI_ISL_859742, EPI_ISL_859743, EPI_ISL_859744, EPI_ISL_859745, EPI_ISL_859746, EPI_ISL_859747, EPI_ISL_859748, EPI_ISL_859749, EPI_ISL_859750, EPI_ISL_859751, EPI_ISL_859752, EPI_ISL_859753, EPI_ISL_859754 | | | |
| see above | BTC, Khalifa University | BTC, Khalifa University | Al Safar et al |
| EPI_ISL_860145 | Keio University School of Medicine | Keio University School of Medicine | Kenjiro Kosaki, Yuka Iwasaki, Hirotsugu Ishizu, Haruhiko Siomi, Kodai Abe |
| EPI_ISL_860548 | St.Olavs hospital/NTNU | IKOM, NTNU | Aleksandr Ianevski, Svein Arne Nordbo, Denis Kainov |
| EPI_ISL_860685 | Respiratory Virus Unit, National Infection Service, Public Health England | COVID-19 Genomics UK (COG-UK) Consortium | PHE Covid Sequencing Team |
| EPI_ISL_860745 | Swiss National Reference Centre for Influenza | Swiss National Reference Centre for Influenza | Ana Rita Gonçalves Cabecinhas, Samuel Cordey, Florian Laubscher,Christoph Grünig, Laurent Kaiser |
| EPI_ISL_862056, EPI_ISL_862057, EPI_ISL_862058, EPI_ISL_862059, EPI_ISL_862060, EPI_ISL_862061, EPI_ISL_862062, EPI_ISL_862063, EPI_ISL_862064, EPI_ISL_862065, EPI_ISL_862066, EPI_ISL_862067, EPI_ISL_862068, EPI_ISL_862069, EPI_ISL_862070, EPI_ISL_862071, EPI_ISL_862072, EPI_ISL_862073, EPI_ISL_862074, EPI_ISL_862082, EPI_ISL_862083, EPI_ISL_862084, EPI_ISL_862085, EPI_ISL_862086, EPI_ISL_862087, EPI_ISL_862088, EPI_ISL_862089, EPI_ISL_862090, EPI_ISL_862091, EPI_ISL_862092, EPI_ISL_862093, EPI_ISL_862094, EPI_ISL_862095, EPI_ISL_862096, EPI_ISL_862097, EPI_ISL_862098, EPI_ISL_862099, EPI_ISL_862100, EPI_ISL_862101, EPI_ISL_862102, EPI_ISL_862103, EPI_ISL_862104, EPI_ISL_862105, EPI_ISL_862106, EPI_ISL_862107, EPI_ISL_862108, EPI_ISL_862109, EPI_ISL_862110, EPI_ISL_862111, EPI_ISL_862112, EPI_ISL_862113, EPI_ISL_862114, EPI_ISL_862115, EPI_ISL_862116, EPI_ISL_862117, EPI_ISL_862118, EPI_ISL_862119, EPI_ISL_862120 | | | |
| see above | Department of Virology and Immunology, University of Helsinki and Helsinki University Hospital, Huslab Finland | Department of Virology, Faculty of Medicine, University of Helsinki, Helsinki, Finland | Teemu Smura, Ravi Kant, Phuoc Truong, Hussein Alburkat, Hanna Liimatainen, Hannimari Kallio-Kokko, Jenni Virtanen, Maija Suvanto, Sari Hannula, Harri Kangas, Pekka Ellonen, Olli Vapalahti |
| EPI_ISL_869242, EPI_ISL_869243, EPI_ISL_869246, EPI_ISL_869248, EPI_ISL_869249, EPI_ISL_869251, EPI_ISL_869253, EPI_ISL_869254, EPI_ISL_869255, EPI_ISL_869256, EPI_ISL_869257, EPI_ISL_869258, EPI_ISL_869259, EPI_ISL_869260, EPI_ISL_872569 | | | |
| see above | UMMC-Health | WHO National Influenza Centre Russian Federation | Andrey Komissarov, Artem Fadeev, Anna Ivanova, Kseniya Komissarova, Dmitry Bazhenov, Mikhail Bakaev,Tatiana Platonova, Daria Danilenko, Ksenia Safina, Elena Nabieva, Georgii Bazykin, Dmitry Lioznov |
| EPI_ISL_876814 | Quest Diagnostics | Quest Diagnostics | Rosenthal,S.H., Gerasimova,A., Kagan,R.M., Anderson, B., Hua, M., Liu Y., Bernstein, L.E., Livingston, K.E., Perez, A., Shalhout, D.F., Shlyakhter, I.A., Owen, R., Tanpaiboon, P., Lacbawan, F. |
| EPI_ISL_878653 | Rady's Childrens Hospital | Andersen lab at Scripps Research | SEARCH Alliance San Diego with Nanda Radamchar, David Dimmock, Linda Luo, Christina Clarke, Kathryn Bouic, Teresa Mueller, Denise Malicki |
| EPI_ISL_882626 | COVID lab, Mymensingh Medical College | Department of Pathology, Bangladesh Agricultural University & Department of Microbiology, Mymensingh Medical College | Afrin, S. Z. Paul, S. K. Parvin, R. |
| EPI_ISL_884306, EPI_ISL_884314, EPI_ISL_884332, EPI_ISL_884347, EPI_ISL_884348, EPI_ISL_884350, EPI_ISL_884351, EPI_ISL_884352, EPI_ISL_884360, EPI_ISL_884362, EPI_ISL_884365, EPI_ISL_884388 | | | |
| see above | Infectious Diseases, Quest Diagnostics | Infectious Diseases, Quest Diagnostics | Rosenthal,S.H., Gerasimova,A., Kagan,R.M., Anderson,B., Bernstein,L.E., Livingston,K.E., Hua,M., Liu,Y., Shalhout,D.F., Owen,R., Lacbawan,F. |
| EPI_ISL_887110 | Institute of Medical Microbiology and Virology, University Hospital Carl Gustav Carus, TU Dresden | DRESDEN-concept Genome Center, CMCB, TU Dresden | Beil J., Brown T., Büttner L., Gscheidel N., Hochauf-Stange K., Klemroth S., Lindemann D., Mehnert G., Petzold A., Reinhardt S., Rost F., Sameith K., Winkler S. |
| EPI_ISL_887153 | Massachusetts General Hospital | Infectious Disease Program, Broad Institute of Harvard and MIT | Lemieux,J.E., Siddle,K.J., Shaw,B., Adams,G., Pierce,V., Turbett,S., Anahtar,M., Branda,J., Slater,D., Harris,J., Lin,A.E., Gladden-Young,A., Lagerborg,K., Rudy,M., DeRuff,K., Carter,A., Normandin,E., Bauer,M., Reilly,S., Tomkins-Tinch,C., Loreth,C., Chaluvadi,S., Neumann,A., Cusick,C., Chapman,S.B., Gnirke,A., Flowers,K., Cerrato,F., Birren,B.W., Gallagher,G., Smole,S., Park,D.J., MacInnis,B.L., Ryan,E., LaRocque,R., Rosenberg,E. and Sabeti,P.C. |
| EPI_ISL_887434 | Instituto Nacional de Saude (INS), Mozambique | KRISP, KZN Research Innovation and Sequencing Platform | Nalia Ismael, Nadia Sitoe, Paulo Arnaldo, Nedio Mabunda, Giandhari J, Pillay S, Tegally H, Wilkinson E, de Oliveira T |
| EPI_ISL_888826 | Hospital | National Reference Center for Viruses of Respiratory Infections, Institut Pasteur, Paris | Marion Barbet, Sylvie Behillil, Méline Bizard, Angela Brisebarre, Camille Capel, Etienne Simon-Lorière, Vincent Enouf, Maud Vanpeene, Sylvie van der Werf, Gisèle Lagathu |
| EPI_ISL_890184 | Seattle Flu Study | Seattle Flu Study | Deborah A. Nickerson, Chris D. Frazar, Jover Lee, Benjamin Pelle, Erica Ryke, Matthew Richardson, Amanda Adler, Elisabeth Brandstetter, Peter D. Han, Kairsten Fay, Misja Ilcisin, Kirsten Lacombe, Thomas R. Sibley, Melissa Truong, Caitlin R. Wolf, Karen Cowgill, Stephanie Schrag, Jeff Duchin, Michael Boeckh, Janet A. Englund, Michael Famulare, Barry R. Lutz, Mark J. Rieder, Lea M. Starita, Matthew Thompson, Helen Y. Chu, Trevor Bedford, Jay Shendure |
| EPI_ISL_891179, EPI_ISL_891191 | DPH, Massachusetts State Public Health Lab | DPH, Massachusetts State Public Health Lab | Lang,A.S., Fink,T., Gallagher,G.R., Smole,S.C. |
| EPI_ISL_891271 | Institute of Biocides and Medical Ecology, Belgarde, Serbia | Virology department Institute of microbiology and immunology Faculty of Medicine University of Belgrade | Banko Ana, Miljanovic Danijela, Milicevic Ognjen, Loncar Ana, Abazovic Dzihan, Despot Dragana |
| EPI_ISL_892604, EPI_ISL_892605, EPI_ISL_892606, EPI_ISL_892607, EPI_ISL_892608, EPI_ISL_892609, EPI_ISL_892610 | Saitama Prefectural Institute of Public Health | Pathogen Genomics Center, National Institute of Infectious Diseases | Tsuyoshi Sekizuka, Kentaro Itokawa, Rina Tanaka, Masanori Hashino, Makoto Kuroda |
| EPI_ISL_893269, EPI_ISL_893270, EPI_ISL_893271 | Pathogen Genomics Center, National Institute of Infectious Diseases | Pathogen Genomics Center, National Institute of Infectious Diseases | Tsuyoshi Sekizuka, Kentaro Itokawa, Rina Tanaka, Masanori Hashino, Makoto Kuroda |
| EPI_ISL_893368, EPI_ISL_893375, EPI_ISL_893376, EPI_ISL_893377, EPI_ISL_893378, EPI_ISL_893379, EPI_ISL_893380, EPI_ISL_893381, EPI_ISL_893382, EPI_ISL_893383, EPI_ISL_893384, EPI_ISL_893385, EPI_ISL_893386, EPI_ISL_893387, EPI_ISL_893388, EPI_ISL_893389, EPI_ISL_893390, EPI_ISL_893391, EPI_ISL_893392, EPI_ISL_893393, EPI_ISL_893394, EPI_ISL_893395, EPI_ISL_893396, EPI_ISL_893397, EPI_ISL_893398, EPI_ISL_893399, EPI_ISL_893400, EPI_ISL_893401, EPI_ISL_893402, EPI_ISL_893403, EPI_ISL_893404, EPI_ISL_893405, EPI_ISL_893406, EPI_ISL_893407, EPI_ISL_893408, EPI_ISL_893420, EPI_ISL_893421, EPI_ISL_893422, EPI_ISL_893423, EPI_ISL_893424, EPI_ISL_893425, EPI_ISL_893426, EPI_ISL_893427, EPI_ISL_893428, EPI_ISL_893429, EPI_ISL_893430, EPI_ISL_893431, EPI_ISL_893432, EPI_ISL_893433, EPI_ISL_893434, EPI_ISL_893435, EPI_ISL_893436, EPI_ISL_893437, EPI_ISL_893438, EPI_ISL_893439, EPI_ISL_893440, EPI_ISL_893441, EPI_ISL_893442, EPI_ISL_893673 | | | |
| see above | Department of Infectious Diseases, Kobe Institute of Health | Department of Infectious Diseases, Kobe Institute of Health | Ryohei Nomoto, Tomotada Iwamoto, Tsuyoshi Sekizuka, Kentaro Itokawa, Rina Tanaka, Masanori Hashino, Makoto Kuroda |
| EPI_ISL_894277, EPI_ISL_894302, EPI_ISL_894465, EPI_ISL_894466, EPI_ISL_894467, EPI_ISL_894468, EPI_ISL_894469, EPI_ISL_894470, EPI_ISL_894471, EPI_ISL_894472, EPI_ISL_894473, EPI_ISL_894474, EPI_ISL_894475, EPI_ISL_894476, EPI_ISL_894477, EPI_ISL_894478, EPI_ISL_894479, EPI_ISL_894480, EPI_ISL_894481, EPI_ISL_894482, EPI_ISL_894483, EPI_ISL_894484, EPI_ISL_894485, EPI_ISL_894486, EPI_ISL_894487, EPI_ISL_894488, EPI_ISL_894489, EPI_ISL_894490, EPI_ISL_894491, EPI_ISL_894492, EPI_ISL_894493, EPI_ISL_894494, EPI_ISL_894495, EPI_ISL_894496, EPI_ISL_894497, EPI_ISL_894498, EPI_ISL_894499, EPI_ISL_894500, EPI_ISL_894501, EPI_ISL_894502, EPI_ISL_894503, EPI_ISL_894504, EPI_ISL_894505, EPI_ISL_894506, EPI_ISL_894507, EPI_ISL_894508, EPI_ISL_894509, EPI_ISL_894510, EPI_ISL_894511, EPI_ISL_894512, EPI_ISL_894513, EPI_ISL_894514, EPI_ISL_894515, EPI_ISL_894516, EPI_ISL_894517, EPI_ISL_894518, EPI_ISL_894519, EPI_ISL_894520, EPI_ISL_894521, EPI_ISL_894522, EPI_ISL_894523, EPI_ISL_894524, EPI_ISL_894525, EPI_ISL_894526, EPI_ISL_894527, EPI_ISL_894528, EPI_ISL_894529, EPI_ISL_894530, EPI_ISL_895096, EPI_ISL_895097, EPI_ISL_895098, EPI_ISL_895099, EPI_ISL_895100, EPI_ISL_895101, EPI_ISL_895102, EPI_ISL_895103, EPI_ISL_895104, EPI_ISL_895105, EPI_ISL_895160, EPI_ISL_895161, EPI_ISL_895162, EPI_ISL_895163, EPI_ISL_895164, EPI_ISL_895165, EPI_ISL_895166, EPI_ISL_895167, EPI_ISL_895168, EPI_ISL_895169, EPI_ISL_895170, EPI_ISL_895171, EPI_ISL_895172, EPI_ISL_895173, EPI_ISL_895174, EPI_ISL_895175, EPI_ISL_895176, EPI_ISL_895177, EPI_ISL_895178, EPI_ISL_895616, EPI_ISL_895617, EPI_ISL_895618, EPI_ISL_895619, EPI_ISL_895620, EPI_ISL_895621, EPI_ISL_895622, EPI_ISL_895623, EPI_ISL_895624, EPI_ISL_895625, EPI_ISL_895626, EPI_ISL_895627, EPI_ISL_895628, EPI_ISL_895629, EPI_ISL_895630, EPI_ISL_895631, EPI_ISL_895632, EPI_ISL_895633, EPI_ISL_895634, EPI_ISL_895635, EPI_ISL_895636, EPI_ISL_895637, EPI_ISL_895638, EPI_ISL_896019 | | | |
| see above | Pathogen Genomics Center, National Institute of Infectious Diseases | Pathogen Genomics Center, National Institute of Infectious Diseases | Tsuyoshi Sekizuka, Kentaro Itokawa, Rina Tanaka, Masanori Hashino, Makoto Kuroda |
| EPI_ISL_896142, EPI_ISL_896147, EPI_ISL_896153, EPI_ISL_896176 | MEPHI, Aix Marseille University | MEPHI, Aix Marseille University | Anthony LEVASSEUR |
| EPI_ISL_896612, EPI_ISL_896613, EPI_ISL_896614, EPI_ISL_896615, EPI_ISL_896616, EPI_ISL_896617, EPI_ISL_896618, EPI_ISL_896619, EPI_ISL_896620, EPI_ISL_896621, EPI_ISL_896622, EPI_ISL_896623, EPI_ISL_896624, EPI_ISL_896625, EPI_ISL_896626, EPI_ISL_896627, EPI_ISL_896628, EPI_ISL_896629, EPI_ISL_896630, EPI_ISL_896631, EPI_ISL_896632, EPI_ISL_896633, EPI_ISL_896634, EPI_ISL_896635, EPI_ISL_896636, EPI_ISL_896637, EPI_ISL_896638, EPI_ISL_896639, EPI_ISL_896640, EPI_ISL_896641, EPI_ISL_896658, EPI_ISL_896698, EPI_ISL_896909, EPI_ISL_896910, EPI_ISL_896911, EPI_ISL_896912, EPI_ISL_896913, EPI_ISL_896914, EPI_ISL_896915, EPI_ISL_896916, EPI_ISL_896917, EPI_ISL_896918, EPI_ISL_896919, EPI_ISL_896920, EPI_ISL_896921, EPI_ISL_896922, EPI_ISL_896923, EPI_ISL_896924, EPI_ISL_896925, EPI_ISL_896926, EPI_ISL_896927, EPI_ISL_896928, EPI_ISL_896929, EPI_ISL_896930, EPI_ISL_896931, EPI_ISL_896932, EPI_ISL_896933, EPI_ISL_896934, EPI_ISL_896935, EPI_ISL_896936, EPI_ISL_896937, EPI_ISL_896938, EPI_ISL_896939, EPI_ISL_897024, EPI_ISL_897025, EPI_ISL_897026, EPI_ISL_897027, EPI_ISL_897028, EPI_ISL_897029, EPI_ISL_897030, EPI_ISL_897031, EPI_ISL_897032, EPI_ISL_897033 | | | |
| see above | Pathogen Genomics Center, National Institute of Infectious Diseases | Pathogen Genomics Center, National Institute of Infectious Diseases | Tsuyoshi Sekizuka, Kentaro Itokawa, Rina Tanaka, Masanori Hashino, Makoto Kuroda |

| | | | |
|---|---|---|---|
| EPI_ISL_897059, EPI_ISL_897060, EPI_ISL_897066 | Niigata Prefectural Institute of Public Health and Environmental Sciences | Pathogen Genomics Center, National Institute of Infectious Diseases | Tsuyoshi Sekizuka, Kentaro Itokawa, Rina Tanaka, Masanori Hashino, Makoto Kuroda |
| EPI_ISL_897081, EPI_ISL_897082, EPI_ISL_897083, EPI_ISL_897084, EPI_ISL_897085, EPI_ISL_897086, EPI_ISL_897087, EPI_ISL_897149, EPI_ISL_897150, EPI_ISL_897273, EPI_ISL_897274, EPI_ISL_897275, EPI_ISL_897276, EPI_ISL_897277, EPI_ISL_897278, EPI_ISL_897279, EPI_ISL_897280, EPI_ISL_897281, EPI_ISL_897282, EPI_ISL_897316, EPI_ISL_897317, EPI_ISL_897318, EPI_ISL_897319, EPI_ISL_897320, EPI_ISL_897321, EPI_ISL_897322, EPI_ISL_897323, EPI_ISL_897324, EPI_ISL_897325, EPI_ISL_897326, EPI_ISL_897327, EPI_ISL_897328, EPI_ISL_897329, EPI_ISL_897330, EPI_ISL_897331, EPI_ISL_897332, EPI_ISL_897333, EPI_ISL_897334, EPI_ISL_897335, EPI_ISL_897336, EPI_ISL_897337, EPI_ISL_897338, EPI_ISL_897339, EPI_ISL_897447, EPI_ISL_897448, EPI_ISL_897449, EPI_ISL_897450, EPI_ISL_897451, EPI_ISL_897452, EPI_ISL_897593, EPI_ISL_897594, EPI_ISL_898033, EPI_ISL_898034, EPI_ISL_898035, EPI_ISL_898036 | | | |
| see above | Pathogen Genomics Center, National Institute of Infectious Diseases | Pathogen Genomics Center, National Institute of Infectious Diseases | Tsuyoshi Sekizuka, Kentaro Itokawa, Rina Tanaka, Masanori Hashino, Makoto Kuroda |
| EPI_ISL_898037, EPI_ISL_898038, EPI_ISL_898039, EPI_ISL_898040, EPI_ISL_898041, EPI_ISL_898042, EPI_ISL_898043, EPI_ISL_898044, EPI_ISL_898045, EPI_ISL_898046, EPI_ISL_898047, EPI_ISL_898048, EPI_ISL_898049, EPI_ISL_898050, EPI_ISL_898051, EPI_ISL_898052, EPI_ISL_898053 | | | |
| see above | Sakai City Institute of Public Health | Pathogen Genomics Center, National Institute of Infectious Diseases | Tsuyoshi Sekizuka, Kentaro Itokawa, Rina Tanaka, Masanori Hashino, Makoto Kuroda |
| EPI_ISL_898057, EPI_ISL_898058, EPI_ISL_898059, EPI_ISL_898060, EPI_ISL_898061, EPI_ISL_898062, EPI_ISL_898063, EPI_ISL_898064, EPI_ISL_898065, EPI_ISL_898066, EPI_ISL_898067, EPI_ISL_898068, EPI_ISL_898069, EPI_ISL_898070, EPI_ISL_898071, EPI_ISL_898072, EPI_ISL_898073, EPI_ISL_898077, EPI_ISL_898078, EPI_ISL_898079, EPI_ISL_898080, EPI_ISL_898081, EPI_ISL_898082, EPI_ISL_898083, EPI_ISL_898084, EPI_ISL_898085, EPI_ISL_898088, EPI_ISL_898134, EPI_ISL_898143, EPI_ISL_898144, EPI_ISL_898145, EPI_ISL_898269, EPI_ISL_898299, EPI_ISL_898300, EPI_ISL_898301, EPI_ISL_898302, EPI_ISL_898303, EPI_ISL_898304, EPI_ISL_898305, EPI_ISL_898306, EPI_ISL_898313, EPI_ISL_898314 | | | |
| see above | Pathogen Genomics Center, National Institute of Infectious Diseases | Pathogen Genomics Center, National Institute of Infectious Diseases | Tsuyoshi Sekizuka, Kentaro Itokawa, Rina Tanaka, Masanori Hashino, Makoto Kuroda |
| EPI_ISL_898321 | Toyama Institute of Health | Pathogen Genomics Center, National Institute of Infectious Diseases | Tsuyoshi Sekizuka, Kentaro Itokawa, Rina Tanaka, Masanori Hashino, Makoto Kuroda |
| EPI_ISL_898326, EPI_ISL_898327, EPI_ISL_898328 | Pathogen Genomics Center, National Institute of Infectious Diseases | Pathogen Genomics Center, National Institute of Infectious Diseases | Tsuyoshi Sekizuka, Kentaro Itokawa, Rina Tanaka, Masanori Hashino, Makoto Kuroda |
| EPI_ISL_898331 | Yamagata Prefectural Institute of Public Health | Pathogen Genomics Center, National Institute of Infectious Diseases | Tsuyoshi Sekizuka, Kentaro Itokawa, Rina Tanaka, Masanori Hashino, Makoto Kuroda |
| EPI_ISL_898539, EPI_ISL_898540, EPI_ISL_898541, EPI_ISL_898542, EPI_ISL_898543, EPI_ISL_898544, EPI_ISL_898545, EPI_ISL_898546, EPI_ISL_898547, EPI_ISL_898548, EPI_ISL_898549, EPI_ISL_898550 | | | |
| see above | Kanagawa Prefectural Institute of Public Health | Pathogen Genomics Center, National Institute of Infectious Diseases | Tsuyoshi Sekizuka, Kentaro Itokawa, Rina Tanaka, Masanori Hashino, Makoto Kuroda |
| EPI_ISL_898552, EPI_ISL_898553, EPI_ISL_898607, EPI_ISL_898608, EPI_ISL_898619, EPI_ISL_898620, EPI_ISL_898621, EPI_ISL_898622, EPI_ISL_898623, EPI_ISL_898624, EPI_ISL_898629, EPI_ISL_898630, EPI_ISL_898631, EPI_ISL_898632, EPI_ISL_898633, EPI_ISL_898634, EPI_ISL_898635, EPI_ISL_898636, EPI_ISL_898696, EPI_ISL_898697, EPI_ISL_898698, EPI_ISL_898699, EPI_ISL_898700, EPI_ISL_898701, EPI_ISL_898702, EPI_ISL_898703, EPI_ISL_898704, EPI_ISL_898705, EPI_ISL_898706, EPI_ISL_898707, EPI_ISL_898708, EPI_ISL_898859, EPI_ISL_898860, EPI_ISL_898861, EPI_ISL_898862, EPI_ISL_898863, EPI_ISL_898864, EPI_ISL_898865, EPI_ISL_898866, EPI_ISL_898867, EPI_ISL_898868, EPI_ISL_898869, EPI_ISL_898870, EPI_ISL_898871, EPI_ISL_898894, EPI_ISL_898895, EPI_ISL_898896, EPI_ISL_898897, EPI_ISL_898898, EPI_ISL_898899, EPI_ISL_898900, EPI_ISL_898901, EPI_ISL_898902, EPI_ISL_898903, EPI_ISL_898904, EPI_ISL_898905, EPI_ISL_898906, EPI_ISL_898907, EPI_ISL_898908, EPI_ISL_898909, EPI_ISL_898910, EPI_ISL_898911, EPI_ISL_898912, EPI_ISL_898913, EPI_ISL_898914 | | | |
| see above | Pathogen Genomics Center, National Institute of Infectious Diseases | Pathogen Genomics Center, National Institute of Infectious Diseases | Tsuyoshi Sekizuka, Kentaro Itokawa, Rina Tanaka, Masanori Hashino, Makoto Kuroda |
| EPI_ISL_900059, EPI_ISL_900074, EPI_ISL_900075, EPI_ISL_900105, EPI_ISL_900127, EPI_ISL_900174, EPI_ISL_900178, EPI_ISL_900188, EPI_ISL_900199, EPI_ISL_900223, EPI_ISL_900225, EPI_ISL_900252, EPI_ISL_900276, EPI_ISL_900326, EPI_ISL_900353, EPI_ISL_900391, EPI_ISL_900406, EPI_ISL_900449, EPI_ISL_900464 | | | |
| see above | MEPHI, Aix Marseille University | MEPHI, Aix Marseille University | Anthony LEVASSEUR |
| EPI_ISL_900760, EPI_ISL_900761, EPI_ISL_900762, EPI_ISL_900763, EPI_ISL_900764, EPI_ISL_900765, EPI_ISL_900766, EPI_ISL_900767, EPI_ISL_900768, EPI_ISL_900769, EPI_ISL_900770, EPI_ISL_900771, EPI_ISL_900776, EPI_ISL_900777, EPI_ISL_900778, EPI_ISL_900779, EPI_ISL_900780, EPI_ISL_900781, EPI_ISL_900782, EPI_ISL_900783, EPI_ISL_900784, EPI_ISL_900785, EPI_ISL_900786, EPI_ISL_900787, EPI_ISL_900788, EPI_ISL_900789, EPI_ISL_900790, EPI_ISL_900791, EPI_ISL_900792, EPI_ISL_900793, EPI_ISL_900794, EPI_ISL_900795, EPI_ISL_900796, EPI_ISL_900797, EPI_ISL_900798, EPI_ISL_900799, EPI_ISL_900800, EPI_ISL_900801, EPI_ISL_900802, EPI_ISL_900803, EPI_ISL_900804, EPI_ISL_900805, EPI_ISL_900806, EPI_ISL_900807, EPI_ISL_900808, EPI_ISL_900809, EPI_ISL_900810, EPI_ISL_900811, EPI_ISL_900812, EPI_ISL_900813, EPI_ISL_900814, EPI_ISL_900815, EPI_ISL_900816, EPI_ISL_900817, EPI_ISL_900818, EPI_ISL_900819, EPI_ISL_900820, EPI_ISL_900821, EPI_ISL_900822, EPI_ISL_900823, EPI_ISL_900824, EPI_ISL_900825, EPI_ISL_900826, EPI_ISL_900827, EPI_ISL_900828, EPI_ISL_900829, EPI_ISL_900830, EPI_ISL_900831, EPI_ISL_900832, EPI_ISL_900833, EPI_ISL_900834, EPI_ISL_900835, EPI_ISL_900836, EPI_ISL_900837, EPI_ISL_900838, EPI_ISL_900839, EPI_ISL_900840, EPI_ISL_900841, EPI_ISL_900842, EPI_ISL_900843, EPI_ISL_900844, EPI_ISL_900845, EPI_ISL_900846, EPI_ISL_900847, EPI_ISL_900848, EPI_ISL_900985, EPI_ISL_900986, EPI_ISL_900987, EPI_ISL_900988, EPI_ISL_900989, EPI_ISL_900990, EPI_ISL_900991, EPI_ISL_900992, EPI_ISL_900993, EPI_ISL_900994, EPI_ISL_900995, EPI_ISL_900996, EPI_ISL_900997, EPI_ISL_900998, EPI_ISL_900999, EPI_ISL_901000, EPI_ISL_901001, EPI_ISL_901002, EPI_ISL_901003, EPI_ISL_901004, EPI_ISL_901005, EPI_ISL_901006, EPI_ISL_901007, EPI_ISL_901008, EPI_ISL_901009, EPI_ISL_901010, EPI_ISL_901011, EPI_ISL_901012, EPI_ISL_901013, EPI_ISL_901014, EPI_ISL_901015, EPI_ISL_901016, EPI_ISL_901017, EPI_ISL_901018, EPI_ISL_901019, EPI_ISL_901020, EPI_ISL_901021, EPI_ISL_901022, EPI_ISL_901023, EPI_ISL_901024, EPI_ISL_901025, EPI_ISL_901026, EPI_ISL_901027, EPI_ISL_901028, EPI_ISL_901029, EPI_ISL_901030, EPI_ISL_901031, EPI_ISL_901032, EPI_ISL_901033, EPI_ISL_901034, EPI_ISL_901035, EPI_ISL_901036, EPI_ISL_901037, EPI_ISL_901038, EPI_ISL_901039, EPI_ISL_901040, EPI_ISL_901041, EPI_ISL_901042, EPI_ISL_901043, EPI_ISL_901044, EPI_ISL_901045, EPI_ISL_901046, EPI_ISL_901047, EPI_ISL_901048, EPI_ISL_901049, EPI_ISL_901050, EPI_ISL_901051, EPI_ISL_901052, EPI_ISL_901053, EPI_ISL_901252, EPI_ISL_901253, EPI_ISL_901254, EPI_ISL_901255, EPI_ISL_901256, EPI_ISL_901257, EPI_ISL_901258, EPI_ISL_901259, EPI_ISL_901260, EPI_ISL_901261, EPI_ISL_901262, EPI_ISL_901263, EPI_ISL_901264, EPI_ISL_901265, EPI_ISL_901266, EPI_ISL_901267, EPI_ISL_901268, EPI_ISL_901269, EPI_ISL_901270, EPI_ISL_901271, EPI_ISL_901306, EPI_ISL_901307, EPI_ISL_901308, EPI_ISL_901309, EPI_ISL_901310, EPI_ISL_901311, EPI_ISL_901312, EPI_ISL_901313, EPI_ISL_901314, EPI_ISL_901315, EPI_ISL_901316, EPI_ISL_901317, EPI_ISL_901318, EPI_ISL_901319, EPI_ISL_901320, EPI_ISL_901321, EPI_ISL_901360, EPI_ISL_901361, EPI_ISL_901362, EPI_ISL_901363, EPI_ISL_901368, EPI_ISL_901369, EPI_ISL_901370, EPI_ISL_901371, EPI_ISL_901372, EPI_ISL_901373, EPI_ISL_901374, EPI_ISL_901375, EPI_ISL_901376, EPI_ISL_901377, EPI_ISL_901378, EPI_ISL_901379, EPI_ISL_901380, EPI_ISL_901381, EPI_ISL_901382, EPI_ISL_901383, EPI_ISL_901384, EPI_ISL_901385, EPI_ISL_901386, EPI_ISL_901387, EPI_ISL_901388, EPI_ISL_901389, EPI_ISL_901390, EPI_ISL_901396, EPI_ISL_901397, EPI_ISL_901398, EPI_ISL_901399, EPI_ISL_901400, EPI_ISL_901401, EPI_ISL_901402, EPI_ISL_901403, EPI_ISL_901404, EPI_ISL_901405, EPI_ISL_901406, EPI_ISL_901407, EPI_ISL_901408, EPI_ISL_901409, EPI_ISL_901410, EPI_ISL_901411, EPI_ISL_901412, EPI_ISL_901413, EPI_ISL_901414, EPI_ISL_901415, EPI_ISL_901416, EPI_ISL_901417, EPI_ISL_901418, EPI_ISL_901419, EPI_ISL_901420, EPI_ISL_901421, EPI_ISL_901422, EPI_ISL_901423, EPI_ISL_901424, EPI_ISL_901425, EPI_ISL_901426, EPI_ISL_901427, EPI_ISL_901428, EPI_ISL_901429, EPI_ISL_901430, EPI_ISL_901431, EPI_ISL_901432, EPI_ISL_901450, EPI_ISL_901451, EPI_ISL_901452, EPI_ISL_901453, EPI_ISL_901454, EPI_ISL_901455, EPI_ISL_901456, EPI_ISL_901457 | | | |
| see above | Pathogen Genomics Center, National Institute of Infectious Diseases | Pathogen Genomics Center, National Institute of Infectious Diseases | Tsuyoshi Sekizuka, Kentaro Itokawa, Rina Tanaka, Masanori Hashino, Makoto Kuroda |
| EPI_ISL_901458, EPI_ISL_901459, EPI_ISL_901460, EPI_ISL_901461, EPI_ISL_901462, EPI_ISL_901463, EPI_ISL_901464, EPI_ISL_901465, EPI_ISL_901466, EPI_ISL_901467, EPI_ISL_901468, EPI_ISL_901469, EPI_ISL_901470, EPI_ISL_901471, EPI_ISL_901472, EPI_ISL_901473, EPI_ISL_901474, EPI_ISL_901475, EPI_ISL_901476, EPI_ISL_901477, EPI_ISL_901478, EPI_ISL_901479, EPI_ISL_901480, EPI_ISL_901481, EPI_ISL_901482, EPI_ISL_901483, EPI_ISL_901484, EPI_ISL_901485, EPI_ISL_901486, EPI_ISL_901487, EPI_ISL_901488, EPI_ISL_901489, EPI_ISL_901490 | | | |
| see above | Gunma Prefectural Institute of Public Health and Environmental Sciences | Pathogen Genomics Center, National Institute of Infectious Diseases | Tsuyoshi Sekizuka, Kentaro Itokawa, Rina Tanaka, Masanori Hashino, Makoto Kuroda |
| EPI_ISL_901491, EPI_ISL_901492, EPI_ISL_901493, EPI_ISL_901494, EPI_ISL_901495, EPI_ISL_901496, EPI_ISL_901497, EPI_ISL_901501, EPI_ISL_901502, EPI_ISL_901503, EPI_ISL_901504, EPI_ISL_901505, EPI_ISL_901506, EPI_ISL_901507, EPI_ISL_901508, EPI_ISL_901509, EPI_ISL_901510, EPI_ISL_901511, EPI_ISL_901512, EPI_ISL_901513, EPI_ISL_901514, EPI_ISL_901515, EPI_ISL_901516, EPI_ISL_901644, EPI_ISL_901645, EPI_ISL_901646, EPI_ISL_901647, EPI_ISL_901648 | | | |
| see above | Pathogen Genomics Center, National Institute of Infectious Diseases | Pathogen Genomics Center, National Institute of Infectious Diseases | Tsuyoshi Sekizuka, Kentaro Itokawa, Rina Tanaka, Masanori Hashino, Makoto Kuroda |
| EPI_ISL_901717, EPI_ISL_901718, EPI_ISL_901719, EPI_ISL_901720, EPI_ISL_901721 | Chiba Prefectural Institute of Public Health | Pathogen Genomics Center, National Institute of Infectious Diseases | Tsuyoshi Sekizuka, Kentaro Itokawa, Rina Tanaka, Masanori Hashino, Makoto Kuroda |
| EPI_ISL_901722, EPI_ISL_901723, EPI_ISL_901724, EPI_ISL_901725, EPI_ISL_901726, EPI_ISL_901727, EPI_ISL_901728, EPI_ISL_901729, EPI_ISL_901730, EPI_ISL_901731, EPI_ISL_901732, EPI_ISL_901733, EPI_ISL_901807, EPI_ISL_901808 | | | |
| see above | Pathogen Genomics Center, National Institute of Infectious Diseases | Pathogen Genomics Center, National Institute of Infectious Diseases | Tsuyoshi Sekizuka, Kentaro Itokawa, Rina Tanaka, Masanori Hashino, Makoto Kuroda |
| EPI_ISL_902024, EPI_ISL_902025, EPI_ISL_902026, EPI_ISL_902027, EPI_ISL_902028, EPI_ISL_902029, EPI_ISL_902030, EPI_ISL_902031, EPI_ISL_902032, EPI_ISL_902033, EPI_ISL_902034, EPI_ISL_902045, EPI_ISL_902046, EPI_ISL_902047, EPI_ISL_902049, EPI_ISL_902050, EPI_ISL_902051, EPI_ISL_902052, EPI_ISL_902053 | | | |
| see above | Department of Infectious Diseases, Kobe Institute of Health | Pathogen Genomics Center, National Institute of Infectious Diseases | Tsuyoshi Sekizuka, Kentaro Itokawa, Rina Tanaka, Masanori Hashino, Makoto Kuroda |
| EPI_ISL_902197, EPI_ISL_902198, EPI_ISL_902199, EPI_ISL_902200, EPI_ISL_902321, EPI_ISL_902322, EPI_ISL_902323, EPI_ISL_902324, EPI_ISL_902325, EPI_ISL_902326, EPI_ISL_902327, EPI_ISL_902328, EPI_ISL_902329, EPI_ISL_902330, EPI_ISL_902331, EPI_ISL_902332, EPI_ISL_902333 | | | |
| see above | Pathogen Genomics Center, National Institute of Infectious Diseases | Pathogen Genomics Center, National Institute of Infectious Diseases | Tsuyoshi Sekizuka, Kentaro Itokawa, Rina Tanaka, Masanori Hashino, Makoto Kuroda |
| EPI_ISL_902344 | Toyama Institute of Health | Pathogen Genomics Center, National Institute of Infectious | Tsuyoshi Sekizuka, Kentaro Itokawa, Rina Tanaka, Masanori Hashino, Makoto Kuroda |

| | | | |
|---|---|---|---|
| | | Diseases | |
| EPI_ISL_902415, EPI_ISL_902416, EPI_ISL_902417, EPI_ISL_902418, EPI_ISL_902419, EPI_ISL_902420, EPI_ISL_902421, EPI_ISL_902422, EPI_ISL_902423, EPI_ISL_902424, EPI_ISL_902425, EPI_ISL_902426, EPI_ISL_902427, EPI_ISL_902428, EPI_ISL_902429, EPI_ISL_902430, EPI_ISL_902431, EPI_ISL_902432, EPI_ISL_902433, EPI_ISL_902434, EPI_ISL_902435, EPI_ISL_902436, EPI_ISL_902437, EPI_ISL_902438, EPI_ISL_902439, EPI_ISL_902440, EPI_ISL_902441, EPI_ISL_902442, EPI_ISL_902444, EPI_ISL_902445, EPI_ISL_902446, EPI_ISL_902447, EPI_ISL_902448, EPI_ISL_902449, EPI_ISL_902450, EPI_ISL_902451, EPI_ISL_902452, EPI_ISL_902453, EPI_ISL_902454, EPI_ISL_902455, EPI_ISL_902456, EPI_ISL_902457, EPI_ISL_902458, EPI_ISL_902459, EPI_ISL_902460, EPI_ISL_902461, EPI_ISL_902462, EPI_ISL_902463, EPI_ISL_902464, EPI_ISL_902465, EPI_ISL_902466, EPI_ISL_902467, EPI_ISL_902468, EPI_ISL_902469, EPI_ISL_902470, EPI_ISL_902471, EPI_ISL_902472, EPI_ISL_902474, EPI_ISL_902500 | | | |
| see above | Pathogen Genomics Center, National Institute of Infectious Diseases | Pathogen Genomics Center, National Institute of Infectious Diseases | Tsuyoshi Sekizuka, Kentaro Itokawa, Rina Tanaka, Masanori Hashino, Makoto Kuroda |
| EPI_ISL_902504, EPI_ISL_902505, EPI_ISL_902506, EPI_ISL_902507, EPI_ISL_902508, EPI_ISL_902509, EPI_ISL_902510, EPI_ISL_902511, EPI_ISL_902512, EPI_ISL_902513, EPI_ISL_902514, EPI_ISL_902515, EPI_ISL_902516, EPI_ISL_902517, EPI_ISL_902518, EPI_ISL_902519, EPI_ISL_902520, EPI_ISL_902521, EPI_ISL_902522, EPI_ISL_902523, EPI_ISL_902524, EPI_ISL_902525, EPI_ISL_902526, EPI_ISL_902527, EPI_ISL_902528, EPI_ISL_902529, EPI_ISL_902530, EPI_ISL_902531, EPI_ISL_902532, EPI_ISL_902533, EPI_ISL_902534, EPI_ISL_902535, EPI_ISL_902536, EPI_ISL_902537, EPI_ISL_902538, EPI_ISL_902539, EPI_ISL_902540, EPI_ISL_902541, EPI_ISL_902542, EPI_ISL_902543, EPI_ISL_902544, EPI_ISL_902545, EPI_ISL_902546, EPI_ISL_902547, EPI_ISL_902548, EPI_ISL_902549, EPI_ISL_902550, EPI_ISL_902551, EPI_ISL_902552 | | | |
| see above | Department of Infectious Diseases, Kobe Institute of Health | Pathogen Genomics Center, National Institute of Infectious Diseases | Tsuyoshi Sekizuka, Kentaro Itokawa, Rina Tanaka, Masanori Hashino, Makoto Kuroda |
| EPI_ISL_902560, EPI_ISL_902561, EPI_ISL_902562, EPI_ISL_902563, EPI_ISL_902564, EPI_ISL_902565, EPI_ISL_902566, EPI_ISL_902567, EPI_ISL_902568, EPI_ISL_902569, EPI_ISL_902570, EPI_ISL_902571, EPI_ISL_902572, EPI_ISL_902573, EPI_ISL_902574, EPI_ISL_902578, EPI_ISL_902682 | | | |
| see above | Pathogen Genomics Center, National Institute of Infectious Diseases | Pathogen Genomics Center, National Institute of Infectious Diseases | Tsuyoshi Sekizuka, Kentaro Itokawa, Rina Tanaka, Masanori Hashino, Makoto Kuroda |
| EPI_ISL_902919 | Tanjungpura University Hospital | Tanjungpura University Hospital | Andriani; Mahyarudin; Virhan Novianry ; Delima Fajar Liana; Sofi Siti Shofiyah; Puji Astuti P; Muhammad Ibnu Kahtan; Ambar Rialita; Eka Ardiani Putri EA; Wiwik Windarti ; Helmi Sastriawan; Arif Wicaksono |
| EPI_ISL_904936, EPI_ISL_904938, EPI_ISL_904943, EPI_ISL_904944 | Vilnius University Hospital Santaros Klinikos, Vilnius University | Institute of Biotechnology, Life Sciences Center, Vilnius University | Emilija Vasiliunaite, Milda Norkiene, Albertas Timinskas, Alma Gedvilaite, Aurelija Zvirbliene, Daniel Naumovas, Laimonas Griskevicius |
| EPI_ISL_906054 | Tilia Laboratories s.r.o. | Tilia Laboratories s.r.o. | Sona Pekova, MD, PhD. |
| EPI_ISL_906055 | Tilia Laboratories s.r.o. | Tilia Laboratories s.r.o. | Sona Pekova MD, PhD. |
| EPI_ISL_906059 | Tilia Laboratories s.r.o. | Tilia Laboratories s.r.o. | Sona Pekova, MD, PhD. |
| EPI_ISL_906469, EPI_ISL_906470, EPI_ISL_906471, EPI_ISL_906472, EPI_ISL_906473 | Pathogen Genomics Center, National Institute of Infectious Diseases | Pathogen Genomics Center, National Institute of Infectious Diseases | Tsuyoshi Sekizuka, Kentaro Itokawa, Rina Tanaka, Masanori Hashino, Makoto Kuroda |
| EPI_ISL_906479, EPI_ISL_906480, EPI_ISL_906481, EPI_ISL_906482, EPI_ISL_906483, EPI_ISL_906484, EPI_ISL_906485, EPI_ISL_906486, EPI_ISL_906487, EPI_ISL_906488, EPI_ISL_906489, EPI_ISL_906490, EPI_ISL_906491, EPI_ISL_906492, EPI_ISL_906493, EPI_ISL_906494, EPI_ISL_906495, EPI_ISL_906496, EPI_ISL_906497, EPI_ISL_906498, EPI_ISL_906499, EPI_ISL_906500, EPI_ISL_906501, EPI_ISL_906502, EPI_ISL_906503, EPI_ISL_906504, EPI_ISL_906505 | | | |
| see above | Sakai City Institute of Public Health | Pathogen Genomics Center, National Institute of Infectious Diseases | Tsuyoshi Sekizuka, Kentaro Itokawa, Rina Tanaka, Masanori Hashino, Makoto Kuroda |
| EPI_ISL_906506, EPI_ISL_906520, EPI_ISL_906521 | Pathogen Genomics Center, National Institute of Infectious Diseases | Pathogen Genomics Center, National Institute of Infectious Diseases | Tsuyoshi Sekizuka, Kentaro Itokawa, Rina Tanaka, Masanori Hashino, Makoto Kuroda |
| EPI_ISL_906711 | Hematology Laboratory, Section of Molecular Diagnostics, University Clinical Centre, Medical University of Gdansk | Laboratory of Recombinant Vaccines | Lukasz Rabalski, Maciej Kosinski, Anna Piotrowska-Mietelska,Izabela Szczygielska, Boguslaw Szewczyk, Krystyna Bienkowska-Szewczyk |
| EPI_ISL_907075 | Department of Biology, University of Basrah | Department of Biology, University of Basrah | Abu-Ali,H.M. and Al-Badran,I.F. |
| EPI_ISL_910252, EPI_ISL_910253, EPI_ISL_910254, EPI_ISL_910255, EPI_ISL_910256, EPI_ISL_910257, EPI_ISL_910258, EPI_ISL_910259, EPI_ISL_910260, EPI_ISL_910261, EPI_ISL_910262, EPI_ISL_910263, EPI_ISL_910264, EPI_ISL_910265, EPI_ISL_910266, EPI_ISL_910267, EPI_ISL_910268, EPI_ISL_910269, EPI_ISL_910270, EPI_ISL_910271, EPI_ISL_910272, EPI_ISL_910273, EPI_ISL_910274, EPI_ISL_910275, EPI_ISL_910276, EPI_ISL_910277, EPI_ISL_910278, EPI_ISL_910279, EPI_ISL_910280, EPI_ISL_910281, EPI_ISL_910282, EPI_ISL_910283, EPI_ISL_910284, EPI_ISL_910285, EPI_ISL_910286, EPI_ISL_910287, EPI_ISL_910288, EPI_ISL_910289, EPI_ISL_910290, EPI_ISL_910291, EPI_ISL_910292, EPI_ISL_910293 | | | |
| see above | CSIR-Centre for Cellular and Molecular Biology | CSIR-Centre for Cellular and Molecular Biology | Payel Mukherjee,Pratheusa Maccha,Namami Gaur,Lamuk Zaveri,Tulasi Nagabandi,Purushotham Vodnala,Blessy B John,Viswagithe S L,B Himasri,Sofia Banu,Priya Singh,Archana Bharadwaj Siva,Karthik Bharadwaj Tallapaka,Rakesh K Mishra,Divya Tej Sowpati |
| EPI_ISL_912193, EPI_ISL_912194 | California Institute of Technology | Chan-Zuckerberg Biohub | CZB Cliahub Consortium |
| EPI_ISL_913092 | Labor Mustafa Wien | Center for Virology | Jeremy V. Camp, Irene Goerzer, Monika Redlberger-Fritz, Stephan W. Aberle |
| EPI_ISL_913094 | Center for Virology | Center for Virology | Jeremy V. Camp, Irene Goerzer, Monika Redlberger-Fritz, Stephan W. Aberle |
| EPI_ISL_915399, EPI_ISL_915400, EPI_ISL_915402 | Keio University School of Medicine | Keio University School of Medicine | Kenjiro Kosaki, Yuka Iwasaki, Hirotsugu Ishizu, Haruhiko Siomi, Kodai Abe |
| EPI_ISL_918378 | Ibaraki Prefectural Institute of Public Health | Ibaraki Prefectural Institute of Public Health | Keiko Goto, Tsuyoshi Sekizuka, Kentaro Itokawa, Rina Tanaka, Masanori Hashino, Makoto Kuroda |
| EPI_ISL_924077 | Quadram Institute Bioscience | COVID-19 Genomics UK (COG-UK) Consortium | Dave J. Baker, Gemma L. Kay, Alp Aydin, Thanh Le-Viet, Steven Rudder, Ana P. Tedim, Anastasia Kolyva, Maria Diaz, Leonardo de Oliveira Martins, Nabil-Fareed Alikhan, Lizzie Meadows, Rachael Stanley, Ngozi Elumogo, Muhammed Yasir, Nicholas M. Thomson, Alexander J Trotter, Rachel Gilroy, Samuel Bloomfield, Claire Stuart, Andrew Bell, Reenesh Prakash, Samir Dervisevic, Alison E. Mather, John Wain, Mark Webber, Andrew J. Page, Justin O'Grady |
| EPI_ISL_925585 | Public Health Ontario Laboratory | Public Health Ontario Laboratory | Vanessa G Allen, Philip Banh, Yao Chen, Richard de Borja, Alireza Eshaghi, Nahuel Fittipaldi, Christine Frantz, Jonathan B Gubbay, Jennifer L Guthrie, Lawrence Heisler, Esha Joshi, Michael Laszloffy, Aimin Li, Michael CY Li, Dean Maxwell, Sandeep Nagra, Samir N Patel, Jared Simpson, Karthikeyan Sivaraman, Ashleigh Sullivan, Yogi Sundaravadanam, Sarah Teatero, Matthew Watson, Andre Villegas, Sandra Zittermann |
| EPI_ISL_931473, EPI_ISL_931474, EPI_ISL_931475, EPI_ISL_931476, EPI_ISL_931477, EPI_ISL_931478, EPI_ISL_931479, EPI_ISL_931480, EPI_ISL_931481, EPI_ISL_931482, EPI_ISL_931483, EPI_ISL_931484, EPI_ISL_931485 | | | |
| see above | City of Milwaukee Health Department Laboratory | City of Milwaukee Health Department Laboratory | Sanjib Bhattacharyya |
| EPI_ISL_935862, EPI_ISL_935864, EPI_ISL_935865, EPI_ISL_935871, EPI_ISL_935872, EPI_ISL_935873 | Cadham Provincial laboratory | National Microbiology Laboratory (NML) | Anna Majer, Shari Tyson, Grace Seo, Philip Mabon, Elsie Grudeski, Rhiannon Huzarewich, Russell Mandes, Anneliese Landgraff, Jennifer Tanner, Natalie Knox, Morag Graham, Gary Van Domselaar, Paul Van Caeseele, Jared Bullard, David Alexander, Kerry Dust, Nathalie Bastien, Yan Li, Timothy Booth, Darian Hole, Madison Chapel, Kirsten Biggar, CanCOGeN's metadata curation team, Public Health Agency of Canada CanCOGeN team |
| EPI_ISL_936849, EPI_ISL_936850, EPI_ISL_936851 | Northwestern Memorial Hospital | Ozer Lab | Ramon Lorenzo-Redondo, Lacy M. Simons, Chad J. Achenbach, Lawrence J. Jennings, Michael G. Ison, Judd F. Hultquist, Egon A. Ozer |
| EPI_ISL_937037, EPI_ISL_937098 | Quest Diagnostics | Quest Diagnostics | Rosenthal,S.H., Gerasimova,A., Kagan,R.M., Anderson, B., Livingston, K.E., Hua, M., Liu Y., Shalhout, D.F., Owen, R., Lacbawan, F. |
| EPI_ISL_940174, EPI_ISL_940188, EPI_ISL_940189, EPI_ISL_940241, EPI_ISL_940245, EPI_ISL_940266, EPI_ISL_940267, EPI_ISL_940271 | Hôpital Bichat Claude Bernard, Laboratoire de Virologie | IAME UMR1137 Inserm, Université de Paris, Hôpital Bichat | Antoine Bridier-Nahmias, Amélie Recoing, Quentin Le Hingrat, Lena Daniel, Siham Hamri, Gilles Collin, Alexandre Storto, Mélanie Bertine, Charlotte Charpentier, Nadhira Houhou-Fidouh, Diane Descamps, Benoit Visseaux |
| EPI_ISL_940960, EPI_ISL_940961, EPI_ISL_940983, EPI_ISL_940986, EPI_ISL_940987, EPI_ISL_942008 | Centers for Disease Control and Prevention, Dengue Branch | Centers for Disease Control and Prevention, Dengue Branch | Gilberto A. Santiago, Glenda Gonzalez, Betzabel Flores, Keyla Charriez, Gabriela Paz-Bailey, Jorge L. Munoz-Jordan |
| EPI_ISL_943974, EPI_ISL_943975, EPI_ISL_943976, EPI_ISL_943977 | LACEN do Estado de Tocantins | Instituto Adolfo Lutz, Interdiciplinary Procedures Center, Strategic Laboratory | Claudio Tavares Sacchi, Claudia Regina Gonçalves, Erica Valessa Ramos Gomes, Karoline Rodrigues Campos |
| EPI_ISL_949231 | Departamento de Microbiología, CDB, Hospital Clínic, Barcelona | SeqCOVID-SPAIN consortium/IBV(CSIC) | Andrea Vergara, Mikel Martínez, Elisa Rubio, Jéssica Navero, Aida Peiró and SeqCOVID-SPAIN consortium |

| | | | |
|---|---|---|---|
| EPI_ISL_956332 | Department of Biology, University of Basrah | Department of Biology, University of Basrah | Abu-Ali,H.F. and Al-Badran,A.I. |
| EPI_ISL_959897, EPI_ISL_959899, EPI_ISL_959904, EPI_ISL_959908, EPI_ISL_959909, EPI_ISL_959920, EPI_ISL_959921, EPI_ISL_959922, EPI_ISL_959923, EPI_ISL_959924, EPI_ISL_959925, EPI_ISL_959926, EPI_ISL_959927, EPI_ISL_959928, EPI_ISL_959929, EPI_ISL_959930, EPI_ISL_959931, EPI_ISL_959932, EPI_ISL_959933, EPI_ISL_959934, EPI_ISL_959935, EPI_ISL_959936, EPI_ISL_959937, EPI_ISL_959938, EPI_ISL_959939, EPI_ISL_959940, EPI_ISL_959941, EPI_ISL_959942, EPI_ISL_959943, EPI_ISL_959944, EPI_ISL_959945, EPI_ISL_959946, EPI_ISL_959947, EPI_ISL_959948, EPI_ISL_959949, EPI_ISL_959950, EPI_ISL_959951, EPI_ISL_959952, EPI_ISL_959953, EPI_ISL_959954, EPI_ISL_959955, EPI_ISL_959956, EPI_ISL_959957 | | | |
| see above | University Medical Center Hamburg Eppendorf | Heinrich Pette Institute, Leibniz Institute for Experimental Virology | Alexis Robitaille, Thomas Günther, Johannes Knobloch, Martin Aepfelbacher, Nicole Fischer, Adam Grundhoff |
| EPI_ISL_960148 | Mitchells Plain Hospital wc MPH | National Health Laboratory Service/UCT | Arash Iranzadeh, Deelan Doolabh, Lynn Tyers, Bruna Galvao, Innocent Mudau, Marvin Hsiao, Kruger Marais, Diana Hardie, Stephen Korsman, Carolyn Williamson |
| EPI_ISL_960149 | Great Brak River Clinic wc GBC | National Health Laboratory Service/UCT | Arash Iranzadeh, Deelan Doolabh, Lynn Tyers, Bruna Galvao, Innocent Mudau, Marvin Hsiao, Kruger Marais, Diana Hardie, Stephen Korsman, Carolyn Williamson |
| EPI_ISL_973760, EPI_ISL_973764, EPI_ISL_973767, EPI_ISL_973769, EPI_ISL_973771, EPI_ISL_973775, EPI_ISL_973776, EPI_ISL_973779, EPI_ISL_973781, EPI_ISL_973783, EPI_ISL_973785, EPI_ISL_973788, EPI_ISL_973791, EPI_ISL_973794, EPI_ISL_973795, EPI_ISL_973797, EPI_ISL_973800, EPI_ISL_973802, EPI_ISL_973805, EPI_ISL_973807, EPI_ISL_973810, EPI_ISL_973813, EPI_ISL_973815, EPI_ISL_973818, EPI_ISL_973821, EPI_ISL_973823, EPI_ISL_973826, EPI_ISL_973828, EPI_ISL_973830, EPI_ISL_973832, EPI_ISL_973834, EPI_ISL_973837, EPI_ISL_973838, EPI_ISL_973839, EPI_ISL_973840, EPI_ISL_973841, EPI_ISL_973842, EPI_ISL_973843, EPI_ISL_973844, EPI_ISL_973845, EPI_ISL_973846, EPI_ISL_973847, EPI_ISL_973848, EPI_ISL_973849, EPI_ISL_973850, EPI_ISL_973851, EPI_ISL_973852, EPI_ISL_973853, EPI_ISL_973854, EPI_ISL_973855, EPI_ISL_973856, EPI_ISL_973857, EPI_ISL_973858, EPI_ISL_973859, EPI_ISL_973860, EPI_ISL_973861, EPI_ISL_973862, EPI_ISL_973863, EPI_ISL_973864, EPI_ISL_973865, EPI_ISL_973866, EPI_ISL_973867, EPI_ISL_973868, EPI_ISL_973869, EPI_ISL_973870, EPI_ISL_973871, EPI_ISL_973872, EPI_ISL_973873, EPI_ISL_973874, EPI_ISL_973875, EPI_ISL_973876, EPI_ISL_973877, EPI_ISL_973878, EPI_ISL_973879, EPI_ISL_973880, EPI_ISL_973881, EPI_ISL_973882, EPI_ISL_973883, EPI_ISL_973884, EPI_ISL_973885, EPI_ISL_973886, EPI_ISL_973887, EPI_ISL_973888, EPI_ISL_973889, EPI_ISL_973890, EPI_ISL_973891, EPI_ISL_973892, EPI_ISL_973893, EPI_ISL_973894, EPI_ISL_973895, EPI_ISL_973896, EPI_ISL_973897, EPI_ISL_973898, EPI_ISL_973899, EPI_ISL_973900, EPI_ISL_973901, EPI_ISL_973902, EPI_ISL_973903, EPI_ISL_973904, EPI_ISL_973905, EPI_ISL_973906, EPI_ISL_973907, EPI_ISL_973908, EPI_ISL_973909, EPI_ISL_973910, EPI_ISL_973911, EPI_ISL_973912, EPI_ISL_973913, EPI_ISL_973914, EPI_ISL_973915, EPI_ISL_973916, EPI_ISL_973917, EPI_ISL_973918, EPI_ISL_973919, EPI_ISL_973920, EPI_ISL_973921, EPI_ISL_973922, EPI_ISL_973923, EPI_ISL_973924, EPI_ISL_973925, EPI_ISL_973926, EPI_ISL_973927, EPI_ISL_973928, EPI_ISL_973929, EPI_ISL_973930, EPI_ISL_973931, EPI_ISL_973932, EPI_ISL_973933, EPI_ISL_973934, EPI_ISL_973935, EPI_ISL_973936, EPI_ISL_973937, EPI_ISL_973938, EPI_ISL_973939, EPI_ISL_973940, EPI_ISL_973941, EPI_ISL_973942, EPI_ISL_973943, EPI_ISL_973944, EPI_ISL_973945, EPI_ISL_973946, EPI_ISL_973947, EPI_ISL_973948, EPI_ISL_973949, EPI_ISL_973950, EPI_ISL_973951, EPI_ISL_973952, EPI_ISL_973953, EPI_ISL_973954, EPI_ISL_973955, EPI_ISL_973956, EPI_ISL_973957, EPI_ISL_973958, EPI_ISL_973959, EPI_ISL_973960, EPI_ISL_973961, EPI_ISL_973962, EPI_ISL_973963, EPI_ISL_973964, EPI_ISL_973965, EPI_ISL_973966, EPI_ISL_973967, EPI_ISL_973968, EPI_ISL_973969, EPI_ISL_973970, EPI_ISL_973971, EPI_ISL_973972, EPI_ISL_973973, EPI_ISL_973974, EPI_ISL_973975, EPI_ISL_973976, EPI_ISL_973977, EPI_ISL_973978, EPI_ISL_973979, EPI_ISL_973980, EPI_ISL_973981, EPI_ISL_973982, EPI_ISL_973983, EPI_ISL_973984, EPI_ISL_973985, EPI_ISL_973986, EPI_ISL_973987, EPI_ISL_973988, EPI_ISL_973989, EPI_ISL_973990, EPI_ISL_973991, EPI_ISL_973992, EPI_ISL_973993, EPI_ISL_973994, EPI_ISL_973995, EPI_ISL_973996, EPI_ISL_973997, EPI_ISL_973998, EPI_ISL_973999, EPI_ISL_974000, EPI_ISL_974001, EPI_ISL_974002, EPI_ISL_974003, EPI_ISL_974004, EPI_ISL_974005, EPI_ISL_974006, EPI_ISL_974007, EPI_ISL_974008, EPI_ISL_974009, EPI_ISL_974010, EPI_ISL_974011, EPI_ISL_974012, EPI_ISL_974013, EPI_ISL_974014, EPI_ISL_974015, EPI_ISL_974016, EPI_ISL_974017 | | | |
| see above | BCCDC Public Health Laboratory | BCCDC Public Health Laboratory | Prystajecky Natalie, Linda Hoang, Dan Fornika, John Tyson, Shannon Russell, Kim Macdonald, Kimia Kamelian, Ana Pacagnella, Corrinne Ng, Loretta Janz, Robert Azana Terry Snutch, Mel Krajden |
| EPI_ISL_977476, EPI_ISL_977477 | Instituto Adolfo Lutz Central | Instituto Adolfo Lutz, Interdiciplinary Procedures Center, Strategic Laboratory | Claudio Tavares Sacchi, Claudia Regina Gonçalves, Erica Valessa Ramos Gomes, Karoline Rodrigues Campos |
