## Supplementary material for "Detection and characterization of the SARS-CoV-2 lineage B.1.526 in New York": Supp. Table 3 GISAID Acknowledgment part 2: gisaid_hcov-19_acknowledgement_table_2021_02_12_23-10.pdf

We gratefully acknowledge the following Authors from the Originating laboratories responsible for obtaining the specimens, as well as the Submitting laboratories where the genome data were generated and shared via GISAID, on which this research is based.

All Submitters of data may be contacted directly via www.gisaid.org

Authors are sorted alphabetically.

| Accession ID | Originating Laboratory | Submitting Laboratory | Authors |
|---|---|---|---|
| EPI_ISL_576558, EPI_ISL_576559, EPI_ISL_576560, EPI_ISL_576561, EPI_ISL_576562, EPI_ISL_576563, EPI_ISL_576564 | UW Virology Lab | UW Virology Lab | Pavitra Roychoudhury, Hong Xie, Lasata Shrestha, Amin Addetia, Victoria M Rachleff, Meei-Li Huang, Keith R Jerome, Alexander Greninger |
| EPI_ISL_577098, EPI_ISL_577099, EPI_ISL_577103, EPI_ISL_577104 | Quadram Institute Bioscience | COVID-19 Genomics UK (COG-UK) Consortium | Dave J. Baker, Gemma L. Kay, Alp Aydin, Thanh Le-Viet, Steven Rudder, Ana P. Tedim, Anastasia Kolyva, Maria Diaz, Leonardo de Oliveira Martins, Nabil-Fareed Alikhan, Lizzie Meadows, Rachael Stanley, Ngozi Elumogo, Muhammed Yasir, Nicholas M. Thomson, Alexander J Trotter, Rachel Gilroy, Samuel Bloomfield, Claire Stuart, Andrew Bell, Reenesh Prakash, Samir Dervisevic, Alison E. Mather, John Wain, Mark Webber, Andrew J. Page, Justin O'Grady |
| EPI_ISL_577124, EPI_ISL_577125, EPI_ISL_577126, EPI_ISL_577127, EPI_ISL_577128, EPI_ISL_577129, EPI_ISL_577130, EPI_ISL_577131, EPI_ISL_577132, EPI_ISL_577133, EPI_ISL_577134, EPI_ISL_577135, EPI_ISL_577136, EPI_ISL_577137, EPI_ISL_577138, EPI_ISL_577139, EPI_ISL_577140, EPI_ISL_577141, EPI_ISL_577142, EPI_ISL_577143, EPI_ISL_577144, EPI_ISL_577145, EPI_ISL_577146, EPI_ISL_577147, EPI_ISL_577148, EPI_ISL_577149, EPI_ISL_577150, EPI_ISL_577151, EPI_ISL_577152, EPI_ISL_577153, EPI_ISL_577154, EPI_ISL_577155, EPI_ISL_577156, EPI_ISL_577174, EPI_ISL_577175, EPI_ISL_577176, EPI_ISL_577177, EPI_ISL_577178, EPI_ISL_577179, EPI_ISL_577180, EPI_ISL_577181, EPI_ISL_577182, EPI_ISL_577183, EPI_ISL_577184, EPI_ISL_577185, EPI_ISL_577186, EPI_ISL_577187, EPI_ISL_577188, EPI_ISL_577189, EPI_ISL_577190, EPI_ISL_577191, EPI_ISL_577192 | | | |
| see above | Queens Medical Centre, Clinical Microbiology Department / DeepSeq Nottingham | COVID-19 Genomics UK (COG-UK) Consortium | Gemma Clark, Wendy Smith, Manjinder Khakh, Vicki M Fleming, Michelle M Lister, Hannah Howson-Wells, Jonathan Ball, Patrick McClure, Joseph Chappell, Theocharis Tsoleridis, Nadine Holmes, Matthew Carlisle, Christopher Moore, Fei Sang, Johnny Debebe, Victoria Wright, Matthew Loose |
| EPI_ISL_577597 | Area of Virology, Serology and Virology Division (SAViD), New South Wales Health Pathology Randwick | Area of Virology, Serology and Virology Division (SAViD), New South Wales Health Pathology Randwick | Rawlinson, W., Bull, R., Deveson, I. |
| EPI_ISL_577604 | Area of Virology, Serology and Virology Division (SAViD), New South Wales Health Pathology Randwick | Area of Virology, Serology and Virology Division (SAViD), New South Wales Health Pathology Randwick | Rawlinson, W., Deveson, I., Bull, R., Van Hal, S. |
| EPI_ISL_578701, EPI_ISL_578702 | Molecular Pathology Division, Department of Pathology, Hong Kong Sanatorium & Hospital | Molecular Pathology Division, Department of Pathology, Hong Kong Sanatorium & Hospital | Chun Hang AU, Wai Sing CHAN, Ho Yin LAM, Dona N. HO, Simon Y.M. LAM, Jonpaul S.T. ZEE, Tsun Leung CHAN, Edmond S.K. MA |
| EPI_ISL_579116, EPI_ISL_579117, EPI_ISL_582017 | LabPLUS | Institute of Environmental Science and Research (ESR) | Xiaoyun Ren, Matt Storey, Nikki Freed, Muhammad Faisal, Jing Wang, Hermes Perez, Anja Werno, Antje van der Linden, Arlo Upton, Chris Mansell, David Hammer, Dragana Drinkovic, Gary McAuliffe, Hana Sofia Andersson, James Ussher, Jill Sherwood, Josh Freeman, Julia Howard, Juliet Elvy, Mary DeAlmeida, Matt Blakiston, Matthew Rogers, Max Bloomfield, Michael Addidle, Michelle Balm, Sally Roberts, Sarah Jefferies, Sharmini Muttaiyah, Susan Morpeth, Susan Taylor, Timothy Blackmore, Vani Sathyendran, Veronica Playle, Virginia Hope, Erasmus Smit, Lauren Jelly, Olin Silander, Joep de Ligt |
| EPI_ISL_582019, EPI_ISL_582020, EPI_ISL_582021 | PathLab Bay of Plenty | Institute of Environmental Science and Research (ESR) | Xiaoyun Ren, Matt Storey, Nikki Freed, Muhammad Faisal, Jing Wang, Hermes Perez, Anja Werno, Antje van der Linden, Arlo Upton, Chris Mansell, David Hammer, Dragana Drinkovic, Gary McAuliffe, Hana Sofia Andersson, James Ussher, Jill Sherwood, Josh Freeman, Julia Howard, Juliet Elvy, Mary DeAlmeida, Matt Blakiston, Matthew Rogers, Max Bloomfield, Michael Addidle, Michelle Balm, Sally Roberts, Sarah Jefferies, Sharmini Muttaiyah, Susan Morpeth, Susan Taylor, Timothy Blackmore, Vani Sathyendran, Veronica Playle, Virginia Hope, Erasmus Smit, Lauren Jelly, Olin Silander, Joep de Ligt |
| EPI_ISL_583533, EPI_ISL_583534 | Michigan Department of Health and Human Services, Bureau of Laboratories | Michigan Department of Health and Human Services, Bureau of Laboratories | Blankenship HM, Riner D, Soehnlen MK |
| EPI_ISL_583975, EPI_ISL_583976, EPI_ISL_583977, EPI_ISL_583978, EPI_ISL_583979, EPI_ISL_583980, EPI_ISL_583981, EPI_ISL_583988, EPI_ISL_583989, EPI_ISL_583992 | Respiratory Virus Unit, Microbiology Services Colindale, Public Health England | Respiratory Virus Unit, Microbiology Services Colindale, Public Health England | PHE Covid Sequencing Team |
| EPI_ISL_584291, EPI_ISL_584299, EPI_ISL_584300, EPI_ISL_584301, EPI_ISL_584302, EPI_ISL_584304, EPI_ISL_584305, EPI_ISL_584306, EPI_ISL_584307, EPI_ISL_584309, EPI_ISL_584310, EPI_ISL_584311, EPI_ISL_584312, EPI_ISL_584313, EPI_ISL_584322, EPI_ISL_584323 | | | |
| see above | Department of Pathology, University of Cambridge | COVID-19 Genomics UK (COG-UK) Consortium | Aminu S. Jahun, Yasmin Chaudhry, Grant Hall, Iliana Georgana, Myra Hosmillo, Martin D. Curran, Malte Pinckert, Surendra Parmar, Ian Goodfellow |
| EPI_ISL_584706, EPI_ISL_584707, EPI_ISL_584708, EPI_ISL_584709, EPI_ISL_584710, EPI_ISL_584711, EPI_ISL_584712, EPI_ISL_584713, EPI_ISL_584714, EPI_ISL_584715 | Northumbria University / South Tees Hospitals NHS Foundation Trust / North Cumbria Integrated Care NHS Foundation Trust / North Tees and Hartlepool NHS Foundation Trust / Newcastle Hospitals NHS Foundation Trust | COVID-19 Genomics UK (COG-UK) Consortium | Darren L Smith,Andrew Nelson,Matthew Bashton,Greg R Young,Joshua Loh,John Allan,Mohammad A Tariq,Giles S Holt,Gary Black,Wen C Yew,Lynn Dover,Paul Baker,Steve Liggett,Sarah Essex,Jane Greenaway,Debra Padgett,Clive Graham,Garren Scott,Edward Barton,Emma Swindells,Brendan Payne,Jennifer Collins,Yusri Taha,Gary Eltringham |
| EPI_ISL_584721, EPI_ISL_584724, EPI_ISL_584725, EPI_ISL_584726, EPI_ISL_584727, EPI_ISL_584732, EPI_ISL_584735, EPI_ISL_584737, EPI_ISL_584738, EPI_ISL_584740, EPI_ISL_584741, EPI_ISL_584742, EPI_ISL_584743, EPI_ISL_584745, EPI_ISL_584746, EPI_ISL_584747 | | | |
| see above | Quadram Institute Bioscience | COVID-19 Genomics UK (COG-UK) Consortium | Dave J. Baker, Gemma L. Kay, Alp Aydin, Thanh Le-Viet, Steven Rudder, Ana P. Tedim, Anastasia Kolyva, Maria Diaz, Leonardo de Oliveira Martins, Nabil-Fareed Alikhan, Lizzie Meadows, Rachael Stanley, Ngozi Elumogo, Muhammed Yasir, Nicholas M. Thomson, Alexander J Trotter, Rachel Gilroy, Samuel Bloomfield, Claire Stuart, Andrew Bell, Reenesh Prakash, Samir Dervisevic, Alison E. Mather, John Wain, Mark Webber, Andrew J. Page, Justin O'Grady |
| EPI_ISL_584883, EPI_ISL_584884, EPI_ISL_584885, EPI_ISL_584886, EPI_ISL_584887, EPI_ISL_584888, EPI_ISL_584889, EPI_ISL_584890, EPI_ISL_584891, EPI_ISL_584892, EPI_ISL_584893, EPI_ISL_584894, EPI_ISL_584895, EPI_ISL_584896, EPI_ISL_584897, EPI_ISL_584900, EPI_ISL_584903, EPI_ISL_584904, EPI_ISL_584905 | | | |
| see above | University of Exeter | COVID-19 Genomics UK (COG-UK) Consortium | Ben Temperton,Aaron Jeffries,Michelle Michelsen,Joanna Warwick-Dugdale,Audrey Farbos,Robyn Manley,Stephen Michell,Jane Masoli |
| EPI_ISL_584953, EPI_ISL_584954, EPI_ISL_584955, EPI_ISL_584957, EPI_ISL_584958, EPI_ISL_584959, EPI_ISL_584960, EPI_ISL_584974, EPI_ISL_584975, EPI_ISL_584976, EPI_ISL_584977, EPI_ISL_584978, EPI_ISL_584979, EPI_ISL_584980, EPI_ISL_585000, EPI_ISL_585002, EPI_ISL_585003, EPI_ISL_585004, EPI_ISL_585005 | | | |
| see above | Quadram Institute Bioscience | COVID-19 Genomics UK (COG-UK) Consortium | Dave J. Baker, Gemma L. Kay, Alp Aydin, Thanh Le-Viet, Steven Rudder, Ana P. Tedim, Anastasia Kolyva, Maria Diaz, Leonardo de Oliveira Martins, Nabil-Fareed Alikhan, Lizzie Meadows, Rachael Stanley, Ngozi Elumogo, Muhammed Yasir, Nicholas M. Thomson, Alexander J Trotter, Rachel Gilroy, Samuel Bloomfield, Claire Stuart, Andrew Bell, Reenesh Prakash, Samir Dervisevic, Alison E. Mather, John Wain, Mark Webber, Andrew J. Page, Justin O'Grady |
| EPI_ISL_585009, EPI_ISL_585010, EPI_ISL_585012, EPI_ISL_585016, EPI_ISL_585018, EPI_ISL_585023, EPI_ISL_585027, EPI_ISL_585028, EPI_ISL_585029, EPI_ISL_585033, EPI_ISL_585042, EPI_ISL_585046, EPI_ISL_585047, EPI_ISL_585050, EPI_ISL_585065, EPI_ISL_585067, EPI_ISL_585069, EPI_ISL_585074, EPI_ISL_585075, EPI_ISL_585079, EPI_ISL_585081, EPI_ISL_585084, EPI_ISL_585091 | | | |
| see above | Virology Department, Sheffield Teaching Hospitals NHS Foundation Trust/Department of Infection, Immunity and Cardiovascular Disease, The Medical School, University of Sheffield | COVID-19 Genomics UK (COG-UK) Consortium | Thushan de Silva, Matthew Parker, Nikki Smith, Adri Angyal, Rebecca Brown, Luke Green, Rachel Tucker, Paul Parsons, Danielle Groves, Katie Johnson, Laura Carrilero, Alex Keeley, Dave Partridge, Matthew Wyles, Benjamin Lindsey, Mehmet Yavuz, Mohammad Raza, Cariad Evans |
| EPI_ISL_585474, EPI_ISL_585475, EPI_ISL_585476, EPI_ISL_585477, EPI_ISL_585478, EPI_ISL_585479, EPI_ISL_585480, EPI_ISL_585482, EPI_ISL_585483, EPI_ISL_585489, EPI_ISL_585490, EPI_ISL_585491, EPI_ISL_585494, EPI_ISL_585495, EPI_ISL_585498, EPI_ISL_585499 | | | |
| see above | Virology Department, Royal Infirmary of Edinburgh, NHS Lothian / School of Biological Sciences, University of | COVID-19 Genomics UK (COG-UK) Consortium | McHugh M, Dewar R, Rooke S, Gallagher M, Balcaza C, O'Toole Á, Scher E, Hill V, McCrone JT, Colquhoun R, Yu X, Jackson B, Rambaut A, Williams TC, Templeton K |

| | | | |
|---|---|---|---|
| | Edinburgh / Institute of Genetics and Molecular Medicine, University of Edinburgh | | |
| EPI_ISL_585533, EPI_ISL_585534, EPI_ISL_585535, EPI_ISL_585536, EPI_ISL_585537, EPI_ISL_585538, EPI_ISL_585539, EPI_ISL_585540, EPI_ISL_585541, EPI_ISL_585542, EPI_ISL_585543, EPI_ISL_585544, EPI_ISL_585545, EPI_ISL_585546, EPI_ISL_585547, EPI_ISL_585548, EPI_ISL_585549, EPI_ISL_585550, EPI_ISL_585551, EPI_ISL_585552, EPI_ISL_585553, EPI_ISL_585554, EPI_ISL_585555, EPI_ISL_585556, EPI_ISL_585557, EPI_ISL_585558, EPI_ISL_585559, EPI_ISL_585560, EPI_ISL_585561, EPI_ISL_585562, EPI_ISL_585563, EPI_ISL_585564, EPI_ISL_585565, EPI_ISL_585566, EPI_ISL_585567, EPI_ISL_585568, EPI_ISL_585569, EPI_ISL_585570, EPI_ISL_585571, EPI_ISL_585572, EPI_ISL_585573, EPI_ISL_585574, EPI_ISL_585575, EPI_ISL_585576, EPI_ISL_585577, EPI_ISL_585578, EPI_ISL_585579, EPI_ISL_585580, EPI_ISL_585581, EPI_ISL_585582 | | | |
| see above | University of Exeter | COVID-19 Genomics UK (COG-UK) Consortium | Ben Temperton,Aaron Jeffries,Michelle Michelsen,Joanna Warwick-Dugdale,Audrey Farbos,Robyn Manley,Stephen Michell,Jane Masoli |
| EPI_ISL_585598, EPI_ISL_585599, EPI_ISL_585600, EPI_ISL_585601, EPI_ISL_585602, EPI_ISL_585603, EPI_ISL_585604, EPI_ISL_585605, EPI_ISL_585609, EPI_ISL_585610, EPI_ISL_585611, EPI_ISL_585612, EPI_ISL_585613, EPI_ISL_585614, EPI_ISL_585615, EPI_ISL_585616, EPI_ISL_585617, EPI_ISL_585618 | | | |
| see above | Quadram Institute Bioscience | COVID-19 Genomics UK (COG-UK) Consortium | Dave J. Baker, Gemma L. Kay, Alp Aydin, Thanh Le-Viet, Steven Rudder, Ana P. Tedim, Anastasia Kolyva, Maria Diaz, Leonardo de Oliveira Martins, Nabil-Fareed Alikhan, Lizzie Meadows, Rachael Stanley, Ngozi Elumogo, Muhammed Yasir, Nicholas M. Thomson, Alexander J Trotter, Rachel Gilroy, Samuel Bloomfield, Claire Stuart, Andrew Bell, Reenesh Prakash, Samir Dervisevic, Alison E. Mather, John Wain, Mark Webber, Andrew J. Page, Justin O'Grady |
| EPI_ISL_585630, EPI_ISL_585632, EPI_ISL_585636, EPI_ISL_585644, EPI_ISL_585648, EPI_ISL_585652, EPI_ISL_585656, EPI_ISL_585659, EPI_ISL_585662, EPI_ISL_585670, EPI_ISL_585671, EPI_ISL_585673, EPI_ISL_585680, EPI_ISL_585682, EPI_ISL_585688, EPI_ISL_585693, EPI_ISL_585695, EPI_ISL_585696, EPI_ISL_585697, EPI_ISL_585701, EPI_ISL_585704, EPI_ISL_585706, EPI_ISL_585717, EPI_ISL_585724, EPI_ISL_585727, EPI_ISL_585731, EPI_ISL_585734, EPI_ISL_585737, EPI_ISL_585740, EPI_ISL_585742, EPI_ISL_585743, EPI_ISL_585744, EPI_ISL_585745, EPI_ISL_585748, EPI_ISL_585753, EPI_ISL_585764, EPI_ISL_585766, EPI_ISL_585767, EPI_ISL_585768, EPI_ISL_585769, EPI_ISL_585770, EPI_ISL_585771, EPI_ISL_585772, EPI_ISL_585773, EPI_ISL_585774, EPI_ISL_585775, EPI_ISL_585777, EPI_ISL_585778, EPI_ISL_585780, EPI_ISL_585784, EPI_ISL_585785, EPI_ISL_585786, EPI_ISL_585787, EPI_ISL_585788, EPI_ISL_585789, EPI_ISL_585790, EPI_ISL_585792, EPI_ISL_585793, EPI_ISL_585795, EPI_ISL_585799, EPI_ISL_585802, EPI_ISL_585803, EPI_ISL_585805, EPI_ISL_585808, EPI_ISL_585809, EPI_ISL_585810, EPI_ISL_585811, EPI_ISL_585812, EPI_ISL_585813, EPI_ISL_585814, EPI_ISL_585815, EPI_ISL_585816, EPI_ISL_585817, EPI_ISL_585818, EPI_ISL_585819, EPI_ISL_585820, EPI_ISL_585821, EPI_ISL_585822, EPI_ISL_585823, EPI_ISL_585824, EPI_ISL_585826, EPI_ISL_585827, EPI_ISL_585828, EPI_ISL_585829, EPI_ISL_585831, EPI_ISL_585832, EPI_ISL_585836, EPI_ISL_585837, EPI_ISL_585838, EPI_ISL_585839, EPI_ISL_585840, EPI_ISL_585841, EPI_ISL_585842, EPI_ISL_585843, EPI_ISL_585844, EPI_ISL_585845, EPI_ISL_585846, EPI_ISL_585847, EPI_ISL_585848, EPI_ISL_585849, EPI_ISL_585850, EPI_ISL_585851, EPI_ISL_585852, EPI_ISL_585853, EPI_ISL_585854, EPI_ISL_585855, EPI_ISL_585856, EPI_ISL_585858, EPI_ISL_585859, EPI_ISL_585860, EPI_ISL_585861, EPI_ISL_585863, EPI_ISL_585865, EPI_ISL_585866, EPI_ISL_585868, EPI_ISL_585869, EPI_ISL_585870, EPI_ISL_585871, EPI_ISL_585872, EPI_ISL_585873, EPI_ISL_585874, EPI_ISL_585875, EPI_ISL_585877, EPI_ISL_585880, EPI_ISL_585881, EPI_ISL_585882, EPI_ISL_585883, EPI_ISL_585884, EPI_ISL_585885, EPI_ISL_585889, EPI_ISL_585890, EPI_ISL_585891, EPI_ISL_585894, EPI_ISL_585896, EPI_ISL_585897, EPI_ISL_585898, EPI_ISL_585901, EPI_ISL_585902, EPI_ISL_585906, EPI_ISL_585907, EPI_ISL_585910, EPI_ISL_585911, EPI_ISL_585912, EPI_ISL_585913, EPI_ISL_585914, EPI_ISL_585915, EPI_ISL_585916, EPI_ISL_585917, EPI_ISL_585921, EPI_ISL_585922, EPI_ISL_585923, EPI_ISL_585924, EPI_ISL_585925, EPI_ISL_585927, EPI_ISL_585928, EPI_ISL_585932, EPI_ISL_585933, EPI_ISL_585934, EPI_ISL_585935, EPI_ISL_585936, EPI_ISL_585939, EPI_ISL_585941, EPI_ISL_585943, EPI_ISL_585944, EPI_ISL_585945, EPI_ISL_585946, EPI_ISL_585947, EPI_ISL_585948, EPI_ISL_585949, EPI_ISL_585950, EPI_ISL_585951, EPI_ISL_585952, EPI_ISL_585953, EPI_ISL_585954, EPI_ISL_585956, EPI_ISL_585957, EPI_ISL_585958, EPI_ISL_585959, EPI_ISL_585960, EPI_ISL_585961, EPI_ISL_585963, EPI_ISL_585965, EPI_ISL_585966, EPI_ISL_585967, EPI_ISL_585968, EPI_ISL_585969, EPI_ISL_585970, EPI_ISL_585971, EPI_ISL_585972, EPI_ISL_585973, EPI_ISL_585974, EPI_ISL_585976, EPI_ISL_585979, EPI_ISL_585981, EPI_ISL_585982, EPI_ISL_585983, EPI_ISL_585984, EPI_ISL_585987, EPI_ISL_585989, EPI_ISL_585991, EPI_ISL_585992, EPI_ISL_585993, EPI_ISL_585994, EPI_ISL_585995, EPI_ISL_585996, EPI_ISL_585997, EPI_ISL_585999, EPI_ISL_586000, EPI_ISL_586001, EPI_ISL_586002, EPI_ISL_586003, EPI_ISL_586004, EPI_ISL_586006, EPI_ISL_586007, EPI_ISL_586008, EPI_ISL_586009, EPI_ISL_586010, EPI_ISL_586015, EPI_ISL_586016, EPI_ISL_586018, EPI_ISL_586019, EPI_ISL_586021, EPI_ISL_586022, EPI_ISL_586031, EPI_ISL_586033, EPI_ISL_586034, EPI_ISL_586037, EPI_ISL_586043, EPI_ISL_586048, EPI_ISL_586052, EPI_ISL_586053, EPI_ISL_586054, EPI_ISL_586055, EPI_ISL_586063, EPI_ISL_586065, EPI_ISL_586070, EPI_ISL_586079, EPI_ISL_586089, EPI_ISL_586091, EPI_ISL_586094, EPI_ISL_586096, EPI_ISL_586097, EPI_ISL_586099, EPI_ISL_586100, EPI_ISL_586108, EPI_ISL_586114, EPI_ISL_586121, EPI_ISL_586122, EPI_ISL_586133, EPI_ISL_586142, EPI_ISL_586145, EPI_ISL_586147, EPI_ISL_586154, EPI_ISL_586155, EPI_ISL_586156, EPI_ISL_586159, EPI_ISL_586163, EPI_ISL_586169, EPI_ISL_586171, EPI_ISL_586180, EPI_ISL_586185, EPI_ISL_586193, EPI_ISL_586195, EPI_ISL_586199, EPI_ISL_586200, EPI_ISL_586202, EPI_ISL_586208, EPI_ISL_586221, EPI_ISL_586227, EPI_ISL_586230, EPI_ISL_586232, EPI_ISL_586235, EPI_ISL_586237 | | | |
| see above | Wales Specialist Virology Centre Sequencing lab: Pathogen Genomics Unit | COVID-19 Genomics UK (COG-UK) Consortium | Catherine Moore, Johnathan Evans, Laura Gifford, Malorie Perry, Simon Cottrell, Angela Marchbank, Alec Birchley, Alexander Adams, Amy Gaskin, Bree Gatica-Wilcox, Jason Coombes, Joel Southgate, Lauren Gilbert, Lee Graham, Nicole Pacchiarini, Sara Kumziene-Summerhayes, Sarah Taylor, Sophie Jones, Sara Rey, Matthew Bull, Joanne Watkins, Sally Corden, Tom Connor |
| EPI_ISL_586572, EPI_ISL_586573, EPI_ISL_586574, EPI_ISL_586575, EPI_ISL_586576, EPI_ISL_586577, EPI_ISL_586578, EPI_ISL_586579, EPI_ISL_586580, EPI_ISL_586581, EPI_ISL_586582, EPI_ISL_586583, EPI_ISL_586584, EPI_ISL_586585, EPI_ISL_586586, EPI_ISL_586587, EPI_ISL_586588, EPI_ISL_586589, EPI_ISL_586590, EPI_ISL_586591, EPI_ISL_586592, EPI_ISL_586593, EPI_ISL_586594, EPI_ISL_586595, EPI_ISL_586596, EPI_ISL_586597, EPI_ISL_586598, EPI_ISL_586599, EPI_ISL_586600, EPI_ISL_586601, EPI_ISL_586602, EPI_ISL_586603, EPI_ISL_586604, EPI_ISL_586605, EPI_ISL_586606, EPI_ISL_586607, EPI_ISL_586608, EPI_ISL_586609, EPI_ISL_586610, EPI_ISL_586611 | | | |
| see above | Lighthouse Lab in Alderley Park | Wellcome Sanger Institute for the COVID-19 Genomics UK (COG-UK) consortium | Jacquelyn Wynn, Mairead Hyland, The Lighthouse Lab in Alderley Park and Alex Alderton, Roberto Amato, Sonia Goncalves, Ewan Harrison, David K. Jackson, Ian Johnston, Dominic Kwiatkowski, Cordelia Langford, John Sillitoe on behalf of the Wellcome Sanger Institute COVID-19 Surveillance Team (http://www.sanger.ac.uk/covid-team) |
| EPI_ISL_586612 | Lighthouse Lab in Alderley Park | Wellcome Sanger Institute for the COVID-19 Genomics UK (COG-UK) Consortium | Jacquelyn Wynn, Mairead Hyland, The Lighthouse Lab in Alderley Park and Alex Alderton, Roberto Amato, Sonia Goncalves, Ewan Harrison, David K. Jackson, Ian Johnston, Dominic Kwiatkowski, Cordelia Langford, John Sillitoe on behalf of the Wellcome Sanger Institute COVID-19 Surveillance Team (http://www.sanger.ac.uk/covid-team) |
| EPI_ISL_586613, EPI_ISL_586614, EPI_ISL_586615, EPI_ISL_586616 | Lighthouse Lab in Alderley Park | Wellcome Sanger Institute for the COVID-19 Genomics UK (COG-UK) consortium | Jacquelyn Wynn, Mairead Hyland, The Lighthouse Lab in Alderley Park and Alex Alderton, Roberto Amato, Sonia Goncalves, Ewan Harrison, David K. Jackson, Ian Johnston, Dominic Kwiatkowski, Cordelia Langford, John Sillitoe on behalf of the Wellcome Sanger Institute COVID-19 Surveillance Team (http://www.sanger.ac.uk/covid-team) |
| EPI_ISL_586617 | Lighthouse Lab in Alderley Park | Wellcome Sanger Institute for the COVID-19 Genomics UK (COG-UK) Consortium | Jacquelyn Wynn, Mairead Hyland, The Lighthouse Lab in Alderley Park and Alex Alderton, Roberto Amato, Sonia Goncalves, Ewan Harrison, David K. Jackson, Ian Johnston, Dominic Kwiatkowski, Cordelia Langford, John Sillitoe on behalf of the Wellcome Sanger Institute COVID-19 Surveillance Team (http://www.sanger.ac.uk/covid-team) |
| EPI_ISL_586618, EPI_ISL_586619, EPI_ISL_586620 | Lighthouse Lab in Alderley Park | Wellcome Sanger Institute for the COVID-19 Genomics UK (COG-UK) consortium | Jacquelyn Wynn, Mairead Hyland, The Lighthouse Lab in Alderley Park and Alex Alderton, Roberto Amato, Sonia Goncalves, Ewan Harrison, David K. Jackson, Ian Johnston, Dominic Kwiatkowski, Cordelia Langford, John Sillitoe on behalf of the Wellcome Sanger Institute COVID-19 Surveillance Team (http://www.sanger.ac.uk/covid-team) |
| EPI_ISL_586621 | Lighthouse Lab in Alderley Park | Wellcome Sanger Institute for the COVID-19 Genomics UK (COG-UK) Consortium | Jacquelyn Wynn, Mairead Hyland, The Lighthouse Lab in Alderley Park and Alex Alderton, Roberto Amato, Sonia Goncalves, Ewan Harrison, David K. Jackson, Ian Johnston, Dominic Kwiatkowski, Cordelia Langford, John Sillitoe on behalf of the Wellcome Sanger Institute COVID-19 Surveillance Team (http://www.sanger.ac.uk/covid-team) |
| EPI_ISL_586622, EPI_ISL_586623, EPI_ISL_586624, EPI_ISL_586625, EPI_ISL_586626, EPI_ISL_586627, EPI_ISL_586628, EPI_ISL_586629, EPI_ISL_586630, EPI_ISL_586631, EPI_ISL_586632, EPI_ISL_586633, EPI_ISL_586634, EPI_ISL_586635, EPI_ISL_586636, EPI_ISL_586637, EPI_ISL_586638, EPI_ISL_586639, EPI_ISL_586640, EPI_ISL_586641, EPI_ISL_586642, EPI_ISL_586643, EPI_ISL_586644, EPI_ISL_586645, EPI_ISL_586646, EPI_ISL_586647, EPI_ISL_586648, EPI_ISL_586649, EPI_ISL_586650, EPI_ISL_586651, EPI_ISL_586652, EPI_ISL_586653, EPI_ISL_586654, EPI_ISL_586655, EPI_ISL_586656, EPI_ISL_586657, EPI_ISL_586658, EPI_ISL_586659, EPI_ISL_586660, EPI_ISL_586661, EPI_ISL_586662, EPI_ISL_586663, EPI_ISL_586664, EPI_ISL_586665, EPI_ISL_586666, EPI_ISL_586667, EPI_ISL_586668, EPI_ISL_586669, EPI_ISL_586670, EPI_ISL_586671, EPI_ISL_586672, EPI_ISL_586673, EPI_ISL_586674, EPI_ISL_586675, EPI_ISL_586676, EPI_ISL_586677, EPI_ISL_586678, EPI_ISL_586679, EPI_ISL_586680, EPI_ISL_586681, EPI_ISL_586682, EPI_ISL_586683, EPI_ISL_586684, EPI_ISL_586685, EPI_ISL_586686, EPI_ISL_586687, EPI_ISL_586688, EPI_ISL_586689, EPI_ISL_586690, EPI_ISL_586691, EPI_ISL_586692, EPI_ISL_586693, EPI_ISL_586694, EPI_ISL_586695, EPI_ISL_586696, EPI_ISL_586697, EPI_ISL_586698, EPI_ISL_586699, EPI_ISL_586700, EPI_ISL_586701, EPI_ISL_586702, EPI_ISL_586703, EPI_ISL_586704, EPI_ISL_586705, EPI_ISL_586706, EPI_ISL_586707, EPI_ISL_586708 | | | |
| see above | Lighthouse Lab in Alderley Park | Wellcome Sanger Institute for the COVID-19 Genomics UK (COG-UK) consortium | Jacquelyn Wynn, Mairead Hyland, The Lighthouse Lab in Alderley Park and Alex Alderton, Roberto Amato, Sonia Goncalves, Ewan Harrison, David K. Jackson, Ian Johnston, Dominic Kwiatkowski, Cordelia Langford, John Sillitoe on behalf of the Wellcome Sanger Institute COVID-19 Surveillance Team (http://www.sanger.ac.uk/covid-team) |
| EPI_ISL_586709 | Lighthouse Lab in Alderley Park | Wellcome Sanger Institute for the COVID-19 Genomics UK (COG-UK) Consortium | Jacquelyn Wynn, Mairead Hyland, The Lighthouse Lab in Alderley Park and Alex Alderton, Roberto Amato, Sonia Goncalves, Ewan Harrison, David K. Jackson, Ian Johnston, Dominic Kwiatkowski, Cordelia Langford, John Sillitoe on behalf of the Wellcome Sanger Institute COVID-19 Surveillance Team (http://www.sanger.ac.uk/covid-team) |
| EPI_ISL_586710, EPI_ISL_586711, EPI_ISL_586712, EPI_ISL_586713, EPI_ISL_586714, EPI_ISL_586715, EPI_ISL_586716, EPI_ISL_586717, EPI_ISL_586718, EPI_ISL_586719, EPI_ISL_586720, EPI_ISL_586721, EPI_ISL_586722, EPI_ISL_586723, EPI_ISL_586724, EPI_ISL_586725, EPI_ISL_586726, EPI_ISL_586727, EPI_ISL_586728, EPI_ISL_586729, EPI_ISL_586730, EPI_ISL_586731, EPI_ISL_586732, EPI_ISL_586733, EPI_ISL_586734, EPI_ISL_586735, EPI_ISL_586736, EPI_ISL_586737, EPI_ISL_586738, EPI_ISL_586739, EPI_ISL_586740, EPI_ISL_586741, EPI_ISL_586742, EPI_ISL_586743, EPI_ISL_586744, EPI_ISL_586745, EPI_ISL_586746, EPI_ISL_586747, EPI_ISL_586748, EPI_ISL_586749, EPI_ISL_586750, EPI_ISL_586751, EPI_ISL_586752, EPI_ISL_586753, EPI_ISL_586754, EPI_ISL_586755, EPI_ISL_586756, EPI_ISL_586757, EPI_ISL_586758, EPI_ISL_586759, EPI_ISL_586760, EPI_ISL_586761, EPI_ISL_586762, EPI_ISL_586763, EPI_ISL_586764, EPI_ISL_586765, EPI_ISL_586766, EPI_ISL_586767, EPI_ISL_586768, EPI_ISL_586769, EPI_ISL_586770, EPI_ISL_586771, EPI_ISL_586772, EPI_ISL_586773, EPI_ISL_586774, EPI_ISL_586775, EPI_ISL_586776, EPI_ISL_586777, EPI_ISL_586778, EPI_ISL_586779, EPI_ISL_586780, EPI_ISL_586781, EPI_ISL_586782, EPI_ISL_586783, EPI_ISL_586784, EPI_ISL_586785, EPI_ISL_586786, EPI_ISL_586787, EPI_ISL_586788, EPI_ISL_586789, EPI_ISL_586790, EPI_ISL_586791, EPI_ISL_586792, EPI_ISL_586793, EPI_ISL_586794, EPI_ISL_586795, EPI_ISL_586796, EPI_ISL_586797, EPI_ISL_586798, EPI_ISL_586799, EPI_ISL_586800, EPI_ISL_586801, EPI_ISL_586802, EPI_ISL_586803, EPI_ISL_586804, EPI_ISL_586805, EPI_ISL_586806, EPI_ISL_586807, EPI_ISL_586808, EPI_ISL_586809, EPI_ISL_586810, EPI_ISL_586811, EPI_ISL_586812, EPI_ISL_586813, EPI_ISL_586814, EPI_ISL_586815, EPI_ISL_586816, EPI_ISL_586817, EPI_ISL_586818, EPI_ISL_586819, EPI_ISL_586820, EPI_ISL_586821, EPI_ISL_586822, EPI_ISL_586823, EPI_ISL_586824, EPI_ISL_586825, EPI_ISL_586826, EPI_ISL_586827, EPI_ISL_586828, EPI_ISL_586829, EPI_ISL_586830, EPI_ISL_586831, EPI_ISL_586832, EPI_ISL_586833, EPI_ISL_586834, EPI_ISL_586835, EPI_ISL_586836, EPI_ISL_586837, EPI_ISL_586838, EPI_ISL_586839, EPI_ISL_586840, EPI_ISL_586841, EPI_ISL_586842, EPI_ISL_586843, EPI_ISL_586844, EPI_ISL_586845, EPI_ISL_586846, EPI_ISL_586847, EPI_ISL_586848, EPI_ISL_586849, EPI_ISL_586850, EPI_ISL_586851, EPI_ISL_586852, EPI_ISL_586853, EPI_ISL_586854, EPI_ISL_586855, EPI_ISL_586856, EPI_ISL_586857, EPI_ISL_586858, EPI_ISL_586859, EPI_ISL_586860, EPI_ISL_586861, EPI_ISL_586862, EPI_ISL_586863, EPI_ISL_586864, EPI_ISL_586865, EPI_ISL_586866, EPI_ISL_586867, EPI_ISL_586868, EPI_ISL_586869, EPI_ISL_586870, EPI_ISL_586871, EPI_ISL_586872, EPI_ISL_586873, EPI_ISL_586874, EPI_ISL_586875, EPI_ISL_586876, EPI_ISL_586877, EPI_ISL_586878, EPI_ISL_586879, EPI_ISL_586880, EPI_ISL_586881, EPI_ISL_586882, EPI_ISL_586883, EPI_ISL_586884, EPI_ISL_586885, EPI_ISL_586886, EPI_ISL_586887, EPI_ISL_586888, EPI_ISL_586889, EPI_ISL_586890, EPI_ISL_586891, EPI_ISL_586892, EPI_ISL_586893, EPI_ISL_586894, EPI_ISL_586895, EPI_ISL_586896, EPI_ISL_586897, EPI_ISL_586898, EPI_ISL_586899, EPI_ISL_586900, EPI_ISL_586901, EPI_ISL_586902, EPI_ISL_586903, EPI_ISL_586904, EPI_ISL_586905, EPI_ISL_586906 | | | |
| see above | Lighthouse Lab in Alderley Park | Wellcome Sanger Institute for the COVID-19 Genomics UK (COG-UK) consortium | Jacquelyn Wynn, Mairead Hyland, The Lighthouse Lab in Alderley Park and Alex Alderton, Roberto Amato, Sonia Goncalves, Ewan Harrison, David K. Jackson, Ian Johnston, Dominic Kwiatkowski, Cordelia Langford, John Sillitoe on behalf of the Wellcome Sanger Institute COVID-19 Surveillance Team (http://www.sanger.ac.uk/covid-team) |
| EPI_ISL_586907 | Lighthouse Lab in Alderley Park | Wellcome Sanger Institute for the COVID-19 Genomics UK (COG-UK) Consortium | Jacquelyn Wynn, Mairead Hyland, The Lighthouse Lab in Alderley Park and Alex Alderton, Roberto Amato, Sonia Goncalves, Ewan Harrison, David K. Jackson, Ian Johnston, Dominic Kwiatkowski, Cordelia Langford, John Sillitoe on behalf of the Wellcome Sanger Institute COVID-19 Surveillance Team |

| | | | |
|---|---|---|---|
| | | | (http://www.sanger.ac.uk/covid-team) |
| EPI_ISL_586908, EPI_ISL_586909, EPI_ISL_586910, EPI_ISL_586911, EPI_ISL_586912, EPI_ISL_586913, EPI_ISL_586914, EPI_ISL_586915, EPI_ISL_586916, EPI_ISL_586917, EPI_ISL_586918, EPI_ISL_586919, EPI_ISL_586920, EPI_ISL_586921, EPI_ISL_586922, EPI_ISL_586923, EPI_ISL_586924, EPI_ISL_586925, EPI_ISL_586926, EPI_ISL_586927, EPI_ISL_586928, EPI_ISL_586929, EPI_ISL_586930, EPI_ISL_586931, EPI_ISL_586932, EPI_ISL_586933, EPI_ISL_586934, EPI_ISL_586935, EPI_ISL_586936, EPI_ISL_586937, EPI_ISL_586938, EPI_ISL_586939, EPI_ISL_586940, EPI_ISL_586941, EPI_ISL_586942, EPI_ISL_586943, EPI_ISL_586944, EPI_ISL_586945, EPI_ISL_586946, EPI_ISL_586947, EPI_ISL_586948, EPI_ISL_586949, EPI_ISL_586950, EPI_ISL_586951, EPI_ISL_586952, EPI_ISL_586953, EPI_ISL_586954, EPI_ISL_586955, EPI_ISL_586956, EPI_ISL_586957, EPI_ISL_586958, EPI_ISL_586959, EPI_ISL_586960, EPI_ISL_586961, EPI_ISL_586962, EPI_ISL_586963, EPI_ISL_586964, EPI_ISL_586965, EPI_ISL_586966, EPI_ISL_586967, EPI_ISL_586968, EPI_ISL_586969, EPI_ISL_586970, EPI_ISL_586971, EPI_ISL_586972, EPI_ISL_586973, EPI_ISL_586974, EPI_ISL_586975, EPI_ISL_586976, EPI_ISL_586977, EPI_ISL_586978, EPI_ISL_586979, EPI_ISL_586980, EPI_ISL_586981, EPI_ISL_586982, EPI_ISL_586983, EPI_ISL_586984, EPI_ISL_586985, EPI_ISL_586986, EPI_ISL_586987, EPI_ISL_586988, EPI_ISL_586989, EPI_ISL_586990, EPI_ISL_586991, EPI_ISL_586992, EPI_ISL_586993, EPI_ISL_586994, EPI_ISL_586995, EPI_ISL_586996, EPI_ISL_586997, EPI_ISL_586998, EPI_ISL_586999, EPI_ISL_587000, EPI_ISL_587001, EPI_ISL_587002, EPI_ISL_587003, EPI_ISL_587004, EPI_ISL_587005, EPI_ISL_587006, EPI_ISL_587007, EPI_ISL_587008, EPI_ISL_587009, EPI_ISL_587010, EPI_ISL_587011, EPI_ISL_587012, EPI_ISL_587013, EPI_ISL_587014, EPI_ISL_587015, EPI_ISL_587016, EPI_ISL_587017, EPI_ISL_587018, EPI_ISL_587019, EPI_ISL_587020, EPI_ISL_587021, EPI_ISL_587022, EPI_ISL_587023, EPI_ISL_587024, EPI_ISL_587025, EPI_ISL_587026, EPI_ISL_587027, EPI_ISL_587028, EPI_ISL_587029, EPI_ISL_587030, EPI_ISL_587031, EPI_ISL_587032, EPI_ISL_587033, EPI_ISL_587034, EPI_ISL_587035, EPI_ISL_587036, EPI_ISL_587037, EPI_ISL_587038, EPI_ISL_587039, EPI_ISL_587040, EPI_ISL_587041, EPI_ISL_587042, EPI_ISL_587043, EPI_ISL_587044, EPI_ISL_587045, EPI_ISL_587046, EPI_ISL_587047, EPI_ISL_587048, EPI_ISL_587049, EPI_ISL_587050, EPI_ISL_587051, EPI_ISL_587052, EPI_ISL_587053, EPI_ISL_587054, EPI_ISL_587055, EPI_ISL_587056, EPI_ISL_587057, EPI_ISL_587058, EPI_ISL_587059, EPI_ISL_587060, EPI_ISL_587061, EPI_ISL_587062, EPI_ISL_587063, EPI_ISL_587064, EPI_ISL_587065, EPI_ISL_587066, EPI_ISL_587067, EPI_ISL_587068, EPI_ISL_587069, EPI_ISL_587070, EPI_ISL_587071 | | | |
| see above | Lighthouse Lab in Alderley Park | Wellcome Sanger Institute for the COVID-19 Genomics UK (COG-UK) consortium | Jacquelyn Wynn, Mairead Hyland, The Lighthouse Lab in Alderley Park and Alex Alderton, Roberto Amato, Sonia Goncalves, Ewan Harrison, David K. Jackson, Ian Johnston, Dominic Kwiatkowski, Cordelia Langford, John Sillitoe on behalf of the Wellcome Sanger Institute COVID-19 Surveillance Team (http://www.sanger.ac.uk/covid-team) |
| EPI_ISL_587072, EPI_ISL_587073, EPI_ISL_587074, EPI_ISL_587075, EPI_ISL_587076, EPI_ISL_587077, EPI_ISL_587078, EPI_ISL_587079, EPI_ISL_587080, EPI_ISL_587081, EPI_ISL_587082, EPI_ISL_587083, EPI_ISL_587084 | | | |
| see above | Lighthouse Lab in Alderley Park | Wellcome Sanger Institute for the COVID-19 Genomics UK (COG-UK) consortium | Jacquelyn Wynn, Mairead Hyland, The Lighthouse Lab in Alderley Park and Alex Alderton, Roberto Amato, Sonia Goncalves, Ewan Harrison, David K. Jackson, Ian Johnston, Dominic Kwiatkowski, Cordelia Langford, John Sillitoe on behalf of the Wellcome Sanger Institute COVID-19 Surveillance Team |
| EPI_ISL_587085 | Lighthouse Lab in Alderley Park | Wellcome Sanger Institute for the COVID-19 Genomics UK (COG-UK) Consortium | Jacquelyn Wynn, Mairead Hyland, The Lighthouse Lab in Alderley Park and Alex Alderton, Roberto Amato, Sonia Goncalves, Ewan Harrison, David K. Jackson, Ian Johnston, Dominic Kwiatkowski, Cordelia Langford, John Sillitoe on behalf of the Wellcome Sanger Institute COVID-19 Surveillance Team |
| EPI_ISL_587086, EPI_ISL_587087, EPI_ISL_587088, EPI_ISL_587089, EPI_ISL_587090, EPI_ISL_587091, EPI_ISL_587092, EPI_ISL_587093, EPI_ISL_587094, EPI_ISL_587095, EPI_ISL_587096, EPI_ISL_587097, EPI_ISL_587098, EPI_ISL_587099, EPI_ISL_587100, EPI_ISL_587101, EPI_ISL_587102, EPI_ISL_587103, EPI_ISL_587104, EPI_ISL_587105, EPI_ISL_587106, EPI_ISL_587107, EPI_ISL_587108, EPI_ISL_587109, EPI_ISL_587110, EPI_ISL_587111, EPI_ISL_587112, EPI_ISL_587113, EPI_ISL_587114, EPI_ISL_587115, EPI_ISL_587116, EPI_ISL_587117, EPI_ISL_587118, EPI_ISL_587119, EPI_ISL_587120, EPI_ISL_587121, EPI_ISL_587122, EPI_ISL_587123, EPI_ISL_587124, EPI_ISL_587125, EPI_ISL_587126, EPI_ISL_587127, EPI_ISL_587128, EPI_ISL_587129, EPI_ISL_587130, EPI_ISL_587131, EPI_ISL_587132, EPI_ISL_587133, EPI_ISL_587134, EPI_ISL_587135, EPI_ISL_587136, EPI_ISL_587137, EPI_ISL_587138, EPI_ISL_587139, EPI_ISL_587140, EPI_ISL_587141, EPI_ISL_587142, EPI_ISL_587143, EPI_ISL_587144, EPI_ISL_587145, EPI_ISL_587146, EPI_ISL_587147, EPI_ISL_587148, EPI_ISL_587149, EPI_ISL_587150, EPI_ISL_587151, EPI_ISL_587152, EPI_ISL_587153, EPI_ISL_587154, EPI_ISL_587155, EPI_ISL_587156, EPI_ISL_587157, EPI_ISL_587158, EPI_ISL_587159, EPI_ISL_587160, EPI_ISL_587161, EPI_ISL_587162, EPI_ISL_587163, EPI_ISL_587164, EPI_ISL_587165, EPI_ISL_587166, EPI_ISL_587167, EPI_ISL_587168, EPI_ISL_587169, EPI_ISL_587170, EPI_ISL_587171, EPI_ISL_587172, EPI_ISL_587173, EPI_ISL_587174, EPI_ISL_587175, EPI_ISL_587176, EPI_ISL_587177, EPI_ISL_587178, EPI_ISL_587179, EPI_ISL_587180, EPI_ISL_587181, EPI_ISL_587182, EPI_ISL_587183, EPI_ISL_587184, EPI_ISL_587185, EPI_ISL_587186, EPI_ISL_587187, EPI_ISL_587188, EPI_ISL_587189, EPI_ISL_587190, EPI_ISL_587191, EPI_ISL_587192, EPI_ISL_587193, EPI_ISL_587194, EPI_ISL_587195, EPI_ISL_587196, EPI_ISL_587197, EPI_ISL_587198, EPI_ISL_587199, EPI_ISL_587200, EPI_ISL_587201, EPI_ISL_587202, EPI_ISL_587203, EPI_ISL_587204, EPI_ISL_587205, EPI_ISL_587206, EPI_ISL_587207, EPI_ISL_587208, EPI_ISL_587209, EPI_ISL_587210, EPI_ISL_587211, EPI_ISL_587212, EPI_ISL_587213, EPI_ISL_587214, EPI_ISL_587215, EPI_ISL_587216, EPI_ISL_587217, EPI_ISL_587218, EPI_ISL_587219, EPI_ISL_587220, EPI_ISL_587221, EPI_ISL_587222, EPI_ISL_587223, EPI_ISL_587224, EPI_ISL_587225, EPI_ISL_587226, EPI_ISL_587227, EPI_ISL_587228, EPI_ISL_587229, EPI_ISL_587230, EPI_ISL_587231, EPI_ISL_587232, EPI_ISL_587233, EPI_ISL_587234, EPI_ISL_587235, EPI_ISL_587236, EPI_ISL_587237, EPI_ISL_587238, EPI_ISL_587239, EPI_ISL_587240, EPI_ISL_587241, EPI_ISL_587242, EPI_ISL_587243, EPI_ISL_587244, EPI_ISL_587245, EPI_ISL_587246, EPI_ISL_587247, EPI_ISL_587248, EPI_ISL_587249, EPI_ISL_587250, EPI_ISL_587251, EPI_ISL_587252, EPI_ISL_587253, EPI_ISL_587254, EPI_ISL_587255, EPI_ISL_587256, EPI_ISL_587257, EPI_ISL_587258, EPI_ISL_587259, EPI_ISL_587260, EPI_ISL_587261, EPI_ISL_587262, EPI_ISL_587263, EPI_ISL_587264 | | | |
| see above | Lighthouse Lab in Alderley Park | Wellcome Sanger Institute for the COVID-19 Genomics UK (COG-UK) consortium | Jacquelyn Wynn, Mairead Hyland, The Lighthouse Lab in Alderley Park and Alex Alderton, Roberto Amato, Sonia Goncalves, Ewan Harrison, David K. Jackson, Ian Johnston, Dominic Kwiatkowski, Cordelia Langford, John Sillitoe on behalf of the Wellcome Sanger Institute COVID-19 Surveillance Team |
| EPI_ISL_587265 | Lighthouse Lab in Alderley Park | Wellcome Sanger Institute for the COVID-19 Genomics UK (COG-UK) Consortium | Jacquelyn Wynn, Mairead Hyland, The Lighthouse Lab in Alderley Park and Alex Alderton, Roberto Amato, Sonia Goncalves, Ewan Harrison, David K. Jackson, Ian Johnston, Dominic Kwiatkowski, Cordelia Langford, John Sillitoe on behalf of the Wellcome Sanger Institute COVID-19 Surveillance Team |
| EPI_ISL_587266, EPI_ISL_587267, EPI_ISL_587268, EPI_ISL_587269, EPI_ISL_587270, EPI_ISL_587271, EPI_ISL_587272, EPI_ISL_587273, EPI_ISL_587274, EPI_ISL_587275, EPI_ISL_587276, EPI_ISL_587277, EPI_ISL_587278, EPI_ISL_587279, EPI_ISL_587280, EPI_ISL_587281, EPI_ISL_587282, EPI_ISL_587283, EPI_ISL_587284, EPI_ISL_587285, EPI_ISL_587286, EPI_ISL_587287, EPI_ISL_587288, EPI_ISL_587289, EPI_ISL_587290, EPI_ISL_587291, EPI_ISL_587292, EPI_ISL_587293, EPI_ISL_587294, EPI_ISL_587295, EPI_ISL_587296, EPI_ISL_587297, EPI_ISL_587298, EPI_ISL_587299, EPI_ISL_587300, EPI_ISL_587301, EPI_ISL_587302, EPI_ISL_587715, EPI_ISL_587716, EPI_ISL_587717, EPI_ISL_587719, EPI_ISL_587723, EPI_ISL_587725, EPI_ISL_587728, EPI_ISL_587732, EPI_ISL_587733, EPI_ISL_587735, EPI_ISL_587736, EPI_ISL_587737, EPI_ISL_587739, EPI_ISL_587740, EPI_ISL_587742, EPI_ISL_587743, EPI_ISL_587745, EPI_ISL_587747, EPI_ISL_587749, EPI_ISL_587752, EPI_ISL_587756, EPI_ISL_587757, EPI_ISL_587759, EPI_ISL_587767, EPI_ISL_587771, EPI_ISL_587780, EPI_ISL_587783, EPI_ISL_587790, EPI_ISL_587795, EPI_ISL_587798, EPI_ISL_587802, EPI_ISL_587804, EPI_ISL_587805, EPI_ISL_587810, EPI_ISL_587811, EPI_ISL_587812, EPI_ISL_587813, EPI_ISL_587815, EPI_ISL_587817, EPI_ISL_587821, EPI_ISL_587823, EPI_ISL_587826, EPI_ISL_587827, EPI_ISL_587831, EPI_ISL_587832, EPI_ISL_587836, EPI_ISL_587839, EPI_ISL_587841, EPI_ISL_587842, EPI_ISL_587845, EPI_ISL_587848, EPI_ISL_587850, EPI_ISL_587851, EPI_ISL_587854, EPI_ISL_587855, EPI_ISL_587857, EPI_ISL_587858, EPI_ISL_587863, EPI_ISL_587868, EPI_ISL_587871, EPI_ISL_587872, EPI_ISL_587877, EPI_ISL_587878, EPI_ISL_587879, EPI_ISL_587880, EPI_ISL_587881, EPI_ISL_587882, EPI_ISL_587889, EPI_ISL_587892, EPI_ISL_587893, EPI_ISL_587894, EPI_ISL_587895, EPI_ISL_587896, EPI_ISL_587903, EPI_ISL_587909, EPI_ISL_587914, EPI_ISL_587915, EPI_ISL_587918, EPI_ISL_587919, EPI_ISL_587921, EPI_ISL_587925, EPI_ISL_587926, EPI_ISL_587936, EPI_ISL_587939, EPI_ISL_587940, EPI_ISL_587942, EPI_ISL_587943, EPI_ISL_587945, EPI_ISL_587948, EPI_ISL_587949, EPI_ISL_587954, EPI_ISL_587955, EPI_ISL_587956, EPI_ISL_587959, EPI_ISL_587962, EPI_ISL_587963, EPI_ISL_587964, EPI_ISL_587968, EPI_ISL_587969, EPI_ISL_587970, EPI_ISL_587974, EPI_ISL_587975, EPI_ISL_587979, EPI_ISL_587980, EPI_ISL_587982, EPI_ISL_587983, EPI_ISL_587985, EPI_ISL_587988, EPI_ISL_587989, EPI_ISL_587991, EPI_ISL_587994, EPI_ISL_587995, EPI_ISL_588003, EPI_ISL_588005, EPI_ISL_588006, EPI_ISL_588008, EPI_ISL_588016, EPI_ISL_588020, EPI_ISL_588023, EPI_ISL_588027, EPI_ISL_588030, EPI_ISL_588033, EPI_ISL_588035, EPI_ISL_588039, EPI_ISL_588041, EPI_ISL_588043, EPI_ISL_588047, EPI_ISL_588050, EPI_ISL_588052, EPI_ISL_588053, EPI_ISL_588056, EPI_ISL_588058 | | | |
| see above | Lighthouse Lab in Alderley Park | Wellcome Sanger Institute for the COVID-19 Genomics UK (COG-UK) consortium | Jacquelyn Wynn, Mairead Hyland, The Lighthouse Lab in Alderley Park and Alex Alderton, Roberto Amato, Sonia Goncalves, Ewan Harrison, David K. Jackson, Ian Johnston, Dominic Kwiatkowski, Cordelia Langford, John Sillitoe on behalf of the Wellcome Sanger Institute COVID-19 Surveillance Team |
| EPI_ISL_590721, EPI_ISL_590741 | University of Michigan Clinical Microbiology Laboratory | Lauring Lab, University of Michigan, Department of Microbiology and Immunology | Valesano |
| EPI_ISL_591488 | Laverty Pathology | NSW Health Pathology - Institute of Clinical Pathology and Medical Research; Westmead Hospital; University of Sydney | CIDM-PH et al. |
| EPI_ISL_591510 | Sydney South West Pathology Service (SSWPS) - Liverpool Hospital - NSW Health Pathology | NSW Health Pathology - Institute of Clinical Pathology and Medical Research; Westmead Hospital; University of Sydney | CIDM-PH et al. |
| EPI_ISL_592567, EPI_ISL_592743, EPI_ISL_592769 | Microbiological Diagnostic Unit - Public Health Laboratory (MDU-PHL) | MDU-PHL | Seemann T., Schultz, M. B., Sait, M., Sherry, N. |
| EPI_ISL_592783 | Victorian Infectious Diseases Reference Laboratory (VIDRL) | VIDRL and MDU-PHL | Caly L., Seemann T., Sait, M., Schultz, M. B., Druce J., Sherry, N. |
| EPI_ISL_593410 | Microbiological Diagnostic Unit - Public Health Laboratory (MDU-PHL) | MDU-PHL | Seemann T., Schultz, M. B., Sait, M., Sherry, N. |
| EPI_ISL_593411, EPI_ISL_593412 | Victorian Infectious Diseases Reference Laboratory (VIDRL) | VIDRL and MDU-PHL | Caly L., Seemann T., Sait, M., Schultz, M. B., Druce J., Sherry, N. |
| EPI_ISL_593427, EPI_ISL_593475 | Microbiological Diagnostic Unit - Public Health Laboratory (MDU-PHL) | MDU-PHL | Seemann T., Schultz, M. B., Sait, M., Sherry, N. |
| EPI_ISL_593745, EPI_ISL_593751 | South Eastern Area Laboratory Services (SEALS) | NSW Health Pathology - Institute of Clinical Pathology and Medical Research; Westmead Hospital; University of Sydney | CIDM-PH et al. |
| EPI_ISL_593810 | Respiratory Virus Unit, Microbiology Services Colindale, Public Health England | Respiratory Virus Unit, Microbiology Services Colindale, Public Health England | PHE Covid Sequencing Team |
| EPI_ISL_593895, EPI_ISL_593896, EPI_ISL_593897, EPI_ISL_593898, EPI_ISL_593899, EPI_ISL_593900 | CHU Purpan - Laboratoire de Virologie - Institut Fédératif de Biologie | CHU Purpan - Laboratoire de Virologie - Institut Fédératif de Biologie | Latour J., Ranger N., Dubois M., Carcenac R., Harter A., Boyer P., Tremeaux P., Izopet J. |
| EPI_ISL_594559, EPI_ISL_594560, EPI_ISL_594561, EPI_ISL_594562, EPI_ISL_594563, EPI_ISL_594564, EPI_ISL_594565, EPI_ISL_594566, EPI_ISL_594567, EPI_ISL_594568, EPI_ISL_594569, EPI_ISL_594570, EPI_ISL_594571, EPI_ISL_594572, EPI_ISL_594573, EPI_ISL_594574, EPI_ISL_594575, EPI_ISL_594576, EPI_ISL_594577, EPI_ISL_594578, EPI_ISL_594579, EPI_ISL_594580, EPI_ISL_594581, EPI_ISL_594582, EPI_ISL_594583, EPI_ISL_594584, EPI_ISL_594585, EPI_ISL_594586, EPI_ISL_594587, EPI_ISL_594588, EPI_ISL_594589, EPI_ISL_594590, EPI_ISL_594591, EPI_ISL_594592, EPI_ISL_594593, EPI_ISL_594594, EPI_ISL_594595, EPI_ISL_594596, EPI_ISL_594599, EPI_ISL_594600, EPI_ISL_594607, EPI_ISL_594614 | | | |
| see above | University of Birmingham | COVID-19 Genomics UK (COG-UK) Consortium | Institute of Microbiology, University of Birmingham: Claire McMurray, Joanne Stockton, Samuel Nicholls, Radoslaw Poplawski, Will Rowe, Josh Quick, Nicholas Loman. University of Birmingham Testing Laboratory: Celina M Whalley, Andrew Bosworth, Charlotte Poxon, Kasun Wanigasooriya, Oliver Pickles, Mike Kidd, Alex Richter, Andrew D Beggs PHE Heartlands Lab: Husam Osman, Andrew Bosworth. Queen Elizabeth Hospital: Anna Casey |
| EPI_ISL_594645, EPI_ISL_594646, EPI_ISL_594647, EPI_ISL_594648, EPI_ISL_594649, EPI_ISL_594650, EPI_ISL_594651, EPI_ISL_594652, EPI_ISL_594653, EPI_ISL_594654, EPI_ISL_594656, EPI_ISL_594657, EPI_ISL_594658, EPI_ISL_594660, EPI_ISL_594661, EPI_ISL_594662, EPI_ISL_594663, EPI_ISL_594664, EPI_ISL_594665, EPI_ISL_594666, EPI_ISL_594668, EPI_ISL_594669, EPI_ISL_594670, EPI_ISL_594672, EPI_ISL_594673, EPI_ISL_594674, EPI_ISL_594675, EPI_ISL_594676, EPI_ISL_594677, EPI_ISL_594678, EPI_ISL_594679, EPI_ISL_594682, EPI_ISL_594683, EPI_ISL_594684, EPI_ISL_594685, EPI_ISL_594686, EPI_ISL_594687, EPI_ISL_594688, EPI_ISL_594689, EPI_ISL_594690, EPI_ISL_594693, EPI_ISL_594694, EPI_ISL_594695, EPI_ISL_594696, EPI_ISL_594697, EPI_ISL_594698, EPI_ISL_594699, EPI_ISL_594701, EPI_ISL_594702, EPI_ISL_594704, EPI_ISL_594705, EPI_ISL_594706, EPI_ISL_594707, EPI_ISL_594708, | | | |

| | | | |
|---|---|---|---|
| EPI_ISL_594709, EPI_ISL_594711, EPI_ISL_594712, EPI_ISL_594715, EPI_ISL_594716, EPI_ISL_594717, EPI_ISL_594718, EPI_ISL_594719, EPI_ISL_594720, EPI_ISL_594722, EPI_ISL_594723, EPI_ISL_594724, EPI_ISL_594725, EPI_ISL_594726, EPI_ISL_594727, EPI_ISL_594728, EPI_ISL_594729, EPI_ISL_594730, EPI_ISL_594731, EPI_ISL_594732, EPI_ISL_594734, EPI_ISL_594735, EPI_ISL_594736, EPI_ISL_594737, EPI_ISL_594738, EPI_ISL_594739, EPI_ISL_594740, EPI_ISL_594741 | | | |
| see above | West of Scotland Specialist Virology Centre, NHSGGC / MRC-University of Glasgow Centre for Virus Research | COVID-19 Genomics UK (COG-UK) Consortium | Ana da Silva Filipe, Natasha Johnson, Kathy Smollett, Daniel Mair, Stephen Carmichael, Lily Tong, Jenna Nichols, Elihu Aranday-Cortes, Kyriaki Nomikou; Sarah McDonald, Marc Niebel, Patawee Asamaphan; Richard Orton, Joseph Hughes, Sreenu Vattipally, David L Robertson; Alasdair MacLean, Rory Gunson; Kathy Li, Igor Starinskij, Natasha Jesudason, Rajiv Shah, James Shepherd, Antonia Ho, Emma Thomson |
| EPI_ISL_594828, EPI_ISL_594829, EPI_ISL_594830, EPI_ISL_594831 | Virology Department, Royal Infirmary of Edinburgh, NHS Lothian / School of Biological Sciences, University of Edinburgh / Institute of Genetics and Molecular Medicine, University of Edinburgh | COVID-19 Genomics UK (COG-UK) Consortium | McHugh M, Dewar R, Rooke S, Gallagher M, Balcaza C, O'Toole Á, Scher E, Hill V, McCrone JT, Colquhoun R, Yu X, Jackson B, Rambaut A, Williams TC, Templeton K |
| EPI_ISL_594850, EPI_ISL_594852, EPI_ISL_594857, EPI_ISL_594858 | Liverpool Clinical Laboratories | COVID-19 Genomics UK (COG-UK) Consortium | Sam Haldenby, Anita Lucaci, Steve Paterson, Julian Hiscox, Alistair Darby, M Almsaud, A Alrezaihi, Muhannad Alruwaili, Stuart D Armstrong, Jones Benjamin, Eleanor G Bentley, Anu Chawla, Jordan J Clark, Angela Cowell, Richard Eccles, Isabel García-Dorival, Matthew Gemmell, Alessandro Gerada, PKF Gilmore, Richard Gregory, Ximeng Han, Catherine Hartley, Margaret Hughes, Miren Iturriza-Gomara, James Johnson, L Luu, Jenifer Manson, Charlotte Nelson, Elaine O'Toole, Cassie Olateju, Rebekah Penrice-Randal , Lucille Rainbow, N.P Randle, Trevor Ian Robinson, Parul Sharma, Ghada T Shawli, James P Stewart, Neil Swainston, Ecaterina Vamos, Joanne Watts, Mark Whitehead |
| EPI_ISL_594865, EPI_ISL_594866, EPI_ISL_594869, EPI_ISL_594870, EPI_ISL_594872, EPI_ISL_594873, EPI_ISL_594874, EPI_ISL_594875, EPI_ISL_594876, EPI_ISL_594880, EPI_ISL_594881, EPI_ISL_594884, EPI_ISL_594888 | | | |
| see above | University College London, Great Ormond Street Hospital for Children NHS Foundation Trust, Imperial College Healthcare NHS Trust | COVID-19 Genomics UK (COG-UK) Consortium | Sergi Castellano, Rachel Williams, Mark Kristiansen, Paola Resende Silva, Sunando Roy, Tony Brooks, Helena Tutill, Paola Niola, Patricia Dyal, Charlotte Williams, Leysa Forrest, Yasmin Panchbhaya, Jacqueline Findlay, Samuel Weeks, Julianne Brown, Kathryn Harris, Paul Randell, James Price, Alison Holmes, Judith Breuer |
| EPI_ISL_595046, EPI_ISL_595047, EPI_ISL_595048, EPI_ISL_595049, EPI_ISL_595050, EPI_ISL_595051, EPI_ISL_595052, EPI_ISL_595053, EPI_ISL_595054, EPI_ISL_595055, EPI_ISL_595056, EPI_ISL_595057, EPI_ISL_595058, EPI_ISL_595059, EPI_ISL_595060, EPI_ISL_595061, EPI_ISL_595062, EPI_ISL_595063, EPI_ISL_595064, EPI_ISL_595065, EPI_ISL_595066, EPI_ISL_595067, EPI_ISL_595068, EPI_ISL_595069, EPI_ISL_595070, EPI_ISL_595071, EPI_ISL_595072, EPI_ISL_595073, EPI_ISL_595074, EPI_ISL_595075, EPI_ISL_595076, EPI_ISL_595077, EPI_ISL_595078, EPI_ISL_595079, EPI_ISL_595080, EPI_ISL_595081, EPI_ISL_595082, EPI_ISL_595083, EPI_ISL_595084, EPI_ISL_595085, EPI_ISL_595086, EPI_ISL_595087, EPI_ISL_595088, EPI_ISL_595089, EPI_ISL_595090, EPI_ISL_595091, EPI_ISL_595092, EPI_ISL_595093, EPI_ISL_595094, EPI_ISL_595099 | | | |
| see above | Queens Medical Centre, Clinical Microbiology Department / DeepSeq Nottingham | COVID-19 Genomics UK (COG-UK) Consortium | Gemma Clark, Wendy Smith, Manjinder Khakh, Vicki M Fleming, Michelle M Lister, Hannah Howson-Wells, Jonathan Ball, Patrick McClure, Joseph Chappell, Theocharis Tsoleridis, Nadine Holmes, Matthew Carlisle, Christopher Moore, Fei Sang, Johnny Debebe, Victoria Wright, Matthew Loose |
| EPI_ISL_595100, EPI_ISL_595101, EPI_ISL_595102, EPI_ISL_595103, EPI_ISL_595104, EPI_ISL_595105, EPI_ISL_595106, EPI_ISL_595107, EPI_ISL_595108, EPI_ISL_595109, EPI_ISL_595110, EPI_ISL_595111, EPI_ISL_595112, EPI_ISL_595113, EPI_ISL_595114 | | | |
| see above | Centre for Enzyme Innovation, University of Portsmouth / Translational Research Laboratory, Portsmouth Hospitals NHS Trust | COVID-19 Genomics UK (COG-UK) Consortium | Angela Beckett,Yann Bourgeois,Garry Scarlett,Sharon Glaysher,Scott Elliott,Kelly Bicknell,Robert Impey,Allyson Lloyd,Sarah Wyllie,Ethan Butcher,Anoop Chauhan,Samuel Robson |
| EPI_ISL_595160, EPI_ISL_595166 | Quadram Institute Bioscience | COVID-19 Genomics UK (COG-UK) Consortium | Dave J. Baker, Gemma L. Kay, Alp Aydin, Thanh Le-Viet, Steven Rudder, Ana P. Tedim, Anastasia Kolyva, Maria Diaz, Leonardo de Oliveira Martins, Nabil-Fareed Alikhan, Lizzie Meadows, Rachael Stanley, Ngozi Elumogo, Muhammed Yasir, Nicholas M. Thomson, Alexander J Trotter, Rachel Gilroy, Samuel Bloomfield, Claire Stuart, Andrew Bell, Reenesh Prakash, Samir Dervisevic, Alison E. Mather, John Wain, Mark Webber, Andrew J. Page, Justin O'Grady |
| EPI_ISL_595386, EPI_ISL_595406, EPI_ISL_595436, EPI_ISL_595461, EPI_ISL_595469, EPI_ISL_595470, EPI_ISL_595479, EPI_ISL_595481, EPI_ISL_595492, EPI_ISL_595503, EPI_ISL_595512, EPI_ISL_595523, EPI_ISL_595524, EPI_ISL_595558, EPI_ISL_595575, EPI_ISL_595598, EPI_ISL_595608 | | | |
| see above | Wales Specialist Virology Centre Sequencing lab: Pathogen Genomics Unit | COVID-19 Genomics UK (COG-UK) Consortium | Catherine Moore, Johnathan Evans, Laura Gifford, Malorie Perry, Simon Cottrell, Angela Marchbank, Alec Birchley, Alexander Adams, Amy Gaskin, Bree Gatica-Wilcox, Jason Coombes, Joel Southgate, Lauren Gilbert, Lee Graham, Nicole Pacchiarini, Sara Kumziene-Summerhayes, Sarah Taylor, Sophie Jones, Sara Rey, Matthew Bull, Joanne Watkins, Sally Corden, Tom Connor |
| EPI_ISL_595800, EPI_ISL_595801 | Liverpool Clinical Laboratories | COVID-19 Genomics UK (COG-UK) Consortium | Sam Haldenby, Anita Lucaci, Steve Paterson, Julian Hiscox, Alistair Darby, M Almsaud, A Alrezaihi, Muhannad Alruwaili, Stuart D Armstrong, Jones Benjamin, Eleanor G Bentley, Anu Chawla, Jordan J Clark, Angela Cowell, Richard Eccles, Isabel García-Dorival, Matthew Gemmell, Alessandro Gerada, PKF Gilmore, Richard Gregory, Ximeng Han, Catherine Hartley, Margaret Hughes, Miren Iturriza-Gomara, James Johnson, L Luu, Jenifer Manson, Charlotte Nelson, Elaine O'Toole, Cassie Olateju, Rebekah Penrice-Randal , Lucille Rainbow, N.P Randle, Trevor Ian Robinson, Parul Sharma, Ghada T Shawli, James P Stewart, Neil Swainston, Ecaterina Vamos, Joanne Watts, Mark Whitehead |
| EPI_ISL_595802, EPI_ISL_595804, EPI_ISL_595805, EPI_ISL_595806, EPI_ISL_595807, EPI_ISL_595808, EPI_ISL_595809, EPI_ISL_595810, EPI_ISL_595811, EPI_ISL_595812, EPI_ISL_595819, EPI_ISL_595820, EPI_ISL_595821 | | | |
| see above | Virology Department, Sheffield Teaching Hospitals NHS Foundation Trust/Department of Infection, Immunity and Cardiovascular Disease, The Medical School, University of Sheffield | COVID-19 Genomics UK (COG-UK) Consortium | Thushan de Silva, Matthew Parker, Nikki Smith, Adri Angyal, Rebecca Brown, Luke Green, Rachel Tucker, Paul Parsons, Danielle Groves, Katie Johnson, Laura Carrilero, Alex Keeley, Dave Partridge, Matthew Wyles, Benjamin Lindsey, Mehmet Yavuz, Mohammad Raza, Cariad Evans |
| EPI_ISL_595822 | Liverpool Clinical Laboratories | COVID-19 Genomics UK (COG-UK) Consortium | Sam Haldenby, Anita Lucaci, Steve Paterson, Julian Hiscox, Alistair Darby, M Almsaud, A Alrezaihi, Muhannad Alruwaili, Stuart D Armstrong, Jones Benjamin, Eleanor G Bentley, Anu Chawla, Jordan J Clark, Angela Cowell, Richard Eccles, Isabel García-Dorival, Matthew Gemmell, Alessandro Gerada, PKF Gilmore, Richard Gregory, Ximeng Han, Catherine Hartley, Margaret Hughes, Miren Iturriza-Gomara, James Johnson, L Luu, Jenifer Manson, Charlotte Nelson, Elaine O'Toole, Cassie Olateju, Rebekah Penrice-Randal , Lucille Rainbow, N.P Randle, Trevor Ian Robinson, Parul Sharma, Ghada T Shawli, James P Stewart, Neil Swainston, Ecaterina Vamos, Joanne Watts, Mark Whitehead |
| EPI_ISL_595828, EPI_ISL_595829, EPI_ISL_595830, EPI_ISL_595832, EPI_ISL_595833, EPI_ISL_595834, EPI_ISL_595835, EPI_ISL_595836 | Virology Department, Sheffield Teaching Hospitals NHS Foundation Trust/Department of Infection, Immunity and Cardiovascular Disease, The Medical School, University of Sheffield | COVID-19 Genomics UK (COG-UK) Consortium | Thushan de Silva, Matthew Parker, Nikki Smith, Adri Angyal, Rebecca Brown, Luke Green, Rachel Tucker, Paul Parsons, Danielle Groves, Katie Johnson, Laura Carrilero, Alex Keeley, Dave Partridge, Matthew Wyles, Benjamin Lindsey, Mehmet Yavuz, Mohammad Raza, Cariad Evans |
| EPI_ISL_595848, EPI_ISL_595849, EPI_ISL_595850, EPI_ISL_595851, EPI_ISL_595852, EPI_ISL_595853, EPI_ISL_595854, EPI_ISL_595855, EPI_ISL_595856, EPI_ISL_595857, EPI_ISL_595858, EPI_ISL_595859, EPI_ISL_595860, EPI_ISL_595861, EPI_ISL_595862, EPI_ISL_595863, EPI_ISL_595864, EPI_ISL_595865, EPI_ISL_595866, EPI_ISL_595867, EPI_ISL_595868, EPI_ISL_595869, EPI_ISL_595870, EPI_ISL_595871, EPI_ISL_595872, EPI_ISL_595873, EPI_ISL_595874, EPI_ISL_595875, EPI_ISL_595876, EPI_ISL_595877, EPI_ISL_595878, EPI_ISL_595879, EPI_ISL_595880, EPI_ISL_595881, EPI_ISL_595882, EPI_ISL_595883, EPI_ISL_595884, EPI_ISL_595885, EPI_ISL_595886, EPI_ISL_595887, EPI_ISL_595888, EPI_ISL_595889, EPI_ISL_595890, EPI_ISL_595891, EPI_ISL_595892, EPI_ISL_595893, EPI_ISL_595894, EPI_ISL_595895, EPI_ISL_595896, EPI_ISL_595897, EPI_ISL_595898, EPI_ISL_595899, EPI_ISL_595900, EPI_ISL_595901, EPI_ISL_595902, EPI_ISL_595903, EPI_ISL_595904, EPI_ISL_595905, EPI_ISL_595906, EPI_ISL_595907, EPI_ISL_595908, EPI_ISL_595909, EPI_ISL_595910, EPI_ISL_595911, EPI_ISL_595912, EPI_ISL_595913, EPI_ISL_595914, EPI_ISL_595915, EPI_ISL_595916, EPI_ISL_595917, EPI_ISL_595918, EPI_ISL_595919, EPI_ISL_595920, EPI_ISL_595921, EPI_ISL_595922, EPI_ISL_595923, EPI_ISL_595924, EPI_ISL_595937, EPI_ISL_595938, EPI_ISL_595954, EPI_ISL_595955, EPI_ISL_595956, EPI_ISL_595958, EPI_ISL_595961, EPI_ISL_595962, EPI_ISL_595963, EPI_ISL_595964, EPI_ISL_595972, EPI_ISL_596012, EPI_ISL_596021, EPI_ISL_596022, EPI_ISL_596023, EPI_ISL_596024, EPI_ISL_596025, EPI_ISL_596026, EPI_ISL_596027, EPI_ISL_596028, EPI_ISL_596029, EPI_ISL_596030, EPI_ISL_596031, EPI_ISL_596049, EPI_ISL_596050, EPI_ISL_596051, EPI_ISL_596052, EPI_ISL_596053, EPI_ISL_596054, EPI_ISL_596055, EPI_ISL_596056, EPI_ISL_596057, EPI_ISL_596058, EPI_ISL_596059, EPI_ISL_596060, EPI_ISL_596061, EPI_ISL_596062, EPI_ISL_596063, EPI_ISL_596064, EPI_ISL_596065, EPI_ISL_596066, EPI_ISL_596067, EPI_ISL_596068, EPI_ISL_596069, EPI_ISL_596070, EPI_ISL_596071, EPI_ISL_596072, EPI_ISL_596073, EPI_ISL_596074, EPI_ISL_596075, EPI_ISL_596076, EPI_ISL_596077, EPI_ISL_596078, EPI_ISL_596079, EPI_ISL_596080, EPI_ISL_596081, EPI_ISL_596082, EPI_ISL_596083, EPI_ISL_596084, EPI_ISL_596085, EPI_ISL_596086, EPI_ISL_596087, EPI_ISL_596088, EPI_ISL_596089, EPI_ISL_596090, EPI_ISL_596091, EPI_ISL_596092, EPI_ISL_596093, EPI_ISL_596094, EPI_ISL_596095, EPI_ISL_596096, EPI_ISL_596097, EPI_ISL_596098, EPI_ISL_596107, EPI_ISL_596108, EPI_ISL_596109, EPI_ISL_596110, EPI_ISL_596111, EPI_ISL_596112, EPI_ISL_596113, EPI_ISL_596114, EPI_ISL_596115 | | | |
| see above | Quadram Institute Bioscience | COVID-19 Genomics UK (COG-UK) Consortium | Dave J. Baker, Gemma L. Kay, Alp Aydin, Thanh Le-Viet, Steven Rudder, Ana P. Tedim, Anastasia Kolyva, Maria Diaz, Leonardo de Oliveira Martins, Nabil-Fareed Alikhan, Lizzie Meadows, Rachael Stanley, Ngozi Elumogo, Muhammed Yasir, Nicholas M. Thomson, Alexander J Trotter, Rachel Gilroy, Samuel Bloomfield, Claire Stuart, Andrew Bell, Reenesh Prakash, Samir Dervisevic, Alison E. Mather, John Wain, Mark Webber, Andrew J. Page, Justin O'Grady |
| EPI_ISL_596191, EPI_ISL_596195, EPI_ISL_596196, EPI_ISL_596197, EPI_ISL_596198, EPI_ISL_596200, EPI_ISL_596204, EPI_ISL_596206, EPI_ISL_596210, EPI_ISL_596212, EPI_ISL_596219, EPI_ISL_596225, EPI_ISL_596227 | | | |
| see above | Virology Department, Sheffield Teaching Hospitals NHS Foundation Trust/Department of Infection, Immunity and Cardiovascular Disease, The Medical School, University of Sheffield | COVID-19 Genomics UK (COG-UK) Consortium | Thushan de Silva, Matthew Parker, Nikki Smith, Adri Angyal, Rebecca Brown, Luke Green, Rachel Tucker, Paul Parsons, Danielle Groves, Katie Johnson, Laura Carrilero, Alex Keeley, Dave Partridge, Matthew Wyles, Benjamin Lindsey, Mehmet Yavuz, Mohammad Raza, Cariad Evans |
| EPI_ISL_596428, EPI_ISL_596429, EPI_ISL_596430, EPI_ISL_596431, | Seattle Flu Study | Seattle Flu Study | Deborah A. Nickerson, Chris D. Frazar, Jover Lee, Benjamin Pelle, Matthew Richardson, Amanda Adler, Elisabeth Brandstetter, Peter D. Han, Kairsten Fay, Misja Ilcisin, Kirsten Lacombe, Thomas R. Sibley, Melissa Truong, Caitlin R. Wolf, Michael Boeckh, Janet A. Englund, Michael Famulare, Barry R. Lutz, Mark J. Rieder, Lea M. Starita, Matthew Thompson, Jay Shendure, Trevor Bedford, Helen Y. Chu |

| | | | |
|---|---|---|---|
| EPI_ISL_596432, EPI_ISL_596433, EPI_ISL_596434, EPI_ISL_596435, EPI_ISL_596436 | | | |
| EPI_ISL_596437, EPI_ISL_596438, EPI_ISL_596439 | Seattle Flu Study | Seattle Flu Study | Deborah A. Nickerson, Chris D. Frazar, Jover Lee, Benjamin Pelle, Matthew Richardson, Amanda Adler, Elisabeth Brandstetter, Peter D. Han, Kairsten Fay, Misja Ilcisin, Kirsten Lacombe, Thomas R. Sibley, Melissa Truong, Caitlin R. Wolf, Karen Cowgill, Stephanie Schrag, Jeff Duchin, Michael Boeckh, Janet A. Englund, Michael Famulare, Barry R. Lutz, Mark J. Rieder, Lea M. Starita, Matthew Thompson, Helen Y. Chu, Trevor Bedford, Jay Shendure |
| EPI_ISL_596440, EPI_ISL_596441, EPI_ISL_596442, EPI_ISL_596443, EPI_ISL_596444, EPI_ISL_596445 | Seattle Flu Study | Seattle Flu Study | Deborah A. Nickerson, Chris D. Frazar, Jover Lee, Benjamin Pelle, Matthew Richardson, Amanda Adler, Elisabeth Brandstetter, Peter D. Han, Kairsten Fay, Misja Ilcisin, Kirsten Lacombe, Thomas R. Sibley, Melissa Truong, Caitlin R. Wolf, Michael Boeckh, Janet A. Englund, Michael Famulare, Barry R. Lutz, Mark J. Rieder, Lea M. Starita, Matthew Thompson, Jay Shendure, Trevor Bedford, Helen Y. Chu |
| EPI_ISL_596446 | Seattle Flu Study | Seattle Flu Study | Deborah A. Nickerson, Chris D. Frazar, Jover Lee, Benjamin Pelle, Matthew Richardson, Amanda Adler, Elisabeth Brandstetter, Peter D. Han, Kairsten Fay, Misja Ilcisin, Kirsten Lacombe, Thomas R. Sibley, Melissa Truong, Caitlin R. Wolf, Karen Cowgill, Stephanie Schrag, Jeff Duchin, Michael Boeckh, Janet A. Englund, Michael Famulare, Barry R. Lutz, Mark J. Rieder, Lea M. Starita, Matthew Thompson, Helen Y. Chu, Trevor Bedford, Jay Shendure |
| EPI_ISL_596456, EPI_ISL_596457, EPI_ISL_596458, EPI_ISL_596459, EPI_ISL_596460, EPI_ISL_596461, EPI_ISL_596462, EPI_ISL_596464, EPI_ISL_596466, EPI_ISL_596467, EPI_ISL_596469, EPI_ISL_596471, EPI_ISL_596473, EPI_ISL_596478, EPI_ISL_596480, EPI_ISL_596482, EPI_ISL_596484, EPI_ISL_596496 | | | |
| see above | National Public Health Laboratory, National Centre for Infectious Diseases | National Public Health Laboratory, National Centre for Infectious Diseases | Tze Minn Mak, Sophie Octavia, Zhenyang Zhou, Lin Cui, Raymond Tzer Pin Lin |
| EPI_ISL_596570, EPI_ISL_596571, EPI_ISL_596573, EPI_ISL_596574, EPI_ISL_596575, EPI_ISL_596576, EPI_ISL_596577, EPI_ISL_596578, EPI_ISL_596579, EPI_ISL_596580, EPI_ISL_596582, EPI_ISL_596583, EPI_ISL_596585, EPI_ISL_596586, EPI_ISL_596587, EPI_ISL_596588, EPI_ISL_596589, EPI_ISL_596590, EPI_ISL_596591, EPI_ISL_596592, EPI_ISL_596593, EPI_ISL_596595, EPI_ISL_596596, EPI_ISL_596597, EPI_ISL_596598, EPI_ISL_596599, EPI_ISL_596602, EPI_ISL_596603, EPI_ISL_596604, EPI_ISL_596605, EPI_ISL_596606, EPI_ISL_596607, EPI_ISL_596608, EPI_ISL_596610 | | | |
| see above | University of Michigan Clinical Microbiology Laboratory | Lauring Lab, University of Michigan, Department of Microbiology and Immunology | Valesano |
| EPI_ISL_596821, EPI_ISL_596842 | PathWest Laboratory Medicine WA | PathWest Laboratory Medicine WA Microbial Surveillance Unit | PathWest Laboratory Medicine WA Microbial Surveillance Unit |
| EPI_ISL_596951, EPI_ISL_597002, EPI_ISL_597135, EPI_ISL_597144, EPI_ISL_597453 | Lighthouse Lab in Cambridge | Wellcome Sanger Institute for the COVID-19 Genomics UK (COG-UK) consortium | Rob Howes, The Lighthouse Lab in Cambridge and Alex Alderton, Roberto Amato, Sonia Goncalves, Ewan Harrison, David K. Jackson, Ian Johnston, Dominic Kwiatkowski, Cordelia Langford, John Sillitoe on behalf of the Wellcome Sanger Institute COVID-19 Surveillance Team (http://www.sanger.ac.uk/covid-team) |
| EPI_ISL_597740, EPI_ISL_597859, EPI_ISL_598038, EPI_ISL_598043, EPI_ISL_598115 | Lighthouse Lab in Alderley Park | Wellcome Sanger Institute for the COVID-19 Genomics UK (COG-UK) consortium | Jacquelyn Wynn, Mairead Hyland, The Lighthouse Lab in Alderley Park and Alex Alderton, Roberto Amato, Sonia Goncalves, Ewan Harrison, David K. Jackson, Ian Johnston, Dominic Kwiatkowski, Cordelia Langford, John Sillitoe on behalf of the Wellcome Sanger Institute COVID-19 Surveillance Team (http://www.sanger.ac.uk/covid-team) |
| EPI_ISL_598249 | Lighthouse Lab in Milton Keynes | Wellcome Sanger Institute for the COVID-19 Genomics UK (COG-UK) Consortium | The Lighthouse Lab in Milton Keynes and Alex Alderton, Roberto Amato, Sonia Goncalves, Ewan Harrison, David K. Jackson, Ian Johnston, Dominic Kwiatkowski, Cordelia Langford, John Sillitoe on behalf of the Wellcome Sanger Institute COVID-19 Surveillance Team (http://www.sanger.ac.uk/covid-team) |
| EPI_ISL_598250, EPI_ISL_598251, EPI_ISL_598252, EPI_ISL_598253, EPI_ISL_598254, EPI_ISL_598255, EPI_ISL_598256, EPI_ISL_598257, EPI_ISL_598258, EPI_ISL_598259, EPI_ISL_598260, EPI_ISL_598261, EPI_ISL_598262, EPI_ISL_598263, EPI_ISL_598264, EPI_ISL_598265, EPI_ISL_598266, EPI_ISL_598267, EPI_ISL_598268, EPI_ISL_598269, EPI_ISL_598270, EPI_ISL_598271, EPI_ISL_598272, EPI_ISL_598273, EPI_ISL_598274, EPI_ISL_598275, EPI_ISL_598276, EPI_ISL_598277, EPI_ISL_598278, EPI_ISL_598279, EPI_ISL_598280, EPI_ISL_598281, EPI_ISL_598282, EPI_ISL_598283, EPI_ISL_598284, EPI_ISL_598285, EPI_ISL_598286, EPI_ISL_598287, EPI_ISL_598288, EPI_ISL_598289, EPI_ISL_598290, EPI_ISL_598291, EPI_ISL_598292, EPI_ISL_598293, EPI_ISL_598294, EPI_ISL_598295, EPI_ISL_598296, EPI_ISL_598297, EPI_ISL_598298, EPI_ISL_598299, EPI_ISL_598300, EPI_ISL_598301, EPI_ISL_598302, EPI_ISL_598303, EPI_ISL_598304 | | | |
| see above | Lighthouse Lab in Milton Keynes | Wellcome Sanger Institute for the COVID-19 Genomics UK (COG-UK) consortium | The Lighthouse Lab in Milton Keynes and Alex Alderton, Roberto Amato, Sonia Goncalves, Ewan Harrison, David K. Jackson, Ian Johnston, Dominic Kwiatkowski, Cordelia Langford, John Sillitoe on behalf of the Wellcome Sanger Institute COVID-19 Surveillance Team (http://www.sanger.ac.uk/covid-team) |
| EPI_ISL_598305 | Lighthouse Lab in Milton Keynes | Wellcome Sanger Institute for the COVID-19 Genomics UK (COG-UK) Consortium | The Lighthouse Lab in Milton Keynes and Alex Alderton, Roberto Amato, Sonia Goncalves, Ewan Harrison, David K. Jackson, Ian Johnston, Dominic Kwiatkowski, Cordelia Langford, John Sillitoe on behalf of the Wellcome Sanger Institute COVID-19 Surveillance Team (http://www.sanger.ac.uk/covid-team) |
| EPI_ISL_598306, EPI_ISL_598307, EPI_ISL_598308, EPI_ISL_598309, EPI_ISL_598310, EPI_ISL_598311, EPI_ISL_598312, EPI_ISL_598313, EPI_ISL_598314, EPI_ISL_598315, EPI_ISL_598316, EPI_ISL_598317, EPI_ISL_598318, EPI_ISL_598319, EPI_ISL_598320, EPI_ISL_598321, EPI_ISL_598322, EPI_ISL_598323, EPI_ISL_598324, EPI_ISL_598325, EPI_ISL_598326, EPI_ISL_598327, EPI_ISL_598328, EPI_ISL_598329 | | | |
| see above | Lighthouse Lab in Milton Keynes | Wellcome Sanger Institute for the COVID-19 Genomics UK (COG-UK) consortium | The Lighthouse Lab in Milton Keynes and Alex Alderton, Roberto Amato, Sonia Goncalves, Ewan Harrison, David K. Jackson, Ian Johnston, Dominic Kwiatkowski, Cordelia Langford, John Sillitoe on behalf of the Wellcome Sanger Institute COVID-19 Surveillance Team (http://www.sanger.ac.uk/covid-team) |
| EPI_ISL_598330, EPI_ISL_598331 | Lighthouse Lab in Milton Keynes | Wellcome Sanger Institute for the COVID-19 Genomics UK (COG-UK) Consortium | The Lighthouse Lab in Milton Keynes and Alex Alderton, Roberto Amato, Sonia Goncalves, Ewan Harrison, David K. Jackson, Ian Johnston, Dominic Kwiatkowski, Cordelia Langford, John Sillitoe on behalf of the Wellcome Sanger Institute COVID-19 Surveillance Team (http://www.sanger.ac.uk/covid-team) |
| EPI_ISL_598332, EPI_ISL_598333, EPI_ISL_598334, EPI_ISL_598335, EPI_ISL_598336, EPI_ISL_598337, EPI_ISL_598338, EPI_ISL_598339, EPI_ISL_598340, EPI_ISL_598341, EPI_ISL_598342, EPI_ISL_598343, EPI_ISL_598344, EPI_ISL_598345, EPI_ISL_598346, EPI_ISL_598347, EPI_ISL_598348, EPI_ISL_598349, EPI_ISL_598350, EPI_ISL_598351, EPI_ISL_598352, EPI_ISL_598353, EPI_ISL_598354, EPI_ISL_598355, EPI_ISL_598356, EPI_ISL_598357, EPI_ISL_598358, EPI_ISL_598359, EPI_ISL_598360 | | | |
| see above | Lighthouse Lab in Milton Keynes | Wellcome Sanger Institute for the COVID-19 Genomics UK (COG-UK) consortium | The Lighthouse Lab in Milton Keynes and Alex Alderton, Roberto Amato, Sonia Goncalves, Ewan Harrison, David K. Jackson, Ian Johnston, Dominic Kwiatkowski, Cordelia Langford, John Sillitoe on behalf of the Wellcome Sanger Institute COVID-19 Surveillance Team (http://www.sanger.ac.uk/covid-team) |
| EPI_ISL_598361 | Lighthouse Lab in Milton Keynes | Wellcome Sanger Institute for the COVID-19 Genomics UK (COG-UK) Consortium | The Lighthouse Lab in Milton Keynes and Alex Alderton, Roberto Amato, Sonia Goncalves, Ewan Harrison, David K. Jackson, Ian Johnston, Dominic Kwiatkowski, Cordelia Langford, John Sillitoe on behalf of the Wellcome Sanger Institute COVID-19 Surveillance Team (http://www.sanger.ac.uk/covid-team) |
| EPI_ISL_598362, EPI_ISL_598363, EPI_ISL_598364, EPI_ISL_598365, EPI_ISL_598366, EPI_ISL_598367, EPI_ISL_598368, EPI_ISL_598369, EPI_ISL_598370, EPI_ISL_598371, EPI_ISL_598372, EPI_ISL_598373, EPI_ISL_598374, EPI_ISL_598375, EPI_ISL_598376, EPI_ISL_598377, EPI_ISL_598378, EPI_ISL_598379, EPI_ISL_598380, EPI_ISL_598381, EPI_ISL_598382, EPI_ISL_598383, EPI_ISL_598384, EPI_ISL_598385, EPI_ISL_598386, EPI_ISL_598387, EPI_ISL_598388, EPI_ISL_598389, EPI_ISL_598390, EPI_ISL_598391, EPI_ISL_598392, EPI_ISL_598393, EPI_ISL_598394, EPI_ISL_598395, EPI_ISL_598396, EPI_ISL_598397, EPI_ISL_598398, EPI_ISL_598399, EPI_ISL_598400, EPI_ISL_598401, EPI_ISL_598402, EPI_ISL_598403, EPI_ISL_598404, EPI_ISL_598405, EPI_ISL_598406, EPI_ISL_598407 | | | |
| see above | Lighthouse Lab in Milton Keynes | Wellcome Sanger Institute for the COVID-19 Genomics UK (COG-UK) consortium | The Lighthouse Lab in Milton Keynes and Alex Alderton, Roberto Amato, Sonia Goncalves, Ewan Harrison, David K. Jackson, Ian Johnston, Dominic Kwiatkowski, Cordelia Langford, John Sillitoe on behalf of the Wellcome Sanger Institute COVID-19 Surveillance Team (http://www.sanger.ac.uk/covid-team) |
| EPI_ISL_598408 | Lighthouse Lab in Milton Keynes | Wellcome Sanger Institute for the COVID-19 Genomics UK (COG-UK) Consortium | The Lighthouse Lab in Milton Keynes and Alex Alderton, Roberto Amato, Sonia Goncalves, Ewan Harrison, David K. Jackson, Ian Johnston, Dominic Kwiatkowski, Cordelia Langford, John Sillitoe on behalf of the Wellcome Sanger Institute COVID-19 Surveillance Team (http://www.sanger.ac.uk/covid-team) |
| EPI_ISL_598409, EPI_ISL_598410, EPI_ISL_598411, EPI_ISL_598412, EPI_ISL_598413, EPI_ISL_598414, EPI_ISL_598415, EPI_ISL_598416, EPI_ISL_598417, EPI_ISL_598418, EPI_ISL_598419, EPI_ISL_598420, EPI_ISL_598421, EPI_ISL_598422, EPI_ISL_598423, EPI_ISL_598424, EPI_ISL_598425, EPI_ISL_598426, EPI_ISL_598427, EPI_ISL_598428, EPI_ISL_598429, EPI_ISL_598430, EPI_ISL_598431, EPI_ISL_598432 | | | |
| see above | Lighthouse Lab in Milton Keynes | Wellcome Sanger Institute for the COVID-19 Genomics UK (COG-UK) consortium | The Lighthouse Lab in Milton Keynes and Alex Alderton, Roberto Amato, Sonia Goncalves, Ewan Harrison, David K. Jackson, Ian Johnston, Dominic Kwiatkowski, Cordelia Langford, John Sillitoe on behalf of the Wellcome Sanger Institute COVID-19 Surveillance Team (http://www.sanger.ac.uk/covid-team) |
| EPI_ISL_598433 | Lighthouse Lab in Milton Keynes | Wellcome Sanger Institute for the COVID-19 Genomics UK (COG-UK) Consortium | The Lighthouse Lab in Milton Keynes and Alex Alderton, Roberto Amato, Sonia Goncalves, Ewan Harrison, David K. Jackson, Ian Johnston, Dominic Kwiatkowski, Cordelia Langford, John Sillitoe on behalf of the Wellcome Sanger Institute COVID-19 Surveillance Team (http://www.sanger.ac.uk/covid-team) |
| EPI_ISL_598434, EPI_ISL_598435 | Lighthouse Lab in Milton Keynes | Wellcome Sanger Institute for the COVID-19 Genomics UK (COG-UK) consortium | The Lighthouse Lab in Milton Keynes and Alex Alderton, Roberto Amato, Sonia Goncalves, Ewan Harrison, David K. Jackson, Ian Johnston, Dominic Kwiatkowski, Cordelia Langford, John Sillitoe on behalf of the Wellcome Sanger Institute COVID-19 Surveillance Team (http://www.sanger.ac.uk/covid-team) |
| EPI_ISL_598436 | Lighthouse Lab in Milton Keynes | Wellcome Sanger Institute for the COVID-19 Genomics UK (COG-UK) Consortium | The Lighthouse Lab in Milton Keynes and Alex Alderton, Roberto Amato, Sonia Goncalves, Ewan Harrison, David K. Jackson, Ian Johnston, Dominic Kwiatkowski, Cordelia Langford, John Sillitoe on behalf of the Wellcome Sanger Institute COVID-19 Surveillance Team (http://www.sanger.ac.uk/covid-team) |
| EPI_ISL_598437, EPI_ISL_598438, EPI_ISL_598439, EPI_ISL_598440, EPI_ISL_598441, EPI_ISL_598442, EPI_ISL_598443, EPI_ISL_598444, EPI_ISL_598445, EPI_ISL_598446, EPI_ISL_598447, EPI_ISL_598448, EPI_ISL_598449, EPI_ISL_598450, EPI_ISL_598451, EPI_ISL_598452, EPI_ISL_598453, EPI_ISL_598454, EPI_ISL_598455, EPI_ISL_598456, EPI_ISL_598457, EPI_ISL_598458, EPI_ISL_598459, EPI_ISL_598460, EPI_ISL_598461, EPI_ISL_598462, EPI_ISL_598463, EPI_ISL_598464, EPI_ISL_598465, EPI_ISL_598466, EPI_ISL_598467, EPI_ISL_598468, EPI_ISL_598469, EPI_ISL_598470, EPI_ISL_598471, EPI_ISL_598472, EPI_ISL_598473, EPI_ISL_598474, EPI_ISL_598475, EPI_ISL_598476, EPI_ISL_598477, EPI_ISL_598478, EPI_ISL_598479, EPI_ISL_598480, EPI_ISL_598481, EPI_ISL_598482, EPI_ISL_598483, EPI_ISL_598484, EPI_ISL_598485, EPI_ISL_598486, EPI_ISL_598487, EPI_ISL_598488, EPI_ISL_598489, EPI_ISL_598490, EPI_ISL_598491, EPI_ISL_598492, EPI_ISL_598493, EPI_ISL_598494, EPI_ISL_598495, EPI_ISL_598496, EPI_ISL_598497, EPI_ISL_598498, EPI_ISL_598499, EPI_ISL_598500, EPI_ISL_598501, EPI_ISL_598502, EPI_ISL_598503, EPI_ISL_598504, EPI_ISL_598505, EPI_ISL_598506, EPI_ISL_598507, EPI_ISL_598508, EPI_ISL_598509, EPI_ISL_598510, EPI_ISL_598511, EPI_ISL_598512, EPI_ISL_598513, EPI_ISL_598514, EPI_ISL_598515, EPI_ISL_598516, EPI_ISL_598517, EPI_ISL_598518, EPI_ISL_598519, EPI_ISL_598520, EPI_ISL_598521, EPI_ISL_598522, EPI_ISL_598523, EPI_ISL_598524, EPI_ISL_598525, EPI_ISL_598526, EPI_ISL_598527, EPI_ISL_598528, EPI_ISL_598529, EPI_ISL_598530, EPI_ISL_598531, EPI_ISL_598532, EPI_ISL_598533, EPI_ISL_598534, EPI_ISL_598535, EPI_ISL_598536, EPI_ISL_598537, EPI_ISL_598538, EPI_ISL_598539, EPI_ISL_598541, EPI_ISL_598542, EPI_ISL_598543, EPI_ISL_598544, EPI_ISL_598545, EPI_ISL_598546, EPI_ISL_598547, EPI_ISL_598548, EPI_ISL_598549, EPI_ISL_598550, EPI_ISL_598551, EPI_ISL_598552, EPI_ISL_598553, EPI_ISL_598554, EPI_ISL_598555, EPI_ISL_598556, EPI_ISL_598557, EPI_ISL_598558, EPI_ISL_598559, EPI_ISL_598560, EPI_ISL_598561, EPI_ISL_598562, EPI_ISL_598563, | | | |

| | | | |
|---|---|---|---|
| EPI_ISL_598564, EPI_ISL_598565, EPI_ISL_598566, EPI_ISL_598568, EPI_ISL_598569, EPI_ISL_598571, EPI_ISL_598572, EPI_ISL_598573, EPI_ISL_598574, EPI_ISL_598575, EPI_ISL_598576, EPI_ISL_598577, EPI_ISL_598578, EPI_ISL_598579, EPI_ISL_598580, EPI_ISL_598581, EPI_ISL_598582, EPI_ISL_598583, EPI_ISL_598584, EPI_ISL_598585, EPI_ISL_598586, EPI_ISL_598587, EPI_ISL_598588, EPI_ISL_598589, EPI_ISL_598590, EPI_ISL_598591, EPI_ISL_598592, EPI_ISL_598593, EPI_ISL_598594, EPI_ISL_598595, EPI_ISL_598596, EPI_ISL_598597, EPI_ISL_598598, EPI_ISL_598599, EPI_ISL_598600, EPI_ISL_598601, EPI_ISL_598602, EPI_ISL_598603, EPI_ISL_598604, EPI_ISL_598605 | | | |
| see above | Lighthouse Lab in Milton Keynes | Wellcome Sanger Institute for the COVID-19 Genomics UK (COG-UK) consortium | The Lighthouse Lab in Milton Keynes and Alex Alderton, Roberto Amato, Sonia Goncalves, Ewan Harrison, David K. Jackson, Ian Johnston, Dominic Kwiatkowski, Cordelia Langford, John Sillitoe on behalf of the Wellcome Sanger Institute COVID-19 Surveillance Team (http://www.sanger.ac.uk/covid-team) |
| EPI_ISL_598606 | Lighthouse Lab in Milton Keynes | Wellcome Sanger Institute for the COVID-19 Genomics UK (COG-UK) Consortium | The Lighthouse Lab in Milton Keynes and Alex Alderton, Roberto Amato, Sonia Goncalves, Ewan Harrison, David K. Jackson, Ian Johnston, Dominic Kwiatkowski, Cordelia Langford, John Sillitoe on behalf of the Wellcome Sanger Institute COVID-19 Surveillance Team (http://www.sanger.ac.uk/covid-team) |
| EPI_ISL_598607, EPI_ISL_598608, EPI_ISL_598609, EPI_ISL_598610, EPI_ISL_598611, EPI_ISL_598612, EPI_ISL_598613, EPI_ISL_598614, EPI_ISL_598615, EPI_ISL_598616, EPI_ISL_598617, EPI_ISL_598618, EPI_ISL_598619, EPI_ISL_598620, EPI_ISL_598621, EPI_ISL_598622, EPI_ISL_598623, EPI_ISL_598624, EPI_ISL_598625, EPI_ISL_598626, EPI_ISL_598627, EPI_ISL_598628, EPI_ISL_598629, EPI_ISL_598630, EPI_ISL_598632, EPI_ISL_598633, EPI_ISL_598635, EPI_ISL_598636 | | | |
| see above | Lighthouse Lab in Milton Keynes | Wellcome Sanger Institute for the COVID-19 Genomics UK (COG-UK) consortium | The Lighthouse Lab in Milton Keynes and Alex Alderton, Roberto Amato, Sonia Goncalves, Ewan Harrison, David K. Jackson, Ian Johnston, Dominic Kwiatkowski, Cordelia Langford, John Sillitoe on behalf of the Wellcome Sanger Institute COVID-19 Surveillance Team (http://www.sanger.ac.uk/covid-team) |
| EPI_ISL_598638 | Lighthouse Lab in Milton Keynes | Wellcome Sanger Institute for the COVID-19 Genomics UK (COG-UK) Consortium | The Lighthouse Lab in Milton Keynes and Alex Alderton, Roberto Amato, Sonia Goncalves, Ewan Harrison, David K. Jackson, Ian Johnston, Dominic Kwiatkowski, Cordelia Langford, John Sillitoe on behalf of the Wellcome Sanger Institute COVID-19 Surveillance Team (http://www.sanger.ac.uk/covid-team) |
| EPI_ISL_598639, EPI_ISL_598640, EPI_ISL_598641, EPI_ISL_598642, EPI_ISL_598643, EPI_ISL_598644, EPI_ISL_598645, EPI_ISL_598646, EPI_ISL_598647, EPI_ISL_598648, EPI_ISL_598649, EPI_ISL_598650, EPI_ISL_598651, EPI_ISL_598652, EPI_ISL_598653, EPI_ISL_598654, EPI_ISL_598655, EPI_ISL_598656, EPI_ISL_598657, EPI_ISL_598658, EPI_ISL_598659, EPI_ISL_598660, EPI_ISL_598661, EPI_ISL_598662, EPI_ISL_598663, EPI_ISL_598664, EPI_ISL_598665, EPI_ISL_598666 | | | |
| see above | Lighthouse Lab in Milton Keynes | Wellcome Sanger Institute for the COVID-19 Genomics UK (COG-UK) consortium | The Lighthouse Lab in Milton Keynes and Alex Alderton, Roberto Amato, Sonia Goncalves, Ewan Harrison, David K. Jackson, Ian Johnston, Dominic Kwiatkowski, Cordelia Langford, John Sillitoe on behalf of the Wellcome Sanger Institute COVID-19 Surveillance Team (http://www.sanger.ac.uk/covid-team) |
| EPI_ISL_598667 | Lighthouse Lab in Milton Keynes | Wellcome Sanger Institute for the COVID-19 Genomics UK (COG-UK) Consortium | The Lighthouse Lab in Milton Keynes and Alex Alderton, Roberto Amato, Sonia Goncalves, Ewan Harrison, David K. Jackson, Ian Johnston, Dominic Kwiatkowski, Cordelia Langford, John Sillitoe on behalf of the Wellcome Sanger Institute COVID-19 Surveillance Team (http://www.sanger.ac.uk/covid-team) |
| EPI_ISL_598668, EPI_ISL_598669, EPI_ISL_598670, EPI_ISL_598671, EPI_ISL_598672, EPI_ISL_598673, EPI_ISL_598674, EPI_ISL_598675, EPI_ISL_598676, EPI_ISL_598677, EPI_ISL_598678, EPI_ISL_598679, EPI_ISL_598680, EPI_ISL_598681, EPI_ISL_598682, EPI_ISL_598683, EPI_ISL_598684, EPI_ISL_598685, EPI_ISL_598686, EPI_ISL_598687, EPI_ISL_598688, EPI_ISL_598689, EPI_ISL_598690, EPI_ISL_598691, EPI_ISL_598692, EPI_ISL_598693, EPI_ISL_598694, EPI_ISL_598695, EPI_ISL_598696, EPI_ISL_598697, EPI_ISL_598698, EPI_ISL_598699, EPI_ISL_598700, EPI_ISL_598701, EPI_ISL_598702, EPI_ISL_598703, EPI_ISL_598704, EPI_ISL_598705, EPI_ISL_598706, EPI_ISL_598707, EPI_ISL_598708, EPI_ISL_598709, EPI_ISL_598710, EPI_ISL_598711, EPI_ISL_598712, EPI_ISL_598713, EPI_ISL_598714, EPI_ISL_598715, EPI_ISL_598716, EPI_ISL_598717, EPI_ISL_598718, EPI_ISL_598719, EPI_ISL_598720, EPI_ISL_598721, EPI_ISL_598722, EPI_ISL_598723, EPI_ISL_598724, EPI_ISL_598725, EPI_ISL_598726, EPI_ISL_598727, EPI_ISL_598728, EPI_ISL_598729, EPI_ISL_598730, EPI_ISL_598731, EPI_ISL_598732, EPI_ISL_598733, EPI_ISL_598734, EPI_ISL_598735, EPI_ISL_598736, EPI_ISL_598738, EPI_ISL_598739, EPI_ISL_598740, EPI_ISL_598741, EPI_ISL_598742, EPI_ISL_598743, EPI_ISL_598744, EPI_ISL_598745, EPI_ISL_598746, EPI_ISL_598747, EPI_ISL_598748, EPI_ISL_598749, EPI_ISL_598750, EPI_ISL_598751, EPI_ISL_598752, EPI_ISL_598753, EPI_ISL_598754, EPI_ISL_598756, EPI_ISL_598757, EPI_ISL_598758, EPI_ISL_598759, EPI_ISL_598760, EPI_ISL_598761, EPI_ISL_598762, EPI_ISL_598763, EPI_ISL_598764, EPI_ISL_598765, EPI_ISL_598766, EPI_ISL_598767, EPI_ISL_598768, EPI_ISL_598769, EPI_ISL_598770, EPI_ISL_598771, EPI_ISL_598772, EPI_ISL_598773, EPI_ISL_598774, EPI_ISL_598775, EPI_ISL_598776, EPI_ISL_598777, EPI_ISL_598778, EPI_ISL_598779, EPI_ISL_598780, EPI_ISL_598781, EPI_ISL_598782, EPI_ISL_598783, EPI_ISL_598784, EPI_ISL_598785, EPI_ISL_598786, EPI_ISL_598787, EPI_ISL_598788, EPI_ISL_598789, EPI_ISL_598790, EPI_ISL_598791, EPI_ISL_598792, EPI_ISL_598793, EPI_ISL_598794, EPI_ISL_598795, EPI_ISL_598796, EPI_ISL_598797, EPI_ISL_598798, EPI_ISL_598801, EPI_ISL_598802, EPI_ISL_598803, EPI_ISL_598804, EPI_ISL_598805, EPI_ISL_598806, EPI_ISL_598807, EPI_ISL_598808, EPI_ISL_598809, EPI_ISL_598810, EPI_ISL_598811, EPI_ISL_598812, EPI_ISL_598813, EPI_ISL_598814, EPI_ISL_598815, EPI_ISL_598816, EPI_ISL_598817, EPI_ISL_598818, EPI_ISL_598819, EPI_ISL_598820 | | | |
| see above | Lighthouse Lab in Milton Keynes | Wellcome Sanger Institute for the COVID-19 Genomics UK (COG-UK) consortium | The Lighthouse Lab in Milton Keynes and Alex Alderton, Roberto Amato, Sonia Goncalves, Ewan Harrison, David K. Jackson, Ian Johnston, Dominic Kwiatkowski, Cordelia Langford, John Sillitoe on behalf of the Wellcome Sanger Institute COVID-19 Surveillance Team (http://www.sanger.ac.uk/covid-team) |
| EPI_ISL_598821 | Lighthouse Lab in Milton Keynes | Wellcome Sanger Institute for the COVID-19 Genomics UK (COG-UK) Consortium | The Lighthouse Lab in Milton Keynes and Alex Alderton, Roberto Amato, Sonia Goncalves, Ewan Harrison, David K. Jackson, Ian Johnston, Dominic Kwiatkowski, Cordelia Langford, John Sillitoe on behalf of the Wellcome Sanger Institute COVID-19 Surveillance Team (http://www.sanger.ac.uk/covid-team) |
| EPI_ISL_598822, EPI_ISL_598823 | Lighthouse Lab in Milton Keynes | Wellcome Sanger Institute for the COVID-19 Genomics UK (COG-UK) consortium | The Lighthouse Lab in Milton Keynes and Alex Alderton, Roberto Amato, Sonia Goncalves, Ewan Harrison, David K. Jackson, Ian Johnston, Dominic Kwiatkowski, Cordelia Langford, John Sillitoe on behalf of the Wellcome Sanger Institute COVID-19 Surveillance Team (http://www.sanger.ac.uk/covid-team) |
| EPI_ISL_598824, EPI_ISL_598825, EPI_ISL_598826, EPI_ISL_598827, EPI_ISL_598828, EPI_ISL_598829, EPI_ISL_598830 | Lighthouse Lab in Alderley Park | Wellcome Sanger Institute for the COVID-19 Genomics UK (COG-UK) consortium | Jacquelyn Wynn, Mairead Hyland, The Lighthouse Lab in Alderley Park and Alex Alderton, Roberto Amato, Sonia Goncalves, Ewan Harrison, David K. Jackson, Ian Johnston, Dominic Kwiatkowski, Cordelia Langford, John Sillitoe on behalf of the Wellcome Sanger Institute COVID-19 Surveillance Team (http://www.sanger.ac.uk/covid-team) |
| EPI_ISL_599138, EPI_ISL_599140, EPI_ISL_599145, EPI_ISL_599146, EPI_ISL_599148, EPI_ISL_599149, EPI_ISL_599154, EPI_ISL_599160, EPI_ISL_599161, EPI_ISL_599162, EPI_ISL_599165 | | | |
| see above | Lighthouse Lab in Glasgow | Wellcome Sanger Institute for the COVID-19 Genomics UK (COG-UK) consortium | Harper VanSteenhouse, Yumi Kasai, David Gray, Carol Clugston, Anna Dominiczak and Alex Alderton, Roberto Amato, Sonia Goncalves, Ewan Harrison, David K. Jackson, Ian Johnston, Dominic Kwiatkowski, Cordelia Langford, John Sillitoe on behalf of the Wellcome Sanger Institute COVID-19 Surveillance Team (http://www.sanger.ac.uk/covid-team) |
| EPI_ISL_599166 | Lighthouse Lab in Glasgow | Wellcome Sanger Institute for the COVID-19 Genomics UK (COG-UK) Consortium | Harper VanSteenhouse, Yumi Kasai, David Gray, Carol Clugston, Anna Dominiczak and Alex Alderton, Roberto Amato, Sonia Goncalves, Ewan Harrison, David K. Jackson, Ian Johnston, Dominic Kwiatkowski, Cordelia Langford, John Sillitoe on behalf of the Wellcome Sanger Institute COVID-19 Surveillance Team |
| EPI_ISL_599169, EPI_ISL_599170, EPI_ISL_599171, EPI_ISL_599173, EPI_ISL_599175, EPI_ISL_599176, EPI_ISL_599177, EPI_ISL_599178, EPI_ISL_599179, EPI_ISL_599181, EPI_ISL_599187, EPI_ISL_599188, EPI_ISL_599189, EPI_ISL_599191 | | | |
| see above | Lighthouse Lab in Glasgow | Wellcome Sanger Institute for the COVID-19 Genomics UK (COG-UK) consortium | Harper VanSteenhouse, Yumi Kasai, David Gray, Carol Clugston, Anna Dominiczak and Alex Alderton, Roberto Amato, Sonia Goncalves, Ewan Harrison, David K. Jackson, Ian Johnston, Dominic Kwiatkowski, Cordelia Langford, John Sillitoe on behalf of the Wellcome Sanger Institute COVID-19 Surveillance Team (http://www.sanger.ac.uk/covid-team) |
| EPI_ISL_599194 | Lighthouse Lab in Glasgow | Wellcome Sanger Institute for the COVID-19 Genomics UK (COG-UK) Consortium | Harper VanSteenhouse, Yumi Kasai, David Gray, Carol Clugston, Anna Dominiczak and Alex Alderton, Roberto Amato, Sonia Goncalves, Ewan Harrison, David K. Jackson, Ian Johnston, Dominic Kwiatkowski, Cordelia Langford, John Sillitoe on behalf of the Wellcome Sanger Institute COVID-19 Surveillance Team |
| EPI_ISL_599196, EPI_ISL_599197, EPI_ISL_599199 | Lighthouse Lab in Glasgow | Wellcome Sanger Institute for the COVID-19 Genomics UK (COG-UK) consortium | Harper VanSteenhouse, Yumi Kasai, David Gray, Carol Clugston, Anna Dominiczak and Alex Alderton, Roberto Amato, Sonia Goncalves, Ewan Harrison, David K. Jackson, Ian Johnston, Dominic Kwiatkowski, Cordelia Langford, John Sillitoe on behalf of the Wellcome Sanger Institute COVID-19 Surveillance Team (http://www.sanger.ac.uk/covid-team) |
| EPI_ISL_599201 | Lighthouse Lab in Glasgow | Wellcome Sanger Institute for the COVID-19 Genomics UK (COG-UK) Consortium | Harper VanSteenhouse, Yumi Kasai, David Gray, Carol Clugston, Anna Dominiczak and Alex Alderton, Roberto Amato, Sonia Goncalves, Ewan Harrison, David K. Jackson, Ian Johnston, Dominic Kwiatkowski, Cordelia Langford, John Sillitoe on behalf of the Wellcome Sanger Institute COVID-19 Surveillance Team |
| EPI_ISL_599202, EPI_ISL_599203, EPI_ISL_599204, EPI_ISL_599206, EPI_ISL_599207, EPI_ISL_599209, EPI_ISL_599210, EPI_ISL_599211, EPI_ISL_599212, EPI_ISL_599214, EPI_ISL_599215, EPI_ISL_599216, EPI_ISL_599219, EPI_ISL_599220, EPI_ISL_599222, EPI_ISL_599223, EPI_ISL_599225, EPI_ISL_599227, EPI_ISL_599228, EPI_ISL_599231, EPI_ISL_599234, EPI_ISL_599235, EPI_ISL_599236, EPI_ISL_599237, EPI_ISL_599238, EPI_ISL_599239, EPI_ISL_599241, EPI_ISL_599246, EPI_ISL_599248, EPI_ISL_599249, EPI_ISL_599250, EPI_ISL_599253, EPI_ISL_599255, EPI_ISL_599256, EPI_ISL_599257, EPI_ISL_599262, EPI_ISL_599266, EPI_ISL_599267, EPI_ISL_599268, EPI_ISL_599269, EPI_ISL_599271, EPI_ISL_599276, EPI_ISL_599277, EPI_ISL_599279, EPI_ISL_599280, EPI_ISL_599286, EPI_ISL_599288, EPI_ISL_599291, EPI_ISL_599293, EPI_ISL_599294, EPI_ISL_599295, EPI_ISL_599299, EPI_ISL_599306, EPI_ISL_599307, EPI_ISL_599310, EPI_ISL_599311, EPI_ISL_599316, EPI_ISL_599318, EPI_ISL_599320, EPI_ISL_599321, EPI_ISL_599323, EPI_ISL_599325, EPI_ISL_599326, EPI_ISL_599328, EPI_ISL_599330, EPI_ISL_599331, EPI_ISL_599332, EPI_ISL_599333, EPI_ISL_599339, EPI_ISL_599340, EPI_ISL_599341, EPI_ISL_599342, EPI_ISL_599343, EPI_ISL_599344, EPI_ISL_599345, EPI_ISL_599347, EPI_ISL_599352, EPI_ISL_599353, EPI_ISL_599354, EPI_ISL_599361, EPI_ISL_599362, EPI_ISL_599366, EPI_ISL_599369, EPI_ISL_599372, EPI_ISL_599374, EPI_ISL_599376, EPI_ISL_599377, EPI_ISL_599378, EPI_ISL_599379, EPI_ISL_599384, EPI_ISL_599385, EPI_ISL_599390, EPI_ISL_599393, EPI_ISL_599394, EPI_ISL_599397, EPI_ISL_599398, EPI_ISL_599399, EPI_ISL_599401, EPI_ISL_599403, EPI_ISL_599404, EPI_ISL_599408, EPI_ISL_599409, EPI_ISL_599410, EPI_ISL_599411, EPI_ISL_599412, EPI_ISL_599413, EPI_ISL_599414, EPI_ISL_599418, EPI_ISL_599419, EPI_ISL_599420, EPI_ISL_599426, EPI_ISL_599429, EPI_ISL_599431, EPI_ISL_599435, EPI_ISL_599436, EPI_ISL_599437, EPI_ISL_599438, EPI_ISL_599440, EPI_ISL_599444, EPI_ISL_599445, EPI_ISL_599447, EPI_ISL_599449, EPI_ISL_599450, EPI_ISL_599452, EPI_ISL_599457, EPI_ISL_599459, EPI_ISL_599463, EPI_ISL_599465, EPI_ISL_599466, EPI_ISL_599468, EPI_ISL_599470, EPI_ISL_599471, EPI_ISL_599472, EPI_ISL_599474, EPI_ISL_599475, EPI_ISL_599477, EPI_ISL_599480, EPI_ISL_599481, EPI_ISL_599482, EPI_ISL_600129, EPI_ISL_600131, EPI_ISL_600132, EPI_ISL_600137, EPI_ISL_600138, EPI_ISL_600140, EPI_ISL_600141, EPI_ISL_600142, EPI_ISL_600143, EPI_ISL_600145, EPI_ISL_600146, EPI_ISL_600149, EPI_ISL_600153, EPI_ISL_600154, EPI_ISL_600155, EPI_ISL_600156, EPI_ISL_600162, EPI_ISL_600164, EPI_ISL_600165, EPI_ISL_600168, EPI_ISL_600169, EPI_ISL_600170, EPI_ISL_600171, EPI_ISL_600173, EPI_ISL_600174, EPI_ISL_600175, EPI_ISL_600176, EPI_ISL_600178, EPI_ISL_600181, EPI_ISL_600183, EPI_ISL_600184, EPI_ISL_600186, EPI_ISL_600187, EPI_ISL_600188, EPI_ISL_600189, EPI_ISL_600190, EPI_ISL_600191, EPI_ISL_600195, EPI_ISL_600198, EPI_ISL_600199, EPI_ISL_600202, EPI_ISL_600203, EPI_ISL_600206, EPI_ISL_600207, EPI_ISL_600208, EPI_ISL_600211, EPI_ISL_600213, EPI_ISL_600214, EPI_ISL_600216, EPI_ISL_600218, EPI_ISL_600219, EPI_ISL_600220 | | | |
| see above | Lighthouse Lab in Glasgow | Wellcome Sanger Institute for the COVID-19 Genomics UK (COG-UK) consortium | Harper VanSteenhouse, Yumi Kasai, David Gray, Carol Clugston, Anna Dominiczak and Alex Alderton, Roberto Amato, Sonia Goncalves, Ewan Harrison, David K. Jackson, Ian Johnston, Dominic Kwiatkowski, Cordelia Langford, John Sillitoe on behalf of the Wellcome Sanger Institute COVID-19 Surveillance Team (http://www.sanger.ac.uk/covid-team) |
| EPI_ISL_600221 | Lighthouse Lab in Glasgow | Wellcome Sanger Institute for the COVID-19 Genomics UK (COG-UK) Consortium | Harper VanSteenhouse, Yumi Kasai, David Gray, Carol Clugston, Anna Dominiczak and Alex Alderton, Roberto Amato, Sonia Goncalves, Ewan Harrison, David K. Jackson, Ian Johnston, Dominic Kwiatkowski, Cordelia Langford, John Sillitoe on behalf of the Wellcome Sanger Institute COVID-19 Surveillance Team |

| | | | |
|---|---|---|---|
| EPI_ISL_600222, EPI_ISL_600226, EPI_ISL_600227, EPI_ISL_600228, EPI_ISL_600229, EPI_ISL_600230, EPI_ISL_600234, EPI_ISL_600235, EPI_ISL_600236, EPI_ISL_600237, EPI_ISL_600242, EPI_ISL_600244, EPI_ISL_600245, EPI_ISL_600246, EPI_ISL_600247, EPI_ISL_600248, EPI_ISL_600251, EPI_ISL_600253, EPI_ISL_600255, EPI_ISL_600256, EPI_ISL_600259, EPI_ISL_600260, EPI_ISL_600261, EPI_ISL_600264, EPI_ISL_600265, EPI_ISL_600268, EPI_ISL_600270, EPI_ISL_600274, EPI_ISL_600275, EPI_ISL_600276, EPI_ISL_600277, EPI_ISL_600278, EPI_ISL_600280, EPI_ISL_600282, EPI_ISL_600284, EPI_ISL_600288, EPI_ISL_600289, EPI_ISL_600291, EPI_ISL_600294, EPI_ISL_600295, EPI_ISL_600296, EPI_ISL_600297, EPI_ISL_600300, EPI_ISL_600301, EPI_ISL_600302, EPI_ISL_600305, EPI_ISL_600307, EPI_ISL_600308, EPI_ISL_600310, EPI_ISL_600312, EPI_ISL_600316, EPI_ISL_600318, EPI_ISL_600319, EPI_ISL_600320, EPI_ISL_600321, EPI_ISL_600322, EPI_ISL_600323, EPI_ISL_600324, EPI_ISL_600326, EPI_ISL_600327, EPI_ISL_600328, EPI_ISL_600329, EPI_ISL_600330, EPI_ISL_600331, EPI_ISL_600333, EPI_ISL_600335, EPI_ISL_600336, EPI_ISL_600337, EPI_ISL_600339, EPI_ISL_600340, EPI_ISL_600341, EPI_ISL_600343, EPI_ISL_600346, EPI_ISL_600347, EPI_ISL_600348, EPI_ISL_600349, EPI_ISL_600350, EPI_ISL_600351, EPI_ISL_600353, EPI_ISL_600354, EPI_ISL_600358, EPI_ISL_600360, EPI_ISL_600362, EPI_ISL_600363, EPI_ISL_600364, EPI_ISL_600365, EPI_ISL_600367, EPI_ISL_600368, EPI_ISL_600370, EPI_ISL_600371, EPI_ISL_600372, EPI_ISL_600373, EPI_ISL_600374, EPI_ISL_600376, EPI_ISL_600377, EPI_ISL_600379, EPI_ISL_600382, EPI_ISL_600383, EPI_ISL_600384, EPI_ISL_600385, EPI_ISL_600386, EPI_ISL_600387, EPI_ISL_600388, EPI_ISL_600390, EPI_ISL_600392, EPI_ISL_600394, EPI_ISL_600395, EPI_ISL_600397, EPI_ISL_600398, EPI_ISL_600399, EPI_ISL_600400, EPI_ISL_600401, EPI_ISL_600403, EPI_ISL_600405, EPI_ISL_600408, EPI_ISL_600409, EPI_ISL_600410, EPI_ISL_600411, EPI_ISL_600412, EPI_ISL_600418, EPI_ISL_600420, EPI_ISL_600423, EPI_ISL_600425, EPI_ISL_600426, EPI_ISL_600450, EPI_ISL_600453, EPI_ISL_600457, EPI_ISL_600458, EPI_ISL_600460, EPI_ISL_600462, EPI_ISL_600464, EPI_ISL_600465, EPI_ISL_600467, EPI_ISL_600469, EPI_ISL_600470, EPI_ISL_600474, EPI_ISL_600481, EPI_ISL_600482, EPI_ISL_600483, EPI_ISL_600486, EPI_ISL_600488, EPI_ISL_600490, EPI_ISL_600491, EPI_ISL_600493, EPI_ISL_600494, EPI_ISL_600497, EPI_ISL_600499, EPI_ISL_600506, EPI_ISL_600508, EPI_ISL_600511, EPI_ISL_600513, EPI_ISL_600514, EPI_ISL_600516, EPI_ISL_600518 | | | |
| see above | Lighthouse Lab in Glasgow | Wellcome Sanger Institute for the COVID-19 Genomics UK (COG-UK) consortium | Harper VanSteenhouse, Yumi Kasai, David Gray, Carol Clugston, Anna Dominiczak and Alex Alderton, Roberto Amato, Sonia Goncalves, Ewan Harrison, David K. Jackson, Ian Johnston, Dominic Kwiatkowski, Cordelia Langford, John Sillitoe on behalf of the Wellcome Sanger Institute COVID-19 Surveillance Team (http://www.sanger.ac.uk/covid-team) |
| EPI_ISL_600526, EPI_ISL_600531, EPI_ISL_600534 | Lighthouse Lab in Alderley Park | Wellcome Sanger Institute for the COVID-19 Genomics UK (COG-UK) consortium | Jacquelyn Wynn, Mairead Hyland, The Lighthouse Lab in Alderley Park and Alex Alderton, Roberto Amato, Sonia Goncalves, Ewan Harrison, David K. Jackson, Ian Johnston, Dominic Kwiatkowski, Cordelia Langford, John Sillitoe on behalf of the Wellcome Sanger Institute COVID-19 Surveillance Team (http://www.sanger.ac.uk/covid-team) |
| EPI_ISL_602264, EPI_ISL_602265, EPI_ISL_602266, EPI_ISL_602267, EPI_ISL_602268, EPI_ISL_602269, EPI_ISL_602270, EPI_ISL_602271, EPI_ISL_602272, EPI_ISL_602273, EPI_ISL_602274, EPI_ISL_602275, EPI_ISL_602276, EPI_ISL_602277 | | | |
| see above | MD PHL | MD PHL | Maryland Department of Health Laboratories Administration |
| EPI_ISL_602564 | Centre for Dengue Research, Department of Immunology and Molecular Medicine | Centre for Dengue Research | Chandima Jeewandara, Deshni Jayathilaka, Dinuka Ariyaratne, Diyanath Ranasinghe, Laksiri Gomes, Ananda Wijewickrama, Malika Karunaratne, Gathsaurie Neelika Malavige |
| EPI_ISL_602570, EPI_ISL_602571 | Centre for Dengue Research, Department of Immunology and Molecular Medicine | Centre for Dengue Research, Department of Immunology and Molecular Medicine | Chandima Jeewandara, Deshni Jayathilaka, Dinuka Ariyaratne, Laksiri Gomes, Diyanath Ranasinghe, Ananda Wijewickrama, Malika Karunaratne, Gathsaurie Neelika Malavige |
| EPI_ISL_602578 | SA Pathology | SA Pathology | Lex Leong, Julien Soubrier, Chuan Kok Lim, Song Gao, Mark Turra, Karin Kassahn, Ivan Bastian, Geoff Higgins |
| EPI_ISL_602867, EPI_ISL_602868, EPI_ISL_602869, EPI_ISL_602870, EPI_ISL_602871, EPI_ISL_602872, EPI_ISL_602873, EPI_ISL_602874, EPI_ISL_602875, EPI_ISL_602876, EPI_ISL_602877, EPI_ISL_602878, EPI_ISL_602879, EPI_ISL_602880, EPI_ISL_602881, EPI_ISL_602882, EPI_ISL_602883, EPI_ISL_602884, EPI_ISL_602886, EPI_ISL_602887, EPI_ISL_602888, EPI_ISL_602889, EPI_ISL_602890, EPI_ISL_602891, EPI_ISL_602892, EPI_ISL_602894, EPI_ISL_602895, EPI_ISL_602900, EPI_ISL_602901, EPI_ISL_602902, EPI_ISL_602903, EPI_ISL_602904, EPI_ISL_602905, EPI_ISL_602906, EPI_ISL_602907, EPI_ISL_602908, EPI_ISL_602909, EPI_ISL_602910, EPI_ISL_602911 | | | |
| see above | NHLS-IALCH | KRISP, KZN Research Innovation and Sequencing Platform | Giandhari J, Pillay S, Lessells R, Mdlalose K, York D, Khan S, Tegally H, Wilkinson E, de Oliveira T |
| EPI_ISL_602985, EPI_ISL_602986, EPI_ISL_602987, EPI_ISL_602988, EPI_ISL_602989, EPI_ISL_602990, EPI_ISL_602991 | Minnesota Department of Health, Public Health Laboratory | Minnesota Department of Health, Public Health Laboratory | Matt Plumb, Jacob Garfin, Alexandra Lorentz, and Xiong Wang |
| EPI_ISL_603005, EPI_ISL_603014, EPI_ISL_603016 | Essentia Health-St. Mary's Medical Center | Minnesota Department of Health, Public Health Laboratory | Matt Plumb, Jacob Garfin, Alexandra Lorentz, and Xiong Wang |
| EPI_ISL_603246, EPI_ISL_603247 | National Institute of Laboratory Medicine and Referral Center | Genomic Research Lab, BCSIR | Md. Saddam Hossain, Abu Sayeed Mohammad Mahmud, Mohammad Samir Uzzaman, Eshrar Osman, Md. Ahashan Habib, Shahina Akter, Tanjina Akhter Banu, Md. Murshed Hasan Sarkar, Barna Goswami, Iffat Jahan, Tasnim Nafisa, Md. Maruf Ahmed Molla, Mahmuda Yeasmin, Asish Kumar Ghosh, A. K. M. Shamsuzzaman, Monira Parveen, Md. Masum Hossain Arif, Md. Salim Khan |
| EPI_ISL_603248 | Brotman Baty Institute for Precision Medicine | Brotman Baty Institute for Precision Medicine | Trevor Bedford, Jennifer K. Logue, Peter D. Han, Caitlin R. Wolf, Chris D. Frazar, Benjamin Pelle, Mark J. Rieder, Deborah A. Nickerson, Christina M. Lockwood, Lea M. Starita, Helen Y. Chu, Jay Shendure |
| EPI_ISL_603249, EPI_ISL_603250 | National Institute of Laboratory Medicine and Referral Center | Genomic Research Lab, BCSIR | Md. Ahashan Habib, Abu Sayeed Mohammad Mahmud, Mohammad Samir Uzzaman, Eshrar Osman, Shahina Akter, Tanjina Akhter Banu, Md. Murshed Hasan Sarkar, Barna Goswami, Iffat Jahan, Md. Saddam Hossain, Tasnim Nafisa, Md. Maruf Ahmed Molla, Mahmuda Yeasmin, Asish Kumar Ghosh, A. K. M. Shamsuzzaman, Monira Parveen, Md. Masum Hossain Arif, Md. Salim Khan |
| EPI_ISL_609219, EPI_ISL_609220, EPI_ISL_609221, EPI_ISL_609222, EPI_ISL_609223, EPI_ISL_609224, EPI_ISL_609225, EPI_ISL_609226, EPI_ISL_609227, EPI_ISL_609228, EPI_ISL_609229 | | | |
| see above | Lighthouse Lab in Milton Keynes | Wellcome Sanger Institute for the COVID-19 Genomics UK (COG-UK) consortium | The Lighthouse Lab in Milton Keynes and Alex Alderton, Roberto Amato, Sonia Goncalves, Ewan Harrison, David K. Jackson, Ian Johnston, Dominic Kwiatkowski, Cordelia Langford, John Sillitoe on behalf of the Wellcome Sanger Institute COVID-19 Surveillance Team |
| EPI_ISL_609230 | Lighthouse Lab in Milton Keynes | Wellcome Sanger Institute for the COVID-19 Genomics UK (COG-UK) Consortium | The Lighthouse Lab in Milton Keynes and Alex Alderton, Roberto Amato, Sonia Goncalves, Ewan Harrison, David K. Jackson, Ian Johnston, Dominic Kwiatkowski, Cordelia Langford, John Sillitoe on behalf of the Wellcome Sanger Institute COVID-19 Surveillance Team |
| EPI_ISL_609231, EPI_ISL_609232, EPI_ISL_609233, EPI_ISL_609234, EPI_ISL_609235, EPI_ISL_609236, EPI_ISL_609237, EPI_ISL_609238, EPI_ISL_609239, EPI_ISL_609240, EPI_ISL_609241, EPI_ISL_609242, EPI_ISL_609243, EPI_ISL_609244, EPI_ISL_609245, EPI_ISL_609246, EPI_ISL_609247, EPI_ISL_609248, EPI_ISL_609249, EPI_ISL_609250, EPI_ISL_609251, EPI_ISL_609252, EPI_ISL_609253, EPI_ISL_609254, EPI_ISL_609255, EPI_ISL_609256, EPI_ISL_609257, EPI_ISL_609258, EPI_ISL_609259, EPI_ISL_609260, EPI_ISL_609261, EPI_ISL_609262, EPI_ISL_609263, EPI_ISL_609264, EPI_ISL_609265, EPI_ISL_609266 | | | |
| see above | Lighthouse Lab in Milton Keynes | Wellcome Sanger Institute for the COVID-19 Genomics UK (COG-UK) consortium | The Lighthouse Lab in Milton Keynes and Alex Alderton, Roberto Amato, Sonia Goncalves, Ewan Harrison, David K. Jackson, Ian Johnston, Dominic Kwiatkowski, Cordelia Langford, John Sillitoe on behalf of the Wellcome Sanger Institute COVID-19 Surveillance Team |
| EPI_ISL_609267 | Lighthouse Lab in Milton Keynes | Wellcome Sanger Institute for the COVID-19 Genomics UK (COG-UK) Consortium | The Lighthouse Lab in Milton Keynes and Alex Alderton, Roberto Amato, Sonia Goncalves, Ewan Harrison, David K. Jackson, Ian Johnston, Dominic Kwiatkowski, Cordelia Langford, John Sillitoe on behalf of the Wellcome Sanger Institute COVID-19 Surveillance Team |
| EPI_ISL_609268, EPI_ISL_609269, EPI_ISL_609270, EPI_ISL_609271, EPI_ISL_609272, EPI_ISL_609273, EPI_ISL_609274, EPI_ISL_609275, EPI_ISL_609276, EPI_ISL_609277, EPI_ISL_609278, EPI_ISL_609279, EPI_ISL_609280, EPI_ISL_609281, EPI_ISL_609282, EPI_ISL_609283, EPI_ISL_609284, EPI_ISL_609285, EPI_ISL_609286, EPI_ISL_609287, EPI_ISL_609288, EPI_ISL_609289, EPI_ISL_609290, EPI_ISL_609291, EPI_ISL_609292, EPI_ISL_609293, EPI_ISL_609294, EPI_ISL_609295, EPI_ISL_609296, EPI_ISL_609298, EPI_ISL_609299, EPI_ISL_609300, EPI_ISL_609301 | | | |
| see above | Lighthouse Lab in Milton Keynes | Wellcome Sanger Institute for the COVID-19 Genomics UK (COG-UK) consortium | The Lighthouse Lab in Milton Keynes and Alex Alderton, Roberto Amato, Sonia Goncalves, Ewan Harrison, David K. Jackson, Ian Johnston, Dominic Kwiatkowski, Cordelia Langford, John Sillitoe on behalf of the Wellcome Sanger Institute COVID-19 Surveillance Team |
| EPI_ISL_609302 | Lighthouse Lab in Milton Keynes | Wellcome Sanger Institute for the COVID-19 Genomics UK (COG-UK) Consortium | The Lighthouse Lab in Milton Keynes and Alex Alderton, Roberto Amato, Sonia Goncalves, Ewan Harrison, David K. Jackson, Ian Johnston, Dominic Kwiatkowski, Cordelia Langford, John Sillitoe on behalf of the Wellcome Sanger Institute COVID-19 Surveillance Team |
| EPI_ISL_609303, EPI_ISL_609304, EPI_ISL_609305, EPI_ISL_609306, EPI_ISL_609307 | Lighthouse Lab in Milton Keynes | Wellcome Sanger Institute for the COVID-19 Genomics UK (COG-UK) consortium | The Lighthouse Lab in Milton Keynes and Alex Alderton, Roberto Amato, Sonia Goncalves, Ewan Harrison, David K. Jackson, Ian Johnston, Dominic Kwiatkowski, Cordelia Langford, John Sillitoe on behalf of the Wellcome Sanger Institute COVID-19 Surveillance Team |
| EPI_ISL_609308, EPI_ISL_609312, EPI_ISL_609313, EPI_ISL_609314, EPI_ISL_609315, EPI_ISL_609316, EPI_ISL_609318, EPI_ISL_609320, EPI_ISL_609322, EPI_ISL_609323, EPI_ISL_609324, EPI_ISL_609326, EPI_ISL_609327, EPI_ISL_609328, EPI_ISL_609329, EPI_ISL_609330, EPI_ISL_609332, EPI_ISL_609333, EPI_ISL_609334, EPI_ISL_609335, EPI_ISL_609337, EPI_ISL_609338, EPI_ISL_609339, EPI_ISL_609340, EPI_ISL_609342, EPI_ISL_609343, EPI_ISL_609346, EPI_ISL_609347, EPI_ISL_609348, EPI_ISL_609350, EPI_ISL_609351, EPI_ISL_609353, EPI_ISL_609355, EPI_ISL_609356, EPI_ISL_609357, EPI_ISL_609358, EPI_ISL_609360, EPI_ISL_609363, EPI_ISL_609364, EPI_ISL_609365, EPI_ISL_609366, EPI_ISL_609367, EPI_ISL_609368, EPI_ISL_609369, EPI_ISL_609370, EPI_ISL_609371, EPI_ISL_609373, EPI_ISL_609374, EPI_ISL_609376, EPI_ISL_609377 | | | |
| see above | Lighthouse Lab in Cambridge | Wellcome Sanger Institute for the COVID-19 Genomics UK (COG-UK) consortium | Rob Howes, The Lighthouse Lab in Cambridge and Alex Alderton, Roberto Amato, Sonia Goncalves, Ewan Harrison, David K. Jackson, Ian Johnston, Dominic Kwiatkowski, Cordelia Langford, John Sillitoe on behalf of the Wellcome Sanger Institute COVID-19 Surveillance Team |
| EPI_ISL_609378 | Lighthouse Lab in Cambridge | Wellcome Sanger Institute for the COVID-19 Genomics UK (COG-UK) Consortium | Rob Howes, The Lighthouse Lab in Cambridge and Alex Alderton, Roberto Amato, Sonia Goncalves, Ewan Harrison, David K. Jackson, Ian Johnston, Dominic Kwiatkowski, Cordelia Langford, John Sillitoe on behalf of the Wellcome Sanger Institute COVID-19 Surveillance Team |
| EPI_ISL_609379, EPI_ISL_609380, EPI_ISL_609381, EPI_ISL_609382, EPI_ISL_609383, EPI_ISL_609385, EPI_ISL_609386, EPI_ISL_609389, EPI_ISL_609390 | Lighthouse Lab in Cambridge | Wellcome Sanger Institute for the COVID-19 Genomics UK (COG-UK) consortium | Rob Howes, The Lighthouse Lab in Cambridge and Alex Alderton, Roberto Amato, Sonia Goncalves, Ewan Harrison, David K. Jackson, Ian Johnston, Dominic Kwiatkowski, Cordelia Langford, John Sillitoe on behalf of the Wellcome Sanger Institute COVID-19 Surveillance Team |
| EPI_ISL_609393 | Lighthouse Lab in Cambridge | Wellcome Sanger Institute for the COVID-19 Genomics UK | Rob Howes, The Lighthouse Lab in Cambridge and Alex Alderton, Roberto Amato, Sonia Goncalves, Ewan Harrison, David K. Jackson, Ian Johnston, Dominic |

| | | | |
|---|---|---|---|
| | | (COG-UK) Consortium | Kwiatkowski, Cordelia Langford, John Sillitoe on behalf of the Wellcome Sanger Institute COVID-19 Surveillance Team |
| EPI_ISL_609394, EPI_ISL_609396, EPI_ISL_609398, EPI_ISL_609401, EPI_ISL_609403, EPI_ISL_609404, EPI_ISL_609405, EPI_ISL_609406, EPI_ISL_609407, EPI_ISL_609409, EPI_ISL_609411, EPI_ISL_609412, EPI_ISL_609413, EPI_ISL_609415, EPI_ISL_609416, EPI_ISL_609417, EPI_ISL_609418, EPI_ISL_609419, EPI_ISL_609420, EPI_ISL_609422, EPI_ISL_609426, EPI_ISL_609427, EPI_ISL_609428, EPI_ISL_609429, EPI_ISL_609430, EPI_ISL_609433, EPI_ISL_609434, EPI_ISL_609440, EPI_ISL_609441, EPI_ISL_609442, EPI_ISL_609444, EPI_ISL_609446, EPI_ISL_609448, EPI_ISL_609449 | | | |
| see above | Lighthouse Lab in Cambridge | Wellcome Sanger Institute for the COVID-19 Genomics UK (COG-UK) consortium | Rob Howes, The Lighthouse Lab in Cambridge and Alex Alderton, Roberto Amato, Sonia Goncalves, Ewan Harrison, David K. Jackson, Ian Johnston, Dominic Kwiatkowski, Cordelia Langford, John Sillitoe on behalf of the Wellcome Sanger Institute COVID-19 Surveillance Team |
| EPI_ISL_609450 | Lighthouse Lab in Cambridge | Wellcome Sanger Institute for the COVID-19 Genomics UK (COG-UK) Consortium | Rob Howes, The Lighthouse Lab in Cambridge and Alex Alderton, Roberto Amato, Sonia Goncalves, Ewan Harrison, David K. Jackson, Ian Johnston, Dominic Kwiatkowski, Cordelia Langford, John Sillitoe on behalf of the Wellcome Sanger Institute COVID-19 Surveillance Team |
| EPI_ISL_609452, EPI_ISL_609454, EPI_ISL_609455, EPI_ISL_609456, EPI_ISL_609458, EPI_ISL_609459, EPI_ISL_609463, EPI_ISL_609464, EPI_ISL_609465, EPI_ISL_609466, EPI_ISL_609467, EPI_ISL_609468, EPI_ISL_609469 | | | |
| see above | Lighthouse Lab in Cambridge | Wellcome Sanger Institute for the COVID-19 Genomics UK (COG-UK) consortium | Rob Howes, The Lighthouse Lab in Cambridge and Alex Alderton, Roberto Amato, Sonia Goncalves, Ewan Harrison, David K. Jackson, Ian Johnston, Dominic Kwiatkowski, Cordelia Langford, John Sillitoe on behalf of the Wellcome Sanger Institute COVID-19 Surveillance Team |
| EPI_ISL_609472 | Lighthouse Lab in Cambridge | Wellcome Sanger Institute for the COVID-19 Genomics UK (COG-UK) Consortium | Rob Howes, The Lighthouse Lab in Cambridge and Alex Alderton, Roberto Amato, Sonia Goncalves, Ewan Harrison, David K. Jackson, Ian Johnston, Dominic Kwiatkowski, Cordelia Langford, John Sillitoe on behalf of the Wellcome Sanger Institute COVID-19 Surveillance Team |
| EPI_ISL_609473, EPI_ISL_609474, EPI_ISL_609475, EPI_ISL_609476, EPI_ISL_609477, EPI_ISL_609478, EPI_ISL_609480, EPI_ISL_609481, EPI_ISL_609482, EPI_ISL_609483, EPI_ISL_609485, EPI_ISL_609486, EPI_ISL_609489, EPI_ISL_609490, EPI_ISL_609493, EPI_ISL_609494, EPI_ISL_609495, EPI_ISL_609498, EPI_ISL_609499, EPI_ISL_609500, EPI_ISL_609501, EPI_ISL_609502, EPI_ISL_609503, EPI_ISL_609505, EPI_ISL_609507, EPI_ISL_609509, EPI_ISL_609510, EPI_ISL_609511, EPI_ISL_609513, EPI_ISL_609515, EPI_ISL_609516, EPI_ISL_609517, EPI_ISL_609518, EPI_ISL_609520, EPI_ISL_609521, EPI_ISL_609523, EPI_ISL_609524, EPI_ISL_609525, EPI_ISL_609526 | | | |
| see above | Lighthouse Lab in Cambridge | Wellcome Sanger Institute for the COVID-19 Genomics UK (COG-UK) consortium | Rob Howes, The Lighthouse Lab in Cambridge and Alex Alderton, Roberto Amato, Sonia Goncalves, Ewan Harrison, David K. Jackson, Ian Johnston, Dominic Kwiatkowski, Cordelia Langford, John Sillitoe on behalf of the Wellcome Sanger Institute COVID-19 Surveillance Team |
| EPI_ISL_609527 | Lighthouse Lab in Cambridge | Wellcome Sanger Institute for the COVID-19 Genomics UK (COG-UK) Consortium | Rob Howes, The Lighthouse Lab in Cambridge and Alex Alderton, Roberto Amato, Sonia Goncalves, Ewan Harrison, David K. Jackson, Ian Johnston, Dominic Kwiatkowski, Cordelia Langford, John Sillitoe on behalf of the Wellcome Sanger Institute COVID-19 Surveillance Team |
| EPI_ISL_609528, EPI_ISL_609529, EPI_ISL_609530, EPI_ISL_609532, EPI_ISL_609533 | Lighthouse Lab in Cambridge | Wellcome Sanger Institute for the COVID-19 Genomics UK (COG-UK) consortium | Rob Howes, The Lighthouse Lab in Cambridge and Alex Alderton, Roberto Amato, Sonia Goncalves, Ewan Harrison, David K. Jackson, Ian Johnston, Dominic Kwiatkowski, Cordelia Langford, John Sillitoe on behalf of the Wellcome Sanger Institute COVID-19 Surveillance Team |
| EPI_ISL_609534 | Lighthouse Lab in Cambridge | Wellcome Sanger Institute for the COVID-19 Genomics UK (COG-UK) Consortium | Rob Howes, The Lighthouse Lab in Cambridge and Alex Alderton, Roberto Amato, Sonia Goncalves, Ewan Harrison, David K. Jackson, Ian Johnston, Dominic Kwiatkowski, Cordelia Langford, John Sillitoe on behalf of the Wellcome Sanger Institute COVID-19 Surveillance Team |
| EPI_ISL_609535, EPI_ISL_609536, EPI_ISL_609537, EPI_ISL_609541, EPI_ISL_609542, EPI_ISL_609543, EPI_ISL_609544, EPI_ISL_609545, EPI_ISL_609546, EPI_ISL_609547, EPI_ISL_609548, EPI_ISL_609549, EPI_ISL_609550, EPI_ISL_609553, EPI_ISL_609554, EPI_ISL_609555, EPI_ISL_609556, EPI_ISL_609557, EPI_ISL_609558, EPI_ISL_609559, EPI_ISL_609560, EPI_ISL_609561, EPI_ISL_609562, EPI_ISL_609563, EPI_ISL_609564, EPI_ISL_609565, EPI_ISL_609566, EPI_ISL_609567, EPI_ISL_609569, EPI_ISL_609570, EPI_ISL_609571, EPI_ISL_609572, EPI_ISL_609573, EPI_ISL_609574, EPI_ISL_609576, EPI_ISL_609578, EPI_ISL_609579, EPI_ISL_609580, EPI_ISL_609581, EPI_ISL_609582, EPI_ISL_609583, EPI_ISL_609584, EPI_ISL_609585, EPI_ISL_609586, EPI_ISL_609587, EPI_ISL_609588, EPI_ISL_609589, EPI_ISL_609590, EPI_ISL_609591, EPI_ISL_609592, EPI_ISL_609593, EPI_ISL_609594, EPI_ISL_609595, EPI_ISL_609597, EPI_ISL_609598, EPI_ISL_609599 | | | |
| see above | Lighthouse Lab in Cambridge | Wellcome Sanger Institute for the COVID-19 Genomics UK (COG-UK) consortium | Rob Howes, The Lighthouse Lab in Cambridge and Alex Alderton, Roberto Amato, Sonia Goncalves, Ewan Harrison, David K. Jackson, Ian Johnston, Dominic Kwiatkowski, Cordelia Langford, John Sillitoe on behalf of the Wellcome Sanger Institute COVID-19 Surveillance Team |
| EPI_ISL_609600 | Lighthouse Lab in Cambridge | Wellcome Sanger Institute for the COVID-19 Genomics UK (COG-UK) Consortium | Rob Howes, The Lighthouse Lab in Cambridge and Alex Alderton, Roberto Amato, Sonia Goncalves, Ewan Harrison, David K. Jackson, Ian Johnston, Dominic Kwiatkowski, Cordelia Langford, John Sillitoe on behalf of the Wellcome Sanger Institute COVID-19 Surveillance Team |
| EPI_ISL_609601, EPI_ISL_609602, EPI_ISL_609603, EPI_ISL_609604, EPI_ISL_609606, EPI_ISL_609607, EPI_ISL_609608, EPI_ISL_609609, EPI_ISL_609610, EPI_ISL_609611, EPI_ISL_609612, EPI_ISL_609613, EPI_ISL_609614, EPI_ISL_609615, EPI_ISL_609616, EPI_ISL_609617, EPI_ISL_609618, EPI_ISL_609619, EPI_ISL_609620, EPI_ISL_609621, EPI_ISL_609622, EPI_ISL_609623, EPI_ISL_609624, EPI_ISL_609625, EPI_ISL_609626, EPI_ISL_609627, EPI_ISL_609628, EPI_ISL_609630, EPI_ISL_609631, EPI_ISL_609632, EPI_ISL_609633, EPI_ISL_609634, EPI_ISL_609637, EPI_ISL_609639, EPI_ISL_609640, EPI_ISL_609641, EPI_ISL_609642, EPI_ISL_609644, EPI_ISL_609645, EPI_ISL_609646, EPI_ISL_609647, EPI_ISL_609648, EPI_ISL_609649, EPI_ISL_609650, EPI_ISL_609651, EPI_ISL_609652, EPI_ISL_609653, EPI_ISL_609654, EPI_ISL_609655, EPI_ISL_609656, EPI_ISL_609657, EPI_ISL_609658, EPI_ISL_609659, EPI_ISL_609660, EPI_ISL_609662, EPI_ISL_609663, EPI_ISL_609664, EPI_ISL_609665, EPI_ISL_609666, EPI_ISL_609668, EPI_ISL_609669, EPI_ISL_609670, EPI_ISL_609671, EPI_ISL_609672, EPI_ISL_609673, EPI_ISL_609674, EPI_ISL_609675, EPI_ISL_609676, EPI_ISL_609677, EPI_ISL_609678, EPI_ISL_609680, EPI_ISL_609681, EPI_ISL_609683, EPI_ISL_609684, EPI_ISL_609685, EPI_ISL_609686, EPI_ISL_609687, EPI_ISL_609688, EPI_ISL_609689, EPI_ISL_609690, EPI_ISL_609691, EPI_ISL_609692, EPI_ISL_609694, EPI_ISL_609695, EPI_ISL_609696, EPI_ISL_609697, EPI_ISL_609698, EPI_ISL_609700, EPI_ISL_609701, EPI_ISL_609702, EPI_ISL_609703, EPI_ISL_609704, EPI_ISL_609705, EPI_ISL_609706, EPI_ISL_609707, EPI_ISL_609708, EPI_ISL_609709, EPI_ISL_609710, EPI_ISL_609712, EPI_ISL_609713, EPI_ISL_609715, EPI_ISL_609716, EPI_ISL_609717, EPI_ISL_609718, EPI_ISL_609719, EPI_ISL_609721, EPI_ISL_609722, EPI_ISL_609723, EPI_ISL_609724, EPI_ISL_609725, EPI_ISL_609726, EPI_ISL_609727, EPI_ISL_609728, EPI_ISL_609729, EPI_ISL_609730, EPI_ISL_609731, EPI_ISL_609732, EPI_ISL_609733, EPI_ISL_609735, EPI_ISL_609736, EPI_ISL_609738, EPI_ISL_609739, EPI_ISL_609740, EPI_ISL_609741, EPI_ISL_609742, EPI_ISL_609743, EPI_ISL_609744, EPI_ISL_609745, EPI_ISL_609748, EPI_ISL_609750, EPI_ISL_609751, EPI_ISL_609752, EPI_ISL_609753, EPI_ISL_609754, EPI_ISL_609755, EPI_ISL_609756, EPI_ISL_609757, EPI_ISL_609758, EPI_ISL_609759, EPI_ISL_609760, EPI_ISL_609761, EPI_ISL_609762, EPI_ISL_609764, EPI_ISL_609765, EPI_ISL_609766, EPI_ISL_609767, EPI_ISL_609768, EPI_ISL_609769, EPI_ISL_609770, EPI_ISL_609771, EPI_ISL_609772, EPI_ISL_609773, EPI_ISL_609774, EPI_ISL_609775, EPI_ISL_609776, EPI_ISL_609777, EPI_ISL_609778, EPI_ISL_609779, EPI_ISL_609780, EPI_ISL_609781, EPI_ISL_609782, EPI_ISL_609783, EPI_ISL_609784, EPI_ISL_609787, EPI_ISL_609788, EPI_ISL_609790, EPI_ISL_609793, EPI_ISL_609794, EPI_ISL_609795, EPI_ISL_609796, EPI_ISL_609797, EPI_ISL_609798, EPI_ISL_609799, EPI_ISL_609800 | | | |
| see above | Lighthouse Lab in Cambridge | Wellcome Sanger Institute for the COVID-19 Genomics UK (COG-UK) consortium | Rob Howes, The Lighthouse Lab in Cambridge and Alex Alderton, Roberto Amato, Sonia Goncalves, Ewan Harrison, David K. Jackson, Ian Johnston, Dominic Kwiatkowski, Cordelia Langford, John Sillitoe on behalf of the Wellcome Sanger Institute COVID-19 Surveillance Team |
| EPI_ISL_609811, EPI_ISL_609822 | Unity Health Toronto | Ontario Institute for Cancer Research | Ramzi Fattouh, Larissa M. Matukas, Yan Chen,Mark Downing, Trina Otterman, Karel Boissinot, Wai Sum Siu, Zhi Cui, Le Luu, Samira Mubareka, TIBDN, Ilinca Lungu, Bernard Lam, Jeremy Johns, Paul Krzyzanowski, Richard de Borja, Felicia Vincelli, Philip Zuzarte, Jared T. Simpson |
| EPI_ISL_609830, EPI_ISL_609831, EPI_ISL_609832, EPI_ISL_609833, EPI_ISL_609834, EPI_ISL_609835, EPI_ISL_609836, EPI_ISL_609837, EPI_ISL_609838, EPI_ISL_609840, EPI_ISL_609841, EPI_ISL_609842, EPI_ISL_609843, EPI_ISL_609844, EPI_ISL_609845, EPI_ISL_609846, EPI_ISL_609847, EPI_ISL_609848, EPI_ISL_609849, EPI_ISL_609850, EPI_ISL_609851, EPI_ISL_609852 | | | |
| see above | Respiratory Virus Unit, Microbiology Services Colindale, Public Health England | Respiratory Virus Unit, Microbiology Services Colindale, Public Health England | PHE Covid Sequencing Team |
| EPI_ISL_610199, EPI_ISL_610200, EPI_ISL_610214, EPI_ISL_610215, EPI_ISL_610222 | Department of Health Technology and Informatics, The Hong Kong Polytechnic University | Department of Health Technology and Informatics, The Hong Kong Polytechnic University | Siu,G.K.-H., Lee,L.-K., Leung,K.S.-S., Leung,J.S.-L., Ng,T.T.-L., Chan,C.T.-M., Tam,K.K.-G., Lao,H.-Y., Wu,A.K.-L., Yau,M.C.-Y., Lai,Y.W.-M., Fung,K.S.-C., Chau,S.K.-Y., Wong,B.K.-C., To,W.-K., Luk,K., Ho,A.Y.-M., Que,T.-L., Yip,K.-T., Yam,W.C., Shum,D.H.-K., Yip,S.P. |
| EPI_ISL_610726 | Lighthouse Lab in Cambridge | Wellcome Sanger Institute for the COVID-19 Genomics UK (COG-UK) consortium | Rob Howes, The Lighthouse Lab in Cambridge and Alex Alderton, Roberto Amato, Sonia Goncalves, Ewan Harrison, David K. Jackson, Ian Johnston, Dominic Kwiatkowski, Cordelia Langford, John Sillitoe on behalf of the Wellcome Sanger Institute COVID-19 Surveillance Team |
| EPI_ISL_611522 | Liverpool Clinical Laboratories | COVID-19 Genomics UK (COG-UK) Consortium | Sam Haldenby, Anita Lucaci, Steve Paterson, Julian Hiscox, Alistair Darby, M Almsaud, A Alrezaihi, Muhannad Alruwaili, Stuart D Armstrong, Jones Benjamin, Eleanor G Bentley, Anu Chawla, Jordan J Clark, Angela Cowell, Richard Eccles, Isabel García-Dorival, Matthew Gemmell, Alessandro Gerada, PKF Gilmore, Richard Gregory, Ximeng Han, Catherine Hartley, Margaret Hughes, Miren Iturriza-Gomara, James Johnson, L Luu, Jenifer Manson, Charlotte Nelson, Elaine O'Toole, Cassie Olateju, Rebekah Penrice-Randal , Lucille Rainbow, N.P Randle, Trevor Ian Robinson, Parul Sharma, Ghada T Shawli, James P Stewart, Neil Swainston, Ecaterina Vamos, Joanne Watts, Mark Whitehead |
| EPI_ISL_611524 | Northumbria University / South Tees Hospitals NHS Foundation Trust / North Cumbria Integrated Care NHS Foundation Trust / North Tees and Hartlepool NHS Foundation Trust / Newcastle Hospitals NHS Foundation Trust | COVID-19 Genomics UK (COG-UK) Consortium | Darren L Smith,Andrew Nelson,Matthew Bashton,Greg R Young,Joshua Loh,John Allan,Mohammad A Tariq,Giles S Holt,Gary Black,Wen C Yew,Lynn Dover,Paul Baker,Steve Liggett,Sarah Essex,Jane Greenaway,Debra Padgett,Clive Graham,Garren Scott,Edward Barton,Emma Swindells,Brendan Payne,Jennifer Collins,Yusri Taha,Gary Eltringham |
| EPI_ISL_611536 | Virology Department, Sheffield Teaching Hospitals NHS Foundation Trust/Department of Infection, Immunity and Cardiovascular Disease, The Medical School, University of Sheffield | COVID-19 Genomics UK (COG-UK) Consortium | Thushan de Silva, Matthew Parker, Nikki Smith, Adri Angyal, Rebecca Brown, Luke Green, Rachel Tucker, Paul Parsons, Danielle Groves, Katie Johnson, Laura Carrilero, Alex Keeley, Dave Partridge, Matthew Wyles, Benjamin Lindsey, Mehmet Yavuz, Mohammad Raza, Cariad Evans |
| EPI_ISL_611568 | Liverpool Clinical Laboratories | COVID-19 Genomics UK (COG-UK) Consortium | Sam Haldenby, Anita Lucaci, Steve Paterson, Julian Hiscox, Alistair Darby, M Almsaud, A Alrezaihi, Muhannad Alruwaili, Stuart D Armstrong, Jones Benjamin, Eleanor G Bentley, Anu Chawla, Jordan J Clark, Angela Cowell, Richard Eccles, Isabel García-Dorival, Matthew Gemmell, Alessandro Gerada, PKF Gilmore, Richard Gregory, Ximeng Han, Catherine Hartley, Margaret Hughes, Miren Iturriza-Gomara, James Johnson, L Luu, Jenifer Manson, Charlotte Nelson, Elaine O'Toole, Cassie Olateju, Rebekah Penrice-Randal , Lucille Rainbow, N.P Randle, Trevor Ian Robinson, Parul Sharma, Ghada T Shawli, James P Stewart, Neil Swainston, Ecaterina Vamos, Joanne Watts, Mark Whitehead |

| | | | |
|---|---|---|---|
| EPI_ISL_611574, EPI_ISL_611576 | Northumbria University / South Tees Hospitals NHS Foundation Trust / North Cumbria Integrated Care NHS Foundation Trust / North Tees and Hartlepool NHS Foundation Trust / Newcastle Hospitals NHS Foundation Trust | COVID-19 Genomics UK (COG-UK) Consortium | Darren L Smith,Andrew Nelson,Matthew Bashton,Greg R Young,Joshua Loh,John Allan,Mohammad A Tariq,Giles S Holt,Gary Black,Wen C Yew,Lynn Dover,Paul Baker,Steve Liggett,Sarah Essex,Jane Greenaway,Debra Padgett,Clive Graham,Garren Scott,Edward Barton,Emma Swindells,Brendan Payne,Jennifer Collins,Yusri Taha,Gary Eltringham |
| EPI_ISL_611580 | Liverpool Clinical Laboratories | COVID-19 Genomics UK (COG-UK) Consortium | Sam Haldenby, Anita Lucaci, Steve Paterson, Julian Hiscox, Alistair Darby, M Almsaud, A Alrezaihi, Muhannad Alruwaili, Stuart D Armstrong, Jones Benjamin, Eleanor G Bentley, Anu Chawla, Jordan J Clark, Angela Cowell, Richard Eccles, Isabel García-Dorival, Matthew Gemmell, Alessandro Gerada, PKF Gilmore, Richard Gregory, Ximeng Han, Catherine Hartley, Margaret Hughes, Miren Iturriza-Gomara, James Johnson, L Luu, Jenifer Manson, Charlotte Nelson, Elaine O'Toole, Cassie Olateju, Rebekah Penrice-Randal , Lucille Rainbow, N.P Randle, Trevor Ian Robinson, Parul Sharma, Ghada T Shawli, James P Stewart, Neil Swainston, Ecaterina Vamos, Joanne Watts, Mark Whitehead |
| EPI_ISL_611591 | Northumbria University / South Tees Hospitals NHS Foundation Trust / North Cumbria Integrated Care NHS Foundation Trust / North Tees and Hartlepool NHS Foundation Trust / Newcastle Hospitals NHS Foundation Trust | COVID-19 Genomics UK (COG-UK) Consortium | Darren L Smith,Andrew Nelson,Matthew Bashton,Greg R Young,Joshua Loh,John Allan,Mohammad A Tariq,Giles S Holt,Gary Black,Wen C Yew,Lynn Dover,Paul Baker,Steve Liggett,Sarah Essex,Jane Greenaway,Debra Padgett,Clive Graham,Garren Scott,Edward Barton,Emma Swindells,Brendan Payne,Jennifer Collins,Yusri Taha,Gary Eltringham |
| EPI_ISL_611600 | Liverpool Clinical Laboratories | COVID-19 Genomics UK (COG-UK) Consortium | Sam Haldenby, Anita Lucaci, Steve Paterson, Julian Hiscox, Alistair Darby, M Almsaud, A Alrezaihi, Muhannad Alruwaili, Stuart D Armstrong, Jones Benjamin, Eleanor G Bentley, Anu Chawla, Jordan J Clark, Angela Cowell, Richard Eccles, Isabel García-Dorival, Matthew Gemmell, Alessandro Gerada, PKF Gilmore, Richard Gregory, Ximeng Han, Catherine Hartley, Margaret Hughes, Miren Iturriza-Gomara, James Johnson, L Luu, Jenifer Manson, Charlotte Nelson, Elaine O'Toole, Cassie Olateju, Rebekah Penrice-Randal , Lucille Rainbow, N.P Randle, Trevor Ian Robinson, Parul Sharma, Ghada T Shawli, James P Stewart, Neil Swainston, Ecaterina Vamos, Joanne Watts, Mark Whitehead |
| EPI_ISL_611643, EPI_ISL_611645, EPI_ISL_611657 | Northumbria University / South Tees Hospitals NHS Foundation Trust / North Cumbria Integrated Care NHS Foundation Trust / North Tees and Hartlepool NHS Foundation Trust / Newcastle Hospitals NHS Foundation Trust | COVID-19 Genomics UK (COG-UK) Consortium | Darren L Smith,Andrew Nelson,Matthew Bashton,Greg R Young,Joshua Loh,John Allan,Mohammad A Tariq,Giles S Holt,Gary Black,Wen C Yew,Lynn Dover,Paul Baker,Steve Liggett,Sarah Essex,Jane Greenaway,Debra Padgett,Clive Graham,Garren Scott,Edward Barton,Emma Swindells,Brendan Payne,Jennifer Collins,Yusri Taha,Gary Eltringham |
| EPI_ISL_611667 | Liverpool Clinical Laboratories | COVID-19 Genomics UK (COG-UK) Consortium | Sam Haldenby, Anita Lucaci, Steve Paterson, Julian Hiscox, Alistair Darby, M Almsaud, A Alrezaihi, Muhannad Alruwaili, Stuart D Armstrong, Jones Benjamin, Eleanor G Bentley, Anu Chawla, Jordan J Clark, Angela Cowell, Richard Eccles, Isabel García-Dorival, Matthew Gemmell, Alessandro Gerada, PKF Gilmore, Richard Gregory, Ximeng Han, Catherine Hartley, Margaret Hughes, Miren Iturriza-Gomara, James Johnson, L Luu, Jenifer Manson, Charlotte Nelson, Elaine O'Toole, Cassie Olateju, Rebekah Penrice-Randal , Lucille Rainbow, N.P Randle, Trevor Ian Robinson, Parul Sharma, Ghada T Shawli, James P Stewart, Neil Swainston, Ecaterina Vamos, Joanne Watts, Mark Whitehead |
| EPI_ISL_611668 | Northumbria University / South Tees Hospitals NHS Foundation Trust / North Cumbria Integrated Care NHS Foundation Trust / North Tees and Hartlepool NHS Foundation Trust / Newcastle Hospitals NHS Foundation Trust | COVID-19 Genomics UK (COG-UK) Consortium | Darren L Smith,Andrew Nelson,Matthew Bashton,Greg R Young,Joshua Loh,John Allan,Mohammad A Tariq,Giles S Holt,Gary Black,Wen C Yew,Lynn Dover,Paul Baker,Steve Liggett,Sarah Essex,Jane Greenaway,Debra Padgett,Clive Graham,Garren Scott,Edward Barton,Emma Swindells,Brendan Payne,Jennifer Collins,Yusri Taha,Gary Eltringham |
| EPI_ISL_611683 | Wales Specialist Virology Centre Sequencing lab: Pathogen Genomics Unit | COVID-19 Genomics UK (COG-UK) Consortium | Catherine Moore, Johnathan Evans, Laura Gifford, Malorie Perry, Simon Cottrell, Angela Marchbank, Alec Birchley, Alexander Adams, Amy Gaskin, Bree Gatica-Wilcox, Jason Coombes, Joel Southgate, Lauren Gilbert, Lee Graham, Nicole Pacchiarini, Sara Kumziene-Summerhayes, Sarah Taylor, Sophie Jones, Sara Rey, Matthew Bull, Joanne Watkins, Sally Corden, Tom Connor |
| EPI_ISL_611685 | Virology Department, Royal Infirmary of Edinburgh, NHS Lothian / School of Biological Sciences, University of Edinburgh / Institute of Genetics and Molecular Medicine, University of Edinburgh | COVID-19 Genomics UK (COG-UK) Consortium | McHugh M, Dewar R, Rooke S, Gallagher M, Balcaza C, O'Toole Á, Scher E, Hill V, McCrone JT, Colquhoun R, Yu X, Jackson B, Rambaut A, Williams TC, Templeton K |
| EPI_ISL_611689 | Centre for Enzyme Innovation, University of Portsmouth / Translational Research Laboratory, Portsmouth Hospitals NHS Trust | COVID-19 Genomics UK (COG-UK) Consortium | Angela Beckett,Yann Bourgeois,Garry Scarlett,Sharon Glaysher,Scott Elliott,Kelly Bicknell,Robert Impey,Allyson Lloyd,Sarah Wyllie,Ethan Butcher,Anoop Chauhan,Samuel Robson |
| EPI_ISL_611707 | Virology Department, Royal Infirmary of Edinburgh, NHS Lothian / School of Biological Sciences, University of Edinburgh / Institute of Genetics and Molecular Medicine, University of Edinburgh | COVID-19 Genomics UK (COG-UK) Consortium | McHugh M, Dewar R, Rooke S, Gallagher M, Balcaza C, O'Toole Á, Scher E, Hill V, McCrone JT, Colquhoun R, Yu X, Jackson B, Rambaut A, Williams TC, Templeton K |
| EPI_ISL_611729, EPI_ISL_611731 | Liverpool Clinical Laboratories | COVID-19 Genomics UK (COG-UK) Consortium | Sam Haldenby, Anita Lucaci, Steve Paterson, Julian Hiscox, Alistair Darby, M Almsaud, A Alrezaihi, Muhannad Alruwaili, Stuart D Armstrong, Jones Benjamin, Eleanor G Bentley, Anu Chawla, Jordan J Clark, Angela Cowell, Richard Eccles, Isabel García-Dorival, Matthew Gemmell, Alessandro Gerada, PKF Gilmore, Richard Gregory, Ximeng Han, Catherine Hartley, Margaret Hughes, Miren Iturriza-Gomara, James Johnson, L Luu, Jenifer Manson, Charlotte Nelson, Elaine O'Toole, Cassie Olateju, Rebekah Penrice-Randal , Lucille Rainbow, N.P Randle, Trevor Ian Robinson, Parul Sharma, Ghada T Shawli, James P Stewart, Neil Swainston, Ecaterina Vamos, Joanne Watts, Mark Whitehead |
| EPI_ISL_611733 | Virology Department, Sheffield Teaching Hospitals NHS Foundation Trust/Department of Infection, Immunity and Cardiovascular Disease, The Medical School, University of Sheffield | COVID-19 Genomics UK (COG-UK) Consortium | Thushan de Silva, Matthew Parker, Nikki Smith, Adri Angyal, Rebecca Brown, Luke Green, Rachel Tucker, Paul Parsons, Danielle Groves, Katie Johnson, Laura Carrilero, Alex Keeley, Dave Partridge, Matthew Wyles, Benjamin Lindsey, Mehmet Yavuz, Mohammad Raza, Cariad Evans |
| EPI_ISL_611756, EPI_ISL_611757, EPI_ISL_611758, EPI_ISL_611759, EPI_ISL_611760, EPI_ISL_611767, EPI_ISL_611768 | Northumbria University / South Tees Hospitals NHS Foundation Trust / North Cumbria Integrated Care NHS Foundation Trust / North Tees and Hartlepool NHS Foundation Trust / Newcastle Hospitals NHS Foundation Trust | COVID-19 Genomics UK (COG-UK) Consortium | Darren L Smith,Andrew Nelson,Matthew Bashton,Greg R Young,Joshua Loh,John Allan,Mohammad A Tariq,Giles S Holt,Gary Black,Wen C Yew,Lynn Dover,Paul Baker,Steve Liggett,Sarah Essex,Jane Greenaway,Debra Padgett,Clive Graham,Garren Scott,Edward Barton,Emma Swindells,Brendan Payne,Jennifer Collins,Yusri Taha,Gary Eltringham |
| EPI_ISL_611789 | Virology Department, Sheffield Teaching Hospitals NHS Foundation Trust/Department of Infection, Immunity and Cardiovascular Disease, The Medical School, University of Sheffield | COVID-19 Genomics UK (COG-UK) Consortium | Thushan de Silva, Matthew Parker, Nikki Smith, Adri Angyal, Rebecca Brown, Luke Green, Rachel Tucker, Paul Parsons, Danielle Groves, Katie Johnson, Laura Carrilero, Alex Keeley, Dave Partridge, Matthew Wyles, Benjamin Lindsey, Mehmet Yavuz, Mohammad Raza, Cariad Evans |
| EPI_ISL_611832, EPI_ISL_611834 | Wales Specialist Virology Centre Sequencing lab: Pathogen Genomics Unit | COVID-19 Genomics UK (COG-UK) Consortium | Catherine Moore, Johnathan Evans, Laura Gifford, Malorie Perry, Simon Cottrell, Angela Marchbank, Alec Birchley, Alexander Adams, Amy Gaskin, Bree Gatica-Wilcox, Jason Coombes, Joel Southgate, Lauren Gilbert, Lee Graham, Nicole Pacchiarini, Sara Kumziene-Summerhayes, Sarah Taylor, Sophie Jones, Sara Rey, Matthew Bull, Joanne Watkins, Sally Corden, Tom Connor |
| EPI_ISL_611836 | University of Exeter | COVID-19 Genomics UK (COG-UK) Consortium | Ben Temperton,Aaron Jeffries,Michelle Michelsen,Joanna Warwick-Dugdale,Audrey Farbos,Robyn Manley,Stephen Michell,Jane Masoli |
| EPI_ISL_611839, EPI_ISL_611864, EPI_ISL_611868 | Northumbria University / South Tees Hospitals NHS Foundation Trust / North Cumbria Integrated Care NHS Foundation Trust / North Tees and Hartlepool NHS Foundation Trust / Newcastle Hospitals NHS Foundation Trust | COVID-19 Genomics UK (COG-UK) Consortium | Darren L Smith,Andrew Nelson,Matthew Bashton,Greg R Young,Joshua Loh,John Allan,Mohammad A Tariq,Giles S Holt,Gary Black,Wen C Yew,Lynn Dover,Paul Baker,Steve Liggett,Sarah Essex,Jane Greenaway,Debra Padgett,Clive Graham,Garren Scott,Edward Barton,Emma Swindells,Brendan Payne,Jennifer Collins,Yusri Taha,Gary Eltringham |
| EPI_ISL_611885, EPI_ISL_611908, EPI_ISL_611910 | Liverpool Clinical Laboratories | COVID-19 Genomics UK (COG-UK) Consortium | Sam Haldenby, Anita Lucaci, Steve Paterson, Julian Hiscox, Alistair Darby, M Almsaud, A Alrezaihi, Muhannad Alruwaili, Stuart D Armstrong, Jones Benjamin, Eleanor G Bentley, Anu Chawla, Jordan J Clark, Angela Cowell, Richard Eccles, Isabel García-Dorival, Matthew Gemmell, Alessandro Gerada, PKF Gilmore, |

| | | | |
|---|---|---|---|
| | | | Richard Gregory, Ximeng Han, Catherine Hartley, Margaret Hughes, Miren Iturriza-Gomara, James Johnson, L Luu, Jenifer Manson, Charlotte Nelson, Elaine O'Toole, Cassie Olateju, Rebekah Penrice-Randal , Lucille Rainbow, N.P Randle, Trevor Ian Robinson, Parul Sharma, Ghada T Shawli, James P Stewart, Neil Swainston, Ecaterina Vamos, Joanne Watts, Mark Whitehead |
| EPI_ISL_611930 | Northumbria University / South Tees Hospitals NHS Foundation Trust / North Cumbria Integrated Care NHS Foundation Trust / North Tees and Hartlepool NHS Foundation Trust / Newcastle Hospitals NHS Foundation Trust | COVID-19 Genomics UK (COG-UK) Consortium | Darren L Smith,Andrew Nelson,Matthew Bashton,Greg R Young,Joshua Loh,John Allan,Mohammad A Tariq,Giles S Holt,Gary Black,Wen C Yew,Lynn Dover,Paul Baker,Steve Liggett,Sarah Essex,Jane Greenaway,Debra Padgett,Clive Graham,Garren Scott,Edward Barton,Emma Swindells,Brendan Payne,Jennifer Collins,Yusri Taha,Gary Eltringham |
| EPI_ISL_611939 | University of Exeter | COVID-19 Genomics UK (COG-UK) Consortium | Ben Temperton,Aaron Jeffries,Michelle Michelsen,Joanna Warwick-Dugdale,Audrey Farbos,Robyn Manley,Stephen Michell,Jane Masoli |
| EPI_ISL_611941, EPI_ISL_611942, EPI_ISL_611943, EPI_ISL_611944, EPI_ISL_611945, EPI_ISL_611946, EPI_ISL_611947, EPI_ISL_611948, EPI_ISL_611949, EPI_ISL_611950, EPI_ISL_611951 | | | |
| see above | Northumbria University / South Tees Hospitals NHS Foundation Trust / North Cumbria Integrated Care NHS Foundation Trust / North Tees and Hartlepool NHS Foundation Trust / Newcastle Hospitals NHS Foundation Trust | COVID-19 Genomics UK (COG-UK) Consortium | Darren L Smith,Andrew Nelson,Matthew Bashton,Greg R Young,Joshua Loh,John Allan,Mohammad A Tariq,Giles S Holt,Gary Black,Wen C Yew,Lynn Dover,Paul Baker,Steve Liggett,Sarah Essex,Jane Greenaway,Debra Padgett,Clive Graham,Garren Scott,Edward Barton,Emma Swindells,Brendan Payne,Jennifer Collins,Yusri Taha,Gary Eltringham |
| EPI_ISL_611957, EPI_ISL_611958, EPI_ISL_611962 | Centre for Enzyme Innovation, University of Portsmouth / Translational Research Laboratory, Portsmouth Hospitals NHS Trust | COVID-19 Genomics UK (COG-UK) Consortium | Angela Beckett,Yann Bourgeois,Garry Scarlett,Sharon Glaysher,Scott Elliott,Kelly Bicknell,Robert Impey,Allyson Lloyd,Sarah Wyllie,Ethan Butcher,Anoop Chauhan,Samuel Robson |
| EPI_ISL_611968 | Wales Specialist Virology Centre Sequencing lab: Pathogen Genomics Unit | COVID-19 Genomics UK (COG-UK) Consortium | Catherine Moore, Johnathan Evans, Laura Gifford, Malorie Perry, Simon Cottrell, Angela Marchbank, Alec Birchley, Alexander Adams, Amy Gaskin, Bree Gatica-Wilcox, Jason Coombes, Joel Southgate, Lauren Gilbert, Lee Graham, Nicole Pacchiarini, Sara Kumziene-Summerhayes, Sarah Taylor, Sophie Jones, Sara Rey, Matthew Bull, Joanne Watkins, Sally Corden, Tom Connor |
| EPI_ISL_611980, EPI_ISL_612008, EPI_ISL_612030 | Northumbria University / South Tees Hospitals NHS Foundation Trust / North Cumbria Integrated Care NHS Foundation Trust / North Tees and Hartlepool NHS Foundation Trust / Newcastle Hospitals NHS Foundation Trust | COVID-19 Genomics UK (COG-UK) Consortium | Darren L Smith,Andrew Nelson,Matthew Bashton,Greg R Young,Joshua Loh,John Allan,Mohammad A Tariq,Giles S Holt,Gary Black,Wen C Yew,Lynn Dover,Paul Baker,Steve Liggett,Sarah Essex,Jane Greenaway,Debra Padgett,Clive Graham,Garren Scott,Edward Barton,Emma Swindells,Brendan Payne,Jennifer Collins,Yusri Taha,Gary Eltringham |
| EPI_ISL_612053 | Wales Specialist Virology Centre Sequencing lab: Pathogen Genomics Unit | COVID-19 Genomics UK (COG-UK) Consortium | Catherine Moore, Johnathan Evans, Laura Gifford, Malorie Perry, Simon Cottrell, Angela Marchbank, Alec Birchley, Alexander Adams, Amy Gaskin, Bree Gatica-Wilcox, Jason Coombes, Joel Southgate, Lauren Gilbert, Lee Graham, Nicole Pacchiarini, Sara Kumziene-Summerhayes, Sarah Taylor, Sophie Jones, Sara Rey, Matthew Bull, Joanne Watkins, Sally Corden, Tom Connor |
| EPI_ISL_612064, EPI_ISL_612088 | Liverpool Clinical Laboratories | COVID-19 Genomics UK (COG-UK) Consortium | Sam Haldenby, Anita Lucaci, Steve Paterson, Julian Hiscox, Alistair Darby, M Almsaud, A Alrezaihi, Muhannad Alruwaili, Stuart D Armstrong, Jones Benjamin, Eleanor G Bentley, Anu Chawla, Jordan J Clark, Angela Cowell, Richard Eccles, Isabel García-Dorival, Matthew Gemmell, Alessandro Gerada, PKF Gilmore, Richard Gregory, Ximeng Han, Catherine Hartley, Margaret Hughes, Miren Iturriza-Gomara, James Johnson, L Luu, Jenifer Manson, Charlotte Nelson, Elaine O'Toole, Cassie Olateju, Rebekah Penrice-Randal , Lucille Rainbow, N.P Randle, Trevor Ian Robinson, Parul Sharma, Ghada T Shawli, James P Stewart, Neil Swainston, Ecaterina Vamos, Joanne Watts, Mark Whitehead |
| EPI_ISL_612097, EPI_ISL_612100 | Northumbria University / South Tees Hospitals NHS Foundation Trust / North Cumbria Integrated Care NHS Foundation Trust / North Tees and Hartlepool NHS Foundation Trust / Newcastle Hospitals NHS Foundation Trust | COVID-19 Genomics UK (COG-UK) Consortium | Darren L Smith,Andrew Nelson,Matthew Bashton,Greg R Young,Joshua Loh,John Allan,Mohammad A Tariq,Giles S Holt,Gary Black,Wen C Yew,Lynn Dover,Paul Baker,Steve Liggett,Sarah Essex,Jane Greenaway,Debra Padgett,Clive Graham,Garren Scott,Edward Barton,Emma Swindells,Brendan Payne,Jennifer Collins,Yusri Taha,Gary Eltringham |
| EPI_ISL_612103 | University College London, Great Ormond Street Hospital for Children NHS Foundation Trust, Imperial College Healthcare NHS Trust | COVID-19 Genomics UK (COG-UK) Consortium | Sergi Castellano, Rachel Williams, Mark Kristiansen, Paola Resende Silva, Sunando Roy, Tony Brooks, Helena Tutill, Paola Niola, Patricia Dyal, Charlotte Williams, Leysa Forrest, Yasmin Panchbhaya, Jacqueline Findlay, Samuel Weeks, Julianne Brown, Kathryn Harris, Paul Randell, James Price, Alison Holmes, Judith Breuer |
| EPI_ISL_612104, EPI_ISL_612110 | Northumbria University / South Tees Hospitals NHS Foundation Trust / North Cumbria Integrated Care NHS Foundation Trust / North Tees and Hartlepool NHS Foundation Trust / Newcastle Hospitals NHS Foundation Trust | COVID-19 Genomics UK (COG-UK) Consortium | Darren L Smith,Andrew Nelson,Matthew Bashton,Greg R Young,Joshua Loh,John Allan,Mohammad A Tariq,Giles S Holt,Gary Black,Wen C Yew,Lynn Dover,Paul Baker,Steve Liggett,Sarah Essex,Jane Greenaway,Debra Padgett,Clive Graham,Garren Scott,Edward Barton,Emma Swindells,Brendan Payne,Jennifer Collins,Yusri Taha,Gary Eltringham |
| EPI_ISL_612112 | Liverpool Clinical Laboratories | COVID-19 Genomics UK (COG-UK) Consortium | Sam Haldenby, Anita Lucaci, Steve Paterson, Julian Hiscox, Alistair Darby, M Almsaud, A Alrezaihi, Muhannad Alruwaili, Stuart D Armstrong, Jones Benjamin, Eleanor G Bentley, Anu Chawla, Jordan J Clark, Angela Cowell, Richard Eccles, Isabel García-Dorival, Matthew Gemmell, Alessandro Gerada, PKF Gilmore, Richard Gregory, Ximeng Han, Catherine Hartley, Margaret Hughes, Miren Iturriza-Gomara, James Johnson, L Luu, Jenifer Manson, Charlotte Nelson, Elaine O'Toole, Cassie Olateju, Rebekah Penrice-Randal , Lucille Rainbow, N.P Randle, Trevor Ian Robinson, Parul Sharma, Ghada T Shawli, James P Stewart, Neil Swainston, Ecaterina Vamos, Joanne Watts, Mark Whitehead |
| EPI_ISL_612117 | Wales Specialist Virology Centre Sequencing lab: Pathogen Genomics Unit | COVID-19 Genomics UK (COG-UK) Consortium | Catherine Moore, Johnathan Evans, Laura Gifford, Malorie Perry, Simon Cottrell, Angela Marchbank, Alec Birchley, Alexander Adams, Amy Gaskin, Bree Gatica-Wilcox, Jason Coombes, Joel Southgate, Lauren Gilbert, Lee Graham, Nicole Pacchiarini, Sara Kumziene-Summerhayes, Sarah Taylor, Sophie Jones, Sara Rey, Matthew Bull, Joanne Watkins, Sally Corden, Tom Connor |
| EPI_ISL_612118 | University of Exeter | COVID-19 Genomics UK (COG-UK) Consortium | Ben Temperton,Aaron Jeffries,Michelle Michelsen,Joanna Warwick-Dugdale,Audrey Farbos,Robyn Manley,Stephen Michell,Jane Masoli |
| EPI_ISL_612122 | Wales Specialist Virology Centre Sequencing lab: Pathogen Genomics Unit | COVID-19 Genomics UK (COG-UK) Consortium | Catherine Moore, Johnathan Evans, Laura Gifford, Malorie Perry, Simon Cottrell, Angela Marchbank, Alec Birchley, Alexander Adams, Amy Gaskin, Bree Gatica-Wilcox, Jason Coombes, Joel Southgate, Lauren Gilbert, Lee Graham, Nicole Pacchiarini, Sara Kumziene-Summerhayes, Sarah Taylor, Sophie Jones, Sara Rey, Matthew Bull, Joanne Watkins, Sally Corden, Tom Connor |
| EPI_ISL_612126 | Virology Department, Royal Infirmary of Edinburgh, NHS Lothian / School of Biological Sciences, University of Edinburgh / Institute of Genetics and Molecular Medicine, University of Edinburgh | COVID-19 Genomics UK (COG-UK) Consortium | McHugh M, Dewar R, Rooke S, Gallagher M, Balcaza C, O'Toole Á, Scher E, Hill V, McCrone JT, Colquhoun R, Yu X, Jackson B, Rambaut A, Williams TC, Templeton K |
| EPI_ISL_612130 | University of Exeter | COVID-19 Genomics UK (COG-UK) Consortium | Ben Temperton,Aaron Jeffries,Michelle Michelsen,Joanna Warwick-Dugdale,Audrey Farbos,Robyn Manley,Stephen Michell,Jane Masoli |
| EPI_ISL_612134 | Northumbria University / South Tees Hospitals NHS Foundation Trust / North Cumbria Integrated Care NHS Foundation Trust / North Tees and Hartlepool NHS Foundation Trust / Newcastle Hospitals NHS Foundation Trust | COVID-19 Genomics UK (COG-UK) Consortium | Darren L Smith,Andrew Nelson,Matthew Bashton,Greg R Young,Joshua Loh,John Allan,Mohammad A Tariq,Giles S Holt,Gary Black,Wen C Yew,Lynn Dover,Paul Baker,Steve Liggett,Sarah Essex,Jane Greenaway,Debra Padgett,Clive Graham,Garren Scott,Edward Barton,Emma Swindells,Brendan Payne,Jennifer Collins,Yusri Taha,Gary Eltringham |
| EPI_ISL_612145, EPI_ISL_612184 | Wales Specialist Virology Centre Sequencing lab: Pathogen Genomics Unit | COVID-19 Genomics UK (COG-UK) Consortium | Catherine Moore, Johnathan Evans, Laura Gifford, Malorie Perry, Simon Cottrell, Angela Marchbank, Alec Birchley, Alexander Adams, Amy Gaskin, Bree Gatica-Wilcox, Jason Coombes, Joel Southgate, Lauren Gilbert, Lee Graham, Nicole Pacchiarini, Sara Kumziene-Summerhayes, Sarah Taylor, Sophie Jones, Sara Rey, Matthew Bull, Joanne Watkins, Sally Corden, Tom Connor |
| EPI_ISL_612198 | University College London, Great Ormond Street Hospital for Children NHS Foundation Trust, Imperial College Healthcare NHS Trust | COVID-19 Genomics UK (COG-UK) Consortium | Sergi Castellano, Rachel Williams, Mark Kristiansen, Paola Resende Silva, Sunando Roy, Tony Brooks, Helena Tutill, Paola Niola, Patricia Dyal, Charlotte Williams, Leysa Forrest, Yasmin Panchbhaya, Jacqueline Findlay, Samuel Weeks, Julianne Brown, Kathryn Harris, Paul Randell, James Price, Alison Holmes, Judith Breuer |
| EPI_ISL_612200, EPI_ISL_612201 | Wales Specialist Virology Centre Sequencing lab: Pathogen | COVID-19 Genomics UK (COG-UK) Consortium | Catherine Moore, Johnathan Evans, Laura Gifford, Malorie Perry, Simon Cottrell, Angela Marchbank, Alec Birchley, Alexander Adams, Amy Gaskin, Bree |

| | | | |
|---|---|---|---|
| | Genomics Unit | | Gatica-Wilcox, Jason Coombes, Joel Southgate, Lauren Gilbert, Lee Graham, Nicole Pacchiarini, Sara Kumziene-Summerhayes, Sarah Taylor, Sophie Jones, Sara Rey, Matthew Bull, Joanne Watkins, Sally Corden, Tom Connor |
| EPI_ISL_612230 | University of Birmingham | COVID-19 Genomics UK (COG-UK) Consortium | Institute of Microbiology, University of Birmingham: Claire McMurray, Joanne Stockton, Samuel Nicholls, Radoslaw Poplawski, Will Rowe, Josh Quick, Nicholas Loman. University of Birmingham Testing Laboratory: Celina M Whalley, Andrew Bosworth, Charlotte Poxon, Kasun Wanigasooriya, Oliver Pickles, Mike Kidd, Alex Richter, Andrew D Beggs PHE Heartlands Lab: Husam Osman, Andrew Bosworth. Queen Elizabeth Hospital: Anna Casey |
| EPI_ISL_612265, EPI_ISL_612267 | Department of Pathology, University of Cambridge | COVID-19 Genomics UK (COG-UK) Consortium | Aminu S. Jahun, Yasmin Chaudhry, Grant Hall, Iliana Georgana, Myra Hosmillo, Martin D. Curran, Malte Pinckert, Surendra Parmar, Ian Goodfellow |
| EPI_ISL_612326, EPI_ISL_612327, EPI_ISL_612328, EPI_ISL_612329, EPI_ISL_612330, EPI_ISL_612331, EPI_ISL_612332, EPI_ISL_612333, EPI_ISL_612334, EPI_ISL_612335, EPI_ISL_612336, EPI_ISL_612337, EPI_ISL_612347 | | | |
| see above | Virology Department, Royal Infirmary of Edinburgh, NHS Lothian / School of Biological Sciences, University of Edinburgh / Institute of Genetics and Molecular Medicine, University of Edinburgh | COVID-19 Genomics UK (COG-UK) Consortium | McHugh M, Dewar R, Rooke S, Gallagher M, Balcaza C, O'Toole Á, Scher E, Hill V, McCrone JT, Colquhoun R, Yu X, Jackson B, Rambaut A, Williams TC, Templeton K |
| EPI_ISL_612380, EPI_ISL_612381, EPI_ISL_612382, EPI_ISL_612383, EPI_ISL_612384, EPI_ISL_612385, EPI_ISL_612386, EPI_ISL_612387, EPI_ISL_612388, EPI_ISL_612389, EPI_ISL_612390, EPI_ISL_612391, EPI_ISL_612392, EPI_ISL_612393 | | | |
| see above | University of Exeter | COVID-19 Genomics UK (COG-UK) Consortium | Ben Temperton,Aaron Jeffries,Michelle Michelsen,Joanna Warwick-Dugdale,Audrey Farbos,Robyn Manley,Stephen Michell,Jane Masoli |
| EPI_ISL_612434 | Liverpool Clinical Laboratories | COVID-19 Genomics UK (COG-UK) Consortium | Sam Haldenby, Anita Lucaci, Steve Paterson, Julian Hiscox, Alistair Darby, M Almsaud, A Alrezaihi, Muhannad Alruwaili, Stuart D Armstrong, Jones Benjamin, Eleanor G Bentley, Anu Chawla, Jordan J Clark, Angela Cowell, Richard Eccles, Isabel García-Dorival, Matthew Gemmell, Alessandro Gerada, PKF Gilmore, Richard Gregory, Ximeng Han, Catherine Hartley, Margaret Hughes, Miren Iturriza-Gomara, James Johnson, L Luu, Jenifer Manson, Charlotte Nelson, Elaine O'Toole, Cassie Olateju, Rebekah Penrice-Randal , Lucille Rainbow, N.P Randle, Trevor Ian Robinson, Parul Sharma, Ghada T Shawli, James P Stewart, Neil Swainston, Ecaterina Vamos, Joanne Watts, Mark Whitehead |
| EPI_ISL_612457 | University College London, Great Ormond Street Hospital for Children NHS Foundation Trust, Imperial College Healthcare NHS Trust | COVID-19 Genomics UK (COG-UK) Consortium | Sergi Castellano, Rachel Williams, Mark Kristiansen, Paola Resende Silva, Sunando Roy, Tony Brooks, Helena Tutill, Paola Niola, Patricia Dyal, Charlotte Williams, Leysa Forrest, Yasmin Panchbhaya, Jacqueline Findlay, Samuel Weeks, Julianne Brown, Kathryn Harris, Paul Randell, James Price, Alison Holmes, Judith Breuer |
| EPI_ISL_612488, EPI_ISL_612489, EPI_ISL_612490, EPI_ISL_612491, EPI_ISL_612492, EPI_ISL_612501, EPI_ISL_612502, EPI_ISL_612503, EPI_ISL_612504, EPI_ISL_612505, EPI_ISL_612506, EPI_ISL_612507, EPI_ISL_612508, EPI_ISL_612509, EPI_ISL_612510, EPI_ISL_612511, EPI_ISL_612512, EPI_ISL_612513, EPI_ISL_612514, EPI_ISL_612515, EPI_ISL_612516, EPI_ISL_612517, EPI_ISL_612518, EPI_ISL_612519, EPI_ISL_612520, EPI_ISL_612521, EPI_ISL_612522 | | | |
| see above | Wales Specialist Virology Centre Sequencing lab: Pathogen Genomics Unit | COVID-19 Genomics UK (COG-UK) Consortium | Catherine Moore, Johnathan Evans, Laura Gifford, Malorie Perry, Simon Cottrell, Angela Marchbank, Alec Birchley, Alexander Adams, Amy Gaskin, Bree Gatica-Wilcox, Jason Coombes, Joel Southgate, Lauren Gilbert, Lee Graham, Nicole Pacchiarini, Sara Kumziene-Summerhayes, Sarah Taylor, Sophie Jones, Sara Rey, Matthew Bull, Joanne Watkins, Sally Corden, Tom Connor |
| EPI_ISL_612523, EPI_ISL_612524, EPI_ISL_612525, EPI_ISL_612526, EPI_ISL_612527, EPI_ISL_612528, EPI_ISL_612529, EPI_ISL_612530, EPI_ISL_612531, EPI_ISL_612532, EPI_ISL_612533, EPI_ISL_612534, EPI_ISL_612535, EPI_ISL_612536, EPI_ISL_612537, EPI_ISL_612538, EPI_ISL_612539, EPI_ISL_612540, EPI_ISL_612541, EPI_ISL_612542, EPI_ISL_612543, EPI_ISL_612544, EPI_ISL_612545, EPI_ISL_612557, EPI_ISL_612564, EPI_ISL_612565 | | | |
| see above | Northumbria University / South Tees Hospitals NHS Foundation Trust / North Cumbria Integrated Care NHS Foundation Trust / North Tees and Hartlepool NHS Foundation Trust / Newcastle Hospitals NHS Foundation Trust | COVID-19 Genomics UK (COG-UK) Consortium | Darren L Smith,Andrew Nelson,Matthew Bashton,Greg R Young,Joshua Loh,John Allan,Mohammad A Tariq,Giles S Holt,Gary Black,Wen C Yew,Lynn Dover,Paul Baker,Steve Liggett,Sarah Essex,Jane Greenaway,Debra Padgett,Clive Graham,Garren Scott,Edward Barton,Emma Swindells,Brendan Payne,Jennifer Collins,Yusri Taha,Gary Eltringham |
| EPI_ISL_612608, EPI_ISL_612636, EPI_ISL_612639, EPI_ISL_612643, EPI_ISL_612646, EPI_ISL_612650, EPI_ISL_612651, EPI_ISL_612662, EPI_ISL_612666, EPI_ISL_612684, EPI_ISL_612688, EPI_ISL_612693, EPI_ISL_612700, EPI_ISL_612704, EPI_ISL_612707, EPI_ISL_612708, EPI_ISL_612716, EPI_ISL_612721, EPI_ISL_612738, EPI_ISL_612746, EPI_ISL_612753, EPI_ISL_612765, EPI_ISL_612768, EPI_ISL_612769, EPI_ISL_612770, EPI_ISL_612773, EPI_ISL_612787, EPI_ISL_612789, EPI_ISL_612794, EPI_ISL_612795, EPI_ISL_612809, EPI_ISL_612816, EPI_ISL_612818, EPI_ISL_612829, EPI_ISL_612836, EPI_ISL_612867, EPI_ISL_612905, EPI_ISL_612923, EPI_ISL_612928, EPI_ISL_612944, EPI_ISL_613100, EPI_ISL_613114, EPI_ISL_613136, EPI_ISL_613168 | | | |
| see above | Wales Specialist Virology Centre Sequencing lab: Pathogen Genomics Unit | COVID-19 Genomics UK (COG-UK) Consortium | Catherine Moore, Johnathan Evans, Laura Gifford, Malorie Perry, Simon Cottrell, Angela Marchbank, Alec Birchley, Alexander Adams, Amy Gaskin, Bree Gatica-Wilcox, Jason Coombes, Joel Southgate, Lauren Gilbert, Lee Graham, Nicole Pacchiarini, Sara Kumziene-Summerhayes, Sarah Taylor, Sophie Jones, Sara Rey, Matthew Bull, Joanne Watkins, Sally Corden, Tom Connor |
| EPI_ISL_613362, EPI_ISL_613364, EPI_ISL_613372, EPI_ISL_613380, EPI_ISL_613389 | Virology Department, Sheffield Teaching Hospitals NHS Foundation Trust/Department of Infection, Immunity and Cardiovascular Disease, The Medical School, University of Sheffield | COVID-19 Genomics UK (COG-UK) Consortium | Thushan de Silva, Matthew Parker, Nikki Smith, Adri Angyal, Rebecca Brown, Luke Green, Rachel Tucker, Paul Parsons, Danielle Groves, Katie Johnson, Laura Carrilero, Alex Keeley, Dave Partridge, Matthew Wyles, Benjamin Lindsey, Mehmet Yavuz, Mohammad Raza, Cariad Evans |
| EPI_ISL_613493, EPI_ISL_613495 | Public Health Laboratory - Infectious Disease Lab, Minnesota Department of Health Infectious Disease Laboratory Submission Group | Minnesota Department of Health, Public Health Laboratory | Plumb,M., Garfin,J., Lorentz,A., Wang,X. |
| EPI_ISL_613713, EPI_ISL_613725, EPI_ISL_613729, EPI_ISL_613740, EPI_ISL_613748, EPI_ISL_613749, EPI_ISL_613750, EPI_ISL_613751, EPI_ISL_613761, EPI_ISL_613766, EPI_ISL_613774, EPI_ISL_613785, EPI_ISL_613793, EPI_ISL_613795, EPI_ISL_613799, EPI_ISL_613801, EPI_ISL_613828, EPI_ISL_613856, EPI_ISL_613857, EPI_ISL_613858, EPI_ISL_613859, EPI_ISL_613860, EPI_ISL_613861, EPI_ISL_613862, EPI_ISL_613863, EPI_ISL_613864, EPI_ISL_613865, EPI_ISL_613866, EPI_ISL_613867, EPI_ISL_613868, EPI_ISL_613869, EPI_ISL_613870, EPI_ISL_613871, EPI_ISL_613872, EPI_ISL_613873 | | | |
| see above | Florida Bureau of Public Health Laboratories | Florida Bureau of Public Health Laboratories | Sarah Schmedes, Jason Blanton |
| EPI_ISL_614063, EPI_ISL_614064, EPI_ISL_614066, EPI_ISL_614067, EPI_ISL_614068, EPI_ISL_614069, EPI_ISL_614089, EPI_ISL_614090, EPI_ISL_614091, EPI_ISL_614092, EPI_ISL_614150, EPI_ISL_614151, EPI_ISL_614152, EPI_ISL_614153, EPI_ISL_614154 | | | |
| see above | Virginia DCLS | Virginia DCLS | Virginia DCLS |
| EPI_ISL_614159, EPI_ISL_614163, EPI_ISL_614167, EPI_ISL_614174, EPI_ISL_614176, EPI_ISL_614180, EPI_ISL_614190, EPI_ISL_614191, EPI_ISL_614196, EPI_ISL_614198, EPI_ISL_614206, EPI_ISL_614218, EPI_ISL_614234 | | | |
| see above | Michigan Department of Health and Human Services, Bureau of Laboratories | Michigan Department of Health and Human Services, Bureau of Laboratories | Blankenship HM, Riner D, Soehnlen MK |
| EPI_ISL_614282, EPI_ISL_614283 | General practitioner | National Reference Center for Viruses of Respiratory Infections, Institut Pasteur, Paris | Marion Barbet, Sylvie Behillil, Méline Bizard, Angela Brisebarre, Camille Capel, Etienne Simon-Lorière, Vincent Enouf, Maud Vanpeene, Sylvie van der Werf |
| EPI_ISL_615163, EPI_ISL_615164 | Texas Department of State Health Services | Texas Department of State Health Services | Rashmi Tuladhar, Bonnie Oh, Jenny Zhang, Maliha Rahman, Anita Pokharel, Myong Koag, Chung Wang, Rachel Lee, Grace Kubin, Mayela Pedrueza |
| EPI_ISL_619406, EPI_ISL_619407, EPI_ISL_619408, EPI_ISL_619409, EPI_ISL_619410, EPI_ISL_619411, EPI_ISL_619412, EPI_ISL_619413, EPI_ISL_619414, EPI_ISL_619415, EPI_ISL_619416, EPI_ISL_619417, EPI_ISL_619418, EPI_ISL_619419, EPI_ISL_619420, EPI_ISL_619421, EPI_ISL_619422, EPI_ISL_619423, EPI_ISL_619424, EPI_ISL_619425, EPI_ISL_619426, EPI_ISL_619427, EPI_ISL_619428, EPI_ISL_619429, EPI_ISL_619430, EPI_ISL_619431, EPI_ISL_619432, EPI_ISL_619433, EPI_ISL_619434, EPI_ISL_619435, EPI_ISL_619436, EPI_ISL_619437, EPI_ISL_619438, EPI_ISL_619439, EPI_ISL_619440, EPI_ISL_619441, EPI_ISL_619442, EPI_ISL_619443, EPI_ISL_619444, EPI_ISL_619445, EPI_ISL_619446, EPI_ISL_619447, EPI_ISL_619448, EPI_ISL_619449, EPI_ISL_619450, EPI_ISL_619451, EPI_ISL_619452, EPI_ISL_619453, EPI_ISL_619454, EPI_ISL_619984, EPI_ISL_619985, EPI_ISL_619986, EPI_ISL_619987, EPI_ISL_619988, EPI_ISL_619989, EPI_ISL_619990, EPI_ISL_619991, EPI_ISL_619992, EPI_ISL_619993, EPI_ISL_619994, EPI_ISL_619995, EPI_ISL_619996, EPI_ISL_619997, EPI_ISL_619998, EPI_ISL_619999, EPI_ISL_620000, EPI_ISL_620001, EPI_ISL_620002, EPI_ISL_620003, EPI_ISL_620004, EPI_ISL_620005, EPI_ISL_620006, EPI_ISL_620007, EPI_ISL_620008, EPI_ISL_620009, EPI_ISL_620010, EPI_ISL_620011, EPI_ISL_620012, EPI_ISL_620013, EPI_ISL_620014, EPI_ISL_620015, EPI_ISL_620016, EPI_ISL_620017, EPI_ISL_620018, EPI_ISL_620019, EPI_ISL_620020, EPI_ISL_620021, EPI_ISL_620022, EPI_ISL_620023, EPI_ISL_620024, EPI_ISL_620025, EPI_ISL_620026, EPI_ISL_620027, EPI_ISL_620028, EPI_ISL_620029, EPI_ISL_620030, EPI_ISL_620031, EPI_ISL_620032, EPI_ISL_620033, EPI_ISL_620034, EPI_ISL_620035, EPI_ISL_620036, EPI_ISL_620037, EPI_ISL_620038, EPI_ISL_620039, EPI_ISL_620040, EPI_ISL_620041, EPI_ISL_620042, EPI_ISL_620043, EPI_ISL_620044, EPI_ISL_620045, EPI_ISL_620046, EPI_ISL_620047, EPI_ISL_620048, EPI_ISL_620049, EPI_ISL_620050, EPI_ISL_620051, EPI_ISL_620052, EPI_ISL_620053, EPI_ISL_620054, EPI_ISL_620055, EPI_ISL_620056, EPI_ISL_620057, EPI_ISL_620058, EPI_ISL_620059, EPI_ISL_620060, EPI_ISL_620061, EPI_ISL_620062, EPI_ISL_620063, EPI_ISL_620064, EPI_ISL_620065, EPI_ISL_620066, EPI_ISL_620067, EPI_ISL_620068, EPI_ISL_620069, EPI_ISL_620070, EPI_ISL_620071, EPI_ISL_620072, EPI_ISL_620073, EPI_ISL_620074, EPI_ISL_620075, EPI_ISL_620076, EPI_ISL_620077, EPI_ISL_620078, EPI_ISL_620079, EPI_ISL_620080, EPI_ISL_620081, EPI_ISL_620082, EPI_ISL_620083, EPI_ISL_620084, EPI_ISL_620085, EPI_ISL_620086, EPI_ISL_620087, EPI_ISL_620088, EPI_ISL_620089, EPI_ISL_620090, EPI_ISL_620091, EPI_ISL_620092, EPI_ISL_620093, EPI_ISL_620094, EPI_ISL_620095, EPI_ISL_620096, EPI_ISL_620097, EPI_ISL_620098, EPI_ISL_620099, EPI_ISL_620100, EPI_ISL_620101, EPI_ISL_620102, EPI_ISL_620103, EPI_ISL_620104, EPI_ISL_620105, EPI_ISL_620106, EPI_ISL_620107, EPI_ISL_620108, EPI_ISL_620109, EPI_ISL_620110, EPI_ISL_620111, EPI_ISL_620112, EPI_ISL_620113, EPI_ISL_620114, EPI_ISL_620115, EPI_ISL_620116, EPI_ISL_620117, EPI_ISL_620118, EPI_ISL_620119, EPI_ISL_620120, EPI_ISL_620121, EPI_ISL_620122, EPI_ISL_620123, EPI_ISL_620124, EPI_ISL_620125, EPI_ISL_620126, EPI_ISL_620148, EPI_ISL_620149, EPI_ISL_620150, EPI_ISL_620151, EPI_ISL_620152, EPI_ISL_620153, EPI_ISL_620154, EPI_ISL_620155, EPI_ISL_620156, EPI_ISL_620157, EPI_ISL_620158, EPI_ISL_620159, EPI_ISL_620160, EPI_ISL_620161, EPI_ISL_620162, EPI_ISL_620163, EPI_ISL_620164, EPI_ISL_620165, EPI_ISL_620166, EPI_ISL_620167, EPI_ISL_620168, EPI_ISL_620169, EPI_ISL_620170, EPI_ISL_620171, EPI_ISL_620172, EPI_ISL_620173, EPI_ISL_620174, EPI_ISL_620175, EPI_ISL_620176, EPI_ISL_620177, EPI_ISL_620178, EPI_ISL_620179, EPI_ISL_620180, EPI_ISL_620181, EPI_ISL_620182, EPI_ISL_620183, EPI_ISL_620184, EPI_ISL_620185, EPI_ISL_620186, EPI_ISL_620187, EPI_ISL_620188, EPI_ISL_620189, EPI_ISL_620190, EPI_ISL_620191, EPI_ISL_620192, EPI_ISL_620193, EPI_ISL_620194, EPI_ISL_620195, EPI_ISL_620196, EPI_ISL_620197, EPI_ISL_620198, EPI_ISL_620199, EPI_ISL_620200, EPI_ISL_620201, EPI_ISL_620202, EPI_ISL_620203, EPI_ISL_620204, EPI_ISL_620205, EPI_ISL_620206, EPI_ISL_620207, EPI_ISL_620208, EPI_ISL_620209, EPI_ISL_620210, EPI_ISL_620211, EPI_ISL_620212, EPI_ISL_620213, EPI_ISL_620214, EPI_ISL_620215, EPI_ISL_620216, EPI_ISL_620217, EPI_ISL_620218, EPI_ISL_620219, EPI_ISL_620220, EPI_ISL_620221, EPI_ISL_620222, EPI_ISL_620223, EPI_ISL_620224, EPI_ISL_620225, EPI_ISL_620226, EPI_ISL_620227, EPI_ISL_620228, EPI_ISL_620229, EPI_ISL_620230, EPI_ISL_620231, EPI_ISL_620232, EPI_ISL_620233, EPI_ISL_620234, EPI_ISL_620235, EPI_ISL_620236, EPI_ISL_620237, EPI_ISL_620238, EPI_ISL_620239, EPI_ISL_620240, EPI_ISL_620241, EPI_ISL_620242, EPI_ISL_620243, EPI_ISL_620244, EPI_ISL_620245, EPI_ISL_620246, EPI_ISL_620247, EPI_ISL_620248, EPI_ISL_620249, EPI_ISL_620250, EPI_ISL_620251, EPI_ISL_620252, EPI_ISL_620253, EPI_ISL_620254, EPI_ISL_620255, EPI_ISL_620256, EPI_ISL_620257, EPI_ISL_620258, EPI_ISL_620259, EPI_ISL_620260, EPI_ISL_620261, EPI_ISL_620262, EPI_ISL_620263, EPI_ISL_620264, EPI_ISL_620265, EPI_ISL_620266, EPI_ISL_620267, EPI_ISL_620268, EPI_ISL_620269, EPI_ISL_620270, EPI_ISL_620271, EPI_ISL_620272, EPI_ISL_620273, EPI_ISL_620274, EPI_ISL_620275, EPI_ISL_620276, EPI_ISL_620277, EPI_ISL_620278, EPI_ISL_620279, | | | |

| | | | |
|---|---|---|---|
| EPI_ISL_620280, EPI_ISL_620281, EPI_ISL_620282, EPI_ISL_620283, EPI_ISL_620284, EPI_ISL_620285, EPI_ISL_620287, EPI_ISL_620288, EPI_ISL_620289, EPI_ISL_620290, EPI_ISL_620291, EPI_ISL_620292, EPI_ISL_620293, EPI_ISL_620294, EPI_ISL_620295, EPI_ISL_620296, EPI_ISL_620297, EPI_ISL_620298, EPI_ISL_620299, EPI_ISL_620300, EPI_ISL_620301, EPI_ISL_620380, EPI_ISL_620381, EPI_ISL_620382, EPI_ISL_620383, EPI_ISL_620384, EPI_ISL_620385, EPI_ISL_620386, EPI_ISL_620387, EPI_ISL_620388, EPI_ISL_620389, EPI_ISL_620390, EPI_ISL_620391, EPI_ISL_620392, EPI_ISL_620393, EPI_ISL_620394, EPI_ISL_620395, EPI_ISL_620396, EPI_ISL_620397, EPI_ISL_620398, EPI_ISL_620399, EPI_ISL_620400, EPI_ISL_620401, EPI_ISL_620402, EPI_ISL_620403, EPI_ISL_620404, EPI_ISL_620405, EPI_ISL_622332, EPI_ISL_622334, EPI_ISL_622335, EPI_ISL_622336, EPI_ISL_622337, EPI_ISL_622338, EPI_ISL_622341, EPI_ISL_622365, EPI_ISL_622366, EPI_ISL_622367, EPI_ISL_622368, EPI_ISL_622397, EPI_ISL_622398 | | | |
| see above | Department of Virus and Microbiological Special Diagnostics, Statens Serum Institut, Denmark | Albertsen lab, Department of Chemistry and Bioscience, Aalborg University, Denmark | Danish Covid-19 Genome Consortia |
| EPI_ISL_622802 | Middlemore Hospital | Institute of Environmental Science and Research (ESR) | Xiaoyun Ren, Matt Storey, Nikki Freed, Muhammad Faisal, Jing Wang, Hermes Perez, Anja Werno, Antje van der Linden, Arlo Upton, Chris Mansell, David Hammer, Dragana Drinkovic, Gary McAuliffe, Hana Sofia Andersson, James Ussher, Jill Sherwood, Josh Freeman, Julia Howard, Juliet Elvy, Mary DeAlmeida, Matt Blakiston, Matthew Rogers, Max Bloomfield, Michael Addidle, Michelle Balm, Sally Roberts, Sarah Jefferies, Sharmini Muttaiyah, Susan Morpeth, Susan Taylor, Timothy Blackmore, Vani Sathyendran, Veronica Playle, Virginia Hope, Erasmus Smit, Lauren Jelly, Olin Silander, Joep de Ligt |
| EPI_ISL_622803, EPI_ISL_622807 | LabPLUS | Institute of Environmental Science and Research (ESR) | Xiaoyun Ren, Matt Storey, Nikki Freed, Muhammad Faisal, Jing Wang, Hermes Perez, Anja Werno, Antje van der Linden, Arlo Upton, Chris Mansell, David Hammer, Dragana Drinkovic, Gary McAuliffe, Hana Sofia Andersson, James Ussher, Jill Sherwood, Josh Freeman, Julia Howard, Juliet Elvy, Mary DeAlmeida, Matt Blakiston, Matthew Rogers, Max Bloomfield, Michael Addidle, Michelle Balm, Sally Roberts, Sarah Jefferies, Sharmini Muttaiyah, Susan Morpeth, Susan Taylor, Timothy Blackmore, Vani Sathyendran, Veronica Playle, Virginia Hope, Erasmus Smit, Lauren Jelly, Olin Silander, Joep de Ligt |
| EPI_ISL_625437, EPI_ISL_625438, EPI_ISL_625439, EPI_ISL_625440, EPI_ISL_625444, EPI_ISL_625445, EPI_ISL_625446, EPI_ISL_625447, EPI_ISL_625448, EPI_ISL_625449, EPI_ISL_625450, EPI_ISL_625451, EPI_ISL_625452, EPI_ISL_625453 | | | |
| see above | Lighthouse Lab in Cambridge | Wellcome Sanger Institute for the COVID-19 Genomics UK (COG-UK) consortium | Rob Howes, The Lighthouse Lab in Cambridge and Alex Alderton, Roberto Amato, Sonia Goncalves, Ewan Harrison, David K. Jackson, Ian Johnston, Dominic Kwiatkowski, Cordelia Langford, John Sillitoe on behalf of the Wellcome Sanger Institute COVID-19 Surveillance Team (http://www.sanger.ac.uk/covid-team) |
| EPI_ISL_625600, EPI_ISL_625602, EPI_ISL_625603, EPI_ISL_625604, EPI_ISL_625605, EPI_ISL_625606, EPI_ISL_625607, EPI_ISL_625608, EPI_ISL_625609 | Santa Clara County Public Health Laboratory | Chan-Zuckerberg Biohub | CZB Cliahub Consortium |
| EPI_ISL_625614, EPI_ISL_625615, EPI_ISL_625616, EPI_ISL_625617, EPI_ISL_625618, EPI_ISL_625619, EPI_ISL_625620 | Alameda County Public Health Lab | Chan-Zuckerberg Biohub | CZB Cliahub Consortium |
| EPI_ISL_625658, EPI_ISL_625659 | Orange County Public Health Lab | Chan-Zuckerberg Biohub | CZB Cliahub Consortium |
| EPI_ISL_625665, EPI_ISL_625666 | UCSF Clinical Microbiology Laboratory | Chan-Zuckerberg Biohub | CZB Cliahub Consortium |
| EPI_ISL_626592, EPI_ISL_626597, EPI_ISL_626598, EPI_ISL_626602, EPI_ISL_626611, EPI_ISL_626616, EPI_ISL_626617, EPI_ISL_626618 | The National Institute of Public Health | State Veterinary Institute Prague | Nagy,A;Jirincova,H;Novakova,L;Trnka,D;Vecerova,J |
| EPI_ISL_626664 | Virology Department, Sheffield Teaching Hospitals NHS Foundation Trust/Department of Infection, Immunity and Cardiovascular Disease, The Medical School, University of Sheffield | COVID-19 Genomics UK (COG-UK) Consortium | Thushan de Silva, Matthew Parker, Nikki Smith, Adri Angyal, Rebecca Brown, Luke Green, Rachel Tucker, Paul Parsons, Danielle Groves, Katie Johnson, Laura Carrilero, Alex Keeley, Dave Partridge, Matthew Wyles, Benjamin Lindsey, Mehmet Yavuz, Mohammad Raza, Cariad Evans |
| EPI_ISL_626668, EPI_ISL_626669 | University of Exeter | COVID-19 Genomics UK (COG-UK) Consortium | Ben Temperton,Aaron Jeffries,Michelle Michelsen,Joanna Warwick-Dugdale,Audrey Farbos,Robyn Manley,Stephen Michell,Jane Masoli |
| EPI_ISL_626679, EPI_ISL_626680 | Virology Department, Sheffield Teaching Hospitals NHS Foundation Trust/Department of Infection, Immunity and Cardiovascular Disease, The Medical School, University of Sheffield | COVID-19 Genomics UK (COG-UK) Consortium | Thushan de Silva, Matthew Parker, Nikki Smith, Adri Angyal, Rebecca Brown, Luke Green, Rachel Tucker, Paul Parsons, Danielle Groves, Katie Johnson, Laura Carrilero, Alex Keeley, Dave Partridge, Matthew Wyles, Benjamin Lindsey, Mehmet Yavuz, Mohammad Raza, Cariad Evans |
| EPI_ISL_626697 | Liverpool Clinical Laboratories | COVID-19 Genomics UK (COG-UK) Consortium | Sam Haldenby, Anita Lucaci, Steve Paterson, Julian Hiscox, Alistair Darby, M Almsaud, A Alrezaihi, Muhannad Alruwaili, Stuart D Armstrong, Jones Benjamin, Eleanor G Bentley, Anu Chawla, Jordan J Clark, Angela Cowell, Richard Eccles, Isabel García-Dorival, Matthew Gemmell, Alessandro Gerada, PKF Gilmore, Richard Gregory, Ximeng Han, Catherine Hartley, Margaret Hughes, Miren Iturriza-Gomara, James Johnson, L Luu, Jenifer Manson, Charlotte Nelson, Elaine O'Toole, Cassie Olateju, Rebekah Penrice-Randal , Lucille Rainbow, N.P Randle, Trevor Ian Robinson, Parul Sharma, Ghada T Shawli, James P Stewart, Neil Swainston, Ecaterina Vamos, Joanne Watts, Mark Whitehead |
| EPI_ISL_626731 | University of Exeter | COVID-19 Genomics UK (COG-UK) Consortium | Ben Temperton,Aaron Jeffries,Michelle Michelsen,Joanna Warwick-Dugdale,Audrey Farbos,Robyn Manley,Stephen Michell,Jane Masoli |
| EPI_ISL_626733 | Wales Specialist Virology Centre Sequencing lab: Pathogen Genomics Unit | COVID-19 Genomics UK (COG-UK) Consortium | Catherine Moore, Johnathan Evans, Laura Gifford, Malorie Perry, Simon Cottrell, Angela Marchbank, Alec Birchley, Alexander Adams, Amy Gaskin, Bree Gatica-Wilcox, Jason Coombes, Joel Southgate, Lauren Gilbert, Lee Graham, Nicole Pacchiarini, Sara Kumziene-Summerhayes, Sarah Taylor, Sophie Jones, Sara Rey, Matthew Bull, Joanne Watkins, Sally Corden, Tom Connor |
| EPI_ISL_626734, EPI_ISL_626739, EPI_ISL_626752, EPI_ISL_626757, EPI_ISL_626771, EPI_ISL_626786, EPI_ISL_626791 | Virology Department, Sheffield Teaching Hospitals NHS Foundation Trust/Department of Infection, Immunity and Cardiovascular Disease, The Medical School, University of Sheffield | COVID-19 Genomics UK (COG-UK) Consortium | Thushan de Silva, Matthew Parker, Nikki Smith, Adri Angyal, Rebecca Brown, Luke Green, Rachel Tucker, Paul Parsons, Danielle Groves, Katie Johnson, Laura Carrilero, Alex Keeley, Dave Partridge, Matthew Wyles, Benjamin Lindsey, Mehmet Yavuz, Mohammad Raza, Cariad Evans |
| EPI_ISL_626810 | University of Exeter | COVID-19 Genomics UK (COG-UK) Consortium | Ben Temperton,Aaron Jeffries,Michelle Michelsen,Joanna Warwick-Dugdale,Audrey Farbos,Robyn Manley,Stephen Michell,Jane Masoli |
| EPI_ISL_626821 | Virology Department, Sheffield Teaching Hospitals NHS Foundation Trust/Department of Infection, Immunity and Cardiovascular Disease, The Medical School, University of Sheffield | COVID-19 Genomics UK (COG-UK) Consortium | Thushan de Silva, Matthew Parker, Nikki Smith, Adri Angyal, Rebecca Brown, Luke Green, Rachel Tucker, Paul Parsons, Danielle Groves, Katie Johnson, Laura Carrilero, Alex Keeley, Dave Partridge, Matthew Wyles, Benjamin Lindsey, Mehmet Yavuz, Mohammad Raza, Cariad Evans |
| EPI_ISL_626869, EPI_ISL_626875 | Wales Specialist Virology Centre Sequencing lab: Pathogen Genomics Unit | COVID-19 Genomics UK (COG-UK) Consortium | Catherine Moore, Johnathan Evans, Laura Gifford, Malorie Perry, Simon Cottrell, Angela Marchbank, Alec Birchley, Alexander Adams, Amy Gaskin, Bree Gatica-Wilcox, Jason Coombes, Joel Southgate, Lauren Gilbert, Lee Graham, Nicole Pacchiarini, Sara Kumziene-Summerhayes, Sarah Taylor, Sophie Jones, Sara Rey, Matthew Bull, Joanne Watkins, Sally Corden, Tom Connor |
| EPI_ISL_626890, EPI_ISL_626891 | University of Exeter | COVID-19 Genomics UK (COG-UK) Consortium | Ben Temperton,Aaron Jeffries,Michelle Michelsen,Joanna Warwick-Dugdale,Audrey Farbos,Robyn Manley,Stephen Michell,Jane Masoli |
| EPI_ISL_626901, EPI_ISL_626902, EPI_ISL_626903, EPI_ISL_626915, EPI_ISL_626916, EPI_ISL_626922, EPI_ISL_626923, EPI_ISL_626924, EPI_ISL_626929, EPI_ISL_626932 | Virology Department, Sheffield Teaching Hospitals NHS Foundation Trust/Department of Infection, Immunity and Cardiovascular Disease, The Medical School, University of Sheffield | COVID-19 Genomics UK (COG-UK) Consortium | Thushan de Silva, Matthew Parker, Nikki Smith, Adri Angyal, Rebecca Brown, Luke Green, Rachel Tucker, Paul Parsons, Danielle Groves, Katie Johnson, Laura Carrilero, Alex Keeley, Dave Partridge, Matthew Wyles, Benjamin Lindsey, Mehmet Yavuz, Mohammad Raza, Cariad Evans |
| EPI_ISL_626944 | University of Exeter | COVID-19 Genomics UK (COG-UK) Consortium | Ben Temperton,Aaron Jeffries,Michelle Michelsen,Joanna Warwick-Dugdale,Audrey Farbos,Robyn Manley,Stephen Michell,Jane Masoli |
| EPI_ISL_626970 | West of Scotland Specialist Virology Centre, NHSGGC / MRC-University of Glasgow Centre for Virus Research | COVID-19 Genomics UK (COG-UK) Consortium | Ana da Silva Filipe, Natasha Johnson, Kathy Smollett, Daniel Mair, Stephen Carmichael, Lily Tong, Jenna Nichols, Elihu Aranday-Cortes, Kyriaki Nomikou; Sarah McDonald, Marc Niebel, Patawee Asamaphan; Richard Orton, Joseph Hughes, Sreenu Vattipally, David L Robertson; Alasdair MacLean, Rory Gunson; Kathy Li, Igor Starinskij, Natasha Jesudason, Rajiv Shah, James Shepherd, Antonia Ho, Emma Thomson |
| EPI_ISL_626994 | Liverpool Clinical Laboratories | COVID-19 Genomics UK (COG-UK) Consortium | Sam Haldenby, Anita Lucaci, Steve Paterson, Julian Hiscox, Alistair Darby, M Almsaud, A Alrezaihi, Muhannad Alruwaili, Stuart D Armstrong, Jones Benjamin, Eleanor G Bentley, Anu Chawla, Jordan J Clark, Angela Cowell, Richard Eccles, Isabel García-Dorival, Matthew Gemmell, Alessandro Gerada, PKF Gilmore, Richard Gregory, Ximeng Han, Catherine Hartley, Margaret Hughes, Miren Iturriza-Gomara, James Johnson, L Luu, Jenifer Manson, Charlotte Nelson, Elaine |

| | | | |
|---|---|---|---|
| | | | O'Toole, Cassie Olateju, Rebekah Penrice-Randal , Lucille Rainbow, N.P Randle, Trevor Ian Robinson, Parul Sharma, Ghada T Shawli, James P Stewart, Neil Swainston, Ecaterina Vamos, Joanne Watts, Mark Whitehead |
| EPI_ISL_627000 | Regional Virus Laboratory, Belfast Health and Social Care Trust | COVID-19 Genomics UK (COG-UK) Consortium | Conall McCaughey, James McKenna, Tanya Curran, Susan Feeney, Alison Watt, Ciara Cox, Mairead Connor, Zoltan Molnar, David Simpson, Derek Fairley |
| EPI_ISL_627014 | West of Scotland Specialist Virology Centre, NHSGGC / MRC-University of Glasgow Centre for Virus Research | COVID-19 Genomics UK (COG-UK) Consortium | Ana da Silva Filipe, Natasha Johnson, Kathy Smollett, Daniel Mair, Stephen Carmichael, Lily Tong, Jenna Nichols, Elihu Aranday-Cortes, Kyriaki Nomikou; Sarah McDonald, Marc Niebel, Patawee Asamaphan; Richard Orton, Joseph Hughes, Sreenu Vattipally, David L Robertson; Alasdair MacLean, Rory Gunson; Kathy Li, Igor Starinskij, Natasha Jesudason, Rajiv Shah, James Shepherd, Antonia Ho, Emma Thomson |
| EPI_ISL_627017 | Liverpool Clinical Laboratories | COVID-19 Genomics UK (COG-UK) Consortium | Sam Haldenby, Anita Lucaci, Steve Paterson, Julian Hiscox, Alistair Darby, M Almsaud, A Alrezaihi, Muhannad Alruwaili, Stuart D Armstrong, Jones Benjamin, Eleanor G Bentley, Anu Chawla, Jordan J Clark, Angela Cowell, Richard Eccles, Isabel García-Dorival, Matthew Gemmell, Alessandro Gerada, PKF Gilmore, Richard Gregory, Ximeng Han, Catherine Hartley, Margaret Hughes, Miren Iturriza-Gomara, James Johnson, L Luu, Jenifer Manson, Charlotte Nelson, Elaine O'Toole, Cassie Olateju, Rebekah Penrice-Randal , Lucille Rainbow, N.P Randle, Trevor Ian Robinson, Parul Sharma, Ghada T Shawli, James P Stewart, Neil Swainston, Ecaterina Vamos, Joanne Watts, Mark Whitehead |
| EPI_ISL_627019 | Virology Department, Sheffield Teaching Hospitals NHS Foundation Trust/Department of Infection, Immunity and Cardiovascular Disease, The Medical School, University of Sheffield | COVID-19 Genomics UK (COG-UK) Consortium | Thushan de Silva, Matthew Parker, Nikki Smith, Adri Angyal, Rebecca Brown, Luke Green, Rachel Tucker, Paul Parsons, Danielle Groves, Katie Johnson, Laura Carrilero, Alex Keeley, Dave Partridge, Matthew Wyles, Benjamin Lindsey, Mehmet Yavuz, Mohammad Raza, Cariad Evans |
| EPI_ISL_627038 | Regional Virus Laboratory, Belfast Health and Social Care Trust | COVID-19 Genomics UK (COG-UK) Consortium | Conall McCaughey, James McKenna, Tanya Curran, Susan Feeney, Alison Watt, Ciara Cox, Mairead Connor, Zoltan Molnar, David Simpson, Derek Fairley |
| EPI_ISL_627067 | University of Exeter | COVID-19 Genomics UK (COG-UK) Consortium | Ben Temperton,Aaron Jeffries,Michelle Michelsen,Joanna Warwick-Dugdale,Audrey Farbos,Robyn Manley,Stephen Michell,Jane Masoli |
| EPI_ISL_627079, EPI_ISL_627085 | Oxford Viromics, NDM, University of Oxford; Oxford University Hospitals; Basingstoke and North Hampshire Hospital | COVID-19 Genomics UK (COG-UK) Consortium | Tanya Golubchik, David Bonsall, George Macintyre, Amy Trebes, Mariateresa de Cesare, Catrin Moore, Alex Mobbs, Anita Justice, Robert Shaw, Monique Andersson, Timothy Peto, Emma Wise, Nathan Moore, Jessica Lynch, Nick Cortes, Matilde Mori, Stephen Kidd, David Buck, John Todd, Christophe Fraser |
| EPI_ISL_627097 | Wales Specialist Virology Centre Sequencing lab: Pathogen Genomics Unit | COVID-19 Genomics UK (COG-UK) Consortium | Catherine Moore, Johnathan Evans, Laura Gifford, Malorie Perry, Simon Cottrell, Angela Marchbank, Alec Birchley, Alexander Adams, Amy Gaskin, Bree Gatica-Wilcox, Jason Coombes, Joel Southgate, Lauren Gilbert, Lee Graham, Nicole Pacchiarini, Sara Kumziene-Summerhayes, Sarah Taylor, Sophie Jones, Sara Rey, Matthew Bull, Joanne Watkins, Sally Corden, Tom Connor |
| EPI_ISL_627102 | University of Exeter | COVID-19 Genomics UK (COG-UK) Consortium | Ben Temperton,Aaron Jeffries,Michelle Michelsen,Joanna Warwick-Dugdale,Audrey Farbos,Robyn Manley,Stephen Michell,Jane Masoli |
| EPI_ISL_627104 | University College London, Great Ormond Street Hospital for Children NHS Foundation Trust, Imperial College Healthcare NHS Trust | COVID-19 Genomics UK (COG-UK) Consortium | Sergi Castellano, Rachel Williams, Mark Kristiansen, Paola Resende Silva, Sunando Roy, Tony Brooks, Helena Tutill, Paola Niola, Patricia Dyal, Charlotte Williams, Leysa Forrest, Yasmin Panchbhaya, Jacqueline Findlay, Samuel Weeks, Julianne Brown, Kathryn Harris, Paul Randell, James Price, Alison Holmes, Judith Breuer |
| EPI_ISL_627112 | University of Exeter | COVID-19 Genomics UK (COG-UK) Consortium | Ben Temperton,Aaron Jeffries,Michelle Michelsen,Joanna Warwick-Dugdale,Audrey Farbos,Robyn Manley,Stephen Michell,Jane Masoli |
| EPI_ISL_627117 | West of Scotland Specialist Virology Centre, NHSGGC / MRC-University of Glasgow Centre for Virus Research | COVID-19 Genomics UK (COG-UK) Consortium | Ana da Silva Filipe, Natasha Johnson, Kathy Smollett, Daniel Mair, Stephen Carmichael, Lily Tong, Jenna Nichols, Elihu Aranday-Cortes, Kyriaki Nomikou; Sarah McDonald, Marc Niebel, Patawee Asamaphan; Richard Orton, Joseph Hughes, Sreenu Vattipally, David L Robertson; Alasdair MacLean, Rory Gunson; Kathy Li, Igor Starinskij, Natasha Jesudason, Rajiv Shah, James Shepherd, Antonia Ho, Emma Thomson |
| EPI_ISL_627131, EPI_ISL_627137, EPI_ISL_627152 | Liverpool Clinical Laboratories | COVID-19 Genomics UK (COG-UK) Consortium | Sam Haldenby, Anita Lucaci, Steve Paterson, Julian Hiscox, Alistair Darby, M Almsaud, A Alrezaihi, Muhannad Alruwaili, Stuart D Armstrong, Jones Benjamin, Eleanor G Bentley, Anu Chawla, Jordan J Clark, Angela Cowell, Richard Eccles, Isabel García-Dorival, Matthew Gemmell, Alessandro Gerada, PKF Gilmore, Richard Gregory, Ximeng Han, Catherine Hartley, Margaret Hughes, Miren Iturriza-Gomara, James Johnson, L Luu, Jenifer Manson, Charlotte Nelson, Elaine O'Toole, Cassie Olateju, Rebekah Penrice-Randal , Lucille Rainbow, N.P Randle, Trevor Ian Robinson, Parul Sharma, Ghada T Shawli, James P Stewart, Neil Swainston, Ecaterina Vamos, Joanne Watts, Mark Whitehead |
| EPI_ISL_627158 | Regional Virus Laboratory, Belfast Health and Social Care Trust | COVID-19 Genomics UK (COG-UK) Consortium | Conall McCaughey, James McKenna, Tanya Curran, Susan Feeney, Alison Watt, Ciara Cox, Mairead Connor, Zoltan Molnar, David Simpson, Derek Fairley |
| EPI_ISL_627164 | Virology Department, Sheffield Teaching Hospitals NHS Foundation Trust/Department of Infection, Immunity and Cardiovascular Disease, The Medical School, University of Sheffield | COVID-19 Genomics UK (COG-UK) Consortium | Thushan de Silva, Matthew Parker, Nikki Smith, Adri Angyal, Rebecca Brown, Luke Green, Rachel Tucker, Paul Parsons, Danielle Groves, Katie Johnson, Laura Carrilero, Alex Keeley, Dave Partridge, Matthew Wyles, Benjamin Lindsey, Mehmet Yavuz, Mohammad Raza, Cariad Evans |
| EPI_ISL_627174, EPI_ISL_627177, EPI_ISL_627178, EPI_ISL_627179 | Liverpool Clinical Laboratories | COVID-19 Genomics UK (COG-UK) Consortium | Sam Haldenby, Anita Lucaci, Steve Paterson, Julian Hiscox, Alistair Darby, M Almsaud, A Alrezaihi, Muhannad Alruwaili, Stuart D Armstrong, Jones Benjamin, Eleanor G Bentley, Anu Chawla, Jordan J Clark, Angela Cowell, Richard Eccles, Isabel García-Dorival, Matthew Gemmell, Alessandro Gerada, PKF Gilmore, Richard Gregory, Ximeng Han, Catherine Hartley, Margaret Hughes, Miren Iturriza-Gomara, James Johnson, L Luu, Jenifer Manson, Charlotte Nelson, Elaine O'Toole, Cassie Olateju, Rebekah Penrice-Randal , Lucille Rainbow, N.P Randle, Trevor Ian Robinson, Parul Sharma, Ghada T Shawli, James P Stewart, Neil Swainston, Ecaterina Vamos, Joanne Watts, Mark Whitehead |
| EPI_ISL_627184 | Virology Department, Sheffield Teaching Hospitals NHS Foundation Trust/Department of Infection, Immunity and Cardiovascular Disease, The Medical School, University of Sheffield | COVID-19 Genomics UK (COG-UK) Consortium | Thushan de Silva, Matthew Parker, Nikki Smith, Adri Angyal, Rebecca Brown, Luke Green, Rachel Tucker, Paul Parsons, Danielle Groves, Katie Johnson, Laura Carrilero, Alex Keeley, Dave Partridge, Matthew Wyles, Benjamin Lindsey, Mehmet Yavuz, Mohammad Raza, Cariad Evans |
| EPI_ISL_627188 | Liverpool Clinical Laboratories | COVID-19 Genomics UK (COG-UK) Consortium | Sam Haldenby, Anita Lucaci, Steve Paterson, Julian Hiscox, Alistair Darby, M Almsaud, A Alrezaihi, Muhannad Alruwaili, Stuart D Armstrong, Jones Benjamin, Eleanor G Bentley, Anu Chawla, Jordan J Clark, Angela Cowell, Richard Eccles, Isabel García-Dorival, Matthew Gemmell, Alessandro Gerada, PKF Gilmore, Richard Gregory, Ximeng Han, Catherine Hartley, Margaret Hughes, Miren Iturriza-Gomara, James Johnson, L Luu, Jenifer Manson, Charlotte Nelson, Elaine O'Toole, Cassie Olateju, Rebekah Penrice-Randal , Lucille Rainbow, N.P Randle, Trevor Ian Robinson, Parul Sharma, Ghada T Shawli, James P Stewart, Neil Swainston, Ecaterina Vamos, Joanne Watts, Mark Whitehead |
| EPI_ISL_627219 | Wales Specialist Virology Centre Sequencing lab: Pathogen Genomics Unit | COVID-19 Genomics UK (COG-UK) Consortium | Catherine Moore, Johnathan Evans, Laura Gifford, Malorie Perry, Simon Cottrell, Angela Marchbank, Alec Birchley, Alexander Adams, Amy Gaskin, Bree Gatica-Wilcox, Jason Coombes, Joel Southgate, Lauren Gilbert, Lee Graham, Nicole Pacchiarini, Sara Kumziene-Summerhayes, Sarah Taylor, Sophie Jones, Sara Rey, Matthew Bull, Joanne Watkins, Sally Corden, Tom Connor |
| EPI_ISL_627251 | Oxford Viromics, NDM, University of Oxford; Oxford University Hospitals; Basingstoke and North Hampshire Hospital | COVID-19 Genomics UK (COG-UK) Consortium | Tanya Golubchik, David Bonsall, George Macintyre, Amy Trebes, Mariateresa de Cesare, Catrin Moore, Alex Mobbs, Anita Justice, Robert Shaw, Monique Andersson, Timothy Peto, Emma Wise, Nathan Moore, Jessica Lynch, Nick Cortes, Matilde Mori, Stephen Kidd, David Buck, John Todd, Christophe Fraser |
| EPI_ISL_627252, EPI_ISL_627253 | Virology Department, Sheffield Teaching Hospitals NHS Foundation Trust/Department of Infection, Immunity and Cardiovascular Disease, The Medical School, University of Sheffield | COVID-19 Genomics UK (COG-UK) Consortium | Thushan de Silva, Matthew Parker, Nikki Smith, Adri Angyal, Rebecca Brown, Luke Green, Rachel Tucker, Paul Parsons, Danielle Groves, Katie Johnson, Laura Carrilero, Alex Keeley, Dave Partridge, Matthew Wyles, Benjamin Lindsey, Mehmet Yavuz, Mohammad Raza, Cariad Evans |
| EPI_ISL_627256, EPI_ISL_627258 | Oxford Viromics, NDM, University of Oxford; Oxford University Hospitals; Basingstoke and North Hampshire Hospital | COVID-19 Genomics UK (COG-UK) Consortium | Tanya Golubchik, David Bonsall, George Macintyre, Amy Trebes, Mariateresa de Cesare, Catrin Moore, Alex Mobbs, Anita Justice, Robert Shaw, Monique Andersson, Timothy Peto, Emma Wise, Nathan Moore, Jessica Lynch, Nick Cortes, Matilde Mori, Stephen Kidd, David Buck, John Todd, Christophe Fraser |
| EPI_ISL_627280 | Virology Department, Sheffield Teaching Hospitals NHS Foundation Trust/Department of Infection, Immunity and Cardiovascular Disease, The Medical School, University of Sheffield | COVID-19 Genomics UK (COG-UK) Consortium | Thushan de Silva, Matthew Parker, Nikki Smith, Adri Angyal, Rebecca Brown, Luke Green, Rachel Tucker, Paul Parsons, Danielle Groves, Katie Johnson, Laura Carrilero, Alex Keeley, Dave Partridge, Matthew Wyles, Benjamin Lindsey, Mehmet Yavuz, Mohammad Raza, Cariad Evans |

| | | | |
|---|---|---|---|
| EPI_ISL_627281, EPI_ISL_627284, EPI_ISL_627285, EPI_ISL_627294, EPI_ISL_627305 | Oxford Viromics, NDM, University of Oxford; Oxford University Hospitals; Basingstoke and North Hampshire Hospital | COVID-19 Genomics UK (COG-UK) Consortium | Tanya Golubchik, David Bonsall, George Macintyre, Amy Trebes, Mariateresa de Cesare, Catrin Moore, Alex Mobbs, Anita Justice, Robert Shaw, Monique Andersson, Timothy Peto, Emma Wise, Nathan Moore, Jessica Lynch, Nick Cortes, Matilde Mori, Stephen Kidd, David Buck, John Todd, Christophe Fraser |
| EPI_ISL_627307, EPI_ISL_627308 | Wales Specialist Virology Centre Sequencing lab: Pathogen Genomics Unit | COVID-19 Genomics UK (COG-UK) Consortium | Catherine Moore, Johnathan Evans, Laura Gifford, Malorie Perry, Simon Cottrell, Angela Marchbank, Alec Birchley, Alexander Adams, Amy Gaskin, Bree Gatica-Wilcox, Jason Coombes, Joel Southgate, Lauren Gilbert, Lee Graham, Nicole Pacchiarini, Sara Kumziene-Summerhayes, Sarah Taylor, Sophie Jones, Sara Rey, Matthew Bull, Joanne Watkins, Sally Corden, Tom Connor |
| EPI_ISL_627435 | University of Exeter | COVID-19 Genomics UK (COG-UK) Consortium | Ben Temperton,Aaron Jeffries,Michelle Michelsen,Joanna Warwick-Dugdale,Audrey Farbos,Robyn Manley,Stephen Michell,Jane Masoli |
| EPI_ISL_627472, EPI_ISL_627473, EPI_ISL_627474, EPI_ISL_627475, EPI_ISL_627476, EPI_ISL_627477, EPI_ISL_627478, EPI_ISL_627479, EPI_ISL_627480, EPI_ISL_627481 | University College London, Great Ormond Street Hospital for Children NHS Foundation Trust, Imperial College Healthcare NHS Trust | COVID-19 Genomics UK (COG-UK) Consortium | Sergi Castellano, Rachel Williams, Mark Kristiansen, Paola Resende Silva, Sunando Roy, Tony Brooks, Helena Tutill, Paola Niola, Patricia Dyal, Charlotte Williams, Leysa Forrest, Yasmin Panchbhaya, Jacqueline Findlay, Samuel Weeks, Julianne Brown, Kathryn Harris, Paul Randell, James Price, Alison Holmes, Judith Breuer |
| EPI_ISL_627488, EPI_ISL_627489, EPI_ISL_627490 | Wales Specialist Virology Centre Sequencing lab: Pathogen Genomics Unit | COVID-19 Genomics UK (COG-UK) Consortium | Catherine Moore, Johnathan Evans, Laura Gifford, Malorie Perry, Simon Cottrell, Angela Marchbank, Alec Birchley, Alexander Adams, Amy Gaskin, Bree Gatica-Wilcox, Jason Coombes, Joel Southgate, Lauren Gilbert, Lee Graham, Nicole Pacchiarini, Sara Kumziene-Summerhayes, Sarah Taylor, Sophie Jones, Sara Rey, Matthew Bull, Joanne Watkins, Sally Corden, Tom Connor |
| EPI_ISL_627542, EPI_ISL_627543, EPI_ISL_627545, EPI_ISL_627547, EPI_ISL_627548, EPI_ISL_627549, EPI_ISL_627550, EPI_ISL_627551, EPI_ISL_627552, EPI_ISL_627553 | Regional Virus Laboratory, Belfast Health and Social Care Trust | COVID-19 Genomics UK (COG-UK) Consortium | Conall McCaughey, James McKenna, Tanya Curran, Susan Feeney, Alison Watt, Ciara Cox, Mairead Connor, Zoltan Molnar, David Simpson, Derek Fairley |
| EPI_ISL_627743, EPI_ISL_627759, EPI_ISL_627780, EPI_ISL_627783, EPI_ISL_627799, EPI_ISL_627806, EPI_ISL_627832, EPI_ISL_627872, EPI_ISL_627875, EPI_ISL_627880 | Wales Specialist Virology Centre Sequencing lab: Pathogen Genomics Unit | COVID-19 Genomics UK (COG-UK) Consortium | Catherine Moore, Johnathan Evans, Laura Gifford, Malorie Perry, Simon Cottrell, Angela Marchbank, Alec Birchley, Alexander Adams, Amy Gaskin, Bree Gatica-Wilcox, Jason Coombes, Joel Southgate, Lauren Gilbert, Lee Graham, Nicole Pacchiarini, Sara Kumziene-Summerhayes, Sarah Taylor, Sophie Jones, Sara Rey, Matthew Bull, Joanne Watkins, Sally Corden, Tom Connor |
| EPI_ISL_628349, EPI_ISL_628350, EPI_ISL_628351, EPI_ISL_628352, EPI_ISL_628353, EPI_ISL_628354, EPI_ISL_628355, EPI_ISL_628356, EPI_ISL_628357, EPI_ISL_628359, EPI_ISL_628360, EPI_ISL_628361, EPI_ISL_628362, EPI_ISL_628363, EPI_ISL_628364, EPI_ISL_628365, EPI_ISL_628366, EPI_ISL_628367, EPI_ISL_628368 | | | |
| see above | Oxford Viromics, NDM, University of Oxford; Oxford University Hospitals; Basingstoke and North Hampshire Hospital | COVID-19 Genomics UK (COG-UK) Consortium | Tanya Golubchik, David Bonsall, George Macintyre, Amy Trebes, Mariateresa de Cesare, Catrin Moore, Alex Mobbs, Anita Justice, Robert Shaw, Monique Andersson, Timothy Peto, Emma Wise, Nathan Moore, Jessica Lynch, Nick Cortes, Matilde Mori, Stephen Kidd, David Buck, John Todd, Christophe Fraser |
| EPI_ISL_628369, EPI_ISL_628370, EPI_ISL_628371, EPI_ISL_628372, EPI_ISL_628373, EPI_ISL_628375, EPI_ISL_628376, EPI_ISL_628377, EPI_ISL_628378, EPI_ISL_628379, EPI_ISL_628380, EPI_ISL_628381, EPI_ISL_628382, EPI_ISL_628383, EPI_ISL_628384, EPI_ISL_628385, EPI_ISL_628386, EPI_ISL_628387, EPI_ISL_628388, EPI_ISL_628389, EPI_ISL_628390, EPI_ISL_628391, EPI_ISL_628392, EPI_ISL_628393, EPI_ISL_628394, EPI_ISL_628395, EPI_ISL_628396, EPI_ISL_628401, EPI_ISL_628402, EPI_ISL_628403, EPI_ISL_628404, EPI_ISL_628405, EPI_ISL_628406, EPI_ISL_628407, EPI_ISL_628408, EPI_ISL_628409, EPI_ISL_628410, EPI_ISL_628411, EPI_ISL_628412, EPI_ISL_628413 | | | |
| see above | Virology Department, Sheffield Teaching Hospitals NHS Foundation Trust/Department of Infection, Immunity and Cardiovascular Disease, The Medical School, University of Sheffield | COVID-19 Genomics UK (COG-UK) Consortium | Thushan de Silva, Matthew Parker, Nikki Smith, Adri Angyal, Rebecca Brown, Luke Green, Rachel Tucker, Paul Parsons, Danielle Groves, Katie Johnson, Laura Carrilero, Alex Keeley, Dave Partridge, Matthew Wyles, Benjamin Lindsey, Mehmet Yavuz, Mohammad Raza, Cariad Evans |
| EPI_ISL_628414, EPI_ISL_628415, EPI_ISL_628416, EPI_ISL_628417, EPI_ISL_628418, EPI_ISL_628419, EPI_ISL_628420 | Oxford Viromics, NDM, University of Oxford; Oxford University Hospitals; Basingstoke and North Hampshire Hospital | COVID-19 Genomics UK (COG-UK) Consortium | Tanya Golubchik, David Bonsall, George Macintyre, Amy Trebes, Mariateresa de Cesare, Catrin Moore, Alex Mobbs, Anita Justice, Robert Shaw, Monique Andersson, Timothy Peto, Emma Wise, Nathan Moore, Jessica Lynch, Nick Cortes, Matilde Mori, Stephen Kidd, David Buck, John Todd, Christophe Fraser |
| EPI_ISL_628421 | Virology Department, Sheffield Teaching Hospitals NHS Foundation Trust/Department of Infection, Immunity and Cardiovascular Disease, The Medical School, University of Sheffield | COVID-19 Genomics UK (COG-UK) Consortium | Thushan de Silva, Matthew Parker, Nikki Smith, Adri Angyal, Rebecca Brown, Luke Green, Rachel Tucker, Paul Parsons, Danielle Groves, Katie Johnson, Laura Carrilero, Alex Keeley, Dave Partridge, Matthew Wyles, Benjamin Lindsey, Mehmet Yavuz, Mohammad Raza, Cariad Evans |
| EPI_ISL_628422, EPI_ISL_628423, EPI_ISL_628424, EPI_ISL_628425, EPI_ISL_628426, EPI_ISL_628427, EPI_ISL_628428, EPI_ISL_628429, EPI_ISL_628430, EPI_ISL_628431, EPI_ISL_628432, EPI_ISL_628433, EPI_ISL_628434, EPI_ISL_628435, EPI_ISL_628436, EPI_ISL_628437, EPI_ISL_628438, EPI_ISL_628439, EPI_ISL_628440, EPI_ISL_628453, EPI_ISL_628454, EPI_ISL_628455, EPI_ISL_628456, EPI_ISL_628457, EPI_ISL_628458, EPI_ISL_628459, EPI_ISL_628460, EPI_ISL_628461, EPI_ISL_628462, EPI_ISL_628463, EPI_ISL_628464, EPI_ISL_628485, EPI_ISL_628509, EPI_ISL_628510, EPI_ISL_628518, EPI_ISL_628519, EPI_ISL_628520, EPI_ISL_628521, EPI_ISL_628522, EPI_ISL_628523, EPI_ISL_628524, EPI_ISL_628525, EPI_ISL_628526, EPI_ISL_628527, EPI_ISL_628528, EPI_ISL_628529, EPI_ISL_628530, EPI_ISL_628531, EPI_ISL_628532, EPI_ISL_628533, EPI_ISL_628534, EPI_ISL_628535, EPI_ISL_628536, EPI_ISL_628537, EPI_ISL_628538, EPI_ISL_628539, EPI_ISL_628541, EPI_ISL_628542, EPI_ISL_628544, EPI_ISL_628545, EPI_ISL_628546, EPI_ISL_628547, EPI_ISL_628548, EPI_ISL_628559, EPI_ISL_628560, EPI_ISL_628561, EPI_ISL_628567, EPI_ISL_628568, EPI_ISL_628571, EPI_ISL_628572, EPI_ISL_628573, EPI_ISL_628574, EPI_ISL_628575, EPI_ISL_628576, EPI_ISL_628577, EPI_ISL_628578, EPI_ISL_628579, EPI_ISL_628580, EPI_ISL_628581, EPI_ISL_628582, EPI_ISL_628583, EPI_ISL_628584, EPI_ISL_628585, EPI_ISL_628586, EPI_ISL_628587, EPI_ISL_628588, EPI_ISL_628593, EPI_ISL_628594, EPI_ISL_628595, EPI_ISL_628596, EPI_ISL_628597, EPI_ISL_628598, EPI_ISL_628599, EPI_ISL_628600, EPI_ISL_628601, EPI_ISL_628602, EPI_ISL_628603, EPI_ISL_628604, EPI_ISL_628605, EPI_ISL_628606, EPI_ISL_628607, EPI_ISL_628608, EPI_ISL_628609, EPI_ISL_628610, EPI_ISL_628611, EPI_ISL_628614, EPI_ISL_628615, EPI_ISL_628616, EPI_ISL_628617, EPI_ISL_628618 | | | |
| see above | Oxford Viromics, NDM, University of Oxford; Oxford University Hospitals; Basingstoke and North Hampshire Hospital | COVID-19 Genomics UK (COG-UK) Consortium | Tanya Golubchik, David Bonsall, George Macintyre, Amy Trebes, Mariateresa de Cesare, Catrin Moore, Alex Mobbs, Anita Justice, Robert Shaw, Monique Andersson, Timothy Peto, Emma Wise, Nathan Moore, Jessica Lynch, Nick Cortes, Matilde Mori, Stephen Kidd, David Buck, John Todd, Christophe Fraser |
| EPI_ISL_628936, EPI_ISL_628937, EPI_ISL_628938 | Utah Public Health Laboratory | Utah Public Health Laboratory | Erin Young, Kelly Oakeson |
| EPI_ISL_629003 | South Eastern Area Laboratory Services (SEALS) | NSW Health Pathology - Institute of Clinical Pathology and Medical Research; Westmead Hospital; University of Sydney | CIDM-PH et al. |
| EPI_ISL_629036, EPI_ISL_629037, EPI_ISL_629038, EPI_ISL_629039, EPI_ISL_629040 | Maryland Public Health Laboratory | Maryland Public Health Laboratory | Maryland Department of Health Laboratories Administration |
| EPI_ISL_630971, EPI_ISL_630976, EPI_ISL_630994, EPI_ISL_631023, EPI_ISL_631057, EPI_ISL_631089, EPI_ISL_631111, EPI_ISL_631116, EPI_ISL_631127, EPI_ISL_631128, EPI_ISL_631163, EPI_ISL_631173, EPI_ISL_631175, EPI_ISL_631202, EPI_ISL_631212, EPI_ISL_631222, EPI_ISL_631228, EPI_ISL_631241 | | | |
| see above | Lighthouse Lab in Alderley Park | Wellcome Sanger Institute for the COVID-19 Genomics UK (COG-UK) consortium | Jacquelyn Wynn, Mairead Hyland, The Lighthouse Lab in Alderley Park and Alex Alderton, Roberto Amato, Sonia Goncalves, Ewan Harrison, David K. Jackson, Ian Johnston, Dominic Kwiatkowski, Cordelia Langford, John Sillitoe on behalf of the Wellcome Sanger Institute COVID-19 Surveillance Team |
| EPI_ISL_631304, EPI_ISL_631305, EPI_ISL_631306, EPI_ISL_631307, EPI_ISL_631308, EPI_ISL_631309, EPI_ISL_631310, EPI_ISL_631311, EPI_ISL_631312, EPI_ISL_631313, EPI_ISL_631314, EPI_ISL_631315, EPI_ISL_631316, EPI_ISL_631317, EPI_ISL_631318, EPI_ISL_631319, EPI_ISL_631320 | | | |
| see above | ZOTZ KLIMAS MVZ Düsseldorf-Centrum GbR ÜBAG für Labormedizin, Genetik, Zytologie, Pathologie | Center of Medical Microbiology, Virology, and Hospital Hygiene, University of Duesseldorf | Maximilian Damagnez, Alexander Dilthey, Ashley-Jane Duplessis, Patrick Finzer, Katrin Hoffmann, Torsten Houwaart, Lisanna Hülse, Malte Kohns Vasconcelos, Marek Korencak, Nadine Lübke, Jessica Nicolai, Klaus Pfeffer, Daniel Strelow, Jörg Timm, Andreas Walker, Tobias Wienemann, Rainer Zotz |
| EPI_ISL_631477, EPI_ISL_631478, EPI_ISL_631479 | Wisconsin State Laboratory of Hygiene Communicable Disease Division | Wisconsin State Laboratory of Hygiene Communicable Disease Division | Kelsey R. Florek, Abigail C. Shockey |
| EPI_ISL_632264 | Communicable Disease Laboratory, Public Health Directorate | Communicable Disease Laboratory, Public Health Directorate | AlWasti,H., AlTaif,Z., AlHujairi,Z., AlAbbas,Z. |
| EPI_ISL_632329, EPI_ISL_632333 | Dutch COVID-19 response team | Erasmus Medical Center | Bas Oude Munnink, David Nieuwenhuijse, Reina Sikkema, Claudia Schapendonk, Irina Chestakova, Anne van der Linden, Theo Bestebroer, Stefan van Nieuwkoop, Mark Pronk, Pascal Lexmond, Corien Swaan, Manon Haverkate, Madelief Mollers, Mart Stein, Sandra Kengne Kamga Mobou, Jeroen van Kampen, Jolanda Voermans, Aura Timen, Corine GeurtsvanKessel, Annemiek van der Eijk, Richard Molenkamp, Marion Koopmans, on behalf of the Dutch |

| | | | |
|---|---|---|---|
| | | | national COVID-19 response team. |
| EPI_ISL_632334 | Dutch COVID-19 response team | Erasmus Medical Center | OH consortium |
| EPI_ISL_632336, EPI_ISL_632337, EPI_ISL_632344, EPI_ISL_632368, EPI_ISL_632369, EPI_ISL_632371, EPI_ISL_632373, EPI_ISL_632375, EPI_ISL_632397, EPI_ISL_632414, EPI_ISL_632423 | | | |
| see above | Dutch COVID-19 response team | Erasmus Medical Center | Bas Oude Munnink, David Nieuwenhuijse, Reina Sikkema, Claudia Schapendonk, Irina Chestakova, Anne van der Linden, Theo Bestebroer, Stefan van Nieuwkoop, Mark Pronk, Pascal Lexmond, Corien Swaan, Manon Haverkate, Madelief Mollers, Mart Stein, Sandra Kengne Kamga Mobou, Jeroen van Kampen, Jolanda Voermans, Aura Timen, Corine GeurtsvanKessel, Annemiek van der Eijk, Richard Molenkamp, Marion Koopmans, on behalf of the Dutch national COVID-19 response team. |
| EPI_ISL_632434, EPI_ISL_632435, EPI_ISL_632436, EPI_ISL_632437, EPI_ISL_632438 | Dutch COVID-19 response team | Erasmus Medical Center | OH consortium |
| EPI_ISL_632446, EPI_ISL_632480, EPI_ISL_632481 | Dutch COVID-19 response team | Erasmus Medical Center | Bas Oude Munnink, David Nieuwenhuijse, Reina Sikkema, Claudia Schapendonk, Irina Chestakova, Anne van der Linden, Theo Bestebroer, Stefan van Nieuwkoop, Mark Pronk, Pascal Lexmond, Corien Swaan, Manon Haverkate, Madelief Mollers, Mart Stein, Sandra Kengne Kamga Mobou, Jeroen van Kampen, Jolanda Voermans, Aura Timen, Corine GeurtsvanKessel, Annemiek van der Eijk, Richard Molenkamp, Marion Koopmans, on behalf of the Dutch national COVID-19 response team. |
| EPI_ISL_632483, EPI_ISL_632484, EPI_ISL_632485 | Dutch COVID-19 response team | Erasmus Medical Center | OH consortium |
| EPI_ISL_632507, EPI_ISL_632508, EPI_ISL_632512, EPI_ISL_632514, EPI_ISL_632515, EPI_ISL_632516, EPI_ISL_632517, EPI_ISL_632518, EPI_ISL_632519, EPI_ISL_632536, EPI_ISL_632537, EPI_ISL_632538, EPI_ISL_632539, EPI_ISL_632540, EPI_ISL_632541, EPI_ISL_632542, EPI_ISL_632559, EPI_ISL_632560, EPI_ISL_632561, EPI_ISL_632562, EPI_ISL_632563, EPI_ISL_632567, EPI_ISL_632584, EPI_ISL_632585, EPI_ISL_632586, EPI_ISL_632594, EPI_ISL_632595, EPI_ISL_632600, EPI_ISL_632601, EPI_ISL_632602, EPI_ISL_632607, EPI_ISL_632609, EPI_ISL_632641, EPI_ISL_632642, EPI_ISL_632643, EPI_ISL_632644, EPI_ISL_632645, EPI_ISL_632646, EPI_ISL_632647, EPI_ISL_632648, EPI_ISL_632649, EPI_ISL_632650, EPI_ISL_632651, EPI_ISL_632652, EPI_ISL_632653, EPI_ISL_632654, EPI_ISL_632658, EPI_ISL_632659, EPI_ISL_632666, EPI_ISL_632669, EPI_ISL_632672, EPI_ISL_632696, EPI_ISL_632699, EPI_ISL_632700, EPI_ISL_632719, EPI_ISL_632734, EPI_ISL_632755, EPI_ISL_632786 | | | |
| see above | Dutch COVID-19 response team | Erasmus Medical Center | Bas Oude Munnink, David Nieuwenhuijse, Reina Sikkema, Claudia Schapendonk, Irina Chestakova, Anne van der Linden, Theo Bestebroer, Stefan van Nieuwkoop, Mark Pronk, Pascal Lexmond, Corien Swaan, Manon Haverkate, Madelief Mollers, Mart Stein, Sandra Kengne Kamga Mobou, Jeroen van Kampen, Jolanda Voermans, Aura Timen, Corine GeurtsvanKessel, Annemiek van der Eijk, Richard Molenkamp, Marion Koopmans, on behalf of the Dutch national COVID-19 response team. |
| EPI_ISL_632904 | Communicable Disease Laboratory, Public Health Directorate | Communicable Disease Laboratory, Public Health Directorate | AlTaif,Z., AlHujairi,Z., AlWasti,H., AlAbbas,Z. |
| EPI_ISL_632992, EPI_ISL_632993 | DOHMH Corona | New York City Public Health Laboratory | Jade Wang, et al. |
| EPI_ISL_632996, EPI_ISL_632997, EPI_ISL_632998 | DOHMH Crown Heights | New York City Public Health Laboratory | Jade Wang, et al. |
| EPI_ISL_633025 | DOHMH Chelsea | New York City Public Health Laboratory | Jade Wang, et al. |
| EPI_ISL_633026 | DOHMH PHL | New York City Public Health Laboratory | Jade Wang, et al. |
| EPI_ISL_633027, EPI_ISL_633028, EPI_ISL_633029, EPI_ISL_633030 | DOHMH Morrisania | New York City Public Health Laboratory | Jade Wang, et al. |
| EPI_ISL_633031 | DOHMH Chelsea | New York City Public Health Laboratory | Jade Wang, et al. |
| EPI_ISL_634885 | Lab voor klinische biologie | Onderzoeksgroep Virologie | Laurens Lambrechts, Nick Vereecke, Marthe Pauwels, Bruno Verhasselt, Linos Vandekerckhove, Hans Nauwynck, Sebastiaan Theuns |
| EPI_ISL_634886, EPI_ISL_634887, EPI_ISL_634888, EPI_ISL_634889 | Lab voor klinische biologie | Onderzoeksgroep Virologie | Nick Vereecke, Laurens Lambrechts, Marthe Pauwels, Bruno Verhasselt, Linos Vandekerckhove, Hans Nauwynck, Sebastiaan Theuns |
| EPI_ISL_634925, EPI_ISL_634926, EPI_ISL_634927, EPI_ISL_634928, EPI_ISL_634929, EPI_ISL_634930, EPI_ISL_634931, EPI_ISL_634932, EPI_ISL_634933 | Utah Public Health Laboratory | Utah Public Health Laboratory | Erin L. Young, Kelly F. Oakeson |
| EPI_ISL_635063 | Foerde Hospital, Department of Microbiology | Norwegian Institute of Public Health, Department of Virology | Kathrine Stene-Johansen, Kamilla Heddeland Instefjord, Hilde Elshaug, Marie Paulsen Madsen, Rasmus Riis Kopperud, Hilde Vollan, Karoline Bragstad, Olav Hungnes |
| EPI_ISL_635089, EPI_ISL_635091 | Haukeland University Hospital, Department of Medical Microbiology | Norwegian Institute of Public Health, Department of Virology | Kathrine Stene-Johansen, Kamilla Heddeland Instefjord, Hilde Elshaug, Marie Paulsen Madsen, Rasmus Riis Kopperud, Hilde Vollan, Karoline Bragstad, Olav Hungnes |
| EPI_ISL_635095 | Oslo University Hospital, Department of Medical Microbiology | Norwegian Institute of Public Health, Department of Virology | Kathrine Stene-Johansen, Kamilla Heddeland Instefjord, Hilde Elshaug, Marie Paulsen Madsen, Rasmus Riis Kopperud, Hilde Vollan, Karoline Bragstad, Olav Hungnes |
| EPI_ISL_635096 | Department of Medical Microbiology, St. Olavs hospital | Norwegian Institute of Public Health, Department of Virology | Kathrine Stene-Johansen, Kamilla Heddeland Instefjord, Hilde Elshaug, Marie Paulsen Madsen, Rasmus Riis Kopperud, Hilde Vollan, Karoline Bragstad, Olav Hungnes |
| EPI_ISL_635097 | Hospital of Southern Norway - Kristiansand, Department of Medical Microbiology | Norwegian Institute of Public Health, Department of Virology | Kathrine Stene-Johansen, Kamilla Heddeland Instefjord, Hilde Elshaug, Marie Paulsen Madsen, Rasmus Riis Kopperud, Hilde Vollan, Karoline Bragstad, Olav Hungnes |
| EPI_ISL_635124, EPI_ISL_635125 | Innlandet Hospital Trust, Division Lillehammer, Department for Medical Microbiology | Norwegian Institute of Public Health, Department of Virology | Kathrine Stene-Johansen, Kamilla Heddeland Instefjord, Hilde Elshaug, Marie Paulsen Madsen, Rasmus Riis Kopperud, Hilde Vollan, Karoline Bragstad, Olav Hungnes |
| EPI_ISL_635128 | Haukeland University Hospital, Dept. of Microbiology | Norwegian Institute of Public Health, Department of Virology | Kathrine Stene-Johansen, Kamilla Heddeland Instefjord, Hilde Elshaug, Marie Paulsen Madsen, Rasmus Riis Kopperud, Hilde Vollan, Karoline Bragstad, Olav Hungnes |
| EPI_ISL_635129 | Oslo University Hospital, Department of Medical Microbiology | Norwegian Institute of Public Health, Department of Virology | Kathrine Stene-Johansen, Kamilla Heddeland Instefjord, Hilde Elshaug, Marie Paulsen Madsen, Rasmus Riis Kopperud, Hilde Vollan, Karoline Bragstad, Olav Hungnes |
| EPI_ISL_635134 | Medical Microbiology Unit, Department for Laboratory Medicine, Drammen Hospital, Vestre Viken Health Trust, | Norwegian Institute of Public Health, Department of Virology | Kathrine Stene-Johansen, Kamilla Heddeland Instefjord, Hilde Elshaug, Marie Paulsen Madsen, Rasmus Riis Kopperud, Hilde Vollan, Karoline Bragstad, Olav Hungnes |
| EPI_ISL_635138 | Hospital of Southern Norway - Kristiansand, Department of Medical Microbiology | Norwegian Institute of Public Health, Department of Virology | Kathrine Stene-Johansen, Kamilla Heddeland Instefjord, Hilde Elshaug, Marie Paulsen Madsen, Rasmus Riis Kopperud, Hilde Vollan, Karoline Bragstad, Olav Hungnes |
| EPI_ISL_635144 | Dept. of Medical Microbiology, Stavanger University Hospital, Helse Stavanger HF | Norwegian Institute of Public Health, Department of Virology | Kathrine Stene-Johansen, Kamilla Heddeland Instefjord, Hilde Elshaug, Marie Paulsen Madsen, Rasmus Riis Kopperud, Hilde Vollan, Karoline Bragstad, Olav Hungnes |
| EPI_ISL_635154 | Unilabs Laboratory Medicine | Norwegian Institute of Public Health, Department of Virology | Kathrine Stene-Johansen, Kamilla Heddeland Instefjord, Hilde Elshaug, Marie Paulsen Madsen, Rasmus Riis Kopperud, Hilde Vollan, Karoline Bragstad, Olav Hungnes |
| EPI_ISL_635159 | Medical Microbiology Unit, Department for Laboratory Medicine, Drammen Hospital, Vestre Viken Health Trust, | Norwegian Institute of Public Health, Department of Virology | Kathrine Stene-Johansen, Kamilla Heddeland Instefjord, Hilde Elshaug, Marie Paulsen Madsen, Rasmus Riis Kopperud, Hilde Vollan, Karoline Bragstad, Olav Hungnes |
| EPI_ISL_635172 | Oslo University Hospital, Department of Medical Microbiology | Norwegian Institute of Public Health, Department of Virology | Kathrine Stene-Johansen, Kamilla Heddeland Instefjord, Hilde Elshaug, Marie Paulsen Madsen, Rasmus Riis Kopperud, Hilde Vollan, Karoline Bragstad, Olav Hungnes |
| EPI_ISL_635173, EPI_ISL_635176, | Foerde Hospital, Department of Microbiology | Norwegian Institute of Public Health, Department of Virology | Kathrine Stene-Johansen, Kamilla Heddeland Instefjord, Hilde Elshaug, Marie Paulsen Madsen, Rasmus Riis Kopperud, Hilde Vollan, Karoline Bragstad, Olav |

| | | | |
|---|---|---|---|
| EPI_ISL_635177 | | | Hungnes |
| EPI_ISL_635183 | Innlandet Hospital Trust, Division Lillehammer, Department for Medical Microbiology | Norwegian Institute of Public Health, Department of Virology | Kathrine Stene-Johansen, Kamilla Heddeland Instefjord, Hilde Elshaug, Marie Paulsen Madsen, Rasmus Riis Kopperud, Hilde Vollan, Karoline Bragstad, Olav Hungnes |
| EPI_ISL_635190 | University Hospital of Northern Norway, Department for Microbiology and Infectious Disease Control | Norwegian Institute of Public Health, Department of Virology | Kathrine Stene-Johansen, Kamilla Heddeland Instefjord, Hilde Elshaug, Marie Paulsen Madsen, Rasmus Riis Kopperud, Hilde Vollan, Karoline Bragstad, Olav Hungnes |
| EPI_ISL_635197 | Oslo University Hospital, Department of Medical Microbiology | Norwegian Institute of Public Health, Department of Virology | Kathrine Stene-Johansen, Kamilla Heddeland Instefjord, Hilde Elshaug, Marie Paulsen Madsen, Rasmus Riis Kopperud, Hilde Vollan, Karoline Bragstad, Olav Hungnes |
| EPI_ISL_636329, EPI_ISL_636330, EPI_ISL_636331, EPI_ISL_636332, EPI_ISL_636333, EPI_ISL_636334, EPI_ISL_636335, EPI_ISL_636336, EPI_ISL_636337, EPI_ISL_636338, EPI_ISL_636339, EPI_ISL_636340, EPI_ISL_636341, EPI_ISL_636342, EPI_ISL_636343, EPI_ISL_636344, EPI_ISL_636345, EPI_ISL_636346, EPI_ISL_636347, EPI_ISL_636348, EPI_ISL_636349, EPI_ISL_636350, EPI_ISL_636351, EPI_ISL_636352, EPI_ISL_636353, EPI_ISL_636354, EPI_ISL_636355, EPI_ISL_636356, EPI_ISL_636357, EPI_ISL_636358, EPI_ISL_636359, EPI_ISL_636360, EPI_ISL_636361, EPI_ISL_636362, EPI_ISL_636363, EPI_ISL_636364, EPI_ISL_636365, EPI_ISL_636366, EPI_ISL_636367, EPI_ISL_636368, EPI_ISL_636369, EPI_ISL_636370, EPI_ISL_636371, EPI_ISL_636372, EPI_ISL_636373, EPI_ISL_636374, EPI_ISL_636375, EPI_ISL_636376, EPI_ISL_636377, EPI_ISL_636378, EPI_ISL_636379, EPI_ISL_636380, EPI_ISL_636381, EPI_ISL_636382, EPI_ISL_636383, EPI_ISL_636384, EPI_ISL_636385, EPI_ISL_636386, EPI_ISL_636387, EPI_ISL_636388, EPI_ISL_636389, EPI_ISL_636390, EPI_ISL_636391, EPI_ISL_636392, EPI_ISL_636393, EPI_ISL_636394, EPI_ISL_636395, EPI_ISL_636396, EPI_ISL_636397, EPI_ISL_636398, EPI_ISL_636399, EPI_ISL_636400, EPI_ISL_636401, EPI_ISL_636402, EPI_ISL_636403, EPI_ISL_636404, EPI_ISL_636405, EPI_ISL_636406, EPI_ISL_636407, EPI_ISL_636408, EPI_ISL_636409, EPI_ISL_636410, EPI_ISL_636411, EPI_ISL_636412, EPI_ISL_636413, EPI_ISL_636414, EPI_ISL_636415, EPI_ISL_636416, EPI_ISL_636417, EPI_ISL_636418, EPI_ISL_636419, EPI_ISL_636420, EPI_ISL_636421, EPI_ISL_636422, EPI_ISL_636423, EPI_ISL_636424, EPI_ISL_636425, EPI_ISL_636426, EPI_ISL_636427, EPI_ISL_636428, EPI_ISL_636429, EPI_ISL_636430, EPI_ISL_636431, EPI_ISL_636432, EPI_ISL_636433, EPI_ISL_636434, EPI_ISL_636435, EPI_ISL_636436, EPI_ISL_636437, EPI_ISL_636438, EPI_ISL_636439, EPI_ISL_636440, EPI_ISL_636441, EPI_ISL_636442, EPI_ISL_636443, EPI_ISL_636444, EPI_ISL_636445, EPI_ISL_636446, EPI_ISL_636447, EPI_ISL_636448, EPI_ISL_636449, EPI_ISL_636450, EPI_ISL_636451, EPI_ISL_636452, EPI_ISL_636453, EPI_ISL_636454, EPI_ISL_636455, EPI_ISL_636456, EPI_ISL_636457, EPI_ISL_636458 | | | |
| see above | Biolab Diagnostic Laboratories | Andersen lab at Scripps Research | Issa Abu-Dayyeh, Ahmad Tibi, Lama Hussein, Lina Mohammad, Zein Naber, Amid Abdelnour with SEARCH Alliance San Diego |
| EPI_ISL_636580 | Dutch COVID-19 response team | National Institute for Public Health and the Environment (RIVM) | Adam Meijer, Harry Vennema, Jeroen Cremer, Sharon van den Brink, Bas van der Veer, AnneMarie van den Brandt, Florian Zwagemaker, Dennis Schmitz, Chantal Reusken, on behalf of the national COVID-19 response team |
| EPI_ISL_636702 | Respiratory Virus Unit, Microbiology Services Colindale, Public Health England | Respiratory Virus Unit, Microbiology Services Colindale, Public Health England | PHE Covid Sequencing Team |
| EPI_ISL_637018 | Department of Infectious Diseases and Immunology, National Hospital Organization Nagoya Medical Center | Clinical Research Center, National Hospital Organization Nagoya Medical Center | Yoshihiro Nakata, Hirotaka Ode, Mai Kubota, Masakazu Matsuda, Kazuhiro Matsuoka, Miho Nakasuji, Mikiko Mori, Mayumi Imahashi, Yoshiyuki Yokomaku, Yasumasa Iwatani |
| EPI_ISL_637305 | Centre for Enzyme Innovation, University of Portsmouth / Translational Research Laboratory, Portsmouth Hospitals NHS Trust | COVID-19 Genomics UK (COG-UK) Consortium | Angela Beckett,Yann Bourgeois,Garry Scarlett,Sharon Glaysher,Scott Elliott,Kelly Bicknell,Robert Impey,Allyson Lloyd,Sarah Wyllie,Ethan Butcher,Anoop Chauhan,Samuel Robson |
| EPI_ISL_637346, EPI_ISL_637347 | Wales Specialist Virology Centre Sequencing lab: Pathogen Genomics Unit | COVID-19 Genomics UK (COG-UK) Consortium | Catherine Moore, Johnathan Evans, Laura Gifford, Malorie Perry, Simon Cottrell, Angela Marchbank, Alec Birchley, Alexander Adams, Amy Gaskin, Bree Gatica-Wilcox, Jason Coombes, Joel Southgate, Lauren Gilbert, Lee Graham, Nicole Pacchiarini, Sara Kumziene-Summerhayes, Sarah Taylor, Sophie Jones, Sara Rey, Matthew Bull, Joanne Watkins, Sally Corden, Tom Connor |
| EPI_ISL_637369, EPI_ISL_637370, EPI_ISL_637371, EPI_ISL_637372 | Centre for Enzyme Innovation, University of Portsmouth / Translational Research Laboratory, Portsmouth Hospitals NHS Trust | COVID-19 Genomics UK (COG-UK) Consortium | Angela Beckett,Yann Bourgeois,Garry Scarlett,Sharon Glaysher,Scott Elliott,Kelly Bicknell,Robert Impey,Allyson Lloyd,Sarah Wyllie,Ethan Butcher,Anoop Chauhan,Samuel Robson |
| EPI_ISL_637398 | Department of Pathology, University of Cambridge | COVID-19 Genomics UK (COG-UK) Consortium | Aminu S. Jahun, Yasmin Chaudhry, Grant Hall, Iliana Georgana, Myra Hosmillo, Martin D. Curran, Malte Pinckert, Surendra Parmar, Ian Goodfellow |
| EPI_ISL_637400, EPI_ISL_637437 | Wales Specialist Virology Centre Sequencing lab: Pathogen Genomics Unit | COVID-19 Genomics UK (COG-UK) Consortium | Catherine Moore, Johnathan Evans, Laura Gifford, Malorie Perry, Simon Cottrell, Angela Marchbank, Alec Birchley, Alexander Adams, Amy Gaskin, Bree Gatica-Wilcox, Jason Coombes, Joel Southgate, Lauren Gilbert, Lee Graham, Nicole Pacchiarini, Sara Kumziene-Summerhayes, Sarah Taylor, Sophie Jones, Sara Rey, Matthew Bull, Joanne Watkins, Sally Corden, Tom Connor |
| EPI_ISL_637453 | Centre for Enzyme Innovation, University of Portsmouth / Translational Research Laboratory, Portsmouth Hospitals NHS Trust | COVID-19 Genomics UK (COG-UK) Consortium | Angela Beckett,Yann Bourgeois,Garry Scarlett,Sharon Glaysher,Scott Elliott,Kelly Bicknell,Robert Impey,Allyson Lloyd,Sarah Wyllie,Ethan Butcher,Anoop Chauhan,Samuel Robson |
| EPI_ISL_637458 | Wales Specialist Virology Centre Sequencing lab: Pathogen Genomics Unit | COVID-19 Genomics UK (COG-UK) Consortium | Catherine Moore, Johnathan Evans, Laura Gifford, Malorie Perry, Simon Cottrell, Angela Marchbank, Alec Birchley, Alexander Adams, Amy Gaskin, Bree Gatica-Wilcox, Jason Coombes, Joel Southgate, Lauren Gilbert, Lee Graham, Nicole Pacchiarini, Sara Kumziene-Summerhayes, Sarah Taylor, Sophie Jones, Sara Rey, Matthew Bull, Joanne Watkins, Sally Corden, Tom Connor |
| EPI_ISL_637496, EPI_ISL_637497 | Centre for Enzyme Innovation, University of Portsmouth / Translational Research Laboratory, Portsmouth Hospitals NHS Trust | COVID-19 Genomics UK (COG-UK) Consortium | Angela Beckett,Yann Bourgeois,Garry Scarlett,Sharon Glaysher,Scott Elliott,Kelly Bicknell,Robert Impey,Allyson Lloyd,Sarah Wyllie,Ethan Butcher,Anoop Chauhan,Samuel Robson |
| EPI_ISL_637519 | Department of Pathology, University of Cambridge | COVID-19 Genomics UK (COG-UK) Consortium | Aminu S. Jahun, Yasmin Chaudhry, Grant Hall, Iliana Georgana, Myra Hosmillo, Martin D. Curran, Malte Pinckert, Surendra Parmar, Ian Goodfellow |
| EPI_ISL_637524, EPI_ISL_637537, EPI_ISL_637596 | Wales Specialist Virology Centre Sequencing lab: Pathogen Genomics Unit | COVID-19 Genomics UK (COG-UK) Consortium | Catherine Moore, Johnathan Evans, Laura Gifford, Malorie Perry, Simon Cottrell, Angela Marchbank, Alec Birchley, Alexander Adams, Amy Gaskin, Bree Gatica-Wilcox, Jason Coombes, Joel Southgate, Lauren Gilbert, Lee Graham, Nicole Pacchiarini, Sara Kumziene-Summerhayes, Sarah Taylor, Sophie Jones, Sara Rey, Matthew Bull, Joanne Watkins, Sally Corden, Tom Connor |
| EPI_ISL_637610, EPI_ISL_637659 | Centre for Enzyme Innovation, University of Portsmouth / Translational Research Laboratory, Portsmouth Hospitals NHS Trust | COVID-19 Genomics UK (COG-UK) Consortium | Angela Beckett,Yann Bourgeois,Garry Scarlett,Sharon Glaysher,Scott Elliott,Kelly Bicknell,Robert Impey,Allyson Lloyd,Sarah Wyllie,Ethan Butcher,Anoop Chauhan,Samuel Robson |
| EPI_ISL_637667 | Wales Specialist Virology Centre Sequencing lab: Pathogen Genomics Unit | COVID-19 Genomics UK (COG-UK) Consortium | Catherine Moore, Johnathan Evans, Laura Gifford, Malorie Perry, Simon Cottrell, Angela Marchbank, Alec Birchley, Alexander Adams, Amy Gaskin, Bree Gatica-Wilcox, Jason Coombes, Joel Southgate, Lauren Gilbert, Lee Graham, Nicole Pacchiarini, Sara Kumziene-Summerhayes, Sarah Taylor, Sophie Jones, Sara Rey, Matthew Bull, Joanne Watkins, Sally Corden, Tom Connor |
| EPI_ISL_637668 | Department of Pathology, University of Cambridge | COVID-19 Genomics UK (COG-UK) Consortium | Aminu S. Jahun, Yasmin Chaudhry, Grant Hall, Iliana Georgana, Myra Hosmillo, Martin D. Curran, Malte Pinckert, Surendra Parmar, Ian Goodfellow |
| EPI_ISL_637782, EPI_ISL_637783 | Centre for Enzyme Innovation, University of Portsmouth / Translational Research Laboratory, Portsmouth Hospitals NHS Trust | COVID-19 Genomics UK (COG-UK) Consortium | Angela Beckett,Yann Bourgeois,Garry Scarlett,Sharon Glaysher,Scott Elliott,Kelly Bicknell,Robert Impey,Allyson Lloyd,Sarah Wyllie,Ethan Butcher,Anoop Chauhan,Samuel Robson |
| EPI_ISL_637794 | Northumbria University / South Tees Hospitals NHS Foundation Trust / North Cumbria Integrated Care NHS Foundation Trust / North Tees and Hartlepool NHS Foundation Trust / Newcastle Hospitals NHS Foundation Trust | COVID-19 Genomics UK (COG-UK) Consortium | Darren L Smith,Andrew Nelson,Matthew Bashton,Greg R Young,Joshua Loh,John Allan,Mohammad A Tariq,Giles S Holt,Gary Black,Wen C Yew,Lynn Dover,Paul Baker,Steve Liggett,Sarah Essex,Jane Greenaway,Debra Padgett,Clive Graham,Garren Scott,Edward Barton,Emma Swindells,Brendan Payne,Jennifer Collins,Yusri Taha,Gary Eltringham |
| EPI_ISL_637810, EPI_ISL_637811, EPI_ISL_637812, EPI_ISL_637813, EPI_ISL_637814, EPI_ISL_637815, EPI_ISL_637816, EPI_ISL_637817, EPI_ISL_637818, EPI_ISL_637819, EPI_ISL_637820, EPI_ISL_637821 | | | |
| see above | Department of Pathology, University of Cambridge | COVID-19 Genomics UK (COG-UK) Consortium | Aminu S. Jahun, Yasmin Chaudhry, Grant Hall, Iliana Georgana, Myra Hosmillo, Martin D. Curran, Malte Pinckert, Surendra Parmar, Ian Goodfellow |
| EPI_ISL_637899 | Wales Specialist Virology Centre Sequencing lab: Pathogen Genomics Unit | COVID-19 Genomics UK (COG-UK) Consortium | Catherine Moore, Johnathan Evans, Laura Gifford, Malorie Perry, Simon Cottrell, Angela Marchbank, Alec Birchley, Alexander Adams, Amy Gaskin, Bree Gatica-Wilcox, Jason Coombes, Joel Southgate, Lauren Gilbert, Lee Graham, Nicole Pacchiarini, Sara Kumziene-Summerhayes, Sarah Taylor, Sophie Jones, Sara Rey, Matthew Bull, Joanne Watkins, Sally Corden, Tom Connor |
| EPI_ISL_637902 | Northumbria University / South Tees Hospitals NHS Foundation Trust / North Cumbria Integrated Care NHS Foundation Trust / North Tees and Hartlepool NHS Foundation Trust / Newcastle Hospitals NHS Foundation Trust | COVID-19 Genomics UK (COG-UK) Consortium | Darren L Smith,Andrew Nelson,Matthew Bashton,Greg R Young,Joshua Loh,John Allan,Mohammad A Tariq,Giles S Holt,Gary Black,Wen C Yew,Lynn Dover,Paul Baker,Steve Liggett,Sarah Essex,Jane Greenaway,Debra Padgett,Clive Graham,Garren Scott,Edward Barton,Emma Swindells,Brendan Payne,Jennifer Collins,Yusri Taha,Gary Eltringham |

| | | | |
|---|---|---|---|
| EPI_ISL_637932 | Centre for Enzyme Innovation, University of Portsmouth / Translational Research Laboratory, Portsmouth Hospitals NHS Trust | COVID-19 Genomics UK (COG-UK) Consortium | Angela Beckett,Yann Bourgeois,Garry Scarlett,Sharon Glaysher,Scott Elliott,Kelly Bicknell,Robert Impey,Allyson Lloyd,Sarah Wyllie,Ethan Butcher,Anoop Chauhan,Samuel Robson |
| EPI_ISL_637994 | Northumbria University / South Tees Hospitals NHS Foundation Trust / North Cumbria Integrated Care NHS Foundation Trust / North Tees and Hartlepool NHS Foundation Trust / Newcastle Hospitals NHS Foundation Trust | COVID-19 Genomics UK (COG-UK) Consortium | Darren L Smith,Andrew Nelson,Matthew Bashton,Greg R Young,Joshua Loh,John Allan,Mohammad A Tariq,Giles S Holt,Gary Black,Wen C Yew,Lynn Dover,Paul Baker,Steve Liggett,Sarah Essex,Jane Greenaway,Debra Padgett,Clive Graham,Garren Scott,Edward Barton,Emma Swindells,Brendan Payne,Jennifer Collins,Yusri Taha,Gary Eltringham |
| EPI_ISL_637997 | Wales Specialist Virology Centre Sequencing lab: Pathogen Genomics Unit | COVID-19 Genomics UK (COG-UK) Consortium | Catherine Moore, Johnathan Evans, Laura Gifford, Malorie Perry, Simon Cottrell, Angela Marchbank, Alec Birchley, Alexander Adams, Amy Gaskin, Bree Gatica-Wilcox, Jason Coombes, Joel Southgate, Lauren Gilbert, Lee Graham, Nicole Pacchiarini, Sara Kumziene-Summerhayes, Sarah Taylor, Sophie Jones, Sara Rey, Matthew Bull, Joanne Watkins, Sally Corden, Tom Connor |
| EPI_ISL_638002 | Centre for Enzyme Innovation, University of Portsmouth / Translational Research Laboratory, Portsmouth Hospitals NHS Trust | COVID-19 Genomics UK (COG-UK) Consortium | Angela Beckett,Yann Bourgeois,Garry Scarlett,Sharon Glaysher,Scott Elliott,Kelly Bicknell,Robert Impey,Allyson Lloyd,Sarah Wyllie,Ethan Butcher,Anoop Chauhan,Samuel Robson |
| EPI_ISL_638017 | Virology Department, Sheffield Teaching Hospitals NHS Foundation Trust/Department of Infection, Immunity and Cardiovascular Disease, The Medical School, University of Sheffield | COVID-19 Genomics UK (COG-UK) Consortium | Thushan de Silva, Matthew Parker, Nikki Smith, Adri Angyal, Rebecca Brown, Luke Green, Rachel Tucker, Paul Parsons, Danielle Groves, Katie Johnson, Laura Carrilero, Alex Keeley, Dave Partridge, Matthew Wyles, Benjamin Lindsey, Mehmet Yavuz, Mohammad Raza, Cariad Evans |
| EPI_ISL_638041, EPI_ISL_638084, EPI_ISL_638085 | Wales Specialist Virology Centre Sequencing lab: Pathogen Genomics Unit | COVID-19 Genomics UK (COG-UK) Consortium | Catherine Moore, Johnathan Evans, Laura Gifford, Malorie Perry, Simon Cottrell, Angela Marchbank, Alec Birchley, Alexander Adams, Amy Gaskin, Bree Gatica-Wilcox, Jason Coombes, Joel Southgate, Lauren Gilbert, Lee Graham, Nicole Pacchiarini, Sara Kumziene-Summerhayes, Sarah Taylor, Sophie Jones, Sara Rey, Matthew Bull, Joanne Watkins, Sally Corden, Tom Connor |
| EPI_ISL_638169 | Department of Pathology, University of Cambridge | COVID-19 Genomics UK (COG-UK) Consortium | Aminu S. Jahun, Yasmin Chaudhry, Grant Hall, Iliana Georgana, Myra Hosmillo, Martin D. Curran, Malte Pinckert, Surendra Parmar, Ian Goodfellow |
| EPI_ISL_638455, EPI_ISL_638456, EPI_ISL_638457, EPI_ISL_638458 | Centre for Enzyme Innovation, University of Portsmouth / Translational Research Laboratory, Portsmouth Hospitals NHS Trust | COVID-19 Genomics UK (COG-UK) Consortium | Angela Beckett,Yann Bourgeois,Garry Scarlett,Sharon Glaysher,Scott Elliott,Kelly Bicknell,Robert Impey,Allyson Lloyd,Sarah Wyllie,Ethan Butcher,Anoop Chauhan,Samuel Robson |
| EPI_ISL_638560, EPI_ISL_638563 | Northumbria University / South Tees Hospitals NHS Foundation Trust / North Cumbria Integrated Care NHS Foundation Trust / North Tees and Hartlepool NHS Foundation Trust / Newcastle Hospitals NHS Foundation Trust | COVID-19 Genomics UK (COG-UK) Consortium | Darren L Smith,Andrew Nelson,Matthew Bashton,Greg R Young,Joshua Loh,John Allan,Mohammad A Tariq,Giles S Holt,Gary Black,Wen C Yew,Lynn Dover,Paul Baker,Steve Liggett,Sarah Essex,Jane Greenaway,Debra Padgett,Clive Graham,Garren Scott,Edward Barton,Emma Swindells,Brendan Payne,Jennifer Collins,Yusri Taha,Gary Eltringham |
| EPI_ISL_638682, EPI_ISL_638683, EPI_ISL_638684, EPI_ISL_638685, EPI_ISL_638686, EPI_ISL_638687, EPI_ISL_638688, EPI_ISL_638689, EPI_ISL_638690, EPI_ISL_638691, EPI_ISL_638692, EPI_ISL_638693, EPI_ISL_638694, EPI_ISL_638695, EPI_ISL_638696, EPI_ISL_638697, EPI_ISL_638698, EPI_ISL_638699, EPI_ISL_638700, EPI_ISL_638701, EPI_ISL_638702, EPI_ISL_638703, EPI_ISL_638704, EPI_ISL_638705, EPI_ISL_638706, EPI_ISL_638707, EPI_ISL_638708, EPI_ISL_638709, EPI_ISL_638710, EPI_ISL_638711, EPI_ISL_638712, EPI_ISL_638713, EPI_ISL_638714, EPI_ISL_638715, EPI_ISL_638716, EPI_ISL_638717, EPI_ISL_638718, EPI_ISL_638719, EPI_ISL_638720, EPI_ISL_638721, EPI_ISL_638722, EPI_ISL_638723, EPI_ISL_638724, EPI_ISL_638725, EPI_ISL_638726, EPI_ISL_638727, EPI_ISL_638728, EPI_ISL_638729, EPI_ISL_638730 | | | |
| see above | Wales Specialist Virology Centre Sequencing lab: Pathogen Genomics Unit | COVID-19 Genomics UK (COG-UK) Consortium | Catherine Moore, Johnathan Evans, Laura Gifford, Malorie Perry, Simon Cottrell, Angela Marchbank, Alec Birchley, Alexander Adams, Amy Gaskin, Bree Gatica-Wilcox, Jason Coombes, Joel Southgate, Lauren Gilbert, Lee Graham, Nicole Pacchiarini, Sara Kumziene-Summerhayes, Sarah Taylor, Sophie Jones, Sara Rey, Matthew Bull, Joanne Watkins, Sally Corden, Tom Connor |
| EPI_ISL_638731, EPI_ISL_638732, EPI_ISL_638733, EPI_ISL_638734 | Department of Pathology, University of Cambridge | COVID-19 Genomics UK (COG-UK) Consortium | Aminu S. Jahun, Yasmin Chaudhry, Grant Hall, Iliana Georgana, Myra Hosmillo, Martin D. Curran, Malte Pinckert, Surendra Parmar, Ian Goodfellow |
| EPI_ISL_638928, EPI_ISL_638929, EPI_ISL_638930, EPI_ISL_638931, EPI_ISL_638932, EPI_ISL_638933, EPI_ISL_638934, EPI_ISL_638935 | Wales Specialist Virology Centre Sequencing lab: Pathogen Genomics Unit | COVID-19 Genomics UK (COG-UK) Consortium | Catherine Moore, Johnathan Evans, Laura Gifford, Malorie Perry, Simon Cottrell, Angela Marchbank, Alec Birchley, Alexander Adams, Amy Gaskin, Bree Gatica-Wilcox, Jason Coombes, Joel Southgate, Lauren Gilbert, Lee Graham, Nicole Pacchiarini, Sara Kumziene-Summerhayes, Sarah Taylor, Sophie Jones, Sara Rey, Matthew Bull, Joanne Watkins, Sally Corden, Tom Connor |
| EPI_ISL_639007 | Centre for Enzyme Innovation, University of Portsmouth / Translational Research Laboratory, Portsmouth Hospitals NHS Trust | COVID-19 Genomics UK (COG-UK) Consortium | Angela Beckett,Yann Bourgeois,Garry Scarlett,Sharon Glaysher,Scott Elliott,Kelly Bicknell,Robert Impey,Allyson Lloyd,Sarah Wyllie,Ethan Butcher,Anoop Chauhan,Samuel Robson |
| EPI_ISL_639626 | Wales Specialist Virology Centre Sequencing lab: Pathogen Genomics Unit | COVID-19 Genomics UK (COG-UK) Consortium | Catherine Moore, Johnathan Evans, Laura Gifford, Malorie Perry, Simon Cottrell, Angela Marchbank, Alec Birchley, Alexander Adams, Amy Gaskin, Bree Gatica-Wilcox, Jason Coombes, Joel Southgate, Lauren Gilbert, Lee Graham, Nicole Pacchiarini, Sara Kumziene-Summerhayes, Sarah Taylor, Sophie Jones, Sara Rey, Matthew Bull, Joanne Watkins, Sally Corden, Tom Connor |
| EPI_ISL_639669, EPI_ISL_639671, EPI_ISL_639684, EPI_ISL_639688, EPI_ISL_639689, EPI_ISL_639692 | E. Gulbja Laboratorija | Latvian Biomedical Research and Study Centre | Ivars Silamielis, Kaspars Megnis, Monta Ustinova, ikita Zrelovs, Vita Rovte, Mikus Gavars, Dmitrijs Perminovs, Uga Dumpis, Jnis Kloviš |
| EPI_ISL_639866, EPI_ISL_639867, EPI_ISL_639868, EPI_ISL_639869, EPI_ISL_639870, EPI_ISL_639871, EPI_ISL_639872, EPI_ISL_639873, EPI_ISL_639874, EPI_ISL_639875, EPI_ISL_639876, EPI_ISL_639877, EPI_ISL_639878, EPI_ISL_639879, EPI_ISL_639880, EPI_ISL_639881, EPI_ISL_639882, EPI_ISL_639883, EPI_ISL_639884, EPI_ISL_639886, EPI_ISL_639887, EPI_ISL_639888, EPI_ISL_639889, EPI_ISL_639890, EPI_ISL_639891, EPI_ISL_639904 | | | |
| see above | National Virus Reference Laboratory | National Virus Reference Laboratory | Michael Carr, Gabriel Gonzalez, Jonathan Dean, Daniel Hare, Cillian F De Gascun |
| EPI_ISL_639974, EPI_ISL_639975, EPI_ISL_639987 | CNR Virus des Infections Respiratoires - France SUD | CNR Virus des Infections Respiratoires - France SUD | Antonin Bal, Gregory Destras, Gwendolyne Burfin, Hadrien Règue, Alexandre Gaymard, Maude Bouscambert-Duchamp, Florence Morfin-Sherpa, Martine Valette, Bruno Lina, Laurence Josset |
| EPI_ISL_640087 | Victoria Hospital wc VHW | NHLS/UCT | Arash Iranzadeh, Deelan Doolabh, Lynn Tyers, Bruna Galvao, Innocent Mudau, Marvin Hsiao, Kruger Marais, Diana Hardie, Stephen Korsman, Carolyn Williamson |
| EPI_ISL_640337, EPI_ISL_640519, EPI_ISL_640588 | Victorian Infectious Diseases Reference Laboratory (VIDRL) | VIDRL and MDU-PHL | Caly L., Seemann T., Sait, M.L., Schultz M.B., Druce J., Sherry, N.L. |
| EPI_ISL_640641, EPI_ISL_640682, EPI_ISL_640754 | Microbiological Diagnostic Unit - Public Health Laboratory (MDU-PHL) | MDU-PHL | Seemann T., Schultz M.B., Sait, M.L., Sherry, N.L. |
| EPI_ISL_640784 | Victorian Infectious Diseases Reference Laboratory (VIDRL) | VIDRL and MDU-PHL | Caly L., Seemann T., Sait, M.L., Schultz M.B., Druce J., Sherry, N.L. |
| EPI_ISL_641220 | Microbiological Diagnostic Unit - Public Health Laboratory (MDU-PHL) | MDU-PHL | Seemann T., Schultz M.B., Sait, M.L., Sherry, N.L. |
| EPI_ISL_641273 | Victorian Infectious Diseases Reference Laboratory (VIDRL) | VIDRL and MDU-PHL | Caly L., Seemann T., Sait, M.L., Schultz M.B., Druce J., Sherry, N.L. |
| EPI_ISL_641318 | Environmental and Global Health, University of Florida | Environmental and Global Health, University of Florida | Nannu-Shankar,S., Witanachchi,C.T., Alam,M.M., Loeb,J.C., Stephenson,C.J., Fan,Z.H., Wu,C.-Y., Lednicky,J.A. |
| EPI_ISL_641444, EPI_ISL_641445, EPI_ISL_641499, EPI_ISL_641500, EPI_ISL_641502 | Department of Virus and Microbiological Special Diagnostics, Statens Serum Institut, Copenhagen, Denmark | Albertsen lab, Department of Chemistry and Bioscience, Aalborg University, Denmark | Thomas Bruun Rasmussen, Jannik Fonager, Morten Rasmussen |
| EPI_ISL_641603, EPI_ISL_641604 | Department of Clinical Microbiology | GIGA Medical Genomics | Keith Durkin, Maria Artesi, Sébastien Bontems, Raphaël Boreux, Bouchra Boujemla, Cécile Meex, Pierrette Melin, Marie-Pierre Hayette, Vincent Bours |
| EPI_ISL_644378, EPI_ISL_644528, EPI_ISL_644529, EPI_ISL_644530, | MEPHI, Aix Marseille University | MEPHI, Aix Marseille University | Anthony LEVASSEUR |

| | | | |
|---|---|---|---|
| EPI_ISL_644531, EPI_ISL_644533, EPI_ISL_644534, EPI_ISL_644535 | | | |
| EPI_ISL_644893 | Virginia DCLS | Virginia DCLS | Virginia DCLS |
| EPI_ISL_644943 | Essentia Health-St. Mary's Medical Center | Minnesota Department of Health, Public Health Laboratory | Matt Plumb, Jacob Garfin, Alexandra Lorentz, and Xiong Wang |
| EPI_ISL_647213 | Lighthouse Lab in Cambridge | Wellcome Sanger Institute for the COVID-19 Genomics UK (COG-UK) Consortium | Rob Howes, The Lighthouse Lab in Cambridge and Alex Alderton, Roberto Amato, Sonia Goncalves, Ewan Harrison, David K. Jackson, Ian Johnston, Dominic Kwiatkowski, Cordelia Langford, John Sillitoe on behalf of the Wellcome Sanger Institute COVID-19 Surveillance Team |
| EPI_ISL_648135, EPI_ISL_648147, EPI_ISL_648149, EPI_ISL_648150, EPI_ISL_648156, EPI_ISL_648185, EPI_ISL_648186, EPI_ISL_648188, EPI_ISL_648194 | The Public Health Agency of Sweden | The Public Health Agency of Sweden | Anna-Malin Linde, Maria Lind Karlberg, Mattias Haukland, Reza Advani, Olov Svartstrom, Oskar Karlsson Lindsjo, Sandra Broddesson, Petra Edquist, Mia Brytting, Anna Risberg, Karin Tegmark-Wisell |
| EPI_ISL_648504, EPI_ISL_648505, EPI_ISL_648506, EPI_ISL_648507, EPI_ISL_648508 | Orange County Public Health Lab | Chan-Zuckerberg Biohub | CZB Cliahub Consortium |
| EPI_ISL_648539, EPI_ISL_648540, EPI_ISL_648541 | Tulare County Public Health Lab | Chan-Zuckerberg Biohub | CZB Cliahub Consortium |
| EPI_ISL_648888, EPI_ISL_648892, EPI_ISL_648894, EPI_ISL_648900, EPI_ISL_648904, EPI_ISL_648914, EPI_ISL_648915, EPI_ISL_648924, EPI_ISL_648926, EPI_ISL_648929, EPI_ISL_648930, EPI_ISL_648932, EPI_ISL_648935, EPI_ISL_648937, EPI_ISL_648943, EPI_ISL_648949, EPI_ISL_648954, EPI_ISL_648962, EPI_ISL_648965, EPI_ISL_648967, EPI_ISL_648969, EPI_ISL_648971, EPI_ISL_648976, EPI_ISL_648978, EPI_ISL_648979, EPI_ISL_649056 | | | |
| see above | San Diego County Public Health Laboratory | Andersen lab at Scripps Research | SEARCH Alliance San Diego with Tracy Basler, Jovan Shephard, Brett Austin |
| EPI_ISL_649389 | Lighthouse Lab in Alderley Park | Wellcome Sanger Institute for the COVID-19 Genomics UK (COG-UK) Consortium | Jacquelyn Wynn, Mairead Hyland, The Lighthouse Lab in Alderley Park and Alex Alderton, Roberto Amato, Sonia Goncalves, Ewan Harrison, David K. Jackson, Ian Johnston, Dominic Kwiatkowski, Cordelia Langford, John Sillitoe on behalf of the Wellcome Sanger Institute COVID-19 Surveillance Team (http://www.sanger.ac.uk/covid-team) |
| EPI_ISL_650168, EPI_ISL_650221 | Centre for Enzyme Innovation, University of Portsmouth / Translational Research Laboratory, Portsmouth Hospitals NHS Trust | COVID-19 Genomics UK (COG-UK) Consortium | Angela Beckett,Yann Bourgeois,Garry Scarlett,Sharon Glaysher,Scott Elliott,Kelly Bicknell,Robert Impey,Allyson Lloyd,Sarah Wyllie,Ethan Butcher,Anoop Chauhan,Samuel Robson |
| EPI_ISL_650241 | Northumbria University / South Tees Hospitals NHS Foundation Trust / North Cumbria Integrated Care NHS Foundation Trust / North Tees and Hartlepool NHS Foundation Trust / Newcastle Hospitals NHS Foundation Trust | COVID-19 Genomics UK (COG-UK) Consortium | Darren L Smith,Andrew Nelson,Matthew Bashton,Greg R Young,Joshua Loh,John Allan,Mohammad A Tariq,Giles S Holt,Gary Black,Wen C Yew,Lynn Dover,Paul Baker,Steve Liggett,Sarah Essex,Jane Greenaway,Debra Padgett,Clive Graham,Garren Scott,Edward Barton,Emma Swindells,Brendan Payne,Jennifer Collins,Yusri Taha,Gary Eltringham |
| EPI_ISL_650244, EPI_ISL_650283 | Virology Department, Sheffield Teaching Hospitals NHS Foundation Trust/Department of Infection, Immunity and Cardiovascular Disease, The Medical School, University of Sheffield | COVID-19 Genomics UK (COG-UK) Consortium | Thushan de Silva, Matthew Parker, Nikki Smith, Adri Angyal, Rebecca Brown, Luke Green, Rachel Tucker, Paul Parsons, Danielle Groves, Katie Johnson, Laura Carrilero, Alex Keeley, Dave Partridge, Matthew Wyles, Benjamin Lindsey, Mehmet Yavuz, Mohammad Raza, Cariad Evans |
| EPI_ISL_650292 | West of Scotland Specialist Virology Centre, NHSGGC / MRC-University of Glasgow Centre for Virus Research | COVID-19 Genomics UK (COG-UK) Consortium | Ana da Silva Filipe, Natasha Johnson, Kathy Smollett, Daniel Mair, Stephen Carmichael, Alice Broos, Lily Tong, Jenna Nichols, Kyriaki Nomikou; Sarah McDonald; Richard Orton, Joseph Hughes, Sreenu Vattipally, David L Robertson; Alasdair MacLean, Rory Gunson; Sharif Shaaban, Matthew Holden; Rachel Blacow, Guy Mollett, Kathy Li, James Shepherd, Antonia Ho, Emma Thomson |
| EPI_ISL_650348 | Northumbria University / South Tees Hospitals NHS Foundation Trust / North Cumbria Integrated Care NHS Foundation Trust / North Tees and Hartlepool NHS Foundation Trust / Newcastle Hospitals NHS Foundation Trust | COVID-19 Genomics UK (COG-UK) Consortium | Darren L Smith,Andrew Nelson,Matthew Bashton,Greg R Young,Joshua Loh,John Allan,Mohammad A Tariq,Giles S Holt,Gary Black,Wen C Yew,Lynn Dover,Paul Baker,Steve Liggett,Sarah Essex,Jane Greenaway,Debra Padgett,Clive Graham,Garren Scott,Edward Barton,Emma Swindells,Brendan Payne,Jennifer Collins,Yusri Taha,Gary Eltringham |
| EPI_ISL_650372 | Virology Department, Sheffield Teaching Hospitals NHS Foundation Trust/Department of Infection, Immunity and Cardiovascular Disease, The Medical School, University of Sheffield | COVID-19 Genomics UK (COG-UK) Consortium | Thushan de Silva, Matthew Parker, Nikki Smith, Adri Angyal, Rebecca Brown, Luke Green, Rachel Tucker, Paul Parsons, Danielle Groves, Katie Johnson, Laura Carrilero, Alex Keeley, Dave Partridge, Matthew Wyles, Benjamin Lindsey, Mehmet Yavuz, Mohammad Raza, Cariad Evans |
| EPI_ISL_650401 | Centre for Enzyme Innovation, University of Portsmouth / Translational Research Laboratory, Portsmouth Hospitals NHS Trust | COVID-19 Genomics UK (COG-UK) Consortium | Angela Beckett,Yann Bourgeois,Garry Scarlett,Sharon Glaysher,Scott Elliott,Kelly Bicknell,Robert Impey,Allyson Lloyd,Sarah Wyllie,Ethan Butcher,Anoop Chauhan,Samuel Robson |
| EPI_ISL_650490, EPI_ISL_650497, EPI_ISL_650517, EPI_ISL_650518, EPI_ISL_650519, EPI_ISL_650523 | West of Scotland Specialist Virology Centre, NHSGGC / MRC-University of Glasgow Centre for Virus Research | COVID-19 Genomics UK (COG-UK) Consortium | Ana da Silva Filipe, Natasha Johnson, Kathy Smollett, Daniel Mair, Stephen Carmichael, Alice Broos, Lily Tong, Jenna Nichols, Kyriaki Nomikou; Sarah McDonald; Richard Orton, Joseph Hughes, Sreenu Vattipally, David L Robertson; Alasdair MacLean, Rory Gunson; Sharif Shaaban, Matthew Holden; Rachel Blacow, Guy Mollett, Kathy Li, James Shepherd, Antonia Ho, Emma Thomson |
| EPI_ISL_650566, EPI_ISL_650567, EPI_ISL_650568, EPI_ISL_650569, EPI_ISL_650570, EPI_ISL_650571, EPI_ISL_650584, EPI_ISL_650585 | Centre for Enzyme Innovation, University of Portsmouth / Translational Research Laboratory, Portsmouth Hospitals NHS Trust | COVID-19 Genomics UK (COG-UK) Consortium | Angela Beckett,Yann Bourgeois,Garry Scarlett,Sharon Glaysher,Scott Elliott,Kelly Bicknell,Robert Impey,Allyson Lloyd,Sarah Wyllie,Ethan Butcher,Anoop Chauhan,Samuel Robson |
| EPI_ISL_650640 | Northumbria University / South Tees Hospitals NHS Foundation Trust / North Cumbria Integrated Care NHS Foundation Trust / North Tees and Hartlepool NHS Foundation Trust / Newcastle Hospitals NHS Foundation Trust | COVID-19 Genomics UK (COG-UK) Consortium | Darren L Smith,Andrew Nelson,Matthew Bashton,Greg R Young,Joshua Loh,John Allan,Mohammad A Tariq,Giles S Holt,Gary Black,Wen C Yew,Lynn Dover,Paul Baker,Steve Liggett,Sarah Essex,Jane Greenaway,Debra Padgett,Clive Graham,Garren Scott,Edward Barton,Emma Swindells,Brendan Payne,Jennifer Collins,Yusri Taha,Gary Eltringham |
| EPI_ISL_650663 | Virology Department, Sheffield Teaching Hospitals NHS Foundation Trust/Department of Infection, Immunity and Cardiovascular Disease, The Medical School, University of Sheffield | COVID-19 Genomics UK (COG-UK) Consortium | Thushan de Silva, Matthew Parker, Nikki Smith, Adri Angyal, Rebecca Brown, Luke Green, Rachel Tucker, Paul Parsons, Danielle Groves, Katie Johnson, Laura Carrilero, Alex Keeley, Dave Partridge, Matthew Wyles, Benjamin Lindsey, Mehmet Yavuz, Mohammad Raza, Cariad Evans |
| EPI_ISL_650711 | Centre for Enzyme Innovation, University of Portsmouth / Translational Research Laboratory, Portsmouth Hospitals NHS Trust | COVID-19 Genomics UK (COG-UK) Consortium | Angela Beckett,Yann Bourgeois,Garry Scarlett,Sharon Glaysher,Scott Elliott,Kelly Bicknell,Robert Impey,Allyson Lloyd,Sarah Wyllie,Ethan Butcher,Anoop Chauhan,Samuel Robson |
| EPI_ISL_650853, EPI_ISL_650927 | West of Scotland Specialist Virology Centre, NHSGGC / MRC-University of Glasgow Centre for Virus Research | COVID-19 Genomics UK (COG-UK) Consortium | Ana da Silva Filipe, Natasha Johnson, Kathy Smollett, Daniel Mair, Stephen Carmichael, Alice Broos, Lily Tong, Jenna Nichols, Kyriaki Nomikou; Sarah McDonald; Richard Orton, Joseph Hughes, Sreenu Vattipally, David L Robertson; Alasdair MacLean, Rory Gunson; Sharif Shaaban, Matthew Holden; Rachel Blacow, Guy Mollett, Kathy Li, James Shepherd, Antonia Ho, Emma Thomson |
| EPI_ISL_650947, EPI_ISL_650949, EPI_ISL_650964, EPI_ISL_650965, EPI_ISL_650966 | Centre for Enzyme Innovation, University of Portsmouth / Translational Research Laboratory, Portsmouth Hospitals NHS Trust | COVID-19 Genomics UK (COG-UK) Consortium | Angela Beckett,Yann Bourgeois,Garry Scarlett,Sharon Glaysher,Scott Elliott,Kelly Bicknell,Robert Impey,Allyson Lloyd,Sarah Wyllie,Ethan Butcher,Anoop Chauhan,Samuel Robson |
| EPI_ISL_651062 | Northumbria University / South Tees Hospitals NHS | COVID-19 Genomics UK (COG-UK) Consortium | Darren L Smith,Andrew Nelson,Matthew Bashton,Greg R Young,Joshua Loh,John Allan,Mohammad A Tariq,Giles S Holt,Gary Black,Wen C Yew,Lynn |

| | | | |
|---|---|---|---|
| | Foundation Trust / North Cumbria Integrated Care NHS Foundation Trust / North Tees and Hartlepool NHS Foundation Trust / Newcastle Hospitals NHS Foundation Trust | | Dover,Paul Baker,Steve Liggett,Sarah Essex,Jane Greenaway,Debra Padgett,Clive Graham,Garren Scott,Edward Barton,Emma Swindells,Brendan Payne,Jennifer Collins,Yusri Taha,Gary Eltringham |
| EPI_ISL_651099 | Centre for Enzyme Innovation, University of Portsmouth / Translational Research Laboratory, Portsmouth Hospitals NHS Trust | COVID-19 Genomics UK (COG-UK) Consortium | Angela Beckett,Yann Bourgeois,Garry Scarlett,Sharon Glaysher,Scott Elliott,Kelly Bicknell,Robert Impey,Allyson Lloyd,Sarah Wyllie,Ethan Butcher,Anoop Chauhan,Samuel Robson |
| EPI_ISL_651117 | Northumbria University / South Tees Hospitals NHS Foundation Trust / North Cumbria Integrated Care NHS Foundation Trust / North Tees and Hartlepool NHS Foundation Trust / Newcastle Hospitals NHS Foundation Trust | COVID-19 Genomics UK (COG-UK) Consortium | Darren L Smith,Andrew Nelson,Matthew Bashton,Greg R Young,Joshua Loh,John Allan,Mohammad A Tariq,Giles S Holt,Gary Black,Wen C Yew,Lynn Dover,Paul Baker,Steve Liggett,Sarah Essex,Jane Greenaway,Debra Padgett,Clive Graham,Garren Scott,Edward Barton,Emma Swindells,Brendan Payne,Jennifer Collins,Yusri Taha,Gary Eltringham |
| EPI_ISL_651243 | Virology Department, Sheffield Teaching Hospitals NHS Foundation Trust/Department of Infection, Immunity and Cardiovascular Disease, The Medical School, University of Sheffield | COVID-19 Genomics UK (COG-UK) Consortium | Thushan de Silva, Matthew Parker, Nikki Smith, Adri Angyal, Rebecca Brown, Luke Green, Rachel Tucker, Paul Parsons, Danielle Groves, Katie Johnson, Laura Carrilero, Alex Keeley, Dave Partridge, Matthew Wyles, Benjamin Lindsey, Mehmet Yavuz, Mohammad Raza, Cariad Evans |
| EPI_ISL_651330 | West of Scotland Specialist Virology Centre, NHSGGC / MRC-University of Glasgow Centre for Virus Research | COVID-19 Genomics UK (COG-UK) Consortium | Ana da Silva Filipe, Natasha Johnson, Kathy Smollett, Daniel Mair, Stephen Carmichael, Alice Broos, Lily Tong, Jenna Nichols, Kyriaki Nomikou; Sarah McDonald; Richard Orton, Joseph Hughes, Sreenu Vattipally, David L Robertson; Alasdair MacLean, Rory Gunson; Sharif Shaaban, Matthew Holden; Rachel Blacow, Guy Mollett, Kathy Li, James Shepherd, Antonia Ho, Emma Thomson |
| EPI_ISL_651345 | Northumbria University / South Tees Hospitals NHS Foundation Trust / North Cumbria Integrated Care NHS Foundation Trust / North Tees and Hartlepool NHS Foundation Trust / Newcastle Hospitals NHS Foundation Trust | COVID-19 Genomics UK (COG-UK) Consortium | Darren L Smith,Andrew Nelson,Matthew Bashton,Greg R Young,Joshua Loh,John Allan,Mohammad A Tariq,Giles S Holt,Gary Black,Wen C Yew,Lynn Dover,Paul Baker,Steve Liggett,Sarah Essex,Jane Greenaway,Debra Padgett,Clive Graham,Garren Scott,Edward Barton,Emma Swindells,Brendan Payne,Jennifer Collins,Yusri Taha,Gary Eltringham |
| EPI_ISL_651733, EPI_ISL_651734, EPI_ISL_651735, EPI_ISL_651736, EPI_ISL_651737, EPI_ISL_651738, EPI_ISL_651786, EPI_ISL_651787, EPI_ISL_651788, EPI_ISL_651789, EPI_ISL_651955 | | | |
| see above | West of Scotland Specialist Virology Centre, NHSGGC / MRC-University of Glasgow Centre for Virus Research | COVID-19 Genomics UK (COG-UK) Consortium | Ana da Silva Filipe, Natasha Johnson, Kathy Smollett, Daniel Mair, Stephen Carmichael, Alice Broos, Lily Tong, Jenna Nichols, Kyriaki Nomikou; Sarah McDonald; Richard Orton, Joseph Hughes, Sreenu Vattipally, David L Robertson; Alasdair MacLean, Rory Gunson; Sharif Shaaban, Matthew Holden; Rachel Blacow, Guy Mollett, Kathy Li, James Shepherd, Antonia Ho, Emma Thomson |
| EPI_ISL_651957 | Virology Department, Royal Infirmary of Edinburgh, NHS Lothian / School of Biological Sciences, University of Edinburgh / Institute of Genetics and Molecular Medicine, University of Edinburgh | COVID-19 Genomics UK (COG-UK) Consortium | McHugh M, Dewar R, Rooke S, Gallagher M, Balcaza C, O'Toole Á, Scher E, Hill V, McCrone JT, Colquhoun R, Yu X, Jackson B, Rambaut A, Williams TC, Templeton K |
| EPI_ISL_652103, EPI_ISL_652104, EPI_ISL_652105, EPI_ISL_652106, EPI_ISL_652107, EPI_ISL_652108, EPI_ISL_652109, EPI_ISL_652110 | Centre for Enzyme Innovation, University of Portsmouth / Translational Research Laboratory, Portsmouth Hospitals NHS Trust | COVID-19 Genomics UK (COG-UK) Consortium | Angela Beckett,Yann Bourgeois,Garry Scarlett,Sharon Glaysher,Scott Elliott,Kelly Bicknell,Robert Impey,Allyson Lloyd,Sarah Wyllie,Ethan Butcher,Anoop Chauhan,Samuel Robson |
| EPI_ISL_652343 | Northumbria University / South Tees Hospitals NHS Foundation Trust / North Cumbria Integrated Care NHS Foundation Trust / North Tees and Hartlepool NHS Foundation Trust / Newcastle Hospitals NHS Foundation Trust | COVID-19 Genomics UK (COG-UK) Consortium | Darren L Smith,Andrew Nelson,Matthew Bashton,Greg R Young,Joshua Loh,John Allan,Mohammad A Tariq,Giles S Holt,Gary Black,Wen C Yew,Lynn Dover,Paul Baker,Steve Liggett,Sarah Essex,Jane Greenaway,Debra Padgett,Clive Graham,Garren Scott,Edward Barton,Emma Swindells,Brendan Payne,Jennifer Collins,Yusri Taha,Gary Eltringham |
| EPI_ISL_653071, EPI_ISL_653074, EPI_ISL_653086, EPI_ISL_653087, EPI_ISL_653093, EPI_ISL_653097 | Virology Department, Sheffield Teaching Hospitals NHS Foundation Trust/Department of Infection, Immunity and Cardiovascular Disease, The Medical School, University of Sheffield | COVID-19 Genomics UK (COG-UK) Consortium | Thushan de Silva, Matthew Parker, Nikki Smith, Adri Angyal, Rebecca Brown, Luke Green, Rachel Tucker, Paul Parsons, Danielle Groves, Katie Johnson, Laura Carrilero, Alex Keeley, Dave Partridge, Matthew Wyles, Benjamin Lindsey, Mehmet Yavuz, Mohammad Raza, Cariad Evans |
| EPI_ISL_654168, EPI_ISL_654231, EPI_ISL_654232, EPI_ISL_654233, EPI_ISL_654234, EPI_ISL_654235, EPI_ISL_654236, EPI_ISL_654237, EPI_ISL_654238, EPI_ISL_654239, EPI_ISL_654240, EPI_ISL_654284, EPI_ISL_654356 | | | |
| see above | Hospital General Universitario Gregorio Marañón | SeqCOVID-SPAIN consortium/IBV(CSIC) | Darío García de Viedma, Laura Pérez-Lago, Marta Herranz, Jon Sicilia, Julia Suárez, Pilar Catalán, Patricia Muñoz and SeqCOVID-SPAIN consortium |
| EPI_ISL_654408, EPI_ISL_654434, EPI_ISL_654455, EPI_ISL_654456 | Servicio de Microbiología, Hospital Miguel Servet, Zaragoza | SeqCOVID-SPAIN consortium/IBV(CSIC) | Antonio Rezusta López, Alexander Tristancho Baró, Ana Milagro, Yolanda Gracia Grataloup, Nieves Martínez Cameo and SeqCOVID-SPAIN consortium |
| EPI_ISL_654504, EPI_ISL_654505 | Klinisk mikrobiologi Vasternorrland | The Public Health Agency of Sweden | Anna-Malin Linde, Maria Lind Karlberg, Mattias Haukland, Reza Advani, Olov Svartstrom, Oskar Karlsson Lindsjo, Sandra Broddesson, Petra Edquist, Mia Brytting, Anna Risberg, Karin Tegmark-Wisell |
| EPI_ISL_654508 | The Public Health Agency of Sweden | The Public Health Agency of Sweden | Anna-Malin Linde, Maria Lind Karlberg, Mattias Haukland, Reza Advani, Olov Svartstrom, Oskar Karlsson Lindsjo, Sandra Broddesson, Petra Edquist, Mia Brytting, Anna Risberg, Karin Tegmark-Wisell |
| EPI_ISL_654509, EPI_ISL_654511, EPI_ISL_654514, EPI_ISL_654518, EPI_ISL_654522, EPI_ISL_654526, EPI_ISL_654527, EPI_ISL_654528, EPI_ISL_654534, EPI_ISL_654537 | Servicio de Microbiología, Laboratori Clínic Metropolitana Nord. Hospital Universitari Germans Trias i Pujol. Institut d'Investigació en Ciències de la Salut Germans Trias i Pujol (IGTP) | SeqCOVID-SPAIN consortium/IBV(CSIC) | Elisa Martró, Antoni E. Bordoy, Anna Not, Adrián Antuori, Anabel Fernández, Nona Romaní and SeqCOVID-SPAIN consortium |
| EPI_ISL_654543, EPI_ISL_654555, EPI_ISL_654556, EPI_ISL_654557, EPI_ISL_654559, EPI_ISL_654560, EPI_ISL_654561, EPI_ISL_654562, EPI_ISL_654563, EPI_ISL_654564 | Servicio de Microbiología. Hospital Universitario Donostia. OSI Donostialdea. Área de Enfermedades Infecciosas, Grupo de Infección Respiratoria y Resistencia Antimicrobiana. Instituto de Investigación Sanitaria Biodonostia | SeqCOVID-SPAIN consortium/IBV(CSIC) | Gustavo Cilla Eguiluz, Milagrosa Montes Ros, Luis Piñeiro Vázquez, Ane Sorrarain, Jose Maria Marimón and SeqCOVID-SPAIN consortium |
| EPI_ISL_660137 | Hamadi | National Health Laboratory Service (NHLS), Tygerberg | Susan Engelbrecht, Draper C, Davis M-A, Siegfried N, Williamson C, Hsiao M, Kayla Delaney, Bronwyn Kleinhans, Houriiyah Tegally, Eduan Wilkindon, Gert van Zyl, Wolfgang Preiser, Tulio de Oliveira |
| EPI_ISL_660184, EPI_ISL_660185 | NHLS-IALCH | KRISP, KZN Research Innovation and Sequencing Platform | Gazy I, Sigal A, Karim F, Cele S, Giandhari J, Pillay S, Tegally H, Wilkinson E, de Oliveira T |
| EPI_ISL_660532 | Institute of Microbiology, Universidad San Francisco de Quito | Institute of Microbiology, Universidad San Francisco de Quito | Sully Márquez, Belén Prado-Vivar, Juan José Guadalupe, Monica Becerra-Wong, Bernardo Gutiérrez, Nabih Dahik, Verónica Barragán, Patricio Rojas-Silva, Gabriel Trueba, Michelle Grunauer, Paúl Cárdenas |
| EPI_ISL_660533, EPI_ISL_660534 | Institute of Microbiology, Universidad San Francisco de Quito | Institute of Microbiology, Universidad San Francisco de Quito | Sully Márquez, Belén Prado-Vivar, Juan José Guadalupe, Monica Becerra-Wong, Bernardo Gutiérrez, Nabih Dahik, Freddy Iza, Verónica Barragán, Patricio Rojas-Silva, Gabriel Trueba, Michelle Grunauer, Paúl Cárdenas |
| EPI_ISL_660539 | Institute of Microbiology, Universidad San Francisco de Quito | Institute of Microbiology, Universidad San Francisco de Quito | Sully Márquez, Belén Prado-Vivar, Juan José Guadalupe, Monica Becerra-Wong, Bernardo Gutiérrez, Nabih Dahik, Verónica Barragán, Patricio Rojas-Silva, Gabriel Trueba, Michelle Grunauer, Paúl Cárdenas |
| EPI_ISL_660545 | Laboratory Medicine | Department of Laboratory Medicine, Lin-Kou Chang Gung Memorial Hospital, Taoyuan, Taiwan | Kuo-Chien Tsao, Yu-Nong Gong, Shu-Li Yang, Yi-Chun Liu, Chung-Guei Huang, Mei-Jen Hsiao, Po-Wei Huang, Cheng-Ta Yang, Cheng-Hsun Chiu, Peng-Nien Huang, Kuo-Ming Lee, Guang-Wu Chen, Shin-Ru Shih |
| EPI_ISL_660875, EPI_ISL_660876, | Gundersen Clinical Microbiology Laboratory | Kabara Cancer Research Institute | Craig S. Richmond, Paraic A. Kenny |

| | | | |
|---|---|---|---|
| EPI_ISL_660877, EPI_ISL_660878, EPI_ISL_660879, EPI_ISL_660880, EPI_ISL_660881 | | | |
| EPI_ISL_660882, EPI_ISL_660883, EPI_ISL_660884, EPI_ISL_660885, EPI_ISL_660886, EPI_ISL_660887, EPI_ISL_660888, EPI_ISL_660889, EPI_ISL_660890, EPI_ISL_660891 | Gundersen Molecular Diagnostics Laboratory | Kabara Cancer Research Institute | Craig S. Richmond, Paraic A. Kenny |
| EPI_ISL_660892, EPI_ISL_660893 | Gundersen Clinical Microbiology Laboratory | Kabara Cancer Research Institute | Craig S. Richmond, Paraic A. Kenny |
| EPI_ISL_660894, EPI_ISL_660895, EPI_ISL_660896, EPI_ISL_660897, EPI_ISL_660898, EPI_ISL_660899, EPI_ISL_660900, EPI_ISL_660901 | Gundersen Molecular Diagnostics Laboratory | Kabara Cancer Research Institute | Craig S. Richmond, Paraic A. Kenny |
| EPI_ISL_661280 | Klinisk mikrobiologi | The Public Health Agency of Sweden | Department of Microbiology, The Public Health Agency of Sweden |
| EPI_ISL_664352, EPI_ISL_664479, EPI_ISL_665186, EPI_ISL_665187, EPI_ISL_665241 | University College London Hospital | COVID-19 Genomics UK (COG-UK) Consortium | Judith Heaney, Matthew Byott, Catherine Houlihan, Dan Frampton, Stuart Kirk, Moira Spyer and Eleni Nastouli |
| EPI_ISL_665296 | Quadram Institute Bioscience | COVID-19 Genomics UK (COG-UK) Consortium | Dave J. Baker, Gemma L. Kay, Alp Aydin, Thanh Le-Viet, Steven Rudder, Ana P. Tedim, Anastasia Kolyva, Maria Diaz, Leonardo de Oliveira Martins, Nabil-Fareed Alikhan, Lizzie Meadows, Rachael Stanley, Ngozi Elumogo, Muhammed Yasir, Nicholas M. Thomson, Alexander J Trotter, Rachel Gilroy, Samuel Bloomfield, Claire Stuart, Andrew Bell, Reenesh Prakash, Samir Dervisevic, Alison E. Mather, John Wain, Mark Webber, Andrew J. Page, Justin O'Grady |
| EPI_ISL_665349 | Northumbria University / South Tees Hospitals NHS Foundation Trust / North Cumbria Integrated Care NHS Foundation Trust / North Tees and Hartlepool NHS Foundation Trust / Newcastle Hospitals NHS Foundation Trust | COVID-19 Genomics UK (COG-UK) Consortium | Darren L Smith,Andrew Nelson,Matthew Bashton,Greg R Young,Joshua Loh,John Allan,Mohammad A Tariq,Giles S Holt,Gary Black,Wen C Yew,Lynn Dover,Paul Baker,Steve Liggett,Sarah Essex,Jane Greenaway,Debra Padgett,Clive Graham,Garren Scott,Edward Barton,Emma Swindells,Brendan Payne,Jennifer Collins,Yusri Taha,Gary Eltringham |
| EPI_ISL_665391 | Centre for Enzyme Innovation, University of Portsmouth / Translational Research Laboratory, Portsmouth Hospitals NHS Trust | COVID-19 Genomics UK (COG-UK) Consortium | Angela Beckett,Yann Bourgeois,Garry Scarlett,Sharon Glaysher,Scott Elliott,Kelly Bicknell,Robert Impey,Allyson Lloyd,Sarah Wyllie,Ethan Butcher,Anoop Chauhan,Samuel Robson |
| EPI_ISL_665505 | Quadram Institute Bioscience | COVID-19 Genomics UK (COG-UK) Consortium | Dave J. Baker, Gemma L. Kay, Alp Aydin, Thanh Le-Viet, Steven Rudder, Ana P. Tedim, Anastasia Kolyva, Maria Diaz, Leonardo de Oliveira Martins, Nabil-Fareed Alikhan, Lizzie Meadows, Rachael Stanley, Ngozi Elumogo, Muhammed Yasir, Nicholas M. Thomson, Alexander J Trotter, Rachel Gilroy, Samuel Bloomfield, Claire Stuart, Andrew Bell, Reenesh Prakash, Samir Dervisevic, Alison E. Mather, John Wain, Mark Webber, Andrew J. Page, Justin O'Grady |
| EPI_ISL_665540 | Northumbria University / South Tees Hospitals NHS Foundation Trust / North Cumbria Integrated Care NHS Foundation Trust / North Tees and Hartlepool NHS Foundation Trust / Newcastle Hospitals NHS Foundation Trust | COVID-19 Genomics UK (COG-UK) Consortium | Darren L Smith,Andrew Nelson,Matthew Bashton,Greg R Young,Joshua Loh,John Allan,Mohammad A Tariq,Giles S Holt,Gary Black,Wen C Yew,Lynn Dover,Paul Baker,Steve Liggett,Sarah Essex,Jane Greenaway,Debra Padgett,Clive Graham,Garren Scott,Edward Barton,Emma Swindells,Brendan Payne,Jennifer Collins,Yusri Taha,Gary Eltringham |
| EPI_ISL_665652 | Centre for Enzyme Innovation, University of Portsmouth / Translational Research Laboratory, Portsmouth Hospitals NHS Trust | COVID-19 Genomics UK (COG-UK) Consortium | Angela Beckett,Yann Bourgeois,Garry Scarlett,Sharon Glaysher,Scott Elliott,Kelly Bicknell,Robert Impey,Allyson Lloyd,Sarah Wyllie,Ethan Butcher,Anoop Chauhan,Samuel Robson |
| EPI_ISL_665656, EPI_ISL_665671 | Quadram Institute Bioscience | COVID-19 Genomics UK (COG-UK) Consortium | Dave J. Baker, Gemma L. Kay, Alp Aydin, Thanh Le-Viet, Steven Rudder, Ana P. Tedim, Anastasia Kolyva, Maria Diaz, Leonardo de Oliveira Martins, Nabil-Fareed Alikhan, Lizzie Meadows, Rachael Stanley, Ngozi Elumogo, Muhammed Yasir, Nicholas M. Thomson, Alexander J Trotter, Rachel Gilroy, Samuel Bloomfield, Claire Stuart, Andrew Bell, Reenesh Prakash, Samir Dervisevic, Alison E. Mather, John Wain, Mark Webber, Andrew J. Page, Justin O'Grady |
| EPI_ISL_665680, EPI_ISL_665744, EPI_ISL_665787 | Northumbria University / South Tees Hospitals NHS Foundation Trust / North Cumbria Integrated Care NHS Foundation Trust / North Tees and Hartlepool NHS Foundation Trust / Newcastle Hospitals NHS Foundation Trust | COVID-19 Genomics UK (COG-UK) Consortium | Darren L Smith,Andrew Nelson,Matthew Bashton,Greg R Young,Joshua Loh,John Allan,Mohammad A Tariq,Giles S Holt,Gary Black,Wen C Yew,Lynn Dover,Paul Baker,Steve Liggett,Sarah Essex,Jane Greenaway,Debra Padgett,Clive Graham,Garren Scott,Edward Barton,Emma Swindells,Brendan Payne,Jennifer Collins,Yusri Taha,Gary Eltringham |
| EPI_ISL_665849 | Centre for Enzyme Innovation, University of Portsmouth / Translational Research Laboratory, Portsmouth Hospitals NHS Trust | COVID-19 Genomics UK (COG-UK) Consortium | Angela Beckett,Yann Bourgeois,Garry Scarlett,Sharon Glaysher,Scott Elliott,Kelly Bicknell,Robert Impey,Allyson Lloyd,Sarah Wyllie,Ethan Butcher,Anoop Chauhan,Samuel Robson |
| EPI_ISL_665868, EPI_ISL_665869, EPI_ISL_665870, EPI_ISL_665871, EPI_ISL_665872 | Northumbria University / South Tees Hospitals NHS Foundation Trust / North Cumbria Integrated Care NHS Foundation Trust / North Tees and Hartlepool NHS Foundation Trust / Newcastle Hospitals NHS Foundation Trust | COVID-19 Genomics UK (COG-UK) Consortium | Darren L Smith,Andrew Nelson,Matthew Bashton,Greg R Young,Joshua Loh,John Allan,Mohammad A Tariq,Giles S Holt,Gary Black,Wen C Yew,Lynn Dover,Paul Baker,Steve Liggett,Sarah Essex,Jane Greenaway,Debra Padgett,Clive Graham,Garren Scott,Edward Barton,Emma Swindells,Brendan Payne,Jennifer Collins,Yusri Taha,Gary Eltringham |
| EPI_ISL_665940 | Queens Medical Centre, Clinical Microbiology Department / DeepSeq Nottingham | COVID-19 Genomics UK (COG-UK) Consortium | Gemma Clark, Wendy Smith, Manjinder Khakh, Vicki M Fleming, Michelle M Lister, Hannah Howson-Wells, Jonathan Ball, Patrick McClure, Joseph Chappell, Theocharis Tsoleridis, Nadine Holmes, Matthew Carlisle, Christopher Moore, Fei Sang, Johnny Debebe, Victoria Wright, Matthew Loose |
| EPI_ISL_665957, EPI_ISL_665958, EPI_ISL_665959, EPI_ISL_665960, EPI_ISL_665961, EPI_ISL_665963 | Quadram Institute Bioscience | COVID-19 Genomics UK (COG-UK) Consortium | Dave J. Baker, Gemma L. Kay, Alp Aydin, Thanh Le-Viet, Steven Rudder, Ana P. Tedim, Anastasia Kolyva, Maria Diaz, Leonardo de Oliveira Martins, Nabil-Fareed Alikhan, Lizzie Meadows, Rachael Stanley, Ngozi Elumogo, Muhammed Yasir, Nicholas M. Thomson, Alexander J Trotter, Rachel Gilroy, Samuel Bloomfield, Claire Stuart, Andrew Bell, Reenesh Prakash, Samir Dervisevic, Alison E. Mather, John Wain, Mark Webber, Andrew J. Page, Justin O'Grady |
| EPI_ISL_666622 | Environmental and Global Health, University of Florida | Environmental and Global Health, University of Florida | Stephenson,C.J., Elbadry,M.A., Loeb,J.C., Silva,L.O., Morris,J.G., Lednicky,J.A. |
| EPI_ISL_667483, EPI_ISL_667485, EPI_ISL_667486, EPI_ISL_667487, EPI_ISL_667488, EPI_ISL_667489, EPI_ISL_667490, EPI_ISL_667491, EPI_ISL_667492, EPI_ISL_667493, EPI_ISL_667494, EPI_ISL_667495, EPI_ISL_667496, EPI_ISL_667497, EPI_ISL_667498, EPI_ISL_667499, EPI_ISL_667500, EPI_ISL_667501, EPI_ISL_667502 | | | |
| see above | OHSU Lab Services Molecular Microbiology Lab | Oregon SARS-CoV-2 Genome Sequencing Center | Brendan L. O'Connell, Ruth V. Nichols, Sally Grindstaff, Alec J. Hirsch, Donna Hansel, Guang Fan, Daniel N. Streblow, William B. Messer, Andrew C. Adey, Benjamin N. Bimber, Brian J. O'Roak |
| EPI_ISL_668406 | Vestfold Hospital, Toensberg Department of Microbiology | Norwegian Institute of Public Health, Department of Virology | Kathrine Stene-Johansen, Kamilla Heddeland Instefjord, Hilde Elshaug, Marie Paulsen Madsen, Rasmus Riis Kopperud, Hilde Vollan, Karoline Bragstad, Olav Hungnes |
| EPI_ISL_670516, EPI_ISL_670650, EPI_ISL_671230 | Department of Virus and Microbiological Special Diagnostics, Statens Serum Institut, Copenhagen, Denmark | Albertsen Lab, Department of Chemistry and Bioscience, Aalborg University, Denmark | Danish Covid-19 Genome Consortium |
| EPI_ISL_671455, EPI_ISL_671456, EPI_ISL_671457, EPI_ISL_671458, EPI_ISL_671459, EPI_ISL_671460, EPI_ISL_671461, EPI_ISL_671462, EPI_ISL_671463, EPI_ISL_671464, EPI_ISL_671465, EPI_ISL_671466, EPI_ISL_671467, EPI_ISL_671468, EPI_ISL_671469, EPI_ISL_671470, EPI_ISL_671471, EPI_ISL_671472, EPI_ISL_671473, EPI_ISL_671474, EPI_ISL_671475, EPI_ISL_671476, EPI_ISL_671477, EPI_ISL_671478, EPI_ISL_671479, EPI_ISL_671480, EPI_ISL_671481, EPI_ISL_671482, EPI_ISL_671483, EPI_ISL_671484 | | | |

| | | | |
|---|---|---|---|
| see above | University of Debrecen, Department of Medical Microbiology | National Laboratory of Virology, Szentágothai Research Centre | Endre Gábor Tóth, Balázs Somogyi, Brigitta Zana, Eszter Csoma, Ferenc Jakab, Gábor Kemenesi |
| EPI_ISL_671654, EPI_ISL_671656, EPI_ISL_671657, EPI_ISL_671658, EPI_ISL_671659, EPI_ISL_671665, EPI_ISL_671666, EPI_ISL_671667, EPI_ISL_671669 | Unity Health Toronto | Ontario Institute for Cancer Research | Ramzi Fattouh, Larissa M. Matukas, Yan Chen,Mark Downing, Trina Otterman, Karel Boissinot, Wai Sum Siu, Zhi Cui, Le Luu, Samira Mubareka, TIBDN, Ilinca Lungu, Bernard Lam, Jeremy Johns, Paul Krzyzanowski, Richard de Borja, Felicia Vincelli, Philip Zuzarte, Jared T. Simpson |
| EPI_ISL_671803 | Hospital Clínico Universitario Lozano Blesa de Zaragoza (España) | SeqCOVID-SPAIN consortium/IBV(CSIC) | Rafael Benito, Sonia Algarate, Jessica Bueno and SeqCOVID-SPAIN consortium |
| EPI_ISL_671868, EPI_ISL_671869, EPI_ISL_671870, EPI_ISL_671871, EPI_ISL_671911, EPI_ISL_671912, EPI_ISL_671913, EPI_ISL_671914, EPI_ISL_671915, EPI_ISL_671916, EPI_ISL_671928, EPI_ISL_671930 | | | |
| see above | National Virus Reference Laboratory | National Virus Reference Laboratory | Michael Carr, Gabriel Gonzalez, Jonathan Dean, Daniel Hare, Cillian F De Gascun |
| EPI_ISL_671944, EPI_ISL_671945, EPI_ISL_671965 | CHU Purpan - Laboratoire de Virologie - Institut Fédératif de Biologie | CHU Purpan - Laboratoire de Virologie - Institut Fédératif de Biologie | Latour J., Ranger N., Dubois M., Carcenac R., Harter A., Boyer P., Tremeaux P., Izopet J. |
| EPI_ISL_672031, EPI_ISL_672043, EPI_ISL_672044, EPI_ISL_672070, EPI_ISL_672080 | Santa Clara County Public Health Laboratory | Chan-Zuckerberg Biohub | CZB Cliahub Consortium |
| EPI_ISL_672110 | Orange County Public Health Lab | Chan-Zuckerberg Biohub | CZB Cliahub Consortium |
| EPI_ISL_672284, EPI_ISL_672285, EPI_ISL_672286, EPI_ISL_672287, EPI_ISL_672288, EPI_ISL_672289, EPI_ISL_672290, EPI_ISL_672291 | Santa Clara County Public Health Laboratory | Chan-Zuckerberg Biohub | CZB Cliahub Consortium |
| EPI_ISL_672374 | Orange County Public Health Lab | Chan-Zuckerberg Biohub | CZB Cliahub Consortium |
| EPI_ISL_676494 | Klinisk mikrobiologi | The Public Health Agency of Sweden | Department of Microbiology, The Public Health Agency of Sweden |
| EPI_ISL_676534 | Kalmar klinisk mikrobiologi | The Public Health Agency of Sweden | Department of Microbiology, The Public Health Agency of Sweden |
| EPI_ISL_677335, EPI_ISL_677336, EPI_ISL_677337, EPI_ISL_677338, EPI_ISL_677340, EPI_ISL_677341, EPI_ISL_677342, EPI_ISL_677347, EPI_ISL_677348, EPI_ISL_677349, EPI_ISL_677351, EPI_ISL_677422, EPI_ISL_677424, EPI_ISL_677425, EPI_ISL_677433, EPI_ISL_677447, EPI_ISL_677449, EPI_ISL_677452, EPI_ISL_677455, EPI_ISL_677459, EPI_ISL_677461, EPI_ISL_677513, EPI_ISL_677519, EPI_ISL_677521, EPI_ISL_677524, EPI_ISL_677537, EPI_ISL_677630, EPI_ISL_677631, EPI_ISL_677658 | | | |
| see above | University of Wisconsin-Madison AIDS Vaccine Research Laboratories | University of Wisconsin-Madison AIDS Vaccine Research Laboratories | Gage Moreno, Katarina Braun, et al. AIDS Vaccine Research Laboratories |
| EPI_ISL_677770, EPI_ISL_677771, EPI_ISL_677775, EPI_ISL_677776, EPI_ISL_677778, EPI_ISL_677779, EPI_ISL_677780, EPI_ISL_677781, EPI_ISL_677783, EPI_ISL_677784, EPI_ISL_677785, EPI_ISL_677786, EPI_ISL_677787, EPI_ISL_677788, EPI_ISL_677791, EPI_ISL_677792, EPI_ISL_677793, EPI_ISL_677794, EPI_ISL_677796, EPI_ISL_677800, EPI_ISL_677806, EPI_ISL_677807, EPI_ISL_677808, EPI_ISL_677809, EPI_ISL_677810, EPI_ISL_677811, EPI_ISL_677812, EPI_ISL_677813, EPI_ISL_677814, EPI_ISL_677816, EPI_ISL_677817, EPI_ISL_677819, EPI_ISL_678290, EPI_ISL_678291 | | | |
| see above | University of Szeged, Institute of Clinical Microbiology | National Laboratory of Virology, Szentágothai Research Centre | Endre Gábor Tóth, Balázs Somogyi, Brigitta, Gabriella Terhes, Ferenc Jakab, Gábor Kemenesi |
| EPI_ISL_678349, EPI_ISL_678350, EPI_ISL_678352, EPI_ISL_678353, EPI_ISL_678362, EPI_ISL_678363, EPI_ISL_678369, EPI_ISL_678370 | Area of Virology, Serology and Virology Division (SAViD), New South Wales Health Pathology Randwick | Virology Research Laboratory; Area of Virology, Serology and Virology Division (SAViD), New South Wales Health Pathology Randwick | Foster, C.; Au, J.; Ruiz Silva, M.; Deveson, I.; Bull, R.; Van Hal, S.; Rawlinson, W. |
| EPI_ISL_679412 | University College London Hospital | COVID-19 Genomics UK (COG-UK) Consortium | Judith Heaney, Matthew Byott, Catherine Houlihan, Dan Frampton, Stuart Kirk, Moira Spyer and Eleni Nastouli |
| EPI_ISL_679555, EPI_ISL_679556, EPI_ISL_679560, EPI_ISL_679563, EPI_ISL_679568 | University College London, Great Ormond Street Hospital for Children NHS Foundation Trust, Imperial College Healthcare NHS Trust | COVID-19 Genomics UK (COG-UK) Consortium | Sergi Castellano, Rachel Williams, Mark Kristiansen, Paola Resende Silva, Sunando Roy, Tony Brooks, Helena Tutill, Paola Niola, Patricia Dyal, Charlotte Williams, Leysa Forrest, Yasmin Panchbhaya, Jacqueline Findlay, Samuel Weeks, Julianne Brown, Kathryn Harris, Paul Randell, James Price, Alison Holmes, Judith Breuer |
| EPI_ISL_680290, EPI_ISL_680291, EPI_ISL_680296, EPI_ISL_680297, EPI_ISL_680316, EPI_ISL_680317, EPI_ISL_680318, EPI_ISL_680319, EPI_ISL_680320, EPI_ISL_680321, EPI_ISL_680322, EPI_ISL_680323, EPI_ISL_680324, EPI_ISL_680325, EPI_ISL_680326, EPI_ISL_680327, EPI_ISL_680328, EPI_ISL_680436, EPI_ISL_680437, EPI_ISL_680438, EPI_ISL_680439 | | | |
| see above | Regional Virus Laboratory, Belfast Health and Social Care Trust | COVID-19 Genomics UK (COG-UK) Consortium | Conall McCaughey, James McKenna, Tanya Curran, Susan Feeney, Alison Watt, Ciara Cox, Mairead Connor, Zoltan Molnar, David Simpson, Derek Fairley |
| EPI_ISL_680469, EPI_ISL_680471 | Virology Department, Royal Infirmary of Edinburgh, NHS Lothian / School of Biological Sciences, University of Edinburgh / Institute of Genetics and Molecular Medicine, University of Edinburgh | COVID-19 Genomics UK (COG-UK) Consortium | McHugh M, Dewar R, Rooke S, Gallagher M, Balcaza C, O'Toole Á, Scher E, Hill V, McCrone JT, Colquhoun R, Yu X, Jackson B, Rambaut A, Williams TC, Templeton K |
| EPI_ISL_680556, EPI_ISL_680557, EPI_ISL_680558, EPI_ISL_680559, EPI_ISL_680560, EPI_ISL_680561 | Virology Department, Sheffield Teaching Hospitals NHS Foundation Trust/Department of Infection, Immunity and Cardiovascular Disease, The Medical School, University of Sheffield | COVID-19 Genomics UK (COG-UK) Consortium | Thushan de Silva, Matthew Parker, Nikki Smith, Adri Angyal, Rebecca Brown, Luke Green, Rachel Tucker, Paul Parsons, Danielle Groves, Katie Johnson, Laura Carrilero, Alex Keeley, Dave Partridge, Matthew Wyles, Benjamin Lindsey, Mehmet Yavuz, Mohammad Raza, Cariad Evans |
| EPI_ISL_680612, EPI_ISL_680614 | Wales Specialist Virology Centre Sequencing lab: Pathogen Genomics Unit | COVID-19 Genomics UK (COG-UK) Consortium | Catherine Moore, Johnathan Evans, Laura Gifford, Malorie Perry, Simon Cottrell, Angela Marchbank, Alec Birchley, Alexander Adams, Amy Gaskin, Bree Gatica-Wilcox, Jason Coombes, Joel Southgate, Lauren Gilbert, Lee Graham, Nicole Pacchiarini, Sara Kumziene-Summerhayes, Sarah Taylor, Sophie Jones, Sara Rey, Matthew Bull, Joanne Watkins, Sally Corden, Tom Connor |
| EPI_ISL_681937, EPI_ISL_682030, EPI_ISL_682031, EPI_ISL_682032 | UPMC Clinical Microbiology Laboratory | Microbial Genomic Epidemiology Laboratory, University of Pittsburgh | Mustapha M. Mustapha, Jane W. Marsh, Dan Snyder, Marissa P. Griffith, Stephanie L. Mitchell, Vatsala R. Srinivasa, Kady D. Waggle, Chinelo Ezeonwuku, Vaughn S. Cooper, Lee H. Harrison |
| EPI_ISL_682339 | NHLS Universitas Academic | UFS Virology | PA Bester, MM Nyaga, P Nthiga, MT Mogotsi, D Goedhals, T de Oliveira |
| EPI_ISL_691669, EPI_ISL_691670, EPI_ISL_691671, EPI_ISL_691672, EPI_ISL_691673 | Servicio de Microbiología, Hospital Universitario Son Espases | SeqCOVID-SPAIN consortium/IBV(CSIC) | Carla López-Causapé, Jordi Reina, Antonio Oliver and SeqCOVID-SPAIN consortium |
| EPI_ISL_692902, EPI_ISL_692905 | Massachusetts State Public Health Laboratory | Massachusetts State Public Health Laboratory | Andrew Lang, Timelia Fink, Glen Gallagher, Sandra Smole |
| EPI_ISL_693482, EPI_ISL_693483 | Central Public Health Laboratory | National Public Health Laboratory, National Centre for Infectious Diseases | Tze Minn Mak, Sophie Octavia, Zhenyang Zhou, Esorom Daoni, Theresa Palou, Lin Cui, Raymond Tzer Pin Lin |
| EPI_ISL_693709, EPI_ISL_693752 | Delaware Public Health Laboratory | Delaware Public Health Laboratory | Gregory Hovan |
| EPI_ISL_705443, EPI_ISL_705464, EPI_ISL_705465, EPI_ISL_705466, EPI_ISL_705499, EPI_ISL_705500 | Oxford Viromics, NDM, University of Oxford; Oxford University Hospitals; Basingstoke and North Hampshire Hospital | COVID-19 Genomics UK (COG-UK) Consortium | Tanya Golubchik, David Bonsall, George Macintyre, Amy Trebes, Mariateresa de Cesare, Catrin Moore, Alex Mobbs, Anita Justice, Robert Shaw, Monique Andersson, Timothy Peto, Emma Wise, Nathan Moore, Jessica Lynch, Nick Cortes, Matilde Mori, Stephen Kidd, David Buck, John Todd, Christophe Fraser |
| EPI_ISL_705761 | Virology Department, Royal Infirmary of Edinburgh, NHS Lothian / School of Biological Sciences, University of Edinburgh / Institute of Genetics and Molecular Medicine, University of Edinburgh | COVID-19 Genomics UK (COG-UK) Consortium | McHugh M, Dewar R, Rooke S, Gallagher M, Balcaza C, O'Toole Á, Scher E, Hill V, McCrone JT, Colquhoun R, Yu X, Jackson B, Rambaut A, Williams TC, Templeton K |

| | | | |
|---|---|---|---|
| EPI_ISL_706891, EPI_ISL_706892, EPI_ISL_706893, EPI_ISL_706894, EPI_ISL_706895, EPI_ISL_706896, EPI_ISL_706897, EPI_ISL_706898, EPI_ISL_706899, EPI_ISL_706900, EPI_ISL_706901, EPI_ISL_706902, EPI_ISL_706903, EPI_ISL_706904, EPI_ISL_706905, EPI_ISL_706906, EPI_ISL_706907, EPI_ISL_706908, EPI_ISL_706909, EPI_ISL_706910, EPI_ISL_706911, EPI_ISL_706912, EPI_ISL_706913, EPI_ISL_706914, EPI_ISL_706915, EPI_ISL_706916, EPI_ISL_706917, EPI_ISL_706918, EPI_ISL_706919, EPI_ISL_706920, EPI_ISL_706921, EPI_ISL_706922, EPI_ISL_706923, EPI_ISL_706924, EPI_ISL_706925, EPI_ISL_706926, EPI_ISL_706927, EPI_ISL_706928, EPI_ISL_706929 | | | |
| see above | Oxford Viromics, NDM, University of Oxford; Oxford University Hospitals; Basingstoke and North Hampshire Hospital | COVID-19 Genomics UK (COG-UK) Consortium | Tanya Golubchik, David Bonsall, George Macintyre, Amy Trebes, Mariateresa de Cesare, Catrin Moore, Alex Mobbs, Anita Justice, Robert Shaw, Monique Andersson, Timothy Peto, Emma Wise, Nathan Moore, Jessica Lynch, Nick Cortes, Matilde Mori, Stephen Kidd, David Buck, John Todd, Christophe Fraser |
| EPI_ISL_707692 | Medical Research Center, Faculty of Medicine, Syarif Hidayatullah State Islamic University Jakarta | Medical Research Center, Faculty of Medicine, Syarif Hidayatullah State Islamic University Jakarta | Chris Adhiyanto, Laifa Hendarmin, Erike Suwarsono, Zeti Harriyati, Rini Puspitaningrum, Ferania Mela, Dennis Nurjadi |
| EPI_ISL_707695 | Los Angeles County Public Health Laboratory | Los Angeles County Public Health Laboratory | P. Hemarajata et al. |
| EPI_ISL_708405, EPI_ISL_708409, EPI_ISL_708413, EPI_ISL_708415, EPI_ISL_708419, EPI_ISL_708421 | Delaware Public Health Lab | Delaware Public Health Lab | Gregory Hovan |
| EPI_ISL_708507, EPI_ISL_708729 | Los Angeles County Public Health Laboratory | Los Angeles County Public Health Laboratory | P. Hemarajata et al. |
| EPI_ISL_708813 | World Medical Hospital | National Institute of Health, Department of Medical Sciences, Ministry of Public Health, Thailand | Pilailuk Okada; Siripaporn Phuygun; Thanutsapa Thanadachakul; Sittiporn Parnmen; Pakorn Piromtong; Warawan Wongboot; Sunthareeya Waicharoen; Malinee Chittaganpitch |
| EPI_ISL_708816, EPI_ISL_708818 | Urban Institute for Disease Prevention and Control | National Institute of Health, Department of Medical Sciences, Ministry of Public Health, Thailand | Pilailuk Okada; Siripaporn Phuygun; Thanutsapa Thanadachakul; Sittiporn Parnmen; Pakorn Piromtong; Warawan Wongboot; Sunthareeya Waicharoen; Malinee Chittaganpitch |
| EPI_ISL_710571, EPI_ISL_710572, EPI_ISL_710573, EPI_ISL_710574 | University Hospital Dubrava | Ruer Boškovic Institute; Forensic Science Centre Ivan Vueti; University of Zagreb Faculty of Science | Robert Beluži, Marina Korolija, Ana Livun, Vjekoslav Tomai, Dunja Glavaš, Maja Kuzman, Paula Štancl, Lucija Markulin, Lucija Basi, Antonela Blažekovi, Fran Boroveki, Lidija Cvetko-Krajinovi, Ivana elap, Fuad osovi, Mirjana Domazet-Lošo, Tomislav Domazet-Lošo, Valentina umlijan-Combaj, Kristina Gotovac Jerei, Jasna Kašman, Vladimir Krajinovi, Danilo Licastro, Boris Maek, Željka Maak Šafranko, Gordana Maravi Vlahoviek, Senica Pejša, Josipa Skelin, Ivan Šamija, Mario Štefanovi, Sanja Tadinac, Katarina Marija Tupek, Petra Vrabec, Rosa Karli, Kristian Vlahoviek |
| EPI_ISL_718236, EPI_ISL_718241 | Hospital | National Reference Center for Viruses of Respiratory Infections, Institut Pasteur, Paris | Marion Barbet, Sylvie Behillil, Méline Bizard, Angela Brisebarre, Camille Capel, Etienne Simon-Lorière, Vincent Enouf, Maud Vanpeene, Sylvie van der Werf, Gisèle Lagathu |
| EPI_ISL_722219, EPI_ISL_722221, EPI_ISL_722222, EPI_ISL_722223, EPI_ISL_722228, EPI_ISL_722229, EPI_ISL_722230, EPI_ISL_722231, EPI_ISL_722232, EPI_ISL_722245, EPI_ISL_722246, EPI_ISL_722248, EPI_ISL_722249, EPI_ISL_722250, EPI_ISL_722251, EPI_ISL_722252, EPI_ISL_722253, EPI_ISL_722254, EPI_ISL_722264, EPI_ISL_722265, EPI_ISL_722266, EPI_ISL_722269 | | | |
| see above | Servicio de Microbiología, Hospital Miguel Servet, Zaragoza | SeqCOVID-SPAIN consortium/IBV(CSIC) | Antonio Rezusta López, Alexander Tristancho Baró, Ana Milagro, Yolanda Gracia Grataloup, Nieves Martínez Cameo and SeqCOVID-SPAIN consortium |
| EPI_ISL_722330, EPI_ISL_722349, EPI_ISL_722425, EPI_ISL_722445, EPI_ISL_722517, EPI_ISL_722570, EPI_ISL_722591, EPI_ISL_722592, EPI_ISL_722593, EPI_ISL_722594, EPI_ISL_722595, EPI_ISL_722596 | | | |
| see above | Dutch COVID-19 response team | Erasmus Medical Center | Bas Oude Munnink, Reina Sikkema, David Nieuwenhuijse, Irina Chestakova, Anne van der Linden, Marjan Boter, Emmanuelle Munger, Corine GeurtsvanKessel, Annemiek van der Eijk, Richard Molenkamp, Marion Koopmans, on behalf of the Dutch national COVID-19 response team. |
| EPI_ISL_723492, EPI_ISL_723520 | Virginia Division of Consolidated Laboratory Services (DCLS) | Virginia Division of Consolidated Laboratory Services (DCLS) | Virginia DCLS |
| EPI_ISL_724979 | Northumbria University / South Tees Hospitals NHS Foundation Trust / North Cumbria Integrated Care NHS Foundation Trust / North Tees and Hartlepool NHS Foundation Trust / Newcastle Hospitals NHS Foundation Trust | COVID-19 Genomics UK (COG-UK) Consortium | Darren L Smith,Andrew Nelson,Matthew Bashton,Greg R Young,Joshua Loh,John Allan,Mohammad A Tariq,Giles S Holt,Gary Black,Wen C Yew,Lynn Dover,Paul Baker,Steve Liggett,Sarah Essex,Jane Greenaway,Debra Padgett,Clive Graham,Garren Scott,Edward Barton,Emma Swindells,Brendan Payne,Jennifer Collins,Yusri Taha,Gary Eltringham |
| EPI_ISL_728014, EPI_ISL_728015, EPI_ISL_728088, EPI_ISL_728128 | University of Wisconsin-Madison AIDS Vaccine Research Laboratories | University of Wisconsin-Madison AIDS Vaccine Research Laboratories | Gage Moreno, Katarina Braun, et al. AIDS Vaccine Research Laboratories |
| EPI_ISL_728160, EPI_ISL_728164 | Institute for Medical Research, Infectious Disease Research Centre, National Institutes of Health, Ministry of Health Malaysia | Institute for Medical Research, Infectious Disease Research Centre, National Institutes of Health, Ministry of Health Malaysia | Suppiah J, Kamel K, Mohd-Zawawi Z, Thayan R |
| EPI_ISL_728756 | Dutch COVID-19 response team | National Institute for Public Health and the Environment (RIVM) | Adam Meijer, Harry Vennema, Jeroen Cremer, Sharon van den Brink, Bas van der Veer, AnneMarie van den Brandt, Florian Zwagemaker, Dennis Schmitz, Chantal Reusken, on behalf of the national COVID-19 response team |
| EPI_ISL_729394, EPI_ISL_729395, EPI_ISL_729396, EPI_ISL_729612, EPI_ISL_729613, EPI_ISL_729614, EPI_ISL_729617, EPI_ISL_729618 | A. Krumbholz, Labor Dr. Krause und Kollegen MVZ GmbH, Kiel | Charité Universitätsmedizin Berlin, Institut für Virologie | Victor M Corman, Barbara Mühlemann, Jörn Beheim-Schwarzbach, Talitha Veith, Julia Schneider, Terry Jones, Christian Drosten |
| EPI_ISL_729982 | Nigeria Centre for Disease Control (NCDC) | African Centre of Excellence for Genomics of Infectious Diseases (ACEGID), Redeemer's University, Ede, Osun State, Nigeria | Oluniyi P.E. et al |
| EPI_ISL_730090, EPI_ISL_730099, EPI_ISL_730122, EPI_ISL_730125 | San Diego County Public Health Laboratory | Andersen lab at Scripps Research | SEARCH Alliance San Diego with Tracy Basler, Jovan Shephard, Brett Austin |
| EPI_ISL_730230, EPI_ISL_730231, EPI_ISL_730233, EPI_ISL_730234, EPI_ISL_730235, EPI_ISL_730236, EPI_ISL_730237, EPI_ISL_730238, EPI_ISL_730239, EPI_ISL_730243, EPI_ISL_730244, EPI_ISL_730245, EPI_ISL_730246, EPI_ISL_730247, EPI_ISL_730250, EPI_ISL_730251, EPI_ISL_730252, EPI_ISL_730256, EPI_ISL_730257, EPI_ISL_730258, EPI_ISL_730259, EPI_ISL_730260, EPI_ISL_730264, EPI_ISL_730265, EPI_ISL_730266, EPI_ISL_730267, EPI_ISL_730271, EPI_ISL_730272, EPI_ISL_730273, EPI_ISL_730274, EPI_ISL_730280, EPI_ISL_730281, EPI_ISL_730282, EPI_ISL_730283, EPI_ISL_730287, EPI_ISL_730288, EPI_ISL_730289, EPI_ISL_730290, EPI_ISL_730429, EPI_ISL_730430, EPI_ISL_730436, EPI_ISL_730443, EPI_ISL_730448, EPI_ISL_730455, EPI_ISL_730466, EPI_ISL_730474, EPI_ISL_730479, EPI_ISL_730480, EPI_ISL_730482, EPI_ISL_730483, EPI_ISL_730484, EPI_ISL_730486, EPI_ISL_730487, EPI_ISL_730491, EPI_ISL_730492, EPI_ISL_730494, EPI_ISL_730495, EPI_ISL_730499, EPI_ISL_730500, EPI_ISL_730503, EPI_ISL_730504, EPI_ISL_730509, EPI_ISL_730510, EPI_ISL_730515 | | | |
| see above | Biolab Diagnostic Laboratories | Andersen lab at Scripps Research | Issa Abu-Dayyeh, Ahmad Tibi, Lama Hussein, Lina Mohammad, Zein Naber, Amid Abdelnour with SEARCH Alliance San Diego |
| EPI_ISL_730570 | Gazi University Faculty of Medicine, Medical Virology Laboratory | Gazi University Faculty of Medicine, Medical Virology Laboratory | Erdem ahin, Hager Muftah, Selin Yiit, Shaknoza Sarzhanova, Özlem Güzel Tunçcan, Murat Dizbay, Il Fidan, Kayhan Çalar, Gülendam Bozday |
| EPI_ISL_730583, EPI_ISL_730584 | Hong Kong Sanatorium & Hospital | Hong Kong Department of Health | Mak Gannon C.K., Lam Edman T.K., Chan Rickjason C.W., Tsang Dominic N.C. |
| EPI_ISL_730606, EPI_ISL_730607, EPI_ISL_730608 | Tuen Mun Hospital | Hong Kong Department of Health | Mak Gannon C.K., Lam Edman T.K., Chan Rickjason C.W., Tsang Dominic N.C. |
| EPI_ISL_730619 | Yan Chai Hospital | Hong Kong Department of Health | Mak Gannon C.K., Lam Edman T.K., Chan Rickjason C.W., Tsang Dominic N.C. |
| EPI_ISL_733083, EPI_ISL_733084, EPI_ISL_733085, EPI_ISL_733086, EPI_ISL_733087, EPI_ISL_733088, EPI_ISL_733089, EPI_ISL_733090, EPI_ISL_733091, EPI_ISL_733092, EPI_ISL_733104, EPI_ISL_733110, EPI_ISL_733119, EPI_ISL_733120, EPI_ISL_733123, EPI_ISL_733137, EPI_ISL_733138, EPI_ISL_733140, EPI_ISL_733142, EPI_ISL_733144, EPI_ISL_733146, EPI_ISL_733150 | | | |
| see above | HELIX LLC | WHO National Influenza Centre Russian Federation | Andrey Komissarov, Artem Fadeev, Anna Ivanova, Kseniya Komissarova, Dmitry Bazhenov, Daria Danilenko, Ksenia Safina, Elena Nabieva, Georgii Bazykin, Dmitry Lioznov |
| EPI_ISL_733243, EPI_ISL_733244, EPI_ISL_733245, EPI_ISL_733246, EPI_ISL_733264, EPI_ISL_733265, EPI_ISL_733266, EPI_ISL_733267, EPI_ISL_733268, EPI_ISL_733293 | WHO National Influenza Centre Russian Federation | WHO National Influenza Centre Russian Federation | Andrey Komissarov, Artem Fadeev, Anna Ivanova, Kseniya Komissarova, Dmitry Bazhenov, Daria Danilenko, Ksenia Safina, Elena Nabieva, Georgii Bazykin, Dmitry Lioznov |
| EPI_ISL_733413, EPI_ISL_733414, | UMMC-Health | WHO National Influenza Centre Russian Federation | Andrey Komissarov, Artem Fadeev, Anna Ivanova, Kseniya Komissarova, Dmitry Bazhenov, Tatiana Platonova, Daria Danilenko, Ksenia Safina, Elena |

| | | | |
|---|---|---|---|
| EPI_ISL_733415, EPI_ISL_733416 | | | Nabieva, Georgii Bazykin, Dmitry Lioznov |
| EPI_ISL_733580, EPI_ISL_733581, EPI_ISL_733583 | Respiratory Virus Unit, National Infection Service, Public Health England | COVID-19 Genomics UK (COG-UK) Consortium | PHE Covid Sequencing Team |
| EPI_ISL_734294, EPI_ISL_734314, EPI_ISL_734315, EPI_ISL_734316, EPI_ISL_734317, EPI_ISL_734318, EPI_ISL_734319, EPI_ISL_734320, EPI_ISL_734321, EPI_ISL_734322, EPI_ISL_734323, EPI_ISL_734324, EPI_ISL_734325, EPI_ISL_734326 | | | |
| see above | Wadsworth Center, New York State Department.of Health | Wadsworth Center, New York State Department.of Health | Kirsten St. George, Daryl M. Lamson, Alexis Russel, Jonathan Plitnick, Navjot Singh, John Kelly, Sara Griesemer, Erasmus Schneider, Erica Lasek-Nesselquist |
| EPI_ISL_734487, EPI_ISL_734488, EPI_ISL_734497, EPI_ISL_735044, EPI_ISL_735045, EPI_ISL_735046, EPI_ISL_735047, EPI_ISL_735048, EPI_ISL_735049, EPI_ISL_735050, EPI_ISL_735051, EPI_ISL_735052, EPI_ISL_735053, EPI_ISL_735054, EPI_ISL_735055, EPI_ISL_735056, EPI_ISL_735057, EPI_ISL_735058, EPI_ISL_735059, EPI_ISL_735060, EPI_ISL_735061, EPI_ISL_735062, EPI_ISL_735063, EPI_ISL_735064, EPI_ISL_735065, EPI_ISL_735066, EPI_ISL_735067, EPI_ISL_735068, EPI_ISL_735069, EPI_ISL_735070, EPI_ISL_735071, EPI_ISL_735072, EPI_ISL_735073, EPI_ISL_735074, EPI_ISL_738240 | | | |
| see above | UZ Leuven, National Reference Laboratory for Coronaviruses, Laboratory Medicine, Leuven, Belgium | KU Leuven, Rega Institute, Clinical and Epidemiological Virology | Tony Wawina-Bokalanga, Joan Marti-Carerras, Bert Vanmechelen, Piet Maes |
| EPI_ISL_739366, EPI_ISL_739571 | Humboldt County Public Health Laboratory | Chan-Zuckerberg Biohub | CZB Cliahub Consortium |
| EPI_ISL_739756, EPI_ISL_739944, EPI_ISL_740003, EPI_ISL_740334 | Laboratoire national de santé, Microbiology, Virology | Laboratoire national de santé, Microbiology, Microbial Genomics Platform | Anke Wienecke-Baldacchino, Catherine Ragimbeau,Jessica Tapp, Fatu Djabi, Lise Pignon, Raoul Salmon, Tamir Abdelrahman |
| EPI_ISL_741348 | Centre for Enzyme Innovation, University of Portsmouth / Translational Research Laboratory, Portsmouth Hospitals NHS Trust | COVID-19 Genomics UK (COG-UK) Consortium | Angela Beckett,Yann Bourgeois,Garry Scarlett,Sharon Glaysher,Scott Elliott,Kelly Bicknell,Robert Impey,Allyson Lloyd,Sarah Wyllie,Ethan Butcher,Anoop Chauhan,Samuel Robson |
| EPI_ISL_741760, EPI_ISL_741762, EPI_ISL_741763, EPI_ISL_741764, EPI_ISL_741765 | Oxford Viromics, NDM, University of Oxford; Oxford University Hospitals; Basingstoke and North Hampshire Hospital | COVID-19 Genomics UK (COG-UK) Consortium | Tanya Golubchik, David Bonsall, George Macintyre, Amy Trebes, Mariateresa de Cesare, Catrin Moore, Alex Mobbs, Anita Justice, Robert Shaw, Monique Andersson, Timothy Peto, Emma Wise, Nathan Moore, Jessica Lynch, Nick Cortes, Matilde Mori, Stephen Kidd, David Buck, John Todd, Christophe Fraser |
| EPI_ISL_742260 | Centre for Enzyme Innovation, University of Portsmouth / Translational Research Laboratory, Portsmouth Hospitals NHS Trust | COVID-19 Genomics UK (COG-UK) Consortium | Angela Beckett,Yann Bourgeois,Garry Scarlett,Sharon Glaysher,Scott Elliott,Kelly Bicknell,Robert Impey,Allyson Lloyd,Sarah Wyllie,Ethan Butcher,Anoop Chauhan,Samuel Robson |
| EPI_ISL_744228, EPI_ISL_744253, EPI_ISL_744600, EPI_ISL_744993 | Laboratoire national de santé, Microbiology, Virology | Laboratoire national de santé, Microbiology, Microbial Genomics Platform | Anke Wienecke-Baldacchino, Catherine Ragimbeau,Jessica Tapp, Fatu Djabi, Lise Pignon, Raoul Salmon, Tamir Abdelrahman |
| EPI_ISL_745033 | West Java Health Laboratory | West Java Health Laboratory | Azzania Fibriani, Ema Rahmawati, Ryan Bayusantika Ristandi, Rifky Waluyajati Rachman, Cut Nur Cinthia Alamanda, Isak Solihin, Rini Robiani, Miftahul Faridl, Karimatu Khoirunnisa |
| EPI_ISL_745581, EPI_ISL_745592, EPI_ISL_745593, EPI_ISL_745623, EPI_ISL_745640, EPI_ISL_746095, EPI_ISL_746099, EPI_ISL_746102, EPI_ISL_746103, EPI_ISL_746106, EPI_ISL_746117, EPI_ISL_746118, EPI_ISL_746131, EPI_ISL_746139, EPI_ISL_746147, EPI_ISL_746162, EPI_ISL_746170, EPI_ISL_746178, EPI_ISL_746184, EPI_ISL_746224, EPI_ISL_746310 | | | |
| see above | Ginkgo Bioworks Clinical Laboratory | Utah Public Health Laboratory | Erin L. Young, Kelly Oakeson, Tara Gallagher, Michael T. Pyne, E. Susan Slechta, Melanie A. Mallory, Jeffrey B. Stevenson, Salika M. Shakir, David R. Hillyard, Malaika McKenzie-Bennett, James McGann, Jim Griffin, Keith Robison, Alex Plocik, Becky Schilling, Martha Pierson, Rebecca Littlefield, Michelle Spencer, Birgitte Simen |
| EPI_ISL_746479, EPI_ISL_746538, EPI_ISL_746539 | Genetica Molecular and Subdepartamento de Virologia ISP Chile | Instituto de Salud Publica de Chile | Javier Tognarelli, Barbara Parra, Loredana Arata, Jaime Lagos, Gisselle Barra, Patricia Bustos, Rodrigo Fasce, Andres Castillo, Jorge Fernandez |
| EPI_ISL_747041 | Respiratory Viruses Branch, Centers for Disease Control and Prevention | Respiratory Viruses Branch, Centers for Disease Control and Prevention | Queen,K., Li,Y., Tao,Y., Uehara,A., Montmayeur,A., Paden,C.R., Cook,P.W., Marine,R., Sheth,M., Wang,H., Lee,J., Tong,S. |
| EPI_ISL_747239 | Kuningan Publlic Health Office | West Java Health Laboratory; School of Life Sciences and Technology, Institut Teknologi Bandung | Azzania Fibriani, Ema Rahmawati, Ryan Bayusantika Ristandi, Rifky Waluyajati Rachman, Cut Nur Cinthia Alamanda, Isak Solihin, Rini Robiani, Miftahul Faridl, Karimatu Khoirunnisa |
| EPI_ISL_747322, EPI_ISL_747327, EPI_ISL_747332, EPI_ISL_747333, EPI_ISL_747356, EPI_ISL_747357, EPI_ISL_747358, EPI_ISL_747359, EPI_ISL_747360, EPI_ISL_747361, EPI_ISL_747362, EPI_ISL_747363, EPI_ISL_747364, EPI_ISL_747365 | | | |
| see above | Division of Emerging Infectious Diseases, Bureau of Infectious Diseases Diagnosis Control, Korea Disease Control and Prevention Agency | Division of Emerging Infectious Diseases, Bureau of Infectious Diseases Diagnosis Control, Korea Disease Control and Prevention Agency | Ae Kyung Park, Il-Hwan Kim, Heui Man Kim, Jeong-Min Kim, Namjoo Lee, Chaeyoung Lee, Sang Hee Woo, Eun-Jin Kim |
| EPI_ISL_748131 | Department of Virus and Microbiological Special Diagnostics, Statens Serum Institut, Copenhagen, Denmark | Albertsen Lab, Department of Chemistry and Bioscience, Aalborg University, Denmark | Danish Covid-19 Genome Consortium |
| EPI_ISL_751495 | CHU Purpan - Laboratoire de Virologie - Institut Fédératif de Biologie | CHU Purpan - Laboratoire de Virologie - Institut Fédératif de Biologie | Latour J., Ranger N., Dubois M., Carcenac R., Harter A., Boyer P., Tremeaux P., Izopet J. |
| EPI_ISL_752978, EPI_ISL_752979, EPI_ISL_752980, EPI_ISL_752981, EPI_ISL_752982, EPI_ISL_752983, EPI_ISL_752986, EPI_ISL_753002, EPI_ISL_753003, EPI_ISL_753009, EPI_ISL_753031, EPI_ISL_753124, EPI_ISL_753125, EPI_ISL_753126, EPI_ISL_753127, EPI_ISL_753128 | | | |
| see above | State Laboratories Division, Hawaii State Department of Health | State Laboratories Division, Hawaii State Department of Health | Pamela O'Brien, Sabrina Diemert, Drew Kuwazaki, Razvan Sultana, Edward Desmond |
| EPI_ISL_753710, EPI_ISL_753716, EPI_ISL_753737, EPI_ISL_753745, EPI_ISL_753758, EPI_ISL_753762, EPI_ISL_753773, EPI_ISL_753801, EPI_ISL_753802, EPI_ISL_753803, EPI_ISL_753815, EPI_ISL_753861, EPI_ISL_753862, EPI_ISL_753863, EPI_ISL_753864, EPI_ISL_753865, EPI_ISL_753866, EPI_ISL_753867, EPI_ISL_753868, EPI_ISL_753878, EPI_ISL_753880, EPI_ISL_753881, EPI_ISL_753889, EPI_ISL_753911, EPI_ISL_754184, EPI_ISL_754188, EPI_ISL_754189 | | | |
| see above | Charité Universitätsmedizin Berlin, Institut für Virologie/Labor Berlin | Charité Universitätsmedizin Berlin, Institut für Virologie | Victor M Corman, Jörn Beheim-Schwarzbach, Barbara Mühlemann, Julia Schneider, Talitha Veith, Terry Jones, Christian Drosten |
| EPI_ISL_754533, EPI_ISL_754534, EPI_ISL_754535, EPI_ISL_754536, EPI_ISL_754537, EPI_ISL_754538, EPI_ISL_754539 | Wadsworth Center, New York State Department.of Health | Wadsworth Center, New York State Department.of Health | Kirsten St. George, Daryl M. Lamson, Alexis Russel, Matthew Shudt, Melissa A Leisner, Jonathan Plitnick, Navjot Singh, John Kelly, Sara Griesemer, Erasmus Schneider, Erica Lasek-Nesselquist |
| EPI_ISL_754628 | University of Wisconsin-Madison AIDS Vaccine Research Laboratories | University of Wisconsin-Madison AIDS Vaccine Research Laboratories | Gage Moreno, Katarina Braun, et al. AIDS Vaccine Research Laboratories |
| EPI_ISL_754916, EPI_ISL_754990 | California Department of Public Health | California Department of Public Health | CDPH IDLB COVIDNet |
| EPI_ISL_755420, EPI_ISL_755421, EPI_ISL_755422, EPI_ISL_755423, EPI_ISL_755424, EPI_ISL_755425, EPI_ISL_755426, EPI_ISL_755427, EPI_ISL_755428, EPI_ISL_755429, EPI_ISL_755430, EPI_ISL_755431, EPI_ISL_755432, EPI_ISL_755433 | | | |
| see above | Maine Health and Environmental Testing Laboratory | Tewhey Lab, The Jackson Laboratory | Matluk,N., Dewey,H., Iosue,F., Barter,M., Lynch,R., Munger,H. and Tewhey,R. |
| EPI_ISL_756045, EPI_ISL_756046, EPI_ISL_756047, EPI_ISL_756048, EPI_ISL_756049, EPI_ISL_756050, EPI_ISL_756051, EPI_ISL_756052, EPI_ISL_756053, EPI_ISL_756054, EPI_ISL_756055, EPI_ISL_756056, EPI_ISL_756057, EPI_ISL_756058, EPI_ISL_756059, EPI_ISL_756060, EPI_ISL_756061, EPI_ISL_756062, EPI_ISL_756063, EPI_ISL_756064, EPI_ISL_756065, EPI_ISL_756066, EPI_ISL_756067, EPI_ISL_756068, EPI_ISL_756069, EPI_ISL_756070, EPI_ISL_756071, EPI_ISL_756072, EPI_ISL_756073, EPI_ISL_756074, EPI_ISL_756075, EPI_ISL_756076, EPI_ISL_756077, EPI_ISL_756078, EPI_ISL_756079, EPI_ISL_756080, EPI_ISL_756081, EPI_ISL_756082, EPI_ISL_756083, EPI_ISL_756084, EPI_ISL_756085, EPI_ISL_756086, EPI_ISL_756087, EPI_ISL_756090, EPI_ISL_756091, EPI_ISL_756092, EPI_ISL_756093, EPI_ISL_756094, EPI_ISL_756095, EPI_ISL_756096, EPI_ISL_756097, EPI_ISL_756098, EPI_ISL_756099, EPI_ISL_756100, EPI_ISL_756101, EPI_ISL_756102, EPI_ISL_756103, EPI_ISL_756104, EPI_ISL_756105, EPI_ISL_756106, EPI_ISL_756107, EPI_ISL_756108, EPI_ISL_756109, EPI_ISL_756110, EPI_ISL_756111, EPI_ISL_756112, EPI_ISL_756113, EPI_ISL_756114, EPI_ISL_756115, EPI_ISL_756116, EPI_ISL_756117, EPI_ISL_756118, EPI_ISL_756119, EPI_ISL_756120, EPI_ISL_756121, EPI_ISL_756122, EPI_ISL_756123, EPI_ISL_756124, EPI_ISL_756125 | | | |
| see above | Department of Virology and Immunology, University of Helsinki and Helsinki University Hospital, Huslab Finland | Department of Virology, Faculty of Medicine, University of Helsinki, Helsinki, Finland | Teemu Smura, Ravi Kant, Phuoc Truong, Hussein Alburkat, Hannimari Kallio-Kokko, Jenni Virtanen, Maija Suvanto, Sari Hannula, Harri Kangas, Pekka Ellonen, Olli Vapalahti |
| EPI_ISL_759968 | RSI Darus Syifa' | Institute of Tropical Disease, Universitas Airlangga | Maria I Lusida, Krisnoadi Rahardjo, Aldise M Nastri, Jezzy R Dewantari, Rima R Prasetya, HM Faiz, Gatot Soegiarto, Laksmi Wulandari, Resti Yudhawati, Soetjipto, Yasuko Mori, Kazufumi Shimizu |
| EPI_ISL_760001, EPI_ISL_760002 | Division of Emerging Infectious Diseases, Bureau of | Division of Emerging Infectious Diseases, Bureau of | Ae Kyung Park, Il-Hwan Kim, Heui Man Kim, Jeong-Min Kim, Namjoo Lee, Chaeyoung Lee, Sang Hee Woo, Eun-Jin Kim |

| | | | |
|---|---|---|---|
| | Infectious Diseases Diagnosis Control, Korea Disease Control and Prevention Agency | Infectious Diseases Diagnosis Control, Korea Disease Control and Prevention Agency | |
| EPI_ISL_760039, EPI_ISL_760040, EPI_ISL_760041, EPI_ISL_760042, EPI_ISL_760045, EPI_ISL_760046, EPI_ISL_760047 | Hong Kong Department of Health | School of Public Health, The University of Hong Kong | Daniel Chu, Haogao Gu, Pavithra Krishnan, Daisy Ng, Gigi Liu, Carrie Wan, Malik Peiris, Leo Poon |
| EPI_ISL_760115, EPI_ISL_760152, EPI_ISL_760205, EPI_ISL_760225 | Division of Emerging Infectious Diseases, Bureau of Infectious Diseases Diagnosis Control, Korea Disease Control and Prevention Agency | Division of Emerging Infectious Diseases, Bureau of Infectious Diseases Diagnosis Control, Korea Disease Control and Prevention Agency | Ae Kyung Park, Il-Hwan Kim, Heui Man Kim, Jeong-Min Kim, Namjoo Lee, Chaeyoung Lee, Sang Hee Woo, Eun-Jin Kim |
| EPI_ISL_763149, EPI_ISL_763230 | Dutch COVID-19 response team | Erasmus Medical Center | Bas Oude Munnink, Reina Sikkema, David Nieuwenhuijse, Irina Chestakova, Anne van der Linden, Marjan Boter, Emmanuelle Munger, Corine GeurtsvanKessel, Annemiek van der Eijk, Richard Molenkamp, Marion Koopmans, on behalf of the Dutch national COVID-19 response team. |
| EPI_ISL_765750 | Massachusetts General Hospital | Infectious Disease Program, Broad Institute of Harvard and MIT | Lemieux,J.E., Siddle,K.J., Shaw,B., Adams,G., Pierce,V., Turbett,S., Anahtar,M., Branda,J., Slater,D., Harris,J., Lin,A.E., Gladden-Young,A., Lagerborg,K., Rudy,M., DeRuff,K., Carter,A., Normandin,E., Bauer,M., Reilly,S., Tomkins-Tinch,C., Loreth,C., Chaluvadi,S., Neumann,A., Cusick,C., Chapman,S.B., Gnirke,A., Flowers,K., Cerrato,F., Birren,B.W., Gallagher,G., Smole,S., Park,D.J., MacInnis,B.L., Ryan,E., LaRocque,R., Rosenberg,E. and Sabeti,P.C. |
| EPI_ISL_765943, EPI_ISL_765944, EPI_ISL_765953, EPI_ISL_765954, EPI_ISL_765955 | Worobey Lab, Department of Ecology and Evolutionary Biology, University of Arizona | Worobey Lab, Department of Ecology and Evolutionary Biology, University of Arizona | Brendan Larsen, Grace Quirk, Thomas Watts, David Baltrus, Michael Worobey |
| EPI_ISL_766611, EPI_ISL_766613 | Klinisk mikrobiologi | The Public Health Agency of Sweden | Department of Microbiology, The Public Health Agency of Sweden |
| EPI_ISL_768774 | AIID | Irish Coronavirus Sequencing Consortium - National Virus Reference Laboratory | Michael Carr, Gabriel Gonzalez, Alejandro Abner Garcia Leon, Patrick Mallon |
| EPI_ISL_770000 | Area De Salud La Cruz | Inciensa, Instituto Costarricense de Investigación y Enseñanza en Nutrición y Salud | Francisco Duarte, Hebleen Porras, Claudio Soto-Garita, Estela Cordero, Adriana Godínez, Melany Calderón & Mariel López |
| EPI_ISL_770004 | Clinica Biblica | Inciensa, Instituto Costarricense de Investigación y Enseñanza en Nutrición y Salud | Francisco Duarte, Hebleen Porras, Claudio Soto-Garita, Estela Cordero, Adriana Godínez, Melany Calderón & Karla Gutiérrez-González |
| EPI_ISL_770656, EPI_ISL_770659, EPI_ISL_770664 | UMMC-Health | WHO National Influenza Centre Russian Federation | Andrey Komissarov, Artem Fadeev, Anna Ivanova, Kseniya Komissarova, Dmitry Bazhenov, Tatiana Platonova, Daria Danilenko, Ksenia Safina, Elena Nabieva, Georgii Bazykin, Dmitry Lioznov |
| EPI_ISL_770731, EPI_ISL_770732, EPI_ISL_770733, EPI_ISL_770772, EPI_ISL_770773, EPI_ISL_770774 | ZOTZ KLIMAS MVZ Düsseldorf-Centrum GbR ÜBAG für Labormedizin, Genetik, Zytologie, Pathologie | Center of Medical Microbiology, Virology, and Hospital Hygiene, University of Duesseldorf | Maximilian Damagnez, Alexander Dilthey, Ashley-Jane Duplessis, Patrick Finzer, Katrin Hoffmann, Torsten Houwaart, Lisanna Hülse, Malte Kohns Vasconcelos, Marek Korencak, Nadine Lübke, Jessica Nicolai, Klaus Pfeffer, Daniel Strelow, Jörg Timm, Andreas Walker, Tobias Wienemann, Rainer Zotz |
| EPI_ISL_775215 | Gonoshasthya-RNA Molecular Research Center | Gonoshasthya-RNA Molecular Research Center | Nihad Adnan, Mohd. Raeed Jamiruddin, Md. Ahsanul Haq, Mohib Ullah Khondoker, Nafisa Azmuda, Firoz Ahmed, Shahana Sharmin, Salma Akter, Taslin Jahan Mou, Mahfuza Marzan, Sayeda Moriam Liza, Nowshin Jahan, Tamanna Ali, Maha Jamiruddin, Mousumi Chaity, Shahad Saif Khandker, Mumtarin Jannat Oishee |
| EPI_ISL_775279 | Furst Medical Laboratory | Norwegian Institute of Public Health, Department of Virology | Kathrine Stene-Johansen, Kamilla Heddeland Instefjord, Hilde Elshaug, Atiya R Ali,Marie Paulsen Madsen, Rasmus Riis Kopperud, Hilde Vollan, Karoline Bragstad, Olav Hungnes |
| EPI_ISL_775309 | Medical Microbiology Unit, Department for Laboratory Medicine, Drammen Hospital, Vestre Viken Health Trust, | Norwegian Institute of Public Health, Department of Virology | Kathrine Stene-Johansen, Kamilla Heddeland Instefjord, Hilde Elshaug, Atiya R Ali,Marie Paulsen Madsen, Rasmus Riis Kopperud, Hilde Vollan, Karoline Bragstad, Olav Hungnes |
| EPI_ISL_775317 | Oslo University Hospital, Department of Medical Microbiology | Norwegian Institute of Public Health, Department of Virology | Kathrine Stene-Johansen, Kamilla Heddeland Instefjord, Hilde Elshaug, Atiya R Ali,Marie Paulsen Madsen, Rasmus Riis Kopperud, Hilde Vollan, Karoline Bragstad, Olav Hungnes |
| EPI_ISL_775440 | Innlandet Hospital Trust, Division Lillehammer, Department for Medical Microbiology | Norwegian Institute of Public Health, Department of Virology | Kathrine Stene-Johansen, Kamilla Heddeland Instefjord, Hilde Elshaug, Atiya R Ali,Marie Paulsen Madsen, Rasmus Riis Kopperud, Hilde Vollan, Karoline Bragstad, Olav Hungnes |
| EPI_ISL_775445, EPI_ISL_775446 | University Hospital of Northern Norway, Department for Microbiology and Infectious Disease Control | Norwegian Institute of Public Health, Department of Virology | Kathrine Stene-Johansen, Kamilla Heddeland Instefjord, Hilde Elshaug, Atiya R Ali,Marie Paulsen Madsen, Rasmus Riis Kopperud, Hilde Vollan, Karoline Bragstad, Olav Hungnes |
| EPI_ISL_775469, EPI_ISL_775470, EPI_ISL_775471 | Unilabs Laboratory Medicine | Norwegian Institute of Public Health, Department of Virology | Kathrine Stene-Johansen, Kamilla Heddeland Instefjord, Hilde Elshaug, Atiya R Ali,Marie Paulsen Madsen, Rasmus Riis Kopperud, Hilde Vollan, Karoline Bragstad, Olav Hungnes |
| EPI_ISL_778844 | ZOTZ KLIMAS MVZ Düsseldorf-Centrum GbR ÜBAG für Labormedizin, Genetik, Zytologie, Pathologie | Center of Medical Microbiology, Virology, and Hospital Hygiene, University of Duesseldorf | Maximilian Damagnez, Alexander Dilthey, Ashley-Jane Duplessis, Patrick Finzer, Katrin Hoffmann, Torsten Houwaart, Lisanna Hülse, Malte Kohns Vasconcelos, Marek Korencak, Nadine Lübke, Jessica Nicolai, Klaus Pfeffer, Daniel Strelow, Jörg Timm, Andreas Walker, Tobias Wienemann, Rainer Zotz |
| EPI_ISL_779732, EPI_ISL_779733, EPI_ISL_779736, EPI_ISL_779740, EPI_ISL_779753, EPI_ISL_779774, EPI_ISL_779775, EPI_ISL_779776, EPI_ISL_779777, EPI_ISL_779778, EPI_ISL_779779 | | | |
| see above | Servicio de Microbiología, Hospital Miguel Servet, Zaragoza | SeqCOVID-SPAIN consortium/IBV(CSIC) | Antonio Rezusta López, Alexander Tristancho Baró, Ana Milagro, Yolanda Gracia Grataloup, Nieves Martínez Cameo and SeqCOVID-SPAIN consortium |
| EPI_ISL_780068 | Hospital General Universitario Gregorio Marañón | SeqCOVID-SPAIN consortium/IBV(CSIC) | Darío García de Viedma, Laura Pérez-Lago, Marta Herranz, Jon Sicilia, Julia Suárez, Pilar Catalán, Patricia Muñoz and SeqCOVID-SPAIN consortium |
| EPI_ISL_787392, EPI_ISL_787393, EPI_ISL_787394, EPI_ISL_787395, EPI_ISL_787396, EPI_ISL_787397, EPI_ISL_787398, EPI_ISL_787399, EPI_ISL_787400, EPI_ISL_787404, EPI_ISL_787405, EPI_ISL_787406, EPI_ISL_787407, EPI_ISL_787408, EPI_ISL_787409, EPI_ISL_787410, EPI_ISL_787411, EPI_ISL_787415, EPI_ISL_787416, EPI_ISL_787417, EPI_ISL_787418, EPI_ISL_787419, EPI_ISL_787420, EPI_ISL_787421, EPI_ISL_787422, EPI_ISL_787423, EPI_ISL_787427, EPI_ISL_787428, EPI_ISL_787429, EPI_ISL_787430, EPI_ISL_787431, EPI_ISL_787432, EPI_ISL_787433, EPI_ISL_787434, EPI_ISL_787435, EPI_ISL_787439, EPI_ISL_787440, EPI_ISL_787441, EPI_ISL_787442, EPI_ISL_787443, EPI_ISL_787444, EPI_ISL_787445, EPI_ISL_787446, EPI_ISL_787447, EPI_ISL_787451, EPI_ISL_787452, EPI_ISL_787453, EPI_ISL_787454, EPI_ISL_787455, EPI_ISL_787456, EPI_ISL_787457, EPI_ISL_787458, EPI_ISL_787459, EPI_ISL_787463, EPI_ISL_787464, EPI_ISL_787465, EPI_ISL_787466, EPI_ISL_787467, EPI_ISL_787468, EPI_ISL_787469, EPI_ISL_787470, EPI_ISL_787471, EPI_ISL_787475, EPI_ISL_787476, EPI_ISL_787477, EPI_ISL_787478, EPI_ISL_787479, EPI_ISL_787480, EPI_ISL_787481, EPI_ISL_787482, EPI_ISL_787483, EPI_ISL_787484, EPI_ISL_787485, EPI_ISL_787486, EPI_ISL_787487, EPI_ISL_787488, EPI_ISL_787489, EPI_ISL_787490, EPI_ISL_787491, EPI_ISL_787492, EPI_ISL_787493, EPI_ISL_787494, EPI_ISL_787495, EPI_ISL_787496, EPI_ISL_787497, EPI_ISL_787498, EPI_ISL_787499, EPI_ISL_787500, EPI_ISL_787501, EPI_ISL_787502, EPI_ISL_787503, EPI_ISL_787504, EPI_ISL_787505, EPI_ISL_787506, EPI_ISL_787507, EPI_ISL_787508, EPI_ISL_787509, EPI_ISL_787510, EPI_ISL_787511, EPI_ISL_787512, EPI_ISL_787513, EPI_ISL_787514, EPI_ISL_787515, EPI_ISL_787516, EPI_ISL_787517, EPI_ISL_787518, EPI_ISL_787519, EPI_ISL_787520, EPI_ISL_787521, EPI_ISL_787522, EPI_ISL_787523, EPI_ISL_787524, EPI_ISL_787525, EPI_ISL_787526, EPI_ISL_787527, EPI_ISL_787528, EPI_ISL_787529, EPI_ISL_787530, EPI_ISL_787531, EPI_ISL_787532, EPI_ISL_787533, EPI_ISL_787534, EPI_ISL_787535, EPI_ISL_787536, EPI_ISL_787537, EPI_ISL_787538, EPI_ISL_787539, EPI_ISL_787540, EPI_ISL_787541, EPI_ISL_787542, EPI_ISL_787543, EPI_ISL_787544, EPI_ISL_787545, EPI_ISL_787546, EPI_ISL_787547, EPI_ISL_787548, EPI_ISL_787549, EPI_ISL_787550, EPI_ISL_787551, EPI_ISL_787552, EPI_ISL_787553, EPI_ISL_787554, EPI_ISL_787555, EPI_ISL_787556, EPI_ISL_787557, EPI_ISL_787558, EPI_ISL_787559, EPI_ISL_787560, EPI_ISL_787561, EPI_ISL_787562, EPI_ISL_787563, EPI_ISL_787564, EPI_ISL_787565, EPI_ISL_787566, EPI_ISL_787567, EPI_ISL_787568, EPI_ISL_787569, EPI_ISL_787570, EPI_ISL_787571, EPI_ISL_787572, EPI_ISL_787573, EPI_ISL_787574, EPI_ISL_787575, EPI_ISL_787576, EPI_ISL_787577, EPI_ISL_787578, EPI_ISL_787579, EPI_ISL_787580, EPI_ISL_787581, EPI_ISL_787582, EPI_ISL_787583, EPI_ISL_787584, EPI_ISL_787585, EPI_ISL_787586, EPI_ISL_787587, EPI_ISL_787588, EPI_ISL_787589, EPI_ISL_787590, EPI_ISL_787591, EPI_ISL_787592, EPI_ISL_787593, EPI_ISL_787594, EPI_ISL_787595, EPI_ISL_787596, EPI_ISL_787597, EPI_ISL_787598, EPI_ISL_787599, EPI_ISL_787600, EPI_ISL_787601, EPI_ISL_787602, EPI_ISL_787603, EPI_ISL_787604, EPI_ISL_787605, EPI_ISL_787606, EPI_ISL_787607, EPI_ISL_787608, EPI_ISL_787609, EPI_ISL_787610, EPI_ISL_787611, EPI_ISL_787612, EPI_ISL_787613, EPI_ISL_787614, EPI_ISL_787615, EPI_ISL_787616, EPI_ISL_787617, EPI_ISL_787618, EPI_ISL_787619, EPI_ISL_787620, EPI_ISL_787621, EPI_ISL_787622, EPI_ISL_787623, EPI_ISL_787624, EPI_ISL_787625, EPI_ISL_787626, EPI_ISL_787627, EPI_ISL_787628, EPI_ISL_787629, EPI_ISL_787630, EPI_ISL_787631, EPI_ISL_787632, EPI_ISL_787633, EPI_ISL_787634, EPI_ISL_787636, EPI_ISL_787637, EPI_ISL_787638, EPI_ISL_787639, EPI_ISL_787640, EPI_ISL_787641, EPI_ISL_787642, EPI_ISL_787643, EPI_ISL_787644, EPI_ISL_787645, EPI_ISL_787646, EPI_ISL_787647, EPI_ISL_787648, EPI_ISL_787649, EPI_ISL_787650, EPI_ISL_787651, EPI_ISL_787652, EPI_ISL_787653, EPI_ISL_787654, EPI_ISL_787655, EPI_ISL_787656, EPI_ISL_787657, EPI_ISL_787658, EPI_ISL_787659, EPI_ISL_787660, EPI_ISL_787661, EPI_ISL_787662, EPI_ISL_787663, EPI_ISL_787664, EPI_ISL_787665, EPI_ISL_787666, EPI_ISL_787667, EPI_ISL_787668, EPI_ISL_787669, EPI_ISL_787670, EPI_ISL_787671, EPI_ISL_787672, EPI_ISL_787673, EPI_ISL_787674, EPI_ISL_787675, EPI_ISL_787676, EPI_ISL_787677, EPI_ISL_787688, EPI_ISL_787689, EPI_ISL_787700, EPI_ISL_787701, EPI_ISL_787712, EPI_ISL_787713, EPI_ISL_787724, EPI_ISL_787736, EPI_ISL_787748, EPI_ISL_787760 | | | |
| see above | Houston Methodist Hospital | Houston Methodist Hospital | S. Wesley Long, Randall J. Olsen, Paul A. Christensen, David W. Bernard, James J. Davis, Maulik Shukla, Marcus Nguyen, Matthew Ojeda Saavedra, Prasanti Yerramilli, Layne Pruitt, Sishir Subedi, Heather Hendrickson, and James M. Musser |
| EPI_ISL_790928, EPI_ISL_790929, EPI_ISL_790943 | Dutch COVID-19 response team | National Institute for Public Health and the Environment (RIVM) | Adam Meijer, Harry Vennema, Jeroen Cremer, Sharon van den Brink, Bas van der Veer, AnneMarie van den Brandt, Florian Zwagemaker, Dennis Schmitz, Chantal Reusken, on behalf of the national COVID-19 response team |
| EPI_ISL_791989 | RSUD Waluyo Jati Kota Kraksaan, Jawa Timur | National Institute of Health Research and Development | Subangkit;Pawestri,HA;Ikawati,HD;Nugraha,AA;Puspa,KD;Muhashonah,I;Pangesti,KNA;Soekarso,T;Puspandari,N;Setiawaty,V |
| EPI_ISL_796051, EPI_ISL_796052, EPI_ISL_796053, EPI_ISL_796059, EPI_ISL_796060 | Institute of Virology, University of Cologne | Institute of Virology, University of Cologne | Saleta Sierra, Gibran Rubio, Zevanya Tessalonica, Dominik Aschenmeier, Eva Heger, Elena Knops, Rolf Kaiser, Maximilian Damagnez, Andreas Walker, Jörg Timm, Alexander Dilthey, Martin Däumer, Alex Thielen |

| | | | |
|---|---|---|---|
| EPI_ISL_801389 | Laboratorio de Ecologia de Doencas Transmissiveis na Amazonia, Instituto Leonidas e Maria Deane - Fiocruz Amazonia | Laboratorio de Ecologia de Doencas Transmissiveis na Amazonia, Instituto Leonidas e Maria Deane - Fiocruz Amazonia | Valdinete Nascimento, Victor Souza, André Corado, Fernanda Nascimento, George Silva, Ágatha Costa, Debora Duarte, Luciana Gonçalves, Maria Júlia Brandão, Michele Jesus, Felipe Naveca |
| EPI_ISL_801404 | Dutch COVID-19 response team | Erasmus Medical Center | Bas Oude Munnink, Reina Sikkema, David Nieuwenhuijse, Irina Chestakova, Anne van der Linden, Marjan Boter, Emmanuelle Munger, Corine GeurtsvanKessel, Annemiek van der Eijk, Richard Molenkamp, Marion Koopmans, on behalf of the Dutch national COVID-19 response team. |
| EPI_ISL_801869, EPI_ISL_801882, EPI_ISL_801884, EPI_ISL_801890, EPI_ISL_801919, EPI_ISL_802326, EPI_ISL_802327, EPI_ISL_802328, EPI_ISL_802329, EPI_ISL_802330, EPI_ISL_802331, EPI_ISL_802332, EPI_ISL_802333, EPI_ISL_802334, EPI_ISL_802335, EPI_ISL_802336, EPI_ISL_802337, EPI_ISL_802338, EPI_ISL_802339, EPI_ISL_802340, EPI_ISL_802341, EPI_ISL_802342, EPI_ISL_802343, EPI_ISL_802344, EPI_ISL_802345, EPI_ISL_802346, EPI_ISL_802347, EPI_ISL_802348, EPI_ISL_802349, EPI_ISL_802350, EPI_ISL_802351, EPI_ISL_802352, EPI_ISL_802353, EPI_ISL_802354, EPI_ISL_802355, EPI_ISL_802356 | | | |
| see above | MSHS Clinical Microbiology Laboratories | MSHS Pathogen Surveillance Program | Ana S. Gonzalez-Reiche, Hala Alshammary, Mitchell J. Sullivan, Brianne Ciferri, Ajay Obla, Angela Amoako, Mahmoud Awawda, Elena Hirsch, Ashley S. Salimbangon, Levy Sominsky, Katherine Beach, Kayla Russo, Charles Gleason, Shelcie Fabre, Giulio Kleiner, Zenab Khan, Bremy Alburquerque, Adriana van de Guchte, Komal Srivastava, Matthew M. Hernandez, Jayeeta Dutta, Denise Jurczyszak, Emily Ferreri, Rachel Chernet, Nancy Francoeur, Betsaida Salom Melo, Irina Oussenko, Gintaras Deikus, Juan Soto, Shwetha Hara Sridhar, Ying-Chih Wang, Kathryn Twyman, Andrew Kasarskis, Deena R. Altman, Robert Sebra, Adolfo García-Sastre, Marta Luksza, Gopi Patel, Sarah Schaefer, Melissa Gitman, Michael D. Nowak, Alberto Paniz-Mondolfi, Emilia Mia Sordillo, Viviana Simon, Harm van Bakel |
| EPI_ISL_802545, EPI_ISL_802546, EPI_ISL_802547, EPI_ISL_802548, EPI_ISL_802549, EPI_ISL_802550, EPI_ISL_802551, EPI_ISL_802552, EPI_ISL_802553, EPI_ISL_802554, EPI_ISL_802555, EPI_ISL_802556, EPI_ISL_802557, EPI_ISL_802558, EPI_ISL_802559, EPI_ISL_802560, EPI_ISL_802561, EPI_ISL_802562, EPI_ISL_802563, EPI_ISL_802564, EPI_ISL_802565, EPI_ISL_802566 | | | |
| see above | Molecular Microbiology and Food Research Laboratory (MMFRLAB) - Universidad San Sebastián | Facultad de Ciencias de la Vida, UNAB | Dayán Sanhueza, Macarena Bastías, César Echeverría, Sebastián Wolter, Waldo Díaz, Jorge Olivares, Eduardo Castro, Claudio Meneses |
| EPI_ISL_803196, EPI_ISL_803197, EPI_ISL_803198, EPI_ISL_803199, EPI_ISL_803200, EPI_ISL_803201, EPI_ISL_803202, EPI_ISL_803203, EPI_ISL_803204, EPI_ISL_803205, EPI_ISL_803206, EPI_ISL_803207, EPI_ISL_803208, EPI_ISL_803209, EPI_ISL_803210, EPI_ISL_803211, EPI_ISL_803212, EPI_ISL_803213, EPI_ISL_803214, EPI_ISL_803215, EPI_ISL_803255, EPI_ISL_803256, EPI_ISL_803257, EPI_ISL_803258, EPI_ISL_803261, EPI_ISL_803262, EPI_ISL_803263, EPI_ISL_803264, EPI_ISL_803265, EPI_ISL_803267, EPI_ISL_803268, EPI_ISL_803272, EPI_ISL_803273, EPI_ISL_803274, EPI_ISL_803275, EPI_ISL_803276, EPI_ISL_803277, EPI_ISL_803278, EPI_ISL_803279, EPI_ISL_803280, EPI_ISL_803281, EPI_ISL_803282, EPI_ISL_803283, EPI_ISL_803284, EPI_ISL_803285, EPI_ISL_803286, EPI_ISL_803287, EPI_ISL_803288, EPI_ISL_803354, EPI_ISL_803355, EPI_ISL_803356, EPI_ISL_803357, EPI_ISL_803358, EPI_ISL_803359, EPI_ISL_803378, EPI_ISL_803379, EPI_ISL_803380, EPI_ISL_803381, EPI_ISL_803577, EPI_ISL_803578, EPI_ISL_803579, EPI_ISL_803580, EPI_ISL_803581, EPI_ISL_803582, EPI_ISL_803583, EPI_ISL_803584, EPI_ISL_803585, EPI_ISL_803598, EPI_ISL_803599, EPI_ISL_803600, EPI_ISL_803602, EPI_ISL_803627, EPI_ISL_803630, EPI_ISL_803631, EPI_ISL_803632, EPI_ISL_803633, EPI_ISL_803634, EPI_ISL_803635, EPI_ISL_803636, EPI_ISL_803637, EPI_ISL_803639, EPI_ISL_803640, EPI_ISL_803641, EPI_ISL_803642, EPI_ISL_803643, EPI_ISL_803644, EPI_ISL_803646, EPI_ISL_803673, EPI_ISL_803674, EPI_ISL_803727, EPI_ISL_803728, EPI_ISL_803729, EPI_ISL_803730, EPI_ISL_803731, EPI_ISL_803732, EPI_ISL_803733, EPI_ISL_803756, EPI_ISL_803757, EPI_ISL_803758, EPI_ISL_803759, EPI_ISL_803828 | | | |
| see above | Wisconsin State Laboratory of Hygiene Communicable Disease Division | Wisconsin State Laboratory of Hygiene Communicable Disease Division | Kelsey R. Florek, Abigail C. Shockey |
| EPI_ISL_804488 | MEPHI, Aix Marseille University | MEPHI, Aix Marseille University | Anthony LEVASSEUR |
| EPI_ISL_813660, EPI_ISL_813662, EPI_ISL_813664, EPI_ISL_813665, EPI_ISL_813666, EPI_ISL_813667, EPI_ISL_813668, EPI_ISL_813669, EPI_ISL_813670, EPI_ISL_813672, EPI_ISL_813673, EPI_ISL_813674, EPI_ISL_813675, EPI_ISL_813676 | | | |
| see above | Liverpool Clinical Laboratories | COVID-19 Genomics UK (COG-UK) Consortium | Sam Haldenby, Anita Lucaci, Steve Paterson, Julian Hiscox, Alistair Darby, M Almsaud, A Alrezaihi, Muhannad Alruwaili, Stuart D Armstrong, Jones Benjamin, Eleanor G Bentley, Anu Chawla, Jordan J Clark, Angela Cowell, Richard Eccles, Isabel García-Dorival, Matthew Gemmell, Alessandro Gerada, PKF Gilmore, Richard Gregory, Ximeng Han, Catherine Hartley, Margaret Hughes, Miren Iturriza-Gomara, James Johnson, L Luu, Jenifer Manson, Charlotte Nelson, Elaine O'Toole, Cassie Olateju, Rebekah Penrice-Randal , Lucille Rainbow, N.P Randle, Trevor Ian Robinson, Parul Sharma, Ghada T Shawli, James P Stewart, Neil Swainston, Ecaterina Vamos, Joanne Watts, Mark Whitehead |
| EPI_ISL_822309 | Lighthouse Lab in Milton Keynes | Wellcome Sanger Institute for the COVID-19 Genomics UK (COG-UK) Consortium | The Lighthouse Lab in Milton Keynes and Alex Alderton, Roberto Amato, Sonia Goncalves, Ewan Harrison, David K. Jackson, Ian Johnston, Dominic Kwiatkowski, Cordelia Langford, John Sillitoe on behalf of the Wellcome Sanger Institute COVID-19 Surveillance Team |
| EPI_ISL_822310, EPI_ISL_822311, EPI_ISL_822312, EPI_ISL_822313, EPI_ISL_822314, EPI_ISL_822315, EPI_ISL_822316 | Lighthouse Lab in Cambridge | Wellcome Sanger Institute for the COVID-19 Genomics UK (COG-UK) Consortium | Rob Howes, The Lighthouse Lab in Cambridge and Alex Alderton, Roberto Amato, Sonia Goncalves, Ewan Harrison, David K. Jackson, Ian Johnston, Dominic Kwiatkowski, Cordelia Langford, John Sillitoe on behalf of the Wellcome Sanger Institute COVID-19 Surveillance Team |
| EPI_ISL_822317 | Lighthouse Lab in Alderley Park | Wellcome Sanger Institute for the COVID-19 Genomics UK (COG-UK) Consortium | Jacquelyn Wynn, Mairead Hyland, The Lighthouse Lab in Alderley Park and Alex Alderton, Roberto Amato, Sonia Goncalves, Ewan Harrison, David K. Jackson, Ian Johnston, Dominic Kwiatkowski, Cordelia Langford, John Sillitoe on behalf of the Wellcome Sanger Institute COVID-19 Surveillance Team |
| EPI_ISL_824158 | Dutch COVID-19 response team | National Institute for Public Health and the Environment (RIVM) | Adam Meijer, Harry Vennema, Jeroen Cremer, Sharon van den Brink, Bas van der Veer, AnneMarie van den Brandt, Florian Zwagemaker, Dennis Schmitz, Chantal Reusken, on behalf of the national COVID-19 response team |
| EPI_ISL_824503, EPI_ISL_824504, EPI_ISL_824505, EPI_ISL_824506, EPI_ISL_824507, EPI_ISL_824508 | Hospital Universitari Vall d'Hebron - Vall d'Hebron Institut de Rercerca | Hospital Universitari Vall d'Hebron | Cristina Andrés, Maria Piñana, Josep F Abril, Damir Garcia-Cehic, Ariadna Rando, Juliana Esperalba, Maria Gema Codina, Carla Castillo, Maria Carmen Martin, Tomàs Pumarola, Josep Quer, Andrés Antón |
| EPI_ISL_825497, EPI_ISL_825576 | Respiratory Virus Unit, National Infection Service, Public Health England | COVID-19 Genomics UK (COG-UK) Consortium | PHE Covid Sequencing Team |
| EPI_ISL_825632 | Nigeria Centre For Disease Control | National reference Laboratory, NCDC, Gaduwa, Abuja | Dr Ndodo Nnaemeka, Olusola Anuoluwapo Akanbi, Chimaobi Chukwu, Dr Omoare Adesuyi, Esebanmen Grace, Anthony Ahumibe, Naidoo Dhamari, Nwando Mba, Dr Chikwe Ihekweazu |
| EPI_ISL_826677, EPI_ISL_826678, EPI_ISL_826681, EPI_ISL_826683 | The National University Hospital of Iceland | deCODE genetics | Daniel F Gudbjartsson; Agnar Helgason; Hakon Jonsson; Olafur T Magnusson; Pall Melsted; Gudmundur L Norddahl; Jona Saemundsdottir; Asgeir Sigurdsson; Patrick Sulem; Arna B Agustsdottir; Hannes Eggertsson; Berglind Eiriksdottir; Run Fridriksdottir; Elisabet E Gardarsdottir; Gudmundur Georgsson; Olafia S Gretarsdottir; Kjartan R Gudmundsson; Thora R Gunnarsdottir; Arnaldur Gylfason; Hilma Holm; Brynjar O Jensson; Aslaug Jonasdottir; Kamilla S Josefsdottir; Thordur Kristjansson; Droplaug N Magnusdottir; Solvi Rognvaldsson; Louise le Roux; Gudrun Sigmundsdottir; Gardar Sveinbjornsson; Kristin E Sveinsdottir; Maney Sveinsdottir; Emil A Thorarensen; Bjarni Thorbjornsson; Gisli Masson; Ingileif Jonsdottir; Alma Moller; Thorolfur Gudnason; Karl G Kristinsson; Unnur Thorsteinsdottir; Kari Stefansson |
| EPI_ISL_826685, EPI_ISL_826686, EPI_ISL_826687, EPI_ISL_826688, EPI_ISL_826689, EPI_ISL_826690, EPI_ISL_826691, EPI_ISL_826692, EPI_ISL_826694, EPI_ISL_826724, EPI_ISL_826725, EPI_ISL_826732, EPI_ISL_826733, EPI_ISL_826734, EPI_ISL_826735, EPI_ISL_826736, EPI_ISL_826737, EPI_ISL_826738, EPI_ISL_826739, EPI_ISL_826851, EPI_ISL_826868, EPI_ISL_826869, EPI_ISL_826870, EPI_ISL_826872, EPI_ISL_826873, EPI_ISL_826874, EPI_ISL_826875, EPI_ISL_826877, EPI_ISL_826878, EPI_ISL_827018, EPI_ISL_827019, EPI_ISL_827020, EPI_ISL_827021, EPI_ISL_827022, EPI_ISL_827023, EPI_ISL_827024, EPI_ISL_827025, EPI_ISL_827028, EPI_ISL_827029 | | | |
| see above | deCODE genetics | deCODE genetics | Daniel F Gudbjartsson; Agnar Helgason; Hakon Jonsson; Olafur T Magnusson; Pall Melsted; Gudmundur L Norddahl; Jona Saemundsdottir; Asgeir Sigurdsson; Patrick Sulem; Arna B Agustsdottir; Hannes Eggertsson; Berglind Eiriksdottir; Run Fridriksdottir; Elisabet E Gardarsdottir; Gudmundur Georgsson; Olafia S Gretarsdottir; Kjartan R Gudmundsson; Thora R Gunnarsdottir; Arnaldur Gylfason; Hilma Holm; Brynjar O Jensson; Aslaug Jonasdottir; Kamilla S Josefsdottir; Thordur Kristjansson; Droplaug N Magnusdottir; Solvi Rognvaldsson; Louise le Roux; Gudrun Sigmundsdottir; Gardar Sveinbjornsson; Kristin E Sveinsdottir; Maney Sveinsdottir; Emil A Thorarensen; Bjarni Thorbjornsson; Gisli Masson; Ingileif Jonsdottir; Alma Moller; Thorolfur Gudnason; Karl G Kristinsson; Unnur Thorsteinsdottir; Kari Stefansson |
| EPI_ISL_827036, EPI_ISL_827037, EPI_ISL_827040 | The National University Hospital of Iceland | deCODE genetics | Daniel F Gudbjartsson; Agnar Helgason; Hakon Jonsson; Olafur T Magnusson; Pall Melsted; Gudmundur L Norddahl; Jona Saemundsdottir; Asgeir Sigurdsson; Patrick Sulem; Arna B Agustsdottir; Hannes Eggertsson; Berglind Eiriksdottir; Run Fridriksdottir; Elisabet E Gardarsdottir; Gudmundur Georgsson; Olafia S Gretarsdottir; Kjartan R Gudmundsson; Thora R Gunnarsdottir; Arnaldur Gylfason; Hilma Holm; Brynjar O Jensson; Aslaug Jonasdottir; Kamilla S Josefsdottir; Thordur Kristjansson; Droplaug N Magnusdottir; Solvi Rognvaldsson; Louise le Roux; Gudrun Sigmundsdottir; Gardar Sveinbjornsson; Kristin E Sveinsdottir; Maney Sveinsdottir; Emil A Thorarensen; Bjarni Thorbjornsson; Gisli Masson; Ingileif Jonsdottir; Alma Moller; Thorolfur Gudnason; Karl G Kristinsson; Unnur Thorsteinsdottir; Kari Stefansson |
| EPI_ISL_827131, EPI_ISL_827132, EPI_ISL_827133, EPI_ISL_827136, EPI_ISL_827137, EPI_ISL_827138, EPI_ISL_827139, EPI_ISL_827294, EPI_ISL_827295, EPI_ISL_827296, EPI_ISL_827297, EPI_ISL_827298, EPI_ISL_827299, EPI_ISL_827300, EPI_ISL_827301, EPI_ISL_827302, EPI_ISL_827303 | | | |
| see above | deCODE genetics | deCODE genetics | Daniel F Gudbjartsson; Agnar Helgason; Hakon Jonsson; Olafur T Magnusson; Pall Melsted; Gudmundur L Norddahl; Jona Saemundsdottir; Asgeir Sigurdsson; Patrick Sulem; Arna B Agustsdottir; Hannes Eggertsson; Berglind Eiriksdottir; Run Fridriksdottir; Elisabet E Gardarsdottir; Gudmundur Georgsson; Olafia S Gretarsdottir; Kjartan R Gudmundsson; Thora R Gunnarsdottir; Arnaldur Gylfason; Hilma Holm; Brynjar O Jensson; Aslaug Jonasdottir; Kamilla S |

| | | | |
|---|---|---|---|
| | | | Josefsdottir; Thordur Kristjansson; Droplaug N Magnusdottir; Solvi Rognvaldsson; Louise le Roux; Gudrun Sigmundsdottir; Gardar Sveinbjornsson; Kristin E Sveinsdottir; Maney Sveinsdottir; Emil A Thorarensen; Bjarni Thorbjornsson; Gisli Masson; Ingileif Jonsdottir; Alma Moller; Thorolfur Gudnason; Karl G Kristinsson; Unnur Thorsteinsdottir; Kari Stefansson |
| EPI_ISL_827306, EPI_ISL_827330, EPI_ISL_827333, EPI_ISL_827337 | The National University Hospital of Iceland | deCODE genetics | Daniel F Gudbjartsson; Agnar Helgason; Hakon Jonsson; Olafur T Magnusson; Pall Melsted; Gudmundur L Norddahl; Jona Saemundsdottir; Asgeir Sigurdsson; Patrick Sulem; Arna B Agustsdottir; Hannes Eggertsson; Berglind Eiriksdottir; Run Fridriksdottir; Elisabet E Gardarsdottir; Gudmundur Georgsson; Olafia S Gretarsdottir; Kjartan R Gudmundsson; Thora R Gunnarsdottir; Arnaldur Gylfason; Hilma Holm; Brynjar O Jensson; Aslaug Jonasdottir; Kamilla S Josefsdottir; Thordur Kristjansson; Droplaug N Magnusdottir; Solvi Rognvaldsson; Louise le Roux; Gudrun Sigmundsdottir; Gardar Sveinbjornsson; Kristin E Sveinsdottir; Maney Sveinsdottir; Emil A Thorarensen; Bjarni Thorbjornsson; Gisli Masson; Ingileif Jonsdottir; Alma Moller; Thorolfur Gudnason; Karl G Kristinsson; Unnur Thorsteinsdottir; Kari Stefansson |
| EPI_ISL_827366, EPI_ISL_827403, EPI_ISL_827404, EPI_ISL_827405, EPI_ISL_827406, EPI_ISL_827407, EPI_ISL_827408, EPI_ISL_827409, EPI_ISL_827410, EPI_ISL_827411, EPI_ISL_827412, EPI_ISL_827415, EPI_ISL_827429, EPI_ISL_827585 | | | |
| see above | deCODE genetics | deCODE genetics | Daniel F Gudbjartsson; Agnar Helgason; Hakon Jonsson; Olafur T Magnusson; Pall Melsted; Gudmundur L Norddahl; Jona Saemundsdottir; Asgeir Sigurdsson; Patrick Sulem; Arna B Agustsdottir; Hannes Eggertsson; Berglind Eiriksdottir; Run Fridriksdottir; Elisabet E Gardarsdottir; Gudmundur Georgsson; Olafia S Gretarsdottir; Kjartan R Gudmundsson; Thora R Gunnarsdottir; Arnaldur Gylfason; Hilma Holm; Brynjar O Jensson; Aslaug Jonasdottir; Kamilla S Josefsdottir; Thordur Kristjansson; Droplaug N Magnusdottir; Solvi Rognvaldsson; Louise le Roux; Gudrun Sigmundsdottir; Gardar Sveinbjornsson; Kristin E Sveinsdottir; Maney Sveinsdottir; Emil A Thorarensen; Bjarni Thorbjornsson; Gisli Masson; Ingileif Jonsdottir; Alma Moller; Thorolfur Gudnason; Karl G Kristinsson; Unnur Thorsteinsdottir; Kari Stefansson |
| EPI_ISL_827586 | The National University Hospital of Iceland | deCODE genetics | Daniel F Gudbjartsson; Agnar Helgason; Hakon Jonsson; Olafur T Magnusson; Pall Melsted; Gudmundur L Norddahl; Jona Saemundsdottir; Asgeir Sigurdsson; Patrick Sulem; Arna B Agustsdottir; Hannes Eggertsson; Berglind Eiriksdottir; Run Fridriksdottir; Elisabet E Gardarsdottir; Gudmundur Georgsson; Olafia S Gretarsdottir; Kjartan R Gudmundsson; Thora R Gunnarsdottir; Arnaldur Gylfason; Hilma Holm; Brynjar O Jensson; Aslaug Jonasdottir; Kamilla S Josefsdottir; Thordur Kristjansson; Droplaug N Magnusdottir; Solvi Rognvaldsson; Louise le Roux; Gudrun Sigmundsdottir; Gardar Sveinbjornsson; Kristin E Sveinsdottir; Maney Sveinsdottir; Emil A Thorarensen; Bjarni Thorbjornsson; Gisli Masson; Ingileif Jonsdottir; Alma Moller; Thorolfur Gudnason; Karl G Kristinsson; Unnur Thorsteinsdottir; Kari Stefansson |
| EPI_ISL_827591, EPI_ISL_827592, EPI_ISL_827593, EPI_ISL_827594, EPI_ISL_827595, EPI_ISL_827596, EPI_ISL_827597, EPI_ISL_827598, EPI_ISL_827623, EPI_ISL_827626 | deCODE genetics | deCODE genetics | Daniel F Gudbjartsson; Agnar Helgason; Hakon Jonsson; Olafur T Magnusson; Pall Melsted; Gudmundur L Norddahl; Jona Saemundsdottir; Asgeir Sigurdsson; Patrick Sulem; Arna B Agustsdottir; Hannes Eggertsson; Berglind Eiriksdottir; Run Fridriksdottir; Elisabet E Gardarsdottir; Gudmundur Georgsson; Olafia S Gretarsdottir; Kjartan R Gudmundsson; Thora R Gunnarsdottir; Arnaldur Gylfason; Hilma Holm; Brynjar O Jensson; Aslaug Jonasdottir; Kamilla S Josefsdottir; Thordur Kristjansson; Droplaug N Magnusdottir; Solvi Rognvaldsson; Louise le Roux; Gudrun Sigmundsdottir; Gardar Sveinbjornsson; Kristin E Sveinsdottir; Maney Sveinsdottir; Emil A Thorarensen; Bjarni Thorbjornsson; Gisli Masson; Ingileif Jonsdottir; Alma Moller; Thorolfur Gudnason; Karl G Kristinsson; Unnur Thorsteinsdottir; Kari Stefansson |
| EPI_ISL_827644 | The National University Hospital of Iceland | deCODE genetics | Daniel F Gudbjartsson; Agnar Helgason; Hakon Jonsson; Olafur T Magnusson; Pall Melsted; Gudmundur L Norddahl; Jona Saemundsdottir; Asgeir Sigurdsson; Patrick Sulem; Arna B Agustsdottir; Hannes Eggertsson; Berglind Eiriksdottir; Run Fridriksdottir; Elisabet E Gardarsdottir; Gudmundur Georgsson; Olafia S Gretarsdottir; Kjartan R Gudmundsson; Thora R Gunnarsdottir; Arnaldur Gylfason; Hilma Holm; Brynjar O Jensson; Aslaug Jonasdottir; Kamilla S Josefsdottir; Thordur Kristjansson; Droplaug N Magnusdottir; Solvi Rognvaldsson; Louise le Roux; Gudrun Sigmundsdottir; Gardar Sveinbjornsson; Kristin E Sveinsdottir; Maney Sveinsdottir; Emil A Thorarensen; Bjarni Thorbjornsson; Gisli Masson; Ingileif Jonsdottir; Alma Moller; Thorolfur Gudnason; Karl G Kristinsson; Unnur Thorsteinsdottir; Kari Stefansson |
| EPI_ISL_827648, EPI_ISL_827649, EPI_ISL_827652, EPI_ISL_827653, EPI_ISL_827654, EPI_ISL_827655, EPI_ISL_827656, EPI_ISL_827657, EPI_ISL_827658, EPI_ISL_827659, EPI_ISL_827664, EPI_ISL_827666, EPI_ISL_827669 | | | |
| see above | deCODE genetics | deCODE genetics | Daniel F Gudbjartsson; Agnar Helgason; Hakon Jonsson; Olafur T Magnusson; Pall Melsted; Gudmundur L Norddahl; Jona Saemundsdottir; Asgeir Sigurdsson; Patrick Sulem; Arna B Agustsdottir; Hannes Eggertsson; Berglind Eiriksdottir; Run Fridriksdottir; Elisabet E Gardarsdottir; Gudmundur Georgsson; Olafia S Gretarsdottir; Kjartan R Gudmundsson; Thora R Gunnarsdottir; Arnaldur Gylfason; Hilma Holm; Brynjar O Jensson; Aslaug Jonasdottir; Kamilla S Josefsdottir; Thordur Kristjansson; Droplaug N Magnusdottir; Solvi Rognvaldsson; Louise le Roux; Gudrun Sigmundsdottir; Gardar Sveinbjornsson; Kristin E Sveinsdottir; Maney Sveinsdottir; Emil A Thorarensen; Bjarni Thorbjornsson; Gisli Masson; Ingileif Jonsdottir; Alma Moller; Thorolfur Gudnason; Karl G Kristinsson; Unnur Thorsteinsdottir; Kari Stefansson |
| EPI_ISL_827693, EPI_ISL_827695 | The National University Hospital of Iceland | deCODE genetics | Daniel F Gudbjartsson; Agnar Helgason; Hakon Jonsson; Olafur T Magnusson; Pall Melsted; Gudmundur L Norddahl; Jona Saemundsdottir; Asgeir Sigurdsson; Patrick Sulem; Arna B Agustsdottir; Hannes Eggertsson; Berglind Eiriksdottir; Run Fridriksdottir; Elisabet E Gardarsdottir; Gudmundur Georgsson; Olafia S Gretarsdottir; Kjartan R Gudmundsson; Thora R Gunnarsdottir; Arnaldur Gylfason; Hilma Holm; Brynjar O Jensson; Aslaug Jonasdottir; Kamilla S Josefsdottir; Thordur Kristjansson; Droplaug N Magnusdottir; Solvi Rognvaldsson; Louise le Roux; Gudrun Sigmundsdottir; Gardar Sveinbjornsson; Kristin E Sveinsdottir; Maney Sveinsdottir; Emil A Thorarensen; Bjarni Thorbjornsson; Gisli Masson; Ingileif Jonsdottir; Alma Moller; Thorolfur Gudnason; Karl G Kristinsson; Unnur Thorsteinsdottir; Kari Stefansson |
| EPI_ISL_827744, EPI_ISL_827745, EPI_ISL_827747, EPI_ISL_827748, EPI_ISL_827749, EPI_ISL_827750, EPI_ISL_827751, EPI_ISL_827752 | deCODE genetics | deCODE genetics | Daniel F Gudbjartsson; Agnar Helgason; Hakon Jonsson; Olafur T Magnusson; Pall Melsted; Gudmundur L Norddahl; Jona Saemundsdottir; Asgeir Sigurdsson; Patrick Sulem; Arna B Agustsdottir; Hannes Eggertsson; Berglind Eiriksdottir; Run Fridriksdottir; Elisabet E Gardarsdottir; Gudmundur Georgsson; Olafia S Gretarsdottir; Kjartan R Gudmundsson; Thora R Gunnarsdottir; Arnaldur Gylfason; Hilma Holm; Brynjar O Jensson; Aslaug Jonasdottir; Kamilla S Josefsdottir; Thordur Kristjansson; Droplaug N Magnusdottir; Solvi Rognvaldsson; Louise le Roux; Gudrun Sigmundsdottir; Gardar Sveinbjornsson; Kristin E Sveinsdottir; Maney Sveinsdottir; Emil A Thorarensen; Bjarni Thorbjornsson; Gisli Masson; Ingileif Jonsdottir; Alma Moller; Thorolfur Gudnason; Karl G Kristinsson; Unnur Thorsteinsdottir; Kari Stefansson |
| EPI_ISL_827801, EPI_ISL_827895, EPI_ISL_827902, EPI_ISL_827903, EPI_ISL_827904, EPI_ISL_827905, EPI_ISL_827907, EPI_ISL_827908, EPI_ISL_827909 | The National University Hospital of Iceland | deCODE genetics | Daniel F Gudbjartsson; Agnar Helgason; Hakon Jonsson; Olafur T Magnusson; Pall Melsted; Gudmundur L Norddahl; Jona Saemundsdottir; Asgeir Sigurdsson; Patrick Sulem; Arna B Agustsdottir; Hannes Eggertsson; Berglind Eiriksdottir; Run Fridriksdottir; Elisabet E Gardarsdottir; Gudmundur Georgsson; Olafia S Gretarsdottir; Kjartan R Gudmundsson; Thora R Gunnarsdottir; Arnaldur Gylfason; Hilma Holm; Brynjar O Jensson; Aslaug Jonasdottir; Kamilla S Josefsdottir; Thordur Kristjansson; Droplaug N Magnusdottir; Solvi Rognvaldsson; Louise le Roux; Gudrun Sigmundsdottir; Gardar Sveinbjornsson; Kristin E Sveinsdottir; Maney Sveinsdottir; Emil A Thorarensen; Bjarni Thorbjornsson; Gisli Masson; Ingileif Jonsdottir; Alma Moller; Thorolfur Gudnason; Karl G Kristinsson; Unnur Thorsteinsdottir; Kari Stefansson |
| EPI_ISL_827948, EPI_ISL_827949, EPI_ISL_827954, EPI_ISL_827955, EPI_ISL_827956, EPI_ISL_827957, EPI_ISL_827958, EPI_ISL_827959, EPI_ISL_827960, EPI_ISL_827961 | deCODE genetics | deCODE genetics | Daniel F Gudbjartsson; Agnar Helgason; Hakon Jonsson; Olafur T Magnusson; Pall Melsted; Gudmundur L Norddahl; Jona Saemundsdottir; Asgeir Sigurdsson; Patrick Sulem; Arna B Agustsdottir; Hannes Eggertsson; Berglind Eiriksdottir; Run Fridriksdottir; Elisabet E Gardarsdottir; Gudmundur Georgsson; Olafia S Gretarsdottir; Kjartan R Gudmundsson; Thora R Gunnarsdottir; Arnaldur Gylfason; Hilma Holm; Brynjar O Jensson; Aslaug Jonasdottir; Kamilla S Josefsdottir; Thordur Kristjansson; Droplaug N Magnusdottir; Solvi Rognvaldsson; Louise le Roux; Gudrun Sigmundsdottir; Gardar Sveinbjornsson; Kristin E Sveinsdottir; Maney Sveinsdottir; Emil A Thorarensen; Bjarni Thorbjornsson; Gisli Masson; Ingileif Jonsdottir; Alma Moller; Thorolfur Gudnason; Karl G Kristinsson; Unnur Thorsteinsdottir; Kari Stefansson |
| EPI_ISL_828029, EPI_ISL_828030, EPI_ISL_828033 | The National University Hospital of Iceland | deCODE genetics | Daniel F Gudbjartsson; Agnar Helgason; Hakon Jonsson; Olafur T Magnusson; Pall Melsted; Gudmundur L Norddahl; Jona Saemundsdottir; Asgeir Sigurdsson; Patrick Sulem; Arna B Agustsdottir; Hannes Eggertsson; Berglind Eiriksdottir; Run Fridriksdottir; Elisabet E Gardarsdottir; Gudmundur Georgsson; Olafia S Gretarsdottir; Kjartan R Gudmundsson; Thora R Gunnarsdottir; Arnaldur Gylfason; Hilma Holm; Brynjar O Jensson; Aslaug Jonasdottir; Kamilla S Josefsdottir; Thordur Kristjansson; Droplaug N Magnusdottir; Solvi Rognvaldsson; Louise le Roux; Gudrun Sigmundsdottir; Gardar Sveinbjornsson; Kristin E Sveinsdottir; Maney Sveinsdottir; Emil A Thorarensen; Bjarni Thorbjornsson; Gisli Masson; Ingileif Jonsdottir; Alma Moller; Thorolfur Gudnason; Karl G Kristinsson; Unnur Thorsteinsdottir; Kari Stefansson |
| EPI_ISL_828034 | deCODE genetics | deCODE genetics | Daniel F Gudbjartsson; Agnar Helgason; Hakon Jonsson; Olafur T Magnusson; Pall Melsted; Gudmundur L Norddahl; Jona Saemundsdottir; Asgeir Sigurdsson; Patrick Sulem; Arna B Agustsdottir; Hannes Eggertsson; Berglind Eiriksdottir; Run Fridriksdottir; Elisabet E Gardarsdottir; Gudmundur Georgsson; Olafia S Gretarsdottir; Kjartan R Gudmundsson; Thora R Gunnarsdottir; Arnaldur Gylfason; Hilma Holm; Brynjar O Jensson; Aslaug Jonasdottir; Kamilla S Josefsdottir; Thordur Kristjansson; Droplaug N Magnusdottir; Solvi Rognvaldsson; Louise le Roux; Gudrun Sigmundsdottir; Gardar Sveinbjornsson; Kristin E Sveinsdottir; Maney Sveinsdottir; Emil A Thorarensen; Bjarni Thorbjornsson; Gisli Masson; Ingileif Jonsdottir; Alma Moller; Thorolfur Gudnason; Karl G Kristinsson; Unnur Thorsteinsdottir; Kari Stefansson |
| EPI_ISL_828037, EPI_ISL_828038 | The National University Hospital of Iceland | deCODE genetics | Daniel F Gudbjartsson; Agnar Helgason; Hakon Jonsson; Olafur T Magnusson; Pall Melsted; Gudmundur L Norddahl; Jona Saemundsdottir; Asgeir Sigurdsson; Patrick Sulem; Arna B Agustsdottir; Hannes Eggertsson; Berglind Eiriksdottir; Run Fridriksdottir; Elisabet E Gardarsdottir; Gudmundur Georgsson; Olafia S Gretarsdottir; Kjartan R Gudmundsson; Thora R Gunnarsdottir; Arnaldur Gylfason; Hilma Holm; Brynjar O Jensson; Aslaug Jonasdottir; Kamilla S |

| | | | |
|---|---|---|---|
| | | | Josefsdottir; Thordur Kristjansson; Droplaug N Magnusdottir; Solvi Rognvaldsson; Louise le Roux; Gudrun Sigmundsdottir; Gardar Sveinbjornsson; Kristin E Sveinsdottir; Maney Sveinsdottir; Emil A Thorarensen; Bjarni Thorbjornsson; Gisli Masson; Ingileif Jonsdottir; Alma Moller; Thorolfur Gudnason; Karl G Kristinsson; Unnur Thorsteinsdottir; Kari Stefansson |
| EPI_ISL_828085, EPI_ISL_828086, EPI_ISL_828095, EPI_ISL_828109, EPI_ISL_828113, EPI_ISL_828124, EPI_ISL_828145, EPI_ISL_828164, EPI_ISL_828178, EPI_ISL_828192, EPI_ISL_828206, EPI_ISL_828230, EPI_ISL_828232, EPI_ISL_828234, EPI_ISL_828235, EPI_ISL_828236, EPI_ISL_828306, EPI_ISL_828310, EPI_ISL_828312, EPI_ISL_828313, EPI_ISL_828314, EPI_ISL_828315, EPI_ISL_828317, EPI_ISL_828349, EPI_ISL_828350, EPI_ISL_828353, EPI_ISL_828354, EPI_ISL_828355, EPI_ISL_828356, EPI_ISL_828357, EPI_ISL_828358, EPI_ISL_828620, EPI_ISL_828621, EPI_ISL_828622, EPI_ISL_828623, EPI_ISL_828624, EPI_ISL_828625, EPI_ISL_828626, EPI_ISL_828627, EPI_ISL_828628, EPI_ISL_828629, EPI_ISL_828690, EPI_ISL_828692, EPI_ISL_828693, EPI_ISL_828699, EPI_ISL_828713, EPI_ISL_828725, EPI_ISL_828742, EPI_ISL_828743, EPI_ISL_828744, EPI_ISL_828745, EPI_ISL_828746, EPI_ISL_828747, EPI_ISL_828748, EPI_ISL_828749, EPI_ISL_828750, EPI_ISL_828751, EPI_ISL_828850, EPI_ISL_828851, EPI_ISL_828852, EPI_ISL_828853, EPI_ISL_828854, EPI_ISL_828855, EPI_ISL_828856, EPI_ISL_828857, EPI_ISL_828858, EPI_ISL_828859, EPI_ISL_828972, EPI_ISL_828973, EPI_ISL_828979, EPI_ISL_828980, EPI_ISL_828981, EPI_ISL_828982, EPI_ISL_828983, EPI_ISL_828984, EPI_ISL_828985, EPI_ISL_828986, EPI_ISL_828987, EPI_ISL_828988, EPI_ISL_828989, EPI_ISL_828990, EPI_ISL_828991, EPI_ISL_829221, EPI_ISL_829240, EPI_ISL_829242, EPI_ISL_829243, EPI_ISL_829245, EPI_ISL_829246, EPI_ISL_829247, EPI_ISL_829248, EPI_ISL_829251, EPI_ISL_829252, EPI_ISL_829253, EPI_ISL_829254, EPI_ISL_829262, EPI_ISL_829263, EPI_ISL_829264, EPI_ISL_829265, EPI_ISL_829266, EPI_ISL_829267, EPI_ISL_829268, EPI_ISL_829282, EPI_ISL_829283 | | | |
| see above | deCODE genetics | deCODE genetics | Daniel F Gudbjartsson; Agnar Helgason; Hakon Jonsson; Olafur T Magnusson; Pall Melsted; Gudmundur L Norddahl; Jona Saemundsdottir; Asgeir Sigurdsson; Patrick Sulem; Arna B Agustsdottir; Hannes Eggertsson; Berglind Eiriksdottir; Run Fridriksdottir; Elisabet E Gardarsdottir; Gudmundur Georgsson; Olafia S Gretarsdottir; Kjartan R Gudmundsson; Thora R Gunnarsdottir; Arnaldur Gylfason; Hilma Holm; Brynjar O Jensson; Aslaug Jonasdottir; Kamilla S Josefsdottir; Thordur Kristjansson; Droplaug N Magnusdottir; Solvi Rognvaldsson; Louise le Roux; Gudrun Sigmundsdottir; Gardar Sveinbjornsson; Kristin E Sveinsdottir; Maney Sveinsdottir; Emil A Thorarensen; Bjarni Thorbjornsson; Gisli Masson; Ingileif Jonsdottir; Alma Moller; Thorolfur Gudnason; Karl G Kristinsson; Unnur Thorsteinsdottir; Kari Stefansson |
| EPI_ISL_829369, EPI_ISL_829370, EPI_ISL_829371, EPI_ISL_829372 | The National University Hospital of Iceland | deCODE genetics | Daniel F Gudbjartsson; Agnar Helgason; Hakon Jonsson; Olafur T Magnusson; Pall Melsted; Gudmundur L Norddahl; Jona Saemundsdottir; Asgeir Sigurdsson; Patrick Sulem; Arna B Agustsdottir; Hannes Eggertsson; Berglind Eiriksdottir; Run Fridriksdottir; Elisabet E Gardarsdottir; Gudmundur Georgsson; Olafia S Gretarsdottir; Kjartan R Gudmundsson; Thora R Gunnarsdottir; Arnaldur Gylfason; Hilma Holm; Brynjar O Jensson; Aslaug Jonasdottir; Kamilla S Josefsdottir; Thordur Kristjansson; Droplaug N Magnusdottir; Solvi Rognvaldsson; Louise le Roux; Gudrun Sigmundsdottir; Gardar Sveinbjornsson; Kristin E Sveinsdottir; Maney Sveinsdottir; Emil A Thorarensen; Bjarni Thorbjornsson; Gisli Masson; Ingileif Jonsdottir; Alma Moller; Thorolfur Gudnason; Karl G Kristinsson; Unnur Thorsteinsdottir; Kari Stefansson |
| EPI_ISL_829463, EPI_ISL_829480, EPI_ISL_829481, EPI_ISL_829482, EPI_ISL_829483, EPI_ISL_829484, EPI_ISL_829485, EPI_ISL_829486, EPI_ISL_829487, EPI_ISL_829488, EPI_ISL_829489, EPI_ISL_829490, EPI_ISL_829491, EPI_ISL_829492, EPI_ISL_829493, EPI_ISL_829494, EPI_ISL_829495, EPI_ISL_829496, EPI_ISL_829497, EPI_ISL_829498, EPI_ISL_829499, EPI_ISL_829500, EPI_ISL_829501, EPI_ISL_829502, EPI_ISL_829503, EPI_ISL_829504, EPI_ISL_829505, EPI_ISL_829619, EPI_ISL_829620, EPI_ISL_829621 | | | |
| see above | deCODE genetics | deCODE genetics | Daniel F Gudbjartsson; Agnar Helgason; Hakon Jonsson; Olafur T Magnusson; Pall Melsted; Gudmundur L Norddahl; Jona Saemundsdottir; Asgeir Sigurdsson; Patrick Sulem; Arna B Agustsdottir; Hannes Eggertsson; Berglind Eiriksdottir; Run Fridriksdottir; Elisabet E Gardarsdottir; Gudmundur Georgsson; Olafia S Gretarsdottir; Kjartan R Gudmundsson; Thora R Gunnarsdottir; Arnaldur Gylfason; Hilma Holm; Brynjar O Jensson; Aslaug Jonasdottir; Kamilla S Josefsdottir; Thordur Kristjansson; Droplaug N Magnusdottir; Solvi Rognvaldsson; Louise le Roux; Gudrun Sigmundsdottir; Gardar Sveinbjornsson; Kristin E Sveinsdottir; Maney Sveinsdottir; Emil A Thorarensen; Bjarni Thorbjornsson; Gisli Masson; Ingileif Jonsdottir; Alma Moller; Thorolfur Gudnason; Karl G Kristinsson; Unnur Thorsteinsdottir; Kari Stefansson |
| EPI_ISL_829622, EPI_ISL_829623, EPI_ISL_829624, EPI_ISL_829625 | The National University Hospital of Iceland | deCODE genetics | Daniel F Gudbjartsson; Agnar Helgason; Hakon Jonsson; Olafur T Magnusson; Pall Melsted; Gudmundur L Norddahl; Jona Saemundsdottir; Asgeir Sigurdsson; Patrick Sulem; Arna B Agustsdottir; Hannes Eggertsson; Berglind Eiriksdottir; Run Fridriksdottir; Elisabet E Gardarsdottir; Gudmundur Georgsson; Olafia S Gretarsdottir; Kjartan R Gudmundsson; Thora R Gunnarsdottir; Arnaldur Gylfason; Hilma Holm; Brynjar O Jensson; Aslaug Jonasdottir; Kamilla S Josefsdottir; Thordur Kristjansson; Droplaug N Magnusdottir; Solvi Rognvaldsson; Louise le Roux; Gudrun Sigmundsdottir; Gardar Sveinbjornsson; Kristin E Sveinsdottir; Maney Sveinsdottir; Emil A Thorarensen; Bjarni Thorbjornsson; Gisli Masson; Ingileif Jonsdottir; Alma Moller; Thorolfur Gudnason; Karl G Kristinsson; Unnur Thorsteinsdottir; Kari Stefansson |
| EPI_ISL_829747, EPI_ISL_829748, EPI_ISL_829749, EPI_ISL_829750, EPI_ISL_829751, EPI_ISL_829752, EPI_ISL_829753, EPI_ISL_829754, EPI_ISL_829756, EPI_ISL_829757, EPI_ISL_829760, EPI_ISL_829763, EPI_ISL_829764 | | | |
| see above | deCODE genetics | deCODE genetics | Daniel F Gudbjartsson; Agnar Helgason; Hakon Jonsson; Olafur T Magnusson; Pall Melsted; Gudmundur L Norddahl; Jona Saemundsdottir; Asgeir Sigurdsson; Patrick Sulem; Arna B Agustsdottir; Hannes Eggertsson; Berglind Eiriksdottir; Run Fridriksdottir; Elisabet E Gardarsdottir; Gudmundur Georgsson; Olafia S Gretarsdottir; Kjartan R Gudmundsson; Thora R Gunnarsdottir; Arnaldur Gylfason; Hilma Holm; Brynjar O Jensson; Aslaug Jonasdottir; Kamilla S Josefsdottir; Thordur Kristjansson; Droplaug N Magnusdottir; Solvi Rognvaldsson; Louise le Roux; Gudrun Sigmundsdottir; Gardar Sveinbjornsson; Kristin E Sveinsdottir; Maney Sveinsdottir; Emil A Thorarensen; Bjarni Thorbjornsson; Gisli Masson; Ingileif Jonsdottir; Alma Moller; Thorolfur Gudnason; Karl G Kristinsson; Unnur Thorsteinsdottir; Kari Stefansson |
| EPI_ISL_829798, EPI_ISL_829799, EPI_ISL_829800, EPI_ISL_829801, EPI_ISL_829802, EPI_ISL_829881, EPI_ISL_829882 | The National University Hospital of Iceland | deCODE genetics | Daniel F Gudbjartsson; Agnar Helgason; Hakon Jonsson; Olafur T Magnusson; Pall Melsted; Gudmundur L Norddahl; Jona Saemundsdottir; Asgeir Sigurdsson; Patrick Sulem; Arna B Agustsdottir; Hannes Eggertsson; Berglind Eiriksdottir; Run Fridriksdottir; Elisabet E Gardarsdottir; Gudmundur Georgsson; Olafia S Gretarsdottir; Kjartan R Gudmundsson; Thora R Gunnarsdottir; Arnaldur Gylfason; Hilma Holm; Brynjar O Jensson; Aslaug Jonasdottir; Kamilla S Josefsdottir; Thordur Kristjansson; Droplaug N Magnusdottir; Solvi Rognvaldsson; Louise le Roux; Gudrun Sigmundsdottir; Gardar Sveinbjornsson; Kristin E Sveinsdottir; Maney Sveinsdottir; Emil A Thorarensen; Bjarni Thorbjornsson; Gisli Masson; Ingileif Jonsdottir; Alma Moller; Thorolfur Gudnason; Karl G Kristinsson; Unnur Thorsteinsdottir; Kari Stefansson |
| EPI_ISL_829990, EPI_ISL_829992, EPI_ISL_829993, EPI_ISL_829994, EPI_ISL_829996, EPI_ISL_829997, EPI_ISL_829998, EPI_ISL_830000, EPI_ISL_830001, EPI_ISL_830002, EPI_ISL_830003, EPI_ISL_830004, EPI_ISL_830055, EPI_ISL_830056, EPI_ISL_830057, EPI_ISL_830058, EPI_ISL_830059, EPI_ISL_830060, EPI_ISL_830179, EPI_ISL_830180, EPI_ISL_830181, EPI_ISL_830182, EPI_ISL_830195, EPI_ISL_830224, EPI_ISL_830233, EPI_ISL_830247, EPI_ISL_830258 | | | |
| see above | deCODE genetics | deCODE genetics | Daniel F Gudbjartsson; Agnar Helgason; Hakon Jonsson; Olafur T Magnusson; Pall Melsted; Gudmundur L Norddahl; Jona Saemundsdottir; Asgeir Sigurdsson; Patrick Sulem; Arna B Agustsdottir; Hannes Eggertsson; Berglind Eiriksdottir; Run Fridriksdottir; Elisabet E Gardarsdottir; Gudmundur Georgsson; Olafia S Gretarsdottir; Kjartan R Gudmundsson; Thora R Gunnarsdottir; Arnaldur Gylfason; Hilma Holm; Brynjar O Jensson; Aslaug Jonasdottir; Kamilla S Josefsdottir; Thordur Kristjansson; Droplaug N Magnusdottir; Solvi Rognvaldsson; Louise le Roux; Gudrun Sigmundsdottir; Gardar Sveinbjornsson; Kristin E Sveinsdottir; Maney Sveinsdottir; Emil A Thorarensen; Bjarni Thorbjornsson; Gisli Masson; Ingileif Jonsdottir; Alma Moller; Thorolfur Gudnason; Karl G Kristinsson; Unnur Thorsteinsdottir; Kari Stefansson |
| EPI_ISL_830307, EPI_ISL_830311 | The National University Hospital of Iceland | deCODE genetics | Daniel F Gudbjartsson; Agnar Helgason; Hakon Jonsson; Olafur T Magnusson; Pall Melsted; Gudmundur L Norddahl; Jona Saemundsdottir; Asgeir Sigurdsson; Patrick Sulem; Arna B Agustsdottir; Hannes Eggertsson; Berglind Eiriksdottir; Run Fridriksdottir; Elisabet E Gardarsdottir; Gudmundur Georgsson; Olafia S Gretarsdottir; Kjartan R Gudmundsson; Thora R Gunnarsdottir; Arnaldur Gylfason; Hilma Holm; Brynjar O Jensson; Aslaug Jonasdottir; Kamilla S Josefsdottir; Thordur Kristjansson; Droplaug N Magnusdottir; Solvi Rognvaldsson; Louise le Roux; Gudrun Sigmundsdottir; Gardar Sveinbjornsson; Kristin E Sveinsdottir; Maney Sveinsdottir; Emil A Thorarensen; Bjarni Thorbjornsson; Gisli Masson; Ingileif Jonsdottir; Alma Moller; Thorolfur Gudnason; Karl G Kristinsson; Unnur Thorsteinsdottir; Kari Stefansson |
| EPI_ISL_830322, EPI_ISL_830324, EPI_ISL_830330, EPI_ISL_830332, EPI_ISL_830336, EPI_ISL_830339, EPI_ISL_830341, EPI_ISL_830343, EPI_ISL_830345, EPI_ISL_830347, EPI_ISL_830394, EPI_ISL_830395, EPI_ISL_830399, EPI_ISL_830400, EPI_ISL_830401, EPI_ISL_830402, EPI_ISL_830403, EPI_ISL_830404, EPI_ISL_830406, EPI_ISL_830507, EPI_ISL_830508, EPI_ISL_830511, EPI_ISL_830512, EPI_ISL_830513, EPI_ISL_830514, EPI_ISL_830515, EPI_ISL_830516, EPI_ISL_830517, EPI_ISL_830518 | | | |
| see above | deCODE genetics | deCODE genetics | Daniel F Gudbjartsson; Agnar Helgason; Hakon Jonsson; Olafur T Magnusson; Pall Melsted; Gudmundur L Norddahl; Jona Saemundsdottir; Asgeir Sigurdsson; Patrick Sulem; Arna B Agustsdottir; Hannes Eggertsson; Berglind Eiriksdottir; Run Fridriksdottir; Elisabet E Gardarsdottir; Gudmundur Georgsson; Olafia S Gretarsdottir; Kjartan R Gudmundsson; Thora R Gunnarsdottir; Arnaldur Gylfason; Hilma Holm; Brynjar O Jensson; Aslaug Jonasdottir; Kamilla S Josefsdottir; Thordur Kristjansson; Droplaug N Magnusdottir; Solvi Rognvaldsson; Louise le Roux; Gudrun Sigmundsdottir; Gardar Sveinbjornsson; Kristin E Sveinsdottir; Maney Sveinsdottir; Emil A Thorarensen; Bjarni Thorbjornsson; Gisli Masson; Ingileif Jonsdottir; Alma Moller; Thorolfur Gudnason; Karl G Kristinsson; Unnur Thorsteinsdottir; Kari Stefansson |
| EPI_ISL_830731, EPI_ISL_830735, EPI_ISL_830745, EPI_ISL_830754, EPI_ISL_830755, EPI_ISL_830769, EPI_ISL_830770, EPI_ISL_830828, EPI_ISL_830829, EPI_ISL_830830, EPI_ISL_830831, EPI_ISL_830832, EPI_ISL_830833, EPI_ISL_830834, EPI_ISL_830835, EPI_ISL_830836, EPI_ISL_830837, EPI_ISL_830838, EPI_ISL_830839, EPI_ISL_830840, EPI_ISL_830841, EPI_ISL_830842, EPI_ISL_830843, EPI_ISL_830844, EPI_ISL_830845, EPI_ISL_830846, EPI_ISL_830847, EPI_ISL_830848, EPI_ISL_830849, EPI_ISL_830850, EPI_ISL_831002, EPI_ISL_831017 | | | |
| see above | University Hospital Basel, Clinical Virology | University Hospital Basel, Clinical Bacteriology | Tim Roloff, Madlen Stange, Helena MB Seth-Smith, Alfredo Mari, Karoline Leuzinger, Julia Bielicki, Manuel Battegay, Hans Hirsch, Adrian Egli |
| EPI_ISL_831034, EPI_ISL_831035, EPI_ISL_831095, EPI_ISL_831096, EPI_ISL_831097, EPI_ISL_831190, EPI_ISL_831191, EPI_ISL_831192, EPI_ISL_831193, EPI_ISL_831194, EPI_ISL_831195, EPI_ISL_831196, EPI_ISL_831197, EPI_ISL_831198, EPI_ISL_831199 | | | |
| see above | Hospital Universitario La Paz (Madrid) | SeqCOVID-SPAIN consortium/IBV(CSIC) | María Rodríguez-Tejedor, Elias Dahdouh, Fernando Lázaro-Perona, Jesús Mingorance and SeqCOVID-SPAIN consortium |
| EPI_ISL_831690, EPI_ISL_831692, EPI_ISL_831723, EPI_ISL_831764, EPI_ISL_831767, EPI_ISL_831768, | United States Air Force School of Aerospace Medicine | United States Air Force School of Aerospace Medicine | Anthony Fries, Jennifer Meyer, William Gruner, Amanda Javorina, Sarah Purves, Clarise Starr, Elizabeth Macias |

| | | | |
|---|---|---|---|
| EPI_ISL_831769 | | | |
| EPI_ISL_832414, EPI_ISL_832415, EPI_ISL_832416, EPI_ISL_832417, EPI_ISL_832418, EPI_ISL_832419, EPI_ISL_832420, EPI_ISL_832421, EPI_ISL_832424, EPI_ISL_832430, EPI_ISL_832431 | | | |
| see above | OHSU Lab Services Molecular Microbiology Lab | Oregon SARS-CoV-2 Genome Sequencing Center | Brendan L. O'Connell, Ruth V. Nichols, Sally Grindstaff, Alec J. Hirsch, Donna Hansel, Guang Fan, Daniel N. Streblow, William B. Messer, Andrew C. Adey, Benjamin N. Bimber, Brian J. O'Roak |
| EPI_ISL_842101, EPI_ISL_842102, EPI_ISL_842103, EPI_ISL_842104, EPI_ISL_842105, EPI_ISL_842106 | Quadram Institute Bioscience | COVID-19 Genomics UK (COG-UK) Consortium | Dave J. Baker, Gemma L. Kay, Alp Aydin, Thanh Le-Viet, Steven Rudder, Ana P. Tedim, Anastasia Kolyva, Maria Diaz, Leonardo de Oliveira Martins, Nabil-Fareed Alikhan, Lizzie Meadows, Rachael Stanley, Ngozi Elumogo, Muhammed Yasir, Nicholas M. Thomson, Alexander J Trotter, Rachel Gilroy, Samuel Bloomfield, Claire Stuart, Andrew Bell, Reenesh Prakash, Samir Dervisevic, Alison E. Mather, John Wain, Mark Webber, Andrew J. Page, Justin O'Grady |
| EPI_ISL_842107 | Oxford Viromics, NDM, University of Oxford; Oxford University Hospitals; Basingstoke and North Hampshire Hospital | COVID-19 Genomics UK (COG-UK) Consortium | Tanya Golubchik, David Bonsall, George Macintyre, Amy Trebes, Mariateresa de Cesare, Catrin Moore, Alex Mobbs, Anita Justice, Robert Shaw, Monique Andersson, Timothy Peto, Emma Wise, Nathan Moore, Jessica Lynch, Nick Cortes, Matilde Mori, Stephen Kidd, David Buck, John Todd, Christophe Fraser |
| EPI_ISL_842109 | Quadram Institute Bioscience | COVID-19 Genomics UK (COG-UK) Consortium | Dave J. Baker, Gemma L. Kay, Alp Aydin, Thanh Le-Viet, Steven Rudder, Ana P. Tedim, Anastasia Kolyva, Maria Diaz, Leonardo de Oliveira Martins, Nabil-Fareed Alikhan, Lizzie Meadows, Rachael Stanley, Ngozi Elumogo, Muhammed Yasir, Nicholas M. Thomson, Alexander J Trotter, Rachel Gilroy, Samuel Bloomfield, Claire Stuart, Andrew Bell, Reenesh Prakash, Samir Dervisevic, Alison E. Mather, John Wain, Mark Webber, Andrew J. Page, Justin O'Grady |
| EPI_ISL_842110 | Department of Pathology, University of Cambridge | COVID-19 Genomics UK (COG-UK) Consortium | Aminu S. Jahun, Yasmin Chaudhry, Grant Hall, Iliana Georgana, Myra Hosmillo, Martin D. Curran, Malte Pinckert, Surendra Parmar, Ian Goodfellow |
| EPI_ISL_842111, EPI_ISL_842112, EPI_ISL_842113 | Oxford Viromics, NDM, University of Oxford; Oxford University Hospitals; Basingstoke and North Hampshire Hospital | COVID-19 Genomics UK (COG-UK) Consortium | Tanya Golubchik, David Bonsall, George Macintyre, Amy Trebes, Mariateresa de Cesare, Catrin Moore, Alex Mobbs, Anita Justice, Robert Shaw, Monique Andersson, Timothy Peto, Emma Wise, Nathan Moore, Jessica Lynch, Nick Cortes, Matilde Mori, Stephen Kidd, David Buck, John Todd, Christophe Fraser |
| EPI_ISL_842114, EPI_ISL_842115 | Quadram Institute Bioscience | COVID-19 Genomics UK (COG-UK) Consortium | Dave J. Baker, Gemma L. Kay, Alp Aydin, Thanh Le-Viet, Steven Rudder, Ana P. Tedim, Anastasia Kolyva, Maria Diaz, Leonardo de Oliveira Martins, Nabil-Fareed Alikhan, Lizzie Meadows, Rachael Stanley, Ngozi Elumogo, Muhammed Yasir, Nicholas M. Thomson, Alexander J Trotter, Rachel Gilroy, Samuel Bloomfield, Claire Stuart, Andrew Bell, Reenesh Prakash, Samir Dervisevic, Alison E. Mather, John Wain, Mark Webber, Andrew J. Page, Justin O'Grady |
| EPI_ISL_842116, EPI_ISL_842117, EPI_ISL_842118 | Oxford Viromics, NDM, University of Oxford; Oxford University Hospitals; Basingstoke and North Hampshire Hospital | COVID-19 Genomics UK (COG-UK) Consortium | Tanya Golubchik, David Bonsall, George Macintyre, Amy Trebes, Mariateresa de Cesare, Catrin Moore, Alex Mobbs, Anita Justice, Robert Shaw, Monique Andersson, Timothy Peto, Emma Wise, Nathan Moore, Jessica Lynch, Nick Cortes, Matilde Mori, Stephen Kidd, David Buck, John Todd, Christophe Fraser |
| EPI_ISL_842646, EPI_ISL_842647, EPI_ISL_842648, EPI_ISL_842649 | University Hospital Basel, Clinical Virology | University Hospital Basel, Clinical Bacteriology | Tim Roloff, Madlen Stange, Helena MB Seth-Smith, Alfredo Mari, Karoline Leuzinger, Julia Bielicki, Manuel Battegay, Hans Hirsch, Adrian Egli |
| EPI_ISL_848079, EPI_ISL_848080, EPI_ISL_848082 | Laboratory of Clinical Research on Dermatozoonoses in Domestic Animals, Evandro Chagas National Institute of Infectious Diseases, Oswaldo Cruz Foundation (Fiocruz), Rio de Janeiro, RJ, Brazil. | Laboratory of Respiratory Viruses and Measles, Oswaldo Cruz Institute, FIOCRUZ | Guilherme Amaral Calvet, Michelle Fernanda Borges da Silva, Anielle de Pina Costa, Ezequias Batista Martins, Isabella Campos Vargas de Moraes, Lusiele Guaraldo, Patrícia Brasil, Sandro Antônio Pereira, Rodrigo Caldas Menezes, Isabella Dib Ferreira Gremião, Lucas Oliveira Keidel, Shanna Araujo dos Santos, Artur Augusto Velho Mendes Junior, Renato Orsini Ornellas, Maria Ogrzewalska, PaolaCristina Resende, Alex Pauvolid-Corrêa, Fernando do Couto Motta, Alice Sampaio Barreto da Rocha, Thiago C. Souza, Marilda Mendonça Siqueira |
| EPI_ISL_848320, EPI_ISL_848321, EPI_ISL_848322, EPI_ISL_848323, EPI_ISL_848324, EPI_ISL_848325, EPI_ISL_848326, EPI_ISL_848495, EPI_ISL_848496, EPI_ISL_848497 | Illinois Department of Public Health | Gagnon Lab, Southern Illinois University | Keith Gagnon |
| EPI_ISL_848960, EPI_ISL_848961, EPI_ISL_848962, EPI_ISL_848963, EPI_ISL_848964, EPI_ISL_848965, EPI_ISL_848966, EPI_ISL_848967, EPI_ISL_848968, EPI_ISL_848969, EPI_ISL_848970, EPI_ISL_848971, EPI_ISL_848972, EPI_ISL_848973, EPI_ISL_848974, EPI_ISL_848975, EPI_ISL_848976, EPI_ISL_848977, EPI_ISL_848978, EPI_ISL_848979, EPI_ISL_848980, EPI_ISL_848981, EPI_ISL_848982, EPI_ISL_848983, EPI_ISL_848984, EPI_ISL_848985, EPI_ISL_848986, EPI_ISL_848987, EPI_ISL_848988, EPI_ISL_848989, EPI_ISL_848990, EPI_ISL_848991, EPI_ISL_848992, EPI_ISL_848993, EPI_ISL_848994, EPI_ISL_848995, EPI_ISL_848996, EPI_ISL_848997, EPI_ISL_848998, EPI_ISL_848999, EPI_ISL_849000, EPI_ISL_849001, EPI_ISL_849002, EPI_ISL_849003, EPI_ISL_849004, EPI_ISL_849005, EPI_ISL_849006, EPI_ISL_849007, EPI_ISL_849008, EPI_ISL_849009, EPI_ISL_849010 | | | |
| see above | Florida Bureau of Public Health Laboratories | Florida Bureau of Public Health Laboratories | Sarah Schmedes, Jason Blanton |
| EPI_ISL_849479, EPI_ISL_849480 | Seattle Flu Study | Seattle Flu Study | Deborah A. Nickerson, Chris D. Frazar, Jover Lee, Benjamin Pelle, Matthew Richardson, Amanda Adler, Elisabeth Brandstetter, Peter D. Han, Kairsten Fay, Misja Ilcisin, Kirsten Lacombe, Thomas R. Sibley, Melissa Truong, Caitlin R. Wolf, Karen Cowgill, Stephanie Schrag, Jeff Duchin, Michael Boeckh, Janet A. Englund, Michael Famulare, Barry R. Lutz, Mark J. Rieder, Lea M. Starita, Matthew Thompson, Helen Y. Chu, Trevor Bedford, Jay Shendure |
| EPI_ISL_849481, EPI_ISL_849482 | Seattle Flu Study | Seattle Flu Study | Deborah A. Nickerson, Chris D. Frazar, Jover Lee, Benjamin Pelle, Matthew Richardson, Amanda Adler, Elisabeth Brandstetter, Peter D. Han, Kairsten Fay, Misja Ilcisin, Kirsten Lacombe, Thomas R. Sibley, Melissa Truong, Caitlin R. Wolf, Michael Boeckh, Janet A. Englund, Michael Famulare, Barry R. Lutz, Mark J. Rieder, Lea M. Starita, Matthew Thompson, Jay Shendure, Trevor Bedford, Helen Y. Chu |
| EPI_ISL_849486, EPI_ISL_849593, EPI_ISL_849596 | Seattle Flu Study | Seattle Flu Study | Deborah A. Nickerson, Chris D. Frazar, Jover Lee, Benjamin Pelle, Matthew Richardson, Amanda Adler, Elisabeth Brandstetter, Peter D. Han, Kairsten Fay, Misja Ilcisin, Kirsten Lacombe, Thomas R. Sibley, Melissa Truong, Caitlin R. Wolf, Karen Cowgill, Stephanie Schrag, Jeff Duchin, Michael Boeckh, Janet A. Englund, Michael Famulare, Barry R. Lutz, Mark J. Rieder, Lea M. Starita, Matthew Thompson, Helen Y. Chu, Trevor Bedford, Jay Shendure |
| EPI_ISL_853295, EPI_ISL_853303, EPI_ISL_853305 | UPMC Clinical Microbiology Laboratory | Microbial Genome Sequencing Center; Microbial Genomic Epidemiology Laboratory | Mustapha M. Mustapha, Jane W. Marsh, Dan Snyder, Marissa P. Griffith, Stephanie L. Mitchell, Vatsala R. Srinivasa, Kady D. Waggle, Chinelo Ezeonwuku, Vaughn S. Cooper, Lee H. Harrison |
| EPI_ISL_854230 | Department of Microbiology, University Innsbruck | Bergthaler laboratory, CeMM Research Center for Molecular Medicine of the Austrian Academy of Sciences | Lukas Endler, Alexandra Popa, Benedikt Agerer, Jakob-Wendelin Genger, Alexander Lercher, Anna Schedl, Thomas Penz, Michael Schuster, Jan Laine, Martin Senekowitsch, Christoph Bock, Andreas Bergthaler |
| EPI_ISL_857366, EPI_ISL_857367, EPI_ISL_857368, EPI_ISL_857369, EPI_ISL_857370, EPI_ISL_857371, EPI_ISL_857372, EPI_ISL_857373, EPI_ISL_857374, EPI_ISL_857375, EPI_ISL_857376, EPI_ISL_857377, EPI_ISL_857378, EPI_ISL_857379, EPI_ISL_857380, EPI_ISL_857381, EPI_ISL_857382, EPI_ISL_857383, EPI_ISL_857384, EPI_ISL_857385, EPI_ISL_857386, EPI_ISL_857387, EPI_ISL_857388, EPI_ISL_857389 | | | |
| see above | Texas Department of State Health Services (TXDSHS) | Texas Department of State Health Services (TXDSHS) | Anita Pokharel, Bonnie Oh, James Daniel Bonser, Rashmi Tuladhar, Mayela Pedrueza, Jenny Zhang, Maliha Rahman, Myong Koag, Chung Wang, Rachel Lee, Grace Kubin |
| EPI_ISL_859793 | BTC, Khalifa University | BTC, Khalifa University | Al Safar et al |
| EPI_ISL_860146 | Keio University School of Medicine | Keio University School of Medicine | Kenjiro Kosaki, Yuka Iwasaki, Hirotsugu Ishizu, Haruhiko Siomi, Kodai Abe |
| EPI_ISL_867939, EPI_ISL_867940 | Centre for Enzyme Innovation, University of Portsmouth / Translational Research Laboratory, Portsmouth Hospitals NHS Trust | COVID-19 Genomics UK (COG-UK) Consortium | Angela Beckett,Yann Bourgeois,Garry Scarlett,Sharon Glaysher,Scott Elliott,Kelly Bicknell,Robert Impey,Allyson Lloyd,Sarah Wyllie,Ethan Butcher,Anoop Chauhan,Samuel Robson |
| EPI_ISL_869244, EPI_ISL_869245, EPI_ISL_869252, EPI_ISL_869261 | UMMC-Health | WHO National Influenza Centre Russian Federation | Andrey Komissarov, Artem Fadeev, Anna Ivanova, Kseniya Komissarova, Dmitry Bazhenov, Mikhail Bakaev,Tatiana Platonova, Daria Danilenko, Ksenia Safina, Elena Nabieva, Georgii Bazykin, Dmitry Lioznov |
| EPI_ISL_871817 | Eurofins Diatherix | Hudsonalpha Genome Sequencing Center | Jane Grimwood, Melissa Williams, Lori H. Handley, Joshua Stough, Leslie Malone, Stefan Brzezinski, Ada Stewart, Teresa Jones, Jenell Webber, John T. Lovell, Jennifer Cart, and Jeremy Schmutz |
| EPI_ISL_872350, EPI_ISL_872351, EPI_ISL_872352, EPI_ISL_872353, EPI_ISL_872354, EPI_ISL_872355, EPI_ISL_872356, EPI_ISL_872357, EPI_ISL_872358, EPI_ISL_872359, EPI_ISL_872360, EPI_ISL_872361 | | | |
| see above | Texas Department of State Health Services (TXDSHS) | Texas Department of State Health Services (TXDSHS) | Bonnie Oh, Anita Pokharel, James Daniel Bonser, Myong Koag, Chung Wang, Rachel Lee, Grace Kubin, Rashmi Tuladhar, Mayela Pedrueza, Maliha Rahman, Jenny Zhang |
| EPI_ISL_872395, EPI_ISL_872396, EPI_ISL_872397, EPI_ISL_872398, EPI_ISL_872399, EPI_ISL_872400, EPI_ISL_872401, EPI_ISL_872402, EPI_ISL_872403, EPI_ISL_872404, EPI_ISL_872405, EPI_ISL_872406, EPI_ISL_872407, EPI_ISL_872408, EPI_ISL_872409, EPI_ISL_872410, EPI_ISL_872411, EPI_ISL_872412, EPI_ISL_872413, EPI_ISL_872414, EPI_ISL_872415, EPI_ISL_872416, EPI_ISL_872417, EPI_ISL_872418, EPI_ISL_872419, EPI_ISL_872420, EPI_ISL_872421, EPI_ISL_872422, EPI_ISL_872423, EPI_ISL_872424, EPI_ISL_872425, EPI_ISL_872426, EPI_ISL_872427, EPI_ISL_872428, EPI_ISL_872429, EPI_ISL_872430, EPI_ISL_872431, EPI_ISL_872432, EPI_ISL_872433, EPI_ISL_872434, EPI_ISL_872435, EPI_ISL_872436, EPI_ISL_872437, EPI_ISL_872438, EPI_ISL_872439, EPI_ISL_872440, EPI_ISL_872441, EPI_ISL_872442, EPI_ISL_872443, EPI_ISL_872444 | | | |

| | | | |
|---|---|---|---|
| see above | Eurofins Diatherix | Hudsonalpha Genome Sequencing Center | Jane Grimwood, Melissa Williams, Lori H. Handley, Joshua Stough, Leslie Malone, Stefan Brzezinski, Ada Stewart, Teresa Jones, Jenell Webber, John Lovell, Jennifer Cart, and Jeremy Schmutz |
| EPI_ISL_876879 | Quest Diagnostics | Quest Diagnostics | Rosenthal,S.H., Gerasimova,A., Kagan,R.M., Anderson, B., Hua, M., Liu Y., Bernstein, L.E., Livingston, K.E., Perez, A., Shalhout, D.F., Shlyakhter, I.A., Owen, R., Tanpaiboon, P., Lacbawan, F. |
| EPI_ISL_877424 | Institute of Microbiology and Immunology, Faculty of Medicine, University of Ljubljana | Institute of Microbiology and Immunology, Faculty of Medicine, University of Ljubljana | Samo Zakotnik, Tomaž Mark Zorec, Matic Brvar, Miša Korva, Mario Poljak, Tatjana Avši - Županc |
| EPI_ISL_882622 | COVID lab, Mymensingh Medical College | Department of Pathology, Bangladesh Agricultural University & Department of Microbiology, Mymensingh Medical College | Afrin, S. Z. Paul, S. K. Parvin, R. |
| EPI_ISL_882680, EPI_ISL_882712, EPI_ISL_882742, EPI_ISL_882747 | 1.AO Universitaria 'S. Giovanni di Dio e Ruggi D'Aragona, Scuola Medica Salernitana' Hospital / 2.UOC di Virologia e Microbiologia, Università della Campania 'L. Vanvitelli' / 3.AO Universitaria 'Federico II' Napoli Hospital / 4.AORN 'San Giuseppe Moscati' Avellino Hospital / 5.AO 'San Pio - presidio G. Rummo' Benevento Hospital / 6.AO 'Sant'Anna e San Sebastiano' Caserta Hospital / 7.PO 'Maria Santissima Addolorata' Eboli Hospital / 8.Biogem Istituto di Ricerche Genetiche | 1. Genome Research Center for Health (CRGS) / 2. Laboratory of Molecular Medicine and Genomics(LMMGe) / 3. Center for Research in Pure and Applied Mathematics (CRMPA) | Giorgio Giurato, Francesca Rizzo, Alessandro Weisz, Gianluigi Franci, Giovanni Nassa, Pasquale Pagliano, Roberta Tarallo, Elena Alexandrova, Ylenia D'Agostino, Carlo Ferravante, Jessica Lamberti, Viola Melone, Domenico Memoli, Valeria Mirici Cappa, Domenico Palumbo, Giovanni Pecoraro, Assunta Sellitto, Oriana Strianese, Ilaria Terenzi, Giuseppe Fenza, Aniello Gentile, Antonello Saccomanno, Sonia Amabile, Teresa Rocco, Annamaria Salvati, Emilia Vaccaro, Massimiliano Galdiero, Michele Cennamo, Giuseppe Portella, Maria Grazia Foti, Mariarosaria Ingino, Maria Landi, Maurizio Fumi, Vincenzo Rocco, Rita Greco, Vittoria Letizia, Arnolfo Petruzziello, Maddalena Schioppa, Gregorio Goffredi, Francesca Marciano, Michele Caraglia, Alessia Cossu, Marianna Scrima |
| EPI_ISL_884305, EPI_ISL_884333, EPI_ISL_884356, EPI_ISL_884363, EPI_ISL_884364, EPI_ISL_884389, EPI_ISL_884390, EPI_ISL_884391, EPI_ISL_884435, EPI_ISL_884436, EPI_ISL_884439, EPI_ISL_884440 | | | |
| see above | Infectious Diseases, Quest Diagnostics | Infectious Diseases, Quest Diagnostics | Rosenthal,S.H., Gerasimova,A., Kagan,R.M., Anderson,B., Bernstein,L.E., Livingston,K.E., Hua,M., Liu,Y., Shalhout,D.F., Owen,R., Lacbawan,F. |
| EPI_ISL_887112, EPI_ISL_887113, EPI_ISL_887114 | Institute of Medical Microbiology and Virology, University Hospital Carl Gustav Carus, TU Dresden | DRESDEN-concept Genome Center, CMCB, TU Dresden | Beil J., Brown T., Büttner L., Gscheidel N., Hochauf-Stange K., Klemroth S., Lindemann D., Mehnert G., Petzold A., Reinhardt S., Rost F., Sameith K., Winkler S. |
| EPI_ISL_888704, EPI_ISL_888713, EPI_ISL_888720, EPI_ISL_888731, EPI_ISL_888732, EPI_ISL_888737 | KU Leuven, Rega Institute, Clinical and Epidemiological Virology | KU Leuven, Rega Institute, Clinical and Epidemiological Virology | Tony Wawina-Bokalanga, Bert Vanmechelen, Joan Marti-Carerras, Piet Maes |
| EPI_ISL_889398, EPI_ISL_889399, EPI_ISL_889400, EPI_ISL_889401, EPI_ISL_889402, EPI_ISL_889403, EPI_ISL_889404, EPI_ISL_889405, EPI_ISL_889406, EPI_ISL_889407, EPI_ISL_889408, EPI_ISL_889409, EPI_ISL_889410, EPI_ISL_889411, EPI_ISL_889412, EPI_ISL_889413, EPI_ISL_889414, EPI_ISL_889415, EPI_ISL_889416, EPI_ISL_889417, EPI_ISL_889418, EPI_ISL_889419, EPI_ISL_889420, EPI_ISL_889421, EPI_ISL_889422, EPI_ISL_889423, EPI_ISL_889424, EPI_ISL_889425, EPI_ISL_889426, EPI_ISL_889427, EPI_ISL_889428, EPI_ISL_889429, EPI_ISL_889430, EPI_ISL_889431, EPI_ISL_889432, EPI_ISL_889433, EPI_ISL_889434, EPI_ISL_889435, EPI_ISL_889436, EPI_ISL_889437, EPI_ISL_889438, EPI_ISL_889439 | | | |
| see above | Olomouc University Hospital | Institute of Applied Biotechnologies a.s. | Petr Klempt, Ondej Brzo, Martin Kašný, Kateina Kvapilová, Marián Hajdúch, Petr Kvapil |
| EPI_ISL_890191 | Gonoshasthaya-RNA Research Center, Gonoshasthaya-RNA Molecular Diagnostics and Research Center | Gonoshasthaya-RNA Research Center, Gonoshasthaya-RNA Molecular Diagnostics and Research Center | Jamiruddin,M.R., Khondoker,M.U., Sharif,N., Azmuda,N., Ahmed,M.F., Sharmin,S., Akter,S., Mou,T.J., Marzan,M., Liza,S.M., Nahar,S., Jahan,N., Ali,T., Khandker,S.S., Jamiruddin,M., Haq,M.A., Adnan,N., Chaity,M., Oishee,M. |
| EPI_ISL_891218 | Olomouc University Hospital | Institute of Applied Biotechnologies a.s. | Petr Klempt, Ondej Brzo, Martin Kašný, Kateina Kvapilová, Marián Hajdúch, Petr Kvapil |
| EPI_ISL_891257 | COVID lab, Mymensingh Medical College | Department of Pathology, Bangladesh Agricultural University and Department of Microbiology, Mymensingh Medical College, | Afrin, S. Z. Paul, S. K. Parvin, R. |
| EPI_ISL_891261 | COVID lab, Mymensingh Medical College | Department of Pathology, Bangladesh Agricultural University and Department of Microbiology, Mymensingh Medical College | Afrin, S. Z. Paul, S. K. Parvin, R. |
| EPI_ISL_891262 | COVID lab, Mymensingh Medical College | Department of Pathology, Bangladesh Agricultural University and Department of Microbiology, Mymensingh Medical College | Afrin, S. Z. Paul, S. K. Parvin, R. |
| EPI_ISL_891268, EPI_ISL_891270 | Institute of Biocides and Medical Ecology, Belgarde, Serbia | Virology department Institute of microbiology and immunology Faculty of Medicine University of Belgrade | Banko Ana, Miljanovic Danijela, Milicevic Ognjen, Loncar Ana, Abazovic Dzihan, Despot Dragana |
| EPI_ISL_900063, EPI_ISL_900124, EPI_ISL_900151, EPI_ISL_900167, EPI_ISL_900220, EPI_ISL_900260, EPI_ISL_900266, EPI_ISL_900279, EPI_ISL_900297, EPI_ISL_900310, EPI_ISL_900358, EPI_ISL_900401, EPI_ISL_900423, EPI_ISL_900425, EPI_ISL_900452, EPI_ISL_900480 | | | |
| see above | MEPHI, Aix Marseille University | MEPHI, Aix Marseille University | Anthony LEVASSEUR |
| EPI_ISL_904426 | Dutch COVID-19 response team | Erasmus Medical Center | Bas Oude Munnink, Reina Sikkema, David Nieuwenhuijse, Irina Chestakova, Anne van der Linden, Marjan Boter, Emmanuelle Munger, Corine GeurtsvanKessel, Annemiek van der Eijk, Richard Molenkamp, Marion Koopmans, on behalf of the Dutch national COVID-19 response team. |
| EPI_ISL_904939 | Vilnius University Hospital Santaros Klinikos, Vilnius University | Institute of Biotechnology, Life Sciences Center, Vilnius University | Emilija Vasiliunaite, Milda Norkiene, Albertas Timinskas, Alma Gedvilaite, Aurelija Zvirbliene, Daniel Naumovas, Laimonas Griskevicius |
| EPI_ISL_909654 | General Hospital - Tetovo | Research Center for Genetic Engineering and Biotechnology "Georgi D. Efremov" , Macedonian Academy of Sciences and Arts | RCGEB - MASA |
| EPI_ISL_909736 | General Hospital - Struga | Research Center for Genetic Engineering and Biotechnology "Georgi D. Efremov" , Macedonian Academy of Sciences and Arts | RCGEB - MASA |
| EPI_ISL_909738 | General Hospital - Bitola | Research Center for Genetic Engineering and Biotechnology "Georgi D. Efremov" , Macedonian Academ | RCGEB - MASA |
| EPI_ISL_909739 | General Hospital - Prilep | Research Center for Genetic Engineering and Biotechnology "Georgi D. Efremov" , Macedonian Academy of Sciences and Arts | RCGEB - MASA |
| EPI_ISL_909740 | General Hospital - Ohrid | Research Center for Genetic Engineering and Biotechnology "Georgi D. Efremov" , Macedonian Academy of Sciences and Arts | RCGEB - MASA |
| EPI_ISL_909742 | General Hospital - Shtip | Research Center for Genetic Engineering and Biotechnology "Georgi D. Efremov" , Macedonian Academy of Sciences and Arts | RCGEB - MASA |
| EPI_ISL_911662, EPI_ISL_911663 | Texas Department of State Health Services (TXDSHS) | Texas Department of State Health Services (TXDSHS) | Bonnie Oh, Anita Pokharel, James Daniel Bonser, Myong Koag, Chung Wang, Rachel Lee, Grace Kubin, Rashmi Tuladhar, Mayela Pedrueza, Maliha Rahman, Jenny Zhang |
| EPI_ISL_913066, EPI_ISL_913069 | Center for Virology | Center for Virology | Jeremy V. Camp, Irene Goerzer, Monika Redlberger-Fritz, Stephan W. Aberle |
| EPI_ISL_913477, EPI_ISL_913478 | Klinisk mikrobiologi | The Public Health Agency of Sweden | Anna-Malin Linde, Maria Lind Karlberg, Carlo Berg, Oskar Karlsson Lindsjo, Sofia Stamouli, Reza Advani, Mattias Haukland, Petra Holmstrom, Noura Walai, Petra Edquist, Mia Brytting, Anna Risberg, Karin Tegmark-Wisell |
| EPI_ISL_915401, EPI_ISL_915403 | Keio University School of Medicine | Keio University School of Medicine | Kenjiro Kosaki, Yuka Iwasaki, Hirotsugu Ishizu, Haruhiko Siomi, Kodai Abe |

| | | | |
|---|---|---|---|
| EPI_ISL_924418 | Virology Department, Sheffield Teaching Hospitals NHS Foundation Trust/Department of Infection, Immunity and Cardiovascular Disease, The Medical School, University of Sheffield | COVID-19 Genomics UK (COG-UK) Consortium | Thushan de Silva, Matthew Parker, Nikki Smith, Adri Angyal, Rebecca Brown, Luke Green, Rachel Tucker, Paul Parsons, Danielle Groves, Katie Johnson, Laura Carrilero, Alex Keeley, Dave Partridge, Matthew Wyles, Benjamin Lindsey, Mehmet Yavuz, Mohammad Raza, Cariad Evans |
| EPI_ISL_935875 | Cadham Provincial laboratory | National Microbiology Laboratory (NML) | Anna Majer, Shari Tyson, Grace Seo, Philip Mabon, Elsie Grudeski, Rhiannon Huzarewich, Russell Mandes, Anneliese Landgraff, Jennifer Tanner, Natalie Knox, Morag Graham, Gary Van Domselaar, Paul Van Caeseele, Jared Bullard, David Alexander, Kerry Dust, Nathalie Bastien, Yan Li, Timothy Booth, Darian Hole, Madison Chapel, Kirsten Biggar, CanCOGeN's metadata curation team, Public Health Agency of Canada CanCOGeN team |
| EPI_ISL_936852, EPI_ISL_936853, EPI_ISL_936854 | Northwestern Memorial Hospital | Ozer Lab | Ramon Lorenzo-Redondo, Lacy M. Simons, Chad J. Achenbach, Lawrence J. Jennings, Michael G. Ison, Judd F. Hultquist, Egon A. Ozer |
| EPI_ISL_937033, EPI_ISL_937093, EPI_ISL_937101, EPI_ISL_937114, EPI_ISL_937115 | Quest Diagnostics | Quest Diagnostics | Rosenthal,S.H., Gerasimova,A., Kagan,R.M., Anderson, B., Livingston, K.E., Hua, M., Liu Y., Shalhout, D.F., Owen, R., Lacbawan, F. |
| EPI_ISL_939656, EPI_ISL_939657, EPI_ISL_939658 | National Institute for Communicable Diseases,National Health Laboratory Services, Gauteng, South Africa | National Institute for Communicable Diseases of the National Health Laboratory Service | Amoako DG, Mohale T, Ntuli N, Mahlangu B, Allam M, Ismail A, Bhiman JN |
| EPI_ISL_940186, EPI_ISL_940190, EPI_ISL_940197, EPI_ISL_940269, EPI_ISL_940270, EPI_ISL_940276, EPI_ISL_940279, EPI_ISL_940283, EPI_ISL_940284, EPI_ISL_940285, EPI_ISL_940286, EPI_ISL_940291, EPI_ISL_940491 | | | |
| see above | Hôpital Bichat Claude Bernard, Laboratoire de Virologie | IAME UMR1137 Inserm, Université de Paris, Hôpital Bichat | Antoine Bridier-Nahmias, Amélie Recoing, Quentin Le Hingrat, Lena Daniel, Siham Hamri, Gilles Collin, Alexandre Storto, Mélanie Bertine, Charlotte Charpentier, Nadhira Houhou-Fidouh, Diane Descamps, Benoit Visseaux |
| EPI_ISL_940984, EPI_ISL_940985, EPI_ISL_940988, EPI_ISL_940989, EPI_ISL_940990, EPI_ISL_940991, EPI_ISL_940992, EPI_ISL_940993 | Centers for Disease Control and Prevention, Dengue Branch | Centers for Disease Control and Prevention, Dengue Branch | Gilberto A. Santiago, Glenda Gonzalez, Betzabel Flores, Keyla Charriez, Gabriela Paz-Bailey, Jorge L. Munoz-Jordan |
| EPI_ISL_941178 | Servicio de Microbiología. Hospital Clínico Universitario de Valencia | SeqCOVID-SPAIN consortium/IBV(CSIC) | David Navarro Ortega, Eliseo Albert Vicent, Ignacio Torres and SeqCOVID-SPAIN consortium |
| EPI_ISL_952942 | Oxford Viromics, NDM, University of Oxford; Oxford University Hospitals; Basingstoke and North Hampshire Hospital | COVID-19 Genomics UK (COG-UK) Consortium | Tanya Golubchik, David Bonsall, George Macintyre, Amy Trebes, Mariateresa de Cesare, Catrin Moore, Alex Mobbs, Anita Justice, Robert Shaw, Monique Andersson, Timothy Peto, Emma Wise, Nathan Moore, Jessica Lynch, Nick Cortes, Matilde Mori, Stephen Kidd, David Buck, John Todd, Christophe Fraser |
| EPI_ISL_952944, EPI_ISL_952945, EPI_ISL_952946, EPI_ISL_952947, EPI_ISL_952949 | Department of Pathology, University of Cambridge | COVID-19 Genomics UK (COG-UK) Consortium | Aminu S. Jahun, Yasmin Chaudhry, Iliana Georgana, Myra Hosmillo, Rhys Izu, Martin D. Curran, Surendra Parmar, Ian Goodfellow |
| EPI_ISL_954183 | 1.AO Universitaria 'S. Giovanni di Dio e Ruggi D'Aragona, Scuola Medica Salernitana' Hospital / 2.UOC di Virologia e Microbiologia, Università della Campania 'L. Vanvitelli' / 3.AO Universitaria 'Federico II' Napoli Hospital / 4.AORN 'San Giuseppe Moscati' Avellino Hospital / 5.AO 'San Pio - presidio G. Rummo' Benevento Hospital / 6.AO 'Sant'Anna e San Sebastiano' Caserta Hospital / 7.PO 'Maria Santissima Addolorata' Eboli Hospital / 8.Biogem Istituto di Ricerche Genetiche | 1. Genome Research Center for Health (CRGS) / 2. Laboratory of Molecular Medicine and Genomics(LMMGe) / 3. Center for Research in Pure and Applied Mathematics (CRMPA) | Giorgio Giurato, Francesca Rizzo, Alessandro Weisz, Gianluigi Franci, Giovanni Nassa, Pasquale Pagliano, Roberta Tarallo, Elena Alexandrova, Ylenia D'Agostino, Carlo Ferravante, Jessica Lamberti, Viola Melone, Domenico Memoli, Valeria Mirici Cappa, Domenico Palumbo, Giovanni Pecoraro, Assunta Sellitto, Oriana Strianese, Ilaria Terenzi, Giuseppe Fenza, Aniello Gentile, Antonello Saccomanno, Sonia Amabile, Teresa Rocco, Annamaria Salvati, Emilia Vaccaro, Massimiliano Galdiero, Michele Cennamo, Giuseppe Portella, Maria Grazia Foti, Mariarosaria Ingino, Maria Landi, Maurizio Fumi, Vincenzo Rocco, Rita Greco, Vittoria Letizia, Arnolfo Petruzziello, Maddalena Schioppa, Gregorio Goffredi, Francesca Marciano, Michele Caraglia, Alessia Cossu, Marianna Scrima, Edmondo Adorisio, Morena D'Avenia, Michela Iacobellis, Rosanna Piluscio, Giorgio Dirani, Vittorio Sambri, Simona Semprini, Silvia Zanoli, Francesco Curcio, Stefania Marzinotto, Andreina Baj, Fausto Sessa. |
| EPI_ISL_955150 | University of Sarajevo, Veterinary Faculty, Laboratory for Molecular Diagnostic and Research Laboratory | University of Sarajevo, Veterinary Faculty, Laboratory for Molecular Diagnostic and Research Laboratory | Goleti T., Goleti Š., Softi A., Ali-Šeho A., Hodži A., Šabi E., Jaži A., Nicevi M., Terzi I. |
| EPI_ISL_955158, EPI_ISL_955159, EPI_ISL_955160, EPI_ISL_955161, EPI_ISL_955162, EPI_ISL_955163 | Institute for Biocides and Medical Ecology | Institute of microbiology and Immunology, Faculty of Medicine, University of Belgrade | Knezevic,A., Jankovic,M., Vidanovic,D., Milicevic,O., Tesovic,B., Sekler,M., Jovanovic,T. |
| EPI_ISL_956301, EPI_ISL_956306 | Laboratorio Gencore- Universidad de los Andes | Instituto Nacional de Salud- Dirección de Investigación en Salud Pública, Universidad de los Andes- Gencore | Katherine Laiton-Donato, Diego A. Álvarez-Díaz, Carlos Franco-Muñoz, Mauricio Pacheco-Montealegre, Hector Alejandro Ruiz-Moreno, Maria T. Herrera-Sepúlveda, Diego Andrés Prada, Jhonnatan Reales-González, Sheryll Corchuelo, Julian Naizaque, Gerardo Santamaría, Magdalena Wiesner, Martha Lucia Ospina Martinez, Marcela Mercado-Reyes |
| EPI_ISL_966324 | Kentucky State Public Health Lab | Kentucky State Public Health Lab | Stephanie Lunn, Karim George, Joshua Tobias, William Grooms, Vaneet Arora, Matthew Johnson, Rachel Zinner, Rhonda Lucas |
| EPI_ISL_974018, EPI_ISL_974019, EPI_ISL_974020, EPI_ISL_974021, EPI_ISL_974022, EPI_ISL_974023, EPI_ISL_974024, EPI_ISL_974025, EPI_ISL_974026, EPI_ISL_974027, EPI_ISL_974028, EPI_ISL_974029, EPI_ISL_974030, EPI_ISL_974031, EPI_ISL_974032, EPI_ISL_974033, EPI_ISL_974034, EPI_ISL_974035, EPI_ISL_974036, EPI_ISL_974037, EPI_ISL_974038, EPI_ISL_974039, EPI_ISL_974040, EPI_ISL_974041, EPI_ISL_974042, EPI_ISL_974043, EPI_ISL_974044, EPI_ISL_974045, EPI_ISL_974046, EPI_ISL_974047, EPI_ISL_974048, EPI_ISL_974049, EPI_ISL_974050, EPI_ISL_974051, EPI_ISL_974052, EPI_ISL_974053, EPI_ISL_974054, EPI_ISL_974055, EPI_ISL_974056, EPI_ISL_974057, EPI_ISL_974058, EPI_ISL_974059, EPI_ISL_974060, EPI_ISL_974061, EPI_ISL_974062, EPI_ISL_974063, EPI_ISL_974064, EPI_ISL_974065, EPI_ISL_974066, EPI_ISL_974067, EPI_ISL_974068, EPI_ISL_974069, EPI_ISL_974070, EPI_ISL_974071, EPI_ISL_974072, EPI_ISL_974073, EPI_ISL_974074, EPI_ISL_974075, EPI_ISL_974076, EPI_ISL_974077, EPI_ISL_974078, EPI_ISL_974079, EPI_ISL_974080, EPI_ISL_974081, EPI_ISL_974082, EPI_ISL_974083, EPI_ISL_974084, EPI_ISL_974085, EPI_ISL_974086, EPI_ISL_974087, EPI_ISL_974088, EPI_ISL_974089, EPI_ISL_974090, EPI_ISL_974091, EPI_ISL_974092, EPI_ISL_974093, EPI_ISL_974094, EPI_ISL_974095, EPI_ISL_974096, EPI_ISL_974097, EPI_ISL_974098, EPI_ISL_974099, EPI_ISL_974100, EPI_ISL_974101, EPI_ISL_974102, EPI_ISL_974103, EPI_ISL_974104, EPI_ISL_974105, EPI_ISL_974106, EPI_ISL_974107, EPI_ISL_974108, EPI_ISL_974109, EPI_ISL_974110, EPI_ISL_974111, EPI_ISL_974112, EPI_ISL_974113, EPI_ISL_974114, EPI_ISL_974115, EPI_ISL_974116, EPI_ISL_974117, EPI_ISL_974118, EPI_ISL_974119, EPI_ISL_974120, EPI_ISL_974121, EPI_ISL_974122, EPI_ISL_974123, EPI_ISL_974124, EPI_ISL_974125, EPI_ISL_974126, EPI_ISL_974127, EPI_ISL_974128, EPI_ISL_974129, EPI_ISL_974130, EPI_ISL_974131, EPI_ISL_974132, EPI_ISL_974133, EPI_ISL_974134, EPI_ISL_974135, EPI_ISL_974136, EPI_ISL_974137, EPI_ISL_974138, EPI_ISL_974139, EPI_ISL_974140, EPI_ISL_974141, EPI_ISL_974142, EPI_ISL_974143, EPI_ISL_974144, EPI_ISL_974145, EPI_ISL_974146, EPI_ISL_974147, EPI_ISL_974148, EPI_ISL_974149, EPI_ISL_974151, EPI_ISL_974152, EPI_ISL_974153, EPI_ISL_974154, EPI_ISL_974155, EPI_ISL_974156, EPI_ISL_974157 | | | |
| see above | BCCDC Public Health Laboratory | BCCDC Public Health Laboratory | Prystajecky Natalie, Linda Hoang, Dan Fornika, John Tyson, Shannon Russell, Kim Macdonald, Kimia Kamelian, Ana Pacagnella, Corrinne Ng, Loretta Janz, Robert Azana Terry Snutch, Mel Krajden |
