## Supplementary material for "Detection and characterization of the SARS-CoV-2 lineage B.1.526 in New York": Supp. Table 3 GISAID Acknowledgment part 2: gisaid_hcov-19_acknowledgement_table_2021_02_12_23-11.pdf

We gratefully acknowledge the following Authors from the Originating laboratories responsible for obtaining the specimens, as well as the Submitting laboratories where the genome data were generated and shared via GISAID, on which this research is based.

All Submitters of data may be contacted directly via www.gisaid.org

Authors are sorted alphabetically.

| Accession ID | Originating Laboratory | Submitting Laboratory | Authors |
| --- | --- | --- | --- |
| EPI_ISL_577157, EPI_ISL_577158, EPI_ISL_577159, EPI_ISL_577160, EPI_ISL_577161, EPI_ISL_577162, EPI_ISL_577163, EPI_ISL_577164, EPI_ISL_577165, EPI_ISL_577166, EPI_ISL_577167, EPI_ISL_577168, EPI_ISL_577169, EPI_ISL_577170, EPI_ISL_577171, EPI_ISL_577172, EPI_ISL_577173 | | | |
| see above | Queens Medical Centre, Clinical Microbiology Department / DeepSeq Nottingham | COVID-19 Genomics UK (COG-UK) Consortium | Gemma Clark, Wendy Smith, Manjinder Khakh, Vicki M Fleming, Michelle M Lister, Hannah Howson-Wells, Jonathan Ball, Patrick McClure, Joseph Chappell, Theocharis Tsoleridis, Nadine Holmes, Matthew Carlisle, Christopher Moore, Fei Sang, Johnny Debebe, Victoria Wright, Matthew Loose |
| EPI_ISL_582018 | PathLab Bay of Plenty | Institute of Environmental Science and Research (ESR) | Xiaoyun Ren, Matt Storey, Nikki Freed, Muhammad Faisal, Jing Wang, Hermes Perez, Anja Werno, Antje van der Linden, Arlo Upton, Chris Mansell, David Hammer, Dragana Drinkovic, Gary McAuliffe, Hana Sofia Andersson, James Ussher, Jill Sherwood, Josh Freeman, Julia Howard, Juliet Elvy, Mary DeAlmeida, Matt Blakiston, Matthew Rogers, Max Bloomfield, Michael Addidle, Michelle Balm, Sally Roberts, Sarah Jefferies, Sharmini Muttaiyah, Susan Morpeth, Susan Taylor, Timothy Blackmore, Vani Sathyendran, Veronica Playle, Virginia Hope, Erasmus Smit, Lauren Jelly, Olin Silander, Joep de Ligt |
| EPI_ISL_582022, EPI_ISL_582023 | Pathlab Lakes | Institute of Environmental Science and Research (ESR) | Xiaoyun Ren, Matt Storey, Nikki Freed, Muhammad Faisal, Jing Wang, Hermes Perez, Anja Werno, Antje van der Linden, Arlo Upton, Chris Mansell, David Hammer, Dragana Drinkovic, Gary McAuliffe, Hana Sofia Andersson, James Ussher, Jill Sherwood, Josh Freeman, Julia Howard, Juliet Elvy, Mary DeAlmeida, Matt Blakiston, Matthew Rogers, Max Bloomfield, Michael Addidle, Michelle Balm, Sally Roberts, Sarah Jefferies, Sharmini Muttaiyah, Susan Morpeth, Susan Taylor, Timothy Blackmore, Vani Sathyendran, Veronica Playle, Virginia Hope, Erasmus Smit, Lauren Jelly, Olin Silander, Joep de Ligt |
| EPI_ISL_584314, EPI_ISL_584316, EPI_ISL_584317, EPI_ISL_584318, EPI_ISL_584321 | Department of Pathology, University of Cambridge | COVID-19 Genomics UK (COG-UK) Consortium | Aminu S. Jahun, Yasmin Chaudhry, Grant Hall, Iliana Georgana, Myra Hosmillo, Martin D. Curran, Malte Pinckert, Surendra Parmar, Ian Goodfellow |
| EPI_ISL_584717, EPI_ISL_584718, EPI_ISL_584729, EPI_ISL_584730, EPI_ISL_584731, EPI_ISL_584733, EPI_ISL_584734, EPI_ISL_584736, EPI_ISL_584739, EPI_ISL_584744, EPI_ISL_584748, EPI_ISL_584749 | | | |
| see above | Quadram Institute Bioscience | COVID-19 Genomics UK (COG-UK) Consortium | Dave J. Baker, Gemma L. Kay, Alp Aydin, Thanh Le-Viet, Steven Rudder, Ana P. Tedim, Anastasia Kolyva, Maria Diaz, Leonardo de Oliveira Martins, Nabil-Fareed Alikhan, Lizzie Meadows, Rachael Stanley, Ngozi Elumogo, Muhammed Yasir, Nicholas M. Thomson, Alexander J Trotter, Rachel Gilroy, Samuel Bloomfield, Claire Stuart, Andrew Bell, Reenesh Prakash, Samir Dervisevic, Alison E. Mather, John Wain, Mark Webber, Andrew J. Page, Justin O'Grady |
| EPI_ISL_584750, EPI_ISL_584751, EPI_ISL_584752, EPI_ISL_584753, EPI_ISL_584754, EPI_ISL_584755, EPI_ISL_584756, EPI_ISL_584757, EPI_ISL_584758, EPI_ISL_584759, EPI_ISL_584760, EPI_ISL_584761, EPI_ISL_584762, EPI_ISL_584763, EPI_ISL_584764, EPI_ISL_584765, EPI_ISL_584766, EPI_ISL_584767, EPI_ISL_584768, EPI_ISL_584769, EPI_ISL_584770, EPI_ISL_584771, EPI_ISL_584772, EPI_ISL_584773, EPI_ISL_584774, EPI_ISL_584775, EPI_ISL_584776, EPI_ISL_584777, EPI_ISL_584778, EPI_ISL_584779, EPI_ISL_584780, EPI_ISL_584781, EPI_ISL_584782, EPI_ISL_584783, EPI_ISL_584784, EPI_ISL_584785, EPI_ISL_584786, EPI_ISL_584787, EPI_ISL_584793, EPI_ISL_584794, EPI_ISL_584795, EPI_ISL_584796, EPI_ISL_584797, EPI_ISL_584798, EPI_ISL_584799, EPI_ISL_584800, EPI_ISL_584801, EPI_ISL_584802, EPI_ISL_584803, EPI_ISL_584804, EPI_ISL_584805, EPI_ISL_584806, EPI_ISL_584807, EPI_ISL_584808 | | | |
| see above | Queens Medical Centre, Clinical Microbiology Department / DeepSeq Nottingham | COVID-19 Genomics UK (COG-UK) Consortium | Gemma Clark, Wendy Smith, Manjinder Khakh, Vicki M Fleming, Michelle M Lister, Hannah Howson-Wells, Jonathan Ball, Patrick McClure, Joseph Chappell, Theocharis Tsoleridis, Nadine Holmes, Matthew Carlisle, Christopher Moore, Fei Sang, Johnny Debebe, Victoria Wright, Matthew Loose |
| EPI_ISL_585481, EPI_ISL_585484, EPI_ISL_585485, EPI_ISL_585486, EPI_ISL_585487, EPI_ISL_585500, EPI_ISL_585501 | Virology Department, Royal Infirmary of Edinburgh, NHS Lothian / School of Biological Sciences, University of Edinburgh / Institute of Genetics and Molecular Medicine, University of Edinburgh | COVID-19 Genomics UK (COG-UK) Consortium | McHugh M, Dewar R, Rooke S, Gallagher M, Balcaza C, O'Toole Á, Scher E, Hill V, McCrone JT, Colquhoun R, Yu X, Jackson B, Rambaut A, Williams TC, Templeton K |
| EPI_ISL_585776, EPI_ISL_585905, EPI_ISL_585909, EPI_ISL_586014, EPI_ISL_586025, EPI_ISL_586026, EPI_ISL_586027, EPI_ISL_586028, EPI_ISL_586029, EPI_ISL_586030, EPI_ISL_586032, EPI_ISL_586035, EPI_ISL_586036, EPI_ISL_586038, EPI_ISL_586039, EPI_ISL_586040, EPI_ISL_586041, EPI_ISL_586042, EPI_ISL_586044, EPI_ISL_586045, EPI_ISL_586046, EPI_ISL_586047, EPI_ISL_586049, EPI_ISL_586050, EPI_ISL_586051, EPI_ISL_586056, EPI_ISL_586058, EPI_ISL_586059, EPI_ISL_586060, EPI_ISL_586061, EPI_ISL_586062, EPI_ISL_586064, EPI_ISL_586066, EPI_ISL_586067, EPI_ISL_586068, EPI_ISL_586069, EPI_ISL_586071, EPI_ISL_586072, EPI_ISL_586073, EPI_ISL_586075, EPI_ISL_586076, EPI_ISL_586077, EPI_ISL_586081, EPI_ISL_586082, EPI_ISL_586083, EPI_ISL_586084, EPI_ISL_586085, EPI_ISL_586086, EPI_ISL_586087, EPI_ISL_586088, EPI_ISL_586090, EPI_ISL_586093, EPI_ISL_586095, EPI_ISL_586098, EPI_ISL_586101, EPI_ISL_586102, EPI_ISL_586103, EPI_ISL_586104, EPI_ISL_586105, EPI_ISL_586106, EPI_ISL_586107, EPI_ISL_586109, EPI_ISL_586110, EPI_ISL_586111, EPI_ISL_586112, EPI_ISL_586113, EPI_ISL_586115, EPI_ISL_586116, EPI_ISL_586117, EPI_ISL_586118, EPI_ISL_586119, EPI_ISL_586120, EPI_ISL_586123, EPI_ISL_586124, EPI_ISL_586125, EPI_ISL_586126, EPI_ISL_586127, EPI_ISL_586129, EPI_ISL_586130, EPI_ISL_586132, EPI_ISL_586134, EPI_ISL_586136, EPI_ISL_586137, EPI_ISL_586138, EPI_ISL_586139, EPI_ISL_586141, EPI_ISL_586143, EPI_ISL_586144, EPI_ISL_586146, EPI_ISL_586148, EPI_ISL_586149, EPI_ISL_586150, EPI_ISL_586151, EPI_ISL_586152, EPI_ISL_586153, EPI_ISL_586157, EPI_ISL_586158, EPI_ISL_586160, EPI_ISL_586161, EPI_ISL_586165, EPI_ISL_586166, EPI_ISL_586168, EPI_ISL_586172, EPI_ISL_586173, EPI_ISL_586174, EPI_ISL_586175, EPI_ISL_586176, EPI_ISL_586177, EPI_ISL_586178, EPI_ISL_586179, EPI_ISL_586181, EPI_ISL_586182, EPI_ISL_586183, EPI_ISL_586184, EPI_ISL_586186, EPI_ISL_586187, EPI_ISL_586189, EPI_ISL_586190, EPI_ISL_586191, EPI_ISL_586192, EPI_ISL_586194, EPI_ISL_586196, EPI_ISL_586197, EPI_ISL_586198, EPI_ISL_586201, EPI_ISL_586203, EPI_ISL_586204, EPI_ISL_586205, EPI_ISL_586206, EPI_ISL_586207, EPI_ISL_586209, EPI_ISL_586210, EPI_ISL_586211, EPI_ISL_586214, EPI_ISL_586215, EPI_ISL_586216, EPI_ISL_586217, EPI_ISL_586218, EPI_ISL_586219, EPI_ISL_586220, EPI_ISL_586222, EPI_ISL_586223, EPI_ISL_586224, EPI_ISL_586226, EPI_ISL_586228, EPI_ISL_586229, EPI_ISL_586231, EPI_ISL_586233, EPI_ISL_586234, EPI_ISL_586236, EPI_ISL_586238, EPI_ISL_586239 | | | |
| see above | Wales Specialist Virology Centre Sequencing lab: Pathogen Genomics Unit | COVID-19 Genomics UK (COG-UK) Consortium | Catherine Moore, Johnathan Evans, Laura Gifford, Malorie Perry, Simon Cottrell, Angela Marchbank, Alec Birchley, Alexander Adams, Amy Gaskin, Bree Gatica-Wilcox, Jason Coombes, Joel Southgate, Lauren Gilbert, Lee Graham, Nicole Pacchiarini, Sara Kumziene-Summerhayes, Sarah Taylor, Sophie Jones, Sara Rey, Matthew Bull, Joanne Watkins, Sally Corden, Tom Connor |
| EPI_ISL_586571 | Area of Virology, Serology and Virology Division (SAViD), New South Wales Health Pathology Randwick | Area of Virology, Serology and Virology Division (SAViD), New South Wales Health Pathology Randwick | Rawlinson, W., Deveson, I., Bull, R. |
| EPI_ISL_590724 | University of Michigan Clinical Microbiology Laboratory | Lauring Lab, University of Michigan, Department of Microbiology and Immunology | Valesano |
| EPI_ISL_591280 | National Institute for Viral Disease Control and Prevention, China CDC | National Institute for Viral Disease Control and Prevention, China CDC | Huilai Ma, Zhaoguo Wang, Xiang Zhao, Jun Han, Yong Zhang, Hong Wang, Cao Chen, Ji Wang, Jingdong Song, Yao Meng, Yuchao Wu, Zhixiao Chen, Dayan Wang, Ruqin Gao, George F.Gao, Wenbo Xu |
| EPI_ISL_591485 | Histopath | NSW Health Pathology - Institute of Clinical Pathology and Medical Research; Westmead Hospital; University of Sydney | CIDM-PH et al. |
| EPI_ISL_591489, EPI_ISL_591490, EPI_ISL_591491 | Laverty Pathology | NSW Health Pathology - Institute of Clinical Pathology and Medical Research; Westmead Hospital; University of Sydney | CIDM-PH et al. |
| EPI_ISL_591492 | Medlab Pathology | NSW Health Pathology - Institute of Clinical Pathology and Medical Research; Westmead Hospital; University of Sydney | CIDM-PH et al. |
| EPI_ISL_591497, EPI_ISL_591498, EPI_ISL_591499, EPI_ISL_591500, EPI_ISL_591501 | Pathology West - NSW Health Pathology | NSW Health Pathology - Institute of Clinical Pathology and Medical Research; Westmead Hospital; University of Sydney | CIDM-PH et al. |
| EPI_ISL_591505, EPI_ISL_591506 | St Vincent's Pathology (SydPath) | NSW Health Pathology - Institute of Clinical Pathology and Medical Research; Westmead Hospital; University of Sydney | CIDM-PH et al. |
| EPI_ISL_591511, EPI_ISL_591512, EPI_ISL_591513, EPI_ISL_591514, EPI_ISL_591515 | Sydney South West Pathology Service (SSWPS) - Liverpool Hospital - NSW Health Pathology | NSW Health Pathology - Institute of Clinical Pathology and Medical Research; Westmead Hospital; University of Sydney | CIDM-PH et al. |
| EPI_ISL_594609, EPI_ISL_594610, EPI_ISL_594611, EPI_ISL_594612, EPI_ISL_594613, EPI_ISL_594615, EPI_ISL_594616 | University of Birmingham | COVID-19 Genomics UK (COG-UK) Consortium | Institute of Microbiology, University of Birmingham: Claire McMurray, Joanne Stockton, Samuel Nicholls, Radoslaw Poplawski, Will Rowe, Josh Quick, Nicholas Loman. University of Birmingham Testing Laboratory: Celina M Whalley, Andrew Bosworth, Charlotte Poxon, Kasun Wanigasooriya, Oliver Pickles, Mike Kidd, Alex Richter, Andrew D Beggs PHE Heartlands Lab: Husam Osman, Andrew Bosworth. Queen Elizabeth Hospital: Anna Casey |
| EPI_ISL_594680, EPI_ISL_594691, EPI_ISL_594703, EPI_ISL_594742, EPI_ISL_594743, EPI_ISL_594744, EPI_ISL_594745, EPI_ISL_594746, EPI_ISL_594747, EPI_ISL_594748, EPI_ISL_594749, EPI_ISL_594750, EPI_ISL_594751, EPI_ISL_594752, EPI_ISL_594753, EPI_ISL_594754, EPI_ISL_594755, EPI_ISL_594756, | | | |

| | | | |
|---|---|---|---|
| EPI_ISL_594757, EPI_ISL_594758, EPI_ISL_594759, EPI_ISL_594760, EPI_ISL_594761, EPI_ISL_594762, EPI_ISL_594763, EPI_ISL_594764, EPI_ISL_594765, EPI_ISL_594766, EPI_ISL_594767, EPI_ISL_594768, EPI_ISL_594769, EPI_ISL_594770, EPI_ISL_594771, EPI_ISL_594772, EPI_ISL_594773, EPI_ISL_594774, EPI_ISL_594775, EPI_ISL_594776, EPI_ISL_594777, EPI_ISL_594778, EPI_ISL_594779, EPI_ISL_594780, EPI_ISL_594781, EPI_ISL_594782, EPI_ISL_594783, EPI_ISL_594784, EPI_ISL_594785, EPI_ISL_594786, EPI_ISL_594787, EPI_ISL_594788, EPI_ISL_594789, EPI_ISL_594790, EPI_ISL_594791, EPI_ISL_594792, EPI_ISL_594793, EPI_ISL_594794, EPI_ISL_594795, EPI_ISL_594796, EPI_ISL_594797, EPI_ISL_594798, EPI_ISL_594799, EPI_ISL_594800, EPI_ISL_594801, EPI_ISL_594802, EPI_ISL_594803, EPI_ISL_594804, EPI_ISL_594805, EPI_ISL_594806, EPI_ISL_594807, EPI_ISL_594808, EPI_ISL_594809, EPI_ISL_594810, EPI_ISL_594811, EPI_ISL_594812, EPI_ISL_594813, EPI_ISL_594814, EPI_ISL_594815 | | | |
| see above | West of Scotland Specialist Virology Centre, NHSGGC / MRC-University of Glasgow Centre for Virus Research | COVID-19 Genomics UK (COG-UK) Consortium | Ana da Silva Filipe, Natasha Johnson, Kathy Smollett, Daniel Mair, Stephen Carmichael, Lily Tong, Jenna Nichols, Elihu Aranday-Cortes, Kyriaki Nomikou; Sarah McDonald, Marc Niebel, Patawee Asamaphan; Richard Orton, Joseph Hughes, Sreenu Vattipally, David L Robertson; Alasdair MacLean, Rory Gunson; Kathy Li, Igor Starinskij, Natasha Jesudason, Rajiv Shah, James Shepherd, Antonia Ho, Emma Thomson |
| EPI_ISL_594832, EPI_ISL_594833, EPI_ISL_594834, EPI_ISL_594835, EPI_ISL_594836, EPI_ISL_594837, EPI_ISL_594838, EPI_ISL_594839, EPI_ISL_594840, EPI_ISL_594841, EPI_ISL_594842, EPI_ISL_594843, EPI_ISL_594844, EPI_ISL_594845 | | | |
| see above | Virology Department, Royal Infirmary of Edinburgh, NHS Lothian / School of Biological Sciences, University of Edinburgh / Institute of Genetics and Molecular Medicine, University of Edinburgh | COVID-19 Genomics UK (COG-UK) Consortium | McHugh M, Dewar R, Rooke S, Gallagher M, Balcaza C, O'Toole Á, Scher E, Hill V, McCrone JT, Colquhoun R, Yu X, Jackson B, Rambaut A, Williams TC, Templeton K |
| EPI_ISL_594877, EPI_ISL_594878, EPI_ISL_594879, EPI_ISL_594882, EPI_ISL_594883, EPI_ISL_594885, EPI_ISL_594886, EPI_ISL_594887 | University College London, Great Ormond Street Hospital for Children NHS Foundation Trust, Imperial College Healthcare NHS Trust | COVID-19 Genomics UK (COG-UK) Consortium | Sergi Castellano, Rachel Williams, Mark Kristiansen, Paola Resende Silva, Sunando Roy, Tony Brooks, Helena Tutill, Paola Niola, Patricia Dyal, Charlotte Williams, Leysa Forrest, Yasmin Panchbhaya, Jacqueline Findlay, Samuel Weeks, Julianne Brown, Kathryn Harris, Paul Randell, James Price, Alison Holmes, Judith Breuer |
| EPI_ISL_595115, EPI_ISL_595116, EPI_ISL_595117, EPI_ISL_595118, EPI_ISL_595119, EPI_ISL_595120, EPI_ISL_595121, EPI_ISL_595122, EPI_ISL_595123, EPI_ISL_595124, EPI_ISL_595125, EPI_ISL_595126, EPI_ISL_595127, EPI_ISL_595128, EPI_ISL_595129, EPI_ISL_595130, EPI_ISL_595131, EPI_ISL_595132, EPI_ISL_595133, EPI_ISL_595134, EPI_ISL_595135, EPI_ISL_595136, EPI_ISL_595137, EPI_ISL_595138, EPI_ISL_595139, EPI_ISL_595140, EPI_ISL_595141, EPI_ISL_595142, EPI_ISL_595143, EPI_ISL_595144, EPI_ISL_595145, EPI_ISL_595146, EPI_ISL_595147, EPI_ISL_595148, EPI_ISL_595149, EPI_ISL_595150, EPI_ISL_595151, EPI_ISL_595152, EPI_ISL_595153, EPI_ISL_595154, EPI_ISL_595155, EPI_ISL_595156, EPI_ISL_595157, EPI_ISL_595158, EPI_ISL_595159, EPI_ISL_595161, EPI_ISL_595162, EPI_ISL_595163, EPI_ISL_595164, EPI_ISL_595165, EPI_ISL_595167, EPI_ISL_595168, EPI_ISL_595169, EPI_ISL_595170, EPI_ISL_595171, EPI_ISL_595172, EPI_ISL_595173, EPI_ISL_595174, EPI_ISL_595175, EPI_ISL_595176, EPI_ISL_595177, EPI_ISL_595178, EPI_ISL_595179, EPI_ISL_595187, EPI_ISL_595190, EPI_ISL_595200 | | | |
| see above | Quadram Institute Bioscience | COVID-19 Genomics UK (COG-UK) Consortium | Dave J. Baker, Gemma L. Kay, Alp Aydin, Thanh Le-Viet, Steven Rudder, Ana P. Tedim, Anastasia Kolyva, Maria Diaz, Leonardo de Oliveira Martins, Nabil-Fareed Alikhan, Lizzie Meadows, Rachael Stanley, Ngozi Elumogo, Muhammed Yasir, Nicholas M. Thomson, Alexander J Trotter, Rachel Gilroy, Samuel Bloomfield, Claire Stuart, Andrew Bell, Reenesh Prakash, Samir Dervisevic, Alison E. Mather, John Wain, Mark Webber, Andrew J. Page, Justin O'Grady |
| EPI_ISL_595232, EPI_ISL_595233, EPI_ISL_595234, EPI_ISL_595235, EPI_ISL_595236 | Queens Medical Centre, Clinical Microbiology Department / DeepSeq Nottingham | COVID-19 Genomics UK (COG-UK) Consortium | Gemma Clark, Wendy Smith, Manjinder Khakh, Vicki M Fleming, Michelle M Lister, Hannah Howson-Wells, Jonathan Ball, Patrick McClure, Joseph Chappell, Theocharis Tsoleridis, Nadine Holmes, Matthew Carlisle, Christopher Moore, Fei Sang, Johnny Debebe, Victoria Wright, Matthew Loose |
| EPI_ISL_595329, EPI_ISL_595330, EPI_ISL_595332, EPI_ISL_595334, EPI_ISL_595335, EPI_ISL_595336, EPI_ISL_595337, EPI_ISL_595338, EPI_ISL_595339, EPI_ISL_595340, EPI_ISL_595341, EPI_ISL_595343, EPI_ISL_595345, EPI_ISL_595346, EPI_ISL_595347, EPI_ISL_595348, EPI_ISL_595351, EPI_ISL_595352, EPI_ISL_595353, EPI_ISL_595354, EPI_ISL_595355, EPI_ISL_595356, EPI_ISL_595357, EPI_ISL_595358, EPI_ISL_595360, EPI_ISL_595361, EPI_ISL_595362, EPI_ISL_595363, EPI_ISL_595364, EPI_ISL_595365, EPI_ISL_595366, EPI_ISL_595367, EPI_ISL_595368, EPI_ISL_595370, EPI_ISL_595372, EPI_ISL_595373, EPI_ISL_595374, EPI_ISL_595375, EPI_ISL_595378, EPI_ISL_595379, EPI_ISL_595383, EPI_ISL_595384, EPI_ISL_595385, EPI_ISL_595387, EPI_ISL_595389, EPI_ISL_595390, EPI_ISL_595393, EPI_ISL_595394, EPI_ISL_595395, EPI_ISL_595396, EPI_ISL_595398, EPI_ISL_595399, EPI_ISL_595400, EPI_ISL_595410, EPI_ISL_595411, EPI_ISL_595415, EPI_ISL_595416, EPI_ISL_595418, EPI_ISL_595419, EPI_ISL_595420, EPI_ISL_595421, EPI_ISL_595422, EPI_ISL_595423, EPI_ISL_595429, EPI_ISL_595431, EPI_ISL_595432, EPI_ISL_595433, EPI_ISL_595434, EPI_ISL_595437, EPI_ISL_595438, EPI_ISL_595439, EPI_ISL_595440, EPI_ISL_595442, EPI_ISL_595443, EPI_ISL_595446, EPI_ISL_595447, EPI_ISL_595449, EPI_ISL_595450, EPI_ISL_595451, EPI_ISL_595454, EPI_ISL_595458, EPI_ISL_595460, EPI_ISL_595462, EPI_ISL_595463, EPI_ISL_595464, EPI_ISL_595465, EPI_ISL_595466, EPI_ISL_595467, EPI_ISL_595472, EPI_ISL_595474, EPI_ISL_595477, EPI_ISL_595480, EPI_ISL_595482, EPI_ISL_595486, EPI_ISL_595487, EPI_ISL_595488, EPI_ISL_595489, EPI_ISL_595490, EPI_ISL_595499, EPI_ISL_595502, EPI_ISL_595507, EPI_ISL_595508, EPI_ISL_595509, EPI_ISL_595511, EPI_ISL_595513, EPI_ISL_595515, EPI_ISL_595516, EPI_ISL_595517, EPI_ISL_595519, EPI_ISL_595521, EPI_ISL_595522, EPI_ISL_595528, EPI_ISL_595531, EPI_ISL_595535, EPI_ISL_595536, EPI_ISL_595537, EPI_ISL_595540, EPI_ISL_595541, EPI_ISL_595542, EPI_ISL_595543, EPI_ISL_595544, EPI_ISL_595545, EPI_ISL_595546, EPI_ISL_595547, EPI_ISL_595548, EPI_ISL_595549, EPI_ISL_595552, EPI_ISL_595553, EPI_ISL_595557, EPI_ISL_595561, EPI_ISL_595563, EPI_ISL_595566, EPI_ISL_595567, EPI_ISL_595568, EPI_ISL_595569, EPI_ISL_595574, EPI_ISL_595578, EPI_ISL_595581, EPI_ISL_595584, EPI_ISL_595585, EPI_ISL_595588, EPI_ISL_595589, EPI_ISL_595590, EPI_ISL_595593, EPI_ISL_595594, EPI_ISL_595595, EPI_ISL_595596, EPI_ISL_595597, EPI_ISL_595599, EPI_ISL_595600, EPI_ISL_595601, EPI_ISL_595602, EPI_ISL_595603, EPI_ISL_595604, EPI_ISL_595605, EPI_ISL_595606, EPI_ISL_595609, EPI_ISL_595611, EPI_ISL_595614, EPI_ISL_595615, EPI_ISL_595616 | | | |
| see above | Wales Specialist Virology Centre Sequencing lab: Pathogen Genomics Unit | COVID-19 Genomics UK (COG-UK) Consortium | Catherine Moore, Johnathan Evans, Laura Gifford, Malorie Perry, Simon Cottrell, Angela Marchbank, Alec Birchley, Alexander Adams, Amy Gaskin, Bree Gatica-Wilcox, Jason Coombes, Joel Southgate, Lauren Gilbert, Lee Graham, Nicole Pacchiarini, Sara Kumziene-Summerhayes, Sarah Taylor, Sophie Jones, Sara Rey, Matthew Bull, Joanne Watkins, Sally Corden, Tom Connor |
| EPI_ISL_595925, EPI_ISL_595926, EPI_ISL_595927, EPI_ISL_595928, EPI_ISL_595929, EPI_ISL_595930, EPI_ISL_595931, EPI_ISL_595932, EPI_ISL_595933, EPI_ISL_595934, EPI_ISL_595935, EPI_ISL_595936, EPI_ISL_595939, EPI_ISL_595940, EPI_ISL_595941, EPI_ISL_595942, EPI_ISL_595943, EPI_ISL_595944, EPI_ISL_595945, EPI_ISL_595946, EPI_ISL_595947, EPI_ISL_595948, EPI_ISL_595949, EPI_ISL_595950, EPI_ISL_595951, EPI_ISL_595952, EPI_ISL_595953, EPI_ISL_595957, EPI_ISL_595959, EPI_ISL_595960, EPI_ISL_595965, EPI_ISL_595966, EPI_ISL_595967, EPI_ISL_595968, EPI_ISL_595969, EPI_ISL_595970, EPI_ISL_595971, EPI_ISL_595973, EPI_ISL_595974, EPI_ISL_595975, EPI_ISL_595976, EPI_ISL_595977, EPI_ISL_595978, EPI_ISL_595979, EPI_ISL_595980, EPI_ISL_595981, EPI_ISL_595982, EPI_ISL_595983, EPI_ISL_595984, EPI_ISL_595985, EPI_ISL_595986, EPI_ISL_595987, EPI_ISL_595988, EPI_ISL_595989, EPI_ISL_595990, EPI_ISL_595991, EPI_ISL_595992, EPI_ISL_595993, EPI_ISL_595994, EPI_ISL_595995, EPI_ISL_595996, EPI_ISL_595997, EPI_ISL_595998, EPI_ISL_595999, EPI_ISL_596000, EPI_ISL_596001, EPI_ISL_596002, EPI_ISL_596003, EPI_ISL_596004, EPI_ISL_596005, EPI_ISL_596006, EPI_ISL_596007, EPI_ISL_596008, EPI_ISL_596009, EPI_ISL_596010, EPI_ISL_596011, EPI_ISL_596013, EPI_ISL_596014, EPI_ISL_596015, EPI_ISL_596016, EPI_ISL_596017, EPI_ISL_596018, EPI_ISL_596019, EPI_ISL_596020, EPI_ISL_596032, EPI_ISL_596033, EPI_ISL_596034, EPI_ISL_596035, EPI_ISL_596036, EPI_ISL_596037, EPI_ISL_596038, EPI_ISL_596039, EPI_ISL_596040, EPI_ISL_596041, EPI_ISL_596042, EPI_ISL_596043, EPI_ISL_596044, EPI_ISL_596045, EPI_ISL_596046, EPI_ISL_596047, EPI_ISL_596048, EPI_ISL_596099, EPI_ISL_596100, EPI_ISL_596101, EPI_ISL_596102, EPI_ISL_596103, EPI_ISL_596104, EPI_ISL_596105, EPI_ISL_596106, EPI_ISL_596116, EPI_ISL_596117, EPI_ISL_596118, EPI_ISL_596119, EPI_ISL_596120, EPI_ISL_596121, EPI_ISL_596122, EPI_ISL_596123, EPI_ISL_596124, EPI_ISL_596125, EPI_ISL_596126, EPI_ISL_596127, EPI_ISL_596129, EPI_ISL_596130, EPI_ISL_596131, EPI_ISL_596132, EPI_ISL_596133, EPI_ISL_596134, EPI_ISL_596135, EPI_ISL_596136, EPI_ISL_596137, EPI_ISL_596138, EPI_ISL_596139, EPI_ISL_596140, EPI_ISL_596141, EPI_ISL_596142, EPI_ISL_596143, EPI_ISL_596144, EPI_ISL_596145, EPI_ISL_596146, EPI_ISL_596147, EPI_ISL_596148, EPI_ISL_596149, EPI_ISL_596150, EPI_ISL_596151, EPI_ISL_596152, EPI_ISL_596153, EPI_ISL_596154, EPI_ISL_596155, EPI_ISL_596156, EPI_ISL_596157, EPI_ISL_596158, EPI_ISL_596159, EPI_ISL_596160, EPI_ISL_596161, EPI_ISL_596162, EPI_ISL_596163, EPI_ISL_596164, EPI_ISL_596165, EPI_ISL_596166, EPI_ISL_596167, EPI_ISL_596168, EPI_ISL_596169, EPI_ISL_596170, EPI_ISL_596171, EPI_ISL_596172, EPI_ISL_596173, EPI_ISL_596174, EPI_ISL_596175, EPI_ISL_596176, EPI_ISL_596177, EPI_ISL_596178, EPI_ISL_596179, EPI_ISL_596180, EPI_ISL_596181, EPI_ISL_596182, EPI_ISL_596184, EPI_ISL_596185, EPI_ISL_596186, EPI_ISL_596187, EPI_ISL_596188, EPI_ISL_596189, EPI_ISL_596190 | | | |
| see above | Quadram Institute Bioscience | COVID-19 Genomics UK (COG-UK) Consortium | Dave J. Baker, Gemma L. Kay, Alp Aydin, Thanh Le-Viet, Steven Rudder, Ana P. Tedim, Anastasia Kolyva, Maria Diaz, Leonardo de Oliveira Martins, Nabil-Fareed Alikhan, Lizzie Meadows, Rachael Stanley, Ngozi Elumogo, Muhammed Yasir, Nicholas M. Thomson, Alexander J Trotter, Rachel Gilroy, Samuel Bloomfield, Claire Stuart, Andrew Bell, Reenesh Prakash, Samir Dervisevic, Alison E. Mather, John Wain, Mark Webber, Andrew J. Page, Justin O'Grady |
| EPI_ISL_596192, EPI_ISL_596193, EPI_ISL_596194, EPI_ISL_596199, EPI_ISL_596203, EPI_ISL_596207, EPI_ISL_596208, EPI_ISL_596211, EPI_ISL_596214, EPI_ISL_596216, EPI_ISL_596221, EPI_ISL_596222, EPI_ISL_596223, EPI_ISL_596224, EPI_ISL_596226 | | | |
| see above | Virology Department, Sheffield Teaching Hospitals NHS Foundation Trust/Department of Infection, Immunity and Cardiovascular Disease, The Medical School, University of Sheffield | COVID-19 Genomics UK (COG-UK) Consortium | Thushan de Silva, Matthew Parker, Nikki Smith, Adri Angyal, Rebecca Brown, Luke Green, Rachel Tucker, Paul Parsons, Danielle Groves, Katie Johnson, Laura Carrilero, Alex Keeley, Dave Partridge, Matthew Wyles, Benjamin Lindsey, Mehmet Yavuz, Mohammad Raza, Cariad Evans |
| EPI_ISL_596483, EPI_ISL_596485, EPI_ISL_596490, EPI_ISL_596492, EPI_ISL_596497 | National Public Health Laboratory, National Centre for Infectious Diseases | National Public Health Laboratory, National Centre for Infectious Diseases | Tze Minn Mak, Sophie Octavia, Zhenyang Zhou, Lin Cui, Raymond Tzer Pin Lin |
| EPI_ISL_596584, EPI_ISL_596594, EPI_ISL_596600, EPI_ISL_596601, EPI_ISL_596609, EPI_ISL_596611, EPI_ISL_596612, EPI_ISL_596613, EPI_ISL_596614, EPI_ISL_596615, EPI_ISL_596616, EPI_ISL_596617, EPI_ISL_596618, EPI_ISL_596619, EPI_ISL_596620, EPI_ISL_596621, EPI_ISL_596622, EPI_ISL_596623 | | | |
| see above | University of Michigan Clinical Microbiology Laboratory | Lauring Lab, University of Michigan, Department of Microbiology and Immunology | Valesano |
| EPI_ISL_596819 | PathWest Laboratory Medicine WA | PathWest Laboratory Medicine WA Microbial Surveillance Unit | PathWest Laboratory Medicine WA Microbial Surveillance Unit |
| EPI_ISL_596928, EPI_ISL_596929, EPI_ISL_596930, EPI_ISL_596931, EPI_ISL_596932, EPI_ISL_596933, EPI_ISL_596934, EPI_ISL_596935, EPI_ISL_596936, EPI_ISL_596937, EPI_ISL_596938, EPI_ISL_596939, EPI_ISL_596940, EPI_ISL_596941, EPI_ISL_596942, EPI_ISL_596943, EPI_ISL_596944, EPI_ISL_596945, EPI_ISL_596946, EPI_ISL_596947, EPI_ISL_596948, EPI_ISL_596949, EPI_ISL_596950, EPI_ISL_596952, EPI_ISL_596953, EPI_ISL_596954, EPI_ISL_596955, EPI_ISL_596956, EPI_ISL_596957, EPI_ISL_596958, EPI_ISL_596959, EPI_ISL_596960, EPI_ISL_596961, EPI_ISL_596962, EPI_ISL_596963, EPI_ISL_596964, EPI_ISL_596965, EPI_ISL_596966, EPI_ISL_596967, EPI_ISL_596968 | | | |
| see above | Lighthouse Lab in Cambridge | Wellcome Sanger Institute for the COVID-19 Genomics UK (COG-UK) consortium | Rob Howes, The Lighthouse Lab in Cambridge and Alex Alderton, Roberto Amato, Sonia Goncalves, Ewan Harrison, David K. Jackson, Ian Johnston, Dominic Kwiatkowski, Cordelia Langford, John Sillitoe on behalf of the Wellcome Sanger Institute COVID-19 Surveillance Team (http://www.sanger.ac.uk/covid-team) |
| EPI_ISL_596969 | Lighthouse Lab in Cambridge | Wellcome Sanger Institute for the COVID-19 Genomics UK (COG-UK) Consortium | Rob Howes, The Lighthouse Lab in Cambridge and Alex Alderton, Roberto Amato, Sonia Goncalves, Ewan Harrison, David K. Jackson, Ian Johnston, Dominic Kwiatkowski, Cordelia Langford, John Sillitoe on behalf of the Wellcome Sanger Institute COVID-19 Surveillance Team |
| EPI_ISL_596970, EPI_ISL_596971, EPI_ISL_596972, EPI_ISL_596973, EPI_ISL_596974, EPI_ISL_596975, EPI_ISL_596976, EPI_ISL_596977, EPI_ISL_596978, EPI_ISL_596979, EPI_ISL_596980, EPI_ISL_596981, EPI_ISL_596982, EPI_ISL_596983, EPI_ISL_596984, EPI_ISL_596985, EPI_ISL_596986, EPI_ISL_596987, EPI_ISL_596988, EPI_ISL_596989, EPI_ISL_596990, EPI_ISL_596991, EPI_ISL_596992, EPI_ISL_596993, EPI_ISL_596994, EPI_ISL_596995, EPI_ISL_596996, EPI_ISL_596997, EPI_ISL_596998, EPI_ISL_596999, EPI_ISL_597000, EPI_ISL_597001, EPI_ISL_597003, EPI_ISL_597004, EPI_ISL_597005, EPI_ISL_597006, EPI_ISL_597007, EPI_ISL_597008, EPI_ISL_597009, EPI_ISL_597010, EPI_ISL_597011, EPI_ISL_597012, EPI_ISL_597013, EPI_ISL_597014, EPI_ISL_597015, EPI_ISL_597016, EPI_ISL_597017, EPI_ISL_597018, EPI_ISL_597019, EPI_ISL_597020, EPI_ISL_597021, EPI_ISL_597022, EPI_ISL_597023, EPI_ISL_597024, | | | |

| | | | |
|---|---|---|---|
| EPI_ISL_597025, EPI_ISL_597026, EPI_ISL_597027, EPI_ISL_597028, EPI_ISL_597029, EPI_ISL_597030, EPI_ISL_597031, EPI_ISL_597032, EPI_ISL_597033, EPI_ISL_597034, EPI_ISL_597035, EPI_ISL_597036, EPI_ISL_597037, EPI_ISL_597038, EPI_ISL_597039, EPI_ISL_597040, EPI_ISL_597041, EPI_ISL_597042, EPI_ISL_597043, EPI_ISL_597044, EPI_ISL_597045, EPI_ISL_597046, EPI_ISL_597047, EPI_ISL_597048, EPI_ISL_597049, EPI_ISL_597050, EPI_ISL_597051, EPI_ISL_597052, EPI_ISL_597053, EPI_ISL_597054, EPI_ISL_597055, EPI_ISL_597056, EPI_ISL_597057, EPI_ISL_597058, EPI_ISL_597059, EPI_ISL_597060, EPI_ISL_597061, EPI_ISL_597062, EPI_ISL_597063, EPI_ISL_597064, EPI_ISL_597065, EPI_ISL_597066, EPI_ISL_597067, EPI_ISL_597068, EPI_ISL_597069, EPI_ISL_597070, EPI_ISL_597071, EPI_ISL_597072, EPI_ISL_597073 | | | |
| see above | Lighthouse Lab in Cambridge | Wellcome Sanger Institute for the COVID-19 Genomics UK (COG-UK) consortium | Rob Howes, The Lighthouse Lab in Cambridge and Alex Alderton, Roberto Amato, Sonia Goncalves, Ewan Harrison, David K. Jackson, Ian Johnston, Dominic Kwiatkowski, Cordelia Langford, John Sillitoe on behalf of the Wellcome Sanger Institute COVID-19 Surveillance Team (http://www.sanger.ac.uk/covid-team) |
| EPI_ISL_597074 | Lighthouse Lab in Cambridge | Wellcome Sanger Institute for the COVID-19 Genomics UK (COG-UK) Consortium | Rob Howes, The Lighthouse Lab in Cambridge and Alex Alderton, Roberto Amato, Sonia Goncalves, Ewan Harrison, David K. Jackson, Ian Johnston, Dominic Kwiatkowski, Cordelia Langford, John Sillitoe on behalf of the Wellcome Sanger Institute COVID-19 Surveillance Team |
| EPI_ISL_597075, EPI_ISL_597076, EPI_ISL_597077, EPI_ISL_597078, EPI_ISL_597079, EPI_ISL_597080, EPI_ISL_597081, EPI_ISL_597082, EPI_ISL_597083, EPI_ISL_597084, EPI_ISL_597085, EPI_ISL_597086, EPI_ISL_597087, EPI_ISL_597088, EPI_ISL_597089, EPI_ISL_597090, EPI_ISL_597091, EPI_ISL_597092, EPI_ISL_597093, EPI_ISL_597094, EPI_ISL_597095, EPI_ISL_597096, EPI_ISL_597097, EPI_ISL_597098 | | | |
| see above | Lighthouse Lab in Cambridge | Wellcome Sanger Institute for the COVID-19 Genomics UK (COG-UK) consortium | Rob Howes, The Lighthouse Lab in Cambridge and Alex Alderton, Roberto Amato, Sonia Goncalves, Ewan Harrison, David K. Jackson, Ian Johnston, Dominic Kwiatkowski, Cordelia Langford, John Sillitoe on behalf of the Wellcome Sanger Institute COVID-19 Surveillance Team (http://www.sanger.ac.uk/covid-team) |
| EPI_ISL_597099 | Lighthouse Lab in Cambridge | Wellcome Sanger Institute for the COVID-19 Genomics UK (COG-UK) Consortium | Rob Howes, The Lighthouse Lab in Cambridge and Alex Alderton, Roberto Amato, Sonia Goncalves, Ewan Harrison, David K. Jackson, Ian Johnston, Dominic Kwiatkowski, Cordelia Langford, John Sillitoe on behalf of the Wellcome Sanger Institute COVID-19 Surveillance Team |
| EPI_ISL_597100, EPI_ISL_597101, EPI_ISL_597102, EPI_ISL_597103, EPI_ISL_597104, EPI_ISL_597105, EPI_ISL_597106, EPI_ISL_597107, EPI_ISL_597108, EPI_ISL_597109, EPI_ISL_597110, EPI_ISL_597111, EPI_ISL_597112, EPI_ISL_597113, EPI_ISL_597114, EPI_ISL_597115, EPI_ISL_597116, EPI_ISL_597117, EPI_ISL_597118, EPI_ISL_597119, EPI_ISL_597120, EPI_ISL_597121, EPI_ISL_597122, EPI_ISL_597123, EPI_ISL_597124, EPI_ISL_597125, EPI_ISL_597126, EPI_ISL_597127, EPI_ISL_597128, EPI_ISL_597129, EPI_ISL_597130, EPI_ISL_597131, EPI_ISL_597132, EPI_ISL_597133, EPI_ISL_597134, EPI_ISL_597136, EPI_ISL_597137, EPI_ISL_597138, EPI_ISL_597139, EPI_ISL_597140, EPI_ISL_597141, EPI_ISL_597142, EPI_ISL_597143, EPI_ISL_597145, EPI_ISL_597146, EPI_ISL_597147, EPI_ISL_597148, EPI_ISL_597149, EPI_ISL_597150, EPI_ISL_597151, EPI_ISL_597152, EPI_ISL_597153, EPI_ISL_597154, EPI_ISL_597155, EPI_ISL_597156, EPI_ISL_597157, EPI_ISL_597158, EPI_ISL_597159, EPI_ISL_597160, EPI_ISL_597161, EPI_ISL_597162, EPI_ISL_597163, EPI_ISL_597164, EPI_ISL_597165, EPI_ISL_597166, EPI_ISL_597167, EPI_ISL_597168, EPI_ISL_597169, EPI_ISL_597170, EPI_ISL_597171, EPI_ISL_597172, EPI_ISL_597173, EPI_ISL_597174, EPI_ISL_597175, EPI_ISL_597176, EPI_ISL_597177, EPI_ISL_597178, EPI_ISL_597179, EPI_ISL_597180, EPI_ISL_597181, EPI_ISL_597182, EPI_ISL_597183, EPI_ISL_597184, EPI_ISL_597185, EPI_ISL_597186, EPI_ISL_597187, EPI_ISL_597188 | | | |
| see above | Lighthouse Lab in Cambridge | Wellcome Sanger Institute for the COVID-19 Genomics UK (COG-UK) consortium | Rob Howes, The Lighthouse Lab in Cambridge and Alex Alderton, Roberto Amato, Sonia Goncalves, Ewan Harrison, David K. Jackson, Ian Johnston, Dominic Kwiatkowski, Cordelia Langford, John Sillitoe on behalf of the Wellcome Sanger Institute COVID-19 Surveillance Team (http://www.sanger.ac.uk/covid-team) |
| EPI_ISL_597189 | Lighthouse Lab in Cambridge | Wellcome Sanger Institute for the COVID-19 Genomics UK (COG-UK) Consortium | Rob Howes, The Lighthouse Lab in Cambridge and Alex Alderton, Roberto Amato, Sonia Goncalves, Ewan Harrison, David K. Jackson, Ian Johnston, Dominic Kwiatkowski, Cordelia Langford, John Sillitoe on behalf of the Wellcome Sanger Institute COVID-19 Surveillance Team |
| EPI_ISL_597190, EPI_ISL_597191, EPI_ISL_597192, EPI_ISL_597193, EPI_ISL_597194, EPI_ISL_597195, EPI_ISL_597196, EPI_ISL_597197, EPI_ISL_597198, EPI_ISL_597199, EPI_ISL_597200, EPI_ISL_597201, EPI_ISL_597202, EPI_ISL_597203, EPI_ISL_597204, EPI_ISL_597205, EPI_ISL_597206, EPI_ISL_597207, EPI_ISL_597208, EPI_ISL_597209, EPI_ISL_597210, EPI_ISL_597211, EPI_ISL_597212, EPI_ISL_597213, EPI_ISL_597214, EPI_ISL_597215, EPI_ISL_597216, EPI_ISL_597217, EPI_ISL_597218, EPI_ISL_597219, EPI_ISL_597220, EPI_ISL_597221, EPI_ISL_597222, EPI_ISL_597223, EPI_ISL_597224, EPI_ISL_597225, EPI_ISL_597226, EPI_ISL_597227, EPI_ISL_597228, EPI_ISL_597229, EPI_ISL_597230, EPI_ISL_597231, EPI_ISL_597232, EPI_ISL_597233 | | | |
| see above | Lighthouse Lab in Cambridge | Wellcome Sanger Institute for the COVID-19 Genomics UK (COG-UK) consortium | Rob Howes, The Lighthouse Lab in Cambridge and Alex Alderton, Roberto Amato, Sonia Goncalves, Ewan Harrison, David K. Jackson, Ian Johnston, Dominic Kwiatkowski, Cordelia Langford, John Sillitoe on behalf of the Wellcome Sanger Institute COVID-19 Surveillance Team (http://www.sanger.ac.uk/covid-team) |
| EPI_ISL_597234, EPI_ISL_597235, EPI_ISL_597236 | Lighthouse Lab in Glasgow | Wellcome Sanger Institute for the COVID-19 Genomics UK (COG-UK) consortium | Harper VanSteenhouse, Yumi Kasai, David Gray, Carol Clugston, Anna Dominiczak and Alex Alderton, Roberto Amato, Sonia Goncalves, Ewan Harrison, David K. Jackson, Ian Johnston, Dominic Kwiatkowski, Cordelia Langford, John Sillitoe on behalf of the Wellcome Sanger Institute COVID-19 Surveillance Team (http://www.sanger.ac.uk/covid-team) |
| EPI_ISL_597237, EPI_ISL_597238, EPI_ISL_597239 | Lighthouse Lab in Cambridge | Wellcome Sanger Institute for the COVID-19 Genomics UK (COG-UK) consortium | Rob Howes, The Lighthouse Lab in Cambridge and Alex Alderton, Roberto Amato, Sonia Goncalves, Ewan Harrison, David K. Jackson, Ian Johnston, Dominic Kwiatkowski, Cordelia Langford, John Sillitoe on behalf of the Wellcome Sanger Institute COVID-19 Surveillance Team (http://www.sanger.ac.uk/covid-team) |
| EPI_ISL_597240 | Lighthouse Lab in Glasgow | Wellcome Sanger Institute for the COVID-19 Genomics UK (COG-UK) consortium | Harper VanSteenhouse, Yumi Kasai, David Gray, Carol Clugston, Anna Dominiczak and Alex Alderton, Roberto Amato, Sonia Goncalves, Ewan Harrison, David K. Jackson, Ian Johnston, Dominic Kwiatkowski, Cordelia Langford, John Sillitoe on behalf of the Wellcome Sanger Institute COVID-19 Surveillance Team (http://www.sanger.ac.uk/covid-team) |
| EPI_ISL_597241 | Lighthouse Lab in Cambridge | Wellcome Sanger Institute for the COVID-19 Genomics UK (COG-UK) consortium | Rob Howes, The Lighthouse Lab in Cambridge and Alex Alderton, Roberto Amato, Sonia Goncalves, Ewan Harrison, David K. Jackson, Ian Johnston, Dominic Kwiatkowski, Cordelia Langford, John Sillitoe on behalf of the Wellcome Sanger Institute COVID-19 Surveillance Team (http://www.sanger.ac.uk/covid-team) |
| EPI_ISL_597242 | Lighthouse Lab in Glasgow | Wellcome Sanger Institute for the COVID-19 Genomics UK (COG-UK) consortium | Harper VanSteenhouse, Yumi Kasai, David Gray, Carol Clugston, Anna Dominiczak and Alex Alderton, Roberto Amato, Sonia Goncalves, Ewan Harrison, David K. Jackson, Ian Johnston, Dominic Kwiatkowski, Cordelia Langford, John Sillitoe on behalf of the Wellcome Sanger Institute COVID-19 Surveillance Team (http://www.sanger.ac.uk/covid-team) |
| EPI_ISL_597243 | Lighthouse Lab in Cambridge | Wellcome Sanger Institute for the COVID-19 Genomics UK (COG-UK) consortium | Rob Howes, The Lighthouse Lab in Cambridge and Alex Alderton, Roberto Amato, Sonia Goncalves, Ewan Harrison, David K. Jackson, Ian Johnston, Dominic Kwiatkowski, Cordelia Langford, John Sillitoe on behalf of the Wellcome Sanger Institute COVID-19 Surveillance Team (http://www.sanger.ac.uk/covid-team) |
| EPI_ISL_597244, EPI_ISL_597245 | Lighthouse Lab in Glasgow | Wellcome Sanger Institute for the COVID-19 Genomics UK (COG-UK) consortium | Harper VanSteenhouse, Yumi Kasai, David Gray, Carol Clugston, Anna Dominiczak and Alex Alderton, Roberto Amato, Sonia Goncalves, Ewan Harrison, David K. Jackson, Ian Johnston, Dominic Kwiatkowski, Cordelia Langford, John Sillitoe on behalf of the Wellcome Sanger Institute COVID-19 Surveillance Team (http://www.sanger.ac.uk/covid-team) |
| EPI_ISL_597246, EPI_ISL_597247, EPI_ISL_597248 | Lighthouse Lab in Cambridge | Wellcome Sanger Institute for the COVID-19 Genomics UK (COG-UK) consortium | Rob Howes, The Lighthouse Lab in Cambridge and Alex Alderton, Roberto Amato, Sonia Goncalves, Ewan Harrison, David K. Jackson, Ian Johnston, Dominic Kwiatkowski, Cordelia Langford, John Sillitoe on behalf of the Wellcome Sanger Institute COVID-19 Surveillance Team (http://www.sanger.ac.uk/covid-team) |
| EPI_ISL_597249 | Lighthouse Lab in Glasgow | Wellcome Sanger Institute for the COVID-19 Genomics UK (COG-UK) consortium | Harper VanSteenhouse, Yumi Kasai, David Gray, Carol Clugston, Anna Dominiczak and Alex Alderton, Roberto Amato, Sonia Goncalves, Ewan Harrison, David K. Jackson, Ian Johnston, Dominic Kwiatkowski, Cordelia Langford, John Sillitoe on behalf of the Wellcome Sanger Institute COVID-19 Surveillance Team (http://www.sanger.ac.uk/covid-team) |
| EPI_ISL_597250, EPI_ISL_597251 | Lighthouse Lab in Cambridge | Wellcome Sanger Institute for the COVID-19 Genomics UK (COG-UK) consortium | Rob Howes, The Lighthouse Lab in Cambridge and Alex Alderton, Roberto Amato, Sonia Goncalves, Ewan Harrison, David K. Jackson, Ian Johnston, Dominic Kwiatkowski, Cordelia Langford, John Sillitoe on behalf of the Wellcome Sanger Institute COVID-19 Surveillance Team (http://www.sanger.ac.uk/covid-team) |
| EPI_ISL_597252 | Lighthouse Lab in Glasgow | Wellcome Sanger Institute for the COVID-19 Genomics UK (COG-UK) consortium | Harper VanSteenhouse, Yumi Kasai, David Gray, Carol Clugston, Anna Dominiczak and Alex Alderton, Roberto Amato, Sonia Goncalves, Ewan Harrison, David K. Jackson, Ian Johnston, Dominic Kwiatkowski, Cordelia Langford, John Sillitoe on behalf of the Wellcome Sanger Institute COVID-19 Surveillance Team (http://www.sanger.ac.uk/covid-team) |
| EPI_ISL_597254, EPI_ISL_597255, EPI_ISL_597256, EPI_ISL_597257, EPI_ISL_597258, EPI_ISL_597259 | Lighthouse Lab in Cambridge | Wellcome Sanger Institute for the COVID-19 Genomics UK (COG-UK) consortium | Rob Howes, The Lighthouse Lab in Cambridge and Alex Alderton, Roberto Amato, Sonia Goncalves, Ewan Harrison, David K. Jackson, Ian Johnston, Dominic Kwiatkowski, Cordelia Langford, John Sillitoe on behalf of the Wellcome Sanger Institute COVID-19 Surveillance Team (http://www.sanger.ac.uk/covid-team) |
| EPI_ISL_597260, EPI_ISL_597261 | Lighthouse Lab in Glasgow | Wellcome Sanger Institute for the COVID-19 Genomics UK (COG-UK) consortium | Harper VanSteenhouse, Yumi Kasai, David Gray, Carol Clugston, Anna Dominiczak and Alex Alderton, Roberto Amato, Sonia Goncalves, Ewan Harrison, David K. Jackson, Ian Johnston, Dominic Kwiatkowski, Cordelia Langford, John Sillitoe on behalf of the Wellcome Sanger Institute COVID-19 Surveillance Team (http://www.sanger.ac.uk/covid-team) |
| EPI_ISL_597262 | Lighthouse Lab in Cambridge | Wellcome Sanger Institute for the COVID-19 Genomics UK (COG-UK) consortium | Rob Howes, The Lighthouse Lab in Cambridge and Alex Alderton, Roberto Amato, Sonia Goncalves, Ewan Harrison, David K. Jackson, Ian Johnston, Dominic Kwiatkowski, Cordelia Langford, John Sillitoe on behalf of the Wellcome Sanger Institute COVID-19 Surveillance Team (http://www.sanger.ac.uk/covid-team) |
| EPI_ISL_597263 | Lighthouse Lab in Glasgow | Wellcome Sanger Institute for the COVID-19 Genomics UK (COG-UK) consortium | Harper VanSteenhouse, Yumi Kasai, David Gray, Carol Clugston, Anna Dominiczak and Alex Alderton, Roberto Amato, Sonia Goncalves, Ewan Harrison, David K. Jackson, Ian Johnston, Dominic Kwiatkowski, Cordelia Langford, John Sillitoe on behalf of the Wellcome Sanger Institute COVID-19 Surveillance Team (http://www.sanger.ac.uk/covid-team) |
| EPI_ISL_597265, EPI_ISL_597266 | Lighthouse Lab in Cambridge | Wellcome Sanger Institute for the COVID-19 Genomics UK (COG-UK) consortium | Rob Howes, The Lighthouse Lab in Cambridge and Alex Alderton, Roberto Amato, Sonia Goncalves, Ewan Harrison, David K. Jackson, Ian Johnston, Dominic Kwiatkowski, Cordelia Langford, John Sillitoe on behalf of the Wellcome Sanger Institute COVID-19 Surveillance Team (http://www.sanger.ac.uk/covid-team) |
| EPI_ISL_597267, EPI_ISL_597268 | Lighthouse Lab in Glasgow | Wellcome Sanger Institute for the COVID-19 Genomics UK (COG-UK) consortium | Harper VanSteenhouse, Yumi Kasai, David Gray, Carol Clugston, Anna Dominiczak and Alex Alderton, Roberto Amato, Sonia Goncalves, Ewan Harrison, David K. Jackson, Ian Johnston, Dominic Kwiatkowski, Cordelia Langford, John Sillitoe on behalf of the Wellcome Sanger Institute COVID-19 Surveillance Team (http://www.sanger.ac.uk/covid-team) |
| EPI_ISL_597269, EPI_ISL_597270, EPI_ISL_597271 | Lighthouse Lab in Cambridge | Wellcome Sanger Institute for the COVID-19 Genomics UK (COG-UK) consortium | Rob Howes, The Lighthouse Lab in Cambridge and Alex Alderton, Roberto Amato, Sonia Goncalves, Ewan Harrison, David K. Jackson, Ian Johnston, Dominic Kwiatkowski, Cordelia Langford, John Sillitoe on behalf of the Wellcome Sanger Institute COVID-19 Surveillance Team (http://www.sanger.ac.uk/covid-team) |

| | | | |
|---|---|---|---|
| EPI_ISL_597272 | Lighthouse Lab in Glasgow | Wellcome Sanger Institute for the COVID-19 Genomics UK (COG-UK) consortium | Harper VanSteenhouse, Yumi Kasai, David Gray, Carol Clugston, Anna Dominiczak and Alex Alderton, Roberto Amato, Sonia Goncalves, Ewan Harrison, David K. Jackson, Ian Johnston, Dominic Kwiatkowski, Cordelia Langford, John Sillitoe on behalf of the Wellcome Sanger Institute COVID-19 Surveillance Team (http://www.sanger.ac.uk/covid-team) |
| EPI_ISL_597273, EPI_ISL_597274 | Lighthouse Lab in Cambridge | Wellcome Sanger Institute for the COVID-19 Genomics UK (COG-UK) consortium | Rob Howes, The Lighthouse Lab in Cambridge and Alex Alderton, Roberto Amato, Sonia Goncalves, Ewan Harrison, David K. Jackson, Ian Johnston, Dominic Kwiatkowski, Cordelia Langford, John Sillitoe on behalf of the Wellcome Sanger Institute COVID-19 Surveillance Team (http://www.sanger.ac.uk/covid-team) |
| EPI_ISL_597275, EPI_ISL_597276, EPI_ISL_597277, EPI_ISL_597278 | Lighthouse Lab in Glasgow | Wellcome Sanger Institute for the COVID-19 Genomics UK (COG-UK) consortium | Harper VanSteenhouse, Yumi Kasai, David Gray, Carol Clugston, Anna Dominiczak and Alex Alderton, Roberto Amato, Sonia Goncalves, Ewan Harrison, David K. Jackson, Ian Johnston, Dominic Kwiatkowski, Cordelia Langford, John Sillitoe on behalf of the Wellcome Sanger Institute COVID-19 Surveillance Team (http://www.sanger.ac.uk/covid-team) |
| EPI_ISL_597279 | Lighthouse Lab in Cambridge | Wellcome Sanger Institute for the COVID-19 Genomics UK (COG-UK) consortium | Rob Howes, The Lighthouse Lab in Cambridge and Alex Alderton, Roberto Amato, Sonia Goncalves, Ewan Harrison, David K. Jackson, Ian Johnston, Dominic Kwiatkowski, Cordelia Langford, John Sillitoe on behalf of the Wellcome Sanger Institute COVID-19 Surveillance Team (http://www.sanger.ac.uk/covid-team) |
| EPI_ISL_597280, EPI_ISL_597281 | Lighthouse Lab in Glasgow | Wellcome Sanger Institute for the COVID-19 Genomics UK (COG-UK) consortium | Harper VanSteenhouse, Yumi Kasai, David Gray, Carol Clugston, Anna Dominiczak and Alex Alderton, Roberto Amato, Sonia Goncalves, Ewan Harrison, David K. Jackson, Ian Johnston, Dominic Kwiatkowski, Cordelia Langford, John Sillitoe on behalf of the Wellcome Sanger Institute COVID-19 Surveillance Team (http://www.sanger.ac.uk/covid-team) |
| EPI_ISL_597282, EPI_ISL_597283, EPI_ISL_597284, EPI_ISL_597285, EPI_ISL_597286, EPI_ISL_597287 | Lighthouse Lab in Cambridge | Wellcome Sanger Institute for the COVID-19 Genomics UK (COG-UK) consortium | Rob Howes, The Lighthouse Lab in Cambridge and Alex Alderton, Roberto Amato, Sonia Goncalves, Ewan Harrison, David K. Jackson, Ian Johnston, Dominic Kwiatkowski, Cordelia Langford, John Sillitoe on behalf of the Wellcome Sanger Institute COVID-19 Surveillance Team (http://www.sanger.ac.uk/covid-team) |
| EPI_ISL_597288 | Lighthouse Lab in Glasgow | Wellcome Sanger Institute for the COVID-19 Genomics UK (COG-UK) consortium | Harper VanSteenhouse, Yumi Kasai, David Gray, Carol Clugston, Anna Dominiczak and Alex Alderton, Roberto Amato, Sonia Goncalves, Ewan Harrison, David K. Jackson, Ian Johnston, Dominic Kwiatkowski, Cordelia Langford, John Sillitoe on behalf of the Wellcome Sanger Institute COVID-19 Surveillance Team (http://www.sanger.ac.uk/covid-team) |
| EPI_ISL_597289 | Lighthouse Lab in Cambridge | Wellcome Sanger Institute for the COVID-19 Genomics UK (COG-UK) consortium | Rob Howes, The Lighthouse Lab in Cambridge and Alex Alderton, Roberto Amato, Sonia Goncalves, Ewan Harrison, David K. Jackson, Ian Johnston, Dominic Kwiatkowski, Cordelia Langford, John Sillitoe on behalf of the Wellcome Sanger Institute COVID-19 Surveillance Team (http://www.sanger.ac.uk/covid-team) |
| EPI_ISL_597290, EPI_ISL_597291, EPI_ISL_597292 | Lighthouse Lab in Glasgow | Wellcome Sanger Institute for the COVID-19 Genomics UK (COG-UK) consortium | Harper VanSteenhouse, Yumi Kasai, David Gray, Carol Clugston, Anna Dominiczak and Alex Alderton, Roberto Amato, Sonia Goncalves, Ewan Harrison, David K. Jackson, Ian Johnston, Dominic Kwiatkowski, Cordelia Langford, John Sillitoe on behalf of the Wellcome Sanger Institute COVID-19 Surveillance Team (http://www.sanger.ac.uk/covid-team) |
| EPI_ISL_597293, EPI_ISL_597294, EPI_ISL_597295 | Lighthouse Lab in Cambridge | Wellcome Sanger Institute for the COVID-19 Genomics UK (COG-UK) consortium | Rob Howes, The Lighthouse Lab in Cambridge and Alex Alderton, Roberto Amato, Sonia Goncalves, Ewan Harrison, David K. Jackson, Ian Johnston, Dominic Kwiatkowski, Cordelia Langford, John Sillitoe on behalf of the Wellcome Sanger Institute COVID-19 Surveillance Team (http://www.sanger.ac.uk/covid-team) |
| EPI_ISL_597296 | Lighthouse Lab in Glasgow | Wellcome Sanger Institute for the COVID-19 Genomics UK (COG-UK) consortium | Harper VanSteenhouse, Yumi Kasai, David Gray, Carol Clugston, Anna Dominiczak and Alex Alderton, Roberto Amato, Sonia Goncalves, Ewan Harrison, David K. Jackson, Ian Johnston, Dominic Kwiatkowski, Cordelia Langford, John Sillitoe on behalf of the Wellcome Sanger Institute COVID-19 Surveillance Team (http://www.sanger.ac.uk/covid-team) |
| EPI_ISL_597297 | Lighthouse Lab in Cambridge | Wellcome Sanger Institute for the COVID-19 Genomics UK (COG-UK) consortium | Rob Howes, The Lighthouse Lab in Cambridge and Alex Alderton, Roberto Amato, Sonia Goncalves, Ewan Harrison, David K. Jackson, Ian Johnston, Dominic Kwiatkowski, Cordelia Langford, John Sillitoe on behalf of the Wellcome Sanger Institute COVID-19 Surveillance Team (http://www.sanger.ac.uk/covid-team) |
| EPI_ISL_597298, EPI_ISL_597299 | Lighthouse Lab in Glasgow | Wellcome Sanger Institute for the COVID-19 Genomics UK (COG-UK) consortium | Harper VanSteenhouse, Yumi Kasai, David Gray, Carol Clugston, Anna Dominiczak and Alex Alderton, Roberto Amato, Sonia Goncalves, Ewan Harrison, David K. Jackson, Ian Johnston, Dominic Kwiatkowski, Cordelia Langford, John Sillitoe on behalf of the Wellcome Sanger Institute COVID-19 Surveillance Team (http://www.sanger.ac.uk/covid-team) |
| EPI_ISL_597300, EPI_ISL_597301 | Lighthouse Lab in Cambridge | Wellcome Sanger Institute for the COVID-19 Genomics UK (COG-UK) consortium | Rob Howes, The Lighthouse Lab in Cambridge and Alex Alderton, Roberto Amato, Sonia Goncalves, Ewan Harrison, David K. Jackson, Ian Johnston, Dominic Kwiatkowski, Cordelia Langford, John Sillitoe on behalf of the Wellcome Sanger Institute COVID-19 Surveillance Team (http://www.sanger.ac.uk/covid-team) |
| EPI_ISL_597302, EPI_ISL_597303 | Lighthouse Lab in Glasgow | Wellcome Sanger Institute for the COVID-19 Genomics UK (COG-UK) consortium | Harper VanSteenhouse, Yumi Kasai, David Gray, Carol Clugston, Anna Dominiczak and Alex Alderton, Roberto Amato, Sonia Goncalves, Ewan Harrison, David K. Jackson, Ian Johnston, Dominic Kwiatkowski, Cordelia Langford, John Sillitoe on behalf of the Wellcome Sanger Institute COVID-19 Surveillance Team (http://www.sanger.ac.uk/covid-team) |
| EPI_ISL_597304, EPI_ISL_597305, EPI_ISL_597306, EPI_ISL_597307, EPI_ISL_597308, EPI_ISL_597309, EPI_ISL_597310 | Lighthouse Lab in Cambridge | Wellcome Sanger Institute for the COVID-19 Genomics UK (COG-UK) consortium | Rob Howes, The Lighthouse Lab in Cambridge and Alex Alderton, Roberto Amato, Sonia Goncalves, Ewan Harrison, David K. Jackson, Ian Johnston, Dominic Kwiatkowski, Cordelia Langford, John Sillitoe on behalf of the Wellcome Sanger Institute COVID-19 Surveillance Team (http://www.sanger.ac.uk/covid-team) |
| EPI_ISL_597311, EPI_ISL_597312, EPI_ISL_597313, EPI_ISL_597314, EPI_ISL_597315 | Lighthouse Lab in Glasgow | Wellcome Sanger Institute for the COVID-19 Genomics UK (COG-UK) consortium | Harper VanSteenhouse, Yumi Kasai, David Gray, Carol Clugston, Anna Dominiczak and Alex Alderton, Roberto Amato, Sonia Goncalves, Ewan Harrison, David K. Jackson, Ian Johnston, Dominic Kwiatkowski, Cordelia Langford, John Sillitoe on behalf of the Wellcome Sanger Institute COVID-19 Surveillance Team (http://www.sanger.ac.uk/covid-team) |
| EPI_ISL_597316, EPI_ISL_597317 | Lighthouse Lab in Cambridge | Wellcome Sanger Institute for the COVID-19 Genomics UK (COG-UK) consortium | Rob Howes, The Lighthouse Lab in Cambridge and Alex Alderton, Roberto Amato, Sonia Goncalves, Ewan Harrison, David K. Jackson, Ian Johnston, Dominic Kwiatkowski, Cordelia Langford, John Sillitoe on behalf of the Wellcome Sanger Institute COVID-19 Surveillance Team (http://www.sanger.ac.uk/covid-team) |
| EPI_ISL_597318 | Lighthouse Lab in Glasgow | Wellcome Sanger Institute for the COVID-19 Genomics UK (COG-UK) consortium | Harper VanSteenhouse, Yumi Kasai, David Gray, Carol Clugston, Anna Dominiczak and Alex Alderton, Roberto Amato, Sonia Goncalves, Ewan Harrison, David K. Jackson, Ian Johnston, Dominic Kwiatkowski, Cordelia Langford, John Sillitoe on behalf of the Wellcome Sanger Institute COVID-19 Surveillance Team (http://www.sanger.ac.uk/covid-team) |
| EPI_ISL_597319, EPI_ISL_597320, EPI_ISL_597321 | Lighthouse Lab in Cambridge | Wellcome Sanger Institute for the COVID-19 Genomics UK (COG-UK) consortium | Rob Howes, The Lighthouse Lab in Cambridge and Alex Alderton, Roberto Amato, Sonia Goncalves, Ewan Harrison, David K. Jackson, Ian Johnston, Dominic Kwiatkowski, Cordelia Langford, John Sillitoe on behalf of the Wellcome Sanger Institute COVID-19 Surveillance Team (http://www.sanger.ac.uk/covid-team) |
| EPI_ISL_597322 | Lighthouse Lab in Glasgow | Wellcome Sanger Institute for the COVID-19 Genomics UK (COG-UK) consortium | Harper VanSteenhouse, Yumi Kasai, David Gray, Carol Clugston, Anna Dominiczak and Alex Alderton, Roberto Amato, Sonia Goncalves, Ewan Harrison, David K. Jackson, Ian Johnston, Dominic Kwiatkowski, Cordelia Langford, John Sillitoe on behalf of the Wellcome Sanger Institute COVID-19 Surveillance Team (http://www.sanger.ac.uk/covid-team) |
| EPI_ISL_597323, EPI_ISL_597324, EPI_ISL_597325 | Lighthouse Lab in Cambridge | Wellcome Sanger Institute for the COVID-19 Genomics UK (COG-UK) consortium | Rob Howes, The Lighthouse Lab in Cambridge and Alex Alderton, Roberto Amato, Sonia Goncalves, Ewan Harrison, David K. Jackson, Ian Johnston, Dominic Kwiatkowski, Cordelia Langford, John Sillitoe on behalf of the Wellcome Sanger Institute COVID-19 Surveillance Team (http://www.sanger.ac.uk/covid-team) |
| EPI_ISL_597326 | Lighthouse Lab in Glasgow | Wellcome Sanger Institute for the COVID-19 Genomics UK (COG-UK) consortium | Harper VanSteenhouse, Yumi Kasai, David Gray, Carol Clugston, Anna Dominiczak and Alex Alderton, Roberto Amato, Sonia Goncalves, Ewan Harrison, David K. Jackson, Ian Johnston, Dominic Kwiatkowski, Cordelia Langford, John Sillitoe on behalf of the Wellcome Sanger Institute COVID-19 Surveillance Team (http://www.sanger.ac.uk/covid-team) |
| EPI_ISL_597327 | Lighthouse Lab in Cambridge | Wellcome Sanger Institute for the COVID-19 Genomics UK (COG-UK) consortium | Rob Howes, The Lighthouse Lab in Cambridge and Alex Alderton, Roberto Amato, Sonia Goncalves, Ewan Harrison, David K. Jackson, Ian Johnston, Dominic Kwiatkowski, Cordelia Langford, John Sillitoe on behalf of the Wellcome Sanger Institute COVID-19 Surveillance Team (http://www.sanger.ac.uk/covid-team) |
| EPI_ISL_597328, EPI_ISL_597329, EPI_ISL_597330 | Lighthouse Lab in Glasgow | Wellcome Sanger Institute for the COVID-19 Genomics UK (COG-UK) consortium | Harper VanSteenhouse, Yumi Kasai, David Gray, Carol Clugston, Anna Dominiczak and Alex Alderton, Roberto Amato, Sonia Goncalves, Ewan Harrison, David K. Jackson, Ian Johnston, Dominic Kwiatkowski, Cordelia Langford, John Sillitoe on behalf of the Wellcome Sanger Institute COVID-19 Surveillance Team (http://www.sanger.ac.uk/covid-team) |
| EPI_ISL_597331 | Lighthouse Lab in Cambridge | Wellcome Sanger Institute for the COVID-19 Genomics UK (COG-UK) consortium | Rob Howes, The Lighthouse Lab in Cambridge and Alex Alderton, Roberto Amato, Sonia Goncalves, Ewan Harrison, David K. Jackson, Ian Johnston, Dominic Kwiatkowski, Cordelia Langford, John Sillitoe on behalf of the Wellcome Sanger Institute COVID-19 Surveillance Team (http://www.sanger.ac.uk/covid-team) |
| EPI_ISL_597332, EPI_ISL_597333, EPI_ISL_597334 | Lighthouse Lab in Glasgow | Wellcome Sanger Institute for the COVID-19 Genomics UK (COG-UK) consortium | Harper VanSteenhouse, Yumi Kasai, David Gray, Carol Clugston, Anna Dominiczak and Alex Alderton, Roberto Amato, Sonia Goncalves, Ewan Harrison, David K. Jackson, Ian Johnston, Dominic Kwiatkowski, Cordelia Langford, John Sillitoe on behalf of the Wellcome Sanger Institute COVID-19 Surveillance Team (http://www.sanger.ac.uk/covid-team) |
| EPI_ISL_597335 | Lighthouse Lab in Cambridge | Wellcome Sanger Institute for the COVID-19 Genomics UK (COG-UK) consortium | Rob Howes, The Lighthouse Lab in Cambridge and Alex Alderton, Roberto Amato, Sonia Goncalves, Ewan Harrison, David K. Jackson, Ian Johnston, Dominic Kwiatkowski, Cordelia Langford, John Sillitoe on behalf of the Wellcome Sanger Institute COVID-19 Surveillance Team (http://www.sanger.ac.uk/covid-team) |

| | | | |
|---|---|---|---|
| EPI_ISL_597336 | Lighthouse Lab in Glasgow | Wellcome Sanger Institute for the COVID-19 Genomics UK (COG-UK) consortium | Harper VanSteenhouse, Yumi Kasai, David Gray, Carol Clugston, Anna Dominiczak and Alex Alderton, Roberto Amato, Sonia Goncalves, Ewan Harrison, David K. Jackson, Ian Johnston, Dominic Kwiatkowski, Cordelia Langford, John Sillitoe on behalf of the Wellcome Sanger Institute COVID-19 Surveillance Team (http://www.sanger.ac.uk/covid-team) |
| EPI_ISL_597337 | Lighthouse Lab in Cambridge | Wellcome Sanger Institute for the COVID-19 Genomics UK (COG-UK) consortium | Rob Howes, The Lighthouse Lab in Cambridge and Alex Alderton, Roberto Amato, Sonia Goncalves, Ewan Harrison, David K. Jackson, Ian Johnston, Dominic Kwiatkowski, Cordelia Langford, John Sillitoe on behalf of the Wellcome Sanger Institute COVID-19 Surveillance Team (http://www.sanger.ac.uk/covid-team) |
| EPI_ISL_597338, EPI_ISL_597339 | Lighthouse Lab in Glasgow | Wellcome Sanger Institute for the COVID-19 Genomics UK (COG-UK) consortium | Harper VanSteenhouse, Yumi Kasai, David Gray, Carol Clugston, Anna Dominiczak and Alex Alderton, Roberto Amato, Sonia Goncalves, Ewan Harrison, David K. Jackson, Ian Johnston, Dominic Kwiatkowski, Cordelia Langford, John Sillitoe on behalf of the Wellcome Sanger Institute COVID-19 Surveillance Team (http://www.sanger.ac.uk/covid-team) |
| EPI_ISL_597340, EPI_ISL_597341, EPI_ISL_597342, EPI_ISL_597343, EPI_ISL_597344, EPI_ISL_597345, EPI_ISL_597346 | Lighthouse Lab in Cambridge | Wellcome Sanger Institute for the COVID-19 Genomics UK (COG-UK) consortium | Rob Howes, The Lighthouse Lab in Cambridge and Alex Alderton, Roberto Amato, Sonia Goncalves, Ewan Harrison, David K. Jackson, Ian Johnston, Dominic Kwiatkowski, Cordelia Langford, John Sillitoe on behalf of the Wellcome Sanger Institute COVID-19 Surveillance Team (http://www.sanger.ac.uk/covid-team) |
| EPI_ISL_597347, EPI_ISL_597348 | Lighthouse Lab in Glasgow | Wellcome Sanger Institute for the COVID-19 Genomics UK (COG-UK) consortium | Harper VanSteenhouse, Yumi Kasai, David Gray, Carol Clugston, Anna Dominiczak and Alex Alderton, Roberto Amato, Sonia Goncalves, Ewan Harrison, David K. Jackson, Ian Johnston, Dominic Kwiatkowski, Cordelia Langford, John Sillitoe on behalf of the Wellcome Sanger Institute COVID-19 Surveillance Team (http://www.sanger.ac.uk/covid-team) |
| EPI_ISL_597349, EPI_ISL_597350 | Lighthouse Lab in Cambridge | Wellcome Sanger Institute for the COVID-19 Genomics UK (COG-UK) consortium | Rob Howes, The Lighthouse Lab in Cambridge and Alex Alderton, Roberto Amato, Sonia Goncalves, Ewan Harrison, David K. Jackson, Ian Johnston, Dominic Kwiatkowski, Cordelia Langford, John Sillitoe on behalf of the Wellcome Sanger Institute COVID-19 Surveillance Team (http://www.sanger.ac.uk/covid-team) |
| EPI_ISL_597351, EPI_ISL_597352, EPI_ISL_597353, EPI_ISL_597355, EPI_ISL_597356, EPI_ISL_597357 | Lighthouse Lab in Glasgow | Wellcome Sanger Institute for the COVID-19 Genomics UK (COG-UK) consortium | Harper VanSteenhouse, Yumi Kasai, David Gray, Carol Clugston, Anna Dominiczak and Alex Alderton, Roberto Amato, Sonia Goncalves, Ewan Harrison, David K. Jackson, Ian Johnston, Dominic Kwiatkowski, Cordelia Langford, John Sillitoe on behalf of the Wellcome Sanger Institute COVID-19 Surveillance Team (http://www.sanger.ac.uk/covid-team) |
| EPI_ISL_597358 | Lighthouse Lab in Cambridge | Wellcome Sanger Institute for the COVID-19 Genomics UK (COG-UK) consortium | Rob Howes, The Lighthouse Lab in Cambridge and Alex Alderton, Roberto Amato, Sonia Goncalves, Ewan Harrison, David K. Jackson, Ian Johnston, Dominic Kwiatkowski, Cordelia Langford, John Sillitoe on behalf of the Wellcome Sanger Institute COVID-19 Surveillance Team (http://www.sanger.ac.uk/covid-team) |
| EPI_ISL_597359, EPI_ISL_597360 | Lighthouse Lab in Glasgow | Wellcome Sanger Institute for the COVID-19 Genomics UK (COG-UK) consortium | Harper VanSteenhouse, Yumi Kasai, David Gray, Carol Clugston, Anna Dominiczak and Alex Alderton, Roberto Amato, Sonia Goncalves, Ewan Harrison, David K. Jackson, Ian Johnston, Dominic Kwiatkowski, Cordelia Langford, John Sillitoe on behalf of the Wellcome Sanger Institute COVID-19 Surveillance Team (http://www.sanger.ac.uk/covid-team) |
| EPI_ISL_597361 | Lighthouse Lab in Cambridge | Wellcome Sanger Institute for the COVID-19 Genomics UK (COG-UK) consortium | Rob Howes, The Lighthouse Lab in Cambridge and Alex Alderton, Roberto Amato, Sonia Goncalves, Ewan Harrison, David K. Jackson, Ian Johnston, Dominic Kwiatkowski, Cordelia Langford, John Sillitoe on behalf of the Wellcome Sanger Institute COVID-19 Surveillance Team (http://www.sanger.ac.uk/covid-team) |
| EPI_ISL_597362, EPI_ISL_597363, EPI_ISL_597364 | Lighthouse Lab in Glasgow | Wellcome Sanger Institute for the COVID-19 Genomics UK (COG-UK) consortium | Harper VanSteenhouse, Yumi Kasai, David Gray, Carol Clugston, Anna Dominiczak and Alex Alderton, Roberto Amato, Sonia Goncalves, Ewan Harrison, David K. Jackson, Ian Johnston, Dominic Kwiatkowski, Cordelia Langford, John Sillitoe on behalf of the Wellcome Sanger Institute COVID-19 Surveillance Team (http://www.sanger.ac.uk/covid-team) |
| EPI_ISL_597365, EPI_ISL_597366, EPI_ISL_597367, EPI_ISL_597368 | Lighthouse Lab in Cambridge | Wellcome Sanger Institute for the COVID-19 Genomics UK (COG-UK) consortium | Rob Howes, The Lighthouse Lab in Cambridge and Alex Alderton, Roberto Amato, Sonia Goncalves, Ewan Harrison, David K. Jackson, Ian Johnston, Dominic Kwiatkowski, Cordelia Langford, John Sillitoe on behalf of the Wellcome Sanger Institute COVID-19 Surveillance Team (http://www.sanger.ac.uk/covid-team) |
| EPI_ISL_597369, EPI_ISL_597370 | Lighthouse Lab in Glasgow | Wellcome Sanger Institute for the COVID-19 Genomics UK (COG-UK) consortium | Harper VanSteenhouse, Yumi Kasai, David Gray, Carol Clugston, Anna Dominiczak and Alex Alderton, Roberto Amato, Sonia Goncalves, Ewan Harrison, David K. Jackson, Ian Johnston, Dominic Kwiatkowski, Cordelia Langford, John Sillitoe on behalf of the Wellcome Sanger Institute COVID-19 Surveillance Team (http://www.sanger.ac.uk/covid-team) |
| EPI_ISL_597371, EPI_ISL_597372, EPI_ISL_597373, EPI_ISL_597374, EPI_ISL_597375 | Lighthouse Lab in Cambridge | Wellcome Sanger Institute for the COVID-19 Genomics UK (COG-UK) consortium | Rob Howes, The Lighthouse Lab in Cambridge and Alex Alderton, Roberto Amato, Sonia Goncalves, Ewan Harrison, David K. Jackson, Ian Johnston, Dominic Kwiatkowski, Cordelia Langford, John Sillitoe on behalf of the Wellcome Sanger Institute COVID-19 Surveillance Team (http://www.sanger.ac.uk/covid-team) |
| EPI_ISL_597376 | Lighthouse Lab in Glasgow | Wellcome Sanger Institute for the COVID-19 Genomics UK (COG-UK) consortium | Harper VanSteenhouse, Yumi Kasai, David Gray, Carol Clugston, Anna Dominiczak and Alex Alderton, Roberto Amato, Sonia Goncalves, Ewan Harrison, David K. Jackson, Ian Johnston, Dominic Kwiatkowski, Cordelia Langford, John Sillitoe on behalf of the Wellcome Sanger Institute COVID-19 Surveillance Team (http://www.sanger.ac.uk/covid-team) |
| EPI_ISL_597377, EPI_ISL_597378 | Lighthouse Lab in Cambridge | Wellcome Sanger Institute for the COVID-19 Genomics UK (COG-UK) consortium | Rob Howes, The Lighthouse Lab in Cambridge and Alex Alderton, Roberto Amato, Sonia Goncalves, Ewan Harrison, David K. Jackson, Ian Johnston, Dominic Kwiatkowski, Cordelia Langford, John Sillitoe on behalf of the Wellcome Sanger Institute COVID-19 Surveillance Team (http://www.sanger.ac.uk/covid-team) |
| EPI_ISL_597379 | Lighthouse Lab in Glasgow | Wellcome Sanger Institute for the COVID-19 Genomics UK (COG-UK) consortium | Harper VanSteenhouse, Yumi Kasai, David Gray, Carol Clugston, Anna Dominiczak and Alex Alderton, Roberto Amato, Sonia Goncalves, Ewan Harrison, David K. Jackson, Ian Johnston, Dominic Kwiatkowski, Cordelia Langford, John Sillitoe on behalf of the Wellcome Sanger Institute COVID-19 Surveillance Team (http://www.sanger.ac.uk/covid-team) |
| EPI_ISL_597380, EPI_ISL_597381 | Lighthouse Lab in Cambridge | Wellcome Sanger Institute for the COVID-19 Genomics UK (COG-UK) consortium | Rob Howes, The Lighthouse Lab in Cambridge and Alex Alderton, Roberto Amato, Sonia Goncalves, Ewan Harrison, David K. Jackson, Ian Johnston, Dominic Kwiatkowski, Cordelia Langford, John Sillitoe on behalf of the Wellcome Sanger Institute COVID-19 Surveillance Team (http://www.sanger.ac.uk/covid-team) |
| EPI_ISL_597382, EPI_ISL_597383, EPI_ISL_597384, EPI_ISL_597385, EPI_ISL_597386, EPI_ISL_597387 | Lighthouse Lab in Glasgow | Wellcome Sanger Institute for the COVID-19 Genomics UK (COG-UK) consortium | Harper VanSteenhouse, Yumi Kasai, David Gray, Carol Clugston, Anna Dominiczak and Alex Alderton, Roberto Amato, Sonia Goncalves, Ewan Harrison, David K. Jackson, Ian Johnston, Dominic Kwiatkowski, Cordelia Langford, John Sillitoe on behalf of the Wellcome Sanger Institute COVID-19 Surveillance Team (http://www.sanger.ac.uk/covid-team) |
| EPI_ISL_597388 | Lighthouse Lab in Cambridge | Wellcome Sanger Institute for the COVID-19 Genomics UK (COG-UK) consortium | Rob Howes, The Lighthouse Lab in Cambridge and Alex Alderton, Roberto Amato, Sonia Goncalves, Ewan Harrison, David K. Jackson, Ian Johnston, Dominic Kwiatkowski, Cordelia Langford, John Sillitoe on behalf of the Wellcome Sanger Institute COVID-19 Surveillance Team (http://www.sanger.ac.uk/covid-team) |
| EPI_ISL_597389 | Lighthouse Lab in Glasgow | Wellcome Sanger Institute for the COVID-19 Genomics UK (COG-UK) consortium | Harper VanSteenhouse, Yumi Kasai, David Gray, Carol Clugston, Anna Dominiczak and Alex Alderton, Roberto Amato, Sonia Goncalves, Ewan Harrison, David K. Jackson, Ian Johnston, Dominic Kwiatkowski, Cordelia Langford, John Sillitoe on behalf of the Wellcome Sanger Institute COVID-19 Surveillance Team (http://www.sanger.ac.uk/covid-team) |
| EPI_ISL_597390 | Lighthouse Lab in Cambridge | Wellcome Sanger Institute for the COVID-19 Genomics UK (COG-UK) consortium | Rob Howes, The Lighthouse Lab in Cambridge and Alex Alderton, Roberto Amato, Sonia Goncalves, Ewan Harrison, David K. Jackson, Ian Johnston, Dominic Kwiatkowski, Cordelia Langford, John Sillitoe on behalf of the Wellcome Sanger Institute COVID-19 Surveillance Team (http://www.sanger.ac.uk/covid-team) |
| EPI_ISL_597391 | Lighthouse Lab in Glasgow | Wellcome Sanger Institute for the COVID-19 Genomics UK (COG-UK) consortium | Harper VanSteenhouse, Yumi Kasai, David Gray, Carol Clugston, Anna Dominiczak and Alex Alderton, Roberto Amato, Sonia Goncalves, Ewan Harrison, David K. Jackson, Ian Johnston, Dominic Kwiatkowski, Cordelia Langford, John Sillitoe on behalf of the Wellcome Sanger Institute COVID-19 Surveillance Team (http://www.sanger.ac.uk/covid-team) |
| EPI_ISL_597392 | Lighthouse Lab in Cambridge | Wellcome Sanger Institute for the COVID-19 Genomics UK (COG-UK) consortium | Rob Howes, The Lighthouse Lab in Cambridge and Alex Alderton, Roberto Amato, Sonia Goncalves, Ewan Harrison, David K. Jackson, Ian Johnston, Dominic Kwiatkowski, Cordelia Langford, John Sillitoe on behalf of the Wellcome Sanger Institute COVID-19 Surveillance Team (http://www.sanger.ac.uk/covid-team) |
| EPI_ISL_597393, EPI_ISL_597394, EPI_ISL_597395 | Lighthouse Lab in Glasgow | Wellcome Sanger Institute for the COVID-19 Genomics UK (COG-UK) consortium | Harper VanSteenhouse, Yumi Kasai, David Gray, Carol Clugston, Anna Dominiczak and Alex Alderton, Roberto Amato, Sonia Goncalves, Ewan Harrison, David K. Jackson, Ian Johnston, Dominic Kwiatkowski, Cordelia Langford, John Sillitoe on behalf of the Wellcome Sanger Institute COVID-19 Surveillance Team (http://www.sanger.ac.uk/covid-team) |
| EPI_ISL_597396, EPI_ISL_597397, EPI_ISL_597398, EPI_ISL_597399 | Lighthouse Lab in Cambridge | Wellcome Sanger Institute for the COVID-19 Genomics UK (COG-UK) consortium | Rob Howes, The Lighthouse Lab in Cambridge and Alex Alderton, Roberto Amato, Sonia Goncalves, Ewan Harrison, David K. Jackson, Ian Johnston, Dominic Kwiatkowski, Cordelia Langford, John Sillitoe on behalf of the Wellcome Sanger Institute COVID-19 Surveillance Team (http://www.sanger.ac.uk/covid-team) |
| EPI_ISL_597400 | Lighthouse Lab in Glasgow | Wellcome Sanger Institute for the COVID-19 Genomics UK (COG-UK) consortium | Harper VanSteenhouse, Yumi Kasai, David Gray, Carol Clugston, Anna Dominiczak and Alex Alderton, Roberto Amato, Sonia Goncalves, Ewan Harrison, David K. Jackson, Ian Johnston, Dominic Kwiatkowski, Cordelia Langford, John Sillitoe on behalf of the Wellcome Sanger Institute COVID-19 Surveillance Team (http://www.sanger.ac.uk/covid-team) |
| EPI_ISL_597401, EPI_ISL_597402, EPI_ISL_597403 | Lighthouse Lab in Cambridge | Wellcome Sanger Institute for the COVID-19 Genomics UK (COG-UK) consortium | Rob Howes, The Lighthouse Lab in Cambridge and Alex Alderton, Roberto Amato, Sonia Goncalves, Ewan Harrison, David K. Jackson, Ian Johnston, Dominic Kwiatkowski, Cordelia Langford, John Sillitoe on behalf of the Wellcome Sanger Institute COVID-19 Surveillance Team (http://www.sanger.ac.uk/covid-team) |

| | | | |
|---|---|---|---|
| EPI_ISL_597404 | Lighthouse Lab in Glasgow | Wellcome Sanger Institute for the COVID-19 Genomics UK (COG-UK) consortium | Harper VanSteenhouse, Yumi Kasai, David Gray, Carol Clugston, Anna Dominiczak and Alex Alderton, Roberto Amato, Sonia Goncalves, Ewan Harrison, David K. Jackson, Ian Johnston, Dominic Kwiatkowski, Cordelia Langford, John Sillitoe on behalf of the Wellcome Sanger Institute COVID-19 Surveillance Team (http://www.sanger.ac.uk/covid-team) |
| EPI_ISL_597405 | Lighthouse Lab in Cambridge | Wellcome Sanger Institute for the COVID-19 Genomics UK (COG-UK) consortium | Rob Howes, The Lighthouse Lab in Cambridge and Alex Alderton, Roberto Amato, Sonia Goncalves, Ewan Harrison, David K. Jackson, Ian Johnston, Dominic Kwiatkowski, Cordelia Langford, John Sillitoe on behalf of the Wellcome Sanger Institute COVID-19 Surveillance Team (http://www.sanger.ac.uk/covid-team) |
| EPI_ISL_597406, EPI_ISL_597407, EPI_ISL_597408, EPI_ISL_597409, EPI_ISL_597410, EPI_ISL_597411 | Lighthouse Lab in Glasgow | Wellcome Sanger Institute for the COVID-19 Genomics UK (COG-UK) consortium | Harper VanSteenhouse, Yumi Kasai, David Gray, Carol Clugston, Anna Dominiczak and Alex Alderton, Roberto Amato, Sonia Goncalves, Ewan Harrison, David K. Jackson, Ian Johnston, Dominic Kwiatkowski, Cordelia Langford, John Sillitoe on behalf of the Wellcome Sanger Institute COVID-19 Surveillance Team (http://www.sanger.ac.uk/covid-team) |
| EPI_ISL_597412 | Lighthouse Lab in Cambridge | Wellcome Sanger Institute for the COVID-19 Genomics UK (COG-UK) consortium | Rob Howes, The Lighthouse Lab in Cambridge and Alex Alderton, Roberto Amato, Sonia Goncalves, Ewan Harrison, David K. Jackson, Ian Johnston, Dominic Kwiatkowski, Cordelia Langford, John Sillitoe on behalf of the Wellcome Sanger Institute COVID-19 Surveillance Team (http://www.sanger.ac.uk/covid-team) |
| EPI_ISL_597413 | Lighthouse Lab in Glasgow | Wellcome Sanger Institute for the COVID-19 Genomics UK (COG-UK) consortium | Harper VanSteenhouse, Yumi Kasai, David Gray, Carol Clugston, Anna Dominiczak and Alex Alderton, Roberto Amato, Sonia Goncalves, Ewan Harrison, David K. Jackson, Ian Johnston, Dominic Kwiatkowski, Cordelia Langford, John Sillitoe on behalf of the Wellcome Sanger Institute COVID-19 Surveillance Team (http://www.sanger.ac.uk/covid-team) |
| EPI_ISL_597414 | Lighthouse Lab in Cambridge | Wellcome Sanger Institute for the COVID-19 Genomics UK (COG-UK) consortium | Rob Howes, The Lighthouse Lab in Cambridge and Alex Alderton, Roberto Amato, Sonia Goncalves, Ewan Harrison, David K. Jackson, Ian Johnston, Dominic Kwiatkowski, Cordelia Langford, John Sillitoe on behalf of the Wellcome Sanger Institute COVID-19 Surveillance Team (http://www.sanger.ac.uk/covid-team) |
| EPI_ISL_597415, EPI_ISL_597416, EPI_ISL_597417 | Lighthouse Lab in Glasgow | Wellcome Sanger Institute for the COVID-19 Genomics UK (COG-UK) consortium | Harper VanSteenhouse, Yumi Kasai, David Gray, Carol Clugston, Anna Dominiczak and Alex Alderton, Roberto Amato, Sonia Goncalves, Ewan Harrison, David K. Jackson, Ian Johnston, Dominic Kwiatkowski, Cordelia Langford, John Sillitoe on behalf of the Wellcome Sanger Institute COVID-19 Surveillance Team (http://www.sanger.ac.uk/covid-team) |
| EPI_ISL_597418 | Lighthouse Lab in Cambridge | Wellcome Sanger Institute for the COVID-19 Genomics UK (COG-UK) consortium | Rob Howes, The Lighthouse Lab in Cambridge and Alex Alderton, Roberto Amato, Sonia Goncalves, Ewan Harrison, David K. Jackson, Ian Johnston, Dominic Kwiatkowski, Cordelia Langford, John Sillitoe on behalf of the Wellcome Sanger Institute COVID-19 Surveillance Team (http://www.sanger.ac.uk/covid-team) |
| EPI_ISL_597419, EPI_ISL_597420, EPI_ISL_597421, EPI_ISL_597422, EPI_ISL_597423 | Lighthouse Lab in Glasgow | Wellcome Sanger Institute for the COVID-19 Genomics UK (COG-UK) consortium | Harper VanSteenhouse, Yumi Kasai, David Gray, Carol Clugston, Anna Dominiczak and Alex Alderton, Roberto Amato, Sonia Goncalves, Ewan Harrison, David K. Jackson, Ian Johnston, Dominic Kwiatkowski, Cordelia Langford, John Sillitoe on behalf of the Wellcome Sanger Institute COVID-19 Surveillance Team (http://www.sanger.ac.uk/covid-team) |
| EPI_ISL_597424, EPI_ISL_597425, EPI_ISL_597426, EPI_ISL_597427, EPI_ISL_597428 | Lighthouse Lab in Cambridge | Wellcome Sanger Institute for the COVID-19 Genomics UK (COG-UK) consortium | Rob Howes, The Lighthouse Lab in Cambridge and Alex Alderton, Roberto Amato, Sonia Goncalves, Ewan Harrison, David K. Jackson, Ian Johnston, Dominic Kwiatkowski, Cordelia Langford, John Sillitoe on behalf of the Wellcome Sanger Institute COVID-19 Surveillance Team (http://www.sanger.ac.uk/covid-team) |
| EPI_ISL_597429, EPI_ISL_597430, EPI_ISL_597431 | Lighthouse Lab in Glasgow | Wellcome Sanger Institute for the COVID-19 Genomics UK (COG-UK) consortium | Harper VanSteenhouse, Yumi Kasai, David Gray, Carol Clugston, Anna Dominiczak and Alex Alderton, Roberto Amato, Sonia Goncalves, Ewan Harrison, David K. Jackson, Ian Johnston, Dominic Kwiatkowski, Cordelia Langford, John Sillitoe on behalf of the Wellcome Sanger Institute COVID-19 Surveillance Team (http://www.sanger.ac.uk/covid-team) |
| EPI_ISL_597432 | Lighthouse Lab in Cambridge | Wellcome Sanger Institute for the COVID-19 Genomics UK (COG-UK) consortium | Rob Howes, The Lighthouse Lab in Cambridge and Alex Alderton, Roberto Amato, Sonia Goncalves, Ewan Harrison, David K. Jackson, Ian Johnston, Dominic Kwiatkowski, Cordelia Langford, John Sillitoe on behalf of the Wellcome Sanger Institute COVID-19 Surveillance Team (http://www.sanger.ac.uk/covid-team) |
| EPI_ISL_597433, EPI_ISL_597434 | Lighthouse Lab in Glasgow | Wellcome Sanger Institute for the COVID-19 Genomics UK (COG-UK) consortium | Harper VanSteenhouse, Yumi Kasai, David Gray, Carol Clugston, Anna Dominiczak and Alex Alderton, Roberto Amato, Sonia Goncalves, Ewan Harrison, David K. Jackson, Ian Johnston, Dominic Kwiatkowski, Cordelia Langford, John Sillitoe on behalf of the Wellcome Sanger Institute COVID-19 Surveillance Team (http://www.sanger.ac.uk/covid-team) |
| EPI_ISL_597435 | Lighthouse Lab in Cambridge | Wellcome Sanger Institute for the COVID-19 Genomics UK (COG-UK) consortium | Rob Howes, The Lighthouse Lab in Cambridge and Alex Alderton, Roberto Amato, Sonia Goncalves, Ewan Harrison, David K. Jackson, Ian Johnston, Dominic Kwiatkowski, Cordelia Langford, John Sillitoe on behalf of the Wellcome Sanger Institute COVID-19 Surveillance Team (http://www.sanger.ac.uk/covid-team) |
| EPI_ISL_597436 | Lighthouse Lab in Glasgow | Wellcome Sanger Institute for the COVID-19 Genomics UK (COG-UK) consortium | Harper VanSteenhouse, Yumi Kasai, David Gray, Carol Clugston, Anna Dominiczak and Alex Alderton, Roberto Amato, Sonia Goncalves, Ewan Harrison, David K. Jackson, Ian Johnston, Dominic Kwiatkowski, Cordelia Langford, John Sillitoe on behalf of the Wellcome Sanger Institute COVID-19 Surveillance Team (http://www.sanger.ac.uk/covid-team) |
| EPI_ISL_597437, EPI_ISL_597438 | Lighthouse Lab in Cambridge | Wellcome Sanger Institute for the COVID-19 Genomics UK (COG-UK) consortium | Rob Howes, The Lighthouse Lab in Cambridge and Alex Alderton, Roberto Amato, Sonia Goncalves, Ewan Harrison, David K. Jackson, Ian Johnston, Dominic Kwiatkowski, Cordelia Langford, John Sillitoe on behalf of the Wellcome Sanger Institute COVID-19 Surveillance Team (http://www.sanger.ac.uk/covid-team) |
| EPI_ISL_597439, EPI_ISL_597440 | Lighthouse Lab in Glasgow | Wellcome Sanger Institute for the COVID-19 Genomics UK (COG-UK) consortium | Harper VanSteenhouse, Yumi Kasai, David Gray, Carol Clugston, Anna Dominiczak and Alex Alderton, Roberto Amato, Sonia Goncalves, Ewan Harrison, David K. Jackson, Ian Johnston, Dominic Kwiatkowski, Cordelia Langford, John Sillitoe on behalf of the Wellcome Sanger Institute COVID-19 Surveillance Team (http://www.sanger.ac.uk/covid-team) |
| EPI_ISL_597441 | Lighthouse Lab in Cambridge | Wellcome Sanger Institute for the COVID-19 Genomics UK (COG-UK) consortium | Rob Howes, The Lighthouse Lab in Cambridge and Alex Alderton, Roberto Amato, Sonia Goncalves, Ewan Harrison, David K. Jackson, Ian Johnston, Dominic Kwiatkowski, Cordelia Langford, John Sillitoe on behalf of the Wellcome Sanger Institute COVID-19 Surveillance Team (http://www.sanger.ac.uk/covid-team) |
| EPI_ISL_597442, EPI_ISL_597443, EPI_ISL_597444 | Lighthouse Lab in Glasgow | Wellcome Sanger Institute for the COVID-19 Genomics UK (COG-UK) consortium | Harper VanSteenhouse, Yumi Kasai, David Gray, Carol Clugston, Anna Dominiczak and Alex Alderton, Roberto Amato, Sonia Goncalves, Ewan Harrison, David K. Jackson, Ian Johnston, Dominic Kwiatkowski, Cordelia Langford, John Sillitoe on behalf of the Wellcome Sanger Institute COVID-19 Surveillance Team (http://www.sanger.ac.uk/covid-team) |
| EPI_ISL_597445, EPI_ISL_597446 | Lighthouse Lab in Cambridge | Wellcome Sanger Institute for the COVID-19 Genomics UK (COG-UK) consortium | Rob Howes, The Lighthouse Lab in Cambridge and Alex Alderton, Roberto Amato, Sonia Goncalves, Ewan Harrison, David K. Jackson, Ian Johnston, Dominic Kwiatkowski, Cordelia Langford, John Sillitoe on behalf of the Wellcome Sanger Institute COVID-19 Surveillance Team (http://www.sanger.ac.uk/covid-team) |
| EPI_ISL_597447, EPI_ISL_597448 | Lighthouse Lab in Glasgow | Wellcome Sanger Institute for the COVID-19 Genomics UK (COG-UK) consortium | Harper VanSteenhouse, Yumi Kasai, David Gray, Carol Clugston, Anna Dominiczak and Alex Alderton, Roberto Amato, Sonia Goncalves, Ewan Harrison, David K. Jackson, Ian Johnston, Dominic Kwiatkowski, Cordelia Langford, John Sillitoe on behalf of the Wellcome Sanger Institute COVID-19 Surveillance Team (http://www.sanger.ac.uk/covid-team) |
| EPI_ISL_597449, EPI_ISL_597450, EPI_ISL_597451 | Lighthouse Lab in Cambridge | Wellcome Sanger Institute for the COVID-19 Genomics UK (COG-UK) consortium | Rob Howes, The Lighthouse Lab in Cambridge and Alex Alderton, Roberto Amato, Sonia Goncalves, Ewan Harrison, David K. Jackson, Ian Johnston, Dominic Kwiatkowski, Cordelia Langford, John Sillitoe on behalf of the Wellcome Sanger Institute COVID-19 Surveillance Team (http://www.sanger.ac.uk/covid-team) |
| EPI_ISL_597452, EPI_ISL_597454, EPI_ISL_597455, EPI_ISL_597456, EPI_ISL_597457 | Lighthouse Lab in Glasgow | Wellcome Sanger Institute for the COVID-19 Genomics UK (COG-UK) consortium | Harper VanSteenhouse, Yumi Kasai, David Gray, Carol Clugston, Anna Dominiczak and Alex Alderton, Roberto Amato, Sonia Goncalves, Ewan Harrison, David K. Jackson, Ian Johnston, Dominic Kwiatkowski, Cordelia Langford, John Sillitoe on behalf of the Wellcome Sanger Institute COVID-19 Surveillance Team (http://www.sanger.ac.uk/covid-team) |
| EPI_ISL_597458, EPI_ISL_597459, EPI_ISL_597460, EPI_ISL_597461, EPI_ISL_597462 | Lighthouse Lab in Cambridge | Wellcome Sanger Institute for the COVID-19 Genomics UK (COG-UK) consortium | Rob Howes, The Lighthouse Lab in Cambridge and Alex Alderton, Roberto Amato, Sonia Goncalves, Ewan Harrison, David K. Jackson, Ian Johnston, Dominic Kwiatkowski, Cordelia Langford, John Sillitoe on behalf of the Wellcome Sanger Institute COVID-19 Surveillance Team (http://www.sanger.ac.uk/covid-team) |
| EPI_ISL_597463 | Lighthouse Lab in Glasgow | Wellcome Sanger Institute for the COVID-19 Genomics UK (COG-UK) consortium | Harper VanSteenhouse, Yumi Kasai, David Gray, Carol Clugston, Anna Dominiczak and Alex Alderton, Roberto Amato, Sonia Goncalves, Ewan Harrison, David K. Jackson, Ian Johnston, Dominic Kwiatkowski, Cordelia Langford, John Sillitoe on behalf of the Wellcome Sanger Institute COVID-19 Surveillance Team (http://www.sanger.ac.uk/covid-team) |
| EPI_ISL_597464 | Lighthouse Lab in Cambridge | Wellcome Sanger Institute for the COVID-19 Genomics UK (COG-UK) consortium | Rob Howes, The Lighthouse Lab in Cambridge and Alex Alderton, Roberto Amato, Sonia Goncalves, Ewan Harrison, David K. Jackson, Ian Johnston, Dominic Kwiatkowski, Cordelia Langford, John Sillitoe on behalf of the Wellcome Sanger Institute COVID-19 Surveillance Team (http://www.sanger.ac.uk/covid-team) |
| EPI_ISL_597465 | Lighthouse Lab in Glasgow | Wellcome Sanger Institute for the COVID-19 Genomics UK (COG-UK) consortium | Harper VanSteenhouse, Yumi Kasai, David Gray, Carol Clugston, Anna Dominiczak and Alex Alderton, Roberto Amato, Sonia Goncalves, Ewan Harrison, David K. Jackson, Ian Johnston, Dominic Kwiatkowski, Cordelia Langford, John Sillitoe on behalf of the Wellcome Sanger Institute COVID-19 Surveillance Team (http://www.sanger.ac.uk/covid-team) |
| EPI_ISL_597466, EPI_ISL_597467, EPI_ISL_597468, EPI_ISL_597469, EPI_ISL_597470 | Lighthouse Lab in Cambridge | Wellcome Sanger Institute for the COVID-19 Genomics UK (COG-UK) consortium | Rob Howes, The Lighthouse Lab in Cambridge and Alex Alderton, Roberto Amato, Sonia Goncalves, Ewan Harrison, David K. Jackson, Ian Johnston, Dominic Kwiatkowski, Cordelia Langford, John Sillitoe on behalf of the Wellcome Sanger Institute COVID-19 Surveillance Team (http://www.sanger.ac.uk/covid-team) |

| | | | |
|---|---|---|---|
| EPI_ISL_597471 | Lighthouse Lab in Glasgow | Wellcome Sanger Institute for the COVID-19 Genomics UK (COG-UK) consortium | Harper VanSteenhouse, Yumi Kasai, David Gray, Carol Clugston, Anna Dominiczak and Alex Alderton, Roberto Amato, Sonia Goncalves, Ewan Harrison, David K. Jackson, Ian Johnston, Dominic Kwiatkowski, Cordelia Langford, John Sillitoe on behalf of the Wellcome Sanger Institute COVID-19 Surveillance Team (http://www.sanger.ac.uk/covid-team) |
| EPI_ISL_597472, EPI_ISL_597473 | Lighthouse Lab in Cambridge | Wellcome Sanger Institute for the COVID-19 Genomics UK (COG-UK) consortium | Rob Howes, The Lighthouse Lab in Cambridge and Alex Alderton, Roberto Amato, Sonia Goncalves, Ewan Harrison, David K. Jackson, Ian Johnston, Dominic Kwiatkowski, Cordelia Langford, John Sillitoe on behalf of the Wellcome Sanger Institute COVID-19 Surveillance Team (http://www.sanger.ac.uk/covid-team) |
| EPI_ISL_597474, EPI_ISL_597475 | Lighthouse Lab in Glasgow | Wellcome Sanger Institute for the COVID-19 Genomics UK (COG-UK) consortium | Harper VanSteenhouse, Yumi Kasai, David Gray, Carol Clugston, Anna Dominiczak and Alex Alderton, Roberto Amato, Sonia Goncalves, Ewan Harrison, David K. Jackson, Ian Johnston, Dominic Kwiatkowski, Cordelia Langford, John Sillitoe on behalf of the Wellcome Sanger Institute COVID-19 Surveillance Team (http://www.sanger.ac.uk/covid-team) |
| EPI_ISL_597476, EPI_ISL_597477, EPI_ISL_597478, EPI_ISL_597479, EPI_ISL_597480 | Lighthouse Lab in Cambridge | Wellcome Sanger Institute for the COVID-19 Genomics UK (COG-UK) consortium | Rob Howes, The Lighthouse Lab in Cambridge and Alex Alderton, Roberto Amato, Sonia Goncalves, Ewan Harrison, David K. Jackson, Ian Johnston, Dominic Kwiatkowski, Cordelia Langford, John Sillitoe on behalf of the Wellcome Sanger Institute COVID-19 Surveillance Team (http://www.sanger.ac.uk/covid-team) |
| EPI_ISL_597481, EPI_ISL_597482 | Lighthouse Lab in Glasgow | Wellcome Sanger Institute for the COVID-19 Genomics UK (COG-UK) consortium | Harper VanSteenhouse, Yumi Kasai, David Gray, Carol Clugston, Anna Dominiczak and Alex Alderton, Roberto Amato, Sonia Goncalves, Ewan Harrison, David K. Jackson, Ian Johnston, Dominic Kwiatkowski, Cordelia Langford, John Sillitoe on behalf of the Wellcome Sanger Institute COVID-19 Surveillance Team (http://www.sanger.ac.uk/covid-team) |
| EPI_ISL_597483 | Lighthouse Lab in Cambridge | Wellcome Sanger Institute for the COVID-19 Genomics UK (COG-UK) consortium | Rob Howes, The Lighthouse Lab in Cambridge and Alex Alderton, Roberto Amato, Sonia Goncalves, Ewan Harrison, David K. Jackson, Ian Johnston, Dominic Kwiatkowski, Cordelia Langford, John Sillitoe on behalf of the Wellcome Sanger Institute COVID-19 Surveillance Team (http://www.sanger.ac.uk/covid-team) |
| EPI_ISL_597484, EPI_ISL_597485, EPI_ISL_597486 | Lighthouse Lab in Glasgow | Wellcome Sanger Institute for the COVID-19 Genomics UK (COG-UK) consortium | Harper VanSteenhouse, Yumi Kasai, David Gray, Carol Clugston, Anna Dominiczak and Alex Alderton, Roberto Amato, Sonia Goncalves, Ewan Harrison, David K. Jackson, Ian Johnston, Dominic Kwiatkowski, Cordelia Langford, John Sillitoe on behalf of the Wellcome Sanger Institute COVID-19 Surveillance Team (http://www.sanger.ac.uk/covid-team) |
| EPI_ISL_597487, EPI_ISL_597488, EPI_ISL_597489, EPI_ISL_597490 | Lighthouse Lab in Cambridge | Wellcome Sanger Institute for the COVID-19 Genomics UK (COG-UK) consortium | Rob Howes, The Lighthouse Lab in Cambridge and Alex Alderton, Roberto Amato, Sonia Goncalves, Ewan Harrison, David K. Jackson, Ian Johnston, Dominic Kwiatkowski, Cordelia Langford, John Sillitoe on behalf of the Wellcome Sanger Institute COVID-19 Surveillance Team (http://www.sanger.ac.uk/covid-team) |
| EPI_ISL_597491, EPI_ISL_597492, EPI_ISL_597493 | Lighthouse Lab in Glasgow | Wellcome Sanger Institute for the COVID-19 Genomics UK (COG-UK) consortium | Harper VanSteenhouse, Yumi Kasai, David Gray, Carol Clugston, Anna Dominiczak and Alex Alderton, Roberto Amato, Sonia Goncalves, Ewan Harrison, David K. Jackson, Ian Johnston, Dominic Kwiatkowski, Cordelia Langford, John Sillitoe on behalf of the Wellcome Sanger Institute COVID-19 Surveillance Team (http://www.sanger.ac.uk/covid-team) |
| EPI_ISL_597494, EPI_ISL_597495 | Lighthouse Lab in Cambridge | Wellcome Sanger Institute for the COVID-19 Genomics UK (COG-UK) consortium | Rob Howes, The Lighthouse Lab in Cambridge and Alex Alderton, Roberto Amato, Sonia Goncalves, Ewan Harrison, David K. Jackson, Ian Johnston, Dominic Kwiatkowski, Cordelia Langford, John Sillitoe on behalf of the Wellcome Sanger Institute COVID-19 Surveillance Team (http://www.sanger.ac.uk/covid-team) |
| EPI_ISL_597496 | Lighthouse Lab in Glasgow | Wellcome Sanger Institute for the COVID-19 Genomics UK (COG-UK) consortium | Harper VanSteenhouse, Yumi Kasai, David Gray, Carol Clugston, Anna Dominiczak and Alex Alderton, Roberto Amato, Sonia Goncalves, Ewan Harrison, David K. Jackson, Ian Johnston, Dominic Kwiatkowski, Cordelia Langford, John Sillitoe on behalf of the Wellcome Sanger Institute COVID-19 Surveillance Team (http://www.sanger.ac.uk/covid-team) |
| EPI_ISL_597497 | Lighthouse Lab in Cambridge | Wellcome Sanger Institute for the COVID-19 Genomics UK (COG-UK) consortium | Rob Howes, The Lighthouse Lab in Cambridge and Alex Alderton, Roberto Amato, Sonia Goncalves, Ewan Harrison, David K. Jackson, Ian Johnston, Dominic Kwiatkowski, Cordelia Langford, John Sillitoe on behalf of the Wellcome Sanger Institute COVID-19 Surveillance Team (http://www.sanger.ac.uk/covid-team) |
| EPI_ISL_597498, EPI_ISL_597499, EPI_ISL_597500, EPI_ISL_597501 | Lighthouse Lab in Glasgow | Wellcome Sanger Institute for the COVID-19 Genomics UK (COG-UK) consortium | Harper VanSteenhouse, Yumi Kasai, David Gray, Carol Clugston, Anna Dominiczak and Alex Alderton, Roberto Amato, Sonia Goncalves, Ewan Harrison, David K. Jackson, Ian Johnston, Dominic Kwiatkowski, Cordelia Langford, John Sillitoe on behalf of the Wellcome Sanger Institute COVID-19 Surveillance Team (http://www.sanger.ac.uk/covid-team) |
| EPI_ISL_597502 | Lighthouse Lab in Cambridge | Wellcome Sanger Institute for the COVID-19 Genomics UK (COG-UK) consortium | Rob Howes, The Lighthouse Lab in Cambridge and Alex Alderton, Roberto Amato, Sonia Goncalves, Ewan Harrison, David K. Jackson, Ian Johnston, Dominic Kwiatkowski, Cordelia Langford, John Sillitoe on behalf of the Wellcome Sanger Institute COVID-19 Surveillance Team (http://www.sanger.ac.uk/covid-team) |
| EPI_ISL_597503 | Lighthouse Lab in Glasgow | Wellcome Sanger Institute for the COVID-19 Genomics UK (COG-UK) consortium | Harper VanSteenhouse, Yumi Kasai, David Gray, Carol Clugston, Anna Dominiczak and Alex Alderton, Roberto Amato, Sonia Goncalves, Ewan Harrison, David K. Jackson, Ian Johnston, Dominic Kwiatkowski, Cordelia Langford, John Sillitoe on behalf of the Wellcome Sanger Institute COVID-19 Surveillance Team (http://www.sanger.ac.uk/covid-team) |
| EPI_ISL_597504, EPI_ISL_597505 | Lighthouse Lab in Cambridge | Wellcome Sanger Institute for the COVID-19 Genomics UK (COG-UK) consortium | Rob Howes, The Lighthouse Lab in Cambridge and Alex Alderton, Roberto Amato, Sonia Goncalves, Ewan Harrison, David K. Jackson, Ian Johnston, Dominic Kwiatkowski, Cordelia Langford, John Sillitoe on behalf of the Wellcome Sanger Institute COVID-19 Surveillance Team (http://www.sanger.ac.uk/covid-team) |
| EPI_ISL_597506 | Lighthouse Lab in Glasgow | Wellcome Sanger Institute for the COVID-19 Genomics UK (COG-UK) consortium | Harper VanSteenhouse, Yumi Kasai, David Gray, Carol Clugston, Anna Dominiczak and Alex Alderton, Roberto Amato, Sonia Goncalves, Ewan Harrison, David K. Jackson, Ian Johnston, Dominic Kwiatkowski, Cordelia Langford, John Sillitoe on behalf of the Wellcome Sanger Institute COVID-19 Surveillance Team (http://www.sanger.ac.uk/covid-team) |
| EPI_ISL_597507, EPI_ISL_597508 | Lighthouse Lab in Cambridge | Wellcome Sanger Institute for the COVID-19 Genomics UK (COG-UK) consortium | Rob Howes, The Lighthouse Lab in Cambridge and Alex Alderton, Roberto Amato, Sonia Goncalves, Ewan Harrison, David K. Jackson, Ian Johnston, Dominic Kwiatkowski, Cordelia Langford, John Sillitoe on behalf of the Wellcome Sanger Institute COVID-19 Surveillance Team (http://www.sanger.ac.uk/covid-team) |
| EPI_ISL_597509 | Lighthouse Lab in Glasgow | Wellcome Sanger Institute for the COVID-19 Genomics UK (COG-UK) consortium | Harper VanSteenhouse, Yumi Kasai, David Gray, Carol Clugston, Anna Dominiczak and Alex Alderton, Roberto Amato, Sonia Goncalves, Ewan Harrison, David K. Jackson, Ian Johnston, Dominic Kwiatkowski, Cordelia Langford, John Sillitoe on behalf of the Wellcome Sanger Institute COVID-19 Surveillance Team (http://www.sanger.ac.uk/covid-team) |
| EPI_ISL_597510 | Lighthouse Lab in Cambridge | Wellcome Sanger Institute for the COVID-19 Genomics UK (COG-UK) consortium | Rob Howes, The Lighthouse Lab in Cambridge and Alex Alderton, Roberto Amato, Sonia Goncalves, Ewan Harrison, David K. Jackson, Ian Johnston, Dominic Kwiatkowski, Cordelia Langford, John Sillitoe on behalf of the Wellcome Sanger Institute COVID-19 Surveillance Team (http://www.sanger.ac.uk/covid-team) |
| EPI_ISL_597511 | Lighthouse Lab in Glasgow | Wellcome Sanger Institute for the COVID-19 Genomics UK (COG-UK) consortium | Harper VanSteenhouse, Yumi Kasai, David Gray, Carol Clugston, Anna Dominiczak and Alex Alderton, Roberto Amato, Sonia Goncalves, Ewan Harrison, David K. Jackson, Ian Johnston, Dominic Kwiatkowski, Cordelia Langford, John Sillitoe on behalf of the Wellcome Sanger Institute COVID-19 Surveillance Team (http://www.sanger.ac.uk/covid-team) |
| EPI_ISL_597512 | Lighthouse Lab in Cambridge | Wellcome Sanger Institute for the COVID-19 Genomics UK (COG-UK) consortium | Rob Howes, The Lighthouse Lab in Cambridge and Alex Alderton, Roberto Amato, Sonia Goncalves, Ewan Harrison, David K. Jackson, Ian Johnston, Dominic Kwiatkowski, Cordelia Langford, John Sillitoe on behalf of the Wellcome Sanger Institute COVID-19 Surveillance Team (http://www.sanger.ac.uk/covid-team) |
| EPI_ISL_597513, EPI_ISL_597514, EPI_ISL_597515, EPI_ISL_597516 | Lighthouse Lab in Glasgow | Wellcome Sanger Institute for the COVID-19 Genomics UK (COG-UK) consortium | Harper VanSteenhouse, Yumi Kasai, David Gray, Carol Clugston, Anna Dominiczak and Alex Alderton, Roberto Amato, Sonia Goncalves, Ewan Harrison, David K. Jackson, Ian Johnston, Dominic Kwiatkowski, Cordelia Langford, John Sillitoe on behalf of the Wellcome Sanger Institute COVID-19 Surveillance Team (http://www.sanger.ac.uk/covid-team) |
| EPI_ISL_597517 | Lighthouse Lab in Cambridge | Wellcome Sanger Institute for the COVID-19 Genomics UK (COG-UK) consortium | Rob Howes, The Lighthouse Lab in Cambridge and Alex Alderton, Roberto Amato, Sonia Goncalves, Ewan Harrison, David K. Jackson, Ian Johnston, Dominic Kwiatkowski, Cordelia Langford, John Sillitoe on behalf of the Wellcome Sanger Institute COVID-19 Surveillance Team (http://www.sanger.ac.uk/covid-team) |
| EPI_ISL_597518 | Lighthouse Lab in Glasgow | Wellcome Sanger Institute for the COVID-19 Genomics UK (COG-UK) consortium | Harper VanSteenhouse, Yumi Kasai, David Gray, Carol Clugston, Anna Dominiczak and Alex Alderton, Roberto Amato, Sonia Goncalves, Ewan Harrison, David K. Jackson, Ian Johnston, Dominic Kwiatkowski, Cordelia Langford, John Sillitoe on behalf of the Wellcome Sanger Institute COVID-19 Surveillance Team (http://www.sanger.ac.uk/covid-team) |
| EPI_ISL_597519, EPI_ISL_597520 | Lighthouse Lab in Cambridge | Wellcome Sanger Institute for the COVID-19 Genomics UK (COG-UK) consortium | Rob Howes, The Lighthouse Lab in Cambridge and Alex Alderton, Roberto Amato, Sonia Goncalves, Ewan Harrison, David K. Jackson, Ian Johnston, Dominic Kwiatkowski, Cordelia Langford, John Sillitoe on behalf of the Wellcome Sanger Institute COVID-19 Surveillance Team (http://www.sanger.ac.uk/covid-team) |
| EPI_ISL_597521, EPI_ISL_597522 | Lighthouse Lab in Glasgow | Wellcome Sanger Institute for the COVID-19 Genomics UK (COG-UK) consortium | Harper VanSteenhouse, Yumi Kasai, David Gray, Carol Clugston, Anna Dominiczak and Alex Alderton, Roberto Amato, Sonia Goncalves, Ewan Harrison, David K. Jackson, Ian Johnston, Dominic Kwiatkowski, Cordelia Langford, John Sillitoe on behalf of the Wellcome Sanger Institute COVID-19 Surveillance Team (http://www.sanger.ac.uk/covid-team) |

EPI_ISL_597523, EPI_ISL_597524, EPI_ISL_597525, EPI_ISL_597526, EPI_ISL_597527, EPI_ISL_597528, EPI_ISL_597529, EPI_ISL_597530, EPI_ISL_597531, EPI_ISL_597532, EPI_ISL_597533, EPI_ISL_597534, EPI_ISL_597535, EPI_ISL_597536, EPI_ISL_597537, EPI_ISL_597538, EPI_ISL_597539, EPI_ISL_597540, EPI_ISL_597541, EPI_ISL_597542, EPI_ISL_597543, EPI_ISL_597544, EPI_ISL_597545, EPI_ISL_597546, EPI_ISL_597547, EPI_ISL_597548, EPI_ISL_597549, EPI_ISL_597550, EPI_ISL_597551, EPI_ISL_597552, EPI_ISL_597553, EPI_ISL_597554, EPI_ISL_597555, EPI_ISL_597556, EPI_ISL_597557, EPI_ISL_597558, EPI_ISL_597559, EPI_ISL_597560, EPI_ISL_597561, EPI_ISL_597562, EPI_ISL_597563, EPI_ISL_597564, EPI_ISL_597565, EPI_ISL_597566, EPI_ISL_597567, EPI_ISL_597568, EPI_ISL_597569, EPI_ISL_597570, EPI_ISL_597571, EPI_ISL_597572, EPI_ISL_597573, EPI_ISL_597574, EPI_ISL_597575, EPI_ISL_597576, EPI_ISL_597577, EPI_ISL_597578, EPI_ISL_597579, EPI_ISL_597580, EPI_ISL_597581, EPI_ISL_597582, EPI_ISL_597583, EPI_ISL_597584, EPI_ISL_597585, EPI_ISL_597586, EPI_ISL_597587, EPI_ISL_597588, EPI_ISL_597589, EPI_ISL_597590, EPI_ISL_597591, EPI_ISL_597592, EPI_ISL_597593, EPI_ISL_597594, EPI_ISL_597595, EPI_ISL_597596, EPI_ISL_597597, EPI_ISL_597598, EPI_ISL_597599, EPI_ISL_597600, EPI_ISL_597601, EPI_ISL_597602, EPI_ISL_597603, EPI_ISL_597604, EPI_ISL_597605, EPI_ISL_597606, EPI_ISL_597607, EPI_ISL_597608, EPI_ISL_597609, EPI_ISL_597610, EPI_ISL_597611, EPI_ISL_597612,

| | | | |
|---|---|---|---|
| EPI_ISL_597613, EPI_ISL_597614, EPI_ISL_597615, EPI_ISL_597616, EPI_ISL_597617, EPI_ISL_597618, EPI_ISL_597619, EPI_ISL_597620, EPI_ISL_597621, EPI_ISL_597622, EPI_ISL_597623, EPI_ISL_597624, EPI_ISL_597625, EPI_ISL_597626, EPI_ISL_597627, EPI_ISL_597628, EPI_ISL_597629, EPI_ISL_597630, EPI_ISL_597631, EPI_ISL_597632 | | | |
| see above | Lighthouse Lab in Alderley Park | Wellcome Sanger Institute for the COVID-19 Genomics UK (COG-UK) consortium | Jacquelyn Wynn, Mairead Hyland, The Lighthouse Lab in Alderley Park and Alex Alderton, Roberto Amato, Sonia Goncalves, Ewan Harrison, David K. Jackson, Ian Johnston, Dominic Kwiatkowski, Cordelia Langford, John Sillitoe on behalf of the Wellcome Sanger Institute COVID-19 Surveillance Team (http://www.sanger.ac.uk/covid-team) |
| EPI_ISL_597633 | Lighthouse Lab in Alderley Park | Wellcome Sanger Institute for the COVID-19 Genomics UK (COG-UK) Consortium | Jacquelyn Wynn, Mairead Hyland, The Lighthouse Lab in Alderley Park and Alex Alderton, Roberto Amato, Sonia Goncalves, Ewan Harrison, David K. Jackson, Ian Johnston, Dominic Kwiatkowski, Cordelia Langford, John Sillitoe on behalf of the Wellcome Sanger Institute COVID-19 Surveillance Team (http://www.sanger.ac.uk/covid-team) |
| EPI_ISL_597634, EPI_ISL_597635, EPI_ISL_597636, EPI_ISL_597637, EPI_ISL_597638, EPI_ISL_597639, EPI_ISL_597640, EPI_ISL_597641, EPI_ISL_597642, EPI_ISL_597643, EPI_ISL_597644, EPI_ISL_597645, EPI_ISL_597646, EPI_ISL_597647, EPI_ISL_597648, EPI_ISL_597649, EPI_ISL_597650, EPI_ISL_597651, EPI_ISL_597652, EPI_ISL_597653, EPI_ISL_597654, EPI_ISL_597655, EPI_ISL_597656, EPI_ISL_597657, EPI_ISL_597658, EPI_ISL_597659, EPI_ISL_597660, EPI_ISL_597661, EPI_ISL_597662, EPI_ISL_597663, EPI_ISL_597664, EPI_ISL_597665, EPI_ISL_597666, EPI_ISL_597667, EPI_ISL_597668, EPI_ISL_597669, EPI_ISL_597670, EPI_ISL_597671, EPI_ISL_597672, EPI_ISL_597673, EPI_ISL_597674, EPI_ISL_597675, EPI_ISL_597676, EPI_ISL_597677, EPI_ISL_597678, EPI_ISL_597679, EPI_ISL_597680, EPI_ISL_597681, EPI_ISL_597682, EPI_ISL_597683, EPI_ISL_597684, EPI_ISL_597685, EPI_ISL_597686, EPI_ISL_597687, EPI_ISL_597688, EPI_ISL_597689, EPI_ISL_597690, EPI_ISL_597691, EPI_ISL_597692, EPI_ISL_597693, EPI_ISL_597694, EPI_ISL_597695, EPI_ISL_597696, EPI_ISL_597697, EPI_ISL_597698, EPI_ISL_597699, EPI_ISL_597700, EPI_ISL_597701, EPI_ISL_597702, EPI_ISL_597703, EPI_ISL_597704, EPI_ISL_597705, EPI_ISL_597706, EPI_ISL_597707, EPI_ISL_597708, EPI_ISL_597709, EPI_ISL_597710, EPI_ISL_597711, EPI_ISL_597712, EPI_ISL_597713, EPI_ISL_597714, EPI_ISL_597715, EPI_ISL_597716, EPI_ISL_597717, EPI_ISL_597718, EPI_ISL_597719, EPI_ISL_597720, EPI_ISL_597721, EPI_ISL_597722, EPI_ISL_597723, EPI_ISL_597724, EPI_ISL_597725, EPI_ISL_597726, EPI_ISL_597727, EPI_ISL_597728, EPI_ISL_597729, EPI_ISL_597730, EPI_ISL_597731, EPI_ISL_597732, EPI_ISL_597733, EPI_ISL_597734, EPI_ISL_597735, EPI_ISL_597736, EPI_ISL_597737, EPI_ISL_597738, EPI_ISL_597739, EPI_ISL_597741, EPI_ISL_597742, EPI_ISL_597743, EPI_ISL_597744, EPI_ISL_597745, EPI_ISL_597746, EPI_ISL_597747, EPI_ISL_597748, EPI_ISL_597749, EPI_ISL_597750, EPI_ISL_597751, EPI_ISL_597752, EPI_ISL_597753, EPI_ISL_597754, EPI_ISL_597755, EPI_ISL_597756, EPI_ISL_597757, EPI_ISL_597758, EPI_ISL_597759, EPI_ISL_597760, EPI_ISL_597761, EPI_ISL_597762, EPI_ISL_597763, EPI_ISL_597764, EPI_ISL_597765, EPI_ISL_597766, EPI_ISL_597767, EPI_ISL_597768, EPI_ISL_597769, EPI_ISL_597770, EPI_ISL_597771, EPI_ISL_597772, EPI_ISL_597773, EPI_ISL_597774, EPI_ISL_597775, EPI_ISL_597776, EPI_ISL_597777, EPI_ISL_597778, EPI_ISL_597779, EPI_ISL_597780, EPI_ISL_597781, EPI_ISL_597782, EPI_ISL_597783, EPI_ISL_597784, EPI_ISL_597785, EPI_ISL_597786, EPI_ISL_597787, EPI_ISL_597788, EPI_ISL_597789, EPI_ISL_597790, EPI_ISL_597791, EPI_ISL_597792, EPI_ISL_597793, EPI_ISL_597794, EPI_ISL_597795, EPI_ISL_597796, EPI_ISL_597797, EPI_ISL_597798, EPI_ISL_597799, EPI_ISL_597800, EPI_ISL_597801, EPI_ISL_597802, EPI_ISL_597803, EPI_ISL_597804, EPI_ISL_597805, EPI_ISL_597806, EPI_ISL_597807, EPI_ISL_597808, EPI_ISL_597809, EPI_ISL_597810, EPI_ISL_597811, EPI_ISL_597812, EPI_ISL_597813, EPI_ISL_597814, EPI_ISL_597815, EPI_ISL_597816, EPI_ISL_597817, EPI_ISL_597818, EPI_ISL_597819, EPI_ISL_597820, EPI_ISL_597821, EPI_ISL_597822, EPI_ISL_597823, EPI_ISL_597824, EPI_ISL_597825, EPI_ISL_597826, EPI_ISL_597827, EPI_ISL_597828, EPI_ISL_597829, EPI_ISL_597830, EPI_ISL_597831, EPI_ISL_597832, EPI_ISL_597833, EPI_ISL_597834, EPI_ISL_597835, EPI_ISL_597836, EPI_ISL_597837, EPI_ISL_597838, EPI_ISL_597839, EPI_ISL_597840, EPI_ISL_597841, EPI_ISL_597842, EPI_ISL_597843, EPI_ISL_597844, EPI_ISL_597845, EPI_ISL_597846, EPI_ISL_597847, EPI_ISL_597848, EPI_ISL_597849, EPI_ISL_597850, EPI_ISL_597851, EPI_ISL_597852, EPI_ISL_597853, EPI_ISL_597854, EPI_ISL_597855, EPI_ISL_597856, EPI_ISL_597857, EPI_ISL_597858, EPI_ISL_597860, EPI_ISL_597861, EPI_ISL_597862, EPI_ISL_597863, EPI_ISL_597864, EPI_ISL_597865, EPI_ISL_597866, EPI_ISL_597867, EPI_ISL_597868, EPI_ISL_597869, EPI_ISL_597870, EPI_ISL_597871, EPI_ISL_597872, EPI_ISL_597873, EPI_ISL_597874, EPI_ISL_597875, EPI_ISL_597876, EPI_ISL_597877, EPI_ISL_597878, EPI_ISL_597879, EPI_ISL_597880, EPI_ISL_597881, EPI_ISL_597882, EPI_ISL_597883, EPI_ISL_597884, EPI_ISL_597885, EPI_ISL_597886, EPI_ISL_597887, EPI_ISL_597888, EPI_ISL_597889, EPI_ISL_597890, EPI_ISL_597891, EPI_ISL_597892, EPI_ISL_597893, EPI_ISL_597894, EPI_ISL_597895, EPI_ISL_597896, EPI_ISL_597897, EPI_ISL_597898, EPI_ISL_597899, EPI_ISL_597900, EPI_ISL_597901, EPI_ISL_597902, EPI_ISL_597903, EPI_ISL_597904, EPI_ISL_597905, EPI_ISL_597906, EPI_ISL_597907, EPI_ISL_597908, EPI_ISL_597909, EPI_ISL_597910, EPI_ISL_597911, EPI_ISL_597912, EPI_ISL_597913, EPI_ISL_597914, EPI_ISL_597915, EPI_ISL_597916, EPI_ISL_597917, EPI_ISL_597918, EPI_ISL_597919, EPI_ISL_597920, EPI_ISL_597921, EPI_ISL_597922, EPI_ISL_597923, EPI_ISL_597924, EPI_ISL_597925, EPI_ISL_597926, EPI_ISL_597927, EPI_ISL_597928, EPI_ISL_597929, EPI_ISL_597930, EPI_ISL_597931, EPI_ISL_597932, EPI_ISL_597933, EPI_ISL_597934, EPI_ISL_597935, EPI_ISL_597936, EPI_ISL_597937, EPI_ISL_597938, EPI_ISL_597939, EPI_ISL_597940, EPI_ISL_597941, EPI_ISL_597942, EPI_ISL_597943, EPI_ISL_597944, EPI_ISL_597945, EPI_ISL_597946, EPI_ISL_597947, EPI_ISL_597948, EPI_ISL_597949, EPI_ISL_597950, EPI_ISL_597951, EPI_ISL_597952, EPI_ISL_597953, EPI_ISL_597954, EPI_ISL_597955, EPI_ISL_597956, EPI_ISL_597957, EPI_ISL_597958, EPI_ISL_597959, EPI_ISL_597960, EPI_ISL_597961, EPI_ISL_597962, EPI_ISL_597963, EPI_ISL_597964, EPI_ISL_597965, EPI_ISL_597966, EPI_ISL_597967, EPI_ISL_597968, EPI_ISL_597969, EPI_ISL_597970, EPI_ISL_597971, EPI_ISL_597972, EPI_ISL_597973, EPI_ISL_597974, EPI_ISL_597975, EPI_ISL_597976, EPI_ISL_597977, EPI_ISL_597978, EPI_ISL_597979, EPI_ISL_597980, EPI_ISL_597981, EPI_ISL_597982, EPI_ISL_597983 | | | |
| see above | Lighthouse Lab in Alderley Park | Wellcome Sanger Institute for the COVID-19 Genomics UK (COG-UK) consortium | Jacquelyn Wynn, Mairead Hyland, The Lighthouse Lab in Alderley Park and Alex Alderton, Roberto Amato, Sonia Goncalves, Ewan Harrison, David K. Jackson, Ian Johnston, Dominic Kwiatkowski, Cordelia Langford, John Sillitoe on behalf of the Wellcome Sanger Institute COVID-19 Surveillance Team (http://www.sanger.ac.uk/covid-team) |
| EPI_ISL_597984 | Lighthouse Lab in Alderley Park | Wellcome Sanger Institute for the COVID-19 Genomics UK (COG-UK) Consortium | Jacquelyn Wynn, Mairead Hyland, The Lighthouse Lab in Alderley Park and Alex Alderton, Roberto Amato, Sonia Goncalves, Ewan Harrison, David K. Jackson, Ian Johnston, Dominic Kwiatkowski, Cordelia Langford, John Sillitoe on behalf of the Wellcome Sanger Institute COVID-19 Surveillance Team (http://www.sanger.ac.uk/covid-team) |
| EPI_ISL_597985, EPI_ISL_597986, EPI_ISL_597987, EPI_ISL_597988, EPI_ISL_597989, EPI_ISL_597990, EPI_ISL_597991, EPI_ISL_597992, EPI_ISL_597993, EPI_ISL_597994, EPI_ISL_597995, EPI_ISL_597996, EPI_ISL_597997, EPI_ISL_597998, EPI_ISL_597999, EPI_ISL_598000, EPI_ISL_598001, EPI_ISL_598002, EPI_ISL_598003, EPI_ISL_598004, EPI_ISL_598005, EPI_ISL_598006, EPI_ISL_598007, EPI_ISL_598008, EPI_ISL_598009, EPI_ISL_598010, EPI_ISL_598011, EPI_ISL_598012, EPI_ISL_598013, EPI_ISL_598014, EPI_ISL_598015, EPI_ISL_598016, EPI_ISL_598017, EPI_ISL_598018, EPI_ISL_598019, EPI_ISL_598020, EPI_ISL_598021, EPI_ISL_598022, EPI_ISL_598023, EPI_ISL_598024, EPI_ISL_598025, EPI_ISL_598026, EPI_ISL_598027, EPI_ISL_598028, EPI_ISL_598029, EPI_ISL_598030, EPI_ISL_598031, EPI_ISL_598032, EPI_ISL_598033, EPI_ISL_598034, EPI_ISL_598035, EPI_ISL_598036, EPI_ISL_598037, EPI_ISL_598039, EPI_ISL_598040, EPI_ISL_598041, EPI_ISL_598042, EPI_ISL_598044, EPI_ISL_598045, EPI_ISL_598046, EPI_ISL_598047, EPI_ISL_598048, EPI_ISL_598049, EPI_ISL_598050, EPI_ISL_598051, EPI_ISL_598052, EPI_ISL_598053, EPI_ISL_598054, EPI_ISL_598055, EPI_ISL_598056, EPI_ISL_598057, EPI_ISL_598058, EPI_ISL_598059, EPI_ISL_598060, EPI_ISL_598061, EPI_ISL_598062, EPI_ISL_598063, EPI_ISL_598064, EPI_ISL_598065, EPI_ISL_598066, EPI_ISL_598067, EPI_ISL_598068, EPI_ISL_598069, EPI_ISL_598070, EPI_ISL_598071, EPI_ISL_598072, EPI_ISL_598073, EPI_ISL_598074, EPI_ISL_598075, EPI_ISL_598076, EPI_ISL_598077, EPI_ISL_598078, EPI_ISL_598079, EPI_ISL_598080, EPI_ISL_598081, EPI_ISL_598082, EPI_ISL_598083, EPI_ISL_598084, EPI_ISL_598085, EPI_ISL_598086, EPI_ISL_598087, EPI_ISL_598088, EPI_ISL_598089, EPI_ISL_598090, EPI_ISL_598091, EPI_ISL_598092, EPI_ISL_598093, EPI_ISL_598094, EPI_ISL_598095, EPI_ISL_598096, EPI_ISL_598097, EPI_ISL_598098, EPI_ISL_598099, EPI_ISL_598100, EPI_ISL_598101, EPI_ISL_598102, EPI_ISL_598103, EPI_ISL_598104, EPI_ISL_598105, EPI_ISL_598106, EPI_ISL_598107, EPI_ISL_598108, EPI_ISL_598109, EPI_ISL_598110, EPI_ISL_598111, EPI_ISL_598112, EPI_ISL_598113, EPI_ISL_598114, EPI_ISL_598116, EPI_ISL_598117, EPI_ISL_598118, EPI_ISL_598119, EPI_ISL_598120, EPI_ISL_598121, EPI_ISL_598122, EPI_ISL_598123, EPI_ISL_598124, EPI_ISL_598125, EPI_ISL_598126, EPI_ISL_598127, EPI_ISL_598128, EPI_ISL_598129, EPI_ISL_598130, EPI_ISL_598131, EPI_ISL_598132, EPI_ISL_598133, EPI_ISL_598134, EPI_ISL_598135, EPI_ISL_598136, EPI_ISL_598137, EPI_ISL_598138, EPI_ISL_598139, EPI_ISL_598140, EPI_ISL_598141, EPI_ISL_598142, EPI_ISL_598143, EPI_ISL_598144, EPI_ISL_598145, EPI_ISL_598146, EPI_ISL_598147, EPI_ISL_598148, EPI_ISL_598149, EPI_ISL_598150, EPI_ISL_598151, EPI_ISL_598152, EPI_ISL_598153, EPI_ISL_598154, EPI_ISL_598155, EPI_ISL_598156, EPI_ISL_598157, EPI_ISL_598158, EPI_ISL_598159, EPI_ISL_598160, EPI_ISL_598161, EPI_ISL_598162 | | | |
| see above | Lighthouse Lab in Alderley Park | Wellcome Sanger Institute for the COVID-19 Genomics UK (COG-UK) consortium | Jacquelyn Wynn, Mairead Hyland, The Lighthouse Lab in Alderley Park and Alex Alderton, Roberto Amato, Sonia Goncalves, Ewan Harrison, David K. Jackson, Ian Johnston, Dominic Kwiatkowski, Cordelia Langford, John Sillitoe on behalf of the Wellcome Sanger Institute COVID-19 Surveillance Team (http://www.sanger.ac.uk/covid-team) |
| EPI_ISL_598163 | Lighthouse Lab in Alderley Park | Wellcome Sanger Institute for the COVID-19 Genomics UK (COG-UK) Consortium | Jacquelyn Wynn, Mairead Hyland, The Lighthouse Lab in Alderley Park and Alex Alderton, Roberto Amato, Sonia Goncalves, Ewan Harrison, David K. Jackson, Ian Johnston, Dominic Kwiatkowski, Cordelia Langford, John Sillitoe on behalf of the Wellcome Sanger Institute COVID-19 Surveillance Team (http://www.sanger.ac.uk/covid-team) |
| EPI_ISL_598164, EPI_ISL_598165, EPI_ISL_598166, EPI_ISL_598167, EPI_ISL_598168, EPI_ISL_598169, EPI_ISL_598170, EPI_ISL_598171, EPI_ISL_598172, EPI_ISL_598173, EPI_ISL_598174, EPI_ISL_598175, EPI_ISL_598176, EPI_ISL_598177, EPI_ISL_598178, EPI_ISL_598179, EPI_ISL_598180, EPI_ISL_598181, EPI_ISL_598182, EPI_ISL_598183, EPI_ISL_598184, EPI_ISL_598185, EPI_ISL_598186, EPI_ISL_598187, EPI_ISL_598188, EPI_ISL_598189, EPI_ISL_598190, EPI_ISL_598191, EPI_ISL_598192, EPI_ISL_598193, EPI_ISL_598194, EPI_ISL_598195, EPI_ISL_598196, EPI_ISL_598197, EPI_ISL_598198, EPI_ISL_598199, EPI_ISL_598200, EPI_ISL_598201, EPI_ISL_598202, EPI_ISL_598203, EPI_ISL_598204, EPI_ISL_598205, EPI_ISL_598206, EPI_ISL_598207, EPI_ISL_598208, EPI_ISL_598209, EPI_ISL_598210, EPI_ISL_598211, EPI_ISL_598212, EPI_ISL_598213, EPI_ISL_598214, EPI_ISL_598215, EPI_ISL_598216, EPI_ISL_598217, EPI_ISL_598218, EPI_ISL_598219, EPI_ISL_598220, EPI_ISL_598221, EPI_ISL_598222, EPI_ISL_598223, EPI_ISL_598224, EPI_ISL_598225, EPI_ISL_598226, EPI_ISL_598227, EPI_ISL_598228, EPI_ISL_598229, EPI_ISL_598230, EPI_ISL_598231, EPI_ISL_598232, EPI_ISL_598233, EPI_ISL_598234, EPI_ISL_598235, EPI_ISL_598236, EPI_ISL_598237, EPI_ISL_598238, EPI_ISL_598239, EPI_ISL_598240, EPI_ISL_598241, EPI_ISL_598242, EPI_ISL_598243, EPI_ISL_598244, EPI_ISL_598245, EPI_ISL_598246, EPI_ISL_598247, EPI_ISL_598248 | | | |
| see above | Lighthouse Lab in Alderley Park | Wellcome Sanger Institute for the COVID-19 Genomics UK (COG-UK) consortium | Jacquelyn Wynn, Mairead Hyland, The Lighthouse Lab in Alderley Park and Alex Alderton, Roberto Amato, Sonia Goncalves, Ewan Harrison, David K. Jackson, Ian Johnston, Dominic Kwiatkowski, Cordelia Langford, John Sillitoe on behalf of the Wellcome Sanger Institute COVID-19 Surveillance Team (http://www.sanger.ac.uk/covid-team) |
| EPI_ISL_598540, EPI_ISL_598567, EPI_ISL_598570, EPI_ISL_598631, EPI_ISL_598634, EPI_ISL_598637, EPI_ISL_598737, EPI_ISL_598755, EPI_ISL_598799, EPI_ISL_598800 | Lighthouse Lab in Milton Keynes | Wellcome Sanger Institute for the COVID-19 Genomics UK (COG-UK) consortium | The Lighthouse Lab in Milton Keynes and Alex Alderton, Roberto Amato, Sonia Goncalves, Ewan Harrison, David K. Jackson, Ian Johnston, Dominic Kwiatkowski, Cordelia Langford, John Sillitoe on behalf of the Wellcome Sanger Institute COVID-19 Surveillance Team (http://www.sanger.ac.uk/covid-team) |
| EPI_ISL_602577, EPI_ISL_602579, EPI_ISL_602580, EPI_ISL_602581 | SA Pathology | SA Pathology | Lex Leong, Julien Soubrier, Chuan Kok Lim, Song Gao, Mark Turra, Karin Kassahn, Ivan Bastian, Geoff Higgins |
| EPI_ISL_602885, EPI_ISL_602893, EPI_ISL_602896, EPI_ISL_602897, EPI_ISL_602898, EPI_ISL_602899 | NHLS-IALCH | KRISP, KZN Research Innovation and Sequencing Platform | Giandhari J, Pillay S, Lessells R, Mdlalose K, York D, Khan S, Tegally H, Wilkinson E, de Oliveira T |
| EPI_ISL_602992, EPI_ISL_602993, EPI_ISL_602994, EPI_ISL_602995, EPI_ISL_602996, EPI_ISL_602997, EPI_ISL_602998, EPI_ISL_602999 | Minnesota Department of Health, Public Health Laboratory | Minnesota Department of Health, Public Health Laboratory | Matt Plumb, Jacob Garfin, Alexandra Lorentz, and Xiong Wang |
| EPI_ISL_603012, EPI_ISL_603013, EPI_ISL_603017 | Essentia Health-St. Mary's Medical Center | Minnesota Department of Health, Public Health Laboratory | Matt Plumb, Jacob Garfin, Alexandra Lorentz, and Xiong Wang |
| EPI_ISL_603188, EPI_ISL_603189, EPI_ISL_603190, EPI_ISL_603191, EPI_ISL_603192, EPI_ISL_603193, EPI_ISL_603194, EPI_ISL_603195, EPI_ISL_603196, EPI_ISL_603197, EPI_ISL_603198, EPI_ISL_603199, EPI_ISL_603200, EPI_ISL_603201, EPI_ISL_603202, EPI_ISL_603203, EPI_ISL_603204, EPI_ISL_603206, EPI_ISL_603212, EPI_ISL_603215 | | | |
| see above | Respiratory Virus Unit, Microbiology Services Colindale, Public Health England | Respiratory Virus Unit, Microbiology Services Colindale, Public Health England | PHE Covid Sequencing Team |

| | | | |
|---|---|---|---|
| EPI_ISL_603222 | National Institute of Laboratory Medicine and Referral Center | Genomic Research Lab, BCSIR | Abu Sayeed Mohammad Mahmud, Mohammad Samir Uzzaman, Eshrar Osman, Md. Ahashan Habib, Shahina Akter, Tanjina Akhter Banu, Md. Murshed Hasan Sarkar, Barna Goswami, Iffat Jahan, Md. Saddam Hossain, Tasnim Nafisa, Md. Maruf Ahmed Molla, Mahmuda Yeasmin, Asish Kumar Ghosh, A. K. M. Shamsuzzaman, Monira Parveen, Md. Masum Hossain Arif, Md. Salim Khan |
| EPI_ISL_603239 | National Institute of Laboratory Medicine and Referral Center | Genomic Research Lab, BCSIR | Shahina Akter, Abu Sayeed Mohammad Mahmud, Mohammad Samir Uzzaman, Eshrar Osman, Md. Ahashan Habib, Tanjina Akhter Banu, Md. Murshed Hasan Sarkar, Barna Goswami, Iffat Jahan, Md. Saddam Hossain, Tasnim Nafisa, Md. Maruf Ahmed Molla, Mahmuda Yeasmin, Asish Kumar Ghosh, A. K. M. Shamsuzzaman, Monira Parveen, Md. Masum Hossain Arif, Md. Salim Khan |
| EPI_ISL_603240 | National Institute of Laboratory Medicine and Referral Center | Genomic Research Lab, BCSIR | Tanjina Akhter Banu, Abu Sayeed Mohammad Mahmud, Mohammad Samir Uzzaman, Eshrar Osman, Md. Ahashan Habib, Shahina Akter, Md. Murshed Hasan Sarkar, Barna Goswami, Iffat Jahan, Md. Saddam Hossain, Tasnim Nafisa, Md. Maruf Ahmed Molla, Mahmuda Yeasmin, Asish Kumar Ghosh, A. K. M. Shamsuzzaman, Monira Parveen, Md. Masum Hossain Arif, Md. Salim Khan |
| EPI_ISL_603242, EPI_ISL_603243 | National Institute of Laboratory Medicine and Referral Center | Genomic Research Lab, BCSIR | Barna Goswami, Abu Sayeed Mohammad Mahmud, Mohammad Samir Uzzaman, Eshrar Osman, Md. Ahashan Habib, Shahina Akter, Tanjina Akhter Banu, Md. Murshed Hasan Sarkar, Iffat Jahan, Md. Saddam Hossain, Tasnim Nafisa, Md. Maruf Ahmed Molla, Mahmuda Yeasmin, Asish Kumar Ghosh, A. K. M. Shamsuzzaman, Monira Parveen, Md. Masum Hossain Arif, Md. Salim Khan |
| EPI_ISL_603244, EPI_ISL_603245 | National Institute of Laboratory Medicine and Referral Center | Genomic Research Lab, BCSIR | Iffat Jahan, Abu Sayeed Mohammad Mahmud, Mohammad Samir Uzzaman, Eshrar Osman, Md. Ahashan Habib, Shahina Akter, Tanjina Akhter Banu, Md. Murshed Hasan Sarkar, Barna Goswami, Md. Saddam Hossain, Tasnim Nafisa, Md. Maruf Ahmed Molla, Mahmuda Yeasmin, Asish Kumar Ghosh, A. K. M. Shamsuzzaman, Monira Parveen, Md. Masum Hossain Arif, Md. Salim Khan |
| EPI_ISL_603423, EPI_ISL_603424, EPI_ISL_603425, EPI_ISL_603426, EPI_ISL_603427, EPI_ISL_603428, EPI_ISL_603429, EPI_ISL_603430, EPI_ISL_603431, EPI_ISL_603432, EPI_ISL_603433, EPI_ISL_603434, EPI_ISL_603435, EPI_ISL_603436, EPI_ISL_603437, EPI_ISL_603438, EPI_ISL_603439, EPI_ISL_603440, EPI_ISL_603441, EPI_ISL_603442, EPI_ISL_603443, EPI_ISL_603444, EPI_ISL_603445, EPI_ISL_603446, EPI_ISL_603447, EPI_ISL_603448, EPI_ISL_603449, EPI_ISL_603450, EPI_ISL_603451, EPI_ISL_603452, EPI_ISL_603453, EPI_ISL_603454, EPI_ISL_603455, EPI_ISL_603456, EPI_ISL_603457, EPI_ISL_603458, EPI_ISL_603459, EPI_ISL_603460, EPI_ISL_603461, EPI_ISL_603462, EPI_ISL_603463, EPI_ISL_603464, EPI_ISL_603465, EPI_ISL_603466, EPI_ISL_603467, EPI_ISL_603468, EPI_ISL_603469, EPI_ISL_603470, EPI_ISL_603471, EPI_ISL_603472, EPI_ISL_603473, EPI_ISL_603474, EPI_ISL_603475, EPI_ISL_603476, EPI_ISL_603477, EPI_ISL_603478, EPI_ISL_603479, EPI_ISL_603480, EPI_ISL_603481, EPI_ISL_603482, EPI_ISL_603483, EPI_ISL_603484, EPI_ISL_603485, EPI_ISL_603486, EPI_ISL_603487, EPI_ISL_603488, EPI_ISL_603489, EPI_ISL_603490, EPI_ISL_603491, EPI_ISL_603492, EPI_ISL_603493, EPI_ISL_603494, EPI_ISL_603495, EPI_ISL_603496, EPI_ISL_603497, EPI_ISL_603498, EPI_ISL_603499, EPI_ISL_603500, EPI_ISL_603501, EPI_ISL_603502, EPI_ISL_603503, EPI_ISL_603504, EPI_ISL_603505, EPI_ISL_603506, EPI_ISL_603507, EPI_ISL_603508, EPI_ISL_603509, EPI_ISL_603510, EPI_ISL_603511, EPI_ISL_603512, EPI_ISL_603513, EPI_ISL_603514, EPI_ISL_603515, EPI_ISL_603516, EPI_ISL_603517, EPI_ISL_603518, EPI_ISL_603519, EPI_ISL_603520, EPI_ISL_603521, EPI_ISL_603522, EPI_ISL_603523, EPI_ISL_603524, EPI_ISL_603525, EPI_ISL_603526, EPI_ISL_603527, EPI_ISL_603528, EPI_ISL_603529, EPI_ISL_603530, EPI_ISL_603531, EPI_ISL_603532, EPI_ISL_603533, EPI_ISL_603534, EPI_ISL_603535, EPI_ISL_603536, EPI_ISL_603537, EPI_ISL_603538, EPI_ISL_603539, EPI_ISL_603540, EPI_ISL_603541, EPI_ISL_603542, EPI_ISL_603543, EPI_ISL_603544, EPI_ISL_603545, EPI_ISL_603546, EPI_ISL_603547, EPI_ISL_603548, EPI_ISL_603549, EPI_ISL_603550, EPI_ISL_603551, EPI_ISL_603552, EPI_ISL_603553, EPI_ISL_603554, EPI_ISL_603555, EPI_ISL_603556, EPI_ISL_603557, EPI_ISL_603558, EPI_ISL_603559, EPI_ISL_603560, EPI_ISL_603561, EPI_ISL_603562, EPI_ISL_603563, EPI_ISL_603564, EPI_ISL_603565, EPI_ISL_603566, EPI_ISL_603567, EPI_ISL_603568, EPI_ISL_603569, EPI_ISL_603570, EPI_ISL_603571, EPI_ISL_603572, EPI_ISL_603573, EPI_ISL_603574, EPI_ISL_603575, EPI_ISL_603576, EPI_ISL_603577, EPI_ISL_603578, EPI_ISL_603579, EPI_ISL_603580, EPI_ISL_603581, EPI_ISL_603582, EPI_ISL_603583, EPI_ISL_603584, EPI_ISL_603585, EPI_ISL_603586, EPI_ISL_603587, EPI_ISL_603588, EPI_ISL_603589, EPI_ISL_603590, EPI_ISL_603591, EPI_ISL_603592, EPI_ISL_603593, EPI_ISL_603594, EPI_ISL_603595, EPI_ISL_603596, EPI_ISL_603597, EPI_ISL_603598, EPI_ISL_603599 | | | |
| see above | Viollier AG | Department of Biosystems Science and Engineering, ETH Zürich | Christian Beisel, Sarah Nadeau, Ivan Topolsky, Pedro Ferreira, Philipp Jablonski, Susana Posada-Céspedes, Tobias Schär, Ina Nissen, Natascha Santacroce, Elodie Burcklen, Christiane Beckmann, Maurice Redondo, Olivier Kobel, Christoph Noppen, Sophie Seidel, Noemie Santamaria de Souza, Niko Beerenwinkel, Tanja Stadler |
| EPI_ISL_605787, EPI_ISL_605788 | NHLS-IALCH | KRISP, KZN Research Innovation and Sequencing Platform | Giandhari J, Pillay S, Lessells R, Mdlalose K, York D, Khan S, Tegally H, Wilkinson E, de Oliveira T |
| EPI_ISL_605837, EPI_ISL_605838, EPI_ISL_605843, EPI_ISL_605859, EPI_ISL_605864, EPI_ISL_605865 | PathWest Laboratory Medicine WA | PathWest Laboratory Medicine WA Microbial Surveillance Unit | PathWest Laboratory Medicine WA Microbial Surveillance Unit |
| EPI_ISL_606296, EPI_ISL_606306, EPI_ISL_606321, EPI_ISL_606328, EPI_ISL_606344, EPI_ISL_606366, EPI_ISL_606386, EPI_ISL_606388, EPI_ISL_606530, EPI_ISL_606532, EPI_ISL_606561, EPI_ISL_606576, EPI_ISL_606584 | | | |
| see above | Lighthouse Lab in Milton Keynes | Wellcome Sanger Institute for the COVID-19 Genomics UK (COG-UK) consortium | The Lighthouse Lab in Milton Keynes and Alex Alderton, Roberto Amato, Sonia Goncalves, Ewan Harrison, David K. Jackson, Ian Johnston, Dominic Kwiatkowski, Cordelia Langford, John Sillitoe on behalf of the Wellcome Sanger Institute COVID-19 Surveillance Team |
| EPI_ISL_606588, EPI_ISL_606608 | Lighthouse Lab in Milton Keynes | Wellcome Sanger Institute for the COVID-19 Genomics UK (COG-UK) Consortium | The Lighthouse Lab in Milton Keynes and Alex Alderton, Roberto Amato, Sonia Goncalves, Ewan Harrison, David K. Jackson, Ian Johnston, Dominic Kwiatkowski, Cordelia Langford, John Sillitoe on behalf of the Wellcome Sanger Institute COVID-19 Surveillance Team |
| EPI_ISL_606611, EPI_ISL_606613, EPI_ISL_606636, EPI_ISL_606660, EPI_ISL_606689, EPI_ISL_606725, EPI_ISL_606727, EPI_ISL_606747, EPI_ISL_606751, EPI_ISL_606761, EPI_ISL_606769, EPI_ISL_606807, EPI_ISL_606822, EPI_ISL_606827, EPI_ISL_606835, EPI_ISL_606863, EPI_ISL_606865, EPI_ISL_606900, EPI_ISL_606902, EPI_ISL_607585 | | | |
| see above | Lighthouse Lab in Milton Keynes | Wellcome Sanger Institute for the COVID-19 Genomics UK (COG-UK) consortium | The Lighthouse Lab in Milton Keynes and Alex Alderton, Roberto Amato, Sonia Goncalves, Ewan Harrison, David K. Jackson, Ian Johnston, Dominic Kwiatkowski, Cordelia Langford, John Sillitoe on behalf of the Wellcome Sanger Institute COVID-19 Surveillance Team |
| EPI_ISL_608347 | Lighthouse Lab in Cambridge | Wellcome Sanger Institute for the COVID-19 Genomics UK (COG-UK) Consortium | Rob Howes, The Lighthouse Lab in Cambridge and Alex Alderton, Roberto Amato, Sonia Goncalves, Ewan Harrison, David K. Jackson, Ian Johnston, Dominic Kwiatkowski, Cordelia Langford, John Sillitoe on behalf of the Wellcome Sanger Institute COVID-19 Surveillance Team |
| EPI_ISL_608348, EPI_ISL_608349 | Lighthouse Lab in Cambridge | Wellcome Sanger Institute for the COVID-19 Genomics UK (COG-UK) consortium | Rob Howes, The Lighthouse Lab in Cambridge and Alex Alderton, Roberto Amato, Sonia Goncalves, Ewan Harrison, David K. Jackson, Ian Johnston, Dominic Kwiatkowski, Cordelia Langford, John Sillitoe on behalf of the Wellcome Sanger Institute COVID-19 Surveillance Team |
| EPI_ISL_608350, EPI_ISL_608351, EPI_ISL_608352 | Lighthouse Lab in Alderley Park | Wellcome Sanger Institute for the COVID-19 Genomics UK (COG-UK) consortium | Jacquelyn Wynn, Mairead Hyland, The Lighthouse Lab in Alderley Park and Alex Alderton, Roberto Amato, Sonia Goncalves, Ewan Harrison, David K. Jackson, Ian Johnston, Dominic Kwiatkowski, Cordelia Langford, John Sillitoe on behalf of the Wellcome Sanger Institute COVID-19 Surveillance Team |
| EPI_ISL_608353, EPI_ISL_608354 | Lighthouse Lab in Cambridge | Wellcome Sanger Institute for the COVID-19 Genomics UK (COG-UK) consortium | Rob Howes, The Lighthouse Lab in Cambridge and Alex Alderton, Roberto Amato, Sonia Goncalves, Ewan Harrison, David K. Jackson, Ian Johnston, Dominic Kwiatkowski, Cordelia Langford, John Sillitoe on behalf of the Wellcome Sanger Institute COVID-19 Surveillance Team |
| EPI_ISL_608355, EPI_ISL_608356 | Lighthouse Lab in Alderley Park | Wellcome Sanger Institute for the COVID-19 Genomics UK (COG-UK) consortium | Jacquelyn Wynn, Mairead Hyland, The Lighthouse Lab in Alderley Park and Alex Alderton, Roberto Amato, Sonia Goncalves, Ewan Harrison, David K. Jackson, Ian Johnston, Dominic Kwiatkowski, Cordelia Langford, John Sillitoe on behalf of the Wellcome Sanger Institute COVID-19 Surveillance Team |
| EPI_ISL_608357 | Lighthouse Lab in Cambridge | Wellcome Sanger Institute for the COVID-19 Genomics UK (COG-UK) consortium | Rob Howes, The Lighthouse Lab in Cambridge and Alex Alderton, Roberto Amato, Sonia Goncalves, Ewan Harrison, David K. Jackson, Ian Johnston, Dominic Kwiatkowski, Cordelia Langford, John Sillitoe on behalf of the Wellcome Sanger Institute COVID-19 Surveillance Team |
| EPI_ISL_608358 | Lighthouse Lab in Alderley Park | Wellcome Sanger Institute for the COVID-19 Genomics UK (COG-UK) consortium | Jacquelyn Wynn, Mairead Hyland, The Lighthouse Lab in Alderley Park and Alex Alderton, Roberto Amato, Sonia Goncalves, Ewan Harrison, David K. Jackson, Ian Johnston, Dominic Kwiatkowski, Cordelia Langford, John Sillitoe on behalf of the Wellcome Sanger Institute COVID-19 Surveillance Team |
| EPI_ISL_608359, EPI_ISL_608360 | Lighthouse Lab in Cambridge | Wellcome Sanger Institute for the COVID-19 Genomics UK (COG-UK) consortium | Rob Howes, The Lighthouse Lab in Cambridge and Alex Alderton, Roberto Amato, Sonia Goncalves, Ewan Harrison, David K. Jackson, Ian Johnston, Dominic Kwiatkowski, Cordelia Langford, John Sillitoe on behalf of the Wellcome Sanger Institute COVID-19 Surveillance Team |
| EPI_ISL_608361, EPI_ISL_608362 | Lighthouse Lab in Alderley Park | Wellcome Sanger Institute for the COVID-19 Genomics UK (COG-UK) consortium | Jacquelyn Wynn, Mairead Hyland, The Lighthouse Lab in Alderley Park and Alex Alderton, Roberto Amato, Sonia Goncalves, Ewan Harrison, David K. Jackson, Ian Johnston, Dominic Kwiatkowski, Cordelia Langford, John Sillitoe on behalf of the Wellcome Sanger Institute COVID-19 Surveillance Team |
| EPI_ISL_608363 | Lighthouse Lab in Cambridge | Wellcome Sanger Institute for the COVID-19 Genomics UK (COG-UK) consortium | Rob Howes, The Lighthouse Lab in Cambridge and Alex Alderton, Roberto Amato, Sonia Goncalves, Ewan Harrison, David K. Jackson, Ian Johnston, Dominic Kwiatkowski, Cordelia Langford, John Sillitoe on behalf of the Wellcome Sanger Institute COVID-19 Surveillance Team |
| EPI_ISL_608364, EPI_ISL_608365, EPI_ISL_608366, EPI_ISL_608367 | Lighthouse Lab in Alderley Park | Wellcome Sanger Institute for the COVID-19 Genomics UK (COG-UK) consortium | Jacquelyn Wynn, Mairead Hyland, The Lighthouse Lab in Alderley Park and Alex Alderton, Roberto Amato, Sonia Goncalves, Ewan Harrison, David K. Jackson, Ian Johnston, Dominic Kwiatkowski, Cordelia Langford, John Sillitoe on behalf of the Wellcome Sanger Institute COVID-19 Surveillance Team |
| EPI_ISL_608368 | Lighthouse Lab in Cambridge | Wellcome Sanger Institute for the COVID-19 Genomics UK (COG-UK) consortium | Rob Howes, The Lighthouse Lab in Cambridge and Alex Alderton, Roberto Amato, Sonia Goncalves, Ewan Harrison, David K. Jackson, Ian Johnston, Dominic Kwiatkowski, Cordelia Langford, John Sillitoe on behalf of the Wellcome Sanger Institute COVID-19 Surveillance Team |
| EPI_ISL_608369 | Lighthouse Lab in Alderley Park | Wellcome Sanger Institute for the COVID-19 Genomics UK (COG-UK) consortium | Jacquelyn Wynn, Mairead Hyland, The Lighthouse Lab in Alderley Park and Alex Alderton, Roberto Amato, Sonia Goncalves, Ewan Harrison, David K. Jackson, Ian Johnston, Dominic Kwiatkowski, Cordelia Langford, John Sillitoe on behalf of the Wellcome Sanger Institute COVID-19 Surveillance Team |
| EPI_ISL_608370 | Lighthouse Lab in Cambridge | Wellcome Sanger Institute for the COVID-19 Genomics UK (COG-UK) consortium | Rob Howes, The Lighthouse Lab in Cambridge and Alex Alderton, Roberto Amato, Sonia Goncalves, Ewan Harrison, David K. Jackson, Ian Johnston, Dominic Kwiatkowski, Cordelia Langford, John Sillitoe on behalf of the Wellcome Sanger Institute COVID-19 Surveillance Team |
| EPI_ISL_608371, EPI_ISL_608372 | Lighthouse Lab in Alderley Park | Wellcome Sanger Institute for the COVID-19 Genomics UK (COG-UK) consortium | Jacquelyn Wynn, Mairead Hyland, The Lighthouse Lab in Alderley Park and Alex Alderton, Roberto Amato, Sonia Goncalves, Ewan Harrison, David K. Jackson, Ian Johnston, Dominic Kwiatkowski, Cordelia Langford, John Sillitoe on behalf of the Wellcome Sanger Institute COVID-19 Surveillance Team |
| EPI_ISL_608373 | Lighthouse Lab in Cambridge | Wellcome Sanger Institute for the COVID-19 Genomics UK (COG-UK) consortium | Rob Howes, The Lighthouse Lab in Cambridge and Alex Alderton, Roberto Amato, Sonia Goncalves, Ewan Harrison, David K. Jackson, Ian Johnston, Dominic Kwiatkowski, Cordelia Langford, John Sillitoe on behalf of the Wellcome Sanger Institute COVID-19 Surveillance Team |

| | | | |
|---|---|---|---|
| EPI_ISL_608374, EPI_ISL_608375, EPI_ISL_608376, EPI_ISL_608377, EPI_ISL_608378, EPI_ISL_608379, EPI_ISL_608380 | Lighthouse Lab in Alderley Park | Wellcome Sanger Institute for the COVID-19 Genomics UK (COG-UK) consortium | Jacquelyn Wynn, Mairead Hyland, The Lighthouse Lab in Alderley Park and Alex Alderton, Roberto Amato, Sonia Goncalves, Ewan Harrison, David K. Jackson, Ian Johnston, Dominic Kwiatkowski, Cordelia Langford, John Sillitoe on behalf of the Wellcome Sanger Institute COVID-19 Surveillance Team |
| EPI_ISL_608381, EPI_ISL_608382 | Lighthouse Lab in Cambridge | Wellcome Sanger Institute for the COVID-19 Genomics UK (COG-UK) consortium | Rob Howes, The Lighthouse Lab in Cambridge and Alex Alderton, Roberto Amato, Sonia Goncalves, Ewan Harrison, David K. Jackson, Ian Johnston, Dominic Kwiatkowski, Cordelia Langford, John Sillitoe on behalf of the Wellcome Sanger Institute COVID-19 Surveillance Team |
| EPI_ISL_608383, EPI_ISL_608384, EPI_ISL_608385, EPI_ISL_608386, EPI_ISL_608387 | Lighthouse Lab in Alderley Park | Wellcome Sanger Institute for the COVID-19 Genomics UK (COG-UK) consortium | Jacquelyn Wynn, Mairead Hyland, The Lighthouse Lab in Alderley Park and Alex Alderton, Roberto Amato, Sonia Goncalves, Ewan Harrison, David K. Jackson, Ian Johnston, Dominic Kwiatkowski, Cordelia Langford, John Sillitoe on behalf of the Wellcome Sanger Institute COVID-19 Surveillance Team |
| EPI_ISL_608388, EPI_ISL_608389 | Lighthouse Lab in Cambridge | Wellcome Sanger Institute for the COVID-19 Genomics UK (COG-UK) consortium | Rob Howes, The Lighthouse Lab in Cambridge and Alex Alderton, Roberto Amato, Sonia Goncalves, Ewan Harrison, David K. Jackson, Ian Johnston, Dominic Kwiatkowski, Cordelia Langford, John Sillitoe on behalf of the Wellcome Sanger Institute COVID-19 Surveillance Team |
| EPI_ISL_608390 | Lighthouse Lab in Alderley Park | Wellcome Sanger Institute for the COVID-19 Genomics UK (COG-UK) consortium | Jacquelyn Wynn, Mairead Hyland, The Lighthouse Lab in Alderley Park and Alex Alderton, Roberto Amato, Sonia Goncalves, Ewan Harrison, David K. Jackson, Ian Johnston, Dominic Kwiatkowski, Cordelia Langford, John Sillitoe on behalf of the Wellcome Sanger Institute COVID-19 Surveillance Team |
| EPI_ISL_608391, EPI_ISL_608392, EPI_ISL_608393, EPI_ISL_608394, EPI_ISL_608395, EPI_ISL_608396, EPI_ISL_608397, EPI_ISL_608398 | Lighthouse Lab in Cambridge | Wellcome Sanger Institute for the COVID-19 Genomics UK (COG-UK) consortium | Rob Howes, The Lighthouse Lab in Cambridge and Alex Alderton, Roberto Amato, Sonia Goncalves, Ewan Harrison, David K. Jackson, Ian Johnston, Dominic Kwiatkowski, Cordelia Langford, John Sillitoe on behalf of the Wellcome Sanger Institute COVID-19 Surveillance Team |
| EPI_ISL_608399 | Lighthouse Lab in Alderley Park | Wellcome Sanger Institute for the COVID-19 Genomics UK (COG-UK) consortium | Jacquelyn Wynn, Mairead Hyland, The Lighthouse Lab in Alderley Park and Alex Alderton, Roberto Amato, Sonia Goncalves, Ewan Harrison, David K. Jackson, Ian Johnston, Dominic Kwiatkowski, Cordelia Langford, John Sillitoe on behalf of the Wellcome Sanger Institute COVID-19 Surveillance Team |
| EPI_ISL_608400 | Lighthouse Lab in Cambridge | Wellcome Sanger Institute for the COVID-19 Genomics UK (COG-UK) consortium | Rob Howes, The Lighthouse Lab in Cambridge and Alex Alderton, Roberto Amato, Sonia Goncalves, Ewan Harrison, David K. Jackson, Ian Johnston, Dominic Kwiatkowski, Cordelia Langford, John Sillitoe on behalf of the Wellcome Sanger Institute COVID-19 Surveillance Team |
| EPI_ISL_608401 | Lighthouse Lab in Alderley Park | Wellcome Sanger Institute for the COVID-19 Genomics UK (COG-UK) consortium | Jacquelyn Wynn, Mairead Hyland, The Lighthouse Lab in Alderley Park and Alex Alderton, Roberto Amato, Sonia Goncalves, Ewan Harrison, David K. Jackson, Ian Johnston, Dominic Kwiatkowski, Cordelia Langford, John Sillitoe on behalf of the Wellcome Sanger Institute COVID-19 Surveillance Team |
| EPI_ISL_608402 | Lighthouse Lab in Cambridge | Wellcome Sanger Institute for the COVID-19 Genomics UK (COG-UK) consortium | Rob Howes, The Lighthouse Lab in Cambridge and Alex Alderton, Roberto Amato, Sonia Goncalves, Ewan Harrison, David K. Jackson, Ian Johnston, Dominic Kwiatkowski, Cordelia Langford, John Sillitoe on behalf of the Wellcome Sanger Institute COVID-19 Surveillance Team |
| EPI_ISL_608403 | Lighthouse Lab in Alderley Park | Wellcome Sanger Institute for the COVID-19 Genomics UK (COG-UK) consortium | Jacquelyn Wynn, Mairead Hyland, The Lighthouse Lab in Alderley Park and Alex Alderton, Roberto Amato, Sonia Goncalves, Ewan Harrison, David K. Jackson, Ian Johnston, Dominic Kwiatkowski, Cordelia Langford, John Sillitoe on behalf of the Wellcome Sanger Institute COVID-19 Surveillance Team |
| EPI_ISL_608404 | Lighthouse Lab in Cambridge | Wellcome Sanger Institute for the COVID-19 Genomics UK (COG-UK) consortium | Rob Howes, The Lighthouse Lab in Cambridge and Alex Alderton, Roberto Amato, Sonia Goncalves, Ewan Harrison, David K. Jackson, Ian Johnston, Dominic Kwiatkowski, Cordelia Langford, John Sillitoe on behalf of the Wellcome Sanger Institute COVID-19 Surveillance Team |
| EPI_ISL_608405, EPI_ISL_608406 | Lighthouse Lab in Alderley Park | Wellcome Sanger Institute for the COVID-19 Genomics UK (COG-UK) consortium | Jacquelyn Wynn, Mairead Hyland, The Lighthouse Lab in Alderley Park and Alex Alderton, Roberto Amato, Sonia Goncalves, Ewan Harrison, David K. Jackson, Ian Johnston, Dominic Kwiatkowski, Cordelia Langford, John Sillitoe on behalf of the Wellcome Sanger Institute COVID-19 Surveillance Team |
| EPI_ISL_608407, EPI_ISL_608408 | Lighthouse Lab in Cambridge | Wellcome Sanger Institute for the COVID-19 Genomics UK (COG-UK) consortium | Rob Howes, The Lighthouse Lab in Cambridge and Alex Alderton, Roberto Amato, Sonia Goncalves, Ewan Harrison, David K. Jackson, Ian Johnston, Dominic Kwiatkowski, Cordelia Langford, John Sillitoe on behalf of the Wellcome Sanger Institute COVID-19 Surveillance Team |
| EPI_ISL_608409 | Lighthouse Lab in Alderley Park | Wellcome Sanger Institute for the COVID-19 Genomics UK (COG-UK) consortium | Jacquelyn Wynn, Mairead Hyland, The Lighthouse Lab in Alderley Park and Alex Alderton, Roberto Amato, Sonia Goncalves, Ewan Harrison, David K. Jackson, Ian Johnston, Dominic Kwiatkowski, Cordelia Langford, John Sillitoe on behalf of the Wellcome Sanger Institute COVID-19 Surveillance Team |
| EPI_ISL_608410, EPI_ISL_608411, EPI_ISL_608412 | Lighthouse Lab in Cambridge | Wellcome Sanger Institute for the COVID-19 Genomics UK (COG-UK) consortium | Rob Howes, The Lighthouse Lab in Cambridge and Alex Alderton, Roberto Amato, Sonia Goncalves, Ewan Harrison, David K. Jackson, Ian Johnston, Dominic Kwiatkowski, Cordelia Langford, John Sillitoe on behalf of the Wellcome Sanger Institute COVID-19 Surveillance Team |
| EPI_ISL_608413, EPI_ISL_608414, EPI_ISL_608415 | Lighthouse Lab in Alderley Park | Wellcome Sanger Institute for the COVID-19 Genomics UK (COG-UK) consortium | Jacquelyn Wynn, Mairead Hyland, The Lighthouse Lab in Alderley Park and Alex Alderton, Roberto Amato, Sonia Goncalves, Ewan Harrison, David K. Jackson, Ian Johnston, Dominic Kwiatkowski, Cordelia Langford, John Sillitoe on behalf of the Wellcome Sanger Institute COVID-19 Surveillance Team |
| EPI_ISL_608416, EPI_ISL_608417 | Lighthouse Lab in Cambridge | Wellcome Sanger Institute for the COVID-19 Genomics UK (COG-UK) consortium | Rob Howes, The Lighthouse Lab in Cambridge and Alex Alderton, Roberto Amato, Sonia Goncalves, Ewan Harrison, David K. Jackson, Ian Johnston, Dominic Kwiatkowski, Cordelia Langford, John Sillitoe on behalf of the Wellcome Sanger Institute COVID-19 Surveillance Team |
| EPI_ISL_608418, EPI_ISL_608419, EPI_ISL_608420, EPI_ISL_608421, EPI_ISL_608422 | Lighthouse Lab in Alderley Park | Wellcome Sanger Institute for the COVID-19 Genomics UK (COG-UK) consortium | Jacquelyn Wynn, Mairead Hyland, The Lighthouse Lab in Alderley Park and Alex Alderton, Roberto Amato, Sonia Goncalves, Ewan Harrison, David K. Jackson, Ian Johnston, Dominic Kwiatkowski, Cordelia Langford, John Sillitoe on behalf of the Wellcome Sanger Institute COVID-19 Surveillance Team |
| EPI_ISL_608423, EPI_ISL_608424 | Lighthouse Lab in Cambridge | Wellcome Sanger Institute for the COVID-19 Genomics UK (COG-UK) consortium | Rob Howes, The Lighthouse Lab in Cambridge and Alex Alderton, Roberto Amato, Sonia Goncalves, Ewan Harrison, David K. Jackson, Ian Johnston, Dominic Kwiatkowski, Cordelia Langford, John Sillitoe on behalf of the Wellcome Sanger Institute COVID-19 Surveillance Team |
| EPI_ISL_608425 | Lighthouse Lab in Alderley Park | Wellcome Sanger Institute for the COVID-19 Genomics UK (COG-UK) consortium | Jacquelyn Wynn, Mairead Hyland, The Lighthouse Lab in Alderley Park and Alex Alderton, Roberto Amato, Sonia Goncalves, Ewan Harrison, David K. Jackson, Ian Johnston, Dominic Kwiatkowski, Cordelia Langford, John Sillitoe on behalf of the Wellcome Sanger Institute COVID-19 Surveillance Team |
| EPI_ISL_608426, EPI_ISL_608427, EPI_ISL_608428, EPI_ISL_608429, EPI_ISL_608430 | Lighthouse Lab in Cambridge | Wellcome Sanger Institute for the COVID-19 Genomics UK (COG-UK) consortium | Rob Howes, The Lighthouse Lab in Cambridge and Alex Alderton, Roberto Amato, Sonia Goncalves, Ewan Harrison, David K. Jackson, Ian Johnston, Dominic Kwiatkowski, Cordelia Langford, John Sillitoe on behalf of the Wellcome Sanger Institute COVID-19 Surveillance Team |
| EPI_ISL_608431 | Lighthouse Lab in Alderley Park | Wellcome Sanger Institute for the COVID-19 Genomics UK (COG-UK) consortium | Jacquelyn Wynn, Mairead Hyland, The Lighthouse Lab in Alderley Park and Alex Alderton, Roberto Amato, Sonia Goncalves, Ewan Harrison, David K. Jackson, Ian Johnston, Dominic Kwiatkowski, Cordelia Langford, John Sillitoe on behalf of the Wellcome Sanger Institute COVID-19 Surveillance Team |
| EPI_ISL_608432, EPI_ISL_608433 | Lighthouse Lab in Cambridge | Wellcome Sanger Institute for the COVID-19 Genomics UK (COG-UK) consortium | Rob Howes, The Lighthouse Lab in Cambridge and Alex Alderton, Roberto Amato, Sonia Goncalves, Ewan Harrison, David K. Jackson, Ian Johnston, Dominic Kwiatkowski, Cordelia Langford, John Sillitoe on behalf of the Wellcome Sanger Institute COVID-19 Surveillance Team |
| EPI_ISL_608434, EPI_ISL_608435 | Lighthouse Lab in Alderley Park | Wellcome Sanger Institute for the COVID-19 Genomics UK (COG-UK) consortium | Jacquelyn Wynn, Mairead Hyland, The Lighthouse Lab in Alderley Park and Alex Alderton, Roberto Amato, Sonia Goncalves, Ewan Harrison, David K. Jackson, Ian Johnston, Dominic Kwiatkowski, Cordelia Langford, John Sillitoe on behalf of the Wellcome Sanger Institute COVID-19 Surveillance Team |
| EPI_ISL_608436, EPI_ISL_608437 | Lighthouse Lab in Cambridge | Wellcome Sanger Institute for the COVID-19 Genomics UK (COG-UK) consortium | Rob Howes, The Lighthouse Lab in Cambridge and Alex Alderton, Roberto Amato, Sonia Goncalves, Ewan Harrison, David K. Jackson, Ian Johnston, Dominic Kwiatkowski, Cordelia Langford, John Sillitoe on behalf of the Wellcome Sanger Institute COVID-19 Surveillance Team |
| EPI_ISL_608438 | Lighthouse Lab in Alderley Park | Wellcome Sanger Institute for the COVID-19 Genomics UK (COG-UK) consortium | Jacquelyn Wynn, Mairead Hyland, The Lighthouse Lab in Alderley Park and Alex Alderton, Roberto Amato, Sonia Goncalves, Ewan Harrison, David K. Jackson, Ian Johnston, Dominic Kwiatkowski, Cordelia Langford, John Sillitoe on behalf of the Wellcome Sanger Institute COVID-19 Surveillance Team |
| EPI_ISL_608439 | Lighthouse Lab in Cambridge | Wellcome Sanger Institute for the COVID-19 Genomics UK (COG-UK) consortium | Rob Howes, The Lighthouse Lab in Cambridge and Alex Alderton, Roberto Amato, Sonia Goncalves, Ewan Harrison, David K. Jackson, Ian Johnston, Dominic Kwiatkowski, Cordelia Langford, John Sillitoe on behalf of the Wellcome Sanger Institute COVID-19 Surveillance Team |
| EPI_ISL_608440, EPI_ISL_608441 | Lighthouse Lab in Alderley Park | Wellcome Sanger Institute for the COVID-19 Genomics UK (COG-UK) consortium | Jacquelyn Wynn, Mairead Hyland, The Lighthouse Lab in Alderley Park and Alex Alderton, Roberto Amato, Sonia Goncalves, Ewan Harrison, David K. Jackson, Ian Johnston, Dominic Kwiatkowski, Cordelia Langford, John Sillitoe on behalf of the Wellcome Sanger Institute COVID-19 Surveillance Team |
| EPI_ISL_608442 | Lighthouse Lab in Cambridge | Wellcome Sanger Institute for the COVID-19 Genomics UK (COG-UK) consortium | Rob Howes, The Lighthouse Lab in Cambridge and Alex Alderton, Roberto Amato, Sonia Goncalves, Ewan Harrison, David K. Jackson, Ian Johnston, Dominic Kwiatkowski, Cordelia Langford, John Sillitoe on behalf of the Wellcome Sanger Institute COVID-19 Surveillance Team |
| EPI_ISL_608443, EPI_ISL_608444 | Lighthouse Lab in Alderley Park | Wellcome Sanger Institute for the COVID-19 Genomics UK (COG-UK) consortium | Jacquelyn Wynn, Mairead Hyland, The Lighthouse Lab in Alderley Park and Alex Alderton, Roberto Amato, Sonia Goncalves, Ewan Harrison, David K. Jackson, Ian Johnston, Dominic Kwiatkowski, Cordelia Langford, John Sillitoe on behalf of the Wellcome Sanger Institute COVID-19 Surveillance Team |
| EPI_ISL_608446, EPI_ISL_608447 | Lighthouse Lab in Cambridge | Wellcome Sanger Institute for the COVID-19 Genomics UK (COG-UK) consortium | Rob Howes, The Lighthouse Lab in Cambridge and Alex Alderton, Roberto Amato, Sonia Goncalves, Ewan Harrison, David K. Jackson, Ian Johnston, Dominic Kwiatkowski, Cordelia Langford, John Sillitoe on behalf of the Wellcome Sanger Institute COVID-19 Surveillance Team |
| EPI_ISL_608448 | Lighthouse Lab in Alderley Park | Wellcome Sanger Institute for the COVID-19 Genomics UK | Jacquelyn Wynn, Mairead Hyland, The Lighthouse Lab in Alderley Park and Alex Alderton, Roberto Amato, Sonia Goncalves, Ewan Harrison, David K. |

| | | | |
|---|---|---|---|
| | | (COG-UK) consortium | Jackson, Ian Johnston, Dominic Kwiatkowski, Cordelia Langford, John Sillitoe on behalf of the Wellcome Sanger Institute COVID-19 Surveillance Team |
| EPI_ISL_608449, EPI_ISL_608450, EPI_ISL_608451 | Lighthouse Lab in Cambridge | Wellcome Sanger Institute for the COVID-19 Genomics UK (COG-UK) consortium | Rob Howes, The Lighthouse Lab in Cambridge and Alex Alderton, Roberto Amato, Sonia Goncalves, Ewan Harrison, David K. Jackson, Ian Johnston, Dominic Kwiatkowski, Cordelia Langford, John Sillitoe on behalf of the Wellcome Sanger Institute COVID-19 Surveillance Team |
| EPI_ISL_608452, EPI_ISL_608453, EPI_ISL_608454 | Lighthouse Lab in Alderley Park | Wellcome Sanger Institute for the COVID-19 Genomics UK (COG-UK) consortium | Jacquelyn Wynn, Mairead Hyland, The Lighthouse Lab in Alderley Park and Alex Alderton, Roberto Amato, Sonia Goncalves, Ewan Harrison, David K. Jackson, Ian Johnston, Dominic Kwiatkowski, Cordelia Langford, John Sillitoe on behalf of the Wellcome Sanger Institute COVID-19 Surveillance Team |
| EPI_ISL_608455 | Lighthouse Lab in Cambridge | Wellcome Sanger Institute for the COVID-19 Genomics UK (COG-UK) Consortium | Rob Howes, The Lighthouse Lab in Cambridge and Alex Alderton, Roberto Amato, Sonia Goncalves, Ewan Harrison, David K. Jackson, Ian Johnston, Dominic Kwiatkowski, Cordelia Langford, John Sillitoe on behalf of the Wellcome Sanger Institute COVID-19 Surveillance Team |
| EPI_ISL_608456 | Lighthouse Lab in Cambridge | Wellcome Sanger Institute for the COVID-19 Genomics UK (COG-UK) consortium | Rob Howes, The Lighthouse Lab in Cambridge and Alex Alderton, Roberto Amato, Sonia Goncalves, Ewan Harrison, David K. Jackson, Ian Johnston, Dominic Kwiatkowski, Cordelia Langford, John Sillitoe on behalf of the Wellcome Sanger Institute COVID-19 Surveillance Team |
| EPI_ISL_608457 | Lighthouse Lab in Alderley Park | Wellcome Sanger Institute for the COVID-19 Genomics UK (COG-UK) consortium | Jacquelyn Wynn, Mairead Hyland, The Lighthouse Lab in Alderley Park and Alex Alderton, Roberto Amato, Sonia Goncalves, Ewan Harrison, David K. Jackson, Ian Johnston, Dominic Kwiatkowski, Cordelia Langford, John Sillitoe on behalf of the Wellcome Sanger Institute COVID-19 Surveillance Team |
| EPI_ISL_608458, EPI_ISL_608459 | Lighthouse Lab in Cambridge | Wellcome Sanger Institute for the COVID-19 Genomics UK (COG-UK) consortium | Rob Howes, The Lighthouse Lab in Cambridge and Alex Alderton, Roberto Amato, Sonia Goncalves, Ewan Harrison, David K. Jackson, Ian Johnston, Dominic Kwiatkowski, Cordelia Langford, John Sillitoe on behalf of the Wellcome Sanger Institute COVID-19 Surveillance Team |
| EPI_ISL_608460, EPI_ISL_608461, EPI_ISL_608462 | Lighthouse Lab in Alderley Park | Wellcome Sanger Institute for the COVID-19 Genomics UK (COG-UK) consortium | Jacquelyn Wynn, Mairead Hyland, The Lighthouse Lab in Alderley Park and Alex Alderton, Roberto Amato, Sonia Goncalves, Ewan Harrison, David K. Jackson, Ian Johnston, Dominic Kwiatkowski, Cordelia Langford, John Sillitoe on behalf of the Wellcome Sanger Institute COVID-19 Surveillance Team |
| EPI_ISL_608463, EPI_ISL_608464 | Lighthouse Lab in Cambridge | Wellcome Sanger Institute for the COVID-19 Genomics UK (COG-UK) consortium | Rob Howes, The Lighthouse Lab in Cambridge and Alex Alderton, Roberto Amato, Sonia Goncalves, Ewan Harrison, David K. Jackson, Ian Johnston, Dominic Kwiatkowski, Cordelia Langford, John Sillitoe on behalf of the Wellcome Sanger Institute COVID-19 Surveillance Team |
| EPI_ISL_608465 | Lighthouse Lab in Alderley Park | Wellcome Sanger Institute for the COVID-19 Genomics UK (COG-UK) consortium | Jacquelyn Wynn, Mairead Hyland, The Lighthouse Lab in Alderley Park and Alex Alderton, Roberto Amato, Sonia Goncalves, Ewan Harrison, David K. Jackson, Ian Johnston, Dominic Kwiatkowski, Cordelia Langford, John Sillitoe on behalf of the Wellcome Sanger Institute COVID-19 Surveillance Team |
| EPI_ISL_608466, EPI_ISL_608467 | Lighthouse Lab in Cambridge | Wellcome Sanger Institute for the COVID-19 Genomics UK (COG-UK) consortium | Rob Howes, The Lighthouse Lab in Cambridge and Alex Alderton, Roberto Amato, Sonia Goncalves, Ewan Harrison, David K. Jackson, Ian Johnston, Dominic Kwiatkowski, Cordelia Langford, John Sillitoe on behalf of the Wellcome Sanger Institute COVID-19 Surveillance Team |
| EPI_ISL_608468, EPI_ISL_608469, EPI_ISL_608470 | Lighthouse Lab in Alderley Park | Wellcome Sanger Institute for the COVID-19 Genomics UK (COG-UK) consortium | Jacquelyn Wynn, Mairead Hyland, The Lighthouse Lab in Alderley Park and Alex Alderton, Roberto Amato, Sonia Goncalves, Ewan Harrison, David K. Jackson, Ian Johnston, Dominic Kwiatkowski, Cordelia Langford, John Sillitoe on behalf of the Wellcome Sanger Institute COVID-19 Surveillance Team |
| EPI_ISL_608471, EPI_ISL_608472 | Lighthouse Lab in Cambridge | Wellcome Sanger Institute for the COVID-19 Genomics UK (COG-UK) consortium | Rob Howes, The Lighthouse Lab in Cambridge and Alex Alderton, Roberto Amato, Sonia Goncalves, Ewan Harrison, David K. Jackson, Ian Johnston, Dominic Kwiatkowski, Cordelia Langford, John Sillitoe on behalf of the Wellcome Sanger Institute COVID-19 Surveillance Team |
| EPI_ISL_608473, EPI_ISL_608474 | Lighthouse Lab in Alderley Park | Wellcome Sanger Institute for the COVID-19 Genomics UK (COG-UK) consortium | Jacquelyn Wynn, Mairead Hyland, The Lighthouse Lab in Alderley Park and Alex Alderton, Roberto Amato, Sonia Goncalves, Ewan Harrison, David K. Jackson, Ian Johnston, Dominic Kwiatkowski, Cordelia Langford, John Sillitoe on behalf of the Wellcome Sanger Institute COVID-19 Surveillance Team |
| EPI_ISL_608475, EPI_ISL_608476, EPI_ISL_608477, EPI_ISL_608478 | Lighthouse Lab in Cambridge | Wellcome Sanger Institute for the COVID-19 Genomics UK (COG-UK) consortium | Rob Howes, The Lighthouse Lab in Cambridge and Alex Alderton, Roberto Amato, Sonia Goncalves, Ewan Harrison, David K. Jackson, Ian Johnston, Dominic Kwiatkowski, Cordelia Langford, John Sillitoe on behalf of the Wellcome Sanger Institute COVID-19 Surveillance Team |
| EPI_ISL_608479 | Lighthouse Lab in Alderley Park | Wellcome Sanger Institute for the COVID-19 Genomics UK (COG-UK) consortium | Jacquelyn Wynn, Mairead Hyland, The Lighthouse Lab in Alderley Park and Alex Alderton, Roberto Amato, Sonia Goncalves, Ewan Harrison, David K. Jackson, Ian Johnston, Dominic Kwiatkowski, Cordelia Langford, John Sillitoe on behalf of the Wellcome Sanger Institute COVID-19 Surveillance Team |
| EPI_ISL_608480 | Lighthouse Lab in Cambridge | Wellcome Sanger Institute for the COVID-19 Genomics UK (COG-UK) consortium | Rob Howes, The Lighthouse Lab in Cambridge and Alex Alderton, Roberto Amato, Sonia Goncalves, Ewan Harrison, David K. Jackson, Ian Johnston, Dominic Kwiatkowski, Cordelia Langford, John Sillitoe on behalf of the Wellcome Sanger Institute COVID-19 Surveillance Team |
| EPI_ISL_608481 | Lighthouse Lab in Alderley Park | Wellcome Sanger Institute for the COVID-19 Genomics UK (COG-UK) consortium | Jacquelyn Wynn, Mairead Hyland, The Lighthouse Lab in Alderley Park and Alex Alderton, Roberto Amato, Sonia Goncalves, Ewan Harrison, David K. Jackson, Ian Johnston, Dominic Kwiatkowski, Cordelia Langford, John Sillitoe on behalf of the Wellcome Sanger Institute COVID-19 Surveillance Team |
| EPI_ISL_608482 | Lighthouse Lab in Cambridge | Wellcome Sanger Institute for the COVID-19 Genomics UK (COG-UK) consortium | Rob Howes, The Lighthouse Lab in Cambridge and Alex Alderton, Roberto Amato, Sonia Goncalves, Ewan Harrison, David K. Jackson, Ian Johnston, Dominic Kwiatkowski, Cordelia Langford, John Sillitoe on behalf of the Wellcome Sanger Institute COVID-19 Surveillance Team |
| EPI_ISL_608483 | Lighthouse Lab in Alderley Park | Wellcome Sanger Institute for the COVID-19 Genomics UK (COG-UK) consortium | Jacquelyn Wynn, Mairead Hyland, The Lighthouse Lab in Alderley Park and Alex Alderton, Roberto Amato, Sonia Goncalves, Ewan Harrison, David K. Jackson, Ian Johnston, Dominic Kwiatkowski, Cordelia Langford, John Sillitoe on behalf of the Wellcome Sanger Institute COVID-19 Surveillance Team |
| EPI_ISL_608484, EPI_ISL_608485, EPI_ISL_608486 | Lighthouse Lab in Cambridge | Wellcome Sanger Institute for the COVID-19 Genomics UK (COG-UK) consortium | Rob Howes, The Lighthouse Lab in Cambridge and Alex Alderton, Roberto Amato, Sonia Goncalves, Ewan Harrison, David K. Jackson, Ian Johnston, Dominic Kwiatkowski, Cordelia Langford, John Sillitoe on behalf of the Wellcome Sanger Institute COVID-19 Surveillance Team |
| EPI_ISL_608487, EPI_ISL_608488, EPI_ISL_608489 | Lighthouse Lab in Alderley Park | Wellcome Sanger Institute for the COVID-19 Genomics UK (COG-UK) consortium | Jacquelyn Wynn, Mairead Hyland, The Lighthouse Lab in Alderley Park and Alex Alderton, Roberto Amato, Sonia Goncalves, Ewan Harrison, David K. Jackson, Ian Johnston, Dominic Kwiatkowski, Cordelia Langford, John Sillitoe on behalf of the Wellcome Sanger Institute COVID-19 Surveillance Team |
| EPI_ISL_608490 | Lighthouse Lab in Cambridge | Wellcome Sanger Institute for the COVID-19 Genomics UK (COG-UK) consortium | Rob Howes, The Lighthouse Lab in Cambridge and Alex Alderton, Roberto Amato, Sonia Goncalves, Ewan Harrison, David K. Jackson, Ian Johnston, Dominic Kwiatkowski, Cordelia Langford, John Sillitoe on behalf of the Wellcome Sanger Institute COVID-19 Surveillance Team |
| EPI_ISL_608491 | Lighthouse Lab in Alderley Park | Wellcome Sanger Institute for the COVID-19 Genomics UK (COG-UK) consortium | Jacquelyn Wynn, Mairead Hyland, The Lighthouse Lab in Alderley Park and Alex Alderton, Roberto Amato, Sonia Goncalves, Ewan Harrison, David K. Jackson, Ian Johnston, Dominic Kwiatkowski, Cordelia Langford, John Sillitoe on behalf of the Wellcome Sanger Institute COVID-19 Surveillance Team |
| EPI_ISL_608492, EPI_ISL_608493, EPI_ISL_608494, EPI_ISL_608495 | Lighthouse Lab in Cambridge | Wellcome Sanger Institute for the COVID-19 Genomics UK (COG-UK) consortium | Rob Howes, The Lighthouse Lab in Cambridge and Alex Alderton, Roberto Amato, Sonia Goncalves, Ewan Harrison, David K. Jackson, Ian Johnston, Dominic Kwiatkowski, Cordelia Langford, John Sillitoe on behalf of the Wellcome Sanger Institute COVID-19 Surveillance Team |
| EPI_ISL_608496 | Lighthouse Lab in Alderley Park | Wellcome Sanger Institute for the COVID-19 Genomics UK (COG-UK) consortium | Jacquelyn Wynn, Mairead Hyland, The Lighthouse Lab in Alderley Park and Alex Alderton, Roberto Amato, Sonia Goncalves, Ewan Harrison, David K. Jackson, Ian Johnston, Dominic Kwiatkowski, Cordelia Langford, John Sillitoe on behalf of the Wellcome Sanger Institute COVID-19 Surveillance Team |
| EPI_ISL_608497, EPI_ISL_608498 | Lighthouse Lab in Cambridge | Wellcome Sanger Institute for the COVID-19 Genomics UK (COG-UK) consortium | Rob Howes, The Lighthouse Lab in Cambridge and Alex Alderton, Roberto Amato, Sonia Goncalves, Ewan Harrison, David K. Jackson, Ian Johnston, Dominic Kwiatkowski, Cordelia Langford, John Sillitoe on behalf of the Wellcome Sanger Institute COVID-19 Surveillance Team |
| EPI_ISL_608499, EPI_ISL_608500 | Lighthouse Lab in Alderley Park | Wellcome Sanger Institute for the COVID-19 Genomics UK (COG-UK) consortium | Jacquelyn Wynn, Mairead Hyland, The Lighthouse Lab in Alderley Park and Alex Alderton, Roberto Amato, Sonia Goncalves, Ewan Harrison, David K. Jackson, Ian Johnston, Dominic Kwiatkowski, Cordelia Langford, John Sillitoe on behalf of the Wellcome Sanger Institute COVID-19 Surveillance Team |
| EPI_ISL_608501 | Lighthouse Lab in Cambridge | Wellcome Sanger Institute for the COVID-19 Genomics UK (COG-UK) consortium | Rob Howes, The Lighthouse Lab in Cambridge and Alex Alderton, Roberto Amato, Sonia Goncalves, Ewan Harrison, David K. Jackson, Ian Johnston, Dominic Kwiatkowski, Cordelia Langford, John Sillitoe on behalf of the Wellcome Sanger Institute COVID-19 Surveillance Team |
| EPI_ISL_608502, EPI_ISL_608503 | Lighthouse Lab in Alderley Park | Wellcome Sanger Institute for the COVID-19 Genomics UK (COG-UK) consortium | Jacquelyn Wynn, Mairead Hyland, The Lighthouse Lab in Alderley Park and Alex Alderton, Roberto Amato, Sonia Goncalves, Ewan Harrison, David K. Jackson, Ian Johnston, Dominic Kwiatkowski, Cordelia Langford, John Sillitoe on behalf of the Wellcome Sanger Institute COVID-19 Surveillance Team |
| EPI_ISL_608504, EPI_ISL_608506, EPI_ISL_608507, EPI_ISL_608508, EPI_ISL_608509, EPI_ISL_608510 | Lighthouse Lab in Cambridge | Wellcome Sanger Institute for the COVID-19 Genomics UK (COG-UK) consortium | Rob Howes, The Lighthouse Lab in Cambridge and Alex Alderton, Roberto Amato, Sonia Goncalves, Ewan Harrison, David K. Jackson, Ian Johnston, Dominic Kwiatkowski, Cordelia Langford, John Sillitoe on behalf of the Wellcome Sanger Institute COVID-19 Surveillance Team |
| EPI_ISL_608511, EPI_ISL_608513 | Lighthouse Lab in Alderley Park | Wellcome Sanger Institute for the COVID-19 Genomics UK (COG-UK) consortium | Jacquelyn Wynn, Mairead Hyland, The Lighthouse Lab in Alderley Park and Alex Alderton, Roberto Amato, Sonia Goncalves, Ewan Harrison, David K. Jackson, Ian Johnston, Dominic Kwiatkowski, Cordelia Langford, John Sillitoe on behalf of the Wellcome Sanger Institute COVID-19 Surveillance Team |
| EPI_ISL_608514, EPI_ISL_608515 | Lighthouse Lab in Cambridge | Wellcome Sanger Institute for the COVID-19 Genomics UK (COG-UK) consortium | Rob Howes, The Lighthouse Lab in Cambridge and Alex Alderton, Roberto Amato, Sonia Goncalves, Ewan Harrison, David K. Jackson, Ian Johnston, Dominic Kwiatkowski, Cordelia Langford, John Sillitoe on behalf of the Wellcome Sanger Institute COVID-19 Surveillance Team |
| EPI_ISL_608516, EPI_ISL_608517, EPI_ISL_608518, EPI_ISL_608519, EPI_ISL_608520 | Lighthouse Lab in Alderley Park | Wellcome Sanger Institute for the COVID-19 Genomics UK (COG-UK) consortium | Jacquelyn Wynn, Mairead Hyland, The Lighthouse Lab in Alderley Park and Alex Alderton, Roberto Amato, Sonia Goncalves, Ewan Harrison, David K. Jackson, Ian Johnston, Dominic Kwiatkowski, Cordelia Langford, John Sillitoe on behalf of the Wellcome Sanger Institute COVID-19 Surveillance Team |
| EPI_ISL_608521, EPI_ISL_608522, EPI_ISL_608523, EPI_ISL_608524, | Lighthouse Lab in Cambridge | Wellcome Sanger Institute for the COVID-19 Genomics UK (COG-UK) consortium | Rob Howes, The Lighthouse Lab in Cambridge and Alex Alderton, Roberto Amato, Sonia Goncalves, Ewan Harrison, David K. Jackson, Ian Johnston, Dominic Kwiatkowski, Cordelia Langford, John Sillitoe on behalf of the Wellcome Sanger Institute COVID-19 Surveillance Team |

| | | | |
|---|---|---|---|
| EPI_ISL_608525 | | | |
| EPI_ISL_608526 | Lighthouse Lab in Alderley Park | Wellcome Sanger Institute for the COVID-19 Genomics UK (COG-UK) consortium | Jacquelyn Wynn, Mairead Hyland, The Lighthouse Lab in Alderley Park and Alex Alderton, Roberto Amato, Sonia Goncalves, Ewan Harrison, David K. Jackson, Ian Johnston, Dominic Kwiatkowski, Cordelia Langford, John Sillitoe on behalf of the Wellcome Sanger Institute COVID-19 Surveillance Team |
| EPI_ISL_608527 | Lighthouse Lab in Cambridge | Wellcome Sanger Institute for the COVID-19 Genomics UK (COG-UK) consortium | Rob Howes, The Lighthouse Lab in Cambridge and Alex Alderton, Roberto Amato, Sonia Goncalves, Ewan Harrison, David K. Jackson, Ian Johnston, Dominic Kwiatkowski, Cordelia Langford, John Sillitoe on behalf of the Wellcome Sanger Institute COVID-19 Surveillance Team |
| EPI_ISL_608528 | Lighthouse Lab in Alderley Park | Wellcome Sanger Institute for the COVID-19 Genomics UK (COG-UK) consortium | Jacquelyn Wynn, Mairead Hyland, The Lighthouse Lab in Alderley Park and Alex Alderton, Roberto Amato, Sonia Goncalves, Ewan Harrison, David K. Jackson, Ian Johnston, Dominic Kwiatkowski, Cordelia Langford, John Sillitoe on behalf of the Wellcome Sanger Institute COVID-19 Surveillance Team |
| EPI_ISL_608529, EPI_ISL_608530, EPI_ISL_608531, EPI_ISL_608532 | Lighthouse Lab in Cambridge | Wellcome Sanger Institute for the COVID-19 Genomics UK (COG-UK) consortium | Rob Howes, The Lighthouse Lab in Cambridge and Alex Alderton, Roberto Amato, Sonia Goncalves, Ewan Harrison, David K. Jackson, Ian Johnston, Dominic Kwiatkowski, Cordelia Langford, John Sillitoe on behalf of the Wellcome Sanger Institute COVID-19 Surveillance Team |
| EPI_ISL_608533, EPI_ISL_608534 | Lighthouse Lab in Alderley Park | Wellcome Sanger Institute for the COVID-19 Genomics UK (COG-UK) consortium | Jacquelyn Wynn, Mairead Hyland, The Lighthouse Lab in Alderley Park and Alex Alderton, Roberto Amato, Sonia Goncalves, Ewan Harrison, David K. Jackson, Ian Johnston, Dominic Kwiatkowski, Cordelia Langford, John Sillitoe on behalf of the Wellcome Sanger Institute COVID-19 Surveillance Team |
| EPI_ISL_608535, EPI_ISL_608536, EPI_ISL_608537, EPI_ISL_608538, EPI_ISL_608540 | Lighthouse Lab in Cambridge | Wellcome Sanger Institute for the COVID-19 Genomics UK (COG-UK) consortium | Rob Howes, The Lighthouse Lab in Cambridge and Alex Alderton, Roberto Amato, Sonia Goncalves, Ewan Harrison, David K. Jackson, Ian Johnston, Dominic Kwiatkowski, Cordelia Langford, John Sillitoe on behalf of the Wellcome Sanger Institute COVID-19 Surveillance Team |
| EPI_ISL_608541, EPI_ISL_608542, EPI_ISL_608543 | Lighthouse Lab in Alderley Park | Wellcome Sanger Institute for the COVID-19 Genomics UK (COG-UK) consortium | Jacquelyn Wynn, Mairead Hyland, The Lighthouse Lab in Alderley Park and Alex Alderton, Roberto Amato, Sonia Goncalves, Ewan Harrison, David K. Jackson, Ian Johnston, Dominic Kwiatkowski, Cordelia Langford, John Sillitoe on behalf of the Wellcome Sanger Institute COVID-19 Surveillance Team |
| EPI_ISL_608544, EPI_ISL_608545, EPI_ISL_608546, EPI_ISL_608547 | Lighthouse Lab in Cambridge | Wellcome Sanger Institute for the COVID-19 Genomics UK (COG-UK) consortium | Rob Howes, The Lighthouse Lab in Cambridge and Alex Alderton, Roberto Amato, Sonia Goncalves, Ewan Harrison, David K. Jackson, Ian Johnston, Dominic Kwiatkowski, Cordelia Langford, John Sillitoe on behalf of the Wellcome Sanger Institute COVID-19 Surveillance Team |
| EPI_ISL_608548, EPI_ISL_608549 | Lighthouse Lab in Alderley Park | Wellcome Sanger Institute for the COVID-19 Genomics UK (COG-UK) consortium | Jacquelyn Wynn, Mairead Hyland, The Lighthouse Lab in Alderley Park and Alex Alderton, Roberto Amato, Sonia Goncalves, Ewan Harrison, David K. Jackson, Ian Johnston, Dominic Kwiatkowski, Cordelia Langford, John Sillitoe on behalf of the Wellcome Sanger Institute COVID-19 Surveillance Team |
| EPI_ISL_608550 | Lighthouse Lab in Cambridge | Wellcome Sanger Institute for the COVID-19 Genomics UK (COG-UK) consortium | Rob Howes, The Lighthouse Lab in Cambridge and Alex Alderton, Roberto Amato, Sonia Goncalves, Ewan Harrison, David K. Jackson, Ian Johnston, Dominic Kwiatkowski, Cordelia Langford, John Sillitoe on behalf of the Wellcome Sanger Institute COVID-19 Surveillance Team |
| EPI_ISL_608551 | Lighthouse Lab in Alderley Park | Wellcome Sanger Institute for the COVID-19 Genomics UK (COG-UK) consortium | Jacquelyn Wynn, Mairead Hyland, The Lighthouse Lab in Alderley Park and Alex Alderton, Roberto Amato, Sonia Goncalves, Ewan Harrison, David K. Jackson, Ian Johnston, Dominic Kwiatkowski, Cordelia Langford, John Sillitoe on behalf of the Wellcome Sanger Institute COVID-19 Surveillance Team |
| EPI_ISL_608552 | Lighthouse Lab in Cambridge | Wellcome Sanger Institute for the COVID-19 Genomics UK (COG-UK) consortium | Rob Howes, The Lighthouse Lab in Cambridge and Alex Alderton, Roberto Amato, Sonia Goncalves, Ewan Harrison, David K. Jackson, Ian Johnston, Dominic Kwiatkowski, Cordelia Langford, John Sillitoe on behalf of the Wellcome Sanger Institute COVID-19 Surveillance Team |
| EPI_ISL_608553, EPI_ISL_608554, EPI_ISL_608555, EPI_ISL_608556 | Lighthouse Lab in Alderley Park | Wellcome Sanger Institute for the COVID-19 Genomics UK (COG-UK) consortium | Jacquelyn Wynn, Mairead Hyland, The Lighthouse Lab in Alderley Park and Alex Alderton, Roberto Amato, Sonia Goncalves, Ewan Harrison, David K. Jackson, Ian Johnston, Dominic Kwiatkowski, Cordelia Langford, John Sillitoe on behalf of the Wellcome Sanger Institute COVID-19 Surveillance Team |
| EPI_ISL_608557, EPI_ISL_608558, EPI_ISL_608559, EPI_ISL_608560, EPI_ISL_608561 | Lighthouse Lab in Cambridge | Wellcome Sanger Institute for the COVID-19 Genomics UK (COG-UK) consortium | Rob Howes, The Lighthouse Lab in Cambridge and Alex Alderton, Roberto Amato, Sonia Goncalves, Ewan Harrison, David K. Jackson, Ian Johnston, Dominic Kwiatkowski, Cordelia Langford, John Sillitoe on behalf of the Wellcome Sanger Institute COVID-19 Surveillance Team |
| EPI_ISL_608562, EPI_ISL_608563, EPI_ISL_608564 | Lighthouse Lab in Alderley Park | Wellcome Sanger Institute for the COVID-19 Genomics UK (COG-UK) consortium | Jacquelyn Wynn, Mairead Hyland, The Lighthouse Lab in Alderley Park and Alex Alderton, Roberto Amato, Sonia Goncalves, Ewan Harrison, David K. Jackson, Ian Johnston, Dominic Kwiatkowski, Cordelia Langford, John Sillitoe on behalf of the Wellcome Sanger Institute COVID-19 Surveillance Team |
| EPI_ISL_608565, EPI_ISL_608566 | Lighthouse Lab in Cambridge | Wellcome Sanger Institute for the COVID-19 Genomics UK (COG-UK) consortium | Rob Howes, The Lighthouse Lab in Cambridge and Alex Alderton, Roberto Amato, Sonia Goncalves, Ewan Harrison, David K. Jackson, Ian Johnston, Dominic Kwiatkowski, Cordelia Langford, John Sillitoe on behalf of the Wellcome Sanger Institute COVID-19 Surveillance Team |
| EPI_ISL_608567 | Lighthouse Lab in Alderley Park | Wellcome Sanger Institute for the COVID-19 Genomics UK (COG-UK) consortium | Jacquelyn Wynn, Mairead Hyland, The Lighthouse Lab in Alderley Park and Alex Alderton, Roberto Amato, Sonia Goncalves, Ewan Harrison, David K. Jackson, Ian Johnston, Dominic Kwiatkowski, Cordelia Langford, John Sillitoe on behalf of the Wellcome Sanger Institute COVID-19 Surveillance Team |
| EPI_ISL_608568, EPI_ISL_608570, EPI_ISL_608571, EPI_ISL_608572, EPI_ISL_608573 | Lighthouse Lab in Cambridge | Wellcome Sanger Institute for the COVID-19 Genomics UK (COG-UK) consortium | Rob Howes, The Lighthouse Lab in Cambridge and Alex Alderton, Roberto Amato, Sonia Goncalves, Ewan Harrison, David K. Jackson, Ian Johnston, Dominic Kwiatkowski, Cordelia Langford, John Sillitoe on behalf of the Wellcome Sanger Institute COVID-19 Surveillance Team |
| EPI_ISL_608574, EPI_ISL_608575 | Lighthouse Lab in Alderley Park | Wellcome Sanger Institute for the COVID-19 Genomics UK (COG-UK) consortium | Jacquelyn Wynn, Mairead Hyland, The Lighthouse Lab in Alderley Park and Alex Alderton, Roberto Amato, Sonia Goncalves, Ewan Harrison, David K. Jackson, Ian Johnston, Dominic Kwiatkowski, Cordelia Langford, John Sillitoe on behalf of the Wellcome Sanger Institute COVID-19 Surveillance Team |
| EPI_ISL_608576, EPI_ISL_608577, EPI_ISL_608578 | Lighthouse Lab in Cambridge | Wellcome Sanger Institute for the COVID-19 Genomics UK (COG-UK) consortium | Rob Howes, The Lighthouse Lab in Cambridge and Alex Alderton, Roberto Amato, Sonia Goncalves, Ewan Harrison, David K. Jackson, Ian Johnston, Dominic Kwiatkowski, Cordelia Langford, John Sillitoe on behalf of the Wellcome Sanger Institute COVID-19 Surveillance Team |
| EPI_ISL_608579, EPI_ISL_608580, EPI_ISL_608581, EPI_ISL_608582, EPI_ISL_608583, EPI_ISL_608584 | Lighthouse Lab in Alderley Park | Wellcome Sanger Institute for the COVID-19 Genomics UK (COG-UK) consortium | Jacquelyn Wynn, Mairead Hyland, The Lighthouse Lab in Alderley Park and Alex Alderton, Roberto Amato, Sonia Goncalves, Ewan Harrison, David K. Jackson, Ian Johnston, Dominic Kwiatkowski, Cordelia Langford, John Sillitoe on behalf of the Wellcome Sanger Institute COVID-19 Surveillance Team |
| EPI_ISL_608585 | Lighthouse Lab in Cambridge | Wellcome Sanger Institute for the COVID-19 Genomics UK (COG-UK) consortium | Rob Howes, The Lighthouse Lab in Cambridge and Alex Alderton, Roberto Amato, Sonia Goncalves, Ewan Harrison, David K. Jackson, Ian Johnston, Dominic Kwiatkowski, Cordelia Langford, John Sillitoe on behalf of the Wellcome Sanger Institute COVID-19 Surveillance Team |
| EPI_ISL_608586, EPI_ISL_608587 | Lighthouse Lab in Alderley Park | Wellcome Sanger Institute for the COVID-19 Genomics UK (COG-UK) consortium | Jacquelyn Wynn, Mairead Hyland, The Lighthouse Lab in Alderley Park and Alex Alderton, Roberto Amato, Sonia Goncalves, Ewan Harrison, David K. Jackson, Ian Johnston, Dominic Kwiatkowski, Cordelia Langford, John Sillitoe on behalf of the Wellcome Sanger Institute COVID-19 Surveillance Team |
| EPI_ISL_608588 | Lighthouse Lab in Cambridge | Wellcome Sanger Institute for the COVID-19 Genomics UK (COG-UK) consortium | Rob Howes, The Lighthouse Lab in Cambridge and Alex Alderton, Roberto Amato, Sonia Goncalves, Ewan Harrison, David K. Jackson, Ian Johnston, Dominic Kwiatkowski, Cordelia Langford, John Sillitoe on behalf of the Wellcome Sanger Institute COVID-19 Surveillance Team |
| EPI_ISL_608589, EPI_ISL_608590, EPI_ISL_608591, EPI_ISL_608592, EPI_ISL_608593, EPI_ISL_608594, EPI_ISL_608595 | Lighthouse Lab in Alderley Park | Wellcome Sanger Institute for the COVID-19 Genomics UK (COG-UK) consortium | Jacquelyn Wynn, Mairead Hyland, The Lighthouse Lab in Alderley Park and Alex Alderton, Roberto Amato, Sonia Goncalves, Ewan Harrison, David K. Jackson, Ian Johnston, Dominic Kwiatkowski, Cordelia Langford, John Sillitoe on behalf of the Wellcome Sanger Institute COVID-19 Surveillance Team |
| EPI_ISL_608596, EPI_ISL_608597, EPI_ISL_608598 | Lighthouse Lab in Cambridge | Wellcome Sanger Institute for the COVID-19 Genomics UK (COG-UK) consortium | Rob Howes, The Lighthouse Lab in Cambridge and Alex Alderton, Roberto Amato, Sonia Goncalves, Ewan Harrison, David K. Jackson, Ian Johnston, Dominic Kwiatkowski, Cordelia Langford, John Sillitoe on behalf of the Wellcome Sanger Institute COVID-19 Surveillance Team |
| EPI_ISL_608599, EPI_ISL_608600, EPI_ISL_608601 | Lighthouse Lab in Alderley Park | Wellcome Sanger Institute for the COVID-19 Genomics UK (COG-UK) consortium | Jacquelyn Wynn, Mairead Hyland, The Lighthouse Lab in Alderley Park and Alex Alderton, Roberto Amato, Sonia Goncalves, Ewan Harrison, David K. Jackson, Ian Johnston, Dominic Kwiatkowski, Cordelia Langford, John Sillitoe on behalf of the Wellcome Sanger Institute COVID-19 Surveillance Team |
| EPI_ISL_608602 | Lighthouse Lab in Cambridge | Wellcome Sanger Institute for the COVID-19 Genomics UK (COG-UK) consortium | Rob Howes, The Lighthouse Lab in Cambridge and Alex Alderton, Roberto Amato, Sonia Goncalves, Ewan Harrison, David K. Jackson, Ian Johnston, Dominic Kwiatkowski, Cordelia Langford, John Sillitoe on behalf of the Wellcome Sanger Institute COVID-19 Surveillance Team |
| EPI_ISL_608603 | Lighthouse Lab in Alderley Park | Wellcome Sanger Institute for the COVID-19 Genomics UK (COG-UK) consortium | Jacquelyn Wynn, Mairead Hyland, The Lighthouse Lab in Alderley Park and Alex Alderton, Roberto Amato, Sonia Goncalves, Ewan Harrison, David K. Jackson, Ian Johnston, Dominic Kwiatkowski, Cordelia Langford, John Sillitoe on behalf of the Wellcome Sanger Institute COVID-19 Surveillance Team |
| EPI_ISL_608604 | Lighthouse Lab in Cambridge | Wellcome Sanger Institute for the COVID-19 Genomics UK (COG-UK) consortium | Rob Howes, The Lighthouse Lab in Cambridge and Alex Alderton, Roberto Amato, Sonia Goncalves, Ewan Harrison, David K. Jackson, Ian Johnston, Dominic Kwiatkowski, Cordelia Langford, John Sillitoe on behalf of the Wellcome Sanger Institute COVID-19 Surveillance Team |
| EPI_ISL_608605, EPI_ISL_608606 | Lighthouse Lab in Alderley Park | Wellcome Sanger Institute for the COVID-19 Genomics UK (COG-UK) consortium | Jacquelyn Wynn, Mairead Hyland, The Lighthouse Lab in Alderley Park and Alex Alderton, Roberto Amato, Sonia Goncalves, Ewan Harrison, David K. Jackson, Ian Johnston, Dominic Kwiatkowski, Cordelia Langford, John Sillitoe on behalf of the Wellcome Sanger Institute COVID-19 Surveillance Team |
| EPI_ISL_608607, EPI_ISL_608608 | Lighthouse Lab in Cambridge | Wellcome Sanger Institute for the COVID-19 Genomics UK (COG-UK) consortium | Rob Howes, The Lighthouse Lab in Cambridge and Alex Alderton, Roberto Amato, Sonia Goncalves, Ewan Harrison, David K. Jackson, Ian Johnston, Dominic Kwiatkowski, Cordelia Langford, John Sillitoe on behalf of the Wellcome Sanger Institute COVID-19 Surveillance Team |

| | | | |
|---|---|---|---|
| EPI_ISL_608609, EPI_ISL_608610 | Lighthouse Lab in Alderley Park | Wellcome Sanger Institute for the COVID-19 Genomics UK (COG-UK) consortium | Jacquelyn Wynn, Mairead Hyland, The Lighthouse Lab in Alderley Park and Alex Alderton, Roberto Amato, Sonia Goncalves, Ewan Harrison, David K. Jackson, Ian Johnston, Dominic Kwiatkowski, Cordelia Langford, John Sillitoe on behalf of the Wellcome Sanger Institute COVID-19 Surveillance Team |
| EPI_ISL_608611, EPI_ISL_608612, EPI_ISL_608613 | Lighthouse Lab in Cambridge | Wellcome Sanger Institute for the COVID-19 Genomics UK (COG-UK) consortium | Rob Howes, The Lighthouse Lab in Cambridge and Alex Alderton, Roberto Amato, Sonia Goncalves, Ewan Harrison, David K. Jackson, Ian Johnston, Dominic Kwiatkowski, Cordelia Langford, John Sillitoe on behalf of the Wellcome Sanger Institute COVID-19 Surveillance Team |
| EPI_ISL_608614 | Lighthouse Lab in Alderley Park | Wellcome Sanger Institute for the COVID-19 Genomics UK (COG-UK) consortium | Jacquelyn Wynn, Mairead Hyland, The Lighthouse Lab in Alderley Park and Alex Alderton, Roberto Amato, Sonia Goncalves, Ewan Harrison, David K. Jackson, Ian Johnston, Dominic Kwiatkowski, Cordelia Langford, John Sillitoe on behalf of the Wellcome Sanger Institute COVID-19 Surveillance Team |
| EPI_ISL_608615, EPI_ISL_608616 | Lighthouse Lab in Cambridge | Wellcome Sanger Institute for the COVID-19 Genomics UK (COG-UK) consortium | Rob Howes, The Lighthouse Lab in Cambridge and Alex Alderton, Roberto Amato, Sonia Goncalves, Ewan Harrison, David K. Jackson, Ian Johnston, Dominic Kwiatkowski, Cordelia Langford, John Sillitoe on behalf of the Wellcome Sanger Institute COVID-19 Surveillance Team |
| EPI_ISL_608617 | Lighthouse Lab in Alderley Park | Wellcome Sanger Institute for the COVID-19 Genomics UK (COG-UK) consortium | Jacquelyn Wynn, Mairead Hyland, The Lighthouse Lab in Alderley Park and Alex Alderton, Roberto Amato, Sonia Goncalves, Ewan Harrison, David K. Jackson, Ian Johnston, Dominic Kwiatkowski, Cordelia Langford, John Sillitoe on behalf of the Wellcome Sanger Institute COVID-19 Surveillance Team |
| EPI_ISL_608618, EPI_ISL_608619, EPI_ISL_608620 | Lighthouse Lab in Cambridge | Wellcome Sanger Institute for the COVID-19 Genomics UK (COG-UK) consortium | Rob Howes, The Lighthouse Lab in Cambridge and Alex Alderton, Roberto Amato, Sonia Goncalves, Ewan Harrison, David K. Jackson, Ian Johnston, Dominic Kwiatkowski, Cordelia Langford, John Sillitoe on behalf of the Wellcome Sanger Institute COVID-19 Surveillance Team |
| EPI_ISL_608621, EPI_ISL_608622, EPI_ISL_608623 | Lighthouse Lab in Alderley Park | Wellcome Sanger Institute for the COVID-19 Genomics UK (COG-UK) consortium | Jacquelyn Wynn, Mairead Hyland, The Lighthouse Lab in Alderley Park and Alex Alderton, Roberto Amato, Sonia Goncalves, Ewan Harrison, David K. Jackson, Ian Johnston, Dominic Kwiatkowski, Cordelia Langford, John Sillitoe on behalf of the Wellcome Sanger Institute COVID-19 Surveillance Team |
| EPI_ISL_608624 | Lighthouse Lab in Cambridge | Wellcome Sanger Institute for the COVID-19 Genomics UK (COG-UK) consortium | Rob Howes, The Lighthouse Lab in Cambridge and Alex Alderton, Roberto Amato, Sonia Goncalves, Ewan Harrison, David K. Jackson, Ian Johnston, Dominic Kwiatkowski, Cordelia Langford, John Sillitoe on behalf of the Wellcome Sanger Institute COVID-19 Surveillance Team |
| EPI_ISL_608625 | Lighthouse Lab in Alderley Park | Wellcome Sanger Institute for the COVID-19 Genomics UK (COG-UK) consortium | Jacquelyn Wynn, Mairead Hyland, The Lighthouse Lab in Alderley Park and Alex Alderton, Roberto Amato, Sonia Goncalves, Ewan Harrison, David K. Jackson, Ian Johnston, Dominic Kwiatkowski, Cordelia Langford, John Sillitoe on behalf of the Wellcome Sanger Institute COVID-19 Surveillance Team |
| EPI_ISL_608626, EPI_ISL_608627, EPI_ISL_608628, EPI_ISL_608629 | Lighthouse Lab in Cambridge | Wellcome Sanger Institute for the COVID-19 Genomics UK (COG-UK) consortium | Rob Howes, The Lighthouse Lab in Cambridge and Alex Alderton, Roberto Amato, Sonia Goncalves, Ewan Harrison, David K. Jackson, Ian Johnston, Dominic Kwiatkowski, Cordelia Langford, John Sillitoe on behalf of the Wellcome Sanger Institute COVID-19 Surveillance Team |
| EPI_ISL_608630 | Lighthouse Lab in Alderley Park | Wellcome Sanger Institute for the COVID-19 Genomics UK (COG-UK) consortium | Jacquelyn Wynn, Mairead Hyland, The Lighthouse Lab in Alderley Park and Alex Alderton, Roberto Amato, Sonia Goncalves, Ewan Harrison, David K. Jackson, Ian Johnston, Dominic Kwiatkowski, Cordelia Langford, John Sillitoe on behalf of the Wellcome Sanger Institute COVID-19 Surveillance Team |
| EPI_ISL_608631, EPI_ISL_608632, EPI_ISL_608633 | Lighthouse Lab in Cambridge | Wellcome Sanger Institute for the COVID-19 Genomics UK (COG-UK) consortium | Rob Howes, The Lighthouse Lab in Cambridge and Alex Alderton, Roberto Amato, Sonia Goncalves, Ewan Harrison, David K. Jackson, Ian Johnston, Dominic Kwiatkowski, Cordelia Langford, John Sillitoe on behalf of the Wellcome Sanger Institute COVID-19 Surveillance Team |
| EPI_ISL_608634, EPI_ISL_608635 | Lighthouse Lab in Alderley Park | Wellcome Sanger Institute for the COVID-19 Genomics UK (COG-UK) consortium | Jacquelyn Wynn, Mairead Hyland, The Lighthouse Lab in Alderley Park and Alex Alderton, Roberto Amato, Sonia Goncalves, Ewan Harrison, David K. Jackson, Ian Johnston, Dominic Kwiatkowski, Cordelia Langford, John Sillitoe on behalf of the Wellcome Sanger Institute COVID-19 Surveillance Team |
| EPI_ISL_608636, EPI_ISL_608637, EPI_ISL_608638 | Lighthouse Lab in Cambridge | Wellcome Sanger Institute for the COVID-19 Genomics UK (COG-UK) consortium | Rob Howes, The Lighthouse Lab in Cambridge and Alex Alderton, Roberto Amato, Sonia Goncalves, Ewan Harrison, David K. Jackson, Ian Johnston, Dominic Kwiatkowski, Cordelia Langford, John Sillitoe on behalf of the Wellcome Sanger Institute COVID-19 Surveillance Team |
| EPI_ISL_608640 | Lighthouse Lab in Alderley Park | Wellcome Sanger Institute for the COVID-19 Genomics UK (COG-UK) consortium | Jacquelyn Wynn, Mairead Hyland, The Lighthouse Lab in Alderley Park and Alex Alderton, Roberto Amato, Sonia Goncalves, Ewan Harrison, David K. Jackson, Ian Johnston, Dominic Kwiatkowski, Cordelia Langford, John Sillitoe on behalf of the Wellcome Sanger Institute COVID-19 Surveillance Team |
| EPI_ISL_608641, EPI_ISL_608642, EPI_ISL_608643, EPI_ISL_608644, EPI_ISL_608645 | Lighthouse Lab in Cambridge | Wellcome Sanger Institute for the COVID-19 Genomics UK (COG-UK) consortium | Rob Howes, The Lighthouse Lab in Cambridge and Alex Alderton, Roberto Amato, Sonia Goncalves, Ewan Harrison, David K. Jackson, Ian Johnston, Dominic Kwiatkowski, Cordelia Langford, John Sillitoe on behalf of the Wellcome Sanger Institute COVID-19 Surveillance Team |
| EPI_ISL_608646, EPI_ISL_608647, EPI_ISL_608648 | Lighthouse Lab in Alderley Park | Wellcome Sanger Institute for the COVID-19 Genomics UK (COG-UK) consortium | Jacquelyn Wynn, Mairead Hyland, The Lighthouse Lab in Alderley Park and Alex Alderton, Roberto Amato, Sonia Goncalves, Ewan Harrison, David K. Jackson, Ian Johnston, Dominic Kwiatkowski, Cordelia Langford, John Sillitoe on behalf of the Wellcome Sanger Institute COVID-19 Surveillance Team |
| EPI_ISL_608650, EPI_ISL_608651 | Lighthouse Lab in Cambridge | Wellcome Sanger Institute for the COVID-19 Genomics UK (COG-UK) consortium | Rob Howes, The Lighthouse Lab in Cambridge and Alex Alderton, Roberto Amato, Sonia Goncalves, Ewan Harrison, David K. Jackson, Ian Johnston, Dominic Kwiatkowski, Cordelia Langford, John Sillitoe on behalf of the Wellcome Sanger Institute COVID-19 Surveillance Team |
| EPI_ISL_608652 | Lighthouse Lab in Alderley Park | Wellcome Sanger Institute for the COVID-19 Genomics UK (COG-UK) consortium | Jacquelyn Wynn, Mairead Hyland, The Lighthouse Lab in Alderley Park and Alex Alderton, Roberto Amato, Sonia Goncalves, Ewan Harrison, David K. Jackson, Ian Johnston, Dominic Kwiatkowski, Cordelia Langford, John Sillitoe on behalf of the Wellcome Sanger Institute COVID-19 Surveillance Team |
| EPI_ISL_608653, EPI_ISL_608654 | Lighthouse Lab in Cambridge | Wellcome Sanger Institute for the COVID-19 Genomics UK (COG-UK) consortium | Rob Howes, The Lighthouse Lab in Cambridge and Alex Alderton, Roberto Amato, Sonia Goncalves, Ewan Harrison, David K. Jackson, Ian Johnston, Dominic Kwiatkowski, Cordelia Langford, John Sillitoe on behalf of the Wellcome Sanger Institute COVID-19 Surveillance Team |
| EPI_ISL_608655 | Lighthouse Lab in Alderley Park | Wellcome Sanger Institute for the COVID-19 Genomics UK (COG-UK) consortium | Jacquelyn Wynn, Mairead Hyland, The Lighthouse Lab in Alderley Park and Alex Alderton, Roberto Amato, Sonia Goncalves, Ewan Harrison, David K. Jackson, Ian Johnston, Dominic Kwiatkowski, Cordelia Langford, John Sillitoe on behalf of the Wellcome Sanger Institute COVID-19 Surveillance Team |
| EPI_ISL_608656 | Lighthouse Lab in Cambridge | Wellcome Sanger Institute for the COVID-19 Genomics UK (COG-UK) consortium | Rob Howes, The Lighthouse Lab in Cambridge and Alex Alderton, Roberto Amato, Sonia Goncalves, Ewan Harrison, David K. Jackson, Ian Johnston, Dominic Kwiatkowski, Cordelia Langford, John Sillitoe on behalf of the Wellcome Sanger Institute COVID-19 Surveillance Team |
| EPI_ISL_608657, EPI_ISL_608659 | Lighthouse Lab in Alderley Park | Wellcome Sanger Institute for the COVID-19 Genomics UK (COG-UK) consortium | Jacquelyn Wynn, Mairead Hyland, The Lighthouse Lab in Alderley Park and Alex Alderton, Roberto Amato, Sonia Goncalves, Ewan Harrison, David K. Jackson, Ian Johnston, Dominic Kwiatkowski, Cordelia Langford, John Sillitoe on behalf of the Wellcome Sanger Institute COVID-19 Surveillance Team |
| EPI_ISL_608660 | Lighthouse Lab in Cambridge | Wellcome Sanger Institute for the COVID-19 Genomics UK (COG-UK) consortium | Rob Howes, The Lighthouse Lab in Cambridge and Alex Alderton, Roberto Amato, Sonia Goncalves, Ewan Harrison, David K. Jackson, Ian Johnston, Dominic Kwiatkowski, Cordelia Langford, John Sillitoe on behalf of the Wellcome Sanger Institute COVID-19 Surveillance Team |
| EPI_ISL_608661, EPI_ISL_608662, EPI_ISL_608663 | Lighthouse Lab in Alderley Park | Wellcome Sanger Institute for the COVID-19 Genomics UK (COG-UK) consortium | Jacquelyn Wynn, Mairead Hyland, The Lighthouse Lab in Alderley Park and Alex Alderton, Roberto Amato, Sonia Goncalves, Ewan Harrison, David K. Jackson, Ian Johnston, Dominic Kwiatkowski, Cordelia Langford, John Sillitoe on behalf of the Wellcome Sanger Institute COVID-19 Surveillance Team |
| EPI_ISL_608664 | Lighthouse Lab in Cambridge | Wellcome Sanger Institute for the COVID-19 Genomics UK (COG-UK) consortium | Rob Howes, The Lighthouse Lab in Cambridge and Alex Alderton, Roberto Amato, Sonia Goncalves, Ewan Harrison, David K. Jackson, Ian Johnston, Dominic Kwiatkowski, Cordelia Langford, John Sillitoe on behalf of the Wellcome Sanger Institute COVID-19 Surveillance Team |
| EPI_ISL_608665 | Lighthouse Lab in Alderley Park | Wellcome Sanger Institute for the COVID-19 Genomics UK (COG-UK) consortium | Jacquelyn Wynn, Mairead Hyland, The Lighthouse Lab in Alderley Park and Alex Alderton, Roberto Amato, Sonia Goncalves, Ewan Harrison, David K. Jackson, Ian Johnston, Dominic Kwiatkowski, Cordelia Langford, John Sillitoe on behalf of the Wellcome Sanger Institute COVID-19 Surveillance Team |
| EPI_ISL_608666 | Lighthouse Lab in Cambridge | Wellcome Sanger Institute for the COVID-19 Genomics UK (COG-UK) consortium | Rob Howes, The Lighthouse Lab in Cambridge and Alex Alderton, Roberto Amato, Sonia Goncalves, Ewan Harrison, David K. Jackson, Ian Johnston, Dominic Kwiatkowski, Cordelia Langford, John Sillitoe on behalf of the Wellcome Sanger Institute COVID-19 Surveillance Team |
| EPI_ISL_608667, EPI_ISL_608668, EPI_ISL_608669, EPI_ISL_608670 | Lighthouse Lab in Alderley Park | Wellcome Sanger Institute for the COVID-19 Genomics UK (COG-UK) consortium | Jacquelyn Wynn, Mairead Hyland, The Lighthouse Lab in Alderley Park and Alex Alderton, Roberto Amato, Sonia Goncalves, Ewan Harrison, David K. Jackson, Ian Johnston, Dominic Kwiatkowski, Cordelia Langford, John Sillitoe on behalf of the Wellcome Sanger Institute COVID-19 Surveillance Team |
| EPI_ISL_608671 | Lighthouse Lab in Cambridge | Wellcome Sanger Institute for the COVID-19 Genomics UK (COG-UK) consortium | Rob Howes, The Lighthouse Lab in Cambridge and Alex Alderton, Roberto Amato, Sonia Goncalves, Ewan Harrison, David K. Jackson, Ian Johnston, Dominic Kwiatkowski, Cordelia Langford, John Sillitoe on behalf of the Wellcome Sanger Institute COVID-19 Surveillance Team |
| EPI_ISL_608672 | Lighthouse Lab in Alderley Park | Wellcome Sanger Institute for the COVID-19 Genomics UK (COG-UK) consortium | Jacquelyn Wynn, Mairead Hyland, The Lighthouse Lab in Alderley Park and Alex Alderton, Roberto Amato, Sonia Goncalves, Ewan Harrison, David K. Jackson, Ian Johnston, Dominic Kwiatkowski, Cordelia Langford, John Sillitoe on behalf of the Wellcome Sanger Institute COVID-19 Surveillance Team |
| EPI_ISL_608673 | Lighthouse Lab in Cambridge | Wellcome Sanger Institute for the COVID-19 Genomics UK (COG-UK) consortium | Rob Howes, The Lighthouse Lab in Cambridge and Alex Alderton, Roberto Amato, Sonia Goncalves, Ewan Harrison, David K. Jackson, Ian Johnston, Dominic Kwiatkowski, Cordelia Langford, John Sillitoe on behalf of the Wellcome Sanger Institute COVID-19 Surveillance Team |
| EPI_ISL_608674, EPI_ISL_608675, EPI_ISL_608676, EPI_ISL_608677 | Lighthouse Lab in Alderley Park | Wellcome Sanger Institute for the COVID-19 Genomics UK (COG-UK) consortium | Jacquelyn Wynn, Mairead Hyland, The Lighthouse Lab in Alderley Park and Alex Alderton, Roberto Amato, Sonia Goncalves, Ewan Harrison, David K. Jackson, Ian Johnston, Dominic Kwiatkowski, Cordelia Langford, John Sillitoe on behalf of the Wellcome Sanger Institute COVID-19 Surveillance Team |
| EPI_ISL_608678, EPI_ISL_608679 | Lighthouse Lab in Cambridge | Wellcome Sanger Institute for the COVID-19 Genomics UK (COG-UK) consortium | Rob Howes, The Lighthouse Lab in Cambridge and Alex Alderton, Roberto Amato, Sonia Goncalves, Ewan Harrison, David K. Jackson, Ian Johnston, Dominic Kwiatkowski, Cordelia Langford, John Sillitoe on behalf of the Wellcome Sanger Institute COVID-19 Surveillance Team |
| EPI_ISL_608680 | Lighthouse Lab in Alderley Park | Wellcome Sanger Institute for the COVID-19 Genomics UK (COG-UK) consortium | Jacquelyn Wynn, Mairead Hyland, The Lighthouse Lab in Alderley Park and Alex Alderton, Roberto Amato, Sonia Goncalves, Ewan Harrison, David K. Jackson, Ian Johnston, Dominic Kwiatkowski, Cordelia Langford, John Sillitoe on behalf of the Wellcome Sanger Institute COVID-19 Surveillance Team |

| | | | |
|---|---|---|---|
| EPI_ISL_608681 | Lighthouse Lab in Cambridge | Wellcome Sanger Institute for the COVID-19 Genomics UK (COG-UK) Consortium | Rob Howes, The Lighthouse Lab in Cambridge and Alex Alderton, Roberto Amato, Sonia Goncalves, Ewan Harrison, David K. Jackson, Ian Johnston, Dominic Kwiatkowski, Cordelia Langford, John Sillitoe on behalf of the Wellcome Sanger Institute COVID-19 Surveillance Team |
| EPI_ISL_608682, EPI_ISL_608683 | Lighthouse Lab in Cambridge | Wellcome Sanger Institute for the COVID-19 Genomics UK (COG-UK) consortium | Rob Howes, The Lighthouse Lab in Cambridge and Alex Alderton, Roberto Amato, Sonia Goncalves, Ewan Harrison, David K. Jackson, Ian Johnston, Dominic Kwiatkowski, Cordelia Langford, John Sillitoe on behalf of the Wellcome Sanger Institute COVID-19 Surveillance Team |
| EPI_ISL_608684, EPI_ISL_608685, EPI_ISL_608686, EPI_ISL_608687 | Lighthouse Lab in Alderley Park | Wellcome Sanger Institute for the COVID-19 Genomics UK (COG-UK) consortium | Jacquelyn Wynn, Mairead Hyland, The Lighthouse Lab in Alderley Park and Alex Alderton, Roberto Amato, Sonia Goncalves, Ewan Harrison, David K. Jackson, Ian Johnston, Dominic Kwiatkowski, Cordelia Langford, John Sillitoe on behalf of the Wellcome Sanger Institute COVID-19 Surveillance Team |
| EPI_ISL_608688, EPI_ISL_608689 | Lighthouse Lab in Cambridge | Wellcome Sanger Institute for the COVID-19 Genomics UK (COG-UK) consortium | Rob Howes, The Lighthouse Lab in Cambridge and Alex Alderton, Roberto Amato, Sonia Goncalves, Ewan Harrison, David K. Jackson, Ian Johnston, Dominic Kwiatkowski, Cordelia Langford, John Sillitoe on behalf of the Wellcome Sanger Institute COVID-19 Surveillance Team |
| EPI_ISL_608690, EPI_ISL_608691, EPI_ISL_608692 | Lighthouse Lab in Alderley Park | Wellcome Sanger Institute for the COVID-19 Genomics UK (COG-UK) consortium | Jacquelyn Wynn, Mairead Hyland, The Lighthouse Lab in Alderley Park and Alex Alderton, Roberto Amato, Sonia Goncalves, Ewan Harrison, David K. Jackson, Ian Johnston, Dominic Kwiatkowski, Cordelia Langford, John Sillitoe on behalf of the Wellcome Sanger Institute COVID-19 Surveillance Team |
| EPI_ISL_608693 | Lighthouse Lab in Cambridge | Wellcome Sanger Institute for the COVID-19 Genomics UK (COG-UK) consortium | Rob Howes, The Lighthouse Lab in Cambridge and Alex Alderton, Roberto Amato, Sonia Goncalves, Ewan Harrison, David K. Jackson, Ian Johnston, Dominic Kwiatkowski, Cordelia Langford, John Sillitoe on behalf of the Wellcome Sanger Institute COVID-19 Surveillance Team |
| EPI_ISL_608694 | Lighthouse Lab in Alderley Park | Wellcome Sanger Institute for the COVID-19 Genomics UK (COG-UK) consortium | Jacquelyn Wynn, Mairead Hyland, The Lighthouse Lab in Alderley Park and Alex Alderton, Roberto Amato, Sonia Goncalves, Ewan Harrison, David K. Jackson, Ian Johnston, Dominic Kwiatkowski, Cordelia Langford, John Sillitoe on behalf of the Wellcome Sanger Institute COVID-19 Surveillance Team |
| EPI_ISL_608695, EPI_ISL_608696 | Lighthouse Lab in Cambridge | Wellcome Sanger Institute for the COVID-19 Genomics UK (COG-UK) consortium | Rob Howes, The Lighthouse Lab in Cambridge and Alex Alderton, Roberto Amato, Sonia Goncalves, Ewan Harrison, David K. Jackson, Ian Johnston, Dominic Kwiatkowski, Cordelia Langford, John Sillitoe on behalf of the Wellcome Sanger Institute COVID-19 Surveillance Team |
| EPI_ISL_608697, EPI_ISL_608698, EPI_ISL_608699, EPI_ISL_608700, EPI_ISL_608701, EPI_ISL_608702, EPI_ISL_608703, EPI_ISL_608704, EPI_ISL_608705, EPI_ISL_608706, EPI_ISL_608707, EPI_ISL_608708, EPI_ISL_608709, EPI_ISL_608710, EPI_ISL_608711, EPI_ISL_608712, EPI_ISL_608713, EPI_ISL_608714, EPI_ISL_608715, EPI_ISL_608716, EPI_ISL_608717, EPI_ISL_608718, EPI_ISL_608719, EPI_ISL_608720, EPI_ISL_608721, EPI_ISL_608722, EPI_ISL_608723, EPI_ISL_608724, EPI_ISL_608725, EPI_ISL_608726, EPI_ISL_608727, EPI_ISL_608728, EPI_ISL_608729, EPI_ISL_608730, EPI_ISL_608731, EPI_ISL_608732, EPI_ISL_608733, EPI_ISL_608734, EPI_ISL_608735, EPI_ISL_608736, EPI_ISL_608737, EPI_ISL_608738, EPI_ISL_608739, EPI_ISL_608740, EPI_ISL_608741, EPI_ISL_608742, EPI_ISL_608743, EPI_ISL_608744, EPI_ISL_608745, EPI_ISL_608746, EPI_ISL_608747 | | | |
| see above | Lighthouse Lab in Milton Keynes | Wellcome Sanger Institute for the COVID-19 Genomics UK (COG-UK) consortium | The Lighthouse Lab in Milton Keynes and Alex Alderton, Roberto Amato, Sonia Goncalves, Ewan Harrison, David K. Jackson, Ian Johnston, Dominic Kwiatkowski, Cordelia Langford, John Sillitoe on behalf of the Wellcome Sanger Institute COVID-19 Surveillance Team |
| EPI_ISL_608748 | Lighthouse Lab in Milton Keynes | Wellcome Sanger Institute for the COVID-19 Genomics UK (COG-UK) Consortium | The Lighthouse Lab in Milton Keynes and Alex Alderton, Roberto Amato, Sonia Goncalves, Ewan Harrison, David K. Jackson, Ian Johnston, Dominic Kwiatkowski, Cordelia Langford, John Sillitoe on behalf of the Wellcome Sanger Institute COVID-19 Surveillance Team |
| EPI_ISL_608749, EPI_ISL_608750, EPI_ISL_608751, EPI_ISL_608752, EPI_ISL_608753, EPI_ISL_608754, EPI_ISL_608755, EPI_ISL_608756, EPI_ISL_608757, EPI_ISL_608758, EPI_ISL_608759, EPI_ISL_608760, EPI_ISL_608761, EPI_ISL_608762, EPI_ISL_608763, EPI_ISL_608764, EPI_ISL_608765, EPI_ISL_608766, EPI_ISL_608767, EPI_ISL_608768, EPI_ISL_608769, EPI_ISL_608770, EPI_ISL_608771, EPI_ISL_608772, EPI_ISL_608773, EPI_ISL_608774, EPI_ISL_608775, EPI_ISL_608776, EPI_ISL_608777, EPI_ISL_608778, EPI_ISL_608779, EPI_ISL_608780, EPI_ISL_608781, EPI_ISL_608782, EPI_ISL_608783, EPI_ISL_608784, EPI_ISL_608785, EPI_ISL_608786, EPI_ISL_608787, EPI_ISL_608788, EPI_ISL_608789, EPI_ISL_608790, EPI_ISL_608791, EPI_ISL_608792, EPI_ISL_608793, EPI_ISL_608794, EPI_ISL_608795, EPI_ISL_608796, EPI_ISL_608797, EPI_ISL_608798, EPI_ISL_608799, EPI_ISL_608800, EPI_ISL_608801, EPI_ISL_608802, EPI_ISL_608803, EPI_ISL_608804, EPI_ISL_608805, EPI_ISL_608806, EPI_ISL_608807, EPI_ISL_608808, EPI_ISL_608809, EPI_ISL_608810, EPI_ISL_608811, EPI_ISL_608812, EPI_ISL_608813, EPI_ISL_608814, EPI_ISL_608815, EPI_ISL_608816, EPI_ISL_608817, EPI_ISL_608818, EPI_ISL_608819, EPI_ISL_608820, EPI_ISL_608821, EPI_ISL_608822, EPI_ISL_608823, EPI_ISL_608824, EPI_ISL_608825, EPI_ISL_608826, EPI_ISL_608827, EPI_ISL_608828, EPI_ISL_608829, EPI_ISL_608830, EPI_ISL_608831, EPI_ISL_608832, EPI_ISL_608833 | | | |
| see above | Lighthouse Lab in Milton Keynes | Wellcome Sanger Institute for the COVID-19 Genomics UK (COG-UK) consortium | The Lighthouse Lab in Milton Keynes and Alex Alderton, Roberto Amato, Sonia Goncalves, Ewan Harrison, David K. Jackson, Ian Johnston, Dominic Kwiatkowski, Cordelia Langford, John Sillitoe on behalf of the Wellcome Sanger Institute COVID-19 Surveillance Team |
| EPI_ISL_608834 | Lighthouse Lab in Milton Keynes | Wellcome Sanger Institute for the COVID-19 Genomics UK (COG-UK) Consortium | The Lighthouse Lab in Milton Keynes and Alex Alderton, Roberto Amato, Sonia Goncalves, Ewan Harrison, David K. Jackson, Ian Johnston, Dominic Kwiatkowski, Cordelia Langford, John Sillitoe on behalf of the Wellcome Sanger Institute COVID-19 Surveillance Team |
| EPI_ISL_608835, EPI_ISL_608836, EPI_ISL_608837, EPI_ISL_608838, EPI_ISL_608839, EPI_ISL_608840, EPI_ISL_608841, EPI_ISL_608842, EPI_ISL_608843, EPI_ISL_608844 | Lighthouse Lab in Milton Keynes | Wellcome Sanger Institute for the COVID-19 Genomics UK (COG-UK) consortium | The Lighthouse Lab in Milton Keynes and Alex Alderton, Roberto Amato, Sonia Goncalves, Ewan Harrison, David K. Jackson, Ian Johnston, Dominic Kwiatkowski, Cordelia Langford, John Sillitoe on behalf of the Wellcome Sanger Institute COVID-19 Surveillance Team |
| EPI_ISL_608845 | Lighthouse Lab in Cambridge | Wellcome Sanger Institute for the COVID-19 Genomics UK (COG-UK) consortium | Rob Howes, The Lighthouse Lab in Cambridge and Alex Alderton, Roberto Amato, Sonia Goncalves, Ewan Harrison, David K. Jackson, Ian Johnston, Dominic Kwiatkowski, Cordelia Langford, John Sillitoe on behalf of the Wellcome Sanger Institute COVID-19 Surveillance Team |
| EPI_ISL_608846 | Lighthouse Lab in Milton Keynes | Wellcome Sanger Institute for the COVID-19 Genomics UK (COG-UK) consortium | The Lighthouse Lab in Milton Keynes and Alex Alderton, Roberto Amato, Sonia Goncalves, Ewan Harrison, David K. Jackson, Ian Johnston, Dominic Kwiatkowski, Cordelia Langford, John Sillitoe on behalf of the Wellcome Sanger Institute COVID-19 Surveillance Team |
| EPI_ISL_608847 | Lighthouse Lab in Alderley Park | Wellcome Sanger Institute for the COVID-19 Genomics UK (COG-UK) consortium | Jacquelyn Wynn, Mairead Hyland, The Lighthouse Lab in Alderley Park and Alex Alderton, Roberto Amato, Sonia Goncalves, Ewan Harrison, David K. Jackson, Ian Johnston, Dominic Kwiatkowski, Cordelia Langford, John Sillitoe on behalf of the Wellcome Sanger Institute COVID-19 Surveillance Team |
| EPI_ISL_608848 | Lighthouse Lab in Milton Keynes | Wellcome Sanger Institute for the COVID-19 Genomics UK (COG-UK) consortium | The Lighthouse Lab in Milton Keynes and Alex Alderton, Roberto Amato, Sonia Goncalves, Ewan Harrison, David K. Jackson, Ian Johnston, Dominic Kwiatkowski, Cordelia Langford, John Sillitoe on behalf of the Wellcome Sanger Institute COVID-19 Surveillance Team |
| EPI_ISL_608849 | Lighthouse Lab in Alderley Park | Wellcome Sanger Institute for the COVID-19 Genomics UK (COG-UK) consortium | Jacquelyn Wynn, Mairead Hyland, The Lighthouse Lab in Alderley Park and Alex Alderton, Roberto Amato, Sonia Goncalves, Ewan Harrison, David K. Jackson, Ian Johnston, Dominic Kwiatkowski, Cordelia Langford, John Sillitoe on behalf of the Wellcome Sanger Institute COVID-19 Surveillance Team |
| EPI_ISL_608850 | Lighthouse Lab in Milton Keynes | Wellcome Sanger Institute for the COVID-19 Genomics UK (COG-UK) consortium | The Lighthouse Lab in Milton Keynes and Alex Alderton, Roberto Amato, Sonia Goncalves, Ewan Harrison, David K. Jackson, Ian Johnston, Dominic Kwiatkowski, Cordelia Langford, John Sillitoe on behalf of the Wellcome Sanger Institute COVID-19 Surveillance Team |
| EPI_ISL_608851 | Lighthouse Lab in Alderley Park | Wellcome Sanger Institute for the COVID-19 Genomics UK (COG-UK) consortium | Jacquelyn Wynn, Mairead Hyland, The Lighthouse Lab in Alderley Park and Alex Alderton, Roberto Amato, Sonia Goncalves, Ewan Harrison, David K. Jackson, Ian Johnston, Dominic Kwiatkowski, Cordelia Langford, John Sillitoe on behalf of the Wellcome Sanger Institute COVID-19 Surveillance Team |
| EPI_ISL_608852 | Lighthouse Lab in Cambridge | Wellcome Sanger Institute for the COVID-19 Genomics UK (COG-UK) consortium | Rob Howes, The Lighthouse Lab in Cambridge and Alex Alderton, Roberto Amato, Sonia Goncalves, Ewan Harrison, David K. Jackson, Ian Johnston, Dominic Kwiatkowski, Cordelia Langford, John Sillitoe on behalf of the Wellcome Sanger Institute COVID-19 Surveillance Team |
| EPI_ISL_608853 | Lighthouse Lab in Alderley Park | Wellcome Sanger Institute for the COVID-19 Genomics UK (COG-UK) consortium | Jacquelyn Wynn, Mairead Hyland, The Lighthouse Lab in Alderley Park and Alex Alderton, Roberto Amato, Sonia Goncalves, Ewan Harrison, David K. Jackson, Ian Johnston, Dominic Kwiatkowski, Cordelia Langford, John Sillitoe on behalf of the Wellcome Sanger Institute COVID-19 Surveillance Team |
| EPI_ISL_608854 | Lighthouse Lab in Milton Keynes | Wellcome Sanger Institute for the COVID-19 Genomics UK (COG-UK) consortium | The Lighthouse Lab in Milton Keynes and Alex Alderton, Roberto Amato, Sonia Goncalves, Ewan Harrison, David K. Jackson, Ian Johnston, Dominic Kwiatkowski, Cordelia Langford, John Sillitoe on behalf of the Wellcome Sanger Institute COVID-19 Surveillance Team |
| EPI_ISL_608855, EPI_ISL_608856 | Lighthouse Lab in Alderley Park | Wellcome Sanger Institute for the COVID-19 Genomics UK (COG-UK) consortium | Jacquelyn Wynn, Mairead Hyland, The Lighthouse Lab in Alderley Park and Alex Alderton, Roberto Amato, Sonia Goncalves, Ewan Harrison, David K. Jackson, Ian Johnston, Dominic Kwiatkowski, Cordelia Langford, John Sillitoe on behalf of the Wellcome Sanger Institute COVID-19 Surveillance Team |
| EPI_ISL_608857 | Lighthouse Lab in Milton Keynes | Wellcome Sanger Institute for the COVID-19 Genomics UK (COG-UK) consortium | The Lighthouse Lab in Milton Keynes and Alex Alderton, Roberto Amato, Sonia Goncalves, Ewan Harrison, David K. Jackson, Ian Johnston, Dominic Kwiatkowski, Cordelia Langford, John Sillitoe on behalf of the Wellcome Sanger Institute COVID-19 Surveillance Team |
| EPI_ISL_608858 | Lighthouse Lab in Alderley Park | Wellcome Sanger Institute for the COVID-19 Genomics UK (COG-UK) consortium | Jacquelyn Wynn, Mairead Hyland, The Lighthouse Lab in Alderley Park and Alex Alderton, Roberto Amato, Sonia Goncalves, Ewan Harrison, David K. Jackson, Ian Johnston, Dominic Kwiatkowski, Cordelia Langford, John Sillitoe on behalf of the Wellcome Sanger Institute COVID-19 Surveillance Team |
| EPI_ISL_608859 | Lighthouse Lab in Milton Keynes | Wellcome Sanger Institute for the COVID-19 Genomics UK (COG-UK) consortium | The Lighthouse Lab in Milton Keynes and Alex Alderton, Roberto Amato, Sonia Goncalves, Ewan Harrison, David K. Jackson, Ian Johnston, Dominic Kwiatkowski, Cordelia Langford, John Sillitoe on behalf of the Wellcome Sanger Institute COVID-19 Surveillance Team |
| EPI_ISL_608860 | Lighthouse Lab in Alderley Park | Wellcome Sanger Institute for the COVID-19 Genomics UK (COG-UK) consortium | Jacquelyn Wynn, Mairead Hyland, The Lighthouse Lab in Alderley Park and Alex Alderton, Roberto Amato, Sonia Goncalves, Ewan Harrison, David K. Jackson, Ian Johnston, Dominic Kwiatkowski, Cordelia Langford, John Sillitoe on behalf of the Wellcome Sanger Institute COVID-19 Surveillance Team |
| EPI_ISL_608861 | Lighthouse Lab in Milton Keynes | Wellcome Sanger Institute for the COVID-19 Genomics UK (COG-UK) consortium | The Lighthouse Lab in Milton Keynes and Alex Alderton, Roberto Amato, Sonia Goncalves, Ewan Harrison, David K. Jackson, Ian Johnston, Dominic Kwiatkowski, Cordelia Langford, John Sillitoe on behalf of the Wellcome Sanger Institute COVID-19 Surveillance Team |
| EPI_ISL_608862, EPI_ISL_608863, EPI_ISL_608864, EPI_ISL_608865, EPI_ISL_608866 | Lighthouse Lab in Alderley Park | Wellcome Sanger Institute for the COVID-19 Genomics UK (COG-UK) consortium | Jacquelyn Wynn, Mairead Hyland, The Lighthouse Lab in Alderley Park and Alex Alderton, Roberto Amato, Sonia Goncalves, Ewan Harrison, David K. Jackson, Ian Johnston, Dominic Kwiatkowski, Cordelia Langford, John Sillitoe on behalf of the Wellcome Sanger Institute COVID-19 Surveillance Team |

| | | | |
|---|---|---|---|
| EPI_ISL_608867, EPI_ISL_608868 | Lighthouse Lab in Cambridge | Wellcome Sanger Institute for the COVID-19 Genomics UK (COG-UK) consortium | Rob Howes, The Lighthouse Lab in Cambridge and Alex Alderton, Roberto Amato, Sonia Goncalves, Ewan Harrison, David K. Jackson, Ian Johnston, Dominic Kwiatkowski, Cordelia Langford, John Sillitoe on behalf of the Wellcome Sanger Institute COVID-19 Surveillance Team |
| EPI_ISL_608869 | Lighthouse Lab in Alderley Park | Wellcome Sanger Institute for the COVID-19 Genomics UK (COG-UK) consortium | Jacquelyn Wynn, Mairead Hyland, The Lighthouse Lab in Alderley Park and Alex Alderton, Roberto Amato, Sonia Goncalves, Ewan Harrison, David K. Jackson, Ian Johnston, Dominic Kwiatkowski, Cordelia Langford, John Sillitoe on behalf of the Wellcome Sanger Institute COVID-19 Surveillance Team |
| EPI_ISL_608870, EPI_ISL_608871 | Lighthouse Lab in Milton Keynes | Wellcome Sanger Institute for the COVID-19 Genomics UK (COG-UK) consortium | The Lighthouse Lab in Milton Keynes and Alex Alderton, Roberto Amato, Sonia Goncalves, Ewan Harrison, David K. Jackson, Ian Johnston, Dominic Kwiatkowski, Cordelia Langford, John Sillitoe on behalf of the Wellcome Sanger Institute COVID-19 Surveillance Team |
| EPI_ISL_608872 | Lighthouse Lab in Alderley Park | Wellcome Sanger Institute for the COVID-19 Genomics UK (COG-UK) consortium | Jacquelyn Wynn, Mairead Hyland, The Lighthouse Lab in Alderley Park and Alex Alderton, Roberto Amato, Sonia Goncalves, Ewan Harrison, David K. Jackson, Ian Johnston, Dominic Kwiatkowski, Cordelia Langford, John Sillitoe on behalf of the Wellcome Sanger Institute COVID-19 Surveillance Team |
| EPI_ISL_608873 | Lighthouse Lab in Cambridge | Wellcome Sanger Institute for the COVID-19 Genomics UK (COG-UK) consortium | Rob Howes, The Lighthouse Lab in Cambridge and Alex Alderton, Roberto Amato, Sonia Goncalves, Ewan Harrison, David K. Jackson, Ian Johnston, Dominic Kwiatkowski, Cordelia Langford, John Sillitoe on behalf of the Wellcome Sanger Institute COVID-19 Surveillance Team |
| EPI_ISL_608874 | Lighthouse Lab in Alderley Park | Wellcome Sanger Institute for the COVID-19 Genomics UK (COG-UK) consortium | Jacquelyn Wynn, Mairead Hyland, The Lighthouse Lab in Alderley Park and Alex Alderton, Roberto Amato, Sonia Goncalves, Ewan Harrison, David K. Jackson, Ian Johnston, Dominic Kwiatkowski, Cordelia Langford, John Sillitoe on behalf of the Wellcome Sanger Institute COVID-19 Surveillance Team |
| EPI_ISL_608875, EPI_ISL_608876 | Lighthouse Lab in Cambridge | Wellcome Sanger Institute for the COVID-19 Genomics UK (COG-UK) consortium | Rob Howes, The Lighthouse Lab in Cambridge and Alex Alderton, Roberto Amato, Sonia Goncalves, Ewan Harrison, David K. Jackson, Ian Johnston, Dominic Kwiatkowski, Cordelia Langford, John Sillitoe on behalf of the Wellcome Sanger Institute COVID-19 Surveillance Team |
| EPI_ISL_608877, EPI_ISL_608878 | Lighthouse Lab in Alderley Park | Wellcome Sanger Institute for the COVID-19 Genomics UK (COG-UK) consortium | Jacquelyn Wynn, Mairead Hyland, The Lighthouse Lab in Alderley Park and Alex Alderton, Roberto Amato, Sonia Goncalves, Ewan Harrison, David K. Jackson, Ian Johnston, Dominic Kwiatkowski, Cordelia Langford, John Sillitoe on behalf of the Wellcome Sanger Institute COVID-19 Surveillance Team |
| EPI_ISL_608879, EPI_ISL_608881 | Lighthouse Lab in Cambridge | Wellcome Sanger Institute for the COVID-19 Genomics UK (COG-UK) consortium | Rob Howes, The Lighthouse Lab in Cambridge and Alex Alderton, Roberto Amato, Sonia Goncalves, Ewan Harrison, David K. Jackson, Ian Johnston, Dominic Kwiatkowski, Cordelia Langford, John Sillitoe on behalf of the Wellcome Sanger Institute COVID-19 Surveillance Team |
| EPI_ISL_608882 | Lighthouse Lab in Milton Keynes | Wellcome Sanger Institute for the COVID-19 Genomics UK (COG-UK) consortium | The Lighthouse Lab in Milton Keynes and Alex Alderton, Roberto Amato, Sonia Goncalves, Ewan Harrison, David K. Jackson, Ian Johnston, Dominic Kwiatkowski, Cordelia Langford, John Sillitoe on behalf of the Wellcome Sanger Institute COVID-19 Surveillance Team |
| EPI_ISL_608883, EPI_ISL_608884, EPI_ISL_608885, EPI_ISL_608886, EPI_ISL_608887, EPI_ISL_608888, EPI_ISL_608889, EPI_ISL_608890, EPI_ISL_608891 | Lighthouse Lab in Alderley Park | Wellcome Sanger Institute for the COVID-19 Genomics UK (COG-UK) consortium | Jacquelyn Wynn, Mairead Hyland, The Lighthouse Lab in Alderley Park and Alex Alderton, Roberto Amato, Sonia Goncalves, Ewan Harrison, David K. Jackson, Ian Johnston, Dominic Kwiatkowski, Cordelia Langford, John Sillitoe on behalf of the Wellcome Sanger Institute COVID-19 Surveillance Team |
| EPI_ISL_608892 | Lighthouse Lab in Cambridge | Wellcome Sanger Institute for the COVID-19 Genomics UK (COG-UK) consortium | Rob Howes, The Lighthouse Lab in Cambridge and Alex Alderton, Roberto Amato, Sonia Goncalves, Ewan Harrison, David K. Jackson, Ian Johnston, Dominic Kwiatkowski, Cordelia Langford, John Sillitoe on behalf of the Wellcome Sanger Institute COVID-19 Surveillance Team |
| EPI_ISL_608893, EPI_ISL_608894 | Lighthouse Lab in Milton Keynes | Wellcome Sanger Institute for the COVID-19 Genomics UK (COG-UK) consortium | The Lighthouse Lab in Milton Keynes and Alex Alderton, Roberto Amato, Sonia Goncalves, Ewan Harrison, David K. Jackson, Ian Johnston, Dominic Kwiatkowski, Cordelia Langford, John Sillitoe on behalf of the Wellcome Sanger Institute COVID-19 Surveillance Team |
| EPI_ISL_608895, EPI_ISL_608896, EPI_ISL_608897, EPI_ISL_608898, EPI_ISL_608899, EPI_ISL_608900 | Lighthouse Lab in Alderley Park | Wellcome Sanger Institute for the COVID-19 Genomics UK (COG-UK) consortium | Jacquelyn Wynn, Mairead Hyland, The Lighthouse Lab in Alderley Park and Alex Alderton, Roberto Amato, Sonia Goncalves, Ewan Harrison, David K. Jackson, Ian Johnston, Dominic Kwiatkowski, Cordelia Langford, John Sillitoe on behalf of the Wellcome Sanger Institute COVID-19 Surveillance Team |
| EPI_ISL_608901 | Lighthouse Lab in Milton Keynes | Wellcome Sanger Institute for the COVID-19 Genomics UK (COG-UK) consortium | The Lighthouse Lab in Milton Keynes and Alex Alderton, Roberto Amato, Sonia Goncalves, Ewan Harrison, David K. Jackson, Ian Johnston, Dominic Kwiatkowski, Cordelia Langford, John Sillitoe on behalf of the Wellcome Sanger Institute COVID-19 Surveillance Team |
| EPI_ISL_608902 | Lighthouse Lab in Cambridge | Wellcome Sanger Institute for the COVID-19 Genomics UK (COG-UK) consortium | Rob Howes, The Lighthouse Lab in Cambridge and Alex Alderton, Roberto Amato, Sonia Goncalves, Ewan Harrison, David K. Jackson, Ian Johnston, Dominic Kwiatkowski, Cordelia Langford, John Sillitoe on behalf of the Wellcome Sanger Institute COVID-19 Surveillance Team |
| EPI_ISL_608903, EPI_ISL_608904 | Lighthouse Lab in Alderley Park | Wellcome Sanger Institute for the COVID-19 Genomics UK (COG-UK) consortium | Jacquelyn Wynn, Mairead Hyland, The Lighthouse Lab in Alderley Park and Alex Alderton, Roberto Amato, Sonia Goncalves, Ewan Harrison, David K. Jackson, Ian Johnston, Dominic Kwiatkowski, Cordelia Langford, John Sillitoe on behalf of the Wellcome Sanger Institute COVID-19 Surveillance Team |
| EPI_ISL_608905 | Lighthouse Lab in Milton Keynes | Wellcome Sanger Institute for the COVID-19 Genomics UK (COG-UK) consortium | The Lighthouse Lab in Milton Keynes and Alex Alderton, Roberto Amato, Sonia Goncalves, Ewan Harrison, David K. Jackson, Ian Johnston, Dominic Kwiatkowski, Cordelia Langford, John Sillitoe on behalf of the Wellcome Sanger Institute COVID-19 Surveillance Team |
| EPI_ISL_608906 | Lighthouse Lab in Cambridge | Wellcome Sanger Institute for the COVID-19 Genomics UK (COG-UK) consortium | Rob Howes, The Lighthouse Lab in Cambridge and Alex Alderton, Roberto Amato, Sonia Goncalves, Ewan Harrison, David K. Jackson, Ian Johnston, Dominic Kwiatkowski, Cordelia Langford, John Sillitoe on behalf of the Wellcome Sanger Institute COVID-19 Surveillance Team |
| EPI_ISL_608907 | Lighthouse Lab in Cambridge | Wellcome Sanger Institute for the COVID-19 Genomics UK (COG-UK) Consortium | Rob Howes, The Lighthouse Lab in Cambridge and Alex Alderton, Roberto Amato, Sonia Goncalves, Ewan Harrison, David K. Jackson, Ian Johnston, Dominic Kwiatkowski, Cordelia Langford, John Sillitoe on behalf of the Wellcome Sanger Institute COVID-19 Surveillance Team |
| EPI_ISL_608908 | Lighthouse Lab in Cambridge | Wellcome Sanger Institute for the COVID-19 Genomics UK (COG-UK) consortium | Rob Howes, The Lighthouse Lab in Cambridge and Alex Alderton, Roberto Amato, Sonia Goncalves, Ewan Harrison, David K. Jackson, Ian Johnston, Dominic Kwiatkowski, Cordelia Langford, John Sillitoe on behalf of the Wellcome Sanger Institute COVID-19 Surveillance Team |
| EPI_ISL_608909 | Lighthouse Lab in Alderley Park | Wellcome Sanger Institute for the COVID-19 Genomics UK (COG-UK) consortium | Jacquelyn Wynn, Mairead Hyland, The Lighthouse Lab in Alderley Park and Alex Alderton, Roberto Amato, Sonia Goncalves, Ewan Harrison, David K. Jackson, Ian Johnston, Dominic Kwiatkowski, Cordelia Langford, John Sillitoe on behalf of the Wellcome Sanger Institute COVID-19 Surveillance Team |
| EPI_ISL_608910 | Lighthouse Lab in Cambridge | Wellcome Sanger Institute for the COVID-19 Genomics UK (COG-UK) consortium | Rob Howes, The Lighthouse Lab in Cambridge and Alex Alderton, Roberto Amato, Sonia Goncalves, Ewan Harrison, David K. Jackson, Ian Johnston, Dominic Kwiatkowski, Cordelia Langford, John Sillitoe on behalf of the Wellcome Sanger Institute COVID-19 Surveillance Team |
| EPI_ISL_608911 | Lighthouse Lab in Alderley Park | Wellcome Sanger Institute for the COVID-19 Genomics UK (COG-UK) consortium | Jacquelyn Wynn, Mairead Hyland, The Lighthouse Lab in Alderley Park and Alex Alderton, Roberto Amato, Sonia Goncalves, Ewan Harrison, David K. Jackson, Ian Johnston, Dominic Kwiatkowski, Cordelia Langford, John Sillitoe on behalf of the Wellcome Sanger Institute COVID-19 Surveillance Team |
| EPI_ISL_608912 | Lighthouse Lab in Cambridge | Wellcome Sanger Institute for the COVID-19 Genomics UK (COG-UK) consortium | Rob Howes, The Lighthouse Lab in Cambridge and Alex Alderton, Roberto Amato, Sonia Goncalves, Ewan Harrison, David K. Jackson, Ian Johnston, Dominic Kwiatkowski, Cordelia Langford, John Sillitoe on behalf of the Wellcome Sanger Institute COVID-19 Surveillance Team |
| EPI_ISL_608913, EPI_ISL_608914, EPI_ISL_608915, EPI_ISL_608916, EPI_ISL_608917 | Lighthouse Lab in Alderley Park | Wellcome Sanger Institute for the COVID-19 Genomics UK (COG-UK) consortium | Jacquelyn Wynn, Mairead Hyland, The Lighthouse Lab in Alderley Park and Alex Alderton, Roberto Amato, Sonia Goncalves, Ewan Harrison, David K. Jackson, Ian Johnston, Dominic Kwiatkowski, Cordelia Langford, John Sillitoe on behalf of the Wellcome Sanger Institute COVID-19 Surveillance Team |
| EPI_ISL_608918 | Lighthouse Lab in Cambridge | Wellcome Sanger Institute for the COVID-19 Genomics UK (COG-UK) consortium | Rob Howes, The Lighthouse Lab in Cambridge and Alex Alderton, Roberto Amato, Sonia Goncalves, Ewan Harrison, David K. Jackson, Ian Johnston, Dominic Kwiatkowski, Cordelia Langford, John Sillitoe on behalf of the Wellcome Sanger Institute COVID-19 Surveillance Team |
| EPI_ISL_608919, EPI_ISL_608920, EPI_ISL_608921, EPI_ISL_608922 | Lighthouse Lab in Alderley Park | Wellcome Sanger Institute for the COVID-19 Genomics UK (COG-UK) consortium | Jacquelyn Wynn, Mairead Hyland, The Lighthouse Lab in Alderley Park and Alex Alderton, Roberto Amato, Sonia Goncalves, Ewan Harrison, David K. Jackson, Ian Johnston, Dominic Kwiatkowski, Cordelia Langford, John Sillitoe on behalf of the Wellcome Sanger Institute COVID-19 Surveillance Team |
| EPI_ISL_608923, EPI_ISL_608924 | Lighthouse Lab in Cambridge | Wellcome Sanger Institute for the COVID-19 Genomics UK (COG-UK) consortium | Rob Howes, The Lighthouse Lab in Cambridge and Alex Alderton, Roberto Amato, Sonia Goncalves, Ewan Harrison, David K. Jackson, Ian Johnston, Dominic Kwiatkowski, Cordelia Langford, John Sillitoe on behalf of the Wellcome Sanger Institute COVID-19 Surveillance Team |
| EPI_ISL_608925, EPI_ISL_608926 | Lighthouse Lab in Alderley Park | Wellcome Sanger Institute for the COVID-19 Genomics UK (COG-UK) consortium | Jacquelyn Wynn, Mairead Hyland, The Lighthouse Lab in Alderley Park and Alex Alderton, Roberto Amato, Sonia Goncalves, Ewan Harrison, David K. Jackson, Ian Johnston, Dominic Kwiatkowski, Cordelia Langford, John Sillitoe on behalf of the Wellcome Sanger Institute COVID-19 Surveillance Team |
| EPI_ISL_608927 | Lighthouse Lab in Cambridge | Wellcome Sanger Institute for the COVID-19 Genomics UK (COG-UK) consortium | Rob Howes, The Lighthouse Lab in Cambridge and Alex Alderton, Roberto Amato, Sonia Goncalves, Ewan Harrison, David K. Jackson, Ian Johnston, Dominic Kwiatkowski, Cordelia Langford, John Sillitoe on behalf of the Wellcome Sanger Institute COVID-19 Surveillance Team |
| EPI_ISL_608928, EPI_ISL_608929, EPI_ISL_608930, EPI_ISL_608931, EPI_ISL_608932, EPI_ISL_608933, EPI_ISL_608934 | Lighthouse Lab in Alderley Park | Wellcome Sanger Institute for the COVID-19 Genomics UK (COG-UK) consortium | Jacquelyn Wynn, Mairead Hyland, The Lighthouse Lab in Alderley Park and Alex Alderton, Roberto Amato, Sonia Goncalves, Ewan Harrison, David K. Jackson, Ian Johnston, Dominic Kwiatkowski, Cordelia Langford, John Sillitoe on behalf of the Wellcome Sanger Institute COVID-19 Surveillance Team |

| | | | |
|---|---|---|---|
| EPI_ISL_608935 | Lighthouse Lab in Cambridge | Wellcome Sanger Institute for the COVID-19 Genomics UK (COG-UK) consortium | Rob Howes, The Lighthouse Lab in Cambridge and Alex Alderton, Roberto Amato, Sonia Goncalves, Ewan Harrison, David K. Jackson, Ian Johnston, Dominic Kwiatkowski, Cordelia Langford, John Sillitoe on behalf of the Wellcome Sanger Institute COVID-19 Surveillance Team |
| EPI_ISL_608936 | Lighthouse Lab in Milton Keynes | Wellcome Sanger Institute for the COVID-19 Genomics UK (COG-UK) consortium | The Lighthouse Lab in Milton Keynes and Alex Alderton, Roberto Amato, Sonia Goncalves, Ewan Harrison, David K. Jackson, Ian Johnston, Dominic Kwiatkowski, Cordelia Langford, John Sillitoe on behalf of the Wellcome Sanger Institute COVID-19 Surveillance Team |
| EPI_ISL_608937 | Lighthouse Lab in Cambridge | Wellcome Sanger Institute for the COVID-19 Genomics UK (COG-UK) consortium | Rob Howes, The Lighthouse Lab in Cambridge and Alex Alderton, Roberto Amato, Sonia Goncalves, Ewan Harrison, David K. Jackson, Ian Johnston, Dominic Kwiatkowski, Cordelia Langford, John Sillitoe on behalf of the Wellcome Sanger Institute COVID-19 Surveillance Team |
| EPI_ISL_608938 | Lighthouse Lab in Alderley Park | Wellcome Sanger Institute for the COVID-19 Genomics UK (COG-UK) consortium | Jacquelyn Wynn, Mairead Hyland, The Lighthouse Lab in Alderley Park and Alex Alderton, Roberto Amato, Sonia Goncalves, Ewan Harrison, David K. Jackson, Ian Johnston, Dominic Kwiatkowski, Cordelia Langford, John Sillitoe on behalf of the Wellcome Sanger Institute COVID-19 Surveillance Team |
| EPI_ISL_608939 | Lighthouse Lab in Cambridge | Wellcome Sanger Institute for the COVID-19 Genomics UK (COG-UK) consortium | Rob Howes, The Lighthouse Lab in Cambridge and Alex Alderton, Roberto Amato, Sonia Goncalves, Ewan Harrison, David K. Jackson, Ian Johnston, Dominic Kwiatkowski, Cordelia Langford, John Sillitoe on behalf of the Wellcome Sanger Institute COVID-19 Surveillance Team |
| EPI_ISL_608940 | Lighthouse Lab in Milton Keynes | Wellcome Sanger Institute for the COVID-19 Genomics UK (COG-UK) consortium | The Lighthouse Lab in Milton Keynes and Alex Alderton, Roberto Amato, Sonia Goncalves, Ewan Harrison, David K. Jackson, Ian Johnston, Dominic Kwiatkowski, Cordelia Langford, John Sillitoe on behalf of the Wellcome Sanger Institute COVID-19 Surveillance Team |
| EPI_ISL_608941, EPI_ISL_608942 | Lighthouse Lab in Alderley Park | Wellcome Sanger Institute for the COVID-19 Genomics UK (COG-UK) consortium | Jacquelyn Wynn, Mairead Hyland, The Lighthouse Lab in Alderley Park and Alex Alderton, Roberto Amato, Sonia Goncalves, Ewan Harrison, David K. Jackson, Ian Johnston, Dominic Kwiatkowski, Cordelia Langford, John Sillitoe on behalf of the Wellcome Sanger Institute COVID-19 Surveillance Team |
| EPI_ISL_608943 | Lighthouse Lab in Cambridge | Wellcome Sanger Institute for the COVID-19 Genomics UK (COG-UK) consortium | Rob Howes, The Lighthouse Lab in Cambridge and Alex Alderton, Roberto Amato, Sonia Goncalves, Ewan Harrison, David K. Jackson, Ian Johnston, Dominic Kwiatkowski, Cordelia Langford, John Sillitoe on behalf of the Wellcome Sanger Institute COVID-19 Surveillance Team |
| EPI_ISL_608944, EPI_ISL_608945, EPI_ISL_608946 | Lighthouse Lab in Alderley Park | Wellcome Sanger Institute for the COVID-19 Genomics UK (COG-UK) consortium | Jacquelyn Wynn, Mairead Hyland, The Lighthouse Lab in Alderley Park and Alex Alderton, Roberto Amato, Sonia Goncalves, Ewan Harrison, David K. Jackson, Ian Johnston, Dominic Kwiatkowski, Cordelia Langford, John Sillitoe on behalf of the Wellcome Sanger Institute COVID-19 Surveillance Team |
| EPI_ISL_608947, EPI_ISL_608949 | Lighthouse Lab in Cambridge | Wellcome Sanger Institute for the COVID-19 Genomics UK (COG-UK) consortium | Rob Howes, The Lighthouse Lab in Cambridge and Alex Alderton, Roberto Amato, Sonia Goncalves, Ewan Harrison, David K. Jackson, Ian Johnston, Dominic Kwiatkowski, Cordelia Langford, John Sillitoe on behalf of the Wellcome Sanger Institute COVID-19 Surveillance Team |
| EPI_ISL_608950 | Lighthouse Lab in Alderley Park | Wellcome Sanger Institute for the COVID-19 Genomics UK (COG-UK) consortium | Jacquelyn Wynn, Mairead Hyland, The Lighthouse Lab in Alderley Park and Alex Alderton, Roberto Amato, Sonia Goncalves, Ewan Harrison, David K. Jackson, Ian Johnston, Dominic Kwiatkowski, Cordelia Langford, John Sillitoe on behalf of the Wellcome Sanger Institute COVID-19 Surveillance Team |
| EPI_ISL_608951 | Lighthouse Lab in Milton Keynes | Wellcome Sanger Institute for the COVID-19 Genomics UK (COG-UK) consortium | The Lighthouse Lab in Milton Keynes and Alex Alderton, Roberto Amato, Sonia Goncalves, Ewan Harrison, David K. Jackson, Ian Johnston, Dominic Kwiatkowski, Cordelia Langford, John Sillitoe on behalf of the Wellcome Sanger Institute COVID-19 Surveillance Team |
| EPI_ISL_608952 | Lighthouse Lab in Alderley Park | Wellcome Sanger Institute for the COVID-19 Genomics UK (COG-UK) consortium | Jacquelyn Wynn, Mairead Hyland, The Lighthouse Lab in Alderley Park and Alex Alderton, Roberto Amato, Sonia Goncalves, Ewan Harrison, David K. Jackson, Ian Johnston, Dominic Kwiatkowski, Cordelia Langford, John Sillitoe on behalf of the Wellcome Sanger Institute COVID-19 Surveillance Team |
| EPI_ISL_608953 | Lighthouse Lab in Cambridge | Wellcome Sanger Institute for the COVID-19 Genomics UK (COG-UK) consortium | Rob Howes, The Lighthouse Lab in Cambridge and Alex Alderton, Roberto Amato, Sonia Goncalves, Ewan Harrison, David K. Jackson, Ian Johnston, Dominic Kwiatkowski, Cordelia Langford, John Sillitoe on behalf of the Wellcome Sanger Institute COVID-19 Surveillance Team |
| EPI_ISL_608954 | Lighthouse Lab in Milton Keynes | Wellcome Sanger Institute for the COVID-19 Genomics UK (COG-UK) consortium | The Lighthouse Lab in Milton Keynes and Alex Alderton, Roberto Amato, Sonia Goncalves, Ewan Harrison, David K. Jackson, Ian Johnston, Dominic Kwiatkowski, Cordelia Langford, John Sillitoe on behalf of the Wellcome Sanger Institute COVID-19 Surveillance Team |
| EPI_ISL_608955, EPI_ISL_608956, EPI_ISL_608957 | Lighthouse Lab in Alderley Park | Wellcome Sanger Institute for the COVID-19 Genomics UK (COG-UK) consortium | Jacquelyn Wynn, Mairead Hyland, The Lighthouse Lab in Alderley Park and Alex Alderton, Roberto Amato, Sonia Goncalves, Ewan Harrison, David K. Jackson, Ian Johnston, Dominic Kwiatkowski, Cordelia Langford, John Sillitoe on behalf of the Wellcome Sanger Institute COVID-19 Surveillance Team |
| EPI_ISL_608958, EPI_ISL_608959, EPI_ISL_608960 | Lighthouse Lab in Cambridge | Wellcome Sanger Institute for the COVID-19 Genomics UK (COG-UK) consortium | Rob Howes, The Lighthouse Lab in Cambridge and Alex Alderton, Roberto Amato, Sonia Goncalves, Ewan Harrison, David K. Jackson, Ian Johnston, Dominic Kwiatkowski, Cordelia Langford, John Sillitoe on behalf of the Wellcome Sanger Institute COVID-19 Surveillance Team |
| EPI_ISL_608961, EPI_ISL_608962, EPI_ISL_608963, EPI_ISL_608964 | Lighthouse Lab in Alderley Park | Wellcome Sanger Institute for the COVID-19 Genomics UK (COG-UK) consortium | Jacquelyn Wynn, Mairead Hyland, The Lighthouse Lab in Alderley Park and Alex Alderton, Roberto Amato, Sonia Goncalves, Ewan Harrison, David K. Jackson, Ian Johnston, Dominic Kwiatkowski, Cordelia Langford, John Sillitoe on behalf of the Wellcome Sanger Institute COVID-19 Surveillance Team |
| EPI_ISL_608965 | Lighthouse Lab in Cambridge | Wellcome Sanger Institute for the COVID-19 Genomics UK (COG-UK) consortium | Rob Howes, The Lighthouse Lab in Cambridge and Alex Alderton, Roberto Amato, Sonia Goncalves, Ewan Harrison, David K. Jackson, Ian Johnston, Dominic Kwiatkowski, Cordelia Langford, John Sillitoe on behalf of the Wellcome Sanger Institute COVID-19 Surveillance Team |
| EPI_ISL_608966 | Lighthouse Lab in Alderley Park | Wellcome Sanger Institute for the COVID-19 Genomics UK (COG-UK) consortium | Jacquelyn Wynn, Mairead Hyland, The Lighthouse Lab in Alderley Park and Alex Alderton, Roberto Amato, Sonia Goncalves, Ewan Harrison, David K. Jackson, Ian Johnston, Dominic Kwiatkowski, Cordelia Langford, John Sillitoe on behalf of the Wellcome Sanger Institute COVID-19 Surveillance Team |
| EPI_ISL_608967, EPI_ISL_608968, EPI_ISL_608969 | Lighthouse Lab in Cambridge | Wellcome Sanger Institute for the COVID-19 Genomics UK (COG-UK) consortium | Rob Howes, The Lighthouse Lab in Cambridge and Alex Alderton, Roberto Amato, Sonia Goncalves, Ewan Harrison, David K. Jackson, Ian Johnston, Dominic Kwiatkowski, Cordelia Langford, John Sillitoe on behalf of the Wellcome Sanger Institute COVID-19 Surveillance Team |
| EPI_ISL_608970, EPI_ISL_608971, EPI_ISL_608972, EPI_ISL_608973 | Lighthouse Lab in Alderley Park | Wellcome Sanger Institute for the COVID-19 Genomics UK (COG-UK) consortium | Jacquelyn Wynn, Mairead Hyland, The Lighthouse Lab in Alderley Park and Alex Alderton, Roberto Amato, Sonia Goncalves, Ewan Harrison, David K. Jackson, Ian Johnston, Dominic Kwiatkowski, Cordelia Langford, John Sillitoe on behalf of the Wellcome Sanger Institute COVID-19 Surveillance Team |
| EPI_ISL_608974 | Lighthouse Lab in Cambridge | Wellcome Sanger Institute for the COVID-19 Genomics UK (COG-UK) consortium | Rob Howes, The Lighthouse Lab in Cambridge and Alex Alderton, Roberto Amato, Sonia Goncalves, Ewan Harrison, David K. Jackson, Ian Johnston, Dominic Kwiatkowski, Cordelia Langford, John Sillitoe on behalf of the Wellcome Sanger Institute COVID-19 Surveillance Team |
| EPI_ISL_608975 | Lighthouse Lab in Milton Keynes | Wellcome Sanger Institute for the COVID-19 Genomics UK (COG-UK) consortium | The Lighthouse Lab in Milton Keynes and Alex Alderton, Roberto Amato, Sonia Goncalves, Ewan Harrison, David K. Jackson, Ian Johnston, Dominic Kwiatkowski, Cordelia Langford, John Sillitoe on behalf of the Wellcome Sanger Institute COVID-19 Surveillance Team |
| EPI_ISL_608976 | Lighthouse Lab in Alderley Park | Wellcome Sanger Institute for the COVID-19 Genomics UK (COG-UK) consortium | Jacquelyn Wynn, Mairead Hyland, The Lighthouse Lab in Alderley Park and Alex Alderton, Roberto Amato, Sonia Goncalves, Ewan Harrison, David K. Jackson, Ian Johnston, Dominic Kwiatkowski, Cordelia Langford, John Sillitoe on behalf of the Wellcome Sanger Institute COVID-19 Surveillance Team |
| EPI_ISL_608977, EPI_ISL_608978 | Lighthouse Lab in Milton Keynes | Wellcome Sanger Institute for the COVID-19 Genomics UK (COG-UK) consortium | The Lighthouse Lab in Milton Keynes and Alex Alderton, Roberto Amato, Sonia Goncalves, Ewan Harrison, David K. Jackson, Ian Johnston, Dominic Kwiatkowski, Cordelia Langford, John Sillitoe on behalf of the Wellcome Sanger Institute COVID-19 Surveillance Team |
| EPI_ISL_608979, EPI_ISL_608980, EPI_ISL_608981 | Lighthouse Lab in Alderley Park | Wellcome Sanger Institute for the COVID-19 Genomics UK (COG-UK) consortium | Jacquelyn Wynn, Mairead Hyland, The Lighthouse Lab in Alderley Park and Alex Alderton, Roberto Amato, Sonia Goncalves, Ewan Harrison, David K. Jackson, Ian Johnston, Dominic Kwiatkowski, Cordelia Langford, John Sillitoe on behalf of the Wellcome Sanger Institute COVID-19 Surveillance Team |
| EPI_ISL_608982 | Lighthouse Lab in Cambridge | Wellcome Sanger Institute for the COVID-19 Genomics UK (COG-UK) consortium | Rob Howes, The Lighthouse Lab in Cambridge and Alex Alderton, Roberto Amato, Sonia Goncalves, Ewan Harrison, David K. Jackson, Ian Johnston, Dominic Kwiatkowski, Cordelia Langford, John Sillitoe on behalf of the Wellcome Sanger Institute COVID-19 Surveillance Team |
| EPI_ISL_608983, EPI_ISL_608984, EPI_ISL_608985 | Lighthouse Lab in Alderley Park | Wellcome Sanger Institute for the COVID-19 Genomics UK (COG-UK) consortium | Jacquelyn Wynn, Mairead Hyland, The Lighthouse Lab in Alderley Park and Alex Alderton, Roberto Amato, Sonia Goncalves, Ewan Harrison, David K. Jackson, Ian Johnston, Dominic Kwiatkowski, Cordelia Langford, John Sillitoe on behalf of the Wellcome Sanger Institute COVID-19 Surveillance Team |
| EPI_ISL_608986 | Lighthouse Lab in Milton Keynes | Wellcome Sanger Institute for the COVID-19 Genomics UK (COG-UK) consortium | The Lighthouse Lab in Milton Keynes and Alex Alderton, Roberto Amato, Sonia Goncalves, Ewan Harrison, David K. Jackson, Ian Johnston, Dominic Kwiatkowski, Cordelia Langford, John Sillitoe on behalf of the Wellcome Sanger Institute COVID-19 Surveillance Team |
| EPI_ISL_608987 | Lighthouse Lab in Cambridge | Wellcome Sanger Institute for the COVID-19 Genomics UK (COG-UK) consortium | Rob Howes, The Lighthouse Lab in Cambridge and Alex Alderton, Roberto Amato, Sonia Goncalves, Ewan Harrison, David K. Jackson, Ian Johnston, Dominic Kwiatkowski, Cordelia Langford, John Sillitoe on behalf of the Wellcome Sanger Institute COVID-19 Surveillance Team |
| EPI_ISL_608988, EPI_ISL_608989 | Lighthouse Lab in Alderley Park | Wellcome Sanger Institute for the COVID-19 Genomics UK (COG-UK) consortium | Jacquelyn Wynn, Mairead Hyland, The Lighthouse Lab in Alderley Park and Alex Alderton, Roberto Amato, Sonia Goncalves, Ewan Harrison, David K. Jackson, Ian Johnston, Dominic Kwiatkowski, Cordelia Langford, John Sillitoe on behalf of the Wellcome Sanger Institute COVID-19 Surveillance Team |
| EPI_ISL_608990 | Lighthouse Lab in Cambridge | Wellcome Sanger Institute for the COVID-19 Genomics UK (COG-UK) consortium | Rob Howes, The Lighthouse Lab in Cambridge and Alex Alderton, Roberto Amato, Sonia Goncalves, Ewan Harrison, David K. Jackson, Ian Johnston, Dominic Kwiatkowski, Cordelia Langford, John Sillitoe on behalf of the Wellcome Sanger Institute COVID-19 Surveillance Team |
| EPI_ISL_608991, EPI_ISL_608992, EPI_ISL_608993, EPI_ISL_608994, EPI_ISL_608995 | Lighthouse Lab in Alderley Park | Wellcome Sanger Institute for the COVID-19 Genomics UK (COG-UK) consortium | Jacquelyn Wynn, Mairead Hyland, The Lighthouse Lab in Alderley Park and Alex Alderton, Roberto Amato, Sonia Goncalves, Ewan Harrison, David K. Jackson, Ian Johnston, Dominic Kwiatkowski, Cordelia Langford, John Sillitoe on behalf of the Wellcome Sanger Institute COVID-19 Surveillance Team |
| EPI_ISL_608996, EPI_ISL_608997 | Lighthouse Lab in Cambridge | Wellcome Sanger Institute for the COVID-19 Genomics UK (COG-UK) consortium | Rob Howes, The Lighthouse Lab in Cambridge and Alex Alderton, Roberto Amato, Sonia Goncalves, Ewan Harrison, David K. Jackson, Ian Johnston, Dominic Kwiatkowski, Cordelia Langford, John Sillitoe on behalf of the Wellcome Sanger Institute COVID-19 Surveillance Team |

| | | | |
|---|---|---|---|
| EPI_ISL_608998, EPI_ISL_608999, EPI_ISL_609000 | Lighthouse Lab in Alderley Park | Wellcome Sanger Institute for the COVID-19 Genomics UK (COG-UK) consortium | Jacquelyn Wynn, Mairead Hyland, The Lighthouse Lab in Alderley Park and Alex Alderton, Roberto Amato, Sonia Goncalves, Ewan Harrison, David K. Jackson, Ian Johnston, Dominic Kwiatkowski, Cordelia Langford, John Sillitoe on behalf of the Wellcome Sanger Institute COVID-19 Surveillance Team |
| EPI_ISL_609001 | Lighthouse Lab in Milton Keynes | Wellcome Sanger Institute for the COVID-19 Genomics UK (COG-UK) consortium | The Lighthouse Lab in Milton Keynes and Alex Alderton, Roberto Amato, Sonia Goncalves, Ewan Harrison, David K. Jackson, Ian Johnston, Dominic Kwiatkowski, Cordelia Langford, John Sillitoe on behalf of the Wellcome Sanger Institute COVID-19 Surveillance Team |
| EPI_ISL_609002 | Lighthouse Lab in Alderley Park | Wellcome Sanger Institute for the COVID-19 Genomics UK (COG-UK) consortium | Jacquelyn Wynn, Mairead Hyland, The Lighthouse Lab in Alderley Park and Alex Alderton, Roberto Amato, Sonia Goncalves, Ewan Harrison, David K. Jackson, Ian Johnston, Dominic Kwiatkowski, Cordelia Langford, John Sillitoe on behalf of the Wellcome Sanger Institute COVID-19 Surveillance Team |
| EPI_ISL_609003 | Lighthouse Lab in Cambridge | Wellcome Sanger Institute for the COVID-19 Genomics UK (COG-UK) consortium | Rob Howes, The Lighthouse Lab in Cambridge and Alex Alderton, Roberto Amato, Sonia Goncalves, Ewan Harrison, David K. Jackson, Ian Johnston, Dominic Kwiatkowski, Cordelia Langford, John Sillitoe on behalf of the Wellcome Sanger Institute COVID-19 Surveillance Team |
| EPI_ISL_609004 | Lighthouse Lab in Alderley Park | Wellcome Sanger Institute for the COVID-19 Genomics UK (COG-UK) consortium | Jacquelyn Wynn, Mairead Hyland, The Lighthouse Lab in Alderley Park and Alex Alderton, Roberto Amato, Sonia Goncalves, Ewan Harrison, David K. Jackson, Ian Johnston, Dominic Kwiatkowski, Cordelia Langford, John Sillitoe on behalf of the Wellcome Sanger Institute COVID-19 Surveillance Team |
| EPI_ISL_609005 | Lighthouse Lab in Cambridge | Wellcome Sanger Institute for the COVID-19 Genomics UK (COG-UK) consortium | Rob Howes, The Lighthouse Lab in Cambridge and Alex Alderton, Roberto Amato, Sonia Goncalves, Ewan Harrison, David K. Jackson, Ian Johnston, Dominic Kwiatkowski, Cordelia Langford, John Sillitoe on behalf of the Wellcome Sanger Institute COVID-19 Surveillance Team |
| EPI_ISL_609006 | Lighthouse Lab in Milton Keynes | Wellcome Sanger Institute for the COVID-19 Genomics UK (COG-UK) consortium | The Lighthouse Lab in Milton Keynes and Alex Alderton, Roberto Amato, Sonia Goncalves, Ewan Harrison, David K. Jackson, Ian Johnston, Dominic Kwiatkowski, Cordelia Langford, John Sillitoe on behalf of the Wellcome Sanger Institute COVID-19 Surveillance Team |
| EPI_ISL_609007 | Lighthouse Lab in Alderley Park | Wellcome Sanger Institute for the COVID-19 Genomics UK (COG-UK) consortium | Jacquelyn Wynn, Mairead Hyland, The Lighthouse Lab in Alderley Park and Alex Alderton, Roberto Amato, Sonia Goncalves, Ewan Harrison, David K. Jackson, Ian Johnston, Dominic Kwiatkowski, Cordelia Langford, John Sillitoe on behalf of the Wellcome Sanger Institute COVID-19 Surveillance Team |
| EPI_ISL_609008 | Lighthouse Lab in Milton Keynes | Wellcome Sanger Institute for the COVID-19 Genomics UK (COG-UK) consortium | The Lighthouse Lab in Milton Keynes and Alex Alderton, Roberto Amato, Sonia Goncalves, Ewan Harrison, David K. Jackson, Ian Johnston, Dominic Kwiatkowski, Cordelia Langford, John Sillitoe on behalf of the Wellcome Sanger Institute COVID-19 Surveillance Team |
| EPI_ISL_609009, EPI_ISL_609010, EPI_ISL_609011, EPI_ISL_609012, EPI_ISL_609013 | Lighthouse Lab in Alderley Park | Wellcome Sanger Institute for the COVID-19 Genomics UK (COG-UK) consortium | Jacquelyn Wynn, Mairead Hyland, The Lighthouse Lab in Alderley Park and Alex Alderton, Roberto Amato, Sonia Goncalves, Ewan Harrison, David K. Jackson, Ian Johnston, Dominic Kwiatkowski, Cordelia Langford, John Sillitoe on behalf of the Wellcome Sanger Institute COVID-19 Surveillance Team |
| EPI_ISL_609014 | Lighthouse Lab in Cambridge | Wellcome Sanger Institute for the COVID-19 Genomics UK (COG-UK) consortium | Rob Howes, The Lighthouse Lab in Cambridge and Alex Alderton, Roberto Amato, Sonia Goncalves, Ewan Harrison, David K. Jackson, Ian Johnston, Dominic Kwiatkowski, Cordelia Langford, John Sillitoe on behalf of the Wellcome Sanger Institute COVID-19 Surveillance Team |
| EPI_ISL_609015 | Lighthouse Lab in Milton Keynes | Wellcome Sanger Institute for the COVID-19 Genomics UK (COG-UK) consortium | The Lighthouse Lab in Milton Keynes and Alex Alderton, Roberto Amato, Sonia Goncalves, Ewan Harrison, David K. Jackson, Ian Johnston, Dominic Kwiatkowski, Cordelia Langford, John Sillitoe on behalf of the Wellcome Sanger Institute COVID-19 Surveillance Team |
| EPI_ISL_609016 | Lighthouse Lab in Alderley Park | Wellcome Sanger Institute for the COVID-19 Genomics UK (COG-UK) consortium | Jacquelyn Wynn, Mairead Hyland, The Lighthouse Lab in Alderley Park and Alex Alderton, Roberto Amato, Sonia Goncalves, Ewan Harrison, David K. Jackson, Ian Johnston, Dominic Kwiatkowski, Cordelia Langford, John Sillitoe on behalf of the Wellcome Sanger Institute COVID-19 Surveillance Team |
| EPI_ISL_609017 | Lighthouse Lab in Milton Keynes | Wellcome Sanger Institute for the COVID-19 Genomics UK (COG-UK) consortium | The Lighthouse Lab in Milton Keynes and Alex Alderton, Roberto Amato, Sonia Goncalves, Ewan Harrison, David K. Jackson, Ian Johnston, Dominic Kwiatkowski, Cordelia Langford, John Sillitoe on behalf of the Wellcome Sanger Institute COVID-19 Surveillance Team |
| EPI_ISL_609018 | Lighthouse Lab in Cambridge | Wellcome Sanger Institute for the COVID-19 Genomics UK (COG-UK) consortium | Rob Howes, The Lighthouse Lab in Cambridge and Alex Alderton, Roberto Amato, Sonia Goncalves, Ewan Harrison, David K. Jackson, Ian Johnston, Dominic Kwiatkowski, Cordelia Langford, John Sillitoe on behalf of the Wellcome Sanger Institute COVID-19 Surveillance Team |
| EPI_ISL_609019, EPI_ISL_609020 | Lighthouse Lab in Alderley Park | Wellcome Sanger Institute for the COVID-19 Genomics UK (COG-UK) consortium | Jacquelyn Wynn, Mairead Hyland, The Lighthouse Lab in Alderley Park and Alex Alderton, Roberto Amato, Sonia Goncalves, Ewan Harrison, David K. Jackson, Ian Johnston, Dominic Kwiatkowski, Cordelia Langford, John Sillitoe on behalf of the Wellcome Sanger Institute COVID-19 Surveillance Team |
| EPI_ISL_609021 | Lighthouse Lab in Milton Keynes | Wellcome Sanger Institute for the COVID-19 Genomics UK (COG-UK) consortium | The Lighthouse Lab in Milton Keynes and Alex Alderton, Roberto Amato, Sonia Goncalves, Ewan Harrison, David K. Jackson, Ian Johnston, Dominic Kwiatkowski, Cordelia Langford, John Sillitoe on behalf of the Wellcome Sanger Institute COVID-19 Surveillance Team |
| EPI_ISL_609022 | Lighthouse Lab in Alderley Park | Wellcome Sanger Institute for the COVID-19 Genomics UK (COG-UK) consortium | Jacquelyn Wynn, Mairead Hyland, The Lighthouse Lab in Alderley Park and Alex Alderton, Roberto Amato, Sonia Goncalves, Ewan Harrison, David K. Jackson, Ian Johnston, Dominic Kwiatkowski, Cordelia Langford, John Sillitoe on behalf of the Wellcome Sanger Institute COVID-19 Surveillance Team |
| EPI_ISL_609023 | Lighthouse Lab in Cambridge | Wellcome Sanger Institute for the COVID-19 Genomics UK (COG-UK) consortium | Rob Howes, The Lighthouse Lab in Cambridge and Alex Alderton, Roberto Amato, Sonia Goncalves, Ewan Harrison, David K. Jackson, Ian Johnston, Dominic Kwiatkowski, Cordelia Langford, John Sillitoe on behalf of the Wellcome Sanger Institute COVID-19 Surveillance Team |
| EPI_ISL_609024 | Lighthouse Lab in Milton Keynes | Wellcome Sanger Institute for the COVID-19 Genomics UK (COG-UK) consortium | The Lighthouse Lab in Milton Keynes and Alex Alderton, Roberto Amato, Sonia Goncalves, Ewan Harrison, David K. Jackson, Ian Johnston, Dominic Kwiatkowski, Cordelia Langford, John Sillitoe on behalf of the Wellcome Sanger Institute COVID-19 Surveillance Team |
| EPI_ISL_609025 | Lighthouse Lab in Cambridge | Wellcome Sanger Institute for the COVID-19 Genomics UK (COG-UK) consortium | Rob Howes, The Lighthouse Lab in Cambridge and Alex Alderton, Roberto Amato, Sonia Goncalves, Ewan Harrison, David K. Jackson, Ian Johnston, Dominic Kwiatkowski, Cordelia Langford, John Sillitoe on behalf of the Wellcome Sanger Institute COVID-19 Surveillance Team |
| EPI_ISL_609026, EPI_ISL_609027 | Lighthouse Lab in Milton Keynes | Wellcome Sanger Institute for the COVID-19 Genomics UK (COG-UK) consortium | The Lighthouse Lab in Milton Keynes and Alex Alderton, Roberto Amato, Sonia Goncalves, Ewan Harrison, David K. Jackson, Ian Johnston, Dominic Kwiatkowski, Cordelia Langford, John Sillitoe on behalf of the Wellcome Sanger Institute COVID-19 Surveillance Team |
| EPI_ISL_609028 | Lighthouse Lab in Alderley Park | Wellcome Sanger Institute for the COVID-19 Genomics UK (COG-UK) consortium | Jacquelyn Wynn, Mairead Hyland, The Lighthouse Lab in Alderley Park and Alex Alderton, Roberto Amato, Sonia Goncalves, Ewan Harrison, David K. Jackson, Ian Johnston, Dominic Kwiatkowski, Cordelia Langford, John Sillitoe on behalf of the Wellcome Sanger Institute COVID-19 Surveillance Team |
| EPI_ISL_609029, EPI_ISL_609030 | Lighthouse Lab in Milton Keynes | Wellcome Sanger Institute for the COVID-19 Genomics UK (COG-UK) consortium | The Lighthouse Lab in Milton Keynes and Alex Alderton, Roberto Amato, Sonia Goncalves, Ewan Harrison, David K. Jackson, Ian Johnston, Dominic Kwiatkowski, Cordelia Langford, John Sillitoe on behalf of the Wellcome Sanger Institute COVID-19 Surveillance Team |
| EPI_ISL_609031, EPI_ISL_609032 | Lighthouse Lab in Alderley Park | Wellcome Sanger Institute for the COVID-19 Genomics UK (COG-UK) consortium | Jacquelyn Wynn, Mairead Hyland, The Lighthouse Lab in Alderley Park and Alex Alderton, Roberto Amato, Sonia Goncalves, Ewan Harrison, David K. Jackson, Ian Johnston, Dominic Kwiatkowski, Cordelia Langford, John Sillitoe on behalf of the Wellcome Sanger Institute COVID-19 Surveillance Team |
| EPI_ISL_609033 | Lighthouse Lab in Cambridge | Wellcome Sanger Institute for the COVID-19 Genomics UK (COG-UK) consortium | Rob Howes, The Lighthouse Lab in Cambridge and Alex Alderton, Roberto Amato, Sonia Goncalves, Ewan Harrison, David K. Jackson, Ian Johnston, Dominic Kwiatkowski, Cordelia Langford, John Sillitoe on behalf of the Wellcome Sanger Institute COVID-19 Surveillance Team |
| EPI_ISL_609034, EPI_ISL_609035 | Lighthouse Lab in Alderley Park | Wellcome Sanger Institute for the COVID-19 Genomics UK (COG-UK) consortium | Jacquelyn Wynn, Mairead Hyland, The Lighthouse Lab in Alderley Park and Alex Alderton, Roberto Amato, Sonia Goncalves, Ewan Harrison, David K. Jackson, Ian Johnston, Dominic Kwiatkowski, Cordelia Langford, John Sillitoe on behalf of the Wellcome Sanger Institute COVID-19 Surveillance Team |
| EPI_ISL_609036 | Lighthouse Lab in Milton Keynes | Wellcome Sanger Institute for the COVID-19 Genomics UK (COG-UK) consortium | The Lighthouse Lab in Milton Keynes and Alex Alderton, Roberto Amato, Sonia Goncalves, Ewan Harrison, David K. Jackson, Ian Johnston, Dominic Kwiatkowski, Cordelia Langford, John Sillitoe on behalf of the Wellcome Sanger Institute COVID-19 Surveillance Team |
| EPI_ISL_609037 | Lighthouse Lab in Cambridge | Wellcome Sanger Institute for the COVID-19 Genomics UK (COG-UK) consortium | Rob Howes, The Lighthouse Lab in Cambridge and Alex Alderton, Roberto Amato, Sonia Goncalves, Ewan Harrison, David K. Jackson, Ian Johnston, Dominic Kwiatkowski, Cordelia Langford, John Sillitoe on behalf of the Wellcome Sanger Institute COVID-19 Surveillance Team |
| EPI_ISL_609038 | Lighthouse Lab in Milton Keynes | Wellcome Sanger Institute for the COVID-19 Genomics UK (COG-UK) consortium | The Lighthouse Lab in Milton Keynes and Alex Alderton, Roberto Amato, Sonia Goncalves, Ewan Harrison, David K. Jackson, Ian Johnston, Dominic Kwiatkowski, Cordelia Langford, John Sillitoe on behalf of the Wellcome Sanger Institute COVID-19 Surveillance Team |
| EPI_ISL_609039, EPI_ISL_609040, EPI_ISL_609041 | Lighthouse Lab in Alderley Park | Wellcome Sanger Institute for the COVID-19 Genomics UK (COG-UK) consortium | Jacquelyn Wynn, Mairead Hyland, The Lighthouse Lab in Alderley Park and Alex Alderton, Roberto Amato, Sonia Goncalves, Ewan Harrison, David K. Jackson, Ian Johnston, Dominic Kwiatkowski, Cordelia Langford, John Sillitoe on behalf of the Wellcome Sanger Institute COVID-19 Surveillance Team |
| EPI_ISL_609042 | Lighthouse Lab in Milton Keynes | Wellcome Sanger Institute for the COVID-19 Genomics UK (COG-UK) consortium | The Lighthouse Lab in Milton Keynes and Alex Alderton, Roberto Amato, Sonia Goncalves, Ewan Harrison, David K. Jackson, Ian Johnston, Dominic Kwiatkowski, Cordelia Langford, John Sillitoe on behalf of the Wellcome Sanger Institute COVID-19 Surveillance Team |
| EPI_ISL_609043 | Lighthouse Lab in Alderley Park | Wellcome Sanger Institute for the COVID-19 Genomics UK (COG-UK) consortium | Jacquelyn Wynn, Mairead Hyland, The Lighthouse Lab in Alderley Park and Alex Alderton, Roberto Amato, Sonia Goncalves, Ewan Harrison, David K. Jackson, Ian Johnston, Dominic Kwiatkowski, Cordelia Langford, John Sillitoe on behalf of the Wellcome Sanger Institute COVID-19 Surveillance Team |
| EPI_ISL_609044 | Lighthouse Lab in Milton Keynes | Wellcome Sanger Institute for the COVID-19 Genomics UK (COG-UK) consortium | The Lighthouse Lab in Milton Keynes and Alex Alderton, Roberto Amato, Sonia Goncalves, Ewan Harrison, David K. Jackson, Ian Johnston, Dominic Kwiatkowski, Cordelia Langford, John Sillitoe on behalf of the Wellcome Sanger Institute COVID-19 Surveillance Team |
| EPI_ISL_609045 | Lighthouse Lab in Cambridge | Wellcome Sanger Institute for the COVID-19 Genomics UK | Rob Howes, The Lighthouse Lab in Cambridge and Alex Alderton, Roberto Amato, Sonia Goncalves, Ewan Harrison, David K. Jackson, Ian Johnston, Dominic |

| | | | |
|---|---|---|---|
| | | (COG-UK) consortium | Kwiatkowski, Cordelia Langford, John Sillitoe on behalf of the Wellcome Sanger Institute COVID-19 Surveillance Team |
| EPI_ISL_609046 | Lighthouse Lab in Milton Keynes | Wellcome Sanger Institute for the COVID-19 Genomics UK (COG-UK) consortium | The Lighthouse Lab in Milton Keynes and Alex Alderton, Roberto Amato, Sonia Goncalves, Ewan Harrison, David K. Jackson, Ian Johnston, Dominic Kwiatkowski, Cordelia Langford, John Sillitoe on behalf of the Wellcome Sanger Institute COVID-19 Surveillance Team |
| EPI_ISL_609047 | Lighthouse Lab in Alderley Park | Wellcome Sanger Institute for the COVID-19 Genomics UK (COG-UK) consortium | Jacquelyn Wynn, Mairead Hyland, The Lighthouse Lab in Alderley Park and Alex Alderton, Roberto Amato, Sonia Goncalves, Ewan Harrison, David K. Jackson, Ian Johnston, Dominic Kwiatkowski, Cordelia Langford, John Sillitoe on behalf of the Wellcome Sanger Institute COVID-19 Surveillance Team |
| EPI_ISL_609048 | Lighthouse Lab in Cambridge | Wellcome Sanger Institute for the COVID-19 Genomics UK (COG-UK) consortium | Rob Howes, The Lighthouse Lab in Cambridge and Alex Alderton, Roberto Amato, Sonia Goncalves, Ewan Harrison, David K. Jackson, Ian Johnston, Dominic Kwiatkowski, Cordelia Langford, John Sillitoe on behalf of the Wellcome Sanger Institute COVID-19 Surveillance Team |
| EPI_ISL_609049 | Lighthouse Lab in Milton Keynes | Wellcome Sanger Institute for the COVID-19 Genomics UK (COG-UK) consortium | The Lighthouse Lab in Milton Keynes and Alex Alderton, Roberto Amato, Sonia Goncalves, Ewan Harrison, David K. Jackson, Ian Johnston, Dominic Kwiatkowski, Cordelia Langford, John Sillitoe on behalf of the Wellcome Sanger Institute COVID-19 Surveillance Team |
| EPI_ISL_609050 | Lighthouse Lab in Cambridge | Wellcome Sanger Institute for the COVID-19 Genomics UK (COG-UK) consortium | Rob Howes, The Lighthouse Lab in Cambridge and Alex Alderton, Roberto Amato, Sonia Goncalves, Ewan Harrison, David K. Jackson, Ian Johnston, Dominic Kwiatkowski, Cordelia Langford, John Sillitoe on behalf of the Wellcome Sanger Institute COVID-19 Surveillance Team |
| EPI_ISL_609051, EPI_ISL_609052, EPI_ISL_609053 | Lighthouse Lab in Alderley Park | Wellcome Sanger Institute for the COVID-19 Genomics UK (COG-UK) consortium | Jacquelyn Wynn, Mairead Hyland, The Lighthouse Lab in Alderley Park and Alex Alderton, Roberto Amato, Sonia Goncalves, Ewan Harrison, David K. Jackson, Ian Johnston, Dominic Kwiatkowski, Cordelia Langford, John Sillitoe on behalf of the Wellcome Sanger Institute COVID-19 Surveillance Team |
| EPI_ISL_609054 | Lighthouse Lab in Cambridge | Wellcome Sanger Institute for the COVID-19 Genomics UK (COG-UK) consortium | Rob Howes, The Lighthouse Lab in Cambridge and Alex Alderton, Roberto Amato, Sonia Goncalves, Ewan Harrison, David K. Jackson, Ian Johnston, Dominic Kwiatkowski, Cordelia Langford, John Sillitoe on behalf of the Wellcome Sanger Institute COVID-19 Surveillance Team |
| EPI_ISL_609055, EPI_ISL_609056, EPI_ISL_609057 | Lighthouse Lab in Alderley Park | Wellcome Sanger Institute for the COVID-19 Genomics UK (COG-UK) consortium | Jacquelyn Wynn, Mairead Hyland, The Lighthouse Lab in Alderley Park and Alex Alderton, Roberto Amato, Sonia Goncalves, Ewan Harrison, David K. Jackson, Ian Johnston, Dominic Kwiatkowski, Cordelia Langford, John Sillitoe on behalf of the Wellcome Sanger Institute COVID-19 Surveillance Team |
| EPI_ISL_609058, EPI_ISL_609059 | Lighthouse Lab in Milton Keynes | Wellcome Sanger Institute for the COVID-19 Genomics UK (COG-UK) consortium | The Lighthouse Lab in Milton Keynes and Alex Alderton, Roberto Amato, Sonia Goncalves, Ewan Harrison, David K. Jackson, Ian Johnston, Dominic Kwiatkowski, Cordelia Langford, John Sillitoe on behalf of the Wellcome Sanger Institute COVID-19 Surveillance Team |
| EPI_ISL_609060, EPI_ISL_609061 | Lighthouse Lab in Alderley Park | Wellcome Sanger Institute for the COVID-19 Genomics UK (COG-UK) consortium | Jacquelyn Wynn, Mairead Hyland, The Lighthouse Lab in Alderley Park and Alex Alderton, Roberto Amato, Sonia Goncalves, Ewan Harrison, David K. Jackson, Ian Johnston, Dominic Kwiatkowski, Cordelia Langford, John Sillitoe on behalf of the Wellcome Sanger Institute COVID-19 Surveillance Team |
| EPI_ISL_609062, EPI_ISL_609063 | Lighthouse Lab in Milton Keynes | Wellcome Sanger Institute for the COVID-19 Genomics UK (COG-UK) consortium | The Lighthouse Lab in Milton Keynes and Alex Alderton, Roberto Amato, Sonia Goncalves, Ewan Harrison, David K. Jackson, Ian Johnston, Dominic Kwiatkowski, Cordelia Langford, John Sillitoe on behalf of the Wellcome Sanger Institute COVID-19 Surveillance Team |
| EPI_ISL_609064, EPI_ISL_609065 | Lighthouse Lab in Alderley Park | Wellcome Sanger Institute for the COVID-19 Genomics UK (COG-UK) consortium | Jacquelyn Wynn, Mairead Hyland, The Lighthouse Lab in Alderley Park and Alex Alderton, Roberto Amato, Sonia Goncalves, Ewan Harrison, David K. Jackson, Ian Johnston, Dominic Kwiatkowski, Cordelia Langford, John Sillitoe on behalf of the Wellcome Sanger Institute COVID-19 Surveillance Team |
| EPI_ISL_609066 | Lighthouse Lab in Milton Keynes | Wellcome Sanger Institute for the COVID-19 Genomics UK (COG-UK) consortium | The Lighthouse Lab in Milton Keynes and Alex Alderton, Roberto Amato, Sonia Goncalves, Ewan Harrison, David K. Jackson, Ian Johnston, Dominic Kwiatkowski, Cordelia Langford, John Sillitoe on behalf of the Wellcome Sanger Institute COVID-19 Surveillance Team |
| EPI_ISL_609067 | Lighthouse Lab in Cambridge | Wellcome Sanger Institute for the COVID-19 Genomics UK (COG-UK) consortium | Rob Howes, The Lighthouse Lab in Cambridge and Alex Alderton, Roberto Amato, Sonia Goncalves, Ewan Harrison, David K. Jackson, Ian Johnston, Dominic Kwiatkowski, Cordelia Langford, John Sillitoe on behalf of the Wellcome Sanger Institute COVID-19 Surveillance Team |
| EPI_ISL_609068, EPI_ISL_609069, EPI_ISL_609070 | Lighthouse Lab in Alderley Park | Wellcome Sanger Institute for the COVID-19 Genomics UK (COG-UK) consortium | Jacquelyn Wynn, Mairead Hyland, The Lighthouse Lab in Alderley Park and Alex Alderton, Roberto Amato, Sonia Goncalves, Ewan Harrison, David K. Jackson, Ian Johnston, Dominic Kwiatkowski, Cordelia Langford, John Sillitoe on behalf of the Wellcome Sanger Institute COVID-19 Surveillance Team |
| EPI_ISL_609071 | Lighthouse Lab in Milton Keynes | Wellcome Sanger Institute for the COVID-19 Genomics UK (COG-UK) consortium | The Lighthouse Lab in Milton Keynes and Alex Alderton, Roberto Amato, Sonia Goncalves, Ewan Harrison, David K. Jackson, Ian Johnston, Dominic Kwiatkowski, Cordelia Langford, John Sillitoe on behalf of the Wellcome Sanger Institute COVID-19 Surveillance Team |
| EPI_ISL_609072 | Lighthouse Lab in Cambridge | Wellcome Sanger Institute for the COVID-19 Genomics UK (COG-UK) consortium | Rob Howes, The Lighthouse Lab in Cambridge and Alex Alderton, Roberto Amato, Sonia Goncalves, Ewan Harrison, David K. Jackson, Ian Johnston, Dominic Kwiatkowski, Cordelia Langford, John Sillitoe on behalf of the Wellcome Sanger Institute COVID-19 Surveillance Team |
| EPI_ISL_609073, EPI_ISL_609074, EPI_ISL_609075 | Lighthouse Lab in Alderley Park | Wellcome Sanger Institute for the COVID-19 Genomics UK (COG-UK) consortium | Jacquelyn Wynn, Mairead Hyland, The Lighthouse Lab in Alderley Park and Alex Alderton, Roberto Amato, Sonia Goncalves, Ewan Harrison, David K. Jackson, Ian Johnston, Dominic Kwiatkowski, Cordelia Langford, John Sillitoe on behalf of the Wellcome Sanger Institute COVID-19 Surveillance Team |
| EPI_ISL_609076 | Lighthouse Lab in Milton Keynes | Wellcome Sanger Institute for the COVID-19 Genomics UK (COG-UK) consortium | The Lighthouse Lab in Milton Keynes and Alex Alderton, Roberto Amato, Sonia Goncalves, Ewan Harrison, David K. Jackson, Ian Johnston, Dominic Kwiatkowski, Cordelia Langford, John Sillitoe on behalf of the Wellcome Sanger Institute COVID-19 Surveillance Team |
| EPI_ISL_609077 | Lighthouse Lab in Cambridge | Wellcome Sanger Institute for the COVID-19 Genomics UK (COG-UK) consortium | Rob Howes, The Lighthouse Lab in Cambridge and Alex Alderton, Roberto Amato, Sonia Goncalves, Ewan Harrison, David K. Jackson, Ian Johnston, Dominic Kwiatkowski, Cordelia Langford, John Sillitoe on behalf of the Wellcome Sanger Institute COVID-19 Surveillance Team |
| EPI_ISL_609078, EPI_ISL_609079, EPI_ISL_609080 | Lighthouse Lab in Alderley Park | Wellcome Sanger Institute for the COVID-19 Genomics UK (COG-UK) consortium | Jacquelyn Wynn, Mairead Hyland, The Lighthouse Lab in Alderley Park and Alex Alderton, Roberto Amato, Sonia Goncalves, Ewan Harrison, David K. Jackson, Ian Johnston, Dominic Kwiatkowski, Cordelia Langford, John Sillitoe on behalf of the Wellcome Sanger Institute COVID-19 Surveillance Team |
| EPI_ISL_609081, EPI_ISL_609082 | Lighthouse Lab in Cambridge | Wellcome Sanger Institute for the COVID-19 Genomics UK (COG-UK) consortium | Rob Howes, The Lighthouse Lab in Cambridge and Alex Alderton, Roberto Amato, Sonia Goncalves, Ewan Harrison, David K. Jackson, Ian Johnston, Dominic Kwiatkowski, Cordelia Langford, John Sillitoe on behalf of the Wellcome Sanger Institute COVID-19 Surveillance Team |
| EPI_ISL_609083 | Lighthouse Lab in Alderley Park | Wellcome Sanger Institute for the COVID-19 Genomics UK (COG-UK) consortium | Jacquelyn Wynn, Mairead Hyland, The Lighthouse Lab in Alderley Park and Alex Alderton, Roberto Amato, Sonia Goncalves, Ewan Harrison, David K. Jackson, Ian Johnston, Dominic Kwiatkowski, Cordelia Langford, John Sillitoe on behalf of the Wellcome Sanger Institute COVID-19 Surveillance Team |
| EPI_ISL_609084 | Lighthouse Lab in Milton Keynes | Wellcome Sanger Institute for the COVID-19 Genomics UK (COG-UK) consortium | The Lighthouse Lab in Milton Keynes and Alex Alderton, Roberto Amato, Sonia Goncalves, Ewan Harrison, David K. Jackson, Ian Johnston, Dominic Kwiatkowski, Cordelia Langford, John Sillitoe on behalf of the Wellcome Sanger Institute COVID-19 Surveillance Team |
| EPI_ISL_609085 | Lighthouse Lab in Alderley Park | Wellcome Sanger Institute for the COVID-19 Genomics UK (COG-UK) consortium | Jacquelyn Wynn, Mairead Hyland, The Lighthouse Lab in Alderley Park and Alex Alderton, Roberto Amato, Sonia Goncalves, Ewan Harrison, David K. Jackson, Ian Johnston, Dominic Kwiatkowski, Cordelia Langford, John Sillitoe on behalf of the Wellcome Sanger Institute COVID-19 Surveillance Team |
| EPI_ISL_609086 | Lighthouse Lab in Cambridge | Wellcome Sanger Institute for the COVID-19 Genomics UK (COG-UK) consortium | Rob Howes, The Lighthouse Lab in Cambridge and Alex Alderton, Roberto Amato, Sonia Goncalves, Ewan Harrison, David K. Jackson, Ian Johnston, Dominic Kwiatkowski, Cordelia Langford, John Sillitoe on behalf of the Wellcome Sanger Institute COVID-19 Surveillance Team |
| EPI_ISL_609087 | Lighthouse Lab in Alderley Park | Wellcome Sanger Institute for the COVID-19 Genomics UK (COG-UK) consortium | Jacquelyn Wynn, Mairead Hyland, The Lighthouse Lab in Alderley Park and Alex Alderton, Roberto Amato, Sonia Goncalves, Ewan Harrison, David K. Jackson, Ian Johnston, Dominic Kwiatkowski, Cordelia Langford, John Sillitoe on behalf of the Wellcome Sanger Institute COVID-19 Surveillance Team |
| EPI_ISL_609088 | Lighthouse Lab in Cambridge | Wellcome Sanger Institute for the COVID-19 Genomics UK (COG-UK) consortium | Rob Howes, The Lighthouse Lab in Cambridge and Alex Alderton, Roberto Amato, Sonia Goncalves, Ewan Harrison, David K. Jackson, Ian Johnston, Dominic Kwiatkowski, Cordelia Langford, John Sillitoe on behalf of the Wellcome Sanger Institute COVID-19 Surveillance Team |
| EPI_ISL_609089, EPI_ISL_609090 | Lighthouse Lab in Alderley Park | Wellcome Sanger Institute for the COVID-19 Genomics UK (COG-UK) consortium | Jacquelyn Wynn, Mairead Hyland, The Lighthouse Lab in Alderley Park and Alex Alderton, Roberto Amato, Sonia Goncalves, Ewan Harrison, David K. Jackson, Ian Johnston, Dominic Kwiatkowski, Cordelia Langford, John Sillitoe on behalf of the Wellcome Sanger Institute COVID-19 Surveillance Team |
| EPI_ISL_609091 | Lighthouse Lab in Cambridge | Wellcome Sanger Institute for the COVID-19 Genomics UK (COG-UK) consortium | Rob Howes, The Lighthouse Lab in Cambridge and Alex Alderton, Roberto Amato, Sonia Goncalves, Ewan Harrison, David K. Jackson, Ian Johnston, Dominic Kwiatkowski, Cordelia Langford, John Sillitoe on behalf of the Wellcome Sanger Institute COVID-19 Surveillance Team |
| EPI_ISL_609092, EPI_ISL_609093, EPI_ISL_609094, EPI_ISL_609095, EPI_ISL_609096 | Lighthouse Lab in Alderley Park | Wellcome Sanger Institute for the COVID-19 Genomics UK (COG-UK) consortium | Jacquelyn Wynn, Mairead Hyland, The Lighthouse Lab in Alderley Park and Alex Alderton, Roberto Amato, Sonia Goncalves, Ewan Harrison, David K. Jackson, Ian Johnston, Dominic Kwiatkowski, Cordelia Langford, John Sillitoe on behalf of the Wellcome Sanger Institute COVID-19 Surveillance Team |
| EPI_ISL_609097 | Lighthouse Lab in Cambridge | Wellcome Sanger Institute for the COVID-19 Genomics UK (COG-UK) consortium | Rob Howes, The Lighthouse Lab in Cambridge and Alex Alderton, Roberto Amato, Sonia Goncalves, Ewan Harrison, David K. Jackson, Ian Johnston, Dominic Kwiatkowski, Cordelia Langford, John Sillitoe on behalf of the Wellcome Sanger Institute COVID-19 Surveillance Team |
| EPI_ISL_609098 | Lighthouse Lab in Alderley Park | Wellcome Sanger Institute for the COVID-19 Genomics UK (COG-UK) consortium | Jacquelyn Wynn, Mairead Hyland, The Lighthouse Lab in Alderley Park and Alex Alderton, Roberto Amato, Sonia Goncalves, Ewan Harrison, David K. Jackson, Ian Johnston, Dominic Kwiatkowski, Cordelia Langford, John Sillitoe on behalf of the Wellcome Sanger Institute COVID-19 Surveillance Team |
| EPI_ISL_609099, EPI_ISL_609100, EPI_ISL_609101 | Lighthouse Lab in Cambridge | Wellcome Sanger Institute for the COVID-19 Genomics UK (COG-UK) consortium | Rob Howes, The Lighthouse Lab in Cambridge and Alex Alderton, Roberto Amato, Sonia Goncalves, Ewan Harrison, David K. Jackson, Ian Johnston, Dominic Kwiatkowski, Cordelia Langford, John Sillitoe on behalf of the Wellcome Sanger Institute COVID-19 Surveillance Team |

| | | | |
|---|---|---|---|
| EPI_ISL_609102, EPI_ISL_609103, EPI_ISL_609104, EPI_ISL_609105, EPI_ISL_609106, EPI_ISL_609107 | Lighthouse Lab in Alderley Park | Wellcome Sanger Institute for the COVID-19 Genomics UK (COG-UK) consortium | Jacquelyn Wynn, Mairead Hyland, The Lighthouse Lab in Alderley Park and Alex Alderton, Roberto Amato, Sonia Goncalves, Ewan Harrison, David K. Jackson, Ian Johnston, Dominic Kwiatkowski, Cordelia Langford, John Sillitoe on behalf of the Wellcome Sanger Institute COVID-19 Surveillance Team |
| EPI_ISL_609108 | Lighthouse Lab in Milton Keynes | Wellcome Sanger Institute for the COVID-19 Genomics UK (COG-UK) consortium | The Lighthouse Lab in Milton Keynes and Alex Alderton, Roberto Amato, Sonia Goncalves, Ewan Harrison, David K. Jackson, Ian Johnston, Dominic Kwiatkowski, Cordelia Langford, John Sillitoe on behalf of the Wellcome Sanger Institute COVID-19 Surveillance Team |
| EPI_ISL_609109 | Lighthouse Lab in Alderley Park | Wellcome Sanger Institute for the COVID-19 Genomics UK (COG-UK) consortium | Jacquelyn Wynn, Mairead Hyland, The Lighthouse Lab in Alderley Park and Alex Alderton, Roberto Amato, Sonia Goncalves, Ewan Harrison, David K. Jackson, Ian Johnston, Dominic Kwiatkowski, Cordelia Langford, John Sillitoe on behalf of the Wellcome Sanger Institute COVID-19 Surveillance Team |
| EPI_ISL_609110 | Lighthouse Lab in Milton Keynes | Wellcome Sanger Institute for the COVID-19 Genomics UK (COG-UK) consortium | The Lighthouse Lab in Milton Keynes and Alex Alderton, Roberto Amato, Sonia Goncalves, Ewan Harrison, David K. Jackson, Ian Johnston, Dominic Kwiatkowski, Cordelia Langford, John Sillitoe on behalf of the Wellcome Sanger Institute COVID-19 Surveillance Team |
| EPI_ISL_609111 | Lighthouse Lab in Cambridge | Wellcome Sanger Institute for the COVID-19 Genomics UK (COG-UK) consortium | Rob Howes, The Lighthouse Lab in Cambridge and Alex Alderton, Roberto Amato, Sonia Goncalves, Ewan Harrison, David K. Jackson, Ian Johnston, Dominic Kwiatkowski, Cordelia Langford, John Sillitoe on behalf of the Wellcome Sanger Institute COVID-19 Surveillance Team |
| EPI_ISL_609112 | Lighthouse Lab in Alderley Park | Wellcome Sanger Institute for the COVID-19 Genomics UK (COG-UK) consortium | Jacquelyn Wynn, Mairead Hyland, The Lighthouse Lab in Alderley Park and Alex Alderton, Roberto Amato, Sonia Goncalves, Ewan Harrison, David K. Jackson, Ian Johnston, Dominic Kwiatkowski, Cordelia Langford, John Sillitoe on behalf of the Wellcome Sanger Institute COVID-19 Surveillance Team |
| EPI_ISL_609113, EPI_ISL_609114 | Lighthouse Lab in Cambridge | Wellcome Sanger Institute for the COVID-19 Genomics UK (COG-UK) consortium | Rob Howes, The Lighthouse Lab in Cambridge and Alex Alderton, Roberto Amato, Sonia Goncalves, Ewan Harrison, David K. Jackson, Ian Johnston, Dominic Kwiatkowski, Cordelia Langford, John Sillitoe on behalf of the Wellcome Sanger Institute COVID-19 Surveillance Team |
| EPI_ISL_609115, EPI_ISL_609116, EPI_ISL_609117 | Lighthouse Lab in Milton Keynes | Wellcome Sanger Institute for the COVID-19 Genomics UK (COG-UK) consortium | The Lighthouse Lab in Milton Keynes and Alex Alderton, Roberto Amato, Sonia Goncalves, Ewan Harrison, David K. Jackson, Ian Johnston, Dominic Kwiatkowski, Cordelia Langford, John Sillitoe on behalf of the Wellcome Sanger Institute COVID-19 Surveillance Team |
| EPI_ISL_609118, EPI_ISL_609119 | Lighthouse Lab in Cambridge | Wellcome Sanger Institute for the COVID-19 Genomics UK (COG-UK) consortium | Rob Howes, The Lighthouse Lab in Cambridge and Alex Alderton, Roberto Amato, Sonia Goncalves, Ewan Harrison, David K. Jackson, Ian Johnston, Dominic Kwiatkowski, Cordelia Langford, John Sillitoe on behalf of the Wellcome Sanger Institute COVID-19 Surveillance Team |
| EPI_ISL_609120 | Lighthouse Lab in Alderley Park | Wellcome Sanger Institute for the COVID-19 Genomics UK (COG-UK) consortium | Jacquelyn Wynn, Mairead Hyland, The Lighthouse Lab in Alderley Park and Alex Alderton, Roberto Amato, Sonia Goncalves, Ewan Harrison, David K. Jackson, Ian Johnston, Dominic Kwiatkowski, Cordelia Langford, John Sillitoe on behalf of the Wellcome Sanger Institute COVID-19 Surveillance Team |
| EPI_ISL_609121 | Lighthouse Lab in Cambridge | Wellcome Sanger Institute for the COVID-19 Genomics UK (COG-UK) Consortium | Rob Howes, The Lighthouse Lab in Cambridge and Alex Alderton, Roberto Amato, Sonia Goncalves, Ewan Harrison, David K. Jackson, Ian Johnston, Dominic Kwiatkowski, Cordelia Langford, John Sillitoe on behalf of the Wellcome Sanger Institute COVID-19 Surveillance Team |
| EPI_ISL_609122, EPI_ISL_609123 | Lighthouse Lab in Cambridge | Wellcome Sanger Institute for the COVID-19 Genomics UK (COG-UK) consortium | Rob Howes, The Lighthouse Lab in Cambridge and Alex Alderton, Roberto Amato, Sonia Goncalves, Ewan Harrison, David K. Jackson, Ian Johnston, Dominic Kwiatkowski, Cordelia Langford, John Sillitoe on behalf of the Wellcome Sanger Institute COVID-19 Surveillance Team |
| EPI_ISL_609124, EPI_ISL_609125 | Lighthouse Lab in Alderley Park | Wellcome Sanger Institute for the COVID-19 Genomics UK (COG-UK) consortium | Jacquelyn Wynn, Mairead Hyland, The Lighthouse Lab in Alderley Park and Alex Alderton, Roberto Amato, Sonia Goncalves, Ewan Harrison, David K. Jackson, Ian Johnston, Dominic Kwiatkowski, Cordelia Langford, John Sillitoe on behalf of the Wellcome Sanger Institute COVID-19 Surveillance Team |
| EPI_ISL_609126 | Lighthouse Lab in Milton Keynes | Wellcome Sanger Institute for the COVID-19 Genomics UK (COG-UK) consortium | The Lighthouse Lab in Milton Keynes and Alex Alderton, Roberto Amato, Sonia Goncalves, Ewan Harrison, David K. Jackson, Ian Johnston, Dominic Kwiatkowski, Cordelia Langford, John Sillitoe on behalf of the Wellcome Sanger Institute COVID-19 Surveillance Team |
| EPI_ISL_609127 | Lighthouse Lab in Alderley Park | Wellcome Sanger Institute for the COVID-19 Genomics UK (COG-UK) consortium | Jacquelyn Wynn, Mairead Hyland, The Lighthouse Lab in Alderley Park and Alex Alderton, Roberto Amato, Sonia Goncalves, Ewan Harrison, David K. Jackson, Ian Johnston, Dominic Kwiatkowski, Cordelia Langford, John Sillitoe on behalf of the Wellcome Sanger Institute COVID-19 Surveillance Team |
| EPI_ISL_609128 | Lighthouse Lab in Milton Keynes | Wellcome Sanger Institute for the COVID-19 Genomics UK (COG-UK) consortium | The Lighthouse Lab in Milton Keynes and Alex Alderton, Roberto Amato, Sonia Goncalves, Ewan Harrison, David K. Jackson, Ian Johnston, Dominic Kwiatkowski, Cordelia Langford, John Sillitoe on behalf of the Wellcome Sanger Institute COVID-19 Surveillance Team |
| EPI_ISL_609129 | Lighthouse Lab in Cambridge | Wellcome Sanger Institute for the COVID-19 Genomics UK (COG-UK) consortium | Rob Howes, The Lighthouse Lab in Cambridge and Alex Alderton, Roberto Amato, Sonia Goncalves, Ewan Harrison, David K. Jackson, Ian Johnston, Dominic Kwiatkowski, Cordelia Langford, John Sillitoe on behalf of the Wellcome Sanger Institute COVID-19 Surveillance Team |
| EPI_ISL_609130 | Lighthouse Lab in Alderley Park | Wellcome Sanger Institute for the COVID-19 Genomics UK (COG-UK) consortium | Jacquelyn Wynn, Mairead Hyland, The Lighthouse Lab in Alderley Park and Alex Alderton, Roberto Amato, Sonia Goncalves, Ewan Harrison, David K. Jackson, Ian Johnston, Dominic Kwiatkowski, Cordelia Langford, John Sillitoe on behalf of the Wellcome Sanger Institute COVID-19 Surveillance Team |
| EPI_ISL_609131 | Lighthouse Lab in Cambridge | Wellcome Sanger Institute for the COVID-19 Genomics UK (COG-UK) consortium | Rob Howes, The Lighthouse Lab in Cambridge and Alex Alderton, Roberto Amato, Sonia Goncalves, Ewan Harrison, David K. Jackson, Ian Johnston, Dominic Kwiatkowski, Cordelia Langford, John Sillitoe on behalf of the Wellcome Sanger Institute COVID-19 Surveillance Team |
| EPI_ISL_609132, EPI_ISL_609133, EPI_ISL_609134, EPI_ISL_609135 | Lighthouse Lab in Alderley Park | Wellcome Sanger Institute for the COVID-19 Genomics UK (COG-UK) consortium | Jacquelyn Wynn, Mairead Hyland, The Lighthouse Lab in Alderley Park and Alex Alderton, Roberto Amato, Sonia Goncalves, Ewan Harrison, David K. Jackson, Ian Johnston, Dominic Kwiatkowski, Cordelia Langford, John Sillitoe on behalf of the Wellcome Sanger Institute COVID-19 Surveillance Team |
| EPI_ISL_609136, EPI_ISL_609137, EPI_ISL_609138, EPI_ISL_609139, EPI_ISL_609140 | Lighthouse Lab in Milton Keynes | Wellcome Sanger Institute for the COVID-19 Genomics UK (COG-UK) consortium | The Lighthouse Lab in Milton Keynes and Alex Alderton, Roberto Amato, Sonia Goncalves, Ewan Harrison, David K. Jackson, Ian Johnston, Dominic Kwiatkowski, Cordelia Langford, John Sillitoe on behalf of the Wellcome Sanger Institute COVID-19 Surveillance Team |
| EPI_ISL_609141 | Lighthouse Lab in Alderley Park | Wellcome Sanger Institute for the COVID-19 Genomics UK (COG-UK) consortium | Jacquelyn Wynn, Mairead Hyland, The Lighthouse Lab in Alderley Park and Alex Alderton, Roberto Amato, Sonia Goncalves, Ewan Harrison, David K. Jackson, Ian Johnston, Dominic Kwiatkowski, Cordelia Langford, John Sillitoe on behalf of the Wellcome Sanger Institute COVID-19 Surveillance Team |
| EPI_ISL_609142, EPI_ISL_609143 | Lighthouse Lab in Cambridge | Wellcome Sanger Institute for the COVID-19 Genomics UK (COG-UK) consortium | Rob Howes, The Lighthouse Lab in Cambridge and Alex Alderton, Roberto Amato, Sonia Goncalves, Ewan Harrison, David K. Jackson, Ian Johnston, Dominic Kwiatkowski, Cordelia Langford, John Sillitoe on behalf of the Wellcome Sanger Institute COVID-19 Surveillance Team |
| EPI_ISL_609144 | Lighthouse Lab in Alderley Park | Wellcome Sanger Institute for the COVID-19 Genomics UK (COG-UK) consortium | Jacquelyn Wynn, Mairead Hyland, The Lighthouse Lab in Alderley Park and Alex Alderton, Roberto Amato, Sonia Goncalves, Ewan Harrison, David K. Jackson, Ian Johnston, Dominic Kwiatkowski, Cordelia Langford, John Sillitoe on behalf of the Wellcome Sanger Institute COVID-19 Surveillance Team |
| EPI_ISL_609145 | Lighthouse Lab in Cambridge | Wellcome Sanger Institute for the COVID-19 Genomics UK (COG-UK) consortium | Rob Howes, The Lighthouse Lab in Cambridge and Alex Alderton, Roberto Amato, Sonia Goncalves, Ewan Harrison, David K. Jackson, Ian Johnston, Dominic Kwiatkowski, Cordelia Langford, John Sillitoe on behalf of the Wellcome Sanger Institute COVID-19 Surveillance Team |
| EPI_ISL_609146, EPI_ISL_609147, EPI_ISL_609148 | Lighthouse Lab in Alderley Park | Wellcome Sanger Institute for the COVID-19 Genomics UK (COG-UK) consortium | Jacquelyn Wynn, Mairead Hyland, The Lighthouse Lab in Alderley Park and Alex Alderton, Roberto Amato, Sonia Goncalves, Ewan Harrison, David K. Jackson, Ian Johnston, Dominic Kwiatkowski, Cordelia Langford, John Sillitoe on behalf of the Wellcome Sanger Institute COVID-19 Surveillance Team |
| EPI_ISL_609149 | Lighthouse Lab in Milton Keynes | Wellcome Sanger Institute for the COVID-19 Genomics UK (COG-UK) consortium | The Lighthouse Lab in Milton Keynes and Alex Alderton, Roberto Amato, Sonia Goncalves, Ewan Harrison, David K. Jackson, Ian Johnston, Dominic Kwiatkowski, Cordelia Langford, John Sillitoe on behalf of the Wellcome Sanger Institute COVID-19 Surveillance Team |
| EPI_ISL_609150 | Lighthouse Lab in Alderley Park | Wellcome Sanger Institute for the COVID-19 Genomics UK (COG-UK) consortium | Jacquelyn Wynn, Mairead Hyland, The Lighthouse Lab in Alderley Park and Alex Alderton, Roberto Amato, Sonia Goncalves, Ewan Harrison, David K. Jackson, Ian Johnston, Dominic Kwiatkowski, Cordelia Langford, John Sillitoe on behalf of the Wellcome Sanger Institute COVID-19 Surveillance Team |
| EPI_ISL_609151 | Lighthouse Lab in Milton Keynes | Wellcome Sanger Institute for the COVID-19 Genomics UK (COG-UK) consortium | The Lighthouse Lab in Milton Keynes and Alex Alderton, Roberto Amato, Sonia Goncalves, Ewan Harrison, David K. Jackson, Ian Johnston, Dominic Kwiatkowski, Cordelia Langford, John Sillitoe on behalf of the Wellcome Sanger Institute COVID-19 Surveillance Team |
| EPI_ISL_609152, EPI_ISL_609153, EPI_ISL_609154 | Lighthouse Lab in Alderley Park | Wellcome Sanger Institute for the COVID-19 Genomics UK (COG-UK) consortium | Jacquelyn Wynn, Mairead Hyland, The Lighthouse Lab in Alderley Park and Alex Alderton, Roberto Amato, Sonia Goncalves, Ewan Harrison, David K. Jackson, Ian Johnston, Dominic Kwiatkowski, Cordelia Langford, John Sillitoe on behalf of the Wellcome Sanger Institute COVID-19 Surveillance Team |
| EPI_ISL_609155 | Lighthouse Lab in Milton Keynes | Wellcome Sanger Institute for the COVID-19 Genomics UK (COG-UK) consortium | The Lighthouse Lab in Milton Keynes and Alex Alderton, Roberto Amato, Sonia Goncalves, Ewan Harrison, David K. Jackson, Ian Johnston, Dominic Kwiatkowski, Cordelia Langford, John Sillitoe on behalf of the Wellcome Sanger Institute COVID-19 Surveillance Team |
| EPI_ISL_609156 | Lighthouse Lab in Cambridge | Wellcome Sanger Institute for the COVID-19 Genomics UK (COG-UK) consortium | Rob Howes, The Lighthouse Lab in Cambridge and Alex Alderton, Roberto Amato, Sonia Goncalves, Ewan Harrison, David K. Jackson, Ian Johnston, Dominic Kwiatkowski, Cordelia Langford, John Sillitoe on behalf of the Wellcome Sanger Institute COVID-19 Surveillance Team |
| EPI_ISL_609157 | Lighthouse Lab in Alderley Park | Wellcome Sanger Institute for the COVID-19 Genomics UK (COG-UK) consortium | Jacquelyn Wynn, Mairead Hyland, The Lighthouse Lab in Alderley Park and Alex Alderton, Roberto Amato, Sonia Goncalves, Ewan Harrison, David K. Jackson, Ian Johnston, Dominic Kwiatkowski, Cordelia Langford, John Sillitoe on behalf of the Wellcome Sanger Institute COVID-19 Surveillance Team |

EPI_ISL_609158, EPI_ISL_609159, EPI_ISL_609160, EPI_ISL_609161, EPI_ISL_609162, EPI_ISL_609163, EPI_ISL_609164, EPI_ISL_609165, EPI_ISL_609166, EPI_ISL_609167, EPI_ISL_609168, EPI_ISL_609169, EPI_ISL_609170, EPI_ISL_609171, EPI_ISL_609172, EPI_ISL_609173, EPI_ISL_609174, EPI_ISL_609175, EPI_ISL_609176, EPI_ISL_609177

| | | | |
|---|---|---|---|
| see above | Lighthouse Lab in Cambridge | Wellcome Sanger Institute for the COVID-19 Genomics UK | Rob Howes, The Lighthouse Lab in Cambridge and Alex Alderton, Roberto Amato, Sonia Goncalves, Ewan Harrison, David K. Jackson, Ian Johnston, Dominic |

| | | | |
|---|---|---|---|
| | | (COG-UK) consortium | Kwiatkowski, Cordelia Langford, John Sillitoe on behalf of the Wellcome Sanger Institute COVID-19 Surveillance Team |
| EPI_ISL_609178, EPI_ISL_609179 | Lighthouse Lab in Glasgow | Wellcome Sanger Institute for the COVID-19 Genomics UK (COG-UK) consortium | Harper VanSteenhouse, Yumi Kasai, David Gray, Carol Clugston, Anna Dominiczak and Alex Alderton, Roberto Amato, Sonia Goncalves, Ewan Harrison, David K. Jackson, Ian Johnston, Dominic Kwiatkowski, Cordelia Langford, John Sillitoe on behalf of the Wellcome Sanger Institute COVID-19 Surveillance Team |
| EPI_ISL_609180, EPI_ISL_609181 | Lighthouse Lab in Cambridge | Wellcome Sanger Institute for the COVID-19 Genomics UK (COG-UK) consortium | Rob Howes, The Lighthouse Lab in Cambridge and Alex Alderton, Roberto Amato, Sonia Goncalves, Ewan Harrison, David K. Jackson, Ian Johnston, Dominic Kwiatkowski, Cordelia Langford, John Sillitoe on behalf of the Wellcome Sanger Institute COVID-19 Surveillance Team |
| EPI_ISL_609182 | Lighthouse Lab in Glasgow | Wellcome Sanger Institute for the COVID-19 Genomics UK (COG-UK) consortium | Harper VanSteenhouse, Yumi Kasai, David Gray, Carol Clugston, Anna Dominiczak and Alex Alderton, Roberto Amato, Sonia Goncalves, Ewan Harrison, David K. Jackson, Ian Johnston, Dominic Kwiatkowski, Cordelia Langford, John Sillitoe on behalf of the Wellcome Sanger Institute COVID-19 Surveillance Team |
| EPI_ISL_609183 | Lighthouse Lab in Cambridge | Wellcome Sanger Institute for the COVID-19 Genomics UK (COG-UK) consortium | Rob Howes, The Lighthouse Lab in Cambridge and Alex Alderton, Roberto Amato, Sonia Goncalves, Ewan Harrison, David K. Jackson, Ian Johnston, Dominic Kwiatkowski, Cordelia Langford, John Sillitoe on behalf of the Wellcome Sanger Institute COVID-19 Surveillance Team |
| EPI_ISL_609184, EPI_ISL_609185, EPI_ISL_609186 | Lighthouse Lab in Glasgow | Wellcome Sanger Institute for the COVID-19 Genomics UK (COG-UK) consortium | Harper VanSteenhouse, Yumi Kasai, David Gray, Carol Clugston, Anna Dominiczak and Alex Alderton, Roberto Amato, Sonia Goncalves, Ewan Harrison, David K. Jackson, Ian Johnston, Dominic Kwiatkowski, Cordelia Langford, John Sillitoe on behalf of the Wellcome Sanger Institute COVID-19 Surveillance Team |
| EPI_ISL_609187 | Lighthouse Lab in Glasgow | Wellcome Sanger Institute for the COVID-19 Genomics UK (COG-UK) Consortium | Harper VanSteenhouse, Yumi Kasai, David Gray, Carol Clugston, Anna Dominiczak and Alex Alderton, Roberto Amato, Sonia Goncalves, Ewan Harrison, David K. Jackson, Ian Johnston, Dominic Kwiatkowski, Cordelia Langford, John Sillitoe on behalf of the Wellcome Sanger Institute COVID-19 Surveillance Team |
| EPI_ISL_609188 | Lighthouse Lab in Cambridge | Wellcome Sanger Institute for the COVID-19 Genomics UK (COG-UK) consortium | Rob Howes, The Lighthouse Lab in Cambridge and Alex Alderton, Roberto Amato, Sonia Goncalves, Ewan Harrison, David K. Jackson, Ian Johnston, Dominic Kwiatkowski, Cordelia Langford, John Sillitoe on behalf of the Wellcome Sanger Institute COVID-19 Surveillance Team |
| EPI_ISL_609189, EPI_ISL_609190, EPI_ISL_609191 | Lighthouse Lab in Glasgow | Wellcome Sanger Institute for the COVID-19 Genomics UK (COG-UK) consortium | Harper VanSteenhouse, Yumi Kasai, David Gray, Carol Clugston, Anna Dominiczak and Alex Alderton, Roberto Amato, Sonia Goncalves, Ewan Harrison, David K. Jackson, Ian Johnston, Dominic Kwiatkowski, Cordelia Langford, John Sillitoe on behalf of the Wellcome Sanger Institute COVID-19 Surveillance Team |
| EPI_ISL_609192 | Lighthouse Lab in Glasgow | Wellcome Sanger Institute for the COVID-19 Genomics UK (COG-UK) Consortium | Harper VanSteenhouse, Yumi Kasai, David Gray, Carol Clugston, Anna Dominiczak and Alex Alderton, Roberto Amato, Sonia Goncalves, Ewan Harrison, David K. Jackson, Ian Johnston, Dominic Kwiatkowski, Cordelia Langford, John Sillitoe on behalf of the Wellcome Sanger Institute COVID-19 Surveillance Team |
| EPI_ISL_609193 | Lighthouse Lab in Cambridge | Wellcome Sanger Institute for the COVID-19 Genomics UK (COG-UK) consortium | Rob Howes, The Lighthouse Lab in Cambridge and Alex Alderton, Roberto Amato, Sonia Goncalves, Ewan Harrison, David K. Jackson, Ian Johnston, Dominic Kwiatkowski, Cordelia Langford, John Sillitoe on behalf of the Wellcome Sanger Institute COVID-19 Surveillance Team |
| EPI_ISL_609194 | Lighthouse Lab in Glasgow | Wellcome Sanger Institute for the COVID-19 Genomics UK (COG-UK) consortium | Harper VanSteenhouse, Yumi Kasai, David Gray, Carol Clugston, Anna Dominiczak and Alex Alderton, Roberto Amato, Sonia Goncalves, Ewan Harrison, David K. Jackson, Ian Johnston, Dominic Kwiatkowski, Cordelia Langford, John Sillitoe on behalf of the Wellcome Sanger Institute COVID-19 Surveillance Team |
| EPI_ISL_609195, EPI_ISL_609196 | Lighthouse Lab in Cambridge | Wellcome Sanger Institute for the COVID-19 Genomics UK (COG-UK) consortium | Rob Howes, The Lighthouse Lab in Cambridge and Alex Alderton, Roberto Amato, Sonia Goncalves, Ewan Harrison, David K. Jackson, Ian Johnston, Dominic Kwiatkowski, Cordelia Langford, John Sillitoe on behalf of the Wellcome Sanger Institute COVID-19 Surveillance Team |
| EPI_ISL_609197, EPI_ISL_609198, EPI_ISL_609199 | Lighthouse Lab in Glasgow | Wellcome Sanger Institute for the COVID-19 Genomics UK (COG-UK) consortium | Harper VanSteenhouse, Yumi Kasai, David Gray, Carol Clugston, Anna Dominiczak and Alex Alderton, Roberto Amato, Sonia Goncalves, Ewan Harrison, David K. Jackson, Ian Johnston, Dominic Kwiatkowski, Cordelia Langford, John Sillitoe on behalf of the Wellcome Sanger Institute COVID-19 Surveillance Team |
| EPI_ISL_609200, EPI_ISL_609201 | Lighthouse Lab in Cambridge | Wellcome Sanger Institute for the COVID-19 Genomics UK (COG-UK) consortium | Rob Howes, The Lighthouse Lab in Cambridge and Alex Alderton, Roberto Amato, Sonia Goncalves, Ewan Harrison, David K. Jackson, Ian Johnston, Dominic Kwiatkowski, Cordelia Langford, John Sillitoe on behalf of the Wellcome Sanger Institute COVID-19 Surveillance Team |
| EPI_ISL_609202 | Lighthouse Lab in Glasgow | Wellcome Sanger Institute for the COVID-19 Genomics UK (COG-UK) consortium | Harper VanSteenhouse, Yumi Kasai, David Gray, Carol Clugston, Anna Dominiczak and Alex Alderton, Roberto Amato, Sonia Goncalves, Ewan Harrison, David K. Jackson, Ian Johnston, Dominic Kwiatkowski, Cordelia Langford, John Sillitoe on behalf of the Wellcome Sanger Institute COVID-19 Surveillance Team |
| EPI_ISL_609203, EPI_ISL_609204, EPI_ISL_609205 | Lighthouse Lab in Cambridge | Wellcome Sanger Institute for the COVID-19 Genomics UK (COG-UK) consortium | Rob Howes, The Lighthouse Lab in Cambridge and Alex Alderton, Roberto Amato, Sonia Goncalves, Ewan Harrison, David K. Jackson, Ian Johnston, Dominic Kwiatkowski, Cordelia Langford, John Sillitoe on behalf of the Wellcome Sanger Institute COVID-19 Surveillance Team |
| EPI_ISL_609206 | Lighthouse Lab in Glasgow | Wellcome Sanger Institute for the COVID-19 Genomics UK (COG-UK) consortium | Harper VanSteenhouse, Yumi Kasai, David Gray, Carol Clugston, Anna Dominiczak and Alex Alderton, Roberto Amato, Sonia Goncalves, Ewan Harrison, David K. Jackson, Ian Johnston, Dominic Kwiatkowski, Cordelia Langford, John Sillitoe on behalf of the Wellcome Sanger Institute COVID-19 Surveillance Team |
| EPI_ISL_609207, EPI_ISL_609208 | Lighthouse Lab in Alderley Park | Wellcome Sanger Institute for the COVID-19 Genomics UK (COG-UK) consortium | Jacquelyn Wynn, Mairead Hyland, The Lighthouse Lab in Alderley Park and Alex Alderton, Roberto Amato, Sonia Goncalves, Ewan Harrison, David K. Jackson, Ian Johnston, Dominic Kwiatkowski, Cordelia Langford, John Sillitoe on behalf of the Wellcome Sanger Institute COVID-19 Surveillance Team |
| EPI_ISL_609209 | Lighthouse Lab in Alderley Park | Wellcome Sanger Institute for the COVID-19 Genomics UK (COG-UK) Consortium | Jacquelyn Wynn, Mairead Hyland, The Lighthouse Lab in Alderley Park and Alex Alderton, Roberto Amato, Sonia Goncalves, Ewan Harrison, David K. Jackson, Ian Johnston, Dominic Kwiatkowski, Cordelia Langford, John Sillitoe on behalf of the Wellcome Sanger Institute COVID-19 Surveillance Team |
| EPI_ISL_609210, EPI_ISL_609211, EPI_ISL_609212, EPI_ISL_609213, EPI_ISL_609214, EPI_ISL_609215, EPI_ISL_609216, EPI_ISL_609217, EPI_ISL_609218 | Lighthouse Lab in Alderley Park | Wellcome Sanger Institute for the COVID-19 Genomics UK (COG-UK) consortium | Jacquelyn Wynn, Mairead Hyland, The Lighthouse Lab in Alderley Park and Alex Alderton, Roberto Amato, Sonia Goncalves, Ewan Harrison, David K. Jackson, Ian Johnston, Dominic Kwiatkowski, Cordelia Langford, John Sillitoe on behalf of the Wellcome Sanger Institute COVID-19 Surveillance Team |
| EPI_ISL_609297 | Lighthouse Lab in Milton Keynes | Wellcome Sanger Institute for the COVID-19 Genomics UK (COG-UK) consortium | The Lighthouse Lab in Milton Keynes and Alex Alderton, Roberto Amato, Sonia Goncalves, Ewan Harrison, David K. Jackson, Ian Johnston, Dominic Kwiatkowski, Cordelia Langford, John Sillitoe on behalf of the Wellcome Sanger Institute COVID-19 Surveillance Team |
| EPI_ISL_609309, EPI_ISL_609310, EPI_ISL_609311, EPI_ISL_609317, EPI_ISL_609319, EPI_ISL_609321, EPI_ISL_609325, EPI_ISL_609331, EPI_ISL_609336, EPI_ISL_609341, EPI_ISL_609344, EPI_ISL_609345, EPI_ISL_609349, EPI_ISL_609352, EPI_ISL_609354, EPI_ISL_609359, EPI_ISL_609361, EPI_ISL_609362, EPI_ISL_609372, EPI_ISL_609375, EPI_ISL_609384, EPI_ISL_609387, EPI_ISL_609388, EPI_ISL_609391, EPI_ISL_609392, EPI_ISL_609395, EPI_ISL_609397, EPI_ISL_609399, EPI_ISL_609400, EPI_ISL_609402, EPI_ISL_609408, EPI_ISL_609410, EPI_ISL_609414, EPI_ISL_609421, EPI_ISL_609423, EPI_ISL_609424, EPI_ISL_609425, EPI_ISL_609431, EPI_ISL_609432, EPI_ISL_609435, EPI_ISL_609436, EPI_ISL_609437, EPI_ISL_609438, EPI_ISL_609439, EPI_ISL_609443, EPI_ISL_609445, EPI_ISL_609447, EPI_ISL_609451, EPI_ISL_609453, EPI_ISL_609457, EPI_ISL_609460, EPI_ISL_609461, EPI_ISL_609462, EPI_ISL_609470, EPI_ISL_609471, EPI_ISL_609479, EPI_ISL_609484, EPI_ISL_609487, EPI_ISL_609488, EPI_ISL_609491, EPI_ISL_609492, EPI_ISL_609496, EPI_ISL_609497, EPI_ISL_609504, EPI_ISL_609506, EPI_ISL_609508, EPI_ISL_609512, EPI_ISL_609514, EPI_ISL_609519, EPI_ISL_609522, EPI_ISL_609531, EPI_ISL_609538, EPI_ISL_609540, EPI_ISL_609551, EPI_ISL_609552, EPI_ISL_609568, EPI_ISL_609575, EPI_ISL_609577, EPI_ISL_609596, EPI_ISL_609605, EPI_ISL_609629, EPI_ISL_609635, EPI_ISL_609636, EPI_ISL_609638, EPI_ISL_609643, EPI_ISL_609661, EPI_ISL_609667, EPI_ISL_609679, EPI_ISL_609682, EPI_ISL_609693, EPI_ISL_609699, EPI_ISL_609711, EPI_ISL_609714, EPI_ISL_609720, EPI_ISL_609734, EPI_ISL_609737 | | | |
| see above | Lighthouse Lab in Cambridge | Wellcome Sanger Institute for the COVID-19 Genomics UK (COG-UK) consortium | Rob Howes, The Lighthouse Lab in Cambridge and Alex Alderton, Roberto Amato, Sonia Goncalves, Ewan Harrison, David K. Jackson, Ian Johnston, Dominic Kwiatkowski, Cordelia Langford, John Sillitoe on behalf of the Wellcome Sanger Institute COVID-19 Surveillance Team |
| EPI_ISL_609746 | Lighthouse Lab in Cambridge | Wellcome Sanger Institute for the COVID-19 Genomics UK (COG-UK) Consortium | Rob Howes, The Lighthouse Lab in Cambridge and Alex Alderton, Roberto Amato, Sonia Goncalves, Ewan Harrison, David K. Jackson, Ian Johnston, Dominic Kwiatkowski, Cordelia Langford, John Sillitoe on behalf of the Wellcome Sanger Institute COVID-19 Surveillance Team |
| EPI_ISL_609747, EPI_ISL_609749, EPI_ISL_609763, EPI_ISL_609785, EPI_ISL_609786, EPI_ISL_609789, EPI_ISL_609791, EPI_ISL_609792 | Lighthouse Lab in Cambridge | Wellcome Sanger Institute for the COVID-19 Genomics UK (COG-UK) consortium | Rob Howes, The Lighthouse Lab in Cambridge and Alex Alderton, Roberto Amato, Sonia Goncalves, Ewan Harrison, David K. Jackson, Ian Johnston, Dominic Kwiatkowski, Cordelia Langford, John Sillitoe on behalf of the Wellcome Sanger Institute COVID-19 Surveillance Team |
| EPI_ISL_609853, EPI_ISL_609854, EPI_ISL_609855, EPI_ISL_609856, EPI_ISL_609858, EPI_ISL_609859, EPI_ISL_609860, EPI_ISL_609861, EPI_ISL_609862, EPI_ISL_609863, EPI_ISL_609864, EPI_ISL_609865, EPI_ISL_609866, EPI_ISL_609867, EPI_ISL_609868, EPI_ISL_609869, EPI_ISL_609870, EPI_ISL_609871, EPI_ISL_609872, EPI_ISL_609873, EPI_ISL_609874, EPI_ISL_609875, EPI_ISL_609876, EPI_ISL_609877, EPI_ISL_609878, EPI_ISL_609879, EPI_ISL_609880, EPI_ISL_609881, EPI_ISL_609882, EPI_ISL_609883, EPI_ISL_609884, EPI_ISL_609885, EPI_ISL_609886, EPI_ISL_609887, EPI_ISL_609888, EPI_ISL_609889, EPI_ISL_609897, EPI_ISL_609909 | | | |
| see above | Respiratory Virus Unit, Microbiology Services Colindale, | Respiratory Virus Unit, Microbiology Services Colindale, | PHE Covid Sequencing Team |

| | | | |
|---|---|---|---|
| | Public Health England | Public Health England | |
| EPI_ISL_610048, EPI_ISL_610050, EPI_ISL_610053, EPI_ISL_610054, EPI_ISL_610055, EPI_ISL_610056, EPI_ISL_610057, EPI_ISL_610058, EPI_ISL_610059, EPI_ISL_610061, EPI_ISL_610062, EPI_ISL_610063, EPI_ISL_610064, EPI_ISL_610065, EPI_ISL_610066, EPI_ISL_610067, EPI_ISL_610068, EPI_ISL_610069, EPI_ISL_610070, EPI_ISL_610071, EPI_ISL_610072, EPI_ISL_610073, EPI_ISL_610075, EPI_ISL_610076, EPI_ISL_610078, EPI_ISL_610080, EPI_ISL_610082, EPI_ISL_610084, EPI_ISL_610085, EPI_ISL_610087, EPI_ISL_610088, EPI_ISL_610091, EPI_ISL_610092, EPI_ISL_610100 | | | |
| see above | University of Michigan Clinical Microbiology Laboratory | Lauring Lab, University of Michigan, Department of Microbiology and Immunology | Valesano |
| EPI_ISL_610154 | Singapore General Hospital | Department of Microbiology | Nurdyana Abdul Rahman, Kun Lee Lim, Chenhao Li, Sui Sin Goh, Kenneth Xin Long Chan, Kian Sing Chan, Lynette Oon, Kern Rei Chng, Niranjan Nagarajan, Karrie Ko |
| EPI_ISL_610201, EPI_ISL_610202, EPI_ISL_610203, EPI_ISL_610223, EPI_ISL_610224, EPI_ISL_610225 | Department of Health Technology and Informatics, The Hong Kong Polytechnic University | Department of Health Technology and Informatics, The Hong Kong Polytechnic University | Siu,G.K.-H., Lee,L.-K., Leung,K.S.-S., Leung,J.S.-L., Ng,T.T.-L., Chan,C.T.-M., Tam,K.K.-G., Lao,H.-Y., Wu,A.K.-L., Yau,M.C.-Y., Lai,Y.W.-M., Fung,K.S.-C., Chau,S.K.-Y., Wong,B.K.-C., To,W.-K., Luk,K., Ho,A.Y.-M., Que,T.-L., Yip,K.-T., Yam,W.C., Shum,D.H.-K., Yip,S.P. |
| EPI_ISL_610247 | University of Michigan Clinical Microbiology Laboratory | Lauring Lab, University of Michigan, Department of Microbiology and Immunology | Valesano |
| EPI_ISL_610632, EPI_ISL_610633, EPI_ISL_610635, EPI_ISL_610637, EPI_ISL_610638, EPI_ISL_610639, EPI_ISL_610643, EPI_ISL_610645, EPI_ISL_610646, EPI_ISL_610648, EPI_ISL_610650, EPI_ISL_610654, EPI_ISL_610656, EPI_ISL_610657, EPI_ISL_610659, EPI_ISL_610661, EPI_ISL_610669, EPI_ISL_610671, EPI_ISL_610673, EPI_ISL_610678, EPI_ISL_610680, EPI_ISL_610682, EPI_ISL_610688, EPI_ISL_610690, EPI_ISL_610696, EPI_ISL_610700, EPI_ISL_610701, EPI_ISL_610703, EPI_ISL_610704, EPI_ISL_610706, EPI_ISL_610707, EPI_ISL_610711, EPI_ISL_610718, EPI_ISL_610724, EPI_ISL_610731, EPI_ISL_610732, EPI_ISL_610733, EPI_ISL_610735, EPI_ISL_610737, EPI_ISL_610739, EPI_ISL_610742, EPI_ISL_610750, EPI_ISL_610751, EPI_ISL_610752, EPI_ISL_610753, EPI_ISL_610754, EPI_ISL_610755, EPI_ISL_610760, EPI_ISL_610764, EPI_ISL_610765, EPI_ISL_610766, EPI_ISL_610767, EPI_ISL_610773, EPI_ISL_610774, EPI_ISL_610777, EPI_ISL_610778, EPI_ISL_610782, EPI_ISL_610785 | | | |
| see above | Lighthouse Lab in Cambridge | Wellcome Sanger Institute for the COVID-19 Genomics UK (COG-UK) consortium | Rob Howes, The Lighthouse Lab in Cambridge and Alex Alderton, Roberto Amato, Sonia Goncalves, Ewan Harrison, David K. Jackson, Ian Johnston, Dominic Kwiatkowski, Cordelia Langford, John Sillitoe on behalf of the Wellcome Sanger Institute COVID-19 Surveillance Team |
| EPI_ISL_610786, EPI_ISL_610788, EPI_ISL_610792, EPI_ISL_610796, EPI_ISL_610799, EPI_ISL_610801, EPI_ISL_610805, EPI_ISL_610806, EPI_ISL_610810, EPI_ISL_610811, EPI_ISL_610814, EPI_ISL_610815, EPI_ISL_610816, EPI_ISL_610819, EPI_ISL_610820, EPI_ISL_610822, EPI_ISL_610823, EPI_ISL_610826, EPI_ISL_610827, EPI_ISL_610829, EPI_ISL_610830, EPI_ISL_610833, EPI_ISL_610834, EPI_ISL_610836, EPI_ISL_610837, EPI_ISL_610839, EPI_ISL_610842, EPI_ISL_610847, EPI_ISL_610852, EPI_ISL_610853, EPI_ISL_610856, EPI_ISL_610857, EPI_ISL_610861, EPI_ISL_610867, EPI_ISL_610871, EPI_ISL_610875, EPI_ISL_610877, EPI_ISL_610879, EPI_ISL_610881, EPI_ISL_610884, EPI_ISL_610885, EPI_ISL_610887, EPI_ISL_610888, EPI_ISL_610889, EPI_ISL_610890, EPI_ISL_610892, EPI_ISL_610895, EPI_ISL_610896, EPI_ISL_610899, EPI_ISL_610900, EPI_ISL_610905, EPI_ISL_610906, EPI_ISL_610907, EPI_ISL_610913, EPI_ISL_610914, EPI_ISL_610915, EPI_ISL_610916, EPI_ISL_610917, EPI_ISL_610920 | | | |
| see above | Lighthouse Lab in Cambridge | Wellcome Sanger Institute for the COVID-19 Genomics UK (COG-UK) consortium | Rob Howes, The Lighthouse Lab in Cambridge and Alex Alderton, Roberto Amato, Sonia Goncalves, Ewan Harrison, David K. Jackson, Ian Johnston, Dominic Kwiatkowski, Cordelia Langford, John Sillitoe on behalf of the Wellcome Sanger Institute COVID-19 Surveillance Team (http://www.sanger.ac.uk/covid-team) |
| EPI_ISL_611515, EPI_ISL_611516 | Liverpool Clinical Laboratories | COVID-19 Genomics UK (COG-UK) Consortium | Sam Haldenby, Anita Lucaci, Steve Paterson, Julian Hiscox, Alistair Darby, M Almsaud, A Alrezaihi, Muhannad Alruwaili, Stuart D Armstrong, Jones Benjamin, Eleanor G Bentley, Anu Chawla, Jordan J Clark, Angela Cowell, Richard Eccles, Isabel García-Dorival, Matthew Gemmell, Alessandro Gerada, PKF Gilmore, Richard Gregory, Ximeng Han, Catherine Hartley, Margaret Hughes, Miren Iturriza-Gomara, James Johnson, L Luu, Jenifer Manson, Charlotte Nelson, Elaine O'Toole, Cassie Olateju, Rebekah Penrice-Randal , Lucille Rainbow, N.P Randle, Trevor Ian Robinson, Parul Sharma, Ghada T Shawli, James P Stewart, Neil Swainston, Ecaterina Vamos, Joanne Watts, Mark Whitehead |
| EPI_ISL_611541 | Virology Department, Sheffield Teaching Hospitals NHS Foundation Trust/Department of Infection, Immunity and Cardiovascular Disease, The Medical School, University of Sheffield | COVID-19 Genomics UK (COG-UK) Consortium | Thushan de Silva, Matthew Parker, Nikki Smith, Adri Angyal, Rebecca Brown, Luke Green, Rachel Tucker, Paul Parsons, Danielle Groves, Katie Johnson, Laura Carrilero, Alex Keeley, Dave Partridge, Matthew Wyles, Benjamin Lindsey, Mehmet Yavuz, Mohammad Raza, Cariad Evans |
| EPI_ISL_611542, EPI_ISL_611543 | Liverpool Clinical Laboratories | COVID-19 Genomics UK (COG-UK) Consortium | Sam Haldenby, Anita Lucaci, Steve Paterson, Julian Hiscox, Alistair Darby, M Almsaud, A Alrezaihi, Muhannad Alruwaili, Stuart D Armstrong, Jones Benjamin, Eleanor G Bentley, Anu Chawla, Jordan J Clark, Angela Cowell, Richard Eccles, Isabel García-Dorival, Matthew Gemmell, Alessandro Gerada, PKF Gilmore, Richard Gregory, Ximeng Han, Catherine Hartley, Margaret Hughes, Miren Iturriza-Gomara, James Johnson, L Luu, Jenifer Manson, Charlotte Nelson, Elaine O'Toole, Cassie Olateju, Rebekah Penrice-Randal , Lucille Rainbow, N.P Randle, Trevor Ian Robinson, Parul Sharma, Ghada T Shawli, James P Stewart, Neil Swainston, Ecaterina Vamos, Joanne Watts, Mark Whitehead |
| EPI_ISL_611545 | Virology Department, Sheffield Teaching Hospitals NHS Foundation Trust/Department of Infection, Immunity and Cardiovascular Disease, The Medical School, University of Sheffield | COVID-19 Genomics UK (COG-UK) Consortium | Thushan de Silva, Matthew Parker, Nikki Smith, Adri Angyal, Rebecca Brown, Luke Green, Rachel Tucker, Paul Parsons, Danielle Groves, Katie Johnson, Laura Carrilero, Alex Keeley, Dave Partridge, Matthew Wyles, Benjamin Lindsey, Mehmet Yavuz, Mohammad Raza, Cariad Evans |
| EPI_ISL_611550, EPI_ISL_611554, EPI_ISL_611556 | University of Birmingham | COVID-19 Genomics UK (COG-UK) Consortium | Institute of Microbiology, University of Birmingham: Claire McMurray, Joanne Stockton, Samuel Nicholls, Radoslaw Poplawski, Will Rowe, Josh Quick, Nicholas Loman. University of Birmingham Testing Laboratory: Celina M Whalley, Andrew Bosworth, Charlotte Poxon, Kasun Wanigasooriya, Oliver Pickles, Mike Kidd, Alex Richter, Andrew D Beggs PHE Heartlands Lab: Husam Osman, Andrew Bosworth. Queen Elizabeth Hospital: Anna Casey |
| EPI_ISL_611571 | Liverpool Clinical Laboratories | COVID-19 Genomics UK (COG-UK) Consortium | Sam Haldenby, Anita Lucaci, Steve Paterson, Julian Hiscox, Alistair Darby, M Almsaud, A Alrezaihi, Muhannad Alruwaili, Stuart D Armstrong, Jones Benjamin, Eleanor G Bentley, Anu Chawla, Jordan J Clark, Angela Cowell, Richard Eccles, Isabel García-Dorival, Matthew Gemmell, Alessandro Gerada, PKF Gilmore, Richard Gregory, Ximeng Han, Catherine Hartley, Margaret Hughes, Miren Iturriza-Gomara, James Johnson, L Luu, Jenifer Manson, Charlotte Nelson, Elaine O'Toole, Cassie Olateju, Rebekah Penrice-Randal , Lucille Rainbow, N.P Randle, Trevor Ian Robinson, Parul Sharma, Ghada T Shawli, James P Stewart, Neil Swainston, Ecaterina Vamos, Joanne Watts, Mark Whitehead |
| EPI_ISL_611577 | Northumbria University / South Tees Hospitals NHS Foundation Trust / North Cumbria Integrated Care NHS Foundation Trust / North Tees and Hartlepool NHS Foundation Trust / Newcastle Hospitals NHS Foundation Trust | COVID-19 Genomics UK (COG-UK) Consortium | Darren L Smith,Andrew Nelson,Matthew Bashton,Greg R Young,Joshua Loh,John Allan,Mohammad A Tariq,Giles S Holt,Gary Black,Wen C Yew,Lynn Dover,Paul Baker,Steve Liggett,Sarah Essex,Jane Greenaway,Debra Padgett,Clive Graham,Garren Scott,Edward Barton,Emma Swindells,Brendan Payne,Jennifer Collins,Yusri Taha,Gary Eltringham |
| EPI_ISL_611589 | Centre for Enzyme Innovation, University of Portsmouth / Translational Research Laboratory, Portsmouth Hospitals NHS Trust | COVID-19 Genomics UK (COG-UK) Consortium | Angela Beckett,Yann Bourgeois,Garry Scarlett,Sharon Glaysher,Scott Elliott,Kelly Bicknell,Robert Impey,Allyson Lloyd,Sarah Wyllie,Ethan Butcher,Anoop Chauhan,Samuel Robson |
| EPI_ISL_611594 | Virology Department, Sheffield Teaching Hospitals NHS Foundation Trust/Department of Infection, Immunity and Cardiovascular Disease, The Medical School, University of Sheffield | COVID-19 Genomics UK (COG-UK) Consortium | Thushan de Silva, Matthew Parker, Nikki Smith, Adri Angyal, Rebecca Brown, Luke Green, Rachel Tucker, Paul Parsons, Danielle Groves, Katie Johnson, Laura Carrilero, Alex Keeley, Dave Partridge, Matthew Wyles, Benjamin Lindsey, Mehmet Yavuz, Mohammad Raza, Cariad Evans |
| EPI_ISL_611599 | University of Exeter | COVID-19 Genomics UK (COG-UK) Consortium | Ben Temperton,Aaron Jeffries,Michelle Michelsen,Joanna Warwick-Dugdale,Audrey Farbos,Robyn Manley,Stephen Michell,Jane Masoli |
| EPI_ISL_611608, EPI_ISL_611612 | University of Birmingham | COVID-19 Genomics UK (COG-UK) Consortium | Institute of Microbiology, University of Birmingham: Claire McMurray, Joanne Stockton, Samuel Nicholls, Radoslaw Poplawski, Will Rowe, Josh Quick, Nicholas Loman. University of Birmingham Testing Laboratory: Celina M Whalley, Andrew Bosworth, Charlotte Poxon, Kasun Wanigasooriya, Oliver Pickles, Mike Kidd, Alex Richter, Andrew D Beggs PHE Heartlands Lab: Husam Osman, Andrew Bosworth. Queen Elizabeth Hospital: Anna Casey |
| EPI_ISL_611613 | Liverpool Clinical Laboratories | COVID-19 Genomics UK (COG-UK) Consortium | Sam Haldenby, Anita Lucaci, Steve Paterson, Julian Hiscox, Alistair Darby, M Almsaud, A Alrezaihi, Muhannad Alruwaili, Stuart D Armstrong, Jones Benjamin, Eleanor G Bentley, Anu Chawla, Jordan J Clark, Angela Cowell, Richard Eccles, Isabel García-Dorival, Matthew Gemmell, Alessandro Gerada, PKF Gilmore, Richard Gregory, Ximeng Han, Catherine Hartley, Margaret Hughes, Miren Iturriza-Gomara, James Johnson, L Luu, Jenifer Manson, Charlotte Nelson, Elaine O'Toole, Cassie Olateju, Rebekah Penrice-Randal , Lucille Rainbow, N.P Randle, Trevor Ian Robinson, Parul Sharma, Ghada T Shawli, James P Stewart, Neil Swainston, Ecaterina Vamos, Joanne Watts, Mark Whitehead |
| EPI_ISL_611622 | University College London, Great Ormond Street Hospital for Children NHS Foundation Trust, Imperial College Healthcare NHS Trust | COVID-19 Genomics UK (COG-UK) Consortium | Sergi Castellano, Rachel Williams, Mark Kristiansen, Paola Resende Silva, Sunando Roy, Tony Brooks, Helena Tutill, Paola Niola, Patricia Dyal, Charlotte Williams, Leysa Forrest, Yasmin Panchbhaya, Jacqueline Findlay, Samuel Weeks, Julianne Brown, Kathryn Harris, Paul Randell, James Price, Alison Holmes, Judith Breuer |
| EPI_ISL_611630, EPI_ISL_611634, EPI_ISL_611635 | Liverpool Clinical Laboratories | COVID-19 Genomics UK (COG-UK) Consortium | Sam Haldenby, Anita Lucaci, Steve Paterson, Julian Hiscox, Alistair Darby, M Almsaud, A Alrezaihi, Muhannad Alruwaili, Stuart D Armstrong, Jones Benjamin, Eleanor G Bentley, Anu Chawla, Jordan J Clark, Angela Cowell, Richard Eccles, Isabel García-Dorival, Matthew Gemmell, Alessandro Gerada, PKF Gilmore, |

| | | | |
|---|---|---|---|
| | | | Richard Gregory, Ximeng Han, Catherine Hartley, Margaret Hughes, Miren Iturriza-Gomara, James Johnson, L Luu, Jenifer Manson, Charlotte Nelson, Elaine O'Toole, Cassie Olateju, Rebekah Penrice-Randal , Lucille Rainbow, N.P Randle, Trevor Ian Robinson, Parul Sharma, Ghada T Shawli, James P Stewart, Neil Swainston, Ecaterina Vamos, Joanne Watts, Mark Whitehead |
| EPI_ISL_611646, EPI_ISL_611647, EPI_ISL_611648, EPI_ISL_611649 | Northumbria University / South Tees Hospitals NHS Foundation Trust / North Cumbria Integrated Care NHS Foundation Trust / North Tees and Hartlepool NHS Foundation Trust / Newcastle Hospitals NHS Foundation Trust | COVID-19 Genomics UK (COG-UK) Consortium | Darren L Smith,Andrew Nelson,Matthew Bashton,Greg R Young,Joshua Loh,John Allan,Mohammad A Tariq,Giles S Holt,Gary Black,Wen C Yew,Lynn Dover,Paul Baker,Steve Liggett,Sarah Essex,Jane Greenaway,Debra Padgett,Clive Graham,Garren Scott,Edward Barton,Emma Swindells,Brendan Payne,Jennifer Collins,Yusri Taha,Gary Eltringham |
| EPI_ISL_611655 | Virology Department, Sheffield Teaching Hospitals NHS Foundation Trust/Department of Infection, Immunity and Cardiovascular Disease, The Medical School, University of Sheffield | COVID-19 Genomics UK (COG-UK) Consortium | Thushan de Silva, Matthew Parker, Nikki Smith, Adri Angyal, Rebecca Brown, Luke Green, Rachel Tucker, Paul Parsons, Danielle Groves, Katie Johnson, Laura Carrilero, Alex Keeley, Dave Partridge, Matthew Wyles, Benjamin Lindsey, Mehmet Yavuz, Mohammad Raza, Cariad Evans |
| EPI_ISL_611658 | Liverpool Clinical Laboratories | COVID-19 Genomics UK (COG-UK) Consortium | Sam Haldenby, Anita Lucaci, Steve Paterson, Julian Hiscox, Alistair Darby, M Almsaud, A Alrezaihi, Muhannad Alruwaili, Stuart D Armstrong, Jones Benjamin, Eleanor G Bentley, Anu Chawla, Jordan J Clark, Angela Cowell, Richard Eccles, Isabel García-Dorival, Matthew Gemmell, Alessandro Gerada, PKF Gilmore, Richard Gregory, Ximeng Han, Catherine Hartley, Margaret Hughes, Miren Iturriza-Gomara, James Johnson, L Luu, Jenifer Manson, Charlotte Nelson, Elaine O'Toole, Cassie Olateju, Rebekah Penrice-Randal , Lucille Rainbow, N.P Randle, Trevor Ian Robinson, Parul Sharma, Ghada T Shawli, James P Stewart, Neil Swainston, Ecaterina Vamos, Joanne Watts, Mark Whitehead |
| EPI_ISL_611661 | Virology Department, Sheffield Teaching Hospitals NHS Foundation Trust/Department of Infection, Immunity and Cardiovascular Disease, The Medical School, University of Sheffield | COVID-19 Genomics UK (COG-UK) Consortium | Thushan de Silva, Matthew Parker, Nikki Smith, Adri Angyal, Rebecca Brown, Luke Green, Rachel Tucker, Paul Parsons, Danielle Groves, Katie Johnson, Laura Carrilero, Alex Keeley, Dave Partridge, Matthew Wyles, Benjamin Lindsey, Mehmet Yavuz, Mohammad Raza, Cariad Evans |
| EPI_ISL_611663 | Wales Specialist Virology Centre Sequencing lab: Pathogen Genomics Unit | COVID-19 Genomics UK (COG-UK) Consortium | Catherine Moore, Johnathan Evans, Laura Gifford, Malorie Perry, Simon Cottrell, Angela Marchbank, Alec Birchley, Alexander Adams, Amy Gaskin, Bree Gatica-Wilcox, Jason Coombes, Joel Southgate, Lauren Gilbert, Lee Graham, Nicole Pacchiarini, Sara Kumziene-Summerhayes, Sarah Taylor, Sophie Jones, Sara Rey, Matthew Bull, Joanne Watkins, Sally Corden, Tom Connor |
| EPI_ISL_611671 | University of Birmingham | COVID-19 Genomics UK (COG-UK) Consortium | Institute of Microbiology, University of Birmingham: Claire McMurray, Joanne Stockton, Samuel Nicholls, Radoslaw Poplawski, Will Rowe, Josh Quick, Nicholas Loman. University of Birmingham Testing Laboratory: Celina M Whalley, Andrew Bosworth, Charlotte Poxon, Kasun Wanigasooriya, Oliver Pickles, Mike Kidd, Alex Richter, Andrew D Beggs PHE Heartlands Lab: Husam Osman, Andrew Bosworth. Queen Elizabeth Hospital: Anna Casey |
| EPI_ISL_611675 | Liverpool Clinical Laboratories | COVID-19 Genomics UK (COG-UK) Consortium | Sam Haldenby, Anita Lucaci, Steve Paterson, Julian Hiscox, Alistair Darby, M Almsaud, A Alrezaihi, Muhannad Alruwaili, Stuart D Armstrong, Jones Benjamin, Eleanor G Bentley, Anu Chawla, Jordan J Clark, Angela Cowell, Richard Eccles, Isabel García-Dorival, Matthew Gemmell, Alessandro Gerada, PKF Gilmore, Richard Gregory, Ximeng Han, Catherine Hartley, Margaret Hughes, Miren Iturriza-Gomara, James Johnson, L Luu, Jenifer Manson, Charlotte Nelson, Elaine O'Toole, Cassie Olateju, Rebekah Penrice-Randal , Lucille Rainbow, N.P Randle, Trevor Ian Robinson, Parul Sharma, Ghada T Shawli, James P Stewart, Neil Swainston, Ecaterina Vamos, Joanne Watts, Mark Whitehead |
| EPI_ISL_611692, EPI_ISL_611696, EPI_ISL_611700, EPI_ISL_611705 | University of Birmingham | COVID-19 Genomics UK (COG-UK) Consortium | Institute of Microbiology, University of Birmingham: Claire McMurray, Joanne Stockton, Samuel Nicholls, Radoslaw Poplawski, Will Rowe, Josh Quick, Nicholas Loman. University of Birmingham Testing Laboratory: Celina M Whalley, Andrew Bosworth, Charlotte Poxon, Kasun Wanigasooriya, Oliver Pickles, Mike Kidd, Alex Richter, Andrew D Beggs PHE Heartlands Lab: Husam Osman, Andrew Bosworth. Queen Elizabeth Hospital: Anna Casey |
| EPI_ISL_611724 | Liverpool Clinical Laboratories | COVID-19 Genomics UK (COG-UK) Consortium | Sam Haldenby, Anita Lucaci, Steve Paterson, Julian Hiscox, Alistair Darby, M Almsaud, A Alrezaihi, Muhannad Alruwaili, Stuart D Armstrong, Jones Benjamin, Eleanor G Bentley, Anu Chawla, Jordan J Clark, Angela Cowell, Richard Eccles, Isabel García-Dorival, Matthew Gemmell, Alessandro Gerada, PKF Gilmore, Richard Gregory, Ximeng Han, Catherine Hartley, Margaret Hughes, Miren Iturriza-Gomara, James Johnson, L Luu, Jenifer Manson, Charlotte Nelson, Elaine O'Toole, Cassie Olateju, Rebekah Penrice-Randal , Lucille Rainbow, N.P Randle, Trevor Ian Robinson, Parul Sharma, Ghada T Shawli, James P Stewart, Neil Swainston, Ecaterina Vamos, Joanne Watts, Mark Whitehead |
| EPI_ISL_611725 | Wales Specialist Virology Centre Sequencing lab: Pathogen Genomics Unit | COVID-19 Genomics UK (COG-UK) Consortium | Catherine Moore, Johnathan Evans, Laura Gifford, Malorie Perry, Simon Cottrell, Angela Marchbank, Alec Birchley, Alexander Adams, Amy Gaskin, Bree Gatica-Wilcox, Jason Coombes, Joel Southgate, Lauren Gilbert, Lee Graham, Nicole Pacchiarini, Sara Kumziene-Summerhayes, Sarah Taylor, Sophie Jones, Sara Rey, Matthew Bull, Joanne Watkins, Sally Corden, Tom Connor |
| EPI_ISL_611737, EPI_ISL_611741, EPI_ISL_611750 | Liverpool Clinical Laboratories | COVID-19 Genomics UK (COG-UK) Consortium | Sam Haldenby, Anita Lucaci, Steve Paterson, Julian Hiscox, Alistair Darby, M Almsaud, A Alrezaihi, Muhannad Alruwaili, Stuart D Armstrong, Jones Benjamin, Eleanor G Bentley, Anu Chawla, Jordan J Clark, Angela Cowell, Richard Eccles, Isabel García-Dorival, Matthew Gemmell, Alessandro Gerada, PKF Gilmore, Richard Gregory, Ximeng Han, Catherine Hartley, Margaret Hughes, Miren Iturriza-Gomara, James Johnson, L Luu, Jenifer Manson, Charlotte Nelson, Elaine O'Toole, Cassie Olateju, Rebekah Penrice-Randal , Lucille Rainbow, N.P Randle, Trevor Ian Robinson, Parul Sharma, Ghada T Shawli, James P Stewart, Neil Swainston, Ecaterina Vamos, Joanne Watts, Mark Whitehead |
| EPI_ISL_611752 | Northumbria University / South Tees Hospitals NHS Foundation Trust / North Cumbria Integrated Care NHS Foundation Trust / North Tees and Hartlepool NHS Foundation Trust / Newcastle Hospitals NHS Foundation Trust | COVID-19 Genomics UK (COG-UK) Consortium | Darren L Smith,Andrew Nelson,Matthew Bashton,Greg R Young,Joshua Loh,John Allan,Mohammad A Tariq,Giles S Holt,Gary Black,Wen C Yew,Lynn Dover,Paul Baker,Steve Liggett,Sarah Essex,Jane Greenaway,Debra Padgett,Clive Graham,Garren Scott,Edward Barton,Emma Swindells,Brendan Payne,Jennifer Collins,Yusri Taha,Gary Eltringham |
| EPI_ISL_611763 | Centre for Enzyme Innovation, University of Portsmouth / Translational Research Laboratory, Portsmouth Hospitals NHS Trust | COVID-19 Genomics UK (COG-UK) Consortium | Angela Beckett,Yann Bourgeois,Garry Scarlett,Sharon Glaysher,Scott Elliott,Kelly Bicknell,Robert Impey,Allyson Lloyd,Sarah Wyllie,Ethan Butcher,Anoop Chauhan,Samuel Robson |
| EPI_ISL_611764, EPI_ISL_611765 | Liverpool Clinical Laboratories | COVID-19 Genomics UK (COG-UK) Consortium | Sam Haldenby, Anita Lucaci, Steve Paterson, Julian Hiscox, Alistair Darby, M Almsaud, A Alrezaihi, Muhannad Alruwaili, Stuart D Armstrong, Jones Benjamin, Eleanor G Bentley, Anu Chawla, Jordan J Clark, Angela Cowell, Richard Eccles, Isabel García-Dorival, Matthew Gemmell, Alessandro Gerada, PKF Gilmore, Richard Gregory, Ximeng Han, Catherine Hartley, Margaret Hughes, Miren Iturriza-Gomara, James Johnson, L Luu, Jenifer Manson, Charlotte Nelson, Elaine O'Toole, Cassie Olateju, Rebekah Penrice-Randal , Lucille Rainbow, N.P Randle, Trevor Ian Robinson, Parul Sharma, Ghada T Shawli, James P Stewart, Neil Swainston, Ecaterina Vamos, Joanne Watts, Mark Whitehead |
| EPI_ISL_611769, EPI_ISL_611770 | Northumbria University / South Tees Hospitals NHS Foundation Trust / North Cumbria Integrated Care NHS Foundation Trust / North Tees and Hartlepool NHS Foundation Trust / Newcastle Hospitals NHS Foundation Trust | COVID-19 Genomics UK (COG-UK) Consortium | Darren L Smith,Andrew Nelson,Matthew Bashton,Greg R Young,Joshua Loh,John Allan,Mohammad A Tariq,Giles S Holt,Gary Black,Wen C Yew,Lynn Dover,Paul Baker,Steve Liggett,Sarah Essex,Jane Greenaway,Debra Padgett,Clive Graham,Garren Scott,Edward Barton,Emma Swindells,Brendan Payne,Jennifer Collins,Yusri Taha,Gary Eltringham |
| EPI_ISL_611776 | University of Birmingham | COVID-19 Genomics UK (COG-UK) Consortium | Institute of Microbiology, University of Birmingham: Claire McMurray, Joanne Stockton, Samuel Nicholls, Radoslaw Poplawski, Will Rowe, Josh Quick, Nicholas Loman. University of Birmingham Testing Laboratory: Celina M Whalley, Andrew Bosworth, Charlotte Poxon, Kasun Wanigasooriya, Oliver Pickles, Mike Kidd, Alex Richter, Andrew D Beggs PHE Heartlands Lab: Husam Osman, Andrew Bosworth. Queen Elizabeth Hospital: Anna Casey |
| EPI_ISL_611786 | Liverpool Clinical Laboratories | COVID-19 Genomics UK (COG-UK) Consortium | Sam Haldenby, Anita Lucaci, Steve Paterson, Julian Hiscox, Alistair Darby, M Almsaud, A Alrezaihi, Muhannad Alruwaili, Stuart D Armstrong, Jones Benjamin, Eleanor G Bentley, Anu Chawla, Jordan J Clark, Angela Cowell, Richard Eccles, Isabel García-Dorival, Matthew Gemmell, Alessandro Gerada, PKF Gilmore, Richard Gregory, Ximeng Han, Catherine Hartley, Margaret Hughes, Miren Iturriza-Gomara, James Johnson, L Luu, Jenifer Manson, Charlotte Nelson, Elaine O'Toole, Cassie Olateju, Rebekah Penrice-Randal , Lucille Rainbow, N.P Randle, Trevor Ian Robinson, Parul Sharma, Ghada T Shawli, James P Stewart, Neil Swainston, Ecaterina Vamos, Joanne Watts, Mark Whitehead |
| EPI_ISL_611787, EPI_ISL_611795, EPI_ISL_611800, EPI_ISL_611805, EPI_ISL_611806 | Virology Department, Sheffield Teaching Hospitals NHS Foundation Trust/Department of Infection, Immunity and Cardiovascular Disease, The Medical School, University of Sheffield | COVID-19 Genomics UK (COG-UK) Consortium | Thushan de Silva, Matthew Parker, Nikki Smith, Adri Angyal, Rebecca Brown, Luke Green, Rachel Tucker, Paul Parsons, Danielle Groves, Katie Johnson, Laura Carrilero, Alex Keeley, Dave Partridge, Matthew Wyles, Benjamin Lindsey, Mehmet Yavuz, Mohammad Raza, Cariad Evans |

| | | | |
|---|---|---|---|
| EPI_ISL_611812 | Queens Medical Centre, Clinical Microbiology Department / DeepSeq Nottingham | COVID-19 Genomics UK (COG-UK) Consortium | Gemma Clark, Wendy Smith, Manjinder Khakh, Vicki M Fleming, Michelle M Lister, Hannah Howson-Wells, Jonathan Ball, Patrick McClure, Joseph Chappell, Theocharis Tsoleridis, Nadine Holmes, Matthew Carlisle, Christopher Moore, Fei Sang, Johnny Debebe, Victoria Wright, Matthew Loose |
| EPI_ISL_611814 | University of Birmingham | COVID-19 Genomics UK (COG-UK) Consortium | Institute of Microbiology, University of Birmingham: Claire McMurray, Joanne Stockton, Samuel Nicholls, Radoslaw Poplawski, Will Rowe, Josh Quick, Nicholas Loman. University of Birmingham Testing Laboratory: Celina M Whalley, Andrew Bosworth, Charlotte Poxon, Kasun Wanigasooriya, Oliver Pickles, Mike Kidd, Alex Richter, Andrew D Beggs PHE Heartlands Lab: Husam Osman, Andrew Bosworth. Queen Elizabeth Hospital: Anna Casey |
| EPI_ISL_611816 | Virology Department, Sheffield Teaching Hospitals NHS Foundation Trust/Department of Infection, Immunity and Cardiovascular Disease, The Medical School, University of Sheffield | COVID-19 Genomics UK (COG-UK) Consortium | Thushan de Silva, Matthew Parker, Nikki Smith, Adri Angyal, Rebecca Brown, Luke Green, Rachel Tucker, Paul Parsons, Danielle Groves, Katie Johnson, Laura Carrilero, Alex Keeley, Dave Partridge, Matthew Wyles, Benjamin Lindsey, Mehmet Yavuz, Mohammad Raza, Cariad Evans |
| EPI_ISL_611825 | University of Birmingham | COVID-19 Genomics UK (COG-UK) Consortium | Institute of Microbiology, University of Birmingham: Claire McMurray, Joanne Stockton, Samuel Nicholls, Radoslaw Poplawski, Will Rowe, Josh Quick, Nicholas Loman. University of Birmingham Testing Laboratory: Celina M Whalley, Andrew Bosworth, Charlotte Poxon, Kasun Wanigasooriya, Oliver Pickles, Mike Kidd, Alex Richter, Andrew D Beggs PHE Heartlands Lab: Husam Osman, Andrew Bosworth. Queen Elizabeth Hospital: Anna Casey |
| EPI_ISL_611829 | Virology Department, Sheffield Teaching Hospitals NHS Foundation Trust/Department of Infection, Immunity and Cardiovascular Disease, The Medical School, University of Sheffield | COVID-19 Genomics UK (COG-UK) Consortium | Thushan de Silva, Matthew Parker, Nikki Smith, Adri Angyal, Rebecca Brown, Luke Green, Rachel Tucker, Paul Parsons, Danielle Groves, Katie Johnson, Laura Carrilero, Alex Keeley, Dave Partridge, Matthew Wyles, Benjamin Lindsey, Mehmet Yavuz, Mohammad Raza, Cariad Evans |
| EPI_ISL_611835 | Wales Specialist Virology Centre Sequencing lab: Pathogen Genomics Unit | COVID-19 Genomics UK (COG-UK) Consortium | Catherine Moore, Johnathan Evans, Laura Gifford, Malorie Perry, Simon Cottrell, Angela Marchbank, Alec Birchley, Alexander Adams, Amy Gaskin, Bree Gatica-Wilcox, Jason Coombes, Joel Southgate, Lauren Gilbert, Lee Graham, Nicole Pacchiarini, Sara Kumziene-Summerhayes, Sarah Taylor, Sophie Jones, Sara Rey, Matthew Bull, Joanne Watkins, Sally Corden, Tom Connor |
| EPI_ISL_611837 | University of Exeter | COVID-19 Genomics UK (COG-UK) Consortium | Ben Temperton,Aaron Jeffries,Michelle Michelsen,Joanna Warwick-Dugdale,Audrey Farbos,Robyn Manley,Stephen Michell,Jane Masoli |
| EPI_ISL_611841 | Centre for Enzyme Innovation, University of Portsmouth / Translational Research Laboratory, Portsmouth Hospitals NHS Trust | COVID-19 Genomics UK (COG-UK) Consortium | Angela Beckett,Yann Bourgeois,Garry Scarlett,Sharon Glaysher,Scott Elliott,Kelly Bicknell,Robert Impey,Allyson Lloyd,Sarah Wyllie,Ethan Butcher,Anoop Chauhan,Samuel Robson |
| EPI_ISL_611846 | Northumbria University / South Tees Hospitals NHS Foundation Trust / North Cumbria Integrated Care NHS Foundation Trust / North Tees and Hartlepool NHS Foundation Trust / Newcastle Hospitals NHS Foundation Trust | COVID-19 Genomics UK (COG-UK) Consortium | Darren L Smith,Andrew Nelson,Matthew Bashton,Greg R Young,Joshua Loh,John Allan,Mohammad A Tariq,Giles S Holt,Gary Black,Wen C Yew,Lynn Dover,Paul Baker,Steve Liggett,Sarah Essex,Jane Greenaway,Debra Padgett,Clive Graham,Garren Scott,Edward Barton,Emma Swindells,Brendan Payne,Jennifer Collins,Yusri Taha,Gary Eltringham |
| EPI_ISL_611847, EPI_ISL_611855, EPI_ISL_611856, EPI_ISL_611858, EPI_ISL_611859, EPI_ISL_611860 | University of Birmingham | COVID-19 Genomics UK (COG-UK) Consortium | Institute of Microbiology, University of Birmingham: Claire McMurray, Joanne Stockton, Samuel Nicholls, Radoslaw Poplawski, Will Rowe, Josh Quick, Nicholas Loman. University of Birmingham Testing Laboratory: Celina M Whalley, Andrew Bosworth, Charlotte Poxon, Kasun Wanigasooriya, Oliver Pickles, Mike Kidd, Alex Richter, Andrew D Beggs PHE Heartlands Lab: Husam Osman, Andrew Bosworth. Queen Elizabeth Hospital: Anna Casey |
| EPI_ISL_611862 | Virology Department, Sheffield Teaching Hospitals NHS Foundation Trust/Department of Infection, Immunity and Cardiovascular Disease, The Medical School, University of Sheffield | COVID-19 Genomics UK (COG-UK) Consortium | Thushan de Silva, Matthew Parker, Nikki Smith, Adri Angyal, Rebecca Brown, Luke Green, Rachel Tucker, Paul Parsons, Danielle Groves, Katie Johnson, Laura Carrilero, Alex Keeley, Dave Partridge, Matthew Wyles, Benjamin Lindsey, Mehmet Yavuz, Mohammad Raza, Cariad Evans |
| EPI_ISL_611889 | Liverpool Clinical Laboratories | COVID-19 Genomics UK (COG-UK) Consortium | Sam Haldenby, Anita Lucaci, Steve Paterson, Julian Hiscox, Alistair Darby, M Almsaud, A Alrezaihi, Muhannad Alruwaili, Stuart D Armstrong, Jones Benjamin, Eleanor G Bentley, Anu Chawla, Jordan J Clark, Angela Cowell, Richard Eccles, Isabel García-Dorival, Matthew Gemmell, Alessandro Gerada, PKF Gilmore, Richard Gregory, Ximeng Han, Catherine Hartley, Margaret Hughes, Miren Iturriza-Gomara, James Johnson, L Luu, Jenifer Manson, Charlotte Nelson, Elaine O'Toole, Cassie Olateju, Rebekah Penrice-Randal , Lucille Rainbow, N.P Randle, Trevor Ian Robinson, Parul Sharma, Ghada T Shawli, James P Stewart, Neil Swainston, Ecaterina Vamos, Joanne Watts, Mark Whitehead |
| EPI_ISL_611896, EPI_ISL_611900, EPI_ISL_611906 | Wales Specialist Virology Centre Sequencing lab: Pathogen Genomics Unit | COVID-19 Genomics UK (COG-UK) Consortium | Catherine Moore, Johnathan Evans, Laura Gifford, Malorie Perry, Simon Cottrell, Angela Marchbank, Alec Birchley, Alexander Adams, Amy Gaskin, Bree Gatica-Wilcox, Jason Coombes, Joel Southgate, Lauren Gilbert, Lee Graham, Nicole Pacchiarini, Sara Kumziene-Summerhayes, Sarah Taylor, Sophie Jones, Sara Rey, Matthew Bull, Joanne Watkins, Sally Corden, Tom Connor |
| EPI_ISL_611911 | Liverpool Clinical Laboratories | COVID-19 Genomics UK (COG-UK) Consortium | Sam Haldenby, Anita Lucaci, Steve Paterson, Julian Hiscox, Alistair Darby, M Almsaud, A Alrezaihi, Muhannad Alruwaili, Stuart D Armstrong, Jones Benjamin, Eleanor G Bentley, Anu Chawla, Jordan J Clark, Angela Cowell, Richard Eccles, Isabel García-Dorival, Matthew Gemmell, Alessandro Gerada, PKF Gilmore, Richard Gregory, Ximeng Han, Catherine Hartley, Margaret Hughes, Miren Iturriza-Gomara, James Johnson, L Luu, Jenifer Manson, Charlotte Nelson, Elaine O'Toole, Cassie Olateju, Rebekah Penrice-Randal , Lucille Rainbow, N.P Randle, Trevor Ian Robinson, Parul Sharma, Ghada T Shawli, James P Stewart, Neil Swainston, Ecaterina Vamos, Joanne Watts, Mark Whitehead |
| EPI_ISL_611912 | Wales Specialist Virology Centre Sequencing lab: Pathogen Genomics Unit | COVID-19 Genomics UK (COG-UK) Consortium | Catherine Moore, Johnathan Evans, Laura Gifford, Malorie Perry, Simon Cottrell, Angela Marchbank, Alec Birchley, Alexander Adams, Amy Gaskin, Bree Gatica-Wilcox, Jason Coombes, Joel Southgate, Lauren Gilbert, Lee Graham, Nicole Pacchiarini, Sara Kumziene-Summerhayes, Sarah Taylor, Sophie Jones, Sara Rey, Matthew Bull, Joanne Watkins, Sally Corden, Tom Connor |
| EPI_ISL_611914 | Liverpool Clinical Laboratories | COVID-19 Genomics UK (COG-UK) Consortium | Sam Haldenby, Anita Lucaci, Steve Paterson, Julian Hiscox, Alistair Darby, M Almsaud, A Alrezaihi, Muhannad Alruwaili, Stuart D Armstrong, Jones Benjamin, Eleanor G Bentley, Anu Chawla, Jordan J Clark, Angela Cowell, Richard Eccles, Isabel García-Dorival, Matthew Gemmell, Alessandro Gerada, PKF Gilmore, Richard Gregory, Ximeng Han, Catherine Hartley, Margaret Hughes, Miren Iturriza-Gomara, James Johnson, L Luu, Jenifer Manson, Charlotte Nelson, Elaine O'Toole, Cassie Olateju, Rebekah Penrice-Randal , Lucille Rainbow, N.P Randle, Trevor Ian Robinson, Parul Sharma, Ghada T Shawli, James P Stewart, Neil Swainston, Ecaterina Vamos, Joanne Watts, Mark Whitehead |
| EPI_ISL_611916 | Centre for Enzyme Innovation, University of Portsmouth / Translational Research Laboratory, Portsmouth Hospitals NHS Trust | COVID-19 Genomics UK (COG-UK) Consortium | Angela Beckett,Yann Bourgeois,Garry Scarlett,Sharon Glaysher,Scott Elliott,Kelly Bicknell,Robert Impey,Allyson Lloyd,Sarah Wyllie,Ethan Butcher,Anoop Chauhan,Samuel Robson |
| EPI_ISL_611919 | Queens Medical Centre, Clinical Microbiology Department / DeepSeq Nottingham | COVID-19 Genomics UK (COG-UK) Consortium | Gemma Clark, Wendy Smith, Manjinder Khakh, Vicki M Fleming, Michelle M Lister, Hannah Howson-Wells, Jonathan Ball, Patrick McClure, Joseph Chappell, Theocharis Tsoleridis, Nadine Holmes, Matthew Carlisle, Christopher Moore, Fei Sang, Johnny Debebe, Victoria Wright, Matthew Loose |
| EPI_ISL_611921 | Liverpool Clinical Laboratories | COVID-19 Genomics UK (COG-UK) Consortium | Sam Haldenby, Anita Lucaci, Steve Paterson, Julian Hiscox, Alistair Darby, M Almsaud, A Alrezaihi, Muhannad Alruwaili, Stuart D Armstrong, Jones Benjamin, Eleanor G Bentley, Anu Chawla, Jordan J Clark, Angela Cowell, Richard Eccles, Isabel García-Dorival, Matthew Gemmell, Alessandro Gerada, PKF Gilmore, Richard Gregory, Ximeng Han, Catherine Hartley, Margaret Hughes, Miren Iturriza-Gomara, James Johnson, L Luu, Jenifer Manson, Charlotte Nelson, Elaine O'Toole, Cassie Olateju, Rebekah Penrice-Randal , Lucille Rainbow, N.P Randle, Trevor Ian Robinson, Parul Sharma, Ghada T Shawli, James P Stewart, Neil Swainston, Ecaterina Vamos, Joanne Watts, Mark Whitehead |
| EPI_ISL_611927, EPI_ISL_611928 | Centre for Enzyme Innovation, University of Portsmouth / Translational Research Laboratory, Portsmouth Hospitals NHS Trust | COVID-19 Genomics UK (COG-UK) Consortium | Angela Beckett,Yann Bourgeois,Garry Scarlett,Sharon Glaysher,Scott Elliott,Kelly Bicknell,Robert Impey,Allyson Lloyd,Sarah Wyllie,Ethan Butcher,Anoop Chauhan,Samuel Robson |
| EPI_ISL_611940 | University of Exeter | COVID-19 Genomics UK (COG-UK) Consortium | Ben Temperton,Aaron Jeffries,Michelle Michelsen,Joanna Warwick-Dugdale,Audrey Farbos,Robyn Manley,Stephen Michell,Jane Masoli |
| EPI_ISL_611953 | Northumbria University / South Tees Hospitals NHS Foundation Trust / North Cumbria Integrated Care NHS Foundation Trust / North Tees and Hartlepool NHS Foundation Trust / Newcastle Hospitals NHS Foundation Trust | COVID-19 Genomics UK (COG-UK) Consortium | Darren L Smith,Andrew Nelson,Matthew Bashton,Greg R Young,Joshua Loh,John Allan,Mohammad A Tariq,Giles S Holt,Gary Black,Wen C Yew,Lynn Dover,Paul Baker,Steve Liggett,Sarah Essex,Jane Greenaway,Debra Padgett,Clive Graham,Garren Scott,Edward Barton,Emma Swindells,Brendan Payne,Jennifer Collins,Yusri Taha,Gary Eltringham |
| EPI_ISL_611959 | Centre for Enzyme Innovation, University of Portsmouth / | COVID-19 Genomics UK (COG-UK) Consortium | Angela Beckett,Yann Bourgeois,Garry Scarlett,Sharon Glaysher,Scott Elliott,Kelly Bicknell,Robert Impey,Allyson Lloyd,Sarah Wyllie,Ethan Butcher,Anoop |

| | | | |
|---|---|---|---|
| | Translational Research Laboratory, Portsmouth Hospitals NHS Trust | | Chauhan,Samuel Robson |
| EPI_ISL_611963 | Queens Medical Centre, Clinical Microbiology Department / DeepSeq Nottingham | COVID-19 Genomics UK (COG-UK) Consortium | Gemma Clark, Wendy Smith, Manjinder Khakh, Vicki M Fleming, Michelle M Lister, Hannah Howson-Wells, Jonathan Ball, Patrick McClure, Joseph Chappell, Theocharis Tsoleridis, Nadine Holmes, Matthew Carlisle, Christopher Moore, Fei Sang, Johnny Debebe, Victoria Wright, Matthew Loose |
| EPI_ISL_611967 | Department of Pathology, University of Cambridge | COVID-19 Genomics UK (COG-UK) Consortium | Aminu S. Jahun, Yasmin Chaudhry, Grant Hall, Iliana Georgana, Myra Hosmillo, Martin D. Curran, Malte Pinckert, Surendra Parmar, Ian Goodfellow |
| EPI_ISL_611971 | Centre for Enzyme Innovation, University of Portsmouth / Translational Research Laboratory, Portsmouth Hospitals NHS Trust | COVID-19 Genomics UK (COG-UK) Consortium | Angela Beckett,Yann Bourgeois,Garry Scarlett,Sharon Glaysher,Scott Elliott,Kelly Bicknell,Robert Impey,Allyson Lloyd,Sarah Wyllie,Ethan Butcher,Anoop Chauhan,Samuel Robson |
| EPI_ISL_611981, EPI_ISL_611985 | Liverpool Clinical Laboratories | COVID-19 Genomics UK (COG-UK) Consortium | Sam Haldenby, Anita Lucaci, Steve Paterson, Julian Hiscox, Alistair Darby, M Almsaud, A Alrezaihi, Muhannad Alruwaili, Stuart D Armstrong, Jones Benjamin, Eleanor G Bentley, Anu Chawla, Jordan J Clark, Angela Cowell, Richard Eccles, Isabel García-Dorival, Matthew Gemmell, Alessandro Gerada, PKF Gilmore, Richard Gregory, Ximeng Han, Catherine Hartley, Margaret Hughes, Miren Iturriza-Gomara, James Johnson, L Luu, Jenifer Manson, Charlotte Nelson, Elaine O'Toole, Cassie Olateju, Rebekah Penrice-Randal , Lucille Rainbow, N.P Randle, Trevor Ian Robinson, Parul Sharma, Ghada T Shawli, James P Stewart, Neil Swainston, Ecaterina Vamos, Joanne Watts, Mark Whitehead |
| EPI_ISL_611988 | Virology Department, Sheffield Teaching Hospitals NHS Foundation Trust/Department of Infection, Immunity and Cardiovascular Disease, The Medical School, University of Sheffield | COVID-19 Genomics UK (COG-UK) Consortium | Thushan de Silva, Matthew Parker, Nikki Smith, Adri Angyal, Rebecca Brown, Luke Green, Rachel Tucker, Paul Parsons, Danielle Groves, Katie Johnson, Laura Carrilero, Alex Keeley, Dave Partridge, Matthew Wyles, Benjamin Lindsey, Mehmet Yavuz, Mohammad Raza, Cariad Evans |
| EPI_ISL_611989 | Virology Department, Royal Infirmary of Edinburgh, NHS Lothian / School of Biological Sciences, University of Edinburgh / Institute of Genetics and Molecular Medicine, University of Edinburgh | COVID-19 Genomics UK (COG-UK) Consortium | McHugh M, Dewar R, Rooke S, Gallagher M, Balcaza C, O'Toole Á, Scher E, Hill V, McCrone JT, Colquhoun R, Yu X, Jackson B, Rambaut A, Williams TC, Templeton K |
| EPI_ISL_611995, EPI_ISL_611999, EPI_ISL_612003 | Liverpool Clinical Laboratories | COVID-19 Genomics UK (COG-UK) Consortium | Sam Haldenby, Anita Lucaci, Steve Paterson, Julian Hiscox, Alistair Darby, M Almsaud, A Alrezaihi, Muhannad Alruwaili, Stuart D Armstrong, Jones Benjamin, Eleanor G Bentley, Anu Chawla, Jordan J Clark, Angela Cowell, Richard Eccles, Isabel García-Dorival, Matthew Gemmell, Alessandro Gerada, PKF Gilmore, Richard Gregory, Ximeng Han, Catherine Hartley, Margaret Hughes, Miren Iturriza-Gomara, James Johnson, L Luu, Jenifer Manson, Charlotte Nelson, Elaine O'Toole, Cassie Olateju, Rebekah Penrice-Randal , Lucille Rainbow, N.P Randle, Trevor Ian Robinson, Parul Sharma, Ghada T Shawli, James P Stewart, Neil Swainston, Ecaterina Vamos, Joanne Watts, Mark Whitehead |
| EPI_ISL_612007 | Department of Pathology, University of Cambridge | COVID-19 Genomics UK (COG-UK) Consortium | Aminu S. Jahun, Yasmin Chaudhry, Grant Hall, Iliana Georgana, Myra Hosmillo, Martin D. Curran, Malte Pinckert, Surendra Parmar, Ian Goodfellow |
| EPI_ISL_612011 | University of Birmingham | COVID-19 Genomics UK (COG-UK) Consortium | Institute of Microbiology, University of Birmingham: Claire McMurray, Joanne Stockton, Samuel Nicholls, Radoslaw Poplawski, Will Rowe, Josh Quick, Nicholas Loman. University of Birmingham Testing Laboratory: Celina M Whalley, Andrew Bosworth, Charlotte Poxon, Kasun Wanigasooriya, Oliver Pickles, Mike Kidd, Alex Richter, Andrew D Beggs PHE Heartlands Lab: Husam Osman, Andrew Bosworth. Queen Elizabeth Hospital: Anna Casey |
| EPI_ISL_612012 | Wales Specialist Virology Centre Sequencing lab: Pathogen Genomics Unit | COVID-19 Genomics UK (COG-UK) Consortium | Catherine Moore, Johnathan Evans, Laura Gifford, Malorie Perry, Simon Cottrell, Angela Marchbank, Alec Birchley, Alexander Adams, Amy Gaskin, Bree Gatica-Wilcox, Jason Coombes, Joel Southgate, Lauren Gilbert, Lee Graham, Nicole Pacchiarini, Sara Kumziene-Summerhayes, Sarah Taylor, Sophie Jones, Sara Rey, Matthew Bull, Joanne Watkins, Sally Corden, Tom Connor |
| EPI_ISL_612015 | University of Birmingham | COVID-19 Genomics UK (COG-UK) Consortium | Institute of Microbiology, University of Birmingham: Claire McMurray, Joanne Stockton, Samuel Nicholls, Radoslaw Poplawski, Will Rowe, Josh Quick, Nicholas Loman. University of Birmingham Testing Laboratory: Celina M Whalley, Andrew Bosworth, Charlotte Poxon, Kasun Wanigasooriya, Oliver Pickles, Mike Kidd, Alex Richter, Andrew D Beggs PHE Heartlands Lab: Husam Osman, Andrew Bosworth. Queen Elizabeth Hospital: Anna Casey |
| EPI_ISL_612019 | Liverpool Clinical Laboratories | COVID-19 Genomics UK (COG-UK) Consortium | Sam Haldenby, Anita Lucaci, Steve Paterson, Julian Hiscox, Alistair Darby, M Almsaud, A Alrezaihi, Muhannad Alruwaili, Stuart D Armstrong, Jones Benjamin, Eleanor G Bentley, Anu Chawla, Jordan J Clark, Angela Cowell, Richard Eccles, Isabel García-Dorival, Matthew Gemmell, Alessandro Gerada, PKF Gilmore, Richard Gregory, Ximeng Han, Catherine Hartley, Margaret Hughes, Miren Iturriza-Gomara, James Johnson, L Luu, Jenifer Manson, Charlotte Nelson, Elaine O'Toole, Cassie Olateju, Rebekah Penrice-Randal , Lucille Rainbow, N.P Randle, Trevor Ian Robinson, Parul Sharma, Ghada T Shawli, James P Stewart, Neil Swainston, Ecaterina Vamos, Joanne Watts, Mark Whitehead |
| EPI_ISL_612020 | Virology Department, Sheffield Teaching Hospitals NHS Foundation Trust/Department of Infection, Immunity and Cardiovascular Disease, The Medical School, University of Sheffield | COVID-19 Genomics UK (COG-UK) Consortium | Thushan de Silva, Matthew Parker, Nikki Smith, Adri Angyal, Rebecca Brown, Luke Green, Rachel Tucker, Paul Parsons, Danielle Groves, Katie Johnson, Laura Carrilero, Alex Keeley, Dave Partridge, Matthew Wyles, Benjamin Lindsey, Mehmet Yavuz, Mohammad Raza, Cariad Evans |
| EPI_ISL_612021, EPI_ISL_612024 | Liverpool Clinical Laboratories | COVID-19 Genomics UK (COG-UK) Consortium | Sam Haldenby, Anita Lucaci, Steve Paterson, Julian Hiscox, Alistair Darby, M Almsaud, A Alrezaihi, Muhannad Alruwaili, Stuart D Armstrong, Jones Benjamin, Eleanor G Bentley, Anu Chawla, Jordan J Clark, Angela Cowell, Richard Eccles, Isabel García-Dorival, Matthew Gemmell, Alessandro Gerada, PKF Gilmore, Richard Gregory, Ximeng Han, Catherine Hartley, Margaret Hughes, Miren Iturriza-Gomara, James Johnson, L Luu, Jenifer Manson, Charlotte Nelson, Elaine O'Toole, Cassie Olateju, Rebekah Penrice-Randal , Lucille Rainbow, N.P Randle, Trevor Ian Robinson, Parul Sharma, Ghada T Shawli, James P Stewart, Neil Swainston, Ecaterina Vamos, Joanne Watts, Mark Whitehead |
| EPI_ISL_612026, EPI_ISL_612027 | Northumbria University / South Tees Hospitals NHS Foundation Trust / North Cumbria Integrated Care NHS Foundation Trust / North Tees and Hartlepool NHS Foundation Trust / Newcastle Hospitals NHS Foundation Trust | COVID-19 Genomics UK (COG-UK) Consortium | Darren L Smith,Andrew Nelson,Matthew Bashton,Greg R Young,Joshua Loh,John Allan,Mohammad A Tariq,Giles S Holt,Gary Black,Wen C Yew,Lynn Dover,Paul Baker,Steve Liggett,Sarah Essex,Jane Greenaway,Debra Padgett,Clive Graham,Garren Scott,Edward Barton,Emma Swindells,Brendan Payne,Jennifer Collins,Yusri Taha,Gary Eltringham |
| EPI_ISL_612029 | Wales Specialist Virology Centre Sequencing lab: Pathogen Genomics Unit | COVID-19 Genomics UK (COG-UK) Consortium | Catherine Moore, Johnathan Evans, Laura Gifford, Malorie Perry, Simon Cottrell, Angela Marchbank, Alec Birchley, Alexander Adams, Amy Gaskin, Bree Gatica-Wilcox, Jason Coombes, Joel Southgate, Lauren Gilbert, Lee Graham, Nicole Pacchiarini, Sara Kumziene-Summerhayes, Sarah Taylor, Sophie Jones, Sara Rey, Matthew Bull, Joanne Watkins, Sally Corden, Tom Connor |
| EPI_ISL_612032 | Queens Medical Centre, Clinical Microbiology Department / DeepSeq Nottingham | COVID-19 Genomics UK (COG-UK) Consortium | Gemma Clark, Wendy Smith, Manjinder Khakh, Vicki M Fleming, Michelle M Lister, Hannah Howson-Wells, Jonathan Ball, Patrick McClure, Joseph Chappell, Theocharis Tsoleridis, Nadine Holmes, Matthew Carlisle, Christopher Moore, Fei Sang, Johnny Debebe, Victoria Wright, Matthew Loose |
| EPI_ISL_612033 | Virology Department, Sheffield Teaching Hospitals NHS Foundation Trust/Department of Infection, Immunity and Cardiovascular Disease, The Medical School, University of Sheffield | COVID-19 Genomics UK (COG-UK) Consortium | Thushan de Silva, Matthew Parker, Nikki Smith, Adri Angyal, Rebecca Brown, Luke Green, Rachel Tucker, Paul Parsons, Danielle Groves, Katie Johnson, Laura Carrilero, Alex Keeley, Dave Partridge, Matthew Wyles, Benjamin Lindsey, Mehmet Yavuz, Mohammad Raza, Cariad Evans |
| EPI_ISL_612034 | Department of Pathology, University of Cambridge | COVID-19 Genomics UK (COG-UK) Consortium | Aminu S. Jahun, Yasmin Chaudhry, Grant Hall, Iliana Georgana, Myra Hosmillo, Martin D. Curran, Malte Pinckert, Surendra Parmar, Ian Goodfellow |
| EPI_ISL_612035, EPI_ISL_612036 | Liverpool Clinical Laboratories | COVID-19 Genomics UK (COG-UK) Consortium | Sam Haldenby, Anita Lucaci, Steve Paterson, Julian Hiscox, Alistair Darby, M Almsaud, A Alrezaihi, Muhannad Alruwaili, Stuart D Armstrong, Jones Benjamin, Eleanor G Bentley, Anu Chawla, Jordan J Clark, Angela Cowell, Richard Eccles, Isabel García-Dorival, Matthew Gemmell, Alessandro Gerada, PKF Gilmore, Richard Gregory, Ximeng Han, Catherine Hartley, Margaret Hughes, Miren Iturriza-Gomara, James Johnson, L Luu, Jenifer Manson, Charlotte Nelson, Elaine O'Toole, Cassie Olateju, Rebekah Penrice-Randal , Lucille Rainbow, N.P Randle, Trevor Ian Robinson, Parul Sharma, Ghada T Shawli, James P Stewart, Neil Swainston, Ecaterina Vamos, Joanne Watts, Mark Whitehead |
| EPI_ISL_612037 | Virology Department, Royal Infirmary of Edinburgh, NHS Lothian / School of Biological Sciences, University of Edinburgh / Institute of Genetics and Molecular Medicine, University of Edinburgh | COVID-19 Genomics UK (COG-UK) Consortium | McHugh M, Dewar R, Rooke S, Gallagher M, Balcaza C, O'Toole Á, Scher E, Hill V, McCrone JT, Colquhoun R, Yu X, Jackson B, Rambaut A, Williams TC, Templeton K |
| EPI_ISL_612041, EPI_ISL_612043, EPI_ISL_612044 | Liverpool Clinical Laboratories | COVID-19 Genomics UK (COG-UK) Consortium | Sam Haldenby, Anita Lucaci, Steve Paterson, Julian Hiscox, Alistair Darby, M Almsaud, A Alrezaihi, Muhannad Alruwaili, Stuart D Armstrong, Jones Benjamin, Eleanor G Bentley, Anu Chawla, Jordan J Clark, Angela Cowell, Richard Eccles, Isabel García-Dorival, Matthew Gemmell, Alessandro Gerada, PKF Gilmore, |

| | | | |
|---|---|---|---|
| | | | Richard Gregory, Ximeng Han, Catherine Hartley, Margaret Hughes, Miren Iturriza-Gomara, James Johnson, L Luu, Jenifer Manson, Charlotte Nelson, Elaine O'Toole, Cassie Olateju, Rebekah Penrice-Randal , Lucille Rainbow, N.P Randle, Trevor Ian Robinson, Parul Sharma, Ghada T Shawli, James P Stewart, Neil Swainston, Ecaterina Vamos, Joanne Watts, Mark Whitehead |
| EPI_ISL_612047 | University of Birmingham | COVID-19 Genomics UK (COG-UK) Consortium | Institute of Microbiology, University of Birmingham: Claire McMurray, Joanne Stockton, Samuel Nicholls, Radoslaw Poplawski, Will Rowe, Josh Quick, Nicholas Loman. University of Birmingham Testing Laboratory: Celina M Whalley, Andrew Bosworth, Charlotte Poxon, Kasun Wanigasooriya, Oliver Pickles, Mike Kidd, Alex Richter, Andrew D Beggs PHE Heartlands Lab: Husam Osman, Andrew Bosworth. Queen Elizabeth Hospital: Anna Casey |
| EPI_ISL_612048 | Wales Specialist Virology Centre Sequencing lab: Pathogen Genomics Unit | COVID-19 Genomics UK (COG-UK) Consortium | Catherine Moore, Johnathan Evans, Laura Gifford, Malorie Perry, Simon Cottrell, Angela Marchbank, Alec Birchley, Alexander Adams, Amy Gaskin, Bree Gatica-Wilcox, Jason Coombes, Joel Southgate, Lauren Gilbert, Lee Graham, Nicole Pacchiarini, Sara Kumziene-Summerhayes, Sarah Taylor, Sophie Jones, Sara Rey, Matthew Bull, Joanne Watkins, Sally Corden, Tom Connor |
| EPI_ISL_612054 | Northumbria University / South Tees Hospitals NHS Foundation Trust / North Cumbria Integrated Care NHS Foundation Trust / North Tees and Hartlepool NHS Foundation Trust / Newcastle Hospitals NHS Foundation Trust | COVID-19 Genomics UK (COG-UK) Consortium | Darren L Smith,Andrew Nelson,Matthew Bashton,Greg R Young,Joshua Loh,John Allan,Mohammad A Tariq,Giles S Holt,Gary Black,Wen C Yew,Lynn Dover,Paul Baker,Steve Liggett,Sarah Essex,Jane Greenaway,Debra Padgett,Clive Graham,Garren Scott,Edward Barton,Emma Swindells,Brendan Payne,Jennifer Collins,Yusri Taha,Gary Eltringham |
| EPI_ISL_612059, EPI_ISL_612060, EPI_ISL_612061, EPI_ISL_612066 | Liverpool Clinical Laboratories | COVID-19 Genomics UK (COG-UK) Consortium | Sam Haldenby, Anita Lucaci, Steve Paterson, Julian Hiscox, Alistair Darby, M Almsaud, A Alrezaihi, Muhannad Alruwaili, Stuart D Armstrong, Jones Benjamin, Eleanor G Bentley, Anu Chawla, Jordan J Clark, Angela Cowell, Richard Eccles, Isabel García-Dorival, Matthew Gemmell, Alessandro Gerada, PKF Gilmore, Richard Gregory, Ximeng Han, Catherine Hartley, Margaret Hughes, Miren Iturriza-Gomara, James Johnson, L Luu, Jenifer Manson, Charlotte Nelson, Elaine O'Toole, Cassie Olateju, Rebekah Penrice-Randal , Lucille Rainbow, N.P Randle, Trevor Ian Robinson, Parul Sharma, Ghada T Shawli, James P Stewart, Neil Swainston, Ecaterina Vamos, Joanne Watts, Mark Whitehead |
| EPI_ISL_612068 | Centre for Enzyme Innovation, University of Portsmouth / Translational Research Laboratory, Portsmouth Hospitals NHS Trust | COVID-19 Genomics UK (COG-UK) Consortium | Angela Beckett,Yann Bourgeois,Garry Scarlett,Sharon Glaysher,Scott Elliott,Kelly Bicknell,Robert Impey,Allyson Lloyd,Sarah Wyllie,Ethan Butcher,Anoop Chauhan,Samuel Robson |
| EPI_ISL_612072 | Liverpool Clinical Laboratories | COVID-19 Genomics UK (COG-UK) Consortium | Sam Haldenby, Anita Lucaci, Steve Paterson, Julian Hiscox, Alistair Darby, M Almsaud, A Alrezaihi, Muhannad Alruwaili, Stuart D Armstrong, Jones Benjamin, Eleanor G Bentley, Anu Chawla, Jordan J Clark, Angela Cowell, Richard Eccles, Isabel García-Dorival, Matthew Gemmell, Alessandro Gerada, PKF Gilmore, Richard Gregory, Ximeng Han, Catherine Hartley, Margaret Hughes, Miren Iturriza-Gomara, James Johnson, L Luu, Jenifer Manson, Charlotte Nelson, Elaine O'Toole, Cassie Olateju, Rebekah Penrice-Randal , Lucille Rainbow, N.P Randle, Trevor Ian Robinson, Parul Sharma, Ghada T Shawli, James P Stewart, Neil Swainston, Ecaterina Vamos, Joanne Watts, Mark Whitehead |
| EPI_ISL_612073 | University of Birmingham | COVID-19 Genomics UK (COG-UK) Consortium | Institute of Microbiology, University of Birmingham: Claire McMurray, Joanne Stockton, Samuel Nicholls, Radoslaw Poplawski, Will Rowe, Josh Quick, Nicholas Loman. University of Birmingham Testing Laboratory: Celina M Whalley, Andrew Bosworth, Charlotte Poxon, Kasun Wanigasooriya, Oliver Pickles, Mike Kidd, Alex Richter, Andrew D Beggs PHE Heartlands Lab: Husam Osman, Andrew Bosworth. Queen Elizabeth Hospital: Anna Casey |
| EPI_ISL_612080 | Liverpool Clinical Laboratories | COVID-19 Genomics UK (COG-UK) Consortium | Sam Haldenby, Anita Lucaci, Steve Paterson, Julian Hiscox, Alistair Darby, M Almsaud, A Alrezaihi, Muhannad Alruwaili, Stuart D Armstrong, Jones Benjamin, Eleanor G Bentley, Anu Chawla, Jordan J Clark, Angela Cowell, Richard Eccles, Isabel García-Dorival, Matthew Gemmell, Alessandro Gerada, PKF Gilmore, Richard Gregory, Ximeng Han, Catherine Hartley, Margaret Hughes, Miren Iturriza-Gomara, James Johnson, L Luu, Jenifer Manson, Charlotte Nelson, Elaine O'Toole, Cassie Olateju, Rebekah Penrice-Randal , Lucille Rainbow, N.P Randle, Trevor Ian Robinson, Parul Sharma, Ghada T Shawli, James P Stewart, Neil Swainston, Ecaterina Vamos, Joanne Watts, Mark Whitehead |
| EPI_ISL_612082 | Centre for Enzyme Innovation, University of Portsmouth / Translational Research Laboratory, Portsmouth Hospitals NHS Trust | COVID-19 Genomics UK (COG-UK) Consortium | Angela Beckett,Yann Bourgeois,Garry Scarlett,Sharon Glaysher,Scott Elliott,Kelly Bicknell,Robert Impey,Allyson Lloyd,Sarah Wyllie,Ethan Butcher,Anoop Chauhan,Samuel Robson |
| EPI_ISL_612084 | Virology Department, Sheffield Teaching Hospitals NHS Foundation Trust/Department of Infection, Immunity and Cardiovascular Disease, The Medical School, University of Sheffield | COVID-19 Genomics UK (COG-UK) Consortium | Thushan de Silva, Matthew Parker, Nikki Smith, Adri Angyal, Rebecca Brown, Luke Green, Rachel Tucker, Paul Parsons, Danielle Groves, Katie Johnson, Laura Carrilero, Alex Keeley, Dave Partridge, Matthew Wyles, Benjamin Lindsey, Mehmet Yavuz, Mohammad Raza, Cariad Evans |
| EPI_ISL_612085 | Northumbria University / South Tees Hospitals NHS Foundation Trust / North Cumbria Integrated Care NHS Foundation Trust / North Tees and Hartlepool NHS Foundation Trust / Newcastle Hospitals NHS Foundation Trust | COVID-19 Genomics UK (COG-UK) Consortium | Darren L Smith,Andrew Nelson,Matthew Bashton,Greg R Young,Joshua Loh,John Allan,Mohammad A Tariq,Giles S Holt,Gary Black,Wen C Yew,Lynn Dover,Paul Baker,Steve Liggett,Sarah Essex,Jane Greenaway,Debra Padgett,Clive Graham,Garren Scott,Edward Barton,Emma Swindells,Brendan Payne,Jennifer Collins,Yusri Taha,Gary Eltringham |
| EPI_ISL_612091 | Wales Specialist Virology Centre Sequencing lab: Pathogen Genomics Unit | COVID-19 Genomics UK (COG-UK) Consortium | Catherine Moore, Johnathan Evans, Laura Gifford, Malorie Perry, Simon Cottrell, Angela Marchbank, Alec Birchley, Alexander Adams, Amy Gaskin, Bree Gatica-Wilcox, Jason Coombes, Joel Southgate, Lauren Gilbert, Lee Graham, Nicole Pacchiarini, Sara Kumziene-Summerhayes, Sarah Taylor, Sophie Jones, Sara Rey, Matthew Bull, Joanne Watkins, Sally Corden, Tom Connor |
| EPI_ISL_612106 | Department of Pathology, University of Cambridge | COVID-19 Genomics UK (COG-UK) Consortium | Aminu S. Jahun, Yasmin Chaudhry, Grant Hall, Iliana Georgana, Myra Hosmillo, Martin D. Curran, Malte Pinckert, Surendra Parmar, Ian Goodfellow |
| EPI_ISL_612107 | Queens Medical Centre, Clinical Microbiology Department / DeepSeq Nottingham | COVID-19 Genomics UK (COG-UK) Consortium | Gemma Clark, Wendy Smith, Manjinder Khakh, Vicki M Fleming, Michelle M Lister, Hannah Howson-Wells, Jonathan Ball, Patrick McClure, Joseph Chappell, Theocharis Tsoleridis, Nadine Holmes, Matthew Carlisle, Christopher Moore, Fei Sang, Johnny Debebe, Victoria Wright, Matthew Loose |
| EPI_ISL_612111, EPI_ISL_612114 | Liverpool Clinical Laboratories | COVID-19 Genomics UK (COG-UK) Consortium | Sam Haldenby, Anita Lucaci, Steve Paterson, Julian Hiscox, Alistair Darby, M Almsaud, A Alrezaihi, Muhannad Alruwaili, Stuart D Armstrong, Jones Benjamin, Eleanor G Bentley, Anu Chawla, Jordan J Clark, Angela Cowell, Richard Eccles, Isabel García-Dorival, Matthew Gemmell, Alessandro Gerada, PKF Gilmore, Richard Gregory, Ximeng Han, Catherine Hartley, Margaret Hughes, Miren Iturriza-Gomara, James Johnson, L Luu, Jenifer Manson, Charlotte Nelson, Elaine O'Toole, Cassie Olateju, Rebekah Penrice-Randal , Lucille Rainbow, N.P Randle, Trevor Ian Robinson, Parul Sharma, Ghada T Shawli, James P Stewart, Neil Swainston, Ecaterina Vamos, Joanne Watts, Mark Whitehead |
| EPI_ISL_612135, EPI_ISL_612137, EPI_ISL_612138, EPI_ISL_612139, EPI_ISL_612141, EPI_ISL_612142, EPI_ISL_612144, EPI_ISL_612162, EPI_ISL_612163, EPI_ISL_612164, EPI_ISL_612165, EPI_ISL_612166, EPI_ISL_612167, EPI_ISL_612168, EPI_ISL_612169, EPI_ISL_612170, EPI_ISL_612171 | | | |
| see above | Wales Specialist Virology Centre Sequencing lab: Pathogen Genomics Unit | COVID-19 Genomics UK (COG-UK) Consortium | Catherine Moore, Johnathan Evans, Laura Gifford, Malorie Perry, Simon Cottrell, Angela Marchbank, Alec Birchley, Alexander Adams, Amy Gaskin, Bree Gatica-Wilcox, Jason Coombes, Joel Southgate, Lauren Gilbert, Lee Graham, Nicole Pacchiarini, Sara Kumziene-Summerhayes, Sarah Taylor, Sophie Jones, Sara Rey, Matthew Bull, Joanne Watkins, Sally Corden, Tom Connor |
| EPI_ISL_612196 | Virology Department, Sheffield Teaching Hospitals NHS Foundation Trust/Department of Infection, Immunity and Cardiovascular Disease, The Medical School, University of Sheffield | COVID-19 Genomics UK (COG-UK) Consortium | Thushan de Silva, Matthew Parker, Nikki Smith, Adri Angyal, Rebecca Brown, Luke Green, Rachel Tucker, Paul Parsons, Danielle Groves, Katie Johnson, Laura Carrilero, Alex Keeley, Dave Partridge, Matthew Wyles, Benjamin Lindsey, Mehmet Yavuz, Mohammad Raza, Cariad Evans |
| EPI_ISL_612207, EPI_ISL_612208, EPI_ISL_612209, EPI_ISL_612210, EPI_ISL_612211, EPI_ISL_612212, EPI_ISL_612213, EPI_ISL_612214, EPI_ISL_612215, EPI_ISL_612216, EPI_ISL_612217, EPI_ISL_612218, EPI_ISL_612219, EPI_ISL_612220, EPI_ISL_612221, EPI_ISL_612222, EPI_ISL_612223, EPI_ISL_612224, EPI_ISL_612225, EPI_ISL_612226, EPI_ISL_612227, EPI_ISL_612228, EPI_ISL_612229, EPI_ISL_612231, EPI_ISL_612232, EPI_ISL_612233, EPI_ISL_612234, EPI_ISL_612235, EPI_ISL_612236, EPI_ISL_612237, EPI_ISL_612238, EPI_ISL_612239, EPI_ISL_612240, EPI_ISL_612241, EPI_ISL_612242, EPI_ISL_612243, EPI_ISL_612244, EPI_ISL_612245, EPI_ISL_612246, EPI_ISL_612247, EPI_ISL_612248, EPI_ISL_612249, EPI_ISL_612250, EPI_ISL_612251, EPI_ISL_612252, EPI_ISL_612253, EPI_ISL_612254, EPI_ISL_612255, EPI_ISL_612256, EPI_ISL_612257, EPI_ISL_612258, EPI_ISL_612259, EPI_ISL_612260 | | | |
| see above | University of Birmingham | COVID-19 Genomics UK (COG-UK) Consortium | Institute of Microbiology, University of Birmingham: Claire McMurray, Joanne Stockton, Samuel Nicholls, Radoslaw Poplawski, Will Rowe, Josh Quick, Nicholas Loman. University of Birmingham Testing Laboratory: Celina M Whalley, Andrew Bosworth, Charlotte Poxon, Kasun Wanigasooriya, Oliver Pickles, Mike Kidd, Alex Richter, Andrew D Beggs PHE Heartlands Lab: Husam Osman, Andrew Bosworth. Queen Elizabeth Hospital: Anna Casey |
| EPI_ISL_612266, EPI_ISL_612268, EPI_ISL_612269, EPI_ISL_612270, EPI_ISL_612271, EPI_ISL_612272, EPI_ISL_612273, EPI_ISL_612274, EPI_ISL_612275, EPI_ISL_612276, EPI_ISL_612280 | | | |
| see above | Department of Pathology, University of Cambridge | COVID-19 Genomics UK (COG-UK) Consortium | Aminu S. Jahun, Yasmin Chaudhry, Grant Hall, Iliana Georgana, Myra Hosmillo, Martin D. Curran, Malte Pinckert, Surendra Parmar, Ian Goodfellow |
| EPI_ISL_612314, EPI_ISL_612315, EPI_ISL_612316, EPI_ISL_612317, | Virology Department, Royal Infirmary of Edinburgh, NHS Lothian / School of Biological Sciences, University of | COVID-19 Genomics UK (COG-UK) Consortium | McHugh M, Dewar R, Rooke S, Gallagher M, Balcaza C, O'Toole Á, Scher E, Hill V, McCrone JT, Colquhoun R, Yu X, Jackson B, Rambaut A, Williams TC, Templeton K |

| | | | |
|---|---|---|---|
| EPI_ISL_612318, EPI_ISL_612319, EPI_ISL_612338, EPI_ISL_612339, EPI_ISL_612340, EPI_ISL_612344 | Edinburgh / Institute of Genetics and Molecular Medicine, University of Edinburgh | | |
| EPI_ISL_612549, EPI_ISL_612566, EPI_ISL_612567, EPI_ISL_612568, EPI_ISL_612569, EPI_ISL_612570 | Northumbria University / South Tees Hospitals NHS Foundation Trust / North Cumbria Integrated Care NHS Foundation Trust / North Tees and Hartlepool NHS Foundation Trust / Newcastle Hospitals NHS Foundation Trust | COVID-19 Genomics UK (COG-UK) Consortium | Darren L Smith,Andrew Nelson,Matthew Bashton,Greg R Young,Joshua Loh,John Allan,Mohammad A Tariq,Giles S Holt,Gary Black,Wen C Yew,Lynn Dover,Paul Baker,Steve Liggett,Sarah Essex,Jane Greenaway,Debra Padgett,Clive Graham,Garren Scott,Edward Barton,Emma Swindells,Brendan Payne,Jennifer Collins,Yusri Taha,Gary Eltringham |
| EPI_ISL_612607, EPI_ISL_612609, EPI_ISL_612610, EPI_ISL_612612, EPI_ISL_612613, EPI_ISL_612615, EPI_ISL_612616, EPI_ISL_612617, EPI_ISL_612620, EPI_ISL_612621, EPI_ISL_612622, EPI_ISL_612624, EPI_ISL_612625, EPI_ISL_612627, EPI_ISL_612628, EPI_ISL_612629, EPI_ISL_612631, EPI_ISL_612632, EPI_ISL_612635, EPI_ISL_612637, EPI_ISL_612641, EPI_ISL_612642, EPI_ISL_612644, EPI_ISL_612647, EPI_ISL_612656, EPI_ISL_612657, EPI_ISL_612658, EPI_ISL_612661, EPI_ISL_612663, EPI_ISL_612664, EPI_ISL_612665, EPI_ISL_612668, EPI_ISL_612669, EPI_ISL_612670, EPI_ISL_612671, EPI_ISL_612672, EPI_ISL_612678, EPI_ISL_612679, EPI_ISL_612680, EPI_ISL_612681, EPI_ISL_612682, EPI_ISL_612687, EPI_ISL_612690, EPI_ISL_612691, EPI_ISL_612695, EPI_ISL_612696, EPI_ISL_612697, EPI_ISL_612701, EPI_ISL_612702, EPI_ISL_612703, EPI_ISL_612706, EPI_ISL_612709, EPI_ISL_612710, EPI_ISL_612711, EPI_ISL_612712, EPI_ISL_612713, EPI_ISL_612714, EPI_ISL_612715, EPI_ISL_612717, EPI_ISL_612718, EPI_ISL_612719, EPI_ISL_612720, EPI_ISL_612725, EPI_ISL_612726, EPI_ISL_612727, EPI_ISL_612728, EPI_ISL_612731, EPI_ISL_612732, EPI_ISL_612733, EPI_ISL_612737, EPI_ISL_612739, EPI_ISL_612740, EPI_ISL_612745, EPI_ISL_612747, EPI_ISL_612748, EPI_ISL_612749, EPI_ISL_612750, EPI_ISL_612751, EPI_ISL_612752, EPI_ISL_612754, EPI_ISL_612755, EPI_ISL_612756, EPI_ISL_612757, EPI_ISL_612759, EPI_ISL_612760, EPI_ISL_612762, EPI_ISL_612763, EPI_ISL_612764, EPI_ISL_612766, EPI_ISL_612767, EPI_ISL_612771, EPI_ISL_612774, EPI_ISL_612777, EPI_ISL_612779, EPI_ISL_612781, EPI_ISL_612782, EPI_ISL_612784, EPI_ISL_612785, EPI_ISL_612786, EPI_ISL_612788, EPI_ISL_612790, EPI_ISL_612791, EPI_ISL_612793, EPI_ISL_612796, EPI_ISL_612797, EPI_ISL_612798, EPI_ISL_612799, EPI_ISL_612802, EPI_ISL_612803, EPI_ISL_612804, EPI_ISL_612808, EPI_ISL_612811, EPI_ISL_612812, EPI_ISL_612813, EPI_ISL_612814, EPI_ISL_612817, EPI_ISL_612819, EPI_ISL_612821, EPI_ISL_612824, EPI_ISL_612826, EPI_ISL_612828, EPI_ISL_612830, EPI_ISL_612831, EPI_ISL_612833, EPI_ISL_612834, EPI_ISL_612835, EPI_ISL_612838, EPI_ISL_612839, EPI_ISL_612840, EPI_ISL_612850, EPI_ISL_612851, EPI_ISL_612854, EPI_ISL_612858, EPI_ISL_612860, EPI_ISL_612862, EPI_ISL_612863, EPI_ISL_612869, EPI_ISL_612870, EPI_ISL_612881, EPI_ISL_612883, EPI_ISL_612885, EPI_ISL_612887, EPI_ISL_612897, EPI_ISL_612901, EPI_ISL_612907, EPI_ISL_612910, EPI_ISL_612911, EPI_ISL_612913, EPI_ISL_612920, EPI_ISL_612925, EPI_ISL_612935, EPI_ISL_612937, EPI_ISL_612938, EPI_ISL_612939, EPI_ISL_612941, EPI_ISL_612950, EPI_ISL_612959, EPI_ISL_612960, EPI_ISL_612961, EPI_ISL_612964, EPI_ISL_612967, EPI_ISL_612974, EPI_ISL_612977, EPI_ISL_612979, EPI_ISL_612982, EPI_ISL_612983, EPI_ISL_612984, EPI_ISL_612985, EPI_ISL_612989, EPI_ISL_612990, EPI_ISL_612993, EPI_ISL_613051, EPI_ISL_613056, EPI_ISL_613073, EPI_ISL_613074, EPI_ISL_613075, EPI_ISL_613080, EPI_ISL_613096, EPI_ISL_613102, EPI_ISL_613103, EPI_ISL_613112, EPI_ISL_613170, EPI_ISL_613180 | | | |
| see above | Wales Specialist Virology Centre Sequencing lab: Pathogen Genomics Unit | COVID-19 Genomics UK (COG-UK) Consortium | Catherine Moore, Johnathan Evans, Laura Gifford, Malorie Perry, Simon Cottrell, Angela Marchbank, Alec Birchley, Alexander Adams, Amy Gaskin, Bree Gatica-Wilcox, Jason Coombes, Joel Southgate, Lauren Gilbert, Lee Graham, Nicole Pacchiarini, Sara Kumziene-Summerhayes, Sarah Taylor, Sophie Jones, Sara Rey, Matthew Bull, Joanne Watkins, Sally Corden, Tom Connor |
| EPI_ISL_613288, EPI_ISL_613289, EPI_ISL_613290, EPI_ISL_613291, EPI_ISL_613292, EPI_ISL_613293, EPI_ISL_613294, EPI_ISL_613295, EPI_ISL_613302, EPI_ISL_613303, EPI_ISL_613304, EPI_ISL_613305, EPI_ISL_613306 | | | |
| see above | Centre for Enzyme Innovation, University of Portsmouth / Translational Research Laboratory, Portsmouth Hospitals NHS Trust | COVID-19 Genomics UK (COG-UK) Consortium | Angela Beckett,Yann Bourgeois,Garry Scarlett,Sharon Glaysher,Scott Elliott,Kelly Bicknell,Robert Impey,Allyson Lloyd,Sarah Wyllie,Ethan Butcher,Anoop Chauhan,Samuel Robson |
| EPI_ISL_613315, EPI_ISL_613318, EPI_ISL_613329, EPI_ISL_613347 | Queens Medical Centre, Clinical Microbiology Department / DeepSeq Nottingham | COVID-19 Genomics UK (COG-UK) Consortium | Gemma Clark, Wendy Smith, Manjinder Khakh, Vicki M Fleming, Michelle M Lister, Hannah Howson-Wells, Jonathan Ball, Patrick McClure, Joseph Chappell, Theocharis Tsoleridis, Nadine Holmes, Matthew Carlisle, Christopher Moore, Fei Sang, Johnny Debebe, Victoria Wright, Matthew Loose |
| EPI_ISL_613358, EPI_ISL_613360, EPI_ISL_613371, EPI_ISL_613375, EPI_ISL_613377, EPI_ISL_613381, EPI_ISL_613384, EPI_ISL_613386, EPI_ISL_613387, EPI_ISL_613392, EPI_ISL_613393, EPI_ISL_613396, EPI_ISL_613399, EPI_ISL_613403, EPI_ISL_613405, EPI_ISL_613407, EPI_ISL_613408, EPI_ISL_613409 | | | |
| see above | Virology Department, Sheffield Teaching Hospitals NHS Foundation Trust/Department of Infection, Immunity and Cardiovascular Disease, The Medical School, University of Sheffield | COVID-19 Genomics UK (COG-UK) Consortium | Thushan de Silva, Matthew Parker, Nikki Smith, Adri Angyal, Rebecca Brown, Luke Green, Rachel Tucker, Paul Parsons, Danielle Groves, Katie Johnson, Laura Carrilero, Alex Keeley, Dave Partridge, Matthew Wyles, Benjamin Lindsey, Mehmet Yavuz, Mohammad Raza, Cariad Evans |
| EPI_ISL_613494, EPI_ISL_613496, EPI_ISL_613497, EPI_ISL_613498, EPI_ISL_613499, EPI_ISL_613500, EPI_ISL_613501, EPI_ISL_613502 | Public Health Laboratory - Infectious Disease Lab, Minnesota Department of Health Infectious Disease Laboratory Submission Group | Minnesota Department of Health, Public Health Laboratory | Plumb,M., Garfin,J., Lorentz,A., Wang,X. |
| EPI_ISL_613587, EPI_ISL_613588, EPI_ISL_613589, EPI_ISL_613633 | Respiratory Virus Unit, Microbiology Services Colindale, Public Health England | Respiratory Virus Unit, Microbiology Services Colindale, Public Health England | PHE Covid Sequencing Team |
| EPI_ISL_614070, EPI_ISL_614073, EPI_ISL_614074, EPI_ISL_614093, EPI_ISL_614094, EPI_ISL_614096, EPI_ISL_614097, EPI_ISL_614098, EPI_ISL_614099, EPI_ISL_614149 | Virginia DCLS | Virginia DCLS | Virginia DCLS |
| EPI_ISL_614179, EPI_ISL_614181, EPI_ISL_614187, EPI_ISL_614193, EPI_ISL_614199, EPI_ISL_614210, EPI_ISL_614213, EPI_ISL_614216, EPI_ISL_614219, EPI_ISL_614223, EPI_ISL_614225, EPI_ISL_614227, EPI_ISL_614228, EPI_ISL_614229, EPI_ISL_614232 | | | |
| see above | Michigan Department of Health and Human Services, Bureau of Laboratories | Michigan Department of Health and Human Services, Bureau of Laboratories | Blankenship HM, Riner D, Soehnlen MK |
| EPI_ISL_614284, EPI_ISL_614285 | General practitioner | National Reference Center for Viruses of Respiratory Infections, Institut Pasteur, Paris | Marion Barbet, Sylvie Behillil, Méline Bizard, Angela Brisebarre, Camille Capel, Etienne Simon-Lorière, Vincent Enouf, Maud Vanpeene, Sylvie van der Werf |
| EPI_ISL_618406, EPI_ISL_618407, EPI_ISL_618408, EPI_ISL_618409, EPI_ISL_618410, EPI_ISL_618411, EPI_ISL_618412, EPI_ISL_618413, EPI_ISL_618414, EPI_ISL_618415, EPI_ISL_618416, EPI_ISL_618417, EPI_ISL_618418, EPI_ISL_618419, EPI_ISL_618420, EPI_ISL_618421, EPI_ISL_618422, EPI_ISL_618423, EPI_ISL_618424, EPI_ISL_618425, EPI_ISL_618426, EPI_ISL_618427, EPI_ISL_618428, EPI_ISL_618429, EPI_ISL_618430, EPI_ISL_618431, EPI_ISL_618432, EPI_ISL_618433, EPI_ISL_618434, EPI_ISL_618435, EPI_ISL_618436, EPI_ISL_618437, EPI_ISL_618438, EPI_ISL_618439, EPI_ISL_618440, EPI_ISL_618441, EPI_ISL_618442, EPI_ISL_618443, EPI_ISL_618444, EPI_ISL_618445, EPI_ISL_618446, EPI_ISL_618447, EPI_ISL_618448, EPI_ISL_618449, EPI_ISL_618450, EPI_ISL_618451, EPI_ISL_618452, EPI_ISL_618453, EPI_ISL_618454, EPI_ISL_618455, EPI_ISL_618456, EPI_ISL_618457, EPI_ISL_618458, EPI_ISL_618459, EPI_ISL_618460, EPI_ISL_618461, EPI_ISL_618462, EPI_ISL_618463, EPI_ISL_618464, EPI_ISL_618465, EPI_ISL_618466, EPI_ISL_618467, EPI_ISL_618468, EPI_ISL_618469, EPI_ISL_618470, EPI_ISL_618471, EPI_ISL_618472, EPI_ISL_618473, EPI_ISL_618474, EPI_ISL_618475, EPI_ISL_618476, EPI_ISL_618477, EPI_ISL_618478, EPI_ISL_618480, EPI_ISL_618481, EPI_ISL_618482, EPI_ISL_618483, EPI_ISL_618484, EPI_ISL_618485, EPI_ISL_618486, EPI_ISL_618487, EPI_ISL_618488, EPI_ISL_618489, EPI_ISL_618490, EPI_ISL_618491, EPI_ISL_618492, EPI_ISL_618493, EPI_ISL_618494, EPI_ISL_618495, EPI_ISL_618497, EPI_ISL_618498, EPI_ISL_618906, EPI_ISL_618907, EPI_ISL_618908, EPI_ISL_618909, EPI_ISL_618910, EPI_ISL_618911, EPI_ISL_618912, EPI_ISL_618913, EPI_ISL_618914, EPI_ISL_618915, EPI_ISL_618916, EPI_ISL_618917, EPI_ISL_618918, EPI_ISL_618919, EPI_ISL_618920, EPI_ISL_618921, EPI_ISL_618922, EPI_ISL_618923, EPI_ISL_618924, EPI_ISL_618925, EPI_ISL_618926, EPI_ISL_618927, EPI_ISL_618928, EPI_ISL_618929, EPI_ISL_618930, EPI_ISL_618931, EPI_ISL_618932, EPI_ISL_618933, EPI_ISL_618934, EPI_ISL_618935, EPI_ISL_618936, EPI_ISL_618937, EPI_ISL_618938, EPI_ISL_618939, EPI_ISL_618940, EPI_ISL_618941, EPI_ISL_618942, EPI_ISL_618943, EPI_ISL_618944, EPI_ISL_618945, EPI_ISL_618946, EPI_ISL_618947, EPI_ISL_618948, EPI_ISL_618949, EPI_ISL_618950, EPI_ISL_618951, EPI_ISL_618952, EPI_ISL_618953, EPI_ISL_618954, EPI_ISL_618955, EPI_ISL_618956, EPI_ISL_618957, EPI_ISL_618958, EPI_ISL_618959, EPI_ISL_618960, EPI_ISL_618961, EPI_ISL_618963, EPI_ISL_618964, EPI_ISL_618966, EPI_ISL_618967, EPI_ISL_618968, EPI_ISL_618970, EPI_ISL_618971, EPI_ISL_618973, EPI_ISL_618974, EPI_ISL_618975, EPI_ISL_618976, EPI_ISL_618977, EPI_ISL_618981, EPI_ISL_618982, EPI_ISL_618983, EPI_ISL_618984, EPI_ISL_618985, EPI_ISL_618986, EPI_ISL_618987, EPI_ISL_618988, EPI_ISL_618989, EPI_ISL_618990, EPI_ISL_618991, EPI_ISL_618992, EPI_ISL_618993, EPI_ISL_618994, EPI_ISL_618995, EPI_ISL_618996, EPI_ISL_618997, EPI_ISL_618998, EPI_ISL_618999, EPI_ISL_619000, EPI_ISL_619001, EPI_ISL_619002, EPI_ISL_619003, EPI_ISL_619004, EPI_ISL_619005, EPI_ISL_619006, EPI_ISL_619007, EPI_ISL_619008, EPI_ISL_619009, EPI_ISL_619010, EPI_ISL_619011, EPI_ISL_619012, EPI_ISL_619013, EPI_ISL_619014, EPI_ISL_619015, EPI_ISL_619016, EPI_ISL_619019, EPI_ISL_619020, EPI_ISL_619021, EPI_ISL_619022, EPI_ISL_619023, EPI_ISL_619024, EPI_ISL_619025, EPI_ISL_619026, EPI_ISL_619027, EPI_ISL_619028, EPI_ISL_619029, EPI_ISL_619030, EPI_ISL_619031, EPI_ISL_619032, EPI_ISL_619033, EPI_ISL_619034, EPI_ISL_619035, EPI_ISL_619036, EPI_ISL_619037, EPI_ISL_619038, EPI_ISL_619039, EPI_ISL_619040, EPI_ISL_619041, EPI_ISL_619042, EPI_ISL_619043, EPI_ISL_619044, EPI_ISL_619045, EPI_ISL_619046, EPI_ISL_619047, EPI_ISL_619048, EPI_ISL_619049, EPI_ISL_619050, EPI_ISL_619051, EPI_ISL_619052, EPI_ISL_619053, EPI_ISL_619054, EPI_ISL_619055, EPI_ISL_619056, EPI_ISL_619057, EPI_ISL_619058, EPI_ISL_619059, EPI_ISL_619060, EPI_ISL_619061, EPI_ISL_619062, EPI_ISL_619064, EPI_ISL_619065, EPI_ISL_619066, EPI_ISL_619067, EPI_ISL_619068, EPI_ISL_619069, EPI_ISL_619070, EPI_ISL_619071, EPI_ISL_619072, EPI_ISL_619073, EPI_ISL_619074, EPI_ISL_619075, EPI_ISL_619076, EPI_ISL_619077, EPI_ISL_619078, EPI_ISL_619079, EPI_ISL_619080, EPI_ISL_619081, EPI_ISL_619082, EPI_ISL_619083, EPI_ISL_619084, EPI_ISL_619085, EPI_ISL_619086, EPI_ISL_619087, EPI_ISL_619088, EPI_ISL_619089, EPI_ISL_619090, EPI_ISL_619091, EPI_ISL_619092, EPI_ISL_619093, EPI_ISL_619094, EPI_ISL_619095, EPI_ISL_619096, EPI_ISL_619097, EPI_ISL_619098, EPI_ISL_619099, EPI_ISL_619100, EPI_ISL_619101, EPI_ISL_619102, EPI_ISL_619103, EPI_ISL_619104, EPI_ISL_619105, EPI_ISL_619106, EPI_ISL_619107, EPI_ISL_619108, EPI_ISL_619109, EPI_ISL_619110, EPI_ISL_619111, EPI_ISL_619112, EPI_ISL_619113, EPI_ISL_619114, EPI_ISL_619115, EPI_ISL_619116, EPI_ISL_619117, EPI_ISL_619118, EPI_ISL_619119, EPI_ISL_619120, EPI_ISL_619121, EPI_ISL_619122, EPI_ISL_619123, EPI_ISL_619124, EPI_ISL_619125, EPI_ISL_619126, EPI_ISL_619127, EPI_ISL_619128, EPI_ISL_619129, EPI_ISL_619130, EPI_ISL_619131, EPI_ISL_619132, EPI_ISL_619134, EPI_ISL_619135, EPI_ISL_619136, EPI_ISL_619137, EPI_ISL_619138, EPI_ISL_619139, EPI_ISL_619140, EPI_ISL_619142, EPI_ISL_619143, EPI_ISL_619144, EPI_ISL_619146, EPI_ISL_619147, EPI_ISL_619148, EPI_ISL_619149, EPI_ISL_619150, EPI_ISL_619152, EPI_ISL_619153, EPI_ISL_619154, EPI_ISL_619155, EPI_ISL_619156, EPI_ISL_619157, EPI_ISL_619158, EPI_ISL_619159, EPI_ISL_619160, EPI_ISL_619161, EPI_ISL_619162, EPI_ISL_619164, EPI_ISL_619165, EPI_ISL_619166, EPI_ISL_619167, EPI_ISL_619168, EPI_ISL_619169, EPI_ISL_619171, EPI_ISL_619172, EPI_ISL_619173, EPI_ISL_619174, EPI_ISL_619175, EPI_ISL_619177, EPI_ISL_619178, EPI_ISL_619179, EPI_ISL_619180, EPI_ISL_619181, EPI_ISL_619182, EPI_ISL_619183, EPI_ISL_619184, EPI_ISL_619186, EPI_ISL_619187, EPI_ISL_619188, EPI_ISL_619189, EPI_ISL_619190, EPI_ISL_619191, EPI_ISL_619192, EPI_ISL_619193, EPI_ISL_619194, EPI_ISL_619195, EPI_ISL_619196, EPI_ISL_619197, EPI_ISL_619198, EPI_ISL_619199, EPI_ISL_619200, EPI_ISL_619201, EPI_ISL_619202, EPI_ISL_619203, EPI_ISL_619204, EPI_ISL_619205, EPI_ISL_619206, EPI_ISL_619207, EPI_ISL_619208, EPI_ISL_619209, EPI_ISL_619210, EPI_ISL_619212, EPI_ISL_619213, EPI_ISL_619214, EPI_ISL_619215, EPI_ISL_619216, EPI_ISL_619217, EPI_ISL_619218, EPI_ISL_619219, EPI_ISL_619220, EPI_ISL_619221, EPI_ISL_619222, EPI_ISL_619223, EPI_ISL_619224, EPI_ISL_619225, EPI_ISL_619227, EPI_ISL_619228, EPI_ISL_619229, EPI_ISL_619230, EPI_ISL_619231, EPI_ISL_619232, EPI_ISL_619234, EPI_ISL_619235, EPI_ISL_619236, EPI_ISL_619237, EPI_ISL_619238, EPI_ISL_619239, EPI_ISL_619240, EPI_ISL_619241, EPI_ISL_619242, EPI_ISL_619243, EPI_ISL_619244, EPI_ISL_619245, EPI_ISL_619246, EPI_ISL_619247, EPI_ISL_619248, EPI_ISL_619249, EPI_ISL_619250, EPI_ISL_619251, EPI_ISL_619252, EPI_ISL_619253, EPI_ISL_619254, EPI_ISL_619255, EPI_ISL_619257, EPI_ISL_619258, EPI_ISL_619259, EPI_ISL_619274, EPI_ISL_619275, EPI_ISL_619276, EPI_ISL_619277, EPI_ISL_619278, EPI_ISL_619279, EPI_ISL_619280, EPI_ISL_619281, EPI_ISL_619282, EPI_ISL_619283, EPI_ISL_619284, EPI_ISL_619285, EPI_ISL_619286, EPI_ISL_619287, EPI_ISL_619288, EPI_ISL_619289, EPI_ISL_619290, EPI_ISL_619291, EPI_ISL_619293, EPI_ISL_619294, EPI_ISL_619295, EPI_ISL_619296, EPI_ISL_619297, EPI_ISL_619298, EPI_ISL_619299, EPI_ISL_619300, EPI_ISL_619301, EPI_ISL_619302, EPI_ISL_619303, EPI_ISL_619304, EPI_ISL_619305, EPI_ISL_619306, EPI_ISL_619307, EPI_ISL_619308, EPI_ISL_619309, EPI_ISL_619310, EPI_ISL_619311, EPI_ISL_619312, EPI_ISL_619313, EPI_ISL_619314, EPI_ISL_619315, EPI_ISL_619316, EPI_ISL_619317, EPI_ISL_619319, EPI_ISL_619320, EPI_ISL_619322, EPI_ISL_619323, EPI_ISL_619324, EPI_ISL_619325, EPI_ISL_619326, EPI_ISL_619327, EPI_ISL_619329, EPI_ISL_619330, EPI_ISL_619331, EPI_ISL_619333, EPI_ISL_619334, EPI_ISL_619335, EPI_ISL_619336, EPI_ISL_619338, EPI_ISL_619339, EPI_ISL_619340, EPI_ISL_619341, EPI_ISL_619342, EPI_ISL_619343, EPI_ISL_619345, EPI_ISL_619346, EPI_ISL_619347, EPI_ISL_619348, EPI_ISL_619349, EPI_ISL_619350, EPI_ISL_619351, EPI_ISL_619352, EPI_ISL_619353, EPI_ISL_619354, EPI_ISL_619355, EPI_ISL_619356, EPI_ISL_619357, EPI_ISL_619358, EPI_ISL_619359, EPI_ISL_619360, EPI_ISL_619361, EPI_ISL_619362, EPI_ISL_619363, EPI_ISL_619364, EPI_ISL_619365, EPI_ISL_619366, EPI_ISL_619367, EPI_ISL_619368, EPI_ISL_619369, EPI_ISL_619370, EPI_ISL_619371, EPI_ISL_619372, EPI_ISL_619373, EPI_ISL_619374, EPI_ISL_619375, EPI_ISL_619376, EPI_ISL_619377, EPI_ISL_619379, EPI_ISL_619380, EPI_ISL_619381, EPI_ISL_619382, EPI_ISL_619383, EPI_ISL_619385, EPI_ISL_619386, EPI_ISL_619387, EPI_ISL_619388, EPI_ISL_619390, EPI_ISL_619391, EPI_ISL_619392, EPI_ISL_619393, EPI_ISL_619394, EPI_ISL_619395, EPI_ISL_619396, EPI_ISL_619397, EPI_ISL_619399, EPI_ISL_619400, EPI_ISL_619401, EPI_ISL_619402, EPI_ISL_619403, EPI_ISL_619404, EPI_ISL_619457, EPI_ISL_619458, EPI_ISL_619459, EPI_ISL_619460, EPI_ISL_619461, EPI_ISL_619462, EPI_ISL_619463, EPI_ISL_619464, EPI_ISL_619465, EPI_ISL_619466, EPI_ISL_619467, EPI_ISL_619468, EPI_ISL_619469, EPI_ISL_619470, EPI_ISL_619471, | | | |

| | | | |
|---|---|---|---|
| EPI_ISL_619472, EPI_ISL_619473, EPI_ISL_619474, EPI_ISL_619475, EPI_ISL_619476, EPI_ISL_619477, EPI_ISL_619478, EPI_ISL_619479, EPI_ISL_619480, EPI_ISL_619481, EPI_ISL_619482, EPI_ISL_619483, EPI_ISL_619484, EPI_ISL_619485, EPI_ISL_619486, EPI_ISL_619487, EPI_ISL_619488, EPI_ISL_619489, EPI_ISL_619490, EPI_ISL_619491, EPI_ISL_619493, EPI_ISL_619494, EPI_ISL_619495, EPI_ISL_619496, EPI_ISL_619497, EPI_ISL_619498, EPI_ISL_619499, EPI_ISL_619500, EPI_ISL_619501, EPI_ISL_619502, EPI_ISL_619503, EPI_ISL_619504, EPI_ISL_619505, EPI_ISL_619506, EPI_ISL_619507, EPI_ISL_619508, EPI_ISL_619509, EPI_ISL_619510, EPI_ISL_619511, EPI_ISL_619512, EPI_ISL_619513, EPI_ISL_619514, EPI_ISL_619515, EPI_ISL_619516, EPI_ISL_619517, EPI_ISL_619518, EPI_ISL_619519, EPI_ISL_619520, EPI_ISL_619521, EPI_ISL_619522, EPI_ISL_619523, EPI_ISL_619524, EPI_ISL_619525, EPI_ISL_619526, EPI_ISL_619527, EPI_ISL_619528, EPI_ISL_619529, EPI_ISL_619530, EPI_ISL_619531, EPI_ISL_619532, EPI_ISL_619533, EPI_ISL_619535, EPI_ISL_619536, EPI_ISL_619537, EPI_ISL_619538, EPI_ISL_619539, EPI_ISL_619540, EPI_ISL_619541, EPI_ISL_619542, EPI_ISL_619543, EPI_ISL_619544, EPI_ISL_619545, EPI_ISL_619546, EPI_ISL_619547, EPI_ISL_619548, EPI_ISL_619549, EPI_ISL_619550, EPI_ISL_619551, EPI_ISL_619552, EPI_ISL_619553, EPI_ISL_619554, EPI_ISL_619555, EPI_ISL_619557, EPI_ISL_619558, EPI_ISL_619559, EPI_ISL_619560, EPI_ISL_619561, EPI_ISL_619562, EPI_ISL_619563, EPI_ISL_619564, EPI_ISL_619565, EPI_ISL_619566, EPI_ISL_619567, EPI_ISL_619568, EPI_ISL_619569, EPI_ISL_619570, EPI_ISL_619571, EPI_ISL_619572, EPI_ISL_619573, EPI_ISL_619574, EPI_ISL_619575, EPI_ISL_619576, EPI_ISL_619577, EPI_ISL_619578, EPI_ISL_619579, EPI_ISL_619580, EPI_ISL_619581, EPI_ISL_619582, EPI_ISL_619583, EPI_ISL_619584, EPI_ISL_619585, EPI_ISL_619586, EPI_ISL_619587, EPI_ISL_619588, EPI_ISL_619589, EPI_ISL_619590, EPI_ISL_619591, EPI_ISL_619592, EPI_ISL_619593, EPI_ISL_619594, EPI_ISL_619595, EPI_ISL_619596, EPI_ISL_619597, EPI_ISL_619598, EPI_ISL_619599, EPI_ISL_619600, EPI_ISL_619601, EPI_ISL_619602, EPI_ISL_619603, EPI_ISL_619604, EPI_ISL_619605, EPI_ISL_619606, EPI_ISL_619607, EPI_ISL_619608, EPI_ISL_619609, EPI_ISL_619610, EPI_ISL_619611, EPI_ISL_619612, EPI_ISL_619613, EPI_ISL_619614, EPI_ISL_619615, EPI_ISL_619616, EPI_ISL_619617, EPI_ISL_619618, EPI_ISL_619619, EPI_ISL_619620, EPI_ISL_619621, EPI_ISL_619622, EPI_ISL_619623, EPI_ISL_619624, EPI_ISL_619625, EPI_ISL_619626, EPI_ISL_619627, EPI_ISL_619628, EPI_ISL_619629, EPI_ISL_619630, EPI_ISL_619631, EPI_ISL_619632, EPI_ISL_619633, EPI_ISL_619634, EPI_ISL_619635, EPI_ISL_619636, EPI_ISL_619637, EPI_ISL_619638, EPI_ISL_619639, EPI_ISL_619640, EPI_ISL_619641, EPI_ISL_619642, EPI_ISL_619643, EPI_ISL_619644, EPI_ISL_619645, EPI_ISL_619646, EPI_ISL_619647, EPI_ISL_619648, EPI_ISL_619649, EPI_ISL_619650, EPI_ISL_619651, EPI_ISL_619652, EPI_ISL_619653, EPI_ISL_619654, EPI_ISL_619655, EPI_ISL_619656, EPI_ISL_619658, EPI_ISL_619659, EPI_ISL_619660, EPI_ISL_619661, EPI_ISL_619662, EPI_ISL_619663, EPI_ISL_619664, EPI_ISL_619665, EPI_ISL_619666, EPI_ISL_619667, EPI_ISL_619668, EPI_ISL_619669, EPI_ISL_619670, EPI_ISL_619671, EPI_ISL_619672, EPI_ISL_619673, EPI_ISL_619674, EPI_ISL_619675, EPI_ISL_619676, EPI_ISL_619677, EPI_ISL_619678, EPI_ISL_619679, EPI_ISL_619680, EPI_ISL_619681, EPI_ISL_619682, EPI_ISL_619683, EPI_ISL_619684, EPI_ISL_619685, EPI_ISL_619686, EPI_ISL_619687, EPI_ISL_619688, EPI_ISL_619689, EPI_ISL_619690, EPI_ISL_619691, EPI_ISL_619692, EPI_ISL_619693, EPI_ISL_619694, EPI_ISL_619695, EPI_ISL_619696, EPI_ISL_619697, EPI_ISL_619698, EPI_ISL_619699, EPI_ISL_619700, EPI_ISL_619701, EPI_ISL_619702, EPI_ISL_619703, EPI_ISL_619704, EPI_ISL_619705, EPI_ISL_619706, EPI_ISL_619707, EPI_ISL_619708, EPI_ISL_619709, EPI_ISL_619710, EPI_ISL_619711, EPI_ISL_619712, EPI_ISL_619713, EPI_ISL_619714, EPI_ISL_619715, EPI_ISL_619716, EPI_ISL_619717, EPI_ISL_619718, EPI_ISL_619719, EPI_ISL_619720, EPI_ISL_619721, EPI_ISL_619722, EPI_ISL_619723, EPI_ISL_619724, EPI_ISL_619725, EPI_ISL_619726, EPI_ISL_619727, EPI_ISL_619728, EPI_ISL_619729, EPI_ISL_619730, EPI_ISL_619731, EPI_ISL_619732, EPI_ISL_619733, EPI_ISL_619734, EPI_ISL_619735, EPI_ISL_619736, EPI_ISL_619737, EPI_ISL_619738, EPI_ISL_619739, EPI_ISL_619740, EPI_ISL_619741, EPI_ISL_619742, EPI_ISL_619743, EPI_ISL_619744, EPI_ISL_619745, EPI_ISL_619746, EPI_ISL_619747, EPI_ISL_619748, EPI_ISL_619749, EPI_ISL_619750, EPI_ISL_619752, EPI_ISL_619753, EPI_ISL_619754, EPI_ISL_619755, EPI_ISL_619756, EPI_ISL_619757, EPI_ISL_619758, EPI_ISL_619759, EPI_ISL_619760, EPI_ISL_619761, EPI_ISL_619762, EPI_ISL_619763, EPI_ISL_619764, EPI_ISL_619765, EPI_ISL_619766, EPI_ISL_619767, EPI_ISL_619768, EPI_ISL_619769, EPI_ISL_619770, EPI_ISL_619771, EPI_ISL_619772, EPI_ISL_619773, EPI_ISL_619774, EPI_ISL_619775, EPI_ISL_619776, EPI_ISL_619777, EPI_ISL_619778, EPI_ISL_619779, EPI_ISL_619780, EPI_ISL_619781, EPI_ISL_619782, EPI_ISL_619783, EPI_ISL_619784, EPI_ISL_619785, EPI_ISL_619786, EPI_ISL_619787, EPI_ISL_619788, EPI_ISL_619789, EPI_ISL_619790, EPI_ISL_619791, EPI_ISL_619792, EPI_ISL_619793, EPI_ISL_619794, EPI_ISL_619795, EPI_ISL_619796, EPI_ISL_619797, EPI_ISL_619798, EPI_ISL_619799, EPI_ISL_619800, EPI_ISL_619801, EPI_ISL_619802, EPI_ISL_619803, EPI_ISL_619804, EPI_ISL_619805, EPI_ISL_619806, EPI_ISL_619807, EPI_ISL_619808, EPI_ISL_619809, EPI_ISL_619810, EPI_ISL_619811, EPI_ISL_619812, EPI_ISL_619813, EPI_ISL_619814, EPI_ISL_619815, EPI_ISL_619816, EPI_ISL_619817, EPI_ISL_619818, EPI_ISL_619819, EPI_ISL_619820, EPI_ISL_619821, EPI_ISL_619822, EPI_ISL_619823, EPI_ISL_619824, EPI_ISL_619825, EPI_ISL_619826, EPI_ISL_619827, EPI_ISL_619828, EPI_ISL_619829, EPI_ISL_619830, EPI_ISL_619831, EPI_ISL_619832, EPI_ISL_619833, EPI_ISL_619834, EPI_ISL_619835, EPI_ISL_619836, EPI_ISL_619837, EPI_ISL_619838, EPI_ISL_619839, EPI_ISL_619841, EPI_ISL_619842, EPI_ISL_619843, EPI_ISL_619844, EPI_ISL_619845, EPI_ISL_619846, EPI_ISL_619847, EPI_ISL_619848, EPI_ISL_619849, EPI_ISL_619850, EPI_ISL_619851, EPI_ISL_619852, EPI_ISL_619853, EPI_ISL_619854, EPI_ISL_619855, EPI_ISL_619856, EPI_ISL_619857, EPI_ISL_619858, EPI_ISL_619859, EPI_ISL_619860, EPI_ISL_619861, EPI_ISL_619862, EPI_ISL_619863, EPI_ISL_619864, EPI_ISL_619865, EPI_ISL_619866, EPI_ISL_619867, EPI_ISL_619868, EPI_ISL_619869, EPI_ISL_619870, EPI_ISL_619871, EPI_ISL_619872, EPI_ISL_619873, EPI_ISL_619874, EPI_ISL_619875, EPI_ISL_619876, EPI_ISL_619877, EPI_ISL_619878, EPI_ISL_619879, EPI_ISL_619880, EPI_ISL_619881, EPI_ISL_619882, EPI_ISL_619883, EPI_ISL_619884, EPI_ISL_619886, EPI_ISL_619887, EPI_ISL_619888, EPI_ISL_619889, EPI_ISL_619890, EPI_ISL_619891, EPI_ISL_619892, EPI_ISL_619893, EPI_ISL_619894, EPI_ISL_619896, EPI_ISL_619897, EPI_ISL_619898, EPI_ISL_619899, EPI_ISL_619900, EPI_ISL_619901, EPI_ISL_619902, EPI_ISL_619903, EPI_ISL_619904, EPI_ISL_620303, EPI_ISL_620304, EPI_ISL_620305, EPI_ISL_620306, EPI_ISL_620307, EPI_ISL_620308, EPI_ISL_620309, EPI_ISL_620310, EPI_ISL_620311, EPI_ISL_620312, EPI_ISL_620313, EPI_ISL_620314, EPI_ISL_620315, EPI_ISL_620316, EPI_ISL_620317, EPI_ISL_620318, EPI_ISL_620319, EPI_ISL_620320, EPI_ISL_620321, EPI_ISL_620322, EPI_ISL_620323, EPI_ISL_620324, EPI_ISL_620325, EPI_ISL_620326, EPI_ISL_620327, EPI_ISL_620328, EPI_ISL_620329, EPI_ISL_620330, EPI_ISL_620331, EPI_ISL_620332, EPI_ISL_620333, EPI_ISL_620334, EPI_ISL_620335, EPI_ISL_620336, EPI_ISL_620337, EPI_ISL_620338, EPI_ISL_620339, EPI_ISL_620340, EPI_ISL_620341, EPI_ISL_620342, EPI_ISL_620343, EPI_ISL_620344, EPI_ISL_620345, EPI_ISL_620346, EPI_ISL_620347, EPI_ISL_620348, EPI_ISL_620349, EPI_ISL_620350, EPI_ISL_620351, EPI_ISL_620352, EPI_ISL_620353, EPI_ISL_620354, EPI_ISL_620355, EPI_ISL_620356, EPI_ISL_620357, EPI_ISL_620358, EPI_ISL_620359, EPI_ISL_620360, EPI_ISL_620361, EPI_ISL_620362, EPI_ISL_620363, EPI_ISL_620364, EPI_ISL_620365, EPI_ISL_620366, EPI_ISL_620367, EPI_ISL_620368, EPI_ISL_620369, EPI_ISL_620370, EPI_ISL_620371, EPI_ISL_620372, EPI_ISL_620373, EPI_ISL_620374, EPI_ISL_620375, EPI_ISL_620376, EPI_ISL_620377, EPI_ISL_620378, EPI_ISL_620379 | | | |
| see above | Department of Virus and Microbiological Special Diagnostics, Statens Serum Institut, Denmark | Albertsen lab, Department of Chemistry and Bioscience, Aalborg University, Denmark | Danish Covid-19 Genome Consortia |
| EPI_ISL_622785 | Waikato Hospital | Institute of Environmental Science and Research (ESR) | Xiaoyun Ren, Matt Storey, Nikki Freed, Muhammad Faisal, Jing Wang, Hermes Perez, Anja Werno, Antje van der Linden, Arlo Upton, Chris Mansell, David Hammer, Dragana Drinkovic, Gary McAuliffe, Hana Sofia Andersson, James Ussher, Jill Sherwood, Josh Freeman, Julia Howard, Juliet Elvy, Mary DeAlmeida, Matt Blakiston, Matthew Rogers, Max Bloomfield, Michael Addidle, Michelle Balm, Sally Roberts, Sarah Jefferies, Sharmini Muttaiyah, Susan Morpeth, Susan Taylor, Timothy Blackmore, Vani Sathyendran, Veronica Playle, Virginia Hope, Erasmus Smit, Lauren Jelly, Olin Silander, Joep de Ligt |
| EPI_ISL_622843, EPI_ISL_622846, EPI_ISL_622848, EPI_ISL_622851, EPI_ISL_622872 | Respiratory Virus Unit, Microbiology Services Colindale, Public Health England | Respiratory Virus Unit, Microbiology Services Colindale, Public Health England | PHE Covid Sequencing Team |
| EPI_ISL_625281, EPI_ISL_625282, EPI_ISL_625284, EPI_ISL_625285, EPI_ISL_625286 | Lighthouse Lab in Cambridge | Wellcome Sanger Institute for the COVID-19 Genomics UK (COG-UK) consortium | Rob Howes, The Lighthouse Lab in Cambridge and Alex Alderton, Roberto Amato, Sonia Goncalves, Ewan Harrison, David K. Jackson, Ian Johnston, Dominic Kwiatkowski, Cordelia Langford, John Sillitoe on behalf of the Wellcome Sanger Institute COVID-19 Surveillance Team (http://www.sanger.ac.uk/covid-team) |
| EPI_ISL_625289 | Lighthouse Lab in Cambridge | Wellcome Sanger Institute for the COVID-19 Genomics UK (COG-UK) Consortium | Rob Howes, The Lighthouse Lab in Cambridge and Alex Alderton, Roberto Amato, Sonia Goncalves, Ewan Harrison, David K. Jackson, Ian Johnston, Dominic Kwiatkowski, Cordelia Langford, John Sillitoe on behalf of the Wellcome Sanger Institute COVID-19 Surveillance Team (http://www.sanger.ac.uk/covid-team) |
| EPI_ISL_625290, EPI_ISL_625292, EPI_ISL_625293, EPI_ISL_625297, EPI_ISL_625298, EPI_ISL_625299, EPI_ISL_625300, EPI_ISL_625301, EPI_ISL_625302, EPI_ISL_625303, EPI_ISL_625307, EPI_ISL_625308, EPI_ISL_625309, EPI_ISL_625313, EPI_ISL_625314 | | | |
| see above | Lighthouse Lab in Cambridge | Wellcome Sanger Institute for the COVID-19 Genomics UK (COG-UK) consortium | Rob Howes, The Lighthouse Lab in Cambridge and Alex Alderton, Roberto Amato, Sonia Goncalves, Ewan Harrison, David K. Jackson, Ian Johnston, Dominic Kwiatkowski, Cordelia Langford, John Sillitoe on behalf of the Wellcome Sanger Institute COVID-19 Surveillance Team (http://www.sanger.ac.uk/covid-team) |
| EPI_ISL_625405, EPI_ISL_625406, EPI_ISL_625407, EPI_ISL_625408, EPI_ISL_625409, EPI_ISL_625410, EPI_ISL_625411, EPI_ISL_625412, EPI_ISL_625413, EPI_ISL_625414, EPI_ISL_625415, EPI_ISL_625416, EPI_ISL_625417, EPI_ISL_625418, EPI_ISL_625419, EPI_ISL_625420, EPI_ISL_625421 | | | |
| see above | Lighthouse Lab in Milton Keynes | Wellcome Sanger Institute for the COVID-19 Genomics UK (COG-UK) consortium | The Lighthouse Lab in Milton Keynes and Alex Alderton, Roberto Amato, Sonia Goncalves, Ewan Harrison, David K. Jackson, Ian Johnston, Dominic Kwiatkowski, Cordelia Langford, John Sillitoe on behalf of the Wellcome Sanger Institute COVID-19 Surveillance Team (http://www.sanger.ac.uk/covid-team) |
| EPI_ISL_625422, EPI_ISL_625423 | Lighthouse Lab in Alderley Park | Wellcome Sanger Institute for the COVID-19 Genomics UK (COG-UK) consortium | Jacquelyn Wynn, Mairead Hyland, The Lighthouse Lab in Alderley Park and Alex Alderton, Roberto Amato, Sonia Goncalves, Ewan Harrison, David K. Jackson, Ian Johnston, Dominic Kwiatkowski, Cordelia Langford, John Sillitoe on behalf of the Wellcome Sanger Institute COVID-19 Surveillance Team (http://www.sanger.ac.uk/covid-team) |
| EPI_ISL_625424 | Lighthouse Lab in Milton Keynes | Wellcome Sanger Institute for the COVID-19 Genomics UK (COG-UK) consortium | The Lighthouse Lab in Milton Keynes and Alex Alderton, Roberto Amato, Sonia Goncalves, Ewan Harrison, David K. Jackson, Ian Johnston, Dominic Kwiatkowski, Cordelia Langford, John Sillitoe on behalf of the Wellcome Sanger Institute COVID-19 Surveillance Team (http://www.sanger.ac.uk/covid-team) |
| EPI_ISL_625425 | Lighthouse Lab in Alderley Park | Wellcome Sanger Institute for the COVID-19 Genomics UK (COG-UK) consortium | Jacquelyn Wynn, Mairead Hyland, The Lighthouse Lab in Alderley Park and Alex Alderton, Roberto Amato, Sonia Goncalves, Ewan Harrison, David K. Jackson, Ian Johnston, Dominic Kwiatkowski, Cordelia Langford, John Sillitoe on behalf of the Wellcome Sanger Institute COVID-19 Surveillance Team (http://www.sanger.ac.uk/covid-team) |
| EPI_ISL_625426 | Lighthouse Lab in Milton Keynes | Wellcome Sanger Institute for the COVID-19 Genomics UK (COG-UK) consortium | The Lighthouse Lab in Milton Keynes and Alex Alderton, Roberto Amato, Sonia Goncalves, Ewan Harrison, David K. Jackson, Ian Johnston, Dominic Kwiatkowski, Cordelia Langford, John Sillitoe on behalf of the Wellcome Sanger Institute COVID-19 Surveillance Team (http://www.sanger.ac.uk/covid-team) |
| EPI_ISL_625427 | Lighthouse Lab in Alderley Park | Wellcome Sanger Institute for the COVID-19 Genomics UK (COG-UK) consortium | Jacquelyn Wynn, Mairead Hyland, The Lighthouse Lab in Alderley Park and Alex Alderton, Roberto Amato, Sonia Goncalves, Ewan Harrison, David K. Jackson, Ian Johnston, Dominic Kwiatkowski, Cordelia Langford, John Sillitoe on behalf of the Wellcome Sanger Institute COVID-19 Surveillance Team (http://www.sanger.ac.uk/covid-team) |
| EPI_ISL_625428 | Lighthouse Lab in Milton Keynes | Wellcome Sanger Institute for the COVID-19 Genomics UK (COG-UK) consortium | The Lighthouse Lab in Milton Keynes and Alex Alderton, Roberto Amato, Sonia Goncalves, Ewan Harrison, David K. Jackson, Ian Johnston, Dominic Kwiatkowski, Cordelia Langford, John Sillitoe on behalf of the Wellcome Sanger Institute COVID-19 Surveillance Team (http://www.sanger.ac.uk/covid-team) |
| EPI_ISL_625429 | Lighthouse Lab in Alderley Park | Wellcome Sanger Institute for the COVID-19 Genomics UK (COG-UK) consortium | Jacquelyn Wynn, Mairead Hyland, The Lighthouse Lab in Alderley Park and Alex Alderton, Roberto Amato, Sonia Goncalves, Ewan Harrison, David K. Jackson, Ian Johnston, Dominic Kwiatkowski, Cordelia Langford, John Sillitoe on behalf of the Wellcome Sanger Institute COVID-19 Surveillance Team (http://www.sanger.ac.uk/covid-team) |
| EPI_ISL_625430 | Lighthouse Lab in Milton Keynes | Wellcome Sanger Institute for the COVID-19 Genomics UK (COG-UK) consortium | The Lighthouse Lab in Milton Keynes and Alex Alderton, Roberto Amato, Sonia Goncalves, Ewan Harrison, David K. Jackson, Ian Johnston, Dominic Kwiatkowski, Cordelia Langford, John Sillitoe on behalf of the Wellcome Sanger Institute COVID-19 Surveillance Team (http://www.sanger.ac.uk/covid-team) |
| EPI_ISL_625431 | Lighthouse Lab in Alderley Park | Wellcome Sanger Institute for the COVID-19 Genomics UK (COG-UK) consortium | Jacquelyn Wynn, Mairead Hyland, The Lighthouse Lab in Alderley Park and Alex Alderton, Roberto Amato, Sonia Goncalves, Ewan Harrison, David K. Jackson, Ian Johnston, Dominic Kwiatkowski, Cordelia Langford, John Sillitoe on behalf of the Wellcome Sanger Institute COVID-19 Surveillance Team (http://www.sanger.ac.uk/covid-team) |
| EPI_ISL_625432 | Lighthouse Lab in Cambridge | Wellcome Sanger Institute for the COVID-19 Genomics UK (COG-UK) consortium | Rob Howes, The Lighthouse Lab in Cambridge and Alex Alderton, Roberto Amato, Sonia Goncalves, Ewan Harrison, David K. Jackson, Ian Johnston, Dominic Kwiatkowski, Cordelia Langford, John Sillitoe on behalf of the Wellcome Sanger Institute COVID-19 Surveillance Team (http://www.sanger.ac.uk/covid-team) |
| EPI_ISL_625433, EPI_ISL_625434, EPI_ISL_625435, EPI_ISL_625436 | Lighthouse Lab in Alderley Park | Wellcome Sanger Institute for the COVID-19 Genomics UK (COG-UK) consortium | Jacquelyn Wynn, Mairead Hyland, The Lighthouse Lab in Alderley Park and Alex Alderton, Roberto Amato, Sonia Goncalves, Ewan Harrison, David K. Jackson, Ian Johnston, Dominic Kwiatkowski, Cordelia Langford, John Sillitoe on behalf of the Wellcome Sanger Institute COVID-19 Surveillance Team (http://www.sanger.ac.uk/covid-team) |
| EPI_ISL_625441, EPI_ISL_625442, | Lighthouse Lab in Cambridge | Wellcome Sanger Institute for the COVID-19 Genomics UK | Rob Howes, The Lighthouse Lab in Cambridge and Alex Alderton, Roberto Amato, Sonia Goncalves, Ewan Harrison, David K. Jackson, Ian Johnston, Dominic |

| | | | |
|---|---|---|---|
| EPI_ISL_625443 | | (COG-UK) consortium | Kwiatkowski, Cordelia Langford, John Sillitoe on behalf of the Wellcome Sanger Institute COVID-19 Surveillance Team (http://www.sanger.ac.uk/covid-team) |
| EPI_ISL_625610 | Santa Clara County Public Health Laboratory | Chan-Zuckerberg Biohub | CZB Cliahub Consortium |
| EPI_ISL_625661, EPI_ISL_625662, EPI_ISL_625663, EPI_ISL_625664 | UCSF Clinical Microbiology Laboratory | Chan-Zuckerberg Biohub | CZB Cliahub Consortium |
| EPI_ISL_625678, EPI_ISL_625679 | Laboratory of Molecular Medicine, University of Magallanes | Centro Asistencial Docente y de Investigacion, Universidad de Magallanes | Jorge Gonzalez, Jacqueline Aldridge, Diego Alvarez, Marcelo Navarrete |
| EPI_ISL_626594, EPI_ISL_626595, EPI_ISL_626596, EPI_ISL_626603, EPI_ISL_626604, EPI_ISL_626605, EPI_ISL_626606, EPI_ISL_626607, EPI_ISL_626608, EPI_ISL_626609, EPI_ISL_626610, EPI_ISL_626612, EPI_ISL_626613, EPI_ISL_626614, EPI_ISL_626615, EPI_ISL_626619, EPI_ISL_626620, EPI_ISL_626621, EPI_ISL_626622 | | | |
| see above | The National Institute of Public Health | State Veterinary Institute Prague | Nagy,A;Jirincova,H;Novakova,L;Trnka,D;Vecerova,J |
| EPI_ISL_626661 | Liverpool Clinical Laboratories | COVID-19 Genomics UK (COG-UK) Consortium | Sam Haldenby, Anita Lucaci, Steve Paterson, Julian Hiscox, Alistair Darby, M Almsaud, A Alrezaihi, Muhannad Alruwaili, Stuart D Armstrong, Jones Benjamin, Eleanor G Bentley, Anu Chawla, Jordan J Clark, Angela Cowell, Richard Eccles, Isabel García-Dorival, Matthew Gemmell, Alessandro Gerada, PKF Gilmore, Richard Gregory, Ximeng Han, Catherine Hartley, Margaret Hughes, Miren Iturriza-Gomara, James Johnson, L Luu, Jenifer Manson, Charlotte Nelson, Elaine O'Toole, Cassie Olateju, Rebekah Penrice-Randal , Lucille Rainbow, N.P Randle, Trevor Ian Robinson, Parul Sharma, Ghada T Shawli, James P Stewart, Neil Swainston, Ecaterina Vamos, Joanne Watts, Mark Whitehead |
| EPI_ISL_626662 | University of Exeter | COVID-19 Genomics UK (COG-UK) Consortium | Ben Temperton,Aaron Jeffries,Michelle Michelsen,Joanna Warwick-Dugdale,Audrey Farbos,Robyn Manley,Stephen Michell,Jane Masoli |
| EPI_ISL_626665 | Liverpool Clinical Laboratories | COVID-19 Genomics UK (COG-UK) Consortium | Sam Haldenby, Anita Lucaci, Steve Paterson, Julian Hiscox, Alistair Darby, M Almsaud, A Alrezaihi, Muhannad Alruwaili, Stuart D Armstrong, Jones Benjamin, Eleanor G Bentley, Anu Chawla, Jordan J Clark, Angela Cowell, Richard Eccles, Isabel García-Dorival, Matthew Gemmell, Alessandro Gerada, PKF Gilmore, Richard Gregory, Ximeng Han, Catherine Hartley, Margaret Hughes, Miren Iturriza-Gomara, James Johnson, L Luu, Jenifer Manson, Charlotte Nelson, Elaine O'Toole, Cassie Olateju, Rebekah Penrice-Randal , Lucille Rainbow, N.P Randle, Trevor Ian Robinson, Parul Sharma, Ghada T Shawli, James P Stewart, Neil Swainston, Ecaterina Vamos, Joanne Watts, Mark Whitehead |
| EPI_ISL_626671, EPI_ISL_626673, EPI_ISL_626676, EPI_ISL_626682 | University of Exeter | COVID-19 Genomics UK (COG-UK) Consortium | Ben Temperton,Aaron Jeffries,Michelle Michelsen,Joanna Warwick-Dugdale,Audrey Farbos,Robyn Manley,Stephen Michell,Jane Masoli |
| EPI_ISL_626709 | Liverpool Clinical Laboratories | COVID-19 Genomics UK (COG-UK) Consortium | Sam Haldenby, Anita Lucaci, Steve Paterson, Julian Hiscox, Alistair Darby, M Almsaud, A Alrezaihi, Muhannad Alruwaili, Stuart D Armstrong, Jones Benjamin, Eleanor G Bentley, Anu Chawla, Jordan J Clark, Angela Cowell, Richard Eccles, Isabel García-Dorival, Matthew Gemmell, Alessandro Gerada, PKF Gilmore, Richard Gregory, Ximeng Han, Catherine Hartley, Margaret Hughes, Miren Iturriza-Gomara, James Johnson, L Luu, Jenifer Manson, Charlotte Nelson, Elaine O'Toole, Cassie Olateju, Rebekah Penrice-Randal , Lucille Rainbow, N.P Randle, Trevor Ian Robinson, Parul Sharma, Ghada T Shawli, James P Stewart, Neil Swainston, Ecaterina Vamos, Joanne Watts, Mark Whitehead |
| EPI_ISL_626710 | Virology Department, Sheffield Teaching Hospitals NHS Foundation Trust/Department of Infection, Immunity and Cardiovascular Disease, The Medical School, University of Sheffield | COVID-19 Genomics UK (COG-UK) Consortium | Thushan de Silva, Matthew Parker, Nikki Smith, Adri Angyal, Rebecca Brown, Luke Green, Rachel Tucker, Paul Parsons, Danielle Groves, Katie Johnson, Laura Carrilero, Alex Keeley, Dave Partridge, Matthew Wyles, Benjamin Lindsey, Mehmet Yavuz, Mohammad Raza, Cariad Evans |
| EPI_ISL_626712, EPI_ISL_626714 | Liverpool Clinical Laboratories | COVID-19 Genomics UK (COG-UK) Consortium | Sam Haldenby, Anita Lucaci, Steve Paterson, Julian Hiscox, Alistair Darby, M Almsaud, A Alrezaihi, Muhannad Alruwaili, Stuart D Armstrong, Jones Benjamin, Eleanor G Bentley, Anu Chawla, Jordan J Clark, Angela Cowell, Richard Eccles, Isabel García-Dorival, Matthew Gemmell, Alessandro Gerada, PKF Gilmore, Richard Gregory, Ximeng Han, Catherine Hartley, Margaret Hughes, Miren Iturriza-Gomara, James Johnson, L Luu, Jenifer Manson, Charlotte Nelson, Elaine O'Toole, Cassie Olateju, Rebekah Penrice-Randal , Lucille Rainbow, N.P Randle, Trevor Ian Robinson, Parul Sharma, Ghada T Shawli, James P Stewart, Neil Swainston, Ecaterina Vamos, Joanne Watts, Mark Whitehead |
| EPI_ISL_626715, EPI_ISL_626716, EPI_ISL_626717, EPI_ISL_626718, EPI_ISL_626719, EPI_ISL_626720, EPI_ISL_626721, EPI_ISL_626722, EPI_ISL_626723, EPI_ISL_626724, EPI_ISL_626725, EPI_ISL_626729 | | | |
| see above | University of Exeter | COVID-19 Genomics UK (COG-UK) Consortium | Ben Temperton,Aaron Jeffries,Michelle Michelsen,Joanna Warwick-Dugdale,Audrey Farbos,Robyn Manley,Stephen Michell,Jane Masoli |
| EPI_ISL_626735 | Virology Department, Sheffield Teaching Hospitals NHS Foundation Trust/Department of Infection, Immunity and Cardiovascular Disease, The Medical School, University of Sheffield | COVID-19 Genomics UK (COG-UK) Consortium | Thushan de Silva, Matthew Parker, Nikki Smith, Adri Angyal, Rebecca Brown, Luke Green, Rachel Tucker, Paul Parsons, Danielle Groves, Katie Johnson, Laura Carrilero, Alex Keeley, Dave Partridge, Matthew Wyles, Benjamin Lindsey, Mehmet Yavuz, Mohammad Raza, Cariad Evans |
| EPI_ISL_626742 | Liverpool Clinical Laboratories | COVID-19 Genomics UK (COG-UK) Consortium | Sam Haldenby, Anita Lucaci, Steve Paterson, Julian Hiscox, Alistair Darby, M Almsaud, A Alrezaihi, Muhannad Alruwaili, Stuart D Armstrong, Jones Benjamin, Eleanor G Bentley, Anu Chawla, Jordan J Clark, Angela Cowell, Richard Eccles, Isabel García-Dorival, Matthew Gemmell, Alessandro Gerada, PKF Gilmore, Richard Gregory, Ximeng Han, Catherine Hartley, Margaret Hughes, Miren Iturriza-Gomara, James Johnson, L Luu, Jenifer Manson, Charlotte Nelson, Elaine O'Toole, Cassie Olateju, Rebekah Penrice-Randal , Lucille Rainbow, N.P Randle, Trevor Ian Robinson, Parul Sharma, Ghada T Shawli, James P Stewart, Neil Swainston, Ecaterina Vamos, Joanne Watts, Mark Whitehead |
| EPI_ISL_626743, EPI_ISL_626747, EPI_ISL_626754 | Virology Department, Sheffield Teaching Hospitals NHS Foundation Trust/Department of Infection, Immunity and Cardiovascular Disease, The Medical School, University of Sheffield | COVID-19 Genomics UK (COG-UK) Consortium | Thushan de Silva, Matthew Parker, Nikki Smith, Adri Angyal, Rebecca Brown, Luke Green, Rachel Tucker, Paul Parsons, Danielle Groves, Katie Johnson, Laura Carrilero, Alex Keeley, Dave Partridge, Matthew Wyles, Benjamin Lindsey, Mehmet Yavuz, Mohammad Raza, Cariad Evans |
| EPI_ISL_626756, EPI_ISL_626759 | Quadram Institute Bioscience | COVID-19 Genomics UK (COG-UK) Consortium | Dave J. Baker, Gemma L. Kay, Alp Aydin, Thanh Le-Viet, Steven Rudder, Ana P. Tedim, Anastasia Kolyva, Maria Diaz, Leonardo de Oliveira Martins, Nabil-Fareed Alikhan, Lizzie Meadows, Rachael Stanley, Ngozi Elumogo, Muhammed Yasir, Nicholas M. Thomson, Alexander J Trotter, Rachel Gilroy, Samuel Bloomfield, Claire Stuart, Andrew Bell, Reenesh Prakash, Samir Dervisevic, Alison E. Mather, John Wain, Mark Webber, Andrew J. Page, Justin O'Grady |
| EPI_ISL_626766 | Liverpool Clinical Laboratories | COVID-19 Genomics UK (COG-UK) Consortium | Sam Haldenby, Anita Lucaci, Steve Paterson, Julian Hiscox, Alistair Darby, M Almsaud, A Alrezaihi, Muhannad Alruwaili, Stuart D Armstrong, Jones Benjamin, Eleanor G Bentley, Anu Chawla, Jordan J Clark, Angela Cowell, Richard Eccles, Isabel García-Dorival, Matthew Gemmell, Alessandro Gerada, PKF Gilmore, Richard Gregory, Ximeng Han, Catherine Hartley, Margaret Hughes, Miren Iturriza-Gomara, James Johnson, L Luu, Jenifer Manson, Charlotte Nelson, Elaine O'Toole, Cassie Olateju, Rebekah Penrice-Randal , Lucille Rainbow, N.P Randle, Trevor Ian Robinson, Parul Sharma, Ghada T Shawli, James P Stewart, Neil Swainston, Ecaterina Vamos, Joanne Watts, Mark Whitehead |
| EPI_ISL_626772, EPI_ISL_626773, EPI_ISL_626774, EPI_ISL_626775 | University of Exeter | COVID-19 Genomics UK (COG-UK) Consortium | Ben Temperton,Aaron Jeffries,Michelle Michelsen,Joanna Warwick-Dugdale,Audrey Farbos,Robyn Manley,Stephen Michell,Jane Masoli |
| EPI_ISL_626780 | Wales Specialist Virology Centre Sequencing lab: Pathogen Genomics Unit | COVID-19 Genomics UK (COG-UK) Consortium | Catherine Moore, Johnathan Evans, Laura Gifford, Malorie Perry, Simon Cottrell, Angela Marchbank, Alec Birchley, Alexander Adams, Amy Gaskin, Bree Gatica-Wilcox, Jason Coombes, Joel Southgate, Lauren Gilbert, Lee Graham, Nicole Pacchiarini, Sara Kumziene-Summerhayes, Sarah Taylor, Sophie Jones, Sara Rey, Matthew Bull, Joanne Watkins, Sally Corden, Tom Connor |
| EPI_ISL_626781 | Virology Department, Sheffield Teaching Hospitals NHS Foundation Trust/Department of Infection, Immunity and Cardiovascular Disease, The Medical School, University of Sheffield | COVID-19 Genomics UK (COG-UK) Consortium | Thushan de Silva, Matthew Parker, Nikki Smith, Adri Angyal, Rebecca Brown, Luke Green, Rachel Tucker, Paul Parsons, Danielle Groves, Katie Johnson, Laura Carrilero, Alex Keeley, Dave Partridge, Matthew Wyles, Benjamin Lindsey, Mehmet Yavuz, Mohammad Raza, Cariad Evans |
| EPI_ISL_626785 | University of Exeter | COVID-19 Genomics UK (COG-UK) Consortium | Ben Temperton,Aaron Jeffries,Michelle Michelsen,Joanna Warwick-Dugdale,Audrey Farbos,Robyn Manley,Stephen Michell,Jane Masoli |
| EPI_ISL_626789 | Liverpool Clinical Laboratories | COVID-19 Genomics UK (COG-UK) Consortium | Sam Haldenby, Anita Lucaci, Steve Paterson, Julian Hiscox, Alistair Darby, M Almsaud, A Alrezaihi, Muhannad Alruwaili, Stuart D Armstrong, Jones Benjamin, Eleanor G Bentley, Anu Chawla, Jordan J Clark, Angela Cowell, Richard Eccles, Isabel García-Dorival, Matthew Gemmell, Alessandro Gerada, PKF Gilmore, Richard Gregory, Ximeng Han, Catherine Hartley, Margaret Hughes, Miren Iturriza-Gomara, James Johnson, L Luu, Jenifer Manson, Charlotte Nelson, Elaine O'Toole, Cassie Olateju, Rebekah Penrice-Randal , Lucille Rainbow, N.P Randle, Trevor Ian Robinson, Parul Sharma, Ghada T Shawli, James P Stewart, Neil Swainston, Ecaterina Vamos, Joanne Watts, Mark Whitehead |
| EPI_ISL_626793, EPI_ISL_626798, | University of Exeter | COVID-19 Genomics UK (COG-UK) Consortium | Ben Temperton,Aaron Jeffries,Michelle Michelsen,Joanna Warwick-Dugdale,Audrey Farbos,Robyn Manley,Stephen Michell,Jane Masoli |

| | | | |
|---|---|---|---|
| EPI_ISL_626799, EPI_ISL_626800, EPI_ISL_626804 | | | |
| EPI_ISL_626808 | Quadram Institute Bioscience | COVID-19 Genomics UK (COG-UK) Consortium | Dave J. Baker, Gemma L. Kay, Alp Aydin, Thanh Le-Viet, Steven Rudder, Ana P. Tedim, Anastasia Kolyva, Maria Diaz, Leonardo de Oliveira Martins, Nabil-Fareed Alikhan, Lizzie Meadows, Rachael Stanley, Ngozi Elumogo, Muhammed Yasir, Nicholas M. Thomson, Alexander J Trotter, Rachel Gilroy, Samuel Bloomfield, Claire Stuart, Andrew Bell, Reenesh Prakash, Samir Dervisevic, Alison E. Mather, John Wain, Mark Webber, Andrew J. Page, Justin O'Grady |
| EPI_ISL_626809 | Virology Department, Sheffield Teaching Hospitals NHS Foundation Trust/Department of Infection, Immunity and Cardiovascular Disease, The Medical School, University of Sheffield | COVID-19 Genomics UK (COG-UK) Consortium | Thushan de Silva, Matthew Parker, Nikki Smith, Adri Angyal, Rebecca Brown, Luke Green, Rachel Tucker, Paul Parsons, Danielle Groves, Katie Johnson, Laura Carrilero, Alex Keeley, Dave Partridge, Matthew Wyles, Benjamin Lindsey, Mehmet Yavuz, Mohammad Raza, Cariad Evans |
| EPI_ISL_626815 | Liverpool Clinical Laboratories | COVID-19 Genomics UK (COG-UK) Consortium | Sam Haldenby, Anita Lucaci, Steve Paterson, Julian Hiscox, Alistair Darby, M Almsaud, A Alrezaihi, Muhannad Alruwaili, Stuart D Armstrong, Jones Benjamin, Eleanor G Bentley, Anu Chawla, Jordan J Clark, Angela Cowell, Richard Eccles, Isabel García-Dorival, Matthew Gemmell, Alessandro Gerada, PKF Gilmore, Richard Gregory, Ximeng Han, Catherine Hartley, Margaret Hughes, Miren Iturriza-Gomara, James Johnson, L Luu, Jenifer Manson, Charlotte Nelson, Elaine O'Toole, Cassie Olateju, Rebekah Penrice-Randal , Lucille Rainbow, N.P Randle, Trevor Ian Robinson, Parul Sharma, Ghada T Shawli, James P Stewart, Neil Swainston, Ecaterina Vamos, Joanne Watts, Mark Whitehead |
| EPI_ISL_626817, EPI_ISL_626823, EPI_ISL_626824, EPI_ISL_626827 | Virology Department, Sheffield Teaching Hospitals NHS Foundation Trust/Department of Infection, Immunity and Cardiovascular Disease, The Medical School, University of Sheffield | COVID-19 Genomics UK (COG-UK) Consortium | Thushan de Silva, Matthew Parker, Nikki Smith, Adri Angyal, Rebecca Brown, Luke Green, Rachel Tucker, Paul Parsons, Danielle Groves, Katie Johnson, Laura Carrilero, Alex Keeley, Dave Partridge, Matthew Wyles, Benjamin Lindsey, Mehmet Yavuz, Mohammad Raza, Cariad Evans |
| EPI_ISL_626828 | Regional Virus Laboratory, Belfast Health and Social Care Trust | COVID-19 Genomics UK (COG-UK) Consortium | Conall McCaughey, James McKenna, Tanya Curran, Susan Feeney, Alison Watt, Ciara Cox, Mairead Connor, Zoltan Molnar, David Simpson, Derek Fairley |
| EPI_ISL_626830 | Virology Department, Sheffield Teaching Hospitals NHS Foundation Trust/Department of Infection, Immunity and Cardiovascular Disease, The Medical School, University of Sheffield | COVID-19 Genomics UK (COG-UK) Consortium | Thushan de Silva, Matthew Parker, Nikki Smith, Adri Angyal, Rebecca Brown, Luke Green, Rachel Tucker, Paul Parsons, Danielle Groves, Katie Johnson, Laura Carrilero, Alex Keeley, Dave Partridge, Matthew Wyles, Benjamin Lindsey, Mehmet Yavuz, Mohammad Raza, Cariad Evans |
| EPI_ISL_626850 | Wales Specialist Virology Centre Sequencing lab: Pathogen Genomics Unit | COVID-19 Genomics UK (COG-UK) Consortium | Catherine Moore, Johnathan Evans, Laura Gifford, Malorie Perry, Simon Cottrell, Angela Marchbank, Alec Birchley, Alexander Adams, Amy Gaskin, Bree Gatica-Wilcox, Jason Coombes, Joel Southgate, Lauren Gilbert, Lee Graham, Nicole Pacchiarini, Sara Kumziene-Summerhayes, Sarah Taylor, Sophie Jones, Sara Rey, Matthew Bull, Joanne Watkins, Sally Corden, Tom Connor |
| EPI_ISL_626852 | Virology Department, Sheffield Teaching Hospitals NHS Foundation Trust/Department of Infection, Immunity and Cardiovascular Disease, The Medical School, University of Sheffield | COVID-19 Genomics UK (COG-UK) Consortium | Thushan de Silva, Matthew Parker, Nikki Smith, Adri Angyal, Rebecca Brown, Luke Green, Rachel Tucker, Paul Parsons, Danielle Groves, Katie Johnson, Laura Carrilero, Alex Keeley, Dave Partridge, Matthew Wyles, Benjamin Lindsey, Mehmet Yavuz, Mohammad Raza, Cariad Evans |
| EPI_ISL_626859 | Regional Virus Laboratory, Belfast Health and Social Care Trust | COVID-19 Genomics UK (COG-UK) Consortium | Conall McCaughey, James McKenna, Tanya Curran, Susan Feeney, Alison Watt, Ciara Cox, Mairead Connor, Zoltan Molnar, David Simpson, Derek Fairley |
| EPI_ISL_626882, EPI_ISL_626883, EPI_ISL_626884, EPI_ISL_626885, EPI_ISL_626886, EPI_ISL_626887, EPI_ISL_626888 | University of Exeter | COVID-19 Genomics UK (COG-UK) Consortium | Ben Temperton,Aaron Jeffries,Michelle Michelsen,Joanna Warwick-Dugdale,Audrey Farbos,Robyn Manley,Stephen Michell,Jane Masoli |
| EPI_ISL_626893 | Quadram Institute Bioscience | COVID-19 Genomics UK (COG-UK) Consortium | Dave J. Baker, Gemma L. Kay, Alp Aydin, Thanh Le-Viet, Steven Rudder, Ana P. Tedim, Anastasia Kolyva, Maria Diaz, Leonardo de Oliveira Martins, Nabil-Fareed Alikhan, Lizzie Meadows, Rachael Stanley, Ngozi Elumogo, Muhammed Yasir, Nicholas M. Thomson, Alexander J Trotter, Rachel Gilroy, Samuel Bloomfield, Claire Stuart, Andrew Bell, Reenesh Prakash, Samir Dervisevic, Alison E. Mather, John Wain, Mark Webber, Andrew J. Page, Justin O'Grady |
| EPI_ISL_626896, EPI_ISL_626898, EPI_ISL_626899, EPI_ISL_626900 | Virology Department, Sheffield Teaching Hospitals NHS Foundation Trust/Department of Infection, Immunity and Cardiovascular Disease, The Medical School, University of Sheffield | COVID-19 Genomics UK (COG-UK) Consortium | Thushan de Silva, Matthew Parker, Nikki Smith, Adri Angyal, Rebecca Brown, Luke Green, Rachel Tucker, Paul Parsons, Danielle Groves, Katie Johnson, Laura Carrilero, Alex Keeley, Dave Partridge, Matthew Wyles, Benjamin Lindsey, Mehmet Yavuz, Mohammad Raza, Cariad Evans |
| EPI_ISL_626905 | Liverpool Clinical Laboratories | COVID-19 Genomics UK (COG-UK) Consortium | Sam Haldenby, Anita Lucaci, Steve Paterson, Julian Hiscox, Alistair Darby, M Almsaud, A Alrezaihi, Muhannad Alruwaili, Stuart D Armstrong, Jones Benjamin, Eleanor G Bentley, Anu Chawla, Jordan J Clark, Angela Cowell, Richard Eccles, Isabel García-Dorival, Matthew Gemmell, Alessandro Gerada, PKF Gilmore, Richard Gregory, Ximeng Han, Catherine Hartley, Margaret Hughes, Miren Iturriza-Gomara, James Johnson, L Luu, Jenifer Manson, Charlotte Nelson, Elaine O'Toole, Cassie Olateju, Rebekah Penrice-Randal , Lucille Rainbow, N.P Randle, Trevor Ian Robinson, Parul Sharma, Ghada T Shawli, James P Stewart, Neil Swainston, Ecaterina Vamos, Joanne Watts, Mark Whitehead |
| EPI_ISL_626906, EPI_ISL_626907, EPI_ISL_626908, EPI_ISL_626909 | Virology Department, Sheffield Teaching Hospitals NHS Foundation Trust/Department of Infection, Immunity and Cardiovascular Disease, The Medical School, University of Sheffield | COVID-19 Genomics UK (COG-UK) Consortium | Thushan de Silva, Matthew Parker, Nikki Smith, Adri Angyal, Rebecca Brown, Luke Green, Rachel Tucker, Paul Parsons, Danielle Groves, Katie Johnson, Laura Carrilero, Alex Keeley, Dave Partridge, Matthew Wyles, Benjamin Lindsey, Mehmet Yavuz, Mohammad Raza, Cariad Evans |
| EPI_ISL_626910 | University of Exeter | COVID-19 Genomics UK (COG-UK) Consortium | Ben Temperton,Aaron Jeffries,Michelle Michelsen,Joanna Warwick-Dugdale,Audrey Farbos,Robyn Manley,Stephen Michell,Jane Masoli |
| EPI_ISL_626911, EPI_ISL_626912, EPI_ISL_626913, EPI_ISL_626918 | Virology Department, Sheffield Teaching Hospitals NHS Foundation Trust/Department of Infection, Immunity and Cardiovascular Disease, The Medical School, University of Sheffield | COVID-19 Genomics UK (COG-UK) Consortium | Thushan de Silva, Matthew Parker, Nikki Smith, Adri Angyal, Rebecca Brown, Luke Green, Rachel Tucker, Paul Parsons, Danielle Groves, Katie Johnson, Laura Carrilero, Alex Keeley, Dave Partridge, Matthew Wyles, Benjamin Lindsey, Mehmet Yavuz, Mohammad Raza, Cariad Evans |
| EPI_ISL_626925 | University of Exeter | COVID-19 Genomics UK (COG-UK) Consortium | Ben Temperton,Aaron Jeffries,Michelle Michelsen,Joanna Warwick-Dugdale,Audrey Farbos,Robyn Manley,Stephen Michell,Jane Masoli |
| EPI_ISL_626933, EPI_ISL_626934 | West of Scotland Specialist Virology Centre, NHSGGC / MRC-University of Glasgow Centre for Virus Research | COVID-19 Genomics UK (COG-UK) Consortium | Ana da Silva Filipe, Natasha Johnson, Kathy Smollett, Daniel Mair, Stephen Carmichael, Lily Tong, Jenna Nichols, Elihu Aranday-Cortes, Kyriaki Nomikou; Sarah McDonald, Marc Niebel, Patawee Asamaphan; Richard Orton, Joseph Hughes, Sreenu Vattipally, David L Robertson; Alasdair MacLean, Rory Gunson; Kathy Li, Igor Starinskij, Natasha Jesudason, Rajiv Shah, James Shepherd, Antonia Ho, Emma Thomson |
| EPI_ISL_626938 | Virology Department, Sheffield Teaching Hospitals NHS Foundation Trust/Department of Infection, Immunity and Cardiovascular Disease, The Medical School, University of Sheffield | COVID-19 Genomics UK (COG-UK) Consortium | Thushan de Silva, Matthew Parker, Nikki Smith, Adri Angyal, Rebecca Brown, Luke Green, Rachel Tucker, Paul Parsons, Danielle Groves, Katie Johnson, Laura Carrilero, Alex Keeley, Dave Partridge, Matthew Wyles, Benjamin Lindsey, Mehmet Yavuz, Mohammad Raza, Cariad Evans |
| EPI_ISL_626939, EPI_ISL_626940 | University of Exeter | COVID-19 Genomics UK (COG-UK) Consortium | Ben Temperton,Aaron Jeffries,Michelle Michelsen,Joanna Warwick-Dugdale,Audrey Farbos,Robyn Manley,Stephen Michell,Jane Masoli |
| EPI_ISL_626947, EPI_ISL_626948, EPI_ISL_626949, EPI_ISL_626950, EPI_ISL_626951 | Virology Department, Sheffield Teaching Hospitals NHS Foundation Trust/Department of Infection, Immunity and Cardiovascular Disease, The Medical School, University of Sheffield | COVID-19 Genomics UK (COG-UK) Consortium | Thushan de Silva, Matthew Parker, Nikki Smith, Adri Angyal, Rebecca Brown, Luke Green, Rachel Tucker, Paul Parsons, Danielle Groves, Katie Johnson, Laura Carrilero, Alex Keeley, Dave Partridge, Matthew Wyles, Benjamin Lindsey, Mehmet Yavuz, Mohammad Raza, Cariad Evans |
| EPI_ISL_626953 | Regional Virus Laboratory, Belfast Health and Social Care Trust | COVID-19 Genomics UK (COG-UK) Consortium | Conall McCaughey, James McKenna, Tanya Curran, Susan Feeney, Alison Watt, Ciara Cox, Mairead Connor, Zoltan Molnar, David Simpson, Derek Fairley |
| EPI_ISL_627004 | Liverpool Clinical Laboratories | COVID-19 Genomics UK (COG-UK) Consortium | Sam Haldenby, Anita Lucaci, Steve Paterson, Julian Hiscox, Alistair Darby, M Almsaud, A Alrezaihi, Muhannad Alruwaili, Stuart D Armstrong, Jones Benjamin, Eleanor G Bentley, Anu Chawla, Jordan J Clark, Angela Cowell, Richard Eccles, Isabel García-Dorival, Matthew Gemmell, Alessandro Gerada, PKF Gilmore, Richard Gregory, Ximeng Han, Catherine Hartley, Margaret Hughes, Miren Iturriza-Gomara, James Johnson, L Luu, Jenifer Manson, Charlotte Nelson, Elaine |

| | | | |
|---|---|---|---|
| | | | O'Toole, Cassie Olateju, Rebekah Penrice-Randal , Lucille Rainbow, N.P Randle, Trevor Ian Robinson, Parul Sharma, Ghada T Shawli, James P Stewart, Neil Swainston, Ecaterina Vamos, Joanne Watts, Mark Whitehead |
| EPI_ISL_627008 | Regional Virus Laboratory, Belfast Health and Social Care Trust | COVID-19 Genomics UK (COG-UK) Consortium | Conall McCaughey, James McKenna, Tanya Curran, Susan Feeney, Alison Watt, Ciara Cox, Mairead Connor, Zoltan Molnar, David Simpson, Derek Fairley |
| EPI_ISL_627012 | Wales Specialist Virology Centre Sequencing lab: Pathogen Genomics Unit | COVID-19 Genomics UK (COG-UK) Consortium | Catherine Moore, Johnathan Evans, Laura Gifford, Malorie Perry, Simon Cottrell, Angela Marchbank, Alec Birchley, Alexander Adams, Amy Gaskin, Bree Gatica-Wilcox, Jason Coombes, Joel Southgate, Lauren Gilbert, Lee Graham, Nicole Pacchiarini, Sara Kumziene-Summerhayes, Sarah Taylor, Sophie Jones, Sara Rey, Matthew Bull, Joanne Watkins, Sally Corden, Tom Connor |
| EPI_ISL_627020 | Regional Virus Laboratory, Belfast Health and Social Care Trust | COVID-19 Genomics UK (COG-UK) Consortium | Conall McCaughey, James McKenna, Tanya Curran, Susan Feeney, Alison Watt, Ciara Cox, Mairead Connor, Zoltan Molnar, David Simpson, Derek Fairley |
| EPI_ISL_627023 | Wales Specialist Virology Centre Sequencing lab: Pathogen Genomics Unit | COVID-19 Genomics UK (COG-UK) Consortium | Catherine Moore, Johnathan Evans, Laura Gifford, Malorie Perry, Simon Cottrell, Angela Marchbank, Alec Birchley, Alexander Adams, Amy Gaskin, Bree Gatica-Wilcox, Jason Coombes, Joel Southgate, Lauren Gilbert, Lee Graham, Nicole Pacchiarini, Sara Kumziene-Summerhayes, Sarah Taylor, Sophie Jones, Sara Rey, Matthew Bull, Joanne Watkins, Sally Corden, Tom Connor |
| EPI_ISL_627031 | University of Exeter | COVID-19 Genomics UK (COG-UK) Consortium | Ben Temperton,Aaron Jeffries,Michelle Michelsen,Joanna Warwick-Dugdale,Audrey Farbos,Robyn Manley,Stephen Michell,Jane Masoli |
| EPI_ISL_627046 | Regional Virus Laboratory, Belfast Health and Social Care Trust | COVID-19 Genomics UK (COG-UK) Consortium | Conall McCaughey, James McKenna, Tanya Curran, Susan Feeney, Alison Watt, Ciara Cox, Mairead Connor, Zoltan Molnar, David Simpson, Derek Fairley |
| EPI_ISL_627047 | Virology Department, Sheffield Teaching Hospitals NHS Foundation Trust/Department of Infection, Immunity and Cardiovascular Disease, The Medical School, University of Sheffield | COVID-19 Genomics UK (COG-UK) Consortium | Thushan de Silva, Matthew Parker, Nikki Smith, Adri Angyal, Rebecca Brown, Luke Green, Rachel Tucker, Paul Parsons, Danielle Groves, Katie Johnson, Laura Carrilero, Alex Keeley, Dave Partridge, Matthew Wyles, Benjamin Lindsey, Mehmet Yavuz, Mohammad Raza, Cariad Evans |
| EPI_ISL_627050 | Wales Specialist Virology Centre Sequencing lab: Pathogen Genomics Unit | COVID-19 Genomics UK (COG-UK) Consortium | Catherine Moore, Johnathan Evans, Laura Gifford, Malorie Perry, Simon Cottrell, Angela Marchbank, Alec Birchley, Alexander Adams, Amy Gaskin, Bree Gatica-Wilcox, Jason Coombes, Joel Southgate, Lauren Gilbert, Lee Graham, Nicole Pacchiarini, Sara Kumziene-Summerhayes, Sarah Taylor, Sophie Jones, Sara Rey, Matthew Bull, Joanne Watkins, Sally Corden, Tom Connor |
| EPI_ISL_627058, EPI_ISL_627059, EPI_ISL_627060, EPI_ISL_627061, EPI_ISL_627062, EPI_ISL_627065 | University of Exeter | COVID-19 Genomics UK (COG-UK) Consortium | Ben Temperton,Aaron Jeffries,Michelle Michelsen,Joanna Warwick-Dugdale,Audrey Farbos,Robyn Manley,Stephen Michell,Jane Masoli |
| EPI_ISL_627070 | Regional Virus Laboratory, Belfast Health and Social Care Trust | COVID-19 Genomics UK (COG-UK) Consortium | Conall McCaughey, James McKenna, Tanya Curran, Susan Feeney, Alison Watt, Ciara Cox, Mairead Connor, Zoltan Molnar, David Simpson, Derek Fairley |
| EPI_ISL_627080 | Quadram Institute Bioscience | COVID-19 Genomics UK (COG-UK) Consortium | Dave J. Baker, Gemma L. Kay, Alp Aydin, Thanh Le-Viet, Steven Rudder, Ana P. Tedim, Anastasia Kolyva, Maria Diaz, Leonardo de Oliveira Martins, Nabil-Fareed Alikhan, Lizzie Meadows, Rachael Stanley, Ngozi Elumogo, Muhammed Yasir, Nicholas M. Thomson, Alexander J Trotter, Rachel Gilroy, Samuel Bloomfield, Claire Stuart, Andrew Bell, Reenesh Prakash, Samir Dervisevic, Alison E. Mather, John Wain, Mark Webber, Andrew J. Page, Justin O'Grady |
| EPI_ISL_627103 | Virology Department, Sheffield Teaching Hospitals NHS Foundation Trust/Department of Infection, Immunity and Cardiovascular Disease, The Medical School, University of Sheffield | COVID-19 Genomics UK (COG-UK) Consortium | Thushan de Silva, Matthew Parker, Nikki Smith, Adri Angyal, Rebecca Brown, Luke Green, Rachel Tucker, Paul Parsons, Danielle Groves, Katie Johnson, Laura Carrilero, Alex Keeley, Dave Partridge, Matthew Wyles, Benjamin Lindsey, Mehmet Yavuz, Mohammad Raza, Cariad Evans |
| EPI_ISL_627108, EPI_ISL_627110 | University College London, Great Ormond Street Hospital for Children NHS Foundation Trust, Imperial College Healthcare NHS Trust | COVID-19 Genomics UK (COG-UK) Consortium | Sergi Castellano, Rachel Williams, Mark Kristiansen, Paola Resende Silva, Sunando Roy, Tony Brooks, Helena Tutill, Paola Niola, Patricia Dyal, Charlotte Williams, Leysa Forrest, Yasmin Panchbhaya, Jacqueline Findlay, Samuel Weeks, Julianne Brown, Kathryn Harris, Paul Randell, James Price, Alison Holmes, Judith Breuer |
| EPI_ISL_627115 | Wales Specialist Virology Centre Sequencing lab: Pathogen Genomics Unit | COVID-19 Genomics UK (COG-UK) Consortium | Catherine Moore, Johnathan Evans, Laura Gifford, Malorie Perry, Simon Cottrell, Angela Marchbank, Alec Birchley, Alexander Adams, Amy Gaskin, Bree Gatica-Wilcox, Jason Coombes, Joel Southgate, Lauren Gilbert, Lee Graham, Nicole Pacchiarini, Sara Kumziene-Summerhayes, Sarah Taylor, Sophie Jones, Sara Rey, Matthew Bull, Joanne Watkins, Sally Corden, Tom Connor |
| EPI_ISL_627118 | University of Exeter | COVID-19 Genomics UK (COG-UK) Consortium | Ben Temperton,Aaron Jeffries,Michelle Michelsen,Joanna Warwick-Dugdale,Audrey Farbos,Robyn Manley,Stephen Michell,Jane Masoli |
| EPI_ISL_627123, EPI_ISL_627127 | Liverpool Clinical Laboratories | COVID-19 Genomics UK (COG-UK) Consortium | Sam Haldenby, Anita Lucaci, Steve Paterson, Julian Hiscox, Alistair Darby, M Almsaud, A Alrezaihi, Muhannad Alruwaili, Stuart D Armstrong, Jones Benjamin, Eleanor G Bentley, Anu Chawla, Jordan J Clark, Angela Cowell, Richard Eccles, Isabel García-Dorival, Matthew Gemmell, Alessandro Gerada, PKF Gilmore, Richard Gregory, Ximeng Han, Catherine Hartley, Margaret Hughes, Miren Iturriza-Gomara, James Johnson, L Luu, Jenifer Manson, Charlotte Nelson, Elaine O'Toole, Cassie Olateju, Rebekah Penrice-Randal , Lucille Rainbow, N.P Randle, Trevor Ian Robinson, Parul Sharma, Ghada T Shawli, James P Stewart, Neil Swainston, Ecaterina Vamos, Joanne Watts, Mark Whitehead |
| EPI_ISL_627130 | Virology Department, Sheffield Teaching Hospitals NHS Foundation Trust/Department of Infection, Immunity and Cardiovascular Disease, The Medical School, University of Sheffield | COVID-19 Genomics UK (COG-UK) Consortium | Thushan de Silva, Matthew Parker, Nikki Smith, Adri Angyal, Rebecca Brown, Luke Green, Rachel Tucker, Paul Parsons, Danielle Groves, Katie Johnson, Laura Carrilero, Alex Keeley, Dave Partridge, Matthew Wyles, Benjamin Lindsey, Mehmet Yavuz, Mohammad Raza, Cariad Evans |
| EPI_ISL_627135, EPI_ISL_627136, EPI_ISL_627139 | Liverpool Clinical Laboratories | COVID-19 Genomics UK (COG-UK) Consortium | Sam Haldenby, Anita Lucaci, Steve Paterson, Julian Hiscox, Alistair Darby, M Almsaud, A Alrezaihi, Muhannad Alruwaili, Stuart D Armstrong, Jones Benjamin, Eleanor G Bentley, Anu Chawla, Jordan J Clark, Angela Cowell, Richard Eccles, Isabel García-Dorival, Matthew Gemmell, Alessandro Gerada, PKF Gilmore, Richard Gregory, Ximeng Han, Catherine Hartley, Margaret Hughes, Miren Iturriza-Gomara, James Johnson, L Luu, Jenifer Manson, Charlotte Nelson, Elaine O'Toole, Cassie Olateju, Rebekah Penrice-Randal , Lucille Rainbow, N.P Randle, Trevor Ian Robinson, Parul Sharma, Ghada T Shawli, James P Stewart, Neil Swainston, Ecaterina Vamos, Joanne Watts, Mark Whitehead |
| EPI_ISL_627143 | University of Exeter | COVID-19 Genomics UK (COG-UK) Consortium | Ben Temperton,Aaron Jeffries,Michelle Michelsen,Joanna Warwick-Dugdale,Audrey Farbos,Robyn Manley,Stephen Michell,Jane Masoli |
| EPI_ISL_627148 | Quadram Institute Bioscience | COVID-19 Genomics UK (COG-UK) Consortium | Dave J. Baker, Gemma L. Kay, Alp Aydin, Thanh Le-Viet, Steven Rudder, Ana P. Tedim, Anastasia Kolyva, Maria Diaz, Leonardo de Oliveira Martins, Nabil-Fareed Alikhan, Lizzie Meadows, Rachael Stanley, Ngozi Elumogo, Muhammed Yasir, Nicholas M. Thomson, Alexander J Trotter, Rachel Gilroy, Samuel Bloomfield, Claire Stuart, Andrew Bell, Reenesh Prakash, Samir Dervisevic, Alison E. Mather, John Wain, Mark Webber, Andrew J. Page, Justin O'Grady |
| EPI_ISL_627149 | University of Exeter | COVID-19 Genomics UK (COG-UK) Consortium | Ben Temperton,Aaron Jeffries,Michelle Michelsen,Joanna Warwick-Dugdale,Audrey Farbos,Robyn Manley,Stephen Michell,Jane Masoli |
| EPI_ISL_627153 | Liverpool Clinical Laboratories | COVID-19 Genomics UK (COG-UK) Consortium | Sam Haldenby, Anita Lucaci, Steve Paterson, Julian Hiscox, Alistair Darby, M Almsaud, A Alrezaihi, Muhannad Alruwaili, Stuart D Armstrong, Jones Benjamin, Eleanor G Bentley, Anu Chawla, Jordan J Clark, Angela Cowell, Richard Eccles, Isabel García-Dorival, Matthew Gemmell, Alessandro Gerada, PKF Gilmore, Richard Gregory, Ximeng Han, Catherine Hartley, Margaret Hughes, Miren Iturriza-Gomara, James Johnson, L Luu, Jenifer Manson, Charlotte Nelson, Elaine O'Toole, Cassie Olateju, Rebekah Penrice-Randal , Lucille Rainbow, N.P Randle, Trevor Ian Robinson, Parul Sharma, Ghada T Shawli, James P Stewart, Neil Swainston, Ecaterina Vamos, Joanne Watts, Mark Whitehead |
| EPI_ISL_627155 | Wales Specialist Virology Centre Sequencing lab: Pathogen Genomics Unit | COVID-19 Genomics UK (COG-UK) Consortium | Catherine Moore, Johnathan Evans, Laura Gifford, Malorie Perry, Simon Cottrell, Angela Marchbank, Alec Birchley, Alexander Adams, Amy Gaskin, Bree Gatica-Wilcox, Jason Coombes, Joel Southgate, Lauren Gilbert, Lee Graham, Nicole Pacchiarini, Sara Kumziene-Summerhayes, Sarah Taylor, Sophie Jones, Sara Rey, Matthew Bull, Joanne Watkins, Sally Corden, Tom Connor |
| EPI_ISL_627159 | University of Exeter | COVID-19 Genomics UK (COG-UK) Consortium | Ben Temperton,Aaron Jeffries,Michelle Michelsen,Joanna Warwick-Dugdale,Audrey Farbos,Robyn Manley,Stephen Michell,Jane Masoli |
| EPI_ISL_627162 | Liverpool Clinical Laboratories | COVID-19 Genomics UK (COG-UK) Consortium | Sam Haldenby, Anita Lucaci, Steve Paterson, Julian Hiscox, Alistair Darby, M Almsaud, A Alrezaihi, Muhannad Alruwaili, Stuart D Armstrong, Jones Benjamin, Eleanor G Bentley, Anu Chawla, Jordan J Clark, Angela Cowell, Richard Eccles, Isabel García-Dorival, Matthew Gemmell, Alessandro Gerada, PKF Gilmore, Richard Gregory, Ximeng Han, Catherine Hartley, Margaret Hughes, Miren Iturriza-Gomara, James Johnson, L Luu, Jenifer Manson, Charlotte Nelson, Elaine O'Toole, Cassie Olateju, Rebekah Penrice-Randal , Lucille Rainbow, N.P Randle, Trevor Ian Robinson, Parul Sharma, Ghada T Shawli, James P Stewart, Neil Swainston, Ecaterina Vamos, Joanne Watts, Mark Whitehead |

| | | | |
|---|---|---|---|
| EPI_ISL_627166 | University of Exeter | COVID-19 Genomics UK (COG-UK) Consortium | Ben Temperton,Aaron Jeffries,Michelle Michelsen,Joanna Warwick-Dugdale,Audrey Farbos,Robyn Manley,Stephen Michell,Jane Masoli |
| EPI_ISL_627170 | Quadram Institute Bioscience | COVID-19 Genomics UK (COG-UK) Consortium | Dave J. Baker, Gemma L. Kay, Alp Aydin, Thanh Le-Viet, Steven Rudder, Ana P. Tedim, Anastasia Kolyva, Maria Diaz, Leonardo de Oliveira Martins, Nabil-Fareed Alikhan, Lizzie Meadows, Rachael Stanley, Ngozi Elumogo, Muhammed Yasir, Nicholas M. Thomson, Alexander J Trotter, Rachel Gilroy, Samuel Bloomfield, Claire Stuart, Andrew Bell, Reenesh Prakash, Samir Dervisevic, Alison E. Mather, John Wain, Mark Webber, Andrew J. Page, Justin O'Grady |
| EPI_ISL_627176, EPI_ISL_627183 | Liverpool Clinical Laboratories | COVID-19 Genomics UK (COG-UK) Consortium | Sam Haldenby, Anita Lucaci, Steve Paterson, Julian Hiscox, Alistair Darby, M Almsaud, A Alrezaihi, Muhannad Alruwaili, Stuart D Armstrong, Jones Benjamin, Eleanor G Bentley, Anu Chawla, Jordan J Clark, Angela Cowell, Richard Eccles, Isabel García-Dorival, Matthew Gemmell, Alessandro Gerada, PKF Gilmore, Richard Gregory, Ximeng Han, Catherine Hartley, Margaret Hughes, Miren Iturriza-Gomara, James Johnson, L Luu, Jenifer Manson, Charlotte Nelson, Elaine O'Toole, Cassie Olateju, Rebekah Penrice-Randal , Lucille Rainbow, N.P Randle, Trevor Ian Robinson, Parul Sharma, Ghada T Shawli, James P Stewart, Neil Swainston, Ecaterina Vamos, Joanne Watts, Mark Whitehead |
| EPI_ISL_627189 | University of Exeter | COVID-19 Genomics UK (COG-UK) Consortium | Ben Temperton,Aaron Jeffries,Michelle Michelsen,Joanna Warwick-Dugdale,Audrey Farbos,Robyn Manley,Stephen Michell,Jane Masoli |
| EPI_ISL_627191 | Virology Department, Sheffield Teaching Hospitals NHS Foundation Trust/Department of Infection, Immunity and Cardiovascular Disease, The Medical School, University of Sheffield | COVID-19 Genomics UK (COG-UK) Consortium | Thushan de Silva, Matthew Parker, Nikki Smith, Adri Angyal, Rebecca Brown, Luke Green, Rachel Tucker, Paul Parsons, Danielle Groves, Katie Johnson, Laura Carrilero, Alex Keeley, Dave Partridge, Matthew Wyles, Benjamin Lindsey, Mehmet Yavuz, Mohammad Raza, Cariad Evans |
| EPI_ISL_627196 | University of Exeter | COVID-19 Genomics UK (COG-UK) Consortium | Ben Temperton,Aaron Jeffries,Michelle Michelsen,Joanna Warwick-Dugdale,Audrey Farbos,Robyn Manley,Stephen Michell,Jane Masoli |
| EPI_ISL_627197 | Wales Specialist Virology Centre Sequencing lab: Pathogen Genomics Unit | COVID-19 Genomics UK (COG-UK) Consortium | Catherine Moore, Johnathan Evans, Laura Gifford, Malorie Perry, Simon Cottrell, Angela Marchbank, Alec Birchley, Alexander Adams, Amy Gaskin, Bree Gatica-Wilcox, Jason Coombes, Joel Southgate, Lauren Gilbert, Lee Graham, Nicole Pacchiarini, Sara Kumziene-Summerhayes, Sarah Taylor, Sophie Jones, Sara Rey, Matthew Bull, Joanne Watkins, Sally Corden, Tom Connor |
| EPI_ISL_627199 | Regional Virus Laboratory, Belfast Health and Social Care Trust | COVID-19 Genomics UK (COG-UK) Consortium | Conall McCaughey, James McKenna, Tanya Curran, Susan Feeney, Alison Watt, Ciara Cox, Mairead Connor, Zoltan Molnar, David Simpson, Derek Fairley |
| EPI_ISL_627250 | Oxford Viromics, NDM, University of Oxford; Oxford University Hospitals; Basingstoke and North Hampshire Hospital | COVID-19 Genomics UK (COG-UK) Consortium | Tanya Golubchik, David Bonsall, George Macintyre, Amy Trebes, Mariateresa de Cesare, Catrin Moore, Alex Mobbs, Anita Justice, Robert Shaw, Monique Andersson, Timothy Peto, Emma Wise, Nathan Moore, Jessica Lynch, Nick Cortes, Matilde Mori, Stephen Kidd, David Buck, John Todd, Christophe Fraser |
| EPI_ISL_627254 | Virology Department, Sheffield Teaching Hospitals NHS Foundation Trust/Department of Infection, Immunity and Cardiovascular Disease, The Medical School, University of Sheffield | COVID-19 Genomics UK (COG-UK) Consortium | Thushan de Silva, Matthew Parker, Nikki Smith, Adri Angyal, Rebecca Brown, Luke Green, Rachel Tucker, Paul Parsons, Danielle Groves, Katie Johnson, Laura Carrilero, Alex Keeley, Dave Partridge, Matthew Wyles, Benjamin Lindsey, Mehmet Yavuz, Mohammad Raza, Cariad Evans |
| EPI_ISL_627255, EPI_ISL_627257, EPI_ISL_627259, EPI_ISL_627261, EPI_ISL_627262, EPI_ISL_627263 | Oxford Viromics, NDM, University of Oxford; Oxford University Hospitals; Basingstoke and North Hampshire Hospital | COVID-19 Genomics UK (COG-UK) Consortium | Tanya Golubchik, David Bonsall, George Macintyre, Amy Trebes, Mariateresa de Cesare, Catrin Moore, Alex Mobbs, Anita Justice, Robert Shaw, Monique Andersson, Timothy Peto, Emma Wise, Nathan Moore, Jessica Lynch, Nick Cortes, Matilde Mori, Stephen Kidd, David Buck, John Todd, Christophe Fraser |
| EPI_ISL_627264 | Virology Department, Sheffield Teaching Hospitals NHS Foundation Trust/Department of Infection, Immunity and Cardiovascular Disease, The Medical School, University of Sheffield | COVID-19 Genomics UK (COG-UK) Consortium | Thushan de Silva, Matthew Parker, Nikki Smith, Adri Angyal, Rebecca Brown, Luke Green, Rachel Tucker, Paul Parsons, Danielle Groves, Katie Johnson, Laura Carrilero, Alex Keeley, Dave Partridge, Matthew Wyles, Benjamin Lindsey, Mehmet Yavuz, Mohammad Raza, Cariad Evans |
| EPI_ISL_627266, EPI_ISL_627267, EPI_ISL_627269 | Wales Specialist Virology Centre Sequencing lab: Pathogen Genomics Unit | COVID-19 Genomics UK (COG-UK) Consortium | Catherine Moore, Johnathan Evans, Laura Gifford, Malorie Perry, Simon Cottrell, Angela Marchbank, Alec Birchley, Alexander Adams, Amy Gaskin, Bree Gatica-Wilcox, Jason Coombes, Joel Southgate, Lauren Gilbert, Lee Graham, Nicole Pacchiarini, Sara Kumziene-Summerhayes, Sarah Taylor, Sophie Jones, Sara Rey, Matthew Bull, Joanne Watkins, Sally Corden, Tom Connor |
| EPI_ISL_627282, EPI_ISL_627283, EPI_ISL_627286, EPI_ISL_627287, EPI_ISL_627288, EPI_ISL_627289, EPI_ISL_627290 | Oxford Viromics, NDM, University of Oxford; Oxford University Hospitals; Basingstoke and North Hampshire Hospital | COVID-19 Genomics UK (COG-UK) Consortium | Tanya Golubchik, David Bonsall, George Macintyre, Amy Trebes, Mariateresa de Cesare, Catrin Moore, Alex Mobbs, Anita Justice, Robert Shaw, Monique Andersson, Timothy Peto, Emma Wise, Nathan Moore, Jessica Lynch, Nick Cortes, Matilde Mori, Stephen Kidd, David Buck, John Todd, Christophe Fraser |
| EPI_ISL_627291 | University of Exeter | COVID-19 Genomics UK (COG-UK) Consortium | Ben Temperton,Aaron Jeffries,Michelle Michelsen,Joanna Warwick-Dugdale,Audrey Farbos,Robyn Manley,Stephen Michell,Jane Masoli |
| EPI_ISL_627292, EPI_ISL_627297 | Oxford Viromics, NDM, University of Oxford; Oxford University Hospitals; Basingstoke and North Hampshire Hospital | COVID-19 Genomics UK (COG-UK) Consortium | Tanya Golubchik, David Bonsall, George Macintyre, Amy Trebes, Mariateresa de Cesare, Catrin Moore, Alex Mobbs, Anita Justice, Robert Shaw, Monique Andersson, Timothy Peto, Emma Wise, Nathan Moore, Jessica Lynch, Nick Cortes, Matilde Mori, Stephen Kidd, David Buck, John Todd, Christophe Fraser |
| EPI_ISL_627298 | Virology Department, Sheffield Teaching Hospitals NHS Foundation Trust/Department of Infection, Immunity and Cardiovascular Disease, The Medical School, University of Sheffield | COVID-19 Genomics UK (COG-UK) Consortium | Thushan de Silva, Matthew Parker, Nikki Smith, Adri Angyal, Rebecca Brown, Luke Green, Rachel Tucker, Paul Parsons, Danielle Groves, Katie Johnson, Laura Carrilero, Alex Keeley, Dave Partridge, Matthew Wyles, Benjamin Lindsey, Mehmet Yavuz, Mohammad Raza, Cariad Evans |
| EPI_ISL_627299, EPI_ISL_627302, EPI_ISL_627303 | Oxford Viromics, NDM, University of Oxford; Oxford University Hospitals; Basingstoke and North Hampshire Hospital | COVID-19 Genomics UK (COG-UK) Consortium | Tanya Golubchik, David Bonsall, George Macintyre, Amy Trebes, Mariateresa de Cesare, Catrin Moore, Alex Mobbs, Anita Justice, Robert Shaw, Monique Andersson, Timothy Peto, Emma Wise, Nathan Moore, Jessica Lynch, Nick Cortes, Matilde Mori, Stephen Kidd, David Buck, John Todd, Christophe Fraser |
| EPI_ISL_627306 | Wales Specialist Virology Centre Sequencing lab: Pathogen Genomics Unit | COVID-19 Genomics UK (COG-UK) Consortium | Catherine Moore, Johnathan Evans, Laura Gifford, Malorie Perry, Simon Cottrell, Angela Marchbank, Alec Birchley, Alexander Adams, Amy Gaskin, Bree Gatica-Wilcox, Jason Coombes, Joel Southgate, Lauren Gilbert, Lee Graham, Nicole Pacchiarini, Sara Kumziene-Summerhayes, Sarah Taylor, Sophie Jones, Sara Rey, Matthew Bull, Joanne Watkins, Sally Corden, Tom Connor |
| EPI_ISL_627313 | Virology Department, Sheffield Teaching Hospitals NHS Foundation Trust/Department of Infection, Immunity and Cardiovascular Disease, The Medical School, University of Sheffield | COVID-19 Genomics UK (COG-UK) Consortium | Thushan de Silva, Matthew Parker, Nikki Smith, Adri Angyal, Rebecca Brown, Luke Green, Rachel Tucker, Paul Parsons, Danielle Groves, Katie Johnson, Laura Carrilero, Alex Keeley, Dave Partridge, Matthew Wyles, Benjamin Lindsey, Mehmet Yavuz, Mohammad Raza, Cariad Evans |
| EPI_ISL_627314, EPI_ISL_627315, EPI_ISL_627316, EPI_ISL_627317, EPI_ISL_627318, EPI_ISL_627319, EPI_ISL_627320, EPI_ISL_627321, EPI_ISL_627322, EPI_ISL_627323, EPI_ISL_627324, EPI_ISL_627325, EPI_ISL_627326, EPI_ISL_627327, EPI_ISL_627328, EPI_ISL_627429 | | | |
| see above | University of Exeter | COVID-19 Genomics UK (COG-UK) Consortium | Ben Temperton,Aaron Jeffries,Michelle Michelsen,Joanna Warwick-Dugdale,Audrey Farbos,Robyn Manley,Stephen Michell,Jane Masoli |
| EPI_ISL_627445, EPI_ISL_627446 | Liverpool Clinical Laboratories | COVID-19 Genomics UK (COG-UK) Consortium | Sam Haldenby, Anita Lucaci, Steve Paterson, Julian Hiscox, Alistair Darby, M Almsaud, A Alrezaihi, Muhannad Alruwaili, Stuart D Armstrong, Jones Benjamin, Eleanor G Bentley, Anu Chawla, Jordan J Clark, Angela Cowell, Richard Eccles, Isabel García-Dorival, Matthew Gemmell, Alessandro Gerada, PKF Gilmore, Richard Gregory, Ximeng Han, Catherine Hartley, Margaret Hughes, Miren Iturriza-Gomara, James Johnson, L Luu, Jenifer Manson, Charlotte Nelson, Elaine O'Toole, Cassie Olateju, Rebekah Penrice-Randal , Lucille Rainbow, N.P Randle, Trevor Ian Robinson, Parul Sharma, Ghada T Shawli, James P Stewart, Neil Swainston, Ecaterina Vamos, Joanne Watts, Mark Whitehead |
| EPI_ISL_627492, EPI_ISL_627493, EPI_ISL_627494, EPI_ISL_627495, EPI_ISL_627496, EPI_ISL_627497, EPI_ISL_627498, EPI_ISL_627499, EPI_ISL_627500, EPI_ISL_627501, EPI_ISL_627502, EPI_ISL_627503, EPI_ISL_627504, EPI_ISL_627505, EPI_ISL_627506, EPI_ISL_627507, EPI_ISL_627508, EPI_ISL_627509, EPI_ISL_627510, EPI_ISL_627511, EPI_ISL_627512, EPI_ISL_627513, EPI_ISL_627514, EPI_ISL_627515, EPI_ISL_627516, EPI_ISL_627517 | | | |
| see above | Virology Department, Sheffield Teaching Hospitals NHS Foundation Trust/Department of Infection, Immunity and Cardiovascular Disease, The Medical School, University of Sheffield | COVID-19 Genomics UK (COG-UK) Consortium | Thushan de Silva, Matthew Parker, Nikki Smith, Adri Angyal, Rebecca Brown, Luke Green, Rachel Tucker, Paul Parsons, Danielle Groves, Katie Johnson, Laura Carrilero, Alex Keeley, Dave Partridge, Matthew Wyles, Benjamin Lindsey, Mehmet Yavuz, Mohammad Raza, Cariad Evans |
| EPI_ISL_627518, EPI_ISL_627519, EPI_ISL_627520, EPI_ISL_627521, EPI_ISL_627522, EPI_ISL_627523, EPI_ISL_627524, EPI_ISL_627525, EPI_ISL_627526, EPI_ISL_627527, EPI_ISL_627528, EPI_ISL_627529, EPI_ISL_627530, EPI_ISL_627531, EPI_ISL_627532, EPI_ISL_627533, EPI_ISL_627534, EPI_ISL_627535, EPI_ISL_627536, EPI_ISL_627537, EPI_ISL_627538, EPI_ISL_627539, EPI_ISL_627540, EPI_ISL_627541, EPI_ISL_627544, EPI_ISL_627546, EPI_ISL_627554, EPI_ISL_627555, EPI_ISL_627556, EPI_ISL_627557, EPI_ISL_627558, EPI_ISL_627559, EPI_ISL_627560, EPI_ISL_627561, EPI_ISL_627562, EPI_ISL_627563, EPI_ISL_627564, EPI_ISL_627565, EPI_ISL_627566, EPI_ISL_627567, EPI_ISL_627568, EPI_ISL_627569 | | | |

| | | | |
|---|---|---|---|
| see above | Regional Virus Laboratory, Belfast Health and Social Care Trust | COVID-19 Genomics UK (COG-UK) Consortium | Conall McCaughey, James McKenna, Tanya Curran, Susan Feeney, Alison Watt, Ciara Cox, Mairead Connor, Zoltan Molnar, David Simpson, Derek Fairley |
| EPI_ISL_627586, EPI_ISL_627603, EPI_ISL_627607, EPI_ISL_627612, EPI_ISL_627613, EPI_ISL_627615 | Quadram Institute Bioscience | COVID-19 Genomics UK (COG-UK) Consortium | Dave J. Baker, Gemma L. Kay, Alp Aydin, Thanh Le-Viet, Steven Rudder, Ana P. Tedim, Anastasia Kolyva, Maria Diaz, Leonardo de Oliveira Martins, Nabil-Fareed Alikhan, Lizzie Meadows, Rachael Stanley, Ngozi Elumogo, Muhammed Yasir, Nicholas M. Thomson, Alexander J Trotter, Rachel Gilroy, Samuel Bloomfield, Claire Stuart, Andrew Bell, Reenesh Prakash, Samir Dervisevic, Alison E. Mather, John Wain, Mark Webber, Andrew J. Page, Justin O'Grady |
| EPI_ISL_627735, EPI_ISL_627741, EPI_ISL_627750, EPI_ISL_627770, EPI_ISL_627784, EPI_ISL_627793, EPI_ISL_627802, EPI_ISL_627803, EPI_ISL_627807, EPI_ISL_627818, EPI_ISL_627825, EPI_ISL_627828, EPI_ISL_627829, EPI_ISL_627833, EPI_ISL_627834, EPI_ISL_627842, EPI_ISL_627848, EPI_ISL_627849, EPI_ISL_627854, EPI_ISL_627855, EPI_ISL_627877, EPI_ISL_627890, EPI_ISL_627891, EPI_ISL_627892, EPI_ISL_627904, EPI_ISL_627905, EPI_ISL_627906, EPI_ISL_627909, EPI_ISL_627910, EPI_ISL_628026 | | | |
| see above | Wales Specialist Virology Centre Sequencing lab: Pathogen Genomics Unit | COVID-19 Genomics UK (COG-UK) Consortium | Catherine Moore, Johnathan Evans, Laura Gifford, Malorie Perry, Simon Cottrell, Angela Marchbank, Alec Birchley, Alexander Adams, Amy Gaskin, Bree Gatica-Wilcox, Jason Coombes, Joel Southgate, Lauren Gilbert, Lee Graham, Nicole Pacchiarini, Sara Kumziene-Summerhayes, Sarah Taylor, Sophie Jones, Sara Rey, Matthew Bull, Joanne Watkins, Sally Corden, Tom Connor |
| EPI_ISL_628334, EPI_ISL_628335, EPI_ISL_628336, EPI_ISL_628337, EPI_ISL_628338, EPI_ISL_628339, EPI_ISL_628340, EPI_ISL_628341, EPI_ISL_628342, EPI_ISL_628343, EPI_ISL_628344, EPI_ISL_628345, EPI_ISL_628346, EPI_ISL_628347, EPI_ISL_628348, EPI_ISL_628358 | | | |
| see above | Oxford Viromics, NDM, University of Oxford; Oxford University Hospitals; Basingstoke and North Hampshire Hospital | COVID-19 Genomics UK (COG-UK) Consortium | Tanya Golubchik, David Bonsall, George Macintyre, Amy Trebes, Mariateresa de Cesare, Catrin Moore, Alex Mobbs, Anita Justice, Robert Shaw, Monique Andersson, Timothy Peto, Emma Wise, Nathan Moore, Jessica Lynch, Nick Cortes, Matilde Mori, Stephen Kidd, David Buck, John Todd, Christophe Fraser |
| EPI_ISL_628397, EPI_ISL_628398, EPI_ISL_628399, EPI_ISL_628400 | Virology Department, Sheffield Teaching Hospitals NHS Foundation Trust/Department of Infection, Immunity and Cardiovascular Disease, The Medical School, University of Sheffield | COVID-19 Genomics UK (COG-UK) Consortium | Thushan de Silva, Matthew Parker, Nikki Smith, Adri Angyal, Rebecca Brown, Luke Green, Rachel Tucker, Paul Parsons, Danielle Groves, Katie Johnson, Laura Carrilero, Alex Keeley, Dave Partridge, Matthew Wyles, Benjamin Lindsey, Mehmet Yavuz, Mohammad Raza, Cariad Evans |
| EPI_ISL_628441, EPI_ISL_628442, EPI_ISL_628443, EPI_ISL_628444, EPI_ISL_628445, EPI_ISL_628446, EPI_ISL_628447, EPI_ISL_628448, EPI_ISL_628449, EPI_ISL_628450, EPI_ISL_628451, EPI_ISL_628452, EPI_ISL_628465, EPI_ISL_628466, EPI_ISL_628467, EPI_ISL_628468, EPI_ISL_628469, EPI_ISL_628470, EPI_ISL_628471, EPI_ISL_628472, EPI_ISL_628473, EPI_ISL_628474, EPI_ISL_628475, EPI_ISL_628476, EPI_ISL_628477, EPI_ISL_628478, EPI_ISL_628479, EPI_ISL_628480, EPI_ISL_628481, EPI_ISL_628482, EPI_ISL_628483, EPI_ISL_628484, EPI_ISL_628486, EPI_ISL_628487, EPI_ISL_628488, EPI_ISL_628489, EPI_ISL_628490, EPI_ISL_628491, EPI_ISL_628492, EPI_ISL_628493, EPI_ISL_628494, EPI_ISL_628495, EPI_ISL_628496, EPI_ISL_628497, EPI_ISL_628498, EPI_ISL_628499, EPI_ISL_628500, EPI_ISL_628501, EPI_ISL_628502, EPI_ISL_628503, EPI_ISL_628504, EPI_ISL_628505, EPI_ISL_628506, EPI_ISL_628507, EPI_ISL_628508, EPI_ISL_628511, EPI_ISL_628512, EPI_ISL_628513, EPI_ISL_628514, EPI_ISL_628515, EPI_ISL_628516, EPI_ISL_628517, EPI_ISL_628540, EPI_ISL_628549, EPI_ISL_628550, EPI_ISL_628551, EPI_ISL_628552, EPI_ISL_628553, EPI_ISL_628554, EPI_ISL_628555, EPI_ISL_628556, EPI_ISL_628557, EPI_ISL_628558, EPI_ISL_628562, EPI_ISL_628563, EPI_ISL_628564, EPI_ISL_628565, EPI_ISL_628566, EPI_ISL_628569, EPI_ISL_628570, EPI_ISL_628589, EPI_ISL_628590, EPI_ISL_628591, EPI_ISL_628592, EPI_ISL_628612, EPI_ISL_628613, EPI_ISL_628619, EPI_ISL_628620, EPI_ISL_628621, EPI_ISL_628622, EPI_ISL_628623, EPI_ISL_628624, EPI_ISL_628625, EPI_ISL_628626, EPI_ISL_628627, EPI_ISL_628634, EPI_ISL_628635, EPI_ISL_628636, EPI_ISL_628637, EPI_ISL_628638, EPI_ISL_628639, EPI_ISL_628640, EPI_ISL_628641, EPI_ISL_628642, EPI_ISL_628643, EPI_ISL_628644, EPI_ISL_628645, EPI_ISL_628647, EPI_ISL_628653, EPI_ISL_628654, EPI_ISL_628655, EPI_ISL_628656, EPI_ISL_628657, EPI_ISL_628658, EPI_ISL_628659, EPI_ISL_628660, EPI_ISL_628661, EPI_ISL_628662, EPI_ISL_628663, EPI_ISL_628664, EPI_ISL_628665, EPI_ISL_628666, EPI_ISL_628667, EPI_ISL_628668, EPI_ISL_628669, EPI_ISL_628670, EPI_ISL_628671, EPI_ISL_628672, EPI_ISL_628673, EPI_ISL_628674, EPI_ISL_628675, EPI_ISL_628676, EPI_ISL_628677, EPI_ISL_628678, EPI_ISL_628679, EPI_ISL_628680, EPI_ISL_628681, EPI_ISL_628682, EPI_ISL_628683, EPI_ISL_628684, EPI_ISL_628685, EPI_ISL_628686, EPI_ISL_628687, EPI_ISL_628688, EPI_ISL_628689, EPI_ISL_628690, EPI_ISL_628691, EPI_ISL_628692, EPI_ISL_628693, EPI_ISL_628694, EPI_ISL_628695, EPI_ISL_628696, EPI_ISL_628697, EPI_ISL_628698, EPI_ISL_628699, EPI_ISL_628700 | | | |
| see above | Oxford Viromics, NDM, University of Oxford; Oxford University Hospitals; Basingstoke and North Hampshire Hospital | COVID-19 Genomics UK (COG-UK) Consortium | Tanya Golubchik, David Bonsall, George Macintyre, Amy Trebes, Mariateresa de Cesare, Catrin Moore, Alex Mobbs, Anita Justice, Robert Shaw, Monique Andersson, Timothy Peto, Emma Wise, Nathan Moore, Jessica Lynch, Nick Cortes, Matilde Mori, Stephen Kidd, David Buck, John Todd, Christophe Fraser |
| EPI_ISL_628932, EPI_ISL_628943, EPI_ISL_628944, EPI_ISL_628945, EPI_ISL_628950, EPI_ISL_628951, EPI_ISL_628952, EPI_ISL_628953, EPI_ISL_628954, EPI_ISL_628955, EPI_ISL_628957, EPI_ISL_628958, EPI_ISL_628959, EPI_ISL_628961 | | | |
| see above | Utah Public Health Laboratory | Utah Public Health Laboratory | Erin Young, Kelly Oakeson |
| EPI_ISL_629004, EPI_ISL_629005, EPI_ISL_629006 | South Eastern Area Laboratory Services (SEALS) | NSW Health Pathology - Institute of Clinical Pathology and Medical Research; Westmead Hospital; University of Sydney | CIDM-PH et al. |
| EPI_ISL_629064, EPI_ISL_629065, EPI_ISL_629068, EPI_ISL_629069, EPI_ISL_629076 | Maryland Public Health Laboratory | Maryland Public Health Laboratory | Maryland Department of Health Laboratories Administration |
| EPI_ISL_630348, EPI_ISL_630349, EPI_ISL_630350, EPI_ISL_630351, EPI_ISL_630352, EPI_ISL_630353, EPI_ISL_630354, EPI_ISL_630355, EPI_ISL_630356, EPI_ISL_630357, EPI_ISL_630358, EPI_ISL_630359, EPI_ISL_630360, EPI_ISL_630361, EPI_ISL_630362, EPI_ISL_630363, EPI_ISL_630364, EPI_ISL_630365, EPI_ISL_630366, EPI_ISL_630367, EPI_ISL_630368, EPI_ISL_630369, EPI_ISL_630370, EPI_ISL_630371, EPI_ISL_630372, EPI_ISL_630373, EPI_ISL_630374, EPI_ISL_630375, EPI_ISL_630376, EPI_ISL_630377, EPI_ISL_630378, EPI_ISL_630379, EPI_ISL_630380, EPI_ISL_630381, EPI_ISL_630382, EPI_ISL_630383, EPI_ISL_630384, EPI_ISL_630385, EPI_ISL_630386, EPI_ISL_630387, EPI_ISL_630388, EPI_ISL_630389, EPI_ISL_630390, EPI_ISL_630391, EPI_ISL_630392, EPI_ISL_630393, EPI_ISL_630394, EPI_ISL_630395, EPI_ISL_630396, EPI_ISL_630397, EPI_ISL_630398, EPI_ISL_630399, EPI_ISL_630400, EPI_ISL_630401, EPI_ISL_630402, EPI_ISL_630403, EPI_ISL_630404, EPI_ISL_630405, EPI_ISL_630406, EPI_ISL_630407, EPI_ISL_630408, EPI_ISL_630409, EPI_ISL_630410, EPI_ISL_630411, EPI_ISL_630412, EPI_ISL_630413, EPI_ISL_630414, EPI_ISL_630415, EPI_ISL_630416, EPI_ISL_630417, EPI_ISL_630418, EPI_ISL_630419, EPI_ISL_630420, EPI_ISL_630421, EPI_ISL_630422, EPI_ISL_630423, EPI_ISL_630424, EPI_ISL_630425, EPI_ISL_630426, EPI_ISL_630427, EPI_ISL_630428, EPI_ISL_630429, EPI_ISL_630430, EPI_ISL_630431, EPI_ISL_630432, EPI_ISL_630433, EPI_ISL_630434, EPI_ISL_630435, EPI_ISL_630436, EPI_ISL_630437, EPI_ISL_630438, EPI_ISL_630439, EPI_ISL_630440, EPI_ISL_630441, EPI_ISL_630442, EPI_ISL_630443, EPI_ISL_630444, EPI_ISL_630445, EPI_ISL_630446, EPI_ISL_630447, EPI_ISL_630448, EPI_ISL_630449, EPI_ISL_630450, EPI_ISL_630451, EPI_ISL_630452, EPI_ISL_630453, EPI_ISL_630454, EPI_ISL_630455, EPI_ISL_630456, EPI_ISL_630457, EPI_ISL_630458, EPI_ISL_630459, EPI_ISL_630460, EPI_ISL_630461, EPI_ISL_630462, EPI_ISL_630463, EPI_ISL_630464, EPI_ISL_630465, EPI_ISL_630466, EPI_ISL_630467, EPI_ISL_630468, EPI_ISL_630469, EPI_ISL_630470, EPI_ISL_630471, EPI_ISL_630472, EPI_ISL_630473 | | | |
| see above | Lighthouse Lab in Alderley Park | Wellcome Sanger Institute for the COVID-19 Genomics UK (COG-UK) consortium | Jacquelyn Wynn, Mairead Hyland, The Lighthouse Lab in Alderley Park and Alex Alderton, Roberto Amato, Sonia Goncalves, Ewan Harrison, David K. Jackson, Ian Johnston, Dominic Kwiatkowski, Cordelia Langford, John Sillitoe on behalf of the Wellcome Sanger Institute COVID-19 Surveillance Team |
| EPI_ISL_630474 | Lighthouse Lab in Alderley Park | Wellcome Sanger Institute for the COVID-19 Genomics UK (COG-UK) Consortium | Jacquelyn Wynn, Mairead Hyland, The Lighthouse Lab in Alderley Park and Alex Alderton, Roberto Amato, Sonia Goncalves, Ewan Harrison, David K. Jackson, Ian Johnston, Dominic Kwiatkowski, Cordelia Langford, John Sillitoe on behalf of the Wellcome Sanger Institute COVID-19 Surveillance Team |
| EPI_ISL_630475, EPI_ISL_630476, EPI_ISL_630477, EPI_ISL_630478, EPI_ISL_630479, EPI_ISL_630480, EPI_ISL_630481, EPI_ISL_630482, EPI_ISL_630483, EPI_ISL_630484, EPI_ISL_630485, EPI_ISL_630486, EPI_ISL_630487, EPI_ISL_630488, EPI_ISL_630489, EPI_ISL_630490, EPI_ISL_630491, EPI_ISL_630492, EPI_ISL_630493, EPI_ISL_630494, EPI_ISL_630495, EPI_ISL_630496, EPI_ISL_630497, EPI_ISL_630498, EPI_ISL_630499, EPI_ISL_630500, EPI_ISL_630501, EPI_ISL_630502, EPI_ISL_630503, EPI_ISL_630504, EPI_ISL_630505, EPI_ISL_630506, EPI_ISL_630507, EPI_ISL_630508, EPI_ISL_630509, EPI_ISL_630510, EPI_ISL_630511, EPI_ISL_630512, EPI_ISL_630513, EPI_ISL_630514, EPI_ISL_630515, EPI_ISL_630516, EPI_ISL_630517, EPI_ISL_630518, EPI_ISL_630519, EPI_ISL_630520, EPI_ISL_630521, EPI_ISL_630522, EPI_ISL_630523, EPI_ISL_630524, EPI_ISL_630525, EPI_ISL_630526, EPI_ISL_630527, EPI_ISL_630528, EPI_ISL_630529, EPI_ISL_630530, EPI_ISL_630531, EPI_ISL_630532 | | | |
| see above | Lighthouse Lab in Alderley Park | Wellcome Sanger Institute for the COVID-19 Genomics UK (COG-UK) consortium | Jacquelyn Wynn, Mairead Hyland, The Lighthouse Lab in Alderley Park and Alex Alderton, Roberto Amato, Sonia Goncalves, Ewan Harrison, David K. Jackson, Ian Johnston, Dominic Kwiatkowski, Cordelia Langford, John Sillitoe on behalf of the Wellcome Sanger Institute COVID-19 Surveillance Team |
| EPI_ISL_630533 | Lighthouse Lab in Alderley Park | Wellcome Sanger Institute for the COVID-19 Genomics UK (COG-UK) Consortium | Jacquelyn Wynn, Mairead Hyland, The Lighthouse Lab in Alderley Park and Alex Alderton, Roberto Amato, Sonia Goncalves, Ewan Harrison, David K. Jackson, Ian Johnston, Dominic Kwiatkowski, Cordelia Langford, John Sillitoe on behalf of the Wellcome Sanger Institute COVID-19 Surveillance Team |
| EPI_ISL_630534, EPI_ISL_630535, EPI_ISL_630536, EPI_ISL_630537, EPI_ISL_630538, EPI_ISL_630539, EPI_ISL_630540, EPI_ISL_630541, EPI_ISL_630542, EPI_ISL_630543, EPI_ISL_630544, EPI_ISL_630545, EPI_ISL_630546 | | | |
| see above | Lighthouse Lab in Alderley Park | Wellcome Sanger Institute for the COVID-19 Genomics UK (COG-UK) consortium | Jacquelyn Wynn, Mairead Hyland, The Lighthouse Lab in Alderley Park and Alex Alderton, Roberto Amato, Sonia Goncalves, Ewan Harrison, David K. Jackson, Ian Johnston, Dominic Kwiatkowski, Cordelia Langford, John Sillitoe on behalf of the Wellcome Sanger Institute COVID-19 Surveillance Team |
| EPI_ISL_630547 | Lighthouse Lab in Alderley Park | Wellcome Sanger Institute for the COVID-19 Genomics UK (COG-UK) Consortium | Jacquelyn Wynn, Mairead Hyland, The Lighthouse Lab in Alderley Park and Alex Alderton, Roberto Amato, Sonia Goncalves, Ewan Harrison, David K. Jackson, Ian Johnston, Dominic Kwiatkowski, Cordelia Langford, John Sillitoe on behalf of the Wellcome Sanger Institute COVID-19 Surveillance Team |
| EPI_ISL_630548, EPI_ISL_630549, EPI_ISL_630550, EPI_ISL_630551, EPI_ISL_630552, EPI_ISL_630553, EPI_ISL_630554, EPI_ISL_630555, EPI_ISL_630556, EPI_ISL_630557, EPI_ISL_630558, EPI_ISL_630559, EPI_ISL_630560, EPI_ISL_630561, EPI_ISL_630562, EPI_ISL_630563, EPI_ISL_630564, EPI_ISL_630565, EPI_ISL_630566, EPI_ISL_630567, EPI_ISL_630568, EPI_ISL_630569, EPI_ISL_630570, EPI_ISL_630571, EPI_ISL_630572, EPI_ISL_630573, EPI_ISL_630574, EPI_ISL_630575, EPI_ISL_630576, EPI_ISL_630577, EPI_ISL_630578, EPI_ISL_630579, EPI_ISL_630580, EPI_ISL_630581, EPI_ISL_630582, EPI_ISL_630583, EPI_ISL_630584, EPI_ISL_630585, EPI_ISL_630586, EPI_ISL_630587, EPI_ISL_630588, EPI_ISL_630589, EPI_ISL_630590, EPI_ISL_630591, EPI_ISL_630592, EPI_ISL_630593, EPI_ISL_630594, EPI_ISL_630595, EPI_ISL_630596, EPI_ISL_630597, EPI_ISL_630598, EPI_ISL_630599, EPI_ISL_630600, EPI_ISL_630601, EPI_ISL_630602, EPI_ISL_630603, EPI_ISL_630604, EPI_ISL_630605, EPI_ISL_630606, EPI_ISL_630607, EPI_ISL_630608, EPI_ISL_630609, EPI_ISL_630610, EPI_ISL_630611, EPI_ISL_630612, EPI_ISL_630613, EPI_ISL_630614, EPI_ISL_630615, EPI_ISL_630616, EPI_ISL_630617, EPI_ISL_630618, EPI_ISL_630619, EPI_ISL_630620, EPI_ISL_630621, EPI_ISL_630622, EPI_ISL_630623, EPI_ISL_630624, EPI_ISL_630625, EPI_ISL_630626, EPI_ISL_630627, EPI_ISL_630628, EPI_ISL_630629, EPI_ISL_630630, EPI_ISL_630631, EPI_ISL_630632, EPI_ISL_630633, EPI_ISL_630634, EPI_ISL_630635, EPI_ISL_630636, EPI_ISL_630637, EPI_ISL_630638, EPI_ISL_630639, EPI_ISL_630640, EPI_ISL_630641, EPI_ISL_630642, EPI_ISL_630643, EPI_ISL_630644, EPI_ISL_630645, EPI_ISL_630646, EPI_ISL_630647, EPI_ISL_630648, EPI_ISL_630649, EPI_ISL_630650, EPI_ISL_630651, EPI_ISL_630652, EPI_ISL_630653, EPI_ISL_630654, EPI_ISL_630655, EPI_ISL_630656, EPI_ISL_630657, EPI_ISL_630658, EPI_ISL_630659, EPI_ISL_630660, EPI_ISL_630661, EPI_ISL_630662, EPI_ISL_630663, EPI_ISL_630664, EPI_ISL_630665, EPI_ISL_630666, EPI_ISL_630667, EPI_ISL_630668, EPI_ISL_630669, EPI_ISL_630670, EPI_ISL_630671, EPI_ISL_630672, EPI_ISL_630673, EPI_ISL_630674, EPI_ISL_630675, EPI_ISL_630676, EPI_ISL_630677, EPI_ISL_630678, EPI_ISL_630679, EPI_ISL_630680, EPI_ISL_630681, EPI_ISL_630682, EPI_ISL_630683, EPI_ISL_630684, EPI_ISL_630685, EPI_ISL_630686, EPI_ISL_630687, EPI_ISL_630688, EPI_ISL_630689, EPI_ISL_630690, EPI_ISL_630691, EPI_ISL_630692, EPI_ISL_630693, EPI_ISL_630694, EPI_ISL_630695, EPI_ISL_630696, EPI_ISL_630697, EPI_ISL_630698, EPI_ISL_630699, EPI_ISL_630700, EPI_ISL_630701, EPI_ISL_630975, EPI_ISL_630982, EPI_ISL_630985, EPI_ISL_630992, EPI_ISL_631011, EPI_ISL_631031, EPI_ISL_631044, EPI_ISL_631050, EPI_ISL_631059, EPI_ISL_631060, EPI_ISL_631074, EPI_ISL_631084, EPI_ISL_631096, EPI_ISL_631113, EPI_ISL_631165, EPI_ISL_631184, EPI_ISL_631190, EPI_ISL_631231, EPI_ISL_631269 | | | |
| see above | Lighthouse Lab in Alderley Park | Wellcome Sanger Institute for the COVID-19 Genomics UK (COG-UK) consortium | Jacquelyn Wynn, Mairead Hyland, The Lighthouse Lab in Alderley Park and Alex Alderton, Roberto Amato, Sonia Goncalves, Ewan Harrison, David K. Jackson, Ian Johnston, Dominic Kwiatkowski, Cordelia Langford, John Sillitoe on behalf of the Wellcome Sanger Institute COVID-19 Surveillance Team |
| EPI_ISL_632364, EPI_ISL_632370, EPI_ISL_632372, EPI_ISL_632379, | Dutch COVID-19 response team | Erasmus Medical Center | Bas Oude Munnink, David Nieuwenhuijse, Reina Sikkema, Claudia Schapendonk, Irina Chestakova, Anne van der Linden, Theo Bestebroer, Stefan van Nieuwkoop, Mark Pronk, Pascal Lexmond, Corien Swaan, Manon Haverkate, Madelief Mollers, Mart Stein, Sandra Kengne Kamga Mobou, Jeroen van |

| | | | |
|---|---|---|---|
| EPI_ISL_632401, EPI_ISL_632408, EPI_ISL_632411, EPI_ISL_632418, EPI_ISL_632440, EPI_ISL_632445 | | | Kampen, Jolanda Voermans, Aura Timen, Corine GeurtsvanKessel, Annemiek van der Eijk, Richard Molenkamp, Marion Koopmans, on behalf of the Dutch national COVID-19 response team. |
| EPI_ISL_632492 | Dutch COVID-19 response team | Erasmus Medical Center | OH consortium |
| EPI_ISL_632564, EPI_ISL_632565, EPI_ISL_632566, EPI_ISL_632573, EPI_ISL_632574, EPI_ISL_632575, EPI_ISL_632576, EPI_ISL_632587, EPI_ISL_632603, EPI_ISL_632604, EPI_ISL_632605, EPI_ISL_632606, EPI_ISL_632608, EPI_ISL_632610, EPI_ISL_632611, EPI_ISL_632623, EPI_ISL_632624, EPI_ISL_632625, EPI_ISL_632626, EPI_ISL_632627, EPI_ISL_632628, EPI_ISL_632629, EPI_ISL_632630, EPI_ISL_632631, EPI_ISL_632632, EPI_ISL_632633, EPI_ISL_632634, EPI_ISL_632635, EPI_ISL_632636, EPI_ISL_632637, EPI_ISL_632638, EPI_ISL_632639, EPI_ISL_632640, EPI_ISL_632655, EPI_ISL_632660, EPI_ISL_632661, EPI_ISL_632662, EPI_ISL_632663, EPI_ISL_632667, EPI_ISL_632668, EPI_ISL_632673, EPI_ISL_632674, EPI_ISL_632698, EPI_ISL_632702, EPI_ISL_632703, EPI_ISL_632707, EPI_ISL_632708, EPI_ISL_632709, EPI_ISL_632714, EPI_ISL_632735, EPI_ISL_632777 | | | |
| see above | Dutch COVID-19 response team | Erasmus Medical Center | Bas Oude Munnink, David Nieuwenhuijse, Reina Sikkema, Claudia Schapendonk, Irina Chestakova, Anne van der Linden, Theo Bestebroer, Stefan van Nieuwkoop, Mark Pronk, Pascal Lexmond, Corien Swaan, Manon Haverkate, Madelief Mollers, Mart Stein, Sandra Kengne Kamga Mobou, Jeroen van Kampen, Jolanda Voermans, Aura Timen, Corine GeurtsvanKessel, Annemiek van der Eijk, Richard Molenkamp, Marion Koopmans, on behalf of the Dutch national COVID-19 response team. |
| EPI_ISL_632796 | Respiratory Virus Unit, Microbiology Services Colindale, Public Health England | Respiratory Virus Unit, Microbiology Services Colindale, Public Health England | PHE Covid Sequencing Team |
| EPI_ISL_633008 | DOHMH Jamaica | New York City Public Health Laboratory | Jade Wang, et al. |
| EPI_ISL_633032 | DOHMH Morrisania | New York City Public Health Laboratory | Jade Wang, et al. |
| EPI_ISL_633033 | DOHMH Chelsea | New York City Public Health Laboratory | Jade Wang, et al. |
| EPI_ISL_633034 | DOHMH Riverside | New York City Public Health Laboratory | Jade Wang, et al. |
| EPI_ISL_633056 | DOHMH Central Harlem | New York City Public Health Laboratory | Jade Wang, et al. |
| EPI_ISL_633057, EPI_ISL_633058 | DOHMH Morrisania | New York City Public Health Laboratory | Jade Wang, et al. |
| EPI_ISL_633059 | DOHMH Corona | New York City Public Health Laboratory | Jade Wang, et al. |
| EPI_ISL_633060 | DOHMH Jamaica | New York City Public Health Laboratory | Jade Wang, et al. |
| EPI_ISL_633065 | DOHMH Morrisania | New York City Public Health Laboratory | Jade Wang, et al. |
| EPI_ISL_633066 | DOHMH Chelsea | New York City Public Health Laboratory | Jade Wang, et al. |
| EPI_ISL_634754, EPI_ISL_634755, EPI_ISL_634756 | Lighthouse Lab in Alderley Park | Wellcome Sanger Institute for the COVID-19 Genomics UK (COG-UK) consortium | Jacquelyn Wynn, Mairead Hyland, The Lighthouse Lab in Alderley Park and Alex Alderton, Roberto Amato, Sonia Goncalves, Ewan Harrison, David K. Jackson, Ian Johnston, Dominic Kwiatkowski, Cordelia Langford, John Sillitoe on behalf of the Wellcome Sanger Institute COVID-19 Surveillance Team (http://www.sanger.ac.uk/covid-team) |
| EPI_ISL_634838 | Minnesota Department of Health, Public Health Laboratory | Minnesota Department of Health, Public Health Laboratory | Matt Plumb, Jacob Garfin, Alexandra Lorentz, and Xiong Wang |
| EPI_ISL_634882 | Lab voor klinische biologie | Onderzoeksgroep Virologie | Laurens Lambrechts, Nick Vereecke, Marthe Pauwels, Bruno Verhasselt, Linos Vandekerckhove, Hans Nauwynck, Sebastiaan Theuns |
| EPI_ISL_634892 | Lab voor klinische biologie | Onderzoeksgroep Virologie | Nick Vereecke, Laurens Lambrechts, Marthe Pauwels, Bruno Verhasselt, Linos Vandekerckhove, Hans Nauwynck, Sebastiaan Theuns |
| EPI_ISL_634924 | Utah Public Health Laboratory | Utah Public Health Laboratory | Erin L. Young, Kelly F. Oakeson |
| EPI_ISL_634993, EPI_ISL_634994, EPI_ISL_634995, EPI_ISL_634996, EPI_ISL_634997, EPI_ISL_634998, EPI_ISL_634999, EPI_ISL_635000, EPI_ISL_635002, EPI_ISL_635005, EPI_ISL_635006, EPI_ISL_635007, EPI_ISL_635008, EPI_ISL_635010, EPI_ISL_635011, EPI_ISL_635012, EPI_ISL_635013, EPI_ISL_635014 | | | |
| see above | National Health Laboratory Service - Inkosi Albert Luthuli Central Hospital (NHLS-IALCH) | KRISP, KZN Research Innovation and Sequencing Platform | Giandhari J, Pillay S, Lessells R, Mdlalose K, York D, Khan S, Tegally H, Wilkinson E, de Oliveira T |
| EPI_ISL_635071, EPI_ISL_635072 | University Hospital of Northern Norway, Department for Microbiology and Infectious Disease Control | Norwegian Institute of Public Health, Department of Virology | Kathrine Stene-Johansen, Kamilla Heddeland Instefjord, Hilde Elshaug, Marie Paulsen Madsen, Rasmus Riis Kopperud, Hilde Vollan, Karoline Bragstad, Olav Hungnes |
| EPI_ISL_635088 | Medical Microbiology Unit, Department for Laboratory Medicine, Drammen Hospital, Vestre Viken Health Trust, | Norwegian Institute of Public Health, Department of Virology | Kathrine Stene-Johansen, Kamilla Heddeland Instefjord, Hilde Elshaug, Marie Paulsen Madsen, Rasmus Riis Kopperud, Hilde Vollan, Karoline Bragstad, Olav Hungnes |
| EPI_ISL_635139, EPI_ISL_635140 | Vestfold Hospital, Toensberg Department of Microbiology | Norwegian Institute of Public Health, Department of Virology | Kathrine Stene-Johansen, Kamilla Heddeland Instefjord, Hilde Elshaug, Marie Paulsen Madsen, Rasmus Riis Kopperud, Hilde Vollan, Karoline Bragstad, Olav Hungnes |
| EPI_ISL_635141 | University Hospital of Northern Norway, Department for Microbiology and Infectious Disease Control | Norwegian Institute of Public Health, Department of Virology | Kathrine Stene-Johansen, Kamilla Heddeland Instefjord, Hilde Elshaug, Marie Paulsen Madsen, Rasmus Riis Kopperud, Hilde Vollan, Karoline Bragstad, Olav Hungnes |
| EPI_ISL_635142 | Nordland Hospital - Bodo, Laboratory Department, Molecular Biology Unit | Norwegian Institute of Public Health, Department of Virology | Kathrine Stene-Johansen, Kamilla Heddeland Instefjord, Hilde Elshaug, Marie Paulsen Madsen, Rasmus Riis Kopperud, Hilde Vollan, Karoline Bragstad, Olav Hungnes |
| EPI_ISL_635143 | Hospital of Southern Norway - Kristiansand, Department of Medical Microbiology | Norwegian Institute of Public Health, Department of Virology | Kathrine Stene-Johansen, Kamilla Heddeland Instefjord, Hilde Elshaug, Marie Paulsen Madsen, Rasmus Riis Kopperud, Hilde Vollan, Karoline Bragstad, Olav Hungnes |
| EPI_ISL_635145 | Nordland Hospital - Bodo, Laboratory Department, Molecular Biology Unit | Norwegian Institute of Public Health, Department of Virology | Kathrine Stene-Johansen, Kamilla Heddeland Instefjord, Hilde Elshaug, Marie Paulsen Madsen, Rasmus Riis Kopperud, Hilde Vollan, Karoline Bragstad, Olav Hungnes |
| EPI_ISL_635153 | Unilabs Laboratory Medicine | Norwegian Institute of Public Health, Department of Virology | Kathrine Stene-Johansen, Kamilla Heddeland Instefjord, Hilde Elshaug, Marie Paulsen Madsen, Rasmus Riis Kopperud, Hilde Vollan, Karoline Bragstad, Olav Hungnes |
| EPI_ISL_635160 | University Hospital of Northern Norway, Department for Microbiology and Infectious Disease Control | Norwegian Institute of Public Health, Department of Virology | Kathrine Stene-Johansen, Kamilla Heddeland Instefjord, Hilde Elshaug, Marie Paulsen Madsen, Rasmus Riis Kopperud, Hilde Vollan, Karoline Bragstad, Olav Hungnes |
| EPI_ISL_635162 | Unilabs Laboratory Medicine | Norwegian Institute of Public Health, Department of Virology | Kathrine Stene-Johansen, Kamilla Heddeland Instefjord, Hilde Elshaug, Marie Paulsen Madsen, Rasmus Riis Kopperud, Hilde Vollan, Karoline Bragstad, Olav Hungnes |
| EPI_ISL_635163 | University Hospital of Northern Norway, Department for Microbiology and Infectious Disease Control | Norwegian Institute of Public Health, Department of Virology | Kathrine Stene-Johansen, Kamilla Heddeland Instefjord, Hilde Elshaug, Marie Paulsen Madsen, Rasmus Riis Kopperud, Hilde Vollan, Karoline Bragstad, Olav Hungnes |
| EPI_ISL_635166 | Dept. of Medical Microbiology, Stavanger University Hospital, Helse Stavanger HF | Norwegian Institute of Public Health, Department of Virology | Kathrine Stene-Johansen, Kamilla Heddeland Instefjord, Hilde Elshaug, Marie Paulsen Madsen, Rasmus Riis Kopperud, Hilde Vollan, Karoline Bragstad, Olav Hungnes |
| EPI_ISL_635167, EPI_ISL_635170 | Foerde Hospital, Department of Microbiology | Norwegian Institute of Public Health, Department of Virology | Kathrine Stene-Johansen, Kamilla Heddeland Instefjord, Hilde Elshaug, Marie Paulsen Madsen, Rasmus Riis Kopperud, Hilde Vollan, Karoline Bragstad, Olav Hungnes |
| EPI_ISL_635171 | Vestfold Hospital, Toensberg Department of Microbiology | Norwegian Institute of Public Health, Department of Virology | Kathrine Stene-Johansen, Kamilla Heddeland Instefjord, Hilde Elshaug, Marie Paulsen Madsen, Rasmus Riis Kopperud, Hilde Vollan, Karoline Bragstad, Olav Hungnes |
| EPI_ISL_635182 | Nordland Hospital - Bodo, Laboratory Department, Molecular Biology Unit | Norwegian Institute of Public Health, Department of Virology | Kathrine Stene-Johansen, Kamilla Heddeland Instefjord, Hilde Elshaug, Marie Paulsen Madsen, Rasmus Riis Kopperud, Hilde Vollan, Karoline Bragstad, Olav Hungnes |
| EPI_ISL_635189 | Hospital of Southern Norway - Kristiansand, Department of Medical Microbiology | Norwegian Institute of Public Health, Department of Virology | Kathrine Stene-Johansen, Kamilla Heddeland Instefjord, Hilde Elshaug, Marie Paulsen Madsen, Rasmus Riis Kopperud, Hilde Vollan, Karoline Bragstad, Olav Hungnes |
| EPI_ISL_635191 | Unilabs Laboratory Medicine | Norwegian Institute of Public Health, Department of Virology | Kathrine Stene-Johansen, Kamilla Heddeland Instefjord, Hilde Elshaug, Marie Paulsen Madsen, Rasmus Riis Kopperud, Hilde Vollan, Karoline Bragstad, Olav Hungnes |
| EPI_ISL_635196 | Dept. of Medical Microbiology, Stavanger University Hospital, | Norwegian Institute of Public Health, Department of Virology | Kathrine Stene-Johansen, Kamilla Heddeland Instefjord, Hilde Elshaug, Marie Paulsen Madsen, Rasmus Riis Kopperud, Hilde Vollan, Karoline Bragstad, Olav |

| | | | |
|---|---|---|---|
| | Helse Stavanger HF | | Hungnes |
| EPI_ISL_636536, EPI_ISL_636538, EPI_ISL_636581, EPI_ISL_636582, EPI_ISL_636584 | Dutch COVID-19 response team | National Institute for Public Health and the Environment (RIVM) | Adam Meijer, Harry Vennema, Jeroen Cremer, Sharon van den Brink, Bas van der Veer, AnneMarie van den Brandt, Florian Zwagemaker, Dennis Schmitz, Chantal Reusken, on behalf of the national COVID-19 response team |
| EPI_ISL_636690, EPI_ISL_636691, EPI_ISL_636692, EPI_ISL_636693, EPI_ISL_636694, EPI_ISL_636695, EPI_ISL_636696, EPI_ISL_636697, EPI_ISL_636698, EPI_ISL_636699, EPI_ISL_636700, EPI_ISL_636701, EPI_ISL_636703 | | | |
| see above | Respiratory Virus Unit, Microbiology Services Colindale, Public Health England | Respiratory Virus Unit, Microbiology Services Colindale, Public Health England | PHE Covid Sequencing Team |
| EPI_ISL_637019 | Department of Infectious Diseases and Immunology, National Hospital Organization Nagoya Medical Center | Clinical Research Center, National Hospital Organization Nagoya Medical Center | Yoshihiro Nakata, Hirotaka Ode, Mai Kubota, Masakazu Matsuda, Kazuhiro Matsuoka, Miho Nakasuji, Mikiko Mori, Mayumi Imahashi, Yoshiyuki Yokomaku, Yasumasa Iwatani |
| EPI_ISL_637114, EPI_ISL_637115, EPI_ISL_637116, EPI_ISL_637117 | Respiratory Virus Unit, Microbiology Services Colindale, Public Health England | COVID-19 Genomics UK (COG-UK) Consortium | PHE Covid Sequencing Team |
| EPI_ISL_637260, EPI_ISL_637264, EPI_ISL_637269, EPI_ISL_637270, EPI_ISL_637271, EPI_ISL_637272, EPI_ISL_637276, EPI_ISL_637287, EPI_ISL_637304, EPI_ISL_637308, EPI_ISL_637309, EPI_ISL_637310 | | | |
| see above | Department of Pathology, University of Cambridge | COVID-19 Genomics UK (COG-UK) Consortium | Aminu S. Jahun, Yasmin Chaudhry, Grant Hall, Iliana Georgana, Myra Hosmillo, Martin D. Curran, Malte Pinckert, Surendra Parmar, Ian Goodfellow |
| EPI_ISL_637312 | Northumbria University / South Tees Hospitals NHS Foundation Trust / North Cumbria Integrated Care NHS Foundation Trust / North Tees and Hartlepool NHS Foundation Trust / Newcastle Hospitals NHS Foundation Trust | COVID-19 Genomics UK (COG-UK) Consortium | Darren L Smith,Andrew Nelson,Matthew Bashton,Greg R Young,Joshua Loh,John Allan,Mohammad A Tariq,Giles S Holt,Gary Black,Wen C Yew,Lynn Dover,Paul Baker,Steve Liggett,Sarah Essex,Jane Greenaway,Debra Padgett,Clive Graham,Garren Scott,Edward Barton,Emma Swindells,Brendan Payne,Jennifer Collins,Yusri Taha,Gary Eltringham |
| EPI_ISL_637314, EPI_ISL_637315, EPI_ISL_637316, EPI_ISL_637317, EPI_ISL_637318, EPI_ISL_637331, EPI_ISL_637348, EPI_ISL_637353, EPI_ISL_637356, EPI_ISL_637360, EPI_ISL_637362, EPI_ISL_637363, EPI_ISL_637364, EPI_ISL_637365, EPI_ISL_637366, EPI_ISL_637367, EPI_ISL_637368 | | | |
| see above | Department of Pathology, University of Cambridge | COVID-19 Genomics UK (COG-UK) Consortium | Aminu S. Jahun, Yasmin Chaudhry, Grant Hall, Iliana Georgana, Myra Hosmillo, Martin D. Curran, Malte Pinckert, Surendra Parmar, Ian Goodfellow |
| EPI_ISL_637376 | Northumbria University / South Tees Hospitals NHS Foundation Trust / North Cumbria Integrated Care NHS Foundation Trust / North Tees and Hartlepool NHS Foundation Trust / Newcastle Hospitals NHS Foundation Trust | COVID-19 Genomics UK (COG-UK) Consortium | Darren L Smith,Andrew Nelson,Matthew Bashton,Greg R Young,Joshua Loh,John Allan,Mohammad A Tariq,Giles S Holt,Gary Black,Wen C Yew,Lynn Dover,Paul Baker,Steve Liggett,Sarah Essex,Jane Greenaway,Debra Padgett,Clive Graham,Garren Scott,Edward Barton,Emma Swindells,Brendan Payne,Jennifer Collins,Yusri Taha,Gary Eltringham |
| EPI_ISL_637384, EPI_ISL_637386, EPI_ISL_637391, EPI_ISL_637393, EPI_ISL_637396, EPI_ISL_637397, EPI_ISL_637403, EPI_ISL_637407, EPI_ISL_637408, EPI_ISL_637424, EPI_ISL_637451, EPI_ISL_637457, EPI_ISL_637461, EPI_ISL_637462 | | | |
| see above | Department of Pathology, University of Cambridge | COVID-19 Genomics UK (COG-UK) Consortium | Aminu S. Jahun, Yasmin Chaudhry, Grant Hall, Iliana Georgana, Myra Hosmillo, Martin D. Curran, Malte Pinckert, Surendra Parmar, Ian Goodfellow |
| EPI_ISL_637463 | Northumbria University / South Tees Hospitals NHS Foundation Trust / North Cumbria Integrated Care NHS Foundation Trust / North Tees and Hartlepool NHS Foundation Trust / Newcastle Hospitals NHS Foundation Trust | COVID-19 Genomics UK (COG-UK) Consortium | Darren L Smith,Andrew Nelson,Matthew Bashton,Greg R Young,Joshua Loh,John Allan,Mohammad A Tariq,Giles S Holt,Gary Black,Wen C Yew,Lynn Dover,Paul Baker,Steve Liggett,Sarah Essex,Jane Greenaway,Debra Padgett,Clive Graham,Garren Scott,Edward Barton,Emma Swindells,Brendan Payne,Jennifer Collins,Yusri Taha,Gary Eltringham |
| EPI_ISL_637465, EPI_ISL_637475, EPI_ISL_637476, EPI_ISL_637477, EPI_ISL_637478, EPI_ISL_637479, EPI_ISL_637480, EPI_ISL_637481, EPI_ISL_637482, EPI_ISL_637483, EPI_ISL_637484, EPI_ISL_637485, EPI_ISL_637486, EPI_ISL_637487, EPI_ISL_637488, EPI_ISL_637489, EPI_ISL_637490, EPI_ISL_637500, EPI_ISL_637501, EPI_ISL_637504, EPI_ISL_637507, EPI_ISL_637513, EPI_ISL_637514, EPI_ISL_637515, EPI_ISL_637516, EPI_ISL_637517, EPI_ISL_637518 | | | |
| see above | Department of Pathology, University of Cambridge | COVID-19 Genomics UK (COG-UK) Consortium | Aminu S. Jahun, Yasmin Chaudhry, Grant Hall, Iliana Georgana, Myra Hosmillo, Martin D. Curran, Malte Pinckert, Surendra Parmar, Ian Goodfellow |
| EPI_ISL_637521 | Northumbria University / South Tees Hospitals NHS Foundation Trust / North Cumbria Integrated Care NHS Foundation Trust / North Tees and Hartlepool NHS Foundation Trust / Newcastle Hospitals NHS Foundation Trust | COVID-19 Genomics UK (COG-UK) Consortium | Darren L Smith,Andrew Nelson,Matthew Bashton,Greg R Young,Joshua Loh,John Allan,Mohammad A Tariq,Giles S Holt,Gary Black,Wen C Yew,Lynn Dover,Paul Baker,Steve Liggett,Sarah Essex,Jane Greenaway,Debra Padgett,Clive Graham,Garren Scott,Edward Barton,Emma Swindells,Brendan Payne,Jennifer Collins,Yusri Taha,Gary Eltringham |
| EPI_ISL_637532, EPI_ISL_637534, EPI_ISL_637548, EPI_ISL_637552 | Department of Pathology, University of Cambridge | COVID-19 Genomics UK (COG-UK) Consortium | Aminu S. Jahun, Yasmin Chaudhry, Grant Hall, Iliana Georgana, Myra Hosmillo, Martin D. Curran, Malte Pinckert, Surendra Parmar, Ian Goodfellow |
| EPI_ISL_637559 | Virology Department, Sheffield Teaching Hospitals NHS Foundation Trust/Department of Infection, Immunity and Cardiovascular Disease, The Medical School, University of Sheffield | COVID-19 Genomics UK (COG-UK) Consortium | Thushan de Silva, Matthew Parker, Nikki Smith, Adri Angyal, Rebecca Brown, Luke Green, Rachel Tucker, Paul Parsons, Danielle Groves, Katie Johnson, Laura Carrilero, Alex Keeley, Dave Partridge, Matthew Wyles, Benjamin Lindsey, Mehmet Yavuz, Mohammad Raza, Cariad Evans |
| EPI_ISL_637581 | Northumbria University / South Tees Hospitals NHS Foundation Trust / North Cumbria Integrated Care NHS Foundation Trust / North Tees and Hartlepool NHS Foundation Trust / Newcastle Hospitals NHS Foundation Trust | COVID-19 Genomics UK (COG-UK) Consortium | Darren L Smith,Andrew Nelson,Matthew Bashton,Greg R Young,Joshua Loh,John Allan,Mohammad A Tariq,Giles S Holt,Gary Black,Wen C Yew,Lynn Dover,Paul Baker,Steve Liggett,Sarah Essex,Jane Greenaway,Debra Padgett,Clive Graham,Garren Scott,Edward Barton,Emma Swindells,Brendan Payne,Jennifer Collins,Yusri Taha,Gary Eltringham |
| EPI_ISL_637583, EPI_ISL_637628, EPI_ISL_637629, EPI_ISL_637630, EPI_ISL_637631, EPI_ISL_637648, EPI_ISL_637649, EPI_ISL_637650, EPI_ISL_637651, EPI_ISL_637652, EPI_ISL_637653, EPI_ISL_637654, EPI_ISL_637655, EPI_ISL_637656, EPI_ISL_637657, EPI_ISL_637658, EPI_ISL_637672, EPI_ISL_637673, EPI_ISL_637676, EPI_ISL_637677, EPI_ISL_637678, EPI_ISL_637680, EPI_ISL_637681, EPI_ISL_637682, EPI_ISL_637683, EPI_ISL_637684, EPI_ISL_637756, EPI_ISL_637757, EPI_ISL_637758, EPI_ISL_637759, EPI_ISL_637760, EPI_ISL_637761, EPI_ISL_637762, EPI_ISL_637763, EPI_ISL_637764, EPI_ISL_637765, EPI_ISL_637766, EPI_ISL_637767, EPI_ISL_637768, EPI_ISL_637769, EPI_ISL_637770, EPI_ISL_637771, EPI_ISL_637772, EPI_ISL_637773, EPI_ISL_637774, EPI_ISL_637775, EPI_ISL_637776, EPI_ISL_637777, EPI_ISL_637778, EPI_ISL_637779 | | | |
| see above | Department of Pathology, University of Cambridge | COVID-19 Genomics UK (COG-UK) Consortium | Aminu S. Jahun, Yasmin Chaudhry, Grant Hall, Iliana Georgana, Myra Hosmillo, Martin D. Curran, Malte Pinckert, Surendra Parmar, Ian Goodfellow |
| EPI_ISL_637784, EPI_ISL_637797 | Northumbria University / South Tees Hospitals NHS Foundation Trust / North Cumbria Integrated Care NHS Foundation Trust / North Tees and Hartlepool NHS Foundation Trust / Newcastle Hospitals NHS Foundation Trust | COVID-19 Genomics UK (COG-UK) Consortium | Darren L Smith,Andrew Nelson,Matthew Bashton,Greg R Young,Joshua Loh,John Allan,Mohammad A Tariq,Giles S Holt,Gary Black,Wen C Yew,Lynn Dover,Paul Baker,Steve Liggett,Sarah Essex,Jane Greenaway,Debra Padgett,Clive Graham,Garren Scott,Edward Barton,Emma Swindells,Brendan Payne,Jennifer Collins,Yusri Taha,Gary Eltringham |
| EPI_ISL_637822, EPI_ISL_637823, EPI_ISL_637824, EPI_ISL_637825, EPI_ISL_637826, EPI_ISL_637827, EPI_ISL_637828, EPI_ISL_637829, EPI_ISL_637830, EPI_ISL_637831, EPI_ISL_637832, EPI_ISL_637833, EPI_ISL_637834, EPI_ISL_637835, EPI_ISL_637836, EPI_ISL_637837, EPI_ISL_637838, EPI_ISL_637839, EPI_ISL_637840, EPI_ISL_637841, EPI_ISL_637842, EPI_ISL_637843, EPI_ISL_637844, EPI_ISL_637845, EPI_ISL_637846, EPI_ISL_637847, EPI_ISL_637848, EPI_ISL_637849, EPI_ISL_637850, EPI_ISL_637851, EPI_ISL_637852 | | | |
| see above | Department of Pathology, University of Cambridge | COVID-19 Genomics UK (COG-UK) Consortium | Aminu S. Jahun, Yasmin Chaudhry, Grant Hall, Iliana Georgana, Myra Hosmillo, Martin D. Curran, Malte Pinckert, Surendra Parmar, Ian Goodfellow |
| EPI_ISL_637855, EPI_ISL_637867 | Oxford Viromics, NDM, University of Oxford; Oxford University Hospitals; Basingstoke and North Hampshire Hospital | COVID-19 Genomics UK (COG-UK) Consortium | Tanya Golubchik, David Bonsall, George Macintyre, Amy Trebes, Mariateresa de Cesare, Catrin Moore, Alex Mobbs, Anita Justice, Robert Shaw, Monique Andersson, Timothy Peto, Emma Wise, Nathan Moore, Jessica Lynch, Nick Cortes, Matilde Mori, Stephen Kidd, David Buck, John Todd, Christophe Fraser |
| EPI_ISL_637873, EPI_ISL_637877, EPI_ISL_637890, EPI_ISL_637906, EPI_ISL_637938, EPI_ISL_637940, EPI_ISL_637963, EPI_ISL_637967, EPI_ISL_637977, EPI_ISL_637980, EPI_ISL_637982, EPI_ISL_637983 | | | |
| see above | Department of Pathology, University of Cambridge | COVID-19 Genomics UK (COG-UK) Consortium | Aminu S. Jahun, Yasmin Chaudhry, Grant Hall, Iliana Georgana, Myra Hosmillo, Martin D. Curran, Malte Pinckert, Surendra Parmar, Ian Goodfellow |
| EPI_ISL_637996 | Northumbria University / South Tees Hospitals NHS Foundation Trust / North Cumbria Integrated Care NHS Foundation Trust / North Tees and Hartlepool NHS Foundation Trust / Newcastle Hospitals NHS Foundation Trust | COVID-19 Genomics UK (COG-UK) Consortium | Darren L Smith,Andrew Nelson,Matthew Bashton,Greg R Young,Joshua Loh,John Allan,Mohammad A Tariq,Giles S Holt,Gary Black,Wen C Yew,Lynn Dover,Paul Baker,Steve Liggett,Sarah Essex,Jane Greenaway,Debra Padgett,Clive Graham,Garren Scott,Edward Barton,Emma Swindells,Brendan Payne,Jennifer Collins,Yusri Taha,Gary Eltringham |

| | | | |
|---|---|---|---|
| EPI_ISL_638000, EPI_ISL_638008 | Department of Pathology, University of Cambridge | COVID-19 Genomics UK (COG-UK) Consortium | Aminu S. Jahun, Yasmin Chaudhry, Grant Hall, Iliana Georgana, Myra Hosmillo, Martin D. Curran, Malte Pinckert, Surendra Parmar, Ian Goodfellow |
| EPI_ISL_638010 | Northumbria University / South Tees Hospitals NHS Foundation Trust / North Cumbria Integrated Care NHS Foundation Trust / North Tees and Hartlepool NHS Foundation Trust / Newcastle Hospitals NHS Foundation Trust | COVID-19 Genomics UK (COG-UK) Consortium | Darren L Smith,Andrew Nelson,Matthew Bashton,Greg R Young,Joshua Loh,John Allan,Mohammad A Tariq,Giles S Holt,Gary Black,Wen C Yew,Lynn Dover,Paul Baker,Steve Liggett,Sarah Essex,Jane Greenaway,Debra Padgett,Clive Graham,Garren Scott,Edward Barton,Emma Swindells,Brendan Payne,Jennifer Collins,Yusri Taha,Gary Eltringham |
| EPI_ISL_638013, EPI_ISL_638014, EPI_ISL_638018, EPI_ISL_638036, EPI_ISL_638072 | Department of Pathology, University of Cambridge | COVID-19 Genomics UK (COG-UK) Consortium | Aminu S. Jahun, Yasmin Chaudhry, Grant Hall, Iliana Georgana, Myra Hosmillo, Martin D. Curran, Malte Pinckert, Surendra Parmar, Ian Goodfellow |
| EPI_ISL_638086, EPI_ISL_638087, EPI_ISL_638145 | Oxford Viromics, NDM, University of Oxford; Oxford University Hospitals; Basingstoke and North Hampshire Hospital | COVID-19 Genomics UK (COG-UK) Consortium | Tanya Golubchik, David Bonsall, George Macintyre, Amy Trebes, Mariateresa de Cesare, Catrin Moore, Alex Mobbs, Anita Justice, Robert Shaw, Monique Andersson, Timothy Peto, Emma Wise, Nathan Moore, Jessica Lynch, Nick Cortes, Matilde Mori, Stephen Kidd, David Buck, John Todd, Christophe Fraser |
| EPI_ISL_638146 | Department of Pathology, University of Cambridge | COVID-19 Genomics UK (COG-UK) Consortium | Aminu S. Jahun, Yasmin Chaudhry, Grant Hall, Iliana Georgana, Myra Hosmillo, Martin D. Curran, Malte Pinckert, Surendra Parmar, Ian Goodfellow |
| EPI_ISL_638153 | Wales Specialist Virology Centre Sequencing lab: Pathogen Genomics Unit | COVID-19 Genomics UK (COG-UK) Consortium | Catherine Moore, Johnathan Evans, Laura Gifford, Malorie Perry, Simon Cottrell, Angela Marchbank, Alec Birchley, Alexander Adams, Amy Gaskin, Bree Gatica-Wilcox, Jason Coombes, Joel Southgate, Lauren Gilbert, Lee Graham, Nicole Pacchiarini, Sara Kumziene-Summerhayes, Sarah Taylor, Sophie Jones, Sara Rey, Matthew Bull, Joanne Watkins, Sally Corden, Tom Connor |
| EPI_ISL_638207 | Oxford Viromics, NDM, University of Oxford; Oxford University Hospitals; Basingstoke and North Hampshire Hospital | COVID-19 Genomics UK (COG-UK) Consortium | Tanya Golubchik, David Bonsall, George Macintyre, Amy Trebes, Mariateresa de Cesare, Catrin Moore, Alex Mobbs, Anita Justice, Robert Shaw, Monique Andersson, Timothy Peto, Emma Wise, Nathan Moore, Jessica Lynch, Nick Cortes, Matilde Mori, Stephen Kidd, David Buck, John Todd, Christophe Fraser |
| EPI_ISL_638444, EPI_ISL_638445, EPI_ISL_638446, EPI_ISL_638447, EPI_ISL_638448 | Department of Pathology, University of Cambridge | COVID-19 Genomics UK (COG-UK) Consortium | Aminu S. Jahun, Yasmin Chaudhry, Grant Hall, Iliana Georgana, Myra Hosmillo, Martin D. Curran, Malte Pinckert, Surendra Parmar, Ian Goodfellow |
| EPI_ISL_638482 | Oxford Viromics, NDM, University of Oxford; Oxford University Hospitals; Basingstoke and North Hampshire Hospital | COVID-19 Genomics UK (COG-UK) Consortium | Tanya Golubchik, David Bonsall, George Macintyre, Amy Trebes, Mariateresa de Cesare, Catrin Moore, Alex Mobbs, Anita Justice, Robert Shaw, Monique Andersson, Timothy Peto, Emma Wise, Nathan Moore, Jessica Lynch, Nick Cortes, Matilde Mori, Stephen Kidd, David Buck, John Todd, Christophe Fraser |
| EPI_ISL_638561, EPI_ISL_638564, EPI_ISL_638565, EPI_ISL_638566, EPI_ISL_638567, EPI_ISL_638568, EPI_ISL_638569, EPI_ISL_638570, EPI_ISL_638578, EPI_ISL_638579 | Northumbria University / South Tees Hospitals NHS Foundation Trust / North Cumbria Integrated Care NHS Foundation Trust / North Tees and Hartlepool NHS Foundation Trust / Newcastle Hospitals NHS Foundation Trust | COVID-19 Genomics UK (COG-UK) Consortium | Darren L Smith,Andrew Nelson,Matthew Bashton,Greg R Young,Joshua Loh,John Allan,Mohammad A Tariq,Giles S Holt,Gary Black,Wen C Yew,Lynn Dover,Paul Baker,Steve Liggett,Sarah Essex,Jane Greenaway,Debra Padgett,Clive Graham,Garren Scott,Edward Barton,Emma Swindells,Brendan Payne,Jennifer Collins,Yusri Taha,Gary Eltringham |
| EPI_ISL_638584 | Queens Medical Centre, Clinical Microbiology Department / DeepSeq Nottingham | COVID-19 Genomics UK (COG-UK) Consortium | Gemma Clark, Wendy Smith, Manjinder Khakh, Vicki M Fleming, Michelle M Lister, Hannah Howson-Wells, Jonathan Ball, Patrick McClure, Joseph Chappell, Theocharis Tsoleridis, Nadine Holmes, Matthew Carlisle, Christopher Moore, Fei Sang, Johnny Debebe, Victoria Wright, Matthew Loose |
| EPI_ISL_638735 | Department of Pathology, University of Cambridge | COVID-19 Genomics UK (COG-UK) Consortium | Aminu S. Jahun, Yasmin Chaudhry, Grant Hall, Iliana Georgana, Myra Hosmillo, Martin D. Curran, Malte Pinckert, Surendra Parmar, Ian Goodfellow |
| EPI_ISL_638736, EPI_ISL_638737, EPI_ISL_638738, EPI_ISL_638741, EPI_ISL_638742, EPI_ISL_638743, EPI_ISL_638744, EPI_ISL_638745, EPI_ISL_638749, EPI_ISL_638750, EPI_ISL_638751, EPI_ISL_638752, EPI_ISL_638753, EPI_ISL_638754, EPI_ISL_638755, EPI_ISL_638756, EPI_ISL_638757, EPI_ISL_638758, EPI_ISL_638759, EPI_ISL_638760, EPI_ISL_638762, EPI_ISL_638763, EPI_ISL_638764, EPI_ISL_638766, EPI_ISL_638767, EPI_ISL_638768, EPI_ISL_638769, EPI_ISL_638770, EPI_ISL_638771, EPI_ISL_638776, EPI_ISL_638777, EPI_ISL_638778, EPI_ISL_638779, EPI_ISL_638780, EPI_ISL_638781, EPI_ISL_638782, EPI_ISL_638783, EPI_ISL_638784, EPI_ISL_638785, EPI_ISL_638786, EPI_ISL_638787, EPI_ISL_638788, EPI_ISL_638789, EPI_ISL_638790, EPI_ISL_638791, EPI_ISL_638792, EPI_ISL_638793, EPI_ISL_638794, EPI_ISL_638795, EPI_ISL_638796, EPI_ISL_638797, EPI_ISL_638798, EPI_ISL_638799, EPI_ISL_638805, EPI_ISL_638806, EPI_ISL_638807, EPI_ISL_638808, EPI_ISL_638809 | | | |
| see above | Oxford Viromics, NDM, University of Oxford; Oxford University Hospitals; Basingstoke and North Hampshire Hospital | COVID-19 Genomics UK (COG-UK) Consortium | Tanya Golubchik, David Bonsall, George Macintyre, Amy Trebes, Mariateresa de Cesare, Catrin Moore, Alex Mobbs, Anita Justice, Robert Shaw, Monique Andersson, Timothy Peto, Emma Wise, Nathan Moore, Jessica Lynch, Nick Cortes, Matilde Mori, Stephen Kidd, David Buck, John Todd, Christophe Fraser |
| EPI_ISL_638878, EPI_ISL_638882 | Virology Department, Sheffield Teaching Hospitals NHS Foundation Trust/Department of Infection, Immunity and Cardiovascular Disease, The Medical School, University of Sheffield | COVID-19 Genomics UK (COG-UK) Consortium | Thushan de Silva, Matthew Parker, Nikki Smith, Adri Angyal, Rebecca Brown, Luke Green, Rachel Tucker, Paul Parsons, Danielle Groves, Katie Johnson, Laura Carrilero, Alex Keeley, Dave Partridge, Matthew Wyles, Benjamin Lindsey, Mehmet Yavuz, Mohammad Raza, Cariad Evans |
| EPI_ISL_638936, EPI_ISL_638937, EPI_ISL_638938, EPI_ISL_638939, EPI_ISL_638940, EPI_ISL_638941, EPI_ISL_638942, EPI_ISL_638943 | Department of Pathology, University of Cambridge | COVID-19 Genomics UK (COG-UK) Consortium | Aminu S. Jahun, Yasmin Chaudhry, Grant Hall, Iliana Georgana, Myra Hosmillo, Martin D. Curran, Malte Pinckert, Surendra Parmar, Ian Goodfellow |
| EPI_ISL_638985 | Oxford Viromics, NDM, University of Oxford; Oxford University Hospitals; Basingstoke and North Hampshire Hospital | COVID-19 Genomics UK (COG-UK) Consortium | Tanya Golubchik, David Bonsall, George Macintyre, Amy Trebes, Mariateresa de Cesare, Catrin Moore, Alex Mobbs, Anita Justice, Robert Shaw, Monique Andersson, Timothy Peto, Emma Wise, Nathan Moore, Jessica Lynch, Nick Cortes, Matilde Mori, Stephen Kidd, David Buck, John Todd, Christophe Fraser |
| EPI_ISL_639063 | Wales Specialist Virology Centre Sequencing lab: Pathogen Genomics Unit | COVID-19 Genomics UK (COG-UK) Consortium | Catherine Moore, Johnathan Evans, Laura Gifford, Malorie Perry, Simon Cottrell, Angela Marchbank, Alec Birchley, Alexander Adams, Amy Gaskin, Bree Gatica-Wilcox, Jason Coombes, Joel Southgate, Lauren Gilbert, Lee Graham, Nicole Pacchiarini, Sara Kumziene-Summerhayes, Sarah Taylor, Sophie Jones, Sara Rey, Matthew Bull, Joanne Watkins, Sally Corden, Tom Connor |
| EPI_ISL_639627 | Oxford Viromics, NDM, University of Oxford; Oxford University Hospitals; Basingstoke and North Hampshire Hospital | COVID-19 Genomics UK (COG-UK) Consortium | Tanya Golubchik, David Bonsall, George Macintyre, Amy Trebes, Mariateresa de Cesare, Catrin Moore, Alex Mobbs, Anita Justice, Robert Shaw, Monique Andersson, Timothy Peto, Emma Wise, Nathan Moore, Jessica Lynch, Nick Cortes, Matilde Mori, Stephen Kidd, David Buck, John Todd, Christophe Fraser |
| EPI_ISL_639672 | Centrl Laboratorija | Latvian Biomedical Research and Study Centre | Ivars Silamielis, Kaspars Megnis, Monta Ustinova, ikita Zrelovs, Vita Rovte, Stella Lapia, Jana Oste, Marta Priedte, Uga Dumpis, Jnis Kloviš |
| EPI_ISL_639681, EPI_ISL_639685, EPI_ISL_639687, EPI_ISL_639690, EPI_ISL_639693, EPI_ISL_639694 | E. Gulbja Laboratorija | Latvian Biomedical Research and Study Centre | Ivars Silamielis, Kaspars Megnis, Monta Ustinova, ikita Zrelovs, Vita Rovte, Mikus Gavars, Dmitrijs Perminovs, Uga Dumpis, Jnis Kloviš |
| EPI_ISL_639731 | Laverty Pathology | NSW Health Pathology - Institute of Clinical Pathology and Medical Research; Westmead Hospital; University of Sydney | CIDM-PH et al. |
| EPI_ISL_640088 | False Bay Hospital wc FBH | NHLS/UCT | Arash Iranzadeh, Deelan Doolabh, Lynn Tyers, Bruna Galvao, Innocent Mudau, Marvin Hsiao, Kruger Marais, Diana Hardie, Stephen Korsman, Carolyn Williamson |
| EPI_ISL_640089 | 2 Military Hospital wc MAA | NHLS/UCT | Arash Iranzadeh, Deelan Doolabh, Lynn Tyers, Bruna Galvao, Innocent Mudau, Marvin Hsiao, Kruger Marais, Diana Hardie, Stephen Korsman, Carolyn Williamson |
| EPI_ISL_640090 | False Bay Hospital wc FBH | NHLS/UCT | Arash Iranzadeh, Deelan Doolabh, Lynn Tyers, Bruna Galvao, Innocent Mudau, Marvin Hsiao, Kruger Marais, Diana Hardie, Stephen Korsman, Carolyn Williamson |
| EPI_ISL_640091 | Du Noon CDC wc DNC | NHLS/UCT | Arash Iranzadeh, Deelan Doolabh, Lynn Tyers, Bruna Galvao, Innocent Mudau, Marvin Hsiao, Kruger Marais, Diana Hardie, Stephen Korsman, Carolyn Williamson |
| EPI_ISL_640092, EPI_ISL_640093 | Kensington CDC wc KSC | NHLS/UCT | Arash Iranzadeh, Deelan Doolabh, Lynn Tyers, Bruna Galvao, Innocent Mudau, Marvin Hsiao, Kruger Marais, Diana Hardie, Stephen Korsman, Carolyn Williamson |
| EPI_ISL_640094, EPI_ISL_640095 | Du Noon CDC wc DNC | NHLS/UCT | Arash Iranzadeh, Deelan Doolabh, Lynn Tyers, Bruna Galvao, Innocent Mudau, Marvin Hsiao, Kruger Marais, Diana Hardie, Stephen Korsman, Carolyn |

| | | | |
|---|---|---|---|
| | | | Williamson |
| EPI_ISL_640137, EPI_ISL_640138, EPI_ISL_640139, EPI_ISL_640140 | Groote Schuur Hospital wc GSH | NHLS/UCT | Arash Iranzadeh, Deelan Doolabh, Lynn Tyers, Bruna Galvao, Innocent Mudau, Marvin Hsiao, Kruger Marais, Diana Hardie, Stephen Korsman, Carolyn Williamson |
| EPI_ISL_640141 | Victoria Hospital wc VHW | NHLS/UCT | Arash Iranzadeh, Deelan Doolabh, Lynn Tyers, Bruna Galvao, Innocent Mudau, Marvin Hsiao, Kruger Marais, Diana Hardie, Stephen Korsman, Carolyn Williamson |
| EPI_ISL_640543, EPI_ISL_640600, EPI_ISL_640689, EPI_ISL_640978, EPI_ISL_641041, EPI_ISL_641050 | Microbiological Diagnostic Unit - Public Health Laboratory (MDU-PHL) | MDU-PHL | Seemann T., Schultz M.B., Sait, M.L., Sherry, N.L. |
| EPI_ISL_641088 | Victorian Infectious Diseases Reference Laboratory (VIDRL) | VIDRL and MDU-PHL | Caly L., Seemann T., Sait, M.L., Schultz M.B., Druce J., Sherry, N.L. |
| EPI_ISL_641104, EPI_ISL_641177, EPI_ISL_641277 | Microbiological Diagnostic Unit - Public Health Laboratory (MDU-PHL) | MDU-PHL | Seemann T., Schultz M.B., Sait, M.L., Sherry, N.L. |
| EPI_ISL_641446, EPI_ISL_641501, EPI_ISL_641503, EPI_ISL_641504, EPI_ISL_641505 | Department of Virus and Microbiological Special Diagnostics, Statens Serum Institut, Copenhagen, Denmark | Albertsen lab, Department of Chemistry and Bioscience, Aalborg University, Denmark | Thomas Bruun Rasmussen, Jannik Fonager, Morten Rasmussen |
| EPI_ISL_644118 | Lighthouse Lab in Alderley Park | Wellcome Sanger Institute for the COVID-19 Genomics UK (COG-UK) Consortium | Jacquelyn Wynn, Mairead Hyland, The Lighthouse Lab in Alderley Park and Alex Alderton, Roberto Amato, Sonia Goncalves, Ewan Harrison, David K. Jackson, Ian Johnston, Dominic Kwiatkowski, Cordelia Langford, John Sillitoe on behalf of the Wellcome Sanger Institute COVID-19 Surveillance Team |
| EPI_ISL_644363 | Michigan Department of Health and Human Services, Bureau of Laboratories | Michigan Department of Health and Human Services, Bureau of Laboratories | Blankenship HM, Riner D, Soehnlen MK |
| EPI_ISL_644376, EPI_ISL_644377, EPI_ISL_644385, EPI_ISL_644390, EPI_ISL_644397, EPI_ISL_644399, EPI_ISL_644536, EPI_ISL_644537, EPI_ISL_644538, EPI_ISL_644539, EPI_ISL_644540, EPI_ISL_644541, EPI_ISL_644542, EPI_ISL_644543, EPI_ISL_644544, EPI_ISL_644545, EPI_ISL_644546, EPI_ISL_644547, EPI_ISL_644548, EPI_ISL_644549, EPI_ISL_644550, EPI_ISL_644551, EPI_ISL_644552, EPI_ISL_644553, EPI_ISL_644554, EPI_ISL_644555 | | | |
| see above | MEPHI, Aix Marseille University | MEPHI, Aix Marseille University | Anthony LEVASSEUR |
| EPI_ISL_644882, EPI_ISL_644883, EPI_ISL_644884, EPI_ISL_644885, EPI_ISL_644886, EPI_ISL_644887, EPI_ISL_644888, EPI_ISL_644889, EPI_ISL_644890, EPI_ISL_644891, EPI_ISL_644892, EPI_ISL_644894, EPI_ISL_644895, EPI_ISL_644896, EPI_ISL_644897, EPI_ISL_644898, EPI_ISL_644899, EPI_ISL_644932 | | | |
| see above | Virginia DCLS | Virginia DCLS | Virginia DCLS |
| EPI_ISL_648130 | Uppsala klinisk mikrobiologi | The Public Health Agency of Sweden | Anna-Malin Linde, Maria Lind Karlberg, Mattias Haukland, Reza Advani, Olov Svartstrom, Oskar Karlsson Lindsjo, Sandra Broddesson, Petra Edquist, Mia Brytting, Anna Risberg, Karin Tegmark-Wisell |
| EPI_ISL_648137, EPI_ISL_648187, EPI_ISL_648189, EPI_ISL_648190, EPI_ISL_648191 | The Public Health Agency of Sweden | The Public Health Agency of Sweden | Anna-Malin Linde, Maria Lind Karlberg, Mattias Haukland, Reza Advani, Olov Svartstrom, Oskar Karlsson Lindsjo, Sandra Broddesson, Petra Edquist, Mia Brytting, Anna Risberg, Karin Tegmark-Wisell |
| EPI_ISL_648384, EPI_ISL_648385, EPI_ISL_648386, EPI_ISL_648387, EPI_ISL_648388, EPI_ISL_648389, EPI_ISL_648390, EPI_ISL_648391, EPI_ISL_648392, EPI_ISL_648393, EPI_ISL_648394, EPI_ISL_648395, EPI_ISL_648396, EPI_ISL_648397, EPI_ISL_648398, EPI_ISL_648399, EPI_ISL_648400, EPI_ISL_648401, EPI_ISL_648402, EPI_ISL_648403, EPI_ISL_648404, EPI_ISL_648405, EPI_ISL_648406, EPI_ISL_648407, EPI_ISL_648408, EPI_ISL_648409, EPI_ISL_648410, EPI_ISL_648411, EPI_ISL_648412, EPI_ISL_648413, EPI_ISL_648414, EPI_ISL_648415, EPI_ISL_648416, EPI_ISL_648417, EPI_ISL_648418, EPI_ISL_648419, EPI_ISL_648420, EPI_ISL_648421, EPI_ISL_648422, EPI_ISL_648423, EPI_ISL_648425, EPI_ISL_648426, EPI_ISL_648427, EPI_ISL_648428, EPI_ISL_648429, EPI_ISL_648430, EPI_ISL_648431, EPI_ISL_648432, EPI_ISL_648433, EPI_ISL_648434, EPI_ISL_648435, EPI_ISL_648436, EPI_ISL_648437 | | | |
| see above | Santa Clara County Public Health Laboratory | Chan-Zuckerberg Biohub | CZB Cliahub Consortium |
| EPI_ISL_648509, EPI_ISL_648510, EPI_ISL_648511, EPI_ISL_648512, EPI_ISL_648515 | Orange County Public Health Lab | Chan-Zuckerberg Biohub | CZB Cliahub Consortium |
| EPI_ISL_648908, EPI_ISL_648913, EPI_ISL_648919, EPI_ISL_648925, EPI_ISL_648936, EPI_ISL_648941, EPI_ISL_648942, EPI_ISL_648944, EPI_ISL_648950, EPI_ISL_648951, EPI_ISL_648953, EPI_ISL_648955, EPI_ISL_648956, EPI_ISL_648957, EPI_ISL_648960, EPI_ISL_648968, EPI_ISL_648972, EPI_ISL_648974, EPI_ISL_648977, EPI_ISL_648980, EPI_ISL_649012, EPI_ISL_649029 | | | |
| see above | San Diego County Public Health Laboratory | Andersen lab at Scripps Research | SEARCH Alliance San Diego with Tracy Basler, Jovan Shephard, Brett Austin |
| EPI_ISL_649219, EPI_ISL_649234, EPI_ISL_649261, EPI_ISL_649309, EPI_ISL_649312, EPI_ISL_649321, EPI_ISL_649368 | Lighthouse Lab in Alderley Park | Wellcome Sanger Institute for the COVID-19 Genomics UK (COG-UK) Consortium | Jacquelyn Wynn, Mairead Hyland, The Lighthouse Lab in Alderley Park and Alex Alderton, Roberto Amato, Sonia Goncalves, Ewan Harrison, David K. Jackson, Ian Johnston, Dominic Kwiatkowski, Cordelia Langford, John Sillitoe on behalf of the Wellcome Sanger Institute COVID-19 Surveillance Team (http://www.sanger.ac.uk/covid-team) |
| EPI_ISL_649817 | Respiratory Virus Unit, Microbiology Services Colindale, Public Health England | COVID-19 Genomics UK (COG-UK) Consortium | PHE Covid Sequencing Team |
| EPI_ISL_650394 | Northumbria University / South Tees Hospitals NHS Foundation Trust / North Cumbria Integrated Care NHS Foundation Trust / North Tees and Hartlepool NHS Foundation Trust / Newcastle Hospitals NHS Foundation Trust | COVID-19 Genomics UK (COG-UK) Consortium | Darren L Smith,Andrew Nelson,Matthew Bashton,Greg R Young,Joshua Loh,John Allan,Mohammad A Tariq,Giles S Holt,Gary Black,Wen C Yew,Lynn Dover,Paul Baker,Steve Liggett,Sarah Essex,Jane Greenaway,Debra Padgett,Clive Graham,Garren Scott,Edward Barton,Emma Swindells,Brendan Payne,Jennifer Collins,Yusri Taha,Gary Eltringham |
| EPI_ISL_650712 | West of Scotland Specialist Virology Centre, NHSGGC / MRC-University of Glasgow Centre for Virus Research | COVID-19 Genomics UK (COG-UK) Consortium | Ana da Silva Filipe, Natasha Johnson, Kathy Smollett, Daniel Mair, Stephen Carmichael, Alice Broos, Lily Tong, Jenna Nichols, Kyriaki Nomikou; Sarah McDonald; Richard Orton, Joseph Hughes, Sreenu Vattipally, David L Robertson; Alasdair MacLean, Rory Gunson; Sharif Shaaban, Matthew Holden; Rachel Blacow, Guy Mollett, Kathy Li, James Shepherd, Antonia Ho, Emma Thomson |
| EPI_ISL_650832 | Virology Department, Royal Infirmary of Edinburgh, NHS Lothian / School of Biological Sciences, University of Edinburgh / Institute of Genetics and Molecular Medicine, University of Edinburgh | COVID-19 Genomics UK (COG-UK) Consortium | McHugh M, Dewar R, Rooke S, Gallagher M, Balcaza C, O'Toole Á, Scher E, Hill V, McCrone JT, Colquhoun R, Yu X, Jackson B, Rambaut A, Williams TC, Templeton K |
| EPI_ISL_651063 | Northumbria University / South Tees Hospitals NHS Foundation Trust / North Cumbria Integrated Care NHS Foundation Trust / North Tees and Hartlepool NHS Foundation Trust / Newcastle Hospitals NHS Foundation Trust | COVID-19 Genomics UK (COG-UK) Consortium | Darren L Smith,Andrew Nelson,Matthew Bashton,Greg R Young,Joshua Loh,John Allan,Mohammad A Tariq,Giles S Holt,Gary Black,Wen C Yew,Lynn Dover,Paul Baker,Steve Liggett,Sarah Essex,Jane Greenaway,Debra Padgett,Clive Graham,Garren Scott,Edward Barton,Emma Swindells,Brendan Payne,Jennifer Collins,Yusri Taha,Gary Eltringham |
| EPI_ISL_651067 | Centre for Enzyme Innovation, University of Portsmouth / Translational Research Laboratory, Portsmouth Hospitals NHS Trust | COVID-19 Genomics UK (COG-UK) Consortium | Angela Beckett,Yann Bourgeois,Garry Scarlett,Sharon Glaysher,Scott Elliott,Kelly Bicknell,Robert Impey,Allyson Lloyd,Sarah Wyllie,Ethan Butcher,Anoop Chauhan,Samuel Robson |
| EPI_ISL_651076 | Virology Department, Sheffield Teaching Hospitals NHS Foundation Trust/Department of Infection, Immunity and Cardiovascular Disease, The Medical School, University of Sheffield | COVID-19 Genomics UK (COG-UK) Consortium | Thushan de Silva, Matthew Parker, Nikki Smith, Adri Angyal, Rebecca Brown, Luke Green, Rachel Tucker, Paul Parsons, Danielle Groves, Katie Johnson, Laura Carrilero, Alex Keeley, Dave Partridge, Matthew Wyles, Benjamin Lindsey, Mehmet Yavuz, Mohammad Raza, Cariad Evans |
| EPI_ISL_651493 | West of Scotland Specialist Virology Centre, NHSGGC / MRC-University of Glasgow Centre for Virus Research | COVID-19 Genomics UK (COG-UK) Consortium | Ana da Silva Filipe, Natasha Johnson, Kathy Smollett, Daniel Mair, Stephen Carmichael, Alice Broos, Lily Tong, Jenna Nichols, Kyriaki Nomikou; Sarah McDonald; Richard Orton, Joseph Hughes, Sreenu Vattipally, David L Robertson; Alasdair MacLean, Rory Gunson; Sharif Shaaban, Matthew Holden; Rachel Blacow, Guy Mollett, Kathy Li, James Shepherd, Antonia Ho, Emma Thomson |
| EPI_ISL_651520 | Virology Department, Royal Infirmary of Edinburgh, NHS Lothian / School of Biological Sciences, University of | COVID-19 Genomics UK (COG-UK) Consortium | McHugh M, Dewar R, Rooke S, Gallagher M, Balcaza C, O'Toole Á, Scher E, Hill V, McCrone JT, Colquhoun R, Yu X, Jackson B, Rambaut A, Williams TC, Templeton K |

| | | | |
|---|---|---|---|
| | Edinburgh / Institute of Genetics and Molecular Medicine, University of Edinburgh | | |
| EPI_ISL_651530 | Virology Department, Sheffield Teaching Hospitals NHS Foundation Trust/Department of Infection, Immunity and Cardiovascular Disease, The Medical School, University of Sheffield | COVID-19 Genomics UK (COG-UK) Consortium | Thushan de Silva, Matthew Parker, Nikki Smith, Adri Angyal, Rebecca Brown, Luke Green, Rachel Tucker, Paul Parsons, Danielle Groves, Katie Johnson, Laura Carrilero, Alex Keeley, Dave Partridge, Matthew Wyles, Benjamin Lindsey, Mehmet Yavuz, Mohammad Raza, Cariad Evans |
| EPI_ISL_651739, EPI_ISL_651785, EPI_ISL_651956 | West of Scotland Specialist Virology Centre, NHSGGC / MRC-University of Glasgow Centre for Virus Research | COVID-19 Genomics UK (COG-UK) Consortium | Ana da Silva Filipe, Natasha Johnson, Kathy Smollett, Daniel Mair, Stephen Carmichael, Alice Broos, Lily Tong, Jenna Nichols, Kyriaki Nomikou; Sarah McDonald; Richard Orton, Joseph Hughes, Sreenu Vattipally, David L Robertson; Alasdair MacLean, Rory Gunson; Sharif Shaaban, Matthew Holden; Rachel Blacow, Guy Mollett, Kathy Li, James Shepherd, Antonia Ho, Emma Thomson |
| EPI_ISL_652623 | Centre for Enzyme Innovation, University of Portsmouth / Translational Research Laboratory, Portsmouth Hospitals NHS Trust | COVID-19 Genomics UK (COG-UK) Consortium | Angela Beckett,Yann Bourgeois,Garry Scarlett,Sharon Glaysher,Scott Elliott,Kelly Bicknell,Robert Impey,Allyson Lloyd,Sarah Wyllie,Ethan Butcher,Anoop Chauhan,Samuel Robson |
| EPI_ISL_653063, EPI_ISL_653069, EPI_ISL_653075 | Virology Department, Sheffield Teaching Hospitals NHS Foundation Trust/Department of Infection, Immunity and Cardiovascular Disease, The Medical School, University of Sheffield | COVID-19 Genomics UK (COG-UK) Consortium | Thushan de Silva, Matthew Parker, Nikki Smith, Adri Angyal, Rebecca Brown, Luke Green, Rachel Tucker, Paul Parsons, Danielle Groves, Katie Johnson, Laura Carrilero, Alex Keeley, Dave Partridge, Matthew Wyles, Benjamin Lindsey, Mehmet Yavuz, Mohammad Raza, Cariad Evans |
| EPI_ISL_654179, EPI_ISL_654184, EPI_ISL_654201, EPI_ISL_654241, EPI_ISL_654242, EPI_ISL_654243, EPI_ISL_654244, EPI_ISL_654245, EPI_ISL_654246, EPI_ISL_654247, EPI_ISL_654285, EPI_ISL_654286, EPI_ISL_654287, EPI_ISL_654288, EPI_ISL_654289, EPI_ISL_654290, EPI_ISL_654291, EPI_ISL_654292, EPI_ISL_654293, EPI_ISL_654294, EPI_ISL_654349 | | | |
| see above | Hospital General Universitario Gregorio Marañón | SeqCOVID-SPAIN consortium/IBV(CSIC) | Darío García de Viedma, Laura Pérez-Lago, Marta Herranz, Jon Sicilia, Julia Suárez, Pilar Catalán, Patricia Muñoz and SeqCOVID-SPAIN consortium |
| EPI_ISL_654402, EPI_ISL_654407, EPI_ISL_654409, EPI_ISL_654413, EPI_ISL_654415, EPI_ISL_654436, EPI_ISL_654440 | Servicio de Microbiología, Hospital Miguel Servet, Zaragoza | SeqCOVID-SPAIN consortium/IBV(CSIC) | Antonio Rezusta López, Alexander Tristancho Baró, Ana Milagro, Yolanda Gracia Grataloup, Nieves Martínez Cameo and SeqCOVID-SPAIN consortium |
| EPI_ISL_654516, EPI_ISL_654517, EPI_ISL_654525, EPI_ISL_654529, EPI_ISL_654530, EPI_ISL_654531, EPI_ISL_654532, EPI_ISL_654533, EPI_ISL_654536 | Servicio de Microbiología, Laboratori Clínic Metropolitana Nord. Hospital Universitari Germans Trias i Pujol. Institut d'Investigació en Ciències de la Salut Germans Trias i Pujol (IGTP) | SeqCOVID-SPAIN consortium/IBV(CSIC) | Elisa Martró, Antoni E. Bordoy, Anna Not, Adrián Antuori, Anabel Fernández, Nona Romaní and SeqCOVID-SPAIN consortium |
| EPI_ISL_654539, EPI_ISL_654541, EPI_ISL_654542, EPI_ISL_654546, EPI_ISL_654565, EPI_ISL_654566, EPI_ISL_654567, EPI_ISL_654568, EPI_ISL_654570, EPI_ISL_654571, EPI_ISL_654573 | | | |
| see above | Servicio de Microbiología. Hospital Universitario Donostia. OSI Donostialdea. Área de Enfermedades Infecciosas, Grupo de Infección Respiratoria y Resistencia Antimicrobiana. Instituto de Investigación Sanitaria Biodonostia | SeqCOVID-SPAIN consortium/IBV(CSIC) | Gustavo Cilla Eguiluz, Milagrosa Montes Ros, Luis Piñeiro Vázquez, Ane Sorrarain, Jose Maria Marimón and SeqCOVID-SPAIN consortium |
| EPI_ISL_654699 | Essentia Health-St. Mary's Medical Center | Minnesota Department of Health, Public Health Laboratory | Matt Plumb, Jacob Garfin, Alexandra Lorentz, and Xiong Wang |
| EPI_ISL_660121, EPI_ISL_660122, EPI_ISL_660123, EPI_ISL_660124, EPI_ISL_660125, EPI_ISL_660126, EPI_ISL_660127, EPI_ISL_660128, EPI_ISL_660129 | Ampath | National Health Laboratory Service (NHLS), Tygerberg | Susan Engelbrecht, Draper C, Davis M-A, Siegfried N, Williamson C, Hsiao M, Kayla Delaney, Bronwyn Kleinhans, Houriiyah Tegally, Eduan Wilkindon, Gert van Zyl, Wolfgang Preiser, Tulio de Oliveira |
| EPI_ISL_660130, EPI_ISL_660131, EPI_ISL_660132, EPI_ISL_660133, EPI_ISL_660134, EPI_ISL_660135, EPI_ISL_660136 | Lancet | National Health Laboratory Service (NHLS), Tygerberg | Susan Engelbrecht, Draper C, Davis M-A, Siegfried N, Williamson C, Hsiao M, Kayla Delaney, Bronwyn Kleinhans, Houriiyah Tegally, Eduan Wilkindon, Gert van Zyl, Wolfgang Preiser, Tulio de Oliveira |
| EPI_ISL_660138, EPI_ISL_660139, EPI_ISL_660141, EPI_ISL_660142 | Hamadi | National Health Laboratory Service (NHLS), Tygerberg | Susan Engelbrecht, Draper C, Davis M-A, Siegfried N, Williamson C, Hsiao M, Kayla Delaney, Bronwyn Kleinhans, Houriiyah Tegally, Eduan Wilkindon, Gert van Zyl, Wolfgang Preiser, Tulio de Oliveira |
| EPI_ISL_660143, EPI_ISL_660144, EPI_ISL_660145, EPI_ISL_660146, EPI_ISL_660147, EPI_ISL_660148, EPI_ISL_660149, EPI_ISL_660150, EPI_ISL_660151, EPI_ISL_660152, EPI_ISL_660154, EPI_ISL_660156, EPI_ISL_660158 | | | |
| see above | PathCare | National Health Laboratory Service (NHLS), Tygerberg | Susan Engelbrecht, Draper C, Davis M-A, Siegfried N, Williamson C, Hsiao M, Kayla Delaney, Bronwyn Kleinhans, Houriiyah Tegally, Eduan Wilkindon, Gert van Zyl, Wolfgang Preiser, Tulio de Oliveira |
| EPI_ISL_660468, EPI_ISL_660488, EPI_ISL_660507, EPI_ISL_660511, EPI_ISL_660521 | Laboratoire de Microbiologie CHU Souro Sanou | Centre Muraz | Abdoul-Salam Ouedraogo, Yacouba Sawadogo, Essia Belarbi, Grit Schubert, Fabian Leendertz, Arsène Zongo, Soumeya Ouangraoua, Zekiba Tarnagda, Lassana Sangaré, Halidou Tinto |
| EPI_ISL_660535, EPI_ISL_660536, EPI_ISL_660537, EPI_ISL_660538 | Institute of Microbiology, Universidad San Francisco de Quito | Institute of Microbiology, Universidad San Francisco de Quito | Sully Márquez, Belén Prado-Vivar, Juan José Guadalupe, Monica Becerra-Wong, Bernardo Gutiérrez, Hermelinda Paguay, Alexandra Tino, Jorge Montaño, Verónica Barragán, Patricio Rojas-Silva, Gabriel Trueba, Michelle Grunauer, Paúl Cárdenas |
| EPI_ISL_660902, EPI_ISL_660903, EPI_ISL_660904, EPI_ISL_660905, EPI_ISL_660906, EPI_ISL_660907, EPI_ISL_660908, EPI_ISL_660909, EPI_ISL_660910, EPI_ISL_660911, EPI_ISL_660912, EPI_ISL_660913, EPI_ISL_660914, EPI_ISL_660915, EPI_ISL_660916, EPI_ISL_660917, EPI_ISL_660918, EPI_ISL_660919, EPI_ISL_660920, EPI_ISL_660921, EPI_ISL_660922, EPI_ISL_660923, EPI_ISL_660924, EPI_ISL_660925, EPI_ISL_660926, EPI_ISL_660927, EPI_ISL_660928, EPI_ISL_660929, EPI_ISL_660930, EPI_ISL_660931 | | | |
| see above | Gundersen Molecular Diagnostics Laboratory | Kabara Cancer Research Institute | Craig S. Richmond, Paraic A. Kenny |
| EPI_ISL_664361, EPI_ISL_664367, EPI_ISL_664500, EPI_ISL_664505, EPI_ISL_665193, EPI_ISL_665194, EPI_ISL_665195, EPI_ISL_665196, EPI_ISL_665197, EPI_ISL_665198, EPI_ISL_665242 | | | |
| see above | University College London Hospital | COVID-19 Genomics UK (COG-UK) Consortium | Judith Heaney, Matthew Byott, Catherine Houlihan, Dan Frampton, Stuart Kirk, Moira Spyer and Eleni Nastouli |
| EPI_ISL_665272 | Northumbria University / South Tees Hospitals NHS Foundation Trust / North Cumbria Integrated Care NHS Foundation Trust / North Tees and Hartlepool NHS Foundation Trust / Newcastle Hospitals NHS Foundation Trust | COVID-19 Genomics UK (COG-UK) Consortium | Darren L Smith,Andrew Nelson,Matthew Bashton,Greg R Young,Joshua Loh,John Allan,Mohammad A Tariq,Giles S Holt,Gary Black,Wen C Yew,Lynn Dover,Paul Baker,Steve Liggett,Sarah Essex,Jane Greenaway,Debra Padgett,Clive Graham,Garren Scott,Edward Barton,Emma Swindells,Brendan Payne,Jennifer Collins,Yusri Taha,Gary Eltringham |
| EPI_ISL_665273, EPI_ISL_665276, EPI_ISL_665277, EPI_ISL_665279, EPI_ISL_665282 | University of Exeter | COVID-19 Genomics UK (COG-UK) Consortium | Ben Temperton,Aaron Jeffries,Michelle Michelsen,Joanna Warwick-Dugdale,Audrey Farbos,Robyn Manley,Stephen Michell,Jane Masoli |
| EPI_ISL_665305, EPI_ISL_665312 | Northumbria University / South Tees Hospitals NHS Foundation Trust / North Cumbria Integrated Care NHS Foundation Trust / North Tees and Hartlepool NHS Foundation Trust / Newcastle Hospitals NHS Foundation Trust | COVID-19 Genomics UK (COG-UK) Consortium | Darren L Smith,Andrew Nelson,Matthew Bashton,Greg R Young,Joshua Loh,John Allan,Mohammad A Tariq,Giles S Holt,Gary Black,Wen C Yew,Lynn Dover,Paul Baker,Steve Liggett,Sarah Essex,Jane Greenaway,Debra Padgett,Clive Graham,Garren Scott,Edward Barton,Emma Swindells,Brendan Payne,Jennifer Collins,Yusri Taha,Gary Eltringham |
| EPI_ISL_665324, EPI_ISL_665325, EPI_ISL_665326, EPI_ISL_665332, | University of Exeter | COVID-19 Genomics UK (COG-UK) Consortium | Ben Temperton,Aaron Jeffries,Michelle Michelsen,Joanna Warwick-Dugdale,Audrey Farbos,Robyn Manley,Stephen Michell,Jane Masoli |

| | | | |
|---|---|---|---|
| EPI_ISL_665333 | | | |
| EPI_ISL_665392, EPI_ISL_665393, EPI_ISL_665394, EPI_ISL_665395, EPI_ISL_665396, EPI_ISL_665408, EPI_ISL_665409 | Northumbria University / South Tees Hospitals NHS Foundation Trust / North Cumbria Integrated Care NHS Foundation Trust / North Tees and Hartlepool NHS Foundation Trust / Newcastle Hospitals NHS Foundation Trust | COVID-19 Genomics UK (COG-UK) Consortium | Darren L Smith,Andrew Nelson,Matthew Bashton,Greg R Young,Joshua Loh,John Allan,Mohammad A Tariq,Giles S Holt,Gary Black,Wen C Yew,Lynn Dover,Paul Baker,Steve Liggett,Sarah Essex,Jane Greenaway,Debra Padgett,Clive Graham,Garren Scott,Edward Barton,Emma Swindells,Brendan Payne,Jennifer Collins,Yusri Taha,Gary Eltringham |
| EPI_ISL_665411 | Quadram Institute Bioscience | COVID-19 Genomics UK (COG-UK) Consortium | Dave J. Baker, Gemma L. Kay, Alp Aydin, Thanh Le-Viet, Steven Rudder, Ana P. Tedim, Anastasia Kolyva, Maria Diaz, Leonardo de Oliveira Martins, Nabil-Fareed Alikhan, Lizzie Meadows, Rachael Stanley, Ngozi Elumogo, Muhammed Yasir, Nicholas M. Thomson, Alexander J Trotter, Rachel Gilroy, Samuel Bloomfield, Claire Stuart, Andrew Bell, Reenesh Prakash, Samir Dervisevic, Alison E. Mather, John Wain, Mark Webber, Andrew J. Page, Justin O'Grady |
| EPI_ISL_665417, EPI_ISL_665419, EPI_ISL_665435 | University of Exeter | COVID-19 Genomics UK (COG-UK) Consortium | Ben Temperton,Aaron Jeffries,Michelle Michelsen,Joanna Warwick-Dugdale,Audrey Farbos,Robyn Manley,Stephen Michell,Jane Masoli |
| EPI_ISL_665445 | Quadram Institute Bioscience | COVID-19 Genomics UK (COG-UK) Consortium | Dave J. Baker, Gemma L. Kay, Alp Aydin, Thanh Le-Viet, Steven Rudder, Ana P. Tedim, Anastasia Kolyva, Maria Diaz, Leonardo de Oliveira Martins, Nabil-Fareed Alikhan, Lizzie Meadows, Rachael Stanley, Ngozi Elumogo, Muhammed Yasir, Nicholas M. Thomson, Alexander J Trotter, Rachel Gilroy, Samuel Bloomfield, Claire Stuart, Andrew Bell, Reenesh Prakash, Samir Dervisevic, Alison E. Mather, John Wain, Mark Webber, Andrew J. Page, Justin O'Grady |
| EPI_ISL_665492, EPI_ISL_665493, EPI_ISL_665494, EPI_ISL_665495, EPI_ISL_665506 | Northumbria University / South Tees Hospitals NHS Foundation Trust / North Cumbria Integrated Care NHS Foundation Trust / North Tees and Hartlepool NHS Foundation Trust / Newcastle Hospitals NHS Foundation Trust | COVID-19 Genomics UK (COG-UK) Consortium | Darren L Smith,Andrew Nelson,Matthew Bashton,Greg R Young,Joshua Loh,John Allan,Mohammad A Tariq,Giles S Holt,Gary Black,Wen C Yew,Lynn Dover,Paul Baker,Steve Liggett,Sarah Essex,Jane Greenaway,Debra Padgett,Clive Graham,Garren Scott,Edward Barton,Emma Swindells,Brendan Payne,Jennifer Collins,Yusri Taha,Gary Eltringham |
| EPI_ISL_665514, EPI_ISL_665515, EPI_ISL_665547, EPI_ISL_665548, EPI_ISL_665549, EPI_ISL_665570, EPI_ISL_665571 | University of Exeter | COVID-19 Genomics UK (COG-UK) Consortium | Ben Temperton,Aaron Jeffries,Michelle Michelsen,Joanna Warwick-Dugdale,Audrey Farbos,Robyn Manley,Stephen Michell,Jane Masoli |
| EPI_ISL_665617, EPI_ISL_665621, EPI_ISL_665626 | Northumbria University / South Tees Hospitals NHS Foundation Trust / North Cumbria Integrated Care NHS Foundation Trust / North Tees and Hartlepool NHS Foundation Trust / Newcastle Hospitals NHS Foundation Trust | COVID-19 Genomics UK (COG-UK) Consortium | Darren L Smith,Andrew Nelson,Matthew Bashton,Greg R Young,Joshua Loh,John Allan,Mohammad A Tariq,Giles S Holt,Gary Black,Wen C Yew,Lynn Dover,Paul Baker,Steve Liggett,Sarah Essex,Jane Greenaway,Debra Padgett,Clive Graham,Garren Scott,Edward Barton,Emma Swindells,Brendan Payne,Jennifer Collins,Yusri Taha,Gary Eltringham |
| EPI_ISL_665663, EPI_ISL_665666, EPI_ISL_665675 | University of Exeter | COVID-19 Genomics UK (COG-UK) Consortium | Ben Temperton,Aaron Jeffries,Michelle Michelsen,Joanna Warwick-Dugdale,Audrey Farbos,Robyn Manley,Stephen Michell,Jane Masoli |
| EPI_ISL_665676 | Northumbria University / South Tees Hospitals NHS Foundation Trust / North Cumbria Integrated Care NHS Foundation Trust / North Tees and Hartlepool NHS Foundation Trust / Newcastle Hospitals NHS Foundation Trust | COVID-19 Genomics UK (COG-UK) Consortium | Darren L Smith,Andrew Nelson,Matthew Bashton,Greg R Young,Joshua Loh,John Allan,Mohammad A Tariq,Giles S Holt,Gary Black,Wen C Yew,Lynn Dover,Paul Baker,Steve Liggett,Sarah Essex,Jane Greenaway,Debra Padgett,Clive Graham,Garren Scott,Edward Barton,Emma Swindells,Brendan Payne,Jennifer Collins,Yusri Taha,Gary Eltringham |
| EPI_ISL_665686, EPI_ISL_665694, EPI_ISL_665700 | University of Exeter | COVID-19 Genomics UK (COG-UK) Consortium | Ben Temperton,Aaron Jeffries,Michelle Michelsen,Joanna Warwick-Dugdale,Audrey Farbos,Robyn Manley,Stephen Michell,Jane Masoli |
| EPI_ISL_665701 | Northumbria University / South Tees Hospitals NHS Foundation Trust / North Cumbria Integrated Care NHS Foundation Trust / North Tees and Hartlepool NHS Foundation Trust / Newcastle Hospitals NHS Foundation Trust | COVID-19 Genomics UK (COG-UK) Consortium | Darren L Smith,Andrew Nelson,Matthew Bashton,Greg R Young,Joshua Loh,John Allan,Mohammad A Tariq,Giles S Holt,Gary Black,Wen C Yew,Lynn Dover,Paul Baker,Steve Liggett,Sarah Essex,Jane Greenaway,Debra Padgett,Clive Graham,Garren Scott,Edward Barton,Emma Swindells,Brendan Payne,Jennifer Collins,Yusri Taha,Gary Eltringham |
| EPI_ISL_665702 | Quadram Institute Bioscience | COVID-19 Genomics UK (COG-UK) Consortium | Dave J. Baker, Gemma L. Kay, Alp Aydin, Thanh Le-Viet, Steven Rudder, Ana P. Tedim, Anastasia Kolyva, Maria Diaz, Leonardo de Oliveira Martins, Nabil-Fareed Alikhan, Lizzie Meadows, Rachael Stanley, Ngozi Elumogo, Muhammed Yasir, Nicholas M. Thomson, Alexander J Trotter, Rachel Gilroy, Samuel Bloomfield, Claire Stuart, Andrew Bell, Reenesh Prakash, Samir Dervisevic, Alison E. Mather, John Wain, Mark Webber, Andrew J. Page, Justin O'Grady |
| EPI_ISL_665718 | Northumbria University / South Tees Hospitals NHS Foundation Trust / North Cumbria Integrated Care NHS Foundation Trust / North Tees and Hartlepool NHS Foundation Trust / Newcastle Hospitals NHS Foundation Trust | COVID-19 Genomics UK (COG-UK) Consortium | Darren L Smith,Andrew Nelson,Matthew Bashton,Greg R Young,Joshua Loh,John Allan,Mohammad A Tariq,Giles S Holt,Gary Black,Wen C Yew,Lynn Dover,Paul Baker,Steve Liggett,Sarah Essex,Jane Greenaway,Debra Padgett,Clive Graham,Garren Scott,Edward Barton,Emma Swindells,Brendan Payne,Jennifer Collins,Yusri Taha,Gary Eltringham |
| EPI_ISL_665721, EPI_ISL_665734, EPI_ISL_665738, EPI_ISL_665760 | University of Exeter | COVID-19 Genomics UK (COG-UK) Consortium | Ben Temperton,Aaron Jeffries,Michelle Michelsen,Joanna Warwick-Dugdale,Audrey Farbos,Robyn Manley,Stephen Michell,Jane Masoli |
| EPI_ISL_665850 | Quadram Institute Bioscience | COVID-19 Genomics UK (COG-UK) Consortium | Dave J. Baker, Gemma L. Kay, Alp Aydin, Thanh Le-Viet, Steven Rudder, Ana P. Tedim, Anastasia Kolyva, Maria Diaz, Leonardo de Oliveira Martins, Nabil-Fareed Alikhan, Lizzie Meadows, Rachael Stanley, Ngozi Elumogo, Muhammed Yasir, Nicholas M. Thomson, Alexander J Trotter, Rachel Gilroy, Samuel Bloomfield, Claire Stuart, Andrew Bell, Reenesh Prakash, Samir Dervisevic, Alison E. Mather, John Wain, Mark Webber, Andrew J. Page, Justin O'Grady |
| EPI_ISL_665851, EPI_ISL_665852, EPI_ISL_665853, EPI_ISL_665854, EPI_ISL_665855, EPI_ISL_665856, EPI_ISL_665857, EPI_ISL_665858, EPI_ISL_665859, EPI_ISL_665860, EPI_ISL_665861, EPI_ISL_665862, EPI_ISL_665863 | | | |
| see above | University of Exeter | COVID-19 Genomics UK (COG-UK) Consortium | Ben Temperton,Aaron Jeffries,Michelle Michelsen,Joanna Warwick-Dugdale,Audrey Farbos,Robyn Manley,Stephen Michell,Jane Masoli |
| EPI_ISL_665873, EPI_ISL_665874, EPI_ISL_665875 | Northumbria University / South Tees Hospitals NHS Foundation Trust / North Cumbria Integrated Care NHS Foundation Trust / North Tees and Hartlepool NHS Foundation Trust / Newcastle Hospitals NHS Foundation Trust | COVID-19 Genomics UK (COG-UK) Consortium | Darren L Smith,Andrew Nelson,Matthew Bashton,Greg R Young,Joshua Loh,John Allan,Mohammad A Tariq,Giles S Holt,Gary Black,Wen C Yew,Lynn Dover,Paul Baker,Steve Liggett,Sarah Essex,Jane Greenaway,Debra Padgett,Clive Graham,Garren Scott,Edward Barton,Emma Swindells,Brendan Payne,Jennifer Collins,Yusri Taha,Gary Eltringham |
| EPI_ISL_665962 | Quadram Institute Bioscience | COVID-19 Genomics UK (COG-UK) Consortium | Dave J. Baker, Gemma L. Kay, Alp Aydin, Thanh Le-Viet, Steven Rudder, Ana P. Tedim, Anastasia Kolyva, Maria Diaz, Leonardo de Oliveira Martins, Nabil-Fareed Alikhan, Lizzie Meadows, Rachael Stanley, Ngozi Elumogo, Muhammed Yasir, Nicholas M. Thomson, Alexander J Trotter, Rachel Gilroy, Samuel Bloomfield, Claire Stuart, Andrew Bell, Reenesh Prakash, Samir Dervisevic, Alison E. Mather, John Wain, Mark Webber, Andrew J. Page, Justin O'Grady |
| EPI_ISL_665964, EPI_ISL_665966, EPI_ISL_665967, EPI_ISL_665968, EPI_ISL_665969, EPI_ISL_665970 | Northumbria University / South Tees Hospitals NHS Foundation Trust / North Cumbria Integrated Care NHS Foundation Trust / North Tees and Hartlepool NHS Foundation Trust / Newcastle Hospitals NHS Foundation Trust | COVID-19 Genomics UK (COG-UK) Consortium | Darren L Smith,Andrew Nelson,Matthew Bashton,Greg R Young,Joshua Loh,John Allan,Mohammad A Tariq,Giles S Holt,Gary Black,Wen C Yew,Lynn Dover,Paul Baker,Steve Liggett,Sarah Essex,Jane Greenaway,Debra Padgett,Clive Graham,Garren Scott,Edward Barton,Emma Swindells,Brendan Payne,Jennifer Collins,Yusri Taha,Gary Eltringham |
| EPI_ISL_665971, EPI_ISL_665972, EPI_ISL_665976 | Queens Medical Centre, Clinical Microbiology Department / DeepSeq Nottingham | COVID-19 Genomics UK (COG-UK) Consortium | Gemma Clark, Wendy Smith, Manjinder Khakh, Vicki M Fleming, Michelle M Lister, Hannah Howson-Wells, Jonathan Ball, Patrick McClure, Joseph Chappell, Theocharis Tsoleridis, Nadine Holmes, Matthew Carlisle, Christopher Moore, Fei Sang, Johnny Debebe, Victoria Wright, Matthew Loose |
| EPI_ISL_666591 | Northumbria University / South Tees Hospitals NHS Foundation Trust / North Cumbria Integrated Care NHS | COVID-19 Genomics UK (COG-UK) Consortium | Darren L Smith,Andrew Nelson,Matthew Bashton,Greg R Young,Joshua Loh,John Allan,Mohammad A Tariq,Giles S Holt,Gary Black,Wen C Yew,Lynn Dover,Paul Baker,Steve Liggett,Sarah Essex,Jane Greenaway,Debra Padgett,Clive Graham,Garren Scott,Edward Barton,Emma Swindells,Brendan Payne,Jennifer Collins,Yusri Taha,Gary Eltringham |

| | | | |
|---|---|---|---|
| | Foundation Trust / North Tees and Hartlepool NHS Foundation Trust / Newcastle Hospitals NHS Foundation Trust | | |
| EPI_ISL_666623, EPI_ISL_666624, EPI_ISL_666625, EPI_ISL_666626, EPI_ISL_666627 | ZOTZ KLIMAS MVZ Düsseldorf-Centrum GbR ÜBAG für Labormedizin, Genetik, Zytologie, Pathologie | Center of Medical Microbiology, Virology, and Hospital Hygiene, University of Duesseldorf | Maximilian Damagnez, Alexander Dilthey, Ashley-Jane Duplessis, Patrick Finzer, Katrin Hoffmann, Torsten Houwaart, Lisanna Hülse, Malte Kohns Vasconcelos, Marek Korencak, Nadine Lübke, Jessica Nicolai, Klaus Pfeffer, Daniel Strelow, Jörg Timm, Andreas Walker, Tobias Wienemann, Rainer Zotz |
| EPI_ISL_667503 | OHSU Lab Services Molecular Microbiology Lab | Oregon SARS-CoV-2 Genome Sequencing Center | Brendan L. O'Connell, Ruth V. Nichols, Sally Grindstaff, Alec J. Hirsch, Donna Hansel, Guang Fan, Daniel N. Streblow, William B. Messer, Andrew C. Adey, Benjamin N. Bimber, Brian J. O'Roak |
| EPI_ISL_667796 | South Eastern Area Laboratory Services (SEALS) | NSW Health Pathology - Institute of Clinical Pathology and Medical Research; Westmead Hospital; University of Sydney | CIDM-PH et al. |
| EPI_ISL_668405 | Haukeland University Hospital, Dept. of Microbiology | Norwegian Institute of Public Health, Department of Virology | Kathrine Stene-Johansen, Kamilla Heddeland Instefjord, Hilde Elshaug, Marie Paulsen Madsen, Rasmus Riis Kopperud, Hilde Vollan, Karoline Bragstad, Olav Hungnes |
| EPI_ISL_670458, EPI_ISL_670502, EPI_ISL_670515, EPI_ISL_670647, EPI_ISL_670648, EPI_ISL_670649, EPI_ISL_670684, EPI_ISL_670685, EPI_ISL_670689, EPI_ISL_670697, EPI_ISL_670698, EPI_ISL_670699, EPI_ISL_670724, EPI_ISL_670725, EPI_ISL_670726, EPI_ISL_670727, EPI_ISL_670771, EPI_ISL_671224, EPI_ISL_671225, EPI_ISL_671226, EPI_ISL_671227, EPI_ISL_671228, EPI_ISL_671229 | | | |
| see above | Department of Virus and Microbiological Special Diagnostics, Statens Serum Institut, Copenhagen, Denmark | Albertsen Lab, Department of Chemistry and Bioscience, Aalborg University, Denmark | Danish Covid-19 Genome Consortium |
| EPI_ISL_671416 | National Laboratory of Virology, Szentágothai Research Centre | National Laboratory of Virology, Szentágothai Research Centre | Endre Gábor Tóth, Balázs Somogyi, Brigitta, Ferenc Jakab, Gábor Kemenesi |
| EPI_ISL_671485, EPI_ISL_671486, EPI_ISL_671487, EPI_ISL_671488, EPI_ISL_671489, EPI_ISL_671490, EPI_ISL_671491, EPI_ISL_671492, EPI_ISL_671493, EPI_ISL_671494, EPI_ISL_671495 | | | |
| see above | University of Debrecen, Department of Medical Microbiology | National Laboratory of Virology, Szentágothai Research Centre | Endre Gábor Tóth, Balázs Somogyi, Brigitta Zana, Eszter Csoma, Ferenc Jakab, Gábor Kemenesi |
| EPI_ISL_671834 | Servicio de Microbiología, Laboratori Clínic Metropolitana Nord. Hospital Universitari Germans Trias i Pujol. Institut d'Investigació en Ciències de la Salut Germans Trias i Pujol (IGTP) | SeqCOVID-SPAIN consortium/IBV(CSIC) | Elisa Martró, Antoni E. Bordoy, Anna Not, Adrián Antuori, Anabel Fernández, Nona Romaní, Verónica Saludes, Cristina Casañ and SeqCOVID-SPAIN consortium |
| EPI_ISL_671864, EPI_ISL_671872, EPI_ISL_671873, EPI_ISL_671874, EPI_ISL_671875, EPI_ISL_671876, EPI_ISL_671877, EPI_ISL_671878, EPI_ISL_671879, EPI_ISL_671880, EPI_ISL_671884, EPI_ISL_671885, EPI_ISL_671886, EPI_ISL_671887, EPI_ISL_671888, EPI_ISL_671889, EPI_ISL_671892, EPI_ISL_671902, EPI_ISL_671908, EPI_ISL_671909, EPI_ISL_671917, EPI_ISL_671918, EPI_ISL_671919, EPI_ISL_671920, EPI_ISL_671921, EPI_ISL_671924, EPI_ISL_671925, EPI_ISL_671926, EPI_ISL_671931, EPI_ISL_671934, EPI_ISL_671935, EPI_ISL_671938, EPI_ISL_671939 | | | |
| see above | National Virus Reference Laboratory | National Virus Reference Laboratory | Michael Carr, Gabriel Gonzalez, Jonathan Dean, Daniel Hare, Cillian F De Gascun |
| EPI_ISL_671971 | CHU Purpan - Laboratoire de Virologie - Institut Fédératif de Biologie | CHU Purpan - Laboratoire de Virologie - Institut Fédératif de Biologie | Latour J., Ranger N., Dubois M., Carcenac R., Harter A., Boyer P., Tremeaux P., Izopet J. |
| EPI_ISL_676507 | Klinisk mikrobiologi | The Public Health Agency of Sweden | Department of Microbiology, The Public Health Agency of Sweden |
| EPI_ISL_677263 | Colorado Department of Public Health and Environment | Colorado Department of Puplic Health and Environment | Laura Bankers, Molly Hetherington-Rauth, Shannon Ely, Shannon R. Matzinger, Sarah Elizabeth Totten, Emily A. Travanty |
| EPI_ISL_677345, EPI_ISL_677423, EPI_ISL_677430, EPI_ISL_677441, EPI_ISL_677445, EPI_ISL_677456, EPI_ISL_677457, EPI_ISL_677458, EPI_ISL_677468, EPI_ISL_677469, EPI_ISL_677470, EPI_ISL_677471, EPI_ISL_677473, EPI_ISL_677479, EPI_ISL_677522, EPI_ISL_677523, EPI_ISL_677526, EPI_ISL_677538, EPI_ISL_677554, EPI_ISL_677555, EPI_ISL_677556, EPI_ISL_677577, EPI_ISL_677629, EPI_ISL_677661, EPI_ISL_677662 | | | |
| see above | University of Wisconsin-Madison AIDS Vaccine Research Laboratories | University of Wisconsin-Madison AIDS Vaccine Research Laboratories | Gage Moreno, Katarina Braun, et al. AIDS Vaccine Research Laboratories |
| EPI_ISL_677818, EPI_ISL_677820 | University of Szeged, Institute of Clinical Microbiology | National Laboratory of Virology, Szentágothai Research Centre | Endre Gábor Tóth, Balázs Somogyi, Brigitta, Gabriella Terhes, Ferenc Jakab, Gábor Kemenesi |
| EPI_ISL_678364, EPI_ISL_678365, EPI_ISL_678366, EPI_ISL_678367, EPI_ISL_678368 | Area of Virology, Serology and Virology Division (SAViD), New South Wales Health Pathology Randwick | Virology Research Laboratory; Area of Virology, Serology and Virology Division (SAViD), New South Wales Health Pathology Randwick | Foster, C.; Au, J.; Ruiz Silva, M.; Deveson, I.; Bull, R.; Van Hal, S.; Rawlinson, W. |
| EPI_ISL_678387, EPI_ISL_678388, EPI_ISL_678389, EPI_ISL_678390, EPI_ISL_678391, EPI_ISL_678392, EPI_ISL_678393, EPI_ISL_678394, EPI_ISL_678395, EPI_ISL_678396 | University of Debrecen, Department of Medical Microbiology | National Laboratory of Virology, Szentágothai Research Centre | Endre Gábor Tóth, Balázs Somogyi, Brigitta Zana, Eszter Csoma, Ferenc Jakab, Gábor Kemenesi |
| EPI_ISL_679413, EPI_ISL_679414, EPI_ISL_679415 | University College London Hospital | COVID-19 Genomics UK (COG-UK) Consortium | Judith Heaney, Matthew Byott, Catherine Houlihan, Dan Frampton, Stuart Kirk, Moira Spyer and Eleni Nastouli |
| EPI_ISL_679550, EPI_ISL_679551, EPI_ISL_679552, EPI_ISL_679553, EPI_ISL_679554, EPI_ISL_679557, EPI_ISL_679558, EPI_ISL_679559, EPI_ISL_679561, EPI_ISL_679562, EPI_ISL_679564, EPI_ISL_679565, EPI_ISL_679566, EPI_ISL_679567, EPI_ISL_679569, EPI_ISL_679570, EPI_ISL_679571, EPI_ISL_679572, EPI_ISL_679573, EPI_ISL_679574 | | | |
| see above | University College London, Great Ormond Street Hospital for Children NHS Foundation Trust, Imperial College Healthcare NHS Trust | COVID-19 Genomics UK (COG-UK) Consortium | Sergi Castellano, Rachel Williams, Mark Kristiansen, Paola Resende Silva, Sunando Roy, Tony Brooks, Helena Tutill, Paola Niola, Patricia Dyal, Charlotte Williams, Leysa Forrest, Yasmin Panchbhaya, Jacqueline Findlay, Samuel Weeks, Julianne Brown, Kathryn Harris, Paul Randell, James Price, Alison Holmes, Judith Breuer |
| EPI_ISL_680340, EPI_ISL_680341 | Regional Virus Laboratory, Belfast Health and Social Care Trust | COVID-19 Genomics UK (COG-UK) Consortium | Conall McCaughey, James McKenna, Tanya Curran, Susan Feeney, Alison Watt, Ciara Cox, Mairead Connor, Zoltan Molnar, David Simpson, Derek Fairley |
| EPI_ISL_680470, EPI_ISL_680472, EPI_ISL_680473, EPI_ISL_680474, EPI_ISL_680475, EPI_ISL_680476, EPI_ISL_680482, EPI_ISL_680495, EPI_ISL_680496 | Virology Department, Royal Infirmary of Edinburgh, NHS Lothian / School of Biological Sciences, University of Edinburgh / Institute of Genetics and Molecular Medicine, University of Edinburgh | COVID-19 Genomics UK (COG-UK) Consortium | McHugh M, Dewar R, Rooke S, Gallagher M, Balcaza C, O'Toole Á, Scher E, Hill V, McCrone JT, Colquhoun R, Yu X, Jackson B, Rambaut A, Williams TC, Templeton K |
| EPI_ISL_681306, EPI_ISL_681307, EPI_ISL_681312 | Communicable Disease Laboratory, Public Health Directorate | Communicable Disease Laboratory, Public Health Directorate | Alwasti,H., Altaif,Z., AlHujairi,Z., AlAbbas,Z. |
| EPI_ISL_682033, EPI_ISL_682039, EPI_ISL_682040, EPI_ISL_682051 | UPMC Clinical Microbiology Laboratory | Microbial Genomic Epidemiology Laboratory, University of Pittsburgh | Mustapha M. Mustapha, Jane W. Marsh, Dan Snyder, Marissa P. Griffith, Stephanie L. Mitchell, Vatsala R. Srinivasa, Kady D. Waggle, Chinelo Ezeonwuku, Vaughn S. Cooper, Lee H. Harrison |
| EPI_ISL_682306, EPI_ISL_682307, EPI_ISL_682308, EPI_ISL_682309, EPI_ISL_682314 | Communicable Disease Laboratory, Public Health Directorate | Communicable Disease Laboratory, Public Health Directorate | Alwasti,H., Altaif,Z., AlHujairi,Z., AlAbbas,Z. |
| EPI_ISL_682327, EPI_ISL_682329, EPI_ISL_682341, EPI_ISL_682343, EPI_ISL_682347 | NHLS Universitas Academic | UFS Virology | PA Bester, MM Nyaga, P Nthiga, MT Mogotsi, D Goedhals, T de Oliveira |
| EPI_ISL_683728 | Mayo Clinic & Mayo Clinic Laboratories | Minnesota Department of Health, Public Health Laboratory | Alexandra Lorentz, Jacob Garfin, Matt Plumb, and Xiong Wang |
| EPI_ISL_691685 | Hospital de Leon | Instituto de Salud Carlos III | Iglesias-Caballero, M. Camarero, S. Molinero Calamita, M. González-Esguevillas, M. Pozo, F. Casas, I. Jiménez, P. Jiménez, M. Zaballos, A. Monzón, S. |

| | | | |
|---|---|---|---|
| | | | Varona, S. Juliá, M. Cuesta, I. Vidan, J. |
| EPI_ISL_691731 | Hospital Universitario Puerta de Hierro | Instituto de Salud Carlos III | Iglesias-Caballero, M. Camarero, S. Molinero Calamita, M. González-Esguevillas, M. Pozo, F. Casas, I. Jiménez, P. Jiménez, M. Zaballos, A. Monzón, S. Varona, S. Juliá, M. Cuesta, I. Velasco, A. |
| EPI_ISL_691732 | Hospital de Madrid | Instituto de Salud Carlos III | Iglesias-Caballero, M. Camarero, S. Molinero Calamita, M. González-Esguevillas, M. Pozo, F. Casas, I. Jiménez, P. Jiménez, M. Zaballos, A. Monzón, S. Varona, S. Juliá, M. Cuesta, I. Alhambra, A. |
| EPI_ISL_692847, EPI_ISL_692856, EPI_ISL_692862, EPI_ISL_692863, EPI_ISL_692864, EPI_ISL_692865, EPI_ISL_692868, EPI_ISL_692871 | Massachusetts State Public Health Laboratory | Massachusetts State Public Health Laboratory | Andrew Lang, Timelia Fink, Glen Gallagher, Sandra Smole |
| EPI_ISL_699572, EPI_ISL_699573 | Group of Genetic Engeneering and Biotechnology, Federal Budget Institution of Science 'Central Research Institute of Epidemiology' of The Federal Service on Customers' Rights Protection and Human Well-being Surveillance | Group of Genetic Engeneering and Biotechnology, Federal Budget Institution of Science 'Central Research Institute of Epidemiology' of The Federal Service on Customers' Rights Protection and Human Well-being Surveillance | Cherkashina,A.S., Golubeva,A.G., Soloviova,E.D., Zotova,M.I., Berlina,Y.Y., Valdokhina,A.V., Bulanenko,V.P., Speranskaya,A.S., Tivanova,E.V., Shipulina,O.Y., Akimkin,V.G. |
| EPI_ISL_699643 | Medlab Pathology | NSW Health Pathology - Institute of Clinical Pathology and Medical Research; Westmead Hospital; University of Sydney | CIDM-PH et al. |
| EPI_ISL_700730, EPI_ISL_700733, EPI_ISL_700736 | Texas Department of State Health Services | Texas Department of State Health Services | Rashmi Tuladhar, Bonnie Oh, Jenny Zhang, Maliha Rahman, Anita Pokharel, Myong Koag, Chung Wang, Rachel Lee, Grace Kubin, Mayela Pedrueza, James Daniel Bonser |
| EPI_ISL_702690, EPI_ISL_703179, EPI_ISL_703332 | University College London, Great Ormond Street Hospital for Children NHS Foundation Trust, Imperial College Healthcare NHS Trust | COVID-19 Genomics UK (COG-UK) Consortium | Sergi Castellano, Rachel Williams, Mark Kristiansen, Paola Resende Silva, Sunando Roy, Tony Brooks, Helena Tutill, Paola Niola, Patricia Dyal, Charlotte Williams, Leysa Forrest, Yasmin Panchbhaya, Jacqueline Findlay, Samuel Weeks, Julianne Brown, Kathryn Harris, Paul Randell, James Price, Alison Holmes, Judith Breuer |
| EPI_ISL_703622, EPI_ISL_703782, EPI_ISL_703784 | Virology Department, Royal Infirmary of Edinburgh, NHS Lothian / School of Biological Sciences, University of Edinburgh / Institute of Genetics and Molecular Medicine, University of Edinburgh | COVID-19 Genomics UK (COG-UK) Consortium | McHugh M, Dewar R, Rooke S, Gallagher M, Balcaza C, O'Toole Á, Scher E, Hill V, McCrone JT, Colquhoun R, Yu X, Jackson B, Rambaut A, Williams TC, Templeton K |
| EPI_ISL_704046, EPI_ISL_704084, EPI_ISL_704163, EPI_ISL_704193 | University College London, Great Ormond Street Hospital for Children NHS Foundation Trust, Imperial College Healthcare NHS Trust | COVID-19 Genomics UK (COG-UK) Consortium | Sergi Castellano, Rachel Williams, Mark Kristiansen, Paola Resende Silva, Sunando Roy, Tony Brooks, Helena Tutill, Paola Niola, Patricia Dyal, Charlotte Williams, Leysa Forrest, Yasmin Panchbhaya, Jacqueline Findlay, Samuel Weeks, Julianne Brown, Kathryn Harris, Paul Randell, James Price, Alison Holmes, Judith Breuer |
| EPI_ISL_704594 | Oxford Viromics, NDM, University of Oxford; Oxford University Hospitals; Basingstoke and North Hampshire Hospital | COVID-19 Genomics UK (COG-UK) Consortium | Tanya Golubchik, David Bonsall, George Macintyre, Amy Trebes, Mariateresa de Cesare, Catrin Moore, Alex Mobbs, Anita Justice, Robert Shaw, Monique Andersson, Timothy Peto, Emma Wise, Nathan Moore, Jessica Lynch, Nick Cortes, Matilde Mori, Stephen Kidd, David Buck, John Todd, Christophe Fraser |
| EPI_ISL_705425 | University College London, Great Ormond Street Hospital for Children NHS Foundation Trust, Imperial College Healthcare NHS Trust | COVID-19 Genomics UK (COG-UK) Consortium | Sergi Castellano, Rachel Williams, Mark Kristiansen, Paola Resende Silva, Sunando Roy, Tony Brooks, Helena Tutill, Paola Niola, Patricia Dyal, Charlotte Williams, Leysa Forrest, Yasmin Panchbhaya, Jacqueline Findlay, Samuel Weeks, Julianne Brown, Kathryn Harris, Paul Randell, James Price, Alison Holmes, Judith Breuer |
| EPI_ISL_705444, EPI_ISL_705489 | Oxford Viromics, NDM, University of Oxford; Oxford University Hospitals; Basingstoke and North Hampshire Hospital | COVID-19 Genomics UK (COG-UK) Consortium | Tanya Golubchik, David Bonsall, George Macintyre, Amy Trebes, Mariateresa de Cesare, Catrin Moore, Alex Mobbs, Anita Justice, Robert Shaw, Monique Andersson, Timothy Peto, Emma Wise, Nathan Moore, Jessica Lynch, Nick Cortes, Matilde Mori, Stephen Kidd, David Buck, John Todd, Christophe Fraser |
| EPI_ISL_705763, EPI_ISL_705778 | Virology Department, Royal Infirmary of Edinburgh, NHS Lothian / School of Biological Sciences, University of Edinburgh / Institute of Genetics and Molecular Medicine, University of Edinburgh | COVID-19 Genomics UK (COG-UK) Consortium | McHugh M, Dewar R, Rooke S, Gallagher M, Balcaza C, O'Toole Á, Scher E, Hill V, McCrone JT, Colquhoun R, Yu X, Jackson B, Rambaut A, Williams TC, Templeton K |
| EPI_ISL_706938, EPI_ISL_706939, EPI_ISL_706940, EPI_ISL_706941, EPI_ISL_706942, EPI_ISL_706943, EPI_ISL_706944, EPI_ISL_706945, EPI_ISL_706946, EPI_ISL_706947, EPI_ISL_706948, EPI_ISL_706949, EPI_ISL_706950, EPI_ISL_706951, EPI_ISL_706952, EPI_ISL_706953, EPI_ISL_706954, EPI_ISL_706955, EPI_ISL_706956, EPI_ISL_706957, EPI_ISL_706958, EPI_ISL_706959, EPI_ISL_706960, EPI_ISL_706961, EPI_ISL_706962, EPI_ISL_706963, EPI_ISL_706964, EPI_ISL_706965, EPI_ISL_706966, EPI_ISL_706967, EPI_ISL_706968, EPI_ISL_706969, EPI_ISL_706970, EPI_ISL_706971, EPI_ISL_706972, EPI_ISL_706973 | | | |
| see above | Oxford Viromics, NDM, University of Oxford; Oxford University Hospitals; Basingstoke and North Hampshire Hospital | COVID-19 Genomics UK (COG-UK) Consortium | Tanya Golubchik, David Bonsall, George Macintyre, Amy Trebes, Mariateresa de Cesare, Catrin Moore, Alex Mobbs, Anita Justice, Robert Shaw, Monique Andersson, Timothy Peto, Emma Wise, Nathan Moore, Jessica Lynch, Nick Cortes, Matilde Mori, Stephen Kidd, David Buck, John Todd, Christophe Fraser |
| EPI_ISL_707898 | Area of Virology, Serology and Virology Division (SAViD), New South Wales Health Pathology Randwick | Virology Research Laboratory; Area of Virology, Serology and Virology Division (SAViD), New South Wales Health Pathology Randwick | Foster, C.; Au, J.; Ruiz Silva, M.; Deveson, I.; Bull, R.; Van Hal, S.; Rawlinson, W. |
| EPI_ISL_707970, EPI_ISL_707971, EPI_ISL_707972, EPI_ISL_707973, EPI_ISL_707974, EPI_ISL_707975, EPI_ISL_708010, EPI_ISL_708011, EPI_ISL_708012 | Virology, Universitätsklinikum des Saarlandes | Epigenetics, Saarland University | Kathrin Kattler, Markus Vogelgesang, Stefan Lohse, Sascha Tierling, Sigrun Smola, Jörn Walter |
| EPI_ISL_708382, EPI_ISL_708385, EPI_ISL_708387, EPI_ISL_708391, EPI_ISL_708402, EPI_ISL_708404, EPI_ISL_708416, EPI_ISL_708427, EPI_ISL_708428 | Delaware Public Health Lab | Delaware Public Health Lab | Gregory Hovan |
| EPI_ISL_708483 | Mayo Clinic & Mayo Clinic Laboratories | Minnesota Department of Health, Public Health Laboratory | Alexandra Lorentz, Jacob Garfin, Matt Plumb, and Xiong Wang |
| EPI_ISL_708814 | World Medical Hospital | National Institute of Health, Department of Medical Sciences, Ministry of Public Health, Thailand | Pilailuk Okada; Siripaporn Phuygun; Thanutsapa Thanadachakul; Sittiporn Parnmen; Pakorn Piromtong; Warawan Wongboot; Sunthareeya Waicharoen; Malinee Chittaganpitch |
| EPI_ISL_708817 | Urban Institute for Disease Prevention and Control | National Institute of Health, Department of Medical Sciences, Ministry of Public Health, Thailand | Pilailuk Okada; Siripaporn Phuygun; Thanutsapa Thanadachakul; Sittiporn Parnmen; Pakorn Piromtong; Warawan Wongboot; Sunthareeya Waicharoen; Malinee Chittaganpitch |
| EPI_ISL_708820 | Vajira Hospital | National Institute of Health, Department of Medical Sciences, Ministry of Public Health, Thailand | Pilailuk Okada; Siripaporn Phuygun; Thanutsapa Thanadachakul; Sittiporn Parnmen; Pakorn Piromtong; Warawan Wongboot; Sunthareeya Waicharoen; Malinee Chittaganpitch |
| EPI_ISL_708821 | Synphaet Hospital | National Institute of Health, Department of Medical Sciences, Ministry of Public Health, Thailand | Pilailuk Okada; Siripaporn Phuygun; Thanutsapa Thanadachakul; Sittiporn Parnmen; Pakorn Piromtong; Warawan Wongboot; Sunthareeya Waicharoen; Malinee Chittaganpitch |
| EPI_ISL_709932, EPI_ISL_709933, EPI_ISL_709934, EPI_ISL_709935, EPI_ISL_709942, EPI_ISL_709943, EPI_ISL_709944, EPI_ISL_709945, EPI_ISL_709946 | Lighthouse Lab in Milton Keynes | Wellcome Sanger Institute for the COVID-19 Genomics UK (COG-UK) Consortium | The Lighthouse Lab in Milton Keynes and Alex Alderton, Roberto Amato, Sonia Goncalves, Ewan Harrison, David K. Jackson, Ian Johnston, Dominic Kwiatkowski, Cordelia Langford, John Sillitoe on behalf of the Wellcome Sanger Institute COVID-19 Surveillance Team |
| EPI_ISL_710268, EPI_ISL_710349, EPI_ISL_710350, EPI_ISL_710351, EPI_ISL_710352, EPI_ISL_710353 | Colorado Department of Public Health and Environment | Colorado Department of Puplic Health and Environment | Laura Bankers, Molly C. Hetherington-Rauth, Shannon Ely, Shannon R. Matzinger, Sarah Elizabeth Totten, Emily A. Travanty |

| | | | |
|---|---|---|---|
| EPI_ISL_710418 | Los Angeles County PHL | Los Angeles County PHL | P. Hemarajata et al. |
| EPI_ISL_712071, EPI_ISL_712073, EPI_ISL_712081, EPI_ISL_712093, EPI_ISL_712094 | Port Elizabeth Provincial Hospital, National Health Laboratory Services, Eastern Cape, South Africa | National Institute for Communicable Diseases of the National Health Laboratory Service | Mohale T, Ntuli N, Mahlangu B, Allam M, Ismail A, Bhiman JN |
| EPI_ISL_717793, EPI_ISL_717818, EPI_ISL_717819, EPI_ISL_717874, EPI_ISL_717875, EPI_ISL_717876, EPI_ISL_717925, EPI_ISL_717960 | Laboratorio de Virologia Molecular / UFRJ | Bioinformatics Laboratory / LNCC | Carolina M Voloch, Ronaldo da Silva F Jr, Luiz G P de Almeida, Cynthia C Cardoso, Otavio Bustrolini, Alexandra L Gerber, Ana Paula de C Guimarães, Diana Mariani, Andréa Cony Cavalcanti, Claudia dos Santos Rodrigues, Terezinha M P P Castiñeira, Amilcar Tanuri, Ana Tereza R de Vasconcelos |
| EPI_ISL_718028, EPI_ISL_718030, EPI_ISL_718031, EPI_ISL_718032, EPI_ISL_718033 | ZOTZ KLIMAS MVZ Düsseldorf-Centrum GbR ÜBAG für Labormedizin, Genetik, Zytologie, Pathologie | Center of Medical Microbiology, Virology, and Hospital Hygiene, University of Duesseldorf | Maximilian Damagnez, Alexander Dilthey, Ashley-Jane Duplessis, Patrick Finzer, Katrin Hoffmann, Torsten Houwaart, Lisanna Hülse, Malte Kohns Vasconcelos, Marek Korencak, Nadine Lübke, Jessica Nicolai, Klaus Pfeffer, Daniel Strelow, Jörg Timm, Andreas Walker, Tobias Wienemann, Rainer Zotz |
| EPI_ISL_718210, EPI_ISL_718219 | Ministry of Health Hospitals | Institute of Health and Community Medicine | David Perera, Ooi Mong How, Chua Hock Hin, Tonnii Sia Loong Loong, Wong Jyn Shan, Wong Kiing Aik, Chan Chia Jui |
| EPI_ISL_718237, EPI_ISL_718244, EPI_ISL_718246 | Hospital | National Reference Center for Viruses of Respiratory Infections, Institut Pasteur, Paris | Marion Barbet, Sylvie Behillil, Méline Bizard, Angela Brisebarre, Camille Capel, Etienne Simon-Lorière, Vincent Enouf, Maud Vanpeene, Sylvie van der Werf, Gisèle Lagathu |
| EPI_ISL_718311 | Institute for Medical Research, Infectious Disease Research Centre, National Institutes of Health, Ministry of Health Malaysia | Institute for Medical Research, Infectious Disease Research Centre, National Institutes of Health, Ministry of Health Malaysia | Suppiah J, Kamel K, Mohd-Zawawi Z, Thayan R |
| EPI_ISL_722217, EPI_ISL_722218, EPI_ISL_722220, EPI_ISL_722234, EPI_ISL_722240, EPI_ISL_722270 | Servicio de Microbiología, Hospital Miguel Servet, Zaragoza | SeqCOVID-SPAIN consortium/IBV(CSIC) | Antonio Rezusta López, Alexander Tristancho Baró, Ana Milagro, Yolanda Gracia Grataloup, Nieves Martínez Cameo and SeqCOVID-SPAIN consortium |
| EPI_ISL_722307, EPI_ISL_722335, EPI_ISL_722347, EPI_ISL_722359, EPI_ISL_722360, EPI_ISL_722382, EPI_ISL_722435, EPI_ISL_722446, EPI_ISL_722496, EPI_ISL_722497, EPI_ISL_722498, EPI_ISL_722499 | | | |
| see above | Dutch COVID-19 response team | Erasmus Medical Center | Bas Oude Munnink, Reina Sikkema, David Nieuwenhuijse, Irina Chestakova, Anne van der Linden, Marjan Boter, Emmanuelle Munger, Corine GeurtsvanKessel, Annemiek van der Eijk, Richard Molenkamp, Marion Koopmans, on behalf of the Dutch national COVID-19 response team. |
| EPI_ISL_722893 | Istituto Zooprofilattico Sperimentale della Puglia e della Basilicata | Istituto Zooprofilattico Sperimentale della Puglia e della Basilicata | Parisi A., Bianco A., Capozzi L., Del Sambro L., Manzulli V, Rondinone V., Pace L., Cipolletta D., Galante D. |
| EPI_ISL_723212 | Dutch COVID-19 response team | National Institute for Public Health and the Environment (RIVM) | Adam Meijer, Harry Vennema, Jeroen Cremer, Sharon van den Brink, Bas van der Veer, AnneMarie van den Brandt, Florian Zwagemaker, Dennis Schmitz, Chantal Reusken, on behalf of the national COVID-19 response team |
| EPI_ISL_723470, EPI_ISL_723471, EPI_ISL_723472 | Virginia DCLS | Virginia DCLS | Virginia DCLS |
| EPI_ISL_724966 | Northumbria University / South Tees Hospitals NHS Foundation Trust / North Cumbria Integrated Care NHS Foundation Trust / North Tees and Hartlepool NHS Foundation Trust / Newcastle Hospitals NHS Foundation Trust | COVID-19 Genomics UK (COG-UK) Consortium | Darren L Smith,Andrew Nelson,Matthew Bashton,Greg R Young,Joshua Loh,John Allan,Mohammad A Tariq,Giles S Holt,Gary Black,Wen C Yew,Lynn Dover,Paul Baker,Steve Liggett,Sarah Essex,Jane Greenaway,Debra Padgett,Clive Graham,Garren Scott,Edward Barton,Emma Swindells,Brendan Payne,Jennifer Collins,Yusri Taha,Gary Eltringham |
| EPI_ISL_725159, EPI_ISL_725161, EPI_ISL_725162, EPI_ISL_725163, EPI_ISL_725165, EPI_ISL_725168, EPI_ISL_725169, EPI_ISL_725170, EPI_ISL_725172, EPI_ISL_725173, EPI_ISL_725187, EPI_ISL_725191 | | | |
| see above | Quadram Institute Bioscience | COVID-19 Genomics UK (COG-UK) Consortium | Dave J. Baker, Gemma L. Kay, Alp Aydin, Thanh Le-Viet, Steven Rudder, Ana P. Tedim, Anastasia Kolyva, Maria Diaz, Leonardo de Oliveira Martins, Nabil-Fareed Alikhan, Lizzie Meadows, Rachael Stanley, Ngozi Elumogo, Muhammed Yasir, Nicholas M. Thomson, Alexander J Trotter, Rachel Gilroy, Samuel Bloomfield, Claire Stuart, Andrew Bell, Reenesh Prakash, Samir Dervisevic, Alison E. Mather, John Wain, Mark Webber, Andrew J. Page, Justin O'Grady |
| EPI_ISL_728017, EPI_ISL_728021, EPI_ISL_728023, EPI_ISL_728027, EPI_ISL_728028, EPI_ISL_728029, EPI_ISL_728030, EPI_ISL_728031, EPI_ISL_728032, EPI_ISL_728033, EPI_ISL_728037, EPI_ISL_728077, EPI_ISL_728092 | | | |
| see above | University of Wisconsin-Madison AIDS Vaccine Research Laboratories | University of Wisconsin-Madison AIDS Vaccine Research Laboratories | Gage Moreno, Katarina Braun, et al. AIDS Vaccine Research Laboratories |
| EPI_ISL_728165, EPI_ISL_728166 | Institute for Medical Research, Infectious Disease Research Centre, National Institutes of Health, Ministry of Health Malaysia | Institute for Medical Research, Infectious Disease Research Centre, National Institutes of Health, Ministry of Health Malaysia | Suppiah J, Kamel K, Mohd-Zawawi Z, Thayan R |
| EPI_ISL_728732 | Dutch COVID-19 response team | National Institute for Public Health and the Environment (RIVM) | Adam Meijer, Harry Vennema, Jeroen Cremer, Sharon van den Brink, Bas van der Veer, AnneMarie van den Brandt, Florian Zwagemaker, Dennis Schmitz, Chantal Reusken, on behalf of the national COVID-19 response team |
| EPI_ISL_729379, EPI_ISL_729390, EPI_ISL_729421 | A. Krumbholz, Labor Dr. Krause und Kollegen MVZ GmbH, Kiel | Charité Universitätsmedizin Berlin, Institut für Virologie | Victor M Corman, Barbara Mühlemann, Jörn Beheim-Schwarzbach, Talitha Veith, Julia Schneider, Terry Jones, Christian Drosten |
| EPI_ISL_729737, EPI_ISL_729740 | Grubaugh Lab - Yale School of Public Health | Grubaugh Lab - Yale School of Public Health | Joseph Fauver, Tara Alpert, Anderson Brito, Annie Watkins, Anne Wyllie, Chantal Vogels, Mary Petrone, Chaney Kalinich, Isabel Ott, Arnau Casanovas, Catherine Muenker, Adam Moore, Alice Lu, Maria Tokuyama, Patrick Wong, Peiwen Lu, Saad Omer, Richard Martinello, Allison Nelson, Shelli Farhadian, Akiko Iwasaki, Charlese Dela Cruz, Albert Ko, Nathan Grubaugh |
| EPI_ISL_729766 | Yale Pathology Lab | Grubaugh Lab - Yale School of Public Health | Joseph Fauver, Tara Alpert, Anderson Brito, Annie Watkins, Anne Wyllie, Chantal Vogels, Mary Petrone, Chaney Kalinich, Isabel Ott, Arnau Casanovas, Catherine Muenker, Adam Moore, Alice Lu, Maria Tokuyama, Patrick Wong, Peiwen Lu, Saad Omer, Richard Martinello, Allison Nelson, Shelli Farhadian, Akiko Iwasaki, Charlese Dela Cruz, Albert Ko, Nathan Grubaugh |
| EPI_ISL_729782 | Grubaugh Lab - Yale School of Public Health | Grubaugh Lab - Yale School of Public Health | Joseph Fauver, Tara Alpert, Anderson Brito, Annie Watkins, Anne Wyllie, Chantal Vogels, Mary Petrone, Chaney Kalinich, Isabel Ott, Arnau Casanovas, Catherine Muenker, Adam Moore, Alice Lu, Maria Tokuyama, Patrick Wong, Peiwen Lu, Saad Omer, Richard Martinello, Allison Nelson, Shelli Farhadian, Akiko Iwasaki, Charlese Dela Cruz, Albert Ko, Nathan Grubaugh |
| EPI_ISL_729876 | Laboratorio de Referencia Nacional de Virus Respiratorios, Instituto Nacional de Salud Peru | Laboratorio de Genómica Microbiana, Universidad Peruana Cayetano Heredia | Pablo Tsukayama, Alejandra Dávila-Barclay, Luis González, Guillermo Salvatierra, Pedro E. Romero, Brenda Ayzanoa, Janet Huancachoque, Pool Marcos, Marco Galarza, Priscila Lope, Nancy Rojas |
| EPI_ISL_729983, EPI_ISL_729984, EPI_ISL_729985 | Nigeria Centre for Disease Control (NCDC) | African Centre of Excellence for Genomics of Infectious Diseases (ACEGID), Redeemer's University, Ede, Osun State, Nigeria | Oluniyi P.E. et al |
| EPI_ISL_730087, EPI_ISL_730095, EPI_ISL_730102, EPI_ISL_730113 | San Diego County Public Health Laboratory | Andersen lab at Scripps Research | SEARCH Alliance San Diego with Tracy Basler, Jovan Shephard, Brett Austin |
| EPI_ISL_730232, EPI_ISL_730240, EPI_ISL_730241, EPI_ISL_730242, EPI_ISL_730248, EPI_ISL_730249, EPI_ISL_730253, EPI_ISL_730254, EPI_ISL_730255, EPI_ISL_730261, EPI_ISL_730262, EPI_ISL_730263, EPI_ISL_730268, EPI_ISL_730269, EPI_ISL_730270, EPI_ISL_730275, EPI_ISL_730276, EPI_ISL_730277, EPI_ISL_730278, EPI_ISL_730279, EPI_ISL_730284, EPI_ISL_730285, EPI_ISL_730286, EPI_ISL_730372, EPI_ISL_730379, EPI_ISL_730380, EPI_ISL_730382, EPI_ISL_730390, EPI_ISL_730391, EPI_ISL_730394, EPI_ISL_730404, EPI_ISL_730411, EPI_ISL_730412, EPI_ISL_730418, EPI_ISL_730419, EPI_ISL_730432, EPI_ISL_730433, EPI_ISL_730437, EPI_ISL_730439, EPI_ISL_730440, EPI_ISL_730441, EPI_ISL_730444, EPI_ISL_730445, EPI_ISL_730449, EPI_ISL_730450, EPI_ISL_730451, EPI_ISL_730456, EPI_ISL_730457, EPI_ISL_730461, EPI_ISL_730462, EPI_ISL_730463, EPI_ISL_730464, EPI_ISL_730467, EPI_ISL_730468, EPI_ISL_730473, EPI_ISL_730475, EPI_ISL_730476, EPI_ISL_730488, EPI_ISL_730493, EPI_ISL_730498, EPI_ISL_730501, EPI_ISL_730505, EPI_ISL_730506, EPI_ISL_730508, EPI_ISL_730511, EPI_ISL_730512, EPI_ISL_730513, EPI_ISL_730516 | | | |
| see above | Biolab Diagnostic Laboratories | Andersen lab at Scripps Research | Issa Abu-Dayyeh, Ahmad Tibi, Lama Hussein, Lina Mohammad, Zein Naber, Amid Abdelnour with SEARCH Alliance San Diego |
| EPI_ISL_730593 | Private medical practitioner | Hong Kong Department of Health | Mak Gannon C.K., Lam Edman T.K., Chan Rickjason C.W., Tsang Dominic N.C. |
| EPI_ISL_730598, EPI_ISL_730599 | Queen Mary Hospital | Hong Kong Department of Health | Mak Gannon C.K., Lam Edman T.K., Chan Rickjason C.W., Tsang Dominic N.C. |
| EPI_ISL_732563 | Bundeswehr Institute of Microbiology | Bundeswehr Institute of Microbiology | Markus Antwerpen, Alexandra Rehn, Mathias Walter, Malena Bestehorn-Willmann, Sabine Zange, Enrico Georgi, Roman Wölfel |

| | | | |
|---|---|---|---|
| EPI_ISL_732658 | Bundeswehr Institute of Microbiology | Bundeswehr Institute of Microbiology | Elham Khatamzas, Markus Antwerpen, Mathias Walter, Alexandra Rehn, Sabine Zange, Enrico Georgi, Michael von Bergwelt-Baildon, Roman Wölfel |
| EPI_ISL_732800, EPI_ISL_732803 | Centro de Investigación Biomédica de La Rioja - Hospital San Pedro Logroño | SeqCOVID-SPAIN consortium/IBV(CSIC) | María de Toro, José Manuel Azcona Gutiérrez, María Pilar Bea Escudero, Miriam Blasco Alberdi and SeqCOVID-SPAIN consortium |
| EPI_ISL_733098, EPI_ISL_733099, EPI_ISL_733100, EPI_ISL_733101, EPI_ISL_733102, EPI_ISL_733103, EPI_ISL_733105, EPI_ISL_733106, EPI_ISL_733107, EPI_ISL_733108, EPI_ISL_733109, EPI_ISL_733111, EPI_ISL_733112, EPI_ISL_733113, EPI_ISL_733114, EPI_ISL_733115, EPI_ISL_733116, EPI_ISL_733117, EPI_ISL_733118, EPI_ISL_733121, EPI_ISL_733122, EPI_ISL_733124, EPI_ISL_733125, EPI_ISL_733126, EPI_ISL_733127, EPI_ISL_733128, EPI_ISL_733129, EPI_ISL_733130, EPI_ISL_733131, EPI_ISL_733132, EPI_ISL_733133, EPI_ISL_733134, EPI_ISL_733135, EPI_ISL_733136, EPI_ISL_733139, EPI_ISL_733141, EPI_ISL_733143, EPI_ISL_733145, EPI_ISL_733147, EPI_ISL_733148, EPI_ISL_733149, EPI_ISL_733151, EPI_ISL_733152, EPI_ISL_733153 | | | |
| see above | HELIX LLC | WHO National Influenza Centre Russian Federation | Andrey Komissarov, Artem Fadeev, Anna Ivanova, Kseniya Komissarova, Dmitry Bazhenov, Daria Danilenko, Ksenia Safina, Elena Nabieva, Georgii Bazykin, Dmitry Lioznov |
| EPI_ISL_733247, EPI_ISL_733269, EPI_ISL_733270, EPI_ISL_733271 | WHO National Influenza Centre Russian Federation | WHO National Influenza Centre Russian Federation | Andrey Komissarov, Artem Fadeev, Anna Ivanova, Kseniya Komissarova, Dmitry Bazhenov, Daria Danilenko, Ksenia Safina, Elena Nabieva, Georgii Bazykin, Dmitry Lioznov |
| EPI_ISL_733419, EPI_ISL_733420, EPI_ISL_733421, EPI_ISL_733424, EPI_ISL_733425, EPI_ISL_733430, EPI_ISL_733435, EPI_ISL_733436, EPI_ISL_733437, EPI_ISL_733439, EPI_ISL_733440, EPI_ISL_733442, EPI_ISL_733444, EPI_ISL_733445, EPI_ISL_733447, EPI_ISL_733448, EPI_ISL_733454, EPI_ISL_733457, EPI_ISL_733458 | | | |
| see above | HELIX LLC | WHO National Influenza Centre Russian Federation | Andrey Komissarov, Artem Fadeev, Anna Ivanova, Kseniya Komissarova, Dmitry Bazhenov, Daria Danilenko, Ksenia Safina, Elena Nabieva, Georgii Bazykin, Dmitry Lioznov |
| EPI_ISL_733584, EPI_ISL_733586, EPI_ISL_733587, EPI_ISL_733588 | Respiratory Virus Unit, National Infection Service, Public Health England | COVID-19 Genomics UK (COG-UK) Consortium | PHE Covid Sequencing Team |
| EPI_ISL_734168 | CHRU Pontchaillou - Laboratoire de Virologie | National Reference Center for Viruses of Respiratory Infections, Institut Pasteur, Paris | Marion Barbet, Sylvie Behillil, Méline Bizard, Angela Brisebarre, Camille Capel, Etienne Simon-Lorière, Vincent Enouf, Maud Vanpeene, Sylvie van der Werf, Gisèle Lagathu |
| EPI_ISL_734257 | Virginia Division of Consolidated Laboratory Services (DCLS) | Virginia Division of Consolidated Laboratory Services (DCLS) | Virginia DCLS |
| EPI_ISL_734306, EPI_ISL_734327, EPI_ISL_734328, EPI_ISL_734329, EPI_ISL_734330, EPI_ISL_734331, EPI_ISL_734332, EPI_ISL_734333, EPI_ISL_734334, EPI_ISL_734335, EPI_ISL_734336, EPI_ISL_734337, EPI_ISL_734338, EPI_ISL_734339, EPI_ISL_734340, EPI_ISL_734341, EPI_ISL_734342, EPI_ISL_734343, EPI_ISL_734344, EPI_ISL_734345, EPI_ISL_734346, EPI_ISL_734347, EPI_ISL_734348, EPI_ISL_734350 | | | |
| see above | Wadsworth Center, New York State Department.of Health | Wadsworth Center, New York State Department.of Health | Kirsten St. George, Daryl M. Lamson, Alexis Russel, Jonathan Plitnick, Navjot Singh, John Kelly, Sara Griesemer, Erasmus Schneider, Erica Lasek-Nesselquist |
| EPI_ISL_734447, EPI_ISL_734448 | Masonic Medical Research Institute | Wadsworth Center, New York State Department.of Health | Kirsten St. George, Nathan Tucker, Ryan D. Pfeiffer, Daryl M. Lamson, Alexis Russel, Jonathan Plitnick, Navjot Singh, John Kelly, Sara Griesemer, Erasmus Schneider, Erica Lasek-Nesselquist |
| EPI_ISL_735075, EPI_ISL_735076, EPI_ISL_735077, EPI_ISL_735078, EPI_ISL_735079, EPI_ISL_735080, EPI_ISL_735081, EPI_ISL_735082, EPI_ISL_735083, EPI_ISL_735084, EPI_ISL_735085, EPI_ISL_735086, EPI_ISL_735087, EPI_ISL_735088, EPI_ISL_735089, EPI_ISL_735090, EPI_ISL_735091, EPI_ISL_735092, EPI_ISL_735093, EPI_ISL_735094 | | | |
| see above | UZ Leuven, National Reference Laboratory for Coronaviruses, Laboratory Medicine, Leuven, Belgium | KU Leuven, Rega Institute, Clinical and Epidemiological Virology | Tony Wawina-Bokalanga, Joan Marti-Carerras, Bert Vanmechelen, Piet Maes |
| EPI_ISL_735250, EPI_ISL_735251, EPI_ISL_735252, EPI_ISL_735253, EPI_ISL_735258 | Group of Genetic Engeneering and Biotechnology, Federal Budget Institution of Science 'Central Research Institute of Epidemiology' of The Federal Service on Customers' Rights Protection and Human Well-being Surveillance | Group of Genetic Engeneering and Biotechnology, Federal Budget Institution of Science 'Central Research Institute of Epidemiology' of The Federal Service on Customers' Rights Protection and Human Well-being Surveillance | Cherkashina,A.S., Golubeva,A.G., Solovyova,E.D., Valdokhina,A.V., Bulanenko,V.P., Zotova,M.I., Berlina,Y.Y., Speranskaya,A.S., Tivanova,E.V., Shipulina,O.Y., Akimkin,V.G. |
| EPI_ISL_735454, EPI_ISL_735455, EPI_ISL_735456, EPI_ISL_735458, EPI_ISL_735464, EPI_ISL_735465, EPI_ISL_735467, EPI_ISL_735468, EPI_ISL_735470, EPI_ISL_735472, EPI_ISL_735473, EPI_ISL_735474 | | | |
| see above | UW Virology Lab | UW Virology Lab | Pavitra Roychoudhury, Hong Xie, Lasata Shrestha, Meei-Li Huang, Keith R Jerome, Alexander Greninger |
| EPI_ISL_737037, EPI_ISL_737038, EPI_ISL_737039, EPI_ISL_737040, EPI_ISL_737041, EPI_ISL_737042, EPI_ISL_737043, EPI_ISL_737044, EPI_ISL_737045, EPI_ISL_737046, EPI_ISL_737047, EPI_ISL_737048, EPI_ISL_737049, EPI_ISL_737050, EPI_ISL_737051, EPI_ISL_737052, EPI_ISL_737053, EPI_ISL_737054, EPI_ISL_737055, EPI_ISL_737056, EPI_ISL_737057, EPI_ISL_737058, EPI_ISL_737059, EPI_ISL_737060, EPI_ISL_737070, EPI_ISL_737071, EPI_ISL_737072, EPI_ISL_737073, EPI_ISL_737074, EPI_ISL_737075, EPI_ISL_737076, EPI_ISL_737077, EPI_ISL_737078, EPI_ISL_737079, EPI_ISL_737080 | | | |
| see above | Department of Virology and Immunology, University of Helsinki and Helsinki University Hospital, Huslab Finland | Department of Virology, Faculty of Medicine, University of Helsinki, Helsinki, Finland | Teemu Smura, Ravi Kant, Phuoc Truong, Hussein Alburkat, Hannimari Kallio-Kokko, Jenni Virtanen, Maija Suvanto, Sari Hannula, Harri Kangas, Pekka Ellonen, Olli Vapalahti |
| EPI_ISL_737573, EPI_ISL_737595, EPI_ISL_737608, EPI_ISL_737628, EPI_ISL_737643, EPI_ISL_737682, EPI_ISL_737683, EPI_ISL_737684, EPI_ISL_737685, EPI_ISL_737686, EPI_ISL_737687, EPI_ISL_737688, EPI_ISL_737689, EPI_ISL_737690, EPI_ISL_737691, EPI_ISL_737751, EPI_ISL_737752, EPI_ISL_737753, EPI_ISL_737754, EPI_ISL_737755, EPI_ISL_737756, EPI_ISL_737757, EPI_ISL_737784, EPI_ISL_737822, EPI_ISL_737823, EPI_ISL_737824, EPI_ISL_737825, EPI_ISL_737826, EPI_ISL_737827, EPI_ISL_737828, EPI_ISL_737829, EPI_ISL_737830, EPI_ISL_737831, EPI_ISL_737832, EPI_ISL_737833, EPI_ISL_737834, EPI_ISL_737835, EPI_ISL_737836, EPI_ISL_737837, EPI_ISL_737838, EPI_ISL_737879, EPI_ISL_737880, EPI_ISL_737881, EPI_ISL_737882, EPI_ISL_737883, EPI_ISL_737884, EPI_ISL_737885, EPI_ISL_737886, EPI_ISL_737887, EPI_ISL_737888, EPI_ISL_737889, EPI_ISL_737890, EPI_ISL_737891, EPI_ISL_737899 | | | |
| see above | Viollier AG | Department of Biosystems Science and Engineering, ETH Zürich | Chaoran Chen, Sarah Nadeau, Ivan Topolsky, Emmanouil Dermitzakis, Keith Harshman, Ioannis Xenarios, Henri Pegeot, Lorenzo Cerutti, Deborah Penet, Philipp Jablonski, Lara Fuhrmann, David Dreifuss, Katharina Jahn, Christiane Beckmann, Maurice Redondo, Olivier Kobel, Christoph Noppen, Sophie Seidel, Noemie Santamaria de Souza, Niko Beerenwinkel, Tanja Stadler |
| EPI_ISL_738498 | UZ Leuven, National Reference Laboratory for Coronaviruses, Laboratory Medicine, Leuven, Belgium | KU Leuven, Rega Institute, Clinical and Epidemiological Virology | Tony Wawina-Bokalanga, Joan Marti-Carerras, Bert Vanmechelen, Piet Maes |
| EPI_ISL_738544 | UCSF Clinical Microbiology Laboratory | Chan-Zuckerberg Biohub | CZB Cliahub Consortium |
| EPI_ISL_738988 | Humboldt County Public Health Laboratory | Chan-Zuckerberg Biohub | CZB Cliahub Consortium |
| EPI_ISL_739203 | UCSF Clinical Microbiology Laboratory | Chan-Zuckerberg Biohub | CZB Cliahub Consortium |
| EPI_ISL_739311, EPI_ISL_739533 | Humboldt County Public Health Laboratory | Chan-Zuckerberg Biohub | CZB Cliahub Consortium |
| EPI_ISL_739678, EPI_ISL_739681, EPI_ISL_739682, EPI_ISL_739683 | Instituto Nacional de Salud, Bogotá, Colombia | Instituto Nacional de Salud, Bogotá, Colombia | Katherine Laiton-Donato, Diego A. Álvarez-Díaz, Carlos Franco-Muñoz, Mauricio Pacheco-Montealegre, Jonathan Reales, Diego Andrés Prada, Sheryl Corchuelo, Magdalena Weisner, Martha Lucia Ospina Martinez, Marcela Mercado-Reyes |
| EPI_ISL_739728, EPI_ISL_739826, EPI_ISL_739828, EPI_ISL_740068, EPI_ISL_740120, EPI_ISL_740131, EPI_ISL_740200, EPI_ISL_740396, EPI_ISL_740401 | Laboratoire national de santé, Microbiology, Virology | Laboratoire national de santé, Microbiology, Microbial Genomics Platform | Anke Wienecke-Baldacchino, Catherine Ragimbeau,Jessica Tapp, Fatu Djabi, Lise Pignon, Raoul Salmon, Tamir Abdelrahman |
| EPI_ISL_741761, EPI_ISL_741766, EPI_ISL_741767, EPI_ISL_741768, EPI_ISL_741769 | Oxford Viromics, NDM, University of Oxford; Oxford University Hospitals; Basingstoke and North Hampshire Hospital | COVID-19 Genomics UK (COG-UK) Consortium | Tanya Golubchik, David Bonsall, George Macintyre, Amy Trebes, Mariateresa de Cesare, Catrin Moore, Alex Mobbs, Anita Justice, Robert Shaw, Monique Andersson, Timothy Peto, Emma Wise, Nathan Moore, Jessica Lynch, Nick Cortes, Matilde Mori, Stephen Kidd, David Buck, John Todd, Christophe Fraser |
| EPI_ISL_744138, EPI_ISL_744234, EPI_ISL_744410, EPI_ISL_744507, EPI_ISL_744515, EPI_ISL_744532, EPI_ISL_744582, EPI_ISL_744633, EPI_ISL_744688, EPI_ISL_744782, EPI_ISL_744903, EPI_ISL_744948 | | | |
| see above | Laboratoire national de santé, Microbiology, Virology | Laboratoire national de santé, Microbiology, Microbial Genomics Platform | Anke Wienecke-Baldacchino, Catherine Ragimbeau,Jessica Tapp, Fatu Djabi, Lise Pignon, Raoul Salmon, Tamir Abdelrahman |
| EPI_ISL_745578, EPI_ISL_745626, EPI_ISL_745650, EPI_ISL_746097, EPI_ISL_746114, EPI_ISL_746136, EPI_ISL_746204, EPI_ISL_746211, EPI_ISL_746237, EPI_ISL_746263, EPI_ISL_746264, EPI_ISL_746311 | | | |
| see above | Ginkgo Bioworks Clinical Laboratory | Utah Public Health Laboratory | Erin L. Young, Kelly Oakeson, Tara Gallagher, Michael T. Pyne, E. Susan Slechta, Melanie A. Mallory, Jeffrey B. Stevenson, Salika M. Shakir, David R. Hillyard, Malaika McKenzie-Bennett, James McGann, Jim Griffin, Keith Robison, Alex Plocik, Becky Schilling, Martha Pierson, Rebecca Littlefield, Michelle Spencer, Birgitte Simen |
| EPI_ISL_746513, EPI_ISL_746514, EPI_ISL_746515, EPI_ISL_746516, EPI_ISL_746530, EPI_ISL_746531, EPI_ISL_746540, EPI_ISL_746541, EPI_ISL_746542, EPI_ISL_746757, EPI_ISL_746758, EPI_ISL_746759, EPI_ISL_746760, EPI_ISL_746761, EPI_ISL_746762, EPI_ISL_746763, EPI_ISL_746764 | | | |
| see above | Genetica Molecular and Subdepartamento de Virologia ISP | Instituto de Salud Publica de Chile | Javier Tognarelli, Barbara Parra, Loredana Arata, Jaime Lagos, Gisselle Barra, Patricia Bustos, Rodrigo Fasce, Andres Castillo, Jorge Fernandez |

| | | | |
|---|---|---|---|
| | Chile | | |
| EPI_ISL_746882 | Utah Public Health Laboratory | Utah Public Health Laboratory | Erin Young, Kelly Oakeson, Tara Gallagher |
| EPI_ISL_747320, EPI_ISL_747321, EPI_ISL_747324, EPI_ISL_747325, EPI_ISL_747366, EPI_ISL_747367, EPI_ISL_747368 | Division of Emerging Infectious Diseases, Bureau of Infectious Diseases Diagnosis Control, Korea Disease Control and Prevention Agency | Division of Emerging Infectious Diseases, Bureau of Infectious Diseases Diagnosis Control, Korea Disease Control and Prevention Agency | Ae Kyung Park, Il-Hwan Kim, Heui Man Kim, Jeong-Min Kim, Namjoo Lee, Chaeyoung Lee, Sang Hee Woo, Eun-Jin Kim |
| EPI_ISL_747520 | ZOTZ KLIMAS MVZ Düsseldorf-Centrum GbR ÜBAG für Labormedizin, Genetik, Zytologie, Pathologie | Center of Medical Microbiology, Virology, and Hospital Hygiene, University of Duesseldorf | Maximilian Damagnez, Alexander Dilthey, Ashley-Jane Duplessis, Patrick Finzer, Katrin Hoffmann, Torsten Houwaart, Lisanna Hülse, Malte Kohns Vasconcelos, Marek Korencak, Nadine Lübke, Jessica Nicolai, Klaus Pfeffer, Daniel Strelow, Jörg Timm, Andreas Walker, Tobias Wienemann, Rainer Zotz |
| EPI_ISL_747749, EPI_ISL_748129, EPI_ISL_748130 | Department of Virus and Microbiological Special Diagnostics, Statens Serum Institut, Copenhagen, Denmark | Albertsen Lab, Department of Chemistry and Bioscience, Aalborg University, Denmark | Danish Covid-19 Genome Consortium |
| EPI_ISL_751480, EPI_ISL_751486, EPI_ISL_751496 | CHU Purpan - Laboratoire de Virologie - Institut Fédératif de Biologie | CHU Purpan - Laboratoire de Virologie - Institut Fédératif de Biologie | Latour J., Ranger N., Dubois M., Carcenac R., Harter A., Boyer P., Tremeaux P., Izopet J. |
| EPI_ISL_752984, EPI_ISL_752985, EPI_ISL_752987, EPI_ISL_752988, EPI_ISL_752989, EPI_ISL_752990, EPI_ISL_752991, EPI_ISL_752992, EPI_ISL_752993, EPI_ISL_752994, EPI_ISL_752995, EPI_ISL_753004, EPI_ISL_753005, EPI_ISL_753006, EPI_ISL_753007, EPI_ISL_753008, EPI_ISL_753010, EPI_ISL_753011, EPI_ISL_753012, EPI_ISL_753013, EPI_ISL_753014, EPI_ISL_753015, EPI_ISL_753016, EPI_ISL_753032, EPI_ISL_753033, EPI_ISL_753034, EPI_ISL_753035, EPI_ISL_753036, EPI_ISL_753037 | | | |
| see above | State Laboratories Division, Hawaii State Department of Health | State Laboratories Division, Hawaii State Department of Health | Pamela O'Brien, Sabrina Diemert, Drew Kuwazaki, Razvan Sultana, Edward Desmond |
| EPI_ISL_753707, EPI_ISL_753717, EPI_ISL_753779, EPI_ISL_753780, EPI_ISL_753781, EPI_ISL_753782, EPI_ISL_753783, EPI_ISL_753784, EPI_ISL_753789, EPI_ISL_753794, EPI_ISL_753795, EPI_ISL_753816, EPI_ISL_753817, EPI_ISL_753818, EPI_ISL_753859, EPI_ISL_753869, EPI_ISL_753870, EPI_ISL_753871, EPI_ISL_753872, EPI_ISL_753873, EPI_ISL_753874, EPI_ISL_753875, EPI_ISL_753882, EPI_ISL_753883, EPI_ISL_753890, EPI_ISL_753903, EPI_ISL_753904, EPI_ISL_753905, EPI_ISL_753906, EPI_ISL_753908, EPI_ISL_753913, EPI_ISL_753914, EPI_ISL_753915, EPI_ISL_754017 | | | |
| see above | Charité Universitätsmedizin Berlin, Institut für Virologie/Labor Berlin | Charité Universitätsmedizin Berlin, Institut für Virologie | Victor M Corman, Jörn Beheim-Schwarzbach, Barbara Mühlemann, Julia Schneider, Talitha Veith, Terry Jones, Christian Drosten |
| EPI_ISL_754138 | CHU Purpan - Laboratoire de Virologie - Institut Fédératif de Biologie | CHU Purpan - Laboratoire de Virologie - Institut Fédératif de Biologie | Latour J., Ranger N., Dubois M., Carcenac R., Harter A., Boyer P., Tremeaux P., Izopet J. |
| EPI_ISL_754183 | Wyoming Public Health Laboratory | Wyoming Public Health Laborator | Noah Hull, Taylor Fearing, Channing Weber, Ashley Norberg, Bailey Bowcutt, and Wanda Manley |
| EPI_ISL_754192, EPI_ISL_754193 | Charité Universitätsmedizin Berlin, Institut für Virologie/Labor Berlin | Charité Universitätsmedizin Berlin, Institut für Virologie | Victor M Corman, Jörn Beheim-Schwarzbach, Barbara Mühlemann, Julia Schneider, Talitha Veith, Terry Jones, Christian Drosten |
| EPI_ISL_754217, EPI_ISL_754218, EPI_ISL_754219, EPI_ISL_754220, EPI_ISL_754221, EPI_ISL_754222, EPI_ISL_754223, EPI_ISL_754224, EPI_ISL_754225, EPI_ISL_754226, EPI_ISL_754227, EPI_ISL_754228 | | | |
| see above | Wyoming Public Health Laboratory | Wyoming Public Health Laboratory | Noah Hull, Taylor Fearing, Channing Weber, Ashley Norberg, Bailey Bowcutt, and Wanda Manley |
| EPI_ISL_754235 | The Republican Research and Practical Center for Epidemiology and Microbiology (RRPCEM) | WHO National Influenza Centre Russian Federation | Elena Gasich, Kirill Bulda, Anatoly Krasko, Andrey Komissarov, Artem Fadeev, Anna Ivanova, Kseniya Komissarova, Dmitry Bazhenov, Daria Danilenko, Ksenia Safina, Elena Nabieva, Georgii Bazykin, Dmitry Lioznov |
| EPI_ISL_755065, EPI_ISL_755066 | California Department of Public Health | California Department of Public Health | CDPH IDLB COVIDNet |
| EPI_ISL_755210, EPI_ISL_755211, EPI_ISL_755212 | Biolab Diagnostic Laboratories | Andersen lab at Scripps Research | Issa Abu-Dayyeh, Ahmad Tibi, Lama Hussein, Lina Mohammad, Zein Naber, Amid Abdelnour with SEARCH Alliance San Diego |
| EPI_ISL_755810 | Toronto Invasive Bacterial Diseases Network | McMaster University | Allison McGeer, Patryk Aftanas, Hooman Derakhshani, Angel Li, Kuganya Nirmalarajah, Emily Panousis, Ahmed Draia, Jalees Nasir, Michael Surette, Samira Mubareka, Andrew G. McArthur |
| EPI_ISL_755943, EPI_ISL_755944, EPI_ISL_755945, EPI_ISL_755946, EPI_ISL_755947, EPI_ISL_755948, EPI_ISL_755949, EPI_ISL_755950, EPI_ISL_755951, EPI_ISL_755952, EPI_ISL_755953, EPI_ISL_755954, EPI_ISL_755955, EPI_ISL_755956, EPI_ISL_755957, EPI_ISL_755958, EPI_ISL_755959, EPI_ISL_755960, EPI_ISL_755961, EPI_ISL_755962, EPI_ISL_755963, EPI_ISL_755964, EPI_ISL_755965, EPI_ISL_755970, EPI_ISL_755971, EPI_ISL_755972, EPI_ISL_755973, EPI_ISL_755974, EPI_ISL_755975, EPI_ISL_755976, EPI_ISL_755977, EPI_ISL_755978, EPI_ISL_755979, EPI_ISL_755980, EPI_ISL_755981, EPI_ISL_755982, EPI_ISL_755983, EPI_ISL_755984, EPI_ISL_755985, EPI_ISL_755986, EPI_ISL_755987, EPI_ISL_755988, EPI_ISL_755989, EPI_ISL_755990, EPI_ISL_755991, EPI_ISL_755992, EPI_ISL_756014, EPI_ISL_756015, EPI_ISL_756016, EPI_ISL_756017, EPI_ISL_756018, EPI_ISL_756019, EPI_ISL_756020, EPI_ISL_756021, EPI_ISL_756022, EPI_ISL_756023, EPI_ISL_756024, EPI_ISL_756025, EPI_ISL_756026, EPI_ISL_756027, EPI_ISL_756028, EPI_ISL_756029, EPI_ISL_756030, EPI_ISL_756031, EPI_ISL_756032, EPI_ISL_756033, EPI_ISL_756034, EPI_ISL_756035, EPI_ISL_756036, EPI_ISL_756037, EPI_ISL_756038, EPI_ISL_756039, EPI_ISL_756040, EPI_ISL_756041, EPI_ISL_756042, EPI_ISL_756043, EPI_ISL_756044 | | | |
| see above | Department of Virology and Immunology, University of Helsinki and Helsinki University Hospital, Huslab Finland | Department of Virology, Faculty of Medicine, University of Helsinki, Helsinki, Finland | Teemu Smura, Ravi Kant, Phuoc Truong, Hussein Alburkat, Hannimari Kallio-Kokko, Jenni Virtanen, Maija Suvanto, Sari Hannula, Harri Kangas, Pekka Ellonen, Olli Vapalahti |
| EPI_ISL_759705 | RSUD Bung Karno, Surakarta | Universitas Sebelas Maret (UNS); Rumah Sakit Universitas Sebelas Maret (RS-UNS), Surakarta; National Institute of Health Research and Development, Indonesian Ministry of Health , Jakarta). | Betty Suryawati, Yulia Sari, Hartono, Maryani, Afif Avicena Gufron, Hana Apsari Pawestri, Kartika, Dewi Puspa, Hartanti Dian Ika, Arie Ardiansyah Nugraha, Vivi Setiawaty. |
| EPI_ISL_759970, EPI_ISL_759971, EPI_ISL_759972 | Department of Medical Microbiology, St. Olavs hospital | Norwegian Institute of Public Health, Department of Virology | Kathrine Stene-Johansen, Kamilla Heddeland Instefjord, Hilde Elshaug, Marie Paulsen Madsen, Rasmus Riis Kopperud, Hilde Vollan, Karoline Bragstad, Olav Hungnes |
| EPI_ISL_759991, EPI_ISL_759995, EPI_ISL_759996, EPI_ISL_759997, EPI_ISL_759998, EPI_ISL_759999, EPI_ISL_760000 | Division of Emerging Infectious Diseases, Bureau of Infectious Diseases Diagnosis Control, Korea Disease Control and Prevention Agency | Division of Emerging Infectious Diseases, Bureau of Infectious Diseases Diagnosis Control, Korea Disease Control and Prevention Agency | Ae Kyung Park, Il-Hwan Kim, Heui Man Kim, Jeong-Min Kim, Namjoo Lee, Chaeyoung Lee, Sang Hee Woo, Eun-Jin Kim |
| EPI_ISL_760049, EPI_ISL_760050, EPI_ISL_760051, EPI_ISL_760052, EPI_ISL_760053, EPI_ISL_760054, EPI_ISL_760055, EPI_ISL_760056 | Hong Kong Department of Health | School of Public Health, The University of Hong Kong | Daniel Chu, Haogao Gu, Pavithra Krishnan, Daisy Ng, Gigi Liu, Carrie Wan, Malik Peiris, Leo Poon |
| EPI_ISL_760062, EPI_ISL_760063, EPI_ISL_760116 | Division of Emerging Infectious Diseases, Bureau of Infectious Diseases Diagnosis Control, Korea Disease Control and Prevention Agency | Division of Emerging Infectious Diseases, Bureau of Infectious Diseases Diagnosis Control, Korea Disease Control and Prevention Agency | Ae Kyung Park, Il-Hwan Kim, Heui Man Kim, Jeong-Min Kim, Namjoo Lee, Chaeyoung Lee, Sang Hee Woo, Eun-Jin Kim |
| EPI_ISL_763063 | RS Panti Waluyo, Surakarta | Universitas Sebelas Maret (UNS); Rumah Sakit UNS (RS-UNS); National Institute of Health Research and Development, Indonesian Ministry of Health. | Betty Suryawati, Yulia Sari, Hartono, Maryani, Afif A Ghufron, Revi G H Novika, Hana A Pawestri, Kartika, Dewi Puspa, Hartanti Dian Ika, Arie A Nugraha, Vivi Setiawaty. |
| EPI_ISL_763066 | Puskesmas Grogol, Sukoharjo | Universitas Sebelas Maret (UNS); Rumah Sakit UNS (RS-UNS); National Institute of Health Research and Development, Indonesian Ministry of Health. | Yulia Sari, Hartono, Maryani, Betty Suryawati, Afif A Ghufron, Revi G H Novika, Hana A Pawestri, Kartika, Dewi Puspa, Hartanti Dian Ika, Arie A Nugraha, Vivi Setiawaty. |
| EPI_ISL_765751, EPI_ISL_765752, EPI_ISL_765753, EPI_ISL_765754, EPI_ISL_765755 | Massachusetts General Hospital | Infectious Disease Program, Broad Institute of Harvard and MIT | Lemieux,J.E., Siddle,K.J., Shaw,B., Adams,G., Pierce,V., Turbett,S., Anahtar,M., Branda,J., Slater,D., Harris,J., Lin,A.E., Gladden-Young,A., Lagerborg,K., Rudy,M., DeRuff,K., Carter,A., Normandin,E., Bauer,M., Reilly,S., Tomkins-Tinch,C., Loreth,C., Chaluvadi,S., Neumann,A., Cusick,C., Chapman,S.B., Gnirke,A., Flowers,K., Cerrato,F., Birren,B.W., Gallagher,G., Smole,S., Park,D.J., MacInnis,B.L., Ryan,E., LaRocque,R., Rosenberg,E. and Sabeti,P.C. |
| EPI_ISL_765952 | Worobey Lab, Department of Ecology and Evolutionary Biology, University of Arizona | Worobey Lab, Department of Ecology and Evolutionary Biology, University of Arizona | Brendan Larsen, Grace Quirk, Thomas Watts, David Baltrus, Michael Worobey |
| EPI_ISL_766882 | Wyoming Public Health Laboratory | Wyoming Public Health Laboratory | Noah Hull, Taylor Fearing, Lynette Gumbleton, Channing Weber, Ashley Norberg, Bailey Bowcutt, and Wanda Manley |
| EPI_ISL_768755, EPI_ISL_768775, EPI_ISL_768776, EPI_ISL_768777, | AIID | Irish Coronavirus Sequencing Consortium - National Virus Reference Laboratory | Michael Carr, Gabriel Gonzalez, Alejandro Abner Garcia Leon, Patrick Mallon |

| | | | |
|---|---|---|---|
| EPI_ISL_768778, EPI_ISL_768779 | | | |
| EPI_ISL_770001, EPI_ISL_770002 | Area De Salud La Cruz | Inciensa, Instituto Costarricense de Investigación y Enseñanza en Nutrición y Salud | Francisco Duarte, Hebleen Porras, Claudio Soto-Garita, Estela Cordero, Adriana Godínez, Melany Calderón & Mariel López |
| EPI_ISL_770065, EPI_ISL_770079, EPI_ISL_770080, EPI_ISL_770081, EPI_ISL_770082, EPI_ISL_770083, EPI_ISL_770084, EPI_ISL_770085 | Wyoming Public Health Laboratory | Wyoming Public Health Laboratory | Noah Hull, Taylor Fearing, Lynette Gumbleton, Channing Weber, Ashley Norberg, Bailey Bowcutt, and Wanda Manley |
| EPI_ISL_770734 | ZOTZ KLIMAS MVZ Düsseldorf-Centrum GbR ÜBAG für Labormedizin, Genetik, Zytologie, Pathologie | Center of Medical Microbiology, Virology, and Hospital Hygiene, University of Duesseldorf | Maximilian Damagnez, Alexander Dilthey, Ashley-Jane Duplessis, Patrick Finzer, Katrin Hoffmann, Torsten Houwaart, Lisanna Hülse, Malte Kohns Vasconcelos, Marek Korencak, Nadine Lübke, Jessica Nicolai, Klaus Pfeffer, Daniel Strelow, Jörg Timm, Andreas Walker, Tobias Wienemann, Rainer Zotz |
| EPI_ISL_770826 | Eastern Ontario Regional Laboratory Association | McMaster University | Leanne Mortimer, Hooman Derakhshani, Emily Panousis, Ahmed Draia, Jalees Nasir, Robert Slinger, Andrew G. McArthur |
| EPI_ISL_771371 | CardioVir EA-4684, University of Reims Champagne Ardenne | CardioVir EA-4684, University of Reims Champagne Ardenne | Callon,D., N'Guyen,Y., Lebreil,A.-L., Fornes,P. and Andreoletti,L. |
| EPI_ISL_775311 | Medical Microbiology Unit, Department for Laboratory Medicine, Drammen Hospital, Vestre Viken Health Trust, | Norwegian Institute of Public Health, Department of Virology | Kathrine Stene-Johansen, Kamilla Heddeland Instefjord, Hilde Elshaug, Atiya R Ali,Marie Paulsen Madsen, Rasmus Riis Kopperud, Hilde Vollan, Karoline Bragstad, Olav Hungnes |
| EPI_ISL_775312, EPI_ISL_775313 | Nordland Hospital - Bodo, Laboratory Department, Molecular Biology Unit | Norwegian Institute of Public Health, Department of Virology | Kathrine Stene-Johansen, Kamilla Heddeland Instefjord, Hilde Elshaug, Atiya R Ali,Marie Paulsen Madsen, Rasmus Riis Kopperud, Hilde Vollan, Karoline Bragstad, Olav Hungnes |
| EPI_ISL_775316, EPI_ISL_775318 | Oslo University Hospital, Department of Medical Microbiology | Norwegian Institute of Public Health, Department of Virology | Kathrine Stene-Johansen, Kamilla Heddeland Instefjord, Hilde Elshaug, Atiya R Ali,Marie Paulsen Madsen, Rasmus Riis Kopperud, Hilde Vollan, Karoline Bragstad, Olav Hungnes |
| EPI_ISL_775322 | Ostfold Hospital Trust - Kalnes, Centre for Laboratory Medicine, Section for gene technology and infection serology | Norwegian Institute of Public Health, Department of Virology | Kathrine Stene-Johansen, Kamilla Heddeland Instefjord, Hilde Elshaug, Atiya R Ali,Marie Paulsen Madsen, Rasmus Riis Kopperud, Hilde Vollan, Karoline Bragstad, Olav Hungnes |
| EPI_ISL_775447, EPI_ISL_775448 | University Hospital of Northern Norway, Department for Microbiology and Infectious Disease Control | Norwegian Institute of Public Health, Department of Virology | Kathrine Stene-Johansen, Kamilla Heddeland Instefjord, Hilde Elshaug, Atiya R Ali,Marie Paulsen Madsen, Rasmus Riis Kopperud, Hilde Vollan, Karoline Bragstad, Olav Hungnes |
| EPI_ISL_775449 | Vestfold Hospital, Toensberg Department of Microbiology | Norwegian Institute of Public Health, Department of Virology | Kathrine Stene-Johansen, Kamilla Heddeland Instefjord, Hilde Elshaug, Atiya R Ali,Marie Paulsen Madsen, Rasmus Riis Kopperud, Hilde Vollan, Karoline Bragstad, Olav Hungnes |
| EPI_ISL_775459 | Norwegian Institute of Public Health, Department of Virology | Norwegian Institute of Public Health, Department of Virology | Kathrine Stene-Johansen, Kamilla Heddeland Instefjord, Hilde Elshaug, Atiya R Ali,Marie Paulsen Madsen, Rasmus Riis Kopperud, Hilde Vollan, Karoline Bragstad, Olav Hungnes |
| EPI_ISL_775472, EPI_ISL_775473 | Vestfold Hospital, Toensberg Department of Microbiology | Norwegian Institute of Public Health, Department of Virology | Kathrine Stene-Johansen, Kamilla Heddeland Instefjord, Hilde Elshaug, Atiya R Ali,Marie Paulsen Madsen, Rasmus Riis Kopperud, Hilde Vollan, Karoline Bragstad, Olav Hungnes |
| EPI_ISL_775476, EPI_ISL_775477 | Unilabs Laboratory Medicine | Norwegian Institute of Public Health, Department of Virology | Kathrine Stene-Johansen, Kamilla Heddeland Instefjord, Hilde Elshaug, Atiya R Ali,Marie Paulsen Madsen, Rasmus Riis Kopperud, Hilde Vollan, Karoline Bragstad, Olav Hungnes |
| EPI_ISL_776661, EPI_ISL_776694 | UW Virology Lab | UW Virology Lab | Pavitra Roychoudhury, Hong Xie, Lasata Shrestha, Meei-Li Huang, Keith R Jerome, Alexander Greninger |
| EPI_ISL_779929 | Center of Medical Microbiology, Virology, and Hospital Hygiene, University of Duesseldorf | Center of Medical Microbiology, Virology, and Hospital Hygiene, University of Duesseldorf | Maximilian Damagnez, Alexander Dilthey, Ashley-Jane Duplessis, Torsten Houwaart, Lisanna Hülse, Malte Kohns Vasconcelos, Nadine Lübke, Jessica Nicolai, Klaus Pfeffer, Daniel Strelow, Teresa Tamayo, Jörg Timm, Andreas Walker, Tobias Wienemann |
| EPI_ISL_780028 | Hospital General Universitario Gregorio Marañón | SeqCOVID-SPAIN consortium/IBV(CSIC) | Darío García de Viedma, Laura Pérez-Lago, Marta Herranz, Jon Sicilia, Julia Suárez, Pilar Catalán, Patricia Muñoz and SeqCOVID-SPAIN consortium |
| EPI_ISL_786198, EPI_ISL_786199, EPI_ISL_786204, EPI_ISL_786205, EPI_ISL_786206, EPI_ISL_786207, EPI_ISL_786208, EPI_ISL_786209, EPI_ISL_786210, EPI_ISL_786211, EPI_ISL_786216, EPI_ISL_786217, EPI_ISL_786218, EPI_ISL_786219, EPI_ISL_786220, EPI_ISL_786221, EPI_ISL_786222, EPI_ISL_786223, EPI_ISL_786228, EPI_ISL_786229, EPI_ISL_786230, EPI_ISL_786231, EPI_ISL_786232, EPI_ISL_786233, EPI_ISL_786234, EPI_ISL_786235, EPI_ISL_786240, EPI_ISL_786241, EPI_ISL_786242, EPI_ISL_786243, EPI_ISL_786244, EPI_ISL_786245, EPI_ISL_786246, EPI_ISL_786247, EPI_ISL_786251, EPI_ISL_786252, EPI_ISL_786253, EPI_ISL_786254, EPI_ISL_786255, EPI_ISL_786256, EPI_ISL_786257, EPI_ISL_786258, EPI_ISL_786259, EPI_ISL_786260, EPI_ISL_786261, EPI_ISL_786262, EPI_ISL_786263, EPI_ISL_786264, EPI_ISL_786272, EPI_ISL_786273, EPI_ISL_786274, EPI_ISL_786275, EPI_ISL_786276, EPI_ISL_786284, EPI_ISL_786285, EPI_ISL_786286, EPI_ISL_786287, EPI_ISL_786296, EPI_ISL_786297, EPI_ISL_786298, EPI_ISL_786304, EPI_ISL_786307, EPI_ISL_786308, EPI_ISL_786309, EPI_ISL_786310, EPI_ISL_786313, EPI_ISL_786319, EPI_ISL_786320, EPI_ISL_786321, EPI_ISL_786322, EPI_ISL_786331, EPI_ISL_786332, EPI_ISL_786333, EPI_ISL_786334, EPI_ISL_786343, EPI_ISL_786344, EPI_ISL_786345, EPI_ISL_786346, EPI_ISL_786349, EPI_ISL_786391, EPI_ISL_787635, EPI_ISL_787678, EPI_ISL_787679, EPI_ISL_787680, EPI_ISL_787681, EPI_ISL_787682, EPI_ISL_787683, EPI_ISL_787684, EPI_ISL_787685, EPI_ISL_787686, EPI_ISL_787687, EPI_ISL_787690, EPI_ISL_787691, EPI_ISL_787692, EPI_ISL_787693, EPI_ISL_787694, EPI_ISL_787695, EPI_ISL_787696, EPI_ISL_787697, EPI_ISL_787698, EPI_ISL_787699, EPI_ISL_787702, EPI_ISL_787703, EPI_ISL_787704, EPI_ISL_787705, EPI_ISL_787706, EPI_ISL_787707, EPI_ISL_787708, EPI_ISL_787709, EPI_ISL_787710, EPI_ISL_787711, EPI_ISL_787714, EPI_ISL_787715, EPI_ISL_787716, EPI_ISL_787717, EPI_ISL_787718, EPI_ISL_787719, EPI_ISL_787720, EPI_ISL_787721, EPI_ISL_787722, EPI_ISL_787723, EPI_ISL_787725, EPI_ISL_787726, EPI_ISL_787727, EPI_ISL_787728, EPI_ISL_787729, EPI_ISL_787730, EPI_ISL_787731, EPI_ISL_787732, EPI_ISL_787733, EPI_ISL_787734, EPI_ISL_787735, EPI_ISL_787737, EPI_ISL_787738, EPI_ISL_787739, EPI_ISL_787740, EPI_ISL_787741, EPI_ISL_787742, EPI_ISL_787743, EPI_ISL_787744, EPI_ISL_787745, EPI_ISL_787746, EPI_ISL_787747, EPI_ISL_787749, EPI_ISL_787750, EPI_ISL_787751, EPI_ISL_787752, EPI_ISL_787753, EPI_ISL_787754, EPI_ISL_787755, EPI_ISL_787756, EPI_ISL_787757, EPI_ISL_787758, EPI_ISL_787759, EPI_ISL_787761, EPI_ISL_787762, EPI_ISL_787763, EPI_ISL_787764, EPI_ISL_787765, EPI_ISL_787766, EPI_ISL_787767, EPI_ISL_787768, EPI_ISL_787769, EPI_ISL_787770, EPI_ISL_787771, EPI_ISL_787772, EPI_ISL_787773, EPI_ISL_787774, EPI_ISL_787775, EPI_ISL_787776, EPI_ISL_787777, EPI_ISL_787778, EPI_ISL_787779, EPI_ISL_787780, EPI_ISL_787781, EPI_ISL_787782, EPI_ISL_787783, EPI_ISL_787784, EPI_ISL_787785, EPI_ISL_787786, EPI_ISL_787787, EPI_ISL_787788, EPI_ISL_787789, EPI_ISL_787790, EPI_ISL_787791, EPI_ISL_787792, EPI_ISL_787793, EPI_ISL_787794, EPI_ISL_787795, EPI_ISL_787796, EPI_ISL_787797, EPI_ISL_787798, EPI_ISL_787799, EPI_ISL_787800, EPI_ISL_787801, EPI_ISL_787802, EPI_ISL_787803, EPI_ISL_787804, EPI_ISL_787805, EPI_ISL_787806, EPI_ISL_787807, EPI_ISL_787808, EPI_ISL_787809, EPI_ISL_787810, EPI_ISL_787811, EPI_ISL_787812, EPI_ISL_787813, EPI_ISL_787814, EPI_ISL_787815, EPI_ISL_787816, EPI_ISL_787817, EPI_ISL_787818, EPI_ISL_787819, EPI_ISL_787820, EPI_ISL_787821, EPI_ISL_787822, EPI_ISL_787823, EPI_ISL_787824, EPI_ISL_787825, EPI_ISL_787826, EPI_ISL_787827, EPI_ISL_787828, EPI_ISL_787829, EPI_ISL_787830, EPI_ISL_787831, EPI_ISL_787832, EPI_ISL_787833, EPI_ISL_787834, EPI_ISL_787835, EPI_ISL_787836, EPI_ISL_787837, EPI_ISL_787838, EPI_ISL_787839, EPI_ISL_787840, EPI_ISL_787841, EPI_ISL_787842, EPI_ISL_787843, EPI_ISL_787844, EPI_ISL_787845, EPI_ISL_787846, EPI_ISL_787847, EPI_ISL_787848, EPI_ISL_787849, EPI_ISL_787850, EPI_ISL_787851, EPI_ISL_787852, EPI_ISL_787853, EPI_ISL_787854, EPI_ISL_787855, EPI_ISL_787856, EPI_ISL_787857, EPI_ISL_787858, EPI_ISL_787859, EPI_ISL_787860, EPI_ISL_787861, EPI_ISL_787862, EPI_ISL_787863, EPI_ISL_787864, EPI_ISL_787865, EPI_ISL_787866, EPI_ISL_787867, EPI_ISL_787868, EPI_ISL_787869, EPI_ISL_787870, EPI_ISL_787871, EPI_ISL_787872, EPI_ISL_787873, EPI_ISL_787874, EPI_ISL_787875, EPI_ISL_787876, EPI_ISL_787877, EPI_ISL_787878, EPI_ISL_787879, EPI_ISL_787880, EPI_ISL_787881, EPI_ISL_787882, EPI_ISL_787883, EPI_ISL_787884, EPI_ISL_787885, EPI_ISL_787886, EPI_ISL_787887, EPI_ISL_787888, EPI_ISL_787889, EPI_ISL_787890, EPI_ISL_787891, EPI_ISL_787892, EPI_ISL_787893, EPI_ISL_787894, EPI_ISL_787895, EPI_ISL_787896, EPI_ISL_787897, EPI_ISL_787898, EPI_ISL_787899, EPI_ISL_787900, EPI_ISL_787901, EPI_ISL_787902, EPI_ISL_787903, EPI_ISL_787904, EPI_ISL_787905, EPI_ISL_787906, EPI_ISL_787907, EPI_ISL_787908, EPI_ISL_787909, EPI_ISL_787910, EPI_ISL_787911, EPI_ISL_787912, EPI_ISL_787913, EPI_ISL_787914, EPI_ISL_787915, EPI_ISL_787916, EPI_ISL_787917, EPI_ISL_787918, EPI_ISL_787919, EPI_ISL_787920, EPI_ISL_787921, EPI_ISL_787922, EPI_ISL_787923, EPI_ISL_787924, EPI_ISL_787925, EPI_ISL_787926, EPI_ISL_787927, EPI_ISL_787928, EPI_ISL_787929, EPI_ISL_787930, EPI_ISL_787931, EPI_ISL_787932, EPI_ISL_787933, EPI_ISL_787934, EPI_ISL_787935, EPI_ISL_787936, EPI_ISL_787937, EPI_ISL_787938, EPI_ISL_787939, EPI_ISL_787940, EPI_ISL_787941, EPI_ISL_787942, EPI_ISL_787943, EPI_ISL_787944, EPI_ISL_787945, EPI_ISL_787946, EPI_ISL_787947, EPI_ISL_787948, EPI_ISL_787949, EPI_ISL_787950, EPI_ISL_787951, EPI_ISL_787952, EPI_ISL_787953, EPI_ISL_787954, EPI_ISL_787955, EPI_ISL_787956, EPI_ISL_787957, EPI_ISL_787958, EPI_ISL_787959, EPI_ISL_787960, EPI_ISL_787961, EPI_ISL_787962, EPI_ISL_787963, EPI_ISL_787968, EPI_ISL_787969, EPI_ISL_787970, EPI_ISL_787971, EPI_ISL_787972, EPI_ISL_787973, EPI_ISL_787974, EPI_ISL_787975, EPI_ISL_787980, EPI_ISL_787981, EPI_ISL_787982, EPI_ISL_787983, EPI_ISL_787984, EPI_ISL_787985, EPI_ISL_787986, EPI_ISL_787987, EPI_ISL_787992, EPI_ISL_787993, EPI_ISL_787994, EPI_ISL_787995, EPI_ISL_787996, EPI_ISL_787997 | | | |
| see above | Houston Methodist Hospital | Houston Methodist Hospital | S. Wesley Long, Randall J. Olsen, Paul A. Christensen, David W. Bernard, James J. Davis, Maulik Shukla, Marcus Nguyen, Matthew Ojeda Saavedra, Prasanti Yerramilli, Layne Pruitt, Sishir Subedi, Heather Hendrickson, and James M. Musser |
| EPI_ISL_791986 | RSUD Morotai Maluku Utara | National Institute of Health Research and Development | Subangkit;Pawestri,HA;Ikawati,HD;Nugraha,AA;Puspa,KD;Denggo,SD;Pangesti,KNA;Soekarso,T;Puspandari,N;Setiawaty,V |
| EPI_ISL_792021, EPI_ISL_792022 | Plateforme COVID IDF | National Reference Center for Viruses of Respiratory Infections, Institut Pasteur, Paris | Marion Barbet, Sylvie Behillil, Méline Bizard, Angela Brisebarre, Camille Capel, Etienne Simon-Lorière, Vincent Enouf, Maud Vanpeene, Sylvie van der Werf,Jacques Fourgeaud |
| EPI_ISL_792622, EPI_ISL_792623 | LACEN-PB | Laboratory of Respiratory Viruses and Measles, Oswaldo Cruz Institute, FIOCRUZ | Paola Resende, Luciana Appolinario, Fernando Motta, Anna Carolina Paixao, Ana Carolina Mendonca, João Felipe Bezerra, Romero Henrique Teixeira de Vasconcelos, Dalane Loudal Florentino Teixeira, Thiago Franco de Oliveira Carneiro, Marilda Siqueira |
| EPI_ISL_796061 | Institute of Virology, University of Cologne | Institute of Virology, University of Cologne | Saleta Sierra, Gibran Rubio, Zevanya Tessalonica, Dominik Aschenmeier, Eva Heger, Elena Knops, Rolf Kaiser, Maximilian Damagnez, Andreas Walker, Jörg Timm, Alexander Dilthey, Martin Däumer, Alex Thielen |
| EPI_ISL_801876, EPI_ISL_801901, EPI_ISL_801939, EPI_ISL_801955, EPI_ISL_801973, EPI_ISL_801974, EPI_ISL_801975, EPI_ISL_801979, EPI_ISL_802357, EPI_ISL_802358, EPI_ISL_802359, EPI_ISL_802360, EPI_ISL_802361, EPI_ISL_802362, EPI_ISL_802363, EPI_ISL_802364, EPI_ISL_802365, EPI_ISL_802366, EPI_ISL_802367, EPI_ISL_802368, EPI_ISL_802369, EPI_ISL_802370, EPI_ISL_802371, EPI_ISL_802372, EPI_ISL_802373, EPI_ISL_802374, EPI_ISL_802375, EPI_ISL_802376, EPI_ISL_802377, EPI_ISL_802378, EPI_ISL_802379, EPI_ISL_802380, EPI_ISL_802381, EPI_ISL_802382, EPI_ISL_802383, EPI_ISL_802384, EPI_ISL_802385, EPI_ISL_802386, EPI_ISL_802387, EPI_ISL_802388, EPI_ISL_802389, EPI_ISL_802390, EPI_ISL_802391, EPI_ISL_802392, EPI_ISL_802393, EPI_ISL_802394, EPI_ISL_802395, EPI_ISL_802396, EPI_ISL_802397, EPI_ISL_802398, EPI_ISL_802399, EPI_ISL_802400, EPI_ISL_802401, EPI_ISL_802402, EPI_ISL_802403, EPI_ISL_802404, EPI_ISL_802405, EPI_ISL_802406, EPI_ISL_802407, EPI_ISL_802408, EPI_ISL_802409, EPI_ISL_802410, EPI_ISL_802411, EPI_ISL_802412, EPI_ISL_802413, EPI_ISL_802414, EPI_ISL_802415, EPI_ISL_802416 | | | |
| see above | MSHS Clinical Microbiology Laboratories | MSHS Pathogen Surveillance Program | Ana S. Gonzalez-Reiche, Hala Alshammary, Mitchell J. Sullivan, Brianne Ciferri, Ajay Obla, Angela Amoako, Mahmoud Awawda, Elena Hirsch, Ashley S. Salimbangon, Levy Sominsky, Katherine Beach, Kayla Russo, Charles Gleason, Shelcie Fabre, Giulio Kleiner, Zenab Khan, Bremy Alburquerque, Adriana van de Guchte, Komal Srivastava, Matthew M. Hernandez, Jayeeta Dutta, Denise Jurczyszak, Emily Ferreri, Rachel Chernet, Nancy Francoeur, Betsaida Salom Melo, Irina Oussenko, Gintaras Deikus, Juan Soto, Shwetha Hara Sridhar, Ying-Chih Wang, Kathryn Twyman, Andrew Kasarskis, Deena R. Altman, Robert Sebra, Adolfo García-Sastre, Marta Luksza, Gopi Patel, Sarah Schaefer, Melissa Gitman, Michael D. Nowak, Alberto Paniz-Mondolfi, Emilia Mia Sordillo, Viviana Simon, Harm van Bakel |

| | | | |
|---|---|---|---|
| EPI_ISL_802572 | UMN Advanced Research and Diagnostics | Minnesota Department of Health, Public Health Laboratory | Alexandra Lorentz, Jacob Garfin, Matt Plumb, and Xiong Wang |
| EPI_ISL_803254, EPI_ISL_803259, EPI_ISL_803260, EPI_ISL_803266, EPI_ISL_803289, EPI_ISL_803290, EPI_ISL_803291, EPI_ISL_803292, EPI_ISL_803293, EPI_ISL_803294, EPI_ISL_803295, EPI_ISL_803296, EPI_ISL_803297, EPI_ISL_803298, EPI_ISL_803299, EPI_ISL_803360, EPI_ISL_803361, EPI_ISL_803362, EPI_ISL_803363, EPI_ISL_803364, EPI_ISL_803365, EPI_ISL_803366, EPI_ISL_803367, EPI_ISL_803368, EPI_ISL_803382, EPI_ISL_803383, EPI_ISL_803384, EPI_ISL_803385, EPI_ISL_803386, EPI_ISL_803387, EPI_ISL_803388, EPI_ISL_803389, EPI_ISL_803390, EPI_ISL_803391, EPI_ISL_803398, EPI_ISL_803645, EPI_ISL_803647, EPI_ISL_803648, EPI_ISL_803649, EPI_ISL_803650, EPI_ISL_803651, EPI_ISL_803652, EPI_ISL_803653, EPI_ISL_803654, EPI_ISL_803655, EPI_ISL_803656, EPI_ISL_803657, EPI_ISL_803675, EPI_ISL_803676, EPI_ISL_803677, EPI_ISL_803678, EPI_ISL_803679, EPI_ISL_803680, EPI_ISL_803681, EPI_ISL_803682, EPI_ISL_803683, EPI_ISL_803684, EPI_ISL_803685, EPI_ISL_803686, EPI_ISL_803687, EPI_ISL_803734, EPI_ISL_803735, EPI_ISL_803736, EPI_ISL_803737, EPI_ISL_803738, EPI_ISL_803760, EPI_ISL_803761, EPI_ISL_803762, EPI_ISL_803763, EPI_ISL_803764, EPI_ISL_803765, EPI_ISL_803766, EPI_ISL_803767, EPI_ISL_803778, EPI_ISL_803789, EPI_ISL_803790 | | | |
| see above | Wisconsin State Laboratory of Hygiene Communicable Disease Division | Wisconsin State Laboratory of Hygiene Communicable Disease Division | Kelsey R. Florek, Abigail C. Shockey |
| EPI_ISL_804264 | Respiratory Virus Unit, National Infection Service, Public Health England | COVID-19 Genomics UK (COG-UK) Consortium | PHE Covid Sequencing Team |
| EPI_ISL_804468, EPI_ISL_804473, EPI_ISL_804477, EPI_ISL_804489, EPI_ISL_804490, EPI_ISL_804491, EPI_ISL_804492, EPI_ISL_804498, EPI_ISL_804499, EPI_ISL_804500, EPI_ISL_804535, EPI_ISL_804536 | | | |
| see above | MEPHI, Aix Marseille University | MEPHI, Aix Marseille University | Anthony LEVASSEUR |
| EPI_ISL_813657, EPI_ISL_813658, EPI_ISL_813659, EPI_ISL_813663, EPI_ISL_813671 | Liverpool Clinical Laboratories | COVID-19 Genomics UK (COG-UK) Consortium | Sam Haldenby, Anita Lucaci, Steve Paterson, Julian Hiscox, Alistair Darby, M Almsaud, A Alrezaihi, Muhannad Alruwaili, Stuart D Armstrong, Jones Benjamin, Eleanor G Bentley, Anu Chawla, Jordan J Clark, Angela Cowell, Richard Eccles, Isabel García-Dorival, Matthew Gemmell, Alessandro Gerada, PKF Gilmore, Richard Gregory, Ximeng Han, Catherine Hartley, Margaret Hughes, Miren Iturriza-Gomara, James Johnson, L Luu, Jenifer Manson, Charlotte Nelson, Elaine O'Toole, Cassie Olateju, Rebekah Penrice-Randal , Lucille Rainbow, N.P Randle, Trevor Ian Robinson, Parul Sharma, Ghada T Shawli, James P Stewart, Neil Swainston, Ecaterina Vamos, Joanne Watts, Mark Whitehead |
| EPI_ISL_816362 | Virology Department, Sheffield Teaching Hospitals NHS Foundation Trust/Department of Infection, Immunity and Cardiovascular Disease, The Medical School, University of Sheffield | COVID-19 Genomics UK (COG-UK) Consortium | Thushan de Silva, Matthew Parker, Nikki Smith, Adri Angyal, Rebecca Brown, Luke Green, Rachel Tucker, Paul Parsons, Danielle Groves, Katie Johnson, Laura Carrilero, Alex Keeley, Dave Partridge, Matthew Wyles, Benjamin Lindsey, Mehmet Yavuz, Mohammad Raza, Cariad Evans |
| EPI_ISL_819448 | Northumbria University / South Tees Hospitals NHS Foundation Trust / North Cumbria Integrated Care NHS Foundation Trust / North Tees and Hartlepool NHS Foundation Trust / Newcastle Hospitals NHS Foundation Trust | COVID-19 Genomics UK (COG-UK) Consortium | Darren L Smith,Andrew Nelson,Matthew Bashton,Greg R Young,Joshua Loh,John Allan,Mohammad A Tariq,Giles S Holt,Gary Black,Wen C Yew,Lynn Dover,Paul Baker,Steve Liggett,Sarah Essex,Jane Greenaway,Debra Padgett,Clive Graham,Garren Scott,Edward Barton,Emma Swindells,Brendan Payne,Jennifer Collins,Yusri Taha,Gary Eltringham |
| EPI_ISL_824024 | Dutch COVID-19 response team | National Institute for Public Health and the Environment (RIVM) | Adam Meijer, Harry Vennema, Jeroen Cremer, Sharon van den Brink, Bas van der Veer, AnneMarie van den Brandt, Florian Zwagemaker, Dennis Schmitz, Chantal Reusken, on behalf of the national COVID-19 response team |
| EPI_ISL_824509, EPI_ISL_824510 | Hospital Universitari Vall d'Hebron - Vall d'Hebron Institut de Rercerca | Hospital Universitari Vall d'Hebron | Cristina Andrés, Maria Piñana, Josep F Abril, Damir Garcia-Cehic, Ariadna Rando, Juliana Esperalba, Maria Gema Codina, Carla Castillo, Maria Carmen Martin, Tomàs Pumarola, Josep Quer, Andrés Antón |
| EPI_ISL_825577, EPI_ISL_825578, EPI_ISL_825579 | Respiratory Virus Unit, National Infection Service, Public Health England | COVID-19 Genomics UK (COG-UK) Consortium | PHE Covid Sequencing Team |
| EPI_ISL_826741, EPI_ISL_826748, EPI_ISL_826753, EPI_ISL_826754, EPI_ISL_826755, EPI_ISL_826756, EPI_ISL_826757, EPI_ISL_826758, EPI_ISL_826759, EPI_ISL_826760, EPI_ISL_826777, EPI_ISL_826778, EPI_ISL_826779, EPI_ISL_826780, EPI_ISL_826781, EPI_ISL_826782, EPI_ISL_826783, EPI_ISL_826784, EPI_ISL_826785, EPI_ISL_826786, EPI_ISL_826787 | | | |
| see above | deCODE genetics | deCODE genetics | Daniel F Gudbjartsson; Agnar Helgason; Hakon Jonsson; Olafur T Magnusson; Pall Melsted; Gudmundur L Norddahl; Jona Saemundsdottir; Asgeir Sigurdsson; Patrick Sulem; Arna B Agustsdottir; Hannes Eggertsson; Berglind Eiriksdottir; Run Fridriksdottir; Elisabet E Gardarsdottir; Gudmundur Georgsson; Olafia S Gretarsdottir; Kjartan R Gudmundsson; Thora R Gunnarsdottir; Arnaldur Gylfason; Hilma Holm; Brynjar O Jensson; Aslaug Jonasdottir; Kamilla S Josefsdottir; Thordur Kristjansson; Droplaug N Magnusdottir; Solvi Rognvaldsson; Louise le Roux; Gudrun Sigmundsdottir; Gardar Sveinbjornsson; Kristin E Sveinsdottir; Maney Sveinsdottir; Emil A Thorarensen; Bjarni Thorbjornsson; Gisli Masson; Ingileif Jonsdottir; Alma Moller; Thorolfur Gudnason; Karl G Kristinsson; Unnur Thorsteinsdottir; Kari Stefansson |
| EPI_ISL_826801 | The National University Hospital of Iceland | deCODE genetics | Daniel F Gudbjartsson; Agnar Helgason; Hakon Jonsson; Olafur T Magnusson; Pall Melsted; Gudmundur L Norddahl; Jona Saemundsdottir; Asgeir Sigurdsson; Patrick Sulem; Arna B Agustsdottir; Hannes Eggertsson; Berglind Eiriksdottir; Run Fridriksdottir; Elisabet E Gardarsdottir; Gudmundur Georgsson; Olafia S Gretarsdottir; Kjartan R Gudmundsson; Thora R Gunnarsdottir; Arnaldur Gylfason; Hilma Holm; Brynjar O Jensson; Aslaug Jonasdottir; Kamilla S Josefsdottir; Thordur Kristjansson; Droplaug N Magnusdottir; Solvi Rognvaldsson; Louise le Roux; Gudrun Sigmundsdottir; Gardar Sveinbjornsson; Kristin E Sveinsdottir; Maney Sveinsdottir; Emil A Thorarensen; Bjarni Thorbjornsson; Gisli Masson; Ingileif Jonsdottir; Alma Moller; Thorolfur Gudnason; Karl G Kristinsson; Unnur Thorsteinsdottir; Kari Stefansson |
| EPI_ISL_826831, EPI_ISL_826833, EPI_ISL_826835, EPI_ISL_826837, EPI_ISL_826838, EPI_ISL_826839, EPI_ISL_826840, EPI_ISL_826845, EPI_ISL_826846 | deCODE genetics | deCODE genetics | Daniel F Gudbjartsson; Agnar Helgason; Hakon Jonsson; Olafur T Magnusson; Pall Melsted; Gudmundur L Norddahl; Jona Saemundsdottir; Asgeir Sigurdsson; Patrick Sulem; Arna B Agustsdottir; Hannes Eggertsson; Berglind Eiriksdottir; Run Fridriksdottir; Elisabet E Gardarsdottir; Gudmundur Georgsson; Olafia S Gretarsdottir; Kjartan R Gudmundsson; Thora R Gunnarsdottir; Arnaldur Gylfason; Hilma Holm; Brynjar O Jensson; Aslaug Jonasdottir; Kamilla S Josefsdottir; Thordur Kristjansson; Droplaug N Magnusdottir; Solvi Rognvaldsson; Louise le Roux; Gudrun Sigmundsdottir; Gardar Sveinbjornsson; Kristin E Sveinsdottir; Maney Sveinsdottir; Emil A Thorarensen; Bjarni Thorbjornsson; Gisli Masson; Ingileif Jonsdottir; Alma Moller; Thorolfur Gudnason; Karl G Kristinsson; Unnur Thorsteinsdottir; Kari Stefansson |
| EPI_ISL_826862 | The National University Hospital of Iceland | deCODE genetics | Daniel F Gudbjartsson; Agnar Helgason; Hakon Jonsson; Olafur T Magnusson; Pall Melsted; Gudmundur L Norddahl; Jona Saemundsdottir; Asgeir Sigurdsson; Patrick Sulem; Arna B Agustsdottir; Hannes Eggertsson; Berglind Eiriksdottir; Run Fridriksdottir; Elisabet E Gardarsdottir; Gudmundur Georgsson; Olafia S Gretarsdottir; Kjartan R Gudmundsson; Thora R Gunnarsdottir; Arnaldur Gylfason; Hilma Holm; Brynjar O Jensson; Aslaug Jonasdottir; Kamilla S Josefsdottir; Thordur Kristjansson; Droplaug N Magnusdottir; Solvi Rognvaldsson; Louise le Roux; Gudrun Sigmundsdottir; Gardar Sveinbjornsson; Kristin E Sveinsdottir; Maney Sveinsdottir; Emil A Thorarensen; Bjarni Thorbjornsson; Gisli Masson; Ingileif Jonsdottir; Alma Moller; Thorolfur Gudnason; Karl G Kristinsson; Unnur Thorsteinsdottir; Kari Stefansson |
| EPI_ISL_826865, EPI_ISL_826866 | deCODE genetics | deCODE genetics | Daniel F Gudbjartsson; Agnar Helgason; Hakon Jonsson; Olafur T Magnusson; Pall Melsted; Gudmundur L Norddahl; Jona Saemundsdottir; Asgeir Sigurdsson; Patrick Sulem; Arna B Agustsdottir; Hannes Eggertsson; Berglind Eiriksdottir; Run Fridriksdottir; Elisabet E Gardarsdottir; Gudmundur Georgsson; Olafia S Gretarsdottir; Kjartan R Gudmundsson; Thora R Gunnarsdottir; Arnaldur Gylfason; Hilma Holm; Brynjar O Jensson; Aslaug Jonasdottir; Kamilla S Josefsdottir; Thordur Kristjansson; Droplaug N Magnusdottir; Solvi Rognvaldsson; Louise le Roux; Gudrun Sigmundsdottir; Gardar Sveinbjornsson; Kristin E Sveinsdottir; Maney Sveinsdottir; Emil A Thorarensen; Bjarni Thorbjornsson; Gisli Masson; Ingileif Jonsdottir; Alma Moller; Thorolfur Gudnason; Karl G Kristinsson; Unnur Thorsteinsdottir; Kari Stefansson |
| EPI_ISL_826981, EPI_ISL_826994 | The National University Hospital of Iceland | deCODE genetics | Daniel F Gudbjartsson; Agnar Helgason; Hakon Jonsson; Olafur T Magnusson; Pall Melsted; Gudmundur L Norddahl; Jona Saemundsdottir; Asgeir Sigurdsson; Patrick Sulem; Arna B Agustsdottir; Hannes Eggertsson; Berglind Eiriksdottir; Run Fridriksdottir; Elisabet E Gardarsdottir; Gudmundur Georgsson; Olafia S Gretarsdottir; Kjartan R Gudmundsson; Thora R Gunnarsdottir; Arnaldur Gylfason; Hilma Holm; Brynjar O Jensson; Aslaug Jonasdottir; Kamilla S Josefsdottir; Thordur Kristjansson; Droplaug N Magnusdottir; Solvi Rognvaldsson; Louise le Roux; Gudrun Sigmundsdottir; Gardar Sveinbjornsson; Kristin E Sveinsdottir; Maney Sveinsdottir; Emil A Thorarensen; Bjarni Thorbjornsson; Gisli Masson; Ingileif Jonsdottir; Alma Moller; Thorolfur Gudnason; Karl G Kristinsson; Unnur Thorsteinsdottir; Kari Stefansson |
| EPI_ISL_827002, EPI_ISL_827003, EPI_ISL_827004, EPI_ISL_827005, EPI_ISL_827007, EPI_ISL_827008, EPI_ISL_827010 | deCODE genetics | deCODE genetics | Daniel F Gudbjartsson; Agnar Helgason; Hakon Jonsson; Olafur T Magnusson; Pall Melsted; Gudmundur L Norddahl; Jona Saemundsdottir; Asgeir Sigurdsson; Patrick Sulem; Arna B Agustsdottir; Hannes Eggertsson; Berglind Eiriksdottir; Run Fridriksdottir; Elisabet E Gardarsdottir; Gudmundur Georgsson; Olafia S Gretarsdottir; Kjartan R Gudmundsson; Thora R Gunnarsdottir; Arnaldur Gylfason; Hilma Holm; Brynjar O Jensson; Aslaug Jonasdottir; Kamilla S Josefsdottir; Thordur Kristjansson; Droplaug N Magnusdottir; Solvi Rognvaldsson; Louise le Roux; Gudrun Sigmundsdottir; Gardar Sveinbjornsson; Kristin E Sveinsdottir; Maney Sveinsdottir; Emil A Thorarensen; Bjarni Thorbjornsson; Gisli Masson; Ingileif Jonsdottir; Alma Moller; Thorolfur Gudnason; Karl G Kristinsson; Unnur Thorsteinsdottir; Kari Stefansson |
| EPI_ISL_827048, EPI_ISL_827077 | The National University Hospital of Iceland | deCODE genetics | Daniel F Gudbjartsson; Agnar Helgason; Hakon Jonsson; Olafur T Magnusson; Pall Melsted; Gudmundur L Norddahl; Jona Saemundsdottir; Asgeir |

| | | | |
|---|---|---|---|
| | | | Sigurdsson; Patrick Sulem; Arna B Agustsdottir; Hannes Eggertsson; Berglind Eiriksdottir; Run Fridriksdottir; Elisabet E Gardarsdottir; Gudmundur Georgsson; Olafia S Gretarsdottir; Kjartan R Gudmundsson; Thora R Gunnarsdottir; Arnaldur Gylfason; Hilma Holm; Brynjar O Jensson; Aslaug Jonasdottir; Kamilla S Josefsdottir; Thordur Kristjansson; Droplaug N Magnusdottir; Solvi Rognvaldsson; Louise le Roux; Gudrun Sigmundsdottir; Gardar Sveinbjornsson; Kristin E Sveinsdottir; Maney Sveinsdottir; Emil A Thorarensen; Bjarni Thorbjornsson; Gisli Masson; Ingileif Jonsdottir; Alma Moller; Thorolfur Gudnason; Karl G Kristinsson; Unnur Thorsteinsdottir; Kari Stefansson |
| EPI_ISL_827091, EPI_ISL_827092, EPI_ISL_827093, EPI_ISL_827094, EPI_ISL_827095, EPI_ISL_827096, EPI_ISL_827097, EPI_ISL_827098, EPI_ISL_827103, EPI_ISL_827104, EPI_ISL_827193, EPI_ISL_827194, EPI_ISL_827196, EPI_ISL_827214, EPI_ISL_827215, EPI_ISL_827216, EPI_ISL_827255 | | | |
| see above | deCODE genetics | deCODE genetics | Daniel F Gudbjartsson; Agnar Helgason; Hakon Jonsson; Olafur T Magnusson; Pall Melsted; Gudmundur L Norddahl; Jona Saemundsdottir; Asgeir Sigurdsson; Patrick Sulem; Arna B Agustsdottir; Hannes Eggertsson; Berglind Eiriksdottir; Run Fridriksdottir; Elisabet E Gardarsdottir; Gudmundur Georgsson; Olafia S Gretarsdottir; Kjartan R Gudmundsson; Thora R Gunnarsdottir; Arnaldur Gylfason; Hilma Holm; Brynjar O Jensson; Aslaug Jonasdottir; Kamilla S Josefsdottir; Thordur Kristjansson; Droplaug N Magnusdottir; Solvi Rognvaldsson; Louise le Roux; Gudrun Sigmundsdottir; Gardar Sveinbjornsson; Kristin E Sveinsdottir; Maney Sveinsdottir; Emil A Thorarensen; Bjarni Thorbjornsson; Gisli Masson; Ingileif Jonsdottir; Alma Moller; Thorolfur Gudnason; Karl G Kristinsson; Unnur Thorsteinsdottir; Kari Stefansson |
| EPI_ISL_827258, EPI_ISL_827309, EPI_ISL_827310, EPI_ISL_827331, EPI_ISL_827334, EPI_ISL_827335 | The National University Hospital of Iceland | deCODE genetics | Daniel F Gudbjartsson; Agnar Helgason; Hakon Jonsson; Olafur T Magnusson; Pall Melsted; Gudmundur L Norddahl; Jona Saemundsdottir; Asgeir Sigurdsson; Patrick Sulem; Arna B Agustsdottir; Hannes Eggertsson; Berglind Eiriksdottir; Run Fridriksdottir; Elisabet E Gardarsdottir; Gudmundur Georgsson; Olafia S Gretarsdottir; Kjartan R Gudmundsson; Thora R Gunnarsdottir; Arnaldur Gylfason; Hilma Holm; Brynjar O Jensson; Aslaug Jonasdottir; Kamilla S Josefsdottir; Thordur Kristjansson; Droplaug N Magnusdottir; Solvi Rognvaldsson; Louise le Roux; Gudrun Sigmundsdottir; Gardar Sveinbjornsson; Kristin E Sveinsdottir; Maney Sveinsdottir; Emil A Thorarensen; Bjarni Thorbjornsson; Gisli Masson; Ingileif Jonsdottir; Alma Moller; Thorolfur Gudnason; Karl G Kristinsson; Unnur Thorsteinsdottir; Kari Stefansson |
| EPI_ISL_827465, EPI_ISL_827572, EPI_ISL_827578, EPI_ISL_827580, EPI_ISL_827622 | deCODE genetics | deCODE genetics | Daniel F Gudbjartsson; Agnar Helgason; Hakon Jonsson; Olafur T Magnusson; Pall Melsted; Gudmundur L Norddahl; Jona Saemundsdottir; Asgeir Sigurdsson; Patrick Sulem; Arna B Agustsdottir; Hannes Eggertsson; Berglind Eiriksdottir; Run Fridriksdottir; Elisabet E Gardarsdottir; Gudmundur Georgsson; Olafia S Gretarsdottir; Kjartan R Gudmundsson; Thora R Gunnarsdottir; Arnaldur Gylfason; Hilma Holm; Brynjar O Jensson; Aslaug Jonasdottir; Kamilla S Josefsdottir; Thordur Kristjansson; Droplaug N Magnusdottir; Solvi Rognvaldsson; Louise le Roux; Gudrun Sigmundsdottir; Gardar Sveinbjornsson; Kristin E Sveinsdottir; Maney Sveinsdottir; Emil A Thorarensen; Bjarni Thorbjornsson; Gisli Masson; Ingileif Jonsdottir; Alma Moller; Thorolfur Gudnason; Karl G Kristinsson; Unnur Thorsteinsdottir; Kari Stefansson |
| EPI_ISL_827696, EPI_ISL_827697 | The National University Hospital of Iceland | deCODE genetics | Daniel F Gudbjartsson; Agnar Helgason; Hakon Jonsson; Olafur T Magnusson; Pall Melsted; Gudmundur L Norddahl; Jona Saemundsdottir; Asgeir Sigurdsson; Patrick Sulem; Arna B Agustsdottir; Hannes Eggertsson; Berglind Eiriksdottir; Run Fridriksdottir; Elisabet E Gardarsdottir; Gudmundur Georgsson; Olafia S Gretarsdottir; Kjartan R Gudmundsson; Thora R Gunnarsdottir; Arnaldur Gylfason; Hilma Holm; Brynjar O Jensson; Aslaug Jonasdottir; Kamilla S Josefsdottir; Thordur Kristjansson; Droplaug N Magnusdottir; Solvi Rognvaldsson; Louise le Roux; Gudrun Sigmundsdottir; Gardar Sveinbjornsson; Kristin E Sveinsdottir; Maney Sveinsdottir; Emil A Thorarensen; Bjarni Thorbjornsson; Gisli Masson; Ingileif Jonsdottir; Alma Moller; Thorolfur Gudnason; Karl G Kristinsson; Unnur Thorsteinsdottir; Kari Stefansson |
| EPI_ISL_827704, EPI_ISL_827705, EPI_ISL_827706, EPI_ISL_827707, EPI_ISL_827708, EPI_ISL_827709, EPI_ISL_827710, EPI_ISL_827711, EPI_ISL_827712, EPI_ISL_827786, EPI_ISL_827809, EPI_ISL_827814, EPI_ISL_827815, EPI_ISL_827817, EPI_ISL_827819, EPI_ISL_827928, EPI_ISL_827936 | | | |
| see above | deCODE genetics | deCODE genetics | Daniel F Gudbjartsson; Agnar Helgason; Hakon Jonsson; Olafur T Magnusson; Pall Melsted; Gudmundur L Norddahl; Jona Saemundsdottir; Asgeir Sigurdsson; Patrick Sulem; Arna B Agustsdottir; Hannes Eggertsson; Berglind Eiriksdottir; Run Fridriksdottir; Elisabet E Gardarsdottir; Gudmundur Georgsson; Olafia S Gretarsdottir; Kjartan R Gudmundsson; Thora R Gunnarsdottir; Arnaldur Gylfason; Hilma Holm; Brynjar O Jensson; Aslaug Jonasdottir; Kamilla S Josefsdottir; Thordur Kristjansson; Droplaug N Magnusdottir; Solvi Rognvaldsson; Louise le Roux; Gudrun Sigmundsdottir; Gardar Sveinbjornsson; Kristin E Sveinsdottir; Maney Sveinsdottir; Emil A Thorarensen; Bjarni Thorbjornsson; Gisli Masson; Ingileif Jonsdottir; Alma Moller; Thorolfur Gudnason; Karl G Kristinsson; Unnur Thorsteinsdottir; Kari Stefansson |
| EPI_ISL_827938, EPI_ISL_827939, EPI_ISL_827940 | The National University Hospital of Iceland | deCODE genetics | Daniel F Gudbjartsson; Agnar Helgason; Hakon Jonsson; Olafur T Magnusson; Pall Melsted; Gudmundur L Norddahl; Jona Saemundsdottir; Asgeir Sigurdsson; Patrick Sulem; Arna B Agustsdottir; Hannes Eggertsson; Berglind Eiriksdottir; Run Fridriksdottir; Elisabet E Gardarsdottir; Gudmundur Georgsson; Olafia S Gretarsdottir; Kjartan R Gudmundsson; Thora R Gunnarsdottir; Arnaldur Gylfason; Hilma Holm; Brynjar O Jensson; Aslaug Jonasdottir; Kamilla S Josefsdottir; Thordur Kristjansson; Droplaug N Magnusdottir; Solvi Rognvaldsson; Louise le Roux; Gudrun Sigmundsdottir; Gardar Sveinbjornsson; Kristin E Sveinsdottir; Maney Sveinsdottir; Emil A Thorarensen; Bjarni Thorbjornsson; Gisli Masson; Ingileif Jonsdottir; Alma Moller; Thorolfur Gudnason; Karl G Kristinsson; Unnur Thorsteinsdottir; Kari Stefansson |
| EPI_ISL_827966, EPI_ISL_827973, EPI_ISL_827989, EPI_ISL_827995, EPI_ISL_827996, EPI_ISL_828001, EPI_ISL_828002, EPI_ISL_828003, EPI_ISL_828004, EPI_ISL_828005, EPI_ISL_828006, EPI_ISL_828007, EPI_ISL_828108, EPI_ISL_828114, EPI_ISL_828115, EPI_ISL_828116, EPI_ISL_828117, EPI_ISL_828118, EPI_ISL_828119, EPI_ISL_828120, EPI_ISL_828125, EPI_ISL_828126, EPI_ISL_828127, EPI_ISL_828128, EPI_ISL_828129, EPI_ISL_828130, EPI_ISL_828131, EPI_ISL_828132, EPI_ISL_828133, EPI_ISL_828135, EPI_ISL_828136 | | | |
| see above | deCODE genetics | deCODE genetics | Daniel F Gudbjartsson; Agnar Helgason; Hakon Jonsson; Olafur T Magnusson; Pall Melsted; Gudmundur L Norddahl; Jona Saemundsdottir; Asgeir Sigurdsson; Patrick Sulem; Arna B Agustsdottir; Hannes Eggertsson; Berglind Eiriksdottir; Run Fridriksdottir; Elisabet E Gardarsdottir; Gudmundur Georgsson; Olafia S Gretarsdottir; Kjartan R Gudmundsson; Thora R Gunnarsdottir; Arnaldur Gylfason; Hilma Holm; Brynjar O Jensson; Aslaug Jonasdottir; Kamilla S Josefsdottir; Thordur Kristjansson; Droplaug N Magnusdottir; Solvi Rognvaldsson; Louise le Roux; Gudrun Sigmundsdottir; Gardar Sveinbjornsson; Kristin E Sveinsdottir; Maney Sveinsdottir; Emil A Thorarensen; Bjarni Thorbjornsson; Gisli Masson; Ingileif Jonsdottir; Alma Moller; Thorolfur Gudnason; Karl G Kristinsson; Unnur Thorsteinsdottir; Kari Stefansson |
| EPI_ISL_828190 | The National University Hospital of Iceland | deCODE genetics | Daniel F Gudbjartsson; Agnar Helgason; Hakon Jonsson; Olafur T Magnusson; Pall Melsted; Gudmundur L Norddahl; Jona Saemundsdottir; Asgeir Sigurdsson; Patrick Sulem; Arna B Agustsdottir; Hannes Eggertsson; Berglind Eiriksdottir; Run Fridriksdottir; Elisabet E Gardarsdottir; Gudmundur Georgsson; Olafia S Gretarsdottir; Kjartan R Gudmundsson; Thora R Gunnarsdottir; Arnaldur Gylfason; Hilma Holm; Brynjar O Jensson; Aslaug Jonasdottir; Kamilla S Josefsdottir; Thordur Kristjansson; Droplaug N Magnusdottir; Solvi Rognvaldsson; Louise le Roux; Gudrun Sigmundsdottir; Gardar Sveinbjornsson; Kristin E Sveinsdottir; Maney Sveinsdottir; Emil A Thorarensen; Bjarni Thorbjornsson; Gisli Masson; Ingileif Jonsdottir; Alma Moller; Thorolfur Gudnason; Karl G Kristinsson; Unnur Thorsteinsdottir; Kari Stefansson |
| EPI_ISL_828222 | deCODE genetics | deCODE genetics | Daniel F Gudbjartsson; Agnar Helgason; Hakon Jonsson; Olafur T Magnusson; Pall Melsted; Gudmundur L Norddahl; Jona Saemundsdottir; Asgeir Sigurdsson; Patrick Sulem; Arna B Agustsdottir; Hannes Eggertsson; Berglind Eiriksdottir; Run Fridriksdottir; Elisabet E Gardarsdottir; Gudmundur Georgsson; Olafia S Gretarsdottir; Kjartan R Gudmundsson; Thora R Gunnarsdottir; Arnaldur Gylfason; Hilma Holm; Brynjar O Jensson; Aslaug Jonasdottir; Kamilla S Josefsdottir; Thordur Kristjansson; Droplaug N Magnusdottir; Solvi Rognvaldsson; Louise le Roux; Gudrun Sigmundsdottir; Gardar Sveinbjornsson; Kristin E Sveinsdottir; Maney Sveinsdottir; Emil A Thorarensen; Bjarni Thorbjornsson; Gisli Masson; Ingileif Jonsdottir; Alma Moller; Thorolfur Gudnason; Karl G Kristinsson; Unnur Thorsteinsdottir; Kari Stefansson |
| EPI_ISL_828224, EPI_ISL_828225, EPI_ISL_828226, EPI_ISL_828227 | The National University Hospital of Iceland | deCODE genetics | Daniel F Gudbjartsson; Agnar Helgason; Hakon Jonsson; Olafur T Magnusson; Pall Melsted; Gudmundur L Norddahl; Jona Saemundsdottir; Asgeir Sigurdsson; Patrick Sulem; Arna B Agustsdottir; Hannes Eggertsson; Berglind Eiriksdottir; Run Fridriksdottir; Elisabet E Gardarsdottir; Gudmundur Georgsson; Olafia S Gretarsdottir; Kjartan R Gudmundsson; Thora R Gunnarsdottir; Arnaldur Gylfason; Hilma Holm; Brynjar O Jensson; Aslaug Jonasdottir; Kamilla S Josefsdottir; Thordur Kristjansson; Droplaug N Magnusdottir; Solvi Rognvaldsson; Louise le Roux; Gudrun Sigmundsdottir; Gardar Sveinbjornsson; Kristin E Sveinsdottir; Maney Sveinsdottir; Emil A Thorarensen; Bjarni Thorbjornsson; Gisli Masson; Ingileif Jonsdottir; Alma Moller; Thorolfur Gudnason; Karl G Kristinsson; Unnur Thorsteinsdottir; Kari Stefansson |
| EPI_ISL_828272, EPI_ISL_828275, EPI_ISL_828276, EPI_ISL_828277, EPI_ISL_828278, EPI_ISL_828279, EPI_ISL_828281, EPI_ISL_828282 | deCODE genetics | deCODE genetics | Daniel F Gudbjartsson; Agnar Helgason; Hakon Jonsson; Olafur T Magnusson; Pall Melsted; Gudmundur L Norddahl; Jona Saemundsdottir; Asgeir Sigurdsson; Patrick Sulem; Arna B Agustsdottir; Hannes Eggertsson; Berglind Eiriksdottir; Run Fridriksdottir; Elisabet E Gardarsdottir; Gudmundur Georgsson; Olafia S Gretarsdottir; Kjartan R Gudmundsson; Thora R Gunnarsdottir; Arnaldur Gylfason; Hilma Holm; Brynjar O Jensson; Aslaug Jonasdottir; Kamilla S Josefsdottir; Thordur Kristjansson; Droplaug N Magnusdottir; Solvi Rognvaldsson; Louise le Roux; Gudrun Sigmundsdottir; Gardar Sveinbjornsson; Kristin E Sveinsdottir; Maney Sveinsdottir; Emil A Thorarensen; Bjarni Thorbjornsson; Gisli Masson; Ingileif Jonsdottir; Alma Moller; Thorolfur Gudnason; Karl G Kristinsson; Unnur Thorsteinsdottir; Kari Stefansson |
| EPI_ISL_828283 | The National University Hospital of Iceland | deCODE genetics | Daniel F Gudbjartsson; Agnar Helgason; Hakon Jonsson; Olafur T Magnusson; Pall Melsted; Gudmundur L Norddahl; Jona Saemundsdottir; Asgeir Sigurdsson; Patrick Sulem; Arna B Agustsdottir; Hannes Eggertsson; Berglind Eiriksdottir; Run Fridriksdottir; Elisabet E Gardarsdottir; Gudmundur Georgsson; Olafia S Gretarsdottir; Kjartan R Gudmundsson; Thora R Gunnarsdottir; Arnaldur Gylfason; Hilma Holm; Brynjar O Jensson; Aslaug Jonasdottir; Kamilla S |

| | | | |
|---|---|---|---|
| | | | Josefsdottir; Thordur Kristjansson; Droplaug N Magnusdottir; Solvi Rognvaldsson; Louise le Roux; Gudrun Sigmundsdottir; Gardar Sveinbjornsson; Kristin E Sveinsdottir; Maney Sveinsdottir; Emil A Thorarensen; Bjarni Thorbjornsson; Gisli Masson; Ingileif Jonsdottir; Alma Moller; Thorolfur Gudnason; Karl G Kristinsson; Unnur Thorsteinsdottir; Kari Stefansson |
| EPI_ISL_828387, EPI_ISL_828388, EPI_ISL_828389, EPI_ISL_828390, EPI_ISL_828392, EPI_ISL_828393, EPI_ISL_828394, EPI_ISL_828395, EPI_ISL_828396, EPI_ISL_828438, EPI_ISL_828439, EPI_ISL_828440, EPI_ISL_828442, EPI_ISL_828443, EPI_ISL_828444, EPI_ISL_828445, EPI_ISL_828446, EPI_ISL_828447, EPI_ISL_828510, EPI_ISL_828511, EPI_ISL_828513, EPI_ISL_828514, EPI_ISL_828515, EPI_ISL_828516, EPI_ISL_828604, EPI_ISL_828605, EPI_ISL_828606, EPI_ISL_828607, EPI_ISL_828608, EPI_ISL_828609, EPI_ISL_828610, EPI_ISL_828611, EPI_ISL_828612, EPI_ISL_828613, EPI_ISL_828645, EPI_ISL_828647, EPI_ISL_828648, EPI_ISL_828649, EPI_ISL_828650, EPI_ISL_828651, EPI_ISL_828652, EPI_ISL_828665, EPI_ISL_828666, EPI_ISL_828667, EPI_ISL_828668, EPI_ISL_828669, EPI_ISL_828670, EPI_ISL_828671, EPI_ISL_828672, EPI_ISL_828673, EPI_ISL_828674, EPI_ISL_828680, EPI_ISL_828731, EPI_ISL_828732, EPI_ISL_828734, EPI_ISL_828735, EPI_ISL_828736, EPI_ISL_828737, EPI_ISL_828738, EPI_ISL_828739, EPI_ISL_828740 | | | |
| see above | deCODE genetics | deCODE genetics | Daniel F Gudbjartsson; Agnar Helgason; Hakon Jonsson; Olafur T Magnusson; Pall Melsted; Gudmundur L Norddahl; Jona Saemundsdottir; Asgeir Sigurdsson; Patrick Sulem; Arna B Agustsdottir; Hannes Eggertsson; Berglind Eiriksdottir; Run Fridriksdottir; Elisabet E Gardarsdottir; Gudmundur Georgsson; Olafia S Gretarsdottir; Kjartan R Gudmundsson; Thora R Gunnarsdottir; Arnaldur Gylfason; Hilma Holm; Brynjar O Jensson; Aslaug Jonasdottir; Kamilla S Josefsdottir; Thordur Kristjansson; Droplaug N Magnusdottir; Solvi Rognvaldsson; Louise le Roux; Gudrun Sigmundsdottir; Gardar Sveinbjornsson; Kristin E Sveinsdottir; Maney Sveinsdottir; Emil A Thorarensen; Bjarni Thorbjornsson; Gisli Masson; Ingileif Jonsdottir; Alma Moller; Thorolfur Gudnason; Karl G Kristinsson; Unnur Thorsteinsdottir; Kari Stefansson |
| EPI_ISL_828759, EPI_ISL_828761, EPI_ISL_828762 | The National University Hospital of Iceland | deCODE genetics | Daniel F Gudbjartsson; Agnar Helgason; Hakon Jonsson; Olafur T Magnusson; Pall Melsted; Gudmundur L Norddahl; Jona Saemundsdottir; Asgeir Sigurdsson; Patrick Sulem; Arna B Agustsdottir; Hannes Eggertsson; Berglind Eiriksdottir; Run Fridriksdottir; Elisabet E Gardarsdottir; Gudmundur Georgsson; Olafia S Gretarsdottir; Kjartan R Gudmundsson; Thora R Gunnarsdottir; Arnaldur Gylfason; Hilma Holm; Brynjar O Jensson; Aslaug Jonasdottir; Kamilla S Josefsdottir; Thordur Kristjansson; Droplaug N Magnusdottir; Solvi Rognvaldsson; Louise le Roux; Gudrun Sigmundsdottir; Gardar Sveinbjornsson; Kristin E Sveinsdottir; Maney Sveinsdottir; Emil A Thorarensen; Bjarni Thorbjornsson; Gisli Masson; Ingileif Jonsdottir; Alma Moller; Thorolfur Gudnason; Karl G Kristinsson; Unnur Thorsteinsdottir; Kari Stefansson |
| EPI_ISL_828773 | deCODE genetics | deCODE genetics | Daniel F Gudbjartsson; Agnar Helgason; Hakon Jonsson; Olafur T Magnusson; Pall Melsted; Gudmundur L Norddahl; Jona Saemundsdottir; Asgeir Sigurdsson; Patrick Sulem; Arna B Agustsdottir; Hannes Eggertsson; Berglind Eiriksdottir; Run Fridriksdottir; Elisabet E Gardarsdottir; Gudmundur Georgsson; Olafia S Gretarsdottir; Kjartan R Gudmundsson; Thora R Gunnarsdottir; Arnaldur Gylfason; Hilma Holm; Brynjar O Jensson; Aslaug Jonasdottir; Kamilla S Josefsdottir; Thordur Kristjansson; Droplaug N Magnusdottir; Solvi Rognvaldsson; Louise le Roux; Gudrun Sigmundsdottir; Gardar Sveinbjornsson; Kristin E Sveinsdottir; Maney Sveinsdottir; Emil A Thorarensen; Bjarni Thorbjornsson; Gisli Masson; Ingileif Jonsdottir; Alma Moller; Thorolfur Gudnason; Karl G Kristinsson; Unnur Thorsteinsdottir; Kari Stefansson |
| EPI_ISL_828774 | The National University Hospital of Iceland | deCODE genetics | Daniel F Gudbjartsson; Agnar Helgason; Hakon Jonsson; Olafur T Magnusson; Pall Melsted; Gudmundur L Norddahl; Jona Saemundsdottir; Asgeir Sigurdsson; Patrick Sulem; Arna B Agustsdottir; Hannes Eggertsson; Berglind Eiriksdottir; Run Fridriksdottir; Elisabet E Gardarsdottir; Gudmundur Georgsson; Olafia S Gretarsdottir; Kjartan R Gudmundsson; Thora R Gunnarsdottir; Arnaldur Gylfason; Hilma Holm; Brynjar O Jensson; Aslaug Jonasdottir; Kamilla S Josefsdottir; Thordur Kristjansson; Droplaug N Magnusdottir; Solvi Rognvaldsson; Louise le Roux; Gudrun Sigmundsdottir; Gardar Sveinbjornsson; Kristin E Sveinsdottir; Maney Sveinsdottir; Emil A Thorarensen; Bjarni Thorbjornsson; Gisli Masson; Ingileif Jonsdottir; Alma Moller; Thorolfur Gudnason; Karl G Kristinsson; Unnur Thorsteinsdottir; Kari Stefansson |
| EPI_ISL_828813, EPI_ISL_828814, EPI_ISL_828815, EPI_ISL_828816, EPI_ISL_828817, EPI_ISL_828818, EPI_ISL_828819, EPI_ISL_828820, EPI_ISL_828821, EPI_ISL_828822, EPI_ISL_828834, EPI_ISL_828835, EPI_ISL_828837, EPI_ISL_828838, EPI_ISL_828839, EPI_ISL_828840, EPI_ISL_828841, EPI_ISL_828848, EPI_ISL_828849, EPI_ISL_828874, EPI_ISL_828875, EPI_ISL_828877, EPI_ISL_828878, EPI_ISL_828880, EPI_ISL_828883, EPI_ISL_828886, EPI_ISL_828887, EPI_ISL_828891, EPI_ISL_828905, EPI_ISL_828906, EPI_ISL_828907, EPI_ISL_828908, EPI_ISL_828909, EPI_ISL_828910, EPI_ISL_828911, EPI_ISL_828912, EPI_ISL_828913, EPI_ISL_828938, EPI_ISL_828951, EPI_ISL_828952, EPI_ISL_828953, EPI_ISL_828954, EPI_ISL_828955, EPI_ISL_828956, EPI_ISL_828957, EPI_ISL_828958, EPI_ISL_828960, EPI_ISL_828961, EPI_ISL_829015, EPI_ISL_829082, EPI_ISL_829083, EPI_ISL_829084, EPI_ISL_829085, EPI_ISL_829086, EPI_ISL_829087, EPI_ISL_829088, EPI_ISL_829089, EPI_ISL_829092, EPI_ISL_829093, EPI_ISL_829107, EPI_ISL_829120, EPI_ISL_829121, EPI_ISL_829125, EPI_ISL_829126, EPI_ISL_829127, EPI_ISL_829128, EPI_ISL_829129, EPI_ISL_829130, EPI_ISL_829131 | | | |
| see above | deCODE genetics | deCODE genetics | Daniel F Gudbjartsson; Agnar Helgason; Hakon Jonsson; Olafur T Magnusson; Pall Melsted; Gudmundur L Norddahl; Jona Saemundsdottir; Asgeir Sigurdsson; Patrick Sulem; Arna B Agustsdottir; Hannes Eggertsson; Berglind Eiriksdottir; Run Fridriksdottir; Elisabet E Gardarsdottir; Gudmundur Georgsson; Olafia S Gretarsdottir; Kjartan R Gudmundsson; Thora R Gunnarsdottir; Arnaldur Gylfason; Hilma Holm; Brynjar O Jensson; Aslaug Jonasdottir; Kamilla S Josefsdottir; Thordur Kristjansson; Droplaug N Magnusdottir; Solvi Rognvaldsson; Louise le Roux; Gudrun Sigmundsdottir; Gardar Sveinbjornsson; Kristin E Sveinsdottir; Maney Sveinsdottir; Emil A Thorarensen; Bjarni Thorbjornsson; Gisli Masson; Ingileif Jonsdottir; Alma Moller; Thorolfur Gudnason; Karl G Kristinsson; Unnur Thorsteinsdottir; Kari Stefansson |
| EPI_ISL_829154, EPI_ISL_829155, EPI_ISL_829156 | The National University Hospital of Iceland | deCODE genetics | Daniel F Gudbjartsson; Agnar Helgason; Hakon Jonsson; Olafur T Magnusson; Pall Melsted; Gudmundur L Norddahl; Jona Saemundsdottir; Asgeir Sigurdsson; Patrick Sulem; Arna B Agustsdottir; Hannes Eggertsson; Berglind Eiriksdottir; Run Fridriksdottir; Elisabet E Gardarsdottir; Gudmundur Georgsson; Olafia S Gretarsdottir; Kjartan R Gudmundsson; Thora R Gunnarsdottir; Arnaldur Gylfason; Hilma Holm; Brynjar O Jensson; Aslaug Jonasdottir; Kamilla S Josefsdottir; Thordur Kristjansson; Droplaug N Magnusdottir; Solvi Rognvaldsson; Louise le Roux; Gudrun Sigmundsdottir; Gardar Sveinbjornsson; Kristin E Sveinsdottir; Maney Sveinsdottir; Emil A Thorarensen; Bjarni Thorbjornsson; Gisli Masson; Ingileif Jonsdottir; Alma Moller; Thorolfur Gudnason; Karl G Kristinsson; Unnur Thorsteinsdottir; Kari Stefansson |
| EPI_ISL_829158, EPI_ISL_829159, EPI_ISL_829160, EPI_ISL_829161, EPI_ISL_829162, EPI_ISL_829183, EPI_ISL_829217, EPI_ISL_829225, EPI_ISL_829227, EPI_ISL_829228, EPI_ISL_829229, EPI_ISL_829230, EPI_ISL_829231, EPI_ISL_829233, EPI_ISL_829234, EPI_ISL_829327, EPI_ISL_829328, EPI_ISL_829336, EPI_ISL_829337, EPI_ISL_829338, EPI_ISL_829339, EPI_ISL_829383, EPI_ISL_829384, EPI_ISL_829385, EPI_ISL_829386, EPI_ISL_829387, EPI_ISL_829389, EPI_ISL_829392, EPI_ISL_829393, EPI_ISL_829435, EPI_ISL_829436, EPI_ISL_829439, EPI_ISL_829440, EPI_ISL_829441, EPI_ISL_829442, EPI_ISL_829443, EPI_ISL_829444, EPI_ISL_829445, EPI_ISL_829557, EPI_ISL_829559, EPI_ISL_829560, EPI_ISL_829561, EPI_ISL_829562, EPI_ISL_829563, EPI_ISL_829564, EPI_ISL_829571, EPI_ISL_829572, EPI_ISL_829605, EPI_ISL_829606, EPI_ISL_829609, EPI_ISL_829610, EPI_ISL_829611, EPI_ISL_829612, EPI_ISL_829613, EPI_ISL_829614, EPI_ISL_829615, EPI_ISL_829616, EPI_ISL_829630, EPI_ISL_829631, EPI_ISL_829632, EPI_ISL_829633, EPI_ISL_829637, EPI_ISL_829634, EPI_ISL_829635, EPI_ISL_829636, EPI_ISL_829637, EPI_ISL_829638, EPI_ISL_829639, EPI_ISL_829685, EPI_ISL_829687, EPI_ISL_829688, EPI_ISL_829689, EPI_ISL_829690, EPI_ISL_829691, EPI_ISL_829692, EPI_ISL_829699, EPI_ISL_829700, EPI_ISL_829814, EPI_ISL_829815, EPI_ISL_829817, EPI_ISL_829818, EPI_ISL_829819, EPI_ISL_829820, EPI_ISL_829824, EPI_ISL_829825 | | | |
| see above | deCODE genetics | deCODE genetics | Daniel F Gudbjartsson; Agnar Helgason; Hakon Jonsson; Olafur T Magnusson; Pall Melsted; Gudmundur L Norddahl; Jona Saemundsdottir; Asgeir Sigurdsson; Patrick Sulem; Arna B Agustsdottir; Hannes Eggertsson; Berglind Eiriksdottir; Run Fridriksdottir; Elisabet E Gardarsdottir; Gudmundur Georgsson; Olafia S Gretarsdottir; Kjartan R Gudmundsson; Thora R Gunnarsdottir; Arnaldur Gylfason; Hilma Holm; Brynjar O Jensson; Aslaug Jonasdottir; Kamilla S Josefsdottir; Thordur Kristjansson; Droplaug N Magnusdottir; Solvi Rognvaldsson; Louise le Roux; Gudrun Sigmundsdottir; Gardar Sveinbjornsson; Kristin E Sveinsdottir; Maney Sveinsdottir; Emil A Thorarensen; Bjarni Thorbjornsson; Gisli Masson; Ingileif Jonsdottir; Alma Moller; Thorolfur Gudnason; Karl G Kristinsson; Unnur Thorsteinsdottir; Kari Stefansson |
| EPI_ISL_829937, EPI_ISL_829940 | The National University Hospital of Iceland | deCODE genetics | Daniel F Gudbjartsson; Agnar Helgason; Hakon Jonsson; Olafur T Magnusson; Pall Melsted; Gudmundur L Norddahl; Jona Saemundsdottir; Asgeir Sigurdsson; Patrick Sulem; Arna B Agustsdottir; Hannes Eggertsson; Berglind Eiriksdottir; Run Fridriksdottir; Elisabet E Gardarsdottir; Gudmundur Georgsson; Olafia S Gretarsdottir; Kjartan R Gudmundsson; Thora R Gunnarsdottir; Arnaldur Gylfason; Hilma Holm; Brynjar O Jensson; Aslaug Jonasdottir; Kamilla S Josefsdottir; Thordur Kristjansson; Droplaug N Magnusdottir; Solvi Rognvaldsson; Louise le Roux; Gudrun Sigmundsdottir; Gardar Sveinbjornsson; Kristin E Sveinsdottir; Maney Sveinsdottir; Emil A Thorarensen; Bjarni Thorbjornsson; Gisli Masson; Ingileif Jonsdottir; Alma Moller; Thorolfur Gudnason; Karl G Kristinsson; Unnur Thorsteinsdottir; Kari Stefansson |
| EPI_ISL_829979, EPI_ISL_830012, EPI_ISL_830016, EPI_ISL_830017, EPI_ISL_830018, EPI_ISL_830020, EPI_ISL_830021, EPI_ISL_830022, EPI_ISL_830023, EPI_ISL_830024, EPI_ISL_830026, EPI_ISL_830027, EPI_ISL_830028, EPI_ISL_830029, EPI_ISL_830030, EPI_ISL_830031 | | | |
| see above | deCODE genetics | deCODE genetics | Daniel F Gudbjartsson; Agnar Helgason; Hakon Jonsson; Olafur T Magnusson; Pall Melsted; Gudmundur L Norddahl; Jona Saemundsdottir; Asgeir Sigurdsson; Patrick Sulem; Arna B Agustsdottir; Hannes Eggertsson; Berglind Eiriksdottir; Run Fridriksdottir; Elisabet E Gardarsdottir; Gudmundur Georgsson; Olafia S Gretarsdottir; Kjartan R Gudmundsson; Thora R Gunnarsdottir; Arnaldur Gylfason; Hilma Holm; Brynjar O Jensson; Aslaug Jonasdottir; Kamilla S Josefsdottir; Thordur Kristjansson; Droplaug N Magnusdottir; Solvi Rognvaldsson; Louise le Roux; Gudrun Sigmundsdottir; Gardar Sveinbjornsson; Kristin E Sveinsdottir; Maney Sveinsdottir; Emil A Thorarensen; Bjarni Thorbjornsson; Gisli Masson; Ingileif Jonsdottir; Alma Moller; Thorolfur Gudnason; Karl G Kristinsson; Unnur Thorsteinsdottir; Kari Stefansson |
| EPI_ISL_830033, EPI_ISL_830034 | The National University Hospital of Iceland | deCODE genetics | Daniel F Gudbjartsson; Agnar Helgason; Hakon Jonsson; Olafur T Magnusson; Pall Melsted; Gudmundur L Norddahl; Jona Saemundsdottir; Asgeir Sigurdsson; Patrick Sulem; Arna B Agustsdottir; Hannes Eggertsson; Berglind Eiriksdottir; Run Fridriksdottir; Elisabet E Gardarsdottir; Gudmundur Georgsson; Olafia S Gretarsdottir; Kjartan R Gudmundsson; Thora R Gunnarsdottir; Arnaldur Gylfason; Hilma Holm; Brynjar O Jensson; Aslaug Jonasdottir; Kamilla S Josefsdottir; Thordur Kristjansson; Droplaug N Magnusdottir; Solvi Rognvaldsson; Louise le Roux; Gudrun Sigmundsdottir; Gardar Sveinbjornsson; Kristin E Sveinsdottir; Maney Sveinsdottir; Emil A Thorarensen; Bjarni Thorbjornsson; Gisli Masson; Ingileif Jonsdottir; Alma Moller; Thorolfur Gudnason; Karl G Kristinsson; Unnur Thorsteinsdottir; Kari Stefansson |
| EPI_ISL_830035, EPI_ISL_830036, EPI_ISL_830041, EPI_ISL_830042, EPI_ISL_830073, EPI_ISL_830074, EPI_ISL_830075, EPI_ISL_830076, EPI_ISL_830077, EPI_ISL_830078, EPI_ISL_830079, EPI_ISL_830080, EPI_ISL_830081, EPI_ISL_830082 | | | |
| see above | deCODE genetics | deCODE genetics | Daniel F Gudbjartsson; Agnar Helgason; Hakon Jonsson; Olafur T Magnusson; Pall Melsted; Gudmundur L Norddahl; Jona Saemundsdottir; Asgeir |

| | | | |
|---|---|---|---|
| | | | Sigurdsson; Patrick Sulem; Arna B Agustsdottir; Hannes Eggertsson; Berglind Eiriksdottir; Run Fridriksdottir; Elisabet E Gardarsdottir; Gudmundur Georgsson; Olafia S Gretarsdottir; Kjartan R Gudmundsson; Thora R Gunnarsdottir; Arnaldur Gylfason; Hilma Holm; Brynjar O Jensson; Aslaug Jonasdottir; Kamilla S Josefsdottir; Thordur Kristjansson; Droplaug N Magnusdottir; Solvi Rognvaldsson; Louise le Roux; Gudrun Sigmundsdottir; Gardar Sveinbjornsson; Kristin E Sveinsdottir; Maney Sveinsdottir; Emil A Thorarensen; Bjarni Thorbjornsson; Gisli Masson; Ingileif Jonsdottir; Alma Moller; Thorolfur Gudnason; Karl G Kristinsson; Unnur Thorsteinsdottir; Kari Stefansson |
| EPI_ISL_830147, EPI_ISL_830152 | The National University Hospital of Iceland | deCODE genetics | Daniel F Gudbjartsson; Agnar Helgason; Hakon Jonsson; Olafur T Magnusson; Pall Melsted; Gudmundur L Norddahl; Jona Saemundsdottir; Asgeir Sigurdsson; Patrick Sulem; Arna B Agustsdottir; Hannes Eggertsson; Berglind Eiriksdottir; Run Fridriksdottir; Elisabet E Gardarsdottir; Gudmundur Georgsson; Olafia S Gretarsdottir; Kjartan R Gudmundsson; Thora R Gunnarsdottir; Arnaldur Gylfason; Hilma Holm; Brynjar O Jensson; Aslaug Jonasdottir; Kamilla S Josefsdottir; Thordur Kristjansson; Droplaug N Magnusdottir; Solvi Rognvaldsson; Louise le Roux; Gudrun Sigmundsdottir; Gardar Sveinbjornsson; Kristin E Sveinsdottir; Maney Sveinsdottir; Emil A Thorarensen; Bjarni Thorbjornsson; Gisli Masson; Ingileif Jonsdottir; Alma Moller; Thorolfur Gudnason; Karl G Kristinsson; Unnur Thorsteinsdottir; Kari Stefansson |
| EPI_ISL_830166, EPI_ISL_830170, EPI_ISL_830171, EPI_ISL_830172, EPI_ISL_830173, EPI_ISL_830350, EPI_ISL_830352, EPI_ISL_830353, EPI_ISL_830357, EPI_ISL_830358 | deCODE genetics | deCODE genetics | Daniel F Gudbjartsson; Agnar Helgason; Hakon Jonsson; Olafur T Magnusson; Pall Melsted; Gudmundur L Norddahl; Jona Saemundsdottir; Asgeir Sigurdsson; Patrick Sulem; Arna B Agustsdottir; Hannes Eggertsson; Berglind Eiriksdottir; Run Fridriksdottir; Elisabet E Gardarsdottir; Gudmundur Georgsson; Olafia S Gretarsdottir; Kjartan R Gudmundsson; Thora R Gunnarsdottir; Arnaldur Gylfason; Hilma Holm; Brynjar O Jensson; Aslaug Jonasdottir; Kamilla S Josefsdottir; Thordur Kristjansson; Droplaug N Magnusdottir; Solvi Rognvaldsson; Louise le Roux; Gudrun Sigmundsdottir; Gardar Sveinbjornsson; Kristin E Sveinsdottir; Maney Sveinsdottir; Emil A Thorarensen; Bjarni Thorbjornsson; Gisli Masson; Ingileif Jonsdottir; Alma Moller; Thorolfur Gudnason; Karl G Kristinsson; Unnur Thorsteinsdottir; Kari Stefansson |
| EPI_ISL_830359, EPI_ISL_830361, EPI_ISL_830362, EPI_ISL_830363, EPI_ISL_830364, EPI_ISL_830365 | The National University Hospital of Iceland | deCODE genetics | Daniel F Gudbjartsson; Agnar Helgason; Hakon Jonsson; Olafur T Magnusson; Pall Melsted; Gudmundur L Norddahl; Jona Saemundsdottir; Asgeir Sigurdsson; Patrick Sulem; Arna B Agustsdottir; Hannes Eggertsson; Berglind Eiriksdottir; Run Fridriksdottir; Elisabet E Gardarsdottir; Gudmundur Georgsson; Olafia S Gretarsdottir; Kjartan R Gudmundsson; Thora R Gunnarsdottir; Arnaldur Gylfason; Hilma Holm; Brynjar O Jensson; Aslaug Jonasdottir; Kamilla S Josefsdottir; Thordur Kristjansson; Droplaug N Magnusdottir; Solvi Rognvaldsson; Louise le Roux; Gudrun Sigmundsdottir; Gardar Sveinbjornsson; Kristin E Sveinsdottir; Maney Sveinsdottir; Emil A Thorarensen; Bjarni Thorbjornsson; Gisli Masson; Ingileif Jonsdottir; Alma Moller; Thorolfur Gudnason; Karl G Kristinsson; Unnur Thorsteinsdottir; Kari Stefansson |
| EPI_ISL_830369, EPI_ISL_830370, EPI_ISL_830372, EPI_ISL_830373, EPI_ISL_830374, EPI_ISL_830375, EPI_ISL_830376, EPI_ISL_830426, EPI_ISL_830427, EPI_ISL_830430, EPI_ISL_830431, EPI_ISL_830432, EPI_ISL_830433, EPI_ISL_830434, EPI_ISL_830435, EPI_ISL_830436, EPI_ISL_830437, EPI_ISL_830439, EPI_ISL_830440, EPI_ISL_830441, EPI_ISL_830442, EPI_ISL_830443, EPI_ISL_830444, EPI_ISL_830446, EPI_ISL_830447 | | | |
| see above | deCODE genetics | deCODE genetics | Daniel F Gudbjartsson; Agnar Helgason; Hakon Jonsson; Olafur T Magnusson; Pall Melsted; Gudmundur L Norddahl; Jona Saemundsdottir; Asgeir Sigurdsson; Patrick Sulem; Arna B Agustsdottir; Hannes Eggertsson; Berglind Eiriksdottir; Run Fridriksdottir; Elisabet E Gardarsdottir; Gudmundur Georgsson; Olafia S Gretarsdottir; Kjartan R Gudmundsson; Thora R Gunnarsdottir; Arnaldur Gylfason; Hilma Holm; Brynjar O Jensson; Aslaug Jonasdottir; Kamilla S Josefsdottir; Thordur Kristjansson; Droplaug N Magnusdottir; Solvi Rognvaldsson; Louise le Roux; Gudrun Sigmundsdottir; Gardar Sveinbjornsson; Kristin E Sveinsdottir; Maney Sveinsdottir; Emil A Thorarensen; Bjarni Thorbjornsson; Gisli Masson; Ingileif Jonsdottir; Alma Moller; Thorolfur Gudnason; Karl G Kristinsson; Unnur Thorsteinsdottir; Kari Stefansson |
| EPI_ISL_830487 | The National University Hospital of Iceland | deCODE genetics | Daniel F Gudbjartsson; Agnar Helgason; Hakon Jonsson; Olafur T Magnusson; Pall Melsted; Gudmundur L Norddahl; Jona Saemundsdottir; Asgeir Sigurdsson; Patrick Sulem; Arna B Agustsdottir; Hannes Eggertsson; Berglind Eiriksdottir; Run Fridriksdottir; Elisabet E Gardarsdottir; Gudmundur Georgsson; Olafia S Gretarsdottir; Kjartan R Gudmundsson; Thora R Gunnarsdottir; Arnaldur Gylfason; Hilma Holm; Brynjar O Jensson; Aslaug Jonasdottir; Kamilla S Josefsdottir; Thordur Kristjansson; Droplaug N Magnusdottir; Solvi Rognvaldsson; Louise le Roux; Gudrun Sigmundsdottir; Gardar Sveinbjornsson; Kristin E Sveinsdottir; Maney Sveinsdottir; Emil A Thorarensen; Bjarni Thorbjornsson; Gisli Masson; Ingileif Jonsdottir; Alma Moller; Thorolfur Gudnason; Karl G Kristinsson; Unnur Thorsteinsdottir; Kari Stefansson |
| EPI_ISL_830534, EPI_ISL_830536, EPI_ISL_830547 | deCODE genetics | deCODE genetics | Daniel F Gudbjartsson; Agnar Helgason; Hakon Jonsson; Olafur T Magnusson; Pall Melsted; Gudmundur L Norddahl; Jona Saemundsdottir; Asgeir Sigurdsson; Patrick Sulem; Arna B Agustsdottir; Hannes Eggertsson; Berglind Eiriksdottir; Run Fridriksdottir; Elisabet E Gardarsdottir; Gudmundur Georgsson; Olafia S Gretarsdottir; Kjartan R Gudmundsson; Thora R Gunnarsdottir; Arnaldur Gylfason; Hilma Holm; Brynjar O Jensson; Aslaug Jonasdottir; Kamilla S Josefsdottir; Thordur Kristjansson; Droplaug N Magnusdottir; Solvi Rognvaldsson; Louise le Roux; Gudrun Sigmundsdottir; Gardar Sveinbjornsson; Kristin E Sveinsdottir; Maney Sveinsdottir; Emil A Thorarensen; Bjarni Thorbjornsson; Gisli Masson; Ingileif Jonsdottir; Alma Moller; Thorolfur Gudnason; Karl G Kristinsson; Unnur Thorsteinsdottir; Kari Stefansson |
| EPI_ISL_830727, EPI_ISL_830730, EPI_ISL_830746, EPI_ISL_830747, EPI_ISL_830748, EPI_ISL_830749, EPI_ISL_830756, EPI_ISL_830757, EPI_ISL_830761, EPI_ISL_830763, EPI_ISL_830768, EPI_ISL_830771, EPI_ISL_830851, EPI_ISL_830852, EPI_ISL_830853, EPI_ISL_830854, EPI_ISL_830855, EPI_ISL_830856, EPI_ISL_830857, EPI_ISL_830858, EPI_ISL_830859, EPI_ISL_830860, EPI_ISL_830861, EPI_ISL_830862, EPI_ISL_830863, EPI_ISL_830864, EPI_ISL_830865, EPI_ISL_830866, EPI_ISL_830867, EPI_ISL_830868, EPI_ISL_830869, EPI_ISL_830870, EPI_ISL_830871, EPI_ISL_830872, EPI_ISL_830873, EPI_ISL_830874, EPI_ISL_830875, EPI_ISL_830876, EPI_ISL_830877, EPI_ISL_830878, EPI_ISL_830879, EPI_ISL_830880, EPI_ISL_830881, EPI_ISL_830882, EPI_ISL_830883, EPI_ISL_830884, EPI_ISL_830885, EPI_ISL_830886, EPI_ISL_830887, EPI_ISL_830888, EPI_ISL_830889, EPI_ISL_830890, EPI_ISL_830891, EPI_ISL_830892, EPI_ISL_830893, EPI_ISL_830894, EPI_ISL_830895, EPI_ISL_830896, EPI_ISL_830897, EPI_ISL_830898, EPI_ISL_830899, EPI_ISL_830900, EPI_ISL_830901, EPI_ISL_830902, EPI_ISL_830903, EPI_ISL_830904, EPI_ISL_830905, EPI_ISL_830906, EPI_ISL_830907, EPI_ISL_830908, EPI_ISL_830909, EPI_ISL_830979, EPI_ISL_830984, EPI_ISL_830990, EPI_ISL_830991, EPI_ISL_831001, EPI_ISL_831013, EPI_ISL_831015, EPI_ISL_831018, EPI_ISL_831019 | | | |
| see above | University Hospital Basel, Clinical Virology | University Hospital Basel, Clinical Bacteriology | Tim Roloff, Madlen Stange, Helena MB Seth-Smith, Alfredo Mari, Karoline Leuzinger, Julia Bielicki, Manuel Battegay, Hans Hirsch, Adrian Egli |
| EPI_ISL_831098, EPI_ISL_831099 | Hospital Universitario La Paz (Madrid) | SeqCOVID-SPAIN consortium/IBV(CSIC) | María Rodríguez-Tejedor, Elias Dahdouh, Fernando Lázaro-Perona, Jesús Mingorance and SeqCOVID-SPAIN consortium |
| EPI_ISL_831765, EPI_ISL_831766 | United States Air Force School of Aerospace Medicine | United States Air Force School of Aerospace Medicine | Anthony Fries, Jennifer Meyer, William Gruner, Amanda Javorina, Sarah Purves, Clarise Starr, Elizabeth Macias |
| EPI_ISL_832422, EPI_ISL_832423, EPI_ISL_832425, EPI_ISL_832426, EPI_ISL_832427, EPI_ISL_832428, EPI_ISL_832429, EPI_ISL_832432, EPI_ISL_832433, EPI_ISL_832434, EPI_ISL_832435, EPI_ISL_832436, EPI_ISL_832437, EPI_ISL_832438, EPI_ISL_832439, EPI_ISL_832440, EPI_ISL_832441, EPI_ISL_832442, EPI_ISL_832443, EPI_ISL_832444, EPI_ISL_832445, EPI_ISL_832446, EPI_ISL_832447, EPI_ISL_832448, EPI_ISL_832449, EPI_ISL_832450, EPI_ISL_832451, EPI_ISL_832452, EPI_ISL_832453, EPI_ISL_832454, EPI_ISL_832455, EPI_ISL_832456, EPI_ISL_832457, EPI_ISL_832458, EPI_ISL_832459, EPI_ISL_832460, EPI_ISL_832461, EPI_ISL_832462, EPI_ISL_832463, EPI_ISL_832464, EPI_ISL_832465, EPI_ISL_832466, EPI_ISL_832467 | | | |
| see above | OHSU Lab Services Molecular Microbiology Lab | Oregon SARS-CoV-2 Genome Sequencing Center | Brendan L. O'Connell, Ruth V. Nichols, Sally Grindstaff, Alec J. Hirsch, Donna Hansel, Guang Fan, Daniel N. Streblow, William B. Messer, Andrew C. Adey, Benjamin N. Bimber, Brian J. O'Roak |
| EPI_ISL_833039 | RS TK II 02.05.01 dr. AK Gani Palembang | National Institute of Health Research and Development | Subangkit;Pawestri,HA;Ikawati,HD;Nugraha,AA;Puspa,KD;Pangesti,KNA;Soekarso,T;Tantula,A;Wiradharma,RP;Puspandari,N;Setiawaty,V |
| EPI_ISL_833335, EPI_ISL_833338 | Siniloan Rural Health Unit | Research Institute for Tropical Medicine | Hannah Leah Morito, Othoniel Jan Onza, John Leonard Chan, Ma Angelica Tujan, Francisco Gerardo Polotan, Inez Andrea Medado, Kirstyn Brunker, Edelwisa Mercado, Daria Manalo, Catalino Demetria |
| EPI_ISL_842119, EPI_ISL_842120, EPI_ISL_842122, EPI_ISL_842123, EPI_ISL_842124, EPI_ISL_842125, EPI_ISL_842126, EPI_ISL_842127, EPI_ISL_842128, EPI_ISL_842129, EPI_ISL_842130, EPI_ISL_842131, EPI_ISL_842132, EPI_ISL_842133, EPI_ISL_842134, EPI_ISL_842135, EPI_ISL_842136, EPI_ISL_842138, EPI_ISL_842140, EPI_ISL_842143, EPI_ISL_842144, EPI_ISL_842145, EPI_ISL_842147, EPI_ISL_842148, EPI_ISL_842149, EPI_ISL_842150, EPI_ISL_842151, EPI_ISL_842152, EPI_ISL_842154, EPI_ISL_842155, EPI_ISL_842156, EPI_ISL_842157, EPI_ISL_842158, EPI_ISL_842159, EPI_ISL_842160, EPI_ISL_842162, EPI_ISL_842163, EPI_ISL_842165, EPI_ISL_842166, EPI_ISL_842167, EPI_ISL_842168, EPI_ISL_842169, EPI_ISL_842170, EPI_ISL_842171 | | | |
| see above | Oxford Viromics, NDM, University of Oxford; Oxford University Hospitals; Basingstoke and North Hampshire Hospital | COVID-19 Genomics UK (COG-UK) Consortium | Tanya Golubchik, David Bonsall, George Macintyre, Amy Trebes, Mariateresa de Cesare, Catrin Moore, Alex Mobbs, Anita Justice, Robert Shaw, Monique Andersson, Timothy Peto, Emma Wise, Nathan Moore, Jessica Lynch, Nick Cortes, Matilde Mori, Stephen Kidd, David Buck, John Todd, Christophe Fraser |
| EPI_ISL_842650 | University Hospital Basel, Clinical Virology | University Hospital Basel, Clinical Bacteriology | Tim Roloff, Madlen Stange, Helena MB Seth-Smith, Alfredo Mari, Karoline Leuzinger, Julia Bielicki, Manuel Battegay, Hans Hirsch, Adrian Egli |
| EPI_ISL_845664, EPI_ISL_845678, EPI_ISL_845679 | Quest Diagnostics | Quest Diagnostics | Rosenthal,S.H., Gerasimova,A., Kagan,R.M., Anderson, B., Bernstein, L.E., Livingston, K.E., Hua, M., Liu Y., Shalhout, D.F., Shlyakhter, I.A., Owen, R., Lacbawan, F. |
| EPI_ISL_848204, EPI_ISL_848230, EPI_ISL_848247, EPI_ISL_848252, EPI_ISL_848253, EPI_ISL_848254, EPI_ISL_848327, EPI_ISL_848328, EPI_ISL_848329, EPI_ISL_848330, EPI_ISL_848331, EPI_ISL_848332, EPI_ISL_848333, EPI_ISL_848334, EPI_ISL_848335, EPI_ISL_848336, EPI_ISL_848337, EPI_ISL_848338, EPI_ISL_848339, EPI_ISL_848340, EPI_ISL_848341, EPI_ISL_848342, EPI_ISL_848343, EPI_ISL_848344, EPI_ISL_848345, EPI_ISL_848346, EPI_ISL_848347, EPI_ISL_848348, EPI_ISL_848349, EPI_ISL_848350, EPI_ISL_848351, EPI_ISL_848352, EPI_ISL_848353, EPI_ISL_848354, EPI_ISL_848355, EPI_ISL_848440, EPI_ISL_848453, EPI_ISL_848463, EPI_ISL_848498, EPI_ISL_848499, EPI_ISL_848500, EPI_ISL_848501, EPI_ISL_848502, EPI_ISL_848503, EPI_ISL_848504, EPI_ISL_848505, EPI_ISL_848506, EPI_ISL_848507, EPI_ISL_848508, EPI_ISL_848509 | | | |
| see above | Illinois Department of Public Health | Gagnon Lab, Southern Illinois University | Keith Gagnon |
| EPI_ISL_848951, EPI_ISL_848952, EPI_ISL_848953, EPI_ISL_848954, | Florida Bureau of Public Health Laboratories | Florida Bureau of Public Health Laboratories | Sarah Schmedes, Jason Blanton |

| | | | |
|---|---|---|---|
| EPI_ISL_848955, EPI_ISL_848956, EPI_ISL_848957, EPI_ISL_848958, EPI_ISL_848959 | | | |
| EPI_ISL_849373, EPI_ISL_849374, EPI_ISL_849375, EPI_ISL_849376, EPI_ISL_849377, EPI_ISL_849378, EPI_ISL_849379, EPI_ISL_849380, EPI_ISL_849381, EPI_ISL_849382 | Servicio Virosis Respiratorias-Departamento Virología-INEI | Instituto Nacional Enfermedades Infecciosas C.G.Malbran | Baumeister E., Avaro M., Benedetti E., Russo M., Dattero ME, Pontoriero A., Cisterna D., Molina V., Perandones C., Tuduri E., Lorenzo F., Poklepovich T., Campos J. |
| EPI_ISL_849424, EPI_ISL_849484, EPI_ISL_849485 | Seattle Flu Study | Seattle Flu Study | Deborah A. Nickerson, Chris D. Frazar, Jover Lee, Benjamin Pelle, Matthew Richardson, Amanda Adler, Elisabeth Brandstetter, Peter D. Han, Kairsten Fay, Misja Ilcisin, Kirsten Lacombe, Thomas R. Sibley, Melissa Truong, Caitlin R. Wolf, Michael Boeckh, Janet A. Englund, Michael Famulare, Barry R. Lutz, Mark J. Rieder, Lea M. Starita, Matthew Thompson, Jay Shendure, Trevor Bedford, Helen Y. Chu |
| EPI_ISL_849487, EPI_ISL_849488 | Seattle Flu Study | Seattle Flu Study | Deborah A. Nickerson, Chris D. Frazar, Jover Lee, Benjamin Pelle, Matthew Richardson, Amanda Adler, Elisabeth Brandstetter, Peter D. Han, Kairsten Fay, Misja Ilcisin, Kirsten Lacombe, Thomas R. Sibley, Melissa Truong, Caitlin R. Wolf, Karen Cowgill, Stephanie Schrag, Jeff Duchin, Michael Boeckh, Janet A. Englund, Michael Famulare, Barry R. Lutz, Mark J. Rieder, Lea M. Starita, Matthew Thompson, Helen Y. Chu, Trevor Bedford, Jay Shendure |
| EPI_ISL_849489, EPI_ISL_849594, EPI_ISL_849595, EPI_ISL_849597, EPI_ISL_849598 | Seattle Flu Study | Seattle Flu Study | Deborah A. Nickerson, Chris D. Frazar, Jover Lee, Benjamin Pelle, Matthew Richardson, Amanda Adler, Elisabeth Brandstetter, Peter D. Han, Kairsten Fay, Misja Ilcisin, Kirsten Lacombe, Thomas R. Sibley, Melissa Truong, Caitlin R. Wolf, Michael Boeckh, Janet A. Englund, Michael Famulare, Barry R. Lutz, Mark J. Rieder, Lea M. Starita, Matthew Thompson, Jay Shendure, Trevor Bedford, Helen Y. Chu |
| EPI_ISL_852654 | Institute of Virology, Medical Center, University of Freiburg, Freiburg, Germany | Institute of Virology, Clinial Virus Genomics, Medical Center, University of Freiburg, Freiburg, Germany | Jonas Fuchs, Lisa Kern, Sandra Reuter, Hajo Grundmann, Marcus Panning |
| EPI_ISL_853302, EPI_ISL_853304, EPI_ISL_853307, EPI_ISL_853323, EPI_ISL_853352 | UPMC Clinical Microbiology Laboratory | Microbial Genome Sequencing Center; Microbial Genomic Epidemiology Laboratory | Mustapha M. Mustapha, Jane W. Marsh, Dan Snyder, Marissa P. Griffith, Stephanie L. Mitchell, Vatsala R. Srinivasa, Kady D. Waggle, Chinelo Ezeonwuku, Vaughn S. Cooper, Lee H. Harrison |
| EPI_ISL_853784, EPI_ISL_853856, EPI_ISL_853857, EPI_ISL_853858, EPI_ISL_853859, EPI_ISL_853860, EPI_ISL_853861, EPI_ISL_853862, EPI_ISL_853863, EPI_ISL_853864, EPI_ISL_853865, EPI_ISL_853915, EPI_ISL_853916, EPI_ISL_854227 | | | |
| see above | Institute for Laboratory Diagnostics and Microbiology, Klinikum Klagenfurt am Worthersee | Bergthaler laboratory, CeMM Research Center for Molecular Medicine of the Austrian Academy of Sciences | Lukas Endler, Alexandra Popa, Benedikt Agerer, Jakob-Wendelin Genger, Alexander Lercher, Anna Schedl, Thomas Penz, Michael Schuster, Jan Laine, Martin Senekowitsch, Christoph Bock, Andreas Bergthaler |
| EPI_ISL_854236 | Department of Microbiology, University Innsbruck | Bergthaler laboratory, CeMM Research Center for Molecular Medicine of the Austrian Academy of Sciences | Lukas Endler, Alexandra Popa, Benedikt Agerer, Jakob-Wendelin Genger, Alexander Lercher, Anna Schedl, Thomas Penz, Michael Schuster, Jan Laine, Martin Senekowitsch, Christoph Bock, Andreas Bergthaler |
| EPI_ISL_855497, EPI_ISL_855501, EPI_ISL_855513 | KEMRI-Wellcome Trust Research Programme/KEMRI-CGMR-C Kilifi | KEMRI-Wellcome Trust Research Programme/KEMRI-CGMR-C Kilifi | Githinji et al |
| EPI_ISL_859792, EPI_ISL_859794, EPI_ISL_859795, EPI_ISL_859796 | BTC, Khalifa University | BTC, Khalifa University | Al Safar et al |
| EPI_ISL_860119, EPI_ISL_860120, EPI_ISL_860121, EPI_ISL_860124, EPI_ISL_860125, EPI_ISL_860126, EPI_ISL_860127, EPI_ISL_860128, EPI_ISL_860129, EPI_ISL_860147 | Keio University School of Medicine | Keio University School of Medicine | Kenjiro Kosaki, Yuka Iwasaki, Hirotsugu Ishizu, Haruhiko Siomi, Kodai Abe |
| EPI_ISL_860689 | Respiratory Virus Unit, National Infection Service, Public Health England | COVID-19 Genomics UK (COG-UK) Consortium | PHE Covid Sequencing Team |
| EPI_ISL_860746 | Swiss National Reference Centre for Influenza | Swiss National Reference Centre for Influenza | Ana Rita Gonçalves Cabecinhas, Samuel Cordey, Florian Laubscher,Christoph Grünig, Laurent Kaiser |
| EPI_ISL_861717, EPI_ISL_861729 | Department of Medical Microbiology, Hospital Pengajar Universiti Putra Malaysia | Malaysia Genome Institute | Mohd Noor Mat Isa, Syafinaz Amin-Nordin, Irni Suhayu Sapian, Hui-Yee Chee, Yusuf Muhammad Noor, Nurhezreen Md Iqbal, Enizza Kasim, Siti Noraini Othman, Mohd Faizal Abu Bakar, Shamsidar Sopie, Azrin Ahmad, Narcisse Joseph, Muhammad MI, Avisha Richards, Nor Zahrin Hasran, Nor Azfa Johari |
| EPI_ISL_867936, EPI_ISL_867937, EPI_ISL_867938, EPI_ISL_867954, EPI_ISL_867958, EPI_ISL_867959, EPI_ISL_867960, EPI_ISL_867961, EPI_ISL_867971, EPI_ISL_867972, EPI_ISL_867973, EPI_ISL_867974, EPI_ISL_867976, EPI_ISL_867977, EPI_ISL_867978, EPI_ISL_867979, EPI_ISL_867980, EPI_ISL_867981 | | | |
| see above | Centre for Enzyme Innovation, University of Portsmouth / Translational Research Laboratory, Portsmouth Hospitals NHS Trust | COVID-19 Genomics UK (COG-UK) Consortium | Angela Beckett,Yann Bourgeois,Garry Scarlett,Sharon Glaysher,Scott Elliott,Kelly Bicknell,Robert Impey,Allyson Lloyd,Sarah Wyllie,Ethan Butcher,Anoop Chauhan,Samuel Robson |
| EPI_ISL_868797, EPI_ISL_868821, EPI_ISL_868823, EPI_ISL_868824, EPI_ISL_868825, EPI_ISL_868826, EPI_ISL_868827, EPI_ISL_868828, EPI_ISL_868833, EPI_ISL_868835, EPI_ISL_868836, EPI_ISL_868839, EPI_ISL_868840 | | | |
| see above | Bioinformatics and Biostatistics Lab, Advanced Sequencing Facility | COVID-19 Genomics UK (COG-UK) Consortium | Aengus Stewart,Jerome Nicod,Chelsea Sawyer,Laura Cubitt,Harshil Patel,Margaret Crawford |
| EPI_ISL_871994 | Hospital de Madrid | Instituto de Salud Carlos III | Iglesias-Caballero, M. Camarero, S. Molinero Calamita, M. González-Esguevillas, M. Pozo, F. Casas, I. Jiménez, P. Jiménez, M. Zaballos, A. Monzón, S. Varona, S. Juliá, M. Cuesta, I. Alhambra, A. |
| EPI_ISL_872362 | Texas Department of State Health Services (TXDSHS) | Texas Department of State Health Services (TXDSHS) | Bonnie Oh, Anita Pokharel, James Daniel Bonser, Myong Koag, Chung Wang, Rachel Lee, Grace Kubin, Rashmi Tuladhar, Mayela Pedrueza, Maliha Rahman, Jenny Zhang |
| EPI_ISL_872445, EPI_ISL_872446, EPI_ISL_872447, EPI_ISL_872448, EPI_ISL_872449, EPI_ISL_872450, EPI_ISL_872451, EPI_ISL_872452, EPI_ISL_872453, EPI_ISL_872454, EPI_ISL_872455, EPI_ISL_872456, EPI_ISL_872457 | | | |
| see above | Eurofins Diatherix | Hudsonalpha Genome Sequencing Center | Jane Grimwood, Melissa Williams, Lori H. Handley, Joshua Stough, Leslie Malone, Stefan Brzezinski, Ada Stewart, Teresa Jones, Jenell Webber, John Lovell, Jennifer Cart, and Jeremy Schmutz |
| EPI_ISL_875538 | Institute of Virology, Biomedical Research Center of the Slovak Academy of Sciences, Bratislava | Faculty of Natural Sciences, Comenius University, Bratislava | Viktória abanová, Kristína Boršová, Broa Brejová, Viktória Hodorová, Sabina Fumaová Havlíková, Juraj Kopáek, Martina Liková, ubomíra Lukáiková, Martina Neboháová, Monika Sláviková, Tomáš Vina, Jozef Nosek, Boris Klempa |
| EPI_ISL_876036 | Montefiore Medical Center | Albert Einstein College of Medicine, Dept. of Microbiology & Immunology, Chandran lab | J. Maximilian Fels, Saad Khan, Ryan Forster, Karin A. Skalina, Surksha Sirichand, Amy S. Fox, Aviv Bergman, William B. Mitchell, Lucia R. Wolgast, Wendy Szymczak, Robert H. Bortz III, M. Eugenia Dieterle, Catalina Florez, Denise Haslwanter, Rohit K. Jangra, Ethan Laudermilch, Ariel S. Wirchnianski, Jason Barnhill, David L. Goldman, Hnin Khine, D. Yitzchak Goldstein, Johanna P. Daily, Kartik Chandran, Libusha Kelly |
| EPI_ISL_876572 | Florida Bureau of Public Health Laboratories | Florida Bureau of Public Health Laboratories | Sarah Schmedes, Jason Blanton |
| EPI_ISL_876877, EPI_ISL_876878 | Quest Diagnostics | Quest Diagnostics | Rosenthal,S.H., Gerasimova,A., Kagan,R.M., Anderson, B., Hua, M., Liu Y., Bernstein, L.E., Livingston, K.E., Perez, A., Shalhout, D.F., Shlyakhter, I.A., Owen, R., Tanpaiboon, P., Lacbawan, F. |
| EPI_ISL_877431 | Institute of Microbiology and Immunology, Faculty of Medicine, University of Ljubljana | Institute of Microbiology and Immunology, Faculty of Medicine, University of Ljubljana | Samo Zakotnik, Tomaž Mark Zorec, Matic Brvar, Miša Korva, Mario Poljak, Tatjana Avši - Županc |
| EPI_ISL_877433 | National laboratory of health, environment and food Celje | Institute of Microbiology and Immunology, Faculty of Medicine, University of Ljubljana | Samo Zakotnik, Tomaž Mark Zorec, Matic Brvar, Miša Korva, Mario Poljak, Tatjana Avši - Županc |
| EPI_ISL_877764 | COVID-19 lab, MMC Department of Microbiology, Mymensingh Medical college | Department of Pathology, Bangladesh Agricultural University & Department of Microbiology, Mymensingh Medical College | Afrin, S. Z. Paul, S. K. Parvin, R. |
| EPI_ISL_878560 | Robert Garry lab | Andersen lab at Scripps Research | Allison Smither, Gilberto Sabino-Santos, Patricia Snarski, Lilia Melnik, Antoinette Bell, Kaylynn Genemaras, Arnaud Drouin, Dahlene Fusco, Robert Garry with SEARCH Alliance San Diego |
| EPI_ISL_878637, EPI_ISL_878640, | Rady's Childrens Hospital | Andersen lab at Scripps Research | SEARCH Alliance San Diego with Nanda Radamchar, David Dimmock, Linda Luo, Christina Clarke, Kathryn Bouic, Teresa Mueller, Denise Malicki |

| | | | |
|---|---|---|---|
| EPI_ISL_878643, EPI_ISL_878645, EPI_ISL_878664, EPI_ISL_878677 | | | |
| EPI_ISL_882677, EPI_ISL_882678, EPI_ISL_882683, EPI_ISL_882713, EPI_ISL_882714, EPI_ISL_882715, EPI_ISL_882716, EPI_ISL_882739, EPI_ISL_882755, EPI_ISL_882756, EPI_ISL_882757, EPI_ISL_882760, EPI_ISL_882762 | | | |
| see above | 1.AO Universitaria 'S. Giovanni di Dio e Ruggi D'Aragona, Scuola Medica Salernitana' Hospital / 2.UOC di Virologia e Microbiologia, Università della Campania 'L. Vanvitelli' / 3.AO Universitaria 'Federico II' Napoli Hospital / 4.AORN 'San Giuseppe Moscati' Avellino Hospital / 5.AO 'San Pio - presidio G. Rummo' Benevento Hospital / 6.AO 'Sant'Anna e San Sebastiano' Caserta Hospital / 7.PO 'Maria Santissima Addolorata' Eboli Hospital / 8.Biogem Istituto di Ricerche Genetiche | 1. Genome Research Center for Health (CRGS) / 2. Laboratory of Molecular Medicine and Genomics(LMMGe) / 3. Center for Research in Pure and Applied Mathematics (CRMPA) | Giorgio Giurato, Francesca Rizzo, Alessandro Weisz, Gianluigi Franci, Giovanni Nassa, Pasquale Pagliano, Roberta Tarallo, Elena Alexandrova, Ylenia D'Agostino, Carlo Ferravante, Jessica Lamberti, Viola Melone, Domenico Memoli, Valeria Mirici Cappa, Domenico Palumbo, Giovanni Pecoraro, Assunta Sellitto, Oriana Strianese, Ilaria Terenzi, Giuseppe Fenza, Aniello Gentile, Antonello Saccomanno, Sonia Amabile, Teresa Rocco, Annamaria Salvati, Emilia Vaccaro, Massimiliano Galdiero, Michele Cennamo, Giuseppe Portella, Maria Grazia Foti, Mariarosaria Ingino, Maria Landi, Maurizio Fumi, Vincenzo Rocco, Rita Greco, Vittoria Letizia, Arnolfo Petruzziello, Maddalena Schioppa, Gregorio Goffredi, Francesca Marciano, Michele Caraglia, Alessia Cossu, Marianna Scrima |
| EPI_ISL_884437, EPI_ISL_884438 | Infectious Diseases, Quest Diagnostics | Infectious Diseases, Quest Diagnostics | Rosenthal,S.H., Gerasimova,A., Kagan,R.M., Anderson,B., Bernstein,L.E., Livingston,K.E., Hua,M., Liu,Y., Shalhout,D.F., Owen,R., Lacbawan,F. |
| EPI_ISL_887154, EPI_ISL_887155 | Massachusetts General Hospital | Infectious Disease Program, Broad Institute of Harvard and MIT | Lemieux,J.E., Siddle,K.J., Shaw,B., Adams,G., Pierce,V., Turbett,S., Anahtar,M., Branda,J., Slater,D., Harris,J., Lin,A.E., Gladden-Young,A., Lagerborg,K., Rudy,M., DeRuff,K., Carter,A., Normandin,E., Bauer,M., Reilly,S., Tomkins-Tinch,C., Loreth,C., Chaluvadi,S., Neumann,A., Cusick,C., Chapman,S.B., Gnirke,A., Flowers,K., Cerrato,F., Birren,B.W., Gallagher,G., Smole,S., Park,D.J., MacInnis,B.L., Ryan,E., LaRocque,R., Rosenberg,E. and Sabeti,P.C. |
| EPI_ISL_888702, EPI_ISL_888703, EPI_ISL_888707, EPI_ISL_888708, EPI_ISL_888709, EPI_ISL_888711, EPI_ISL_888712, EPI_ISL_888716, EPI_ISL_888717, EPI_ISL_888718, EPI_ISL_888721, EPI_ISL_888722, EPI_ISL_888723, EPI_ISL_888725, EPI_ISL_888726, EPI_ISL_888727, EPI_ISL_888728, EPI_ISL_888730, EPI_ISL_888733, EPI_ISL_888736, EPI_ISL_888738 | | | |
| see above | KU Leuven, Rega Institute, Clinical and Epidemiological Virology | KU Leuven, Rega Institute, Clinical and Epidemiological Virology | Tony Wawina-Bokalanga, Bert Vanmechelen, Joan Marti-Carerras, Piet Maes |
| EPI_ISL_891182 | DPH, Massachusetts State Public Health Lab | DPH, Massachusetts State Public Health Lab | Lang,A.S., Fink,T., Gallagher,G.R., Smole,S.C. |
| EPI_ISL_892225 | Lighthouse Lab in Cambridge | Wellcome Sanger Institute for the COVID-19 Genomics UK (COG-UK) Consortium | Rob Howes, The Lighthouse Lab in Cambridge and Alex Alderton, Roberto Amato, Sonia Goncalves, Ewan Harrison, David K. Jackson, Ian Johnston, Dominic Kwiatkowski, Cordelia Langford, John Sillitoe on behalf of the Wellcome Sanger Institute COVID-19 Surveillance Team |
| EPI_ISL_892226 | Lighthouse Lab in Alderley Park | Wellcome Sanger Institute for the COVID-19 Genomics UK (COG-UK) Consortium | Jacquelyn Wynn, Mairead Hyland, The Lighthouse Lab in Alderley Park and Alex Alderton, Roberto Amato, Sonia Goncalves, Ewan Harrison, David K. Jackson, Ian Johnston, Dominic Kwiatkowski, Cordelia Langford, John Sillitoe on behalf of the Wellcome Sanger Institute COVID-19 Surveillance Team |
| EPI_ISL_896122, EPI_ISL_896132, EPI_ISL_896137, EPI_ISL_896139, EPI_ISL_900055, EPI_ISL_900056, EPI_ISL_900073, EPI_ISL_900143, EPI_ISL_900169, EPI_ISL_900196, EPI_ISL_900203, EPI_ISL_900211, EPI_ISL_900227, EPI_ISL_900259, EPI_ISL_900269, EPI_ISL_900271, EPI_ISL_900284, EPI_ISL_900289, EPI_ISL_900307, EPI_ISL_900320, EPI_ISL_900364, EPI_ISL_900372, EPI_ISL_900393, EPI_ISL_900400, EPI_ISL_900412, EPI_ISL_900413, EPI_ISL_900414, EPI_ISL_900429, EPI_ISL_900482, EPI_ISL_900486, EPI_ISL_900487, EPI_ISL_900490 | | | |
| see above | MEPHI, Aix Marseille University | MEPHI, Aix Marseille University | Anthony LEVASSEUR |
| EPI_ISL_904171, EPI_ISL_904229, EPI_ISL_904427, EPI_ISL_904428, EPI_ISL_904429, EPI_ISL_904430 | Dutch COVID-19 response team | Erasmus Medical Center | Bas Oude Munnink, Reina Sikkema, David Nieuwenhuijse, Irina Chestakova, Anne van der Linden, Marjan Boter, Emmanuelle Munger, Corine GeurtsvanKessel, Annemiek van der Eijk, Richard Molenkamp, Marion Koopmans, on behalf of the Dutch national COVID-19 response team. |
| EPI_ISL_906117 | Institute of Microbiology and Immunology, Faculty of Medicine, University of Ljubljana | Institute of Microbiology and Immunology, Faculty of Medicine, University of Ljubljana | Samo Zakotnik, Tomaž Mark Zorec, Matic Brvar, Miša Korva, Mario Poljak, Tatjana Avši - Županc |
| EPI_ISL_909744 | Institute for Lung Diseases in Children - Skopje | Research Center for Genetic Engineering and Biotechnology "Georgi D. Efremov" , Macedonian Academy of Sciences and Arts | RCGEB - MASA |
| EPI_ISL_913070, EPI_ISL_913074, EPI_ISL_913080, EPI_ISL_913081 | Center for Virology | Center for Virology | Jeremy V. Camp, Irene Goerzer, Monika Redlberger-Fritz, Stephan W. Aberle |
| EPI_ISL_913326 | Klinisk mikrobiologi | The Public Health Agency of Sweden | Anna-Malin Linde, Maria Lind Karlberg, Carlo Berg, Oskar Karlsson Lindsjo, Sofia Stamouli, Reza Advani, Mattias Haukland, Petra Holmstrom, Noura Walai, Petra Edquist, Mia Brytting, Anna Risberg, Karin Tegmark-Wisell |
| EPI_ISL_915404 | Keio University School of Medicine | Keio University School of Medicine | Kenjiro Kosaki, Yuka Iwasaki, Hirotsugu Ishizu, Haruhiko Siomi, Kodai Abe |
| EPI_ISL_918125 | Lighthouse Lab in Alderley Park | Wellcome Sanger Institute for the COVID-19 Genomics UK (COG-UK) Consortium | Jacquelyn Wynn, Mairead Hyland, The Lighthouse Lab in Alderley Park and Alex Alderton, Roberto Amato, Sonia Goncalves, Ewan Harrison, David K. Jackson, Ian Johnston, Dominic Kwiatkowski, Cordelia Langford, John Sillitoe on behalf of the Wellcome Sanger Institute COVID-19 Surveillance Team |
| EPI_ISL_924078 | Oxford Viromics, NDM, University of Oxford; Oxford University Hospitals; Basingstoke and North Hampshire Hospital | COVID-19 Genomics UK (COG-UK) Consortium | Tanya Golubchik, David Bonsall, George Macintyre, Amy Trebes, Mariateresa de Cesare, Catrin Moore, Alex Mobbs, Anita Justice, Robert Shaw, Monique Andersson, Timothy Peto, Emma Wise, Nathan Moore, Jessica Lynch, Nick Cortes, Matilde Mori, Stephen Kidd, David Buck, John Todd, Christophe Fraser |
| EPI_ISL_924341, EPI_ISL_924426 | Virology Department, Sheffield Teaching Hospitals NHS Foundation Trust/Department of Infection, Immunity and Cardiovascular Disease, The Medical School, University of Sheffield | COVID-19 Genomics UK (COG-UK) Consortium | Thushan de Silva, Matthew Parker, Nikki Smith, Adri Angyal, Rebecca Brown, Luke Green, Rachel Tucker, Paul Parsons, Danielle Groves, Katie Johnson, Laura Carrilero, Alex Keeley, Dave Partridge, Matthew Wyles, Benjamin Lindsey, Mehmet Yavuz, Mohammad Raza, Cariad Evans |
| EPI_ISL_925097, EPI_ISL_925102, EPI_ISL_925119, EPI_ISL_925124 | 1.AO Universitaria 'S. Giovanni di Dio e Ruggi D'Aragona, Scuola Medica Salernitana' Hospital / 2.UOC di Virologia e Microbiologia, Università della Campania 'L. Vanvitelli' / 3.AO Universitaria 'Federico II' Napoli Hospital / 4.AORN 'San Giuseppe Moscati' Avellino Hospital / 5.AO 'San Pio - presidio G. Rummo' Benevento Hospital / 6.AO 'Sant'Anna e San Sebastiano' Caserta Hospital / 7.PO 'Maria Santissima Addolorata' Eboli Hospital / 8.Biogem Istituto di Ricerche Genetiche | 1. Genome Research Center for Health (CRGS) / 2. Laboratory of Molecular Medicine and Genomics(LMMGe) / 3. Center for Research in Pure and Applied Mathematics (CRMPA) | Giorgio Giurato, Francesca Rizzo, Alessandro Weisz, Gianluigi Franci, Giovanni Nassa, Pasquale Pagliano, Roberta Tarallo, Elena Alexandrova, Ylenia D'Agostino, Carlo Ferravante, Jessica Lamberti, Viola Melone, Domenico Memoli, Valeria Mirici Cappa, Domenico Palumbo, Giovanni Pecoraro, Assunta Sellitto, Oriana Strianese, Ilaria Terenzi, Giuseppe Fenza, Aniello Gentile, Antonello Saccomanno, Sonia Amabile, Teresa Rocco, Annamaria Salvati, Emilia Vaccaro, Massimiliano Galdiero, Michele Cennamo, Giuseppe Portella, Maria Grazia Foti, Mariarosaria Ingino, Maria Landi, Maurizio Fumi, Vincenzo Rocco, Rita Greco, Vittoria Letizia, Arnolfo Petruzziello, Maddalena Schioppa, Gregorio Goffredi, Francesca Marciano, Michele Caraglia, Alessia Cossu, Marianna Scrima |
| EPI_ISL_925440, EPI_ISL_925472, EPI_ISL_925473, EPI_ISL_925474, EPI_ISL_925475 | Department of Clinical Microbiology | GIGA Medical Genomics | Keith Durkin, Maria Artesi, Sébastien Bontems, Raphaël Boreux, Bouchra Boujemla, Cécile Meex, Pierrette Melin, Marie-Pierre Hayette, Vincent Bours |
| EPI_ISL_935129, EPI_ISL_935130, EPI_ISL_935131 | 1.AO Universitaria 'S. Giovanni di Dio e Ruggi D'Aragona, Scuola Medica Salernitana' Hospital / 2.UOC di Virologia e Microbiologia, Università della Campania 'L. Vanvitelli' / 3.AO Universitaria 'Federico II' Napoli Hospital / 4.AORN 'San Giuseppe Moscati' Avellino Hospital / 5.AO 'San Pio - presidio G. Rummo' Benevento Hospital / 6.AO 'Sant'Anna e San Sebastiano' Caserta Hospital / 7.PO 'Maria Santissima Addolorata' Eboli Hospital / 8.Biogem Istituto di Ricerche Genetiche | 1. Genome Research Center for Health (CRGS) / 2. Laboratory of Molecular Medicine and Genomics(LMMGe) / 3. Center for Research in Pure and Applied Mathematics (CRMPA) | Giorgio Giurato (Corresponding Author), Francesca Rizzo (Corresponding Author), Alessandro Weisz (Corresponding Author), Gianluigi Franci, Giovanni Nassa, Pasquale Pagliano, Roberta Tarallo, Elena Alexandrova, Ylenia D'Agostino, Carlo Ferravante, Jessica Lamberti, Viola Melone, Domenico Memoli, Valeria Mirici Cappa, Domenico Palumbo, Giovanni Pecoraro, Assunta Sellitto, Oriana Strianese, Ilaria Terenzi, Giuseppe Fenza, Aniello Gentile, Antonello Saccomanno, Sonia Amabile, Teresa Rocco, Annamaria Salvati, Emilia Vaccaro, Massimiliano Galdiero, Michele Cennamo, Giuseppe Portella, Maria Grazia Foti, Mariarosaria Ingino, Maria Landi, Maurizio Fumi, Vincenzo Rocco, Rita Greco, Vittoria Letizia, Arnolfo Petruzziello, Maddalena Schioppa, Gregorio Goffredi, Francesca Marciano, Michele Caraglia, Alessia Cossu, Marianna Scrima |
| EPI_ISL_936496 | Institute for Medical Research, Infectious Disease Research Centre, National Institutes of Health, Ministry of Health Malaysia | Institute for Medical Research, Infectious Disease Research Centre, National Institutes of Health, Ministry of Health Malaysia | Suppiah J, Kamel K, Azizan MA, Thayan R |
| EPI_ISL_936583, EPI_ISL_936584, | Northwestern Memorial Hospital | Ozer Lab | Ramon Lorenzo-Redondo, Lacy M. Simons, Chad J. Achenbach, Lawrence J. Jennings, Michael G. Ison, Judd F. Hultquist, Egon A. Ozer |

| | | | |
|---|---|---|---|
| EPI_ISL_936585, EPI_ISL_936586, EPI_ISL_936587, EPI_ISL_936855, EPI_ISL_936856, EPI_ISL_936857 | | | |
| EPI_ISL_937054, EPI_ISL_937055, EPI_ISL_937060, EPI_ISL_937061, EPI_ISL_937071, EPI_ISL_937081, EPI_ISL_937082, EPI_ISL_937095, EPI_ISL_937102, EPI_ISL_937103, EPI_ISL_937104, EPI_ISL_937109, EPI_ISL_937116, EPI_ISL_937117 | | | |
| see above | Quest Diagnostics | Quest Diagnostics | Rosenthal,S.H., Gerasimova,A., Kagan,R.M., Anderson, B., Livingston, K.E., Hua, M., Liu Y., Shalhout, D.F., Owen, R., Lacbawan, F. |
| EPI_ISL_940976, EPI_ISL_940977, EPI_ISL_940978, EPI_ISL_940979, EPI_ISL_940981, EPI_ISL_940982 | Centers for Disease Control and Prevention, Dengue Branch | Centers for Disease Control and Prevention, Dengue Branch | Gilberto A. Santiago, Glenda Gonzalez, Betzabel Flores, Keyla Charriez, Gabriela Paz-Bailey, Jorge L. Munoz-Jordan |
| EPI_ISL_941175, EPI_ISL_941176, EPI_ISL_941177, EPI_ISL_941179, EPI_ISL_941180, EPI_ISL_941181, EPI_ISL_941182, EPI_ISL_941183, EPI_ISL_941184 | Servicio de Microbiología. Hospital Clínico Universitario de Valencia | SeqCOVID-SPAIN consortium/IBV(CSIC) | David Navarro Ortega, Eliseo Albert Vicent, Ignacio Torres and SeqCOVID-SPAIN consortium |
| EPI_ISL_943978, EPI_ISL_943979, EPI_ISL_943980 | LACEN do Estado de Tocantins | Instituto Adolfo Lutz, Interdiciplinary Procedures Center, Strategic Laboratory | Claudio Tavares Sacchi, Claudia Regina Gonçalves, Erica Valessa Ramos Gomes, Karoline Rodrigues Campos |
| EPI_ISL_949205, EPI_ISL_949216 | Departamento de Microbiología, CDB, Hospital Clínic, Barcelona | SeqCOVID-SPAIN consortium/IBV(CSIC) | Andrea Vergara, Mikel Martínez, Elisa Rubio, Jéssica Navero, Aida Peiró and SeqCOVID-SPAIN consortium |
| EPI_ISL_952943, EPI_ISL_952948 | Department of Pathology, University of Cambridge | COVID-19 Genomics UK (COG-UK) Consortium | Aminu S. Jahun, Yasmin Chaudhry, Iliana Georgana, Myra Hosmillo, Rhys Izu, Martin D. Curran, Surendra Parmar, Ian Goodfellow |
| EPI_ISL_954184, EPI_ISL_954185 | 1.AO Universitaria 'S. Giovanni di Dio e Ruggi D'Aragona, Scuola Medica Salernitana' Hospital / 2.UOC di Virologia e Microbiologia, Università della Campania 'L. Vanvitelli' / 3.AO Universitaria 'Federico II' Napoli Hospital / 4.AORN 'San Giuseppe Moscati' Avellino Hospital / 5.AO 'San Pio - presidio G. Rummo' Benevento Hospital / 6.AO 'Sant'Anna e San Sebastiano' Caserta Hospital / 7.PO 'Maria Santissima Addolorata' Eboli Hospital / 8.Biogem Istituto di Ricerche Genetiche | 1. Genome Research Center for Health (CRGS) / 2. Laboratory of Molecular Medicine and Genomics(LMMGe) / 3. Center for Research in Pure and Applied Mathematics (CRMPA) | Giorgio Giurato, Francesca Rizzo, Alessandro Weisz, Gianluigi Franci, Giovanni Nassa, Pasquale Pagliano, Roberta Tarallo, Elena Alexandrova, Ylenia D'Agostino, Carlo Ferravante, Jessica Lamberti, Viola Melone, Domenico Memoli, Valeria Mirici Cappa, Domenico Palumbo, Giovanni Pecoraro, Assunta Sellitto, Oriana Strianese, Ilaria Terenzi, Giuseppe Fenza, Aniello Gentile, Antonello Saccomanno, Sonia Amabile, Teresa Rocco, Annamaria Salvati, Emilia Vaccaro, Massimiliano Galdiero, Michele Cennamo, Giuseppe Portella, Maria Grazia Foti, Mariarosaria Ingino, Maria Landi, Maurizio Fumi, Vincenzo Rocco, Rita Greco, Vittoria Letizia, Arnolfo Petruzziello, Maddalena Schioppa, Gregorio Goffredi, Francesca Marciano, Michele Caraglia, Alessia Cossu, Marianna Scrima, Edmondo Adorisio, Morena D'Avenia, Michela Iacobellis, Rosanna Piluscio, Giorgio Dirani, Vittorio Sambri, Simona Semprini, Silvia Zanoli, Francesco Curcio, Stefania Marzinotto, Andreina Baj, Fausto Sessa. |
| EPI_ISL_959504 | Division of Emerging Infectious Diseases, Bureau of Infectious Diseases Diagnosis Control, Korea Disease Control and Prevention Agency | Division of Emerging Infectious Diseases, Bureau of Infectious Diseases Diagnosis Control, Korea Disease Control and Prevention Agency | Ae Kyung Park, Il-Hwan Kim, Heui Man Kim, Jeong-Min Kim, Namjoo Lee, Chae Young Lee, Sang Hee Woo, Eun-Jin Kim |
| EPI_ISL_960141 | Beaufort West Hospital wc BWH | National Health Laboratory Service/UCT | Arash Iranzadeh, Deelan Doolabh, Lynn Tyers, Bruna Galvao, Innocent Mudau, Marvin Hsiao, Kruger Marais, Diana Hardie, Stephen Korsman, Carolyn Williamson |
| EPI_ISL_960142, EPI_ISL_960143 | Groote Schuur Hospital wc GSH | National Health Laboratory Service/UCT | Arash Iranzadeh, Deelan Doolabh, Lynn Tyers, Bruna Galvao, Innocent Mudau, Marvin Hsiao, Kruger Marais, Diana Hardie, Stephen Korsman, Carolyn Williamson |
| EPI_ISL_960144 | Oudtshoorn Hospital wc OUD | National Health Laboratory Service/UCT | Arash Iranzadeh, Deelan Doolabh, Lynn Tyers, Bruna Galvao, Innocent Mudau, Marvin Hsiao, Kruger Marais, Diana Hardie, Stephen Korsman, Carolyn Williamson |
| EPI_ISL_960145 | Groote Schuur Hospital wc GSH | National Health Laboratory Service/UCT | Arash Iranzadeh, Deelan Doolabh, Lynn Tyers, Bruna Galvao, Innocent Mudau, Marvin Hsiao, Kruger Marais, Diana Hardie, Stephen Korsman, Carolyn Williamson |
| EPI_ISL_960146, EPI_ISL_960147 | Oudtshoorn Hospital wc OUD | National Health Laboratory Service/UCT | Arash Iranzadeh, Deelan Doolabh, Lynn Tyers, Bruna Galvao, Innocent Mudau, Marvin Hsiao, Kruger Marais, Diana Hardie, Stephen Korsman, Carolyn Williamson |
| EPI_ISL_960339, EPI_ISL_960342 | University of Wisconsin-Madison AIDS Vaccine Research Laboratories | University of Wisconsin-Madison AIDS Vaccine Research Laboratories | Gage Moreno, Katarina Braun, et al. AIDS Vaccine Research Laboratories |
| EPI_ISL_961921, EPI_ISL_961945, EPI_ISL_961946, EPI_ISL_961975, EPI_ISL_961976 | Illinois Department of Public Health | Gagnon Lab, Southern Illinois University | Keith Gagnon |
| EPI_ISL_965624, EPI_ISL_965718 | Dutch COVID-19 response team | Medical Microbiology, Maastricht University Medical Centre | Jozef Dingemans*, Brian van der Veer*, Erik Beuken, Carmen Reumkens, Lieke van Alphen, Christian Hoebe, Paul Savelkoul |
| EPI_ISL_966325, EPI_ISL_966326 | Kentucky State Public Health Lab | Kentucky State Public Health Lab | Stephanie Lunn, Karim George, Joshua Tobias, William Grooms, Vaneet Arora, Matthew Johnson, Rachel Zinner, Rhonda Lucas |
| EPI_ISL_968186 | Clinical Molecular Microbiology Laboratory, UNC Hospital | Dirk Dittmer | Justin T. Landis , Razia Moorad , Brent A. Eason, Melissa B. Miller, Linda Pluta, Dirk Dittmer, Angelica Juarez, Cecilia Thompson, Shawn Hawken, Cameroon Grant, Evelyn Hoffman, Patricio Cano, Jason Wong, Carolina Caro-Vegas, Ryan McNamara, Blossom Damania. |
| EPI_ISL_968881, EPI_ISL_968882 | KEMRI-Wellcome Trust Research Programme/KEMRI-CGMR-C Kilifi | KEMRI-Wellcome Trust Research Programme/KEMRI-CGMR-C Kilifi | Githinji et al |
| EPI_ISL_974158, EPI_ISL_974159, EPI_ISL_974160, EPI_ISL_974161, EPI_ISL_974162, EPI_ISL_974163, EPI_ISL_974164, EPI_ISL_974165, EPI_ISL_974166, EPI_ISL_974167, EPI_ISL_974168, EPI_ISL_974169, EPI_ISL_974170, EPI_ISL_974171, EPI_ISL_974172, EPI_ISL_974173, EPI_ISL_974174, EPI_ISL_974175, EPI_ISL_974176, EPI_ISL_974177, EPI_ISL_974178, EPI_ISL_974179, EPI_ISL_974180, EPI_ISL_974181, EPI_ISL_974182, EPI_ISL_974183, EPI_ISL_974184, EPI_ISL_974185, EPI_ISL_974186, EPI_ISL_974187, EPI_ISL_974188, EPI_ISL_974189, EPI_ISL_974190, EPI_ISL_974191, EPI_ISL_974192, EPI_ISL_974193, EPI_ISL_974194, EPI_ISL_974195, EPI_ISL_974196, EPI_ISL_974197, EPI_ISL_974198, EPI_ISL_974199, EPI_ISL_974200, EPI_ISL_974201, EPI_ISL_974202, EPI_ISL_974203, EPI_ISL_974204, EPI_ISL_974205, EPI_ISL_974206, EPI_ISL_974207, EPI_ISL_974208, EPI_ISL_974209, EPI_ISL_974210, EPI_ISL_974211, EPI_ISL_974212, EPI_ISL_974213, EPI_ISL_974214, EPI_ISL_974215, EPI_ISL_974216, EPI_ISL_974217, EPI_ISL_974218, EPI_ISL_974219, EPI_ISL_974220, EPI_ISL_974221, EPI_ISL_974222, EPI_ISL_974223, EPI_ISL_974224, EPI_ISL_974225, EPI_ISL_974226, EPI_ISL_974227, EPI_ISL_974228, EPI_ISL_974229, EPI_ISL_974230, EPI_ISL_974231, EPI_ISL_974232, EPI_ISL_974233, EPI_ISL_974234, EPI_ISL_974235, EPI_ISL_974236, EPI_ISL_974237, EPI_ISL_974238, EPI_ISL_974239, EPI_ISL_974240, EPI_ISL_974241, EPI_ISL_974242, EPI_ISL_974243, EPI_ISL_974244, EPI_ISL_974245, EPI_ISL_974246, EPI_ISL_974247, EPI_ISL_974248, EPI_ISL_974249, EPI_ISL_974250, EPI_ISL_974251, EPI_ISL_974252, EPI_ISL_974253, EPI_ISL_974254, EPI_ISL_974255, EPI_ISL_974256, EPI_ISL_974257, EPI_ISL_974258, EPI_ISL_974259, EPI_ISL_974260, EPI_ISL_974261, EPI_ISL_974262, EPI_ISL_974263, EPI_ISL_974264, EPI_ISL_974265, EPI_ISL_974266, EPI_ISL_974267, EPI_ISL_974268, EPI_ISL_974269, EPI_ISL_974270, EPI_ISL_974271, EPI_ISL_974272, EPI_ISL_974273, EPI_ISL_974274, EPI_ISL_974275, EPI_ISL_974276, EPI_ISL_974277, EPI_ISL_974278, EPI_ISL_974279, EPI_ISL_974280, EPI_ISL_974281, EPI_ISL_974282, EPI_ISL_974283 | | | |
| see above | BCCDC Public Health Laboratory | BCCDC Public Health Laboratory | Prystajecky Natalie, Linda Hoang, Dan Fornika, John Tyson, Shannon Russell, Kim Macdonald, Kimia Kamelian, Ana Pacagnella, Corrinne Ng, Loretta Janz, Robert Azana Terry Snutch, Mel Krajden |
| EPI_ISL_977585 | Department of Medical Microbiology, Hospital Pengajar Universiti Putra Malaysia | Malaysia Genome Institute | Mohd Noor Mat Isa, Syafinaz Amin-Nordin, Irni Suhayu Sapian, Hui-Yee Chee, Yusuf Muhammad Noor, Nurhezreen Md Iqbal, Enizza Kasim, Siti Noraini Othman, Mohd Faizal Abu Bakar, Shamsidar Sopie, Azrin Ahmad, Narcisse Joseph, Muhammad MI, Avisha Richards, Nor Zahrin Hasran, Nor Azfa Johari |
| EPI_ISL_977911, EPI_ISL_977913, EPI_ISL_977914, EPI_ISL_977917 | Chiu Laboratory, University of California, San Francisco | Chiu Laboratory, University of California, San Francisco | Charles Chiu, Xianding (Wayne) Deng, Candace Wang, Venice Servellita, Jill Hacker, Debra Wadford |
| EPI_ISL_982229, EPI_ISL_982230 | MEPHI, Aix Marseille University | MEPHI, Aix Marseille University | Anthony LEVASSEUR |
