## Supplementary material for "Detection and characterization of the SARS-CoV-2 lineage B.1.526 in New York": Supp. Table 3 GISAID Acknowledgment part 2: gisaid_hcov-19_acknowledgement_table_2021_02_12_23.pdf

We gratefully acknowledge the following Authors from the Originating laboratories responsible for obtaining the specimens, as well as the Submitting laboratories where the genome data were generated and shared via GISAID, on which this research is based.

All Submitters of data may be contacted directly via www.gisaid.org

Authors are sorted alphabetically.

| Accession ID | Originating Laboratory | Submitting Laboratory | Authors |
|---|---|---|---|
| EPI_ISL_493607 | Queens Medical Centre, Clinical Microbiology Department / DeepSeq Nottingham | COVID-19 Genomics UK (COG-UK) Consortium | Gemma Clark, Wendy Smith, Manjinder Khakh, Vicki M Fleming, Michelle M Lister, Hannah Howson-Wells, Jonathan Ball, Patrick McClure, Joseph Chappell, Theocharis Tsoleridis, Nadine Holmes, Matthew Carlisle, Christopher Moore, Fei Sang, Johnny Debebe, Victoria Wright, Matthew Loose |
| EPI_ISL_493684, EPI_ISL_493724 | Virology Department, Sheffield Teaching Hospitals NHS Foundation Trust/Department of Infection, Immunity and Cardiovascular Disease, The Medical School, University of Sheffield | COVID-19 Genomics UK (COG-UK) Consortium | Thushan de Silva, Matthew Parker, Nikki Smith, Adri Angyal, Rebecca Brown, Luke Green, Rachel Tucker, Paul Parsons, Danielle Groves, Katie Johnson, Laura Carrilero, Alex Keeley, Dave Partridge, Matthew Wyles, Benjamin Lindsey, Mehmet Yavuz, Mohammad Raza, Cariad Evans |
| EPI_ISL_497778, EPI_ISL_497785, EPI_ISL_497790, EPI_ISL_497794, EPI_ISL_497800, EPI_ISL_497801, EPI_ISL_497815, EPI_ISL_497838, EPI_ISL_497849, EPI_ISL_497853, EPI_ISL_497857, EPI_ISL_497867, EPI_ISL_497868, EPI_ISL_497869 | | | |
| see above | Department of Microbiology, The University of Hong Kong | Department of Microbiology, The University of Hong Kong | Kelvin K.W. To, Kwok-Yung Yuen |
| EPI_ISL_497950 | Shaoxing CDC | Zhejiang Provincial Center for Disease Control and Prevention | Yin Chen, Yanjun Zhang, Haiyan Mao, Junhang Pan, Xiuyu Lou, Yi Sun, Hao Yan, Zhen Li, Wen Shi |
| EPI_ISL_498069, EPI_ISL_498070, EPI_ISL_498071, EPI_ISL_498072, EPI_ISL_498073, EPI_ISL_498074, EPI_ISL_498075, EPI_ISL_498076, EPI_ISL_498077, EPI_ISL_498079, EPI_ISL_498080, EPI_ISL_498082, EPI_ISL_498083, EPI_ISL_498084, EPI_ISL_498085, EPI_ISL_498086, EPI_ISL_498087, EPI_ISL_498088, EPI_ISL_498089, EPI_ISL_498090, EPI_ISL_498091, EPI_ISL_498092, EPI_ISL_498093, EPI_ISL_498094, EPI_ISL_498095, EPI_ISL_498096, EPI_ISL_498097, EPI_ISL_498098, EPI_ISL_498099, EPI_ISL_498100, EPI_ISL_498101, EPI_ISL_498102, EPI_ISL_498103, EPI_ISL_498104, EPI_ISL_498105, EPI_ISL_498106, EPI_ISL_498107, EPI_ISL_498109, EPI_ISL_498110, EPI_ISL_498111, EPI_ISL_498112, EPI_ISL_498113 | | | |
| see above | NHLS-IALCH | KRISP, KZN Research Innovation and Sequencing Platform | Giandhari J, Pillay S, Lessells R, Chimukangara B, Mdlalose K, York D, Khan S, Tegally H, Wilkinson E, de Oliveira T |
| EPI_ISL_498268 | National Institute of Laboratory Medicine and Referral Center | Genomic Research Lab, BCSIR | Barna Goswami, Abu Sayeed Mohammad Mahmud, Mohammad Samir Uzzaman, Eshrar Osman, Md. Ahasan Habib, Shahina Akter, Tanjina Akhter Banu, Md. Murshed Hasan Sarkar, Iffat Jahan, Md. Saddam Hossain, Tasnim Nafisa, Md. Maruf Ahmed Molla, Mahmuda Yeasmin, Asish Kumar Ghosh, A. K. M. Shamsuzzaman, Sheikh Md. Selim Al Din, Utpal Chandra Ray, Salek Ahmed Sajib, Md. Salim Khan |
| EPI_ISL_498272, EPI_ISL_498273 | National Institute of Laboratory Medicine and Referral Center | Genomic Research Lab, BCSIR | Iffat Jahan, Abu Sayeed Mohammad Mahmud, Mohammad Samir Uzzaman, Eshrar Osman, Md. Ahasan Habib, Shahina Akter, Tanjina Akhter Banu, Md. Murshed Hasan Sarkar, Barna Goswami, Md. Saddam Hossain, Tasnim Nafisa, Md. Maruf Ahmed Molla, Mahmuda Yeasmin, Asish Kumar Ghosh, A. K. M. Shamsuzzaman, Sheikh Md. Selim Al Din, Utpal Chandra Ray, Salek Ahmed Sajib, Md. Salim Khan |
| EPI_ISL_498274, EPI_ISL_498417 | National Institute of Laboratory Medicine and Referral Center | Genomic Research Lab, BCSIR | Tasnim Nafisa, Abu Sayeed Mohammad Mahmud, Mohammad Samir Uzzaman, Eshrar Osman, Md. Ahasan Habib, Shahina Akter, Tanjina Akhter Banu, Md. Murshed Hasan Sarkar, Barna Goswami, Iffat Jahan, Md. Saddam Hossain, Md. Maruf Ahmed Molla, Mahmuda Yeasmin, Asish Kumar Ghosh, A. K. M. Shamsuzzaman, Sheikh Md. Selim Al Din, Utpal Chandra Ray, Salek Ahmed Sajib, Md. Salim Khan |
| EPI_ISL_498418, EPI_ISL_498419 | National Institute of Laboratory Medicine and Referral Center | Genomic Research Lab, BCSIR | Md. Maruf Ahmed Molla, Abu Sayeed Mohammad Mahmud, Mohammad Samir Uzzaman, Eshrar Osman, Md. Ahasan Habib, Shahina Akter, Tanjina Akhter Banu, Md. Murshed Hasan Sarkar, Barna Goswami, Iffat Jahan, Md. Saddam Hossain, Tasnim Nafisa, Mahmuda Yeasmin, Asish Kumar Ghosh, A. K. M. Shamsuzzaman, Sheikh Md. Selim Al Din, Utpal Chandra Ray, Salek Ahmed Sajib, Md. Salim Khan |
| EPI_ISL_498466, EPI_ISL_498467 | National Institute of Laboratory Medicine and Referral Center | Genomic Research Lab, BCSIR | Mahmuda Yeasmin, Abu Sayeed Mohammad Mahmud, Mohammad Samir Uzzaman, Eshrar Osman, Md. Ahasan Habib, Shahina Akter, Tanjina Akhter Banu, Md. Murshed Hasan Sarkar, Barna Goswami, Iffat Jahan, Md. Saddam Hossain, Tasnim Nafisa, Md. Maruf Ahmed Molla, Asish Kumar Ghosh, A. K. M. Shamsuzzaman, Sheikh Md. Selim Al Din, Utpal Chandra Ray, Salek Ahmed Sajib, Md. Salim Khan |
| EPI_ISL_498549, EPI_ISL_498550 | National Institute of Laboratory Medicine and Referral Center | Genomic Research Lab, BCSIR | Asish Kumar Ghosh, Abu Sayeed Mohammad Mahmud, Mohammad Samir Uzzaman, Eshrar Osman, Md. Ahasan Habib, Shahina Akter, Tanjina Akhter Banu, Md. Murshed Hasan Sarkar, Barna Goswami, Iffat Jahan, Md. Saddam Hossain, Tasnim Nafisa, Md. Maruf Ahmed Molla, Mahmuda Yeasmin, A. K. M. Shamsuzzaman, Sheikh Md. Selim Al Din, Utpal Chandra Ray, Salek Ahmed Sajib, Md. Salim Khan |
| EPI_ISL_498583, EPI_ISL_498586, EPI_ISL_498589, EPI_ISL_498592, EPI_ISL_498593, EPI_ISL_498594, EPI_ISL_498596, EPI_ISL_498599, EPI_ISL_498608, EPI_ISL_498609, EPI_ISL_498610, EPI_ISL_498611, EPI_ISL_498612, EPI_ISL_498613, EPI_ISL_498614, EPI_ISL_498615, EPI_ISL_498616, EPI_ISL_498617, EPI_ISL_498618 | | | |
| see above | National Public Health Laboratory, National Centre for Infectious Diseases | National Public Health Laboratory, National Centre for Infectious Diseases | Mak TM, Octavia S, Zhou Z, Chavatte JM, Cui L, Lin RTP |
| EPI_ISL_498691, EPI_ISL_498692, EPI_ISL_498693, EPI_ISL_498694 | National Institute for Viral Disease Control and Prevention, China CDC | National Institute for Viral Disease Control and Prevention, China CDC | Xiang Zhao,LingLing Mao,Yao Meng,Zhixiao Chen,Yuchao Wu,Yong ZhangBo ZhijianJianqun Zhang,Yang Song,Dayan Wang,WenQing YaoWenbo Xu |
| EPI_ISL_498764 | Sydney South West Pathology Service (SSWPS) - Liverpool Hospital - NSW Health Pathology | NSW Health Pathology - Institute of Clinical Pathology and Medical Research; Westmead Hospital; University of Sydney | CIDM-PH et al. |
| EPI_ISL_498766, EPI_ISL_498767 | Pathology West - NSW Health Pathology | NSW Health Pathology - Institute of Clinical Pathology and Medical Research; Westmead Hospital; University of Sydney | CIDM-PH et al. |
| EPI_ISL_498768 | South Eastern Area Laboratory Services (SEALS) | NSW Health Pathology - Institute of Clinical Pathology and Medical Research; Westmead Hospital; University of Sydney | CIDM-PH et al. |
| EPI_ISL_498769, EPI_ISL_498770, EPI_ISL_498771, EPI_ISL_498772 | Laverty Pathology | NSW Health Pathology - Institute of Clinical Pathology and Medical Research; Westmead Hospital; University of Sydney | CIDM-PH et al. |
| EPI_ISL_498773 | Douglass Hanly Moir Pathology | NSW Health Pathology - Institute of Clinical Pathology and Medical Research; Westmead Hospital; University of Sydney | CIDM-PH et al. |
| EPI_ISL_498774, EPI_ISL_498775 | Histopath | NSW Health Pathology - Institute of Clinical Pathology and Medical Research; Westmead Hospital; University of Sydney | CIDM-PH et al. |
| EPI_ISL_498776, EPI_ISL_498777 | Pathology West - NSW Health Pathology | NSW Health Pathology - Institute of Clinical Pathology and Medical Research; Westmead Hospital; University of Sydney | CIDM-PH et al. |
| EPI_ISL_498778 | Sydney South West Pathology Service (SSWPS) - Liverpool Hospital - NSW Health Pathology | NSW Health Pathology - Institute of Clinical Pathology and Medical Research; Westmead Hospital; University of Sydney | CIDM-PH et al. |
| EPI_ISL_498779, EPI_ISL_498780 | Pathology West - NSW Health Pathology | NSW Health Pathology - Institute of Clinical Pathology and Medical Research; Westmead Hospital; University of Sydney | CIDM-PH et al. |
| EPI_ISL_498781, EPI_ISL_498782 | Sydney South West Pathology Service (SSWPS) - Liverpool Hospital - NSW Health Pathology | NSW Health Pathology - Institute of Clinical Pathology and Medical Research; Westmead Hospital; University of Sydney | CIDM-PH et al. |
| EPI_ISL_498784, EPI_ISL_498785, EPI_ISL_498786, EPI_ISL_498787 | National Institute of Laboratory Medicine and Referral Center | Genomic Research Lab, BCSIR | Md. Saddam Hossain, Abu Sayeed Mohammad Mahmud, Mohammad Samir Uzzaman, Eshrar Osman, Md. Ahasan Habib, Shahina Akter, Tanjina Akhter Banu, Md. Murshed Hasan Sarkar, Barna Goswami, Iffat Jahan, Tasnim Nafisa, Md. Maruf Ahmed Molla, Mahmuda Yeasmin, Asish Kumar Ghosh, A. K. M. Shamsuzzaman, Sheikh Md. Selim Al Din, Utpal Chandra Ray, Salek Ahmed Sajib, Md. Salim Khan |
| EPI_ISL_498797, EPI_ISL_498800, | National Institute of Laboratory Medicine and Referral Center | Genomic Research Lab, BCSIR | Md. Murshed Hasan Sarkar, Abu Sayeed Mohammad Mahmud, Mohammad Samir Uzzaman, Eshrar Osman, Md. Ahasan Habib, Shahina Akter, Tanjina |

| | | | |
|---|---|---|---|
| EPI_ISL_498801, EPI_ISL_498802, EPI_ISL_498803, EPI_ISL_498804, EPI_ISL_498805, EPI_ISL_498806 | | | Akhter Banu, Barna Goswami, Iffat Jahan, Md. Saddam Hossain, Tasnim Nafisa, Md. Maruf Ahmed Molla, Mahmuda Yeasmin, Asish Kumar Ghosh, A. K. M. Shamsuzzaman, Sheikh Md. Selim Al Din, Utpal Chandra Ray, Salek Ahmed Sajib, Md. Salim Khan |
| EPI_ISL_498809, EPI_ISL_498811, EPI_ISL_498814, EPI_ISL_498815, EPI_ISL_498816, EPI_ISL_498817, EPI_ISL_498818, EPI_ISL_498830, EPI_ISL_498892 | National Institute of Laboratory Medicine and Referral Center | Genomic Research Lab, BCSIR | Abu Sayeed Mohammad Mahmud, Mohammad Samir Uzzaman, Eshrar Osman, Md. Ahasan Habib, Shahina Akter, Tanjina Akhter Banu, Md. Murshed Hasan Sarkar, Barna Goswami, Iffat Jahan, Md. Saddam Hossain, Tasnim Nafisa, Md. Maruf Ahmed Molla, Mahmuda Yeasmin, Asish Kumar Ghosh, A. K. M. Shamsuzzaman, Sheikh Md. Selim Al Din, Utpal Chandra Ray, Salek Ahmed Sajib, Md. Salim Khan |
| EPI_ISL_499266, EPI_ISL_499267, EPI_ISL_499268, EPI_ISL_499269 | Queens Medical Centre, Clinical Microbiology Department / DeepSeq Nottingham | COVID-19 Genomics UK (COG-UK) Consortium | Gemma Clark, Wendy Smith, Manjinder Khakh, Vicki M Fleming, Michelle M Lister, Hannah Howson-Wells, Jonathan Ball, Patrick McClure, Joseph Chappell, Theocharis Tsoleridis, Nadine Holmes, Matthew Carlisle, Christopher Moore, Fei Sang, Johnny Debebe, Victoria Wright, Matthew Loose |
| EPI_ISL_499354 | West of Scotland Specialist Virology Centre, NHSGGC / MRC-University of Glasgow Centre for Virus Research | COVID-19 Genomics UK (COG-UK) Consortium | Ana da Silva Filipe, Natasha Johnson, Kathy Smollett, Daniel Mair, Stephen Carmichael, Lily Tong, Jenna Nichols, Elihu Aranday-Cortes, Kirstyn Brunker, Yasmin Parr, Alice Broos, Kyriaki Nomikou; Sarah McDonald, Marc Niebel, Patawee Asamaphan; Richard Orton, Joseph Hughes, Sreenu Vattipally, David L Robertson; Alasdair MacLean, Rory Gunson; Kathy Li, Natasha Jesudason, Rajiv Shah, James Shepherd, Antonia Ho, Emma Thomson |
| EPI_ISL_499361, EPI_ISL_499381, EPI_ISL_499383, EPI_ISL_499386, EPI_ISL_499400, EPI_ISL_499407, EPI_ISL_499410, EPI_ISL_499431 | Wales Specialist Virology Centre Sequencing lab: Pathogen Genomics Unit | COVID-19 Genomics UK (COG-UK) Consortium | Catherine Moore, Johnathan Evans, Laura Gifford, Malorie Perry, Simon Cottrell, Angela Marchbank, Alec Birchley, Alexander Adams, Amy Gaskin, Bree Gatica-Wilcox, Jason Coombes, Joel Southgate, Lauren Gilbert, Lee Graham, Nicole Pacchiarini, Sara Kumziene-Summerhayes, Sarah Taylor, Sophie Jones, Sara Rey, Matthew Bull, Joanne Watkins, Sally Corden, Tom Connor |
| EPI_ISL_499808, EPI_ISL_499809 | Queens Medical Centre, Clinical Microbiology Department / DeepSeq Nottingham | COVID-19 Genomics UK (COG-UK) Consortium | Gemma Clark, Wendy Smith, Manjinder Khakh, Vicki M Fleming, Michelle M Lister, Hannah Howson-Wells, Jonathan Ball, Patrick McClure, Joseph Chappell, Theocharis Tsoleridis, Nadine Holmes, Matthew Carlisle, Christopher Moore, Fei Sang, Johnny Debebe, Victoria Wright, Matthew Loose |
| EPI_ISL_499905, EPI_ISL_499969 | Department of Pathology, University of Cambridge | COVID-19 Genomics UK (COG-UK) Consortium | Luke W Meredith, M. Estée Török, Myra Hosmillo, William L. Hamilton, Martin D. Curran, Theresa Feltwell, Grant Hall, Anna Yakovleva, Fahad A Khokhar, Charlotte J. Houldcroft, Laura G Caller, Aminu S. Jahun, Sarah L. Caddy, Yasmin Chaudhry, Malte Pinckert, Ian Goodfellow |
| EPI_ISL_500572 | Singapore General Hospital | Department of Microbiology | Nurdyana Abdul Rahman, Kun Lee Lim, Chenhao Li, Kian Sing Chan, Lynette Oon, Kern Rei Chng, Niranjan Nagarajan, Karrie Ko |
| EPI_ISL_500713, EPI_ISL_500714, EPI_ISL_500715 | Respiratory Virus Unit, Microbiology Services Colindale, Public Health England | Respiratory Virus Unit, Microbiology Services Colindale, Public Health England | PHE Covid Sequencing Team |
| EPI_ISL_500796 | Akershus University Hospital, Department for Microbiology and Infectious Disease Control | Norwegian Institute of Public Health, Department of Virology | Kathrine Stene-Johansen, Kamilla Heddeland Instefjord, Hilde Elshaug, Rasmus Riis Kopperud, Karoline Bragstad, Olav Hungnes |
| EPI_ISL_500797, EPI_ISL_500798, EPI_ISL_500799 | Hospital of Southern Norway - Kristiansand, Department of Medical Microbiology | Norwegian Institute of Public Health, Department of Virology | Kathrine Stene-Johansen, Kamilla Heddeland Instefjord, Hilde Elshaug, Rasmus Riis Kopperud, Karoline Bragstad, Olav Hungnes |
| EPI_ISL_501271, EPI_ISL_501272, EPI_ISL_501273, EPI_ISL_501274 | National Virus Reference Laboratory | National Virus Reference Laboratory | Michael Carr, Gabriel Gonzalez, Jonathan Dean, Suzie Coughlan, Cillian F De Gascun |
| EPI_ISL_501823, EPI_ISL_501829, EPI_ISL_501833, EPI_ISL_501839, EPI_ISL_501849, EPI_ISL_501894 | E. Gulbja Laboratorija | Latvian Biomedical Research and Study Centre | Ivars Silamielis, Kaspars Megnis, Monta Ustinova, ikita Zrelovs, Vita Rovte, Mikus Gavars, Dmitrijs Perminovs, Uga Dumpis, Jnis Kloviš |
| EPI_ISL_501895 | Centrl laboratorija | Latvian Biomedical Research and Study Centre | Ivars Silamielis, Kaspars Megnis, Monta Ustinova, ikita Zrelovs, Vita Rovte, Stella Lapia, Jana Oste, Marta Priedte, Uga Dumpis, Jnis Kloviš |
| EPI_ISL_504137, EPI_ISL_504176, EPI_ISL_504177, EPI_ISL_504178, EPI_ISL_504179, EPI_ISL_504183, EPI_ISL_504184 | National Institute of Laboratory Medicine and Referral Center | Genomic Research Lab, BCSIR | Abu Sayeed Mohammad Mahmud, Mohammad Samir Uzzaman, Eshrar Osman, Md. Ahasan Habib, Shahina Akter, Tanjina Akhter Banu, Md. Murshed Hasan Sarkar, Barna Goswami, Iffat Jahan, Md. Saddam Hossain, Tarannum Taznin, Tasnim Nafisa, Md. Maruf Ahmed Molla, Mahmuda Yeasmin, Asish Kumar Ghosh, A. K. M. Shamsuzzaman, Sheikh Md. Selim Al Din, Utpal Chandra Ray, Salek Ahmed Sajib, Md. Salim Khan |
| EPI_ISL_507130 | Quadram Institute Bioscience | COVID-19 Genomics UK (COG-UK) Consortium | Dave J. Baker, Gemma L. Kay, Alp Aydin, Thanh Le-Viet, Steven Rudder, Ana P. Tedim, Anastasia Kolyva, Maria Diaz, Leonardo de Oliveira Martins, Nabil-Fareed Alikhan, Lizzie Meadows, Rachael Stanley, Ngozi Elumogo, Muhammed Yasir, Nicholas M. Thomson, Alexander J Trotter, Rachel Gilroy, Samuel Bloomfield, Claire Stuart, Andrew Bell, Reenesh Prakash, Samir Dervisevic, Alison E. Mather, John Wain, Mark Webber, Andrew J. Page, Justin O'Grady |
| EPI_ISL_507164, EPI_ISL_507167, EPI_ISL_507173, EPI_ISL_507174, EPI_ISL_507175, EPI_ISL_507176 | Virology Department, Sheffield Teaching Hospitals NHS Foundation Trust/Department of Infection, Immunity and Cardiovascular Disease, The Medical School, University of Sheffield | COVID-19 Genomics UK (COG-UK) Consortium | Thushan de Silva, Matthew Parker, Nikki Smith, Adri Angyal, Rebecca Brown, Luke Green, Rachel Tucker, Paul Parsons, Danielle Groves, Katie Johnson, Laura Carrilero, Alex Keeley, Dave Partridge, Matthew Wyles, Benjamin Lindsey, Mehmet Yavuz, Mohammad Raza, Cariad Evans |
| EPI_ISL_508134 | SA Pathology | SA Pathology | Lex Leong, Chuan Kok Lim, Mark Turra, Ivan Bastian, Geoff Higgins |
| EPI_ISL_509154, EPI_ISL_509155, EPI_ISL_509156, EPI_ISL_509157, EPI_ISL_509158, EPI_ISL_509159, EPI_ISL_509160, EPI_ISL_509161, EPI_ISL_509162, EPI_ISL_509163, EPI_ISL_509164, EPI_ISL_509165, EPI_ISL_509166, EPI_ISL_509167, EPI_ISL_509168, EPI_ISL_509169, EPI_ISL_509170, EPI_ISL_509171, EPI_ISL_509172, EPI_ISL_509173, EPI_ISL_509174, EPI_ISL_509175, EPI_ISL_509176, EPI_ISL_509177, EPI_ISL_509178, EPI_ISL_509179, EPI_ISL_509180, EPI_ISL_509181, EPI_ISL_509182, EPI_ISL_509183, EPI_ISL_509184, EPI_ISL_509185, EPI_ISL_509186, EPI_ISL_509187, EPI_ISL_509188, EPI_ISL_509189, EPI_ISL_509190, EPI_ISL_509191, EPI_ISL_509192, EPI_ISL_509193, EPI_ISL_509194, EPI_ISL_509195, EPI_ISL_509196, EPI_ISL_509197, EPI_ISL_509198, EPI_ISL_509199, EPI_ISL_509200, EPI_ISL_509201, EPI_ISL_509202, EPI_ISL_509203, EPI_ISL_509204, EPI_ISL_509205, EPI_ISL_509206, EPI_ISL_509207, EPI_ISL_509208, EPI_ISL_509209, EPI_ISL_509210, EPI_ISL_509211, EPI_ISL_509212, EPI_ISL_509213, EPI_ISL_509214 | | | |
| see above | OHSU Lab Services Molecular Microbiology Lab | Oregon SARS-CoV-2 Genome Sequencing Center | Brendan L. O'Connell, Ruth V. Nichols, Sally B. Grindstaff, Alec J. Hirsch, Guang Fan, Daniel N. Streblow, William B. Messer, Andrew C. Adey, Benjamin N. Bimber, Brian J. O'Roak |
| EPI_ISL_509223, EPI_ISL_509224, EPI_ISL_509225, EPI_ISL_509226, EPI_ISL_509227, EPI_ISL_509228, EPI_ISL_509230, EPI_ISL_509231, EPI_ISL_509232, EPI_ISL_509233, EPI_ISL_509234, EPI_ISL_509236, EPI_ISL_509237, EPI_ISL_509238, EPI_ISL_509239, EPI_ISL_509240, EPI_ISL_509241, EPI_ISL_509242, EPI_ISL_509243, EPI_ISL_509244, EPI_ISL_509245, EPI_ISL_509246, EPI_ISL_509250, EPI_ISL_509252, EPI_ISL_509254, EPI_ISL_509260, EPI_ISL_509265, EPI_ISL_509277, EPI_ISL_509279, EPI_ISL_509286, EPI_ISL_509287, EPI_ISL_509288, EPI_ISL_509289, EPI_ISL_509290, EPI_ISL_509291, EPI_ISL_509293, EPI_ISL_509294, EPI_ISL_509295, EPI_ISL_509296, EPI_ISL_509297, EPI_ISL_509298, EPI_ISL_509299, EPI_ISL_509300, EPI_ISL_509301, EPI_ISL_509302, EPI_ISL_509303, EPI_ISL_509304, EPI_ISL_509306, EPI_ISL_509307, EPI_ISL_509308, EPI_ISL_509309, EPI_ISL_509310, EPI_ISL_509312, EPI_ISL_509313, EPI_ISL_509324, EPI_ISL_509325, EPI_ISL_509326, EPI_ISL_509327, EPI_ISL_509328, EPI_ISL_509335, EPI_ISL_509338, EPI_ISL_509339, EPI_ISL_509340, EPI_ISL_509341, EPI_ISL_509343, EPI_ISL_509345, EPI_ISL_509346, EPI_ISL_509348, EPI_ISL_509352, EPI_ISL_509353, EPI_ISL_509355, EPI_ISL_509356, EPI_ISL_509357, EPI_ISL_509358, EPI_ISL_509359, EPI_ISL_509360, EPI_ISL_509361, EPI_ISL_509362, EPI_ISL_509363, EPI_ISL_509364, EPI_ISL_509365, EPI_ISL_509366, EPI_ISL_509367, EPI_ISL_509368, EPI_ISL_509369, EPI_ISL_509370, EPI_ISL_509371 | | | |
| see above | NHLS-IALCH | KRISP, KZN Research Innovation and Sequencing Platform | Giandhari J, Pillay S, Lessells R, Mdlalose K, York D, Tegally H, Wilkinson E, de Oliveira T |
| EPI_ISL_509379, EPI_ISL_509380, EPI_ISL_509381, EPI_ISL_509382, EPI_ISL_509383, EPI_ISL_509384, EPI_ISL_509385, EPI_ISL_509386, EPI_ISL_509387, EPI_ISL_509388, EPI_ISL_509389 | | | |
| see above | Singapore General Hospital | Department of Microbiology | Nurdyana Abdul Rahman, Kun Lee Lim, Chenhao Li, Kian Sing Chan, Lynette Oon, Kern Rei Chng, Niranjan Nagarajan, Karrie Ko |
| EPI_ISL_509430, EPI_ISL_509431, EPI_ISL_509432, EPI_ISL_509433, EPI_ISL_509435 | Centro de Desenvolvimento Tecnologico em Saude, Fundacao Oswaldo Cruz | Centro de Desenvolvimento Tecnologico em Saude, Fundacao Oswaldo Cruz | Souza,T.M., Fintelman-Rodrigues,N., De Paula,A.D., Saraiva,F.B., Ferreira,M.A., Sacramento,C.Q., Medeiros,M.A. |
| EPI_ISL_509522, EPI_ISL_509523 | Area of Virology, Serology and Virology Division (SAViD), New South Wales Health Pathology Randwick | Area of Virology, Serology and Virology Division (SAViD), New South Wales Health Pathology Randwick | Rawlinson, W. |
| EPI_ISL_509776, EPI_ISL_509777, EPI_ISL_509778, EPI_ISL_509779, EPI_ISL_509780, EPI_ISL_509781, EPI_ISL_509782, EPI_ISL_509783, EPI_ISL_509784, EPI_ISL_509785, EPI_ISL_509786, EPI_ISL_509787, EPI_ISL_509788, EPI_ISL_509789, EPI_ISL_509790, EPI_ISL_509791, EPI_ISL_509792, EPI_ISL_509793, EPI_ISL_509794, EPI_ISL_509795, EPI_ISL_509796, EPI_ISL_509797, EPI_ISL_509798 | | | |
| see above | Florida Bureau of Public Health Laboratories | Florida Bureau of Public Health Laboratories | Sarah Schmedes, Jason Blanton |
| EPI_ISL_509801, EPI_ISL_509803, EPI_ISL_509804, EPI_ISL_509805, EPI_ISL_509806, EPI_ISL_509807, EPI_ISL_509809, EPI_ISL_509810, EPI_ISL_509811, EPI_ISL_509812, EPI_ISL_509813, EPI_ISL_509814, EPI_ISL_509817, EPI_ISL_509864, EPI_ISL_509865, EPI_ISL_509866, EPI_ISL_509875, EPI_ISL_509876, EPI_ISL_509877, EPI_ISL_509878, EPI_ISL_509879, EPI_ISL_509880, EPI_ISL_509881, EPI_ISL_509882, EPI_ISL_509883, EPI_ISL_509884, EPI_ISL_509885, EPI_ISL_509887, EPI_ISL_509888, EPI_ISL_509889, EPI_ISL_509890, EPI_ISL_509891, EPI_ISL_509892, EPI_ISL_509893, EPI_ISL_509895, EPI_ISL_509896, | | | |

| | | | |
|---|---|---|---|
| EPI_ISL_509897, EPI_ISL_509898, EPI_ISL_509899, EPI_ISL_509900, EPI_ISL_509901, EPI_ISL_509902, EPI_ISL_509903, EPI_ISL_509904, EPI_ISL_509905, EPI_ISL_509907, EPI_ISL_509909, EPI_ISL_509913, EPI_ISL_509917, EPI_ISL_509918, EPI_ISL_509920, EPI_ISL_509921, EPI_ISL_509922, EPI_ISL_509923, EPI_ISL_509925, EPI_ISL_509927, EPI_ISL_509930, EPI_ISL_509931, EPI_ISL_509932, EPI_ISL_509971, EPI_ISL_509972, EPI_ISL_509973, EPI_ISL_509974, EPI_ISL_509975, EPI_ISL_509976, EPI_ISL_509977, EPI_ISL_509978, EPI_ISL_509979, EPI_ISL_509980, EPI_ISL_509981, EPI_ISL_509982, EPI_ISL_509983, EPI_ISL_509984, EPI_ISL_509985, EPI_ISL_509986, EPI_ISL_509987, EPI_ISL_509988, EPI_ISL_509989, EPI_ISL_509990, EPI_ISL_509991, EPI_ISL_509992, EPI_ISL_509993 | | | |
| see above | University of Wisconsin-Madison AIDS Vaccine Research Laboratories | University of Wisconsin-Madison AIDS Vaccine Research Laboratories | Gage Moreno, Katarina Braun, et al. AIDS Vaccine Research Laboratories |
| EPI_ISL_510442, EPI_ISL_510443, EPI_ISL_510445, EPI_ISL_510447, EPI_ISL_510460 | Hospital Universitario Virgen de las Nieves de Granada-SAS | SeqCOVID-SPAIN consortium/IBV(CSIC) | Mercedes Pérez Ruiz, Sara Sanbonmatsu Gámez, Irene Pedrosa Corral, José M. Navarro-Marí and SeqCOVID-SPAIN consortium |
| EPI_ISL_510536 | Centro de Desenvolvimento Tecnologico em Saude, Fundacao Oswaldo Cruz | Centro de Desenvolvimento Tecnologico em Saude, Fundacao Oswaldo Cruz | Souza,T.M., Fintelman-Rodrigues,N., De Paula,A.D., Saraiva,F.B., Ferreira,M.A., Sacramento,C.Q. and Medeiros,M.A. |
| EPI_ISL_510538, EPI_ISL_510539, EPI_ISL_510540 | Department of Microbiology, The University of Hong Kong | Department of Microbiology, The University of Hong Kong | Kelvin K.W. To, Kwok-Yung Yuen |
| EPI_ISL_510541 | Centro de Desenvolvimento Tecnologico em Saude, Fundacao Oswaldo Cruz | Centro de Desenvolvimento Tecnologico em Saude, Fundacao Oswaldo Cruz | Souza,T.M., Fintelman-Rodrigues,N., De Paula,A.D., Saraiva,F.B., Ferreira,M.A., Sacramento,C.Q. and Medeiros,M.A. |
| EPI_ISL_510698, EPI_ISL_510699, EPI_ISL_510700, EPI_ISL_510701, EPI_ISL_510702, EPI_ISL_510703, EPI_ISL_510704, EPI_ISL_510705, EPI_ISL_510706, EPI_ISL_510707, EPI_ISL_510708, EPI_ISL_510709, EPI_ISL_510710, EPI_ISL_510711, EPI_ISL_510712, EPI_ISL_510713, EPI_ISL_510714, EPI_ISL_510715, EPI_ISL_510716, EPI_ISL_510717, EPI_ISL_510718, EPI_ISL_510719, EPI_ISL_510720, EPI_ISL_510721, EPI_ISL_510722, EPI_ISL_510723, EPI_ISL_510724, EPI_ISL_510725, EPI_ISL_510726, EPI_ISL_510727, EPI_ISL_510728, EPI_ISL_510729, EPI_ISL_510735, EPI_ISL_510736, EPI_ISL_510737, EPI_ISL_510738, EPI_ISL_510739, EPI_ISL_510740, EPI_ISL_510741, EPI_ISL_510742, EPI_ISL_510743, EPI_ISL_510744, EPI_ISL_510745, EPI_ISL_510746, EPI_ISL_510747, EPI_ISL_510748, EPI_ISL_510749, EPI_ISL_510750, EPI_ISL_510751, EPI_ISL_510752, EPI_ISL_510753, EPI_ISL_510754, EPI_ISL_510755, EPI_ISL_510756, EPI_ISL_510757, EPI_ISL_510758, EPI_ISL_510759, EPI_ISL_510760, EPI_ISL_510761, EPI_ISL_510762, EPI_ISL_510763, EPI_ISL_510764, EPI_ISL_510765, EPI_ISL_510766, EPI_ISL_510767, EPI_ISL_510768, EPI_ISL_510769, EPI_ISL_510770, EPI_ISL_510771, EPI_ISL_510772, EPI_ISL_510773, EPI_ISL_510774, EPI_ISL_510775, EPI_ISL_510776, EPI_ISL_510777, EPI_ISL_510778, EPI_ISL_510785, EPI_ISL_510786, EPI_ISL_510787, EPI_ISL_510788, EPI_ISL_510789, EPI_ISL_510790, EPI_ISL_510791, EPI_ISL_510792, EPI_ISL_510793, EPI_ISL_510794, EPI_ISL_510795, EPI_ISL_510796, EPI_ISL_510797, EPI_ISL_510798, EPI_ISL_510799, EPI_ISL_510800, EPI_ISL_510801, EPI_ISL_510802, EPI_ISL_510803, EPI_ISL_510804, EPI_ISL_510805 | | | |
| see above | Viollier AG | Department of Biosystems Science and Engineering, ETH Zürich | Christian Beisel, Sarah Nadeau, Ivan Topolsky, Pedro Ferreira, Philipp Jablonski, Susana Posada-Céspedes, Tobias Schär, Ina Nissen, Natascha Santacroce, Elodie Burcklen, Christiane Beckmann, Maurice Redondo, Olivier Kobel, Christoph Noppen, Sophie Seidel, Noemie Santamaria de Souza, Niko Beerenwinkel, Tanja Stadler |
| EPI_ISL_510878, EPI_ISL_510879, EPI_ISL_510880, EPI_ISL_510881, EPI_ISL_510882, EPI_ISL_510883, EPI_ISL_510884, EPI_ISL_510885, EPI_ISL_510886, EPI_ISL_510887, EPI_ISL_510888 | | | |
| see above | New Mexico Department of Health Scientific Laboratory | New Mexico Department of Health Scientific Laboratory | Ellie Johnson, Anastacia Griego-Fisher |
| EPI_ISL_511878 | Santosa Hospital Bandung Kopo | Laboratorium Kesehatan Provinsi Jawa Barat; Molecular Genetics Laboratory-Faculty of Medicine-Universitas Padjadjaran; School of Life Sciences and Technology & School of Pharmacy-Institut Teknologi Bandung | Ema Rahmawati, Marselina Irasonia Tan, Yunia Sribudiani, Catur Riani, Azzania Fibriani, Husna Nugrahapraja, Tarwadi, Hesti Lina Wiraswati, Lia Faridah, Savira Ekawardhani, Ryan Bayusantika Ristandi, Rifky Waluyajati Rachman, Cut Nur Cinthia Alamanda, Hammam Riza, Soni Solistia Wirawan, Agung Eru Wibowo |
| EPI_ISL_511984, EPI_ISL_511985, EPI_ISL_511986, EPI_ISL_511987, EPI_ISL_511988, EPI_ISL_511990, EPI_ISL_511991, EPI_ISL_511992 | Viollier AG | Department of Biosystems Science and Engineering, ETH Zürich | Christian Beisel, Sarah Nadeau, Ivan Topolsky, Pedro Ferreira, Philipp Jablonski, Susana Posada-Céspedes, Tobias Schär, Ina Nissen, Natascha Santacroce, Elodie Burcklen, Christiane Beckmann, Maurice Redondo, Olivier Kobel, Christoph Noppen, Sophie Seidel, Noemie Santamaria de Souza, Niko Beerenwinkel, Tanja Stadler |
| EPI_ISL_512087, EPI_ISL_512088, EPI_ISL_512090, EPI_ISL_512091, EPI_ISL_512101, EPI_ISL_512102 | National Virus Reference Laboratory | National Virus Reference Laboratory | Michael Carr, Gabriel Gonzalez, Jonathan Dean, Aditi Chaturvedi, Suzie Coughlan, Cillian F De Gascun |
| EPI_ISL_512335 | Department of Pathology, University of Cambridge | COVID-19 Genomics UK (COG-UK) Consortium | Luke W Meredith, M. Estée Török, Myra Hosmillo, William L. Hamilton, Martin D. Curran, Theresa Feltwell, Grant Hall, Anna Yakovleva, Fahad A Khokhar, Charlotte J. Houldcroft, Laura G Caller, Aminu S. Jahun, Sarah L. Caddy, Yasmin Chaudhry, Malte Pinckert, Ian Goodfellow |
| EPI_ISL_512350, EPI_ISL_512351, EPI_ISL_512352, EPI_ISL_512353, EPI_ISL_512354, EPI_ISL_512355, EPI_ISL_512356, EPI_ISL_512357, EPI_ISL_512358, EPI_ISL_512359, EPI_ISL_512360, EPI_ISL_512361, EPI_ISL_512362 | | | |
| see above | Northumbria University / South Tees Hospitals NHS Foundation Trust / North Cumbria Integrated Care NHS Foundation Trust / North Tees and Hartlepool NHS Foundation Trust / Newcastle Hospitals NHS Foundation Trust | COVID-19 Genomics UK (COG-UK) Consortium | Darren L Smith,Andrew Nelson,Matthew Bashton,Greg R Young,Joshua Loh,John Allan,Mohammad A Tariq,Giles S Holt,Gary Black,Wen C Yew,Lynn Dover,Paul Baker,Steve Liggett,Sarah Essex,Jane Greenaway,Debra Padgett,Clive Graham,Garren Scott,Edward Barton,Emma Swindells,Brendan Payne,Jennifer Collins,Yusri Taha,Gary Eltringham |
| EPI_ISL_512373 | Quadram Institute Bioscience | COVID-19 Genomics UK (COG-UK) Consortium | Dave J. Baker, Gemma L. Kay, Alp Aydin, Thanh Le-Viet, Steven Rudder, Ana P. Tedim, Anastasia Kolyva, Maria Diaz, Leonardo de Oliveira Martins, Nabil-Fareed Alikhan, Lizzie Meadows, Rachael Stanley, Ngozi Elumogo, Muhammed Yasir, Nicholas M. Thomson, Alexander J Trotter, Rachel Gilroy, Samuel Bloomfield, Claire Stuart, Andrew Bell, Reenesh Prakash, Samir Dervisevic, Alison E. Mather, John Wain, Mark Webber, Andrew J. Page, Justin O'Grady |
| EPI_ISL_512380 | Queens Medical Centre, Clinical Microbiology Department / DeepSeq Nottingham | COVID-19 Genomics UK (COG-UK) Consortium | Gemma Clark, Wendy Smith, Manjinder Khakh, Vicki M Fleming, Michelle M Lister, Hannah Howson-Wells, Jonathan Ball, Patrick McClure, Joseph Chappell, Theocharis Tsoleridis, Nadine Holmes, Matthew Carlisle, Christopher Moore, Fei Sang, Johnny Debebe, Victoria Wright, Matthew Loose |
| EPI_ISL_512481, EPI_ISL_512484, EPI_ISL_512485, EPI_ISL_512486, EPI_ISL_512488, EPI_ISL_512489, EPI_ISL_512490, EPI_ISL_512494, EPI_ISL_512496, EPI_ISL_512501, EPI_ISL_512503, EPI_ISL_512504, EPI_ISL_512506, EPI_ISL_512510, EPI_ISL_512511, EPI_ISL_512512, EPI_ISL_512515, EPI_ISL_512516, EPI_ISL_512517, EPI_ISL_512522, EPI_ISL_512523, EPI_ISL_512524, EPI_ISL_512525, EPI_ISL_512527, EPI_ISL_512531, EPI_ISL_512532, EPI_ISL_512533, EPI_ISL_512534, EPI_ISL_512535, EPI_ISL_512537, EPI_ISL_512538, EPI_ISL_512539, EPI_ISL_512540, EPI_ISL_512541, EPI_ISL_512543, EPI_ISL_512546 | | | |
| see above | Wales Specialist Virology Centre Sequencing lab: Pathogen Genomics Unit | COVID-19 Genomics UK (COG-UK) Consortium | Catherine Moore, Johnathan Evans, Laura Gifford, Malorie Perry, Simon Cottrell, Angela Marchbank, Alec Birchley, Alexander Adams, Amy Gaskin, Bree Gatica-Wilcox, Jason Coombes, Joel Southgate, Lauren Gilbert, Lee Graham, Nicole Pacchiarini, Sara Kumziene-Summerhayes, Sarah Taylor, Sophie Jones, Sara Rey, Matthew Bull, Joanne Watkins, Sally Corden, Tom Connor |
| EPI_ISL_512618 | National Laboratory for Influenza/Virology reference laboratory, Public Health Center of the Ministry of Health of Ukraine | Respiratory Virus Unit, Microbiology Services Colindale, Public Health England | PHE Covid Sequencing Team, Dr. Iryna Demchyshyna |
| EPI_ISL_512645 | E. Gulbja Laboratorija | Latvian Biomedical Research and Study Centre | Ivars Silamielis, Kaspars Megnis, Monta Ustinova, ikita Zrelovs, Vita Rovte, Mikus Gavars, Dmitrijs Perminovs, Uga Dumpis, Jnis Kloviš |
| EPI_ISL_512648, EPI_ISL_512649, EPI_ISL_512650 | Latvijas Infektoloijas centrs | Latvian Biomedical Research and Study Centre | Ivars Silamielis, Kaspars Megnis, Monta Ustinova, ikita Zrelovs, Vita Rovte, Jeena Storoženko, Tatjana Kolupajeva, Oksana Savicka, Uga Dumpis, Jnis Kloviš |
| EPI_ISL_512731, EPI_ISL_512732, EPI_ISL_512733, EPI_ISL_512734, EPI_ISL_512735, EPI_ISL_512765 | PathWest Laboratory Medicine WA | PathWest Laboratory Medicine WA Microbial Surveillance Unit | PathWest Laboratory Medicine WA Microbial Surveillance Unit |
| EPI_ISL_512832, EPI_ISL_512834, EPI_ISL_512835, EPI_ISL_512836 | National Public Health Laboratory, National Centre for Infectious Diseases | National Public Health Laboratory, National Centre for Infectious Diseases | Mak TM, Octavia S, Zhou Z, Chavatte JM, Cui L, Lin RTP |
| EPI_ISL_513318, EPI_ISL_513319, EPI_ISL_513320 | Histopath | NSW Health Pathology - Institute of Clinical Pathology and Medical Research; Westmead Hospital; University of Sydney | CIDM-PH et al. |
| EPI_ISL_513321, EPI_ISL_513322 | Pathology West - NSW Health Pathology | NSW Health Pathology - Institute of Clinical Pathology and Medical Research; Westmead Hospital; University of Sydney | CIDM-PH et al. |
| EPI_ISL_513323 | Sydney South West Pathology Service (SSWPS) - Liverpool Hospital - NSW Health Pathology | NSW Health Pathology - Institute of Clinical Pathology and Medical Research; Westmead Hospital; University of Sydney | CIDM-PH et al. |
| EPI_ISL_513324, EPI_ISL_513325, | Pathology West - NSW Health Pathology | NSW Health Pathology - Institute of Clinical Pathology and | CIDM-PH et al. |

| | | | |
|---|---|---|---|
| EPI_ISL_513326 | | Medical Research; Westmead Hospital; University of Sydney | |
| EPI_ISL_513327, EPI_ISL_513328, EPI_ISL_513329 | Sydney South West Pathology Service (SSWPS) - Liverpool Hospital - NSW Health Pathology | NSW Health Pathology - Institute of Clinical Pathology and Medical Research; Westmead Hospital; University of Sydney | CIDM-PH et al. |
| EPI_ISL_513330 | St Vincent's Pathology (SydPath) | NSW Health Pathology - Institute of Clinical Pathology and Medical Research; Westmead Hospital; University of Sydney | CIDM-PH et al. |
| EPI_ISL_513331 | Australian Clinical Labs | NSW Health Pathology - Institute of Clinical Pathology and Medical Research; Westmead Hospital; University of Sydney | CIDM-PH et al. |
| EPI_ISL_513332 | Pathology West - NSW Health Pathology | NSW Health Pathology - Institute of Clinical Pathology and Medical Research; Westmead Hospital; University of Sydney | CIDM-PH et al. |
| EPI_ISL_513333, EPI_ISL_513334 | Sydney South West Pathology Service (SSWPS) - Liverpool Hospital - NSW Health Pathology | NSW Health Pathology - Institute of Clinical Pathology and Medical Research; Westmead Hospital; University of Sydney | CIDM-PH et al. |
| EPI_ISL_513335 | Pathology North - Hunter - NSW Health Pathology | NSW Health Pathology - Institute of Clinical Pathology and Medical Research; Westmead Hospital; University of Sydney | CIDM-PH et al. |
| EPI_ISL_513336, EPI_ISL_513337 | Sydney South West Pathology Service (SSWPS) - Liverpool Hospital - NSW Health Pathology | NSW Health Pathology - Institute of Clinical Pathology and Medical Research; Westmead Hospital; University of Sydney | CIDM-PH et al. |
| EPI_ISL_513338, EPI_ISL_513339 | Pathology West - NSW Health Pathology | NSW Health Pathology - Institute of Clinical Pathology and Medical Research; Westmead Hospital; University of Sydney | CIDM-PH et al. |
| EPI_ISL_513340 | Austech Medical Laboratories | NSW Health Pathology - Institute of Clinical Pathology and Medical Research; Westmead Hospital; University of Sydney | CIDM-PH et al. |
| EPI_ISL_513341 | Medlab Pathology | NSW Health Pathology - Institute of Clinical Pathology and Medical Research; Westmead Hospital; University of Sydney | CIDM-PH et al. |
| EPI_ISL_513342 | Pathology West - NSW Health Pathology | NSW Health Pathology - Institute of Clinical Pathology and Medical Research; Westmead Hospital; University of Sydney | CIDM-PH et al. |
| EPI_ISL_513347 | St Vincent's Pathology (SydPath) | NSW Health Pathology - Institute of Clinical Pathology and Medical Research; Westmead Hospital; University of Sydney | CIDM-PH et al. |
| EPI_ISL_513389 | Douglas Hanly Moir | NSW Health Pathology - Institute of Clinical Pathology and Medical Research; Westmead Hospital; University of Sydney | CIDM-PH et al. |
| EPI_ISL_513396, EPI_ISL_513397, EPI_ISL_513398, EPI_ISL_513399 | St Vincent's Pathology (SydPath) | NSW Health Pathology - Institute of Clinical Pathology and Medical Research; Westmead Hospital; University of Sydney | CIDM-PH et al. |
| EPI_ISL_513403, EPI_ISL_513404, EPI_ISL_513405, EPI_ISL_513406, EPI_ISL_513407 | Laverty Pathology | NSW Health Pathology - Institute of Clinical Pathology and Medical Research; Westmead Hospital; University of Sydney | CIDM-PH et al. |
| EPI_ISL_513584 | PHE South West Regional Laboratory, National Infection Service | Wellcome Sanger Institute for the COVID-19 Genomics UK (COG-UK) consortium | Stephanie Hutchings, Hannah Pymont, Dr Peter Muir, Barry Vipond, Rich Hopes; and Alex Alderton, Roberto Amato, Sonia Goncalves, Ewan Harrison, David K. Jackson, Ian Johnston, Dominic Kwiatkowski, Cordelia Langford, John Sillitoe on behalf of the Wellcome Sanger Institute COVID-19 Surveillance Team (http://www.sanger.ac.uk/covid-team) |
| EPI_ISL_514147, EPI_ISL_514148, EPI_ISL_514223, EPI_ISL_514224, EPI_ISL_514225 | Florida Bureau of Public Health Laboratories | Florida Bureau of Public Health Laboratories | Sarah Schmedes, Jason Blanton |
| EPI_ISL_514228, EPI_ISL_514229, EPI_ISL_514230 | National Institute of Laboratory Medicine and Referral Center | Genomic Research Lab, BCSIR | Barna Goswami, Abu Sayeed Mohammad Mahmud, Mohammad Samir Uzzaman, Eshrar Osman, Md. Ahashan Habib, Shahina Akter, Tanjina Akhter Banu, Md. Murshed Hasan Sarkar, Iffat Jahan, Md. Saddam Hossain, Tasnim Nafisa, Md. Maruf Ahmed Molla, Mahmuda Yeasmin, Asish Kumar Ghosh, A. K. M. Shamsuzzaman, Sheikh Md. Selim Al Din, Utpal Chandra Ray, Salek Ahmed Sajib, Md. Salim Khan |
| EPI_ISL_514231, EPI_ISL_514232 | National Institute of Laboratory Medicine and Referral Center | Genomic Research Lab, BCSIR | Shahina Akter, Abu Sayeed Mohammad Mahmud, Mohammad Samir Uzzaman, Eshrar Osman, Md. Ahashan Habib, Tanjina Akhter Banu, Md. Murshed Hasan Sarkar, Barna Goswami, Iffat Jahan, Md. Saddam Hossain, Tasnim Nafisa, Md. Maruf Ahmed Molla, Mahmuda Yeasmin, Asish Kumar Ghosh, A. K. M. Shamsuzzaman, Sheikh Md. Selim Al Din, Utpal Chandra Ray, Salek Ahmed Sajib, Md. Salim Khan |
| EPI_ISL_514238, EPI_ISL_514239, EPI_ISL_514240, EPI_ISL_514241 | National Institute of Laboratory Medicine and Referral Center | Genomic Research Lab, BCSIR | Md. Saddam Hossain, Abu Sayeed Mohammad Mahmud, Mohammad Samir Uzzaman, Eshrar Osman, Md. Ahashan Habib, Shahina Akter, Tanjina Akhter Banu, Md. Murshed Hasan Sarkar, Barna Goswami, Iffat Jahan, Tasnim Nafisa, Md. Maruf Ahmed Molla, Mahmuda Yeasmin, Asish Kumar Ghosh, A. K. M. Shamsuzzaman, Sheikh Md. Selim Al Din, Utpal Chandra Ray, Salek Ahmed Sajib, Md. Salim Khan |
| EPI_ISL_514242, EPI_ISL_514243, EPI_ISL_514244 | National Institute of Laboratory Medicine and Referral Center | Genomic Research Lab, BCSIR | Tanjina Akhter Banu, Abu Sayeed Mohammad Mahmud, Mohammad Samir Uzzaman, Eshrar Osman, Md. Ahashan Habib, Shahina Akter, Md. Murshed Hasan Sarkar, Barna Goswami, Iffat Jahan, Md. Saddam Hossain, Tasnim Nafisa, Md. Maruf Ahmed Molla, Mahmuda Yeasmin, Asish Kumar Ghosh, A. K. M. Shamsuzzaman, Sheikh Md. Selim Al Din, Utpal Chandra Ray, Salek Ahmed Sajib, Md. Salim Khan |
| EPI_ISL_514245, EPI_ISL_514246, EPI_ISL_514247 | National Institute of Laboratory Medicine and Referral Center | Genomic Research Lab, BCSIR | Iffat Jahan, Abu Sayeed Mohammad Mahmud, Mohammad Samir Uzzaman, Eshrar Osman, Md. Ahashan Habib, Shahina Akter, Tanjina Akhter Banu, Md. Murshed Hasan Sarkar, Barna Goswami, Md. Saddam Hossain, Tasnim Nafisa, Md. Maruf Ahmed Molla, Mahmuda Yeasmin, Asish Kumar Ghosh, A. K. M. Shamsuzzaman, Sheikh Md. Selim Al Din, Utpal Chandra Ray, Salek Ahmed Sajib, Md. Salim Khan |
| EPI_ISL_514251, EPI_ISL_514252 | National Institute of Laboratory Medicine and Referral Center | Genomic Research Lab, BCSIR | Abu Sayeed Mohammad Mahmud, Mohammad Samir Uzzaman, Eshrar Osman, Md. Ahashan Habib, Shahina Akter, Tanjina Akhter Banu, Md. Murshed Hasan Sarkar, Barna Goswami, Iffat Jahan, Md. Saddam Hossain, Tasnim Nafisa, Md. Maruf Ahmed Molla, Mahmuda Yeasmin, Asish Kumar Ghosh, A. K. M. Shamsuzzaman, Sheikh Md. Selim Al Din, Utpal Chandra Ray, Salek Ahmed Sajib, Md. Salim Khan |
| EPI_ISL_514253 | Advanced Biotechnology Laboratory | Genomic Research Lab, BCSIR | Abu Sayeed Mohammad Mahmud, Mohammad Samir Uzzaman, Eshrar Osman, Hossain Uddin Shekhar, M. Aftab Uddin, Md. Bayejid Hosen, Eunus Ali, Md. Ahashan Habib, Shahina Akter, Tanjina Akhter Banu, Md. Murshed Hasan Sarkar, Barna Goswami, Iffat Jahan, Md. Saddam Hossain, Utpal Chandra Ray, Salek Ahmed Sajib, Md. Salim Khan |
| EPI_ISL_514267, EPI_ISL_514268, EPI_ISL_514269, EPI_ISL_514270, EPI_ISL_514271, EPI_ISL_514272, EPI_ISL_514273, EPI_ISL_514274 | Israel Central Virology laboratory | Israel Central Virology laboratory | Neta Zuckerman, Efrat Dahan Bucris, Oran Erster, Ella Mendelson, Michal Mandelboim |
| EPI_ISL_514275 | Israeli Central Virology laboratory | Israel Central Virology laboratory | Neta Zuckerman, Efrat Dahan Bucris, Oran Erster, Ella Mendelson, Michal Mandelboim |
| EPI_ISL_514276, EPI_ISL_514277, EPI_ISL_514278, EPI_ISL_514279, EPI_ISL_514280, EPI_ISL_514281, EPI_ISL_514282, EPI_ISL_514283, EPI_ISL_514284, EPI_ISL_514285, EPI_ISL_514286, EPI_ISL_514287, EPI_ISL_514288, EPI_ISL_514289, EPI_ISL_514290, EPI_ISL_514291, EPI_ISL_514292, EPI_ISL_514293, EPI_ISL_514294, EPI_ISL_514295, EPI_ISL_514296, EPI_ISL_514297, EPI_ISL_514298, EPI_ISL_514299, EPI_ISL_514300, EPI_ISL_514301, EPI_ISL_514302, EPI_ISL_514303, EPI_ISL_514304, EPI_ISL_514307 | | | |
| see above | Israel Central Virology laboratory | Israel Central Virology laboratory | Neta Zuckerman, Efrat Dahan Bucris, Oran Erster, Ella Mendelson, Michal Mandelboim |
| EPI_ISL_514355, EPI_ISL_514356, EPI_ISL_514357, EPI_ISL_514358, EPI_ISL_514359, EPI_ISL_514360, EPI_ISL_514361, EPI_ISL_514362, EPI_ISL_514363, EPI_ISL_514364, EPI_ISL_514365, EPI_ISL_514366, EPI_ISL_514367, EPI_ISL_514368, EPI_ISL_514369, EPI_ISL_514370, EPI_ISL_514371, EPI_ISL_514372, EPI_ISL_514373, EPI_ISL_514374, EPI_ISL_514375, EPI_ISL_514376, EPI_ISL_514377, EPI_ISL_514378, EPI_ISL_514379, EPI_ISL_514380, EPI_ISL_514381, EPI_ISL_514382, EPI_ISL_514383, EPI_ISL_514384, EPI_ISL_514385, EPI_ISL_514386, EPI_ISL_514387, EPI_ISL_514388, EPI_ISL_514389, EPI_ISL_514390, EPI_ISL_514391, EPI_ISL_514392, EPI_ISL_514393, EPI_ISL_514394, EPI_ISL_514395, EPI_ISL_514396, EPI_ISL_514397, EPI_ISL_514398, EPI_ISL_514399, EPI_ISL_514400, EPI_ISL_514401, EPI_ISL_514402, EPI_ISL_514403, EPI_ISL_514404, EPI_ISL_514405, EPI_ISL_514406, EPI_ISL_514407, EPI_ISL_514408, EPI_ISL_514409, EPI_ISL_514410, EPI_ISL_514411, EPI_ISL_514412, EPI_ISL_514413, EPI_ISL_514414, EPI_ISL_514415, EPI_ISL_514416, EPI_ISL_514417, EPI_ISL_514419, EPI_ISL_514423 | | | |
| see above | National Institute for Communicable Diseases of the National Health Laboratory Service | National Institute for Communicable Diseases of the National Health Laboratory Service | Allam M, Ismail A, Khumalo Z, Kwenda S, Mtshali P, Mnyameni F, Mohale T, Bhiman JN |

| | | | |
|---|---|---|---|
| EPI_ISL_514425, EPI_ISL_514427, EPI_ISL_514428, EPI_ISL_514429 | Laboratory Diagnostic, Veterinary Specialized Institute Kraljevo | Laboratory Diagnostic, Veterinary Specialized Institute Kraljevo | Vidanovic,D., Tesovic,B., Knezevic,A., Jankovic,M., Sekler,M., Banovic Djeri,B., Volkening,J., Afonso,C., Petrovic,T. |
| EPI_ISL_514434 | NSTU COVID-19 Diagnostic Center, | NSU Genome Research Institute (NGRI), North South University | Dr. Muhammad Maqsud Hossain, Aura Rahman, Prof. Firoz Ahmed, Tahrima Huq, Abdus Sadique, Jahidul Alam, Md Aminul Islam, Prof. Md. Didar-Ul-Alam, Prof. Kazi Nadim Hasan, Prof. Abdul Khaleque, Prof, Hasan Mahmud Reza |
| EPI_ISL_514440 | NSTU COVID-19 Diagnostic Center | NSU Genome Research Institute (NGRI), North South University | Dr. Muhammad Maqsud Hossain, Aura Rahman, Prof. Firoz Ahmed, Tahrima Huq, Abdus Sadique, Tamanna Afroze, Jahidul Alam, Md Aminul Islam, Prof. Md. Didar-Ul-Alam, Prof. Kazi Nadim Hasan, Prof. Abdul Khaleque, Prof, Hasan Mahmud Reza |
| EPI_ISL_514580, EPI_ISL_514613, EPI_ISL_514614, EPI_ISL_514615 | NSTU COVID-19 Diagnostic Center | NSU Genome Research Institute (NGRI), North South University | Dr. Muhammad Maqsud Hossain, Aura Rahman, Prof. Firoz Ahmed, Tahrima Huq, Abdus Sadique, Jahidul Alam, Tamanna Afroze, Md Aminul Islam, Prof. Md. Didar-Ul-Alam, Prof. Kazi Nadim Hasan, Prof. Abdul Khaleque, Prof, Hasan Mahmud Reza |
| EPI_ISL_515049, EPI_ISL_515050, EPI_ISL_515051, EPI_ISL_515052, EPI_ISL_515053, EPI_ISL_515054 | Division of Viral Diseases, Center for Laboratory Control of Infectious Diseases, Korea Centers for Diseases Control and Prevention | Division of Viral Diseases, Center for Laboratory Control of Infectious Diseases, Korea Centers for Diseases Control and Prevention | Jeong-Min Kim, Yoon-Seok Chung, Namjoo Lee, Sang Hee Woo, Hye-Jun Jo, Heui Man Kim, Jun-Sub Kim, Myung Guk Han |
| EPI_ISL_515055, EPI_ISL_515056, EPI_ISL_515057, EPI_ISL_515059, EPI_ISL_515060, EPI_ISL_515061, EPI_ISL_515062, EPI_ISL_515063, EPI_ISL_515064, EPI_ISL_515066, EPI_ISL_515067, EPI_ISL_515068, EPI_ISL_515069, EPI_ISL_515070, EPI_ISL_515071, EPI_ISL_515072, EPI_ISL_515073, EPI_ISL_515075, EPI_ISL_515076, EPI_ISL_515077, EPI_ISL_515078, EPI_ISL_515079 | | | |
| see above | Department of Clinical Microbiology | GIGA Medical Genomics | Keith Durkin, Maria Artesi, Sebastien Bontems, Raphael Boreux, Cecile Meex, Axelle Chaslain, Celine Fombellida-Lopez, Pierrette Melin, Marie-Pierre Hayette, Vincent Bours. |
| EPI_ISL_515466 | Discovery DNA | Discovery DNA | Dustin Hittel, Marina Kerr, Leo Dimnik, Desmond Koo, Alice Li, Aneal Khan |
| EPI_ISL_515569, EPI_ISL_515570, EPI_ISL_515571, EPI_ISL_515589, EPI_ISL_515592, EPI_ISL_515595, EPI_ISL_515596, EPI_ISL_515597, EPI_ISL_515600, EPI_ISL_515602, EPI_ISL_515603, EPI_ISL_515604, EPI_ISL_515613, EPI_ISL_515614, EPI_ISL_515615, EPI_ISL_515616, EPI_ISL_515617, EPI_ISL_515618, EPI_ISL_515619, EPI_ISL_515620, EPI_ISL_515621, EPI_ISL_515622, EPI_ISL_515623, EPI_ISL_515624, EPI_ISL_515625, EPI_ISL_515626, EPI_ISL_515627, EPI_ISL_515628, EPI_ISL_515629, EPI_ISL_515630, EPI_ISL_515631, EPI_ISL_515632, EPI_ISL_515633, EPI_ISL_515634, EPI_ISL_515635, EPI_ISL_515636, EPI_ISL_515637, EPI_ISL_515638, EPI_ISL_515639, EPI_ISL_515640, EPI_ISL_515641, EPI_ISL_515642, EPI_ISL_515643, EPI_ISL_515644, EPI_ISL_515645, EPI_ISL_515646, EPI_ISL_515647, EPI_ISL_515648, EPI_ISL_515649, EPI_ISL_515650, EPI_ISL_515651, EPI_ISL_515652, EPI_ISL_515653, EPI_ISL_515654, EPI_ISL_515655, EPI_ISL_515656, EPI_ISL_515657, EPI_ISL_515658, EPI_ISL_515659, EPI_ISL_515660, EPI_ISL_515661, EPI_ISL_515662, EPI_ISL_515663, EPI_ISL_515664, EPI_ISL_515665, EPI_ISL_515666, EPI_ISL_515667, EPI_ISL_515669, EPI_ISL_515670, EPI_ISL_515673, EPI_ISL_515674, EPI_ISL_515676, EPI_ISL_515677, EPI_ISL_515678, EPI_ISL_515682, EPI_ISL_515684, EPI_ISL_515687, EPI_ISL_515688, EPI_ISL_515689, EPI_ISL_515690, EPI_ISL_515691, EPI_ISL_515692, EPI_ISL_515693, EPI_ISL_515694, EPI_ISL_515695, EPI_ISL_515696, EPI_ISL_515697, EPI_ISL_515698, EPI_ISL_515700, EPI_ISL_515701, EPI_ISL_515702, EPI_ISL_515703, EPI_ISL_515704, EPI_ISL_515705, EPI_ISL_515706, EPI_ISL_515708, EPI_ISL_515711, EPI_ISL_515712, EPI_ISL_515713, EPI_ISL_515714, EPI_ISL_515715, EPI_ISL_515717, EPI_ISL_515720, EPI_ISL_515722, EPI_ISL_515723, EPI_ISL_515724, EPI_ISL_515725, EPI_ISL_515726, EPI_ISL_515727, EPI_ISL_515728, EPI_ISL_515729, EPI_ISL_515730, EPI_ISL_515731, EPI_ISL_515732, EPI_ISL_515733, EPI_ISL_515734, EPI_ISL_515735, EPI_ISL_515736, EPI_ISL_515737, EPI_ISL_515738, EPI_ISL_515739, EPI_ISL_515740, EPI_ISL_515741, EPI_ISL_515742, EPI_ISL_515743, EPI_ISL_515744, EPI_ISL_515745, EPI_ISL_515746, EPI_ISL_515747, EPI_ISL_515748, EPI_ISL_515749, EPI_ISL_515750, EPI_ISL_515751, EPI_ISL_515752, EPI_ISL_515753, EPI_ISL_515754, EPI_ISL_515756 | | | |
| see above | NHLS-IALCH | KRISP, KZN Research Innovation and Sequencing Platform | Giandhari J, Pillay S, Lessells R, Mdlalose K, York D, Khan S, Tegally H, Wilkinson E, de Oliveira T |
| EPI_ISL_515800, EPI_ISL_515801, EPI_ISL_515802, EPI_ISL_515803, EPI_ISL_515804, EPI_ISL_515805, EPI_ISL_515806, EPI_ISL_515807, EPI_ISL_515808, EPI_ISL_515809, EPI_ISL_515810, EPI_ISL_515811, EPI_ISL_515812, EPI_ISL_515813, EPI_ISL_515814, EPI_ISL_515815, EPI_ISL_515816, EPI_ISL_515817, EPI_ISL_515818, EPI_ISL_515819, EPI_ISL_515820, EPI_ISL_515821, EPI_ISL_515822, EPI_ISL_515823, EPI_ISL_515829, EPI_ISL_515830, EPI_ISL_515831, EPI_ISL_515840, EPI_ISL_515841, EPI_ISL_515842, EPI_ISL_515843, EPI_ISL_515844, EPI_ISL_515845, EPI_ISL_515846, EPI_ISL_515847, EPI_ISL_515848, EPI_ISL_515849, EPI_ISL_515850, EPI_ISL_515851, EPI_ISL_515852, EPI_ISL_515853, EPI_ISL_515854, EPI_ISL_515855, EPI_ISL_515856, EPI_ISL_515857, EPI_ISL_515858, EPI_ISL_515859, EPI_ISL_515860, EPI_ISL_515861, EPI_ISL_515862, EPI_ISL_515863, EPI_ISL_515865, EPI_ISL_515866, EPI_ISL_515867, EPI_ISL_515868, EPI_ISL_515870, EPI_ISL_515872, EPI_ISL_515874, EPI_ISL_515875, EPI_ISL_515876, EPI_ISL_515877, EPI_ISL_515878, EPI_ISL_515879, EPI_ISL_515880, EPI_ISL_515881, EPI_ISL_515882, EPI_ISL_515883, EPI_ISL_515884, EPI_ISL_515885, EPI_ISL_515886, EPI_ISL_515887, EPI_ISL_515888, EPI_ISL_515889, EPI_ISL_515890, EPI_ISL_515891, EPI_ISL_515892, EPI_ISL_515893 | | | |
| see above | Medical Disagnostics Services (MDS) | KRISP, KZN Research Innovation and Sequencing Platform | Giandhari J, Pillay S, Lessells R, ChimukangaraB, Mdlalose K, York D, Khan S, Tegally H, Wilkinson E, de Oliveira T |
| EPI_ISL_516198, EPI_ISL_516199, EPI_ISL_516200, EPI_ISL_516203, EPI_ISL_516204, EPI_ISL_516205, EPI_ISL_516206, EPI_ISL_516207, EPI_ISL_516208, EPI_ISL_516209, EPI_ISL_516210, EPI_ISL_516381, EPI_ISL_516385, EPI_ISL_516388, EPI_ISL_516390, EPI_ISL_516392, EPI_ISL_516393, EPI_ISL_516394, EPI_ISL_516395, EPI_ISL_516397, EPI_ISL_516399, EPI_ISL_516403 | | | |
| see above | Michigan Department of Health and Human Services, Bureau of Laboratories | Michigan Department of Health and Human Services, Bureau of Laboratories | Blankenship HM, Riner D, Soehnlen MK |
| EPI_ISL_516429, EPI_ISL_516430, EPI_ISL_516431 | Clinical Hospital - Shtip | Research Center for Genetic Engineering and Biotechnology "Georgi D. Efremov" , Macedonian Academy of Sciences and Arts | RCGEB - MASA |
| EPI_ISL_516432, EPI_ISL_516433, EPI_ISL_516434, EPI_ISL_516435, EPI_ISL_516436, EPI_ISL_516437, EPI_ISL_516438, EPI_ISL_516439, EPI_ISL_516440, EPI_ISL_516441, EPI_ISL_516442, EPI_ISL_516443, EPI_ISL_516444, EPI_ISL_516445 | | | |
| see above | University of Wisconsin-Madison AIDS Vaccine Research Laboratories | University of Wisconsin-Madison AIDS Vaccine Research Laboratories | Gage Moreno, Katarina Braun, et al. AIDS Vaccine Research Laboratories |
| EPI_ISL_516648 | Institute of Microbiology, Universidad San Francisco de Quito | Institute of Microbiology, Universidad San Francisco de Quito | Juan José Guadalupe, Monica Becerra-Wong, Prado-Vivar, Sully Márquez, Bernardo Gutiérrez, Nabih Dahik, Carlos Mena, Ligia Briceño, Verónica Barragán, Patricio Rojas-Silva, Gabriel Trueba, Michelle Grunauer, Paúl Cárdenas |
| EPI_ISL_516649 | Institute of Microbiology, Universidad San Francisco de Quito | Institute of Microbiology, Universidad San Francisco de Quito | Juan José Guadalupe, Monica Becerra-Wong, Bernardo Gutiérrez, Nabih Dahik, Carlos Mena, Ligia Briceño, Verónica Barragán, Patricio Rojas-Silva, Gabriel Trueba, Michelle Grunauer, Paúl Cárdenas |
| EPI_ISL_516650 | Institute of Microbiology, Universidad San Francisco de Quito | Institute of Microbiology, Universidad San Francisco de Quito | Prado-Vivar, Sully Márquez, Juan José Guadalupe, Monica Becerra-Wong, Bernardo Gutiérrez, Verónica Barragán, Patricio Rojas-Silva, Gabriel Trueba, Michelle Grunauer, Paúl Cárdenas |
| EPI_ISL_516724, EPI_ISL_516725, EPI_ISL_516726, EPI_ISL_516727, EPI_ISL_516728, EPI_ISL_516729, EPI_ISL_516730, EPI_ISL_516731, EPI_ISL_516732, EPI_ISL_516733, EPI_ISL_516734 | | | |
| see above | UCSF Clinical Microbiology Laboratory | Chan-Zuckerberg Biohub | CZB Cliahub Consortium |
| EPI_ISL_516772, EPI_ISL_516775, EPI_ISL_516777, EPI_ISL_516778, EPI_ISL_516779, EPI_ISL_516780, EPI_ISL_516781, EPI_ISL_516782 | van Bakel Laboratory, Genetics and Genomics Sciences, Icahn School of Medicine at Mount Sinai | van Bakel Laboratory, Genetics and Genomics Sciences, Icahn School of Medicine at Mount Sinai | Andrew G. Letizia, Irene Ramos, Ajay Obla, Carl Goforth, Dawn Weir, Yongchao Ge, Marcas M. Bamman, Jayeeta Dutta, Ethan Ellis, Luis Estrella, Mary-Catherine George, Ana S. Gonzalez-Reiche, Darnell Graham, Adriana van de Guchte, Ramiro Gutierrez, Franca Jones, Aspasia Kalomoiri, Rhonda Lizewski, Stephen Lizewski, Jan Marayag, Nada Marjanovic, Eugene V. Millar, Venugopalan Nair, German Nudelman, Edgar Nunez, Brian Pike, James Regeimbal, Stas Rirak , Ernesto Santa Ana, Rachel S. Gelernter Sealfon, Robert Sebra, Mark Simons, Alessandra Soares-Schanoski, Michael Termini, Sindhu Vangeti, Carlos Williams, Harm van Bakel, Stuart C. Sealfon |
| EPI_ISL_516783, EPI_ISL_516784, EPI_ISL_516785 | Center for Laboratory Control of Infectious Diseases, Korea Centers for Diseases Control and Prevention | Center for Laboratory Control of Infectious Diseases, Korea Centers for Diseases Control and Prevention | Junyoung Kim, Ae Kyung Park, Eunkyung Shin, Jin Sun No, Jeong-Min Kim, Yoon-Seok Chung, Heui Man Kim, Myung Guk Han |
| EPI_ISL_516925, EPI_ISL_516926, EPI_ISL_516927, EPI_ISL_516928, EPI_ISL_516929, EPI_ISL_516930, EPI_ISL_516931, EPI_ISL_516932, EPI_ISL_516933 | Department for Molecular Diagnostics, Centre for Medical Microbiology, Institute of Public Health of Montenegro | Charite Universitatsmedizin Berlin, Institut fur Virologie | Victor M Corman, Terry Jones, Jörn Beheim-Schwarzbach, Barbara Muehlemann, Talitha Veith, Julia Schneider, Marija Govedarica and Danijela Vujoševi, Christian Drosten |
| EPI_ISL_517371, EPI_ISL_517372 | Liverpool Clinical Laboratories | COVID-19 Genomics UK (COG-UK) Consortium | Sam Haldenby, Anita Lucaci, Steve Paterson, Julian Hiscox, Alistair Darby, M Almsaud, A Alrezaihi, Muhannad Alruwaili, Stuart D Armstrong, Jones Benjamin, Eleanor G Bentley, Anu Chawla, Jordan J Clark, Angela Cowell, Richard Eccles, Isabel García-Dorival, Matthew Gemmell, Alessandro Gerada, PKF Gilmore, Richard Gregory, Ximeng Han, Catherine Hartley, Margaret Hughes, Miren Iturriza-Gomara, James Johnson, L Luu, Jenifer Manson, Charlotte Nelson, Elaine O'Toole, Cassie Olateju, Rebekah Penrice-Randal , Lucille Rainbow, N.P Randle, Trevor Ian Robinson, Parul Sharma, Ghada T Shawli, James P Stewart, Neil Swainston, Ecaterina Vamos, Joanne Watts, Mark Whitehead |
| EPI_ISL_517546, EPI_ISL_517549, EPI_ISL_517551, EPI_ISL_517554, EPI_ISL_517559, EPI_ISL_517564, EPI_ISL_517565, EPI_ISL_517569, EPI_ISL_517573, EPI_ISL_517578 | Virology Department, Sheffield Teaching Hospitals NHS Foundation Trust/Department of Infection, Immunity and Cardiovascular Disease, The Medical School, University of Sheffield | COVID-19 Genomics UK (COG-UK) Consortium | Thushan de Silva, Matthew Parker, Nikki Smith, Adri Angyal, Rebecca Brown, Luke Green, Rachel Tucker, Paul Parsons, Danielle Groves, Katie Johnson, Laura Carrilero, Alex Keeley, Dave Partridge, Matthew Wyles, Benjamin Lindsey, Mehmet Yavuz, Mohammad Raza, Cariad Evans |
| EPI_ISL_517591 | Wales Specialist Virology Centre Sequencing lab: Pathogen Genomics Unit | COVID-19 Genomics UK (COG-UK) Consortium | Catherine Moore, Johnathan Evans, Laura Gifford, Malorie Perry, Simon Cottrell, Angela Marchbank, Alec Birchley, Alexander Adams, Amy Gaskin, Bree Gatica-Wilcox, Jason Coombes, Joel Southgate, Lauren Gilbert, Lee Graham, Nicole Pacchiarini, Sara Kumziene-Summerhayes, Sarah Taylor, Sophie Jones, |

| | | | |
|---|---|---|---|
| | | | Sara Rey, Matthew Bull, Joanne Watkins, Sally Corden, Tom Connor |
| EPI_ISL_517658, EPI_ISL_517659, EPI_ISL_517660 | Academic Hospital Paramaribo | Erasmus Medical Center | Bas Oude Munnink, Dion Gajadin, Ed Ijzerman, Emmanuelle Munger, Gary Gummels, Ingrid Krishnadath, Lycke Woittiez, Marion Koopmans, Mireille Van de Veer, Princes Wongsowidjojo, Radjesh Ori, Rohma Banwari, Stephen Vreden |
| EPI_ISL_517812, EPI_ISL_517813, EPI_ISL_517818, EPI_ISL_517820, EPI_ISL_517821, EPI_ISL_517822, EPI_ISL_517823, EPI_ISL_517824, EPI_ISL_517825, EPI_ISL_517826, EPI_ISL_517827, EPI_ISL_517828, EPI_ISL_517829, EPI_ISL_517830, EPI_ISL_517831, EPI_ISL_517832, EPI_ISL_517833, EPI_ISL_517834, EPI_ISL_517835, EPI_ISL_517836, EPI_ISL_517837, EPI_ISL_517838, EPI_ISL_517893, EPI_ISL_517894, EPI_ISL_517896, EPI_ISL_517897, EPI_ISL_517898, EPI_ISL_517900, EPI_ISL_517901, EPI_ISL_517902, EPI_ISL_517903, EPI_ISL_517904, EPI_ISL_517905, EPI_ISL_517906, EPI_ISL_517907, EPI_ISL_517908, EPI_ISL_517909, EPI_ISL_517910, EPI_ISL_517911, EPI_ISL_517912, EPI_ISL_517913, EPI_ISL_517914, EPI_ISL_517915, EPI_ISL_517916, EPI_ISL_517917, EPI_ISL_517918, EPI_ISL_517919, EPI_ISL_517920, EPI_ISL_517921, EPI_ISL_517924, EPI_ISL_517925, EPI_ISL_517926, EPI_ISL_517927, EPI_ISL_517948, EPI_ISL_517949, EPI_ISL_517950, EPI_ISL_517951, EPI_ISL_517952, EPI_ISL_517953, EPI_ISL_517954, EPI_ISL_517955, EPI_ISL_517956, EPI_ISL_517957 | | | |
| see above | Florida Bureau of Public Health Laboratories | Florida Bureau of Public Health Laboratories | Sarah Schmedes, Jason Blanton |
| EPI_ISL_518004, EPI_ISL_518005, EPI_ISL_518006, EPI_ISL_518007, EPI_ISL_518008, EPI_ISL_518009, EPI_ISL_518010, EPI_ISL_518011 | Singapore General Hospital | Department of Microbiology | Nurdyana Abdul Rahman, Kun Lee Lim, Chenhao Li, Kian Sing Chan, Lynette Oon, Kern Rei Chng, Niranjan Nagarajan, Karrie Ko |
| EPI_ISL_518037, EPI_ISL_518038, EPI_ISL_518039, EPI_ISL_518040, EPI_ISL_518041, EPI_ISL_518046, EPI_ISL_518047, EPI_ISL_518051 | NHLS-IALCH | KRISP, KZN Research Innovation and Sequencing Platform | Giandhari J, Pillay S, Lessells R, Mdlalose K, York D, Khan S, Tegally H, Wilkinson E, de Oliveira T |
| EPI_ISL_518064, EPI_ISL_518065, EPI_ISL_518077, EPI_ISL_518078 | Microbiological Diagnostic Unit - Public Health Laboratory (MDU-PHL) | MDU-PHL | Seemann T., Schultz M., Sait, M., Sherry, N. |
| EPI_ISL_518081, EPI_ISL_518082, EPI_ISL_518083, EPI_ISL_518084, EPI_ISL_518085, EPI_ISL_518086, EPI_ISL_518087, EPI_ISL_518088, EPI_ISL_518089, EPI_ISL_518090, EPI_ISL_518091, EPI_ISL_518092, EPI_ISL_518098, EPI_ISL_518099, EPI_ISL_518100, EPI_ISL_518101, EPI_ISL_518102, EPI_ISL_518103, EPI_ISL_518104, EPI_ISL_518105, EPI_ISL_518106, EPI_ISL_518107, EPI_ISL_518108, EPI_ISL_518109, EPI_ISL_518111, EPI_ISL_518112, EPI_ISL_518113, EPI_ISL_518114, EPI_ISL_518115, EPI_ISL_518116, EPI_ISL_518117, EPI_ISL_518119, EPI_ISL_518120, EPI_ISL_518122, EPI_ISL_518123, EPI_ISL_518124, EPI_ISL_518125, EPI_ISL_518126, EPI_ISL_518127, EPI_ISL_518128, EPI_ISL_518129, EPI_ISL_518131, EPI_ISL_518132, EPI_ISL_518133, EPI_ISL_518134, EPI_ISL_518135, EPI_ISL_518136, EPI_ISL_518137, EPI_ISL_518138, EPI_ISL_518139, EPI_ISL_518140, EPI_ISL_518141, EPI_ISL_518142, EPI_ISL_518143, EPI_ISL_518144, EPI_ISL_518145, EPI_ISL_518146, EPI_ISL_518147, EPI_ISL_518149, EPI_ISL_518150, EPI_ISL_518151, EPI_ISL_518152, EPI_ISL_518153, EPI_ISL_518154, EPI_ISL_518155, EPI_ISL_518156, EPI_ISL_518157, EPI_ISL_518158, EPI_ISL_518159, EPI_ISL_518160, EPI_ISL_518161, EPI_ISL_518162, EPI_ISL_518163, EPI_ISL_518164, EPI_ISL_518165, EPI_ISL_518166, EPI_ISL_518167, EPI_ISL_518168, EPI_ISL_518169, EPI_ISL_518176, EPI_ISL_518177, EPI_ISL_518178, EPI_ISL_518179, EPI_ISL_518180, EPI_ISL_518181, EPI_ISL_518182, EPI_ISL_518183, EPI_ISL_518184, EPI_ISL_518185, EPI_ISL_518186, EPI_ISL_518188, EPI_ISL_518189, EPI_ISL_518190, EPI_ISL_518191, EPI_ISL_518192, EPI_ISL_518193, EPI_ISL_518194, EPI_ISL_518198, EPI_ISL_518199, EPI_ISL_518210, EPI_ISL_518212, EPI_ISL_518218, EPI_ISL_518219, EPI_ISL_518220, EPI_ISL_518221, EPI_ISL_518223, EPI_ISL_518225, EPI_ISL_518226, EPI_ISL_518227, EPI_ISL_518228, EPI_ISL_518229, EPI_ISL_518230 | | | |
| see above | Victorian Infectious Diseases Reference Laboratory (VIDRL) | VIDRL and MDU-PHL | Caly L., Seemann T., Sait, M., Schultz M., Druce J., Sherry, N. |
| EPI_ISL_518233, EPI_ISL_518235 | Microbiological Diagnostic Unit - Public Health Laboratory (MDU-PHL) | MDU-PHL | Seemann T., Schultz M., Sait, M., Sherry, N. |
| EPI_ISL_518239, EPI_ISL_518240, EPI_ISL_518245, EPI_ISL_518246, EPI_ISL_518247, EPI_ISL_518248, EPI_ISL_518249 | Victorian Infectious Diseases Reference Laboratory (VIDRL) | VIDRL and MDU-PHL | Caly L., Seemann T., Sait, M., Schultz M., Druce J., Sherry, N. |
| EPI_ISL_518269, EPI_ISL_518270, EPI_ISL_518271, EPI_ISL_518275, EPI_ISL_518276, EPI_ISL_518277, EPI_ISL_518278, EPI_ISL_518279, EPI_ISL_518280, EPI_ISL_518281, EPI_ISL_518282, EPI_ISL_518283, EPI_ISL_518288, EPI_ISL_518289, EPI_ISL_518292, EPI_ISL_518293, EPI_ISL_518316, EPI_ISL_518320, EPI_ISL_518322, EPI_ISL_518323, EPI_ISL_518327, EPI_ISL_518328, EPI_ISL_518329, EPI_ISL_518330, EPI_ISL_518331, EPI_ISL_518332, EPI_ISL_518333, EPI_ISL_518334, EPI_ISL_518335, EPI_ISL_518336, EPI_ISL_518337, EPI_ISL_518339, EPI_ISL_518340, EPI_ISL_518341, EPI_ISL_518342, EPI_ISL_518343, EPI_ISL_518344, EPI_ISL_518345, EPI_ISL_518346, EPI_ISL_518347, EPI_ISL_518348, EPI_ISL_518349, EPI_ISL_518350, EPI_ISL_518351, EPI_ISL_518352, EPI_ISL_518353, EPI_ISL_518354, EPI_ISL_518355, EPI_ISL_518356, EPI_ISL_518357, EPI_ISL_518358, EPI_ISL_518359, EPI_ISL_518360, EPI_ISL_518361, EPI_ISL_518362, EPI_ISL_518363, EPI_ISL_518364, EPI_ISL_518365, EPI_ISL_518366, EPI_ISL_518367, EPI_ISL_518368, EPI_ISL_518369, EPI_ISL_518370, EPI_ISL_518371, EPI_ISL_518372, EPI_ISL_518373, EPI_ISL_518374, EPI_ISL_518375, EPI_ISL_518376, EPI_ISL_518377, EPI_ISL_518378, EPI_ISL_518379, EPI_ISL_518380, EPI_ISL_518383, EPI_ISL_518384, EPI_ISL_518385, EPI_ISL_518387 | | | |
| see above | Microbiological Diagnostic Unit - Public Health Laboratory (MDU-PHL) | MDU-PHL | Seemann T., Schultz M., Sait, M., Sherry, N. |
| EPI_ISL_518388, EPI_ISL_518389, EPI_ISL_518390, EPI_ISL_518391, EPI_ISL_518392, EPI_ISL_518393, EPI_ISL_518394, EPI_ISL_518395, EPI_ISL_518396, EPI_ISL_518397, EPI_ISL_518398, EPI_ISL_518399, EPI_ISL_518400, EPI_ISL_518401 | | | |
| see above | Victorian Infectious Diseases Reference Laboratory (VIDRL) | VIDRL and MDU-PHL | Caly L., Seemann T., Sait, M., Schultz M., Druce J., Sherry, N. |
| EPI_ISL_518402, EPI_ISL_518403, EPI_ISL_518404, EPI_ISL_518405, EPI_ISL_518406, EPI_ISL_518407, EPI_ISL_518408, EPI_ISL_518409, EPI_ISL_518410, EPI_ISL_518411, EPI_ISL_518412, EPI_ISL_518413, EPI_ISL_518415, EPI_ISL_518416, EPI_ISL_518417, EPI_ISL_518418, EPI_ISL_518419, EPI_ISL_518420, EPI_ISL_518421, EPI_ISL_518422, EPI_ISL_518423 | | | |
| see above | Microbiological Diagnostic Unit - Public Health Laboratory (MDU-PHL) | MDU-PHL | Seemann T., Schultz M., Sait, M., Sherry, N. |
| EPI_ISL_518424, EPI_ISL_518425, EPI_ISL_518426, EPI_ISL_518427, EPI_ISL_518428, EPI_ISL_518429, EPI_ISL_518430, EPI_ISL_518431, EPI_ISL_518432, EPI_ISL_518433, EPI_ISL_518434, EPI_ISL_518435, EPI_ISL_518436, EPI_ISL_518437, EPI_ISL_518438, EPI_ISL_518439, EPI_ISL_518440, EPI_ISL_518441, EPI_ISL_518442, EPI_ISL_518443, EPI_ISL_518444, EPI_ISL_518445, EPI_ISL_518446, EPI_ISL_518447, EPI_ISL_518448, EPI_ISL_518449, EPI_ISL_518450, EPI_ISL_518451, EPI_ISL_518452, EPI_ISL_518453, EPI_ISL_518454, EPI_ISL_518455, EPI_ISL_518456, EPI_ISL_518457, EPI_ISL_518458, EPI_ISL_518459, EPI_ISL_518460, EPI_ISL_518461, EPI_ISL_518462, EPI_ISL_518463, EPI_ISL_518464, EPI_ISL_518465, EPI_ISL_518466, EPI_ISL_518467, EPI_ISL_518468, EPI_ISL_518469, EPI_ISL_518470, EPI_ISL_518471 | | | |
| see above | Victorian Infectious Diseases Reference Laboratory (VIDRL) | VIDRL and MDU-PHL | Caly L., Seemann T., Sait, M., Schultz M., Druce J., Sherry, N. |
| EPI_ISL_518474, EPI_ISL_518475, EPI_ISL_518476, EPI_ISL_518477, EPI_ISL_518478, EPI_ISL_518479, EPI_ISL_518491, EPI_ISL_518492, EPI_ISL_518493, EPI_ISL_518494, EPI_ISL_518495, EPI_ISL_518496, EPI_ISL_518497, EPI_ISL_518498, EPI_ISL_518499, EPI_ISL_518500, EPI_ISL_518501, EPI_ISL_518502, EPI_ISL_518503, EPI_ISL_518504, EPI_ISL_518505, EPI_ISL_518510, EPI_ISL_518511, EPI_ISL_518512, EPI_ISL_518513, EPI_ISL_518514, EPI_ISL_518515, EPI_ISL_518516, EPI_ISL_518517, EPI_ISL_518518, EPI_ISL_518519, EPI_ISL_518520, EPI_ISL_518521, EPI_ISL_518522, EPI_ISL_518523, EPI_ISL_518524, EPI_ISL_518525, EPI_ISL_518526, EPI_ISL_518527, EPI_ISL_518528, EPI_ISL_518529, EPI_ISL_518530, EPI_ISL_518531, EPI_ISL_518532, EPI_ISL_518533, EPI_ISL_518534, EPI_ISL_518535, EPI_ISL_518536, EPI_ISL_518537, EPI_ISL_518538, EPI_ISL_518539, EPI_ISL_518540, EPI_ISL_518541, EPI_ISL_518542, EPI_ISL_518543, EPI_ISL_518544, EPI_ISL_518545, EPI_ISL_518546, EPI_ISL_518547, EPI_ISL_518549, EPI_ISL_518550, EPI_ISL_518551, EPI_ISL_518552, EPI_ISL_518553, EPI_ISL_518554, EPI_ISL_518555, EPI_ISL_518556, EPI_ISL_518557, EPI_ISL_518559, EPI_ISL_518560, EPI_ISL_518561, EPI_ISL_518562, EPI_ISL_518563, EPI_ISL_518564, EPI_ISL_518565, EPI_ISL_518566, EPI_ISL_518567, EPI_ISL_518568, EPI_ISL_518569, EPI_ISL_518570, EPI_ISL_518571, EPI_ISL_518572, EPI_ISL_518573, EPI_ISL_518574, EPI_ISL_518575, EPI_ISL_518576, EPI_ISL_518577, EPI_ISL_518578, EPI_ISL_518579, EPI_ISL_518580, EPI_ISL_518581, EPI_ISL_518582, EPI_ISL_518583, EPI_ISL_518584, EPI_ISL_518585, EPI_ISL_518586, EPI_ISL_518588, EPI_ISL_518589, EPI_ISL_518590, EPI_ISL_518592, EPI_ISL_518593, EPI_ISL_518594, EPI_ISL_518595, EPI_ISL_518596, EPI_ISL_518597, EPI_ISL_518598, EPI_ISL_518600, EPI_ISL_518601, EPI_ISL_518602, EPI_ISL_518603, EPI_ISL_518604, EPI_ISL_518605, EPI_ISL_518606, EPI_ISL_518607, EPI_ISL_518608, EPI_ISL_518609, EPI_ISL_518610, EPI_ISL_518611, EPI_ISL_518613, EPI_ISL_518614, EPI_ISL_518619, EPI_ISL_518622, EPI_ISL_518623, EPI_ISL_518624, EPI_ISL_518626, EPI_ISL_518627, EPI_ISL_518628, EPI_ISL_518629, EPI_ISL_518630, EPI_ISL_518631, EPI_ISL_518632, EPI_ISL_518633, EPI_ISL_518634, EPI_ISL_518635, EPI_ISL_518636, EPI_ISL_518637, EPI_ISL_518638, EPI_ISL_518639, EPI_ISL_518640, EPI_ISL_518641, EPI_ISL_518642, EPI_ISL_518643, EPI_ISL_518644, EPI_ISL_518645, EPI_ISL_518646, EPI_ISL_518647, EPI_ISL_518648, EPI_ISL_518649, EPI_ISL_518650 | | | |
| see above | Microbiological Diagnostic Unit - Public Health Laboratory (MDU-PHL) | MDU-PHL | Seemann T., Schultz M., Sait, M., Sherry, N. |
| EPI_ISL_518651 | Microbiological Diagnostic Unit - Public Health Laboratory (MDU-PHL) | Microbiological Diagnostic Unit - Public Health Laboratory (MDU-PHL) | Seemann T., Schultz M., Sait, M., Sherry, N. |
| EPI_ISL_518652, EPI_ISL_518653, EPI_ISL_518654, EPI_ISL_518655, EPI_ISL_518656, EPI_ISL_518657, EPI_ISL_518658, EPI_ISL_518659, EPI_ISL_518661, EPI_ISL_518662, EPI_ISL_518663, EPI_ISL_518664, EPI_ISL_518665, EPI_ISL_518666, EPI_ISL_518667, EPI_ISL_518669, EPI_ISL_518670, EPI_ISL_518671, EPI_ISL_518672, EPI_ISL_518673, EPI_ISL_518674, EPI_ISL_518675, EPI_ISL_518676, EPI_ISL_518677, EPI_ISL_518678, EPI_ISL_518679, EPI_ISL_518680, EPI_ISL_518681, EPI_ISL_518682, EPI_ISL_518683, EPI_ISL_518684, EPI_ISL_518685, EPI_ISL_518686, EPI_ISL_518687, EPI_ISL_518688, EPI_ISL_518689, EPI_ISL_518690, EPI_ISL_518691, EPI_ISL_518692, EPI_ISL_518693, EPI_ISL_518694, EPI_ISL_518695, EPI_ISL_518696, EPI_ISL_518697, EPI_ISL_518698, EPI_ISL_518699, EPI_ISL_518700, EPI_ISL_518701, EPI_ISL_518703, EPI_ISL_518704, EPI_ISL_518705, EPI_ISL_518706, EPI_ISL_518707, EPI_ISL_518708, EPI_ISL_518711, EPI_ISL_518712, EPI_ISL_518713, EPI_ISL_518714, EPI_ISL_518715, EPI_ISL_518716, EPI_ISL_518717, EPI_ISL_518718, EPI_ISL_518719, EPI_ISL_518720, EPI_ISL_518721, EPI_ISL_518722 | | | |
| see above | Microbiological Diagnostic Unit - Public Health Laboratory (MDU-PHL) | MDU-PHL | Seemann T., Schultz M., Sait, M., Sherry, N. |
| EPI_ISL_518723 | Victorian Infectious Diseases Reference Laboratory (VIDRL) | VIDRL and MDU-PHL | Caly L., Seemann T., Sait, M., Schultz M., Druce J., Sherry, N. |
| EPI_ISL_518725, EPI_ISL_518726, EPI_ISL_518727, EPI_ISL_518728, EPI_ISL_518729, EPI_ISL_518732, EPI_ISL_518733, EPI_ISL_518735, EPI_ISL_518736, EPI_ISL_518745, EPI_ISL_518746, EPI_ISL_518747, EPI_ISL_518748, EPI_ISL_518749, EPI_ISL_518750, EPI_ISL_518751, EPI_ISL_518752, EPI_ISL_518753, EPI_ISL_518754, EPI_ISL_518755, EPI_ISL_518756, EPI_ISL_518757, EPI_ISL_518758, EPI_ISL_518759, EPI_ISL_518760, EPI_ISL_518761, EPI_ISL_518762, EPI_ISL_518763, EPI_ISL_518764, EPI_ISL_518765, EPI_ISL_518766, EPI_ISL_518767, EPI_ISL_518768, EPI_ISL_518769, EPI_ISL_518770, EPI_ISL_518771, EPI_ISL_518772, EPI_ISL_518773, EPI_ISL_518789, EPI_ISL_518791 | | | |
| see above | Microbiological Diagnostic Unit - Public Health Laboratory (MDU-PHL) | MDU-PHL | Seemann T., Schultz M., Sait, M., Sherry, N. |

EPI_ISL_518799, EPI_ISL_518800, EPI_ISL_518801, EPI_ISL_518802, EPI_ISL_518803, EPI_ISL_518804, EPI_ISL_518805, EPI_ISL_518806, EPI_ISL_518807, EPI_ISL_518808, EPI_ISL_518809, EPI_ISL_518810, EPI_ISL_518811, EPI_ISL_518812, EPI_ISL_518813, EPI_ISL_518814, EPI_ISL_518815, EPI_ISL_518816, EPI_ISL_518817

| | | | |
|---|---|---|---|
| see above | Academic Hospital Paramaribo | Erasmus Medical Center | Bas Oude Munnink, Dion Gajadin, Ed Ijzerman, Emmanuelle Munger, Gary Gummels, Ingrid Krishnadath, Lycke Woittiez, Marion Koopmans, Mireille Van de Veer, Princes Wongsowidjojo, Radjesh Ori, Rohma Banwari, Stephen Vreden |

EPI_ISL_518901, EPI_ISL_518902, EPI_ISL_518904, EPI_ISL_518906, EPI_ISL_518907, EPI_ISL_518908, EPI_ISL_518909, EPI_ISL_518910, EPI_ISL_518911, EPI_ISL_518912, EPI_ISL_518913, EPI_ISL_518914, EPI_ISL_518915, EPI_ISL_518917, EPI_ISL_518918, EPI_ISL_518919, EPI_ISL_518920, EPI_ISL_518921, EPI_ISL_518922, EPI_ISL_518923, EPI_ISL_518924, EPI_ISL_518925, EPI_ISL_518926, EPI_ISL_518927, EPI_ISL_518928, EPI_ISL_518929, EPI_ISL_518930, EPI_ISL_518931, EPI_ISL_518933, EPI_ISL_518939, EPI_ISL_518940, EPI_ISL_518941, EPI_ISL_518942, EPI_ISL_518943, EPI_ISL_518944, EPI_ISL_518945, EPI_ISL_518946, EPI_ISL_518947, EPI_ISL_518948, EPI_ISL_518949, EPI_ISL_518950, EPI_ISL_518951, EPI_ISL_518952, EPI_ISL_518953, EPI_ISL_518954, EPI_ISL_518955, EPI_ISL_518956, EPI_ISL_518957, EPI_ISL_518958, EPI_ISL_518959, EPI_ISL_518960, EPI_ISL_518961, EPI_ISL_518962, EPI_ISL_518963, EPI_ISL_518964, EPI_ISL_518965, EPI_ISL_518966, EPI_ISL_518967, EPI_ISL_518968, EPI_ISL_518969, EPI_ISL_518970, EPI_ISL_518971, EPI_ISL_518972, EPI_ISL_518973, EPI_ISL_518975

| | | | |
|---|---|---|---|
| see above | Microbiological Diagnostic Unit - Public Health Laboratory (MDU-PHL) | MDU-PHL | Seemann T., Schultz M., Sait, M., Sherry, N. |
| EPI_ISL_518976 | Victorian Infectious Diseases Reference Laboratory (VIDRL) | VIDRL and MDU-PHL | Caly L., Seemann T., Sait, M., Schultz M., Druce J., Sherry, N. |
| EPI_ISL_518977, EPI_ISL_518978, EPI_ISL_518979, EPI_ISL_518980 | Microbiological Diagnostic Unit - Public Health Laboratory (MDU-PHL) | MDU-PHL | Seemann T., Schultz M., Sait, M., Sherry, N. |

EPI_ISL_518981, EPI_ISL_518982, EPI_ISL_518983, EPI_ISL_518984, EPI_ISL_518985, EPI_ISL_518986, EPI_ISL_518987, EPI_ISL_518988, EPI_ISL_518989, EPI_ISL_518990, EPI_ISL_518991, EPI_ISL_518992, EPI_ISL_518993, EPI_ISL_518994, EPI_ISL_518995, EPI_ISL_518996, EPI_ISL_518997, EPI_ISL_518998, EPI_ISL_518999, EPI_ISL_519000, EPI_ISL_519001, EPI_ISL_519002, EPI_ISL_519003, EPI_ISL_519004, EPI_ISL_519005, EPI_ISL_519006, EPI_ISL_519007, EPI_ISL_519008, EPI_ISL_519009, EPI_ISL_519010, EPI_ISL_519011, EPI_ISL_519012, EPI_ISL_519013, EPI_ISL_519014, EPI_ISL_519015, EPI_ISL_519016, EPI_ISL_519018, EPI_ISL_519019, EPI_ISL_519020, EPI_ISL_519021, EPI_ISL_519022, EPI_ISL_519029, EPI_ISL_519030, EPI_ISL_519031, EPI_ISL_519032, EPI_ISL_519033, EPI_ISL_519034, EPI_ISL_519035, EPI_ISL_519036, EPI_ISL_519037, EPI_ISL_519038, EPI_ISL_519039, EPI_ISL_519040, EPI_ISL_519041, EPI_ISL_519042, EPI_ISL_519043, EPI_ISL_519044, EPI_ISL_519045, EPI_ISL_519046, EPI_ISL_519048, EPI_ISL_519049, EPI_ISL_519050, EPI_ISL_519051

| | | | |
|---|---|---|---|
| see above | Victorian Infectious Diseases Reference Laboratory (VIDRL) | VIDRL and MDU-PHL | Caly L., Seemann T., Sait, M., Schultz M., Druce J., Sherry, N. |

EPI_ISL_519052, EPI_ISL_519054, EPI_ISL_519055, EPI_ISL_519056, EPI_ISL_519057, EPI_ISL_519059, EPI_ISL_519060, EPI_ISL_519061, EPI_ISL_519062, EPI_ISL_519063, EPI_ISL_519064, EPI_ISL_519065, EPI_ISL_519066, EPI_ISL_519067, EPI_ISL_519068, EPI_ISL_519069, EPI_ISL_519070, EPI_ISL_519071, EPI_ISL_519072, EPI_ISL_519073, EPI_ISL_519074, EPI_ISL_519075, EPI_ISL_519076, EPI_ISL_519077, EPI_ISL_519078, EPI_ISL_519079, EPI_ISL_519080, EPI_ISL_519081, EPI_ISL_519082, EPI_ISL_519083, EPI_ISL_519084, EPI_ISL_519085, EPI_ISL_519086, EPI_ISL_519087, EPI_ISL_519088, EPI_ISL_519089, EPI_ISL_519090, EPI_ISL_519091, EPI_ISL_519093, EPI_ISL_519094, EPI_ISL_519095, EPI_ISL_519096, EPI_ISL_519097, EPI_ISL_519098, EPI_ISL_519099, EPI_ISL_519100, EPI_ISL_519101, EPI_ISL_519102, EPI_ISL_519104, EPI_ISL_519105, EPI_ISL_519106, EPI_ISL_519107, EPI_ISL_519108, EPI_ISL_519109, EPI_ISL_519110, EPI_ISL_519111, EPI_ISL_519112, EPI_ISL_519113, EPI_ISL_519114, EPI_ISL_519115, EPI_ISL_519116, EPI_ISL_519117, EPI_ISL_519118, EPI_ISL_519119, EPI_ISL_519120, EPI_ISL_519121, EPI_ISL_519122, EPI_ISL_519123, EPI_ISL_519124, EPI_ISL_519125, EPI_ISL_519126, EPI_ISL_519127, EPI_ISL_519128, EPI_ISL_519129, EPI_ISL_519130, EPI_ISL_519131, EPI_ISL_519132, EPI_ISL_519133, EPI_ISL_519134, EPI_ISL_519135, EPI_ISL_519136, EPI_ISL_519137, EPI_ISL_519138, EPI_ISL_519139, EPI_ISL_519140, EPI_ISL_519141, EPI_ISL_519142, EPI_ISL_519143, EPI_ISL_519144, EPI_ISL_519145, EPI_ISL_519146, EPI_ISL_519147, EPI_ISL_519148, EPI_ISL_519149, EPI_ISL_519150, EPI_ISL_519151, EPI_ISL_519152, EPI_ISL_519153, EPI_ISL_519154, EPI_ISL_519155, EPI_ISL_519156, EPI_ISL_519157, EPI_ISL_519158, EPI_ISL_519159, EPI_ISL_519160, EPI_ISL_519161, EPI_ISL_519162, EPI_ISL_519163, EPI_ISL_519164, EPI_ISL_519165, EPI_ISL_519166, EPI_ISL_519168, EPI_ISL_519169, EPI_ISL_519170, EPI_ISL_519171, EPI_ISL_519173, EPI_ISL_519174, EPI_ISL_519175, EPI_ISL_519176, EPI_ISL_519177, EPI_ISL_519178, EPI_ISL_519179, EPI_ISL_519180, EPI_ISL_519181, EPI_ISL_519182, EPI_ISL_519183, EPI_ISL_519184, EPI_ISL_519185, EPI_ISL_519186, EPI_ISL_519187, EPI_ISL_519188, EPI_ISL_519189, EPI_ISL_519190, EPI_ISL_519191, EPI_ISL_519192, EPI_ISL_519193, EPI_ISL_519194, EPI_ISL_519195, EPI_ISL_519196, EPI_ISL_519197, EPI_ISL_519198, EPI_ISL_519199, EPI_ISL_519200, EPI_ISL_519201, EPI_ISL_519202, EPI_ISL_519203, EPI_ISL_519204, EPI_ISL_519205, EPI_ISL_519206, EPI_ISL_519207, EPI_ISL_519208, EPI_ISL_519209, EPI_ISL_519210, EPI_ISL_519211, EPI_ISL_519212, EPI_ISL_519213, EPI_ISL_519214, EPI_ISL_519215, EPI_ISL_519216, EPI_ISL_519217, EPI_ISL_519218, EPI_ISL_519219, EPI_ISL_519220, EPI_ISL_519221, EPI_ISL_519222, EPI_ISL_519223, EPI_ISL_519226, EPI_ISL_519228, EPI_ISL_519230, EPI_ISL_519231, EPI_ISL_519232, EPI_ISL_519233, EPI_ISL_519234, EPI_ISL_519235, EPI_ISL_519236, EPI_ISL_519238, EPI_ISL_519239, EPI_ISL_519240, EPI_ISL_519241, EPI_ISL_519242, EPI_ISL_519243, EPI_ISL_519244, EPI_ISL_519245, EPI_ISL_519246, EPI_ISL_519247, EPI_ISL_519248, EPI_ISL_519249, EPI_ISL_519250, EPI_ISL_519251, EPI_ISL_519252, EPI_ISL_519253, EPI_ISL_519254, EPI_ISL_519255

| | | | |
|---|---|---|---|
| see above | Microbiological Diagnostic Unit - Public Health Laboratory (MDU-PHL) | MDU-PHL | Seemann T., Schultz M., Sait, M., Sherry, N. |

EPI_ISL_519256, EPI_ISL_519257, EPI_ISL_519258, EPI_ISL_519259, EPI_ISL_519260, EPI_ISL_519261, EPI_ISL_519262, EPI_ISL_519263, EPI_ISL_519264, EPI_ISL_519265, EPI_ISL_519266, EPI_ISL_519267, EPI_ISL_519268, EPI_ISL_519269, EPI_ISL_519270, EPI_ISL_519271, EPI_ISL_519272, EPI_ISL_519273, EPI_ISL_519274, EPI_ISL_519275, EPI_ISL_519276, EPI_ISL_519277, EPI_ISL_519278, EPI_ISL_519279, EPI_ISL_519280, EPI_ISL_519281, EPI_ISL_519282, EPI_ISL_519283, EPI_ISL_519284, EPI_ISL_519285, EPI_ISL_519286, EPI_ISL_519287, EPI_ISL_519288, EPI_ISL_519289, EPI_ISL_519290, EPI_ISL_519291, EPI_ISL_519292, EPI_ISL_519293, EPI_ISL_519294, EPI_ISL_519295, EPI_ISL_519296, EPI_ISL_519297, EPI_ISL_519298, EPI_ISL_519299, EPI_ISL_519300, EPI_ISL_519301, EPI_ISL_519302, EPI_ISL_519303, EPI_ISL_519304, EPI_ISL_519305, EPI_ISL_519306, EPI_ISL_519307, EPI_ISL_519308, EPI_ISL_519309, EPI_ISL_519310, EPI_ISL_519311, EPI_ISL_519312, EPI_ISL_519313, EPI_ISL_519314, EPI_ISL_519315, EPI_ISL_519316, EPI_ISL_519317, EPI_ISL_519318, EPI_ISL_519319, EPI_ISL_519320, EPI_ISL_519321, EPI_ISL_519324, EPI_ISL_519325, EPI_ISL_519326

| | | | |
|---|---|---|---|
| see above | Victorian Infectious Diseases Reference Laboratory (VIDRL) | VIDRL and MDU-PHL | Caly L., Seemann T., Sait, M., Schultz M., Druce J., Sherry, N. |

EPI_ISL_519327, EPI_ISL_519328, EPI_ISL_519329, EPI_ISL_519330, EPI_ISL_519333, EPI_ISL_519334, EPI_ISL_519335, EPI_ISL_519336, EPI_ISL_519339, EPI_ISL_519340, EPI_ISL_519341, EPI_ISL_519342, EPI_ISL_519343, EPI_ISL_519344, EPI_ISL_519345, EPI_ISL_519346, EPI_ISL_519347, EPI_ISL_519348, EPI_ISL_519349, EPI_ISL_519350, EPI_ISL_519351, EPI_ISL_519352, EPI_ISL_519353, EPI_ISL_519354, EPI_ISL_519355, EPI_ISL_519356, EPI_ISL_519357, EPI_ISL_519359, EPI_ISL_519360, EPI_ISL_519363, EPI_ISL_519364, EPI_ISL_519366, EPI_ISL_519368, EPI_ISL_519371, EPI_ISL_519372, EPI_ISL_519373, EPI_ISL_519374, EPI_ISL_519375, EPI_ISL_519376, EPI_ISL_519377, EPI_ISL_519378, EPI_ISL_519379, EPI_ISL_519380, EPI_ISL_519381, EPI_ISL_519382, EPI_ISL_519383, EPI_ISL_519384, EPI_ISL_519387, EPI_ISL_519391, EPI_ISL_519393, EPI_ISL_519396, EPI_ISL_519413, EPI_ISL_519414, EPI_ISL_519415, EPI_ISL_519416, EPI_ISL_519418, EPI_ISL_519422, EPI_ISL_519435, EPI_ISL_519442, EPI_ISL_519445, EPI_ISL_519450, EPI_ISL_519455, EPI_ISL_519456, EPI_ISL_519457, EPI_ISL_519458, EPI_ISL_519460, EPI_ISL_519461, EPI_ISL_519462, EPI_ISL_519463, EPI_ISL_519464, EPI_ISL_519465, EPI_ISL_519466, EPI_ISL_519467, EPI_ISL_519468, EPI_ISL_519469, EPI_ISL_519470, EPI_ISL_519471, EPI_ISL_519472, EPI_ISL_519473, EPI_ISL_519474, EPI_ISL_519475, EPI_ISL_519476, EPI_ISL_519477, EPI_ISL_519478, EPI_ISL_519479, EPI_ISL_519480, EPI_ISL_519481, EPI_ISL_519482, EPI_ISL_519483, EPI_ISL_519484, EPI_ISL_519485, EPI_ISL_519486, EPI_ISL_519487, EPI_ISL_519488, EPI_ISL_519489, EPI_ISL_519490, EPI_ISL_519491, EPI_ISL_519492, EPI_ISL_519493, EPI_ISL_519494, EPI_ISL_519495, EPI_ISL_519496, EPI_ISL_519498, EPI_ISL_519501, EPI_ISL_519503, EPI_ISL_519504, EPI_ISL_519505, EPI_ISL_519506, EPI_ISL_519511, EPI_ISL_519514, EPI_ISL_519515, EPI_ISL_519516, EPI_ISL_519517, EPI_ISL_519518

| | | | |
|---|---|---|---|
| see above | Microbiological Diagnostic Unit - Public Health Laboratory (MDU-PHL) | MDU-PHL | Seemann T., Schultz M., Sait, M., Sherry, N. |
| EPI_ISL_519520, EPI_ISL_519521 | Victorian Infectious Diseases Reference Laboratory (VIDRL) | VIDRL and MDU-PHL | Caly L., Seemann T., Sait, M., Schultz M., Druce J., Sherry, N. |

EPI_ISL_519523, EPI_ISL_519524, EPI_ISL_519525, EPI_ISL_519526, EPI_ISL_519527, EPI_ISL_519528, EPI_ISL_519529, EPI_ISL_519530, EPI_ISL_519531, EPI_ISL_519532, EPI_ISL_519533, EPI_ISL_519534, EPI_ISL_519535, EPI_ISL_519536, EPI_ISL_519537, EPI_ISL_519538, EPI_ISL_519539, EPI_ISL_519540, EPI_ISL_519541, EPI_ISL_519542, EPI_ISL_519543, EPI_ISL_519547, EPI_ISL_519548, EPI_ISL_519549, EPI_ISL_519550, EPI_ISL_519551, EPI_ISL_519552, EPI_ISL_519553, EPI_ISL_519554, EPI_ISL_519555, EPI_ISL_519556, EPI_ISL_519557, EPI_ISL_519558, EPI_ISL_519559, EPI_ISL_519561, EPI_ISL_519562, EPI_ISL_519563, EPI_ISL_519565, EPI_ISL_519566, EPI_ISL_519567, EPI_ISL_519568, EPI_ISL_519569, EPI_ISL_519571, EPI_ISL_519572, EPI_ISL_519573, EPI_ISL_519574, EPI_ISL_519575, EPI_ISL_519576, EPI_ISL_519577, EPI_ISL_519578, EPI_ISL_519579, EPI_ISL_519580, EPI_ISL_519581, EPI_ISL_519582, EPI_ISL_519583, EPI_ISL_519584, EPI_ISL_519585, EPI_ISL_519586, EPI_ISL_519587, EPI_ISL_519588, EPI_ISL_519589, EPI_ISL_519590, EPI_ISL_519591, EPI_ISL_519592, EPI_ISL_519593, EPI_ISL_519594, EPI_ISL_519595, EPI_ISL_519596, EPI_ISL_519597, EPI_ISL_519598, EPI_ISL_519599, EPI_ISL_519600, EPI_ISL_519602, EPI_ISL_519603, EPI_ISL_519604, EPI_ISL_519605, EPI_ISL_519607, EPI_ISL_519608, EPI_ISL_519609, EPI_ISL_519610, EPI_ISL_519611, EPI_ISL_519612, EPI_ISL_519613, EPI_ISL_519614, EPI_ISL_519615, EPI_ISL_519616, EPI_ISL_519617, EPI_ISL_519618, EPI_ISL_519619, EPI_ISL_519620, EPI_ISL_519621, EPI_ISL_519622, EPI_ISL_519623, EPI_ISL_519624, EPI_ISL_519625, EPI_ISL_519626, EPI_ISL_519627, EPI_ISL_519628, EPI_ISL_519629, EPI_ISL_519630, EPI_ISL_519631, EPI_ISL_519632, EPI_ISL_519633, EPI_ISL_519634, EPI_ISL_519635, EPI_ISL_519636, EPI_ISL_519637, EPI_ISL_519638, EPI_ISL_519639, EPI_ISL_519640, EPI_ISL_519641, EPI_ISL_519642, EPI_ISL_519643, EPI_ISL_519645, EPI_ISL_519646, EPI_ISL_519647, EPI_ISL_519648, EPI_ISL_519649, EPI_ISL_519651, EPI_ISL_519652, EPI_ISL_519653, EPI_ISL_519654, EPI_ISL_519655, EPI_ISL_519656, EPI_ISL_519657, EPI_ISL_519658, EPI_ISL_519659, EPI_ISL_519660, EPI_ISL_519661, EPI_ISL_519662, EPI_ISL_519663, EPI_ISL_519664, EPI_ISL_519665, EPI_ISL_519666, EPI_ISL_519667, EPI_ISL_519668, EPI_ISL_519669, EPI_ISL_519670, EPI_ISL_519671, EPI_ISL_519672, EPI_ISL_519673, EPI_ISL_519674, EPI_ISL_519675, EPI_ISL_519676, EPI_ISL_519677, EPI_ISL_519678, EPI_ISL_519679, EPI_ISL_519680, EPI_ISL_519681, EPI_ISL_519682, EPI_ISL_519683, EPI_ISL_519684, EPI_ISL_519685, EPI_ISL_519686, EPI_ISL_519687, EPI_ISL_519688, EPI_ISL_519689, EPI_ISL_519690, EPI_ISL_519691, EPI_ISL_519692, EPI_ISL_519693, EPI_ISL_519694, EPI_ISL_519695, EPI_ISL_519696, EPI_ISL_519697, EPI_ISL_519698, EPI_ISL_519699, EPI_ISL_519700, EPI_ISL_519701, EPI_ISL_519702, EPI_ISL_519703, EPI_ISL_519704, EPI_ISL_519705, EPI_ISL_519706, EPI_ISL_519707, EPI_ISL_519708, EPI_ISL_519709, EPI_ISL_519710, EPI_ISL_519711, EPI_ISL_519712, EPI_ISL_519713, EPI_ISL_519714, EPI_ISL_519715, EPI_ISL_519716, EPI_ISL_519717, EPI_ISL_519718, EPI_ISL_519719, EPI_ISL_519720, EPI_ISL_519721, EPI_ISL_519722, EPI_ISL_519723, EPI_ISL_519724, EPI_ISL_519725, EPI_ISL_519726, EPI_ISL_519727, EPI_ISL_519728, EPI_ISL_519729, EPI_ISL_519730, EPI_ISL_519731, EPI_ISL_519732, EPI_ISL_519733, EPI_ISL_519734, EPI_ISL_519735, EPI_ISL_519736, EPI_ISL_519737, EPI_ISL_519738, EPI_ISL_519739, EPI_ISL_519740, EPI_ISL_519741, EPI_ISL_519742, EPI_ISL_519743, EPI_ISL_519744, EPI_ISL_519745, EPI_ISL_519746, EPI_ISL_519747, EPI_ISL_519748, EPI_ISL_519749, EPI_ISL_519750, EPI_ISL_519751, EPI_ISL_519752, EPI_ISL_519753, EPI_ISL_519754, EPI_ISL_519755, EPI_ISL_519756, EPI_ISL_519757, EPI_ISL_519758, EPI_ISL_519759, EPI_ISL_519760, EPI_ISL_519761, EPI_ISL_519762, EPI_ISL_519763, EPI_ISL_519764, EPI_ISL_519765, EPI_ISL_519766, EPI_ISL_519767, EPI_ISL_519770, EPI_ISL_519771, EPI_ISL_519773, EPI_ISL_519780, EPI_ISL_519781, EPI_ISL_519782, EPI_ISL_519783, EPI_ISL_519784, EPI_ISL_519785, EPI_ISL_519786, EPI_ISL_519787, EPI_ISL_519788, EPI_ISL_519789, EPI_ISL_519790, EPI_ISL_519791, EPI_ISL_519792, EPI_ISL_519793, EPI_ISL_519794, EPI_ISL_519795, EPI_ISL_519797, EPI_ISL_519798, EPI_ISL_519799, EPI_ISL_519800, EPI_ISL_519801, EPI_ISL_519802, EPI_ISL_519803, EPI_ISL_519804, EPI_ISL_519805, EPI_ISL_519806, EPI_ISL_519807, EPI_ISL_519808, EPI_ISL_519810, EPI_ISL_519811, EPI_ISL_519812, EPI_ISL_519813, EPI_ISL_519815, EPI_ISL_519816, EPI_ISL_519817, EPI_ISL_519818, EPI_ISL_519819, EPI_ISL_519820, EPI_ISL_519822, EPI_ISL_519823, EPI_ISL_519824, EPI_ISL_519825, EPI_ISL_519826, EPI_ISL_519827, EPI_ISL_519828, EPI_ISL_519829, EPI_ISL_519830, EPI_ISL_519831, EPI_ISL_519832, EPI_ISL_519833, EPI_ISL_519834, EPI_ISL_519835, EPI_ISL_519836, EPI_ISL_519837, EPI_ISL_519838, EPI_ISL_519839, EPI_ISL_519840, EPI_ISL_519841, EPI_ISL_519842, EPI_ISL_519843, EPI_ISL_519844, EPI_ISL_519845, EPI_ISL_519846, EPI_ISL_519847, EPI_ISL_519848, EPI_ISL_519849, EPI_ISL_519850, EPI_ISL_519851, EPI_ISL_519852, EPI_ISL_519853, EPI_ISL_519854, EPI_ISL_519855, EPI_ISL_519856, EPI_ISL_519857, EPI_ISL_519858, EPI_ISL_519859, EPI_ISL_519860, EPI_ISL_519861, EPI_ISL_519862, EPI_ISL_519863, EPI_ISL_519864, EPI_ISL_519865, EPI_ISL_519866, EPI_ISL_519867, EPI_ISL_519868, EPI_ISL_519869, EPI_ISL_519870, EPI_ISL_519871, EPI_ISL_519872, EPI_ISL_519873, EPI_ISL_519874, EPI_ISL_519875, EPI_ISL_519876, EPI_ISL_519877, EPI_ISL_519878, EPI_ISL_519879, EPI_ISL_519880, EPI_ISL_519881, EPI_ISL_519882, EPI_ISL_519883, EPI_ISL_519884, EPI_ISL_519885, EPI_ISL_519886, EPI_ISL_519887, EPI_ISL_519888, EPI_ISL_519889, EPI_ISL_519890, EPI_ISL_519891, EPI_ISL_519892, EPI_ISL_519893, EPI_ISL_519894

| | | | |
|---|---|---|---|
| see above | Microbiological Diagnostic Unit - Public Health Laboratory (MDU-PHL) | MDU-PHL | Seemann T., Schultz M., Sait, M., Sherry, N. |
| EPI_ISL_519895 | Microbiological Diagnostic Unit - Public Health Laboratory (MDU-PHL) | Microbiological Diagnostic Unit - Public Health Laboratory (MDU-PHL) | Seemann T., Schultz M., Sait, M., Sherry, N. |

EPI_ISL_519896, EPI_ISL_519897, EPI_ISL_519898, EPI_ISL_519899, EPI_ISL_519900, EPI_ISL_519901, EPI_ISL_519902, EPI_ISL_519903, EPI_ISL_519904, EPI_ISL_519905, EPI_ISL_519906, EPI_ISL_519907, EPI_ISL_519908, EPI_ISL_519909, EPI_ISL_519910, EPI_ISL_519911, EPI_ISL_519913, EPI_ISL_519914, EPI_ISL_519915, EPI_ISL_519916, EPI_ISL_519917, EPI_ISL_519918, EPI_ISL_519919, EPI_ISL_519920, EPI_ISL_519921, EPI_ISL_519922, EPI_ISL_519923, EPI_ISL_519924, EPI_ISL_519925, EPI_ISL_519926, EPI_ISL_519927, EPI_ISL_519928, EPI_ISL_519929, EPI_ISL_519930, EPI_ISL_519931, EPI_ISL_519932, EPI_ISL_519933, EPI_ISL_519934, EPI_ISL_519935, EPI_ISL_519936, EPI_ISL_519937, EPI_ISL_519938, EPI_ISL_519939, EPI_ISL_519940, EPI_ISL_519941, EPI_ISL_519942, EPI_ISL_519943, EPI_ISL_519944, EPI_ISL_519945, EPI_ISL_519946

| | | | |
|---|---|---|---|
| see above | Microbiological Diagnostic Unit - Public Health Laboratory (MDU-PHL) | MDU-PHL | Seemann T., Schultz M., Sait, M., Sherry, N. |
| EPI_ISL_519948, EPI_ISL_519949, EPI_ISL_519950 | Victorian Infectious Diseases Reference Laboratory (VIDRL) | VIDRL and MDU-PHL | Caly L., Seemann T., Sait, M., Schultz M., Druce J., Sherry, N. |

| | | | |
|---|---|---|---|
| EPI_ISL_519951, EPI_ISL_519952, EPI_ISL_519953, EPI_ISL_519954, EPI_ISL_519955, EPI_ISL_519956, EPI_ISL_519957, EPI_ISL_519958, EPI_ISL_519963, EPI_ISL_519964, EPI_ISL_519965, EPI_ISL_519966, EPI_ISL_519967, EPI_ISL_519968, EPI_ISL_519969, EPI_ISL_519970, EPI_ISL_519971, EPI_ISL_519972, EPI_ISL_519973, EPI_ISL_519974, EPI_ISL_519975, EPI_ISL_519976, EPI_ISL_519977, EPI_ISL_519978, EPI_ISL_519979, EPI_ISL_519980, EPI_ISL_519983, EPI_ISL_519984, EPI_ISL_519985, EPI_ISL_519986, EPI_ISL_519987, EPI_ISL_519988, EPI_ISL_519989, EPI_ISL_519990, EPI_ISL_519992, EPI_ISL_519993, EPI_ISL_519994, EPI_ISL_519995, EPI_ISL_519996, EPI_ISL_519997, EPI_ISL_519998, EPI_ISL_519999, EPI_ISL_520000, EPI_ISL_520001, EPI_ISL_520002, EPI_ISL_520003, EPI_ISL_520004, EPI_ISL_520005, EPI_ISL_520007, EPI_ISL_520008, EPI_ISL_520009, EPI_ISL_520010, EPI_ISL_520016, EPI_ISL_520017, EPI_ISL_520018, EPI_ISL_520019, EPI_ISL_520020, EPI_ISL_520021, EPI_ISL_520022, EPI_ISL_520023, EPI_ISL_520024, EPI_ISL_520025, EPI_ISL_520026, EPI_ISL_520027, EPI_ISL_520028, EPI_ISL_520029, EPI_ISL_520030, EPI_ISL_520031, EPI_ISL_520041, EPI_ISL_520042, EPI_ISL_520043, EPI_ISL_520044, EPI_ISL_520045, EPI_ISL_520046, EPI_ISL_520047, EPI_ISL_520048, EPI_ISL_520049, EPI_ISL_520050, EPI_ISL_520051, EPI_ISL_520052, EPI_ISL_520053, EPI_ISL_520054, EPI_ISL_520055, EPI_ISL_520057, EPI_ISL_520058, EPI_ISL_520059, EPI_ISL_520061, EPI_ISL_520062, EPI_ISL_520063, EPI_ISL_520066, EPI_ISL_520067, EPI_ISL_520068, EPI_ISL_520069, EPI_ISL_520070, EPI_ISL_520072, EPI_ISL_520073, EPI_ISL_520074, EPI_ISL_520075, EPI_ISL_520077, EPI_ISL_520079, EPI_ISL_520080, EPI_ISL_520081, EPI_ISL_520082, EPI_ISL_520083, EPI_ISL_520085, EPI_ISL_520086, EPI_ISL_520087, EPI_ISL_520089, EPI_ISL_520090, EPI_ISL_520091, EPI_ISL_520092, EPI_ISL_520093, EPI_ISL_520094, EPI_ISL_520095, EPI_ISL_520096, EPI_ISL_520097, EPI_ISL_520098, EPI_ISL_520099, EPI_ISL_520100, EPI_ISL_520101, EPI_ISL_520102, EPI_ISL_520103, EPI_ISL_520104, EPI_ISL_520105, EPI_ISL_520106, EPI_ISL_520107, EPI_ISL_520108, EPI_ISL_520109, EPI_ISL_520110, EPI_ISL_520111, EPI_ISL_520112, EPI_ISL_520113, EPI_ISL_520114, EPI_ISL_520115, EPI_ISL_520116, EPI_ISL_520117, EPI_ISL_520118, EPI_ISL_520119, EPI_ISL_520121, EPI_ISL_520122, EPI_ISL_520123, EPI_ISL_520124, EPI_ISL_520125, EPI_ISL_520126, EPI_ISL_520127, EPI_ISL_520128, EPI_ISL_520129, EPI_ISL_520130, EPI_ISL_520131, EPI_ISL_520132, EPI_ISL_520133, EPI_ISL_520134, EPI_ISL_520135, EPI_ISL_520136, EPI_ISL_520137, EPI_ISL_520138, EPI_ISL_520139, EPI_ISL_520140, EPI_ISL_520141, EPI_ISL_520142, EPI_ISL_520143, EPI_ISL_520144, EPI_ISL_520145, EPI_ISL_520146, EPI_ISL_520147, EPI_ISL_520148, EPI_ISL_520149, EPI_ISL_520150, EPI_ISL_520151, EPI_ISL_520152, EPI_ISL_520153, EPI_ISL_520154, EPI_ISL_520155, EPI_ISL_520156, EPI_ISL_520157, EPI_ISL_520158, EPI_ISL_520159, EPI_ISL_520160, EPI_ISL_520161, EPI_ISL_520162, EPI_ISL_520163, EPI_ISL_520164, EPI_ISL_520166, EPI_ISL_520167, EPI_ISL_520168 | | | |
| see above | Microbiological Diagnostic Unit - Public Health Laboratory (MDU-PHL) | MDU-PHL | Seemann T., Schultz M., Sait, M., Sherry, N. |
| EPI_ISL_520181, EPI_ISL_520182, EPI_ISL_520183, EPI_ISL_520184, EPI_ISL_520185, EPI_ISL_520186, EPI_ISL_520187, EPI_ISL_520188, EPI_ISL_520189, EPI_ISL_520190, EPI_ISL_520191, EPI_ISL_520192, EPI_ISL_520193, EPI_ISL_520194, EPI_ISL_520195, EPI_ISL_520196, EPI_ISL_520197, EPI_ISL_520198, EPI_ISL_520199, EPI_ISL_520200, EPI_ISL_520202, EPI_ISL_520203, EPI_ISL_520204, EPI_ISL_520205, EPI_ISL_520206, EPI_ISL_520207, EPI_ISL_520208, EPI_ISL_520209, EPI_ISL_520210, EPI_ISL_520211, EPI_ISL_520212, EPI_ISL_520213, EPI_ISL_520214 | | | |
| see above | Victorian Infectious Diseases Reference Laboratory (VIDRL) | VIDRL and MDU-PHL | Caly L., Seemann T., Sait, M., Schultz M., Druce J., Sherry, N. |
| EPI_ISL_520215, EPI_ISL_520216, EPI_ISL_520217, EPI_ISL_520218, EPI_ISL_520219, EPI_ISL_520220, EPI_ISL_520221, EPI_ISL_520222, EPI_ISL_520223, EPI_ISL_520224, EPI_ISL_520225, EPI_ISL_520226, EPI_ISL_520227, EPI_ISL_520228, EPI_ISL_520229, EPI_ISL_520230, EPI_ISL_520231, EPI_ISL_520233, EPI_ISL_520234, EPI_ISL_520235, EPI_ISL_520236, EPI_ISL_520237, EPI_ISL_520238, EPI_ISL_520239, EPI_ISL_520240, EPI_ISL_520241, EPI_ISL_520242, EPI_ISL_520243, EPI_ISL_520244, EPI_ISL_520245, EPI_ISL_520246, EPI_ISL_520247, EPI_ISL_520248, EPI_ISL_520392, EPI_ISL_520393, EPI_ISL_520400, EPI_ISL_520409, EPI_ISL_520413, EPI_ISL_520414, EPI_ISL_520415, EPI_ISL_520417, EPI_ISL_520418, EPI_ISL_520419, EPI_ISL_520423, EPI_ISL_520424, EPI_ISL_520425, EPI_ISL_520426, EPI_ISL_520427, EPI_ISL_520428, EPI_ISL_520429, EPI_ISL_520430, EPI_ISL_520431, EPI_ISL_520432, EPI_ISL_520433, EPI_ISL_520437, EPI_ISL_520438, EPI_ISL_520439, EPI_ISL_520440, EPI_ISL_520441, EPI_ISL_520442, EPI_ISL_520443, EPI_ISL_520444, EPI_ISL_520445, EPI_ISL_520446, EPI_ISL_520449, EPI_ISL_520450, EPI_ISL_520451, EPI_ISL_520452, EPI_ISL_520453, EPI_ISL_520454, EPI_ISL_520455, EPI_ISL_520457, EPI_ISL_520458, EPI_ISL_520459, EPI_ISL_520460, EPI_ISL_520461, EPI_ISL_520464, EPI_ISL_520465, EPI_ISL_520466, EPI_ISL_520467, EPI_ISL_520468, EPI_ISL_520470, EPI_ISL_520471, EPI_ISL_520472, EPI_ISL_520473, EPI_ISL_520474, EPI_ISL_520475, EPI_ISL_520477, EPI_ISL_520481, EPI_ISL_520482, EPI_ISL_520485, EPI_ISL_520487, EPI_ISL_520488, EPI_ISL_520489, EPI_ISL_520492, EPI_ISL_520493, EPI_ISL_520495, EPI_ISL_520496, EPI_ISL_520497, EPI_ISL_520498, EPI_ISL_520500, EPI_ISL_520501, EPI_ISL_520502, EPI_ISL_520503, EPI_ISL_520504, EPI_ISL_520505, EPI_ISL_520507, EPI_ISL_520508, EPI_ISL_520509, EPI_ISL_520510, EPI_ISL_520511, EPI_ISL_520512, EPI_ISL_520513, EPI_ISL_520514, EPI_ISL_520516, EPI_ISL_520517, EPI_ISL_520518, EPI_ISL_520519, EPI_ISL_520520, EPI_ISL_520521, EPI_ISL_520522, EPI_ISL_520523, EPI_ISL_520524, EPI_ISL_520525, EPI_ISL_520526, EPI_ISL_520527, EPI_ISL_520528, EPI_ISL_520529, EPI_ISL_520530, EPI_ISL_520531, EPI_ISL_520532, EPI_ISL_520533, EPI_ISL_520543, EPI_ISL_520544, EPI_ISL_520545, EPI_ISL_520546, EPI_ISL_520547, EPI_ISL_520548, EPI_ISL_520549, EPI_ISL_520552, EPI_ISL_520553, EPI_ISL_520554, EPI_ISL_520555, EPI_ISL_520556, EPI_ISL_520557, EPI_ISL_520558, EPI_ISL_520559, EPI_ISL_520560, EPI_ISL_520561, EPI_ISL_520563, EPI_ISL_520564, EPI_ISL_520565, EPI_ISL_520567, EPI_ISL_520568, EPI_ISL_520573, EPI_ISL_520574, EPI_ISL_520575, EPI_ISL_520576, EPI_ISL_520577, EPI_ISL_520578, EPI_ISL_520579, EPI_ISL_520580, EPI_ISL_520581, EPI_ISL_520582, EPI_ISL_520583, EPI_ISL_520584, EPI_ISL_520585, EPI_ISL_520586, EPI_ISL_520587, EPI_ISL_520588, EPI_ISL_520589, EPI_ISL_520590, EPI_ISL_520591, EPI_ISL_520592, EPI_ISL_520593, EPI_ISL_520594, EPI_ISL_520595, EPI_ISL_520596, EPI_ISL_520597, EPI_ISL_520598, EPI_ISL_520599, EPI_ISL_520600, EPI_ISL_520601, EPI_ISL_520602, EPI_ISL_520603 | | | |
| see above | Microbiological Diagnostic Unit - Public Health Laboratory (MDU-PHL) | MDU-PHL | Seemann T., Schultz M., Sait, M., Sherry, N. |
| EPI_ISL_520604, EPI_ISL_520605 | Microbiological Diagnostic Unit - Public Health Laboratory (MDU-PHL) | Microbiological Diagnostic Unit - Public Health Laboratory (MDU-PHL) | Seemann T., Schultz M., Sait, M., Sherry, N. |
| EPI_ISL_520606, EPI_ISL_520607, EPI_ISL_520608, EPI_ISL_520609, EPI_ISL_520610, EPI_ISL_520611, EPI_ISL_520612, EPI_ISL_520613, EPI_ISL_520614, EPI_ISL_520620, EPI_ISL_520621, EPI_ISL_520624, EPI_ISL_520625, EPI_ISL_520626, EPI_ISL_520627, EPI_ISL_520628, EPI_ISL_520629, EPI_ISL_520630, EPI_ISL_520635, EPI_ISL_520636, EPI_ISL_520637, EPI_ISL_520638, EPI_ISL_520639, EPI_ISL_520640, EPI_ISL_520641, EPI_ISL_520642, EPI_ISL_520643 | | | |
| see above | Microbiological Diagnostic Unit - Public Health Laboratory (MDU-PHL) | MDU-PHL | Seemann T., Schultz M., Sait, M., Sherry, N. |
| EPI_ISL_520644 | Microbiological Diagnostic Unit - Public Health Laboratory (MDU-PHL) | Microbiological Diagnostic Unit - Public Health Laboratory (MDU-PHL) | Seemann T., Schultz M., Sait, M., Sherry, N. |
| EPI_ISL_520645, EPI_ISL_520647, EPI_ISL_520648, EPI_ISL_520649, EPI_ISL_520650, EPI_ISL_520651, EPI_ISL_520652, EPI_ISL_520653, EPI_ISL_520654, EPI_ISL_520655, EPI_ISL_520656, EPI_ISL_520657, EPI_ISL_520658, EPI_ISL_520659, EPI_ISL_520660, EPI_ISL_520661, EPI_ISL_520662, EPI_ISL_520845, EPI_ISL_520847, EPI_ISL_520850, EPI_ISL_520853, EPI_ISL_520854, EPI_ISL_520857, EPI_ISL_520860, EPI_ISL_520863, EPI_ISL_520929, EPI_ISL_520935, EPI_ISL_520936, EPI_ISL_520937, EPI_ISL_520938, EPI_ISL_520939, EPI_ISL_520940, EPI_ISL_520941, EPI_ISL_520942, EPI_ISL_520943, EPI_ISL_520944, EPI_ISL_520983, EPI_ISL_520986, EPI_ISL_520989, EPI_ISL_520990, EPI_ISL_520995, EPI_ISL_521078, EPI_ISL_521079, EPI_ISL_521080, EPI_ISL_521082, EPI_ISL_521090, EPI_ISL_521092, EPI_ISL_521093, EPI_ISL_521094, EPI_ISL_521097, EPI_ISL_521098, EPI_ISL_521099, EPI_ISL_521100, EPI_ISL_521101, EPI_ISL_521154, EPI_ISL_521156, EPI_ISL_521157, EPI_ISL_521158, EPI_ISL_521159, EPI_ISL_521160, EPI_ISL_521161, EPI_ISL_521162, EPI_ISL_521163, EPI_ISL_521165, EPI_ISL_521195, EPI_ISL_521196, EPI_ISL_521202, EPI_ISL_521203, EPI_ISL_521204, EPI_ISL_521205, EPI_ISL_521206, EPI_ISL_521209, EPI_ISL_521212, EPI_ISL_521230, EPI_ISL_521239, EPI_ISL_521248, EPI_ISL_521260, EPI_ISL_521261, EPI_ISL_521262, EPI_ISL_521263, EPI_ISL_521264, EPI_ISL_521266, EPI_ISL_521267, EPI_ISL_521268, EPI_ISL_521269 | | | |
| see above | Microbiological Diagnostic Unit - Public Health Laboratory (MDU-PHL) | MDU-PHL | Seemann T., Schultz M., Sait, M., Sherry, N. |
| EPI_ISL_521272, EPI_ISL_521273, EPI_ISL_521275, EPI_ISL_521277 | Victorian Infectious Diseases Reference Laboratory (VIDRL) | VIDRL and MDU-PHL | Caly L., Seemann T., Sait, M., Schultz M., Druce J., Sherry, N. |
| EPI_ISL_521302 | Microbiological Diagnostic Unit - Public Health Laboratory (MDU-PHL) | MDU-PHL | Seemann T., Schultz M., Sait, M., Sherry, N. |
| EPI_ISL_521315, EPI_ISL_521316, EPI_ISL_521317, EPI_ISL_521318, EPI_ISL_521319, EPI_ISL_521320, EPI_ISL_521321, EPI_ISL_521322, EPI_ISL_521323, EPI_ISL_521324, EPI_ISL_521325, EPI_ISL_521326, EPI_ISL_521327, EPI_ISL_521328, EPI_ISL_521329, EPI_ISL_521330, EPI_ISL_521331, EPI_ISL_521332, EPI_ISL_521333, EPI_ISL_521334, EPI_ISL_521335, EPI_ISL_521336, EPI_ISL_521337, EPI_ISL_521338, EPI_ISL_521339, EPI_ISL_521340, EPI_ISL_521341, EPI_ISL_521342, EPI_ISL_521343, EPI_ISL_521344, EPI_ISL_521345, EPI_ISL_521346, EPI_ISL_521347, EPI_ISL_521348, EPI_ISL_521349, EPI_ISL_521350, EPI_ISL_521351, EPI_ISL_521352, EPI_ISL_521353, EPI_ISL_521354, EPI_ISL_521355, EPI_ISL_521356, EPI_ISL_521357, EPI_ISL_521358, EPI_ISL_521359, EPI_ISL_521360, EPI_ISL_521361, EPI_ISL_521362, EPI_ISL_521363, EPI_ISL_521364, EPI_ISL_521365, EPI_ISL_521366, EPI_ISL_521367, EPI_ISL_521368, EPI_ISL_521369, EPI_ISL_521370, EPI_ISL_521371, EPI_ISL_521372, EPI_ISL_521373, EPI_ISL_521374, EPI_ISL_521375, EPI_ISL_521376, EPI_ISL_521377, EPI_ISL_521378, EPI_ISL_521379, EPI_ISL_521380, EPI_ISL_521381, EPI_ISL_521382, EPI_ISL_521383, EPI_ISL_521384, EPI_ISL_521385, EPI_ISL_521386, EPI_ISL_521387, EPI_ISL_521388, EPI_ISL_521389, EPI_ISL_521390, EPI_ISL_521391, EPI_ISL_521392, EPI_ISL_521393, EPI_ISL_521394, EPI_ISL_521395, EPI_ISL_521396, EPI_ISL_521397, EPI_ISL_521398, EPI_ISL_521399, EPI_ISL_521400, EPI_ISL_521401, EPI_ISL_521402, EPI_ISL_521403, EPI_ISL_521404, EPI_ISL_521405, EPI_ISL_521406, EPI_ISL_521407, EPI_ISL_521408, EPI_ISL_521410, EPI_ISL_521411, EPI_ISL_521412, EPI_ISL_521413, EPI_ISL_521414, EPI_ISL_521415, EPI_ISL_521416, EPI_ISL_521417, EPI_ISL_521418, EPI_ISL_521419, EPI_ISL_521420, EPI_ISL_521421, EPI_ISL_521422, EPI_ISL_521423, EPI_ISL_521424, EPI_ISL_521425, EPI_ISL_521426, EPI_ISL_521427, EPI_ISL_521428, EPI_ISL_521429, EPI_ISL_521430, EPI_ISL_521431, EPI_ISL_521432, EPI_ISL_521433, EPI_ISL_521434, EPI_ISL_521435, EPI_ISL_521436, EPI_ISL_521437, EPI_ISL_521438, EPI_ISL_521439, EPI_ISL_521440, EPI_ISL_521441, EPI_ISL_521442, EPI_ISL_521443, EPI_ISL_521444, EPI_ISL_521445, EPI_ISL_521446, EPI_ISL_521447, EPI_ISL_521448, EPI_ISL_521449 | | | |
| see above | Victorian Infectious Diseases Reference Laboratory (VIDRL) | VIDRL and MDU-PHL | Caly L., Seemann T., Sait, M., Schultz M., Druce J., Sherry, N. |
| EPI_ISL_521774, EPI_ISL_521789, EPI_ISL_521791, EPI_ISL_521797, EPI_ISL_521823, EPI_ISL_521825, EPI_ISL_521827, EPI_ISL_521836 | Microbiological Diagnostic Unit - Public Health Laboratory (MDU-PHL) | MDU-PHL | Seemann T., Schultz M., Sait, M., Sherry, N. |
| EPI_ISL_522491, EPI_ISL_522492, EPI_ISL_522493, EPI_ISL_522494, EPI_ISL_522495, EPI_ISL_522496, EPI_ISL_522497 | Center for Laboratory Control of Infectious Diseases, Korea Centers for Diseases Control and Prevention | Center for Laboratory Control of Infectious Diseases, Korea Centers for Diseases Control and Prevention | Junyoung Kim, Ae Kyung Park, Eunkyung Shin, Jin Sun No, Jeong-Min Kim, Yoon-Seok Chung, Heui Man Kim, Myung Guk Han |
| EPI_ISL_522498, EPI_ISL_522499, EPI_ISL_522500, EPI_ISL_522501 | Division of Viral Diseases, Center for Laboratory Control of Infectious Diseases, Korea Centers for Diseases Control and Prevention | Division of Viral Diseases, Center for Laboratory Control of Infectious Diseases, Korea Centers for Diseases Control and Prevention | Jeong-Min Kim, Yoon-Seok Chung, Namjoo Lee, Sang Hee Woo, Hye-Jun Jo, Heui Man Kim, Jun-Sub Kim, Myung Guk Han |
| EPI_ISL_522502 | Center for Laboratory Control of Infectious Diseases, Korea Centers for Diseases Control and Prevention | Center for Laboratory Control of Infectious Diseases, Korea Centers for Diseases Control and Prevention | Junyoung Kim, Ae Kyung Park, Eunkyung Shin, Jin Sun No, Jeong-Min Kim, Yoon-Seok Chung, Heui Man Kim, Myung Guk Han |
| EPI_ISL_522503, EPI_ISL_522504 | Division of Viral Diseases, Center for Laboratory Control of Infectious Diseases, Korea Centers for Diseases Control and Prevention | Division of Viral Diseases, Center for Laboratory Control of Infectious Diseases, Korea Centers for Diseases Control and Prevention | Jeong-Min Kim, Yoon-Seok Chung, Namjoo Lee, Sang Hee Woo, Hye-Jun Jo, Heui Man Kim, Jun-Sub Kim, Myung Guk Han |
| EPI_ISL_522505 | Center for Laboratory Control of Infectious Diseases, Korea Centers for Diseases Control and Prevention | Center for Laboratory Control of Infectious Diseases, Korea Centers for Diseases Control and Prevention | Junyoung Kim, Ae Kyung Park, Eunkyung Shin, Jin Sun No, Jeong-Min Kim, Yoon-Seok Chung, Heui Man Kim, Myung Guk Han |
| EPI_ISL_522506, EPI_ISL_522507 | Division of Viral Diseases, Center for Laboratory Control of | Division of Viral Diseases, Center for Laboratory Control of | Jeong-Min Kim, Yoon-Seok Chung, Namjoo Lee, Sang Hee Woo, Hye-Jun Jo, Heui Man Kim, Jun-Sub Kim, Myung Guk Han |

| | | | |
|---|---|---|---|
| | Infectious Diseases, Korea Centers for Diseases Control and Prevention | Infectious Diseases, Korea Centers for Diseases Control and Prevention | |
| EPI_ISL_522508 | Center for Laboratory Control of Infectious Diseases, Korea Centers for Diseases Control and Prevention | Center for Laboratory Control of Infectious Diseases, Korea Centers for Diseases Control and Prevention | Junyoung Kim, Ae Kyung Park, Eunkyung Shin, Jin Sun No, Jeong-Min Kim, Yoon-Seok Chung, Heui Man Kim, Myung Guk Han |
| EPI_ISL_522509 | Division of Viral Diseases, Center for Laboratory Control of Infectious Diseases, Korea Centers for Diseases Control and Prevention | Division of Viral Diseases, Center for Laboratory Control of Infectious Diseases, Korea Centers for Diseases Control and Prevention | Jeong-Min Kim, Yoon-Seok Chung, Namjoo Lee, Sang Hee Woo, Hye-Jun Jo, Heui Man Kim, Jun-Sub Kim, Myung Guk Han |
| EPI_ISL_522510 | Center for Laboratory Control of Infectious Diseases, Korea Centers for Diseases Control and Prevention | Center for Laboratory Control of Infectious Diseases, Korea Centers for Diseases Control and Prevention | Junyoung Kim, Ae Kyung Park, Eunkyung Shin, Jin Sun No, Jeong-Min Kim, Yoon-Seok Chung, Heui Man Kim, Myung Guk Han |
| EPI_ISL_522511, EPI_ISL_522512, EPI_ISL_522513, EPI_ISL_522514 | Division of Viral Diseases, Center for Laboratory Control of Infectious Diseases, Korea Centers for Diseases Control and Prevention | Division of Viral Diseases, Center for Laboratory Control of Infectious Diseases, Korea Centers for Diseases Control and Prevention | Jeong-Min Kim, Yoon-Seok Chung, Namjoo Lee, Sang Hee Woo, Hye-Jun Jo, Heui Man Kim, Jun-Sub Kim, Myung Guk Han |
| EPI_ISL_522515, EPI_ISL_522516 | Center for Laboratory Control of Infectious Diseases, Korea Centers for Diseases Control and Prevention | Center for Laboratory Control of Infectious Diseases, Korea Centers for Diseases Control and Prevention | Junyoung Kim, Ae Kyung Park, Eunkyung Shin, Jin Sun No, Jeong-Min Kim, Yoon-Seok Chung, Heui Man Kim, Myung Guk Han |
| EPI_ISL_522517 | Division of Viral Diseases, Center for Laboratory Control of Infectious Diseases, Korea Centers for Diseases Control and Prevention | Division of Viral Diseases, Center for Laboratory Control of Infectious Diseases, Korea Centers for Diseases Control and Prevention | Jeong-Min Kim, Yoon-Seok Chung, Namjoo Lee, Sang Hee Woo, Hye-Jun Jo, Heui Man Kim, Jun-Sub Kim, Myung Guk Han |
| EPI_ISL_522518 | Center for Laboratory Control of Infectious Diseases, Korea Centers for Diseases Control and Prevention | Center for Laboratory Control of Infectious Diseases, Korea Centers for Diseases Control and Prevention | Junyoung Kim, Ae Kyung Park, Eunkyung Shin, Jin Sun No, Jeong-Min Kim, Yoon-Seok Chung, Heui Man Kim, Myung Guk Han |
| EPI_ISL_522519, EPI_ISL_522520, EPI_ISL_522521, EPI_ISL_522522, EPI_ISL_522523, EPI_ISL_522524, EPI_ISL_522525, EPI_ISL_522526, EPI_ISL_522527, EPI_ISL_522528, EPI_ISL_522529, EPI_ISL_522530, EPI_ISL_522531, EPI_ISL_522532, EPI_ISL_522533, EPI_ISL_522534, EPI_ISL_522535, EPI_ISL_522536, EPI_ISL_522537, EPI_ISL_522538, EPI_ISL_522539, EPI_ISL_522540, EPI_ISL_522541, EPI_ISL_522542, EPI_ISL_522543, EPI_ISL_522544 | | | |
| see above | Division of Viral Diseases, Center for Laboratory Control of Infectious Diseases, Korea Centers for Diseases Control and Prevention | Division of Viral Diseases, Center for Laboratory Control of Infectious Diseases, Korea Centers for Diseases Control and Prevention | Jeong-Min Kim, Yoon-Seok Chung, Namjoo Lee, Sang Hee Woo, Hye-Jun Jo, Heui Man Kim, Jun-Sub Kim, Myung Guk Han |
| EPI_ISL_522705 | Royal Hobart Hospital Microbiology Department | MDU-PHL | Cooley L., van Haeften R., Seemann T., Sait M., Schultz, M.B., Sherry N. |
| EPI_ISL_522810, EPI_ISL_522811, EPI_ISL_522812, EPI_ISL_522813, EPI_ISL_522814, EPI_ISL_522815, EPI_ISL_522816 | Virginia DCLS | Virginia DCLS | Virginia DCLS |
| EPI_ISL_523074, EPI_ISL_523076, EPI_ISL_523077, EPI_ISL_523078, EPI_ISL_523079, EPI_ISL_523080, EPI_ISL_523081, EPI_ISL_523082 | Dutch COVID-19 response team | Erasmus Medical Center | OH consortium |
| EPI_ISL_523349, EPI_ISL_523534, EPI_ISL_523535, EPI_ISL_523685, EPI_ISL_523686, EPI_ISL_523688, EPI_ISL_523689, EPI_ISL_523690, EPI_ISL_523691, EPI_ISL_523731, EPI_ISL_523732, EPI_ISL_523733, EPI_ISL_523740, EPI_ISL_523744 | | | |
| see above | Dutch COVID-19 response team | Erasmus Medical Center | Bas Oude Munnink, David Nieuwenhuijse, Reina Sikkema, Claudia Schapendonk, Irina Chestakova, Anne van der Linden, Theo Bestebroer, Stefan van Nieuwkoop, Mark Pronk, Pascal Lexmond, Corien Swaan, Manon Haverkate, Madelief Mollers, Mart Stein, Sandra Kengne Kamga Mobou, Jeroen van Kampen, Jolanda Voermans, Aura Timen, Corine GeurtsvanKessel, Annemiek van der Eijk, Richard Molenkamp, Marion Koopmans, on behalf of the Dutch national COVID-19 response team. |
| EPI_ISL_523832, EPI_ISL_523833, EPI_ISL_523834 | Tuen Mun Hospital | Hong Kong Department of Health | Mak Gannon C.K., Lam Edman T.K., Chan Rickjason C.W., Tsang Dominic N.C. |
| EPI_ISL_523836 | Queen Elizabeth Hospital | Hong Kong Department of Health | Mak Gannon C.K., Lam Edman T.K., Chan Rickjason C.W., Tsang Dominic N.C. |
| EPI_ISL_523838 | Princess Margaret Hospital | Hong Kong Department of Health | Mak Gannon C.K., Lam Edman T.K., Chan Rickjason C.W., Tsang Dominic N.C. |
| EPI_ISL_523839 | Tseung Kwan O Hospital | Hong Kong Department of Health | Mak Gannon C.K., Lam Edman T.K., Chan Rickjason C.W., Tsang Dominic N.C. |
| EPI_ISL_523840 | Our Lady of Maryknoll Hospital | Hong Kong Department of Health | Mak Gannon C.K., Lam Edman T.K., Chan Rickjason C.W., Tsang Dominic N.C. |
| EPI_ISL_523841 | Queen Elizabeth Hospital | Hong Kong Department of Health | Mak Gannon C.K., Lam Edman T.K., Chan Rickjason C.W., Tsang Dominic N.C. |
| EPI_ISL_523842 | Kwong Wah Hospital | Hong Kong Department of Health | Mak Gannon C.K., Lam Edman T.K., Chan Rickjason C.W., Tsang Dominic N.C. |
| EPI_ISL_523843, EPI_ISL_523844, EPI_ISL_523845, EPI_ISL_523846, EPI_ISL_523847 | Queen Elizabeth Hospital | Hong Kong Department of Health | Mak Gannon C.K., Lam Edman T.K., Chan Rickjason C.W., Tsang Dominic N.C. |
| EPI_ISL_523848, EPI_ISL_523849 | United Christian Hospital | Hong Kong Department of Health | Mak Gannon C.K., Lam Edman T.K., Chan Rickjason C.W., Tsang Dominic N.C. |
| EPI_ISL_523850, EPI_ISL_523851, EPI_ISL_523852 | Tseung Kwan O Hospital | Hong Kong Department of Health | Mak Gannon C.K., Lam Edman T.K., Chan Rickjason C.W., Tsang Dominic N.C. |
| EPI_ISL_523853 | Hong Kong Baptist Hospital | Hong Kong Department of Health | Mak Gannon C.K., Lam Edman T.K., Chan Rickjason C.W., Tsang Dominic N.C. |
| EPI_ISL_523854 | Alice Ho Miu Ling Nethersole Hospital | Hong Kong Department of Health | Mak Gannon C.K., Lam Edman T.K., Chan Rickjason C.W., Tsang Dominic N.C. |
| EPI_ISL_523855 | Kwong Wah Hospital | Hong Kong Department of Health | Mak Gannon C.K., Lam Edman T.K., Chan Rickjason C.W., Tsang Dominic N.C. |
| EPI_ISL_524426 | Egyptian National Cancer Institute (ENCI) | Egyptian National Cancer Institute (ENCI) | Zekri, Abdel Rahman N., Amer,K.E., Ahmed,O.S., Soliman,H.K., Ali,M.A., Hassan,W.A., Mahmoud,A.A., Khattab,A.A., Hafez,M.M., Abouelhoda, Mohamed |
| EPI_ISL_524427 | Egyptian National Cancer Institute (ENCI) | Egyptian National Cancer Institute (ENCI) | Zekri, Abdel Rahman N, Amer,K.E., Ahmed,O.S., Soliman,H.K., Hafez,M.M., Bahnassy,A.A., Abdelhamid,W., Gad,A., Ali,M., Hassan,W., Samir,M., Raouf,A., Hamdy,M.S., Soliman,M.S., Elsissy,M.H., Elkhateeb,S.M., Ezzelarab,M.H., Abouelhoda, Mohamed |
| EPI_ISL_524735 | Government General Hospital, Jam Khambhaliya, Devbhoomi Dwarka | Gujarat Biotechnology Research Centre | Harish Matani, Janvi Raval, Zarna Patel, Monika Gandhi, Pinal Trivedi, Maharshi Pandya, Nidhi Patel, Nitin Savaliya, Raghawendra Kumar, Dinesh Kumar, Zuber Saiyed, Komal Patel, Labdhi Pandya, Afzal Ansari, Nikha Trivedi, Apurvasinh Puvar, R D Dixit, A M Kadri, Harsh Bakshi, Chaitanya Joshi, Madhvi Joshi |
| EPI_ISL_524736 | Government General Hospital, Jam Khambhaliya, Devbhoomi Dwarka | Gujarat Biotechnology Research Centre | Janvi Raval, Zarna Patel, Monika Gandhi, Pinal Trivedi, Maharshi Pandya, Nidhi Patel, Nitin Savaliya, Raghawendra Kumar, Dinesh Kumar, Zuber Saiyed, Komal Patel, Labdhi Pandya, Afzal Ansari, Nikha Trivedi, Harish Matani, Apurvasinh Puvar, R D Dixit, A M Kadri, Harsh Bakshi, Chaitanya Joshi, Madhvi Joshi |
| EPI_ISL_524746 | Sir M P Shah Government Medical College, Jamnagar | Gujarat Biotechnology Research Centre | Binita Aring, Afzal Ansari, Nikha Trivedi, Apurvasinh Puvar, Janvi Raval, Zarna Patel, Monika Gandhi, Pinal Trivedi, Maharshi Pandya, Nidhi Patel, Nitin Savaliya, Raghawendra Kumar, Dinesh Kumar, Zuber Saiyed, Komal Patel, Labdhi Pandya, R D Dixit, A M Kadri, Harsh Bakshi, Chaitanya Joshi, Madhvi Joshi |
| EPI_ISL_524747 | Sir M P Shah Government Medical College, Jamnagar | Gujarat Biotechnology Research Centre | Afzal Ansari, Nikha Trivedi, Binita Aring, Apurvasinh Puvar, Janvi Raval, Zarna Patel, Monika Gandhi, Pinal Trivedi, Maharshi Pandya, Nidhi Patel, Nitin Savaliya, Raghawendra Kumar, Dinesh Kumar, Zuber Saiyed, Komal Patel, Labdhi Pandya, R D Dixit, A M Kadri, Harsh Bakshi, Chaitanya Joshi, Madhvi Joshi |
| EPI_ISL_524748 | Sir M P Shah Government Medical College, Jamnagar | Gujarat Biotechnology Research Centre | Nikha Trivedi, Apurvasinh Puvar, Janvi Raval, Binita Aring, Zarna Patel, Monika Gandhi, Pinal Trivedi, Maharshi Pandya, Nidhi Patel, Nitin Savaliya, Raghawendra Kumar, Dinesh Kumar, Zuber Saiyed, Komal Patel, Labdhi Pandya, Afzal Ansari, R D Dixit, A M Kadri, Harsh Bakshi, Chaitanya Joshi, Madhvi Joshi |

| | | | |
|---|---|---|---|
| EPI_ISL_524749 | Sir M P Shah Government Medical College, Jamnagar | Gujarat Biotechnology Research Centre | Apurvasinh Puvar, Janvi Raval, Zarna Patel, Monika Gandhi, Pinal Trivedi, Maharshi Pandya, Nidhi Patel, Nitin Savaliya, Raghawendra Kumar, Dinesh Kumar, Zuber Saiyed, Komal Patel, Labdhi Pandya, Afzal Ansari, Nikha Trivedi, Binita Aring, R D Dixit, A M Kadri, Harsh Bakshi, Chaitanya Joshi, Madhvi Joshi |
| EPI_ISL_524750 | Sir M P Shah Government Medical College, Jamnagar | Gujarat Biotechnology Research Centre | Janvi Raval, Zarna Patel, Monika Gandhi, Pinal Trivedi, Maharshi Pandya, Nidhi Patel, Nitin Savaliya, Raghawendra Kumar, Dinesh Kumar, Zuber Saiyed, Komal Patel, Labdhi Pandya, Afzal Ansari, Nikha Trivedi, Binita Aring, Apurvasinh Puvar, R D Dixit, A M Kadri, Harsh Bakshi, Chaitanya Joshi, Madhvi Joshi |
| EPI_ISL_524751 | Sir M P Shah Government Medical College, Jamnagar | Gujarat Biotechnology Research Centre | Zarna Patel, Monika Gandhi, Pinal Trivedi, Maharshi Pandya, Nidhi Patel, Nitin Savaliya, Raghawendra Kumar, Dinesh Kumar, Zuber Saiyed, Komal Patel, Labdhi Pandya, Afzal Ansari, Nikha Trivedi, Binita Aring, Apurvasinh Puvar, Janvi Raval, R D Dixit, A M Kadri, Harsh Bakshi, Chaitanya Joshi, Madhvi Joshi |
| EPI_ISL_524883, EPI_ISL_524884, EPI_ISL_524885, EPI_ISL_524886, EPI_ISL_524887, EPI_ISL_524888, EPI_ISL_524889, EPI_ISL_524890, EPI_ISL_524891 | MD PHL | MD PHL | Maryland Department of Health Laboratories Administration |
| EPI_ISL_525118 | Maryland Department of Health Laboratory | Maryland Department of Health Laboratory | Maryland Department of Health Laboratories Administration |
| EPI_ISL_525193, EPI_ISL_525194, EPI_ISL_525195, EPI_ISL_525196, EPI_ISL_525197, EPI_ISL_525198, EPI_ISL_525199 | Virginia DCLS | Virginia DCLS | Virginia DCLS |
| EPI_ISL_525486, EPI_ISL_525488, EPI_ISL_525489 | Centre for Dengue Research | Centre for Dengue Research | Chandima Jeewandara, Deshni Jayathilaka, Dinuka Ariyaratne, Laksiri Gomes, Diyanath Ranasinghe, Ananda Wijewickrama, Malika Karunaratne, Eranga Narangoda, Damayanthi Idampitiya, Gathsaurie Neelika Malavige |
| EPI_ISL_525554 | Istituto Zooprofilattico Sperimentale Puglia e Basilicata; Dipartimento di Bioscienze, Biotecnologie e Biofarmaceutica dell'Università degli Studi di Bari "A.Moro"; Istituto di Biomembrane. Bioenergetica e Biotecnologie Molecolari del Consiglio Nazionale delle Ricerche di Bari | Beaconlab (Bioinformatics, Evolution and Comparative Genomics lab), Dept of Biosciences, University on Milan | Parisi A.,Pesole G., Manzari C., Chiara M |
| EPI_ISL_525720, EPI_ISL_525721, EPI_ISL_525722, EPI_ISL_525723, EPI_ISL_525724, EPI_ISL_525725, EPI_ISL_525726, EPI_ISL_525727 | Seattle Flu Study | Seattle Flu Study | Deborah A. Nickerson, Chris D. Frazar, Jover Lee, Benjamin Pelle, Matthew Richardson, Amanda Adler, Elisabeth Brandstetter, Peter D. Han, Kairsten Fay, Misja Ilcisin, Kirsten Lacombe, Thomas R. Sibley, Melissa Truong, Caitlin R. Wolf, Karen Cowgill, Stephanie Schrag, Jeff Duchin, Michael Boeckh, Janet A. Englund, Michael Famulare, Barry R. Lutz, Mark J. Rieder, Lea M. Starita, Matthew Thompson, Helen Y. Chu, Trevor Bedford, Jay Shendure |
| EPI_ISL_525728 | Seattle Flu Study | Seattle Flu Study | Deborah A. Nickerson, Chris D. Frazar, Jover Lee, Benjamin Pelle, Matthew Richardson, Amanda Adler, Elisabeth Brandstetter, Peter D. Han, Kairsten Fay, Misja Ilcisin, Kirsten Lacombe, Thomas R. Sibley, Melissa Truong, Caitlin R. Wolf, Michael Boeckh, Janet A. Englund, Michael Famulare, Barry R. Lutz, Mark J. Rieder, Lea M. Starita, Matthew Thompson, Jay Shendure, Trevor Bedford, Helen Y. Chu |
| EPI_ISL_525729, EPI_ISL_525730, EPI_ISL_525731, EPI_ISL_525732, EPI_ISL_525733, EPI_ISL_525734, EPI_ISL_525735, EPI_ISL_525736, EPI_ISL_525737, EPI_ISL_525738, EPI_ISL_525739, EPI_ISL_525740 | | | |
| see above | Seattle Flu Study | Seattle Flu Study | Deborah A. Nickerson, Chris D. Frazar, Jover Lee, Benjamin Pelle, Matthew Richardson, Amanda Adler, Elisabeth Brandstetter, Peter D. Han, Kairsten Fay, Misja Ilcisin, Kirsten Lacombe, Thomas R. Sibley, Melissa Truong, Caitlin R. Wolf, Karen Cowgill, Stephanie Schrag, Jeff Duchin, Michael Boeckh, Janet A. Englund, Michael Famulare, Barry R. Lutz, Mark J. Rieder, Lea M. Starita, Matthew Thompson, Helen Y. Chu, Trevor Bedford, Jay Shendure |
| EPI_ISL_525802, EPI_ISL_525971, EPI_ISL_525972, EPI_ISL_525973, EPI_ISL_525974, EPI_ISL_525975, EPI_ISL_525976, EPI_ISL_525977, EPI_ISL_525978, EPI_ISL_525979, EPI_ISL_525980, EPI_ISL_525981, EPI_ISL_525982, EPI_ISL_525983, EPI_ISL_525984, EPI_ISL_525985, EPI_ISL_525986, EPI_ISL_525987, EPI_ISL_525988, EPI_ISL_525989, EPI_ISL_525990, EPI_ISL_525991, EPI_ISL_525992, EPI_ISL_525993, EPI_ISL_525994, EPI_ISL_525995, EPI_ISL_525996, EPI_ISL_525997, EPI_ISL_525998, EPI_ISL_525999, EPI_ISL_526000, EPI_ISL_526001, EPI_ISL_526002, EPI_ISL_526003, EPI_ISL_526086, EPI_ISL_526103, EPI_ISL_526104, EPI_ISL_526105, EPI_ISL_526106 | | | |
| see above | OHSU Lab Services Molecular Microbiology Lab | Oregon SARS-CoV-2 Genome Sequencing Center | Brendan L. O'Connell, Ruth V. Nichols, Alec J. Hirsch, Guang Fan, Daniel N. Streblow, William B. Messer, Andrew C. Adey, Benjamin N. Bimber, Brian J. O'Roak |
| EPI_ISL_526123 | St Vincent's Pathology (SydPath) | NSW Health Pathology - Institute of Clinical Pathology and Medical Research; Westmead Hospital; University of Sydney | CIDM-PH et al. |
| EPI_ISL_526142 | South Eastern Area Laboratory Services (SEALS) | NSW Health Pathology - Institute of Clinical Pathology and Medical Research; Westmead Hospital; University of Sydney | CIDM-PH et al. |
| EPI_ISL_526143 | Laverty Pathology | NSW Health Pathology - Institute of Clinical Pathology and Medical Research; Westmead Hospital; University of Sydney | CIDM-PH et al. |
| EPI_ISL_526191 | South Eastern Area Laboratory Services (SEALS) | NSW Health Pathology - Institute of Clinical Pathology and Medical Research; Westmead Hospital; University of Sydney | CIDM-PH et al. |
| EPI_ISL_526287, EPI_ISL_526288, EPI_ISL_526289, EPI_ISL_526290, EPI_ISL_526291, EPI_ISL_526292, EPI_ISL_526293, EPI_ISL_526294, EPI_ISL_526295, EPI_ISL_526296, EPI_ISL_526297, EPI_ISL_526298, EPI_ISL_526299, EPI_ISL_526300, EPI_ISL_526301, EPI_ISL_526302, EPI_ISL_526303, EPI_ISL_526304, EPI_ISL_526305, EPI_ISL_526306, EPI_ISL_526307, EPI_ISL_526308, EPI_ISL_526309, EPI_ISL_526310, EPI_ISL_526311, EPI_ISL_526312, EPI_ISL_526313, EPI_ISL_526314, EPI_ISL_526315, EPI_ISL_526316, EPI_ISL_526317, EPI_ISL_526318, EPI_ISL_526319, EPI_ISL_526320, EPI_ISL_526321, EPI_ISL_526322, EPI_ISL_526323, EPI_ISL_526324, EPI_ISL_526325, EPI_ISL_526326, EPI_ISL_526327, EPI_ISL_526328, EPI_ISL_526329, EPI_ISL_526330, EPI_ISL_526331, EPI_ISL_526332, EPI_ISL_526333, EPI_ISL_526334 | | | |
| see above | University of Birmingham | COVID-19 Genomics UK (COG-UK) Consortium | Institute of Microbiology, University of Birmingham: Claire McMurray, Joanne Stockton, Samuel Nicholls, Radoslaw Poplawski, Will Rowe, Josh Quick, Nicholas Loman. University of Birmingham Testing Laboratory: Celina M Whalley, Andrew Bosworth, Charlotte Poxon, Kasun Wanigasooriya, Oliver Pickles, Mike Kidd, Alex Richter, Andrew D Beggs PHE Heartlands Lab: Husam Osman, Andrew Bosworth. Queen Elizabeth Hospital: Anna Casey |
| EPI_ISL_526367, EPI_ISL_526369, EPI_ISL_526371, EPI_ISL_526373, EPI_ISL_526375, EPI_ISL_526376, EPI_ISL_526378, EPI_ISL_526380, EPI_ISL_526381, EPI_ISL_526384, EPI_ISL_526385, EPI_ISL_526386, EPI_ISL_526387, EPI_ISL_526388, EPI_ISL_526391, EPI_ISL_526392 | | | |
| see above | Liverpool Clinical Laboratories | COVID-19 Genomics UK (COG-UK) Consortium | Sam Haldenby, Anita Lucaci, Steve Paterson, Julian Hiscox, Alistair Darby, M Almsaud, A Alrezaihi, Muhannad Alruwaili, Stuart D Armstrong, Jones Benjamin, Eleanor G Bentley, Anu Chawla, Jordan J Clark, Angela Cowell, Richard Eccles, Isabel García-Dorival, Matthew Gemmell, Alessandro Gerada, PKF Gilmore, Richard Gregory, Ximeng Han, Catherine Hartley, Margaret Hughes, Miren Iturriza-Gomara, James Johnson, L Luu, Jenifer Manson, Charlotte Nelson, Elaine O'Toole, Cassie Olateju, Rebekah Penrice-Randal , Lucille Rainbow, N.P Randle, Trevor Ian Robinson, Parul Sharma, Ghada T Shawli, James P Stewart, Neil Swainston, Ecaterina Vamos, Joanne Watts, Mark Whitehead |
| EPI_ISL_526447 | Virology Department, Sheffield Teaching Hospitals NHS Foundation Trust/Department of Infection, Immunity and Cardiovascular Disease, The Medical School, University of Sheffield | COVID-19 Genomics UK (COG-UK) Consortium | Thushan de Silva, Matthew Parker, Nikki Smith, Adri Angyal, Rebecca Brown, Luke Green, Rachel Tucker, Paul Parsons, Danielle Groves, Katie Johnson, Laura Carrilero, Alex Keeley, Dave Partridge, Matthew Wyles, Benjamin Lindsey, Mehmet Yavuz, Mohammad Raza, Cariad Evans |
| EPI_ISL_526551 | OHSU Lab Services Molecular Microbiology Lab | Oregon SARS-CoV-2 Genome Sequencing Center | Brendan L. O'Connell, Ruth V. Nichols, Alec J. Hirsch, Guang Fan, Daniel N. Streblow, William B. Messer, Andrew C. Adey, Benjamin N. Bimber, Brian J. O'Roak |
| EPI_ISL_526700, EPI_ISL_526701, EPI_ISL_526702, EPI_ISL_526703, EPI_ISL_526704, EPI_ISL_526705, EPI_ISL_526706 | Division of Viral Diseases, Center for Laboratory Control of Infectious Diseases, Korea Centers for Diseases Control and Prevention | Division of Viral Diseases, Center for Laboratory Control of Infectious Diseases, Korea Centers for Diseases Control and Prevention | Jeong-Min Kim, Yoon-Seok Chung, Namjoo Lee, Sang Hee Woo, Hye-Jun Jo, Heui Man Kim, Jun-Sub Kim, Myung Guk Han |
| EPI_ISL_526855, EPI_ISL_526856 | Virginia DCLS | Virginia DCLS | Virginia DCLS |
| EPI_ISL_526958, EPI_ISL_526960, EPI_ISL_526961, EPI_ISL_526973 | Instituto Nacional de Salud, Bogotá, Colombia | Instituto Nacional de Salud, Bogotá, Colombia | Katherine Laiton-Donato, Diego A. Álvarez-Díaz, Carlos Franco-Muñoz, Mauricio Pacheco-Montealegre, Jonathan Reales, Diego Andrés Prada, Jose A. Usme-Ciro, Zulma M. Cucunubá, Christian Julian VillabonaArenas, Liz Villabona-Arenas, Sussy Echeverria, Astrid C. Flórez, Carolina Ferro, Diana Marcela Walteros-Acero, Franklin Prieto, Carlos Andrés Durán, Martha Lucia Ospina Martinez, Marcela Mercado-Reyes |

| | | | |
|---|---|---|---|
| EPI_ISL_527036 | Area of Virology, Serology and Virology Division (SAViD), New South Wales Health Pathology Randwick | Area of Virology, Serology and Virology Division (SAViD), New South Wales Health Pathology Randwick | Rawlinson, W. |
| EPI_ISL_527584, EPI_ISL_527585, EPI_ISL_527586, EPI_ISL_527587, EPI_ISL_527588, EPI_ISL_527589, EPI_ISL_527590, EPI_ISL_527591, EPI_ISL_527592, EPI_ISL_527593, EPI_ISL_527594, EPI_ISL_527595, EPI_ISL_527596, EPI_ISL_527597, EPI_ISL_527598, EPI_ISL_527599, EPI_ISL_527600 | | | |
| see above | Minnesota Department of Health, Public Health Laboratory | Minnesota Department of Health, Public Health Laboratory | Matt Plumb, Jacob Garfin, and Xiong Wang |
| EPI_ISL_527632, EPI_ISL_527633, EPI_ISL_527634, EPI_ISL_527635, EPI_ISL_527636, EPI_ISL_527637, EPI_ISL_527638, EPI_ISL_527639, EPI_ISL_527640, EPI_ISL_527641, EPI_ISL_527642, EPI_ISL_527643, EPI_ISL_527644, EPI_ISL_527645, EPI_ISL_527646, EPI_ISL_527647, EPI_ISL_527648, EPI_ISL_527649, EPI_ISL_527650 | | | |
| see above | AR Dept. of Health-Public Health Lab | Pathogen Discovery, Respiratory Viruses Branch, Division of Viral Diseases, Centers for Disease Control and Prevention | Ying Tao, Jing Zhang, Yan Li, Krista Queen, Anna Uehara, Clinton Paden, Haibin Wang, Suxiang Tong |
| EPI_ISL_527666 | IN State Department of Health Laboratory Services | Pathogen Discovery, Respiratory Viruses Branch, Division of Viral Diseases, Centers for Disease Control and Prevention | Krista Queen, Brian Lynch, Yan Li, Anna Montmayeur, Jing Zhang, Ying Tao, Anna Uehara, Rachel Marine, Clinton R. Paden, Haibin Wang, Suxiang Tong |
| EPI_ISL_527755 | Centro Nacional De Rehabilitacion Humberto Araya Rojas (Cenare) | Inciensa, Instituto Costarricense de Investigación y Enseñanza en Nutrición y Salud | Francisco Duarte, Hebleen Porras, Claudio Soto-Garita, Estela Cordero, Adriana Godinez & Melany Calderon |
| EPI_ISL_528420, EPI_ISL_528422, EPI_ISL_528424 | TNMC & BYL NAIR CH. HOSPITAL | Institute of Genomics and Integrative Biology - Council of Scientific and Industrial Research | Rajesh Pandey, Jayanthi Shastri, Akshay Kanakan, Vivekanand A, Janani Srinivasa Vasudevan, Ranjeet Maurya, Sachee Agrawal, Nirjhar Chatterjee, Swapneil Parikh, Manish Pathak, Subrat Thanapati, Jasmina Savak, Suresh Poojari, Mahesh Sangar, Amol Borse, Shweta Kawankar, Vasil Nachan, Mayuresh Vishwanathan, Shruthi Sachidanandan, Shrutika Pophale, Utkarsha Yelve |
| EPI_ISL_528489, EPI_ISL_528490, EPI_ISL_528491, EPI_ISL_528492, EPI_ISL_528493, EPI_ISL_528494, EPI_ISL_528495, EPI_ISL_528496, EPI_ISL_528497, EPI_ISL_528498, EPI_ISL_528499, EPI_ISL_528500, EPI_ISL_528501, EPI_ISL_528502, EPI_ISL_528503, EPI_ISL_528504, EPI_ISL_528505, EPI_ISL_528506, EPI_ISL_528507, EPI_ISL_528508, EPI_ISL_528510, EPI_ISL_528511, EPI_ISL_528513, EPI_ISL_528514, EPI_ISL_528515 | | | |
| see above | Alaska State Virology Laboratory | Alaska State Virology Laboratory | Chen J et al with Pathogenomics group Dagdag R, Redlinger M, Milton E, George W, Kovalenko A, Drown DM, Bortz E |
| EPI_ISL_528746 | Immanuel Hospital | Molecular Genetics Laboratory-Faculty of Medicine-Universitas Padjadjaran; School of Life Sciences and Technology & School of Pharmacy-Institut Teknologi Bandung; Laboratorium Kesehatan Provinsi Jawa Barat | Yunia Sribudiani,Tri Hanggono Achmad, Mas Rizky A.A. Syamsunarno, Fensi Amalina, Catur Riani, Azzania Fibriani, Husna Nugrahapraja, Marselina Irasonia Tan, Tarwadi, Ema Rahmawati, Savira Ekawardhani, Hesti Lina Wiraswati, Ryan Bayusantika Ristandi, Rifky Waluyajati Rachman, Cut Nur Cinthia Alamanda, Lia Faridah, Hammam Riza, Sony Solistia Wirawan, Agung Eru Wibowo, Irvan Faizal |
| EPI_ISL_528813 | Department of Medicine, Gandhi hospital, Hyderabad | CSIR-Centre for Cellular and Molecular Biology | Vinayasekhar Aedula,Thrilok Chander Bingi, Rajarao Mesipogu, Shagufta Khan, Lamuk Zaveri, Namami Gaur, Sakshi Shambhavi, Nikhil Hajirnis, M Soujanya Reddy, Pratheusa Maccha,Tulasi Nagabandi, Purushotham Vodnala, Payel Mukherjee, Sofia Banu, Priya Singh, Onkar Kulkarni, Dhiviya Vedagiri, Divya Gupta, Vishal Sah, Santosh Kumar Kuncha, Krishnan Harinivas Harshan, Archana Bharadwaj Siva, Karthik Bharadwaj Tallapaka,Umesh Kumar, Unis Ahmad Bhat, Ajay Sarawagi, Priyanka Pant, Rajkanwar Nathawat, Rakesh K Mishra, Divya Tej Sowpati |
| EPI_ISL_528823 | CSIR-Centre for Cellular and Molecular Biology | CSIR-Centre for Cellular and Molecular Biology | Shagufta Khan, Lamuk Zaveri, Namami Gaur, Sakshi Shambhavi, Nikhil Hajirnis, M Soujanya Reddy, Pratheusa Maccha, Tulasi Nagabandi, Purushotham Vodnala, Payel Mukherjee, Sofia Banu, Priya Singh, Onkar Kulkarni, Dhiviya Vedagiri, Divya Gupta, Vishal Sah, Santosh Kumar Kuncha, Krishnan Harinivas Harshan, Archana Bharadwaj Siva, Karthik Bharadwaj Tallapaka, Renu Sudhakar, Somesh Gorde, Gangumala Srinivas Reddy, Sujoy Deb, Swati Bayyana, Rakesh K Mishra, Divya Tej Sowpati |
| EPI_ISL_528824 | CSIR-Centre for Cellular and Molecular Biology | CSIR-Centre for Cellular and Molecular Biology | Tulasi Nagabandi, Namami Gaur, Sakshi Shambhavi, Lamuk Zaveri, Shagufta Khan, Nikhil Hajirnis, M Soujanya Reddy, Pratheusa Maccha, Purushotham Vodnala, Payel Mukherjee, Sofia Banu, Priya Singh, Onkar Kulkarni, Dhiviya Vedagiri, Divya Gupta, Vishal Sah, Santosh Kumar Kuncha, Krishnan Harinivas Harshan, Archana Bharadwaj Siva, Karthik Bharadwaj Tallapaka,G. Aditya Kumar, Koushick Sivakumar, Pooja Ramesh Gupta, Rajan Kumar Jha, Shraddha Vijay Lahoti, Rakesh K Mishra, Divya Tej Sowpati |
| EPI_ISL_528825, EPI_ISL_528826 | CSIR-Centre for Cellular and Molecular Biology | CSIR-Centre for Cellular and Molecular Biology | Shagufta Khan, Lamuk Zaveri, Namami Gaur, Sakshi Shambhavi, Nikhil Hajirnis, M Soujanya Reddy, Pratheusa Maccha, Tulasi Nagabandi, Purushotham Vodnala, Payel Mukherjee, Sofia Banu, Priya Singh, Onkar Kulkarni, Dhiviya Vedagiri, Divya Gupta, Vishal Sah, Santosh Kumar Kuncha, Krishnan Harinivas Harshan, Archana Bharadwaj Siva, Karthik Bharadwaj Tallapaka, Renu Sudhakar, Somesh Gorde, Gangumala Srinivas Reddy, Sujoy Deb, Swati Bayyana, Rakesh K Mishra, Divya Tej Sowpati |
| EPI_ISL_528827 | CSIR-Centre for Cellular and Molecular Biology | CSIR-Centre for Cellular and Molecular Biology | Namami Gaur, Sakshi Shambhavi, Lamuk Zaveri, Shagufta Khan, Nikhil Hajirnis, M Soujanya Reddy, Pratheusa Maccha, Tulasi Nagabandi, Purushotham Vodnala, Payel Mukherjee, Sofia Banu, Priya Singh, Onkar Kulkarni, Dhiviya Vedagiri, Divya Gupta, Vishal Sah, Santosh Kumar Kuncha, Krishnan Harinivas Harshan, Archana Bharadwaj Siva, Karthik Bharadwaj Tallapaka, Zeba Rizvi, Zuberwasim Sayyad, Kakade Aishwarya Arun, Amrutha H C, Ananga Ghosh, Rakesh K Mishra, Divya Tej Sowpati |
| EPI_ISL_528828 | CSIR-Centre for Cellular and Molecular Biology | CSIR-Centre for Cellular and Molecular Biology | Sofia Banu, Payel Mukherjee, Priya Singh,Onkar Kulkarni, Dhiviya Vedagiri, Divya Gupta, Vishal Sah, Santosh Kumar Kuncha, Krishnan Harinivas Harshan, Archana Bharadwaj Siva, Karthik Bharadwaj Tallapaka, Shagufta Khan, Lamuk Zaveri, Namami Gaur, Sakshi Shambhavi, Nikhil Hajirnis, M Soujanya Reddy, Pratheusa Maccha,Tulasi Nagabandi, Purushotham Vodnala, Deepak Kumar, Devi Prasad Vijayashankar, Disha Nanda, Divya Das, Jotin Gogoi, Manish Bhattacharjee, Rakesh K Mishra, Divya Tej Sowpati |
| EPI_ISL_528829 | CSIR-Centre for Cellular and Molecular Biology | CSIR-Centre for Cellular and Molecular Biology | Sakshi Shambhavi, Lamuk Zaveri, Shagufta Khan, Namami Gaur, Nikhil Hajirnis, M Soujanya Reddy, Pratheusa Maccha, Tulasi Nagabandi, Purushotham Vodnala, Payel Mukherjee, Sofia Banu, Priya Singh, Onkar Kulkarni, Dhiviya Vedagiri, Divya Gupta, Vishal Sah, Santosh Kumar Kuncha, Krishnan Harinivas Harshan, Archana Bharadwaj Siva, Karthik Bharadwaj Tallapaka, Deepak Kumar, Devi Prasad Vijayashankar, Disha Nanda, Divya Das, Jotin Gogoi, Manish Bhattacharjee, Rakesh K Mishra, Divya Tej Sowpati |
| EPI_ISL_528830 | CSIR-Centre for Cellular and Molecular Biology | CSIR-Centre for Cellular and Molecular Biology | Tulasi Nagabandi, Namami Gaur, Sakshi Shambhavi, Lamuk Zaveri, Shagufta Khan, Nikhil Hajirnis, M Soujanya Reddy, Pratheusa Maccha, Purushotham Vodnala, Payel Mukherjee, Sofia Banu, Priya Singh, Onkar Kulkarni, Dhiviya Vedagiri, Divya Gupta, Vishal Sah, Santosh Kumar Kuncha, Krishnan Harinivas Harshan, Archana Bharadwaj Siva, Karthik Bharadwaj Tallapaka,G. Aditya Kumar, Koushick Sivakumar, Pooja Ramesh Gupta, Rajan Kumar Jha, Shraddha Vijay Lahoti, Rakesh K Mishra, Divya Tej Sowpati |
| EPI_ISL_528831 | CSIR-Centre for Cellular and Molecular Biology | CSIR-Centre for Cellular and Molecular Biology | Lamuk Zaveri, Shagufta Khan, Namami Gaur, Sakshi Shambhavi, Nikhil Hajirnis, M Soujanya Reddy, Pratheusa Maccha, Tulasi Nagabandi, Purushotham Vodnala, Payel Mukherjee, Sofia Banu, Priya Singh, Onkar Kulkarni, Dhiviya Vedagiri, Divya Gupta, Vishal Sah, Santosh Kumar Kuncha, Krishnan Harinivas Harshan, Archana Bharadwaj Siva, Karthik Bharadwaj Tallapaka, Renu Sudhakar, Somesh Gorde, Gangumala Srinivas Reddy, Sujoy Deb, Swati Bayyana, Rakesh K Mishra, Divya Tej Sowpati |
| EPI_ISL_528832 | CSIR-Centre for Cellular and Molecular Biology | CSIR-Centre for Cellular and Molecular Biology | Sofia Banu, Payel Mukherjee, Priya Singh,Onkar Kulkarni, Dhiviya Vedagiri, Divya Gupta, Vishal Sah, Santosh Kumar Kuncha, Krishnan Harinivas Harshan, Archana Bharadwaj Siva, Karthik Bharadwaj Tallapaka, Shagufta Khan, Lamuk Zaveri, Namami Gaur, Sakshi Shambhavi, Nikhil Hajirnis, M Soujanya Reddy, Pratheusa Maccha,Tulasi Nagabandi, Purushotham Vodnala, Deepak Kumar, Devi Prasad Vijayashankar, Disha Nanda, Divya Das, Jotin Gogoi, Manish Bhattacharjee, Rakesh K Mishra, Divya Tej Sowpati |
| EPI_ISL_528833 | CSIR-Centre for Cellular and Molecular Biology | CSIR-Centre for Cellular and Molecular Biology | Lamuk Zaveri, Shagufta Khan, Namami Gaur, Sakshi Shambhavi, Nikhil Hajirnis, M Soujanya Reddy, Pratheusa Maccha, Tulasi Nagabandi, Purushotham Vodnala, Payel Mukherjee, Sofia Banu, Priya Singh, Onkar Kulkarni, Dhiviya Vedagiri, Divya Gupta, Vishal Sah, Santosh Kumar Kuncha, Krishnan Harinivas Harshan, Archana Bharadwaj Siva, Karthik Bharadwaj Tallapaka, Renu Sudhakar, Somesh Gorde, Gangumala Srinivas Reddy, Sujoy Deb, Swati Bayyana, Rakesh K Mishra, Divya Tej Sowpati |
| EPI_ISL_528834 | CSIR-Centre for Cellular and Molecular Biology | CSIR-Centre for Cellular and Molecular Biology | Pratheusa Maccha, Sakshi Shambhavi, Lamuk Zaveri, Shagufta Khan, Namami Gaur, Nikhil Hajirnis, M Soujanya Reddy, Tulasi Nagabandi, Purushotham Vodnala, Payel Mukherjee, Sofia Banu, Priya Singh,Onkar Kulkarni, Dhiviya Vedagiri, Divya Gupta, Vishal Sah, Santosh Kumar Kuncha, Krishnan Harinivas Harshan, Archana Bharadwaj Siva, Karthik Bharadwaj Tallapaka, G. Aditya Kumar, Koushick Sivakumar,Disha Nanda, Divya Das, Jotin Gogoi, Manish Bhattacharjee, Ravi Prasad Mukku, Rakesh K Mishra, Divya Tej Sowpati |
| EPI_ISL_528835 | CSIR-Centre for Cellular and Molecular Biology | CSIR-Centre for Cellular and Molecular Biology | M Soujanya Reddy, Nikhil Hajirnis, Pratheusa Maccha, Namami Gaur, Sakshi Shambhavi, Lamuk Zaveri, Shagufta Khan, Tulasi Nagabandi, Purushotham Vodnala, Payel Mukherjee, Sofia Banu, Priya Singh, Onkar Kulkarni, Dhiviya Vedagiri, Divya Gupta, Vishal Sah, Santosh Kumar Kuncha, Krishnan Harinivas Harshan, Archana Bharadwaj Siva, Karthik Bharadwaj Tallapaka, Zeba Rizvi, Zuberwasim Sayyad, Kakade Aishwarya Arun, Amrutha H C, Ananga Ghosh, Rakesh K Mishra, Divya Tej Sowpati |
| EPI_ISL_528836 | CSIR-Centre for Cellular and Molecular Biology | CSIR-Centre for Cellular and Molecular Biology | Tulasi Nagabandi, Namami Gaur, Sakshi Shambhavi, Lamuk Zaveri, Shagufta Khan, Nikhil Hajirnis, M Soujanya Reddy, Pratheusa Maccha, Purushotham |

| | | | |
|---|---|---|---|
| | | | Vodnala, Payel Mukherjee, Sofia Banu, Priya Singh, Onkar Kulkarni, Dhiviya Vedagiri, Divya Gupta, Vishal Sah, Santosh Kumar Kuncha, Krishnan Harinivas Harshan, Archana Bharadwaj Siva, Karthik Bharadwaj Tallapaka,G. Aditya Kumar, Koushick Sivakumar, Pooja Ramesh Gupta, Rajan Kumar Jha, Shraddha Vijay Lahoti, Rakesh K Mishra, Divya Tej Sowpati |
| EPI_ISL_528837 | CSIR-Centre for Cellular and Molecular Biology | CSIR-Centre for Cellular and Molecular Biology | Sakshi Shambhavi, Lamuk Zaveri, Shagufta Khan, Namami Gaur, Nikhil Hajirnis, M Soujanya Reddy, Pratheusa Maccha, Tulasi Nagabandi, Purushotham Vodnala, Payel Mukherjee, Sofia Banu, Priya Singh, Onkar Kulkarni, Dhiviya Vedagiri, Divya Gupta, Vishal Sah, Santosh Kumar Kuncha, Krishnan Harinivas Harshan, Archana Bharadwaj Siva, Karthik Bharadwaj Tallapaka, Deepak Kumar, Devi Prasad Vijayashankar, Disha Nanda, Divya Das, Jotin Gogoi, Manish Bhattacharjee, Rakesh K Mishra, Divya Tej Sowpati |
| EPI_ISL_528838 | CSIR-Centre for Cellular and Molecular Biology | CSIR-Centre for Cellular and Molecular Biology | Payel Mukherjee, Sofia Banu, Priya Singh, Onkar Kulkarni, Dhiviya Vedagiri, Divya Gupta, Vishal Sah, Santosh Kumar Kuncha, Krishnan Harinivas Harshan, Archana Bharadwaj Siva, Karthik Bharadwaj Tallapaka, Shagufta Khan, Lamuk Zaveri, Nikhil Hajirnis, M Soujanya Reddy, Pratheusa Maccha, Namami Gaur, Sakshi Shambhavi, Tulasi Nagabandi, Purushotham Vodnala, Rakesh K Mishra, Sonu Uday, Sudipta Mondal, Annapoorna P Karthyayani, Debabrata Jana, Debrya Saha, Divya Tej Sowpati |
| EPI_ISL_528839 | CSIR-Centre for Cellular and Molecular Biology | CSIR-Centre for Cellular and Molecular Biology | Pratheusa Maccha, Sakshi Shambhavi, Lamuk Zaveri, Shagufta Khan, Namami Gaur, Nikhil Hajirnis, M Soujanya Reddy, Tulasi Nagabandi, Purushotham Vodnala, Payel Mukherjee, Sofia Banu, Priya Singh,Onkar Kulkarni, Dhiviya Vedagiri, Divya Gupta, Vishal Sah, Santosh Kumar Kuncha, Krishnan Harinivas Harshan, Archana Bharadwaj Siva, Karthik Bharadwaj Tallapaka, G. Aditya Kumar, Koushick Sivakumar,Disha Nanda, Divya Das, Jotin Gogoi, Manish Bhattacharjee, Ravi Prasad Mukku, Rakesh K Mishra, Divya Tej Sowpati |
| EPI_ISL_528840 | CSIR-Centre for Cellular and Molecular Biology | CSIR-Centre for Cellular and Molecular Biology | Tulasi Nagabandi, Namami Gaur, Sakshi Shambhavi, Lamuk Zaveri, Shagufta Khan, Nikhil Hajirnis, M Soujanya Reddy, Pratheusa Maccha, Purushotham Vodnala, Payel Mukherjee, Sofia Banu, Priya Singh, Onkar Kulkarni, Dhiviya Vedagiri, Divya Gupta, Vishal Sah, Santosh Kumar Kuncha, Krishnan Harinivas Harshan, Archana Bharadwaj Siva, Karthik Bharadwaj Tallapaka,G. Aditya Kumar, Koushick Sivakumar, Pooja Ramesh Gupta, Rajan Kumar Jha, Shraddha Vijay Lahoti, Rakesh K Mishra, Divya Tej Sowpati |
| EPI_ISL_528841, EPI_ISL_528842 | CSIR-Centre for Cellular and Molecular Biology | CSIR-Centre for Cellular and Molecular Biology | Payel Mukherjee, Sofia Banu, Priya Singh, Onkar Kulkarni, Dhiviya Vedagiri, Divya Gupta, Vishal Sah, Santosh Kumar Kuncha, Krishnan Harinivas Harshan, Archana Bharadwaj Siva, Karthik Bharadwaj Tallapaka, Shagufta Khan, Lamuk Zaveri, Nikhil Hajirnis, M Soujanya Reddy, Pratheusa Maccha, Namami Gaur, Sakshi Shambhavi, Tulasi Nagabandi, Purushotham Vodnala, Rakesh K Mishra, Sonu Uday, Sudipta Mondal, Annapoorna P Karthyayani, Debabrata Jana, Debrya Saha, Divya Tej Sowpati |
| EPI_ISL_528843, EPI_ISL_528844 | CSIR-Centre for Cellular and Molecular Biology | CSIR-Centre for Cellular and Molecular Biology | Namami Gaur, Sakshi Shambhavi, Lamuk Zaveri, Shagufta Khan, Nikhil Hajirnis, M Soujanya Reddy, Pratheusa Maccha, Tulasi Nagabandi, Purushotham Vodnala, Payel Mukherjee, Sofia Banu, Priya Singh, Onkar Kulkarni, Dhiviya Vedagiri, Divya Gupta, Vishal Sah, Santosh Kumar Kuncha, Krishnan Harinivas Harshan, Archana Bharadwaj Siva, Karthik Bharadwaj Tallapaka, Zeba Rizvi, Zuberwasim Sayyad, Kakade Aishwarya Arun, Amrutha H C, Ananga Ghosh, Rakesh K Mishra, Divya Tej Sowpati |
| EPI_ISL_528845 | CSIR-Centre for Cellular and Molecular Biology | CSIR-Centre for Cellular and Molecular Biology | Payel Mukherjee, Sofia Banu, Priya Singh, Onkar Kulkarni, Dhiviya Vedagiri, Divya Gupta, Vishal Sah, Santosh Kumar Kuncha, Krishnan Harinivas Harshan, Archana Bharadwaj Siva, Karthik Bharadwaj Tallapaka, Shagufta Khan, Lamuk Zaveri, Nikhil Hajirnis, M Soujanya Reddy, Pratheusa Maccha, Namami Gaur, Sakshi Shambhavi, Tulasi Nagabandi, Purushotham Vodnala, Rakesh K Mishra, Sonu Uday, Sudipta Mondal, Annapoorna P Karthyayani, Debabrata Jana, Debrya Saha, Divya Tej Sowpati |
| EPI_ISL_528846 | CSIR-Centre for Cellular and Molecular Biology | CSIR-Centre for Cellular and Molecular Biology | Namami Gaur, Sakshi Shambhavi, Lamuk Zaveri, Shagufta Khan, Nikhil Hajirnis, M Soujanya Reddy, Pratheusa Maccha, Tulasi Nagabandi, Purushotham Vodnala, Payel Mukherjee, Sofia Banu, Priya Singh, Onkar Kulkarni, Dhiviya Vedagiri, Divya Gupta, Vishal Sah, Santosh Kumar Kuncha, Krishnan Harinivas Harshan, Archana Bharadwaj Siva, Karthik Bharadwaj Tallapaka, Zeba Rizvi, Zuberwasim Sayyad, Kakade Aishwarya Arun, Amrutha H C, Ananga Ghosh, Rakesh K Mishra, Divya Tej Sowpati |
| EPI_ISL_528847 | CSIR-Centre for Cellular and Molecular Biology | CSIR-Centre for Cellular and Molecular Biology | Shagufta Khan, Lamuk Zaveri, Namami Gaur, Sakshi Shambhavi, Nikhil Hajirnis, M Soujanya Reddy, Pratheusa Maccha, Tulasi Nagabandi, Purushotham Vodnala, Payel Mukherjee, Sofia Banu, Priya Singh, Onkar Kulkarni, Dhiviya Vedagiri, Divya Gupta, Vishal Sah, Santosh Kumar Kuncha, Krishnan Harinivas Harshan, Archana Bharadwaj Siva, Karthik Bharadwaj Tallapaka, Renu Sudhakar, Somesh Gorde, Gangumala Srinivas Reddy, Sujoy Deb, Swati Bayyana, Rakesh K Mishra, Divya Tej Sowpati |
| EPI_ISL_528848 | CSIR-Centre for Cellular and Molecular Biology | CSIR-Centre for Cellular and Molecular Biology | Sofia Banu, Payel Mukherjee, Priya Singh,Onkar Kulkarni, Dhiviya Vedagiri, Divya Gupta, Vishal Sah, Santosh Kumar Kuncha, Krishnan Harinivas Harshan, Archana Bharadwaj Siva, Karthik Bharadwaj Tallapaka, Shagufta Khan, Lamuk Zaveri, Namami Gaur, Sakshi Shambhavi, Nikhil Hajirnis, M Soujanya Reddy, Pratheusa Maccha,Tulasi Nagabandi, Purushotham Vodnala, Deepak Kumar, Devi Prasad Vijayashankar, Disha Nanda, Divya Das, Jotin Gogoi, Manish Bhattacharjee, Rakesh K Mishra, Divya Tej Sowpati |
| EPI_ISL_528849 | CSIR-Centre for Cellular and Molecular Biology | CSIR-Centre for Cellular and Molecular Biology | Nikhil Hajirnis, M Soujanya Reddy, Pratheusa Maccha, Lamuk Zaveri, Shagufta Khan, Namami Gaur, Sakshi Shambhavi, Tulasi Nagabandi, Purushotham Vodnala, Payel Mukherjee, Sofia Banu, Priya Singh, Onkar Kulkarni, Dhiviya Vedagiri, Divya Gupta, Vishal Sah, Santosh Kumar Kuncha, Krishnan Harinivas Harshan, Archana Bharadwaj Siva, Karthik Bharadwaj Tallapaka,Zeba Rizvi, Zuberwasim Sayyad, Kakade Aishwarya Arun, Amrutha H C, Ananga Ghosh, Rakesh K Mishra, Divya Tej Sowpati |
| EPI_ISL_528850 | CSIR-Centre for Cellular and Molecular Biology | CSIR-Centre for Cellular and Molecular Biology | Lamuk Zaveri, Shagufta Khan, Namami Gaur, Sakshi Shambhavi, Nikhil Hajirnis, M Soujanya Reddy, Pratheusa Maccha, Tulasi Nagabandi, Purushotham Vodnala, Payel Mukherjee, Sofia Banu, Priya Singh, Onkar Kulkarni, Dhiviya Vedagiri, Divya Gupta, Vishal Sah, Santosh Kumar Kuncha, Krishnan Harinivas Harshan, Archana Bharadwaj Siva, Karthik Bharadwaj Tallapaka, Renu Sudhakar, Somesh Gorde, Gangumala Srinivas Reddy, Sujoy Deb, Swati Bayyana, Rakesh K Mishra, Divya Tej Sowpati |
| EPI_ISL_528851 | CSIR-Centre for Cellular and Molecular Biology | CSIR-Centre for Cellular and Molecular Biology | Payel Mukherjee, Sofia Banu, Priya Singh, Onkar Kulkarni, Dhiviya Vedagiri, Divya Gupta, Vishal Sah, Santosh Kumar Kuncha, Krishnan Harinivas Harshan, Archana Bharadwaj Siva, Karthik Bharadwaj Tallapaka, Shagufta Khan, Lamuk Zaveri, Nikhil Hajirnis, M Soujanya Reddy, Pratheusa Maccha, Namami Gaur, Sakshi Shambhavi, Tulasi Nagabandi, Purushotham Vodnala, Rakesh K Mishra, Sonu Uday, Sudipta Mondal, Annapoorna P Karthyayani, Debabrata Jana, Debrya Saha, Divya Tej Sowpati |
| EPI_ISL_528852 | CSIR-Centre for Cellular and Molecular Biology | CSIR-Centre for Cellular and Molecular Biology | Namami Gaur, Sakshi Shambhavi, Lamuk Zaveri, Shagufta Khan, Nikhil Hajirnis, M Soujanya Reddy, Pratheusa Maccha, Tulasi Nagabandi, Purushotham Vodnala, Payel Mukherjee, Sofia Banu, Priya Singh, Onkar Kulkarni, Dhiviya Vedagiri, Divya Gupta, Vishal Sah, Santosh Kumar Kuncha, Krishnan Harinivas Harshan, Archana Bharadwaj Siva, Karthik Bharadwaj Tallapaka, Zeba Rizvi, Zuberwasim Sayyad, Kakade Aishwarya Arun, Amrutha H C, Ananga Ghosh, Rakesh K Mishra, Divya Tej Sowpati |
| EPI_ISL_528853 | CSIR-Centre for Cellular and Molecular Biology | CSIR-Centre for Cellular and Molecular Biology | Tulasi Nagabandi, Namami Gaur, Sakshi Shambhavi, Lamuk Zaveri, Shagufta Khan, Nikhil Hajirnis, M Soujanya Reddy, Pratheusa Maccha, Purushotham Vodnala, Payel Mukherjee, Sofia Banu, Priya Singh, Onkar Kulkarni, Dhiviya Vedagiri, Divya Gupta, Vishal Sah, Santosh Kumar Kuncha, Krishnan Harinivas Harshan, Archana Bharadwaj Siva, Karthik Bharadwaj Tallapaka,G. Aditya Kumar, Koushick Sivakumar, Pooja Ramesh Gupta, Rajan Kumar Jha, Shraddha Vijay Lahoti, Rakesh K Mishra, Divya Tej Sowpati |
| EPI_ISL_528854 | CSIR-Centre for Cellular and Molecular Biology | CSIR-Centre for Cellular and Molecular Biology | Payel Mukherjee, Sofia Banu, Priya Singh, Onkar Kulkarni, Dhiviya Vedagiri, Divya Gupta, Vishal Sah, Santosh Kumar Kuncha, Krishnan Harinivas Harshan, Archana Bharadwaj Siva, Karthik Bharadwaj Tallapaka, Shagufta Khan, Lamuk Zaveri, Nikhil Hajirnis, M Soujanya Reddy, Pratheusa Maccha, Namami Gaur, Sakshi Shambhavi, Tulasi Nagabandi, Purushotham Vodnala, Rakesh K Mishra, Sonu Uday, Sudipta Mondal, Annapoorna P Karthyayani, Debabrata Jana, Debrya Saha, Divya Tej Sowpati |
| EPI_ISL_528855 | CSIR-Centre for Cellular and Molecular Biology | CSIR-Centre for Cellular and Molecular Biology | Nikhil Hajirnis, M Soujanya Reddy, Pratheusa Maccha, Lamuk Zaveri, Shagufta Khan, Namami Gaur, Sakshi Shambhavi, Tulasi Nagabandi, Purushotham Vodnala, Payel Mukherjee, Sofia Banu, Priya Singh, Onkar Kulkarni, Dhiviya Vedagiri, Divya Gupta, Vishal Sah, Santosh Kumar Kuncha, Krishnan Harinivas Harshan, Archana Bharadwaj Siva, Karthik Bharadwaj Tallapaka,Zeba Rizvi, Zuberwasim Sayyad, Kakade Aishwarya Arun, Amrutha H C, Ananga Ghosh, Rakesh K Mishra, Divya Tej Sowpati |
| EPI_ISL_528856 | CSIR-Centre for Cellular and Molecular Biology | CSIR-Centre for Cellular and Molecular Biology | Pratheusa Maccha, Sakshi Shambhavi, Lamuk Zaveri, Shagufta Khan, Namami Gaur, Nikhil Hajirnis, M Soujanya Reddy, Tulasi Nagabandi, Purushotham Vodnala, Payel Mukherjee, Sofia Banu, Priya Singh,Onkar Kulkarni, Dhiviya Vedagiri, Divya Gupta, Vishal Sah, Santosh Kumar Kuncha, Krishnan Harinivas Harshan, Archana Bharadwaj Siva, Karthik Bharadwaj Tallapaka, G. Aditya Kumar, Koushick Sivakumar,Disha Nanda, Divya Das, Jotin Gogoi, Manish Bhattacharjee, Ravi Prasad Mukku, Rakesh K Mishra, Divya Tej Sowpati |
| EPI_ISL_528857 | CSIR-Centre for Cellular and Molecular Biology | CSIR-Centre for Cellular and Molecular Biology | Nikhil Hajirnis, M Soujanya Reddy, Pratheusa Maccha, Lamuk Zaveri, Shagufta Khan, Namami Gaur, Sakshi Shambhavi, Tulasi Nagabandi, Purushotham Vodnala, Payel Mukherjee, Sofia Banu, Priya Singh, Onkar Kulkarni, Dhiviya Vedagiri, Divya Gupta, Vishal Sah, Santosh Kumar Kuncha, Krishnan Harinivas Harshan, Archana Bharadwaj Siva, Karthik Bharadwaj Tallapaka,Zeba Rizvi, Zuberwasim Sayyad, Kakade Aishwarya Arun, Amrutha H C, Ananga Ghosh, |

| | | | |
|---|---|---|---|
| | | | Rakesh K Mishra, Divya Tej Sowpati |
| EPI_ISL_528858, EPI_ISL_528859 | CSIR-Centre for Cellular and Molecular Biology | CSIR-Centre for Cellular and Molecular Biology | Pratheusa Maccha, Sakshi Shambhavi, Lamuk Zaveri, Shagufta Khan, Namami Gaur, Nikhil Hajirnis, M Soujanya Reddy, Tulasi Nagabandi, Purushotham Vodnala, Payel Mukherjee, Sofia Banu, Priya Singh,Onkar Kulkarni, Dhiviya Vedagiri, Divya Gupta, Vishal Sah, Santosh Kumar Kuncha, Krishnan Harinivas Harshan, Archana Bharadwaj Siva, Karthik Bharadwaj Tallapaka, G. Aditya Kumar, Koushick Sivakumar,Disha Nanda, Divya Das, Jotin Gogoi, Manish Bhattacharjee, Ravi Prasad Mukku, Rakesh K Mishra, Divya Tej Sowpati |
| EPI_ISL_528860 | CSIR-Centre for Cellular and Molecular Biology | CSIR-Centre for Cellular and Molecular Biology | Tulasi Nagabandi, Namami Gaur, Sakshi Shambhavi, Lamuk Zaveri, Shagufta Khan, Nikhil Hajirnis, M Soujanya Reddy, Pratheusa Maccha, Purushotham Vodnala, Payel Mukherjee, Sofia Banu, Priya Singh, Onkar Kulkarni, Dhiviya Vedagiri, Divya Gupta, Vishal Sah, Santosh Kumar Kuncha, Krishnan Harinivas Harshan, Archana Bharadwaj Siva, Karthik Bharadwaj Tallapaka,G. Aditya Kumar, Koushick Sivakumar, Pooja Ramesh Gupta, Rajan Kumar Jha, Shraddha Vijay Lahoti, Rakesh K Mishra, Divya Tej Sowpati |
| EPI_ISL_528861 | CSIR-Centre for Cellular and Molecular Biology | CSIR-Centre for Cellular and Molecular Biology | Sakshi Shambhavi, Lamuk Zaveri, Shagufta Khan, Namami Gaur, Nikhil Hajirnis, M Soujanya Reddy, Pratheusa Maccha, Tulasi Nagabandi, Purushotham Vodnala, Payel Mukherjee, Sofia Banu, Priya Singh, Onkar Kulkarni, Dhiviya Vedagiri, Divya Gupta, Vishal Sah, Santosh Kumar Kuncha, Krishnan Harinivas Harshan, Archana Bharadwaj Siva, Karthik Bharadwaj Tallapaka, Deepak Kumar, Devi Prasad Vijayashankar, Disha Nanda, Divya Das, Jotin Gogoi, Manish Bhattacharjee, Rakesh K Mishra, Divya Tej Sowpati |
| EPI_ISL_528862 | CSIR-Centre for Cellular and Molecular Biology | CSIR-Centre for Cellular and Molecular Biology | Lamuk Zaveri, Shagufta Khan, Namami Gaur, Sakshi Shambhavi, Nikhil Hajirnis, M Soujanya Reddy, Pratheusa Maccha, Tulasi Nagabandi, Purushotham Vodnala, Payel Mukherjee, Sofia Banu, Priya Singh, Onkar Kulkarni, Dhiviya Vedagiri, Divya Gupta, Vishal Sah, Santosh Kumar Kuncha, Krishnan Harinivas Harshan, Archana Bharadwaj Siva, Karthik Bharadwaj Tallapaka, Renu Sudhakar, Somesh Gorde, Gangumala Srinivas Reddy, Sujoy Deb, Swati Bayyana, Rakesh K Mishra, Divya Tej Sowpati |
| EPI_ISL_528863 | CSIR-Centre for Cellular and Molecular Biology | CSIR-Centre for Cellular and Molecular Biology | Payel Mukherjee, Sofia Banu, Priya Singh, Onkar Kulkarni, Dhiviya Vedagiri, Divya Gupta, Vishal Sah, Santosh Kumar Kuncha, Krishnan Harinivas Harshan, Archana Bharadwaj Siva, Karthik Bharadwaj Tallapaka, Shagufta Khan, Lamuk Zaveri, Nikhil Hajirnis, M Soujanya Reddy, Pratheusa Maccha, Namami Gaur, Sakshi Shambhavi, Tulasi Nagabandi, Purushotham Vodnala, Rakesh K Mishra, Sonu Uday, Sudipta Mondal, Annapoorna P Karthyayani, Debabrata Jana, Debrya Saha, Divya Tej Sowpati |
| EPI_ISL_528864 | CSIR-Centre for Cellular and Molecular Biology | CSIR-Centre for Cellular and Molecular Biology | Sofia Banu, Payel Mukherjee, Priya Singh,Onkar Kulkarni, Dhiviya Vedagiri, Divya Gupta, Vishal Sah, Santosh Kumar Kuncha, Krishnan Harinivas Harshan, Archana Bharadwaj Siva, Karthik Bharadwaj Tallapaka, Shagufta Khan, Lamuk Zaveri, Namami Gaur, Sakshi Shambhavi, Nikhil Hajirnis, M Soujanya Reddy, Pratheusa Maccha,Tulasi Nagabandi, Purushotham Vodnala, Deepak Kumar, Devi Prasad Vijayashankar, Disha Nanda, Divya Das, Jotin Gogoi, Manish Bhattacharjee, Rakesh K Mishra, Divya Tej Sowpati |
| EPI_ISL_528865 | CSIR-Centre for Cellular and Molecular Biology | CSIR-Centre for Cellular and Molecular Biology | Shagufta Khan, Lamuk Zaveri, Namami Gaur, Sakshi Shambhavi, Nikhil Hajirnis, M Soujanya Reddy, Pratheusa Maccha, Tulasi Nagabandi, Purushotham Vodnala, Payel Mukherjee, Sofia Banu, Priya Singh, Onkar Kulkarni, Dhiviya Vedagiri, Divya Gupta, Vishal Sah, Santosh Kumar Kuncha, Krishnan Harinivas Harshan, Archana Bharadwaj Siva, Karthik Bharadwaj Tallapaka, Renu Sudhakar, Somesh Gorde, Gangumala Srinivas Reddy, Sujoy Deb, Swati Bayyana, Rakesh K Mishra, Divya Tej Sowpati |
| EPI_ISL_528866 | CSIR-Centre for Cellular and Molecular Biology | CSIR-Centre for Cellular and Molecular Biology | Sakshi Shambhavi, Lamuk Zaveri, Shagufta Khan, Namami Gaur, Nikhil Hajirnis, M Soujanya Reddy, Pratheusa Maccha, Tulasi Nagabandi, Purushotham Vodnala, Payel Mukherjee, Sofia Banu, Priya Singh, Onkar Kulkarni, Dhiviya Vedagiri, Divya Gupta, Vishal Sah, Santosh Kumar Kuncha, Krishnan Harinivas Harshan, Archana Bharadwaj Siva, Karthik Bharadwaj Tallapaka, Deepak Kumar, Devi Prasad Vijayashankar, Disha Nanda, Divya Das, Jotin Gogoi, Manish Bhattacharjee, Rakesh K Mishra, Divya Tej Sowpati |
| EPI_ISL_528867 | CSIR-Centre for Cellular and Molecular Biology | CSIR-Centre for Cellular and Molecular Biology | M Soujanya Reddy, Nikhil Hajirnis, Pratheusa Maccha, Namami Gaur, Sakshi Shambhavi, Lamuk Zaveri, Shagufta Khan, Tulasi Nagabandi, Purushotham Vodnala, Payel Mukherjee, Sofia Banu, Priya Singh, Onkar Kulkarni, Dhiviya Vedagiri, Divya Gupta, Vishal Sah, Santosh Kumar Kuncha, Krishnan Harinivas Harshan, Archana Bharadwaj Siva, Karthik Bharadwaj Tallapaka, Zeba Rizvi, Zuberwasim Sayyad, Kakade Aishwarya Arun, Amrutha H C, Ananga Ghosh, Rakesh K Mishra, Divya Tej Sowpati |
| EPI_ISL_528868, EPI_ISL_528869 | CSIR-Centre for Cellular and Molecular Biology | CSIR-Centre for Cellular and Molecular Biology | Shagufta Khan, Lamuk Zaveri, Namami Gaur, Sakshi Shambhavi, Nikhil Hajirnis, M Soujanya Reddy, Pratheusa Maccha, Tulasi Nagabandi, Purushotham Vodnala, Payel Mukherjee, Sofia Banu, Priya Singh, Onkar Kulkarni, Dhiviya Vedagiri, Divya Gupta, Vishal Sah, Santosh Kumar Kuncha, Krishnan Harinivas Harshan, Archana Bharadwaj Siva, Karthik Bharadwaj Tallapaka, Renu Sudhakar, Somesh Gorde, Gangumala Srinivas Reddy, Sujoy Deb, Swati Bayyana, Rakesh K Mishra, Divya Tej Sowpati |
| EPI_ISL_529014, EPI_ISL_529015 | Ospedale "Ss. Annunziata" | Istituto Zooprofilattico Sperimentale dell'Abruzzo e Molise "G.Caporale" | Lorusso A, Marcacci M, Di Domenico M, Curini V, Ancora M, Cammà C, Rinaldi A, Mangone I, Di Pasquale A, Puglia I, Savini G. |
| EPI_ISL_529139 | Centro de Desenvolvimento Tecnologico em Saude, Fundacao Oswaldo Cruz | Centro de Desenvolvimento Tecnologico em Saude, Fundacao Oswaldo Cruz | Souza,T.M., Fintelman-Rodrigues,N., De Paula,A.D., Saraiva,F.B., Ferreira,M.A., Sacramento,C.Q., Medeiros,M.A. |
| EPI_ISL_529140 | Centro de Desenvolvimento Tecnologico em Saude, Fundacao Oswaldo Cruz | Centro de Desenvolvimento Tecnologico em Saude, Fundacao Oswaldo Cruz | Souza,T.M., Fintelman-Rodrigues,N., De Paula,A.D., Saraiva,F.B., Ferreira,M.A., Sacramento,C.Q., Medeiros,M.A. |
| EPI_ISL_529141, EPI_ISL_529142, EPI_ISL_529143, EPI_ISL_529144, EPI_ISL_529145 | Egyptian National Cancer Institute (ENCI) | Egyptian National Cancer Institute (ENCI) | Zekri, Abdel Rahman N., Amer,K.E., Ahmed,O.S., Soliman,H.K., Ali,M.A., Hassan,W.A., Mahmoud,A.A., Khattab,A.A., Hafez,M.M., Abouelhoda, Mohamed |
| EPI_ISL_529237 | University of Birmingham | COVID-19 Genomics UK (COG-UK) Consortium | Institute of Microbiology, University of Birmingham: Claire McMurray, Joanne Stockton, Samuel Nicholls, Radoslaw Poplawski, Will Rowe, Josh Quick, Nicholas Loman. University of Birmingham Testing Laboratory: Celina M Whalley, Andrew Bosworth, Charlotte Poxon, Kasun Wanigasooriya, Oliver Pickles, Mike Kidd, Alex Richter, Andrew D Beggs PHE Heartlands Lab: Husam Osman, Andrew Bosworth. Queen Elizabeth Hospital: Anna Casey |
| EPI_ISL_529276 | Queens Medical Centre, Clinical Microbiology Department / DeepSeq Nottingham | COVID-19 Genomics UK (COG-UK) Consortium | Gemma Clark, Wendy Smith, Manjinder Khakh, Vicki M Fleming, Michelle M Lister, Hannah Howson-Wells, Jonathan Ball, Patrick McClure, Joseph Chappell, Theocharis Tsoleridis, Nadine Holmes, Matthew Carlisle, Christopher Moore, Fei Sang, Johnny Debebe, Victoria Wright, Matthew Loose |
| EPI_ISL_529281 | Quadram Institute Bioscience | COVID-19 Genomics UK (COG-UK) Consortium | Dave J. Baker, Gemma L. Kay, Alp Aydin, Thanh Le-Viet, Steven Rudder, Ana P. Tedim, Anastasia Kolyva, Maria Diaz, Leonardo de Oliveira Martins, Nabil-Fareed Alikhan, Lizzie Meadows, Rachael Stanley, Ngozi Elumogo, Muhammed Yasir, Nicholas M. Thomson, Alexander J Trotter, Rachel Gilroy, Samuel Bloomfield, Claire Stuart, Andrew Bell, Reenesh Prakash, Samir Dervisevic, Alison E. Mather, John Wain, Mark Webber, Andrew J. Page, Justin O'Grady |
| EPI_ISL_529306 | Queens Medical Centre, Clinical Microbiology Department / DeepSeq Nottingham | COVID-19 Genomics UK (COG-UK) Consortium | Gemma Clark, Wendy Smith, Manjinder Khakh, Vicki M Fleming, Michelle M Lister, Hannah Howson-Wells, Jonathan Ball, Patrick McClure, Joseph Chappell, Theocharis Tsoleridis, Nadine Holmes, Matthew Carlisle, Christopher Moore, Fei Sang, Johnny Debebe, Victoria Wright, Matthew Loose |
| EPI_ISL_529312, EPI_ISL_529320, EPI_ISL_529322, EPI_ISL_529323, EPI_ISL_529324, EPI_ISL_529329, EPI_ISL_529360, EPI_ISL_529365 | University of Birmingham | COVID-19 Genomics UK (COG-UK) Consortium | Institute of Microbiology, University of Birmingham: Claire McMurray, Joanne Stockton, Samuel Nicholls, Radoslaw Poplawski, Will Rowe, Josh Quick, Nicholas Loman. University of Birmingham Testing Laboratory: Celina M Whalley, Andrew Bosworth, Charlotte Poxon, Kasun Wanigasooriya, Oliver Pickles, Mike Kidd, Alex Richter, Andrew D Beggs PHE Heartlands Lab: Husam Osman, Andrew Bosworth. Queen Elizabeth Hospital: Anna Casey |
| EPI_ISL_529378 | Quadram Institute Bioscience | COVID-19 Genomics UK (COG-UK) Consortium | Dave J. Baker, Gemma L. Kay, Alp Aydin, Thanh Le-Viet, Steven Rudder, Ana P. Tedim, Anastasia Kolyva, Maria Diaz, Leonardo de Oliveira Martins, Nabil-Fareed Alikhan, Lizzie Meadows, Rachael Stanley, Ngozi Elumogo, Muhammed Yasir, Nicholas M. Thomson, Alexander J Trotter, Rachel Gilroy, Samuel Bloomfield, Claire Stuart, Andrew Bell, Reenesh Prakash, Samir Dervisevic, Alison E. Mather, John Wain, Mark Webber, Andrew J. Page, Justin O'Grady |
| EPI_ISL_529387 | Queens Medical Centre, Clinical Microbiology Department / DeepSeq Nottingham | COVID-19 Genomics UK (COG-UK) Consortium | Gemma Clark, Wendy Smith, Manjinder Khakh, Vicki M Fleming, Michelle M Lister, Hannah Howson-Wells, Jonathan Ball, Patrick McClure, Joseph Chappell, Theocharis Tsoleridis, Nadine Holmes, Matthew Carlisle, Christopher Moore, Fei Sang, Johnny Debebe, Victoria Wright, Matthew Loose |
| EPI_ISL_529389 | Quadram Institute Bioscience | COVID-19 Genomics UK (COG-UK) Consortium | Dave J. Baker, Gemma L. Kay, Alp Aydin, Thanh Le-Viet, Steven Rudder, Ana P. Tedim, Anastasia Kolyva, Maria Diaz, Leonardo de Oliveira Martins, Nabil-Fareed Alikhan, Lizzie Meadows, Rachael Stanley, Ngozi Elumogo, Muhammed Yasir, Nicholas M. Thomson, Alexander J Trotter, Rachel Gilroy, Samuel Bloomfield, Claire Stuart, Andrew Bell, Reenesh Prakash, Samir Dervisevic, Alison E. Mather, John Wain, Mark Webber, Andrew J. Page, Justin O'Grady |
| EPI_ISL_529391 | University of Birmingham | COVID-19 Genomics UK (COG-UK) Consortium | Institute of Microbiology, University of Birmingham: Claire McMurray, Joanne Stockton, Samuel Nicholls, Radoslaw Poplawski, Will Rowe, Josh Quick, |

| | | | |
|---|---|---|---|
| | | | Nicholas Loman. University of Birmingham Testing Laboratory: Celina M Whalley, Andrew Bosworth, Charlotte Poxon, Kasun Wanigasooriya, Oliver Pickles, Mike Kidd, Alex Richter, Andrew D Beggs PHE Heartlands Lab: Husam Osman, Andrew Bosworth. Queen Elizabeth Hospital: Anna Casey |
| EPI_ISL_529412 | Queens Medical Centre, Clinical Microbiology Department / DeepSeq Nottingham | COVID-19 Genomics UK (COG-UK) Consortium | Gemma Clark, Wendy Smith, Manjinder Khakh, Vicki M Fleming, Michelle M Lister, Hannah Howson-Wells, Jonathan Ball, Patrick McClure, Joseph Chappell, Theocharis Tsoleridis, Nadine Holmes, Matthew Carlisle, Christopher Moore, Fei Sang, Johnny Debebe, Victoria Wright, Matthew Loose |
| EPI_ISL_529421 | University of Birmingham | COVID-19 Genomics UK (COG-UK) Consortium | Institute of Microbiology, University of Birmingham: Claire McMurray, Joanne Stockton, Samuel Nicholls, Radoslaw Poplawski, Will Rowe, Josh Quick, Nicholas Loman. University of Birmingham Testing Laboratory: Celina M Whalley, Andrew Bosworth, Charlotte Poxon, Kasun Wanigasooriya, Oliver Pickles, Mike Kidd, Alex Richter, Andrew D Beggs PHE Heartlands Lab: Husam Osman, Andrew Bosworth. Queen Elizabeth Hospital: Anna Casey |
| EPI_ISL_529423 | Queens Medical Centre, Clinical Microbiology Department / DeepSeq Nottingham | COVID-19 Genomics UK (COG-UK) Consortium | Gemma Clark, Wendy Smith, Manjinder Khakh, Vicki M Fleming, Michelle M Lister, Hannah Howson-Wells, Jonathan Ball, Patrick McClure, Joseph Chappell, Theocharis Tsoleridis, Nadine Holmes, Matthew Carlisle, Christopher Moore, Fei Sang, Johnny Debebe, Victoria Wright, Matthew Loose |
| EPI_ISL_529424, EPI_ISL_529439, EPI_ISL_529440, EPI_ISL_529468, EPI_ISL_529470, EPI_ISL_529471, EPI_ISL_529475 | University of Birmingham | COVID-19 Genomics UK (COG-UK) Consortium | Institute of Microbiology, University of Birmingham: Claire McMurray, Joanne Stockton, Samuel Nicholls, Radoslaw Poplawski, Will Rowe, Josh Quick, Nicholas Loman. University of Birmingham Testing Laboratory: Celina M Whalley, Andrew Bosworth, Charlotte Poxon, Kasun Wanigasooriya, Oliver Pickles, Mike Kidd, Alex Richter, Andrew D Beggs PHE Heartlands Lab: Husam Osman, Andrew Bosworth. Queen Elizabeth Hospital: Anna Casey |
| EPI_ISL_529479 | Queens Medical Centre, Clinical Microbiology Department / DeepSeq Nottingham | COVID-19 Genomics UK (COG-UK) Consortium | Gemma Clark, Wendy Smith, Manjinder Khakh, Vicki M Fleming, Michelle M Lister, Hannah Howson-Wells, Jonathan Ball, Patrick McClure, Joseph Chappell, Theocharis Tsoleridis, Nadine Holmes, Matthew Carlisle, Christopher Moore, Fei Sang, Johnny Debebe, Victoria Wright, Matthew Loose |
| EPI_ISL_529513 | Liverpool Clinical Laboratories | COVID-19 Genomics UK (COG-UK) Consortium | Sam Haldenby, Anita Lucaci, Steve Paterson, Julian Hiscox, Alistair Darby, M Almsaud, A Alrezaihi, Muhannad Alruwaili, Stuart D Armstrong, Jones Benjamin, Eleanor G Bentley, Anu Chawla, Jordan J Clark, Angela Cowell, Richard Eccles, Isabel García-Dorival, Matthew Gemmell, Alessandro Gerada, PKF Gilmore, Richard Gregory, Ximeng Han, Catherine Hartley, Margaret Hughes, Miren Iturriza-Gomara, James Johnson, L Luu, Jenifer Manson, Charlotte Nelson, Elaine O'Toole, Cassie Olateju, Rebekah Penrice-Randal , Lucille Rainbow, N.P Randle, Trevor Ian Robinson, Parul Sharma, Ghada T Shawli, James P Stewart, Neil Swainston, Ecaterina Vamos, Joanne Watts, Mark Whitehead |
| EPI_ISL_529553, EPI_ISL_529554, EPI_ISL_529555, EPI_ISL_529556 | Queens Medical Centre, Clinical Microbiology Department / DeepSeq Nottingham | COVID-19 Genomics UK (COG-UK) Consortium | Gemma Clark, Wendy Smith, Manjinder Khakh, Vicki M Fleming, Michelle M Lister, Hannah Howson-Wells, Jonathan Ball, Patrick McClure, Joseph Chappell, Theocharis Tsoleridis, Nadine Holmes, Matthew Carlisle, Christopher Moore, Fei Sang, Johnny Debebe, Victoria Wright, Matthew Loose |
| EPI_ISL_529557, EPI_ISL_529558, EPI_ISL_529559, EPI_ISL_529570, EPI_ISL_529571 | Quadram Institute Bioscience | COVID-19 Genomics UK (COG-UK) Consortium | Dave J. Baker, Gemma L. Kay, Alp Aydin, Thanh Le-Viet, Steven Rudder, Ana P. Tedim, Anastasia Kolyva, Maria Diaz, Leonardo de Oliveira Martins, Nabil-Fareed Alikhan, Lizzie Meadows, Rachael Stanley, Ngozi Elumogo, Muhammed Yasir, Nicholas M. Thomson, Alexander J Trotter, Rachel Gilroy, Samuel Bloomfield, Claire Stuart, Andrew Bell, Reenesh Prakash, Samir Dervisevic, Alison E. Mather, John Wain, Mark Webber, Andrew J. Page, Justin O'Grady |
| EPI_ISL_529590, EPI_ISL_529591, EPI_ISL_529592, EPI_ISL_529593, EPI_ISL_529594, EPI_ISL_529595, EPI_ISL_529663, EPI_ISL_529664, EPI_ISL_529665, EPI_ISL_529666 | University of Birmingham | COVID-19 Genomics UK (COG-UK) Consortium | Institute of Microbiology, University of Birmingham: Claire McMurray, Joanne Stockton, Samuel Nicholls, Radoslaw Poplawski, Will Rowe, Josh Quick, Nicholas Loman. University of Birmingham Testing Laboratory: Celina M Whalley, Andrew Bosworth, Charlotte Poxon, Kasun Wanigasooriya, Oliver Pickles, Mike Kidd, Alex Richter, Andrew D Beggs PHE Heartlands Lab: Husam Osman, Andrew Bosworth. Queen Elizabeth Hospital: Anna Casey |
| EPI_ISL_529718 | Santo Borromeus Hospital | Laboratorium Kesehatan Provinsi Jawa Barat; School of Life Sciences and Technology & School of Pharmacy-Institut Teknologi Bandung; Molecular Genetics Laboratory-Faculty of Medicine-Universitas Padjadjaran | Ema Rahmawati, Marselina Irasonia Tan, Yunia Sribudiani, Catur Riani, Azzania Fibriani, Husna Nugrahapraja, Tarwadi, Savira Ekawardhani, Hesti Lina Wiraswati, Ryan Bayusantika Ristandi, Rifky Waluyajati Rachman, Cut Nur Cinthia Alamanda, Lia Faridah, Gusti Ayu Prani Pradani, Adelina Khristiani Rahayu, Hammam Riza, Sony Solistia Wirawan, Agung Eru Wibowo, Irvan Faizal |
| EPI_ISL_529719 | Sumedang Regional General Hospital | Laboratorium Kesehatan Provinsi Jawa Barat; School of Life Sciences and Technology & School of Pharmacy-Institut Teknologi Bandung; Molecular Genetics Laboratory-Faculty of Medicine-Universitas Padjadjaran | Ema Rahmawati, Marselina Irasonia Tan, Yunia Sribudiani, Catur Riani, Azzania Fibriani, Husna Nugrahapraja, Tarwadi, Savira Ekawardhani, Hesti Lina Wiraswati, Ryan Bayusantika Ristandi, Rifky Waluyajati Rachman, Cut Nur Cinthia Alamanda, Lia Faridah, Gusti Ayu Prani Pradani, Adelina Khristiani Rahayu, Hammam Riza, Sony Solistia Wirawan, Agung Eru Wibowo, Irvan Faizal |
| EPI_ISL_529804, EPI_ISL_529834, EPI_ISL_529835, EPI_ISL_529836, EPI_ISL_529837, EPI_ISL_529838, EPI_ISL_529839, EPI_ISL_529840, EPI_ISL_529841, EPI_ISL_529842, EPI_ISL_529871 | | | |
| see above | Michigan Department of Health and Human Services, Bureau of Laboratories | Michigan Department of Health and Human Services, Bureau of Laboratories | Blankenship HM, Riner D, Soehnlen MK |
| EPI_ISL_529963 | Universitas Airlangga Hospital | Institute of Tropical Disease, Universitas Airlangga | Rima R Prasetya, Krisnoadi Rahardjo, Aldise M Nastri, Jezzy R Dewantari, Nasronudin, Gatot Soegiarto, Laksmi Wulandari, Retno A Setyoningrum, Resti Yudhawati, Yohko K Shimizu, Mitsuhiro Nishimura, Yasuko Mori, Soetjipto, Kazufumi Shimizu, Maria I Lusida |
| EPI_ISL_530126 | Seattle Flu Study | Seattle Flu Study | Deborah A. Nickerson, Chris D. Frazar, Jover Lee, Benjamin Pelle, Matthew Richardson, Amanda Adler, Elisabeth Brandstetter, Peter D. Han, Kairsten Fay, Misja Ilcisin, Kirsten Lacombe, Thomas R. Sibley, Melissa Truong, Caitlin R. Wolf, Karen Cowgill, Stephanie Schrag, Jeff Duchin, Michael Boeckh, Janet A. Englund, Michael Famulare, Barry R. Lutz, Mark J. Rieder, Lea M. Starita, Matthew Thompson, Helen Y. Chu, Trevor Bedford, Jay Shendure |
| EPI_ISL_530252, EPI_ISL_530253 | Queensland Health Forensic and Scientific Services, Public Health Virology | Public Health Virology Laboratory, Forensic and Scientific Services, Queensland Health | Son Nguyen et al |
| EPI_ISL_532542, EPI_ISL_532551, EPI_ISL_532557, EPI_ISL_532558, EPI_ISL_532560, EPI_ISL_532562, EPI_ISL_532567, EPI_ISL_532568, EPI_ISL_532578, EPI_ISL_532579 | Lighthouse Lab in Glasgow | Wellcome Sanger Institute for the COVID-19 Genomics UK (COG-UK) consortium | Harper VanSteenhouse, Yumi Kasai, David Gray, Carol Clugston, Anna Dominiczak and Alex Alderton, Roberto Amato, Sonia Goncalves, Ewan Harrison, David K. Jackson, Ian Johnston, Dominic Kwiatkowski, Cordelia Langford, John Sillitoe |
| EPI_ISL_532582 | NHSGGC West of Scotland Specialist Virology Centre / MRC-University of Glasgow Centre for Virus Research | Wellcome Sanger Institute for the COVID-19 Genomics UK (COG-UK) consortium | Ana da Silva Filipe, Natasha Johnson, Kathy Smollett, Daniel Mair, Stephen Carmichael, Lily Tong, Jenna Nichols, Elihu Aranday-Cortes, Kirstyn Brunker, Yasmin Parr, Kyriaki Nomikou; Sarah McDonald, Marc Niebel, Patawee Asamaphan; Richard Orton, Joseph Hughes, Sreenu Vattipally, David L Robertson; Alasdair MacLean, Rory Gunson; Kathy Li, Natasha Jesudason, Rajiv Shah, James Shepherd, Antonia Ho, Alice Broos, Emma Thomson and Alex Alderton, Roberto Amato, Sonia Goncalves, Ewan Harrison, David K. Jackson, Ian Johnston, Dominic Kwiatkowski, Cordelia Langford, John Sillitoe |
| EPI_ISL_532589, EPI_ISL_532595, EPI_ISL_532617, EPI_ISL_532624, EPI_ISL_532627, EPI_ISL_532651, EPI_ISL_532652 | Lighthouse Lab in Glasgow | Wellcome Sanger Institute for the COVID-19 Genomics UK (COG-UK) consortium | Harper VanSteenhouse, Yumi Kasai, David Gray, Carol Clugston, Anna Dominiczak and Alex Alderton, Roberto Amato, Sonia Goncalves, Ewan Harrison, David K. Jackson, Ian Johnston, Dominic Kwiatkowski, Cordelia Langford, John Sillitoe |
| EPI_ISL_532659 | Virology Department, Royal Infirmary of Edinburgh, NHS Lothian / School of Biological Sciences, University of Edinburgh | Wellcome Sanger Institute for the COVID-19 Genomics UK (COG-UK) consortium | McHugh M, Dewar R, Rooke S, O'Toole Á, Scher E, Hill V, McCrone JT, Colquhoun R, Yu X, Jackson B, Rambaut A, Templeton K and Alex Alderton, Roberto Amato, Sonia Goncalves, Ewan Harrison, David K. Jackson, Ian Johnston, Dominic Kwiatkowski, Cordelia Langford, John Sillitoe |
| EPI_ISL_532661, EPI_ISL_532666, EPI_ISL_532667, EPI_ISL_532669, EPI_ISL_532677 | Lighthouse Lab in Glasgow | Wellcome Sanger Institute for the COVID-19 Genomics UK (COG-UK) consortium | Harper VanSteenhouse, Yumi Kasai, David Gray, Carol Clugston, Anna Dominiczak and Alex Alderton, Roberto Amato, Sonia Goncalves, Ewan Harrison, David K. Jackson, Ian Johnston, Dominic Kwiatkowski, Cordelia Langford, John Sillitoe |
| EPI_ISL_532683, EPI_ISL_532689 | Virology Department, Royal Infirmary of Edinburgh, NHS Lothian / School of Biological Sciences, University of Edinburgh | Wellcome Sanger Institute for the COVID-19 Genomics UK (COG-UK) consortium | McHugh M, Dewar R, Rooke S, O'Toole Á, Scher E, Hill V, McCrone JT, Colquhoun R, Yu X, Jackson B, Rambaut A, Templeton K and Alex Alderton, Roberto Amato, Sonia Goncalves, Ewan Harrison, David K. Jackson, Ian Johnston, Dominic Kwiatkowski, Cordelia Langford, John Sillitoe |
| EPI_ISL_532691, EPI_ISL_532693 | Lighthouse Lab in Glasgow | Wellcome Sanger Institute for the COVID-19 Genomics UK (COG-UK) consortium | Harper VanSteenhouse, Yumi Kasai, David Gray, Carol Clugston, Anna Dominiczak and Alex Alderton, Roberto Amato, Sonia Goncalves, Ewan Harrison, David K. Jackson, Ian Johnston, Dominic Kwiatkowski, Cordelia Langford, John Sillitoe |
| EPI_ISL_532705 | Virology Department, Royal Infirmary of Edinburgh, NHS Lothian / School of Biological Sciences, University of | Wellcome Sanger Institute for the COVID-19 Genomics UK (COG-UK) consortium | McHugh M, Dewar R, Rooke S, O'Toole Á, Scher E, Hill V, McCrone JT, Colquhoun R, Yu X, Jackson B, Rambaut A, Templeton K and Alex Alderton, Roberto Amato, Sonia Goncalves, Ewan Harrison, David K. Jackson, Ian Johnston, Dominic Kwiatkowski, Cordelia Langford, John Sillitoe |

| | | | |
|---|---|---|---|
| | Edinburgh | | |
| EPI_ISL_532706, EPI_ISL_532716, EPI_ISL_532719, EPI_ISL_532728, EPI_ISL_532730, EPI_ISL_532745, EPI_ISL_532761, EPI_ISL_532788, EPI_ISL_532806, EPI_ISL_532821, EPI_ISL_532836, EPI_ISL_532837, EPI_ISL_532839, EPI_ISL_532840, EPI_ISL_532841, EPI_ISL_532842, EPI_ISL_532843, EPI_ISL_532845, EPI_ISL_532846, EPI_ISL_532847, EPI_ISL_532848 | | | |
| see above | Lighthouse Lab in Glasgow | Wellcome Sanger Institute for the COVID-19 Genomics UK (COG-UK) consortium | Harper VanSteenhouse, Yumi Kasai, David Gray, Carol Clugston, Anna Dominiczak and Alex Alderton, Roberto Amato, Sonia Goncalves, Ewan Harrison, David K. Jackson, Ian Johnston, Dominic Kwiatkowski, Cordelia Langford, John Sillitoe |
| EPI_ISL_532851 | Lighthouse Lab in Glasgow | Wellcome Sanger Institute for the COVID-19 Genomics UK (COG-UK) Consortium | Harper VanSteenhouse, Yumi Kasai, David Gray, Carol Clugston, Anna Dominiczak and Alex Alderton, Roberto Amato, Sonia Goncalves, Ewan Harrison, David K. Jackson, Ian Johnston, Dominic Kwiatkowski, Cordelia Langford, John Sillitoe on behalf of the Wellcome Sanger Institute COVID-19 Surveillance Team |
| EPI_ISL_532852, EPI_ISL_532853, EPI_ISL_532856, EPI_ISL_532857, EPI_ISL_532859, EPI_ISL_532860, EPI_ISL_532861, EPI_ISL_532862, EPI_ISL_532863 | Lighthouse Lab in Glasgow | Wellcome Sanger Institute for the COVID-19 Genomics UK (COG-UK) consortium | Harper VanSteenhouse, Yumi Kasai, David Gray, Carol Clugston, Anna Dominiczak and Alex Alderton, Roberto Amato, Sonia Goncalves, Ewan Harrison, David K. Jackson, Ian Johnston, Dominic Kwiatkowski, Cordelia Langford, John Sillitoe |
| EPI_ISL_532864, EPI_ISL_532865 | NHSGGC West of Scotland Specialist Virology Centre / MRC-University of Glasgow Centre for Virus Research | Wellcome Sanger Institute for the COVID-19 Genomics UK (COG-UK) consortium | Ana da Silva Filipe, Natasha Johnson, Kathy Smollett, Daniel Mair, Stephen Carmichael, Lily Tong, Jenna Nichols, Elihu Aranday-Cortes, Kirstyn Brunker, Yasmin Parr, Kyriaki Nomikou; Sarah McDonald, Marc Niebel, Patawee Asamaphan; Richard Orton, Joseph Hughes, Sreenu Vattipally, David L Robertson; Alasdair MacLean, Rory Gunson; Kathy Li, Natasha Jesudason, Rajiv Shah, James Shepherd, Antonia Ho, Alice Broos, Emma Thomson and Alex Alderton, Roberto Amato, Sonia Goncalves, Ewan Harrison, David K. Jackson, Ian Johnston, Dominic Kwiatkowski, Cordelia Langford, John Sillitoe |
| EPI_ISL_532866, EPI_ISL_532867, EPI_ISL_532868, EPI_ISL_532869, EPI_ISL_532870, EPI_ISL_532871, EPI_ISL_532872 | Lighthouse Lab in Glasgow | Wellcome Sanger Institute for the COVID-19 Genomics UK (COG-UK) consortium | Harper VanSteenhouse, Yumi Kasai, David Gray, Carol Clugston, Anna Dominiczak and Alex Alderton, Roberto Amato, Sonia Goncalves, Ewan Harrison, David K. Jackson, Ian Johnston, Dominic Kwiatkowski, Cordelia Langford, John Sillitoe |
| EPI_ISL_532874, EPI_ISL_532875, EPI_ISL_532876 | NHSGGC West of Scotland Specialist Virology Centre / MRC-University of Glasgow Centre for Virus Research | Wellcome Sanger Institute for the COVID-19 Genomics UK (COG-UK) consortium | Ana da Silva Filipe, Natasha Johnson, Kathy Smollett, Daniel Mair, Stephen Carmichael, Lily Tong, Jenna Nichols, Elihu Aranday-Cortes, Kirstyn Brunker, Yasmin Parr, Kyriaki Nomikou; Sarah McDonald, Marc Niebel, Patawee Asamaphan; Richard Orton, Joseph Hughes, Sreenu Vattipally, David L Robertson; Alasdair MacLean, Rory Gunson; Kathy Li, Natasha Jesudason, Rajiv Shah, James Shepherd, Antonia Ho, Alice Broos, Emma Thomson and Alex Alderton, Roberto Amato, Sonia Goncalves, Ewan Harrison, David K. Jackson, Ian Johnston, Dominic Kwiatkowski, Cordelia Langford, John Sillitoe |
| EPI_ISL_532877, EPI_ISL_532879, EPI_ISL_532880, EPI_ISL_532881, EPI_ISL_532882, EPI_ISL_532883, EPI_ISL_532884 | Lighthouse Lab in Glasgow | Wellcome Sanger Institute for the COVID-19 Genomics UK (COG-UK) consortium | Harper VanSteenhouse, Yumi Kasai, David Gray, Carol Clugston, Anna Dominiczak and Alex Alderton, Roberto Amato, Sonia Goncalves, Ewan Harrison, David K. Jackson, Ian Johnston, Dominic Kwiatkowski, Cordelia Langford, John Sillitoe |
| EPI_ISL_532886 | NHSGGC West of Scotland Specialist Virology Centre / MRC-University of Glasgow Centre for Virus Research | Wellcome Sanger Institute for the COVID-19 Genomics UK (COG-UK) consortium | Ana da Silva Filipe, Natasha Johnson, Kathy Smollett, Daniel Mair, Stephen Carmichael, Lily Tong, Jenna Nichols, Elihu Aranday-Cortes, Kirstyn Brunker, Yasmin Parr, Kyriaki Nomikou; Sarah McDonald, Marc Niebel, Patawee Asamaphan; Richard Orton, Joseph Hughes, Sreenu Vattipally, David L Robertson; Alasdair MacLean, Rory Gunson; Kathy Li, Natasha Jesudason, Rajiv Shah, James Shepherd, Antonia Ho, Alice Broos, Emma Thomson and Alex Alderton, Roberto Amato, Sonia Goncalves, Ewan Harrison, David K. Jackson, Ian Johnston, Dominic Kwiatkowski, Cordelia Langford, John Sillitoe |
| EPI_ISL_532887, EPI_ISL_532888, EPI_ISL_532889, EPI_ISL_532890, EPI_ISL_532892, EPI_ISL_532893 | Lighthouse Lab in Glasgow | Wellcome Sanger Institute for the COVID-19 Genomics UK (COG-UK) consortium | Harper VanSteenhouse, Yumi Kasai, David Gray, Carol Clugston, Anna Dominiczak and Alex Alderton, Roberto Amato, Sonia Goncalves, Ewan Harrison, David K. Jackson, Ian Johnston, Dominic Kwiatkowski, Cordelia Langford, John Sillitoe |
| EPI_ISL_532894 | NHSGGC West of Scotland Specialist Virology Centre / MRC-University of Glasgow Centre for Virus Research | Wellcome Sanger Institute for the COVID-19 Genomics UK (COG-UK) consortium | Ana da Silva Filipe, Natasha Johnson, Kathy Smollett, Daniel Mair, Stephen Carmichael, Lily Tong, Jenna Nichols, Elihu Aranday-Cortes, Kirstyn Brunker, Yasmin Parr, Kyriaki Nomikou; Sarah McDonald, Marc Niebel, Patawee Asamaphan; Richard Orton, Joseph Hughes, Sreenu Vattipally, David L Robertson; Alasdair MacLean, Rory Gunson; Kathy Li, Natasha Jesudason, Rajiv Shah, James Shepherd, Antonia Ho, Alice Broos, Emma Thomson and Alex Alderton, Roberto Amato, Sonia Goncalves, Ewan Harrison, David K. Jackson, Ian Johnston, Dominic Kwiatkowski, Cordelia Langford, John Sillitoe |
| EPI_ISL_532895, EPI_ISL_532897, EPI_ISL_532898, EPI_ISL_532899, EPI_ISL_532900, EPI_ISL_532901, EPI_ISL_532902, EPI_ISL_532904 | Lighthouse Lab in Glasgow | Wellcome Sanger Institute for the COVID-19 Genomics UK (COG-UK) consortium | Harper VanSteenhouse, Yumi Kasai, David Gray, Carol Clugston, Anna Dominiczak and Alex Alderton, Roberto Amato, Sonia Goncalves, Ewan Harrison, David K. Jackson, Ian Johnston, Dominic Kwiatkowski, Cordelia Langford, John Sillitoe |
| EPI_ISL_532905, EPI_ISL_532906 | NHSGGC West of Scotland Specialist Virology Centre / MRC-University of Glasgow Centre for Virus Research | Wellcome Sanger Institute for the COVID-19 Genomics UK (COG-UK) consortium | Ana da Silva Filipe, Natasha Johnson, Kathy Smollett, Daniel Mair, Stephen Carmichael, Lily Tong, Jenna Nichols, Elihu Aranday-Cortes, Kirstyn Brunker, Yasmin Parr, Kyriaki Nomikou; Sarah McDonald, Marc Niebel, Patawee Asamaphan; Richard Orton, Joseph Hughes, Sreenu Vattipally, David L Robertson; Alasdair MacLean, Rory Gunson; Kathy Li, Natasha Jesudason, Rajiv Shah, James Shepherd, Antonia Ho, Alice Broos, Emma Thomson and Alex Alderton, Roberto Amato, Sonia Goncalves, Ewan Harrison, David K. Jackson, Ian Johnston, Dominic Kwiatkowski, Cordelia Langford, John Sillitoe |
| EPI_ISL_532907, EPI_ISL_532908, EPI_ISL_532910, EPI_ISL_532911, EPI_ISL_532914 | Lighthouse Lab in Glasgow | Wellcome Sanger Institute for the COVID-19 Genomics UK (COG-UK) consortium | Harper VanSteenhouse, Yumi Kasai, David Gray, Carol Clugston, Anna Dominiczak and Alex Alderton, Roberto Amato, Sonia Goncalves, Ewan Harrison, David K. Jackson, Ian Johnston, Dominic Kwiatkowski, Cordelia Langford, John Sillitoe |
| EPI_ISL_532915 | NHSGGC West of Scotland Specialist Virology Centre / MRC-University of Glasgow Centre for Virus Research | Wellcome Sanger Institute for the COVID-19 Genomics UK (COG-UK) consortium | Ana da Silva Filipe, Natasha Johnson, Kathy Smollett, Daniel Mair, Stephen Carmichael, Lily Tong, Jenna Nichols, Elihu Aranday-Cortes, Kirstyn Brunker, Yasmin Parr, Kyriaki Nomikou; Sarah McDonald, Marc Niebel, Patawee Asamaphan; Richard Orton, Joseph Hughes, Sreenu Vattipally, David L Robertson; Alasdair MacLean, Rory Gunson; Kathy Li, Natasha Jesudason, Rajiv Shah, James Shepherd, Antonia Ho, Alice Broos, Emma Thomson and Alex Alderton, Roberto Amato, Sonia Goncalves, Ewan Harrison, David K. Jackson, Ian Johnston, Dominic Kwiatkowski, Cordelia Langford, John Sillitoe |
| EPI_ISL_532916, EPI_ISL_532917, EPI_ISL_532918, EPI_ISL_532919, EPI_ISL_532920, EPI_ISL_532921, EPI_ISL_532922, EPI_ISL_532923, EPI_ISL_532925 | Lighthouse Lab in Glasgow | Wellcome Sanger Institute for the COVID-19 Genomics UK (COG-UK) consortium | Harper VanSteenhouse, Yumi Kasai, David Gray, Carol Clugston, Anna Dominiczak and Alex Alderton, Roberto Amato, Sonia Goncalves, Ewan Harrison, David K. Jackson, Ian Johnston, Dominic Kwiatkowski, Cordelia Langford, John Sillitoe |
| EPI_ISL_532926 | Virology Department, Royal Infirmary of Edinburgh, NHS Lothian / School of Biological Sciences, University of Edinburgh | Wellcome Sanger Institute for the COVID-19 Genomics UK (COG-UK) consortium | McHugh M, Dewar R, Rooke S, O'Toole Á, Scher E, Hill V, McCrone JT, Colquhoun R, Yu X, Jackson B, Rambaut A, Templeton K and Alex Alderton, Roberto Amato, Sonia Goncalves, Ewan Harrison, David K. Jackson, Ian Johnston, Dominic Kwiatkowski, Cordelia Langford, John Sillitoe |
| EPI_ISL_532927, EPI_ISL_532928, EPI_ISL_532929, EPI_ISL_532930, EPI_ISL_532931 | Lighthouse Lab in Glasgow | Wellcome Sanger Institute for the COVID-19 Genomics UK (COG-UK) consortium | Harper VanSteenhouse, Yumi Kasai, David Gray, Carol Clugston, Anna Dominiczak and Alex Alderton, Roberto Amato, Sonia Goncalves, Ewan Harrison, David K. Jackson, Ian Johnston, Dominic Kwiatkowski, Cordelia Langford, John Sillitoe |
| EPI_ISL_532932, EPI_ISL_532933 | NHSGGC West of Scotland Specialist Virology Centre / MRC-University of Glasgow Centre for Virus Research | Wellcome Sanger Institute for the COVID-19 Genomics UK (COG-UK) consortium | Ana da Silva Filipe, Natasha Johnson, Kathy Smollett, Daniel Mair, Stephen Carmichael, Lily Tong, Jenna Nichols, Elihu Aranday-Cortes, Kirstyn Brunker, Yasmin Parr, Kyriaki Nomikou; Sarah McDonald, Marc Niebel, Patawee Asamaphan; Richard Orton, Joseph Hughes, Sreenu Vattipally, David L Robertson; Alasdair MacLean, Rory Gunson; Kathy Li, Natasha Jesudason, Rajiv Shah, James Shepherd, Antonia Ho, Alice Broos, Emma Thomson and Alex Alderton, Roberto Amato, Sonia Goncalves, Ewan Harrison, David K. Jackson, Ian Johnston, Dominic Kwiatkowski, Cordelia Langford, John Sillitoe |
| EPI_ISL_532934, EPI_ISL_532935, EPI_ISL_532936, EPI_ISL_532937, EPI_ISL_532939, EPI_ISL_532940, EPI_ISL_532941, EPI_ISL_532942, EPI_ISL_532943, EPI_ISL_532944, EPI_ISL_532946, EPI_ISL_532947 | | | |
| see above | Lighthouse Lab in Glasgow | Wellcome Sanger Institute for the COVID-19 Genomics UK (COG-UK) consortium | Harper VanSteenhouse, Yumi Kasai, David Gray, Carol Clugston, Anna Dominiczak and Alex Alderton, Roberto Amato, Sonia Goncalves, Ewan Harrison, David K. Jackson, Ian Johnston, Dominic Kwiatkowski, Cordelia Langford, John Sillitoe |
| EPI_ISL_532948 | NHSGGC West of Scotland Specialist Virology Centre / MRC-University of Glasgow Centre for Virus Research | Wellcome Sanger Institute for the COVID-19 Genomics UK (COG-UK) consortium | Ana da Silva Filipe, Natasha Johnson, Kathy Smollett, Daniel Mair, Stephen Carmichael, Lily Tong, Jenna Nichols, Elihu Aranday-Cortes, Kirstyn Brunker, Yasmin Parr, Kyriaki Nomikou; Sarah McDonald, Marc Niebel, Patawee Asamaphan; Richard Orton, Joseph Hughes, Sreenu Vattipally, David L Robertson; Alasdair MacLean, Rory Gunson; Kathy Li, Natasha Jesudason, Rajiv Shah, James Shepherd, Antonia Ho, Alice Broos, Emma Thomson and Alex Alderton, |

| | | | |
|---|---|---|---|
| | | | Roberto Amato, Sonia Goncalves, Ewan Harrison, David K. Jackson, Ian Johnston, Dominic Kwiatkowski, Cordelia Langford, John Sillitoe |
| EPI_ISL_532949, EPI_ISL_532950, EPI_ISL_532951, EPI_ISL_532954, EPI_ISL_532955, EPI_ISL_532956, EPI_ISL_532957, EPI_ISL_532958, EPI_ISL_532959, EPI_ISL_532960, EPI_ISL_532961, EPI_ISL_532963, EPI_ISL_532964, EPI_ISL_532965, EPI_ISL_532966, EPI_ISL_532967, EPI_ISL_532968, EPI_ISL_532969, EPI_ISL_532970, EPI_ISL_532971, EPI_ISL_532972, EPI_ISL_532973, EPI_ISL_532974, EPI_ISL_532975, EPI_ISL_532976, EPI_ISL_532977 | | | |
| see above | Lighthouse Lab in Glasgow | Wellcome Sanger Institute for the COVID-19 Genomics UK (COG-UK) consortium | Harper VanSteenhouse, Yumi Kasai, David Gray, Carol Clugston, Anna Dominiczak and Alex Alderton, Roberto Amato, Sonia Goncalves, Ewan Harrison, David K. Jackson, Ian Johnston, Dominic Kwiatkowski, Cordelia Langford, John Sillitoe |
| EPI_ISL_532978 | Virology Department, Royal Infirmary of Edinburgh, NHS Lothian / School of Biological Sciences, University of Edinburgh | Wellcome Sanger Institute for the COVID-19 Genomics UK (COG-UK) consortium | McHugh M, Dewar R, Rooke S, O'Toole Á, Scher E, Hill V, McCrone JT, Colquhoun R, Yu X, Jackson B, Rambaut A, Templeton K and Alex Alderton, Roberto Amato, Sonia Goncalves, Ewan Harrison, David K. Jackson, Ian Johnston, Dominic Kwiatkowski, Cordelia Langford, John Sillitoe |
| EPI_ISL_532979, EPI_ISL_532980, EPI_ISL_532981, EPI_ISL_532982 | Lighthouse Lab in Glasgow | Wellcome Sanger Institute for the COVID-19 Genomics UK (COG-UK) consortium | Harper VanSteenhouse, Yumi Kasai, David Gray, Carol Clugston, Anna Dominiczak and Alex Alderton, Roberto Amato, Sonia Goncalves, Ewan Harrison, David K. Jackson, Ian Johnston, Dominic Kwiatkowski, Cordelia Langford, John Sillitoe |
| EPI_ISL_532983 | Virology Department, Royal Infirmary of Edinburgh, NHS Lothian / School of Biological Sciences, University of Edinburgh | Wellcome Sanger Institute for the COVID-19 Genomics UK (COG-UK) consortium | McHugh M, Dewar R, Rooke S, O'Toole Á, Scher E, Hill V, McCrone JT, Colquhoun R, Yu X, Jackson B, Rambaut A, Templeton K and Alex Alderton, Roberto Amato, Sonia Goncalves, Ewan Harrison, David K. Jackson, Ian Johnston, Dominic Kwiatkowski, Cordelia Langford, John Sillitoe |
| EPI_ISL_532984, EPI_ISL_532985, EPI_ISL_532986, EPI_ISL_532987 | Lighthouse Lab in Glasgow | Wellcome Sanger Institute for the COVID-19 Genomics UK (COG-UK) consortium | Harper VanSteenhouse, Yumi Kasai, David Gray, Carol Clugston, Anna Dominiczak and Alex Alderton, Roberto Amato, Sonia Goncalves, Ewan Harrison, David K. Jackson, Ian Johnston, Dominic Kwiatkowski, Cordelia Langford, John Sillitoe |
| EPI_ISL_532988 | NHSGGC West of Scotland Specialist Virology Centre / MRC-University of Glasgow Centre for Virus Research | Wellcome Sanger Institute for the COVID-19 Genomics UK (COG-UK) consortium | Ana da Silva Filipe, Natasha Johnson, Kathy Smollett, Daniel Mair, Stephen Carmichael, Lily Tong, Jenna Nichols, Elihu Aranday-Cortes, Kirstyn Brunker, Yasmin Parr, Kyriaki Nomikou; Sarah McDonald, Marc Niebel, Patawee Asamaphan; Richard Orton, Joseph Hughes, Sreenu Vattipally, David L Robertson; Alasdair MacLean, Rory Gunson; Kathy Li, Natasha Jesudason, Rajiv Shah, James Shepherd, Antonia Ho, Alice Broos, Emma Thomson and Alex Alderton, Roberto Amato, Sonia Goncalves, Ewan Harrison, David K. Jackson, Ian Johnston, Dominic Kwiatkowski, Cordelia Langford, John Sillitoe |
| EPI_ISL_532989 | Lighthouse Lab in Glasgow | Wellcome Sanger Institute for the COVID-19 Genomics UK (COG-UK) consortium | Harper VanSteenhouse, Yumi Kasai, David Gray, Carol Clugston, Anna Dominiczak and Alex Alderton, Roberto Amato, Sonia Goncalves, Ewan Harrison, David K. Jackson, Ian Johnston, Dominic Kwiatkowski, Cordelia Langford, John Sillitoe |
| EPI_ISL_532990 | Virology Department, Royal Infirmary of Edinburgh, NHS Lothian / School of Biological Sciences, University of Edinburgh | Wellcome Sanger Institute for the COVID-19 Genomics UK (COG-UK) consortium | McHugh M, Dewar R, Rooke S, O'Toole Á, Scher E, Hill V, McCrone JT, Colquhoun R, Yu X, Jackson B, Rambaut A, Templeton K and Alex Alderton, Roberto Amato, Sonia Goncalves, Ewan Harrison, David K. Jackson, Ian Johnston, Dominic Kwiatkowski, Cordelia Langford, John Sillitoe |
| EPI_ISL_532991 | Lighthouse Lab in Glasgow | Wellcome Sanger Institute for the COVID-19 Genomics UK (COG-UK) consortium | Harper VanSteenhouse, Yumi Kasai, David Gray, Carol Clugston, Anna Dominiczak and Alex Alderton, Roberto Amato, Sonia Goncalves, Ewan Harrison, David K. Jackson, Ian Johnston, Dominic Kwiatkowski, Cordelia Langford, John Sillitoe |
| EPI_ISL_532992 | Virology Department, Royal Infirmary of Edinburgh, NHS Lothian / School of Biological Sciences, University of Edinburgh | Wellcome Sanger Institute for the COVID-19 Genomics UK (COG-UK) consortium | McHugh M, Dewar R, Rooke S, O'Toole Á, Scher E, Hill V, McCrone JT, Colquhoun R, Yu X, Jackson B, Rambaut A, Templeton K and Alex Alderton, Roberto Amato, Sonia Goncalves, Ewan Harrison, David K. Jackson, Ian Johnston, Dominic Kwiatkowski, Cordelia Langford, John Sillitoe |
| EPI_ISL_532993 | Lighthouse Lab in Glasgow | Wellcome Sanger Institute for the COVID-19 Genomics UK (COG-UK) consortium | Harper VanSteenhouse, Yumi Kasai, David Gray, Carol Clugston, Anna Dominiczak and Alex Alderton, Roberto Amato, Sonia Goncalves, Ewan Harrison, David K. Jackson, Ian Johnston, Dominic Kwiatkowski, Cordelia Langford, John Sillitoe |
| EPI_ISL_532994 | Virology Department, Royal Infirmary of Edinburgh, NHS Lothian / School of Biological Sciences, University of Edinburgh | Wellcome Sanger Institute for the COVID-19 Genomics UK (COG-UK) consortium | McHugh M, Dewar R, Rooke S, O'Toole Á, Scher E, Hill V, McCrone JT, Colquhoun R, Yu X, Jackson B, Rambaut A, Templeton K and Alex Alderton, Roberto Amato, Sonia Goncalves, Ewan Harrison, David K. Jackson, Ian Johnston, Dominic Kwiatkowski, Cordelia Langford, John Sillitoe |
| EPI_ISL_532995, EPI_ISL_532996, EPI_ISL_532997, EPI_ISL_532998, EPI_ISL_532999 | Lighthouse Lab in Glasgow | Wellcome Sanger Institute for the COVID-19 Genomics UK (COG-UK) consortium | Harper VanSteenhouse, Yumi Kasai, David Gray, Carol Clugston, Anna Dominiczak and Alex Alderton, Roberto Amato, Sonia Goncalves, Ewan Harrison, David K. Jackson, Ian Johnston, Dominic Kwiatkowski, Cordelia Langford, John Sillitoe |
| EPI_ISL_533000 | NHSGGC West of Scotland Specialist Virology Centre / MRC-University of Glasgow Centre for Virus Research | Wellcome Sanger Institute for the COVID-19 Genomics UK (COG-UK) consortium | Ana da Silva Filipe, Natasha Johnson, Kathy Smollett, Daniel Mair, Stephen Carmichael, Lily Tong, Jenna Nichols, Elihu Aranday-Cortes, Kirstyn Brunker, Yasmin Parr, Kyriaki Nomikou; Sarah McDonald, Marc Niebel, Patawee Asamaphan; Richard Orton, Joseph Hughes, Sreenu Vattipally, David L Robertson; Alasdair MacLean, Rory Gunson; Kathy Li, Natasha Jesudason, Rajiv Shah, James Shepherd, Antonia Ho, Alice Broos, Emma Thomson and Alex Alderton, Roberto Amato, Sonia Goncalves, Ewan Harrison, David K. Jackson, Ian Johnston, Dominic Kwiatkowski, Cordelia Langford, John Sillitoe |
| EPI_ISL_533001, EPI_ISL_533002, EPI_ISL_533003, EPI_ISL_533004, EPI_ISL_533005, EPI_ISL_533006, EPI_ISL_533007, EPI_ISL_533008, EPI_ISL_533009, EPI_ISL_533011, EPI_ISL_533012, EPI_ISL_533013, EPI_ISL_533014, EPI_ISL_533015 | | | |
| see above | Lighthouse Lab in Glasgow | Wellcome Sanger Institute for the COVID-19 Genomics UK (COG-UK) consortium | Harper VanSteenhouse, Yumi Kasai, David Gray, Carol Clugston, Anna Dominiczak and Alex Alderton, Roberto Amato, Sonia Goncalves, Ewan Harrison, David K. Jackson, Ian Johnston, Dominic Kwiatkowski, Cordelia Langford, John Sillitoe |
| EPI_ISL_533016 | NHSGGC West of Scotland Specialist Virology Centre / MRC-University of Glasgow Centre for Virus Research | Wellcome Sanger Institute for the COVID-19 Genomics UK (COG-UK) consortium | Ana da Silva Filipe, Natasha Johnson, Kathy Smollett, Daniel Mair, Stephen Carmichael, Lily Tong, Jenna Nichols, Elihu Aranday-Cortes, Kirstyn Brunker, Yasmin Parr, Kyriaki Nomikou; Sarah McDonald, Marc Niebel, Patawee Asamaphan; Richard Orton, Joseph Hughes, Sreenu Vattipally, David L Robertson; Alasdair MacLean, Rory Gunson; Kathy Li, Natasha Jesudason, Rajiv Shah, James Shepherd, Antonia Ho, Alice Broos, Emma Thomson and Alex Alderton, Roberto Amato, Sonia Goncalves, Ewan Harrison, David K. Jackson, Ian Johnston, Dominic Kwiatkowski, Cordelia Langford, John Sillitoe |
| EPI_ISL_533017, EPI_ISL_533018, EPI_ISL_533019, EPI_ISL_533020, EPI_ISL_533021, EPI_ISL_533022, EPI_ISL_533023, EPI_ISL_533024, EPI_ISL_533025, EPI_ISL_533026 | Lighthouse Lab in Glasgow | Wellcome Sanger Institute for the COVID-19 Genomics UK (COG-UK) consortium | Harper VanSteenhouse, Yumi Kasai, David Gray, Carol Clugston, Anna Dominiczak and Alex Alderton, Roberto Amato, Sonia Goncalves, Ewan Harrison, David K. Jackson, Ian Johnston, Dominic Kwiatkowski, Cordelia Langford, John Sillitoe |
| EPI_ISL_533027 | NHSGGC West of Scotland Specialist Virology Centre / MRC-University of Glasgow Centre for Virus Research | Wellcome Sanger Institute for the COVID-19 Genomics UK (COG-UK) consortium | Ana da Silva Filipe, Natasha Johnson, Kathy Smollett, Daniel Mair, Stephen Carmichael, Lily Tong, Jenna Nichols, Elihu Aranday-Cortes, Kirstyn Brunker, Yasmin Parr, Kyriaki Nomikou; Sarah McDonald, Marc Niebel, Patawee Asamaphan; Richard Orton, Joseph Hughes, Sreenu Vattipally, David L Robertson; Alasdair MacLean, Rory Gunson; Kathy Li, Natasha Jesudason, Rajiv Shah, James Shepherd, Antonia Ho, Alice Broos, Emma Thomson and Alex Alderton, Roberto Amato, Sonia Goncalves, Ewan Harrison, David K. Jackson, Ian Johnston, Dominic Kwiatkowski, Cordelia Langford, John Sillitoe |
| EPI_ISL_533028, EPI_ISL_533029, EPI_ISL_533030, EPI_ISL_533031, EPI_ISL_533033, EPI_ISL_533034, EPI_ISL_533035, EPI_ISL_533036, EPI_ISL_533037, EPI_ISL_533038, EPI_ISL_533039, EPI_ISL_533040, EPI_ISL_533041, EPI_ISL_533043, EPI_ISL_533044, EPI_ISL_533045, EPI_ISL_533046, EPI_ISL_533047, EPI_ISL_533048, EPI_ISL_533049, EPI_ISL_533050, EPI_ISL_533051, EPI_ISL_533052, EPI_ISL_533053, EPI_ISL_533054, EPI_ISL_533055, EPI_ISL_533056, EPI_ISL_533057, EPI_ISL_533058, EPI_ISL_533060, EPI_ISL_533061, EPI_ISL_533062, EPI_ISL_533063, EPI_ISL_533064, EPI_ISL_533065, EPI_ISL_533066, EPI_ISL_533067, EPI_ISL_533068, EPI_ISL_533069, EPI_ISL_533070, EPI_ISL_533071 | | | |
| see above | Lighthouse Lab in Glasgow | Wellcome Sanger Institute for the COVID-19 Genomics UK (COG-UK) consortium | Harper VanSteenhouse, Yumi Kasai, David Gray, Carol Clugston, Anna Dominiczak and Alex Alderton, Roberto Amato, Sonia Goncalves, Ewan Harrison, David K. Jackson, Ian Johnston, Dominic Kwiatkowski, Cordelia Langford, John Sillitoe |
| EPI_ISL_533072 | NHSGGC West of Scotland Specialist Virology Centre / MRC-University of Glasgow Centre for Virus Research | Wellcome Sanger Institute for the COVID-19 Genomics UK (COG-UK) consortium | Ana da Silva Filipe, Natasha Johnson, Kathy Smollett, Daniel Mair, Stephen Carmichael, Lily Tong, Jenna Nichols, Elihu Aranday-Cortes, Kirstyn Brunker, Yasmin Parr, Kyriaki Nomikou; Sarah McDonald, Marc Niebel, Patawee Asamaphan; Richard Orton, Joseph Hughes, Sreenu Vattipally, David L Robertson; Alasdair MacLean, Rory Gunson; Kathy Li, Natasha Jesudason, Rajiv Shah, James Shepherd, Antonia Ho, Alice Broos, Emma Thomson and Alex Alderton, Roberto Amato, Sonia Goncalves, Ewan Harrison, David K. Jackson, Ian Johnston, Dominic Kwiatkowski, Cordelia Langford, John Sillitoe |
| EPI_ISL_533073, EPI_ISL_533074, EPI_ISL_533075, EPI_ISL_533076, EPI_ISL_533077, EPI_ISL_533078, EPI_ISL_533079, EPI_ISL_533080, EPI_ISL_533081, EPI_ISL_533082, EPI_ISL_533083, EPI_ISL_533085 | | | |
| see above | Lighthouse Lab in Glasgow | Wellcome Sanger Institute for the COVID-19 Genomics UK (COG-UK) consortium | Harper VanSteenhouse, Yumi Kasai, David Gray, Carol Clugston, Anna Dominiczak and Alex Alderton, Roberto Amato, Sonia Goncalves, Ewan Harrison, David K. Jackson, Ian Johnston, Dominic Kwiatkowski, Cordelia Langford, John Sillitoe |
| EPI_ISL_533086 | Lighthouse Lab in Glasgow | Wellcome Sanger Institute for the COVID-19 Genomics UK (COG-UK) Consortium | Harper VanSteenhouse, Yumi Kasai, David Gray, Carol Clugston, Anna Dominiczak and Alex Alderton, Roberto Amato, Sonia Goncalves, Ewan Harrison, David K. Jackson, Ian Johnston, Dominic Kwiatkowski, Cordelia Langford, John Sillitoe on behalf of the Wellcome Sanger Institute COVID-19 Surveillance Team |
| EPI_ISL_533087, EPI_ISL_533088, EPI_ISL_533089, EPI_ISL_533090, EPI_ISL_533091, EPI_ISL_533092, EPI_ISL_533093, EPI_ISL_533094, EPI_ISL_533095, EPI_ISL_533096, EPI_ISL_533097, EPI_ISL_533098, EPI_ISL_533100, EPI_ISL_533101, EPI_ISL_533102, EPI_ISL_533103, EPI_ISL_533105, EPI_ISL_533106, EPI_ISL_533107, EPI_ISL_533108, EPI_ISL_533109, EPI_ISL_533110, EPI_ISL_533111, EPI_ISL_533112, EPI_ISL_533114, EPI_ISL_533115, EPI_ISL_533116, EPI_ISL_533117, EPI_ISL_533118, EPI_ISL_533119, EPI_ISL_533120, EPI_ISL_533121, EPI_ISL_533122, EPI_ISL_533123, EPI_ISL_533124, EPI_ISL_533125, | | | |

| | | | |
|---|---|---|---|
| EPI_ISL_533126, EPI_ISL_533127, EPI_ISL_533128, EPI_ISL_533129, EPI_ISL_533130, EPI_ISL_533131, EPI_ISL_533132, EPI_ISL_533133, EPI_ISL_533134, EPI_ISL_533135, EPI_ISL_533136, EPI_ISL_533137, EPI_ISL_533138, EPI_ISL_533139, EPI_ISL_533140, EPI_ISL_533141, EPI_ISL_533142, EPI_ISL_533143, EPI_ISL_533144, EPI_ISL_533145, EPI_ISL_533146, EPI_ISL_533147, EPI_ISL_533148, EPI_ISL_533149, EPI_ISL_533151, EPI_ISL_533152, EPI_ISL_533153, EPI_ISL_533154, EPI_ISL_533155, EPI_ISL_533156, EPI_ISL_533157, EPI_ISL_533158, EPI_ISL_533159, EPI_ISL_533160, EPI_ISL_533161, EPI_ISL_533162, EPI_ISL_533163, EPI_ISL_533164, EPI_ISL_533165, EPI_ISL_533166, EPI_ISL_533167, EPI_ISL_533168, EPI_ISL_533169, EPI_ISL_533170, EPI_ISL_533171, EPI_ISL_533172, EPI_ISL_533173, EPI_ISL_533174, EPI_ISL_533175, EPI_ISL_533176, EPI_ISL_533177, EPI_ISL_533178, EPI_ISL_533179, EPI_ISL_533180, EPI_ISL_533181, EPI_ISL_533182, EPI_ISL_533183, EPI_ISL_533184, EPI_ISL_533185, EPI_ISL_533186, EPI_ISL_533187, EPI_ISL_533188, EPI_ISL_533189, EPI_ISL_533190, EPI_ISL_533191, EPI_ISL_533192, EPI_ISL_533193, EPI_ISL_533194, EPI_ISL_533195, EPI_ISL_533196, EPI_ISL_533197, EPI_ISL_533198, EPI_ISL_533199, EPI_ISL_533200, EPI_ISL_533201, EPI_ISL_533202, EPI_ISL_533203, EPI_ISL_533204, EPI_ISL_533205, EPI_ISL_533206, EPI_ISL_533207, EPI_ISL_533208, EPI_ISL_533209, EPI_ISL_533210, EPI_ISL_533211, EPI_ISL_533212, EPI_ISL_533213, EPI_ISL_533214, EPI_ISL_533216, EPI_ISL_533217, EPI_ISL_533218, EPI_ISL_533219, EPI_ISL_533220, EPI_ISL_533221, EPI_ISL_533222, EPI_ISL_533223, EPI_ISL_533224 | | | |
| see above | Lighthouse Lab in Glasgow | Wellcome Sanger Institute for the COVID-19 Genomics UK (COG-UK) consortium | Harper VanSteenhouse, Yumi Kasai, David Gray, Carol Clugston, Anna Dominiczak and Alex Alderton, Roberto Amato, Sonia Goncalves, Ewan Harrison, David K. Jackson, Ian Johnston, Dominic Kwiatkowski, Cordelia Langford, John Sillitoe |
| EPI_ISL_534224 | Universitetssjukhuset i Linkoping | The Public Health Agency of Sweden | Anna-Malin Linde, Maria Lind Karlberg, Mattias Haukland, Reza Advani, Olov Svartstrom, Oskar Karlsson Lindsjo, Sandra Broddesson, Petra Edquist, Mia Brytting, Anna Risberg, Karin Tegmark-Wisell |
| EPI_ISL_534225, EPI_ISL_534226, EPI_ISL_534227, EPI_ISL_534228, EPI_ISL_534229 | Skanes universitetssjukhus Lund | The Public Health Agency of Sweden | Anna-Malin Linde, Maria Lind Karlberg, Mattias Haukland, Reza Advani, Olov Svartstrom, Oskar Karlsson Lindsjo, Sandra Broddesson, Petra Edquist, Mia Brytting, Anna Risberg, Karin Tegmark-Wisell |
| EPI_ISL_534240 | Norra Alvsborgs lanssjukhus | The Public Health Agency of Sweden | Anna-Malin Linde, Maria Lind Karlberg, Mattias Haukland, Reza Advani, Olov Svartstrom, Oskar Karlsson Lindsjo, Sandra Broddesson, Petra Edquist, Mia Brytting, Anna Risberg, Karin Tegmark-Wisell |
| EPI_ISL_534244 | Sundsvalls sjukhus | The Public Health Agency of Sweden | Anna-Malin Linde, Maria Lind Karlberg, Mattias Haukland, Reza Advani, Olov Svartstrom, Oskar Karlsson Lindsjo, Sandra Broddesson, Petra Edquist, Mia Brytting, Anna Risberg, Karin Tegmark-Wisell |
| EPI_ISL_534245 | Kliniskt mikrobiologiska laboratoriet | The Public Health Agency of Sweden | Anna-Malin Linde, Maria Lind Karlberg, Mattias Haukland, Reza Advani, Olov Svartstrom, Oskar Karlsson Lindsjo, Sandra Broddesson, Petra Edquist, Mia Brytting, Anna Risberg, Karin Tegmark-Wisell |
| EPI_ISL_534337, EPI_ISL_534338, EPI_ISL_534339, EPI_ISL_534340, EPI_ISL_534341, EPI_ISL_534342, EPI_ISL_534343, EPI_ISL_534344, EPI_ISL_534345 | Molecular diagnostic laboratory of Federal Budget Institution of Science "Central Research Institute of Epidemiology" of The Federal Service on Customers' Rights Protection and Human Well-being Surveillance | Group of Genomics and Postgenomic Technologies of Central Research Institute of Epidemilology | Speranskaya AS, Kaptelova VV, Samoilov AE, Valdokhina AV, Bulanenko VP, Bukharina A.Y., Tivanova EV, Shipulina OY, Akimkin VG |
| EPI_ISL_534719 | Respiratory Virus Unit, Microbiology Services Colindale, Public Health England | Respiratory Virus Unit, Microbiology Services Colindale, Public Health England | PHE Covid Sequencing Team |
| EPI_ISL_535040 | Centre for Enzyme Innovation, University of Portsmouth / Translational Research Laboratory, Portsmouth Hospitals NHS Trust | COVID-19 Genomics UK (COG-UK) Consortium | Angela Beckett,Yann Bourgeois,Garry Scarlett,Sharon Glaysher,Scott Elliott,Kelly Bicknell,Robert Impey,Allyson Lloyd,Sarah Wyllie,Ethan Butcher,Anoop Chauhan,Samuel Robson |
| EPI_ISL_535297, EPI_ISL_535298, EPI_ISL_535299, EPI_ISL_535300, EPI_ISL_535301, EPI_ISL_535302, EPI_ISL_535303 | New Mexico Department of Health Scientific Laboratory | New Mexico Department of Health Scientific Laboratory | Ellie Johnson, Anastacia Griego-Fisher, D'Eldra Malone |
| EPI_ISL_535657 | AR Dept. of Health-Public Health Lab | Pathogen Discovery, Respiratory Viruses Branch, Division of Viral Diseases, Centers for Disease Control and Prevention | Yan Li, Jing Zhang, Ying Tao, Krista Queen, Brian Lynch, Anna Uehara, Clinton R. Paden, Rachel Marine, Haibin Wang, Suxiang Tong |
| EPI_ISL_535658, EPI_ISL_535659 | AR Dept. of Health-Public Health Lab | Pathogen Discovery, Respiratory Viruses Branch, Division of Viral Diseases, Centers for Disease Control and Prevention | Brian Lynch, Yan Li, Jing Zhang, Ying Tao, Krista Queen, Anna Uehara, Clinton R. Paden, Rachel Marine, Haibin Wang, Suxiang Tong |
| EPI_ISL_535660 | AR Dept. of Health-Public Health Lab | Pathogen Discovery, Respiratory Viruses Branch, Division of Viral Diseases, Centers for Disease Control and Prevention | Yan Li, Jing Zhang, Ying Tao, Krista Queen, Brian Lynch, Anna Uehara, Clinton R. Paden, Rachel Marine, Haibin Wang, Suxiang Tong |
| EPI_ISL_535661 | AR Dept. of Health-Public Health Lab | Pathogen Discovery, Respiratory Viruses Branch, Division of Viral Diseases, Centers for Disease Control and Prevention | Brian Lynch, Yan Li, Jing Zhang, Ying Tao, Krista Queen, Anna Uehara, Clinton R. Paden, Rachel Marine, Haibin Wang, Suxiang Tong |
| EPI_ISL_536787 | University of Wisconsin-Madison AIDS Vaccine Research Laboratories | University of Wisconsin-Madison AIDS Vaccine Research Laboratories | Gage Moreno, Katarina Braun, et al. AIDS Vaccine Research Laboratories |
| EPI_ISL_537380 | Complejo Hospitalario Universitario de Vigo | SeqCOVID-SPAIN consortium/IBV(CSIC) | Benito Regueiro and SeqCOVID-SPAIN consortium |
| EPI_ISL_539803 | Queen Mary Hospital | Hong Kong Department of Health | Alan K.L. Tsang, Peter C.W. Yip, Edman T.K. Lam, Rickjason C.W. Chan, Dominic N.C. Tsang |
| EPI_ISL_539809, EPI_ISL_539810, EPI_ISL_539811, EPI_ISL_539812 | Asiaworld Expo Command Post | Hong Kong Department of Health | Alan K.L. Tsang, Peter C.W. Yip, Edman T.K. Lam, Rickjason C.W. Chan, Dominic N.C. Tsang |
| EPI_ISL_539813 | Prince of Wales Hospital | Hong Kong Department of Health | Alan K.L. Tsang, Peter C.W. Yip, Edman T.K. Lam, Rickjason C.W. Chan, Dominic N.C. Tsang |
| EPI_ISL_539814 | Tuen Mun Hospital | Hong Kong Department of Health | Alan K.L. Tsang, Peter C.W. Yip, Edman T.K. Lam, Rickjason C.W. Chan, Dominic N.C. Tsang |
| EPI_ISL_539815 | Queen Elizabeth Hospital | Hong Kong Department of Health | Alan K.L. Tsang, Peter C.W. Yip, Edman T.K. Lam, Rickjason C.W. Chan, Dominic N.C. Tsang |
| EPI_ISL_539884 | Omtanken Grimmered | The Public Health Agency of Sweden | Anna-Malin Linde, Maria Lind Karlberg, Oskar Karlsson Lindsjo, Olov Svartstrom, Mattias Haukland, Reza Advani, Sandra Broddesson, Anna Risberg, Theresa Enkirch, Mia Brytting, Karin Tegmark-Wisell |
| EPI_ISL_541079, EPI_ISL_541080 | Hospital de la Santa Creu i Sant Pau. Servicio de Microbiología | SeqCOVID-SPAIN consortium/Institute of Biomedicine of Valencia, IBV-CSIC | Ferran Navarro, Núria Rabella, Elisenda Miró and SeqCOVID-SPAIN consortium |
| EPI_ISL_541096, EPI_ISL_541097, EPI_ISL_541098, EPI_ISL_541099, EPI_ISL_541100, EPI_ISL_541101, EPI_ISL_541102, EPI_ISL_541103, EPI_ISL_541104, EPI_ISL_541105, EPI_ISL_541106, EPI_ISL_541107, EPI_ISL_541108, EPI_ISL_541109, EPI_ISL_541110, EPI_ISL_541111, EPI_ISL_541112, EPI_ISL_541113, EPI_ISL_541114, EPI_ISL_541115, EPI_ISL_541116, EPI_ISL_541117, EPI_ISL_541118, EPI_ISL_541119, EPI_ISL_541120, EPI_ISL_541121, EPI_ISL_541122, EPI_ISL_541123, EPI_ISL_541124, EPI_ISL_541125, EPI_ISL_541126, EPI_ISL_541127, EPI_ISL_541128, EPI_ISL_541129 | | | |
| see above | Servicio de Microbiología. Hospital Universitario Donostia. OSI Donostialdea. Área de Enfermedades Infecciosas, Grupo de Infección Respiratoria y Resistencia Antimicrobiana. Instituto de Investigación Sanitaria Biodonostia | SeqCOVID-SPAIN consortium/Institute of Biomedicine of Valencia, IBV-CSIC | Gustavo Cilla, Milagrosa Montes, Luis Piñeiro, Jose Maria Marimón and SeqCOVID-SPAIN consortium |
| EPI_ISL_541175, EPI_ISL_541176, EPI_ISL_541177, EPI_ISL_541179, EPI_ISL_541251, EPI_ISL_541252, EPI_ISL_541253, EPI_ISL_541254, EPI_ISL_541255, EPI_ISL_541256, EPI_ISL_541257, EPI_ISL_541258, EPI_ISL_541259, EPI_ISL_541260, EPI_ISL_541261, EPI_ISL_541262, EPI_ISL_541263, EPI_ISL_541264, EPI_ISL_541265, EPI_ISL_541266, EPI_ISL_541267, EPI_ISL_541268, EPI_ISL_541269, EPI_ISL_541270, EPI_ISL_541271, EPI_ISL_541272, EPI_ISL_541273, EPI_ISL_541274, EPI_ISL_541275, EPI_ISL_541276, EPI_ISL_541277, EPI_ISL_541278, EPI_ISL_541279, EPI_ISL_541280, EPI_ISL_541281, EPI_ISL_541282, EPI_ISL_541283, EPI_ISL_541284, EPI_ISL_541285, EPI_ISL_541286, EPI_ISL_541287, EPI_ISL_541288, EPI_ISL_541289, EPI_ISL_541290, EPI_ISL_541291, EPI_ISL_541292, EPI_ISL_541293, EPI_ISL_541294, EPI_ISL_541295, EPI_ISL_541296, EPI_ISL_541297, EPI_ISL_541298, EPI_ISL_541299, EPI_ISL_541300, EPI_ISL_541301, EPI_ISL_541302, EPI_ISL_541303, EPI_ISL_541304, EPI_ISL_541305, EPI_ISL_541306, EPI_ISL_541307, EPI_ISL_541308, EPI_ISL_541309, EPI_ISL_541310, EPI_ISL_541311, EPI_ISL_541312, EPI_ISL_541313, EPI_ISL_541314, EPI_ISL_541315, EPI_ISL_541316, EPI_ISL_541317, EPI_ISL_541318, EPI_ISL_541319, EPI_ISL_541320, EPI_ISL_541321, EPI_ISL_541322, EPI_ISL_541323, EPI_ISL_541324, EPI_ISL_541325, EPI_ISL_541326, EPI_ISL_541327, EPI_ISL_541328, EPI_ISL_541329, EPI_ISL_541330 | | | |
| see above | Florida Bureau of Public Health Laboratories, Florida Department of Health | Florida Bureau of Public Health Laboratories, Florida Department of Health | Schmedes,S., Blanton,J. |
| EPI_ISL_541650, EPI_ISL_541652, EPI_ISL_541653, EPI_ISL_541654, EPI_ISL_541660, EPI_ISL_541662 | Laboratory Diagnostic, Veterinary Specialized Institute Kraljevo | Laboratory Diagnostic, Veterinary Specialized Institute Kraljevo | Vidanovic,D., Tesovic,B., Knezevic,A., Jovanovic,T., Jankovic,M., Sekler,M., Banovic Djeri,B., Volkening,J., Afonso,C., Petrovic,T. |
| EPI_ISL_541684, EPI_ISL_541685, EPI_ISL_541686, EPI_ISL_541687, EPI_ISL_541688, EPI_ISL_541689, EPI_ISL_541690, EPI_ISL_541691, EPI_ISL_541692, EPI_ISL_541693, EPI_ISL_541694 | | | |
| see above | National Institute of Virology, NIV Influenza | National Institute of Virology, NIV Influenza | Potdar V |

| | | | |
|---|---|---|---|
| EPI_ISL_541886, EPI_ISL_541887, EPI_ISL_541888, EPI_ISL_541889, EPI_ISL_541890 | Hospital General Universitario Gregorio Marañón | SeqCOVID-SPAIN consortium/IBV(CSIC) | Laura Pérez-Lago, Marta Herranz, Jon Sicilia, Julia Suárez, Pilar Catalán, Patricia Muñoz, Darío García de Viedma and SeqCOVID-SPAIN consortium |
| EPI_ISL_542036, EPI_ISL_542037, EPI_ISL_542038 | New Mexico Department of Health Scientific Laboratory | New Mexico Department of Health Scientific Laboratory | Ellie Johnson, Anastacia Griego-Fisher, D'Eldra Malone |
| EPI_ISL_542990 | TriCore Reference Laboratories | Center for Global Health, University of New Mexico Health Sciences Center | Daryl Domman, Kurt Schwalm, Twila Kunde, Joseph Hicks, Michael Edwards, Darrell Dinwiddie |
| EPI_ISL_544957 | Laverty Pathology | NSW Health Pathology - Institute of Clinical Pathology and Medical Research; Westmead Hospital; University of Sydney | CIDM-PH et al. |
| EPI_ISL_546436 | The National Institute of Public Health | State Veterinary Institute Prague | Nagy,A; Jirincova,H; Novakova,L; Trnka,D; Vecerova,J |
| EPI_ISL_547434, EPI_ISL_547437, EPI_ISL_547444 | Microbiology, Department of Pathology, St. Bernard's Hospital, Gibraltar Health Authority | Respiratory Virus Unit, Microbiology Services Colindale, Public Health England | PHE Covid Sequencing Team, Dr Nicholas Cortes (Gibraltar), Charlotte Gillborn-Jones (Gibraltar) |
| EPI_ISL_547478, EPI_ISL_547513 | Dutch COVID-19 response team | National Institute for Public Health and the Environment (RIVM) | Adam Meijer, Harry Vennema, Jeroen Cremer, Sharon van den Brink, Bas van der Veer, AnneMarie van den Brandt, Florian Zwagemaker, Dennis Schmitz, Chantal Reusken, on behalf of the national COVID-19 response team |
| EPI_ISL_547608, EPI_ISL_547609, EPI_ISL_547610 | Gundersen Clinical Microbiology Laboratory | Kabara Cancer Research Institute | Craig S. Richmond, Paraic A. Kenny |
| EPI_ISL_547611, EPI_ISL_547612, EPI_ISL_547613, EPI_ISL_547614, EPI_ISL_547615, EPI_ISL_547616, EPI_ISL_547617, EPI_ISL_547618, EPI_ISL_547619, EPI_ISL_547620, EPI_ISL_547621, EPI_ISL_547622, EPI_ISL_547623, EPI_ISL_547624, EPI_ISL_547625, EPI_ISL_547626, EPI_ISL_547627, EPI_ISL_547628, EPI_ISL_547629, EPI_ISL_547630, EPI_ISL_547631, EPI_ISL_547632, EPI_ISL_547633, EPI_ISL_547634, EPI_ISL_547635, EPI_ISL_547636, EPI_ISL_547637 | | | |
| see above | Gundersen Molecular Diagnostics Laboratory | Kabara Cancer Research Institute | Craig S. Richmond, Paraic A. Kenny |
| EPI_ISL_547666, EPI_ISL_547667 | Gundersen Clinical Microbiology Laboratory | Kabara Cancer Research Institute | Craig S. Richmond, Paraic A. Kenny |
| EPI_ISL_548140 | Canterbury Health Laboratories | Institute of Environmental Science and Research (ESR) | Xiaoyun Ren, Matt Storey, Nikki Freed, Muhammad Faisal, Jing Wang, Hermes Perez, Anja Werno, Antje van der Linden, Arlo Upton, Chris Mansell, David Hammer, Dragana Drinkovic, Gary McAuliffe, Hana Sofia Andersson, James Ussher, Jill Sherwood, Josh Freeman, Julia Howard, Juliet Elvy, Mary DeAlmeida, Matt Blakiston, Matthew Rogers, Max Bloomfield, Michael Addidle, Michelle Balm, Sally Roberts, Sarah Jefferies, Sharmini Muttaiyah, Susan Morpeth, Susan Taylor, Timothy Blackmore, Vani Sathyendran, Veronica Playle, Virginia Hope, Erasmus Smit, Lauren Jelly, Olin Silander, Joep de Ligt |
| EPI_ISL_548264 | County of Santa Clara Public Health Department | Chan-Zuckerberg Biohub | CZB Cliahub Consortium |
| EPI_ISL_548265 | County of San Luis Obispo Public Health Laboratory | Chan-Zuckerberg Biohub | CZB Cliahub Consortium |
| EPI_ISL_548266, EPI_ISL_548267 | County of Santa Clara Public Health Department | Chan-Zuckerberg Biohub | CZB Cliahub Consortium |
| EPI_ISL_548268, EPI_ISL_548269 | County of San Luis Obispo Public Health Laboratory | Chan-Zuckerberg Biohub | CZB Cliahub Consortium |
| EPI_ISL_548270, EPI_ISL_548271, EPI_ISL_548273, EPI_ISL_548274, EPI_ISL_548275 | Orange County Public Health Laboratory | Chan-Zuckerberg Biohub | CZB Cliahub Consortium |
| EPI_ISL_548277 | County of Santa Clara Public Health Department | Chan-Zuckerberg Biohub | CZB Cliahub Consortium |
| EPI_ISL_548278 | Orange County Public Health Laboratory | Chan-Zuckerberg Biohub | CZB Cliahub Consortium |
| EPI_ISL_548279 | County of Santa Clara Public Health Department | Chan-Zuckerberg Biohub | CZB Cliahub Consortium |
| EPI_ISL_548280, EPI_ISL_548281 | County of San Luis Obispo Public Health Laboratory | Chan-Zuckerberg Biohub | CZB Cliahub Consortium |
| EPI_ISL_548282 | County of Santa Clara Public Health Department | Chan-Zuckerberg Biohub | CZB Cliahub Consortium |
| EPI_ISL_548283 | Orange County Public Health Laboratory | Chan-Zuckerberg Biohub | CZB Cliahub Consortium |
| EPI_ISL_548284 | County of San Luis Obispo Public Health Laboratory | Chan-Zuckerberg Biohub | CZB Cliahub Consortium |
| EPI_ISL_548285 | Orange County Public Health Laboratory | Chan-Zuckerberg Biohub | CZB Cliahub Consortium |
| EPI_ISL_548286 | County of San Luis Obispo Public Health Laboratory | Chan-Zuckerberg Biohub | CZB Cliahub Consortium |
| EPI_ISL_548287 | Orange County Public Health Laboratory | Chan-Zuckerberg Biohub | CZB Cliahub Consortium |
| EPI_ISL_548288 | County of San Luis Obispo Public Health Laboratory | Chan-Zuckerberg Biohub | CZB Cliahub Consortium |
| EPI_ISL_548289 | Orange County Public Health Laboratory | Chan-Zuckerberg Biohub | CZB Cliahub Consortium |
| EPI_ISL_548290 | County of Santa Clara Public Health Department | Chan-Zuckerberg Biohub | CZB Cliahub Consortium |
| EPI_ISL_548291 | Orange County Public Health Laboratory | Chan-Zuckerberg Biohub | CZB Cliahub Consortium |
| EPI_ISL_548292 | County of San Luis Obispo Public Health Laboratory | Chan-Zuckerberg Biohub | CZB Cliahub Consortium |
| EPI_ISL_548293 | Orange County Public Health Laboratory | Chan-Zuckerberg Biohub | CZB Cliahub Consortium |
| EPI_ISL_548294, EPI_ISL_548295 | County of San Luis Obispo Public Health Laboratory | Chan-Zuckerberg Biohub | CZB Cliahub Consortium |
| EPI_ISL_548296, EPI_ISL_548297, EPI_ISL_548298 | Orange County Public Health Laboratory | Chan-Zuckerberg Biohub | CZB Cliahub Consortium |
| EPI_ISL_548299 | County of Santa Clara Public Health Department | Chan-Zuckerberg Biohub | CZB Cliahub Consortium |
| EPI_ISL_548300, EPI_ISL_548301 | Orange County Public Health Laboratory | Chan-Zuckerberg Biohub | CZB Cliahub Consortium |
| EPI_ISL_548302, EPI_ISL_548303 | County of Santa Clara Public Health Department | Chan-Zuckerberg Biohub | CZB Cliahub Consortium |
| EPI_ISL_548304 | County of San Luis Obispo Public Health Laboratory | Chan-Zuckerberg Biohub | CZB Cliahub Consortium |
| EPI_ISL_548306 | Orange County Public Health Laboratory | Chan-Zuckerberg Biohub | CZB Cliahub Consortium |
| EPI_ISL_548307 | County of Santa Clara Public Health Department | Chan-Zuckerberg Biohub | CZB Cliahub Consortium |
| EPI_ISL_548308, EPI_ISL_548309 | Orange County Public Health Laboratory | Chan-Zuckerberg Biohub | CZB Cliahub Consortium |
| EPI_ISL_548310 | County of Santa Clara Public Health Department | Chan-Zuckerberg Biohub | CZB Cliahub Consortium |
| EPI_ISL_548311 | County of San Luis Obispo Public Health Laboratory | Chan-Zuckerberg Biohub | CZB Cliahub Consortium |
| EPI_ISL_548312 | Orange County Public Health Laboratory | Chan-Zuckerberg Biohub | CZB Cliahub Consortium |
| EPI_ISL_548314 | County of Santa Clara Public Health Department | Chan-Zuckerberg Biohub | CZB Cliahub Consortium |
| EPI_ISL_548315 | Orange County Public Health Laboratory | Chan-Zuckerberg Biohub | CZB Cliahub Consortium |
| EPI_ISL_548316 | County of San Luis Obispo Public Health Laboratory | Chan-Zuckerberg Biohub | CZB Cliahub Consortium |
| EPI_ISL_548318 | Orange County Public Health Laboratory | Chan-Zuckerberg Biohub | CZB Cliahub Consortium |

| | | | |
|---|---|---|---|
| EPI_ISL_548319, EPI_ISL_548320, EPI_ISL_548321 | County of Santa Clara Public Health Department | Chan-Zuckerberg Biohub | CZB Cliahub Consortium |
| EPI_ISL_548322 | County of San Luis Obispo Public Health Laboratory | Chan-Zuckerberg Biohub | CZB Cliahub Consortium |
| EPI_ISL_548323 | Orange County Public Health Laboratory | Chan-Zuckerberg Biohub | CZB Cliahub Consortium |
| EPI_ISL_548324 | County of Santa Clara Public Health Department | Chan-Zuckerberg Biohub | CZB Cliahub Consortium |
| EPI_ISL_548325 | County of San Luis Obispo Public Health Laboratory | Chan-Zuckerberg Biohub | CZB Cliahub Consortium |
| EPI_ISL_548326 | Orange County Public Health Laboratory | Chan-Zuckerberg Biohub | CZB Cliahub Consortium |
| EPI_ISL_548327 | County of Santa Clara Public Health Department | Chan-Zuckerberg Biohub | CZB Cliahub Consortium |
| EPI_ISL_548328 | Orange County Public Health Laboratory | Chan-Zuckerberg Biohub | CZB Cliahub Consortium |
| EPI_ISL_548329, EPI_ISL_548330 | County of San Luis Obispo Public Health Laboratory | Chan-Zuckerberg Biohub | CZB Cliahub Consortium |
| EPI_ISL_548331 | County of Santa Clara Public Health Department | Chan-Zuckerberg Biohub | CZB Cliahub Consortium |
| EPI_ISL_548332 | Orange County Public Health Laboratory | Chan-Zuckerberg Biohub | CZB Cliahub Consortium |
| EPI_ISL_548333 | County of Santa Clara Public Health Department | Chan-Zuckerberg Biohub | CZB Cliahub Consortium |
| EPI_ISL_548334 | Orange County Public Health Laboratory | Chan-Zuckerberg Biohub | CZB Cliahub Consortium |
| EPI_ISL_548335, EPI_ISL_548336 | County of San Luis Obispo Public Health Laboratory | Chan-Zuckerberg Biohub | CZB Cliahub Consortium |
| EPI_ISL_548345 | County of Santa Clara Public Health Department | Chan-Zuckerberg Biohub | CZB Cliahub Consortium |
| EPI_ISL_548363 | Tuolumne County Public Health | Chan-Zuckerberg Biohub | CZB Cliahub Consortium |
| EPI_ISL_548371 | County of Santa Clara Public Health Department | Chan-Zuckerberg Biohub | CZB Cliahub Consortium |
| EPI_ISL_548373 | Orange County Public Health Laboratory | Chan-Zuckerberg Biohub | CZB Cliahub Consortium |
| EPI_ISL_548374 | University of California, Davis | Chan-Zuckerberg Biohub | CZB Cliahub Consortium |
| EPI_ISL_548377 | Orange County Public Health Laboratory | Chan-Zuckerberg Biohub | CZB Cliahub Consortium |
| EPI_ISL_548379 | County of Santa Clara Public Health Department | Chan-Zuckerberg Biohub | CZB Cliahub Consortium |
| EPI_ISL_548380 | University of California, Davis | Chan-Zuckerberg Biohub | CZB Cliahub Consortium |
| EPI_ISL_548384, EPI_ISL_548385 | County of Santa Clara Public Health Department | Chan-Zuckerberg Biohub | CZB Cliahub Consortium |
| EPI_ISL_548386 | Orange County Public Health Laboratory | Chan-Zuckerberg Biohub | CZB Cliahub Consortium |
| EPI_ISL_548388, EPI_ISL_548392, EPI_ISL_548394, EPI_ISL_548395, EPI_ISL_548396 | County of Santa Clara Public Health Department | Chan-Zuckerberg Biohub | CZB Cliahub Consortium |
| EPI_ISL_548397 | University of California, Davis | Chan-Zuckerberg Biohub | CZB Cliahub Consortium |
| EPI_ISL_548399, EPI_ISL_548401, EPI_ISL_548403 | County of Santa Clara Public Health Department | Chan-Zuckerberg Biohub | CZB Cliahub Consortium |
| EPI_ISL_548404, EPI_ISL_548408 | Orange County Public Health Laboratory | Chan-Zuckerberg Biohub | CZB Cliahub Consortium |
| EPI_ISL_548414 | University of California, Davis | Chan-Zuckerberg Biohub | CZB Cliahub Consortium |
| EPI_ISL_548415 | County of Santa Clara Public Health Department | Chan-Zuckerberg Biohub | CZB Cliahub Consortium |
| EPI_ISL_548416, EPI_ISL_548418 | Orange County Public Health Laboratory | Chan-Zuckerberg Biohub | CZB Cliahub Consortium |
| EPI_ISL_548419, EPI_ISL_548421, EPI_ISL_548422, EPI_ISL_548423, EPI_ISL_548426, EPI_ISL_548429, EPI_ISL_548433 | County of Santa Clara Public Health Department | Chan-Zuckerberg Biohub | CZB Cliahub Consortium |
| EPI_ISL_548435 | University of California, Davis | Chan-Zuckerberg Biohub | CZB Cliahub Consortium |
| EPI_ISL_548437 | County of Santa Clara Public Health Department | Chan-Zuckerberg Biohub | CZB Cliahub Consortium |
| EPI_ISL_548438, EPI_ISL_548439, EPI_ISL_548444 | Orange County Public Health Laboratory | Chan-Zuckerberg Biohub | CZB Cliahub Consortium |
| EPI_ISL_548446, EPI_ISL_548448 | County of Santa Clara Public Health Department | Chan-Zuckerberg Biohub | CZB Cliahub Consortium |
| EPI_ISL_548449 | Orange County Public Health Laboratory | Chan-Zuckerberg Biohub | CZB Cliahub Consortium |
| EPI_ISL_548451 | County of Santa Clara Public Health Department | Chan-Zuckerberg Biohub | CZB Cliahub Consortium |
| EPI_ISL_548460, EPI_ISL_548464 | Orange County Public Health Laboratory | Chan-Zuckerberg Biohub | CZB Cliahub Consortium |
| EPI_ISL_548466, EPI_ISL_548468, EPI_ISL_548472 | County of Santa Clara Public Health Department | Chan-Zuckerberg Biohub | CZB Cliahub Consortium |
| EPI_ISL_548474, EPI_ISL_548477 | University of California, Davis | Chan-Zuckerberg Biohub | CZB Cliahub Consortium |
| EPI_ISL_548478 | County of Santa Clara Public Health Department | Chan-Zuckerberg Biohub | CZB Cliahub Consortium |
| EPI_ISL_548479 | University of California, Davis | Chan-Zuckerberg Biohub | CZB Cliahub Consortium |
| EPI_ISL_548482 | County of Santa Clara Public Health Department | Chan-Zuckerberg Biohub | CZB Cliahub Consortium |
| EPI_ISL_548486 | University of California, Davis | Chan-Zuckerberg Biohub | CZB Cliahub Consortium |
| EPI_ISL_548494, EPI_ISL_548496, EPI_ISL_548497, EPI_ISL_548498, EPI_ISL_548500, EPI_ISL_548503, EPI_ISL_548509, EPI_ISL_548511, EPI_ISL_548512, EPI_ISL_548514, EPI_ISL_548518, EPI_ISL_548523, EPI_ISL_548525, EPI_ISL_548526, EPI_ISL_548529, EPI_ISL_548530, EPI_ISL_548533, EPI_ISL_548536, EPI_ISL_548543 | | | |
| see above | County of Santa Clara Public Health Department | Chan-Zuckerberg Biohub | CZB Cliahub Consortium |
| EPI_ISL_548545 | Contra Costa Public Health Lab | Chan-Zuckerberg Biohub | CZB Cliahub Consortium |
| EPI_ISL_548546, EPI_ISL_548547, EPI_ISL_548550, EPI_ISL_548551, EPI_ISL_548553, EPI_ISL_548559, EPI_ISL_548562, EPI_ISL_548563 | County of Santa Clara Public Health Department | Chan-Zuckerberg Biohub | CZB Cliahub Consortium |
| EPI_ISL_548567 | Contra Costa Public Health Lab | Chan-Zuckerberg Biohub | CZB Cliahub Consortium |

| | | | |
|---|---|---|---|
| EPI_ISL_548570, EPI_ISL_548576, EPI_ISL_548578 | County of Santa Clara Public Health Department | Chan-Zuckerberg Biohub | CZB Cliahub Consortium |
| EPI_ISL_548603, EPI_ISL_548631, EPI_ISL_548644, EPI_ISL_548663, EPI_ISL_548671 | County of San Luis Obispo Public Health Laboratory | Chan-Zuckerberg Biohub | CZB Cliahub Consortium |
| EPI_ISL_549093, EPI_ISL_549097, EPI_ISL_549102, EPI_ISL_549109, EPI_ISL_549114, EPI_ISL_549116 | Ostfold Hospital Trust - Kalnes, Centre for Laboratory Medicine, Section for gene technology and infection serology | Norwegian Institute of Public Health, Department of Virology | Kathrine Stene-Johansen, Kamilla Heddeland Instefjord, Hilde Elshaug, Rasmus Riis Kopperud, Hilde Synnøve Vollan, Karoline Bragstad, Olav Hungnes |
| EPI_ISL_549132, EPI_ISL_549168 | Furst Medical Laboratory | Norwegian Institute of Public Health, Department of Virology | Kathrine Stene-Johansen, Kamilla Heddeland Instefjord, Hilde Elshaug, Rasmus Riis Kopperud, Hilde Synnøve Vollan, Karoline Bragstad, Olav Hungnes |
| EPI_ISL_549169, EPI_ISL_549170, EPI_ISL_549171 | Akershus University Hospital, Department for Microbiology and Infectious Disease Control | Norwegian Institute of Public Health, Department of Virology | Kathrine Stene-Johansen, Kamilla Heddeland Instefjord, Hilde Elshaug, Rasmus Riis Kopperud, Hilde Synnøve Vollan, Karoline Bragstad, Olav Hungnes |
| EPI_ISL_549172 | Unilabs Laboratory Medicine | Norwegian Institute of Public Health, Department of Virology | Kathrine Stene-Johansen, Kamilla Heddeland Instefjord, Hilde Elshaug, Rasmus Riis Kopperud, Hilde Synnøve Vollan, Karoline Bragstad, Olav Hungnes |
| EPI_ISL_550442, EPI_ISL_552664 | Lighthouse Lab in Milton Keynes | Wellcome Sanger Institute for the COVID-19 Genomics UK (COG-UK) consortium | The Lighthouse Lab in Milton Keynes and Alex Alderton, Roberto Amato, Sonia Goncalves, Ewan Harrison, David K. Jackson, Ian Johnston, Dominic Kwiatkowski, Cordelia Langford, John Sillitoe on behalf of the Wellcome Sanger Institute COVID-19 Surveillance Team (http://www.sanger.ac.uk/covid-team) |
| EPI_ISL_555844, EPI_ISL_556517, EPI_ISL_556518, EPI_ISL_556519, EPI_ISL_556520, EPI_ISL_556521, EPI_ISL_556522, EPI_ISL_556523, EPI_ISL_556524, EPI_ISL_556525, EPI_ISL_556526, EPI_ISL_556527, EPI_ISL_556529, EPI_ISL_556533, EPI_ISL_556534, EPI_ISL_556536, EPI_ISL_556537, EPI_ISL_556538, EPI_ISL_556539, EPI_ISL_556540, EPI_ISL_556541, EPI_ISL_556542, EPI_ISL_556543, EPI_ISL_556544, EPI_ISL_556545, EPI_ISL_556547, EPI_ISL_556548, EPI_ISL_556549, EPI_ISL_556550, EPI_ISL_556551, EPI_ISL_556552, EPI_ISL_556553, EPI_ISL_556554, EPI_ISL_556555, EPI_ISL_556556, EPI_ISL_556558, EPI_ISL_556559, EPI_ISL_556560, EPI_ISL_556561, EPI_ISL_556562, EPI_ISL_556564, EPI_ISL_556566, EPI_ISL_556567, EPI_ISL_556568, EPI_ISL_556569, EPI_ISL_556570, EPI_ISL_556571, EPI_ISL_556572, EPI_ISL_556573, EPI_ISL_556574, EPI_ISL_556639, EPI_ISL_556640, EPI_ISL_556641, EPI_ISL_556642, EPI_ISL_556643 | | | |
| see above | Lighthouse Lab in Cambridge | Wellcome Sanger Institute for the COVID-19 Genomics UK (COG-UK) consortium | Rob Howes, The Lighthouse Lab in Cambridge and Alex Alderton, Roberto Amato, Sonia Goncalves, Ewan Harrison, David K. Jackson, Ian Johnston, Dominic Kwiatkowski, Cordelia Langford, John Sillitoe on behalf of the Wellcome Sanger Institute COVID-19 Surveillance Team |
| EPI_ISL_556644 | Lighthouse Lab in Cambridge | Wellcome Sanger Institute for the COVID-19 Genomics UK (COG-UK) consortium | Rob Howes, The Lighthouse Lab in Cambridge and Alex Alderton, Roberto Amato, Sonia Goncalves, Ewan Harrison, David K. Jackson, Ian Johnston, Dominic Kwiatkowski, Cordelia Langford, John Sillitoe on behalf of the Wellcome Sanger Institute COVID-19 Surveillance Team (http://www.sanger.ac.uk/covid-team) |
| EPI_ISL_556645, EPI_ISL_556646, EPI_ISL_556647, EPI_ISL_556648, EPI_ISL_556649, EPI_ISL_556651, EPI_ISL_556652, EPI_ISL_556653, EPI_ISL_556654, EPI_ISL_556655, EPI_ISL_556656, EPI_ISL_556657, EPI_ISL_556658, EPI_ISL_556659, EPI_ISL_556660, EPI_ISL_556661, EPI_ISL_556662, EPI_ISL_556663, EPI_ISL_556664, EPI_ISL_556665, EPI_ISL_556666, EPI_ISL_556667, EPI_ISL_556668, EPI_ISL_556669, EPI_ISL_556670, EPI_ISL_556671, EPI_ISL_556672, EPI_ISL_556673, EPI_ISL_556674, EPI_ISL_556675, EPI_ISL_556676, EPI_ISL_556677, EPI_ISL_556678, EPI_ISL_556679, EPI_ISL_556680, EPI_ISL_556681 | | | |
| see above | Lighthouse Lab in Cambridge | Wellcome Sanger Institute for the COVID-19 Genomics UK (COG-UK) consortium | Rob Howes, The Lighthouse Lab in Cambridge and Alex Alderton, Roberto Amato, Sonia Goncalves, Ewan Harrison, David K. Jackson, Ian Johnston, Dominic Kwiatkowski, Cordelia Langford, John Sillitoe on behalf of the Wellcome Sanger Institute COVID-19 Surveillance Team |
| EPI_ISL_556682 | Lighthouse Lab in Cambridge | Wellcome Sanger Institute for the COVID-19 Genomics UK (COG-UK) consortium | Rob Howes, The Lighthouse Lab in Cambridge and Alex Alderton, Roberto Amato, Sonia Goncalves, Ewan Harrison, David K. Jackson, Ian Johnston, Dominic Kwiatkowski, Cordelia Langford, John Sillitoe on behalf of the Wellcome Sanger Institute COVID-19 Surveillance Team (http://www.sanger.ac.uk/covid-team) |
| EPI_ISL_556683, EPI_ISL_556684, EPI_ISL_556685, EPI_ISL_556686, EPI_ISL_556687, EPI_ISL_556688, EPI_ISL_556689, EPI_ISL_556690, EPI_ISL_556691, EPI_ISL_556692, EPI_ISL_556693, EPI_ISL_556694, EPI_ISL_556695, EPI_ISL_556696, EPI_ISL_556697 | | | |
| see above | Lighthouse Lab in Cambridge | Wellcome Sanger Institute for the COVID-19 Genomics UK (COG-UK) consortium | Rob Howes, The Lighthouse Lab in Cambridge and Alex Alderton, Roberto Amato, Sonia Goncalves, Ewan Harrison, David K. Jackson, Ian Johnston, Dominic Kwiatkowski, Cordelia Langford, John Sillitoe on behalf of the Wellcome Sanger Institute COVID-19 Surveillance Team |
| EPI_ISL_556698 | Lighthouse Lab in Cambridge | Wellcome Sanger Institute for the COVID-19 Genomics UK (COG-UK) Consortium | Rob Howes, The Lighthouse Lab in Cambridge and Alex Alderton, Roberto Amato, Sonia Goncalves, Ewan Harrison, David K. Jackson, Ian Johnston, Dominic Kwiatkowski, Cordelia Langford, John Sillitoe on behalf of the Wellcome Sanger Institute COVID-19 Surveillance Team |
| EPI_ISL_556699, EPI_ISL_556700, EPI_ISL_556701, EPI_ISL_556702, EPI_ISL_556703, EPI_ISL_556704, EPI_ISL_556705, EPI_ISL_556706, EPI_ISL_556707, EPI_ISL_556708, EPI_ISL_556709, EPI_ISL_556710, EPI_ISL_556711, EPI_ISL_556712, EPI_ISL_556713, EPI_ISL_556714, EPI_ISL_556715, EPI_ISL_556716, EPI_ISL_556717, EPI_ISL_556718, EPI_ISL_556719, EPI_ISL_556720, EPI_ISL_556721, EPI_ISL_556722, EPI_ISL_556723, EPI_ISL_556724, EPI_ISL_556725, EPI_ISL_556726, EPI_ISL_556727, EPI_ISL_556728, EPI_ISL_556729, EPI_ISL_556730, EPI_ISL_556731, EPI_ISL_556732, EPI_ISL_556733, EPI_ISL_556734, EPI_ISL_556735, EPI_ISL_556736, EPI_ISL_556737, EPI_ISL_556738, EPI_ISL_556739, EPI_ISL_556740, EPI_ISL_556741, EPI_ISL_556742, EPI_ISL_556743, EPI_ISL_556744, EPI_ISL_556745, EPI_ISL_556746, EPI_ISL_556747, EPI_ISL_556748, EPI_ISL_556749, EPI_ISL_556750 | | | |
| see above | Lighthouse Lab in Cambridge | Wellcome Sanger Institute for the COVID-19 Genomics UK (COG-UK) consortium | Rob Howes, The Lighthouse Lab in Cambridge and Alex Alderton, Roberto Amato, Sonia Goncalves, Ewan Harrison, David K. Jackson, Ian Johnston, Dominic Kwiatkowski, Cordelia Langford, John Sillitoe on behalf of the Wellcome Sanger Institute COVID-19 Surveillance Team |
| EPI_ISL_556751 | Lighthouse Lab in Cambridge | Wellcome Sanger Institute for the COVID-19 Genomics UK (COG-UK) consortium | Rob Howes, The Lighthouse Lab in Cambridge and Alex Alderton, Roberto Amato, Sonia Goncalves, Ewan Harrison, David K. Jackson, Ian Johnston, Dominic Kwiatkowski, Cordelia Langford, John Sillitoe on behalf of the Wellcome Sanger Institute COVID-19 Surveillance Team (http://www.sanger.ac.uk/covid-team) |
| EPI_ISL_556752, EPI_ISL_556753, EPI_ISL_556754, EPI_ISL_556755, EPI_ISL_556756, EPI_ISL_556757, EPI_ISL_556758, EPI_ISL_556759, EPI_ISL_556760 | Lighthouse Lab in Cambridge | Wellcome Sanger Institute for the COVID-19 Genomics UK (COG-UK) consortium | Rob Howes, The Lighthouse Lab in Cambridge and Alex Alderton, Roberto Amato, Sonia Goncalves, Ewan Harrison, David K. Jackson, Ian Johnston, Dominic Kwiatkowski, Cordelia Langford, John Sillitoe on behalf of the Wellcome Sanger Institute COVID-19 Surveillance Team |
| EPI_ISL_556761 | Lighthouse Lab in Cambridge | Wellcome Sanger Institute for the COVID-19 Genomics UK (COG-UK) consortium | Rob Howes, The Lighthouse Lab in Cambridge and Alex Alderton, Roberto Amato, Sonia Goncalves, Ewan Harrison, David K. Jackson, Ian Johnston, Dominic Kwiatkowski, Cordelia Langford, John Sillitoe on behalf of the Wellcome Sanger Institute COVID-19 Surveillance Team (http://www.sanger.ac.uk/covid-team) |
| EPI_ISL_556762, EPI_ISL_556763, EPI_ISL_556764, EPI_ISL_556765, EPI_ISL_556766, EPI_ISL_556767, EPI_ISL_556768, EPI_ISL_556769 | Lighthouse Lab in Cambridge | Wellcome Sanger Institute for the COVID-19 Genomics UK (COG-UK) consortium | Rob Howes, The Lighthouse Lab in Cambridge and Alex Alderton, Roberto Amato, Sonia Goncalves, Ewan Harrison, David K. Jackson, Ian Johnston, Dominic Kwiatkowski, Cordelia Langford, John Sillitoe on behalf of the Wellcome Sanger Institute COVID-19 Surveillance Team |
| EPI_ISL_556770 | Lighthouse Lab in Cambridge | Wellcome Sanger Institute for the COVID-19 Genomics UK (COG-UK) Consortium | Rob Howes, The Lighthouse Lab in Cambridge and Alex Alderton, Roberto Amato, Sonia Goncalves, Ewan Harrison, David K. Jackson, Ian Johnston, Dominic Kwiatkowski, Cordelia Langford, John Sillitoe on behalf of the Wellcome Sanger Institute COVID-19 Surveillance Team |
| EPI_ISL_556771, EPI_ISL_556772, EPI_ISL_556773, EPI_ISL_556774, EPI_ISL_556775, EPI_ISL_556776, EPI_ISL_556777, EPI_ISL_556778, EPI_ISL_556779, EPI_ISL_556780, EPI_ISL_556781, EPI_ISL_556783, EPI_ISL_556784, EPI_ISL_556785, EPI_ISL_556786 | | | |
| see above | Lighthouse Lab in Cambridge | Wellcome Sanger Institute for the COVID-19 Genomics UK (COG-UK) consortium | Rob Howes, The Lighthouse Lab in Cambridge and Alex Alderton, Roberto Amato, Sonia Goncalves, Ewan Harrison, David K. Jackson, Ian Johnston, Dominic Kwiatkowski, Cordelia Langford, John Sillitoe on behalf of the Wellcome Sanger Institute COVID-19 Surveillance Team |
| EPI_ISL_556787 | Lighthouse Lab in Cambridge | Wellcome Sanger Institute for the COVID-19 Genomics UK (COG-UK) consortium | Rob Howes, The Lighthouse Lab in Cambridge and Alex Alderton, Roberto Amato, Sonia Goncalves, Ewan Harrison, David K. Jackson, Ian Johnston, Dominic Kwiatkowski, Cordelia Langford, John Sillitoe on behalf of the Wellcome Sanger Institute COVID-19 Surveillance Team (http://www.sanger.ac.uk/covid-team) |
| EPI_ISL_556788, EPI_ISL_556789, EPI_ISL_556790, EPI_ISL_556791, EPI_ISL_556792, EPI_ISL_556793, EPI_ISL_556794, EPI_ISL_556795, EPI_ISL_556796, EPI_ISL_556797, EPI_ISL_556798, EPI_ISL_556799, EPI_ISL_556800, EPI_ISL_556801, EPI_ISL_556802, EPI_ISL_556803, EPI_ISL_556804, EPI_ISL_556805, EPI_ISL_556806, EPI_ISL_556807, EPI_ISL_556808, EPI_ISL_556809, EPI_ISL_556810, EPI_ISL_556811, EPI_ISL_556812, EPI_ISL_556813, EPI_ISL_556814, EPI_ISL_556815, EPI_ISL_556816, EPI_ISL_556817, EPI_ISL_556818, EPI_ISL_556819, EPI_ISL_556820, EPI_ISL_556821, EPI_ISL_556822, EPI_ISL_556823, EPI_ISL_556824, EPI_ISL_556826, EPI_ISL_556827, EPI_ISL_556830, EPI_ISL_556831, EPI_ISL_556832, EPI_ISL_556834, EPI_ISL_556837, EPI_ISL_556838, EPI_ISL_556840, EPI_ISL_556842, EPI_ISL_556844, EPI_ISL_556846, EPI_ISL_556847, EPI_ISL_556848, EPI_ISL_556852, EPI_ISL_556853, EPI_ISL_556854, EPI_ISL_556857, EPI_ISL_556863, EPI_ISL_556864, EPI_ISL_556870, EPI_ISL_556871, EPI_ISL_556872, EPI_ISL_556876, EPI_ISL_556877, EPI_ISL_556880, EPI_ISL_556881 | | | |
| see above | Lighthouse Lab in Cambridge | Wellcome Sanger Institute for the COVID-19 Genomics UK (COG-UK) consortium | Rob Howes, The Lighthouse Lab in Cambridge and Alex Alderton, Roberto Amato, Sonia Goncalves, Ewan Harrison, David K. Jackson, Ian Johnston, Dominic Kwiatkowski, Cordelia Langford, John Sillitoe on behalf of the Wellcome Sanger Institute COVID-19 Surveillance Team |
| EPI_ISL_556883 | Lighthouse Lab in Cambridge | Wellcome Sanger Institute for the COVID-19 Genomics UK (COG-UK) consortium | Rob Howes, The Lighthouse Lab in Cambridge and Alex Alderton, Roberto Amato, Sonia Goncalves, Ewan Harrison, David K. Jackson, Ian Johnston, Dominic Kwiatkowski, Cordelia Langford, John Sillitoe on behalf of the Wellcome Sanger Institute COVID-19 Surveillance Team (http://www.sanger.ac.uk/covid-team) |
| EPI_ISL_556884, EPI_ISL_556886, | Lighthouse Lab in Cambridge | Wellcome Sanger Institute for the COVID-19 Genomics UK | Rob Howes, The Lighthouse Lab in Cambridge and Alex Alderton, Roberto Amato, Sonia Goncalves, Ewan Harrison, David K. Jackson, Ian Johnston, |

| | | | |
|---|---|---|---|
| EPI_ISL_556888, EPI_ISL_556889, EPI_ISL_556891, EPI_ISL_556892 | | (COG-UK) consortium | Dominic Kwiatkowski, Cordelia Langford, John Sillitoe on behalf of the Wellcome Sanger Institute COVID-19 Surveillance Team |
| EPI_ISL_556894 | Lighthouse Lab in Cambridge | Wellcome Sanger Institute for the COVID-19 Genomics UK (COG-UK) consortium | Rob Howes, The Lighthouse Lab in Cambridge and Alex Alderton, Roberto Amato, Sonia Goncalves, Ewan Harrison, David K. Jackson, Ian Johnston, Dominic Kwiatkowski, Cordelia Langford, John Sillitoe on behalf of the Wellcome Sanger Institute COVID-19 Surveillance Team (http://www.sanger.ac.uk/covid-team) |
| EPI_ISL_556895, EPI_ISL_556897, EPI_ISL_556901, EPI_ISL_556902, EPI_ISL_556903, EPI_ISL_556906, EPI_ISL_556909, EPI_ISL_556912, EPI_ISL_556914, EPI_ISL_556915, EPI_ISL_556916, EPI_ISL_556917 | | | |
| see above | Lighthouse Lab in Cambridge | Wellcome Sanger Institute for the COVID-19 Genomics UK (COG-UK) consortium | Rob Howes, The Lighthouse Lab in Cambridge and Alex Alderton, Roberto Amato, Sonia Goncalves, Ewan Harrison, David K. Jackson, Ian Johnston, Dominic Kwiatkowski, Cordelia Langford, John Sillitoe on behalf of the Wellcome Sanger Institute COVID-19 Surveillance Team |
| EPI_ISL_556918 | Lighthouse Lab in Cambridge | Wellcome Sanger Institute for the COVID-19 Genomics UK (COG-UK) consortium | Rob Howes, The Lighthouse Lab in Cambridge and Alex Alderton, Roberto Amato, Sonia Goncalves, Ewan Harrison, David K. Jackson, Ian Johnston, Dominic Kwiatkowski, Cordelia Langford, John Sillitoe on behalf of the Wellcome Sanger Institute COVID-19 Surveillance Team (http://www.sanger.ac.uk/covid-team) |
| EPI_ISL_556919, EPI_ISL_556920, EPI_ISL_556921, EPI_ISL_556922, EPI_ISL_556923, EPI_ISL_556924, EPI_ISL_556925, EPI_ISL_556926, EPI_ISL_556927, EPI_ISL_556928 | Lighthouse Lab in Cambridge | Wellcome Sanger Institute for the COVID-19 Genomics UK (COG-UK) consortium | Rob Howes, The Lighthouse Lab in Cambridge and Alex Alderton, Roberto Amato, Sonia Goncalves, Ewan Harrison, David K. Jackson, Ian Johnston, Dominic Kwiatkowski, Cordelia Langford, John Sillitoe on behalf of the Wellcome Sanger Institute COVID-19 Surveillance Team |
| EPI_ISL_556929 | Lighthouse Lab in Cambridge | Wellcome Sanger Institute for the COVID-19 Genomics UK (COG-UK) consortium | Rob Howes, The Lighthouse Lab in Cambridge and Alex Alderton, Roberto Amato, Sonia Goncalves, Ewan Harrison, David K. Jackson, Ian Johnston, Dominic Kwiatkowski, Cordelia Langford, John Sillitoe on behalf of the Wellcome Sanger Institute COVID-19 Surveillance Team (http://www.sanger.ac.uk/covid-team) |
| EPI_ISL_556930, EPI_ISL_556931, EPI_ISL_556932, EPI_ISL_556933, EPI_ISL_556934, EPI_ISL_556935, EPI_ISL_556936, EPI_ISL_556937, EPI_ISL_556938, EPI_ISL_556939 | Lighthouse Lab in Cambridge | Wellcome Sanger Institute for the COVID-19 Genomics UK (COG-UK) consortium | Rob Howes, The Lighthouse Lab in Cambridge and Alex Alderton, Roberto Amato, Sonia Goncalves, Ewan Harrison, David K. Jackson, Ian Johnston, Dominic Kwiatkowski, Cordelia Langford, John Sillitoe on behalf of the Wellcome Sanger Institute COVID-19 Surveillance Team |
| EPI_ISL_556940 | Lighthouse Lab in Cambridge | Wellcome Sanger Institute for the COVID-19 Genomics UK (COG-UK) Consortium | Rob Howes, The Lighthouse Lab in Cambridge and Alex Alderton, Roberto Amato, Sonia Goncalves, Ewan Harrison, David K. Jackson, Ian Johnston, Dominic Kwiatkowski, Cordelia Langford, John Sillitoe on behalf of the Wellcome Sanger Institute COVID-19 Surveillance Team |
| EPI_ISL_556941, EPI_ISL_556942, EPI_ISL_556943, EPI_ISL_556944, EPI_ISL_556945, EPI_ISL_556946, EPI_ISL_556947, EPI_ISL_556948, EPI_ISL_556949, EPI_ISL_556950, EPI_ISL_556951, EPI_ISL_556952, EPI_ISL_556953, EPI_ISL_556954, EPI_ISL_556955, EPI_ISL_556956, EPI_ISL_556957, EPI_ISL_556958, EPI_ISL_556959, EPI_ISL_556961, EPI_ISL_556962 | | | |
| see above | Lighthouse Lab in Cambridge | Wellcome Sanger Institute for the COVID-19 Genomics UK (COG-UK) consortium | Rob Howes, The Lighthouse Lab in Cambridge and Alex Alderton, Roberto Amato, Sonia Goncalves, Ewan Harrison, David K. Jackson, Ian Johnston, Dominic Kwiatkowski, Cordelia Langford, John Sillitoe on behalf of the Wellcome Sanger Institute COVID-19 Surveillance Team |
| EPI_ISL_556963 | Lighthouse Lab in Cambridge | Wellcome Sanger Institute for the COVID-19 Genomics UK (COG-UK) consortium | Rob Howes, The Lighthouse Lab in Cambridge and Alex Alderton, Roberto Amato, Sonia Goncalves, Ewan Harrison, David K. Jackson, Ian Johnston, Dominic Kwiatkowski, Cordelia Langford, John Sillitoe on behalf of the Wellcome Sanger Institute COVID-19 Surveillance Team (http://www.sanger.ac.uk/covid-team) |
| EPI_ISL_556964, EPI_ISL_556965, EPI_ISL_556966, EPI_ISL_556967, EPI_ISL_556968, EPI_ISL_556969, EPI_ISL_556970, EPI_ISL_556971, EPI_ISL_556972, EPI_ISL_556973, EPI_ISL_556974, EPI_ISL_556975, EPI_ISL_556976, EPI_ISL_556977, EPI_ISL_556978, EPI_ISL_556979, EPI_ISL_556980, EPI_ISL_556981, EPI_ISL_556982, EPI_ISL_556983 | | | |
| see above | Lighthouse Lab in Cambridge | Wellcome Sanger Institute for the COVID-19 Genomics UK (COG-UK) consortium | Rob Howes, The Lighthouse Lab in Cambridge and Alex Alderton, Roberto Amato, Sonia Goncalves, Ewan Harrison, David K. Jackson, Ian Johnston, Dominic Kwiatkowski, Cordelia Langford, John Sillitoe on behalf of the Wellcome Sanger Institute COVID-19 Surveillance Team |
| EPI_ISL_556984 | Lighthouse Lab in Cambridge | Wellcome Sanger Institute for the COVID-19 Genomics UK (COG-UK) consortium | Rob Howes, The Lighthouse Lab in Cambridge and Alex Alderton, Roberto Amato, Sonia Goncalves, Ewan Harrison, David K. Jackson, Ian Johnston, Dominic Kwiatkowski, Cordelia Langford, John Sillitoe on behalf of the Wellcome Sanger Institute COVID-19 Surveillance Team (http://www.sanger.ac.uk/covid-team) |
| EPI_ISL_557166, EPI_ISL_557167, EPI_ISL_557168, EPI_ISL_557169, EPI_ISL_557170, EPI_ISL_557171, EPI_ISL_557172, EPI_ISL_557173, EPI_ISL_557174, EPI_ISL_557175, EPI_ISL_557176, EPI_ISL_557177, EPI_ISL_557178, EPI_ISL_557179, EPI_ISL_557180, EPI_ISL_557181, EPI_ISL_557182, EPI_ISL_557183, EPI_ISL_557184, EPI_ISL_557185 | | | |
| see above | Lighthouse Lab in Milton Keynes | Wellcome Sanger Institute for the COVID-19 Genomics UK (COG-UK) consortium | The Lighthouse Lab in Milton Keynes and Alex Alderton, Roberto Amato, Sonia Goncalves, Ewan Harrison, David K. Jackson, Ian Johnston, Dominic Kwiatkowski, Cordelia Langford, John Sillitoe on behalf of the Wellcome Sanger Institute COVID-19 Surveillance Team (http://www.sanger.ac.uk/covid-team) |
| EPI_ISL_557186 | Lighthouse Lab in Milton Keynes | Wellcome Sanger Institute for the COVID-19 Genomics UK (COG-UK) Consortium | The Lighthouse Lab in Milton Keynes and Alex Alderton, Roberto Amato, Sonia Goncalves, Ewan Harrison, David K. Jackson, Ian Johnston, Dominic Kwiatkowski, Cordelia Langford, John Sillitoe on behalf of the Wellcome Sanger Institute COVID-19 Surveillance Team |
| EPI_ISL_557187, EPI_ISL_557188 | Lighthouse Lab in Milton Keynes | Wellcome Sanger Institute for the COVID-19 Genomics UK (COG-UK) consortium | The Lighthouse Lab in Milton Keynes and Alex Alderton, Roberto Amato, Sonia Goncalves, Ewan Harrison, David K. Jackson, Ian Johnston, Dominic Kwiatkowski, Cordelia Langford, John Sillitoe on behalf of the Wellcome Sanger Institute COVID-19 Surveillance Team (http://www.sanger.ac.uk/covid-team) |
| EPI_ISL_557189 | Lighthouse Lab in Milton Keynes | Wellcome Sanger Institute for the COVID-19 Genomics UK (COG-UK) Consortium | The Lighthouse Lab in Milton Keynes and Alex Alderton, Roberto Amato, Sonia Goncalves, Ewan Harrison, David K. Jackson, Ian Johnston, Dominic Kwiatkowski, Cordelia Langford, John Sillitoe on behalf of the Wellcome Sanger Institute COVID-19 Surveillance Team |
| EPI_ISL_557190, EPI_ISL_557191, EPI_ISL_557192, EPI_ISL_557193, EPI_ISL_557194, EPI_ISL_557195, EPI_ISL_557196, EPI_ISL_557197, EPI_ISL_557198, EPI_ISL_557199, EPI_ISL_557200, EPI_ISL_557201, EPI_ISL_557202, EPI_ISL_557203, EPI_ISL_557204, EPI_ISL_557205, EPI_ISL_557206, EPI_ISL_557207, EPI_ISL_557208, EPI_ISL_557209, EPI_ISL_557210, EPI_ISL_557211, EPI_ISL_557212 | | | |
| see above | Lighthouse Lab in Milton Keynes | Wellcome Sanger Institute for the COVID-19 Genomics UK (COG-UK) consortium | The Lighthouse Lab in Milton Keynes and Alex Alderton, Roberto Amato, Sonia Goncalves, Ewan Harrison, David K. Jackson, Ian Johnston, Dominic Kwiatkowski, Cordelia Langford, John Sillitoe on behalf of the Wellcome Sanger Institute COVID-19 Surveillance Team (http://www.sanger.ac.uk/covid-team) |
| EPI_ISL_557213 | Lighthouse Lab in Milton Keynes | Wellcome Sanger Institute for the COVID-19 Genomics UK (COG-UK) Consortium | The Lighthouse Lab in Milton Keynes and Alex Alderton, Roberto Amato, Sonia Goncalves, Ewan Harrison, David K. Jackson, Ian Johnston, Dominic Kwiatkowski, Cordelia Langford, John Sillitoe on behalf of the Wellcome Sanger Institute COVID-19 Surveillance Team |
| EPI_ISL_557214, EPI_ISL_557215, EPI_ISL_557216, EPI_ISL_557217, EPI_ISL_557218, EPI_ISL_557219, EPI_ISL_557220, EPI_ISL_557221, EPI_ISL_557222, EPI_ISL_557223, EPI_ISL_557224, EPI_ISL_557225, EPI_ISL_557226, EPI_ISL_557227, EPI_ISL_557228, EPI_ISL_557229, EPI_ISL_557230, EPI_ISL_557231, EPI_ISL_557232, EPI_ISL_557233, EPI_ISL_557234, EPI_ISL_557235, EPI_ISL_557236, EPI_ISL_557237 | | | |
| see above | Lighthouse Lab in Milton Keynes | Wellcome Sanger Institute for the COVID-19 Genomics UK (COG-UK) consortium | The Lighthouse Lab in Milton Keynes and Alex Alderton, Roberto Amato, Sonia Goncalves, Ewan Harrison, David K. Jackson, Ian Johnston, Dominic Kwiatkowski, Cordelia Langford, John Sillitoe on behalf of the Wellcome Sanger Institute COVID-19 Surveillance Team (http://www.sanger.ac.uk/covid-team) |
| EPI_ISL_557238 | Lighthouse Lab in Milton Keynes | Wellcome Sanger Institute for the COVID-19 Genomics UK (COG-UK) Consortium | The Lighthouse Lab in Milton Keynes and Alex Alderton, Roberto Amato, Sonia Goncalves, Ewan Harrison, David K. Jackson, Ian Johnston, Dominic Kwiatkowski, Cordelia Langford, John Sillitoe on behalf of the Wellcome Sanger Institute COVID-19 Surveillance Team |
| EPI_ISL_557239, EPI_ISL_557240, EPI_ISL_557241, EPI_ISL_557242, EPI_ISL_557243, EPI_ISL_557244, EPI_ISL_557245, EPI_ISL_557246, EPI_ISL_557247, EPI_ISL_557248, EPI_ISL_557249, EPI_ISL_557250, EPI_ISL_557251, EPI_ISL_557252, EPI_ISL_557253, EPI_ISL_557254, EPI_ISL_557255, EPI_ISL_557256, EPI_ISL_557257, EPI_ISL_557258, EPI_ISL_557259, EPI_ISL_557260, EPI_ISL_557261, EPI_ISL_557262, EPI_ISL_557263, EPI_ISL_557264, EPI_ISL_557265, EPI_ISL_557266, EPI_ISL_557267, EPI_ISL_557268, EPI_ISL_557269, EPI_ISL_557270, EPI_ISL_557271, EPI_ISL_557272, EPI_ISL_557273, EPI_ISL_557274, EPI_ISL_557275, EPI_ISL_557276, EPI_ISL_557277, EPI_ISL_557278, EPI_ISL_557279, EPI_ISL_557280, EPI_ISL_557281, EPI_ISL_557282, EPI_ISL_557283, EPI_ISL_557284, EPI_ISL_557285, EPI_ISL_557286, EPI_ISL_557287, EPI_ISL_557288, EPI_ISL_557289, EPI_ISL_557290, EPI_ISL_557291, EPI_ISL_557292, EPI_ISL_557293, EPI_ISL_557294, EPI_ISL_557295, EPI_ISL_557296, EPI_ISL_557297, EPI_ISL_557298, EPI_ISL_557299, EPI_ISL_557300, EPI_ISL_557301, EPI_ISL_557302, EPI_ISL_557303, EPI_ISL_557304, EPI_ISL_557305, EPI_ISL_557306, EPI_ISL_557307, EPI_ISL_557308, EPI_ISL_557309, EPI_ISL_557310, EPI_ISL_557311, EPI_ISL_557312, EPI_ISL_557313, EPI_ISL_557314, EPI_ISL_557315, EPI_ISL_557316, EPI_ISL_557317, EPI_ISL_557318, EPI_ISL_557319, EPI_ISL_557320, EPI_ISL_557321, EPI_ISL_557322, EPI_ISL_557323, EPI_ISL_557324, EPI_ISL_557325, EPI_ISL_557326, EPI_ISL_557327, EPI_ISL_557328, EPI_ISL_557329, EPI_ISL_557330, EPI_ISL_557331, EPI_ISL_557332, EPI_ISL_557333, EPI_ISL_557334, EPI_ISL_557335, EPI_ISL_557336 | | | |
| see above | Lighthouse Lab in Milton Keynes | Wellcome Sanger Institute for the COVID-19 Genomics UK (COG-UK) consortium | The Lighthouse Lab in Milton Keynes and Alex Alderton, Roberto Amato, Sonia Goncalves, Ewan Harrison, David K. Jackson, Ian Johnston, Dominic Kwiatkowski, Cordelia Langford, John Sillitoe on behalf of the Wellcome Sanger Institute COVID-19 Surveillance Team (http://www.sanger.ac.uk/covid-team) |
| EPI_ISL_557337 | Lighthouse Lab in Milton Keynes | Wellcome Sanger Institute for the COVID-19 Genomics UK (COG-UK) consortium | The Lighthouse Lab in Alderley Park and Alex Alderton, Roberto Amato, Sonia Goncalves, Ewan Harrison, David K. Jackson, Ian Johnston, Dominic Kwiatkowski, Cordelia Langford, John Sillitoe on behalf of the Wellcome Sanger Institute COVID-19 Surveillance Team |
| EPI_ISL_557338, EPI_ISL_557339, EPI_ISL_557340, EPI_ISL_557341, EPI_ISL_557342, EPI_ISL_557343, EPI_ISL_557344, EPI_ISL_557345, EPI_ISL_557346, EPI_ISL_557347, EPI_ISL_557348, EPI_ISL_557349, EPI_ISL_557350, EPI_ISL_557352, EPI_ISL_557353, EPI_ISL_557358, EPI_ISL_557360, EPI_ISL_557364, | | | |

| | | | |
|---|---|---|---|
| EPI_ISL_557365, EPI_ISL_557372, EPI_ISL_557377, EPI_ISL_557382, EPI_ISL_557384, EPI_ISL_557387, EPI_ISL_557391, EPI_ISL_557393, EPI_ISL_557394, EPI_ISL_557396, EPI_ISL_557397, EPI_ISL_557400, EPI_ISL_557402, EPI_ISL_557413, EPI_ISL_557416, EPI_ISL_557420, EPI_ISL_557421, EPI_ISL_557422, EPI_ISL_557424, EPI_ISL_557427, EPI_ISL_557429, EPI_ISL_557431, EPI_ISL_557432, EPI_ISL_557441, EPI_ISL_557442, EPI_ISL_557443, EPI_ISL_557448, EPI_ISL_557450, EPI_ISL_557451, EPI_ISL_557455, EPI_ISL_557457 | | | |
| see above | Lighthouse Lab in Milton Keynes | Wellcome Sanger Institute for the COVID-19 Genomics UK (COG-UK) consortium | The Lighthouse Lab in Milton Keynes and Alex Alderton, Roberto Amato, Sonia Goncalves, Ewan Harrison, David K. Jackson, Ian Johnston, Dominic Kwiatkowski, Cordelia Langford, John Sillitoe on behalf of the Wellcome Sanger Institute COVID-19 Surveillance Team (http://www.sanger.ac.uk/covid-team) |
| EPI_ISL_557459 | Lighthouse Lab in Milton Keynes | Wellcome Sanger Institute for the COVID-19 Genomics UK (COG-UK) consortium | The Lighthouse Lab in Milton Keynes and Alex Alderton, Roberto Amato, Sonia Goncalves, Ewan Harrison, David K. Jackson, Ian Johnston, Dominic Kwiatkowski, Cordelia Langford, John Sillitoe on behalf of the Wellcome Sanger Institute COVID-19 Surveillance Team |
| EPI_ISL_557460, EPI_ISL_557462, EPI_ISL_557463, EPI_ISL_557464, EPI_ISL_557470, EPI_ISL_557472, EPI_ISL_557473, EPI_ISL_557475, EPI_ISL_557477, EPI_ISL_557479, EPI_ISL_557480, EPI_ISL_557483, EPI_ISL_557486, EPI_ISL_557489, EPI_ISL_557490, EPI_ISL_557494, EPI_ISL_557495, EPI_ISL_557498, EPI_ISL_557501 | | | |
| see above | Lighthouse Lab in Milton Keynes | Wellcome Sanger Institute for the COVID-19 Genomics UK (COG-UK) consortium | The Lighthouse Lab in Milton Keynes and Alex Alderton, Roberto Amato, Sonia Goncalves, Ewan Harrison, David K. Jackson, Ian Johnston, Dominic Kwiatkowski, Cordelia Langford, John Sillitoe on behalf of the Wellcome Sanger Institute COVID-19 Surveillance Team (http://www.sanger.ac.uk/covid-team) |
| EPI_ISL_557504 | Lighthouse Lab in Milton Keynes | Wellcome Sanger Institute for the COVID-19 Genomics UK (COG-UK) Consortium | The Lighthouse Lab in Milton Keynes and Alex Alderton, Roberto Amato, Sonia Goncalves, Ewan Harrison, David K. Jackson, Ian Johnston, Dominic Kwiatkowski, Cordelia Langford, John Sillitoe on behalf of the Wellcome Sanger Institute COVID-19 Surveillance Team |
| EPI_ISL_557506, EPI_ISL_557508, EPI_ISL_557509, EPI_ISL_557510, EPI_ISL_557514, EPI_ISL_557515, EPI_ISL_557517, EPI_ISL_557519, EPI_ISL_557520, EPI_ISL_557524, EPI_ISL_557525, EPI_ISL_557529, EPI_ISL_557534, EPI_ISL_557538, EPI_ISL_557542, EPI_ISL_557547, EPI_ISL_557548, EPI_ISL_557552, EPI_ISL_557782 | | | |
| see above | Lighthouse Lab in Milton Keynes | Wellcome Sanger Institute for the COVID-19 Genomics UK (COG-UK) consortium | The Lighthouse Lab in Milton Keynes and Alex Alderton, Roberto Amato, Sonia Goncalves, Ewan Harrison, David K. Jackson, Ian Johnston, Dominic Kwiatkowski, Cordelia Langford, John Sillitoe on behalf of the Wellcome Sanger Institute COVID-19 Surveillance Team (http://www.sanger.ac.uk/covid-team) |
| EPI_ISL_560127 | Wales Specialist Virology Centre Sequencing lab: Pathogen Genomics Unit | COVID-19 Genomics UK (COG-UK) Consortium | Catherine Moore, Johnathan Evans, Laura Gifford, Malorie Perry, Simon Cottrell, Angela Marchbank, Alec Birchley, Alexander Adams, Amy Gaskin, Bree Gatica-Wilcox, Jason Coombes, Joel Southgate, Lauren Gilbert, Lee Graham, Nicole Pacchiarini, Sara Kumziene-Summerhayes, Sarah Taylor, Sophie Jones, Sara Rey, Matthew Bull, Joanne Watkins, Sally Corden, Tom Connor |
| EPI_ISL_560311, EPI_ISL_560312, EPI_ISL_560313, EPI_ISL_560314, EPI_ISL_560315, EPI_ISL_560316 | UMMC-Health | WHO National Influenza Centre Russian Federation | Andrey Komissarov, Artem Fadeev, Anna Ivanova, Tatiana Platonova, Daria Danilenko |
| EPI_ISL_560321 | CMS, Roorkee | CSIR-Institute of Microbial Technology | Kanika Bansal, Sanjeet Kumar, Anu Singh, Debarghya Ghose, Amandeep Kaur, Rajesh Kumar Mishra, Poushali Chakraborty, Harsh Goar, Navin Baid, Ashwani Kumar, Dipak Dutta, Sanjeev Khosla, Prabhu B. Patil |
| EPI_ISL_560643, EPI_ISL_560645, EPI_ISL_560646 | Hospital | National Reference Center for Viruses of Respiratory Infections, Institut Pasteur, Paris | Sylvie Behillil, Fabiana Gambaro, Etienne Simon-Lorière, Vincent Enouf, Maud Vanpeene, Sylvie van der Werf |
| EPI_ISL_560977, EPI_ISL_560978, EPI_ISL_560979 | Universitetssjukhuset i Linkoping | The Public Health Agency of Sweden | Anna-Malin Linde, Maria Lind Karlberg, Mattias Haukland, Reza Advani, Olov Svartstrom, Oskar Karlsson Lindsjo, Sandra Broddesson, Petra Edquist, Mia Brytting, Anna Risberg, Karin Tegmark-Wisell |
| EPI_ISL_560980 | Skanes universitetssjukhus Lund | The Public Health Agency of Sweden | Anna-Malin Linde, Maria Lind Karlberg, Mattias Haukland, Reza Advani, Olov Svartstrom, Oskar Karlsson Lindsjo, Sandra Broddesson, Petra Edquist, Mia Brytting, Anna Risberg, Karin Tegmark-Wisell |
| EPI_ISL_560986 | Sundsvalls sjukhus | The Public Health Agency of Sweden | Anna-Malin Linde, Maria Lind Karlberg, Mattias Haukland, Reza Advani, Olov Svartstrom, Oskar Karlsson Lindsjo, Sandra Broddesson, Petra Edquist, Mia Brytting, Anna Risberg, Karin Tegmark-Wisell |
| EPI_ISL_560987 | Kliniskt mikrobiologiska laboratoriet | The Public Health Agency of Sweden | Anna-Malin Linde, Maria Lind Karlberg, Mattias Haukland, Reza Advani, Olov Svartstrom, Oskar Karlsson Lindsjo, Sandra Broddesson, Petra Edquist, Mia Brytting, Anna Risberg, Karin Tegmark-Wisell |
| EPI_ISL_561105, EPI_ISL_561106, EPI_ISL_561107, EPI_ISL_561108, EPI_ISL_561109, EPI_ISL_561110, EPI_ISL_561111, EPI_ISL_561112, EPI_ISL_561113, EPI_ISL_561114, EPI_ISL_561115, EPI_ISL_561116, EPI_ISL_561117, EPI_ISL_561118, EPI_ISL_561119, EPI_ISL_561120, EPI_ISL_561121, EPI_ISL_561122, EPI_ISL_561123, EPI_ISL_561124, EPI_ISL_561125, EPI_ISL_561126, EPI_ISL_561127, EPI_ISL_561128, EPI_ISL_561129, EPI_ISL_561130, EPI_ISL_561131, EPI_ISL_561132, EPI_ISL_561133, EPI_ISL_561134, EPI_ISL_561135, EPI_ISL_561136, EPI_ISL_561137, EPI_ISL_561138, EPI_ISL_561139, EPI_ISL_561140, EPI_ISL_561141, EPI_ISL_561142, EPI_ISL_561143, EPI_ISL_561144, EPI_ISL_561145, EPI_ISL_561146, EPI_ISL_561147, EPI_ISL_561148, EPI_ISL_561149, EPI_ISL_561238, EPI_ISL_561240, EPI_ISL_561241, EPI_ISL_561249, EPI_ISL_561252, EPI_ISL_561253, EPI_ISL_561255, EPI_ISL_561256, EPI_ISL_561258, EPI_ISL_561259, EPI_ISL_561260, EPI_ISL_561261, EPI_ISL_561262, EPI_ISL_561263, EPI_ISL_561265, EPI_ISL_561267, EPI_ISL_561269, EPI_ISL_561272, EPI_ISL_561274, EPI_ISL_561276, EPI_ISL_561310 | | | |
| see above | MRCG at LSHTM Genomics lab | MRCG at LSHTM Genomics lab | Abdul Karim sesay, Abdoulie Kanteh, Jarra Manneh, Mariama Kujabi, Bakary Sanyang |
| EPI_ISL_561388 | Victorian Infectious Diseases Reference Laboratory (VIDRL) | VIDRL and MDU-PHL | Caly, L., Seemann, T., Sait, M., Schultz, M. B., Druce J., Sherry, N. |
| EPI_ISL_561415, EPI_ISL_561423 | Microbiological Diagnostic Unit - Public Health Laboratory (MDU-PHL) | MDU-PHL | Seemann, T., Schultz M. B., Sait, M., Sherry, N. |
| EPI_ISL_561425, EPI_ISL_561434, EPI_ISL_561453 | Victorian Infectious Diseases Reference Laboratory (VIDRL) | VIDRL and MDU-PHL | Caly, L., Seemann, T., Sait, M., Schultz, M. B., Druce J., Sherry, N. |
| EPI_ISL_561456 | Microbiological Diagnostic Unit - Public Health Laboratory (MDU-PHL) | MDU-PHL | Seemann, T., Schultz M. B., Sait, M., Sherry, N. |
| EPI_ISL_561458, EPI_ISL_561491 | Victorian Infectious Diseases Reference Laboratory (VIDRL) | VIDRL and MDU-PHL | Caly, L., Seemann, T., Sait, M., Schultz, M. B., Druce J., Sherry, N. |
| EPI_ISL_561509, EPI_ISL_561510, EPI_ISL_561528, EPI_ISL_561542, EPI_ISL_561568, EPI_ISL_561569 | Microbiological Diagnostic Unit - Public Health Laboratory (MDU-PHL) | MDU-PHL | Seemann, T., Schultz M. B., Sait, M., Sherry, N. |
| EPI_ISL_561584 | Victorian Infectious Diseases Reference Laboratory (VIDRL) | VIDRL and MDU-PHL | Caly, L., Seemann, T., Sait, M., Schultz, M. B., Druce J., Sherry, N. |
| EPI_ISL_561605, EPI_ISL_561644 | Microbiological Diagnostic Unit - Public Health Laboratory (MDU-PHL) | MDU-PHL | Seemann, T., Schultz M. B., Sait, M., Sherry, N. |
| EPI_ISL_561686 | Victorian Infectious Diseases Reference Laboratory (VIDRL) | VIDRL and MDU-PHL | Caly, L., Seemann, T., Sait, M., Schultz, M. B., Druce J., Sherry, N. |
| EPI_ISL_561692 | Microbiological Diagnostic Unit - Public Health Laboratory (MDU-PHL) | MDU-PHL | Seemann, T., Schultz M. B., Sait, M., Sherry, N. |
| EPI_ISL_561741 | Victorian Infectious Diseases Reference Laboratory (VIDRL) | VIDRL and MDU-PHL | Caly, L., Seemann, T., Sait, M., Schultz, M. B., Druce J., Sherry, N. |
| EPI_ISL_561757, EPI_ISL_561758 | Microbiological Diagnostic Unit - Public Health Laboratory (MDU-PHL) | MDU-PHL | Seemann, T., Schultz M. B., Sait, M., Sherry, N. |
| EPI_ISL_561765 | Victorian Infectious Diseases Reference Laboratory (VIDRL) | VIDRL and MDU-PHL | Caly, L., Seemann, T., Sait, M., Schultz, M. B., Druce J., Sherry, N. |
| EPI_ISL_561774, EPI_ISL_561790 | Microbiological Diagnostic Unit - Public Health Laboratory (MDU-PHL) | MDU-PHL | Seemann, T., Schultz M. B., Sait, M., Sherry, N. |
| EPI_ISL_561791 | Victorian Infectious Diseases Reference Laboratory (VIDRL) | VIDRL and MDU-PHL | Caly, L., Seemann, T., Sait, M., Schultz, M. B., Druce J., Sherry, N. |
| EPI_ISL_561811, EPI_ISL_561817, EPI_ISL_561821, EPI_ISL_561841, EPI_ISL_561850 | Microbiological Diagnostic Unit - Public Health Laboratory (MDU-PHL) | MDU-PHL | Seemann, T., Schultz M. B., Sait, M., Sherry, N. |
| EPI_ISL_561863, EPI_ISL_561866 | Victorian Infectious Diseases Reference Laboratory (VIDRL) | VIDRL and MDU-PHL | Caly, L., Seemann, T., Sait, M., Schultz, M. B., Druce J., Sherry, N. |
| EPI_ISL_561867, EPI_ISL_561873, EPI_ISL_561883, EPI_ISL_561919, EPI_ISL_561930, EPI_ISL_561933, EPI_ISL_561939, EPI_ISL_561947, EPI_ISL_561952, EPI_ISL_561964, EPI_ISL_561973, EPI_ISL_561996, EPI_ISL_562015 | | | |
| see above | Microbiological Diagnostic Unit - Public Health Laboratory (MDU-PHL) | MDU-PHL | Seemann, T., Schultz M. B., Sait, M., Sherry, N. |

| | | | |
|---|---|---|---|
| EPI_ISL_562033 | Victorian Infectious Diseases Reference Laboratory (VIDRL) | VIDRL and MDU-PHL | Caly, L., Seemann, T., Sait, M., Schultz, M. B., Druce J., Sherry, N. |
| EPI_ISL_562035, EPI_ISL_562044, EPI_ISL_562049, EPI_ISL_562050 | Microbiological Diagnostic Unit - Public Health Laboratory (MDU-PHL) | MDU-PHL | Seemann, T., Schultz M. B., Sait, M., Sherry, N. |
| EPI_ISL_562067 | Victorian Infectious Diseases Reference Laboratory (VIDRL) | VIDRL and MDU-PHL | Caly, L., Seemann, T., Sait, M., Schultz, M. B., Druce J., Sherry, N. |
| EPI_ISL_562091, EPI_ISL_562099, EPI_ISL_562112, EPI_ISL_562126, EPI_ISL_562171 | Microbiological Diagnostic Unit - Public Health Laboratory (MDU-PHL) | MDU-PHL | Seemann, T., Schultz M. B., Sait, M., Sherry, N. |
| EPI_ISL_562179 | Victorian Infectious Diseases Reference Laboratory (VIDRL) | VIDRL and MDU-PHL | Caly, L., Seemann, T., Sait, M., Schultz, M. B., Druce J., Sherry, N. |
| EPI_ISL_562205, EPI_ISL_562223 | Microbiological Diagnostic Unit - Public Health Laboratory (MDU-PHL) | MDU-PHL | Seemann, T., Schultz M. B., Sait, M., Sherry, N. |
| EPI_ISL_562224, EPI_ISL_562228, EPI_ISL_562237 | Victorian Infectious Diseases Reference Laboratory (VIDRL) | VIDRL and MDU-PHL | Caly, L., Seemann, T., Sait, M., Schultz, M. B., Druce J., Sherry, N. |
| EPI_ISL_562240 | Microbiological Diagnostic Unit - Public Health Laboratory (MDU-PHL) | MDU-PHL | Seemann, T., Schultz M. B., Sait, M., Sherry, N. |
| EPI_ISL_562244 | Victorian Infectious Diseases Reference Laboratory (VIDRL) | VIDRL and MDU-PHL | Caly, L., Seemann, T., Sait, M., Schultz, M. B., Druce J., Sherry, N. |
| EPI_ISL_562266, EPI_ISL_562268, EPI_ISL_562271, EPI_ISL_562281 | Microbiological Diagnostic Unit - Public Health Laboratory (MDU-PHL) | MDU-PHL | Seemann, T., Schultz M. B., Sait, M., Sherry, N. |
| EPI_ISL_562282, EPI_ISL_562294, EPI_ISL_562322 | Victorian Infectious Diseases Reference Laboratory (VIDRL) | VIDRL and MDU-PHL | Caly, L., Seemann, T., Sait, M., Schultz, M. B., Druce J., Sherry, N. |
| EPI_ISL_562327, EPI_ISL_562337 | Microbiological Diagnostic Unit - Public Health Laboratory (MDU-PHL) | MDU-PHL | Seemann, T., Schultz M. B., Sait, M., Sherry, N. |
| EPI_ISL_562356 | Victorian Infectious Diseases Reference Laboratory (VIDRL) | VIDRL and MDU-PHL | Caly, L., Seemann, T., Sait, M., Schultz, M. B., Druce J., Sherry, N. |
| EPI_ISL_562360 | Microbiological Diagnostic Unit - Public Health Laboratory (MDU-PHL) | MDU-PHL | Seemann, T., Schultz M. B., Sait, M., Sherry, N. |
| EPI_ISL_562376 | Victorian Infectious Diseases Reference Laboratory (VIDRL) | VIDRL and MDU-PHL | Caly, L., Seemann, T., Sait, M., Schultz, M. B., Druce J., Sherry, N. |
| EPI_ISL_562387, EPI_ISL_562398 | Microbiological Diagnostic Unit - Public Health Laboratory (MDU-PHL) | MDU-PHL | Seemann, T., Schultz M. B., Sait, M., Sherry, N. |
| EPI_ISL_562406, EPI_ISL_562412 | Victorian Infectious Diseases Reference Laboratory (VIDRL) | VIDRL and MDU-PHL | Caly, L., Seemann, T., Sait, M., Schultz, M. B., Druce J., Sherry, N. |
| EPI_ISL_562439, EPI_ISL_562528 | Microbiological Diagnostic Unit - Public Health Laboratory (MDU-PHL) | MDU-PHL | Seemann, T., Schultz M. B., Sait, M., Sherry, N. |
| EPI_ISL_562536 | Victorian Infectious Diseases Reference Laboratory (VIDRL) | VIDRL and MDU-PHL | Caly, L., Seemann, T., Sait, M., Schultz, M. B., Druce J., Sherry, N. |
| EPI_ISL_562544, EPI_ISL_562553, EPI_ISL_562578, EPI_ISL_562580, EPI_ISL_562584, EPI_ISL_562667, EPI_ISL_562704 | Microbiological Diagnostic Unit - Public Health Laboratory (MDU-PHL) | MDU-PHL | Seemann, T., Schultz M. B., Sait, M., Sherry, N. |
| EPI_ISL_562754 | Victorian Infectious Diseases Reference Laboratory (VIDRL) | VIDRL and MDU-PHL | Caly, L., Seemann, T., Sait, M., Schultz, M. B., Druce J., Sherry, N. |
| EPI_ISL_562757, EPI_ISL_562759, EPI_ISL_562760, EPI_ISL_562919, EPI_ISL_562921, EPI_ISL_562922, EPI_ISL_562923, EPI_ISL_562927, EPI_ISL_562928, EPI_ISL_562931, EPI_ISL_562947, EPI_ISL_562949, EPI_ISL_562956, EPI_ISL_562964 | | | |
| see above | Microbiological Diagnostic Unit - Public Health Laboratory (MDU-PHL) | MDU-PHL | Seemann, T., Schultz M. B., Sait, M., Sherry, N. |
| EPI_ISL_563133 | Victorian Infectious Diseases Reference Laboratory (VIDRL) | VIDRL and MDU-PHL | Caly, L., Seemann, T., Sait, M., Schultz, M. B., Druce J., Sherry, N. |
| EPI_ISL_563265, EPI_ISL_563312, EPI_ISL_563313, EPI_ISL_563314, EPI_ISL_563315, EPI_ISL_563316, EPI_ISL_563317, EPI_ISL_563318, EPI_ISL_563319, EPI_ISL_563320, EPI_ISL_563321, EPI_ISL_563328, EPI_ISL_563425, EPI_ISL_563428, EPI_ISL_563429, EPI_ISL_563434, EPI_ISL_563438, EPI_ISL_563439, EPI_ISL_563440, EPI_ISL_563448, EPI_ISL_563452, EPI_ISL_563456, EPI_ISL_563458, EPI_ISL_563460, EPI_ISL_563461, EPI_ISL_563462, EPI_ISL_563464, EPI_ISL_563468, EPI_ISL_563470, EPI_ISL_563472, EPI_ISL_563475, EPI_ISL_563477, EPI_ISL_563478, EPI_ISL_563481, EPI_ISL_563482, EPI_ISL_563483, EPI_ISL_563484, EPI_ISL_563485, EPI_ISL_563487, EPI_ISL_563488, EPI_ISL_563489, EPI_ISL_563490, EPI_ISL_563492, EPI_ISL_563493, EPI_ISL_563494, EPI_ISL_563495, EPI_ISL_563501, EPI_ISL_563502, EPI_ISL_563504, EPI_ISL_563512, EPI_ISL_563515, EPI_ISL_563520, EPI_ISL_563523, EPI_ISL_563525, EPI_ISL_563527, EPI_ISL_563528, EPI_ISL_563529, EPI_ISL_563531, EPI_ISL_563533, EPI_ISL_563537, EPI_ISL_563830, EPI_ISL_563886, EPI_ISL_563887, EPI_ISL_563889, EPI_ISL_563890, EPI_ISL_563894, EPI_ISL_563896, EPI_ISL_563900, EPI_ISL_563903, EPI_ISL_563904, EPI_ISL_563906, EPI_ISL_563908, EPI_ISL_563911, EPI_ISL_563912, EPI_ISL_563914, EPI_ISL_563915, EPI_ISL_563917, EPI_ISL_563919, EPI_ISL_563920, EPI_ISL_563922, EPI_ISL_563926, EPI_ISL_563930, EPI_ISL_563934, EPI_ISL_563989, EPI_ISL_563990 | | | |
| see above | Microbiological Diagnostic Unit - Public Health Laboratory (MDU-PHL) | MDU-PHL | Seemann, T., Schultz M. B., Sait, M., Sherry, N. |
| EPI_ISL_563997, EPI_ISL_563998, EPI_ISL_563999, EPI_ISL_564000, EPI_ISL_564001, EPI_ISL_564002, EPI_ISL_564055, EPI_ISL_564056, EPI_ISL_564057, EPI_ISL_564058, EPI_ISL_564059, EPI_ISL_564060, EPI_ISL_564061, EPI_ISL_564062, EPI_ISL_564063, EPI_ISL_564064, EPI_ISL_564065, EPI_ISL_564066, EPI_ISL_564067, EPI_ISL_564068, EPI_ISL_564069, EPI_ISL_564070, EPI_ISL_564071, EPI_ISL_564072, EPI_ISL_564073, EPI_ISL_564074, EPI_ISL_564076 | | | |
| see above | Victorian Infectious Diseases Reference Laboratory (VIDRL) | VIDRL and MDU-PHL | Caly, L., Seemann, T., Sait, M., Schultz, M. B., Druce J., Sherry, N. |
| EPI_ISL_564115, EPI_ISL_564116, EPI_ISL_564117, EPI_ISL_564118, EPI_ISL_564120, EPI_ISL_564121, EPI_ISL_564122, EPI_ISL_564123, EPI_ISL_564124, EPI_ISL_564125, EPI_ISL_564126, EPI_ISL_564127, EPI_ISL_564128, EPI_ISL_564129, EPI_ISL_564131, EPI_ISL_564132, EPI_ISL_564133, EPI_ISL_564134, EPI_ISL_564135, EPI_ISL_564139, EPI_ISL_564140, EPI_ISL_564141, EPI_ISL_564142, EPI_ISL_564143, EPI_ISL_564144, EPI_ISL_564146, EPI_ISL_564147, EPI_ISL_564149, EPI_ISL_564150, EPI_ISL_564151, EPI_ISL_564152, EPI_ISL_564155, EPI_ISL_564156, EPI_ISL_564158, EPI_ISL_564159, EPI_ISL_564160, EPI_ISL_564161, EPI_ISL_564163, EPI_ISL_564178 | | | |
| see above | Microbiological Diagnostic Unit - Public Health Laboratory (MDU-PHL) | MDU-PHL | Seemann, T., Schultz M. B., Sait, M., Sherry, N. |
| EPI_ISL_564181, EPI_ISL_564203, EPI_ISL_564204, EPI_ISL_564205, EPI_ISL_564206, EPI_ISL_564207, EPI_ISL_564208, EPI_ISL_564210, EPI_ISL_564211, EPI_ISL_564212, EPI_ISL_564213, EPI_ISL_564214 | | | |
| see above | Victorian Infectious Diseases Reference Laboratory (VIDRL) | VIDRL and MDU-PHL | Caly, L., Seemann, T., Sait, M., Schultz, M. B., Druce J., Sherry, N. |
| EPI_ISL_564273, EPI_ISL_564275, EPI_ISL_564278, EPI_ISL_564282, EPI_ISL_564283, EPI_ISL_564287, EPI_ISL_564295 | Microbiological Diagnostic Unit - Public Health Laboratory (MDU-PHL) | MDU-PHL | Seemann, T., Schultz M. B., Sait, M., Sherry, N. |
| EPI_ISL_564330, EPI_ISL_564331, EPI_ISL_564332, EPI_ISL_564333, EPI_ISL_564334, EPI_ISL_564358, EPI_ISL_564359, EPI_ISL_564360, EPI_ISL_564361, EPI_ISL_564364 | Victorian Infectious Diseases Reference Laboratory (VIDRL) | VIDRL and MDU-PHL | Caly, L., Seemann, T., Sait, M., Schultz, M. B., Druce J., Sherry, N. |
| EPI_ISL_564417, EPI_ISL_564421, EPI_ISL_564428, EPI_ISL_564431, EPI_ISL_564436, EPI_ISL_564439, EPI_ISL_564440, EPI_ISL_564451, EPI_ISL_564453, EPI_ISL_564471, EPI_ISL_564473, EPI_ISL_564480, EPI_ISL_564503, EPI_ISL_564504, EPI_ISL_564505, EPI_ISL_564506, EPI_ISL_564507, EPI_ISL_564514, EPI_ISL_564519, EPI_ISL_564529, EPI_ISL_564530, EPI_ISL_564531, EPI_ISL_564532, EPI_ISL_564533, EPI_ISL_564534, EPI_ISL_564538, EPI_ISL_564543, EPI_ISL_564545, EPI_ISL_564565, EPI_ISL_564566, EPI_ISL_564567, EPI_ISL_564682, EPI_ISL_565008, EPI_ISL_565109 | | | |
| see above | Microbiological Diagnostic Unit - Public Health Laboratory (MDU-PHL) | MDU-PHL | Seemann, T., Schultz M. B., Sait, M., Sherry, N. |

| | | | |
|---|---|---|---|
| EPI_ISL_565141, EPI_ISL_565142, EPI_ISL_565143 | Victorian Infectious Diseases Reference Laboratory (VIDRL) | VIDRL and MDU-PHL | Caly, L., Seemann, T., Sait, M., Schultz, M. B., Druce J., Sherry, N. |
| EPI_ISL_565227, EPI_ISL_565230, EPI_ISL_565240, EPI_ISL_565242, EPI_ISL_565247, EPI_ISL_565256, EPI_ISL_565257, EPI_ISL_565265, EPI_ISL_565268, EPI_ISL_565269, EPI_ISL_565270, EPI_ISL_565272, EPI_ISL_565273, EPI_ISL_565274, EPI_ISL_565275, EPI_ISL_565276, EPI_ISL_565280, EPI_ISL_565282, EPI_ISL_565700, EPI_ISL_565701, EPI_ISL_565702, EPI_ISL_565703, EPI_ISL_565705, EPI_ISL_565706, EPI_ISL_565716, EPI_ISL_565717, EPI_ISL_565722, EPI_ISL_565724, EPI_ISL_565726, EPI_ISL_565727, EPI_ISL_565728, EPI_ISL_565747, EPI_ISL_565761, EPI_ISL_565788, EPI_ISL_565789, EPI_ISL_565796, EPI_ISL_565800, EPI_ISL_565804, EPI_ISL_565812, EPI_ISL_565818 | | | |
| see above | Microbiological Diagnostic Unit - Public Health Laboratory (MDU-PHL) | MDU-PHL | Seemann, T., Schultz M. B., Sait, M., Sherry, N. |
| EPI_ISL_565850, EPI_ISL_565851, EPI_ISL_565852, EPI_ISL_565853, EPI_ISL_565854, EPI_ISL_565855 | Michigan Department of Health and Human Services, Bureau of Laboratories | Michigan Department of Health and Human Services, Bureau of Laboratories | Blankenship HM, Riner D, Soehnlen MK |
| EPI_ISL_566063 | Respiratory Virus Unit, Microbiology Services Colindale, Public Health England | Respiratory Virus Unit, Microbiology Services Colindale, Public Health England | PHE Covid Sequencing Team |
| EPI_ISL_568472, EPI_ISL_568473 | Lighthouse Lab in Cambridge | Wellcome Sanger Institute for the COVID-19 Genomics UK (COG-UK) consortium | Rob Howes, The Lighthouse Lab in Cambridge and Alex Alderton, Roberto Amato, Sonia Goncalves, Ewan Harrison, David K. Jackson, Ian Johnston, Dominic Kwiatkowski, Cordelia Langford, John Sillitoe on behalf of the Wellcome Sanger Institute COVID-19 Surveillance Team |
| EPI_ISL_568564, EPI_ISL_568565, EPI_ISL_568566, EPI_ISL_568567, EPI_ISL_568568, EPI_ISL_568569 | Department of Infectious Diseases and Immunology, National Hospital Organization Nagoya Medical Center | Clinical Research Center, National Hospital Organization Nagoya Medical Center | Yoshihiro Nakata, Hirotaka Ode, Mai Kubota, Masakazu Matsuda, Kazuhiro Matsuoka, Nakasuji Miho, Mikiko Mori, Mayumi Imahashi, Yoshiyuki Yokomaku, Yasumasa Iwatani |
| EPI_ISL_568612 | Florida Bureau of Public Health Laboratories | Florida Bureau of Public Health Laboratories | Sarah Schmedes, Jason Blanton |
| EPI_ISL_569007, EPI_ISL_569008, EPI_ISL_569009, EPI_ISL_569010, EPI_ISL_569011, EPI_ISL_569012, EPI_ISL_569013, EPI_ISL_569014, EPI_ISL_569015, EPI_ISL_569016, EPI_ISL_569017, EPI_ISL_569018, EPI_ISL_569019, EPI_ISL_569020, EPI_ISL_569021, EPI_ISL_569022 | | | |
| see above | MEPHI, Aix Marseille University | MEPHI, Aix Marseille University | Anthony LEVASSEUR |
| EPI_ISL_569615 | Quick Care Huron | South Dakota Public Health Laboratory | Matt Plumb, Jacob Garfin, Xiong Wang, and Chris Carlson |
| EPI_ISL_569616 | Aurora County Clinic | South Dakota Public Health Laboratory | Matt Plumb, Jacob Garfin, Xiong Wang, and Chris Carlson |
| EPI_ISL_569784, EPI_ISL_569785, EPI_ISL_569786, EPI_ISL_569787, EPI_ISL_569788, EPI_ISL_569789, EPI_ISL_569790, EPI_ISL_569807, EPI_ISL_569813, EPI_ISL_569823, EPI_ISL_569824, EPI_ISL_569830, EPI_ISL_569831, EPI_ISL_569838, EPI_ISL_569839, EPI_ISL_569844, EPI_ISL_569846, EPI_ISL_569847 | | | |
| see above | Omsk Research Institute of Natural Focal Infections | WHO National Influenza Centre Russian Federation | Artem Fadeev, Ekaterina Gradoboeva, Ekaterina Savkina, Daria Nashatyreva, Elena Poleshchuk, Aleksei Vasilenko, Valery Yakimenko, Andrey Komissarov |
| EPI_ISL_569920, EPI_ISL_569946 | Innovative Genomics Institute, UC Berkeley | Innovative Genomics Institute, UC Berkeley | Stacia Wyman, Haridha Shivram, Phil Frankino, Liana Lareau, Shana McDevitt, Justin Choi |
| EPI_ISL_569982, EPI_ISL_569983, EPI_ISL_569984, EPI_ISL_569987 | Unity Health Toronto | Ontario Institute for Cancer Research | Ramzi Fattouh, Larissa M. Matukas, Yan Chen,Mark Downing, Trina Otterman, Karel Boissinot, Wai Sum Siu, Zhi Cui, Le Luu, Samira Mubareka, TIBDN, Ilinca Lungu, Bernard Lam, Jeremy Johns, Paul Krzyzanowski, Richard de Borja, Felicia Vincelli, Philip Zuzarte, Jared T. Simpson |
| EPI_ISL_572238, EPI_ISL_572239, EPI_ISL_572240, EPI_ISL_572241, EPI_ISL_572242, EPI_ISL_572243, EPI_ISL_572244, EPI_ISL_572245, EPI_ISL_572248, EPI_ISL_572256 | Virginia DCLS | Virginia DCLS | Virginia DCLS |
| EPI_ISL_572491 | Northumbria University / South Tees Hospitals NHS Foundation Trust / North Cumbria Integrated Care NHS Foundation Trust / North Tees and Hartlepool NHS Foundation Trust / Newcastle Hospitals NHS Foundation Trust | COVID-19 Genomics UK (COG-UK) Consortium | Darren L Smith,Andrew Nelson,Matthew Bashton,Greg R Young,Joshua Loh,John Allan,Mohammad A Tariq,Giles S Holt,Gary Black,Wen C Yew,Lynn Dover,Paul Baker,Steve Liggett,Sarah Essex,Jane Greenaway,Debra Padgett,Clive Graham,Garren Scott,Edward Barton,Emma Swindells,Brendan Payne,Jennifer Collins,Yusri Taha,Gary Eltringham |
| EPI_ISL_572540 | Wales Specialist Virology Centre Sequencing lab: Pathogen Genomics Unit | COVID-19 Genomics UK (COG-UK) Consortium | Catherine Moore, Johnathan Evans, Laura Gifford, Malorie Perry, Simon Cottrell, Angela Marchbank, Alec Birchley, Alexander Adams, Amy Gaskin, Bree Gatica-Wilcox, Jason Coombes, Joel Southgate, Lauren Gilbert, Lee Graham, Nicole Pacchiarini, Sara Kumziene-Summerhayes, Sarah Taylor, Sophie Jones, Sara Rey, Matthew Bull, Joanne Watkins, Sally Corden, Tom Connor |
| EPI_ISL_573313, EPI_ISL_573324, EPI_ISL_573325, EPI_ISL_573326, EPI_ISL_573327, EPI_ISL_573328, EPI_ISL_573329, EPI_ISL_573330, EPI_ISL_573331, EPI_ISL_573332, EPI_ISL_573333 | | | |
| see above | Northumbria University / South Tees Hospitals NHS Foundation Trust / North Cumbria Integrated Care NHS Foundation Trust / North Tees and Hartlepool NHS Foundation Trust / Newcastle Hospitals NHS Foundation Trust | COVID-19 Genomics UK (COG-UK) Consortium | Darren L Smith,Andrew Nelson,Matthew Bashton,Greg R Young,Joshua Loh,John Allan,Mohammad A Tariq,Giles S Holt,Gary Black,Wen C Yew,Lynn Dover,Paul Baker,Steve Liggett,Sarah Essex,Jane Greenaway,Debra Padgett,Clive Graham,Garren Scott,Edward Barton,Emma Swindells,Brendan Payne,Jennifer Collins,Yusri Taha,Gary Eltringham |
| EPI_ISL_574317, EPI_ISL_574318, EPI_ISL_574319 | LSUHS Emerging Viral Threat Laboratory | Microbial Genome Sequencing Center | Rona S. Scott, Jeremy P. Kamil, Maarten Van Diest, Malgorzata Bienkowska-Haba, Katarzyna Zwolinska, Andrew D. Yurochko, Christopher G. Kevil, Martin J. Sapp, Daniel J. Snyder, Vaughn S. Cooper, John A. Vanchiere |
| EPI_ISL_574547, EPI_ISL_574548, EPI_ISL_574549 | Microbiology Division, South Carolina Department of Health and Environmental Control | Microbiology Division, South Carolina Department of Health and Environmental Control | Flores,H. |
| EPI_ISL_576196, EPI_ISL_576197 | AR Dept. of Health-Public Health Lab | Pathogen Discovery, Respiratory Viruses Branch, Division of Viral Diseases, Centers for Disease Control and Prevention | Ying Tao, Jing Zhang, Brian Lynch, Yan Li, Krista Queen, Anna Uehara, Clinton R. Paden, Peter Cook, Haibin Wang, Suxiang Tong |
| EPI_ISL_576257 | Instituto de Diagnostico y Referencia Epidemiologicos (INDRE) | Instituto de Diagnostico y Referencia Epidemiologicos (INDRE) | Gisela Barrera-Badillo , Abril Rodriguez-Maldonado, Claudia Wong-Arambula , Natividad Cruz-Ortiz, Tatiana Nunez-Garcia, Dayanira Arellano-Suarez, Fabiola Garces-Ayala, Edgar Mendieta-Condado, Lucia Hernandez-Rivas, Irma Lopez-Martinez, Ernesto Ramirez-Gonzalez. |
| EPI_ISL_576292, EPI_ISL_576293, EPI_ISL_576294, EPI_ISL_576295, EPI_ISL_576296, EPI_ISL_576297, EPI_ISL_576298, EPI_ISL_576299, EPI_ISL_576300, EPI_ISL_576301, EPI_ISL_576302, EPI_ISL_576303, EPI_ISL_576304, EPI_ISL_576305, EPI_ISL_576306, EPI_ISL_576307, EPI_ISL_576308, EPI_ISL_576309, EPI_ISL_576310, EPI_ISL_576311, EPI_ISL_576312, EPI_ISL_576313, EPI_ISL_576314, EPI_ISL_576315, EPI_ISL_576316, EPI_ISL_576317, EPI_ISL_576318, EPI_ISL_576319, EPI_ISL_576320, EPI_ISL_576321, EPI_ISL_576322, EPI_ISL_576323, EPI_ISL_576324, EPI_ISL_576325 | | | |
| see above | Texas Department of State Health Services | Texas Department of State Health Services | Rashmi Tuladhar, Bonnie Oh, Jenny Zhang, Maliha Rahman, Anita Pokharel, Myong Koag, Chun Wang, Rachel Lee, Grace Kubin |
| EPI_ISL_576327, EPI_ISL_576328, EPI_ISL_576329, EPI_ISL_576330, EPI_ISL_576331, EPI_ISL_576332, EPI_ISL_576333, EPI_ISL_576334, EPI_ISL_576335 | Texas Department of State Health Services | Texas Department of State Health Services | Rashmi Tuladhar, Bonnie Oh, Mayela Pedrueza, Jenny Zhang, Maliha Rahman, Anita Pokharel, Myong Koag, Chun Wang, Rachel Lee, Grace Kubin |
| EPI_ISL_576371, EPI_ISL_576372, EPI_ISL_576373 | Cancer Biology Department, National Cancer Institute | Cancer Biology Department, National Cancer Institute | Zekri,A.N., Soliman,H.K., Ahmed,O.S., Hafez,M.M., Hamdy,M.S., Abouelhoda,M. |
| EPI_ISL_576389, EPI_ISL_576390, EPI_ISL_576391, EPI_ISL_576392, EPI_ISL_576393 | Genome Centre | Genome Centre | Selina Akter Pravas Chandra Roy Amina Ferdaus manami Habiba Ibnat A. S. M. Rubayet Ul Alam Shireen Nigar Iqbal Kabir Jahid and M. Anwar Hossain |
| EPI_ISL_576518, EPI_ISL_576519, EPI_ISL_576521, EPI_ISL_576523, EPI_ISL_576525, EPI_ISL_576527, EPI_ISL_576532, EPI_ISL_576536, EPI_ISL_576538, EPI_ISL_576540, EPI_ISL_576545, EPI_ISL_576550 | | | |
| see above | Innovative Genomics Institute, UC Berkeley | Innovative Genomics Institute, UC Berkeley | Stacia Wyman, Haridha Shivram, Phil Frankino, Liana Lareau, Shana McDevitt, Justin Choi |
| EPI_ISL_577724, EPI_ISL_577725, EPI_ISL_577726 | NIV Influenza | NIV Influenza | Potdar V |

| | | | |
|---|---|---|---|
| EPI_ISL_578188 | Hospital Virgen de las Nieves | Instituto de Salud Carlos III | Iglesias-Caballero, M. Molinero Calamita, M. González-Esguevillas, M. Camarero, S. Pozo, F. Casas, I. Jiménez, P. Jiménez, M. Zaballos, A. Monzón, S. Varona, S. Juliá, M. Cuesta, I, J.M Navarro |
| EPI_ISL_581428 | Lighthouse Lab in Milton Keynes | Wellcome Sanger Institute for the COVID-19 Genomics UK (COG-UK) consortium | The Lighthouse Lab in Milton Keynes and Alex Alderton, Roberto Amato, Sonia Goncalves, Ewan Harrison, David K. Jackson, Ian Johnston, Dominic Kwiatkowski, Cordelia Langford, John Sillitoe on behalf of the Wellcome Sanger Institute COVID-19 Surveillance Team |
| EPI_ISL_581542, EPI_ISL_581543, EPI_ISL_581544, EPI_ISL_581545, EPI_ISL_581546, EPI_ISL_581547, EPI_ISL_581548, EPI_ISL_581549, EPI_ISL_581550, EPI_ISL_581551, EPI_ISL_581552, EPI_ISL_581553, EPI_ISL_581554, EPI_ISL_581555, EPI_ISL_581556, EPI_ISL_581557 | | | |
| see above | Virginia DCLS | Virginia DCLS | Virginia DCLS |
| EPI_ISL_581884, EPI_ISL_581885, EPI_ISL_581886, EPI_ISL_581887, EPI_ISL_581888, EPI_ISL_581889, EPI_ISL_581900, EPI_ISL_581901, EPI_ISL_581902, EPI_ISL_581905, EPI_ISL_581906, EPI_ISL_581909, EPI_ISL_581910, EPI_ISL_581911, EPI_ISL_581912, EPI_ISL_581913 | | | |
| see above | University Hospital Basel, Clinical Virology | University Hospital Basel, Clinical Bacteriology | Madlen Stange, Alfredo Mari, Tim Roloff, Helena MB Seth-Smith, Michael Schweitzer, Myrta Brunner, Karoline Leuzinger, Kirstine K. Soegaard, Alexander Gensch, Sarah Tschudin-Sutter, Simon Fuchs, Julia Bielicki, Hans Pargger, Martin Siegemund, Christian Nickel, Roland Bingisser, Michael Osthoff, Stefano Bassetti, Rita Schneider-Sliwa, Manuel Battegay, Hans Hirsch, Adrian Egli |
| EPI_ISL_582320, EPI_ISL_582321, EPI_ISL_582322, EPI_ISL_582323, EPI_ISL_582324, EPI_ISL_582356, EPI_ISL_582357, EPI_ISL_582358, EPI_ISL_582363, EPI_ISL_582364, EPI_ISL_582365, EPI_ISL_582367, EPI_ISL_582368, EPI_ISL_582373, EPI_ISL_582374, EPI_ISL_582375, EPI_ISL_582376, EPI_ISL_582381, EPI_ISL_582382, EPI_ISL_582383, EPI_ISL_582384, EPI_ISL_582389, EPI_ISL_582390, EPI_ISL_582391, EPI_ISL_582392, EPI_ISL_582395, EPI_ISL_582396, EPI_ISL_582397, EPI_ISL_582402, EPI_ISL_582403, EPI_ISL_582404, EPI_ISL_582409, EPI_ISL_582410, EPI_ISL_582411, EPI_ISL_582423, EPI_ISL_582463, EPI_ISL_582464, EPI_ISL_582465, EPI_ISL_582466, EPI_ISL_582468, EPI_ISL_582474, EPI_ISL_582475, EPI_ISL_582476, EPI_ISL_582477, EPI_ISL_582486, EPI_ISL_582487, EPI_ISL_582505 | | | |
| see above | Cadham Provincial Laboratory | National Microbiology Laboratory (NML) | Anna Majer, Shari Tyson, Grace Seo, Philip Mabon, Elsie Grudeski, Rhiannon Huzarewich, Russell Mandes, Anneliese Landgraff, Jennifer Tanner, Natalie Knox, Morag Graham, Gary Van Domselaar, Paul Van Caeseele, Jared Bullard, David Alexander, Kerry Dust, Nathalie Bastien, Yan Li, Timothy Booth, Darian Hole, Madison Chapel, CanCOGeN's metadata curation team, Public Health Agency of Canada CanCOGeN team |
| EPI_ISL_582530 | Veterinary Specialized Institute "Kraljevo", Serbia | Veterinary Specialized Institute "Kraljevo", Serbia | Vidanovic,D., Tesovic,B., Knezevic,A., Jovanovic,T., Jankovic,M., Sekler,M., Banovic Djeri,B., Petrovic,T., Volkening,J., Afonso,C. |
| EPI_ISL_582780 | Uppsala klinisk mikrobiologi | The Public Health Agency of Sweden | Anna-Malin Linde, Maria Lind Karlberg, Mattias Haukland, Reza Advani, Olov Svartstrom, Oskar Karlsson Lindsjo, Sandra Broddesson, Petra Edquist, Mia Brytting, Anna Risberg, Karin Tegmark-Wisell |
| EPI_ISL_582880, EPI_ISL_582881, EPI_ISL_582882, EPI_ISL_582883, EPI_ISL_582884, EPI_ISL_582885, EPI_ISL_582886 | County of Santa Clara Public Health Department | Chan-Zuckerberg Biohub | CZB Cliahub Consortium |
| EPI_ISL_582959, EPI_ISL_582960, EPI_ISL_582961, EPI_ISL_582962, EPI_ISL_582963, EPI_ISL_582964, EPI_ISL_582965, EPI_ISL_582966, EPI_ISL_582967, EPI_ISL_582968 | San Luis Obispo Public Health Department | Chan-Zuckerberg Biohub | CZB Cliahub Consortium |
| EPI_ISL_583070, EPI_ISL_583071, EPI_ISL_583072, EPI_ISL_583073, EPI_ISL_583074, EPI_ISL_583075, EPI_ISL_583076, EPI_ISL_583077, EPI_ISL_583078, EPI_ISL_583079, EPI_ISL_583080, EPI_ISL_583081, EPI_ISL_583082, EPI_ISL_583083, EPI_ISL_583084, EPI_ISL_583085, EPI_ISL_583086, EPI_ISL_583087 | | | |
| see above | Humboldt County Public Health Laboratory | Chan-Zuckerberg Biohub | CZB Cliahub Consortium |
| EPI_ISL_583210, EPI_ISL_583211, EPI_ISL_583212, EPI_ISL_583213, EPI_ISL_583214, EPI_ISL_583215 | UCSF Clinical Microbiology Laboratory | Chan-Zuckerberg Biohub | CZB Cliahub Consortium |
| EPI_ISL_583452, EPI_ISL_583453, EPI_ISL_583454, EPI_ISL_583471, EPI_ISL_583473, EPI_ISL_583477 | Garcia-Sastre Laboratory, Department of Microbiology, Icahn School of Medicine at Mount Sinai | van Bakel Laboratory, Genetics and Genomics Sciences, Icahn School of Medicine at Mount Sinai | Teresa Aydillo, Ana S. Gonzalez-Reiche, Sadaf Aslam, Adriana van de Guchte, Zenab Khan, Ajay Obla, Jayeeta Dutta, Harm van Bakel, Judith Aberg, Adolfo García-Sastre, Gunjan Shah, Tobias Hohl, Genovefa Papanicolaou, Miguel-Angel Perales, Kent Sepkowitz, Ngolela Esther Babady, and Mini Kamboj |
| EPI_ISL_583885, EPI_ISL_583886 | Austrian Agency for Health and Food Safety (AGES) | Bergthaler laboratory, CeMM Research Center for Molecular Medicine of the Austrian Academy of Sciences | Alexandra Popa, Benedikt Agerer, Henrique Colaco, Lukas Endler, Jakob-Wendelin Genger, Alexander Lercher, Mark Smyth, Thomas Penz, Michael Schuster, Jan Laine, Martin Senekowitsch, Judith Aberle, Stephan Aberle, Peter Hufnagl, Daniela Schmid, Franz Allerberger, Elisabeth Puchhammer-Stoeckl, Manfred Nairz, Guenter Weiss, Gregor Hörmann, Kinga Rigler-Hohenwarter, Rainer Gattringer, Wegene Borena, Dorothee von Laer, Gernot Walder, Peter Obrist, Christian Paar, Sabine Sussitz-Rack, Gunther Vogl, Adi Steinrigl, Christoph Bock, Andreas Bergthaler |
| EPI_ISL_586531 | General Hospital,Chhotaudepur | Gujarat Biotechnology Research Centre | Yogesh Parmar, Arti Thakur, Dharak Pandya, Maharshi Pandya, Nidhi Patel, Nitin Savaliya, Raghawendra Kumar, Dinesh Kumar, Zuber Saiyed, Komal Patel, Labdhi Pandya, Afzal Ansari, Nikha Trivedi, Apurvasinh Puvar, Janvi Raval, Zarna Patel, Monika Gandhi, Pinal Trivedi, R D Dixit, A M Kadri, Harsh Bakshi, Chaitanya Joshi, Madhvi Joshi |
| EPI_ISL_586532 | General Hospital,Chhotaudepur | Gujarat Biotechnology Research Centre | Arti Thakur, Dharak Pandya, Maharshi Pandya, Nidhi Patel, Nitin Savaliya, Raghawendra Kumar, Dinesh Kumar, Zuber Saiyed, Komal Patel, Labdhi Pandya, Afzal Ansari, Nikha Trivedi, Apurvasinh Puvar, Janvi Raval, Zarna Patel, Monika Gandhi, Pinal Trivedi, Yogesh Parmar, R D Dixit, A M Kadri, Harsh Bakshi, Chaitanya Joshi, Madhvi Joshi |
| EPI_ISL_586533 | General Hospital,Chhotaudepur | Gujarat Biotechnology Research Centre | Dharak Pandya, Maharshi Pandya, Nidhi Patel, Nitin Savaliya, Raghawendra Kumar, Dinesh Kumar, Zuber Saiyed, Komal Patel, Labdhi Pandya, Afzal Ansari, Nikha Trivedi, Apurvasinh Puvar, Janvi Raval, Zarna Patel, Monika Gandhi, Pinal Trivedi, Yogesh Parmar, Arti Thakur, R D Dixit, A M Kadri, Harsh Bakshi, Chaitanya Joshi, Madhvi Joshi |
| EPI_ISL_586534 | General Hospital,Chhotaudepur | Gujarat Biotechnology Research Centre | Maharshi Pandya, Nidhi Patel, Nitin Savaliya, Raghawendra Kumar, Dinesh Kumar, Zuber Saiyed, Komal Patel, Labdhi Pandya, Afzal Ansari, Nikha Trivedi, Apurvasinh Puvar, Janvi Raval, Zarna Patel, Monika Gandhi, Pinal Trivedi, Yogesh Parmar, Arti Thakur, Dharak Pandya, R D Dixit, A M Kadri, Harsh Bakshi, Chaitanya Joshi, Madhvi Joshi |
| EPI_ISL_586535 | General Hospital,Chhotaudepur | Gujarat Biotechnology Research Centre | Nidhi Patel, Nitin Savaliya, Raghawendra Kumar, Dinesh Kumar, Zuber Saiyed, Komal Patel, Labdhi Pandya, Afzal Ansari, Nikha Trivedi, Apurvasinh Puvar, Janvi Raval, Zarna Patel, Monika Gandhi, Pinal Trivedi, Yogesh Parmar, Arti Thakur, Dharak Pandya, Maharshi Pandya, R D Dixit, A M Kadri, Harsh Bakshi, Chaitanya Joshi, Madhvi Joshi |
| EPI_ISL_586536 | Zydus Medical Hospital,Dahod | Gujarat Biotechnology Research Centre | Kalpesh Labana, Mohit Desai, Nitin Savaliya, Raghawendra Kumar, Dinesh Kumar, Zuber Saiyed, Komal Patel, Labdhi Pandya, Afzal Ansari, Nikha Trivedi, Apurvasinh Puvar, Janvi Raval, Zarna Patel, Monika Gandhi, Pinal Trivedi, Maharshi Pandya, Nidhi Patel, R D Dixit, A M Kadri, Harsh Bakshi, Chaitanya Joshi, Madhvi Joshi |
| EPI_ISL_586537 | Zydus Medical Hospital,Dahod | Gujarat Biotechnology Research Centre | Mohit Desai, Nitin Savaliya, Raghawendra Kumar, Dinesh Kumar, Zuber Saiyed, Komal Patel, Labdhi Pandya, Afzal Ansari, Nikha Trivedi, Apurvasinh Puvar, Janvi Raval, Zarna Patel, Monika Gandhi, Pinal Trivedi, Maharshi Pandya, Nidhi Patel, Kalpesh Labana, R D Dixit, A M Kadri, Harsh Bakshi, Chaitanya Joshi, Madhvi Joshi |
| EPI_ISL_586538 | Zydus Medical Hospital,Dahod | Gujarat Biotechnology Research Centre | Nitin Savaliya, Raghawendra Kumar, Dinesh Kumar, Zuber Saiyed, Komal Patel, Labdhi Pandya, Afzal Ansari, Nikha Trivedi, Apurvasinh Puvar, Janvi Raval, Zarna Patel, Monika Gandhi, Pinal Trivedi, Maharshi Pandya, Nidhi Patel, Kalpesh Labana, Mohit Desai, R D Dixit, A M Kadri, Harsh Bakshi, Chaitanya Joshi, Madhvi Joshi |
| EPI_ISL_586539 | Zydus Medical Hospital,Dahod | Gujarat Biotechnology Research Centre | Raghawendra Kumar, Dinesh Kumar, Zuber Saiyed, Komal Patel, Labdhi Pandya, Afzal Ansari, Nikha Trivedi, Apurvasinh Puvar, Janvi Raval, Zarna Patel, Monika Gandhi, Pinal Trivedi, Maharshi Pandya, Nidhi Patel, Kalpesh Labana, Mohit Desai, Nitin Savaliya, R D Dixit, A M Kadri, Harsh Bakshi, Chaitanya Joshi, Madhvi Joshi |
| EPI_ISL_586540 | Zydus Medical Hospital,Dahod | Gujarat Biotechnology Research Centre | Dinesh Kumar, Zuber Saiyed, Komal Patel, Labdhi Pandya, Afzal Ansari, Nikha Trivedi, Apurvasinh Puvar, Janvi Raval, Zarna Patel, Monika Gandhi, Pinal Trivedi, Maharshi Pandya, Nidhi Patel, Kalpesh Labana, Mohit Desai, Nitin Savaliya, Raghawendra Kumar, R D Dixit, A M Kadri, Harsh Bakshi, Chaitanya Joshi, Madhvi Joshi |
| EPI_ISL_586541 | Zydus Medical Hospital,Dahod | Gujarat Biotechnology Research Centre | Zuber Saiyed, Komal Patel, Labdhi Pandya, Afzal Ansari, Nikha Trivedi, Apurvasinh Puvar, Janvi Raval, Zarna Patel, Monika Gandhi, Pinal Trivedi, Maharshi Pandya, Nidhi Patel, Kalpesh Labana, Mohit Desai, Nitin Savaliya, Raghawendra Kumar, Dinesh Kumar, R D Dixit, A M Kadri, Harsh Bakshi, Chaitanya Joshi, Madhvi Joshi |

| | | | |
|---|---|---|---|
| EPI_ISL_586542 | Government General Hospital, Jam Khambhaliya, Devbhoomi Dwarka | Gujarat Biotechnology Research Centre | Komal Patel, Labdhi Pandya, Afzal Ansari, Nikha Trivedi, Harish Matani, Apurvasinh Puvar, Janvi Raval, Zarna Patel, Monika Gandhi, Pinal Trivedi, Maharshi Pandya, Nidhi Patel, Nitin Savaliya, Raghawendra Kumar, Dinesh Kumar, Zuber Saiyed, R D Dixit, A M Kadri, Harsh Bakshi, Chaitanya Joshi, Madhvi Joshi |
| EPI_ISL_586543 | Government General Hospital, Jam Khambhaliya, Devbhoomi Dwarka | Gujarat Biotechnology Research Centre | Labdhi Pandya, Afzal Ansari, Nikha Trivedi, Harish Matani, Apurvasinh Puvar, Janvi Raval, Zarna Patel, Monika Gandhi, Pinal Trivedi, Maharshi Pandya, Nidhi Patel, Nitin Savaliya, Raghawendra Kumar, Dinesh Kumar, Zuber Saiyed, Komal Patel, R D Dixit, A M Kadri, Harsh Bakshi, Chaitanya Joshi, Madhvi Joshi |
| EPI_ISL_586544 | Government General Hospital, Jam Khambhaliya, Devbhoomi Dwarka | Gujarat Biotechnology Research Centre | Afzal Ansari, Nikha Trivedi, Harish Matani, Apurvasinh Puvar, Janvi Raval, Zarna Patel, Monika Gandhi, Pinal Trivedi, Maharshi Pandya, Nidhi Patel, Nitin Savaliya, Raghawendra Kumar, Dinesh Kumar, Zuber Saiyed, Komal Patel, Labdhi Pandya, R D Dixit, A M Kadri, Harsh Bakshi, Chaitanya Joshi, Madhvi Joshi |
| EPI_ISL_586545 | Government General Hospital, Jam Khambhaliya, Devbhoomi Dwarka | Gujarat Biotechnology Research Centre | Nikha Trivedi, Harish Matani, Apurvasinh Puvar, Janvi Raval, Zarna Patel, Monika Gandhi, Pinal Trivedi, Maharshi Pandya, Nidhi Patel, Nitin Savaliya, Raghawendra Kumar, Dinesh Kumar, Zuber Saiyed, Komal Patel, Labdhi Pandya, Afzal Ansari, R D Dixit, A M Kadri, Harsh Bakshi, Chaitanya Joshi, Madhvi Joshi |
| EPI_ISL_586546 | General Hospital,Godhra | Gujarat Biotechnology Research Centre | Neela Katara, Pinal Gandhi, Sangeeta Kumar, Apurvasinh Puvar, Janvi Raval, Zarna Patel, Monika Gandhi, Pinal Trivedi, Maharshi Pandya, Nidhi Patel, Nitin Savaliya, Raghawendra Kumar, Dinesh Kumar, Zuber Saiyed, Komal Patel, Labdhi Pandya, Afzal Ansari, Nikha Trivedi, R D Dixit, A M Kadri, Harsh Bakshi, Chaitanya Joshi, Madhvi Joshi |
| EPI_ISL_586547 | General Hospital,Godhra | Gujarat Biotechnology Research Centre | Pinal Gandhi, Sangeeta Kumar, Apurvasinh Puvar, Janvi Raval, Zarna Patel, Monika Gandhi, Pinal Trivedi, Maharshi Pandya, Nidhi Patel, Nitin Savaliya, Raghawendra Kumar, Dinesh Kumar, Zuber Saiyed, Komal Patel, Labdhi Pandya, Afzal Ansari, Nikha Trivedi, Neela Katara, R D Dixit, A M Kadri, Harsh Bakshi, Chaitanya Joshi, Madhvi Joshi |
| EPI_ISL_586548 | General Hospital,Godhra | Gujarat Biotechnology Research Centre | Sangeeta Kumar, Apurvasinh Puvar, Janvi Raval, Zarna Patel, Monika Gandhi, Pinal Trivedi, Maharshi Pandya, Nidhi Patel, Nitin Savaliya, Raghawendra Kumar, Dinesh Kumar, Zuber Saiyed, Komal Patel, Labdhi Pandya, Afzal Ansari, Nikha Trivedi, Neela Katara, Pinal Gandhi, R D Dixit, A M Kadri, Harsh Bakshi, Chaitanya Joshi, Madhvi Joshi |
| EPI_ISL_586549 | CCC,Veraval | Gujarat Biotechnology Research Centre | Jignesh Parmar, Jitendra Bamrotia, Apurvasinh Puvar, Janvi Raval, Zarna Patel, Monika Gandhi, Pinal Trivedi, Maharshi Pandya, Nidhi Patel, Nitin Savaliya, Raghawendra Kumar, Dinesh Kumar, Zuber Saiyed, Komal Patel, Labdhi Pandya, Afzal Ansari, Nikha Trivedi, R D Dixit, A M Kadri, Harsh Bakshi, Chaitanya Joshi, Madhvi Joshi |
| EPI_ISL_586550 | CCC,Veraval | Gujarat Biotechnology Research Centre | Jitendra Bamrotia, Apurvasinh Puvar, Janvi Raval, Zarna Patel, Monika Gandhi, Pinal Trivedi, Maharshi Pandya, Nidhi Patel, Nitin Savaliya, Raghawendra Kumar, Dinesh Kumar, Zuber Saiyed, Komal Patel, Labdhi Pandya, Afzal Ansari, Nikha Trivedi, Jignesh Parmar, R D Dixit, A M Kadri, Harsh Bakshi, Chaitanya Joshi, Madhvi Joshi |
| EPI_ISL_586551 | Government Hospital,Veraval | Gujarat Biotechnology Research Centre | Monika Gandhi, Pinal Trivedi, Maharshi Pandya, Nidhi Patel, Nitin Savaliya, Raghawendra Kumar, Dinesh Kumar, Zuber Saiyed, Komal Patel, Labdhi Pandya, Afzal Ansari, Nikha Trivedi, Jignesh Parmar, Jitendra Bamrotia, Apurvasinh Puvar, Janvi Raval, Zarna Patel, R D Dixit, A M Kadri, Harsh Bakshi, Chaitanya Joshi, Madhvi Joshi |
| EPI_ISL_586552 | Government Hospital,Veraval | Gujarat Biotechnology Research Centre | Pinal Trivedi, Maharshi Pandya, Nidhi Patel, Nitin Savaliya, Raghawendra Kumar, Dinesh Kumar, Zuber Saiyed, Komal Patel, Labdhi Pandya, Afzal Ansari, Nikha Trivedi, Jignesh Parmar, Jitendra Bamrotia, Apurvasinh Puvar, Janvi Raval, Zarna Patel, Monika Gandhi, R D Dixit, A M Kadri, Harsh Bakshi, Chaitanya Joshi, Madhvi Joshi |
| EPI_ISL_586553 | Government Hospital,Veraval | Gujarat Biotechnology Research Centre | Maharshi Pandya, Nidhi Patel, Nitin Savaliya, Raghawendra Kumar, Dinesh Kumar, Zuber Saiyed, Komal Patel, Labdhi Pandya, Afzal Ansari, Nikha Trivedi, Jignesh Parmar, Jitendra Bamrotia, Apurvasinh Puvar, Janvi Raval, Zarna Patel, Monika Gandhi, Pinal Trivedi, R D Dixit, A M Kadri, Harsh Bakshi, Chaitanya Joshi, Madhvi Joshi |
| EPI_ISL_586554 | Government Hospital,Veraval | Gujarat Biotechnology Research Centre | Nidhi Patel, Nitin Savaliya, Raghawendra Kumar, Dinesh Kumar, Zuber Saiyed, Komal Patel, Labdhi Pandya, Afzal Ansari, Nikha Trivedi, Jignesh Parmar, Jitendra Bamrotia, Apurvasinh Puvar, Janvi Raval, Zarna Patel, Monika Gandhi, Pinal Trivedi, Maharshi Pandya, R D Dixit, A M Kadri, Harsh Bakshi, Chaitanya Joshi, Madhvi Joshi |
| EPI_ISL_586555 | CCC,Veraval | Gujarat Biotechnology Research Centre | Nitin Savaliya, Raghawendra Kumar, Dinesh Kumar, Zuber Saiyed, Komal Patel, Labdhi Pandya, Afzal Ansari, Nikha Trivedi, Jignesh Parmar, Jitendra Bamrotia, Apurvasinh Puvar, Janvi Raval, Zarna Patel, Monika Gandhi, Pinal Trivedi, Maharshi Pandya, Nidhi Patel, R D Dixit, A M Kadri, Harsh Bakshi, Chaitanya Joshi, Madhvi Joshi |
| EPI_ISL_586556 | CCC,Veraval | Gujarat Biotechnology Research Centre | Raghawendra Kumar, Dinesh Kumar, Zuber Saiyed, Komal Patel, Labdhi Pandya, Afzal Ansari, Nikha Trivedi, Jignesh Parmar, Jitendra Bamrotia, Apurvasinh Puvar, Janvi Raval, Zarna Patel, Monika Gandhi, Pinal Trivedi, Maharshi Pandya, Nidhi Patel, Nitin Savaliya, R D Dixit, A M Kadri, Harsh Bakshi, Chaitanya Joshi, Madhvi Joshi |
| EPI_ISL_586557 | CCC,Veraval | Gujarat Biotechnology Research Centre | Dinesh Kumar, Zuber Saiyed, Komal Patel, Labdhi Pandya, Afzal Ansari, Nikha Trivedi, Jignesh Parmar, Jitendra Bamrotia, Apurvasinh Puvar, Janvi Raval, Zarna Patel, Monika Gandhi, Pinal Trivedi, Maharshi Pandya, Nidhi Patel, Nitin Savaliya, Raghawendra Kumar, R D Dixit, A M Kadri, Harsh Bakshi, Chaitanya Joshi, Madhvi Joshi |
| EPI_ISL_586558 | CCC,Veraval | Gujarat Biotechnology Research Centre | Zuber Saiyed, Komal Patel, Labdhi Pandya, Afzal Ansari, Nikha Trivedi, Jignesh Parmar, Jitendra Bamrotia, Apurvasinh Puvar, Janvi Raval, Zarna Patel, Monika Gandhi, Pinal Trivedi, Maharshi Pandya, Nidhi Patel, Nitin Savaliya, Raghawendra Kumar, Dinesh Kumar, R D Dixit, A M Kadri, Harsh Bakshi, Chaitanya Joshi, Madhvi Joshi |
| EPI_ISL_586559 | CCC,Veraval | Gujarat Biotechnology Research Centre | Komal Patel, Labdhi Pandya, Afzal Ansari, Nikha Trivedi, Jignesh Parmar, Jitendra Bamrotia, Apurvasinh Puvar, Janvi Raval, Zarna Patel, Monika Gandhi, Pinal Trivedi, Maharshi Pandya, Nidhi Patel, Nitin Savaliya, Raghawendra Kumar, Dinesh Kumar, Zuber Saiyed, R D Dixit, A M Kadri, Harsh Bakshi, Chaitanya Joshi, Madhvi Joshi |
| EPI_ISL_586560 | CCC,Veraval | Gujarat Biotechnology Research Centre | Labdhi Pandya, Afzal Ansari, Nikha Trivedi, Jignesh Parmar, Jitendra Bamrotia, Apurvasinh Puvar, Janvi Raval, Zarna Patel, Monika Gandhi, Pinal Trivedi, Maharshi Pandya, Nidhi Patel, Nitin Savaliya, Raghawendra Kumar, Dinesh Kumar, Zuber Saiyed, Komal Patel, R D Dixit, A M Kadri, Harsh Bakshi, Chaitanya Joshi, Madhvi Joshi |
| EPI_ISL_586561 | CCC,Veraval | Gujarat Biotechnology Research Centre | Afzal Ansari, Nikha Trivedi, Jignesh Parmar, Jitendra Bamrotia, Apurvasinh Puvar, Janvi Raval, Zarna Patel, Monika Gandhi, Pinal Trivedi, Maharshi Pandya, Nidhi Patel, Nitin Savaliya, Raghawendra Kumar, Dinesh Kumar, Zuber Saiyed, Komal Patel, Labdhi Pandya, R D Dixit, A M Kadri, Harsh Bakshi, Chaitanya Joshi, Madhvi Joshi |
| EPI_ISL_586562 | GMERS GMERS Medical College and Hospital, Junagadh | Gujarat Biotechnology Research Centre | Nikha Trivedi, Bhavesh Bagda , Ravi Dedaniya, Apurvasinh Puvar, Janvi Raval, Zarna Patel, Monika Gandhi, Pinal Trivedi, Maharshi Pandya, Nidhi Patel, Nitin Savaliya, Raghawendra Kumar, Dinesh Kumar, Zuber Saiyed, Komal Patel, Labdhi Pandya, Afzal Ansari, R D Dixit, A M Kadri, Harsh Bakshi, Chaitanya Joshi, Madhvi Joshi |
| EPI_ISL_586563 | GMERS GMERS Medical College and Hospital, Junagadh | Gujarat Biotechnology Research Centre | Bhavesh Bagda , Ravi Dedaniya, Apurvasinh Puvar, Janvi Raval, Zarna Patel, Monika Gandhi, Pinal Trivedi, Maharshi Pandya, Nidhi Patel, Nitin Savaliya, Raghawendra Kumar, Dinesh Kumar, Zuber Saiyed, Komal Patel, Labdhi Pandya, Afzal Ansari, Nikha Trivedi, R D Dixit, A M Kadri, Harsh Bakshi, Chaitanya Joshi, Madhvi Joshi |
| EPI_ISL_586564 | GMERS GMERS Medical College and Hospital, Junagadh | Gujarat Biotechnology Research Centre | Ravi Dedaniya, Apurvasinh Puvar, Janvi Raval, Zarna Patel, Monika Gandhi, Pinal Trivedi, Maharshi Pandya, Nidhi Patel, Nitin Savaliya, Raghawendra Kumar, Dinesh Kumar, Zuber Saiyed, Komal Patel, Labdhi Pandya, Afzal Ansari, Nikha Trivedi, Bhavesh Bagda , R D Dixit, A M Kadri, Harsh Bakshi, Chaitanya Joshi, Madhvi Joshi |
| EPI_ISL_586565 | GMERS GMERS Medical College and Hospital, Junagadh | Gujarat Biotechnology Research Centre | Apurvasinh Puvar, Janvi Raval, Zarna Patel, Monika Gandhi, Pinal Trivedi, Maharshi Pandya, Nidhi Patel, Nitin Savaliya, Raghawendra Kumar, Dinesh Kumar, Zuber Saiyed, Komal Patel, Labdhi Pandya, Afzal Ansari, Nikha Trivedi, Bhavesh Bagda , Ravi Dedaniya, R D Dixit, A M Kadri, Harsh Bakshi, Chaitanya Joshi, Madhvi Joshi |
| EPI_ISL_586566 | GMERS GMERS Medical College and Hospital, Junagadh | Gujarat Biotechnology Research Centre | Janvi Raval, Zarna Patel, Monika Gandhi, Pinal Trivedi, Maharshi Pandya, Nidhi Patel, Nitin Savaliya, Raghawendra Kumar, Dinesh Kumar, Zuber Saiyed, Komal Patel, Labdhi Pandya, Afzal Ansari, Nikha Trivedi, Bhavesh Bagda , Ravi Dedaniya, Apurvasinh Puvar, R D Dixit, A M Kadri, Harsh Bakshi, Chaitanya Joshi, Madhvi Joshi |
| EPI_ISL_590690 | CCC,Veraval | Gujarat Biotechnology Research Centre | Apurvasinh Puvar, Janvi Raval, Zarna Patel, Monika Gandhi, Pinal Trivedi, Maharshi Pandya, Nidhi Patel, Nitin Savaliya, Raghawendra Kumar, Dinesh Kumar, Zuber Saiyed, Komal Patel, Labdhi Pandya, Afzal Ansari, Nikha Trivedi, Jignesh Parmar, Jitendra Bamrotia, R D Dixit, A M Kadri, Harsh Bakshi, |

| | | | |
|---|---|---|---|
| | | | Chaitanya Joshi, Madhvi Joshi |
| EPI_ISL_590691 | Government Hospital,Veraval | Gujarat Biotechnology Research Centre | Janvi Raval, Zarna Patel, Monika Gandhi, Pinal Trivedi, Maharshi Pandya, Nidhi Patel, Nitin Savaliya, Raghawendra Kumar, Dinesh Kumar, Zuber Saiyed, Komal Patel, Labdhi Pandya, Afzal Ansari, Nikha Trivedi, Jignesh Parmar, Jitendra Bamrotia, Apurvasinh Puvar, R D Dixit, A M Kadri, Harsh Bakshi, Chaitanya Joshi, Madhvi Joshi |
| EPI_ISL_590692 | Government Hospital,Veraval | Gujarat Biotechnology Research Centre | Zarna Patel, Monika Gandhi, Pinal Trivedi, Maharshi Pandya, Nidhi Patel, Nitin Savaliya, Raghawendra Kumar, Dinesh Kumar, Zuber Saiyed, Komal Patel, Labdhi Pandya, Afzal Ansari, Nikha Trivedi, Jignesh Parmar, Jitendra Bamrotia, Apurvasinh Puvar, Janvi Raval, R D Dixit, A M Kadri, Harsh Bakshi, Chaitanya Joshi, Madhvi Joshi |
| EPI_ISL_591054, EPI_ISL_591055, EPI_ISL_591056 | Virus Ecology, Rocky Mountain Laboratories, National Institutes of Health | Virus Ecology, Rocky Mountain Laboratories, National Institutes of Health | Van Doremalen,N., Holbrook,M.G., Barbian,K.D., Bushmaker,C., Bushmaker,T., Martens,C.A., Munster,V.J. |
| EPI_ISL_591494, EPI_ISL_591495 | Pathology West - NSW Health Pathology | NSW Health Pathology - Institute of Clinical Pathology and Medical Research; Westmead Hospital; University of Sydney | CIDM-PH et al. |
| EPI_ISL_591503 | South Eastern Area Laboratory Services (SEALS) | NSW Health Pathology - Institute of Clinical Pathology and Medical Research; Westmead Hospital; University of Sydney | CIDM-PH et al. |
| EPI_ISL_591508 | Sydney South West Pathology Service (SSWPS) - Liverpool Hospital - NSW Health Pathology | NSW Health Pathology - Institute of Clinical Pathology and Medical Research; Westmead Hospital; University of Sydney | CIDM-PH et al. |
| EPI_ISL_591563, EPI_ISL_591628, EPI_ISL_591662, EPI_ISL_591683, EPI_ISL_591764, EPI_ISL_591811, EPI_ISL_591905, EPI_ISL_591914, EPI_ISL_591916, EPI_ISL_591921, EPI_ISL_591922, EPI_ISL_591934, EPI_ISL_591935 | | | |
| see above | Microbiological Diagnostic Unit - Public Health Laboratory (MDU-PHL) | MDU-PHL | Seemann T., Schultz, M. B., Sait, M., Sherry, N. |
| EPI_ISL_592160, EPI_ISL_592188 | Victorian Infectious Diseases Reference Laboratory (VIDRL) | VIDRL and MDU-PHL | Caly L., Seemann T., Sait, M., Schultz, M. B., Druce J., Sherry, N. |
| EPI_ISL_592295, EPI_ISL_592296, EPI_ISL_592297, EPI_ISL_592298, EPI_ISL_592299, EPI_ISL_592300, EPI_ISL_592301, EPI_ISL_592302, EPI_ISL_592303, EPI_ISL_592304, EPI_ISL_592305, EPI_ISL_592306, EPI_ISL_592510, EPI_ISL_592667, EPI_ISL_592740, EPI_ISL_593017, EPI_ISL_593124 | | | |
| see above | Microbiological Diagnostic Unit - Public Health Laboratory (MDU-PHL) | MDU-PHL | Seemann T., Schultz, M. B., Sait, M., Sherry, N. |
| EPI_ISL_593646 | 4Cyte Pathology | NSW Health Pathology - Institute of Clinical Pathology and Medical Research; Westmead Hospital; University of Sydney | CIDM-PH et al. |
| EPI_ISL_593649 | Austech Medical Laboratories | NSW Health Pathology - Institute of Clinical Pathology and Medical Research; Westmead Hospital; University of Sydney | CIDM-PH et al. |
| EPI_ISL_593650 | Douglass Hanly Moir Pathology | NSW Health Pathology - Institute of Clinical Pathology and Medical Research; Westmead Hospital; University of Sydney | CIDM-PH et al. |
| EPI_ISL_593655 | Pathology North - Hunter - NSW Health Pathology | NSW Health Pathology - Institute of Clinical Pathology and Medical Research; Westmead Hospital; University of Sydney | CIDM-PH et al. |
| EPI_ISL_593669, EPI_ISL_593670 | Pathology West - NSW Health Pathology | NSW Health Pathology - Institute of Clinical Pathology and Medical Research; Westmead Hospital; University of Sydney | CIDM-PH et al. |
| EPI_ISL_593704, EPI_ISL_593705, EPI_ISL_593709 | South Eastern Area Laboratory Services (SEALS) | NSW Health Pathology - Institute of Clinical Pathology and Medical Research; Westmead Hospital; University of Sydney | CIDM-PH et al. |
| EPI_ISL_593868, EPI_ISL_593869, EPI_ISL_593870, EPI_ISL_593871, EPI_ISL_593872, EPI_ISL_593873, EPI_ISL_593883 | CHU Purpan - Laboratoire de Virologie - Institut Fédératif de Biologie | CHU Purpan - Laboratoire de Virologie - Institut Fédératif de Biologie | Latour J., Ranger N., Dubois M., Carcenac R., Harter A., Boyer P., Tremeaux P., Izopet J. |
| EPI_ISL_594138, EPI_ISL_594146 | MDU-PHL, The Peter Doherty Institute for Infection and Immunity | MDU-PHL, The Peter Doherty Institute for Infection and Immunity | Caly,L., Seemann,T., Sait,M.L., Schultz,M.B., Druce,J., Sherry,N.L. |
| EPI_ISL_594159, EPI_ISL_594160, EPI_ISL_594161 | Israel Institute for Biological Research | Israel Institute for Biological Research | Galia Zaide, Inbar Cohen-Gihon, Ofir Israeli, Dana Stein, Shay Weiss, Orly Laskar, Yoav Gal, Libby Weiss, Emanuelle Mamroud, Adi Beth-Din and Anat Zvi |
| EPI_ISL_596264 | WHO National Influenza Centre Russian Federation | WHO National Influenza Centre Russian Federation | Andrey Komissarov, Artem Fadeev, Anna Ivanova, Mariia Sergeeva, Kseniya Komissarova, Dmitry Bazhenov, Daria Danilenko |
| EPI_ISL_596500, EPI_ISL_596501, EPI_ISL_596507, EPI_ISL_596514, EPI_ISL_596522, EPI_ISL_596533, EPI_ISL_596555, EPI_ISL_596564, EPI_ISL_596565, EPI_ISL_596567 | Palestinian Ministry of Health | Molecular Genetics Lab | Nouar Qutob, Zaidoun Salah, Damien Richard, Hisham Darwish, Husam Sallam, Issa Shtayeh, Osama Najjar, Mahmoud Ruzayqat, Dana Najjar, Francois Balloux, Lucy van Dorp |
| EPI_ISL_596708, EPI_ISL_596712, EPI_ISL_596717, EPI_ISL_596719, EPI_ISL_596720, EPI_ISL_596722, EPI_ISL_596727, EPI_ISL_596732, EPI_ISL_596735, EPI_ISL_596753, EPI_ISL_596759, EPI_ISL_596760, EPI_ISL_596783, EPI_ISL_596784, EPI_ISL_596785, EPI_ISL_596790, EPI_ISL_596793, EPI_ISL_596799, EPI_ISL_596802, EPI_ISL_596807, EPI_ISL_596818 | | | |
| see above | PathWest Laboratory Medicine WA | PathWest Laboratory Medicine WA Microbial Surveillance Unit | PathWest Laboratory Medicine WA Microbial Surveillance Unit |
| EPI_ISL_602245, EPI_ISL_602246, EPI_ISL_602247, EPI_ISL_602248, EPI_ISL_602249, EPI_ISL_602250, EPI_ISL_602251, EPI_ISL_602252, EPI_ISL_602253, EPI_ISL_602254, EPI_ISL_602255 | | | |
| see above | Texas Department of State Health Services | Texas Department of State Health Services | Rashmi Tuladhar, Bonnie Oh, Jenny Zhang, Maliha Rahman, Anita Pokharel, Myong Koag, Chung Wang, Rachel Lee, Grace Kubin, Mayela Pedrueza |
| EPI_ISL_602582, EPI_ISL_602583 | Private medical practitioner | Hong Kong Department of Health | Mak Gannon C.K., Lam Edman T.K., Chan Rickjason C.W., Tsang Dominic N.C. |
| EPI_ISL_602584 | Queen Mary Hospital | Hong Kong Department of Health | Mak Gannon C.K., Lam Edman T.K., Chan Rickjason C.W., Tsang Dominic N.C. |
| EPI_ISL_602585 | Private medical practitioner | Hong Kong Department of Health | Mak Gannon C.K., Lam Edman T.K., Chan Rickjason C.W., Tsang Dominic N.C. |
| EPI_ISL_602586 | Queen Elizabeth Hospital | Hong Kong Department of Health | Mak Gannon C.K., Lam Edman T.K., Chan Rickjason C.W., Tsang Dominic N.C. |
| EPI_ISL_602587 | Private medical practitioner | Hong Kong Department of Health | Mak Gannon C.K., Lam Edman T.K., Chan Rickjason C.W., Tsang Dominic N.C. |
| EPI_ISL_602588 | United Christian Hospital | Hong Kong Department of Health | Mak Gannon C.K., Lam Edman T.K., Chan Rickjason C.W., Tsang Dominic N.C. |
| EPI_ISL_602589 | Queen Elizabeth Hospital | Hong Kong Department of Health | Mak Gannon C.K., Lam Edman T.K., Chan Rickjason C.W., Tsang Dominic N.C. |
| EPI_ISL_602590 | Caritas Medical Centre | Hong Kong Department of Health | Mak Gannon C.K., Lam Edman T.K., Chan Rickjason C.W., Tsang Dominic N.C. |
| EPI_ISL_602591 | Queen Elizabeth Hospital | Hong Kong Department of Health | Mak Gannon C.K., Lam Edman T.K., Chan Rickjason C.W., Tsang Dominic N.C. |
| EPI_ISL_602592 | United Christian Hospital | Hong Kong Department of Health | Mak Gannon C.K., Lam Edman T.K., Chan Rickjason C.W., Tsang Dominic N.C. |
| EPI_ISL_602593 | Queen Mary Hospital | Hong Kong Department of Health | Mak Gannon C.K., Lam Edman T.K., Chan Rickjason C.W., Tsang Dominic N.C. |
| EPI_ISL_602594 | Prince of Wales Hospital | Hong Kong Department of Health | Mak Gannon C.K., Lam Edman T.K., Chan Rickjason C.W., Tsang Dominic N.C. |
| EPI_ISL_602595 | Pamela Youde Nethersole Eastern Hospital | Hong Kong Department of Health | Mak Gannon C.K., Lam Edman T.K., Chan Rickjason C.W., Tsang Dominic N.C. |
| EPI_ISL_602597 | Private medical practitioner | Hong Kong Department of Health | Mak Gannon C.K., Lam Edman T.K., Chan Rickjason C.W., Tsang Dominic N.C. |

| | | | |
|---|---|---|---|
| EPI_ISL_602600 | United Christian Hospital | Hong Kong Department of Health | Mak Gannon C.K., Lam Edman T.K., Chan Rickjason C.W., Tsang Dominic N.C. |
| EPI_ISL_602625 | AHRI-Sigal | KRISP, KZN Research Innovation and Sequencing Platform | Gazy I, Sigl A, Karim F, Cele S, Giandhari J, Pillay S, Tegally H, Wilkinson E, de Oliveira T |
| EPI_ISL_603021 | Pronto Socorro Dr. Conrado Cesarino Nuvolini | Instituto Adolfo Lutz, Interdiciplinary Procedures Center, Strategic Laboratory | Claudio Tavares Sacchi, Claudia Regina Gonçalves, Erica Valessa Ramos Gomes, Karoline Rodrigues Campos |
| EPI_ISL_603022 | Departamento de Vigilância à Saúde | Instituto Adolfo Lutz, Interdiciplinary Procedures Center, Strategic Laboratory | Claudio Tavares Sacchi, Claudia Regina Gonçalves, Erica Valessa Ramos Gomes, Karoline Rodrigues Campos |
| EPI_ISL_603030 | Hospital Domingos Leonardo Ceravolo Presidente Prudente | Instituto Adolfo Lutz, Interdiciplinary Procedures Center, Strategic Laboratory | Claudio Tavares Sacchi, Claudia Regina Gonçalves, Erica Valessa Ramos Gomes, Karoline Rodrigues Campos |
| EPI_ISL_603031 | Santa Casa de Presidente Epitácio | Instituto Adolfo Lutz, Interdiciplinary Procedures Center, Strategic Laboratory | Claudio Tavares Sacchi, Claudia Regina Gonçalves, Erica Valessa Ramos Gomes, Karoline Rodrigues Campos |
| EPI_ISL_603032 | Santa Casa da Misericórdia de Presidente Prudente | Instituto Adolfo Lutz, Interdiciplinary Procedures Center, Strategic Laboratory | Claudio Tavares Sacchi, Claudia Regina Gonçalves, Erica Valessa Ramos Gomes, Karoline Rodrigues Campos |
| EPI_ISL_603033 | Vigilancia Epidemiologica de São Bernardo do Campo | Instituto Adolfo Lutz, Interdiciplinary Procedures Center, Strategic Laboratory | Claudio Tavares Sacchi, Claudia Regina Gonçalves, Erica Valessa Ramos Gomes, Karoline Rodrigues Campos |
| EPI_ISL_603034 | Departamento de Vigilância à Saúde | Instituto Adolfo Lutz, Interdiciplinary Procedures Center, Strategic Laboratory | Claudio Tavares Sacchi, Claudia Regina Gonçalves, Erica Valessa Ramos Gomes, Karoline Rodrigues Campos |
| EPI_ISL_603037 | Hospital Geral de Pedreira | Instituto Adolfo Lutz, Interdiciplinary Procedures Center, Strategic Laboratory | Claudio Tavares Sacchi, Claudia Regina Gonçalves, Erica Valessa Ramos Gomes, Karoline Rodrigues Campos |
| EPI_ISL_603038 | Santa Casa de Misericordia de Araçatuba | Instituto Adolfo Lutz, Interdiciplinary Procedures Center, Strategic Laboratory | Claudio Tavares Sacchi, Claudia Regina Gonçalves, Erica Valessa Ramos Gomes, Karoline Rodrigues Campos |
| EPI_ISL_603039 | Hospital Municipal Mário Gatti | Instituto Adolfo Lutz, Interdiciplinary Procedures Center, Strategic Laboratory | Claudio Tavares Sacchi, Claudia Regina Gonçalves, Erica Valessa Ramos Gomes, Karoline Rodrigues Campos |
| EPI_ISL_605156, EPI_ISL_605157, EPI_ISL_605158, EPI_ISL_605159, EPI_ISL_605160, EPI_ISL_605161, EPI_ISL_605162, EPI_ISL_605163, EPI_ISL_605164, EPI_ISL_605165, EPI_ISL_605166, EPI_ISL_605167, EPI_ISL_605168, EPI_ISL_605169, EPI_ISL_605170, EPI_ISL_605171, EPI_ISL_605172, EPI_ISL_605173, EPI_ISL_605174, EPI_ISL_605175, EPI_ISL_605176, EPI_ISL_605177, EPI_ISL_605178, EPI_ISL_605179, EPI_ISL_605180, EPI_ISL_605181, EPI_ISL_605182, EPI_ISL_605183, EPI_ISL_605184, EPI_ISL_605185, EPI_ISL_605186, EPI_ISL_605187, EPI_ISL_605188, EPI_ISL_605189, EPI_ISL_605190, EPI_ISL_605191, EPI_ISL_605192, EPI_ISL_605193, EPI_ISL_605194, EPI_ISL_605195, EPI_ISL_605196, EPI_ISL_605197, EPI_ISL_605198, EPI_ISL_605199, EPI_ISL_605200, EPI_ISL_605201, EPI_ISL_605202, EPI_ISL_605203, EPI_ISL_605204, EPI_ISL_605205, EPI_ISL_605206, EPI_ISL_605207, EPI_ISL_605208, EPI_ISL_605209, EPI_ISL_605210, EPI_ISL_605211, EPI_ISL_605212, EPI_ISL_605213, EPI_ISL_605214, EPI_ISL_605215, EPI_ISL_605216, EPI_ISL_605217, EPI_ISL_605218, EPI_ISL_605219, EPI_ISL_605220, EPI_ISL_605221, EPI_ISL_605222, EPI_ISL_605223, EPI_ISL_605224, EPI_ISL_605225, EPI_ISL_605226, EPI_ISL_605227, EPI_ISL_605228, EPI_ISL_605229, EPI_ISL_605230, EPI_ISL_605231, EPI_ISL_605232, EPI_ISL_605233, EPI_ISL_605234, EPI_ISL_605235, EPI_ISL_605236, EPI_ISL_605237, EPI_ISL_605238, EPI_ISL_605239, EPI_ISL_605240, EPI_ISL_605241, EPI_ISL_605242, EPI_ISL_605243, EPI_ISL_605244, EPI_ISL_605245, EPI_ISL_605246, EPI_ISL_605247, EPI_ISL_605248, EPI_ISL_605249, EPI_ISL_605250, EPI_ISL_605251, EPI_ISL_605252, EPI_ISL_605253, EPI_ISL_605254, EPI_ISL_605255, EPI_ISL_605256, EPI_ISL_605257, EPI_ISL_605258, EPI_ISL_605259, EPI_ISL_605260, EPI_ISL_605261, EPI_ISL_605262, EPI_ISL_605263, EPI_ISL_605264, EPI_ISL_605265, EPI_ISL_605266, EPI_ISL_605267, EPI_ISL_605268, EPI_ISL_605269, EPI_ISL_605270, EPI_ISL_605271, EPI_ISL_605272, EPI_ISL_605273, EPI_ISL_605274, EPI_ISL_605275, EPI_ISL_605276, EPI_ISL_605277, EPI_ISL_605278, EPI_ISL_605279, EPI_ISL_605280, EPI_ISL_605281, EPI_ISL_605282, EPI_ISL_605283, EPI_ISL_605284, EPI_ISL_605285, EPI_ISL_605286, EPI_ISL_605287, EPI_ISL_605288, EPI_ISL_605289, EPI_ISL_605290, EPI_ISL_605291, EPI_ISL_605292, EPI_ISL_605293, EPI_ISL_605294, EPI_ISL_605295, EPI_ISL_605296, EPI_ISL_605297, EPI_ISL_605298, EPI_ISL_605299, EPI_ISL_605300, EPI_ISL_605301, EPI_ISL_605302, EPI_ISL_605303, EPI_ISL_605304, EPI_ISL_605305, EPI_ISL_605306, EPI_ISL_605307, EPI_ISL_605308, EPI_ISL_605309, EPI_ISL_605310, EPI_ISL_605311, EPI_ISL_605312, EPI_ISL_605313, EPI_ISL_605314, EPI_ISL_605315, EPI_ISL_605316, EPI_ISL_605317, EPI_ISL_605318, EPI_ISL_605319 | | | |
| see above | Utah Public Health Laboratory | Utah Public Health Laboratory | Erin L. Young, Kelly Oakeson, Tara Gallagher, Michael T. Pyne, E. Susan Slechta, Melanie A. Mallory, Jeffrey B. Stevenson, Salika M. Shakir, David R. Hillyard |
| EPI_ISL_610182, EPI_ISL_610183, EPI_ISL_610184, EPI_ISL_610188, EPI_ISL_610189, EPI_ISL_610190, EPI_ISL_610191, EPI_ISL_610192 | Department of Health Technology and Informatics, The Hong Kong Polytechnic University | Department of Health Technology and Informatics, The Hong Kong Polytechnic University | Siu,G.K.-H., Lee,L.-K., Leung,K.S.-S., Leung,J.S.-L., Ng,T.T.-L., Chan,C.T.-M., Tam,K.K.-G., Lao,H.-Y., Wu,A.K.-L., Yau,M.C.-Y., Lai,Y.W.-M., Fung,K.S.-C., Chau,S.K.-Y., Wong,B.K.-C., To,W.-K., Luk,K., Ho,A.Y.-M., Que,T.-L., Yip,K.-T., Yam,W.C., Shum,D.H.-K., Yip,S.P. |
| EPI_ISL_610227, EPI_ISL_610228, EPI_ISL_610229, EPI_ISL_610230, EPI_ISL_610231, EPI_ISL_610232, EPI_ISL_610233, EPI_ISL_610234, EPI_ISL_610235, EPI_ISL_610236 | Molecular diagnostic laboratory of Federal Budget Institution of Science "Central Research Institute of Epidemiology" of The Federal Service on Customers' Rights Protection and Human Well-being Surveillance | Group of Genomics and Postgenomic Technologies of Central Research Institute of Epidemilology | Samoilov AE, Kaptelova VV, Valdokhina AV, Bulanenko VP, Speranskaya AS, Tivanova EV, Shipulina OY, Akimkin VG |
| EPI_ISL_611977 | Liverpool Clinical Laboratories | COVID-19 Genomics UK (COG-UK) Consortium | Sam Haldenby, Anita Lucaci, Steve Paterson, Julian Hiscox, Alistair Darby, M Almsaud, A Alrezaihi, Muhannad Alruwaili, Stuart D Armstrong, Jones Benjamin, Eleanor G Bentley, Anu Chawla, Jordan J Clark, Angela Cowell, Richard Eccles, Isabel García-Dorival, Matthew Gemmell, Alessandro Gerada, PKF Gilmore, Richard Gregory, Ximeng Han, Catherine Hartley, Margaret Hughes, Miren Iturriza-Gomara, James Johnson, L Luu, Jenifer Manson, Charlotte Nelson, Elaine O'Toole, Cassie Olateju, Rebekah Penrice-Randal , Lucille Rainbow, N.P Randle, Trevor Ian Robinson, Parul Sharma, Ghada T Shawli, James P Stewart, Neil Swainston, Ecaterina Vamos, Joanne Watts, Mark Whitehead |
| EPI_ISL_613712, EPI_ISL_613715, EPI_ISL_613716, EPI_ISL_613726, EPI_ISL_613727, EPI_ISL_613730, EPI_ISL_613732, EPI_ISL_613735, EPI_ISL_613739, EPI_ISL_613742, EPI_ISL_613745, EPI_ISL_613759, EPI_ISL_613763, EPI_ISL_613764, EPI_ISL_613768, EPI_ISL_613769, EPI_ISL_613770, EPI_ISL_613771, EPI_ISL_613772, EPI_ISL_613773, EPI_ISL_613777, EPI_ISL_613787, EPI_ISL_613791, EPI_ISL_613805, EPI_ISL_613807, EPI_ISL_613808, EPI_ISL_613809, EPI_ISL_613816, EPI_ISL_613826, EPI_ISL_613829, EPI_ISL_613831, EPI_ISL_613832, EPI_ISL_613833, EPI_ISL_613834, EPI_ISL_613835, EPI_ISL_613836, EPI_ISL_613838, EPI_ISL_613949 | | | |
| see above | Florida Bureau of Public Health Laboratories | Florida Bureau of Public Health Laboratories | Sarah Schmedes, Jason Blanton |
| EPI_ISL_614377, EPI_ISL_614378, EPI_ISL_614379, EPI_ISL_614380 | Molecular diagnostic unit for viral haemorrhagic fevers and emerging viruses, Bouaké CHU Laboratory | Project group Epidemology of Highly Pathogenic Microorganisms, Robert Koch-Institute | Chantal Akoua-Koffi, Diané Bamourou, Etilé Anoh, Essia Belarbi, Safiatou Karidioula, Grit Schubert, Adjaratou Traoré, Soundélé Maïté, Monemo Pacome, Coulibaly Mbegnan, Bamba Fatoumata Touré, Kra Ouffoué, Fabian Leendertz |
| EPI_ISL_618178, EPI_ISL_618179, EPI_ISL_618180, EPI_ISL_618181, EPI_ISL_618182, EPI_ISL_618183, EPI_ISL_618184, EPI_ISL_618191, EPI_ISL_618192, EPI_ISL_618193, EPI_ISL_618194, EPI_ISL_618195, EPI_ISL_618199, EPI_ISL_618200, EPI_ISL_618201, EPI_ISL_618202, EPI_ISL_618203, EPI_ISL_618204, EPI_ISL_618205, EPI_ISL_618206, EPI_ISL_618207, EPI_ISL_618208, EPI_ISL_618209, EPI_ISL_618210, EPI_ISL_618211, EPI_ISL_618212, EPI_ISL_618213, EPI_ISL_618214, EPI_ISL_618215, EPI_ISL_618216, EPI_ISL_618217, EPI_ISL_618218, EPI_ISL_618219, EPI_ISL_618220, EPI_ISL_618221, EPI_ISL_618222, EPI_ISL_618223, EPI_ISL_618224, EPI_ISL_618225, EPI_ISL_618226, EPI_ISL_618227, EPI_ISL_618228, EPI_ISL_618229, EPI_ISL_618232, EPI_ISL_618240, EPI_ISL_618241, EPI_ISL_618245, EPI_ISL_618246, EPI_ISL_618250, EPI_ISL_618251, EPI_ISL_618258, EPI_ISL_618259, EPI_ISL_618260, EPI_ISL_618261, EPI_ISL_618262, EPI_ISL_618270, EPI_ISL_618271, EPI_ISL_618274, EPI_ISL_618275, EPI_ISL_618276, EPI_ISL_618277, EPI_ISL_618279, EPI_ISL_618282, EPI_ISL_618283, EPI_ISL_618284, EPI_ISL_618285, EPI_ISL_618286, EPI_ISL_618287, EPI_ISL_618288, EPI_ISL_618290, EPI_ISL_618291, EPI_ISL_618292, EPI_ISL_618293, EPI_ISL_618295, EPI_ISL_618296, EPI_ISL_618297, EPI_ISL_618298, EPI_ISL_618299, EPI_ISL_618300, EPI_ISL_618301, EPI_ISL_618302, EPI_ISL_618303, EPI_ISL_618304, EPI_ISL_618305, EPI_ISL_618306, EPI_ISL_618307, EPI_ISL_618308, EPI_ISL_618309, EPI_ISL_618310, EPI_ISL_618311, EPI_ISL_618312, EPI_ISL_618315, EPI_ISL_618316, EPI_ISL_618317, EPI_ISL_618318, EPI_ISL_618319, EPI_ISL_618320, EPI_ISL_618321, EPI_ISL_618322, EPI_ISL_618323, EPI_ISL_618324, EPI_ISL_618325, EPI_ISL_618326, EPI_ISL_618327, EPI_ISL_618328, EPI_ISL_618329, EPI_ISL_618330, EPI_ISL_618331, EPI_ISL_618332, EPI_ISL_618333, EPI_ISL_618334, EPI_ISL_618336, EPI_ISL_618345, EPI_ISL_618357, EPI_ISL_618359, EPI_ISL_618360, EPI_ISL_618361, EPI_ISL_618362, EPI_ISL_618372, EPI_ISL_618373, EPI_ISL_618374, EPI_ISL_618380, EPI_ISL_618402, EPI_ISL_618403, EPI_ISL_618404, EPI_ISL_622571, EPI_ISL_622572, EPI_ISL_622573, EPI_ISL_622599, EPI_ISL_622600, EPI_ISL_622603, EPI_ISL_622604, EPI_ISL_622605, EPI_ISL_622606, EPI_ISL_622607, EPI_ISL_622608, EPI_ISL_622609, EPI_ISL_622610, EPI_ISL_622611, EPI_ISL_622612, EPI_ISL_622614 | | | |
| see above | Department of Virus and Microbiological Special Diagnostics, Statens Serum Institut, Denmark | Albertsen lab, Department of Chemistry and Bioscience, Aalborg University, Denmark | Danish Covid-19 Genome Consortia |
| EPI_ISL_622903, EPI_ISL_622907, EPI_ISL_622908, EPI_ISL_622911, EPI_ISL_622912 | National Institute for Communicable Diseases of the National Health Laboratory Service | National Institute for Communicable Diseases of the National Health Laboratory Service | Allam M, Ismail A, Khumalo Z, Kwenda S, Mtshali P, Mnyameni F, Mohale T, Subramoney K, Bhiman JN |
| EPI_ISL_625474, EPI_ISL_625476, EPI_ISL_625477 | Child Health Research Foundation | Child Health Research Foundation | Senjuti Saha, Md Saiful Islam Sajib, Nikkon Sarkar, Syed Muktadir Al Sium, Afroza Akter Tanni, Roly Malaker, Arif Mohammad Tanmoy, Md Hafizur Rahman, Samir K Saha |
| EPI_ISL_625547, EPI_ISL_625548, EPI_ISL_625549 | County of San Luis Obispo Public Health Laboratory | Chan-Zuckerberg Biohub | CZB Cliahub Consortium |
| EPI_ISL_625550, EPI_ISL_625551, EPI_ISL_625552, EPI_ISL_625553, | UCSF Clinical Microbiology Laboratory | Chan-Zuckerberg Biohub | CZB Cliahub Consortium |

| | | | |
|---|---|---|---|
| EPI_ISL_625554, EPI_ISL_625555, EPI_ISL_625556, EPI_ISL_625557, EPI_ISL_625558 | | | |
| EPI_ISL_625573, EPI_ISL_625574, EPI_ISL_625575, EPI_ISL_625576, EPI_ISL_625577, EPI_ISL_625578 | NaN | Chan-Zuckerberg Biohub | CZB Cliahub Consortium |
| EPI_ISL_625624 | County of San Luis Obispo Public Health Laboratory | Chan-Zuckerberg Biohub | CZB Cliahub Consortium |
| EPI_ISL_626528, EPI_ISL_626529, EPI_ISL_626530, EPI_ISL_626531, EPI_ISL_626532, EPI_ISL_626533, EPI_ISL_626534, EPI_ISL_626535, EPI_ISL_626536, EPI_ISL_626537, EPI_ISL_626538, EPI_ISL_626539 | | | |
| see above | Northwestern Memorial Hospital | Ozer Lab | Ramon Lorenzo-Redondo, Hannah H. Nam, Scott C. Roberts, Lacy M. Simons, Chad J. Achenbach, Lawrence J. Jennings, Chao Qi, Alan R. Hauser, Michael G. Ison, Judd F. Hultquist, Egon A. Ozer |
| EPI_ISL_626550, EPI_ISL_626552 | Laboratorio de Biología Molecular, Facultad de Medicina, Universidad de Atacama, Copiapo, Chile/ FONDAP CRG, Universidad Andrés Bello, Santiago, Chile | Center for Mathematical Modeling and Center for Genome Regulation. Santiago, Chile | Echeverría C, Manríquez R, Bastias M, Sanhueza D, Travisany D, Allende ML, Maass A, González M, Montecino, M, Orellana A, Castro E, Meneses C. |
| EPI_ISL_628750, EPI_ISL_628753, EPI_ISL_628759 | UHAS COVID-19 Lab | UHAS COVID-19 Lab | Kwabena O. Duedu, Jones Gyamfi, Reuben Ayivor-Djanie, John O. Gyapong and the UHAS COVID-19 Lab Team |
| EPI_ISL_631398 | Wisconsin State Laboratory of Hygiene Communicable Disease Division | Wisconsin State Laboratory of Hygiene Communicable Disease Division | Kelsey R. Florek, Abigail C. Shockey |
| EPI_ISL_632287 | Environmental and Global Health, University of Florida | Environmental and Global Health, University of Florida | Alam,M.M., Lauzardo,M., Elbadry,M.A., Stephenson,C.J., Gibson,J.C., Morris,J.G. and Lednicky,J.A. |
| EPI_ISL_632963 | NYC Department Of Health And Mental Hygiene | New York City Public Health Laboratory | Jade Wang, et al. |
| EPI_ISL_635573, EPI_ISL_635574, EPI_ISL_635575 | Centro de Diagnstico COVID-19 UABC Tijuana | Andersen lab at Scripps Research | SEARCH Alliance San Diego with Idanya Rubí Serafín Higuera, Manuel Sánchez Alavez, Jorge Luis Jiménez Niebla, Germán Ibarra, Jonathan Vincent Baena, Oscar Efrén Zazueta Fierro |
| EPI_ISL_635580, EPI_ISL_635582, EPI_ISL_635583, EPI_ISL_635584, EPI_ISL_635585, EPI_ISL_635587, EPI_ISL_635588, EPI_ISL_635589, EPI_ISL_635592, EPI_ISL_635595, EPI_ISL_635596, EPI_ISL_635597, EPI_ISL_635601, EPI_ISL_635602, EPI_ISL_635603, EPI_ISL_635604, EPI_ISL_635605, EPI_ISL_635606, EPI_ISL_635607, EPI_ISL_635608, EPI_ISL_635610, EPI_ISL_635611, EPI_ISL_635612, EPI_ISL_635614, EPI_ISL_635616, EPI_ISL_635618, EPI_ISL_635619, EPI_ISL_635620, EPI_ISL_635621, EPI_ISL_635624, EPI_ISL_635625, EPI_ISL_635626, EPI_ISL_635627, EPI_ISL_635632, EPI_ISL_635633, EPI_ISL_635634, EPI_ISL_635635, EPI_ISL_635636, EPI_ISL_635637, EPI_ISL_635638, EPI_ISL_635639, EPI_ISL_635640, EPI_ISL_635641, EPI_ISL_635642, EPI_ISL_635643, EPI_ISL_635644, EPI_ISL_635645, EPI_ISL_635646, EPI_ISL_635647, EPI_ISL_635648, EPI_ISL_635649, EPI_ISL_635650, EPI_ISL_635651, EPI_ISL_635652, EPI_ISL_635653, EPI_ISL_635654, EPI_ISL_635655, EPI_ISL_635656, EPI_ISL_635657, EPI_ISL_635658, EPI_ISL_635659, EPI_ISL_635660, EPI_ISL_635661, EPI_ISL_635662, EPI_ISL_635663, EPI_ISL_635664, EPI_ISL_635665, EPI_ISL_635666, EPI_ISL_635667, EPI_ISL_635668, EPI_ISL_635669, EPI_ISL_635670, EPI_ISL_635671, EPI_ISL_635672, EPI_ISL_635673, EPI_ISL_635674, EPI_ISL_635675, EPI_ISL_635676, EPI_ISL_635677, EPI_ISL_635678, EPI_ISL_635679, EPI_ISL_635686, EPI_ISL_635689, EPI_ISL_635690, EPI_ISL_635691, EPI_ISL_635694, EPI_ISL_635695, EPI_ISL_635699, EPI_ISL_635700, EPI_ISL_635702, EPI_ISL_635722, EPI_ISL_635724, EPI_ISL_635725, EPI_ISL_635726, EPI_ISL_635727, EPI_ISL_635728, EPI_ISL_635730, EPI_ISL_635732, EPI_ISL_635734, EPI_ISL_635736, EPI_ISL_635738, EPI_ISL_635739, EPI_ISL_635740, EPI_ISL_635741, EPI_ISL_635742, EPI_ISL_635743, EPI_ISL_635744, EPI_ISL_635745, EPI_ISL_635746, EPI_ISL_635747, EPI_ISL_635748, EPI_ISL_635783, EPI_ISL_635784, EPI_ISL_635785, EPI_ISL_635786, EPI_ISL_635787, EPI_ISL_635788, EPI_ISL_635845, EPI_ISL_635846, EPI_ISL_635848, EPI_ISL_635849, EPI_ISL_635850, EPI_ISL_635851, EPI_ISL_635852, EPI_ISL_635853, EPI_ISL_635855, EPI_ISL_635856, EPI_ISL_635857, EPI_ISL_635858, EPI_ISL_635872, EPI_ISL_635874, EPI_ISL_635883, EPI_ISL_635889, EPI_ISL_635890, EPI_ISL_635912, EPI_ISL_635920, EPI_ISL_635921, EPI_ISL_635923, EPI_ISL_635931, EPI_ISL_636045, EPI_ISL_636058, EPI_ISL_636064, EPI_ISL_636065, EPI_ISL_636232, EPI_ISL_636239, EPI_ISL_636256 | | | |
| see above | San Diego County Public Health Laboratory | Andersen lab at Scripps Research | SEARCH Alliance San Diego with Tracy Basler, Jovan Shephard, Brett Austin |
| EPI_ISL_636604, EPI_ISL_636839, EPI_ISL_636840 | Lithuanian University of Health Sciences Hospital, Department of Laboratory Medicine | Lithuanian University of Health Sciences, Molecular cardiology lab. | Lukas Zemaitis, Ingrida Olendrait, Arnoldas Pautienius, Kamile Tamusauskaite, Dovydas Gecys, Laura Pareckaite, Vaiva Lesauskaite, Astra Vitkauskiene |
| EPI_ISL_636976, EPI_ISL_636977, EPI_ISL_636978 | HP Pemba | KRISP, KZN Research Innovation and Sequencing Platform | Ismael N, Giandhari J, Pillay S, Tegally H, Wilkinson E, de Oliveira T, Nadia Sitoe, Paulo Arnaldo, Nedio Mabunda |
| EPI_ISL_636980 | CS Xai Xai | KRISP, KZN Research Innovation and Sequencing Platform | Ismael N, Giandhari J, Pillay S, Tegally H, Wilkinson E, de Oliveira T, Nadia Sitoe, Paulo Arnaldo, Nedio Mabunda |
| EPI_ISL_636995, EPI_ISL_636996 | Department of Infectious Diseases and Immunology, National Hospital Organization Nagoya Medical Center | Clinical Research Center, National Hospital Organization Nagoya Medical Center | Yoshihiro Nakata, Hirotaka Ode, Mai Kubota, Masakazu Matsuda, Kazuhiro Matsuoka, Miho Nakasuji, Mikiko Mori, Mayumi Imahashi, Yoshiyuki Yokomaku, Yasumasa Iwatani |
| EPI_ISL_639739 | Centre of Nanotechnologies, INCD IMT-Bucuresti (National Institute for Research and Development in Microtechnologies - Bucharest) | Centre of Nanotechnologies, INCD IMT-Bucuresti (National Institute for Research and Development in Microtechnologies - Bucharest) | Salceanu,A., Gogianu,L. and Baisan,M. |
| EPI_ISL_640064 | Ethembeni Clinic [Prieska] | NHLS/UCT | Arash Iranzadeh, Deelan Doolabh, Lynn Tyers, Bruna Galvao, Innocent Mudau, Marvin Hsiao, Kruger Marais, Diana Hardie, Stephen Korsman, Carolyn Williamson |
| EPI_ISL_640074 | Mamre CDC wc MRC | NHLS/UCT | Arash Iranzadeh, Deelan Doolabh, Lynn Tyers, Bruna Galvao, Innocent Mudau, Marvin Hsiao, Kruger Marais, Diana Hardie, Stephen Korsman, Carolyn Williamson |
| EPI_ISL_640344, EPI_ISL_640347, EPI_ISL_640383, EPI_ISL_640480, EPI_ISL_640502, EPI_ISL_640507, EPI_ISL_640529, EPI_ISL_640538, EPI_ISL_640623, EPI_ISL_640624, EPI_ISL_640639, EPI_ISL_640680, EPI_ISL_640725, EPI_ISL_640811, EPI_ISL_640835, EPI_ISL_640867, EPI_ISL_640869, EPI_ISL_640873, EPI_ISL_640890, EPI_ISL_640894, EPI_ISL_640908, EPI_ISL_640933, EPI_ISL_640991, EPI_ISL_641000, EPI_ISL_641003, EPI_ISL_641033, EPI_ISL_641060, EPI_ISL_641086, EPI_ISL_641091, EPI_ISL_641107, EPI_ISL_641110, EPI_ISL_641112, EPI_ISL_641181, EPI_ISL_641216, EPI_ISL_641294 | | | |
| see above | Microbiological Diagnostic Unit - Public Health Laboratory (MDU-PHL) | MDU-PHL | Seemann T., Schultz M.B., Sait, M.L., Sherry, N.L. |
| EPI_ISL_644565, EPI_ISL_644566, EPI_ISL_644567, EPI_ISL_644569, EPI_ISL_644571, EPI_ISL_644573 | Veterinary Specialized Institute "Kraljevo", Serbia | Veterinary Specialized Institute "Kraljevo", Serbia | Vidanovic,D., Tesovic,B., Knezevic,A., Jovanovic,T., Jankovic,M., Sekler,M., Banovic Djeri,B., Petrovic,T., Volkening,J., Afonso,C. |
| EPI_ISL_644961, EPI_ISL_644962, EPI_ISL_644963, EPI_ISL_644964, EPI_ISL_644965, EPI_ISL_644966, EPI_ISL_644967, EPI_ISL_644968, EPI_ISL_644969 | Department of Infectious Diseases, Keio University School of Medicine, Tokyo, Japan | Center for Medical Genetics, Keio University School of Medicine, Tokyo, Japan | Kenjiro Kosaki, Yuka Iwasaki, Hirotsugu Ishizu, Haruhiko Siomi, Kodai Abe |
| EPI_ISL_648134, EPI_ISL_648138, EPI_ISL_648139 | The Public Health Agency of Sweden | The Public Health Agency of Sweden | Anna-Malin Linde, Maria Lind Karlberg, Mattias Haukland, Reza Advani, Olov Svartstrom, Oskar Karlsson Lindsjo, Sandra Broddesson, Petra Edquist, Mia Brytting, Anna Risberg, Karin Tegmark-Wisell |
| EPI_ISL_648209, EPI_ISL_648210, EPI_ISL_648211, EPI_ISL_648212, EPI_ISL_648213, EPI_ISL_648214, EPI_ISL_648215, EPI_ISL_648216 | INBIRS-UBA | Laboratorio Mixto de Biotecnología Acuática (LMBA) | Joaquín Ezpeleta, Ignacio García Labari, Victoria Posner, Vanina Villanova, Pablo Casal, Sofía Lavista Llanos, Federico Remes Lenicov, Ana Paletta, Flavio Spetale, Agustina Cerri, Silvana Spinelli, Elisa Bolatti, Diego Chouhy, María Re, Gastón Viarengo, Ana Cavatorta, Julian Acosta, Javier Murillo, Laura Angelone, Leandro Ciappina, Pilar Bulacio, Adriana Giri, Silvia Arranz, Elizabeth Tapia |
| EPI_ISL_648217 | SILAB | Laboratorio Mixto de Biotecnología Acuática (LMBA) | Joaquín Ezpeleta, Ignacio García Labari, Victoria Posner, Vanina Villanova, Pablo Casal, Sofía Lavista Llanos, Federico Remes Lenicov, Ana Paletta, Flavio Spetale, Agustina Cerri, Silvana Spinelli, Elisa Bolatti, Diego Chouhy, María Re, Gastón Viarengo, Ana Cavatorta, Julian Acosta, Javier Murillo, Laura Angelone, Leandro Ciappina, Pilar Bulacio, Adriana Giri, Silvia Arranz, Elizabeth Tapia |
| EPI_ISL_648218 | INBIRS-UBA | Laboratorio Mixto de Biotecnología Acuática (LMBA) | Joaquín Ezpeleta, Ignacio García Labari, Victoria Posner, Vanina Villanova, Pablo Casal, Sofía Lavista Llanos, Federico Remes Lenicov, Ana Paletta, Flavio Spetale, Agustina Cerri, Silvana Spinelli, Elisa Bolatti, Diego Chouhy, María Re, Gastón Viarengo, Ana Cavatorta, Julian Acosta, Javier Murillo, Laura Angelone, Leandro Ciappina, Pilar Bulacio, Adriana Giri, Silvia Arranz, Elizabeth Tapia |
| EPI_ISL_648327, EPI_ISL_648328, EPI_ISL_648329, EPI_ISL_648330, EPI_ISL_648376, EPI_ISL_648377 | Laboratorio de Investigaciones de Baney | University Hospital Basel, Clinical Bacteriology | Carlos Cortes, Claudia Daubenberger, Adrian Egli, Guillermo Garcia, Salome Hosch, Bonifacio Manguire Nlavo, Alfredo Mari, Maximilian Mpina, Elizabeth Nyakarungu, Diosdado Odjama Nseng Ada, Mitoha Ondo O Ayekaba, Tim Roloff, Tobias Schindler, Helena Seth-Smith, Madlen Stange, Philip Wonder Phiri |
| EPI_ISL_648677 | INBIRS-UBA | Laboratorio Mixto de Biotecnología Acuática (LMBA) | Joaquín Ezpeleta, Ignacio García Labari, Victoria Posner, Vanina Villanova, Pablo Casal, Sofía Lavista Llanos, Federico Remes Lenicov, Ana Paletta, Flavio |

| | | | |
|---|---|---|---|
| | | | Spetale, Agustina Cerri, Silvana Spinelli, Elisa Bolatti, Diego Chouhy, María Re, Gastón Viarengo, Ana Cavatorta, Julian Acosta, Javier Murillo, Laura Angelone, Leandro Ciappina, Pilar Bulacio, Adriana Giri, Silvia Arranz, Elizabeth Tapia |
| EPI_ISL_648680, EPI_ISL_648681, EPI_ISL_648735, EPI_ISL_648736, EPI_ISL_648737, EPI_ISL_648738, EPI_ISL_648739, EPI_ISL_648740 | Department of Laboratory Medicine, Tan Tock Seng Hospital | Department of Laboratory Medicine, Tan Tock Seng Hospital | Chen YYC, Zair X, Lim JX, Li C, Tang WY, Maurer-Stroh S, Barkham TMS, Nagarajan N, Sessions OM |
| EPI_ISL_648982, EPI_ISL_648984, EPI_ISL_648986, EPI_ISL_648987, EPI_ISL_648991, EPI_ISL_648992, EPI_ISL_648995, EPI_ISL_648996, EPI_ISL_648997, EPI_ISL_648998, EPI_ISL_648999, EPI_ISL_649004 | | | |
| see above | San Diego County Public Health Laboratory | Andersen lab at Scripps Research | SEARCH Alliance San Diego with Tracy Basler, Jovan Shephard, Brett Austin |
| EPI_ISL_649152 | Infectious diseases, Chan Zuckerberg Biohub | Infectious diseases, Chan Zuckerberg Biohub | Bhatt,K.D. |
| EPI_ISL_649167, EPI_ISL_649168, EPI_ISL_649169 | Laboratorio de Investigaciones de Baney | University Hospital Basel, Clinical Bacteriology | Carlos Cortes, Claudia Daubenberger, Adrian Egli, Guillermo Garcia, Salome Hosch, Bonifacio Manguire Nlavo, Alfredo Mari, Maximilian Mpina, Elizabeth Nyakarungu, Diosdado Odjama Nseng Ada, Mitoha Ondo O Ayekaba, Tim Roloff, Tobias Schindler, Helena Seth-Smith, Madlen Stange, Philip Wonder Phiri |
| EPI_ISL_653188, EPI_ISL_653189, EPI_ISL_653190, EPI_ISL_653191, EPI_ISL_653192, EPI_ISL_653193, EPI_ISL_653194, EPI_ISL_653195, EPI_ISL_653196, EPI_ISL_653197, EPI_ISL_653209, EPI_ISL_653223, EPI_ISL_653224, EPI_ISL_653225, EPI_ISL_653230, EPI_ISL_653239, EPI_ISL_653240, EPI_ISL_653241, EPI_ISL_653242, EPI_ISL_653243, EPI_ISL_653244, EPI_ISL_653251, EPI_ISL_653252, EPI_ISL_653342, EPI_ISL_653343, EPI_ISL_653344 | | | |
| see above | Florida Bureau of Public Health Laboratories | Florida Bureau of Public Health Laboratories | Sarah Schmedes, Jason Blanton |
| EPI_ISL_653345, EPI_ISL_653346, EPI_ISL_653347, EPI_ISL_653348, EPI_ISL_653349, EPI_ISL_653350, EPI_ISL_653351, EPI_ISL_653352, EPI_ISL_653353, EPI_ISL_653354, EPI_ISL_653355, EPI_ISL_653356, EPI_ISL_653357, EPI_ISL_653358, EPI_ISL_653359, EPI_ISL_653360, EPI_ISL_653361, EPI_ISL_653362, EPI_ISL_653363, EPI_ISL_653364, EPI_ISL_653365, EPI_ISL_653366, EPI_ISL_653367, EPI_ISL_653368, EPI_ISL_653369 | | | |
| see above | LSUHS Emerging Viral Threat Laboratory | Microbial Genome Sequencing Center | Maarten Van Diest, Jeremy P. Kamil, Rona S. Scott, Malgorzata Bienkowska-Haba, Katarzyna Zwolinska, Andrew D. Yurochko, Christopher G. Kevil, Martin J. Sapp, Daniel J. Snyder, Vaughn S. Cooper, John A. Vanchiere |
| EPI_ISL_653370, EPI_ISL_653371, EPI_ISL_653372, EPI_ISL_653373, EPI_ISL_653374, EPI_ISL_653375, EPI_ISL_653376, EPI_ISL_653377, EPI_ISL_653378, EPI_ISL_653379, EPI_ISL_653380, EPI_ISL_653381, EPI_ISL_653382, EPI_ISL_653383, EPI_ISL_653384, EPI_ISL_653385, EPI_ISL_653386, EPI_ISL_653387, EPI_ISL_653388, EPI_ISL_653389, EPI_ISL_653390, EPI_ISL_653391, EPI_ISL_653392, EPI_ISL_653393, EPI_ISL_653394 | | | |
| see above | LSUHS Emerging Viral Threat Laboratory | Microbial Genome Sequencing Center | Rona S. Scott, Jeremy P. Kamil, Maarten Van Diest, Malgorzata Bienkowska-Haba, Katarzyna Zwolinska, Andrew D. Yurochko, Christopher G. Kevil, Martin J. Sapp, Daniel J. Snyder, Vaughn S. Cooper, John A. Vanchiere |
| EPI_ISL_653395, EPI_ISL_653396, EPI_ISL_653397, EPI_ISL_653398, EPI_ISL_653399, EPI_ISL_653400, EPI_ISL_653401, EPI_ISL_653402, EPI_ISL_653403, EPI_ISL_653404, EPI_ISL_653405, EPI_ISL_653406, EPI_ISL_653407, EPI_ISL_653408, EPI_ISL_653409, EPI_ISL_653410, EPI_ISL_653411, EPI_ISL_653412, EPI_ISL_653413, EPI_ISL_653414, EPI_ISL_653415, EPI_ISL_653416, EPI_ISL_653417, EPI_ISL_653418, EPI_ISL_653419, EPI_ISL_653420, EPI_ISL_653421, EPI_ISL_653422, EPI_ISL_653423, EPI_ISL_653424, EPI_ISL_653425, EPI_ISL_653426, EPI_ISL_653427, EPI_ISL_653428, EPI_ISL_653429, EPI_ISL_653430, EPI_ISL_653431, EPI_ISL_653432, EPI_ISL_653433, EPI_ISL_653434, EPI_ISL_653435, EPI_ISL_653436, EPI_ISL_653437, EPI_ISL_653438, EPI_ISL_653439, EPI_ISL_653440, EPI_ISL_653441, EPI_ISL_653442, EPI_ISL_653443, EPI_ISL_653444, EPI_ISL_653445, EPI_ISL_653446, EPI_ISL_653447, EPI_ISL_653448, EPI_ISL_653449, EPI_ISL_653450, EPI_ISL_653451, EPI_ISL_653452, EPI_ISL_653453, EPI_ISL_653454, EPI_ISL_653455, EPI_ISL_653456, EPI_ISL_653457, EPI_ISL_653458, EPI_ISL_653459, EPI_ISL_653460, EPI_ISL_653461, EPI_ISL_653462, EPI_ISL_653463, EPI_ISL_653464, EPI_ISL_653465, EPI_ISL_653466, EPI_ISL_653467, EPI_ISL_653468, EPI_ISL_653469, EPI_ISL_653470, EPI_ISL_653471, EPI_ISL_653472, EPI_ISL_653473, EPI_ISL_653474 | | | |
| see above | LSUHS Emerging Viral Threat Laboratory | Microbial Genome Sequencing Center | Jeremy P. Kamil, Rona S. Scott, Maarten Van Diest, Malgorzata Bienkowska-Haba, Katarzyna Zwolinska, Andrew D. Yurochko, Christopher G. Kevil, Martin J. Sapp, Daniel J. Snyder, Vaughn S. Cooper, John A. Vanchiere |
| EPI_ISL_654191 | Hospital General Universitario Gregorio Marañón | SeqCOVID-SPAIN consortium/IBV(CSIC) | Darío García de Viedma, Laura Pérez-Lago, Marta Herranz, Jon Sicilia, Julia Suárez, Pilar Catalán, Patricia Muñoz and SeqCOVID-SPAIN consortium |
| EPI_ISL_654498 | The Public Health Agency of Sweden | The Public Health Agency of Sweden | Anna-Malin Linde, Maria Lind Karlberg, Mattias Haukland, Reza Advani, Olov Svartstrom, Oskar Karlsson Lindsjo, Sandra Broddesson, Petra Edquist, Mia Brytting, Anna Risberg, Karin Tegmark-Wisell |
| EPI_ISL_654943 | Orebro klinisk mikrobiologi | The Public Health Agency of Sweden | Anna-Malin Linde, Maria Lind Karlberg, Mattias Haukland, Reza Advani, Olov Svartstrom, Oskar Karlsson Lindsjo, Sandra Broddesson, Petra Edquist, Mia Brytting, Anna Risberg, Karin Tegmark-Wisell |
| EPI_ISL_654944 | Klinisk mikrobiologi | The Public Health Agency of Sweden | Anna-Malin Linde, Maria Lind Karlberg, Mattias Haukland, Reza Advani, Olov Svartstrom, Oskar Karlsson Lindsjo, Sandra Broddesson, Petra Edquist, Mia Brytting, Anna Risberg, Karin Tegmark-Wisell |
| EPI_ISL_654955 | Skovde/Unilabs | The Public Health Agency of Sweden | Anna-Malin Linde, Maria Lind Karlberg, Mattias Haukland, Reza Advani, Olov Svartstrom, Oskar Karlsson Lindsjo, Sandra Broddesson, Petra Edquist, Mia Brytting, Anna Risberg, Karin Tegmark-Wisell |
| EPI_ISL_658899 | Instituto de Diagnostico y Referencia Epidemiologicos (INDRE) | Instituto de Diagnostico y Referencia Epidemiologicos (INDRE) | Gisela Barrera-Badillo , Abril Rodriguez-Maldonado, Claudia Wong-Arambula , Natividad Cruz-Ortiz, Tatiana Nunez-Garcia, Dayanira Arellano-Suarez, Fabiola Garces-Ayala, Lucia Hernandez-Rivas, Irma Lopez-Martinez, Ernesto Ramirez-Gonzalez. |
| EPI_ISL_660069 | Zurita & Zurita Laboratorios | Zurita & Zurita Laboratorios | Gabriela Sevillano Camilo Zurita-Salinas Karen Loaiza David Ortega-Paredes Jeannete Zurita |
| EPI_ISL_660170, EPI_ISL_660172 | NHLS-IALCH | KRISP, KZN Research Innovation and Sequencing Platform | Gazy I, Sigal A, Karim F, Cele S, Giandhari J, Pillay S, Tegally H, Wilkinson E, de Oliveira T |
| EPI_ISL_660433, EPI_ISL_660434, EPI_ISL_660435, EPI_ISL_660436 | Molecular diagnostic laboratory of Federal Budget Institution of Science "Central Research Institute of Epidemiology" of The Federal Service on Customers' Rights Protection and Human Well-being Surveillance | Group of Genomics and Postgenomic Technologies of Central Research Institute of Epidemilology | Samoilov AE, Kaptelova VV, Valdokhina AV, Bulanenko VP, Speranskaya AS, Tivanova EV, Shipulina OY, Akimkin VG |
| EPI_ISL_660446 | Laboratoire de Microbiologie CHU Souro Sanou | Centre Muraz | Abdoul-Salam Ouedraogo, Yacouba Sawadogo, Essia Belarbi, Grit Schubert, Fabian Leendertz, Arsène Zongo, Soumeya Ouangraoua, Zekiba Tarnagda, Lassana Sangaré, Halidou Tinto |
| EPI_ISL_661190, EPI_ISL_661194 | Scientific Veterinary Institute Novi Sad | Veterinary Specialized Institute "Kraljevo", Serbia | Vidanovic,D., Tesovic,B., Knezevic,A., Jovanovic,T., Jankovic,M., Sekler,M., Banovic Djeri,B., Petrovic,T., Volkening,J., Afonso,C. |
| EPI_ISL_661273 | Stockholm_StGoran | The Public Health Agency of Sweden | Department of Microbiology, The Public Health Agency of Sweden |
| EPI_ISL_663318, EPI_ISL_663352, EPI_ISL_663367, EPI_ISL_663439, EPI_ISL_663440, EPI_ISL_663469, EPI_ISL_663478, EPI_ISL_663483, EPI_ISL_663484, EPI_ISL_663512, EPI_ISL_663545, EPI_ISL_663683, EPI_ISL_663915, EPI_ISL_663921, EPI_ISL_663933, EPI_ISL_663934, EPI_ISL_663936, EPI_ISL_663939, EPI_ISL_663945, EPI_ISL_663951, EPI_ISL_663956, EPI_ISL_663957, EPI_ISL_663958 | | | |
| see above | Microbiological Diagnostic Unit - Public Health Laboratory (MDU-PHL) | MDU-PHL | Seemann T., Schultz M.B., Sait, M.L., Sherry, N.L. |
| EPI_ISL_666609 | LSUHS Emerging Viral Threat Laboratory | Microbial Genome Sequencing Center | Jeremy P. Kamil, Rona S. Scott, Maarten Van Diest, Malgorzata Bienkowska-Haba, Katarzyna Zwolinska, Andrew D. Yurochko, Christopher G. Kevil, Martin J. Sapp, Daniel J. Snyder, Vaughn S. Cooper, John A. Vanchiere |
| EPI_ISL_666814 | Florida Bureau of Public Health Laboratories | Florida Bureau of Public Health Laboratories | Sarah Schmedes, Jason Blanton |
| EPI_ISL_667014, EPI_ISL_667016, EPI_ISL_667017, EPI_ISL_667018, EPI_ISL_667021, EPI_ISL_667022, EPI_ISL_667023, EPI_ISL_667024, EPI_ISL_667029, EPI_ISL_667033 | San Diego County Public Health Laboratory | Andersen lab at Scripps Research | SEARCH Alliance San Diego with Tracy Basler, Jovan Shephard, Brett Austin |
| EPI_ISL_671847, EPI_ISL_671848, EPI_ISL_671849, EPI_ISL_671850, EPI_ISL_671851, EPI_ISL_671852, EPI_ISL_671853, EPI_ISL_671854, EPI_ISL_671855, EPI_ISL_671856, EPI_ISL_671857, EPI_ISL_671858 | | | |
| see above | Servicio de Microbiología, Laboratori Clínic Metropolitana Nord. Hospital Universitari Germans Trias i Pujol. Institut d'Investigació en Ciències de la Salut Germans Trias i Pujol (IGTP) | SeqCOVID-SPAIN consortium/IBV(CSIC) | Elisa Martró, Antoni E. Bordoy, Anna Not, Adrián Antuori, Anabel Fernández, Nona Romaní, Verónica Saludes, Cristina Casañ and SeqCOVID-SPAIN consortium |
| EPI_ISL_672015 | UCSF Clinical Microbiology Laboratory | Chan-Zuckerberg Biohub | CZB Cliahub Consortium |
| EPI_ISL_672055 | The Ashley Laboratory, Stanford University | Chan-Zuckerberg Biohub | CZB Cliahub Consortium |
| EPI_ISL_672068 | County of San Luis Obispo Public Health Laboratory | Chan-Zuckerberg Biohub | CZB Cliahub Consortium |

| | | | |
|---|---|---|---|
| EPI_ISL_672077 | San Francisco Public Health Laboratory | Chan-Zuckerberg Biohub | CZB Cliahub Consortium |
| EPI_ISL_672087 | County of San Luis Obispo Public Health Laboratory | Chan-Zuckerberg Biohub | CZB Cliahub Consortium |
| EPI_ISL_672185, EPI_ISL_672186, EPI_ISL_672187, EPI_ISL_672227, EPI_ISL_672234 | The Ashley Laboratory, Stanford University | Chan-Zuckerberg Biohub | CZB Cliahub Consortium |
| EPI_ISL_672354, EPI_ISL_672355, EPI_ISL_672356, EPI_ISL_672357, EPI_ISL_672358, EPI_ISL_672359 | County of San Luis Obispo Public Health Laboratory | Chan-Zuckerberg Biohub | CZB Cliahub Consortium |
| EPI_ISL_672375, EPI_ISL_672376, EPI_ISL_672377, EPI_ISL_672379, EPI_ISL_672384 | San Francisco Public Health Laboratory | Chan-Zuckerberg Biohub | CZB Cliahub Consortium |
| EPI_ISL_672386, EPI_ISL_672387, EPI_ISL_672389, EPI_ISL_672390, EPI_ISL_672391, EPI_ISL_672392 | UCSF Clinical Microbiology Laboratory | Chan-Zuckerberg Biohub | CZB Cliahub Consortium |
| EPI_ISL_672502, EPI_ISL_672503 | Madera County Department of Public Health | Chan-Zuckerberg Biohub | CZB Cliahub Consortium |
| EPI_ISL_672629, EPI_ISL_672630 | PathWest Laboratory Medicine WA | PathWest Laboratory Medicine WA Microbial Surveillance Unit | PathWest Laboratory Medicine WA Microbial Surveillance Unit |
| EPI_ISL_676529 | Uppsala klinisk mikrobiologi | The Public Health Agency of Sweden | Department of Microbiology, The Public Health Agency of Sweden |
| EPI_ISL_676575 | Scientific Veterinary Institute Novi Sad | Veterinary Specialized Institute "Kraljevo", Serbia | Vidanovic,D., Tesovic,B., Knezevic,A., Jovanovic,T., Jankovic,M., Sekler,M., Banovic Djeri,B., Petrovic,T., Volkening,J., Afonso,C. |
| EPI_ISL_676663 | Masonic Medical Research Institute | Wadsworth Center, New York State Department.of Health | Nathan Tucker, Kirsten St. George, Daryl M. Lamson, Alexis Russel, Jonathan Plitnick, Navjot Singh, John Kelly, Sara Griesemer, Erasmus Schneider, Erica Lasek-Nesselquist |
| EPI_ISL_676676 | Wadsworth Center, New York State Department.of Health | Wadsworth Center, New York State Department.of Health | Kirsten St. George, Daryl M. Lamson, Alexis Russel, Jonathan Plitnick, Navjot Singh, John Kelly, Sara Griesemer, Erasmus Schneider, Erica Lasek-Nesselquist |
| EPI_ISL_676726 | Masonic Medical Research Institute | Wadsworth Center, New York State Department.of Health | Nathan Tucker, Kirsten St. George, Daryl M. Lamson, Alexis Russel, Jonathan Plitnick, Navjot Singh, John Kelly, Sara Griesemer, Erasmus Schneider, Erica Lasek-Nesselquist |
| EPI_ISL_676993 | Wadsworth Center, New York State Department.of Health | Wadsworth Center, New York State Department.of Health | Kirsten St. George, Daryl M. Lamson, Alexis Russel, Jonathan Plitnick, Navjot Singh, John Kelly, Sara Griesemer, Erasmus Schneider, Erica Lasek-Nesselquist |
| EPI_ISL_677024, EPI_ISL_677071, EPI_ISL_677072, EPI_ISL_677073, EPI_ISL_677074, EPI_ISL_677075 | Masonic Medical Research Institute | Wadsworth Center, New York State Department.of Health | Nathan Tucker, Kirsten St. George, Daryl M. Lamson, Alexis Russel, Jonathan Plitnick, Navjot Singh, John Kelly, Sara Griesemer, Erasmus Schneider, Erica Lasek-Nesselquist |
| EPI_ISL_677352, EPI_ISL_677467 | University of Wisconsin-Madison AIDS Vaccine Research Laboratories | University of Wisconsin-Madison AIDS Vaccine Research Laboratories | Gage Moreno, Katarina Braun, et al. AIDS Vaccine Research Laboratories |
| EPI_ISL_677668 | Wadsworth Center, New York State Department.of Health | Wadsworth Center, New York State Department.of Health | Kirsten St. George, Daryl M. Lamson, Alexis Russel, Jonathan Plitnick, Navjot Singh, John Kelly, Sara Griesemer, Erasmus Schneider, Erica Lasek-Nesselquist |
| EPI_ISL_677717 | Clinical Hospital - Shtip | Research Center for Genetic Engineering and Biotechnology "Georgi D. Efremov" , Macedonian Academy of Sciences and Arts | RCGEB - MASA |
| EPI_ISL_677718 | Clinical Hospital - Bitola | Research Center for Genetic Engineering and Biotechnology "Georgi D. Efremov" , Macedonian Academy of Sciences and Arts | RCGEB - MASA |
| EPI_ISL_677824, EPI_ISL_677890, EPI_ISL_677897, EPI_ISL_677900, EPI_ISL_677902 | Innovative Genomics Institute, UC Berkeley | Innovative Genomics Institute, UC Berkeley | Stacia Wyman, Haridha Shivram, Phil Frankino, Liana Lareau, Shana McDevitt, Justin Choi |
| EPI_ISL_678185 | Pathogen Genomics Lab King Abdullah University of Science and Technology(KAUST) | Pathogen Genomics Lab King Abdullah University of Science and Technology(KAUST) | Sara Mfarrej, Raushan Nugmanova, Olga Douvropoulou, Raeece Naeem, Sharif Hala, Luke Esau, Amanda Ooi, Awad Al-Omari, Samer Salih, Abbas Al Mutair, Arnab Pain |
| EPI_ISL_678220 | Pathogen Genomics Lab King Abdullah University of Science and Technology(KAUST) | Pathogen Genomics Lab King Abdullah University of Science and Technology(KAUST) | Sara Mfarrej, Sharif Hala, Luke Esau, Amanda Ooi, Raeece Naeem, Awad Al-Omari, Samer Salih, Abbas Al Mutair, Arnab Pain |
| EPI_ISL_678221, EPI_ISL_678222, EPI_ISL_678223 | Pathogen Genomics Lab King Abdullah University of Science and Technology(KAUST) | Pathogen Genomics Lab King Abdullah University of Science and Technology(KAUST) | Sara Mfarrej, Olga Douvropoulou, Raushan Nugmanova, Raeece Naeem, Sharif Hala, Awad Al-Omari, Samer Salih, Abbas Al Mutair, Arnab Pain |
| EPI_ISL_678224 | Pathogen Genomics Lab King Abdullah University of Science and Technology(KAUST) | Pathogen Genomics Lab King Abdullah University of Science and Technology(KAUST) | Sara Mfarrej, Sharif Hala, Luke Esau, Amanda Ooi, Raeece Naeem, Awad Al-Omari, Samer Salih, Abbas Al Mutair, Arnab Pain |
| EPI_ISL_678225 | Pathogen Genomics Lab King Abdullah University of Science and Technology(KAUST) | Pathogen Genomics Lab King Abdullah University of Science and Technology(KAUST) | Sara Mfarrej, Olga Douvropoulou, Raushan Nugmanova, Raeece Naeem, Sharif Hala, Awad Al-Omari, Samer Salih, Abbas Al Mutair, Arnab Pain |
| EPI_ISL_678286 | Mikrobiologie, RARI | Mikrobiologie, RARI | Krasnov,Y.M., Naryshkina,E.A., Guseva,N.P., Sosedova,E.A., Fedorov,A.V., Badanin,D.V., Sharapova,N.A., Portenko,S.A., Shcherbakova,S.A., Kutyrev,V.V. |
| EPI_ISL_678313, EPI_ISL_678317, EPI_ISL_678318, EPI_ISL_678321, EPI_ISL_678325, EPI_ISL_678327, EPI_ISL_678328, EPI_ISL_678329, EPI_ISL_678330, EPI_ISL_678331, EPI_ISL_678332 | | | |
| see above | Area of Virology, Serology and Virology Division (SAViD), New South Wales Health Pathology Randwick | Virology Research Laboratory; Area of Virology, Serology and Virology Division (SAViD), New South Wales Health Pathology Randwick | Foster, C.; Au, J.; Ruiz Silva, M.; Deveson, I.; Bull, R.; Van Hal, S.; Rawlinson, W. |
| EPI_ISL_678493 | Veterinary Specialized Institute "Sabac", Serbia | Veterinary Specialized Institute "Kraljevo", Serbia | Vidanovic,D., Tesovic,B., Knezevic,A., Jovanovic,T., Jankovic,M., Sekler,M., Banovic Djeri,B., Petrovic,T., Mrkovacki, S., Volkening,J., Afonso,C. |
| EPI_ISL_681694 | Molecular Medicine Laboratory, University of Magallanes | Centro Asistencial Docente y de Investigacion, Universidad de Magallanes | Jorge González, Jacqueline Aldridge, Diego Alvarez, Marco Montes de Oca, Hermy Alvarez, Roberto Uribe-Paredes, Marcelo Navarrete |
| EPI_ISL_681838 | Molecular diagnostic unit for viral haemorrhagic fevers and emerging viruses, Bouaké CHU Laboratory | Project group Epidemology of Highly Pathogenic Microorganisms, Robert Koch-Institute | Chantal Akoua-Koffi, Diané Bamourou, Etilé Anoh, Essia Belarbi, Safiatou Karidioula, Grit Schubert, Adjaratou Traoré, Soundélé Maïté, Monemo Pacome, Coulibaly Mbegnan, Bamba Fatoumata Touré, Kra Ouffoué, Fabian Leendertz |
| EPI_ISL_681930, EPI_ISL_682019, EPI_ISL_682021, EPI_ISL_682022, EPI_ISL_682025 | UPMC Clinical Microbiology Laboratory | Microbial Genomic Epidemiology Laboratory, University of Pittsburgh | Mustapha M. Mustapha, Jane W. Marsh, Dan Snyder, Marissa P. Griffith, Stephanie L. Mitchell, Vatsala R. Srinivasa, Kady D. Waggle, Chinelo Ezeonwuku, Vaughn S. Cooper, Lee H. Harrison |
| EPI_ISL_682058 | Molecular diagnostic unit for viral haemorrhagic fevers and emerging viruses, Bouaké CHU Laboratory | Project group Epidemology of Highly Pathogenic Microorganisms, Robert Koch-Institute | Chantal Akoua-Koffi, Diané Bamourou, Etilé Anoh, Essia Belarbi, Safiatou Karidioula, Grit Schubert, Adjaratou Traoré, Soundélé Maïté, Monemo Pacome, Coulibaly Mbegnan, Bamba Fatoumata Touré, Kra Ouffoué, Fabian Leendertz |
| EPI_ISL_682241 | HOSPITAL SAN JUAN DE DIOS | Inciensa, Instituto Costarricense de Investigación y Enseñanza en Nutrición y Salud | Francisco Duarte, Hebleen Porras, Claudio Soto-Garita, Estela Cordero, Adriana Godinez & Melany Calderon |
| EPI_ISL_682242 | HOSPITAL SAN VICENTE DE PAUL | Inciensa, Instituto Costarricense de Investigación y | Francisco Duarte, Hebleen Porras, Claudio Soto-Garita, Estela Cordero, Adriana Godinez, Melany Calderon & Mariel López |

| | | | |
|---|---|---|---|
| | | Enseñanza en Nutrición y Salud | |
| EPI_ISL_682243 | HOSPITAL SAN JUAN DE DIOS | Inciensa, Instituto Costarricense de Investigación y Enseñanza en Nutrición y Salud | Francisco Duarte, Hebleen Porras, Claudio Soto-Garita, Estela Cordero, Adriana Godinez & Melany Calderon |
| EPI_ISL_682244 | AREA DE SALUD LA CRUZ | Inciensa, Instituto Costarricense de Investigación y Enseñanza en Nutrición y Salud | Francisco Duarte, Hebleen Porras, Claudio Soto-Garita, Estela Cordero, Adriana Godinez, Melany Calderon & Mariel López |
| EPI_ISL_682245, EPI_ISL_682246 | HOSPITAL SAN JUAN DE DIOS | Inciensa, Instituto Costarricense de Investigación y Enseñanza en Nutrición y Salud | Francisco Duarte, Hebleen Porras, Claudio Soto-Garita, Estela Cordero, Adriana Godinez & Melany Calderon |
| EPI_ISL_682247 | AREA DE SALUD CIUDAD QUESADA | Inciensa, Instituto Costarricense de Investigación y Enseñanza en Nutrición y Salud | Francisco Duarte, Hebleen Porras, Claudio Soto-Garita, Estela Cordero, Adriana Godinez & Melany Calderon |
| EPI_ISL_682248 | AREA DE SALUD CIUDAD QUESADA | Inciensa, Instituto Costarricense de Investigación y Enseñanza en Nutrición y Salud | Francisco Duarte, Hebleen Porras, Claudio Soto-Garita, Estela Cordero, Adriana Godinez, Melany Calderon & Mariel López |
| EPI_ISL_682249 | AREA DE SALUD LA UNION | Inciensa, Instituto Costarricense de Investigación y Enseñanza en Nutrición y Salud | Francisco Duarte, Hebleen Porras, Claudio Soto-Garita, Estela Cordero, Adriana Godinez, Melany Calderon & Mariel López |
| EPI_ISL_682250, EPI_ISL_682251, EPI_ISL_682252 | AREA DE SALUD ESCAZU (COOPESANA) | Inciensa, Instituto Costarricense de Investigación y Enseñanza en Nutrición y Salud | Francisco Duarte, Hebleen Porras, Claudio Soto-Garita, Estela Cordero, Adriana Godinez & Melany Calderon |
| EPI_ISL_682253, EPI_ISL_682254 | HOSPITAL SAN JUAN DE DIOS | Inciensa, Instituto Costarricense de Investigación y Enseñanza en Nutrición y Salud | Francisco Duarte, Hebleen Porras, Claudio Soto-Garita, Estela Cordero, Adriana Godinez & Melany Calderon |
| EPI_ISL_682255 | HOSPITAL SAN VICENTE DE PAUL | Inciensa, Instituto Costarricense de Investigación y Enseñanza en Nutrición y Salud | Francisco Duarte, Hebleen Porras, Claudio Soto-Garita, Estela Cordero, Adriana Godinez & Melany Calderon |
| EPI_ISL_682256 | AREA DE SALUD CORREDORES | Inciensa, Instituto Costarricense de Investigación y Enseñanza en Nutrición y Salud | Francisco Duarte, Hebleen Porras, Claudio Soto-Garita, Estela Cordero, Adriana Godinez, Melany Calderon & Mariel López |
| EPI_ISL_683622 | Servicio de Microbiología, Laboratori Clínic Metropolitana Nord. Hospital Universitari Germans Trias i Pujol. Institut d'Investigació en Ciències de la Salut Germans Trias i Pujol (IGTP) | SeqCOVID-SPAIN consortium/IBV(CSIC) | Elisa Martró, Antoni E. Bordoy, Anna Not, Adrián Antuori, Anabel Fernández, Nona Romaní, Verónica Saludes, Cristina Casañ and SeqCOVID-SPAIN consortium |
| EPI_ISL_691680 | Servicio de Microbiología, Hospital Universitario Son Espases | SeqCOVID-SPAIN consortium/IBV(CSIC) | Carla López-Causapé, Jordi Reina, Antonio Oliver and SeqCOVID-SPAIN consortium |
| EPI_ISL_693214 | Unidade de Pronto Atendimento Central de Caraguatatuba | Instituto Adolfo Lutz, Interdiciplinary Procedures Center, Strategic Laboratory | Claudio Tavares Sacchi, Claudia Regina Gonçalves, Erica Valessa Ramos Gomes, Karoline Rodrigues Campos |
| EPI_ISL_693471, EPI_ISL_693472, EPI_ISL_693474, EPI_ISL_693475, EPI_ISL_693476 | Central Public Health Laboratory | National Public Health Laboratory, National Centre for Infectious Diseases | Tze Minn Mak, Sophie Octavia, Zhenyang Zhou, Esorom Daoni, Theresa Palou, Lin Cui, Raymond Tzer Pin Lin |
| EPI_ISL_693532, EPI_ISL_693626, EPI_ISL_693627, EPI_ISL_693628, EPI_ISL_693629 | Instituto Nacional de Saude (INSA) | Instituto Nacional de Saude (INSA) | Borges et al |
| EPI_ISL_695231, EPI_ISL_695232, EPI_ISL_695233, EPI_ISL_695234, EPI_ISL_695235, EPI_ISL_695236, EPI_ISL_695237, EPI_ISL_695238, EPI_ISL_695239, EPI_ISL_695240, EPI_ISL_695241, EPI_ISL_695242, EPI_ISL_695243, EPI_ISL_695244, EPI_ISL_695245, EPI_ISL_695246, EPI_ISL_695247, EPI_ISL_695248, EPI_ISL_695249, EPI_ISL_695250, EPI_ISL_695259, EPI_ISL_695260, EPI_ISL_695261, EPI_ISL_695262, EPI_ISL_695263, EPI_ISL_695265, EPI_ISL_695266, EPI_ISL_695267, EPI_ISL_695268, EPI_ISL_695269, EPI_ISL_695270, EPI_ISL_695271, EPI_ISL_695272, EPI_ISL_695273, EPI_ISL_695274, EPI_ISL_695275, EPI_ISL_695276, EPI_ISL_695277, EPI_ISL_695278, EPI_ISL_695279, EPI_ISL_695280, EPI_ISL_695281, EPI_ISL_695282, EPI_ISL_695283, EPI_ISL_695284, EPI_ISL_695285, EPI_ISL_695286, EPI_ISL_695287, EPI_ISL_695288, EPI_ISL_695289, EPI_ISL_695290, EPI_ISL_695291, EPI_ISL_695292, EPI_ISL_695293, EPI_ISL_695294, EPI_ISL_695295, EPI_ISL_695296, EPI_ISL_695297, EPI_ISL_695298, EPI_ISL_695299, EPI_ISL_695300, EPI_ISL_695301, EPI_ISL_695302, EPI_ISL_695303, EPI_ISL_695304, EPI_ISL_695305, EPI_ISL_695306, EPI_ISL_695307, EPI_ISL_695308, EPI_ISL_695309, EPI_ISL_695310, EPI_ISL_695311, EPI_ISL_695312, EPI_ISL_695313, EPI_ISL_695314, EPI_ISL_695315, EPI_ISL_695372, EPI_ISL_695373, EPI_ISL_695374, EPI_ISL_695375, EPI_ISL_695376, EPI_ISL_695377, EPI_ISL_695378, EPI_ISL_695379, EPI_ISL_695380, EPI_ISL_695381, EPI_ISL_695382, EPI_ISL_695383, EPI_ISL_695384, EPI_ISL_695385, EPI_ISL_695386, EPI_ISL_695387, EPI_ISL_695388, EPI_ISL_695389, EPI_ISL_695390, EPI_ISL_695391, EPI_ISL_695392 | | | |
| see above | TGen North | TGen North | Jolene Bowers, Megan Folkerts, Chris French, Hayley Yaglom, Ashlyn Pfeiffer, Darrin Lemmer, Dave Engelthaler, The Arizona COVID Genomics Union (ACGU) |
| EPI_ISL_695593, EPI_ISL_695594, EPI_ISL_695595, EPI_ISL_695596, EPI_ISL_695597, EPI_ISL_695598, EPI_ISL_695599, EPI_ISL_695600, EPI_ISL_695601, EPI_ISL_695602, EPI_ISL_695603, EPI_ISL_695604, EPI_ISL_695605, EPI_ISL_695606, EPI_ISL_695607, EPI_ISL_695608, EPI_ISL_695609, EPI_ISL_695610, EPI_ISL_695611, EPI_ISL_695612, EPI_ISL_695613, EPI_ISL_695642, EPI_ISL_695643, EPI_ISL_695644, EPI_ISL_695645, EPI_ISL_695646, EPI_ISL_695647, EPI_ISL_695648, EPI_ISL_695649, EPI_ISL_695650, EPI_ISL_695651, EPI_ISL_695652, EPI_ISL_695653, EPI_ISL_695654, EPI_ISL_695655, EPI_ISL_695665, EPI_ISL_695666, EPI_ISL_695667, EPI_ISL_695668, EPI_ISL_695669, EPI_ISL_695670, EPI_ISL_695671, EPI_ISL_695672, EPI_ISL_695673, EPI_ISL_695674, EPI_ISL_695675, EPI_ISL_695676, EPI_ISL_695677, EPI_ISL_695678, EPI_ISL_695679, EPI_ISL_695680, EPI_ISL_695681, EPI_ISL_695682, EPI_ISL_695683, EPI_ISL_695684, EPI_ISL_695685, EPI_ISL_695686, EPI_ISL_695687, EPI_ISL_695688, EPI_ISL_695689, EPI_ISL_695690, EPI_ISL_695691, EPI_ISL_695692, EPI_ISL_695693, EPI_ISL_695694, EPI_ISL_695695, EPI_ISL_695696, EPI_ISL_695697, EPI_ISL_695698, EPI_ISL_695699, EPI_ISL_695700 | | | |
| see above | AZ SPHL, Arizona Department of Health Services | TGen North | Jolene Bowers, Megan Folkerts, Chris French, Hayley Yaglom, Ashlyn Pfeiffer, Darrin Lemmer, Dave Engelthaler, The Arizona COVID Genomics Union (ACGU) |
| EPI_ISL_697783 | Institute of Microbiology, Universidad San Francisco de Quito | Institute of Microbiology, Universidad San Francisco de Quito | Belén Prado-Vivar, Sully Márquez, Juan José Guadalupe, Monica Becerra-Wong, Bernardo Gutiérrez, Jonathan Araujo, Verónica Barragán, Patricio Rojas-Silva, Gabriel Trueba, Michelle Grunauer, Paúl Cárdenas |
| EPI_ISL_697797 | Centro de Investigaciones, Universidad de Especialidades Espíritu Santo | Institute of Microbiology, Universidad San Francisco de Quito | Derly Andrade, Juan Carlos Fernandez, Belén Prado-Vivar, Sully Márquez, Juan José Guadalupe, Monica Becerra-Wong, Bernardo Gutiérrez, Gabriel Morey, Ruben Armas, Jose Pedro Barberan, Fernando Espinoza, Edith Lopez, Verónica Barragán, Patricio Rojas-Silva, Gabriel Trueba, Michelle Grunauer, Paúl Cárdenas |
| EPI_ISL_699507 | Diagnostic Virology Laboratory, USDA National Veterinary Services Laboratories | Diagnostic Virology Laboratory, USDA National Veterinary Services Laboratories | Hamer,S.A., Pauvolid-Correa,A., Zecca,I.B., Davila,E., Auckland,L.D., Roundy,C.M., Tang,W., Torchetti,M., Killian,M.L., Jenkins-Moore,M., Akpalu,Y., Ghai,R.R., Spengler,J., Barton Behravesh,C., Fischer,R.S., Hamer,G.L., Franzen,K.M., Love,E.R. |
| EPI_ISL_699574, EPI_ISL_699575, EPI_ISL_699576, EPI_ISL_699577 | Group of Genetic Engeneering and Biotechnology, Federal Budget Institution of Science 'Central Research Institute of Epidemiology' of The Federal Service on Customers' Rights Protection and Human Well-being Surveillance | Group of Genetic Engeneering and Biotechnology, Federal Budget Institution of Science 'Central Research Institute of Epidemiology' of The Federal Service on Customers' Rights Protection and Human Well-being Surveillance | Cherkashina,A.S., Golubeva,A.G., Solovyova,E.D., Zotova,M.I., Berlina,Y.Y., Valdokhina,A.V., Bulanenko,V.P., Speranskaya,A.S., Tivanova,E.V., Shipulina,O.Y., Akimkin,V.G. |
| EPI_ISL_699930, EPI_ISL_699931, EPI_ISL_699932, EPI_ISL_699933, EPI_ISL_699934, EPI_ISL_699935, EPI_ISL_699936, EPI_ISL_699937, EPI_ISL_699938, EPI_ISL_699939, EPI_ISL_699940, EPI_ISL_699941, EPI_ISL_699942, EPI_ISL_699943, EPI_ISL_699944, EPI_ISL_699945, EPI_ISL_699946, EPI_ISL_699947, EPI_ISL_699948, EPI_ISL_699949, EPI_ISL_699950, EPI_ISL_699951, EPI_ISL_699952, EPI_ISL_699953, EPI_ISL_699954, EPI_ISL_699955, EPI_ISL_699956, EPI_ISL_699957, EPI_ISL_699958, EPI_ISL_699959, EPI_ISL_699960, EPI_ISL_699961, EPI_ISL_699962, EPI_ISL_699963, EPI_ISL_699964, EPI_ISL_699965, EPI_ISL_699966, EPI_ISL_699967, EPI_ISL_699968, EPI_ISL_699969, EPI_ISL_699970, EPI_ISL_699971, EPI_ISL_699972, EPI_ISL_699973, EPI_ISL_699974, EPI_ISL_699975, EPI_ISL_699976, EPI_ISL_699977, EPI_ISL_699978, EPI_ISL_699979, EPI_ISL_699980, EPI_ISL_699981, EPI_ISL_699982, EPI_ISL_699983, EPI_ISL_699984, EPI_ISL_699985, EPI_ISL_699986, EPI_ISL_699987, EPI_ISL_699988, EPI_ISL_699989, EPI_ISL_699990, EPI_ISL_699991, EPI_ISL_699992, EPI_ISL_699993, EPI_ISL_699994, EPI_ISL_699995, EPI_ISL_699996, EPI_ISL_699997, EPI_ISL_699998, EPI_ISL_699999, EPI_ISL_700000, EPI_ISL_700001, EPI_ISL_700002, EPI_ISL_700003, EPI_ISL_700004 | | | |
| see above | Hematopathology Laboratory, ACTREC, TMC | Hematopathology Laboratory, ACTREC, TMC | Hematopathology Laboratory, ACTREC |
| EPI_ISL_707779 | Rwanda National Reference Laboratory | Rwanda National Reference Laboratory | Enatha Mukantwari,Jeanne d'Arc UMuringa |
| EPI_ISL_707780 | Rwanda National Reference Laboratory | Rwanda National Reference Laboratory | Enatha Mukantwari,Jeanne d'Arc Umuringa |
| EPI_ISL_707783 | Rwanda National Reference Laboratory | Rwanda National Reference Laboratory | ENatha MUkantwari,Jeanne d'Arc Mukantwari |
| EPI_ISL_708730, EPI_ISL_708731, EPI_ISL_708732, EPI_ISL_708733, EPI_ISL_708734 | Los Angeles County Public Health Laboratory | Los Angeles County Public Health Laboratory | P. Hemarajata et al. |
| EPI_ISL_710214, EPI_ISL_710216, | Colorado Department of Public Health and Environment | Colorado Department of Puplic Health and Environment | Laura Bankers, Molly C. Hetherington-Rauth, Shannon Ely, Shannon R. Matzinger, Sarah Elizabeth Totten, Emily A. Travanty |

| | | | |
|---|---|---|---|
| EPI_ISL_710292, EPI_ISL_710293, EPI_ISL_710318 | | | |
| EPI_ISL_710479 | Group of Genetic Engineering and Biotechnology, Federal Budget Institution of Science 'Central Research Institute of Epidemiology' of The Federal Service on Customers' Rights Protection and Human Well-being Surveillance | Group of Genetic Engineering and Biotechnology, Federal Budget Institution of Science 'Central Research Institute of Epidemiology' of The Federal Service on Customers' Rights Protection and Human Well-being Surveillance | Cherkashina,A.S., Golubeva,A.G., Soloviova,E.D., Zotova,M.I., Berlina,Y.Y., Valdokhina,A.V., Bulanenko,V.P., Speranskaya,A.S.,Tivanova,E.V., Shipulina,O.Y. and Akimkin,V.G. |
| EPI_ISL_710587 | Omtanken Grimmered | The Public Health Agency of Sweden | Department of Microbiology, The Public Health Agency of Sweden |
| EPI_ISL_710595 | Klinisk mikrobiologi | The Public Health Agency of Sweden | Department of Microbiology, The Public Health Agency of Sweden |
| EPI_ISL_717767 | UW Virology Lab | UW Virology Lab | Pavitra Roychoudhury, Hong Xie, Lasata Shrestha, Michelle Lin, Meei-Li Huang, Keith R Jerome, Alexander Greninger |
| EPI_ISL_717907 | LACEN RJ - Noel Nutels | Bioinformatics Laboratory / LNCC | Carolina M Voloch, Ronaldo da Silva F Jr, Luiz G P de Almeida, Cynthia C Cardoso, Otavio Bustrolini, Alexandra L Gerber, Ana Paula de C Guimarães, Diana Mariani, Andréa Cony Cavalcanti, Claudia dos Santos Rodrigues, Terezinha M P P Castiñeira, Amilcar Tanuri, Ana Tereza R de Vasconcelos |
| EPI_ISL_718146 | Ministry of Health Hospitals | Institute of Health and Community Medicine | David Perera, Ooi Mong How, Chua Hock Hin, Tonnii Sia Loong Loong, Wong Jyn Shan, Wong Kiing Aik, Chan Chia Jui |
| EPI_ISL_721629, EPI_ISL_721630 | Armed Forces Medical College | National Centre For Cell Science | Dhiraj Paul, Kunal Jani, Radha Chauhan, Janesh Kumar, Vasudevan Seshadri, Girdhari Lal, Rajesh Karyakarte, Suvarna Joshi, Murlidhar Tambe, Sourav Sen, Santosh Karade, Kavita Bala Anand, Shelinder Pal Singh Shergill, Rajiv Mohan Gupta, Manoj Kumar Bhat, Arvind Sahu, Yogesh S Shouche |
| EPI_ISL_721647, EPI_ISL_721648, EPI_ISL_721649, EPI_ISL_721650, EPI_ISL_721651, EPI_ISL_721653, EPI_ISL_722180 | National Centre For Cell Science | National Centre For Cell Science | Dhiraj Paul, Kunal Jani, Radha Chauhan, Janesh Kumar, Vasudevan Seshadri, Girdhari Lal, Rajesh Karyakarte, Suvarna Joshi, Murlidhar Tambe, Sourav Sen, Santosh Karade, Kavita Bala Anand, Shelinder Pal Singh Shergill, Rajiv Mohan Gupta, Manoj Kumar Bhat, Arvind Sahu, Yogesh S Shouche |
| EPI_ISL_722197 | Armed Forces Medical College | National Centre For Cell Science | Dhiraj Paul, Kunal Jani, Radha Chauhan, Janesh Kumar, Vasudevan Seshadri, Girdhari Lal, Rajesh Karyakarte, Suvarna Joshi, Murlidhar Tambe, Sourav Sen, Santosh Karade, Kavita Bala Anand, Shelinder Pal Singh Shergill, Rajiv Mohan Gupta, Manoj Kumar Bhat, Arvind Sahu, Yogesh S Shouche |
| EPI_ISL_722387, EPI_ISL_722801, EPI_ISL_722802, EPI_ISL_722803, EPI_ISL_722804, EPI_ISL_722805, EPI_ISL_722806, EPI_ISL_722807, EPI_ISL_722808, EPI_ISL_722809 | Dutch COVID-19 response team | Erasmus Medical Center | Bas Oude Munnink, Reina Sikkema, David Nieuwenhuijse, Irina Chestakova, Anne van der Linden, Marjan Boter, Emmanuelle Munger, Corine GeurtsvanKessel, Annemiek van der Eijk, Richard Molenkamp, Marion Koopmans, on behalf of the Dutch national COVID-19 response team. |
| EPI_ISL_722855, EPI_ISL_722856, EPI_ISL_722857, EPI_ISL_722858, EPI_ISL_722872 | Dipartimento di Scienze Biomediche e Oncologia Umana - Azienda Ospedaliero Universitaria Consorziale Policlinico | Istituto Zooprofilattico Sperimentale della Puglia e della Basilicata | Parisi A., Bianco A., Capozzi L., Del Sambro L., Chironna M., Loconsole D. |
| EPI_ISL_723050, EPI_ISL_723051, EPI_ISL_723052 | Hematopathology Laboratory, ACTREC, TMC | Hematopathology Laboratory, ACTREC, TMC | Hematopathology Laboratory, ACTREC |
| EPI_ISL_728254, EPI_ISL_728255, EPI_ISL_728256, EPI_ISL_728269, EPI_ISL_728271, EPI_ISL_728327 | B.J. Govt. Medical College | National Centre For Cell Science | Dhiraj Paul, Kunal Jani, Radha Chauhan, Janesh Kumar, Vasudevan Seshadri, Girdhari Lal, Rajesh Karyakarte, Suvarna Joshi, Murlidhar Tambe, Sourav Sen, Santosh Karade, Kavita Bala Anand, Shelinder Pal Singh Shergill, Rajiv Mohan Gupta, Manoj Kumar Bhat, Arvind Sahu, Yogesh S Shouche |
| EPI_ISL_729469, EPI_ISL_729470, EPI_ISL_729564, EPI_ISL_729565, EPI_ISL_729566, EPI_ISL_729567, EPI_ISL_729568, EPI_ISL_729569, EPI_ISL_729570, EPI_ISL_729571 | A. Krumbholz, Labor Dr. Krause und Kollegen MVZ GmbH, Kiel | Charité Universitätsmedizin Berlin, Institut für Virologie | Victor M Corman, Barbara Mühlemann, Jörn Beheim-Schwarzbach, Talitha Veith, Julia Schneider, Terry Jones, Christian Drosten |
| EPI_ISL_729801, EPI_ISL_729844, EPI_ISL_729850, EPI_ISL_729856, EPI_ISL_729857 | Laboratorio Central de Saude Publica do Estado do Rio Grande do Sul (LACEN-RS) | Laboratory of Respiratory Viruses and Measles, Oswaldo Cruz Institute, FIOCRUZ | Paola Resende, Luciana Appolinario, Fernando Motta, Anna Carolina Paixão, Ana Carolina Mendonça, Tatiana Schaffer Gregianini, Marilda Tereza Mar da Rosa, Marilda Siqueira |
| EPI_ISL_729956, EPI_ISL_729957, EPI_ISL_729990, EPI_ISL_729991, EPI_ISL_729992, EPI_ISL_729993, EPI_ISL_730046, EPI_ISL_730047, EPI_ISL_730048 | Nigeria Centre for Disease Control (NCDC) | African Centre of Excellence for Genomics of Infectious Diseases (ACEGID), Redeemer's University, Ede, Osun State, Nigeria | Oluniyi P.E. et al |
| EPI_ISL_730084 | Yale Clinical Virology Laboratory | Grubaugh Lab - Yale School of Public Health | Joseph Fauver, Tara Alpert, Anderson Brito, Annie Watkins, Anne Wyllie, Chantal Vogels, Mary Petrone, Chaney Kalinich, Isabel Ott, Arnau Casanovas, Catherine Muenker, Adam Moore, Alice Lu, Maria Tokuyama, Patrick Wong, Peiwen Lu, Saad Omer, Richard Martinello, Allison Nelson, Shelli Farhadian, Akiko Iwasaki, Charlese Dela Cruz, Albert Ko, Nathan Grubaugh |
| EPI_ISL_730206, EPI_ISL_730207, EPI_ISL_730208 | Genomica Lab Molecular, M©xico | Andersen lab at Scripps Research | SEARCH Alliance San Diego with Jonathan Gonzalez Garcia, Jose Roman Chavez Mendez, Jose Horacio Reyna Verdugo, Martin Gonzalez Ibarra, Luis Alberto Rangel Gonzalez |
| EPI_ISL_732775, EPI_ISL_732776, EPI_ISL_732777, EPI_ISL_732806 | Centro de Investigación Biomédica de La Rioja - Hospital San Pedro Logroño | SeqCOVID-SPAIN consortium/IBV(CSIC) | María de Toro, José Manuel Azcona Gutiérrez, María Pilar Bea Escudero, Miriam Blasco Alberdi and SeqCOVID-SPAIN consortium |
| EPI_ISL_732993 | UMMC-Health | WHO National Influenza Centre Russian Federation | Andrey Komissarov, Artem Fadeev, Anna Ivanova, Kseniya Komissarova, Dmitry Bazhenov, Tatiana Platonova, Daria Danilenko, Ksenia Safina, Elena Nabieva, Georgii Bazykin, Dmitry Lioznov |
| EPI_ISL_733500 | 1-Laboratory of Microbiology, National Reference Lab, Charles Nicolle Hospital; 2-University of Tunis ElManar, Faculty of Medicine of Tunis, LR99ES09, Tunis, Tunisia | 1-Clinical and Experimental Pharmacology Lab, LR16SP02, National Center of Pharmacovigilance, University of Tunis El Manar, Tunis, Tunisia. 2-Neurodegenerative diseases and psychiatric troubles, LR18SP03, Razi Hospital, University of Tunis El Manar, Tunis, Tunisia. 3- Ministry of Health, National Observatory of New and Emerging Diseases, 1006, Tunis, Tunisia | Ilhem Boutiba-Ben Boubaker, Sameh Trabelsi, Nissaf Ben Alaya, Maher Kharrat, Alia Ben Kahla, Jalila Ben Khelil, Salma Abid, Sana Ferjani, Mouna Ben Sassi, Mouna Safer, Guedi Ali Barreh, Habiba Ben Romdhane, Souissi Amira, Sarra Chamman, Hanen El Jebari, Asma Ferjani, Gaies Emna, Riadh Daghfous, Riadh Gouider. |
| EPI_ISL_734494, EPI_ISL_734796, EPI_ISL_734797, EPI_ISL_734801, EPI_ISL_734802, EPI_ISL_734803, EPI_ISL_734804, EPI_ISL_734805, EPI_ISL_734806, EPI_ISL_734807, EPI_ISL_734808, EPI_ISL_734809, EPI_ISL_734810, EPI_ISL_734811, EPI_ISL_734812, EPI_ISL_734813, EPI_ISL_734814, EPI_ISL_734815, EPI_ISL_734816, EPI_ISL_734817, EPI_ISL_734818, EPI_ISL_734819, EPI_ISL_734820, EPI_ISL_734821, EPI_ISL_734822, EPI_ISL_734823, EPI_ISL_734824 | | | |
| see above | UZ Leuven, National Reference Laboratory for Coronaviruses, Laboratory Medicine, Leuven, Belgium | KU Leuven, Rega Institute, Clinical and Epidemiological Virology | Tony Wawina-Bokalanga, Joan Marti-Carerras, Bert Vanmechelen, Piet Maes |
| EPI_ISL_735254, EPI_ISL_735255, EPI_ISL_735256 | Group of Genetic Engeneering and Biotechnology, Federal Budget Institution of Science 'Central Research Institute of Epidemiology' of The Federal Service on Customers' Rights Protection and Human Well-being Surveillance | Group of Genetic Engeneering and Biotechnology, Federal Budget Institution of Science 'Central Research Institute of Epidemiology' of The Federal Service on Customers' Rights Protection and Human Well-being Surveillance | Cherkashina,A.S., Golubeva,A.G., Solovyova,E.D., Valdokhina,A.V., Bulanenko,V.P., Zotova,M.I., Berlina,Y.Y., Speranskaya,A.S., Tivanova,E.V., Shipulina,O.Y., Akimkin,V.G. |
| EPI_ISL_735378, EPI_ISL_735379, EPI_ISL_735380, EPI_ISL_735381, EPI_ISL_735382, EPI_ISL_735383 | Los Angeles County Public Health Laboratory | Los Angeles County Public Health Laboratory | P. Hemarajata et al. |
| EPI_ISL_737939, EPI_ISL_737972, | Uganda Central Public Health Lab and Uganda Virus | MRC/UVRI & LSHTM Uganda Research Unit | Matthew Cotten, Dan Lule Bugembe, My V.T. Phan, Pontiano Kaleebu et al. |

| | | | |
|---|---|---|---|
| EPI_ISL_737973, EPI_ISL_737977, EPI_ISL_737978, EPI_ISL_737979 | Research Institute | | |
| EPI_ISL_738222 | UZ Leuven, National Reference Laboratory for Coronaviruses, Laboratory Medicine, Leuven, Belgium | KU Leuven, Rega Institute, Clinical and Epidemiological Virology | Tony Wawina-Bokalanga, Joan Marti-Carerras, Bert Vanmechelen, Piet Maes |
| EPI_ISL_738535, EPI_ISL_738622, EPI_ISL_738983, EPI_ISL_739083, EPI_ISL_739084, EPI_ISL_739134, EPI_ISL_739200, EPI_ISL_739349, EPI_ISL_739422, EPI_ISL_739516, EPI_ISL_739598, EPI_ISL_739622, EPI_ISL_739656 | | | |
| see above | Alameda County Public Health Lab | Chan-Zuckerberg Biohub | CZB Cliahub Consortium |
| EPI_ISL_739792, EPI_ISL_739848, EPI_ISL_739995, EPI_ISL_740032, EPI_ISL_740127, EPI_ISL_740132, EPI_ISL_740186, EPI_ISL_740204, EPI_ISL_740314, EPI_ISL_744147, EPI_ISL_744486, EPI_ISL_744561, EPI_ISL_744696, EPI_ISL_744711, EPI_ISL_744720, EPI_ISL_744811 | | | |
| see above | Laboratoire national de santé, Microbiology, Virology | Laboratoire national de santé, Microbiology, Microbial Genomics Platform | Anke Wienecke-Baldacchino, Catherine Ragimbeau, Tamir Abdelrahman, Jessica Tapp, Fatu Djabi |
| EPI_ISL_745776, EPI_ISL_745783, EPI_ISL_745792, EPI_ISL_745799 | Ginkgo Bioworks Clinical Laboratory | Utah Public Health Laboratory | Erin L. Young, Kelly Oakeson, Tara Gallagher, Michael T. Pyne, E. Susan Slechta, Melanie A. Mallory, Jeffrey B. Stevenson, Salika M. Shakir, David R. Hillyard, Malaika McKenzie-Bennett, James McGann, Jim Griffin, Keith Robison, Alex Plocik, Becky Schilling, Martha Pierson, Rebecca Littlefield, Michelle Spencer, Birgitte Simen |
| EPI_ISL_746464 | Utah Public Health Laboratory | Utah Public Health Laboratory | Erin Young, Kelly Oakeson, Tara Gallagher |
| EPI_ISL_746485, EPI_ISL_746497, EPI_ISL_746594, EPI_ISL_746595, EPI_ISL_746596, EPI_ISL_746597, EPI_ISL_746598, EPI_ISL_746599, EPI_ISL_746600, EPI_ISL_746601, EPI_ISL_746602, EPI_ISL_746603, EPI_ISL_746604, EPI_ISL_746605, EPI_ISL_746606, EPI_ISL_746607, EPI_ISL_746608, EPI_ISL_746609, EPI_ISL_746610, EPI_ISL_746611, EPI_ISL_746612, EPI_ISL_746613, EPI_ISL_746614, EPI_ISL_746615, EPI_ISL_746616, EPI_ISL_746617, EPI_ISL_746618, EPI_ISL_746619, EPI_ISL_746620, EPI_ISL_746621, EPI_ISL_746622, EPI_ISL_746623, EPI_ISL_746624, EPI_ISL_746627 | | | |
| see above | Genetica Molecular and Subdepartamento de Virologia ISP Chile | Instituto de Salud Publica de Chile | Javier Tognarelli, Barbara Parra, Loredana Arata, Jaime Lagos, Gisselle Barra, Patricia Bustos, Rodrigo Fasce, Andres Castillo, Jorge Fernandez |
| EPI_ISL_747419 | Division of Emerging Infectious Diseases, Bureau of Infectious Diseases Diagnosis Control, Korea Disease Control and Prevention Agency | Division of Emerging Infectious Diseases, Bureau of Infectious Diseases Diagnosis Control, Korea Disease Control and Prevention Agency | Ae Kyung Park, Il-Hwan Kim, Heui Man Kim, Jeong-Min Kim, Namjoo Lee, Chaeyoung Lee, Sang Hee Woo, Eun-Jin Kim |
| EPI_ISL_750175 | CENUR Este-Sede Rocha-UdelaR | Institut Pasteur de Montevideo | Daiana Mir, Natalia Rego, Paola Cristina Resende, Fernando Lopez-Tort, Tamara Fernandez-Calero, Veronica Noya, Mariana Brandes, Tania Possi, Mailen Arleo, Natalia Reyes, Matias Victoria, Andres Lizasoain, Matias Castells, Leticia Maya, Matías Salvo, Tatiana Schäffer Gregianini, Marilda Tereza Mar da Rosa, Letícia Garay Martins, Cecilia Alonso, Yasser Vega, Cecilia Salazar, Ignacio Ferrés, Pablo Smircich, Jose Sotelo, Ighor Arantes, Luciana Appolinario, Ana Carolina Mendonça, Maria Jose Benitez-Galeano, Martín Graña, Camila Simoes, Fernando Motta, Marilda Mendonça Siqueira, Gonzalo Bello, Rodney Colina, Lucia Spangenberg |
| EPI_ISL_750177, EPI_ISL_750178 | Sanatorio Americano | Institut Pasteur de Montevideo | Daiana Mir, Natalia Rego, Paola Cristina Resende, Fernando Lopez-Tort, Tamara Fernandez-Calero, Veronica Noya, Mariana Brandes, Tania Possi, Mailen Arleo, Natalia Reyes, Matias Victoria, Andres Lizasoain, Matias Castells, Leticia Maya, Matías Salvo, Tatiana Schäffer Gregianini, Marilda Tereza Mar da Rosa, Letícia Garay Martins, Cecilia Alonso, Yasser Vega, Cecilia Salazar, Ignacio Ferrés, Pablo Smircich, Jose Sotelo, Ighor Arantes, Luciana Appolinario, Ana Carolina Mendonça, Maria Jose Benitez-Galeano, Martín Graña, Camila Simoes, Fernando Motta, Marilda Mendonça Siqueira, Gonzalo Bello, Rodney Colina, Lucia Spangenberg |
| EPI_ISL_751185, EPI_ISL_751186, EPI_ISL_751187, EPI_ISL_751188 | CENUR Litoral Norte - UdelaR, Salto, Uruguay | Institut Pasteur de Montevideo | Daiana Mir, Natalia Rego, Paola Cristina Resende, Fernando Lopez-Tort, Tamara Fernandez-Calero, Veronica Noya, Mariana Brandes, Tania Possi, Mailen Arleo, Natalia Reyes, Matias Victoria, Andres Lizasoain, Matias Castells, Leticia Maya, Matías Salvo, Tatiana Schäffer Gregianini, Marilda Tereza Mar da Rosa, Letícia Garay Martins, Cecilia Alonso, Yasser Vega, Cecilia Salazar, Ignacio Ferrés, Pablo Smircich, Jose Sotelo, Ighor Arantes, Luciana Appolinario, Ana Carolina Mendonça, Maria Jose Benitez-Galeano, Martín Graña, Camila Simoes, Fernando Motta, Marilda Mendonça Siqueira, Gonzalo Bello, Rodney Colina, Lucia Spangenberg |
| EPI_ISL_751201 | Laboratorio DILAVE/MGAP-INIA-UdelaR -Tacuarembó | Institut Pasteur de Montevideo | Daiana Mir, Natalia Rego, Paola Cristina Resende, Fernando Lopez-Tort, Tamara Fernandez-Calero, Veronica Noya, Mariana Brandes, Tania Possi, Mailen Arleo, Natalia Reyes, Matias Victoria, Andres Lizasoain, Matias Castells, Leticia Maya, Matías Salvo, Tatiana Schäffer Gregianini, Marilda Tereza Mar da Rosa, Letícia Garay Martins, Cecilia Alonso, Yasser Vega, Cecilia Salazar, Ignacio Ferrés, Pablo Smircich, Jose Sotelo, Ighor Arantes, Luciana Appolinario, Ana Carolina Mendonça, Maria Jose Benitez-Galeano, Martín Graña, Camila Simoes, Fernando Motta, Marilda Mendonça Siqueira, Gonzalo Bello, Rodney Colina, Lucia Spangenberg |
| EPI_ISL_751210 | Pathogen Genomics Lab King Abdullah University of Science and Technology(KAUST) | Pathogen Genomics Lab King Abdullah University of Science and Technology(KAUST) | Sara Mfarrej, Raushan Nugmanova, Olga Douvropoulou, Sharif Hala, Raeece Naeem, Fadwa Alofi, Asim Khogeer, Afrah Alsomali, Jumana Taha, Abdulaziz Alahmadi, Kahled Algithami, Anwar Hashem, Naif Almontashiri, Arnab Pain |
| EPI_ISL_751214 | Pathogen Genomics Lab King Abdullah University of Science and Technology(KAUST) | Pathogen Genomics Lab King Abdullah University of Science and Technology(KAUST) | Sara Mfarrej, Raushan Nugmanova, Olga Douvropoulou, Sharif Hala, Raeece Naeem, Fadwa Alofi, Asim Khogeer, Afrah Alsomali, Jumana Taha, Abdulaziz Alahmadi, Kahled Algithami, Anwar Hashem, Naif Almontashiri, Arnab Pain |
| EPI_ISL_751226 | Pathogen Genomics Lab King Abdullah University of Science and Technology(KAUST) | Pathogen Genomics Lab King Abdullah University of Science and Technology(KAUST) | Sara Mfarrej, Raushan Nugmanova, Olga Douvropoulou, Sharif Hala, Raeece Naeem, Fadwa Alofi, Asim Khogeer, Afrah Alsomali, Jumana Taha, Abdulaziz Alahmadi, Kahled Algithami, Anwar Hashem, Naif Almontashiri, Arnab Pain |
| EPI_ISL_751228 | Pathogen Genomics Lab King Abdullah University of Science and Technology(KAUST) | Pathogen Genomics Lab King Abdullah University of Science and Technology(KAUST) | Sara Mfarrej, Raushan Nugmanova, Olga Douvropoulou, Sharif Hala, Raeece Naeem, Fadwa Alofi, Asim Khogeer, Afrah Alsomali, Jumana Taha, Abdulaziz Alahmadi, Kahled Algithami, Anwar Hashem, Naif Almontashiri, Arnab Pain |
| EPI_ISL_751234 | Pathogen Genomics Lab King Abdullah University of Science and Technology(KAUST) | Pathogen Genomics Lab King Abdullah University of Science and Technology(KAUST) | Sara Mfarrej, Olga Douvropoulou, Raushan Nugmanova, Raeece Naeem, Sharif Hala, Fadwa Alofi, Asim Khogeer, Afrah Alsomali, Jumana Taha, Abdulaziz Alahmadi, Kahled Algithami, Anwar Hashem, Naif Almontashiri, Arnab Pain |
| EPI_ISL_751500 | Pathogen Genomics Lab King Abdullah University of Science and Technology(KAUST) | Pathogen Genomics Lab King Abdullah University of Science and Technology(KAUST) | Sara Mfarrej, Raushan Nugmanova, Olga Douvropoulou, Sharif Hala, Raeece Naeem, Fadwa Alofi, Asim Khogeer, Afrah Alsomali, Jumana Taha, Abdulaziz Alahmadi, Kahled Algithami, Anwar Hashem, Naif Almontashiri, Arnab Pain |
| EPI_ISL_751509, EPI_ISL_751510, EPI_ISL_751519, EPI_ISL_751524, EPI_ISL_751526, EPI_ISL_751527, EPI_ISL_751533, EPI_ISL_751534, EPI_ISL_751536, EPI_ISL_751539, EPI_ISL_751541 | | | |
| see above | Texas Department of State Health Services | Texas Department of State Health Services | Rashmi Tuladhar, Bonnie Oh, Jenny Zhang, Maliha Rahman, Anita Pokharel, Myong Koag, Chung Wang, Rachel Lee, Grace Kubin, Mayela Pedrueza, James Daniel Bonser |
| EPI_ISL_752701, EPI_ISL_752702, EPI_ISL_752703, EPI_ISL_752709, EPI_ISL_752712, EPI_ISL_752713, EPI_ISL_752714, EPI_ISL_752715, EPI_ISL_752717, EPI_ISL_752718, EPI_ISL_752726, EPI_ISL_752727, EPI_ISL_752728, EPI_ISL_752729, EPI_ISL_752730 | | | |
| see above | State Laboratories Division, Hawaii State Department of Health | State Laboratories Division, Hawaii State Department of Health | Pamela O'Brien, Sabrina Diemert, Drew Kuwazaki, Razvan Sultana, Edward Desmond |
| EPI_ISL_753778, EPI_ISL_753981, EPI_ISL_753982, EPI_ISL_753983, EPI_ISL_753984, EPI_ISL_753985, EPI_ISL_753992, EPI_ISL_754039 | Charité Universitätsmedizin Berlin, Institut für Virologie/Labor Berlin | Charité Universitätsmedizin Berlin, Institut für Virologie | Victor M Corman, Jörn Beheim-Schwarzbach, Barbara Mühlemann, Julia Schneider, Talitha Veith, Terry Jones, Christian Drosten |
| EPI_ISL_754239 | Santosa Hospital Bandung Central | "School of Life Sciences and Technology & School of Pharmacy-Institut Teknologi Bandung; Molecular Genetics Laboratory-Faculty of Medicine-Universitas Padjadjaran; Laboratorium Kesehatan Provinsi Jawa Barat" | Marselina Irasonia Tan, Yunia Sribudiani, Catur Riani, Azzania Fibriani, Husna Nugrahapraja, Tarwadi, Ema Rahmawati, Savira Ekawardhani, Hesti Lina Wiraswati, Ryan Bayusantika Ristandi, Rifky Waluyajati Rachman, Cut Nur Cinthia Alamanda, Lia Faridah, Miftahul Faridl, Karimatu Khoirunnisa, Hammam Riza, Soni Solistia Wirawan, Agung Eru Wibowo, Irvan Faizal |
| EPI_ISL_754391 | Genetica Molecular and Subdepartamento de Virologia ISP Chile | Insituto de Salud Publica de Chile | Javier Tognarelli, Barbara Parra, Loredana Arata, Jaime Lagos, Gisselle Barra, Patricia Bustos, Rodrigo Fasce, Andres Castillo, Jorge Fernandez |
| EPI_ISL_754857, EPI_ISL_754858, EPI_ISL_754859, EPI_ISL_754860, EPI_ISL_754861, EPI_ISL_754862, EPI_ISL_754863, EPI_ISL_754865, EPI_ISL_754866, EPI_ISL_754867, EPI_ISL_754868, EPI_ISL_754869, EPI_ISL_754870, EPI_ISL_754871, EPI_ISL_754872, EPI_ISL_754873, EPI_ISL_754874, EPI_ISL_754875, EPI_ISL_754876, EPI_ISL_754877, EPI_ISL_754879, EPI_ISL_754880, EPI_ISL_754882, EPI_ISL_754883, EPI_ISL_754884, EPI_ISL_754886, EPI_ISL_754887, EPI_ISL_754888, EPI_ISL_754890, EPI_ISL_754891, EPI_ISL_754892, EPI_ISL_754896, EPI_ISL_754898, EPI_ISL_754900 | | | |
| see above | Innovative Genomics Institute, UC Berkeley | Innovative Genomics Institute, UC Berkeley | Stacia Wyman, Haridha Shivram, Phil Frankino, Liana Lareau, Shana McDevitt, Justin Choi |
| EPI_ISL_754920, EPI_ISL_754944, EPI_ISL_754945, EPI_ISL_754947, EPI_ISL_754950, EPI_ISL_754961, EPI_ISL_754968, EPI_ISL_754998, EPI_ISL_755004, EPI_ISL_755025, EPI_ISL_755028, EPI_ISL_755043 | | | |

| | | | |
|---|---|---|---|
| see above | California Department of Public Health | California Department of Public Health | CDPH IDLB COVIDNet |
| EPI_ISL_756312, EPI_ISL_756313, EPI_ISL_756314, EPI_ISL_756316, EPI_ISL_756317, EPI_ISL_756318, EPI_ISL_756319, EPI_ISL_756320, EPI_ISL_756321, EPI_ISL_756322, EPI_ISL_756324, EPI_ISL_756325, EPI_ISL_756326, EPI_ISL_756327, EPI_ISL_756328, EPI_ISL_756329 | | | |
| see above | Innovative Genomics Institute, UC Berkeley | Innovative Genomics Institute, UC Berkeley | Stacia Wyman, Haridha Shivram, Phil Frankino, Liana Lareau, Shana McDevitt, Justin Choi |
| EPI_ISL_759960 | Surabaya Public Health Office | Institute of Tropical Disease, Universitas Airlangga | Aldise M Nastri, Jezzy R Dewantari, Rima R Prasetya, Krisnoadi Rahardjo, Febria Rachmanita, Gatot Soegiarto, Laksmi Wulandari, Resti Yudhawati, Soetjipto, Yasuko Mori, Maria I Lusida, Kazufumi Shimizu |
| EPI_ISL_760123, EPI_ISL_760124, EPI_ISL_760125, EPI_ISL_760126, EPI_ISL_760127, EPI_ISL_760128 | Division of Emerging Infectious Diseases, Bureau of Infectious Diseases Diagnosis Control, Korea Disease Control and Prevention Agency | Division of Emerging Infectious Diseases, Bureau of Infectious Diseases Diagnosis Control, Korea Disease Control and Prevention Agency | Ae Kyung Park, Il-Hwan Kim, Heui Man Kim, Jeong-Min Kim, Namjoo Lee, Chaeyoung Lee, Sang Hee Woo, Eun-Jin Kim |
| EPI_ISL_765672, EPI_ISL_765673, EPI_ISL_765674, EPI_ISL_765675, EPI_ISL_765676, EPI_ISL_765677, EPI_ISL_765678, EPI_ISL_765679, EPI_ISL_765682 | Massachusetts General Hospital | Infectious Disease Program, Broad Institute of Harvard and MIT | Lemieux,J.E., Siddle,K.J., Shaw,B., Adams,G., Pierce,V., Turbett,S., Anahtar,M., Branda,J., Slater,D., Harris,J., Lin,A.E., Gladden-Young,A., Lagerborg,K., Rudy,M., DeRuff,K., Carter,A., Normandin,E., Bauer,M., Reilly,S., Tomkins-Tinch,C., Loreth,C., Chaluvadi,S., Neumann,A., Cusick,C., Chapman,S.B., Gnirke,A., Flowers,K., Cerrato,F., Birren,B.W., Gallagher,G., Smole,S., Park,D.J., MacInnis,B.L., Ryan,E., LaRocque,R., Rosenberg,E. and Sabeti,P.C. |
| EPI_ISL_765996, EPI_ISL_765997 | USC Clinical Lab | Los Angeles County PHL | P. Hemarajata et al. |
| EPI_ISL_766647, EPI_ISL_766649, EPI_ISL_766663, EPI_ISL_766668, EPI_ISL_766673, EPI_ISL_766682, EPI_ISL_766686 | Texas Department of State Health Services | Texas Department of State Health Services | Rashmi Tuladhar, Bonnie Oh, Jenny Zhang, Maliha Rahman, Anita Pokharel, Myong Koag, Chung Wang, Rachel Lee, Grace Kubin, Mayela Pedrueza, James Daniel Bonser |
| EPI_ISL_766864 | NIC Viral Respiratory Unit - Institut Pasteur of Algeria | National Reference Center for Viruses of Respiratory Infections, Institut Pasteur, Paris | Mélanie Albert, Marion Barbet, Sylvie Behillil, Méline Bizard, Angela Brisebarre, Flora Donati, Etienne Simon-Lorière, Vincent Enouf, Maud Vanpeene, Sylvie van der Werf, Fawzi Derrar |
| EPI_ISL_768741 | Child Health Research Foundation | Child Health Research Foundation | Senjuti Saha, Afroza Akter Tanni, Roly Malaker, Sharmistha Goswami, Syed Muktadir Al Sium, Arif Mohammad Tanmoy, Md Hafizur Rahman, Samir K Saha |
| EPI_ISL_769907, EPI_ISL_769908, EPI_ISL_769909, EPI_ISL_769910, EPI_ISL_769911, EPI_ISL_769912, EPI_ISL_769913, EPI_ISL_769914, EPI_ISL_769915 | Wyoming Public Health Laboratory | Wyoming Public Health Laboratory | Noah Hull, Taylor Fearing, Lynette Gumbleton, Channing Weber, Ashley Norberg, Bailey Bowcutt, and Wanda Manley |
| EPI_ISL_770009 | Area De Salud Escazu (Coopesana) | Inciensa, Instituto Costarricense de Investigación y Enseñanza en Nutrición y Salud | Francisco Duarte, Hebleen Porras, Claudio Soto-Garita, Estela Cordero, Adriana Godínez & Melany Calderón |
| EPI_ISL_770125, EPI_ISL_770126, EPI_ISL_770127 | Wyoming Public Health Laboratory | Wyoming Public Health Laboratory | Noah Hull, Taylor Fearing, Lynette Gumbleton, Channing Weber, Ashley Norberg, Bailey Bowcutt, and Wanda Manley |
| EPI_ISL_771219, EPI_ISL_771230, EPI_ISL_771231, EPI_ISL_771232, EPI_ISL_771233 | Colorado Department of Public Health and Environment | Colorado Department of Puplic Health and Environment | Laura Bankers, Molly C. Hetherington-Rauth, Diana Ir, Shannon Ely, Shannon R. Matzinger, Sarah Elizabeth Totten, Emily A. Travanty |
| EPI_ISL_774971, EPI_ISL_774972, EPI_ISL_774973, EPI_ISL_774974, EPI_ISL_774975, EPI_ISL_774978, EPI_ISL_774980, EPI_ISL_774981, EPI_ISL_774982, EPI_ISL_774983, EPI_ISL_774986, EPI_ISL_774989, EPI_ISL_774991, EPI_ISL_774993, EPI_ISL_775000, EPI_ISL_775004, EPI_ISL_775005, EPI_ISL_775006, EPI_ISL_775010, EPI_ISL_775012, EPI_ISL_775013, EPI_ISL_775014, EPI_ISL_775015, EPI_ISL_775016, EPI_ISL_775017, EPI_ISL_775018 | | | |
| see above | Designated Reference Institute for Chemical Measurements (DRICM) | DNA SOLUTION LTD. | Md. Imran Khan, Kazi Nadim Hasan, Abu Sufian, Jannatun Naima, Abdul Khaleque, Mizanur Rahman, MSM Chowdhury, Hasan Ul Haider, Mamudul Hasan Razu, Mala Khan, Mohammad Fazle Alam Rabbi |
| EPI_ISL_776658, EPI_ISL_776673, EPI_ISL_776681, EPI_ISL_776682, EPI_ISL_776684, EPI_ISL_776686 | UW Virology Lab | UW Virology Lab | Pavitra Roychoudhury, Hong Xie, Lasata Shrestha, Meei-Li Huang, Keith R Jerome, Alexander Greninger |
| EPI_ISL_776772, EPI_ISL_776870 | Hospital General Universitario Gregorio Marañón | Hospital General Universitario Gregorio Marañón | Sergio Buenestado Serrano, Pedro Sola Campoy, Laura Perez-Lago, Pilar Catalán, Patricia Muñoz, Darío García de Viedma |
| EPI_ISL_779442, EPI_ISL_779443, EPI_ISL_779447, EPI_ISL_779453, EPI_ISL_779455, EPI_ISL_779457, EPI_ISL_779458, EPI_ISL_779459, EPI_ISL_779460, EPI_ISL_779461, EPI_ISL_779462, EPI_ISL_779464, EPI_ISL_779466, EPI_ISL_779467, EPI_ISL_779468, EPI_ISL_779469, EPI_ISL_779470, EPI_ISL_779471, EPI_ISL_779476, EPI_ISL_779477, EPI_ISL_779478, EPI_ISL_779479, EPI_ISL_779480, EPI_ISL_779481, EPI_ISL_779482, EPI_ISL_779483, EPI_ISL_779484 | | | |
| see above | Microbiological Diagnostic Unit - Public Health Laboratory (MDU-PHL) | MDU-PHL | Seemann T., Sait, M.L., Sherry, N.L. |
| EPI_ISL_785706, EPI_ISL_789378, EPI_ISL_789390 | Houston Methodist Hospital | Houston Methodist Hospital | S. Wesley Long, Randall J. Olsen, Paul A. Christensen, David W. Bernard, James J. Davis, Maulik Shukla, Marcus Nguyen, Matthew Ojeda Saavedra, Prasanti Yerramilli, Layne Pruitt, Sishir Subedi, Heather Hendrickson, and James M. Musser |
| EPI_ISL_791982 | RSUP DR KARIADI Semarang | National Institute of Health Research and Development | Ikawati,HD;Subangkit;Pawestri,HA;Nugraha,AA;Puspa,KD;Puspitasari,I;Pangesti,KNA;Soekarso,T;Puspandari,N;Setiawaty,V |
| EPI_ISL_791987 | RSUD dr. Loekmono Hadi Kudus | National Institute of Health Research and Development | Puspa,KD;Subangkit;Pawestri,HA;Ikawati,HD;Nugraha,AA;Khoiriyah,S;Pangesti,KNA;Soekarso,T;Puspandari,N;Setiawaty,V |
| EPI_ISL_802137, EPI_ISL_802138, EPI_ISL_802139 | MSHS Clinical Microbiology Laboratories | MSHS Pathogen Surveillance Program | Ana S. Gonzalez-Reiche, Hala Alshammary, Mitchell J. Sullivan, Brianne Ciferri, Ajay Obla, Angela Amoako, Mahmoud Awawda, Elena Hirsch, Ashley S. Salimbangon, Levy Sominsky, Katherine Beach, Kayla Russo, Charles Gleason, Shelcie Fabre, Giulio Kleiner, Zenab Khan, Bremy Alburquerque, Adriana van de Guchte, Komal Srivastava, Matthew M. Hernandez, Jayeeta Dutta, Denise Jurczyszak, Emily Ferreri, Rachel Chernet, Nancy Francoeur, Betsaida Salom Melo, Irina Oussenko, Gintaras Deikus, Juan Soto, Shwetha Hara Sridhar, Ying-Chih Wang, Kathryn Twyman, Andrew Kasarskis, Deena R. Altman, Robert Sebra, Adolfo García-Sastre, Marta Luksza, Gopi Patel, Sarah Schaefer, Melissa Gitman, Michael D. Nowak, Alberto Paniz-Mondolfi, Emilia Mia Sordillo, Viviana Simon, Harm van Bakel |
| EPI_ISL_803107 | COVID-19 National Reference Laboratory, Pasteur Institute of Iran | COVID-19 National Reference Laboratory, Pasteur Institute of Iran | Jalali,T., Pouriayevali,M.H., Shoja,Z., Ahmadi,Z., Sadjadi,M.,TavakoliRad,M., Azadmanjiri,S., Fereydouni,Z., Mohebali,M.,Zarei,Z., Hassanpour,G., Zainali,M., Gouya,M.M., Khayatzadeh,S.,Parsaei,M., Sarafraz,N., Hassanzadeh,M., Azarm,A., Karami,C.,Sasani,F., Heidari,Z. and Salehi-Vaziri,M. |
| EPI_ISL_803722 | Wisconsin State Laboratory of Hygiene Communicable Disease Division | Wisconsin State Laboratory of Hygiene Communicable Disease Division | Kelsey R. Florek, Abigail C. Shockey |
| EPI_ISL_803870 | National Institute of Laboratory Medicine and Referral Center | Genomic Research Lab, BCSIR | Md. Murshed Hasan Sarkar,Mohammad Samir Uzzaman, Eshrar Osman, Md. Ahashan Habib, Shahina Akter, Tanjina Akhter Banu, Abu Sayeed Mohammad Mahmud,Barna Goswami, Iffat Jahan, Md. Saddam Hossain, Tasnim Nafisa, Md. Maruf Ahmed Molla, Mahmuda Yeasmin, Asish Kumar Ghosh, A. K. M. Shamsuzzaman, Monira Parveen, Md. Masum Hossain Arif, Md. Salim Khan |
| EPI_ISL_804925, EPI_ISL_804926, EPI_ISL_804927, EPI_ISL_804928, EPI_ISL_804929, EPI_ISL_804930, EPI_ISL_804931, EPI_ISL_804932 | DC Public Health Lab/ Dept. of Forensic Sciences | DC Public Health Lab/ Dept. of Forensic Sciences | Scott Nguyen, Elizabeth Zelaya, Connie Maza, Monica Mann, Brittany Hamilton, David Payne, Jocelyn Hauser |
| EPI_ISL_806564, EPI_ISL_806565, EPI_ISL_806567, EPI_ISL_806568, EPI_ISL_806569, EPI_ISL_806570, EPI_ISL_806591, EPI_ISL_806636, EPI_ISL_806637, EPI_ISL_806697 | KEMRI-Wellcome Trust Research Programme/KEMRI-CGMR-C Kilifi | KEMRI-Wellcome Trust Research Programme/KEMRI-CGMR-C Kilifi | Githinji et al |
| EPI_ISL_810990, EPI_ISL_810992, EPI_ISL_810993, EPI_ISL_810994, EPI_ISL_810995, EPI_ISL_810996, EPI_ISL_810997, EPI_ISL_810998, EPI_ISL_810999, EPI_ISL_811000, EPI_ISL_811001 | | | |
| see above | MRCG at LSHTM Genomics lab | MRCG at LSHTM Genomics lab | Abdul Karim sesay, Abdoulie Kanteh, Jarra Manneh, Mariama Kujabi, Bakary Sanyang |

| | | | |
|---|---|---|---|
| EPI_ISL_811157, EPI_ISL_811159, EPI_ISL_811163, EPI_ISL_811166, EPI_ISL_811167, EPI_ISL_811171, EPI_ISL_811172, EPI_ISL_811173, EPI_ISL_811174, EPI_ISL_811176, EPI_ISL_811177, EPI_ISL_811178, EPI_ISL_811179, EPI_ISL_811181, EPI_ISL_811183, EPI_ISL_811184, EPI_ISL_811186, EPI_ISL_811187, EPI_ISL_811188 | | | |
| see above | Dharwad | CSIR Institute of Genomics and Integrative Biology | Dr. Shivarudrapp B Bhairappanavar, Rahul Bhoyar, Mohammed Imran, Mohit Divakar, Disha Sharma, Dr. Vijay A Yenagi, Dr. Suresh B Arakera, Dr. Amit Ugargol, Dr. Rgavendra B Nayak, Bani Jolly, Abhinav Jain, Paras Sehgal, Gyan Ranjan, Vinod Scaria, Sridhar Sivasubbu |
| EPI_ISL_812156, EPI_ISL_812157, EPI_ISL_812158, EPI_ISL_812159, EPI_ISL_812160 | GA Department of Public Health Laboratory | Pathogen Discovery, Respiratory Viruses Branch, Division of Viral Diseases, Centers for Disease Control and Prevention | Yan Li, Ying Tao, Anna Montmayeur, Jing Zhang, Brian Lynch, Krista Queen, Anna Uehara, Rachel Marine, Peter Cook, Clinton R. Paden, Haibin Wang, Suxiang Tong |
| EPI_ISL_812527, EPI_ISL_812528, EPI_ISL_812529, EPI_ISL_812530, EPI_ISL_812531, EPI_ISL_812532, EPI_ISL_812533, EPI_ISL_812534, EPI_ISL_812535, EPI_ISL_812536, EPI_ISL_812539, EPI_ISL_812541, EPI_ISL_812542, EPI_ISL_812543, EPI_ISL_812544, EPI_ISL_812545, EPI_ISL_812546, EPI_ISL_812547, EPI_ISL_812548, EPI_ISL_812549, EPI_ISL_812550, EPI_ISL_812551, EPI_ISL_812553, EPI_ISL_812556, EPI_ISL_812557, EPI_ISL_812558, EPI_ISL_812559 | | | |
| see above | United States Air Force School of Aerospace Medicine | United States Air Force School of Aerospace Medicine | Anthony Fries, Jennifer Meyer, Amanda Javorina, Sarah Purves, William Gruner, Clarise Starr, Elizabeth Macias |
| EPI_ISL_812808, EPI_ISL_812809, EPI_ISL_812811, EPI_ISL_812816, EPI_ISL_812824, EPI_ISL_812825, EPI_ISL_812827, EPI_ISL_812832, EPI_ISL_812845, EPI_ISL_812849, EPI_ISL_812859, EPI_ISL_812861, EPI_ISL_812864, EPI_ISL_812871 | | | |
| see above | Genomics Program, Children Cancer Hospital | Genomics Program, Children Cancer Hospital | Hatem,A., Hadad,A., Abouelnaga,S., Amer,K., Salah,H., Farawyla,H., Halafawy,A., Mansour,T., shalaby,L., Hassan,W., Soliman,M., Gomaa,C., Hassan,R., Soliman,S., Monuir,G., Hammad,M., Hussein,S., Abdo,I., Jalal,D., El-Zayat,M., El-Shaqnqery,H., Diab,A., Bakry,U., Samir,O., Magdeldin,S., Sayed,A. |
| EPI_ISL_815256, EPI_ISL_815260, EPI_ISL_815261, EPI_ISL_815262, EPI_ISL_815271, EPI_ISL_815281, EPI_ISL_815282, EPI_ISL_815297, EPI_ISL_815321, EPI_ISL_815324, EPI_ISL_815334, EPI_ISL_815352, EPI_ISL_815357, EPI_ISL_815364, EPI_ISL_815366, EPI_ISL_815388, EPI_ISL_815392, EPI_ISL_815393, EPI_ISL_815394, EPI_ISL_815395, EPI_ISL_815396 | | | |
| see above | Centogene | Centogene | Peter Bauer, Krishna Kumar Kandaswamy, Vivi Hue-Trang Lieu |
| EPI_ISL_824409, EPI_ISL_824412, EPI_ISL_824413, EPI_ISL_824415, EPI_ISL_824429, EPI_ISL_824430, EPI_ISL_824431, EPI_ISL_824432, EPI_ISL_824433, EPI_ISL_824434, EPI_ISL_824435, EPI_ISL_824436, EPI_ISL_824437, EPI_ISL_824438, EPI_ISL_824439, EPI_ISL_824440, EPI_ISL_824441, EPI_ISL_824442, EPI_ISL_824443, EPI_ISL_824444, EPI_ISL_824445, EPI_ISL_824446, EPI_ISL_824447, EPI_ISL_824448, EPI_ISL_824449, EPI_ISL_824450, EPI_ISL_824451, EPI_ISL_824452 | | | |
| see above | Hospital Universitari Vall d'Hebron - Vall d'Hebron Institut de Recerca | Hospital Universitari Vall d'Hebron | Cristina Andrés, Maria Piñana, Josep F Abril, Damir Garcia-Cehic, Ariadna Rando, Juliana Esperalba, Maria Gema Codina, Carla Castillo, Maria Carmen Martin, Tomàs Pumarola, Josep Quer, Andrés Antón |
| EPI_ISL_825650, EPI_ISL_825665, EPI_ISL_825666, EPI_ISL_825667, EPI_ISL_825668, EPI_ISL_825669, EPI_ISL_825670, EPI_ISL_825671, EPI_ISL_825672, EPI_ISL_825673, EPI_ISL_825674, EPI_ISL_825675, EPI_ISL_825676, EPI_ISL_825677, EPI_ISL_825678, EPI_ISL_825679, EPI_ISL_825680, EPI_ISL_825681, EPI_ISL_825682, EPI_ISL_825683, EPI_ISL_825684, EPI_ISL_825685, EPI_ISL_825686, EPI_ISL_825936, EPI_ISL_825937, EPI_ISL_825938, EPI_ISL_825939, EPI_ISL_825941, EPI_ISL_826118, EPI_ISL_826119, EPI_ISL_826120, EPI_ISL_826121, EPI_ISL_826122 | | | |
| see above | Laboratoire de santé publique du Québec | Laboratoire de santé publique du Québec | Sandrine Moreira, Ioannis Ragoussis, Guillaume Bourque, Jesse Shapiro, Mark Lathrop and Michel Roger on behalf of the CoVSeQ research group (http://covseq.ca/researchgroup) |
| EPI_ISL_830209, EPI_ISL_830213 | The National University Hospital of Iceland | deCODE genetics | Daniel F Gudbjartsson; Agnar Helgason; Hakon Jonsson; Olafur T Magnusson; Pall Melsted; Gudmundur L Norddahl; Jona Saemundsdottir; Asgeir Sigurdsson; Patrick Sulem; Arna B Agustsdottir; Hannes Eggertsson; Berglind Eiriksdottir; Run Fridriksdottir; Elisabet E Gardarsdottir; Gudmundur Georgsson; Olafia S Gretarsdottir; Kjartan R Gudmundsson; Thora R Gunnarsdottir; Arnaldur Gylfason; Hilma Holm; Brynjar O Jensson; Aslaug Jonasdottir; Kamilla S Josefsdottir; Thordur Kristjansson; Droplaug N Magnusdottir; Solvi Rognvaldsson; Louise le Roux; Gudrun Sigmundsdottir; Gardar Sveinbjornsson; Kristin E Sveinsdottir; Maney Sveinsdottir; Emil A Thorarensen; Bjarni Thorbjornsson; Gisli Masson; Ingileif Jonsdottir; Alma Moller; Thorolfur Gudnason; Karl G Kristinsson; Unnur Thorsteinsdottir; Kari Stefansson |
| EPI_ISL_831688 | Laboratório de Microbiologia Molecular - Universidade FEEVALE | Universidade Federal de Ciências da Saúde de Porto Alegre | Vinícius Bonetti Franceschi, Amanda de Menezes Mayer, Gabriel Dickin Caldana, Carla Andretta Moreira Neves, Patrícia Aline Gröhs Ferrareze, Gabriela Bettella Cybis, Ricardo Ariel Zimerman, Lívia Kmetzsch, Fernando Rosado Spilki, Claudia Elizabeth Thompson |
| EPI_ISL_831918 | New Mexico Department of Health Scientific Laboratory | New Mexico Department of Health Scientific Laboratory | Ellie Johnson, Anastacia Griego-Fisher, D'eldra Malone |
| EPI_ISL_832023, EPI_ISL_832024, EPI_ISL_832025, EPI_ISL_832026, EPI_ISL_832027, EPI_ISL_832028, EPI_ISL_832029, EPI_ISL_832030, EPI_ISL_832031 | Wyoming Public Health Laboratory | Wyoming Public Health Laboratory | Noah Hull, Taylor Fearing, Lynette Gumbleton, Channing Weber, Ashley Norberg, Bailey Bowcutt, and Wanda Manley |
| EPI_ISL_832406, EPI_ISL_832407, EPI_ISL_832408, EPI_ISL_832409, EPI_ISL_832410, EPI_ISL_832411 | OHSU Lab Services Molecular Microbiology Lab | Oregon SARS-CoV-2 Genome Sequencing Center | Brendan L. O'Connell, Ruth V. Nichols, Sally Grindstaff, Alec J. Hirsch, Donna Hansel, Guang Fan, Daniel N. Streblow, William B. Messer, Andrew C. Adey, Benjamin N. Bimber, Brian J. O'Roak |
| EPI_ISL_833181 | National Institute of Laboratory Medicine and Referral Center | Genomic Research Lab, BCSIR | Abu Sayeed Mohammad Mahmud, Mohammad Samir Uzzaman, Eshrar Osman, Md. Ahashan Habib, Shahina Akter, Tanjina Akhtar Banu,Md. Murshed Hasan Sarkar, Barna Goswami, Iffat Jahan, Md. Saddam Hossain, Tasnim Nafisa, Md. Maruf Ahmed Molla, Mahmuda Yeasmin, Asish Kumar Ghosh, A. K. M. Shamsuzzaman, Monira Parveen, Md. Masum Hossain Arif, Md. Salim Khan |
| EPI_ISL_833196, EPI_ISL_833198 | Hôpital Bichat Claude Bernard, Laboratoire de Virologie | IAME UMR1137 Inserm, Université de Paris, Hôpital Bichat | Antoine Bridier, Amélie Recoing, Quentin Le Hingrat, Lena Daniel, Siham Hamri, Gilles Collin, Alexandre Storto, Mélanie Bertine, Charlotte Charpentier, Nadhira Houhou-Fidouh, Diane Descamps, Benoit Visseaux |
| EPI_ISL_833493 | RS PMI Bogor, Indonesia | Biosafety Level-3 Laboratory, Indonesian Institute of Sciences (LIPI) | Isa Nuryana, Ade Andriani, Anik Budhi Dharmayanthi, Syam Budi Iryanto, Andri Wardiana, Anggia Prasetyoputri, Ahmad Fathoni, Indriawati, Nurulliawaty, Ratih Asmana Ningrum |
| EPI_ISL_833506 | RS PMI, Bogor, Indonesia | Biosafety Level-3 Laboratory, Indonesian Institute of Sciences (LIPI) | Isa Nuryana, Ade Andriani, Ahmad Fathoni, Anik Budhi Dharmayanthi, Syam Budi Iryanto, Andri Wardiana, Anggia Prasetyoputri, Satrio Haryo Benowo, Gilang Akbariani, Ratih Asmana Ningrum |
| EPI_ISL_833508 | Klinik Apotek Dein, Jakarta, Indonesia | Biosafety Level-3 Laboratory, Indonesian Institute of Sciences (LIPI) | Andri Wardiana, Anggia Prasetyoputri, Anik Budhi Dharmayanthi, Syam Budi Iryanto, Isa Nuryana, Ade Andriani, Ahmad Fathoni, Ratih Asmana Ningrum |
| EPI_ISL_833516 | Veterinary Specialized Instute "Nis" | Veterinary Specialized Institute "Kraljevo", Serbia | Vidanovic,D., Tesovic,B., Manic,M., Petrovic,M.,Knezevic,A., Jovanovic,T., Jankovic,M., Sekler,M., Banovic Djeri,B., Petrovic,T., Volkening,J., Afonso,C. |
| EPI_ISL_837556 | Centro Nacional de Enfermedades Tropicales (CENETROP) | Laboratory of Respiratory Viruses and Measles, Oswaldo Cruz Institute, FIOCRUZ | Paola Resende, Roxana Loayza, Cinthia Avila, Luciana Appolinario, Fernando Motta, Anna Carolina Paixao, Ana Carolina Mendonca, Marilda Siqueira |
| EPI_ISL_837586, EPI_ISL_837587, EPI_ISL_837588, EPI_ISL_837589 | Laboratorio Nacional de Salud | Laboratory of Respiratory Viruses and Measles, Oswaldo Cruz Institute, FIOCRUZ | Paola Resende, Cesar Roberto Conde Pereira, Claudia Estrada, Luciana Appolinario, Fernando Motta, Anna Carolina Paixao, Ana Carolina Mendonca, Marilda Siqueira |
| EPI_ISL_837763, EPI_ISL_837764, EPI_ISL_837765, EPI_ISL_837766, EPI_ISL_837767, EPI_ISL_837768, EPI_ISL_837769, EPI_ISL_837770, EPI_ISL_837771, EPI_ISL_837772, EPI_ISL_837773, EPI_ISL_837774, EPI_ISL_837775, EPI_ISL_837776, EPI_ISL_837777, EPI_ISL_837778, EPI_ISL_837779, EPI_ISL_837780 | | | |
| see above | Instituto Nacional de Enfermedades Respiratorias (INER) | Instituto Nacional de Enfermedades Respiratorias (INER) | Celia Boukadida, Margarita Matías-Florentino, Alma Rincón-Rubio, Hector Esteban Paz-Juárez, Olivia Briceño, Edgar Sevilla-Reyes, Fidencio Mejía-Nepomuceno, Mario Mújica-Sánchez, Eduardo Becerril-Vargas, José Arturo Martínez-Orozco, Alejandra Hernández-Terán, Jorge Salas-Hernández, Santiago Ávila-Ríos, Joel Armando Vázquez-Pérez |
| EPI_ISL_845807 | National Institute of Laboratory Medicine and Referral Center | Genomic Research Lab, BCSIR | Md. Murshed Hasan Sarkar,Mohammad Samir Uzzaman, Eshrar Osman,Md. Ahashan Habib, Shahina Akter, Tanjina Akhtar Banu,Abu Sayeed Mohammad Mahmud,Barna Goswami, Iffat Jahan, Md. Saddam Hossain, Tasnim Nafisa, Md. Maruf Ahmed Molla, Mahmuda Yeasmin, Asish Kumar Ghosh, A. K. M. Shamsuzzaman, Md. Salim Khan |
| EPI_ISL_848073, EPI_ISL_848074, EPI_ISL_848075 | Laboratory of Clinical Research on Dermatozoonoses in Domestic Animals, Evandro Chagas National Institute of Infectious Diseases, Oswaldo Cruz Foundation (Fiocruz), Rio de Janeiro, RJ, Brazil. | Laboratory of Respiratory Viruses and Measles, Oswaldo Cruz Institute, FIOCRUZ | Guilherme Amaral Calvet, Michelle Fernanda Borges da Silva, Anielle de Pina Costa, Ezequias Batista Martins, Isabella Campos Vargas de Moraes, Lusiele Guaraldo, Patrícia Brasil, Sandro Antônio Pereira, Rodrigo Caldas Menezes, Isabella Dib Ferreira Gremião, Lucas Oliveira Keidel, Shanna Araujo dos Santos, Artur Augusto Velho Mendes Junior, Renato Orsini Ornellas, Maria Ogrzewalska, PaolaCristina Resende, Alex Pauvolid-Corrêa, Fernando do Couto Motta, Alice Sampaio Barreto da Rocha, Thiago C. Souza, Marilda Mendonça Siqueira |
| EPI_ISL_848297, EPI_ISL_848298, EPI_ISL_848299, EPI_ISL_848484, EPI_ISL_848485 | Illinois Department of Public Health | Gagnon Lab, Southern Illinois University | Keith Gagnon |
| EPI_ISL_848561, EPI_ISL_848609 | Evandro Chagas Institute | Evandro Chagas Institute | Santos, M.C.; Silva, A.M.; Junior, W.D.C.; Barbagelata, L.S.; Ferreira, J.A.; Sousa, E.M.A.; da Silva, P.S.; Pinheiro, K.C.; L.C.; Sousa Junior, E.C. |

| | | | |
|---|---|---|---|
| EPI_ISL_849650, EPI_ISL_849651, EPI_ISL_849652 | Servizio di igiene epidemiologia e sanità pubblica (SIESP)-Chieti | Istituto Zooprofilattico Sperimentale dell'Abruzzo e Molise "G.Caporale" | Lorusso A, Marcacci M, Di Domenico M, Curini V, Ancora M, Cammà C, Rinaldi A, Mangone I, Di Pasquale A, Puglia I, Savini G. |
| EPI_ISL_849655 | Servizio di igiene e sanità pubblica (SIESP)-Teramo | Istituto Zooprofilattico Sperimentale dell'Abruzzo e Molise "G.Caporale" | Lorusso A, Marcacci M, Di Domenico M, Curini V, Ancora M, Cammà C, Rinaldi A, Mangone I, Di Pasquale A, Puglia I, Savini G. |
| EPI_ISL_849663 | Servizio di igiene epidemiologia e sanità pubblica (SIESP)-Chieti | Istituto Zooprofilattico Sperimentale dell'Abruzzo e Molise "G.Caporale" | Lorusso A, Marcacci M, Di Domenico M, Curini V, Ancora M, Cammà C, Rinaldi A, Mangone I, Di Pasquale A, Puglia I, Savini G. |
| EPI_ISL_849931, EPI_ISL_849932, EPI_ISL_849947 | UC Davis- Department of Pathology and Laboratory Medicine | Chan-Zuckerberg Biohub | CZB Cliahub Consortium |
| EPI_ISL_850230, EPI_ISL_850231, EPI_ISL_850232, EPI_ISL_850233, EPI_ISL_850234, EPI_ISL_850235, EPI_ISL_850236, EPI_ISL_850237, EPI_ISL_850238, EPI_ISL_850239 | Division of Emerging Infectious Diseases, Bureau of Infectious Diseases Diagnosis Control, Korea Disease Control and Prevention Agency | Division of Emerging Infectious Diseases, Bureau of Infectious Diseases Diagnosis Control, Korea Disease Control and Prevention Agency | Ae Kyung Park, Il-Hwan Kim, Heui Man Kim, Jeong-Min Kim, Namjoo Lee, Chaeyoung Lee, Sang Hee Woo, Eun-Jin Kim |
| EPI_ISL_850951 | National Institute for Viral Disease Control and Prevention, China CDC | National Institute for Viral Disease Control and Prevention, China CDC | Xiang Zhao, Yenan Feng, Zhixiao Chen, Yao Meng, Yuchao Wu, Yang Song, Ji Wang, Kai Nie, Yong Zhang, Yanhai Wang, Weimin Zhou, Wenjie Tan, Jun Han, Shiwen Wang, Wenbo Xu, Cao Chen, Dayan Wang |
| EPI_ISL_851050 | Gandhi Medical College and Hospital | CSIR-Centre for Cellular and Molecular Biology | Nagamani Kammili, Winnie Thomas, Madhavi Latha Manolla, Shailaja VV, Sudhamadhuri Devara, Vanisree Rajoli, Archana GJ, Sushma Rajyalakshmi Gudiseva, Sunitha Pakalapati, Manisha Rani, Amrithesh Kumar, Raja Rao Mesepogu, Vinay Shekar Reddy, Thrilok Chander Bingi, Sofia Banu, Divya Tej Sowpati |
| EPI_ISL_851051 | Gandhi Medical College and Hospital | CSIR-Centre for Cellular and Molecular Biology | Nagamani Kammili, Madhavi Latha Manolla, Winnie Thomas, Shailaja VV, Sudhamadhuri Devara, Vanisree Rajoli, Archana GJ,Sushma Rajyalakshmi Gudiseva, Sunitha Pakalapati, Manisha Rani, Amrithesh Kumar, Raja Rao Mesepogu, Vinay Shekar Reddy, Thrilok Chander Bingi, Sofia Banu, Divya Tej Sowpati |
| EPI_ISL_853287, EPI_ISL_853289, EPI_ISL_853291, EPI_ISL_853292 | UPMC Clinical Microbiology Laboratory | Microbial Genome Sequencing Center; Microbial Genomic Epidemiology Laboratory | Mustapha M. Mustapha, Jane W. Marsh, Dan Snyder, Marissa P. Griffith, Stephanie L. Mitchell, Vatsala R. Srinivasa, Kady D. Waggle, Chinelo Ezeonwuku, Vaughn S. Cooper, Lee H. Harrison |
| EPI_ISL_853751, EPI_ISL_853836, EPI_ISL_853837, EPI_ISL_853838, EPI_ISL_853839, EPI_ISL_853840, EPI_ISL_853841, EPI_ISL_853887, EPI_ISL_853888, EPI_ISL_853889, EPI_ISL_853890, EPI_ISL_853891, EPI_ISL_853959 | | | |
| see above | Center for Virology, Medical University of Vienna | Bergthaler laboratory, CeMM Research Center for Molecular Medicine of the Austrian Academy of Sciences | Lukas Endler, Alexandra Popa, Benedikt Agerer, Jakob-Wendelin Genger, Alexander Lercher, Anna Schedl, Thomas Penz, Michael Schuster, Jan Laine, Martin Senekowitsch, Christoph Bock, Andreas Bergthaler |
| EPI_ISL_860552 | Ohio Department of Health Laboratory | Ohio Department of Health Laboratory | Holmes, Jennifer; Eric Brandt, Keoni Omura, Glen McGillivary, Caitlin McDonnell, Kirtana Ramadugu, Erica Leasure, Kelsey Florek, Heather Blankenship, Quanta Brown, and Tammy Bannerman |
| EPI_ISL_861875 | LATE - Laboratório de Técnicas Especiais - Hospital Israelita Albert Einstein | LATE - Laboratório de Técnicas Especiais - Hospital Israelita Albert Einstein | Deyvid Amgarten, Fernanda de Mello Malta, Raquel Riyuzo, Ana Paula Moreira Salles, Pedro Henrique Sebe Rodrigues, João Renato Rebello Pinho |
| EPI_ISL_865894 | University College London, Great Ormond Street Hospital for Children NHS Foundation Trust, Imperial College Healthcare NHS Trust | COVID-19 Genomics UK (COG-UK) Consortium | Sergi Castellano, Rachel Williams, Mark Kristiansen, Paola Resende Silva, Sunando Roy, Tony Brooks, Helena Tutill, Paola Niola, Patricia Dyal, Charlotte Williams, Leysa Forrest, Yasmin Panchbhaya, Jacqueline Findlay, Samuel Weeks, Julianne Brown, Kathryn Harris, Paul Randell, James Price, Alison Holmes, Judith Breuer |
| EPI_ISL_876528, EPI_ISL_876577, EPI_ISL_876578, EPI_ISL_876579, EPI_ISL_876580, EPI_ISL_876581, EPI_ISL_876582, EPI_ISL_876583, EPI_ISL_876584, EPI_ISL_876585, EPI_ISL_876586, EPI_ISL_876587 | | | |
| see above | Florida Bureau of Public Health Laboratories | Florida Bureau of Public Health Laboratories | Sarah Schmedes, Jason Blanton |
| EPI_ISL_876954, EPI_ISL_876955, EPI_ISL_876956, EPI_ISL_876957 | Quest Diagnostics | Quest Diagnostics | Rosenthal,S.H., Gerasimova,A., Kagan,R.M., Anderson, B., Hua, M., Liu Y., Bernstein, L.E., Livingston, K.E., Perez, A., Shalhout, D.F., Shlyakhter, I.A., Owen, R., Tanpaiboon, P., Lacbawan, F. |
| EPI_ISL_877648, EPI_ISL_877649, EPI_ISL_877650, EPI_ISL_877651, EPI_ISL_877652, EPI_ISL_877653 | Clinical Molecular Microbiology Laboratory, UNC Hospital | Dirk Dittmer | Razia Moorad , Justin T. Landis , Brent A. Eason, Melissa B. Miller, Linda Pluta, Dirk Dittmer, Angelica Juarez, Cecilia Thompson , Cameroon Grant, Evelyn Hoffman, Patricio Cano, Jason Wong, Carolina Caro-Vegas, Blossom Damania. |
| EPI_ISL_882634 | COVID lab, Mymensingh Medical College | Department of Pathology, Bangladesh Agricultural University & Department of Microbiology, Mymensingh Medical College | Afrin, S. Z. Paul, S. K. Parvin, R. |
| EPI_ISL_884327, EPI_ISL_884345, EPI_ISL_884404, EPI_ISL_884416, EPI_ISL_884429, EPI_ISL_884432 | Infectious Diseases, Quest Diagnostics | Infectious Diseases, Quest Diagnostics | Rosenthal,S.H., Gerasimova,A., Kagan,R.M., Anderson,B., Bernstein,L.E., Livingston,K.E., Hua,M., Liu,Y., Shalhout,D.F., Owen,R., Lacbawan,F. |
| EPI_ISL_884530 | Molecular Microbiology & Immunology, University of Missouri | Molecular Microbiology & Immunology, University of Missouri | Tang,C.Y., Li,T., Hang,J., Lidl,G.M., Wan,X.-F. |
| EPI_ISL_884826, EPI_ISL_884827, EPI_ISL_884828, EPI_ISL_884829, EPI_ISL_884830, EPI_ISL_884831, EPI_ISL_884832, EPI_ISL_884833, EPI_ISL_884834, EPI_ISL_884836, EPI_ISL_884837, EPI_ISL_884838, EPI_ISL_884844, EPI_ISL_884845, EPI_ISL_884846, EPI_ISL_884847, EPI_ISL_884848, EPI_ISL_884849, EPI_ISL_884850, EPI_ISL_884851 | | | |
| see above | Department of Biochemistry, Cell and Molecular Biology, West African Centre for Cell Biology of Infectious Pathogens (WACCBIP), University of Ghana | Department of Biochemistry, Cell and Molecular Biology, West African Centre for Cell Biology of Infectious Pathogens (WACCBIP), University of Ghana | Ngoi,J.M., Tei-Maya,F., Morang'a,C.M., Magnussen,V., Amuzu,D.S., Mohammed,A., Tapela,K., Kibinge,N., Diallo,A.B., Kumi-Ansah,F., Odoom,T., Boakye,O.D., Amoako,E., Abass,A.-K., Quashie,P., Amenga-Etego,L.N., Akoriyea,S.K., Awandare,G.A., Bediako,Y. |
| EPI_ISL_887145, EPI_ISL_887146 | Massachusetts General Hospital | Infectious Disease Program, Broad Institute of Harvard and MIT | Lemieux,J.E., Siddle,K.J., Shaw,B., Adams,G., Pierce,V., Turbett,S., Anahtar,M., Branda,J., Slater,D., Harris,J., Lin,A.E., Gladden-Young,A., Lagerborg,K., Rudy,M., DeRuff,K., Carter,A., Normandin,E., Bauer,M., Reilly,S., Tomkins-Tinch,C., Loreth,C., Chaluvadi,S., Neumann,A., Cusick,C., Chapman,S.B., Gnirke,A., Flowers,K., Cerrato,F., Birren,B.W., Gallagher,G., Smole,S., Park,D.J., MacInnis,B.L., Ryan,E., LaRocque,R., Rosenberg,E. and Sabeti,P.C. |
| EPI_ISL_889334, EPI_ISL_889335, EPI_ISL_889340, EPI_ISL_889341, EPI_ISL_889342, EPI_ISL_889343, EPI_ISL_889344 | The University Hospital Brno | Institute of Applied Biotechnologies a.s. | Petr Klempt, Ondej Brzo, Martin Kašný, Kateina Kvapilová, Martina Lengerová, Petr Kvapil |
| EPI_ISL_889361, EPI_ISL_889362, EPI_ISL_889363 | Motol University Hospital | Institute of Applied Biotechnologies a.s. | Petr Klempt, Ondej Brzo, Martin Kašný, Kateina Kvapilová, Pavel Devínek, Petr Kvapil |
| EPI_ISL_890110 | Laboratoire de santé publique du Québec | Laboratoire de santé publique du Québec | Sandrine Moreira, Ioannis Ragoussis, Guillaume Bourque, Jesse Shapiro, Mark Lathrop and Michel Roger on behalf of the CoVSeQ research group |
| EPI_ISL_891223, EPI_ISL_891227, EPI_ISL_891230 | The Oncology Institute "Prof. Dr. Ion Chiricuta" Cluj Napoca | "Stefan cel Mare" University Metagenomics Lab | Lobiuc Andrei, Gheorghita Roxana |
| EPI_ISL_896135, EPI_ISL_900134, EPI_ISL_900156, EPI_ISL_900179, EPI_ISL_900200, EPI_ISL_900208, EPI_ISL_900335, EPI_ISL_900376, EPI_ISL_900473 | MEPHI, Aix Marseille University | MEPHI, Aix Marseille University | Anthony LEVASSEUR |
| EPI_ISL_900703, EPI_ISL_900704, EPI_ISL_900705, EPI_ISL_900706, EPI_ISL_900707, EPI_ISL_900708, EPI_ISL_900709, EPI_ISL_900710, EPI_ISL_900711, EPI_ISL_900727, EPI_ISL_900728, EPI_ISL_900729, EPI_ISL_900730, EPI_ISL_900731, EPI_ISL_900732, EPI_ISL_900733 | | | |
| see above | Bozeman Health Deaconess Hospital | Wiedenheft lab, Montana State University | Artem Nemudryi, Anna Nemudraia, Tanner Wiegand, Joseph Nichols, Deann T. Snyder, Jodi F. Hedges, Calvin Cicha, Helen Lee, Karl K. Vanderwood, Diane Bimczok, Mark A. Jutila and Blake Wiedenheft |
| EPI_ISL_902744 | Hospital Universitari Germans Trias i Pujol (HUGTiP) / | IrsiCaixa - Can Ruti CovidSeq | Fundació irsiCaixa. Hospital Universitari Germans Trias i Pujol(HUGTiP), 2a planta, maternal Ctra Canyet s/n, Badalona Marta Massanella, Ester Ballana, |

| | | | |
|---|---|---|---|
| | Fundació Lluita contra la SIDA (FLSida) | | Lidia Ruiz, Nuria Izquierdo, Jorge Carrillo, Roger Paredes, Julia Blanco, Joaquim Segalés, Bonaventura Clotet |
| EPI_ISL_910306, EPI_ISL_910307, EPI_ISL_910308, EPI_ISL_910309, EPI_ISL_910310, EPI_ISL_910311, EPI_ISL_910312, EPI_ISL_910313 | CSIR-Centre for Cellular and Molecular Biology | CSIR-Centre for Cellular and Molecular Biology | Payel Mukherjee,Pratheusa Maccha,Namami Gaur,Lamuk Zaveri,Tulasi Nagabandi,Purushotham Vodnala,Blessy B John,Viswagithe S L,B Himasri,Sofia Banu,Priya Singh,Archana Bharadwaj Siva,Karthik Bharadwaj Tallapaka,Rakesh K Mishra,Divya Tej Sowpati |
| EPI_ISL_912537 | NHLS Universitas Academic | UFS Virology | PA Bester, MM Nyaga, P Nthiga, MT Mogotsi, D Goedhals, T de Oliveira |
| EPI_ISL_913911, EPI_ISL_913912, EPI_ISL_913917, EPI_ISL_913919, EPI_ISL_913969 | Instituto de Diagnostico y Referencia Epidemiologicos INDRE_RNLSP | Instituto de Diagnostico y Referencia Epidemiologicos (INDRE) | Claudia Wong-Arambula, Abril Rodriguez-Maldonado, Fabiola Garces-Ayala, Adnan Araiza-Rodriguez, David Fragoso-Fonseca, Sergio Rangel-Guerrero, Mayra Jimenez-Morales, Nancy Munoz-Hernandez, Natividad Cruz-Ortiz, Tatiana Nunez-Garcia, Gisela Barrera-Badillo, Lucia Hernandez-Rivas, Irma Lopez-Martinez, Ernesto Ramirez-Gonzalez. |
| EPI_ISL_914533, EPI_ISL_914534, EPI_ISL_914535, EPI_ISL_914536, EPI_ISL_914537, EPI_ISL_914538, EPI_ISL_914539, EPI_ISL_914540, EPI_ISL_914543, EPI_ISL_914544, EPI_ISL_914545, EPI_ISL_914546, EPI_ISL_914547, EPI_ISL_914548, EPI_ISL_914549, EPI_ISL_914550, EPI_ISL_914551, EPI_ISL_914552, EPI_ISL_914553, EPI_ISL_914554, EPI_ISL_914555, EPI_ISL_914556, EPI_ISL_914557, EPI_ISL_914558, EPI_ISL_914559, EPI_ISL_914560, EPI_ISL_914561, EPI_ISL_914562, EPI_ISL_914563, EPI_ISL_914564, EPI_ISL_914565, EPI_ISL_914566, EPI_ISL_914568, EPI_ISL_914569, EPI_ISL_914570, EPI_ISL_914571, EPI_ISL_914572 | | | |
| see above | TGen North | TGen North | "Jolene Bowers, Megan Folkerts, Chris French, Hayley Yaglom, Ashlyn Pfeiffer, Darrin Lemmer, Dave Engelthaler, The Arizona COVID Genomics Union (ACGU)" |
| EPI_ISL_930854, EPI_ISL_930855 | Central Laboratory of Public Health of Rio Grande do Sul(Lacen_RS) | State Center for Health Surveillance of the Health Department of the State of Rio Grande do Sul(CEVS_SES-RS) | Barcellos R, Campos A, Dornelles C, Godinho F, Gonzalez A, Gregianini T, Molina C, Salvato R, Schaurich A, |
| EPI_ISL_933650, EPI_ISL_933651, EPI_ISL_933652 | Toronto Invasive Bacterial Diseases Network | McMaster University | Allison McGeer, Patryk Aftanas, Hooman Derakhshani, Angel Li, Kuganya Nirmalarajah, Emily Panousis, Ahmed Draia, Jalees Nasir, Michael Surette, Samira Mubareka, Andrew G. McArthur |
| EPI_ISL_936380, EPI_ISL_936381, EPI_ISL_936382, EPI_ISL_936383, EPI_ISL_936384, EPI_ISL_936385, EPI_ISL_936386 | Genetica y Virologia, Facultad de Ciencias | Genetica y Virologia, Facultad de Ciencias | Panzera,Y., Ramos,N., Frabasile,S., Calleros,L., Marandino,A., Tomas,G., Techera,C., Grecco,S., Fuques,E., Goni,N., Ramas,V., Coppola,L., Chiparelli,H., Sorhouet,C., Mogdasy,C., Arbiza,J., Delfraro,A., Perez,R. |
| EPI_ISL_936574, EPI_ISL_936575, EPI_ISL_936576, EPI_ISL_936577, EPI_ISL_936578, EPI_ISL_936579, EPI_ISL_936580, EPI_ISL_936581 | Northwestern Memorial Hospital | Ozer Lab | Ramon Lorenzo-Redondo, Lacy M. Simons, Chad J. Achenbach, Lawrence J. Jennings, Michael G. Ison, Judd F. Hultquist, Egon A. Ozer |
| EPI_ISL_940396, EPI_ISL_940402, EPI_ISL_940403, EPI_ISL_940541 | Hôpital Bichat Claude Bernard, Laboratoire de Virologie | IAME UMR1137 Inserm, Université de Paris, Hôpital Bichat | Antoine Bridier-Nahmias, Amélie Recoing, Quentin Le Hingrat, Lena Daniel, Siham Hamri, Gilles Collin, Alexandre Storto, Mélanie Bertine, Charlotte Charpentier, Nadhira Houhou-Fidouh, Diane Descamps, Benoit Visseaux |
| EPI_ISL_940902, EPI_ISL_940906, EPI_ISL_940909, EPI_ISL_940910, EPI_ISL_940911, EPI_ISL_940913, EPI_ISL_940914, EPI_ISL_940915, EPI_ISL_940916, EPI_ISL_940917, EPI_ISL_940918, EPI_ISL_940919, EPI_ISL_940928 | | | |
| see above | Centers for Disease Control and Prevention, Dengue Branch | Centers for Disease Control and Prevention, Dengue Branch | Gilberto A. Santiago, Glenda Gonzalez, Betzabel Flores, Keyla Charriez, Gabriela Paz-Bailey, Jorge L. Munoz-Jordan |
| EPI_ISL_943576, EPI_ISL_943577, EPI_ISL_943578, EPI_ISL_943579, EPI_ISL_943580 | Lacen_RS | State Center for Health Surveillance. Rio Grande do Sul State Secretary of Health | Aline Campos, Amanda da Silva, Anelise Schaurich, Claudia Dornelles, Cynthia Molina, Fernanda Godinho, Lara Crescente, Leticia Garay, Regina Barcellos, Richard Salvato, Tatiana Gregianini, Vagner Fonseca |
| EPI_ISL_953413 | Laboratorio de Investigaciones de Baney | "Swiss Tropical and Public Health Institute" | "Carlos Cortes, Claudia Daubenberger, Guillermo Garcia, Salome Hosch, Bonifacio Manguire Nlavo, Maximilian Mpina, Elizabeth Nyakarungu, Diosdado Odjama Nseng Ada, Mitoha Ondo O Ayekaba, Tobias Schindler, Philip Wonder Phiri" |
| EPI_ISL_954883 | Colorado Department of Public Health and Environment | Colorado Department of Puplic Health and Environment | Laura Bankers, Molly C. Hetherington-Rauth, Diana Ir, Shannon Ely, Shannon R. Matzinger, Sarah Elizabeth Totten, Emily A. Travanty |
| EPI_ISL_955142 | University of Sarajevo, Veterinary Faculty, Laboratory for Molecular Diagnostic and Research Laboratory | University of Sarajevo, Veterinary Faculty, Laboratory for Molecular Diagnostic and Research Laboratory | Goleti T., Goleti Š., Softi A., Ali-Šeho A., Šabi E., Jaži A., Nicevi M., Hodži A., Terzi I. |
| EPI_ISL_956277 | Paru Hospital | Institute of Tropical Disease, Universitas Airlangga | Krisnoadi Rahardjo, Aldise M Nastri, Jezzy R Dewantari, Rima R Prasetya, Dyah Retno, Gatot Soegiarto, Laksmi Wulandari, Resti Yudhawati, Soetjipto, Yasuko Mori, Maria I Lusida, Kazufumi Shimizu |
| EPI_ISL_956310 | RSUD Dr. M. Soewandhie | Institute of Tropical Disease, Universitas Airlangga | Aldise M Nastri, Jezzy R Dewantari, Rima R Prasetya, Krisnoadi Rahardjo, Febria Rachmanita, Gatot Soegiarto, Laksmi Wulandari, Resti Yudhawati, Yasuko Mori, Soetjipto, Kazufumi Shimizu, Maria I Lusida |
| EPI_ISL_956311 | Islamic Hospital | Institute of Tropical Disease, Universitas Airlangga | Kazufumi Shimizu, Krisnoadi Rahardjo, Aldise M Nastri, Jezzy R Dewantari, Rima R Prasetya, Dodo Anondo, Gatot Soegiarto, Laksmi Wulandari, Resti Yudhawati, Yasuko Mori, Soetjipto, Maria I Lusida |
| EPI_ISL_960153 | Heideveld Emergency Centre | National Health Laboratory Service/UCT | Arash Iranzadeh, Deelan Doolabh, Lynn Tyers, Bruna Galvao, Innocent Mudau, Marvin Hsiao, Kruger Marais, Diana Hardie, Stephen Korsman, Carolyn Williamson |
| EPI_ISL_961766, EPI_ISL_961767, EPI_ISL_961768, EPI_ISL_961769, EPI_ISL_961770, EPI_ISL_961771, EPI_ISL_961772, EPI_ISL_961773 | Laboratorio de Infectología, Servicio de Infectología, Hospital Universitario Dr. José Eleuterio González - Universidad Autónoma de Nuevo León | Laboratorio de Infectología Molecular, Departamento de Bioquímica y Medicina Molecular,Facultad de Medicina - Universidad Autónoma de Nuevo León | Kame A. Galán-Huerta, María F. Herrera-Saldivar, Natalia Martínez-Acuña, Sonia A. Lozano-Sepúlveda, Daniel Arellanos-Soto, Ana M. Rivas-Estilla, Paola Bocanegra-Ibarias, Samantha M. Flores-Treviño, Elvira Garza-González, Eduardo Perez-Alba, Laura Nuzzolo-Shihadeh, Adrian Camacho-Ortiz |
| EPI_ISL_964887 | Hospital Cutral Co | Laboratorio Central Mg. Luis Alfredo Pianciola on behalf of 'Proyecto Argentino Interinstitucional de genomica de SARS-CoV-2' (PAIS Consortium) | L Pianciola, M Mazzeo, C Ziehm, C Pintos, M Fernandez, J Ousset, M Nabaes, M Viegas. |
| EPI_ISL_964900 | Hospital Senillosa | Laboratorio Central Mg. Luis Alfredo Pianciola on behalf of 'Proyecto Argentino Interinstitucional de genomica de SARS-CoV-2' (PAIS Consortium) | L Pianciola, M Mazzeo, C Ziehm, C Pintos, M Fernandez, J Ousset, M Nabaes, M Viegas. |
| EPI_ISL_964902 | Cs. San Lorenzo Norte | Laboratorio Central Mg. Luis Alfredo Pianciola on behalf of 'Proyecto Argentino Interinstitucional de genomica de SARS-CoV-2' (PAIS Consortium) | L Pianciola, M Mazzeo, C Ziehm, C Pintos, M Fernandez, J Ousset, M Nabaes, M Viegas. |
| EPI_ISL_964903 | Hospital Centenario | Laboratorio Central Mg. Luis Alfredo Pianciola on behalf of 'Proyecto Argentino Interinstitucional de genomica de SARS-CoV-2' (PAIS Consortium) | L Pianciola, M Mazzeo, C Ziehm, C Pintos, M Fernandez, J Ousset, M Nabaes, M Viegas. |
| EPI_ISL_965030, EPI_ISL_965032, EPI_ISL_965034 | Ohio Department of Health Laboratory | Ohio Department of Health Laboratory | Holmes, Jennifer; Eric Brandt, Keoni Omura, Glen McGillivary, Caitlin McDonnell, Jade Mowery, Stephanie Mccracken, Tyler Payne, Kirtana Ramadugu, Erica Leasure, Brent Lee, Kelsey Florek, Heather Blankenship, Quanta Brown, and Tammy Bannerman |
| EPI_ISL_968149, EPI_ISL_968159, EPI_ISL_968160, EPI_ISL_968161, EPI_ISL_968162, EPI_ISL_968163 | Clinical Molecular Microbiology Laboratory, UNC Hospital | Dirk Dittmer | Justin T. Landis , Razia Moorad , Brent A. Eason, Melissa B. Miller, Linda Pluta, Dirk Dittmer, Angelica Juarez, Cecilia Thompson, Shawn Hawken, Cameroon Grant, Evelyn Hoffman, Patricio Cano, Jason Wong, Carolina Caro-Vegas, Ryan McNamara, Blossom Damania. |
| EPI_ISL_968317, EPI_ISL_968318, EPI_ISL_968319, EPI_ISL_968320, EPI_ISL_968321, EPI_ISL_968322, EPI_ISL_968323, EPI_ISL_968324, EPI_ISL_968325, EPI_ISL_968326, EPI_ISL_968327, EPI_ISL_968328, EPI_ISL_968329, EPI_ISL_968330, EPI_ISL_968331, EPI_ISL_968332, EPI_ISL_968333, EPI_ISL_968334, EPI_ISL_968335, EPI_ISL_968336, EPI_ISL_968337, EPI_ISL_968338, EPI_ISL_968339, EPI_ISL_968340, EPI_ISL_968341, EPI_ISL_968342, EPI_ISL_968343, EPI_ISL_968344, EPI_ISL_968345, EPI_ISL_968346, EPI_ISL_968347, EPI_ISL_968348, EPI_ISL_968349, EPI_ISL_968350, EPI_ISL_968351, EPI_ISL_968352, EPI_ISL_968353, EPI_ISL_968354, EPI_ISL_968355, EPI_ISL_968356, EPI_ISL_968357, EPI_ISL_968358, EPI_ISL_968359, EPI_ISL_968360, EPI_ISL_968361, EPI_ISL_968362 | | | |
| see above | BCCDC Public Health Laboratory | BCCDC Public Health Laboratory | Prystajecky Natalie, Linda Hoang, Dan Fornika, John Tyson, Shannon Russell, Kim Macdonald, Kimia Kamelian, Ana Pacagnella, Corrinne Ng, Loretta Janz, Robert Azana Terry Snutch, Mel Krajden |

EPI_ISL_977251, EPI_ISL_977252, EPI_ISL_977261, EPI_ISL_977262, EPI_ISL_977263, EPI_ISL_977264, EPI_ISL_977266, EPI_ISL_977267, EPI_ISL_977273, EPI_ISL_977275, EPI_ISL_977282, EPI_ISL_977295, EPI_ISL_977296, EPI_ISL_977297, EPI_ISL_977299, EPI_ISL_977303, EPI_ISL_977310, EPI_ISL_977315, EPI_ISL_977316, EPI_ISL_977317, EPI_ISL_977416, EPI_ISL_977418, EPI_ISL_977419, EPI_ISL_977420, EPI_ISL_977423, EPI_ISL_977424, EPI_ISL_977425, EPI_ISL_977426, EPI_ISL_977427, EPI_ISL_977428, EPI_ISL_977429, EPI_ISL_977430, EPI_ISL_977432, EPI_ISL_977433, EPI_ISL_977434, EPI_ISL_977435, EPI_ISL_977436, EPI_ISL_977437, EPI_ISL_977438, EPI_ISL_977439, EPI_ISL_977440, EPI_ISL_977445, EPI_ISL_977446, EPI_ISL_977447, EPI_ISL_977448, EPI_ISL_977449, EPI_ISL_977450, EPI_ISL_977452, EPI_ISL_977453, EPI_ISL_977454, EPI_ISL_977455, EPI_ISL_977456, EPI_ISL_977457

| | | | |
|---|---|---|---|
| see above | University of Zambia, School of Veterinary Medicine | UNZAVET and PATH | Mulenga Mwenda-Chimfwembe, Ngonda Saasa, Daniel Bridges |
| EPI_ISL_981034 | Hospital Dr. Francisco López Lima | Laboratorio Central Mg. Luis Alfredo Pianciola on behalf of 'Proyecto Argentino Interinstitucional de genomica de SARS-CoV-2' (PAIS Consortium) | L Pianciola, M Mazzeo, C Ziehm, C Pintos, M Fernandez, J Ousset, M Nabaes, M Viegas. |
| EPI_ISL_981040 | Hospital Privado Regional | Laboratorio Central Mg. Luis Alfredo Pianciola on behalf of 'Proyecto Argentino Interinstitucional de genomica de SARS-CoV-2' (PAIS Consortium) | L Pianciola, M Mazzeo, C Ziehm, C Pintos, M Fernandez, J Ousset, M Nabaes, M Viegas. |
| EPI_ISL_981043, EPI_ISL_981044 | Hospital Bariloche | Laboratorio Central Mg. Luis Alfredo Pianciola on behalf of 'Proyecto Argentino Interinstitucional de genomica de SARS-CoV-2' (PAIS Consortium) | L Pianciola, M Mazzeo, C Ziehm, C Pintos, M Fernandez, J Ousset, M Nabaes, M Viegas. |
