## Supplementary material for "Detection and characterization of the SARS-CoV-2 lineage B.1.526 in New York": Supp. Table 3 GISAID Acknowledgment part 2: gisaid_hcov-19_acknowledgement_table_2021_02_13_00-3.pdf

We gratefully acknowledge the following Authors from the Originating laboratories responsible for obtaining the specimens, as well as the Submitting laboratories where the genome data were generated and shared via GISAID, on which this research is based.

All Submitters of data may be contacted directly via www.gisaid.org

Authors are sorted alphabetically.

| Accession ID | Originating Laboratory | Submitting Laboratory | Authors |
|---|---|---|---|
| EPI_ISL_611659, EPI_ISL_611662, EPI_ISL_611791, EPI_ISL_611956, EPI_ISL_612591, EPI_ISL_612592, EPI_ISL_612593, EPI_ISL_612594, EPI_ISL_612595, EPI_ISL_612596, EPI_ISL_612597, EPI_ISL_612598, EPI_ISL_612599, EPI_ISL_612600, EPI_ISL_612601, EPI_ISL_612602, EPI_ISL_612603, EPI_ISL_612604, EPI_ISL_612605, EPI_ISL_612606 | | | |
| see above | Queens Medical Centre, Clinical Microbiology Department / DeepSeq Nottingham | COVID-19 Genomics UK (COG-UK) Consortium | Gemma Clark, Wendy Smith, Manjinder Khakh, Vicki M Fleming, Michelle M Lister, Hannah Howson-Wells, Jonathan Ball, Patrick McClure, Joseph Chappell, Theocharis Tsoleridis, Nadine Holmes, Matthew Carlisle, Christopher Moore, Fei Sang, Johnny Debebe, Victoria Wright, Matthew Loose |
| EPI_ISL_612857 | Wales Specialist Virology Centre Sequencing lab: Pathogen Genomics Unit | COVID-19 Genomics UK (COG-UK) Consortium | Catherine Moore, Johnathan Evans, Laura Gifford, Malorie Perry, Simon Cottrell, Angela Marchbank, Alec Birchley, Alexander Adams, Amy Gaskin, Bree Gatica-Wilcox, Jason Coombes, Joel Southgate, Lauren Gilbert, Lee Graham, Nicole Pacchiarini, Sara Kumziene-Summerhayes, Sarah Taylor, Sophie Jones, Sara Rey, Matthew Bull, Joanne Watkins, Sally Corden, Tom Connor |
| EPI_ISL_613566, EPI_ISL_613567, EPI_ISL_613570, EPI_ISL_613636, EPI_ISL_613637, EPI_ISL_613638, EPI_ISL_613639, EPI_ISL_613640, EPI_ISL_613641, EPI_ISL_613642, EPI_ISL_613643, EPI_ISL_613644, EPI_ISL_613645, EPI_ISL_613646, EPI_ISL_613647, EPI_ISL_613648, EPI_ISL_613649, EPI_ISL_613650, EPI_ISL_613651, EPI_ISL_613652, EPI_ISL_613653, EPI_ISL_613654, EPI_ISL_613655, EPI_ISL_613656, EPI_ISL_613657, EPI_ISL_613658, EPI_ISL_613659, EPI_ISL_613660, EPI_ISL_613661, EPI_ISL_613662, EPI_ISL_613663, EPI_ISL_613664, EPI_ISL_613665, EPI_ISL_613666, EPI_ISL_613667, EPI_ISL_613668, EPI_ISL_613669, EPI_ISL_613670, EPI_ISL_613671, EPI_ISL_613672, EPI_ISL_613673, EPI_ISL_613674, EPI_ISL_613675, EPI_ISL_613676, EPI_ISL_613677, EPI_ISL_613678, EPI_ISL_613679, EPI_ISL_613680, EPI_ISL_613681, EPI_ISL_613682, EPI_ISL_613683, EPI_ISL_613684, EPI_ISL_613685, EPI_ISL_613686, EPI_ISL_613687, EPI_ISL_613688, EPI_ISL_613689, EPI_ISL_613690, EPI_ISL_613691, EPI_ISL_613692, EPI_ISL_613693, EPI_ISL_613694, EPI_ISL_613695, EPI_ISL_613696, EPI_ISL_613697, EPI_ISL_613698, EPI_ISL_613699, EPI_ISL_613700, EPI_ISL_613701, EPI_ISL_613702, EPI_ISL_613703, EPI_ISL_613704, EPI_ISL_613705, EPI_ISL_613958, EPI_ISL_613959, EPI_ISL_613960, EPI_ISL_614981, EPI_ISL_614982, EPI_ISL_614983, EPI_ISL_614984, EPI_ISL_614985, EPI_ISL_614986, EPI_ISL_614987, EPI_ISL_614988, EPI_ISL_614989, EPI_ISL_614990, EPI_ISL_614991, EPI_ISL_614992, EPI_ISL_614993, EPI_ISL_614994, EPI_ISL_614995, EPI_ISL_614996, EPI_ISL_614997, EPI_ISL_614998, EPI_ISL_614999, EPI_ISL_615000, EPI_ISL_615001, EPI_ISL_615002, EPI_ISL_615003, EPI_ISL_615004, EPI_ISL_615005, EPI_ISL_615006, EPI_ISL_615007, EPI_ISL_615008, EPI_ISL_615009, EPI_ISL_615010, EPI_ISL_615011, EPI_ISL_615012, EPI_ISL_615013, EPI_ISL_615014, EPI_ISL_615015, EPI_ISL_615016, EPI_ISL_615017, EPI_ISL_615018, EPI_ISL_615019, EPI_ISL_615020, EPI_ISL_615021, EPI_ISL_615022, EPI_ISL_615023, EPI_ISL_615024, EPI_ISL_615025, EPI_ISL_615026, EPI_ISL_615027, EPI_ISL_615028, EPI_ISL_615029, EPI_ISL_615030, EPI_ISL_615031, EPI_ISL_615032, EPI_ISL_615033, EPI_ISL_615034, EPI_ISL_615035, EPI_ISL_615036, EPI_ISL_615037, EPI_ISL_615038, EPI_ISL_615039, EPI_ISL_615040, EPI_ISL_615041, EPI_ISL_615042, EPI_ISL_615043, EPI_ISL_615044, EPI_ISL_615045, EPI_ISL_615046, EPI_ISL_615047, EPI_ISL_615048, EPI_ISL_615049, EPI_ISL_615050, EPI_ISL_615051, EPI_ISL_615052, EPI_ISL_615053, EPI_ISL_615054, EPI_ISL_615055, EPI_ISL_615056, EPI_ISL_615057, EPI_ISL_615058, EPI_ISL_615059, EPI_ISL_615060, EPI_ISL_615061, EPI_ISL_615062 | | | |
| see above | Microbiology, Department of Pathology, St. Bernard's Hospital, Gibraltar Health Authority | Respiratory Virus Unit, Microbiology Services Colindale, Public Health England | PHE Covid Sequencing Team, Dr Nicholas Cortes (Gibraltar), Charlotte Gillborn-Jones (Gibraltar) |
| EPI_ISL_616906, EPI_ISL_616907, EPI_ISL_616908, EPI_ISL_616909, EPI_ISL_616910, EPI_ISL_616911, EPI_ISL_616912, EPI_ISL_616913, EPI_ISL_616914, EPI_ISL_616915, EPI_ISL_616916, EPI_ISL_616917, EPI_ISL_616918, EPI_ISL_616919, EPI_ISL_616920, EPI_ISL_616921, EPI_ISL_616922, EPI_ISL_616925, EPI_ISL_616926, EPI_ISL_616927, EPI_ISL_616928, EPI_ISL_616929, EPI_ISL_616930, EPI_ISL_616931, EPI_ISL_616932, EPI_ISL_616933, EPI_ISL_616934, EPI_ISL_616935, EPI_ISL_616936, EPI_ISL_616937, EPI_ISL_616938, EPI_ISL_616939, EPI_ISL_616940, EPI_ISL_616941, EPI_ISL_616942, EPI_ISL_616943, EPI_ISL_616944, EPI_ISL_616945, EPI_ISL_616946, EPI_ISL_616947, EPI_ISL_616948, EPI_ISL_616949, EPI_ISL_616950, EPI_ISL_616951, EPI_ISL_616952, EPI_ISL_616953, EPI_ISL_616954, EPI_ISL_616955, EPI_ISL_616956, EPI_ISL_616957, EPI_ISL_616958, EPI_ISL_616959, EPI_ISL_616960, EPI_ISL_616961, EPI_ISL_616962, EPI_ISL_616963, EPI_ISL_616964, EPI_ISL_616965, EPI_ISL_616966, EPI_ISL_616967, EPI_ISL_616968, EPI_ISL_616969, EPI_ISL_616970, EPI_ISL_616971, EPI_ISL_616972, EPI_ISL_616973, EPI_ISL_616974, EPI_ISL_616976, EPI_ISL_616977, EPI_ISL_616978, EPI_ISL_616979, EPI_ISL_616980, EPI_ISL_616981, EPI_ISL_616982, EPI_ISL_616983, EPI_ISL_616984, EPI_ISL_616985, EPI_ISL_616986, EPI_ISL_616987, EPI_ISL_616988, EPI_ISL_616989, EPI_ISL_616990, EPI_ISL_616991, EPI_ISL_616992, EPI_ISL_616993, EPI_ISL_616994, EPI_ISL_616995, EPI_ISL_616996, EPI_ISL_616997, EPI_ISL_616998, EPI_ISL_616999, EPI_ISL_617000, EPI_ISL_617001, EPI_ISL_617002, EPI_ISL_617003, EPI_ISL_617004, EPI_ISL_617005, EPI_ISL_617006, EPI_ISL_617007, EPI_ISL_617008, EPI_ISL_617009, EPI_ISL_617010, EPI_ISL_617011, EPI_ISL_617012, EPI_ISL_617013, EPI_ISL_617014, EPI_ISL_617015, EPI_ISL_617016, EPI_ISL_617017, EPI_ISL_617018, EPI_ISL_617019, EPI_ISL_617020, EPI_ISL_617021, EPI_ISL_617022, EPI_ISL_617023, EPI_ISL_617024, EPI_ISL_617025, EPI_ISL_617027, EPI_ISL_617028, EPI_ISL_617029, EPI_ISL_617030, EPI_ISL_617031, EPI_ISL_617032, EPI_ISL_617033, EPI_ISL_617034, EPI_ISL_617035, EPI_ISL_617036, EPI_ISL_617037, EPI_ISL_617038, EPI_ISL_617039, EPI_ISL_617040, EPI_ISL_617041, EPI_ISL_617042, EPI_ISL_617043, EPI_ISL_617044, EPI_ISL_617045, EPI_ISL_617046, EPI_ISL_617047, EPI_ISL_617048, EPI_ISL_617049, EPI_ISL_617050, EPI_ISL_617051, EPI_ISL_617052, EPI_ISL_617053, EPI_ISL_617054, EPI_ISL_617055, EPI_ISL_617056, EPI_ISL_617057, EPI_ISL_617058, EPI_ISL_617059, EPI_ISL_617060, EPI_ISL_617061, EPI_ISL_617062, EPI_ISL_617063, EPI_ISL_617064, EPI_ISL_617065, EPI_ISL_617066, EPI_ISL_617067, EPI_ISL_617068, EPI_ISL_617069, EPI_ISL_617070, EPI_ISL_617071, EPI_ISL_617072, EPI_ISL_617073, EPI_ISL_617074, EPI_ISL_617075, EPI_ISL_617076, EPI_ISL_617077, EPI_ISL_617078, EPI_ISL_617079, EPI_ISL_617080, EPI_ISL_617081, EPI_ISL_617082, EPI_ISL_617083, EPI_ISL_617084, EPI_ISL_617085, EPI_ISL_617086, EPI_ISL_617087, EPI_ISL_617088, EPI_ISL_617089, EPI_ISL_617090, EPI_ISL_617091, EPI_ISL_617092, EPI_ISL_617093, EPI_ISL_617094, EPI_ISL_617095, EPI_ISL_617096, EPI_ISL_617097, EPI_ISL_617098, EPI_ISL_617099, EPI_ISL_617100, EPI_ISL_617101, EPI_ISL_617102, EPI_ISL_617103, EPI_ISL_617104, EPI_ISL_617105, EPI_ISL_617106, EPI_ISL_617108, EPI_ISL_617109, EPI_ISL_617110, EPI_ISL_617111, EPI_ISL_617112, EPI_ISL_617113, EPI_ISL_617114, EPI_ISL_617115, EPI_ISL_617116, EPI_ISL_617117, EPI_ISL_617118, EPI_ISL_617119, EPI_ISL_617120, EPI_ISL_617121, EPI_ISL_617122, EPI_ISL_617123, EPI_ISL_617124, EPI_ISL_617125, EPI_ISL_617126, EPI_ISL_617127, EPI_ISL_617128, EPI_ISL_617129, EPI_ISL_617130, EPI_ISL_617131, EPI_ISL_617132, EPI_ISL_617133, EPI_ISL_617134, EPI_ISL_617135, EPI_ISL_617136, EPI_ISL_617137, EPI_ISL_617138, EPI_ISL_617139, EPI_ISL_617140, EPI_ISL_617141, EPI_ISL_617142, EPI_ISL_617143, EPI_ISL_617144, EPI_ISL_617145, EPI_ISL_617146, EPI_ISL_617147, EPI_ISL_617148, EPI_ISL_617149, EPI_ISL_617150, EPI_ISL_617151, EPI_ISL_617152, EPI_ISL_617153, EPI_ISL_617154, EPI_ISL_617155, EPI_ISL_617156, EPI_ISL_617157, EPI_ISL_617158, EPI_ISL_617159, EPI_ISL_617160, EPI_ISL_617161, EPI_ISL_617162, EPI_ISL_617163, EPI_ISL_617164, EPI_ISL_617165, EPI_ISL_617166, EPI_ISL_617167, EPI_ISL_617168, EPI_ISL_617169, EPI_ISL_617170, EPI_ISL_617171, EPI_ISL_617172, EPI_ISL_617173, EPI_ISL_617174, EPI_ISL_617175, EPI_ISL_617176, EPI_ISL_617177, EPI_ISL_617178, EPI_ISL_617179, EPI_ISL_617180, EPI_ISL_617181, EPI_ISL_617182, EPI_ISL_617183, EPI_ISL_617184, EPI_ISL_617185, EPI_ISL_617186, EPI_ISL_617187, EPI_ISL_617188, EPI_ISL_617189, EPI_ISL_617190, EPI_ISL_617191, EPI_ISL_617192, EPI_ISL_617193, EPI_ISL_617194, EPI_ISL_617195, EPI_ISL_617196, EPI_ISL_617197, EPI_ISL_617198, EPI_ISL_617199, EPI_ISL_617200, EPI_ISL_617201, EPI_ISL_617202, EPI_ISL_617203, EPI_ISL_617204, EPI_ISL_617205, EPI_ISL_617206, EPI_ISL_617207, EPI_ISL_617208, EPI_ISL_617209, EPI_ISL_617210, EPI_ISL_617211, EPI_ISL_617212, EPI_ISL_617213, EPI_ISL_617214, EPI_ISL_617215, EPI_ISL_617216, EPI_ISL_617217, EPI_ISL_617218, EPI_ISL_617219, EPI_ISL_617220, EPI_ISL_617221, EPI_ISL_617222, EPI_ISL_617223, EPI_ISL_617224, EPI_ISL_617225, EPI_ISL_617226, EPI_ISL_617227, EPI_ISL_617228, EPI_ISL_617229, EPI_ISL_617230, EPI_ISL_617231, EPI_ISL_617232, EPI_ISL_617233, EPI_ISL_617234, EPI_ISL_617235, EPI_ISL_617236, EPI_ISL_617237, EPI_ISL_617238, EPI_ISL_617239, EPI_ISL_617240, EPI_ISL_617241, EPI_ISL_617242, EPI_ISL_617243, EPI_ISL_617244, EPI_ISL_617245, EPI_ISL_617246, EPI_ISL_617247, EPI_ISL_617248, EPI_ISL_617249, EPI_ISL_617250, EPI_ISL_617252, EPI_ISL_617253, EPI_ISL_617254, EPI_ISL_617255, EPI_ISL_617256, EPI_ISL_617258, EPI_ISL_617259, EPI_ISL_617260, EPI_ISL_617261, EPI_ISL_617262, EPI_ISL_617263, EPI_ISL_617264, EPI_ISL_617265, EPI_ISL_617266, EPI_ISL_617267, EPI_ISL_617268, EPI_ISL_617269, EPI_ISL_617270, EPI_ISL_617271, EPI_ISL_617272, EPI_ISL_617273, EPI_ISL_617274, EPI_ISL_617275, EPI_ISL_617276, EPI_ISL_617277, EPI_ISL_617278, EPI_ISL_617279, EPI_ISL_617280, EPI_ISL_617281, EPI_ISL_617282, EPI_ISL_617283, EPI_ISL_617285, EPI_ISL_617286, EPI_ISL_617287, EPI_ISL_617288, EPI_ISL_617289, EPI_ISL_617290, EPI_ISL_617291, EPI_ISL_617292, EPI_ISL_617293, EPI_ISL_617294, EPI_ISL_617295, EPI_ISL_617296, EPI_ISL_617297, EPI_ISL_617298, EPI_ISL_617299, EPI_ISL_617300, EPI_ISL_617301, EPI_ISL_617302, EPI_ISL_617303, EPI_ISL_617304, EPI_ISL_617305, EPI_ISL_617306, EPI_ISL_617307, EPI_ISL_617308, EPI_ISL_617309, EPI_ISL_617310, EPI_ISL_617311, EPI_ISL_617312, EPI_ISL_617313, EPI_ISL_617314, EPI_ISL_617315, EPI_ISL_617316, EPI_ISL_617317, EPI_ISL_617318, EPI_ISL_617319, EPI_ISL_617320, EPI_ISL_617321, EPI_ISL_617322, EPI_ISL_617323, EPI_ISL_617324, EPI_ISL_617325, EPI_ISL_617326, EPI_ISL_617327, EPI_ISL_617328, EPI_ISL_617329, EPI_ISL_617330, EPI_ISL_617331, EPI_ISL_617332, EPI_ISL_617333, EPI_ISL_617334, EPI_ISL_617335, EPI_ISL_617336, EPI_ISL_617337, EPI_ISL_617338, EPI_ISL_617339, EPI_ISL_617340, EPI_ISL_617341, EPI_ISL_617342, EPI_ISL_617343, EPI_ISL_617344, EPI_ISL_617345, EPI_ISL_617346, EPI_ISL_617347, EPI_ISL_617348, EPI_ISL_617349, EPI_ISL_617350, EPI_ISL_617351, EPI_ISL_617352, EPI_ISL_617353, EPI_ISL_617354, EPI_ISL_617355, EPI_ISL_617356, EPI_ISL_617357, EPI_ISL_617358, EPI_ISL_617359, EPI_ISL_617360, EPI_ISL_617361, EPI_ISL_617362, EPI_ISL_617363, EPI_ISL_617364, EPI_ISL_617365, EPI_ISL_617366, EPI_ISL_617367, EPI_ISL_617368, EPI_ISL_617369, EPI_ISL_617370, EPI_ISL_617371, EPI_ISL_617372, EPI_ISL_617373, EPI_ISL_617374, EPI_ISL_617375, EPI_ISL_617376, EPI_ISL_617377, EPI_ISL_617378, EPI_ISL_617379, EPI_ISL_617380, EPI_ISL_617381, EPI_ISL_617385, EPI_ISL_617386, EPI_ISL_617387, EPI_ISL_617388, EPI_ISL_617389, EPI_ISL_617390, EPI_ISL_617391, EPI_ISL_617392, EPI_ISL_617393, EPI_ISL_617394, EPI_ISL_617395, EPI_ISL_617396, EPI_ISL_617397, EPI_ISL_617398, EPI_ISL_617399, EPI_ISL_617400, EPI_ISL_617401, EPI_ISL_617402, EPI_ISL_617403, EPI_ISL_617404, EPI_ISL_617405, EPI_ISL_617906, EPI_ISL_617907, EPI_ISL_617908, EPI_ISL_617909, EPI_ISL_617910, EPI_ISL_617911, EPI_ISL_617912, EPI_ISL_617913, EPI_ISL_617914, EPI_ISL_617915, EPI_ISL_617916, EPI_ISL_617917, EPI_ISL_617918, EPI_ISL_617919, EPI_ISL_617920, EPI_ISL_617921, EPI_ISL_617922, EPI_ISL_617923, EPI_ISL_617924, EPI_ISL_617925, EPI_ISL_617926, EPI_ISL_617927, EPI_ISL_617928, EPI_ISL_617929, EPI_ISL_617930, EPI_ISL_617931, EPI_ISL_617932, EPI_ISL_617933, EPI_ISL_617934, EPI_ISL_617935, EPI_ISL_617936, EPI_ISL_617937, EPI_ISL_617938, EPI_ISL_618809, EPI_ISL_618810, EPI_ISL_618811, EPI_ISL_618812, EPI_ISL_618814, EPI_ISL_618815, EPI_ISL_618816, EPI_ISL_618817, EPI_ISL_618818, EPI_ISL_618819, EPI_ISL_618820, EPI_ISL_618821, EPI_ISL_618823, EPI_ISL_618824, EPI_ISL_618825, EPI_ISL_618826, EPI_ISL_618827, EPI_ISL_618828, EPI_ISL_618829, EPI_ISL_618831, EPI_ISL_618832, EPI_ISL_618834, EPI_ISL_618835, EPI_ISL_618836, EPI_ISL_618838, EPI_ISL_618839, EPI_ISL_618840, EPI_ISL_618841, EPI_ISL_618842, EPI_ISL_618843, EPI_ISL_618845, EPI_ISL_618848, EPI_ISL_618849, EPI_ISL_618850, EPI_ISL_618851, EPI_ISL_618853, EPI_ISL_618854, EPI_ISL_618856, EPI_ISL_618858, EPI_ISL_618859, EPI_ISL_618860, EPI_ISL_618861, EPI_ISL_618862, EPI_ISL_618863, EPI_ISL_618864, EPI_ISL_618865, EPI_ISL_618866, EPI_ISL_618867, EPI_ISL_618868, EPI_ISL_618869, EPI_ISL_618870, EPI_ISL_618871, EPI_ISL_618873, EPI_ISL_618874, EPI_ISL_618875, EPI_ISL_618876, EPI_ISL_618877, EPI_ISL_618878, EPI_ISL_618879, EPI_ISL_618880, EPI_ISL_618881, EPI_ISL_618882, EPI_ISL_618883, EPI_ISL_618884, EPI_ISL_618885, EPI_ISL_618886, EPI_ISL_618887, EPI_ISL_618888, EPI_ISL_618889, EPI_ISL_618890, EPI_ISL_618891, EPI_ISL_618892, EPI_ISL_618893, EPI_ISL_618894, EPI_ISL_618895, EPI_ISL_618896, EPI_ISL_618897, EPI_ISL_618898, EPI_ISL_618899, EPI_ISL_618900, EPI_ISL_618901, EPI_ISL_618902, EPI_ISL_618903, EPI_ISL_618904, EPI_ISL_618905 | | | |
| see above | Department of Virus and Microbiological Special Diagnostics, Statens Serum Institut, Denmark | Albertsen lab, Department of Chemistry and Bioscience, Aalborg University, Denmark | Danish Covid-19 Genome Consortia |
| EPI_ISL_622762, EPI_ISL_622763, EPI_ISL_622764, EPI_ISL_622765, EPI_ISL_622766 | SA Pathology | SA Pathology | Lex Leong, Julien Soubrier, Chuan Kok Lim, Song Gao, Mark Turra, Karin Kassahn, Ivan Bastian, Geoff Higgins |
| EPI_ISL_622773 | LabPLUS | Institute of Environmental Science and Research (ESR) | Xiaoyun Ren, Matt Storey, Nikki Freed, Muhammad Faisal, Jing Wang, Hermes Perez, Anja Werno, Antje van der Linden, Arlo Upton, Chris Mansell, David Hammer, Dragana Drinkovic, Gary McAuliffe, Hana Sofia Andersson, James Ussher, Jill Sherwood, Josh Freeman, Julia Howard, Juliet Elvy, Mary DeAlmeida, Matt Blakiston, Matthew Rogers, Max Bloomfield, Michael Addidle, Michelle Balm, Sally Roberts, Sarah Jefferies, Sharmini Muttaiyah, Susan Morpeth, Susan Taylor, Timothy Blackmore, Vani Sathyendran, Veronica Playle, Virginia Hope, Erasmus Smit, Lauren Jelly, Olin Silander, Joep de Ligt |
| EPI_ISL_622783, EPI_ISL_622787, EPI_ISL_622817 | Middlemore Hospital | Institute of Environmental Science and Research (ESR) | Xiaoyun Ren, Matt Storey, Nikki Freed, Muhammad Faisal, Jing Wang, Hermes Perez, Anja Werno, Antje van der Linden, Arlo Upton, Chris Mansell, David Hammer, Dragana Drinkovic, Gary McAuliffe, Hana Sofia Andersson, James Ussher, Jill Sherwood, Josh Freeman, Julia Howard, Juliet Elvy, Mary DeAlmeida, Matt Blakiston, Matthew Rogers, Max Bloomfield, Michael Addidle, Michelle Balm, Sally Roberts, Sarah Jefferies, Sharmini Muttaiyah, Susan Morpeth, Susan Taylor, Timothy Blackmore, Vani Sathyendran, Veronica Playle, Virginia Hope, Erasmus Smit, Lauren Jelly, Olin Silander, Joep de Ligt |
| EPI_ISL_622821, EPI_ISL_622822, EPI_ISL_622823 | LabPLUS | Institute of Environmental Science and Research (ESR) | Xiaoyun Ren, Matt Storey, Nikki Freed, Muhammad Faisal, Jing Wang, Hermes Perez, Anja Werno, Antje van der Linden, Arlo Upton, Chris Mansell, David Hammer, Dragana Drinkovic, Gary McAuliffe, Hana Sofia Andersson, James Ussher, Jill Sherwood, Josh Freeman, Julia Howard, Juliet Elvy, Mary DeAlmeida, Matt Blakiston, Matthew Rogers, Max Bloomfield, Michael Addidle, Michelle Balm, Sally Roberts, Sarah Jefferies, Sharmini Muttaiyah, Susan Morpeth, Susan Taylor, Timothy Blackmore, Vani Sathyendran, Veronica Playle, Virginia Hope, Erasmus Smit, Lauren Jelly, Olin Silander, Joep de Ligt |
| EPI_ISL_622830 | Canterbury Health Laboratories | Institute of Environmental Science and Research (ESR) | Xiaoyun Ren, Matt Storey, Nikki Freed, Muhammad Faisal, Jing Wang, Hermes Perez, Anja Werno, Antje van der Linden, Arlo Upton, Chris Mansell, David |

| | | | |
|---|---|---|---|
| | | | Hammer, Dragana Drinkovic, Gary McAuliffe, Hana Sofia Andersson, James Ussher, Jill Sherwood, Josh Freeman, Julia Howard, Juliet Elvy, Mary DeAlmeida, Matt Blakiston, Matthew Rogers, Max Bloomfield, Michael Addidle, Michelle Balm, Sally Roberts, Sarah Jefferies, Sharmini Muttaiyah, Susan Morpeth, Susan Taylor, Timothy Blackmore, Vani Sathyendran, Veronica Playle, Virginia Hope, Erasmus Smit, Lauren Jelly, Olin Silander, Joep de Ligt |
| EPI_ISL_622832 | Middlemore Hospital | Institute of Environmental Science and Research (ESR) | Xiaoyun Ren, Matt Storey, Nikki Freed, Muhammad Faisal, Jing Wang, Hermes Perez, Anja Werno, Antje van der Linden, Arlo Upton, Chris Mansell, David Hammer, Dragana Drinkovic, Gary McAuliffe, Hana Sofia Andersson, James Ussher, Jill Sherwood, Josh Freeman, Julia Howard, Juliet Elvy, Mary DeAlmeida, Matt Blakiston, Matthew Rogers, Max Bloomfield, Michael Addidle, Michelle Balm, Sally Roberts, Sarah Jefferies, Sharmini Muttaiyah, Susan Morpeth, Susan Taylor, Timothy Blackmore, Vani Sathyendran, Veronica Playle, Virginia Hope, Erasmus Smit, Lauren Jelly, Olin Silander, Joep de Ligt |
| EPI_ISL_622878 | Respiratory Virus Unit, Microbiology Services Colindale, Public Health England | Respiratory Virus Unit, Microbiology Services Colindale, Public Health England | PHE Covid Sequencing Team |
| EPI_ISL_625684, EPI_ISL_625685, EPI_ISL_625686, EPI_ISL_625687, EPI_ISL_625688, EPI_ISL_625689, EPI_ISL_625690, EPI_ISL_625691, EPI_ISL_625692, EPI_ISL_625693, EPI_ISL_625694, EPI_ISL_625696, EPI_ISL_625697, EPI_ISL_625699, EPI_ISL_625700, EPI_ISL_625701, EPI_ISL_625702, EPI_ISL_625703, EPI_ISL_625704, EPI_ISL_625705, EPI_ISL_625707, EPI_ISL_625708, EPI_ISL_625710, EPI_ISL_625711, EPI_ISL_625712, EPI_ISL_625713, EPI_ISL_625714, EPI_ISL_625715, EPI_ISL_625716, EPI_ISL_625717, EPI_ISL_625718, EPI_ISL_625719, EPI_ISL_625720, EPI_ISL_625721, EPI_ISL_625722, EPI_ISL_625723, EPI_ISL_625724, EPI_ISL_625725, EPI_ISL_625726, EPI_ISL_625727, EPI_ISL_625728, EPI_ISL_625729, EPI_ISL_625730, EPI_ISL_625731, EPI_ISL_625732, EPI_ISL_625733, EPI_ISL_625734, EPI_ISL_625735, EPI_ISL_625736, EPI_ISL_625737, EPI_ISL_625738, EPI_ISL_625739, EPI_ISL_625740, EPI_ISL_625741, EPI_ISL_625743, EPI_ISL_625744, EPI_ISL_625745, EPI_ISL_625746, EPI_ISL_625747, EPI_ISL_625748, EPI_ISL_625749, EPI_ISL_625750, EPI_ISL_625751, EPI_ISL_625752, EPI_ISL_625753, EPI_ISL_625754, EPI_ISL_625755, EPI_ISL_625756, EPI_ISL_625757, EPI_ISL_625758, EPI_ISL_625759, EPI_ISL_625760, EPI_ISL_625761, EPI_ISL_625762, EPI_ISL_625763, EPI_ISL_625764, EPI_ISL_625765, EPI_ISL_625766, EPI_ISL_625767, EPI_ISL_625768, EPI_ISL_625769, EPI_ISL_625770, EPI_ISL_625771, EPI_ISL_625772, EPI_ISL_625773, EPI_ISL_625774, EPI_ISL_625775, EPI_ISL_625776, EPI_ISL_625777, EPI_ISL_625778, EPI_ISL_625779, EPI_ISL_625780, EPI_ISL_625781, EPI_ISL_625782, EPI_ISL_625783, EPI_ISL_625784, EPI_ISL_625785, EPI_ISL_625786, EPI_ISL_625787, EPI_ISL_625788, EPI_ISL_625789, EPI_ISL_625790, EPI_ISL_625791, EPI_ISL_625792, EPI_ISL_625793, EPI_ISL_625794, EPI_ISL_625795, EPI_ISL_625796, EPI_ISL_625797, EPI_ISL_625798, EPI_ISL_625799, EPI_ISL_625800, EPI_ISL_625801, EPI_ISL_625802, EPI_ISL_625803, EPI_ISL_625804, EPI_ISL_625805, EPI_ISL_625806, EPI_ISL_625807, EPI_ISL_625808, EPI_ISL_625809, EPI_ISL_625810, EPI_ISL_625811, EPI_ISL_625812, EPI_ISL_625813, EPI_ISL_625814, EPI_ISL_625815, EPI_ISL_625816, EPI_ISL_625817, EPI_ISL_625818, EPI_ISL_625819, EPI_ISL_625820, EPI_ISL_625821, EPI_ISL_625824, EPI_ISL_625825, EPI_ISL_625826, EPI_ISL_625827, EPI_ISL_625828, EPI_ISL_625829, EPI_ISL_625830, EPI_ISL_625831, EPI_ISL_625832, EPI_ISL_625833, EPI_ISL_625834, EPI_ISL_625835, EPI_ISL_625836, EPI_ISL_625837, EPI_ISL_625838, EPI_ISL_625839, EPI_ISL_625840, EPI_ISL_625841, EPI_ISL_625842, EPI_ISL_625843, EPI_ISL_625844, EPI_ISL_625845, EPI_ISL_625846, EPI_ISL_625847, EPI_ISL_625848, EPI_ISL_625849, EPI_ISL_625850, EPI_ISL_625851, EPI_ISL_625852, EPI_ISL_625853, EPI_ISL_625854, EPI_ISL_625855, EPI_ISL_625856, EPI_ISL_625857, EPI_ISL_625858, EPI_ISL_625859, EPI_ISL_625860, EPI_ISL_625861, EPI_ISL_625862, EPI_ISL_625863, EPI_ISL_625865, EPI_ISL_625866, EPI_ISL_625867, EPI_ISL_625868, EPI_ISL_625869, EPI_ISL_625870, EPI_ISL_625871, EPI_ISL_625872, EPI_ISL_625873, EPI_ISL_625874, EPI_ISL_625875, EPI_ISL_625876, EPI_ISL_625877, EPI_ISL_625878, EPI_ISL_625879, EPI_ISL_625880, EPI_ISL_625881, EPI_ISL_625882, EPI_ISL_625884, EPI_ISL_625885, EPI_ISL_625886, EPI_ISL_625887, EPI_ISL_625888, EPI_ISL_625889, EPI_ISL_625891, EPI_ISL_625892, EPI_ISL_625893, EPI_ISL_625894, EPI_ISL_625895, EPI_ISL_625896, EPI_ISL_625897, EPI_ISL_625898, EPI_ISL_625899, EPI_ISL_625900, EPI_ISL_625901, EPI_ISL_625902, EPI_ISL_625903, EPI_ISL_625904, EPI_ISL_625905, EPI_ISL_625906, EPI_ISL_625907, EPI_ISL_625908, EPI_ISL_625909, EPI_ISL_625910, EPI_ISL_625911, EPI_ISL_625912, EPI_ISL_625913, EPI_ISL_625914, EPI_ISL_625915, EPI_ISL_625916, EPI_ISL_625917, EPI_ISL_625918, EPI_ISL_625919, EPI_ISL_625920, EPI_ISL_625921, EPI_ISL_625922, EPI_ISL_625923, EPI_ISL_625924, EPI_ISL_625925, EPI_ISL_625926, EPI_ISL_625927, EPI_ISL_625928, EPI_ISL_625929, EPI_ISL_625930, EPI_ISL_625931, EPI_ISL_625932, EPI_ISL_625934, EPI_ISL_625935, EPI_ISL_625936, EPI_ISL_625937, EPI_ISL_625938, EPI_ISL_625939, EPI_ISL_625940, EPI_ISL_625941, EPI_ISL_625942, EPI_ISL_625943, EPI_ISL_625944, EPI_ISL_625945, EPI_ISL_625946, EPI_ISL_625947, EPI_ISL_625948, EPI_ISL_625949, EPI_ISL_625950, EPI_ISL_625951, EPI_ISL_625952, EPI_ISL_625953, EPI_ISL_625954, EPI_ISL_625955, EPI_ISL_625956, EPI_ISL_625957, EPI_ISL_625958, EPI_ISL_625959, EPI_ISL_625961, EPI_ISL_625962, EPI_ISL_625963, EPI_ISL_625964, EPI_ISL_625965, EPI_ISL_625966, EPI_ISL_625967, EPI_ISL_625968, EPI_ISL_625969, EPI_ISL_625970, EPI_ISL_625971, EPI_ISL_625974, EPI_ISL_625975, EPI_ISL_625976, EPI_ISL_625977, EPI_ISL_625978, EPI_ISL_625979, EPI_ISL_625980, EPI_ISL_625981, EPI_ISL_625982, EPI_ISL_625983, EPI_ISL_625984, EPI_ISL_625985, EPI_ISL_625986, EPI_ISL_625987, EPI_ISL_625988, EPI_ISL_625989, EPI_ISL_625990, EPI_ISL_625991, EPI_ISL_625992, EPI_ISL_625993, EPI_ISL_625994, EPI_ISL_625995, EPI_ISL_625996, EPI_ISL_625997, EPI_ISL_625999, EPI_ISL_626000, EPI_ISL_626001, EPI_ISL_626002, EPI_ISL_626003, EPI_ISL_626004, EPI_ISL_626005, EPI_ISL_626006, EPI_ISL_626007, EPI_ISL_626008, EPI_ISL_626009, EPI_ISL_626011, EPI_ISL_626012, EPI_ISL_626013, EPI_ISL_626014, EPI_ISL_626015, EPI_ISL_626016, EPI_ISL_626017, EPI_ISL_626018, EPI_ISL_626019, EPI_ISL_626020, EPI_ISL_626021, EPI_ISL_626022, EPI_ISL_626023, EPI_ISL_626024, EPI_ISL_626025, EPI_ISL_626026, EPI_ISL_626027, EPI_ISL_626028, EPI_ISL_626029, EPI_ISL_626030, EPI_ISL_626031, EPI_ISL_626032, EPI_ISL_626033, EPI_ISL_626034, EPI_ISL_626035, EPI_ISL_626036, EPI_ISL_626037, EPI_ISL_626038, EPI_ISL_626039, EPI_ISL_626040, EPI_ISL_626041, EPI_ISL_626042, EPI_ISL_626043, EPI_ISL_626044, EPI_ISL_626045, EPI_ISL_626046, EPI_ISL_626047, EPI_ISL_626048, EPI_ISL_626049, EPI_ISL_626050, EPI_ISL_626051, EPI_ISL_626052, EPI_ISL_626053, EPI_ISL_626054, EPI_ISL_626055, EPI_ISL_626056, EPI_ISL_626057, EPI_ISL_626058, EPI_ISL_626059, EPI_ISL_626060, EPI_ISL_626061, EPI_ISL_626062, EPI_ISL_626063, EPI_ISL_626064, EPI_ISL_626065, EPI_ISL_626066, EPI_ISL_626067, EPI_ISL_626068, EPI_ISL_626069, EPI_ISL_626070, EPI_ISL_626071, EPI_ISL_626072, EPI_ISL_626073, EPI_ISL_626074, EPI_ISL_626075, EPI_ISL_626076, EPI_ISL_626077, EPI_ISL_626078, EPI_ISL_626079, EPI_ISL_626080, EPI_ISL_626081, EPI_ISL_626082, EPI_ISL_626083, EPI_ISL_626084, EPI_ISL_626085, EPI_ISL_626086, EPI_ISL_626087, EPI_ISL_626088, EPI_ISL_626089, EPI_ISL_626090, EPI_ISL_626091, EPI_ISL_626092, EPI_ISL_626093, EPI_ISL_626094, EPI_ISL_626095, EPI_ISL_626096, EPI_ISL_626097, EPI_ISL_626098, EPI_ISL_626099, EPI_ISL_626100, EPI_ISL_626101, EPI_ISL_626102, EPI_ISL_626103, EPI_ISL_626104, EPI_ISL_626105, EPI_ISL_626106, EPI_ISL_626107, EPI_ISL_626108, EPI_ISL_626109, EPI_ISL_626110, EPI_ISL_626111, EPI_ISL_626112, EPI_ISL_626113, EPI_ISL_626114, EPI_ISL_626115, EPI_ISL_626116, EPI_ISL_626117, EPI_ISL_626118, EPI_ISL_626119, EPI_ISL_626120, EPI_ISL_626121, EPI_ISL_626122, EPI_ISL_626123, EPI_ISL_626125, EPI_ISL_626126, EPI_ISL_626128, EPI_ISL_626130, EPI_ISL_626131, EPI_ISL_626135, EPI_ISL_626136, EPI_ISL_626140, EPI_ISL_626141, EPI_ISL_626142, EPI_ISL_626143, EPI_ISL_626145, EPI_ISL_626147, EPI_ISL_626150, EPI_ISL_626151, EPI_ISL_626152, EPI_ISL_626153, EPI_ISL_626154, EPI_ISL_626155, EPI_ISL_626156, EPI_ISL_626158, EPI_ISL_626159, EPI_ISL_626160, EPI_ISL_626161, EPI_ISL_626163, EPI_ISL_626165, EPI_ISL_626166, EPI_ISL_626167, EPI_ISL_626168, EPI_ISL_626169, EPI_ISL_626170, EPI_ISL_626171, EPI_ISL_626172, EPI_ISL_626173, EPI_ISL_626174, EPI_ISL_626175, EPI_ISL_626176, EPI_ISL_626177, EPI_ISL_626178, EPI_ISL_626179, EPI_ISL_626180, EPI_ISL_626181, EPI_ISL_626182, EPI_ISL_626183, EPI_ISL_626184, EPI_ISL_626185, EPI_ISL_626186, EPI_ISL_626187, EPI_ISL_626188, EPI_ISL_626189, EPI_ISL_626190, EPI_ISL_626191, EPI_ISL_626192, EPI_ISL_626193, EPI_ISL_626194, EPI_ISL_626195, EPI_ISL_626196, EPI_ISL_626197, EPI_ISL_626198, EPI_ISL_626199, EPI_ISL_626200, EPI_ISL_626201, EPI_ISL_626202, EPI_ISL_626203, EPI_ISL_626204, EPI_ISL_626205, EPI_ISL_626206, EPI_ISL_626207, EPI_ISL_626208, EPI_ISL_626209, EPI_ISL_626210 | | | |
| see above | Department of Virus and Microbiological Special Diagnostics, Statens Serum Institut, Denmark | Albertsen lab, Department of Chemistry and Bioscience, Aalborg University, Denmark | Danish Covid-19 Genome Consortia |
| EPI_ISL_626355 | LabPLUS | Institute of Environmental Science and Research (ESR) | Xiaoyun Ren, Matt Storey, Nikki Freed, Muhammad Faisal, Jing Wang, Hermes Perez, Anja Werno, Antje van der Linden, Arlo Upton, Chris Mansell, David Hammer, Dragana Drinkovic, Gary McAuliffe, Hana Sofia Andersson, James Ussher, Jill Sherwood, Josh Freeman, Julia Howard, Juliet Elvy, Mary DeAlmeida, Matt Blakiston, Matthew Rogers, Max Bloomfield, Michael Addidle, Michelle Balm, Sally Roberts, Sarah Jefferies, Sharmini Muttaiyah, Susan Morpeth, Susan Taylor, Timothy Blackmore, Vani Sathyendran, Veronica Playle, Virginia Hope, Erasmus Smit, Lauren Jelly, Olin Silander, Joep de Ligt |
| EPI_ISL_626628, EPI_ISL_626629, EPI_ISL_626633, EPI_ISL_626636, EPI_ISL_626637, EPI_ISL_626639, EPI_ISL_626640 | National Public Health Laboratory, National Centre for Infectious Diseases | National Public Health Laboratory, National Centre for Infectious Diseases | Tze Minn Mak, Sophie Octavia, Zhenyang Zhou, Lin Cui, Raymond Tzer Pin Lin |
| EPI_ISL_626659, EPI_ISL_626688 | Queens Medical Centre, Clinical Microbiology Department / DeepSeq Nottingham | COVID-19 Genomics UK (COG-UK) Consortium | Gemma Clark, Wendy Smith, Manjinder Khakh, Vicki M Fleming, Michelle M Lister, Hannah Howson-Wells, Jonathan Ball, Patrick McClure, Joseph Chappell, Theocharis Tsoleridis, Nadine Holmes, Matthew Carlisle, Christopher Moore, Fei Sang, Johnny Debebe, Victoria Wright, Matthew Loose |
| EPI_ISL_626689 | Centre for Enzyme Innovation, University of Portsmouth / Translational Research Laboratory, Portsmouth Hospitals NHS Trust | COVID-19 Genomics UK (COG-UK) Consortium | Angela Beckett,Yann Bourgeois,Garry Scarlett,Sharon Glaysher,Scott Elliott,Kelly Bicknell,Robert Impey,Allyson Lloyd,Sarah Wyllie,Ethan Butcher,Anoop Chauhan,Samuel Robson |
| EPI_ISL_626695 | Queens Medical Centre, Clinical Microbiology Department / DeepSeq Nottingham | COVID-19 Genomics UK (COG-UK) Consortium | Gemma Clark, Wendy Smith, Manjinder Khakh, Vicki M Fleming, Michelle M Lister, Hannah Howson-Wells, Jonathan Ball, Patrick McClure, Joseph Chappell, Theocharis Tsoleridis, Nadine Holmes, Matthew Carlisle, Christopher Moore, Fei Sang, Johnny Debebe, Victoria Wright, Matthew Loose |
| EPI_ISL_626761, EPI_ISL_626763 | Wales Specialist Virology Centre Sequencing lab: Pathogen Genomics Unit | COVID-19 Genomics UK (COG-UK) Consortium | Catherine Moore, Johnathan Evans, Laura Gifford, Malorie Perry, Simon Cottrell, Angela Marchbank, Alec Birchley, Alexander Adams, Amy Gaskin, Bree Gatica-Wilcox, Jason Coombes, Joel Southgate, Lauren Gilbert, Lee Graham, Nicole Pacchiarini, Sara Kumziene-Summerhayes, Sarah Taylor, Sophie Jones, Sara Rey, Matthew Bull, Joanne Watkins, Sally Corden, Tom Connor |
| EPI_ISL_626782 | Quadram Institute Bioscience | COVID-19 Genomics UK (COG-UK) Consortium | Dave J. Baker, Gemma L. Kay, Alp Aydin, Thanh Le-Viet, Steven Rudder, Ana P. Tedim, Anastasia Kolyva, Maria Diaz, Leonardo de Oliveira Martins, Nabil-Fareed Alikhan, Lizzie Meadows, Rachael Stanley, Ngozi Elumogo, Muhammed Yasir, Nicholas M. Thomson, Alexander J Trotter, Rachel Gilroy, Samuel Bloomfield, Claire Stuart, Andrew Bell, Reenesh Prakash, Samir Dervisevic, Alison E. Mather, John Wain, Mark Webber, Andrew J. Page, Justin O'Grady |
| EPI_ISL_626816, EPI_ISL_626834 | Queens Medical Centre, Clinical Microbiology Department / DeepSeq Nottingham | COVID-19 Genomics UK (COG-UK) Consortium | Gemma Clark, Wendy Smith, Manjinder Khakh, Vicki M Fleming, Michelle M Lister, Hannah Howson-Wells, Jonathan Ball, Patrick McClure, Joseph Chappell, Theocharis Tsoleridis, Nadine Holmes, Matthew Carlisle, Christopher Moore, Fei Sang, Johnny Debebe, Victoria Wright, Matthew Loose |
| EPI_ISL_626879 | Centre for Enzyme Innovation, University of Portsmouth / Translational Research Laboratory, Portsmouth Hospitals NHS Trust | COVID-19 Genomics UK (COG-UK) Consortium | Angela Beckett,Yann Bourgeois,Garry Scarlett,Sharon Glaysher,Scott Elliott,Kelly Bicknell,Robert Impey,Allyson Lloyd,Sarah Wyllie,Ethan Butcher,Anoop Chauhan,Samuel Robson |
| EPI_ISL_626919 | Queens Medical Centre, Clinical Microbiology Department / DeepSeq Nottingham | COVID-19 Genomics UK (COG-UK) Consortium | Gemma Clark, Wendy Smith, Manjinder Khakh, Vicki M Fleming, Michelle M Lister, Hannah Howson-Wells, Jonathan Ball, Patrick McClure, Joseph Chappell, Theocharis Tsoleridis, Nadine Holmes, Matthew Carlisle, Christopher Moore, Fei Sang, Johnny Debebe, Victoria Wright, Matthew Loose |
| EPI_ISL_626928, EPI_ISL_627001, EPI_ISL_627007, EPI_ISL_627013 | Wales Specialist Virology Centre Sequencing lab: Pathogen Genomics Unit | COVID-19 Genomics UK (COG-UK) Consortium | Catherine Moore, Johnathan Evans, Laura Gifford, Malorie Perry, Simon Cottrell, Angela Marchbank, Alec Birchley, Alexander Adams, Amy Gaskin, Bree Gatica-Wilcox, Jason Coombes, Joel Southgate, Lauren Gilbert, Lee Graham, Nicole Pacchiarini, Sara Kumziene-Summerhayes, Sarah Taylor, Sophie Jones, Sara Rey, Matthew Bull, Joanne Watkins, Sally Corden, Tom Connor |
| EPI_ISL_627053 | Queens Medical Centre, Clinical Microbiology Department / DeepSeq Nottingham | COVID-19 Genomics UK (COG-UK) Consortium | Gemma Clark, Wendy Smith, Manjinder Khakh, Vicki M Fleming, Michelle M Lister, Hannah Howson-Wells, Jonathan Ball, Patrick McClure, Joseph Chappell, Theocharis Tsoleridis, Nadine Holmes, Matthew Carlisle, Christopher Moore, Fei Sang, Johnny Debebe, Victoria Wright, Matthew Loose |
| EPI_ISL_627056 | Centre for Enzyme Innovation, University of Portsmouth / Translational Research Laboratory, Portsmouth Hospitals NHS Trust | COVID-19 Genomics UK (COG-UK) Consortium | Angela Beckett,Yann Bourgeois,Garry Scarlett,Sharon Glaysher,Scott Elliott,Kelly Bicknell,Robert Impey,Allyson Lloyd,Sarah Wyllie,Ethan Butcher,Anoop Chauhan,Samuel Robson |
| EPI_ISL_627071, EPI_ISL_627073 | Quadram Institute Bioscience | COVID-19 Genomics UK (COG-UK) Consortium | Dave J. Baker, Gemma L. Kay, Alp Aydin, Thanh Le-Viet, Steven Rudder, Ana P. Tedim, Anastasia Kolyva, Maria Diaz, Leonardo de Oliveira Martins, Nabil-Fareed Alikhan, Lizzie Meadows, Rachael Stanley, Ngozi Elumogo, Muhammed Yasir, Nicholas M. Thomson, Alexander J Trotter, Rachel Gilroy, |

| | | | |
|---|---|---|---|
| | | | Samuel Bloomfield, Claire Stuart, Andrew Bell, Reenesh Prakash, Samir Dervisevic, Alison E. Mather, John Wain, Mark Webber, Andrew J. Page, Justin O'Grady |
| EPI_ISL_627087, EPI_ISL_627140 | Wales Specialist Virology Centre Sequencing lab: Pathogen Genomics Unit | COVID-19 Genomics UK (COG-UK) Consortium | Catherine Moore, Johnathan Evans, Laura Gifford, Malorie Perry, Simon Cottrell, Angela Marchbank, Alec Birchley, Alexander Adams, Amy Gaskin, Bree Gatica-Wilcox, Jason Coombes, Joel Southgate, Lauren Gilbert, Lee Graham, Nicole Pacchiarini, Sara Kumziene-Summerhayes, Sarah Taylor, Sophie Jones, Sara Rey, Matthew Bull, Joanne Watkins, Sally Corden, Tom Connor |
| EPI_ISL_627147 | Queens Medical Centre, Clinical Microbiology Department / DeepSeq Nottingham | COVID-19 Genomics UK (COG-UK) Consortium | Gemma Clark, Wendy Smith, Manjinder Khakh, Vicki M Fleming, Michelle M Lister, Hannah Howson-Wells, Jonathan Ball, Patrick McClure, Joseph Chappell, Theocharis Tsoleridis, Nadine Holmes, Matthew Carlisle, Christopher Moore, Fei Sang, Johnny Debebe, Victoria Wright, Matthew Loose |
| EPI_ISL_627151 | Quadram Institute Bioscience | COVID-19 Genomics UK (COG-UK) Consortium | Dave J. Baker, Gemma L. Kay, Alp Aydin, Thanh Le-Viet, Steven Rudder, Ana P. Tedim, Anastasia Kolyva, Maria Diaz, Leonardo de Oliveira Martins, Nabil-Fareed Alikhan, Lizzie Meadows, Rachael Stanley, Ngozi Elumogo, Muhammed Yasir, Nicholas M. Thomson, Alexander J Trotter, Rachel Gilroy, Samuel Bloomfield, Claire Stuart, Andrew Bell, Reenesh Prakash, Samir Dervisevic, Alison E. Mather, John Wain, Mark Webber, Andrew J. Page, Justin O'Grady |
| EPI_ISL_627167, EPI_ISL_627205, EPI_ISL_627207, EPI_ISL_627237, EPI_ISL_627242, EPI_ISL_627244, EPI_ISL_627248, EPI_ISL_627249 | Wales Specialist Virology Centre Sequencing lab: Pathogen Genomics Unit | COVID-19 Genomics UK (COG-UK) Consortium | Catherine Moore, Johnathan Evans, Laura Gifford, Malorie Perry, Simon Cottrell, Angela Marchbank, Alec Birchley, Alexander Adams, Amy Gaskin, Bree Gatica-Wilcox, Jason Coombes, Joel Southgate, Lauren Gilbert, Lee Graham, Nicole Pacchiarini, Sara Kumziene-Summerhayes, Sarah Taylor, Sophie Jones, Sara Rey, Matthew Bull, Joanne Watkins, Sally Corden, Tom Connor |
| EPI_ISL_627309 | Queens Medical Centre, Clinical Microbiology Department / DeepSeq Nottingham | COVID-19 Genomics UK (COG-UK) Consortium | Gemma Clark, Wendy Smith, Manjinder Khakh, Vicki M Fleming, Michelle M Lister, Hannah Howson-Wells, Jonathan Ball, Patrick McClure, Joseph Chappell, Theocharis Tsoleridis, Nadine Holmes, Matthew Carlisle, Christopher Moore, Fei Sang, Johnny Debebe, Victoria Wright, Matthew Loose |
| EPI_ISL_627659 | Quadram Institute Bioscience | COVID-19 Genomics UK (COG-UK) Consortium | Dave J. Baker, Gemma L. Kay, Alp Aydin, Thanh Le-Viet, Steven Rudder, Ana P. Tedim, Anastasia Kolyva, Maria Diaz, Leonardo de Oliveira Martins, Nabil-Fareed Alikhan, Lizzie Meadows, Rachael Stanley, Ngozi Elumogo, Muhammed Yasir, Nicholas M. Thomson, Alexander J Trotter, Rachel Gilroy, Samuel Bloomfield, Claire Stuart, Andrew Bell, Reenesh Prakash, Samir Dervisevic, Alison E. Mather, John Wain, Mark Webber, Andrew J. Page, Justin O'Grady |
| EPI_ISL_627674, EPI_ISL_627675, EPI_ISL_627676, EPI_ISL_627677, EPI_ISL_627678, EPI_ISL_627679, EPI_ISL_627680, EPI_ISL_627681, EPI_ISL_627682, EPI_ISL_627683, EPI_ISL_627684, EPI_ISL_627685, EPI_ISL_627686, EPI_ISL_627687, EPI_ISL_627688, EPI_ISL_627689, EPI_ISL_627690, EPI_ISL_627691, EPI_ISL_627692, EPI_ISL_627693, EPI_ISL_627694, EPI_ISL_627695, EPI_ISL_627696, EPI_ISL_627697, EPI_ISL_627698, EPI_ISL_627699 | | | |
| see above | Queens Medical Centre, Clinical Microbiology Department / DeepSeq Nottingham | COVID-19 Genomics UK (COG-UK) Consortium | Gemma Clark, Wendy Smith, Manjinder Khakh, Vicki M Fleming, Michelle M Lister, Hannah Howson-Wells, Jonathan Ball, Patrick McClure, Joseph Chappell, Theocharis Tsoleridis, Nadine Holmes, Matthew Carlisle, Christopher Moore, Fei Sang, Johnny Debebe, Victoria Wright, Matthew Loose |
| EPI_ISL_628149, EPI_ISL_628159, EPI_ISL_628164, EPI_ISL_628167, EPI_ISL_628177, EPI_ISL_628179, EPI_ISL_628183, EPI_ISL_628187, EPI_ISL_628198, EPI_ISL_628202, EPI_ISL_628206, EPI_ISL_628207, EPI_ISL_628212, EPI_ISL_628219, EPI_ISL_628230, EPI_ISL_628234, EPI_ISL_628236, EPI_ISL_628237, EPI_ISL_628240, EPI_ISL_628243, EPI_ISL_628246, EPI_ISL_628258, EPI_ISL_628260, EPI_ISL_628264, EPI_ISL_628268, EPI_ISL_628271, EPI_ISL_628272, EPI_ISL_628273, EPI_ISL_628288, EPI_ISL_628298, EPI_ISL_628304, EPI_ISL_628307, EPI_ISL_628308 | | | |
| see above | Wales Specialist Virology Centre Sequencing lab: Pathogen Genomics Unit | COVID-19 Genomics UK (COG-UK) Consortium | Catherine Moore, Johnathan Evans, Laura Gifford, Malorie Perry, Simon Cottrell, Angela Marchbank, Alec Birchley, Alexander Adams, Amy Gaskin, Bree Gatica-Wilcox, Jason Coombes, Joel Southgate, Lauren Gilbert, Lee Graham, Nicole Pacchiarini, Sara Kumziene-Summerhayes, Sarah Taylor, Sophie Jones, Sara Rey, Matthew Bull, Joanne Watkins, Sally Corden, Tom Connor |
| EPI_ISL_628321, EPI_ISL_628322, EPI_ISL_628323, EPI_ISL_628324, EPI_ISL_628327, EPI_ISL_628330, EPI_ISL_628331 | Centre for Enzyme Innovation, University of Portsmouth / Translational Research Laboratory, Portsmouth Hospitals NHS Trust | COVID-19 Genomics UK (COG-UK) Consortium | Angela Beckett,Yann Bourgeois,Garry Scarlett,Sharon Glaysher,Scott Elliott,Kelly Bicknell,Robert Impey,Allyson Lloyd,Sarah Wyllie,Ethan Butcher,Anoop Chauhan,Samuel Robson |
| EPI_ISL_629008 | Laverty Pathology | NSW Health Pathology - Institute of Clinical Pathology and Medical Research; Westmead Hospital; University of Sydney | CIDM-PH et al. |
| EPI_ISL_629756, EPI_ISL_629757, EPI_ISL_629762, EPI_ISL_629766, EPI_ISL_629767, EPI_ISL_629769, EPI_ISL_629771, EPI_ISL_629773, EPI_ISL_629776, EPI_ISL_629778, EPI_ISL_629786, EPI_ISL_629787, EPI_ISL_629792, EPI_ISL_629793, EPI_ISL_629794, EPI_ISL_629795, EPI_ISL_629796, EPI_ISL_629798, EPI_ISL_629799, EPI_ISL_629800, EPI_ISL_629802, EPI_ISL_629807, EPI_ISL_629808, EPI_ISL_629809, EPI_ISL_629810, EPI_ISL_629811, EPI_ISL_629813, EPI_ISL_629815, EPI_ISL_629817, EPI_ISL_629819, EPI_ISL_629820, EPI_ISL_629822, EPI_ISL_629823, EPI_ISL_629824, EPI_ISL_629825, EPI_ISL_629827, EPI_ISL_629830, EPI_ISL_629831, EPI_ISL_629834, EPI_ISL_629835, EPI_ISL_629836, EPI_ISL_629837, EPI_ISL_629840, EPI_ISL_629841, EPI_ISL_629842, EPI_ISL_629843, EPI_ISL_629849, EPI_ISL_629850, EPI_ISL_629852, EPI_ISL_629853, EPI_ISL_629854, EPI_ISL_629855, EPI_ISL_629856, EPI_ISL_629858, EPI_ISL_629860, EPI_ISL_629871, EPI_ISL_629872, EPI_ISL_629874, EPI_ISL_629875, EPI_ISL_629876, EPI_ISL_629877, EPI_ISL_629878, EPI_ISL_629880, EPI_ISL_629881, EPI_ISL_629886, EPI_ISL_629888, EPI_ISL_629889, EPI_ISL_629890, EPI_ISL_629892, EPI_ISL_629893, EPI_ISL_629894, EPI_ISL_629895, EPI_ISL_629897, EPI_ISL_629898, EPI_ISL_629899, EPI_ISL_629901, EPI_ISL_629903, EPI_ISL_629904, EPI_ISL_629905, EPI_ISL_629906, EPI_ISL_629907, EPI_ISL_629908, EPI_ISL_629910, EPI_ISL_629912, EPI_ISL_629914, EPI_ISL_629915, EPI_ISL_629916, EPI_ISL_629917, EPI_ISL_629918, EPI_ISL_629919, EPI_ISL_629921, EPI_ISL_629922, EPI_ISL_629923, EPI_ISL_629924, EPI_ISL_629925, EPI_ISL_629926, EPI_ISL_629927, EPI_ISL_629929, EPI_ISL_629931, EPI_ISL_629935, EPI_ISL_629938 | | | |
| see above | Lighthouse Lab in Glasgow | Wellcome Sanger Institute for the COVID-19 Genomics UK (COG-UK) consortium | Harper VanSteenhouse, Yumi Kasai, David Gray, Carol Clugston, Anna Dominiczak and Alex Alderton, Roberto Amato, Sonia Goncalves, Ewan Harrison, David K. Jackson, Ian Johnston, Dominic Kwiatkowski, Cordelia Langford, John Sillitoe on behalf of the Wellcome Sanger Institute COVID-19 Surveillance Team |
| EPI_ISL_629940 | Lighthouse Lab in Glasgow | Wellcome Sanger Institute for the COVID-19 Genomics UK (COG-UK) Consortium | Harper VanSteenhouse, Yumi Kasai, David Gray, Carol Clugston, Anna Dominiczak and Alex Alderton, Roberto Amato, Sonia Goncalves, Ewan Harrison, David K. Jackson, Ian Johnston, Dominic Kwiatkowski, Cordelia Langford, John Sillitoe on behalf of the Wellcome Sanger Institute COVID-19 Surveillance Team |
| EPI_ISL_629941, EPI_ISL_629942, EPI_ISL_629943, EPI_ISL_629945, EPI_ISL_629946, EPI_ISL_629949, EPI_ISL_629950, EPI_ISL_629954, EPI_ISL_629955, EPI_ISL_629957, EPI_ISL_629959, EPI_ISL_629961, EPI_ISL_629963, EPI_ISL_629965, EPI_ISL_629966, EPI_ISL_629969, EPI_ISL_629971, EPI_ISL_629972, EPI_ISL_629973, EPI_ISL_629974, EPI_ISL_629976, EPI_ISL_629977, EPI_ISL_629980, EPI_ISL_629982, EPI_ISL_629983, EPI_ISL_629988, EPI_ISL_629990, EPI_ISL_629991, EPI_ISL_629992, EPI_ISL_629993, EPI_ISL_629994, EPI_ISL_629995, EPI_ISL_629996, EPI_ISL_629998, EPI_ISL_630000, EPI_ISL_630001, EPI_ISL_630004 | | | |
| see above | Lighthouse Lab in Glasgow | Wellcome Sanger Institute for the COVID-19 Genomics UK (COG-UK) consortium | Harper VanSteenhouse, Yumi Kasai, David Gray, Carol Clugston, Anna Dominiczak and Alex Alderton, Roberto Amato, Sonia Goncalves, Ewan Harrison, David K. Jackson, Ian Johnston, Dominic Kwiatkowski, Cordelia Langford, John Sillitoe on behalf of the Wellcome Sanger Institute COVID-19 Surveillance Team |
| EPI_ISL_631182 | Lighthouse Lab in Milton Keynes | Wellcome Sanger Institute for the COVID-19 Genomics UK (COG-UK) consortium | The Lighthouse Lab in Milton Keynes and Alex Alderton, Roberto Amato, Sonia Goncalves, Ewan Harrison, David K. Jackson, Ian Johnston, Dominic Kwiatkowski, Cordelia Langford, John Sillitoe on behalf of the Wellcome Sanger Institute COVID-19 Surveillance Team |
| EPI_ISL_632325, EPI_ISL_632331, EPI_ISL_632335, EPI_ISL_632354, EPI_ISL_632725, EPI_ISL_632738, EPI_ISL_632756, EPI_ISL_632757, EPI_ISL_632758, EPI_ISL_632764, EPI_ISL_632765, EPI_ISL_632768, EPI_ISL_632770, EPI_ISL_632790, EPI_ISL_632791, EPI_ISL_632792, EPI_ISL_632795 | | | |
| see above | Dutch COVID-19 response team | Erasmus Medical Center | Bas Oude Munnink, David Nieuwenhuijse, Reina Sikkema, Claudia Schapendonk, Irina Chestakova, Anne van der Linden, Theo Bestebroer, Stefan van Nieuwkoop, Mark Pronk, Pascal Lexmond, Corien Swaan, Manon Haverkate, Madelief Mollers, Mart Stein, Sandra Kengne Kamga Mobou, Jeroen van Kampen, Jolanda Voermans, Aura Timen, Corine GeurtsvanKessel, Annemiek van der Eijk, Richard Molenkamp, Marion Koopmans, on behalf of the Dutch national COVID-19 response team. |
| EPI_ISL_632797, EPI_ISL_632798, EPI_ISL_632799, EPI_ISL_632800, EPI_ISL_632801, EPI_ISL_632802, EPI_ISL_632803, EPI_ISL_632804, EPI_ISL_632805, EPI_ISL_632806, EPI_ISL_632807, EPI_ISL_632808, EPI_ISL_632809, EPI_ISL_632810, EPI_ISL_632811, EPI_ISL_632812, EPI_ISL_632813, EPI_ISL_632814, EPI_ISL_632815, EPI_ISL_632816, EPI_ISL_632817, EPI_ISL_632818, EPI_ISL_632819, EPI_ISL_632820, EPI_ISL_632821, EPI_ISL_632822, EPI_ISL_632846 | | | |
| see above | Microbiology, Department of Pathology, St. Bernard's Hospital, Gibraltar Health Authority | Respiratory Virus Unit, Microbiology Services Colindale, Public Health England | PHE Covid Sequencing Team, Dr Nicholas Cortes (Gibraltar), Charlotte Gillborn-Jones (Gibraltar) |
| EPI_ISL_633102 | Lighthouse Lab in Cambridge | Wellcome Sanger Institute for the COVID-19 Genomics UK (COG-UK) consortium | Rob Howes, The Lighthouse Lab in Cambridge and Alex Alderton, Roberto Amato, Sonia Goncalves, Ewan Harrison, David K. Jackson, Ian Johnston, Dominic Kwiatkowski, Cordelia Langford, John Sillitoe on behalf of the Wellcome Sanger Institute COVID-19 Surveillance Team |
| EPI_ISL_633103 | Lighthouse Lab in Glasgow | Wellcome Sanger Institute for the COVID-19 Genomics UK (COG-UK) consortium | Harper VanSteenhouse, Yumi Kasai, David Gray, Carol Clugston, Anna Dominiczak and Alex Alderton, Roberto Amato, Sonia Goncalves, Ewan Harrison, David K. Jackson, Ian Johnston, Dominic Kwiatkowski, Cordelia Langford, John Sillitoe on behalf of the Wellcome Sanger Institute COVID-19 Surveillance Team |
| EPI_ISL_633104, EPI_ISL_633107, EPI_ISL_633110 | Lighthouse Lab in Cambridge | Wellcome Sanger Institute for the COVID-19 Genomics UK (COG-UK) consortium | Rob Howes, The Lighthouse Lab in Cambridge and Alex Alderton, Roberto Amato, Sonia Goncalves, Ewan Harrison, David K. Jackson, Ian Johnston, Dominic Kwiatkowski, Cordelia Langford, John Sillitoe on behalf of the Wellcome Sanger Institute COVID-19 Surveillance Team |
| EPI_ISL_633111, EPI_ISL_633112, EPI_ISL_633113, EPI_ISL_633115 | Lighthouse Lab in Glasgow | Wellcome Sanger Institute for the COVID-19 Genomics UK (COG-UK) consortium | Harper VanSteenhouse, Yumi Kasai, David Gray, Carol Clugston, Anna Dominiczak and Alex Alderton, Roberto Amato, Sonia Goncalves, Ewan Harrison, David K. Jackson, Ian Johnston, Dominic Kwiatkowski, Cordelia Langford, John Sillitoe on behalf of the Wellcome Sanger Institute COVID-19 Surveillance |

| | | | |
|---|---|---|---|
| | | | Team |
| EPI_ISL_633116 | Lighthouse Lab in Cambridge | Wellcome Sanger Institute for the COVID-19 Genomics UK (COG-UK) consortium | Rob Howes, The Lighthouse Lab in Cambridge and Alex Alderton, Roberto Amato, Sonia Goncalves, Ewan Harrison, David K. Jackson, Ian Johnston, Dominic Kwiatkowski, Cordelia Langford, John Sillitoe on behalf of the Wellcome Sanger Institute COVID-19 Surveillance Team |
| EPI_ISL_633118 | Lighthouse Lab in Glasgow | Wellcome Sanger Institute for the COVID-19 Genomics UK (COG-UK) Consortium | Harper VanSteenhouse, Yumi Kasai, David Gray, Carol Clugston, Anna Dominiczak and Alex Alderton, Roberto Amato, Sonia Goncalves, Ewan Harrison, David K. Jackson, Ian Johnston, Dominic Kwiatkowski, Cordelia Langford, John Sillitoe on behalf of the Wellcome Sanger Institute COVID-19 Surveillance Team |
| EPI_ISL_633119 | Lighthouse Lab in Glasgow | Wellcome Sanger Institute for the COVID-19 Genomics UK (COG-UK) consortium | Harper VanSteenhouse, Yumi Kasai, David Gray, Carol Clugston, Anna Dominiczak and Alex Alderton, Roberto Amato, Sonia Goncalves, Ewan Harrison, David K. Jackson, Ian Johnston, Dominic Kwiatkowski, Cordelia Langford, John Sillitoe on behalf of the Wellcome Sanger Institute COVID-19 Surveillance Team |
| EPI_ISL_633120, EPI_ISL_633122, EPI_ISL_633123 | Lighthouse Lab in Cambridge | Wellcome Sanger Institute for the COVID-19 Genomics UK (COG-UK) consortium | Rob Howes, The Lighthouse Lab in Cambridge and Alex Alderton, Roberto Amato, Sonia Goncalves, Ewan Harrison, David K. Jackson, Ian Johnston, Dominic Kwiatkowski, Cordelia Langford, John Sillitoe on behalf of the Wellcome Sanger Institute COVID-19 Surveillance Team |
| EPI_ISL_633125 | Lighthouse Lab in Glasgow | Wellcome Sanger Institute for the COVID-19 Genomics UK (COG-UK) consortium | Harper VanSteenhouse, Yumi Kasai, David Gray, Carol Clugston, Anna Dominiczak and Alex Alderton, Roberto Amato, Sonia Goncalves, Ewan Harrison, David K. Jackson, Ian Johnston, Dominic Kwiatkowski, Cordelia Langford, John Sillitoe on behalf of the Wellcome Sanger Institute COVID-19 Surveillance Team |
| EPI_ISL_633126, EPI_ISL_633127, EPI_ISL_633128, EPI_ISL_633131 | Lighthouse Lab in Cambridge | Wellcome Sanger Institute for the COVID-19 Genomics UK (COG-UK) consortium | Rob Howes, The Lighthouse Lab in Cambridge and Alex Alderton, Roberto Amato, Sonia Goncalves, Ewan Harrison, David K. Jackson, Ian Johnston, Dominic Kwiatkowski, Cordelia Langford, John Sillitoe on behalf of the Wellcome Sanger Institute COVID-19 Surveillance Team |
| EPI_ISL_633132, EPI_ISL_633133, EPI_ISL_633134 | Lighthouse Lab in Glasgow | Wellcome Sanger Institute for the COVID-19 Genomics UK (COG-UK) consortium | Harper VanSteenhouse, Yumi Kasai, David Gray, Carol Clugston, Anna Dominiczak and Alex Alderton, Roberto Amato, Sonia Goncalves, Ewan Harrison, David K. Jackson, Ian Johnston, Dominic Kwiatkowski, Cordelia Langford, John Sillitoe on behalf of the Wellcome Sanger Institute COVID-19 Surveillance Team |
| EPI_ISL_633135 | Lighthouse Lab in Cambridge | Wellcome Sanger Institute for the COVID-19 Genomics UK (COG-UK) consortium | Rob Howes, The Lighthouse Lab in Cambridge and Alex Alderton, Roberto Amato, Sonia Goncalves, Ewan Harrison, David K. Jackson, Ian Johnston, Dominic Kwiatkowski, Cordelia Langford, John Sillitoe on behalf of the Wellcome Sanger Institute COVID-19 Surveillance Team |
| EPI_ISL_633137, EPI_ISL_633138, EPI_ISL_633139 | Lighthouse Lab in Glasgow | Wellcome Sanger Institute for the COVID-19 Genomics UK (COG-UK) consortium | Harper VanSteenhouse, Yumi Kasai, David Gray, Carol Clugston, Anna Dominiczak and Alex Alderton, Roberto Amato, Sonia Goncalves, Ewan Harrison, David K. Jackson, Ian Johnston, Dominic Kwiatkowski, Cordelia Langford, John Sillitoe on behalf of the Wellcome Sanger Institute COVID-19 Surveillance Team |
| EPI_ISL_633141 | Lighthouse Lab in Cambridge | Wellcome Sanger Institute for the COVID-19 Genomics UK (COG-UK) consortium | Rob Howes, The Lighthouse Lab in Cambridge and Alex Alderton, Roberto Amato, Sonia Goncalves, Ewan Harrison, David K. Jackson, Ian Johnston, Dominic Kwiatkowski, Cordelia Langford, John Sillitoe on behalf of the Wellcome Sanger Institute COVID-19 Surveillance Team |
| EPI_ISL_633143, EPI_ISL_633144, EPI_ISL_633145, EPI_ISL_633146 | Lighthouse Lab in Glasgow | Wellcome Sanger Institute for the COVID-19 Genomics UK (COG-UK) consortium | Harper VanSteenhouse, Yumi Kasai, David Gray, Carol Clugston, Anna Dominiczak and Alex Alderton, Roberto Amato, Sonia Goncalves, Ewan Harrison, David K. Jackson, Ian Johnston, Dominic Kwiatkowski, Cordelia Langford, John Sillitoe on behalf of the Wellcome Sanger Institute COVID-19 Surveillance Team |
| EPI_ISL_633147, EPI_ISL_633148, EPI_ISL_633149, EPI_ISL_633150 | Lighthouse Lab in Cambridge | Wellcome Sanger Institute for the COVID-19 Genomics UK (COG-UK) consortium | Rob Howes, The Lighthouse Lab in Cambridge and Alex Alderton, Roberto Amato, Sonia Goncalves, Ewan Harrison, David K. Jackson, Ian Johnston, Dominic Kwiatkowski, Cordelia Langford, John Sillitoe on behalf of the Wellcome Sanger Institute COVID-19 Surveillance Team |
| EPI_ISL_633151, EPI_ISL_633152 | Lighthouse Lab in Glasgow | Wellcome Sanger Institute for the COVID-19 Genomics UK (COG-UK) consortium | Harper VanSteenhouse, Yumi Kasai, David Gray, Carol Clugston, Anna Dominiczak and Alex Alderton, Roberto Amato, Sonia Goncalves, Ewan Harrison, David K. Jackson, Ian Johnston, Dominic Kwiatkowski, Cordelia Langford, John Sillitoe on behalf of the Wellcome Sanger Institute COVID-19 Surveillance Team |
| EPI_ISL_633153 | Lighthouse Lab in Cambridge | Wellcome Sanger Institute for the COVID-19 Genomics UK (COG-UK) consortium | Rob Howes, The Lighthouse Lab in Cambridge and Alex Alderton, Roberto Amato, Sonia Goncalves, Ewan Harrison, David K. Jackson, Ian Johnston, Dominic Kwiatkowski, Cordelia Langford, John Sillitoe on behalf of the Wellcome Sanger Institute COVID-19 Surveillance Team |
| EPI_ISL_633154 | Lighthouse Lab in Glasgow | Wellcome Sanger Institute for the COVID-19 Genomics UK (COG-UK) consortium | Harper VanSteenhouse, Yumi Kasai, David Gray, Carol Clugston, Anna Dominiczak and Alex Alderton, Roberto Amato, Sonia Goncalves, Ewan Harrison, David K. Jackson, Ian Johnston, Dominic Kwiatkowski, Cordelia Langford, John Sillitoe on behalf of the Wellcome Sanger Institute COVID-19 Surveillance Team |
| EPI_ISL_633155, EPI_ISL_633156 | Lighthouse Lab in Cambridge | Wellcome Sanger Institute for the COVID-19 Genomics UK (COG-UK) consortium | Rob Howes, The Lighthouse Lab in Cambridge and Alex Alderton, Roberto Amato, Sonia Goncalves, Ewan Harrison, David K. Jackson, Ian Johnston, Dominic Kwiatkowski, Cordelia Langford, John Sillitoe on behalf of the Wellcome Sanger Institute COVID-19 Surveillance Team |
| EPI_ISL_633158, EPI_ISL_633160 | Lighthouse Lab in Glasgow | Wellcome Sanger Institute for the COVID-19 Genomics UK (COG-UK) consortium | Harper VanSteenhouse, Yumi Kasai, David Gray, Carol Clugston, Anna Dominiczak and Alex Alderton, Roberto Amato, Sonia Goncalves, Ewan Harrison, David K. Jackson, Ian Johnston, Dominic Kwiatkowski, Cordelia Langford, John Sillitoe on behalf of the Wellcome Sanger Institute COVID-19 Surveillance Team |
| EPI_ISL_633161 | Lighthouse Lab in Cambridge | Wellcome Sanger Institute for the COVID-19 Genomics UK (COG-UK) consortium | Rob Howes, The Lighthouse Lab in Cambridge and Alex Alderton, Roberto Amato, Sonia Goncalves, Ewan Harrison, David K. Jackson, Ian Johnston, Dominic Kwiatkowski, Cordelia Langford, John Sillitoe on behalf of the Wellcome Sanger Institute COVID-19 Surveillance Team |
| EPI_ISL_633162 | Lighthouse Lab in Glasgow | Wellcome Sanger Institute for the COVID-19 Genomics UK (COG-UK) consortium | Harper VanSteenhouse, Yumi Kasai, David Gray, Carol Clugston, Anna Dominiczak and Alex Alderton, Roberto Amato, Sonia Goncalves, Ewan Harrison, David K. Jackson, Ian Johnston, Dominic Kwiatkowski, Cordelia Langford, John Sillitoe on behalf of the Wellcome Sanger Institute COVID-19 Surveillance Team |
| EPI_ISL_633163, EPI_ISL_633164 | Lighthouse Lab in Cambridge | Wellcome Sanger Institute for the COVID-19 Genomics UK (COG-UK) consortium | Rob Howes, The Lighthouse Lab in Cambridge and Alex Alderton, Roberto Amato, Sonia Goncalves, Ewan Harrison, David K. Jackson, Ian Johnston, Dominic Kwiatkowski, Cordelia Langford, John Sillitoe on behalf of the Wellcome Sanger Institute COVID-19 Surveillance Team |
| EPI_ISL_633165 | Lighthouse Lab in Cambridge | Wellcome Sanger Institute for the COVID-19 Genomics UK (COG-UK) Consortium | Rob Howes, The Lighthouse Lab in Cambridge and Alex Alderton, Roberto Amato, Sonia Goncalves, Ewan Harrison, David K. Jackson, Ian Johnston, Dominic Kwiatkowski, Cordelia Langford, John Sillitoe on behalf of the Wellcome Sanger Institute COVID-19 Surveillance Team |
| EPI_ISL_633166, EPI_ISL_633167, EPI_ISL_633168, EPI_ISL_633169 | Lighthouse Lab in Glasgow | Wellcome Sanger Institute for the COVID-19 Genomics UK (COG-UK) consortium | Harper VanSteenhouse, Yumi Kasai, David Gray, Carol Clugston, Anna Dominiczak and Alex Alderton, Roberto Amato, Sonia Goncalves, Ewan Harrison, David K. Jackson, Ian Johnston, Dominic Kwiatkowski, Cordelia Langford, John Sillitoe on behalf of the Wellcome Sanger Institute COVID-19 Surveillance Team |
| EPI_ISL_633170 | Lighthouse Lab in Cambridge | Wellcome Sanger Institute for the COVID-19 Genomics UK (COG-UK) consortium | Rob Howes, The Lighthouse Lab in Cambridge and Alex Alderton, Roberto Amato, Sonia Goncalves, Ewan Harrison, David K. Jackson, Ian Johnston, Dominic Kwiatkowski, Cordelia Langford, John Sillitoe on behalf of the Wellcome Sanger Institute COVID-19 Surveillance Team |
| EPI_ISL_633171 | Lighthouse Lab in Glasgow | Wellcome Sanger Institute for the COVID-19 Genomics UK (COG-UK) consortium | Harper VanSteenhouse, Yumi Kasai, David Gray, Carol Clugston, Anna Dominiczak and Alex Alderton, Roberto Amato, Sonia Goncalves, Ewan Harrison, David K. Jackson, Ian Johnston, Dominic Kwiatkowski, Cordelia Langford, John Sillitoe on behalf of the Wellcome Sanger Institute COVID-19 Surveillance Team |
| EPI_ISL_633173 | Lighthouse Lab in Cambridge | Wellcome Sanger Institute for the COVID-19 Genomics UK (COG-UK) consortium | Rob Howes, The Lighthouse Lab in Cambridge and Alex Alderton, Roberto Amato, Sonia Goncalves, Ewan Harrison, David K. Jackson, Ian Johnston, Dominic Kwiatkowski, Cordelia Langford, John Sillitoe on behalf of the Wellcome Sanger Institute COVID-19 Surveillance Team |
| EPI_ISL_633174, EPI_ISL_633175 | Lighthouse Lab in Glasgow | Wellcome Sanger Institute for the COVID-19 Genomics UK (COG-UK) consortium | Harper VanSteenhouse, Yumi Kasai, David Gray, Carol Clugston, Anna Dominiczak and Alex Alderton, Roberto Amato, Sonia Goncalves, Ewan Harrison, David K. Jackson, Ian Johnston, Dominic Kwiatkowski, Cordelia Langford, John Sillitoe on behalf of the Wellcome Sanger Institute COVID-19 Surveillance Team |
| EPI_ISL_633176 | Lighthouse Lab in Cambridge | Wellcome Sanger Institute for the COVID-19 Genomics UK (COG-UK) consortium | Rob Howes, The Lighthouse Lab in Cambridge and Alex Alderton, Roberto Amato, Sonia Goncalves, Ewan Harrison, David K. Jackson, Ian Johnston, Dominic Kwiatkowski, Cordelia Langford, John Sillitoe on behalf of the Wellcome Sanger Institute COVID-19 Surveillance Team |
| EPI_ISL_633178, EPI_ISL_633179, EPI_ISL_633180, EPI_ISL_633181, EPI_ISL_633182 | Lighthouse Lab in Glasgow | Wellcome Sanger Institute for the COVID-19 Genomics UK (COG-UK) consortium | Harper VanSteenhouse, Yumi Kasai, David Gray, Carol Clugston, Anna Dominiczak and Alex Alderton, Roberto Amato, Sonia Goncalves, Ewan Harrison, David K. Jackson, Ian Johnston, Dominic Kwiatkowski, Cordelia Langford, John Sillitoe on behalf of the Wellcome Sanger Institute COVID-19 Surveillance Team |
| EPI_ISL_633183 | Lighthouse Lab in Cambridge | Wellcome Sanger Institute for the COVID-19 Genomics UK (COG-UK) consortium | Rob Howes, The Lighthouse Lab in Cambridge and Alex Alderton, Roberto Amato, Sonia Goncalves, Ewan Harrison, David K. Jackson, Ian Johnston, Dominic Kwiatkowski, Cordelia Langford, John Sillitoe on behalf of the Wellcome Sanger Institute COVID-19 Surveillance Team |

| | | | |
|---|---|---|---|
| EPI_ISL_633184, EPI_ISL_633185 | Lighthouse Lab in Glasgow | Wellcome Sanger Institute for the COVID-19 Genomics UK (COG-UK) consortium | Harper VanSteenhouse, Yumi Kasai, David Gray, Carol Clugston, Anna Dominiczak and Alex Alderton, Roberto Amato, Sonia Goncalves, Ewan Harrison, David K. Jackson, Ian Johnston, Dominic Kwiatkowski, Cordelia Langford, John Sillitoe on behalf of the Wellcome Sanger Institute COVID-19 Surveillance Team |
| EPI_ISL_633186, EPI_ISL_633187, EPI_ISL_633188, EPI_ISL_633189 | Lighthouse Lab in Cambridge | Wellcome Sanger Institute for the COVID-19 Genomics UK (COG-UK) consortium | Rob Howes, The Lighthouse Lab in Cambridge and Alex Alderton, Roberto Amato, Sonia Goncalves, Ewan Harrison, David K. Jackson, Ian Johnston, Dominic Kwiatkowski, Cordelia Langford, John Sillitoe on behalf of the Wellcome Sanger Institute COVID-19 Surveillance Team |
| EPI_ISL_633190, EPI_ISL_633191 | Lighthouse Lab in Glasgow | Wellcome Sanger Institute for the COVID-19 Genomics UK (COG-UK) consortium | Harper VanSteenhouse, Yumi Kasai, David Gray, Carol Clugston, Anna Dominiczak and Alex Alderton, Roberto Amato, Sonia Goncalves, Ewan Harrison, David K. Jackson, Ian Johnston, Dominic Kwiatkowski, Cordelia Langford, John Sillitoe on behalf of the Wellcome Sanger Institute COVID-19 Surveillance Team |
| EPI_ISL_633192 | Lighthouse Lab in Cambridge | Wellcome Sanger Institute for the COVID-19 Genomics UK (COG-UK) consortium | Rob Howes, The Lighthouse Lab in Cambridge and Alex Alderton, Roberto Amato, Sonia Goncalves, Ewan Harrison, David K. Jackson, Ian Johnston, Dominic Kwiatkowski, Cordelia Langford, John Sillitoe on behalf of the Wellcome Sanger Institute COVID-19 Surveillance Team |
| EPI_ISL_633193, EPI_ISL_633195, EPI_ISL_633196 | Lighthouse Lab in Glasgow | Wellcome Sanger Institute for the COVID-19 Genomics UK (COG-UK) consortium | Harper VanSteenhouse, Yumi Kasai, David Gray, Carol Clugston, Anna Dominiczak and Alex Alderton, Roberto Amato, Sonia Goncalves, Ewan Harrison, David K. Jackson, Ian Johnston, Dominic Kwiatkowski, Cordelia Langford, John Sillitoe on behalf of the Wellcome Sanger Institute COVID-19 Surveillance Team |
| EPI_ISL_633197, EPI_ISL_633198 | Lighthouse Lab in Cambridge | Wellcome Sanger Institute for the COVID-19 Genomics UK (COG-UK) consortium | Rob Howes, The Lighthouse Lab in Cambridge and Alex Alderton, Roberto Amato, Sonia Goncalves, Ewan Harrison, David K. Jackson, Ian Johnston, Dominic Kwiatkowski, Cordelia Langford, John Sillitoe on behalf of the Wellcome Sanger Institute COVID-19 Surveillance Team |
| EPI_ISL_633201, EPI_ISL_633202, EPI_ISL_633203, EPI_ISL_633204 | Lighthouse Lab in Glasgow | Wellcome Sanger Institute for the COVID-19 Genomics UK (COG-UK) consortium | Harper VanSteenhouse, Yumi Kasai, David Gray, Carol Clugston, Anna Dominiczak and Alex Alderton, Roberto Amato, Sonia Goncalves, Ewan Harrison, David K. Jackson, Ian Johnston, Dominic Kwiatkowski, Cordelia Langford, John Sillitoe on behalf of the Wellcome Sanger Institute COVID-19 Surveillance Team |
| EPI_ISL_633205 | Lighthouse Lab in Cambridge | Wellcome Sanger Institute for the COVID-19 Genomics UK (COG-UK) consortium | Rob Howes, The Lighthouse Lab in Cambridge and Alex Alderton, Roberto Amato, Sonia Goncalves, Ewan Harrison, David K. Jackson, Ian Johnston, Dominic Kwiatkowski, Cordelia Langford, John Sillitoe on behalf of the Wellcome Sanger Institute COVID-19 Surveillance Team |
| EPI_ISL_633206, EPI_ISL_633207 | Lighthouse Lab in Glasgow | Wellcome Sanger Institute for the COVID-19 Genomics UK (COG-UK) consortium | Harper VanSteenhouse, Yumi Kasai, David Gray, Carol Clugston, Anna Dominiczak and Alex Alderton, Roberto Amato, Sonia Goncalves, Ewan Harrison, David K. Jackson, Ian Johnston, Dominic Kwiatkowski, Cordelia Langford, John Sillitoe on behalf of the Wellcome Sanger Institute COVID-19 Surveillance Team |
| EPI_ISL_633208 | Lighthouse Lab in Cambridge | Wellcome Sanger Institute for the COVID-19 Genomics UK (COG-UK) consortium | Rob Howes, The Lighthouse Lab in Cambridge and Alex Alderton, Roberto Amato, Sonia Goncalves, Ewan Harrison, David K. Jackson, Ian Johnston, Dominic Kwiatkowski, Cordelia Langford, John Sillitoe on behalf of the Wellcome Sanger Institute COVID-19 Surveillance Team |
| EPI_ISL_633209, EPI_ISL_633210 | Lighthouse Lab in Glasgow | Wellcome Sanger Institute for the COVID-19 Genomics UK (COG-UK) consortium | Harper VanSteenhouse, Yumi Kasai, David Gray, Carol Clugston, Anna Dominiczak and Alex Alderton, Roberto Amato, Sonia Goncalves, Ewan Harrison, David K. Jackson, Ian Johnston, Dominic Kwiatkowski, Cordelia Langford, John Sillitoe on behalf of the Wellcome Sanger Institute COVID-19 Surveillance Team |
| EPI_ISL_633212 | Lighthouse Lab in Cambridge | Wellcome Sanger Institute for the COVID-19 Genomics UK (COG-UK) consortium | Rob Howes, The Lighthouse Lab in Cambridge and Alex Alderton, Roberto Amato, Sonia Goncalves, Ewan Harrison, David K. Jackson, Ian Johnston, Dominic Kwiatkowski, Cordelia Langford, John Sillitoe on behalf of the Wellcome Sanger Institute COVID-19 Surveillance Team |
| EPI_ISL_633213 | Lighthouse Lab in Glasgow | Wellcome Sanger Institute for the COVID-19 Genomics UK (COG-UK) consortium | Harper VanSteenhouse, Yumi Kasai, David Gray, Carol Clugston, Anna Dominiczak and Alex Alderton, Roberto Amato, Sonia Goncalves, Ewan Harrison, David K. Jackson, Ian Johnston, Dominic Kwiatkowski, Cordelia Langford, John Sillitoe on behalf of the Wellcome Sanger Institute COVID-19 Surveillance Team |
| EPI_ISL_633214, EPI_ISL_633216, EPI_ISL_633217, EPI_ISL_633218, EPI_ISL_633219 | Lighthouse Lab in Cambridge | Wellcome Sanger Institute for the COVID-19 Genomics UK (COG-UK) consortium | Rob Howes, The Lighthouse Lab in Cambridge and Alex Alderton, Roberto Amato, Sonia Goncalves, Ewan Harrison, David K. Jackson, Ian Johnston, Dominic Kwiatkowski, Cordelia Langford, John Sillitoe on behalf of the Wellcome Sanger Institute COVID-19 Surveillance Team |
| EPI_ISL_633222 | Lighthouse Lab in Glasgow | Wellcome Sanger Institute for the COVID-19 Genomics UK (COG-UK) consortium | Harper VanSteenhouse, Yumi Kasai, David Gray, Carol Clugston, Anna Dominiczak and Alex Alderton, Roberto Amato, Sonia Goncalves, Ewan Harrison, David K. Jackson, Ian Johnston, Dominic Kwiatkowski, Cordelia Langford, John Sillitoe on behalf of the Wellcome Sanger Institute COVID-19 Surveillance Team |
| EPI_ISL_633223, EPI_ISL_633224 | Lighthouse Lab in Cambridge | Wellcome Sanger Institute for the COVID-19 Genomics UK (COG-UK) consortium | Rob Howes, The Lighthouse Lab in Cambridge and Alex Alderton, Roberto Amato, Sonia Goncalves, Ewan Harrison, David K. Jackson, Ian Johnston, Dominic Kwiatkowski, Cordelia Langford, John Sillitoe on behalf of the Wellcome Sanger Institute COVID-19 Surveillance Team |
| EPI_ISL_633226, EPI_ISL_633228 | Lighthouse Lab in Glasgow | Wellcome Sanger Institute for the COVID-19 Genomics UK (COG-UK) consortium | Harper VanSteenhouse, Yumi Kasai, David Gray, Carol Clugston, Anna Dominiczak and Alex Alderton, Roberto Amato, Sonia Goncalves, Ewan Harrison, David K. Jackson, Ian Johnston, Dominic Kwiatkowski, Cordelia Langford, John Sillitoe on behalf of the Wellcome Sanger Institute COVID-19 Surveillance Team |
| EPI_ISL_633229 | Lighthouse Lab in Cambridge | Wellcome Sanger Institute for the COVID-19 Genomics UK (COG-UK) consortium | Rob Howes, The Lighthouse Lab in Cambridge and Alex Alderton, Roberto Amato, Sonia Goncalves, Ewan Harrison, David K. Jackson, Ian Johnston, Dominic Kwiatkowski, Cordelia Langford, John Sillitoe on behalf of the Wellcome Sanger Institute COVID-19 Surveillance Team |
| EPI_ISL_633230 | Lighthouse Lab in Glasgow | Wellcome Sanger Institute for the COVID-19 Genomics UK (COG-UK) consortium | Harper VanSteenhouse, Yumi Kasai, David Gray, Carol Clugston, Anna Dominiczak and Alex Alderton, Roberto Amato, Sonia Goncalves, Ewan Harrison, David K. Jackson, Ian Johnston, Dominic Kwiatkowski, Cordelia Langford, John Sillitoe on behalf of the Wellcome Sanger Institute COVID-19 Surveillance Team |
| EPI_ISL_633231 | Lighthouse Lab in Cambridge | Wellcome Sanger Institute for the COVID-19 Genomics UK (COG-UK) consortium | Rob Howes, The Lighthouse Lab in Cambridge and Alex Alderton, Roberto Amato, Sonia Goncalves, Ewan Harrison, David K. Jackson, Ian Johnston, Dominic Kwiatkowski, Cordelia Langford, John Sillitoe on behalf of the Wellcome Sanger Institute COVID-19 Surveillance Team |
| EPI_ISL_633232, EPI_ISL_633233 | Lighthouse Lab in Glasgow | Wellcome Sanger Institute for the COVID-19 Genomics UK (COG-UK) consortium | Harper VanSteenhouse, Yumi Kasai, David Gray, Carol Clugston, Anna Dominiczak and Alex Alderton, Roberto Amato, Sonia Goncalves, Ewan Harrison, David K. Jackson, Ian Johnston, Dominic Kwiatkowski, Cordelia Langford, John Sillitoe on behalf of the Wellcome Sanger Institute COVID-19 Surveillance Team |
| EPI_ISL_633235, EPI_ISL_633236 | Lighthouse Lab in Cambridge | Wellcome Sanger Institute for the COVID-19 Genomics UK (COG-UK) consortium | Rob Howes, The Lighthouse Lab in Cambridge and Alex Alderton, Roberto Amato, Sonia Goncalves, Ewan Harrison, David K. Jackson, Ian Johnston, Dominic Kwiatkowski, Cordelia Langford, John Sillitoe on behalf of the Wellcome Sanger Institute COVID-19 Surveillance Team |
| EPI_ISL_633237 | Lighthouse Lab in Glasgow | Wellcome Sanger Institute for the COVID-19 Genomics UK (COG-UK) consortium | Harper VanSteenhouse, Yumi Kasai, David Gray, Carol Clugston, Anna Dominiczak and Alex Alderton, Roberto Amato, Sonia Goncalves, Ewan Harrison, David K. Jackson, Ian Johnston, Dominic Kwiatkowski, Cordelia Langford, John Sillitoe on behalf of the Wellcome Sanger Institute COVID-19 Surveillance Team |
| EPI_ISL_633238, EPI_ISL_633239 | Lighthouse Lab in Cambridge | Wellcome Sanger Institute for the COVID-19 Genomics UK (COG-UK) consortium | Rob Howes, The Lighthouse Lab in Cambridge and Alex Alderton, Roberto Amato, Sonia Goncalves, Ewan Harrison, David K. Jackson, Ian Johnston, Dominic Kwiatkowski, Cordelia Langford, John Sillitoe on behalf of the Wellcome Sanger Institute COVID-19 Surveillance Team |
| EPI_ISL_633240, EPI_ISL_633241 | Lighthouse Lab in Glasgow | Wellcome Sanger Institute for the COVID-19 Genomics UK (COG-UK) consortium | Harper VanSteenhouse, Yumi Kasai, David Gray, Carol Clugston, Anna Dominiczak and Alex Alderton, Roberto Amato, Sonia Goncalves, Ewan Harrison, David K. Jackson, Ian Johnston, Dominic Kwiatkowski, Cordelia Langford, John Sillitoe on behalf of the Wellcome Sanger Institute COVID-19 Surveillance Team |
| EPI_ISL_633242 | Lighthouse Lab in Cambridge | Wellcome Sanger Institute for the COVID-19 Genomics UK (COG-UK) consortium | Rob Howes, The Lighthouse Lab in Cambridge and Alex Alderton, Roberto Amato, Sonia Goncalves, Ewan Harrison, David K. Jackson, Ian Johnston, Dominic Kwiatkowski, Cordelia Langford, John Sillitoe on behalf of the Wellcome Sanger Institute COVID-19 Surveillance Team |
| EPI_ISL_633243, EPI_ISL_633244 | Lighthouse Lab in Glasgow | Wellcome Sanger Institute for the COVID-19 Genomics UK (COG-UK) consortium | Harper VanSteenhouse, Yumi Kasai, David Gray, Carol Clugston, Anna Dominiczak and Alex Alderton, Roberto Amato, Sonia Goncalves, Ewan Harrison, David K. Jackson, Ian Johnston, Dominic Kwiatkowski, Cordelia Langford, John Sillitoe on behalf of the Wellcome Sanger Institute COVID-19 Surveillance Team |
| EPI_ISL_633245, EPI_ISL_633246, EPI_ISL_633248, EPI_ISL_633249 | Lighthouse Lab in Cambridge | Wellcome Sanger Institute for the COVID-19 Genomics UK (COG-UK) consortium | Rob Howes, The Lighthouse Lab in Cambridge and Alex Alderton, Roberto Amato, Sonia Goncalves, Ewan Harrison, David K. Jackson, Ian Johnston, Dominic Kwiatkowski, Cordelia Langford, John Sillitoe on behalf of the Wellcome Sanger Institute COVID-19 Surveillance Team |
| EPI_ISL_633251 | Lighthouse Lab in Glasgow | Wellcome Sanger Institute for the COVID-19 Genomics UK (COG-UK) consortium | Harper VanSteenhouse, Yumi Kasai, David Gray, Carol Clugston, Anna Dominiczak and Alex Alderton, Roberto Amato, Sonia Goncalves, Ewan Harrison, David K. Jackson, Ian Johnston, Dominic Kwiatkowski, Cordelia Langford, John Sillitoe on behalf of the Wellcome Sanger Institute COVID-19 Surveillance |

| | | | |
|---|---|---|---|
| | | | Team |
| EPI_ISL_633252, EPI_ISL_633253, EPI_ISL_633254, EPI_ISL_633255 | Lighthouse Lab in Cambridge | Wellcome Sanger Institute for the COVID-19 Genomics UK (COG-UK) consortium | Rob Howes, The Lighthouse Lab in Cambridge and Alex Alderton, Roberto Amato, Sonia Goncalves, Ewan Harrison, David K. Jackson, Ian Johnston, Dominic Kwiatkowski, Cordelia Langford, John Sillitoe on behalf of the Wellcome Sanger Institute COVID-19 Surveillance Team |
| EPI_ISL_633256, EPI_ISL_633259 | Lighthouse Lab in Glasgow | Wellcome Sanger Institute for the COVID-19 Genomics UK (COG-UK) consortium | Harper VanSteenhouse, Yumi Kasai, David Gray, Carol Clugston, Anna Dominiczak and Alex Alderton, Roberto Amato, Sonia Goncalves, Ewan Harrison, David K. Jackson, Ian Johnston, Dominic Kwiatkowski, Cordelia Langford, John Sillitoe on behalf of the Wellcome Sanger Institute COVID-19 Surveillance Team |
| EPI_ISL_633260 | Lighthouse Lab in Cambridge | Wellcome Sanger Institute for the COVID-19 Genomics UK (COG-UK) consortium | Rob Howes, The Lighthouse Lab in Cambridge and Alex Alderton, Roberto Amato, Sonia Goncalves, Ewan Harrison, David K. Jackson, Ian Johnston, Dominic Kwiatkowski, Cordelia Langford, John Sillitoe on behalf of the Wellcome Sanger Institute COVID-19 Surveillance Team |
| EPI_ISL_633261, EPI_ISL_633262 | Lighthouse Lab in Glasgow | Wellcome Sanger Institute for the COVID-19 Genomics UK (COG-UK) consortium | Harper VanSteenhouse, Yumi Kasai, David Gray, Carol Clugston, Anna Dominiczak and Alex Alderton, Roberto Amato, Sonia Goncalves, Ewan Harrison, David K. Jackson, Ian Johnston, Dominic Kwiatkowski, Cordelia Langford, John Sillitoe on behalf of the Wellcome Sanger Institute COVID-19 Surveillance Team |
| EPI_ISL_633263, EPI_ISL_633264, EPI_ISL_633265, EPI_ISL_633266 | Lighthouse Lab in Cambridge | Wellcome Sanger Institute for the COVID-19 Genomics UK (COG-UK) consortium | Rob Howes, The Lighthouse Lab in Cambridge and Alex Alderton, Roberto Amato, Sonia Goncalves, Ewan Harrison, David K. Jackson, Ian Johnston, Dominic Kwiatkowski, Cordelia Langford, John Sillitoe on behalf of the Wellcome Sanger Institute COVID-19 Surveillance Team |
| EPI_ISL_633268, EPI_ISL_633269 | Lighthouse Lab in Glasgow | Wellcome Sanger Institute for the COVID-19 Genomics UK (COG-UK) consortium | Harper VanSteenhouse, Yumi Kasai, David Gray, Carol Clugston, Anna Dominiczak and Alex Alderton, Roberto Amato, Sonia Goncalves, Ewan Harrison, David K. Jackson, Ian Johnston, Dominic Kwiatkowski, Cordelia Langford, John Sillitoe on behalf of the Wellcome Sanger Institute COVID-19 Surveillance Team |
| EPI_ISL_633270 | Lighthouse Lab in Cambridge | Wellcome Sanger Institute for the COVID-19 Genomics UK (COG-UK) consortium | Rob Howes, The Lighthouse Lab in Cambridge and Alex Alderton, Roberto Amato, Sonia Goncalves, Ewan Harrison, David K. Jackson, Ian Johnston, Dominic Kwiatkowski, Cordelia Langford, John Sillitoe on behalf of the Wellcome Sanger Institute COVID-19 Surveillance Team |
| EPI_ISL_633271, EPI_ISL_633272 | Lighthouse Lab in Glasgow | Wellcome Sanger Institute for the COVID-19 Genomics UK (COG-UK) consortium | Harper VanSteenhouse, Yumi Kasai, David Gray, Carol Clugston, Anna Dominiczak and Alex Alderton, Roberto Amato, Sonia Goncalves, Ewan Harrison, David K. Jackson, Ian Johnston, Dominic Kwiatkowski, Cordelia Langford, John Sillitoe on behalf of the Wellcome Sanger Institute COVID-19 Surveillance Team |
| EPI_ISL_633273 | Lighthouse Lab in Cambridge | Wellcome Sanger Institute for the COVID-19 Genomics UK (COG-UK) consortium | Rob Howes, The Lighthouse Lab in Cambridge and Alex Alderton, Roberto Amato, Sonia Goncalves, Ewan Harrison, David K. Jackson, Ian Johnston, Dominic Kwiatkowski, Cordelia Langford, John Sillitoe on behalf of the Wellcome Sanger Institute COVID-19 Surveillance Team |
| EPI_ISL_633274 | Lighthouse Lab in Glasgow | Wellcome Sanger Institute for the COVID-19 Genomics UK (COG-UK) consortium | Harper VanSteenhouse, Yumi Kasai, David Gray, Carol Clugston, Anna Dominiczak and Alex Alderton, Roberto Amato, Sonia Goncalves, Ewan Harrison, David K. Jackson, Ian Johnston, Dominic Kwiatkowski, Cordelia Langford, John Sillitoe on behalf of the Wellcome Sanger Institute COVID-19 Surveillance Team |
| EPI_ISL_633275 | Lighthouse Lab in Cambridge | Wellcome Sanger Institute for the COVID-19 Genomics UK (COG-UK) consortium | Rob Howes, The Lighthouse Lab in Cambridge and Alex Alderton, Roberto Amato, Sonia Goncalves, Ewan Harrison, David K. Jackson, Ian Johnston, Dominic Kwiatkowski, Cordelia Langford, John Sillitoe on behalf of the Wellcome Sanger Institute COVID-19 Surveillance Team |
| EPI_ISL_633276 | Lighthouse Lab in Glasgow | Wellcome Sanger Institute for the COVID-19 Genomics UK (COG-UK) consortium | Harper VanSteenhouse, Yumi Kasai, David Gray, Carol Clugston, Anna Dominiczak and Alex Alderton, Roberto Amato, Sonia Goncalves, Ewan Harrison, David K. Jackson, Ian Johnston, Dominic Kwiatkowski, Cordelia Langford, John Sillitoe on behalf of the Wellcome Sanger Institute COVID-19 Surveillance Team |
| EPI_ISL_633277, EPI_ISL_633278 | Lighthouse Lab in Cambridge | Wellcome Sanger Institute for the COVID-19 Genomics UK (COG-UK) consortium | Rob Howes, The Lighthouse Lab in Cambridge and Alex Alderton, Roberto Amato, Sonia Goncalves, Ewan Harrison, David K. Jackson, Ian Johnston, Dominic Kwiatkowski, Cordelia Langford, John Sillitoe on behalf of the Wellcome Sanger Institute COVID-19 Surveillance Team |
| EPI_ISL_633279, EPI_ISL_633280 | Lighthouse Lab in Glasgow | Wellcome Sanger Institute for the COVID-19 Genomics UK (COG-UK) consortium | Harper VanSteenhouse, Yumi Kasai, David Gray, Carol Clugston, Anna Dominiczak and Alex Alderton, Roberto Amato, Sonia Goncalves, Ewan Harrison, David K. Jackson, Ian Johnston, Dominic Kwiatkowski, Cordelia Langford, John Sillitoe on behalf of the Wellcome Sanger Institute COVID-19 Surveillance Team |
| EPI_ISL_633282 | Lighthouse Lab in Glasgow | Wellcome Sanger Institute for the COVID-19 Genomics UK (COG-UK) Consortium | Harper VanSteenhouse, Yumi Kasai, David Gray, Carol Clugston, Anna Dominiczak and Alex Alderton, Roberto Amato, Sonia Goncalves, Ewan Harrison, David K. Jackson, Ian Johnston, Dominic Kwiatkowski, Cordelia Langford, John Sillitoe on behalf of the Wellcome Sanger Institute COVID-19 Surveillance Team |
| EPI_ISL_633283 | Lighthouse Lab in Glasgow | Wellcome Sanger Institute for the COVID-19 Genomics UK (COG-UK) consortium | Harper VanSteenhouse, Yumi Kasai, David Gray, Carol Clugston, Anna Dominiczak and Alex Alderton, Roberto Amato, Sonia Goncalves, Ewan Harrison, David K. Jackson, Ian Johnston, Dominic Kwiatkowski, Cordelia Langford, John Sillitoe on behalf of the Wellcome Sanger Institute COVID-19 Surveillance Team |
| EPI_ISL_633284 | Lighthouse Lab in Cambridge | Wellcome Sanger Institute for the COVID-19 Genomics UK (COG-UK) consortium | Rob Howes, The Lighthouse Lab in Cambridge and Alex Alderton, Roberto Amato, Sonia Goncalves, Ewan Harrison, David K. Jackson, Ian Johnston, Dominic Kwiatkowski, Cordelia Langford, John Sillitoe on behalf of the Wellcome Sanger Institute COVID-19 Surveillance Team |
| EPI_ISL_633285 | Lighthouse Lab in Glasgow | Wellcome Sanger Institute for the COVID-19 Genomics UK (COG-UK) consortium | Harper VanSteenhouse, Yumi Kasai, David Gray, Carol Clugston, Anna Dominiczak and Alex Alderton, Roberto Amato, Sonia Goncalves, Ewan Harrison, David K. Jackson, Ian Johnston, Dominic Kwiatkowski, Cordelia Langford, John Sillitoe on behalf of the Wellcome Sanger Institute COVID-19 Surveillance Team |
| EPI_ISL_633286 | Lighthouse Lab in Cambridge | Wellcome Sanger Institute for the COVID-19 Genomics UK (COG-UK) consortium | Rob Howes, The Lighthouse Lab in Cambridge and Alex Alderton, Roberto Amato, Sonia Goncalves, Ewan Harrison, David K. Jackson, Ian Johnston, Dominic Kwiatkowski, Cordelia Langford, John Sillitoe on behalf of the Wellcome Sanger Institute COVID-19 Surveillance Team |
| EPI_ISL_633287 | Lighthouse Lab in Glasgow | Wellcome Sanger Institute for the COVID-19 Genomics UK (COG-UK) consortium | Harper VanSteenhouse, Yumi Kasai, David Gray, Carol Clugston, Anna Dominiczak and Alex Alderton, Roberto Amato, Sonia Goncalves, Ewan Harrison, David K. Jackson, Ian Johnston, Dominic Kwiatkowski, Cordelia Langford, John Sillitoe on behalf of the Wellcome Sanger Institute COVID-19 Surveillance Team |
| EPI_ISL_633288 | Lighthouse Lab in Cambridge | Wellcome Sanger Institute for the COVID-19 Genomics UK (COG-UK) consortium | Rob Howes, The Lighthouse Lab in Cambridge and Alex Alderton, Roberto Amato, Sonia Goncalves, Ewan Harrison, David K. Jackson, Ian Johnston, Dominic Kwiatkowski, Cordelia Langford, John Sillitoe on behalf of the Wellcome Sanger Institute COVID-19 Surveillance Team |
| EPI_ISL_633289, EPI_ISL_633290, EPI_ISL_633293 | Lighthouse Lab in Glasgow | Wellcome Sanger Institute for the COVID-19 Genomics UK (COG-UK) consortium | Harper VanSteenhouse, Yumi Kasai, David Gray, Carol Clugston, Anna Dominiczak and Alex Alderton, Roberto Amato, Sonia Goncalves, Ewan Harrison, David K. Jackson, Ian Johnston, Dominic Kwiatkowski, Cordelia Langford, John Sillitoe on behalf of the Wellcome Sanger Institute COVID-19 Surveillance Team |
| EPI_ISL_633294, EPI_ISL_633295 | Lighthouse Lab in Cambridge | Wellcome Sanger Institute for the COVID-19 Genomics UK (COG-UK) consortium | Rob Howes, The Lighthouse Lab in Cambridge and Alex Alderton, Roberto Amato, Sonia Goncalves, Ewan Harrison, David K. Jackson, Ian Johnston, Dominic Kwiatkowski, Cordelia Langford, John Sillitoe on behalf of the Wellcome Sanger Institute COVID-19 Surveillance Team |
| EPI_ISL_633296 | Lighthouse Lab in Glasgow | Wellcome Sanger Institute for the COVID-19 Genomics UK (COG-UK) consortium | Harper VanSteenhouse, Yumi Kasai, David Gray, Carol Clugston, Anna Dominiczak and Alex Alderton, Roberto Amato, Sonia Goncalves, Ewan Harrison, David K. Jackson, Ian Johnston, Dominic Kwiatkowski, Cordelia Langford, John Sillitoe on behalf of the Wellcome Sanger Institute COVID-19 Surveillance Team |
| EPI_ISL_633297, EPI_ISL_633298, EPI_ISL_633299 | Lighthouse Lab in Cambridge | Wellcome Sanger Institute for the COVID-19 Genomics UK (COG-UK) consortium | Rob Howes, The Lighthouse Lab in Cambridge and Alex Alderton, Roberto Amato, Sonia Goncalves, Ewan Harrison, David K. Jackson, Ian Johnston, Dominic Kwiatkowski, Cordelia Langford, John Sillitoe on behalf of the Wellcome Sanger Institute COVID-19 Surveillance Team |
| EPI_ISL_633301, EPI_ISL_633303, EPI_ISL_633304, EPI_ISL_633305, EPI_ISL_633306, EPI_ISL_633307 | Lighthouse Lab in Glasgow | Wellcome Sanger Institute for the COVID-19 Genomics UK (COG-UK) consortium | Harper VanSteenhouse, Yumi Kasai, David Gray, Carol Clugston, Anna Dominiczak and Alex Alderton, Roberto Amato, Sonia Goncalves, Ewan Harrison, David K. Jackson, Ian Johnston, Dominic Kwiatkowski, Cordelia Langford, John Sillitoe on behalf of the Wellcome Sanger Institute COVID-19 Surveillance Team |
| EPI_ISL_633308 | Lighthouse Lab in Cambridge | Wellcome Sanger Institute for the COVID-19 Genomics UK (COG-UK) consortium | Rob Howes, The Lighthouse Lab in Cambridge and Alex Alderton, Roberto Amato, Sonia Goncalves, Ewan Harrison, David K. Jackson, Ian Johnston, Dominic Kwiatkowski, Cordelia Langford, John Sillitoe on behalf of the Wellcome Sanger Institute COVID-19 Surveillance Team |
| EPI_ISL_633309, EPI_ISL_633310 | Lighthouse Lab in Glasgow | Wellcome Sanger Institute for the COVID-19 Genomics UK (COG-UK) consortium | Harper VanSteenhouse, Yumi Kasai, David Gray, Carol Clugston, Anna Dominiczak and Alex Alderton, Roberto Amato, Sonia Goncalves, Ewan Harrison, David K. Jackson, Ian Johnston, Dominic Kwiatkowski, Cordelia Langford, John Sillitoe on behalf of the Wellcome Sanger Institute COVID-19 Surveillance Team |
| EPI_ISL_633311 | Lighthouse Lab in Cambridge | Wellcome Sanger Institute for the COVID-19 Genomics UK | Rob Howes, The Lighthouse Lab in Cambridge and Alex Alderton, Roberto Amato, Sonia Goncalves, Ewan Harrison, David K. Jackson, Ian Johnston, |

| | | | |
|---|---|---|---|
| | | (COG-UK) consortium | Dominic Kwiatkowski, Cordelia Langford, John Sillitoe on behalf of the Wellcome Sanger Institute COVID-19 Surveillance Team |
| EPI_ISL_633312 | Lighthouse Lab in Glasgow | Wellcome Sanger Institute for the COVID-19 Genomics UK (COG-UK) consortium | Harper VanSteenhouse, Yumi Kasai, David Gray, Carol Clugston, Anna Dominiczak and Alex Alderton, Roberto Amato, Sonia Goncalves, Ewan Harrison, David K. Jackson, Ian Johnston, Dominic Kwiatkowski, Cordelia Langford, John Sillitoe on behalf of the Wellcome Sanger Institute COVID-19 Surveillance Team |
| EPI_ISL_633314, EPI_ISL_633315 | Lighthouse Lab in Cambridge | Wellcome Sanger Institute for the COVID-19 Genomics UK (COG-UK) consortium | Rob Howes, The Lighthouse Lab in Cambridge and Alex Alderton, Roberto Amato, Sonia Goncalves, Ewan Harrison, David K. Jackson, Ian Johnston, Dominic Kwiatkowski, Cordelia Langford, John Sillitoe on behalf of the Wellcome Sanger Institute COVID-19 Surveillance Team |
| EPI_ISL_633316, EPI_ISL_633317, EPI_ISL_633318 | Lighthouse Lab in Glasgow | Wellcome Sanger Institute for the COVID-19 Genomics UK (COG-UK) consortium | Harper VanSteenhouse, Yumi Kasai, David Gray, Carol Clugston, Anna Dominiczak and Alex Alderton, Roberto Amato, Sonia Goncalves, Ewan Harrison, David K. Jackson, Ian Johnston, Dominic Kwiatkowski, Cordelia Langford, John Sillitoe on behalf of the Wellcome Sanger Institute COVID-19 Surveillance Team |
| EPI_ISL_633319 | Lighthouse Lab in Cambridge | Wellcome Sanger Institute for the COVID-19 Genomics UK (COG-UK) consortium | Rob Howes, The Lighthouse Lab in Cambridge and Alex Alderton, Roberto Amato, Sonia Goncalves, Ewan Harrison, David K. Jackson, Ian Johnston, Dominic Kwiatkowski, Cordelia Langford, John Sillitoe on behalf of the Wellcome Sanger Institute COVID-19 Surveillance Team |
| EPI_ISL_633320 | Lighthouse Lab in Glasgow | Wellcome Sanger Institute for the COVID-19 Genomics UK (COG-UK) consortium | Harper VanSteenhouse, Yumi Kasai, David Gray, Carol Clugston, Anna Dominiczak and Alex Alderton, Roberto Amato, Sonia Goncalves, Ewan Harrison, David K. Jackson, Ian Johnston, Dominic Kwiatkowski, Cordelia Langford, John Sillitoe on behalf of the Wellcome Sanger Institute COVID-19 Surveillance Team |
| EPI_ISL_633321, EPI_ISL_633322 | Lighthouse Lab in Cambridge | Wellcome Sanger Institute for the COVID-19 Genomics UK (COG-UK) consortium | Rob Howes, The Lighthouse Lab in Cambridge and Alex Alderton, Roberto Amato, Sonia Goncalves, Ewan Harrison, David K. Jackson, Ian Johnston, Dominic Kwiatkowski, Cordelia Langford, John Sillitoe on behalf of the Wellcome Sanger Institute COVID-19 Surveillance Team |
| EPI_ISL_633323, EPI_ISL_633326, EPI_ISL_633327, EPI_ISL_633328 | Lighthouse Lab in Glasgow | Wellcome Sanger Institute for the COVID-19 Genomics UK (COG-UK) consortium | Harper VanSteenhouse, Yumi Kasai, David Gray, Carol Clugston, Anna Dominiczak and Alex Alderton, Roberto Amato, Sonia Goncalves, Ewan Harrison, David K. Jackson, Ian Johnston, Dominic Kwiatkowski, Cordelia Langford, John Sillitoe on behalf of the Wellcome Sanger Institute COVID-19 Surveillance Team |
| EPI_ISL_633329, EPI_ISL_633330 | Lighthouse Lab in Cambridge | Wellcome Sanger Institute for the COVID-19 Genomics UK (COG-UK) consortium | Rob Howes, The Lighthouse Lab in Cambridge and Alex Alderton, Roberto Amato, Sonia Goncalves, Ewan Harrison, David K. Jackson, Ian Johnston, Dominic Kwiatkowski, Cordelia Langford, John Sillitoe on behalf of the Wellcome Sanger Institute COVID-19 Surveillance Team |
| EPI_ISL_633331, EPI_ISL_633332 | Lighthouse Lab in Glasgow | Wellcome Sanger Institute for the COVID-19 Genomics UK (COG-UK) consortium | Harper VanSteenhouse, Yumi Kasai, David Gray, Carol Clugston, Anna Dominiczak and Alex Alderton, Roberto Amato, Sonia Goncalves, Ewan Harrison, David K. Jackson, Ian Johnston, Dominic Kwiatkowski, Cordelia Langford, John Sillitoe on behalf of the Wellcome Sanger Institute COVID-19 Surveillance Team |
| EPI_ISL_633333, EPI_ISL_633334 | Lighthouse Lab in Cambridge | Wellcome Sanger Institute for the COVID-19 Genomics UK (COG-UK) consortium | Rob Howes, The Lighthouse Lab in Cambridge and Alex Alderton, Roberto Amato, Sonia Goncalves, Ewan Harrison, David K. Jackson, Ian Johnston, Dominic Kwiatkowski, Cordelia Langford, John Sillitoe on behalf of the Wellcome Sanger Institute COVID-19 Surveillance Team |
| EPI_ISL_633335, EPI_ISL_633338, EPI_ISL_633339 | Lighthouse Lab in Glasgow | Wellcome Sanger Institute for the COVID-19 Genomics UK (COG-UK) consortium | Harper VanSteenhouse, Yumi Kasai, David Gray, Carol Clugston, Anna Dominiczak and Alex Alderton, Roberto Amato, Sonia Goncalves, Ewan Harrison, David K. Jackson, Ian Johnston, Dominic Kwiatkowski, Cordelia Langford, John Sillitoe on behalf of the Wellcome Sanger Institute COVID-19 Surveillance Team |
| EPI_ISL_633340 | Lighthouse Lab in Cambridge | Wellcome Sanger Institute for the COVID-19 Genomics UK (COG-UK) consortium | Rob Howes, The Lighthouse Lab in Cambridge and Alex Alderton, Roberto Amato, Sonia Goncalves, Ewan Harrison, David K. Jackson, Ian Johnston, Dominic Kwiatkowski, Cordelia Langford, John Sillitoe on behalf of the Wellcome Sanger Institute COVID-19 Surveillance Team |
| EPI_ISL_633342, EPI_ISL_633343 | Lighthouse Lab in Glasgow | Wellcome Sanger Institute for the COVID-19 Genomics UK (COG-UK) consortium | Harper VanSteenhouse, Yumi Kasai, David Gray, Carol Clugston, Anna Dominiczak and Alex Alderton, Roberto Amato, Sonia Goncalves, Ewan Harrison, David K. Jackson, Ian Johnston, Dominic Kwiatkowski, Cordelia Langford, John Sillitoe on behalf of the Wellcome Sanger Institute COVID-19 Surveillance Team |
| EPI_ISL_633345, EPI_ISL_633346, EPI_ISL_633347 | Lighthouse Lab in Cambridge | Wellcome Sanger Institute for the COVID-19 Genomics UK (COG-UK) consortium | Rob Howes, The Lighthouse Lab in Cambridge and Alex Alderton, Roberto Amato, Sonia Goncalves, Ewan Harrison, David K. Jackson, Ian Johnston, Dominic Kwiatkowski, Cordelia Langford, John Sillitoe on behalf of the Wellcome Sanger Institute COVID-19 Surveillance Team |
| EPI_ISL_633348, EPI_ISL_633351, EPI_ISL_633352, EPI_ISL_633354, EPI_ISL_633355 | Lighthouse Lab in Glasgow | Wellcome Sanger Institute for the COVID-19 Genomics UK (COG-UK) consortium | Harper VanSteenhouse, Yumi Kasai, David Gray, Carol Clugston, Anna Dominiczak and Alex Alderton, Roberto Amato, Sonia Goncalves, Ewan Harrison, David K. Jackson, Ian Johnston, Dominic Kwiatkowski, Cordelia Langford, John Sillitoe on behalf of the Wellcome Sanger Institute COVID-19 Surveillance Team |
| EPI_ISL_633357 | Lighthouse Lab in Cambridge | Wellcome Sanger Institute for the COVID-19 Genomics UK (COG-UK) consortium | Rob Howes, The Lighthouse Lab in Cambridge and Alex Alderton, Roberto Amato, Sonia Goncalves, Ewan Harrison, David K. Jackson, Ian Johnston, Dominic Kwiatkowski, Cordelia Langford, John Sillitoe on behalf of the Wellcome Sanger Institute COVID-19 Surveillance Team |
| EPI_ISL_633358 | Lighthouse Lab in Glasgow | Wellcome Sanger Institute for the COVID-19 Genomics UK (COG-UK) consortium | Harper VanSteenhouse, Yumi Kasai, David Gray, Carol Clugston, Anna Dominiczak and Alex Alderton, Roberto Amato, Sonia Goncalves, Ewan Harrison, David K. Jackson, Ian Johnston, Dominic Kwiatkowski, Cordelia Langford, John Sillitoe on behalf of the Wellcome Sanger Institute COVID-19 Surveillance Team |
| EPI_ISL_633360 | Lighthouse Lab in Cambridge | Wellcome Sanger Institute for the COVID-19 Genomics UK (COG-UK) consortium | Rob Howes, The Lighthouse Lab in Cambridge and Alex Alderton, Roberto Amato, Sonia Goncalves, Ewan Harrison, David K. Jackson, Ian Johnston, Dominic Kwiatkowski, Cordelia Langford, John Sillitoe on behalf of the Wellcome Sanger Institute COVID-19 Surveillance Team |
| EPI_ISL_633361 | Lighthouse Lab in Glasgow | Wellcome Sanger Institute for the COVID-19 Genomics UK (COG-UK) Consortium | Harper VanSteenhouse, Yumi Kasai, David Gray, Carol Clugston, Anna Dominiczak and Alex Alderton, Roberto Amato, Sonia Goncalves, Ewan Harrison, David K. Jackson, Ian Johnston, Dominic Kwiatkowski, Cordelia Langford, John Sillitoe on behalf of the Wellcome Sanger Institute COVID-19 Surveillance Team |
| EPI_ISL_633362 | Lighthouse Lab in Glasgow | Wellcome Sanger Institute for the COVID-19 Genomics UK (COG-UK) consortium | Harper VanSteenhouse, Yumi Kasai, David Gray, Carol Clugston, Anna Dominiczak and Alex Alderton, Roberto Amato, Sonia Goncalves, Ewan Harrison, David K. Jackson, Ian Johnston, Dominic Kwiatkowski, Cordelia Langford, John Sillitoe on behalf of the Wellcome Sanger Institute COVID-19 Surveillance Team |
| EPI_ISL_633363 | Lighthouse Lab in Cambridge | Wellcome Sanger Institute for the COVID-19 Genomics UK (COG-UK) consortium | Rob Howes, The Lighthouse Lab in Cambridge and Alex Alderton, Roberto Amato, Sonia Goncalves, Ewan Harrison, David K. Jackson, Ian Johnston, Dominic Kwiatkowski, Cordelia Langford, John Sillitoe on behalf of the Wellcome Sanger Institute COVID-19 Surveillance Team |
| EPI_ISL_633364 | Lighthouse Lab in Glasgow | Wellcome Sanger Institute for the COVID-19 Genomics UK (COG-UK) consortium | Harper VanSteenhouse, Yumi Kasai, David Gray, Carol Clugston, Anna Dominiczak and Alex Alderton, Roberto Amato, Sonia Goncalves, Ewan Harrison, David K. Jackson, Ian Johnston, Dominic Kwiatkowski, Cordelia Langford, John Sillitoe on behalf of the Wellcome Sanger Institute COVID-19 Surveillance Team |
| EPI_ISL_633365, EPI_ISL_633366 | Lighthouse Lab in Cambridge | Wellcome Sanger Institute for the COVID-19 Genomics UK (COG-UK) consortium | Rob Howes, The Lighthouse Lab in Cambridge and Alex Alderton, Roberto Amato, Sonia Goncalves, Ewan Harrison, David K. Jackson, Ian Johnston, Dominic Kwiatkowski, Cordelia Langford, John Sillitoe on behalf of the Wellcome Sanger Institute COVID-19 Surveillance Team |
| EPI_ISL_633367, EPI_ISL_633368 | Lighthouse Lab in Glasgow | Wellcome Sanger Institute for the COVID-19 Genomics UK (COG-UK) consortium | Harper VanSteenhouse, Yumi Kasai, David Gray, Carol Clugston, Anna Dominiczak and Alex Alderton, Roberto Amato, Sonia Goncalves, Ewan Harrison, David K. Jackson, Ian Johnston, Dominic Kwiatkowski, Cordelia Langford, John Sillitoe on behalf of the Wellcome Sanger Institute COVID-19 Surveillance Team |
| EPI_ISL_633369, EPI_ISL_633370 | Lighthouse Lab in Cambridge | Wellcome Sanger Institute for the COVID-19 Genomics UK (COG-UK) consortium | Rob Howes, The Lighthouse Lab in Cambridge and Alex Alderton, Roberto Amato, Sonia Goncalves, Ewan Harrison, David K. Jackson, Ian Johnston, Dominic Kwiatkowski, Cordelia Langford, John Sillitoe on behalf of the Wellcome Sanger Institute COVID-19 Surveillance Team |
| EPI_ISL_633371 | Lighthouse Lab in Glasgow | Wellcome Sanger Institute for the COVID-19 Genomics UK (COG-UK) consortium | Harper VanSteenhouse, Yumi Kasai, David Gray, Carol Clugston, Anna Dominiczak and Alex Alderton, Roberto Amato, Sonia Goncalves, Ewan Harrison, David K. Jackson, Ian Johnston, Dominic Kwiatkowski, Cordelia Langford, John Sillitoe on behalf of the Wellcome Sanger Institute COVID-19 Surveillance Team |
| EPI_ISL_633372, EPI_ISL_633374, EPI_ISL_633375 | Lighthouse Lab in Cambridge | Wellcome Sanger Institute for the COVID-19 Genomics UK (COG-UK) consortium | Rob Howes, The Lighthouse Lab in Cambridge and Alex Alderton, Roberto Amato, Sonia Goncalves, Ewan Harrison, David K. Jackson, Ian Johnston, Dominic Kwiatkowski, Cordelia Langford, John Sillitoe on behalf of the Wellcome Sanger Institute COVID-19 Surveillance Team |
| EPI_ISL_633376, EPI_ISL_633377 | Lighthouse Lab in Glasgow | Wellcome Sanger Institute for the COVID-19 Genomics UK (COG-UK) consortium | Harper VanSteenhouse, Yumi Kasai, David Gray, Carol Clugston, Anna Dominiczak and Alex Alderton, Roberto Amato, Sonia Goncalves, Ewan Harrison, David K. Jackson, Ian Johnston, Dominic Kwiatkowski, Cordelia Langford, John Sillitoe on behalf of the Wellcome Sanger Institute COVID-19 Surveillance Team |
| EPI_ISL_633378 | Lighthouse Lab in Cambridge | Wellcome Sanger Institute for the COVID-19 Genomics UK | Rob Howes, The Lighthouse Lab in Cambridge and Alex Alderton, Roberto Amato, Sonia Goncalves, Ewan Harrison, David K. Jackson, Ian Johnston, |

| | | | |
|---|---|---|---|
| | | (COG-UK) consortium | Dominic Kwiatkowski, Cordelia Langford, John Sillitoe on behalf of the Wellcome Sanger Institute COVID-19 Surveillance Team |
| EPI_ISL_633379, EPI_ISL_633381, EPI_ISL_633382 | Lighthouse Lab in Glasgow | Wellcome Sanger Institute for the COVID-19 Genomics UK (COG-UK) consortium | Harper VanSteenhouse, Yumi Kasai, David Gray, Carol Clugston, Anna Dominiczak and Alex Alderton, Roberto Amato, Sonia Goncalves, Ewan Harrison, David K. Jackson, Ian Johnston, Dominic Kwiatkowski, Cordelia Langford, John Sillitoe on behalf of the Wellcome Sanger Institute COVID-19 Surveillance Team |
| EPI_ISL_633383 | Lighthouse Lab in Cambridge | Wellcome Sanger Institute for the COVID-19 Genomics UK (COG-UK) consortium | Rob Howes, The Lighthouse Lab in Cambridge and Alex Alderton, Roberto Amato, Sonia Goncalves, Ewan Harrison, David K. Jackson, Ian Johnston, Dominic Kwiatkowski, Cordelia Langford, John Sillitoe on behalf of the Wellcome Sanger Institute COVID-19 Surveillance Team |
| EPI_ISL_633384, EPI_ISL_633386 | Lighthouse Lab in Glasgow | Wellcome Sanger Institute for the COVID-19 Genomics UK (COG-UK) consortium | Harper VanSteenhouse, Yumi Kasai, David Gray, Carol Clugston, Anna Dominiczak and Alex Alderton, Roberto Amato, Sonia Goncalves, Ewan Harrison, David K. Jackson, Ian Johnston, Dominic Kwiatkowski, Cordelia Langford, John Sillitoe on behalf of the Wellcome Sanger Institute COVID-19 Surveillance Team |
| EPI_ISL_633390 | Lighthouse Lab in Cambridge | Wellcome Sanger Institute for the COVID-19 Genomics UK (COG-UK) consortium | Rob Howes, The Lighthouse Lab in Cambridge and Alex Alderton, Roberto Amato, Sonia Goncalves, Ewan Harrison, David K. Jackson, Ian Johnston, Dominic Kwiatkowski, Cordelia Langford, John Sillitoe on behalf of the Wellcome Sanger Institute COVID-19 Surveillance Team |
| EPI_ISL_633391, EPI_ISL_633392 | Lighthouse Lab in Glasgow | Wellcome Sanger Institute for the COVID-19 Genomics UK (COG-UK) consortium | Harper VanSteenhouse, Yumi Kasai, David Gray, Carol Clugston, Anna Dominiczak and Alex Alderton, Roberto Amato, Sonia Goncalves, Ewan Harrison, David K. Jackson, Ian Johnston, Dominic Kwiatkowski, Cordelia Langford, John Sillitoe on behalf of the Wellcome Sanger Institute COVID-19 Surveillance Team |
| EPI_ISL_633393 | Lighthouse Lab in Cambridge | Wellcome Sanger Institute for the COVID-19 Genomics UK (COG-UK) consortium | Rob Howes, The Lighthouse Lab in Cambridge and Alex Alderton, Roberto Amato, Sonia Goncalves, Ewan Harrison, David K. Jackson, Ian Johnston, Dominic Kwiatkowski, Cordelia Langford, John Sillitoe on behalf of the Wellcome Sanger Institute COVID-19 Surveillance Team |
| EPI_ISL_633394 | Lighthouse Lab in Glasgow | Wellcome Sanger Institute for the COVID-19 Genomics UK (COG-UK) consortium | Harper VanSteenhouse, Yumi Kasai, David Gray, Carol Clugston, Anna Dominiczak and Alex Alderton, Roberto Amato, Sonia Goncalves, Ewan Harrison, David K. Jackson, Ian Johnston, Dominic Kwiatkowski, Cordelia Langford, John Sillitoe on behalf of the Wellcome Sanger Institute COVID-19 Surveillance Team |
| EPI_ISL_633395 | Lighthouse Lab in Cambridge | Wellcome Sanger Institute for the COVID-19 Genomics UK (COG-UK) consortium | Rob Howes, The Lighthouse Lab in Cambridge and Alex Alderton, Roberto Amato, Sonia Goncalves, Ewan Harrison, David K. Jackson, Ian Johnston, Dominic Kwiatkowski, Cordelia Langford, John Sillitoe on behalf of the Wellcome Sanger Institute COVID-19 Surveillance Team |
| EPI_ISL_633396 | Lighthouse Lab in Glasgow | Wellcome Sanger Institute for the COVID-19 Genomics UK (COG-UK) consortium | Harper VanSteenhouse, Yumi Kasai, David Gray, Carol Clugston, Anna Dominiczak and Alex Alderton, Roberto Amato, Sonia Goncalves, Ewan Harrison, David K. Jackson, Ian Johnston, Dominic Kwiatkowski, Cordelia Langford, John Sillitoe on behalf of the Wellcome Sanger Institute COVID-19 Surveillance Team |
| EPI_ISL_633399 | Lighthouse Lab in Cambridge | Wellcome Sanger Institute for the COVID-19 Genomics UK (COG-UK) consortium | Rob Howes, The Lighthouse Lab in Cambridge and Alex Alderton, Roberto Amato, Sonia Goncalves, Ewan Harrison, David K. Jackson, Ian Johnston, Dominic Kwiatkowski, Cordelia Langford, John Sillitoe on behalf of the Wellcome Sanger Institute COVID-19 Surveillance Team |
| EPI_ISL_633400, EPI_ISL_633401, EPI_ISL_633402 | Lighthouse Lab in Glasgow | Wellcome Sanger Institute for the COVID-19 Genomics UK (COG-UK) consortium | Harper VanSteenhouse, Yumi Kasai, David Gray, Carol Clugston, Anna Dominiczak and Alex Alderton, Roberto Amato, Sonia Goncalves, Ewan Harrison, David K. Jackson, Ian Johnston, Dominic Kwiatkowski, Cordelia Langford, John Sillitoe on behalf of the Wellcome Sanger Institute COVID-19 Surveillance Team |
| EPI_ISL_633403 | Lighthouse Lab in Cambridge | Wellcome Sanger Institute for the COVID-19 Genomics UK (COG-UK) consortium | Rob Howes, The Lighthouse Lab in Cambridge and Alex Alderton, Roberto Amato, Sonia Goncalves, Ewan Harrison, David K. Jackson, Ian Johnston, Dominic Kwiatkowski, Cordelia Langford, John Sillitoe on behalf of the Wellcome Sanger Institute COVID-19 Surveillance Team |
| EPI_ISL_633404 | Lighthouse Lab in Glasgow | Wellcome Sanger Institute for the COVID-19 Genomics UK (COG-UK) consortium | Harper VanSteenhouse, Yumi Kasai, David Gray, Carol Clugston, Anna Dominiczak and Alex Alderton, Roberto Amato, Sonia Goncalves, Ewan Harrison, David K. Jackson, Ian Johnston, Dominic Kwiatkowski, Cordelia Langford, John Sillitoe on behalf of the Wellcome Sanger Institute COVID-19 Surveillance Team |
| EPI_ISL_633405 | Lighthouse Lab in Cambridge | Wellcome Sanger Institute for the COVID-19 Genomics UK (COG-UK) consortium | Rob Howes, The Lighthouse Lab in Cambridge and Alex Alderton, Roberto Amato, Sonia Goncalves, Ewan Harrison, David K. Jackson, Ian Johnston, Dominic Kwiatkowski, Cordelia Langford, John Sillitoe on behalf of the Wellcome Sanger Institute COVID-19 Surveillance Team |
| EPI_ISL_633406 | Lighthouse Lab in Glasgow | Wellcome Sanger Institute for the COVID-19 Genomics UK (COG-UK) consortium | Harper VanSteenhouse, Yumi Kasai, David Gray, Carol Clugston, Anna Dominiczak and Alex Alderton, Roberto Amato, Sonia Goncalves, Ewan Harrison, David K. Jackson, Ian Johnston, Dominic Kwiatkowski, Cordelia Langford, John Sillitoe on behalf of the Wellcome Sanger Institute COVID-19 Surveillance Team |
| EPI_ISL_633407, EPI_ISL_633408 | Lighthouse Lab in Cambridge | Wellcome Sanger Institute for the COVID-19 Genomics UK (COG-UK) consortium | Rob Howes, The Lighthouse Lab in Cambridge and Alex Alderton, Roberto Amato, Sonia Goncalves, Ewan Harrison, David K. Jackson, Ian Johnston, Dominic Kwiatkowski, Cordelia Langford, John Sillitoe on behalf of the Wellcome Sanger Institute COVID-19 Surveillance Team |
| EPI_ISL_633409, EPI_ISL_633410, EPI_ISL_633411, EPI_ISL_633412, EPI_ISL_633413, EPI_ISL_633414, EPI_ISL_633415, EPI_ISL_633416, EPI_ISL_633417, EPI_ISL_633418, EPI_ISL_633419, EPI_ISL_633420, EPI_ISL_633421, EPI_ISL_633422, EPI_ISL_633423, EPI_ISL_633424, EPI_ISL_633425, EPI_ISL_633426, EPI_ISL_633427, EPI_ISL_633428, EPI_ISL_633429, EPI_ISL_633430, EPI_ISL_633431, EPI_ISL_633432, EPI_ISL_633433, EPI_ISL_633434, EPI_ISL_633435, EPI_ISL_633436, EPI_ISL_633438, EPI_ISL_633440, EPI_ISL_633442, EPI_ISL_633443, EPI_ISL_633444, EPI_ISL_633445, EPI_ISL_633446, EPI_ISL_633447, EPI_ISL_633448, EPI_ISL_633449, EPI_ISL_633450, EPI_ISL_633451, EPI_ISL_633452, EPI_ISL_633453, EPI_ISL_633454, EPI_ISL_633455, EPI_ISL_633456, EPI_ISL_633457, EPI_ISL_633458, EPI_ISL_633459, EPI_ISL_633460, EPI_ISL_633461, EPI_ISL_633462, EPI_ISL_633463, EPI_ISL_633464, EPI_ISL_633465, EPI_ISL_633466, EPI_ISL_633467, EPI_ISL_633468, EPI_ISL_633469, EPI_ISL_633470, EPI_ISL_633471, EPI_ISL_633472, EPI_ISL_633473, EPI_ISL_633474, EPI_ISL_633475, EPI_ISL_633476, EPI_ISL_633477, EPI_ISL_633478, EPI_ISL_633479, EPI_ISL_633480, EPI_ISL_633481, EPI_ISL_633482, EPI_ISL_633483, EPI_ISL_633484, EPI_ISL_633485, EPI_ISL_633486, EPI_ISL_633487, EPI_ISL_633488, EPI_ISL_633489, EPI_ISL_633490, EPI_ISL_633491 | | | |
| see above | Lighthouse Lab in Glasgow | Wellcome Sanger Institute for the COVID-19 Genomics UK (COG-UK) consortium | Harper VanSteenhouse, Yumi Kasai, David Gray, Carol Clugston, Anna Dominiczak and Alex Alderton, Roberto Amato, Sonia Goncalves, Ewan Harrison, David K. Jackson, Ian Johnston, Dominic Kwiatkowski, Cordelia Langford, John Sillitoe on behalf of the Wellcome Sanger Institute COVID-19 Surveillance Team |
| EPI_ISL_633492 | Lighthouse Lab in Glasgow | Wellcome Sanger Institute for the COVID-19 Genomics UK (COG-UK) Consortium | Harper VanSteenhouse, Yumi Kasai, David Gray, Carol Clugston, Anna Dominiczak and Alex Alderton, Roberto Amato, Sonia Goncalves, Ewan Harrison, David K. Jackson, Ian Johnston, Dominic Kwiatkowski, Cordelia Langford, John Sillitoe on behalf of the Wellcome Sanger Institute COVID-19 Surveillance Team |
| EPI_ISL_633493, EPI_ISL_633494, EPI_ISL_633495, EPI_ISL_633496, EPI_ISL_633497, EPI_ISL_633498, EPI_ISL_633500, EPI_ISL_633501, EPI_ISL_633502, EPI_ISL_633504, EPI_ISL_633505, EPI_ISL_633506, EPI_ISL_633508, EPI_ISL_633509, EPI_ISL_633510, EPI_ISL_633511, EPI_ISL_633512, EPI_ISL_633513, EPI_ISL_633514, EPI_ISL_633515, EPI_ISL_633516, EPI_ISL_633517, EPI_ISL_633518, EPI_ISL_633519, EPI_ISL_633520, EPI_ISL_633521, EPI_ISL_633523, EPI_ISL_633524, EPI_ISL_633525, EPI_ISL_633526, EPI_ISL_633527, EPI_ISL_633528, EPI_ISL_633529, EPI_ISL_633530, EPI_ISL_633531, EPI_ISL_633532, EPI_ISL_633533, EPI_ISL_633534, EPI_ISL_633535, EPI_ISL_633536, EPI_ISL_633537, EPI_ISL_633538, EPI_ISL_633539, EPI_ISL_633540, EPI_ISL_633541, EPI_ISL_633542, EPI_ISL_633544, EPI_ISL_633545, EPI_ISL_633546, EPI_ISL_633548, EPI_ISL_633549, EPI_ISL_633550, EPI_ISL_633552, EPI_ISL_633554, EPI_ISL_633556, EPI_ISL_633557, EPI_ISL_633558, EPI_ISL_633559, EPI_ISL_633560, EPI_ISL_633561, EPI_ISL_633562, EPI_ISL_633563, EPI_ISL_633564, EPI_ISL_633565, EPI_ISL_633566, EPI_ISL_633567, EPI_ISL_633568, EPI_ISL_633569, EPI_ISL_633570, EPI_ISL_633571, EPI_ISL_633572, EPI_ISL_633573, EPI_ISL_633574, EPI_ISL_633575, EPI_ISL_633576 | | | |
| see above | Lighthouse Lab in Glasgow | Wellcome Sanger Institute for the COVID-19 Genomics UK (COG-UK) consortium | Harper VanSteenhouse, Yumi Kasai, David Gray, Carol Clugston, Anna Dominiczak and Alex Alderton, Roberto Amato, Sonia Goncalves, Ewan Harrison, David K. Jackson, Ian Johnston, Dominic Kwiatkowski, Cordelia Langford, John Sillitoe on behalf of the Wellcome Sanger Institute COVID-19 Surveillance Team |
| EPI_ISL_633577 | Lighthouse Lab in Glasgow | Wellcome Sanger Institute for the COVID-19 Genomics UK (COG-UK) Consortium | Harper VanSteenhouse, Yumi Kasai, David Gray, Carol Clugston, Anna Dominiczak and Alex Alderton, Roberto Amato, Sonia Goncalves, Ewan Harrison, David K. Jackson, Ian Johnston, Dominic Kwiatkowski, Cordelia Langford, John Sillitoe on behalf of the Wellcome Sanger Institute COVID-19 Surveillance Team |
| EPI_ISL_633578, EPI_ISL_633579, EPI_ISL_633580, EPI_ISL_633581, EPI_ISL_633582, EPI_ISL_633583, EPI_ISL_633584, EPI_ISL_633585, EPI_ISL_633586, EPI_ISL_633587, EPI_ISL_633588, EPI_ISL_633589, EPI_ISL_633590, EPI_ISL_633591, EPI_ISL_633592, EPI_ISL_633593, EPI_ISL_633594, EPI_ISL_633596, EPI_ISL_633597, EPI_ISL_633598, EPI_ISL_633599, EPI_ISL_633601, EPI_ISL_633602, EPI_ISL_633603, EPI_ISL_633605, EPI_ISL_633606, EPI_ISL_633607, EPI_ISL_633608, EPI_ISL_633609, EPI_ISL_633611, EPI_ISL_633612, EPI_ISL_633613, EPI_ISL_633614, EPI_ISL_633615, EPI_ISL_633616, EPI_ISL_633617, EPI_ISL_633618, EPI_ISL_633619, EPI_ISL_633620, EPI_ISL_633621, EPI_ISL_633622, EPI_ISL_633623, EPI_ISL_633624, EPI_ISL_633625, EPI_ISL_633626, EPI_ISL_633627, EPI_ISL_633628, EPI_ISL_633629, EPI_ISL_633631, EPI_ISL_633632, EPI_ISL_633633, EPI_ISL_633634, EPI_ISL_633635, EPI_ISL_633636, EPI_ISL_633637, EPI_ISL_633638, EPI_ISL_633639, EPI_ISL_633640, EPI_ISL_633641, EPI_ISL_633642, EPI_ISL_633643, EPI_ISL_633644, EPI_ISL_633645, EPI_ISL_633646, EPI_ISL_633647, EPI_ISL_633648, EPI_ISL_633649, EPI_ISL_633650, EPI_ISL_633651, EPI_ISL_633652 | | | |
| see above | Lighthouse Lab in Glasgow | Wellcome Sanger Institute for the COVID-19 Genomics UK (COG-UK) consortium | Harper VanSteenhouse, Yumi Kasai, David Gray, Carol Clugston, Anna Dominiczak and Alex Alderton, Roberto Amato, Sonia Goncalves, Ewan Harrison, David K. Jackson, Ian Johnston, Dominic Kwiatkowski, Cordelia Langford, John Sillitoe on behalf of the Wellcome Sanger Institute COVID-19 Surveillance Team |
| EPI_ISL_633653 | Lighthouse Lab in Glasgow | Wellcome Sanger Institute for the COVID-19 Genomics UK (COG-UK) Consortium | Harper VanSteenhouse, Yumi Kasai, David Gray, Carol Clugston, Anna Dominiczak and Alex Alderton, Roberto Amato, Sonia Goncalves, Ewan Harrison, David K. Jackson, Ian Johnston, Dominic Kwiatkowski, Cordelia Langford, John Sillitoe on behalf of the Wellcome Sanger Institute COVID-19 Surveillance Team |
| EPI_ISL_633654, EPI_ISL_633656, EPI_ISL_633657, EPI_ISL_633659, EPI_ISL_633660, EPI_ISL_633661, EPI_ISL_633662, EPI_ISL_633663, EPI_ISL_633664, EPI_ISL_633665, EPI_ISL_633666, EPI_ISL_633667, EPI_ISL_633668, EPI_ISL_633669, EPI_ISL_633670, EPI_ISL_633671, EPI_ISL_633672, EPI_ISL_633673, | | | |

| | | | |
|---|---|---|---|
| EPI_ISL_633674, EPI_ISL_633675, EPI_ISL_633677, EPI_ISL_633678, EPI_ISL_633679, EPI_ISL_633680, EPI_ISL_633681, EPI_ISL_633682, EPI_ISL_633684, EPI_ISL_633685, EPI_ISL_633686, EPI_ISL_633687, EPI_ISL_633688, EPI_ISL_633689, EPI_ISL_633690, EPI_ISL_633691, EPI_ISL_633692, EPI_ISL_633693, EPI_ISL_633696, EPI_ISL_633697, EPI_ISL_633701, EPI_ISL_633702, EPI_ISL_633703, EPI_ISL_633704, EPI_ISL_633708, EPI_ISL_633709, EPI_ISL_633710, EPI_ISL_633711, EPI_ISL_633713, EPI_ISL_633714, EPI_ISL_633715, EPI_ISL_633717, EPI_ISL_633718, EPI_ISL_633719, EPI_ISL_633720, EPI_ISL_633721, EPI_ISL_633722, EPI_ISL_633724, EPI_ISL_633725, EPI_ISL_633726, EPI_ISL_633727, EPI_ISL_633728, EPI_ISL_633730, EPI_ISL_633732, EPI_ISL_633733, EPI_ISL_633734, EPI_ISL_633735, EPI_ISL_633737, EPI_ISL_633739, EPI_ISL_633740, EPI_ISL_633741, EPI_ISL_633742, EPI_ISL_633743, EPI_ISL_633745, EPI_ISL_633746, EPI_ISL_633747, EPI_ISL_633748, EPI_ISL_633749, EPI_ISL_633750, EPI_ISL_633751, EPI_ISL_633752, EPI_ISL_633753, EPI_ISL_633754, EPI_ISL_633755, EPI_ISL_633759 | | | |
| see above | Lighthouse Lab in Glasgow | Wellcome Sanger Institute for the COVID-19 Genomics UK (COG-UK) consortium | Harper VanSteenhouse, Yumi Kasai, David Gray, Carol Clugston, Anna Dominiczak and Alex Alderton, Roberto Amato, Sonia Goncalves, Ewan Harrison, David K. Jackson, Ian Johnston, Dominic Kwiatkowski, Cordelia Langford, John Sillitoe on behalf of the Wellcome Sanger Institute COVID-19 Surveillance Team |
| EPI_ISL_633760 | Lighthouse Lab in Glasgow | Wellcome Sanger Institute for the COVID-19 Genomics UK (COG-UK) Consortium | Harper VanSteenhouse, Yumi Kasai, David Gray, Carol Clugston, Anna Dominiczak and Alex Alderton, Roberto Amato, Sonia Goncalves, Ewan Harrison, David K. Jackson, Ian Johnston, Dominic Kwiatkowski, Cordelia Langford, John Sillitoe on behalf of the Wellcome Sanger Institute COVID-19 Surveillance Team |
| EPI_ISL_633761, EPI_ISL_633762, EPI_ISL_633763, EPI_ISL_633768, EPI_ISL_633769, EPI_ISL_633770, EPI_ISL_633775, EPI_ISL_633776, EPI_ISL_633777, EPI_ISL_633778, EPI_ISL_633779, EPI_ISL_633780, EPI_ISL_633781 | | | |
| see above | Lighthouse Lab in Glasgow | Wellcome Sanger Institute for the COVID-19 Genomics UK (COG-UK) consortium | Harper VanSteenhouse, Yumi Kasai, David Gray, Carol Clugston, Anna Dominiczak and Alex Alderton, Roberto Amato, Sonia Goncalves, Ewan Harrison, David K. Jackson, Ian Johnston, Dominic Kwiatkowski, Cordelia Langford, John Sillitoe on behalf of the Wellcome Sanger Institute COVID-19 Surveillance Team |
| EPI_ISL_633782 | Lighthouse Lab in Glasgow | Wellcome Sanger Institute for the COVID-19 Genomics UK (COG-UK) Consortium | Harper VanSteenhouse, Yumi Kasai, David Gray, Carol Clugston, Anna Dominiczak and Alex Alderton, Roberto Amato, Sonia Goncalves, Ewan Harrison, David K. Jackson, Ian Johnston, Dominic Kwiatkowski, Cordelia Langford, John Sillitoe on behalf of the Wellcome Sanger Institute COVID-19 Surveillance Team |
| EPI_ISL_633783, EPI_ISL_633784, EPI_ISL_633785, EPI_ISL_633786, EPI_ISL_633789, EPI_ISL_633790, EPI_ISL_633792, EPI_ISL_633793, EPI_ISL_633794, EPI_ISL_633795, EPI_ISL_633796, EPI_ISL_633797, EPI_ISL_633801, EPI_ISL_633802, EPI_ISL_633803, EPI_ISL_633805, EPI_ISL_633806, EPI_ISL_633808, EPI_ISL_633809, EPI_ISL_633810, EPI_ISL_633811, EPI_ISL_633812, EPI_ISL_633813, EPI_ISL_633814, EPI_ISL_633818, EPI_ISL_633819, EPI_ISL_633822, EPI_ISL_633823, EPI_ISL_633824, EPI_ISL_633825, EPI_ISL_633829, EPI_ISL_633830, EPI_ISL_633831, EPI_ISL_633832, EPI_ISL_633834, EPI_ISL_633836, EPI_ISL_633838, EPI_ISL_633840, EPI_ISL_633843, EPI_ISL_633844, EPI_ISL_633848, EPI_ISL_633849, EPI_ISL_633850, EPI_ISL_633851, EPI_ISL_633852, EPI_ISL_633854, EPI_ISL_633855, EPI_ISL_633856, EPI_ISL_633857, EPI_ISL_633858, EPI_ISL_633859, EPI_ISL_633860, EPI_ISL_633861, EPI_ISL_633862, EPI_ISL_633863, EPI_ISL_633864, EPI_ISL_633868, EPI_ISL_633869, EPI_ISL_633870, EPI_ISL_633871, EPI_ISL_633873, EPI_ISL_633874, EPI_ISL_633875, EPI_ISL_633878, EPI_ISL_633879, EPI_ISL_633880, EPI_ISL_633881, EPI_ISL_633883, EPI_ISL_633885, EPI_ISL_633887, EPI_ISL_633888, EPI_ISL_633889, EPI_ISL_633890, EPI_ISL_633892, EPI_ISL_633894, EPI_ISL_633895, EPI_ISL_633897, EPI_ISL_633899, EPI_ISL_633900, EPI_ISL_633901, EPI_ISL_633902, EPI_ISL_633903, EPI_ISL_633904, EPI_ISL_633905, EPI_ISL_633906, EPI_ISL_633907, EPI_ISL_633909, EPI_ISL_633911, EPI_ISL_633912, EPI_ISL_633913, EPI_ISL_633915, EPI_ISL_633916, EPI_ISL_633917, EPI_ISL_633919, EPI_ISL_633920, EPI_ISL_633922, EPI_ISL_633923, EPI_ISL_633925, EPI_ISL_633926, EPI_ISL_633927, EPI_ISL_633928, EPI_ISL_633931, EPI_ISL_633932, EPI_ISL_633933, EPI_ISL_633934, EPI_ISL_633935, EPI_ISL_633936, EPI_ISL_633938, EPI_ISL_633941, EPI_ISL_633942, EPI_ISL_633943, EPI_ISL_633944, EPI_ISL_633945, EPI_ISL_633946, EPI_ISL_633949, EPI_ISL_633950, EPI_ISL_633951, EPI_ISL_633952, EPI_ISL_633953, EPI_ISL_633954, EPI_ISL_633956, EPI_ISL_633957, EPI_ISL_633959, EPI_ISL_633960, EPI_ISL_633961, EPI_ISL_633963, EPI_ISL_633964, EPI_ISL_633965, EPI_ISL_633966, EPI_ISL_633968, EPI_ISL_633969, EPI_ISL_633970, EPI_ISL_633971, EPI_ISL_633972, EPI_ISL_633973, EPI_ISL_633974, EPI_ISL_633975, EPI_ISL_633976, EPI_ISL_633977, EPI_ISL_633978, EPI_ISL_633981, EPI_ISL_633982, EPI_ISL_633983, EPI_ISL_633986, EPI_ISL_633987, EPI_ISL_633988, EPI_ISL_633989, EPI_ISL_633991, EPI_ISL_633992, EPI_ISL_633993, EPI_ISL_633995, EPI_ISL_633998, EPI_ISL_633999, EPI_ISL_634000, EPI_ISL_634001, EPI_ISL_634002, EPI_ISL_634004, EPI_ISL_634005, EPI_ISL_634006, EPI_ISL_634007, EPI_ISL_634009, EPI_ISL_634010, EPI_ISL_634011, EPI_ISL_634012, EPI_ISL_634013, EPI_ISL_634014, EPI_ISL_634015, EPI_ISL_634016, EPI_ISL_634017, EPI_ISL_634018, EPI_ISL_634020, EPI_ISL_634021, EPI_ISL_634022, EPI_ISL_634028, EPI_ISL_634029, EPI_ISL_634030, EPI_ISL_634031, EPI_ISL_634032, EPI_ISL_634033, EPI_ISL_634034, EPI_ISL_634035, EPI_ISL_634037, EPI_ISL_634038, EPI_ISL_634039, EPI_ISL_634040, EPI_ISL_634041, EPI_ISL_634042, EPI_ISL_634043, EPI_ISL_634044, EPI_ISL_634046, EPI_ISL_634047, EPI_ISL_634048, EPI_ISL_634049, EPI_ISL_634050, EPI_ISL_634051, EPI_ISL_634052, EPI_ISL_634053, EPI_ISL_634054, EPI_ISL_634055, EPI_ISL_634057, EPI_ISL_634058, EPI_ISL_634059, EPI_ISL_634060, EPI_ISL_634061, EPI_ISL_634063, EPI_ISL_634065, EPI_ISL_634067, EPI_ISL_634068, EPI_ISL_634069, EPI_ISL_634070, EPI_ISL_634071, EPI_ISL_634072, EPI_ISL_634074, EPI_ISL_634075, EPI_ISL_634076, EPI_ISL_634077, EPI_ISL_634078, EPI_ISL_634079, EPI_ISL_634080, EPI_ISL_634081, EPI_ISL_634082, EPI_ISL_634083, EPI_ISL_634084, EPI_ISL_634087, EPI_ISL_634088, EPI_ISL_634089, EPI_ISL_634092, EPI_ISL_634096, EPI_ISL_634097, EPI_ISL_634098, EPI_ISL_634099, EPI_ISL_634100, EPI_ISL_634101, EPI_ISL_634102, EPI_ISL_634104, EPI_ISL_634105, EPI_ISL_634107, EPI_ISL_634108, EPI_ISL_634109, EPI_ISL_634110, EPI_ISL_634113, EPI_ISL_634114, EPI_ISL_634115, EPI_ISL_634116, EPI_ISL_634117, EPI_ISL_634118, EPI_ISL_634119, EPI_ISL_634121, EPI_ISL_634122, EPI_ISL_634123, EPI_ISL_634124, EPI_ISL_634125, EPI_ISL_634127, EPI_ISL_634128, EPI_ISL_634129, EPI_ISL_634130, EPI_ISL_634131, EPI_ISL_634132, EPI_ISL_634134, EPI_ISL_634135, EPI_ISL_634136, EPI_ISL_634137, EPI_ISL_634138, EPI_ISL_634139, EPI_ISL_634140, EPI_ISL_634141, EPI_ISL_634142, EPI_ISL_634145, EPI_ISL_634148, EPI_ISL_634150, EPI_ISL_634152, EPI_ISL_634153, EPI_ISL_634154 | | | |
| see above | Lighthouse Lab in Glasgow | Wellcome Sanger Institute for the COVID-19 Genomics UK (COG-UK) consortium | Harper VanSteenhouse, Yumi Kasai, David Gray, Carol Clugston, Anna Dominiczak and Alex Alderton, Roberto Amato, Sonia Goncalves, Ewan Harrison, David K. Jackson, Ian Johnston, Dominic Kwiatkowski, Cordelia Langford, John Sillitoe on behalf of the Wellcome Sanger Institute COVID-19 Surveillance Team |
| EPI_ISL_634156 | Lighthouse Lab in Glasgow | Wellcome Sanger Institute for the COVID-19 Genomics UK (COG-UK) Consortium | Harper VanSteenhouse, Yumi Kasai, David Gray, Carol Clugston, Anna Dominiczak and Alex Alderton, Roberto Amato, Sonia Goncalves, Ewan Harrison, David K. Jackson, Ian Johnston, Dominic Kwiatkowski, Cordelia Langford, John Sillitoe on behalf of the Wellcome Sanger Institute COVID-19 Surveillance Team |
| EPI_ISL_634158, EPI_ISL_634159, EPI_ISL_634160, EPI_ISL_634161, EPI_ISL_634163, EPI_ISL_634164, EPI_ISL_634165, EPI_ISL_634166, EPI_ISL_634168, EPI_ISL_634169, EPI_ISL_634170, EPI_ISL_634171, EPI_ISL_634174, EPI_ISL_634175, EPI_ISL_634177, EPI_ISL_634178, EPI_ISL_634180, EPI_ISL_634181, EPI_ISL_634182, EPI_ISL_634183, EPI_ISL_634184, EPI_ISL_634186, EPI_ISL_634187, EPI_ISL_634188, EPI_ISL_634189, EPI_ISL_634190, EPI_ISL_634191, EPI_ISL_634194, EPI_ISL_634195, EPI_ISL_634196, EPI_ISL_634198, EPI_ISL_634199, EPI_ISL_634200, EPI_ISL_634201, EPI_ISL_634205, EPI_ISL_634206, EPI_ISL_634208, EPI_ISL_634210, EPI_ISL_634211, EPI_ISL_634212, EPI_ISL_634215, EPI_ISL_634216, EPI_ISL_634217, EPI_ISL_634218, EPI_ISL_634219, EPI_ISL_634220, EPI_ISL_634221, EPI_ISL_634223, EPI_ISL_634225, EPI_ISL_634226, EPI_ISL_634227, EPI_ISL_634229, EPI_ISL_634230, EPI_ISL_634231, EPI_ISL_634232, EPI_ISL_634233, EPI_ISL_634234, EPI_ISL_634235, EPI_ISL_634237, EPI_ISL_634238, EPI_ISL_634239, EPI_ISL_634240, EPI_ISL_634241, EPI_ISL_634242, EPI_ISL_634243, EPI_ISL_634244, EPI_ISL_634246, EPI_ISL_634247, EPI_ISL_634248, EPI_ISL_634250, EPI_ISL_634251, EPI_ISL_634253, EPI_ISL_634254, EPI_ISL_634255, EPI_ISL_634520, EPI_ISL_634521, EPI_ISL_634522, EPI_ISL_634523, EPI_ISL_634525, EPI_ISL_634526, EPI_ISL_634527, EPI_ISL_634529, EPI_ISL_634531, EPI_ISL_634533, EPI_ISL_634534, EPI_ISL_634535, EPI_ISL_634536, EPI_ISL_634537, EPI_ISL_634538, EPI_ISL_634539, EPI_ISL_634540, EPI_ISL_634542, EPI_ISL_634543, EPI_ISL_634544, EPI_ISL_634546, EPI_ISL_634548, EPI_ISL_634549, EPI_ISL_634550, EPI_ISL_634551, EPI_ISL_634552, EPI_ISL_634553, EPI_ISL_634554, EPI_ISL_634555, EPI_ISL_634556, EPI_ISL_634557, EPI_ISL_634558, EPI_ISL_634560, EPI_ISL_634561, EPI_ISL_634562, EPI_ISL_634563, EPI_ISL_634564, EPI_ISL_634565, EPI_ISL_634566, EPI_ISL_634568, EPI_ISL_634569, EPI_ISL_634570, EPI_ISL_634571, EPI_ISL_634572, EPI_ISL_634573, EPI_ISL_634574, EPI_ISL_634575, EPI_ISL_634576, EPI_ISL_634577, EPI_ISL_634578, EPI_ISL_634579, EPI_ISL_634580, EPI_ISL_634581, EPI_ISL_634584, EPI_ISL_634585, EPI_ISL_634586, EPI_ISL_634587, EPI_ISL_634588, EPI_ISL_634589, EPI_ISL_634590, EPI_ISL_634591, EPI_ISL_634592, EPI_ISL_634593, EPI_ISL_634594, EPI_ISL_634595, EPI_ISL_634596, EPI_ISL_634597, EPI_ISL_634599 | | | |
| see above | Lighthouse Lab in Glasgow | Wellcome Sanger Institute for the COVID-19 Genomics UK (COG-UK) consortium | Harper VanSteenhouse, Yumi Kasai, David Gray, Carol Clugston, Anna Dominiczak and Alex Alderton, Roberto Amato, Sonia Goncalves, Ewan Harrison, David K. Jackson, Ian Johnston, Dominic Kwiatkowski, Cordelia Langford, John Sillitoe on behalf of the Wellcome Sanger Institute COVID-19 Surveillance Team |
| EPI_ISL_634600, EPI_ISL_634601, EPI_ISL_634602, EPI_ISL_634603, EPI_ISL_634604, EPI_ISL_634605, EPI_ISL_634606, EPI_ISL_634607, EPI_ISL_634608, EPI_ISL_634609, EPI_ISL_634610, EPI_ISL_634611, EPI_ISL_634613, EPI_ISL_634614, EPI_ISL_634615, EPI_ISL_634616, EPI_ISL_634617, EPI_ISL_634618, EPI_ISL_634620, EPI_ISL_634621, EPI_ISL_634623, EPI_ISL_634624, EPI_ISL_634625, EPI_ISL_634626, EPI_ISL_634627, EPI_ISL_634628, EPI_ISL_634629, EPI_ISL_634630, EPI_ISL_634633, EPI_ISL_634634, EPI_ISL_634635, EPI_ISL_634636, EPI_ISL_634637, EPI_ISL_634638, EPI_ISL_634639, EPI_ISL_634641, EPI_ISL_634642, EPI_ISL_634643, EPI_ISL_634644, EPI_ISL_634645, EPI_ISL_634646, EPI_ISL_634647, EPI_ISL_634648, EPI_ISL_634649, EPI_ISL_634650, EPI_ISL_634651, EPI_ISL_634652, EPI_ISL_634653, EPI_ISL_634655, EPI_ISL_634656, EPI_ISL_634657, EPI_ISL_634659, EPI_ISL_634660, EPI_ISL_634661, EPI_ISL_634662, EPI_ISL_634663, EPI_ISL_634664, EPI_ISL_634665, EPI_ISL_634666, EPI_ISL_634667, EPI_ISL_634669, EPI_ISL_634670, EPI_ISL_634672, EPI_ISL_634673, EPI_ISL_634674, EPI_ISL_634675, EPI_ISL_634676, EPI_ISL_634677 | | | |
| see above | Lighthouse Lab in Glasgow | Wellcome Sanger Institute for the COVID-19 Genomics UK (COG-UK) consortium | Harper VanSteenhouse, Yumi Kasai, David Gray, Carol Clugston, Anna Dominiczak and Alex Alderton, Roberto Amato, Sonia Goncalves, Ewan Harrison, David K. Jackson, Ian Johnston, Dominic Kwiatkowski, Cordelia Langford, John Sillitoe on behalf of the Wellcome Sanger Institute COVID-19 Surveillance Team (http://www.sanger.ac.uk/covid-team) |
| EPI_ISL_634678 | Lighthouse Lab in Glasgow | Wellcome Sanger Institute for the COVID-19 Genomics UK (COG-UK) Consortium | Harper VanSteenhouse, Yumi Kasai, David Gray, Carol Clugston, Anna Dominiczak and Alex Alderton, Roberto Amato, Sonia Goncalves, Ewan Harrison, David K. Jackson, Ian Johnston, Dominic Kwiatkowski, Cordelia Langford, John Sillitoe on behalf of the Wellcome Sanger Institute COVID-19 Surveillance Team (http://www.sanger.ac.uk/covid-team) |
| EPI_ISL_634679, EPI_ISL_634680, EPI_ISL_634681, EPI_ISL_634682, EPI_ISL_634684, EPI_ISL_634685, EPI_ISL_634686, EPI_ISL_634687, EPI_ISL_634688, EPI_ISL_634690, EPI_ISL_634691, EPI_ISL_634692, EPI_ISL_634693, EPI_ISL_634694, EPI_ISL_634695, EPI_ISL_634696, EPI_ISL_634698, EPI_ISL_634699, EPI_ISL_634700, EPI_ISL_634701, EPI_ISL_634702, EPI_ISL_634703, EPI_ISL_634704, EPI_ISL_634705, EPI_ISL_634706, EPI_ISL_634707, EPI_ISL_634708, EPI_ISL_634709, EPI_ISL_634710, EPI_ISL_634711, EPI_ISL_634712, EPI_ISL_634714, EPI_ISL_634715, EPI_ISL_634716, EPI_ISL_634717, EPI_ISL_634719, EPI_ISL_634720, EPI_ISL_634721, EPI_ISL_634722, EPI_ISL_634723, EPI_ISL_634724, EPI_ISL_634725, EPI_ISL_634726, EPI_ISL_634727, EPI_ISL_634728, EPI_ISL_634729, EPI_ISL_634730, EPI_ISL_634731, EPI_ISL_634732, EPI_ISL_634733, EPI_ISL_634734, EPI_ISL_634735, EPI_ISL_634737, EPI_ISL_634738, EPI_ISL_634740, EPI_ISL_634741, EPI_ISL_634742, EPI_ISL_634743, EPI_ISL_634746, EPI_ISL_634747, EPI_ISL_634748, EPI_ISL_634749, EPI_ISL_634751, EPI_ISL_634753 | | | |
| see above | Lighthouse Lab in Glasgow | Wellcome Sanger Institute for the COVID-19 Genomics UK (COG-UK) consortium | Harper VanSteenhouse, Yumi Kasai, David Gray, Carol Clugston, Anna Dominiczak and Alex Alderton, Roberto Amato, Sonia Goncalves, Ewan Harrison, David K. Jackson, Ian Johnston, Dominic Kwiatkowski, Cordelia Langford, John Sillitoe on behalf of the Wellcome Sanger Institute COVID-19 Surveillance Team (http://www.sanger.ac.uk/covid-team) |
| EPI_ISL_634858, EPI_ISL_634859, EPI_ISL_634860, EPI_ISL_634861, EPI_ISL_634862, EPI_ISL_634863, EPI_ISL_634866, EPI_ISL_634867, EPI_ISL_634868, EPI_ISL_634869, EPI_ISL_634870, EPI_ISL_634871, EPI_ISL_634872, EPI_ISL_634873, EPI_ISL_634874, EPI_ISL_634875 | | | |
| see above | Minnesota Department of Health, Public Health Laboratory | Minnesota Department of Health, Public Health Laboratory | Matt Plumb, Jacob Garfin, Alexandra Lorentz, and Xiong Wang |
| EPI_ISL_635074, EPI_ISL_635075, EPI_ISL_635076, EPI_ISL_635077, EPI_ISL_635078, EPI_ISL_635079 | Norwegian Institute of Public Health, Department of Virology | Norwegian Institute of Public Health, Department of Virology | Kathrine Stene-Johansen, Kamilla Heddeland Instefjord, Hilde Elshaug, Marie Paulsen Madsen, Rasmus Riis Kopperud, Hilde Vollan, Karoline Bragstad, Olav Hungnes |
| EPI_ISL_635080, EPI_ISL_635081, EPI_ISL_635082, EPI_ISL_635084 | Haukeland University Hospital, Department of Medical Microbiology | Norwegian Institute of Public Health, Department of Virology | Kathrine Stene-Johansen, Kamilla Heddeland Instefjord, Hilde Elshaug, Marie Paulsen Madsen, Rasmus Riis Kopperud, Hilde Vollan, Karoline Bragstad, Olav Hungnes |
| EPI_ISL_635086, EPI_ISL_635087 | Furst Medical Laboratory | Norwegian Institute of Public Health, Department of Virology | Kathrine Stene-Johansen, Kamilla Heddeland Instefjord, Hilde Elshaug, Marie Paulsen Madsen, Rasmus Riis Kopperud, Hilde Vollan, Karoline Bragstad, Olav Hungnes |
| EPI_ISL_636605, EPI_ISL_636606, EPI_ISL_636608, EPI_ISL_636609, EPI_ISL_636611, EPI_ISL_636612, EPI_ISL_636613, EPI_ISL_636614, EPI_ISL_636616, EPI_ISL_636618, EPI_ISL_636619, EPI_ISL_636620, EPI_ISL_636621, EPI_ISL_636622, EPI_ISL_636623, EPI_ISL_636624, EPI_ISL_636625, EPI_ISL_636626, | | | |

| | | | |
|---|---|---|---|
| EPI_ISL_636628, EPI_ISL_636629, EPI_ISL_636630, EPI_ISL_636631, EPI_ISL_636632, EPI_ISL_636633, EPI_ISL_636634, EPI_ISL_636635, EPI_ISL_636636, EPI_ISL_636637, EPI_ISL_636638, EPI_ISL_636639, EPI_ISL_636640, EPI_ISL_636641, EPI_ISL_636642, EPI_ISL_636643, EPI_ISL_636644, EPI_ISL_636645, EPI_ISL_636646, EPI_ISL_636647, EPI_ISL_636648, EPI_ISL_636650, EPI_ISL_636651, EPI_ISL_636652, EPI_ISL_636653, EPI_ISL_636654, EPI_ISL_636655, EPI_ISL_636656, EPI_ISL_636658, EPI_ISL_636659, EPI_ISL_636660, EPI_ISL_636661, EPI_ISL_636662, EPI_ISL_636663, EPI_ISL_636664, EPI_ISL_636665, EPI_ISL_636666, EPI_ISL_636667, EPI_ISL_636668, EPI_ISL_636669, EPI_ISL_636670, EPI_ISL_636671, EPI_ISL_636672, EPI_ISL_636673, EPI_ISL_636674, EPI_ISL_636675, EPI_ISL_636676, EPI_ISL_636677, EPI_ISL_636678, EPI_ISL_636679, EPI_ISL_636680, EPI_ISL_636681, EPI_ISL_636682, EPI_ISL_636683, EPI_ISL_636684, EPI_ISL_636685 | | | |
| see above | Department of Clinical Microbiology | GIGA Medical Genomics | Keith Durkin, Maria Artesi, Sébastien Bontems, Raphaël Boreux, Bouchra Boujemla, Cécile Meex, Pierrette Melin, Marie-Pierre Hayette, Vincent Bours |
| EPI_ISL_636732, EPI_ISL_636733 | Respiratory Virus Unit, Microbiology Services Colindale, Public Health England | Respiratory Virus Unit, Microbiology Services Colindale, Public Health England | PHE Covid Sequencing Team |
| EPI_ISL_637021 | Department of Infectious Diseases and Immunology, National Hospital Organization Nagoya Medical Center | Clinical Research Center, National Hospital Organization Nagoya Medical Center | Yoshihiro Nakata, Hirotaka Ode, Mai Kubota, Masakazu Matsuda, Kazuhiro Matsuoka, Miho Nakasuji, Mikiko Mori, Mayumi Imahashi, Yoshiyuki Yokomaku, Yasumasa Iwatani |
| EPI_ISL_637150, EPI_ISL_637151, EPI_ISL_637152, EPI_ISL_637158, EPI_ISL_637159, EPI_ISL_637160, EPI_ISL_637161, EPI_ISL_637162, EPI_ISL_637163, EPI_ISL_637164, EPI_ISL_637165, EPI_ISL_637166, EPI_ISL_637171, EPI_ISL_637173, EPI_ISL_637175, EPI_ISL_637178, EPI_ISL_637179 | | | |
| see above | Respiratory Virus Unit, Microbiology Services Colindale, Public Health England | COVID-19 Genomics UK (COG-UK) Consortium | PHE Covid Sequencing Team |
| EPI_ISL_637274 | Liverpool Clinical Laboratories | COVID-19 Genomics UK (COG-UK) Consortium | Sam Haldenby, Anita Lucaci, Steve Paterson, Julian Hiscox, Alistair Darby, M Almsaud, A Alrezaihi, Muhannad Alruwaili, Stuart D Armstrong, Jones Benjamin, Eleanor G Bentley, Anu Chawla, Jordan J Clark, Angela Cowell, Richard Eccles, Isabel García-Dorival, Matthew Gemmell, Alessandro Gerada, PKF Gilmore, Richard Gregory, Ximeng Han, Catherine Hartley, Margaret Hughes, Miren Iturriza-Gomara, James Johnson, L Luu, Jenifer Manson, Charlotte Nelson, Elaine O'Toole, Cassie Olateju, Rebekah Penrice-Randal , Lucille Rainbow, N.P Randle, Trevor Ian Robinson, Parul Sharma, Ghada T Shawli, James P Stewart, Neil Swainston, Ecaterina Vamos, Joanne Watts, Mark Whitehead |
| EPI_ISL_637280 | Northumbria University / South Tees Hospitals NHS Foundation Trust / North Cumbria Integrated Care NHS Foundation Trust / North Tees and Hartlepool NHS Foundation Trust / Newcastle Hospitals NHS Foundation Trust | COVID-19 Genomics UK (COG-UK) Consortium | Darren L Smith,Andrew Nelson,Matthew Bashton,Greg R Young,Joshua Loh,John Allan,Mohammad A Tariq,Giles S Holt,Gary Black,Wen C Yew,Lynn Dover,Paul Baker,Steve Liggett,Sarah Essex,Jane Greenaway,Debra Padgett,Clive Graham,Garren Scott,Edward Barton,Emma Swindells,Brendan Payne,Jennifer Collins,Yusri Taha,Gary Eltringham |
| EPI_ISL_637288 | Wales Specialist Virology Centre Sequencing lab: Pathogen Genomics Unit | COVID-19 Genomics UK (COG-UK) Consortium | Catherine Moore, Johnathan Evans, Laura Gifford, Malorie Perry, Simon Cottrell, Angela Marchbank, Alec Birchley, Alexander Adams, Amy Gaskin, Bree Gatica-Wilcox, Jason Coombes, Joel Southgate, Lauren Gilbert, Lee Graham, Nicole Pacchiarini, Sara Kumziene-Summerhayes, Sarah Taylor, Sophie Jones, Sara Rey, Matthew Bull, Joanne Watkins, Sally Corden, Tom Connor |
| EPI_ISL_637291 | Virology Department, Sheffield Teaching Hospitals NHS Foundation Trust/Department of Infection, Immunity and Cardiovascular Disease, The Medical School, University of Sheffield | COVID-19 Genomics UK (COG-UK) Consortium | Thushan de Silva, Matthew Parker, Nikki Smith, Adri Angyal, Rebecca Brown, Luke Green, Rachel Tucker, Paul Parsons, Danielle Groves, Katie Johnson, Laura Carrilero, Alex Keeley, Dave Partridge, Matthew Wyles, Benjamin Lindsey, Mehmet Yavuz, Mohammad Raza, Cariad Evans |
| EPI_ISL_637296 | Liverpool Clinical Laboratories | COVID-19 Genomics UK (COG-UK) Consortium | Sam Haldenby, Anita Lucaci, Steve Paterson, Julian Hiscox, Alistair Darby, M Almsaud, A Alrezaihi, Muhannad Alruwaili, Stuart D Armstrong, Jones Benjamin, Eleanor G Bentley, Anu Chawla, Jordan J Clark, Angela Cowell, Richard Eccles, Isabel García-Dorival, Matthew Gemmell, Alessandro Gerada, PKF Gilmore, Richard Gregory, Ximeng Han, Catherine Hartley, Margaret Hughes, Miren Iturriza-Gomara, James Johnson, L Luu, Jenifer Manson, Charlotte Nelson, Elaine O'Toole, Cassie Olateju, Rebekah Penrice-Randal , Lucille Rainbow, N.P Randle, Trevor Ian Robinson, Parul Sharma, Ghada T Shawli, James P Stewart, Neil Swainston, Ecaterina Vamos, Joanne Watts, Mark Whitehead |
| EPI_ISL_637297 | Virology Department, Sheffield Teaching Hospitals NHS Foundation Trust/Department of Infection, Immunity and Cardiovascular Disease, The Medical School, University of Sheffield | COVID-19 Genomics UK (COG-UK) Consortium | Thushan de Silva, Matthew Parker, Nikki Smith, Adri Angyal, Rebecca Brown, Luke Green, Rachel Tucker, Paul Parsons, Danielle Groves, Katie Johnson, Laura Carrilero, Alex Keeley, Dave Partridge, Matthew Wyles, Benjamin Lindsey, Mehmet Yavuz, Mohammad Raza, Cariad Evans |
| EPI_ISL_637313 | Liverpool Clinical Laboratories | COVID-19 Genomics UK (COG-UK) Consortium | Sam Haldenby, Anita Lucaci, Steve Paterson, Julian Hiscox, Alistair Darby, M Almsaud, A Alrezaihi, Muhannad Alruwaili, Stuart D Armstrong, Jones Benjamin, Eleanor G Bentley, Anu Chawla, Jordan J Clark, Angela Cowell, Richard Eccles, Isabel García-Dorival, Matthew Gemmell, Alessandro Gerada, PKF Gilmore, Richard Gregory, Ximeng Han, Catherine Hartley, Margaret Hughes, Miren Iturriza-Gomara, James Johnson, L Luu, Jenifer Manson, Charlotte Nelson, Elaine O'Toole, Cassie Olateju, Rebekah Penrice-Randal , Lucille Rainbow, N.P Randle, Trevor Ian Robinson, Parul Sharma, Ghada T Shawli, James P Stewart, Neil Swainston, Ecaterina Vamos, Joanne Watts, Mark Whitehead |
| EPI_ISL_637330 | Wales Specialist Virology Centre Sequencing lab: Pathogen Genomics Unit | COVID-19 Genomics UK (COG-UK) Consortium | Catherine Moore, Johnathan Evans, Laura Gifford, Malorie Perry, Simon Cottrell, Angela Marchbank, Alec Birchley, Alexander Adams, Amy Gaskin, Bree Gatica-Wilcox, Jason Coombes, Joel Southgate, Lauren Gilbert, Lee Graham, Nicole Pacchiarini, Sara Kumziene-Summerhayes, Sarah Taylor, Sophie Jones, Sara Rey, Matthew Bull, Joanne Watkins, Sally Corden, Tom Connor |
| EPI_ISL_637337 | Liverpool Clinical Laboratories | COVID-19 Genomics UK (COG-UK) Consortium | Sam Haldenby, Anita Lucaci, Steve Paterson, Julian Hiscox, Alistair Darby, M Almsaud, A Alrezaihi, Muhannad Alruwaili, Stuart D Armstrong, Jones Benjamin, Eleanor G Bentley, Anu Chawla, Jordan J Clark, Angela Cowell, Richard Eccles, Isabel García-Dorival, Matthew Gemmell, Alessandro Gerada, PKF Gilmore, Richard Gregory, Ximeng Han, Catherine Hartley, Margaret Hughes, Miren Iturriza-Gomara, James Johnson, L Luu, Jenifer Manson, Charlotte Nelson, Elaine O'Toole, Cassie Olateju, Rebekah Penrice-Randal , Lucille Rainbow, N.P Randle, Trevor Ian Robinson, Parul Sharma, Ghada T Shawli, James P Stewart, Neil Swainston, Ecaterina Vamos, Joanne Watts, Mark Whitehead |
| EPI_ISL_637359 | Quadram Institute Bioscience | COVID-19 Genomics UK (COG-UK) Consortium | Dave J. Baker, Gemma L. Kay, Alp Aydin, Thanh Le-Viet, Steven Rudder, Ana P. Tedim, Anastasia Kolyva, Maria Diaz, Leonardo de Oliveira Martins, Nabil-Fareed Alikhan, Lizzie Meadows, Rachael Stanley, Ngozi Elumogo, Muhammed Yasir, Nicholas M. Thomson, Alexander J Trotter, Rachel Gilroy, Samuel Bloomfield, Claire Stuart, Andrew Bell, Reenesh Prakash, Samir Dervisevic, Alison E. Mather, John Wain, Mark Webber, Andrew J. Page, Justin O'Grady |
| EPI_ISL_637375 | Northumbria University / South Tees Hospitals NHS Foundation Trust / North Cumbria Integrated Care NHS Foundation Trust / North Tees and Hartlepool NHS Foundation Trust / Newcastle Hospitals NHS Foundation Trust | COVID-19 Genomics UK (COG-UK) Consortium | Darren L Smith,Andrew Nelson,Matthew Bashton,Greg R Young,Joshua Loh,John Allan,Mohammad A Tariq,Giles S Holt,Gary Black,Wen C Yew,Lynn Dover,Paul Baker,Steve Liggett,Sarah Essex,Jane Greenaway,Debra Padgett,Clive Graham,Garren Scott,Edward Barton,Emma Swindells,Brendan Payne,Jennifer Collins,Yusri Taha,Gary Eltringham |
| EPI_ISL_637385, EPI_ISL_637412, EPI_ISL_637422 | Wales Specialist Virology Centre Sequencing lab: Pathogen Genomics Unit | COVID-19 Genomics UK (COG-UK) Consortium | Catherine Moore, Johnathan Evans, Laura Gifford, Malorie Perry, Simon Cottrell, Angela Marchbank, Alec Birchley, Alexander Adams, Amy Gaskin, Bree Gatica-Wilcox, Jason Coombes, Joel Southgate, Lauren Gilbert, Lee Graham, Nicole Pacchiarini, Sara Kumziene-Summerhayes, Sarah Taylor, Sophie Jones, Sara Rey, Matthew Bull, Joanne Watkins, Sally Corden, Tom Connor |
| EPI_ISL_637446 | University College London, Great Ormond Street Hospital for Children NHS Foundation Trust, Imperial College Healthcare NHS Trust | COVID-19 Genomics UK (COG-UK) Consortium | Sergi Castellano, Rachel Williams, Mark Kristiansen, Paola Resende Silva, Sunando Roy, Tony Brooks, Helena Tutill, Paola Niola, Patricia Dyal, Charlotte Williams, Leysa Forrest, Yasmin Panchbhaya, Jacqueline Findlay, Samuel Weeks, Julianne Brown, Kathryn Harris, Paul Randell, James Price, Alison Holmes, Judith Breuer |
| EPI_ISL_637447 | Wales Specialist Virology Centre Sequencing lab: Pathogen Genomics Unit | COVID-19 Genomics UK (COG-UK) Consortium | Catherine Moore, Johnathan Evans, Laura Gifford, Malorie Perry, Simon Cottrell, Angela Marchbank, Alec Birchley, Alexander Adams, Amy Gaskin, Bree Gatica-Wilcox, Jason Coombes, Joel Southgate, Lauren Gilbert, Lee Graham, Nicole Pacchiarini, Sara Kumziene-Summerhayes, Sarah Taylor, Sophie Jones, Sara Rey, Matthew Bull, Joanne Watkins, Sally Corden, Tom Connor |
| EPI_ISL_637470 | Department of Pathology, University of Cambridge | COVID-19 Genomics UK (COG-UK) Consortium | Aminu S. Jahun, Yasmin Chaudhry, Grant Hall, Iliana Georgana, Myra Hosmillo, Martin D. Curran, Malte Pinckert, Surendra Parmar, Ian Goodfellow |
| EPI_ISL_637522, EPI_ISL_637523 | Northumbria University / South Tees Hospitals NHS Foundation Trust / North Cumbria Integrated Care NHS Foundation Trust / North Tees and Hartlepool NHS Foundation Trust / Newcastle Hospitals NHS Foundation Trust | COVID-19 Genomics UK (COG-UK) Consortium | Darren L Smith,Andrew Nelson,Matthew Bashton,Greg R Young,Joshua Loh,John Allan,Mohammad A Tariq,Giles S Holt,Gary Black,Wen C Yew,Lynn Dover,Paul Baker,Steve Liggett,Sarah Essex,Jane Greenaway,Debra Padgett,Clive Graham,Garren Scott,Edward Barton,Emma Swindells,Brendan Payne,Jennifer Collins,Yusri Taha,Gary Eltringham |
| EPI_ISL_637543 | Wales Specialist Virology Centre Sequencing lab: Pathogen Genomics Unit | COVID-19 Genomics UK (COG-UK) Consortium | Catherine Moore, Johnathan Evans, Laura Gifford, Malorie Perry, Simon Cottrell, Angela Marchbank, Alec Birchley, Alexander Adams, Amy Gaskin, Bree Gatica-Wilcox, Jason Coombes, Joel Southgate, Lauren Gilbert, Lee Graham, Nicole Pacchiarini, Sara Kumziene-Summerhayes, Sarah Taylor, Sophie Jones, Sara Rey, Matthew Bull, Joanne Watkins, Sally Corden, Tom Connor |
| EPI_ISL_637550 | Quadram Institute Bioscience | COVID-19 Genomics UK (COG-UK) Consortium | Dave J. Baker, Gemma L. Kay, Alp Aydin, Thanh Le-Viet, Steven Rudder, Ana P. Tedim, Anastasia Kolyva, Maria Diaz, Leonardo de Oliveira Martins, |

| | | | |
|---|---|---|---|
| | | | Nabil-Fareed Alikhan, Lizzie Meadows, Rachael Stanley, Ngozi Elumogo, Muhammed Yasir, Nicholas M. Thomson, Alexander J Trotter, Rachel Gilroy, Samuel Bloomfield, Claire Stuart, Andrew Bell, Reenesh Prakash, Samir Dervisevic, Alison E. Mather, John Wain, Mark Webber, Andrew J. Page, Justin O'Grady |
| EPI_ISL_637556 | Wales Specialist Virology Centre Sequencing lab: Pathogen Genomics Unit | COVID-19 Genomics UK (COG-UK) Consortium | Catherine Moore, Johnathan Evans, Laura Gifford, Malorie Perry, Simon Cottrell, Angela Marchbank, Alec Birchley, Alexander Adams, Amy Gaskin, Bree Gatica-Wilcox, Jason Coombes, Joel Southgate, Lauren Gilbert, Lee Graham, Nicole Pacchiarini, Sara Kumziene-Summerhayes, Sarah Taylor, Sophie Jones, Sara Rey, Matthew Bull, Joanne Watkins, Sally Corden, Tom Connor |
| EPI_ISL_637584 | Virology Department, Sheffield Teaching Hospitals NHS Foundation Trust/Department of Infection, Immunity and Cardiovascular Disease, The Medical School, University of Sheffield | COVID-19 Genomics UK (COG-UK) Consortium | Thushan de Silva, Matthew Parker, Nikki Smith, Adri Angyal, Rebecca Brown, Luke Green, Rachel Tucker, Paul Parsons, Danielle Groves, Katie Johnson, Laura Carrilero, Alex Keeley, Dave Partridge, Matthew Wyles, Benjamin Lindsey, Mehmet Yavuz, Mohammad Raza, Cariad Evans |
| EPI_ISL_637589 | Quadram Institute Bioscience | COVID-19 Genomics UK (COG-UK) Consortium | Dave J. Baker, Gemma L. Kay, Alp Aydin, Thanh Le-Viet, Steven Rudder, Ana P. Tedim, Anastasia Kolyva, Maria Diaz, Leonardo de Oliveira Martins, Nabil-Fareed Alikhan, Lizzie Meadows, Rachael Stanley, Ngozi Elumogo, Muhammed Yasir, Nicholas M. Thomson, Alexander J Trotter, Rachel Gilroy, Samuel Bloomfield, Claire Stuart, Andrew Bell, Reenesh Prakash, Samir Dervisevic, Alison E. Mather, John Wain, Mark Webber, Andrew J. Page, Justin O'Grady |
| EPI_ISL_637597, EPI_ISL_637603, EPI_ISL_637604 | Wales Specialist Virology Centre Sequencing lab: Pathogen Genomics Unit | COVID-19 Genomics UK (COG-UK) Consortium | Catherine Moore, Johnathan Evans, Laura Gifford, Malorie Perry, Simon Cottrell, Angela Marchbank, Alec Birchley, Alexander Adams, Amy Gaskin, Bree Gatica-Wilcox, Jason Coombes, Joel Southgate, Lauren Gilbert, Lee Graham, Nicole Pacchiarini, Sara Kumziene-Summerhayes, Sarah Taylor, Sophie Jones, Sara Rey, Matthew Bull, Joanne Watkins, Sally Corden, Tom Connor |
| EPI_ISL_637605 | Northumbria University / South Tees Hospitals NHS Foundation Trust / North Cumbria Integrated Care NHS Foundation Trust / North Tees and Hartlepool NHS Foundation Trust / Newcastle Hospitals NHS Foundation Trust | COVID-19 Genomics UK (COG-UK) Consortium | Darren L Smith,Andrew Nelson,Matthew Bashton,Greg R Young,Joshua Loh,John Allan,Mohammad A Tariq,Giles S Holt,Gary Black,Wen C Yew,Lynn Dover,Paul Baker,Steve Liggett,Sarah Essex,Jane Greenaway,Debra Padgett,Clive Graham,Garren Scott,Edward Barton,Emma Swindells,Brendan Payne,Jennifer Collins,Yusri Taha,Gary Eltringham |
| EPI_ISL_637607, EPI_ISL_637611, EPI_ISL_637612, EPI_ISL_637614, EPI_ISL_637622, EPI_ISL_637627 | Wales Specialist Virology Centre Sequencing lab: Pathogen Genomics Unit | COVID-19 Genomics UK (COG-UK) Consortium | Catherine Moore, Johnathan Evans, Laura Gifford, Malorie Perry, Simon Cottrell, Angela Marchbank, Alec Birchley, Alexander Adams, Amy Gaskin, Bree Gatica-Wilcox, Jason Coombes, Joel Southgate, Lauren Gilbert, Lee Graham, Nicole Pacchiarini, Sara Kumziene-Summerhayes, Sarah Taylor, Sophie Jones, Sara Rey, Matthew Bull, Joanne Watkins, Sally Corden, Tom Connor |
| EPI_ISL_637791, EPI_ISL_637792 | Northumbria University / South Tees Hospitals NHS Foundation Trust / North Cumbria Integrated Care NHS Foundation Trust / North Tees and Hartlepool NHS Foundation Trust / Newcastle Hospitals NHS Foundation Trust | COVID-19 Genomics UK (COG-UK) Consortium | Darren L Smith,Andrew Nelson,Matthew Bashton,Greg R Young,Joshua Loh,John Allan,Mohammad A Tariq,Giles S Holt,Gary Black,Wen C Yew,Lynn Dover,Paul Baker,Steve Liggett,Sarah Essex,Jane Greenaway,Debra Padgett,Clive Graham,Garren Scott,Edward Barton,Emma Swindells,Brendan Payne,Jennifer Collins,Yusri Taha,Gary Eltringham |
| EPI_ISL_637857, EPI_ISL_637864 | Quadram Institute Bioscience | COVID-19 Genomics UK (COG-UK) Consortium | Dave J. Baker, Gemma L. Kay, Alp Aydin, Thanh Le-Viet, Steven Rudder, Ana P. Tedim, Anastasia Kolyva, Maria Diaz, Leonardo de Oliveira Martins, Nabil-Fareed Alikhan, Lizzie Meadows, Rachael Stanley, Ngozi Elumogo, Muhammed Yasir, Nicholas M. Thomson, Alexander J Trotter, Rachel Gilroy, Samuel Bloomfield, Claire Stuart, Andrew Bell, Reenesh Prakash, Samir Dervisevic, Alison E. Mather, John Wain, Mark Webber, Andrew J. Page, Justin O'Grady |
| EPI_ISL_637870, EPI_ISL_637884, EPI_ISL_637885, EPI_ISL_637886, EPI_ISL_637896, EPI_ISL_637900, EPI_ISL_637904 | Wales Specialist Virology Centre Sequencing lab: Pathogen Genomics Unit | COVID-19 Genomics UK (COG-UK) Consortium | Catherine Moore, Johnathan Evans, Laura Gifford, Malorie Perry, Simon Cottrell, Angela Marchbank, Alec Birchley, Alexander Adams, Amy Gaskin, Bree Gatica-Wilcox, Jason Coombes, Joel Southgate, Lauren Gilbert, Lee Graham, Nicole Pacchiarini, Sara Kumziene-Summerhayes, Sarah Taylor, Sophie Jones, Sara Rey, Matthew Bull, Joanne Watkins, Sally Corden, Tom Connor |
| EPI_ISL_637908 | Liverpool Clinical Laboratories | COVID-19 Genomics UK (COG-UK) Consortium | Sam Haldenby, Anita Lucaci, Steve Paterson, Julian Hiscox, Alistair Darby, M Almsaud, A Alrezaihi, Muhannad Alruwaili, Stuart D Armstrong, Jones Benjamin, Eleanor G Bentley, Anu Chawla, Jordan J Clark, Angela Cowell, Richard Eccles, Isabel García-Dorival, Matthew Gemmell, Alessandro Gerada, PKF Gilmore, Richard Gregory, Ximeng Han, Catherine Hartley, Margaret Hughes, Miren Iturriza-Gomara, James Johnson, L Luu, Jenifer Manson, Charlotte Nelson, Elaine O'Toole, Cassie Olateju, Rebekah Penrice-Randal , Lucille Rainbow, N.P Randle, Trevor Ian Robinson, Parul Sharma, Ghada T Shawli, James P Stewart, Neil Swainston, Ecaterina Vamos, Joanne Watts, Mark Whitehead |
| EPI_ISL_637912 | University College London, Great Ormond Street Hospital for Children NHS Foundation Trust, Imperial College Healthcare NHS Trust | COVID-19 Genomics UK (COG-UK) Consortium | Sergi Castellano, Rachel Williams, Mark Kristiansen, Paola Resende Silva, Sunando Roy, Tony Brooks, Helena Tutill, Paola Niola, Patricia Dyal, Charlotte Williams, Leysa Forrest, Yasmin Panchbhaya, Jacqueline Findlay, Samuel Weeks, Julianne Brown, Kathryn Harris, Paul Randell, James Price, Alison Holmes, Judith Breuer |
| EPI_ISL_637924, EPI_ISL_637927 | Wales Specialist Virology Centre Sequencing lab: Pathogen Genomics Unit | COVID-19 Genomics UK (COG-UK) Consortium | Catherine Moore, Johnathan Evans, Laura Gifford, Malorie Perry, Simon Cottrell, Angela Marchbank, Alec Birchley, Alexander Adams, Amy Gaskin, Bree Gatica-Wilcox, Jason Coombes, Joel Southgate, Lauren Gilbert, Lee Graham, Nicole Pacchiarini, Sara Kumziene-Summerhayes, Sarah Taylor, Sophie Jones, Sara Rey, Matthew Bull, Joanne Watkins, Sally Corden, Tom Connor |
| EPI_ISL_637929, EPI_ISL_637934 | Liverpool Clinical Laboratories | COVID-19 Genomics UK (COG-UK) Consortium | Sam Haldenby, Anita Lucaci, Steve Paterson, Julian Hiscox, Alistair Darby, M Almsaud, A Alrezaihi, Muhannad Alruwaili, Stuart D Armstrong, Jones Benjamin, Eleanor G Bentley, Anu Chawla, Jordan J Clark, Angela Cowell, Richard Eccles, Isabel García-Dorival, Matthew Gemmell, Alessandro Gerada, PKF Gilmore, Richard Gregory, Ximeng Han, Catherine Hartley, Margaret Hughes, Miren Iturriza-Gomara, James Johnson, L Luu, Jenifer Manson, Charlotte Nelson, Elaine O'Toole, Cassie Olateju, Rebekah Penrice-Randal , Lucille Rainbow, N.P Randle, Trevor Ian Robinson, Parul Sharma, Ghada T Shawli, James P Stewart, Neil Swainston, Ecaterina Vamos, Joanne Watts, Mark Whitehead |
| EPI_ISL_637937 | University College London, Great Ormond Street Hospital for Children NHS Foundation Trust, Imperial College Healthcare NHS Trust | COVID-19 Genomics UK (COG-UK) Consortium | Sergi Castellano, Rachel Williams, Mark Kristiansen, Paola Resende Silva, Sunando Roy, Tony Brooks, Helena Tutill, Paola Niola, Patricia Dyal, Charlotte Williams, Leysa Forrest, Yasmin Panchbhaya, Jacqueline Findlay, Samuel Weeks, Julianne Brown, Kathryn Harris, Paul Randell, James Price, Alison Holmes, Judith Breuer |
| EPI_ISL_637952, EPI_ISL_637956, EPI_ISL_637958 | Wales Specialist Virology Centre Sequencing lab: Pathogen Genomics Unit | COVID-19 Genomics UK (COG-UK) Consortium | Catherine Moore, Johnathan Evans, Laura Gifford, Malorie Perry, Simon Cottrell, Angela Marchbank, Alec Birchley, Alexander Adams, Amy Gaskin, Bree Gatica-Wilcox, Jason Coombes, Joel Southgate, Lauren Gilbert, Lee Graham, Nicole Pacchiarini, Sara Kumziene-Summerhayes, Sarah Taylor, Sophie Jones, Sara Rey, Matthew Bull, Joanne Watkins, Sally Corden, Tom Connor |
| EPI_ISL_637959, EPI_ISL_637960, EPI_ISL_637972 | Liverpool Clinical Laboratories | COVID-19 Genomics UK (COG-UK) Consortium | Sam Haldenby, Anita Lucaci, Steve Paterson, Julian Hiscox, Alistair Darby, M Almsaud, A Alrezaihi, Muhannad Alruwaili, Stuart D Armstrong, Jones Benjamin, Eleanor G Bentley, Anu Chawla, Jordan J Clark, Angela Cowell, Richard Eccles, Isabel García-Dorival, Matthew Gemmell, Alessandro Gerada, PKF Gilmore, Richard Gregory, Ximeng Han, Catherine Hartley, Margaret Hughes, Miren Iturriza-Gomara, James Johnson, L Luu, Jenifer Manson, Charlotte Nelson, Elaine O'Toole, Cassie Olateju, Rebekah Penrice-Randal , Lucille Rainbow, N.P Randle, Trevor Ian Robinson, Parul Sharma, Ghada T Shawli, James P Stewart, Neil Swainston, Ecaterina Vamos, Joanne Watts, Mark Whitehead |
| EPI_ISL_637985 | Virology Department, Sheffield Teaching Hospitals NHS Foundation Trust/Department of Infection, Immunity and Cardiovascular Disease, The Medical School, University of Sheffield | COVID-19 Genomics UK (COG-UK) Consortium | Thushan de Silva, Matthew Parker, Nikki Smith, Adri Angyal, Rebecca Brown, Luke Green, Rachel Tucker, Paul Parsons, Danielle Groves, Katie Johnson, Laura Carrilero, Alex Keeley, Dave Partridge, Matthew Wyles, Benjamin Lindsey, Mehmet Yavuz, Mohammad Raza, Cariad Evans |
| EPI_ISL_637990 | Liverpool Clinical Laboratories | COVID-19 Genomics UK (COG-UK) Consortium | Sam Haldenby, Anita Lucaci, Steve Paterson, Julian Hiscox, Alistair Darby, M Almsaud, A Alrezaihi, Muhannad Alruwaili, Stuart D Armstrong, Jones Benjamin, Eleanor G Bentley, Anu Chawla, Jordan J Clark, Angela Cowell, Richard Eccles, Isabel García-Dorival, Matthew Gemmell, Alessandro Gerada, PKF Gilmore, Richard Gregory, Ximeng Han, Catherine Hartley, Margaret Hughes, Miren Iturriza-Gomara, James Johnson, L Luu, Jenifer Manson, Charlotte Nelson, Elaine O'Toole, Cassie Olateju, Rebekah Penrice-Randal , Lucille Rainbow, N.P Randle, Trevor Ian Robinson, Parul Sharma, Ghada T Shawli, James P Stewart, Neil Swainston, Ecaterina Vamos, Joanne Watts, Mark Whitehead |
| EPI_ISL_637998 | Wales Specialist Virology Centre Sequencing lab: Pathogen Genomics Unit | COVID-19 Genomics UK (COG-UK) Consortium | Catherine Moore, Johnathan Evans, Laura Gifford, Malorie Perry, Simon Cottrell, Angela Marchbank, Alec Birchley, Alexander Adams, Amy Gaskin, Bree Gatica-Wilcox, Jason Coombes, Joel Southgate, Lauren Gilbert, Lee Graham, Nicole Pacchiarini, Sara Kumziene-Summerhayes, Sarah Taylor, Sophie Jones, Sara Rey, Matthew Bull, Joanne Watkins, Sally Corden, Tom Connor |
| EPI_ISL_637999 | University College London, Great Ormond Street Hospital for Children NHS Foundation Trust, Imperial College Healthcare | COVID-19 Genomics UK (COG-UK) Consortium | Sergi Castellano, Rachel Williams, Mark Kristiansen, Paola Resende Silva, Sunando Roy, Tony Brooks, Helena Tutill, Paola Niola, Patricia Dyal, Charlotte Williams, Leysa Forrest, Yasmin Panchbhaya, Jacqueline Findlay, Samuel Weeks, Julianne Brown, Kathryn Harris, Paul Randell, James Price, Alison |

| | | | |
|---|---|---|---|
| | NHS Trust | | Holmes, Judith Breuer |
| EPI_ISL_638015, EPI_ISL_638021 | Liverpool Clinical Laboratories | COVID-19 Genomics UK (COG-UK) Consortium | Sam Haldenby, Anita Lucaci, Steve Paterson, Julian Hiscox, Alistair Darby, M Almsaud, A Alrezaihi, Muhannad Alruwaili, Stuart D Armstrong, Jones Benjamin, Eleanor G Bentley, Anu Chawla, Jordan J Clark, Angela Cowell, Richard Eccles, Isabel García-Dorival, Matthew Gemmell, Alessandro Gerada, PKF Gilmore, Richard Gregory, Ximeng Han, Catherine Hartley, Margaret Hughes, Miren Iturriza-Gomara, James Johnson, L Luu, Jenifer Manson, Charlotte Nelson, Elaine O'Toole, Cassie Olateju, Rebekah Penrice-Randal , Lucille Rainbow, N.P Randle, Trevor Ian Robinson, Parul Sharma, Ghada T Shawli, James P Stewart, Neil Swainston, Ecaterina Vamos, Joanne Watts, Mark Whitehead |
| EPI_ISL_638022 | University College London, Great Ormond Street Hospital for Children NHS Foundation Trust, Imperial College Healthcare NHS Trust | COVID-19 Genomics UK (COG-UK) Consortium | Sergi Castellano, Rachel Williams, Mark Kristiansen, Paola Resende Silva, Sunando Roy, Tony Brooks, Helena Tutill, Paola Niola, Patricia Dyal, Charlotte Williams, Leysa Forrest, Yasmin Panchbhaya, Jacqueline Findlay, Samuel Weeks, Julianne Brown, Kathryn Harris, Paul Randell, James Price, Alison Holmes, Judith Breuer |
| EPI_ISL_638024 | Virology Department, Sheffield Teaching Hospitals NHS Foundation Trust/Department of Infection, Immunity and Cardiovascular Disease, The Medical School, University of Sheffield | COVID-19 Genomics UK (COG-UK) Consortium | Thushan de Silva, Matthew Parker, Nikki Smith, Adri Angyal, Rebecca Brown, Luke Green, Rachel Tucker, Paul Parsons, Danielle Groves, Katie Johnson, Laura Carrilero, Alex Keeley, Dave Partridge, Matthew Wyles, Benjamin Lindsey, Mehmet Yavuz, Mohammad Raza, Cariad Evans |
| EPI_ISL_638028 | Wales Specialist Virology Centre Sequencing lab: Pathogen Genomics Unit | COVID-19 Genomics UK (COG-UK) Consortium | Catherine Moore, Johnathan Evans, Laura Gifford, Malorie Perry, Simon Cottrell, Angela Marchbank, Alec Birchley, Alexander Adams, Amy Gaskin, Bree Gatica-Wilcox, Jason Coombes, Joel Southgate, Lauren Gilbert, Lee Graham, Nicole Pacchiarini, Sara Kumziene-Summerhayes, Sarah Taylor, Sophie Jones, Sara Rey, Matthew Bull, Joanne Watkins, Sally Corden, Tom Connor |
| EPI_ISL_638032 | University College London, Great Ormond Street Hospital for Children NHS Foundation Trust, Imperial College Healthcare NHS Trust | COVID-19 Genomics UK (COG-UK) Consortium | Sergi Castellano, Rachel Williams, Mark Kristiansen, Paola Resende Silva, Sunando Roy, Tony Brooks, Helena Tutill, Paola Niola, Patricia Dyal, Charlotte Williams, Leysa Forrest, Yasmin Panchbhaya, Jacqueline Findlay, Samuel Weeks, Julianne Brown, Kathryn Harris, Paul Randell, James Price, Alison Holmes, Judith Breuer |
| EPI_ISL_638034, EPI_ISL_638042, EPI_ISL_638043, EPI_ISL_638045, EPI_ISL_638048, EPI_ISL_638049, EPI_ISL_638052, EPI_ISL_638053, EPI_ISL_638054, EPI_ISL_638055, EPI_ISL_638056, EPI_ISL_638057, EPI_ISL_638093, EPI_ISL_638094, EPI_ISL_638095, EPI_ISL_638097, EPI_ISL_638100, EPI_ISL_638102, EPI_ISL_638103, EPI_ISL_638105, EPI_ISL_638108, EPI_ISL_638109, EPI_ISL_638110, EPI_ISL_638111, EPI_ISL_638114, EPI_ISL_638115, EPI_ISL_638116, EPI_ISL_638119, EPI_ISL_638124, EPI_ISL_638150, EPI_ISL_638155, EPI_ISL_638158, EPI_ISL_638160, EPI_ISL_638161 | | | |
| see above | Wales Specialist Virology Centre Sequencing lab: Pathogen Genomics Unit | COVID-19 Genomics UK (COG-UK) Consortium | Catherine Moore, Johnathan Evans, Laura Gifford, Malorie Perry, Simon Cottrell, Angela Marchbank, Alec Birchley, Alexander Adams, Amy Gaskin, Bree Gatica-Wilcox, Jason Coombes, Joel Southgate, Lauren Gilbert, Lee Graham, Nicole Pacchiarini, Sara Kumziene-Summerhayes, Sarah Taylor, Sophie Jones, Sara Rey, Matthew Bull, Joanne Watkins, Sally Corden, Tom Connor |
| EPI_ISL_638364, EPI_ISL_638369 | Department of Pathology, University of Cambridge | COVID-19 Genomics UK (COG-UK) Consortium | Aminu S. Jahun, Yasmin Chaudhry, Grant Hall, Iliana Georgana, Myra Hosmillo, Martin D. Curran, Malte Pinckert, Surendra Parmar, Ian Goodfellow |
| EPI_ISL_638581 | Quadram Institute Bioscience | COVID-19 Genomics UK (COG-UK) Consortium | Dave J. Baker, Gemma L. Kay, Alp Aydin, Thanh Le-Viet, Steven Rudder, Ana P. Tedim, Anastasia Kolyva, Maria Diaz, Leonardo de Oliveira Martins, Nabil-Fareed Alikhan, Lizzie Meadows, Rachael Stanley, Ngozi Elumogo, Muhammed Yasir, Nicholas M. Thomson, Alexander J Trotter, Rachel Gilroy, Samuel Bloomfield, Claire Stuart, Andrew Bell, Reenesh Prakash, Samir Dervisevic, Alison E. Mather, John Wain, Mark Webber, Andrew J. Page, Justin O'Grady |
| EPI_ISL_638879, EPI_ISL_638883, EPI_ISL_638885, EPI_ISL_638886 | Virology Department, Sheffield Teaching Hospitals NHS Foundation Trust/Department of Infection, Immunity and Cardiovascular Disease, The Medical School, University of Sheffield | COVID-19 Genomics UK (COG-UK) Consortium | Thushan de Silva, Matthew Parker, Nikki Smith, Adri Angyal, Rebecca Brown, Luke Green, Rachel Tucker, Paul Parsons, Danielle Groves, Katie Johnson, Laura Carrilero, Alex Keeley, Dave Partridge, Matthew Wyles, Benjamin Lindsey, Mehmet Yavuz, Mohammad Raza, Cariad Evans |
| EPI_ISL_639015, EPI_ISL_639016, EPI_ISL_639019, EPI_ISL_639020, EPI_ISL_639022, EPI_ISL_639024, EPI_ISL_639026, EPI_ISL_639028, EPI_ISL_639029, EPI_ISL_639033, EPI_ISL_639038, EPI_ISL_639039, EPI_ISL_639042, EPI_ISL_639043, EPI_ISL_639044, EPI_ISL_639047, EPI_ISL_639051, EPI_ISL_639054, EPI_ISL_639055, EPI_ISL_639058, EPI_ISL_639061, EPI_ISL_639062, EPI_ISL_639066, EPI_ISL_639068, EPI_ISL_639069, EPI_ISL_639070, EPI_ISL_639074, EPI_ISL_639075, EPI_ISL_639076, EPI_ISL_639078, EPI_ISL_639079, EPI_ISL_639084, EPI_ISL_639089, EPI_ISL_639091, EPI_ISL_639092, EPI_ISL_639093, EPI_ISL_639096, EPI_ISL_639098, EPI_ISL_639102, EPI_ISL_639103, EPI_ISL_639106, EPI_ISL_639109, EPI_ISL_639110, EPI_ISL_639111, EPI_ISL_639112, EPI_ISL_639113, EPI_ISL_639114, EPI_ISL_639116, EPI_ISL_639118, EPI_ISL_639120, EPI_ISL_639123, EPI_ISL_639126, EPI_ISL_639128, EPI_ISL_639131, EPI_ISL_639132, EPI_ISL_639133, EPI_ISL_639137, EPI_ISL_639138, EPI_ISL_639139, EPI_ISL_639140, EPI_ISL_639142, EPI_ISL_639144, EPI_ISL_639145, EPI_ISL_639152, EPI_ISL_639154, EPI_ISL_639156, EPI_ISL_639159, EPI_ISL_639164, EPI_ISL_639167, EPI_ISL_639171, EPI_ISL_639178, EPI_ISL_639179, EPI_ISL_639182, EPI_ISL_639185, EPI_ISL_639186, EPI_ISL_639189, EPI_ISL_639190, EPI_ISL_639191, EPI_ISL_639192, EPI_ISL_639194, EPI_ISL_639196, EPI_ISL_639198, EPI_ISL_639200, EPI_ISL_639201, EPI_ISL_639202, EPI_ISL_639208, EPI_ISL_639209, EPI_ISL_639210, EPI_ISL_639214, EPI_ISL_639215, EPI_ISL_639216, EPI_ISL_639218, EPI_ISL_639219, EPI_ISL_639220, EPI_ISL_639221, EPI_ISL_639223, EPI_ISL_639224, EPI_ISL_639226, EPI_ISL_639228, EPI_ISL_639229, EPI_ISL_639230, EPI_ISL_639232, EPI_ISL_639237, EPI_ISL_639241, EPI_ISL_639242, EPI_ISL_639243, EPI_ISL_639245, EPI_ISL_639246, EPI_ISL_639247, EPI_ISL_639248, EPI_ISL_639250, EPI_ISL_639251, EPI_ISL_639255, EPI_ISL_639256, EPI_ISL_639257, EPI_ISL_639260, EPI_ISL_639263, EPI_ISL_639264, EPI_ISL_639266, EPI_ISL_639268, EPI_ISL_639270, EPI_ISL_639272, EPI_ISL_639274, EPI_ISL_639276, EPI_ISL_639278, EPI_ISL_639279, EPI_ISL_639283, EPI_ISL_639284, EPI_ISL_639285, EPI_ISL_639286, EPI_ISL_639287, EPI_ISL_639289, EPI_ISL_639290, EPI_ISL_639291, EPI_ISL_639292, EPI_ISL_639293, EPI_ISL_639294, EPI_ISL_639295, EPI_ISL_639297, EPI_ISL_639298, EPI_ISL_639299, EPI_ISL_639300, EPI_ISL_639301, EPI_ISL_639302, EPI_ISL_639303, EPI_ISL_639304, EPI_ISL_639305, EPI_ISL_639306, EPI_ISL_639307, EPI_ISL_639309, EPI_ISL_639311, EPI_ISL_639312, EPI_ISL_639313, EPI_ISL_639314, EPI_ISL_639315, EPI_ISL_639316, EPI_ISL_639317, EPI_ISL_639318, EPI_ISL_639320, EPI_ISL_639323, EPI_ISL_639324, EPI_ISL_639329, EPI_ISL_639331, EPI_ISL_639333, EPI_ISL_639335, EPI_ISL_639336, EPI_ISL_639337, EPI_ISL_639338, EPI_ISL_639340, EPI_ISL_639341, EPI_ISL_639342, EPI_ISL_639343, EPI_ISL_639346, EPI_ISL_639347, EPI_ISL_639348, EPI_ISL_639349, EPI_ISL_639350, EPI_ISL_639352, EPI_ISL_639353, EPI_ISL_639354, EPI_ISL_639355, EPI_ISL_639356, EPI_ISL_639357, EPI_ISL_639361, EPI_ISL_639364, EPI_ISL_639366, EPI_ISL_639367, EPI_ISL_639368, EPI_ISL_639370, EPI_ISL_639371, EPI_ISL_639372, EPI_ISL_639373, EPI_ISL_639374, EPI_ISL_639375, EPI_ISL_639376, EPI_ISL_639377, EPI_ISL_639378, EPI_ISL_639380, EPI_ISL_639382, EPI_ISL_639383, EPI_ISL_639385, EPI_ISL_639386, EPI_ISL_639387, EPI_ISL_639389, EPI_ISL_639391, EPI_ISL_639392, EPI_ISL_639393, EPI_ISL_639394, EPI_ISL_639400, EPI_ISL_639401, EPI_ISL_639406, EPI_ISL_639408, EPI_ISL_639409, EPI_ISL_639411, EPI_ISL_639412, EPI_ISL_639415, EPI_ISL_639417, EPI_ISL_639419, EPI_ISL_639420, EPI_ISL_639423, EPI_ISL_639430, EPI_ISL_639435, EPI_ISL_639439, EPI_ISL_639442, EPI_ISL_639444, EPI_ISL_639451, EPI_ISL_639456, EPI_ISL_639465, EPI_ISL_639466, EPI_ISL_639471, EPI_ISL_639472, EPI_ISL_639480, EPI_ISL_639487, EPI_ISL_639488, EPI_ISL_639493, EPI_ISL_639502, EPI_ISL_639505, EPI_ISL_639513, EPI_ISL_639517, EPI_ISL_639519, EPI_ISL_639520, EPI_ISL_639524, EPI_ISL_639533, EPI_ISL_639538, EPI_ISL_639543, EPI_ISL_639546, EPI_ISL_639552, EPI_ISL_639555, EPI_ISL_639559, EPI_ISL_639564, EPI_ISL_639577, EPI_ISL_639579, EPI_ISL_639585, EPI_ISL_639590, EPI_ISL_639594, EPI_ISL_639595, EPI_ISL_639598, EPI_ISL_639600, EPI_ISL_639604, EPI_ISL_639608, EPI_ISL_639611, EPI_ISL_639619, EPI_ISL_639622 | | | |
| see above | Wales Specialist Virology Centre Sequencing lab: Pathogen Genomics Unit | COVID-19 Genomics UK (COG-UK) Consortium | Catherine Moore, Johnathan Evans, Laura Gifford, Malorie Perry, Simon Cottrell, Angela Marchbank, Alec Birchley, Alexander Adams, Amy Gaskin, Bree Gatica-Wilcox, Jason Coombes, Joel Southgate, Lauren Gilbert, Lee Graham, Nicole Pacchiarini, Sara Kumziene-Summerhayes, Sarah Taylor, Sophie Jones, Sara Rey, Matthew Bull, Joanne Watkins, Sally Corden, Tom Connor |
| EPI_ISL_639738 | South Eastern Area Laboratory Services (SEALS) | NSW Health Pathology - Institute of Clinical Pathology and Medical Research; Westmead Hospital; University of Sydney | CIDM-PH et al. |
| EPI_ISL_639819 | unknown | Public Health Virology Laboratory, Forensic and Scientific Services (PHV-FSS) | Son Nguyen et al. |
| EPI_ISL_639989, EPI_ISL_639990, EPI_ISL_639996, EPI_ISL_640001, EPI_ISL_640005, EPI_ISL_640007, EPI_ISL_640010, EPI_ISL_640011 | CNR Virus des Infections Respiratoires - France SUD | CNR Virus des Infections Respiratoires - France SUD | Antonin Bal, Gregory Destras, Gwendolyne Burfin, Hadrien Règue, Alexandre Gaymard, Maude Bouscambert-Duchamp, Florence Morfin-Sherpa, Martine Valette, Bruno Lina, Laurence Josset |
| EPI_ISL_641397, EPI_ISL_641398, EPI_ISL_641399, EPI_ISL_641400, EPI_ISL_641401, EPI_ISL_641402, EPI_ISL_641403, EPI_ISL_641414, EPI_ISL_641415, EPI_ISL_641416 | Department of Virus and Microbiological Special Diagnostics, Statens Serum Institut, Copenhagen, Denmark | Albertsen lab, Department of Chemistry and Bioscience, Aalborg University, Denmark | Thomas Bruun Rasmussen, Jannik Fonager, Morten Rasmussen |
| EPI_ISL_643322, EPI_ISL_643324, EPI_ISL_643325, EPI_ISL_643329, EPI_ISL_643330, EPI_ISL_643331, EPI_ISL_643334, EPI_ISL_643337, EPI_ISL_643338, EPI_ISL_643339, EPI_ISL_643341, EPI_ISL_643349, EPI_ISL_643350, EPI_ISL_643351, EPI_ISL_643352, EPI_ISL_643353, EPI_ISL_643355, EPI_ISL_643356, EPI_ISL_643364, EPI_ISL_643365, EPI_ISL_643368, EPI_ISL_643370, EPI_ISL_643372, EPI_ISL_643374, EPI_ISL_643375, EPI_ISL_643376, EPI_ISL_643378, EPI_ISL_643380, EPI_ISL_643382, EPI_ISL_643383, EPI_ISL_643386, EPI_ISL_643389, EPI_ISL_643391, EPI_ISL_643392, EPI_ISL_643393, EPI_ISL_643396, EPI_ISL_643399, EPI_ISL_643403, EPI_ISL_643408, EPI_ISL_643411, EPI_ISL_643412, EPI_ISL_643413, EPI_ISL_643414, EPI_ISL_643415, EPI_ISL_643416, EPI_ISL_643417, EPI_ISL_643418, EPI_ISL_643421, EPI_ISL_643423, EPI_ISL_643424, EPI_ISL_643428, EPI_ISL_643430, EPI_ISL_643432, EPI_ISL_643434, EPI_ISL_643435, EPI_ISL_643442, EPI_ISL_643446, EPI_ISL_643447, EPI_ISL_643448, EPI_ISL_643449, EPI_ISL_643450, EPI_ISL_643451, EPI_ISL_643454, EPI_ISL_643458, EPI_ISL_643459, EPI_ISL_643461, EPI_ISL_643462, EPI_ISL_643467, EPI_ISL_643470, EPI_ISL_643471, EPI_ISL_643472, EPI_ISL_643474, EPI_ISL_643475, EPI_ISL_643477, EPI_ISL_643478, EPI_ISL_643480, EPI_ISL_643481, EPI_ISL_643482, EPI_ISL_643484, EPI_ISL_643485, EPI_ISL_643488, EPI_ISL_643489, EPI_ISL_643491, EPI_ISL_643492, EPI_ISL_643494, EPI_ISL_643495, EPI_ISL_643498, EPI_ISL_643499, EPI_ISL_643504, EPI_ISL_643505, EPI_ISL_643506, EPI_ISL_643509, EPI_ISL_643510, EPI_ISL_643512, EPI_ISL_643513, EPI_ISL_643514, EPI_ISL_643516, EPI_ISL_643517, EPI_ISL_643518, EPI_ISL_643519, EPI_ISL_643520, EPI_ISL_643521, EPI_ISL_643526, EPI_ISL_643527, EPI_ISL_643528, EPI_ISL_643531, EPI_ISL_643533, EPI_ISL_643534, EPI_ISL_643536, EPI_ISL_643539, EPI_ISL_643540, EPI_ISL_643542, EPI_ISL_643543, EPI_ISL_643544, EPI_ISL_643545, EPI_ISL_643546, EPI_ISL_643548, EPI_ISL_643551, EPI_ISL_643553, EPI_ISL_643554, EPI_ISL_643557, EPI_ISL_643565, EPI_ISL_643568, EPI_ISL_643572, EPI_ISL_643575, EPI_ISL_643576, EPI_ISL_643577, EPI_ISL_643581, EPI_ISL_643584, EPI_ISL_643586, EPI_ISL_643588, EPI_ISL_643590, EPI_ISL_643593, EPI_ISL_643594, EPI_ISL_643595, EPI_ISL_643598, EPI_ISL_643600, EPI_ISL_643602, EPI_ISL_643603, EPI_ISL_643605, EPI_ISL_643609, EPI_ISL_643610, EPI_ISL_643612, EPI_ISL_643613, EPI_ISL_643615, EPI_ISL_643616, EPI_ISL_643617, EPI_ISL_643626, EPI_ISL_643628, EPI_ISL_643629, EPI_ISL_643630, EPI_ISL_643631, EPI_ISL_643632, EPI_ISL_643633, EPI_ISL_643634, EPI_ISL_643635, EPI_ISL_643636, EPI_ISL_643637, EPI_ISL_643638, EPI_ISL_643640, EPI_ISL_643641, EPI_ISL_643642, EPI_ISL_643643, EPI_ISL_643644, EPI_ISL_643646, EPI_ISL_643650, EPI_ISL_643651, EPI_ISL_643653, EPI_ISL_643654, EPI_ISL_643655, EPI_ISL_643656, EPI_ISL_643658, EPI_ISL_643662, EPI_ISL_643663, EPI_ISL_643664, EPI_ISL_643668, EPI_ISL_643669, EPI_ISL_643671, EPI_ISL_643672, EPI_ISL_643675, EPI_ISL_643676, EPI_ISL_643677, EPI_ISL_643678, EPI_ISL_643679, EPI_ISL_643681, EPI_ISL_643682, EPI_ISL_643684, EPI_ISL_643686, EPI_ISL_643689, EPI_ISL_643690, EPI_ISL_643691, EPI_ISL_643692, EPI_ISL_643693, EPI_ISL_643694, EPI_ISL_643695, EPI_ISL_643696, EPI_ISL_643699, EPI_ISL_643700, EPI_ISL_643703, EPI_ISL_643705, EPI_ISL_643706, EPI_ISL_643708, EPI_ISL_643710, EPI_ISL_643711, EPI_ISL_643712, EPI_ISL_643713, EPI_ISL_643714, EPI_ISL_643715, EPI_ISL_643718, EPI_ISL_643720, EPI_ISL_643721, EPI_ISL_643725, EPI_ISL_643729, EPI_ISL_643730, EPI_ISL_643731, EPI_ISL_643732, EPI_ISL_643733, EPI_ISL_643734, EPI_ISL_643735, EPI_ISL_643737, EPI_ISL_643740, EPI_ISL_643741, EPI_ISL_643742, EPI_ISL_643744, EPI_ISL_643745, EPI_ISL_643746, EPI_ISL_643748, EPI_ISL_643750, EPI_ISL_643751, EPI_ISL_643753, EPI_ISL_643754, EPI_ISL_643755, EPI_ISL_643757, EPI_ISL_643760, EPI_ISL_643763, EPI_ISL_643764, EPI_ISL_643768, EPI_ISL_643769, EPI_ISL_643770, EPI_ISL_643771, EPI_ISL_643772, EPI_ISL_643773, EPI_ISL_643776, EPI_ISL_643777, EPI_ISL_643780, EPI_ISL_643783, EPI_ISL_643784, EPI_ISL_643785, EPI_ISL_643786, EPI_ISL_643787, EPI_ISL_643788, EPI_ISL_643790, EPI_ISL_643791, EPI_ISL_643793, EPI_ISL_643794, EPI_ISL_643796, EPI_ISL_643798, EPI_ISL_643799, EPI_ISL_643800, EPI_ISL_643801, EPI_ISL_643802, EPI_ISL_643803, EPI_ISL_643806, EPI_ISL_643807, EPI_ISL_643811, EPI_ISL_643814, EPI_ISL_643815, EPI_ISL_643816, EPI_ISL_643818, EPI_ISL_643827, | | | |

| | | | |
|---|---|---|---|
| EPI_ISL_643829, EPI_ISL_643830, EPI_ISL_643832, EPI_ISL_643833, EPI_ISL_643834, EPI_ISL_643835, EPI_ISL_643836, EPI_ISL_643838, EPI_ISL_643839, EPI_ISL_643841, EPI_ISL_643842, EPI_ISL_643844, EPI_ISL_643845, EPI_ISL_643846, EPI_ISL_643847, EPI_ISL_643848, EPI_ISL_643849, EPI_ISL_643850, EPI_ISL_643851, EPI_ISL_643852, EPI_ISL_643853, EPI_ISL_643854, EPI_ISL_643855, EPI_ISL_643856, EPI_ISL_643857, EPI_ISL_643858, EPI_ISL_643864, EPI_ISL_643865, EPI_ISL_643867, EPI_ISL_643869, EPI_ISL_643871, EPI_ISL_643872, EPI_ISL_643873, EPI_ISL_643874, EPI_ISL_643875, EPI_ISL_643878, EPI_ISL_643881, EPI_ISL_643882, EPI_ISL_643884, EPI_ISL_643887, EPI_ISL_643889, EPI_ISL_643895, EPI_ISL_643899, EPI_ISL_643901, EPI_ISL_643902, EPI_ISL_643903, EPI_ISL_643904, EPI_ISL_643905, EPI_ISL_643906, EPI_ISL_643907, EPI_ISL_643908, EPI_ISL_643909, EPI_ISL_643911, EPI_ISL_643912, EPI_ISL_643914, EPI_ISL_643915, EPI_ISL_643917, EPI_ISL_643919, EPI_ISL_643920, EPI_ISL_643922, EPI_ISL_643923, EPI_ISL_643925, EPI_ISL_643927, EPI_ISL_643928, EPI_ISL_643929, EPI_ISL_643930, EPI_ISL_643931, EPI_ISL_643932, EPI_ISL_643933, EPI_ISL_643937, EPI_ISL_643939, EPI_ISL_643940, EPI_ISL_643943, EPI_ISL_643944, EPI_ISL_643945, EPI_ISL_643948, EPI_ISL_643949, EPI_ISL_643950, EPI_ISL_643954, EPI_ISL_643955, EPI_ISL_643956, EPI_ISL_643958, EPI_ISL_643960, EPI_ISL_643962, EPI_ISL_643964, EPI_ISL_643965, EPI_ISL_643966, EPI_ISL_643967, EPI_ISL_643968, EPI_ISL_643969, EPI_ISL_643971, EPI_ISL_643972, EPI_ISL_643974, EPI_ISL_643976, EPI_ISL_643977, EPI_ISL_643978, EPI_ISL_643979, EPI_ISL_643981 | | | |
| see above | Lighthouse Lab in Cambridge | Wellcome Sanger Institute for the COVID-19 Genomics UK (COG-UK) Consortium | Rob Howes, The Lighthouse Lab in Cambridge and Alex Alderton, Roberto Amato, Sonia Goncalves, Ewan Harrison, David K. Jackson, Ian Johnston, Dominic Kwiatkowski, Cordelia Langford, John Sillitoe on behalf of the Wellcome Sanger Institute COVID-19 Surveillance Team |
| EPI_ISL_643992, EPI_ISL_643993, EPI_ISL_643994, EPI_ISL_643995, EPI_ISL_643996, EPI_ISL_643997, EPI_ISL_643998, EPI_ISL_643999, EPI_ISL_644000, EPI_ISL_644002, EPI_ISL_644004, EPI_ISL_644006, EPI_ISL_644007, EPI_ISL_644008, EPI_ISL_644009, EPI_ISL_644010, EPI_ISL_644011, EPI_ISL_644012, EPI_ISL_644013, EPI_ISL_644014, EPI_ISL_644015, EPI_ISL_644016, EPI_ISL_644017, EPI_ISL_644018, EPI_ISL_644019, EPI_ISL_644020, EPI_ISL_644021, EPI_ISL_644022, EPI_ISL_644023, EPI_ISL_644024, EPI_ISL_644025, EPI_ISL_644026, EPI_ISL_644027, EPI_ISL_644028, EPI_ISL_644029, EPI_ISL_644030, EPI_ISL_644032, EPI_ISL_644033, EPI_ISL_644034, EPI_ISL_644035, EPI_ISL_644036, EPI_ISL_644037, EPI_ISL_644038, EPI_ISL_644039, EPI_ISL_644040, EPI_ISL_644043, EPI_ISL_644044, EPI_ISL_644045, EPI_ISL_644047, EPI_ISL_644048, EPI_ISL_644049, EPI_ISL_644051 | | | |
| see above | Lighthouse Lab in Glasgow | Wellcome Sanger Institute for the COVID-19 Genomics UK (COG-UK) Consortium | Harper VanSteenhouse, Yumi Kasai, David Gray, Carol Clugston, Anna Dominiczak and Alex Alderton, Roberto Amato, Sonia Goncalves, Ewan Harrison, David K. Jackson, Ian Johnston, Dominic Kwiatkowski, Cordelia Langford, John Sillitoe on behalf of the Wellcome Sanger Institute COVID-19 Surveillance Team |
| EPI_ISL_644347, EPI_ISL_644349, EPI_ISL_644350, EPI_ISL_644351, EPI_ISL_644353, EPI_ISL_644356, EPI_ISL_644358, EPI_ISL_644359, EPI_ISL_644360, EPI_ISL_644361, EPI_ISL_644362, EPI_ISL_644364, EPI_ISL_644365, EPI_ISL_644366, EPI_ISL_644367, EPI_ISL_644370, EPI_ISL_644371, EPI_ISL_644374 | | | |
| see above | Michigan Department of Health and Human Services, Bureau of Laboratories | Michigan Department of Health and Human Services, Bureau of Laboratories | Blankenship HM, Riner D, Soehnlen MK |
| EPI_ISL_644933 | Essentia Health-St. Mary's Medical Center | Minnesota Department of Health, Public Health Laboratory | Matt Plumb, Jacob Garfin, Alexandra Lorentz, and Xiong Wang |
| EPI_ISL_644941 | Allina Health Laboratory | Minnesota Department of Health, Public Health Laboratory | Matt Plumb, Jacob Garfin, Alexandra Lorentz, and Xiong Wang |
| EPI_ISL_645115, EPI_ISL_645119 | National Public Health Laboratory, National Centre for Infectious Diseases | National Public Health Laboratory, National Centre for Infectious Diseases | Tze Minn Mak, Sophie Octavia, Zhenyang Zhou, Lin Cui, Raymond Tzer Pin Lin |
| EPI_ISL_645266, EPI_ISL_645362, EPI_ISL_645368 | Lighthouse Lab in Alderley Park | Wellcome Sanger Institute for the COVID-19 Genomics UK (COG-UK) Consortium | Jacquelyn Wynn, Mairead Hyland, The Lighthouse Lab in Alderley Park and Alex Alderton, Roberto Amato, Sonia Goncalves, Ewan Harrison, David K. Jackson, Ian Johnston, Dominic Kwiatkowski, Cordelia Langford, John Sillitoe on behalf of the Wellcome Sanger Institute COVID-19 Surveillance Team |
| EPI_ISL_647343 | Lighthouse Lab in Cambridge | Wellcome Sanger Institute for the COVID-19 Genomics UK (COG-UK) Consortium | Rob Howes, The Lighthouse Lab in Cambridge and Alex Alderton, Roberto Amato, Sonia Goncalves, Ewan Harrison, David K. Jackson, Ian Johnston, Dominic Kwiatkowski, Cordelia Langford, John Sillitoe on behalf of the Wellcome Sanger Institute COVID-19 Surveillance Team |
| EPI_ISL_647873, EPI_ISL_647874, EPI_ISL_647875, EPI_ISL_647876, EPI_ISL_647877, EPI_ISL_647878, EPI_ISL_647879, EPI_ISL_647880, EPI_ISL_647881, EPI_ISL_647882, EPI_ISL_647883, EPI_ISL_647884, EPI_ISL_647885, EPI_ISL_647886, EPI_ISL_647887, EPI_ISL_647888, EPI_ISL_647890, EPI_ISL_647891, EPI_ISL_647892, EPI_ISL_647894, EPI_ISL_647896, EPI_ISL_647897, EPI_ISL_647898, EPI_ISL_647899 | | | |
| see above | Lighthouse Lab in Glasgow | Wellcome Sanger Institute for the COVID-19 Genomics UK (COG-UK) Consortium | Harper VanSteenhouse, Yumi Kasai, David Gray, Carol Clugston, Anna Dominiczak and Alex Alderton, Roberto Amato, Sonia Goncalves, Ewan Harrison, David K. Jackson, Ian Johnston, Dominic Kwiatkowski, Cordelia Langford, John Sillitoe on behalf of the Wellcome Sanger Institute COVID-19 Surveillance Team |
| EPI_ISL_647900, EPI_ISL_647901 | Lighthouse Lab in Cambridge | Wellcome Sanger Institute for the COVID-19 Genomics UK (COG-UK) Consortium | Rob Howes, The Lighthouse Lab in Cambridge and Alex Alderton, Roberto Amato, Sonia Goncalves, Ewan Harrison, David K. Jackson, Ian Johnston, Dominic Kwiatkowski, Cordelia Langford, John Sillitoe on behalf of the Wellcome Sanger Institute COVID-19 Surveillance Team |
| EPI_ISL_647902, EPI_ISL_647903, EPI_ISL_647904, EPI_ISL_647905 | Lighthouse Lab in Glasgow | Wellcome Sanger Institute for the COVID-19 Genomics UK (COG-UK) Consortium | Harper VanSteenhouse, Yumi Kasai, David Gray, Carol Clugston, Anna Dominiczak and Alex Alderton, Roberto Amato, Sonia Goncalves, Ewan Harrison, David K. Jackson, Ian Johnston, Dominic Kwiatkowski, Cordelia Langford, John Sillitoe on behalf of the Wellcome Sanger Institute COVID-19 Surveillance Team |
| EPI_ISL_647906 | Lighthouse Lab in Cambridge | Wellcome Sanger Institute for the COVID-19 Genomics UK (COG-UK) Consortium | Rob Howes, The Lighthouse Lab in Cambridge and Alex Alderton, Roberto Amato, Sonia Goncalves, Ewan Harrison, David K. Jackson, Ian Johnston, Dominic Kwiatkowski, Cordelia Langford, John Sillitoe on behalf of the Wellcome Sanger Institute COVID-19 Surveillance Team |
| EPI_ISL_647907 | Lighthouse Lab in Glasgow | Wellcome Sanger Institute for the COVID-19 Genomics UK (COG-UK) Consortium | Harper VanSteenhouse, Yumi Kasai, David Gray, Carol Clugston, Anna Dominiczak and Alex Alderton, Roberto Amato, Sonia Goncalves, Ewan Harrison, David K. Jackson, Ian Johnston, Dominic Kwiatkowski, Cordelia Langford, John Sillitoe on behalf of the Wellcome Sanger Institute COVID-19 Surveillance Team |
| EPI_ISL_647908, EPI_ISL_647909 | Lighthouse Lab in Cambridge | Wellcome Sanger Institute for the COVID-19 Genomics UK (COG-UK) Consortium | Rob Howes, The Lighthouse Lab in Cambridge and Alex Alderton, Roberto Amato, Sonia Goncalves, Ewan Harrison, David K. Jackson, Ian Johnston, Dominic Kwiatkowski, Cordelia Langford, John Sillitoe on behalf of the Wellcome Sanger Institute COVID-19 Surveillance Team |
| EPI_ISL_647910, EPI_ISL_647911, EPI_ISL_647912 | Lighthouse Lab in Glasgow | Wellcome Sanger Institute for the COVID-19 Genomics UK (COG-UK) Consortium | Harper VanSteenhouse, Yumi Kasai, David Gray, Carol Clugston, Anna Dominiczak and Alex Alderton, Roberto Amato, Sonia Goncalves, Ewan Harrison, David K. Jackson, Ian Johnston, Dominic Kwiatkowski, Cordelia Langford, John Sillitoe on behalf of the Wellcome Sanger Institute COVID-19 Surveillance Team |
| EPI_ISL_647913 | Lighthouse Lab in Cambridge | Wellcome Sanger Institute for the COVID-19 Genomics UK (COG-UK) Consortium | Rob Howes, The Lighthouse Lab in Cambridge and Alex Alderton, Roberto Amato, Sonia Goncalves, Ewan Harrison, David K. Jackson, Ian Johnston, Dominic Kwiatkowski, Cordelia Langford, John Sillitoe on behalf of the Wellcome Sanger Institute COVID-19 Surveillance Team |
| EPI_ISL_647914, EPI_ISL_647915 | Lighthouse Lab in Glasgow | Wellcome Sanger Institute for the COVID-19 Genomics UK (COG-UK) Consortium | Harper VanSteenhouse, Yumi Kasai, David Gray, Carol Clugston, Anna Dominiczak and Alex Alderton, Roberto Amato, Sonia Goncalves, Ewan Harrison, David K. Jackson, Ian Johnston, Dominic Kwiatkowski, Cordelia Langford, John Sillitoe on behalf of the Wellcome Sanger Institute COVID-19 Surveillance Team |
| EPI_ISL_647916 | Lighthouse Lab in Cambridge | Wellcome Sanger Institute for the COVID-19 Genomics UK (COG-UK) Consortium | Rob Howes, The Lighthouse Lab in Cambridge and Alex Alderton, Roberto Amato, Sonia Goncalves, Ewan Harrison, David K. Jackson, Ian Johnston, Dominic Kwiatkowski, Cordelia Langford, John Sillitoe on behalf of the Wellcome Sanger Institute COVID-19 Surveillance Team |
| EPI_ISL_647917, EPI_ISL_647918, EPI_ISL_647919, EPI_ISL_647920, EPI_ISL_647922, EPI_ISL_647923, EPI_ISL_647924, EPI_ISL_647925, EPI_ISL_647927, EPI_ISL_647928, EPI_ISL_647929, EPI_ISL_647930, EPI_ISL_647931, EPI_ISL_647932, EPI_ISL_647934, EPI_ISL_647935, EPI_ISL_647937, EPI_ISL_647938, EPI_ISL_647939, EPI_ISL_647940, EPI_ISL_647941, EPI_ISL_647942, EPI_ISL_647943, EPI_ISL_647944, EPI_ISL_647945, EPI_ISL_647947, EPI_ISL_647948, EPI_ISL_647949, EPI_ISL_647950, EPI_ISL_647951, EPI_ISL_647953, EPI_ISL_647955, EPI_ISL_647958, EPI_ISL_647960, EPI_ISL_647961, EPI_ISL_647963, EPI_ISL_647967, EPI_ISL_647968 | | | |
| see above | Lighthouse Lab in Glasgow | Wellcome Sanger Institute for the COVID-19 Genomics UK (COG-UK) Consortium | Harper VanSteenhouse, Yumi Kasai, David Gray, Carol Clugston, Anna Dominiczak and Alex Alderton, Roberto Amato, Sonia Goncalves, Ewan Harrison, David K. Jackson, Ian Johnston, Dominic Kwiatkowski, Cordelia Langford, John Sillitoe on behalf of the Wellcome Sanger Institute COVID-19 Surveillance Team |
| EPI_ISL_648129 | The Public Health Agency of Sweden | The Public Health Agency of Sweden | Anna-Malin Linde, Maria Lind Karlberg, Mattias Haukland, Reza Advani, Olov Svartstrom, Oskar Karlsson Lindsjo, Sandra Broddesson, Petra Edquist, Mia Brytting, Anna Risberg, Karin Tegmark-Wisell |
| EPI_ISL_648159, EPI_ISL_648160, EPI_ISL_648161 | Sundsvall | The Public Health Agency of Sweden | Anna-Malin Linde, Maria Lind Karlberg, Mattias Haukland, Reza Advani, Olov Svartstrom, Oskar Karlsson Lindsjo, Sandra Broddesson, Petra Edquist, Mia Brytting, Anna Risberg, Karin Tegmark-Wisell |
| EPI_ISL_648166 | Eskilstuna | The Public Health Agency of Sweden | Anna-Malin Linde, Maria Lind Karlberg, Mattias Haukland, Reza Advani, Olov Svartstrom, Oskar Karlsson Lindsjo, Sandra Broddesson, Petra Edquist, Mia Brytting, Anna Risberg, Karin Tegmark-Wisell |
| EPI_ISL_648200, EPI_ISL_648201, EPI_ISL_648202, EPI_ISL_648203 | Linkoping | The Public Health Agency of Sweden | Anna-Malin Linde, Maria Lind Karlberg, Mattias Haukland, Reza Advani, Olov Svartstrom, Oskar Karlsson Lindsjo, Sandra Broddesson, Petra Edquist, Mia Brytting, Anna Risberg, Karin Tegmark-Wisell |
| EPI_ISL_648207, EPI_ISL_648208 | Orebro | The Public Health Agency of Sweden | Anna-Malin Linde, Maria Lind Karlberg, Mattias Haukland, Reza Advani, Olov Svartstrom, Oskar Karlsson Lindsjo, Sandra Broddesson, Petra Edquist, Mia Brytting, Anna Risberg, Karin Tegmark-Wisell |
| EPI_ISL_648301, EPI_ISL_648302 | MD PHL | MD PHL | Maryland Department of Health Laboratories Administration |
| EPI_ISL_649007, EPI_ISL_649009, EPI_ISL_649011, EPI_ISL_649013, EPI_ISL_649015, EPI_ISL_649020, EPI_ISL_649022, EPI_ISL_649025 | San Diego County Public Health Laboratory | Andersen lab at Scripps Research | SEARCH Alliance San Diego with Tracy Basler, Jovan Shephard, Brett Austin |

| | | | |
|---|---|---|---|
| EPI_ISL_649228, EPI_ISL_649252, EPI_ISL_649305, EPI_ISL_649354, EPI_ISL_649432 | Lighthouse Lab in Alderley Park | Wellcome Sanger Institute for the COVID-19 Genomics UK (COG-UK) Consortium | Jacquelyn Wynn, Mairead Hyland, The Lighthouse Lab in Alderley Park and Alex Alderton, Roberto Amato, Sonia Goncalves, Ewan Harrison, David K. Jackson, Ian Johnston, Dominic Kwiatkowski, Cordelia Langford, John Sillitoe on behalf of the Wellcome Sanger Institute COVID-19 Surveillance Team (http://www.sanger.ac.uk/covid-team) |
| EPI_ISL_649625, EPI_ISL_649626, EPI_ISL_649627, EPI_ISL_649628, EPI_ISL_649630, EPI_ISL_649631, EPI_ISL_649633 | Lighthouse Lab in Cambridge | Wellcome Sanger Institute for the COVID-19 Genomics UK (COG-UK) Consortium | Rob Howes, The Lighthouse Lab in Cambridge and Alex Alderton, Roberto Amato, Sonia Goncalves, Ewan Harrison, David K. Jackson, Ian Johnston, Dominic Kwiatkowski, Cordelia Langford, John Sillitoe on behalf of the Wellcome Sanger Institute COVID-19 Surveillance Team (http://www.sanger.ac.uk/covid-team) |
| EPI_ISL_649634, EPI_ISL_649635, EPI_ISL_649636, EPI_ISL_649637, EPI_ISL_649638, EPI_ISL_649639, EPI_ISL_649641, EPI_ISL_649642, EPI_ISL_649643, EPI_ISL_649644, EPI_ISL_649645, EPI_ISL_649646, EPI_ISL_649647, EPI_ISL_649648, EPI_ISL_649649, EPI_ISL_649651, EPI_ISL_649652, EPI_ISL_649653, EPI_ISL_649654, EPI_ISL_649655, EPI_ISL_649656, EPI_ISL_649657, EPI_ISL_649658, EPI_ISL_649659, EPI_ISL_649660, EPI_ISL_649661, EPI_ISL_649662, EPI_ISL_649663, EPI_ISL_649664, EPI_ISL_649665, EPI_ISL_649666, EPI_ISL_649667, EPI_ISL_649668, EPI_ISL_649669, EPI_ISL_649670, EPI_ISL_649671, EPI_ISL_649672, EPI_ISL_649673, EPI_ISL_649674, EPI_ISL_649676, EPI_ISL_649677, EPI_ISL_649678, EPI_ISL_649679, EPI_ISL_649680, EPI_ISL_649681, EPI_ISL_649682, EPI_ISL_649683, EPI_ISL_649684, EPI_ISL_649685, EPI_ISL_649686, EPI_ISL_649687, EPI_ISL_649688, EPI_ISL_649689, EPI_ISL_649690, EPI_ISL_649691, EPI_ISL_649693, EPI_ISL_649694, EPI_ISL_649695, EPI_ISL_649696, EPI_ISL_649697, EPI_ISL_649699, EPI_ISL_649700, EPI_ISL_649702, EPI_ISL_649703, EPI_ISL_649704, EPI_ISL_649705, EPI_ISL_649706, EPI_ISL_649707, EPI_ISL_649708, EPI_ISL_649709, EPI_ISL_649712, EPI_ISL_649713, EPI_ISL_649714, EPI_ISL_649715, EPI_ISL_649716, EPI_ISL_649717, EPI_ISL_649718, EPI_ISL_649720, EPI_ISL_649721, EPI_ISL_649722, EPI_ISL_649723, EPI_ISL_649726, EPI_ISL_649727, EPI_ISL_649728, EPI_ISL_649729, EPI_ISL_649730, EPI_ISL_649731, EPI_ISL_649732, EPI_ISL_649733, EPI_ISL_649734, EPI_ISL_649735, EPI_ISL_649736, EPI_ISL_649737, EPI_ISL_649738, EPI_ISL_649739, EPI_ISL_649740, EPI_ISL_649741, EPI_ISL_649742, EPI_ISL_649743, EPI_ISL_649744, EPI_ISL_649745, EPI_ISL_649746, EPI_ISL_649747, EPI_ISL_649748, EPI_ISL_649749, EPI_ISL_649750, EPI_ISL_649751, EPI_ISL_649752, EPI_ISL_649753, EPI_ISL_649754, EPI_ISL_649755, EPI_ISL_649756, EPI_ISL_649757, EPI_ISL_649758, EPI_ISL_649759, EPI_ISL_649760, EPI_ISL_649761, EPI_ISL_649762, EPI_ISL_649763, EPI_ISL_649764, EPI_ISL_649765, EPI_ISL_649766, EPI_ISL_649767, EPI_ISL_649768, EPI_ISL_649769, EPI_ISL_649770, EPI_ISL_649771, EPI_ISL_649772, EPI_ISL_649773, EPI_ISL_649774, EPI_ISL_649775, EPI_ISL_649776, EPI_ISL_649778, EPI_ISL_649780, EPI_ISL_649781, EPI_ISL_649782, EPI_ISL_649783 | | | |
| see above | Lighthouse Lab in Glasgow | Wellcome Sanger Institute for the COVID-19 Genomics UK (COG-UK) Consortium | Harper VanSteenhouse, Yumi Kasai, David Gray, Carol Clugston, Anna Dominiczak and Alex Alderton, Roberto Amato, Sonia Goncalves, Ewan Harrison, David K. Jackson, Ian Johnston, Dominic Kwiatkowski, Cordelia Langford, John Sillitoe on behalf of the Wellcome Sanger Institute COVID-19 Surveillance Team (http://www.sanger.ac.uk/covid-team) |
| EPI_ISL_650000, EPI_ISL_650019, EPI_ISL_650020, EPI_ISL_650021, EPI_ISL_650022, EPI_ISL_650023, EPI_ISL_650024, EPI_ISL_650025, EPI_ISL_650026, EPI_ISL_650027, EPI_ISL_650028, EPI_ISL_650029, EPI_ISL_650030, EPI_ISL_650031, EPI_ISL_650032, EPI_ISL_650033, EPI_ISL_650034, EPI_ISL_650035, EPI_ISL_650036, EPI_ISL_650037, EPI_ISL_650038, EPI_ISL_650039, EPI_ISL_650040, EPI_ISL_650041, EPI_ISL_650042, EPI_ISL_650043, EPI_ISL_650044, EPI_ISL_650045, EPI_ISL_650046, EPI_ISL_650047, EPI_ISL_650048, EPI_ISL_650049, EPI_ISL_650050, EPI_ISL_650051, EPI_ISL_650052, EPI_ISL_650053, EPI_ISL_650054, EPI_ISL_650055, EPI_ISL_650056, EPI_ISL_650057, EPI_ISL_650058, EPI_ISL_650059, EPI_ISL_650060, EPI_ISL_650061, EPI_ISL_650062, EPI_ISL_650063, EPI_ISL_650064, EPI_ISL_650065, EPI_ISL_650066, EPI_ISL_650067, EPI_ISL_650068, EPI_ISL_650069, EPI_ISL_650070, EPI_ISL_650071, EPI_ISL_650072, EPI_ISL_650073, EPI_ISL_650074, EPI_ISL_650075, EPI_ISL_650076, EPI_ISL_650077, EPI_ISL_650078, EPI_ISL_650079, EPI_ISL_650080, EPI_ISL_650081, EPI_ISL_650082, EPI_ISL_650083, EPI_ISL_650084, EPI_ISL_650085, EPI_ISL_650086, EPI_ISL_650087, EPI_ISL_650088, EPI_ISL_650089, EPI_ISL_650090, EPI_ISL_650091, EPI_ISL_650092, EPI_ISL_650093, EPI_ISL_650094, EPI_ISL_650095, EPI_ISL_650096, EPI_ISL_650097, EPI_ISL_650098, EPI_ISL_650099, EPI_ISL_650100, EPI_ISL_650101, EPI_ISL_650102, EPI_ISL_650103, EPI_ISL_650104, EPI_ISL_650106, EPI_ISL_650109, EPI_ISL_650110, EPI_ISL_650111 | | | |
| see above | University of Michigan Clinical Microbiology Laboratory | Lauring Lab, University of Michigan, Department of Microbiology and Immunology | Valesano |
| EPI_ISL_650117 | Quadram Institute Bioscience | COVID-19 Genomics UK (COG-UK) Consortium | Dave J. Baker, Gemma L. Kay, Alp Aydin, Thanh Le-Viet, Steven Rudder, Ana P. Tedim, Anastasia Kolyva, Maria Diaz, Leonardo de Oliveira Martins, Nabil-Fareed Alikhan, Lizzie Meadows, Rachael Stanley, Ngozi Elumogo, Muhammed Yasir, Nicholas M. Thomson, Alexander J Trotter, Rachel Gilroy, Samuel Bloomfield, Claire Stuart, Andrew Bell, Reenesh Prakash, Samir Dervisevic, Alison E. Mather, John Wain, Mark Webber, Andrew J. Page, Justin O'Grady |
| EPI_ISL_650122 | Department of Pathology, University of Cambridge | COVID-19 Genomics UK (COG-UK) Consortium | Aminu S. Jahun, Yasmin Chaudhry, Grant Hall, Iliana Georgana, Myra Hosmillo, Martin D. Curran, Malte Pinckert, Surendra Parmar, Ian Goodfellow |
| EPI_ISL_650128 | Quadram Institute Bioscience | COVID-19 Genomics UK (COG-UK) Consortium | Dave J. Baker, Gemma L. Kay, Alp Aydin, Thanh Le-Viet, Steven Rudder, Ana P. Tedim, Anastasia Kolyva, Maria Diaz, Leonardo de Oliveira Martins, Nabil-Fareed Alikhan, Lizzie Meadows, Rachael Stanley, Ngozi Elumogo, Muhammed Yasir, Nicholas M. Thomson, Alexander J Trotter, Rachel Gilroy, Samuel Bloomfield, Claire Stuart, Andrew Bell, Reenesh Prakash, Samir Dervisevic, Alison E. Mather, John Wain, Mark Webber, Andrew J. Page, Justin O'Grady |
| EPI_ISL_650130 | Department of Pathology, University of Cambridge | COVID-19 Genomics UK (COG-UK) Consortium | Aminu S. Jahun, Yasmin Chaudhry, Grant Hall, Iliana Georgana, Myra Hosmillo, Martin D. Curran, Malte Pinckert, Surendra Parmar, Ian Goodfellow |
| EPI_ISL_650171 | University of Birmingham | COVID-19 Genomics UK (COG-UK) Consortium | Institute of Microbiology, University of Birmingham: Claire McMurray, Joanne Stockton, Samuel Nicholls, Radoslaw Poplawski, Will Rowe, Josh Quick, Nicholas Loman. University of Birmingham Testing Laboratory: Celina M Whalley, Andrew Bosworth, Charlotte Poxon, Kasun Wanigasooriya, Oliver Pickles, Mike Kidd, Alex Richter, Andrew D Beggs PHE Heartlands Lab: Husam Osman, Andrew Bosworth. Queen Elizabeth Hospital: Anna Casey |
| EPI_ISL_650173 | Virology Department, Royal Infirmary of Edinburgh, NHS Lothian / School of Biological Sciences, University of Edinburgh / Institute of Genetics and Molecular Medicine, University of Edinburgh | COVID-19 Genomics UK (COG-UK) Consortium | McHugh M, Dewar R, Rooke S, Gallagher M, Balcaza C, O'Toole Á, Scher E, Hill V, McCrone JT, Colquhoun R, Yu X, Jackson B, Rambaut A, Williams TC, Templeton K |
| EPI_ISL_650176 | University of Birmingham | COVID-19 Genomics UK (COG-UK) Consortium | Institute of Microbiology, University of Birmingham: Claire McMurray, Joanne Stockton, Samuel Nicholls, Radoslaw Poplawski, Will Rowe, Josh Quick, Nicholas Loman. University of Birmingham Testing Laboratory: Celina M Whalley, Andrew Bosworth, Charlotte Poxon, Kasun Wanigasooriya, Oliver Pickles, Mike Kidd, Alex Richter, Andrew D Beggs PHE Heartlands Lab: Husam Osman, Andrew Bosworth. Queen Elizabeth Hospital: Anna Casey |
| EPI_ISL_650190, EPI_ISL_650197 | Quadram Institute Bioscience | COVID-19 Genomics UK (COG-UK) Consortium | Dave J. Baker, Gemma L. Kay, Alp Aydin, Thanh Le-Viet, Steven Rudder, Ana P. Tedim, Anastasia Kolyva, Maria Diaz, Leonardo de Oliveira Martins, Nabil-Fareed Alikhan, Lizzie Meadows, Rachael Stanley, Ngozi Elumogo, Muhammed Yasir, Nicholas M. Thomson, Alexander J Trotter, Rachel Gilroy, Samuel Bloomfield, Claire Stuart, Andrew Bell, Reenesh Prakash, Samir Dervisevic, Alison E. Mather, John Wain, Mark Webber, Andrew J. Page, Justin O'Grady |
| EPI_ISL_650206 | Wales Specialist Virology Centre Sequencing lab: Pathogen Genomics Unit | COVID-19 Genomics UK (COG-UK) Consortium | Catherine Moore, Johnathan Evans, Laura Gifford, Malorie Perry, Simon Cottrell, Angela Marchbank, Alec Birchley, Alexander Adams, Amy Gaskin, Bree Gatica-Wilcox, Jason Coombes, Joel Southgate, Lauren Gilbert, Lee Graham, Nicole Pacchiarini, Sara Kumziene-Summerhayes, Sarah Taylor, Sophie Jones, Sara Rey, Matthew Bull, Joanne Watkins, Sally Corden, Tom Connor |
| EPI_ISL_650214 | Quadram Institute Bioscience | COVID-19 Genomics UK (COG-UK) Consortium | Dave J. Baker, Gemma L. Kay, Alp Aydin, Thanh Le-Viet, Steven Rudder, Ana P. Tedim, Anastasia Kolyva, Maria Diaz, Leonardo de Oliveira Martins, Nabil-Fareed Alikhan, Lizzie Meadows, Rachael Stanley, Ngozi Elumogo, Muhammed Yasir, Nicholas M. Thomson, Alexander J Trotter, Rachel Gilroy, Samuel Bloomfield, Claire Stuart, Andrew Bell, Reenesh Prakash, Samir Dervisevic, Alison E. Mather, John Wain, Mark Webber, Andrew J. Page, Justin O'Grady |
| EPI_ISL_650217, EPI_ISL_650240 | Department of Pathology, University of Cambridge | COVID-19 Genomics UK (COG-UK) Consortium | Aminu S. Jahun, Yasmin Chaudhry, Grant Hall, Iliana Georgana, Myra Hosmillo, Martin D. Curran, Malte Pinckert, Surendra Parmar, Ian Goodfellow |
| EPI_ISL_650248 | Centre for Enzyme Innovation, University of Portsmouth / Translational Research Laboratory, Portsmouth Hospitals NHS Trust | COVID-19 Genomics UK (COG-UK) Consortium | Angela Beckett,Yann Bourgeois,Garry Scarlett,Sharon Glaysher,Scott Elliott,Kelly Bicknell,Robert Impey,Allyson Lloyd,Sarah Wyllie,Ethan Butcher,Anoop Chauhan,Samuel Robson |
| EPI_ISL_650257 | Liverpool Clinical Laboratories | COVID-19 Genomics UK (COG-UK) Consortium | Sam Haldenby, Anita Lucaci, Steve Paterson, Julian Hiscox, Alistair Darby, M Almsaud, A Alrezaihi, Muhannad Alruwaili, Stuart D Armstrong, Jones Benjamin, Eleanor G Bentley, Anu Chawla, Jordan J Clark, Angela Cowell, Richard Eccles, Isabel García-Dorival, Matthew Gemmell, Alessandro Gerada, PKF Gilmore, Richard Gregory, Ximeng Han, Catherine Hartley, Margaret Hughes, Miren Iturriza-Gomara, James Johnson, L Luu, Jenifer Manson, Charlotte Nelson, Elaine O'Toole, Cassie Olateju, Rebekah Penrice-Randal , Lucille Rainbow, N.P Randle, Trevor Ian Robinson, Parul Sharma, Ghada T Shawli, James P Stewart, Neil Swainston, Ecaterina Vamos, Joanne Watts, Mark Whitehead |
| EPI_ISL_650264 | Virology Department, Royal Infirmary of Edinburgh, NHS Lothian / School of Biological Sciences, University of Edinburgh / Institute of Genetics and Molecular Medicine, University of Edinburgh | COVID-19 Genomics UK (COG-UK) Consortium | McHugh M, Dewar R, Rooke S, Gallagher M, Balcaza C, O'Toole Á, Scher E, Hill V, McCrone JT, Colquhoun R, Yu X, Jackson B, Rambaut A, Williams TC, Templeton K |
| EPI_ISL_650265 | Department of Pathology, University of Cambridge | COVID-19 Genomics UK (COG-UK) Consortium | Aminu S. Jahun, Yasmin Chaudhry, Grant Hall, Iliana Georgana, Myra Hosmillo, Martin D. Curran, Malte Pinckert, Surendra Parmar, Ian Goodfellow |
| EPI_ISL_650268 | Liverpool Clinical Laboratories | COVID-19 Genomics UK (COG-UK) Consortium | Sam Haldenby, Anita Lucaci, Steve Paterson, Julian Hiscox, Alistair Darby, M Almsaud, A Alrezaihi, Muhannad Alruwaili, Stuart D Armstrong, Jones Benjamin, Eleanor G Bentley, Anu Chawla, Jordan J Clark, Angela Cowell, Richard Eccles, Isabel García-Dorival, Matthew Gemmell, Alessandro Gerada, PKF Gilmore, Richard Gregory, Ximeng Han, Catherine Hartley, Margaret Hughes, Miren Iturriza-Gomara, James Johnson, L Luu, Jenifer Manson, Charlotte Nelson, Elaine O'Toole, Cassie Olateju, Rebekah Penrice-Randal , Lucille Rainbow, N.P Randle, Trevor Ian Robinson, Parul Sharma, Ghada T Shawli, James P Stewart, Neil Swainston, Ecaterina Vamos, Joanne Watts, Mark Whitehead |

| | | | |
|---|---|---|---|
| EPI_ISL_650280, EPI_ISL_650286 | Virology Department, Royal Infirmary of Edinburgh, NHS Lothian / School of Biological Sciences, University of Edinburgh / Institute of Genetics and Molecular Medicine, University of Edinburgh | COVID-19 Genomics UK (COG-UK) Consortium | McHugh M, Dewar R, Rooke S, Gallagher M, Balcaza C, O'Toole Á, Scher E, Hill V, McCrone JT, Colquhoun R, Yu X, Jackson B, Rambaut A, Williams TC, Templeton K |
| EPI_ISL_650293, EPI_ISL_650297 | West of Scotland Specialist Virology Centre, NHSGGC / MRC-University of Glasgow Centre for Virus Research | COVID-19 Genomics UK (COG-UK) Consortium | Ana da Silva Filipe, Natasha Johnson, Kathy Smollett, Daniel Mair, Stephen Carmichael, Alice Broos, Lily Tong, Jenna Nichols, Kyriaki Nomikou; Sarah McDonald; Richard Orton, Joseph Hughes, Sreenu Vattipally, David L Robertson; Alasdair MacLean, Rory Gunson; Sharif Shaaban, Matthew Holden; Rachel Blacow, Guy Mollett, Kathy Li, James Shepherd, Antonia Ho, Emma Thomson |
| EPI_ISL_650309, EPI_ISL_650319, EPI_ISL_650322, EPI_ISL_650325 | Quadram Institute Bioscience | COVID-19 Genomics UK (COG-UK) Consortium | Dave J. Baker, Gemma L. Kay, Alp Aydin, Thanh Le-Viet, Steven Rudder, Ana P. Tedim, Anastasia Kolyva, Maria Diaz, Leonardo de Oliveira Martins, Nabil-Fareed Alikhan, Lizzie Meadows, Rachael Stanley, Ngozi Elumogo, Muhammed Yasir, Nicholas M. Thomson, Alexander J Trotter, Rachel Gilroy, Samuel Bloomfield, Claire Stuart, Andrew Bell, Reenesh Prakash, Samir Dervisevic, Alison E. Mather, John Wain, Mark Webber, Andrew J. Page, Justin O'Grady |
| EPI_ISL_650328 | West of Scotland Specialist Virology Centre, NHSGGC / MRC-University of Glasgow Centre for Virus Research | COVID-19 Genomics UK (COG-UK) Consortium | Ana da Silva Filipe, Natasha Johnson, Kathy Smollett, Daniel Mair, Stephen Carmichael, Alice Broos, Lily Tong, Jenna Nichols, Kyriaki Nomikou; Sarah McDonald; Richard Orton, Joseph Hughes, Sreenu Vattipally, David L Robertson; Alasdair MacLean, Rory Gunson; Sharif Shaaban, Matthew Holden; Rachel Blacow, Guy Mollett, Kathy Li, James Shepherd, Antonia Ho, Emma Thomson |
| EPI_ISL_650330 | Quadram Institute Bioscience | COVID-19 Genomics UK (COG-UK) Consortium | Dave J. Baker, Gemma L. Kay, Alp Aydin, Thanh Le-Viet, Steven Rudder, Ana P. Tedim, Anastasia Kolyva, Maria Diaz, Leonardo de Oliveira Martins, Nabil-Fareed Alikhan, Lizzie Meadows, Rachael Stanley, Ngozi Elumogo, Muhammed Yasir, Nicholas M. Thomson, Alexander J Trotter, Rachel Gilroy, Samuel Bloomfield, Claire Stuart, Andrew Bell, Reenesh Prakash, Samir Dervisevic, Alison E. Mather, John Wain, Mark Webber, Andrew J. Page, Justin O'Grady |
| EPI_ISL_650334 | Regional Virus Laboratory, Belfast Health and Social Care Trust | COVID-19 Genomics UK (COG-UK) Consortium | Conall McCaughey, James McKenna, Tanya Curran, Susan Feeney, Alison Watt, Ciara Cox, Mairead Connor, Zoltan Molnar, David Simpson, Derek Fairley |
| EPI_ISL_650339 | Wales Specialist Virology Centre Sequencing lab: Pathogen Genomics Unit | COVID-19 Genomics UK (COG-UK) Consortium | Catherine Moore, Johnathan Evans, Laura Gifford, Malorie Perry, Simon Cottrell, Angela Marchbank, Alec Birchley, Alexander Adams, Amy Gaskin, Bree Gatica-Wilcox, Jason Coombes, Joel Southgate, Lauren Gilbert, Lee Graham, Nicole Pacchiarini, Sara Kumziene-Summerhayes, Sarah Taylor, Sophie Jones, Sara Rey, Matthew Bull, Joanne Watkins, Sally Corden, Tom Connor |
| EPI_ISL_650375 | West of Scotland Specialist Virology Centre, NHSGGC / MRC-University of Glasgow Centre for Virus Research | COVID-19 Genomics UK (COG-UK) Consortium | Ana da Silva Filipe, Natasha Johnson, Kathy Smollett, Daniel Mair, Stephen Carmichael, Alice Broos, Lily Tong, Jenna Nichols, Kyriaki Nomikou; Sarah McDonald; Richard Orton, Joseph Hughes, Sreenu Vattipally, David L Robertson; Alasdair MacLean, Rory Gunson; Sharif Shaaban, Matthew Holden; Rachel Blacow, Guy Mollett, Kathy Li, James Shepherd, Antonia Ho, Emma Thomson |
| EPI_ISL_650379 | Centre for Enzyme Innovation, University of Portsmouth / Translational Research Laboratory, Portsmouth Hospitals NHS Trust | COVID-19 Genomics UK (COG-UK) Consortium | Angela Beckett,Yann Bourgeois,Garry Scarlett,Sharon Glaysher,Scott Elliott,Kelly Bicknell,Robert Impey,Allyson Lloyd,Sarah Wyllie,Ethan Butcher,Anoop Chauhan,Samuel Robson |
| EPI_ISL_650400 | Department of Pathology, University of Cambridge | COVID-19 Genomics UK (COG-UK) Consortium | Aminu S. Jahun, Yasmin Chaudhry, Grant Hall, Iliana Georgana, Myra Hosmillo, Martin D. Curran, Malte Pinckert, Surendra Parmar, Ian Goodfellow |
| EPI_ISL_650402 | Quadram Institute Bioscience | COVID-19 Genomics UK (COG-UK) Consortium | Dave J. Baker, Gemma L. Kay, Alp Aydin, Thanh Le-Viet, Steven Rudder, Ana P. Tedim, Anastasia Kolyva, Maria Diaz, Leonardo de Oliveira Martins, Nabil-Fareed Alikhan, Lizzie Meadows, Rachael Stanley, Ngozi Elumogo, Muhammed Yasir, Nicholas M. Thomson, Alexander J Trotter, Rachel Gilroy, Samuel Bloomfield, Claire Stuart, Andrew Bell, Reenesh Prakash, Samir Dervisevic, Alison E. Mather, John Wain, Mark Webber, Andrew J. Page, Justin O'Grady |
| EPI_ISL_650412 | West of Scotland Specialist Virology Centre, NHSGGC / MRC-University of Glasgow Centre for Virus Research | COVID-19 Genomics UK (COG-UK) Consortium | Ana da Silva Filipe, Natasha Johnson, Kathy Smollett, Daniel Mair, Stephen Carmichael, Alice Broos, Lily Tong, Jenna Nichols, Kyriaki Nomikou; Sarah McDonald; Richard Orton, Joseph Hughes, Sreenu Vattipally, David L Robertson; Alasdair MacLean, Rory Gunson; Sharif Shaaban, Matthew Holden; Rachel Blacow, Guy Mollett, Kathy Li, James Shepherd, Antonia Ho, Emma Thomson |
| EPI_ISL_650413, EPI_ISL_650417 | Quadram Institute Bioscience | COVID-19 Genomics UK (COG-UK) Consortium | Dave J. Baker, Gemma L. Kay, Alp Aydin, Thanh Le-Viet, Steven Rudder, Ana P. Tedim, Anastasia Kolyva, Maria Diaz, Leonardo de Oliveira Martins, Nabil-Fareed Alikhan, Lizzie Meadows, Rachael Stanley, Ngozi Elumogo, Muhammed Yasir, Nicholas M. Thomson, Alexander J Trotter, Rachel Gilroy, Samuel Bloomfield, Claire Stuart, Andrew Bell, Reenesh Prakash, Samir Dervisevic, Alison E. Mather, John Wain, Mark Webber, Andrew J. Page, Justin O'Grady |
| EPI_ISL_650423 | Virology Department, Sheffield Teaching Hospitals NHS Foundation Trust/Department of Infection, Immunity and Cardiovascular Disease, The Medical School, University of Sheffield | COVID-19 Genomics UK (COG-UK) Consortium | Thushan de Silva, Matthew Parker, Nikki Smith, Adri Angyal, Rebecca Brown, Luke Green, Rachel Tucker, Paul Parsons, Danielle Groves, Katie Johnson, Laura Carrilero, Alex Keeley, Dave Partridge, Matthew Wyles, Benjamin Lindsey, Mehmet Yavuz, Mohammad Raza, Cariad Evans |
| EPI_ISL_650426, EPI_ISL_650427, EPI_ISL_650435 | Virology Department, Royal Infirmary of Edinburgh, NHS Lothian / School of Biological Sciences, University of Edinburgh / Institute of Genetics and Molecular Medicine, University of Edinburgh | COVID-19 Genomics UK (COG-UK) Consortium | McHugh M, Dewar R, Rooke S, Gallagher M, Balcaza C, O'Toole Á, Scher E, Hill V, McCrone JT, Colquhoun R, Yu X, Jackson B, Rambaut A, Williams TC, Templeton K |
| EPI_ISL_650438 | West of Scotland Specialist Virology Centre, NHSGGC / MRC-University of Glasgow Centre for Virus Research | COVID-19 Genomics UK (COG-UK) Consortium | Ana da Silva Filipe, Natasha Johnson, Kathy Smollett, Daniel Mair, Stephen Carmichael, Alice Broos, Lily Tong, Jenna Nichols, Kyriaki Nomikou; Sarah McDonald; Richard Orton, Joseph Hughes, Sreenu Vattipally, David L Robertson; Alasdair MacLean, Rory Gunson; Sharif Shaaban, Matthew Holden; Rachel Blacow, Guy Mollett, Kathy Li, James Shepherd, Antonia Ho, Emma Thomson |
| EPI_ISL_650446 | Quadram Institute Bioscience | COVID-19 Genomics UK (COG-UK) Consortium | Dave J. Baker, Gemma L. Kay, Alp Aydin, Thanh Le-Viet, Steven Rudder, Ana P. Tedim, Anastasia Kolyva, Maria Diaz, Leonardo de Oliveira Martins, Nabil-Fareed Alikhan, Lizzie Meadows, Rachael Stanley, Ngozi Elumogo, Muhammed Yasir, Nicholas M. Thomson, Alexander J Trotter, Rachel Gilroy, Samuel Bloomfield, Claire Stuart, Andrew Bell, Reenesh Prakash, Samir Dervisevic, Alison E. Mather, John Wain, Mark Webber, Andrew J. Page, Justin O'Grady |
| EPI_ISL_650456 | Virology Department, Royal Infirmary of Edinburgh, NHS Lothian / School of Biological Sciences, University of Edinburgh / Institute of Genetics and Molecular Medicine, University of Edinburgh | COVID-19 Genomics UK (COG-UK) Consortium | McHugh M, Dewar R, Rooke S, Gallagher M, Balcaza C, O'Toole Á, Scher E, Hill V, McCrone JT, Colquhoun R, Yu X, Jackson B, Rambaut A, Williams TC, Templeton K |
| EPI_ISL_650477, EPI_ISL_650483, EPI_ISL_650484 | West of Scotland Specialist Virology Centre, NHSGGC / MRC-University of Glasgow Centre for Virus Research | COVID-19 Genomics UK (COG-UK) Consortium | Ana da Silva Filipe, Natasha Johnson, Kathy Smollett, Daniel Mair, Stephen Carmichael, Alice Broos, Lily Tong, Jenna Nichols, Kyriaki Nomikou; Sarah McDonald; Richard Orton, Joseph Hughes, Sreenu Vattipally, David L Robertson; Alasdair MacLean, Rory Gunson; Sharif Shaaban, Matthew Holden; Rachel Blacow, Guy Mollett, Kathy Li, James Shepherd, Antonia Ho, Emma Thomson |
| EPI_ISL_650488 | Virology Department, Sheffield Teaching Hospitals NHS Foundation Trust/Department of Infection, Immunity and Cardiovascular Disease, The Medical School, University of Sheffield | COVID-19 Genomics UK (COG-UK) Consortium | Thushan de Silva, Matthew Parker, Nikki Smith, Adri Angyal, Rebecca Brown, Luke Green, Rachel Tucker, Paul Parsons, Danielle Groves, Katie Johnson, Laura Carrilero, Alex Keeley, Dave Partridge, Matthew Wyles, Benjamin Lindsey, Mehmet Yavuz, Mohammad Raza, Cariad Evans |
| EPI_ISL_650494 | West of Scotland Specialist Virology Centre, NHSGGC / MRC-University of Glasgow Centre for Virus Research | COVID-19 Genomics UK (COG-UK) Consortium | Ana da Silva Filipe, Natasha Johnson, Kathy Smollett, Daniel Mair, Stephen Carmichael, Alice Broos, Lily Tong, Jenna Nichols, Kyriaki Nomikou; Sarah McDonald; Richard Orton, Joseph Hughes, Sreenu Vattipally, David L Robertson; Alasdair MacLean, Rory Gunson; Sharif Shaaban, Matthew Holden; Rachel Blacow, Guy Mollett, Kathy Li, James Shepherd, Antonia Ho, Emma Thomson |
| EPI_ISL_650496 | Liverpool Clinical Laboratories | COVID-19 Genomics UK (COG-UK) Consortium | Sam Haldenby, Anita Lucaci, Steve Paterson, Julian Hiscox, Alistair Darby, M Almsaud, A Alrezaihi, Muhannad Alruwaili, Stuart D Armstrong, Jones Benjamin, Eleanor G Bentley, Anu Chawla, Jordan J Clark, Angela Cowell, Richard Eccles, Isabel García-Dorival, Matthew Gemmell, Alessandro Gerada, PKF Gilmore, Richard Gregory, Ximeng Han, Catherine Hartley, Margaret Hughes, Miren Iturriza-Gomara, James Johnson, L Luu, Jenifer Manson, Charlotte Nelson, Elaine O'Toole, Cassie Olateju, Rebekah Penrice-Randal , Lucille Rainbow, N.P Randle, Trevor Ian Robinson, Parul Sharma, Ghada T Shawli, James P Stewart, Neil Swainston, Ecaterina Vamos, Joanne Watts, Mark Whitehead |
| EPI_ISL_650508, EPI_ISL_650510, | West of Scotland Specialist Virology Centre, NHSGGC / | COVID-19 Genomics UK (COG-UK) Consortium | Ana da Silva Filipe, Natasha Johnson, Kathy Smollett, Daniel Mair, Stephen Carmichael, Alice Broos, Lily Tong, Jenna Nichols, Kyriaki Nomikou; Sarah |

| | | | |
|---|---|---|---|
| EPI_ISL_650521, EPI_ISL_650525 | MRC-University of Glasgow Centre for Virus Research | | McDonald; Richard Orton, Joseph Hughes, Sreenu Vattipally, David L Robertson; Alasdair MacLean, Rory Gunson; Sharif Shaaban, Matthew Holden; Rachel Blacow, Guy Mollett, Kathy Li, James Shepherd, Antonia Ho, Emma Thomson |
| EPI_ISL_650528, EPI_ISL_650545 | Quadram Institute Bioscience | COVID-19 Genomics UK (COG-UK) Consortium | Dave J. Baker, Gemma L. Kay, Alp Aydin, Thanh Le-Viet, Steven Rudder, Ana P. Tedim, Anastasia Kolyva, Maria Diaz, Leonardo de Oliveira Martins, Nabil-Fareed Alikhan, Lizzie Meadows, Rachael Stanley, Ngozi Elumogo, Muhammed Yasir, Nicholas M. Thomson, Alexander J Trotter, Rachel Gilroy, Samuel Bloomfield, Claire Stuart, Andrew Bell, Reenesh Prakash, Samir Dervisevic, Alison E. Mather, John Wain, Mark Webber, Andrew J. Page, Justin O'Grady |
| EPI_ISL_650562, EPI_ISL_650596 | Department of Pathology, University of Cambridge | COVID-19 Genomics UK (COG-UK) Consortium | Aminu S. Jahun, Yasmin Chaudhry, Grant Hall, Iliana Georgana, Myra Hosmillo, Martin D. Curran, Malte Pinckert, Surendra Parmar, Ian Goodfellow |
| EPI_ISL_650601, EPI_ISL_650602, EPI_ISL_650603, EPI_ISL_650604, EPI_ISL_650605 | Centre for Enzyme Innovation, University of Portsmouth / Translational Research Laboratory, Portsmouth Hospitals NHS Trust | COVID-19 Genomics UK (COG-UK) Consortium | Angela Beckett,Yann Bourgeois,Garry Scarlett,Sharon Glaysher,Scott Elliott,Kelly Bicknell,Robert Impey,Allyson Lloyd,Sarah Wyllie,Ethan Butcher,Anoop Chauhan,Samuel Robson |
| EPI_ISL_650618 | Virology Department, Royal Infirmary of Edinburgh, NHS Lothian / School of Biological Sciences, University of Edinburgh / Institute of Genetics and Molecular Medicine, University of Edinburgh | COVID-19 Genomics UK (COG-UK) Consortium | McHugh M, Dewar R, Rooke S, Gallagher M, Balcaza C, O'Toole Á, Scher E, Hill V, McCrone JT, Colquhoun R, Yu X, Jackson B, Rambaut A, Williams TC, Templeton K |
| EPI_ISL_650633 | Department of Pathology, University of Cambridge | COVID-19 Genomics UK (COG-UK) Consortium | Aminu S. Jahun, Yasmin Chaudhry, Grant Hall, Iliana Georgana, Myra Hosmillo, Martin D. Curran, Malte Pinckert, Surendra Parmar, Ian Goodfellow |
| EPI_ISL_650660 | Wales Specialist Virology Centre Sequencing lab: Pathogen Genomics Unit | COVID-19 Genomics UK (COG-UK) Consortium | Catherine Moore, Johnathan Evans, Laura Gifford, Malorie Perry, Simon Cottrell, Angela Marchbank, Alec Birchley, Alexander Adams, Amy Gaskin, Bree Gatica-Wilcox, Jason Coombes, Joel Southgate, Lauren Gilbert, Lee Graham, Nicole Pacchiarini, Sara Kumziene-Summerhayes, Sarah Taylor, Sophie Jones, Sara Rey, Matthew Bull, Joanne Watkins, Sally Corden, Tom Connor |
| EPI_ISL_650661, EPI_ISL_650666 | Department of Pathology, University of Cambridge | COVID-19 Genomics UK (COG-UK) Consortium | Aminu S. Jahun, Yasmin Chaudhry, Grant Hall, Iliana Georgana, Myra Hosmillo, Martin D. Curran, Malte Pinckert, Surendra Parmar, Ian Goodfellow |
| EPI_ISL_650674, EPI_ISL_650678, EPI_ISL_650679 | Virology Department, Sheffield Teaching Hospitals NHS Foundation Trust/Department of Infection, Immunity and Cardiovascular Disease, The Medical School, University of Sheffield | COVID-19 Genomics UK (COG-UK) Consortium | Thushan de Silva, Matthew Parker, Nikki Smith, Adri Angyal, Rebecca Brown, Luke Green, Rachel Tucker, Paul Parsons, Danielle Groves, Katie Johnson, Laura Carrilero, Alex Keeley, Dave Partridge, Matthew Wyles, Benjamin Lindsey, Mehmet Yavuz, Mohammad Raza, Cariad Evans |
| EPI_ISL_650681 | Quadram Institute Bioscience | COVID-19 Genomics UK (COG-UK) Consortium | Dave J. Baker, Gemma L. Kay, Alp Aydin, Thanh Le-Viet, Steven Rudder, Ana P. Tedim, Anastasia Kolyva, Maria Diaz, Leonardo de Oliveira Martins, Nabil-Fareed Alikhan, Lizzie Meadows, Rachael Stanley, Ngozi Elumogo, Muhammed Yasir, Nicholas M. Thomson, Alexander J Trotter, Rachel Gilroy, Samuel Bloomfield, Claire Stuart, Andrew Bell, Reenesh Prakash, Samir Dervisevic, Alison E. Mather, John Wain, Mark Webber, Andrew J. Page, Justin O'Grady |
| EPI_ISL_650682 | Department of Pathology, University of Cambridge | COVID-19 Genomics UK (COG-UK) Consortium | Aminu S. Jahun, Yasmin Chaudhry, Grant Hall, Iliana Georgana, Myra Hosmillo, Martin D. Curran, Malte Pinckert, Surendra Parmar, Ian Goodfellow |
| EPI_ISL_650683 | Quadram Institute Bioscience | COVID-19 Genomics UK (COG-UK) Consortium | Dave J. Baker, Gemma L. Kay, Alp Aydin, Thanh Le-Viet, Steven Rudder, Ana P. Tedim, Anastasia Kolyva, Maria Diaz, Leonardo de Oliveira Martins, Nabil-Fareed Alikhan, Lizzie Meadows, Rachael Stanley, Ngozi Elumogo, Muhammed Yasir, Nicholas M. Thomson, Alexander J Trotter, Rachel Gilroy, Samuel Bloomfield, Claire Stuart, Andrew Bell, Reenesh Prakash, Samir Dervisevic, Alison E. Mather, John Wain, Mark Webber, Andrew J. Page, Justin O'Grady |
| EPI_ISL_650695 | Wales Specialist Virology Centre Sequencing lab: Pathogen Genomics Unit | COVID-19 Genomics UK (COG-UK) Consortium | Catherine Moore, Johnathan Evans, Laura Gifford, Malorie Perry, Simon Cottrell, Angela Marchbank, Alec Birchley, Alexander Adams, Amy Gaskin, Bree Gatica-Wilcox, Jason Coombes, Joel Southgate, Lauren Gilbert, Lee Graham, Nicole Pacchiarini, Sara Kumziene-Summerhayes, Sarah Taylor, Sophie Jones, Sara Rey, Matthew Bull, Joanne Watkins, Sally Corden, Tom Connor |
| EPI_ISL_650705, EPI_ISL_650706 | Virology Department, Sheffield Teaching Hospitals NHS Foundation Trust/Department of Infection, Immunity and Cardiovascular Disease, The Medical School, University of Sheffield | COVID-19 Genomics UK (COG-UK) Consortium | Thushan de Silva, Matthew Parker, Nikki Smith, Adri Angyal, Rebecca Brown, Luke Green, Rachel Tucker, Paul Parsons, Danielle Groves, Katie Johnson, Laura Carrilero, Alex Keeley, Dave Partridge, Matthew Wyles, Benjamin Lindsey, Mehmet Yavuz, Mohammad Raza, Cariad Evans |
| EPI_ISL_650734, EPI_ISL_650739, EPI_ISL_650746, EPI_ISL_650747, EPI_ISL_650748, EPI_ISL_650749 | West of Scotland Specialist Virology Centre, NHSGGC / MRC-University of Glasgow Centre for Virus Research | COVID-19 Genomics UK (COG-UK) Consortium | Ana da Silva Filipe, Natasha Johnson, Kathy Smollett, Daniel Mair, Stephen Carmichael, Alice Broos, Lily Tong, Jenna Nichols, Kyriaki Nomikou; Sarah McDonald; Richard Orton, Joseph Hughes, Sreenu Vattipally, David L Robertson; Alasdair MacLean, Rory Gunson; Sharif Shaaban, Matthew Holden; Rachel Blacow, Guy Mollett, Kathy Li, James Shepherd, Antonia Ho, Emma Thomson |
| EPI_ISL_650756 | Quadram Institute Bioscience | COVID-19 Genomics UK (COG-UK) Consortium | Dave J. Baker, Gemma L. Kay, Alp Aydin, Thanh Le-Viet, Steven Rudder, Ana P. Tedim, Anastasia Kolyva, Maria Diaz, Leonardo de Oliveira Martins, Nabil-Fareed Alikhan, Lizzie Meadows, Rachael Stanley, Ngozi Elumogo, Muhammed Yasir, Nicholas M. Thomson, Alexander J Trotter, Rachel Gilroy, Samuel Bloomfield, Claire Stuart, Andrew Bell, Reenesh Prakash, Samir Dervisevic, Alison E. Mather, John Wain, Mark Webber, Andrew J. Page, Justin O'Grady |
| EPI_ISL_650757 | Virology Department, Royal Infirmary of Edinburgh, NHS Lothian / School of Biological Sciences, University of Edinburgh / Institute of Genetics and Molecular Medicine, University of Edinburgh | COVID-19 Genomics UK (COG-UK) Consortium | McHugh M, Dewar R, Rooke S, Gallagher M, Balcaza C, O'Toole Á, Scher E, Hill V, McCrone JT, Colquhoun R, Yu X, Jackson B, Rambaut A, Williams TC, Templeton K |
| EPI_ISL_650758 | Quadram Institute Bioscience | COVID-19 Genomics UK (COG-UK) Consortium | Dave J. Baker, Gemma L. Kay, Alp Aydin, Thanh Le-Viet, Steven Rudder, Ana P. Tedim, Anastasia Kolyva, Maria Diaz, Leonardo de Oliveira Martins, Nabil-Fareed Alikhan, Lizzie Meadows, Rachael Stanley, Ngozi Elumogo, Muhammed Yasir, Nicholas M. Thomson, Alexander J Trotter, Rachel Gilroy, Samuel Bloomfield, Claire Stuart, Andrew Bell, Reenesh Prakash, Samir Dervisevic, Alison E. Mather, John Wain, Mark Webber, Andrew J. Page, Justin O'Grady |
| EPI_ISL_650759 | West of Scotland Specialist Virology Centre, NHSGGC / MRC-University of Glasgow Centre for Virus Research | COVID-19 Genomics UK (COG-UK) Consortium | Ana da Silva Filipe, Natasha Johnson, Kathy Smollett, Daniel Mair, Stephen Carmichael, Alice Broos, Lily Tong, Jenna Nichols, Kyriaki Nomikou; Sarah McDonald; Richard Orton, Joseph Hughes, Sreenu Vattipally, David L Robertson; Alasdair MacLean, Rory Gunson; Sharif Shaaban, Matthew Holden; Rachel Blacow, Guy Mollett, Kathy Li, James Shepherd, Antonia Ho, Emma Thomson |
| EPI_ISL_650761, EPI_ISL_650765 | Quadram Institute Bioscience | COVID-19 Genomics UK (COG-UK) Consortium | Dave J. Baker, Gemma L. Kay, Alp Aydin, Thanh Le-Viet, Steven Rudder, Ana P. Tedim, Anastasia Kolyva, Maria Diaz, Leonardo de Oliveira Martins, Nabil-Fareed Alikhan, Lizzie Meadows, Rachael Stanley, Ngozi Elumogo, Muhammed Yasir, Nicholas M. Thomson, Alexander J Trotter, Rachel Gilroy, Samuel Bloomfield, Claire Stuart, Andrew Bell, Reenesh Prakash, Samir Dervisevic, Alison E. Mather, John Wain, Mark Webber, Andrew J. Page, Justin O'Grady |
| EPI_ISL_650768 | Department of Pathology, University of Cambridge | COVID-19 Genomics UK (COG-UK) Consortium | Aminu S. Jahun, Yasmin Chaudhry, Grant Hall, Iliana Georgana, Myra Hosmillo, Martin D. Curran, Malte Pinckert, Surendra Parmar, Ian Goodfellow |
| EPI_ISL_650777, EPI_ISL_650778, EPI_ISL_650779 | Virology Department, Royal Infirmary of Edinburgh, NHS Lothian / School of Biological Sciences, University of Edinburgh / Institute of Genetics and Molecular Medicine, University of Edinburgh | COVID-19 Genomics UK (COG-UK) Consortium | McHugh M, Dewar R, Rooke S, Gallagher M, Balcaza C, O'Toole Á, Scher E, Hill V, McCrone JT, Colquhoun R, Yu X, Jackson B, Rambaut A, Williams TC, Templeton K |
| EPI_ISL_650791 | University of Birmingham | COVID-19 Genomics UK (COG-UK) Consortium | Institute of Microbiology, University of Birmingham: Claire McMurray, Joanne Stockton, Samuel Nicholls, Radoslaw Poplawski, Will Rowe, Josh Quick, Nicholas Loman. University of Birmingham Testing Laboratory: Celina M Whalley, Andrew Bosworth, Charlotte Poxon, Kasun Wanigasooriya, Oliver Pickles, Mike Kidd, Alex Richter, Andrew D Beggs PHE Heartlands Lab: Husam Osman, Andrew Bosworth. Queen Elizabeth Hospital: Anna Casey |
| EPI_ISL_650802, EPI_ISL_650804 | West of Scotland Specialist Virology Centre, NHSGGC / MRC-University of Glasgow Centre for Virus Research | COVID-19 Genomics UK (COG-UK) Consortium | Ana da Silva Filipe, Natasha Johnson, Kathy Smollett, Daniel Mair, Stephen Carmichael, Alice Broos, Lily Tong, Jenna Nichols, Kyriaki Nomikou; Sarah McDonald; Richard Orton, Joseph Hughes, Sreenu Vattipally, David L Robertson; Alasdair MacLean, Rory Gunson; Sharif Shaaban, Matthew Holden; Rachel Blacow, Guy Mollett, Kathy Li, James Shepherd, Antonia Ho, Emma Thomson |
| EPI_ISL_650807 | Virology Department, Sheffield Teaching Hospitals NHS Foundation Trust/Department of Infection, Immunity and Cardiovascular Disease, The Medical School, University of | COVID-19 Genomics UK (COG-UK) Consortium | Thushan de Silva, Matthew Parker, Nikki Smith, Adri Angyal, Rebecca Brown, Luke Green, Rachel Tucker, Paul Parsons, Danielle Groves, Katie Johnson, Laura Carrilero, Alex Keeley, Dave Partridge, Matthew Wyles, Benjamin Lindsey, Mehmet Yavuz, Mohammad Raza, Cariad Evans |

| | | | |
|---|---|---|---|
| | Sheffield | | |
| EPI_ISL_650812, EPI_ISL_650813 | Virology Department, Royal Infirmary of Edinburgh, NHS Lothian / School of Biological Sciences, University of Edinburgh / Institute of Genetics and Molecular Medicine, University of Edinburgh | COVID-19 Genomics UK (COG-UK) Consortium | McHugh M, Dewar R, Rooke S, Gallagher M, Balcaza C, O'Toole Á, Scher E, Hill V, McCrone JT, Colquhoun R, Yu X, Jackson B, Rambaut A, Williams TC, Templeton K |
| EPI_ISL_650817 | Quadram Institute Bioscience | COVID-19 Genomics UK (COG-UK) Consortium | Dave J. Baker, Gemma L. Kay, Alp Aydin, Thanh Le-Viet, Steven Rudder, Ana P. Tedim, Anastasia Kolyva, Maria Diaz, Leonardo de Oliveira Martins, Nabil-Fareed Alikhan, Lizzie Meadows, Rachael Stanley, Ngozi Elumogo, Muhammed Yasir, Nicholas M. Thomson, Alexander J Trotter, Rachel Gilroy, Samuel Bloomfield, Claire Stuart, Andrew Bell, Reenesh Prakash, Samir Dervisevic, Alison E. Mather, John Wain, Mark Webber, Andrew J. Page, Justin O'Grady |
| EPI_ISL_650822, EPI_ISL_650823 | University of Birmingham | COVID-19 Genomics UK (COG-UK) Consortium | Institute of Microbiology, University of Birmingham: Claire McMurray, Joanne Stockton, Samuel Nicholls, Radoslaw Poplawski, Will Rowe, Josh Quick, Nicholas Loman. University of Birmingham Testing Laboratory: Celina M Whalley, Andrew Bosworth, Charlotte Poxon, Kasun Wanigasooriya, Oliver Pickles, Mike Kidd, Alex Richter, Andrew D Beggs PHE Heartlands Lab: Husam Osman, Andrew Bosworth. Queen Elizabeth Hospital: Anna Casey |
| EPI_ISL_650828 | Quadram Institute Bioscience | COVID-19 Genomics UK (COG-UK) Consortium | Dave J. Baker, Gemma L. Kay, Alp Aydin, Thanh Le-Viet, Steven Rudder, Ana P. Tedim, Anastasia Kolyva, Maria Diaz, Leonardo de Oliveira Martins, Nabil-Fareed Alikhan, Lizzie Meadows, Rachael Stanley, Ngozi Elumogo, Muhammed Yasir, Nicholas M. Thomson, Alexander J Trotter, Rachel Gilroy, Samuel Bloomfield, Claire Stuart, Andrew Bell, Reenesh Prakash, Samir Dervisevic, Alison E. Mather, John Wain, Mark Webber, Andrew J. Page, Justin O'Grady |
| EPI_ISL_650834 | Virology Department, Royal Infirmary of Edinburgh, NHS Lothian / School of Biological Sciences, University of Edinburgh / Institute of Genetics and Molecular Medicine, University of Edinburgh | COVID-19 Genomics UK (COG-UK) Consortium | McHugh M, Dewar R, Rooke S, Gallagher M, Balcaza C, O'Toole Á, Scher E, Hill V, McCrone JT, Colquhoun R, Yu X, Jackson B, Rambaut A, Williams TC, Templeton K |
| EPI_ISL_650838, EPI_ISL_650839, EPI_ISL_650842, EPI_ISL_650845, EPI_ISL_650864, EPI_ISL_650865, EPI_ISL_650872, EPI_ISL_650873, EPI_ISL_650874, EPI_ISL_650875 | West of Scotland Specialist Virology Centre, NHSGGC / MRC-University of Glasgow Centre for Virus Research | COVID-19 Genomics UK (COG-UK) Consortium | Ana da Silva Filipe, Natasha Johnson, Kathy Smollett, Daniel Mair, Stephen Carmichael, Alice Broos, Lily Tong, Jenna Nichols, Kyriaki Nomikou; Sarah McDonald; Richard Orton, Joseph Hughes, Sreenu Vattipally, David L Robertson; Alasdair MacLean, Rory Gunson; Sharif Shaaban, Matthew Holden; Rachel Blacow, Guy Mollett, Kathy Li, James Shepherd, Antonia Ho, Emma Thomson |
| EPI_ISL_650894, EPI_ISL_650895 | Quadram Institute Bioscience | COVID-19 Genomics UK (COG-UK) Consortium | Dave J. Baker, Gemma L. Kay, Alp Aydin, Thanh Le-Viet, Steven Rudder, Ana P. Tedim, Anastasia Kolyva, Maria Diaz, Leonardo de Oliveira Martins, Nabil-Fareed Alikhan, Lizzie Meadows, Rachael Stanley, Ngozi Elumogo, Muhammed Yasir, Nicholas M. Thomson, Alexander J Trotter, Rachel Gilroy, Samuel Bloomfield, Claire Stuart, Andrew Bell, Reenesh Prakash, Samir Dervisevic, Alison E. Mather, John Wain, Mark Webber, Andrew J. Page, Justin O'Grady |
| EPI_ISL_650896, EPI_ISL_650914 | Wales Specialist Virology Centre Sequencing lab: Pathogen Genomics Unit | COVID-19 Genomics UK (COG-UK) Consortium | Catherine Moore, Johnathan Evans, Laura Gifford, Malorie Perry, Simon Cottrell, Angela Marchbank, Alec Birchley, Alexander Adams, Amy Gaskin, Bree Gatica-Wilcox, Jason Coombes, Joel Southgate, Lauren Gilbert, Lee Graham, Nicole Pacchiarini, Sara Kumziene-Summerhayes, Sarah Taylor, Sophie Jones, Sara Rey, Matthew Bull, Joanne Watkins, Sally Corden, Tom Connor |
| EPI_ISL_650931, EPI_ISL_650944, EPI_ISL_650953, EPI_ISL_650954, EPI_ISL_650955, EPI_ISL_650974, EPI_ISL_650975, EPI_ISL_650976, EPI_ISL_650977 | Quadram Institute Bioscience | COVID-19 Genomics UK (COG-UK) Consortium | Dave J. Baker, Gemma L. Kay, Alp Aydin, Thanh Le-Viet, Steven Rudder, Ana P. Tedim, Anastasia Kolyva, Maria Diaz, Leonardo de Oliveira Martins, Nabil-Fareed Alikhan, Lizzie Meadows, Rachael Stanley, Ngozi Elumogo, Muhammed Yasir, Nicholas M. Thomson, Alexander J Trotter, Rachel Gilroy, Samuel Bloomfield, Claire Stuart, Andrew Bell, Reenesh Prakash, Samir Dervisevic, Alison E. Mather, John Wain, Mark Webber, Andrew J. Page, Justin O'Grady |
| EPI_ISL_650978 | Department of Pathology, University of Cambridge | COVID-19 Genomics UK (COG-UK) Consortium | Aminu S. Jahun, Yasmin Chaudhry, Grant Hall, Iliana Georgana, Myra Hosmillo, Martin D. Curran, Malte Pinckert, Surendra Parmar, Ian Goodfellow |
| EPI_ISL_650995 | University of Birmingham | COVID-19 Genomics UK (COG-UK) Consortium | Institute of Microbiology, University of Birmingham: Claire McMurray, Joanne Stockton, Samuel Nicholls, Radoslaw Poplawski, Will Rowe, Josh Quick, Nicholas Loman. University of Birmingham Testing Laboratory: Celina M Whalley, Andrew Bosworth, Charlotte Poxon, Kasun Wanigasooriya, Oliver Pickles, Mike Kidd, Alex Richter, Andrew D Beggs PHE Heartlands Lab: Husam Osman, Andrew Bosworth. Queen Elizabeth Hospital: Anna Casey |
| EPI_ISL_651001 | Department of Pathology, University of Cambridge | COVID-19 Genomics UK (COG-UK) Consortium | Aminu S. Jahun, Yasmin Chaudhry, Grant Hall, Iliana Georgana, Myra Hosmillo, Martin D. Curran, Malte Pinckert, Surendra Parmar, Ian Goodfellow |
| EPI_ISL_651012 | Virology Department, Royal Infirmary of Edinburgh, NHS Lothian / School of Biological Sciences, University of Edinburgh / Institute of Genetics and Molecular Medicine, University of Edinburgh | COVID-19 Genomics UK (COG-UK) Consortium | McHugh M, Dewar R, Rooke S, Gallagher M, Balcaza C, O'Toole Á, Scher E, Hill V, McCrone JT, Colquhoun R, Yu X, Jackson B, Rambaut A, Williams TC, Templeton K |
| EPI_ISL_651048 | Quadram Institute Bioscience | COVID-19 Genomics UK (COG-UK) Consortium | Dave J. Baker, Gemma L. Kay, Alp Aydin, Thanh Le-Viet, Steven Rudder, Ana P. Tedim, Anastasia Kolyva, Maria Diaz, Leonardo de Oliveira Martins, Nabil-Fareed Alikhan, Lizzie Meadows, Rachael Stanley, Ngozi Elumogo, Muhammed Yasir, Nicholas M. Thomson, Alexander J Trotter, Rachel Gilroy, Samuel Bloomfield, Claire Stuart, Andrew Bell, Reenesh Prakash, Samir Dervisevic, Alison E. Mather, John Wain, Mark Webber, Andrew J. Page, Justin O'Grady |
| EPI_ISL_651049, EPI_ISL_651050, EPI_ISL_651051, EPI_ISL_651052 | Department of Pathology, University of Cambridge | COVID-19 Genomics UK (COG-UK) Consortium | Aminu S. Jahun, Yasmin Chaudhry, Grant Hall, Iliana Georgana, Myra Hosmillo, Martin D. Curran, Malte Pinckert, Surendra Parmar, Ian Goodfellow |
| EPI_ISL_651068, EPI_ISL_651069 | Centre for Enzyme Innovation, University of Portsmouth / Translational Research Laboratory, Portsmouth Hospitals NHS Trust | COVID-19 Genomics UK (COG-UK) Consortium | Angela Beckett,Yann Bourgeois,Garry Scarlett,Sharon Glaysher,Scott Elliott,Kelly Bicknell,Robert Impey,Allyson Lloyd,Sarah Wyllie,Ethan Butcher,Anoop Chauhan,Samuel Robson |
| EPI_ISL_651079 | Quadram Institute Bioscience | COVID-19 Genomics UK (COG-UK) Consortium | Dave J. Baker, Gemma L. Kay, Alp Aydin, Thanh Le-Viet, Steven Rudder, Ana P. Tedim, Anastasia Kolyva, Maria Diaz, Leonardo de Oliveira Martins, Nabil-Fareed Alikhan, Lizzie Meadows, Rachael Stanley, Ngozi Elumogo, Muhammed Yasir, Nicholas M. Thomson, Alexander J Trotter, Rachel Gilroy, Samuel Bloomfield, Claire Stuart, Andrew Bell, Reenesh Prakash, Samir Dervisevic, Alison E. Mather, John Wain, Mark Webber, Andrew J. Page, Justin O'Grady |
| EPI_ISL_651108 | Wales Specialist Virology Centre Sequencing lab: Pathogen Genomics Unit | COVID-19 Genomics UK (COG-UK) Consortium | Catherine Moore, Johnathan Evans, Laura Gifford, Malorie Perry, Simon Cottrell, Angela Marchbank, Alec Birchley, Alexander Adams, Amy Gaskin, Bree Gatica-Wilcox, Jason Coombes, Joel Southgate, Lauren Gilbert, Lee Graham, Nicole Pacchiarini, Sara Kumziene-Summerhayes, Sarah Taylor, Sophie Jones, Sara Rey, Matthew Bull, Joanne Watkins, Sally Corden, Tom Connor |
| EPI_ISL_651119 | Quadram Institute Bioscience | COVID-19 Genomics UK (COG-UK) Consortium | Dave J. Baker, Gemma L. Kay, Alp Aydin, Thanh Le-Viet, Steven Rudder, Ana P. Tedim, Anastasia Kolyva, Maria Diaz, Leonardo de Oliveira Martins, Nabil-Fareed Alikhan, Lizzie Meadows, Rachael Stanley, Ngozi Elumogo, Muhammed Yasir, Nicholas M. Thomson, Alexander J Trotter, Rachel Gilroy, Samuel Bloomfield, Claire Stuart, Andrew Bell, Reenesh Prakash, Samir Dervisevic, Alison E. Mather, John Wain, Mark Webber, Andrew J. Page, Justin O'Grady |
| EPI_ISL_651121 | Virology Department, Sheffield Teaching Hospitals NHS Foundation Trust/Department of Infection, Immunity and Cardiovascular Disease, The Medical School, University of Sheffield | COVID-19 Genomics UK (COG-UK) Consortium | Thushan de Silva, Matthew Parker, Nikki Smith, Adri Angyal, Rebecca Brown, Luke Green, Rachel Tucker, Paul Parsons, Danielle Groves, Katie Johnson, Laura Carrilero, Alex Keeley, Dave Partridge, Matthew Wyles, Benjamin Lindsey, Mehmet Yavuz, Mohammad Raza, Cariad Evans |
| EPI_ISL_651155 | Liverpool Clinical Laboratories | COVID-19 Genomics UK (COG-UK) Consortium | Sam Haldenby, Anita Lucaci, Steve Paterson, Julian Hiscox, Alistair Darby, M Almsaud, A Alrezaihi, Muhannad Alruwaili, Stuart D Armstrong, Jones Benjamin, Eleanor G Bentley, Anu Chawla, Jordan J Clark, Angela Cowell, Richard Eccles, Isabel García-Dorival, Matthew Gemmell, Alessandro Gerada, PKF Gilmore, Richard Gregory, Ximeng Han, Catherine Hartley, Margaret Hughes, Miren Iturriza-Gomara, James Johnson, L Luu, Jenifer Manson, Charlotte Nelson, Elaine O'Toole, Cassie Olateju, Rebekah Penrice-Randal , Lucille Rainbow, N.P Randle, Trevor Ian Robinson, Parul Sharma, Ghada T Shawli, James P Stewart, Neil Swainston, Ecaterina Vamos, Joanne Watts, Mark Whitehead |
| EPI_ISL_651156 | Quadram Institute Bioscience | COVID-19 Genomics UK (COG-UK) Consortium | Dave J. Baker, Gemma L. Kay, Alp Aydin, Thanh Le-Viet, Steven Rudder, Ana P. Tedim, Anastasia Kolyva, Maria Diaz, Leonardo de Oliveira Martins, |

| | | | |
|---|---|---|---|
| | | | Nabil-Fareed Alikhan, Lizzie Meadows, Rachael Stanley, Ngozi Elumogo, Muhammed Yasir, Nicholas M. Thomson, Alexander J Trotter, Rachel Gilroy, Samuel Bloomfield, Claire Stuart, Andrew Bell, Reenesh Prakash, Samir Dervisevic, Alison E. Mather, John Wain, Mark Webber, Andrew J. Page, Justin O'Grady |
| EPI_ISL_651160 | West of Scotland Specialist Virology Centre, NHSGGC / MRC-University of Glasgow Centre for Virus Research | COVID-19 Genomics UK (COG-UK) Consortium | Ana da Silva Filipe, Natasha Johnson, Kathy Smollett, Daniel Mair, Stephen Carmichael, Alice Broos, Lily Tong, Jenna Nichols, Kyriaki Nomikou; Sarah McDonald; Richard Orton, Joseph Hughes, Sreenu Vattipally, David L Robertson; Alasdair MacLean, Rory Gunson; Sharif Shaaban, Matthew Holden; Rachel Blacow, Guy Mollett, Kathy Li, James Shepherd, Antonia Ho, Emma Thomson |
| EPI_ISL_651167, EPI_ISL_651168 | Quadram Institute Bioscience | COVID-19 Genomics UK (COG-UK) Consortium | Dave J. Baker, Gemma L. Kay, Alp Aydin, Thanh Le-Viet, Steven Rudder, Ana P. Tedim, Anastasia Kolyva, Maria Diaz, Leonardo de Oliveira Martins, Nabil-Fareed Alikhan, Lizzie Meadows, Rachael Stanley, Ngozi Elumogo, Muhammed Yasir, Nicholas M. Thomson, Alexander J Trotter, Rachel Gilroy, Samuel Bloomfield, Claire Stuart, Andrew Bell, Reenesh Prakash, Samir Dervisevic, Alison E. Mather, John Wain, Mark Webber, Andrew J. Page, Justin O'Grady |
| EPI_ISL_651169 | Virology Department, Royal Infirmary of Edinburgh, NHS Lothian / School of Biological Sciences, University of Edinburgh / Institute of Genetics and Molecular Medicine, University of Edinburgh | COVID-19 Genomics UK (COG-UK) Consortium | McHugh M, Dewar R, Rooke S, Gallagher M, Balcaza C, O'Toole Á, Scher E, Hill V, McCrone JT, Colquhoun R, Yu X, Jackson B, Rambaut A, Williams TC, Templeton K |
| EPI_ISL_651170, EPI_ISL_651175 | Liverpool Clinical Laboratories | COVID-19 Genomics UK (COG-UK) Consortium | Sam Haldenby, Anita Lucaci, Steve Paterson, Julian Hiscox, Alistair Darby, M Almsaud, A Alrezaihi, Muhannad Alruwaili, Stuart D Armstrong, Jones Benjamin, Eleanor G Bentley, Anu Chawla, Jordan J Clark, Angela Cowell, Richard Eccles, Isabel García-Dorival, Matthew Gemmell, Alessandro Gerada, PKF Gilmore, Richard Gregory, Ximeng Han, Catherine Hartley, Margaret Hughes, Miren Iturriza-Gomara, James Johnson, L Luu, Jenifer Manson, Charlotte Nelson, Elaine O'Toole, Cassie Olateju, Rebekah Penrice-Randal , Lucille Rainbow, N.P Randle, Trevor Ian Robinson, Parul Sharma, Ghada T Shawli, James P Stewart, Neil Swainston, Ecaterina Vamos, Joanne Watts, Mark Whitehead |
| EPI_ISL_651176 | Department of Pathology, University of Cambridge | COVID-19 Genomics UK (COG-UK) Consortium | Aminu S. Jahun, Yasmin Chaudhry, Grant Hall, Iliana Georgana, Myra Hosmillo, Martin D. Curran, Malte Pinckert, Surendra Parmar, Ian Goodfellow |
| EPI_ISL_651178, EPI_ISL_651181 | Liverpool Clinical Laboratories | COVID-19 Genomics UK (COG-UK) Consortium | Sam Haldenby, Anita Lucaci, Steve Paterson, Julian Hiscox, Alistair Darby, M Almsaud, A Alrezaihi, Muhannad Alruwaili, Stuart D Armstrong, Jones Benjamin, Eleanor G Bentley, Anu Chawla, Jordan J Clark, Angela Cowell, Richard Eccles, Isabel García-Dorival, Matthew Gemmell, Alessandro Gerada, PKF Gilmore, Richard Gregory, Ximeng Han, Catherine Hartley, Margaret Hughes, Miren Iturriza-Gomara, James Johnson, L Luu, Jenifer Manson, Charlotte Nelson, Elaine O'Toole, Cassie Olateju, Rebekah Penrice-Randal , Lucille Rainbow, N.P Randle, Trevor Ian Robinson, Parul Sharma, Ghada T Shawli, James P Stewart, Neil Swainston, Ecaterina Vamos, Joanne Watts, Mark Whitehead |
| EPI_ISL_651187 | Centre for Enzyme Innovation, University of Portsmouth / Translational Research Laboratory, Portsmouth Hospitals NHS Trust | COVID-19 Genomics UK (COG-UK) Consortium | Angela Beckett,Yann Bourgeois,Garry Scarlett,Sharon Glaysher,Scott Elliott,Kelly Bicknell,Robert Impey,Allyson Lloyd,Sarah Wyllie,Ethan Butcher,Anoop Chauhan,Samuel Robson |
| EPI_ISL_651188 | Virology Department, Royal Infirmary of Edinburgh, NHS Lothian / School of Biological Sciences, University of Edinburgh / Institute of Genetics and Molecular Medicine, University of Edinburgh | COVID-19 Genomics UK (COG-UK) Consortium | McHugh M, Dewar R, Rooke S, Gallagher M, Balcaza C, O'Toole Á, Scher E, Hill V, McCrone JT, Colquhoun R, Yu X, Jackson B, Rambaut A, Williams TC, Templeton K |
| EPI_ISL_651193 | University of Birmingham | COVID-19 Genomics UK (COG-UK) Consortium | Institute of Microbiology, University of Birmingham: Claire McMurray, Joanne Stockton, Samuel Nicholls, Radoslaw Poplawski, Will Rowe, Josh Quick, Nicholas Loman. University of Birmingham Testing Laboratory: Celina M Whalley, Andrew Bosworth, Charlotte Poxon, Kasun Wanigasooriya, Oliver Pickles, Mike Kidd, Alex Richter, Andrew D Beggs PHE Heartlands Lab: Husam Osman, Andrew Bosworth. Queen Elizabeth Hospital: Anna Casey |
| EPI_ISL_651201 | University College London, Great Ormond Street Hospital for Children NHS Foundation Trust, Imperial College Healthcare NHS Trust | COVID-19 Genomics UK (COG-UK) Consortium | Sergi Castellano, Rachel Williams, Mark Kristiansen, Paola Resende Silva, Sunando Roy, Tony Brooks, Helena Tutill, Paola Niola, Patricia Dyal, Charlotte Williams, Leysa Forrest, Yasmin Panchbhaya, Jacqueline Findlay, Samuel Weeks, Julianne Brown, Kathryn Harris, Paul Randell, James Price, Alison Holmes, Judith Breuer |
| EPI_ISL_651207 | Wales Specialist Virology Centre Sequencing lab: Pathogen Genomics Unit | COVID-19 Genomics UK (COG-UK) Consortium | Catherine Moore, Johnathan Evans, Laura Gifford, Malorie Perry, Simon Cottrell, Angela Marchbank, Alec Birchley, Alexander Adams, Amy Gaskin, Bree Gatica-Wilcox, Jason Coombes, Joel Southgate, Lauren Gilbert, Lee Graham, Nicole Pacchiarini, Sara Kumziene-Summerhayes, Sarah Taylor, Sophie Jones, Sara Rey, Matthew Bull, Joanne Watkins, Sally Corden, Tom Connor |
| EPI_ISL_651208 | Liverpool Clinical Laboratories | COVID-19 Genomics UK (COG-UK) Consortium | Sam Haldenby, Anita Lucaci, Steve Paterson, Julian Hiscox, Alistair Darby, M Almsaud, A Alrezaihi, Muhannad Alruwaili, Stuart D Armstrong, Jones Benjamin, Eleanor G Bentley, Anu Chawla, Jordan J Clark, Angela Cowell, Richard Eccles, Isabel García-Dorival, Matthew Gemmell, Alessandro Gerada, PKF Gilmore, Richard Gregory, Ximeng Han, Catherine Hartley, Margaret Hughes, Miren Iturriza-Gomara, James Johnson, L Luu, Jenifer Manson, Charlotte Nelson, Elaine O'Toole, Cassie Olateju, Rebekah Penrice-Randal , Lucille Rainbow, N.P Randle, Trevor Ian Robinson, Parul Sharma, Ghada T Shawli, James P Stewart, Neil Swainston, Ecaterina Vamos, Joanne Watts, Mark Whitehead |
| EPI_ISL_651209 | Quadram Institute Bioscience | COVID-19 Genomics UK (COG-UK) Consortium | Dave J. Baker, Gemma L. Kay, Alp Aydin, Thanh Le-Viet, Steven Rudder, Ana P. Tedim, Anastasia Kolyva, Maria Diaz, Leonardo de Oliveira Martins, Nabil-Fareed Alikhan, Lizzie Meadows, Rachael Stanley, Ngozi Elumogo, Muhammed Yasir, Nicholas M. Thomson, Alexander J Trotter, Rachel Gilroy, Samuel Bloomfield, Claire Stuart, Andrew Bell, Reenesh Prakash, Samir Dervisevic, Alison E. Mather, John Wain, Mark Webber, Andrew J. Page, Justin O'Grady |
| EPI_ISL_651213 | Liverpool Clinical Laboratories | COVID-19 Genomics UK (COG-UK) Consortium | Sam Haldenby, Anita Lucaci, Steve Paterson, Julian Hiscox, Alistair Darby, M Almsaud, A Alrezaihi, Muhannad Alruwaili, Stuart D Armstrong, Jones Benjamin, Eleanor G Bentley, Anu Chawla, Jordan J Clark, Angela Cowell, Richard Eccles, Isabel García-Dorival, Matthew Gemmell, Alessandro Gerada, PKF Gilmore, Richard Gregory, Ximeng Han, Catherine Hartley, Margaret Hughes, Miren Iturriza-Gomara, James Johnson, L Luu, Jenifer Manson, Charlotte Nelson, Elaine O'Toole, Cassie Olateju, Rebekah Penrice-Randal , Lucille Rainbow, N.P Randle, Trevor Ian Robinson, Parul Sharma, Ghada T Shawli, James P Stewart, Neil Swainston, Ecaterina Vamos, Joanne Watts, Mark Whitehead |
| EPI_ISL_651216, EPI_ISL_651226 | Quadram Institute Bioscience | COVID-19 Genomics UK (COG-UK) Consortium | Dave J. Baker, Gemma L. Kay, Alp Aydin, Thanh Le-Viet, Steven Rudder, Ana P. Tedim, Anastasia Kolyva, Maria Diaz, Leonardo de Oliveira Martins, Nabil-Fareed Alikhan, Lizzie Meadows, Rachael Stanley, Ngozi Elumogo, Muhammed Yasir, Nicholas M. Thomson, Alexander J Trotter, Rachel Gilroy, Samuel Bloomfield, Claire Stuart, Andrew Bell, Reenesh Prakash, Samir Dervisevic, Alison E. Mather, John Wain, Mark Webber, Andrew J. Page, Justin O'Grady |
| EPI_ISL_651240 | University College London, Great Ormond Street Hospital for Children NHS Foundation Trust, Imperial College Healthcare NHS Trust | COVID-19 Genomics UK (COG-UK) Consortium | Sergi Castellano, Rachel Williams, Mark Kristiansen, Paola Resende Silva, Sunando Roy, Tony Brooks, Helena Tutill, Paola Niola, Patricia Dyal, Charlotte Williams, Leysa Forrest, Yasmin Panchbhaya, Jacqueline Findlay, Samuel Weeks, Julianne Brown, Kathryn Harris, Paul Randell, James Price, Alison Holmes, Judith Breuer |
| EPI_ISL_651250, EPI_ISL_651252 | Quadram Institute Bioscience | COVID-19 Genomics UK (COG-UK) Consortium | Dave J. Baker, Gemma L. Kay, Alp Aydin, Thanh Le-Viet, Steven Rudder, Ana P. Tedim, Anastasia Kolyva, Maria Diaz, Leonardo de Oliveira Martins, Nabil-Fareed Alikhan, Lizzie Meadows, Rachael Stanley, Ngozi Elumogo, Muhammed Yasir, Nicholas M. Thomson, Alexander J Trotter, Rachel Gilroy, Samuel Bloomfield, Claire Stuart, Andrew Bell, Reenesh Prakash, Samir Dervisevic, Alison E. Mather, John Wain, Mark Webber, Andrew J. Page, Justin O'Grady |
| EPI_ISL_651253 | University College London, Great Ormond Street Hospital for Children NHS Foundation Trust, Imperial College Healthcare NHS Trust | COVID-19 Genomics UK (COG-UK) Consortium | Sergi Castellano, Rachel Williams, Mark Kristiansen, Paola Resende Silva, Sunando Roy, Tony Brooks, Helena Tutill, Paola Niola, Patricia Dyal, Charlotte Williams, Leysa Forrest, Yasmin Panchbhaya, Jacqueline Findlay, Samuel Weeks, Julianne Brown, Kathryn Harris, Paul Randell, James Price, Alison Holmes, Judith Breuer |
| EPI_ISL_651268 | Quadram Institute Bioscience | COVID-19 Genomics UK (COG-UK) Consortium | Dave J. Baker, Gemma L. Kay, Alp Aydin, Thanh Le-Viet, Steven Rudder, Ana P. Tedim, Anastasia Kolyva, Maria Diaz, Leonardo de Oliveira Martins, Nabil-Fareed Alikhan, Lizzie Meadows, Rachael Stanley, Ngozi Elumogo, Muhammed Yasir, Nicholas M. Thomson, Alexander J Trotter, Rachel Gilroy, Samuel Bloomfield, Claire Stuart, Andrew Bell, Reenesh Prakash, Samir Dervisevic, Alison E. Mather, John Wain, Mark Webber, Andrew J. Page, Justin O'Grady |
| EPI_ISL_651272 | Liverpool Clinical Laboratories | COVID-19 Genomics UK (COG-UK) Consortium | Sam Haldenby, Anita Lucaci, Steve Paterson, Julian Hiscox, Alistair Darby, M Almsaud, A Alrezaihi, Muhannad Alruwaili, Stuart D Armstrong, Jones Benjamin, Eleanor G Bentley, Anu Chawla, Jordan J Clark, Angela Cowell, Richard Eccles, Isabel García-Dorival, Matthew Gemmell, Alessandro Gerada, |

| | | | |
|---|---|---|---|
| | | | PKF Gilmore, Richard Gregory, Ximeng Han, Catherine Hartley, Margaret Hughes, Miren Iturriza-Gomara, James Johnson, L Luu, Jenifer Manson, Charlotte Nelson, Elaine O'Toole, Cassie Olateju, Rebekah Penrice-Randal , Lucille Rainbow, N.P Randle, Trevor Ian Robinson, Parul Sharma, Ghada T Shawli, James P Stewart, Neil Swainston, Ecaterina Vamos, Joanne Watts, Mark Whitehead |
| EPI_ISL_651277 | Centre for Enzyme Innovation, University of Portsmouth / Translational Research Laboratory, Portsmouth Hospitals NHS Trust | COVID-19 Genomics UK (COG-UK) Consortium | Angela Beckett,Yann Bourgeois,Garry Scarlett,Sharon Glaysher,Scott Elliott,Kelly Bicknell,Robert Impey,Allyson Lloyd,Sarah Wyllie,Ethan Butcher,Anoop Chauhan,Samuel Robson |
| EPI_ISL_651287 | University of Birmingham | COVID-19 Genomics UK (COG-UK) Consortium | Institute of Microbiology, University of Birmingham: Claire McMurray, Joanne Stockton, Samuel Nicholls, Radoslaw Poplawski, Will Rowe, Josh Quick, Nicholas Loman. University of Birmingham Testing Laboratory: Celina M Whalley, Andrew Bosworth, Charlotte Poxon, Kasun Wanigasooriya, Oliver Pickles, Mike Kidd, Alex Richter, Andrew D Beggs PHE Heartlands Lab: Husam Osman, Andrew Bosworth. Queen Elizabeth Hospital: Anna Casey |
| EPI_ISL_651288 | Liverpool Clinical Laboratories | COVID-19 Genomics UK (COG-UK) Consortium | Sam Haldenby, Anita Lucaci, Steve Paterson, Julian Hiscox, Alistair Darby, M Almsaud, A Alrezaihi, Muhannad Alruwaili, Stuart D Armstrong, Jones Benjamin, Eleanor G Bentley, Anu Chawla, Jordan J Clark, Angela Cowell, Richard Eccles, Isabel García-Dorival, Matthew Gemmell, Alessandro Gerada, PKF Gilmore, Richard Gregory, Ximeng Han, Catherine Hartley, Margaret Hughes, Miren Iturriza-Gomara, James Johnson, L Luu, Jenifer Manson, Charlotte Nelson, Elaine O'Toole, Cassie Olateju, Rebekah Penrice-Randal , Lucille Rainbow, N.P Randle, Trevor Ian Robinson, Parul Sharma, Ghada T Shawli, James P Stewart, Neil Swainston, Ecaterina Vamos, Joanne Watts, Mark Whitehead |
| EPI_ISL_651291 | Virology Department, Sheffield Teaching Hospitals NHS Foundation Trust/Department of Infection, Immunity and Cardiovascular Disease, The Medical School, University of Sheffield | COVID-19 Genomics UK (COG-UK) Consortium | Thushan de Silva, Matthew Parker, Nikki Smith, Adri Angyal, Rebecca Brown, Luke Green, Rachel Tucker, Paul Parsons, Danielle Groves, Katie Johnson, Laura Carrilero, Alex Keeley, Dave Partridge, Matthew Wyles, Benjamin Lindsey, Mehmet Yavuz, Mohammad Raza, Cariad Evans |
| EPI_ISL_651298 | Quadram Institute Bioscience | COVID-19 Genomics UK (COG-UK) Consortium | Dave J. Baker, Gemma L. Kay, Alp Aydin, Thanh Le-Viet, Steven Rudder, Ana P. Tedim, Anastasia Kolyva, Maria Diaz, Leonardo de Oliveira Martins, Nabil-Fareed Alikhan, Lizzie Meadows, Rachael Stanley, Ngozi Elumogo, Muhammed Yasir, Nicholas M. Thomson, Alexander J Trotter, Rachel Gilroy, Samuel Bloomfield, Claire Stuart, Andrew Bell, Reenesh Prakash, Samir Dervisevic, Alison E. Mather, John Wain, Mark Webber, Andrew J. Page, Justin O'Grady |
| EPI_ISL_651299 | Virology Department, Royal Infirmary of Edinburgh, NHS Lothian / School of Biological Sciences, University of Edinburgh / Institute of Genetics and Molecular Medicine, University of Edinburgh | COVID-19 Genomics UK (COG-UK) Consortium | McHugh M, Dewar R, Rooke S, Gallagher M, Balcaza C, O'Toole Á, Scher E, Hill V, McCrone JT, Colquhoun R, Yu X, Jackson B, Rambaut A, Williams TC, Templeton K |
| EPI_ISL_651306, EPI_ISL_651312 | Liverpool Clinical Laboratories | COVID-19 Genomics UK (COG-UK) Consortium | Sam Haldenby, Anita Lucaci, Steve Paterson, Julian Hiscox, Alistair Darby, M Almsaud, A Alrezaihi, Muhannad Alruwaili, Stuart D Armstrong, Jones Benjamin, Eleanor G Bentley, Anu Chawla, Jordan J Clark, Angela Cowell, Richard Eccles, Isabel García-Dorival, Matthew Gemmell, Alessandro Gerada, PKF Gilmore, Richard Gregory, Ximeng Han, Catherine Hartley, Margaret Hughes, Miren Iturriza-Gomara, James Johnson, L Luu, Jenifer Manson, Charlotte Nelson, Elaine O'Toole, Cassie Olateju, Rebekah Penrice-Randal , Lucille Rainbow, N.P Randle, Trevor Ian Robinson, Parul Sharma, Ghada T Shawli, James P Stewart, Neil Swainston, Ecaterina Vamos, Joanne Watts, Mark Whitehead |
| EPI_ISL_651324 | Quadram Institute Bioscience | COVID-19 Genomics UK (COG-UK) Consortium | Dave J. Baker, Gemma L. Kay, Alp Aydin, Thanh Le-Viet, Steven Rudder, Ana P. Tedim, Anastasia Kolyva, Maria Diaz, Leonardo de Oliveira Martins, Nabil-Fareed Alikhan, Lizzie Meadows, Rachael Stanley, Ngozi Elumogo, Muhammed Yasir, Nicholas M. Thomson, Alexander J Trotter, Rachel Gilroy, Samuel Bloomfield, Claire Stuart, Andrew Bell, Reenesh Prakash, Samir Dervisevic, Alison E. Mather, John Wain, Mark Webber, Andrew J. Page, Justin O'Grady |
| EPI_ISL_651328 | Liverpool Clinical Laboratories | COVID-19 Genomics UK (COG-UK) Consortium | Sam Haldenby, Anita Lucaci, Steve Paterson, Julian Hiscox, Alistair Darby, M Almsaud, A Alrezaihi, Muhannad Alruwaili, Stuart D Armstrong, Jones Benjamin, Eleanor G Bentley, Anu Chawla, Jordan J Clark, Angela Cowell, Richard Eccles, Isabel García-Dorival, Matthew Gemmell, Alessandro Gerada, PKF Gilmore, Richard Gregory, Ximeng Han, Catherine Hartley, Margaret Hughes, Miren Iturriza-Gomara, James Johnson, L Luu, Jenifer Manson, Charlotte Nelson, Elaine O'Toole, Cassie Olateju, Rebekah Penrice-Randal , Lucille Rainbow, N.P Randle, Trevor Ian Robinson, Parul Sharma, Ghada T Shawli, James P Stewart, Neil Swainston, Ecaterina Vamos, Joanne Watts, Mark Whitehead |
| EPI_ISL_651337, EPI_ISL_651338, EPI_ISL_651349, EPI_ISL_651351 | Quadram Institute Bioscience | COVID-19 Genomics UK (COG-UK) Consortium | Dave J. Baker, Gemma L. Kay, Alp Aydin, Thanh Le-Viet, Steven Rudder, Ana P. Tedim, Anastasia Kolyva, Maria Diaz, Leonardo de Oliveira Martins, Nabil-Fareed Alikhan, Lizzie Meadows, Rachael Stanley, Ngozi Elumogo, Muhammed Yasir, Nicholas M. Thomson, Alexander J Trotter, Rachel Gilroy, Samuel Bloomfield, Claire Stuart, Andrew Bell, Reenesh Prakash, Samir Dervisevic, Alison E. Mather, John Wain, Mark Webber, Andrew J. Page, Justin O'Grady |
| EPI_ISL_651374 | Virology Department, Sheffield Teaching Hospitals NHS Foundation Trust/Department of Infection, Immunity and Cardiovascular Disease, The Medical School, University of Sheffield | COVID-19 Genomics UK (COG-UK) Consortium | Thushan de Silva, Matthew Parker, Nikki Smith, Adri Angyal, Rebecca Brown, Luke Green, Rachel Tucker, Paul Parsons, Danielle Groves, Katie Johnson, Laura Carrilero, Alex Keeley, Dave Partridge, Matthew Wyles, Benjamin Lindsey, Mehmet Yavuz, Mohammad Raza, Cariad Evans |
| EPI_ISL_651376 | Liverpool Clinical Laboratories | COVID-19 Genomics UK (COG-UK) Consortium | Sam Haldenby, Anita Lucaci, Steve Paterson, Julian Hiscox, Alistair Darby, M Almsaud, A Alrezaihi, Muhannad Alruwaili, Stuart D Armstrong, Jones Benjamin, Eleanor G Bentley, Anu Chawla, Jordan J Clark, Angela Cowell, Richard Eccles, Isabel García-Dorival, Matthew Gemmell, Alessandro Gerada, PKF Gilmore, Richard Gregory, Ximeng Han, Catherine Hartley, Margaret Hughes, Miren Iturriza-Gomara, James Johnson, L Luu, Jenifer Manson, Charlotte Nelson, Elaine O'Toole, Cassie Olateju, Rebekah Penrice-Randal , Lucille Rainbow, N.P Randle, Trevor Ian Robinson, Parul Sharma, Ghada T Shawli, James P Stewart, Neil Swainston, Ecaterina Vamos, Joanne Watts, Mark Whitehead |
| EPI_ISL_651385 | Virology Department, Royal Infirmary of Edinburgh, NHS Lothian / School of Biological Sciences, University of Edinburgh / Institute of Genetics and Molecular Medicine, University of Edinburgh | COVID-19 Genomics UK (COG-UK) Consortium | McHugh M, Dewar R, Rooke S, Gallagher M, Balcaza C, O'Toole Á, Scher E, Hill V, McCrone JT, Colquhoun R, Yu X, Jackson B, Rambaut A, Williams TC, Templeton K |
| EPI_ISL_651400 | Liverpool Clinical Laboratories | COVID-19 Genomics UK (COG-UK) Consortium | Sam Haldenby, Anita Lucaci, Steve Paterson, Julian Hiscox, Alistair Darby, M Almsaud, A Alrezaihi, Muhannad Alruwaili, Stuart D Armstrong, Jones Benjamin, Eleanor G Bentley, Anu Chawla, Jordan J Clark, Angela Cowell, Richard Eccles, Isabel García-Dorival, Matthew Gemmell, Alessandro Gerada, PKF Gilmore, Richard Gregory, Ximeng Han, Catherine Hartley, Margaret Hughes, Miren Iturriza-Gomara, James Johnson, L Luu, Jenifer Manson, Charlotte Nelson, Elaine O'Toole, Cassie Olateju, Rebekah Penrice-Randal , Lucille Rainbow, N.P Randle, Trevor Ian Robinson, Parul Sharma, Ghada T Shawli, James P Stewart, Neil Swainston, Ecaterina Vamos, Joanne Watts, Mark Whitehead |
| EPI_ISL_651402 | West of Scotland Specialist Virology Centre, NHSGGC / MRC-University of Glasgow Centre for Virus Research | COVID-19 Genomics UK (COG-UK) Consortium | Ana da Silva Filipe, Natasha Johnson, Kathy Smollett, Daniel Mair, Stephen Carmichael, Alice Broos, Lily Tong, Jenna Nichols, Kyriaki Nomikou; Sarah McDonald; Richard Orton, Joseph Hughes, Sreenu Vattipally, David L Robertson; Alasdair MacLean, Rory Gunson; Sharif Shaaban, Matthew Holden; Rachel Blacow, Guy Mollett, Kathy Li, James Shepherd, Antonia Ho, Emma Thomson |
| EPI_ISL_651405 | University of Birmingham | COVID-19 Genomics UK (COG-UK) Consortium | Institute of Microbiology, University of Birmingham: Claire McMurray, Joanne Stockton, Samuel Nicholls, Radoslaw Poplawski, Will Rowe, Josh Quick, Nicholas Loman. University of Birmingham Testing Laboratory: Celina M Whalley, Andrew Bosworth, Charlotte Poxon, Kasun Wanigasooriya, Oliver Pickles, Mike Kidd, Alex Richter, Andrew D Beggs PHE Heartlands Lab: Husam Osman, Andrew Bosworth. Queen Elizabeth Hospital: Anna Casey |
| EPI_ISL_651406 | West of Scotland Specialist Virology Centre, NHSGGC / MRC-University of Glasgow Centre for Virus Research | COVID-19 Genomics UK (COG-UK) Consortium | Ana da Silva Filipe, Natasha Johnson, Kathy Smollett, Daniel Mair, Stephen Carmichael, Alice Broos, Lily Tong, Jenna Nichols, Kyriaki Nomikou; Sarah McDonald; Richard Orton, Joseph Hughes, Sreenu Vattipally, David L Robertson; Alasdair MacLean, Rory Gunson; Sharif Shaaban, Matthew Holden; Rachel Blacow, Guy Mollett, Kathy Li, James Shepherd, Antonia Ho, Emma Thomson |
| EPI_ISL_651411, EPI_ISL_651412 | Quadram Institute Bioscience | COVID-19 Genomics UK (COG-UK) Consortium | Dave J. Baker, Gemma L. Kay, Alp Aydin, Thanh Le-Viet, Steven Rudder, Ana P. Tedim, Anastasia Kolyva, Maria Diaz, Leonardo de Oliveira Martins, Nabil-Fareed Alikhan, Lizzie Meadows, Rachael Stanley, Ngozi Elumogo, Muhammed Yasir, Nicholas M. Thomson, Alexander J Trotter, Rachel Gilroy, Samuel Bloomfield, Claire Stuart, Andrew Bell, Reenesh Prakash, Samir Dervisevic, Alison E. Mather, John Wain, Mark Webber, Andrew J. Page, Justin O'Grady |
| EPI_ISL_651413 | University of Birmingham | COVID-19 Genomics UK (COG-UK) Consortium | Institute of Microbiology, University of Birmingham: Claire McMurray, Joanne Stockton, Samuel Nicholls, Radoslaw Poplawski, Will Rowe, Josh Quick, Nicholas Loman. University of Birmingham Testing Laboratory: Celina M Whalley, Andrew Bosworth, Charlotte Poxon, Kasun Wanigasooriya, Oliver Pickles, Mike Kidd, Alex Richter, Andrew D Beggs PHE Heartlands Lab: Husam Osman, Andrew Bosworth. Queen Elizabeth Hospital: Anna Casey |

| | | | |
|---|---|---|---|
| EPI_ISL_651420 | Wales Specialist Virology Centre Sequencing lab: Pathogen Genomics Unit | COVID-19 Genomics UK (COG-UK) Consortium | Catherine Moore, Johnathan Evans, Laura Gifford, Malorie Perry, Simon Cottrell, Angela Marchbank, Alec Birchley, Alexander Adams, Amy Gaskin, Bree Gatica-Wilcox, Jason Coombes, Joel Southgate, Lauren Gilbert, Lee Graham, Nicole Pacchiarini, Sara Kumziene-Summerhayes, Sarah Taylor, Sophie Jones, Sara Rey, Matthew Bull, Joanne Watkins, Sally Corden, Tom Connor |
| EPI_ISL_651425, EPI_ISL_651426 | Virology Department, Royal Infirmary of Edinburgh, NHS Lothian / School of Biological Sciences, University of Edinburgh / Institute of Genetics and Molecular Medicine, University of Edinburgh | COVID-19 Genomics UK (COG-UK) Consortium | McHugh M, Dewar R, Rooke S, Gallagher M, Balcaza C, O'Toole Á, Scher E, Hill V, McCrone JT, Colquhoun R, Yu X, Jackson B, Rambaut A, Williams TC, Templeton K |
| EPI_ISL_651438 | Quadram Institute Bioscience | COVID-19 Genomics UK (COG-UK) Consortium | Dave J. Baker, Gemma L. Kay, Alp Aydin, Thanh Le-Viet, Steven Rudder, Ana P. Tedim, Anastasia Kolyva, Maria Diaz, Leonardo de Oliveira Martins, Nabil-Fareed Alikhan, Lizzie Meadows, Rachael Stanley, Ngozi Elumogo, Muhammed Yasir, Nicholas M. Thomson, Alexander J Trotter, Rachel Gilroy, Samuel Bloomfield, Claire Stuart, Andrew Bell, Reenesh Prakash, Samir Dervisevic, Alison E. Mather, John Wain, Mark Webber, Andrew J. Page, Justin O'Grady |
| EPI_ISL_651440, EPI_ISL_651441, EPI_ISL_651442, EPI_ISL_651443, EPI_ISL_651444, EPI_ISL_651445, EPI_ISL_651446, EPI_ISL_651447, EPI_ISL_651449, EPI_ISL_651451, EPI_ISL_651455, EPI_ISL_651457, EPI_ISL_651465 | | | |
| see above | Wales Specialist Virology Centre Sequencing lab: Pathogen Genomics Unit | COVID-19 Genomics UK (COG-UK) Consortium | Catherine Moore, Johnathan Evans, Laura Gifford, Malorie Perry, Simon Cottrell, Angela Marchbank, Alec Birchley, Alexander Adams, Amy Gaskin, Bree Gatica-Wilcox, Jason Coombes, Joel Southgate, Lauren Gilbert, Lee Graham, Nicole Pacchiarini, Sara Kumziene-Summerhayes, Sarah Taylor, Sophie Jones, Sara Rey, Matthew Bull, Joanne Watkins, Sally Corden, Tom Connor |
| EPI_ISL_651488, EPI_ISL_651489, EPI_ISL_651490, EPI_ISL_651492 | Department of Pathology, University of Cambridge | COVID-19 Genomics UK (COG-UK) Consortium | Aminu S. Jahun, Yasmin Chaudhry, Grant Hall, Iliana Georgana, Myra Hosmillo, Martin D. Curran, Malte Pinckert, Surendra Parmar, Ian Goodfellow |
| EPI_ISL_651495 | West of Scotland Specialist Virology Centre, NHSGGC / MRC-University of Glasgow Centre for Virus Research | COVID-19 Genomics UK (COG-UK) Consortium | Ana da Silva Filipe, Natasha Johnson, Kathy Smollett, Daniel Mair, Stephen Carmichael, Alice Broos, Lily Tong, Jenna Nichols, Kyriaki Nomikou; Sarah McDonald; Richard Orton, Joseph Hughes, Sreenu Vattipally, David L Robertson; Alasdair MacLean, Rory Gunson; Sharif Shaaban, Matthew Holden; Rachel Blacow, Guy Mollett, Kathy Li, James Shepherd, Antonia Ho, Emma Thomson |
| EPI_ISL_651503 | Wales Specialist Virology Centre Sequencing lab: Pathogen Genomics Unit | COVID-19 Genomics UK (COG-UK) Consortium | Catherine Moore, Johnathan Evans, Laura Gifford, Malorie Perry, Simon Cottrell, Angela Marchbank, Alec Birchley, Alexander Adams, Amy Gaskin, Bree Gatica-Wilcox, Jason Coombes, Joel Southgate, Lauren Gilbert, Lee Graham, Nicole Pacchiarini, Sara Kumziene-Summerhayes, Sarah Taylor, Sophie Jones, Sara Rey, Matthew Bull, Joanne Watkins, Sally Corden, Tom Connor |
| EPI_ISL_651512, EPI_ISL_651516 | Quadram Institute Bioscience | COVID-19 Genomics UK (COG-UK) Consortium | Dave J. Baker, Gemma L. Kay, Alp Aydin, Thanh Le-Viet, Steven Rudder, Ana P. Tedim, Anastasia Kolyva, Maria Diaz, Leonardo de Oliveira Martins, Nabil-Fareed Alikhan, Lizzie Meadows, Rachael Stanley, Ngozi Elumogo, Muhammed Yasir, Nicholas M. Thomson, Alexander J Trotter, Rachel Gilroy, Samuel Bloomfield, Claire Stuart, Andrew Bell, Reenesh Prakash, Samir Dervisevic, Alison E. Mather, John Wain, Mark Webber, Andrew J. Page, Justin O'Grady |
| EPI_ISL_651519 | University of Birmingham | COVID-19 Genomics UK (COG-UK) Consortium | Institute of Microbiology, University of Birmingham: Claire McMurray, Joanne Stockton, Samuel Nicholls, Radoslaw Poplawski, Will Rowe, Josh Quick, Nicholas Loman. University of Birmingham Testing Laboratory: Celina M Whalley, Andrew Bosworth, Charlotte Poxon, Kasun Wanigasooriya, Oliver Pickles, Mike Kidd, Alex Richter, Andrew D Beggs PHE Heartlands Lab: Husam Osman, Andrew Bosworth. Queen Elizabeth Hospital: Anna Casey |
| EPI_ISL_651526 | Quadram Institute Bioscience | COVID-19 Genomics UK (COG-UK) Consortium | Dave J. Baker, Gemma L. Kay, Alp Aydin, Thanh Le-Viet, Steven Rudder, Ana P. Tedim, Anastasia Kolyva, Maria Diaz, Leonardo de Oliveira Martins, Nabil-Fareed Alikhan, Lizzie Meadows, Rachael Stanley, Ngozi Elumogo, Muhammed Yasir, Nicholas M. Thomson, Alexander J Trotter, Rachel Gilroy, Samuel Bloomfield, Claire Stuart, Andrew Bell, Reenesh Prakash, Samir Dervisevic, Alison E. Mather, John Wain, Mark Webber, Andrew J. Page, Justin O'Grady |
| EPI_ISL_651531, EPI_ISL_651532, EPI_ISL_651533, EPI_ISL_651534, EPI_ISL_651535, EPI_ISL_651536, EPI_ISL_651537 | West of Scotland Specialist Virology Centre, NHSGGC / MRC-University of Glasgow Centre for Virus Research | COVID-19 Genomics UK (COG-UK) Consortium | Ana da Silva Filipe, Natasha Johnson, Kathy Smollett, Daniel Mair, Stephen Carmichael, Alice Broos, Lily Tong, Jenna Nichols, Kyriaki Nomikou; Sarah McDonald; Richard Orton, Joseph Hughes, Sreenu Vattipally, David L Robertson; Alasdair MacLean, Rory Gunson; Sharif Shaaban, Matthew Holden; Rachel Blacow, Guy Mollett, Kathy Li, James Shepherd, Antonia Ho, Emma Thomson |
| EPI_ISL_651600, EPI_ISL_651601, EPI_ISL_651603, EPI_ISL_651604, EPI_ISL_651605, EPI_ISL_651606, EPI_ISL_651607, EPI_ISL_651608, EPI_ISL_651609, EPI_ISL_651619, EPI_ISL_651620, EPI_ISL_651621 | | | |
| see above | University of Birmingham | COVID-19 Genomics UK (COG-UK) Consortium | Institute of Microbiology, University of Birmingham: Claire McMurray, Joanne Stockton, Samuel Nicholls, Radoslaw Poplawski, Will Rowe, Josh Quick, Nicholas Loman. University of Birmingham Testing Laboratory: Celina M Whalley, Andrew Bosworth, Charlotte Poxon, Kasun Wanigasooriya, Oliver Pickles, Mike Kidd, Alex Richter, Andrew D Beggs PHE Heartlands Lab: Husam Osman, Andrew Bosworth. Queen Elizabeth Hospital: Anna Casey |
| EPI_ISL_651653, EPI_ISL_651656, EPI_ISL_651659, EPI_ISL_651660, EPI_ISL_651663, EPI_ISL_651664, EPI_ISL_651665, EPI_ISL_651668, EPI_ISL_651671, EPI_ISL_651673, EPI_ISL_651674, EPI_ISL_651675, EPI_ISL_651676, EPI_ISL_651677, EPI_ISL_651681 | | | |
| see above | Department of Pathology, University of Cambridge | COVID-19 Genomics UK (COG-UK) Consortium | Aminu S. Jahun, Yasmin Chaudhry, Grant Hall, Iliana Georgana, Myra Hosmillo, Martin D. Curran, Malte Pinckert, Surendra Parmar, Ian Goodfellow |
| EPI_ISL_651740, EPI_ISL_651767, EPI_ISL_651769, EPI_ISL_651770, EPI_ISL_651771, EPI_ISL_651772, EPI_ISL_651773, EPI_ISL_651774, EPI_ISL_651775, EPI_ISL_651776, EPI_ISL_651777, EPI_ISL_651778, EPI_ISL_651779, EPI_ISL_651832, EPI_ISL_651833, EPI_ISL_651834, EPI_ISL_651835, EPI_ISL_651836, EPI_ISL_651837, EPI_ISL_651838, EPI_ISL_651839, EPI_ISL_651840, EPI_ISL_651841, EPI_ISL_651842, EPI_ISL_651843, EPI_ISL_651844, EPI_ISL_651845, EPI_ISL_651846, EPI_ISL_651847, EPI_ISL_651848, EPI_ISL_651849, EPI_ISL_651874, EPI_ISL_651875, EPI_ISL_651876, EPI_ISL_651877, EPI_ISL_651878, EPI_ISL_651879, EPI_ISL_651880, EPI_ISL_651881, EPI_ISL_651882 | | | |
| see above | West of Scotland Specialist Virology Centre, NHSGGC / MRC-University of Glasgow Centre for Virus Research | COVID-19 Genomics UK (COG-UK) Consortium | Ana da Silva Filipe, Natasha Johnson, Kathy Smollett, Daniel Mair, Stephen Carmichael, Alice Broos, Lily Tong, Jenna Nichols, Kyriaki Nomikou; Sarah McDonald; Richard Orton, Joseph Hughes, Sreenu Vattipally, David L Robertson; Alasdair MacLean, Rory Gunson; Sharif Shaaban, Matthew Holden; Rachel Blacow, Guy Mollett, Kathy Li, James Shepherd, Antonia Ho, Emma Thomson |
| EPI_ISL_651992, EPI_ISL_651994, EPI_ISL_651995, EPI_ISL_651996, EPI_ISL_652000, EPI_ISL_652001, EPI_ISL_652002, EPI_ISL_652003, EPI_ISL_652004, EPI_ISL_652005, EPI_ISL_652006, EPI_ISL_652007, EPI_ISL_652008, EPI_ISL_652009, EPI_ISL_652010, EPI_ISL_652011, EPI_ISL_652012, EPI_ISL_652013, EPI_ISL_652014, EPI_ISL_652015, EPI_ISL_652016, EPI_ISL_652017, EPI_ISL_652018, EPI_ISL_652019, EPI_ISL_652020, EPI_ISL_652021, EPI_ISL_652022, EPI_ISL_652023, EPI_ISL_652041, EPI_ISL_652064 | | | |
| see above | Virology Department, Royal Infirmary of Edinburgh, NHS Lothian / School of Biological Sciences, University of Edinburgh / Institute of Genetics and Molecular Medicine, University of Edinburgh | COVID-19 Genomics UK (COG-UK) Consortium | McHugh M, Dewar R, Rooke S, Gallagher M, Balcaza C, O'Toole Á, Scher E, Hill V, McCrone JT, Colquhoun R, Yu X, Jackson B, Rambaut A, Williams TC, Templeton K |
| EPI_ISL_652241, EPI_ISL_652242, EPI_ISL_652243, EPI_ISL_652244, EPI_ISL_652245, EPI_ISL_652246, EPI_ISL_652247, EPI_ISL_652248, EPI_ISL_652249, EPI_ISL_652250, EPI_ISL_652251, EPI_ISL_652252, EPI_ISL_652253, EPI_ISL_652254, EPI_ISL_652255, EPI_ISL_652256, EPI_ISL_652257, EPI_ISL_652258, EPI_ISL_652259, EPI_ISL_652260, EPI_ISL_652261, EPI_ISL_652262, EPI_ISL_652263, EPI_ISL_652264, EPI_ISL_652265, EPI_ISL_652266, EPI_ISL_652267, EPI_ISL_652268, EPI_ISL_652269, EPI_ISL_652270, EPI_ISL_652271, EPI_ISL_652272, EPI_ISL_652273, EPI_ISL_652274, EPI_ISL_652275, EPI_ISL_652276, EPI_ISL_652277, EPI_ISL_652278, EPI_ISL_652279, EPI_ISL_652280, EPI_ISL_652281, EPI_ISL_652282, EPI_ISL_652283, EPI_ISL_652284, EPI_ISL_652285 | | | |
| see above | Quadram Institute Bioscience | COVID-19 Genomics UK (COG-UK) Consortium | Dave J. Baker, Gemma L. Kay, Alp Aydin, Thanh Le-Viet, Steven Rudder, Ana P. Tedim, Anastasia Kolyva, Maria Diaz, Leonardo de Oliveira Martins, Nabil-Fareed Alikhan, Lizzie Meadows, Rachael Stanley, Ngozi Elumogo, Muhammed Yasir, Nicholas M. Thomson, Alexander J Trotter, Rachel Gilroy, Samuel Bloomfield, Claire Stuart, Andrew Bell, Reenesh Prakash, Samir Dervisevic, Alison E. Mather, John Wain, Mark Webber, Andrew J. Page, Justin O'Grady |
| EPI_ISL_652286, EPI_ISL_652287, EPI_ISL_652288 | Department of Pathology, University of Cambridge | COVID-19 Genomics UK (COG-UK) Consortium | Aminu S. Jahun, Yasmin Chaudhry, Grant Hall, Iliana Georgana, Myra Hosmillo, Martin D. Curran, Malte Pinckert, Surendra Parmar, Ian Goodfellow |
| EPI_ISL_652338, EPI_ISL_652340 | Northumbria University / South Tees Hospitals NHS Foundation Trust / North Cumbria Integrated Care NHS Foundation Trust / North Tees and Hartlepool NHS Foundation Trust / Newcastle Hospitals NHS Foundation Trust | COVID-19 Genomics UK (COG-UK) Consortium | Darren L Smith,Andrew Nelson,Matthew Bashton,Greg R Young,Joshua Loh,John Allan,Mohammad A Tariq,Giles S Holt,Gary Black,Wen C Yew,Lynn Dover,Paul Baker,Steve Liggett,Sarah Essex,Jane Greenaway,Debra Padgett,Clive Graham,Garren Scott,Edward Barton,Emma Swindells,Brendan Payne,Jennifer Collins,Yusri Taha,Gary Eltringham |
| EPI_ISL_652417, EPI_ISL_652419, EPI_ISL_652420, EPI_ISL_652421, EPI_ISL_652422, EPI_ISL_652423, EPI_ISL_652425, EPI_ISL_652426, EPI_ISL_652428, EPI_ISL_652429, EPI_ISL_652430, EPI_ISL_652431, EPI_ISL_652432, EPI_ISL_652433, EPI_ISL_652435, EPI_ISL_652438, EPI_ISL_652439, EPI_ISL_652441, EPI_ISL_652443, EPI_ISL_652444, EPI_ISL_652446, EPI_ISL_652447, EPI_ISL_652448, EPI_ISL_652450, EPI_ISL_652451, EPI_ISL_652452, EPI_ISL_652453, EPI_ISL_652455, EPI_ISL_652457, EPI_ISL_652458, EPI_ISL_652460, EPI_ISL_652461, EPI_ISL_652462, EPI_ISL_652463, EPI_ISL_652465, EPI_ISL_652466, EPI_ISL_652470, EPI_ISL_652472, EPI_ISL_652473, EPI_ISL_652474, EPI_ISL_652475, EPI_ISL_652476, EPI_ISL_652478, EPI_ISL_652479, EPI_ISL_652480, EPI_ISL_652481, EPI_ISL_652482, EPI_ISL_652486, EPI_ISL_652487, EPI_ISL_652488, EPI_ISL_652489, EPI_ISL_652503, EPI_ISL_652507, EPI_ISL_652512, EPI_ISL_652522, EPI_ISL_652523, EPI_ISL_652527, EPI_ISL_652536, EPI_ISL_652543, EPI_ISL_652546, EPI_ISL_652553, EPI_ISL_652555, EPI_ISL_652557, EPI_ISL_652568, EPI_ISL_652571, EPI_ISL_652574, EPI_ISL_652580, EPI_ISL_652591, EPI_ISL_652592, EPI_ISL_652594, EPI_ISL_652596, EPI_ISL_652598, EPI_ISL_652603, EPI_ISL_652606, EPI_ISL_652607, EPI_ISL_652608, EPI_ISL_652609, EPI_ISL_652613 | | | |
| see above | Wales Specialist Virology Centre Sequencing lab: Pathogen | COVID-19 Genomics UK (COG-UK) Consortium | Catherine Moore, Johnathan Evans, Laura Gifford, Malorie Perry, Simon Cottrell, Angela Marchbank, Alec Birchley, Alexander Adams, Amy Gaskin, Bree |

| | | | |
|---|---|---|---|
| | Genomics Unit | | Gatica-Wilcox, Jason Coombes, Joel Southgate, Lauren Gilbert, Lee Graham, Nicole Pacchiarini, Sara Kumziene-Summerhayes, Sarah Taylor, Sophie Jones, Sara Rey, Matthew Bull, Joanne Watkins, Sally Corden, Tom Connor |
| EPI_ISL_652626 | Centre for Enzyme Innovation, University of Portsmouth / Translational Research Laboratory, Portsmouth Hospitals NHS Trust | COVID-19 Genomics UK (COG-UK) Consortium | Angela Beckett,Yann Bourgeois,Garry Scarlett,Sharon Glaysher,Scott Elliott,Kelly Bicknell,Robert Impey,Allyson Lloyd,Sarah Wyllie,Ethan Butcher,Anoop Chauhan,Samuel Robson |
| EPI_ISL_653046, EPI_ISL_653047, EPI_ISL_653048, EPI_ISL_653049, EPI_ISL_653050, EPI_ISL_653051, EPI_ISL_653052, EPI_ISL_653053, EPI_ISL_653054, EPI_ISL_653055, EPI_ISL_653056, EPI_ISL_653057, EPI_ISL_653058 | | | |
| see above | Quadram Institute Bioscience | COVID-19 Genomics UK (COG-UK) Consortium | Dave J. Baker, Gemma L. Kay, Alp Aydin, Thanh Le-Viet, Steven Rudder, Ana P. Tedim, Anastasia Kolyva, Maria Diaz, Leonardo de Oliveira Martins, Nabil-Fareed Alikhan, Lizzie Meadows, Rachael Stanley, Ngozi Elumogo, Muhammed Yasir, Nicholas M. Thomson, Alexander J Trotter, Rachel Gilroy, Samuel Bloomfield, Claire Stuart, Andrew Bell, Reenesh Prakash, Samir Dervisevic, Alison E. Mather, John Wain, Mark Webber, Andrew J. Page, Justin O'Grady |
| EPI_ISL_653059, EPI_ISL_653060, EPI_ISL_653062, EPI_ISL_653065, EPI_ISL_653066, EPI_ISL_653073, EPI_ISL_653083, EPI_ISL_653085, EPI_ISL_653088, EPI_ISL_653090, EPI_ISL_653094, EPI_ISL_653095, EPI_ISL_653096 | | | |
| see above | Virology Department, Sheffield Teaching Hospitals NHS Foundation Trust/Department of Infection, Immunity and Cardiovascular Disease, The Medical School, University of Sheffield | COVID-19 Genomics UK (COG-UK) Consortium | Thushan de Silva, Matthew Parker, Nikki Smith, Adri Angyal, Rebecca Brown, Luke Green, Rachel Tucker, Paul Parsons, Danielle Groves, Katie Johnson, Laura Carrilero, Alex Keeley, Dave Partridge, Matthew Wyles, Benjamin Lindsey, Mehmet Yavuz, Mohammad Raza, Cariad Evans |
| EPI_ISL_653821 | I.R.C.C.S. "S. De Bellis" - Ente Ospedaliero | Istituto Zooprofilattico Sperimentale della Puglia e della Basilicata | Parisi A., Bianco A., Capozzi L., Del Sambro L., Lippolis A., Notarnicola M., Cipolletta D., Galante D. |
| EPI_ISL_653933, EPI_ISL_653935, EPI_ISL_653936, EPI_ISL_653937, EPI_ISL_653941, EPI_ISL_653942, EPI_ISL_653945, EPI_ISL_653946, EPI_ISL_653949, EPI_ISL_653951, EPI_ISL_653952, EPI_ISL_653955, EPI_ISL_653956, EPI_ISL_653961, EPI_ISL_653962, EPI_ISL_653965, EPI_ISL_653966, EPI_ISL_653967, EPI_ISL_653968, EPI_ISL_653969, EPI_ISL_653970, EPI_ISL_653971, EPI_ISL_653972, EPI_ISL_653973, EPI_ISL_653974, EPI_ISL_653976, EPI_ISL_653977, EPI_ISL_653978, EPI_ISL_653979, EPI_ISL_653980, EPI_ISL_653981, EPI_ISL_653982, EPI_ISL_653983, EPI_ISL_653984, EPI_ISL_653985, EPI_ISL_653986, EPI_ISL_653987, EPI_ISL_653988, EPI_ISL_653989, EPI_ISL_653990, EPI_ISL_653991, EPI_ISL_653993, EPI_ISL_653994, EPI_ISL_653995, EPI_ISL_653996, EPI_ISL_653997, EPI_ISL_653998, EPI_ISL_653999, EPI_ISL_654000, EPI_ISL_654001, EPI_ISL_654005, EPI_ISL_654007, EPI_ISL_654010 | | | |
| see above | Respiratory Virus Unit, Microbiology Services Colindale, Public Health England | COVID-19 Genomics UK (COG-UK) Consortium | PHE Covid Sequencing Team |
| EPI_ISL_654020 | Laboratory of Microbiology, National Reference Lab, Charles Nicolle Hospital; 2-University of Tunis ElManar, Faculty of Medicine of Tunis, LR99ES09, Tunis, Tunisia | 1-Clinical and Experimental Pharmacology Lab, LR16SP02, National Center of Pharmacovigilance, University of Tunis El Manar, Tunis, Tunisia. 2-Neurodegenerative diseases and psychiatric troubles, LR18SP03, Razi Hospital, University of Tunis El Manar, Tunis, Tunisia. 3- Ministry of Health, National Observatory of New and Emerging Diseases, 1006, Tunis, Tunisia | Ilhem Boutiba-Ben Boubaker, Sameh Trabelsi, Nissaf Ben Alaya, Maher Kharrat, Alia Ben Kahla, Jalila Ben Khelil, Salma Abid, Sana Ferjani, Mouna Ben Sassi, Mouna Safer, Imen Mkada, Imen Kacem, Gaies Emna, Soumaya Rammeh, Riadh Daghfous, Riadh Gouider. |
| EPI_ISL_654328, EPI_ISL_654334, EPI_ISL_654344, EPI_ISL_654347, EPI_ISL_654350, EPI_ISL_654352, EPI_ISL_654355, EPI_ISL_654371, EPI_ISL_654372, EPI_ISL_654373, EPI_ISL_654374, EPI_ISL_654375, EPI_ISL_654376, EPI_ISL_654377, EPI_ISL_654378, EPI_ISL_654379, EPI_ISL_654380, EPI_ISL_654381, EPI_ISL_654382, EPI_ISL_654383, EPI_ISL_654384 | | | |
| see above | Hospital General Universitario Gregorio Marañón | SeqCOVID-SPAIN consortium/IBV(CSIC) | Darío García de Viedma, Laura Pérez-Lago, Marta Herranz, Jon Sicilia, Julia Suárez, Pilar Catalán, Patricia Muñoz and SeqCOVID-SPAIN consortium |
| EPI_ISL_654502, EPI_ISL_654503 | The Public Health Agency of Sweden | The Public Health Agency of Sweden | Anna-Malin Linde, Maria Lind Karlberg, Mattias Haukland, Reza Advani, Olov Svartstrom, Oskar Karlsson Lindsjo, Sandra Broddesson, Petra Edquist, Mia Brytting, Anna Risberg, Karin Tegmark-Wisell |
| EPI_ISL_654548, EPI_ISL_654558, EPI_ISL_654569, EPI_ISL_654572, EPI_ISL_654586, EPI_ISL_654595, EPI_ISL_654596 | Servicio de Microbiología. Hospital Universitario Donostia. OSI Donostialdea. Área de Enfermedades Infecciosas, Grupo de Infección Respiratoria y Resistencia Antimicrobiana. Instituto de Investigación Sanitaria Biodonostia | SeqCOVID-SPAIN consortium/IBV(CSIC) | Gustavo Cilla Eguiluz, Milagrosa Montes Ros, Luis Piñeiro Vázquez, Ane Sorrarain, Jose Maria Marimón and SeqCOVID-SPAIN consortium |
| EPI_ISL_660072, EPI_ISL_660073, EPI_ISL_660074, EPI_ISL_660075, EPI_ISL_660076, EPI_ISL_660078, EPI_ISL_660079, EPI_ISL_660080, EPI_ISL_660081, EPI_ISL_660082, EPI_ISL_660083, EPI_ISL_660084, EPI_ISL_660085, EPI_ISL_660099, EPI_ISL_660100, EPI_ISL_660101, EPI_ISL_660102, EPI_ISL_660104, EPI_ISL_660105 | | | |
| see above | MD PHL | MD PHL | Maryland Department of Health Laboratories Administration |
| EPI_ISL_660186, EPI_ISL_660187, EPI_ISL_660188, EPI_ISL_660189, EPI_ISL_660192, EPI_ISL_660193, EPI_ISL_660194, EPI_ISL_660195, EPI_ISL_660196, EPI_ISL_660197, EPI_ISL_660201, EPI_ISL_660203, EPI_ISL_660205, EPI_ISL_660211, EPI_ISL_660212, EPI_ISL_660213, EPI_ISL_660214, EPI_ISL_660215, EPI_ISL_660216, EPI_ISL_660217, EPI_ISL_660219, EPI_ISL_660220, EPI_ISL_660242, EPI_ISL_660243, EPI_ISL_660244, EPI_ISL_660245, EPI_ISL_660246, EPI_ISL_660247, EPI_ISL_660248, EPI_ISL_660249, EPI_ISL_660250, EPI_ISL_660251 | | | |
| see above | NHLS-IALCH | KRISP, KZN Research Innovation and Sequencing Platform | Giandhari J, Pillay S, Lessells R, Mdlalose K, York D, Khan S, Tegally H, Wilkinson E, de Oliveira T |
| EPI_ISL_660323 | Hospital | National Reference Center for Viruses of Respiratory Infections, Institut Pasteur, Paris | Marion Barbet, Sylvie Behillil, Méline Bizard, Angela Brisebarre, Camille Capel, Etienne Simon-Lorière, Vincent Enouf, Maud Vanpeene, Sylvie van der Werf, Alexandra Ducancelle |
| EPI_ISL_660471, EPI_ISL_660472 | Laboratoire de Microbiologie CHU Souro Sanou | Centre Muraz | Abdoul-Salam Ouedraogo, Yacouba Sawadogo, Essia Belarbi, Grit Schubert, Fabian Leendertz, Arsène Zongo, Soumeya Ouangraoua, Zekiba Tarnagda, Lassana Sangaré, Halidou Tinto |
| EPI_ISL_660602, EPI_ISL_660603, EPI_ISL_660604, EPI_ISL_660752, EPI_ISL_660753, EPI_ISL_660756, EPI_ISL_660761, EPI_ISL_660762 | Respiratory Virus Unit, Microbiology Services Colindale, Public Health England | COVID-19 Genomics UK (COG-UK) Consortium | PHE Covid Sequencing Team |
| EPI_ISL_660988, EPI_ISL_660989, EPI_ISL_660990, EPI_ISL_660991, EPI_ISL_660992, EPI_ISL_660993, EPI_ISL_660994, EPI_ISL_660995, EPI_ISL_660996, EPI_ISL_660997, EPI_ISL_660998, EPI_ISL_660999, EPI_ISL_661000, EPI_ISL_661001, EPI_ISL_661002, EPI_ISL_661003, EPI_ISL_661004, EPI_ISL_661005, EPI_ISL_661006, EPI_ISL_661007, EPI_ISL_661008, EPI_ISL_661009, EPI_ISL_661010, EPI_ISL_661011, EPI_ISL_661012, EPI_ISL_661013, EPI_ISL_661014, EPI_ISL_661015, EPI_ISL_661016, EPI_ISL_661017, EPI_ISL_661018, EPI_ISL_661019, EPI_ISL_661020, EPI_ISL_661021, EPI_ISL_661022, EPI_ISL_661023, EPI_ISL_661024, EPI_ISL_661025, EPI_ISL_661026 | | | |
| see above | Gundersen Molecular Diagnostics Laboratory | Kabara Cancer Research Institute | Craig S. Richmond, Paraic A. Kenny |
| EPI_ISL_661289 | Klinisk mikrobiologi | The Public Health Agency of Sweden | Department of Microbiology, The Public Health Agency of Sweden |
| EPI_ISL_661291 | Mikrobiologen | The Public Health Agency of Sweden | Department of Microbiology, The Public Health Agency of Sweden |
| EPI_ISL_661295, EPI_ISL_661296, EPI_ISL_661298 | Unilabs | The Public Health Agency of Sweden | Department of Microbiology, The Public Health Agency of Sweden |
| EPI_ISL_661304, EPI_ISL_661305, EPI_ISL_661306, EPI_ISL_661307, EPI_ISL_661308, EPI_ISL_661309, EPI_ISL_661310, EPI_ISL_661311 | CSIR-Indian Institute of Chemical Biology, MEDICA Supercpecialty Hospital Kolkata | CSIR-Indian Institute of Chemical Biology, MEDICA Supercpecialty Hospital Kolkata | Sujay Krishna Maity, Priyanka Mallick, Debaleena Bhowmik, Abhishake Lahiri, Dr. Aviral Roy, Dr. Soumen Saha, Dr. Arpita Ghosh Mitra, Dr. Rajesh Pandey, Dr. Sandip Paul, Dr. Partha Chakrabarti, Dr. Saikat Chakrabarti |
| EPI_ISL_662655, EPI_ISL_662658, EPI_ISL_662660, EPI_ISL_662661, EPI_ISL_662664, EPI_ISL_662665, EPI_ISL_662666, EPI_ISL_662667, EPI_ISL_662669, EPI_ISL_662670, EPI_ISL_662671, EPI_ISL_662672, EPI_ISL_662674, EPI_ISL_662676, EPI_ISL_662678, EPI_ISL_662679, EPI_ISL_662680, EPI_ISL_662681, EPI_ISL_662682, EPI_ISL_662684, EPI_ISL_662685, EPI_ISL_662686, EPI_ISL_662688, EPI_ISL_662689, EPI_ISL_662690, EPI_ISL_662692, EPI_ISL_662694, EPI_ISL_662696, EPI_ISL_662697, EPI_ISL_662698, EPI_ISL_662699, EPI_ISL_662701, EPI_ISL_662703, EPI_ISL_662704, EPI_ISL_662705, EPI_ISL_662706, EPI_ISL_662707, EPI_ISL_662709, EPI_ISL_662710, EPI_ISL_662713, EPI_ISL_662714, EPI_ISL_662715, EPI_ISL_662716, EPI_ISL_662718, EPI_ISL_662719, EPI_ISL_662720, EPI_ISL_662721, EPI_ISL_662724, EPI_ISL_662725, EPI_ISL_662726, EPI_ISL_662728, EPI_ISL_662729, EPI_ISL_662732, EPI_ISL_662736, EPI_ISL_662740, EPI_ISL_662741, EPI_ISL_662742, EPI_ISL_662745, EPI_ISL_662746, EPI_ISL_662747, EPI_ISL_662748, EPI_ISL_662749, EPI_ISL_662751, EPI_ISL_662752, EPI_ISL_662754, EPI_ISL_662755, EPI_ISL_662756, EPI_ISL_662757, EPI_ISL_662758, EPI_ISL_662759, EPI_ISL_662760, EPI_ISL_662761, EPI_ISL_662763, EPI_ISL_662764, EPI_ISL_662766, EPI_ISL_662767, EPI_ISL_662768, EPI_ISL_662770, EPI_ISL_662771, EPI_ISL_662772, EPI_ISL_662773, EPI_ISL_662774, EPI_ISL_662776, EPI_ISL_662777, EPI_ISL_662778, EPI_ISL_662779, EPI_ISL_662781, EPI_ISL_662783, EPI_ISL_662785, EPI_ISL_662786, EPI_ISL_662787, EPI_ISL_662788, EPI_ISL_662789, EPI_ISL_662790, EPI_ISL_662791, EPI_ISL_662792, EPI_ISL_662794, EPI_ISL_662795, EPI_ISL_662796, EPI_ISL_662797, EPI_ISL_662798, EPI_ISL_662799, EPI_ISL_662801, EPI_ISL_662805, EPI_ISL_662806, EPI_ISL_662808, EPI_ISL_662809, EPI_ISL_662811, EPI_ISL_662812, EPI_ISL_662813, EPI_ISL_662815, EPI_ISL_662816, EPI_ISL_662817, EPI_ISL_662818, EPI_ISL_662820, EPI_ISL_662821, EPI_ISL_662822, EPI_ISL_662824, EPI_ISL_662829, EPI_ISL_662830, EPI_ISL_662832, EPI_ISL_662834, EPI_ISL_662835, EPI_ISL_662836, EPI_ISL_662839, EPI_ISL_662841, EPI_ISL_662842, EPI_ISL_662844, EPI_ISL_662845, EPI_ISL_662846, EPI_ISL_662847, EPI_ISL_662849, EPI_ISL_662853, EPI_ISL_662854, EPI_ISL_662855, EPI_ISL_662856, EPI_ISL_662857, EPI_ISL_662858, EPI_ISL_662859, EPI_ISL_662860, EPI_ISL_662861, EPI_ISL_662864, EPI_ISL_662865, EPI_ISL_662867, EPI_ISL_662868, EPI_ISL_662869, EPI_ISL_662871, EPI_ISL_662872, EPI_ISL_662873, EPI_ISL_662875, EPI_ISL_662876, EPI_ISL_662877, EPI_ISL_662878, EPI_ISL_662879, EPI_ISL_662880, EPI_ISL_662882, EPI_ISL_662884, EPI_ISL_662886, EPI_ISL_662887, EPI_ISL_662888, EPI_ISL_662889, EPI_ISL_663134 | | | |

| | | | |
|---|---|---|---|
| see above | Lighthouse Lab in Glasgow | Wellcome Sanger Institute for the COVID-19 Genomics UK (COG-UK) Consortium | Harper VanSteenhouse, Yumi Kasai, David Gray, Carol Clugston, Anna Dominiczak and Alex Alderton, Roberto Amato, Sonia Goncalves, Ewan Harrison, David K. Jackson, Ian Johnston, Dominic Kwiatkowski, Cordelia Langford, John Sillitoe on behalf of the Wellcome Sanger Institute COVID-19 Surveillance Team |
| EPI_ISL_664111 | Liverpool Clinical Laboratories | COVID-19 Genomics UK (COG-UK) Consortium | Sam Haldenby, Anita Lucaci, Steve Paterson, Julian Hiscox, Alistair Darby, M Almsaud, A Alrezaihi, Muhannad Alruwaili, Stuart D Armstrong, Jones Benjamin, Eleanor G Bentley, Anu Chawla, Jordan J Clark, Angela Cowell, Richard Eccles, Isabel García-Dorival, Matthew Gemmell, Alessandro Gerada, PKF Gilmore, Richard Gregory, Ximeng Han, Catherine Hartley, Margaret Hughes, Miren Iturriza-Gomara, James Johnson, L Luu, Jenifer Manson, Charlotte Nelson, Elaine O'Toole, Cassie Olateju, Rebekah Penrice-Randal , Lucille Rainbow, N.P Randle, Trevor Ian Robinson, Parul Sharma, Ghada T Shawli, James P Stewart, Neil Swainston, Ecaterina Vamos, Joanne Watts, Mark Whitehead |
| EPI_ISL_664123, EPI_ISL_664124, EPI_ISL_664127 | University of Exeter | COVID-19 Genomics UK (COG-UK) Consortium | Ben Temperton,Aaron Jeffries,Michelle Michelsen,Joanna Warwick-Dugdale,Audrey Farbos,Robyn Manley,Stephen Michell,Jane Masoli |
| EPI_ISL_664134, EPI_ISL_664138, EPI_ISL_664142 | Liverpool Clinical Laboratories | COVID-19 Genomics UK (COG-UK) Consortium | Sam Haldenby, Anita Lucaci, Steve Paterson, Julian Hiscox, Alistair Darby, M Almsaud, A Alrezaihi, Muhannad Alruwaili, Stuart D Armstrong, Jones Benjamin, Eleanor G Bentley, Anu Chawla, Jordan J Clark, Angela Cowell, Richard Eccles, Isabel García-Dorival, Matthew Gemmell, Alessandro Gerada, PKF Gilmore, Richard Gregory, Ximeng Han, Catherine Hartley, Margaret Hughes, Miren Iturriza-Gomara, James Johnson, L Luu, Jenifer Manson, Charlotte Nelson, Elaine O'Toole, Cassie Olateju, Rebekah Penrice-Randal , Lucille Rainbow, N.P Randle, Trevor Ian Robinson, Parul Sharma, Ghada T Shawli, James P Stewart, Neil Swainston, Ecaterina Vamos, Joanne Watts, Mark Whitehead |
| EPI_ISL_664147 | University College London Hospital | COVID-19 Genomics UK (COG-UK) Consortium | Judith Heaney, Matthew Byott, Catherine Houlihan, Dan Frampton, Stuart Kirk, Moira Spyer and Eleni Nastouli |
| EPI_ISL_664164, EPI_ISL_664203 | Liverpool Clinical Laboratories | COVID-19 Genomics UK (COG-UK) Consortium | Sam Haldenby, Anita Lucaci, Steve Paterson, Julian Hiscox, Alistair Darby, M Almsaud, A Alrezaihi, Muhannad Alruwaili, Stuart D Armstrong, Jones Benjamin, Eleanor G Bentley, Anu Chawla, Jordan J Clark, Angela Cowell, Richard Eccles, Isabel García-Dorival, Matthew Gemmell, Alessandro Gerada, PKF Gilmore, Richard Gregory, Ximeng Han, Catherine Hartley, Margaret Hughes, Miren Iturriza-Gomara, James Johnson, L Luu, Jenifer Manson, Charlotte Nelson, Elaine O'Toole, Cassie Olateju, Rebekah Penrice-Randal , Lucille Rainbow, N.P Randle, Trevor Ian Robinson, Parul Sharma, Ghada T Shawli, James P Stewart, Neil Swainston, Ecaterina Vamos, Joanne Watts, Mark Whitehead |
| EPI_ISL_664207, EPI_ISL_664208, EPI_ISL_664218 | University of Exeter | COVID-19 Genomics UK (COG-UK) Consortium | Ben Temperton,Aaron Jeffries,Michelle Michelsen,Joanna Warwick-Dugdale,Audrey Farbos,Robyn Manley,Stephen Michell,Jane Masoli |
| EPI_ISL_664219, EPI_ISL_664229, EPI_ISL_664231, EPI_ISL_664232, EPI_ISL_664278, EPI_ISL_664301 | Liverpool Clinical Laboratories | COVID-19 Genomics UK (COG-UK) Consortium | Sam Haldenby, Anita Lucaci, Steve Paterson, Julian Hiscox, Alistair Darby, M Almsaud, A Alrezaihi, Muhannad Alruwaili, Stuart D Armstrong, Jones Benjamin, Eleanor G Bentley, Anu Chawla, Jordan J Clark, Angela Cowell, Richard Eccles, Isabel García-Dorival, Matthew Gemmell, Alessandro Gerada, PKF Gilmore, Richard Gregory, Ximeng Han, Catherine Hartley, Margaret Hughes, Miren Iturriza-Gomara, James Johnson, L Luu, Jenifer Manson, Charlotte Nelson, Elaine O'Toole, Cassie Olateju, Rebekah Penrice-Randal , Lucille Rainbow, N.P Randle, Trevor Ian Robinson, Parul Sharma, Ghada T Shawli, James P Stewart, Neil Swainston, Ecaterina Vamos, Joanne Watts, Mark Whitehead |
| EPI_ISL_664318, EPI_ISL_664319, EPI_ISL_664328, EPI_ISL_664330, EPI_ISL_664331 | University of Exeter | COVID-19 Genomics UK (COG-UK) Consortium | Ben Temperton,Aaron Jeffries,Michelle Michelsen,Joanna Warwick-Dugdale,Audrey Farbos,Robyn Manley,Stephen Michell,Jane Masoli |
| EPI_ISL_664333, EPI_ISL_664336, EPI_ISL_664338, EPI_ISL_664345, EPI_ISL_664348, EPI_ISL_664349, EPI_ISL_664355 | Liverpool Clinical Laboratories | COVID-19 Genomics UK (COG-UK) Consortium | Sam Haldenby, Anita Lucaci, Steve Paterson, Julian Hiscox, Alistair Darby, M Almsaud, A Alrezaihi, Muhannad Alruwaili, Stuart D Armstrong, Jones Benjamin, Eleanor G Bentley, Anu Chawla, Jordan J Clark, Angela Cowell, Richard Eccles, Isabel García-Dorival, Matthew Gemmell, Alessandro Gerada, PKF Gilmore, Richard Gregory, Ximeng Han, Catherine Hartley, Margaret Hughes, Miren Iturriza-Gomara, James Johnson, L Luu, Jenifer Manson, Charlotte Nelson, Elaine O'Toole, Cassie Olateju, Rebekah Penrice-Randal , Lucille Rainbow, N.P Randle, Trevor Ian Robinson, Parul Sharma, Ghada T Shawli, James P Stewart, Neil Swainston, Ecaterina Vamos, Joanne Watts, Mark Whitehead |
| EPI_ISL_664356 | University College London Hospital | COVID-19 Genomics UK (COG-UK) Consortium | Judith Heaney, Matthew Byott, Catherine Houlihan, Dan Frampton, Stuart Kirk, Moira Spyer and Eleni Nastouli |
| EPI_ISL_664362, EPI_ISL_664364, EPI_ISL_664424 | Liverpool Clinical Laboratories | COVID-19 Genomics UK (COG-UK) Consortium | Sam Haldenby, Anita Lucaci, Steve Paterson, Julian Hiscox, Alistair Darby, M Almsaud, A Alrezaihi, Muhannad Alruwaili, Stuart D Armstrong, Jones Benjamin, Eleanor G Bentley, Anu Chawla, Jordan J Clark, Angela Cowell, Richard Eccles, Isabel García-Dorival, Matthew Gemmell, Alessandro Gerada, PKF Gilmore, Richard Gregory, Ximeng Han, Catherine Hartley, Margaret Hughes, Miren Iturriza-Gomara, James Johnson, L Luu, Jenifer Manson, Charlotte Nelson, Elaine O'Toole, Cassie Olateju, Rebekah Penrice-Randal , Lucille Rainbow, N.P Randle, Trevor Ian Robinson, Parul Sharma, Ghada T Shawli, James P Stewart, Neil Swainston, Ecaterina Vamos, Joanne Watts, Mark Whitehead |
| EPI_ISL_664439 | University of Exeter | COVID-19 Genomics UK (COG-UK) Consortium | Ben Temperton,Aaron Jeffries,Michelle Michelsen,Joanna Warwick-Dugdale,Audrey Farbos,Robyn Manley,Stephen Michell,Jane Masoli |
| EPI_ISL_664443, EPI_ISL_664444, EPI_ISL_664445, EPI_ISL_664448 | Liverpool Clinical Laboratories | COVID-19 Genomics UK (COG-UK) Consortium | Sam Haldenby, Anita Lucaci, Steve Paterson, Julian Hiscox, Alistair Darby, M Almsaud, A Alrezaihi, Muhannad Alruwaili, Stuart D Armstrong, Jones Benjamin, Eleanor G Bentley, Anu Chawla, Jordan J Clark, Angela Cowell, Richard Eccles, Isabel García-Dorival, Matthew Gemmell, Alessandro Gerada, PKF Gilmore, Richard Gregory, Ximeng Han, Catherine Hartley, Margaret Hughes, Miren Iturriza-Gomara, James Johnson, L Luu, Jenifer Manson, Charlotte Nelson, Elaine O'Toole, Cassie Olateju, Rebekah Penrice-Randal , Lucille Rainbow, N.P Randle, Trevor Ian Robinson, Parul Sharma, Ghada T Shawli, James P Stewart, Neil Swainston, Ecaterina Vamos, Joanne Watts, Mark Whitehead |
| EPI_ISL_664465 | University of Exeter | COVID-19 Genomics UK (COG-UK) Consortium | Ben Temperton,Aaron Jeffries,Michelle Michelsen,Joanna Warwick-Dugdale,Audrey Farbos,Robyn Manley,Stephen Michell,Jane Masoli |
| EPI_ISL_664468, EPI_ISL_664469 | Liverpool Clinical Laboratories | COVID-19 Genomics UK (COG-UK) Consortium | Sam Haldenby, Anita Lucaci, Steve Paterson, Julian Hiscox, Alistair Darby, M Almsaud, A Alrezaihi, Muhannad Alruwaili, Stuart D Armstrong, Jones Benjamin, Eleanor G Bentley, Anu Chawla, Jordan J Clark, Angela Cowell, Richard Eccles, Isabel García-Dorival, Matthew Gemmell, Alessandro Gerada, PKF Gilmore, Richard Gregory, Ximeng Han, Catherine Hartley, Margaret Hughes, Miren Iturriza-Gomara, James Johnson, L Luu, Jenifer Manson, Charlotte Nelson, Elaine O'Toole, Cassie Olateju, Rebekah Penrice-Randal , Lucille Rainbow, N.P Randle, Trevor Ian Robinson, Parul Sharma, Ghada T Shawli, James P Stewart, Neil Swainston, Ecaterina Vamos, Joanne Watts, Mark Whitehead |
| EPI_ISL_664474 | University of Exeter | COVID-19 Genomics UK (COG-UK) Consortium | Ben Temperton,Aaron Jeffries,Michelle Michelsen,Joanna Warwick-Dugdale,Audrey Farbos,Robyn Manley,Stephen Michell,Jane Masoli |
| EPI_ISL_664475, EPI_ISL_664486, EPI_ISL_664489 | Liverpool Clinical Laboratories | COVID-19 Genomics UK (COG-UK) Consortium | Sam Haldenby, Anita Lucaci, Steve Paterson, Julian Hiscox, Alistair Darby, M Almsaud, A Alrezaihi, Muhannad Alruwaili, Stuart D Armstrong, Jones Benjamin, Eleanor G Bentley, Anu Chawla, Jordan J Clark, Angela Cowell, Richard Eccles, Isabel García-Dorival, Matthew Gemmell, Alessandro Gerada, PKF Gilmore, Richard Gregory, Ximeng Han, Catherine Hartley, Margaret Hughes, Miren Iturriza-Gomara, James Johnson, L Luu, Jenifer Manson, Charlotte Nelson, Elaine O'Toole, Cassie Olateju, Rebekah Penrice-Randal , Lucille Rainbow, N.P Randle, Trevor Ian Robinson, Parul Sharma, Ghada T Shawli, James P Stewart, Neil Swainston, Ecaterina Vamos, Joanne Watts, Mark Whitehead |
| EPI_ISL_664491 | University of Exeter | COVID-19 Genomics UK (COG-UK) Consortium | Ben Temperton,Aaron Jeffries,Michelle Michelsen,Joanna Warwick-Dugdale,Audrey Farbos,Robyn Manley,Stephen Michell,Jane Masoli |
| EPI_ISL_664501, EPI_ISL_664513 | Liverpool Clinical Laboratories | COVID-19 Genomics UK (COG-UK) Consortium | Sam Haldenby, Anita Lucaci, Steve Paterson, Julian Hiscox, Alistair Darby, M Almsaud, A Alrezaihi, Muhannad Alruwaili, Stuart D Armstrong, Jones Benjamin, Eleanor G Bentley, Anu Chawla, Jordan J Clark, Angela Cowell, Richard Eccles, Isabel García-Dorival, Matthew Gemmell, Alessandro Gerada, PKF Gilmore, Richard Gregory, Ximeng Han, Catherine Hartley, Margaret Hughes, Miren Iturriza-Gomara, James Johnson, L Luu, Jenifer Manson, Charlotte Nelson, Elaine O'Toole, Cassie Olateju, Rebekah Penrice-Randal , Lucille Rainbow, N.P Randle, Trevor Ian Robinson, Parul Sharma, Ghada T Shawli, James P Stewart, Neil Swainston, Ecaterina Vamos, Joanne Watts, Mark Whitehead |
| EPI_ISL_664517 | University College London Hospital | COVID-19 Genomics UK (COG-UK) Consortium | Judith Heaney, Matthew Byott, Catherine Houlihan, Dan Frampton, Stuart Kirk, Moira Spyer and Eleni Nastouli |
| EPI_ISL_664519, EPI_ISL_664523, EPI_ISL_664524 | Liverpool Clinical Laboratories | COVID-19 Genomics UK (COG-UK) Consortium | Sam Haldenby, Anita Lucaci, Steve Paterson, Julian Hiscox, Alistair Darby, M Almsaud, A Alrezaihi, Muhannad Alruwaili, Stuart D Armstrong, Jones Benjamin, Eleanor G Bentley, Anu Chawla, Jordan J Clark, Angela Cowell, Richard Eccles, Isabel García-Dorival, Matthew Gemmell, Alessandro Gerada, PKF Gilmore, Richard Gregory, Ximeng Han, Catherine Hartley, Margaret Hughes, Miren Iturriza-Gomara, James Johnson, L Luu, Jenifer Manson, Charlotte Nelson, Elaine O'Toole, Cassie Olateju, Rebekah Penrice-Randal , Lucille Rainbow, N.P Randle, Trevor Ian Robinson, Parul Sharma, Ghada T Shawli, James P Stewart, Neil Swainston, Ecaterina Vamos, Joanne Watts, Mark Whitehead |
| EPI_ISL_664603, EPI_ISL_664612 | University of Exeter | COVID-19 Genomics UK (COG-UK) Consortium | Ben Temperton,Aaron Jeffries,Michelle Michelsen,Joanna Warwick-Dugdale,Audrey Farbos,Robyn Manley,Stephen Michell,Jane Masoli |
| EPI_ISL_664632, EPI_ISL_664649, EPI_ISL_664652, EPI_ISL_664653, EPI_ISL_664664, EPI_ISL_664665, | Liverpool Clinical Laboratories | COVID-19 Genomics UK (COG-UK) Consortium | Sam Haldenby, Anita Lucaci, Steve Paterson, Julian Hiscox, Alistair Darby, M Almsaud, A Alrezaihi, Muhannad Alruwaili, Stuart D Armstrong, Jones Benjamin, Eleanor G Bentley, Anu Chawla, Jordan J Clark, Angela Cowell, Richard Eccles, Isabel García-Dorival, Matthew Gemmell, Alessandro Gerada, PKF Gilmore, Richard Gregory, Ximeng Han, Catherine Hartley, Margaret Hughes, Miren Iturriza-Gomara, James Johnson, L Luu, Jenifer Manson, |

| | | | |
|---|---|---|---|
| EPI_ISL_664667, EPI_ISL_664668, EPI_ISL_664670 | | | Charlotte Nelson, Elaine O'Toole, Cassie Olateju, Rebekah Penrice-Randal , Lucille Rainbow, N.P Randle, Trevor Ian Robinson, Parul Sharma, Ghada T Shawli, James P Stewart, Neil Swainston, Ecaterina Vamos, Joanne Watts, Mark Whitehead |
| EPI_ISL_664683, EPI_ISL_664686, EPI_ISL_664691 | University of Exeter | COVID-19 Genomics UK (COG-UK) Consortium | Ben Temperton,Aaron Jeffries,Michelle Michelsen,Joanna Warwick-Dugdale,Audrey Farbos,Robyn Manley,Stephen Michell,Jane Masoli |
| EPI_ISL_664693, EPI_ISL_664695, EPI_ISL_664705 | Liverpool Clinical Laboratories | COVID-19 Genomics UK (COG-UK) Consortium | Sam Haldenby, Anita Lucaci, Steve Paterson, Julian Hiscox, Alistair Darby, M Almsaud, A Alrezaihi, Muhannad Alruwaili, Stuart D Armstrong, Jones Benjamin, Eleanor G Bentley, Anu Chawla, Jordan J Clark, Angela Cowell, Richard Eccles, Isabel García-Dorival, Matthew Gemmell, Alessandro Gerada, PKF Gilmore, Richard Gregory, Ximeng Han, Catherine Hartley, Margaret Hughes, Miren Iturriza-Gomara, James Johnson, L Luu, Jenifer Manson, Charlotte Nelson, Elaine O'Toole, Cassie Olateju, Rebekah Penrice-Randal , Lucille Rainbow, N.P Randle, Trevor Ian Robinson, Parul Sharma, Ghada T Shawli, James P Stewart, Neil Swainston, Ecaterina Vamos, Joanne Watts, Mark Whitehead |
| EPI_ISL_664711 | University College London Hospital | COVID-19 Genomics UK (COG-UK) Consortium | Judith Heaney, Matthew Byott, Catherine Houlihan, Dan Frampton, Stuart Kirk, Moira Spyer and Eleni Nastouli |
| EPI_ISL_664716 | University of Exeter | COVID-19 Genomics UK (COG-UK) Consortium | Ben Temperton,Aaron Jeffries,Michelle Michelsen,Joanna Warwick-Dugdale,Audrey Farbos,Robyn Manley,Stephen Michell,Jane Masoli |
| EPI_ISL_664726, EPI_ISL_664732, EPI_ISL_664733 | Liverpool Clinical Laboratories | COVID-19 Genomics UK (COG-UK) Consortium | Sam Haldenby, Anita Lucaci, Steve Paterson, Julian Hiscox, Alistair Darby, M Almsaud, A Alrezaihi, Muhannad Alruwaili, Stuart D Armstrong, Jones Benjamin, Eleanor G Bentley, Anu Chawla, Jordan J Clark, Angela Cowell, Richard Eccles, Isabel García-Dorival, Matthew Gemmell, Alessandro Gerada, PKF Gilmore, Richard Gregory, Ximeng Han, Catherine Hartley, Margaret Hughes, Miren Iturriza-Gomara, James Johnson, L Luu, Jenifer Manson, Charlotte Nelson, Elaine O'Toole, Cassie Olateju, Rebekah Penrice-Randal , Lucille Rainbow, N.P Randle, Trevor Ian Robinson, Parul Sharma, Ghada T Shawli, James P Stewart, Neil Swainston, Ecaterina Vamos, Joanne Watts, Mark Whitehead |
| EPI_ISL_664754, EPI_ISL_664839 | University College London Hospital | COVID-19 Genomics UK (COG-UK) Consortium | Judith Heaney, Matthew Byott, Catherine Houlihan, Dan Frampton, Stuart Kirk, Moira Spyer and Eleni Nastouli |
| EPI_ISL_665048, EPI_ISL_665049, EPI_ISL_665050, EPI_ISL_665052, EPI_ISL_665053, EPI_ISL_665054, EPI_ISL_665055, EPI_ISL_665056, EPI_ISL_665057, EPI_ISL_665058, EPI_ISL_665059, EPI_ISL_665064, EPI_ISL_665065 | | | |
| see above | University of Exeter | COVID-19 Genomics UK (COG-UK) Consortium | Ben Temperton,Aaron Jeffries,Michelle Michelsen,Joanna Warwick-Dugdale,Audrey Farbos,Robyn Manley,Stephen Michell,Jane Masoli |
| EPI_ISL_665128, EPI_ISL_665129, EPI_ISL_665130, EPI_ISL_665131, EPI_ISL_665132, EPI_ISL_665133, EPI_ISL_665134, EPI_ISL_665135, EPI_ISL_665136, EPI_ISL_665137, EPI_ISL_665138, EPI_ISL_665139 | | | |
| see above | Liverpool Clinical Laboratories | COVID-19 Genomics UK (COG-UK) Consortium | Sam Haldenby, Anita Lucaci, Steve Paterson, Julian Hiscox, Alistair Darby, M Almsaud, A Alrezaihi, Muhannad Alruwaili, Stuart D Armstrong, Jones Benjamin, Eleanor G Bentley, Anu Chawla, Jordan J Clark, Angela Cowell, Richard Eccles, Isabel García-Dorival, Matthew Gemmell, Alessandro Gerada, PKF Gilmore, Richard Gregory, Ximeng Han, Catherine Hartley, Margaret Hughes, Miren Iturriza-Gomara, James Johnson, L Luu, Jenifer Manson, Charlotte Nelson, Elaine O'Toole, Cassie Olateju, Rebekah Penrice-Randal , Lucille Rainbow, N.P Randle, Trevor Ian Robinson, Parul Sharma, Ghada T Shawli, James P Stewart, Neil Swainston, Ecaterina Vamos, Joanne Watts, Mark Whitehead |
| EPI_ISL_665229, EPI_ISL_665230, EPI_ISL_665231, EPI_ISL_665232, EPI_ISL_665233, EPI_ISL_665234, EPI_ISL_665235, EPI_ISL_665236, EPI_ISL_665237, EPI_ISL_665238 | University College London Hospital | COVID-19 Genomics UK (COG-UK) Consortium | Judith Heaney, Matthew Byott, Catherine Houlihan, Dan Frampton, Stuart Kirk, Moira Spyer and Eleni Nastouli |
| EPI_ISL_665261 | Northumbria University / South Tees Hospitals NHS Foundation Trust / North Cumbria Integrated Care NHS Foundation Trust / North Tees and Hartlepool NHS Foundation Trust / Newcastle Hospitals NHS Foundation Trust | COVID-19 Genomics UK (COG-UK) Consortium | Darren L Smith,Andrew Nelson,Matthew Bashton,Greg R Young,Joshua Loh,John Allan,Mohammad A Tariq,Giles S Holt,Gary Black,Wen C Yew,Lynn Dover,Paul Baker,Steve Liggett,Sarah Essex,Jane Greenaway,Debra Padgett,Clive Graham,Garren Scott,Edward Barton,Emma Swindells,Brendan Payne,Jennifer Collins,Yusri Taha,Gary Eltringham |
| EPI_ISL_665269 | Quadram Institute Bioscience | COVID-19 Genomics UK (COG-UK) Consortium | Dave J. Baker, Gemma L. Kay, Alp Aydin, Thanh Le-Viet, Steven Rudder, Ana P. Tedim, Anastasia Kolyva, Maria Diaz, Leonardo de Oliveira Martins, Nabil-Fareed Alikhan, Lizzie Meadows, Rachael Stanley, Ngozi Elumogo, Muhammed Yasir, Nicholas M. Thomson, Alexander J Trotter, Rachel Gilroy, Samuel Bloomfield, Claire Stuart, Andrew Bell, Reenesh Prakash, Samir Dervisevic, Alison E. Mather, John Wain, Mark Webber, Andrew J. Page, Justin O'Grady |
| EPI_ISL_665271, EPI_ISL_665307, EPI_ISL_665318 | Northumbria University / South Tees Hospitals NHS Foundation Trust / North Cumbria Integrated Care NHS Foundation Trust / North Tees and Hartlepool NHS Foundation Trust / Newcastle Hospitals NHS Foundation Trust | COVID-19 Genomics UK (COG-UK) Consortium | Darren L Smith,Andrew Nelson,Matthew Bashton,Greg R Young,Joshua Loh,John Allan,Mohammad A Tariq,Giles S Holt,Gary Black,Wen C Yew,Lynn Dover,Paul Baker,Steve Liggett,Sarah Essex,Jane Greenaway,Debra Padgett,Clive Graham,Garren Scott,Edward Barton,Emma Swindells,Brendan Payne,Jennifer Collins,Yusri Taha,Gary Eltringham |
| EPI_ISL_665382 | University College London Hospital | COVID-19 Genomics UK (COG-UK) Consortium | Judith Heaney, Matthew Byott, Catherine Houlihan, Dan Frampton, Stuart Kirk, Moira Spyer and Eleni Nastouli |
| EPI_ISL_665390, EPI_ISL_665399, EPI_ISL_665400, EPI_ISL_665401, EPI_ISL_665410 | Northumbria University / South Tees Hospitals NHS Foundation Trust / North Cumbria Integrated Care NHS Foundation Trust / North Tees and Hartlepool NHS Foundation Trust / Newcastle Hospitals NHS Foundation Trust | COVID-19 Genomics UK (COG-UK) Consortium | Darren L Smith,Andrew Nelson,Matthew Bashton,Greg R Young,Joshua Loh,John Allan,Mohammad A Tariq,Giles S Holt,Gary Black,Wen C Yew,Lynn Dover,Paul Baker,Steve Liggett,Sarah Essex,Jane Greenaway,Debra Padgett,Clive Graham,Garren Scott,Edward Barton,Emma Swindells,Brendan Payne,Jennifer Collins,Yusri Taha,Gary Eltringham |
| EPI_ISL_665487, EPI_ISL_665488 | Centre for Enzyme Innovation, University of Portsmouth / Translational Research Laboratory, Portsmouth Hospitals NHS Trust | COVID-19 Genomics UK (COG-UK) Consortium | Angela Beckett,Yann Bourgeois,Garry Scarlett,Sharon Glaysher,Scott Elliott,Kelly Bicknell,Robert Impey,Allyson Lloyd,Sarah Wyllie,Ethan Butcher,Anoop Chauhan,Samuel Robson |
| EPI_ISL_665496, EPI_ISL_665497, EPI_ISL_665498, EPI_ISL_665503, EPI_ISL_665509, EPI_ISL_665629, EPI_ISL_665651 | Northumbria University / South Tees Hospitals NHS Foundation Trust / North Cumbria Integrated Care NHS Foundation Trust / North Tees and Hartlepool NHS Foundation Trust / Newcastle Hospitals NHS Foundation Trust | COVID-19 Genomics UK (COG-UK) Consortium | Darren L Smith,Andrew Nelson,Matthew Bashton,Greg R Young,Joshua Loh,John Allan,Mohammad A Tariq,Giles S Holt,Gary Black,Wen C Yew,Lynn Dover,Paul Baker,Steve Liggett,Sarah Essex,Jane Greenaway,Debra Padgett,Clive Graham,Garren Scott,Edward Barton,Emma Swindells,Brendan Payne,Jennifer Collins,Yusri Taha,Gary Eltringham |
| EPI_ISL_665653 | Wales Specialist Virology Centre Sequencing lab: Pathogen Genomics Unit | COVID-19 Genomics UK (COG-UK) Consortium | Catherine Moore, Johnathan Evans, Laura Gifford, Malorie Perry, Simon Cottrell, Angela Marchbank, Alec Birchley, Alexander Adams, Amy Gaskin, Bree Gatica-Wilcox, Jason Coombes, Joel Southgate, Lauren Gilbert, Lee Graham, Nicole Pacchiarini, Sara Kumziene-Summerhayes, Sarah Taylor, Sophie Jones, Sara Rey, Matthew Bull, Joanne Watkins, Sally Corden, Tom Connor |
| EPI_ISL_665679, EPI_ISL_665687 | Northumbria University / South Tees Hospitals NHS Foundation Trust / North Cumbria Integrated Care NHS Foundation Trust / North Tees and Hartlepool NHS Foundation Trust / Newcastle Hospitals NHS Foundation Trust | COVID-19 Genomics UK (COG-UK) Consortium | Darren L Smith,Andrew Nelson,Matthew Bashton,Greg R Young,Joshua Loh,John Allan,Mohammad A Tariq,Giles S Holt,Gary Black,Wen C Yew,Lynn Dover,Paul Baker,Steve Liggett,Sarah Essex,Jane Greenaway,Debra Padgett,Clive Graham,Garren Scott,Edward Barton,Emma Swindells,Brendan Payne,Jennifer Collins,Yusri Taha,Gary Eltringham |
| EPI_ISL_665711 | University College London, Great Ormond Street Hospital for Children NHS Foundation Trust, Imperial College Healthcare NHS Trust | COVID-19 Genomics UK (COG-UK) Consortium | Sergi Castellano, Rachel Williams, Mark Kristiansen, Paola Resende Silva, Sunando Roy, Tony Brooks, Helena Tutill, Paola Niola, Patricia Dyal, Charlotte Williams, Leysa Forrest, Yasmin Panchbhaya, Jacqueline Findlay, Samuel Weeks, Julianne Brown, Kathryn Harris, Paul Randell, James Price, Alison Holmes, Judith Breuer |
| EPI_ISL_665717, EPI_ISL_665736, EPI_ISL_665742 | Northumbria University / South Tees Hospitals NHS Foundation Trust / North Cumbria Integrated Care NHS Foundation Trust / North Tees and Hartlepool NHS Foundation Trust / Newcastle Hospitals NHS Foundation Trust | COVID-19 Genomics UK (COG-UK) Consortium | Darren L Smith,Andrew Nelson,Matthew Bashton,Greg R Young,Joshua Loh,John Allan,Mohammad A Tariq,Giles S Holt,Gary Black,Wen C Yew,Lynn Dover,Paul Baker,Steve Liggett,Sarah Essex,Jane Greenaway,Debra Padgett,Clive Graham,Garren Scott,Edward Barton,Emma Swindells,Brendan Payne,Jennifer Collins,Yusri Taha,Gary Eltringham |
| EPI_ISL_665795 | Wales Specialist Virology Centre Sequencing lab: Pathogen Genomics Unit | COVID-19 Genomics UK (COG-UK) Consortium | Catherine Moore, Johnathan Evans, Laura Gifford, Malorie Perry, Simon Cottrell, Angela Marchbank, Alec Birchley, Alexander Adams, Amy Gaskin, Bree Gatica-Wilcox, Jason Coombes, Joel Southgate, Lauren Gilbert, Lee Graham, Nicole Pacchiarini, Sara Kumziene-Summerhayes, Sarah Taylor, Sophie Jones, Sara Rey, Matthew Bull, Joanne Watkins, Sally Corden, Tom Connor |
| EPI_ISL_665866, EPI_ISL_665867, EPI_ISL_665892 | Quadram Institute Bioscience | COVID-19 Genomics UK (COG-UK) Consortium | Dave J. Baker, Gemma L. Kay, Alp Aydin, Thanh Le-Viet, Steven Rudder, Ana P. Tedim, Anastasia Kolyva, Maria Diaz, Leonardo de Oliveira Martins, Nabil-Fareed Alikhan, Lizzie Meadows, Rachael Stanley, Ngozi Elumogo, Muhammed Yasir, Nicholas M. Thomson, Alexander J Trotter, Rachel Gilroy, Samuel Bloomfield, Claire Stuart, Andrew Bell, Reenesh Prakash, Samir Dervisevic, Alison E. Mather, John Wain, Mark Webber, Andrew J. Page, Justin O'Grady |

| | | | |
|---|---|---|---|
| EPI_ISL_666003, EPI_ISL_666004, EPI_ISL_666005, EPI_ISL_666006, EPI_ISL_666008, EPI_ISL_666010, EPI_ISL_666011, EPI_ISL_666012, EPI_ISL_666013, EPI_ISL_666014, EPI_ISL_666015, EPI_ISL_666016, EPI_ISL_666017, EPI_ISL_666101, EPI_ISL_666102, EPI_ISL_666103, EPI_ISL_666104, EPI_ISL_666105, EPI_ISL_666106 | | | |
| see above | Northumbria University / South Tees Hospitals NHS Foundation Trust / North Cumbria Integrated Care NHS Foundation Trust / North Tees and Hartlepool NHS Foundation Trust / Newcastle Hospitals NHS Foundation Trust | COVID-19 Genomics UK (COG-UK) Consortium | Darren L Smith,Andrew Nelson,Matthew Bashton,Greg R Young,Joshua Loh,John Allan,Mohammad A Tariq,Giles S Holt,Gary Black,Wen C Yew,Lynn Dover,Paul Baker,Steve Liggett,Sarah Essex,Jane Greenaway,Debra Padgett,Clive Graham,Garren Scott,Edward Barton,Emma Swindells,Brendan Payne,Jennifer Collins,Yusri Taha,Gary Eltringham |
| EPI_ISL_666113, EPI_ISL_666120, EPI_ISL_666131, EPI_ISL_666144, EPI_ISL_666151, EPI_ISL_666153, EPI_ISL_666161, EPI_ISL_666182, EPI_ISL_666210, EPI_ISL_666220, EPI_ISL_666222, EPI_ISL_666224, EPI_ISL_666226, EPI_ISL_666241, EPI_ISL_666244, EPI_ISL_666245, EPI_ISL_666252, EPI_ISL_666258, EPI_ISL_666273, EPI_ISL_666276, EPI_ISL_666277, EPI_ISL_666286, EPI_ISL_666305, EPI_ISL_666315, EPI_ISL_666318, EPI_ISL_666319, EPI_ISL_666322, EPI_ISL_666329, EPI_ISL_666350, EPI_ISL_666352, EPI_ISL_666355, EPI_ISL_666361 | | | |
| see above | Wales Specialist Virology Centre Sequencing lab: Pathogen Genomics Unit | COVID-19 Genomics UK (COG-UK) Consortium | Catherine Moore, Johnathan Evans, Laura Gifford, Malorie Perry, Simon Cottrell, Angela Marchbank, Alec Birchley, Alexander Adams, Amy Gaskin, Bree Gatica-Wilcox, Jason Coombes, Joel Southgate, Lauren Gilbert, Lee Graham, Nicole Pacchiarini, Sara Kumziene-Summerhayes, Sarah Taylor, Sophie Jones, Sara Rey, Matthew Bull, Joanne Watkins, Sally Corden, Tom Connor |
| EPI_ISL_666592 | Northumbria University / South Tees Hospitals NHS Foundation Trust / North Cumbria Integrated Care NHS Foundation Trust / North Tees and Hartlepool NHS Foundation Trust / Newcastle Hospitals NHS Foundation Trust | COVID-19 Genomics UK (COG-UK) Consortium | Darren L Smith,Andrew Nelson,Matthew Bashton,Greg R Young,Joshua Loh,John Allan,Mohammad A Tariq,Giles S Holt,Gary Black,Wen C Yew,Lynn Dover,Paul Baker,Steve Liggett,Sarah Essex,Jane Greenaway,Debra Padgett,Clive Graham,Garren Scott,Edward Barton,Emma Swindells,Brendan Payne,Jennifer Collins,Yusri Taha,Gary Eltringham |
| EPI_ISL_666740 | Respiratory Virus Unit, Microbiology Services Colindale, Public Health England | COVID-19 Genomics UK (COG-UK) Consortium | PHE Covid Sequencing Team |
| EPI_ISL_666872, EPI_ISL_666873, EPI_ISL_666874, EPI_ISL_666875, EPI_ISL_666876, EPI_ISL_666877, EPI_ISL_666878, EPI_ISL_666879, EPI_ISL_666880, EPI_ISL_666881, EPI_ISL_666882, EPI_ISL_666883, EPI_ISL_666884, EPI_ISL_666885 | | | |
| see above | Department of Clinical Microbiology | GIGA Medical Genomics | Keith Durkin, Maria Artesi, Justine Defêche, Gilles Darcis, Michel Moutschen, Sébastien Bontems, Raphaël Boreux, Bouchra Boujemla, Cécile Meex, Pierrette Melin, Marie-Pierre Hayette, Vincent Bours |
| EPI_ISL_666892, EPI_ISL_666897, EPI_ISL_666901, EPI_ISL_666904, EPI_ISL_666908, EPI_ISL_666914, EPI_ISL_666915, EPI_ISL_666916, EPI_ISL_666925, EPI_ISL_666938, EPI_ISL_666939, EPI_ISL_666950, EPI_ISL_666956, EPI_ISL_666960 | | | |
| see above | Michigan Department of Health and Human Services, Bureau of Laboratories | Michigan Department of Health and Human Services, Bureau of Laboratories | Blankenship HM, Riner D, Soehnlen MK |
| EPI_ISL_666965, EPI_ISL_666968, EPI_ISL_666969, EPI_ISL_666970, EPI_ISL_666975, EPI_ISL_666976, EPI_ISL_666983, EPI_ISL_666991, EPI_ISL_667062 | San Diego County Public Health Laboratory | Andersen lab at Scripps Research | SEARCH Alliance San Diego with Tracy Basler, Jovan Shephard, Brett Austin |
| EPI_ISL_667230, EPI_ISL_667231, EPI_ISL_667512, EPI_ISL_667513, EPI_ISL_667514, EPI_ISL_667515, EPI_ISL_667523, EPI_ISL_667524, EPI_ISL_667525, EPI_ISL_667548 | OHSU Lab Services Molecular Microbiology Lab | Oregon SARS-CoV-2 Genome Sequencing Center | Brendan L. O'Connell, Ruth V. Nichols, Sally Grindstaff, Alec J. Hirsch, Donna Hansel, Guang Fan, Daniel N. Streblow, William B. Messer, Andrew C. Adey, Benjamin N. Bimber, Brian J. O'Roak |
| EPI_ISL_667778 | Sydney South West Pathology Service (SSWPS) - Royal Prince Alfred Hospital - NSW Health Pathology | NSW Health Pathology - Institute of Clinical Pathology and Medical Research; Westmead Hospital; University of Sydney | CIDM-PH et al. |
| EPI_ISL_667783, EPI_ISL_667789 | South Eastern Area Laboratory Services (SEALS) | NSW Health Pathology - Institute of Clinical Pathology and Medical Research; Westmead Hospital; University of Sydney | CIDM-PH et al. |
| EPI_ISL_668410, EPI_ISL_668411 | Norwegian Institute of Public Health, Department of Virology | Norwegian Institute of Public Health, Department of Virology | Kathrine Stene-Johansen, Kamilla Heddeland Instefjord, Hilde Elshaug, Marie Paulsen Madsen, Rasmus Riis Kopperud, Hilde Vollan, Karoline Bragstad, Olav Hungnes |
| EPI_ISL_668424 | University Hospital of Northern Norway, Department for Microbiology and Infectious Disease Control | Norwegian Institute of Public Health, Department of Virology | Kathrine Stene-Johansen, Kamilla Heddeland Instefjord, Hilde Elshaug, Marie Paulsen Madsen, Rasmus Riis Kopperud, Hilde Vollan, Karoline Bragstad, Olav Hungnes |
| EPI_ISL_668428 | Hospital of Southern Norway - Kristiansand, Department of Medical Microbiology | Norwegian Institute of Public Health, Department of Virology | Kathrine Stene-Johansen, Kamilla Heddeland Instefjord, Hilde Elshaug, Marie Paulsen Madsen, Rasmus Riis Kopperud, Hilde Vollan, Karoline Bragstad, Olav Hungnes |
| EPI_ISL_668435, EPI_ISL_668436, EPI_ISL_668437 | Norwegian Institute of Public Health, Department of Virology | Norwegian Institute of Public Health, Department of Virology | Kathrine Stene-Johansen, Kamilla Heddeland Instefjord, Hilde Elshaug, Marie Paulsen Madsen, Rasmus Riis Kopperud, Hilde Vollan, Karoline Bragstad, Olav Hungnes |
| EPI_ISL_670460, EPI_ISL_670471, EPI_ISL_670472, EPI_ISL_670473, EPI_ISL_670478, EPI_ISL_670482, EPI_ISL_670484, EPI_ISL_670485, EPI_ISL_670489, EPI_ISL_670490, EPI_ISL_670493, EPI_ISL_670494, EPI_ISL_670495, EPI_ISL_670496, EPI_ISL_670498, EPI_ISL_670500, EPI_ISL_670510, EPI_ISL_670511, EPI_ISL_670512, EPI_ISL_670513, EPI_ISL_670514, EPI_ISL_670518, EPI_ISL_670522, EPI_ISL_670524, EPI_ISL_670525, EPI_ISL_670526, EPI_ISL_670527, EPI_ISL_670528, EPI_ISL_670530, EPI_ISL_670531, EPI_ISL_670534, EPI_ISL_670535, EPI_ISL_670538, EPI_ISL_670540, EPI_ISL_670541, EPI_ISL_670544, EPI_ISL_670545, EPI_ISL_670546, EPI_ISL_670547, EPI_ISL_670549, EPI_ISL_670556, EPI_ISL_670557, EPI_ISL_670569, EPI_ISL_670570, EPI_ISL_670571, EPI_ISL_670572, EPI_ISL_670573, EPI_ISL_670574, EPI_ISL_670575, EPI_ISL_670626, EPI_ISL_670627, EPI_ISL_670628, EPI_ISL_670629, EPI_ISL_670630, EPI_ISL_670631, EPI_ISL_670632, EPI_ISL_670633, EPI_ISL_670634, EPI_ISL_670635, EPI_ISL_670636, EPI_ISL_670637, EPI_ISL_670638, EPI_ISL_670639, EPI_ISL_670640, EPI_ISL_670641, EPI_ISL_670642, EPI_ISL_670643, EPI_ISL_670670, EPI_ISL_670671, EPI_ISL_670675, EPI_ISL_670681, EPI_ISL_670682, EPI_ISL_670683, EPI_ISL_670688, EPI_ISL_670694, EPI_ISL_670695, EPI_ISL_670696, EPI_ISL_670714, EPI_ISL_670715, EPI_ISL_670716, EPI_ISL_670717, EPI_ISL_670718, EPI_ISL_670719, EPI_ISL_670720, EPI_ISL_670721, EPI_ISL_670722, EPI_ISL_670723, EPI_ISL_670730, EPI_ISL_670741, EPI_ISL_670742, EPI_ISL_670743, EPI_ISL_670744, EPI_ISL_670754, EPI_ISL_670755, EPI_ISL_670756, EPI_ISL_670757, EPI_ISL_670758, EPI_ISL_670759, EPI_ISL_670760, EPI_ISL_670761, EPI_ISL_670762, EPI_ISL_670763, EPI_ISL_670764, EPI_ISL_670765, EPI_ISL_670766, EPI_ISL_670769, EPI_ISL_670770, EPI_ISL_671040, EPI_ISL_671041, EPI_ISL_671042, EPI_ISL_671043, EPI_ISL_671044, EPI_ISL_671045, EPI_ISL_671046, EPI_ISL_671047, EPI_ISL_671048, EPI_ISL_671049, EPI_ISL_671050, EPI_ISL_671051, EPI_ISL_671052, EPI_ISL_671053, EPI_ISL_671054, EPI_ISL_671055, EPI_ISL_671056, EPI_ISL_671057, EPI_ISL_671058, EPI_ISL_671059, EPI_ISL_671060, EPI_ISL_671061, EPI_ISL_671062, EPI_ISL_671063, EPI_ISL_671064, EPI_ISL_671065, EPI_ISL_671066, EPI_ISL_671067, EPI_ISL_671068, EPI_ISL_671069, EPI_ISL_671070, EPI_ISL_671071, EPI_ISL_671072, EPI_ISL_671073, EPI_ISL_671074, EPI_ISL_671075, EPI_ISL_671076, EPI_ISL_671077, EPI_ISL_671078, EPI_ISL_671079, EPI_ISL_671080, EPI_ISL_671081, EPI_ISL_671082, EPI_ISL_671083, EPI_ISL_671084, EPI_ISL_671085, EPI_ISL_671086, EPI_ISL_671087, EPI_ISL_671088, EPI_ISL_671089, EPI_ISL_671090, EPI_ISL_671091, EPI_ISL_671092, EPI_ISL_671093, EPI_ISL_671094, EPI_ISL_671095, EPI_ISL_671096, EPI_ISL_671097, EPI_ISL_671098, EPI_ISL_671099, EPI_ISL_671100, EPI_ISL_671101, EPI_ISL_671102, EPI_ISL_671103, EPI_ISL_671104, EPI_ISL_671105, EPI_ISL_671106, EPI_ISL_671107, EPI_ISL_671108, EPI_ISL_671109, EPI_ISL_671110, EPI_ISL_671111, EPI_ISL_671112, EPI_ISL_671113, EPI_ISL_671114, EPI_ISL_671115, EPI_ISL_671116, EPI_ISL_671117, EPI_ISL_671118, EPI_ISL_671119, EPI_ISL_671120, EPI_ISL_671121, EPI_ISL_671122, EPI_ISL_671123, EPI_ISL_671124, EPI_ISL_671125, EPI_ISL_671126, EPI_ISL_671127, EPI_ISL_671128, EPI_ISL_671129, EPI_ISL_671130, EPI_ISL_671131, EPI_ISL_671132, EPI_ISL_671133, EPI_ISL_671134, EPI_ISL_671135, EPI_ISL_671136, EPI_ISL_671137, EPI_ISL_671138, EPI_ISL_671139, EPI_ISL_671140, EPI_ISL_671141, EPI_ISL_671142, EPI_ISL_671143, EPI_ISL_671144, EPI_ISL_671145, EPI_ISL_671146, EPI_ISL_671147, EPI_ISL_671148, EPI_ISL_671149, EPI_ISL_671150, EPI_ISL_671151, EPI_ISL_671152, EPI_ISL_671153, EPI_ISL_671154, EPI_ISL_671155, EPI_ISL_671156, EPI_ISL_671157, EPI_ISL_671158, EPI_ISL_671159, EPI_ISL_671160, EPI_ISL_671161, EPI_ISL_671162, EPI_ISL_671163, EPI_ISL_671164, EPI_ISL_671165, EPI_ISL_671166, EPI_ISL_671167, EPI_ISL_671168, EPI_ISL_671169, EPI_ISL_671170, EPI_ISL_671171, EPI_ISL_671172, EPI_ISL_671173, EPI_ISL_671174, EPI_ISL_671175, EPI_ISL_671176, EPI_ISL_671177, EPI_ISL_671178, EPI_ISL_671179, EPI_ISL_671180, EPI_ISL_671181, EPI_ISL_671182, EPI_ISL_671183, EPI_ISL_671184, EPI_ISL_671185, EPI_ISL_671186, EPI_ISL_671187, EPI_ISL_671188, EPI_ISL_671189, EPI_ISL_671190, EPI_ISL_671191, EPI_ISL_671192, EPI_ISL_671193, EPI_ISL_671194, EPI_ISL_671195, EPI_ISL_671196, EPI_ISL_671197, EPI_ISL_671198, EPI_ISL_671199, EPI_ISL_671200, EPI_ISL_671201, EPI_ISL_671202, EPI_ISL_671203, EPI_ISL_671204, EPI_ISL_671205, EPI_ISL_671206, EPI_ISL_671207, EPI_ISL_671208, EPI_ISL_671209, EPI_ISL_671210, EPI_ISL_671211, EPI_ISL_671212, EPI_ISL_671213, EPI_ISL_671214, EPI_ISL_671215, EPI_ISL_671216, EPI_ISL_671217, EPI_ISL_671218 | | | |
| see above | Department of Virus and Microbiological Special Diagnostics, Statens Serum Institut, Copenhagen, Denmark | Albertsen Lab, Department of Chemistry and Bioscience, Aalborg University, Denmark | Danish Covid-19 Genome Consortium |
| EPI_ISL_671709 | DOHMH Riverside | New York City Public Health Laboratory | Jade Wang, et al. |
| EPI_ISL_671710, EPI_ISL_671711 | DOHMH Corona | New York City Public Health Laboratory | Jade Wang, et al. |
| EPI_ISL_671712 | DOHMH Jamaica | New York City Public Health Laboratory | Jade Wang, et al. |
| EPI_ISL_671713 | DOHMH Chelsea | New York City Public Health Laboratory | Jade Wang, et al. |
| EPI_ISL_671714 | DOHMH Jamaica | New York City Public Health Laboratory | Jade Wang, et al. |
| EPI_ISL_671715, EPI_ISL_671716 | DOHMH PHL | New York City Public Health Laboratory | Jade Wang, et al. |
| EPI_ISL_671717, EPI_ISL_671718, EPI_ISL_671719, EPI_ISL_671720 | DOHMH Corona | New York City Public Health Laboratory | Jade Wang, et al. |
| EPI_ISL_671730 | DOHMH Jamaica | New York City Public Health Laboratory | Jade Wang, et al. |
| EPI_ISL_671731, EPI_ISL_671732 | DOHMH Corona | New York City Public Health Laboratory | Jade Wang, et al. |

| | | | |
|---|---|---|---|
| EPI_ISL_671733 | DOHMH Riverside | New York City Public Health Laboratory | Jade Wang, et al. |
| EPI_ISL_671734 | DOHMH PHL | New York City Public Health Laboratory | Jade Wang, et al. |
| EPI_ISL_671735 | DOHMH Jamaica | New York City Public Health Laboratory | Jade Wang, et al. |
| EPI_ISL_671736 | DOHMH Morrisania | New York City Public Health Laboratory | Jade Wang, et al. |
| EPI_ISL_671737 | DOHMH PHL | New York City Public Health Laboratory | Jade Wang, et al. |
| EPI_ISL_671738 | DOHMH Fort Greene | New York City Public Health Laboratory | Jade Wang, et al. |
| EPI_ISL_671739 | DOHMH Jamaica | New York City Public Health Laboratory | Jade Wang, et al. |
| EPI_ISL_671740, EPI_ISL_671741, EPI_ISL_671742 | DOHMH Morrisania | New York City Public Health Laboratory | Jade Wang, et al. |
| EPI_ISL_671743 | DOHMH Jamaica | New York City Public Health Laboratory | Jade Wang, et al. |
| EPI_ISL_671744 | DOHMH PHL | New York City Public Health Laboratory | Jade Wang, et al. |
| EPI_ISL_671745 | DOHMH Riverside | New York City Public Health Laboratory | Jade Wang, et al. |
| EPI_ISL_671746, EPI_ISL_671747 | DOHMH Corona | New York City Public Health Laboratory | Jade Wang, et al. |
| EPI_ISL_671748, EPI_ISL_671749, EPI_ISL_671750 | DOHMH Riverside | New York City Public Health Laboratory | Jade Wang, et al. |
| EPI_ISL_671751, EPI_ISL_671752, EPI_ISL_671753, EPI_ISL_671754, EPI_ISL_671755, EPI_ISL_671756, EPI_ISL_671757, EPI_ISL_671758 | DOHMH Morrisania | New York City Public Health Laboratory | Jade Wang, et al. |
| EPI_ISL_671835, EPI_ISL_671836, EPI_ISL_671837, EPI_ISL_671838, EPI_ISL_671839, EPI_ISL_671840, EPI_ISL_671841, EPI_ISL_671842, EPI_ISL_671843 | Servicio de Microbiología, Laboratori Clínic Metropolitana Nord. Hospital Universitari Germans Trias i Pujol. Institut d'Investigació en Ciències de la Salut Germans Trias i Pujol (IGTP) | SeqCOVID-SPAIN consortium/IBV(CSIC) | Elisa Martró, Antoni E. Bordoy, Anna Not, Adrián Antuori, Anabel Fernández, Nona Romaní, Verónica Saludes, Cristina Casañ and SeqCOVID-SPAIN consortium |
| EPI_ISL_672019, EPI_ISL_672022, EPI_ISL_672037, EPI_ISL_672041, EPI_ISL_672046 | Madera County Department of Public Health | Chan-Zuckerberg Biohub | CZB Cliahub Consortium |
| EPI_ISL_672052 | Santa Clara County Public Health Laboratory | Chan-Zuckerberg Biohub | CZB Cliahub Consortium |
| EPI_ISL_672059 | Madera County Department of Public Health | Chan-Zuckerberg Biohub | CZB Cliahub Consortium |
| EPI_ISL_672065, EPI_ISL_672073 | Santa Clara County Public Health Laboratory | Chan-Zuckerberg Biohub | CZB Cliahub Consortium |
| EPI_ISL_672081 | Madera County Department of Public Health | Chan-Zuckerberg Biohub | CZB Cliahub Consortium |
| EPI_ISL_672085 | Santa Clara County Public Health Laboratory | Chan-Zuckerberg Biohub | CZB Cliahub Consortium |
| EPI_ISL_672098, EPI_ISL_672099 | Madera County Department of Public Health | Chan-Zuckerberg Biohub | CZB Cliahub Consortium |
| EPI_ISL_672300, EPI_ISL_672301, EPI_ISL_672302, EPI_ISL_672303, EPI_ISL_672304 | Santa Clara County Public Health Laboratory | Chan-Zuckerberg Biohub | CZB Cliahub Consortium |
| EPI_ISL_672394, EPI_ISL_672395, EPI_ISL_672396, EPI_ISL_672397, EPI_ISL_672398, EPI_ISL_672399, EPI_ISL_672400, EPI_ISL_672401, EPI_ISL_672402, EPI_ISL_672403, EPI_ISL_672404, EPI_ISL_672405, EPI_ISL_672406, EPI_ISL_672407, EPI_ISL_672408, EPI_ISL_672409, EPI_ISL_672410, EPI_ISL_672411, EPI_ISL_672412, EPI_ISL_672413, EPI_ISL_672414, EPI_ISL_672415, EPI_ISL_672416, EPI_ISL_672417, EPI_ISL_672418, EPI_ISL_672419, EPI_ISL_672420, EPI_ISL_672421, EPI_ISL_672422 | | | |
| see above | Madera County Department of Public Health | Chan-Zuckerberg Biohub | CZB Cliahub Consortium |
| EPI_ISL_672426, EPI_ISL_672427 | UCSF Clinical Microbiology Laboratory | Chan-Zuckerberg Biohub | CZB Cliahub Consortium |
| EPI_ISL_672429, EPI_ISL_672430, EPI_ISL_672431, EPI_ISL_672432, EPI_ISL_672433 | Santa Clara County Public Health Laboratory | Chan-Zuckerberg Biohub | CZB Cliahub Consortium |
| EPI_ISL_672477, EPI_ISL_672478, EPI_ISL_672479, EPI_ISL_672480 | Orange County Public Health Lab | Chan-Zuckerberg Biohub | CZB Cliahub Consortium |
| EPI_ISL_672507, EPI_ISL_672508, EPI_ISL_672509, EPI_ISL_672510, EPI_ISL_672511, EPI_ISL_672512, EPI_ISL_672513, EPI_ISL_672514, EPI_ISL_672515, EPI_ISL_672516, EPI_ISL_672517, EPI_ISL_672518, EPI_ISL_672519, EPI_ISL_672520, EPI_ISL_672521 | | | |
| see above | Madera County Department of Public Health | Chan-Zuckerberg Biohub | CZB Cliahub Consortium |
| EPI_ISL_672912, EPI_ISL_672916, EPI_ISL_672917, EPI_ISL_672918, EPI_ISL_672919, EPI_ISL_672922, EPI_ISL_672923, EPI_ISL_672924, EPI_ISL_672925, EPI_ISL_672926, EPI_ISL_672927, EPI_ISL_672934, EPI_ISL_672935, EPI_ISL_672939, EPI_ISL_672943, EPI_ISL_672944, EPI_ISL_672945, EPI_ISL_672947, EPI_ISL_672949, EPI_ISL_672951, EPI_ISL_672956, EPI_ISL_672959, EPI_ISL_672961, EPI_ISL_672963, EPI_ISL_672964, EPI_ISL_672965, EPI_ISL_672968, EPI_ISL_672971, EPI_ISL_672972, EPI_ISL_672974, EPI_ISL_672977, EPI_ISL_672978, EPI_ISL_672980, EPI_ISL_672983, EPI_ISL_672984, EPI_ISL_672985, EPI_ISL_672987, EPI_ISL_672989, EPI_ISL_672990, EPI_ISL_672991, EPI_ISL_672992, EPI_ISL_672993, EPI_ISL_672998, EPI_ISL_672999, EPI_ISL_673001, EPI_ISL_673004, EPI_ISL_673005, EPI_ISL_673006, EPI_ISL_673007, EPI_ISL_673009, EPI_ISL_673011, EPI_ISL_673012, EPI_ISL_673015, EPI_ISL_673016, EPI_ISL_673017, EPI_ISL_673018, EPI_ISL_673019, EPI_ISL_673022, EPI_ISL_673023, EPI_ISL_673025, EPI_ISL_673027, EPI_ISL_673028, EPI_ISL_673029, EPI_ISL_673032, EPI_ISL_673037, EPI_ISL_673039, EPI_ISL_673040, EPI_ISL_673041, EPI_ISL_673048, EPI_ISL_673050, EPI_ISL_673051, EPI_ISL_673053, EPI_ISL_673054, EPI_ISL_673055, EPI_ISL_673056, EPI_ISL_673057, EPI_ISL_673059, EPI_ISL_673061, EPI_ISL_673062, EPI_ISL_673065, EPI_ISL_673067, EPI_ISL_673069, EPI_ISL_673070, EPI_ISL_673071, EPI_ISL_673073, EPI_ISL_673074, EPI_ISL_673075, EPI_ISL_673076, EPI_ISL_673077, EPI_ISL_673078, EPI_ISL_673080, EPI_ISL_673082, EPI_ISL_673083, EPI_ISL_673085, EPI_ISL_673086, EPI_ISL_673090, EPI_ISL_673092, EPI_ISL_673098, EPI_ISL_673099, EPI_ISL_673100, EPI_ISL_673101, EPI_ISL_673102, EPI_ISL_673103, EPI_ISL_673104, EPI_ISL_673105, EPI_ISL_673106, EPI_ISL_673110, EPI_ISL_673113, EPI_ISL_673115, EPI_ISL_673116, EPI_ISL_673118, EPI_ISL_673122, EPI_ISL_673125, EPI_ISL_673126, EPI_ISL_673130, EPI_ISL_673131, EPI_ISL_673132, EPI_ISL_673133, EPI_ISL_673137, EPI_ISL_673138, EPI_ISL_673141, EPI_ISL_673145, EPI_ISL_673146, EPI_ISL_673149, EPI_ISL_673152, EPI_ISL_673155, EPI_ISL_673157, EPI_ISL_673159, EPI_ISL_673160, EPI_ISL_673163, EPI_ISL_673165, EPI_ISL_673172, EPI_ISL_673174, EPI_ISL_673176, EPI_ISL_673177, EPI_ISL_673178, EPI_ISL_673179, EPI_ISL_673181, EPI_ISL_673182, EPI_ISL_673183, EPI_ISL_673184, EPI_ISL_673185, EPI_ISL_673186, EPI_ISL_673187, EPI_ISL_673189, EPI_ISL_673190, EPI_ISL_673191, EPI_ISL_673192, EPI_ISL_673194, EPI_ISL_673195, EPI_ISL_673197, EPI_ISL_673198, EPI_ISL_673199, EPI_ISL_673203, EPI_ISL_673205, EPI_ISL_673209, EPI_ISL_673211, EPI_ISL_673212, EPI_ISL_673216, EPI_ISL_673219, EPI_ISL_673220, EPI_ISL_673222, EPI_ISL_673223, EPI_ISL_673228, EPI_ISL_673233, EPI_ISL_673234 | | | |
| see above | Lighthouse Lab in Cambridge | Wellcome Sanger Institute for the COVID-19 Genomics UK (COG-UK) Consortium | Rob Howes, The Lighthouse Lab in Cambridge and Alex Alderton, Roberto Amato, Sonia Goncalves, Ewan Harrison, David K. Jackson, Ian Johnston, Dominic Kwiatkowski, Cordelia Langford, John Sillitoe on behalf of the Wellcome Sanger Institute COVID-19 Surveillance Team |
| EPI_ISL_676444, EPI_ISL_676450, EPI_ISL_676477, EPI_ISL_676486 | Lighthouse Lab in Glasgow | Wellcome Sanger Institute for the COVID-19 Genomics UK (COG-UK) Consortium | Harper VanSteenhouse, Yumi Kasai, David Gray, Carol Clugston, Anna Dominiczak and Alex Alderton, Roberto Amato, Sonia Goncalves, Ewan Harrison, David K. Jackson, Ian Johnston, Dominic Kwiatkowski, Cordelia Langford, John Sillitoe on behalf of the Wellcome Sanger Institute COVID-19 Surveillance Team |
| EPI_ISL_676496 | Klinisk mikrobiologi | The Public Health Agency of Sweden | Department of Microbiology, The Public Health Agency of Sweden |
| EPI_ISL_676516, EPI_ISL_676517, EPI_ISL_676535, EPI_ISL_676536, EPI_ISL_676537, EPI_ISL_676538 | Laboratoriemedicin, Klinisk mikrobiologi | The Public Health Agency of Sweden | Department of Microbiology, The Public Health Agency of Sweden |
| EPI_ISL_677147, EPI_ISL_677148, EPI_ISL_677149, EPI_ISL_677150, EPI_ISL_677151, EPI_ISL_677152, EPI_ISL_677153, EPI_ISL_677154, EPI_ISL_677155, EPI_ISL_677156, EPI_ISL_677157, EPI_ISL_677158, EPI_ISL_677159, EPI_ISL_677160, EPI_ISL_677161, EPI_ISL_677162, EPI_ISL_677163, EPI_ISL_677217, EPI_ISL_677218, EPI_ISL_677219, EPI_ISL_677224, EPI_ISL_677225, EPI_ISL_677226, EPI_ISL_677227, EPI_ISL_677228, EPI_ISL_677237 | | | |
| see above | Virginia Division of Consolidated Laboratory Services | Virginia Division of Consolidated Laboratory Services | Virginia DCLS |

| | | | |
|---|---|---|---|
| EPI_ISL_677321, EPI_ISL_677322, EPI_ISL_677323, EPI_ISL_677324, EPI_ISL_677325, EPI_ISL_677326, EPI_ISL_677327, EPI_ISL_677328, EPI_ISL_677329, EPI_ISL_677501, EPI_ISL_677502, EPI_ISL_677578, EPI_ISL_677579, EPI_ISL_677580, EPI_ISL_677581, EPI_ISL_677594, EPI_ISL_677608, EPI_ISL_677609, EPI_ISL_677610, EPI_ISL_677611, EPI_ISL_677612, EPI_ISL_677613, EPI_ISL_677616, EPI_ISL_677617, EPI_ISL_677618, EPI_ISL_677628 | | | |
| see above | University of Wisconsin-Madison AIDS Vaccine Research Laboratories | University of Wisconsin-Madison AIDS Vaccine Research Laboratories | Gage Moreno, Katarina Braun, et al. AIDS Vaccine Research Laboratories |
| EPI_ISL_678379 | Area of Virology, Serology and Virology Division (SAViD), New South Wales Health Pathology Randwick | Virology Research Laboratory; Area of Virology, Serology and Virology Division (SAViD), New South Wales Health Pathology Randwick | Foster, C.; Au, J.; Ruiz Silva, M.; Deveson, I.; Bull, R.; Van Hal, S.; Rawlinson, W. |
| EPI_ISL_678600, EPI_ISL_678604, EPI_ISL_678606 | NHLS-IALCH | KRISP, KZN Research Innovation and Sequencing Platform | Giandhari J, Pillay S, Lessells R, ChimukangaraB, Mdlalose K, York D, Khan S, Tegally H, Wilkinson E, de Oliveira T |
| EPI_ISL_678662 | Respiratory Virus Unit, Microbiology Services Colindale, Public Health England | COVID-19 Genomics UK (COG-UK) Consortium | PHE Covid Sequencing Team |
| EPI_ISL_679133 | University of Birmingham | COVID-19 Genomics UK (COG-UK) Consortium | Institute of Microbiology, University of Birmingham: Claire McMurray, Joanne Stockton, Samuel Nicholls, Radoslaw Poplawski, Will Rowe, Josh Quick, Nicholas Loman. University of Birmingham Testing Laboratory: Celina M Whalley, Andrew Bosworth, Charlotte Poxon, Kasun Wanigasooriya, Oliver Pickles, Mike Kidd, Alex Richter, Andrew D Beggs PHE Heartlands Lab: Husam Osman, Andrew Bosworth. Queen Elizabeth Hospital: Anna Casey |
| EPI_ISL_679390, EPI_ISL_679391 | Liverpool Clinical Laboratories | COVID-19 Genomics UK (COG-UK) Consortium | Sam Haldenby, Anita Lucaci, Steve Paterson, Julian Hiscox, Alistair Darby, M Almsaud, A Alrezaihi, Muhannad Alruwaili, Stuart D Armstrong, Jones Benjamin, Eleanor G Bentley, Anu Chawla, Jordan J Clark, Angela Cowell, Richard Eccles, Isabel García-Dorival, Matthew Gemmell, Alessandro Gerada, PKF Gilmore, Richard Gregory, Ximeng Han, Catherine Hartley, Margaret Hughes, Miren Iturriza-Gomara, James Johnson, L Luu, Jenifer Manson, Charlotte Nelson, Elaine O'Toole, Cassie Olateju, Rebekah Penrice-Randal , Lucille Rainbow, N.P Randle, Trevor Ian Robinson, Parul Sharma, Ghada T Shawli, James P Stewart, Neil Swainston, Ecaterina Vamos, Joanne Watts, Mark Whitehead |
| EPI_ISL_679457, EPI_ISL_679466 | University College London, Great Ormond Street Hospital for Children NHS Foundation Trust, Imperial College Healthcare NHS Trust | COVID-19 Genomics UK (COG-UK) Consortium | Sergi Castellano, Rachel Williams, Mark Kristiansen, Paola Resende Silva, Sunando Roy, Tony Brooks, Helena Tutill, Paola Niola, Patricia Dyal, Charlotte Williams, Leysa Forrest, Yasmin Panchbhaya, Jacqueline Findlay, Samuel Weeks, Julianne Brown, Kathryn Harris, Paul Randell, James Price, Alison Holmes, Judith Breuer |
| EPI_ISL_679727, EPI_ISL_679728 | Oxford Viromics, NDM, University of Oxford; Oxford University Hospitals; Basingstoke and North Hampshire Hospital | COVID-19 Genomics UK (COG-UK) Consortium | Tanya Golubchik, David Bonsall, George Macintyre, Amy Trebes, Mariateresa de Cesare, Catrin Moore, Alex Mobbs, Anita Justice, Robert Shaw, Monique Andersson, Timothy Peto, Emma Wise, Nathan Moore, Jessica Lynch, Nick Cortes, Matilde Mori, Stephen Kidd, David Buck, John Todd, Christophe Fraser |
| EPI_ISL_679926 | Queens Medical Centre, Clinical Microbiology Department / DeepSeq Nottingham | COVID-19 Genomics UK (COG-UK) Consortium | Gemma Clark, Wendy Smith, Manjinder Khakh, Vicki M Fleming, Michelle M Lister, Hannah Howson-Wells, Jonathan Ball, Patrick McClure, Joseph Chappell, Theocharis Tsoleridis, Nadine Holmes, Matthew Carlisle, Christopher Moore, Fei Sang, Johnny Debebe, Victoria Wright, Matthew Loose |
| EPI_ISL_680442, EPI_ISL_680443, EPI_ISL_680444 | West of Scotland Specialist Virology Centre, NHSGGC / MRC-University of Glasgow Centre for Virus Research | COVID-19 Genomics UK (COG-UK) Consortium | Ana da Silva Filipe, Natasha Johnson, Kathy Smollett, Daniel Mair, Stephen Carmichael, Alice Broos, Lily Tong, Jenna Nichols, Kyriaki Nomikou; Sarah McDonald; Richard Orton, Joseph Hughes, Sreenu Vattipally, David L Robertson; Alasdair MacLean, Rory Gunson; Sharif Shaaban, Matthew Holden; Rachel Blacow, Guy Mollett, Kathy Li, James Shepherd, Antonia Ho, Emma Thomson |
| EPI_ISL_680504, EPI_ISL_680505, EPI_ISL_680506, EPI_ISL_680507, EPI_ISL_680513, EPI_ISL_680514, EPI_ISL_680522, EPI_ISL_680523 | Virology Department, Royal Infirmary of Edinburgh, NHS Lothian / School of Biological Sciences, University of Edinburgh / Institute of Genetics and Molecular Medicine, University of Edinburgh | COVID-19 Genomics UK (COG-UK) Consortium | McHugh M, Dewar R, Rooke S, Gallagher M, Balcaza C, O'Toole Á, Scher E, Hill V, McCrone JT, Colquhoun R, Yu X, Jackson B, Rambaut A, Williams TC, Templeton K |
| EPI_ISL_680620, EPI_ISL_680621 | Wales Specialist Virology Centre Sequencing lab: Pathogen Genomics Unit | COVID-19 Genomics UK (COG-UK) Consortium | Catherine Moore, Johnathan Evans, Laura Gifford, Malorie Perry, Simon Cottrell, Angela Marchbank, Alec Birchley, Alexander Adams, Amy Gaskin, Bree Gatica-Wilcox, Jason Coombes, Joel Southgate, Lauren Gilbert, Lee Graham, Nicole Pacchiarini, Sara Kumziene-Summerhayes, Sarah Taylor, Sophie Jones, Sara Rey, Matthew Bull, Joanne Watkins, Sally Corden, Tom Connor |
| EPI_ISL_681264, EPI_ISL_681265, EPI_ISL_681266 | Department of Clinical Microbiology | GIGA Medical Genomics | Keith Durkin, Maria Artesi, Sébastien Bontems, Raphaël Boreux, Bouchra Boujemla, Cécile Meex, Pierrette Melin, Marie-Pierre Hayette, Vincent Bours |
| EPI_ISL_681267 | Department of Clinical Microbiology | GIGA Medical Genomics | Keith Durkin, Maria Artesi, Justine Defêche, Gilles Darcis, Michel Moutschen, Sébastien Bontems, Raphaël Boreux, Bouchra Boujemla, Cécile Meex, Pierrette Melin, Marie-Pierre Hayette, Vincent Bours |
| EPI_ISL_681310 | Communicable Disease Laboratory, Public Health Directorate | Communicable Disease Laboratory, Public Health Directorate | Alwasti,H., Altaif,Z., AlHujairi,Z., AlAbbas,Z. |
| EPI_ISL_681656 | Lighthouse Lab in Cambridge | Wellcome Sanger Institute for the COVID-19 Genomics UK (COG-UK) Consortium | Rob Howes, The Lighthouse Lab in Cambridge and Alex Alderton, Roberto Amato, Sonia Goncalves, Ewan Harrison, David K. Jackson, Ian Johnston, Dominic Kwiatkowski, Cordelia Langford, John Sillitoe on behalf of the Wellcome Sanger Institute COVID-19 Surveillance Team |
| EPI_ISL_681800, EPI_ISL_681801, EPI_ISL_681802, EPI_ISL_681803, EPI_ISL_681804, EPI_ISL_681805, EPI_ISL_681806, EPI_ISL_681807, EPI_ISL_681808, EPI_ISL_681809, EPI_ISL_681810, EPI_ISL_681811, EPI_ISL_681812, EPI_ISL_681813, EPI_ISL_681814, EPI_ISL_681815, EPI_ISL_681816, EPI_ISL_681817, EPI_ISL_681822, EPI_ISL_681823, EPI_ISL_681824, EPI_ISL_681825, EPI_ISL_681826, EPI_ISL_681827, EPI_ISL_681828 | | | |
| see above | GA Department of Public Health Laboratory | Pathogen Discovery, Respiratory Viruses Branch, Division of Viral Diseases, Centers for Disease Control and Prevention | Yan Li, Ying Tao, Jing Zhang, Krista Queen, Anna Uehara, Peter Cook, Clinton R. Paden, Haibin Wang, Suxiang Tong |
| EPI_ISL_682079, EPI_ISL_682082, EPI_ISL_682083, EPI_ISL_682084, EPI_ISL_682085, EPI_ISL_682086, EPI_ISL_682087, EPI_ISL_682088, EPI_ISL_682089, EPI_ISL_682090, EPI_ISL_682091, EPI_ISL_682092, EPI_ISL_682093, EPI_ISL_682096, EPI_ISL_682097, EPI_ISL_682098, EPI_ISL_682099, EPI_ISL_682101, EPI_ISL_682102, EPI_ISL_682112, EPI_ISL_682114, EPI_ISL_682115, EPI_ISL_682119, EPI_ISL_682120, EPI_ISL_682126, EPI_ISL_682130 | | | |
| see above | University of Michigan Clinical Microbiology Laboratory | Lauring Lab, University of Michigan, Department of Microbiology and Immunology | Valesano |
| EPI_ISL_682301 | Communicable Disease Laboratory, Public Health Directorate | Communicable Disease Laboratory, Public Health Directorate | Alwasti,H., Altaif,Z., AlHujairi,Z., AlAbbas,Z. |
| EPI_ISL_682983, EPI_ISL_682984, EPI_ISL_682985, EPI_ISL_682986, EPI_ISL_682987, EPI_ISL_682988, EPI_ISL_682989, EPI_ISL_682990, EPI_ISL_682991, EPI_ISL_682992, EPI_ISL_682993, EPI_ISL_682994, EPI_ISL_682995, EPI_ISL_682996, EPI_ISL_682997, EPI_ISL_682998, EPI_ISL_682999, EPI_ISL_683000, EPI_ISL_683001, EPI_ISL_683181, EPI_ISL_683182, EPI_ISL_683183, EPI_ISL_683184, EPI_ISL_683185, EPI_ISL_683186, EPI_ISL_683187, EPI_ISL_683188, EPI_ISL_683189, EPI_ISL_683190, EPI_ISL_683191, EPI_ISL_683192, EPI_ISL_683193, EPI_ISL_683194, EPI_ISL_683195, EPI_ISL_683196, EPI_ISL_683197, EPI_ISL_683198, EPI_ISL_683199, EPI_ISL_683200, EPI_ISL_683201, EPI_ISL_683202, EPI_ISL_683203 | | | |
| see above | Department of Virus and Microbiological Special Diagnostics, Statens Serum Institut, Copenhagen, Denmark | Albertsen Lab, Department of Chemistry and Bioscience, Aalborg University, Denmark | Danish Covid-19 Genome Consortium |
| EPI_ISL_683704 | Essentia Health-St. Mary's Medical Center | Minnesota Department of Health, Public Health Laboratory | Alexandra Lorentz, Jacob Garfin, Matt Plumb, and Xiong Wang |
| EPI_ISL_683714, EPI_ISL_683715, EPI_ISL_683719, EPI_ISL_683720, EPI_ISL_683721 | Mayo Clinic & Mayo Clinic Laboratories | Minnesota Department of Health, Public Health Laboratory | Alexandra Lorentz, Jacob Garfin, Matt Plumb, and Xiong Wang |
| EPI_ISL_683771 | DOHMH Fort Greene | New York City Public Health Laboratory | Jade Wang, et al. |
| EPI_ISL_683885 | DOHMH Jamaica | New York City Public Health Laboratory | Jade Wang, et al. |
| EPI_ISL_692806, EPI_ISL_692807, EPI_ISL_692813, EPI_ISL_692815, EPI_ISL_692816, EPI_ISL_692820 | Massachusetts State Public Health Laboratory | Massachusetts State Public Health Laboratory | Andrew Lang, Timelia Fink, Glen Gallagher, Sandra Smole |
| EPI_ISL_693660, EPI_ISL_693664, EPI_ISL_693670, EPI_ISL_693673, EPI_ISL_693681, EPI_ISL_693683, EPI_ISL_693687 | The National Institute of Public Health | State Veterinary Institute Prague | Nagy,A;Jirincova,H;Trnka,D;Vecerova,J |
| EPI_ISL_700424 | Thembalethu CDC wc THC | NHLS/UCT | Houriiyah Tegally, Arash Iranzadeh, Deelan Doolabh, Lynn Tyers, Bruna Galvao, Innocent Mudau, Marvin Hsiao, Kruger Marais, Diana Hardie, Stephen Korsman, Carolyn Williamson |

| | | | |
|---|---|---|---|
| EPI_ISL_700435 | Pacaltsdorp Clinic wc PAC | NHLS/UCT | Houriiyah Tegally, Arash Iranzadeh, Deelan Doolabh, Lynn Tyers, Bruna Galvao, Innocent Mudau, Marvin Hsiao, Kruger Marais, Diana Hardie, Stephen Korsman, Carolyn Williamson |
| EPI_ISL_700464 | Khayelethu Clinic wc KLC | NHLS/UCT | Houriiyah Tegally, Arash Iranzadeh, Deelan Doolabh, Lynn Tyers, Bruna Galvao, Innocent Mudau, Marvin Hsiao, Kruger Marais, Diana Hardie, Stephen Korsman, Carolyn Williamson |
| EPI_ISL_700468 | Khayelethu Clinic wc KLC | NHLS/UCT | Arash Iranzadeh, Deelan Doolabh, Lynn Tyers, Bruna Galvao, Innocent Mudau, Marvin Hsiao, Kruger Marais, Diana Hardie, Stephen Korsman, Carolyn Williamson |
| EPI_ISL_700497 | George Hospital wc GRH | NHLS/UCT | Arash Iranzadeh, Deelan Doolabh, Lynn Tyers, Bruna Galvao, Innocent Mudau, Marvin Hsiao, Kruger Marais, Diana Hardie, Stephen Korsman, Carolyn Williamson |
| EPI_ISL_700498 | Thembalethu CDC wc THC | NHLS/UCT | Houriiyah Tegally, Arash Iranzadeh, Deelan Doolabh, Lynn Tyers, Bruna Galvao, Innocent Mudau, Marvin Hsiao, Kruger Marais, Diana Hardie, Stephen Korsman, Carolyn Williamson |
| EPI_ISL_700533 | Conville CDC wc CVC | NHLS/UCT | Arash Iranzadeh, Deelan Doolabh, Lynn Tyers, Bruna Galvao, Innocent Mudau, Marvin Hsiao, Kruger Marais, Diana Hardie, Stephen Korsman, Carolyn Williamson |
| EPI_ISL_700539 | Conville CDC wc CVC | NHLS/UCT | Houriiyah Tegally, Arash Iranzadeh, Deelan Doolabh, Lynn Tyers, Bruna Galvao, Innocent Mudau, Marvin Hsiao, Kruger Marais, Diana Hardie, Stephen Korsman, Carolyn Williamson |
| EPI_ISL_700540 | Khayelethu Clinic wc KLC | NHLS/UCT | Houriiyah Tegally, Arash Iranzadeh, Deelan Doolabh, Lynn Tyers, Bruna Galvao, Innocent Mudau, Marvin Hsiao, Kruger Marais, Diana Hardie, Stephen Korsman, Carolyn Williamson |
| EPI_ISL_700563, EPI_ISL_700590 | Conville CDC wc CVC | NHLS/UCT | Houriiyah Tegally, Arash Iranzadeh, Deelan Doolabh, Lynn Tyers, Bruna Galvao, Innocent Mudau, Marvin Hsiao, Kruger Marais, Diana Hardie, Stephen Korsman, Carolyn Williamson |
| EPI_ISL_702505 | Northumbria University / South Tees Hospitals NHS Foundation Trust / North Cumbria Integrated Care NHS Foundation Trust / North Tees and Hartlepool NHS Foundation Trust / Newcastle Hospitals NHS Foundation Trust | COVID-19 Genomics UK (COG-UK) Consortium | Darren L Smith,Andrew Nelson,Matthew Bashton,Greg R Young,Joshua Loh,John Allan,Mohammad A Tariq,Giles S Holt,Gary Black,Wen C Yew,Lynn Dover,Paul Baker,Steve Liggett,Sarah Essex,Jane Greenaway,Debra Padgett,Clive Graham,Garren Scott,Edward Barton,Emma Swindells,Brendan Payne,Jennifer Collins,Yusri Taha,Gary Eltringham |
| EPI_ISL_702890, EPI_ISL_703227, EPI_ISL_703299, EPI_ISL_703346, EPI_ISL_703627 | Wales Specialist Virology Centre Sequencing lab: Pathogen Genomics Unit | COVID-19 Genomics UK (COG-UK) Consortium | Catherine Moore, Johnathan Evans, Laura Gifford, Malorie Perry, Simon Cottrell, Angela Marchbank, Alec Birchley, Alexander Adams, Amy Gaskin, Bree Gatica-Wilcox, Jason Coombes, Joel Southgate, Lauren Gilbert, Lee Graham, Nicole Pacchiarini, Sara Kumziene-Summerhayes, Sarah Taylor, Sophie Jones, Sara Rey, Matthew Bull, Joanne Watkins, Sally Corden, Tom Connor |
| EPI_ISL_703755, EPI_ISL_703771 | West of Scotland Specialist Virology Centre, NHSGGC / MRC-University of Glasgow Centre for Virus Research | COVID-19 Genomics UK (COG-UK) Consortium | Ana da Silva Filipe, Natasha Johnson, Kathy Smollett, Daniel Mair, Stephen Carmichael, Alice Broos, Lily Tong, Jenna Nichols, Kyriaki Nomikou; Sarah McDonald; Richard Orton, Joseph Hughes, Sreenu Vattipally, David L Robertson; Alasdair MacLean, Rory Gunson; Sharif Shaaban, Matthew Holden; Rachel Blacow, Guy Mollett, Kathy Li, James Shepherd, Antonia Ho, Emma Thomson |
| EPI_ISL_703787 | Virology Department, Royal Infirmary of Edinburgh, NHS Lothian / School of Biological Sciences, University of Edinburgh / Institute of Genetics and Molecular Medicine, University of Edinburgh | COVID-19 Genomics UK (COG-UK) Consortium | McHugh M, Dewar R, Rooke S, Gallagher M, Balcaza C, O'Toole Á, Scher E, Hill V, McCrone JT, Colquhoun R, Yu X, Jackson B, Rambaut A, Williams TC, Templeton K |
| EPI_ISL_703855, EPI_ISL_703902 | Wales Specialist Virology Centre Sequencing lab: Pathogen Genomics Unit | COVID-19 Genomics UK (COG-UK) Consortium | Catherine Moore, Johnathan Evans, Laura Gifford, Malorie Perry, Simon Cottrell, Angela Marchbank, Alec Birchley, Alexander Adams, Amy Gaskin, Bree Gatica-Wilcox, Jason Coombes, Joel Southgate, Lauren Gilbert, Lee Graham, Nicole Pacchiarini, Sara Kumziene-Summerhayes, Sarah Taylor, Sophie Jones, Sara Rey, Matthew Bull, Joanne Watkins, Sally Corden, Tom Connor |
| EPI_ISL_704120 | Northumbria University / South Tees Hospitals NHS Foundation Trust / North Cumbria Integrated Care NHS Foundation Trust / North Tees and Hartlepool NHS Foundation Trust / Newcastle Hospitals NHS Foundation Trust | COVID-19 Genomics UK (COG-UK) Consortium | Darren L Smith,Andrew Nelson,Matthew Bashton,Greg R Young,Joshua Loh,John Allan,Mohammad A Tariq,Giles S Holt,Gary Black,Wen C Yew,Lynn Dover,Paul Baker,Steve Liggett,Sarah Essex,Jane Greenaway,Debra Padgett,Clive Graham,Garren Scott,Edward Barton,Emma Swindells,Brendan Payne,Jennifer Collins,Yusri Taha,Gary Eltringham |
| EPI_ISL_704171, EPI_ISL_704320 | Wales Specialist Virology Centre Sequencing lab: Pathogen Genomics Unit | COVID-19 Genomics UK (COG-UK) Consortium | Catherine Moore, Johnathan Evans, Laura Gifford, Malorie Perry, Simon Cottrell, Angela Marchbank, Alec Birchley, Alexander Adams, Amy Gaskin, Bree Gatica-Wilcox, Jason Coombes, Joel Southgate, Lauren Gilbert, Lee Graham, Nicole Pacchiarini, Sara Kumziene-Summerhayes, Sarah Taylor, Sophie Jones, Sara Rey, Matthew Bull, Joanne Watkins, Sally Corden, Tom Connor |
| EPI_ISL_704470, EPI_ISL_704473 | Department of Pathology, University of Cambridge | COVID-19 Genomics UK (COG-UK) Consortium | Aminu S. Jahun, Yasmin Chaudhry, Grant Hall, Iliana Georgana, Myra Hosmillo, Martin D. Curran, Malte Pinckert, Surendra Parmar, Ian Goodfellow |
| EPI_ISL_704581 | Queens Medical Centre, Clinical Microbiology Department / DeepSeq Nottingham | COVID-19 Genomics UK (COG-UK) Consortium | Gemma Clark, Wendy Smith, Manjinder Khakh, Vicki M Fleming, Michelle M Lister, Hannah Howson-Wells, Jonathan Ball, Patrick McClure, Joseph Chappell, Theocharis Tsoleridis, Nadine Holmes, Matthew Carlisle, Christopher Moore, Fei Sang, Johnny Debebe, Victoria Wright, Matthew Loose |
| EPI_ISL_705355 | Northumbria University / South Tees Hospitals NHS Foundation Trust / North Cumbria Integrated Care NHS Foundation Trust / North Tees and Hartlepool NHS Foundation Trust / Newcastle Hospitals NHS Foundation Trust | COVID-19 Genomics UK (COG-UK) Consortium | Darren L Smith,Andrew Nelson,Matthew Bashton,Greg R Young,Joshua Loh,John Allan,Mohammad A Tariq,Giles S Holt,Gary Black,Wen C Yew,Lynn Dover,Paul Baker,Steve Liggett,Sarah Essex,Jane Greenaway,Debra Padgett,Clive Graham,Garren Scott,Edward Barton,Emma Swindells,Brendan Payne,Jennifer Collins,Yusri Taha,Gary Eltringham |
| EPI_ISL_705459, EPI_ISL_705461 | Wales Specialist Virology Centre Sequencing lab: Pathogen Genomics Unit | COVID-19 Genomics UK (COG-UK) Consortium | Catherine Moore, Johnathan Evans, Laura Gifford, Malorie Perry, Simon Cottrell, Angela Marchbank, Alec Birchley, Alexander Adams, Amy Gaskin, Bree Gatica-Wilcox, Jason Coombes, Joel Southgate, Lauren Gilbert, Lee Graham, Nicole Pacchiarini, Sara Kumziene-Summerhayes, Sarah Taylor, Sophie Jones, Sara Rey, Matthew Bull, Joanne Watkins, Sally Corden, Tom Connor |
| EPI_ISL_705502, EPI_ISL_705544, EPI_ISL_705546, EPI_ISL_705547, EPI_ISL_705548, EPI_ISL_705550 | Department of Pathology, University of Cambridge | COVID-19 Genomics UK (COG-UK) Consortium | Aminu S. Jahun, Yasmin Chaudhry, Grant Hall, Iliana Georgana, Myra Hosmillo, Martin D. Curran, Malte Pinckert, Surendra Parmar, Ian Goodfellow |
| EPI_ISL_706471, EPI_ISL_706472, EPI_ISL_706666, EPI_ISL_706667, EPI_ISL_706668, EPI_ISL_706669, EPI_ISL_706670, EPI_ISL_706671, EPI_ISL_706672, EPI_ISL_706673, EPI_ISL_706674 | | | |
| see above | Wales Specialist Virology Centre Sequencing lab: Pathogen Genomics Unit | COVID-19 Genomics UK (COG-UK) Consortium | Catherine Moore, Johnathan Evans, Laura Gifford, Malorie Perry, Simon Cottrell, Angela Marchbank, Alec Birchley, Alexander Adams, Amy Gaskin, Bree Gatica-Wilcox, Jason Coombes, Joel Southgate, Lauren Gilbert, Lee Graham, Nicole Pacchiarini, Sara Kumziene-Summerhayes, Sarah Taylor, Sophie Jones, Sara Rey, Matthew Bull, Joanne Watkins, Sally Corden, Tom Connor |
| EPI_ISL_707900, EPI_ISL_707901, EPI_ISL_707902 | Area of Virology, Serology and Virology Division (SAViD), New South Wales Health Pathology Randwick | Virology Research Laboratory; Area of Virology, Serology and Virology Division (SAViD), New South Wales Health Pathology Randwick | Foster, C.; Au, J.; Ruiz Silva, M.; Deveson, I.; Bull, R.; Van Hal, S.; Rawlinson, W. |
| EPI_ISL_708035 | Furst Medical Laboratory | Norwegian Institute of Public Health, Department of Virology | Kathrine Stene-Johansen, Kamilla Heddeland Instefjord, Hilde Elshaug, Marie Paulsen Madsen, Rasmus Riis Kopperud, Hilde Vollan, Karoline Bragstad, Olav Hungnes |
| EPI_ISL_708036, EPI_ISL_708037, EPI_ISL_708038 | Innlandet Hospital Trust, Division Lillehammer, Department for Medical Microbiology | Norwegian Institute of Public Health, Department of Virology | Kathrine Stene-Johansen, Kamilla Heddeland Instefjord, Hilde Elshaug, Marie Paulsen Madsen, Rasmus Riis Kopperud, Hilde Vollan, Karoline Bragstad, Olav Hungnes |
| EPI_ISL_708041 | Medical Microbiology Unit, Department for Laboratory Medicine, Drammen Hospital, Vestre Viken Health Trust, | Norwegian Institute of Public Health, Department of Virology | Kathrine Stene-Johansen, Kamilla Heddeland Instefjord, Hilde Elshaug, Marie Paulsen Madsen, Rasmus Riis Kopperud, Hilde Vollan, Karoline Bragstad, Olav Hungnes |
| EPI_ISL_708042 | Department of Medical Microbiology, St. Olavs hospital | Norwegian Institute of Public Health, Department of Virology | Kathrine Stene-Johansen, Kamilla Heddeland Instefjord, Hilde Elshaug, Marie Paulsen Madsen, Rasmus Riis Kopperud, Hilde Vollan, Karoline Bragstad, Olav Hungnes |
| EPI_ISL_708046, EPI_ISL_708047, EPI_ISL_708048 | Unilabs Laboratory Medicine | Norwegian Institute of Public Health, Department of Virology | Kathrine Stene-Johansen, Kamilla Heddeland Instefjord, Hilde Elshaug, Marie Paulsen Madsen, Rasmus Riis Kopperud, Hilde Vollan, Karoline Bragstad, Olav Hungnes |
| EPI_ISL_708049, EPI_ISL_708051 | Norwegian Institute of Public Health, Department of Virology | Norwegian Institute of Public Health, Department of Virology | Kathrine Stene-Johansen, Kamilla Heddeland Instefjord, Hilde Elshaug, Marie Paulsen Madsen, Rasmus Riis Kopperud, Hilde Vollan, Karoline Bragstad, |

| | | | |
|---|---|---|---|
| | | | Olav Hungnes |
| EPI_ISL_708052, EPI_ISL_708053, EPI_ISL_708054 | Oslo University Hospital, Department of Medical Microbiology | Norwegian Institute of Public Health, Department of Virology | Kathrine Stene-Johansen, Kamilla Heddeland Instefjord, Hilde Elshaug, Marie Paulsen Madsen, Rasmus Riis Kopperud, Hilde Vollan, Karoline Bragstad, Olav Hungnes |
| EPI_ISL_708069, EPI_ISL_708070, EPI_ISL_708071, EPI_ISL_708072 | University Hospital of Northern Norway, Department for Microbiology and Infectious Disease Control | Norwegian Institute of Public Health, Department of Virology | Kathrine Stene-Johansen, Kamilla Heddeland Instefjord, Hilde Elshaug, Marie Paulsen Madsen, Rasmus Riis Kopperud, Hilde Vollan, Karoline Bragstad, Olav Hungnes |
| EPI_ISL_708073, EPI_ISL_708074 | Foerde Hospital, Department of Microbiology | Norwegian Institute of Public Health, Department of Virology | Kathrine Stene-Johansen, Kamilla Heddeland Instefjord, Hilde Elshaug, Marie Paulsen Madsen, Rasmus Riis Kopperud, Hilde Vollan, Karoline Bragstad, Olav Hungnes |
| EPI_ISL_708079, EPI_ISL_708082, EPI_ISL_708083, EPI_ISL_708084, EPI_ISL_708085, EPI_ISL_708091, EPI_ISL_708095 | University Hospital of Northern Norway, Department for Microbiology and Infectious Disease Control | Norwegian Institute of Public Health, Department of Virology | Kathrine Stene-Johansen, Kamilla Heddeland Instefjord, Hilde Elshaug, Marie Paulsen Madsen, Rasmus Riis Kopperud, Hilde Vollan, Karoline Bragstad, Olav Hungnes |
| EPI_ISL_708098, EPI_ISL_708099, EPI_ISL_708100 | Nordland Hospital - Bodo, Laboratory Department, Molecular Biology Unit | Norwegian Institute of Public Health, Department of Virology | Kathrine Stene-Johansen, Kamilla Heddeland Instefjord, Hilde Elshaug, Marie Paulsen Madsen, Rasmus Riis Kopperud, Hilde Vollan, Karoline Bragstad, Olav Hungnes |
| EPI_ISL_708154, EPI_ISL_708155, EPI_ISL_708156, EPI_ISL_708157, EPI_ISL_708158, EPI_ISL_708159, EPI_ISL_708160, EPI_ISL_708161, EPI_ISL_708162, EPI_ISL_708165, EPI_ISL_708166, EPI_ISL_708167, EPI_ISL_708168 | | | |
| see above | Vestfold Hospital, Toensberg Department of Microbiology | Norwegian Institute of Public Health, Department of Virology | Kathrine Stene-Johansen, Kamilla Heddeland Instefjord, Hilde Elshaug, Marie Paulsen Madsen, Rasmus Riis Kopperud, Hilde Vollan, Karoline Bragstad, Olav Hungnes |
| EPI_ISL_708196 | Medical Microbiology Unit, Department for Laboratory Medicine, Drammen Hospital, Vestre Viken Health Trust, | Norwegian Institute of Public Health, Department of Virology | Kathrine Stene-Johansen, Kamilla Heddeland Instefjord, Hilde Elshaug, Marie Paulsen Madsen, Rasmus Riis Kopperud, Hilde Vollan, Karoline Bragstad, Olav Hungnes |
| EPI_ISL_708197, EPI_ISL_708198 | University Hospital of Northern Norway, Department for Microbiology and Infectious Disease Control | Norwegian Institute of Public Health, Department of Virology | Kathrine Stene-Johansen, Kamilla Heddeland Instefjord, Hilde Elshaug, Marie Paulsen Madsen, Rasmus Riis Kopperud, Hilde Vollan, Karoline Bragstad, Olav Hungnes |
| EPI_ISL_708308 | University Hospitals of Geneva, Laboratory of Virology | University Hospitals of Geneva, Laboratory of Virology | Cordey Samuel, Laubscher Florian |
| EPI_ISL_708410, EPI_ISL_708437, EPI_ISL_708438, EPI_ISL_708442, EPI_ISL_708443, EPI_ISL_708447, EPI_ISL_708449, EPI_ISL_708451 | Delaware Public Health Lab | Delaware Public Health Lab | Gregory Hovan |
| EPI_ISL_708545, EPI_ISL_708564, EPI_ISL_708577, EPI_ISL_708584 | Michigan Department of Health and Human Services, Bureau of Laboratories | Michigan Department of Health and Human Services, Bureau of Laboratories | Blankenship HM, Riner D, Soehnlen MK |
| EPI_ISL_708776 | PathWest Laboratory Medicine WA | PathWest Laboratory Medicine WA Microbial Surveillance Unit | PathWest Laboratory Medicine WA Microbial Surveillance Unit |
| EPI_ISL_708811 | Regional medical sciences center 2 Phitsanulok | National Institute of Health, Department of Medical Sciences, Ministry of Public Health, Thailand | Pilailuk Okada; Siripaporn Phuygun; Thanutsapa Thanadachakul; Sittiporn Parnmen; Pakorn Piromtong; Warawan Wongboot; Sunthareeya Waicharoen; Malinee Chittaganpitch |
| EPI_ISL_710165, EPI_ISL_710173, EPI_ISL_710240, EPI_ISL_710285, EPI_ISL_710286, EPI_ISL_710287, EPI_ISL_710296, EPI_ISL_710360, EPI_ISL_710361, EPI_ISL_710362, EPI_ISL_710363, EPI_ISL_710364, EPI_ISL_710365, EPI_ISL_710366, EPI_ISL_710367, EPI_ISL_710368 | | | |
| see above | Colorado Department of Public Health and Environment | Colorado Department of Puplic Health and Environment | Laura Bankers, Molly C. Hetherington-Rauth, Shannon Ely, Shannon R. Matzinger, Sarah Elizabeth Totten, Emily A. Travanty |
| EPI_ISL_710419, EPI_ISL_710422 | Los Angeles County PHL | Los Angeles County PHL | P. Hemarajata et al. |
| EPI_ISL_710542 | National Institute for Infectious Diseases, INMI, "L. Spallanzani" IRCCS | National Institute for Infectious Diseases, INMI, "L. Spallanzani" IRCCS | C.E.M Gruber, B Bartolini, M Rueca, F Messina, E Giombini, A Di Caro, MR Capobianchi |
| EPI_ISL_710543 | National Institute for Infectious Diseases, INMI, "L. Spallanzani" IRCCS | National Institute for Infectious Diseases, INMI, "L. Spallanzani" IRCCS | B Bartolini, C.E.M Gruber, M Rueca, F Messina, E Giombini, MR Capobianchi, A Di Caro |
| EPI_ISL_710544 | National Institute for Infectious Diseases, INMI, "L. Spallanzani" IRCCS | National Institute for Infectious Diseases, INMI, "L. Spallanzani" IRCCS | M Rueca, B Bartolini, C.E.M Gruber, F Messina, E Giombini, A Di Caro, MR Capobianchi |
| EPI_ISL_710545 | National Institute for Infectious Diseases, INMI, "L. Spallanzani" IRCCS | National Institute for Infectious Diseases, INMI, "L. Spallanzani" IRCCS | F Messina, E Giombini, M Rueca, B Bartolini, C.E.M Gruber, MR Capobianchi, A Di Caro |
| EPI_ISL_710611 | Ektorps Vardcentral | The Public Health Agency of Sweden | Department of Microbiology, The Public Health Agency of Sweden |
| EPI_ISL_710836 | Lighthouse Lab in Cambridge | Wellcome Sanger Institute for the COVID-19 Genomics UK (COG-UK) Consortium | Rob Howes, The Lighthouse Lab in Cambridge and Alex Alderton, Roberto Amato, Sonia Goncalves, Ewan Harrison, David K. Jackson, Ian Johnston, Dominic Kwiatkowski, Cordelia Langford, John Sillitoe on behalf of the Wellcome Sanger Institute COVID-19 Surveillance Team |
| EPI_ISL_711048, EPI_ISL_711049, EPI_ISL_711050, EPI_ISL_711051, EPI_ISL_711052, EPI_ISL_711053, EPI_ISL_711054 | Lighthouse Lab in Glasgow | Wellcome Sanger Institute for the COVID-19 Genomics UK (COG-UK) Consortium | Harper VanSteenhouse, Yumi Kasai, David Gray, Carol Clugston, Anna Dominiczak and Alex Alderton, Roberto Amato, Sonia Goncalves, Ewan Harrison, David K. Jackson, Ian Johnston, Dominic Kwiatkowski, Cordelia Langford, John Sillitoe on behalf of the Wellcome Sanger Institute COVID-19 Surveillance Team |
| EPI_ISL_712079, EPI_ISL_712086, EPI_ISL_712089, EPI_ISL_712091 | Port Elizabeth Provincial Hospital, National Health Laboratory Services, Eastern Cape, South Africa | National Institute for Communicable Diseases of the National Health Laboratory Service | Mohale T, Ntuli N, Mahlangu B, Allam M, Ismail A, Bhiman JN |
| EPI_ISL_714419, EPI_ISL_714420, EPI_ISL_714421, EPI_ISL_714422 | Department of Virus and Microbiological Special Diagnostics, Statens Serum Institut, Copenhagen, Denmark | Albertsen Lab, Department of Chemistry and Bioscience, Aalborg University, Denmark | Danish Covid-19 Genome Consortium |
| EPI_ISL_717629 | Lab voor klinische biologie | Onderzoeksgroep Virologie | Nick Vereecke, Laurens Lambrechts, Marthe Pauwels, Bruno Verhasselt, Linos Vandekerckhove, Hans Nauwynck, Sebastiaan Theuns |
| EPI_ISL_717820, EPI_ISL_717821, EPI_ISL_717822, EPI_ISL_717823, EPI_ISL_717824, EPI_ISL_717877, EPI_ISL_717878, EPI_ISL_717879, EPI_ISL_717880, EPI_ISL_717881, EPI_ISL_717882, EPI_ISL_717883, EPI_ISL_717884, EPI_ISL_717885, EPI_ISL_717886, EPI_ISL_717887, EPI_ISL_717888, EPI_ISL_717920, EPI_ISL_717921, EPI_ISL_717922, EPI_ISL_717924, EPI_ISL_717926, EPI_ISL_717927, EPI_ISL_717928, EPI_ISL_717929, EPI_ISL_717930, EPI_ISL_717931, EPI_ISL_717932, EPI_ISL_717933, EPI_ISL_717934, EPI_ISL_717935, EPI_ISL_717936, EPI_ISL_717937, EPI_ISL_717938, EPI_ISL_717939 | | | |
| see above | Laboratorio de Virologia Molecular / UFRJ | Bioinformatics Laboratory / LNCC | Carolina M Voloch, Ronaldo da Silva F Jr, Luiz G P de Almeida, Cynthia C Cardoso, Otavio Bustrolini, Alexandra L Gerber, Ana Paula de C Guimarães, Diana Mariani, Andréa Cony Cavalcanti, Claudia dos Santos Rodrigues, Terezinha M P P Castiñeira, Amilcar Tanuri, Ana Tereza R de Vasconcelos |
| EPI_ISL_717991, EPI_ISL_717992, EPI_ISL_717993 | Lab voor klinische biologie | Onderzoeksgroep Virologie | Laurens Lambrechts, Nick Vereecke, Marthe Pauwels, Bruno Verhasselt, Linos Vandekerckhove, Hans Nauwynck, Sebastiaan Theuns |
| EPI_ISL_718169, EPI_ISL_718170 | Ministry of Health Hospitals | Institute of Health and Community Medicine | David Perera, Ooi Mong How, Chua Hock Hin, Tonnii Sia Loong Loong, Wong Jyn Shan, Wong Kiing Aik, Chan Chia Jui |
| EPI_ISL_718174, EPI_ISL_718182, EPI_ISL_718197, EPI_ISL_718198 | Borneo Medical Centre | Institute of Health and Community Medicine | David Perera, Ooi Mong How, Chua Hock Hin, Tonnii Sia Loong Loong, Wong Jyn Shan, Wong Kiing Aik, Chan Chia Jui |
| EPI_ISL_718206 | Ministry of Health Hospitals | Institute of Health and Community Medicine | David Perera, Ooi Mong How, Chua Hock Hin, Tonnii Sia Loong Loong, Wong Jyn Shan, Wong Kiing Aik, Chan Chia Jui |
| EPI_ISL_718228, EPI_ISL_718249 | Hospital | National Reference Center for Viruses of Respiratory Infections, Institut Pasteur, Paris | Marion Barbet, Sylvie Behillil, Méline Bizard, Angela Brisebarre, Camille Capel, Etienne Simon-Lorière, Vincent Enouf, Maud Vanpeene, Sylvie van der Werf, Gisèle Lagathu |
| EPI_ISL_721672, EPI_ISL_721676, EPI_ISL_721680, EPI_ISL_721683, EPI_ISL_721685, EPI_ISL_721687, EPI_ISL_721688, EPI_ISL_721697, EPI_ISL_721698, EPI_ISL_721700, EPI_ISL_721703, EPI_ISL_721704, EPI_ISL_721705, EPI_ISL_721706, EPI_ISL_721707, EPI_ISL_721708, EPI_ISL_721709, EPI_ISL_721710, EPI_ISL_721737, EPI_ISL_721738, EPI_ISL_721739, EPI_ISL_721742, EPI_ISL_721743, EPI_ISL_721744, EPI_ISL_721745, EPI_ISL_721746, EPI_ISL_721747, EPI_ISL_721748, EPI_ISL_721749, EPI_ISL_721750, EPI_ISL_721751, EPI_ISL_721752, EPI_ISL_721753, EPI_ISL_721754, EPI_ISL_721755, EPI_ISL_721756, EPI_ISL_721773, EPI_ISL_721774, EPI_ISL_721775, EPI_ISL_721776, EPI_ISL_721777, EPI_ISL_721806, EPI_ISL_721807, EPI_ISL_721808, EPI_ISL_721809, EPI_ISL_721810, EPI_ISL_721811, EPI_ISL_721812, EPI_ISL_721815, EPI_ISL_721816, EPI_ISL_721817, EPI_ISL_721823, EPI_ISL_721824, EPI_ISL_721825, EPI_ISL_721854, EPI_ISL_721855, EPI_ISL_721856, EPI_ISL_721857, EPI_ISL_721867, EPI_ISL_721868, EPI_ISL_721897, EPI_ISL_721898, EPI_ISL_721899, EPI_ISL_721900, EPI_ISL_721913, EPI_ISL_721914, EPI_ISL_721919, EPI_ISL_721920, EPI_ISL_721921, EPI_ISL_721929, EPI_ISL_721930, EPI_ISL_721931, EPI_ISL_721932, EPI_ISL_721933, EPI_ISL_721934, EPI_ISL_721935, EPI_ISL_721936, EPI_ISL_721937, EPI_ISL_721938, EPI_ISL_721939 | | | |

| | | | |
|---|---|---|---|
| see above | Viollier AG | Department of Biosystems Science and Engineering, ETH Zürich | Christian Beisel |
| EPI_ISL_722283, EPI_ISL_722324, EPI_ISL_722343, EPI_ISL_722344, EPI_ISL_722380, EPI_ISL_722399, EPI_ISL_722400, EPI_ISL_722444, EPI_ISL_722549, EPI_ISL_722550, EPI_ISL_722551, EPI_ISL_722552, EPI_ISL_722553, EPI_ISL_722554, EPI_ISL_722555, EPI_ISL_722556, EPI_ISL_722557, EPI_ISL_722558, EPI_ISL_722559, EPI_ISL_722560, EPI_ISL_722561, EPI_ISL_722562, EPI_ISL_722563, EPI_ISL_722564, EPI_ISL_722565, EPI_ISL_722566, EPI_ISL_722567, EPI_ISL_722568, EPI_ISL_722569 | | | |
| see above | Dutch COVID-19 response team | Erasmus Medical Center | Bas Oude Munnink, Reina Sikkema, David Nieuwenhuijse, Irina Chestakova, Anne van der Linden, Marjan Boter, Emmanuelle Munger, Corine GeurtsvanKessel, Annemiek van der Eijk, Richard Molenkamp, Marion Koopmans, on behalf of the Dutch national COVID-19 response team. |
| EPI_ISL_722873, EPI_ISL_722874, EPI_ISL_722875, EPI_ISL_722876, EPI_ISL_722877, EPI_ISL_722883, EPI_ISL_722887, EPI_ISL_722901, EPI_ISL_722902, EPI_ISL_722903, EPI_ISL_722905, EPI_ISL_722906, EPI_ISL_722913, EPI_ISL_722914 | | | |
| see above | Istituto Zooprofilattico Sperimentale della Puglia e della Basilicata | Istituto Zooprofilattico Sperimentale della Puglia e della Basilicata | Parisi A., Bianco A., Capozzi L., Del Sambro L., Manzulli V, Rondinone V., Pace L., Cipolletta D., Galante D. |
| EPI_ISL_724964, EPI_ISL_724965, EPI_ISL_724974 | Northumbria University / South Tees Hospitals NHS Foundation Trust / North Cumbria Integrated Care NHS Foundation Trust / North Tees and Hartlepool NHS Foundation Trust / Newcastle Hospitals NHS Foundation Trust | COVID-19 Genomics UK (COG-UK) Consortium | Darren L Smith,Andrew Nelson,Matthew Bashton,Gregory R Young,Joshua Loh,John Allan,Mohammad A Tariq,Giles S Holt,Gary Black,Wen C Yew,Lynn Dover,Paul Baker,Steve Liggett,Sarah Essex,Jane Greenaway,Debra Padgett,Clive Graham,Garren Scott,Edward Barton,Emma Swindells,Brendan Payne,Jennifer Collins,Yusri Taha,Gary Eltringham |
| EPI_ISL_725264, EPI_ISL_725274, EPI_ISL_725287, EPI_ISL_725289 | Quadram Institute Bioscience | COVID-19 Genomics UK (COG-UK) Consortium | Dave J. Baker, Gemma L. Kay, Alp Aydin, Thanh Le-Viet, Steven Rudder, Ana P. Tedim, Anastasia Kolyva, Maria Diaz, Leonardo de Oliveira Martins, Nabil-Fareed Alikhan, Lizzie Meadows, Rachael Stanley, Ngozi Elumogo, Muhammed Yasir, Nicholas M. Thomson, Alexander J Trotter, Rachel Gilroy, Samuel Bloomfield, Claire Stuart, Andrew Bell, Reenesh Prakash, Samir Dervisevic, Alison E. Mather, John Wain, Mark Webber, Andrew J. Page, Justin O'Grady |
| EPI_ISL_725999 | Wales Specialist Virology Centre Sequencing lab: Pathogen Genomics Unit | COVID-19 Genomics UK (COG-UK) Consortium | Catherine Moore, Johnathan Evans, Laura Gifford, Malorie Perry, Simon Cottrell, Angela Marchbank, Alec Birchley, Alexander Adams, Amy Gaskin, Bree Gatica-Wilcox, Jason Coombes, Joel Southgate, Lauren Gilbert, Lee Graham, Nicole Pacchiarini, Sara Kumziene-Summerhayes, Sarah Taylor, Sophie Jones, Sara Rey, Matthew Bull, Joanne Watkins, Sally Corden, Tom Connor |
| EPI_ISL_728035, EPI_ISL_728036, EPI_ISL_728052, EPI_ISL_728146, EPI_ISL_728147, EPI_ISL_728148, EPI_ISL_728149, EPI_ISL_728150, EPI_ISL_728151, EPI_ISL_728152 | University of Wisconsin-Madison AIDS Vaccine Research Laboratories | University of Wisconsin-Madison AIDS Vaccine Research Laboratories | Gage Moreno, Katarina Braun, et al. AIDS Vaccine Research Laboratories |
| EPI_ISL_728668, EPI_ISL_728669, EPI_ISL_728670, EPI_ISL_728671, EPI_ISL_728672, EPI_ISL_728673, EPI_ISL_728704, EPI_ISL_728705, EPI_ISL_728706, EPI_ISL_728707 | Dutch COVID-19 response team | National Institute for Public Health and the Environment (RIVM) | Adam Meijer, Harry Vennema, Jeroen Cremer, Sharon van den Brink, Bas van der Veer, AnneMarie van den Brandt, Florian Zwagemaker, Dennis Schmitz, Chantal Reusken, on behalf of the national COVID-19 response team |
| EPI_ISL_730054, EPI_ISL_730056, EPI_ISL_730057 | Connecticut Department of Health | Grubaugh Lab - Yale School of Public Health | Joseph Fauver, Tara Alpert, Anderson Brito, Annie Watkins, Anne Wyllie, Chantal Vogels, Mary Petrone, Chaney Kalinich, Isabel Ott, Arnau Casanovas, Catherine Muenker, Adam Moore, Alice Lu, Maria Tokuyama, Patrick Wong, Peiwen Lu, Saad Omer, Richard Martinello, Allison Nelson, Shelli Farhadian, Akiko Iwasaki, Charlese Dela Cruz, Albert Ko, Nathan Grubaugh |
| EPI_ISL_730094, EPI_ISL_730100, EPI_ISL_730108, EPI_ISL_730112, EPI_ISL_730124, EPI_ISL_730163, EPI_ISL_730165, EPI_ISL_730172, EPI_ISL_730173, EPI_ISL_730180, EPI_ISL_730185, EPI_ISL_730192 | | | |
| see above | San Diego County Public Health Laboratory | Andersen lab at Scripps Research | SEARCH Alliance San Diego with Tracy Basler, Jovan Shephard, Brett Austin |
| EPI_ISL_730581 | Home Quarantine Taskforce | Hong Kong Department of Health | Mak Gannon C.K., Lam Edman T.K., Chan Rickjason C.W., Tsang Dominic N.C. |
| EPI_ISL_730594, EPI_ISL_730595 | Queen Elizabeth Hospital | Hong Kong Department of Health | Mak Gannon C.K., Lam Edman T.K., Chan Rickjason C.W., Tsang Dominic N.C. |
| EPI_ISL_732406, EPI_ISL_732454 | National Virus Reference Laboratory | National Virus Reference Laboratory | Michael Carr, Gabriel Gonzalez, Jonathan Dean, Daniel Hare, Cillian F De Gascun |
| EPI_ISL_732699, EPI_ISL_732700, EPI_ISL_732701, EPI_ISL_732702 | CNR Virus des Infections Respiratoires - France SUD | CNR Virus des Infections Respiratoires - France SUD | Antonin Bal, Gregory Destras, Claudia Gonzalez, Gwendolyne Burfin, Quentin Semanas, Martine Valette, Bruno Lina, Laurence Josset |
| EPI_ISL_732786, EPI_ISL_732793, EPI_ISL_732805 | Centro de Investigación Biomédica de La Rioja - Hospital San Pedro Logroño | SeqCOVID-SPAIN consortium/IBV(CSIC) | María de Toro, José Manuel Azcona Gutiérrez, María Pilar Bea Escudero, Miriam Blasco Alberdi and SeqCOVID-SPAIN consortium |
| EPI_ISL_733184 | WHO National Influenza Centre Russian Federation | WHO National Influenza Centre Russian Federation | Andrey Komissarov, Artem Fadeev, Anna Ivanova, Kseniya Komissarova, Dmitry Bazhenov, Daria Danilenko, Ksenia Safina, Elena Nabieva, Georgii Bazykin, Dmitry Lioznov |
| EPI_ISL_733238 | UMMC-Health | WHO National Influenza Centre Russian Federation | Andrey Komissarov, Artem Fadeev, Anna Ivanova, Kseniya Komissarova, Dmitry Bazhenov, Tatiana Platonova, Daria Danilenko, Ksenia Safina, Elena Nabieva, Georgii Bazykin, Dmitry Lioznov |
| EPI_ISL_733259 | WHO National Influenza Centre Russian Federation | WHO National Influenza Centre Russian Federation | Andrey Komissarov, Artem Fadeev, Anna Ivanova, Kseniya Komissarova, Dmitry Bazhenov, Daria Danilenko, Ksenia Safina, Elena Nabieva, Georgii Bazykin, Dmitry Lioznov |
| EPI_ISL_733397 | HELIX LLC | WHO National Influenza Centre Russian Federation | Andrey Komissarov, Artem Fadeev, Anna Ivanova, Kseniya Komissarova, Dmitry Bazhenov, Daria Danilenko, Ksenia Safina, Elena Nabieva, Georgii Bazykin, Dmitry Lioznov |
| EPI_ISL_733503, EPI_ISL_733504, EPI_ISL_733505, EPI_ISL_733520, EPI_ISL_733521, EPI_ISL_733522, EPI_ISL_733523 | ZOTZ KLIMAS MVZ Düsseldorf-Centrum GbR ÜBAG für Labormedizin, Genetik, Zytologie, Pathologie | Center of Medical Microbiology, Virology, and Hospital Hygiene, University of Duesseldorf | Maximilian Damagnez, Alexander Dilthey, Ashley-Jane Duplessis, Patrick Finzer, Katrin Hoffmann, Torsten Houwaart, Lisanna Hülse, Malte Kohns Vasconcelos, Marek Korencak, Nadine Lübke, Jessica Nicolai, Klaus Pfeffer, Daniel Strelow, Jörg Timm, Andreas Walker, Tobias Wienemann, Rainer Zotz |
| EPI_ISL_734232, EPI_ISL_734233, EPI_ISL_734234, EPI_ISL_734235, EPI_ISL_734236, EPI_ISL_734237, EPI_ISL_734238, EPI_ISL_734239, EPI_ISL_734240, EPI_ISL_734241, EPI_ISL_734242, EPI_ISL_734243, EPI_ISL_734244, EPI_ISL_734270, EPI_ISL_734271, EPI_ISL_734272, EPI_ISL_734273, EPI_ISL_734274, EPI_ISL_734275, EPI_ISL_734276 | | | |
| see above | Virginia Division of Consolidated Laboratory Services (DCLS) | Virginia Division of Consolidated Laboratory Services (DCLS) | Virginia DCLS |
| EPI_ISL_734406, EPI_ISL_734407, EPI_ISL_734408, EPI_ISL_734409, EPI_ISL_734411, EPI_ISL_734412 | Wadsworth Center, New York State Department.of Health | Wadsworth Center, New York State Department.of Health | Kirsten St. George, Daryl M. Lamson, Alexis Russel, Jonathan Plitnick, Navjot Singh, John Kelly, Sara Griesemer, Erasmus Schneider, Erica Lasek-Nesselquist |
| EPI_ISL_734462, EPI_ISL_734463, EPI_ISL_734464, EPI_ISL_734465, EPI_ISL_734466, EPI_ISL_734467, EPI_ISL_734468, EPI_ISL_734469 | Masonic Medical Research Institute | Wadsworth Center, New York State Department.of Health | Kirsten St. George, Nathan Tucker, Ryan D. Pfeiffer, Daryl M. Lamson, Alexis Russel, Jonathan Plitnick, Navjot Singh, John Kelly, Sara Griesemer, Erasmus Schneider, Erica Lasek-Nesselquist |
| EPI_ISL_736777, EPI_ISL_736778, EPI_ISL_736779, EPI_ISL_736780, EPI_ISL_736781, EPI_ISL_736788, EPI_ISL_736789, EPI_ISL_736790, EPI_ISL_736791, EPI_ISL_736792, EPI_ISL_736793, EPI_ISL_736794, EPI_ISL_736795, EPI_ISL_736796, EPI_ISL_736797, EPI_ISL_736798, EPI_ISL_736799, EPI_ISL_736800, EPI_ISL_736801, EPI_ISL_736802, EPI_ISL_736803, EPI_ISL_736804, EPI_ISL_736805, EPI_ISL_736806, EPI_ISL_736807, EPI_ISL_736808, EPI_ISL_736809, EPI_ISL_736810, EPI_ISL_736811, EPI_ISL_736812, EPI_ISL_736813, EPI_ISL_736814, EPI_ISL_736815, EPI_ISL_736816, EPI_ISL_736817, EPI_ISL_736818, EPI_ISL_736819, EPI_ISL_736820, EPI_ISL_736821, EPI_ISL_736822, EPI_ISL_736823, EPI_ISL_736824, EPI_ISL_736825, EPI_ISL_736826, EPI_ISL_736827, EPI_ISL_736828, EPI_ISL_736829, EPI_ISL_736830, EPI_ISL_736831, EPI_ISL_736832, EPI_ISL_736833, EPI_ISL_736834, EPI_ISL_736835, EPI_ISL_736836, EPI_ISL_736837, EPI_ISL_736838, EPI_ISL_736839, EPI_ISL_736840, EPI_ISL_736841, EPI_ISL_736842, EPI_ISL_736843, EPI_ISL_736844, EPI_ISL_736845, EPI_ISL_736846, EPI_ISL_736847, EPI_ISL_736848, EPI_ISL_736849, EPI_ISL_736850, EPI_ISL_736851, EPI_ISL_736852, EPI_ISL_736853, EPI_ISL_736854, EPI_ISL_736855, EPI_ISL_736856, EPI_ISL_736857, EPI_ISL_736858, EPI_ISL_736859, EPI_ISL_736860, EPI_ISL_736861, EPI_ISL_736862, EPI_ISL_736863, EPI_ISL_736864, EPI_ISL_736865, EPI_ISL_736866, EPI_ISL_736867, EPI_ISL_736868, EPI_ISL_736869, EPI_ISL_736870, EPI_ISL_736871, EPI_ISL_736872, EPI_ISL_736873, EPI_ISL_736874, EPI_ISL_736875, EPI_ISL_736876, EPI_ISL_736877, EPI_ISL_736878, EPI_ISL_736879, EPI_ISL_736880, EPI_ISL_736881, EPI_ISL_736882, EPI_ISL_736883, EPI_ISL_736884, EPI_ISL_736885, EPI_ISL_736886, EPI_ISL_736887, EPI_ISL_736888, EPI_ISL_736889 | | | |
| see above | Istituto Zooprofilattico Sperimentale del Mezzogiorno | TIGEM | Patrizia Annunziata, Andrea Ballabio, Valentina Bouche, Davide Cacchiarelli (CorrespAuthor), Pellegrino Cerino, Chiara Colantuono, Lucio Di Filippo, Antonio Grimaldi, Antonio Limone, Gabriella Loconte, Anna Manfredi, Francesco Panariello, Biancamaria Pierri, Marcello Salvi, Lucia Vassallo |
| EPI_ISL_737002, EPI_ISL_737003, EPI_ISL_737004, EPI_ISL_737005, EPI_ISL_737006, EPI_ISL_737007, EPI_ISL_737008, EPI_ISL_737009, EPI_ISL_737010, EPI_ISL_737011, EPI_ISL_737012, EPI_ISL_737013, EPI_ISL_737029, EPI_ISL_737033 | | | |

| | | | |
|---|---|---|---|
| see above | The National Institute of Public Health | State Veterinary Institute Prague | Nagy,A;Jirincova,H;Trnka,D;Vecerova,J |
| EPI_ISL_738754, EPI_ISL_738836 | Napa-Solano-Yolo- Marin County (NSYM) Public Health Laboratories | Chan-Zuckerberg Biohub | CZB Cliahub Consortium |
| EPI_ISL_738951, EPI_ISL_739148 | Humboldt County Public Health Laboratory | Chan-Zuckerberg Biohub | CZB Cliahub Consortium |
| EPI_ISL_739315, EPI_ISL_739325 | Napa-Solano-Yolo- Marin County (NSYM) Public Health Laboratories | Chan-Zuckerberg Biohub | CZB Cliahub Consortium |
| EPI_ISL_739402 | Santa Clara County Public Health Laboratory | Chan-Zuckerberg Biohub | CZB Cliahub Consortium |
| EPI_ISL_739474, EPI_ISL_739628 | Napa-Solano-Yolo- Marin County (NSYM) Public Health Laboratories | Chan-Zuckerberg Biohub | CZB Cliahub Consortium |
| EPI_ISL_739639 | Tulare County Public Health Lab | Chan-Zuckerberg Biohub | CZB Cliahub Consortium |
| EPI_ISL_739702, EPI_ISL_739704, EPI_ISL_739715, EPI_ISL_739745, EPI_ISL_739748, EPI_ISL_739751, EPI_ISL_739754, EPI_ISL_739765, EPI_ISL_739767, EPI_ISL_739770, EPI_ISL_739777, EPI_ISL_739819, EPI_ISL_739832, EPI_ISL_739853, EPI_ISL_739861, EPI_ISL_739864, EPI_ISL_739866, EPI_ISL_739881, EPI_ISL_739882, EPI_ISL_739896, EPI_ISL_739909, EPI_ISL_739911, EPI_ISL_739914, EPI_ISL_739922, EPI_ISL_739926, EPI_ISL_739946, EPI_ISL_739968, EPI_ISL_739981, EPI_ISL_739985, EPI_ISL_739993, EPI_ISL_740001, EPI_ISL_740010, EPI_ISL_740020, EPI_ISL_740034, EPI_ISL_740042, EPI_ISL_740043, EPI_ISL_740046, EPI_ISL_740050, EPI_ISL_740054, EPI_ISL_740078, EPI_ISL_740082, EPI_ISL_740099, EPI_ISL_740117, EPI_ISL_740153, EPI_ISL_740154, EPI_ISL_740159, EPI_ISL_740168, EPI_ISL_740170, EPI_ISL_740173, EPI_ISL_740180, EPI_ISL_740181, EPI_ISL_740192, EPI_ISL_740196, EPI_ISL_740197, EPI_ISL_740206, EPI_ISL_740209, EPI_ISL_740232, EPI_ISL_740233, EPI_ISL_740243, EPI_ISL_740247, EPI_ISL_740260, EPI_ISL_740265, EPI_ISL_740275, EPI_ISL_740292, EPI_ISL_740300, EPI_ISL_740305, EPI_ISL_740310, EPI_ISL_740324, EPI_ISL_740329, EPI_ISL_740337, EPI_ISL_740346, EPI_ISL_740352, EPI_ISL_740356, EPI_ISL_740366, EPI_ISL_740373, EPI_ISL_740384, EPI_ISL_740388, EPI_ISL_740407, EPI_ISL_740414, EPI_ISL_740442, EPI_ISL_740443, EPI_ISL_740461, EPI_ISL_740468, EPI_ISL_740470, EPI_ISL_740474, EPI_ISL_740482, EPI_ISL_740497, EPI_ISL_740513, EPI_ISL_740525, EPI_ISL_740526, EPI_ISL_740537, EPI_ISL_740542 | | | |
| see above | Laboratoire national de santé, Microbiology, Virology | Laboratoire national de santé, Microbiology, Microbial Genomics Platform | Anke Wienecke-Baldacchino, Catherine Ragimbeau,Jessica Tapp, Fatu Djabi, Lise Pignon, Raoul Salmon, Tamir Abdelrahman |
| EPI_ISL_742283, EPI_ISL_742284, EPI_ISL_742285, EPI_ISL_742302, EPI_ISL_742303, EPI_ISL_742304, EPI_ISL_742305, EPI_ISL_742306, EPI_ISL_742307 | Wales Specialist Virology Centre Sequencing lab: Pathogen Genomics Unit | COVID-19 Genomics UK (COG-UK) Consortium | Catherine Moore, Johnathan Evans, Laura Gifford, Malorie Perry, Simon Cottrell, Angela Marchbank, Alec Birchley, Alexander Adams, Amy Gaskin, Bree Gatica-Wilcox, Jason Coombes, Joel Southgate, Lauren Gilbert, Lee Graham, Nicole Pacchiarini, Sara Kumziene-Summerhayes, Sarah Taylor, Sophie Jones, Sara Rey, Matthew Bull, Joanne Watkins, Sally Corden, Tom Connor |
| EPI_ISL_744140, EPI_ISL_744156, EPI_ISL_744159, EPI_ISL_744167, EPI_ISL_744192, EPI_ISL_744200, EPI_ISL_744204, EPI_ISL_744205, EPI_ISL_744216, EPI_ISL_744238, EPI_ISL_744241, EPI_ISL_744252, EPI_ISL_744278, EPI_ISL_744285, EPI_ISL_744289, EPI_ISL_744306, EPI_ISL_744307, EPI_ISL_744323, EPI_ISL_744339, EPI_ISL_744344, EPI_ISL_744362, EPI_ISL_744372, EPI_ISL_744375, EPI_ISL_744384, EPI_ISL_744394, EPI_ISL_744408, EPI_ISL_744413, EPI_ISL_744436, EPI_ISL_744441, EPI_ISL_744459, EPI_ISL_744487, EPI_ISL_744503, EPI_ISL_744509, EPI_ISL_744524, EPI_ISL_744546, EPI_ISL_744548, EPI_ISL_744563, EPI_ISL_744569, EPI_ISL_744583, EPI_ISL_744594, EPI_ISL_744607, EPI_ISL_744610, EPI_ISL_744613, EPI_ISL_744619, EPI_ISL_744628, EPI_ISL_744639, EPI_ISL_744640, EPI_ISL_744642, EPI_ISL_744651, EPI_ISL_744655, EPI_ISL_744658, EPI_ISL_744662, EPI_ISL_744664, EPI_ISL_744669, EPI_ISL_744685, EPI_ISL_744694, EPI_ISL_744705, EPI_ISL_744710, EPI_ISL_744713, EPI_ISL_744718, EPI_ISL_744722, EPI_ISL_744726, EPI_ISL_744731, EPI_ISL_744735, EPI_ISL_744736, EPI_ISL_744744, EPI_ISL_744757, EPI_ISL_744759, EPI_ISL_744770, EPI_ISL_744778, EPI_ISL_744799, EPI_ISL_744815, EPI_ISL_744855, EPI_ISL_744860, EPI_ISL_744862, EPI_ISL_744875, EPI_ISL_744892, EPI_ISL_744901, EPI_ISL_744911, EPI_ISL_744919, EPI_ISL_744929, EPI_ISL_744932, EPI_ISL_744934, EPI_ISL_744939, EPI_ISL_744945, EPI_ISL_744956, EPI_ISL_744960, EPI_ISL_744974, EPI_ISL_744992, EPI_ISL_744998, EPI_ISL_745007, EPI_ISL_745020 | | | |
| see above | Laboratoire national de santé, Microbiology, Virology | Laboratoire national de santé, Microbiology, Microbial Genomics Platform | Anke Wienecke-Baldacchino, Catherine Ragimbeau,Jessica Tapp, Fatu Djabi, Lise Pignon, Raoul Salmon, Tamir Abdelrahman |
| EPI_ISL_745494, EPI_ISL_745513, EPI_ISL_745517, EPI_ISL_745523, EPI_ISL_745542, EPI_ISL_745612, EPI_ISL_745654, EPI_ISL_745658, EPI_ISL_746207, EPI_ISL_746208, EPI_ISL_746214, EPI_ISL_746220, EPI_ISL_746221, EPI_ISL_746222, EPI_ISL_746228, EPI_ISL_746229, EPI_ISL_746233, EPI_ISL_746235, EPI_ISL_746241, EPI_ISL_746242, EPI_ISL_746246, EPI_ISL_746247, EPI_ISL_746248, EPI_ISL_746249, EPI_ISL_746255, EPI_ISL_746256 | | | |
| see above | Ginkgo Bioworks Clinical Laboratory | Utah Public Health Laboratory | Erin L. Young, Kelly Oakeson, Tara Gallagher, Michael T. Pyne, E. Susan Slechta, Melanie A. Mallory, Jeffrey B. Stevenson, Salika M. Shakir, David R. Hillyard, Malaika McKenzie-Bennett, James McGann, Jim Griffin, Keith Robison, Alex Plocik, Becky Schilling, Martha Pierson, Rebecca Littlefield, Michelle Spencer, Birgitte Simen |
| EPI_ISL_747035, EPI_ISL_747043, EPI_ISL_747064, EPI_ISL_747065, EPI_ISL_747066, EPI_ISL_747068, EPI_ISL_747118, EPI_ISL_747133, EPI_ISL_747146, EPI_ISL_747147, EPI_ISL_747148 | | | |
| see above | Respiratory Viruses Branch, Centers for Disease Control and Prevention | Respiratory Viruses Branch, Centers for Disease Control and Prevention | Queen,K., Li,Y., Tao,Y., Uehara,A., Montmayeur,A., Paden,C.R., Cook,P.W., Marine,R., Sheth,M., Wang,H., Lee,J., Tong,S. |
| EPI_ISL_747413 | Division of Emerging Infectious Diseases, Bureau of Infectious Diseases Diagnosis Control, Korea Disease Control and Prevention Agency | Division of Emerging Infectious Diseases, Bureau of Infectious Diseases Diagnosis Control, Korea Disease Control and Prevention Agency | Ae Kyung Park, Il-Hwan Kim, Heui Man Kim, Jeong-Min Kim, Namjoo Lee, Chaeyoung Lee, Sang Hee Woo, Eun-Jin Kim |
| EPI_ISL_747651, EPI_ISL_747747, EPI_ISL_748124 | Department of Virus and Microbiological Special Diagnostics, Statens Serum Institut, Copenhagen, Denmark | Albertsen Lab, Department of Chemistry and Bioscience, Aalborg University, Denmark | Danish Covid-19 Genome Consortium |
| EPI_ISL_751483, EPI_ISL_751484, EPI_ISL_751485 | CHU Purpan - Laboratoire de Virologie - Institut Fédératif de Biologie | CHU Purpan - Laboratoire de Virologie - Institut Fédératif de Biologie | Latour J., Ranger N., Dubois M., Carcenac R., Harter A., Boyer P., Tremeaux P., Izopet J. |
| EPI_ISL_751563, EPI_ISL_751583 | NE Public Health Laboratory | Genomics and Discovery, Respiratory Viruses Branch, Division of Viral Diseases, Centers for Disease Control and Prevention | Krista Queen, Yan Li, Ying Tao, Jing Zhang, Anna Uehara, Anna Montmayeur, Clinton R. Paden, Peter W. Cook,Rachel Marine, Mili Sheth, Haibin Wang, Justin Lee, Suxiang Tong |
| EPI_ISL_751605 | DE Public Health Laboratory | Genomics and Discovery, Respiratory Viruses Branch, Division of Viral Diseases, Centers for Disease Control and Prevention | Krista Queen, Yan Li, Ying Tao, Jing Zhang, Anna Uehara, Anna Montmayeur, Clinton R. Paden, Peter W. Cook,Rachel Marine, Mili Sheth, Haibin Wang, Justin Lee, Suxiang Tong |
| EPI_ISL_751608 | NE Public Health Laboratory | Genomics and Discovery, Respiratory Viruses Branch, Division of Viral Diseases, Centers for Disease Control and Prevention | Krista Queen, Yan Li, Ying Tao, Jing Zhang, Anna Uehara, Anna Montmayeur, Clinton R. Paden, Peter W. Cook,Rachel Marine, Mili Sheth, Haibin Wang, Justin Lee, Suxiang Tong |
| EPI_ISL_751621, EPI_ISL_751623, EPI_ISL_751627, EPI_ISL_751631 | DE Public Health Laboratory | Genomics and Discovery, Respiratory Viruses Branch, Division of Viral Diseases, Centers for Disease Control and Prevention | Krista Queen, Yan Li, Ying Tao, Jing Zhang, Anna Uehara, Anna Montmayeur, Clinton R. Paden, Peter W. Cook,Rachel Marine, Mili Sheth, Haibin Wang, Justin Lee, Suxiang Tong |
| EPI_ISL_751682 | NYSDOH Wadsworth Center, Virology Lab | Genomics and Discovery, Respiratory Viruses Branch, Division of Viral Diseases, Centers for Disease Control and Prevention | Krista Queen, Yan Li, Ying Tao, Jing Zhang, Anna Uehara, Anna Montmayeur, Clinton R. Paden, Peter W. Cook,Rachel Marine, Mili Sheth, Haibin Wang, Justin Lee, Suxiang Tong |
| EPI_ISL_753073, EPI_ISL_753080, EPI_ISL_753108, EPI_ISL_753109 | State Laboratories Division, Hawaii State Department of Health | State Laboratories Division, Hawaii State Department of Health | Pamela O'Brien, Sabrina Diemert, Drew Kuwazaki, Razvan Sultana, Edward Desmond |
| EPI_ISL_754137 | CHU Purpan - Laboratoire de Virologie - Institut Fédératif de Biologie | CHU Purpan - Laboratoire de Virologie - Institut Fédératif de Biologie | Latour J., Ranger N., Dubois M., Carcenac R., Harter A., Boyer P., Tremeaux P., Izopet J. |
| EPI_ISL_754972 | California Department of Public Health | California Department of Public Health | CDPH IDLB COVIDNet |
| EPI_ISL_756277 | State Laboratories Division, Hawaii State Department of Health | State Laboratories Division, Hawaii State Department of Health | Pamela O'Brien, Sabrina Diemert, Drew Kuwazaki, Razvan Sultana, Edward Desmond |
| EPI_ISL_756294 | Center for Biotechnology and Cell Therapy, São Rafael Hospital, Salvador, Brazil | Center for Biotechnology and Cell Therapy, São Rafael Hospital, Salvador, Brazil | Carolina Kymie Vasques Nonaka, Marília Miranda Franco, Tiago Gräf, Ana Verena Almeida Mendes, Renato Santana de Aguiar, Marta Giovanetti, Bruno Solano de Freitas Souza |
| EPI_ISL_756307 | The Caribbean Public Health Agency | Carrington Lab, Department of PreClinical Sciences, Faculty of Medical Sciences, The University of the West Indies | Nikita S. D. Sahadeo, Arianne Brown-Jordan, Sarah Hill, Vernie Ramkissoon, Roshan Parasram, Naresh Nandram, Avery Hinds, Jerome Foster, Stanley Giddings, Karla Georges, Marsha Ivey, Rahul Naidu, Risha Singh, SueMin Nathaniel, Rajini Haraksingh, Jaya Jayaraman, Chinna Chinnadurai, Adesh Ramsubhag, Nuno Faria, Oliver Pybus, Christopher Oura, Gabriel Escobar, Christine V. F. Carrington |

| | | | |
|---|---|---|---|
| EPI_ISL_756361 | The Caribbean Public Health Agency | Carrington Lab, Department of PreClinical Sciences, Faculty of Medical Sciences, The University of the West Indies | Nikita S. D. Sahadeo, Gabriel Escobar, Sarah Hill, Vernie Ramkissoon, Risha Singh, SueMin Nathaniel, Arianne Brown-Jordan, Naresh Nandram, Avery Hinds, Jerome Foster, Stanley Giddings, Karla Georges, Marsha Ivey, Rahul Naidu, , Rajini Haraksingh, Jaya Jayaraman, Chinna Chinnadurai, Adesh Ramsubhag, Nuno Faria, Oliver Pybus, Christopher Oura, Christine V. F. Carrington |
| EPI_ISL_759929, EPI_ISL_759930, EPI_ISL_759931, EPI_ISL_759932, EPI_ISL_759933, EPI_ISL_759934, EPI_ISL_759935, EPI_ISL_759936, EPI_ISL_759937, EPI_ISL_759938, EPI_ISL_759939, EPI_ISL_759940, EPI_ISL_759941, EPI_ISL_759942, EPI_ISL_759943, EPI_ISL_759944 | | | |
| see above | Department of Virology and Immunology, University of Helsinki and Helsinki University Hospital, Huslab Finland | Department of Virology, Faculty of Medicine, University of Helsinki, Helsinki, Finland | Teemu Smura, Ravi Kant, Phuoc Truong, Hussein Alburkat, Hannimari Kallio-Kokko, Jenni Virtanen, Maija Suvanto, Sari Hannula, Harri Kangas, Pekka Ellonen, Olli Vapalahti |
| EPI_ISL_760084, EPI_ISL_760085, EPI_ISL_760086, EPI_ISL_760087, EPI_ISL_760088, EPI_ISL_760089, EPI_ISL_760090, EPI_ISL_760091, EPI_ISL_760092, EPI_ISL_760093, EPI_ISL_760094, EPI_ISL_760095, EPI_ISL_760098, EPI_ISL_760099, EPI_ISL_760100, EPI_ISL_760101, EPI_ISL_760102, EPI_ISL_760103, EPI_ISL_760181, EPI_ISL_760188, EPI_ISL_760201, EPI_ISL_760202, EPI_ISL_760218, EPI_ISL_760221, EPI_ISL_760222 | | | |
| see above | Division of Emerging Infectious Diseases, Bureau of Infectious Diseases Diagnosis Control, Korea Disease Control and Prevention Agency | Division of Emerging Infectious Diseases, Bureau of Infectious Diseases Diagnosis Control, Korea Disease Control and Prevention Agency | Ae Kyung Park, Il-Hwan Kim, Heui Man Kim, Jeong-Min Kim, Namjoo Lee, Chaeyoung Lee, Sang Hee Woo, Eun-Jin Kim |
| EPI_ISL_763069, EPI_ISL_763071, EPI_ISL_763072, EPI_ISL_763073, EPI_ISL_763094, EPI_ISL_763096, EPI_ISL_763097, EPI_ISL_763098, EPI_ISL_763318, EPI_ISL_763320, EPI_ISL_763322, EPI_ISL_763324, EPI_ISL_763325, EPI_ISL_763326, EPI_ISL_763327, EPI_ISL_763328, EPI_ISL_763329, EPI_ISL_763330 | | | |
| see above | Istituto Zooprofilattico Sperimentale dell' Umbria e delle Marche -Togo Rosati | Istituto Superiore di Sanità | Massimo Biagetti , Monica Giammarioli, Luca De Sabato, Gabriele Vaccari, Ilaria Di Bartolo, Giovanni Ianiro |
| EPI_ISL_764402, EPI_ISL_764441, EPI_ISL_764442, EPI_ISL_764443, EPI_ISL_764444 | Regional Virus Laboratory, Belfast Health and Social Care Trust | COVID-19 Genomics UK (COG-UK) Consortium | Conall McCaughey, James McKenna, Tanya Curran, Susan Feeney, Alison Watt, Ciara Cox, Mairead Connor, Zoltan Molnar, David Simpson, Derek Fairley |
| EPI_ISL_765660, EPI_ISL_765772, EPI_ISL_765773, EPI_ISL_765774, EPI_ISL_765776, EPI_ISL_765777 | Massachusetts General Hospital | Infectious Disease Program, Broad Institute of Harvard and MIT | Lemieux,J.E., Siddle,K.J., Shaw,B., Adams,G., Pierce,V., Turbett,S., Anahtar,M., Branda,J., Slater,D., Harris,J., Lin,A.E., Gladden-Young,A., Lagerborg,K., Rudy,M., DeRuff,K., Carter,A., Normandin,E., Bauer,M., Reilly,S., Tomkins-Tinch,C., Loreth,C., Chaluvadi,S., Neumann,A., Cusick,C., Chapman,S.B., Gnirke,A., Flowers,K., Cerrato,F., Birren,B.W., Gallagher,G., Smole,S., Park,D.J., MacInnis,B.L., Ryan,E., LaRocque,R., Rosenberg,E. and Sabeti,P.C. |
| EPI_ISL_765959 | Worobey Lab, Department of Ecology and Evolutionary Biology, University of Arizona | Worobey Lab, Department of Ecology and Evolutionary Biology, University of Arizona | Brendan Larsen, Grace Quirk, Thomas Watts, David Baltrus, Michael Worobey |
| EPI_ISL_766616 | Klinisk mikrobiologi | The Public Health Agency of Sweden | Department of Microbiology, The Public Health Agency of Sweden |
| EPI_ISL_770018, EPI_ISL_770019 | Laboratorio Cenahce | Inciensa, Instituto Costarricense de Investigación y Enseñanza en Nutrición y Salud | Francisco Duarte, Hebleen Porras, Claudio Soto-Garita, Estela Cordero, Adriana Godínez, Melany Calderón & Andrés Feoli-Grant |
| EPI_ISL_775323 | Foerde Hospital, Department of Microbiology | Norwegian Institute of Public Health, Department of Virology | Kathrine Stene-Johansen, Kamilla Heddeland Instefjord, Hilde Elshaug, Atiya R Ali,Marie Paulsen Madsen, Rasmus Riis Kopperud, Hilde Vollan, Karoline Bragstad, Olav Hungnes |
| EPI_ISL_775331 | Vestfold Hospital, Toensberg Department of Microbiology | Norwegian Institute of Public Health, Department of Virology | Kathrine Stene-Johansen, Kamilla Heddeland Instefjord, Hilde Elshaug, Atiya R Ali,Marie Paulsen Madsen, Rasmus Riis Kopperud, Hilde Vollan, Karoline Bragstad, Olav Hungnes |
| EPI_ISL_775344 | Akershus University Hospital, Department for Microbiology and Infectious Disease Control | Norwegian Institute of Public Health, Department of Virology | Kathrine Stene-Johansen, Kamilla Heddeland Instefjord, Hilde Elshaug, Atiya R Ali,Marie Paulsen Madsen, Rasmus Riis Kopperud, Hilde Vollan, Karoline Bragstad, Olav Hungnes |
| EPI_ISL_775452, EPI_ISL_775453, EPI_ISL_775454, EPI_ISL_775456 | University Hospital of Northern Norway, Department for Microbiology and Infectious Disease Control | Norwegian Institute of Public Health, Department of Virology | Kathrine Stene-Johansen, Kamilla Heddeland Instefjord, Hilde Elshaug, Atiya R Ali,Marie Paulsen Madsen, Rasmus Riis Kopperud, Hilde Vollan, Karoline Bragstad, Olav Hungnes |
| EPI_ISL_775482, EPI_ISL_775483 | Vestfold Hospital, Toensberg Department of Microbiology | Norwegian Institute of Public Health, Department of Virology | Kathrine Stene-Johansen, Kamilla Heddeland Instefjord, Hilde Elshaug, Atiya R Ali,Marie Paulsen Madsen, Rasmus Riis Kopperud, Hilde Vollan, Karoline Bragstad, Olav Hungnes |
| EPI_ISL_775532 | Klinisk mikrobiologi, Viruslab | The Public Health Agency of Sweden | Department of Microbiology, The Public Health Agency of Sweden |
| EPI_ISL_776879, EPI_ISL_776880, EPI_ISL_776881, EPI_ISL_776882, EPI_ISL_776883, EPI_ISL_776884, EPI_ISL_776885, EPI_ISL_776886, EPI_ISL_776887, EPI_ISL_776888, EPI_ISL_776889, EPI_ISL_776890, EPI_ISL_776891, EPI_ISL_776892, EPI_ISL_776893, EPI_ISL_776894, EPI_ISL_776895, EPI_ISL_776896, EPI_ISL_776897, EPI_ISL_776898, EPI_ISL_776899, EPI_ISL_776900, EPI_ISL_776901, EPI_ISL_776902, EPI_ISL_776903, EPI_ISL_776904, EPI_ISL_776905, EPI_ISL_776906, EPI_ISL_776907, EPI_ISL_776908, EPI_ISL_776909, EPI_ISL_776910, EPI_ISL_776911, EPI_ISL_776912, EPI_ISL_776913, EPI_ISL_776914, EPI_ISL_776915, EPI_ISL_776916, EPI_ISL_776917, EPI_ISL_776918, EPI_ISL_776919, EPI_ISL_776920, EPI_ISL_776921, EPI_ISL_776922, EPI_ISL_776923, EPI_ISL_776924, EPI_ISL_776925, EPI_ISL_776926, EPI_ISL_776927, EPI_ISL_776928, EPI_ISL_776929, EPI_ISL_776930, EPI_ISL_776931, EPI_ISL_776932, EPI_ISL_776933, EPI_ISL_776934, EPI_ISL_776935, EPI_ISL_776936, EPI_ISL_776937, EPI_ISL_776938, EPI_ISL_776939, EPI_ISL_776940, EPI_ISL_776941, EPI_ISL_776942, EPI_ISL_776943, EPI_ISL_776944, EPI_ISL_776945, EPI_ISL_776946, EPI_ISL_776947, EPI_ISL_776948, EPI_ISL_776949, EPI_ISL_776950, EPI_ISL_776951, EPI_ISL_776952, EPI_ISL_776953, EPI_ISL_776954, EPI_ISL_776955, EPI_ISL_776956, EPI_ISL_776957, EPI_ISL_776958, EPI_ISL_776959 | | | |
| see above | Istituto Zooprofilattico Sperimentale del Mezzogiorno | TIGEM | Patrizia Annunziata, Andrea Ballabio, Valentina Bouche, Davide Cacchiarelli (CorrespAuthor), Pellegrino Cerino, Chiara Colantuono, Lucio Di Filippo, Antonio Grimaldi, Antonio Limone, Gabriella Loconte, Anna Manfredi, Francesco Panariello, Biancamaria Pierri, Marcello Salvi, Lucia Vassallo |
| EPI_ISL_779961, EPI_ISL_780021 | Hospital General Universitario Gregorio Marañón | SeqCOVID-SPAIN consortium/IBV(CSIC) | Darío García de Viedma, Laura Pérez-Lago, Marta Herranz, Jon Sicilia, Julia Suárez, Pilar Catalán, Patricia Muñoz and SeqCOVID-SPAIN consortium |
| EPI_ISL_780402, EPI_ISL_780403 | Bermuda Government Molecular Diagnostics Laboratory (MDL) | Respiratory Virus Unit, National Infection Service, Public Health England | PHE Covid Sequencing Team, Dr Carika Weldon (Bermuda), Dr Ayoola Oyinloye (Bermuda) |
| EPI_ISL_780414 | Hospital General Universitario Gregorio Marañón | SeqCOVID-SPAIN consortium/IBV(CSIC) | Darío García de Viedma, Laura Pérez-Lago, Marta Herranz, Jon Sicilia, Julia Suárez, Pilar Catalán, Patricia Muñoz and SeqCOVID-SPAIN consortium |
| EPI_ISL_785300, EPI_ISL_785301, EPI_ISL_785302, EPI_ISL_785303, EPI_ISL_785304, EPI_ISL_785305, EPI_ISL_785306, EPI_ISL_785307, EPI_ISL_785308, EPI_ISL_785309, EPI_ISL_785312, EPI_ISL_785313, EPI_ISL_785314, EPI_ISL_785315, EPI_ISL_785316, EPI_ISL_785317, EPI_ISL_785318, EPI_ISL_785319, EPI_ISL_785320, EPI_ISL_785321, EPI_ISL_785707, EPI_ISL_785708, EPI_ISL_785710, EPI_ISL_785711, EPI_ISL_785712, EPI_ISL_785713, EPI_ISL_785714, EPI_ISL_785715, EPI_ISL_785716, EPI_ISL_785717, EPI_ISL_785719, EPI_ISL_785720, EPI_ISL_785721, EPI_ISL_785723, EPI_ISL_785724, EPI_ISL_785725, EPI_ISL_785726, EPI_ISL_785727, EPI_ISL_785728, EPI_ISL_785729, EPI_ISL_785730, EPI_ISL_785731, EPI_ISL_785732, EPI_ISL_785733, EPI_ISL_785734, EPI_ISL_785735, EPI_ISL_785736, EPI_ISL_785737, EPI_ISL_785738, EPI_ISL_785739, EPI_ISL_785740, EPI_ISL_785742, EPI_ISL_785743, EPI_ISL_785744, EPI_ISL_785745, EPI_ISL_785746, EPI_ISL_785747, EPI_ISL_785748, EPI_ISL_785749, EPI_ISL_785750, EPI_ISL_785751, EPI_ISL_785752, EPI_ISL_785753, EPI_ISL_785754, EPI_ISL_785755, EPI_ISL_785756, EPI_ISL_785757, EPI_ISL_785758, EPI_ISL_785759, EPI_ISL_785760, EPI_ISL_785761, EPI_ISL_785762, EPI_ISL_785763, EPI_ISL_785764, EPI_ISL_785765, EPI_ISL_785766, EPI_ISL_785767, EPI_ISL_785768, EPI_ISL_785769, EPI_ISL_785772, EPI_ISL_785773, EPI_ISL_785774, EPI_ISL_785775, EPI_ISL_785776, EPI_ISL_785777, EPI_ISL_785778, EPI_ISL_785779, EPI_ISL_785781, EPI_ISL_785782, EPI_ISL_785783, EPI_ISL_785784, EPI_ISL_785785, EPI_ISL_785786, EPI_ISL_785787, EPI_ISL_785788, EPI_ISL_785789, EPI_ISL_785790, EPI_ISL_785791, EPI_ISL_785792, EPI_ISL_785793, EPI_ISL_785794, EPI_ISL_785795, EPI_ISL_785796, EPI_ISL_785797, EPI_ISL_785798, EPI_ISL_785799, EPI_ISL_785800, EPI_ISL_785801, EPI_ISL_785802, EPI_ISL_785803, EPI_ISL_785804, EPI_ISL_785805, EPI_ISL_785806, EPI_ISL_785808, EPI_ISL_785809, EPI_ISL_785810, EPI_ISL_785811, EPI_ISL_785813, EPI_ISL_785814, EPI_ISL_785815, EPI_ISL_785816, EPI_ISL_785817, EPI_ISL_785818, EPI_ISL_785819, EPI_ISL_785822, EPI_ISL_785823, EPI_ISL_785825, EPI_ISL_785826, EPI_ISL_785827, EPI_ISL_785828, EPI_ISL_785829, EPI_ISL_785830, EPI_ISL_785831, EPI_ISL_785833, EPI_ISL_785834, EPI_ISL_785837, EPI_ISL_785838, EPI_ISL_785839, EPI_ISL_785840, EPI_ISL_785841, EPI_ISL_785842, EPI_ISL_785843, EPI_ISL_785844, EPI_ISL_785845, EPI_ISL_785847, EPI_ISL_785850, EPI_ISL_785851, EPI_ISL_785852, EPI_ISL_785853, EPI_ISL_785855, EPI_ISL_785856, EPI_ISL_785857, EPI_ISL_785859, EPI_ISL_785861, EPI_ISL_785863, EPI_ISL_785864, EPI_ISL_785865, EPI_ISL_785866, EPI_ISL_785867, EPI_ISL_785868, EPI_ISL_785872, EPI_ISL_785873, EPI_ISL_785874, EPI_ISL_785875, EPI_ISL_785876, EPI_ISL_785877, EPI_ISL_785878, EPI_ISL_785879, EPI_ISL_785880, EPI_ISL_785881, EPI_ISL_785882, EPI_ISL_785884, EPI_ISL_785885, EPI_ISL_785886, EPI_ISL_785887, EPI_ISL_785888, EPI_ISL_785889, EPI_ISL_785890, EPI_ISL_785891, EPI_ISL_785892, EPI_ISL_785893, EPI_ISL_785895, EPI_ISL_785896, EPI_ISL_785897, EPI_ISL_785898, EPI_ISL_785909, EPI_ISL_785910, EPI_ISL_785915, EPI_ISL_785921, EPI_ISL_785922, EPI_ISL_785933, EPI_ISL_785946, EPI_ISL_785957, EPI_ISL_785958, EPI_ISL_785969, EPI_ISL_785970, EPI_ISL_785980, EPI_ISL_785981, EPI_ISL_785982, EPI_ISL_785992, EPI_ISL_785993, EPI_ISL_785995, EPI_ISL_785996, EPI_ISL_786007, EPI_ISL_786008, EPI_ISL_786019, EPI_ISL_786020, EPI_ISL_786031, EPI_ISL_786043, EPI_ISL_786055, EPI_ISL_786067, EPI_ISL_786079, EPI_ISL_787027, EPI_ISL_787028, EPI_ISL_787029, EPI_ISL_787030, EPI_ISL_787031, EPI_ISL_787032, EPI_ISL_787034, EPI_ISL_787035, EPI_ISL_787036, EPI_ISL_787037, EPI_ISL_787039, EPI_ISL_787040, EPI_ISL_787041, EPI_ISL_787042, EPI_ISL_787043, EPI_ISL_787044, EPI_ISL_787046, EPI_ISL_787047, EPI_ISL_787048, EPI_ISL_787049, EPI_ISL_787051, EPI_ISL_787052, EPI_ISL_787053, EPI_ISL_787054, EPI_ISL_787055, EPI_ISL_787056, EPI_ISL_787058, EPI_ISL_787059, EPI_ISL_787060, EPI_ISL_787061, EPI_ISL_787063, EPI_ISL_787065, EPI_ISL_787066, EPI_ISL_787068, EPI_ISL_787070, EPI_ISL_787071, EPI_ISL_787072, EPI_ISL_787073, EPI_ISL_787075, EPI_ISL_787077, EPI_ISL_787078, EPI_ISL_787079, EPI_ISL_787080, EPI_ISL_787082, EPI_ISL_787083, EPI_ISL_787084, EPI_ISL_787085, EPI_ISL_787088, EPI_ISL_787089, EPI_ISL_787090, EPI_ISL_787092, EPI_ISL_787094, EPI_ISL_787095, EPI_ISL_787096, EPI_ISL_787097 | | | |
| see above | Houston Methodist Hospital | Houston Methodist Hospital | S. Wesley Long, Randall J. Olsen, Paul A. Christensen, David W. Bernard, James J. Davis, Maulik Shukla, Marcus Nguyen, Matthew Ojeda Saavedra, Prasanti Yerramilli, Layne Pruitt, Sishir Subedi, Heather Hendrickson, and James M. Musser |
| EPI_ISL_791352, EPI_ISL_791358 | Johns Hopkins Hospital Department of Pathology | Johns Hopkins Hospital Department of Pathology | C. Paul Morris, Chun Huai Luo, Heba H. Mostafa |
| EPI_ISL_792645 | LACEN-PR | Laboratory of Respiratory Viruses and Measles, Oswaldo Cruz Institute, FIOCRUZ | Paola Resende, Luciana Appolinario, Fernando Motta, Anna Carolina Paixao, Ana Carolina Mendonca, Maria do Carmo Debur, Irina Nastassja Riediger, Marilda Siqueira |
| EPI_ISL_794068, EPI_ISL_794069, EPI_ISL_794070, EPI_ISL_794071, EPI_ISL_794072, EPI_ISL_794073, EPI_ISL_794074, EPI_ISL_794075, EPI_ISL_794076, EPI_ISL_794077 | Wadsworth Center, New York State Department.of Health | Wadsworth Center, New York State Department.of Health | Kirsten St. George, Daryl M. Lamson, Alexis Russel, Matthew Shudt, Melissa A Leisner, Jonathan Plitnick, Navjot Singh, John Kelly, Sara Griesemer, Erasmus Schneider, Erica Lasek-Nesselquist |
| EPI_ISL_796062, EPI_ISL_796063 | Institute of Virology, University of Cologne | Institute of Virology, University of Cologne | Saleta Sierra, Gibran Rubio, Zevanya Tessalonica, Dominik Aschenmeier, Eva Heger, Elena Knops, Rolf Kaiser, Maximilian Damagnez, Andreas Walker, |

| | | | |
|---|---|---|---|
| | | | Jörg Timm, Alexander Dilthey, Martin Däumer, Alex Thielen |
| EPI_ISL_796677 | Dept. of Medical Microbiology, Stavanger University Hospital, Helse Stavanger HF | Norwegian Institute of Public Health, Department of Virology | Kathrine Stene-Johansen, Kamilla Heddeland Instefjord, Hilde Elshaug, Atiya R Ali,Marie Paulsen Madsen, Rasmus Riis Kopperud, Hilde Vollan, Karoline Bragstad, Olav Hungnes |
| EPI_ISL_801393, EPI_ISL_801394 | Laboratorio de Ecologia de Doencas Transmissiveis na Amazonia, Instituto Leonidas e Maria Deane - Fiocruz Amazonia | Laboratorio de Ecologia de Doencas Transmissiveis na Amazonia, Instituto Leonidas e Maria Deane - Fiocruz Amazonia | Valdinete Nascimento, Victor Souza, André Corado, Fernanda Nascimento, George Silva, Ágatha Costa, Debora Duarte, Luciana Gonçalves, Maria Júlia Brandão, Michele Jesus, Felipe Naveca |
| EPI_ISL_801413, EPI_ISL_801487 | Dutch COVID-19 response team | Erasmus Medical Center | Bas Oude Munnink, Reina Sikkema, David Nieuwenhuijse, Irina Chestakova, Anne van der Linden, Marjan Boter, Emmanuelle Munger, Corine GeurtsvanKessel, Annemiek van der Eijk, Richard Molenkamp, Marion Koopmans, on behalf of the Dutch national COVID-19 response team. |
| EPI_ISL_803271, EPI_ISL_803416, EPI_ISL_803417, EPI_ISL_803418, EPI_ISL_803419, EPI_ISL_803420, EPI_ISL_803421, EPI_ISL_803422, EPI_ISL_803423, EPI_ISL_803424, EPI_ISL_803425, EPI_ISL_803461, EPI_ISL_803462, EPI_ISL_803463, EPI_ISL_803464, EPI_ISL_803465, EPI_ISL_803466, EPI_ISL_803467, EPI_ISL_803468, EPI_ISL_803469, EPI_ISL_803470, EPI_ISL_803471 | | | |
| see above | Wisconsin State Laboratory of Hygiene Communicable Disease Division | Wisconsin State Laboratory of Hygiene Communicable Disease Division | Kelsey R. Florek, Abigail C. Shockey |
| EPI_ISL_804471, EPI_ISL_804476, EPI_ISL_804531, EPI_ISL_804532, EPI_ISL_804533, EPI_ISL_804534, EPI_ISL_804562, EPI_ISL_804563, EPI_ISL_804564, EPI_ISL_804565, EPI_ISL_804566, EPI_ISL_804567 | | | |
| see above | MEPHI, Aix Marseille University | MEPHI, Aix Marseille University | Anthony LEVASSEUR |
| EPI_ISL_804664 | Michigan Department of Health and Human Services, Bureau of Laboratories | Michigan Department of Health and Human Services, Bureau of Laboratories | Blankenship HM, Riner D, Soehnlen MK |
| EPI_ISL_812136, EPI_ISL_812140, EPI_ISL_812169, EPI_ISL_812170, EPI_ISL_812171, EPI_ISL_812172, EPI_ISL_812173, EPI_ISL_812174, EPI_ISL_812175, EPI_ISL_812176, EPI_ISL_812177, EPI_ISL_812178, EPI_ISL_812179 | | | |
| see above | GA Department of Public Health Laboratory | Pathogen Discovery, Respiratory Viruses Branch, Division of Viral Diseases, Centers for Disease Control and Prevention | Yan Li, Ying Tao, Anna Montmayeur, Jing Zhang, Brian Lynch, Krista Queen, Anna Uehara, Rachel Marine, Peter Cook, Clinton R. Paden, Haibin Wang, Suxiang Tong |
| EPI_ISL_812973, EPI_ISL_812974, EPI_ISL_812975, EPI_ISL_812976, EPI_ISL_812977, EPI_ISL_812978, EPI_ISL_812979, EPI_ISL_812980, EPI_ISL_812981, EPI_ISL_812982, EPI_ISL_812983, EPI_ISL_812984, EPI_ISL_812985, EPI_ISL_812986, EPI_ISL_812987, EPI_ISL_812988, EPI_ISL_812989, EPI_ISL_812990 | | | |
| see above | University of Birmingham | COVID-19 Genomics UK (COG-UK) Consortium | Institute of Microbiology, University of Birmingham: Claire McMurray, Joanne Stockton, Samuel Nicholls, Radoslaw Poplawski, Will Rowe, Josh Quick, Nicholas Loman. University of Birmingham Testing Laboratory: Celina M Whalley, Andrew Bosworth, Charlotte Poxon, Kasun Wanigasooriya, Oliver Pickles, Mike Kidd, Alex Richter, Andrew D Beggs PHE Heartlands Lab: Husam Osman, Andrew Bosworth. Queen Elizabeth Hospital: Anna Casey |
| EPI_ISL_814681 | Virology Department, Royal Infirmary of Edinburgh, NHS Lothian / School of Biological Sciences, University of Edinburgh / Institute of Genetics and Molecular Medicine, University of Edinburgh | COVID-19 Genomics UK (COG-UK) Consortium | McHugh M, Dewar R, Rooke S, Gallagher M, Balcaza C, O'Toole Á, Scher E, Hill V, McCrone JT, Colquhoun R, Yu X, Jackson B, Rambaut A, Williams TC, Templeton K |
| EPI_ISL_816132, EPI_ISL_816133, EPI_ISL_816134 | Department of Virus and Microbiological Special Diagnostics, Statens Serum Institut, Copenhagen, Denmark | Albertsen Lab, Department of Chemistry and Bioscience, Aalborg University, Denmark | Danish Covid-19 Genome Consortium |
| EPI_ISL_816317 | Virology Department, Sheffield Teaching Hospitals NHS Foundation Trust/Department of Infection, Immunity and Cardiovascular Disease, The Medical School, University of Sheffield | COVID-19 Genomics UK (COG-UK) Consortium | Thushan de Silva, Matthew Parker, Nikki Smith, Adri Angyal, Rebecca Brown, Luke Green, Rachel Tucker, Paul Parsons, Danielle Groves, Katie Johnson, Laura Carrilero, Alex Keeley, Dave Partridge, Matthew Wyles, Benjamin Lindsey, Mehmet Yavuz, Mohammad Raza, Cariad Evans |
| EPI_ISL_819203 | ZOTZ KLIMAS MVZ Düsseldorf-Centrum GbR ÜBAG für Labormedizin, Genetik, Zytologie, Pathologie | Center of Medical Microbiology, Virology, and Hospital Hygiene, University of Duesseldorf | Maximilian Damagnez, Alexander Dilthey, Ashley-Jane Duplessis, Patrick Finzer, Katrin Hoffmann, Torsten Houwaart, Lisanna Hülse, Malte Kohns Vasconcelos, Marek Korencak, Nadine Lübke, Jessica Nicolai, Klaus Pfeffer, Daniel Strelow, Jörg Timm, Andreas Walker, Tobias Wienemann, Rainer Zotz |
| EPI_ISL_819452, EPI_ISL_819453 | Northumbria University / South Tees Hospitals NHS Foundation Trust / North Cumbria Integrated Care NHS Foundation Trust / North Tees and Hartlepool NHS Foundation Trust / Newcastle Hospitals NHS Foundation Trust | COVID-19 Genomics UK (COG-UK) Consortium | Darren L Smith,Andrew Nelson,Matthew Bashton,Greg R Young,Joshua Loh,John Allan,Mohammad A Tariq,Giles S Holt,Gary Black,Wen C Yew,Lynn Dover,Paul Baker,Steve Liggett,Sarah Essex,Jane Greenaway,Debra Padgett,Clive Graham,Garren Scott,Edward Barton,Emma Swindells,Brendan Payne,Jennifer Collins,Yusri Taha,Gary Eltringham |
| EPI_ISL_825339, EPI_ISL_825340, EPI_ISL_825341, EPI_ISL_825342, EPI_ISL_825343, EPI_ISL_825344, EPI_ISL_825345, EPI_ISL_825346 | Hospital Universitari Vall d'Hebron - Vall d'Hebron Institut de Rercerca | Hospital Universitari Vall d'Hebron | Cristina Andrés, Maria Piñana, Josep F Abril, Damir Garcia-Cehic, Ariadna Rando, Juliana Esperalba, Maria Gema Codina, Carla Castillo, Maria Carmen Martin, Tomàs Pumarola, Josep Quer, Andrés Antón |
| EPI_ISL_825589, EPI_ISL_825590 | Respiratory Virus Unit, National Infection Service, Public Health England | COVID-19 Genomics UK (COG-UK) Consortium | PHE Covid Sequencing Team |
| EPI_ISL_826761, EPI_ISL_826769, EPI_ISL_826770, EPI_ISL_826771, EPI_ISL_826772, EPI_ISL_826773, EPI_ISL_826775, EPI_ISL_826776, EPI_ISL_826793, EPI_ISL_826794, EPI_ISL_826795, EPI_ISL_826796, EPI_ISL_826797, EPI_ISL_826799, EPI_ISL_826803, EPI_ISL_826888, EPI_ISL_826917, EPI_ISL_827015, EPI_ISL_827046, EPI_ISL_827058, EPI_ISL_827059, EPI_ISL_827108, EPI_ISL_827111, EPI_ISL_827112, EPI_ISL_827113, EPI_ISL_827114, EPI_ISL_827115, EPI_ISL_827116, EPI_ISL_827117, EPI_ISL_827118, EPI_ISL_827119, EPI_ISL_827144, EPI_ISL_827151, EPI_ISL_827200, EPI_ISL_827209, EPI_ISL_827217, EPI_ISL_827218, EPI_ISL_827219, EPI_ISL_827220, EPI_ISL_827221, EPI_ISL_827222, EPI_ISL_827223, EPI_ISL_827224, EPI_ISL_827225, EPI_ISL_827226, EPI_ISL_827272, EPI_ISL_827320, EPI_ISL_827380, EPI_ISL_827381, EPI_ISL_827384, EPI_ISL_827386, EPI_ISL_827387, EPI_ISL_827388, EPI_ISL_827389, EPI_ISL_827390, EPI_ISL_827417, EPI_ISL_827497, EPI_ISL_827498, EPI_ISL_827500, EPI_ISL_827501, EPI_ISL_827502, EPI_ISL_827503, EPI_ISL_827506, EPI_ISL_827515, EPI_ISL_827516, EPI_ISL_827517, EPI_ISL_827518, EPI_ISL_827519, EPI_ISL_827520, EPI_ISL_827521, EPI_ISL_827525 | | | |
| see above | deCODE genetics | deCODE genetics | Daniel F Gudbjartsson; Agnar Helgason; Hakon Jonsson; Olafur T Magnusson; Pall Melsted; Gudmundur L Norddahl; Jona Saemundsdottir; Asgeir Sigurdsson; Patrick Sulem; Arna B Agustsdottir; Hannes Eggertsson; Berglind Eiriksdottir; Run Fridriksdottir; Elisabet E Gardarsdottir; Gudmundur Georgsson; Olafia S Gretarsdottir; Kjartan R Gudmundsson; Thora R Gunnarsdottir; Arnaldur Gylfason; Hilma Holm; Brynjar O Jensson; Aslaug Jonasdottir; Kamilla S Josefsdottir; Thordur Kristjansson; Droplaug N Magnusdottir; Solvi Rognvaldsson; Louise le Roux; Gudrun Sigmundsdottir; Gardar Sveinbjornsson; Kristin E Sveinsdottir; Maney Sveinsdottir; Emil A Thorarensen; Bjarni Thorbjornsson; Gisli Masson; Ingileif Jonsdottir; Alma Moller; Thorolfur Gudnason; Karl G Kristinsson; Unnur Thorsteinsdottir; Kari Stefansson |
| EPI_ISL_827541, EPI_ISL_827542, EPI_ISL_827543, EPI_ISL_827544 | The National University Hospital of Iceland | deCODE genetics | Daniel F Gudbjartsson; Agnar Helgason; Hakon Jonsson; Olafur T Magnusson; Pall Melsted; Gudmundur L Norddahl; Jona Saemundsdottir; Asgeir Sigurdsson; Patrick Sulem; Arna B Agustsdottir; Hannes Eggertsson; Berglind Eiriksdottir; Run Fridriksdottir; Elisabet E Gardarsdottir; Gudmundur Georgsson; Olafia S Gretarsdottir; Kjartan R Gudmundsson; Thora R Gunnarsdottir; Arnaldur Gylfason; Hilma Holm; Brynjar O Jensson; Aslaug Jonasdottir; Kamilla S Josefsdottir; Thordur Kristjansson; Droplaug N Magnusdottir; Solvi Rognvaldsson; Louise le Roux; Gudrun Sigmundsdottir; Gardar Sveinbjornsson; Kristin E Sveinsdottir; Maney Sveinsdottir; Emil A Thorarensen; Bjarni Thorbjornsson; Gisli Masson; Ingileif Jonsdottir; Alma Moller; Thorolfur Gudnason; Karl G Kristinsson; Unnur Thorsteinsdottir; Kari Stefansson |
| EPI_ISL_827579, EPI_ISL_827601, EPI_ISL_827620, EPI_ISL_827629, EPI_ISL_827647, EPI_ISL_827671, EPI_ISL_827672, EPI_ISL_827673, EPI_ISL_827674, EPI_ISL_827675, EPI_ISL_827677, EPI_ISL_827678, EPI_ISL_827691, EPI_ISL_827698, EPI_ISL_827699, EPI_ISL_827700, EPI_ISL_827701, EPI_ISL_827702, EPI_ISL_827703 | | | |
| see above | deCODE genetics | deCODE genetics | Daniel F Gudbjartsson; Agnar Helgason; Hakon Jonsson; Olafur T Magnusson; Pall Melsted; Gudmundur L Norddahl; Jona Saemundsdottir; Asgeir Sigurdsson; Patrick Sulem; Arna B Agustsdottir; Hannes Eggertsson; Berglind Eiriksdottir; Run Fridriksdottir; Elisabet E Gardarsdottir; Gudmundur Georgsson; Olafia S Gretarsdottir; Kjartan R Gudmundsson; Thora R Gunnarsdottir; Arnaldur Gylfason; Hilma Holm; Brynjar O Jensson; Aslaug Jonasdottir; Kamilla S Josefsdottir; Thordur Kristjansson; Droplaug N Magnusdottir; Solvi Rognvaldsson; Louise le Roux; Gudrun Sigmundsdottir; Gardar Sveinbjornsson; Kristin E Sveinsdottir; Maney Sveinsdottir; Emil A Thorarensen; Bjarni Thorbjornsson; Gisli Masson; Ingileif Jonsdottir; Alma Moller; Thorolfur Gudnason; Karl G Kristinsson; Unnur Thorsteinsdottir; Kari Stefansson |
| EPI_ISL_827736 | The National University Hospital of Iceland | deCODE genetics | Daniel F Gudbjartsson; Agnar Helgason; Hakon Jonsson; Olafur T Magnusson; Pall Melsted; Gudmundur L Norddahl; Jona Saemundsdottir; Asgeir Sigurdsson; Patrick Sulem; Arna B Agustsdottir; Hannes Eggertsson; Berglind Eiriksdottir; Run Fridriksdottir; Elisabet E Gardarsdottir; Gudmundur Georgsson; Olafia S Gretarsdottir; Kjartan R Gudmundsson; Thora R Gunnarsdottir; Arnaldur Gylfason; Hilma Holm; Brynjar O Jensson; Aslaug Jonasdottir; Kamilla S Josefsdottir; Thordur Kristjansson; Droplaug N Magnusdottir; Solvi Rognvaldsson; Louise le Roux; Gudrun Sigmundsdottir; Gardar Sveinbjornsson; Kristin E Sveinsdottir; Maney Sveinsdottir; Emil A Thorarensen; Bjarni Thorbjornsson; Gisli Masson; Ingileif Jonsdottir; Alma Moller; Thorolfur Gudnason; Karl G Kristinsson; Unnur Thorsteinsdottir; Kari Stefansson |

| | | | |
|---|---|---|---|
| EPI_ISL_827847, EPI_ISL_827849, EPI_ISL_827850 | deCODE genetics | deCODE genetics | Daniel F Gudbjartsson; Agnar Helgason; Hakon Jonsson; Olafur T Magnusson; Pall Melsted; Gudmundur L Norddahl; Jona Saemundsdottir; Asgeir Sigurdsson; Patrick Sulem; Arna B Agustsdottir; Hannes Eggertsson; Berglind Eiriksdottir; Run Fridriksdottir; Elisabet E Gardarsdottir; Gudmundur Georgsson; Olafia S Gretarsdottir; Kjartan R Gudmundsson; Thora R Gunnarsdottir; Arnaldur Gylfason; Hilma Holm; Brynjar O Jensson; Aslaug Jonasdottir; Kamilla S Josefsdottir; Thordur Kristjansson; Droplaug N Magnusdottir; Solvi Rognvaldsson; Louise le Roux; Gudrun Sigmundsdottir; Gardar Sveinbjornsson; Kristin E Sveinsdottir; Maney Sveinsdottir; Emil A Thorarensen; Bjarni Thorbjornsson; Gisli Masson; Ingileif Jonsdottir; Alma Moller; Thorolfur Gudnason; Karl G Kristinsson; Unnur Thorsteinsdottir; Kari Stefansson |
| EPI_ISL_827859, EPI_ISL_827860, EPI_ISL_827863, EPI_ISL_827865, EPI_ISL_827866, EPI_ISL_827867, EPI_ISL_827868, EPI_ISL_827869, EPI_ISL_827870 | The National University Hospital of Iceland | deCODE genetics | Daniel F Gudbjartsson; Agnar Helgason; Hakon Jonsson; Olafur T Magnusson; Pall Melsted; Gudmundur L Norddahl; Jona Saemundsdottir; Asgeir Sigurdsson; Patrick Sulem; Arna B Agustsdottir; Hannes Eggertsson; Berglind Eiriksdottir; Run Fridriksdottir; Elisabet E Gardarsdottir; Gudmundur Georgsson; Olafia S Gretarsdottir; Kjartan R Gudmundsson; Thora R Gunnarsdottir; Arnaldur Gylfason; Hilma Holm; Brynjar O Jensson; Aslaug Jonasdottir; Kamilla S Josefsdottir; Thordur Kristjansson; Droplaug N Magnusdottir; Solvi Rognvaldsson; Louise le Roux; Gudrun Sigmundsdottir; Gardar Sveinbjornsson; Kristin E Sveinsdottir; Maney Sveinsdottir; Emil A Thorarensen; Bjarni Thorbjornsson; Gisli Masson; Ingileif Jonsdottir; Alma Moller; Thorolfur Gudnason; Karl G Kristinsson; Unnur Thorsteinsdottir; Kari Stefansson |
| EPI_ISL_827967, EPI_ISL_827968, EPI_ISL_827971, EPI_ISL_827972, EPI_ISL_827975, EPI_ISL_827976, EPI_ISL_827977, EPI_ISL_827978, EPI_ISL_827993, EPI_ISL_827994, EPI_ISL_827997, EPI_ISL_827999, EPI_ISL_828000 | | | |
| see above | deCODE genetics | deCODE genetics | Daniel F Gudbjartsson; Agnar Helgason; Hakon Jonsson; Olafur T Magnusson; Pall Melsted; Gudmundur L Norddahl; Jona Saemundsdottir; Asgeir Sigurdsson; Patrick Sulem; Arna B Agustsdottir; Hannes Eggertsson; Berglind Eiriksdottir; Run Fridriksdottir; Elisabet E Gardarsdottir; Gudmundur Georgsson; Olafia S Gretarsdottir; Kjartan R Gudmundsson; Thora R Gunnarsdottir; Arnaldur Gylfason; Hilma Holm; Brynjar O Jensson; Aslaug Jonasdottir; Kamilla S Josefsdottir; Thordur Kristjansson; Droplaug N Magnusdottir; Solvi Rognvaldsson; Louise le Roux; Gudrun Sigmundsdottir; Gardar Sveinbjornsson; Kristin E Sveinsdottir; Maney Sveinsdottir; Emil A Thorarensen; Bjarni Thorbjornsson; Gisli Masson; Ingileif Jonsdottir; Alma Moller; Thorolfur Gudnason; Karl G Kristinsson; Unnur Thorsteinsdottir; Kari Stefansson |
| EPI_ISL_828023 | The National University Hospital of Iceland | deCODE genetics | Daniel F Gudbjartsson; Agnar Helgason; Hakon Jonsson; Olafur T Magnusson; Pall Melsted; Gudmundur L Norddahl; Jona Saemundsdottir; Asgeir Sigurdsson; Patrick Sulem; Arna B Agustsdottir; Hannes Eggertsson; Berglind Eiriksdottir; Run Fridriksdottir; Elisabet E Gardarsdottir; Gudmundur Georgsson; Olafia S Gretarsdottir; Kjartan R Gudmundsson; Thora R Gunnarsdottir; Arnaldur Gylfason; Hilma Holm; Brynjar O Jensson; Aslaug Jonasdottir; Kamilla S Josefsdottir; Thordur Kristjansson; Droplaug N Magnusdottir; Solvi Rognvaldsson; Louise le Roux; Gudrun Sigmundsdottir; Gardar Sveinbjornsson; Kristin E Sveinsdottir; Maney Sveinsdottir; Emil A Thorarensen; Bjarni Thorbjornsson; Gisli Masson; Ingileif Jonsdottir; Alma Moller; Thorolfur Gudnason; Karl G Kristinsson; Unnur Thorsteinsdottir; Kari Stefansson |
| EPI_ISL_828068 | deCODE genetics | deCODE genetics | Daniel F Gudbjartsson; Agnar Helgason; Hakon Jonsson; Olafur T Magnusson; Pall Melsted; Gudmundur L Norddahl; Jona Saemundsdottir; Asgeir Sigurdsson; Patrick Sulem; Arna B Agustsdottir; Hannes Eggertsson; Berglind Eiriksdottir; Run Fridriksdottir; Elisabet E Gardarsdottir; Gudmundur Georgsson; Olafia S Gretarsdottir; Kjartan R Gudmundsson; Thora R Gunnarsdottir; Arnaldur Gylfason; Hilma Holm; Brynjar O Jensson; Aslaug Jonasdottir; Kamilla S Josefsdottir; Thordur Kristjansson; Droplaug N Magnusdottir; Solvi Rognvaldsson; Louise le Roux; Gudrun Sigmundsdottir; Gardar Sveinbjornsson; Kristin E Sveinsdottir; Maney Sveinsdottir; Emil A Thorarensen; Bjarni Thorbjornsson; Gisli Masson; Ingileif Jonsdottir; Alma Moller; Thorolfur Gudnason; Karl G Kristinsson; Unnur Thorsteinsdottir; Kari Stefansson |
| EPI_ISL_828089, EPI_ISL_828137 | The National University Hospital of Iceland | deCODE genetics | Daniel F Gudbjartsson; Agnar Helgason; Hakon Jonsson; Olafur T Magnusson; Pall Melsted; Gudmundur L Norddahl; Jona Saemundsdottir; Asgeir Sigurdsson; Patrick Sulem; Arna B Agustsdottir; Hannes Eggertsson; Berglind Eiriksdottir; Run Fridriksdottir; Elisabet E Gardarsdottir; Gudmundur Georgsson; Olafia S Gretarsdottir; Kjartan R Gudmundsson; Thora R Gunnarsdottir; Arnaldur Gylfason; Hilma Holm; Brynjar O Jensson; Aslaug Jonasdottir; Kamilla S Josefsdottir; Thordur Kristjansson; Droplaug N Magnusdottir; Solvi Rognvaldsson; Louise le Roux; Gudrun Sigmundsdottir; Gardar Sveinbjornsson; Kristin E Sveinsdottir; Maney Sveinsdottir; Emil A Thorarensen; Bjarni Thorbjornsson; Gisli Masson; Ingileif Jonsdottir; Alma Moller; Thorolfur Gudnason; Karl G Kristinsson; Unnur Thorsteinsdottir; Kari Stefansson |
| EPI_ISL_828161, EPI_ISL_828165, EPI_ISL_828166, EPI_ISL_828167, EPI_ISL_828168, EPI_ISL_828169, EPI_ISL_828170, EPI_ISL_828171 | deCODE genetics | deCODE genetics | Daniel F Gudbjartsson; Agnar Helgason; Hakon Jonsson; Olafur T Magnusson; Pall Melsted; Gudmundur L Norddahl; Jona Saemundsdottir; Asgeir Sigurdsson; Patrick Sulem; Arna B Agustsdottir; Hannes Eggertsson; Berglind Eiriksdottir; Run Fridriksdottir; Elisabet E Gardarsdottir; Gudmundur Georgsson; Olafia S Gretarsdottir; Kjartan R Gudmundsson; Thora R Gunnarsdottir; Arnaldur Gylfason; Hilma Holm; Brynjar O Jensson; Aslaug Jonasdottir; Kamilla S Josefsdottir; Thordur Kristjansson; Droplaug N Magnusdottir; Solvi Rognvaldsson; Louise le Roux; Gudrun Sigmundsdottir; Gardar Sveinbjornsson; Kristin E Sveinsdottir; Maney Sveinsdottir; Emil A Thorarensen; Bjarni Thorbjornsson; Gisli Masson; Ingileif Jonsdottir; Alma Moller; Thorolfur Gudnason; Karl G Kristinsson; Unnur Thorsteinsdottir; Kari Stefansson |
| EPI_ISL_828174, EPI_ISL_828209, EPI_ISL_828214 | The National University Hospital of Iceland | deCODE genetics | Daniel F Gudbjartsson; Agnar Helgason; Hakon Jonsson; Olafur T Magnusson; Pall Melsted; Gudmundur L Norddahl; Jona Saemundsdottir; Asgeir Sigurdsson; Patrick Sulem; Arna B Agustsdottir; Hannes Eggertsson; Berglind Eiriksdottir; Run Fridriksdottir; Elisabet E Gardarsdottir; Gudmundur Georgsson; Olafia S Gretarsdottir; Kjartan R Gudmundsson; Thora R Gunnarsdottir; Arnaldur Gylfason; Hilma Holm; Brynjar O Jensson; Aslaug Jonasdottir; Kamilla S Josefsdottir; Thordur Kristjansson; Droplaug N Magnusdottir; Solvi Rognvaldsson; Louise le Roux; Gudrun Sigmundsdottir; Gardar Sveinbjornsson; Kristin E Sveinsdottir; Maney Sveinsdottir; Emil A Thorarensen; Bjarni Thorbjornsson; Gisli Masson; Ingileif Jonsdottir; Alma Moller; Thorolfur Gudnason; Karl G Kristinsson; Unnur Thorsteinsdottir; Kari Stefansson |
| EPI_ISL_828381, EPI_ISL_828412, EPI_ISL_828415, EPI_ISL_828416, EPI_ISL_828427, EPI_ISL_828428, EPI_ISL_828429, EPI_ISL_828430, EPI_ISL_828431, EPI_ISL_828432, EPI_ISL_828433, EPI_ISL_828434, EPI_ISL_828436, EPI_ISL_828437, EPI_ISL_828525, EPI_ISL_828565, EPI_ISL_828566 | | | |
| see above | deCODE genetics | deCODE genetics | Daniel F Gudbjartsson; Agnar Helgason; Hakon Jonsson; Olafur T Magnusson; Pall Melsted; Gudmundur L Norddahl; Jona Saemundsdottir; Asgeir Sigurdsson; Patrick Sulem; Arna B Agustsdottir; Hannes Eggertsson; Berglind Eiriksdottir; Run Fridriksdottir; Elisabet E Gardarsdottir; Gudmundur Georgsson; Olafia S Gretarsdottir; Kjartan R Gudmundsson; Thora R Gunnarsdottir; Arnaldur Gylfason; Hilma Holm; Brynjar O Jensson; Aslaug Jonasdottir; Kamilla S Josefsdottir; Thordur Kristjansson; Droplaug N Magnusdottir; Solvi Rognvaldsson; Louise le Roux; Gudrun Sigmundsdottir; Gardar Sveinbjornsson; Kristin E Sveinsdottir; Maney Sveinsdottir; Emil A Thorarensen; Bjarni Thorbjornsson; Gisli Masson; Ingileif Jonsdottir; Alma Moller; Thorolfur Gudnason; Karl G Kristinsson; Unnur Thorsteinsdottir; Kari Stefansson |
| EPI_ISL_828679 | The National University Hospital of Iceland | deCODE genetics | Daniel F Gudbjartsson; Agnar Helgason; Hakon Jonsson; Olafur T Magnusson; Pall Melsted; Gudmundur L Norddahl; Jona Saemundsdottir; Asgeir Sigurdsson; Patrick Sulem; Arna B Agustsdottir; Hannes Eggertsson; Berglind Eiriksdottir; Run Fridriksdottir; Elisabet E Gardarsdottir; Gudmundur Georgsson; Olafia S Gretarsdottir; Kjartan R Gudmundsson; Thora R Gunnarsdottir; Arnaldur Gylfason; Hilma Holm; Brynjar O Jensson; Aslaug Jonasdottir; Kamilla S Josefsdottir; Thordur Kristjansson; Droplaug N Magnusdottir; Solvi Rognvaldsson; Louise le Roux; Gudrun Sigmundsdottir; Gardar Sveinbjornsson; Kristin E Sveinsdottir; Maney Sveinsdottir; Emil A Thorarensen; Bjarni Thorbjornsson; Gisli Masson; Ingileif Jonsdottir; Alma Moller; Thorolfur Gudnason; Karl G Kristinsson; Unnur Thorsteinsdottir; Kari Stefansson |
| EPI_ISL_828706, EPI_ISL_828708, EPI_ISL_828711, EPI_ISL_828726 | deCODE genetics | deCODE genetics | Daniel F Gudbjartsson; Agnar Helgason; Hakon Jonsson; Olafur T Magnusson; Pall Melsted; Gudmundur L Norddahl; Jona Saemundsdottir; Asgeir Sigurdsson; Patrick Sulem; Arna B Agustsdottir; Hannes Eggertsson; Berglind Eiriksdottir; Run Fridriksdottir; Elisabet E Gardarsdottir; Gudmundur Georgsson; Olafia S Gretarsdottir; Kjartan R Gudmundsson; Thora R Gunnarsdottir; Arnaldur Gylfason; Hilma Holm; Brynjar O Jensson; Aslaug Jonasdottir; Kamilla S Josefsdottir; Thordur Kristjansson; Droplaug N Magnusdottir; Solvi Rognvaldsson; Louise le Roux; Gudrun Sigmundsdottir; Gardar Sveinbjornsson; Kristin E Sveinsdottir; Maney Sveinsdottir; Emil A Thorarensen; Bjarni Thorbjornsson; Gisli Masson; Ingileif Jonsdottir; Alma Moller; Thorolfur Gudnason; Karl G Kristinsson; Unnur Thorsteinsdottir; Kari Stefansson |
| EPI_ISL_828753, EPI_ISL_828754, EPI_ISL_828755, EPI_ISL_828757, EPI_ISL_828758, EPI_ISL_828780 | The National University Hospital of Iceland | deCODE genetics | Daniel F Gudbjartsson; Agnar Helgason; Hakon Jonsson; Olafur T Magnusson; Pall Melsted; Gudmundur L Norddahl; Jona Saemundsdottir; Asgeir Sigurdsson; Patrick Sulem; Arna B Agustsdottir; Hannes Eggertsson; Berglind Eiriksdottir; Run Fridriksdottir; Elisabet E Gardarsdottir; Gudmundur Georgsson; Olafia S Gretarsdottir; Kjartan R Gudmundsson; Thora R Gunnarsdottir; Arnaldur Gylfason; Hilma Holm; Brynjar O Jensson; Aslaug Jonasdottir; Kamilla S Josefsdottir; Thordur Kristjansson; Droplaug N Magnusdottir; Solvi Rognvaldsson; Louise le Roux; Gudrun Sigmundsdottir; Gardar Sveinbjornsson; Kristin E Sveinsdottir; Maney Sveinsdottir; Emil A Thorarensen; Bjarni Thorbjornsson; Gisli Masson; Ingileif Jonsdottir; Alma Moller; Thorolfur Gudnason; Karl G Kristinsson; Unnur Thorsteinsdottir; Kari Stefansson |
| EPI_ISL_828823, EPI_ISL_828825, EPI_ISL_828915, EPI_ISL_828916, EPI_ISL_828917, EPI_ISL_828919, EPI_ISL_828921, EPI_ISL_828922, EPI_ISL_828925, EPI_ISL_828928, EPI_ISL_828945 | | | |
| see above | deCODE genetics | deCODE genetics | Daniel F Gudbjartsson; Agnar Helgason; Hakon Jonsson; Olafur T Magnusson; Pall Melsted; Gudmundur L Norddahl; Jona Saemundsdottir; Asgeir Sigurdsson; Patrick Sulem; Arna B Agustsdottir; Hannes Eggertsson; Berglind Eiriksdottir; Run Fridriksdottir; Elisabet E Gardarsdottir; Gudmundur Georgsson; Olafia S Gretarsdottir; Kjartan R Gudmundsson; Thora R Gunnarsdottir; Arnaldur Gylfason; Hilma Holm; Brynjar O Jensson; Aslaug |

| | | | |
|---|---|---|---|
| | | | Jonasdottir; Kamilla S Josefsdottir; Thordur Kristjansson; Droplaug N Magnusdottir; Solvi Rognvaldsson; Louise le Roux; Gudrun Sigmundsdottir; Gardar Sveinbjornsson; Kristin E Sveinsdottir; Maney Sveinsdottir; Emil A Thorarensen; Bjarni Thorbjornsson; Gisli Masson; Ingileif Jonsdottir; Alma Moller; Thorolfur Gudnason; Karl G Kristinsson; Unnur Thorsteinsdottir; Kari Stefansson |
| EPI_ISL_828995, EPI_ISL_828997, EPI_ISL_828999, EPI_ISL_829003, EPI_ISL_829004, EPI_ISL_829005, EPI_ISL_829006 | The National University Hospital of Iceland | deCODE genetics | Daniel F Gudbjartsson; Agnar Helgason; Hakon Jonsson; Olafur T Magnusson; Pall Melsted; Gudmundur L Norddahl; Jona Saemundsdottir; Asgeir Sigurdsson; Patrick Sulem; Arna B Agustsdottir; Hannes Eggertsson; Berglind Eiriksdottir; Run Fridriksdottir; Elisabet E Gardarsdottir; Gudmundur Georgsson; Olafia S Gretarsdottir; Kjartan R Gudmundsson; Thora R Gunnarsdottir; Arnaldur Gylfason; Hilma Holm; Brynjar O Jensson; Aslaug Jonasdottir; Kamilla S Josefsdottir; Thordur Kristjansson; Droplaug N Magnusdottir; Solvi Rognvaldsson; Louise le Roux; Gudrun Sigmundsdottir; Gardar Sveinbjornsson; Kristin E Sveinsdottir; Maney Sveinsdottir; Emil A Thorarensen; Bjarni Thorbjornsson; Gisli Masson; Ingileif Jonsdottir; Alma Moller; Thorolfur Gudnason; Karl G Kristinsson; Unnur Thorsteinsdottir; Kari Stefansson |
| EPI_ISL_829012, EPI_ISL_829023, EPI_ISL_829024, EPI_ISL_829027, EPI_ISL_829028, EPI_ISL_829029, EPI_ISL_829036, EPI_ISL_829076 | deCODE genetics | deCODE genetics | Daniel F Gudbjartsson; Agnar Helgason; Hakon Jonsson; Olafur T Magnusson; Pall Melsted; Gudmundur L Norddahl; Jona Saemundsdottir; Asgeir Sigurdsson; Patrick Sulem; Arna B Agustsdottir; Hannes Eggertsson; Berglind Eiriksdottir; Run Fridriksdottir; Elisabet E Gardarsdottir; Gudmundur Georgsson; Olafia S Gretarsdottir; Kjartan R Gudmundsson; Thora R Gunnarsdottir; Arnaldur Gylfason; Hilma Holm; Brynjar O Jensson; Aslaug Jonasdottir; Kamilla S Josefsdottir; Thordur Kristjansson; Droplaug N Magnusdottir; Solvi Rognvaldsson; Louise le Roux; Gudrun Sigmundsdottir; Gardar Sveinbjornsson; Kristin E Sveinsdottir; Maney Sveinsdottir; Emil A Thorarensen; Bjarni Thorbjornsson; Gisli Masson; Ingileif Jonsdottir; Alma Moller; Thorolfur Gudnason; Karl G Kristinsson; Unnur Thorsteinsdottir; Kari Stefansson |
| EPI_ISL_829142, EPI_ISL_829153, EPI_ISL_829166 | The National University Hospital of Iceland | deCODE genetics | Daniel F Gudbjartsson; Agnar Helgason; Hakon Jonsson; Olafur T Magnusson; Pall Melsted; Gudmundur L Norddahl; Jona Saemundsdottir; Asgeir Sigurdsson; Patrick Sulem; Arna B Agustsdottir; Hannes Eggertsson; Berglind Eiriksdottir; Run Fridriksdottir; Elisabet E Gardarsdottir; Gudmundur Georgsson; Olafia S Gretarsdottir; Kjartan R Gudmundsson; Thora R Gunnarsdottir; Arnaldur Gylfason; Hilma Holm; Brynjar O Jensson; Aslaug Jonasdottir; Kamilla S Josefsdottir; Thordur Kristjansson; Droplaug N Magnusdottir; Solvi Rognvaldsson; Louise le Roux; Gudrun Sigmundsdottir; Gardar Sveinbjornsson; Kristin E Sveinsdottir; Maney Sveinsdottir; Emil A Thorarensen; Bjarni Thorbjornsson; Gisli Masson; Ingileif Jonsdottir; Alma Moller; Thorolfur Gudnason; Karl G Kristinsson; Unnur Thorsteinsdottir; Kari Stefansson |
| EPI_ISL_829219 | deCODE genetics | deCODE genetics | Daniel F Gudbjartsson; Agnar Helgason; Hakon Jonsson; Olafur T Magnusson; Pall Melsted; Gudmundur L Norddahl; Jona Saemundsdottir; Asgeir Sigurdsson; Patrick Sulem; Arna B Agustsdottir; Hannes Eggertsson; Berglind Eiriksdottir; Run Fridriksdottir; Elisabet E Gardarsdottir; Gudmundur Georgsson; Olafia S Gretarsdottir; Kjartan R Gudmundsson; Thora R Gunnarsdottir; Arnaldur Gylfason; Hilma Holm; Brynjar O Jensson; Aslaug Jonasdottir; Kamilla S Josefsdottir; Thordur Kristjansson; Droplaug N Magnusdottir; Solvi Rognvaldsson; Louise le Roux; Gudrun Sigmundsdottir; Gardar Sveinbjornsson; Kristin E Sveinsdottir; Maney Sveinsdottir; Emil A Thorarensen; Bjarni Thorbjornsson; Gisli Masson; Ingileif Jonsdottir; Alma Moller; Thorolfur Gudnason; Karl G Kristinsson; Unnur Thorsteinsdottir; Kari Stefansson |
| EPI_ISL_829223 | The National University Hospital of Iceland | deCODE genetics | Daniel F Gudbjartsson; Agnar Helgason; Hakon Jonsson; Olafur T Magnusson; Pall Melsted; Gudmundur L Norddahl; Jona Saemundsdottir; Asgeir Sigurdsson; Patrick Sulem; Arna B Agustsdottir; Hannes Eggertsson; Berglind Eiriksdottir; Run Fridriksdottir; Elisabet E Gardarsdottir; Gudmundur Georgsson; Olafia S Gretarsdottir; Kjartan R Gudmundsson; Thora R Gunnarsdottir; Arnaldur Gylfason; Hilma Holm; Brynjar O Jensson; Aslaug Jonasdottir; Kamilla S Josefsdottir; Thordur Kristjansson; Droplaug N Magnusdottir; Solvi Rognvaldsson; Louise le Roux; Gudrun Sigmundsdottir; Gardar Sveinbjornsson; Kristin E Sveinsdottir; Maney Sveinsdottir; Emil A Thorarensen; Bjarni Thorbjornsson; Gisli Masson; Ingileif Jonsdottir; Alma Moller; Thorolfur Gudnason; Karl G Kristinsson; Unnur Thorsteinsdottir; Kari Stefansson |
| EPI_ISL_829244, EPI_ISL_829271, EPI_ISL_829287, EPI_ISL_829294, EPI_ISL_829297, EPI_ISL_829298, EPI_ISL_829299, EPI_ISL_829300, EPI_ISL_829301, EPI_ISL_829302, EPI_ISL_829303, EPI_ISL_829304, EPI_ISL_829308, EPI_ISL_829309, EPI_ISL_829312 | | | |
| see above | deCODE genetics | deCODE genetics | Daniel F Gudbjartsson; Agnar Helgason; Hakon Jonsson; Olafur T Magnusson; Pall Melsted; Gudmundur L Norddahl; Jona Saemundsdottir; Asgeir Sigurdsson; Patrick Sulem; Arna B Agustsdottir; Hannes Eggertsson; Berglind Eiriksdottir; Run Fridriksdottir; Elisabet E Gardarsdottir; Gudmundur Georgsson; Olafia S Gretarsdottir; Kjartan R Gudmundsson; Thora R Gunnarsdottir; Arnaldur Gylfason; Hilma Holm; Brynjar O Jensson; Aslaug Jonasdottir; Kamilla S Josefsdottir; Thordur Kristjansson; Droplaug N Magnusdottir; Solvi Rognvaldsson; Louise le Roux; Gudrun Sigmundsdottir; Gardar Sveinbjornsson; Kristin E Sveinsdottir; Maney Sveinsdottir; Emil A Thorarensen; Bjarni Thorbjornsson; Gisli Masson; Ingileif Jonsdottir; Alma Moller; Thorolfur Gudnason; Karl G Kristinsson; Unnur Thorsteinsdottir; Kari Stefansson |
| EPI_ISL_829360, EPI_ISL_829464 | The National University Hospital of Iceland | deCODE genetics | Daniel F Gudbjartsson; Agnar Helgason; Hakon Jonsson; Olafur T Magnusson; Pall Melsted; Gudmundur L Norddahl; Jona Saemundsdottir; Asgeir Sigurdsson; Patrick Sulem; Arna B Agustsdottir; Hannes Eggertsson; Berglind Eiriksdottir; Run Fridriksdottir; Elisabet E Gardarsdottir; Gudmundur Georgsson; Olafia S Gretarsdottir; Kjartan R Gudmundsson; Thora R Gunnarsdottir; Arnaldur Gylfason; Hilma Holm; Brynjar O Jensson; Aslaug Jonasdottir; Kamilla S Josefsdottir; Thordur Kristjansson; Droplaug N Magnusdottir; Solvi Rognvaldsson; Louise le Roux; Gudrun Sigmundsdottir; Gardar Sveinbjornsson; Kristin E Sveinsdottir; Maney Sveinsdottir; Emil A Thorarensen; Bjarni Thorbjornsson; Gisli Masson; Ingileif Jonsdottir; Alma Moller; Thorolfur Gudnason; Karl G Kristinsson; Unnur Thorsteinsdottir; Kari Stefansson |
| EPI_ISL_829533, EPI_ISL_829534, EPI_ISL_829537, EPI_ISL_829538, EPI_ISL_829539, EPI_ISL_829540, EPI_ISL_829541, EPI_ISL_829565, EPI_ISL_829592, EPI_ISL_829593 | deCODE genetics | deCODE genetics | Daniel F Gudbjartsson; Agnar Helgason; Hakon Jonsson; Olafur T Magnusson; Pall Melsted; Gudmundur L Norddahl; Jona Saemundsdottir; Asgeir Sigurdsson; Patrick Sulem; Arna B Agustsdottir; Hannes Eggertsson; Berglind Eiriksdottir; Run Fridriksdottir; Elisabet E Gardarsdottir; Gudmundur Georgsson; Olafia S Gretarsdottir; Kjartan R Gudmundsson; Thora R Gunnarsdottir; Arnaldur Gylfason; Hilma Holm; Brynjar O Jensson; Aslaug Jonasdottir; Kamilla S Josefsdottir; Thordur Kristjansson; Droplaug N Magnusdottir; Solvi Rognvaldsson; Louise le Roux; Gudrun Sigmundsdottir; Gardar Sveinbjornsson; Kristin E Sveinsdottir; Maney Sveinsdottir; Emil A Thorarensen; Bjarni Thorbjornsson; Gisli Masson; Ingileif Jonsdottir; Alma Moller; Thorolfur Gudnason; Karl G Kristinsson; Unnur Thorsteinsdottir; Kari Stefansson |
| EPI_ISL_829608 | The National University Hospital of Iceland | deCODE genetics | Daniel F Gudbjartsson; Agnar Helgason; Hakon Jonsson; Olafur T Magnusson; Pall Melsted; Gudmundur L Norddahl; Jona Saemundsdottir; Asgeir Sigurdsson; Patrick Sulem; Arna B Agustsdottir; Hannes Eggertsson; Berglind Eiriksdottir; Run Fridriksdottir; Elisabet E Gardarsdottir; Gudmundur Georgsson; Olafia S Gretarsdottir; Kjartan R Gudmundsson; Thora R Gunnarsdottir; Arnaldur Gylfason; Hilma Holm; Brynjar O Jensson; Aslaug Jonasdottir; Kamilla S Josefsdottir; Thordur Kristjansson; Droplaug N Magnusdottir; Solvi Rognvaldsson; Louise le Roux; Gudrun Sigmundsdottir; Gardar Sveinbjornsson; Kristin E Sveinsdottir; Maney Sveinsdottir; Emil A Thorarensen; Bjarni Thorbjornsson; Gisli Masson; Ingileif Jonsdottir; Alma Moller; Thorolfur Gudnason; Karl G Kristinsson; Unnur Thorsteinsdottir; Kari Stefansson |
| EPI_ISL_829676, EPI_ISL_829722, EPI_ISL_829724, EPI_ISL_829741, EPI_ISL_829746, EPI_ISL_829762, EPI_ISL_829765, EPI_ISL_829776, EPI_ISL_829806, EPI_ISL_829807, EPI_ISL_829808, EPI_ISL_829809, EPI_ISL_829810, EPI_ISL_829812, EPI_ISL_829813, EPI_ISL_829863 | | | |
| see above | deCODE genetics | deCODE genetics | Daniel F Gudbjartsson; Agnar Helgason; Hakon Jonsson; Olafur T Magnusson; Pall Melsted; Gudmundur L Norddahl; Jona Saemundsdottir; Asgeir Sigurdsson; Patrick Sulem; Arna B Agustsdottir; Hannes Eggertsson; Berglind Eiriksdottir; Run Fridriksdottir; Elisabet E Gardarsdottir; Gudmundur Georgsson; Olafia S Gretarsdottir; Kjartan R Gudmundsson; Thora R Gunnarsdottir; Arnaldur Gylfason; Hilma Holm; Brynjar O Jensson; Aslaug Jonasdottir; Kamilla S Josefsdottir; Thordur Kristjansson; Droplaug N Magnusdottir; Solvi Rognvaldsson; Louise le Roux; Gudrun Sigmundsdottir; Gardar Sveinbjornsson; Kristin E Sveinsdottir; Maney Sveinsdottir; Emil A Thorarensen; Bjarni Thorbjornsson; Gisli Masson; Ingileif Jonsdottir; Alma Moller; Thorolfur Gudnason; Karl G Kristinsson; Unnur Thorsteinsdottir; Kari Stefansson |
| EPI_ISL_829864 | The National University Hospital of Iceland | deCODE genetics | Daniel F Gudbjartsson; Agnar Helgason; Hakon Jonsson; Olafur T Magnusson; Pall Melsted; Gudmundur L Norddahl; Jona Saemundsdottir; Asgeir Sigurdsson; Patrick Sulem; Arna B Agustsdottir; Hannes Eggertsson; Berglind Eiriksdottir; Run Fridriksdottir; Elisabet E Gardarsdottir; Gudmundur Georgsson; Olafia S Gretarsdottir; Kjartan R Gudmundsson; Thora R Gunnarsdottir; Arnaldur Gylfason; Hilma Holm; Brynjar O Jensson; Aslaug Jonasdottir; Kamilla S Josefsdottir; Thordur Kristjansson; Droplaug N Magnusdottir; Solvi Rognvaldsson; Louise le Roux; Gudrun Sigmundsdottir; Gardar Sveinbjornsson; Kristin E Sveinsdottir; Maney Sveinsdottir; Emil A Thorarensen; Bjarni Thorbjornsson; Gisli Masson; Ingileif Jonsdottir; Alma Moller; Thorolfur Gudnason; Karl G Kristinsson; Unnur Thorsteinsdottir; Kari Stefansson |
| EPI_ISL_829865, EPI_ISL_829866, EPI_ISL_829867, EPI_ISL_829868, EPI_ISL_829949, EPI_ISL_830063, EPI_ISL_830088, EPI_ISL_830099 | deCODE genetics | deCODE genetics | Daniel F Gudbjartsson; Agnar Helgason; Hakon Jonsson; Olafur T Magnusson; Pall Melsted; Gudmundur L Norddahl; Jona Saemundsdottir; Asgeir Sigurdsson; Patrick Sulem; Arna B Agustsdottir; Hannes Eggertsson; Berglind Eiriksdottir; Run Fridriksdottir; Elisabet E Gardarsdottir; Gudmundur Georgsson; Olafia S Gretarsdottir; Kjartan R Gudmundsson; Thora R Gunnarsdottir; Arnaldur Gylfason; Hilma Holm; Brynjar O Jensson; Aslaug Jonasdottir; Kamilla S Josefsdottir; Thordur Kristjansson; Droplaug N Magnusdottir; Solvi Rognvaldsson; Louise le Roux; Gudrun Sigmundsdottir; Gardar Sveinbjornsson; Kristin E Sveinsdottir; Maney Sveinsdottir; Emil A Thorarensen; Bjarni Thorbjornsson; Gisli Masson; Ingileif Jonsdottir; Alma Moller; Thorolfur Gudnason; Karl G Kristinsson; Unnur Thorsteinsdottir; Kari Stefansson |
| EPI_ISL_830119, EPI_ISL_830127, EPI_ISL_830129, EPI_ISL_830143, EPI_ISL_830144, EPI_ISL_830146 | The National University Hospital of Iceland | deCODE genetics | Daniel F Gudbjartsson; Agnar Helgason; Hakon Jonsson; Olafur T Magnusson; Pall Melsted; Gudmundur L Norddahl; Jona Saemundsdottir; Asgeir Sigurdsson; Patrick Sulem; Arna B Agustsdottir; Hannes Eggertsson; Berglind Eiriksdottir; Run Fridriksdottir; Elisabet E Gardarsdottir; Gudmundur Georgsson; Olafia S Gretarsdottir; Kjartan R Gudmundsson; Thora R Gunnarsdottir; Arnaldur Gylfason; Hilma Holm; Brynjar O Jensson; Aslaug |

| | | | |
|---|---|---|---|
| | | | Jonasdottir; Kamilla S Josefsdottir; Thordur Kristjansson; Droplaug N Magnusdottir; Solvi Rognvaldsson; Louise le Roux; Gudrun Sigmundsdottir; Gardar Sveinbjornsson; Kristin E Sveinsdottir; Maney Sveinsdottir; Emil A Thorarensen; Bjarni Thorbjornsson; Gisli Masson; Ingileif Jonsdottir; Alma Moller; Thorolfur Gudnason; Karl G Kristinsson; Unnur Thorsteinsdottir; Kari Stefansson |
| EPI_ISL_830285, EPI_ISL_830301, EPI_ISL_830303 | deCODE genetics | deCODE genetics | Daniel F Gudbjartsson; Agnar Helgason; Hakon Jonsson; Olafur T Magnusson; Pall Melsted; Gudmundur L Norddahl; Jona Saemundsdottir; Asgeir Sigurdsson; Patrick Sulem; Arna B Agustsdottir; Hannes Eggertsson; Berglind Eiriksdottir; Run Fridriksdottir; Elisabet E Gardarsdottir; Gudmundur Georgsson; Olafia S Gretarsdottir; Kjartan R Gudmundsson; Thora R Gunnarsdottir; Arnaldur Gylfason; Hilma Holm; Brynjar O Jensson; Aslaug Jonasdottir; Kamilla S Josefsdottir; Thordur Kristjansson; Droplaug N Magnusdottir; Solvi Rognvaldsson; Louise le Roux; Gudrun Sigmundsdottir; Gardar Sveinbjornsson; Kristin E Sveinsdottir; Maney Sveinsdottir; Emil A Thorarensen; Bjarni Thorbjornsson; Gisli Masson; Ingileif Jonsdottir; Alma Moller; Thorolfur Gudnason; Karl G Kristinsson; Unnur Thorsteinsdottir; Kari Stefansson |
| EPI_ISL_830328, EPI_ISL_830354 | The National University Hospital of Iceland | deCODE genetics | Daniel F Gudbjartsson; Agnar Helgason; Hakon Jonsson; Olafur T Magnusson; Pall Melsted; Gudmundur L Norddahl; Jona Saemundsdottir; Asgeir Sigurdsson; Patrick Sulem; Arna B Agustsdottir; Hannes Eggertsson; Berglind Eiriksdottir; Run Fridriksdottir; Elisabet E Gardarsdottir; Gudmundur Georgsson; Olafia S Gretarsdottir; Kjartan R Gudmundsson; Thora R Gunnarsdottir; Arnaldur Gylfason; Hilma Holm; Brynjar O Jensson; Aslaug Jonasdottir; Kamilla S Josefsdottir; Thordur Kristjansson; Droplaug N Magnusdottir; Solvi Rognvaldsson; Louise le Roux; Gudrun Sigmundsdottir; Gardar Sveinbjornsson; Kristin E Sveinsdottir; Maney Sveinsdottir; Emil A Thorarensen; Bjarni Thorbjornsson; Gisli Masson; Ingileif Jonsdottir; Alma Moller; Thorolfur Gudnason; Karl G Kristinsson; Unnur Thorsteinsdottir; Kari Stefansson |
| EPI_ISL_830360 | deCODE genetics | deCODE genetics | Daniel F Gudbjartsson; Agnar Helgason; Hakon Jonsson; Olafur T Magnusson; Pall Melsted; Gudmundur L Norddahl; Jona Saemundsdottir; Asgeir Sigurdsson; Patrick Sulem; Arna B Agustsdottir; Hannes Eggertsson; Berglind Eiriksdottir; Run Fridriksdottir; Elisabet E Gardarsdottir; Gudmundur Georgsson; Olafia S Gretarsdottir; Kjartan R Gudmundsson; Thora R Gunnarsdottir; Arnaldur Gylfason; Hilma Holm; Brynjar O Jensson; Aslaug Jonasdottir; Kamilla S Josefsdottir; Thordur Kristjansson; Droplaug N Magnusdottir; Solvi Rognvaldsson; Louise le Roux; Gudrun Sigmundsdottir; Gardar Sveinbjornsson; Kristin E Sveinsdottir; Maney Sveinsdottir; Emil A Thorarensen; Bjarni Thorbjornsson; Gisli Masson; Ingileif Jonsdottir; Alma Moller; Thorolfur Gudnason; Karl G Kristinsson; Unnur Thorsteinsdottir; Kari Stefansson |
| EPI_ISL_830397, EPI_ISL_830407 | The National University Hospital of Iceland | deCODE genetics | Daniel F Gudbjartsson; Agnar Helgason; Hakon Jonsson; Olafur T Magnusson; Pall Melsted; Gudmundur L Norddahl; Jona Saemundsdottir; Asgeir Sigurdsson; Patrick Sulem; Arna B Agustsdottir; Hannes Eggertsson; Berglind Eiriksdottir; Run Fridriksdottir; Elisabet E Gardarsdottir; Gudmundur Georgsson; Olafia S Gretarsdottir; Kjartan R Gudmundsson; Thora R Gunnarsdottir; Arnaldur Gylfason; Hilma Holm; Brynjar O Jensson; Aslaug Jonasdottir; Kamilla S Josefsdottir; Thordur Kristjansson; Droplaug N Magnusdottir; Solvi Rognvaldsson; Louise le Roux; Gudrun Sigmundsdottir; Gardar Sveinbjornsson; Kristin E Sveinsdottir; Maney Sveinsdottir; Emil A Thorarensen; Bjarni Thorbjornsson; Gisli Masson; Ingileif Jonsdottir; Alma Moller; Thorolfur Gudnason; Karl G Kristinsson; Unnur Thorsteinsdottir; Kari Stefansson |
| EPI_ISL_830451, EPI_ISL_830452, EPI_ISL_830455, EPI_ISL_830456, EPI_ISL_830457, EPI_ISL_830458, EPI_ISL_830460, EPI_ISL_830520, EPI_ISL_830533, EPI_ISL_830548, EPI_ISL_830569, EPI_ISL_830570 | | | |
| see above | deCODE genetics | deCODE genetics | Daniel F Gudbjartsson; Agnar Helgason; Hakon Jonsson; Olafur T Magnusson; Pall Melsted; Gudmundur L Norddahl; Jona Saemundsdottir; Asgeir Sigurdsson; Patrick Sulem; Arna B Agustsdottir; Hannes Eggertsson; Berglind Eiriksdottir; Run Fridriksdottir; Elisabet E Gardarsdottir; Gudmundur Georgsson; Olafia S Gretarsdottir; Kjartan R Gudmundsson; Thora R Gunnarsdottir; Arnaldur Gylfason; Hilma Holm; Brynjar O Jensson; Aslaug Jonasdottir; Kamilla S Josefsdottir; Thordur Kristjansson; Droplaug N Magnusdottir; Solvi Rognvaldsson; Louise le Roux; Gudrun Sigmundsdottir; Gardar Sveinbjornsson; Kristin E Sveinsdottir; Maney Sveinsdottir; Emil A Thorarensen; Bjarni Thorbjornsson; Gisli Masson; Ingileif Jonsdottir; Alma Moller; Thorolfur Gudnason; Karl G Kristinsson; Unnur Thorsteinsdottir; Kari Stefansson |
| EPI_ISL_831110, EPI_ISL_831111, EPI_ISL_831112, EPI_ISL_831113, EPI_ISL_831114, EPI_ISL_831228, EPI_ISL_831229 | Hospital Universitario La Paz (Madrid) | SeqCOVID-SPAIN consortium/IBV(CSIC) | María Rodríguez-Tejedor, Elias Dahdouh, Fernando Lázaro-Perona, Jesús Mingorance and SeqCOVID-SPAIN consortium |
| EPI_ISL_831291, EPI_ISL_831293, EPI_ISL_831298, EPI_ISL_831301, EPI_ISL_831303, EPI_ISL_831306, EPI_ISL_831315, EPI_ISL_831316, EPI_ISL_831317, EPI_ISL_831319, EPI_ISL_831320, EPI_ISL_831321, EPI_ISL_831326, EPI_ISL_831328, EPI_ISL_831332 | | | |
| see above | Texas Department of State Health Services | Texas Department of State Health Services | Anita Pokharel, Bonnie Oh, James Daniel Bonser, Rashmi Tuladhar, Mayela Pedrueza, Jenny Zhang, Maliha Rahman, Myong Koag, Chung Wang, Rachel Lee, Grace Kubin |
| EPI_ISL_832569, EPI_ISL_832570, EPI_ISL_832571, EPI_ISL_832572, EPI_ISL_832573, EPI_ISL_832574, EPI_ISL_832575, EPI_ISL_832576, EPI_ISL_832577, EPI_ISL_832578, EPI_ISL_832579, EPI_ISL_832580, EPI_ISL_832581, EPI_ISL_832582, EPI_ISL_832583, EPI_ISL_832584, EPI_ISL_832585, EPI_ISL_832586, EPI_ISL_832587, EPI_ISL_832588, EPI_ISL_832589, EPI_ISL_832590, EPI_ISL_832591, EPI_ISL_832592, EPI_ISL_832593, EPI_ISL_832594, EPI_ISL_832595, EPI_ISL_832596, EPI_ISL_832621, EPI_ISL_832622, EPI_ISL_832623, EPI_ISL_832624, EPI_ISL_832625, EPI_ISL_832626, EPI_ISL_832627, EPI_ISL_832628, EPI_ISL_832629, EPI_ISL_832630, EPI_ISL_832631, EPI_ISL_832632, EPI_ISL_832633, EPI_ISL_832634, EPI_ISL_832635, EPI_ISL_832636, EPI_ISL_832637, EPI_ISL_832638, EPI_ISL_832639, EPI_ISL_832640, EPI_ISL_832641, EPI_ISL_832642, EPI_ISL_832643, EPI_ISL_832644, EPI_ISL_832645, EPI_ISL_832646, EPI_ISL_832647, EPI_ISL_832648, EPI_ISL_832649, EPI_ISL_832650, EPI_ISL_832651, EPI_ISL_832652, EPI_ISL_832653, EPI_ISL_832654, EPI_ISL_832687, EPI_ISL_832688, EPI_ISL_832689, EPI_ISL_832690, EPI_ISL_832691, EPI_ISL_832692 | | | |
| see above | OHSU Lab Services Molecular Microbiology Lab | Oregon SARS-CoV-2 Genome Sequencing Center | Brendan L. O'Connell, Ruth V. Nichols, Sally Grindstaff, Alec J. Hirsch, Donna Hansel, Guang Fan, Daniel N. Streblow, William B. Messer, Andrew C. Adey, Benjamin N. Bimber, Brian J. O'Roak |
| EPI_ISL_833336 | Bureau of Quarantine | Research Institute for Tropical Medicine | Hannah Leah Morito, Othoniel Jan Onza, John Leonard Chan, Ma Angelica Tujan, Francisco Gerardo Polotan, Inez Andrea Medado, Kirstyn Brunker, Edelwisa Mercado, Daria Manalo, Catalino Demetria |
| EPI_ISL_839983, EPI_ISL_839984 | Queens Medical Centre, Clinical Microbiology Department / DeepSeq Nottingham | COVID-19 Genomics UK (COG-UK) Consortium | Gemma Clark, Wendy Smith, Manjinder Khakh, Vicki M Fleming, Michelle M Lister, Hannah Howson-Wells, Jonathan Ball, Patrick McClure, Joseph Chappell, Theocharis Tsoleridis, Nadine Holmes, Matthew Carlisle, Christopher Moore, Fei Sang, Johnny Debebe, Victoria Wright, Matthew Loose |
| EPI_ISL_842865 | Barts Health NHS Trust | COVID-19 Genomics UK (COG-UK) Consortium | CUTINO-MOGUEL, Maria-Teresa; HARRINGTON, David; OWOYEMI, Dola; SHYLINI, Raghavendran; BROAD, Claire; KELE, Beatrix |
| EPI_ISL_845553, EPI_ISL_845554, EPI_ISL_845555 | National Public Health Laboratory, Cameroon | African Centre of Excellence for Genomics of Infectious Diseases (ACEGID), Redeemer's University | Oluniyi P.E. et al |
| EPI_ISL_848232, EPI_ISL_848249, EPI_ISL_848266, EPI_ISL_848378, EPI_ISL_848379, EPI_ISL_848380, EPI_ISL_848381, EPI_ISL_848523 | Illinois Department of Public Health | Gagnon Lab, Southern Illinois University | Keith Gagnon |
| EPI_ISL_848874, EPI_ISL_848875, EPI_ISL_848876 | Florida Bureau of Public Health Laboratories | Florida Bureau of Public Health Laboratories | Sarah Schmedes, Jason Blanton |
| EPI_ISL_849372 | Servicio Virosis Respiratorias-Departamento Virología-INEI | Instituto Nacional Enfermedades Infecciosas C.G.Malbran | Baumeister E., Avaro M., Benedetti E., Russo M., Dattero ME, Pontoriero A., Cisterna D., Molina V., Perandones C., Tuduri E., Lorenzo F., Poklepovich T., Campos J. |
| EPI_ISL_849497, EPI_ISL_849498, EPI_ISL_849606 | Seattle Flu Study | Seattle Flu Study | Deborah A. Nickerson, Chris D. Frazar, Jover Lee, Benjamin Pelle, Matthew Richardson, Amanda Adler, Elisabeth Brandstetter, Peter D. Han, Kairsten Fay, Misja Ilcisin, Kirsten Lacombe, Thomas R. Sibley, Melissa Truong, Caitlin R. Wolf, Michael Boeckh, Janet A. Englund, Michael Famulare, Barry R. Lutz, Mark J. Rieder, Lea M. Starita, Matthew Thompson, Jay Shendure, Trevor Bedford, Helen Y. Chu |
| EPI_ISL_853728 | Department of Microbiology, University Innsbruck | Bergthaler laboratory, CeMM Research Center for Molecular Medicine of the Austrian Academy of Sciences | Lukas Endler, Alexandra Popa, Benedikt Agerer, Jakob-Wendelin Genger, Alexander Lercher, Anna Schedl, Thomas Penz, Michael Schuster, Jan Laine, Martin Senekowitsch, Christoph Bock, Andreas Bergthaler |
| EPI_ISL_853966, EPI_ISL_853967, EPI_ISL_853978, EPI_ISL_853979, EPI_ISL_853983, EPI_ISL_853986, EPI_ISL_853987, EPI_ISL_853993, EPI_ISL_853996, EPI_ISL_853997, EPI_ISL_854002, EPI_ISL_854005, EPI_ISL_854008, EPI_ISL_854009, EPI_ISL_854012, EPI_ISL_854016, EPI_ISL_854019, EPI_ISL_854020, EPI_ISL_854023, EPI_ISL_854025, EPI_ISL_854033, EPI_ISL_854034, EPI_ISL_854039, EPI_ISL_854231, EPI_ISL_854254, EPI_ISL_854277 | | | |
| see above | Austrian Agency for Health and Food Safety (AGES) | Bergthaler laboratory, CeMM Research Center for Molecular Medicine of the Austrian Academy of Sciences | Lukas Endler, Alexandra Popa, Benedikt Agerer, Jakob-Wendelin Genger, Alexander Lercher, Anna Schedl, Thomas Penz, Michael Schuster, Jan Laine, Martin Senekowitsch, Christoph Bock, Andreas Bergthaler |
| EPI_ISL_855502, EPI_ISL_855523, EPI_ISL_855526, EPI_ISL_855533, EPI_ISL_855538 | KEMRI-Wellcome Trust Research Programme/KEMRI-CGMR-C Kilifi | KEMRI-Wellcome Trust Research Programme/KEMRI-CGMR-C Kilifi | Githinji et al |
| EPI_ISL_856790, EPI_ISL_856791, | Servicio Virosis Respiratorias-Departamento Virología-INEI | Instituto Nacional Enfermedades Infecciosas C.G.Malbran | Baumeister E., Avaro M., Benedetti E., Russo M., Dattero ME, Pontoriero A., Cisterna D., Molina V., Perandones C., Tuduri E., Lorenzo F., Poklepovich T., |

| | | | |
|---|---|---|---|
| EPI_ISL_856792, EPI_ISL_856793 | | | Campos J. |
| EPI_ISL_859775, EPI_ISL_859778, EPI_ISL_859779, EPI_ISL_859780, EPI_ISL_859786, EPI_ISL_859787, EPI_ISL_859788, EPI_ISL_859798, EPI_ISL_859799, EPI_ISL_859801, EPI_ISL_859802, EPI_ISL_859803, EPI_ISL_859804, EPI_ISL_859805, EPI_ISL_859806, EPI_ISL_859807, EPI_ISL_859843, EPI_ISL_859844, EPI_ISL_859845 | | | |
| see above | BTC, Khalifa University | BTC, Khalifa University | Al Safar et al |
| EPI_ISL_860131 | Keio University School of Medicine | Keio University School of Medicine | Kenjiro Kosaki, Yuka Iwasaki, Hirotsugu Ishizu, Haruhiko Siomi, Kodai Abe |
| EPI_ISL_860731, EPI_ISL_860740, EPI_ISL_860743, EPI_ISL_860762, EPI_ISL_860763, EPI_ISL_860764, EPI_ISL_860765, EPI_ISL_860766, EPI_ISL_860767, EPI_ISL_860768, EPI_ISL_860769, EPI_ISL_860770, EPI_ISL_860771, EPI_ISL_860772 | | | |
| see above | Swiss National Reference Centre for Influenza | Swiss National Reference Centre for Influenza | Ana Rita Gonçalves Cabecinhas, Samuel Cordey, Florian Laubscher,Christoph Grünig, Laurent Kaiser |
| EPI_ISL_860798, EPI_ISL_860799, EPI_ISL_860800, EPI_ISL_860801, EPI_ISL_860813, EPI_ISL_860815, EPI_ISL_860816, EPI_ISL_860817, EPI_ISL_860818 | WHO/Minsk | Charité Universitätsmedizin Berlin, Institut für Virologie | Victor M Corman, Barbara Mühlemann, Jörn Beheim-Schwarzbach, Talitha Veith, Julia Tesch, Tobias Bleicker, Julia Schneider, Shmialiova Natallia, Sivets Natallia, Terry Jones, Christian Drosten |
| EPI_ISL_861487 | Wyoming Public Health Laboratory | Wyoming Public Health Laboratory | Noah Hull, Taylor Fearing, Lynette Gumbleton, Channing Weber, Ashley Norberg, Bailey Bowcutt, and Wanda Manley |
| EPI_ISL_861721, EPI_ISL_861722 | Department of Medical Microbiology, Hospital Pengajar Universiti Putra Malaysia | Malaysia Genome Institute | Mohd Noor Mat Isa, Syafinaz Amin-Nordin, Irni Suhayu Sapian, Hui-Yee Chee, Yusuf Muhammad Noor, Nurhezreen Md Iqbal, Enizza Kasim, Siti Noraini Othman, Mohd Faizal Abu Bakar, Shamsidar Sopie, Azrin Ahmad, Narcisse Joseph, Muhammad MI, Avisha Richards, Nor Zahrin Hasran, Nor Azfa Johari |
| EPI_ISL_861855 | Hospital General Universitario Gregorio Marañón | SeqCOVID-SPAIN consortium/IBV(CSIC) | Darío García de Viedma, Laura Pérez-Lago, Pedro J Sola-Campoy, Sergio Buenestado-Serrano, Marta Herranz, Victor Manuel de la Cueva, Julia Suárez, Pilar Catalán, Patricia Muñoz and SeqCOVID-SPAIN consortium |
| EPI_ISL_864575 | Institute of Medical Microbiology and Hospital Hygiene | Institute of Medical Microbiology and Hospital Hygiene | Prof. Dr. Achim Kaasch, Aljoscha Tersteegen |
| EPI_ISL_866803 | Quadram Institute Bioscience | COVID-19 Genomics UK (COG-UK) Consortium | Dave J. Baker, Gemma L. Kay, Alp Aydin, Thanh Le-Viet, Steven Rudder, Ana P. Tedim, Anastasia Kolyva, Maria Diaz, Leonardo de Oliveira Martins, Nabil-Fareed Alikhan, Lizzie Meadows, Rachael Stanley, Ngozi Elumogo, Muhammed Yasir, Nicholas M. Thomson, Alexander J Trotter, Rachel Gilroy, Samuel Bloomfield, Claire Stuart, Andrew Bell, Reenesh Prakash, Samir Dervisevic, Alison E. Mather, John Wain, Mark Webber, Andrew J. Page, Justin O'Grady |
| EPI_ISL_867944, EPI_ISL_867965 | Centre for Enzyme Innovation, University of Portsmouth / Translational Research Laboratory, Portsmouth Hospitals NHS Trust | COVID-19 Genomics UK (COG-UK) Consortium | Angela Beckett,Yann Bourgeois,Garry Scarlett,Sharon Glaysher,Scott Elliott,Kelly Bicknell,Robert Impey,Allyson Lloyd,Sarah Wyllie,Ethan Butcher,Anoop Chauhan,Samuel Robson |
| EPI_ISL_868792, EPI_ISL_868806, EPI_ISL_868807, EPI_ISL_868811, EPI_ISL_868819 | Bioinformatics and Biostatistics Lab, Advanced Sequencing Facility | COVID-19 Genomics UK (COG-UK) Consortium | Aengus Stewart,Jerome Nicod,Chelsea Sawyer,Laura Cubitt,Harshil Patel,Margaret Crawford |
| EPI_ISL_871972 | Instituto de Salud Carlos III | Instituto de Salud Carlos III | Iglesias-Caballero, M. Camarero, S. Molinero Calamita, M. González-Esguevillas, M. Pozo, F. Casas, I. Jiménez, P. Jiménez, M. Zaballos, A. Monzón, S. Varona, S. Juliá, M. Cuesta, I. |
| EPI_ISL_872258 | Labo Analyses Med | National Reference Center for Viruses of Respiratory Infections, Institut Pasteur, Paris | Marion Barbet, Sylvie Behillil, Méline Bizard, Angela Brisebarre, Camille Capel, Etienne Simon-Lorière, Vincent Enouf, Maud Vanpeene, Sylvie van der Werf,Girard Sophie |
| EPI_ISL_872681 | Rhode Island Department of Health | Infectious Disease Program, Broad Institute of Harvard and MIT | Lemieux,J.E., Siddle,K.J., Huard,R., King,E., Azevedo,K., Miller,A., Adams,G., Gladden-Young,A., Lagerborg,K., Rudy,M., DeRuff,K., Carter,A., Normandin,E., Bauer,M., Reilly,S., Tomkins-Tinch,C., Loreth,C., Chaluvadi,S., Birren,B.W., Gallagher,G., Smole,S., Park,D.J., MacInnis,B.L., and Sabeti,P.C. |
| EPI_ISL_875570, EPI_ISL_875571, EPI_ISL_875572, EPI_ISL_875573, EPI_ISL_875574, EPI_ISL_875575, EPI_ISL_875576, EPI_ISL_875577, EPI_ISL_875578, EPI_ISL_875579, EPI_ISL_875580, EPI_ISL_875581, EPI_ISL_875582, EPI_ISL_875583, EPI_ISL_875584, EPI_ISL_875585, EPI_ISL_875586, EPI_ISL_875587, EPI_ISL_875588, EPI_ISL_875589, EPI_ISL_875590, EPI_ISL_875591, EPI_ISL_875592, EPI_ISL_875593, EPI_ISL_875594, EPI_ISL_875595, EPI_ISL_875596, EPI_ISL_875597, EPI_ISL_875598, EPI_ISL_875599, EPI_ISL_875600, EPI_ISL_875601, EPI_ISL_875602, EPI_ISL_875603, EPI_ISL_875604, EPI_ISL_875605, EPI_ISL_875606, EPI_ISL_875607, EPI_ISL_875608, EPI_ISL_875609, EPI_ISL_875610, EPI_ISL_875611, EPI_ISL_875612, EPI_ISL_875613, EPI_ISL_875614, EPI_ISL_875615, EPI_ISL_875616, EPI_ISL_875617, EPI_ISL_875618, EPI_ISL_875619, EPI_ISL_875620, EPI_ISL_875621, EPI_ISL_875622, EPI_ISL_875623, EPI_ISL_875624, EPI_ISL_875625, EPI_ISL_875626, EPI_ISL_875627, EPI_ISL_875628, EPI_ISL_875629, EPI_ISL_875630, EPI_ISL_875631, EPI_ISL_875632, EPI_ISL_875633, EPI_ISL_875634, EPI_ISL_875635, EPI_ISL_875636, EPI_ISL_875637, EPI_ISL_875638, EPI_ISL_875639, EPI_ISL_875640, EPI_ISL_875641, EPI_ISL_875642, EPI_ISL_875643, EPI_ISL_875644, EPI_ISL_875645, EPI_ISL_875646, EPI_ISL_875647, EPI_ISL_875648, EPI_ISL_875649, EPI_ISL_875650, EPI_ISL_875651, EPI_ISL_875652, EPI_ISL_875653, EPI_ISL_875654, EPI_ISL_875655, EPI_ISL_875656, EPI_ISL_875657, EPI_ISL_875658, EPI_ISL_875659, EPI_ISL_875875, EPI_ISL_875887, EPI_ISL_875896, EPI_ISL_875897, EPI_ISL_875898, EPI_ISL_875903, EPI_ISL_875904, EPI_ISL_875927, EPI_ISL_875948, EPI_ISL_875954 | | | |
| see above | Eurofins Diatherix | Hudsonalpha Genome Sequencing Center | Jane Grimwood, Melissa Williams, Lori H. Handley, Joshua Stough, Leslie Malone, Stefan Brzezinski, Ada Stewart, Teresa Jones, Jenell Webber, John Lovell, Jennifer Cart, and Jeremy Schmutz |
| EPI_ISL_876860, EPI_ISL_876861, EPI_ISL_876862, EPI_ISL_876863, EPI_ISL_876864, EPI_ISL_876865 | Quest Diagnostics | Quest Diagnostics | Rosenthal,S.H., Gerasimova,A., Kagan,R.M., Anderson, B., Hua, M., Liu Y., Bernstein, L.E., Livingston, K.E., Perez, A., Shalhout, D.F., Shlyakhter, I.A., Owen, R., Tanpaiboon, P., Lacbawan, F. |
| EPI_ISL_877430 | Institute of Microbiology and Immunology, Faculty of Medicine, University of Ljubljana | Institute of Microbiology and Immunology, Faculty of Medicine, University of Ljubljana | Samo Zakotnik, Tomaž Mark Zorec, Matic Brvar, Miša Korva, Mario Poljak, Tatjana Avši - Županc |
| EPI_ISL_878563 | Robert Garry lab | Andersen lab at Scripps Research | Allison Smither, Gilberto Sabino-Santos, Patricia Snarski, Lilia Melnik, Antoinette Bell, Kaylynn Genemaras, Arnaud Drouin, Dahlene Fusco, Robert Garry with SEARCH Alliance San Diego |
| EPI_ISL_878669, EPI_ISL_878679, EPI_ISL_878710 | Rady's Childrens Hospital | Andersen lab at Scripps Research | SEARCH Alliance San Diego with Nanda Radamchar, David Dimmock, Linda Luo, Christina Clarke, Kathryn Bouic, Teresa Mueller, Denise Malicki |
| EPI_ISL_882727, EPI_ISL_882737, EPI_ISL_882738 | 1.AO Universitaria 'S. Giovanni di Dio e Ruggi D'Aragona, Scuola Medica Salernitana' Hospital / 2.UOC di Virologia e Microbiologia, Università della Campania 'L. Vanvitelli' / 3.AO Universitaria 'Federico II' Napoli Hospital / 4.AORN 'San Giuseppe Moscati' Avellino Hospital / 5.AO 'San Pio - presidio G. Rummo' Benevento Hospital / 6.AO 'Sant'Anna e San Sebastiano' Caserta Hospital / 7.PO 'Maria Santissima Addolorata' Eboli Hospital / 8.Biogem Istituto di Ricerche Genetiche | 1. Genome Research Center for Health (CRGS) / 2. Laboratory of Molecular Medicine and Genomics(LMMGe) / 3. Center for Research in Pure and Applied Mathematics (CRMPA) | Giorgio Giurato, Francesca Rizzo, Alessandro Weisz, Gianluigi Franci, Giovanni Nassa, Pasquale Pagliano, Roberta Tarallo, Elena Alexandrova, Ylenia D'Agostino, Carlo Ferravante, Jessica Lamberti, Viola Melone, Domenico Memoli, Valeria Mirici Cappa, Domenico Palumbo, Giovanni Pecoraro, Assunta Sellitto, Oriana Strianese, Ilaria Terenzi, Giuseppe Fenza, Aniello Gentile, Antonello Saccomanno, Sonia Amabile, Teresa Rocco, Annamaria Salvati, Emilia Vaccaro, Massimiliano Galdiero, Michele Cennamo, Giuseppe Portella, Maria Grazia Foti, Mariarosaria Ingino, Maria Landi, Maurizio Fumi, Vincenzo Rocco, Rita Greco, Vittoria Letizia, Arnolfo Petruzziello, Maddalena Schioppa, Gregorio Goffredi, Francesca Marciano, Michele Caraglia, Alessia Cossu, Marianna Scrima |
| EPI_ISL_882960 | Laboratorium Diagnostyi Mikrobiologicznej z Pracowni Prtka Grulicy SPSzW im. Jana Boego w Lublinie | National Institute of Public Health - National Institute of Hygiene | Wokowicz Tomasz, Zacharczuk Katarzyna, Gawor Jan |
| EPI_ISL_884317, EPI_ISL_884321, EPI_ISL_884335, EPI_ISL_884355 | Infectious Diseases, Quest Diagnostics | Infectious Diseases, Quest Diagnostics | Rosenthal,S.H., Gerasimova,A., Kagan,R.M., Anderson,B., Bernstein,L.E., Livingston,K.E., Hua,M., Liu,Y., Shalhout,D.F., Owen,R., Lacbawan,F. |
| EPI_ISL_887157 | Massachusetts General Hospital | Infectious Disease Program, Broad Institute of Harvard and MIT | Lemieux,J.E., Siddle,K.J., Shaw,B., Adams,G., Pierce,V., Turbett,S., Anahtar,M., Branda,J., Slater,D., Harris,J., Lin,A.E., Gladden-Young,A., Lagerborg,K., Rudy,M., DeRuff,K., Carter,A., Normandin,E., Bauer,M., Reilly,S., Tomkins-Tinch,C., Loreth,C., Chaluvadi,S., Neumann,A., Cusick,C., Chapman,S.B., Gnirke,A., Flowers,K., Cerrato,F., Birren,B.W., Gallagher,G., Smole,S., Park,D.J., MacInnis,B.L., Ryan,E., LaRocque,R., Rosenberg,E. and Sabeti,P.C. |
| EPI_ISL_888828, EPI_ISL_888829 | General practitioner office | National Reference Center for Viruses of Respiratory Infections, Institut Pasteur, Paris | Marion Barbet, Sylvie Behillil, Méline Bizard, Angela Brisebarre, Camille Capel, Etienne Simon-Lorière, Vincent Enouf, Maud Vanpeene, Sylvie van der Werf |
| EPI_ISL_888884, EPI_ISL_888885, EPI_ISL_888886, EPI_ISL_888889, EPI_ISL_888890, EPI_ISL_888891, | Michigan Department of Health and Human Services, Bureau of Laboratories | Michigan Department of Health and Human Services, Bureau of Laboratories | Blankenship HM, Riner D, Soehnlen MK |

| | | | |
|---|---|---|---|
| EPI_ISL_888892, EPI_ISL_888893 | | | |
| EPI_ISL_896148 | MEPHI, Aix Marseille University | MEPHI, Aix Marseille University | Anthony LEVASSEUR |
| EPI_ISL_896374 | University of Medicine and Pharmacy of Craiova | "Stefan cel Mare" University Metagenomics Lab | Lobiuc Andrei, Gheorghita Roxana |
| EPI_ISL_900129, EPI_ISL_900181, EPI_ISL_900228, EPI_ISL_900370, EPI_ISL_900483 | MEPHI, Aix Marseille University | MEPHI, Aix Marseille University | Anthony LEVASSEUR |
| EPI_ISL_900570 | Hôpital Paris Saint-Joseph | CNR Virus des Infections Respiratoires - France SUD | Antonin Bal, Gregory Destras, Gwendolyne Burfin, Hadrien Règue, Quentin Semanas, Martine Valette, Bruno Lina, Sylvie Larrat, Laurence Josset |
| EPI_ISL_900573 | Centre Hospitalier de Guéret | CNR Virus des Infections Respiratoires - France SUD | Antonin Bal, Gregory Destras, Gwendolyne Burfin, Hadrien Règue, Quentin Semanas, Martine Valette, Bruno Lina, Sylvie Larrat, Laurence Josset |
| EPI_ISL_904252, EPI_ISL_904452, EPI_ISL_904453, EPI_ISL_904454 | Dutch COVID-19 response team | Erasmus Medical Center | Bas Oude Munnink, Reina Sikkema, David Nieuwenhuijse, Irina Chestakova, Anne van der Linden, Marjan Boter, Emmanuelle Munger, Corine GeurtsvanKessel, Annemiek van der Eijk, Richard Molenkamp, Marion Koopmans, on behalf of the Dutch national COVID-19 response team. |
| EPI_ISL_906119 | Institute of Microbiology and Immunology, Faculty of Medicine, University of Ljubljana | Institute of Microbiology and Immunology, Faculty of Medicine, University of Ljubljana | Samo Zakotnik, Tomaž Mark Zorec, Matic Brvar, Miša Korva, Mario Poljak, Tatjana Avši - Županc |
| EPI_ISL_912482, EPI_ISL_912495, EPI_ISL_912514, EPI_ISL_912518, EPI_ISL_912534 | NHLS Universitas Academic | UFS Virology | PA Bester, MM Nyaga, P Nthiga, MT Mogotsi, D Goedhals, T de Oliveira |
| EPI_ISL_913075, EPI_ISL_913096 | Center for Virology | Center for Virology | Jeremy V. Camp, Irene Goerzer, Monika Redlberger-Fritz, Stephan W. Aberle |
| EPI_ISL_915405, EPI_ISL_915406, EPI_ISL_915407 | Keio University School of Medicine | Keio University School of Medicine | Kenjiro Kosaki, Yuka Iwasaki, Hirotsugu Ishizu, Haruhiko Siomi, Kodai Abe |
| EPI_ISL_924425 | Virology Department, Sheffield Teaching Hospitals NHS Foundation Trust/Department of Infection, Immunity and Cardiovascular Disease, The Medical School, University of Sheffield | COVID-19 Genomics UK (COG-UK) Consortium | Thushan de Silva, Matthew Parker, Nikki Smith, Adri Angyal, Rebecca Brown, Luke Green, Rachel Tucker, Paul Parsons, Danielle Groves, Katie Johnson, Laura Carrilero, Alex Keeley, Dave Partridge, Matthew Wyles, Benjamin Lindsey, Mehmet Yavuz, Mohammad Raza, Cariad Evans |
| EPI_ISL_924608, EPI_ISL_924610, EPI_ISL_924611, EPI_ISL_924618, EPI_ISL_924620, EPI_ISL_925017, EPI_ISL_925019, EPI_ISL_925020, EPI_ISL_925027, EPI_ISL_925029 | Bioinformatics and Biostatistics Lab, Advanced Sequencing Facility | COVID-19 Genomics UK (COG-UK) Consortium | Aengus Stewart,Jerome Nicod,Chelsea Sawyer,Laura Cubitt,Harshil Patel,Margaret Crawford |
| EPI_ISL_925094, EPI_ISL_925095, EPI_ISL_925116, EPI_ISL_925117 | 1.AO Universitaria 'S. Giovanni di Dio e Ruggi D'Aragona, Scuola Medica Salernitana' Hospital / 2.UOC di Virologia e Microbiologia, Università della Campania 'L. Vanvitelli' / 3.AO Universitaria 'Federico II' Napoli Hospital / 4.AORN 'San Giuseppe Moscati' Avellino Hospital / 5.AO 'San Pio - presidio G. Rummo' Benevento Hospital / 6.AO 'Sant'Anna e San Sebastiano' Caserta Hospital / 7.PO 'Maria Santissima Addolorata' Eboli Hospital / 8.Biogem Istituto di Ricerche Genetiche | 1. Genome Research Center for Health (CRGS) / 2. Laboratory of Molecular Medicine and Genomics(LMMGe) / 3. Center for Research in Pure and Applied Mathematics (CRMPA) | Giorgio Giurato, Francesca Rizzo, Alessandro Weisz, Gianluigi Franci, Giovanni Nassa, Pasquale Pagliano, Roberta Tarallo, Elena Alexandrova, Ylenia D'Agostino, Carlo Ferravante, Jessica Lamberti, Viola Melone, Domenico Memoli, Valeria Mirici Cappa, Domenico Palumbo, Giovanni Pecoraro, Assunta Sellitto, Oriana Strianese, Ilaria Terenzi, Giuseppe Fenza, Aniello Gentile, Antonello Saccomanno, Sonia Amabile, Teresa Rocco, Annamaria Salvati, Emilia Vaccaro, Massimiliano Galdiero, Michele Cennamo, Giuseppe Portella, Maria Grazia Foti, Mariarosaria Ingino, Maria Landi, Maurizio Fumi, Vincenzo Rocco, Rita Greco, Vittoria Letizia, Arnolfo Petruzziello, Maddalena Schioppa, Gregorio Goffredi, Francesca Marciano, Michele Caraglia, Alessia Cossu, Marianna Scrima |
| EPI_ISL_925371, EPI_ISL_925372, EPI_ISL_925373, EPI_ISL_925374, EPI_ISL_925375, EPI_ISL_925449, EPI_ISL_925450, EPI_ISL_925451, EPI_ISL_925452, EPI_ISL_925453, EPI_ISL_925454, EPI_ISL_925455, EPI_ISL_925456, EPI_ISL_925457, EPI_ISL_925458, EPI_ISL_925459, EPI_ISL_925460, EPI_ISL_925461, EPI_ISL_925462, EPI_ISL_925490, EPI_ISL_925491, EPI_ISL_925492 | | | |
| see above | Department of Clinical Microbiology | GIGA Medical Genomics | Keith Durkin, Maria Artesi, Sébastien Bontems, Raphaël Boreux, Bouchra Boujemla, Cécile Meex, Pierrette Melin, Marie-Pierre Hayette, Vincent Bours |
| EPI_ISL_930964, EPI_ISL_931174, EPI_ISL_931175, EPI_ISL_931176, EPI_ISL_931177, EPI_ISL_931178, EPI_ISL_931179, EPI_ISL_931180, EPI_ISL_931181, EPI_ISL_931182, EPI_ISL_931183, EPI_ISL_931184, EPI_ISL_931185, EPI_ISL_931186, EPI_ISL_931187, EPI_ISL_931188, EPI_ISL_931190, EPI_ISL_931191, EPI_ISL_931192, EPI_ISL_931193, EPI_ISL_931194, EPI_ISL_931195, EPI_ISL_931196, EPI_ISL_931197, EPI_ISL_931199, EPI_ISL_931200, EPI_ISL_931201, EPI_ISL_931202, EPI_ISL_931203, EPI_ISL_931204, EPI_ISL_931205, EPI_ISL_931206, EPI_ISL_931207, EPI_ISL_931208, EPI_ISL_931209, EPI_ISL_931210, EPI_ISL_931211, EPI_ISL_931212, EPI_ISL_931213, EPI_ISL_931214, EPI_ISL_931215, EPI_ISL_931216, EPI_ISL_931217, EPI_ISL_931218, EPI_ISL_931219, EPI_ISL_931220, EPI_ISL_931221, EPI_ISL_931222, EPI_ISL_931223, EPI_ISL_931224, EPI_ISL_931225, EPI_ISL_931226, EPI_ISL_931227, EPI_ISL_931228, EPI_ISL_931229, EPI_ISL_931230, EPI_ISL_931231, EPI_ISL_931232, EPI_ISL_931233, EPI_ISL_931234, EPI_ISL_931235, EPI_ISL_931236, EPI_ISL_931237, EPI_ISL_931243, EPI_ISL_931244, EPI_ISL_931245, EPI_ISL_931246, EPI_ISL_931247, EPI_ISL_931248, EPI_ISL_931249, EPI_ISL_931250, EPI_ISL_931251, EPI_ISL_931252, EPI_ISL_931253, EPI_ISL_931254, EPI_ISL_931255, EPI_ISL_931256, EPI_ISL_931257, EPI_ISL_931258, EPI_ISL_931259, EPI_ISL_931260, EPI_ISL_931261, EPI_ISL_931262, EPI_ISL_931263, EPI_ISL_931264, EPI_ISL_931265, EPI_ISL_931266, EPI_ISL_931267, EPI_ISL_931268, EPI_ISL_931269, EPI_ISL_931270, EPI_ISL_931271, EPI_ISL_931272, EPI_ISL_931273, EPI_ISL_931274, EPI_ISL_931275, EPI_ISL_931276, EPI_ISL_931277, EPI_ISL_931278, EPI_ISL_931279, EPI_ISL_931280, EPI_ISL_931281, EPI_ISL_931282, EPI_ISL_931283, EPI_ISL_931284, EPI_ISL_931285, EPI_ISL_931286, EPI_ISL_931287, EPI_ISL_931288, EPI_ISL_931289, EPI_ISL_931290, EPI_ISL_931291, EPI_ISL_931292, EPI_ISL_931293, EPI_ISL_931294, EPI_ISL_931295, EPI_ISL_931296, EPI_ISL_931297, EPI_ISL_931298, EPI_ISL_931299, EPI_ISL_931300, EPI_ISL_931301, EPI_ISL_931302, EPI_ISL_931303, EPI_ISL_931304, EPI_ISL_931305, EPI_ISL_931306, EPI_ISL_931307, EPI_ISL_931308, EPI_ISL_931309, EPI_ISL_931310 | | | |
| see above | University Hospital Basel, Clinical Virology | University Hospital Basel, Clinical Bacteriology | Tim Roloff, Madlen Stange, Helena MB Seth-Smith, Alfredo Mari, Karoline Leuzinger, Julia Bielicki, Manuel Battegay, Hans Hirsch, Adrian Egli |
| EPI_ISL_933573 | General Hospital - Kumanovo | Research Center for Genetic Engineering and Biotechnology "Georgi D. Efremov" , Macedonian Academy of Sciences and Arts | RCGEB - MASA |
| EPI_ISL_933574 | General Hospital - Tetovo | Research Center for Genetic Engineering and Biotechnology "Georgi D. Efremov" , Macedonian Academy of Sciences and Arts | RCGEB - MASA |
| EPI_ISL_933577 | Special hospital for orthopedic surgery and traumatology "Sv Erazmo" - Ohrid | Research Center for Genetic Engineering and Biotechnology "Georgi D. Efremov" , Macedonian Academy of Sciences and Arts | RCGEB - MASA |
| EPI_ISL_933578 | General Hospital - Veles | Research Center for Genetic Engineering and Biotechnology "Georgi D. Efremov" , Macedonian Academy of Sciences and Arts | RCGEB - MASA |
| EPI_ISL_933579 | General Hospital - Strumica | Research Center for Genetic Engineering and Biotechnology "Georgi D. Efremov" , Macedonian Academy of Sciences and Arts | RCGEB - MASA |
| EPI_ISL_933602 | General Hospital - Shtip | Research Center for Genetic Engineering and Biotechnology "Georgi D. Efremov" , Macedonian Academy of Sciences and Arts | RCGEB - MASA |
| EPI_ISL_935149 | 1.AO Universitaria 'S. Giovanni di Dio e Ruggi D'Aragona, Scuola Medica Salernitana' Hospital / 2.UOC di Virologia e Microbiologia, Università della Campania 'L. Vanvitelli' / 3.AO Universitaria 'Federico II' Napoli Hospital / 4.AORN 'San Giuseppe Moscati' Avellino Hospital / 5.AO 'San Pio - presidio G. Rummo' Benevento Hospital / 6.AO 'Sant'Anna e San Sebastiano' Caserta Hospital / 7.PO 'Maria Santissima Addolorata' Eboli Hospital / 8.Biogem Istituto di Ricerche Genetiche | 1. Genome Research Center for Health (CRGS) / 2. Laboratory of Molecular Medicine and Genomics(LMMGe) / 3. Center for Research in Pure and Applied Mathematics (CRMPA) | Giorgio Giurato (Corresponding Author), Francesca Rizzo (Corresponding Author), Alessandro Weisz (Corresponding Author), Gianluigi Franci, Giovanni Nassa, Pasquale Pagliano, Roberta Tarallo, Elena Alexandrova, Ylenia D'Agostino, Carlo Ferravante, Jessica Lamberti, Viola Melone, Domenico Memoli, Valeria Mirici Cappa, Domenico Palumbo, Giovanni Pecoraro, Assunta Sellitto, Oriana Strianese, Ilaria Terenzi, Giuseppe Fenza, Aniello Gentile, Antonello Saccomanno, Sonia Amabile, Teresa Rocco, Annamaria Salvati, Emilia Vaccaro, Massimiliano Galdiero, Michele Cennamo, Giuseppe Portella, Maria Grazia Foti, Mariarosaria Ingino, Maria Landi, Maurizio Fumi, Vincenzo Rocco, Rita Greco, Vittoria Letizia, Arnolfo Petruzziello, Maddalena Schioppa, Gregorio Goffredi, Francesca Marciano, Michele Caraglia, Alessia Cossu, Marianna Scrima |

| | | | |
|---|---|---|---|
| EPI_ISL_936495 | Sarawak General Hospital | Institute for Medical Research, Infectious Disease Research Centre, National Institutes of Health, Ministry of Health Malaysia | Suppiah J, Kamel K, Azizan MA, Thayan R |
| EPI_ISL_937046, EPI_ISL_937047 | Quest Diagnostics | Quest Diagnostics | Rosenthal,S.H., Gerasimova,A., Kagan,R.M., Anderson, B., Livingston, K.E., Hua, M., Liu Y., Shalhout, D.F., Owen, R., Lacbawan, F. |
| EPI_ISL_943984, EPI_ISL_943985 | LACEN do Estado de Tocantins | Instituto Adolfo Lutz, Interdiciplinary Procedures Center, Strategic Laboratory | Claudio Tavares Sacchi, Claudia Regina Gonçalves, Erica Valessa Ramos Gomes, Karoline Rodrigues Campos |
| EPI_ISL_944762, EPI_ISL_944763, EPI_ISL_944765, EPI_ISL_944768, EPI_ISL_944769, EPI_ISL_944771, EPI_ISL_944772, EPI_ISL_944773, EPI_ISL_944775, EPI_ISL_944776, EPI_ISL_944781, EPI_ISL_944783, EPI_ISL_944784, EPI_ISL_944788, EPI_ISL_944789, EPI_ISL_944791 | | | |
| see above | CSIR-Indian Institute of Chemical Biology, MEDICA Supercpecialty Hospital Kolkata | CSIR-Indian Institute of Chemical Biology, MEDICA Supercpecialty Hospital Kolkata | Sujay Krishna Maity, Priyanka Mallick, Debaleena Bhowmik, Abhishake Lahiri, Dr. Aviral Roy, Dr. Soumen Saha, Dr. Arpita Ghosh Mitra, Dr. Rajesh Pandey, Dr. Sandip Paul, Dr. Partha Chakrabarti, Dr. Saikat Chakrabarti |
| EPI_ISL_949647, EPI_ISL_949648 | Virology Department, Royal Infirmary of Edinburgh, NHS Lothian / School of Biological Sciences, University of Edinburgh / Institute of Genetics and Molecular Medicine, University of Edinburgh | COVID-19 Genomics UK (COG-UK) Consortium | McHugh M, Dewar R, Rooke S, Gallagher M, Balcaza C, O'Toole Á, Scher E, Hill V, McCrone JT, Colquhoun R, Yu X, Jackson B, Rambaut A, Williams TC, Templeton K |
| EPI_ISL_955151 | University of Sarajevo, Veterinary Faculty, Laboratory for Molecular Diagnostic and Research Laboratory | University of Sarajevo, Veterinary Faculty, Laboratory for Molecular Diagnostic and Research Laboratory | Goleti T., Goleti Š., Softi A., Ali-Šeho A., Terzi I., Šabi E., Jaži A., Nicevi M., Hodži A. |
| EPI_ISL_956317 | Laboratory Medicine | Department of Laboratory Medicine, Lin-Kou Chang Gung Memorial Hospital, Taoyuan, Taiwan | Kuo-Chien Tsao, Yu-Nong Gong, Shu-Li Yang, Yi-Chun Liu, Chung-Guei Huang, Mei-Jen Hsiao, Po-Wei Huang, Cheng-Ta Yang, Cheng-Hsun Chiu, Peng-Nien Huang, Kuo-Ming Lee, Guang-Wu Chen, Shin-Ru Shih |
| EPI_ISL_960127 | Conville CDC wc CVC | National Health Laboratory Service/UCT | Arash Iranzadeh, Deelan Doolabh, Lynn Tyers, Bruna Galvao, Innocent Mudau, Marvin Hsiao, Kruger Marais, Diana Hardie, Stephen Korsman, Carolyn Williamson |
| EPI_ISL_960321, EPI_ISL_960354, EPI_ISL_960391 | University of Wisconsin-Madison AIDS Vaccine Research Laboratories | University of Wisconsin-Madison AIDS Vaccine Research Laboratories | Gage Moreno, Katarina Braun, et al. AIDS Vaccine Research Laboratories |
| EPI_ISL_961909, EPI_ISL_961947, EPI_ISL_961948, EPI_ISL_961959, EPI_ISL_961993, EPI_ISL_962012, EPI_ISL_962155, EPI_ISL_962156, EPI_ISL_962157, EPI_ISL_962178 | Illinois Department of Public Health | Gagnon Lab, Southern Illinois University | Keith Gagnon |
| EPI_ISL_965340, EPI_ISL_965355, EPI_ISL_965365, EPI_ISL_965463, EPI_ISL_965525 | University of Liège COVID-19 testing center | GIGA Medical Genomics | Keith Durkin, Maria Artesi, Bouchra Boujemla, Emmanuel André, Marc Van Ranst, Fabrice Bureau, Laurent Gillet, Wouter Coppieters, Vincent Bours |
| EPI_ISL_965546, EPI_ISL_965567, EPI_ISL_965570, EPI_ISL_965656 | Dutch COVID-19 response team | Medical Microbiology, Maastricht University Medical Centre | Jozef Dingemans*, Brian van der Veer*, Erik Beuken, Carmen Reumkens, Lieke van Alphen, Christian Hoebe, Paul Savelkoul |
| EPI_ISL_968838, EPI_ISL_968896, EPI_ISL_968897, EPI_ISL_968901, EPI_ISL_968902, EPI_ISL_968903, EPI_ISL_968904, EPI_ISL_968984 | KEMRI-Wellcome Trust Research Programme/KEMRI-CGMR-C Kilifi | KEMRI-Wellcome Trust Research Programme/KEMRI-CGMR-C Kilifi | Githinji et al |
| EPI_ISL_974642, EPI_ISL_974643, EPI_ISL_974644, EPI_ISL_974645, EPI_ISL_974646, EPI_ISL_974647, EPI_ISL_974648, EPI_ISL_974649, EPI_ISL_974650, EPI_ISL_974651, EPI_ISL_974652, EPI_ISL_974653, EPI_ISL_974654, EPI_ISL_974655, EPI_ISL_974656, EPI_ISL_974657, EPI_ISL_974658, EPI_ISL_974659, EPI_ISL_974660, EPI_ISL_974661, EPI_ISL_974662, EPI_ISL_974663, EPI_ISL_974664, EPI_ISL_974665, EPI_ISL_974666, EPI_ISL_974667, EPI_ISL_974668, EPI_ISL_974669, EPI_ISL_974670, EPI_ISL_974671, EPI_ISL_974672, EPI_ISL_974673, EPI_ISL_974674, EPI_ISL_974675, EPI_ISL_974676, EPI_ISL_974677, EPI_ISL_974678, EPI_ISL_974679, EPI_ISL_974680, EPI_ISL_974681, EPI_ISL_974682, EPI_ISL_974683, EPI_ISL_974684, EPI_ISL_974685, EPI_ISL_974686, EPI_ISL_974687, EPI_ISL_974688, EPI_ISL_974689, EPI_ISL_974690, EPI_ISL_974691, EPI_ISL_974692, EPI_ISL_974693, EPI_ISL_974694, EPI_ISL_974695, EPI_ISL_974696, EPI_ISL_974697, EPI_ISL_974698, EPI_ISL_974699, EPI_ISL_974700, EPI_ISL_974701, EPI_ISL_974702, EPI_ISL_974703, EPI_ISL_974704, EPI_ISL_974705, EPI_ISL_974706, EPI_ISL_974707, EPI_ISL_974708, EPI_ISL_974709, EPI_ISL_974710, EPI_ISL_974711, EPI_ISL_974712, EPI_ISL_974713, EPI_ISL_974714, EPI_ISL_974715, EPI_ISL_974716, EPI_ISL_974717, EPI_ISL_974718, EPI_ISL_974719, EPI_ISL_974720, EPI_ISL_974721, EPI_ISL_974722, EPI_ISL_974723, EPI_ISL_974724, EPI_ISL_974725, EPI_ISL_974726, EPI_ISL_974727, EPI_ISL_974728, EPI_ISL_974729, EPI_ISL_974730, EPI_ISL_974731, EPI_ISL_974732, EPI_ISL_974733, EPI_ISL_974734, EPI_ISL_974735, EPI_ISL_974736, EPI_ISL_974737, EPI_ISL_974738, EPI_ISL_974739, EPI_ISL_974740, EPI_ISL_974741, EPI_ISL_974742, EPI_ISL_974743, EPI_ISL_974744, EPI_ISL_974746, EPI_ISL_974747, EPI_ISL_974748, EPI_ISL_974749, EPI_ISL_974750, EPI_ISL_974751, EPI_ISL_974752, EPI_ISL_974753, EPI_ISL_974754, EPI_ISL_974755, EPI_ISL_974756, EPI_ISL_974757, EPI_ISL_974758 | | | |
| see above | BCCDC Public Health Laboratory | BCCDC Public Health Laboratory | Prystajecky Natalie, Linda Hoang, Dan Fornika, John Tyson, Shannon Russell, Kim Macdonald, Kimia Kamelian, Ana Pacagnella, Corrinne Ng, Loretta Janz, Robert Azana Terry Snutch, Mel Krajden |
| EPI_ISL_977421 | University of Zambia, School of Veterinary Medicine | UNZAVET and PATH | Mulenga Mwenda-Chimfwembe, Ngonda Saasa, Daniel Bridges |
| EPI_ISL_977482 | Instituto Adolfo Lutz - Regional de Aracatuba | Instituto Adolfo Lutz, Interdiciplinary Procedures Center, Strategic Laboratory | Claudio Tavares Sacchi, Claudia Regina Gonçalves, Erica Valessa Ramos Gomes, Karoline Rodrigues Campos |
| EPI_ISL_977487 | Instituto Adolfo Lutz Central | Instituto Adolfo Lutz, Interdiciplinary Procedures Center, Strategic Laboratory | Claudio Tavares Sacchi, Claudia Regina Gonçalves, Erica Valessa Ramos Gomes, Karoline Rodrigues Campos |
| EPI_ISL_982507, EPI_ISL_982508, EPI_ISL_982509, EPI_ISL_982510 | Kentucky State Public Health Lab | Kentucky State Public Health Lab | Stephanie Lunn, Karim George, Joshua Tobias, William Grooms, Vaneet Arora, Matthew Johnson, Rachel Zinner, Rhonda Lucas |
