## Supplementary material for "Detection and characterization of the SARS-CoV-2 lineage B.1.526 in New York": Supp. Table 3 GISAID Acknowledgment part 2: gisaid_hcov-19_acknowledgement_table_2021_02_13_00-4.pdf

We gratefully acknowledge the following Authors from the Originating laboratories responsible for obtaining the specimens, as well as the Submitting laboratories where the genome data were generated and shared via GISAID, on which this research is based.

All Submitters of data may be contacted directly via www.gisaid.org

Authors are sorted alphabetically.

| Accession ID | Originating Laboratory | Submitting Laboratory | Authors |
|---|---|---|---|
| EPI_ISL_622780 | Canterbury Health Laboratories | Institute of Environmental Science and Research (ESR) | Xiaoyun Ren, Matt Storey, Nikki Freed, Muhammad Faisal, Jing Wang, Hermes Perez, Anja Werno, Antje van der Linden, Arlo Upton, Chris Mansell, David Hammer, Dragana Drinkovic, Gary McAuliffe, Hana Sofia Andersson, James Ussher, Jill Sherwood, Josh Freeman, Julia Howard, Juliet Elvy, Mary DeAlmeida, Matt Blakiston, Matthew Rogers, Max Bloomfield, Michael Addidle, Michelle Balm, Sally Roberts, Sarah Jefferies, Sharmini Muttaiyah, Susan Morpeth, Susan Taylor, Timothy Blackmore, Vani Sathyendran, Veronica Playle, Virginia Hope, Erasmus Smit, Lauren Jelly, Olin Silander, Joep de Ligt |
| EPI_ISL_622792 | Middlemore Hospital | Institute of Environmental Science and Research (ESR) | Xiaoyun Ren, Matt Storey, Nikki Freed, Muhammad Faisal, Jing Wang, Hermes Perez, Anja Werno, Antje van der Linden, Arlo Upton, Chris Mansell, David Hammer, Dragana Drinkovic, Gary McAuliffe, Hana Sofia Andersson, James Ussher, Jill Sherwood, Josh Freeman, Julia Howard, Juliet Elvy, Mary DeAlmeida, Matt Blakiston, Matthew Rogers, Max Bloomfield, Michael Addidle, Michelle Balm, Sally Roberts, Sarah Jefferies, Sharmini Muttaiyah, Susan Morpeth, Susan Taylor, Timothy Blackmore, Vani Sathyendran, Veronica Playle, Virginia Hope, Erasmus Smit, Lauren Jelly, Olin Silander, Joep de Ligt |
| EPI_ISL_622824, EPI_ISL_622828, EPI_ISL_622829 | Canterbury Health Laboratories | Institute of Environmental Science and Research (ESR) | Xiaoyun Ren, Matt Storey, Nikki Freed, Muhammad Faisal, Jing Wang, Hermes Perez, Anja Werno, Antje van der Linden, Arlo Upton, Chris Mansell, David Hammer, Dragana Drinkovic, Gary McAuliffe, Hana Sofia Andersson, James Ussher, Jill Sherwood, Josh Freeman, Julia Howard, Juliet Elvy, Mary DeAlmeida, Matt Blakiston, Matthew Rogers, Max Bloomfield, Michael Addidle, Michelle Balm, Sally Roberts, Sarah Jefferies, Sharmini Muttaiyah, Susan Morpeth, Susan Taylor, Timothy Blackmore, Vani Sathyendran, Veronica Playle, Virginia Hope, Erasmus Smit, Lauren Jelly, Olin Silander, Joep de Ligt |
| EPI_ISL_622831 | Middlemore Hospital | Institute of Environmental Science and Research (ESR) | Xiaoyun Ren, Matt Storey, Nikki Freed, Muhammad Faisal, Jing Wang, Hermes Perez, Anja Werno, Antje van der Linden, Arlo Upton, Chris Mansell, David Hammer, Dragana Drinkovic, Gary McAuliffe, Hana Sofia Andersson, James Ussher, Jill Sherwood, Josh Freeman, Julia Howard, Juliet Elvy, Mary DeAlmeida, Matt Blakiston, Matthew Rogers, Max Bloomfield, Michael Addidle, Michelle Balm, Sally Roberts, Sarah Jefferies, Sharmini Muttaiyah, Susan Morpeth, Susan Taylor, Timothy Blackmore, Vani Sathyendran, Veronica Playle, Virginia Hope, Erasmus Smit, Lauren Jelly, Olin Silander, Joep de Ligt |
| EPI_ISL_626352, EPI_ISL_626353, EPI_ISL_626354 | LabPLUS | Institute of Environmental Science and Research (ESR) | Xiaoyun Ren, Matt Storey, Nikki Freed, Muhammad Faisal, Jing Wang, Hermes Perez, Anja Werno, Antje van der Linden, Arlo Upton, Chris Mansell, David Hammer, Dragana Drinkovic, Gary McAuliffe, Hana Sofia Andersson, James Ussher, Jill Sherwood, Josh Freeman, Julia Howard, Juliet Elvy, Mary DeAlmeida, Matt Blakiston, Matthew Rogers, Max Bloomfield, Michael Addidle, Michelle Balm, Sally Roberts, Sarah Jefferies, Sharmini Muttaiyah, Susan Morpeth, Susan Taylor, Timothy Blackmore, Vani Sathyendran, Veronica Playle, Virginia Hope, Erasmus Smit, Lauren Jelly, Olin Silander, Joep de Ligt |
| EPI_ISL_626641, EPI_ISL_626642, EPI_ISL_626643, EPI_ISL_626644 | National Public Health Laboratory, National Centre for Infectious Diseases | National Public Health Laboratory, National Centre for Infectious Diseases | Tze Minn Mak, Sophie Octavia, Zhenyang Zhou, Lin Cui, Raymond Tzer Pin Lin |
| EPI_ISL_626670, EPI_ISL_626740, EPI_ISL_626750, EPI_ISL_626770, EPI_ISL_627215, EPI_ISL_627310, EPI_ISL_627700, EPI_ISL_627701, EPI_ISL_627702, EPI_ISL_627703, EPI_ISL_627704, EPI_ISL_627705, EPI_ISL_627706, EPI_ISL_627707, EPI_ISL_627708, EPI_ISL_627709, EPI_ISL_627710, EPI_ISL_627716 | | | |
| see above | Queens Medical Centre, Clinical Microbiology Department / DeepSeq Nottingham | COVID-19 Genomics UK (COG-UK) Consortium | Gemma Clark, Wendy Smith, Manjinder Khakh, Vicki M Fleming, Michelle M Lister, Hannah Howson-Wells, Jonathan Ball, Patrick McClure, Joseph Chappell, Theocharis Tsoleridis, Nadine Holmes, Matthew Carlisle, Christopher Moore, Fei Sang, Johnny Debebe, Victoria Wright, Matthew Loose |
| EPI_ISL_629009 | Laverty Pathology | NSW Health Pathology - Institute of Clinical Pathology and Medical Research; Westmead Hospital; University of Sydney | CIDM-PH et al. |
| EPI_ISL_632310 | 1-Laboratory of Microbiology, National Reference Lab, Charles Nicolle Hospital; 2-University of Tunis ElManar, Faculty of Medicine of Tunis, LR99ES09, Tunis, Tunisia | 1-Clinical and Experimental Pharmacology Lab, LR16SP02, National Center of Pharmacovigilance, University of Tunis El Manar, Tunis, Tunisia. 2-Neurodegenerative diseases and psychiatric troubles, LR18SP03, Razi Hospital, University of Tunis El Manar, Tunis, Tunisia. 3- Ministry of Health, National Observatory of New and Emerging Diseases, 1006, Tunis, Tunisia | Ilhem Boutiba-Ben Boubaker, Sameh Trabelsi, Nissaf Ben Alaya, Maher Kharrat, Alia Ben Kahla, Jalila Ben Khelil, Salma Abid, Sana Ferjani, Mouna Ben Sassi, Mouna Safer, Imen Mkada, Imen Kacem, Gaies Emna, Soumaya Rammeh, Riadh Daghfous, Riadh Gouider. |
| EPI_ISL_632398, EPI_ISL_632406, EPI_ISL_632449, EPI_ISL_632766, EPI_ISL_632767, EPI_ISL_632769, EPI_ISL_632793, EPI_ISL_632794 | Dutch COVID-19 response team | Erasmus Medical Center | Bas Oude Munnink, David Nieuwenhuijse, Reina Sikkema, Claudia Schapendonk, Irina Chestakova, Anne van der Linden, Theo Bestebroer, Stefan van Nieuwkoop, Mark Pronk, Pascal Lexmond, Corien Swaan, Manon Haverkate, Madelief Mollers, Mart Stein, Sandra Kengne Kamga Mobou, Jeroen van Kampen, Jolanda Voermans, Aura Timen, Corine GeurtsvanKessel, Annemiek van der Eijk, Richard Molenkamp, Marion Koopmans, on behalf of the Dutch national COVID-19 response team. |
| EPI_ISL_634977 | 1-Laboratory of Microbiology, National Reference Lab, Charles Nicolle Hospital; 2-University of Tunis ElManar, Faculty of Medicine of Tunis, LR99ES09, Tunis, Tunisia | 1-Clinical and Experimental Pharmacology Lab, LR16SP02, National Center of Pharmacovigilance, University of Tunis El Manar, Tunis, Tunisia. 2-Neurodegenerative diseases and psychiatric troubles, LR18SP03, Razi Hospital, University of Tunis El Manar, Tunis, Tunisia. 3- Ministry of Health, National Observatory of New and Emerging Diseases, 1006, Tunis, Tunisia | Ilhem Boutiba-Ben Boubaker, Sameh Trabelsi, Nissaf Ben Alaya, Maher Kharrat, Alia Ben Kahla, Jalila Ben Khelil, Salma Abid, Sana Ferjani, Mouna Ben Sassi, Mouna Safer, Imen Mkada, Imen Kacem, Gaies Emna, Soumaya Rammeh, Riadh Daghfous, Riadh Gouider. |
| EPI_ISL_635083 | Haukeland University Hospital, Department of Medical Microbiology | Norwegian Institute of Public Health, Department of Virology | Kathrine Stene-Johansen, Kamilla Heddeland Instefjord, Hilde Elshaug, Marie Paulsen Madsen, Rasmus Riis Kopperud, Hilde Vollan, Karoline Bragstad, Olav Hungnes |
| EPI_ISL_637168, EPI_ISL_637169, EPI_ISL_637170, EPI_ISL_637172, EPI_ISL_637174, EPI_ISL_637176, EPI_ISL_637177 | Respiratory Virus Unit, Microbiology Services Colindale, Public Health England | COVID-19 Genomics UK (COG-UK) Consortium | PHE Covid Sequencing Team |
| EPI_ISL_637257, EPI_ISL_637259, EPI_ISL_637263 | Liverpool Clinical Laboratories | COVID-19 Genomics UK (COG-UK) Consortium | Sam Haldenby, Anita Lucaci, Steve Paterson, Julian Hiscox, Alistair Darby, M Almsaud, A Alrezaihi, Muhannad Alruwaili, Stuart D Armstrong, Jones Benjamin, Eleanor G Bentley, Anu Chawla, Jordan J Clark, Angela Cowell, Richard Eccles, Isabel García-Dorival, Matthew Gemmell, Alessandro Gerada, PKF Gilmore, Richard Gregory, Ximeng Han, Catherine Hartley, Margaret Hughes, Miren Iturriza-Gomara, James Johnson, L Luu, Jenifer Manson, Charlotte Nelson, Elaine O'Toole, Cassie Olateju, Rebekah Penrice-Randal , Lucille Rainbow, N.P Randle, Trevor Ian Robinson, Parul Sharma, Ghada T Shawli, James P Stewart, Neil Swainston, Ecaterina Vamos, Joanne Watts, Mark Whitehead |
| EPI_ISL_637273 | Virology Department, Sheffield Teaching Hospitals NHS Foundation Trust/Department of Infection, Immunity and Cardiovascular Disease, The Medical School, University of Sheffield | COVID-19 Genomics UK (COG-UK) Consortium | Thushan de Silva, Matthew Parker, Nikki Smith, Adri Angyal, Rebecca Brown, Luke Green, Rachel Tucker, Paul Parsons, Danielle Groves, Katie Johnson, Laura Carrilero, Alex Keeley, Dave Partridge, Matthew Wyles, Benjamin Lindsey, Mehmet Yavuz, Mohammad Raza, Cariad Evans |
| EPI_ISL_637289 | Wales Specialist Virology Centre Sequencing lab: Pathogen Genomics Unit | COVID-19 Genomics UK (COG-UK) Consortium | Catherine Moore, Johnathan Evans, Laura Gifford, Malorie Perry, Simon Cottrell, Angela Marchbank, Alec Birchley, Alexander Adams, Amy Gaskin, Bree Gatica-Wilcox, Jason Coombes, Joel Southgate, Lauren Gilbert, Lee Graham, Nicole Pacchiarini, Sara Kumziene-Summerhayes, Sarah Taylor, Sophie Jones, Sara Rey, Matthew Bull, Joanne Watkins, Sally Corden, Tom Connor |
| EPI_ISL_637300 | Queens Medical Centre, Clinical Microbiology Department / DeepSeq Nottingham | COVID-19 Genomics UK (COG-UK) Consortium | Gemma Clark, Wendy Smith, Manjinder Khakh, Vicki M Fleming, Michelle M Lister, Hannah Howson-Wells, Jonathan Ball, Patrick McClure, Joseph Chappell, Theocharis Tsoleridis, Nadine Holmes, Matthew Carlisle, Christopher Moore, Fei Sang, Johnny Debebe, Victoria Wright, Matthew Loose |
| EPI_ISL_637301 | Quadram Institute Bioscience | COVID-19 Genomics UK (COG-UK) Consortium | Dave J. Baker, Gemma L. Kay, Alp Aydin, Thanh Le-Viet, Steven Rudder, Ana P. Tedim, Anastasia Kolyva, Maria Diaz, Leonardo de Oliveira Martins, |

| | | | |
|---|---|---|---|
| | | | Nabil-Fareed Alikhan, Lizzie Meadows, Rachael Stanley, Ngozi Elumogo, Muhammed Yasir, Nicholas M. Thomson, Alexander J Trotter, Rachel Gilroy, Samuel Bloomfield, Claire Stuart, Andrew Bell, Reenesh Prakash, Samir Dervisevic, Alison E. Mather, John Wain, Mark Webber, Andrew J. Page, Justin O'Grady |
| EPI_ISL_637302, EPI_ISL_637319 | Wales Specialist Virology Centre Sequencing lab: Pathogen Genomics Unit | COVID-19 Genomics UK (COG-UK) Consortium | Catherine Moore, Johnathan Evans, Laura Gifford, Malorie Perry, Simon Cottrell, Angela Marchbank, Alec Birchley, Alexander Adams, Amy Gaskin, Bree Gatica-Wilcox, Jason Coombes, Joel Southgate, Lauren Gilbert, Lee Graham, Nicole Pacchiarini, Sara Kumziene-Summerhayes, Sarah Taylor, Sophie Jones, Sara Rey, Matthew Bull, Joanne Watkins, Sally Corden, Tom Connor |
| EPI_ISL_637336 | Liverpool Clinical Laboratories | COVID-19 Genomics UK (COG-UK) Consortium | Sam Haldenby, Anita Lucaci, Steve Paterson, Julian Hiscox, Alistair Darby, M Almsaud, A Alrezaihi, Muhannad Alruwaili, Stuart D Armstrong, Jones Benjamin, Eleanor G Bentley, Anu Chawla, Jordan J Clark, Angela Cowell, Richard Eccles, Isabel García-Dorival, Matthew Gemmell, Alessandro Gerada, PKF Gilmore, Richard Gregory, Ximeng Han, Catherine Hartley, Margaret Hughes, Miren Iturriza-Gomara, James Johnson, L Luu, Jenifer Manson, Charlotte Nelson, Elaine O'Toole, Cassie Olateju, Rebekah Penrice-Randal , Lucille Rainbow, N.P Randle, Trevor Ian Robinson, Parul Sharma, Ghada T Shawli, James P Stewart, Neil Swainston, Ecaterina Vamos, Joanne Watts, Mark Whitehead |
| EPI_ISL_637343 | Queens Medical Centre, Clinical Microbiology Department / DeepSeq Nottingham | COVID-19 Genomics UK (COG-UK) Consortium | Gemma Clark, Wendy Smith, Manjinder Khakh, Vicki M Fleming, Michelle M Lister, Hannah Howson-Wells, Jonathan Ball, Patrick McClure, Joseph Chappell, Theocharis Tsoleridis, Nadine Holmes, Matthew Carlisle, Christopher Moore, Fei Sang, Johnny Debebe, Victoria Wright, Matthew Loose |
| EPI_ISL_637349 | Wales Specialist Virology Centre Sequencing lab: Pathogen Genomics Unit | COVID-19 Genomics UK (COG-UK) Consortium | Catherine Moore, Johnathan Evans, Laura Gifford, Malorie Perry, Simon Cottrell, Angela Marchbank, Alec Birchley, Alexander Adams, Amy Gaskin, Bree Gatica-Wilcox, Jason Coombes, Joel Southgate, Lauren Gilbert, Lee Graham, Nicole Pacchiarini, Sara Kumziene-Summerhayes, Sarah Taylor, Sophie Jones, Sara Rey, Matthew Bull, Joanne Watkins, Sally Corden, Tom Connor |
| EPI_ISL_637358 | Virology Department, Sheffield Teaching Hospitals NHS Foundation Trust/Department of Infection, Immunity and Cardiovascular Disease, The Medical School, University of Sheffield | COVID-19 Genomics UK (COG-UK) Consortium | Thushan de Silva, Matthew Parker, Nikki Smith, Adri Angyal, Rebecca Brown, Luke Green, Rachel Tucker, Paul Parsons, Danielle Groves, Katie Johnson, Laura Carrilero, Alex Keeley, Dave Partridge, Matthew Wyles, Benjamin Lindsey, Mehmet Yavuz, Mohammad Raza, Cariad Evans |
| EPI_ISL_637380, EPI_ISL_637395 | Wales Specialist Virology Centre Sequencing lab: Pathogen Genomics Unit | COVID-19 Genomics UK (COG-UK) Consortium | Catherine Moore, Johnathan Evans, Laura Gifford, Malorie Perry, Simon Cottrell, Angela Marchbank, Alec Birchley, Alexander Adams, Amy Gaskin, Bree Gatica-Wilcox, Jason Coombes, Joel Southgate, Lauren Gilbert, Lee Graham, Nicole Pacchiarini, Sara Kumziene-Summerhayes, Sarah Taylor, Sophie Jones, Sara Rey, Matthew Bull, Joanne Watkins, Sally Corden, Tom Connor |
| EPI_ISL_637434 | Liverpool Clinical Laboratories | COVID-19 Genomics UK (COG-UK) Consortium | Sam Haldenby, Anita Lucaci, Steve Paterson, Julian Hiscox, Alistair Darby, M Almsaud, A Alrezaihi, Muhannad Alruwaili, Stuart D Armstrong, Jones Benjamin, Eleanor G Bentley, Anu Chawla, Jordan J Clark, Angela Cowell, Richard Eccles, Isabel García-Dorival, Matthew Gemmell, Alessandro Gerada, PKF Gilmore, Richard Gregory, Ximeng Han, Catherine Hartley, Margaret Hughes, Miren Iturriza-Gomara, James Johnson, L Luu, Jenifer Manson, Charlotte Nelson, Elaine O'Toole, Cassie Olateju, Rebekah Penrice-Randal , Lucille Rainbow, N.P Randle, Trevor Ian Robinson, Parul Sharma, Ghada T Shawli, James P Stewart, Neil Swainston, Ecaterina Vamos, Joanne Watts, Mark Whitehead |
| EPI_ISL_637436 | Virology Department, Sheffield Teaching Hospitals NHS Foundation Trust/Department of Infection, Immunity and Cardiovascular Disease, The Medical School, University of Sheffield | COVID-19 Genomics UK (COG-UK) Consortium | Thushan de Silva, Matthew Parker, Nikki Smith, Adri Angyal, Rebecca Brown, Luke Green, Rachel Tucker, Paul Parsons, Danielle Groves, Katie Johnson, Laura Carrilero, Alex Keeley, Dave Partridge, Matthew Wyles, Benjamin Lindsey, Mehmet Yavuz, Mohammad Raza, Cariad Evans |
| EPI_ISL_637442 | Quadram Institute Bioscience | COVID-19 Genomics UK (COG-UK) Consortium | Dave J. Baker, Gemma L. Kay, Alp Aydin, Thanh Le-Viet, Steven Rudder, Ana P. Tedim, Anastasia Kolyva, Maria Diaz, Leonardo de Oliveira Martins, Nabil-Fareed Alikhan, Lizzie Meadows, Rachael Stanley, Ngozi Elumogo, Muhammed Yasir, Nicholas M. Thomson, Alexander J Trotter, Rachel Gilroy, Samuel Bloomfield, Claire Stuart, Andrew Bell, Reenesh Prakash, Samir Dervisevic, Alison E. Mather, John Wain, Mark Webber, Andrew J. Page, Justin O'Grady |
| EPI_ISL_637450, EPI_ISL_637452, EPI_ISL_637525 | Wales Specialist Virology Centre Sequencing lab: Pathogen Genomics Unit | COVID-19 Genomics UK (COG-UK) Consortium | Catherine Moore, Johnathan Evans, Laura Gifford, Malorie Perry, Simon Cottrell, Angela Marchbank, Alec Birchley, Alexander Adams, Amy Gaskin, Bree Gatica-Wilcox, Jason Coombes, Joel Southgate, Lauren Gilbert, Lee Graham, Nicole Pacchiarini, Sara Kumziene-Summerhayes, Sarah Taylor, Sophie Jones, Sara Rey, Matthew Bull, Joanne Watkins, Sally Corden, Tom Connor |
| EPI_ISL_637533 | Virology Department, Sheffield Teaching Hospitals NHS Foundation Trust/Department of Infection, Immunity and Cardiovascular Disease, The Medical School, University of Sheffield | COVID-19 Genomics UK (COG-UK) Consortium | Thushan de Silva, Matthew Parker, Nikki Smith, Adri Angyal, Rebecca Brown, Luke Green, Rachel Tucker, Paul Parsons, Danielle Groves, Katie Johnson, Laura Carrilero, Alex Keeley, Dave Partridge, Matthew Wyles, Benjamin Lindsey, Mehmet Yavuz, Mohammad Raza, Cariad Evans |
| EPI_ISL_637536 | Quadram Institute Bioscience | COVID-19 Genomics UK (COG-UK) Consortium | Dave J. Baker, Gemma L. Kay, Alp Aydin, Thanh Le-Viet, Steven Rudder, Ana P. Tedim, Anastasia Kolyva, Maria Diaz, Leonardo de Oliveira Martins, Nabil-Fareed Alikhan, Lizzie Meadows, Rachael Stanley, Ngozi Elumogo, Muhammed Yasir, Nicholas M. Thomson, Alexander J Trotter, Rachel Gilroy, Samuel Bloomfield, Claire Stuart, Andrew Bell, Reenesh Prakash, Samir Dervisevic, Alison E. Mather, John Wain, Mark Webber, Andrew J. Page, Justin O'Grady |
| EPI_ISL_637542, EPI_ISL_637551 | Wales Specialist Virology Centre Sequencing lab: Pathogen Genomics Unit | COVID-19 Genomics UK (COG-UK) Consortium | Catherine Moore, Johnathan Evans, Laura Gifford, Malorie Perry, Simon Cottrell, Angela Marchbank, Alec Birchley, Alexander Adams, Amy Gaskin, Bree Gatica-Wilcox, Jason Coombes, Joel Southgate, Lauren Gilbert, Lee Graham, Nicole Pacchiarini, Sara Kumziene-Summerhayes, Sarah Taylor, Sophie Jones, Sara Rey, Matthew Bull, Joanne Watkins, Sally Corden, Tom Connor |
| EPI_ISL_637557 | Quadram Institute Bioscience | COVID-19 Genomics UK (COG-UK) Consortium | Dave J. Baker, Gemma L. Kay, Alp Aydin, Thanh Le-Viet, Steven Rudder, Ana P. Tedim, Anastasia Kolyva, Maria Diaz, Leonardo de Oliveira Martins, Nabil-Fareed Alikhan, Lizzie Meadows, Rachael Stanley, Ngozi Elumogo, Muhammed Yasir, Nicholas M. Thomson, Alexander J Trotter, Rachel Gilroy, Samuel Bloomfield, Claire Stuart, Andrew Bell, Reenesh Prakash, Samir Dervisevic, Alison E. Mather, John Wain, Mark Webber, Andrew J. Page, Justin O'Grady |
| EPI_ISL_637585 | Virology Department, Sheffield Teaching Hospitals NHS Foundation Trust/Department of Infection, Immunity and Cardiovascular Disease, The Medical School, University of Sheffield | COVID-19 Genomics UK (COG-UK) Consortium | Thushan de Silva, Matthew Parker, Nikki Smith, Adri Angyal, Rebecca Brown, Luke Green, Rachel Tucker, Paul Parsons, Danielle Groves, Katie Johnson, Laura Carrilero, Alex Keeley, Dave Partridge, Matthew Wyles, Benjamin Lindsey, Mehmet Yavuz, Mohammad Raza, Cariad Evans |
| EPI_ISL_637586, EPI_ISL_637593, EPI_ISL_637600, EPI_ISL_637608, EPI_ISL_637609 | Wales Specialist Virology Centre Sequencing lab: Pathogen Genomics Unit | COVID-19 Genomics UK (COG-UK) Consortium | Catherine Moore, Johnathan Evans, Laura Gifford, Malorie Perry, Simon Cottrell, Angela Marchbank, Alec Birchley, Alexander Adams, Amy Gaskin, Bree Gatica-Wilcox, Jason Coombes, Joel Southgate, Lauren Gilbert, Lee Graham, Nicole Pacchiarini, Sara Kumziene-Summerhayes, Sarah Taylor, Sophie Jones, Sara Rey, Matthew Bull, Joanne Watkins, Sally Corden, Tom Connor |
| EPI_ISL_637617 | Liverpool Clinical Laboratories | COVID-19 Genomics UK (COG-UK) Consortium | Sam Haldenby, Anita Lucaci, Steve Paterson, Julian Hiscox, Alistair Darby, M Almsaud, A Alrezaihi, Muhannad Alruwaili, Stuart D Armstrong, Jones Benjamin, Eleanor G Bentley, Anu Chawla, Jordan J Clark, Angela Cowell, Richard Eccles, Isabel García-Dorival, Matthew Gemmell, Alessandro Gerada, PKF Gilmore, Richard Gregory, Ximeng Han, Catherine Hartley, Margaret Hughes, Miren Iturriza-Gomara, James Johnson, L Luu, Jenifer Manson, Charlotte Nelson, Elaine O'Toole, Cassie Olateju, Rebekah Penrice-Randal , Lucille Rainbow, N.P Randle, Trevor Ian Robinson, Parul Sharma, Ghada T Shawli, James P Stewart, Neil Swainston, Ecaterina Vamos, Joanne Watts, Mark Whitehead |
| EPI_ISL_637626 | Wales Specialist Virology Centre Sequencing lab: Pathogen Genomics Unit | COVID-19 Genomics UK (COG-UK) Consortium | Catherine Moore, Johnathan Evans, Laura Gifford, Malorie Perry, Simon Cottrell, Angela Marchbank, Alec Birchley, Alexander Adams, Amy Gaskin, Bree Gatica-Wilcox, Jason Coombes, Joel Southgate, Lauren Gilbert, Lee Graham, Nicole Pacchiarini, Sara Kumziene-Summerhayes, Sarah Taylor, Sophie Jones, Sara Rey, Matthew Bull, Joanne Watkins, Sally Corden, Tom Connor |
| EPI_ISL_637660 | Queens Medical Centre, Clinical Microbiology Department / DeepSeq Nottingham | COVID-19 Genomics UK (COG-UK) Consortium | Gemma Clark, Wendy Smith, Manjinder Khakh, Vicki M Fleming, Michelle M Lister, Hannah Howson-Wells, Jonathan Ball, Patrick McClure, Joseph Chappell, Theocharis Tsoleridis, Nadine Holmes, Matthew Carlisle, Christopher Moore, Fei Sang, Johnny Debebe, Victoria Wright, Matthew Loose |
| EPI_ISL_637675 | Wales Specialist Virology Centre Sequencing lab: Pathogen Genomics Unit | COVID-19 Genomics UK (COG-UK) Consortium | Catherine Moore, Johnathan Evans, Laura Gifford, Malorie Perry, Simon Cottrell, Angela Marchbank, Alec Birchley, Alexander Adams, Amy Gaskin, Bree Gatica-Wilcox, Jason Coombes, Joel Southgate, Lauren Gilbert, Lee Graham, Nicole Pacchiarini, Sara Kumziene-Summerhayes, Sarah Taylor, Sophie Jones, Sara Rey, Matthew Bull, Joanne Watkins, Sally Corden, Tom Connor |
| EPI_ISL_637865, EPI_ISL_637866 | Quadram Institute Bioscience | COVID-19 Genomics UK (COG-UK) Consortium | Dave J. Baker, Gemma L. Kay, Alp Aydin, Thanh Le-Viet, Steven Rudder, Ana P. Tedim, Anastasia Kolyva, Maria Diaz, Leonardo de Oliveira Martins, Nabil-Fareed Alikhan, Lizzie Meadows, Rachael Stanley, Ngozi Elumogo, Muhammed Yasir, Nicholas M. Thomson, Alexander J Trotter, Rachel Gilroy, Samuel Bloomfield, Claire Stuart, Andrew Bell, Reenesh Prakash, Samir Dervisevic, Alison E. Mather, John Wain, Mark Webber, Andrew J. Page, Justin |

| | | | |
|---|---|---|---|
| | | | O'Grady |
| EPI_ISL_637868, EPI_ISL_637869 | Wales Specialist Virology Centre Sequencing lab: Pathogen Genomics Unit | COVID-19 Genomics UK (COG-UK) Consortium | Catherine Moore, Johnathan Evans, Laura Gifford, Malorie Perry, Simon Cottrell, Angela Marchbank, Alec Birchley, Alexander Adams, Amy Gaskin, Bree Gatica-Wilcox, Jason Coombes, Joel Southgate, Lauren Gilbert, Lee Graham, Nicole Pacchiarini, Sara Kumziene-Summerhayes, Sarah Taylor, Sophie Jones, Sara Rey, Matthew Bull, Joanne Watkins, Sally Corden, Tom Connor |
| EPI_ISL_637872 | Virology Department, Sheffield Teaching Hospitals NHS Foundation Trust/Department of Infection, Immunity and Cardiovascular Disease, The Medical School, University of Sheffield | COVID-19 Genomics UK (COG-UK) Consortium | Thushan de Silva, Matthew Parker, Nikki Smith, Adri Angyal, Rebecca Brown, Luke Green, Rachel Tucker, Paul Parsons, Danielle Groves, Katie Johnson, Laura Carrilero, Alex Keeley, Dave Partridge, Matthew Wyles, Benjamin Lindsey, Mehmet Yavuz, Mohammad Raza, Cariad Evans |
| EPI_ISL_637878 | Quadram Institute Bioscience | COVID-19 Genomics UK (COG-UK) Consortium | Dave J. Baker, Gemma L. Kay, Alp Aydin, Thanh Le-Viet, Steven Rudder, Ana P. Tedim, Anastasia Kolyva, Maria Diaz, Leonardo de Oliveira Martins, Nabil-Fareed Alikhan, Lizzie Meadows, Rachael Stanley, Ngozi Elumogo, Muhammed Yasir, Nicholas M. Thomson, Alexander J Trotter, Rachel Gilroy, Samuel Bloomfield, Claire Stuart, Andrew Bell, Reenesh Prakash, Samir Dervisevic, Alison E. Mather, John Wain, Mark Webber, Andrew J. Page, Justin O'Grady |
| EPI_ISL_637898 | Liverpool Clinical Laboratories | COVID-19 Genomics UK (COG-UK) Consortium | Sam Haldenby, Anita Lucaci, Steve Paterson, Julian Hiscox, Alistair Darby, M Almsaud, A Alrezaihi, Muhannad Alruwaili, Stuart D Armstrong, Jones Benjamin, Eleanor G Bentley, Anu Chawla, Jordan J Clark, Angela Cowell, Richard Eccles, Isabel García-Dorival, Matthew Gemmell, Alessandro Gerada, PKF Gilmore, Richard Gregory, Ximeng Han, Catherine Hartley, Margaret Hughes, Miren Iturriza-Gomara, James Johnson, L Luu, Jenifer Manson, Charlotte Nelson, Elaine O'Toole, Cassie Olateju, Rebekah Penrice-Randal , Lucille Rainbow, N.P Randle, Trevor Ian Robinson, Parul Sharma, Ghada T Shawli, James P Stewart, Neil Swainston, Ecaterina Vamos, Joanne Watts, Mark Whitehead |
| EPI_ISL_637907, EPI_ISL_637909, EPI_ISL_637918 | Wales Specialist Virology Centre Sequencing lab: Pathogen Genomics Unit | COVID-19 Genomics UK (COG-UK) Consortium | Catherine Moore, Johnathan Evans, Laura Gifford, Malorie Perry, Simon Cottrell, Angela Marchbank, Alec Birchley, Alexander Adams, Amy Gaskin, Bree Gatica-Wilcox, Jason Coombes, Joel Southgate, Lauren Gilbert, Lee Graham, Nicole Pacchiarini, Sara Kumziene-Summerhayes, Sarah Taylor, Sophie Jones, Sara Rey, Matthew Bull, Joanne Watkins, Sally Corden, Tom Connor |
| EPI_ISL_637922 | Quadram Institute Bioscience | COVID-19 Genomics UK (COG-UK) Consortium | Dave J. Baker, Gemma L. Kay, Alp Aydin, Thanh Le-Viet, Steven Rudder, Ana P. Tedim, Anastasia Kolyva, Maria Diaz, Leonardo de Oliveira Martins, Nabil-Fareed Alikhan, Lizzie Meadows, Rachael Stanley, Ngozi Elumogo, Muhammed Yasir, Nicholas M. Thomson, Alexander J Trotter, Rachel Gilroy, Samuel Bloomfield, Claire Stuart, Andrew Bell, Reenesh Prakash, Samir Dervisevic, Alison E. Mather, John Wain, Mark Webber, Andrew J. Page, Justin O'Grady |
| EPI_ISL_637945, EPI_ISL_637976 | Liverpool Clinical Laboratories | COVID-19 Genomics UK (COG-UK) Consortium | Sam Haldenby, Anita Lucaci, Steve Paterson, Julian Hiscox, Alistair Darby, M Almsaud, A Alrezaihi, Muhannad Alruwaili, Stuart D Armstrong, Jones Benjamin, Eleanor G Bentley, Anu Chawla, Jordan J Clark, Angela Cowell, Richard Eccles, Isabel García-Dorival, Matthew Gemmell, Alessandro Gerada, PKF Gilmore, Richard Gregory, Ximeng Han, Catherine Hartley, Margaret Hughes, Miren Iturriza-Gomara, James Johnson, L Luu, Jenifer Manson, Charlotte Nelson, Elaine O'Toole, Cassie Olateju, Rebekah Penrice-Randal , Lucille Rainbow, N.P Randle, Trevor Ian Robinson, Parul Sharma, Ghada T Shawli, James P Stewart, Neil Swainston, Ecaterina Vamos, Joanne Watts, Mark Whitehead |
| EPI_ISL_638007 | Wales Specialist Virology Centre Sequencing lab: Pathogen Genomics Unit | COVID-19 Genomics UK (COG-UK) Consortium | Catherine Moore, Johnathan Evans, Laura Gifford, Malorie Perry, Simon Cottrell, Angela Marchbank, Alec Birchley, Alexander Adams, Amy Gaskin, Bree Gatica-Wilcox, Jason Coombes, Joel Southgate, Lauren Gilbert, Lee Graham, Nicole Pacchiarini, Sara Kumziene-Summerhayes, Sarah Taylor, Sophie Jones, Sara Rey, Matthew Bull, Joanne Watkins, Sally Corden, Tom Connor |
| EPI_ISL_638011 | Liverpool Clinical Laboratories | COVID-19 Genomics UK (COG-UK) Consortium | Sam Haldenby, Anita Lucaci, Steve Paterson, Julian Hiscox, Alistair Darby, M Almsaud, A Alrezaihi, Muhannad Alruwaili, Stuart D Armstrong, Jones Benjamin, Eleanor G Bentley, Anu Chawla, Jordan J Clark, Angela Cowell, Richard Eccles, Isabel García-Dorival, Matthew Gemmell, Alessandro Gerada, PKF Gilmore, Richard Gregory, Ximeng Han, Catherine Hartley, Margaret Hughes, Miren Iturriza-Gomara, James Johnson, L Luu, Jenifer Manson, Charlotte Nelson, Elaine O'Toole, Cassie Olateju, Rebekah Penrice-Randal , Lucille Rainbow, N.P Randle, Trevor Ian Robinson, Parul Sharma, Ghada T Shawli, James P Stewart, Neil Swainston, Ecaterina Vamos, Joanne Watts, Mark Whitehead |
| EPI_ISL_638012 | Wales Specialist Virology Centre Sequencing lab: Pathogen Genomics Unit | COVID-19 Genomics UK (COG-UK) Consortium | Catherine Moore, Johnathan Evans, Laura Gifford, Malorie Perry, Simon Cottrell, Angela Marchbank, Alec Birchley, Alexander Adams, Amy Gaskin, Bree Gatica-Wilcox, Jason Coombes, Joel Southgate, Lauren Gilbert, Lee Graham, Nicole Pacchiarini, Sara Kumziene-Summerhayes, Sarah Taylor, Sophie Jones, Sara Rey, Matthew Bull, Joanne Watkins, Sally Corden, Tom Connor |
| EPI_ISL_638025 | Liverpool Clinical Laboratories | COVID-19 Genomics UK (COG-UK) Consortium | Sam Haldenby, Anita Lucaci, Steve Paterson, Julian Hiscox, Alistair Darby, M Almsaud, A Alrezaihi, Muhannad Alruwaili, Stuart D Armstrong, Jones Benjamin, Eleanor G Bentley, Anu Chawla, Jordan J Clark, Angela Cowell, Richard Eccles, Isabel García-Dorival, Matthew Gemmell, Alessandro Gerada, PKF Gilmore, Richard Gregory, Ximeng Han, Catherine Hartley, Margaret Hughes, Miren Iturriza-Gomara, James Johnson, L Luu, Jenifer Manson, Charlotte Nelson, Elaine O'Toole, Cassie Olateju, Rebekah Penrice-Randal , Lucille Rainbow, N.P Randle, Trevor Ian Robinson, Parul Sharma, Ghada T Shawli, James P Stewart, Neil Swainston, Ecaterina Vamos, Joanne Watts, Mark Whitehead |
| EPI_ISL_638033 | Quadram Institute Bioscience | COVID-19 Genomics UK (COG-UK) Consortium | Dave J. Baker, Gemma L. Kay, Alp Aydin, Thanh Le-Viet, Steven Rudder, Ana P. Tedim, Anastasia Kolyva, Maria Diaz, Leonardo de Oliveira Martins, Nabil-Fareed Alikhan, Lizzie Meadows, Rachael Stanley, Ngozi Elumogo, Muhammed Yasir, Nicholas M. Thomson, Alexander J Trotter, Rachel Gilroy, Samuel Bloomfield, Claire Stuart, Andrew Bell, Reenesh Prakash, Samir Dervisevic, Alison E. Mather, John Wain, Mark Webber, Andrew J. Page, Justin O'Grady |
| EPI_ISL_638035, EPI_ISL_638044, EPI_ISL_638046, EPI_ISL_638047, EPI_ISL_638060, EPI_ISL_638091, EPI_ISL_638092, EPI_ISL_638098, EPI_ISL_638101, EPI_ISL_638104, EPI_ISL_638106, EPI_ISL_638107, EPI_ISL_638112, EPI_ISL_638120, EPI_ISL_638121, EPI_ISL_638129, EPI_ISL_638159, EPI_ISL_638172 | | | |
| see above | Wales Specialist Virology Centre Sequencing lab: Pathogen Genomics Unit | COVID-19 Genomics UK (COG-UK) Consortium | Catherine Moore, Johnathan Evans, Laura Gifford, Malorie Perry, Simon Cottrell, Angela Marchbank, Alec Birchley, Alexander Adams, Amy Gaskin, Bree Gatica-Wilcox, Jason Coombes, Joel Southgate, Lauren Gilbert, Lee Graham, Nicole Pacchiarini, Sara Kumziene-Summerhayes, Sarah Taylor, Sophie Jones, Sara Rey, Matthew Bull, Joanne Watkins, Sally Corden, Tom Connor |
| EPI_ISL_638449, EPI_ISL_638450, EPI_ISL_638451, EPI_ISL_638452, EPI_ISL_638453 | Liverpool Clinical Laboratories | COVID-19 Genomics UK (COG-UK) Consortium | Sam Haldenby, Anita Lucaci, Steve Paterson, Julian Hiscox, Alistair Darby, M Almsaud, A Alrezaihi, Muhannad Alruwaili, Stuart D Armstrong, Jones Benjamin, Eleanor G Bentley, Anu Chawla, Jordan J Clark, Angela Cowell, Richard Eccles, Isabel García-Dorival, Matthew Gemmell, Alessandro Gerada, PKF Gilmore, Richard Gregory, Ximeng Han, Catherine Hartley, Margaret Hughes, Miren Iturriza-Gomara, James Johnson, L Luu, Jenifer Manson, Charlotte Nelson, Elaine O'Toole, Cassie Olateju, Rebekah Penrice-Randal , Lucille Rainbow, N.P Randle, Trevor Ian Robinson, Parul Sharma, Ghada T Shawli, James P Stewart, Neil Swainston, Ecaterina Vamos, Joanne Watts, Mark Whitehead |
| EPI_ISL_638582, EPI_ISL_638583 | Quadram Institute Bioscience | COVID-19 Genomics UK (COG-UK) Consortium | Dave J. Baker, Gemma L. Kay, Alp Aydin, Thanh Le-Viet, Steven Rudder, Ana P. Tedim, Anastasia Kolyva, Maria Diaz, Leonardo de Oliveira Martins, Nabil-Fareed Alikhan, Lizzie Meadows, Rachael Stanley, Ngozi Elumogo, Muhammed Yasir, Nicholas M. Thomson, Alexander J Trotter, Rachel Gilroy, Samuel Bloomfield, Claire Stuart, Andrew Bell, Reenesh Prakash, Samir Dervisevic, Alison E. Mather, John Wain, Mark Webber, Andrew J. Page, Justin O'Grady |
| EPI_ISL_638586 | Queens Medical Centre, Clinical Microbiology Department / DeepSeq Nottingham | COVID-19 Genomics UK (COG-UK) Consortium | Gemma Clark, Wendy Smith, Manjinder Khakh, Vicki M Fleming, Michelle M Lister, Hannah Howson-Wells, Jonathan Ball, Patrick McClure, Joseph Chappell, Theocharis Tsoleridis, Nadine Holmes, Matthew Carlisle, Christopher Moore, Fei Sang, Johnny Debebe, Victoria Wright, Matthew Loose |
| EPI_ISL_638877, EPI_ISL_638880, EPI_ISL_638881, EPI_ISL_638884, EPI_ISL_638887, EPI_ISL_638888, EPI_ISL_638889, EPI_ISL_638891, EPI_ISL_638893 | Virology Department, Sheffield Teaching Hospitals NHS Foundation Trust/Department of Infection, Immunity and Cardiovascular Disease, The Medical School, University of Sheffield | COVID-19 Genomics UK (COG-UK) Consortium | Thushan de Silva, Matthew Parker, Nikki Smith, Adri Angyal, Rebecca Brown, Luke Green, Rachel Tucker, Paul Parsons, Danielle Groves, Katie Johnson, Laura Carrilero, Alex Keeley, Dave Partridge, Matthew Wyles, Benjamin Lindsey, Mehmet Yavuz, Mohammad Raza, Cariad Evans |
| EPI_ISL_639011, EPI_ISL_639017, EPI_ISL_639018, EPI_ISL_639021, EPI_ISL_639027, EPI_ISL_639031, EPI_ISL_639032, EPI_ISL_639034, EPI_ISL_639035, EPI_ISL_639037, EPI_ISL_639040, EPI_ISL_639041, EPI_ISL_639045, EPI_ISL_639053, EPI_ISL_639056, EPI_ISL_639057, EPI_ISL_639060, EPI_ISL_639067, EPI_ISL_639077, EPI_ISL_639080, EPI_ISL_639082, EPI_ISL_639085, EPI_ISL_639088, EPI_ISL_639090, EPI_ISL_639105, EPI_ISL_639108, EPI_ISL_639115, EPI_ISL_639119, EPI_ISL_639121, EPI_ISL_639122, EPI_ISL_639125, EPI_ISL_639146, EPI_ISL_639148, EPI_ISL_639155, EPI_ISL_639157, EPI_ISL_639158, EPI_ISL_639160, EPI_ISL_639161, EPI_ISL_639166, EPI_ISL_639173, EPI_ISL_639174, EPI_ISL_639175, EPI_ISL_639176, EPI_ISL_639183, EPI_ISL_639188, EPI_ISL_639193, EPI_ISL_639195, EPI_ISL_639197, EPI_ISL_639199, EPI_ISL_639203, EPI_ISL_639204, EPI_ISL_639205, EPI_ISL_639206, EPI_ISL_639211, EPI_ISL_639212, EPI_ISL_639213, EPI_ISL_639217, EPI_ISL_639222, EPI_ISL_639225, EPI_ISL_639227, EPI_ISL_639233, EPI_ISL_639234, EPI_ISL_639235, EPI_ISL_639238, EPI_ISL_639239, EPI_ISL_639240, EPI_ISL_639244, EPI_ISL_639249, EPI_ISL_639252, EPI_ISL_639253, EPI_ISL_639254, EPI_ISL_639258, EPI_ISL_639259, EPI_ISL_639261, EPI_ISL_639262, EPI_ISL_639267, EPI_ISL_639269, EPI_ISL_639271, EPI_ISL_639273, EPI_ISL_639275, EPI_ISL_639277, EPI_ISL_639280, EPI_ISL_639282, EPI_ISL_639288, EPI_ISL_639296, EPI_ISL_639308, EPI_ISL_639310, EPI_ISL_639319, EPI_ISL_639321, EPI_ISL_639322, EPI_ISL_639325, EPI_ISL_639327, EPI_ISL_639328, EPI_ISL_639330, EPI_ISL_639332, EPI_ISL_639334, EPI_ISL_639339, EPI_ISL_639344, EPI_ISL_639345, EPI_ISL_639351, EPI_ISL_639358, EPI_ISL_639359, EPI_ISL_639360, EPI_ISL_639362, EPI_ISL_639363, EPI_ISL_639365, EPI_ISL_639369, EPI_ISL_639379, EPI_ISL_639381, EPI_ISL_639384, EPI_ISL_639388, EPI_ISL_639390, EPI_ISL_639395, EPI_ISL_639397, EPI_ISL_639398, EPI_ISL_639403, EPI_ISL_639404, EPI_ISL_639405, EPI_ISL_639407, EPI_ISL_639418, EPI_ISL_639421, EPI_ISL_639426, EPI_ISL_639427, EPI_ISL_639428, EPI_ISL_639433, EPI_ISL_639434, EPI_ISL_639441, EPI_ISL_639443, EPI_ISL_639447, EPI_ISL_639448, EPI_ISL_639452, EPI_ISL_639453, EPI_ISL_639460, EPI_ISL_639462, EPI_ISL_639464, EPI_ISL_639467, EPI_ISL_639468, EPI_ISL_639469, EPI_ISL_639475, EPI_ISL_639481, EPI_ISL_639482, EPI_ISL_639484, EPI_ISL_639485, EPI_ISL_639490, EPI_ISL_639492, EPI_ISL_639495, EPI_ISL_639496, EPI_ISL_639498, EPI_ISL_639499, EPI_ISL_639500, EPI_ISL_639504, EPI_ISL_639506, EPI_ISL_639510, EPI_ISL_639515, EPI_ISL_639516, EPI_ISL_639518, EPI_ISL_639526, EPI_ISL_639529, EPI_ISL_639531, EPI_ISL_639532, EPI_ISL_639535, EPI_ISL_639536, | | | |

| | | | |
|---|---|---|---|
| EPI_ISL_639537, EPI_ISL_639540, EPI_ISL_639545, EPI_ISL_639547, EPI_ISL_639548, EPI_ISL_639551, EPI_ISL_639556, EPI_ISL_639558, EPI_ISL_639560, EPI_ISL_639562, EPI_ISL_639565, EPI_ISL_639568, EPI_ISL_639572, EPI_ISL_639574, EPI_ISL_639576, EPI_ISL_639581, EPI_ISL_639586, EPI_ISL_639587, EPI_ISL_639591, EPI_ISL_639596, EPI_ISL_639599, EPI_ISL_639601, EPI_ISL_639602, EPI_ISL_639605, EPI_ISL_639609, EPI_ISL_639610, EPI_ISL_639614, EPI_ISL_639620, EPI_ISL_639621, EPI_ISL_639623 | | | |
| see above | Wales Specialist Virology Centre Sequencing lab: Pathogen Genomics Unit | COVID-19 Genomics UK (COG-UK) Consortium | Catherine Moore, Johnathan Evans, Laura Gifford, Malorie Perry, Simon Cottrell, Angela Marchbank, Alec Birchley, Alexander Adams, Amy Gaskin, Bree Gatica-Wilcox, Jason Coombes, Joel Southgate, Lauren Gilbert, Lee Graham, Nicole Pacchiarini, Sara Kumziene-Summerhayes, Sarah Taylor, Sophie Jones, Sara Rey, Matthew Bull, Joanne Watkins, Sally Corden, Tom Connor |
| EPI_ISL_639822 | Respiratory Virus Unit, Microbiology Services Colindale, Public Health England | COVID-19 Genomics UK (COG-UK) Consortium | PHE Covid Sequencing Team |
| EPI_ISL_639986, EPI_ISL_639988, EPI_ISL_640008 | CNR Virus des Infections Respiratoires - France SUD | CNR Virus des Infections Respiratoires - France SUD | Antonin Bal, Gregory Destras, Gwendolyne Burfin, Hadrien Règue, Alexandre Gaymard, Maude Bouscambert-Duchamp, Florence Morfin-Sherpa, Martine Valette, Bruno Lina, Laurence Josset |
| EPI_ISL_641396 | Department of Virus and Microbiological Special Diagnostics, Statens Serum Institut, Copenhagen, Denmark | Albertsen lab, Department of Chemistry and Bioscience, Aalborg University, Denmark | Thomas Bruun Rasmussen, Jannik Fonager, Morten Rasmussen |
| EPI_ISL_641610, EPI_ISL_641613, EPI_ISL_641628, EPI_ISL_641631, EPI_ISL_641635, EPI_ISL_641649, EPI_ISL_641650, EPI_ISL_641652, EPI_ISL_641667, EPI_ISL_641668, EPI_ISL_641671, EPI_ISL_641685, EPI_ISL_641686, EPI_ISL_641687, EPI_ISL_641696, EPI_ISL_641707, EPI_ISL_641710, EPI_ISL_641711, EPI_ISL_641715, EPI_ISL_641720, EPI_ISL_641733, EPI_ISL_641734, EPI_ISL_641744, EPI_ISL_641745, EPI_ISL_641747, EPI_ISL_641753, EPI_ISL_641756, EPI_ISL_641758, EPI_ISL_641759, EPI_ISL_641761, EPI_ISL_641762, EPI_ISL_641763, EPI_ISL_641769, EPI_ISL_641775, EPI_ISL_641780, EPI_ISL_641782, EPI_ISL_641785, EPI_ISL_641788, EPI_ISL_641795, EPI_ISL_641796, EPI_ISL_641799, EPI_ISL_641802, EPI_ISL_641804, EPI_ISL_641814, EPI_ISL_641815, EPI_ISL_641816, EPI_ISL_641832, EPI_ISL_641833, EPI_ISL_641842, EPI_ISL_641847, EPI_ISL_641850, EPI_ISL_641857, EPI_ISL_641861, EPI_ISL_641864, EPI_ISL_641868, EPI_ISL_641878, EPI_ISL_641879, EPI_ISL_641882, EPI_ISL_641884, EPI_ISL_641888, EPI_ISL_641897, EPI_ISL_641900, EPI_ISL_641901, EPI_ISL_641911, EPI_ISL_641960, EPI_ISL_641979, EPI_ISL_641994, EPI_ISL_642018, EPI_ISL_642038, EPI_ISL_642069, EPI_ISL_642074, EPI_ISL_642155, EPI_ISL_642166, EPI_ISL_642169, EPI_ISL_642170, EPI_ISL_642203, EPI_ISL_642222, EPI_ISL_642243, EPI_ISL_642247, EPI_ISL_642251, EPI_ISL_642284, EPI_ISL_642354, EPI_ISL_642366, EPI_ISL_642379, EPI_ISL_642420, EPI_ISL_642437, EPI_ISL_642439, EPI_ISL_642477, EPI_ISL_642506, EPI_ISL_642508, EPI_ISL_642521, EPI_ISL_642569, EPI_ISL_642581, EPI_ISL_642599, EPI_ISL_642605, EPI_ISL_642634, EPI_ISL_642635, EPI_ISL_642644, EPI_ISL_642655, EPI_ISL_642663, EPI_ISL_642664, EPI_ISL_642684, EPI_ISL_642702, EPI_ISL_642703, EPI_ISL_642729, EPI_ISL_642744 | | | |
| see above | Lighthouse Lab in Glasgow | Wellcome Sanger Institute for the COVID-19 Genomics UK (COG-UK) Consortium | Harper VanSteenhouse, Yumi Kasai, David Gray, Carol Clugston, Anna Dominiczak and Alex Alderton, Roberto Amato, Sonia Goncalves, Ewan Harrison, David K. Jackson, Ian Johnston, Dominic Kwiatkowski, Cordelia Langford, John Sillitoe on behalf of the Wellcome Sanger Institute COVID-19 Surveillance Team |
| EPI_ISL_643315, EPI_ISL_643316, EPI_ISL_643317, EPI_ISL_643318, EPI_ISL_643319, EPI_ISL_643320, EPI_ISL_643321, EPI_ISL_643323, EPI_ISL_643326, EPI_ISL_643327, EPI_ISL_643328, EPI_ISL_643332, EPI_ISL_643333, EPI_ISL_643335, EPI_ISL_643336, EPI_ISL_643340, EPI_ISL_643342, EPI_ISL_643343, EPI_ISL_643344, EPI_ISL_643345, EPI_ISL_643346, EPI_ISL_643347, EPI_ISL_643348, EPI_ISL_643354, EPI_ISL_643357, EPI_ISL_643358, EPI_ISL_643359, EPI_ISL_643360, EPI_ISL_643361, EPI_ISL_643362, EPI_ISL_643363, EPI_ISL_643366, EPI_ISL_643367, EPI_ISL_643369, EPI_ISL_643371, EPI_ISL_643373, EPI_ISL_643377, EPI_ISL_643379, EPI_ISL_643381, EPI_ISL_643384, EPI_ISL_643385, EPI_ISL_643387, EPI_ISL_643388, EPI_ISL_643390, EPI_ISL_643394, EPI_ISL_643395, EPI_ISL_643397, EPI_ISL_643398, EPI_ISL_643400, EPI_ISL_643401, EPI_ISL_643402, EPI_ISL_643404, EPI_ISL_643405, EPI_ISL_643406, EPI_ISL_643407, EPI_ISL_643409, EPI_ISL_643410, EPI_ISL_643419, EPI_ISL_643420, EPI_ISL_643422, EPI_ISL_643425, EPI_ISL_643426, EPI_ISL_643427, EPI_ISL_643429, EPI_ISL_643431, EPI_ISL_643433, EPI_ISL_643436, EPI_ISL_643437, EPI_ISL_643438, EPI_ISL_643439, EPI_ISL_643440, EPI_ISL_643441, EPI_ISL_643443, EPI_ISL_643444, EPI_ISL_643445, EPI_ISL_643452, EPI_ISL_643453, EPI_ISL_643455, EPI_ISL_643456, EPI_ISL_643457, EPI_ISL_643460, EPI_ISL_643463, EPI_ISL_643464, EPI_ISL_643465, EPI_ISL_643466, EPI_ISL_643468, EPI_ISL_643469, EPI_ISL_643473, EPI_ISL_643476, EPI_ISL_643479, EPI_ISL_643483, EPI_ISL_643486, EPI_ISL_643487, EPI_ISL_643490, EPI_ISL_643493, EPI_ISL_643496, EPI_ISL_643497, EPI_ISL_643500, EPI_ISL_643501, EPI_ISL_643502, EPI_ISL_643503, EPI_ISL_643507, EPI_ISL_643508, EPI_ISL_643511, EPI_ISL_643515, EPI_ISL_643522, EPI_ISL_643523, EPI_ISL_643524, EPI_ISL_643525, EPI_ISL_643529, EPI_ISL_643530, EPI_ISL_643532, EPI_ISL_643535, EPI_ISL_643537, EPI_ISL_643538, EPI_ISL_643541, EPI_ISL_643547, EPI_ISL_643549, EPI_ISL_643550, EPI_ISL_643552, EPI_ISL_643555, EPI_ISL_643556, EPI_ISL_643558, EPI_ISL_643559, EPI_ISL_643560, EPI_ISL_643561, EPI_ISL_643562, EPI_ISL_643563, EPI_ISL_643564, EPI_ISL_643566, EPI_ISL_643567, EPI_ISL_643569, EPI_ISL_643570, EPI_ISL_643571, EPI_ISL_643573, EPI_ISL_643574, EPI_ISL_643578, EPI_ISL_643579, EPI_ISL_643580, EPI_ISL_643582, EPI_ISL_643583, EPI_ISL_643585, EPI_ISL_643587, EPI_ISL_643589, EPI_ISL_643591, EPI_ISL_643592, EPI_ISL_643596, EPI_ISL_643597, EPI_ISL_643599, EPI_ISL_643601, EPI_ISL_643604, EPI_ISL_643606, EPI_ISL_643607, EPI_ISL_643608, EPI_ISL_643611, EPI_ISL_643614, EPI_ISL_643618, EPI_ISL_643619, EPI_ISL_643620, EPI_ISL_643621, EPI_ISL_643622, EPI_ISL_643623, EPI_ISL_643624, EPI_ISL_643625, EPI_ISL_643627, EPI_ISL_643639, EPI_ISL_643645, EPI_ISL_643647, EPI_ISL_643648, EPI_ISL_643649, EPI_ISL_643652, EPI_ISL_643657, EPI_ISL_643659, EPI_ISL_643660, EPI_ISL_643661, EPI_ISL_643665, EPI_ISL_643666, EPI_ISL_643667, EPI_ISL_643670, EPI_ISL_643673, EPI_ISL_643674, EPI_ISL_643680, EPI_ISL_643683, EPI_ISL_643685, EPI_ISL_643687, EPI_ISL_643688, EPI_ISL_643697, EPI_ISL_643698, EPI_ISL_643701, EPI_ISL_643702, EPI_ISL_643704, EPI_ISL_643707, EPI_ISL_643709, EPI_ISL_643716, EPI_ISL_643717, EPI_ISL_643719, EPI_ISL_643722, EPI_ISL_643723, EPI_ISL_643724, EPI_ISL_643726, EPI_ISL_643727, EPI_ISL_643728, EPI_ISL_643736, EPI_ISL_643738, EPI_ISL_643739, EPI_ISL_643743, EPI_ISL_643747, EPI_ISL_643749, EPI_ISL_643752, EPI_ISL_643756, EPI_ISL_643758, EPI_ISL_643759, EPI_ISL_643761, EPI_ISL_643762, EPI_ISL_643765, EPI_ISL_643766, EPI_ISL_643767, EPI_ISL_643774, EPI_ISL_643775, EPI_ISL_643778, EPI_ISL_643779, EPI_ISL_643781, EPI_ISL_643782, EPI_ISL_643789, EPI_ISL_643792, EPI_ISL_643795, EPI_ISL_643797, EPI_ISL_643804, EPI_ISL_643805, EPI_ISL_643808, EPI_ISL_643809, EPI_ISL_643810, EPI_ISL_643812, EPI_ISL_643813, EPI_ISL_643817, EPI_ISL_643819, EPI_ISL_643820, EPI_ISL_643821, EPI_ISL_643822, EPI_ISL_643823, EPI_ISL_643824, EPI_ISL_643825, EPI_ISL_643826, EPI_ISL_643828, EPI_ISL_643831, EPI_ISL_643837, EPI_ISL_643840, EPI_ISL_643843, EPI_ISL_643859, EPI_ISL_643860, EPI_ISL_643861, EPI_ISL_643862, EPI_ISL_643863, EPI_ISL_643866, EPI_ISL_643868, EPI_ISL_643870, EPI_ISL_643876, EPI_ISL_643877, EPI_ISL_643879, EPI_ISL_643880, EPI_ISL_643883, EPI_ISL_643885, EPI_ISL_643886, EPI_ISL_643888, EPI_ISL_643890, EPI_ISL_643891, EPI_ISL_643892, EPI_ISL_643893, EPI_ISL_643894, EPI_ISL_643896, EPI_ISL_643897, EPI_ISL_643898, EPI_ISL_643900, EPI_ISL_643910, EPI_ISL_643913, EPI_ISL_643916, EPI_ISL_643918, EPI_ISL_643921, EPI_ISL_643924, EPI_ISL_643926, EPI_ISL_643934, EPI_ISL_643935, EPI_ISL_643936, EPI_ISL_643938, EPI_ISL_643941, EPI_ISL_643942, EPI_ISL_643946, EPI_ISL_643947, EPI_ISL_643951, EPI_ISL_643952, EPI_ISL_643953, EPI_ISL_643957, EPI_ISL_643959, EPI_ISL_643961, EPI_ISL_643963, EPI_ISL_643970, EPI_ISL_643973, EPI_ISL_643975, EPI_ISL_643980, EPI_ISL_643982 | | | |
| see above | Lighthouse Lab in Cambridge | Wellcome Sanger Institute for the COVID-19 Genomics UK (COG-UK) Consortium | Rob Howes, The Lighthouse Lab in Cambridge and Alex Alderton, Roberto Amato, Sonia Goncalves, Ewan Harrison, David K. Jackson, Ian Johnston, Dominic Kwiatkowski, Cordelia Langford, John Sillitoe on behalf of the Wellcome Sanger Institute COVID-19 Surveillance Team |
| EPI_ISL_644611, EPI_ISL_644612, EPI_ISL_644613, EPI_ISL_644616, EPI_ISL_644617, EPI_ISL_644618, EPI_ISL_644619, EPI_ISL_644620, EPI_ISL_644621, EPI_ISL_644622, EPI_ISL_644623, EPI_ISL_644624, EPI_ISL_644625, EPI_ISL_644626 | | | |
| see above | Department of Clinical Microbiology | GIGA Medical Genomics | Keith Durkin, Maria Artesi, Sébastien Bontems, Raphaël Boreux, Bouchra Boujemla, Cécile Meex, Pierrette Melin, Marie-Pierre Hayette, Vincent Bours |
| EPI_ISL_645221, EPI_ISL_645226, EPI_ISL_645241, EPI_ISL_645243, EPI_ISL_645244, EPI_ISL_645263, EPI_ISL_645285, EPI_ISL_645300, EPI_ISL_645308, EPI_ISL_645319, EPI_ISL_645324, EPI_ISL_645327, EPI_ISL_645329, EPI_ISL_645367, EPI_ISL_645369, EPI_ISL_645374, EPI_ISL_645383, EPI_ISL_645391, EPI_ISL_645393, EPI_ISL_645395, EPI_ISL_645399, EPI_ISL_645405, EPI_ISL_645424, EPI_ISL_645441, EPI_ISL_645453, EPI_ISL_645477, EPI_ISL_645513, EPI_ISL_645518, EPI_ISL_645520 | | | |
| see above | Lighthouse Lab in Milton Keynes | Wellcome Sanger Institute for the COVID-19 Genomics UK (COG-UK) Consortium | The Lighthouse Lab in Milton Keynes and Alex Alderton, Roberto Amato, Sonia Goncalves, Ewan Harrison, David K. Jackson, Ian Johnston, Dominic Kwiatkowski, Cordelia Langford, John Sillitoe on behalf of the Wellcome Sanger Institute COVID-19 Surveillance Team |
| EPI_ISL_645530, EPI_ISL_645531, EPI_ISL_645532, EPI_ISL_645533, EPI_ISL_645534, EPI_ISL_645535, EPI_ISL_645536, EPI_ISL_645537, EPI_ISL_645540, EPI_ISL_645541, EPI_ISL_645542, EPI_ISL_645543, EPI_ISL_645544, EPI_ISL_645546, EPI_ISL_645547, EPI_ISL_645548, EPI_ISL_645549, EPI_ISL_645550, EPI_ISL_645552, EPI_ISL_645553, EPI_ISL_645554, EPI_ISL_645555, EPI_ISL_645556, EPI_ISL_645557, EPI_ISL_645558, EPI_ISL_645559, EPI_ISL_645560, EPI_ISL_645561, EPI_ISL_645562, EPI_ISL_645563, EPI_ISL_645564, EPI_ISL_645565, EPI_ISL_645566, EPI_ISL_645567, EPI_ISL_645568, EPI_ISL_645569, EPI_ISL_645570, EPI_ISL_645571, EPI_ISL_645573, EPI_ISL_645574, EPI_ISL_645575, EPI_ISL_645577, EPI_ISL_645578, EPI_ISL_645579, EPI_ISL_645580, EPI_ISL_645581, EPI_ISL_645582, EPI_ISL_645583, EPI_ISL_645585, EPI_ISL_645586, EPI_ISL_645587, EPI_ISL_645588, EPI_ISL_645589, EPI_ISL_645590, EPI_ISL_645591, EPI_ISL_645592, EPI_ISL_645593, EPI_ISL_645594, EPI_ISL_645595, EPI_ISL_645596, EPI_ISL_645597, EPI_ISL_645598, EPI_ISL_645599, EPI_ISL_645600, EPI_ISL_645601, EPI_ISL_645602, EPI_ISL_645603, EPI_ISL_645604, EPI_ISL_645605, EPI_ISL_645606, EPI_ISL_645607, EPI_ISL_645609, EPI_ISL_645611, EPI_ISL_645612, EPI_ISL_645613, EPI_ISL_645614, EPI_ISL_645615, EPI_ISL_645616, EPI_ISL_645617, EPI_ISL_645618, EPI_ISL_645619, EPI_ISL_645620, EPI_ISL_645621, EPI_ISL_645622, EPI_ISL_645624, EPI_ISL_645625, EPI_ISL_645626, EPI_ISL_645627, EPI_ISL_645628, EPI_ISL_645629, EPI_ISL_645630, EPI_ISL_645632, EPI_ISL_645633, EPI_ISL_645634, EPI_ISL_645635, EPI_ISL_645636, EPI_ISL_645637, EPI_ISL_645638, EPI_ISL_645639, EPI_ISL_645640, EPI_ISL_645641, EPI_ISL_645642, EPI_ISL_645643, EPI_ISL_645644, EPI_ISL_645645, EPI_ISL_645646, EPI_ISL_645647, EPI_ISL_645648, EPI_ISL_645649, EPI_ISL_645650, EPI_ISL_645651, EPI_ISL_645652, EPI_ISL_645653, EPI_ISL_645654, EPI_ISL_645655, EPI_ISL_645656, EPI_ISL_645657, EPI_ISL_645658, EPI_ISL_645660, EPI_ISL_645661, EPI_ISL_645662, EPI_ISL_645663, EPI_ISL_645664, EPI_ISL_645666, EPI_ISL_645667, EPI_ISL_645668, EPI_ISL_645669, EPI_ISL_645670, EPI_ISL_645671, EPI_ISL_645672, EPI_ISL_645673, EPI_ISL_645674, EPI_ISL_645675, EPI_ISL_645676, EPI_ISL_645677, EPI_ISL_645679, EPI_ISL_645680, EPI_ISL_645681, EPI_ISL_645682, EPI_ISL_645683, EPI_ISL_645684, EPI_ISL_645685, EPI_ISL_645687, EPI_ISL_645688, EPI_ISL_645689, EPI_ISL_645690, EPI_ISL_645691, EPI_ISL_645693, EPI_ISL_645694, EPI_ISL_645695, EPI_ISL_645696, EPI_ISL_645697, EPI_ISL_645698, EPI_ISL_645699, EPI_ISL_645700, EPI_ISL_645701, EPI_ISL_645702, EPI_ISL_645703, EPI_ISL_645704, EPI_ISL_645705, EPI_ISL_645706, EPI_ISL_645707, EPI_ISL_645708, EPI_ISL_645709, EPI_ISL_645710, EPI_ISL_645711, EPI_ISL_645712, EPI_ISL_645713, EPI_ISL_645714, EPI_ISL_645715, EPI_ISL_645716, EPI_ISL_645717, EPI_ISL_645718, EPI_ISL_645719, EPI_ISL_645720, EPI_ISL_645721, EPI_ISL_645722, EPI_ISL_645723, EPI_ISL_645724, EPI_ISL_645725, EPI_ISL_645726, EPI_ISL_645727, EPI_ISL_645728, EPI_ISL_645729, EPI_ISL_645730, EPI_ISL_645732, EPI_ISL_645733, EPI_ISL_645734, EPI_ISL_645736, EPI_ISL_645737, EPI_ISL_645738, EPI_ISL_645739, EPI_ISL_645740, EPI_ISL_645741, EPI_ISL_645742, EPI_ISL_645743, EPI_ISL_645744, EPI_ISL_645745, EPI_ISL_645746, EPI_ISL_645747, EPI_ISL_645748, EPI_ISL_645749, EPI_ISL_645750, EPI_ISL_645751, EPI_ISL_645752, EPI_ISL_645753, EPI_ISL_645754, EPI_ISL_645756, EPI_ISL_645759, EPI_ISL_645760, EPI_ISL_645761, EPI_ISL_645762, EPI_ISL_645763, EPI_ISL_645764, EPI_ISL_645765, EPI_ISL_645766, EPI_ISL_645767, EPI_ISL_645769, EPI_ISL_645770, EPI_ISL_645771, EPI_ISL_645772, EPI_ISL_645773, EPI_ISL_645774, EPI_ISL_645775, EPI_ISL_645776, EPI_ISL_645777, EPI_ISL_645778, EPI_ISL_645779, EPI_ISL_645780, EPI_ISL_645781, EPI_ISL_645782, EPI_ISL_645783, EPI_ISL_645784, EPI_ISL_645785, EPI_ISL_645786, EPI_ISL_645787, EPI_ISL_645788, EPI_ISL_645789, EPI_ISL_645790, EPI_ISL_645792, EPI_ISL_645793, EPI_ISL_645794, EPI_ISL_645795, EPI_ISL_645797, EPI_ISL_645798, EPI_ISL_645799, EPI_ISL_645800, EPI_ISL_645801, EPI_ISL_645802, EPI_ISL_645803, EPI_ISL_645804, EPI_ISL_645805, EPI_ISL_645806, EPI_ISL_645807, EPI_ISL_645808, EPI_ISL_645809, EPI_ISL_645810, EPI_ISL_645811, EPI_ISL_645814, EPI_ISL_645815, EPI_ISL_645816, EPI_ISL_645817, EPI_ISL_645818, EPI_ISL_645819, EPI_ISL_645820, EPI_ISL_645821, EPI_ISL_645822, EPI_ISL_645823, EPI_ISL_645824, EPI_ISL_645827, EPI_ISL_645830, EPI_ISL_645831, EPI_ISL_645833, EPI_ISL_645834, EPI_ISL_645835, EPI_ISL_645837 | | | |
| see above | Lighthouse Lab in Alderley Park | Wellcome Sanger Institute for the COVID-19 Genomics UK (COG-UK) Consortium | Jacquelyn Wynn, Mairead Hyland, The Lighthouse Lab in Alderley Park and Alex Alderton, Roberto Amato, Sonia Goncalves, Ewan Harrison, David K. Jackson, Ian Johnston, Dominic Kwiatkowski, Cordelia Langford, John Sillitoe on behalf of the Wellcome Sanger Institute COVID-19 Surveillance Team |
| EPI_ISL_645840 | Lighthouse Lab in Cambridge | Wellcome Sanger Institute for the COVID-19 Genomics UK (COG-UK) Consortium | Rob Howes, The Lighthouse Lab in Cambridge and Alex Alderton, Roberto Amato, Sonia Goncalves, Ewan Harrison, David K. Jackson, Ian Johnston, Dominic Kwiatkowski, Cordelia Langford, John Sillitoe on behalf of the Wellcome Sanger Institute COVID-19 Surveillance Team |
| EPI_ISL_645841, EPI_ISL_645842, EPI_ISL_645843, EPI_ISL_645844, EPI_ISL_645845, EPI_ISL_645846, EPI_ISL_645847, EPI_ISL_645848, EPI_ISL_645849, EPI_ISL_645853, EPI_ISL_645854, EPI_ISL_645855, EPI_ISL_645856, EPI_ISL_645857, EPI_ISL_645858, EPI_ISL_645859, EPI_ISL_645860, EPI_ISL_645861, EPI_ISL_645862, EPI_ISL_645863, EPI_ISL_645864, EPI_ISL_645865, EPI_ISL_645867, EPI_ISL_645870, EPI_ISL_645871, EPI_ISL_645872, EPI_ISL_645873, EPI_ISL_645874, EPI_ISL_645876, EPI_ISL_645878, EPI_ISL_645879, EPI_ISL_645883, EPI_ISL_645884 | | | |
| see above | Lighthouse Lab in Alderley Park | Wellcome Sanger Institute for the COVID-19 Genomics UK (COG-UK) Consortium | Jacquelyn Wynn, Mairead Hyland, The Lighthouse Lab in Alderley Park and Alex Alderton, Roberto Amato, Sonia Goncalves, Ewan Harrison, David K. Jackson, Ian Johnston, Dominic Kwiatkowski, Cordelia Langford, John Sillitoe on behalf of the Wellcome Sanger Institute COVID-19 Surveillance Team |
| EPI_ISL_645886 | Lighthouse Lab in Cambridge | Wellcome Sanger Institute for the COVID-19 Genomics UK (COG-UK) Consortium | Rob Howes, The Lighthouse Lab in Cambridge and Alex Alderton, Roberto Amato, Sonia Goncalves, Ewan Harrison, David K. Jackson, Ian Johnston, Dominic Kwiatkowski, Cordelia Langford, John Sillitoe on behalf of the Wellcome Sanger Institute COVID-19 Surveillance Team |
| EPI_ISL_645887, EPI_ISL_645888, EPI_ISL_645890, EPI_ISL_645891, EPI_ISL_645892, EPI_ISL_645893, EPI_ISL_645894, EPI_ISL_645895 | Lighthouse Lab in Alderley Park | Wellcome Sanger Institute for the COVID-19 Genomics UK (COG-UK) Consortium | Jacquelyn Wynn, Mairead Hyland, The Lighthouse Lab in Alderley Park and Alex Alderton, Roberto Amato, Sonia Goncalves, Ewan Harrison, David K. Jackson, Ian Johnston, Dominic Kwiatkowski, Cordelia Langford, John Sillitoe on behalf of the Wellcome Sanger Institute COVID-19 Surveillance Team |
| EPI_ISL_645896, EPI_ISL_645897 | Lighthouse Lab in Cambridge | Wellcome Sanger Institute for the COVID-19 Genomics UK (COG-UK) Consortium | Rob Howes, The Lighthouse Lab in Cambridge and Alex Alderton, Roberto Amato, Sonia Goncalves, Ewan Harrison, David K. Jackson, Ian Johnston, Dominic Kwiatkowski, Cordelia Langford, John Sillitoe on behalf of the Wellcome Sanger Institute COVID-19 Surveillance Team |

| | | | |
|---|---|---|---|
| EPI_ISL_645898, EPI_ISL_645900, EPI_ISL_645901 | Lighthouse Lab in Alderley Park | Wellcome Sanger Institute for the COVID-19 Genomics UK (COG-UK) Consortium | Jacquelyn Wynn, Mairead Hyland, The Lighthouse Lab in Alderley Park and Alex Alderton, Roberto Amato, Sonia Goncalves, Ewan Harrison, David K. Jackson, Ian Johnston, Dominic Kwiatkowski, Cordelia Langford, John Sillitoe on behalf of the Wellcome Sanger Institute COVID-19 Surveillance Team |
| EPI_ISL_645902 | Lighthouse Lab in Cambridge | Wellcome Sanger Institute for the COVID-19 Genomics UK (COG-UK) Consortium | Rob Howes, The Lighthouse Lab in Cambridge and Alex Alderton, Roberto Amato, Sonia Goncalves, Ewan Harrison, David K. Jackson, Ian Johnston, Dominic Kwiatkowski, Cordelia Langford, John Sillitoe on behalf of the Wellcome Sanger Institute COVID-19 Surveillance Team |
| EPI_ISL_645903, EPI_ISL_645904 | Lighthouse Lab in Alderley Park | Wellcome Sanger Institute for the COVID-19 Genomics UK (COG-UK) Consortium | Jacquelyn Wynn, Mairead Hyland, The Lighthouse Lab in Alderley Park and Alex Alderton, Roberto Amato, Sonia Goncalves, Ewan Harrison, David K. Jackson, Ian Johnston, Dominic Kwiatkowski, Cordelia Langford, John Sillitoe on behalf of the Wellcome Sanger Institute COVID-19 Surveillance Team |
| EPI_ISL_645905 | Lighthouse Lab in Cambridge | Wellcome Sanger Institute for the COVID-19 Genomics UK (COG-UK) Consortium | Rob Howes, The Lighthouse Lab in Cambridge and Alex Alderton, Roberto Amato, Sonia Goncalves, Ewan Harrison, David K. Jackson, Ian Johnston, Dominic Kwiatkowski, Cordelia Langford, John Sillitoe on behalf of the Wellcome Sanger Institute COVID-19 Surveillance Team |
| EPI_ISL_645906, EPI_ISL_645907, EPI_ISL_645909, EPI_ISL_645910, EPI_ISL_645911, EPI_ISL_645912, EPI_ISL_645913, EPI_ISL_645914, EPI_ISL_645915 | Lighthouse Lab in Alderley Park | Wellcome Sanger Institute for the COVID-19 Genomics UK (COG-UK) Consortium | Jacquelyn Wynn, Mairead Hyland, The Lighthouse Lab in Alderley Park and Alex Alderton, Roberto Amato, Sonia Goncalves, Ewan Harrison, David K. Jackson, Ian Johnston, Dominic Kwiatkowski, Cordelia Langford, John Sillitoe on behalf of the Wellcome Sanger Institute COVID-19 Surveillance Team |
| EPI_ISL_645916 | Lighthouse Lab in Cambridge | Wellcome Sanger Institute for the COVID-19 Genomics UK (COG-UK) Consortium | Rob Howes, The Lighthouse Lab in Cambridge and Alex Alderton, Roberto Amato, Sonia Goncalves, Ewan Harrison, David K. Jackson, Ian Johnston, Dominic Kwiatkowski, Cordelia Langford, John Sillitoe on behalf of the Wellcome Sanger Institute COVID-19 Surveillance Team |
| EPI_ISL_645919, EPI_ISL_645921, EPI_ISL_645922, EPI_ISL_645924, EPI_ISL_645928, EPI_ISL_645929 | Lighthouse Lab in Alderley Park | Wellcome Sanger Institute for the COVID-19 Genomics UK (COG-UK) Consortium | Jacquelyn Wynn, Mairead Hyland, The Lighthouse Lab in Alderley Park and Alex Alderton, Roberto Amato, Sonia Goncalves, Ewan Harrison, David K. Jackson, Ian Johnston, Dominic Kwiatkowski, Cordelia Langford, John Sillitoe on behalf of the Wellcome Sanger Institute COVID-19 Surveillance Team |
| EPI_ISL_645930 | Lighthouse Lab in Cambridge | Wellcome Sanger Institute for the COVID-19 Genomics UK (COG-UK) Consortium | Rob Howes, The Lighthouse Lab in Cambridge and Alex Alderton, Roberto Amato, Sonia Goncalves, Ewan Harrison, David K. Jackson, Ian Johnston, Dominic Kwiatkowski, Cordelia Langford, John Sillitoe on behalf of the Wellcome Sanger Institute COVID-19 Surveillance Team |
| EPI_ISL_645931, EPI_ISL_645932, EPI_ISL_645933, EPI_ISL_645934, EPI_ISL_645936, EPI_ISL_645937, EPI_ISL_645938, EPI_ISL_645939, EPI_ISL_645940, EPI_ISL_645941, EPI_ISL_645942, EPI_ISL_645943, EPI_ISL_645944 | | | |
| see above | Lighthouse Lab in Alderley Park | Wellcome Sanger Institute for the COVID-19 Genomics UK (COG-UK) Consortium | Jacquelyn Wynn, Mairead Hyland, The Lighthouse Lab in Alderley Park and Alex Alderton, Roberto Amato, Sonia Goncalves, Ewan Harrison, David K. Jackson, Ian Johnston, Dominic Kwiatkowski, Cordelia Langford, John Sillitoe on behalf of the Wellcome Sanger Institute COVID-19 Surveillance Team |
| EPI_ISL_645945 | Lighthouse Lab in Cambridge | Wellcome Sanger Institute for the COVID-19 Genomics UK (COG-UK) Consortium | Rob Howes, The Lighthouse Lab in Cambridge and Alex Alderton, Roberto Amato, Sonia Goncalves, Ewan Harrison, David K. Jackson, Ian Johnston, Dominic Kwiatkowski, Cordelia Langford, John Sillitoe on behalf of the Wellcome Sanger Institute COVID-19 Surveillance Team |
| EPI_ISL_645946, EPI_ISL_645947, EPI_ISL_645948, EPI_ISL_645951, EPI_ISL_645952, EPI_ISL_645953, EPI_ISL_645954, EPI_ISL_645955, EPI_ISL_645956, EPI_ISL_645957 | Lighthouse Lab in Alderley Park | Wellcome Sanger Institute for the COVID-19 Genomics UK (COG-UK) Consortium | Jacquelyn Wynn, Mairead Hyland, The Lighthouse Lab in Alderley Park and Alex Alderton, Roberto Amato, Sonia Goncalves, Ewan Harrison, David K. Jackson, Ian Johnston, Dominic Kwiatkowski, Cordelia Langford, John Sillitoe on behalf of the Wellcome Sanger Institute COVID-19 Surveillance Team |
| EPI_ISL_645959 | Lighthouse Lab in Cambridge | Wellcome Sanger Institute for the COVID-19 Genomics UK (COG-UK) Consortium | Rob Howes, The Lighthouse Lab in Cambridge and Alex Alderton, Roberto Amato, Sonia Goncalves, Ewan Harrison, David K. Jackson, Ian Johnston, Dominic Kwiatkowski, Cordelia Langford, John Sillitoe on behalf of the Wellcome Sanger Institute COVID-19 Surveillance Team |
| EPI_ISL_645960, EPI_ISL_645961, EPI_ISL_645964, EPI_ISL_645965, EPI_ISL_645966, EPI_ISL_645967 | Lighthouse Lab in Alderley Park | Wellcome Sanger Institute for the COVID-19 Genomics UK (COG-UK) Consortium | Jacquelyn Wynn, Mairead Hyland, The Lighthouse Lab in Alderley Park and Alex Alderton, Roberto Amato, Sonia Goncalves, Ewan Harrison, David K. Jackson, Ian Johnston, Dominic Kwiatkowski, Cordelia Langford, John Sillitoe on behalf of the Wellcome Sanger Institute COVID-19 Surveillance Team |
| EPI_ISL_645968 | Lighthouse Lab in Cambridge | Wellcome Sanger Institute for the COVID-19 Genomics UK (COG-UK) Consortium | Rob Howes, The Lighthouse Lab in Cambridge and Alex Alderton, Roberto Amato, Sonia Goncalves, Ewan Harrison, David K. Jackson, Ian Johnston, Dominic Kwiatkowski, Cordelia Langford, John Sillitoe on behalf of the Wellcome Sanger Institute COVID-19 Surveillance Team |
| EPI_ISL_645969, EPI_ISL_645971, EPI_ISL_645972, EPI_ISL_645973, EPI_ISL_645975, EPI_ISL_645976, EPI_ISL_645977, EPI_ISL_645978 | Lighthouse Lab in Alderley Park | Wellcome Sanger Institute for the COVID-19 Genomics UK (COG-UK) Consortium | Jacquelyn Wynn, Mairead Hyland, The Lighthouse Lab in Alderley Park and Alex Alderton, Roberto Amato, Sonia Goncalves, Ewan Harrison, David K. Jackson, Ian Johnston, Dominic Kwiatkowski, Cordelia Langford, John Sillitoe on behalf of the Wellcome Sanger Institute COVID-19 Surveillance Team |
| EPI_ISL_645979 | Lighthouse Lab in Cambridge | Wellcome Sanger Institute for the COVID-19 Genomics UK (COG-UK) Consortium | Rob Howes, The Lighthouse Lab in Cambridge and Alex Alderton, Roberto Amato, Sonia Goncalves, Ewan Harrison, David K. Jackson, Ian Johnston, Dominic Kwiatkowski, Cordelia Langford, John Sillitoe on behalf of the Wellcome Sanger Institute COVID-19 Surveillance Team |
| EPI_ISL_645982, EPI_ISL_645984, EPI_ISL_645985, EPI_ISL_645986 | Lighthouse Lab in Alderley Park | Wellcome Sanger Institute for the COVID-19 Genomics UK (COG-UK) Consortium | Jacquelyn Wynn, Mairead Hyland, The Lighthouse Lab in Alderley Park and Alex Alderton, Roberto Amato, Sonia Goncalves, Ewan Harrison, David K. Jackson, Ian Johnston, Dominic Kwiatkowski, Cordelia Langford, John Sillitoe on behalf of the Wellcome Sanger Institute COVID-19 Surveillance Team |
| EPI_ISL_645987 | Lighthouse Lab in Cambridge | Wellcome Sanger Institute for the COVID-19 Genomics UK (COG-UK) Consortium | Rob Howes, The Lighthouse Lab in Cambridge and Alex Alderton, Roberto Amato, Sonia Goncalves, Ewan Harrison, David K. Jackson, Ian Johnston, Dominic Kwiatkowski, Cordelia Langford, John Sillitoe on behalf of the Wellcome Sanger Institute COVID-19 Surveillance Team |
| EPI_ISL_645988, EPI_ISL_645990 | Lighthouse Lab in Alderley Park | Wellcome Sanger Institute for the COVID-19 Genomics UK (COG-UK) Consortium | Jacquelyn Wynn, Mairead Hyland, The Lighthouse Lab in Alderley Park and Alex Alderton, Roberto Amato, Sonia Goncalves, Ewan Harrison, David K. Jackson, Ian Johnston, Dominic Kwiatkowski, Cordelia Langford, John Sillitoe on behalf of the Wellcome Sanger Institute COVID-19 Surveillance Team |
| EPI_ISL_645994 | Lighthouse Lab in Cambridge | Wellcome Sanger Institute for the COVID-19 Genomics UK (COG-UK) Consortium | Rob Howes, The Lighthouse Lab in Cambridge and Alex Alderton, Roberto Amato, Sonia Goncalves, Ewan Harrison, David K. Jackson, Ian Johnston, Dominic Kwiatkowski, Cordelia Langford, John Sillitoe on behalf of the Wellcome Sanger Institute COVID-19 Surveillance Team |
| EPI_ISL_645996, EPI_ISL_645997, EPI_ISL_645999, EPI_ISL_646000, EPI_ISL_646001, EPI_ISL_646002, EPI_ISL_646003, EPI_ISL_646004, EPI_ISL_646005, EPI_ISL_646006, EPI_ISL_646007, EPI_ISL_646008, EPI_ISL_646010, EPI_ISL_646011, EPI_ISL_646012, EPI_ISL_646013, EPI_ISL_646015, EPI_ISL_646016, EPI_ISL_646019, EPI_ISL_646020, EPI_ISL_646021, EPI_ISL_646023, EPI_ISL_646024, EPI_ISL_646025, EPI_ISL_646026 | | | |
| see above | Lighthouse Lab in Alderley Park | Wellcome Sanger Institute for the COVID-19 Genomics UK (COG-UK) Consortium | Jacquelyn Wynn, Mairead Hyland, The Lighthouse Lab in Alderley Park and Alex Alderton, Roberto Amato, Sonia Goncalves, Ewan Harrison, David K. Jackson, Ian Johnston, Dominic Kwiatkowski, Cordelia Langford, John Sillitoe on behalf of the Wellcome Sanger Institute COVID-19 Surveillance Team |
| EPI_ISL_646027 | Lighthouse Lab in Cambridge | Wellcome Sanger Institute for the COVID-19 Genomics UK (COG-UK) Consortium | Rob Howes, The Lighthouse Lab in Cambridge and Alex Alderton, Roberto Amato, Sonia Goncalves, Ewan Harrison, David K. Jackson, Ian Johnston, Dominic Kwiatkowski, Cordelia Langford, John Sillitoe on behalf of the Wellcome Sanger Institute COVID-19 Surveillance Team |
| EPI_ISL_646033, EPI_ISL_646036, EPI_ISL_646037, EPI_ISL_646038 | Lighthouse Lab in Alderley Park | Wellcome Sanger Institute for the COVID-19 Genomics UK (COG-UK) Consortium | Jacquelyn Wynn, Mairead Hyland, The Lighthouse Lab in Alderley Park and Alex Alderton, Roberto Amato, Sonia Goncalves, Ewan Harrison, David K. Jackson, Ian Johnston, Dominic Kwiatkowski, Cordelia Langford, John Sillitoe on behalf of the Wellcome Sanger Institute COVID-19 Surveillance Team |
| EPI_ISL_646039 | Lighthouse Lab in Cambridge | Wellcome Sanger Institute for the COVID-19 Genomics UK (COG-UK) Consortium | Rob Howes, The Lighthouse Lab in Cambridge and Alex Alderton, Roberto Amato, Sonia Goncalves, Ewan Harrison, David K. Jackson, Ian Johnston, Dominic Kwiatkowski, Cordelia Langford, John Sillitoe on behalf of the Wellcome Sanger Institute COVID-19 Surveillance Team |
| EPI_ISL_646040, EPI_ISL_646042, EPI_ISL_646043, EPI_ISL_646044, EPI_ISL_646045 | Lighthouse Lab in Alderley Park | Wellcome Sanger Institute for the COVID-19 Genomics UK (COG-UK) Consortium | Jacquelyn Wynn, Mairead Hyland, The Lighthouse Lab in Alderley Park and Alex Alderton, Roberto Amato, Sonia Goncalves, Ewan Harrison, David K. Jackson, Ian Johnston, Dominic Kwiatkowski, Cordelia Langford, John Sillitoe on behalf of the Wellcome Sanger Institute COVID-19 Surveillance Team |
| EPI_ISL_646047 | Lighthouse Lab in Cambridge | Wellcome Sanger Institute for the COVID-19 Genomics UK (COG-UK) Consortium | Rob Howes, The Lighthouse Lab in Cambridge and Alex Alderton, Roberto Amato, Sonia Goncalves, Ewan Harrison, David K. Jackson, Ian Johnston, Dominic Kwiatkowski, Cordelia Langford, John Sillitoe on behalf of the Wellcome Sanger Institute COVID-19 Surveillance Team |
| EPI_ISL_646048, EPI_ISL_646049, EPI_ISL_646050, EPI_ISL_646051, EPI_ISL_646053, EPI_ISL_646055, EPI_ISL_646057, EPI_ISL_646059, EPI_ISL_646060, EPI_ISL_646061, EPI_ISL_646063, EPI_ISL_646064, EPI_ISL_646065, EPI_ISL_646066, EPI_ISL_646068 | | | |
| see above | Lighthouse Lab in Alderley Park | Wellcome Sanger Institute for the COVID-19 Genomics UK (COG-UK) Consortium | Jacquelyn Wynn, Mairead Hyland, The Lighthouse Lab in Alderley Park and Alex Alderton, Roberto Amato, Sonia Goncalves, Ewan Harrison, David K. Jackson, Ian Johnston, Dominic Kwiatkowski, Cordelia Langford, John Sillitoe on behalf of the Wellcome Sanger Institute COVID-19 Surveillance Team |
| EPI_ISL_646070 | Lighthouse Lab in Cambridge | Wellcome Sanger Institute for the COVID-19 Genomics UK (COG-UK) Consortium | Rob Howes, The Lighthouse Lab in Cambridge and Alex Alderton, Roberto Amato, Sonia Goncalves, Ewan Harrison, David K. Jackson, Ian Johnston, Dominic Kwiatkowski, Cordelia Langford, John Sillitoe on behalf of the Wellcome Sanger Institute COVID-19 Surveillance Team |
| EPI_ISL_646071, EPI_ISL_646072, | Lighthouse Lab in Alderley Park | Wellcome Sanger Institute for the COVID-19 Genomics UK | Jacquelyn Wynn, Mairead Hyland, The Lighthouse Lab in Alderley Park and Alex Alderton, Roberto Amato, Sonia Goncalves, Ewan Harrison, David K. |

| | | | |
|---|---|---|---|
| EPI_ISL_646073, EPI_ISL_646074, EPI_ISL_646075 | | (COG-UK) Consortium | Jackson, Ian Johnston, Dominic Kwiatkowski, Cordelia Langford, John Sillitoe on behalf of the Wellcome Sanger Institute COVID-19 Surveillance Team |
| EPI_ISL_646076 | Lighthouse Lab in Cambridge | Wellcome Sanger Institute for the COVID-19 Genomics UK (COG-UK) Consortium | Rob Howes, The Lighthouse Lab in Cambridge and Alex Alderton, Roberto Amato, Sonia Goncalves, Ewan Harrison, David K. Jackson, Ian Johnston, Dominic Kwiatkowski, Cordelia Langford, John Sillitoe on behalf of the Wellcome Sanger Institute COVID-19 Surveillance Team |
| EPI_ISL_646077, EPI_ISL_646079, EPI_ISL_646080, EPI_ISL_646081, EPI_ISL_646082 | Lighthouse Lab in Alderley Park | Wellcome Sanger Institute for the COVID-19 Genomics UK (COG-UK) Consortium | Jacquelyn Wynn, Mairead Hyland, The Lighthouse Lab in Alderley Park and Alex Alderton, Roberto Amato, Sonia Goncalves, Ewan Harrison, David K. Jackson, Ian Johnston, Dominic Kwiatkowski, Cordelia Langford, John Sillitoe on behalf of the Wellcome Sanger Institute COVID-19 Surveillance Team |
| EPI_ISL_646086 | Lighthouse Lab in Cambridge | Wellcome Sanger Institute for the COVID-19 Genomics UK (COG-UK) Consortium | Rob Howes, The Lighthouse Lab in Cambridge and Alex Alderton, Roberto Amato, Sonia Goncalves, Ewan Harrison, David K. Jackson, Ian Johnston, Dominic Kwiatkowski, Cordelia Langford, John Sillitoe on behalf of the Wellcome Sanger Institute COVID-19 Surveillance Team |
| EPI_ISL_646087, EPI_ISL_646088, EPI_ISL_646090, EPI_ISL_646092, EPI_ISL_646093, EPI_ISL_646094, EPI_ISL_646095, EPI_ISL_646097, EPI_ISL_646099, EPI_ISL_646100, EPI_ISL_646103, EPI_ISL_646104 | | | |
| see above | Lighthouse Lab in Alderley Park | Wellcome Sanger Institute for the COVID-19 Genomics UK (COG-UK) Consortium | Jacquelyn Wynn, Mairead Hyland, The Lighthouse Lab in Alderley Park and Alex Alderton, Roberto Amato, Sonia Goncalves, Ewan Harrison, David K. Jackson, Ian Johnston, Dominic Kwiatkowski, Cordelia Langford, John Sillitoe on behalf of the Wellcome Sanger Institute COVID-19 Surveillance Team |
| EPI_ISL_646105 | Lighthouse Lab in Cambridge | Wellcome Sanger Institute for the COVID-19 Genomics UK (COG-UK) Consortium | Rob Howes, The Lighthouse Lab in Cambridge and Alex Alderton, Roberto Amato, Sonia Goncalves, Ewan Harrison, David K. Jackson, Ian Johnston, Dominic Kwiatkowski, Cordelia Langford, John Sillitoe on behalf of the Wellcome Sanger Institute COVID-19 Surveillance Team |
| EPI_ISL_646106, EPI_ISL_646107, EPI_ISL_646108, EPI_ISL_646109, EPI_ISL_646110, EPI_ISL_646111, EPI_ISL_646112, EPI_ISL_646113, EPI_ISL_646114, EPI_ISL_646116, EPI_ISL_646117, EPI_ISL_646118, EPI_ISL_646119, EPI_ISL_646120, EPI_ISL_646121, EPI_ISL_646123, EPI_ISL_646124, EPI_ISL_646126, EPI_ISL_646127, EPI_ISL_646130 | | | |
| see above | Lighthouse Lab in Alderley Park | Wellcome Sanger Institute for the COVID-19 Genomics UK (COG-UK) Consortium | Jacquelyn Wynn, Mairead Hyland, The Lighthouse Lab in Alderley Park and Alex Alderton, Roberto Amato, Sonia Goncalves, Ewan Harrison, David K. Jackson, Ian Johnston, Dominic Kwiatkowski, Cordelia Langford, John Sillitoe on behalf of the Wellcome Sanger Institute COVID-19 Surveillance Team |
| EPI_ISL_646131 | Lighthouse Lab in Cambridge | Wellcome Sanger Institute for the COVID-19 Genomics UK (COG-UK) Consortium | Rob Howes, The Lighthouse Lab in Cambridge and Alex Alderton, Roberto Amato, Sonia Goncalves, Ewan Harrison, David K. Jackson, Ian Johnston, Dominic Kwiatkowski, Cordelia Langford, John Sillitoe on behalf of the Wellcome Sanger Institute COVID-19 Surveillance Team |
| EPI_ISL_646132, EPI_ISL_646133, EPI_ISL_646134, EPI_ISL_646137, EPI_ISL_646138 | Lighthouse Lab in Alderley Park | Wellcome Sanger Institute for the COVID-19 Genomics UK (COG-UK) Consortium | Jacquelyn Wynn, Mairead Hyland, The Lighthouse Lab in Alderley Park and Alex Alderton, Roberto Amato, Sonia Goncalves, Ewan Harrison, David K. Jackson, Ian Johnston, Dominic Kwiatkowski, Cordelia Langford, John Sillitoe on behalf of the Wellcome Sanger Institute COVID-19 Surveillance Team |
| EPI_ISL_646153, EPI_ISL_646222, EPI_ISL_646461, EPI_ISL_646503, EPI_ISL_646507, EPI_ISL_646519, EPI_ISL_646527, EPI_ISL_646534, EPI_ISL_646574, EPI_ISL_646579, EPI_ISL_646590, EPI_ISL_646593, EPI_ISL_646602, EPI_ISL_646608, EPI_ISL_646619, EPI_ISL_646632, EPI_ISL_646645, EPI_ISL_646679, EPI_ISL_646704, EPI_ISL_646764, EPI_ISL_646779, EPI_ISL_647073 | | | |
| see above | Lighthouse Lab in Glasgow | Wellcome Sanger Institute for the COVID-19 Genomics UK (COG-UK) Consortium | Harper VanSteenhouse, Yumi Kasai, David Gray, Carol Clugston, Anna Dominiczak and Alex Alderton, Roberto Amato, Sonia Goncalves, Ewan Harrison, David K. Jackson, Ian Johnston, Dominic Kwiatkowski, Cordelia Langford, John Sillitoe on behalf of the Wellcome Sanger Institute COVID-19 Surveillance Team |
| EPI_ISL_647134, EPI_ISL_647135, EPI_ISL_647139, EPI_ISL_647145, EPI_ISL_647147, EPI_ISL_647148, EPI_ISL_647152, EPI_ISL_647153, EPI_ISL_647160, EPI_ISL_647161, EPI_ISL_647165, EPI_ISL_647172, EPI_ISL_647178, EPI_ISL_647180, EPI_ISL_647187, EPI_ISL_647197, EPI_ISL_647199, EPI_ISL_647200, EPI_ISL_647201, EPI_ISL_647205, EPI_ISL_647209, EPI_ISL_647214, EPI_ISL_647216, EPI_ISL_647223, EPI_ISL_647224, EPI_ISL_647230, EPI_ISL_647232, EPI_ISL_647234, EPI_ISL_647240, EPI_ISL_647247, EPI_ISL_647250, EPI_ISL_647251, EPI_ISL_647254, EPI_ISL_647255, EPI_ISL_647256, EPI_ISL_647259, EPI_ISL_647260, EPI_ISL_647263, EPI_ISL_647266, EPI_ISL_647267, EPI_ISL_647269, EPI_ISL_647273, EPI_ISL_647275, EPI_ISL_647279, EPI_ISL_647280, EPI_ISL_647281, EPI_ISL_647282, EPI_ISL_647290, EPI_ISL_647291, EPI_ISL_647292, EPI_ISL_647294, EPI_ISL_647296, EPI_ISL_647308, EPI_ISL_647314, EPI_ISL_647315, EPI_ISL_647321, EPI_ISL_647323, EPI_ISL_647324, EPI_ISL_647328, EPI_ISL_647329, EPI_ISL_647330, EPI_ISL_647332, EPI_ISL_647334, EPI_ISL_647337, EPI_ISL_647339, EPI_ISL_647342, EPI_ISL_647347, EPI_ISL_647354, EPI_ISL_647361, EPI_ISL_647362, EPI_ISL_647363, EPI_ISL_647399, EPI_ISL_647405, EPI_ISL_647411, EPI_ISL_647419, EPI_ISL_647420, EPI_ISL_647425, EPI_ISL_647427, EPI_ISL_647428, EPI_ISL_647429, EPI_ISL_647436, EPI_ISL_647443, EPI_ISL_647445, EPI_ISL_647452, EPI_ISL_647460 | | | |
| see above | Lighthouse Lab in Cambridge | Wellcome Sanger Institute for the COVID-19 Genomics UK (COG-UK) Consortium | Rob Howes, The Lighthouse Lab in Cambridge and Alex Alderton, Roberto Amato, Sonia Goncalves, Ewan Harrison, David K. Jackson, Ian Johnston, Dominic Kwiatkowski, Cordelia Langford, John Sillitoe on behalf of the Wellcome Sanger Institute COVID-19 Surveillance Team |
| EPI_ISL_647468, EPI_ISL_647469, EPI_ISL_647473, EPI_ISL_647474, EPI_ISL_647477 | Lighthouse Lab in Alderley Park | Wellcome Sanger Institute for the COVID-19 Genomics UK (COG-UK) Consortium | Jacquelyn Wynn, Mairead Hyland, The Lighthouse Lab in Alderley Park and Alex Alderton, Roberto Amato, Sonia Goncalves, Ewan Harrison, David K. Jackson, Ian Johnston, Dominic Kwiatkowski, Cordelia Langford, John Sillitoe on behalf of the Wellcome Sanger Institute COVID-19 Surveillance Team |
| EPI_ISL_647478, EPI_ISL_647479, EPI_ISL_647483, EPI_ISL_647485, EPI_ISL_647486, EPI_ISL_647492 | Lighthouse Lab in Cambridge | Wellcome Sanger Institute for the COVID-19 Genomics UK (COG-UK) Consortium | Rob Howes, The Lighthouse Lab in Cambridge and Alex Alderton, Roberto Amato, Sonia Goncalves, Ewan Harrison, David K. Jackson, Ian Johnston, Dominic Kwiatkowski, Cordelia Langford, John Sillitoe on behalf of the Wellcome Sanger Institute COVID-19 Surveillance Team |
| EPI_ISL_647497 | Lighthouse Lab in Alderley Park | Wellcome Sanger Institute for the COVID-19 Genomics UK (COG-UK) Consortium | Jacquelyn Wynn, Mairead Hyland, The Lighthouse Lab in Alderley Park and Alex Alderton, Roberto Amato, Sonia Goncalves, Ewan Harrison, David K. Jackson, Ian Johnston, Dominic Kwiatkowski, Cordelia Langford, John Sillitoe on behalf of the Wellcome Sanger Institute COVID-19 Surveillance Team |
| EPI_ISL_647498 | Lighthouse Lab in Cambridge | Wellcome Sanger Institute for the COVID-19 Genomics UK (COG-UK) Consortium | Rob Howes, The Lighthouse Lab in Cambridge and Alex Alderton, Roberto Amato, Sonia Goncalves, Ewan Harrison, David K. Jackson, Ian Johnston, Dominic Kwiatkowski, Cordelia Langford, John Sillitoe on behalf of the Wellcome Sanger Institute COVID-19 Surveillance Team |
| EPI_ISL_647499, EPI_ISL_647503, EPI_ISL_647506 | Lighthouse Lab in Alderley Park | Wellcome Sanger Institute for the COVID-19 Genomics UK (COG-UK) Consortium | Jacquelyn Wynn, Mairead Hyland, The Lighthouse Lab in Alderley Park and Alex Alderton, Roberto Amato, Sonia Goncalves, Ewan Harrison, David K. Jackson, Ian Johnston, Dominic Kwiatkowski, Cordelia Langford, John Sillitoe on behalf of the Wellcome Sanger Institute COVID-19 Surveillance Team |
| EPI_ISL_647509 | Lighthouse Lab in Cambridge | Wellcome Sanger Institute for the COVID-19 Genomics UK (COG-UK) Consortium | Rob Howes, The Lighthouse Lab in Cambridge and Alex Alderton, Roberto Amato, Sonia Goncalves, Ewan Harrison, David K. Jackson, Ian Johnston, Dominic Kwiatkowski, Cordelia Langford, John Sillitoe on behalf of the Wellcome Sanger Institute COVID-19 Surveillance Team |
| EPI_ISL_647511, EPI_ISL_647513 | Lighthouse Lab in Alderley Park | Wellcome Sanger Institute for the COVID-19 Genomics UK (COG-UK) Consortium | Jacquelyn Wynn, Mairead Hyland, The Lighthouse Lab in Alderley Park and Alex Alderton, Roberto Amato, Sonia Goncalves, Ewan Harrison, David K. Jackson, Ian Johnston, Dominic Kwiatkowski, Cordelia Langford, John Sillitoe on behalf of the Wellcome Sanger Institute COVID-19 Surveillance Team |
| EPI_ISL_647514, EPI_ISL_647515, EPI_ISL_647518, EPI_ISL_647519, EPI_ISL_647522 | Lighthouse Lab in Cambridge | Wellcome Sanger Institute for the COVID-19 Genomics UK (COG-UK) Consortium | Rob Howes, The Lighthouse Lab in Cambridge and Alex Alderton, Roberto Amato, Sonia Goncalves, Ewan Harrison, David K. Jackson, Ian Johnston, Dominic Kwiatkowski, Cordelia Langford, John Sillitoe on behalf of the Wellcome Sanger Institute COVID-19 Surveillance Team |
| EPI_ISL_647526, EPI_ISL_647528 | Lighthouse Lab in Alderley Park | Wellcome Sanger Institute for the COVID-19 Genomics UK (COG-UK) Consortium | Jacquelyn Wynn, Mairead Hyland, The Lighthouse Lab in Alderley Park and Alex Alderton, Roberto Amato, Sonia Goncalves, Ewan Harrison, David K. Jackson, Ian Johnston, Dominic Kwiatkowski, Cordelia Langford, John Sillitoe on behalf of the Wellcome Sanger Institute COVID-19 Surveillance Team |
| EPI_ISL_647531, EPI_ISL_647534 | Lighthouse Lab in Cambridge | Wellcome Sanger Institute for the COVID-19 Genomics UK (COG-UK) Consortium | Rob Howes, The Lighthouse Lab in Cambridge and Alex Alderton, Roberto Amato, Sonia Goncalves, Ewan Harrison, David K. Jackson, Ian Johnston, Dominic Kwiatkowski, Cordelia Langford, John Sillitoe on behalf of the Wellcome Sanger Institute COVID-19 Surveillance Team |
| EPI_ISL_647535, EPI_ISL_647536, EPI_ISL_647541 | Lighthouse Lab in Alderley Park | Wellcome Sanger Institute for the COVID-19 Genomics UK (COG-UK) Consortium | Jacquelyn Wynn, Mairead Hyland, The Lighthouse Lab in Alderley Park and Alex Alderton, Roberto Amato, Sonia Goncalves, Ewan Harrison, David K. Jackson, Ian Johnston, Dominic Kwiatkowski, Cordelia Langford, John Sillitoe on behalf of the Wellcome Sanger Institute COVID-19 Surveillance Team |
| EPI_ISL_647543 | Lighthouse Lab in Cambridge | Wellcome Sanger Institute for the COVID-19 Genomics UK (COG-UK) Consortium | Rob Howes, The Lighthouse Lab in Cambridge and Alex Alderton, Roberto Amato, Sonia Goncalves, Ewan Harrison, David K. Jackson, Ian Johnston, Dominic Kwiatkowski, Cordelia Langford, John Sillitoe on behalf of the Wellcome Sanger Institute COVID-19 Surveillance Team |
| EPI_ISL_647546 | Lighthouse Lab in Alderley Park | Wellcome Sanger Institute for the COVID-19 Genomics UK (COG-UK) Consortium | Jacquelyn Wynn, Mairead Hyland, The Lighthouse Lab in Alderley Park and Alex Alderton, Roberto Amato, Sonia Goncalves, Ewan Harrison, David K. Jackson, Ian Johnston, Dominic Kwiatkowski, Cordelia Langford, John Sillitoe on behalf of the Wellcome Sanger Institute COVID-19 Surveillance Team |
| EPI_ISL_647547 | Lighthouse Lab in Cambridge | Wellcome Sanger Institute for the COVID-19 Genomics UK (COG-UK) Consortium | Rob Howes, The Lighthouse Lab in Cambridge and Alex Alderton, Roberto Amato, Sonia Goncalves, Ewan Harrison, David K. Jackson, Ian Johnston, Dominic Kwiatkowski, Cordelia Langford, John Sillitoe on behalf of the Wellcome Sanger Institute COVID-19 Surveillance Team |
| EPI_ISL_647550, EPI_ISL_647553, EPI_ISL_647558, EPI_ISL_647560 | Lighthouse Lab in Alderley Park | Wellcome Sanger Institute for the COVID-19 Genomics UK (COG-UK) Consortium | Jacquelyn Wynn, Mairead Hyland, The Lighthouse Lab in Alderley Park and Alex Alderton, Roberto Amato, Sonia Goncalves, Ewan Harrison, David K. Jackson, Ian Johnston, Dominic Kwiatkowski, Cordelia Langford, John Sillitoe on behalf of the Wellcome Sanger Institute COVID-19 Surveillance Team |
| EPI_ISL_647562, EPI_ISL_647563, EPI_ISL_647569, EPI_ISL_647570 | Lighthouse Lab in Cambridge | Wellcome Sanger Institute for the COVID-19 Genomics UK (COG-UK) Consortium | Rob Howes, The Lighthouse Lab in Cambridge and Alex Alderton, Roberto Amato, Sonia Goncalves, Ewan Harrison, David K. Jackson, Ian Johnston, Dominic Kwiatkowski, Cordelia Langford, John Sillitoe on behalf of the Wellcome Sanger Institute COVID-19 Surveillance Team |
| EPI_ISL_647572, EPI_ISL_647573, EPI_ISL_647576, EPI_ISL_647579, | Lighthouse Lab in Alderley Park | Wellcome Sanger Institute for the COVID-19 Genomics UK (COG-UK) Consortium | Jacquelyn Wynn, Mairead Hyland, The Lighthouse Lab in Alderley Park and Alex Alderton, Roberto Amato, Sonia Goncalves, Ewan Harrison, David K. Jackson, Ian Johnston, Dominic Kwiatkowski, Cordelia Langford, John Sillitoe on behalf of the Wellcome Sanger Institute COVID-19 Surveillance Team |

| | | | |
|---|---|---|---|
| EPI_ISL_647580, EPI_ISL_647584, EPI_ISL_647591 | | | |
| EPI_ISL_647592 | Lighthouse Lab in Cambridge | Wellcome Sanger Institute for the COVID-19 Genomics UK (COG-UK) Consortium | Rob Howes, The Lighthouse Lab in Cambridge and Alex Alderton, Roberto Amato, Sonia Goncalves, Ewan Harrison, David K. Jackson, Ian Johnston, Dominic Kwiatkowski, Cordelia Langford, John Sillitoe on behalf of the Wellcome Sanger Institute COVID-19 Surveillance Team |
| EPI_ISL_647594 | Lighthouse Lab in Alderley Park | Wellcome Sanger Institute for the COVID-19 Genomics UK (COG-UK) Consortium | Jacquelyn Wynn, Mairead Hyland, The Lighthouse Lab in Alderley Park and Alex Alderton, Roberto Amato, Sonia Goncalves, Ewan Harrison, David K. Jackson, Ian Johnston, Dominic Kwiatkowski, Cordelia Langford, John Sillitoe on behalf of the Wellcome Sanger Institute COVID-19 Surveillance Team |
| EPI_ISL_647595, EPI_ISL_647598, EPI_ISL_647600 | Lighthouse Lab in Cambridge | Wellcome Sanger Institute for the COVID-19 Genomics UK (COG-UK) Consortium | Rob Howes, The Lighthouse Lab in Cambridge and Alex Alderton, Roberto Amato, Sonia Goncalves, Ewan Harrison, David K. Jackson, Ian Johnston, Dominic Kwiatkowski, Cordelia Langford, John Sillitoe on behalf of the Wellcome Sanger Institute COVID-19 Surveillance Team |
| EPI_ISL_647603 | Lighthouse Lab in Alderley Park | Wellcome Sanger Institute for the COVID-19 Genomics UK (COG-UK) Consortium | Jacquelyn Wynn, Mairead Hyland, The Lighthouse Lab in Alderley Park and Alex Alderton, Roberto Amato, Sonia Goncalves, Ewan Harrison, David K. Jackson, Ian Johnston, Dominic Kwiatkowski, Cordelia Langford, John Sillitoe on behalf of the Wellcome Sanger Institute COVID-19 Surveillance Team |
| EPI_ISL_647604, EPI_ISL_647605 | Lighthouse Lab in Cambridge | Wellcome Sanger Institute for the COVID-19 Genomics UK (COG-UK) Consortium | Rob Howes, The Lighthouse Lab in Cambridge and Alex Alderton, Roberto Amato, Sonia Goncalves, Ewan Harrison, David K. Jackson, Ian Johnston, Dominic Kwiatkowski, Cordelia Langford, John Sillitoe on behalf of the Wellcome Sanger Institute COVID-19 Surveillance Team |
| EPI_ISL_647606, EPI_ISL_647608, EPI_ISL_647612 | Lighthouse Lab in Alderley Park | Wellcome Sanger Institute for the COVID-19 Genomics UK (COG-UK) Consortium | Jacquelyn Wynn, Mairead Hyland, The Lighthouse Lab in Alderley Park and Alex Alderton, Roberto Amato, Sonia Goncalves, Ewan Harrison, David K. Jackson, Ian Johnston, Dominic Kwiatkowski, Cordelia Langford, John Sillitoe on behalf of the Wellcome Sanger Institute COVID-19 Surveillance Team |
| EPI_ISL_647614 | Lighthouse Lab in Cambridge | Wellcome Sanger Institute for the COVID-19 Genomics UK (COG-UK) Consortium | Rob Howes, The Lighthouse Lab in Cambridge and Alex Alderton, Roberto Amato, Sonia Goncalves, Ewan Harrison, David K. Jackson, Ian Johnston, Dominic Kwiatkowski, Cordelia Langford, John Sillitoe on behalf of the Wellcome Sanger Institute COVID-19 Surveillance Team |
| EPI_ISL_647615, EPI_ISL_647618 | Lighthouse Lab in Alderley Park | Wellcome Sanger Institute for the COVID-19 Genomics UK (COG-UK) Consortium | Jacquelyn Wynn, Mairead Hyland, The Lighthouse Lab in Alderley Park and Alex Alderton, Roberto Amato, Sonia Goncalves, Ewan Harrison, David K. Jackson, Ian Johnston, Dominic Kwiatkowski, Cordelia Langford, John Sillitoe on behalf of the Wellcome Sanger Institute COVID-19 Surveillance Team |
| EPI_ISL_647619, EPI_ISL_647620 | Lighthouse Lab in Cambridge | Wellcome Sanger Institute for the COVID-19 Genomics UK (COG-UK) Consortium | Rob Howes, The Lighthouse Lab in Cambridge and Alex Alderton, Roberto Amato, Sonia Goncalves, Ewan Harrison, David K. Jackson, Ian Johnston, Dominic Kwiatkowski, Cordelia Langford, John Sillitoe on behalf of the Wellcome Sanger Institute COVID-19 Surveillance Team |
| EPI_ISL_647624, EPI_ISL_647630, EPI_ISL_647632 | Lighthouse Lab in Alderley Park | Wellcome Sanger Institute for the COVID-19 Genomics UK (COG-UK) Consortium | Jacquelyn Wynn, Mairead Hyland, The Lighthouse Lab in Alderley Park and Alex Alderton, Roberto Amato, Sonia Goncalves, Ewan Harrison, David K. Jackson, Ian Johnston, Dominic Kwiatkowski, Cordelia Langford, John Sillitoe on behalf of the Wellcome Sanger Institute COVID-19 Surveillance Team |
| EPI_ISL_647633 | Lighthouse Lab in Cambridge | Wellcome Sanger Institute for the COVID-19 Genomics UK (COG-UK) Consortium | Rob Howes, The Lighthouse Lab in Cambridge and Alex Alderton, Roberto Amato, Sonia Goncalves, Ewan Harrison, David K. Jackson, Ian Johnston, Dominic Kwiatkowski, Cordelia Langford, John Sillitoe on behalf of the Wellcome Sanger Institute COVID-19 Surveillance Team |
| EPI_ISL_647634, EPI_ISL_647635 | Lighthouse Lab in Alderley Park | Wellcome Sanger Institute for the COVID-19 Genomics UK (COG-UK) Consortium | Jacquelyn Wynn, Mairead Hyland, The Lighthouse Lab in Alderley Park and Alex Alderton, Roberto Amato, Sonia Goncalves, Ewan Harrison, David K. Jackson, Ian Johnston, Dominic Kwiatkowski, Cordelia Langford, John Sillitoe on behalf of the Wellcome Sanger Institute COVID-19 Surveillance Team |
| EPI_ISL_647637 | Lighthouse Lab in Cambridge | Wellcome Sanger Institute for the COVID-19 Genomics UK (COG-UK) Consortium | Rob Howes, The Lighthouse Lab in Cambridge and Alex Alderton, Roberto Amato, Sonia Goncalves, Ewan Harrison, David K. Jackson, Ian Johnston, Dominic Kwiatkowski, Cordelia Langford, John Sillitoe on behalf of the Wellcome Sanger Institute COVID-19 Surveillance Team |
| EPI_ISL_647638 | Lighthouse Lab in Alderley Park | Wellcome Sanger Institute for the COVID-19 Genomics UK (COG-UK) Consortium | Jacquelyn Wynn, Mairead Hyland, The Lighthouse Lab in Alderley Park and Alex Alderton, Roberto Amato, Sonia Goncalves, Ewan Harrison, David K. Jackson, Ian Johnston, Dominic Kwiatkowski, Cordelia Langford, John Sillitoe on behalf of the Wellcome Sanger Institute COVID-19 Surveillance Team |
| EPI_ISL_647640, EPI_ISL_647642 | Lighthouse Lab in Cambridge | Wellcome Sanger Institute for the COVID-19 Genomics UK (COG-UK) Consortium | Rob Howes, The Lighthouse Lab in Cambridge and Alex Alderton, Roberto Amato, Sonia Goncalves, Ewan Harrison, David K. Jackson, Ian Johnston, Dominic Kwiatkowski, Cordelia Langford, John Sillitoe on behalf of the Wellcome Sanger Institute COVID-19 Surveillance Team |
| EPI_ISL_647643 | Lighthouse Lab in Alderley Park | Wellcome Sanger Institute for the COVID-19 Genomics UK (COG-UK) Consortium | Jacquelyn Wynn, Mairead Hyland, The Lighthouse Lab in Alderley Park and Alex Alderton, Roberto Amato, Sonia Goncalves, Ewan Harrison, David K. Jackson, Ian Johnston, Dominic Kwiatkowski, Cordelia Langford, John Sillitoe on behalf of the Wellcome Sanger Institute COVID-19 Surveillance Team |
| EPI_ISL_647646, EPI_ISL_647648 | Lighthouse Lab in Cambridge | Wellcome Sanger Institute for the COVID-19 Genomics UK (COG-UK) Consortium | Rob Howes, The Lighthouse Lab in Cambridge and Alex Alderton, Roberto Amato, Sonia Goncalves, Ewan Harrison, David K. Jackson, Ian Johnston, Dominic Kwiatkowski, Cordelia Langford, John Sillitoe on behalf of the Wellcome Sanger Institute COVID-19 Surveillance Team |
| EPI_ISL_647650, EPI_ISL_647655 | Lighthouse Lab in Alderley Park | Wellcome Sanger Institute for the COVID-19 Genomics UK (COG-UK) Consortium | Jacquelyn Wynn, Mairead Hyland, The Lighthouse Lab in Alderley Park and Alex Alderton, Roberto Amato, Sonia Goncalves, Ewan Harrison, David K. Jackson, Ian Johnston, Dominic Kwiatkowski, Cordelia Langford, John Sillitoe on behalf of the Wellcome Sanger Institute COVID-19 Surveillance Team |
| EPI_ISL_647658 | Lighthouse Lab in Cambridge | Wellcome Sanger Institute for the COVID-19 Genomics UK (COG-UK) Consortium | Rob Howes, The Lighthouse Lab in Cambridge and Alex Alderton, Roberto Amato, Sonia Goncalves, Ewan Harrison, David K. Jackson, Ian Johnston, Dominic Kwiatkowski, Cordelia Langford, John Sillitoe on behalf of the Wellcome Sanger Institute COVID-19 Surveillance Team |
| EPI_ISL_647659 | Lighthouse Lab in Alderley Park | Wellcome Sanger Institute for the COVID-19 Genomics UK (COG-UK) Consortium | Jacquelyn Wynn, Mairead Hyland, The Lighthouse Lab in Alderley Park and Alex Alderton, Roberto Amato, Sonia Goncalves, Ewan Harrison, David K. Jackson, Ian Johnston, Dominic Kwiatkowski, Cordelia Langford, John Sillitoe on behalf of the Wellcome Sanger Institute COVID-19 Surveillance Team |
| EPI_ISL_647662 | Lighthouse Lab in Cambridge | Wellcome Sanger Institute for the COVID-19 Genomics UK (COG-UK) Consortium | Rob Howes, The Lighthouse Lab in Cambridge and Alex Alderton, Roberto Amato, Sonia Goncalves, Ewan Harrison, David K. Jackson, Ian Johnston, Dominic Kwiatkowski, Cordelia Langford, John Sillitoe on behalf of the Wellcome Sanger Institute COVID-19 Surveillance Team |
| EPI_ISL_647667, EPI_ISL_647668 | Lighthouse Lab in Alderley Park | Wellcome Sanger Institute for the COVID-19 Genomics UK (COG-UK) Consortium | Jacquelyn Wynn, Mairead Hyland, The Lighthouse Lab in Alderley Park and Alex Alderton, Roberto Amato, Sonia Goncalves, Ewan Harrison, David K. Jackson, Ian Johnston, Dominic Kwiatkowski, Cordelia Langford, John Sillitoe on behalf of the Wellcome Sanger Institute COVID-19 Surveillance Team |
| EPI_ISL_647671 | Lighthouse Lab in Cambridge | Wellcome Sanger Institute for the COVID-19 Genomics UK (COG-UK) Consortium | Rob Howes, The Lighthouse Lab in Cambridge and Alex Alderton, Roberto Amato, Sonia Goncalves, Ewan Harrison, David K. Jackson, Ian Johnston, Dominic Kwiatkowski, Cordelia Langford, John Sillitoe on behalf of the Wellcome Sanger Institute COVID-19 Surveillance Team |
| EPI_ISL_647672, EPI_ISL_647677 | Lighthouse Lab in Alderley Park | Wellcome Sanger Institute for the COVID-19 Genomics UK (COG-UK) Consortium | Jacquelyn Wynn, Mairead Hyland, The Lighthouse Lab in Alderley Park and Alex Alderton, Roberto Amato, Sonia Goncalves, Ewan Harrison, David K. Jackson, Ian Johnston, Dominic Kwiatkowski, Cordelia Langford, John Sillitoe on behalf of the Wellcome Sanger Institute COVID-19 Surveillance Team |
| EPI_ISL_647680 | Lighthouse Lab in Cambridge | Wellcome Sanger Institute for the COVID-19 Genomics UK (COG-UK) Consortium | Rob Howes, The Lighthouse Lab in Cambridge and Alex Alderton, Roberto Amato, Sonia Goncalves, Ewan Harrison, David K. Jackson, Ian Johnston, Dominic Kwiatkowski, Cordelia Langford, John Sillitoe on behalf of the Wellcome Sanger Institute COVID-19 Surveillance Team |
| EPI_ISL_647682, EPI_ISL_647683 | Lighthouse Lab in Alderley Park | Wellcome Sanger Institute for the COVID-19 Genomics UK (COG-UK) Consortium | Jacquelyn Wynn, Mairead Hyland, The Lighthouse Lab in Alderley Park and Alex Alderton, Roberto Amato, Sonia Goncalves, Ewan Harrison, David K. Jackson, Ian Johnston, Dominic Kwiatkowski, Cordelia Langford, John Sillitoe on behalf of the Wellcome Sanger Institute COVID-19 Surveillance Team |
| EPI_ISL_647686, EPI_ISL_647687 | Lighthouse Lab in Cambridge | Wellcome Sanger Institute for the COVID-19 Genomics UK (COG-UK) Consortium | Rob Howes, The Lighthouse Lab in Cambridge and Alex Alderton, Roberto Amato, Sonia Goncalves, Ewan Harrison, David K. Jackson, Ian Johnston, Dominic Kwiatkowski, Cordelia Langford, John Sillitoe on behalf of the Wellcome Sanger Institute COVID-19 Surveillance Team |
| EPI_ISL_647689, EPI_ISL_647693 | Lighthouse Lab in Alderley Park | Wellcome Sanger Institute for the COVID-19 Genomics UK (COG-UK) Consortium | Jacquelyn Wynn, Mairead Hyland, The Lighthouse Lab in Alderley Park and Alex Alderton, Roberto Amato, Sonia Goncalves, Ewan Harrison, David K. Jackson, Ian Johnston, Dominic Kwiatkowski, Cordelia Langford, John Sillitoe on behalf of the Wellcome Sanger Institute COVID-19 Surveillance Team |
| EPI_ISL_647694 | Lighthouse Lab in Cambridge | Wellcome Sanger Institute for the COVID-19 Genomics UK (COG-UK) Consortium | Rob Howes, The Lighthouse Lab in Cambridge and Alex Alderton, Roberto Amato, Sonia Goncalves, Ewan Harrison, David K. Jackson, Ian Johnston, Dominic Kwiatkowski, Cordelia Langford, John Sillitoe on behalf of the Wellcome Sanger Institute COVID-19 Surveillance Team |
| EPI_ISL_647695, EPI_ISL_647698, EPI_ISL_647700, EPI_ISL_647704, EPI_ISL_647705 | Lighthouse Lab in Alderley Park | Wellcome Sanger Institute for the COVID-19 Genomics UK (COG-UK) Consortium | Jacquelyn Wynn, Mairead Hyland, The Lighthouse Lab in Alderley Park and Alex Alderton, Roberto Amato, Sonia Goncalves, Ewan Harrison, David K. Jackson, Ian Johnston, Dominic Kwiatkowski, Cordelia Langford, John Sillitoe on behalf of the Wellcome Sanger Institute COVID-19 Surveillance Team |
| EPI_ISL_647706, EPI_ISL_647707 | Lighthouse Lab in Cambridge | Wellcome Sanger Institute for the COVID-19 Genomics UK (COG-UK) Consortium | Rob Howes, The Lighthouse Lab in Cambridge and Alex Alderton, Roberto Amato, Sonia Goncalves, Ewan Harrison, David K. Jackson, Ian Johnston, Dominic Kwiatkowski, Cordelia Langford, John Sillitoe on behalf of the Wellcome Sanger Institute COVID-19 Surveillance Team |
| EPI_ISL_647708 | Lighthouse Lab in Alderley Park | Wellcome Sanger Institute for the COVID-19 Genomics UK (COG-UK) Consortium | Jacquelyn Wynn, Mairead Hyland, The Lighthouse Lab in Alderley Park and Alex Alderton, Roberto Amato, Sonia Goncalves, Ewan Harrison, David K. Jackson, Ian Johnston, Dominic Kwiatkowski, Cordelia Langford, John Sillitoe on behalf of the Wellcome Sanger Institute COVID-19 Surveillance Team |
| EPI_ISL_647710 | Lighthouse Lab in Cambridge | Wellcome Sanger Institute for the COVID-19 Genomics UK (COG-UK) Consortium | Rob Howes, The Lighthouse Lab in Cambridge and Alex Alderton, Roberto Amato, Sonia Goncalves, Ewan Harrison, David K. Jackson, Ian Johnston, Dominic Kwiatkowski, Cordelia Langford, John Sillitoe on behalf of the Wellcome Sanger Institute COVID-19 Surveillance Team |
| EPI_ISL_647711 | Lighthouse Lab in Alderley Park | Wellcome Sanger Institute for the COVID-19 Genomics UK (COG-UK) Consortium | Jacquelyn Wynn, Mairead Hyland, The Lighthouse Lab in Alderley Park and Alex Alderton, Roberto Amato, Sonia Goncalves, Ewan Harrison, David K. Jackson, Ian Johnston, Dominic Kwiatkowski, Cordelia Langford, John Sillitoe on behalf of the Wellcome Sanger Institute COVID-19 Surveillance Team |

| | | | |
|---|---|---|---|
| EPI_ISL_647713, EPI_ISL_647714, EPI_ISL_647715, EPI_ISL_647716, EPI_ISL_647720, EPI_ISL_647722, EPI_ISL_647724, EPI_ISL_647725 | Lighthouse Lab in Cambridge | Wellcome Sanger Institute for the COVID-19 Genomics UK (COG-UK) Consortium | Rob Howes, The Lighthouse Lab in Cambridge and Alex Alderton, Roberto Amato, Sonia Goncalves, Ewan Harrison, David K. Jackson, Ian Johnston, Dominic Kwiatkowski, Cordelia Langford, John Sillitoe on behalf of the Wellcome Sanger Institute COVID-19 Surveillance Team |
| EPI_ISL_647726 | Lighthouse Lab in Alderley Park | Wellcome Sanger Institute for the COVID-19 Genomics UK (COG-UK) Consortium | Jacquelyn Wynn, Mairead Hyland, The Lighthouse Lab in Alderley Park and Alex Alderton, Roberto Amato, Sonia Goncalves, Ewan Harrison, David K. Jackson, Ian Johnston, Dominic Kwiatkowski, Cordelia Langford, John Sillitoe on behalf of the Wellcome Sanger Institute COVID-19 Surveillance Team |
| EPI_ISL_647727, EPI_ISL_647732 | Lighthouse Lab in Cambridge | Wellcome Sanger Institute for the COVID-19 Genomics UK (COG-UK) Consortium | Rob Howes, The Lighthouse Lab in Cambridge and Alex Alderton, Roberto Amato, Sonia Goncalves, Ewan Harrison, David K. Jackson, Ian Johnston, Dominic Kwiatkowski, Cordelia Langford, John Sillitoe on behalf of the Wellcome Sanger Institute COVID-19 Surveillance Team |
| EPI_ISL_647733 | Lighthouse Lab in Alderley Park | Wellcome Sanger Institute for the COVID-19 Genomics UK (COG-UK) Consortium | Jacquelyn Wynn, Mairead Hyland, The Lighthouse Lab in Alderley Park and Alex Alderton, Roberto Amato, Sonia Goncalves, Ewan Harrison, David K. Jackson, Ian Johnston, Dominic Kwiatkowski, Cordelia Langford, John Sillitoe on behalf of the Wellcome Sanger Institute COVID-19 Surveillance Team |
| EPI_ISL_647738, EPI_ISL_647739 | Lighthouse Lab in Cambridge | Wellcome Sanger Institute for the COVID-19 Genomics UK (COG-UK) Consortium | Rob Howes, The Lighthouse Lab in Cambridge and Alex Alderton, Roberto Amato, Sonia Goncalves, Ewan Harrison, David K. Jackson, Ian Johnston, Dominic Kwiatkowski, Cordelia Langford, John Sillitoe on behalf of the Wellcome Sanger Institute COVID-19 Surveillance Team |
| EPI_ISL_647741 | Lighthouse Lab in Alderley Park | Wellcome Sanger Institute for the COVID-19 Genomics UK (COG-UK) Consortium | Jacquelyn Wynn, Mairead Hyland, The Lighthouse Lab in Alderley Park and Alex Alderton, Roberto Amato, Sonia Goncalves, Ewan Harrison, David K. Jackson, Ian Johnston, Dominic Kwiatkowski, Cordelia Langford, John Sillitoe on behalf of the Wellcome Sanger Institute COVID-19 Surveillance Team |
| EPI_ISL_647748 | Lighthouse Lab in Cambridge | Wellcome Sanger Institute for the COVID-19 Genomics UK (COG-UK) Consortium | Rob Howes, The Lighthouse Lab in Cambridge and Alex Alderton, Roberto Amato, Sonia Goncalves, Ewan Harrison, David K. Jackson, Ian Johnston, Dominic Kwiatkowski, Cordelia Langford, John Sillitoe on behalf of the Wellcome Sanger Institute COVID-19 Surveillance Team |
| EPI_ISL_647750, EPI_ISL_647752 | Lighthouse Lab in Alderley Park | Wellcome Sanger Institute for the COVID-19 Genomics UK (COG-UK) Consortium | Jacquelyn Wynn, Mairead Hyland, The Lighthouse Lab in Alderley Park and Alex Alderton, Roberto Amato, Sonia Goncalves, Ewan Harrison, David K. Jackson, Ian Johnston, Dominic Kwiatkowski, Cordelia Langford, John Sillitoe on behalf of the Wellcome Sanger Institute COVID-19 Surveillance Team |
| EPI_ISL_647755, EPI_ISL_647756, EPI_ISL_647758, EPI_ISL_647760, EPI_ISL_647761 | Lighthouse Lab in Cambridge | Wellcome Sanger Institute for the COVID-19 Genomics UK (COG-UK) Consortium | Rob Howes, The Lighthouse Lab in Cambridge and Alex Alderton, Roberto Amato, Sonia Goncalves, Ewan Harrison, David K. Jackson, Ian Johnston, Dominic Kwiatkowski, Cordelia Langford, John Sillitoe on behalf of the Wellcome Sanger Institute COVID-19 Surveillance Team |
| EPI_ISL_647762 | Lighthouse Lab in Alderley Park | Wellcome Sanger Institute for the COVID-19 Genomics UK (COG-UK) Consortium | Jacquelyn Wynn, Mairead Hyland, The Lighthouse Lab in Alderley Park and Alex Alderton, Roberto Amato, Sonia Goncalves, Ewan Harrison, David K. Jackson, Ian Johnston, Dominic Kwiatkowski, Cordelia Langford, John Sillitoe on behalf of the Wellcome Sanger Institute COVID-19 Surveillance Team |
| EPI_ISL_647764 | Lighthouse Lab in Cambridge | Wellcome Sanger Institute for the COVID-19 Genomics UK (COG-UK) Consortium | Rob Howes, The Lighthouse Lab in Cambridge and Alex Alderton, Roberto Amato, Sonia Goncalves, Ewan Harrison, David K. Jackson, Ian Johnston, Dominic Kwiatkowski, Cordelia Langford, John Sillitoe on behalf of the Wellcome Sanger Institute COVID-19 Surveillance Team |
| EPI_ISL_647771 | Lighthouse Lab in Alderley Park | Wellcome Sanger Institute for the COVID-19 Genomics UK (COG-UK) Consortium | Jacquelyn Wynn, Mairead Hyland, The Lighthouse Lab in Alderley Park and Alex Alderton, Roberto Amato, Sonia Goncalves, Ewan Harrison, David K. Jackson, Ian Johnston, Dominic Kwiatkowski, Cordelia Langford, John Sillitoe on behalf of the Wellcome Sanger Institute COVID-19 Surveillance Team |
| EPI_ISL_647774, EPI_ISL_647775 | Lighthouse Lab in Cambridge | Wellcome Sanger Institute for the COVID-19 Genomics UK (COG-UK) Consortium | Rob Howes, The Lighthouse Lab in Cambridge and Alex Alderton, Roberto Amato, Sonia Goncalves, Ewan Harrison, David K. Jackson, Ian Johnston, Dominic Kwiatkowski, Cordelia Langford, John Sillitoe on behalf of the Wellcome Sanger Institute COVID-19 Surveillance Team |
| EPI_ISL_647776 | Lighthouse Lab in Alderley Park | Wellcome Sanger Institute for the COVID-19 Genomics UK (COG-UK) Consortium | Jacquelyn Wynn, Mairead Hyland, The Lighthouse Lab in Alderley Park and Alex Alderton, Roberto Amato, Sonia Goncalves, Ewan Harrison, David K. Jackson, Ian Johnston, Dominic Kwiatkowski, Cordelia Langford, John Sillitoe on behalf of the Wellcome Sanger Institute COVID-19 Surveillance Team |
| EPI_ISL_647779 | Lighthouse Lab in Cambridge | Wellcome Sanger Institute for the COVID-19 Genomics UK (COG-UK) Consortium | Rob Howes, The Lighthouse Lab in Cambridge and Alex Alderton, Roberto Amato, Sonia Goncalves, Ewan Harrison, David K. Jackson, Ian Johnston, Dominic Kwiatkowski, Cordelia Langford, John Sillitoe on behalf of the Wellcome Sanger Institute COVID-19 Surveillance Team |
| EPI_ISL_647780, EPI_ISL_647786, EPI_ISL_647787 | Lighthouse Lab in Alderley Park | Wellcome Sanger Institute for the COVID-19 Genomics UK (COG-UK) Consortium | Jacquelyn Wynn, Mairead Hyland, The Lighthouse Lab in Alderley Park and Alex Alderton, Roberto Amato, Sonia Goncalves, Ewan Harrison, David K. Jackson, Ian Johnston, Dominic Kwiatkowski, Cordelia Langford, John Sillitoe on behalf of the Wellcome Sanger Institute COVID-19 Surveillance Team |
| EPI_ISL_647789 | Lighthouse Lab in Cambridge | Wellcome Sanger Institute for the COVID-19 Genomics UK (COG-UK) Consortium | Rob Howes, The Lighthouse Lab in Cambridge and Alex Alderton, Roberto Amato, Sonia Goncalves, Ewan Harrison, David K. Jackson, Ian Johnston, Dominic Kwiatkowski, Cordelia Langford, John Sillitoe on behalf of the Wellcome Sanger Institute COVID-19 Surveillance Team |
| EPI_ISL_647792 | Lighthouse Lab in Alderley Park | Wellcome Sanger Institute for the COVID-19 Genomics UK (COG-UK) Consortium | Jacquelyn Wynn, Mairead Hyland, The Lighthouse Lab in Alderley Park and Alex Alderton, Roberto Amato, Sonia Goncalves, Ewan Harrison, David K. Jackson, Ian Johnston, Dominic Kwiatkowski, Cordelia Langford, John Sillitoe on behalf of the Wellcome Sanger Institute COVID-19 Surveillance Team |
| EPI_ISL_647795, EPI_ISL_647796 | Lighthouse Lab in Cambridge | Wellcome Sanger Institute for the COVID-19 Genomics UK (COG-UK) Consortium | Rob Howes, The Lighthouse Lab in Cambridge and Alex Alderton, Roberto Amato, Sonia Goncalves, Ewan Harrison, David K. Jackson, Ian Johnston, Dominic Kwiatkowski, Cordelia Langford, John Sillitoe on behalf of the Wellcome Sanger Institute COVID-19 Surveillance Team |
| EPI_ISL_647798, EPI_ISL_647800, EPI_ISL_647801, EPI_ISL_647804, EPI_ISL_647806, EPI_ISL_647861, EPI_ISL_647863, EPI_ISL_647864, EPI_ISL_647865 | Lighthouse Lab in Glasgow | Wellcome Sanger Institute for the COVID-19 Genomics UK (COG-UK) Consortium | Harper VanSteenhouse, Yumi Kasai, David Gray, Carol Clugston, Anna Dominiczak and Alex Alderton, Roberto Amato, Sonia Goncalves, Ewan Harrison, David K. Jackson, Ian Johnston, Dominic Kwiatkowski, Cordelia Langford, John Sillitoe on behalf of the Wellcome Sanger Institute COVID-19 Surveillance Team |
| EPI_ISL_648151 | Halmstad | The Public Health Agency of Sweden | Anna-Malin Linde, Maria Lind Karlberg, Mattias Haukland, Reza Advani, Olov Svartstrom, Oskar Karlsson Lindsjo, Sandra Broddesson, Petra Edquist, Mia Brytting, Anna Risberg, Karin Tegmark-Wisell |
| EPI_ISL_648157, EPI_ISL_648158 | Sundsvall | The Public Health Agency of Sweden | Anna-Malin Linde, Maria Lind Karlberg, Mattias Haukland, Reza Advani, Olov Svartstrom, Oskar Karlsson Lindsjo, Sandra Broddesson, Petra Edquist, Mia Brytting, Anna Risberg, Karin Tegmark-Wisell |
| EPI_ISL_649018 | San Diego County Public Health Laboratory | Andersen lab at Scripps Research | SEARCH Alliance San Diego with Tracy Basler, Jovan Shephard, Brett Austin |
| EPI_ISL_649058 | University of Michigan Clinical Microbiology Laboratory | Lauring Lab, University of Michigan, Department of Microbiology and Immunology | Valesano |
| EPI_ISL_649483, EPI_ISL_649484, EPI_ISL_649485, EPI_ISL_649486, EPI_ISL_649490, EPI_ISL_649491 | Lighthouse Lab in Alderley Park | Wellcome Sanger Institute for the COVID-19 Genomics UK (COG-UK) Consortium | Jacquelyn Wynn, Mairead Hyland, The Lighthouse Lab in Alderley Park and Alex Alderton, Roberto Amato, Sonia Goncalves, Ewan Harrison, David K. Jackson, Ian Johnston, Dominic Kwiatkowski, Cordelia Langford, John Sillitoe on behalf of the Wellcome Sanger Institute COVID-19 Surveillance Team (http://www.sanger.ac.uk/covid-team) |
| EPI_ISL_649492, EPI_ISL_649493 | Lighthouse Lab in Cambridge | Wellcome Sanger Institute for the COVID-19 Genomics UK (COG-UK) Consortium | Rob Howes, The Lighthouse Lab in Cambridge and Alex Alderton, Roberto Amato, Sonia Goncalves, Ewan Harrison, David K. Jackson, Ian Johnston, Dominic Kwiatkowski, Cordelia Langford, John Sillitoe on behalf of the Wellcome Sanger Institute COVID-19 Surveillance Team (http://www.sanger.ac.uk/covid-team) |
| EPI_ISL_649499, EPI_ISL_649503 | Lighthouse Lab in Glasgow | Wellcome Sanger Institute for the COVID-19 Genomics UK (COG-UK) Consortium | Harper VanSteenhouse, Yumi Kasai, David Gray, Carol Clugston, Anna Dominiczak and Alex Alderton, Roberto Amato, Sonia Goncalves, Ewan Harrison, David K. Jackson, Ian Johnston, Dominic Kwiatkowski, Cordelia Langford, John Sillitoe on behalf of the Wellcome Sanger Institute COVID-19 Surveillance Team (http://www.sanger.ac.uk/covid-team) |
| EPI_ISL_649629, EPI_ISL_649632 | Lighthouse Lab in Cambridge | Wellcome Sanger Institute for the COVID-19 Genomics UK (COG-UK) Consortium | Rob Howes, The Lighthouse Lab in Cambridge and Alex Alderton, Roberto Amato, Sonia Goncalves, Ewan Harrison, David K. Jackson, Ian Johnston, Dominic Kwiatkowski, Cordelia Langford, John Sillitoe on behalf of the Wellcome Sanger Institute COVID-19 Surveillance Team (http://www.sanger.ac.uk/covid-team) |
| EPI_ISL_649890 | Respiratory Virus Unit, Microbiology Services Colindale, Public Health England | COVID-19 Genomics UK (COG-UK) Consortium | PHE Covid Sequencing Team |
| EPI_ISL_650105 | University of Michigan Clinical Microbiology Laboratory | Lauring Lab, University of Michigan, Department of Microbiology and Immunology | Valesano |
| EPI_ISL_650123 | University College London, Great Ormond Street Hospital for Children NHS Foundation Trust, Imperial College Healthcare NHS Trust | COVID-19 Genomics UK (COG-UK) Consortium | Sergi Castellano, Rachel Williams, Mark Kristiansen, Paola Resende Silva, Sunando Roy, Tony Brooks, Helena Tutill, Paola Niola, Patricia Dyal, Charlotte Williams, Leysa Forrest, Yasmin Panchbhaya, Jacqueline Findlay, Samuel Weeks, Julianne Brown, Kathryn Harris, Paul Randell, James Price, Alison Holmes, Judith Breuer |

| | | | |
|---|---|---|---|
| EPI_ISL_650131 | Virology Department, Sheffield Teaching Hospitals NHS Foundation Trust/Department of Infection, Immunity and Cardiovascular Disease, The Medical School, University of Sheffield | COVID-19 Genomics UK (COG-UK) Consortium | Thushan de Silva, Matthew Parker, Nikki Smith, Adri Angyal, Rebecca Brown, Luke Green, Rachel Tucker, Paul Parsons, Danielle Groves, Katie Johnson, Laura Carrilero, Alex Keeley, Dave Partridge, Matthew Wyles, Benjamin Lindsey, Mehmet Yavuz, Mohammad Raza, Cariad Evans |
| EPI_ISL_650135 | Virology Department, Royal Infirmary of Edinburgh, NHS Lothian / School of Biological Sciences, University of Edinburgh / Institute of Genetics and Molecular Medicine, University of Edinburgh | COVID-19 Genomics UK (COG-UK) Consortium | McHugh M, Dewar R, Rooke S, Gallagher M, Balcaza C, O'Toole Á, Scher E, Hill V, McCrone JT, Colquhoun R, Yu X, Jackson B, Rambaut A, Williams TC, Templeton K |
| EPI_ISL_650141 | University of Exeter | COVID-19 Genomics UK (COG-UK) Consortium | Ben Temperton,Aaron Jeffries,Michelle Michelsen,Joanna Warwick-Dugdale,Audrey Farbos,Robyn Manley,Stephen Michell,Jane Masoli |
| EPI_ISL_650143 | Department of Pathology, University of Cambridge | COVID-19 Genomics UK (COG-UK) Consortium | Aminu S. Jahun, Yasmin Chaudhry, Grant Hall, Iliana Georgana, Myra Hosmillo, Martin D. Curran, Malte Pinckert, Surendra Parmar, Ian Goodfellow |
| EPI_ISL_650153, EPI_ISL_650158 | University of Birmingham | COVID-19 Genomics UK (COG-UK) Consortium | Institute of Microbiology, University of Birmingham: Claire McMurray, Joanne Stockton, Samuel Nicholls, Radoslaw Poplawski, Will Rowe, Josh Quick, Nicholas Loman. University of Birmingham Testing Laboratory: Celina M Whalley, Andrew Bosworth, Charlotte Poxon, Kasun Wanigasooriya, Oliver Pickles, Mike Kidd, Alex Richter, Andrew D Beggs PHE Heartlands Lab: Husam Osman, Andrew Bosworth. Queen Elizabeth Hospital: Anna Casey |
| EPI_ISL_650162 | Liverpool Clinical Laboratories | COVID-19 Genomics UK (COG-UK) Consortium | Sam Haldenby, Anita Lucaci, Steve Paterson, Julian Hiscox, Alistair Darby, M Almsaud, A Alrezaihi, Muhannad Alruwaili, Stuart D Armstrong, Jones Benjamin, Eleanor G Bentley, Anu Chawla, Jordan J Clark, Angela Cowell, Richard Eccles, Isabel García-Dorival, Matthew Gemmell, Alessandro Gerada, PKF Gilmore, Richard Gregory, Ximeng Han, Catherine Hartley, Margaret Hughes, Miren Iturriza-Gomara, James Johnson, L Luu, Jenifer Manson, Charlotte Nelson, Elaine O'Toole, Cassie Olateju, Rebekah Penrice-Randal , Lucille Rainbow, N.P Randle, Trevor Ian Robinson, Parul Sharma, Ghada T Shawli, James P Stewart, Neil Swainston, Ecaterina Vamos, Joanne Watts, Mark Whitehead |
| EPI_ISL_650164 | West of Scotland Specialist Virology Centre, NHSGGC / MRC-University of Glasgow Centre for Virus Research | COVID-19 Genomics UK (COG-UK) Consortium | Ana da Silva Filipe, Natasha Johnson, Kathy Smollett, Daniel Mair, Stephen Carmichael, Alice Broos, Lily Tong, Jenna Nichols, Kyriaki Nomikou; Sarah McDonald; Richard Orton, Joseph Hughes, Sreenu Vattipally, David L Robertson; Alasdair MacLean, Rory Gunson; Sharif Shaaban, Matthew Holden; Rachel Blacow, Guy Mollett, Kathy Li, James Shepherd, Antonia Ho, Emma Thomson |
| EPI_ISL_650184 | Liverpool Clinical Laboratories | COVID-19 Genomics UK (COG-UK) Consortium | Sam Haldenby, Anita Lucaci, Steve Paterson, Julian Hiscox, Alistair Darby, M Almsaud, A Alrezaihi, Muhannad Alruwaili, Stuart D Armstrong, Jones Benjamin, Eleanor G Bentley, Anu Chawla, Jordan J Clark, Angela Cowell, Richard Eccles, Isabel García-Dorival, Matthew Gemmell, Alessandro Gerada, PKF Gilmore, Richard Gregory, Ximeng Han, Catherine Hartley, Margaret Hughes, Miren Iturriza-Gomara, James Johnson, L Luu, Jenifer Manson, Charlotte Nelson, Elaine O'Toole, Cassie Olateju, Rebekah Penrice-Randal , Lucille Rainbow, N.P Randle, Trevor Ian Robinson, Parul Sharma, Ghada T Shawli, James P Stewart, Neil Swainston, Ecaterina Vamos, Joanne Watts, Mark Whitehead |
| EPI_ISL_650210 | Virology Department, Sheffield Teaching Hospitals NHS Foundation Trust/Department of Infection, Immunity and Cardiovascular Disease, The Medical School, University of Sheffield | COVID-19 Genomics UK (COG-UK) Consortium | Thushan de Silva, Matthew Parker, Nikki Smith, Adri Angyal, Rebecca Brown, Luke Green, Rachel Tucker, Paul Parsons, Danielle Groves, Katie Johnson, Laura Carrilero, Alex Keeley, Dave Partridge, Matthew Wyles, Benjamin Lindsey, Mehmet Yavuz, Mohammad Raza, Cariad Evans |
| EPI_ISL_650215 | University of Exeter | COVID-19 Genomics UK (COG-UK) Consortium | Ben Temperton,Aaron Jeffries,Michelle Michelsen,Joanna Warwick-Dugdale,Audrey Farbos,Robyn Manley,Stephen Michell,Jane Masoli |
| EPI_ISL_650235, EPI_ISL_650251, EPI_ISL_650252 | University of Birmingham | COVID-19 Genomics UK (COG-UK) Consortium | Institute of Microbiology, University of Birmingham: Claire McMurray, Joanne Stockton, Samuel Nicholls, Radoslaw Poplawski, Will Rowe, Josh Quick, Nicholas Loman. University of Birmingham Testing Laboratory: Celina M Whalley, Andrew Bosworth, Charlotte Poxon, Kasun Wanigasooriya, Oliver Pickles, Mike Kidd, Alex Richter, Andrew D Beggs PHE Heartlands Lab: Husam Osman, Andrew Bosworth. Queen Elizabeth Hospital: Anna Casey |
| EPI_ISL_650269 | Liverpool Clinical Laboratories | COVID-19 Genomics UK (COG-UK) Consortium | Sam Haldenby, Anita Lucaci, Steve Paterson, Julian Hiscox, Alistair Darby, M Almsaud, A Alrezaihi, Muhannad Alruwaili, Stuart D Armstrong, Jones Benjamin, Eleanor G Bentley, Anu Chawla, Jordan J Clark, Angela Cowell, Richard Eccles, Isabel García-Dorival, Matthew Gemmell, Alessandro Gerada, PKF Gilmore, Richard Gregory, Ximeng Han, Catherine Hartley, Margaret Hughes, Miren Iturriza-Gomara, James Johnson, L Luu, Jenifer Manson, Charlotte Nelson, Elaine O'Toole, Cassie Olateju, Rebekah Penrice-Randal , Lucille Rainbow, N.P Randle, Trevor Ian Robinson, Parul Sharma, Ghada T Shawli, James P Stewart, Neil Swainston, Ecaterina Vamos, Joanne Watts, Mark Whitehead |
| EPI_ISL_650273 | Department of Pathology, University of Cambridge | COVID-19 Genomics UK (COG-UK) Consortium | Aminu S. Jahun, Yasmin Chaudhry, Grant Hall, Iliana Georgana, Myra Hosmillo, Martin D. Curran, Malte Pinckert, Surendra Parmar, Ian Goodfellow |
| EPI_ISL_650298 | West of Scotland Specialist Virology Centre, NHSGGC / MRC-University of Glasgow Centre for Virus Research | COVID-19 Genomics UK (COG-UK) Consortium | Ana da Silva Filipe, Natasha Johnson, Kathy Smollett, Daniel Mair, Stephen Carmichael, Alice Broos, Lily Tong, Jenna Nichols, Kyriaki Nomikou; Sarah McDonald; Richard Orton, Joseph Hughes, Sreenu Vattipally, David L Robertson; Alasdair MacLean, Rory Gunson; Sharif Shaaban, Matthew Holden; Rachel Blacow, Guy Mollett, Kathy Li, James Shepherd, Antonia Ho, Emma Thomson |
| EPI_ISL_650299 | Liverpool Clinical Laboratories | COVID-19 Genomics UK (COG-UK) Consortium | Sam Haldenby, Anita Lucaci, Steve Paterson, Julian Hiscox, Alistair Darby, M Almsaud, A Alrezaihi, Muhannad Alruwaili, Stuart D Armstrong, Jones Benjamin, Eleanor G Bentley, Anu Chawla, Jordan J Clark, Angela Cowell, Richard Eccles, Isabel García-Dorival, Matthew Gemmell, Alessandro Gerada, PKF Gilmore, Richard Gregory, Ximeng Han, Catherine Hartley, Margaret Hughes, Miren Iturriza-Gomara, James Johnson, L Luu, Jenifer Manson, Charlotte Nelson, Elaine O'Toole, Cassie Olateju, Rebekah Penrice-Randal , Lucille Rainbow, N.P Randle, Trevor Ian Robinson, Parul Sharma, Ghada T Shawli, James P Stewart, Neil Swainston, Ecaterina Vamos, Joanne Watts, Mark Whitehead |
| EPI_ISL_650310 | West of Scotland Specialist Virology Centre, NHSGGC / MRC-University of Glasgow Centre for Virus Research | COVID-19 Genomics UK (COG-UK) Consortium | Ana da Silva Filipe, Natasha Johnson, Kathy Smollett, Daniel Mair, Stephen Carmichael, Alice Broos, Lily Tong, Jenna Nichols, Kyriaki Nomikou; Sarah McDonald; Richard Orton, Joseph Hughes, Sreenu Vattipally, David L Robertson; Alasdair MacLean, Rory Gunson; Sharif Shaaban, Matthew Holden; Rachel Blacow, Guy Mollett, Kathy Li, James Shepherd, Antonia Ho, Emma Thomson |
| EPI_ISL_650311 | Virology Department, Royal Infirmary of Edinburgh, NHS Lothian / School of Biological Sciences, University of Edinburgh / Institute of Genetics and Molecular Medicine, University of Edinburgh | COVID-19 Genomics UK (COG-UK) Consortium | McHugh M, Dewar R, Rooke S, Gallagher M, Balcaza C, O'Toole Á, Scher E, Hill V, McCrone JT, Colquhoun R, Yu X, Jackson B, Rambaut A, Williams TC, Templeton K |
| EPI_ISL_650316 | West of Scotland Specialist Virology Centre, NHSGGC / MRC-University of Glasgow Centre for Virus Research | COVID-19 Genomics UK (COG-UK) Consortium | Ana da Silva Filipe, Natasha Johnson, Kathy Smollett, Daniel Mair, Stephen Carmichael, Alice Broos, Lily Tong, Jenna Nichols, Kyriaki Nomikou; Sarah McDonald; Richard Orton, Joseph Hughes, Sreenu Vattipally, David L Robertson; Alasdair MacLean, Rory Gunson; Sharif Shaaban, Matthew Holden; Rachel Blacow, Guy Mollett, Kathy Li, James Shepherd, Antonia Ho, Emma Thomson |
| EPI_ISL_650335 | Quadram Institute Bioscience | COVID-19 Genomics UK (COG-UK) Consortium | Dave J. Baker, Gemma L. Kay, Alp Aydin, Thanh Le-Viet, Steven Rudder, Ana P. Tedim, Anastasia Kolyva, Maria Diaz, Leonardo de Oliveira Martins, Nabil-Fareed Alikhan, Lizzie Meadows, Rachael Stanley, Ngozi Elumogo, Muhammed Yasir, Nicholas M. Thomson, Alexander J Trotter, Rachel Gilroy, Samuel Bloomfield, Claire Stuart, Andrew Bell, Reenesh Prakash, Samir Dervisevic, Alison E. Mather, John Wain, Mark Webber, Andrew J. Page, Justin O'Grady |
| EPI_ISL_650354 | University of Birmingham | COVID-19 Genomics UK (COG-UK) Consortium | Institute of Microbiology, University of Birmingham: Claire McMurray, Joanne Stockton, Samuel Nicholls, Radoslaw Poplawski, Will Rowe, Josh Quick, Nicholas Loman. University of Birmingham Testing Laboratory: Celina M Whalley, Andrew Bosworth, Charlotte Poxon, Kasun Wanigasooriya, Oliver Pickles, Mike Kidd, Alex Richter, Andrew D Beggs PHE Heartlands Lab: Husam Osman, Andrew Bosworth. Queen Elizabeth Hospital: Anna Casey |
| EPI_ISL_650357 | University of Exeter | COVID-19 Genomics UK (COG-UK) Consortium | Ben Temperton,Aaron Jeffries,Michelle Michelsen,Joanna Warwick-Dugdale,Audrey Farbos,Robyn Manley,Stephen Michell,Jane Masoli |
| EPI_ISL_650364, EPI_ISL_650365 | Department of Pathology, University of Cambridge | COVID-19 Genomics UK (COG-UK) Consortium | Aminu S. Jahun, Yasmin Chaudhry, Grant Hall, Iliana Georgana, Myra Hosmillo, Martin D. Curran, Malte Pinckert, Surendra Parmar, Ian Goodfellow |
| EPI_ISL_650370 | Virology Department, Sheffield Teaching Hospitals NHS Foundation Trust/Department of Infection, Immunity and Cardiovascular Disease, The Medical School, University of Sheffield | COVID-19 Genomics UK (COG-UK) Consortium | Thushan de Silva, Matthew Parker, Nikki Smith, Adri Angyal, Rebecca Brown, Luke Green, Rachel Tucker, Paul Parsons, Danielle Groves, Katie Johnson, Laura Carrilero, Alex Keeley, Dave Partridge, Matthew Wyles, Benjamin Lindsey, Mehmet Yavuz, Mohammad Raza, Cariad Evans |
| EPI_ISL_650390 | Department of Pathology, University of Cambridge | COVID-19 Genomics UK (COG-UK) Consortium | Aminu S. Jahun, Yasmin Chaudhry, Grant Hall, Iliana Georgana, Myra Hosmillo, Martin D. Curran, Malte Pinckert, Surendra Parmar, Ian Goodfellow |
| EPI_ISL_650424 | West of Scotland Specialist Virology Centre, NHSGGC / MRC-University of Glasgow Centre for Virus Research | COVID-19 Genomics UK (COG-UK) Consortium | Ana da Silva Filipe, Natasha Johnson, Kathy Smollett, Daniel Mair, Stephen Carmichael, Alice Broos, Lily Tong, Jenna Nichols, Kyriaki Nomikou; Sarah McDonald; Richard Orton, Joseph Hughes, Sreenu Vattipally, David L Robertson; Alasdair MacLean, Rory Gunson; Sharif Shaaban, Matthew Holden; Rachel Blacow, Guy Mollett, Kathy Li, James Shepherd, Antonia Ho, Emma Thomson |

| | | | |
|---|---|---|---|
| EPI_ISL_650436, EPI_ISL_650437 | University of Birmingham | COVID-19 Genomics UK (COG-UK) Consortium | Institute of Microbiology, University of Birmingham: Claire McMurray, Joanne Stockton, Samuel Nicholls, Radoslaw Poplawski, Will Rowe, Josh Quick, Nicholas Loman. University of Birmingham Testing Laboratory: Celina M Whalley, Andrew Bosworth, Charlotte Poxon, Kasun Wanigasooriya, Oliver Pickles, Mike Kidd, Alex Richter, Andrew D Beggs PHE Heartlands Lab: Husam Osman, Andrew Bosworth. Queen Elizabeth Hospital: Anna Casey |
| EPI_ISL_650439 | Virology Department, Royal Infirmary of Edinburgh, NHS Lothian / School of Biological Sciences, University of Edinburgh / Institute of Genetics and Molecular Medicine, University of Edinburgh | COVID-19 Genomics UK (COG-UK) Consortium | McHugh M, Dewar R, Rooke S, Gallagher M, Balcaza C, O'Toole Á, Scher E, Hill V, McCrone JT, Colquhoun R, Yu X, Jackson B, Rambaut A, Williams TC, Templeton K |
| EPI_ISL_650447 | Department of Pathology, University of Cambridge | COVID-19 Genomics UK (COG-UK) Consortium | Aminu S. Jahun, Yasmin Chaudhry, Grant Hall, Iliana Georgana, Myra Hosmillo, Martin D. Curran, Malte Pinckert, Surendra Parmar, Ian Goodfellow |
| EPI_ISL_650476 | Quadram Institute Bioscience | COVID-19 Genomics UK (COG-UK) Consortium | Dave J. Baker, Gemma L. Kay, Alp Aydin, Thanh Le-Viet, Steven Rudder, Ana P. Tedim, Anastasia Kolyva, Maria Diaz, Leonardo de Oliveira Martins, Nabil-Fareed Alikhan, Lizzie Meadows, Rachael Stanley, Ngozi Elumogo, Muhammed Yasir, Nicholas M. Thomson, Alexander J Trotter, Rachel Gilroy, Samuel Bloomfield, Claire Stuart, Andrew Bell, Reenesh Prakash, Samir Dervisevic, Alison E. Mather, John Wain, Mark Webber, Andrew J. Page, Justin O'Grady |
| EPI_ISL_650478 | Virology Department, Royal Infirmary of Edinburgh, NHS Lothian / School of Biological Sciences, University of Edinburgh / Institute of Genetics and Molecular Medicine, University of Edinburgh | COVID-19 Genomics UK (COG-UK) Consortium | McHugh M, Dewar R, Rooke S, Gallagher M, Balcaza C, O'Toole Á, Scher E, Hill V, McCrone JT, Colquhoun R, Yu X, Jackson B, Rambaut A, Williams TC, Templeton K |
| EPI_ISL_650481 | Department of Pathology, University of Cambridge | COVID-19 Genomics UK (COG-UK) Consortium | Aminu S. Jahun, Yasmin Chaudhry, Grant Hall, Iliana Georgana, Myra Hosmillo, Martin D. Curran, Malte Pinckert, Surendra Parmar, Ian Goodfellow |
| EPI_ISL_650482 | West of Scotland Specialist Virology Centre, NHSGGC / MRC-University of Glasgow Centre for Virus Research | COVID-19 Genomics UK (COG-UK) Consortium | Ana da Silva Filipe, Natasha Johnson, Kathy Smollett, Daniel Mair, Stephen Carmichael, Alice Broos, Lily Tong, Jenna Nichols, Kyriaki Nomikou; Sarah McDonald; Richard Orton, Joseph Hughes, Sreenu Vattipally, David L Robertson; Alasdair MacLean, Rory Gunson; Sharif Shaaban, Matthew Holden; Rachel Blacow, Guy Mollett, Kathy Li, James Shepherd, Antonia Ho, Emma Thomson |
| EPI_ISL_650495 | Virology Department, Royal Infirmary of Edinburgh, NHS Lothian / School of Biological Sciences, University of Edinburgh / Institute of Genetics and Molecular Medicine, University of Edinburgh | COVID-19 Genomics UK (COG-UK) Consortium | McHugh M, Dewar R, Rooke S, Gallagher M, Balcaza C, O'Toole Á, Scher E, Hill V, McCrone JT, Colquhoun R, Yu X, Jackson B, Rambaut A, Williams TC, Templeton K |
| EPI_ISL_650506, EPI_ISL_650507, EPI_ISL_650511, EPI_ISL_650514, EPI_ISL_650538, EPI_ISL_650550 | West of Scotland Specialist Virology Centre, NHSGGC / MRC-University of Glasgow Centre for Virus Research | COVID-19 Genomics UK (COG-UK) Consortium | Ana da Silva Filipe, Natasha Johnson, Kathy Smollett, Daniel Mair, Stephen Carmichael, Alice Broos, Lily Tong, Jenna Nichols, Kyriaki Nomikou; Sarah McDonald; Richard Orton, Joseph Hughes, Sreenu Vattipally, David L Robertson; Alasdair MacLean, Rory Gunson; Sharif Shaaban, Matthew Holden; Rachel Blacow, Guy Mollett, Kathy Li, James Shepherd, Antonia Ho, Emma Thomson |
| EPI_ISL_650563 | Liverpool Clinical Laboratories | COVID-19 Genomics UK (COG-UK) Consortium | Sam Haldenby, Anita Lucaci, Steve Paterson, Julian Hiscox, Alistair Darby, M Almsaud, A Alrezaihi, Muhannad Alruwaili, Stuart D Armstrong, Jones Benjamin, Eleanor G Bentley, Anu Chawla, Jordan J Clark, Angela Cowell, Richard Eccles, Isabel García-Dorival, Matthew Gemmell, Alessandro Gerada, PKF Gilmore, Richard Gregory, Ximeng Han, Catherine Hartley, Margaret Hughes, Miren Iturriza-Gomara, James Johnson, L Luu, Jenifer Manson, Charlotte Nelson, Elaine O'Toole, Cassie Olateju, Rebekah Penrice-Randal , Lucille Rainbow, N.P Randle, Trevor Ian Robinson, Parul Sharma, Ghada T Shawli, James P Stewart, Neil Swainston, Ecaterina Vamos, Joanne Watts, Mark Whitehead |
| EPI_ISL_650565 | University of Exeter | COVID-19 Genomics UK (COG-UK) Consortium | Ben Temperton,Aaron Jeffries,Michelle Michelsen,Joanna Warwick-Dugdale,Audrey Farbos,Robyn Manley,Stephen Michell,Jane Masoli |
| EPI_ISL_650580 | Department of Pathology, University of Cambridge | COVID-19 Genomics UK (COG-UK) Consortium | Aminu S. Jahun, Yasmin Chaudhry, Grant Hall, Iliana Georgana, Myra Hosmillo, Martin D. Curran, Malte Pinckert, Surendra Parmar, Ian Goodfellow |
| EPI_ISL_650606 | University of Exeter | COVID-19 Genomics UK (COG-UK) Consortium | Ben Temperton,Aaron Jeffries,Michelle Michelsen,Joanna Warwick-Dugdale,Audrey Farbos,Robyn Manley,Stephen Michell,Jane Masoli |
| EPI_ISL_650608 | Centre for Enzyme Innovation, University of Portsmouth / Translational Research Laboratory, Portsmouth Hospitals NHS Trust | COVID-19 Genomics UK (COG-UK) Consortium | Angela Beckett,Yann Bourgeois,Garry Scarlett,Sharon Glaysher,Scott Elliott,Kelly Bicknell,Robert Impey,Allyson Lloyd,Sarah Wyllie,Ethan Butcher,Anoop Chauhan,Samuel Robson |
| EPI_ISL_650614, EPI_ISL_650664 | Department of Pathology, University of Cambridge | COVID-19 Genomics UK (COG-UK) Consortium | Aminu S. Jahun, Yasmin Chaudhry, Grant Hall, Iliana Georgana, Myra Hosmillo, Martin D. Curran, Malte Pinckert, Surendra Parmar, Ian Goodfellow |
| EPI_ISL_650665, EPI_ISL_650668 | University of Birmingham | COVID-19 Genomics UK (COG-UK) Consortium | Institute of Microbiology, University of Birmingham: Claire McMurray, Joanne Stockton, Samuel Nicholls, Radoslaw Poplawski, Will Rowe, Josh Quick, Nicholas Loman. University of Birmingham Testing Laboratory: Celina M Whalley, Andrew Bosworth, Charlotte Poxon, Kasun Wanigasooriya, Oliver Pickles, Mike Kidd, Alex Richter, Andrew D Beggs PHE Heartlands Lab: Husam Osman, Andrew Bosworth. Queen Elizabeth Hospital: Anna Casey |
| EPI_ISL_650691 | Virology Department, Sheffield Teaching Hospitals NHS Foundation Trust/Department of Infection, Immunity and Cardiovascular Disease, The Medical School, University of Sheffield | COVID-19 Genomics UK (COG-UK) Consortium | Thushan de Silva, Matthew Parker, Nikki Smith, Adri Angyal, Rebecca Brown, Luke Green, Rachel Tucker, Paul Parsons, Danielle Groves, Katie Johnson, Laura Carrilero, Alex Keeley, Dave Partridge, Matthew Wyles, Benjamin Lindsey, Mehmet Yavuz, Mohammad Raza, Cariad Evans |
| EPI_ISL_650724 | University of Birmingham | COVID-19 Genomics UK (COG-UK) Consortium | Institute of Microbiology, University of Birmingham: Claire McMurray, Joanne Stockton, Samuel Nicholls, Radoslaw Poplawski, Will Rowe, Josh Quick, Nicholas Loman. University of Birmingham Testing Laboratory: Celina M Whalley, Andrew Bosworth, Charlotte Poxon, Kasun Wanigasooriya, Oliver Pickles, Mike Kidd, Alex Richter, Andrew D Beggs PHE Heartlands Lab: Husam Osman, Andrew Bosworth. Queen Elizabeth Hospital: Anna Casey |
| EPI_ISL_650727 | West of Scotland Specialist Virology Centre, NHSGGC / MRC-University of Glasgow Centre for Virus Research | COVID-19 Genomics UK (COG-UK) Consortium | Ana da Silva Filipe, Natasha Johnson, Kathy Smollett, Daniel Mair, Stephen Carmichael, Alice Broos, Lily Tong, Jenna Nichols, Kyriaki Nomikou; Sarah McDonald; Richard Orton, Joseph Hughes, Sreenu Vattipally, David L Robertson; Alasdair MacLean, Rory Gunson; Sharif Shaaban, Matthew Holden; Rachel Blacow, Guy Mollett, Kathy Li, James Shepherd, Antonia Ho, Emma Thomson |
| EPI_ISL_650729 | University of Birmingham | COVID-19 Genomics UK (COG-UK) Consortium | Institute of Microbiology, University of Birmingham: Claire McMurray, Joanne Stockton, Samuel Nicholls, Radoslaw Poplawski, Will Rowe, Josh Quick, Nicholas Loman. University of Birmingham Testing Laboratory: Celina M Whalley, Andrew Bosworth, Charlotte Poxon, Kasun Wanigasooriya, Oliver Pickles, Mike Kidd, Alex Richter, Andrew D Beggs PHE Heartlands Lab: Husam Osman, Andrew Bosworth. Queen Elizabeth Hospital: Anna Casey |
| EPI_ISL_650733, EPI_ISL_650735 | West of Scotland Specialist Virology Centre, NHSGGC / MRC-University of Glasgow Centre for Virus Research | COVID-19 Genomics UK (COG-UK) Consortium | Ana da Silva Filipe, Natasha Johnson, Kathy Smollett, Daniel Mair, Stephen Carmichael, Alice Broos, Lily Tong, Jenna Nichols, Kyriaki Nomikou; Sarah McDonald; Richard Orton, Joseph Hughes, Sreenu Vattipally, David L Robertson; Alasdair MacLean, Rory Gunson; Sharif Shaaban, Matthew Holden; Rachel Blacow, Guy Mollett, Kathy Li, James Shepherd, Antonia Ho, Emma Thomson |
| EPI_ISL_650744 | University of Birmingham | COVID-19 Genomics UK (COG-UK) Consortium | Institute of Microbiology, University of Birmingham: Claire McMurray, Joanne Stockton, Samuel Nicholls, Radoslaw Poplawski, Will Rowe, Josh Quick, Nicholas Loman. University of Birmingham Testing Laboratory: Celina M Whalley, Andrew Bosworth, Charlotte Poxon, Kasun Wanigasooriya, Oliver Pickles, Mike Kidd, Alex Richter, Andrew D Beggs PHE Heartlands Lab: Husam Osman, Andrew Bosworth. Queen Elizabeth Hospital: Anna Casey |
| EPI_ISL_650750, EPI_ISL_650751, EPI_ISL_650752, EPI_ISL_650753, EPI_ISL_650754, EPI_ISL_650764 | West of Scotland Specialist Virology Centre, NHSGGC / MRC-University of Glasgow Centre for Virus Research | COVID-19 Genomics UK (COG-UK) Consortium | Ana da Silva Filipe, Natasha Johnson, Kathy Smollett, Daniel Mair, Stephen Carmichael, Alice Broos, Lily Tong, Jenna Nichols, Kyriaki Nomikou; Sarah McDonald; Richard Orton, Joseph Hughes, Sreenu Vattipally, David L Robertson; Alasdair MacLean, Rory Gunson; Sharif Shaaban, Matthew Holden; Rachel Blacow, Guy Mollett, Kathy Li, James Shepherd, Antonia Ho, Emma Thomson |
| EPI_ISL_650769, EPI_ISL_650770, EPI_ISL_650771, EPI_ISL_650773 | University of Birmingham | COVID-19 Genomics UK (COG-UK) Consortium | Institute of Microbiology, University of Birmingham: Claire McMurray, Joanne Stockton, Samuel Nicholls, Radoslaw Poplawski, Will Rowe, Josh Quick, Nicholas Loman. University of Birmingham Testing Laboratory: Celina M Whalley, Andrew Bosworth, Charlotte Poxon, Kasun Wanigasooriya, Oliver Pickles, Mike Kidd, Alex Richter, Andrew D Beggs PHE Heartlands Lab: Husam Osman, Andrew Bosworth. Queen Elizabeth Hospital: Anna Casey |
| EPI_ISL_650780 | Virology Department, Royal Infirmary of Edinburgh, NHS Lothian / School of Biological Sciences, University of Edinburgh / Institute of Genetics and Molecular Medicine, University of Edinburgh | COVID-19 Genomics UK (COG-UK) Consortium | McHugh M, Dewar R, Rooke S, Gallagher M, Balcaza C, O'Toole Á, Scher E, Hill V, McCrone JT, Colquhoun R, Yu X, Jackson B, Rambaut A, Williams TC, Templeton K |
| EPI_ISL_650783, EPI_ISL_650784 | University of Exeter | COVID-19 Genomics UK (COG-UK) Consortium | Ben Temperton,Aaron Jeffries,Michelle Michelsen,Joanna Warwick-Dugdale,Audrey Farbos,Robyn Manley,Stephen Michell,Jane Masoli |
| EPI_ISL_650792 | University of Birmingham | COVID-19 Genomics UK (COG-UK) Consortium | Institute of Microbiology, University of Birmingham: Claire McMurray, Joanne Stockton, Samuel Nicholls, Radoslaw Poplawski, Will Rowe, Josh Quick, Nicholas Loman. University of Birmingham Testing Laboratory: Celina M Whalley, Andrew Bosworth, Charlotte Poxon, Kasun Wanigasooriya, Oliver |

| | | | |
|---|---|---|---|
| | | | Pickles, Mike Kidd, Alex Richter, Andrew D Beggs PHE Heartlands Lab: Husam Osman, Andrew Bosworth. Queen Elizabeth Hospital: Anna Casey |
| EPI_ISL_650808 | Virology Department, Sheffield Teaching Hospitals NHS Foundation Trust/Department of Infection, Immunity and Cardiovascular Disease, The Medical School, University of Sheffield | COVID-19 Genomics UK (COG-UK) Consortium | Thushan de Silva, Matthew Parker, Nikki Smith, Adri Angyal, Rebecca Brown, Luke Green, Rachel Tucker, Paul Parsons, Danielle Groves, Katie Johnson, Laura Carrilero, Alex Keeley, Dave Partridge, Matthew Wyles, Benjamin Lindsey, Mehmet Yavuz, Mohammad Raza, Cariad Evans |
| EPI_ISL_650837 | West of Scotland Specialist Virology Centre, NHSGGC / MRC-University of Glasgow Centre for Virus Research | COVID-19 Genomics UK (COG-UK) Consortium | Ana da Silva Filipe, Natasha Johnson, Kathy Smollett, Daniel Mair, Stephen Carmichael, Alice Broos, Lily Tong, Jenna Nichols, Kyriaki Nomikou; Sarah McDonald; Richard Orton, Joseph Hughes, Sreenu Vattipally, David L Robertson; Alasdair MacLean, Rory Gunson; Sharif Shaaban, Matthew Holden; Rachel Blacow, Guy Mollett, Kathy Li, James Shepherd, Antonia Ho, Emma Thomson |
| EPI_ISL_650849 | University of Birmingham | COVID-19 Genomics UK (COG-UK) Consortium | Institute of Microbiology, University of Birmingham: Claire McMurray, Joanne Stockton, Samuel Nicholls, Radoslaw Poplawski, Will Rowe, Josh Quick, Nicholas Loman. University of Birmingham Testing Laboratory: Celina M Whalley, Andrew Bosworth, Charlotte Poxon, Kasun Wanigasooriya, Oliver Pickles, Mike Kidd, Alex Richter, Andrew D Beggs PHE Heartlands Lab: Husam Osman, Andrew Bosworth. Queen Elizabeth Hospital: Anna Casey |
| EPI_ISL_650866, EPI_ISL_650867, EPI_ISL_650868, EPI_ISL_650869 | West of Scotland Specialist Virology Centre, NHSGGC / MRC-University of Glasgow Centre for Virus Research | COVID-19 Genomics UK (COG-UK) Consortium | Ana da Silva Filipe, Natasha Johnson, Kathy Smollett, Daniel Mair, Stephen Carmichael, Alice Broos, Lily Tong, Jenna Nichols, Kyriaki Nomikou; Sarah McDonald; Richard Orton, Joseph Hughes, Sreenu Vattipally, David L Robertson; Alasdair MacLean, Rory Gunson; Sharif Shaaban, Matthew Holden; Rachel Blacow, Guy Mollett, Kathy Li, James Shepherd, Antonia Ho, Emma Thomson |
| EPI_ISL_650885 | Quadram Institute Bioscience | COVID-19 Genomics UK (COG-UK) Consortium | Dave J. Baker, Gemma L. Kay, Alp Aydin, Thanh Le-Viet, Steven Rudder, Ana P. Tedim, Anastasia Kolyva, Maria Diaz, Leonardo de Oliveira Martins, Nabil-Fareed Alikhan, Lizzie Meadows, Rachael Stanley, Ngozi Elumogo, Muhammed Yasir, Nicholas M. Thomson, Alexander J Trotter, Rachel Gilroy, Samuel Bloomfield, Claire Stuart, Andrew Bell, Reenesh Prakash, Samir Dervisevic, Alison E. Mather, John Wain, Mark Webber, Andrew J. Page, Justin O'Grady |
| EPI_ISL_650906 | Wales Specialist Virology Centre Sequencing lab: Pathogen Genomics Unit | COVID-19 Genomics UK (COG-UK) Consortium | Catherine Moore, Johnathan Evans, Laura Gifford, Malorie Perry, Simon Cottrell, Angela Marchbank, Alec Birchley, Alexander Adams, Amy Gaskin, Bree Gatica-Wilcox, Jason Coombes, Joel Southgate, Lauren Gilbert, Lee Graham, Nicole Pacchiarini, Sara Kumziene-Summerhayes, Sarah Taylor, Sophie Jones, Sara Rey, Matthew Bull, Joanne Watkins, Sally Corden, Tom Connor |
| EPI_ISL_650911 | Quadram Institute Bioscience | COVID-19 Genomics UK (COG-UK) Consortium | Dave J. Baker, Gemma L. Kay, Alp Aydin, Thanh Le-Viet, Steven Rudder, Ana P. Tedim, Anastasia Kolyva, Maria Diaz, Leonardo de Oliveira Martins, Nabil-Fareed Alikhan, Lizzie Meadows, Rachael Stanley, Ngozi Elumogo, Muhammed Yasir, Nicholas M. Thomson, Alexander J Trotter, Rachel Gilroy, Samuel Bloomfield, Claire Stuart, Andrew Bell, Reenesh Prakash, Samir Dervisevic, Alison E. Mather, John Wain, Mark Webber, Andrew J. Page, Justin O'Grady |
| EPI_ISL_650916 | Department of Pathology, University of Cambridge | COVID-19 Genomics UK (COG-UK) Consortium | Aminu S. Jahun, Yasmin Chaudhry, Grant Hall, Iliana Georgana, Myra Hosmillo, Martin D. Curran, Malte Pinckert, Surendra Parmar, Ian Goodfellow |
| EPI_ISL_650938, EPI_ISL_650940, EPI_ISL_650958 | Quadram Institute Bioscience | COVID-19 Genomics UK (COG-UK) Consortium | Dave J. Baker, Gemma L. Kay, Alp Aydin, Thanh Le-Viet, Steven Rudder, Ana P. Tedim, Anastasia Kolyva, Maria Diaz, Leonardo de Oliveira Martins, Nabil-Fareed Alikhan, Lizzie Meadows, Rachael Stanley, Ngozi Elumogo, Muhammed Yasir, Nicholas M. Thomson, Alexander J Trotter, Rachel Gilroy, Samuel Bloomfield, Claire Stuart, Andrew Bell, Reenesh Prakash, Samir Dervisevic, Alison E. Mather, John Wain, Mark Webber, Andrew J. Page, Justin O'Grady |
| EPI_ISL_650996 | University of Birmingham | COVID-19 Genomics UK (COG-UK) Consortium | Institute of Microbiology, University of Birmingham: Claire McMurray, Joanne Stockton, Samuel Nicholls, Radoslaw Poplawski, Will Rowe, Josh Quick, Nicholas Loman. University of Birmingham Testing Laboratory: Celina M Whalley, Andrew Bosworth, Charlotte Poxon, Kasun Wanigasooriya, Oliver Pickles, Mike Kidd, Alex Richter, Andrew D Beggs PHE Heartlands Lab: Husam Osman, Andrew Bosworth. Queen Elizabeth Hospital: Anna Casey |
| EPI_ISL_651002 | Department of Pathology, University of Cambridge | COVID-19 Genomics UK (COG-UK) Consortium | Aminu S. Jahun, Yasmin Chaudhry, Grant Hall, Iliana Georgana, Myra Hosmillo, Martin D. Curran, Malte Pinckert, Surendra Parmar, Ian Goodfellow |
| EPI_ISL_651053, EPI_ISL_651055, EPI_ISL_651057, EPI_ISL_651059, EPI_ISL_651083, EPI_ISL_651085 | University of Exeter | COVID-19 Genomics UK (COG-UK) Consortium | Ben Temperton,Aaron Jeffries,Michelle Michelsen,Joanna Warwick-Dugdale,Audrey Farbos,Robyn Manley,Stephen Michell,Jane Masoli |
| EPI_ISL_651123 | Liverpool Clinical Laboratories | COVID-19 Genomics UK (COG-UK) Consortium | Sam Haldenby, Anita Lucaci, Steve Paterson, Julian Hiscox, Alistair Darby, M Almsaud, A Alrezaihi, Muhannad Alruwaili, Stuart D Armstrong, Jones Benjamin, Eleanor G Bentley, Anu Chawla, Jordan J Clark, Angela Cowell, Richard Eccles, Isabel García-Dorival, Matthew Gemmell, Alessandro Gerada, PKF Gilmore, Richard Gregory, Ximeng Han, Catherine Hartley, Margaret Hughes, Miren Iturriza-Gomara, James Johnson, L Luu, Jenifer Manson, Charlotte Nelson, Elaine O'Toole, Cassie Olateju, Rebekah Penrice-Randal , Lucille Rainbow, N.P Randle, Trevor Ian Robinson, Parul Sharma, Ghada T Shawli, James P Stewart, Neil Swainston, Ecaterina Vamos, Joanne Watts, Mark Whitehead |
| EPI_ISL_651127 | University of Birmingham | COVID-19 Genomics UK (COG-UK) Consortium | Institute of Microbiology, University of Birmingham: Claire McMurray, Joanne Stockton, Samuel Nicholls, Radoslaw Poplawski, Will Rowe, Josh Quick, Nicholas Loman. University of Birmingham Testing Laboratory: Celina M Whalley, Andrew Bosworth, Charlotte Poxon, Kasun Wanigasooriya, Oliver Pickles, Mike Kidd, Alex Richter, Andrew D Beggs PHE Heartlands Lab: Husam Osman, Andrew Bosworth. Queen Elizabeth Hospital: Anna Casey |
| EPI_ISL_651133, EPI_ISL_651134 | Quadram Institute Bioscience | COVID-19 Genomics UK (COG-UK) Consortium | Dave J. Baker, Gemma L. Kay, Alp Aydin, Thanh Le-Viet, Steven Rudder, Ana P. Tedim, Anastasia Kolyva, Maria Diaz, Leonardo de Oliveira Martins, Nabil-Fareed Alikhan, Lizzie Meadows, Rachael Stanley, Ngozi Elumogo, Muhammed Yasir, Nicholas M. Thomson, Alexander J Trotter, Rachel Gilroy, Samuel Bloomfield, Claire Stuart, Andrew Bell, Reenesh Prakash, Samir Dervisevic, Alison E. Mather, John Wain, Mark Webber, Andrew J. Page, Justin O'Grady |
| EPI_ISL_651137 | University of Birmingham | COVID-19 Genomics UK (COG-UK) Consortium | Institute of Microbiology, University of Birmingham: Claire McMurray, Joanne Stockton, Samuel Nicholls, Radoslaw Poplawski, Will Rowe, Josh Quick, Nicholas Loman. University of Birmingham Testing Laboratory: Celina M Whalley, Andrew Bosworth, Charlotte Poxon, Kasun Wanigasooriya, Oliver Pickles, Mike Kidd, Alex Richter, Andrew D Beggs PHE Heartlands Lab: Husam Osman, Andrew Bosworth. Queen Elizabeth Hospital: Anna Casey |
| EPI_ISL_651141 | Virology Department, Royal Infirmary of Edinburgh, NHS Lothian / School of Biological Sciences, University of Edinburgh / Institute of Genetics and Molecular Medicine, University of Edinburgh | COVID-19 Genomics UK (COG-UK) Consortium | McHugh M, Dewar R, Rooke S, Gallagher M, Balcaza C, O'Toole Á, Scher E, Hill V, McCrone JT, Colquhoun R, Yu X, Jackson B, Rambaut A, Williams TC, Templeton K |
| EPI_ISL_651142 | Quadram Institute Bioscience | COVID-19 Genomics UK (COG-UK) Consortium | Dave J. Baker, Gemma L. Kay, Alp Aydin, Thanh Le-Viet, Steven Rudder, Ana P. Tedim, Anastasia Kolyva, Maria Diaz, Leonardo de Oliveira Martins, Nabil-Fareed Alikhan, Lizzie Meadows, Rachael Stanley, Ngozi Elumogo, Muhammed Yasir, Nicholas M. Thomson, Alexander J Trotter, Rachel Gilroy, Samuel Bloomfield, Claire Stuart, Andrew Bell, Reenesh Prakash, Samir Dervisevic, Alison E. Mather, John Wain, Mark Webber, Andrew J. Page, Justin O'Grady |
| EPI_ISL_651146 | Liverpool Clinical Laboratories | COVID-19 Genomics UK (COG-UK) Consortium | Sam Haldenby, Anita Lucaci, Steve Paterson, Julian Hiscox, Alistair Darby, M Almsaud, A Alrezaihi, Muhannad Alruwaili, Stuart D Armstrong, Jones Benjamin, Eleanor G Bentley, Anu Chawla, Jordan J Clark, Angela Cowell, Richard Eccles, Isabel García-Dorival, Matthew Gemmell, Alessandro Gerada, PKF Gilmore, Richard Gregory, Ximeng Han, Catherine Hartley, Margaret Hughes, Miren Iturriza-Gomara, James Johnson, L Luu, Jenifer Manson, Charlotte Nelson, Elaine O'Toole, Cassie Olateju, Rebekah Penrice-Randal , Lucille Rainbow, N.P Randle, Trevor Ian Robinson, Parul Sharma, Ghada T Shawli, James P Stewart, Neil Swainston, Ecaterina Vamos, Joanne Watts, Mark Whitehead |
| EPI_ISL_651154 | Wales Specialist Virology Centre Sequencing lab: Pathogen Genomics Unit | COVID-19 Genomics UK (COG-UK) Consortium | Catherine Moore, Johnathan Evans, Laura Gifford, Malorie Perry, Simon Cottrell, Angela Marchbank, Alec Birchley, Alexander Adams, Amy Gaskin, Bree Gatica-Wilcox, Jason Coombes, Joel Southgate, Lauren Gilbert, Lee Graham, Nicole Pacchiarini, Sara Kumziene-Summerhayes, Sarah Taylor, Sophie Jones, Sara Rey, Matthew Bull, Joanne Watkins, Sally Corden, Tom Connor |
| EPI_ISL_651177 | University of Exeter | COVID-19 Genomics UK (COG-UK) Consortium | Ben Temperton,Aaron Jeffries,Michelle Michelsen,Joanna Warwick-Dugdale,Audrey Farbos,Robyn Manley,Stephen Michell,Jane Masoli |
| EPI_ISL_651179 | University College London, Great Ormond Street Hospital for Children NHS Foundation Trust, Imperial College Healthcare NHS Trust | COVID-19 Genomics UK (COG-UK) Consortium | Sergi Castellano, Rachel Williams, Mark Kristiansen, Paola Resende Silva, Sunando Roy, Tony Brooks, Helena Tutill, Paola Niola, Patricia Dyal, Charlotte Williams, Leysa Forrest, Yasmin Panchbhaya, Jacqueline Findlay, Samuel Weeks, Julianne Brown, Kathryn Harris, Paul Randell, James Price, Alison Holmes, Judith Breuer |
| EPI_ISL_651183 | University of Exeter | COVID-19 Genomics UK (COG-UK) Consortium | Ben Temperton,Aaron Jeffries,Michelle Michelsen,Joanna Warwick-Dugdale,Audrey Farbos,Robyn Manley,Stephen Michell,Jane Masoli |
| EPI_ISL_651190 | University of Birmingham | COVID-19 Genomics UK (COG-UK) Consortium | Institute of Microbiology, University of Birmingham: Claire McMurray, Joanne Stockton, Samuel Nicholls, Radoslaw Poplawski, Will Rowe, Josh Quick, Nicholas Loman. University of Birmingham Testing Laboratory: Celina M Whalley, Andrew Bosworth, Charlotte Poxon, Kasun Wanigasooriya, Oliver |

| | | | |
|---|---|---|---|
| | | | Pickles, Mike Kidd, Alex Richter, Andrew D Beggs PHE Heartlands Lab: Husam Osman, Andrew Bosworth. Queen Elizabeth Hospital: Anna Casey |
| EPI_ISL_651197 | Liverpool Clinical Laboratories | COVID-19 Genomics UK (COG-UK) Consortium | Sam Haldenby, Anita Lucaci, Steve Paterson, Julian Hiscox, Alistair Darby, M Almsaud, A Alrezaihi, Muhannad Alruwaili, Stuart D Armstrong, Jones Benjamin, Eleanor G Bentley, Anu Chawla, Jordan J Clark, Angela Cowell, Richard Eccles, Isabel García-Dorival, Matthew Gemmell, Alessandro Gerada, PKF Gilmore, Richard Gregory, Ximeng Han, Catherine Hartley, Margaret Hughes, Miren Iturriza-Gomara, James Johnson, L Luu, Jenifer Manson, Charlotte Nelson, Elaine O'Toole, Cassie Olateju, Rebekah Penrice-Randal , Lucille Rainbow, N.P Randle, Trevor Ian Robinson, Parul Sharma, Ghada T Shawli, James P Stewart, Neil Swainston, Ecaterina Vamos, Joanne Watts, Mark Whitehead |
| EPI_ISL_651224 | Virology Department, Sheffield Teaching Hospitals NHS Foundation Trust/Department of Infection, Immunity and Cardiovascular Disease, The Medical School, University of Sheffield | COVID-19 Genomics UK (COG-UK) Consortium | Thushan de Silva, Matthew Parker, Nikki Smith, Adri Angyal, Rebecca Brown, Luke Green, Rachel Tucker, Paul Parsons, Danielle Groves, Katie Johnson, Laura Carrilero, Alex Keeley, Dave Partridge, Matthew Wyles, Benjamin Lindsey, Mehmet Yavuz, Mohammad Raza, Cariad Evans |
| EPI_ISL_651227 | University of Birmingham | COVID-19 Genomics UK (COG-UK) Consortium | Institute of Microbiology, University of Birmingham: Claire McMurray, Joanne Stockton, Samuel Nicholls, Radoslaw Poplawski, Will Rowe, Josh Quick, Nicholas Loman. University of Birmingham Testing Laboratory: Celina M Whalley, Andrew Bosworth, Charlotte Poxon, Kasun Wanigasooriya, Oliver Pickles, Mike Kidd, Alex Richter, Andrew D Beggs PHE Heartlands Lab: Husam Osman, Andrew Bosworth. Queen Elizabeth Hospital: Anna Casey |
| EPI_ISL_651229, EPI_ISL_651230 | Virology Department, Royal Infirmary of Edinburgh, NHS Lothian / School of Biological Sciences, University of Edinburgh / Institute of Genetics and Molecular Medicine, University of Edinburgh | COVID-19 Genomics UK (COG-UK) Consortium | McHugh M, Dewar R, Rooke S, Gallagher M, Balcaza C, O'Toole Á, Scher E, Hill V, McCrone JT, Colquhoun R, Yu X, Jackson B, Rambaut A, Williams TC, Templeton K |
| EPI_ISL_651233 | Virology Department, Sheffield Teaching Hospitals NHS Foundation Trust/Department of Infection, Immunity and Cardiovascular Disease, The Medical School, University of Sheffield | COVID-19 Genomics UK (COG-UK) Consortium | Thushan de Silva, Matthew Parker, Nikki Smith, Adri Angyal, Rebecca Brown, Luke Green, Rachel Tucker, Paul Parsons, Danielle Groves, Katie Johnson, Laura Carrilero, Alex Keeley, Dave Partridge, Matthew Wyles, Benjamin Lindsey, Mehmet Yavuz, Mohammad Raza, Cariad Evans |
| EPI_ISL_651234 | Liverpool Clinical Laboratories | COVID-19 Genomics UK (COG-UK) Consortium | Sam Haldenby, Anita Lucaci, Steve Paterson, Julian Hiscox, Alistair Darby, M Almsaud, A Alrezaihi, Muhannad Alruwaili, Stuart D Armstrong, Jones Benjamin, Eleanor G Bentley, Anu Chawla, Jordan J Clark, Angela Cowell, Richard Eccles, Isabel García-Dorival, Matthew Gemmell, Alessandro Gerada, PKF Gilmore, Richard Gregory, Ximeng Han, Catherine Hartley, Margaret Hughes, Miren Iturriza-Gomara, James Johnson, L Luu, Jenifer Manson, Charlotte Nelson, Elaine O'Toole, Cassie Olateju, Rebekah Penrice-Randal , Lucille Rainbow, N.P Randle, Trevor Ian Robinson, Parul Sharma, Ghada T Shawli, James P Stewart, Neil Swainston, Ecaterina Vamos, Joanne Watts, Mark Whitehead |
| EPI_ISL_651247, EPI_ISL_651255 | Quadram Institute Bioscience | COVID-19 Genomics UK (COG-UK) Consortium | Dave J. Baker, Gemma L. Kay, Alp Aydin, Thanh Le-Viet, Steven Rudder, Ana P. Tedim, Anastasia Kolyva, Maria Diaz, Leonardo de Oliveira Martins, Nabil-Fareed Alikhan, Lizzie Meadows, Rachael Stanley, Ngozi Elumogo, Muhammed Yasir, Nicholas M. Thomson, Alexander J Trotter, Rachel Gilroy, Samuel Bloomfield, Claire Stuart, Andrew Bell, Reenesh Prakash, Samir Dervisevic, Alison E. Mather, John Wain, Mark Webber, Andrew J. Page, Justin O'Grady |
| EPI_ISL_651265 | Liverpool Clinical Laboratories | COVID-19 Genomics UK (COG-UK) Consortium | Sam Haldenby, Anita Lucaci, Steve Paterson, Julian Hiscox, Alistair Darby, M Almsaud, A Alrezaihi, Muhannad Alruwaili, Stuart D Armstrong, Jones Benjamin, Eleanor G Bentley, Anu Chawla, Jordan J Clark, Angela Cowell, Richard Eccles, Isabel García-Dorival, Matthew Gemmell, Alessandro Gerada, PKF Gilmore, Richard Gregory, Ximeng Han, Catherine Hartley, Margaret Hughes, Miren Iturriza-Gomara, James Johnson, L Luu, Jenifer Manson, Charlotte Nelson, Elaine O'Toole, Cassie Olateju, Rebekah Penrice-Randal , Lucille Rainbow, N.P Randle, Trevor Ian Robinson, Parul Sharma, Ghada T Shawli, James P Stewart, Neil Swainston, Ecaterina Vamos, Joanne Watts, Mark Whitehead |
| EPI_ISL_651269 | Department of Pathology, University of Cambridge | COVID-19 Genomics UK (COG-UK) Consortium | Aminu S. Jahun, Yasmin Chaudhry, Grant Hall, Iliana Georgana, Myra Hosmillo, Martin D. Curran, Malte Pinckert, Surendra Parmar, Ian Goodfellow |
| EPI_ISL_651271, EPI_ISL_651273 | Liverpool Clinical Laboratories | COVID-19 Genomics UK (COG-UK) Consortium | Sam Haldenby, Anita Lucaci, Steve Paterson, Julian Hiscox, Alistair Darby, M Almsaud, A Alrezaihi, Muhannad Alruwaili, Stuart D Armstrong, Jones Benjamin, Eleanor G Bentley, Anu Chawla, Jordan J Clark, Angela Cowell, Richard Eccles, Isabel García-Dorival, Matthew Gemmell, Alessandro Gerada, PKF Gilmore, Richard Gregory, Ximeng Han, Catherine Hartley, Margaret Hughes, Miren Iturriza-Gomara, James Johnson, L Luu, Jenifer Manson, Charlotte Nelson, Elaine O'Toole, Cassie Olateju, Rebekah Penrice-Randal , Lucille Rainbow, N.P Randle, Trevor Ian Robinson, Parul Sharma, Ghada T Shawli, James P Stewart, Neil Swainston, Ecaterina Vamos, Joanne Watts, Mark Whitehead |
| EPI_ISL_651276 | Department of Pathology, University of Cambridge | COVID-19 Genomics UK (COG-UK) Consortium | Aminu S. Jahun, Yasmin Chaudhry, Grant Hall, Iliana Georgana, Myra Hosmillo, Martin D. Curran, Malte Pinckert, Surendra Parmar, Ian Goodfellow |
| EPI_ISL_651290 | University of Exeter | COVID-19 Genomics UK (COG-UK) Consortium | Ben Temperton,Aaron Jeffries,Michelle Michelsen,Joanna Warwick-Dugdale,Audrey Farbos,Robyn Manley,Stephen Michell,Jane Masoli |
| EPI_ISL_651294 | Virology Department, Royal Infirmary of Edinburgh, NHS Lothian / School of Biological Sciences, University of Edinburgh / Institute of Genetics and Molecular Medicine, University of Edinburgh | COVID-19 Genomics UK (COG-UK) Consortium | McHugh M, Dewar R, Rooke S, Gallagher M, Balcaza C, O'Toole Á, Scher E, Hill V, McCrone JT, Colquhoun R, Yu X, Jackson B, Rambaut A, Williams TC, Templeton K |
| EPI_ISL_651303 | West of Scotland Specialist Virology Centre, NHSGGC / MRC-University of Glasgow Centre for Virus Research | COVID-19 Genomics UK (COG-UK) Consortium | Ana da Silva Filipe, Natasha Johnson, Kathy Smollett, Daniel Mair, Stephen Carmichael, Alice Broos, Lily Tong, Jenna Nichols, Kyriaki Nomikou; Sarah McDonald; Richard Orton, Joseph Hughes, Sreenu Vattipally, David L Robertson; Alasdair MacLean, Rory Gunson; Sharif Shaaban, Matthew Holden; Rachel Blacow, Guy Mollett, Kathy Li, James Shepherd, Antonia Ho, Emma Thomson |
| EPI_ISL_651305 | University of Birmingham | COVID-19 Genomics UK (COG-UK) Consortium | Institute of Microbiology, University of Birmingham: Claire McMurray, Joanne Stockton, Samuel Nicholls, Radoslaw Poplawski, Will Rowe, Josh Quick, Nicholas Loman. University of Birmingham Testing Laboratory: Celina M Whalley, Andrew Bosworth, Charlotte Poxon, Kasun Wanigasooriya, Oliver Pickles, Mike Kidd, Alex Richter, Andrew D Beggs PHE Heartlands Lab: Husam Osman, Andrew Bosworth. Queen Elizabeth Hospital: Anna Casey |
| EPI_ISL_651307 | Liverpool Clinical Laboratories | COVID-19 Genomics UK (COG-UK) Consortium | Sam Haldenby, Anita Lucaci, Steve Paterson, Julian Hiscox, Alistair Darby, M Almsaud, A Alrezaihi, Muhannad Alruwaili, Stuart D Armstrong, Jones Benjamin, Eleanor G Bentley, Anu Chawla, Jordan J Clark, Angela Cowell, Richard Eccles, Isabel García-Dorival, Matthew Gemmell, Alessandro Gerada, PKF Gilmore, Richard Gregory, Ximeng Han, Catherine Hartley, Margaret Hughes, Miren Iturriza-Gomara, James Johnson, L Luu, Jenifer Manson, Charlotte Nelson, Elaine O'Toole, Cassie Olateju, Rebekah Penrice-Randal , Lucille Rainbow, N.P Randle, Trevor Ian Robinson, Parul Sharma, Ghada T Shawli, James P Stewart, Neil Swainston, Ecaterina Vamos, Joanne Watts, Mark Whitehead |
| EPI_ISL_651315 | West of Scotland Specialist Virology Centre, NHSGGC / MRC-University of Glasgow Centre for Virus Research | COVID-19 Genomics UK (COG-UK) Consortium | Ana da Silva Filipe, Natasha Johnson, Kathy Smollett, Daniel Mair, Stephen Carmichael, Alice Broos, Lily Tong, Jenna Nichols, Kyriaki Nomikou; Sarah McDonald; Richard Orton, Joseph Hughes, Sreenu Vattipally, David L Robertson; Alasdair MacLean, Rory Gunson; Sharif Shaaban, Matthew Holden; Rachel Blacow, Guy Mollett, Kathy Li, James Shepherd, Antonia Ho, Emma Thomson |
| EPI_ISL_651317 | Quadram Institute Bioscience | COVID-19 Genomics UK (COG-UK) Consortium | Dave J. Baker, Gemma L. Kay, Alp Aydin, Thanh Le-Viet, Steven Rudder, Ana P. Tedim, Anastasia Kolyva, Maria Diaz, Leonardo de Oliveira Martins, Nabil-Fareed Alikhan, Lizzie Meadows, Rachael Stanley, Ngozi Elumogo, Muhammed Yasir, Nicholas M. Thomson, Alexander J Trotter, Rachel Gilroy, Samuel Bloomfield, Claire Stuart, Andrew Bell, Reenesh Prakash, Samir Dervisevic, Alison E. Mather, John Wain, Mark Webber, Andrew J. Page, Justin O'Grady |
| EPI_ISL_651321 | University of Birmingham | COVID-19 Genomics UK (COG-UK) Consortium | Institute of Microbiology, University of Birmingham: Claire McMurray, Joanne Stockton, Samuel Nicholls, Radoslaw Poplawski, Will Rowe, Josh Quick, Nicholas Loman. University of Birmingham Testing Laboratory: Celina M Whalley, Andrew Bosworth, Charlotte Poxon, Kasun Wanigasooriya, Oliver Pickles, Mike Kidd, Alex Richter, Andrew D Beggs PHE Heartlands Lab: Husam Osman, Andrew Bosworth. Queen Elizabeth Hospital: Anna Casey |
| EPI_ISL_651357 | Quadram Institute Bioscience | COVID-19 Genomics UK (COG-UK) Consortium | Dave J. Baker, Gemma L. Kay, Alp Aydin, Thanh Le-Viet, Steven Rudder, Ana P. Tedim, Anastasia Kolyva, Maria Diaz, Leonardo de Oliveira Martins, Nabil-Fareed Alikhan, Lizzie Meadows, Rachael Stanley, Ngozi Elumogo, Muhammed Yasir, Nicholas M. Thomson, Alexander J Trotter, Rachel Gilroy, Samuel Bloomfield, Claire Stuart, Andrew Bell, Reenesh Prakash, Samir Dervisevic, Alison E. Mather, John Wain, Mark Webber, Andrew J. Page, Justin O'Grady |
| EPI_ISL_651367 | Virology Department, Royal Infirmary of Edinburgh, NHS Lothian / School of Biological Sciences, University of Edinburgh / Institute of Genetics and Molecular Medicine, University of Edinburgh | COVID-19 Genomics UK (COG-UK) Consortium | McHugh M, Dewar R, Rooke S, Gallagher M, Balcaza C, O'Toole Á, Scher E, Hill V, McCrone JT, Colquhoun R, Yu X, Jackson B, Rambaut A, Williams TC, Templeton K |

| | | | |
|---|---|---|---|
| EPI_ISL_651378 | University of Birmingham | COVID-19 Genomics UK (COG-UK) Consortium | Institute of Microbiology, University of Birmingham: Claire McMurray, Joanne Stockton, Samuel Nicholls, Radoslaw Poplawski, Will Rowe, Josh Quick, Nicholas Loman. University of Birmingham Testing Laboratory: Celina M Whalley, Andrew Bosworth, Charlotte Poxon, Kasun Wanigasooriya, Oliver Pickles, Mike Kidd, Alex Richter, Andrew D Beggs PHE Heartlands Lab: Husam Osman, Andrew Bosworth. Queen Elizabeth Hospital: Anna Casey |
| EPI_ISL_651394 | Virology Department, Royal Infirmary of Edinburgh, NHS Lothian / School of Biological Sciences, University of Edinburgh / Institute of Genetics and Molecular Medicine, University of Edinburgh | COVID-19 Genomics UK (COG-UK) Consortium | McHugh M, Dewar R, Rooke S, Gallagher M, Balcaza C, O'Toole Á, Scher E, Hill V, McCrone JT, Colquhoun R, Yu X, Jackson B, Rambaut A, Williams TC, Templeton K |
| EPI_ISL_651418, EPI_ISL_651419 | Quadram Institute Bioscience | COVID-19 Genomics UK (COG-UK) Consortium | Dave J. Baker, Gemma L. Kay, Alp Aydin, Thanh Le-Viet, Steven Rudder, Ana P. Tedim, Anastasia Kolyva, Maria Diaz, Leonardo de Oliveira Martins, Nabil-Fareed Alikhan, Lizzie Meadows, Rachael Stanley, Ngozi Elumogo, Muhammed Yasir, Nicholas M. Thomson, Alexander J Trotter, Rachel Gilroy, Samuel Bloomfield, Claire Stuart, Andrew Bell, Reenesh Prakash, Samir Dervisevic, Alison E. Mather, John Wain, Mark Webber, Andrew J. Page, Justin O'Grady |
| EPI_ISL_651423 | Department of Pathology, University of Cambridge | COVID-19 Genomics UK (COG-UK) Consortium | Aminu S. Jahun, Yasmin Chaudhry, Grant Hall, Iliana Georgana, Myra Hosmillo, Martin D. Curran, Malte Pinckert, Surendra Parmar, Ian Goodfellow |
| EPI_ISL_651427 | Virology Department, Royal Infirmary of Edinburgh, NHS Lothian / School of Biological Sciences, University of Edinburgh / Institute of Genetics and Molecular Medicine, University of Edinburgh | COVID-19 Genomics UK (COG-UK) Consortium | McHugh M, Dewar R, Rooke S, Gallagher M, Balcaza C, O'Toole Á, Scher E, Hill V, McCrone JT, Colquhoun R, Yu X, Jackson B, Rambaut A, Williams TC, Templeton K |
| EPI_ISL_651484 | Quadram Institute Bioscience | COVID-19 Genomics UK (COG-UK) Consortium | Dave J. Baker, Gemma L. Kay, Alp Aydin, Thanh Le-Viet, Steven Rudder, Ana P. Tedim, Anastasia Kolyva, Maria Diaz, Leonardo de Oliveira Martins, Nabil-Fareed Alikhan, Lizzie Meadows, Rachael Stanley, Ngozi Elumogo, Muhammed Yasir, Nicholas M. Thomson, Alexander J Trotter, Rachel Gilroy, Samuel Bloomfield, Claire Stuart, Andrew Bell, Reenesh Prakash, Samir Dervisevic, Alison E. Mather, John Wain, Mark Webber, Andrew J. Page, Justin O'Grady |
| EPI_ISL_651502 | Wales Specialist Virology Centre Sequencing lab: Pathogen Genomics Unit | COVID-19 Genomics UK (COG-UK) Consortium | Catherine Moore, Johnathan Evans, Laura Gifford, Malorie Perry, Simon Cottrell, Angela Marchbank, Alec Birchley, Alexander Adams, Amy Gaskin, Bree Gatica-Wilcox, Jason Coombes, Joel Southgate, Lauren Gilbert, Lee Graham, Nicole Pacchiarini, Sara Kumziene-Summerhayes, Sarah Taylor, Sophie Jones, Sara Rey, Matthew Bull, Joanne Watkins, Sally Corden, Tom Connor |
| EPI_ISL_651542, EPI_ISL_651556, EPI_ISL_651557 | Department of Pathology, University of Cambridge | COVID-19 Genomics UK (COG-UK) Consortium | Aminu S. Jahun, Yasmin Chaudhry, Grant Hall, Iliana Georgana, Myra Hosmillo, Martin D. Curran, Malte Pinckert, Surendra Parmar, Ian Goodfellow |
| EPI_ISL_651592, EPI_ISL_651622, EPI_ISL_651623, EPI_ISL_651628 | University of Birmingham | COVID-19 Genomics UK (COG-UK) Consortium | Institute of Microbiology, University of Birmingham: Claire McMurray, Joanne Stockton, Samuel Nicholls, Radoslaw Poplawski, Will Rowe, Josh Quick, Nicholas Loman. University of Birmingham Testing Laboratory: Celina M Whalley, Andrew Bosworth, Charlotte Poxon, Kasun Wanigasooriya, Oliver Pickles, Mike Kidd, Alex Richter, Andrew D Beggs PHE Heartlands Lab: Husam Osman, Andrew Bosworth. Queen Elizabeth Hospital: Anna Casey |
| EPI_ISL_651666, EPI_ISL_651667, EPI_ISL_651669, EPI_ISL_651670, EPI_ISL_651672, EPI_ISL_651679, EPI_ISL_651680, EPI_ISL_651684, EPI_ISL_651685, EPI_ISL_651686 | Department of Pathology, University of Cambridge | COVID-19 Genomics UK (COG-UK) Consortium | Aminu S. Jahun, Yasmin Chaudhry, Grant Hall, Iliana Georgana, Myra Hosmillo, Martin D. Curran, Malte Pinckert, Surendra Parmar, Ian Goodfellow |
| EPI_ISL_651717 | Virology Department, Sheffield Teaching Hospitals NHS Foundation Trust/Department of Infection, Immunity and Cardiovascular Disease, The Medical School, University of Sheffield | COVID-19 Genomics UK (COG-UK) Consortium | Thushan de Silva, Matthew Parker, Nikki Smith, Adri Angyal, Rebecca Brown, Luke Green, Rachel Tucker, Paul Parsons, Danielle Groves, Katie Johnson, Laura Carrilero, Alex Keeley, Dave Partridge, Matthew Wyles, Benjamin Lindsey, Mehmet Yavuz, Mohammad Raza, Cariad Evans |
| EPI_ISL_651811, EPI_ISL_651812, EPI_ISL_651813, EPI_ISL_651821, EPI_ISL_651822, EPI_ISL_651823, EPI_ISL_651824, EPI_ISL_651825, EPI_ISL_651826, EPI_ISL_651827, EPI_ISL_651828, EPI_ISL_651829, EPI_ISL_651850, EPI_ISL_651851, EPI_ISL_651852, EPI_ISL_651853, EPI_ISL_651854, EPI_ISL_651855, EPI_ISL_651856, EPI_ISL_651857, EPI_ISL_651858, EPI_ISL_651859, EPI_ISL_651863, EPI_ISL_651865, EPI_ISL_651866, EPI_ISL_651867, EPI_ISL_651868, EPI_ISL_651873, EPI_ISL_651883 | | | |
| see above | West of Scotland Specialist Virology Centre, NHSGGC / MRC-University of Glasgow Centre for Virus Research | COVID-19 Genomics UK (COG-UK) Consortium | Ana da Silva Filipe, Natasha Johnson, Kathy Smollett, Daniel Mair, Stephen Carmichael, Alice Broos, Lily Tong, Jenna Nichols, Kyriaki Nomikou; Sarah McDonald; Richard Orton, Joseph Hughes, Sreenu Vattipally, David L Robertson; Alasdair MacLean, Rory Gunson; Sharif Shaaban, Matthew Holden; Rachel Blacow, Guy Mollett, Kathy Li, James Shepherd, Antonia Ho, Emma Thomson |
| EPI_ISL_652024, EPI_ISL_652025, EPI_ISL_652026, EPI_ISL_652027, EPI_ISL_652028, EPI_ISL_652029, EPI_ISL_652030, EPI_ISL_652031, EPI_ISL_652032, EPI_ISL_652033, EPI_ISL_652034, EPI_ISL_652035, EPI_ISL_652036, EPI_ISL_652038, EPI_ISL_652039, EPI_ISL_652040, EPI_ISL_652051, EPI_ISL_652057, EPI_ISL_652058, EPI_ISL_652059, EPI_ISL_652060 | | | |
| see above | Virology Department, Royal Infirmary of Edinburgh, NHS Lothian / School of Biological Sciences, University of Edinburgh / Institute of Genetics and Molecular Medicine, University of Edinburgh | COVID-19 Genomics UK (COG-UK) Consortium | McHugh M, Dewar R, Rooke S, Gallagher M, Balcaza C, O'Toole Á, Scher E, Hill V, McCrone JT, Colquhoun R, Yu X, Jackson B, Rambaut A, Williams TC, Templeton K |
| EPI_ISL_652289, EPI_ISL_652295, EPI_ISL_652296, EPI_ISL_652297, EPI_ISL_652298, EPI_ISL_652299, EPI_ISL_652301, EPI_ISL_652303 | University of Exeter | COVID-19 Genomics UK (COG-UK) Consortium | Ben Temperton,Aaron Jeffries,Michelle Michelsen,Joanna Warwick-Dugdale,Audrey Farbos,Robyn Manley,Stephen Michell,Jane Masoli |
| EPI_ISL_652530, EPI_ISL_652556, EPI_ISL_652566, EPI_ISL_652576, EPI_ISL_652601, EPI_ISL_652616 | Wales Specialist Virology Centre Sequencing lab: Pathogen Genomics Unit | COVID-19 Genomics UK (COG-UK) Consortium | Catherine Moore, Johnathan Evans, Laura Gifford, Malorie Perry, Simon Cottrell, Angela Marchbank, Alec Birchley, Alexander Adams, Amy Gaskin, Bree Gatica-Wilcox, Jason Coombes, Joel Southgate, Lauren Gilbert, Lee Graham, Nicole Pacchiarini, Sara Kumziene-Summerhayes, Sarah Taylor, Sophie Jones, Sara Rey, Matthew Bull, Joanne Watkins, Sally Corden, Tom Connor |
| EPI_ISL_653027, EPI_ISL_653028, EPI_ISL_653029, EPI_ISL_653030, EPI_ISL_653031, EPI_ISL_653032, EPI_ISL_653033, EPI_ISL_653034, EPI_ISL_653035, EPI_ISL_653036, EPI_ISL_653037, EPI_ISL_653038, EPI_ISL_653039, EPI_ISL_653040, EPI_ISL_653041, EPI_ISL_653042, EPI_ISL_653043, EPI_ISL_653044, EPI_ISL_653045 | | | |
| see above | Quadram Institute Bioscience | COVID-19 Genomics UK (COG-UK) Consortium | Dave J. Baker, Gemma L. Kay, Alp Aydin, Thanh Le-Viet, Steven Rudder, Ana P. Tedim, Anastasia Kolyva, Maria Diaz, Leonardo de Oliveira Martins, Nabil-Fareed Alikhan, Lizzie Meadows, Rachael Stanley, Ngozi Elumogo, Muhammed Yasir, Nicholas M. Thomson, Alexander J Trotter, Rachel Gilroy, Samuel Bloomfield, Claire Stuart, Andrew Bell, Reenesh Prakash, Samir Dervisevic, Alison E. Mather, John Wain, Mark Webber, Andrew J. Page, Justin O'Grady |
| EPI_ISL_653067, EPI_ISL_653068, EPI_ISL_653082, EPI_ISL_653091 | Virology Department, Sheffield Teaching Hospitals NHS Foundation Trust/Department of Infection, Immunity and Cardiovascular Disease, The Medical School, University of Sheffield | COVID-19 Genomics UK (COG-UK) Consortium | Thushan de Silva, Matthew Parker, Nikki Smith, Adri Angyal, Rebecca Brown, Luke Green, Rachel Tucker, Paul Parsons, Danielle Groves, Katie Johnson, Laura Carrilero, Alex Keeley, Dave Partridge, Matthew Wyles, Benjamin Lindsey, Mehmet Yavuz, Mohammad Raza, Cariad Evans |
| EPI_ISL_653782 | I.R.C.C.S. "S. De Bellis" - Ente Ospedaliero | Istituto Zooprofilattico Sperimentale della Puglia e della Basilicata | Parisi A., Bianco A., Capozzi L., Del Sambro L., Lippolis A., Notarnicola M., Manzulli V, Rondinone V., Pace L. |
| EPI_ISL_653810 | I.R.C.C.S. "S. De Bellis" - Ente Ospedaliero | Istituto Zooprofilattico Sperimentale della Puglia e della Basilicata | Parisi A., Bianco A., Capozzi L., Del Sambro L., Lippolis A., Notarnicola M., Cipolletta D., Galante D. |
| EPI_ISL_653938, EPI_ISL_653947, EPI_ISL_653953, EPI_ISL_653954, EPI_ISL_653992, EPI_ISL_654002, EPI_ISL_654003, EPI_ISL_654004, EPI_ISL_654006, EPI_ISL_654008, EPI_ISL_654009, EPI_ISL_654011, EPI_ISL_654012, EPI_ISL_654013, EPI_ISL_654014, EPI_ISL_654015 | | | |
| see above | Respiratory Virus Unit, Microbiology Services Colindale, Public Health England | COVID-19 Genomics UK (COG-UK) Consortium | PHE Covid Sequencing Team |
| EPI_ISL_654400 | Hospital General Universitario Gregorio Marañón | SeqCOVID-SPAIN consortium/IBV(CSIC) | Darío García de Viedma, Laura Pérez-Lago, Marta Herranz, Jon Sicilia, Julia Suárez, Pilar Catalán, Patricia Muñoz and SeqCOVID-SPAIN consortium |
| EPI_ISL_654700, EPI_ISL_654701, EPI_ISL_654702, EPI_ISL_654704 | Minnesota Department of Health, Public Health Laboratory | Minnesota Department of Health, Public Health Laboratory | Matt Plumb, Jacob Garfin, Alexandra Lorentz, and Xiong Wang |

| | | | |
|---|---|---|---|
| EPI_ISL_658741, EPI_ISL_658742, EPI_ISL_658743, EPI_ISL_658746, EPI_ISL_658747, EPI_ISL_658748, EPI_ISL_658749, EPI_ISL_658751, EPI_ISL_658757, EPI_ISL_658758, EPI_ISL_658761, EPI_ISL_658762, EPI_ISL_658766, EPI_ISL_658770, EPI_ISL_658771, EPI_ISL_658773, EPI_ISL_658774, EPI_ISL_658776, EPI_ISL_658778, EPI_ISL_658779, EPI_ISL_658782, EPI_ISL_658783, EPI_ISL_658784, EPI_ISL_658785, EPI_ISL_658786, EPI_ISL_658790, EPI_ISL_658792, EPI_ISL_658793, EPI_ISL_658794, EPI_ISL_658796, EPI_ISL_658797, EPI_ISL_658798, EPI_ISL_658799, EPI_ISL_658801, EPI_ISL_658802, EPI_ISL_658805, EPI_ISL_658806, EPI_ISL_658808, EPI_ISL_658810, EPI_ISL_658811, EPI_ISL_658813, EPI_ISL_658814, EPI_ISL_658815, EPI_ISL_658817, EPI_ISL_658818, EPI_ISL_658819, EPI_ISL_658820, EPI_ISL_658822, EPI_ISL_658823, EPI_ISL_658824, EPI_ISL_658825, EPI_ISL_658826, EPI_ISL_658828, EPI_ISL_658830, EPI_ISL_658831, EPI_ISL_658834, EPI_ISL_658835, EPI_ISL_658839, EPI_ISL_658840, EPI_ISL_658841, EPI_ISL_658842, EPI_ISL_658843, EPI_ISL_658844, EPI_ISL_658845, EPI_ISL_658848, EPI_ISL_658849, EPI_ISL_658850, EPI_ISL_658853, EPI_ISL_658854, EPI_ISL_658856, EPI_ISL_658857, EPI_ISL_658858, EPI_ISL_658859, EPI_ISL_658860, EPI_ISL_658861, EPI_ISL_658862, EPI_ISL_658867, EPI_ISL_658869, EPI_ISL_658871, EPI_ISL_658872, EPI_ISL_658874, EPI_ISL_658876, EPI_ISL_658882, EPI_ISL_658884, EPI_ISL_658887, EPI_ISL_658889, EPI_ISL_658890 | | | |
| see above | Lighthouse Lab in Milton Keynes | Wellcome Sanger Institute for the COVID-19 Genomics UK (COG-UK) Consortium | The Lighthouse Lab in Milton Keynes and Alex Alderton, Roberto Amato, Sonia Goncalves, Ewan Harrison, David K. Jackson, Ian Johnston, Dominic Kwiatkowski, Cordelia Langford, John Sillitoe on behalf of the Wellcome Sanger Institute COVID-19 Surveillance Team |
| EPI_ISL_658891 | Instituto de Diagnostico y Referencia Epidemiologicos (INDRE) | Instituto de Diagnostico y Referencia Epidemiologicos (INDRE) | Ernesto Ramirez-Gonzalez, Abril Rodriguez-Maldonado, Claudia Wong-Arambula , Natividad Cruz-Ortiz, Tatiana Nunez-Garcia, Dayanira Arellano-Suarez, Fabiola Garces-Ayala, Lucia Hernandez-Rivas, Irma Lopez-Martinez, Gisela Barrera-Badillo. |
| EPI_ISL_658892, EPI_ISL_658894, EPI_ISL_658895, EPI_ISL_658897, EPI_ISL_658898, EPI_ISL_658902, EPI_ISL_658903, EPI_ISL_658906, EPI_ISL_658907, EPI_ISL_658910, EPI_ISL_658911, EPI_ISL_658915, EPI_ISL_658917, EPI_ISL_658918, EPI_ISL_658920, EPI_ISL_658924, EPI_ISL_658926, EPI_ISL_658927, EPI_ISL_658929, EPI_ISL_658932, EPI_ISL_658934, EPI_ISL_658937, EPI_ISL_658939, EPI_ISL_658940, EPI_ISL_658941, EPI_ISL_658942, EPI_ISL_658943, EPI_ISL_658944, EPI_ISL_658945, EPI_ISL_658946, EPI_ISL_658947, EPI_ISL_658949, EPI_ISL_658950, EPI_ISL_658952, EPI_ISL_658953, EPI_ISL_658956, EPI_ISL_658957, EPI_ISL_658958, EPI_ISL_658960, EPI_ISL_658961, EPI_ISL_658962, EPI_ISL_658964, EPI_ISL_658965, EPI_ISL_658966, EPI_ISL_658967, EPI_ISL_658968, EPI_ISL_658969, EPI_ISL_658972, EPI_ISL_658973, EPI_ISL_658974, EPI_ISL_658975, EPI_ISL_658977, EPI_ISL_658978, EPI_ISL_658980, EPI_ISL_658981, EPI_ISL_658982, EPI_ISL_658983, EPI_ISL_658984, EPI_ISL_658986, EPI_ISL_658987, EPI_ISL_658988, EPI_ISL_658989, EPI_ISL_658990, EPI_ISL_658991, EPI_ISL_658992, EPI_ISL_658993, EPI_ISL_658997, EPI_ISL_658998, EPI_ISL_658999, EPI_ISL_659002, EPI_ISL_659003, EPI_ISL_659004, EPI_ISL_659006, EPI_ISL_659010, EPI_ISL_659012, EPI_ISL_659013, EPI_ISL_659015, EPI_ISL_659016, EPI_ISL_659020, EPI_ISL_659022, EPI_ISL_659024, EPI_ISL_659027, EPI_ISL_659029, EPI_ISL_659030, EPI_ISL_659031, EPI_ISL_659035, EPI_ISL_659036, EPI_ISL_659038, EPI_ISL_659039, EPI_ISL_659042, EPI_ISL_659043, EPI_ISL_659044, EPI_ISL_659046, EPI_ISL_659047, EPI_ISL_659048, EPI_ISL_659049, EPI_ISL_659051, EPI_ISL_659052, EPI_ISL_659055, EPI_ISL_659057, EPI_ISL_659058, EPI_ISL_659059, EPI_ISL_659060, EPI_ISL_659061, EPI_ISL_659062, EPI_ISL_659063, EPI_ISL_659064, EPI_ISL_659065, EPI_ISL_659066, EPI_ISL_659067, EPI_ISL_659068, EPI_ISL_659069, EPI_ISL_659074, EPI_ISL_659075, EPI_ISL_659078, EPI_ISL_659081, EPI_ISL_659085, EPI_ISL_659086, EPI_ISL_659087, EPI_ISL_659089, EPI_ISL_659092, EPI_ISL_659095, EPI_ISL_659096 | | | |
| see above | Lighthouse Lab in Milton Keynes | Wellcome Sanger Institute for the COVID-19 Genomics UK (COG-UK) Consortium | The Lighthouse Lab in Milton Keynes and Alex Alderton, Roberto Amato, Sonia Goncalves, Ewan Harrison, David K. Jackson, Ian Johnston, Dominic Kwiatkowski, Cordelia Langford, John Sillitoe on behalf of the Wellcome Sanger Institute COVID-19 Surveillance Team |
| EPI_ISL_659772 | Lighthouse Lab in Glasgow | Wellcome Sanger Institute for the COVID-19 Genomics UK (COG-UK) Consortium | Harper VanSteenhouse, Yumi Kasai, David Gray, Carol Clugston, Anna Dominiczak and Alex Alderton, Roberto Amato, Sonia Goncalves, Ewan Harrison, David K. Jackson, Ian Johnston, Dominic Kwiatkowski, Cordelia Langford, John Sillitoe on behalf of the Wellcome Sanger Institute COVID-19 Surveillance Team |
| EPI_ISL_660020 | Lighthouse Lab in Glasgow | Wellcome Sanger Institute for the COVID-19 Genomics UK (COG-UK) Consortium | Harper VanSteenhouse, Yumi Kasai, David Gray, Carol Clugston, Anna Dominiczak and Alex Alderton, Roberto Amato, Sonia Goncalves, Ewan Harrison, David K. Jackson, Ian Johnston, Dominic Kwiatkowski, Cordelia Langford, John Sillitoe on behalf of the Wellcome Sanger Institute COVID-19 Surveillance Team (http://www.sanger.ac.uk/covid-team) |
| EPI_ISL_660068 | Instituto de Diagnostico y Referencia Epidemiologicos (INDRE) | Instituto de diagnóstico y Referencia Epidemiologicos (INDRE) Departamento de Virología | Gisela Barrera-Badillo , Abril Rodriguez-Maldonado, Claudia Wong-Arambula , Natividad Cruz-Ortiz, Tatiana Nunez-Garcia, Dayanira Arellano-Suarez, Fabiola Garces-Ayala, Lucia Hernandez-Rivas, Irma Lopez-Martinez, Ernesto Ramirez-Gonzalez. |
| EPI_ISL_660094 | MD PHL | MD PHL | Maryland Department of Health Laboratories Administration |
| EPI_ISL_660223, EPI_ISL_660224, EPI_ISL_660226, EPI_ISL_660227, EPI_ISL_660643, EPI_ISL_660652, EPI_ISL_660657 | NHLS-IALCH | KRISP, KZN Research Innovation and Sequencing Platform | Giandhari J, Pillay S, Lessells R, Mdlalose K, York D, Khan S, Tegally H, Wilkinson E, de Oliveira T |
| EPI_ISL_660754, EPI_ISL_660755, EPI_ISL_660757, EPI_ISL_660758, EPI_ISL_660759, EPI_ISL_660760, EPI_ISL_660763, EPI_ISL_660764, EPI_ISL_660765, EPI_ISL_660766, EPI_ISL_660767, EPI_ISL_660768, EPI_ISL_660769, EPI_ISL_660770, EPI_ISL_660771, EPI_ISL_660772, EPI_ISL_660773, EPI_ISL_660774, EPI_ISL_660775, EPI_ISL_660776, EPI_ISL_660777, EPI_ISL_660778, EPI_ISL_660779, EPI_ISL_660780, EPI_ISL_660781, EPI_ISL_660783 | | | |
| see above | Respiratory Virus Unit, Microbiology Services Colindale, Public Health England | COVID-19 Genomics UK (COG-UK) Consortium | PHE Covid Sequencing Team |
| EPI_ISL_661027, EPI_ISL_661028, EPI_ISL_661029, EPI_ISL_661030, EPI_ISL_661031, EPI_ISL_661032, EPI_ISL_661033, EPI_ISL_661034, EPI_ISL_661035, EPI_ISL_661036, EPI_ISL_661037, EPI_ISL_661038, EPI_ISL_661039, EPI_ISL_661040, EPI_ISL_661041, EPI_ISL_661042, EPI_ISL_661043, EPI_ISL_661044 | | | |
| see above | Gundersen Molecular Diagnostics Laboratory | Kabara Cancer Research Institute | Craig S. Richmond, Paraic A. Kenny |
| EPI_ISL_661292 | Mikrobiologen | The Public Health Agency of Sweden | Department of Microbiology, The Public Health Agency of Sweden |
| EPI_ISL_661293 | Klinisk mikrobiologi | The Public Health Agency of Sweden | Department of Microbiology, The Public Health Agency of Sweden |
| EPI_ISL_661297 | Unilabs | The Public Health Agency of Sweden | Department of Microbiology, The Public Health Agency of Sweden |
| EPI_ISL_662637, EPI_ISL_662639, EPI_ISL_662640, EPI_ISL_662641, EPI_ISL_662644, EPI_ISL_662646 | Lighthouse Lab in Milton Keynes | Wellcome Sanger Institute for the COVID-19 Genomics UK (COG-UK) Consortium | The Lighthouse Lab in Milton Keynes and Alex Alderton, Roberto Amato, Sonia Goncalves, Ewan Harrison, David K. Jackson, Ian Johnston, Dominic Kwiatkowski, Cordelia Langford, John Sillitoe on behalf of the Wellcome Sanger Institute COVID-19 Surveillance Team |
| EPI_ISL_662656, EPI_ISL_662657, EPI_ISL_662659, EPI_ISL_662662, EPI_ISL_662663, EPI_ISL_662668, EPI_ISL_662673, EPI_ISL_662675, EPI_ISL_662677, EPI_ISL_662683, EPI_ISL_662687, EPI_ISL_662691, EPI_ISL_662693, EPI_ISL_662695, EPI_ISL_662700, EPI_ISL_662702, EPI_ISL_662708, EPI_ISL_662711, EPI_ISL_662712, EPI_ISL_662717, EPI_ISL_662722, EPI_ISL_662723, EPI_ISL_662727, EPI_ISL_662730, EPI_ISL_662731, EPI_ISL_662733, EPI_ISL_662734, EPI_ISL_662735, EPI_ISL_662737, EPI_ISL_662738, EPI_ISL_662739, EPI_ISL_662743, EPI_ISL_662744, EPI_ISL_662750, EPI_ISL_662753, EPI_ISL_662762, EPI_ISL_662765, EPI_ISL_662769, EPI_ISL_662775, EPI_ISL_662780, EPI_ISL_662782, EPI_ISL_662784, EPI_ISL_662793, EPI_ISL_662800, EPI_ISL_662802, EPI_ISL_662803, EPI_ISL_662804, EPI_ISL_662807, EPI_ISL_662810, EPI_ISL_662814, EPI_ISL_662819, EPI_ISL_662823, EPI_ISL_662825, EPI_ISL_662826, EPI_ISL_662827, EPI_ISL_662828, EPI_ISL_662831, EPI_ISL_662833, EPI_ISL_662837, EPI_ISL_662838, EPI_ISL_662840, EPI_ISL_662843, EPI_ISL_662848, EPI_ISL_662850, EPI_ISL_662851, EPI_ISL_662852, EPI_ISL_662862, EPI_ISL_662863, EPI_ISL_662866, EPI_ISL_662870, EPI_ISL_662874, EPI_ISL_662881, EPI_ISL_662883, EPI_ISL_662885, EPI_ISL_662890, EPI_ISL_662891, EPI_ISL_662892, EPI_ISL_662893, EPI_ISL_662894, EPI_ISL_662895, EPI_ISL_662896, EPI_ISL_662897, EPI_ISL_662898, EPI_ISL_662899, EPI_ISL_662900, EPI_ISL_662901, EPI_ISL_662902, EPI_ISL_662903, EPI_ISL_662904, EPI_ISL_662905, EPI_ISL_662906, EPI_ISL_662907, EPI_ISL_662908, EPI_ISL_662909, EPI_ISL_662910, EPI_ISL_662911, EPI_ISL_662912, EPI_ISL_662913, EPI_ISL_662914, EPI_ISL_662915, EPI_ISL_662916, EPI_ISL_662917, EPI_ISL_662918, EPI_ISL_662919, EPI_ISL_662920, EPI_ISL_662921, EPI_ISL_662922, EPI_ISL_662923, EPI_ISL_662924, EPI_ISL_662925, EPI_ISL_662926, EPI_ISL_662927, EPI_ISL_662928, EPI_ISL_662929, EPI_ISL_662930, EPI_ISL_662931, EPI_ISL_662932, EPI_ISL_662933, EPI_ISL_662934, EPI_ISL_662935, EPI_ISL_662936, EPI_ISL_662937, EPI_ISL_662938, EPI_ISL_662939, EPI_ISL_662940, EPI_ISL_662941, EPI_ISL_662942, EPI_ISL_662943, EPI_ISL_662944, EPI_ISL_662945, EPI_ISL_662946, EPI_ISL_662947, EPI_ISL_662948, EPI_ISL_662949, EPI_ISL_662950, EPI_ISL_662951, EPI_ISL_662952, EPI_ISL_662953, EPI_ISL_662954, EPI_ISL_662955, EPI_ISL_662956, EPI_ISL_662957, EPI_ISL_662958, EPI_ISL_662959, EPI_ISL_662960, EPI_ISL_662961, EPI_ISL_662962, EPI_ISL_662963, EPI_ISL_662964, EPI_ISL_662965, EPI_ISL_662966, EPI_ISL_662967, EPI_ISL_662968, EPI_ISL_662969, EPI_ISL_662970, EPI_ISL_662971, EPI_ISL_662972, EPI_ISL_662973, EPI_ISL_662974, EPI_ISL_662975, EPI_ISL_662976, EPI_ISL_662977, EPI_ISL_662978, EPI_ISL_662979, EPI_ISL_662980, EPI_ISL_662981, EPI_ISL_662982, EPI_ISL_662983, EPI_ISL_662984, EPI_ISL_662985, EPI_ISL_662986, EPI_ISL_662987, EPI_ISL_662988, EPI_ISL_662989, EPI_ISL_662990, EPI_ISL_662991, EPI_ISL_662992, EPI_ISL_662993, EPI_ISL_662994, EPI_ISL_662995, EPI_ISL_662996, EPI_ISL_662997, EPI_ISL_662998, EPI_ISL_662999, EPI_ISL_663000, EPI_ISL_663001, EPI_ISL_663002, EPI_ISL_663003, EPI_ISL_663004, EPI_ISL_663005, EPI_ISL_663006, EPI_ISL_663007, EPI_ISL_663008, EPI_ISL_663009, EPI_ISL_663010, EPI_ISL_663011, EPI_ISL_663012, EPI_ISL_663013, EPI_ISL_663014, EPI_ISL_663015, EPI_ISL_663016, EPI_ISL_663017, EPI_ISL_663018, EPI_ISL_663019, EPI_ISL_663020, EPI_ISL_663021, EPI_ISL_663022, EPI_ISL_663023, EPI_ISL_663024, EPI_ISL_663025, EPI_ISL_663026, EPI_ISL_663027, EPI_ISL_663028, EPI_ISL_663029, EPI_ISL_663030, EPI_ISL_663031, EPI_ISL_663032, EPI_ISL_663033, EPI_ISL_663034, EPI_ISL_663035, EPI_ISL_663036, EPI_ISL_663037, EPI_ISL_663038, EPI_ISL_663039, EPI_ISL_663040, EPI_ISL_663041, EPI_ISL_663042, EPI_ISL_663043, EPI_ISL_663044, EPI_ISL_663045, EPI_ISL_663046, EPI_ISL_663047, EPI_ISL_663048, EPI_ISL_663049, EPI_ISL_663050, EPI_ISL_663051, EPI_ISL_663052, EPI_ISL_663053, EPI_ISL_663054, EPI_ISL_663055, EPI_ISL_663056, EPI_ISL_663057, EPI_ISL_663058, EPI_ISL_663059, EPI_ISL_663060, EPI_ISL_663061, EPI_ISL_663062, EPI_ISL_663063, EPI_ISL_663064, EPI_ISL_663065, EPI_ISL_663066, EPI_ISL_663067, EPI_ISL_663068, EPI_ISL_663069, EPI_ISL_663070, EPI_ISL_663071, EPI_ISL_663072, EPI_ISL_663073, EPI_ISL_663074, EPI_ISL_663075, EPI_ISL_663076, EPI_ISL_663077, EPI_ISL_663078, EPI_ISL_663079, EPI_ISL_663080, EPI_ISL_663081, EPI_ISL_663082, EPI_ISL_663083, EPI_ISL_663084, EPI_ISL_663085, EPI_ISL_663086, EPI_ISL_663087, EPI_ISL_663088, EPI_ISL_663089, EPI_ISL_663090, EPI_ISL_663091, EPI_ISL_663092, EPI_ISL_663093, EPI_ISL_663094, EPI_ISL_663095, EPI_ISL_663096, EPI_ISL_663097, EPI_ISL_663098, EPI_ISL_663099, EPI_ISL_663100, EPI_ISL_663101, EPI_ISL_663102, EPI_ISL_663103, EPI_ISL_663104, EPI_ISL_663105, EPI_ISL_663106, EPI_ISL_663107, EPI_ISL_663108, EPI_ISL_663109, EPI_ISL_663110, EPI_ISL_663111, EPI_ISL_663112, EPI_ISL_663113, EPI_ISL_663114, EPI_ISL_663115, EPI_ISL_663116, EPI_ISL_663117, EPI_ISL_663118, EPI_ISL_663119, EPI_ISL_663120, EPI_ISL_663121, EPI_ISL_663122, EPI_ISL_663123, EPI_ISL_663124, EPI_ISL_663125, EPI_ISL_663126, EPI_ISL_663127, EPI_ISL_663128, EPI_ISL_663129, EPI_ISL_663130, EPI_ISL_663131, EPI_ISL_663132, EPI_ISL_663133, EPI_ISL_663135, EPI_ISL_663136, EPI_ISL_663137, EPI_ISL_663138, EPI_ISL_663139, EPI_ISL_663140, EPI_ISL_663141, EPI_ISL_663142, EPI_ISL_663143, EPI_ISL_663144, EPI_ISL_663145, EPI_ISL_663146, EPI_ISL_663147, EPI_ISL_663148, EPI_ISL_663149, EPI_ISL_663150, EPI_ISL_663151, EPI_ISL_663152, EPI_ISL_663153, EPI_ISL_663154, EPI_ISL_663155, EPI_ISL_663156, EPI_ISL_663157, EPI_ISL_663158, EPI_ISL_663159, EPI_ISL_663160, EPI_ISL_663161, EPI_ISL_663162, EPI_ISL_663163, EPI_ISL_663164, EPI_ISL_663165, EPI_ISL_663166, EPI_ISL_663167, EPI_ISL_663168, EPI_ISL_663169, EPI_ISL_663170, EPI_ISL_663171, EPI_ISL_663172, EPI_ISL_663173, EPI_ISL_663174, EPI_ISL_663175, EPI_ISL_663176, EPI_ISL_663177, EPI_ISL_663178, EPI_ISL_663179, EPI_ISL_663180, EPI_ISL_663181, EPI_ISL_663182, EPI_ISL_663183, EPI_ISL_663184, EPI_ISL_663185, EPI_ISL_663186, EPI_ISL_663187, EPI_ISL_663188, EPI_ISL_663189, EPI_ISL_663190, EPI_ISL_663191, EPI_ISL_663192, EPI_ISL_663193, EPI_ISL_663194, EPI_ISL_663195, EPI_ISL_663196, EPI_ISL_663197, EPI_ISL_663198, EPI_ISL_663199, EPI_ISL_663200, EPI_ISL_663201, EPI_ISL_663202, EPI_ISL_663203, EPI_ISL_663204, EPI_ISL_663205 | | | |
| see above | Lighthouse Lab in Glasgow | Wellcome Sanger Institute for the COVID-19 Genomics UK (COG-UK) Consortium | Harper VanSteenhouse, Yumi Kasai, David Gray, Carol Clugston, Anna Dominiczak and Alex Alderton, Roberto Amato, Sonia Goncalves, Ewan Harrison, David K. Jackson, Ian Johnston, Dominic Kwiatkowski, Cordelia Langford, John Sillitoe on behalf of the Wellcome Sanger Institute COVID-19 Surveillance Team |
| EPI_ISL_664117, EPI_ISL_664162, EPI_ISL_664205, EPI_ISL_664206 | University of Exeter | COVID-19 Genomics UK (COG-UK) Consortium | Ben Temperton,Aaron Jeffries,Michelle Michelsen,Joanna Warwick-Dugdale,Audrey Farbos,Robyn Manley,Stephen Michell,Jane Masoli |
| EPI_ISL_664228 | University College London Hospital | COVID-19 Genomics UK (COG-UK) Consortium | Judith Heaney, Matthew Byott, Catherine Houlihan, Dan Frampton, Stuart Kirk, Moira Spyer and Eleni Nastouli |
| EPI_ISL_664289 | University of Exeter | COVID-19 Genomics UK (COG-UK) Consortium | Ben Temperton,Aaron Jeffries,Michelle Michelsen,Joanna Warwick-Dugdale,Audrey Farbos,Robyn Manley,Stephen Michell,Jane Masoli |
| EPI_ISL_664302 | Centre for Enzyme Innovation, University of Portsmouth / Translational Research Laboratory, Portsmouth Hospitals NHS Trust | COVID-19 Genomics UK (COG-UK) Consortium | Angela Beckett,Yann Bourgeois,Garry Scarlett,Sharon Glaysher,Scott Elliott,Kelly Bicknell,Robert Impey,Allyson Lloyd,Sarah Wyllie,Ethan Butcher,Anoop Chauhan,Samuel Robson |

| | | | |
|---|---|---|---|
| EPI_ISL_664327 | University of Exeter | COVID-19 Genomics UK (COG-UK) Consortium | Ben Temperton,Aaron Jeffries,Michelle Michelsen,Joanna Warwick-Dugdale,Audrey Farbos,Robyn Manley,Stephen Michell,Jane Masoli |
| EPI_ISL_664343, EPI_ISL_664354 | Liverpool Clinical Laboratories | COVID-19 Genomics UK (COG-UK) Consortium | Sam Haldenby, Anita Lucaci, Steve Paterson, Julian Hiscox, Alistair Darby, M Almsaud, A Alrezaihi, Muhannad Alruwaili, Stuart D Armstrong, Jones Benjamin, Eleanor G Bentley, Anu Chawla, Jordan J Clark, Angela Cowell, Richard Eccles, Isabel García-Dorival, Matthew Gemmell, Alessandro Gerada, PKF Gilmore, Richard Gregory, Ximeng Han, Catherine Hartley, Margaret Hughes, Miren Iturriza-Gomara, James Johnson, L Luu, Jenifer Manson, Charlotte Nelson, Elaine O'Toole, Cassie Olateju, Rebekah Penrice-Randal , Lucille Rainbow, N.P Randle, Trevor Ian Robinson, Parul Sharma, Ghada T Shawli, James P Stewart, Neil Swainston, Ecaterina Vamos, Joanne Watts, Mark Whitehead |
| EPI_ISL_664376, EPI_ISL_664377 | Centre for Enzyme Innovation, University of Portsmouth / Translational Research Laboratory, Portsmouth Hospitals NHS Trust | COVID-19 Genomics UK (COG-UK) Consortium | Angela Beckett,Yann Bourgeois,Garry Scarlett,Sharon Glaysher,Scott Elliott,Kelly Bicknell,Robert Impey,Allyson Lloyd,Sarah Wyllie,Ethan Butcher,Anoop Chauhan,Samuel Robson |
| EPI_ISL_664399, EPI_ISL_664473 | University of Exeter | COVID-19 Genomics UK (COG-UK) Consortium | Ben Temperton,Aaron Jeffries,Michelle Michelsen,Joanna Warwick-Dugdale,Audrey Farbos,Robyn Manley,Stephen Michell,Jane Masoli |
| EPI_ISL_664492, EPI_ISL_664557, EPI_ISL_664558, EPI_ISL_664559 | Centre for Enzyme Innovation, University of Portsmouth / Translational Research Laboratory, Portsmouth Hospitals NHS Trust | COVID-19 Genomics UK (COG-UK) Consortium | Angela Beckett,Yann Bourgeois,Garry Scarlett,Sharon Glaysher,Scott Elliott,Kelly Bicknell,Robert Impey,Allyson Lloyd,Sarah Wyllie,Ethan Butcher,Anoop Chauhan,Samuel Robson |
| EPI_ISL_664592 | University of Exeter | COVID-19 Genomics UK (COG-UK) Consortium | Ben Temperton,Aaron Jeffries,Michelle Michelsen,Joanna Warwick-Dugdale,Audrey Farbos,Robyn Manley,Stephen Michell,Jane Masoli |
| EPI_ISL_664606, EPI_ISL_664623, EPI_ISL_664637 | Liverpool Clinical Laboratories | COVID-19 Genomics UK (COG-UK) Consortium | Sam Haldenby, Anita Lucaci, Steve Paterson, Julian Hiscox, Alistair Darby, M Almsaud, A Alrezaihi, Muhannad Alruwaili, Stuart D Armstrong, Jones Benjamin, Eleanor G Bentley, Anu Chawla, Jordan J Clark, Angela Cowell, Richard Eccles, Isabel García-Dorival, Matthew Gemmell, Alessandro Gerada, PKF Gilmore, Richard Gregory, Ximeng Han, Catherine Hartley, Margaret Hughes, Miren Iturriza-Gomara, James Johnson, L Luu, Jenifer Manson, Charlotte Nelson, Elaine O'Toole, Cassie Olateju, Rebekah Penrice-Randal , Lucille Rainbow, N.P Randle, Trevor Ian Robinson, Parul Sharma, Ghada T Shawli, James P Stewart, Neil Swainston, Ecaterina Vamos, Joanne Watts, Mark Whitehead |
| EPI_ISL_664689 | Virology Department, Sheffield Teaching Hospitals NHS Foundation Trust/Department of Infection, Immunity and Cardiovascular Disease, The Medical School, University of Sheffield | COVID-19 Genomics UK (COG-UK) Consortium | Thushan de Silva, Matthew Parker, Nikki Smith, Adri Angyal, Rebecca Brown, Luke Green, Rachel Tucker, Paul Parsons, Danielle Groves, Katie Johnson, Laura Carrilero, Alex Keeley, Dave Partridge, Matthew Wyles, Benjamin Lindsey, Mehmet Yavuz, Mohammad Raza, Cariad Evans |
| EPI_ISL_664690 | Centre for Enzyme Innovation, University of Portsmouth / Translational Research Laboratory, Portsmouth Hospitals NHS Trust | COVID-19 Genomics UK (COG-UK) Consortium | Angela Beckett,Yann Bourgeois,Garry Scarlett,Sharon Glaysher,Scott Elliott,Kelly Bicknell,Robert Impey,Allyson Lloyd,Sarah Wyllie,Ethan Butcher,Anoop Chauhan,Samuel Robson |
| EPI_ISL_664720 | University of Exeter | COVID-19 Genomics UK (COG-UK) Consortium | Ben Temperton,Aaron Jeffries,Michelle Michelsen,Joanna Warwick-Dugdale,Audrey Farbos,Robyn Manley,Stephen Michell,Jane Masoli |
| EPI_ISL_664752 | Virology Department, Sheffield Teaching Hospitals NHS Foundation Trust/Department of Infection, Immunity and Cardiovascular Disease, The Medical School, University of Sheffield | COVID-19 Genomics UK (COG-UK) Consortium | Thushan de Silva, Matthew Parker, Nikki Smith, Adri Angyal, Rebecca Brown, Luke Green, Rachel Tucker, Paul Parsons, Danielle Groves, Katie Johnson, Laura Carrilero, Alex Keeley, Dave Partridge, Matthew Wyles, Benjamin Lindsey, Mehmet Yavuz, Mohammad Raza, Cariad Evans |
| EPI_ISL_664763, EPI_ISL_664767, EPI_ISL_664841, EPI_ISL_664842, EPI_ISL_664846, EPI_ISL_664848, EPI_ISL_664850, EPI_ISL_664851, EPI_ISL_664852, EPI_ISL_664853, EPI_ISL_664854, EPI_ISL_664855 | | | |
| see above | Centre for Enzyme Innovation, University of Portsmouth / Translational Research Laboratory, Portsmouth Hospitals NHS Trust | COVID-19 Genomics UK (COG-UK) Consortium | Angela Beckett,Yann Bourgeois,Garry Scarlett,Sharon Glaysher,Scott Elliott,Kelly Bicknell,Robert Impey,Allyson Lloyd,Sarah Wyllie,Ethan Butcher,Anoop Chauhan,Samuel Robson |
| EPI_ISL_665046, EPI_ISL_665067 | University of Exeter | COVID-19 Genomics UK (COG-UK) Consortium | Ben Temperton,Aaron Jeffries,Michelle Michelsen,Joanna Warwick-Dugdale,Audrey Farbos,Robyn Manley,Stephen Michell,Jane Masoli |
| EPI_ISL_665140 | Liverpool Clinical Laboratories | COVID-19 Genomics UK (COG-UK) Consortium | Sam Haldenby, Anita Lucaci, Steve Paterson, Julian Hiscox, Alistair Darby, M Almsaud, A Alrezaihi, Muhannad Alruwaili, Stuart D Armstrong, Jones Benjamin, Eleanor G Bentley, Anu Chawla, Jordan J Clark, Angela Cowell, Richard Eccles, Isabel García-Dorival, Matthew Gemmell, Alessandro Gerada, PKF Gilmore, Richard Gregory, Ximeng Han, Catherine Hartley, Margaret Hughes, Miren Iturriza-Gomara, James Johnson, L Luu, Jenifer Manson, Charlotte Nelson, Elaine O'Toole, Cassie Olateju, Rebekah Penrice-Randal , Lucille Rainbow, N.P Randle, Trevor Ian Robinson, Parul Sharma, Ghada T Shawli, James P Stewart, Neil Swainston, Ecaterina Vamos, Joanne Watts, Mark Whitehead |
| EPI_ISL_665239 | University College London Hospital | COVID-19 Genomics UK (COG-UK) Consortium | Judith Heaney, Matthew Byott, Catherine Houlihan, Dan Frampton, Stuart Kirk, Moira Spyer and Eleni Nastouli |
| EPI_ISL_665263 | Quadram Institute Bioscience | COVID-19 Genomics UK (COG-UK) Consortium | Dave J. Baker, Gemma L. Kay, Alp Aydin, Thanh Le-Viet, Steven Rudder, Ana P. Tedim, Anastasia Kolyva, Maria Diaz, Leonardo de Oliveira Martins, Nabil-Fareed Alikhan, Lizzie Meadows, Rachael Stanley, Ngozi Elumogo, Muhammed Yasir, Nicholas M. Thomson, Alexander J Trotter, Rachel Gilroy, Samuel Bloomfield, Claire Stuart, Andrew Bell, Reenesh Prakash, Samir Dervisevic, Alison E. Mather, John Wain, Mark Webber, Andrew J. Page, Justin O'Grady |
| EPI_ISL_665288, EPI_ISL_665302, EPI_ISL_665331 | Wales Specialist Virology Centre Sequencing lab: Pathogen Genomics Unit | COVID-19 Genomics UK (COG-UK) Consortium | Catherine Moore, Johnathan Evans, Laura Gifford, Malorie Perry, Simon Cottrell, Angela Marchbank, Alec Birchley, Alexander Adams, Amy Gaskin, Bree Gatica-Wilcox, Jason Coombes, Joel Southgate, Lauren Gilbert, Lee Graham, Nicole Pacchiarini, Sara Kumziene-Summerhayes, Sarah Taylor, Sophie Jones, Sara Rey, Matthew Bull, Joanne Watkins, Sally Corden, Tom Connor |
| EPI_ISL_665335 | University College London, Great Ormond Street Hospital for Children NHS Foundation Trust, Imperial College Healthcare NHS Trust | COVID-19 Genomics UK (COG-UK) Consortium | Sergi Castellano, Rachel Williams, Mark Kristiansen, Paola Resende Silva, Sunando Roy, Tony Brooks, Helena Tutill, Paola Niola, Patricia Dyal, Charlotte Williams, Leysa Forrest, Yasmin Panchbhaya, Jacqueline Findlay, Samuel Weeks, Julianne Brown, Kathryn Harris, Paul Randell, James Price, Alison Holmes, Judith Breuer |
| EPI_ISL_665337 | Quadram Institute Bioscience | COVID-19 Genomics UK (COG-UK) Consortium | Dave J. Baker, Gemma L. Kay, Alp Aydin, Thanh Le-Viet, Steven Rudder, Ana P. Tedim, Anastasia Kolyva, Maria Diaz, Leonardo de Oliveira Martins, Nabil-Fareed Alikhan, Lizzie Meadows, Rachael Stanley, Ngozi Elumogo, Muhammed Yasir, Nicholas M. Thomson, Alexander J Trotter, Rachel Gilroy, Samuel Bloomfield, Claire Stuart, Andrew Bell, Reenesh Prakash, Samir Dervisevic, Alison E. Mather, John Wain, Mark Webber, Andrew J. Page, Justin O'Grady |
| EPI_ISL_665338, EPI_ISL_665347 | Centre for Enzyme Innovation, University of Portsmouth / Translational Research Laboratory, Portsmouth Hospitals NHS Trust | COVID-19 Genomics UK (COG-UK) Consortium | Angela Beckett,Yann Bourgeois,Garry Scarlett,Sharon Glaysher,Scott Elliott,Kelly Bicknell,Robert Impey,Allyson Lloyd,Sarah Wyllie,Ethan Butcher,Anoop Chauhan,Samuel Robson |
| EPI_ISL_665428 | University College London, Great Ormond Street Hospital for Children NHS Foundation Trust, Imperial College Healthcare NHS Trust | COVID-19 Genomics UK (COG-UK) Consortium | Sergi Castellano, Rachel Williams, Mark Kristiansen, Paola Resende Silva, Sunando Roy, Tony Brooks, Helena Tutill, Paola Niola, Patricia Dyal, Charlotte Williams, Leysa Forrest, Yasmin Panchbhaya, Jacqueline Findlay, Samuel Weeks, Julianne Brown, Kathryn Harris, Paul Randell, James Price, Alison Holmes, Judith Breuer |
| EPI_ISL_665457, EPI_ISL_665458, EPI_ISL_665467 | Quadram Institute Bioscience | COVID-19 Genomics UK (COG-UK) Consortium | Dave J. Baker, Gemma L. Kay, Alp Aydin, Thanh Le-Viet, Steven Rudder, Ana P. Tedim, Anastasia Kolyva, Maria Diaz, Leonardo de Oliveira Martins, Nabil-Fareed Alikhan, Lizzie Meadows, Rachael Stanley, Ngozi Elumogo, Muhammed Yasir, Nicholas M. Thomson, Alexander J Trotter, Rachel Gilroy, Samuel Bloomfield, Claire Stuart, Andrew Bell, Reenesh Prakash, Samir Dervisevic, Alison E. Mather, John Wain, Mark Webber, Andrew J. Page, Justin O'Grady |
| EPI_ISL_665469, EPI_ISL_665474 | Wales Specialist Virology Centre Sequencing lab: Pathogen Genomics Unit | COVID-19 Genomics UK (COG-UK) Consortium | Catherine Moore, Johnathan Evans, Laura Gifford, Malorie Perry, Simon Cottrell, Angela Marchbank, Alec Birchley, Alexander Adams, Amy Gaskin, Bree Gatica-Wilcox, Jason Coombes, Joel Southgate, Lauren Gilbert, Lee Graham, Nicole Pacchiarini, Sara Kumziene-Summerhayes, Sarah Taylor, Sophie Jones, Sara Rey, Matthew Bull, Joanne Watkins, Sally Corden, Tom Connor |
| EPI_ISL_665568 | Quadram Institute Bioscience | COVID-19 Genomics UK (COG-UK) Consortium | Dave J. Baker, Gemma L. Kay, Alp Aydin, Thanh Le-Viet, Steven Rudder, Ana P. Tedim, Anastasia Kolyva, Maria Diaz, Leonardo de Oliveira Martins, Nabil-Fareed Alikhan, Lizzie Meadows, Rachael Stanley, Ngozi Elumogo, Muhammed Yasir, Nicholas M. Thomson, Alexander J Trotter, Rachel Gilroy, Samuel Bloomfield, Claire Stuart, Andrew Bell, Reenesh Prakash, Samir Dervisevic, Alison E. Mather, John Wain, Mark Webber, Andrew J. Page, Justin O'Grady |
| EPI_ISL_665576, EPI_ISL_665588 | Wales Specialist Virology Centre Sequencing lab: Pathogen Genomics Unit | COVID-19 Genomics UK (COG-UK) Consortium | Catherine Moore, Johnathan Evans, Laura Gifford, Malorie Perry, Simon Cottrell, Angela Marchbank, Alec Birchley, Alexander Adams, Amy Gaskin, Bree Gatica-Wilcox, Jason Coombes, Joel Southgate, Lauren Gilbert, Lee Graham, Nicole Pacchiarini, Sara Kumziene-Summerhayes, Sarah Taylor, Sophie |

| | | | |
|---|---|---|---|
| | | | Jones, Sara Rey, Matthew Bull, Joanne Watkins, Sally Corden, Tom Connor |
| EPI_ISL_665657 | Quadram Institute Bioscience | COVID-19 Genomics UK (COG-UK) Consortium | Dave J. Baker, Gemma L. Kay, Alp Aydin, Thanh Le-Viet, Steven Rudder, Ana P. Tedim, Anastasia Kolyva, Maria Diaz, Leonardo de Oliveira Martins, Nabil-Fareed Alikhan, Lizzie Meadows, Rachael Stanley, Ngozi Elumogo, Muhammed Yasir, Nicholas M. Thomson, Alexander J Trotter, Rachel Gilroy, Samuel Bloomfield, Claire Stuart, Andrew Bell, Reenesh Prakash, Samir Dervisevic, Alison E. Mather, John Wain, Mark Webber, Andrew J. Page, Justin O'Grady |
| EPI_ISL_665673 | Northumbria University / South Tees Hospitals NHS Foundation Trust / North Cumbria Integrated Care NHS Foundation Trust / North Tees and Hartlepool NHS Foundation Trust / Newcastle Hospitals NHS Foundation Trust | COVID-19 Genomics UK (COG-UK) Consortium | Darren L Smith,Andrew Nelson,Matthew Bashton,Greg R Young,Joshua Loh,John Allan,Mohammad A Tariq,Giles S Holt,Gary Black,Wen C Yew,Lynn Dover,Paul Baker,Steve Liggett,Sarah Essex,Jane Greenaway,Debra Padgett,Clive Graham,Garren Scott,Edward Barton,Emma Swindells,Brendan Payne,Jennifer Collins,Yusri Taha,Gary Eltringham |
| EPI_ISL_665691 | Wales Specialist Virology Centre Sequencing lab: Pathogen Genomics Unit | COVID-19 Genomics UK (COG-UK) Consortium | Catherine Moore, Johnathan Evans, Laura Gifford, Malorie Perry, Simon Cottrell, Angela Marchbank, Alec Birchley, Alexander Adams, Amy Gaskin, Bree Gatica-Wilcox, Jason Coombes, Joel Southgate, Lauren Gilbert, Lee Graham, Nicole Pacchiarini, Sara Kumziene-Summerhayes, Sarah Taylor, Sophie Jones, Sara Rey, Matthew Bull, Joanne Watkins, Sally Corden, Tom Connor |
| EPI_ISL_665723, EPI_ISL_665726 | Centre for Enzyme Innovation, University of Portsmouth / Translational Research Laboratory, Portsmouth Hospitals NHS Trust | COVID-19 Genomics UK (COG-UK) Consortium | Angela Beckett,Yann Bourgeois,Garry Scarlett,Sharon Glaysher,Scott Elliott,Kelly Bicknell,Robert Impey,Allyson Lloyd,Sarah Wyllie,Ethan Butcher,Anoop Chauhan,Samuel Robson |
| EPI_ISL_665768 | Wales Specialist Virology Centre Sequencing lab: Pathogen Genomics Unit | COVID-19 Genomics UK (COG-UK) Consortium | Catherine Moore, Johnathan Evans, Laura Gifford, Malorie Perry, Simon Cottrell, Angela Marchbank, Alec Birchley, Alexander Adams, Amy Gaskin, Bree Gatica-Wilcox, Jason Coombes, Joel Southgate, Lauren Gilbert, Lee Graham, Nicole Pacchiarini, Sara Kumziene-Summerhayes, Sarah Taylor, Sophie Jones, Sara Rey, Matthew Bull, Joanne Watkins, Sally Corden, Tom Connor |
| EPI_ISL_665877, EPI_ISL_665887, EPI_ISL_665889, EPI_ISL_665891, EPI_ISL_665895 | Quadram Institute Bioscience | COVID-19 Genomics UK (COG-UK) Consortium | Dave J. Baker, Gemma L. Kay, Alp Aydin, Thanh Le-Viet, Steven Rudder, Ana P. Tedim, Anastasia Kolyva, Maria Diaz, Leonardo de Oliveira Martins, Nabil-Fareed Alikhan, Lizzie Meadows, Rachael Stanley, Ngozi Elumogo, Muhammed Yasir, Nicholas M. Thomson, Alexander J Trotter, Rachel Gilroy, Samuel Bloomfield, Claire Stuart, Andrew Bell, Reenesh Prakash, Samir Dervisevic, Alison E. Mather, John Wain, Mark Webber, Andrew J. Page, Justin O'Grady |
| EPI_ISL_666007, EPI_ISL_666009, EPI_ISL_666018, EPI_ISL_666019, EPI_ISL_666020, EPI_ISL_666021 | Northumbria University / South Tees Hospitals NHS Foundation Trust / North Cumbria Integrated Care NHS Foundation Trust / North Tees and Hartlepool NHS Foundation Trust / Newcastle Hospitals NHS Foundation Trust | COVID-19 Genomics UK (COG-UK) Consortium | Darren L Smith,Andrew Nelson,Matthew Bashton,Greg R Young,Joshua Loh,John Allan,Mohammad A Tariq,Giles S Holt,Gary Black,Wen C Yew,Lynn Dover,Paul Baker,Steve Liggett,Sarah Essex,Jane Greenaway,Debra Padgett,Clive Graham,Garren Scott,Edward Barton,Emma Swindells,Brendan Payne,Jennifer Collins,Yusri Taha,Gary Eltringham |
| EPI_ISL_666060 | Virology Department, Royal Infirmary of Edinburgh, NHS Lothian / School of Biological Sciences, University of Edinburgh / Institute of Genetics and Molecular Medicine, University of Edinburgh | COVID-19 Genomics UK (COG-UK) Consortium | McHugh M, Dewar R, Rooke S, Gallagher M, Balcaza C, O'Toole Á, Scher E, Hill V, McCrone JT, Colquhoun R, Yu X, Jackson B, Rambaut A, Williams TC, Templeton K |
| EPI_ISL_666107, EPI_ISL_666108, EPI_ISL_666109 | Centre for Enzyme Innovation, University of Portsmouth / Translational Research Laboratory, Portsmouth Hospitals NHS Trust | COVID-19 Genomics UK (COG-UK) Consortium | Angela Beckett,Yann Bourgeois,Garry Scarlett,Sharon Glaysher,Scott Elliott,Kelly Bicknell,Robert Impey,Allyson Lloyd,Sarah Wyllie,Ethan Butcher,Anoop Chauhan,Samuel Robson |
| EPI_ISL_666112, EPI_ISL_666127, EPI_ISL_666136, EPI_ISL_666147, EPI_ISL_666149, EPI_ISL_666157, EPI_ISL_666165, EPI_ISL_666166, EPI_ISL_666172, EPI_ISL_666177, EPI_ISL_666185, EPI_ISL_666197, EPI_ISL_666204, EPI_ISL_666207, EPI_ISL_666214, EPI_ISL_666218, EPI_ISL_666221, EPI_ISL_666223, EPI_ISL_666227, EPI_ISL_666229, EPI_ISL_666235, EPI_ISL_666253, EPI_ISL_666261, EPI_ISL_666264, EPI_ISL_666265, EPI_ISL_666280, EPI_ISL_666283, EPI_ISL_666291, EPI_ISL_666292, EPI_ISL_666307, EPI_ISL_666321, EPI_ISL_666327, EPI_ISL_666338, EPI_ISL_666343, EPI_ISL_666344, EPI_ISL_666354, EPI_ISL_666358, EPI_ISL_666360, EPI_ISL_666367, EPI_ISL_666379, EPI_ISL_666418, EPI_ISL_666430, EPI_ISL_666470, EPI_ISL_666480, EPI_ISL_666483, EPI_ISL_666509, EPI_ISL_666513, EPI_ISL_666525, EPI_ISL_666565, EPI_ISL_666581 | | | |
| see above | Wales Specialist Virology Centre Sequencing lab: Pathogen Genomics Unit | COVID-19 Genomics UK (COG-UK) Consortium | Catherine Moore, Johnathan Evans, Laura Gifford, Malorie Perry, Simon Cottrell, Angela Marchbank, Alec Birchley, Alexander Adams, Amy Gaskin, Bree Gatica-Wilcox, Jason Coombes, Joel Southgate, Lauren Gilbert, Lee Graham, Nicole Pacchiarini, Sara Kumziene-Summerhayes, Sarah Taylor, Sophie Jones, Sara Rey, Matthew Bull, Joanne Watkins, Sally Corden, Tom Connor |
| EPI_ISL_666726, EPI_ISL_666770 | Respiratory Virus Unit, Microbiology Services Colindale, Public Health England | COVID-19 Genomics UK (COG-UK) Consortium | PHE Covid Sequencing Team |
| EPI_ISL_666903, EPI_ISL_666913, EPI_ISL_666917, EPI_ISL_666920, EPI_ISL_666931, EPI_ISL_666940, EPI_ISL_666944, EPI_ISL_666948, EPI_ISL_666953, EPI_ISL_666958 | Michigan Department of Health and Human Services, Bureau of Laboratories | Michigan Department of Health and Human Services, Bureau of Laboratories | Blankenship HM, Riner D, Soehnlen MK |
| EPI_ISL_666966, EPI_ISL_666967, EPI_ISL_666971, EPI_ISL_666974 | San Diego County Public Health Laboratory | Andersen lab at Scripps Research | SEARCH Alliance San Diego with Tracy Basler, Jovan Shephard, Brett Austin |
| EPI_ISL_667516, EPI_ISL_667517, EPI_ISL_667526, EPI_ISL_667527, EPI_ISL_667528, EPI_ISL_667536 | OHSU Lab Services Molecular Microbiology Lab | Oregon SARS-CoV-2 Genome Sequencing Center | Brendan L. O'Connell, Ruth V. Nichols, Sally Grindstaff, Alec J. Hirsch, Donna Hansel, Guang Fan, Daniel N. Streblow, William B. Messer, Andrew C. Adey, Benjamin N. Bimber, Brian J. O'Roak |
| EPI_ISL_667694, EPI_ISL_667695, EPI_ISL_667696, EPI_ISL_667697, EPI_ISL_667698, EPI_ISL_667699, EPI_ISL_667700, EPI_ISL_667701, EPI_ISL_667702, EPI_ISL_667703, EPI_ISL_667704, EPI_ISL_667705, EPI_ISL_667706, EPI_ISL_667707, EPI_ISL_667708, EPI_ISL_667709, EPI_ISL_667710, EPI_ISL_667711, EPI_ISL_667712, EPI_ISL_667713, EPI_ISL_667714, EPI_ISL_667715, EPI_ISL_667716, EPI_ISL_667717, EPI_ISL_667718, EPI_ISL_667719, EPI_ISL_667720, EPI_ISL_667721, EPI_ISL_667722, EPI_ISL_667723, EPI_ISL_667724, EPI_ISL_667725, EPI_ISL_667726, EPI_ISL_667727, EPI_ISL_667728, EPI_ISL_667729, EPI_ISL_667730, EPI_ISL_667731, EPI_ISL_667732, EPI_ISL_667733, EPI_ISL_667734, EPI_ISL_667740, EPI_ISL_667741, EPI_ISL_667742, EPI_ISL_667743, EPI_ISL_667744, EPI_ISL_667745, EPI_ISL_667746, EPI_ISL_667747, EPI_ISL_667748, EPI_ISL_667749, EPI_ISL_667750, EPI_ISL_667751, EPI_ISL_667752, EPI_ISL_667753, EPI_ISL_667754 | | | |
| see above | Pathogen Genomics Center, National Institute of Infectious Diseases | Pathogen Genomics Center, National Institute of Infectious Diseases | Tsuyoshi Sekizuka, Kentaro Itokawa, Rina Tanaka, Masanori Hashino, Makoto Kuroda |
| EPI_ISL_667780, EPI_ISL_667788, EPI_ISL_667790, EPI_ISL_667793, EPI_ISL_667794, EPI_ISL_667795 | South Eastern Area Laboratory Services (SEALS) | NSW Health Pathology - Institute of Clinical Pathology and Medical Research; Westmead Hospital; University of Sydney | CIDM-PH et al. |
| EPI_ISL_671759 | DOHMH Fort Greene | New York City Public Health Laboratory | Jade Wang, et al. |
| EPI_ISL_671760 | DOHMH Crown Heights | New York City Public Health Laboratory | Jade Wang, et al. |
| EPI_ISL_671761 | DOHMH Corona | New York City Public Health Laboratory | Jade Wang, et al. |
| EPI_ISL_671762 | DOHMH Riverside | New York City Public Health Laboratory | Jade Wang, et al. |
| EPI_ISL_671763 | DOHMH Chelsea | New York City Public Health Laboratory | Jade Wang, et al. |
| EPI_ISL_671866 | National Virus Reference Laboratory | National Virus Reference Laboratory | Michael Carr, Gabriel Gonzalez, Jonathan Dean, Daniel Hare, Cillian F De Gascun |
| EPI_ISL_672072, EPI_ISL_672104, EPI_ISL_672106, EPI_ISL_672107, EPI_ISL_672108 | Santa Clara County Public Health Laboratory | Chan-Zuckerberg Biohub | CZB Cliahub Consortium |
| EPI_ISL_672428 | UCSF Clinical Microbiology Laboratory | Chan-Zuckerberg Biohub | CZB Cliahub Consortium |
| EPI_ISL_672506 | Madera County Department of Public Health | Chan-Zuckerberg Biohub | CZB Cliahub Consortium |
| EPI_ISL_672913, EPI_ISL_672914, EPI_ISL_672915, EPI_ISL_672920, EPI_ISL_672921, EPI_ISL_672928, EPI_ISL_672929, EPI_ISL_672930, EPI_ISL_672931, EPI_ISL_672932, EPI_ISL_672933, EPI_ISL_672936, EPI_ISL_672937, EPI_ISL_672938, EPI_ISL_672940, EPI_ISL_672941, EPI_ISL_672942, EPI_ISL_672946, EPI_ISL_672948, EPI_ISL_672950, EPI_ISL_672952, EPI_ISL_672953, EPI_ISL_672954, EPI_ISL_672955, EPI_ISL_672957, EPI_ISL_672958, EPI_ISL_672960, EPI_ISL_672962, EPI_ISL_672966, EPI_ISL_672967, EPI_ISL_672969, EPI_ISL_672970, EPI_ISL_672973, EPI_ISL_672975, EPI_ISL_672976, EPI_ISL_672979, | | | |

| | | | |
|---|---|---|---|
| EPI_ISL_672981, EPI_ISL_672982, EPI_ISL_672986, EPI_ISL_672988, EPI_ISL_672994, EPI_ISL_672995, EPI_ISL_672996, EPI_ISL_672997, EPI_ISL_673000, EPI_ISL_673002, EPI_ISL_673003, EPI_ISL_673008, EPI_ISL_673010, EPI_ISL_673013, EPI_ISL_673014, EPI_ISL_673020, EPI_ISL_673021, EPI_ISL_673024, EPI_ISL_673026, EPI_ISL_673030, EPI_ISL_673031, EPI_ISL_673033, EPI_ISL_673034, EPI_ISL_673035, EPI_ISL_673036, EPI_ISL_673038, EPI_ISL_673042, EPI_ISL_673043, EPI_ISL_673044, EPI_ISL_673045, EPI_ISL_673046, EPI_ISL_673047, EPI_ISL_673049, EPI_ISL_673052, EPI_ISL_673058, EPI_ISL_673060, EPI_ISL_673063, EPI_ISL_673064, EPI_ISL_673066, EPI_ISL_673068, EPI_ISL_673072, EPI_ISL_673079, EPI_ISL_673081, EPI_ISL_673084, EPI_ISL_673087, EPI_ISL_673088, EPI_ISL_673089, EPI_ISL_673091, EPI_ISL_673093, EPI_ISL_673094, EPI_ISL_673095, EPI_ISL_673096, EPI_ISL_673097, EPI_ISL_673107, EPI_ISL_673108, EPI_ISL_673109, EPI_ISL_673111, EPI_ISL_673112, EPI_ISL_673114, EPI_ISL_673117, EPI_ISL_673119, EPI_ISL_673120, EPI_ISL_673121, EPI_ISL_673123, EPI_ISL_673124, EPI_ISL_673127, EPI_ISL_673128, EPI_ISL_673129, EPI_ISL_673134, EPI_ISL_673135, EPI_ISL_673136, EPI_ISL_673139, EPI_ISL_673140, EPI_ISL_673142, EPI_ISL_673143, EPI_ISL_673144, EPI_ISL_673147, EPI_ISL_673148, EPI_ISL_673150, EPI_ISL_673151, EPI_ISL_673153, EPI_ISL_673154, EPI_ISL_673156, EPI_ISL_673158, EPI_ISL_673161, EPI_ISL_673162, EPI_ISL_673164, EPI_ISL_673166, EPI_ISL_673167, EPI_ISL_673168, EPI_ISL_673169, EPI_ISL_673170, EPI_ISL_673171, EPI_ISL_673173, EPI_ISL_673175, EPI_ISL_673180, EPI_ISL_673188, EPI_ISL_673193, EPI_ISL_673196, EPI_ISL_673200, EPI_ISL_673201, EPI_ISL_673202, EPI_ISL_673204, EPI_ISL_673206, EPI_ISL_673207, EPI_ISL_673208, EPI_ISL_673210, EPI_ISL_673213, EPI_ISL_673214, EPI_ISL_673215, EPI_ISL_673217, EPI_ISL_673218, EPI_ISL_673221, EPI_ISL_673224, EPI_ISL_673225, EPI_ISL_673226, EPI_ISL_673227, EPI_ISL_673229, EPI_ISL_673230, EPI_ISL_673231, EPI_ISL_673232, EPI_ISL_673235 | | | |
| see above | Lighthouse Lab in Cambridge | Wellcome Sanger Institute for the COVID-19 Genomics UK (COG-UK) Consortium | Rob Howes, The Lighthouse Lab in Cambridge and Alex Alderton, Roberto Amato, Sonia Goncalves, Ewan Harrison, David K. Jackson, Ian Johnston, Dominic Kwiatkowski, Cordelia Langford, John Sillitoe on behalf of the Wellcome Sanger Institute COVID-19 Surveillance Team |
| EPI_ISL_676434, EPI_ISL_676441, EPI_ISL_676442, EPI_ISL_676443, EPI_ISL_676445, EPI_ISL_676446, EPI_ISL_676447, EPI_ISL_676448, EPI_ISL_676449, EPI_ISL_676451, EPI_ISL_676452, EPI_ISL_676453, EPI_ISL_676454, EPI_ISL_676455, EPI_ISL_676456, EPI_ISL_676457, EPI_ISL_676458, EPI_ISL_676459, EPI_ISL_676460, EPI_ISL_676461, EPI_ISL_676462, EPI_ISL_676463, EPI_ISL_676464, EPI_ISL_676465, EPI_ISL_676466, EPI_ISL_676467, EPI_ISL_676468, EPI_ISL_676469, EPI_ISL_676470, EPI_ISL_676471, EPI_ISL_676472, EPI_ISL_676473, EPI_ISL_676474, EPI_ISL_676475, EPI_ISL_676476, EPI_ISL_676478, EPI_ISL_676479, EPI_ISL_676480, EPI_ISL_676481, EPI_ISL_676482, EPI_ISL_676483, EPI_ISL_676484, EPI_ISL_676485, EPI_ISL_676487, EPI_ISL_676488 | | | |
| see above | Lighthouse Lab in Glasgow | Wellcome Sanger Institute for the COVID-19 Genomics UK (COG-UK) Consortium | Harper VanSteenhouse, Yumi Kasai, David Gray, Carol Clugston, Anna Dominiczak and Alex Alderton, Roberto Amato, Sonia Goncalves, Ewan Harrison, David K. Jackson, Ian Johnston, Dominic Kwiatkowski, Cordelia Langford, John Sillitoe on behalf of the Wellcome Sanger Institute COVID-19 Surveillance Team |
| EPI_ISL_677319 | Colorado Department of Public Health and Environment | Colorado Department of Puplic Health and Environment | Laura Bankers, Molly Hetherington-Rauth, Shannon Ely, Shannon R. Matzinger, Sarah Elizabeth Totten, Emily A. Travanty |
| EPI_ISL_677330, EPI_ISL_677331, EPI_ISL_677332, EPI_ISL_677333, EPI_ISL_677334, EPI_ISL_677503, EPI_ISL_677504 | University of Wisconsin-Madison AIDS Vaccine Research Laboratories | University of Wisconsin-Madison AIDS Vaccine Research Laboratories | Gage Moreno, Katarina Braun, et al. AIDS Vaccine Research Laboratories |
| EPI_ISL_678375, EPI_ISL_678377 | Area of Virology, Serology and Virology Division (SAViD), New South Wales Health Pathology Randwick | Virology Research Laboratory; Area of Virology, Serology and Virology Division (SAViD), New South Wales Health Pathology Randwick | Foster, C.; Au, J.; Ruiz Silva, M.; Deveson, I.; Bull, R.; Van Hal, S.; Rawlinson, W. |
| EPI_ISL_678601, EPI_ISL_678602, EPI_ISL_678603 | NHLS-IALCH | KRISP, KZN Research Innovation and Sequencing Platform | Giandhari J, Pillay S, Lessells R, ChimukangaraB, Mdlalose K, York D, Khan S, Tegally H, Wilkinson E, de Oliveira T |
| EPI_ISL_678787, EPI_ISL_678788, EPI_ISL_678789, EPI_ISL_678790, EPI_ISL_678791, EPI_ISL_678792, EPI_ISL_678793, EPI_ISL_678794, EPI_ISL_678795, EPI_ISL_678796, EPI_ISL_678797, EPI_ISL_678798, EPI_ISL_678799, EPI_ISL_678800, EPI_ISL_678801, EPI_ISL_678802, EPI_ISL_678803, EPI_ISL_678804, EPI_ISL_678805, EPI_ISL_678806, EPI_ISL_678807, EPI_ISL_678808, EPI_ISL_678809, EPI_ISL_678810, EPI_ISL_678811, EPI_ISL_678812, EPI_ISL_678813, EPI_ISL_678814, EPI_ISL_678815, EPI_ISL_678816, EPI_ISL_678817, EPI_ISL_678818, EPI_ISL_678819, EPI_ISL_678820, EPI_ISL_678821, EPI_ISL_678822, EPI_ISL_678823, EPI_ISL_678824, EPI_ISL_678825, EPI_ISL_678826, EPI_ISL_678827 | | | |
| see above | Respiratory Virus Unit, Microbiology Services Colindale, Public Health England | COVID-19 Genomics UK (COG-UK) Consortium | PHE Covid Sequencing Team |
| EPI_ISL_679130, EPI_ISL_679131 | University of Birmingham | COVID-19 Genomics UK (COG-UK) Consortium | Institute of Microbiology, University of Birmingham: Claire McMurray, Joanne Stockton, Samuel Nicholls, Radoslaw Poplawski, Will Rowe, Josh Quick, Nicholas Loman. University of Birmingham Testing Laboratory: Celina M Whalley, Andrew Bosworth, Charlotte Poxon, Kasun Wanigasooriya, Oliver Pickles, Mike Kidd, Alex Richter, Andrew D Beggs PHE Heartlands Lab: Husam Osman, Andrew Bosworth. Queen Elizabeth Hospital: Anna Casey |
| EPI_ISL_679474, EPI_ISL_679475, EPI_ISL_679476, EPI_ISL_679479 | University College London, Great Ormond Street Hospital for Children NHS Foundation Trust, Imperial College Healthcare NHS Trust | COVID-19 Genomics UK (COG-UK) Consortium | Sergi Castellano, Rachel Williams, Mark Kristiansen, Paola Resende Silva, Sunando Roy, Tony Brooks, Helena Tutill, Paola Niola, Patricia Dyal, Charlotte Williams, Leysa Forrest, Yasmin Panchbhaya, Jacqueline Findlay, Samuel Weeks, Julianne Brown, Kathryn Harris, Paul Randell, James Price, Alison Holmes, Judith Breuer |
| EPI_ISL_679631, EPI_ISL_679634, EPI_ISL_679635, EPI_ISL_679636, EPI_ISL_679637, EPI_ISL_679638, EPI_ISL_679639, EPI_ISL_679640, EPI_ISL_679641, EPI_ISL_679646, EPI_ISL_679647, EPI_ISL_679649, EPI_ISL_679650, EPI_ISL_679654, EPI_ISL_679656, EPI_ISL_679657, EPI_ISL_679660, EPI_ISL_679661, EPI_ISL_679662, EPI_ISL_679663, EPI_ISL_679664, EPI_ISL_679665, EPI_ISL_679667, EPI_ISL_679668, EPI_ISL_679670, EPI_ISL_679672, EPI_ISL_679674, EPI_ISL_679675, EPI_ISL_679676, EPI_ISL_679677, EPI_ISL_679681, EPI_ISL_679686, EPI_ISL_679687, EPI_ISL_679689, EPI_ISL_679691, EPI_ISL_679701, EPI_ISL_679708, EPI_ISL_679710, EPI_ISL_679711, EPI_ISL_679713, EPI_ISL_679715, EPI_ISL_679722, EPI_ISL_679724, EPI_ISL_679725, EPI_ISL_679726, EPI_ISL_679729, EPI_ISL_679730, EPI_ISL_679731, EPI_ISL_679732, EPI_ISL_679733, EPI_ISL_679734, EPI_ISL_679735, EPI_ISL_679736, EPI_ISL_679737, EPI_ISL_679738, EPI_ISL_679739, EPI_ISL_679740, EPI_ISL_679741, EPI_ISL_679742, EPI_ISL_679743, EPI_ISL_679744, EPI_ISL_679745, EPI_ISL_679746, EPI_ISL_679747, EPI_ISL_679748, EPI_ISL_679749, EPI_ISL_679750, EPI_ISL_679751, EPI_ISL_679752, EPI_ISL_679753, EPI_ISL_679754, EPI_ISL_679755, EPI_ISL_679756, EPI_ISL_679757, EPI_ISL_679758, EPI_ISL_679759, EPI_ISL_679760, EPI_ISL_679761, EPI_ISL_679762, EPI_ISL_679763, EPI_ISL_679764, EPI_ISL_679765, EPI_ISL_679768, EPI_ISL_679769, EPI_ISL_679771, EPI_ISL_679772, EPI_ISL_679775, EPI_ISL_679776, EPI_ISL_679778, EPI_ISL_679779, EPI_ISL_679780, EPI_ISL_679781, EPI_ISL_679782, EPI_ISL_679786, EPI_ISL_679789, EPI_ISL_679790, EPI_ISL_679791, EPI_ISL_679792, EPI_ISL_679793, EPI_ISL_679794, EPI_ISL_679795, EPI_ISL_679796, EPI_ISL_679797, EPI_ISL_679798, EPI_ISL_679799, EPI_ISL_679800, EPI_ISL_679801, EPI_ISL_679802, EPI_ISL_679803, EPI_ISL_679804, EPI_ISL_679805, EPI_ISL_679807, EPI_ISL_679808, EPI_ISL_679809, EPI_ISL_679813, EPI_ISL_679815, EPI_ISL_679817, EPI_ISL_679820, EPI_ISL_679821, EPI_ISL_679822, EPI_ISL_679823, EPI_ISL_679824, EPI_ISL_679825, EPI_ISL_679826, EPI_ISL_679827, EPI_ISL_679828, EPI_ISL_679829, EPI_ISL_679830, EPI_ISL_679831, EPI_ISL_679832, EPI_ISL_679833, EPI_ISL_679834, EPI_ISL_679835, EPI_ISL_679836, EPI_ISL_679837, EPI_ISL_679838, EPI_ISL_679839, EPI_ISL_679840, EPI_ISL_679841, EPI_ISL_679842, EPI_ISL_679843, EPI_ISL_679844, EPI_ISL_679845, EPI_ISL_679846, EPI_ISL_679847, EPI_ISL_679848, EPI_ISL_679849, EPI_ISL_679850, EPI_ISL_679851 | | | |
| see above | Oxford Viromics, NDM, University of Oxford; Oxford University Hospitals; Basingstoke and North Hampshire Hospital | COVID-19 Genomics UK (COG-UK) Consortium | Tanya Golubchik, David Bonsall, George Macintyre, Amy Trebes, Mariateresa de Cesare, Catrin Moore, Alex Mobbs, Anita Justice, Robert Shaw, Monique Andersson, Timothy Peto, Emma Wise, Nathan Moore, Jessica Lynch, Nick Cortes, Matilde Mori, Stephen Kidd, David Buck, John Todd, Christophe Fraser |
| EPI_ISL_679927 | Queens Medical Centre, Clinical Microbiology Department / DeepSeq Nottingham | COVID-19 Genomics UK (COG-UK) Consortium | Gemma Clark, Wendy Smith, Manjinder Khakh, Vicki M Fleming, Michelle M Lister, Hannah Howson-Wells, Jonathan Ball, Patrick McClure, Joseph Chappell, Theocharis Tsoleridis, Nadine Holmes, Matthew Carlisle, Christopher Moore, Fei Sang, Johnny Debebe, Victoria Wright, Matthew Loose |
| EPI_ISL_680071 | Department of Pathology, University of Cambridge | COVID-19 Genomics UK (COG-UK) Consortium | Aminu S. Jahun, Yasmin Chaudhry, Grant Hall, Iliana Georgana, Myra Hosmillo, Martin D. Curran, Malte Pinckert, Surendra Parmar, Ian Goodfellow |
| EPI_ISL_680508, EPI_ISL_680509, EPI_ISL_680515, EPI_ISL_680516, EPI_ISL_680517, EPI_ISL_680518, EPI_ISL_680519, EPI_ISL_680520, EPI_ISL_680521, EPI_ISL_680524, EPI_ISL_680525, EPI_ISL_680526, EPI_ISL_680527, EPI_ISL_680528, EPI_ISL_680529, EPI_ISL_680530, EPI_ISL_680531, EPI_ISL_680532, EPI_ISL_680533, EPI_ISL_680534, EPI_ISL_680535, EPI_ISL_680536, EPI_ISL_680537 | | | |
| see above | Virology Department, Royal Infirmary of Edinburgh, NHS Lothian / School of Biological Sciences, University of Edinburgh / Institute of Genetics and Molecular Medicine, University of Edinburgh | COVID-19 Genomics UK (COG-UK) Consortium | McHugh M, Dewar R, Rooke S, Gallagher M, Balcaza C, O'Toole Á, Scher E, Hill V, McCrone JT, Colquhoun R, Yu X, Jackson B, Rambaut A, Williams TC, Templeton K |
| EPI_ISL_680578 | Centre for Enzyme Innovation, University of Portsmouth / Translational Research Laboratory, Portsmouth Hospitals NHS Trust | COVID-19 Genomics UK (COG-UK) Consortium | Angela Beckett,Yann Bourgeois,Garry Scarlett,Sharon Glaysher,Scott Elliott,Kelly Bicknell,Robert Impey,Allyson Lloyd,Sarah Wyllie,Ethan Butcher,Anoop Chauhan,Samuel Robson |
| EPI_ISL_680579 | Oxford Viromics, NDM, University of Oxford; Oxford University Hospitals; Basingstoke and North Hampshire Hospital | COVID-19 Genomics UK (COG-UK) Consortium | Tanya Golubchik, David Bonsall, George Macintyre, Amy Trebes, Mariateresa de Cesare, Catrin Moore, Alex Mobbs, Anita Justice, Robert Shaw, Monique Andersson, Timothy Peto, Emma Wise, Nathan Moore, Jessica Lynch, Nick Cortes, Matilde Mori, Stephen Kidd, David Buck, John Todd, Christophe Fraser |
| EPI_ISL_680582, EPI_ISL_680587, EPI_ISL_680622, EPI_ISL_680623, EPI_ISL_680624, EPI_ISL_680625, EPI_ISL_680626, EPI_ISL_680627, EPI_ISL_680628, EPI_ISL_680629, EPI_ISL_680630, EPI_ISL_680631, EPI_ISL_680632, EPI_ISL_680633, EPI_ISL_680684, EPI_ISL_680714, EPI_ISL_680715, EPI_ISL_680716, EPI_ISL_680717, EPI_ISL_680718, EPI_ISL_680719, EPI_ISL_680720, EPI_ISL_680721, EPI_ISL_680722, EPI_ISL_680723, EPI_ISL_680724, EPI_ISL_680725, EPI_ISL_680726, EPI_ISL_680727, EPI_ISL_680728, EPI_ISL_680729, EPI_ISL_680730, EPI_ISL_680731 | | | |
| see above | Wales Specialist Virology Centre Sequencing lab: Pathogen Genomics Unit | COVID-19 Genomics UK (COG-UK) Consortium | Catherine Moore, Johnathan Evans, Laura Gifford, Malorie Perry, Simon Cottrell, Angela Marchbank, Alec Birchley, Alexander Adams, Amy Gaskin, Bree Gatica-Wilcox, Jason Coombes, Joel Southgate, Lauren Gilbert, Lee Graham, Nicole Pacchiarini, Sara Kumziene-Summerhayes, Sarah Taylor, Sophie Jones, Sara Rey, Matthew Bull, Joanne Watkins, Sally Corden, Tom Connor |
| EPI_ISL_681308, EPI_ISL_681311 | Communicable Disease Laboratory, Public Health Directorate | Communicable Disease Laboratory, Public Health Directorate | Alwasti,H., Altaif,Z., AlHujairi,Z., AlAbbas,Z. |
| EPI_ISL_681323, EPI_ISL_681324 | Respiratory Virus Unit, Microbiology Services Colindale, Public Health England | COVID-19 Genomics UK (COG-UK) Consortium | PHE Covid Sequencing Team |
| EPI_ISL_681657 | Lighthouse Lab in Cambridge | Wellcome Sanger Institute for the COVID-19 Genomics UK (COG-UK) Consortium | Rob Howes, The Lighthouse Lab in Cambridge and Alex Alderton, Roberto Amato, Sonia Goncalves, Ewan Harrison, David K. Jackson, Ian Johnston, Dominic Kwiatkowski, Cordelia Langford, John Sillitoe on behalf of the Wellcome Sanger Institute COVID-19 Surveillance Team |
| EPI_ISL_682080, EPI_ISL_682081, EPI_ISL_682094, EPI_ISL_682095, EPI_ISL_682100, EPI_ISL_682103, EPI_ISL_682104, EPI_ISL_682106, EPI_ISL_682107, EPI_ISL_682109, EPI_ISL_682110, EPI_ISL_682113, EPI_ISL_682116, EPI_ISL_682117, EPI_ISL_682118, EPI_ISL_682121, EPI_ISL_682122, EPI_ISL_682123, EPI_ISL_682127, EPI_ISL_682128, EPI_ISL_682131, EPI_ISL_682132, EPI_ISL_682134, EPI_ISL_682135, EPI_ISL_682136, EPI_ISL_682137, EPI_ISL_682138, EPI_ISL_682139, EPI_ISL_682140, EPI_ISL_682141, EPI_ISL_682142, EPI_ISL_682143, EPI_ISL_682144, EPI_ISL_682145, EPI_ISL_682146, EPI_ISL_682147, EPI_ISL_682148, EPI_ISL_682149, EPI_ISL_682150, EPI_ISL_682151, EPI_ISL_682152, EPI_ISL_682153, EPI_ISL_682154, EPI_ISL_682155, EPI_ISL_682156, EPI_ISL_682157, EPI_ISL_682158, EPI_ISL_682159, EPI_ISL_682160 | | | |

| | | | |
|---|---|---|---|
| see above | University of Michigan Clinical Microbiology Laboratory | Lauring Lab, University of Michigan, Department of Microbiology and Immunology | Valesano |
| EPI_ISL_682278 | Middlemore Hospital | Institute of Environmental Science and Research (ESR) | Xiaoyun Ren, Matt Storey, Nikki Freed, Muhammad Faisal, Jing Wang, Hermes Perez, Anja Werno, Antje van der Linden, Arlo Upton, Chris Mansell, David Hammer, Dragana Drinkovic, Gary McAuliffe, Hana Sofia Andersson, James Ussher, Jill Sherwood, Josh Freeman, Julia Howard, Juliet Elvy, Mary DeAlmeida, Matt Blakiston, Matthew Rogers, Max Bloomfield, Michael Addidle, Michelle Balm, Sally Roberts, Sarah Jefferies, Sharmini Muttaiyah, Susan Morpeth, Susan Taylor, Timothy Blackmore, Vani Sathyendran, Veronica Playle, Virginia Hope, Erasmus Smit, Lauren Jelly, Olin Silander, Joep de Ligt |
| EPI_ISL_682299, EPI_ISL_682302 | Communicable Disease Laboratory, Public Health Directorate | Communicable Disease Laboratory, Public Health Directorate | Alwasti,H., Altaif,Z., AlHujairi,Z., AlAbbas,Z. |
| EPI_ISL_683087, EPI_ISL_683088, EPI_ISL_683089, EPI_ISL_683090, EPI_ISL_683091 | Department of Virus and Microbiological Special Diagnostics, Statens Serum Institut, Copenhagen, Denmark | Albertsen Lab, Department of Chemistry and Bioscience, Aalborg University, Denmark | Danish Covid-19 Genome Consortium |
| EPI_ISL_683400, EPI_ISL_683401 | CNR Virus des Infections Respiratoires - France SUD | CNR Virus des Infections Respiratoires - France SUD | Antonin Bal, Gregory Destras, Gwendolyne Burfin, Quentin Semanas, Martine Valette, Bruno Lina, Laurence Josset |
| EPI_ISL_683646 | Servicio de Microbiología, Laboratori Clínic Metropolitana Nord. Hospital Universitari Germans Trias i Pujol. Institut d'Investigació en Ciències de la Salut Germans Trias i Pujol (IGTP) | SeqCOVID-SPAIN consortium/IBV(CSIC) | Elisa Martró, Antoni E. Bordoy, Anna Not, Adrián Antuori, Anabel Fernández, Nona Romaní, Verónica Saludes, Cristina Casañ and SeqCOVID-SPAIN consortium |
| EPI_ISL_683696, EPI_ISL_683702, EPI_ISL_683703, EPI_ISL_683705, EPI_ISL_683706, EPI_ISL_683707, EPI_ISL_683708, EPI_ISL_683710, EPI_ISL_683712, EPI_ISL_683713 | Essentia Health-St. Mary's Medical Center | Minnesota Department of Health, Public Health Laboratory | Alexandra Lorentz, Jacob Garfin, Matt Plumb, and Xiong Wang |
| EPI_ISL_683716, EPI_ISL_683717, EPI_ISL_683718 | Mayo Clinic & Mayo Clinic Laboratories | Minnesota Department of Health, Public Health Laboratory | Alexandra Lorentz, Jacob Garfin, Matt Plumb, and Xiong Wang |
| EPI_ISL_683744, EPI_ISL_683745, EPI_ISL_683746, EPI_ISL_683747, EPI_ISL_683748, EPI_ISL_683749, EPI_ISL_683750 | Minnesota Department of Health, Public Health Laboratory | Minnesota Department of Health, Public Health Laboratory | Alexandra Lorentz, Jacob Garfin, Matt Plumb, and Xiong Wang |
| EPI_ISL_683772, EPI_ISL_683884, EPI_ISL_683886 | DOHMH Morrisania | New York City Public Health Laboratory | Jade Wang, et al. |
| EPI_ISL_683937 | DOHMH Central Harlem | New York City Public Health Laboratory | Jade Wang, et al. |
| EPI_ISL_683938 | DOHMH Morrisania | New York City Public Health Laboratory | Jade Wang, et al. |
| EPI_ISL_692809, EPI_ISL_692810, EPI_ISL_692811, EPI_ISL_692812, EPI_ISL_692814, EPI_ISL_692817, EPI_ISL_692818, EPI_ISL_692819 | Massachusetts State Public Health Laboratory | Massachusetts State Public Health Laboratory | Andrew Lang, Timelia Fink, Glen Gallagher, Sandra Smole |
| EPI_ISL_693269, EPI_ISL_693297 | unknown | Public Health Virology Laboratory, Forensic and Scientific Services (PHV-FSS) | Son Nguyen et al. |
| EPI_ISL_693667, EPI_ISL_693671, EPI_ISL_693676, EPI_ISL_693680 | The National Institute of Public Health | State Veterinary Institute Prague | Nagy,A;Jirincova,H;Trnka,D;Vecerova,J |
| EPI_ISL_693768 | General practitioner | National Reference Center for Viruses of Respiratory Infections, Institut Pasteur, Paris | Marion Barbet, Sylvie Behillil, Méline Bizard, Angela Brisebarre, Camille Capel, Etienne Simon-Lorière, Vincent Enouf, Maud Vanpeene, Sylvie van der Werf |
| EPI_ISL_693844, EPI_ISL_693845, EPI_ISL_693846, EPI_ISL_693847, EPI_ISL_693848, EPI_ISL_693849, EPI_ISL_693850, EPI_ISL_693851, EPI_ISL_693852, EPI_ISL_693853, EPI_ISL_693854, EPI_ISL_693855, EPI_ISL_693856, EPI_ISL_693857, EPI_ISL_693858, EPI_ISL_693859 | | | |
| see above | Viollier AG | Department of Biosystems Science and Engineering, ETH Zürich | Christian Beisel, Sarah Nadeau, Chaoran Chen, Ivan Topolsky, Pedro Ferreira, Philipp Jablonski, Susana Posada-Céspedes, Tobias Schär, Ina Nissen, Natascha Santacroce, Elodie Burcklen, Christiane Beckmann, Maurice Redondo, Olivier Kobel, Christoph Noppen, Sophie Seidel, Noemie Santamaria de Souza, Niko Beerenwinkel, Tanja Stadler |
| EPI_ISL_700428 | Thembalethu CDC wc THC | NHLS/UCT | Houriiyah Tegally, Arash Iranzadeh, Deelan Doolabh, Lynn Tyers, Bruna Galvao, Innocent Mudau, Marvin Hsiao, Kruger Marais, Diana Hardie, Stephen Korsman, Carolyn Williamson |
| EPI_ISL_702820 | Northumbria University / South Tees Hospitals NHS Foundation Trust / North Cumbria Integrated Care NHS Foundation Trust / North Tees and Hartlepool NHS Foundation Trust / Newcastle Hospitals NHS Foundation Trust | COVID-19 Genomics UK (COG-UK) Consortium | Darren L Smith,Andrew Nelson,Matthew Bashton,Greg R Young,Joshua Loh,John Allan,Mohammad A Tariq,Giles S Holt,Gary Black,Wen C Yew,Lynn Dover,Paul Baker,Steve Liggett,Sarah Essex,Jane Greenaway,Debra Padgett,Clive Graham,Garren Scott,Edward Barton,Emma Swindells,Brendan Payne,Jennifer Collins,Yusri Taha,Gary Eltringham |
| EPI_ISL_702896 | Wales Specialist Virology Centre Sequencing lab: Pathogen Genomics Unit | COVID-19 Genomics UK (COG-UK) Consortium | Catherine Moore, Johnathan Evans, Laura Gifford, Malorie Perry, Simon Cottrell, Angela Marchbank, Alec Birchley, Alexander Adams, Amy Gaskin, Bree Gatica-Wilcox, Jason Coombes, Joel Southgate, Lauren Gilbert, Lee Graham, Nicole Pacchiarini, Sara Kumziene-Summerhayes, Sarah Taylor, Sophie Jones, Sara Rey, Matthew Bull, Joanne Watkins, Sally Corden, Tom Connor |
| EPI_ISL_703002 | Northumbria University / South Tees Hospitals NHS Foundation Trust / North Cumbria Integrated Care NHS Foundation Trust / North Tees and Hartlepool NHS Foundation Trust / Newcastle Hospitals NHS Foundation Trust | COVID-19 Genomics UK (COG-UK) Consortium | Darren L Smith,Andrew Nelson,Matthew Bashton,Greg R Young,Joshua Loh,John Allan,Mohammad A Tariq,Giles S Holt,Gary Black,Wen C Yew,Lynn Dover,Paul Baker,Steve Liggett,Sarah Essex,Jane Greenaway,Debra Padgett,Clive Graham,Garren Scott,Edward Barton,Emma Swindells,Brendan Payne,Jennifer Collins,Yusri Taha,Gary Eltringham |
| EPI_ISL_703642 | Wales Specialist Virology Centre Sequencing lab: Pathogen Genomics Unit | COVID-19 Genomics UK (COG-UK) Consortium | Catherine Moore, Johnathan Evans, Laura Gifford, Malorie Perry, Simon Cottrell, Angela Marchbank, Alec Birchley, Alexander Adams, Amy Gaskin, Bree Gatica-Wilcox, Jason Coombes, Joel Southgate, Lauren Gilbert, Lee Graham, Nicole Pacchiarini, Sara Kumziene-Summerhayes, Sarah Taylor, Sophie Jones, Sara Rey, Matthew Bull, Joanne Watkins, Sally Corden, Tom Connor |
| EPI_ISL_703863 | Virology Department, Royal Infirmary of Edinburgh, NHS Lothian / School of Biological Sciences, University of Edinburgh / Institute of Genetics and Molecular Medicine, University of Edinburgh | COVID-19 Genomics UK (COG-UK) Consortium | McHugh M, Dewar R, Rooke S, Gallagher M, Balcaza C, O'Toole Á, Scher E, Hill V, McCrone JT, Colquhoun R, Yu X, Jackson B, Rambaut A, Williams TC, Templeton K |
| EPI_ISL_704017 | Wales Specialist Virology Centre Sequencing lab: Pathogen Genomics Unit | COVID-19 Genomics UK (COG-UK) Consortium | Catherine Moore, Johnathan Evans, Laura Gifford, Malorie Perry, Simon Cottrell, Angela Marchbank, Alec Birchley, Alexander Adams, Amy Gaskin, Bree Gatica-Wilcox, Jason Coombes, Joel Southgate, Lauren Gilbert, Lee Graham, Nicole Pacchiarini, Sara Kumziene-Summerhayes, Sarah Taylor, Sophie Jones, Sara Rey, Matthew Bull, Joanne Watkins, Sally Corden, Tom Connor |
| EPI_ISL_704089 | West of Scotland Specialist Virology Centre, NHSGGC / MRC-University of Glasgow Centre for Virus Research | COVID-19 Genomics UK (COG-UK) Consortium | Ana da Silva Filipe, Natasha Johnson, Kathy Smollett, Daniel Mair, Stephen Carmichael, Alice Broos, Lily Tong, Jenna Nichols, Kyriaki Nomikou; Sarah McDonald; Richard Orton, Joseph Hughes, Sreenu Vattipally, David L Robertson; Alasdair MacLean, Rory Gunson; Sharif Shaaban, Matthew Holden; Rachel Blacow, Guy Mollett, Kathy Li, James Shepherd, Antonia Ho, Emma Thomson |
| EPI_ISL_704166, EPI_ISL_704579 | Northumbria University / South Tees Hospitals NHS Foundation Trust / North Cumbria Integrated Care NHS Foundation Trust / North Tees and Hartlepool NHS Foundation Trust / Newcastle Hospitals NHS Foundation Trust | COVID-19 Genomics UK (COG-UK) Consortium | Darren L Smith,Andrew Nelson,Matthew Bashton,Greg R Young,Joshua Loh,John Allan,Mohammad A Tariq,Giles S Holt,Gary Black,Wen C Yew,Lynn Dover,Paul Baker,Steve Liggett,Sarah Essex,Jane Greenaway,Debra Padgett,Clive Graham,Garren Scott,Edward Barton,Emma Swindells,Brendan Payne,Jennifer Collins,Yusri Taha,Gary Eltringham |

| | | | |
|---|---|---|---|
| EPI_ISL_704682, EPI_ISL_704685, EPI_ISL_704687, EPI_ISL_704690, EPI_ISL_704698, EPI_ISL_704701, EPI_ISL_704703, EPI_ISL_704817, EPI_ISL_704839, EPI_ISL_704861, EPI_ISL_704864, EPI_ISL_704872, EPI_ISL_704874, EPI_ISL_704877, EPI_ISL_704882 | | | |
| see above | Oxford Viromics, NDM, University of Oxford; Oxford University Hospitals; Basingstoke and North Hampshire Hospital | COVID-19 Genomics UK (COG-UK) Consortium | Tanya Golubchik, David Bonsall, George Macintyre, Amy Trebes, Mariateresa de Cesare, Catrin Moore, Alex Mobbs, Anita Justice, Robert Shaw, Monique Andersson, Timothy Peto, Emma Wise, Nathan Moore, Jessica Lynch, Nick Cortes, Matilde Mori, Stephen Kidd, David Buck, John Todd, Christophe Fraser |
| EPI_ISL_704952 | West of Scotland Specialist Virology Centre, NHSGGC / MRC-University of Glasgow Centre for Virus Research | COVID-19 Genomics UK (COG-UK) Consortium | Ana da Silva Filipe, Natasha Johnson, Kathy Smollett, Daniel Mair, Stephen Carmichael, Alice Broos, Lily Tong, Jenna Nichols, Kyriaki Nomikou; Sarah McDonald; Richard Orton, Joseph Hughes, Sreenu Vattipally, David L Robertson; Alasdair MacLean, Rory Gunson; Sharif Shaaban, Matthew Holden; Rachel Blacow, Guy Mollett, Kathy Li, James Shepherd, Antonia Ho, Emma Thomson |
| EPI_ISL_705105, EPI_ISL_705219 | Northumbria University / South Tees Hospitals NHS Foundation Trust / North Cumbria Integrated Care NHS Foundation Trust / North Tees and Hartlepool NHS Foundation Trust / Newcastle Hospitals NHS Foundation Trust | COVID-19 Genomics UK (COG-UK) Consortium | Darren L Smith,Andrew Nelson,Matthew Bashton,Greg R Young,Joshua Loh,John Allan,Mohammad A Tariq,Giles S Holt,Gary Black,Wen C Yew,Lynn Dover,Paul Baker,Steve Liggett,Sarah Essex,Jane Greenaway,Debra Padgett,Clive Graham,Garren Scott,Edward Barton,Emma Swindells,Brendan Payne,Jennifer Collins,Yusri Taha,Gary Eltringham |
| EPI_ISL_705265 | Wales Specialist Virology Centre Sequencing lab: Pathogen Genomics Unit | COVID-19 Genomics UK (COG-UK) Consortium | Catherine Moore, Johnathan Evans, Laura Gifford, Malorie Perry, Simon Cottrell, Angela Marchbank, Alec Birchley, Alexander Adams, Amy Gaskin, Bree Gatica-Wilcox, Jason Coombes, Joel Southgate, Lauren Gilbert, Lee Graham, Nicole Pacchiarini, Sara Kumziene-Summerhayes, Sarah Taylor, Sophie Jones, Sara Rey, Matthew Bull, Joanne Watkins, Sally Corden, Tom Connor |
| EPI_ISL_705293 | Northumbria University / South Tees Hospitals NHS Foundation Trust / North Cumbria Integrated Care NHS Foundation Trust / North Tees and Hartlepool NHS Foundation Trust / Newcastle Hospitals NHS Foundation Trust | COVID-19 Genomics UK (COG-UK) Consortium | Darren L Smith,Andrew Nelson,Matthew Bashton,Greg R Young,Joshua Loh,John Allan,Mohammad A Tariq,Giles S Holt,Gary Black,Wen C Yew,Lynn Dover,Paul Baker,Steve Liggett,Sarah Essex,Jane Greenaway,Debra Padgett,Clive Graham,Garren Scott,Edward Barton,Emma Swindells,Brendan Payne,Jennifer Collins,Yusri Taha,Gary Eltringham |
| EPI_ISL_705316 | West of Scotland Specialist Virology Centre, NHSGGC / MRC-University of Glasgow Centre for Virus Research | COVID-19 Genomics UK (COG-UK) Consortium | Ana da Silva Filipe, Natasha Johnson, Kathy Smollett, Daniel Mair, Stephen Carmichael, Alice Broos, Lily Tong, Jenna Nichols, Kyriaki Nomikou; Sarah McDonald; Richard Orton, Joseph Hughes, Sreenu Vattipally, David L Robertson; Alasdair MacLean, Rory Gunson; Sharif Shaaban, Matthew Holden; Rachel Blacow, Guy Mollett, Kathy Li, James Shepherd, Antonia Ho, Emma Thomson |
| EPI_ISL_705390 | Wales Specialist Virology Centre Sequencing lab: Pathogen Genomics Unit | COVID-19 Genomics UK (COG-UK) Consortium | Catherine Moore, Johnathan Evans, Laura Gifford, Malorie Perry, Simon Cottrell, Angela Marchbank, Alec Birchley, Alexander Adams, Amy Gaskin, Bree Gatica-Wilcox, Jason Coombes, Joel Southgate, Lauren Gilbert, Lee Graham, Nicole Pacchiarini, Sara Kumziene-Summerhayes, Sarah Taylor, Sophie Jones, Sara Rey, Matthew Bull, Joanne Watkins, Sally Corden, Tom Connor |
| EPI_ISL_705426, EPI_ISL_705427, EPI_ISL_705428, EPI_ISL_705450, EPI_ISL_705451, EPI_ISL_705452, EPI_ISL_705455 | Oxford Viromics, NDM, University of Oxford; Oxford University Hospitals; Basingstoke and North Hampshire Hospital | COVID-19 Genomics UK (COG-UK) Consortium | Tanya Golubchik, David Bonsall, George Macintyre, Amy Trebes, Mariateresa de Cesare, Catrin Moore, Alex Mobbs, Anita Justice, Robert Shaw, Monique Andersson, Timothy Peto, Emma Wise, Nathan Moore, Jessica Lynch, Nick Cortes, Matilde Mori, Stephen Kidd, David Buck, John Todd, Christophe Fraser |
| EPI_ISL_705462 | Wales Specialist Virology Centre Sequencing lab: Pathogen Genomics Unit | COVID-19 Genomics UK (COG-UK) Consortium | Catherine Moore, Johnathan Evans, Laura Gifford, Malorie Perry, Simon Cottrell, Angela Marchbank, Alec Birchley, Alexander Adams, Amy Gaskin, Bree Gatica-Wilcox, Jason Coombes, Joel Southgate, Lauren Gilbert, Lee Graham, Nicole Pacchiarini, Sara Kumziene-Summerhayes, Sarah Taylor, Sophie Jones, Sara Rey, Matthew Bull, Joanne Watkins, Sally Corden, Tom Connor |
| EPI_ISL_705469, EPI_ISL_705475, EPI_ISL_705477, EPI_ISL_705478, EPI_ISL_705479, EPI_ISL_705480, EPI_ISL_705482, EPI_ISL_705483, EPI_ISL_705484, EPI_ISL_705485, EPI_ISL_705493, EPI_ISL_705526 | | | |
| see above | Oxford Viromics, NDM, University of Oxford; Oxford University Hospitals; Basingstoke and North Hampshire Hospital | COVID-19 Genomics UK (COG-UK) Consortium | Tanya Golubchik, David Bonsall, George Macintyre, Amy Trebes, Mariateresa de Cesare, Catrin Moore, Alex Mobbs, Anita Justice, Robert Shaw, Monique Andersson, Timothy Peto, Emma Wise, Nathan Moore, Jessica Lynch, Nick Cortes, Matilde Mori, Stephen Kidd, David Buck, John Todd, Christophe Fraser |
| EPI_ISL_705771, EPI_ISL_705798 | Virology Department, Royal Infirmary of Edinburgh, NHS Lothian / School of Biological Sciences, University of Edinburgh / Institute of Genetics and Molecular Medicine, University of Edinburgh | COVID-19 Genomics UK (COG-UK) Consortium | McHugh M, Dewar R, Rooke S, Gallagher M, Balcaza C, O'Toole Á, Scher E, Hill V, McCrone JT, Colquhoun R, Yu X, Jackson B, Rambaut A, Williams TC, Templeton K |
| EPI_ISL_706013, EPI_ISL_706014, EPI_ISL_706015, EPI_ISL_706016, EPI_ISL_706017, EPI_ISL_706018, EPI_ISL_706019, EPI_ISL_706020, EPI_ISL_706021, EPI_ISL_706022, EPI_ISL_706023, EPI_ISL_706024, EPI_ISL_706028, EPI_ISL_706029, EPI_ISL_706030, EPI_ISL_706031, EPI_ISL_706035, EPI_ISL_706036, EPI_ISL_706039, EPI_ISL_706040, EPI_ISL_706041, EPI_ISL_706042, EPI_ISL_706043, EPI_ISL_706044, EPI_ISL_706045, EPI_ISL_706046, EPI_ISL_706047, EPI_ISL_706048, EPI_ISL_706049, EPI_ISL_706050, EPI_ISL_706051, EPI_ISL_706052, EPI_ISL_706053, EPI_ISL_706054, EPI_ISL_706055, EPI_ISL_706056, EPI_ISL_706057, EPI_ISL_706061, EPI_ISL_706062, EPI_ISL_706064, EPI_ISL_706067, EPI_ISL_706071, EPI_ISL_706074, EPI_ISL_706075, EPI_ISL_706076, EPI_ISL_706077, EPI_ISL_706078, EPI_ISL_706079, EPI_ISL_706080, EPI_ISL_706082, EPI_ISL_706083, EPI_ISL_706084, EPI_ISL_706086, EPI_ISL_706092, EPI_ISL_706094, EPI_ISL_706099, EPI_ISL_706100, EPI_ISL_706103, EPI_ISL_706107, EPI_ISL_706108, EPI_ISL_706109, EPI_ISL_706111, EPI_ISL_706112, EPI_ISL_706113, EPI_ISL_706114, EPI_ISL_706115, EPI_ISL_706116, EPI_ISL_706117, EPI_ISL_706118, EPI_ISL_706119, EPI_ISL_706120, EPI_ISL_706121, EPI_ISL_706122, EPI_ISL_706123, EPI_ISL_706124, EPI_ISL_706125, EPI_ISL_706126, EPI_ISL_706127, EPI_ISL_706129, EPI_ISL_706130, EPI_ISL_706131, EPI_ISL_706132, EPI_ISL_706133, EPI_ISL_706135, EPI_ISL_706137, EPI_ISL_706138, EPI_ISL_706139, EPI_ISL_706140, EPI_ISL_706141, EPI_ISL_706142, EPI_ISL_706143, EPI_ISL_706146, EPI_ISL_706148, EPI_ISL_706151, EPI_ISL_706152, EPI_ISL_706153, EPI_ISL_706154, EPI_ISL_706155, EPI_ISL_706157, EPI_ISL_706159, EPI_ISL_706161, EPI_ISL_706162, EPI_ISL_706164, EPI_ISL_706165, EPI_ISL_706167, EPI_ISL_706171, EPI_ISL_706172, EPI_ISL_706173, EPI_ISL_706174, EPI_ISL_706175, EPI_ISL_706176, EPI_ISL_706177, EPI_ISL_706178, EPI_ISL_706179, EPI_ISL_706180, EPI_ISL_706181, EPI_ISL_706182, EPI_ISL_706183, EPI_ISL_706184, EPI_ISL_706186, EPI_ISL_706187, EPI_ISL_706188, EPI_ISL_706189, EPI_ISL_706190, EPI_ISL_706191, EPI_ISL_706192, EPI_ISL_706193, EPI_ISL_706195, EPI_ISL_706196, EPI_ISL_706197, EPI_ISL_706198, EPI_ISL_706199, EPI_ISL_706200, EPI_ISL_706215, EPI_ISL_706217, EPI_ISL_706219, EPI_ISL_706220, EPI_ISL_706221, EPI_ISL_706224, EPI_ISL_706225, EPI_ISL_706226, EPI_ISL_706227, EPI_ISL_706228, EPI_ISL_706231, EPI_ISL_706232, EPI_ISL_706233, EPI_ISL_706234, EPI_ISL_706235, EPI_ISL_706236, EPI_ISL_706237, EPI_ISL_706238, EPI_ISL_706242, EPI_ISL_706245, EPI_ISL_706246, EPI_ISL_706248, EPI_ISL_706249, EPI_ISL_706250, EPI_ISL_706251, EPI_ISL_706252, EPI_ISL_706253, EPI_ISL_706254, EPI_ISL_706255, EPI_ISL_706256, EPI_ISL_706257, EPI_ISL_706258, EPI_ISL_706259, EPI_ISL_706260, EPI_ISL_706261, EPI_ISL_706262, EPI_ISL_706263, EPI_ISL_706264, EPI_ISL_706265, EPI_ISL_706266, EPI_ISL_706267, EPI_ISL_706268 | | | |
| see above | Oxford Viromics, NDM, University of Oxford; Oxford University Hospitals; Basingstoke and North Hampshire Hospital | COVID-19 Genomics UK (COG-UK) Consortium | Tanya Golubchik, David Bonsall, George Macintyre, Amy Trebes, Mariateresa de Cesare, Catrin Moore, Alex Mobbs, Anita Justice, Robert Shaw, Monique Andersson, Timothy Peto, Emma Wise, Nathan Moore, Jessica Lynch, Nick Cortes, Matilde Mori, Stephen Kidd, David Buck, John Todd, Christophe Fraser |
| EPI_ISL_706276, EPI_ISL_706353 | Northumbria University / South Tees Hospitals NHS Foundation Trust / North Cumbria Integrated Care NHS Foundation Trust / North Tees and Hartlepool NHS Foundation Trust / Newcastle Hospitals NHS Foundation Trust | COVID-19 Genomics UK (COG-UK) Consortium | Darren L Smith,Andrew Nelson,Matthew Bashton,Greg R Young,Joshua Loh,John Allan,Mohammad A Tariq,Giles S Holt,Gary Black,Wen C Yew,Lynn Dover,Paul Baker,Steve Liggett,Sarah Essex,Jane Greenaway,Debra Padgett,Clive Graham,Garren Scott,Edward Barton,Emma Swindells,Brendan Payne,Jennifer Collins,Yusri Taha,Gary Eltringham |
| EPI_ISL_706473, EPI_ISL_706474, EPI_ISL_706475, EPI_ISL_706476, EPI_ISL_706477, EPI_ISL_706478, EPI_ISL_706479, EPI_ISL_706480, EPI_ISL_706481, EPI_ISL_706482 | Wales Specialist Virology Centre Sequencing lab: Pathogen Genomics Unit | COVID-19 Genomics UK (COG-UK) Consortium | Catherine Moore, Johnathan Evans, Laura Gifford, Malorie Perry, Simon Cottrell, Angela Marchbank, Alec Birchley, Alexander Adams, Amy Gaskin, Bree Gatica-Wilcox, Jason Coombes, Joel Southgate, Lauren Gilbert, Lee Graham, Nicole Pacchiarini, Sara Kumziene-Summerhayes, Sarah Taylor, Sophie Jones, Sara Rey, Matthew Bull, Joanne Watkins, Sally Corden, Tom Connor |
| EPI_ISL_708039 | Innlandet Hospital Trust, Division Lillehammer, Department for Medical Microbiology | Norwegian Institute of Public Health, Department of Virology | Kathrine Stene-Johansen, Kamilla Heddeland Instefjord, Hilde Elshaug, Marie Paulsen Madsen, Rasmus Riis Kopperud, Hilde Vollan, Karoline Bragstad, Olav Hungnes |
| EPI_ISL_708050 | Norwegian Institute of Public Health, Department of Virology | Norwegian Institute of Public Health, Department of Virology | Kathrine Stene-Johansen, Kamilla Heddeland Instefjord, Hilde Elshaug, Marie Paulsen Madsen, Rasmus Riis Kopperud, Hilde Vollan, Karoline Bragstad, Olav Hungnes |
| EPI_ISL_708055, EPI_ISL_708056 | Hospital of Southern Norway - Kristiansand, Department of Medical Microbiology | Norwegian Institute of Public Health, Department of Virology | Kathrine Stene-Johansen, Kamilla Heddeland Instefjord, Hilde Elshaug, Marie Paulsen Madsen, Rasmus Riis Kopperud, Hilde Vollan, Karoline Bragstad, Olav Hungnes |
| EPI_ISL_708093 | University Hospital of Northern Norway, Department for Microbiology and Infectious Disease Control | Norwegian Institute of Public Health, Department of Virology | Kathrine Stene-Johansen, Kamilla Heddeland Instefjord, Hilde Elshaug, Marie Paulsen Madsen, Rasmus Riis Kopperud, Hilde Vollan, Karoline Bragstad, Olav Hungnes |
| EPI_ISL_708102, EPI_ISL_708103 | Nordland Hospital - Bodo, Laboratory Department, Molecular Biology Unit | Norwegian Institute of Public Health, Department of Virology | Kathrine Stene-Johansen, Kamilla Heddeland Instefjord, Hilde Elshaug, Marie Paulsen Madsen, Rasmus Riis Kopperud, Hilde Vollan, Karoline Bragstad, Olav Hungnes |
| EPI_ISL_708141 | Norwegian Institute of Public Health, Department of Virology | Norwegian Institute of Public Health, Department of Virology | Kathrine Stene-Johansen, Kamilla Heddeland Instefjord, Hilde Elshaug, Marie Paulsen Madsen, Rasmus Riis Kopperud, Hilde Vollan, Karoline Bragstad, Olav Hungnes |
| EPI_ISL_708243 | Michigan Department of Health and Human Services, Bureau of Laboratories | Michigan Department of Health and Human Services, Bureau of Laboratories | Blankenship HM, Riner D, Soehnlen MK |
| EPI_ISL_708315 | University Hospitals of Geneva, Laboratory of Virology | University Hospitals of Geneva, Laboratory of Virology | Cordey Samuel, Laubscher Florian |
| EPI_ISL_708341, EPI_ISL_708367 | Michigan Department of Health and Human Services, Bureau of Laboratories | Michigan Department of Health and Human Services, Bureau of Laboratories | Blankenship HM, Riner D, Soehnlen MK |

| | | | |
|---|---|---|---|
| EPI_ISL_708379, EPI_ISL_708380 | University Hospitals of Geneva, Laboratory of Virology | University Hospitals of Geneva, Laboratory of Virology | Cordey Samuel, Laubscher Florian |
| EPI_ISL_708450, EPI_ISL_708452 | Delaware Public Health Lab | Delaware Public Health Lab | Gregory Hovan |
| EPI_ISL_708460, EPI_ISL_708461 | Minnesota Department of Health, Public Health Laboratory | Minnesota Department of Health, Public Health Laboratory | Alexandra Lorentz, Jacob Garfin, Matt Plumb, and Xiong Wang |
| EPI_ISL_708550, EPI_ISL_708552, EPI_ISL_708582 | Michigan Department of Health and Human Services, Bureau of Laboratories | Michigan Department of Health and Human Services, Bureau of Laboratories | Blankenship HM, Riner D, Soehnlen MK |
| EPI_ISL_708747, EPI_ISL_708774 | PathWest Laboratory Medicine WA | PathWest Laboratory Medicine WA Microbial Surveillance Unit | PathWest Laboratory Medicine WA Microbial Surveillance Unit |
| EPI_ISL_708819 | Urban Institute for Disease Prevention and Control | National Institute of Health, Department of Medical Sciences, Ministry of Public Health, Thailand | Pilailuk Okada; Siripaporn Phuygun; Thanutsapa Thanadachakul; Sittiporn Parnmen; Pakorn Piromtong; Warawan Wongboot; Sunthareeya Waicharoen; Malinee Chittaganpitch |
| EPI_ISL_708823 | Vibharam Hospital | National Institute of Health, Department of Medical Sciences, Ministry of Public Health, Thailand | Pilailuk Okada; Siripaporn Phuygun; Thanutsapa Thanadachakul; Sittiporn Parnmen; Pakorn Piromtong; Warawan Wongboot; Sunthareeya Waicharoen; Malinee Chittaganpitch |
| EPI_ISL_709903, EPI_ISL_709904, EPI_ISL_709905 | Lighthouse Lab in Milton Keynes | Wellcome Sanger Institute for the COVID-19 Genomics UK (COG-UK) Consortium | The Lighthouse Lab in Milton Keynes and Alex Alderton, Roberto Amato, Sonia Goncalves, Ewan Harrison, David K. Jackson, Ian Johnston, Dominic Kwiatkowski, Cordelia Langford, John Sillitoe on behalf of the Wellcome Sanger Institute COVID-19 Surveillance Team |
| EPI_ISL_710039 | University of Michigan Clinical Microbiology Laboratory | Lauring Lab, University of Michigan, Department of Microbiology and Immunology | Valesano |
| EPI_ISL_710157, EPI_ISL_710241, EPI_ISL_710242, EPI_ISL_710243, EPI_ISL_710244, EPI_ISL_710246, EPI_ISL_710369, EPI_ISL_710370, EPI_ISL_710371, EPI_ISL_710372, EPI_ISL_710373 | | | |
| see above | Colorado Department of Public Health and Environment | Colorado Department of Puplic Health and Environment | Laura Bankers, Molly C. Hetherington-Rauth, Shannon Ely, Shannon R. Matzinger, Sarah Elizabeth Totten, Emily A. Travanty |
| EPI_ISL_710596, EPI_ISL_710598, EPI_ISL_710599, EPI_ISL_710600 | Klinisk mikrobiologi | The Public Health Agency of Sweden | Department of Microbiology, The Public Health Agency of Sweden |
| EPI_ISL_710907 | Lighthouse Lab in Milton Keynes | Wellcome Sanger Institute for the COVID-19 Genomics UK (COG-UK) Consortium | The Lighthouse Lab in Milton Keynes and Alex Alderton, Roberto Amato, Sonia Goncalves, Ewan Harrison, David K. Jackson, Ian Johnston, Dominic Kwiatkowski, Cordelia Langford, John Sillitoe on behalf of the Wellcome Sanger Institute COVID-19 Surveillance Team |
| EPI_ISL_710908, EPI_ISL_710909, EPI_ISL_710910, EPI_ISL_710911, EPI_ISL_710912, EPI_ISL_710913, EPI_ISL_710914, EPI_ISL_710915, EPI_ISL_710917, EPI_ISL_710918, EPI_ISL_710919, EPI_ISL_710920, EPI_ISL_710921, EPI_ISL_710922, EPI_ISL_710923, EPI_ISL_710925, EPI_ISL_710926, EPI_ISL_710927, EPI_ISL_710928, EPI_ISL_710929, EPI_ISL_710931, EPI_ISL_710932, EPI_ISL_710933, EPI_ISL_710934, EPI_ISL_710935, EPI_ISL_710936, EPI_ISL_710937, EPI_ISL_710938, EPI_ISL_710939, EPI_ISL_710940, EPI_ISL_710941, EPI_ISL_710942, EPI_ISL_710943, EPI_ISL_710944, EPI_ISL_710945, EPI_ISL_710946, EPI_ISL_710947, EPI_ISL_710948, EPI_ISL_710949, EPI_ISL_710950, EPI_ISL_710951, EPI_ISL_710952, EPI_ISL_710953, EPI_ISL_710954, EPI_ISL_710955, EPI_ISL_710956, EPI_ISL_710957, EPI_ISL_710958, EPI_ISL_710959, EPI_ISL_710960, EPI_ISL_710961, EPI_ISL_710962, EPI_ISL_710963, EPI_ISL_710964, EPI_ISL_710965, EPI_ISL_710966, EPI_ISL_710967, EPI_ISL_710968, EPI_ISL_710969, EPI_ISL_710970, EPI_ISL_710971, EPI_ISL_710972, EPI_ISL_710973, EPI_ISL_710974, EPI_ISL_710975, EPI_ISL_710976, EPI_ISL_710977, EPI_ISL_710978, EPI_ISL_710979, EPI_ISL_710980, EPI_ISL_710981, EPI_ISL_710982, EPI_ISL_710983, EPI_ISL_710984, EPI_ISL_710985, EPI_ISL_710986, EPI_ISL_710987, EPI_ISL_710988, EPI_ISL_710989, EPI_ISL_710990, EPI_ISL_710991, EPI_ISL_710992, EPI_ISL_710993, EPI_ISL_710994 | | | |
| see above | Lighthouse Lab in Alderley Park | Wellcome Sanger Institute for the COVID-19 Genomics UK (COG-UK) Consortium | Jacquelyn Wynn, Mairead Hyland, The Lighthouse Lab in Alderley Park and Alex Alderton, Roberto Amato, Sonia Goncalves, Ewan Harrison, David K. Jackson, Ian Johnston, Dominic Kwiatkowski, Cordelia Langford, John Sillitoe on behalf of the Wellcome Sanger Institute COVID-19 Surveillance Team |
| EPI_ISL_711014 | Lighthouse Lab in Glasgow | Wellcome Sanger Institute for the COVID-19 Genomics UK (COG-UK) Consortium | Harper VanSteenhouse, Yumi Kasai, David Gray, Carol Clugston, Anna Dominiczak and Alex Alderton, Roberto Amato, Sonia Goncalves, Ewan Harrison, David K. Jackson, Ian Johnston, Dominic Kwiatkowski, Cordelia Langford, John Sillitoe on behalf of the Wellcome Sanger Institute COVID-19 Surveillance Team |
| EPI_ISL_711025, EPI_ISL_711026 | Lighthouse Lab in Cambridge | Wellcome Sanger Institute for the COVID-19 Genomics UK (COG-UK) Consortium | Rob Howes, The Lighthouse Lab in Cambridge and Alex Alderton, Roberto Amato, Sonia Goncalves, Ewan Harrison, David K. Jackson, Ian Johnston, Dominic Kwiatkowski, Cordelia Langford, John Sillitoe on behalf of the Wellcome Sanger Institute COVID-19 Surveillance Team |
| EPI_ISL_711027, EPI_ISL_711028, EPI_ISL_711029, EPI_ISL_711030, EPI_ISL_711031, EPI_ISL_711032, EPI_ISL_711033, EPI_ISL_711034, EPI_ISL_711035, EPI_ISL_711036, EPI_ISL_711037 | | | |
| see above | Lighthouse Lab in Alderley Park | Wellcome Sanger Institute for the COVID-19 Genomics UK (COG-UK) Consortium | Jacquelyn Wynn, Mairead Hyland, The Lighthouse Lab in Alderley Park and Alex Alderton, Roberto Amato, Sonia Goncalves, Ewan Harrison, David K. Jackson, Ian Johnston, Dominic Kwiatkowski, Cordelia Langford, John Sillitoe on behalf of the Wellcome Sanger Institute COVID-19 Surveillance Team |
| EPI_ISL_711038 | Lighthouse Lab in Cambridge | Wellcome Sanger Institute for the COVID-19 Genomics UK (COG-UK) Consortium | Rob Howes, The Lighthouse Lab in Cambridge and Alex Alderton, Roberto Amato, Sonia Goncalves, Ewan Harrison, David K. Jackson, Ian Johnston, Dominic Kwiatkowski, Cordelia Langford, John Sillitoe on behalf of the Wellcome Sanger Institute COVID-19 Surveillance Team |
| EPI_ISL_711039, EPI_ISL_711040, EPI_ISL_711041, EPI_ISL_711042, EPI_ISL_711043, EPI_ISL_711044, EPI_ISL_711045, EPI_ISL_711046, EPI_ISL_711047 | Lighthouse Lab in Alderley Park | Wellcome Sanger Institute for the COVID-19 Genomics UK (COG-UK) Consortium | Jacquelyn Wynn, Mairead Hyland, The Lighthouse Lab in Alderley Park and Alex Alderton, Roberto Amato, Sonia Goncalves, Ewan Harrison, David K. Jackson, Ian Johnston, Dominic Kwiatkowski, Cordelia Langford, John Sillitoe on behalf of the Wellcome Sanger Institute COVID-19 Surveillance Team |
| EPI_ISL_712084, EPI_ISL_712085, EPI_ISL_712090 | Port Elizabeth Provincial Hospital, National Health Laboratory Services, Eastern Cape, South Africa | National Institute for Communicable Diseases of the National Health Laboratory Service | Mohale T, Ntuli N, Mahlangu B, Allam M, Ismail A, Bhiman JN |
| EPI_ISL_717831, EPI_ISL_717889, EPI_ISL_717940, EPI_ISL_717941 | Laboratorio de Virologia Molecular / UFRJ | Bioinformatics Laboratory / LNCC | Carolina M Voloch, Ronaldo da Silva F Jr, Luiz G P de Almeida, Cynthia C Cardoso, Otavio Bustrolini, Alexandra L Gerber, Ana Paula de C Guimarães, Diana Mariani, Andréa Cony Cavalcanti, Claudia dos Santos Rodrigues, Terezinha M P P Castiñeira, Amilcar Tanuri, Ana Tereza R de Vasconcelos |
| EPI_ISL_717994 | Lab voor klinische biologie | Onderzoeksgroep Virologie | Laurens Lambrechts, Nick Vereecke, Marthe Pauwels, Bruno Verhasselt, Linos Vandekerckhove, Hans Nauwynck, Sebastiaan Theuns |
| EPI_ISL_718060, EPI_ISL_718061, EPI_ISL_718062, EPI_ISL_718063, EPI_ISL_718064, EPI_ISL_718065, EPI_ISL_718066, EPI_ISL_718067, EPI_ISL_718068, EPI_ISL_718069, EPI_ISL_718070, EPI_ISL_718071, EPI_ISL_718072, EPI_ISL_718073, EPI_ISL_718074, EPI_ISL_718075, EPI_ISL_718076, EPI_ISL_718077, EPI_ISL_718078, EPI_ISL_718079, EPI_ISL_718080, EPI_ISL_718081, EPI_ISL_718082, EPI_ISL_718083 | | | |
| see above | ZOTZ KLIMAS MVZ Düsseldorf-Centrum GbR ÜBAG für Labormedizin, Genetik, Zytologie, Pathologie | Center of Medical Microbiology, Virology, and Hospital Hygiene, University of Duesseldorf | Maximilian Damagnez, Alexander Dilthey, Ashley-Jane Duplessis, Patrick Finzer, Katrin Hoffmann, Torsten Houwaart, Lisanna Hülse, Malte Kohns Vasconcelos, Marek Korencak, Nadine Lübke, Jessica Nicolai, Klaus Pfeffer, Daniel Strelow, Jörg Timm, Andreas Walker, Tobias Wienemann, Rainer Zotz |
| EPI_ISL_718167 | Borneo Medical Centre | Institute of Health and Community Medicine | David Perera, Ooi Mong How, Chua Hock Hin, Tonnii Sia Loong Loong, Wong Jyn Shan, Wong Kiing Aik, Chan Chia Jui |
| EPI_ISL_718171, EPI_ISL_718177 | Ministry of Health Hospitals | Institute of Health and Community Medicine | David Perera, Ooi Mong How, Chua Hock Hin, Tonnii Sia Loong Loong, Wong Jyn Shan, Wong Kiing Aik, Chan Chia Jui |
| EPI_ISL_718203, EPI_ISL_718205 | Borneo Medical Centre | Institute of Health and Community Medicine | David Perera, Ooi Mong How, Chua Hock Hin, Tonnii Sia Loong Loong, Wong Jyn Shan, Wong Kiing Aik, Chan Chia Jui |
| EPI_ISL_718208, EPI_ISL_718209 | Ministry of Health Hospitals | Institute of Health and Community Medicine | David Perera, Ooi Mong How, Chua Hock Hin, Tonnii Sia Loong Loong, Wong Jyn Shan, Wong Kiing Aik, Chan Chia Jui |
| EPI_ISL_718226, EPI_ISL_718227, EPI_ISL_718247 | Hospital | National Reference Center for Viruses of Respiratory Infections, Institut Pasteur, Paris | Marion Barbet, Sylvie Behillil, Méline Bizard, Angela Brisebarre, Camille Capel, Etienne Simon-Lorière, Vincent Enouf, Maud Vanpeene, Sylvie van der Werf, Gisèle Lagathu |
| EPI_ISL_718314, EPI_ISL_718315 | Institute for Medical Research, Infectious Disease Research Centre, National Institutes of Health, Ministry of Health Malaysia | Institute for Medical Research, Infectious Disease Research Centre, National Institutes of Health, Ministry of Health Malaysia | Suppiah J, Kamel K, Mohd-Zawawi Z, Thayan R |
| EPI_ISL_721535 | Lighthouse Lab in Glasgow | Wellcome Sanger Institute for the COVID-19 Genomics UK (COG-UK) Consortium | Harper VanSteenhouse, Yumi Kasai, David Gray, Carol Clugston, Anna Dominiczak and Alex Alderton, Roberto Amato, Sonia Goncalves, Ewan Harrison, David K. Jackson, Ian Johnston, Dominic Kwiatkowski, Cordelia Langford, John Sillitoe on behalf of the Wellcome Sanger Institute COVID-19 Surveillance Team |
| EPI_ISL_721569, EPI_ISL_721570, EPI_ISL_721571, EPI_ISL_721572, EPI_ISL_721574, EPI_ISL_721576, EPI_ISL_721577, EPI_ISL_721578, EPI_ISL_721579, EPI_ISL_721580, EPI_ISL_721581, EPI_ISL_721582, EPI_ISL_721583, EPI_ISL_721584, EPI_ISL_721585, EPI_ISL_721586, EPI_ISL_721587, EPI_ISL_721588, EPI_ISL_721589, EPI_ISL_721590, EPI_ISL_721591, EPI_ISL_721592, EPI_ISL_721593, EPI_ISL_721594, EPI_ISL_721595, EPI_ISL_721596, EPI_ISL_721597, EPI_ISL_721601, EPI_ISL_721602, EPI_ISL_721603, EPI_ISL_721604, EPI_ISL_721605, EPI_ISL_721606, EPI_ISL_721607, EPI_ISL_721608, EPI_ISL_721609, EPI_ISL_721610, EPI_ISL_721618, EPI_ISL_721619, EPI_ISL_721620, EPI_ISL_721621 | | | |
| see above | Pathogen Genomics Center, National Institute of Infectious Diseases | Pathogen Genomics Center, National Institute of Infectious Diseases | Tsuyoshi Sekizuka, Kentaro Itokawa, Rina Tanaka, Masanori Hashino, Makoto Kuroda |
| EPI_ISL_722374, EPI_ISL_722396, EPI_ISL_722405, EPI_ISL_722412, EPI_ISL_722571, EPI_ISL_722572, EPI_ISL_722573, EPI_ISL_722574, EPI_ISL_722575, EPI_ISL_722576, EPI_ISL_722577, EPI_ISL_722578, EPI_ISL_722579, EPI_ISL_722580, EPI_ISL_722581, EPI_ISL_722582, EPI_ISL_722583, EPI_ISL_722584, EPI_ISL_722585, EPI_ISL_722586, EPI_ISL_722587, EPI_ISL_722588, EPI_ISL_722589, EPI_ISL_722590, EPI_ISL_722597, EPI_ISL_722598, EPI_ISL_722599, EPI_ISL_722600, EPI_ISL_722601, EPI_ISL_722602, EPI_ISL_722603, EPI_ISL_722604, EPI_ISL_722605, EPI_ISL_722606, EPI_ISL_722607 | | | |

| | | | |
|---|---|---|---|
| see above | Dutch COVID-19 response team | Erasmus Medical Center | Bas Oude Munnink, Reina Sikkema, David Nieuwenhuijse, Irina Chestakova, Anne van der Linden, Marjan Boter, Emmanuelle Munger, Corine GeurtsvanKessel, Annemiek van der Eijk, Richard Molenkamp, Marion Koopmans, on behalf of the Dutch national COVID-19 response team. |
| EPI_ISL_723469 | Laboratoire Biolife | Laboratoire de Biotechnologie | Mouna Ouadghiri, Tarik Aanniz, Mohammed Walid Chemao Elfihri, Mohamed Chenaoui, Hanae Dakka, Afaf Alaoui, Otmane Touzani, Bouchra Belfquih, Lahecn Belyamani, Saaid Amzazi and Azeddine Ibrahimi |
| EPI_ISL_723518, EPI_ISL_723519 | Virginia Division of Consolidated Laboratory Services (DCLS) | Virginia Division of Consolidated Laboratory Services (DCLS) | Virginia DCLS |
| EPI_ISL_723521, EPI_ISL_723522, EPI_ISL_723523, EPI_ISL_723524, EPI_ISL_723527, EPI_ISL_723528, EPI_ISL_723529, EPI_ISL_723530, EPI_ISL_723531, EPI_ISL_723532, EPI_ISL_723533, EPI_ISL_723534, EPI_ISL_723535, EPI_ISL_723536, EPI_ISL_723537, EPI_ISL_723538, EPI_ISL_723540, EPI_ISL_723541, EPI_ISL_723542, EPI_ISL_723543, EPI_ISL_723544, EPI_ISL_723545, EPI_ISL_723546, EPI_ISL_723548, EPI_ISL_723549, EPI_ISL_723550, EPI_ISL_723551, EPI_ISL_723552, EPI_ISL_723553, EPI_ISL_723554, EPI_ISL_723555, EPI_ISL_723557 | | | |
| see above | Centre for Enzyme Innovation, University of Portsmouth / Translational Research Laboratory, Portsmouth Hospitals NHS Trust | COVID-19 Genomics UK (COG-UK) Consortium | Angela Beckett,Yann Bourgeois,Garry Scarlett,Sharon Glaysher,Scott Elliott,Kelly Bicknell,Robert Impey,Allyson Lloyd,Sarah Wyllie,Ethan Butcher,Anoop Chauhan,Samuel Robson |
| EPI_ISL_724673, EPI_ISL_724705, EPI_ISL_724719, EPI_ISL_724725 | University College London, Great Ormond Street Hospital for Children NHS Foundation Trust, Imperial College Healthcare NHS Trust | COVID-19 Genomics UK (COG-UK) Consortium | Sergi Castellano, Rachel Williams, Mark Kristiansen, Paola Resende Silva, Sunando Roy, Tony Brooks, Helena Tutill, Paola Niola, Patricia Dyal, Charlotte Williams, Leysa Forrest, Yasmin Panchbhaya, Jacqueline Findlay, Samuel Weeks, Julianne Brown, Kathryn Harris, Paul Randell, James Price, Alison Holmes, Judith Breuer |
| EPI_ISL_726017 | Wales Specialist Virology Centre Sequencing lab: Pathogen Genomics Unit | COVID-19 Genomics UK (COG-UK) Consortium | Catherine Moore, Johnathan Evans, Laura Gifford, Malorie Perry, Simon Cottrell, Angela Marchbank, Alec Birchley, Alexander Adams, Amy Gaskin, Bree Gatica-Wilcox, Jason Coombes, Joel Southgate, Lauren Gilbert, Lee Graham, Nicole Pacchiarini, Sara Kumziene-Summerhayes, Sarah Taylor, Sophie Jones, Sara Rey, Matthew Bull, Joanne Watkins, Sally Corden, Tom Connor |
| EPI_ISL_727990, EPI_ISL_727992, EPI_ISL_727993, EPI_ISL_727994, EPI_ISL_727995 | Virology Department, Sheffield Teaching Hospitals NHS Foundation Trust/Department of Infection, Immunity and Cardiovascular Disease, The Medical School, University of Sheffield | COVID-19 Genomics UK (COG-UK) Consortium | Thushan de Silva, Matthew Parker, Nikki Smith, Adri Angyal, Rebecca Brown, Luke Green, Rachel Tucker, Paul Parsons, Danielle Groves, Katie Johnson, Laura Carrilero, Alex Keeley, Dave Partridge, Matthew Wyles, Benjamin Lindsey, Mehmet Yavuz, Mohammad Raza, Cariad Evans |
| EPI_ISL_728034, EPI_ISL_728053, EPI_ISL_728141, EPI_ISL_728142, EPI_ISL_728143, EPI_ISL_728144, EPI_ISL_728145 | University of Wisconsin-Madison AIDS Vaccine Research Laboratories | University of Wisconsin-Madison AIDS Vaccine Research Laboratories | Gage Moreno, Katarina Braun, et al. AIDS Vaccine Research Laboratories |
| EPI_ISL_728167 | Institute for Medical Research, Infectious Disease Research Centre, National Institutes of Health, Ministry of Health Malaysia | Institute for Medical Research, Infectious Disease Research Centre, National Institutes of Health, Ministry of Health Malaysia | Suppiah J, Kamel K, Mohd-Zawawi Z, Thayan R |
| EPI_ISL_728219 | Laboratoire Biolife | Laboratoire de Biotechnologie | Mouna Ouadghiri, Tarik Aanniz, Mohammed Walid Chemao Elfihri, Mohamed Chenaoui, Hanae Dakka, Afaf Alaoui, Otmane Touzani, Bouchra Belfquih, Lahcen Belyamani, Saaid Amzazi and Azeddine Ibrahimi |
| EPI_ISL_728272, EPI_ISL_728273, EPI_ISL_728274, EPI_ISL_728275, EPI_ISL_728276, EPI_ISL_728277, EPI_ISL_728289, EPI_ISL_728290, EPI_ISL_728291, EPI_ISL_728295, EPI_ISL_728297, EPI_ISL_728301, EPI_ISL_728322, EPI_ISL_728332 | | | |
| see above | Laboratoire Biolife | Laboratoire de Biotechnologie | Mouna Ouadghiri, Tarik Aanniz, Mohammed Walid Chemao Elfihri, Mohamed Chenaoui, Hanae Dakka, Afaf Alaoui, Otmane Touzani, Bouchra Belfquih, Lahcen belyamani, Saaid Amzazi and Azeddine Ibrahimi |
| EPI_ISL_728556 | University of Wisconsin-Madison AIDS Vaccine Research Laboratories | University of Wisconsin-Madison AIDS Vaccine Research Laboratories | Gage Moreno, Katarina Braun, et al. AIDS Vaccine Research Laboratories |
| EPI_ISL_728726 | Dutch COVID-19 response team | National Institute for Public Health and the Environment (RIVM) | Adam Meijer, Harry Vennema, Jeroen Cremer, Sharon van den Brink, Bas van der Veer, AnneMarie van den Brandt, Florian Zwagemaker, Dennis Schmitz, Chantal Reusken, on behalf of the national COVID-19 response team |
| EPI_ISL_728780, EPI_ISL_728784, EPI_ISL_728787, EPI_ISL_728789, EPI_ISL_728796, EPI_ISL_728797, EPI_ISL_728801, EPI_ISL_728803, EPI_ISL_728805, EPI_ISL_728812, EPI_ISL_728814, EPI_ISL_728815, EPI_ISL_728822, EPI_ISL_728823, EPI_ISL_728834, EPI_ISL_728845, EPI_ISL_728846, EPI_ISL_728847, EPI_ISL_728848, EPI_ISL_728849, EPI_ISL_728850, EPI_ISL_728851, EPI_ISL_728864, EPI_ISL_728867, EPI_ISL_728869, EPI_ISL_728870, EPI_ISL_728871, EPI_ISL_728872, EPI_ISL_728873, EPI_ISL_728874, EPI_ISL_728875, EPI_ISL_728931, EPI_ISL_728959, EPI_ISL_728965, EPI_ISL_728966, EPI_ISL_728976, EPI_ISL_728982, EPI_ISL_728983, EPI_ISL_729000, EPI_ISL_729004, EPI_ISL_729022, EPI_ISL_729025, EPI_ISL_729030, EPI_ISL_729190, EPI_ISL_729191 | | | |
| see above | Viollier AG | Department of Biosystems Science and Engineering, ETH Zürich | Chaoran Chen, Sarah Nadeau, Catharine Aquino, Ivan Topolsky, Pedro Ferreira, Philipp Jablonski, Susana Posada-Céspedes, Andreia Cabral de Gouvea, Maria Domenica Moccia, Simon Grüter, Timothy Sykes, Lennart Opitz, Ralph Schlapbach, Christiane Beckmann, Maurice Redondo, Olivier Kobel, Christoph Noppen, Sophie Seidel, Noemie Santamaria de Souza, Niko Beerenwinkel, Tanja Stadler |
| EPI_ISL_729384, EPI_ISL_729424, EPI_ISL_729425, EPI_ISL_729426, EPI_ISL_729621, EPI_ISL_729683 | A. Krumbholz, Labor Dr. Krause und Kollegen MVZ GmbH, Kiel | Charité Universitätsmedizin Berlin, Institut für Virologie | Victor M Corman, Barbara Mühlemann, Jörn Beheim-Schwarzbach, Talitha Veith, Julia Schneider, Terry Jones, Christian Drosten |
| EPI_ISL_730053 | Connecticut Department of Health | Grubaugh Lab - Yale School of Public Health | Joseph Fauver, Tara Alpert, Anderson Brito, Annie Watkins, Anne Wyllie, Chantal Vogels, Mary Petrone, Chaney Kalinich, Isabel Ott, Arnau Casanovas, Catherine Muenker, Adam Moore, Alice Lu, Maria Tokuyama, Patrick Wong, Peiwen Lu, Saad Omer, Richard Martinello, Allison Nelson, Shelli Farhadian, Akiko Iwasaki, Charlese Dela Cruz, Albert Ko, Nathan Grubaugh |
| EPI_ISL_730093, EPI_ISL_730111, EPI_ISL_730117, EPI_ISL_730123 | San Diego County Public Health Laboratory | Andersen lab at Scripps Research | SEARCH Alliance San Diego with Tracy Basler, Jovan Shephard, Brett Austin |
| EPI_ISL_730521, EPI_ISL_730527, EPI_ISL_730533, EPI_ISL_730537, EPI_ISL_730542, EPI_ISL_730547 | Biolab Diagnostic Laboratories | Andersen lab at Scripps Research | Issa Abu-Dayyeh, Ahmad Tibi, Lama Hussein, Lina Mohammad, Zein Naber, Amid Abdelnour with SEARCH Alliance San Diego |
| EPI_ISL_732565, EPI_ISL_732566, EPI_ISL_732568, EPI_ISL_732569, EPI_ISL_732571, EPI_ISL_732573, EPI_ISL_732599, EPI_ISL_732601, EPI_ISL_732602, EPI_ISL_732603, EPI_ISL_732618, EPI_ISL_732625, EPI_ISL_732626, EPI_ISL_732628, EPI_ISL_732632, EPI_ISL_732633, EPI_ISL_732634, EPI_ISL_732636, EPI_ISL_732637, EPI_ISL_732638, EPI_ISL_732639, EPI_ISL_732640, EPI_ISL_732641, EPI_ISL_732642, EPI_ISL_732643, EPI_ISL_732644, EPI_ISL_732645, EPI_ISL_732646, EPI_ISL_732647, EPI_ISL_732648, EPI_ISL_732649, EPI_ISL_732650, EPI_ISL_732651, EPI_ISL_732653 | | | |
| see above | Department of Virology and Immunology, University of Helsinki and Helsinki University Hospital, Huslab Finland | Department of Virology, Faculty of Medicine, University of Helsinki, Helsinki, Finland | Teemu Smura, Ravi Kant, Phuoc Truong, Hussein Alburkat, Hannimari Kallio-Kokko, Jenni Virtanen, Maija Suvanto, Sari Hannula, Harri Kangas, Pekka Ellonen, Olli Vapalahti |
| EPI_ISL_732703 | CNR Virus des Infections Respiratoires - France SUD | CNR Virus des Infections Respiratoires - France SUD | Antonin Bal, Gregory Destras, Claudia Gonzalez, Gwendolyne Burfin, Quentin Semanas, Martine Valette, Bruno Lina, Laurence Josset |
| EPI_ISL_732787, EPI_ISL_732791 | Centro de Investigación Biomédica de La Rioja - Hospital San Pedro Logroño | SeqCOVID-SPAIN consortium/IBV(CSIC) | María de Toro, José Manuel Azcona Gutiérrez, María Pilar Bea Escudero, Miriam Blasco Alberdi and SeqCOVID-SPAIN consortium |
| EPI_ISL_732827, EPI_ISL_732828, EPI_ISL_732829, EPI_ISL_732830, EPI_ISL_732831, EPI_ISL_732832, EPI_ISL_732833, EPI_ISL_732834, EPI_ISL_732835, EPI_ISL_732836, EPI_ISL_732837, EPI_ISL_732838, EPI_ISL_732839, EPI_ISL_732840, EPI_ISL_732841, EPI_ISL_732842, EPI_ISL_732843, EPI_ISL_732844, EPI_ISL_732845, EPI_ISL_732846, EPI_ISL_732847, EPI_ISL_732848, EPI_ISL_732849, EPI_ISL_732850, EPI_ISL_732851, EPI_ISL_732852, EPI_ISL_732853, EPI_ISL_732854, EPI_ISL_732855, EPI_ISL_732856, EPI_ISL_732857, EPI_ISL_732858, EPI_ISL_732859, EPI_ISL_732860, EPI_ISL_732861, EPI_ISL_732862, EPI_ISL_732863, EPI_ISL_732864, EPI_ISL_732865, EPI_ISL_732866, EPI_ISL_732867, EPI_ISL_732868, EPI_ISL_732869, EPI_ISL_732870, EPI_ISL_732871, EPI_ISL_732872, EPI_ISL_732873, EPI_ISL_732874, EPI_ISL_732875, EPI_ISL_732876, EPI_ISL_732877, EPI_ISL_732878, EPI_ISL_732879, EPI_ISL_732880, EPI_ISL_732881, EPI_ISL_732883, EPI_ISL_732884, EPI_ISL_732885, EPI_ISL_732886, EPI_ISL_732887, EPI_ISL_732888, EPI_ISL_732889, EPI_ISL_732890, EPI_ISL_732891, EPI_ISL_732892, EPI_ISL_732893, EPI_ISL_732894, EPI_ISL_732895, EPI_ISL_732896, EPI_ISL_732897, EPI_ISL_732898, EPI_ISL_732899, EPI_ISL_732900, EPI_ISL_732901, EPI_ISL_732902, EPI_ISL_732903, EPI_ISL_732904, EPI_ISL_732905, EPI_ISL_732906, EPI_ISL_732907, EPI_ISL_732908, EPI_ISL_732909, EPI_ISL_732910, EPI_ISL_732911, EPI_ISL_732912, EPI_ISL_732913, EPI_ISL_732914, EPI_ISL_732915, EPI_ISL_732916, EPI_ISL_732917, EPI_ISL_732918, EPI_ISL_732919, EPI_ISL_732920, EPI_ISL_732921, EPI_ISL_732922, EPI_ISL_732923, EPI_ISL_732924, EPI_ISL_732925, EPI_ISL_732926 | | | |
| see above | genXone SA, Molecular Diagnostics Laboratory / NZOZ | genXone SA, Research & Development Laboratory | Maciej Sykulski, Grzegorz Nowicki, Monika Makowska-Woniak, Jakub Grabowski, Natalia Drwska-Matelska, ukasz Krych, Micha Kaszuba |
| EPI_ISL_734218, EPI_ISL_734245, EPI_ISL_734277 | Virginia Division of Consolidated Laboratory Services (DCLS) | Virginia Division of Consolidated Laboratory Services (DCLS) | Virginia DCLS |
| EPI_ISL_734293, EPI_ISL_734410, EPI_ISL_734413 | Wadsworth Center, New York State Department.of Health | Wadsworth Center, New York State Department.of Health | Kirsten St. George, Daryl M. Lamson, Alexis Russel, Jonathan Plitnick, Navjot Singh, John Kelly, Sara Griesemer, Erasmus Schneider, Erica Lasek-Nesselquist |
| EPI_ISL_734470 | Masonic Medical Research Institute | Wadsworth Center, New York State Department.of Health | Kirsten St. George, Nathan Tucker, Ryan D. Pfeiffer, Daryl M. Lamson, Alexis Russel, Jonathan Plitnick, Navjot Singh, John Kelly, Sara Griesemer, Erasmus Schneider, Erica Lasek-Nesselquist |

| | | | |
|---|---|---|---|
| EPI_ISL_737016, EPI_ISL_737028, EPI_ISL_737030, EPI_ISL_737031, EPI_ISL_737032 | The National Institute of Public Health | State Veterinary Institute Prague | Nagy,A;Jirincova,H;Trnka,D;Vecerova,J |
| EPI_ISL_739430 | Humboldt County Public Health Laboratory | Chan-Zuckerberg Biohub | CZB Cliahub Consortium |
| EPI_ISL_739663, EPI_ISL_739666 | Instituto Nacional de Salud, Bogotá, Colombia | Instituto Nacional de Salud, Bogotá, Colombia | Katherine Laiton-Donato, Diego A. Álvarez-Díaz, Carlos Franco-Muñoz, Mauricio Pacheco-Montealegre, Jonathan Reales, Diego Andrés Prada, Sheryl Corchuelo, Magdalena Weisner, Martha Lucia Ospina Martinez, Marcela Mercado-Reyes |
| EPI_ISL_739711, EPI_ISL_739718, EPI_ISL_739734, EPI_ISL_739743, EPI_ISL_739747, EPI_ISL_739791, EPI_ISL_739794, EPI_ISL_739800, EPI_ISL_739802, EPI_ISL_739803, EPI_ISL_739806, EPI_ISL_739822, EPI_ISL_739824, EPI_ISL_739825, EPI_ISL_739827, EPI_ISL_739831, EPI_ISL_739838, EPI_ISL_739886, EPI_ISL_739898, EPI_ISL_739917, EPI_ISL_739920, EPI_ISL_739927, EPI_ISL_739953, EPI_ISL_739955, EPI_ISL_739983, EPI_ISL_739992, EPI_ISL_739996, EPI_ISL_740004, EPI_ISL_740006, EPI_ISL_740008, EPI_ISL_740026, EPI_ISL_740031, EPI_ISL_740036, EPI_ISL_740041, EPI_ISL_740053, EPI_ISL_740055, EPI_ISL_740057, EPI_ISL_740060, EPI_ISL_740076, EPI_ISL_740080, EPI_ISL_740095, EPI_ISL_740101, EPI_ISL_740105, EPI_ISL_740106, EPI_ISL_740114, EPI_ISL_740116, EPI_ISL_740155, EPI_ISL_740169, EPI_ISL_740187, EPI_ISL_740224, EPI_ISL_740244, EPI_ISL_740266, EPI_ISL_740272, EPI_ISL_740298, EPI_ISL_740349, EPI_ISL_740361, EPI_ISL_740382, EPI_ISL_740390, EPI_ISL_740398, EPI_ISL_740399, EPI_ISL_740410, EPI_ISL_740423, EPI_ISL_740445, EPI_ISL_740449, EPI_ISL_740453, EPI_ISL_740455, EPI_ISL_740477, EPI_ISL_740498, EPI_ISL_740507, EPI_ISL_740512, EPI_ISL_740514, EPI_ISL_740516, EPI_ISL_740535 | | | |
| see above | Laboratoire national de santé, Microbiology, Virology | Laboratoire national de santé, Microbiology, Microbial Genomics Platform | Anke Wienecke-Baldacchino, Catherine Ragimbeau,Jessica Tapp, Fatu Djabi, Lise Pignon, Raoul Salmon, Tamir Abdelrahman |
| EPI_ISL_740623, EPI_ISL_740624, EPI_ISL_740625, EPI_ISL_740626, EPI_ISL_740627, EPI_ISL_740628, EPI_ISL_740629, EPI_ISL_740630, EPI_ISL_740631, EPI_ISL_740632, EPI_ISL_740633, EPI_ISL_740634, EPI_ISL_740635, EPI_ISL_740636, EPI_ISL_740637, EPI_ISL_740638, EPI_ISL_740639, EPI_ISL_740640, EPI_ISL_740641, EPI_ISL_740642, EPI_ISL_740643, EPI_ISL_740644, EPI_ISL_740645, EPI_ISL_740646, EPI_ISL_740647, EPI_ISL_740648, EPI_ISL_740649, EPI_ISL_740650, EPI_ISL_740651, EPI_ISL_740652, EPI_ISL_740653, EPI_ISL_740654, EPI_ISL_740656, EPI_ISL_740657, EPI_ISL_740658, EPI_ISL_740659, EPI_ISL_740660, EPI_ISL_740661, EPI_ISL_740662, EPI_ISL_740665, EPI_ISL_740666, EPI_ISL_740667, EPI_ISL_740668, EPI_ISL_740669, EPI_ISL_740670, EPI_ISL_740675, EPI_ISL_740678, EPI_ISL_740679, EPI_ISL_740680, EPI_ISL_740683, EPI_ISL_740685, EPI_ISL_740686, EPI_ISL_740687, EPI_ISL_740688, EPI_ISL_740689, EPI_ISL_740690, EPI_ISL_740691, EPI_ISL_740692, EPI_ISL_740693, EPI_ISL_740694, EPI_ISL_740695, EPI_ISL_740696, EPI_ISL_740697, EPI_ISL_740698, EPI_ISL_740699, EPI_ISL_740700, EPI_ISL_740701, EPI_ISL_740702, EPI_ISL_740703, EPI_ISL_740704, EPI_ISL_740705, EPI_ISL_740706, EPI_ISL_740707, EPI_ISL_740708, EPI_ISL_740709, EPI_ISL_740710, EPI_ISL_740711, EPI_ISL_740712, EPI_ISL_740713, EPI_ISL_740714, EPI_ISL_740715, EPI_ISL_740716, EPI_ISL_740717, EPI_ISL_740718, EPI_ISL_740719, EPI_ISL_740720, EPI_ISL_740721, EPI_ISL_740722, EPI_ISL_740723, EPI_ISL_740724, EPI_ISL_740725, EPI_ISL_740726, EPI_ISL_740727, EPI_ISL_740728, EPI_ISL_740729, EPI_ISL_740730, EPI_ISL_740731, EPI_ISL_740732, EPI_ISL_740733, EPI_ISL_740734, EPI_ISL_740735, EPI_ISL_740736, EPI_ISL_740737, EPI_ISL_740738, EPI_ISL_740739, EPI_ISL_740740, EPI_ISL_740741, EPI_ISL_740742, EPI_ISL_740743, EPI_ISL_740744, EPI_ISL_740745, EPI_ISL_740746, EPI_ISL_740747, EPI_ISL_740748, EPI_ISL_740749, EPI_ISL_740750, EPI_ISL_740751, EPI_ISL_740752, EPI_ISL_740753, EPI_ISL_740754, EPI_ISL_740755, EPI_ISL_740756, EPI_ISL_740757, EPI_ISL_740758, EPI_ISL_740759, EPI_ISL_740760, EPI_ISL_740761, EPI_ISL_740762, EPI_ISL_740763, EPI_ISL_740764, EPI_ISL_740765, EPI_ISL_740766, EPI_ISL_740768, EPI_ISL_740769, EPI_ISL_740770, EPI_ISL_740771, EPI_ISL_740772, EPI_ISL_740773, EPI_ISL_740774, EPI_ISL_740775, EPI_ISL_740776, EPI_ISL_740782, EPI_ISL_740783, EPI_ISL_740790, EPI_ISL_740791, EPI_ISL_740792, EPI_ISL_740793, EPI_ISL_740794, EPI_ISL_740795, EPI_ISL_740796, EPI_ISL_740797, EPI_ISL_740798, EPI_ISL_740799, EPI_ISL_740800, EPI_ISL_740801, EPI_ISL_740802, EPI_ISL_740803, EPI_ISL_740804, EPI_ISL_740805, EPI_ISL_740806, EPI_ISL_740807, EPI_ISL_740808, EPI_ISL_740809, EPI_ISL_740810, EPI_ISL_740811, EPI_ISL_740812, EPI_ISL_740813, EPI_ISL_740814, EPI_ISL_740815, EPI_ISL_740816, EPI_ISL_740817, EPI_ISL_740818, EPI_ISL_740819, EPI_ISL_740820, EPI_ISL_740821, EPI_ISL_740822, EPI_ISL_740823, EPI_ISL_740824, EPI_ISL_740825, EPI_ISL_740826, EPI_ISL_740827, EPI_ISL_740828, EPI_ISL_740830, EPI_ISL_740831, EPI_ISL_740832, EPI_ISL_740833, EPI_ISL_740834, EPI_ISL_740835, EPI_ISL_740836, EPI_ISL_740837, EPI_ISL_740838, EPI_ISL_740839, EPI_ISL_740840, EPI_ISL_740841, EPI_ISL_740842, EPI_ISL_740843, EPI_ISL_740844, EPI_ISL_740845, EPI_ISL_740846, EPI_ISL_740847, EPI_ISL_740848, EPI_ISL_740849, EPI_ISL_740850, EPI_ISL_740851, EPI_ISL_740852, EPI_ISL_740853, EPI_ISL_740854, EPI_ISL_740855, EPI_ISL_740856, EPI_ISL_740857, EPI_ISL_740858, EPI_ISL_740859, EPI_ISL_740860, EPI_ISL_740861, EPI_ISL_740862 | | | |
| see above | BCCDC Public Health Laboratory | BCCDC Public Health Laboratory | Prystajecky Natalie, Linda Hoang, Dan Fornika, Shannon Russell, Kim Macdonald, Kimia Kamelian, John Tyson, Inna Sekirov, Mel Krajden |
| EPI_ISL_741359, EPI_ISL_741360, EPI_ISL_741361 | University of Exeter | COVID-19 Genomics UK (COG-UK) Consortium | Ben Temperton,Aaron Jeffries,Michelle Michelsen,Joanna Warwick-Dugdale,Audrey Farbos,Robyn Manley,Stephen Michell,Jane Masoli |
| EPI_ISL_741362, EPI_ISL_741363, EPI_ISL_741364, EPI_ISL_741365, EPI_ISL_741366, EPI_ISL_741369, EPI_ISL_741371, EPI_ISL_741372, EPI_ISL_741374, EPI_ISL_741375, EPI_ISL_741376, EPI_ISL_741377, EPI_ISL_741378, EPI_ISL_741379, EPI_ISL_741380, EPI_ISL_741381, EPI_ISL_741382, EPI_ISL_741383, EPI_ISL_741389, EPI_ISL_741391, EPI_ISL_741392, EPI_ISL_741395, EPI_ISL_741396, EPI_ISL_741397, EPI_ISL_741398, EPI_ISL_741399, EPI_ISL_741400, EPI_ISL_741401, EPI_ISL_741402, EPI_ISL_741403, EPI_ISL_741407, EPI_ISL_741408, EPI_ISL_741410, EPI_ISL_741414, EPI_ISL_741416, EPI_ISL_741417, EPI_ISL_741419, EPI_ISL_741420 | | | |
| see above | Oxford Viromics, NDM, University of Oxford; Oxford University Hospitals; Basingstoke and North Hampshire Hospital | COVID-19 Genomics UK (COG-UK) Consortium | Tanya Golubchik, David Bonsall, George Macintyre, Amy Trebes, Mariateresa de Cesare, Catrin Moore, Alex Mobbs, Anita Justice, Robert Shaw, Monique Andersson, Timothy Peto, Emma Wise, Nathan Moore, Jessica Lynch, Nick Cortes, Matilde Mori, Stephen Kidd, David Buck, John Todd, Christophe Fraser |
| EPI_ISL_741567 | Quadram Institute Bioscience | COVID-19 Genomics UK (COG-UK) Consortium | Dave J. Baker, Gemma L. Kay, Alp Aydin, Thanh Le-Viet, Steven Rudder, Ana P. Tedim, Anastasia Kolyva, Maria Diaz, Leonardo de Oliveira Martins, Nabil-Fareed Alikhan, Lizzie Meadows, Rachael Stanley, Ngozi Elumogo, Muhammed Yasir, Nicholas M. Thomson, Alexander J Trotter, Rachel Gilroy, Samuel Bloomfield, Claire Stuart, Andrew Bell, Reenesh Prakash, Samir Dervisevic, Alison E. Mather, John Wain, Mark Webber, Andrew J. Page, Justin O'Grady |
| EPI_ISL_742117 | University of Exeter | COVID-19 Genomics UK (COG-UK) Consortium | Ben Temperton,Aaron Jeffries,Michelle Michelsen,Joanna Warwick-Dugdale,Audrey Farbos,Robyn Manley,Stephen Michell,Jane Masoli |
| EPI_ISL_742261 | Oxford Viromics, NDM, University of Oxford; Oxford University Hospitals; Basingstoke and North Hampshire Hospital | COVID-19 Genomics UK (COG-UK) Consortium | Tanya Golubchik, David Bonsall, George Macintyre, Amy Trebes, Mariateresa de Cesare, Catrin Moore, Alex Mobbs, Anita Justice, Robert Shaw, Monique Andersson, Timothy Peto, Emma Wise, Nathan Moore, Jessica Lynch, Nick Cortes, Matilde Mori, Stephen Kidd, David Buck, John Todd, Christophe Fraser |
| EPI_ISL_742300, EPI_ISL_742301, EPI_ISL_742308, EPI_ISL_742309, EPI_ISL_742310, EPI_ISL_742311, EPI_ISL_742312, EPI_ISL_742313, EPI_ISL_742314, EPI_ISL_742315, EPI_ISL_742316 | | | |
| see above | Wales Specialist Virology Centre Sequencing lab: Pathogen Genomics Unit | COVID-19 Genomics UK (COG-UK) Consortium | Catherine Moore, Johnathan Evans, Laura Gifford, Malorie Perry, Simon Cottrell, Angela Marchbank, Alec Birchley, Alexander Adams, Amy Gaskin, Bree Gatica-Wilcox, Jason Coombes, Joel Southgate, Lauren Gilbert, Lee Graham, Nicole Pacchiarini, Sara Kumziene-Summerhayes, Sarah Taylor, Sophie Jones, Sara Rey, Matthew Bull, Joanne Watkins, Sally Corden, Tom Connor |
| EPI_ISL_744136, EPI_ISL_744164, EPI_ISL_744191, EPI_ISL_744193, EPI_ISL_744201, EPI_ISL_744207, EPI_ISL_744215, EPI_ISL_744219, EPI_ISL_744220, EPI_ISL_744232, EPI_ISL_744261, EPI_ISL_744290, EPI_ISL_744324, EPI_ISL_744325, EPI_ISL_744332, EPI_ISL_744373, EPI_ISL_744374, EPI_ISL_744378, EPI_ISL_744381, EPI_ISL_744385, EPI_ISL_744401, EPI_ISL_744402, EPI_ISL_744405, EPI_ISL_744437, EPI_ISL_744467, EPI_ISL_744473, EPI_ISL_744512, EPI_ISL_744529, EPI_ISL_744556, EPI_ISL_744559, EPI_ISL_744604, EPI_ISL_744616, EPI_ISL_744617, EPI_ISL_744625, EPI_ISL_744627, EPI_ISL_744630, EPI_ISL_744649, EPI_ISL_744668, EPI_ISL_744675, EPI_ISL_744683, EPI_ISL_744701, EPI_ISL_744729, EPI_ISL_744733, EPI_ISL_744763, EPI_ISL_744783, EPI_ISL_744789, EPI_ISL_744800, EPI_ISL_744806, EPI_ISL_744818, EPI_ISL_744848, EPI_ISL_744872, EPI_ISL_744876, EPI_ISL_744882, EPI_ISL_744895, EPI_ISL_744902, EPI_ISL_744926, EPI_ISL_744955, EPI_ISL_744964, EPI_ISL_744978, EPI_ISL_744984, EPI_ISL_744988, EPI_ISL_745010, EPI_ISL_745025 | | | |
| see above | Laboratoire national de santé, Microbiology, Virology | Laboratoire national de santé, Microbiology, Microbial Genomics Platform | Anke Wienecke-Baldacchino, Catherine Ragimbeau,Jessica Tapp, Fatu Djabi, Lise Pignon, Raoul Salmon, Tamir Abdelrahman |
| EPI_ISL_747032, EPI_ISL_747033, EPI_ISL_747034, EPI_ISL_747067, EPI_ISL_747099, EPI_ISL_747107, EPI_ISL_747108, EPI_ISL_747109, EPI_ISL_747119, EPI_ISL_747127 | Respiratory Viruses Branch, Centers for Disease Control and Prevention | Respiratory Viruses Branch, Centers for Disease Control and Prevention | Queen,K., Li,Y., Tao,Y., Uehara,A., Montmayeur,A., Paden,C.R., Cook,P.W., Marine,R., Sheth,M., Wang,H., Lee,J., Tong,S. |
| EPI_ISL_751448, EPI_ISL_751490 | CHU Purpan - Laboratoire de Virologie - Institut Fédératif de Biologie | CHU Purpan - Laboratoire de Virologie - Institut Fédératif de Biologie | Latour J., Ranger N., Dubois M., Carcenac R., Harter A., Boyer P., Tremeaux P., Izopet J. |
| EPI_ISL_751548 | NYSDOH Wadsworth Center, Virology Lab | Genomics and Discovery, Respiratory Viruses Branch, Division of Viral Diseases, Centers for Disease Control and Prevention | Krista Queen, Yan Li, Ying Tao, Jing Zhang, Anna Uehara, Anna Montmayeur, Clinton R. Paden, Peter W. Cook,Rachel Marine, Mili Sheth, Haibin Wang, Justin Lee, Suxiang Tong |
| EPI_ISL_751732 | NE Public Health Laboratory | Genomics and Discovery, Respiratory Viruses Branch, Division of Viral Diseases, Centers for Disease Control and Prevention | Krista Queen, Yan Li, Ying Tao, Jing Zhang, Anna Uehara, Anna Montmayeur, Clinton R. Paden, Peter W. Cook,Rachel Marine, Mili Sheth, Haibin Wang, Justin Lee, Suxiang Tong |
| EPI_ISL_753054 | State Laboratories Division, Hawaii State Department of Health | State Laboratories Division, Hawaii State Department of Health | Pamela O'Brien, Sabrina Diemert, Drew Kuwazaki, Razvan Sultana, Edward Desmond |
| EPI_ISL_753324 | Clinical virology Laboratory, Children's Hospital Los Angeles | Center for Personalized Medicine, Children's Hospital Los Angeles | Gai et al |
| EPI_ISL_754626, EPI_ISL_754627 | University of Wisconsin-Madison AIDS Vaccine Research Laboratories | University of Wisconsin-Madison AIDS Vaccine Research Laboratories | Gage Moreno, Katarina Braun, et al. AIDS Vaccine Research Laboratories |
| EPI_ISL_757342 | Department of Virology and Immunology, University of Helsinki and Helsinki University Hospital, Huslab Finland | Department of Virology, Faculty of Medicine, University of Helsinki, Helsinki, Finland | Teemu Smura, Ravi Kant, Phuoc Truong, Hussein Alburkat, Hannimari Kallio-Kokko, Jenni Virtanen, Maija Suvanto, Sari Hannula, Harri Kangas, Pekka Ellonen, Olli Vapalahti |
| EPI_ISL_760096, EPI_ISL_760097, EPI_ISL_760118, EPI_ISL_760122, EPI_ISL_760147, EPI_ISL_760175, EPI_ISL_760176, EPI_ISL_760177, EPI_ISL_760178, EPI_ISL_760179, EPI_ISL_760180, EPI_ISL_760199, EPI_ISL_760200, EPI_ISL_760216, EPI_ISL_760217 | | | |
| see above | Division of Emerging Infectious Diseases, Bureau of Infectious Diseases Diagnosis Control, Korea Disease Control and | Division of Emerging Infectious Diseases, Bureau of Infectious Diseases Diagnosis Control, Korea Disease Control and | Ae Kyung Park, Il-Hwan Kim, Heui Man Kim, Jeong-Min Kim, Namjoo Lee, Chaeyoung Lee, Sang Hee Woo, Eun-Jin Kim |

| | | | |
|---|---|---|---|
| | Prevention Agency | Prevention Agency | |
| EPI_ISL_763232 | Dutch COVID-19 response team | Erasmus Medical Center | Bas Oude Munnink, Reina Sikkema, David Nieuwenhuijse, Irina Chestakova, Anne van der Linden, Marjan Boter, Emmanuelle Munger, Corine GeurtsvanKessel, Annemiek van der Eijk, Richard Molenkamp, Marion Koopmans, on behalf of the Dutch national COVID-19 response team. |
| EPI_ISL_765778, EPI_ISL_765781, EPI_ISL_765782, EPI_ISL_765783, EPI_ISL_765784, EPI_ISL_765785 | Massachusetts General Hospital | Infectious Disease Program, Broad Institute of Harvard and MIT | Lemieux,J.E., Siddle,K.J., Shaw,B., Adams,G., Pierce,V., Turbett,S., Anahtar,M., Branda,J., Slater,D., Harris,J., Lin,A.E., Gladden-Young,A., Lagerborg,K., Rudy,M., DeRuff,K., Carter,A., Normandin,E., Bauer,M., Reilly,S., Tomkins-Tinch,C., Loreth,C., Chaluvadi,S., Neumann,A., Cusick,C., Chapman,S.B., Gnirke,A., Flowers,K., Cerrato,F., Birren,B.W., Gallagher,G., Smole,S., Park,D.J., MacInnis,B.L., Ryan,E., LaRocque,R., Rosenberg,E. and Sabeti,P.C. |
| EPI_ISL_765942, EPI_ISL_765958 | Worobey Lab, Department of Ecology and Evolutionary Biology, University of Arizona | Worobey Lab, Department of Ecology and Evolutionary Biology, University of Arizona | Brendan Larsen, Grace Quirk, Thomas Watts, David Baltrus, Michael Worobey |
| EPI_ISL_766615 | Klinisk mikrobiologi | The Public Health Agency of Sweden | Department of Microbiology, The Public Health Agency of Sweden |
| EPI_ISL_768634, EPI_ISL_768635, EPI_ISL_768636, EPI_ISL_768637, EPI_ISL_768638, EPI_ISL_768639, EPI_ISL_768651, EPI_ISL_768652, EPI_ISL_768653, EPI_ISL_768654, EPI_ISL_768702 | | | |
| see above | Pathogen Genomics Center, National Institute of Infectious Diseases | Pathogen Genomics Center, National Institute of Infectious Diseases | Tsuyoshi Sekizuka, Kentaro Itokawa, Rina Tanaka, Masanori Hashino, Makoto Kuroda |
| EPI_ISL_768753, EPI_ISL_768757, EPI_ISL_768786 | AIID | Irish Coronavirus Sequencing Consortium - National Virus Reference Laboratory | Michael Carr, Gabriel Gonzalez, Alejandro Abner Garcia Leon, Patrick Mallon |
| EPI_ISL_768839 | Laboratoire Biolife | Laboratoire de Biotechnologie | Mouna Ouadghiri, Tarik Aanniz, Mohammed Walid Chemao Elfihri, Mohamed Chenaoui, Hanae Dakka, Afaf Alaoui, Otmane Touzani, Amina Benouda, Bouchra Belfquih, Lahcen belyamani, Saaid Amzazi and Azeddine Ibrahimi |
| EPI_ISL_770029 | Area De Salud Corredores | Inciensa, Instituto Costarricense de Investigación y Enseñanza en Nutrición y Salud | Francisco Duarte, Hebleen Porras, Claudio Soto-Garita, Estela Cordero, Adriana Godínez, Melany Calderón & Mariel López |
| EPI_ISL_770533, EPI_ISL_770534, EPI_ISL_770535, EPI_ISL_770536, EPI_ISL_770537, EPI_ISL_770538, EPI_ISL_770539, EPI_ISL_770540, EPI_ISL_770541 | Private lab | Lithuanian University of Health Sciences, Molecular cardiology lab. | Lukas Zemaitis, Ingrida Olendrait, Arnoldas Pautienius, Kamile Tamusauskaite, Dovydas Gecys, Laura Pareckaite, Vaiva Lesauskaite, Astra Vitkauskiene |
| EPI_ISL_770810 | Essentia Health-St. Mary's Medical Center | Minnesota Department of Health, Public Health Laboratory | Alexandra Lorentz, Jacob Garfin, Matt Plumb, and Xiong Wang |
| EPI_ISL_775021, EPI_ISL_775022, EPI_ISL_775023, EPI_ISL_775024, EPI_ISL_775025, EPI_ISL_775026, EPI_ISL_775027, EPI_ISL_775028, EPI_ISL_775029, EPI_ISL_775030, EPI_ISL_775031, EPI_ISL_775032, EPI_ISL_775033, EPI_ISL_775034, EPI_ISL_775035, EPI_ISL_775036, EPI_ISL_775037, EPI_ISL_775038, EPI_ISL_775039, EPI_ISL_775040, EPI_ISL_775041, EPI_ISL_775042, EPI_ISL_775043, EPI_ISL_775044, EPI_ISL_775045, EPI_ISL_775046, EPI_ISL_775047, EPI_ISL_775048, EPI_ISL_775049, EPI_ISL_775050, EPI_ISL_775051, EPI_ISL_775052, EPI_ISL_775053, EPI_ISL_775054, EPI_ISL_775055, EPI_ISL_775056, EPI_ISL_775057, EPI_ISL_775058, EPI_ISL_775059, EPI_ISL_775060, EPI_ISL_775061, EPI_ISL_775062, EPI_ISL_775063, EPI_ISL_775064, EPI_ISL_775065, EPI_ISL_775066, EPI_ISL_775067, EPI_ISL_775068, EPI_ISL_775069, EPI_ISL_775070, EPI_ISL_775071, EPI_ISL_775072, EPI_ISL_775073, EPI_ISL_775074, EPI_ISL_775075, EPI_ISL_775076, EPI_ISL_775077, EPI_ISL_775078, EPI_ISL_775079, EPI_ISL_775080, EPI_ISL_775081, EPI_ISL_775082, EPI_ISL_775083, EPI_ISL_775084, EPI_ISL_775085, EPI_ISL_775086, EPI_ISL_775087, EPI_ISL_775088, EPI_ISL_775089, EPI_ISL_775090, EPI_ISL_775091, EPI_ISL_775092, EPI_ISL_775093, EPI_ISL_775094, EPI_ISL_775095, EPI_ISL_775096, EPI_ISL_775097, EPI_ISL_775098, EPI_ISL_775099, EPI_ISL_775100, EPI_ISL_775101, EPI_ISL_775102, EPI_ISL_775103, EPI_ISL_775104, EPI_ISL_775105, EPI_ISL_775106, EPI_ISL_775107, EPI_ISL_775108, EPI_ISL_775109, EPI_ISL_775110, EPI_ISL_775111, EPI_ISL_775112, EPI_ISL_775113, EPI_ISL_775114, EPI_ISL_775115, EPI_ISL_775116, EPI_ISL_775117, EPI_ISL_775118, EPI_ISL_775119, EPI_ISL_775120, EPI_ISL_775121, EPI_ISL_775122, EPI_ISL_775123, EPI_ISL_775124, EPI_ISL_775125, EPI_ISL_775126, EPI_ISL_775127, EPI_ISL_775128, EPI_ISL_775129, EPI_ISL_775130, EPI_ISL_775131, EPI_ISL_775132, EPI_ISL_775133, EPI_ISL_775134, EPI_ISL_775135, EPI_ISL_775136, EPI_ISL_775137, EPI_ISL_775138, EPI_ISL_775139, EPI_ISL_775140, EPI_ISL_775141, EPI_ISL_775142, EPI_ISL_775143, EPI_ISL_775144, EPI_ISL_775145, EPI_ISL_775146, EPI_ISL_775147, EPI_ISL_775148, EPI_ISL_775149, EPI_ISL_775150, EPI_ISL_775151, EPI_ISL_775152, EPI_ISL_775153, EPI_ISL_775154, EPI_ISL_775155, EPI_ISL_775156, EPI_ISL_775157, EPI_ISL_775158, EPI_ISL_775159, EPI_ISL_775160, EPI_ISL_775161, EPI_ISL_775162, EPI_ISL_775163, EPI_ISL_775164, EPI_ISL_775165, EPI_ISL_775166, EPI_ISL_775167, EPI_ISL_775168, EPI_ISL_775169, EPI_ISL_775170, EPI_ISL_775171, EPI_ISL_775172, EPI_ISL_775173, EPI_ISL_775174, EPI_ISL_775175, EPI_ISL_775176, EPI_ISL_775177, EPI_ISL_775178, EPI_ISL_775179, EPI_ISL_775180, EPI_ISL_775181, EPI_ISL_775182, EPI_ISL_775183, EPI_ISL_775184, EPI_ISL_775185, EPI_ISL_775186, EPI_ISL_775187, EPI_ISL_775188, EPI_ISL_775189, EPI_ISL_775190, EPI_ISL_775191, EPI_ISL_775192, EPI_ISL_775193, EPI_ISL_775194, EPI_ISL_775195, EPI_ISL_775196, EPI_ISL_775197, EPI_ISL_775198, EPI_ISL_775199, EPI_ISL_775200, EPI_ISL_775201, EPI_ISL_775202, EPI_ISL_775203, EPI_ISL_775204, EPI_ISL_775205, EPI_ISL_775206, EPI_ISL_775207, EPI_ISL_775208, EPI_ISL_775209, EPI_ISL_775210, EPI_ISL_775211, EPI_ISL_775212 | | | |
| see above | Public Health Ontario Laboratory | Public Health Ontario Laboratory | Vanessa G Allen, Philip Banh, Richard de Borja, Yao Chen, Alireza Eshaghi, Nahuel Fittipaldi, Christine Frantz, Jonathan B Gubbay, Jennifer L Guthrie, Lawrence Heisler, Esha Joshi, Michael Laszloffy, Aimin Li, Michael CY Li, Dean Maxwell, Sandeep Nagra, Samir N Patel, Heather Rilkoff, Jared Simpson, Karthikeyan Sivaraman, Yogi Sundaravadanam, Sarah Teatero, Andre Villegas, Sandra Zittermann |
| EPI_ISL_775250, EPI_ISL_775252, EPI_ISL_775264, EPI_ISL_775265 | Laboratoire Biolife | Laboratoire de Biotechnologie | Mouna Ouadghiri, Tarik Aanniz, Mohammed Walid Chemao Elfihri, Mohamed Chenaoui, Hanae Dakka, Afaf Alaoui, Otmane Touzani, Amina Benouda, Bouchra Belfquih, Lahcen belyamani, Saaid Amzazi and Azeddine Ibrahimi |
| EPI_ISL_775327 | Dept. of Medical Microbiology, Stavanger University Hospital, Helse Stavanger HF | Norwegian Institute of Public Health, Department of Virology | Kathrine Stene-Johansen, Kamilla Heddeland Instefjord, Hilde Elshaug, Atiya R Ali,Marie Paulsen Madsen, Rasmus Riis Kopperud, Hilde Vollan, Karoline Bragstad, Olav Hungnes |
| EPI_ISL_775330, EPI_ISL_775382 | Nordland Hospital - Bodo, Laboratory Department, Molecular Biology Unit | Norwegian Institute of Public Health, Department of Virology | Kathrine Stene-Johansen, Kamilla Heddeland Instefjord, Hilde Elshaug, Atiya R Ali,Marie Paulsen Madsen, Rasmus Riis Kopperud, Hilde Vollan, Karoline Bragstad, Olav Hungnes |
| EPI_ISL_775393 | Vestfold Hospital, Toensberg Department of Microbiology | Norwegian Institute of Public Health, Department of Virology | Kathrine Stene-Johansen, Kamilla Heddeland Instefjord, Hilde Elshaug, Atiya R Ali,Marie Paulsen Madsen, Rasmus Riis Kopperud, Hilde Vollan, Karoline Bragstad, Olav Hungnes |
| EPI_ISL_775460, EPI_ISL_775461 | Norwegian Institute of Public Health, Department of Virology | Norwegian Institute of Public Health, Department of Virology | Kathrine Stene-Johansen, Kamilla Heddeland Instefjord, Hilde Elshaug, Atiya R Ali,Marie Paulsen Madsen, Rasmus Riis Kopperud, Hilde Vollan, Karoline Bragstad, Olav Hungnes |
| EPI_ISL_775501, EPI_ISL_775502, EPI_ISL_775503 | Vestfold Hospital, Toensberg Department of Microbiology | Norwegian Institute of Public Health, Department of Virology | Kathrine Stene-Johansen, Kamilla Heddeland Instefjord, Hilde Elshaug, Atiya R Ali,Marie Paulsen Madsen, Rasmus Riis Kopperud, Hilde Vollan, Karoline Bragstad, Olav Hungnes |
| EPI_ISL_778845, EPI_ISL_778846, EPI_ISL_778847, EPI_ISL_778848, EPI_ISL_778849, EPI_ISL_778850, EPI_ISL_778851, EPI_ISL_778852, EPI_ISL_778853, EPI_ISL_778854 | Public Health Ontario Laboratory | Public Health Ontario Laboratory | Vanessa G Allen, Philip Banh, Richard de Borja, Yao Chen, Alireza Eshaghi, Nahuel Fittipaldi, Christine Frantz, Jonathan B Gubbay, Jennifer L Guthrie, Lawrence Heisler, Esha Joshi, Michael Laszloffy, Aimin Li, Michael CY Li, Dean Maxwell, Sandeep Nagra, Samir N Patel, Heather Rilkoff, Jared Simpson, Karthikeyan Sivaraman, Yogi Sundaravadanam, Sarah Teatero, Andre Villegas, Sandra Zittermann |
| EPI_ISL_779228, EPI_ISL_779229, EPI_ISL_779230, EPI_ISL_779231, EPI_ISL_779670, EPI_ISL_779671, EPI_ISL_779672, EPI_ISL_779673, EPI_ISL_779674, EPI_ISL_779675, EPI_ISL_779676, EPI_ISL_779677, EPI_ISL_779678, EPI_ISL_779699, EPI_ISL_779700, EPI_ISL_779701, EPI_ISL_779702 | | | |
| see above | Pathogen Genomics Center, National Institute of Infectious Diseases | Pathogen Genomics Center, National Institute of Infectious Diseases | Tsuyoshi Sekizuka, Kentaro Itokawa, Rina Tanaka, Masanori Hashino, Makoto Kuroda |
| EPI_ISL_780022, EPI_ISL_780025, EPI_ISL_780074 | Hospital General Universitario Gregorio Marañón | SeqCOVID-SPAIN consortium/IBV(CSIC) | Darío García de Viedma, Laura Pérez-Lago, Marta Herranz, Jon Sicilia, Julia Suárez, Pilar Catalán, Patricia Muñoz and SeqCOVID-SPAIN consortium |
| EPI_ISL_780383, EPI_ISL_780385, EPI_ISL_780386, EPI_ISL_780404, EPI_ISL_780405, EPI_ISL_780406, EPI_ISL_780410 | Bermuda Government Molecular Diagnostics Laboratory (MDL) | Respiratory Virus Unit, National Infection Service, Public Health England | PHE Covid Sequencing Team, Dr Carika Weldon (Bermuda), Dr Ayoola Oyinloye (Bermuda) |
| EPI_ISL_785709, EPI_ISL_785722, EPI_ISL_785741, EPI_ISL_785770, EPI_ISL_785771, EPI_ISL_785780, EPI_ISL_785812, EPI_ISL_785820, EPI_ISL_785824, EPI_ISL_785832, EPI_ISL_785836, EPI_ISL_785846, EPI_ISL_785848, EPI_ISL_785849, EPI_ISL_785858, EPI_ISL_785860, EPI_ISL_785862, EPI_ISL_785870, EPI_ISL_785871, EPI_ISL_785883, EPI_ISL_785894, EPI_ISL_785899, EPI_ISL_785900, EPI_ISL_785901, EPI_ISL_785902, EPI_ISL_785903, EPI_ISL_785905, EPI_ISL_785906, EPI_ISL_785907, EPI_ISL_785908, EPI_ISL_785911, EPI_ISL_785912, EPI_ISL_785913, EPI_ISL_785914, EPI_ISL_785916, EPI_ISL_785917, EPI_ISL_785918, EPI_ISL_785919, EPI_ISL_785920, EPI_ISL_785923, EPI_ISL_785924, EPI_ISL_785925, EPI_ISL_785926, EPI_ISL_785927, EPI_ISL_785928, EPI_ISL_785929, EPI_ISL_785930, EPI_ISL_785931, EPI_ISL_785932, EPI_ISL_785935, EPI_ISL_785936, EPI_ISL_785937, EPI_ISL_785938, EPI_ISL_785939, EPI_ISL_785940, EPI_ISL_785941, EPI_ISL_785942, EPI_ISL_785943, EPI_ISL_785944, EPI_ISL_785947, EPI_ISL_785948, EPI_ISL_785949, EPI_ISL_785950, EPI_ISL_785951, EPI_ISL_785952, EPI_ISL_785953, EPI_ISL_785954, EPI_ISL_785955, EPI_ISL_785956, EPI_ISL_785959, EPI_ISL_785960, EPI_ISL_785961, EPI_ISL_785962, EPI_ISL_785963, EPI_ISL_785964, EPI_ISL_785965, EPI_ISL_785966, EPI_ISL_785967, EPI_ISL_785968, EPI_ISL_785971, EPI_ISL_785972, EPI_ISL_785973, EPI_ISL_785974, EPI_ISL_785975, EPI_ISL_785976, EPI_ISL_785977, EPI_ISL_785978, EPI_ISL_785979, EPI_ISL_785983, EPI_ISL_785984, EPI_ISL_785985, EPI_ISL_785986, EPI_ISL_785987, EPI_ISL_785988, EPI_ISL_785989, EPI_ISL_785990, EPI_ISL_785991, EPI_ISL_785997, EPI_ISL_785998, EPI_ISL_785999, EPI_ISL_786000, EPI_ISL_786009, EPI_ISL_786010, EPI_ISL_786011, EPI_ISL_786021, EPI_ISL_786022, EPI_ISL_786023, EPI_ISL_786032, EPI_ISL_786033, EPI_ISL_786034, EPI_ISL_786035, EPI_ISL_786044, EPI_ISL_786045, EPI_ISL_786046, EPI_ISL_786047, EPI_ISL_786056, EPI_ISL_786057, EPI_ISL_786058, EPI_ISL_786059, EPI_ISL_786068, EPI_ISL_786069, EPI_ISL_786070, EPI_ISL_786071, EPI_ISL_786080, EPI_ISL_786081, EPI_ISL_786082, EPI_ISL_786083 | | | |
| see above | Houston Methodist Hospital | Houston Methodist Hospital | S. Wesley Long, Randall J. Olsen, Paul A. Christensen, David W. Bernard, James J. Davis, Maulik Shukla, Marcus Nguyen, Matthew Ojeda Saavedra, Prasanti Yerramilli, Layne Pruitt, Sishir Subedi, Heather Hendrickson, and James M. Musser |

| | | | |
|---|---|---|---|
| EPI_ISL_791338, EPI_ISL_791347, EPI_ISL_791353 | Johns Hopkins Hospital Department of Pathology | Johns Hopkins Hospital Department of Pathology | C. Paul Morris, Chun Huai Luo, Heba H. Mostafa |
| EPI_ISL_794078, EPI_ISL_794079, EPI_ISL_794080, EPI_ISL_794081, EPI_ISL_794082 | Wadsworth Center, New York State Department.of Health | Wadsworth Center, New York State Department.of Health | Kirsten St. George, Daryl M. Lamson, Alexis Russel, Matthew Shudt, Melissa A Leisner, Jonathan Plitnick, Navjot Singh, John Kelly, Sara Griesemer, Erasmus Schneider, Erica Lasek-Nesselquist |
| EPI_ISL_801407, EPI_ISL_801430, EPI_ISL_801488 | Dutch COVID-19 response team | Erasmus Medical Center | Bas Oude Munnink, Reina Sikkema, David Nieuwenhuijse, Irina Chestakova, Anne van der Linden, Marjan Boter, Emmanuelle Munger, Corine GeurtsvanKessel, Annemiek van der Eijk, Richard Molenkamp, Marion Koopmans, on behalf of the Dutch national COVID-19 response team. |
| EPI_ISL_803102 | Respiratory Viruses Branch, Centers for Disease Control and Prevention | Respiratory Viruses Branch, Centers for Disease Control and Prevention | Queen,K., Li,Y., Tao,Y., Uehara,A., Montmayeur,A., Paden,C.R., Cook,P.W., Marine,R., Sheth,M., Wang,H., Lee,J., Tong,S. |
| EPI_ISL_803426, EPI_ISL_803427, EPI_ISL_803472, EPI_ISL_803473, EPI_ISL_803474, EPI_ISL_803475, EPI_ISL_803476, EPI_ISL_803477, EPI_ISL_803478 | Wisconsin State Laboratory of Hygiene Communicable Disease Division | Wisconsin State Laboratory of Hygiene Communicable Disease Division | Kelsey R. Florek, Abigail C. Shockey |
| EPI_ISL_803866 | Institute of Virology, University of Cologne | Institute of Virology, University of Cologne | Saleta Sierra, Gibran Rubio, Zevanya Tessalonica, Dominik Aschenmeier, Eva Heger, Elena Knops, Rolf Kaiser, Maximilian Damagnez, Andreas Walker, Jörg Timm, Alexander Dilthey, Martin Däumer, Alex Thielen |
| EPI_ISL_803879 | Keio University School of Medicine | Keio University School of Medicine | Kenjiro Kosaki, Yuka Iwasaki, Hirotsugu Ishizu, Haruhiko Siomi, Kodai Abe |
| EPI_ISL_803917, EPI_ISL_803918, EPI_ISL_803921, EPI_ISL_803922, EPI_ISL_803923, EPI_ISL_803924, EPI_ISL_803925, EPI_ISL_803926, EPI_ISL_803927, EPI_ISL_803928, EPI_ISL_803929, EPI_ISL_803930, EPI_ISL_803931, EPI_ISL_803932, EPI_ISL_803933, EPI_ISL_803934, EPI_ISL_803935, EPI_ISL_803936, EPI_ISL_803937, EPI_ISL_803938, EPI_ISL_803939 | | | |
| see above | Pathogen Genomics Center, National Institute of Infectious Diseases | Pathogen Genomics Center, National Institute of Infectious Diseases | Tsuyoshi Sekizuka, Kentaro Itokawa, Rina Tanaka, Masanori Hashino, Makoto Kuroda |
| EPI_ISL_804568, EPI_ISL_804569 | MEPHI, Aix Marseille University | MEPHI, Aix Marseille University | Anthony LEVASSEUR |
| EPI_ISL_812133, EPI_ISL_812137, EPI_ISL_812180, EPI_ISL_812181 | GA Department of Public Health Laboratory | Pathogen Discovery, Respiratory Viruses Branch, Division of Viral Diseases, Centers for Disease Control and Prevention | Yan Li, Ying Tao, Anna Montmayeur, Jing Zhang, Brian Lynch, Krista Queen, Anna Uehara, Rachel Marine, Peter Cook, Clinton R. Paden, Haibin Wang, Suxiang Tong |
| EPI_ISL_812991, EPI_ISL_812992, EPI_ISL_812993, EPI_ISL_812994, EPI_ISL_812995, EPI_ISL_812996 | University of Birmingham | COVID-19 Genomics UK (COG-UK) Consortium | Institute of Microbiology, University of Birmingham: Claire McMurray, Joanne Stockton, Samuel Nicholls, Radoslaw Poplawski, Will Rowe, Josh Quick, Nicholas Loman. University of Birmingham Testing Laboratory: Celina M Whalley, Andrew Bosworth, Charlotte Poxon, Kasun Wanigasooriya, Oliver Pickles, Mike Kidd, Alex Richter, Andrew D Beggs PHE Heartlands Lab: Husam Osman, Andrew Bosworth. Queen Elizabeth Hospital: Anna Casey |
| EPI_ISL_814682, EPI_ISL_814683, EPI_ISL_814684, EPI_ISL_814687 | Virology Department, Royal Infirmary of Edinburgh, NHS Lothian / School of Biological Sciences, University of Edinburgh / Institute of Genetics and Molecular Medicine, University of Edinburgh | COVID-19 Genomics UK (COG-UK) Consortium | McHugh M, Dewar R, Rooke S, Gallagher M, Balcaza C, O'Toole Á, Scher E, Hill V, McCrone JT, Colquhoun R, Yu X, Jackson B, Rambaut A, Williams TC, Templeton K |
| EPI_ISL_814836 | Wales Specialist Virology Centre Sequencing lab: Pathogen Genomics Unit | COVID-19 Genomics UK (COG-UK) Consortium | Catherine Moore, Johnathan Evans, Laura Gifford, Malorie Perry, Simon Cottrell, Angela Marchbank, Alec Birchley, Alexander Adams, Amy Gaskin, Bree Gatica-Wilcox, Jason Coombes, Joel Southgate, Lauren Gilbert, Lee Graham, Nicole Pacchiarini, Sara Kumziene-Summerhayes, Sarah Taylor, Sophie Jones, Sara Rey, Matthew Bull, Joanne Watkins, Sally Corden, Tom Connor |
| EPI_ISL_819454, EPI_ISL_819455 | Northumbria University / South Tees Hospitals NHS Foundation Trust / North Cumbria Integrated Care NHS Foundation Trust / North Tees and Hartlepool NHS Foundation Trust / Newcastle Hospitals NHS Foundation Trust | COVID-19 Genomics UK (COG-UK) Consortium | Darren L Smith,Andrew Nelson,Matthew Bashton,Greg R Young,Joshua Loh,John Allan,Mohammad A Tariq,Giles S Holt,Gary Black,Wen C Yew,Lynn Dover,Paul Baker,Steve Liggett,Sarah Essex,Jane Greenaway,Debra Padgett,Clive Graham,Garren Scott,Edward Barton,Emma Swindells,Brendan Payne,Jennifer Collins,Yusri Taha,Gary Eltringham |
| EPI_ISL_825326, EPI_ISL_825347, EPI_ISL_825348 | Hospital Universitari Vall d'Hebron - Vall d'Hebron Institut de Rercerca | Hospital Universitari Vall d'Hebron | Cristina Andrés, Maria Piñana, Josep F Abril, Damir Garcia-Cehic, Ariadna Rando, Juliana Esperalba, Maria Gema Codina, Carla Castillo, Maria Carmen Martin, Tomàs Pumarola, Josep Quer, Andrés Antón |
| EPI_ISL_825591, EPI_ISL_825592 | Respiratory Virus Unit, National Infection Service, Public Health England | COVID-19 Genomics UK (COG-UK) Consortium | PHE Covid Sequencing Team |
| EPI_ISL_826591, EPI_ISL_826669, EPI_ISL_826670 | Montefiore Medical Center | Albert Einstein College of Medicine, Dept. of Microbiology & Immunology, Chandran lab | J. Maximilian Fels, Saad Khan, Ryan Forster, Karin A. Skalina, Surksha Sirichand, Amy S. Fox, Aviv Bergman, William B. Mitchell, Lucia R. Wolgast, Wendy Szymczak, Robert H. Bortz III, M. Eugenia Dieterle, Catalina Florez, Denise Haslwanter, Rohit K. Jangra, Ethan Laudermilch, Ariel S. Wirchnianski, Jason Barnhill, David L. Goldman, Hnin Khine, D. Yitzchak Goldstein, Johanna P. Daily, Kartik Chandran, Libusha Kelly |
| EPI_ISL_826991 | deCODE genetics | deCODE genetics | Daniel F Gudbjartsson; Agnar Helgason; Hakon Jonsson; Olafur T Magnusson; Pall Melsted; Gudmundur L Norddahl; Jona Saemundsdottir; Asgeir Sigurdsson; Patrick Sulem; Arna B Agustsdottir; Hannes Eggertsson; Berglind Eiriksdottir; Run Fridriksdottir; Elisabet E Gardarsdottir; Gudmundur Georgsson; Olafia S Gretarsdottir; Kjartan R Gudmundsson; Thora R Gunnarsdottir; Arnaldur Gylfason; Hilma Holm; Brynjar O Jensson; Aslaug Jonasdottir; Kamilla S Josefsdottir; Thordur Kristjansson; Droplaug N Magnusdottir; Solvi Rognvaldsson; Louise le Roux; Gudrun Sigmundsdottir; Gardar Sveinbjornsson; Kristin E Sveinsdottir; Maney Sveinsdottir; Emil A Thorarensen; Bjarni Thorbjornsson; Gisli Masson; Ingileif Jonsdottir; Alma Moller; Thorolfur Gudnason; Karl G Kristinsson; Unnur Thorsteinsdottir; Kari Stefansson |
| EPI_ISL_827074 | The National University Hospital of Iceland | deCODE genetics | Daniel F Gudbjartsson; Agnar Helgason; Hakon Jonsson; Olafur T Magnusson; Pall Melsted; Gudmundur L Norddahl; Jona Saemundsdottir; Asgeir Sigurdsson; Patrick Sulem; Arna B Agustsdottir; Hannes Eggertsson; Berglind Eiriksdottir; Run Fridriksdottir; Elisabet E Gardarsdottir; Gudmundur Georgsson; Olafia S Gretarsdottir; Kjartan R Gudmundsson; Thora R Gunnarsdottir; Arnaldur Gylfason; Hilma Holm; Brynjar O Jensson; Aslaug Jonasdottir; Kamilla S Josefsdottir; Thordur Kristjansson; Droplaug N Magnusdottir; Solvi Rognvaldsson; Louise le Roux; Gudrun Sigmundsdottir; Gardar Sveinbjornsson; Kristin E Sveinsdottir; Maney Sveinsdottir; Emil A Thorarensen; Bjarni Thorbjornsson; Gisli Masson; Ingileif Jonsdottir; Alma Moller; Thorolfur Gudnason; Karl G Kristinsson; Unnur Thorsteinsdottir; Kari Stefansson |
| EPI_ISL_827278, EPI_ISL_827280, EPI_ISL_827281, EPI_ISL_827282, EPI_ISL_827283, EPI_ISL_827284, EPI_ISL_827285, EPI_ISL_827305, EPI_ISL_827308, EPI_ISL_827311, EPI_ISL_827312, EPI_ISL_827314, EPI_ISL_827315, EPI_ISL_827316, EPI_ISL_827317, EPI_ISL_827318, EPI_ISL_827319, EPI_ISL_827321, EPI_ISL_827370, EPI_ISL_827401, EPI_ISL_827418, EPI_ISL_827627, EPI_ISL_827628, EPI_ISL_827726, EPI_ISL_827728, EPI_ISL_827729, EPI_ISL_827730, EPI_ISL_827731, EPI_ISL_827962, EPI_ISL_828019 | | | |
| see above | deCODE genetics | deCODE genetics | Daniel F Gudbjartsson; Agnar Helgason; Hakon Jonsson; Olafur T Magnusson; Pall Melsted; Gudmundur L Norddahl; Jona Saemundsdottir; Asgeir Sigurdsson; Patrick Sulem; Arna B Agustsdottir; Hannes Eggertsson; Berglind Eiriksdottir; Run Fridriksdottir; Elisabet E Gardarsdottir; Gudmundur Georgsson; Olafia S Gretarsdottir; Kjartan R Gudmundsson; Thora R Gunnarsdottir; Arnaldur Gylfason; Hilma Holm; Brynjar O Jensson; Aslaug Jonasdottir; Kamilla S Josefsdottir; Thordur Kristjansson; Droplaug N Magnusdottir; Solvi Rognvaldsson; Louise le Roux; Gudrun Sigmundsdottir; Gardar Sveinbjornsson; Kristin E Sveinsdottir; Maney Sveinsdottir; Emil A Thorarensen; Bjarni Thorbjornsson; Gisli Masson; Ingileif Jonsdottir; Alma Moller; Thorolfur Gudnason; Karl G Kristinsson; Unnur Thorsteinsdottir; Kari Stefansson |
| EPI_ISL_828020, EPI_ISL_828022, EPI_ISL_828025, EPI_ISL_828027 | The National University Hospital of Iceland | deCODE genetics | Daniel F Gudbjartsson; Agnar Helgason; Hakon Jonsson; Olafur T Magnusson; Pall Melsted; Gudmundur L Norddahl; Jona Saemundsdottir; Asgeir Sigurdsson; Patrick Sulem; Arna B Agustsdottir; Hannes Eggertsson; Berglind Eiriksdottir; Run Fridriksdottir; Elisabet E Gardarsdottir; Gudmundur Georgsson; Olafia S Gretarsdottir; Kjartan R Gudmundsson; Thora R Gunnarsdottir; Arnaldur Gylfason; Hilma Holm; Brynjar O Jensson; Aslaug Jonasdottir; Kamilla S Josefsdottir; Thordur Kristjansson; Droplaug N Magnusdottir; Solvi Rognvaldsson; Louise le Roux; Gudrun Sigmundsdottir; Gardar Sveinbjornsson; Kristin E Sveinsdottir; Maney Sveinsdottir; Emil A Thorarensen; Bjarni Thorbjornsson; Gisli Masson; Ingileif Jonsdottir; Alma Moller; Thorolfur Gudnason; Karl G Kristinsson; Unnur Thorsteinsdottir; Kari Stefansson |
| EPI_ISL_828028, EPI_ISL_828039, EPI_ISL_828047, EPI_ISL_828072, EPI_ISL_828295, EPI_ISL_828296, EPI_ISL_828298, EPI_ISL_828299, EPI_ISL_828301, EPI_ISL_828303, EPI_ISL_828468, EPI_ISL_828496, EPI_ISL_828512, EPI_ISL_828569, EPI_ISL_828570, EPI_ISL_828571, EPI_ISL_828573, EPI_ISL_828575, EPI_ISL_828675, EPI_ISL_828691, EPI_ISL_828717, EPI_ISL_828729, EPI_ISL_828760, EPI_ISL_828787, EPI_ISL_828900 | | | |
| see above | deCODE genetics | deCODE genetics | Daniel F Gudbjartsson; Agnar Helgason; Hakon Jonsson; Olafur T Magnusson; Pall Melsted; Gudmundur L Norddahl; Jona Saemundsdottir; Asgeir Sigurdsson; Patrick Sulem; Arna B Agustsdottir; Hannes Eggertsson; Berglind Eiriksdottir; Run Fridriksdottir; Elisabet E Gardarsdottir; Gudmundur |

| | | | |
|---|---|---|---|
| | | | Georgsson; Olafia S Gretarsdottir; Kjartan R Gudmundsson; Thora R Gunnarsdottir; Arnaldur Gylfason; Hilma Holm; Brynjar O Jensson; Aslaug Jonasdottir; Kamilla S Josefsdottir; Thordur Kristjansson; Droplaug N Magnusdottir; Solvi Rognvaldsson; Louise le Roux; Gudrun Sigmundsdottir; Gardar Sveinbjornsson; Kristin E Sveinsdottir; Maney Sveinsdottir; Emil A Thorarensen; Bjarni Thorbjornsson; Gisli Masson; Ingileif Jonsdottir; Alma Moller; Thorolfur Gudnason; Karl G Kristinsson; Unnur Thorsteinsdottir; Kari Stefansson |
| EPI_ISL_829060 | The National University Hospital of Iceland | deCODE genetics | Daniel F Gudbjartsson; Agnar Helgason; Hakon Jonsson; Olafur T Magnusson; Pall Melsted; Gudmundur L Norddahl; Jona Saemundsdottir; Asgeir Sigurdsson; Patrick Sulem; Arna B Agustsdottir; Hannes Eggertsson; Berglind Eiriksdottir; Run Fridriksdottir; Elisabet E Gardarsdottir; Gudmundur Georgsson; Olafia S Gretarsdottir; Kjartan R Gudmundsson; Thora R Gunnarsdottir; Arnaldur Gylfason; Hilma Holm; Brynjar O Jensson; Aslaug Jonasdottir; Kamilla S Josefsdottir; Thordur Kristjansson; Droplaug N Magnusdottir; Solvi Rognvaldsson; Louise le Roux; Gudrun Sigmundsdottir; Gardar Sveinbjornsson; Kristin E Sveinsdottir; Maney Sveinsdottir; Emil A Thorarensen; Bjarni Thorbjornsson; Gisli Masson; Ingileif Jonsdottir; Alma Moller; Thorolfur Gudnason; Karl G Kristinsson; Unnur Thorsteinsdottir; Kari Stefansson |
| EPI_ISL_829061, EPI_ISL_829063, EPI_ISL_829064, EPI_ISL_829065, EPI_ISL_829067, EPI_ISL_829068, EPI_ISL_829069, EPI_ISL_829070, EPI_ISL_829095, EPI_ISL_829163, EPI_ISL_829187 | | | |
| see above | deCODE genetics | deCODE genetics | Daniel F Gudbjartsson; Agnar Helgason; Hakon Jonsson; Olafur T Magnusson; Pall Melsted; Gudmundur L Norddahl; Jona Saemundsdottir; Asgeir Sigurdsson; Patrick Sulem; Arna B Agustsdottir; Hannes Eggertsson; Berglind Eiriksdottir; Run Fridriksdottir; Elisabet E Gardarsdottir; Gudmundur Georgsson; Olafia S Gretarsdottir; Kjartan R Gudmundsson; Thora R Gunnarsdottir; Arnaldur Gylfason; Hilma Holm; Brynjar O Jensson; Aslaug Jonasdottir; Kamilla S Josefsdottir; Thordur Kristjansson; Droplaug N Magnusdottir; Solvi Rognvaldsson; Louise le Roux; Gudrun Sigmundsdottir; Gardar Sveinbjornsson; Kristin E Sveinsdottir; Maney Sveinsdottir; Emil A Thorarensen; Bjarni Thorbjornsson; Gisli Masson; Ingileif Jonsdottir; Alma Moller; Thorolfur Gudnason; Karl G Kristinsson; Unnur Thorsteinsdottir; Kari Stefansson |
| EPI_ISL_829314, EPI_ISL_829315, EPI_ISL_829317, EPI_ISL_829318 | The National University Hospital of Iceland | deCODE genetics | Daniel F Gudbjartsson; Agnar Helgason; Hakon Jonsson; Olafur T Magnusson; Pall Melsted; Gudmundur L Norddahl; Jona Saemundsdottir; Asgeir Sigurdsson; Patrick Sulem; Arna B Agustsdottir; Hannes Eggertsson; Berglind Eiriksdottir; Run Fridriksdottir; Elisabet E Gardarsdottir; Gudmundur Georgsson; Olafia S Gretarsdottir; Kjartan R Gudmundsson; Thora R Gunnarsdottir; Arnaldur Gylfason; Hilma Holm; Brynjar O Jensson; Aslaug Jonasdottir; Kamilla S Josefsdottir; Thordur Kristjansson; Droplaug N Magnusdottir; Solvi Rognvaldsson; Louise le Roux; Gudrun Sigmundsdottir; Gardar Sveinbjornsson; Kristin E Sveinsdottir; Maney Sveinsdottir; Emil A Thorarensen; Bjarni Thorbjornsson; Gisli Masson; Ingileif Jonsdottir; Alma Moller; Thorolfur Gudnason; Karl G Kristinsson; Unnur Thorsteinsdottir; Kari Stefansson |
| EPI_ISL_829566, EPI_ISL_829569, EPI_ISL_829784, EPI_ISL_829785, EPI_ISL_829786, EPI_ISL_829787, EPI_ISL_829788, EPI_ISL_829789, EPI_ISL_829790, EPI_ISL_829791, EPI_ISL_829792, EPI_ISL_829822, EPI_ISL_829823, EPI_ISL_829826, EPI_ISL_829827, EPI_ISL_829960, EPI_ISL_829961, EPI_ISL_829984, EPI_ISL_829986, EPI_ISL_830006 | | | |
| see above | deCODE genetics | deCODE genetics | Daniel F Gudbjartsson; Agnar Helgason; Hakon Jonsson; Olafur T Magnusson; Pall Melsted; Gudmundur L Norddahl; Jona Saemundsdottir; Asgeir Sigurdsson; Patrick Sulem; Arna B Agustsdottir; Hannes Eggertsson; Berglind Eiriksdottir; Run Fridriksdottir; Elisabet E Gardarsdottir; Gudmundur Georgsson; Olafia S Gretarsdottir; Kjartan R Gudmundsson; Thora R Gunnarsdottir; Arnaldur Gylfason; Hilma Holm; Brynjar O Jensson; Aslaug Jonasdottir; Kamilla S Josefsdottir; Thordur Kristjansson; Droplaug N Magnusdottir; Solvi Rognvaldsson; Louise le Roux; Gudrun Sigmundsdottir; Gardar Sveinbjornsson; Kristin E Sveinsdottir; Maney Sveinsdottir; Emil A Thorarensen; Bjarni Thorbjornsson; Gisli Masson; Ingileif Jonsdottir; Alma Moller; Thorolfur Gudnason; Karl G Kristinsson; Unnur Thorsteinsdottir; Kari Stefansson |
| EPI_ISL_830316, EPI_ISL_830327, EPI_ISL_830537 | The National University Hospital of Iceland | deCODE genetics | Daniel F Gudbjartsson; Agnar Helgason; Hakon Jonsson; Olafur T Magnusson; Pall Melsted; Gudmundur L Norddahl; Jona Saemundsdottir; Asgeir Sigurdsson; Patrick Sulem; Arna B Agustsdottir; Hannes Eggertsson; Berglind Eiriksdottir; Run Fridriksdottir; Elisabet E Gardarsdottir; Gudmundur Georgsson; Olafia S Gretarsdottir; Kjartan R Gudmundsson; Thora R Gunnarsdottir; Arnaldur Gylfason; Hilma Holm; Brynjar O Jensson; Aslaug Jonasdottir; Kamilla S Josefsdottir; Thordur Kristjansson; Droplaug N Magnusdottir; Solvi Rognvaldsson; Louise le Roux; Gudrun Sigmundsdottir; Gardar Sveinbjornsson; Kristin E Sveinsdottir; Maney Sveinsdottir; Emil A Thorarensen; Bjarni Thorbjornsson; Gisli Masson; Ingileif Jonsdottir; Alma Moller; Thorolfur Gudnason; Karl G Kristinsson; Unnur Thorsteinsdottir; Kari Stefansson |
| EPI_ISL_830732 | University Hospital Basel, Clinical Virology | University Hospital Basel, Clinical Bacteriology | Tim Roloff, Madlen Stange, Helena MB Seth-Smith, Alfredo Mari, Karoline Leuzinger, Julia Bielicki, Manuel Battegay, Hans Hirsch, Adrian Egli |
| EPI_ISL_831224, EPI_ISL_831225, EPI_ISL_831226, EPI_ISL_831227 | Hospital Universitario La Paz (Madrid) | SeqCOVID-SPAIN consortium/IBV(CSIC) | María Rodríguez-Tejedor, Elias Dahdouh, Fernando Lázaro-Perona, Jesús Mingorance and SeqCOVID-SPAIN consortium |
| EPI_ISL_831763 | United States Air Force School of Aerospace Medicine | United States Air Force School of Aerospace Medicine | Anthony Fries, Jennifer Meyer, William Gruner, Amanda Javorina, Sarah Purves, Clarise Starr, Elizabeth Macias |
| EPI_ISL_832597, EPI_ISL_832598, EPI_ISL_832599, EPI_ISL_832600, EPI_ISL_832601, EPI_ISL_832602, EPI_ISL_832603, EPI_ISL_832604, EPI_ISL_832605, EPI_ISL_832606, EPI_ISL_832607, EPI_ISL_832608, EPI_ISL_832655, EPI_ISL_832656, EPI_ISL_832657, EPI_ISL_832658, EPI_ISL_832659, EPI_ISL_832660, EPI_ISL_832661, EPI_ISL_832662, EPI_ISL_832663, EPI_ISL_832664, EPI_ISL_832679, EPI_ISL_832680, EPI_ISL_832681, EPI_ISL_832682, EPI_ISL_832683, EPI_ISL_832684, EPI_ISL_832694 | | | |
| see above | OHSU Lab Services Molecular Microbiology Lab | Oregon SARS-CoV-2 Genome Sequencing Center | Brendan L. O'Connell, Ruth V. Nichols, Sally Grindstaff, Alec J. Hirsch, Donna Hansel, Guang Fan, Daniel N. Streblow, William B. Messer, Andrew C. Adey, Benjamin N. Bimber, Brian J. O'Roak |
| EPI_ISL_839985 | Queens Medical Centre, Clinical Microbiology Department / DeepSeq Nottingham | COVID-19 Genomics UK (COG-UK) Consortium | Gemma Clark, Wendy Smith, Manjinder Khakh, Vicki M Fleming, Michelle M Lister, Hannah Howson-Wells, Jonathan Ball, Patrick McClure, Joseph Chappell, Theocharis Tsoleridis, Nadine Holmes, Matthew Carlisle, Christopher Moore, Fei Sang, Johnny Debebe, Victoria Wright, Matthew Loose |
| EPI_ISL_842347, EPI_ISL_842352 | Virology Department, Sheffield Teaching Hospitals NHS Foundation Trust/Department of Infection, Immunity and Cardiovascular Disease, The Medical School, University of Sheffield | COVID-19 Genomics UK (COG-UK) Consortium | Thushan de Silva, Matthew Parker, Nikki Smith, Adri Angyal, Rebecca Brown, Luke Green, Rachel Tucker, Paul Parsons, Danielle Groves, Katie Johnson, Laura Carrilero, Alex Keeley, Dave Partridge, Matthew Wyles, Benjamin Lindsey, Mehmet Yavuz, Mohammad Raza, Cariad Evans |
| EPI_ISL_842867, EPI_ISL_842868 | Barts Health NHS Trust | COVID-19 Genomics UK (COG-UK) Consortium | CUTINO-MOGUEL, Maria-Teresa; HARRINGTON, David; OWOYEMI, Dola; SHYLINI, Raghavendran; BROAD, Claire; KELE, Beatrix |
| EPI_ISL_849441, EPI_ISL_849499, EPI_ISL_849607 | Seattle Flu Study | Seattle Flu Study | Deborah A. Nickerson, Chris D. Frazar, Jover Lee, Benjamin Pelle, Matthew Richardson, Amanda Adler, Elisabeth Brandstetter, Peter D. Han, Kairsten Fay, Misja Ilcisin, Kirsten Lacombe, Thomas R. Sibley, Melissa Truong, Caitlin R. Wolf, Michael Boeckh, Janet A. Englund, Michael Famulare, Barry R. Lutz, Mark J. Rieder, Lea M. Starita, Matthew Thompson, Jay Shendure, Trevor Bedford, Helen Y. Chu |
| EPI_ISL_850950 | National Institute for Viral Disease Control and Prevention, China CDC | National Institute for Viral Disease Control and Prevention, China CDC | Xiang Zhao, Yenan Feng, Zhixiao Chen, Yao Meng, Yuchao Wu, Yang Song, Ji Wang, Kai Nie, Yong Zhang, Yanhai Wang, Weimin Zhou, Wenjie Tan, Jun Han, Shiwen Wang, Wenbo Xu, Cao Chen, Dayan Wang |
| EPI_ISL_851055, EPI_ISL_851056, EPI_ISL_851057, EPI_ISL_851058, EPI_ISL_851059, EPI_ISL_851060, EPI_ISL_851061 | National Food and Veterinary Risk Assessment Institute | Lithuanian University of Health Sciences, Molecular cardiology lab. | Lukas Zemaitis, Arnoldas Pautienius, Ingrida Olendrait, Kamile Tamusauskaite, Dovydas Gecys, Laura Pareckaite, Vaiva Lesauskaite, Astra Vitkauskiene, Arunas Stankevicius, Algirdas Malakauskas |
| EPI_ISL_852983 | Hospital General Universitario Gregorio Marañón | SeqCOVID-SPAIN consortium/IBV(CSIC) | Darío García de Viedma, Laura Pérez-Lago, Pedro J Sola-Campoy, Sergio Buenestado-Serrano, Marta Herranz, Victor Manuel de la Cueva, Julia Suárez, Pilar Catalán, Patricia Muñoz and SeqCOVID-SPAIN consortium |
| EPI_ISL_853316 | UPMC Clinical Microbiology Laboratory | Microbial Genome Sequencing Center; Microbial Genomic Epidemiology Laboratory | Mustapha M. Mustapha, Jane W. Marsh, Dan Snyder, Marissa P. Griffith, Stephanie L. Mitchell, Vatsala R. Srinivasa, Kady D. Waggle, Chinelo Ezeonwuku, Vaughn S. Cooper, Lee H. Harrison |
| EPI_ISL_853936, EPI_ISL_853938, EPI_ISL_853944 | Department of Microbiology, University Innsbruck | Bergthaler laboratory, CeMM Research Center for Molecular Medicine of the Austrian Academy of Sciences | Lukas Endler, Alexandra Popa, Benedikt Agerer, Jakob-Wendelin Genger, Alexander Lercher, Anna Schedl, Thomas Penz, Michael Schuster, Jan Laine, Martin Senekowitsch, Christoph Bock, Andreas Bergthaler |
| EPI_ISL_854613, EPI_ISL_854614, EPI_ISL_854615, EPI_ISL_854616, EPI_ISL_854617, EPI_ISL_854618, EPI_ISL_854619, EPI_ISL_854620, EPI_ISL_854621, EPI_ISL_854622, EPI_ISL_854623, EPI_ISL_854624, EPI_ISL_854715, EPI_ISL_854736 | | | |
| see above | Public Health Ontario Laboratory | Public Health Ontario Laboratory | Vanessa G Allen, Philip Banh, Yao Chen, Richard de Borja, Alireza Eshaghi, Nahuel Fittipaldi, Christine Frantz, Jonathan B Gubbay, Jennifer L Guthrie, Lawrence Heisler, Esha Joshi, Michael Laszloffy, Aimin Li, Michael CY Li, Dean Maxwell, Sandeep Nagra, Samir N Patel, Jared Simpson, Karthikeyan Sivaraman, Ashleigh Sullivan, Yogi Sundaravadanam, Sarah Teatero, Matthew Watson, Andre Villegas, Sandra Zittermann |
| EPI_ISL_855500 | KEMRI-Wellcome Trust Research Programme/KEMRI-CGMR-C Kilifi | KEMRI-Wellcome Trust Research Programme/KEMRI-CGMR-C Kilifi | Githinji et al |
| EPI_ISL_856794, EPI_ISL_856795, EPI_ISL_856796 | Servicio Virosis Respiratorias-Departamento Virología-INEI | Instituto Nacional Enfermedades Infecciosas C.G.Malbran | Baumeister E., Avaro M., Benedetti E., Russo M., Dattero ME, Pontoriero A., Cisterna D., Molina V., Perandones C., Tuduri E., Lorenzo F., Poklepovich T., Campos J. |

| | | | |
|---|---|---|---|
| EPI_ISL_859797, EPI_ISL_859800 | BTC, Khalifa University | BTC, Khalifa University | Al Safar et al |
| EPI_ISL_860160 | Keio University School of Medicine | Keio University School of Medicine | Kenjiro Kosaki, Yuka Iwasaki, Hirotsugu Ishizu, Haruhiko Siomi, Kodai Abe |
| EPI_ISL_860732, EPI_ISL_860773, EPI_ISL_860774, EPI_ISL_860775, EPI_ISL_860776 | Swiss National Reference Centre for Influenza | Swiss National Reference Centre for Influenza | Ana Rita Gonçalves Cabecinhas, Samuel Cordey, Florian Laubscher,Christoph Grünig, Laurent Kaiser |
| EPI_ISL_860814 | WHO/Minsk | Charité Universitätsmedizin Berlin, Institut für Virologie | Victor M Corman, Barbara Mühlemann, Jörn Beheim-Schwarzbach, Talitha Veith, Julia Tesch, Tobias Bleicker, Julia Schneider, Shmialiova Natallia, Sivets Natallia, Terry Jones, Christian Drosten |
| EPI_ISL_861461 | UHAS COVID-19 Lab | UHAS COVID-19 Lab | Kwabena O. Duedu, Jones Gyamfi, Reuben Ayivor-Djanie, John O. Gyapong and the UHAS COVID-19 Lab Team |
| EPI_ISL_861671 | Hospital Municipal Prefeito Waldemar Costa Filho | Instituto Adolfo Lutz, Interdiciplinary Procedures Center, Strategic Laboratory | Claudio Tavares Sacchi, Claudia Regina Gonçalves, Erica Valessa Ramos Gomes, Karoline Rodrigues Campos |
| EPI_ISL_867964 | Centre for Enzyme Innovation, University of Portsmouth / Translational Research Laboratory, Portsmouth Hospitals NHS Trust | COVID-19 Genomics UK (COG-UK) Consortium | Angela Beckett,Yann Bourgeois,Garry Scarlett,Sharon Glaysher,Scott Elliott,Kelly Bicknell,Robert Impey,Allyson Lloyd,Sarah Wyllie,Ethan Butcher,Anoop Chauhan,Samuel Robson |
| EPI_ISL_868794, EPI_ISL_868796, EPI_ISL_868803, EPI_ISL_868812, EPI_ISL_868814 | Bioinformatics and Biostatistics Lab, Advanced Sequencing Facility | COVID-19 Genomics UK (COG-UK) Consortium | Aengus Stewart,Jerome Nicod,Chelsea Sawyer,Laura Cubitt,Harshil Patel,Margaret Crawford |
| EPI_ISL_872745 | Colorado Mesa University | Infectious Disease Program, Broad Institute of Harvard and MIT | Lemieux,J.E., Siddle,K.J., Marshall,J., ONeill,M., Bronson,A., Adams,G., Gladden-Young,A., Lagerborg,K., Rudy,M., DeRuff,K., Carter,A., Normandin,E., Bauer,M., Reilly,S., Tomkins-Tinch,C., Loreth,C., Chaluvadi,S., Birren,B.W., Gallagher,G., Smole,S., Park,D.J., MacInnis,B.L., and Sabeti,P.C. |
| EPI_ISL_875814, EPI_ISL_875815 | Public Health Ontario Laboratory | Public Health Ontario Laboratory | Vanessa G Allen, Philip Banh, Yao Chen, Richard de Borja, Alireza Eshaghi, Nahuel Fittipaldi, Christine Frantz, Jonathan B Gubbay, Jennifer L Guthrie, Lawrence Heisler, Esha Joshi, Michael Laszloffy, Aimin Li, Michael CY Li, Dean Maxwell, Sandeep Nagra, Samir N Patel, Jared Simpson, Karthikeyan Sivaraman, Ashleigh Sullivan, Yogi Sundaravadanam, Sarah Teatero, Matthew Watson, Andre Villegas, Sandra Zittermann |
| EPI_ISL_875876, EPI_ISL_875878, EPI_ISL_875880, EPI_ISL_875881, EPI_ISL_875885, EPI_ISL_875888, EPI_ISL_875889, EPI_ISL_875900, EPI_ISL_875902, EPI_ISL_875905, EPI_ISL_875913, EPI_ISL_875914, EPI_ISL_875916, EPI_ISL_875917, EPI_ISL_875918, EPI_ISL_875919, EPI_ISL_875920, EPI_ISL_875923, EPI_ISL_875925, EPI_ISL_875926, EPI_ISL_875928, EPI_ISL_875929, EPI_ISL_875930, EPI_ISL_875931, EPI_ISL_875932, EPI_ISL_875933, EPI_ISL_875935, EPI_ISL_875936, EPI_ISL_875937, EPI_ISL_875940, EPI_ISL_875941, EPI_ISL_875942, EPI_ISL_875943, EPI_ISL_875944, EPI_ISL_875947, EPI_ISL_875951, EPI_ISL_875955, EPI_ISL_875956, EPI_ISL_875957 | | | |
| see above | Eurofins Diatherix | Hudsonalpha Genome Sequencing Center | Jane Grimwood, Melissa Williams, Lori H. Handley, Joshua Stough, Leslie Malone, Stefan Brzezinski, Ada Stewart, Teresa Jones, Jenell Webber, John Lovell, Jennifer Cart, and Jeremy Schmutz |
| EPI_ISL_876819, EPI_ISL_876829, EPI_ISL_876850, EPI_ISL_876851, EPI_ISL_876852, EPI_ISL_876853, EPI_ISL_876854, EPI_ISL_876855, EPI_ISL_876856, EPI_ISL_876857, EPI_ISL_876858, EPI_ISL_876859 | | | |
| see above | Quest Diagnostics | Quest Diagnostics | Rosenthal,S.H., Gerasimova,A., Kagan,R.M., Anderson, B., Hua, M., Liu Y., Bernstein, L.E., Livingston, K.E., Perez, A., Shalhout, D.F., Shlyakhter, I.A., Owen, R., Tanpaiboon, P., Lacbawan, F. |
| EPI_ISL_877428, EPI_ISL_877429 | Institute of Microbiology and Immunology, Faculty of Medicine, University of Ljubljana | Institute of Microbiology and Immunology, Faculty of Medicine, University of Ljubljana | Samo Zakotnik, Tomaž Mark Zorec, Matic Brvar, Miša Korva, Mario Poljak, Tatjana Avši - Županc |
| EPI_ISL_882687, EPI_ISL_882710, EPI_ISL_882761 | 1.AO Universitaria 'S. Giovanni di Dio e Ruggi D'Aragona, Scuola Medica Salernitana' Hospital / 2.UOC di Virologia e Microbiologia, Università della Campania 'L. Vanvitelli' / 3.AO Universitaria 'Federico II' Napoli Hospital / 4.AORN 'San Giuseppe Moscati' Avellino Hospital / 5.AO 'San Pio - presidio G. Rummo' Benevento Hospital / 6.AO 'Sant'Anna e San Sebastiano' Caserta Hospital / 7.PO 'Maria Santissima Addolorata' Eboli Hospital / 8.Biogem Istituto di Ricerche Genetiche | 1. Genome Research Center for Health (CRGS) / 2. Laboratory of Molecular Medicine and Genomics(LMMGe) / 3. Center for Research in Pure and Applied Mathematics (CRMPA) | Giorgio Giurato, Francesca Rizzo, Alessandro Weisz, Gianluigi Franci, Giovanni Nassa, Pasquale Pagliano, Roberta Tarallo, Elena Alexandrova, Ylenia D'Agostino, Carlo Ferravante, Jessica Lamberti, Viola Melone, Domenico Memoli, Valeria Mirici Cappa, Domenico Palumbo, Giovanni Pecoraro, Assunta Sellitto, Oriana Strianese, Ilaria Terenzi, Giuseppe Fenza, Aniello Gentile, Antonello Saccomanno, Sonia Amabile, Teresa Rocco, Annamaria Salvati, Emilia Vaccaro, Massimiliano Galdiero, Michele Cennamo, Giuseppe Portella, Maria Grazia Foti, Mariarosaria Ingino, Maria Landi, Maurizio Fumi, Vincenzo Rocco, Rita Greco, Vittoria Letizia, Arnolfo Petruzziello, Maddalena Schioppa, Gregorio Goffredi, Francesca Marciano, Michele Caraglia, Alessia Cossu, Marianna Scrima |
| EPI_ISL_884318, EPI_ISL_884319, EPI_ISL_884370 | Infectious Diseases, Quest Diagnostics | Infectious Diseases, Quest Diagnostics | Rosenthal,S.H., Gerasimova,A., Kagan,R.M., Anderson,B., Bernstein,L.E., Livingston,K.E., Hua,M., Liu,Y., Shalhout,D.F., Owen,R., Lacbawan,F. |
| EPI_ISL_885128 | California Institute of Technology | Chan-Zuckerberg Biohub | CZB Cliahub Consortium |
| EPI_ISL_887158 | Massachusetts General Hospital | Infectious Disease Program, Broad Institute of Harvard and MIT | Lemieux,J.E., Siddle,K.J., Shaw,B., Adams,G., Pierce,V., Turbett,S., Anahtar,M., Branda,J., Slater,D., Harris,J., Lin,A.E., Gladden-Young,A., Lagerborg,K., Rudy,M., DeRuff,K., Carter,A., Normandin,E., Bauer,M., Reilly,S., Tomkins-Tinch,C., Loreth,C., Chaluvadi,S., Neumann,A., Cusick,C., Chapman,S.B., Gnirke,A., Flowers,K., Cerrato,F., Birren,B.W., Gallagher,G., Smole,S., Park,D.J., MacInnis,B.L., Ryan,E., LaRocque,R., Rosenberg,E. and Sabeti,P.C. |
| EPI_ISL_892373, EPI_ISL_892374, EPI_ISL_892375, EPI_ISL_892376, EPI_ISL_892377, EPI_ISL_892407, EPI_ISL_892408, EPI_ISL_892409, EPI_ISL_892410, EPI_ISL_892411, EPI_ISL_892412, EPI_ISL_892413, EPI_ISL_892414, EPI_ISL_892566, EPI_ISL_892567, EPI_ISL_892568, EPI_ISL_892569, EPI_ISL_892570, EPI_ISL_892571, EPI_ISL_892572, EPI_ISL_892573, EPI_ISL_892574, EPI_ISL_892575, EPI_ISL_892576, EPI_ISL_892577, EPI_ISL_892578, EPI_ISL_892579, EPI_ISL_892580, EPI_ISL_892581, EPI_ISL_892582, EPI_ISL_892583, EPI_ISL_892584, EPI_ISL_892700, EPI_ISL_893252, EPI_ISL_893253, EPI_ISL_893254, EPI_ISL_893272, EPI_ISL_893273, EPI_ISL_893274, EPI_ISL_893275, EPI_ISL_893276, EPI_ISL_893277, EPI_ISL_893278, EPI_ISL_893279, EPI_ISL_893280 | | | |
| see above | Pathogen Genomics Center, National Institute of Infectious Diseases | Pathogen Genomics Center, National Institute of Infectious Diseases | Tsuyoshi Sekizuka, Kentaro Itokawa, Rina Tanaka, Masanori Hashino, Makoto Kuroda |
| EPI_ISL_893369, EPI_ISL_893370, EPI_ISL_893372, EPI_ISL_893373, EPI_ISL_893374, EPI_ISL_893409, EPI_ISL_893410, EPI_ISL_893411, EPI_ISL_893412, EPI_ISL_893413, EPI_ISL_893414, EPI_ISL_893415, EPI_ISL_893416, EPI_ISL_893417, EPI_ISL_893418, EPI_ISL_893419, EPI_ISL_893443, EPI_ISL_893444, EPI_ISL_893445, EPI_ISL_893446, EPI_ISL_893447, EPI_ISL_893448, EPI_ISL_893449, EPI_ISL_893450, EPI_ISL_893451, EPI_ISL_893452, EPI_ISL_893453, EPI_ISL_893454, EPI_ISL_893455, EPI_ISL_893456, EPI_ISL_893457, EPI_ISL_893458, EPI_ISL_893459, EPI_ISL_893460, EPI_ISL_893461, EPI_ISL_893462, EPI_ISL_893463, EPI_ISL_893464, EPI_ISL_893465, EPI_ISL_893466, EPI_ISL_893467, EPI_ISL_893468, EPI_ISL_893469, EPI_ISL_893470, EPI_ISL_893471, EPI_ISL_893472, EPI_ISL_893473, EPI_ISL_893474, EPI_ISL_893475, EPI_ISL_893476, EPI_ISL_893477, EPI_ISL_893478, EPI_ISL_893479, EPI_ISL_893480, EPI_ISL_893481, EPI_ISL_893482, EPI_ISL_893483, EPI_ISL_893484, EPI_ISL_893485, EPI_ISL_893486, EPI_ISL_893487, EPI_ISL_893488, EPI_ISL_893489, EPI_ISL_893490, EPI_ISL_893491, EPI_ISL_893492, EPI_ISL_893493, EPI_ISL_893494, EPI_ISL_893495, EPI_ISL_893496, EPI_ISL_893497, EPI_ISL_893498, EPI_ISL_893499, EPI_ISL_893500, EPI_ISL_893501, EPI_ISL_893502, EPI_ISL_893503, EPI_ISL_893504, EPI_ISL_893505, EPI_ISL_893506, EPI_ISL_893507, EPI_ISL_893508, EPI_ISL_893509, EPI_ISL_893510, EPI_ISL_893511, EPI_ISL_893512, EPI_ISL_893513, EPI_ISL_893514, EPI_ISL_893515, EPI_ISL_893516, EPI_ISL_893517, EPI_ISL_893518, EPI_ISL_893519, EPI_ISL_893520, EPI_ISL_893521, EPI_ISL_893522, EPI_ISL_893523, EPI_ISL_893524, EPI_ISL_893525, EPI_ISL_893526, EPI_ISL_893527, EPI_ISL_893528, EPI_ISL_893529, EPI_ISL_893530, EPI_ISL_893531, EPI_ISL_893532, EPI_ISL_893533, EPI_ISL_893534, EPI_ISL_893535, EPI_ISL_893536, EPI_ISL_893537, EPI_ISL_893538, EPI_ISL_893539, EPI_ISL_893540, EPI_ISL_893541, EPI_ISL_893542, EPI_ISL_893543, EPI_ISL_893544, EPI_ISL_893545, EPI_ISL_893546, EPI_ISL_893547, EPI_ISL_893548, EPI_ISL_893549, EPI_ISL_893550, EPI_ISL_893551, EPI_ISL_893552, EPI_ISL_893553, EPI_ISL_893554, EPI_ISL_893555, EPI_ISL_893556, EPI_ISL_893557, EPI_ISL_893558, EPI_ISL_893559, EPI_ISL_893560, EPI_ISL_893561, EPI_ISL_893562, EPI_ISL_893563, EPI_ISL_893564, EPI_ISL_893565, EPI_ISL_893566, EPI_ISL_893567, EPI_ISL_893568, EPI_ISL_893569, EPI_ISL_893570, EPI_ISL_893571, EPI_ISL_893572, EPI_ISL_893573, EPI_ISL_893574, EPI_ISL_893575, EPI_ISL_893576, EPI_ISL_893577, EPI_ISL_893578, EPI_ISL_893579, EPI_ISL_893580, EPI_ISL_893581, EPI_ISL_893582, EPI_ISL_893583, EPI_ISL_893584, EPI_ISL_893585, EPI_ISL_893586, EPI_ISL_893587, EPI_ISL_893588, EPI_ISL_893589, EPI_ISL_893590, EPI_ISL_893591, EPI_ISL_893592, EPI_ISL_893593, EPI_ISL_893594, EPI_ISL_893595, EPI_ISL_893596, EPI_ISL_893597, EPI_ISL_893598, EPI_ISL_893599, EPI_ISL_893600, EPI_ISL_893601, EPI_ISL_893602, EPI_ISL_893603, EPI_ISL_893604, EPI_ISL_893605, EPI_ISL_893606, EPI_ISL_893607, EPI_ISL_893674, EPI_ISL_893675, EPI_ISL_893676 | | | |
| see above | Department of Infectious Diseases, Kobe Institute of Health | Department of Infectious Diseases, Kobe Institute of Health | Ryohei Nomoto, Tomotada Iwamoto, Tsuyoshi Sekizuka, Kentaro Itokawa, Rina Tanaka, Masanori Hashino, Makoto Kuroda |
| EPI_ISL_894287 | Fukuoka Institute of Health and Environmental Sciences | Pathogen Genomics Center, National Institute of Infectious Diseases | Tsuyoshi Sekizuka, Kentaro Itokawa, Rina Tanaka, Masanori Hashino, Makoto Kuroda |
| EPI_ISL_894297, EPI_ISL_894300, EPI_ISL_894303, EPI_ISL_894531, EPI_ISL_894532, EPI_ISL_894533, EPI_ISL_894534, EPI_ISL_894535, EPI_ISL_894536, EPI_ISL_894537, EPI_ISL_894538, EPI_ISL_894539, EPI_ISL_894540, EPI_ISL_894541, EPI_ISL_894542, EPI_ISL_894543, EPI_ISL_894544, EPI_ISL_894545, EPI_ISL_894546, EPI_ISL_894547, EPI_ISL_894548, EPI_ISL_894549, EPI_ISL_894550, EPI_ISL_894551, EPI_ISL_894552, EPI_ISL_894553, EPI_ISL_894554, EPI_ISL_894555, EPI_ISL_894556, EPI_ISL_894557, EPI_ISL_894558, EPI_ISL_894559, EPI_ISL_894560, EPI_ISL_894561, EPI_ISL_894562, EPI_ISL_894563, EPI_ISL_894564, EPI_ISL_894565, EPI_ISL_894566, EPI_ISL_894567, EPI_ISL_894568, EPI_ISL_894569, EPI_ISL_894570, EPI_ISL_894571, EPI_ISL_894572, EPI_ISL_894573, EPI_ISL_894574, EPI_ISL_894575, EPI_ISL_894576, EPI_ISL_894577, EPI_ISL_894578, EPI_ISL_894579, EPI_ISL_894580, EPI_ISL_894581, EPI_ISL_894582, EPI_ISL_894583, EPI_ISL_894584, EPI_ISL_894585, EPI_ISL_894586, EPI_ISL_894587, EPI_ISL_894588, EPI_ISL_894589, EPI_ISL_894590, EPI_ISL_894591, EPI_ISL_894592, EPI_ISL_894593, EPI_ISL_894594, EPI_ISL_894595, EPI_ISL_894596, EPI_ISL_894597, EPI_ISL_894598, EPI_ISL_894599, EPI_ISL_894600, EPI_ISL_894601, EPI_ISL_894602, EPI_ISL_894603, EPI_ISL_894604, EPI_ISL_894605, EPI_ISL_894606, EPI_ISL_894607, EPI_ISL_894608, EPI_ISL_894609, EPI_ISL_894610, EPI_ISL_894611, EPI_ISL_894612, EPI_ISL_894613, EPI_ISL_894614, EPI_ISL_894615, EPI_ISL_894616, EPI_ISL_894617, EPI_ISL_894618, EPI_ISL_894619, EPI_ISL_894620, EPI_ISL_894621, EPI_ISL_894622, EPI_ISL_894623 | | | |
| see above | Pathogen Genomics Center, National Institute of Infectious Diseases | Pathogen Genomics Center, National Institute of Infectious Diseases | Tsuyoshi Sekizuka, Kentaro Itokawa, Rina Tanaka, Masanori Hashino, Makoto Kuroda |
| EPI_ISL_894672, EPI_ISL_894673, EPI_ISL_894674, EPI_ISL_894675, EPI_ISL_894676, EPI_ISL_894677, EPI_ISL_894678, EPI_ISL_894679, EPI_ISL_894680, EPI_ISL_894688, EPI_ISL_894689, EPI_ISL_894690, EPI_ISL_894691, EPI_ISL_894692, EPI_ISL_894693, EPI_ISL_894694 | | | |

| | | | |
|---|---|---|---|
| see above | Fukuoka Institute of Health and Environmental Sciences | Pathogen Genomics Center, National Institute of Infectious Diseases | Tsuyoshi Sekizuka, Kentaro Itokawa, Rina Tanaka, Masanori Hashino, Makoto Kuroda |
| EPI_ISL_894817, EPI_ISL_894818, EPI_ISL_894819, EPI_ISL_894893, EPI_ISL_894894, EPI_ISL_894895, EPI_ISL_894896, EPI_ISL_894897, EPI_ISL_894898, EPI_ISL_894899, EPI_ISL_894900, EPI_ISL_894901, EPI_ISL_894902, EPI_ISL_894903, EPI_ISL_894904, EPI_ISL_894905, EPI_ISL_894906, EPI_ISL_894907, EPI_ISL_894908, EPI_ISL_894909, EPI_ISL_894910, EPI_ISL_894911, EPI_ISL_894912, EPI_ISL_894948, EPI_ISL_894949, EPI_ISL_894950, EPI_ISL_894951, EPI_ISL_894952, EPI_ISL_894953, EPI_ISL_894954, EPI_ISL_894955, EPI_ISL_894956, EPI_ISL_894957, EPI_ISL_894958, EPI_ISL_894967, EPI_ISL_894968, EPI_ISL_894969, EPI_ISL_894970, EPI_ISL_894971, EPI_ISL_894972, EPI_ISL_894973, EPI_ISL_894974, EPI_ISL_894975, EPI_ISL_894976, EPI_ISL_894977, EPI_ISL_894978, EPI_ISL_894979, EPI_ISL_894980, EPI_ISL_894981, EPI_ISL_894982, EPI_ISL_894983, EPI_ISL_894984, EPI_ISL_894985, EPI_ISL_894986, EPI_ISL_894987, EPI_ISL_894988, EPI_ISL_894989, EPI_ISL_894990, EPI_ISL_894991, EPI_ISL_894992, EPI_ISL_894993, EPI_ISL_894994, EPI_ISL_894995, EPI_ISL_894996, EPI_ISL_894997, EPI_ISL_894998, EPI_ISL_894999, EPI_ISL_895000, EPI_ISL_895001 | | | |
| see above | Pathogen Genomics Center, National Institute of Infectious Diseases | Pathogen Genomics Center, National Institute of Infectious Diseases | Tsuyoshi Sekizuka, Kentaro Itokawa, Rina Tanaka, Masanori Hashino, Makoto Kuroda |
| EPI_ISL_895002, EPI_ISL_895003, EPI_ISL_895004, EPI_ISL_895005, EPI_ISL_895006, EPI_ISL_895007, EPI_ISL_895008, EPI_ISL_895009, EPI_ISL_895010, EPI_ISL_895011, EPI_ISL_895012 | | | |
| see above | Niigata Prefectural Institute of Public Health and Environmental Sciences | Pathogen Genomics Center, National Institute of Infectious Diseases | Tsuyoshi Sekizuka, Kentaro Itokawa, Rina Tanaka, Masanori Hashino, Makoto Kuroda |
| EPI_ISL_895014, EPI_ISL_895015, EPI_ISL_895016, EPI_ISL_895017, EPI_ISL_895018, EPI_ISL_895019, EPI_ISL_895020, EPI_ISL_895021, EPI_ISL_895022, EPI_ISL_895070, EPI_ISL_895071, EPI_ISL_895072, EPI_ISL_895073, EPI_ISL_895074, EPI_ISL_895075, EPI_ISL_895076, EPI_ISL_895077, EPI_ISL_895078, EPI_ISL_895079, EPI_ISL_895080, EPI_ISL_895081, EPI_ISL_895082, EPI_ISL_895083, EPI_ISL_895084, EPI_ISL_895085, EPI_ISL_895086, EPI_ISL_895087, EPI_ISL_895088, EPI_ISL_895093 | | | |
| see above | Department of Infectious Diseases, Kobe Institute of Health | Pathogen Genomics Center, National Institute of Infectious Diseases | Tsuyoshi Sekizuka, Kentaro Itokawa, Rina Tanaka, Masanori Hashino, Makoto Kuroda |
| EPI_ISL_895106, EPI_ISL_895107, EPI_ISL_895108, EPI_ISL_895109, EPI_ISL_895110, EPI_ISL_895111, EPI_ISL_895112, EPI_ISL_895113, EPI_ISL_895114, EPI_ISL_895115, EPI_ISL_895116, EPI_ISL_895117, EPI_ISL_895118, EPI_ISL_895119, EPI_ISL_895120, EPI_ISL_895121, EPI_ISL_895122, EPI_ISL_895123, EPI_ISL_895124, EPI_ISL_895125, EPI_ISL_895126, EPI_ISL_895127, EPI_ISL_895128, EPI_ISL_895129, EPI_ISL_895130, EPI_ISL_895131, EPI_ISL_895132, EPI_ISL_895133, EPI_ISL_895134, EPI_ISL_895135, EPI_ISL_895136, EPI_ISL_895137, EPI_ISL_895138, EPI_ISL_895139, EPI_ISL_895140, EPI_ISL_895141, EPI_ISL_895142, EPI_ISL_895143, EPI_ISL_895144, EPI_ISL_895145, EPI_ISL_895146, EPI_ISL_895147, EPI_ISL_895148, EPI_ISL_895149, EPI_ISL_895150, EPI_ISL_895179, EPI_ISL_895180, EPI_ISL_895181, EPI_ISL_895182, EPI_ISL_895183, EPI_ISL_895184, EPI_ISL_895185, EPI_ISL_895189, EPI_ISL_895190, EPI_ISL_895191, EPI_ISL_895192, EPI_ISL_895193, EPI_ISL_895194, EPI_ISL_895195, EPI_ISL_895196, EPI_ISL_895197, EPI_ISL_895198, EPI_ISL_895199, EPI_ISL_895200, EPI_ISL_895201, EPI_ISL_895202, EPI_ISL_895203, EPI_ISL_895204, EPI_ISL_895205, EPI_ISL_895206, EPI_ISL_895207, EPI_ISL_895208, EPI_ISL_895209, EPI_ISL_895210, EPI_ISL_895211, EPI_ISL_895212, EPI_ISL_895213, EPI_ISL_895214, EPI_ISL_895215, EPI_ISL_895216, EPI_ISL_895217, EPI_ISL_895218, EPI_ISL_895219, EPI_ISL_895220, EPI_ISL_895221, EPI_ISL_895222, EPI_ISL_895223, EPI_ISL_895224, EPI_ISL_895225, EPI_ISL_895226, EPI_ISL_895227, EPI_ISL_895228, EPI_ISL_895229, EPI_ISL_895230, EPI_ISL_895231, EPI_ISL_895232, EPI_ISL_895233, EPI_ISL_895234, EPI_ISL_895235, EPI_ISL_895236, EPI_ISL_895237, EPI_ISL_895238, EPI_ISL_895239, EPI_ISL_895240, EPI_ISL_895241, EPI_ISL_895242, EPI_ISL_895243, EPI_ISL_895244, EPI_ISL_895245, EPI_ISL_895246, EPI_ISL_895247, EPI_ISL_895248, EPI_ISL_895249, EPI_ISL_895250, EPI_ISL_895251, EPI_ISL_895252, EPI_ISL_895253, EPI_ISL_895254, EPI_ISL_895255, EPI_ISL_895256, EPI_ISL_895257, EPI_ISL_895258, EPI_ISL_895259, EPI_ISL_895260, EPI_ISL_895261, EPI_ISL_895262, EPI_ISL_895263, EPI_ISL_895264, EPI_ISL_895265, EPI_ISL_895396, EPI_ISL_895397, EPI_ISL_895398, EPI_ISL_895399, EPI_ISL_895400, EPI_ISL_895401, EPI_ISL_895402, EPI_ISL_895403, EPI_ISL_895404, EPI_ISL_895405, EPI_ISL_895406, EPI_ISL_895407, EPI_ISL_895408, EPI_ISL_895409, EPI_ISL_895410, EPI_ISL_895411, EPI_ISL_895412, EPI_ISL_895413, EPI_ISL_895414, EPI_ISL_895415, EPI_ISL_895416, EPI_ISL_895417, EPI_ISL_895418, EPI_ISL_895419, EPI_ISL_895420, EPI_ISL_895421, EPI_ISL_895422, EPI_ISL_895423, EPI_ISL_895424, EPI_ISL_895425, EPI_ISL_895426, EPI_ISL_895427, EPI_ISL_895428, EPI_ISL_895429, EPI_ISL_895430, EPI_ISL_895431, EPI_ISL_895432, EPI_ISL_895433, EPI_ISL_895434, EPI_ISL_895435, EPI_ISL_895436, EPI_ISL_895437, EPI_ISL_895438, EPI_ISL_895439, EPI_ISL_895440, EPI_ISL_895441, EPI_ISL_895442, EPI_ISL_895443, EPI_ISL_895444, EPI_ISL_895445, EPI_ISL_895446, EPI_ISL_895447, EPI_ISL_895448, EPI_ISL_895449, EPI_ISL_895450, EPI_ISL_895451, EPI_ISL_895452, EPI_ISL_895453, EPI_ISL_895454, EPI_ISL_895455, EPI_ISL_895456, EPI_ISL_895457, EPI_ISL_895458, EPI_ISL_895459, EPI_ISL_895460, EPI_ISL_895461, EPI_ISL_895462, EPI_ISL_895463, EPI_ISL_895464, EPI_ISL_895465, EPI_ISL_895466, EPI_ISL_895487, EPI_ISL_895490, EPI_ISL_895491, EPI_ISL_895492, EPI_ISL_895493, EPI_ISL_895494, EPI_ISL_895495, EPI_ISL_895496, EPI_ISL_895497, EPI_ISL_895498, EPI_ISL_895499, EPI_ISL_895500, EPI_ISL_895501, EPI_ISL_895502, EPI_ISL_895503, EPI_ISL_895504, EPI_ISL_895505, EPI_ISL_895506, EPI_ISL_895507, EPI_ISL_895518, EPI_ISL_895519, EPI_ISL_895520, EPI_ISL_895521, EPI_ISL_895522, EPI_ISL_895523, EPI_ISL_895524, EPI_ISL_895525, EPI_ISL_895526, EPI_ISL_895527, EPI_ISL_895528, EPI_ISL_895529, EPI_ISL_895530, EPI_ISL_895531, EPI_ISL_895532, EPI_ISL_895536, EPI_ISL_895537, EPI_ISL_895538, EPI_ISL_895539, EPI_ISL_895540, EPI_ISL_895541, EPI_ISL_895542, EPI_ISL_895543, EPI_ISL_895544, EPI_ISL_895545, EPI_ISL_895546, EPI_ISL_895547, EPI_ISL_895548, EPI_ISL_895549, EPI_ISL_895550, EPI_ISL_895551, EPI_ISL_895552, EPI_ISL_895553, EPI_ISL_895554, EPI_ISL_895555, EPI_ISL_895556, EPI_ISL_895557, EPI_ISL_895558, EPI_ISL_895559, EPI_ISL_895560, EPI_ISL_895561, EPI_ISL_895562, EPI_ISL_895563, EPI_ISL_895564, EPI_ISL_895565, EPI_ISL_895566, EPI_ISL_895639, EPI_ISL_895640, EPI_ISL_895641, EPI_ISL_895642, EPI_ISL_895643, EPI_ISL_895644, EPI_ISL_895645, EPI_ISL_895646, EPI_ISL_895647, EPI_ISL_895648, EPI_ISL_895649, EPI_ISL_895650, EPI_ISL_895651, EPI_ISL_895652, EPI_ISL_895653, EPI_ISL_895654, EPI_ISL_895655, EPI_ISL_895656, EPI_ISL_895657 | | | |
| see above | Pathogen Genomics Center, National Institute of Infectious Diseases | Pathogen Genomics Center, National Institute of Infectious Diseases | Tsuyoshi Sekizuka, Kentaro Itokawa, Rina Tanaka, Masanori Hashino, Makoto Kuroda |
| EPI_ISL_895979, EPI_ISL_895980, EPI_ISL_895981, EPI_ISL_895982, EPI_ISL_895983, EPI_ISL_895984, EPI_ISL_895985, EPI_ISL_895986, EPI_ISL_895987, EPI_ISL_895988 | Tokyo Metropolitan Institute of Public Health | Pathogen Genomics Center, National Institute of Infectious Diseases | Tsuyoshi Sekizuka, Kentaro Itokawa, Rina Tanaka, Masanori Hashino, Makoto Kuroda |
| EPI_ISL_896013, EPI_ISL_896014, EPI_ISL_896015, EPI_ISL_896016, EPI_ISL_896017, EPI_ISL_896018, EPI_ISL_896020, EPI_ISL_896021, EPI_ISL_896022, EPI_ISL_896023, EPI_ISL_896024, EPI_ISL_896025, EPI_ISL_896026, EPI_ISL_896027, EPI_ISL_896028, EPI_ISL_896029, EPI_ISL_896030, EPI_ISL_896031, EPI_ISL_896032, EPI_ISL_896033, EPI_ISL_896034, EPI_ISL_896035, EPI_ISL_896036, EPI_ISL_896037, EPI_ISL_896038, EPI_ISL_896039, EPI_ISL_896040, EPI_ISL_896041, EPI_ISL_896042, EPI_ISL_896043, EPI_ISL_896044, EPI_ISL_896045, EPI_ISL_896046, EPI_ISL_896047, EPI_ISL_896048, EPI_ISL_896049, EPI_ISL_896050, EPI_ISL_896051, EPI_ISL_896052, EPI_ISL_896053, EPI_ISL_896054, EPI_ISL_896055, EPI_ISL_896056, EPI_ISL_896057, EPI_ISL_896058, EPI_ISL_896059, EPI_ISL_896060, EPI_ISL_896061, EPI_ISL_896062, EPI_ISL_896063 | | | |
| see above | Pathogen Genomics Center, National Institute of Infectious Diseases | Pathogen Genomics Center, National Institute of Infectious Diseases | Tsuyoshi Sekizuka, Kentaro Itokawa, Rina Tanaka, Masanori Hashino, Makoto Kuroda |
| EPI_ISL_896375 | University of Medicine and Pharmacy of Craiova | "Stefan cel Mare" University Metagenomics Lab | Lobiuc Andrei, Gheorghita Roxana |
| EPI_ISL_896602, EPI_ISL_896603, EPI_ISL_896604, EPI_ISL_896605, EPI_ISL_896606, EPI_ISL_896607, EPI_ISL_896608, EPI_ISL_896609, EPI_ISL_896610, EPI_ISL_896611, EPI_ISL_896642, EPI_ISL_896643, EPI_ISL_896644, EPI_ISL_896645, EPI_ISL_896646, EPI_ISL_896659, EPI_ISL_896660, EPI_ISL_896661, EPI_ISL_896662, EPI_ISL_896663, EPI_ISL_896664, EPI_ISL_896665, EPI_ISL_896666, EPI_ISL_896667, EPI_ISL_896668, EPI_ISL_896669, EPI_ISL_896670, EPI_ISL_896671, EPI_ISL_896672, EPI_ISL_896673, EPI_ISL_896674, EPI_ISL_896675, EPI_ISL_896676, EPI_ISL_896677, EPI_ISL_896678, EPI_ISL_896679, EPI_ISL_896680, EPI_ISL_896681, EPI_ISL_896682, EPI_ISL_896683, EPI_ISL_896684, EPI_ISL_896685, EPI_ISL_896686, EPI_ISL_896687, EPI_ISL_896688, EPI_ISL_896689, EPI_ISL_896690, EPI_ISL_896691, EPI_ISL_896692, EPI_ISL_896699, EPI_ISL_896700, EPI_ISL_896701, EPI_ISL_896702, EPI_ISL_896703, EPI_ISL_896704, EPI_ISL_896705, EPI_ISL_896706, EPI_ISL_896707, EPI_ISL_896708, EPI_ISL_896709, EPI_ISL_896710, EPI_ISL_896711, EPI_ISL_896712, EPI_ISL_896713, EPI_ISL_896714, EPI_ISL_896715, EPI_ISL_896716, EPI_ISL_896717, EPI_ISL_896718, EPI_ISL_896719, EPI_ISL_896720, EPI_ISL_896721, EPI_ISL_896722, EPI_ISL_896723, EPI_ISL_896724, EPI_ISL_896725, EPI_ISL_896726, EPI_ISL_896727, EPI_ISL_896728, EPI_ISL_896729, EPI_ISL_896730, EPI_ISL_896731, EPI_ISL_896732, EPI_ISL_896733, EPI_ISL_896734, EPI_ISL_896735, EPI_ISL_896736, EPI_ISL_896737, EPI_ISL_896738, EPI_ISL_896739, EPI_ISL_896740, EPI_ISL_896741, EPI_ISL_896742, EPI_ISL_896743, EPI_ISL_896744, EPI_ISL_896745, EPI_ISL_896746, EPI_ISL_896747, EPI_ISL_896748, EPI_ISL_896749, EPI_ISL_896750, EPI_ISL_896751, EPI_ISL_896752, EPI_ISL_896753, EPI_ISL_896754, EPI_ISL_896755, EPI_ISL_896756, EPI_ISL_896757, EPI_ISL_896758, EPI_ISL_896759, EPI_ISL_896760, EPI_ISL_896761, EPI_ISL_896762, EPI_ISL_896763 | | | |
| see above | Pathogen Genomics Center, National Institute of Infectious Diseases | Pathogen Genomics Center, National Institute of Infectious Diseases | Tsuyoshi Sekizuka, Kentaro Itokawa, Rina Tanaka, Masanori Hashino, Makoto Kuroda |
| EPI_ISL_896764, EPI_ISL_896765, EPI_ISL_896766, EPI_ISL_896767, EPI_ISL_896768, EPI_ISL_896769, EPI_ISL_896770, EPI_ISL_896771, EPI_ISL_896772, EPI_ISL_896773, EPI_ISL_896774, EPI_ISL_896775, EPI_ISL_896776, EPI_ISL_896777, EPI_ISL_896778, EPI_ISL_896779, EPI_ISL_896780, EPI_ISL_896781, EPI_ISL_896782, EPI_ISL_896783, EPI_ISL_896784, EPI_ISL_896785, EPI_ISL_896786, EPI_ISL_896787, EPI_ISL_896788, EPI_ISL_896789, EPI_ISL_896790, EPI_ISL_896791, EPI_ISL_896792, EPI_ISL_896793, EPI_ISL_896794 | | | |
| see above | Gunma Prefectural Institute of Public Health and Environmental Sciences | Pathogen Genomics Center, National Institute of Infectious Diseases | Tsuyoshi Sekizuka, Kentaro Itokawa, Rina Tanaka, Masanori Hashino, Makoto Kuroda |
| EPI_ISL_896795, EPI_ISL_896796, EPI_ISL_896797, EPI_ISL_896798, EPI_ISL_896799, EPI_ISL_896800, EPI_ISL_896801, EPI_ISL_896802, EPI_ISL_896803, EPI_ISL_896804, EPI_ISL_896805, EPI_ISL_896806, EPI_ISL_896807, EPI_ISL_896808, EPI_ISL_896809, EPI_ISL_896810, EPI_ISL_896811, EPI_ISL_896812, EPI_ISL_896813, EPI_ISL_896814, EPI_ISL_896815, EPI_ISL_896816, EPI_ISL_896817, EPI_ISL_896818, EPI_ISL_896819, EPI_ISL_896820, EPI_ISL_896821, EPI_ISL_896822, EPI_ISL_896823, EPI_ISL_896824, EPI_ISL_896825, EPI_ISL_896826, EPI_ISL_896827, EPI_ISL_896828, EPI_ISL_896829, EPI_ISL_896830, EPI_ISL_896831, EPI_ISL_896832, EPI_ISL_896833, EPI_ISL_896834, EPI_ISL_896835, EPI_ISL_896836, EPI_ISL_896837, EPI_ISL_896838, EPI_ISL_896839, EPI_ISL_896840, EPI_ISL_896841, EPI_ISL_896842, EPI_ISL_896843, EPI_ISL_896844, EPI_ISL_896845, EPI_ISL_896846, EPI_ISL_896847, EPI_ISL_896848, EPI_ISL_896849, EPI_ISL_896850, EPI_ISL_896851, EPI_ISL_896852, EPI_ISL_896853, EPI_ISL_896854 | | | |
| see above | Department of Infectious Diseases, Kobe Institute of Health | Pathogen Genomics Center, National Institute of Infectious Diseases | Tsuyoshi Sekizuka, Kentaro Itokawa, Rina Tanaka, Masanori Hashino, Makoto Kuroda |
| EPI_ISL_896940, EPI_ISL_896941, EPI_ISL_896942, EPI_ISL_896943, EPI_ISL_896944, EPI_ISL_896945, EPI_ISL_896946, EPI_ISL_896947, EPI_ISL_896948, EPI_ISL_896949, EPI_ISL_896950, EPI_ISL_896951, EPI_ISL_896952, EPI_ISL_896953, EPI_ISL_896954, EPI_ISL_896955, EPI_ISL_896956, EPI_ISL_896957, EPI_ISL_896958, EPI_ISL_896959, EPI_ISL_896960, EPI_ISL_896961, EPI_ISL_896962, EPI_ISL_896963, EPI_ISL_896964, EPI_ISL_896965, EPI_ISL_896966, EPI_ISL_896967, EPI_ISL_896968, EPI_ISL_896969, EPI_ISL_896970, EPI_ISL_896971, EPI_ISL_896972, EPI_ISL_896973, EPI_ISL_896974, EPI_ISL_896975, EPI_ISL_896976, EPI_ISL_896977, EPI_ISL_896978, EPI_ISL_896979, EPI_ISL_896980, EPI_ISL_896981, EPI_ISL_896982, EPI_ISL_896983, EPI_ISL_896984, EPI_ISL_896985, EPI_ISL_896986, EPI_ISL_896987, EPI_ISL_896988, EPI_ISL_896989, EPI_ISL_896990, EPI_ISL_896991, EPI_ISL_896992, EPI_ISL_896993, EPI_ISL_896994, EPI_ISL_896995, EPI_ISL_896996, EPI_ISL_896997, EPI_ISL_896998, EPI_ISL_896999, EPI_ISL_897000, EPI_ISL_897001, EPI_ISL_897002, EPI_ISL_897003, EPI_ISL_897004, EPI_ISL_897005, EPI_ISL_897006, EPI_ISL_897007, EPI_ISL_897008, EPI_ISL_897009, EPI_ISL_897010, EPI_ISL_897011, EPI_ISL_897012, EPI_ISL_897013, EPI_ISL_897014, EPI_ISL_897015, EPI_ISL_897016, EPI_ISL_897017, EPI_ISL_897018, EPI_ISL_897019, EPI_ISL_897020, EPI_ISL_897021, EPI_ISL_897022, EPI_ISL_897023, EPI_ISL_897034, EPI_ISL_897035, EPI_ISL_897036, EPI_ISL_897037, EPI_ISL_897038, EPI_ISL_897039, EPI_ISL_897040, EPI_ISL_897041, EPI_ISL_897042, EPI_ISL_897043, EPI_ISL_897044, EPI_ISL_897045, EPI_ISL_897046, EPI_ISL_897047, EPI_ISL_897048, EPI_ISL_897049, EPI_ISL_897050, EPI_ISL_897051, EPI_ISL_897052, EPI_ISL_897053, EPI_ISL_897054, EPI_ISL_897055 | | | |
| see above | Pathogen Genomics Center, National Institute of Infectious Diseases | Pathogen Genomics Center, National Institute of Infectious Diseases | Tsuyoshi Sekizuka, Kentaro Itokawa, Rina Tanaka, Masanori Hashino, Makoto Kuroda |
| EPI_ISL_897061, EPI_ISL_897062, EPI_ISL_897063, EPI_ISL_897064, EPI_ISL_897065, EPI_ISL_897067, EPI_ISL_897068, EPI_ISL_897069, EPI_ISL_897070, EPI_ISL_897071, EPI_ISL_897072, EPI_ISL_897073, EPI_ISL_897074, EPI_ISL_897075, EPI_ISL_897076, EPI_ISL_897077, EPI_ISL_897078, EPI_ISL_897079, EPI_ISL_897080 | | | |
| see above | Niigata Prefectural Institute of Public Health and Environmental Sciences | Pathogen Genomics Center, National Institute of Infectious Diseases | Tsuyoshi Sekizuka, Kentaro Itokawa, Rina Tanaka, Masanori Hashino, Makoto Kuroda |
| EPI_ISL_897088, EPI_ISL_897089, EPI_ISL_897090, EPI_ISL_897091, EPI_ISL_897092, EPI_ISL_897093, EPI_ISL_897094, EPI_ISL_897095, EPI_ISL_897096, EPI_ISL_897098, EPI_ISL_897099, EPI_ISL_897100, EPI_ISL_897101, EPI_ISL_897102, EPI_ISL_897103, EPI_ISL_897104, EPI_ISL_897105, EPI_ISL_897106, | | | |

| | | | |
|---|---|---|---|
| EPI_ISL_897107, EPI_ISL_897108, EPI_ISL_897109, EPI_ISL_897110, EPI_ISL_897111, EPI_ISL_897112, EPI_ISL_897113, EPI_ISL_897114, EPI_ISL_897115, EPI_ISL_897116, EPI_ISL_897117, EPI_ISL_897118, EPI_ISL_897119, EPI_ISL_897120, EPI_ISL_897121, EPI_ISL_897122, EPI_ISL_897123, EPI_ISL_897124, EPI_ISL_897125, EPI_ISL_897126, EPI_ISL_897127, EPI_ISL_897128, EPI_ISL_897129, EPI_ISL_897130, EPI_ISL_897131, EPI_ISL_897132, EPI_ISL_897133, EPI_ISL_897134, EPI_ISL_897139, EPI_ISL_897140, EPI_ISL_897141, EPI_ISL_897142, EPI_ISL_897143, EPI_ISL_897144, EPI_ISL_897145, EPI_ISL_897146, EPI_ISL_897147, EPI_ISL_897148, EPI_ISL_897151, EPI_ISL_897152, EPI_ISL_897153, EPI_ISL_897154, EPI_ISL_897155, EPI_ISL_897156, EPI_ISL_897157, EPI_ISL_897158, EPI_ISL_897159, EPI_ISL_897160, EPI_ISL_897161, EPI_ISL_897162, EPI_ISL_897163, EPI_ISL_897164, EPI_ISL_897165, EPI_ISL_897166, EPI_ISL_897167, EPI_ISL_897168, EPI_ISL_897169, EPI_ISL_897170 | | | |
| see above | Pathogen Genomics Center, National Institute of Infectious Diseases | Pathogen Genomics Center, National Institute of Infectious Diseases | Tsuyoshi Sekizuka, Kentaro Itokawa, Rina Tanaka, Masanori Hashino, Makoto Kuroda |
| EPI_ISL_897171, EPI_ISL_897172, EPI_ISL_897173, EPI_ISL_897174, EPI_ISL_897175, EPI_ISL_897176, EPI_ISL_897177, EPI_ISL_897178, EPI_ISL_897179, EPI_ISL_897180, EPI_ISL_897181, EPI_ISL_897182, EPI_ISL_897183, EPI_ISL_897184, EPI_ISL_897185, EPI_ISL_897186, EPI_ISL_897187, EPI_ISL_897188, EPI_ISL_897189, EPI_ISL_897190, EPI_ISL_897191, EPI_ISL_897192, EPI_ISL_897193, EPI_ISL_897194, EPI_ISL_897195, EPI_ISL_897196, EPI_ISL_897197, EPI_ISL_897198, EPI_ISL_897199, EPI_ISL_897200, EPI_ISL_897201, EPI_ISL_897202, EPI_ISL_897203, EPI_ISL_897204, EPI_ISL_897205 | | | |
| see above | Tokyo Metropolitan Institute of Public Health | Pathogen Genomics Center, National Institute of Infectious Diseases | Tsuyoshi Sekizuka, Kentaro Itokawa, Rina Tanaka, Masanori Hashino, Makoto Kuroda |
| EPI_ISL_897206, EPI_ISL_897207, EPI_ISL_897208, EPI_ISL_897209, EPI_ISL_897210, EPI_ISL_897211, EPI_ISL_897212, EPI_ISL_897213, EPI_ISL_897214, EPI_ISL_897215, EPI_ISL_897216, EPI_ISL_897217, EPI_ISL_897218, EPI_ISL_897219, EPI_ISL_897220, EPI_ISL_897221, EPI_ISL_897222, EPI_ISL_897223, EPI_ISL_897224, EPI_ISL_897225, EPI_ISL_897226, EPI_ISL_897227, EPI_ISL_897228, EPI_ISL_897229, EPI_ISL_897230, EPI_ISL_897231, EPI_ISL_897232, EPI_ISL_897233, EPI_ISL_897234, EPI_ISL_897235, EPI_ISL_897236, EPI_ISL_897237, EPI_ISL_897238, EPI_ISL_897239, EPI_ISL_897240, EPI_ISL_897241, EPI_ISL_897242, EPI_ISL_897243, EPI_ISL_897244, EPI_ISL_897245, EPI_ISL_897246, EPI_ISL_897247, EPI_ISL_897251, EPI_ISL_897283, EPI_ISL_897284, EPI_ISL_897285, EPI_ISL_897286, EPI_ISL_897287, EPI_ISL_897288, EPI_ISL_897289, EPI_ISL_897290, EPI_ISL_897291, EPI_ISL_897292, EPI_ISL_897293, EPI_ISL_897294, EPI_ISL_897295, EPI_ISL_897296, EPI_ISL_897297, EPI_ISL_897298, EPI_ISL_897299, EPI_ISL_897300, EPI_ISL_897301, EPI_ISL_897302, EPI_ISL_897303, EPI_ISL_897304, EPI_ISL_897305, EPI_ISL_897306, EPI_ISL_897307, EPI_ISL_897308, EPI_ISL_897309, EPI_ISL_897310, EPI_ISL_897311, EPI_ISL_897312, EPI_ISL_897340, EPI_ISL_897341, EPI_ISL_897342, EPI_ISL_897343, EPI_ISL_897344, EPI_ISL_897345, EPI_ISL_897346, EPI_ISL_897347, EPI_ISL_897348, EPI_ISL_897349, EPI_ISL_897350, EPI_ISL_897351, EPI_ISL_897352, EPI_ISL_897353, EPI_ISL_897354, EPI_ISL_897355, EPI_ISL_897356, EPI_ISL_897357, EPI_ISL_897358, EPI_ISL_897359, EPI_ISL_897360, EPI_ISL_897361, EPI_ISL_897362, EPI_ISL_897363, EPI_ISL_897364, EPI_ISL_897365, EPI_ISL_897366, EPI_ISL_897367, EPI_ISL_897368, EPI_ISL_897369, EPI_ISL_897370, EPI_ISL_897371, EPI_ISL_897372, EPI_ISL_897373, EPI_ISL_897374, EPI_ISL_897375, EPI_ISL_897376, EPI_ISL_897377, EPI_ISL_897378, EPI_ISL_897379, EPI_ISL_897380, EPI_ISL_897381, EPI_ISL_897382, EPI_ISL_897383, EPI_ISL_897384, EPI_ISL_897385, EPI_ISL_897386, EPI_ISL_897387, EPI_ISL_897388, EPI_ISL_897389, EPI_ISL_897390, EPI_ISL_897391, EPI_ISL_897392, EPI_ISL_897393, EPI_ISL_897394, EPI_ISL_897395, EPI_ISL_897396, EPI_ISL_897397, EPI_ISL_897398, EPI_ISL_897399, EPI_ISL_897400, EPI_ISL_897401, EPI_ISL_897402, EPI_ISL_897403, EPI_ISL_897404, EPI_ISL_897405, EPI_ISL_897406, EPI_ISL_897407, EPI_ISL_897408, EPI_ISL_897409, EPI_ISL_897410, EPI_ISL_897411, EPI_ISL_897412, EPI_ISL_897413, EPI_ISL_897414, EPI_ISL_897415, EPI_ISL_897416, EPI_ISL_897417, EPI_ISL_897418, EPI_ISL_897419, EPI_ISL_897420, EPI_ISL_897421, EPI_ISL_897422, EPI_ISL_897423, EPI_ISL_897424, EPI_ISL_897425, EPI_ISL_897426, EPI_ISL_897427, EPI_ISL_897428, EPI_ISL_897429, EPI_ISL_897430, EPI_ISL_897431, EPI_ISL_897432, EPI_ISL_897433, EPI_ISL_897453, EPI_ISL_897454, EPI_ISL_897455, EPI_ISL_897456, EPI_ISL_897457, EPI_ISL_897458, EPI_ISL_897459, EPI_ISL_897460, EPI_ISL_897461, EPI_ISL_897462, EPI_ISL_897463, EPI_ISL_897464, EPI_ISL_897465, EPI_ISL_897466, EPI_ISL_897467, EPI_ISL_897468, EPI_ISL_897469, EPI_ISL_897470, EPI_ISL_897471, EPI_ISL_897472, EPI_ISL_897473, EPI_ISL_897474, EPI_ISL_897475, EPI_ISL_897476, EPI_ISL_897477, EPI_ISL_897478, EPI_ISL_897479, EPI_ISL_897480, EPI_ISL_897481, EPI_ISL_897482, EPI_ISL_897483, EPI_ISL_897484, EPI_ISL_897485, EPI_ISL_897486, EPI_ISL_897487, EPI_ISL_897488, EPI_ISL_897489, EPI_ISL_897490, EPI_ISL_897491, EPI_ISL_897492, EPI_ISL_897493, EPI_ISL_897494, EPI_ISL_897495, EPI_ISL_897496, EPI_ISL_897497, EPI_ISL_897498, EPI_ISL_897499, EPI_ISL_897500, EPI_ISL_897501, EPI_ISL_897502, EPI_ISL_897503, EPI_ISL_897504, EPI_ISL_897505, EPI_ISL_897506, EPI_ISL_897507, EPI_ISL_897508, EPI_ISL_897509, EPI_ISL_897510, EPI_ISL_897511, EPI_ISL_897512, EPI_ISL_897513, EPI_ISL_897514, EPI_ISL_897515, EPI_ISL_897516, EPI_ISL_897517, EPI_ISL_897518, EPI_ISL_897519, EPI_ISL_897520, EPI_ISL_897521, EPI_ISL_897522, EPI_ISL_897523, EPI_ISL_897524, EPI_ISL_897525, EPI_ISL_897526, EPI_ISL_897527, EPI_ISL_897595, EPI_ISL_897596, EPI_ISL_897597 | | | |
| see above | Pathogen Genomics Center, National Institute of Infectious Diseases | Pathogen Genomics Center, National Institute of Infectious Diseases | Tsuyoshi Sekizuka, Kentaro Itokawa, Rina Tanaka, Masanori Hashino, Makoto Kuroda |
| EPI_ISL_898054, EPI_ISL_898055, EPI_ISL_898056 | Sakai City Institute of Public Health | Pathogen Genomics Center, National Institute of Infectious Diseases | Tsuyoshi Sekizuka, Kentaro Itokawa, Rina Tanaka, Masanori Hashino, Makoto Kuroda |
| EPI_ISL_898074, EPI_ISL_898075, EPI_ISL_898076, EPI_ISL_898086, EPI_ISL_898087, EPI_ISL_898089, EPI_ISL_898090, EPI_ISL_898091, EPI_ISL_898092, EPI_ISL_898093, EPI_ISL_898094, EPI_ISL_898095, EPI_ISL_898096, EPI_ISL_898097, EPI_ISL_898098, EPI_ISL_898099, EPI_ISL_898100, EPI_ISL_898101, EPI_ISL_898102, EPI_ISL_898103, EPI_ISL_898104, EPI_ISL_898105, EPI_ISL_898106, EPI_ISL_898107, EPI_ISL_898108, EPI_ISL_898109, EPI_ISL_898110, EPI_ISL_898111, EPI_ISL_898112, EPI_ISL_898113, EPI_ISL_898114, EPI_ISL_898115, EPI_ISL_898116, EPI_ISL_898117, EPI_ISL_898118, EPI_ISL_898119, EPI_ISL_898120, EPI_ISL_898121, EPI_ISL_898122, EPI_ISL_898123, EPI_ISL_898124, EPI_ISL_898125, EPI_ISL_898126, EPI_ISL_898127, EPI_ISL_898128, EPI_ISL_898129, EPI_ISL_898130, EPI_ISL_898131, EPI_ISL_898132, EPI_ISL_898133, EPI_ISL_898136, EPI_ISL_898137, EPI_ISL_898138, EPI_ISL_898139, EPI_ISL_898140, EPI_ISL_898141, EPI_ISL_898142, EPI_ISL_898146, EPI_ISL_898147, EPI_ISL_898148, EPI_ISL_898149, EPI_ISL_898150, EPI_ISL_898151, EPI_ISL_898152, EPI_ISL_898153, EPI_ISL_898154, EPI_ISL_898155, EPI_ISL_898156, EPI_ISL_898157, EPI_ISL_898158, EPI_ISL_898159, EPI_ISL_898160, EPI_ISL_898161, EPI_ISL_898162, EPI_ISL_898163, EPI_ISL_898164, EPI_ISL_898165, EPI_ISL_898166, EPI_ISL_898167, EPI_ISL_898168, EPI_ISL_898169, EPI_ISL_898170, EPI_ISL_898171, EPI_ISL_898172, EPI_ISL_898173, EPI_ISL_898174, EPI_ISL_898175, EPI_ISL_898176, EPI_ISL_898177, EPI_ISL_898178, EPI_ISL_898179, EPI_ISL_898180, EPI_ISL_898181, EPI_ISL_898182, EPI_ISL_898183, EPI_ISL_898184, EPI_ISL_898185, EPI_ISL_898186, EPI_ISL_898187, EPI_ISL_898188, EPI_ISL_898189, EPI_ISL_898190, EPI_ISL_898191, EPI_ISL_898192, EPI_ISL_898193, EPI_ISL_898194, EPI_ISL_898195, EPI_ISL_898196, EPI_ISL_898197, EPI_ISL_898198, EPI_ISL_898199, EPI_ISL_898200, EPI_ISL_898201, EPI_ISL_898202, EPI_ISL_898203, EPI_ISL_898204, EPI_ISL_898205, EPI_ISL_898206, EPI_ISL_898207, EPI_ISL_898208, EPI_ISL_898209, EPI_ISL_898210, EPI_ISL_898211, EPI_ISL_898212, EPI_ISL_898213, EPI_ISL_898214, EPI_ISL_898215, EPI_ISL_898216, EPI_ISL_898217, EPI_ISL_898218, EPI_ISL_898219, EPI_ISL_898220, EPI_ISL_898221, EPI_ISL_898222, EPI_ISL_898223, EPI_ISL_898224, EPI_ISL_898225, EPI_ISL_898226, EPI_ISL_898227, EPI_ISL_898228, EPI_ISL_898229, EPI_ISL_898230, EPI_ISL_898231, EPI_ISL_898232, EPI_ISL_898233, EPI_ISL_898234, EPI_ISL_898235, EPI_ISL_898236, EPI_ISL_898237, EPI_ISL_898238, EPI_ISL_898239, EPI_ISL_898240, EPI_ISL_898241, EPI_ISL_898242, EPI_ISL_898243, EPI_ISL_898244, EPI_ISL_898245, EPI_ISL_898246, EPI_ISL_898247, EPI_ISL_898248, EPI_ISL_898249, EPI_ISL_898250, EPI_ISL_898251, EPI_ISL_898252, EPI_ISL_898253, EPI_ISL_898254, EPI_ISL_898255, EPI_ISL_898256, EPI_ISL_898257, EPI_ISL_898258, EPI_ISL_898259, EPI_ISL_898260, EPI_ISL_898261, EPI_ISL_898262, EPI_ISL_898263, EPI_ISL_898264, EPI_ISL_898265, EPI_ISL_898266, EPI_ISL_898267, EPI_ISL_898268, EPI_ISL_898270, EPI_ISL_898271, EPI_ISL_898272, EPI_ISL_898273, EPI_ISL_898274, EPI_ISL_898275, EPI_ISL_898307, EPI_ISL_898308, EPI_ISL_898309, EPI_ISL_898310, EPI_ISL_898311, EPI_ISL_898312, EPI_ISL_898315, EPI_ISL_898316, EPI_ISL_898317, EPI_ISL_898318, EPI_ISL_898319, EPI_ISL_898320 | | | |
| see above | Pathogen Genomics Center, National Institute of Infectious Diseases | Pathogen Genomics Center, National Institute of Infectious Diseases | Tsuyoshi Sekizuka, Kentaro Itokawa, Rina Tanaka, Masanori Hashino, Makoto Kuroda |
| EPI_ISL_898322, EPI_ISL_898323, EPI_ISL_898324 | Toyama Institute of Health | Pathogen Genomics Center, National Institute of Infectious Diseases | Tsuyoshi Sekizuka, Kentaro Itokawa, Rina Tanaka, Masanori Hashino, Makoto Kuroda |
| EPI_ISL_898332, EPI_ISL_898333, EPI_ISL_898334, EPI_ISL_898335, EPI_ISL_898336, EPI_ISL_898337 | Yamagata Prefectural Institute of Public Health | Pathogen Genomics Center, National Institute of Infectious Diseases | Tsuyoshi Sekizuka, Kentaro Itokawa, Rina Tanaka, Masanori Hashino, Makoto Kuroda |
| EPI_ISL_898554, EPI_ISL_898555, EPI_ISL_898556, EPI_ISL_898557, EPI_ISL_898558, EPI_ISL_898559, EPI_ISL_898560, EPI_ISL_898561, EPI_ISL_898562, EPI_ISL_898563, EPI_ISL_898564, EPI_ISL_898609, EPI_ISL_898610, EPI_ISL_898611, EPI_ISL_898612, EPI_ISL_898613, EPI_ISL_898614, EPI_ISL_898615, EPI_ISL_898625, EPI_ISL_898626, EPI_ISL_898627, EPI_ISL_898628, EPI_ISL_898637, EPI_ISL_898638, EPI_ISL_898639, EPI_ISL_898640, EPI_ISL_898641, EPI_ISL_898642, EPI_ISL_898643, EPI_ISL_898644, EPI_ISL_898645, EPI_ISL_898646, EPI_ISL_898647, EPI_ISL_898648, EPI_ISL_898649, EPI_ISL_898650, EPI_ISL_898651, EPI_ISL_898652, EPI_ISL_898653, EPI_ISL_898654, EPI_ISL_898655, EPI_ISL_898656, EPI_ISL_898657, EPI_ISL_898658, EPI_ISL_898659, EPI_ISL_898709, EPI_ISL_898710 | | | |
| see above | Pathogen Genomics Center, National Institute of Infectious Diseases | Pathogen Genomics Center, National Institute of Infectious Diseases | Tsuyoshi Sekizuka, Kentaro Itokawa, Rina Tanaka, Masanori Hashino, Makoto Kuroda |
| EPI_ISL_898711, EPI_ISL_898712, EPI_ISL_898713, EPI_ISL_898714, EPI_ISL_898715, EPI_ISL_898716, EPI_ISL_898717, EPI_ISL_898718, EPI_ISL_898719, EPI_ISL_898720, EPI_ISL_898721, EPI_ISL_898722, EPI_ISL_898723, EPI_ISL_898724, EPI_ISL_898725, EPI_ISL_898726, EPI_ISL_898727, EPI_ISL_898728, EPI_ISL_898729, EPI_ISL_898730, EPI_ISL_898731, EPI_ISL_898732, EPI_ISL_898733, EPI_ISL_898734, EPI_ISL_898735, EPI_ISL_898736, EPI_ISL_898737, EPI_ISL_898738, EPI_ISL_898739, EPI_ISL_898740, EPI_ISL_898741, EPI_ISL_898742, EPI_ISL_898743, EPI_ISL_898744, EPI_ISL_898745, EPI_ISL_898746, EPI_ISL_898747, EPI_ISL_898748, EPI_ISL_898749, EPI_ISL_898750, EPI_ISL_898751, EPI_ISL_898752, EPI_ISL_898753, EPI_ISL_898754, EPI_ISL_898755, EPI_ISL_898756, EPI_ISL_898757, EPI_ISL_898758, EPI_ISL_898759, EPI_ISL_898760, EPI_ISL_898761, EPI_ISL_898762, EPI_ISL_898763, EPI_ISL_898764, EPI_ISL_898765, EPI_ISL_898766, EPI_ISL_898767, EPI_ISL_898768, EPI_ISL_898769, EPI_ISL_898770, EPI_ISL_898771, EPI_ISL_898772, EPI_ISL_898773, EPI_ISL_898774, EPI_ISL_898775, EPI_ISL_898776, EPI_ISL_898777, EPI_ISL_898778, EPI_ISL_898779, EPI_ISL_898780, EPI_ISL_898781, EPI_ISL_898782, EPI_ISL_898783, EPI_ISL_898784, EPI_ISL_898785, EPI_ISL_898786, EPI_ISL_898787, EPI_ISL_898788, EPI_ISL_898789, EPI_ISL_898790, EPI_ISL_898791, EPI_ISL_898792, EPI_ISL_898793, EPI_ISL_898794, EPI_ISL_898795, EPI_ISL_898796, EPI_ISL_898797, EPI_ISL_898798, EPI_ISL_898799, EPI_ISL_898800, EPI_ISL_898801, EPI_ISL_898802, EPI_ISL_898803, EPI_ISL_898804, EPI_ISL_898805, EPI_ISL_898806, EPI_ISL_898807, EPI_ISL_898808, EPI_ISL_898809 | | | |
| see above | Department of Infectious Diseases, Kobe Institute of Health | Pathogen Genomics Center, National Institute of Infectious Diseases | Tsuyoshi Sekizuka, Kentaro Itokawa, Rina Tanaka, Masanori Hashino, Makoto Kuroda |
| EPI_ISL_898810, EPI_ISL_898811, EPI_ISL_898812, EPI_ISL_898813, EPI_ISL_898814, EPI_ISL_898815, EPI_ISL_898816, EPI_ISL_898817, EPI_ISL_898818, EPI_ISL_898819, EPI_ISL_898820, EPI_ISL_898821, EPI_ISL_898822, EPI_ISL_898823, EPI_ISL_898824, EPI_ISL_898825, EPI_ISL_898826, EPI_ISL_898827, EPI_ISL_898828, EPI_ISL_898829, EPI_ISL_898830, EPI_ISL_898831, EPI_ISL_898832, EPI_ISL_898833, EPI_ISL_898834, EPI_ISL_898835, EPI_ISL_898836, EPI_ISL_898837, EPI_ISL_898838, EPI_ISL_898839, EPI_ISL_898840, EPI_ISL_898841, EPI_ISL_898842, EPI_ISL_898843, EPI_ISL_898844, EPI_ISL_898845, EPI_ISL_898846, EPI_ISL_898847, EPI_ISL_898848, EPI_ISL_898849, EPI_ISL_898850, EPI_ISL_898851, EPI_ISL_898852, EPI_ISL_898853, EPI_ISL_898854, EPI_ISL_898855, EPI_ISL_898856, EPI_ISL_898857, EPI_ISL_898858, EPI_ISL_898872, EPI_ISL_898873, EPI_ISL_898874, EPI_ISL_898875, EPI_ISL_898876, EPI_ISL_898877, EPI_ISL_898878, EPI_ISL_898879, EPI_ISL_898880, EPI_ISL_898881, EPI_ISL_898882, EPI_ISL_898883, EPI_ISL_898884, EPI_ISL_898885, EPI_ISL_898886, EPI_ISL_898887, EPI_ISL_898888, EPI_ISL_898889, EPI_ISL_898890, EPI_ISL_898891, EPI_ISL_898892, EPI_ISL_898893, EPI_ISL_898915, EPI_ISL_898916, EPI_ISL_898917, EPI_ISL_898918, EPI_ISL_898919, EPI_ISL_898920, EPI_ISL_898921, EPI_ISL_898922, EPI_ISL_898923, EPI_ISL_898924, EPI_ISL_898925, EPI_ISL_898926, EPI_ISL_898927, EPI_ISL_898928, EPI_ISL_898929, EPI_ISL_898930, EPI_ISL_898931, EPI_ISL_898932, EPI_ISL_898933, EPI_ISL_898934, EPI_ISL_898935, EPI_ISL_898936, EPI_ISL_898937, EPI_ISL_898938, EPI_ISL_898939, EPI_ISL_898940, EPI_ISL_898941, EPI_ISL_898942, EPI_ISL_898943, EPI_ISL_898944, EPI_ISL_898945, EPI_ISL_898946, EPI_ISL_898947, EPI_ISL_898948, EPI_ISL_898949, EPI_ISL_898950, EPI_ISL_898951, EPI_ISL_898952, EPI_ISL_898953, EPI_ISL_898954, EPI_ISL_898955, EPI_ISL_898956, EPI_ISL_898957, EPI_ISL_898958, EPI_ISL_898959, EPI_ISL_898960, EPI_ISL_898961, EPI_ISL_898962, EPI_ISL_898963, EPI_ISL_898964, EPI_ISL_898965, EPI_ISL_898966, EPI_ISL_898967, EPI_ISL_898968, EPI_ISL_898969, EPI_ISL_898970, EPI_ISL_898971, EPI_ISL_898972, EPI_ISL_898973, EPI_ISL_898974, EPI_ISL_898975, EPI_ISL_898976, EPI_ISL_898977, EPI_ISL_898978, EPI_ISL_898979, EPI_ISL_898980, EPI_ISL_898981, EPI_ISL_898982, EPI_ISL_898983, EPI_ISL_898984, EPI_ISL_898985, EPI_ISL_898986, EPI_ISL_898987, EPI_ISL_898988, EPI_ISL_898989, EPI_ISL_898990, EPI_ISL_898991, EPI_ISL_898992, EPI_ISL_901433, EPI_ISL_901434, EPI_ISL_901435, EPI_ISL_901436, EPI_ISL_901437, EPI_ISL_901438, EPI_ISL_901439, EPI_ISL_901440, EPI_ISL_901441, EPI_ISL_901442, EPI_ISL_901443, EPI_ISL_901444, EPI_ISL_901445, EPI_ISL_901446, EPI_ISL_901447, EPI_ISL_901448, EPI_ISL_901449, EPI_ISL_901498 | | | |
| see above | Pathogen Genomics Center, National Institute of Infectious Diseases | Pathogen Genomics Center, National Institute of Infectious Diseases | Tsuyoshi Sekizuka, Kentaro Itokawa, Rina Tanaka, Masanori Hashino, Makoto Kuroda |
| EPI_ISL_901499, EPI_ISL_901500 | Tokyo Metropolitan Institute of Public Health | Pathogen Genomics Center, National Institute of Infectious Diseases | Tsuyoshi Sekizuka, Kentaro Itokawa, Rina Tanaka, Masanori Hashino, Makoto Kuroda |
| EPI_ISL_904004 | Scientific Veterinary Institute Novi Sad | Veterinary Specialized Institute "Kraljevo", Serbia | Vidanovic,D., Tesovic,B., Knezevic,A., Jovanovic,T., Jankovic,M., Sekler,M., Banovic Djeri,B., Petrovic,T., Volkening,J., Afonso,C. |
| EPI_ISL_904241, EPI_ISL_904251, EPI_ISL_904282, EPI_ISL_904455, EPI_ISL_904456, EPI_ISL_904457, | Dutch COVID-19 response team | Erasmus Medical Center | Bas Oude Munnink, Reina Sikkema, David Nieuwenhuijse, Irina Chestakova, Anne van der Linden, Marjan Boter, Emmanuelle Munger, Corine GeurtsvanKessel, Annemiek van der Eijk, Richard Molenkamp, Marion Koopmans, on behalf of the Dutch national COVID-19 response team. |

| | | | |
|---|---|---|---|
| EPI_ISL_904458, EPI_ISL_904459, EPI_ISL_904460 | | | |
| EPI_ISL_906516, EPI_ISL_906517, EPI_ISL_906518 | Department of Infectious Diseases, Kobe Institute of Health | Pathogen Genomics Center, National Institute of Infectious Diseases | Tsuyoshi Sekizuka, Kentaro Itokawa, Rina Tanaka, Masanori Hashino, Makoto Kuroda |
| EPI_ISL_906519, EPI_ISL_906522, EPI_ISL_906524, EPI_ISL_906525 | Pathogen Genomics Center, National Institute of Infectious Diseases | Pathogen Genomics Center, National Institute of Infectious Diseases | Tsuyoshi Sekizuka, Kentaro Itokawa, Rina Tanaka, Masanori Hashino, Makoto Kuroda |
| EPI_ISL_907076 | Biology, MCL | Biology, MCL | Seadawy,M.G., Zekri,A.N., Shamel,M.D., Gad,A.F., Elhoseiny,M.F. |
| EPI_ISL_911675 | Tanjungpura University Hospital | Tanjungpura University Hospital | Mahyarudin; Andriani; Virhan Novianry; Delima Fajar Liana ; Sofi Siti Shofiyah; Puji Astuti ; Muhammad Ibnu Kahtan ; Ambar Rialita; Eka Ardiani Putri; Wiwik Windarti; Helmi Sastriawan |
| EPI_ISL_912457, EPI_ISL_912503, EPI_ISL_912533 | NHLS Universitas Academic | UFS Virology | PA Bester, MM Nyaga, P Nthiga, MT Mogotsi, D Goedhals, T de Oliveira |
| EPI_ISL_913091 | Center for Virology | Center for Virology | Jeremy V. Camp, Irene Goerzer, Monika Redlberger-Fritz, Stephan W. Aberle |
| EPI_ISL_913352, EPI_ISL_913353 | Klinisk mikrobiologi | The Public Health Agency of Sweden | Anna-Malin Linde, Maria Lind Karlberg, Carlo Berg, Oskar Karlsson Lindsjo, Sofia Stamouli, Reza Advani, Mattias Haukland, Petra Holmstrom, Noura Walai, Petra Edquist, Mia Brytting, Anna Risberg, Karin Tegmark-Wisell |
| EPI_ISL_914884, EPI_ISL_914886 | Vilnius university hospital Santaros Klinikos, Center of Laboratory Medicine | Vilnius University Hospital Santaros Klinikos | Ingrida Olendraite, Daniel Naumovas, Rimvydas Norvilas, Dovil Ežerskyt, Justinas Šlikas |
| EPI_ISL_918377, EPI_ISL_918380, EPI_ISL_918381, EPI_ISL_918382, EPI_ISL_918383, EPI_ISL_918384, EPI_ISL_918385, EPI_ISL_918386, EPI_ISL_918387, EPI_ISL_918388, EPI_ISL_918389, EPI_ISL_918390, EPI_ISL_918391, EPI_ISL_918392, EPI_ISL_918393, EPI_ISL_918394, EPI_ISL_918395, EPI_ISL_918396, EPI_ISL_918397, EPI_ISL_918398, EPI_ISL_918399, EPI_ISL_918400, EPI_ISL_918401, EPI_ISL_918402, EPI_ISL_918403, EPI_ISL_918404, EPI_ISL_918405, EPI_ISL_918406 | | | |
| see above | Ibaraki Prefectural Institute of Public Health | Ibaraki Prefectural Institute of Public Health | Keiko Goto, Tsuyoshi Sekizuka, Kentaro Itokawa, Rina Tanaka, Masanori Hashino, Makoto Kuroda |
| EPI_ISL_923224 | Centre for Enzyme Innovation, University of Portsmouth / Translational Research Laboratory, Portsmouth Hospitals NHS Trust | COVID-19 Genomics UK (COG-UK) Consortium | Angela Beckett,Salman Goudarzi,Christopher Fearn,Kate Cook,Katie Loveson,Sharon Glaysher,Scott Elliott,Samuel Robson |
| EPI_ISL_925104, EPI_ISL_925126 | 1.AO Universitaria 'S. Giovanni di Dio e Ruggi D'Aragona, Scuola Medica Salernitana' Hospital / 2.UOC di Virologia e Microbiologia, Università della Campania 'L. Vanvitelli' / 3.AO Universitaria 'Federico II' Napoli Hospital / 4.AORN 'San Giuseppe Moscati' Avellino Hospital / 5.AO 'San Pio - presidio G. Rummo' Benevento Hospital / 6.AO 'Sant'Anna e San Sebastiano' Caserta Hospital / 7.PO 'Maria Santissima Addolorata' Eboli Hospital / 8.Biogem Istituto di Ricerche Genetiche | 1. Genome Research Center for Health (CRGS) / 2. Laboratory of Molecular Medicine and Genomics(LMMGe) / 3. Center for Research in Pure and Applied Mathematics (CRMPA) | Giorgio Giurato, Francesca Rizzo, Alessandro Weisz, Gianluigi Franci, Giovanni Nassa, Pasquale Pagliano, Roberta Tarallo, Elena Alexandrova, Ylenia D'Agostino, Carlo Ferravante, Jessica Lamberti, Viola Melone, Domenico Memoli, Valeria Mirici Cappa, Domenico Palumbo, Giovanni Pecoraro, Assunta Sellitto, Oriana Strianese, Ilaria Terenzi, Giuseppe Fenza, Aniello Gentile, Antonello Saccomanno, Sonia Amabile, Teresa Rocco, Annamaria Salvati, Emilia Vaccaro, Massimiliano Galdiero, Michele Cennamo, Giuseppe Portella, Maria Grazia Foti, Mariarosaria Ingino, Maria Landi, Maurizio Fumi, Vincenzo Rocco, Rita Greco, Vittoria Letizia, Arnolfo Petruzziello, Maddalena Schioppa, Gregorio Goffredi, Francesca Marciano, Michele Caraglia, Alessia Cossu, Marianna Scrima |
| EPI_ISL_925147 | Virginia DCLS | Virginia DCLS | Virginia DCLS |
| EPI_ISL_925538, EPI_ISL_925586, EPI_ISL_925587, EPI_ISL_925588, EPI_ISL_925589, EPI_ISL_925590, EPI_ISL_925591, EPI_ISL_925592, EPI_ISL_925593, EPI_ISL_925594, EPI_ISL_925595, EPI_ISL_925596, EPI_ISL_925698, EPI_ISL_925699, EPI_ISL_925700, EPI_ISL_925762 | | | |
| see above | Public Health Ontario Laboratory | Public Health Ontario Laboratory | Vanessa G Allen, Philip Banh, Yao Chen, Richard de Borja, Alireza Eshaghi, Nahuel Fittipaldi, Christine Frantz, Jonathan B Gubbay, Jennifer L Guthrie, Lawrence Heisler, Esha Joshi, Michael Laszloffy, Aimin Li, Michael CY Li, Dean Maxwell, Sandeep Nagra, Samir N Patel, Jared Simpson, Karthikeyan Sivaraman, Ashleigh Sullivan, Yogi Sundaravadanam, Sarah Teatero, Matthew Watson, Andre Villegas, Sandra Zittermann |
| EPI_ISL_931311, EPI_ISL_931312, EPI_ISL_931313, EPI_ISL_931314, EPI_ISL_931315, EPI_ISL_931316, EPI_ISL_931317, EPI_ISL_931318, EPI_ISL_931319, EPI_ISL_931320, EPI_ISL_931321, EPI_ISL_931322, EPI_ISL_931323, EPI_ISL_931324, EPI_ISL_931325, EPI_ISL_931326, EPI_ISL_931327, EPI_ISL_931328, EPI_ISL_931329, EPI_ISL_931330, EPI_ISL_931331, EPI_ISL_931332, EPI_ISL_931333, EPI_ISL_931334, EPI_ISL_931335, EPI_ISL_931336, EPI_ISL_931337, EPI_ISL_931338, EPI_ISL_931339, EPI_ISL_931340, EPI_ISL_931341, EPI_ISL_931376, EPI_ISL_931377, EPI_ISL_931378, EPI_ISL_931379, EPI_ISL_931380, EPI_ISL_931381, EPI_ISL_931382, EPI_ISL_931383, EPI_ISL_931384, EPI_ISL_931385, EPI_ISL_931386, EPI_ISL_931387, EPI_ISL_931388, EPI_ISL_931395, EPI_ISL_931400, EPI_ISL_931401, EPI_ISL_931402, EPI_ISL_931403, EPI_ISL_931404, EPI_ISL_931405, EPI_ISL_931406, EPI_ISL_931407, EPI_ISL_931408, EPI_ISL_931409, EPI_ISL_931410, EPI_ISL_931411, EPI_ISL_931412, EPI_ISL_931413, EPI_ISL_931414, EPI_ISL_931415, EPI_ISL_931416, EPI_ISL_931417, EPI_ISL_931418, EPI_ISL_931419, EPI_ISL_931420, EPI_ISL_931421, EPI_ISL_931422, EPI_ISL_931423, EPI_ISL_931424, EPI_ISL_931425, EPI_ISL_931426, EPI_ISL_931427, EPI_ISL_931428, EPI_ISL_931429, EPI_ISL_931430, EPI_ISL_931431, EPI_ISL_931432, EPI_ISL_931433, EPI_ISL_931434, EPI_ISL_931435, EPI_ISL_931436, EPI_ISL_931437, EPI_ISL_931438, EPI_ISL_931439, EPI_ISL_931440, EPI_ISL_931441, EPI_ISL_931442, EPI_ISL_931443, EPI_ISL_931444, EPI_ISL_931445, EPI_ISL_931446, EPI_ISL_931447 | | | |
| see above | University Hospital Basel, Clinical Virology | University Hospital Basel, Clinical Bacteriology | Tim Roloff, Madlen Stange, Helena MB Seth-Smith, Alfredo Mari, Karoline Leuzinger, Julia Bielicki, Manuel Battegay, Hans Hirsch, Adrian Egli |
| EPI_ISL_934335, EPI_ISL_934336 | Klinisk mikrobiologi | The Public Health Agency of Sweden | Anna-Malin Linde, Maria Lind Karlberg, Carlo Berg, Oskar Karlsson Lindsjo, Sofia Stamouli, Reza Advani, Mattias Haukland, Petra Holmstrom, Noura Walai, Petra Edquist, Mia Brytting, Anna Risberg, Karin Tegmark-Wisell |
| EPI_ISL_936589, EPI_ISL_936590, EPI_ISL_936591, EPI_ISL_936592, EPI_ISL_936593, EPI_ISL_936594, EPI_ISL_936595, EPI_ISL_936596, EPI_ISL_936597, EPI_ISL_936598, EPI_ISL_936599, EPI_ISL_936600 | | | |
| see above | Northwestern Memorial Hospital | Ozer Lab | Ramon Lorenzo-Redondo, Lacy M. Simons, Chad J. Achenbach, Lawrence J. Jennings, Michael G. Ison, Judd F. Hultquist, Egon A. Ozer |
| EPI_ISL_937048 | Quest Diagnostics | Quest Diagnostics | Rosenthal,S.H., Gerasimova,A., Kagan,R.M., Anderson, B., Livingston, K.E., Hua, M., Liu Y., Shalhout, D.F., Owen, R., Lacbawan, F. |
| EPI_ISL_939642 | Department of Infectious Diseases, Kobe Institute of Health | Department of Infectious Diseases, Kobe Institute of Health | Ryohei Nomoto, Tomotada Iwamoto, Tsuyoshi Sekizuka, Kentaro Itokawa, Rina Tanaka, Masanori Hashino, Makoto Kuroda |
| EPI_ISL_940162 | Hôpital Bichat Claude Bernard, Laboratoire de Virologie | IAME UMR1137 Inserm, Université de Paris, Hôpital Bichat | Antoine Bridier-Nahmias, Amélie Recoing, Quentin Le Hingrat, Lena Daniel, Siham Hamri, Gilles Collin, Alexandre Storto, Mélanie Bertine, Charlotte Charpentier, Nadhira Houhou-Fidouh, Diane Descamps, Benoit Visseaux |
| EPI_ISL_949190 | University Hospital Basel, Clinical Virology | University Hospital Basel, Clinical Bacteriology | Tim Roloff, Madlen Stange, Helena MB Seth-Smith, Alfredo Mari, Karoline Leuzinger, Julia Bielicki, Manuel Battegay, Hans Hirsch, Adrian Egli |
| EPI_ISL_949224 | Departamento de Microbiología, CDB, Hospital Clínic, Barcelona | SeqCOVID-SPAIN consortium/IBV(CSIC) | Andrea Vergara, Mikel Martínez, Elisa Rubio, Jéssica Navero, Aida Peiró and SeqCOVID-SPAIN consortium |
| EPI_ISL_949649 | Virology Department, Royal Infirmary of Edinburgh, NHS Lothian / School of Biological Sciences, University of Edinburgh / Institute of Genetics and Molecular Medicine, University of Edinburgh | COVID-19 Genomics UK (COG-UK) Consortium | McHugh M, Dewar R, Rooke S, Gallagher M, Balcaza C, O'Toole Á, Scher E, Hill V, McCrone JT, Colquhoun R, Yu X, Jackson B, Rambaut A, Williams TC, Templeton K |
| EPI_ISL_954189 | 1.AO Universitaria 'S. Giovanni di Dio e Ruggi D'Aragona, Scuola Medica Salernitana' Hospital / 2.UOC di Virologia e Microbiologia, Università della Campania 'L. Vanvitelli' / 3.AO Universitaria 'Federico II' Napoli Hospital / 4.AORN 'San Giuseppe Moscati' Avellino Hospital / 5.AO 'San Pio - presidio G. Rummo' Benevento Hospital / 6.AO 'Sant'Anna e San Sebastiano' Caserta Hospital / 7.PO 'Maria Santissima Addolorata' Eboli Hospital / 8.Biogem Istituto di Ricerche Genetiche | 1. Genome Research Center for Health (CRGS) / 2. Laboratory of Molecular Medicine and Genomics(LMMGe) / 3. Center for Research in Pure and Applied Mathematics (CRMPA) | Giorgio Giurato, Francesca Rizzo, Alessandro Weisz, Gianluigi Franci, Giovanni Nassa, Pasquale Pagliano, Roberta Tarallo, Elena Alexandrova, Ylenia D'Agostino, Carlo Ferravante, Jessica Lamberti, Viola Melone, Domenico Memoli, Valeria Mirici Cappa, Domenico Palumbo, Giovanni Pecoraro, Assunta Sellitto, Oriana Strianese, Ilaria Terenzi, Giuseppe Fenza, Aniello Gentile, Antonello Saccomanno, Sonia Amabile, Teresa Rocco, Annamaria Salvati, Emilia Vaccaro, Massimiliano Galdiero, Michele Cennamo, Giuseppe Portella, Maria Grazia Foti, Mariarosaria Ingino, Maria Landi, Maurizio Fumi, Vincenzo Rocco, Rita Greco, Vittoria Letizia, Arnolfo Petruzziello, Maddalena Schioppa, Gregorio Goffredi, Francesca Marciano, Michele Caraglia, Alessia Cossu, Marianna Scrima, Edmondo Adorisio, Morena D'Avenia, Michela Iacobellis, Rosanna Piluscio, Giorgio Dirani, Vittorio Sambri, Simona Semprini, Silvia Zanoli, Francesco Curcio, Stefania Marzinotto, Andreina Baj, Fausto Sessa. |
| EPI_ISL_956318, EPI_ISL_956320 | Laboratory Medicine | Department of Laboratory Medicine, Lin-Kou Chang Gung Memorial Hospital, Taoyuan, Taiwan | Kuo-Chien Tsao, Yu-Nong Gong, Shu-Li Yang, Yi-Chun Liu, Chung-Guei Huang, Mei-Jen Hsiao, Po-Wei Huang, Cheng-Ta Yang, Cheng-Hsun Chiu, Peng-Nien Huang, Kuo-Ming Lee, Guang-Wu Chen, Shin-Ru Shih |
| EPI_ISL_960125 | Mitchells Plain Hospital wc MPH | National Health Laboratory Service/UCT | Arash Iranzadeh, Deelan Doolabh, Lynn Tyers, Bruna Galvao, Innocent Mudau, Marvin Hsiao, Kruger Marais, Diana Hardie, Stephen Korsman, Carolyn Williamson |
| EPI_ISL_960126 | Groote Schuur Hospital wc GSH | National Health Laboratory Service/UCT | Arash Iranzadeh, Deelan Doolabh, Lynn Tyers, Bruna Galvao, Innocent Mudau, Marvin Hsiao, Kruger Marais, Diana Hardie, Stephen Korsman, Carolyn Williamson |

| | | | |
|---|---|---|---|
| EPI_ISL_960355 | University of Wisconsin-Madison AIDS Vaccine Research Laboratories | University of Wisconsin-Madison AIDS Vaccine Research Laboratories | Gage Moreno, Katarina Braun, et al. AIDS Vaccine Research Laboratories |
| EPI_ISL_962102 | Illinois Department of Public Health | Gagnon Lab, Southern Illinois University | Keith Gagnon |
| EPI_ISL_965339, EPI_ISL_965342, EPI_ISL_965346, EPI_ISL_965352, EPI_ISL_965382, EPI_ISL_965457 | University of Liège COVID-19 testing center | GIGA Medical Genomics | Keith Durkin, Maria Artesi, Bouchra Boujemla, Emmanuel André, Marc Van Ranst, Fabrice Bureau, Laurent Gillet, Wouter Coppieters, Vincent Bours |

EPI_ISL_974759, EPI_ISL_974760, EPI_ISL_974761, EPI_ISL_974762, EPI_ISL_974763, EPI_ISL_974764, EPI_ISL_974765, EPI_ISL_974766, EPI_ISL_974767, EPI_ISL_974768, EPI_ISL_974769, EPI_ISL_974770, EPI_ISL_974771, EPI_ISL_974772, EPI_ISL_974773, EPI_ISL_974774, EPI_ISL_974775, EPI_ISL_974776, EPI_ISL_974777, EPI_ISL_974778, EPI_ISL_974779, EPI_ISL_974780, EPI_ISL_974781, EPI_ISL_974782, EPI_ISL_974783, EPI_ISL_974784, EPI_ISL_974785, EPI_ISL_974786, EPI_ISL_974787, EPI_ISL_974788, EPI_ISL_974789, EPI_ISL_974790, EPI_ISL_974791, EPI_ISL_974792, EPI_ISL_974793, EPI_ISL_974794, EPI_ISL_974795, EPI_ISL_974796, EPI_ISL_974797, EPI_ISL_974798, EPI_ISL_974799, EPI_ISL_974800, EPI_ISL_974801, EPI_ISL_974802, EPI_ISL_974803, EPI_ISL_974804, EPI_ISL_974805, EPI_ISL_974806, EPI_ISL_974807, EPI_ISL_974808, EPI_ISL_974809, EPI_ISL_974810, EPI_ISL_974811, EPI_ISL_974812, EPI_ISL_974813, EPI_ISL_974814, EPI_ISL_974815, EPI_ISL_974816, EPI_ISL_974817, EPI_ISL_974818, EPI_ISL_974819, EPI_ISL_974820, EPI_ISL_974821, EPI_ISL_974822, EPI_ISL_974823, EPI_ISL_974824

| | | | |
|---|---|---|---|
| see above | BCCDC Public Health Laboratory | BCCDC Public Health Laboratory | Prystajecky Natalie, Linda Hoang, Dan Fornika, John Tyson, Shannon Russell, Kim Macdonald, Kimia Kamelian, Ana Pacagnella, Corrinne Ng, Loretta Janz, Robert Azana Terry Snutch, Mel Krajden |
