## Supplementary material for "Detection and characterization of the SARS-CoV-2 lineage B.1.526 in New York": Supp. Table 3 GISAID Acknowledgment part 2: gisaid_hcov-19_acknowledgement_table_2021_02_13_00-5.pdf

We gratefully acknowledge the following Authors from the Originating laboratories responsible for obtaining the specimens, as well as the Submitting laboratories where the genome data were generated and shared via GISAID, on which this research is based.

All Submitters of data may be contacted directly via www.gisaid.org

Authors are sorted alphabetically.

| Accession ID | Originating Laboratory | Submitting Laboratory | Authors |
|---|---|---|---|
| EPI_ISL_622786 | Canterbury Health Laboratories | Institute of Environmental Science and Research (ESR) | Xiaoyun Ren, Matt Storey, Nikki Freed, Muhammad Faisal, Jing Wang, Hermes Perez, Anja Werno, Antje van der Linden, Arlo Upton, Chris Mansell, David Hammer, Dragana Drinkovic, Gary McAuliffe, Hana Sofia Andersson, James Ussher, Jill Sherwood, Josh Freeman, Julia Howard, Juliet Elvy, Mary DeAlmeida, Matt Blakiston, Matthew Rogers, Max Bloomfield, Michael Addidle, Michelle Balm, Sally Roberts, Sarah Jefferies, Sharmini Muttaiyah, Susan Morpeth, Susan Taylor, Timothy Blackmore, Vani Sathyendran, Veronica Playle, Virginia Hope, Erasmus Smit, Lauren Jelly, Olin Silander, Joep de Ligt |
| EPI_ISL_625695, EPI_ISL_625698, EPI_ISL_625706, EPI_ISL_625709, EPI_ISL_625742, EPI_ISL_625822, EPI_ISL_625823, EPI_ISL_625864, EPI_ISL_625883, EPI_ISL_625890, EPI_ISL_626124, EPI_ISL_626127, EPI_ISL_626129, EPI_ISL_626132, EPI_ISL_626133, EPI_ISL_626134, EPI_ISL_626137, EPI_ISL_626138, EPI_ISL_626139, EPI_ISL_626144, EPI_ISL_626146, EPI_ISL_626148, EPI_ISL_626149, EPI_ISL_626157, EPI_ISL_626162, EPI_ISL_626164 | | | |
| see above | Department of Virus and Microbiological Special Diagnostics, Statens Serum Institut, Denmark | Albertsen lab, Department of Chemistry and Bioscience, Aalborg University, Denmark | Danish Covid-19 Genome Consortia |
| EPI_ISL_626356 | LabPLUS | Institute of Environmental Science and Research (ESR) | Xiaoyun Ren, Matt Storey, Nikki Freed, Muhammad Faisal, Jing Wang, Hermes Perez, Anja Werno, Antje van der Linden, Arlo Upton, Chris Mansell, David Hammer, Dragana Drinkovic, Gary McAuliffe, Hana Sofia Andersson, James Ussher, Jill Sherwood, Josh Freeman, Julia Howard, Juliet Elvy, Mary DeAlmeida, Matt Blakiston, Matthew Rogers, Max Bloomfield, Michael Addidle, Michelle Balm, Sally Roberts, Sarah Jefferies, Sharmini Muttaiyah, Susan Morpeth, Susan Taylor, Timothy Blackmore, Vani Sathyendran, Veronica Playle, Virginia Hope, Erasmus Smit, Lauren Jelly, Olin Silander, Joep de Ligt |
| EPI_ISL_626646 | National Public Health Laboratory, National Centre for Infectious Diseases | National Public Health Laboratory, National Centre for Infectious Diseases | Tze Minn Mak, Sophie Octavia, Zhenyang Zhou, Lin Cui, Raymond Tzer Pin Lin |
| EPI_ISL_626811, EPI_ISL_627216, EPI_ISL_627304, EPI_ISL_627711, EPI_ISL_627712, EPI_ISL_627713, EPI_ISL_627714, EPI_ISL_627715, EPI_ISL_627717, EPI_ISL_627718, EPI_ISL_627719, EPI_ISL_627720, EPI_ISL_627721, EPI_ISL_627727, EPI_ISL_627728, EPI_ISL_627729 | | | |
| see above | Queens Medical Centre, Clinical Microbiology Department / DeepSeq Nottingham | COVID-19 Genomics UK (COG-UK) Consortium | Gemma Clark, Wendy Smith, Manjinder Khakh, Vicki M Fleming, Michelle M Lister, Hannah Howson-Wells, Jonathan Ball, Patrick McClure, Joseph Chappell, Theocharis Tsoleridis, Nadine Holmes, Matthew Carlisle, Christopher Moore, Fei Sang, Johnny Debebe, Victoria Wright, Matthew Loose |
| EPI_ISL_635059, EPI_ISL_635060, EPI_ISL_635061, EPI_ISL_635062 | 1-Laboratory of Microbiology, National Reference Lab, Charles Nicolle Hospital; 2-University of Tunis ElManar, Faculty of Medicine of Tunis, LR99ES09, Tunis, Tunisia | 1-Clinical and Experimental Pharmacology Lab, LR16SP02, National Center of Pharmacovigilance, University of Tunis El Manar, Tunis, Tunisia. 2-Neurodegenerative diseases and psychiatric troubles, LR18SP03, Razi Hospital, University of Tunis El Manar, Tunis, Tunisia. 3- Ministry of Health, National Observatory of New and Emerging Diseases, 1006, Tunis, Tunisia | Ilhem Boutiba-Ben Boubaker, Sameh Trabelsi, Nissaf Ben Alaya, Maher Kharrat, Alia Ben Kahla, Jalila Ben Khelil, Salma Abid, Sana Ferjani, Mouna Ben Sassi, Mouna Safer, Imen Mkada, Imen Kacem, Gaies Emna, Soumaya Rammeh, Riadh Daghfous, Riadh Gouider. |
| EPI_ISL_637086 | Middlemore Hospital | Institute of Environmental Science and Research (ESR) | Xiaoyun Ren, Matt Storey, Nikki Freed, Muhammad Faisal, Jing Wang, Hermes Perez, Anja Werno, Antje van der Linden, Arlo Upton, Chris Mansell, David Hammer, Dragana Drinkovic, Gary McAuliffe, Hana Sofia Andersson, James Ussher, Jill Sherwood, Josh Freeman, Julia Howard, Juliet Elvy, Mary DeAlmeida, Matt Blakiston, Matthew Rogers, Max Bloomfield, Michael Addidle, Michelle Balm, Sally Roberts, Sarah Jefferies, Sharmini Muttaiyah, Susan Morpeth, Susan Taylor, Timothy Blackmore, Vani Sathyendran, Veronica Playle, Virginia Hope, Erasmus Smit, Lauren Jelly, Olin Silander, Joep de Ligt |
| EPI_ISL_637180, EPI_ISL_637181, EPI_ISL_637182, EPI_ISL_637183, EPI_ISL_637184 | Respiratory Virus Unit, Microbiology Services Colindale, Public Health England | COVID-19 Genomics UK (COG-UK) Consortium | PHE Covid Sequencing Team |
| EPI_ISL_637284 | Quadram Institute Bioscience | COVID-19 Genomics UK (COG-UK) Consortium | Dave J. Baker, Gemma L. Kay, Alp Aydin, Thanh Le-Viet, Steven Rudder, Ana P. Tedim, Anastasia Kolyva, Maria Diaz, Leonardo de Oliveira Martins, Nabil-Fareed Alikhan, Lizzie Meadows, Rachael Stanley, Ngozi Elumogo, Muhammed Yasir, Nicholas M. Thomson, Alexander J Trotter, Rachel Gilroy, Samuel Bloomfield, Claire Stuart, Andrew Bell, Reenesh Prakash, Samir Dervisevic, Alison E. Mather, John Wain, Mark Webber, Andrew J. Page, Justin O'Grady |
| EPI_ISL_637332, EPI_ISL_637378, EPI_ISL_637379, EPI_ISL_637454, EPI_ISL_637587 | Wales Specialist Virology Centre Sequencing lab: Pathogen Genomics Unit | COVID-19 Genomics UK (COG-UK) Consortium | Catherine Moore, Johnathan Evans, Laura Gifford, Malorie Perry, Simon Cottrell, Angela Marchbank, Alec Birchley, Alexander Adams, Amy Gaskin, Bree Gatica-Wilcox, Jason Coombes, Joel Southgate, Lauren Gilbert, Lee Graham, Nicole Pacchiarini, Sara Kumziene-Summerhayes, Sarah Taylor, Sophie Jones, Sara Rey, Matthew Bull, Joanne Watkins, Sally Corden, Tom Connor |
| EPI_ISL_637588, EPI_ISL_637602 | Quadram Institute Bioscience | COVID-19 Genomics UK (COG-UK) Consortium | Dave J. Baker, Gemma L. Kay, Alp Aydin, Thanh Le-Viet, Steven Rudder, Ana P. Tedim, Anastasia Kolyva, Maria Diaz, Leonardo de Oliveira Martins, Nabil-Fareed Alikhan, Lizzie Meadows, Rachael Stanley, Ngozi Elumogo, Muhammed Yasir, Nicholas M. Thomson, Alexander J Trotter, Rachel Gilroy, Samuel Bloomfield, Claire Stuart, Andrew Bell, Reenesh Prakash, Samir Dervisevic, Alison E. Mather, John Wain, Mark Webber, Andrew J. Page, Justin O'Grady |
| EPI_ISL_637615, EPI_ISL_637915, EPI_ISL_637928, EPI_ISL_637971, EPI_ISL_638058, EPI_ISL_638059, EPI_ISL_638061, EPI_ISL_638062, EPI_ISL_638063, EPI_ISL_638064, EPI_ISL_638113, EPI_ISL_638123, EPI_ISL_638125, EPI_ISL_638128, EPI_ISL_638162, EPI_ISL_638163, EPI_ISL_638206, EPI_ISL_639396, EPI_ISL_639399, EPI_ISL_639402, EPI_ISL_639410, EPI_ISL_639413, EPI_ISL_639414, EPI_ISL_639416, EPI_ISL_639422, EPI_ISL_639424, EPI_ISL_639425, EPI_ISL_639431, EPI_ISL_639432, EPI_ISL_639436, EPI_ISL_639437, EPI_ISL_639438, EPI_ISL_639440, EPI_ISL_639445, EPI_ISL_639449, EPI_ISL_639450, EPI_ISL_639457, EPI_ISL_639459, EPI_ISL_639461, EPI_ISL_639463, EPI_ISL_639473, EPI_ISL_639474, EPI_ISL_639476, EPI_ISL_639477, EPI_ISL_639479, EPI_ISL_639483, EPI_ISL_639486, EPI_ISL_639489, EPI_ISL_639491, EPI_ISL_639494, EPI_ISL_639497, EPI_ISL_639501, EPI_ISL_639503, EPI_ISL_639507, EPI_ISL_639508, EPI_ISL_639509, EPI_ISL_639512, EPI_ISL_639514, EPI_ISL_639521, EPI_ISL_639523, EPI_ISL_639525, EPI_ISL_639527, EPI_ISL_639528, EPI_ISL_639530, EPI_ISL_639534, EPI_ISL_639539, EPI_ISL_639541, EPI_ISL_639542, EPI_ISL_639544, EPI_ISL_639549, EPI_ISL_639553, EPI_ISL_639557, EPI_ISL_639561, EPI_ISL_639563, EPI_ISL_639566, EPI_ISL_639567, EPI_ISL_639569, EPI_ISL_639571, EPI_ISL_639578, EPI_ISL_639580, EPI_ISL_639584, EPI_ISL_639588, EPI_ISL_639592, EPI_ISL_639593, EPI_ISL_639597, EPI_ISL_639603, EPI_ISL_639606, EPI_ISL_639607, EPI_ISL_639612, EPI_ISL_639615, EPI_ISL_639616, EPI_ISL_639617, EPI_ISL_639618 | | | |
| see above | Wales Specialist Virology Centre Sequencing lab: Pathogen Genomics Unit | COVID-19 Genomics UK (COG-UK) Consortium | Catherine Moore, Johnathan Evans, Laura Gifford, Malorie Perry, Simon Cottrell, Angela Marchbank, Alec Birchley, Alexander Adams, Amy Gaskin, Bree Gatica-Wilcox, Jason Coombes, Joel Southgate, Lauren Gilbert, Lee Graham, Nicole Pacchiarini, Sara Kumziene-Summerhayes, Sarah Taylor, Sophie Jones, Sara Rey, Matthew Bull, Joanne Watkins, Sally Corden, Tom Connor |
| EPI_ISL_639733 | Laverty Pathology | NSW Health Pathology - Institute of Clinical Pathology and Medical Research; Westmead Hospital; University of Sydney | CIDM-PH et al. |
| EPI_ISL_639734, EPI_ISL_639735 | Sydney South West Pathology Service (SSWPS) - Liverpool Hospital - NSW Health Pathology | NSW Health Pathology - Institute of Clinical Pathology and Medical Research; Westmead Hospital; University of Sydney | CIDM-PH et al. |
| EPI_ISL_639983 | CNR Virus des Infections Respiratoires - France SUD | CNR Virus des Infections Respiratoires - France SUD | Antonin Bal, Gregory Destras, Gwendolyne Burfin, Hadrien Règue, Alexandre Gaymard, Maude Bouscambert-Duchamp, Florence Morfin-Sherpa, Martine Valette, Bruno Lina, Laurence Josset |
| EPI_ISL_641607, EPI_ISL_641608, EPI_ISL_641609, EPI_ISL_641611, EPI_ISL_641612, EPI_ISL_641614, EPI_ISL_641615, EPI_ISL_641616, EPI_ISL_641617, EPI_ISL_641618, EPI_ISL_641619, EPI_ISL_641620, EPI_ISL_641621, EPI_ISL_641622, EPI_ISL_641623, EPI_ISL_641624, EPI_ISL_641625, EPI_ISL_641626, EPI_ISL_641627, EPI_ISL_641629, EPI_ISL_641630, EPI_ISL_641632, EPI_ISL_641633, EPI_ISL_641634, EPI_ISL_641636, EPI_ISL_641637, EPI_ISL_641638, EPI_ISL_641639, EPI_ISL_641640, EPI_ISL_641641, EPI_ISL_641642, EPI_ISL_641643, EPI_ISL_641644, EPI_ISL_641645, EPI_ISL_641646, EPI_ISL_641647, EPI_ISL_641648, EPI_ISL_641651, EPI_ISL_641653, EPI_ISL_641654, EPI_ISL_641655, EPI_ISL_641656, EPI_ISL_641657, EPI_ISL_641658, EPI_ISL_641659, EPI_ISL_641660, EPI_ISL_641661, EPI_ISL_641662, EPI_ISL_641663, EPI_ISL_641664, EPI_ISL_641665, EPI_ISL_641666, EPI_ISL_641669, EPI_ISL_641670, EPI_ISL_641672, EPI_ISL_641673, EPI_ISL_641674, EPI_ISL_641675, EPI_ISL_641676, EPI_ISL_641677, EPI_ISL_641678, EPI_ISL_641679, EPI_ISL_641680, EPI_ISL_641681, EPI_ISL_641682, EPI_ISL_641683, EPI_ISL_641684, EPI_ISL_641688, EPI_ISL_641689, EPI_ISL_641690, EPI_ISL_641691, EPI_ISL_641692, EPI_ISL_641693, EPI_ISL_641694, EPI_ISL_641695, EPI_ISL_641697, EPI_ISL_641698, EPI_ISL_641699, EPI_ISL_641700, EPI_ISL_641701, EPI_ISL_641702, EPI_ISL_641703, EPI_ISL_641704, EPI_ISL_641705, EPI_ISL_641706, EPI_ISL_641708, EPI_ISL_641709, EPI_ISL_641712, EPI_ISL_641713, EPI_ISL_641714, EPI_ISL_641716, EPI_ISL_641717, EPI_ISL_641718, EPI_ISL_641719, EPI_ISL_641721, EPI_ISL_641722, EPI_ISL_641723, EPI_ISL_641724, EPI_ISL_641725, EPI_ISL_641726, EPI_ISL_641727, EPI_ISL_641728, EPI_ISL_641729, EPI_ISL_641730, EPI_ISL_641731, EPI_ISL_641732, EPI_ISL_641735, EPI_ISL_641736, EPI_ISL_641737, EPI_ISL_641738, EPI_ISL_641739, EPI_ISL_641740, EPI_ISL_641741, EPI_ISL_641742, EPI_ISL_641743, EPI_ISL_641746, EPI_ISL_641748, EPI_ISL_641749, EPI_ISL_641750, EPI_ISL_641751, EPI_ISL_641752, EPI_ISL_641754, EPI_ISL_641755, EPI_ISL_641757, EPI_ISL_641760, EPI_ISL_641764, EPI_ISL_641765, EPI_ISL_641766, EPI_ISL_641767, EPI_ISL_641768, EPI_ISL_641770, EPI_ISL_641771, EPI_ISL_641772, EPI_ISL_641773, EPI_ISL_641774, EPI_ISL_641776, EPI_ISL_641777, EPI_ISL_641778, EPI_ISL_641779, EPI_ISL_641781, EPI_ISL_641783, EPI_ISL_641784, EPI_ISL_641786, EPI_ISL_641787, EPI_ISL_641789, EPI_ISL_641790, EPI_ISL_641791, EPI_ISL_641792, EPI_ISL_641793, EPI_ISL_641794, EPI_ISL_641797, EPI_ISL_641798, EPI_ISL_641800, EPI_ISL_641801, EPI_ISL_641803, EPI_ISL_641805, EPI_ISL_641806, EPI_ISL_641807, EPI_ISL_641808, EPI_ISL_641809, EPI_ISL_641810, EPI_ISL_641811, | | | |

EPI_ISL_641812, EPI_ISL_641813, EPI_ISL_641817, EPI_ISL_641818, EPI_ISL_641819, EPI_ISL_641820, EPI_ISL_641821, EPI_ISL_641822, EPI_ISL_641823, EPI_ISL_641824, EPI_ISL_641825, EPI_ISL_641826, EPI_ISL_641827, EPI_ISL_641828, EPI_ISL_641829, EPI_ISL_641830, EPI_ISL_641831, EPI_ISL_641834, EPI_ISL_641835, EPI_ISL_641836, EPI_ISL_641837, EPI_ISL_641838, EPI_ISL_641839, EPI_ISL_641840, EPI_ISL_641841, EPI_ISL_641843, EPI_ISL_641844, EPI_ISL_641845, EPI_ISL_641846, EPI_ISL_641848, EPI_ISL_641849, EPI_ISL_641851, EPI_ISL_641852, EPI_ISL_641853, EPI_ISL_641854, EPI_ISL_641855, EPI_ISL_641856, EPI_ISL_641858, EPI_ISL_641859, EPI_ISL_641860, EPI_ISL_641862, EPI_ISL_641863, EPI_ISL_641865, EPI_ISL_641866, EPI_ISL_641867, EPI_ISL_641869, EPI_ISL_641870, EPI_ISL_641871, EPI_ISL_641872, EPI_ISL_641873, EPI_ISL_641874, EPI_ISL_641875, EPI_ISL_641876, EPI_ISL_641880, EPI_ISL_641881, EPI_ISL_641883, EPI_ISL_641885, EPI_ISL_641886, EPI_ISL_641887, EPI_ISL_641889, EPI_ISL_641890, EPI_ISL_641891, EPI_ISL_641892, EPI_ISL_641893, EPI_ISL_641894, EPI_ISL_641895, EPI_ISL_641896, EPI_ISL_641898, EPI_ISL_641899, EPI_ISL_641902, EPI_ISL_641903, EPI_ISL_641904, EPI_ISL_641905, EPI_ISL_641906, EPI_ISL_641907, EPI_ISL_641908, EPI_ISL_641909, EPI_ISL_641910, EPI_ISL_641912, EPI_ISL_641913, EPI_ISL_641914, EPI_ISL_641915, EPI_ISL_641916, EPI_ISL_641917, EPI_ISL_641918, EPI_ISL_641919, EPI_ISL_641920, EPI_ISL_641921, EPI_ISL_641922, EPI_ISL_641923, EPI_ISL_641924, EPI_ISL_641925, EPI_ISL_641926, EPI_ISL_641927, EPI_ISL_641928, EPI_ISL_641929, EPI_ISL_641930, EPI_ISL_641931, EPI_ISL_641932, EPI_ISL_641933, EPI_ISL_641934, EPI_ISL_641935, EPI_ISL_641936, EPI_ISL_641937, EPI_ISL_641938, EPI_ISL_641939, EPI_ISL_641940, EPI_ISL_641941, EPI_ISL_641942, EPI_ISL_641943, EPI_ISL_641944, EPI_ISL_641945, EPI_ISL_641946, EPI_ISL_641947, EPI_ISL_641948, EPI_ISL_641949, EPI_ISL_641950, EPI_ISL_641951, EPI_ISL_641952, EPI_ISL_641953, EPI_ISL_641954, EPI_ISL_641955, EPI_ISL_641956, EPI_ISL_641957, EPI_ISL_641958, EPI_ISL_641959, EPI_ISL_641961, EPI_ISL_641962, EPI_ISL_641963, EPI_ISL_641964, EPI_ISL_641965, EPI_ISL_641966, EPI_ISL_641967, EPI_ISL_641968, EPI_ISL_641969, EPI_ISL_641970, EPI_ISL_641971, EPI_ISL_641972, EPI_ISL_641973, EPI_ISL_641974, EPI_ISL_641975, EPI_ISL_641976, EPI_ISL_641977, EPI_ISL_641978, EPI_ISL_641980, EPI_ISL_641981, EPI_ISL_641982, EPI_ISL_641983, EPI_ISL_641984, EPI_ISL_641985, EPI_ISL_641986, EPI_ISL_641987, EPI_ISL_641988, EPI_ISL_641989, EPI_ISL_641990, EPI_ISL_641991, EPI_ISL_641992, EPI_ISL_641993, EPI_ISL_641995, EPI_ISL_641996, EPI_ISL_641997, EPI_ISL_641998, EPI_ISL_641999, EPI_ISL_642000, EPI_ISL_642001, EPI_ISL_642002, EPI_ISL_642003, EPI_ISL_642004, EPI_ISL_642005, EPI_ISL_642006, EPI_ISL_642007, EPI_ISL_642008, EPI_ISL_642009, EPI_ISL_642010, EPI_ISL_642011, EPI_ISL_642012, EPI_ISL_642013, EPI_ISL_642014, EPI_ISL_642015, EPI_ISL_642016, EPI_ISL_642017, EPI_ISL_642019, EPI_ISL_642020, EPI_ISL_642021, EPI_ISL_642022, EPI_ISL_642023, EPI_ISL_642024, EPI_ISL_642025, EPI_ISL_642026, EPI_ISL_642027, EPI_ISL_642028, EPI_ISL_642029, EPI_ISL_642030, EPI_ISL_642031, EPI_ISL_642032, EPI_ISL_642033, EPI_ISL_642034, EPI_ISL_642035, EPI_ISL_642036, EPI_ISL_642037, EPI_ISL_642039, EPI_ISL_642040, EPI_ISL_642041, EPI_ISL_642042, EPI_ISL_642043, EPI_ISL_642044, EPI_ISL_642045, EPI_ISL_642046, EPI_ISL_642047, EPI_ISL_642048, EPI_ISL_642049, EPI_ISL_642050, EPI_ISL_642051, EPI_ISL_642052, EPI_ISL_642053, EPI_ISL_642054, EPI_ISL_642055, EPI_ISL_642056, EPI_ISL_642057, EPI_ISL_642058, EPI_ISL_642059, EPI_ISL_642060, EPI_ISL_642061, EPI_ISL_642062, EPI_ISL_642063, EPI_ISL_642064, EPI_ISL_642065, EPI_ISL_642066, EPI_ISL_642067, EPI_ISL_642068, EPI_ISL_642070, EPI_ISL_642071, EPI_ISL_642072, EPI_ISL_642073, EPI_ISL_642075, EPI_ISL_642076, EPI_ISL_642077, EPI_ISL_642078, EPI_ISL_642079, EPI_ISL_642080, EPI_ISL_642081, EPI_ISL_642082, EPI_ISL_642083, EPI_ISL_642084, EPI_ISL_642085, EPI_ISL_642086, EPI_ISL_642087, EPI_ISL_642088, EPI_ISL_642089, EPI_ISL_642090, EPI_ISL_642091, EPI_ISL_642092, EPI_ISL_642093, EPI_ISL_642094, EPI_ISL_642095, EPI_ISL_642096, EPI_ISL_642097, EPI_ISL_642098, EPI_ISL_642099, EPI_ISL_642100, EPI_ISL_642101, EPI_ISL_642102, EPI_ISL_642103, EPI_ISL_642104, EPI_ISL_642105, EPI_ISL_642106, EPI_ISL_642107, EPI_ISL_642108, EPI_ISL_642109, EPI_ISL_642110, EPI_ISL_642111, EPI_ISL_642112, EPI_ISL_642113, EPI_ISL_642114, EPI_ISL_642115, EPI_ISL_642116, EPI_ISL_642117, EPI_ISL_642118, EPI_ISL_642119, EPI_ISL_642120, EPI_ISL_642121, EPI_ISL_642122, EPI_ISL_642123, EPI_ISL_642124, EPI_ISL_642125, EPI_ISL_642126, EPI_ISL_642127, EPI_ISL_642128, EPI_ISL_642129, EPI_ISL_642130, EPI_ISL_642131, EPI_ISL_642132, EPI_ISL_642133, EPI_ISL_642134, EPI_ISL_642135, EPI_ISL_642136, EPI_ISL_642137, EPI_ISL_642138, EPI_ISL_642139, EPI_ISL_642140, EPI_ISL_642141, EPI_ISL_642142, EPI_ISL_642143, EPI_ISL_642144, EPI_ISL_642145, EPI_ISL_642146, EPI_ISL_642147, EPI_ISL_642148, EPI_ISL_642149, EPI_ISL_642150, EPI_ISL_642151, EPI_ISL_642152, EPI_ISL_642153, EPI_ISL_642154, EPI_ISL_642156, EPI_ISL_642157, EPI_ISL_642158, EPI_ISL_642159, EPI_ISL_642160, EPI_ISL_642161, EPI_ISL_642162, EPI_ISL_642163, EPI_ISL_642164, EPI_ISL_642165, EPI_ISL_642167, EPI_ISL_642168, EPI_ISL_642171, EPI_ISL_642172, EPI_ISL_642173, EPI_ISL_642174, EPI_ISL_642175, EPI_ISL_642176, EPI_ISL_642177, EPI_ISL_642178, EPI_ISL_642179, EPI_ISL_642180, EPI_ISL_642181, EPI_ISL_642182, EPI_ISL_642183, EPI_ISL_642184, EPI_ISL_642185, EPI_ISL_642186, EPI_ISL_642187, EPI_ISL_642188, EPI_ISL_642189, EPI_ISL_642190, EPI_ISL_642191, EPI_ISL_642192, EPI_ISL_642193, EPI_ISL_642194, EPI_ISL_642195, EPI_ISL_642196, EPI_ISL_642197, EPI_ISL_642198, EPI_ISL_642199, EPI_ISL_642200, EPI_ISL_642201, EPI_ISL_642202, EPI_ISL_642204, EPI_ISL_642205, EPI_ISL_642206, EPI_ISL_642207, EPI_ISL_642208, EPI_ISL_642209, EPI_ISL_642210, EPI_ISL_642211, EPI_ISL_642212, EPI_ISL_642213, EPI_ISL_642214, EPI_ISL_642215, EPI_ISL_642216, EPI_ISL_642217, EPI_ISL_642218, EPI_ISL_642219, EPI_ISL_642220, EPI_ISL_642221, EPI_ISL_642223, EPI_ISL_642224, EPI_ISL_642225, EPI_ISL_642226, EPI_ISL_642227, EPI_ISL_642228, EPI_ISL_642229, EPI_ISL_642230, EPI_ISL_642231, EPI_ISL_642232, EPI_ISL_642233, EPI_ISL_642234, EPI_ISL_642235, EPI_ISL_642236, EPI_ISL_642237, EPI_ISL_642238, EPI_ISL_642239, EPI_ISL_642240, EPI_ISL_642241, EPI_ISL_642242, EPI_ISL_642244, EPI_ISL_642245, EPI_ISL_642246, EPI_ISL_642248, EPI_ISL_642249, EPI_ISL_642250, EPI_ISL_642252, EPI_ISL_642253, EPI_ISL_642254, EPI_ISL_642255, EPI_ISL_642256, EPI_ISL_642257, EPI_ISL_642258, EPI_ISL_642259, EPI_ISL_642260, EPI_ISL_642261, EPI_ISL_642262, EPI_ISL_642263, EPI_ISL_642264, EPI_ISL_642265, EPI_ISL_642266, EPI_ISL_642267, EPI_ISL_642268, EPI_ISL_642269, EPI_ISL_642270, EPI_ISL_642271, EPI_ISL_642272, EPI_ISL_642273, EPI_ISL_642274, EPI_ISL_642275, EPI_ISL_642276, EPI_ISL_642277, EPI_ISL_642278, EPI_ISL_642279, EPI_ISL_642280, EPI_ISL_642281, EPI_ISL_642282, EPI_ISL_642283, EPI_ISL_642285, EPI_ISL_642286, EPI_ISL_642287, EPI_ISL_642288, EPI_ISL_642289, EPI_ISL_642290, EPI_ISL_642291, EPI_ISL_642292, EPI_ISL_642293, EPI_ISL_642294, EPI_ISL_642295, EPI_ISL_642296, EPI_ISL_642297, EPI_ISL_642298, EPI_ISL_642299, EPI_ISL_642300, EPI_ISL_642301, EPI_ISL_642302, EPI_ISL_642303, EPI_ISL_642304, EPI_ISL_642305, EPI_ISL_642306, EPI_ISL_642307, EPI_ISL_642308, EPI_ISL_642309, EPI_ISL_642310, EPI_ISL_642311, EPI_ISL_642312, EPI_ISL_642313, EPI_ISL_642314, EPI_ISL_642315, EPI_ISL_642316, EPI_ISL_642317, EPI_ISL_642318, EPI_ISL_642319, EPI_ISL_642320, EPI_ISL_642321, EPI_ISL_642322, EPI_ISL_642323, EPI_ISL_642324, EPI_ISL_642325, EPI_ISL_642326, EPI_ISL_642327, EPI_ISL_642328, EPI_ISL_642329, EPI_ISL_642330, EPI_ISL_642331, EPI_ISL_642332, EPI_ISL_642333, EPI_ISL_642334, EPI_ISL_642335, EPI_ISL_642336, EPI_ISL_642337, EPI_ISL_642338, EPI_ISL_642339, EPI_ISL_642340, EPI_ISL_642341, EPI_ISL_642342, EPI_ISL_642343, EPI_ISL_642344, EPI_ISL_642345, EPI_ISL_642346, EPI_ISL_642347, EPI_ISL_642348, EPI_ISL_642349, EPI_ISL_642350, EPI_ISL_642351, EPI_ISL_642352, EPI_ISL_642353, EPI_ISL_642355, EPI_ISL_642356, EPI_ISL_642357, EPI_ISL_642358, EPI_ISL_642359, EPI_ISL_642360, EPI_ISL_642361, EPI_ISL_642362, EPI_ISL_642363, EPI_ISL_642364, EPI_ISL_642365, EPI_ISL_642367, EPI_ISL_642368, EPI_ISL_642369, EPI_ISL_642370, EPI_ISL_642371, EPI_ISL_642372, EPI_ISL_642373, EPI_ISL_642374, EPI_ISL_642375, EPI_ISL_642376, EPI_ISL_642377, EPI_ISL_642378, EPI_ISL_642380, EPI_ISL_642381, EPI_ISL_642382, EPI_ISL_642383, EPI_ISL_642384, EPI_ISL_642385, EPI_ISL_642386, EPI_ISL_642387, EPI_ISL_642388, EPI_ISL_642389, EPI_ISL_642390, EPI_ISL_642391, EPI_ISL_642392, EPI_ISL_642393, EPI_ISL_642394, EPI_ISL_642395, EPI_ISL_642396, EPI_ISL_642397, EPI_ISL_642398, EPI_ISL_642399, EPI_ISL_642400, EPI_ISL_642401, EPI_ISL_642402, EPI_ISL_642403, EPI_ISL_642404, EPI_ISL_642405, EPI_ISL_642406, EPI_ISL_642407, EPI_ISL_642408, EPI_ISL_642409, EPI_ISL_642410, EPI_ISL_642411, EPI_ISL_642412, EPI_ISL_642413, EPI_ISL_642414, EPI_ISL_642415, EPI_ISL_642416, EPI_ISL_642417, EPI_ISL_642418, EPI_ISL_642419, EPI_ISL_642421, EPI_ISL_642422, EPI_ISL_642423, EPI_ISL_642424, EPI_ISL_642425, EPI_ISL_642426, EPI_ISL_642427, EPI_ISL_642428, EPI_ISL_642429, EPI_ISL_642430, EPI_ISL_642431, EPI_ISL_642432, EPI_ISL_642433, EPI_ISL_642434, EPI_ISL_642435, EPI_ISL_642436, EPI_ISL_642438, EPI_ISL_642440, EPI_ISL_642441, EPI_ISL_642442, EPI_ISL_642443, EPI_ISL_642444, EPI_ISL_642445, EPI_ISL_642446, EPI_ISL_642447, EPI_ISL_642448, EPI_ISL_642449, EPI_ISL_642450, EPI_ISL_642451, EPI_ISL_642452, EPI_ISL_642453, EPI_ISL_642454, EPI_ISL_642455, EPI_ISL_642456, EPI_ISL_642457, EPI_ISL_642458, EPI_ISL_642459, EPI_ISL_642460, EPI_ISL_642461, EPI_ISL_642462, EPI_ISL_642463, EPI_ISL_642464, EPI_ISL_642465, EPI_ISL_642466, EPI_ISL_642467, EPI_ISL_642468, EPI_ISL_642469, EPI_ISL_642470, EPI_ISL_642471, EPI_ISL_642472, EPI_ISL_642473, EPI_ISL_642474, EPI_ISL_642475, EPI_ISL_642476, EPI_ISL_642478, EPI_ISL_642479, EPI_ISL_642480, EPI_ISL_642481, EPI_ISL_642482, EPI_ISL_642483, EPI_ISL_642484, EPI_ISL_642485, EPI_ISL_642486, EPI_ISL_642487, EPI_ISL_642488, EPI_ISL_642489, EPI_ISL_642490, EPI_ISL_642491, EPI_ISL_642492, EPI_ISL_642493, EPI_ISL_642494, EPI_ISL_642495, EPI_ISL_642496, EPI_ISL_642497, EPI_ISL_642498, EPI_ISL_642499, EPI_ISL_642500, EPI_ISL_642501, EPI_ISL_642502, EPI_ISL_642503, EPI_ISL_642504, EPI_ISL_642505, EPI_ISL_642507, EPI_ISL_642509, EPI_ISL_642510, EPI_ISL_642511, EPI_ISL_642512, EPI_ISL_642513, EPI_ISL_642514, EPI_ISL_642515, EPI_ISL_642516, EPI_ISL_642517, EPI_ISL_642518, EPI_ISL_642519, EPI_ISL_642520, EPI_ISL_642522, EPI_ISL_642523, EPI_ISL_642524, EPI_ISL_642525, EPI_ISL_642526, EPI_ISL_642527, EPI_ISL_642528, EPI_ISL_642529, EPI_ISL_642530, EPI_ISL_642531, EPI_ISL_642532, EPI_ISL_642533, EPI_ISL_642534, EPI_ISL_642535, EPI_ISL_642536, EPI_ISL_642537, EPI_ISL_642538, EPI_ISL_642539, EPI_ISL_642540, EPI_ISL_642541, EPI_ISL_642542, EPI_ISL_642543, EPI_ISL_642544, EPI_ISL_642545, EPI_ISL_642546, EPI_ISL_642547, EPI_ISL_642548, EPI_ISL_642549, EPI_ISL_642550, EPI_ISL_642551, EPI_ISL_642552, EPI_ISL_642553, EPI_ISL_642554, EPI_ISL_642555, EPI_ISL_642556, EPI_ISL_642557, EPI_ISL_642558, EPI_ISL_642559, EPI_ISL_642560, EPI_ISL_642561, EPI_ISL_642562, EPI_ISL_642563, EPI_ISL_642564, EPI_ISL_642565, EPI_ISL_642566, EPI_ISL_642567, EPI_ISL_642568, EPI_ISL_642570, EPI_ISL_642571, EPI_ISL_642572, EPI_ISL_642573, EPI_ISL_642574, EPI_ISL_642575, EPI_ISL_642576, EPI_ISL_642577, EPI_ISL_642578, EPI_ISL_642579, EPI_ISL_642580, EPI_ISL_642582, EPI_ISL_642583, EPI_ISL_642584, EPI_ISL_642585, EPI_ISL_642586, EPI_ISL_642587, EPI_ISL_642588, EPI_ISL_642589, EPI_ISL_642590, EPI_ISL_642591, EPI_ISL_642592, EPI_ISL_642593, EPI_ISL_642594, EPI_ISL_642595, EPI_ISL_642596, EPI_ISL_642597, EPI_ISL_642598, EPI_ISL_642600, EPI_ISL_642601, EPI_ISL_642602, EPI_ISL_642603, EPI_ISL_642604, EPI_ISL_642606, EPI_ISL_642607, EPI_ISL_642608, EPI_ISL_642609, EPI_ISL_642610, EPI_ISL_642611, EPI_ISL_642612, EPI_ISL_642613, EPI_ISL_642614, EPI_ISL_642615, EPI_ISL_642616, EPI_ISL_642617, EPI_ISL_642618, EPI_ISL_642619, EPI_ISL_642620, EPI_ISL_642621, EPI_ISL_642622, EPI_ISL_642623, EPI_ISL_642624, EPI_ISL_642625, EPI_ISL_642626, EPI_ISL_642627, EPI_ISL_642628, EPI_ISL_642629, EPI_ISL_642630, EPI_ISL_642631, EPI_ISL_642632, EPI_ISL_642633, EPI_ISL_642636, EPI_ISL_642637, EPI_ISL_642638, EPI_ISL_642639, EPI_ISL_642640, EPI_ISL_642641, EPI_ISL_642642, EPI_ISL_642643, EPI_ISL_642645, EPI_ISL_642646, EPI_ISL_642647, EPI_ISL_642648, EPI_ISL_642649, EPI_ISL_642650, EPI_ISL_642651, EPI_ISL_642652, EPI_ISL_642653, EPI_ISL_642654, EPI_ISL_642656, EPI_ISL_642657, EPI_ISL_642658, EPI_ISL_642659, EPI_ISL_642660, EPI_ISL_642661, EPI_ISL_642662, EPI_ISL_642665, EPI_ISL_642666, EPI_ISL_642667, EPI_ISL_642668, EPI_ISL_642669, EPI_ISL_642670, EPI_ISL_642671, EPI_ISL_642672, EPI_ISL_642673, EPI_ISL_642674, EPI_ISL_642675, EPI_ISL_642676, EPI_ISL_642677, EPI_ISL_642678, EPI_ISL_642679, EPI_ISL_642680, EPI_ISL_642681, EPI_ISL_642682, EPI_ISL_642683, EPI_ISL_642685, EPI_ISL_642686, EPI_ISL_642687, EPI_ISL_642688, EPI_ISL_642689, EPI_ISL_642690, EPI_ISL_642691, EPI_ISL_642692, EPI_ISL_642693, EPI_ISL_642694, EPI_ISL_642695, EPI_ISL_642696, EPI_ISL_642697, EPI_ISL_642698, EPI_ISL_642699, EPI_ISL_642700, EPI_ISL_642701, EPI_ISL_642704, EPI_ISL_642705, EPI_ISL_642706, EPI_ISL_642707, EPI_ISL_642708, EPI_ISL_642709, EPI_ISL_642710, EPI_ISL_642711, EPI_ISL_642712, EPI_ISL_642713, EPI_ISL_642714, EPI_ISL_642715, EPI_ISL_642716, EPI_ISL_642717, EPI_ISL_642718, EPI_ISL_642719, EPI_ISL_642720, EPI_ISL_642721, EPI_ISL_642722, EPI_ISL_642723, EPI_ISL_642724, EPI_ISL_642725, EPI_ISL_642726, EPI_ISL_642727, EPI_ISL_642728, EPI_ISL_642730, EPI_ISL_642731, EPI_ISL_642732, EPI_ISL_642733, EPI_ISL_642734, EPI_ISL_642735, EPI_ISL_642736, EPI_ISL_642737, EPI_ISL_642738, EPI_ISL_642739, EPI_ISL_642740, EPI_ISL_642741, EPI_ISL_642742, EPI_ISL_642743, EPI_ISL_642745, EPI_ISL_642746, EPI_ISL_642747, EPI_ISL_642748, EPI_ISL_642749, EPI_ISL_642750, EPI_ISL_642751, EPI_ISL_642752, EPI_ISL_642753, EPI_ISL_642754, EPI_ISL_642755, EPI_ISL_642756, EPI_ISL_642757, EPI_ISL_642758, EPI_ISL_642759, EPI_ISL_642760

| | | | |
|---|---|---|---|
| see above | Lighthouse Lab in Glasgow | Wellcome Sanger Institute for the COVID-19 Genomics UK (COG-UK) Consortium | Harper VanSteenhouse, Yumi Kasai, David Gray, Carol Clugston, Anna Dominiczak and Alex Alderton, Roberto Amato, Sonia Goncalves, Ewan Harrison, David K. Jackson, Ian Johnston, Dominic Kwiatkowski, Cordelia Langford, John Sillitoe on behalf of the Wellcome Sanger Institute COVID-19 Surveillance Team |
| EPI_ISL_644614, EPI_ISL_644615 | Department of Clinical Microbiology | GIGA Medical Genomics | Keith Durkin, Maria Artesi, Sébastien Bontems, Raphaël Boreux, Bouchra Boujemla, Cécile Meex, Pierrette Melin, Marie-Pierre Hayette, Vincent Bours |
| EPI_ISL_645118 | National Public Health Laboratory, National Centre for Infectious Diseases | National Public Health Laboratory, National Centre for Infectious Diseases | Tze Minn Mak, Sophie Octavia, Zhenyang Zhou, Lin Cui, Raymond Tzer Pin Lin |
| EPI_ISL_645264, EPI_ISL_645296, EPI_ISL_645302 | Lighthouse Lab in Milton Keynes | Wellcome Sanger Institute for the COVID-19 Genomics UK (COG-UK) Consortium | The Lighthouse Lab in Milton Keynes and Alex Alderton, Roberto Amato, Sonia Goncalves, Ewan Harrison, David K. Jackson, Ian Johnston, Dominic Kwiatkowski, Cordelia Langford, John Sillitoe on behalf of the Wellcome Sanger Institute COVID-19 Surveillance Team |
| EPI_ISL_645322 | Lighthouse Lab in Cambridge | Wellcome Sanger Institute for the COVID-19 Genomics UK (COG-UK) Consortium | Rob Howes, The Lighthouse Lab in Cambridge and Alex Alderton, Roberto Amato, Sonia Goncalves, Ewan Harrison, David K. Jackson, Ian Johnston, Dominic Kwiatkowski, Cordelia Langford, John Sillitoe on behalf of the Wellcome Sanger Institute COVID-19 Surveillance Team |
| EPI_ISL_645345, EPI_ISL_645354, EPI_ISL_645360, EPI_ISL_645375, EPI_ISL_645406 | Lighthouse Lab in Milton Keynes | Wellcome Sanger Institute for the COVID-19 Genomics UK (COG-UK) Consortium | The Lighthouse Lab in Milton Keynes and Alex Alderton, Roberto Amato, Sonia Goncalves, Ewan Harrison, David K. Jackson, Ian Johnston, Dominic Kwiatkowski, Cordelia Langford, John Sillitoe on behalf of the Wellcome Sanger Institute COVID-19 Surveillance Team |
| EPI_ISL_645414 | Lighthouse Lab in Cambridge | Wellcome Sanger Institute for the COVID-19 Genomics UK (COG-UK) Consortium | Rob Howes, The Lighthouse Lab in Cambridge and Alex Alderton, Roberto Amato, Sonia Goncalves, Ewan Harrison, David K. Jackson, Ian Johnston, Dominic Kwiatkowski, Cordelia Langford, John Sillitoe on behalf of the Wellcome Sanger Institute COVID-19 Surveillance Team |
| EPI_ISL_645421, EPI_ISL_645438, EPI_ISL_645489 | Lighthouse Lab in Milton Keynes | Wellcome Sanger Institute for the COVID-19 Genomics UK (COG-UK) Consortium | The Lighthouse Lab in Milton Keynes and Alex Alderton, Roberto Amato, Sonia Goncalves, Ewan Harrison, David K. Jackson, Ian Johnston, Dominic Kwiatkowski, Cordelia Langford, John Sillitoe on behalf of the Wellcome Sanger Institute COVID-19 Surveillance Team |

EPI_ISL_645538, EPI_ISL_645539, EPI_ISL_645545, EPI_ISL_645551, EPI_ISL_645572, EPI_ISL_645576, EPI_ISL_645584, EPI_ISL_645608, EPI_ISL_645610, EPI_ISL_645623, EPI_ISL_645631, EPI_ISL_645659, EPI_ISL_645665, EPI_ISL_645678, EPI_ISL_645686, EPI_ISL_645692, EPI_ISL_645731, EPI_ISL_645735, EPI_ISL_645755, EPI_ISL_645757, EPI_ISL_645758, EPI_ISL_645768, EPI_ISL_645791, EPI_ISL_645796, EPI_ISL_645812, EPI_ISL_645813, EPI_ISL_645825

| | | | |
|---|---|---|---|
| see above | Lighthouse Lab in Alderley Park | Wellcome Sanger Institute for the COVID-19 Genomics UK (COG-UK) Consortium | Jacquelyn Wynn, Mairead Hyland, The Lighthouse Lab in Alderley Park and Alex Alderton, Roberto Amato, Sonia Goncalves, Ewan Harrison, David K. Jackson, Ian Johnston, Dominic Kwiatkowski, Cordelia Langford, John Sillitoe on behalf of the Wellcome Sanger Institute COVID-19 Surveillance Team |
| EPI_ISL_645826, EPI_ISL_645828, EPI_ISL_645829, EPI_ISL_645832, EPI_ISL_645836, EPI_ISL_645838 | Lighthouse Lab in Cambridge | Wellcome Sanger Institute for the COVID-19 Genomics UK (COG-UK) Consortium | Rob Howes, The Lighthouse Lab in Cambridge and Alex Alderton, Roberto Amato, Sonia Goncalves, Ewan Harrison, David K. Jackson, Ian Johnston, Dominic Kwiatkowski, Cordelia Langford, John Sillitoe on behalf of the Wellcome Sanger Institute COVID-19 Surveillance Team |
| EPI_ISL_645839 | Lighthouse Lab in Alderley Park | Wellcome Sanger Institute for the COVID-19 Genomics UK (COG-UK) Consortium | Jacquelyn Wynn, Mairead Hyland, The Lighthouse Lab in Alderley Park and Alex Alderton, Roberto Amato, Sonia Goncalves, Ewan Harrison, David K. Jackson, Ian Johnston, Dominic Kwiatkowski, Cordelia Langford, John Sillitoe on behalf of the Wellcome Sanger Institute COVID-19 Surveillance Team |
| EPI_ISL_645850, EPI_ISL_645851, EPI_ISL_645852, EPI_ISL_645866, EPI_ISL_645868, EPI_ISL_645869 | Lighthouse Lab in Cambridge | Wellcome Sanger Institute for the COVID-19 Genomics UK (COG-UK) Consortium | Rob Howes, The Lighthouse Lab in Cambridge and Alex Alderton, Roberto Amato, Sonia Goncalves, Ewan Harrison, David K. Jackson, Ian Johnston, Dominic Kwiatkowski, Cordelia Langford, John Sillitoe on behalf of the Wellcome Sanger Institute COVID-19 Surveillance Team |
| EPI_ISL_645875 | Lighthouse Lab in Alderley Park | Wellcome Sanger Institute for the COVID-19 Genomics UK | Jacquelyn Wynn, Mairead Hyland, The Lighthouse Lab in Alderley Park and Alex Alderton, Roberto Amato, Sonia Goncalves, Ewan Harrison, David K. |

| | | | |
|---|---|---|---|
| | | (COG-UK) Consortium | Jackson, Ian Johnston, Dominic Kwiatkowski, Cordelia Langford, John Sillitoe on behalf of the Wellcome Sanger Institute COVID-19 Surveillance Team |
| EPI_ISL_645877, EPI_ISL_645880 | Lighthouse Lab in Cambridge | Wellcome Sanger Institute for the COVID-19 Genomics UK (COG-UK) Consortium | Rob Howes, The Lighthouse Lab in Cambridge and Alex Alderton, Roberto Amato, Sonia Goncalves, Ewan Harrison, David K. Jackson, Ian Johnston, Dominic Kwiatkowski, Cordelia Langford, John Sillitoe on behalf of the Wellcome Sanger Institute COVID-19 Surveillance Team |
| EPI_ISL_645881 | Lighthouse Lab in Alderley Park | Wellcome Sanger Institute for the COVID-19 Genomics UK (COG-UK) Consortium | Jacquelyn Wynn, Mairead Hyland, The Lighthouse Lab in Alderley Park and Alex Alderton, Roberto Amato, Sonia Goncalves, Ewan Harrison, David K. Jackson, Ian Johnston, Dominic Kwiatkowski, Cordelia Langford, John Sillitoe on behalf of the Wellcome Sanger Institute COVID-19 Surveillance Team |
| EPI_ISL_645882, EPI_ISL_645885 | Lighthouse Lab in Cambridge | Wellcome Sanger Institute for the COVID-19 Genomics UK (COG-UK) Consortium | Rob Howes, The Lighthouse Lab in Cambridge and Alex Alderton, Roberto Amato, Sonia Goncalves, Ewan Harrison, David K. Jackson, Ian Johnston, Dominic Kwiatkowski, Cordelia Langford, John Sillitoe on behalf of the Wellcome Sanger Institute COVID-19 Surveillance Team |
| EPI_ISL_645889 | Lighthouse Lab in Alderley Park | Wellcome Sanger Institute for the COVID-19 Genomics UK (COG-UK) Consortium | Jacquelyn Wynn, Mairead Hyland, The Lighthouse Lab in Alderley Park and Alex Alderton, Roberto Amato, Sonia Goncalves, Ewan Harrison, David K. Jackson, Ian Johnston, Dominic Kwiatkowski, Cordelia Langford, John Sillitoe on behalf of the Wellcome Sanger Institute COVID-19 Surveillance Team |
| EPI_ISL_645899, EPI_ISL_645908, EPI_ISL_645917, EPI_ISL_645918, EPI_ISL_645920, EPI_ISL_645923, EPI_ISL_645925, EPI_ISL_645926, EPI_ISL_645927 | Lighthouse Lab in Cambridge | Wellcome Sanger Institute for the COVID-19 Genomics UK (COG-UK) Consortium | Rob Howes, The Lighthouse Lab in Cambridge and Alex Alderton, Roberto Amato, Sonia Goncalves, Ewan Harrison, David K. Jackson, Ian Johnston, Dominic Kwiatkowski, Cordelia Langford, John Sillitoe on behalf of the Wellcome Sanger Institute COVID-19 Surveillance Team |
| EPI_ISL_645935 | Lighthouse Lab in Alderley Park | Wellcome Sanger Institute for the COVID-19 Genomics UK (COG-UK) Consortium | Jacquelyn Wynn, Mairead Hyland, The Lighthouse Lab in Alderley Park and Alex Alderton, Roberto Amato, Sonia Goncalves, Ewan Harrison, David K. Jackson, Ian Johnston, Dominic Kwiatkowski, Cordelia Langford, John Sillitoe on behalf of the Wellcome Sanger Institute COVID-19 Surveillance Team |
| EPI_ISL_645949 | Lighthouse Lab in Cambridge | Wellcome Sanger Institute for the COVID-19 Genomics UK (COG-UK) Consortium | Rob Howes, The Lighthouse Lab in Cambridge and Alex Alderton, Roberto Amato, Sonia Goncalves, Ewan Harrison, David K. Jackson, Ian Johnston, Dominic Kwiatkowski, Cordelia Langford, John Sillitoe on behalf of the Wellcome Sanger Institute COVID-19 Surveillance Team |
| EPI_ISL_645950 | Lighthouse Lab in Alderley Park | Wellcome Sanger Institute for the COVID-19 Genomics UK (COG-UK) Consortium | Jacquelyn Wynn, Mairead Hyland, The Lighthouse Lab in Alderley Park and Alex Alderton, Roberto Amato, Sonia Goncalves, Ewan Harrison, David K. Jackson, Ian Johnston, Dominic Kwiatkowski, Cordelia Langford, John Sillitoe on behalf of the Wellcome Sanger Institute COVID-19 Surveillance Team |
| EPI_ISL_645958, EPI_ISL_645962 | Lighthouse Lab in Cambridge | Wellcome Sanger Institute for the COVID-19 Genomics UK (COG-UK) Consortium | Rob Howes, The Lighthouse Lab in Cambridge and Alex Alderton, Roberto Amato, Sonia Goncalves, Ewan Harrison, David K. Jackson, Ian Johnston, Dominic Kwiatkowski, Cordelia Langford, John Sillitoe on behalf of the Wellcome Sanger Institute COVID-19 Surveillance Team |
| EPI_ISL_645963 | Lighthouse Lab in Alderley Park | Wellcome Sanger Institute for the COVID-19 Genomics UK (COG-UK) Consortium | Jacquelyn Wynn, Mairead Hyland, The Lighthouse Lab in Alderley Park and Alex Alderton, Roberto Amato, Sonia Goncalves, Ewan Harrison, David K. Jackson, Ian Johnston, Dominic Kwiatkowski, Cordelia Langford, John Sillitoe on behalf of the Wellcome Sanger Institute COVID-19 Surveillance Team |
| EPI_ISL_645970 | Lighthouse Lab in Cambridge | Wellcome Sanger Institute for the COVID-19 Genomics UK (COG-UK) Consortium | Rob Howes, The Lighthouse Lab in Cambridge and Alex Alderton, Roberto Amato, Sonia Goncalves, Ewan Harrison, David K. Jackson, Ian Johnston, Dominic Kwiatkowski, Cordelia Langford, John Sillitoe on behalf of the Wellcome Sanger Institute COVID-19 Surveillance Team |
| EPI_ISL_645974 | Lighthouse Lab in Alderley Park | Wellcome Sanger Institute for the COVID-19 Genomics UK (COG-UK) Consortium | Jacquelyn Wynn, Mairead Hyland, The Lighthouse Lab in Alderley Park and Alex Alderton, Roberto Amato, Sonia Goncalves, Ewan Harrison, David K. Jackson, Ian Johnston, Dominic Kwiatkowski, Cordelia Langford, John Sillitoe on behalf of the Wellcome Sanger Institute COVID-19 Surveillance Team |
| EPI_ISL_645980, EPI_ISL_645981, EPI_ISL_645983, EPI_ISL_645989 | Lighthouse Lab in Cambridge | Wellcome Sanger Institute for the COVID-19 Genomics UK (COG-UK) Consortium | Rob Howes, The Lighthouse Lab in Cambridge and Alex Alderton, Roberto Amato, Sonia Goncalves, Ewan Harrison, David K. Jackson, Ian Johnston, Dominic Kwiatkowski, Cordelia Langford, John Sillitoe on behalf of the Wellcome Sanger Institute COVID-19 Surveillance Team |
| EPI_ISL_645991 | Lighthouse Lab in Alderley Park | Wellcome Sanger Institute for the COVID-19 Genomics UK (COG-UK) Consortium | Jacquelyn Wynn, Mairead Hyland, The Lighthouse Lab in Alderley Park and Alex Alderton, Roberto Amato, Sonia Goncalves, Ewan Harrison, David K. Jackson, Ian Johnston, Dominic Kwiatkowski, Cordelia Langford, John Sillitoe on behalf of the Wellcome Sanger Institute COVID-19 Surveillance Team |
| EPI_ISL_645992, EPI_ISL_645993, EPI_ISL_645995, EPI_ISL_645998, EPI_ISL_646009, EPI_ISL_646014, EPI_ISL_646017, EPI_ISL_646018, EPI_ISL_646022, EPI_ISL_646028 | Lighthouse Lab in Cambridge | Wellcome Sanger Institute for the COVID-19 Genomics UK (COG-UK) Consortium | Rob Howes, The Lighthouse Lab in Cambridge and Alex Alderton, Roberto Amato, Sonia Goncalves, Ewan Harrison, David K. Jackson, Ian Johnston, Dominic Kwiatkowski, Cordelia Langford, John Sillitoe on behalf of the Wellcome Sanger Institute COVID-19 Surveillance Team |
| EPI_ISL_646029 | Lighthouse Lab in Alderley Park | Wellcome Sanger Institute for the COVID-19 Genomics UK (COG-UK) Consortium | Jacquelyn Wynn, Mairead Hyland, The Lighthouse Lab in Alderley Park and Alex Alderton, Roberto Amato, Sonia Goncalves, Ewan Harrison, David K. Jackson, Ian Johnston, Dominic Kwiatkowski, Cordelia Langford, John Sillitoe on behalf of the Wellcome Sanger Institute COVID-19 Surveillance Team |
| EPI_ISL_646030, EPI_ISL_646031, EPI_ISL_646032, EPI_ISL_646034, EPI_ISL_646035, EPI_ISL_646041, EPI_ISL_646046, EPI_ISL_646052 | Lighthouse Lab in Cambridge | Wellcome Sanger Institute for the COVID-19 Genomics UK (COG-UK) Consortium | Rob Howes, The Lighthouse Lab in Cambridge and Alex Alderton, Roberto Amato, Sonia Goncalves, Ewan Harrison, David K. Jackson, Ian Johnston, Dominic Kwiatkowski, Cordelia Langford, John Sillitoe on behalf of the Wellcome Sanger Institute COVID-19 Surveillance Team |
| EPI_ISL_646054 | Lighthouse Lab in Alderley Park | Wellcome Sanger Institute for the COVID-19 Genomics UK (COG-UK) Consortium | Jacquelyn Wynn, Mairead Hyland, The Lighthouse Lab in Alderley Park and Alex Alderton, Roberto Amato, Sonia Goncalves, Ewan Harrison, David K. Jackson, Ian Johnston, Dominic Kwiatkowski, Cordelia Langford, John Sillitoe on behalf of the Wellcome Sanger Institute COVID-19 Surveillance Team |
| EPI_ISL_646056, EPI_ISL_646058 | Lighthouse Lab in Cambridge | Wellcome Sanger Institute for the COVID-19 Genomics UK (COG-UK) Consortium | Rob Howes, The Lighthouse Lab in Cambridge and Alex Alderton, Roberto Amato, Sonia Goncalves, Ewan Harrison, David K. Jackson, Ian Johnston, Dominic Kwiatkowski, Cordelia Langford, John Sillitoe on behalf of the Wellcome Sanger Institute COVID-19 Surveillance Team |
| EPI_ISL_646062 | Lighthouse Lab in Alderley Park | Wellcome Sanger Institute for the COVID-19 Genomics UK (COG-UK) Consortium | Jacquelyn Wynn, Mairead Hyland, The Lighthouse Lab in Alderley Park and Alex Alderton, Roberto Amato, Sonia Goncalves, Ewan Harrison, David K. Jackson, Ian Johnston, Dominic Kwiatkowski, Cordelia Langford, John Sillitoe on behalf of the Wellcome Sanger Institute COVID-19 Surveillance Team |
| EPI_ISL_646067 | Lighthouse Lab in Cambridge | Wellcome Sanger Institute for the COVID-19 Genomics UK (COG-UK) Consortium | Rob Howes, The Lighthouse Lab in Cambridge and Alex Alderton, Roberto Amato, Sonia Goncalves, Ewan Harrison, David K. Jackson, Ian Johnston, Dominic Kwiatkowski, Cordelia Langford, John Sillitoe on behalf of the Wellcome Sanger Institute COVID-19 Surveillance Team |
| EPI_ISL_646069, EPI_ISL_646078 | Lighthouse Lab in Alderley Park | Wellcome Sanger Institute for the COVID-19 Genomics UK (COG-UK) Consortium | Jacquelyn Wynn, Mairead Hyland, The Lighthouse Lab in Alderley Park and Alex Alderton, Roberto Amato, Sonia Goncalves, Ewan Harrison, David K. Jackson, Ian Johnston, Dominic Kwiatkowski, Cordelia Langford, John Sillitoe on behalf of the Wellcome Sanger Institute COVID-19 Surveillance Team |
| EPI_ISL_646083, EPI_ISL_646084, EPI_ISL_646085, EPI_ISL_646089, EPI_ISL_646091 | Lighthouse Lab in Cambridge | Wellcome Sanger Institute for the COVID-19 Genomics UK (COG-UK) Consortium | Rob Howes, The Lighthouse Lab in Cambridge and Alex Alderton, Roberto Amato, Sonia Goncalves, Ewan Harrison, David K. Jackson, Ian Johnston, Dominic Kwiatkowski, Cordelia Langford, John Sillitoe on behalf of the Wellcome Sanger Institute COVID-19 Surveillance Team |
| EPI_ISL_646096, EPI_ISL_646098 | Lighthouse Lab in Alderley Park | Wellcome Sanger Institute for the COVID-19 Genomics UK (COG-UK) Consortium | Jacquelyn Wynn, Mairead Hyland, The Lighthouse Lab in Alderley Park and Alex Alderton, Roberto Amato, Sonia Goncalves, Ewan Harrison, David K. Jackson, Ian Johnston, Dominic Kwiatkowski, Cordelia Langford, John Sillitoe on behalf of the Wellcome Sanger Institute COVID-19 Surveillance Team |
| EPI_ISL_646101, EPI_ISL_646102, EPI_ISL_646115, EPI_ISL_646122 | Lighthouse Lab in Cambridge | Wellcome Sanger Institute for the COVID-19 Genomics UK (COG-UK) Consortium | Rob Howes, The Lighthouse Lab in Cambridge and Alex Alderton, Roberto Amato, Sonia Goncalves, Ewan Harrison, David K. Jackson, Ian Johnston, Dominic Kwiatkowski, Cordelia Langford, John Sillitoe on behalf of the Wellcome Sanger Institute COVID-19 Surveillance Team |
| EPI_ISL_646125 | Lighthouse Lab in Alderley Park | Wellcome Sanger Institute for the COVID-19 Genomics UK (COG-UK) Consortium | Jacquelyn Wynn, Mairead Hyland, The Lighthouse Lab in Alderley Park and Alex Alderton, Roberto Amato, Sonia Goncalves, Ewan Harrison, David K. Jackson, Ian Johnston, Dominic Kwiatkowski, Cordelia Langford, John Sillitoe on behalf of the Wellcome Sanger Institute COVID-19 Surveillance Team |
| EPI_ISL_646128, EPI_ISL_646129, EPI_ISL_646135, EPI_ISL_646136, EPI_ISL_646139 | Lighthouse Lab in Cambridge | Wellcome Sanger Institute for the COVID-19 Genomics UK (COG-UK) Consortium | Rob Howes, The Lighthouse Lab in Cambridge and Alex Alderton, Roberto Amato, Sonia Goncalves, Ewan Harrison, David K. Jackson, Ian Johnston, Dominic Kwiatkowski, Cordelia Langford, John Sillitoe on behalf of the Wellcome Sanger Institute COVID-19 Surveillance Team |
| EPI_ISL_646142, EPI_ISL_646152, EPI_ISL_646154 | Lighthouse Lab in Glasgow | Wellcome Sanger Institute for the COVID-19 Genomics UK (COG-UK) Consortium | Harper VanSteenhouse, Yumi Kasai, David Gray, Carol Clugston, Anna Dominiczak and Alex Alderton, Roberto Amato, Sonia Goncalves, Ewan Harrison, David K. Jackson, Ian Johnston, Dominic Kwiatkowski, Cordelia Langford, John Sillitoe on behalf of the Wellcome Sanger Institute COVID-19 Surveillance Team |
| EPI_ISL_646162 | Lighthouse Lab in Alderley Park | Wellcome Sanger Institute for the COVID-19 Genomics UK (COG-UK) Consortium | Jacquelyn Wynn, Mairead Hyland, The Lighthouse Lab in Alderley Park and Alex Alderton, Roberto Amato, Sonia Goncalves, Ewan Harrison, David K. Jackson, Ian Johnston, Dominic Kwiatkowski, Cordelia Langford, John Sillitoe on behalf of the Wellcome Sanger Institute COVID-19 Surveillance Team |
| EPI_ISL_646164, EPI_ISL_646168, EPI_ISL_646174, EPI_ISL_646179, EPI_ISL_646181 | Lighthouse Lab in Glasgow | Wellcome Sanger Institute for the COVID-19 Genomics UK (COG-UK) Consortium | Harper VanSteenhouse, Yumi Kasai, David Gray, Carol Clugston, Anna Dominiczak and Alex Alderton, Roberto Amato, Sonia Goncalves, Ewan Harrison, David K. Jackson, Ian Johnston, Dominic Kwiatkowski, Cordelia Langford, John Sillitoe on behalf of the Wellcome Sanger Institute COVID-19 Surveillance Team |

| | | | |
|---|---|---|---|
| EPI_ISL_646182 | Lighthouse Lab in Alderley Park | Wellcome Sanger Institute for the COVID-19 Genomics UK (COG-UK) Consortium | Jacquelyn Wynn, Mairead Hyland, The Lighthouse Lab in Alderley Park and Alex Alderton, Roberto Amato, Sonia Goncalves, Ewan Harrison, David K. Jackson, Ian Johnston, Dominic Kwiatkowski, Cordelia Langford, John Sillitoe on behalf of the Wellcome Sanger Institute COVID-19 Surveillance Team |
| EPI_ISL_646186, EPI_ISL_646188 | Lighthouse Lab in Glasgow | Wellcome Sanger Institute for the COVID-19 Genomics UK (COG-UK) Consortium | Harper VanSteenhouse, Yumi Kasai, David Gray, Carol Clugston, Anna Dominiczak and Alex Alderton, Roberto Amato, Sonia Goncalves, Ewan Harrison, David K. Jackson, Ian Johnston, Dominic Kwiatkowski, Cordelia Langford, John Sillitoe on behalf of the Wellcome Sanger Institute COVID-19 Surveillance Team |
| EPI_ISL_646191 | Lighthouse Lab in Alderley Park | Wellcome Sanger Institute for the COVID-19 Genomics UK (COG-UK) Consortium | Jacquelyn Wynn, Mairead Hyland, The Lighthouse Lab in Alderley Park and Alex Alderton, Roberto Amato, Sonia Goncalves, Ewan Harrison, David K. Jackson, Ian Johnston, Dominic Kwiatkowski, Cordelia Langford, John Sillitoe on behalf of the Wellcome Sanger Institute COVID-19 Surveillance Team |
| EPI_ISL_646192, EPI_ISL_646200, EPI_ISL_646225, EPI_ISL_646226, EPI_ISL_646237, EPI_ISL_646241, EPI_ISL_646243, EPI_ISL_646247, EPI_ISL_646254, EPI_ISL_646264, EPI_ISL_646266, EPI_ISL_646282, EPI_ISL_646284, EPI_ISL_646286, EPI_ISL_646292, EPI_ISL_646295, EPI_ISL_646298, EPI_ISL_646300 | | | |
| see above | Lighthouse Lab in Glasgow | Wellcome Sanger Institute for the COVID-19 Genomics UK (COG-UK) Consortium | Harper VanSteenhouse, Yumi Kasai, David Gray, Carol Clugston, Anna Dominiczak and Alex Alderton, Roberto Amato, Sonia Goncalves, Ewan Harrison, David K. Jackson, Ian Johnston, Dominic Kwiatkowski, Cordelia Langford, John Sillitoe on behalf of the Wellcome Sanger Institute COVID-19 Surveillance Team |
| EPI_ISL_646303 | Lighthouse Lab in Alderley Park | Wellcome Sanger Institute for the COVID-19 Genomics UK (COG-UK) Consortium | Jacquelyn Wynn, Mairead Hyland, The Lighthouse Lab in Alderley Park and Alex Alderton, Roberto Amato, Sonia Goncalves, Ewan Harrison, David K. Jackson, Ian Johnston, Dominic Kwiatkowski, Cordelia Langford, John Sillitoe on behalf of the Wellcome Sanger Institute COVID-19 Surveillance Team |
| EPI_ISL_646308, EPI_ISL_646310, EPI_ISL_646316, EPI_ISL_646321, EPI_ISL_646322, EPI_ISL_646330, EPI_ISL_646344, EPI_ISL_646351, EPI_ISL_646355, EPI_ISL_646357, EPI_ISL_646358, EPI_ISL_646359, EPI_ISL_646365, EPI_ISL_646368 | | | |
| see above | Lighthouse Lab in Glasgow | Wellcome Sanger Institute for the COVID-19 Genomics UK (COG-UK) Consortium | Harper VanSteenhouse, Yumi Kasai, David Gray, Carol Clugston, Anna Dominiczak and Alex Alderton, Roberto Amato, Sonia Goncalves, Ewan Harrison, David K. Jackson, Ian Johnston, Dominic Kwiatkowski, Cordelia Langford, John Sillitoe on behalf of the Wellcome Sanger Institute COVID-19 Surveillance Team |
| EPI_ISL_646370 | Lighthouse Lab in Alderley Park | Wellcome Sanger Institute for the COVID-19 Genomics UK (COG-UK) Consortium | Jacquelyn Wynn, Mairead Hyland, The Lighthouse Lab in Alderley Park and Alex Alderton, Roberto Amato, Sonia Goncalves, Ewan Harrison, David K. Jackson, Ian Johnston, Dominic Kwiatkowski, Cordelia Langford, John Sillitoe on behalf of the Wellcome Sanger Institute COVID-19 Surveillance Team |
| EPI_ISL_646371, EPI_ISL_646372, EPI_ISL_646380, EPI_ISL_646386, EPI_ISL_646387, EPI_ISL_646391, EPI_ISL_646394, EPI_ISL_646396, EPI_ISL_646401, EPI_ISL_646405, EPI_ISL_646407, EPI_ISL_646410, EPI_ISL_646416, EPI_ISL_646417, EPI_ISL_646422, EPI_ISL_646423, EPI_ISL_646424, EPI_ISL_646427, EPI_ISL_646436, EPI_ISL_646447, EPI_ISL_646451, EPI_ISL_646453, EPI_ISL_646455, EPI_ISL_646456, EPI_ISL_646457, EPI_ISL_646458, EPI_ISL_646459, EPI_ISL_646460, EPI_ISL_646462, EPI_ISL_646464, EPI_ISL_646465, EPI_ISL_646470, EPI_ISL_646471, EPI_ISL_646472, EPI_ISL_646474, EPI_ISL_646475, EPI_ISL_646480, EPI_ISL_646481, EPI_ISL_646482, EPI_ISL_646483, EPI_ISL_646484, EPI_ISL_646495, EPI_ISL_646496, EPI_ISL_646498, EPI_ISL_646499, EPI_ISL_646501, EPI_ISL_646504, EPI_ISL_646517, EPI_ISL_646523, EPI_ISL_646524, EPI_ISL_646530, EPI_ISL_646531, EPI_ISL_646536, EPI_ISL_646539, EPI_ISL_646547, EPI_ISL_646550, EPI_ISL_646552, EPI_ISL_646553, EPI_ISL_646554, EPI_ISL_646556, EPI_ISL_646562, EPI_ISL_646565, EPI_ISL_646570, EPI_ISL_646575, EPI_ISL_646578, EPI_ISL_646582, EPI_ISL_646584, EPI_ISL_646587, EPI_ISL_646588, EPI_ISL_646589, EPI_ISL_646596, EPI_ISL_646598, EPI_ISL_646604, EPI_ISL_646609, EPI_ISL_646611, EPI_ISL_646615, EPI_ISL_646617, EPI_ISL_646620, EPI_ISL_646622, EPI_ISL_646624, EPI_ISL_646627, EPI_ISL_646629, EPI_ISL_646630, EPI_ISL_646633, EPI_ISL_646634, EPI_ISL_646636, EPI_ISL_646637, EPI_ISL_646641, EPI_ISL_646643, EPI_ISL_646644, EPI_ISL_646646, EPI_ISL_646647, EPI_ISL_646649, EPI_ISL_646650, EPI_ISL_646654, EPI_ISL_646656, EPI_ISL_646657, EPI_ISL_646659, EPI_ISL_646661, EPI_ISL_646663, EPI_ISL_646667, EPI_ISL_646669, EPI_ISL_646670, EPI_ISL_646672, EPI_ISL_646674, EPI_ISL_646677, EPI_ISL_646680, EPI_ISL_646681, EPI_ISL_646682, EPI_ISL_646685, EPI_ISL_646688, EPI_ISL_646690, EPI_ISL_646694, EPI_ISL_646699, EPI_ISL_646700, EPI_ISL_646701, EPI_ISL_646703, EPI_ISL_646705, EPI_ISL_646709, EPI_ISL_646714, EPI_ISL_646717, EPI_ISL_646720, EPI_ISL_646723, EPI_ISL_646725, EPI_ISL_646726, EPI_ISL_646728, EPI_ISL_646729, EPI_ISL_646730, EPI_ISL_646732, EPI_ISL_646733, EPI_ISL_646735, EPI_ISL_646742, EPI_ISL_646745, EPI_ISL_646748, EPI_ISL_646750, EPI_ISL_646754, EPI_ISL_646759, EPI_ISL_646765, EPI_ISL_646766, EPI_ISL_646772, EPI_ISL_646774, EPI_ISL_646775, EPI_ISL_646777, EPI_ISL_646778, EPI_ISL_646780, EPI_ISL_646781, EPI_ISL_646783, EPI_ISL_646785, EPI_ISL_646786, EPI_ISL_646788, EPI_ISL_646792, EPI_ISL_646794, EPI_ISL_646798, EPI_ISL_646810, EPI_ISL_646818, EPI_ISL_646825, EPI_ISL_646826, EPI_ISL_646837, EPI_ISL_646840, EPI_ISL_646850, EPI_ISL_646853, EPI_ISL_646854, EPI_ISL_646868, EPI_ISL_646869, EPI_ISL_646870, EPI_ISL_646872, EPI_ISL_646877, EPI_ISL_646880, EPI_ISL_646885, EPI_ISL_646891, EPI_ISL_646892, EPI_ISL_646902 | | | |
| see above | Lighthouse Lab in Glasgow | Wellcome Sanger Institute for the COVID-19 Genomics UK (COG-UK) Consortium | Harper VanSteenhouse, Yumi Kasai, David Gray, Carol Clugston, Anna Dominiczak and Alex Alderton, Roberto Amato, Sonia Goncalves, Ewan Harrison, David K. Jackson, Ian Johnston, Dominic Kwiatkowski, Cordelia Langford, John Sillitoe on behalf of the Wellcome Sanger Institute COVID-19 Surveillance Team |
| EPI_ISL_646903 | Lighthouse Lab in Alderley Park | Wellcome Sanger Institute for the COVID-19 Genomics UK (COG-UK) Consortium | Jacquelyn Wynn, Mairead Hyland, The Lighthouse Lab in Alderley Park and Alex Alderton, Roberto Amato, Sonia Goncalves, Ewan Harrison, David K. Jackson, Ian Johnston, Dominic Kwiatkowski, Cordelia Langford, John Sillitoe on behalf of the Wellcome Sanger Institute COVID-19 Surveillance Team |
| EPI_ISL_646905, EPI_ISL_646912, EPI_ISL_646915, EPI_ISL_646921, EPI_ISL_646929, EPI_ISL_646936, EPI_ISL_646937, EPI_ISL_646939, EPI_ISL_646948, EPI_ISL_646955, EPI_ISL_646960, EPI_ISL_646964, EPI_ISL_646967, EPI_ISL_646974, EPI_ISL_646975, EPI_ISL_646976, EPI_ISL_646977, EPI_ISL_646983, EPI_ISL_646988, EPI_ISL_646992, EPI_ISL_646996, EPI_ISL_647001, EPI_ISL_647003, EPI_ISL_647006, EPI_ISL_647014, EPI_ISL_647023, EPI_ISL_647031, EPI_ISL_647032, EPI_ISL_647035, EPI_ISL_647038, EPI_ISL_647045, EPI_ISL_647050, EPI_ISL_647053, EPI_ISL_647059, EPI_ISL_647061, EPI_ISL_647067, EPI_ISL_647076, EPI_ISL_647081, EPI_ISL_647087, EPI_ISL_647090, EPI_ISL_647097, EPI_ISL_647102, EPI_ISL_647105, EPI_ISL_647109, EPI_ISL_647114, EPI_ISL_647115, EPI_ISL_647117, EPI_ISL_647121 | | | |
| see above | Lighthouse Lab in Glasgow | Wellcome Sanger Institute for the COVID-19 Genomics UK (COG-UK) Consortium | Harper VanSteenhouse, Yumi Kasai, David Gray, Carol Clugston, Anna Dominiczak and Alex Alderton, Roberto Amato, Sonia Goncalves, Ewan Harrison, David K. Jackson, Ian Johnston, Dominic Kwiatkowski, Cordelia Langford, John Sillitoe on behalf of the Wellcome Sanger Institute COVID-19 Surveillance Team |
| EPI_ISL_647123, EPI_ISL_647124, EPI_ISL_647125, EPI_ISL_647126, EPI_ISL_647127, EPI_ISL_647128, EPI_ISL_647129, EPI_ISL_647130, EPI_ISL_647131, EPI_ISL_647132, EPI_ISL_647133, EPI_ISL_647136, EPI_ISL_647137, EPI_ISL_647138, EPI_ISL_647140, EPI_ISL_647141, EPI_ISL_647142, EPI_ISL_647143, EPI_ISL_647144, EPI_ISL_647146, EPI_ISL_647149, EPI_ISL_647150, EPI_ISL_647151, EPI_ISL_647154, EPI_ISL_647155, EPI_ISL_647156, EPI_ISL_647157, EPI_ISL_647158, EPI_ISL_647159, EPI_ISL_647162, EPI_ISL_647163, EPI_ISL_647164, EPI_ISL_647166, EPI_ISL_647167, EPI_ISL_647168, EPI_ISL_647169, EPI_ISL_647170, EPI_ISL_647171, EPI_ISL_647173, EPI_ISL_647174, EPI_ISL_647175, EPI_ISL_647176, EPI_ISL_647177, EPI_ISL_647179, EPI_ISL_647181, EPI_ISL_647182, EPI_ISL_647183, EPI_ISL_647184, EPI_ISL_647185, EPI_ISL_647186, EPI_ISL_647188, EPI_ISL_647189, EPI_ISL_647190, EPI_ISL_647191, EPI_ISL_647192, EPI_ISL_647193, EPI_ISL_647194, EPI_ISL_647195, EPI_ISL_647196, EPI_ISL_647198, EPI_ISL_647202, EPI_ISL_647203, EPI_ISL_647204, EPI_ISL_647206, EPI_ISL_647207, EPI_ISL_647208, EPI_ISL_647210, EPI_ISL_647211, EPI_ISL_647212, EPI_ISL_647215, EPI_ISL_647217, EPI_ISL_647218, EPI_ISL_647219, EPI_ISL_647220, EPI_ISL_647221, EPI_ISL_647222, EPI_ISL_647225, EPI_ISL_647226, EPI_ISL_647227, EPI_ISL_647228, EPI_ISL_647229, EPI_ISL_647231, EPI_ISL_647233, EPI_ISL_647235, EPI_ISL_647236, EPI_ISL_647237, EPI_ISL_647238, EPI_ISL_647239, EPI_ISL_647241, EPI_ISL_647242, EPI_ISL_647243, EPI_ISL_647244, EPI_ISL_647245, EPI_ISL_647246, EPI_ISL_647248, EPI_ISL_647249, EPI_ISL_647252, EPI_ISL_647253, EPI_ISL_647257, EPI_ISL_647258, EPI_ISL_647261, EPI_ISL_647262, EPI_ISL_647264, EPI_ISL_647265, EPI_ISL_647268, EPI_ISL_647270, EPI_ISL_647271, EPI_ISL_647272, EPI_ISL_647274, EPI_ISL_647276, EPI_ISL_647277, EPI_ISL_647278, EPI_ISL_647283, EPI_ISL_647284, EPI_ISL_647285, EPI_ISL_647286, EPI_ISL_647287, EPI_ISL_647288, EPI_ISL_647289, EPI_ISL_647293, EPI_ISL_647295, EPI_ISL_647297, EPI_ISL_647298, EPI_ISL_647299, EPI_ISL_647300, EPI_ISL_647301, EPI_ISL_647302, EPI_ISL_647303, EPI_ISL_647304, EPI_ISL_647305, EPI_ISL_647306, EPI_ISL_647307, EPI_ISL_647309, EPI_ISL_647310, EPI_ISL_647311, EPI_ISL_647312, EPI_ISL_647313, EPI_ISL_647316, EPI_ISL_647317, EPI_ISL_647318, EPI_ISL_647319, EPI_ISL_647320, EPI_ISL_647322, EPI_ISL_647325, EPI_ISL_647326, EPI_ISL_647327, EPI_ISL_647331, EPI_ISL_647333, EPI_ISL_647335, EPI_ISL_647336, EPI_ISL_647338, EPI_ISL_647340, EPI_ISL_647341, EPI_ISL_647344, EPI_ISL_647345, EPI_ISL_647346, EPI_ISL_647348, EPI_ISL_647349, EPI_ISL_647350, EPI_ISL_647351, EPI_ISL_647352, EPI_ISL_647353, EPI_ISL_647355, EPI_ISL_647356, EPI_ISL_647357, EPI_ISL_647358, EPI_ISL_647359, EPI_ISL_647360, EPI_ISL_647364, EPI_ISL_647365, EPI_ISL_647366, EPI_ISL_647367, EPI_ISL_647368, EPI_ISL_647369, EPI_ISL_647370, EPI_ISL_647371, EPI_ISL_647372, EPI_ISL_647373, EPI_ISL_647374, EPI_ISL_647375, EPI_ISL_647376, EPI_ISL_647377, EPI_ISL_647378, EPI_ISL_647379, EPI_ISL_647380, EPI_ISL_647381, EPI_ISL_647382, EPI_ISL_647383, EPI_ISL_647384, EPI_ISL_647385, EPI_ISL_647386, EPI_ISL_647387, EPI_ISL_647388, EPI_ISL_647389, EPI_ISL_647390, EPI_ISL_647391, EPI_ISL_647392, EPI_ISL_647393, EPI_ISL_647394, EPI_ISL_647395, EPI_ISL_647396, EPI_ISL_647397, EPI_ISL_647398, EPI_ISL_647400, EPI_ISL_647401, EPI_ISL_647402, EPI_ISL_647403, EPI_ISL_647404, EPI_ISL_647406, EPI_ISL_647407, EPI_ISL_647408, EPI_ISL_647409, EPI_ISL_647410, EPI_ISL_647412, EPI_ISL_647413, EPI_ISL_647414, EPI_ISL_647415, EPI_ISL_647416, EPI_ISL_647417, EPI_ISL_647418, EPI_ISL_647421, EPI_ISL_647422, EPI_ISL_647423, EPI_ISL_647424, EPI_ISL_647426, EPI_ISL_647430, EPI_ISL_647431, EPI_ISL_647432, EPI_ISL_647433, EPI_ISL_647434, EPI_ISL_647435, EPI_ISL_647437, EPI_ISL_647438, EPI_ISL_647439, EPI_ISL_647440, EPI_ISL_647441, EPI_ISL_647442, EPI_ISL_647444, EPI_ISL_647446, EPI_ISL_647447, EPI_ISL_647448, EPI_ISL_647449, EPI_ISL_647450, EPI_ISL_647451, EPI_ISL_647453, EPI_ISL_647454, EPI_ISL_647455, EPI_ISL_647456, EPI_ISL_647457, EPI_ISL_647458, EPI_ISL_647459, EPI_ISL_647461, EPI_ISL_647462, EPI_ISL_647463, EPI_ISL_647464, EPI_ISL_647465, EPI_ISL_647466, EPI_ISL_647467, EPI_ISL_647470, EPI_ISL_647471, EPI_ISL_647472, EPI_ISL_647475, EPI_ISL_647476, EPI_ISL_647480, EPI_ISL_647481, EPI_ISL_647482, EPI_ISL_647484, EPI_ISL_647487, EPI_ISL_647488, EPI_ISL_647489, EPI_ISL_647490, EPI_ISL_647491, EPI_ISL_647493, EPI_ISL_647494, EPI_ISL_647495, EPI_ISL_647496, EPI_ISL_647500, EPI_ISL_647501, EPI_ISL_647502, EPI_ISL_647504, EPI_ISL_647505, EPI_ISL_647507, EPI_ISL_647508, EPI_ISL_647510, EPI_ISL_647512, EPI_ISL_647516, EPI_ISL_647517, EPI_ISL_647520, EPI_ISL_647521, EPI_ISL_647523, EPI_ISL_647524, EPI_ISL_647525, EPI_ISL_647527, EPI_ISL_647529, EPI_ISL_647530, EPI_ISL_647532, EPI_ISL_647533, EPI_ISL_647537, EPI_ISL_647538, EPI_ISL_647539, EPI_ISL_647540, EPI_ISL_647542, EPI_ISL_647544, EPI_ISL_647545, EPI_ISL_647548, EPI_ISL_647549, EPI_ISL_647551, EPI_ISL_647552, EPI_ISL_647554, EPI_ISL_647555, EPI_ISL_647556, EPI_ISL_647557, EPI_ISL_647559, EPI_ISL_647561, EPI_ISL_647564, EPI_ISL_647565, EPI_ISL_647566, EPI_ISL_647567, EPI_ISL_647568, EPI_ISL_647571, EPI_ISL_647574, EPI_ISL_647575, EPI_ISL_647577, EPI_ISL_647578, EPI_ISL_647581, EPI_ISL_647582, EPI_ISL_647583, EPI_ISL_647585, EPI_ISL_647586, EPI_ISL_647587, EPI_ISL_647588, EPI_ISL_647589, EPI_ISL_647590, EPI_ISL_647593, EPI_ISL_647596, EPI_ISL_647597, EPI_ISL_647599, EPI_ISL_647601, EPI_ISL_647602, EPI_ISL_647607, EPI_ISL_647609, EPI_ISL_647610, EPI_ISL_647611, EPI_ISL_647613, EPI_ISL_647616, EPI_ISL_647617, EPI_ISL_647621, EPI_ISL_647622, EPI_ISL_647623, EPI_ISL_647625, EPI_ISL_647626, EPI_ISL_647627, EPI_ISL_647628, EPI_ISL_647629, EPI_ISL_647631, EPI_ISL_647636, EPI_ISL_647639, EPI_ISL_647641, EPI_ISL_647644, EPI_ISL_647645, EPI_ISL_647647, EPI_ISL_647649, EPI_ISL_647651, EPI_ISL_647652, EPI_ISL_647653, EPI_ISL_647654, EPI_ISL_647656, EPI_ISL_647657, EPI_ISL_647660, EPI_ISL_647661, EPI_ISL_647663, EPI_ISL_647664, EPI_ISL_647665, EPI_ISL_647666, EPI_ISL_647669, EPI_ISL_647670, EPI_ISL_647673, EPI_ISL_647674, EPI_ISL_647675, EPI_ISL_647676, EPI_ISL_647678, EPI_ISL_647679, EPI_ISL_647681, EPI_ISL_647684, EPI_ISL_647685, EPI_ISL_647688, EPI_ISL_647690, EPI_ISL_647691, EPI_ISL_647692, EPI_ISL_647696, EPI_ISL_647699, EPI_ISL_647701, EPI_ISL_647702, EPI_ISL_647703, EPI_ISL_647709, EPI_ISL_647712, EPI_ISL_647717, EPI_ISL_647718, EPI_ISL_647719, EPI_ISL_647721, EPI_ISL_647723, EPI_ISL_647728, EPI_ISL_647729, EPI_ISL_647730, EPI_ISL_647731, EPI_ISL_647734, EPI_ISL_647735, EPI_ISL_647736, EPI_ISL_647737, EPI_ISL_647740, EPI_ISL_647742, EPI_ISL_647743, EPI_ISL_647744, EPI_ISL_647745, EPI_ISL_647746, EPI_ISL_647747, EPI_ISL_647749, EPI_ISL_647751, EPI_ISL_647753, EPI_ISL_647754, EPI_ISL_647757, EPI_ISL_647759, EPI_ISL_647763, EPI_ISL_647765, EPI_ISL_647766, EPI_ISL_647767, EPI_ISL_647768, EPI_ISL_647769, EPI_ISL_647770, EPI_ISL_647772, EPI_ISL_647773, EPI_ISL_647777, EPI_ISL_647778, EPI_ISL_647781, EPI_ISL_647782, EPI_ISL_647783, EPI_ISL_647784, EPI_ISL_647785, EPI_ISL_647788, EPI_ISL_647790, EPI_ISL_647791, EPI_ISL_647793, EPI_ISL_647794 | | | |
| see above | Lighthouse Lab in Cambridge | Wellcome Sanger Institute for the COVID-19 Genomics UK (COG-UK) Consortium | Rob Howes, The Lighthouse Lab in Cambridge and Alex Alderton, Roberto Amato, Sonia Goncalves, Ewan Harrison, David K. Jackson, Ian Johnston, Dominic Kwiatkowski, Cordelia Langford, John Sillitoe on behalf of the Wellcome Sanger Institute COVID-19 Surveillance Team |
| EPI_ISL_647797, EPI_ISL_647799, EPI_ISL_647802, EPI_ISL_647803, EPI_ISL_647805, EPI_ISL_647862, EPI_ISL_647866, EPI_ISL_647867, EPI_ISL_647868, EPI_ISL_647869, EPI_ISL_647870, EPI_ISL_647871, EPI_ISL_647872 | | | |
| see above | Lighthouse Lab in Glasgow | Wellcome Sanger Institute for the COVID-19 Genomics UK (COG-UK) Consortium | Harper VanSteenhouse, Yumi Kasai, David Gray, Carol Clugston, Anna Dominiczak and Alex Alderton, Roberto Amato, Sonia Goncalves, Ewan Harrison, David K. Jackson, Ian Johnston, Dominic Kwiatkowski, Cordelia Langford, John Sillitoe on behalf of the Wellcome Sanger Institute COVID-19 Surveillance Team |
| EPI_ISL_649494, EPI_ISL_649495, EPI_ISL_649496, EPI_ISL_649497, EPI_ISL_649498, EPI_ISL_649500, EPI_ISL_649501, EPI_ISL_649502, EPI_ISL_649504, EPI_ISL_649505, EPI_ISL_649506, EPI_ISL_649507, EPI_ISL_649508, EPI_ISL_649509, EPI_ISL_649510, EPI_ISL_649511, EPI_ISL_649512, EPI_ISL_649513, EPI_ISL_649514, EPI_ISL_649515, EPI_ISL_649516, EPI_ISL_649517, EPI_ISL_649518, EPI_ISL_649519, EPI_ISL_649520, EPI_ISL_649521, EPI_ISL_649522, EPI_ISL_649523, EPI_ISL_649524, EPI_ISL_649525, EPI_ISL_649526, EPI_ISL_649527, EPI_ISL_649528, EPI_ISL_649529, EPI_ISL_649530, EPI_ISL_649531, EPI_ISL_649532, EPI_ISL_649533, EPI_ISL_649534, EPI_ISL_649535, EPI_ISL_649536, EPI_ISL_649537, EPI_ISL_649538, EPI_ISL_649539, EPI_ISL_649540, EPI_ISL_649541, EPI_ISL_649542, EPI_ISL_649543, EPI_ISL_649544, EPI_ISL_649545, EPI_ISL_649546, EPI_ISL_649547, EPI_ISL_649569, EPI_ISL_649570, EPI_ISL_649571, EPI_ISL_649572, EPI_ISL_649573, EPI_ISL_649574, EPI_ISL_649575, EPI_ISL_649576, EPI_ISL_649577, EPI_ISL_649578, EPI_ISL_649579, EPI_ISL_649580, EPI_ISL_649581, EPI_ISL_649582, EPI_ISL_649583, EPI_ISL_649584, EPI_ISL_649585, EPI_ISL_649586, EPI_ISL_649587, EPI_ISL_649588, EPI_ISL_649589, EPI_ISL_649590, EPI_ISL_649591, EPI_ISL_649592, EPI_ISL_649593, EPI_ISL_649594, EPI_ISL_649595, EPI_ISL_649596, EPI_ISL_649597, EPI_ISL_649598, EPI_ISL_649599, EPI_ISL_649600, EPI_ISL_649601, EPI_ISL_649602, EPI_ISL_649603, EPI_ISL_649604, EPI_ISL_649605, EPI_ISL_649606, EPI_ISL_649607, EPI_ISL_649608, EPI_ISL_649609, EPI_ISL_649610, EPI_ISL_649611, EPI_ISL_649612, EPI_ISL_649613, EPI_ISL_649614, EPI_ISL_649615, EPI_ISL_649616, EPI_ISL_649617, EPI_ISL_649618, EPI_ISL_649619, EPI_ISL_649620, EPI_ISL_649621, EPI_ISL_649622, EPI_ISL_649623, EPI_ISL_649624, EPI_ISL_649710 | | | |

| | | | |
|---|---|---|---|
| see above | Lighthouse Lab in Glasgow | Wellcome Sanger Institute for the COVID-19 Genomics UK (COG-UK) Consortium | Harper VanSteenhouse, Yumi Kasai, David Gray, Carol Clugston, Anna Dominiczak and Alex Alderton, Roberto Amato, Sonia Goncalves, Ewan Harrison, David K. Jackson, Ian Johnston, Dominic Kwiatkowski, Cordelia Langford, John Sillitoe on behalf of the Wellcome Sanger Institute COVID-19 Surveillance Team (http://www.sanger.ac.uk/covid-team) |
| EPI_ISL_649786, EPI_ISL_649787, EPI_ISL_649788, EPI_ISL_649789, EPI_ISL_649790, EPI_ISL_649792, EPI_ISL_649793, EPI_ISL_649794, EPI_ISL_649795, EPI_ISL_649796, EPI_ISL_649797, EPI_ISL_649799, EPI_ISL_649800, EPI_ISL_649801, EPI_ISL_649802, EPI_ISL_649803, EPI_ISL_649806, EPI_ISL_649807 | | | |
| see above | Respiratory Virus Unit, Microbiology Services Colindale, Public Health England | COVID-19 Genomics UK (COG-UK) Consortium | PHE Covid Sequencing Team |
| EPI_ISL_649999, EPI_ISL_650001 | University of Michigan Clinical Microbiology Laboratory | Lauring Lab, University of Michigan, Department of Microbiology and Immunology | Valesano |
| EPI_ISL_650115 | Liverpool Clinical Laboratories | COVID-19 Genomics UK (COG-UK) Consortium | Sam Haldenby, Anita Lucaci, Steve Paterson, Julian Hiscox, Alistair Darby, M Almsaud, A Alrezaihi, Muhannad Alruwaili, Stuart D Armstrong, Jones Benjamin, Eleanor G Bentley, Anu Chawla, Jordan J Clark, Angela Cowell, Richard Eccles, Isabel García-Dorival, Matthew Gemmell, Alessandro Gerada, PKF Gilmore, Richard Gregory, Ximeng Han, Catherine Hartley, Margaret Hughes, Miren Iturriza-Gomara, James Johnson, L Luu, Jenifer Manson, Charlotte Nelson, Elaine O'Toole, Cassie Olateju, Rebekah Penrice-Randal , Lucille Rainbow, N.P Randle, Trevor Ian Robinson, Parul Sharma, Ghada T Shawli, James P Stewart, Neil Swainston, Ecaterina Vamos, Joanne Watts, Mark Whitehead |
| EPI_ISL_650121 | Virology Department, Sheffield Teaching Hospitals NHS Foundation Trust/Department of Infection, Immunity and Cardiovascular Disease, The Medical School, University of Sheffield | COVID-19 Genomics UK (COG-UK) Consortium | Thushan de Silva, Matthew Parker, Nikki Smith, Adri Angyal, Rebecca Brown, Luke Green, Rachel Tucker, Paul Parsons, Danielle Groves, Katie Johnson, Laura Carrilero, Alex Keeley, Dave Partridge, Matthew Wyles, Benjamin Lindsey, Mehmet Yavuz, Mohammad Raza, Cariad Evans |
| EPI_ISL_650139 | Liverpool Clinical Laboratories | COVID-19 Genomics UK (COG-UK) Consortium | Sam Haldenby, Anita Lucaci, Steve Paterson, Julian Hiscox, Alistair Darby, M Almsaud, A Alrezaihi, Muhannad Alruwaili, Stuart D Armstrong, Jones Benjamin, Eleanor G Bentley, Anu Chawla, Jordan J Clark, Angela Cowell, Richard Eccles, Isabel García-Dorival, Matthew Gemmell, Alessandro Gerada, PKF Gilmore, Richard Gregory, Ximeng Han, Catherine Hartley, Margaret Hughes, Miren Iturriza-Gomara, James Johnson, L Luu, Jenifer Manson, Charlotte Nelson, Elaine O'Toole, Cassie Olateju, Rebekah Penrice-Randal , Lucille Rainbow, N.P Randle, Trevor Ian Robinson, Parul Sharma, Ghada T Shawli, James P Stewart, Neil Swainston, Ecaterina Vamos, Joanne Watts, Mark Whitehead |
| EPI_ISL_650146 | West of Scotland Specialist Virology Centre, NHSGGC / MRC-University of Glasgow Centre for Virus Research | COVID-19 Genomics UK (COG-UK) Consortium | Ana da Silva Filipe, Natasha Johnson, Kathy Smollett, Daniel Mair, Stephen Carmichael, Alice Broos, Lily Tong, Jenna Nichols, Kyriaki Nomikou; Sarah McDonald; Richard Orton, Joseph Hughes, Sreenu Vattipally, David L Robertson; Alasdair MacLean, Rory Gunson; Sharif Shaaban, Matthew Holden; Rachel Blacow, Guy Mollett, Kathy Li, James Shepherd, Antonia Ho, Emma Thomson |
| EPI_ISL_650151 | University of Exeter | COVID-19 Genomics UK (COG-UK) Consortium | Ben Temperton,Aaron Jeffries,Michelle Michelsen,Joanna Warwick-Dugdale,Audrey Farbos,Robyn Manley,Stephen Michell,Jane Masoli |
| EPI_ISL_650160, EPI_ISL_650161 | Liverpool Clinical Laboratories | COVID-19 Genomics UK (COG-UK) Consortium | Sam Haldenby, Anita Lucaci, Steve Paterson, Julian Hiscox, Alistair Darby, M Almsaud, A Alrezaihi, Muhannad Alruwaili, Stuart D Armstrong, Jones Benjamin, Eleanor G Bentley, Anu Chawla, Jordan J Clark, Angela Cowell, Richard Eccles, Isabel García-Dorival, Matthew Gemmell, Alessandro Gerada, PKF Gilmore, Richard Gregory, Ximeng Han, Catherine Hartley, Margaret Hughes, Miren Iturriza-Gomara, James Johnson, L Luu, Jenifer Manson, Charlotte Nelson, Elaine O'Toole, Cassie Olateju, Rebekah Penrice-Randal , Lucille Rainbow, N.P Randle, Trevor Ian Robinson, Parul Sharma, Ghada T Shawli, James P Stewart, Neil Swainston, Ecaterina Vamos, Joanne Watts, Mark Whitehead |
| EPI_ISL_650163 | University of Exeter | COVID-19 Genomics UK (COG-UK) Consortium | Ben Temperton,Aaron Jeffries,Michelle Michelsen,Joanna Warwick-Dugdale,Audrey Farbos,Robyn Manley,Stephen Michell,Jane Masoli |
| EPI_ISL_650189 | Liverpool Clinical Laboratories | COVID-19 Genomics UK (COG-UK) Consortium | Sam Haldenby, Anita Lucaci, Steve Paterson, Julian Hiscox, Alistair Darby, M Almsaud, A Alrezaihi, Muhannad Alruwaili, Stuart D Armstrong, Jones Benjamin, Eleanor G Bentley, Anu Chawla, Jordan J Clark, Angela Cowell, Richard Eccles, Isabel García-Dorival, Matthew Gemmell, Alessandro Gerada, PKF Gilmore, Richard Gregory, Ximeng Han, Catherine Hartley, Margaret Hughes, Miren Iturriza-Gomara, James Johnson, L Luu, Jenifer Manson, Charlotte Nelson, Elaine O'Toole, Cassie Olateju, Rebekah Penrice-Randal , Lucille Rainbow, N.P Randle, Trevor Ian Robinson, Parul Sharma, Ghada T Shawli, James P Stewart, Neil Swainston, Ecaterina Vamos, Joanne Watts, Mark Whitehead |
| EPI_ISL_650204 | Wales Specialist Virology Centre Sequencing lab: Pathogen Genomics Unit | COVID-19 Genomics UK (COG-UK) Consortium | Catherine Moore, Johnathan Evans, Laura Gifford, Malorie Perry, Simon Cottrell, Angela Marchbank, Alec Birchley, Alexander Adams, Amy Gaskin, Bree Gatica-Wilcox, Jason Coombes, Joel Southgate, Lauren Gilbert, Lee Graham, Nicole Pacchiarini, Sara Kumziene-Summerhayes, Sarah Taylor, Sophie Jones, Sara Rey, Matthew Bull, Joanne Watkins, Sally Corden, Tom Connor |
| EPI_ISL_650208 | Virology Department, Sheffield Teaching Hospitals NHS Foundation Trust/Department of Infection, Immunity and Cardiovascular Disease, The Medical School, University of Sheffield | COVID-19 Genomics UK (COG-UK) Consortium | Thushan de Silva, Matthew Parker, Nikki Smith, Adri Angyal, Rebecca Brown, Luke Green, Rachel Tucker, Paul Parsons, Danielle Groves, Katie Johnson, Laura Carrilero, Alex Keeley, Dave Partridge, Matthew Wyles, Benjamin Lindsey, Mehmet Yavuz, Mohammad Raza, Cariad Evans |
| EPI_ISL_650219, EPI_ISL_650222, EPI_ISL_650226 | University of Exeter | COVID-19 Genomics UK (COG-UK) Consortium | Ben Temperton,Aaron Jeffries,Michelle Michelsen,Joanna Warwick-Dugdale,Audrey Farbos,Robyn Manley,Stephen Michell,Jane Masoli |
| EPI_ISL_650232 | Department of Pathology, University of Cambridge | COVID-19 Genomics UK (COG-UK) Consortium | Aminu S. Jahun, Yasmin Chaudhry, Grant Hall, Iliana Georgana, Myra Hosmillo, Martin D. Curran, Malte Pinckert, Surendra Parmar, Ian Goodfellow |
| EPI_ISL_650234 | University of Birmingham | COVID-19 Genomics UK (COG-UK) Consortium | Institute of Microbiology, University of Birmingham: Claire McMurray, Joanne Stockton, Samuel Nicholls, Radoslaw Poplawski, Will Rowe, Josh Quick, Nicholas Loman. University of Birmingham Testing Laboratory: Celina M Whalley, Andrew Bosworth, Charlotte Poxon, Kasun Wanigasooriya, Oliver Pickles, Mike Kidd, Alex Richter, Andrew D Beggs PHE Heartlands Lab: Husam Osman, Andrew Bosworth. Queen Elizabeth Hospital: Anna Casey |
| EPI_ISL_650256 | Liverpool Clinical Laboratories | COVID-19 Genomics UK (COG-UK) Consortium | Sam Haldenby, Anita Lucaci, Steve Paterson, Julian Hiscox, Alistair Darby, M Almsaud, A Alrezaihi, Muhannad Alruwaili, Stuart D Armstrong, Jones Benjamin, Eleanor G Bentley, Anu Chawla, Jordan J Clark, Angela Cowell, Richard Eccles, Isabel García-Dorival, Matthew Gemmell, Alessandro Gerada, PKF Gilmore, Richard Gregory, Ximeng Han, Catherine Hartley, Margaret Hughes, Miren Iturriza-Gomara, James Johnson, L Luu, Jenifer Manson, Charlotte Nelson, Elaine O'Toole, Cassie Olateju, Rebekah Penrice-Randal , Lucille Rainbow, N.P Randle, Trevor Ian Robinson, Parul Sharma, Ghada T Shawli, James P Stewart, Neil Swainston, Ecaterina Vamos, Joanne Watts, Mark Whitehead |
| EPI_ISL_650258 | University of Exeter | COVID-19 Genomics UK (COG-UK) Consortium | Ben Temperton,Aaron Jeffries,Michelle Michelsen,Joanna Warwick-Dugdale,Audrey Farbos,Robyn Manley,Stephen Michell,Jane Masoli |
| EPI_ISL_650267 | Virology Department, Royal Infirmary of Edinburgh, NHS Lothian / School of Biological Sciences, University of Edinburgh / Institute of Genetics and Molecular Medicine, University of Edinburgh | COVID-19 Genomics UK (COG-UK) Consortium | McHugh M, Dewar R, Rooke S, Gallagher M, Balcaza C, O'Toole Á, Scher E, Hill V, McCrone JT, Colquhoun R, Yu X, Jackson B, Rambaut A, Williams TC, Templeton K |
| EPI_ISL_650278 | Liverpool Clinical Laboratories | COVID-19 Genomics UK (COG-UK) Consortium | Sam Haldenby, Anita Lucaci, Steve Paterson, Julian Hiscox, Alistair Darby, M Almsaud, A Alrezaihi, Muhannad Alruwaili, Stuart D Armstrong, Jones Benjamin, Eleanor G Bentley, Anu Chawla, Jordan J Clark, Angela Cowell, Richard Eccles, Isabel García-Dorival, Matthew Gemmell, Alessandro Gerada, PKF Gilmore, Richard Gregory, Ximeng Han, Catherine Hartley, Margaret Hughes, Miren Iturriza-Gomara, James Johnson, L Luu, Jenifer Manson, Charlotte Nelson, Elaine O'Toole, Cassie Olateju, Rebekah Penrice-Randal , Lucille Rainbow, N.P Randle, Trevor Ian Robinson, Parul Sharma, Ghada T Shawli, James P Stewart, Neil Swainston, Ecaterina Vamos, Joanne Watts, Mark Whitehead |
| EPI_ISL_650281 | University of Birmingham | COVID-19 Genomics UK (COG-UK) Consortium | Institute of Microbiology, University of Birmingham: Claire McMurray, Joanne Stockton, Samuel Nicholls, Radoslaw Poplawski, Will Rowe, Josh Quick, Nicholas Loman. University of Birmingham Testing Laboratory: Celina M Whalley, Andrew Bosworth, Charlotte Poxon, Kasun Wanigasooriya, Oliver Pickles, Mike Kidd, Alex Richter, Andrew D Beggs PHE Heartlands Lab: Husam Osman, Andrew Bosworth. Queen Elizabeth Hospital: Anna Casey |
| EPI_ISL_650320 | Wales Specialist Virology Centre Sequencing lab: Pathogen Genomics Unit | COVID-19 Genomics UK (COG-UK) Consortium | Catherine Moore, Johnathan Evans, Laura Gifford, Malorie Perry, Simon Cottrell, Angela Marchbank, Alec Birchley, Alexander Adams, Amy Gaskin, Bree Gatica-Wilcox, Jason Coombes, Joel Southgate, Lauren Gilbert, Lee Graham, Nicole Pacchiarini, Sara Kumziene-Summerhayes, Sarah Taylor, Sophie Jones, Sara Rey, Matthew Bull, Joanne Watkins, Sally Corden, Tom Connor |
| EPI_ISL_650323 | Quadram Institute Bioscience | COVID-19 Genomics UK (COG-UK) Consortium | Dave J. Baker, Gemma L. Kay, Alp Aydin, Thanh Le-Viet, Steven Rudder, Ana P. Tedim, Anastasia Kolyva, Maria Diaz, Leonardo de Oliveira Martins, Nabil-Fareed Alikhan, Lizzie Meadows, Rachael Stanley, Ngozi Elumogo, Muhammed Yasir, Nicholas M. Thomson, Alexander J Trotter, Rachel Gilroy, Samuel Bloomfield, Claire Stuart, Andrew Bell, Reenesh Prakash, Samir Dervisevic, Alison E. Mather, John Wain, Mark Webber, Andrew J. Page, Justin O'Grady |

| | | | |
|---|---|---|---|
| EPI_ISL_650349 | Virology Department, Sheffield Teaching Hospitals NHS Foundation Trust/Department of Infection, Immunity and Cardiovascular Disease, The Medical School, University of Sheffield | COVID-19 Genomics UK (COG-UK) Consortium | Thushan de Silva, Matthew Parker, Nikki Smith, Adri Angyal, Rebecca Brown, Luke Green, Rachel Tucker, Paul Parsons, Danielle Groves, Katie Johnson, Laura Carrilero, Alex Keeley, Dave Partridge, Matthew Wyles, Benjamin Lindsey, Mehmet Yavuz, Mohammad Raza, Cariad Evans |
| EPI_ISL_650352 | University of Exeter | COVID-19 Genomics UK (COG-UK) Consortium | Ben Temperton,Aaron Jeffries,Michelle Michelsen,Joanna Warwick-Dugdale,Audrey Farbos,Robyn Manley,Stephen Michell,Jane Masoli |
| EPI_ISL_650353 | Department of Pathology, University of Cambridge | COVID-19 Genomics UK (COG-UK) Consortium | Aminu S. Jahun, Yasmin Chaudhry, Grant Hall, Iliana Georgana, Myra Hosmillo, Martin D. Curran, Malte Pinckert, Surendra Parmar, Ian Goodfellow |
| EPI_ISL_650358, EPI_ISL_650377 | University of Exeter | COVID-19 Genomics UK (COG-UK) Consortium | Ben Temperton,Aaron Jeffries,Michelle Michelsen,Joanna Warwick-Dugdale,Audrey Farbos,Robyn Manley,Stephen Michell,Jane Masoli |
| EPI_ISL_650388 | Centre for Enzyme Innovation, University of Portsmouth / Translational Research Laboratory, Portsmouth Hospitals NHS Trust | COVID-19 Genomics UK (COG-UK) Consortium | Angela Beckett,Yann Bourgeois,Garry Scarlett,Sharon Glaysher,Scott Elliott,Kelly Bicknell,Robert Impey,Allyson Lloyd,Sarah Wyllie,Ethan Butcher,Anoop Chauhan,Samuel Robson |
| EPI_ISL_650395 | Virology Department, Sheffield Teaching Hospitals NHS Foundation Trust/Department of Infection, Immunity and Cardiovascular Disease, The Medical School, University of Sheffield | COVID-19 Genomics UK (COG-UK) Consortium | Thushan de Silva, Matthew Parker, Nikki Smith, Adri Angyal, Rebecca Brown, Luke Green, Rachel Tucker, Paul Parsons, Danielle Groves, Katie Johnson, Laura Carrilero, Alex Keeley, Dave Partridge, Matthew Wyles, Benjamin Lindsey, Mehmet Yavuz, Mohammad Raza, Cariad Evans |
| EPI_ISL_650397 | Department of Pathology, University of Cambridge | COVID-19 Genomics UK (COG-UK) Consortium | Aminu S. Jahun, Yasmin Chaudhry, Grant Hall, Iliana Georgana, Myra Hosmillo, Martin D. Curran, Malte Pinckert, Surendra Parmar, Ian Goodfellow |
| EPI_ISL_650399 | University of Exeter | COVID-19 Genomics UK (COG-UK) Consortium | Ben Temperton,Aaron Jeffries,Michelle Michelsen,Joanna Warwick-Dugdale,Audrey Farbos,Robyn Manley,Stephen Michell,Jane Masoli |
| EPI_ISL_650405 | University of Birmingham | COVID-19 Genomics UK (COG-UK) Consortium | Institute of Microbiology, University of Birmingham: Claire McMurray, Joanne Stockton, Samuel Nicholls, Radoslaw Poplawski, Will Rowe, Josh Quick, Nicholas Loman. University of Birmingham Testing Laboratory: Celina M Whalley, Andrew Bosworth, Charlotte Poxon, Kasun Wanigasooriya, Oliver Pickles, Mike Kidd, Alex Richter, Andrew D Beggs PHE Heartlands Lab: Husam Osman, Andrew Bosworth. Queen Elizabeth Hospital: Anna Casey |
| EPI_ISL_650408 | Virology Department, Sheffield Teaching Hospitals NHS Foundation Trust/Department of Infection, Immunity and Cardiovascular Disease, The Medical School, University of Sheffield | COVID-19 Genomics UK (COG-UK) Consortium | Thushan de Silva, Matthew Parker, Nikki Smith, Adri Angyal, Rebecca Brown, Luke Green, Rachel Tucker, Paul Parsons, Danielle Groves, Katie Johnson, Laura Carrilero, Alex Keeley, Dave Partridge, Matthew Wyles, Benjamin Lindsey, Mehmet Yavuz, Mohammad Raza, Cariad Evans |
| EPI_ISL_650414 | University of Exeter | COVID-19 Genomics UK (COG-UK) Consortium | Ben Temperton,Aaron Jeffries,Michelle Michelsen,Joanna Warwick-Dugdale,Audrey Farbos,Robyn Manley,Stephen Michell,Jane Masoli |
| EPI_ISL_650422 | Virology Department, Sheffield Teaching Hospitals NHS Foundation Trust/Department of Infection, Immunity and Cardiovascular Disease, The Medical School, University of Sheffield | COVID-19 Genomics UK (COG-UK) Consortium | Thushan de Silva, Matthew Parker, Nikki Smith, Adri Angyal, Rebecca Brown, Luke Green, Rachel Tucker, Paul Parsons, Danielle Groves, Katie Johnson, Laura Carrilero, Alex Keeley, Dave Partridge, Matthew Wyles, Benjamin Lindsey, Mehmet Yavuz, Mohammad Raza, Cariad Evans |
| EPI_ISL_650433 | Virology Department, Royal Infirmary of Edinburgh, NHS Lothian / School of Biological Sciences, University of Edinburgh / Institute of Genetics and Molecular Medicine, University of Edinburgh | COVID-19 Genomics UK (COG-UK) Consortium | McHugh M, Dewar R, Rooke S, Gallagher M, Balcaza C, O'Toole Á, Scher E, Hill V, McCrone JT, Colquhoun R, Yu X, Jackson B, Rambaut A, Williams TC, Templeton K |
| EPI_ISL_650453 | Liverpool Clinical Laboratories | COVID-19 Genomics UK (COG-UK) Consortium | Sam Haldenby, Anita Lucaci, Steve Paterson, Julian Hiscox, Alistair Darby, M Almsaud, A Alrezaihi, Muhannad Alruwaili, Stuart D Armstrong, Jones Benjamin, Eleanor G Bentley, Anu Chawla, Jordan J Clark, Angela Cowell, Richard Eccles, Isabel García-Dorival, Matthew Gemmell, Alessandro Gerada, PKF Gilmore, Richard Gregory, Ximeng Han, Catherine Hartley, Margaret Hughes, Miren Iturriza-Gomara, James Johnson, L Luu, Jenifer Manson, Charlotte Nelson, Elaine O'Toole, Cassie Olateju, Rebekah Penrice-Randal , Lucille Rainbow, N.P Randle, Trevor Ian Robinson, Parul Sharma, Ghada T Shawli, James P Stewart, Neil Swainston, Ecaterina Vamos, Joanne Watts, Mark Whitehead |
| EPI_ISL_650501 | Virology Department, Royal Infirmary of Edinburgh, NHS Lothian / School of Biological Sciences, University of Edinburgh / Institute of Genetics and Molecular Medicine, University of Edinburgh | COVID-19 Genomics UK (COG-UK) Consortium | McHugh M, Dewar R, Rooke S, Gallagher M, Balcaza C, O'Toole Á, Scher E, Hill V, McCrone JT, Colquhoun R, Yu X, Jackson B, Rambaut A, Williams TC, Templeton K |
| EPI_ISL_650551 | Wales Specialist Virology Centre Sequencing lab: Pathogen Genomics Unit | COVID-19 Genomics UK (COG-UK) Consortium | Catherine Moore, Johnathan Evans, Laura Gifford, Malorie Perry, Simon Cottrell, Angela Marchbank, Alec Birchley, Alexander Adams, Amy Gaskin, Bree Gatica-Wilcox, Jason Coombes, Joel Southgate, Lauren Gilbert, Lee Graham, Nicole Pacchiarini, Sara Kumziene-Summerhayes, Sarah Taylor, Sophie Jones, Sara Rey, Matthew Bull, Joanne Watkins, Sally Corden, Tom Connor |
| EPI_ISL_650575, EPI_ISL_650576 | Centre for Enzyme Innovation, University of Portsmouth / Translational Research Laboratory, Portsmouth Hospitals NHS Trust | COVID-19 Genomics UK (COG-UK) Consortium | Angela Beckett,Yann Bourgeois,Garry Scarlett,Sharon Glaysher,Scott Elliott,Kelly Bicknell,Robert Impey,Allyson Lloyd,Sarah Wyllie,Ethan Butcher,Anoop Chauhan,Samuel Robson |
| EPI_ISL_650583 | Department of Pathology, University of Cambridge | COVID-19 Genomics UK (COG-UK) Consortium | Aminu S. Jahun, Yasmin Chaudhry, Grant Hall, Iliana Georgana, Myra Hosmillo, Martin D. Curran, Malte Pinckert, Surendra Parmar, Ian Goodfellow |
| EPI_ISL_650597, EPI_ISL_650607, EPI_ISL_650610 | University of Exeter | COVID-19 Genomics UK (COG-UK) Consortium | Ben Temperton,Aaron Jeffries,Michelle Michelsen,Joanna Warwick-Dugdale,Audrey Farbos,Robyn Manley,Stephen Michell,Jane Masoli |
| EPI_ISL_650615, EPI_ISL_650616 | Department of Pathology, University of Cambridge | COVID-19 Genomics UK (COG-UK) Consortium | Aminu S. Jahun, Yasmin Chaudhry, Grant Hall, Iliana Georgana, Myra Hosmillo, Martin D. Curran, Malte Pinckert, Surendra Parmar, Ian Goodfellow |
| EPI_ISL_650634, EPI_ISL_650635, EPI_ISL_650636, EPI_ISL_650637, EPI_ISL_650638, EPI_ISL_650639 | University of Exeter | COVID-19 Genomics UK (COG-UK) Consortium | Ben Temperton,Aaron Jeffries,Michelle Michelsen,Joanna Warwick-Dugdale,Audrey Farbos,Robyn Manley,Stephen Michell,Jane Masoli |
| EPI_ISL_650646, EPI_ISL_650662 | Virology Department, Sheffield Teaching Hospitals NHS Foundation Trust/Department of Infection, Immunity and Cardiovascular Disease, The Medical School, University of Sheffield | COVID-19 Genomics UK (COG-UK) Consortium | Thushan de Silva, Matthew Parker, Nikki Smith, Adri Angyal, Rebecca Brown, Luke Green, Rachel Tucker, Paul Parsons, Danielle Groves, Katie Johnson, Laura Carrilero, Alex Keeley, Dave Partridge, Matthew Wyles, Benjamin Lindsey, Mehmet Yavuz, Mohammad Raza, Cariad Evans |
| EPI_ISL_650667 | University of Exeter | COVID-19 Genomics UK (COG-UK) Consortium | Ben Temperton,Aaron Jeffries,Michelle Michelsen,Joanna Warwick-Dugdale,Audrey Farbos,Robyn Manley,Stephen Michell,Jane Masoli |
| EPI_ISL_650670, EPI_ISL_650689 | Virology Department, Sheffield Teaching Hospitals NHS Foundation Trust/Department of Infection, Immunity and Cardiovascular Disease, The Medical School, University of Sheffield | COVID-19 Genomics UK (COG-UK) Consortium | Thushan de Silva, Matthew Parker, Nikki Smith, Adri Angyal, Rebecca Brown, Luke Green, Rachel Tucker, Paul Parsons, Danielle Groves, Katie Johnson, Laura Carrilero, Alex Keeley, Dave Partridge, Matthew Wyles, Benjamin Lindsey, Mehmet Yavuz, Mohammad Raza, Cariad Evans |
| EPI_ISL_650692 | University of Birmingham | COVID-19 Genomics UK (COG-UK) Consortium | Institute of Microbiology, University of Birmingham: Claire McMurray, Joanne Stockton, Samuel Nicholls, Radoslaw Poplawski, Will Rowe, Josh Quick, Nicholas Loman. University of Birmingham Testing Laboratory: Celina M Whalley, Andrew Bosworth, Charlotte Poxon, Kasun Wanigasooriya, Oliver Pickles, Mike Kidd, Alex Richter, Andrew D Beggs PHE Heartlands Lab: Husam Osman, Andrew Bosworth. Queen Elizabeth Hospital: Anna Casey |
| EPI_ISL_650702 | University of Exeter | COVID-19 Genomics UK (COG-UK) Consortium | Ben Temperton,Aaron Jeffries,Michelle Michelsen,Joanna Warwick-Dugdale,Audrey Farbos,Robyn Manley,Stephen Michell,Jane Masoli |
| EPI_ISL_650709 | West of Scotland Specialist Virology Centre, NHSGGC / MRC-University of Glasgow Centre for Virus Research | COVID-19 Genomics UK (COG-UK) Consortium | Ana da Silva Filipe, Natasha Johnson, Kathy Smollett, Daniel Mair, Stephen Carmichael, Alice Broos, Lily Tong, Jenna Nichols, Kyriaki Nomikou; Sarah McDonald; Richard Orton, Joseph Hughes, Sreenu Vattipally, David L Robertson; Alasdair MacLean, Rory Gunson; Sharif Shaaban, Matthew Holden; Rachel Blacow, Guy Mollett, Kathy Li, James Shepherd, Antonia Ho, Emma Thomson |
| EPI_ISL_650713, EPI_ISL_650715 | Virology Department, Sheffield Teaching Hospitals NHS Foundation Trust/Department of Infection, Immunity and Cardiovascular Disease, The Medical School, University of Sheffield | COVID-19 Genomics UK (COG-UK) Consortium | Thushan de Silva, Matthew Parker, Nikki Smith, Adri Angyal, Rebecca Brown, Luke Green, Rachel Tucker, Paul Parsons, Danielle Groves, Katie Johnson, Laura Carrilero, Alex Keeley, Dave Partridge, Matthew Wyles, Benjamin Lindsey, Mehmet Yavuz, Mohammad Raza, Cariad Evans |

| | | | |
|---|---|---|---|
| EPI_ISL_650740 | University of Birmingham | COVID-19 Genomics UK (COG-UK) Consortium | Institute of Microbiology, University of Birmingham: Claire McMurray, Joanne Stockton, Samuel Nicholls, Radoslaw Poplawski, Will Rowe, Josh Quick, Nicholas Loman. University of Birmingham Testing Laboratory: Celina M Whalley, Andrew Bosworth, Charlotte Poxon, Kasun Wanigasooriya, Oliver Pickles, Mike Kidd, Alex Richter, Andrew D Beggs PHE Heartlands Lab: Husam Osman, Andrew Bosworth. Queen Elizabeth Hospital: Anna Casey |
| EPI_ISL_650774 | Department of Pathology, University of Cambridge | COVID-19 Genomics UK (COG-UK) Consortium | Aminu S. Jahun, Yasmin Chaudhry, Grant Hall, Iliana Georgana, Myra Hosmillo, Martin D. Curran, Malte Pinckert, Surendra Parmar, Ian Goodfellow |
| EPI_ISL_650781 | Virology Department, Royal Infirmary of Edinburgh, NHS Lothian / School of Biological Sciences, University of Edinburgh / Institute of Genetics and Molecular Medicine, University of Edinburgh | COVID-19 Genomics UK (COG-UK) Consortium | McHugh M, Dewar R, Rooke S, Gallagher M, Balcaza C, O'Toole Á, Scher E, Hill V, McCrone JT, Colquhoun R, Yu X, Jackson B, Rambaut A, Williams TC, Templeton K |
| EPI_ISL_650787 | Wales Specialist Virology Centre Sequencing lab: Pathogen Genomics Unit | COVID-19 Genomics UK (COG-UK) Consortium | Catherine Moore, Johnathan Evans, Laura Gifford, Malorie Perry, Simon Cottrell, Angela Marchbank, Alec Birchley, Alexander Adams, Amy Gaskin, Bree Gatica-Wilcox, Jason Coombes, Joel Southgate, Lauren Gilbert, Lee Graham, Nicole Pacchiarini, Sara Kumziene-Summerhayes, Sarah Taylor, Sophie Jones, Sara Rey, Matthew Bull, Joanne Watkins, Sally Corden, Tom Connor |
| EPI_ISL_650789 | West of Scotland Specialist Virology Centre, NHSGGC / MRC-University of Glasgow Centre for Virus Research | COVID-19 Genomics UK (COG-UK) Consortium | Ana da Silva Filipe, Natasha Johnson, Kathy Smollett, Daniel Mair, Stephen Carmichael, Alice Broos, Lily Tong, Jenna Nichols, Kyriaki Nomikou; Sarah McDonald; Richard Orton, Joseph Hughes, Sreenu Vattipally, David L Robertson; Alasdair MacLean, Rory Gunson; Sharif Shaaban, Matthew Holden; Rachel Blacow, Guy Mollett, Kathy Li, James Shepherd, Antonia Ho, Emma Thomson |
| EPI_ISL_650799 | Virology Department, Royal Infirmary of Edinburgh, NHS Lothian / School of Biological Sciences, University of Edinburgh / Institute of Genetics and Molecular Medicine, University of Edinburgh | COVID-19 Genomics UK (COG-UK) Consortium | McHugh M, Dewar R, Rooke S, Gallagher M, Balcaza C, O'Toole Á, Scher E, Hill V, McCrone JT, Colquhoun R, Yu X, Jackson B, Rambaut A, Williams TC, Templeton K |
| EPI_ISL_650805 | West of Scotland Specialist Virology Centre, NHSGGC / MRC-University of Glasgow Centre for Virus Research | COVID-19 Genomics UK (COG-UK) Consortium | Ana da Silva Filipe, Natasha Johnson, Kathy Smollett, Daniel Mair, Stephen Carmichael, Alice Broos, Lily Tong, Jenna Nichols, Kyriaki Nomikou; Sarah McDonald; Richard Orton, Joseph Hughes, Sreenu Vattipally, David L Robertson; Alasdair MacLean, Rory Gunson; Sharif Shaaban, Matthew Holden; Rachel Blacow, Guy Mollett, Kathy Li, James Shepherd, Antonia Ho, Emma Thomson |
| EPI_ISL_650826 | Virology Department, Royal Infirmary of Edinburgh, NHS Lothian / School of Biological Sciences, University of Edinburgh / Institute of Genetics and Molecular Medicine, University of Edinburgh | COVID-19 Genomics UK (COG-UK) Consortium | McHugh M, Dewar R, Rooke S, Gallagher M, Balcaza C, O'Toole Á, Scher E, Hill V, McCrone JT, Colquhoun R, Yu X, Jackson B, Rambaut A, Williams TC, Templeton K |
| EPI_ISL_650870, EPI_ISL_650871 | West of Scotland Specialist Virology Centre, NHSGGC / MRC-University of Glasgow Centre for Virus Research | COVID-19 Genomics UK (COG-UK) Consortium | Ana da Silva Filipe, Natasha Johnson, Kathy Smollett, Daniel Mair, Stephen Carmichael, Alice Broos, Lily Tong, Jenna Nichols, Kyriaki Nomikou; Sarah McDonald; Richard Orton, Joseph Hughes, Sreenu Vattipally, David L Robertson; Alasdair MacLean, Rory Gunson; Sharif Shaaban, Matthew Holden; Rachel Blacow, Guy Mollett, Kathy Li, James Shepherd, Antonia Ho, Emma Thomson |
| EPI_ISL_650884, EPI_ISL_650905, EPI_ISL_650929 | Wales Specialist Virology Centre Sequencing lab: Pathogen Genomics Unit | COVID-19 Genomics UK (COG-UK) Consortium | Catherine Moore, Johnathan Evans, Laura Gifford, Malorie Perry, Simon Cottrell, Angela Marchbank, Alec Birchley, Alexander Adams, Amy Gaskin, Bree Gatica-Wilcox, Jason Coombes, Joel Southgate, Lauren Gilbert, Lee Graham, Nicole Pacchiarini, Sara Kumziene-Summerhayes, Sarah Taylor, Sophie Jones, Sara Rey, Matthew Bull, Joanne Watkins, Sally Corden, Tom Connor |
| EPI_ISL_650941 | Centre for Enzyme Innovation, University of Portsmouth / Translational Research Laboratory, Portsmouth Hospitals NHS Trust | COVID-19 Genomics UK (COG-UK) Consortium | Angela Beckett,Yann Bourgeois,Garry Scarlett,Sharon Glaysher,Scott Elliott,Kelly Bicknell,Robert Impey,Allyson Lloyd,Sarah Wyllie,Ethan Butcher,Anoop Chauhan,Samuel Robson |
| EPI_ISL_650990, EPI_ISL_650991, EPI_ISL_650993 | Department of Pathology, University of Cambridge | COVID-19 Genomics UK (COG-UK) Consortium | Aminu S. Jahun, Yasmin Chaudhry, Grant Hall, Iliana Georgana, Myra Hosmillo, Martin D. Curran, Malte Pinckert, Surendra Parmar, Ian Goodfellow |
| EPI_ISL_650998 | University of Birmingham | COVID-19 Genomics UK (COG-UK) Consortium | Institute of Microbiology, University of Birmingham: Claire McMurray, Joanne Stockton, Samuel Nicholls, Radoslaw Poplawski, Will Rowe, Josh Quick, Nicholas Loman. University of Birmingham Testing Laboratory: Celina M Whalley, Andrew Bosworth, Charlotte Poxon, Kasun Wanigasooriya, Oliver Pickles, Mike Kidd, Alex Richter, Andrew D Beggs PHE Heartlands Lab: Husam Osman, Andrew Bosworth. Queen Elizabeth Hospital: Anna Casey |
| EPI_ISL_651003, EPI_ISL_651004, EPI_ISL_651005, EPI_ISL_651006, EPI_ISL_651007, EPI_ISL_651008 | Department of Pathology, University of Cambridge | COVID-19 Genomics UK (COG-UK) Consortium | Aminu S. Jahun, Yasmin Chaudhry, Grant Hall, Iliana Georgana, Myra Hosmillo, Martin D. Curran, Malte Pinckert, Surendra Parmar, Ian Goodfellow |
| EPI_ISL_651054, EPI_ISL_651056, EPI_ISL_651058 | University of Exeter | COVID-19 Genomics UK (COG-UK) Consortium | Ben Temperton,Aaron Jeffries,Michelle Michelsen,Joanna Warwick-Dugdale,Audrey Farbos,Robyn Manley,Stephen Michell,Jane Masoli |
| EPI_ISL_651078 | Virology Department, Sheffield Teaching Hospitals NHS Foundation Trust/Department of Infection, Immunity and Cardiovascular Disease, The Medical School, University of Sheffield | COVID-19 Genomics UK (COG-UK) Consortium | Thushan de Silva, Matthew Parker, Nikki Smith, Adri Angyal, Rebecca Brown, Luke Green, Rachel Tucker, Paul Parsons, Danielle Groves, Katie Johnson, Laura Carrilero, Alex Keeley, Dave Partridge, Matthew Wyles, Benjamin Lindsey, Mehmet Yavuz, Mohammad Raza, Cariad Evans |
| EPI_ISL_651084 | University of Exeter | COVID-19 Genomics UK (COG-UK) Consortium | Ben Temperton,Aaron Jeffries,Michelle Michelsen,Joanna Warwick-Dugdale,Audrey Farbos,Robyn Manley,Stephen Michell,Jane Masoli |
| EPI_ISL_651109 | Virology Department, Sheffield Teaching Hospitals NHS Foundation Trust/Department of Infection, Immunity and Cardiovascular Disease, The Medical School, University of Sheffield | COVID-19 Genomics UK (COG-UK) Consortium | Thushan de Silva, Matthew Parker, Nikki Smith, Adri Angyal, Rebecca Brown, Luke Green, Rachel Tucker, Paul Parsons, Danielle Groves, Katie Johnson, Laura Carrilero, Alex Keeley, Dave Partridge, Matthew Wyles, Benjamin Lindsey, Mehmet Yavuz, Mohammad Raza, Cariad Evans |
| EPI_ISL_651122 | Department of Pathology, University of Cambridge | COVID-19 Genomics UK (COG-UK) Consortium | Aminu S. Jahun, Yasmin Chaudhry, Grant Hall, Iliana Georgana, Myra Hosmillo, Martin D. Curran, Malte Pinckert, Surendra Parmar, Ian Goodfellow |
| EPI_ISL_651125 | West of Scotland Specialist Virology Centre, NHSGGC / MRC-University of Glasgow Centre for Virus Research | COVID-19 Genomics UK (COG-UK) Consortium | Ana da Silva Filipe, Natasha Johnson, Kathy Smollett, Daniel Mair, Stephen Carmichael, Alice Broos, Lily Tong, Jenna Nichols, Kyriaki Nomikou; Sarah McDonald; Richard Orton, Joseph Hughes, Sreenu Vattipally, David L Robertson; Alasdair MacLean, Rory Gunson; Sharif Shaaban, Matthew Holden; Rachel Blacow, Guy Mollett, Kathy Li, James Shepherd, Antonia Ho, Emma Thomson |
| EPI_ISL_651130 | University of Exeter | COVID-19 Genomics UK (COG-UK) Consortium | Ben Temperton,Aaron Jeffries,Michelle Michelsen,Joanna Warwick-Dugdale,Audrey Farbos,Robyn Manley,Stephen Michell,Jane Masoli |
| EPI_ISL_651136 | Department of Pathology, University of Cambridge | COVID-19 Genomics UK (COG-UK) Consortium | Aminu S. Jahun, Yasmin Chaudhry, Grant Hall, Iliana Georgana, Myra Hosmillo, Martin D. Curran, Malte Pinckert, Surendra Parmar, Ian Goodfellow |
| EPI_ISL_651140 | University of Birmingham | COVID-19 Genomics UK (COG-UK) Consortium | Institute of Microbiology, University of Birmingham: Claire McMurray, Joanne Stockton, Samuel Nicholls, Radoslaw Poplawski, Will Rowe, Josh Quick, Nicholas Loman. University of Birmingham Testing Laboratory: Celina M Whalley, Andrew Bosworth, Charlotte Poxon, Kasun Wanigasooriya, Oliver Pickles, Mike Kidd, Alex Richter, Andrew D Beggs PHE Heartlands Lab: Husam Osman, Andrew Bosworth. Queen Elizabeth Hospital: Anna Casey |
| EPI_ISL_651148 | Liverpool Clinical Laboratories | COVID-19 Genomics UK (COG-UK) Consortium | Sam Haldenby, Anita Lucaci, Steve Paterson, Julian Hiscox, Alistair Darby, M Almsaud, A Alrezaihi, Muhannad Alruwaili, Stuart D Armstrong, Jones Benjamin, Eleanor G Bentley, Anu Chawla, Jordan J Clark, Angela Cowell, Richard Eccles, Isabel García-Dorival, Matthew Gemmell, Alessandro Gerada, PKF Gilmore, Richard Gregory, Ximeng Han, Catherine Hartley, Margaret Hughes, Miren Iturriza-Gomara, James Johnson, L Luu, Jenifer Manson, Charlotte Nelson, Elaine O'Toole, Cassie Olateju, Rebekah Penrice-Randal , Lucille Rainbow, N.P Randle, Trevor Ian Robinson, Parul Sharma, Ghada T Shawli, James P Stewart, Neil Swainston, Ecaterina Vamos, Joanne Watts, Mark Whitehead |
| EPI_ISL_651152 | Virology Department, Sheffield Teaching Hospitals NHS Foundation Trust/Department of Infection, Immunity and Cardiovascular Disease, The Medical School, University of Sheffield | COVID-19 Genomics UK (COG-UK) Consortium | Thushan de Silva, Matthew Parker, Nikki Smith, Adri Angyal, Rebecca Brown, Luke Green, Rachel Tucker, Paul Parsons, Danielle Groves, Katie Johnson, Laura Carrilero, Alex Keeley, Dave Partridge, Matthew Wyles, Benjamin Lindsey, Mehmet Yavuz, Mohammad Raza, Cariad Evans |
| EPI_ISL_651158 | Virology Department, Royal Infirmary of Edinburgh, NHS Lothian / School of Biological Sciences, University of Edinburgh / Institute of Genetics and Molecular Medicine, University of Edinburgh | COVID-19 Genomics UK (COG-UK) Consortium | McHugh M, Dewar R, Rooke S, Gallagher M, Balcaza C, O'Toole Á, Scher E, Hill V, McCrone JT, Colquhoun R, Yu X, Jackson B, Rambaut A, Williams TC, Templeton K |

| | | | |
|---|---|---|---|
| EPI_ISL_651223 | Liverpool Clinical Laboratories | COVID-19 Genomics UK (COG-UK) Consortium | Sam Haldenby, Anita Lucaci, Steve Paterson, Julian Hiscox, Alistair Darby, M Almsaud, A Alrezaihi, Muhannad Alruwaili, Stuart D Armstrong, Jones Benjamin, Eleanor G Bentley, Anu Chawla, Jordan J Clark, Angela Cowell, Richard Eccles, Isabel García-Dorival, Matthew Gemmell, Alessandro Gerada, PKF Gilmore, Richard Gregory, Ximeng Han, Catherine Hartley, Margaret Hughes, Miren Iturriza-Gomara, James Johnson, L Luu, Jenifer Manson, Charlotte Nelson, Elaine O'Toole, Cassie Olateju, Rebekah Penrice-Randal , Lucille Rainbow, N.P Randle, Trevor Ian Robinson, Parul Sharma, Ghada T Shawli, James P Stewart, Neil Swainston, Ecaterina Vamos, Joanne Watts, Mark Whitehead |
| EPI_ISL_651238 | University of Exeter | COVID-19 Genomics UK (COG-UK) Consortium | Ben Temperton,Aaron Jeffries,Michelle Michelsen,Joanna Warwick-Dugdale,Audrey Farbos,Robyn Manley,Stephen Michell,Jane Masoli |
| EPI_ISL_651239 | Liverpool Clinical Laboratories | COVID-19 Genomics UK (COG-UK) Consortium | Sam Haldenby, Anita Lucaci, Steve Paterson, Julian Hiscox, Alistair Darby, M Almsaud, A Alrezaihi, Muhannad Alruwaili, Stuart D Armstrong, Jones Benjamin, Eleanor G Bentley, Anu Chawla, Jordan J Clark, Angela Cowell, Richard Eccles, Isabel García-Dorival, Matthew Gemmell, Alessandro Gerada, PKF Gilmore, Richard Gregory, Ximeng Han, Catherine Hartley, Margaret Hughes, Miren Iturriza-Gomara, James Johnson, L Luu, Jenifer Manson, Charlotte Nelson, Elaine O'Toole, Cassie Olateju, Rebekah Penrice-Randal , Lucille Rainbow, N.P Randle, Trevor Ian Robinson, Parul Sharma, Ghada T Shawli, James P Stewart, Neil Swainston, Ecaterina Vamos, Joanne Watts, Mark Whitehead |
| EPI_ISL_651241 | University of Exeter | COVID-19 Genomics UK (COG-UK) Consortium | Ben Temperton,Aaron Jeffries,Michelle Michelsen,Joanna Warwick-Dugdale,Audrey Farbos,Robyn Manley,Stephen Michell,Jane Masoli |
| EPI_ISL_651248, EPI_ISL_651251 | Liverpool Clinical Laboratories | COVID-19 Genomics UK (COG-UK) Consortium | Sam Haldenby, Anita Lucaci, Steve Paterson, Julian Hiscox, Alistair Darby, M Almsaud, A Alrezaihi, Muhannad Alruwaili, Stuart D Armstrong, Jones Benjamin, Eleanor G Bentley, Anu Chawla, Jordan J Clark, Angela Cowell, Richard Eccles, Isabel García-Dorival, Matthew Gemmell, Alessandro Gerada, PKF Gilmore, Richard Gregory, Ximeng Han, Catherine Hartley, Margaret Hughes, Miren Iturriza-Gomara, James Johnson, L Luu, Jenifer Manson, Charlotte Nelson, Elaine O'Toole, Cassie Olateju, Rebekah Penrice-Randal , Lucille Rainbow, N.P Randle, Trevor Ian Robinson, Parul Sharma, Ghada T Shawli, James P Stewart, Neil Swainston, Ecaterina Vamos, Joanne Watts, Mark Whitehead |
| EPI_ISL_651267 | Quadram Institute Bioscience | COVID-19 Genomics UK (COG-UK) Consortium | Dave J. Baker, Gemma L. Kay, Alp Aydin, Thanh Le-Viet, Steven Rudder, Ana P. Tedim, Anastasia Kolyva, Maria Diaz, Leonardo de Oliveira Martins, Nabil-Fareed Alikhan, Lizzie Meadows, Rachael Stanley, Ngozi Elumogo, Muhammed Yasir, Nicholas M. Thomson, Alexander J Trotter, Rachel Gilroy, Samuel Bloomfield, Claire Stuart, Andrew Bell, Reenesh Prakash, Samir Dervisevic, Alison E. Mather, John Wain, Mark Webber, Andrew J. Page, Justin O'Grady |
| EPI_ISL_651282 | University of Birmingham | COVID-19 Genomics UK (COG-UK) Consortium | Institute of Microbiology, University of Birmingham: Claire McMurray, Joanne Stockton, Samuel Nicholls, Radoslaw Poplawski, Will Rowe, Josh Quick, Nicholas Loman. University of Birmingham Testing Laboratory: Celina M Whalley, Andrew Bosworth, Charlotte Poxon, Kasun Wanigasooriya, Oliver Pickles, Mike Kidd, Alex Richter, Andrew D Beggs PHE Heartlands Lab: Husam Osman, Andrew Bosworth. Queen Elizabeth Hospital: Anna Casey |
| EPI_ISL_651289 | University of Exeter | COVID-19 Genomics UK (COG-UK) Consortium | Ben Temperton,Aaron Jeffries,Michelle Michelsen,Joanna Warwick-Dugdale,Audrey Farbos,Robyn Manley,Stephen Michell,Jane Masoli |
| EPI_ISL_651292 | Department of Pathology, University of Cambridge | COVID-19 Genomics UK (COG-UK) Consortium | Aminu S. Jahun, Yasmin Chaudhry, Grant Hall, Iliana Georgana, Myra Hosmillo, Martin D. Curran, Malte Pinckert, Surendra Parmar, Ian Goodfellow |
| EPI_ISL_651304 | University of Birmingham | COVID-19 Genomics UK (COG-UK) Consortium | Institute of Microbiology, University of Birmingham: Claire McMurray, Joanne Stockton, Samuel Nicholls, Radoslaw Poplawski, Will Rowe, Josh Quick, Nicholas Loman. University of Birmingham Testing Laboratory: Celina M Whalley, Andrew Bosworth, Charlotte Poxon, Kasun Wanigasooriya, Oliver Pickles, Mike Kidd, Alex Richter, Andrew D Beggs PHE Heartlands Lab: Husam Osman, Andrew Bosworth. Queen Elizabeth Hospital: Anna Casey |
| EPI_ISL_651355 | Liverpool Clinical Laboratories | COVID-19 Genomics UK (COG-UK) Consortium | Sam Haldenby, Anita Lucaci, Steve Paterson, Julian Hiscox, Alistair Darby, M Almsaud, A Alrezaihi, Muhannad Alruwaili, Stuart D Armstrong, Jones Benjamin, Eleanor G Bentley, Anu Chawla, Jordan J Clark, Angela Cowell, Richard Eccles, Isabel García-Dorival, Matthew Gemmell, Alessandro Gerada, PKF Gilmore, Richard Gregory, Ximeng Han, Catherine Hartley, Margaret Hughes, Miren Iturriza-Gomara, James Johnson, L Luu, Jenifer Manson, Charlotte Nelson, Elaine O'Toole, Cassie Olateju, Rebekah Penrice-Randal , Lucille Rainbow, N.P Randle, Trevor Ian Robinson, Parul Sharma, Ghada T Shawli, James P Stewart, Neil Swainston, Ecaterina Vamos, Joanne Watts, Mark Whitehead |
| EPI_ISL_651359 | West of Scotland Specialist Virology Centre, NHSGGC / MRC-University of Glasgow Centre for Virus Research | COVID-19 Genomics UK (COG-UK) Consortium | Ana da Silva Filipe, Natasha Johnson, Kathy Smollett, Daniel Mair, Stephen Carmichael, Alice Broos, Lily Tong, Jenna Nichols, Kyriaki Nomikou; Sarah McDonald; Richard Orton, Joseph Hughes, Sreenu Vattipally, David L Robertson; Alasdair MacLean, Rory Gunson; Sharif Shaaban, Matthew Holden; Rachel Blacow, Guy Mollett, Kathy Li, James Shepherd, Antonia Ho, Emma Thomson |
| EPI_ISL_651363 | Department of Pathology, University of Cambridge | COVID-19 Genomics UK (COG-UK) Consortium | Aminu S. Jahun, Yasmin Chaudhry, Grant Hall, Iliana Georgana, Myra Hosmillo, Martin D. Curran, Malte Pinckert, Surendra Parmar, Ian Goodfellow |
| EPI_ISL_651373 | University of Exeter | COVID-19 Genomics UK (COG-UK) Consortium | Ben Temperton,Aaron Jeffries,Michelle Michelsen,Joanna Warwick-Dugdale,Audrey Farbos,Robyn Manley,Stephen Michell,Jane Masoli |
| EPI_ISL_651383 | Department of Pathology, University of Cambridge | COVID-19 Genomics UK (COG-UK) Consortium | Aminu S. Jahun, Yasmin Chaudhry, Grant Hall, Iliana Georgana, Myra Hosmillo, Martin D. Curran, Malte Pinckert, Surendra Parmar, Ian Goodfellow |
| EPI_ISL_651398 | Queens Medical Centre, Clinical Microbiology Department / DeepSeq Nottingham | COVID-19 Genomics UK (COG-UK) Consortium | Gemma Clark, Wendy Smith, Manjinder Khakh, Vicki M Fleming, Michelle M Lister, Hannah Howson-Wells, Jonathan Ball, Patrick McClure, Joseph Chappell, Theocharis Tsoleridis, Nadine Holmes, Matthew Carlisle, Christopher Moore, Fei Sang, Johnny Debebe, Victoria Wright, Matthew Loose |
| EPI_ISL_651414 | Virology Department, Sheffield Teaching Hospitals NHS Foundation Trust/Department of Infection, Immunity and Cardiovascular Disease, The Medical School, University of Sheffield | COVID-19 Genomics UK (COG-UK) Consortium | Thushan de Silva, Matthew Parker, Nikki Smith, Adri Angyal, Rebecca Brown, Luke Green, Rachel Tucker, Paul Parsons, Danielle Groves, Katie Johnson, Laura Carrilero, Alex Keeley, Dave Partridge, Matthew Wyles, Benjamin Lindsey, Mehmet Yavuz, Mohammad Raza, Cariad Evans |
| EPI_ISL_651416 | Wales Specialist Virology Centre Sequencing lab: Pathogen Genomics Unit | COVID-19 Genomics UK (COG-UK) Consortium | Catherine Moore, Johnathan Evans, Laura Gifford, Malorie Perry, Simon Cottrell, Angela Marchbank, Alec Birchley, Alexander Adams, Amy Gaskin, Bree Gatica-Wilcox, Jason Coombes, Joel Southgate, Lauren Gilbert, Lee Graham, Nicole Pacchiarini, Sara Kumziene-Summerhayes, Sarah Taylor, Sophie Jones, Sara Rey, Matthew Bull, Joanne Watkins, Sally Corden, Tom Connor |
| EPI_ISL_651439 | University of Exeter | COVID-19 Genomics UK (COG-UK) Consortium | Ben Temperton,Aaron Jeffries,Michelle Michelsen,Joanna Warwick-Dugdale,Audrey Farbos,Robyn Manley,Stephen Michell,Jane Masoli |
| EPI_ISL_651450, EPI_ISL_651452, EPI_ISL_651453, EPI_ISL_651454, EPI_ISL_651460, EPI_ISL_651461, EPI_ISL_651463, EPI_ISL_651464 | Wales Specialist Virology Centre Sequencing lab: Pathogen Genomics Unit | COVID-19 Genomics UK (COG-UK) Consortium | Catherine Moore, Johnathan Evans, Laura Gifford, Malorie Perry, Simon Cottrell, Angela Marchbank, Alec Birchley, Alexander Adams, Amy Gaskin, Bree Gatica-Wilcox, Jason Coombes, Joel Southgate, Lauren Gilbert, Lee Graham, Nicole Pacchiarini, Sara Kumziene-Summerhayes, Sarah Taylor, Sophie Jones, Sara Rey, Matthew Bull, Joanne Watkins, Sally Corden, Tom Connor |
| EPI_ISL_651486 | University of Birmingham | COVID-19 Genomics UK (COG-UK) Consortium | Institute of Microbiology, University of Birmingham: Claire McMurray, Joanne Stockton, Samuel Nicholls, Radoslaw Poplawski, Will Rowe, Josh Quick, Nicholas Loman. University of Birmingham Testing Laboratory: Celina M Whalley, Andrew Bosworth, Charlotte Poxon, Kasun Wanigasooriya, Oliver Pickles, Mike Kidd, Alex Richter, Andrew D Beggs PHE Heartlands Lab: Husam Osman, Andrew Bosworth. Queen Elizabeth Hospital: Anna Casey |
| EPI_ISL_651514, EPI_ISL_651518 | Virology Department, Sheffield Teaching Hospitals NHS Foundation Trust/Department of Infection, Immunity and Cardiovascular Disease, The Medical School, University of Sheffield | COVID-19 Genomics UK (COG-UK) Consortium | Thushan de Silva, Matthew Parker, Nikki Smith, Adri Angyal, Rebecca Brown, Luke Green, Rachel Tucker, Paul Parsons, Danielle Groves, Katie Johnson, Laura Carrilero, Alex Keeley, Dave Partridge, Matthew Wyles, Benjamin Lindsey, Mehmet Yavuz, Mohammad Raza, Cariad Evans |
| EPI_ISL_651523, EPI_ISL_651524 | Virology Department, Royal Infirmary of Edinburgh, NHS Lothian / School of Biological Sciences, University of Edinburgh / Institute of Genetics and Molecular Medicine, University of Edinburgh | COVID-19 Genomics UK (COG-UK) Consortium | McHugh M, Dewar R, Rooke S, Gallagher M, Balcaza C, O'Toole Á, Scher E, Hill V, McCrone JT, Colquhoun R, Yu X, Jackson B, Rambaut A, Williams TC, Templeton K |
| EPI_ISL_651527 | University of Exeter | COVID-19 Genomics UK (COG-UK) Consortium | Ben Temperton,Aaron Jeffries,Michelle Michelsen,Joanna Warwick-Dugdale,Audrey Farbos,Robyn Manley,Stephen Michell,Jane Masoli |
| EPI_ISL_651546 | Wales Specialist Virology Centre Sequencing lab: Pathogen Genomics Unit | COVID-19 Genomics UK (COG-UK) Consortium | Catherine Moore, Johnathan Evans, Laura Gifford, Malorie Perry, Simon Cottrell, Angela Marchbank, Alec Birchley, Alexander Adams, Amy Gaskin, Bree Gatica-Wilcox, Jason Coombes, Joel Southgate, Lauren Gilbert, Lee Graham, Nicole Pacchiarini, Sara Kumziene-Summerhayes, Sarah Taylor, Sophie Jones, Sara Rey, Matthew Bull, Joanne Watkins, Sally Corden, Tom Connor |
| EPI_ISL_651560, EPI_ISL_651562, EPI_ISL_651564, EPI_ISL_651581 | Department of Pathology, University of Cambridge | COVID-19 Genomics UK (COG-UK) Consortium | Aminu S. Jahun, Yasmin Chaudhry, Grant Hall, Iliana Georgana, Myra Hosmillo, Martin D. Curran, Malte Pinckert, Surendra Parmar, Ian Goodfellow |
| EPI_ISL_651624 | University of Birmingham | COVID-19 Genomics UK (COG-UK) Consortium | Institute of Microbiology, University of Birmingham: Claire McMurray, Joanne Stockton, Samuel Nicholls, Radoslaw Poplawski, Will Rowe, Josh Quick, Nicholas Loman. University of Birmingham Testing Laboratory: Celina M Whalley, Andrew Bosworth, Charlotte Poxon, Kasun Wanigasooriya, Oliver Pickles, Mike Kidd, Alex Richter, Andrew D Beggs PHE Heartlands Lab: Husam Osman, Andrew Bosworth. Queen Elizabeth Hospital: Anna Casey |

| | | | |
|---|---|---|---|
| EPI_ISL_651682, EPI_ISL_651683, EPI_ISL_651687, EPI_ISL_651688, EPI_ISL_651689, EPI_ISL_651690, EPI_ISL_651691, EPI_ISL_651692, EPI_ISL_651693, EPI_ISL_651694, EPI_ISL_651695, EPI_ISL_651696, EPI_ISL_651697, EPI_ISL_651698, EPI_ISL_651703, EPI_ISL_651704, EPI_ISL_651705, EPI_ISL_651706, EPI_ISL_651707 | | | |
| see above | Department of Pathology, University of Cambridge | COVID-19 Genomics UK (COG-UK) Consortium | Aminu S. Jahun, Yasmin Chaudhry, Grant Hall, Iliana Georgana, Myra Hosmillo, Martin D. Curran, Malte Pinckert, Surendra Parmar, Ian Goodfellow |
| EPI_ISL_651715, EPI_ISL_651716, EPI_ISL_651718, EPI_ISL_651719, EPI_ISL_651720, EPI_ISL_651721, EPI_ISL_651722, EPI_ISL_651723, EPI_ISL_651724, EPI_ISL_651725, EPI_ISL_651726, EPI_ISL_651727, EPI_ISL_651728, EPI_ISL_651729, EPI_ISL_651730, EPI_ISL_651731, EPI_ISL_651732 | | | |
| see above | Virology Department, Sheffield Teaching Hospitals NHS Foundation Trust/Department of Infection, Immunity and Cardiovascular Disease, The Medical School, University of Sheffield | COVID-19 Genomics UK (COG-UK) Consortium | Thushan de Silva, Matthew Parker, Nikki Smith, Adri Angyal, Rebecca Brown, Luke Green, Rachel Tucker, Paul Parsons, Danielle Groves, Katie Johnson, Laura Carrilero, Alex Keeley, Dave Partridge, Matthew Wyles, Benjamin Lindsey, Mehmet Yavuz, Mohammad Raza, Cariad Evans |
| EPI_ISL_651830, EPI_ISL_651831, EPI_ISL_651860, EPI_ISL_651861, EPI_ISL_651862, EPI_ISL_651864, EPI_ISL_651869, EPI_ISL_651870, EPI_ISL_651871, EPI_ISL_651872, EPI_ISL_651887, EPI_ISL_651892, EPI_ISL_651946 | | | |
| see above | West of Scotland Specialist Virology Centre, NHSGGC / MRC-University of Glasgow Centre for Virus Research | COVID-19 Genomics UK (COG-UK) Consortium | Ana da Silva Filipe, Natasha Johnson, Kathy Smollett, Daniel Mair, Stephen Carmichael, Alice Broos, Lily Tong, Jenna Nichols, Kyriaki Nomikou; Sarah McDonald; Richard Orton, Joseph Hughes, Sreenu Vattipally, David L Robertson; Alasdair MacLean, Rory Gunson; Sharif Shaaban, Matthew Holden; Rachel Blacow, Guy Mollett, Kathy Li, James Shepherd, Antonia Ho, Emma Thomson |
| EPI_ISL_652037, EPI_ISL_652049, EPI_ISL_652050, EPI_ISL_652052, EPI_ISL_652061, EPI_ISL_652065, EPI_ISL_652066 | Virology Department, Royal Infirmary of Edinburgh, NHS Lothian / School of Biological Sciences, University of Edinburgh / Institute of Genetics and Molecular Medicine, University of Edinburgh | COVID-19 Genomics UK (COG-UK) Consortium | McHugh M, Dewar R, Rooke S, Gallagher M, Balcaza C, O'Toole Á, Scher E, Hill V, McCrone JT, Colquhoun R, Yu X, Jackson B, Rambaut A, Williams TC, Templeton K |
| EPI_ISL_652290, EPI_ISL_652291, EPI_ISL_652292, EPI_ISL_652293, EPI_ISL_652294, EPI_ISL_652300, EPI_ISL_652302 | University of Exeter | COVID-19 Genomics UK (COG-UK) Consortium | Ben Temperton,Aaron Jeffries,Michelle Michelsen,Joanna Warwick-Dugdale,Audrey Farbos,Robyn Manley,Stephen Michell,Jane Masoli |
| EPI_ISL_652362 | Quadram Institute Bioscience | COVID-19 Genomics UK (COG-UK) Consortium | Dave J. Baker, Gemma L. Kay, Alp Aydin, Thanh Le-Viet, Steven Rudder, Ana P. Tedim, Anastasia Kolyva, Maria Diaz, Leonardo de Oliveira Martins, Nabil-Fareed Alikhan, Lizzie Meadows, Rachael Stanley, Ngozi Elumogo, Muhammed Yasir, Nicholas M. Thomson, Alexander J Trotter, Rachel Gilroy, Samuel Bloomfield, Claire Stuart, Andrew Bell, Reenesh Prakash, Samir Dervisevic, Alison E. Mather, John Wain, Mark Webber, Andrew J. Page, Justin O'Grady |
| EPI_ISL_652496, EPI_ISL_652497, EPI_ISL_652500, EPI_ISL_652501, EPI_ISL_652502, EPI_ISL_652508, EPI_ISL_652509, EPI_ISL_652510, EPI_ISL_652511, EPI_ISL_652513, EPI_ISL_652514, EPI_ISL_652516, EPI_ISL_652518, EPI_ISL_652519, EPI_ISL_652520, EPI_ISL_652525, EPI_ISL_652526, EPI_ISL_652528, EPI_ISL_652531, EPI_ISL_652532, EPI_ISL_652538, EPI_ISL_652540, EPI_ISL_652541, EPI_ISL_652542, EPI_ISL_652544, EPI_ISL_652547, EPI_ISL_652549, EPI_ISL_652558, EPI_ISL_652559, EPI_ISL_652563, EPI_ISL_652564, EPI_ISL_652565, EPI_ISL_652567, EPI_ISL_652569, EPI_ISL_652570, EPI_ISL_652577, EPI_ISL_652583, EPI_ISL_652588, EPI_ISL_652589, EPI_ISL_652597, EPI_ISL_652602, EPI_ISL_652604, EPI_ISL_652610, EPI_ISL_652611, EPI_ISL_652612, EPI_ISL_652614, EPI_ISL_652615, EPI_ISL_652618 | | | |
| see above | Wales Specialist Virology Centre Sequencing lab: Pathogen Genomics Unit | COVID-19 Genomics UK (COG-UK) Consortium | Catherine Moore, Johnathan Evans, Laura Gifford, Malorie Perry, Simon Cottrell, Angela Marchbank, Alec Birchley, Alexander Adams, Amy Gaskin, Bree Gatica-Wilcox, Jason Coombes, Joel Southgate, Lauren Gilbert, Lee Graham, Nicole Pacchiarini, Sara Kumziene-Summerhayes, Sarah Taylor, Sophie Jones, Sara Rey, Matthew Bull, Joanne Watkins, Sally Corden, Tom Connor |
| EPI_ISL_653061, EPI_ISL_653064, EPI_ISL_653072, EPI_ISL_653081 | Virology Department, Sheffield Teaching Hospitals NHS Foundation Trust/Department of Infection, Immunity and Cardiovascular Disease, The Medical School, University of Sheffield | COVID-19 Genomics UK (COG-UK) Consortium | Thushan de Silva, Matthew Parker, Nikki Smith, Adri Angyal, Rebecca Brown, Luke Green, Rachel Tucker, Paul Parsons, Danielle Groves, Katie Johnson, Laura Carrilero, Alex Keeley, Dave Partridge, Matthew Wyles, Benjamin Lindsey, Mehmet Yavuz, Mohammad Raza, Cariad Evans |
| EPI_ISL_653765, EPI_ISL_653766, EPI_ISL_653767, EPI_ISL_653768, EPI_ISL_653769, EPI_ISL_653770, EPI_ISL_653771, EPI_ISL_653772, EPI_ISL_653773, EPI_ISL_653774, EPI_ISL_653775, EPI_ISL_653776, EPI_ISL_653777, EPI_ISL_653778, EPI_ISL_653779, EPI_ISL_653780, EPI_ISL_653781, EPI_ISL_653783, EPI_ISL_653787, EPI_ISL_653788, EPI_ISL_653789, EPI_ISL_653790, EPI_ISL_653791, EPI_ISL_653792, EPI_ISL_653793 | | | |
| see above | I.R.C.C.S. "S. De Bellis" - Ente Ospedaliero | Istituto Zooprofilattico Sperimentale della Puglia e della Basilicata | Parisi A., Bianco A., Capozzi L., Del Sambro L., Lippolis A., Notarnicola M., Manzulli V, Rondinone V., Pace L. |
| EPI_ISL_653794, EPI_ISL_653795, EPI_ISL_653796, EPI_ISL_653797, EPI_ISL_653798, EPI_ISL_653799, EPI_ISL_653800, EPI_ISL_653801, EPI_ISL_653804, EPI_ISL_653805, EPI_ISL_653806, EPI_ISL_653807, EPI_ISL_653808, EPI_ISL_653809, EPI_ISL_653811, EPI_ISL_653812 | | | |
| see above | I.R.C.C.S. "S. De Bellis" - Ente Ospedaliero | Istituto Zooprofilattico Sperimentale della Puglia e della Basilicata | Parisi A., Bianco A., Capozzi L., Del Sambro L., Lippolis A., Notarnicola M., Cipolletta D., Galante D. |
| EPI_ISL_654017, EPI_ISL_654018, EPI_ISL_654019 | Laboratory of Microbiology, National Reference Lab, Charles Nicolle Hospital; 2-University of Tunis ElManar, Faculty of Medicine of Tunis, LR99ES09, Tunis, Tunisia | 1-Clinical and Experimental Pharmacology Lab, LR16SP02, National Center of Pharmacovigilance, University of Tunis El Manar, Tunis, Tunisia. 2-Neurodegenerative diseases and psychiatric troubles, LR18SP03, Razi Hospital, University of Tunis El Manar, Tunis, Tunisia. 3- Ministry of Health, National Observatory of New and Emerging Diseases, 1006, Tunis, Tunisia | Ilhem Boutiba-Ben Boubaker, Sameh Trabelsi, Nissaf Ben Alaya, Maher Kharrat, Alia Ben Kahla, Jalila Ben Khelil, Salma Abid, Sana Ferjani, Mouna Ben Sassi, Mouna Safer, Imen Mkada, Imen Kacem, Gaies Emna, Soumaya Rammeh, Riadh Daghfous, Riadh Gouider. |
| EPI_ISL_654703, EPI_ISL_654705, EPI_ISL_654706, EPI_ISL_654707, EPI_ISL_654708, EPI_ISL_654709, EPI_ISL_654710, EPI_ISL_654711, EPI_ISL_654712, EPI_ISL_654713, EPI_ISL_654714, EPI_ISL_654715, EPI_ISL_654716, EPI_ISL_654717, EPI_ISL_654718, EPI_ISL_654719, EPI_ISL_654720, EPI_ISL_654721, EPI_ISL_654722, EPI_ISL_654723, EPI_ISL_654724, EPI_ISL_654727 | | | |
| see above | Minnesota Department of Health, Public Health Laboratory | Minnesota Department of Health, Public Health Laboratory | Matt Plumb, Jacob Garfin, Alexandra Lorentz, and Xiong Wang |
| EPI_ISL_657472, EPI_ISL_657861, EPI_ISL_657991, EPI_ISL_658261, EPI_ISL_658754, EPI_ISL_658755, EPI_ISL_658765, EPI_ISL_658772, EPI_ISL_658777, EPI_ISL_658780, EPI_ISL_658789, EPI_ISL_658791, EPI_ISL_658803, EPI_ISL_658804, EPI_ISL_658812, EPI_ISL_658821, EPI_ISL_658827, EPI_ISL_658832, EPI_ISL_658847, EPI_ISL_658851, EPI_ISL_658855 | | | |
| see above | Lighthouse Lab in Milton Keynes | Wellcome Sanger Institute for the COVID-19 Genomics UK (COG-UK) Consortium | The Lighthouse Lab in Milton Keynes and Alex Alderton, Roberto Amato, Sonia Goncalves, Ewan Harrison, David K. Jackson, Ian Johnston, Dominic Kwiatkowski, Cordelia Langford, John Sillitoe on behalf of the Wellcome Sanger Institute COVID-19 Surveillance Team |
| EPI_ISL_658863, EPI_ISL_658865, EPI_ISL_658868, EPI_ISL_658870 | Instituto de Diagnostico y Referencia Epidemiologicos (INDRE) | Instituto de Diagnostico y Referencia Epidemiologicos (INDRE) | Gisela Barrera-Badillo , Abril Rodriguez-Maldonado, Claudia Wong-Arambula , Natividad Cruz-Ortiz, Tatiana Nunez-Garcia, Dayanira Arellano-Suarez, Fabiola Garces-Ayala, Lucia Hernandez-Rivas, Irma Lopez-Martinez, Ernesto Ramirez-Gonzalez. |
| EPI_ISL_658873, EPI_ISL_658875 | Instituto de Diagnostico y Referencia Epidemiologicos (INDRE) | Instituto de Diagnostico y Referencia Epidemiologicos (INDRE) | Gisela Barrera-Badillo , Abril Rodriguez-Maldonado, Claudia Wong-Arambula , Natividad Cruz-Ortiz, Tatiana Nunez-Garcia, Dayanira Arellano-Suarez, Fabiola Garces-Ayala, Lucia Hernandez-Rivas, Irma Lopez-Martinez, Ernesto Ramirez-Gonzalez. |
| EPI_ISL_658878 | Instituto de Diagnostico y Referencia Epidemiologicos (INDRE) | Instituto de Diagnostico y Referencia Epidemiologicos (INDRE) | Ernesto Ramirez-Gonzalez, Abril Rodriguez-Maldonado, Claudia Wong-Arambula , Natividad Cruz-Ortiz, Tatiana Nunez-Garcia, Dayanira Arellano-Suarez, Fabiola Garces-Ayala, Lucia Hernandez-Rivas, Irma Lopez-Martinez, Gisela Barrera-Badillo. |
| EPI_ISL_658879 | Lighthouse Lab in Milton Keynes | Wellcome Sanger Institute for the COVID-19 Genomics UK (COG-UK) Consortium | The Lighthouse Lab in Milton Keynes and Alex Alderton, Roberto Amato, Sonia Goncalves, Ewan Harrison, David K. Jackson, Ian Johnston, Dominic Kwiatkowski, Cordelia Langford, John Sillitoe on behalf of the Wellcome Sanger Institute COVID-19 Surveillance Team |
| EPI_ISL_658880, EPI_ISL_658883, EPI_ISL_658886, EPI_ISL_658888, EPI_ISL_658893 | Instituto de Diagnostico y Referencia Epidemiologicos (INDRE) | Instituto de Diagnostico y Referencia Epidemiologicos (INDRE) | Ernesto Ramirez-Gonzalez, Abril Rodriguez-Maldonado, Claudia Wong-Arambula , Natividad Cruz-Ortiz, Tatiana Nunez-Garcia, Dayanira Arellano-Suarez, Fabiola Garces-Ayala, Lucia Hernandez-Rivas, Irma Lopez-Martinez, Gisela Barrera-Badillo. |
| EPI_ISL_658908, EPI_ISL_658922, EPI_ISL_658928, EPI_ISL_658935, EPI_ISL_658938, EPI_ISL_658951, EPI_ISL_658955, EPI_ISL_658985, EPI_ISL_658994, EPI_ISL_659001, EPI_ISL_659008, EPI_ISL_659009, EPI_ISL_659014, EPI_ISL_659021, EPI_ISL_659023, EPI_ISL_659026, EPI_ISL_659032, EPI_ISL_659033, EPI_ISL_659034, EPI_ISL_659037, EPI_ISL_659041, EPI_ISL_659056, EPI_ISL_659088 | | | |
| see above | Lighthouse Lab in Milton Keynes | Wellcome Sanger Institute for the COVID-19 Genomics UK (COG-UK) Consortium | The Lighthouse Lab in Milton Keynes and Alex Alderton, Roberto Amato, Sonia Goncalves, Ewan Harrison, David K. Jackson, Ian Johnston, Dominic Kwiatkowski, Cordelia Langford, John Sillitoe on behalf of the Wellcome Sanger Institute COVID-19 Surveillance Team |
| EPI_ISL_659135, EPI_ISL_659163, EPI_ISL_659167, EPI_ISL_659345, EPI_ISL_659395 | Lighthouse Lab in Cambridge | Wellcome Sanger Institute for the COVID-19 Genomics UK (COG-UK) Consortium | Rob Howes, The Lighthouse Lab in Cambridge and Alex Alderton, Roberto Amato, Sonia Goncalves, Ewan Harrison, David K. Jackson, Ian Johnston, Dominic Kwiatkowski, Cordelia Langford, John Sillitoe on behalf of the Wellcome Sanger Institute COVID-19 Surveillance Team |

| | | | |
|---|---|---|---|
| EPI_ISL_659745, EPI_ISL_659746, EPI_ISL_659747, EPI_ISL_659749, EPI_ISL_659751, EPI_ISL_659754, EPI_ISL_659758, EPI_ISL_659766, EPI_ISL_659773, EPI_ISL_659774, EPI_ISL_659775, EPI_ISL_659778, EPI_ISL_659780, EPI_ISL_659781, EPI_ISL_659782, EPI_ISL_659785, EPI_ISL_659786, EPI_ISL_659791, EPI_ISL_659796, EPI_ISL_659797, EPI_ISL_659799, EPI_ISL_659800, EPI_ISL_659803, EPI_ISL_659806, EPI_ISL_659810, EPI_ISL_659813, EPI_ISL_659814, EPI_ISL_659816, EPI_ISL_659817, EPI_ISL_659819, EPI_ISL_659822, EPI_ISL_659829, EPI_ISL_659831, EPI_ISL_659834, EPI_ISL_659838, EPI_ISL_659839, EPI_ISL_659841, EPI_ISL_659844, EPI_ISL_659847, EPI_ISL_659848, EPI_ISL_659849, EPI_ISL_659851, EPI_ISL_659854, EPI_ISL_659855, EPI_ISL_659859, EPI_ISL_659861, EPI_ISL_659862, EPI_ISL_659865, EPI_ISL_659866, EPI_ISL_659870, EPI_ISL_659873, EPI_ISL_659876, EPI_ISL_659878, EPI_ISL_659881, EPI_ISL_659885, EPI_ISL_659886, EPI_ISL_659887, EPI_ISL_659889, EPI_ISL_659893, EPI_ISL_659894, EPI_ISL_659895, EPI_ISL_659898, EPI_ISL_659899, EPI_ISL_659901, EPI_ISL_659902, EPI_ISL_659911, EPI_ISL_659912, EPI_ISL_659914, EPI_ISL_659915, EPI_ISL_659916, EPI_ISL_659917, EPI_ISL_659921, EPI_ISL_659922, EPI_ISL_659923, EPI_ISL_659925, EPI_ISL_659927, EPI_ISL_659929, EPI_ISL_659930, EPI_ISL_659932, EPI_ISL_659933, EPI_ISL_659936, EPI_ISL_659937, EPI_ISL_659938, EPI_ISL_659940, EPI_ISL_659943, EPI_ISL_659945, EPI_ISL_659946, EPI_ISL_659950, EPI_ISL_659952, EPI_ISL_659953, EPI_ISL_659958, EPI_ISL_659959, EPI_ISL_659960, EPI_ISL_659965, EPI_ISL_659969, EPI_ISL_659972, EPI_ISL_659976, EPI_ISL_659978, EPI_ISL_659982 | | | |
| see above | Lighthouse Lab in Glasgow | Wellcome Sanger Institute for the COVID-19 Genomics UK (COG-UK) Consortium | Harper VanSteenhouse, Yumi Kasai, David Gray, Carol Clugston, Anna Dominiczak and Alex Alderton, Roberto Amato, Sonia Goncalves, Ewan Harrison, David K. Jackson, Ian Johnston, Dominic Kwiatkowski, Cordelia Langford, John Sillitoe on behalf of the Wellcome Sanger Institute COVID-19 Surveillance Team |
| EPI_ISL_659985, EPI_ISL_659991, EPI_ISL_659994, EPI_ISL_659996, EPI_ISL_659997, EPI_ISL_660000, EPI_ISL_660002, EPI_ISL_660003, EPI_ISL_660004, EPI_ISL_660005, EPI_ISL_660007, EPI_ISL_660009, EPI_ISL_660010, EPI_ISL_660017, EPI_ISL_660018, EPI_ISL_660021, EPI_ISL_660028, EPI_ISL_660030, EPI_ISL_660032, EPI_ISL_660033, EPI_ISL_660035, EPI_ISL_660039, EPI_ISL_660041, EPI_ISL_660044, EPI_ISL_660045, EPI_ISL_660047, EPI_ISL_660050, EPI_ISL_660051, EPI_ISL_660052, EPI_ISL_660055, EPI_ISL_660056, EPI_ISL_660057, EPI_ISL_660058, EPI_ISL_660059, EPI_ISL_660063, EPI_ISL_660065, EPI_ISL_660066 | | | |
| see above | Lighthouse Lab in Glasgow | Wellcome Sanger Institute for the COVID-19 Genomics UK (COG-UK) Consortium | Harper VanSteenhouse, Yumi Kasai, David Gray, Carol Clugston, Anna Dominiczak and Alex Alderton, Roberto Amato, Sonia Goncalves, Ewan Harrison, David K. Jackson, Ian Johnston, Dominic Kwiatkowski, Cordelia Langford, John Sillitoe on behalf of the Wellcome Sanger Institute COVID-19 Surveillance Team (http://www.sanger.ac.uk/covid-team) |
| EPI_ISL_660086, EPI_ISL_660087, EPI_ISL_660088, EPI_ISL_660089, EPI_ISL_660090, EPI_ISL_660091, EPI_ISL_660092, EPI_ISL_660093, EPI_ISL_660097 | MD PHL | MD PHL | Maryland Department of Health Laboratories Administration |
| EPI_ISL_660225 | NHLS-IALCH | KRISP, KZN Research Innovation and Sequencing Platform | Giandhari J, Pillay S, Lessells R, Mdlalose K, York D, Khan S, Tegally H, Wilkinson E, de Oliveira T |
| EPI_ISL_660324, EPI_ISL_660325 | General practitioner | National Reference Center for Viruses of Respiratory Infections, Institut Pasteur, Paris | Marion Barbet, Sylvie Behillil, Méline Bizard, Angela Brisebarre, Camille Capel, Etienne Simon-Lorière, Vincent Enouf, Maud Vanpeene, Sylvie van der Werf |
| EPI_ISL_660473, EPI_ISL_660519 | Laboratoire de Microbiologie CHU Souro Sanou | Centre Muraz | Abdoul-Salam Ouedraogo, Yacouba Sawadogo, Essia Belarbi, Grit Schubert, Fabian Leendertz, Arsène Zongo, Soumeya Ouangraoua, Zekiba Tarnagda, Lassana Sangaré, Halidou Tinto |
| EPI_ISL_660637, EPI_ISL_660639, EPI_ISL_660642, EPI_ISL_660644, EPI_ISL_660645, EPI_ISL_660650, EPI_ISL_660651, EPI_ISL_660653, EPI_ISL_660654, EPI_ISL_660655, EPI_ISL_660656, EPI_ISL_660659, EPI_ISL_660660, EPI_ISL_660661, EPI_ISL_660663 | | | |
| see above | NHLS-IALCH | KRISP, KZN Research Innovation and Sequencing Platform | Giandhari J, Pillay S, Lessells R, Mdlalose K, York D, Khan S, Tegally H, Wilkinson E, de Oliveira T |
| EPI_ISL_660782, EPI_ISL_660784, EPI_ISL_660785, EPI_ISL_660786, EPI_ISL_660787, EPI_ISL_660788, EPI_ISL_660789, EPI_ISL_660790, EPI_ISL_660791, EPI_ISL_660792, EPI_ISL_660793 | | | |
| see above | Respiratory Virus Unit, Microbiology Services Colindale, Public Health England | COVID-19 Genomics UK (COG-UK) Consortium | PHE Covid Sequencing Team |
| EPI_ISL_661045, EPI_ISL_661046, EPI_ISL_661047, EPI_ISL_661048, EPI_ISL_661049, EPI_ISL_661050, EPI_ISL_661051, EPI_ISL_661052, EPI_ISL_661053, EPI_ISL_661054, EPI_ISL_661055, EPI_ISL_661056, EPI_ISL_661057, EPI_ISL_661058, EPI_ISL_661059, EPI_ISL_661060, EPI_ISL_661061, EPI_ISL_661062, EPI_ISL_661063, EPI_ISL_661064, EPI_ISL_661065, EPI_ISL_661066, EPI_ISL_661067, EPI_ISL_661068, EPI_ISL_661069, EPI_ISL_661070, EPI_ISL_661071, EPI_ISL_661072, EPI_ISL_661073, EPI_ISL_661074, EPI_ISL_661075, EPI_ISL_661076, EPI_ISL_661077, EPI_ISL_661078, EPI_ISL_661079, EPI_ISL_661080 | | | |
| see above | Gundersen Molecular Diagnostics Laboratory | Kabara Cancer Research Institute | Craig S. Richmond, Paraic A. Kenny |
| EPI_ISL_661294 | Unilabs | The Public Health Agency of Sweden | Department of Microbiology, The Public Health Agency of Sweden |
| EPI_ISL_661300 | Laboratoriemedicin, Klinisk mikrobiologi | The Public Health Agency of Sweden | Department of Microbiology, The Public Health Agency of Sweden |
| EPI_ISL_662638, EPI_ISL_662642, EPI_ISL_662643, EPI_ISL_662645 | Lighthouse Lab in Milton Keynes | Wellcome Sanger Institute for the COVID-19 Genomics UK (COG-UK) Consortium | The Lighthouse Lab in Milton Keynes and Alex Alderton, Roberto Amato, Sonia Goncalves, Ewan Harrison, David K. Jackson, Ian Johnston, Dominic Kwiatkowski, Cordelia Langford, John Sillitoe on behalf of the Wellcome Sanger Institute COVID-19 Surveillance Team |
| EPI_ISL_662654 | Lighthouse Lab in Glasgow | Wellcome Sanger Institute for the COVID-19 Genomics UK (COG-UK) Consortium | Harper VanSteenhouse, Yumi Kasai, David Gray, Carol Clugston, Anna Dominiczak and Alex Alderton, Roberto Amato, Sonia Goncalves, Ewan Harrison, David K. Jackson, Ian Johnston, Dominic Kwiatkowski, Cordelia Langford, John Sillitoe on behalf of the Wellcome Sanger Institute COVID-19 Surveillance Team |
| EPI_ISL_664137 | Centre for Enzyme Innovation, University of Portsmouth / Translational Research Laboratory, Portsmouth Hospitals NHS Trust | COVID-19 Genomics UK (COG-UK) Consortium | Angela Beckett,Yann Bourgeois,Garry Scarlett,Sharon Glaysher,Scott Elliott,Kelly Bicknell,Robert Impey,Allyson Lloyd,Sarah Wyllie,Ethan Butcher,Anoop Chauhan,Samuel Robson |
| EPI_ISL_664170 | University of Exeter | COVID-19 Genomics UK (COG-UK) Consortium | Ben Temperton,Aaron Jeffries,Michelle Michelsen,Joanna Warwick-Dugdale,Audrey Farbos,Robyn Manley,Stephen Michell,Jane Masoli |
| EPI_ISL_664191 | Centre for Enzyme Innovation, University of Portsmouth / Translational Research Laboratory, Portsmouth Hospitals NHS Trust | COVID-19 Genomics UK (COG-UK) Consortium | Angela Beckett,Yann Bourgeois,Garry Scarlett,Sharon Glaysher,Scott Elliott,Kelly Bicknell,Robert Impey,Allyson Lloyd,Sarah Wyllie,Ethan Butcher,Anoop Chauhan,Samuel Robson |
| EPI_ISL_664230 | Liverpool Clinical Laboratories | COVID-19 Genomics UK (COG-UK) Consortium | Sam Haldenby, Anita Lucaci, Steve Paterson, Julian Hiscox, Alistair Darby, M Almsaud, A Alrezaihi, Muhannad Alruwaili, Stuart D Armstrong, Jones Benjamin, Eleanor G Bentley, Anu Chawla, Jordan J Clark, Angela Cowell, Richard Eccles, Isabel García-Dorival, Matthew Gemmell, Alessandro Gerada, PKF Gilmore, Richard Gregory, Ximeng Han, Catherine Hartley, Margaret Hughes, Miren Iturriza-Gomara, James Johnson, L Luu, Jenifer Manson, Charlotte Nelson, Elaine O'Toole, Cassie Olateju, Rebekah Penrice-Randal , Lucille Rainbow, N.P Randle, Trevor Ian Robinson, Parul Sharma, Ghada T Shawli, James P Stewart, Neil Swainston, Ecaterina Vamos, Joanne Watts, Mark Whitehead |
| EPI_ISL_664283 | Virology Department, Sheffield Teaching Hospitals NHS Foundation Trust/Department of Infection, Immunity and Cardiovascular Disease, The Medical School, University of Sheffield | COVID-19 Genomics UK (COG-UK) Consortium | Thushan de Silva, Matthew Parker, Nikki Smith, Adri Angyal, Rebecca Brown, Luke Green, Rachel Tucker, Paul Parsons, Danielle Groves, Katie Johnson, Laura Carrilero, Alex Keeley, Dave Partridge, Matthew Wyles, Benjamin Lindsey, Mehmet Yavuz, Mohammad Raza, Cariad Evans |
| EPI_ISL_664295 | University of Exeter | COVID-19 Genomics UK (COG-UK) Consortium | Ben Temperton,Aaron Jeffries,Michelle Michelsen,Joanna Warwick-Dugdale,Audrey Farbos,Robyn Manley,Stephen Michell,Jane Masoli |
| EPI_ISL_664298 | Virology Department, Sheffield Teaching Hospitals NHS Foundation Trust/Department of Infection, Immunity and Cardiovascular Disease, The Medical School, University of Sheffield | COVID-19 Genomics UK (COG-UK) Consortium | Thushan de Silva, Matthew Parker, Nikki Smith, Adri Angyal, Rebecca Brown, Luke Green, Rachel Tucker, Paul Parsons, Danielle Groves, Katie Johnson, Laura Carrilero, Alex Keeley, Dave Partridge, Matthew Wyles, Benjamin Lindsey, Mehmet Yavuz, Mohammad Raza, Cariad Evans |
| EPI_ISL_664339 | Liverpool Clinical Laboratories | COVID-19 Genomics UK (COG-UK) Consortium | Sam Haldenby, Anita Lucaci, Steve Paterson, Julian Hiscox, Alistair Darby, M Almsaud, A Alrezaihi, Muhannad Alruwaili, Stuart D Armstrong, Jones Benjamin, Eleanor G Bentley, Anu Chawla, Jordan J Clark, Angela Cowell, Richard Eccles, Isabel García-Dorival, Matthew Gemmell, Alessandro Gerada, PKF Gilmore, Richard Gregory, Ximeng Han, Catherine Hartley, Margaret Hughes, Miren Iturriza-Gomara, James Johnson, L Luu, Jenifer Manson, Charlotte Nelson, Elaine O'Toole, Cassie Olateju, Rebekah Penrice-Randal , Lucille Rainbow, N.P Randle, Trevor Ian Robinson, Parul Sharma, Ghada T Shawli, James P Stewart, Neil Swainston, Ecaterina Vamos, Joanne Watts, Mark Whitehead |
| EPI_ISL_664359 | University College London Hospital | COVID-19 Genomics UK (COG-UK) Consortium | Judith Heaney, Matthew Byott, Catherine Houlihan, Dan Frampton, Stuart Kirk, Moira Spyer and Eleni Nastouli |
| EPI_ISL_664370 | Virology Department, Sheffield Teaching Hospitals NHS Foundation Trust/Department of Infection, Immunity and Cardiovascular Disease, The Medical School, University of Sheffield | COVID-19 Genomics UK (COG-UK) Consortium | Thushan de Silva, Matthew Parker, Nikki Smith, Adri Angyal, Rebecca Brown, Luke Green, Rachel Tucker, Paul Parsons, Danielle Groves, Katie Johnson, Laura Carrilero, Alex Keeley, Dave Partridge, Matthew Wyles, Benjamin Lindsey, Mehmet Yavuz, Mohammad Raza, Cariad Evans |
| EPI_ISL_664373 | Northumbria University / South Tees Hospitals NHS Foundation Trust / North Cumbria Integrated Care NHS | COVID-19 Genomics UK (COG-UK) Consortium | Darren L Smith,Andrew Nelson,Matthew Bashton,Greg R Young,Joshua Loh,John Allan,Mohammad A Tariq,Giles S Holt,Gary Black,Wen C Yew,Lynn Dover,Paul Baker,Steve Liggett,Sarah Essex,Jane Greenaway,Debra Padgett,Clive Graham,Garren Scott,Edward Barton,Emma Swindells,Brendan |

| | | | |
|---|---|---|---|
| | Foundation Trust / North Tees and Hartlepool NHS Foundation Trust / Newcastle Hospitals NHS Foundation Trust | | Payne,Jennifer Collins,Yusri Taha,Gary Eltringham |
| EPI_ISL_664374, EPI_ISL_664375 | Centre for Enzyme Innovation, University of Portsmouth / Translational Research Laboratory, Portsmouth Hospitals NHS Trust | COVID-19 Genomics UK (COG-UK) Consortium | Angela Beckett,Yann Bourgeois,Garry Scarlett,Sharon Glaysher,Scott Elliott,Kelly Bicknell,Robert Impey,Allyson Lloyd,Sarah Wyllie,Ethan Butcher,Anoop Chauhan,Samuel Robson |
| EPI_ISL_664398 | University of Exeter | COVID-19 Genomics UK (COG-UK) Consortium | Ben Temperton,Aaron Jeffries,Michelle Michelsen,Joanna Warwick-Dugdale,Audrey Farbos,Robyn Manley,Stephen Michell,Jane Masoli |
| EPI_ISL_664425, EPI_ISL_664426, EPI_ISL_664431, EPI_ISL_664437 | Virology Department, Sheffield Teaching Hospitals NHS Foundation Trust/Department of Infection, Immunity and Cardiovascular Disease, The Medical School, University of Sheffield | COVID-19 Genomics UK (COG-UK) Consortium | Thushan de Silva, Matthew Parker, Nikki Smith, Adri Angyal, Rebecca Brown, Luke Green, Rachel Tucker, Paul Parsons, Danielle Groves, Katie Johnson, Laura Carrilero, Alex Keeley, Dave Partridge, Matthew Wyles, Benjamin Lindsey, Mehmet Yavuz, Mohammad Raza, Cariad Evans |
| EPI_ISL_664526, EPI_ISL_664556 | Centre for Enzyme Innovation, University of Portsmouth / Translational Research Laboratory, Portsmouth Hospitals NHS Trust | COVID-19 Genomics UK (COG-UK) Consortium | Angela Beckett,Yann Bourgeois,Garry Scarlett,Sharon Glaysher,Scott Elliott,Kelly Bicknell,Robert Impey,Allyson Lloyd,Sarah Wyllie,Ethan Butcher,Anoop Chauhan,Samuel Robson |
| EPI_ISL_664600, EPI_ISL_664602, EPI_ISL_664604 | Virology Department, Sheffield Teaching Hospitals NHS Foundation Trust/Department of Infection, Immunity and Cardiovascular Disease, The Medical School, University of Sheffield | COVID-19 Genomics UK (COG-UK) Consortium | Thushan de Silva, Matthew Parker, Nikki Smith, Adri Angyal, Rebecca Brown, Luke Green, Rachel Tucker, Paul Parsons, Danielle Groves, Katie Johnson, Laura Carrilero, Alex Keeley, Dave Partridge, Matthew Wyles, Benjamin Lindsey, Mehmet Yavuz, Mohammad Raza, Cariad Evans |
| EPI_ISL_664614 | Liverpool Clinical Laboratories | COVID-19 Genomics UK (COG-UK) Consortium | Sam Haldenby, Anita Lucaci, Steve Paterson, Julian Hiscox, Alistair Darby, M Almsaud, A Alrezaihi, Muhannad Alruwaili, Stuart D Armstrong, Jones Benjamin, Eleanor G Bentley, Anu Chawla, Jordan J Clark, Angela Cowell, Richard Eccles, Isabel García-Dorival, Matthew Gemmell, Alessandro Gerada, PKF Gilmore, Richard Gregory, Ximeng Han, Catherine Hartley, Margaret Hughes, Miren Iturriza-Gomara, James Johnson, L Luu, Jenifer Manson, Charlotte Nelson, Elaine O'Toole, Cassie Olateju, Rebekah Penrice-Randal , Lucille Rainbow, N.P Randle, Trevor Ian Robinson, Parul Sharma, Ghada T Shawli, James P Stewart, Neil Swainston, Ecaterina Vamos, Joanne Watts, Mark Whitehead |
| EPI_ISL_664630 | Virology Department, Sheffield Teaching Hospitals NHS Foundation Trust/Department of Infection, Immunity and Cardiovascular Disease, The Medical School, University of Sheffield | COVID-19 Genomics UK (COG-UK) Consortium | Thushan de Silva, Matthew Parker, Nikki Smith, Adri Angyal, Rebecca Brown, Luke Green, Rachel Tucker, Paul Parsons, Danielle Groves, Katie Johnson, Laura Carrilero, Alex Keeley, Dave Partridge, Matthew Wyles, Benjamin Lindsey, Mehmet Yavuz, Mohammad Raza, Cariad Evans |
| EPI_ISL_664679 | Centre for Enzyme Innovation, University of Portsmouth / Translational Research Laboratory, Portsmouth Hospitals NHS Trust | COVID-19 Genomics UK (COG-UK) Consortium | Angela Beckett,Yann Bourgeois,Garry Scarlett,Sharon Glaysher,Scott Elliott,Kelly Bicknell,Robert Impey,Allyson Lloyd,Sarah Wyllie,Ethan Butcher,Anoop Chauhan,Samuel Robson |
| EPI_ISL_664699 | University of Exeter | COVID-19 Genomics UK (COG-UK) Consortium | Ben Temperton,Aaron Jeffries,Michelle Michelsen,Joanna Warwick-Dugdale,Audrey Farbos,Robyn Manley,Stephen Michell,Jane Masoli |
| EPI_ISL_664713 | Virology Department, Sheffield Teaching Hospitals NHS Foundation Trust/Department of Infection, Immunity and Cardiovascular Disease, The Medical School, University of Sheffield | COVID-19 Genomics UK (COG-UK) Consortium | Thushan de Silva, Matthew Parker, Nikki Smith, Adri Angyal, Rebecca Brown, Luke Green, Rachel Tucker, Paul Parsons, Danielle Groves, Katie Johnson, Laura Carrilero, Alex Keeley, Dave Partridge, Matthew Wyles, Benjamin Lindsey, Mehmet Yavuz, Mohammad Raza, Cariad Evans |
| EPI_ISL_664734 | University College London Hospital | COVID-19 Genomics UK (COG-UK) Consortium | Judith Heaney, Matthew Byott, Catherine Houlihan, Dan Frampton, Stuart Kirk, Moira Spyer and Eleni Nastouli |
| EPI_ISL_664740 | University of Exeter | COVID-19 Genomics UK (COG-UK) Consortium | Ben Temperton,Aaron Jeffries,Michelle Michelsen,Joanna Warwick-Dugdale,Audrey Farbos,Robyn Manley,Stephen Michell,Jane Masoli |
| EPI_ISL_664746 | Centre for Enzyme Innovation, University of Portsmouth / Translational Research Laboratory, Portsmouth Hospitals NHS Trust | COVID-19 Genomics UK (COG-UK) Consortium | Angela Beckett,Yann Bourgeois,Garry Scarlett,Sharon Glaysher,Scott Elliott,Kelly Bicknell,Robert Impey,Allyson Lloyd,Sarah Wyllie,Ethan Butcher,Anoop Chauhan,Samuel Robson |
| EPI_ISL_664755 | University College London Hospital | COVID-19 Genomics UK (COG-UK) Consortium | Judith Heaney, Matthew Byott, Catherine Houlihan, Dan Frampton, Stuart Kirk, Moira Spyer and Eleni Nastouli |
| EPI_ISL_664840, EPI_ISL_664843, EPI_ISL_664844, EPI_ISL_664845, EPI_ISL_664847, EPI_ISL_664849 | Centre for Enzyme Innovation, University of Portsmouth / Translational Research Laboratory, Portsmouth Hospitals NHS Trust | COVID-19 Genomics UK (COG-UK) Consortium | Angela Beckett,Yann Bourgeois,Garry Scarlett,Sharon Glaysher,Scott Elliott,Kelly Bicknell,Robert Impey,Allyson Lloyd,Sarah Wyllie,Ethan Butcher,Anoop Chauhan,Samuel Robson |
| EPI_ISL_665023, EPI_ISL_665024, EPI_ISL_665025, EPI_ISL_665026, EPI_ISL_665027, EPI_ISL_665028, EPI_ISL_665029, EPI_ISL_665030, EPI_ISL_665031, EPI_ISL_665032, EPI_ISL_665033, EPI_ISL_665034, EPI_ISL_665035, EPI_ISL_665036, EPI_ISL_665037, EPI_ISL_665038, EPI_ISL_665039, EPI_ISL_665040, EPI_ISL_665041, EPI_ISL_665042, EPI_ISL_665043, EPI_ISL_665044, EPI_ISL_665045 | | | |
| see above | Virology Department, Sheffield Teaching Hospitals NHS Foundation Trust/Department of Infection, Immunity and Cardiovascular Disease, The Medical School, University of Sheffield | COVID-19 Genomics UK (COG-UK) Consortium | Thushan de Silva, Matthew Parker, Nikki Smith, Adri Angyal, Rebecca Brown, Luke Green, Rachel Tucker, Paul Parsons, Danielle Groves, Katie Johnson, Laura Carrilero, Alex Keeley, Dave Partridge, Matthew Wyles, Benjamin Lindsey, Mehmet Yavuz, Mohammad Raza, Cariad Evans |
| EPI_ISL_665243, EPI_ISL_665244, EPI_ISL_665245, EPI_ISL_665246, EPI_ISL_665247, EPI_ISL_665248, EPI_ISL_665249, EPI_ISL_665250 | University College London Hospital | COVID-19 Genomics UK (COG-UK) Consortium | Judith Heaney, Matthew Byott, Catherine Houlihan, Dan Frampton, Stuart Kirk, Moira Spyer and Eleni Nastouli |
| EPI_ISL_665292 | Quadram Institute Bioscience | COVID-19 Genomics UK (COG-UK) Consortium | Dave J. Baker, Gemma L. Kay, Alp Aydin, Thanh Le-Viet, Steven Rudder, Ana P. Tedim, Anastasia Kolyva, Maria Diaz, Leonardo de Oliveira Martins, Nabil-Fareed Alikhan, Lizzie Meadows, Rachael Stanley, Ngozi Elumogo, Muhammed Yasir, Nicholas M. Thomson, Alexander J Trotter, Rachel Gilroy, Samuel Bloomfield, Claire Stuart, Andrew Bell, Reenesh Prakash, Samir Dervisevic, Alison E. Mather, John Wain, Mark Webber, Andrew J. Page, Justin O'Grady |
| EPI_ISL_665294 | Liverpool Clinical Laboratories | COVID-19 Genomics UK (COG-UK) Consortium | Sam Haldenby, Anita Lucaci, Steve Paterson, Julian Hiscox, Alistair Darby, M Almsaud, A Alrezaihi, Muhannad Alruwaili, Stuart D Armstrong, Jones Benjamin, Eleanor G Bentley, Anu Chawla, Jordan J Clark, Angela Cowell, Richard Eccles, Isabel García-Dorival, Matthew Gemmell, Alessandro Gerada, PKF Gilmore, Richard Gregory, Ximeng Han, Catherine Hartley, Margaret Hughes, Miren Iturriza-Gomara, James Johnson, L Luu, Jenifer Manson, Charlotte Nelson, Elaine O'Toole, Cassie Olateju, Rebekah Penrice-Randal , Lucille Rainbow, N.P Randle, Trevor Ian Robinson, Parul Sharma, Ghada T Shawli, James P Stewart, Neil Swainston, Ecaterina Vamos, Joanne Watts, Mark Whitehead |
| EPI_ISL_665315 | Wales Specialist Virology Centre Sequencing lab: Pathogen Genomics Unit | COVID-19 Genomics UK (COG-UK) Consortium | Catherine Moore, Johnathan Evans, Laura Gifford, Malorie Perry, Simon Cottrell, Angela Marchbank, Alec Birchley, Alexander Adams, Amy Gaskin, Bree Gatica-Wilcox, Jason Coombes, Joel Southgate, Lauren Gilbert, Lee Graham, Nicole Pacchiarini, Sara Kumziene-Summerhayes, Sarah Taylor, Sophie Jones, Sara Rey, Matthew Bull, Joanne Watkins, Sally Corden, Tom Connor |
| EPI_ISL_665351, EPI_ISL_665371, EPI_ISL_665406 | Quadram Institute Bioscience | COVID-19 Genomics UK (COG-UK) Consortium | Dave J. Baker, Gemma L. Kay, Alp Aydin, Thanh Le-Viet, Steven Rudder, Ana P. Tedim, Anastasia Kolyva, Maria Diaz, Leonardo de Oliveira Martins, Nabil-Fareed Alikhan, Lizzie Meadows, Rachael Stanley, Ngozi Elumogo, Muhammed Yasir, Nicholas M. Thomson, Alexander J Trotter, Rachel Gilroy, Samuel Bloomfield, Claire Stuart, Andrew Bell, Reenesh Prakash, Samir Dervisevic, Alison E. Mather, John Wain, Mark Webber, Andrew J. Page, Justin O'Grady |
| EPI_ISL_665414 | Liverpool Clinical Laboratories | COVID-19 Genomics UK (COG-UK) Consortium | Sam Haldenby, Anita Lucaci, Steve Paterson, Julian Hiscox, Alistair Darby, M Almsaud, A Alrezaihi, Muhannad Alruwaili, Stuart D Armstrong, Jones Benjamin, Eleanor G Bentley, Anu Chawla, Jordan J Clark, Angela Cowell, Richard Eccles, Isabel García-Dorival, Matthew Gemmell, Alessandro Gerada, PKF Gilmore, Richard Gregory, Ximeng Han, Catherine Hartley, Margaret Hughes, Miren Iturriza-Gomara, James Johnson, L Luu, Jenifer Manson, Charlotte Nelson, Elaine O'Toole, Cassie Olateju, Rebekah Penrice-Randal , Lucille Rainbow, N.P Randle, Trevor Ian Robinson, Parul Sharma, Ghada T Shawli, James P Stewart, Neil Swainston, Ecaterina Vamos, Joanne Watts, Mark Whitehead |
| EPI_ISL_665454, EPI_ISL_665461 | Quadram Institute Bioscience | COVID-19 Genomics UK (COG-UK) Consortium | Dave J. Baker, Gemma L. Kay, Alp Aydin, Thanh Le-Viet, Steven Rudder, Ana P. Tedim, Anastasia Kolyva, Maria Diaz, Leonardo de Oliveira Martins, |

| | | | |
|---|---|---|---|
| | | | Nabil-Fareed Alikhan, Lizzie Meadows, Rachael Stanley, Ngozi Elumogo, Muhammed Yasir, Nicholas M. Thomson, Alexander J Trotter, Rachel Gilroy, Samuel Bloomfield, Claire Stuart, Andrew Bell, Reenesh Prakash, Samir Dervisevic, Alison E. Mather, John Wain, Mark Webber, Andrew J. Page, Justin O'Grady |
| EPI_ISL_665468 | Wales Specialist Virology Centre Sequencing lab: Pathogen Genomics Unit | COVID-19 Genomics UK (COG-UK) Consortium | Catherine Moore, Johnathan Evans, Laura Gifford, Malorie Perry, Simon Cottrell, Angela Marchbank, Alec Birchley, Alexander Adams, Amy Gaskin, Bree Gatica-Wilcox, Jason Coombes, Joel Southgate, Lauren Gilbert, Lee Graham, Nicole Pacchiarini, Sara Kumziene-Summerhayes, Sarah Taylor, Sophie Jones, Sara Rey, Matthew Bull, Joanne Watkins, Sally Corden, Tom Connor |
| EPI_ISL_665481 | Quadram Institute Bioscience | COVID-19 Genomics UK (COG-UK) Consortium | Dave J. Baker, Gemma L. Kay, Alp Aydin, Thanh Le-Viet, Steven Rudder, Ana P. Tedim, Anastasia Kolyva, Maria Diaz, Leonardo de Oliveira Martins, Nabil-Fareed Alikhan, Lizzie Meadows, Rachael Stanley, Ngozi Elumogo, Muhammed Yasir, Nicholas M. Thomson, Alexander J Trotter, Rachel Gilroy, Samuel Bloomfield, Claire Stuart, Andrew Bell, Reenesh Prakash, Samir Dervisevic, Alison E. Mather, John Wain, Mark Webber, Andrew J. Page, Justin O'Grady |
| EPI_ISL_665504, EPI_ISL_665516, EPI_ISL_665526 | Wales Specialist Virology Centre Sequencing lab: Pathogen Genomics Unit | COVID-19 Genomics UK (COG-UK) Consortium | Catherine Moore, Johnathan Evans, Laura Gifford, Malorie Perry, Simon Cottrell, Angela Marchbank, Alec Birchley, Alexander Adams, Amy Gaskin, Bree Gatica-Wilcox, Jason Coombes, Joel Southgate, Lauren Gilbert, Lee Graham, Nicole Pacchiarini, Sara Kumziene-Summerhayes, Sarah Taylor, Sophie Jones, Sara Rey, Matthew Bull, Joanne Watkins, Sally Corden, Tom Connor |
| EPI_ISL_665533 | Quadram Institute Bioscience | COVID-19 Genomics UK (COG-UK) Consortium | Dave J. Baker, Gemma L. Kay, Alp Aydin, Thanh Le-Viet, Steven Rudder, Ana P. Tedim, Anastasia Kolyva, Maria Diaz, Leonardo de Oliveira Martins, Nabil-Fareed Alikhan, Lizzie Meadows, Rachael Stanley, Ngozi Elumogo, Muhammed Yasir, Nicholas M. Thomson, Alexander J Trotter, Rachel Gilroy, Samuel Bloomfield, Claire Stuart, Andrew Bell, Reenesh Prakash, Samir Dervisevic, Alison E. Mather, John Wain, Mark Webber, Andrew J. Page, Justin O'Grady |
| EPI_ISL_665552 | Virology Department, Royal Infirmary of Edinburgh, NHS Lothian / School of Biological Sciences, University of Edinburgh / Institute of Genetics and Molecular Medicine, University of Edinburgh | COVID-19 Genomics UK (COG-UK) Consortium | McHugh M, Dewar R, Rooke S, Gallagher M, Balcaza C, O'Toole Á, Scher E, Hill V, McCrone JT, Colquhoun R, Yu X, Jackson B, Rambaut A, Williams TC, Templeton K |
| EPI_ISL_665573 | Wales Specialist Virology Centre Sequencing lab: Pathogen Genomics Unit | COVID-19 Genomics UK (COG-UK) Consortium | Catherine Moore, Johnathan Evans, Laura Gifford, Malorie Perry, Simon Cottrell, Angela Marchbank, Alec Birchley, Alexander Adams, Amy Gaskin, Bree Gatica-Wilcox, Jason Coombes, Joel Southgate, Lauren Gilbert, Lee Graham, Nicole Pacchiarini, Sara Kumziene-Summerhayes, Sarah Taylor, Sophie Jones, Sara Rey, Matthew Bull, Joanne Watkins, Sally Corden, Tom Connor |
| EPI_ISL_665581, EPI_ISL_665583 | Quadram Institute Bioscience | COVID-19 Genomics UK (COG-UK) Consortium | Dave J. Baker, Gemma L. Kay, Alp Aydin, Thanh Le-Viet, Steven Rudder, Ana P. Tedim, Anastasia Kolyva, Maria Diaz, Leonardo de Oliveira Martins, Nabil-Fareed Alikhan, Lizzie Meadows, Rachael Stanley, Ngozi Elumogo, Muhammed Yasir, Nicholas M. Thomson, Alexander J Trotter, Rachel Gilroy, Samuel Bloomfield, Claire Stuart, Andrew Bell, Reenesh Prakash, Samir Dervisevic, Alison E. Mather, John Wain, Mark Webber, Andrew J. Page, Justin O'Grady |
| EPI_ISL_665584 | Wales Specialist Virology Centre Sequencing lab: Pathogen Genomics Unit | COVID-19 Genomics UK (COG-UK) Consortium | Catherine Moore, Johnathan Evans, Laura Gifford, Malorie Perry, Simon Cottrell, Angela Marchbank, Alec Birchley, Alexander Adams, Amy Gaskin, Bree Gatica-Wilcox, Jason Coombes, Joel Southgate, Lauren Gilbert, Lee Graham, Nicole Pacchiarini, Sara Kumziene-Summerhayes, Sarah Taylor, Sophie Jones, Sara Rey, Matthew Bull, Joanne Watkins, Sally Corden, Tom Connor |
| EPI_ISL_665591 | Quadram Institute Bioscience | COVID-19 Genomics UK (COG-UK) Consortium | Dave J. Baker, Gemma L. Kay, Alp Aydin, Thanh Le-Viet, Steven Rudder, Ana P. Tedim, Anastasia Kolyva, Maria Diaz, Leonardo de Oliveira Martins, Nabil-Fareed Alikhan, Lizzie Meadows, Rachael Stanley, Ngozi Elumogo, Muhammed Yasir, Nicholas M. Thomson, Alexander J Trotter, Rachel Gilroy, Samuel Bloomfield, Claire Stuart, Andrew Bell, Reenesh Prakash, Samir Dervisevic, Alison E. Mather, John Wain, Mark Webber, Andrew J. Page, Justin O'Grady |
| EPI_ISL_665608, EPI_ISL_665660 | Wales Specialist Virology Centre Sequencing lab: Pathogen Genomics Unit | COVID-19 Genomics UK (COG-UK) Consortium | Catherine Moore, Johnathan Evans, Laura Gifford, Malorie Perry, Simon Cottrell, Angela Marchbank, Alec Birchley, Alexander Adams, Amy Gaskin, Bree Gatica-Wilcox, Jason Coombes, Joel Southgate, Lauren Gilbert, Lee Graham, Nicole Pacchiarini, Sara Kumziene-Summerhayes, Sarah Taylor, Sophie Jones, Sara Rey, Matthew Bull, Joanne Watkins, Sally Corden, Tom Connor |
| EPI_ISL_665662 | Liverpool Clinical Laboratories | COVID-19 Genomics UK (COG-UK) Consortium | Sam Haldenby, Anita Lucaci, Steve Paterson, Julian Hiscox, Alistair Darby, M Almsaud, A Alrezaihi, Muhannad Alruwaili, Stuart D Armstrong, Jones Benjamin, Eleanor G Bentley, Anu Chawla, Jordan J Clark, Angela Cowell, Richard Eccles, Isabel García-Dorival, Matthew Gemmell, Alessandro Gerada, PKF Gilmore, Richard Gregory, Ximeng Han, Catherine Hartley, Margaret Hughes, Miren Iturriza-Gomara, James Johnson, L Luu, Jenifer Manson, Charlotte Nelson, Elaine O'Toole, Cassie Olateju, Rebekah Penrice-Randal , Lucille Rainbow, N.P Randle, Trevor Ian Robinson, Parul Sharma, Ghada T Shawli, James P Stewart, Neil Swainston, Ecaterina Vamos, Joanne Watts, Mark Whitehead |
| EPI_ISL_665672 | Quadram Institute Bioscience | COVID-19 Genomics UK (COG-UK) Consortium | Dave J. Baker, Gemma L. Kay, Alp Aydin, Thanh Le-Viet, Steven Rudder, Ana P. Tedim, Anastasia Kolyva, Maria Diaz, Leonardo de Oliveira Martins, Nabil-Fareed Alikhan, Lizzie Meadows, Rachael Stanley, Ngozi Elumogo, Muhammed Yasir, Nicholas M. Thomson, Alexander J Trotter, Rachel Gilroy, Samuel Bloomfield, Claire Stuart, Andrew Bell, Reenesh Prakash, Samir Dervisevic, Alison E. Mather, John Wain, Mark Webber, Andrew J. Page, Justin O'Grady |
| EPI_ISL_665677 | Wales Specialist Virology Centre Sequencing lab: Pathogen Genomics Unit | COVID-19 Genomics UK (COG-UK) Consortium | Catherine Moore, Johnathan Evans, Laura Gifford, Malorie Perry, Simon Cottrell, Angela Marchbank, Alec Birchley, Alexander Adams, Amy Gaskin, Bree Gatica-Wilcox, Jason Coombes, Joel Southgate, Lauren Gilbert, Lee Graham, Nicole Pacchiarini, Sara Kumziene-Summerhayes, Sarah Taylor, Sophie Jones, Sara Rey, Matthew Bull, Joanne Watkins, Sally Corden, Tom Connor |
| EPI_ISL_665684 | Quadram Institute Bioscience | COVID-19 Genomics UK (COG-UK) Consortium | Dave J. Baker, Gemma L. Kay, Alp Aydin, Thanh Le-Viet, Steven Rudder, Ana P. Tedim, Anastasia Kolyva, Maria Diaz, Leonardo de Oliveira Martins, Nabil-Fareed Alikhan, Lizzie Meadows, Rachael Stanley, Ngozi Elumogo, Muhammed Yasir, Nicholas M. Thomson, Alexander J Trotter, Rachel Gilroy, Samuel Bloomfield, Claire Stuart, Andrew Bell, Reenesh Prakash, Samir Dervisevic, Alison E. Mather, John Wain, Mark Webber, Andrew J. Page, Justin O'Grady |
| EPI_ISL_665703 | University College London, Great Ormond Street Hospital for Children NHS Foundation Trust, Imperial College Healthcare NHS Trust | COVID-19 Genomics UK (COG-UK) Consortium | Sergi Castellano, Rachel Williams, Mark Kristiansen, Paola Resende Silva, Sunando Roy, Tony Brooks, Helena Tutill, Paola Niola, Patricia Dyal, Charlotte Williams, Leysa Forrest, Yasmin Panchbhaya, Jacqueline Findlay, Samuel Weeks, Julianne Brown, Kathryn Harris, Paul Randell, James Price, Alison Holmes, Judith Breuer |
| EPI_ISL_665704, EPI_ISL_665707 | Liverpool Clinical Laboratories | COVID-19 Genomics UK (COG-UK) Consortium | Sam Haldenby, Anita Lucaci, Steve Paterson, Julian Hiscox, Alistair Darby, M Almsaud, A Alrezaihi, Muhannad Alruwaili, Stuart D Armstrong, Jones Benjamin, Eleanor G Bentley, Anu Chawla, Jordan J Clark, Angela Cowell, Richard Eccles, Isabel García-Dorival, Matthew Gemmell, Alessandro Gerada, PKF Gilmore, Richard Gregory, Ximeng Han, Catherine Hartley, Margaret Hughes, Miren Iturriza-Gomara, James Johnson, L Luu, Jenifer Manson, Charlotte Nelson, Elaine O'Toole, Cassie Olateju, Rebekah Penrice-Randal , Lucille Rainbow, N.P Randle, Trevor Ian Robinson, Parul Sharma, Ghada T Shawli, James P Stewart, Neil Swainston, Ecaterina Vamos, Joanne Watts, Mark Whitehead |
| EPI_ISL_665713 | Wales Specialist Virology Centre Sequencing lab: Pathogen Genomics Unit | COVID-19 Genomics UK (COG-UK) Consortium | Catherine Moore, Johnathan Evans, Laura Gifford, Malorie Perry, Simon Cottrell, Angela Marchbank, Alec Birchley, Alexander Adams, Amy Gaskin, Bree Gatica-Wilcox, Jason Coombes, Joel Southgate, Lauren Gilbert, Lee Graham, Nicole Pacchiarini, Sara Kumziene-Summerhayes, Sarah Taylor, Sophie Jones, Sara Rey, Matthew Bull, Joanne Watkins, Sally Corden, Tom Connor |
| EPI_ISL_665720 | University College London, Great Ormond Street Hospital for Children NHS Foundation Trust, Imperial College Healthcare NHS Trust | COVID-19 Genomics UK (COG-UK) Consortium | Sergi Castellano, Rachel Williams, Mark Kristiansen, Paola Resende Silva, Sunando Roy, Tony Brooks, Helena Tutill, Paola Niola, Patricia Dyal, Charlotte Williams, Leysa Forrest, Yasmin Panchbhaya, Jacqueline Findlay, Samuel Weeks, Julianne Brown, Kathryn Harris, Paul Randell, James Price, Alison Holmes, Judith Breuer |
| EPI_ISL_665743 | Wales Specialist Virology Centre Sequencing lab: Pathogen Genomics Unit | COVID-19 Genomics UK (COG-UK) Consortium | Catherine Moore, Johnathan Evans, Laura Gifford, Malorie Perry, Simon Cottrell, Angela Marchbank, Alec Birchley, Alexander Adams, Amy Gaskin, Bree Gatica-Wilcox, Jason Coombes, Joel Southgate, Lauren Gilbert, Lee Graham, Nicole Pacchiarini, Sara Kumziene-Summerhayes, Sarah Taylor, Sophie Jones, Sara Rey, Matthew Bull, Joanne Watkins, Sally Corden, Tom Connor |
| EPI_ISL_665747 | Quadram Institute Bioscience | COVID-19 Genomics UK (COG-UK) Consortium | Dave J. Baker, Gemma L. Kay, Alp Aydin, Thanh Le-Viet, Steven Rudder, Ana P. Tedim, Anastasia Kolyva, Maria Diaz, Leonardo de Oliveira Martins, Nabil-Fareed Alikhan, Lizzie Meadows, Rachael Stanley, Ngozi Elumogo, Muhammed Yasir, Nicholas M. Thomson, Alexander J Trotter, Rachel Gilroy, Samuel Bloomfield, Claire Stuart, Andrew Bell, Reenesh Prakash, Samir Dervisevic, Alison E. Mather, John Wain, Mark Webber, Andrew J. Page, Justin O'Grady |
| EPI_ISL_665756, EPI_ISL_665764, EPI_ISL_665771 | Wales Specialist Virology Centre Sequencing lab: Pathogen Genomics Unit | COVID-19 Genomics UK (COG-UK) Consortium | Catherine Moore, Johnathan Evans, Laura Gifford, Malorie Perry, Simon Cottrell, Angela Marchbank, Alec Birchley, Alexander Adams, Amy Gaskin, Bree Gatica-Wilcox, Jason Coombes, Joel Southgate, Lauren Gilbert, Lee Graham, Nicole Pacchiarini, Sara Kumziene-Summerhayes, Sarah Taylor, Sophie |

| | | | |
|---|---|---|---|
| | | | Jones, Sara Rey, Matthew Bull, Joanne Watkins, Sally Corden, Tom Connor |
| EPI_ISL_665788 | Quadram Institute Bioscience | COVID-19 Genomics UK (COG-UK) Consortium | Dave J. Baker, Gemma L. Kay, Alp Aydin, Thanh Le-Viet, Steven Rudder, Ana P. Tedim, Anastasia Kolyva, Maria Diaz, Leonardo de Oliveira Martins, Nabil-Fareed Alikhan, Lizzie Meadows, Rachael Stanley, Ngozi Elumogo, Muhammed Yasir, Nicholas M. Thomson, Alexander J Trotter, Rachel Gilroy, Samuel Bloomfield, Claire Stuart, Andrew Bell, Reenesh Prakash, Samir Dervisevic, Alison E. Mather, John Wain, Mark Webber, Andrew J. Page, Justin O'Grady |
| EPI_ISL_665794, EPI_ISL_665800 | Wales Specialist Virology Centre Sequencing lab: Pathogen Genomics Unit | COVID-19 Genomics UK (COG-UK) Consortium | Catherine Moore, Johnathan Evans, Laura Gifford, Malorie Perry, Simon Cottrell, Angela Marchbank, Alec Birchley, Alexander Adams, Amy Gaskin, Bree Gatica-Wilcox, Jason Coombes, Joel Southgate, Lauren Gilbert, Lee Graham, Nicole Pacchiarini, Sara Kumziene-Summerhayes, Sarah Taylor, Sophie Jones, Sara Rey, Matthew Bull, Joanne Watkins, Sally Corden, Tom Connor |
| EPI_ISL_665878, EPI_ISL_665884, EPI_ISL_665885, EPI_ISL_665888, EPI_ISL_665890, EPI_ISL_665893, EPI_ISL_665896 | Quadram Institute Bioscience | COVID-19 Genomics UK (COG-UK) Consortium | Dave J. Baker, Gemma L. Kay, Alp Aydin, Thanh Le-Viet, Steven Rudder, Ana P. Tedim, Anastasia Kolyva, Maria Diaz, Leonardo de Oliveira Martins, Nabil-Fareed Alikhan, Lizzie Meadows, Rachael Stanley, Ngozi Elumogo, Muhammed Yasir, Nicholas M. Thomson, Alexander J Trotter, Rachel Gilroy, Samuel Bloomfield, Claire Stuart, Andrew Bell, Reenesh Prakash, Samir Dervisevic, Alison E. Mather, John Wain, Mark Webber, Andrew J. Page, Justin O'Grady |
| EPI_ISL_666022 | Liverpool Clinical Laboratories | COVID-19 Genomics UK (COG-UK) Consortium | Sam Haldenby, Anita Lucaci, Steve Paterson, Julian Hiscox, Alistair Darby, M Almsaud, A Alrezaihi, Muhannad Alruwaili, Stuart D Armstrong, Jones Benjamin, Eleanor G Bentley, Anu Chawla, Jordan J Clark, Angela Cowell, Richard Eccles, Isabel García-Dorival, Matthew Gemmell, Alessandro Gerada, PKF Gilmore, Richard Gregory, Ximeng Han, Catherine Hartley, Margaret Hughes, Miren Iturriza-Gomara, James Johnson, L Luu, Jenifer Manson, Charlotte Nelson, Elaine O'Toole, Cassie Olateju, Rebekah Penrice-Randal , Lucille Rainbow, N.P Randle, Trevor Ian Robinson, Parul Sharma, Ghada T Shawli, James P Stewart, Neil Swainston, Ecaterina Vamos, Joanne Watts, Mark Whitehead |
| EPI_ISL_666061, EPI_ISL_666062, EPI_ISL_666063, EPI_ISL_666066, EPI_ISL_666068, EPI_ISL_666069 | Virology Department, Royal Infirmary of Edinburgh, NHS Lothian / School of Biological Sciences, University of Edinburgh / Institute of Genetics and Molecular Medicine, University of Edinburgh | COVID-19 Genomics UK (COG-UK) Consortium | McHugh M, Dewar R, Rooke S, Gallagher M, Balcaza C, O'Toole Á, Scher E, Hill V, McCrone JT, Colquhoun R, Yu X, Jackson B, Rambaut A, Williams TC, Templeton K |
| EPI_ISL_666111, EPI_ISL_666117, EPI_ISL_666118, EPI_ISL_666119, EPI_ISL_666123, EPI_ISL_666124, EPI_ISL_666128, EPI_ISL_666132, EPI_ISL_666134, EPI_ISL_666141, EPI_ISL_666148, EPI_ISL_666160, EPI_ISL_666162, EPI_ISL_666164, EPI_ISL_666168, EPI_ISL_666171, EPI_ISL_666174, EPI_ISL_666179, EPI_ISL_666180, EPI_ISL_666184, EPI_ISL_666193, EPI_ISL_666194, EPI_ISL_666199, EPI_ISL_666208, EPI_ISL_666209, EPI_ISL_666211, EPI_ISL_666212, EPI_ISL_666230, EPI_ISL_666234, EPI_ISL_666238, EPI_ISL_666247, EPI_ISL_666250, EPI_ISL_666254, EPI_ISL_666266, EPI_ISL_666268, EPI_ISL_666270, EPI_ISL_666285, EPI_ISL_666287, EPI_ISL_666301, EPI_ISL_666310, EPI_ISL_666312, EPI_ISL_666320, EPI_ISL_666328, EPI_ISL_666330, EPI_ISL_666332, EPI_ISL_666340, EPI_ISL_666345, EPI_ISL_666347, EPI_ISL_666349, EPI_ISL_666356, EPI_ISL_666365, EPI_ISL_666371, EPI_ISL_666376, EPI_ISL_666377, EPI_ISL_666383, EPI_ISL_666396, EPI_ISL_666397, EPI_ISL_666399, EPI_ISL_666405, EPI_ISL_666406, EPI_ISL_666407, EPI_ISL_666409, EPI_ISL_666411, EPI_ISL_666417, EPI_ISL_666424, EPI_ISL_666425, EPI_ISL_666437, EPI_ISL_666443, EPI_ISL_666451, EPI_ISL_666452, EPI_ISL_666454, EPI_ISL_666457, EPI_ISL_666458, EPI_ISL_666460, EPI_ISL_666461, EPI_ISL_666465, EPI_ISL_666467, EPI_ISL_666474, EPI_ISL_666476, EPI_ISL_666493, EPI_ISL_666498, EPI_ISL_666499, EPI_ISL_666504, EPI_ISL_666514, EPI_ISL_666516, EPI_ISL_666518, EPI_ISL_666520, EPI_ISL_666532, EPI_ISL_666536, EPI_ISL_666537, EPI_ISL_666539, EPI_ISL_666547, EPI_ISL_666548, EPI_ISL_666551, EPI_ISL_666554, EPI_ISL_666555, EPI_ISL_666580, EPI_ISL_666586, EPI_ISL_666589 | | | |
| see above | Wales Specialist Virology Centre Sequencing lab: Pathogen Genomics Unit | COVID-19 Genomics UK (COG-UK) Consortium | Catherine Moore, Johnathan Evans, Laura Gifford, Malorie Perry, Simon Cottrell, Angela Marchbank, Alec Birchley, Alexander Adams, Amy Gaskin, Bree Gatica-Wilcox, Jason Coombes, Joel Southgate, Lauren Gilbert, Lee Graham, Nicole Pacchiarini, Sara Kumziene-Summerhayes, Sarah Taylor, Sophie Jones, Sara Rey, Matthew Bull, Joanne Watkins, Sally Corden, Tom Connor |
| EPI_ISL_666729, EPI_ISL_666732, EPI_ISL_666733, EPI_ISL_666734, EPI_ISL_666735, EPI_ISL_666736, EPI_ISL_666741 | Respiratory Virus Unit, Microbiology Services Colindale, Public Health England | COVID-19 Genomics UK (COG-UK) Consortium | PHE Covid Sequencing Team |
| EPI_ISL_666792 | Maryland Public Health Laboratory | Maryland Public Health Laboratory | Maryland Department of Health Laboratories Administration |
| EPI_ISL_666894, EPI_ISL_666899, EPI_ISL_666907, EPI_ISL_666909, EPI_ISL_666910, EPI_ISL_666919, EPI_ISL_666924, EPI_ISL_666928, EPI_ISL_666932, EPI_ISL_666933, EPI_ISL_666936, EPI_ISL_666941, EPI_ISL_666945, EPI_ISL_666949, EPI_ISL_666954, EPI_ISL_666957, EPI_ISL_666963 | | | |
| see above | Michigan Department of Health and Human Services, Bureau of Laboratories | Michigan Department of Health and Human Services, Bureau of Laboratories | Blankenship HM, Riner D, Soehnlen MK |
| EPI_ISL_666972, EPI_ISL_666973, EPI_ISL_667042, EPI_ISL_667043, EPI_ISL_667060, EPI_ISL_667061, EPI_ISL_667076, EPI_ISL_667078 | San Diego County Public Health Laboratory | Andersen lab at Scripps Research | SEARCH Alliance San Diego with Tracy Basler, Jovan Shephard, Brett Austin |
| EPI_ISL_667518, EPI_ISL_667529, EPI_ISL_667530, EPI_ISL_667531, EPI_ISL_667532, EPI_ISL_667533, EPI_ISL_667534, EPI_ISL_667535, EPI_ISL_667539, EPI_ISL_667540, EPI_ISL_667550 | | | |
| see above | OHSU Lab Services Molecular Microbiology Lab | Oregon SARS-CoV-2 Genome Sequencing Center | Brendan L. O'Connell, Ruth V. Nichols, Sally Grindstaff, Alec J. Hirsch, Donna Hansel, Guang Fan, Daniel N. Streblow, William B. Messer, Andrew C. Adey, Benjamin N. Bimber, Brian J. O'Roak |
| EPI_ISL_667781, EPI_ISL_667806 | South Eastern Area Laboratory Services (SEALS) | NSW Health Pathology - Institute of Clinical Pathology and Medical Research; Westmead Hospital; University of Sydney | CIDM-PH et al. |
| EPI_ISL_668447, EPI_ISL_668448, EPI_ISL_668453, EPI_ISL_668454, EPI_ISL_668455 | Centre for Dengue Research, Department of Immunology and Molecular Medicine | Centre for Dengue Research, Department of Immunology and Molecular Medicine | Chandima Jeewandara, Deshni Jayathilaka, Dinuka Ariyaratne, Diyanath Ranasinghe, Laksiri Gomes, Gathsaurie Neelika Malavige |
| EPI_ISL_669458, EPI_ISL_669460, EPI_ISL_669461, EPI_ISL_669462, EPI_ISL_669464, EPI_ISL_669466, EPI_ISL_669467, EPI_ISL_669468, EPI_ISL_669469, EPI_ISL_669470, EPI_ISL_669471, EPI_ISL_669472, EPI_ISL_669473, EPI_ISL_669474, EPI_ISL_669475, EPI_ISL_669476, EPI_ISL_669477, EPI_ISL_669478, EPI_ISL_669479, EPI_ISL_669480, EPI_ISL_669481, EPI_ISL_669482, EPI_ISL_669483, EPI_ISL_669484, EPI_ISL_669485, EPI_ISL_669487, EPI_ISL_669488, EPI_ISL_669489, EPI_ISL_669490, EPI_ISL_669491, EPI_ISL_669492, EPI_ISL_669493, EPI_ISL_669494, EPI_ISL_669496, EPI_ISL_669497, EPI_ISL_669498, EPI_ISL_669499, EPI_ISL_669500, EPI_ISL_669501, EPI_ISL_669502, EPI_ISL_669505, EPI_ISL_669506, EPI_ISL_669507, EPI_ISL_669508, EPI_ISL_669509, EPI_ISL_669510, EPI_ISL_669511, EPI_ISL_669512, EPI_ISL_669513, EPI_ISL_669514, EPI_ISL_669515, EPI_ISL_669516, EPI_ISL_669517, EPI_ISL_669518, EPI_ISL_669519, EPI_ISL_669520, EPI_ISL_669522, EPI_ISL_669523, EPI_ISL_669524, EPI_ISL_669526, EPI_ISL_669527, EPI_ISL_669528, EPI_ISL_669529, EPI_ISL_669530, EPI_ISL_669531, EPI_ISL_669532, EPI_ISL_669533, EPI_ISL_669534, EPI_ISL_669535, EPI_ISL_669536, EPI_ISL_669538, EPI_ISL_669539, EPI_ISL_669540, EPI_ISL_669541, EPI_ISL_669542, EPI_ISL_669543, EPI_ISL_669544, EPI_ISL_669545, EPI_ISL_669546, EPI_ISL_669547, EPI_ISL_669549, EPI_ISL_669550, EPI_ISL_669551, EPI_ISL_669552, EPI_ISL_669553, EPI_ISL_669554, EPI_ISL_669555, EPI_ISL_669556, EPI_ISL_669557, EPI_ISL_669558, EPI_ISL_669559, EPI_ISL_669560, EPI_ISL_669561, EPI_ISL_669562, EPI_ISL_669563, EPI_ISL_669565, EPI_ISL_669566, EPI_ISL_669567, EPI_ISL_669568, EPI_ISL_669569, EPI_ISL_669570, EPI_ISL_669571, EPI_ISL_669572, EPI_ISL_669573, EPI_ISL_669574, EPI_ISL_669575, EPI_ISL_669576, EPI_ISL_669577, EPI_ISL_669578, EPI_ISL_669579, EPI_ISL_669580, EPI_ISL_669581, EPI_ISL_669582, EPI_ISL_669584, EPI_ISL_669585, EPI_ISL_669586, EPI_ISL_669587, EPI_ISL_669588, EPI_ISL_669589, EPI_ISL_669590, EPI_ISL_669591, EPI_ISL_669592, EPI_ISL_669593, EPI_ISL_669594, EPI_ISL_669595, EPI_ISL_669596, EPI_ISL_669597, EPI_ISL_669598, EPI_ISL_669599, EPI_ISL_669600, EPI_ISL_669601, EPI_ISL_669602, EPI_ISL_669603, EPI_ISL_669604, EPI_ISL_669605, EPI_ISL_669606, EPI_ISL_669607, EPI_ISL_669608, EPI_ISL_669609, EPI_ISL_669610, EPI_ISL_669611, EPI_ISL_669614, EPI_ISL_669615, EPI_ISL_669616, EPI_ISL_669617, EPI_ISL_669618, EPI_ISL_669619, EPI_ISL_669620, EPI_ISL_669621, EPI_ISL_669622, EPI_ISL_669623, EPI_ISL_669624, EPI_ISL_669625, EPI_ISL_669626, EPI_ISL_669627, EPI_ISL_669628, EPI_ISL_669629, EPI_ISL_669630, EPI_ISL_669631, EPI_ISL_669632, EPI_ISL_669633, EPI_ISL_669634, EPI_ISL_669635, EPI_ISL_669644, EPI_ISL_669645, EPI_ISL_669646, EPI_ISL_669647, EPI_ISL_669648, EPI_ISL_669649, EPI_ISL_669650, EPI_ISL_669651, EPI_ISL_669652, EPI_ISL_669653, EPI_ISL_669654, EPI_ISL_669655, EPI_ISL_669656, EPI_ISL_669657, EPI_ISL_669658, EPI_ISL_669659, EPI_ISL_669660, EPI_ISL_669661, EPI_ISL_669662, EPI_ISL_669663, EPI_ISL_669664, EPI_ISL_669665, EPI_ISL_669666, EPI_ISL_669667, EPI_ISL_669668, EPI_ISL_669669, EPI_ISL_669670, EPI_ISL_669671, EPI_ISL_669672, EPI_ISL_669673, EPI_ISL_669674, EPI_ISL_669675, EPI_ISL_669676, EPI_ISL_669677, EPI_ISL_669678, EPI_ISL_669679, EPI_ISL_669680, EPI_ISL_669681, EPI_ISL_669682, EPI_ISL_669683, EPI_ISL_669684, EPI_ISL_669685, EPI_ISL_669686, EPI_ISL_669687, EPI_ISL_669688, EPI_ISL_669689, EPI_ISL_669690, EPI_ISL_669691, EPI_ISL_669692, EPI_ISL_669693, EPI_ISL_669694, EPI_ISL_669695, EPI_ISL_669696, EPI_ISL_669697, EPI_ISL_669698, EPI_ISL_669699, EPI_ISL_669700, EPI_ISL_669701, EPI_ISL_669702, EPI_ISL_669703, EPI_ISL_669704, EPI_ISL_669705, EPI_ISL_669706, EPI_ISL_669707, EPI_ISL_669708, EPI_ISL_669709, EPI_ISL_669710, EPI_ISL_669711, EPI_ISL_669712, EPI_ISL_669713, EPI_ISL_669714, EPI_ISL_669715, EPI_ISL_669716, EPI_ISL_669717, EPI_ISL_669718, EPI_ISL_669719, EPI_ISL_669720, EPI_ISL_669721, EPI_ISL_669722, EPI_ISL_669723, EPI_ISL_669724, EPI_ISL_669725, EPI_ISL_669726, EPI_ISL_669727, EPI_ISL_669728, EPI_ISL_669729, EPI_ISL_669730, EPI_ISL_669732, EPI_ISL_669736, EPI_ISL_669737, EPI_ISL_669738, EPI_ISL_669739, EPI_ISL_669740, EPI_ISL_669741, EPI_ISL_669742, EPI_ISL_669743, EPI_ISL_669744, EPI_ISL_669745, EPI_ISL_669746, EPI_ISL_669747, EPI_ISL_669748, EPI_ISL_669749, EPI_ISL_669750, EPI_ISL_669751, EPI_ISL_669752, EPI_ISL_669753, EPI_ISL_669754, EPI_ISL_669755, EPI_ISL_669756, EPI_ISL_669758, EPI_ISL_669759, EPI_ISL_669761, EPI_ISL_669762, EPI_ISL_669763, EPI_ISL_669764, EPI_ISL_669765, EPI_ISL_669766, EPI_ISL_669767, EPI_ISL_669768, EPI_ISL_669769, EPI_ISL_669770, EPI_ISL_669771, EPI_ISL_669772, EPI_ISL_669775, EPI_ISL_669776, EPI_ISL_669777, EPI_ISL_669778, EPI_ISL_669779, EPI_ISL_669780, EPI_ISL_669781, EPI_ISL_669782, EPI_ISL_669783, EPI_ISL_669784, EPI_ISL_669788, EPI_ISL_669789, EPI_ISL_669790, EPI_ISL_669791, EPI_ISL_669792, EPI_ISL_669793, EPI_ISL_669794, EPI_ISL_669795, EPI_ISL_669796, EPI_ISL_669797, EPI_ISL_669798, EPI_ISL_669799, EPI_ISL_669800, EPI_ISL_669801, EPI_ISL_669802, EPI_ISL_669803, EPI_ISL_669804, EPI_ISL_669805, EPI_ISL_669806, EPI_ISL_669807, EPI_ISL_669808, EPI_ISL_669809, EPI_ISL_669810, EPI_ISL_669811, EPI_ISL_669812, EPI_ISL_669813, EPI_ISL_669814, EPI_ISL_669815, EPI_ISL_669816, EPI_ISL_669818, EPI_ISL_669819, EPI_ISL_669820, EPI_ISL_669821, EPI_ISL_669822, EPI_ISL_669823, EPI_ISL_669824, EPI_ISL_669825, EPI_ISL_669826, EPI_ISL_669827, EPI_ISL_669828, EPI_ISL_669829, EPI_ISL_669830, EPI_ISL_669831, EPI_ISL_669832, EPI_ISL_669882, EPI_ISL_669883, EPI_ISL_669884, EPI_ISL_669885, EPI_ISL_669886, EPI_ISL_669887, EPI_ISL_669888, EPI_ISL_669889, EPI_ISL_669890, EPI_ISL_669891, EPI_ISL_669892, EPI_ISL_669893, EPI_ISL_669894, EPI_ISL_669895, EPI_ISL_669896, EPI_ISL_669897, EPI_ISL_669898, EPI_ISL_669899, EPI_ISL_669900, EPI_ISL_669901, EPI_ISL_669902, EPI_ISL_669903, EPI_ISL_669904, EPI_ISL_669905, EPI_ISL_669906, EPI_ISL_669907, EPI_ISL_669908, EPI_ISL_669909, EPI_ISL_669910, EPI_ISL_669911, EPI_ISL_669912, EPI_ISL_669913, EPI_ISL_669914, EPI_ISL_669915, EPI_ISL_669916, EPI_ISL_669917, EPI_ISL_669918, EPI_ISL_669919, EPI_ISL_669920, EPI_ISL_669921, EPI_ISL_669922, EPI_ISL_669923, EPI_ISL_669924, EPI_ISL_669925, EPI_ISL_669926, EPI_ISL_669927, EPI_ISL_669928, EPI_ISL_669929, EPI_ISL_669930, EPI_ISL_669931, EPI_ISL_669932, EPI_ISL_669933, EPI_ISL_669934, EPI_ISL_669935, EPI_ISL_669936, EPI_ISL_669937, EPI_ISL_669938, EPI_ISL_669939, EPI_ISL_669940, EPI_ISL_669941, EPI_ISL_669942, EPI_ISL_669943, EPI_ISL_669944, EPI_ISL_669945, EPI_ISL_669946, EPI_ISL_669947, EPI_ISL_669948, EPI_ISL_669949, EPI_ISL_669950, EPI_ISL_669951, EPI_ISL_669952, EPI_ISL_669953, EPI_ISL_669954, EPI_ISL_669955, EPI_ISL_669956, EPI_ISL_669957, EPI_ISL_669958, EPI_ISL_669959, EPI_ISL_669960, EPI_ISL_669961, EPI_ISL_669962, EPI_ISL_669963, EPI_ISL_669964, EPI_ISL_669965, EPI_ISL_669966, EPI_ISL_669967, EPI_ISL_669968, EPI_ISL_669969, EPI_ISL_669970, EPI_ISL_669971, EPI_ISL_669972, EPI_ISL_669973, EPI_ISL_669974, EPI_ISL_669975, EPI_ISL_669976, EPI_ISL_669977, EPI_ISL_669978, EPI_ISL_669979, EPI_ISL_669980, EPI_ISL_669981, EPI_ISL_669982, EPI_ISL_669983, EPI_ISL_669984, EPI_ISL_669985, EPI_ISL_669986, EPI_ISL_669987, EPI_ISL_669988, EPI_ISL_669989, EPI_ISL_669990, EPI_ISL_669991, EPI_ISL_669992, EPI_ISL_669993, EPI_ISL_669994, EPI_ISL_669995, EPI_ISL_669996, EPI_ISL_669997, EPI_ISL_669998, EPI_ISL_669999, EPI_ISL_670000, EPI_ISL_670001, EPI_ISL_670002, EPI_ISL_670003, EPI_ISL_670004, EPI_ISL_670005, EPI_ISL_670006, EPI_ISL_670007, EPI_ISL_670008, EPI_ISL_670009, EPI_ISL_670010, EPI_ISL_670011, EPI_ISL_670012, EPI_ISL_670013, EPI_ISL_670014, EPI_ISL_670015, EPI_ISL_670016, EPI_ISL_670017, EPI_ISL_670018, EPI_ISL_670019, EPI_ISL_670020, EPI_ISL_670021, EPI_ISL_670022, EPI_ISL_670023, EPI_ISL_670024, EPI_ISL_670025, EPI_ISL_670026, EPI_ISL_670027, EPI_ISL_670028, EPI_ISL_670029, EPI_ISL_670030, EPI_ISL_670031, EPI_ISL_670032, EPI_ISL_670033, EPI_ISL_670034, EPI_ISL_670035, EPI_ISL_670036, EPI_ISL_670037, EPI_ISL_670038, EPI_ISL_670039, EPI_ISL_670040, EPI_ISL_670041, EPI_ISL_670042, EPI_ISL_670043, EPI_ISL_670044, EPI_ISL_670045, EPI_ISL_670046, EPI_ISL_670047, EPI_ISL_670048, EPI_ISL_670049, EPI_ISL_670050, EPI_ISL_670051, EPI_ISL_670052, EPI_ISL_670053, EPI_ISL_670054, EPI_ISL_670055, EPI_ISL_670056, EPI_ISL_670057, EPI_ISL_670058, EPI_ISL_670059, EPI_ISL_670060, EPI_ISL_670061, EPI_ISL_670062, EPI_ISL_670063, | | | |

| | | | |
|---|---|---|---|
| EPI_ISL_670064, EPI_ISL_670065, EPI_ISL_670066, EPI_ISL_670067, EPI_ISL_670068, EPI_ISL_670069, EPI_ISL_670070, EPI_ISL_670071, EPI_ISL_670072, EPI_ISL_670073, EPI_ISL_670074, EPI_ISL_670075, EPI_ISL_670076, EPI_ISL_670077, EPI_ISL_670078, EPI_ISL_670079, EPI_ISL_670080, EPI_ISL_670081, EPI_ISL_670082, EPI_ISL_670083, EPI_ISL_670084, EPI_ISL_670085, EPI_ISL_670086, EPI_ISL_670087, EPI_ISL_670088, EPI_ISL_670089, EPI_ISL_670090, EPI_ISL_670091, EPI_ISL_670092, EPI_ISL_670093, EPI_ISL_670094, EPI_ISL_670095, EPI_ISL_670096, EPI_ISL_670097, EPI_ISL_670098, EPI_ISL_670099, EPI_ISL_670100, EPI_ISL_670101, EPI_ISL_670102, EPI_ISL_670103, EPI_ISL_670104, EPI_ISL_670105, EPI_ISL_670106, EPI_ISL_670107, EPI_ISL_670108, EPI_ISL_670109, EPI_ISL_670110, EPI_ISL_670111, EPI_ISL_670112, EPI_ISL_670113, EPI_ISL_670114, EPI_ISL_670115, EPI_ISL_670116, EPI_ISL_670117, EPI_ISL_670118, EPI_ISL_670119, EPI_ISL_670120, EPI_ISL_670121, EPI_ISL_670122, EPI_ISL_670123, EPI_ISL_670124, EPI_ISL_670125, EPI_ISL_670126, EPI_ISL_670127, EPI_ISL_670128, EPI_ISL_670129, EPI_ISL_670130, EPI_ISL_670131, EPI_ISL_670132, EPI_ISL_670133, EPI_ISL_670134, EPI_ISL_670135, EPI_ISL_670136, EPI_ISL_670137, EPI_ISL_670138, EPI_ISL_670139, EPI_ISL_670140, EPI_ISL_670141, EPI_ISL_670142, EPI_ISL_670143, EPI_ISL_670144, EPI_ISL_670145, EPI_ISL_670146, EPI_ISL_670147, EPI_ISL_670148, EPI_ISL_670149, EPI_ISL_670150, EPI_ISL_670151, EPI_ISL_670152, EPI_ISL_670153, EPI_ISL_670154, EPI_ISL_670155, EPI_ISL_670156, EPI_ISL_670157, EPI_ISL_670158, EPI_ISL_670159, EPI_ISL_670160, EPI_ISL_670161, EPI_ISL_670162, EPI_ISL_670163, EPI_ISL_670164, EPI_ISL_670165, EPI_ISL_670166, EPI_ISL_670167, EPI_ISL_670168, EPI_ISL_670169, EPI_ISL_670170, EPI_ISL_670171, EPI_ISL_670172, EPI_ISL_670173, EPI_ISL_670174, EPI_ISL_670175, EPI_ISL_670176, EPI_ISL_670177, EPI_ISL_670178, EPI_ISL_670179, EPI_ISL_670180, EPI_ISL_670181, EPI_ISL_670182, EPI_ISL_670183, EPI_ISL_670184, EPI_ISL_670185, EPI_ISL_670186, EPI_ISL_670187, EPI_ISL_670188, EPI_ISL_670189, EPI_ISL_670190, EPI_ISL_670191, EPI_ISL_670192, EPI_ISL_670193, EPI_ISL_670194, EPI_ISL_670195, EPI_ISL_670196, EPI_ISL_670197, EPI_ISL_670198, EPI_ISL_670199, EPI_ISL_670200, EPI_ISL_670201, EPI_ISL_670202, EPI_ISL_670203, EPI_ISL_670204, EPI_ISL_670205, EPI_ISL_670206, EPI_ISL_670207, EPI_ISL_670208, EPI_ISL_670209, EPI_ISL_670210, EPI_ISL_670211, EPI_ISL_670212, EPI_ISL_670213, EPI_ISL_670214, EPI_ISL_670215, EPI_ISL_670216, EPI_ISL_670217, EPI_ISL_670218, EPI_ISL_670219, EPI_ISL_670220, EPI_ISL_670221, EPI_ISL_670222, EPI_ISL_670223, EPI_ISL_670224, EPI_ISL_670225, EPI_ISL_670226, EPI_ISL_670227, EPI_ISL_670228, EPI_ISL_670229, EPI_ISL_670230, EPI_ISL_670231, EPI_ISL_670232, EPI_ISL_670233, EPI_ISL_670234, EPI_ISL_670235, EPI_ISL_670236, EPI_ISL_670237, EPI_ISL_670238, EPI_ISL_670239, EPI_ISL_670240, EPI_ISL_670241, EPI_ISL_670242, EPI_ISL_670243, EPI_ISL_670244, EPI_ISL_670245, EPI_ISL_670246, EPI_ISL_670247, EPI_ISL_670248, EPI_ISL_670249, EPI_ISL_670250, EPI_ISL_670251, EPI_ISL_670252, EPI_ISL_670253, EPI_ISL_670254, EPI_ISL_670255, EPI_ISL_670256, EPI_ISL_670257, EPI_ISL_670258, EPI_ISL_670259, EPI_ISL_670260, EPI_ISL_670261, EPI_ISL_670262, EPI_ISL_670263, EPI_ISL_670264, EPI_ISL_670265, EPI_ISL_670266, EPI_ISL_670267, EPI_ISL_670268, EPI_ISL_670269, EPI_ISL_670270, EPI_ISL_670271, EPI_ISL_670272, EPI_ISL_670273, EPI_ISL_670274, EPI_ISL_670275, EPI_ISL_670276, EPI_ISL_670277, EPI_ISL_670278, EPI_ISL_670279, EPI_ISL_670280, EPI_ISL_670281, EPI_ISL_670282, EPI_ISL_670283, EPI_ISL_670284, EPI_ISL_670285, EPI_ISL_670286, EPI_ISL_670287, EPI_ISL_670288, EPI_ISL_670289, EPI_ISL_670290, EPI_ISL_670291, EPI_ISL_670292, EPI_ISL_670293, EPI_ISL_670294, EPI_ISL_670295, EPI_ISL_670296, EPI_ISL_670297, EPI_ISL_670298, EPI_ISL_670299, EPI_ISL_670300, EPI_ISL_670301, EPI_ISL_670302, EPI_ISL_670303, EPI_ISL_670304, EPI_ISL_670305, EPI_ISL_670306, EPI_ISL_670307, EPI_ISL_670308, EPI_ISL_670309, EPI_ISL_670310, EPI_ISL_670311, EPI_ISL_670312, EPI_ISL_670313, EPI_ISL_670314, EPI_ISL_670315, EPI_ISL_670316, EPI_ISL_670317, EPI_ISL_670318, EPI_ISL_670319, EPI_ISL_670320, EPI_ISL_670321, EPI_ISL_670322, EPI_ISL_670323, EPI_ISL_670324, EPI_ISL_670325, EPI_ISL_670326, EPI_ISL_670327, EPI_ISL_670328, EPI_ISL_670329, EPI_ISL_670330, EPI_ISL_670331, EPI_ISL_670332, EPI_ISL_670333, EPI_ISL_670334, EPI_ISL_670335, EPI_ISL_670336, EPI_ISL_670337, EPI_ISL_670338, EPI_ISL_670339, EPI_ISL_670340, EPI_ISL_670341, EPI_ISL_670342, EPI_ISL_670343, EPI_ISL_670344, EPI_ISL_670345, EPI_ISL_670346, EPI_ISL_670347, EPI_ISL_670348, EPI_ISL_670349, EPI_ISL_670350, EPI_ISL_670351, EPI_ISL_670352, EPI_ISL_670353, EPI_ISL_670354, EPI_ISL_670355, EPI_ISL_670356, EPI_ISL_670357, EPI_ISL_670358, EPI_ISL_670359, EPI_ISL_670360, EPI_ISL_670361, EPI_ISL_670362, EPI_ISL_670363, EPI_ISL_670364, EPI_ISL_670365, EPI_ISL_670366, EPI_ISL_670367, EPI_ISL_670368, EPI_ISL_670369, EPI_ISL_670370, EPI_ISL_670371, EPI_ISL_670372, EPI_ISL_670373, EPI_ISL_670374, EPI_ISL_670375, EPI_ISL_670376, EPI_ISL_670377, EPI_ISL_670378, EPI_ISL_670379, EPI_ISL_670380, EPI_ISL_670381, EPI_ISL_670382, EPI_ISL_670383, EPI_ISL_670384, EPI_ISL_670385, EPI_ISL_670386, EPI_ISL_670387, EPI_ISL_670388, EPI_ISL_670389, EPI_ISL_670390, EPI_ISL_670391, EPI_ISL_670392, EPI_ISL_670393, EPI_ISL_670394, EPI_ISL_670395, EPI_ISL_670396, EPI_ISL_670397, EPI_ISL_670398, EPI_ISL_670399, EPI_ISL_670400, EPI_ISL_670401, EPI_ISL_670402, EPI_ISL_670403, EPI_ISL_670404, EPI_ISL_670405, EPI_ISL_670406, EPI_ISL_670407, EPI_ISL_670408, EPI_ISL_670409, EPI_ISL_670410, EPI_ISL_670411, EPI_ISL_670412, EPI_ISL_670413, EPI_ISL_670414, EPI_ISL_670415, EPI_ISL_670416, EPI_ISL_670417, EPI_ISL_670418, EPI_ISL_670419, EPI_ISL_670420, EPI_ISL_670421, EPI_ISL_670422, EPI_ISL_670423, EPI_ISL_670424, EPI_ISL_670425, EPI_ISL_670426, EPI_ISL_670427, EPI_ISL_670428, EPI_ISL_670429, EPI_ISL_670430, EPI_ISL_670431, EPI_ISL_670432, EPI_ISL_670433, EPI_ISL_670434, EPI_ISL_670435, EPI_ISL_670436, EPI_ISL_670437, EPI_ISL_670438, EPI_ISL_670439, EPI_ISL_670440, EPI_ISL_670441, EPI_ISL_670442, EPI_ISL_670443, EPI_ISL_670444, EPI_ISL_670445, EPI_ISL_670446, EPI_ISL_670447, EPI_ISL_670448, EPI_ISL_670449, EPI_ISL_670450, EPI_ISL_670451, EPI_ISL_670452, EPI_ISL_670453, EPI_ISL_670454, EPI_ISL_670455, EPI_ISL_670456, EPI_ISL_670457, EPI_ISL_670459, EPI_ISL_670462, EPI_ISL_670463, EPI_ISL_670464, EPI_ISL_670465, EPI_ISL_670466, EPI_ISL_670467, EPI_ISL_670468, EPI_ISL_670469, EPI_ISL_670470, EPI_ISL_670474, EPI_ISL_670475, EPI_ISL_670476, EPI_ISL_670479, EPI_ISL_670480, EPI_ISL_670481, EPI_ISL_670483, EPI_ISL_670486, EPI_ISL_670487, EPI_ISL_670488, EPI_ISL_670492, EPI_ISL_670497, EPI_ISL_670499, EPI_ISL_670503, EPI_ISL_670504, EPI_ISL_670505, EPI_ISL_670506, EPI_ISL_670507, EPI_ISL_670508, EPI_ISL_670509, EPI_ISL_670519, EPI_ISL_670520, EPI_ISL_670521, EPI_ISL_670523, EPI_ISL_670529, EPI_ISL_670532, EPI_ISL_670533, EPI_ISL_670539, EPI_ISL_670542, EPI_ISL_670543, EPI_ISL_670548, EPI_ISL_670550, EPI_ISL_670551, EPI_ISL_670552, EPI_ISL_670553, EPI_ISL_670554, EPI_ISL_670555, EPI_ISL_670558, EPI_ISL_670559, EPI_ISL_670560, EPI_ISL_670561, EPI_ISL_670562, EPI_ISL_670563, EPI_ISL_670564, EPI_ISL_670565, EPI_ISL_670566, EPI_ISL_670567, EPI_ISL_670568, EPI_ISL_670576, EPI_ISL_670577, EPI_ISL_670578, EPI_ISL_670579, EPI_ISL_670580, EPI_ISL_670581, EPI_ISL_670582, EPI_ISL_670583, EPI_ISL_670584, EPI_ISL_670585, EPI_ISL_670586, EPI_ISL_670587, EPI_ISL_670588, EPI_ISL_670589, EPI_ISL_670590, EPI_ISL_670591, EPI_ISL_670592, EPI_ISL_670593, EPI_ISL_670594, EPI_ISL_670595, EPI_ISL_670596, EPI_ISL_670597, EPI_ISL_670598, EPI_ISL_670599, EPI_ISL_670600, EPI_ISL_670601, EPI_ISL_670602, EPI_ISL_670603, EPI_ISL_670604, EPI_ISL_670605, EPI_ISL_670606, EPI_ISL_670607, EPI_ISL_670608, EPI_ISL_670609, EPI_ISL_670610, EPI_ISL_670611, EPI_ISL_670612, EPI_ISL_670613, EPI_ISL_670614, EPI_ISL_670615, EPI_ISL_670616, EPI_ISL_670617, EPI_ISL_670618, EPI_ISL_670619, EPI_ISL_670620, EPI_ISL_670621, EPI_ISL_670622, EPI_ISL_670623, EPI_ISL_670624, EPI_ISL_670625, EPI_ISL_670666, EPI_ISL_670667, EPI_ISL_670668, EPI_ISL_670669, EPI_ISL_670676, EPI_ISL_670677, EPI_ISL_670678, EPI_ISL_670679, EPI_ISL_670680, EPI_ISL_670686, EPI_ISL_670687, EPI_ISL_670690, EPI_ISL_670691, EPI_ISL_670692, EPI_ISL_670693, EPI_ISL_670702, EPI_ISL_670703, EPI_ISL_670704, EPI_ISL_670705, EPI_ISL_670706, EPI_ISL_670707, EPI_ISL_670708, EPI_ISL_670709, EPI_ISL_670710, EPI_ISL_670711, EPI_ISL_670712, EPI_ISL_670713, EPI_ISL_670731, EPI_ISL_670732, EPI_ISL_670733, EPI_ISL_670734, EPI_ISL_670735, EPI_ISL_670736, EPI_ISL_670737, EPI_ISL_670738, EPI_ISL_670739, EPI_ISL_670740, EPI_ISL_670746, EPI_ISL_670747, EPI_ISL_670748, EPI_ISL_670749, EPI_ISL_670750, EPI_ISL_670751, EPI_ISL_670752, EPI_ISL_670753, EPI_ISL_670768, EPI_ISL_670772, EPI_ISL_670773, EPI_ISL_670774, EPI_ISL_670775, EPI_ISL_670776, EPI_ISL_670777, EPI_ISL_670778, EPI_ISL_670779, EPI_ISL_670780, EPI_ISL_670781, EPI_ISL_670782, EPI_ISL_670783, EPI_ISL_670784, EPI_ISL_670785, EPI_ISL_670786, EPI_ISL_670787, EPI_ISL_670788, EPI_ISL_670789, EPI_ISL_670790, EPI_ISL_670791, EPI_ISL_670792, EPI_ISL_670793, EPI_ISL_670794, EPI_ISL_670795, EPI_ISL_670796, EPI_ISL_670797, EPI_ISL_670798, EPI_ISL_670799, EPI_ISL_670800, EPI_ISL_670801, EPI_ISL_670802, EPI_ISL_670803, EPI_ISL_670804, EPI_ISL_670805, EPI_ISL_670806, EPI_ISL_670807, EPI_ISL_670808, EPI_ISL_670809, EPI_ISL_670810, EPI_ISL_670811, EPI_ISL_670812, EPI_ISL_670813, EPI_ISL_670814, EPI_ISL_670815, EPI_ISL_670816, EPI_ISL_670817, EPI_ISL_670818, EPI_ISL_670819, EPI_ISL_670820, EPI_ISL_670821, EPI_ISL_670822, EPI_ISL_670823, EPI_ISL_670824, EPI_ISL_670825, EPI_ISL_670826, EPI_ISL_670827, EPI_ISL_670828, EPI_ISL_670829, EPI_ISL_670830, EPI_ISL_670831, EPI_ISL_670832, EPI_ISL_670833, EPI_ISL_670834, EPI_ISL_670835, EPI_ISL_670836, EPI_ISL_670837, EPI_ISL_670838, EPI_ISL_670839, EPI_ISL_670840, EPI_ISL_670841, EPI_ISL_670842, EPI_ISL_670843, EPI_ISL_670844, EPI_ISL_670845, EPI_ISL_670846, EPI_ISL_670847, EPI_ISL_670848, EPI_ISL_670849, EPI_ISL_670850, EPI_ISL_670851, EPI_ISL_670852, EPI_ISL_670853, EPI_ISL_670854, EPI_ISL_670855, EPI_ISL_670856, EPI_ISL_670857, EPI_ISL_670858, EPI_ISL_670859, EPI_ISL_670860, EPI_ISL_670861, EPI_ISL_670862, EPI_ISL_670863, EPI_ISL_670864, EPI_ISL_670865, EPI_ISL_670866, EPI_ISL_670867, EPI_ISL_670868, EPI_ISL_670869, EPI_ISL_670870, EPI_ISL_670871, EPI_ISL_670872, EPI_ISL_670873, EPI_ISL_670874, EPI_ISL_670875, EPI_ISL_670876, EPI_ISL_670877, EPI_ISL_670878, EPI_ISL_670879, EPI_ISL_670880, EPI_ISL_670881, EPI_ISL_670882, EPI_ISL_670883, EPI_ISL_670884, EPI_ISL_670885, EPI_ISL_670886, EPI_ISL_670887, EPI_ISL_670888, EPI_ISL_670889, EPI_ISL_670890, EPI_ISL_670891, EPI_ISL_670892, EPI_ISL_670893, EPI_ISL_670894, EPI_ISL_670895, EPI_ISL_670896, EPI_ISL_670897, EPI_ISL_670898, EPI_ISL_670899, EPI_ISL_670900, EPI_ISL_670901, EPI_ISL_670902, EPI_ISL_670903, EPI_ISL_670904, EPI_ISL_670905, EPI_ISL_670906, EPI_ISL_670907, EPI_ISL_670908, EPI_ISL_670909, EPI_ISL_670910, EPI_ISL_670911, EPI_ISL_670912, EPI_ISL_670913, EPI_ISL_670914, EPI_ISL_670915, EPI_ISL_670916, EPI_ISL_670917, EPI_ISL_670918, EPI_ISL_670919, EPI_ISL_670920, EPI_ISL_670921, EPI_ISL_670922, EPI_ISL_670923, EPI_ISL_670924, EPI_ISL_670925, EPI_ISL_670926, EPI_ISL_670927, EPI_ISL_670928, EPI_ISL_670929, EPI_ISL_670930, EPI_ISL_670931, EPI_ISL_670932, EPI_ISL_670933, EPI_ISL_670934, EPI_ISL_670935, EPI_ISL_670936, EPI_ISL_670937, EPI_ISL_670938, EPI_ISL_670939, EPI_ISL_670940, EPI_ISL_670941, EPI_ISL_670942, EPI_ISL_670943, EPI_ISL_670944, EPI_ISL_670945, EPI_ISL_670946, EPI_ISL_670947, EPI_ISL_670948, EPI_ISL_670949, EPI_ISL_670950, EPI_ISL_670951, EPI_ISL_670952, EPI_ISL_670953, EPI_ISL_670954, EPI_ISL_670955, EPI_ISL_670956, EPI_ISL_670957, EPI_ISL_670958, EPI_ISL_670959, EPI_ISL_670960, EPI_ISL_670961, EPI_ISL_670962, EPI_ISL_670963, EPI_ISL_670964, EPI_ISL_670965, EPI_ISL_670966, EPI_ISL_670967, EPI_ISL_670968, EPI_ISL_670969, EPI_ISL_670970, EPI_ISL_670971, EPI_ISL_670972, EPI_ISL_670973, EPI_ISL_670974, EPI_ISL_670975, EPI_ISL_670976, EPI_ISL_670977, EPI_ISL_670978, EPI_ISL_670979, EPI_ISL_670980, EPI_ISL_670981, EPI_ISL_670982, EPI_ISL_670983, EPI_ISL_670984, EPI_ISL_670985, EPI_ISL_670986, EPI_ISL_670987, EPI_ISL_670988, EPI_ISL_670989, EPI_ISL_670990, EPI_ISL_670991, EPI_ISL_670992, EPI_ISL_670993, EPI_ISL_670994, EPI_ISL_670995, EPI_ISL_670996, EPI_ISL_670997, EPI_ISL_670998, EPI_ISL_670999, EPI_ISL_671000, EPI_ISL_671001, EPI_ISL_671002, EPI_ISL_671003, EPI_ISL_671004, EPI_ISL_671005, EPI_ISL_671006, EPI_ISL_671007, EPI_ISL_671008, EPI_ISL_671009, EPI_ISL_671010, EPI_ISL_671011, EPI_ISL_671012, EPI_ISL_671013, EPI_ISL_671014, EPI_ISL_671015, EPI_ISL_671016, EPI_ISL_671017, EPI_ISL_671018, EPI_ISL_671019, EPI_ISL_671020, EPI_ISL_671021, EPI_ISL_671022, EPI_ISL_671023, EPI_ISL_671024, EPI_ISL_671025, EPI_ISL_671026, EPI_ISL_671027, EPI_ISL_671028, EPI_ISL_671029, EPI_ISL_671030, EPI_ISL_671031, EPI_ISL_671032, EPI_ISL_671033, EPI_ISL_671034, EPI_ISL_671035, EPI_ISL_671036, EPI_ISL_671037, EPI_ISL_671038, EPI_ISL_671039 | | | |
| see above | Department of Virus and Microbiological Special Diagnostics, Statens Serum Institut, Copenhagen, Denmark | Albertsen Lab, Department of Chemistry and Bioscience, Aalborg University, Denmark | Danish Covid-19 Genome Consortium |
| EPI_ISL_672048, EPI_ISL_672067 | Madera County Department of Public Health | Chan-Zuckerberg Biohub | CZB Cliahub Consortium |
| EPI_ISL_672434, EPI_ISL_672435, EPI_ISL_672436, EPI_ISL_672437, EPI_ISL_672438 | Santa Clara County Public Health Laboratory | Chan-Zuckerberg Biohub | CZB Cliahub Consortium |
| EPI_ISL_672481, EPI_ISL_672482, EPI_ISL_672483, EPI_ISL_672484, EPI_ISL_672485, EPI_ISL_672486, EPI_ISL_672487 | Orange County Public Health Lab | Chan-Zuckerberg Biohub | CZB Cliahub Consortium |
| EPI_ISL_672504, EPI_ISL_672505 | Madera County Department of Public Health | Chan-Zuckerberg Biohub | CZB Cliahub Consortium |
| EPI_ISL_676395, EPI_ISL_676396, EPI_ISL_676397, EPI_ISL_676398, EPI_ISL_676401, EPI_ISL_676402, EPI_ISL_676403, EPI_ISL_676404, EPI_ISL_676429, EPI_ISL_676430, EPI_ISL_676431, EPI_ISL_676432, EPI_ISL_676433, EPI_ISL_676435, EPI_ISL_676436, EPI_ISL_676437, EPI_ISL_676438, EPI_ISL_676439, EPI_ISL_676440 | | | |
| see above | Lighthouse Lab in Glasgow | Wellcome Sanger Institute for the COVID-19 Genomics UK (COG-UK) Consortium | Harper VanSteenhouse, Yumi Kasai, David Gray, Carol Clugston, Anna Dominiczak and Alex Alderton, Roberto Amato, Sonia Goncalves, Ewan Harrison, David K. Jackson, Ian Johnston, Dominic Kwiatkowski, Cordelia Langford, John Sillitoe on behalf of the Wellcome Sanger Institute COVID-19 Surveillance Team |
| EPI_ISL_676499 | Klinisk Mikrobiologi | The Public Health Agency of Sweden | Department of Microbiology, The Public Health Agency of Sweden |
| EPI_ISL_676539, EPI_ISL_676540 | Laboratoriemedicin, Klinisk mikrobiologi | The Public Health Agency of Sweden | Department of Microbiology, The Public Health Agency of Sweden |
| EPI_ISL_677498, EPI_ISL_677499, EPI_ISL_677500, EPI_ISL_677632, EPI_ISL_677633 | University of Wisconsin-Madison AIDS Vaccine Research Laboratories | University of Wisconsin-Madison AIDS Vaccine Research Laboratories | Gage Moreno, Katarina Braun, et al. AIDS Vaccine Research Laboratories |
| EPI_ISL_678376 | Area of Virology, Serology and Virology Division (SAViD), New South Wales Health Pathology Randwick | Virology Research Laboratory; Area of Virology, Serology and Virology Division (SAViD), New South Wales Health Pathology Randwick | Foster, C.; Au, J.; Ruiz Silva, M.; Deveson, I.; Bull, R.; Van Hal, S.; Rawlinson, W. |
| EPI_ISL_678599 | NHLS-IALCH | KRISP, KZN Research Innovation and Sequencing Platform | Giandhari J, Pillay S, Lessells R, ChimukangaraB, Mdlalose K, York D, Khan S, Tegally H, Wilkinson E, de Oliveira T |
| EPI_ISL_678644, EPI_ISL_678645, EPI_ISL_678646, EPI_ISL_678647, EPI_ISL_678648, EPI_ISL_678649, EPI_ISL_678651, EPI_ISL_678652, EPI_ISL_678653, EPI_ISL_678654, EPI_ISL_678655, EPI_ISL_678656, EPI_ISL_678657, EPI_ISL_678659, EPI_ISL_678661, EPI_ISL_678663, EPI_ISL_678664, EPI_ISL_678665, EPI_ISL_678667, EPI_ISL_678669, EPI_ISL_678671, EPI_ISL_678676, EPI_ISL_678680 | | | |
| see above | Respiratory Virus Unit, Microbiology Services Colindale, Public Health England | COVID-19 Genomics UK (COG-UK) Consortium | PHE Covid Sequencing Team |
| EPI_ISL_678831 | Centre for Enzyme Innovation, University of Portsmouth / Translational Research Laboratory, Portsmouth Hospitals NHS Trust | COVID-19 Genomics UK (COG-UK) Consortium | Angela Beckett,Yann Bourgeois,Garry Scarlett,Sharon Glaysher,Scott Elliott,Kelly Bicknell,Robert Impey,Allyson Lloyd,Sarah Wyllie,Ethan Butcher,Anoop Chauhan,Samuel Robson |
| EPI_ISL_679140 | Department of Pathology, University of Cambridge | COVID-19 Genomics UK (COG-UK) Consortium | Aminu S. Jahun, Yasmin Chaudhry, Grant Hall, Iliana Georgana, Myra Hosmillo, Martin D. Curran, Malte Pinckert, Surendra Parmar, Ian Goodfellow |

| | | | |
|---|---|---|---|
| EPI_ISL_679450, EPI_ISL_679458, EPI_ISL_679459, EPI_ISL_679460, EPI_ISL_679461, EPI_ISL_679462, EPI_ISL_679463, EPI_ISL_679464, EPI_ISL_679465, EPI_ISL_679467, EPI_ISL_679468, EPI_ISL_679469, EPI_ISL_679470, EPI_ISL_679471, EPI_ISL_679472, EPI_ISL_679473, EPI_ISL_679477 | | | |
| see above | University College London, Great Ormond Street Hospital for Children NHS Foundation Trust, Imperial College Healthcare NHS Trust | COVID-19 Genomics UK (COG-UK) Consortium | Sergi Castellano, Rachel Williams, Mark Kristiansen, Paola Resende Silva, Sunando Roy, Tony Brooks, Helena Tutill, Paola Niola, Patricia Dyal, Charlotte Williams, Leysa Forrest, Yasmin Panchbhaya, Jacqueline Findlay, Samuel Weeks, Julianne Brown, Kathryn Harris, Paul Randell, James Price, Alison Holmes, Judith Breuer |
| EPI_ISL_679629, EPI_ISL_679630, EPI_ISL_679632, EPI_ISL_679633, EPI_ISL_679642, EPI_ISL_679643, EPI_ISL_679644, EPI_ISL_679645, EPI_ISL_679648, EPI_ISL_679651, EPI_ISL_679652, EPI_ISL_679653, EPI_ISL_679655, EPI_ISL_679658, EPI_ISL_679659, EPI_ISL_679666, EPI_ISL_679669, EPI_ISL_679671, EPI_ISL_679673, EPI_ISL_679678, EPI_ISL_679679, EPI_ISL_679680, EPI_ISL_679682, EPI_ISL_679683, EPI_ISL_679684, EPI_ISL_679685, EPI_ISL_679688, EPI_ISL_679690, EPI_ISL_679692, EPI_ISL_679693, EPI_ISL_679694, EPI_ISL_679695, EPI_ISL_679696, EPI_ISL_679697, EPI_ISL_679698, EPI_ISL_679699, EPI_ISL_679700, EPI_ISL_679702, EPI_ISL_679703, EPI_ISL_679704, EPI_ISL_679705, EPI_ISL_679706, EPI_ISL_679707, EPI_ISL_679709, EPI_ISL_679712, EPI_ISL_679714, EPI_ISL_679716, EPI_ISL_679717, EPI_ISL_679718, EPI_ISL_679719, EPI_ISL_679720, EPI_ISL_679721, EPI_ISL_679723, EPI_ISL_679766, EPI_ISL_679767, EPI_ISL_679770, EPI_ISL_679773, EPI_ISL_679774, EPI_ISL_679777, EPI_ISL_679783, EPI_ISL_679784, EPI_ISL_679785, EPI_ISL_679787, EPI_ISL_679788, EPI_ISL_679806, EPI_ISL_679810, EPI_ISL_679811, EPI_ISL_679812, EPI_ISL_679814, EPI_ISL_679816, EPI_ISL_679818, EPI_ISL_679819 | | | |
| see above | Oxford Viromics, NDM, University of Oxford; Oxford University Hospitals; Basingstoke and North Hampshire Hospital | COVID-19 Genomics UK (COG-UK) Consortium | Tanya Golubchik, David Bonsall, George Macintyre, Amy Trebes, Mariateresa de Cesare, Catrin Moore, Alex Mobbs, Anita Justice, Robert Shaw, Monique Andersson, Timothy Peto, Emma Wise, Nathan Moore, Jessica Lynch, Nick Cortes, Matilde Mori, Stephen Kidd, David Buck, John Todd, Christophe Fraser |
| EPI_ISL_679905 | Virology Department, Sheffield Teaching Hospitals NHS Foundation Trust/Department of Infection, Immunity and Cardiovascular Disease, The Medical School, University of Sheffield | COVID-19 Genomics UK (COG-UK) Consortium | Thushan de Silva, Matthew Parker, Nikki Smith, Adri Angyal, Rebecca Brown, Luke Green, Rachel Tucker, Paul Parsons, Danielle Groves, Katie Johnson, Laura Carrilero, Alex Keeley, Dave Partridge, Matthew Wyles, Benjamin Lindsey, Mehmet Yavuz, Mohammad Raza, Cariad Evans |
| EPI_ISL_679928, EPI_ISL_679929, EPI_ISL_679930 | Queens Medical Centre, Clinical Microbiology Department / DeepSeq Nottingham | COVID-19 Genomics UK (COG-UK) Consortium | Gemma Clark, Wendy Smith, Manjinder Khakh, Vicki M Fleming, Michelle M Lister, Hannah Howson-Wells, Jonathan Ball, Patrick McClure, Joseph Chappell, Theocharis Tsoleridis, Nadine Holmes, Matthew Carlisle, Christopher Moore, Fei Sang, Johnny Debebe, Victoria Wright, Matthew Loose |
| EPI_ISL_680049, EPI_ISL_680055, EPI_ISL_680059, EPI_ISL_680068, EPI_ISL_680069, EPI_ISL_680070, EPI_ISL_680073, EPI_ISL_680074, EPI_ISL_680075, EPI_ISL_680076, EPI_ISL_680077 | | | |
| see above | Department of Pathology, University of Cambridge | COVID-19 Genomics UK (COG-UK) Consortium | Aminu S. Jahun, Yasmin Chaudhry, Grant Hall, Iliana Georgana, Myra Hosmillo, Martin D. Curran, Malte Pinckert, Surendra Parmar, Ian Goodfellow |
| EPI_ISL_680131 | Virology Department, Sheffield Teaching Hospitals NHS Foundation Trust/Department of Infection, Immunity and Cardiovascular Disease, The Medical School, University of Sheffield | COVID-19 Genomics UK (COG-UK) Consortium | Thushan de Silva, Matthew Parker, Nikki Smith, Adri Angyal, Rebecca Brown, Luke Green, Rachel Tucker, Paul Parsons, Danielle Groves, Katie Johnson, Laura Carrilero, Alex Keeley, Dave Partridge, Matthew Wyles, Benjamin Lindsey, Mehmet Yavuz, Mohammad Raza, Cariad Evans |
| EPI_ISL_680544, EPI_ISL_680545 | Virology Department, Royal Infirmary of Edinburgh, NHS Lothian / School of Biological Sciences, University of Edinburgh / Institute of Genetics and Molecular Medicine, University of Edinburgh | COVID-19 Genomics UK (COG-UK) Consortium | McHugh M, Dewar R, Rooke S, Gallagher M, Balcaza C, O'Toole Á, Scher E, Hill V, McCrone JT, Colquhoun R, Yu X, Jackson B, Rambaut A, Williams TC, Templeton K |
| EPI_ISL_680571, EPI_ISL_680572, EPI_ISL_680573 | Department of Pathology, University of Cambridge | COVID-19 Genomics UK (COG-UK) Consortium | Aminu S. Jahun, Yasmin Chaudhry, Grant Hall, Iliana Georgana, Myra Hosmillo, Martin D. Curran, Malte Pinckert, Surendra Parmar, Ian Goodfellow |
| EPI_ISL_680585, EPI_ISL_680590, EPI_ISL_680685, EPI_ISL_680732, EPI_ISL_680733, EPI_ISL_680734, EPI_ISL_680735, EPI_ISL_680736, EPI_ISL_680737, EPI_ISL_680738, EPI_ISL_680740, EPI_ISL_680961, EPI_ISL_680962, EPI_ISL_680963, EPI_ISL_680964, EPI_ISL_680965, EPI_ISL_680966, EPI_ISL_680967, EPI_ISL_681028, EPI_ISL_681029, EPI_ISL_681030, EPI_ISL_681031, EPI_ISL_681032, EPI_ISL_681033, EPI_ISL_681034 | | | |
| see above | Wales Specialist Virology Centre Sequencing lab: Pathogen Genomics Unit | COVID-19 Genomics UK (COG-UK) Consortium | Catherine Moore, Johnathan Evans, Laura Gifford, Malorie Perry, Simon Cottrell, Angela Marchbank, Alec Birchley, Alexander Adams, Amy Gaskin, Bree Gatica-Wilcox, Jason Coombes, Joel Southgate, Lauren Gilbert, Lee Graham, Nicole Pacchiarini, Sara Kumziene-Summerhayes, Sarah Taylor, Sophie Jones, Sara Rey, Matthew Bull, Joanne Watkins, Sally Corden, Tom Connor |
| EPI_ISL_681260 | Department of Pathology, University of Cambridge | COVID-19 Genomics UK (COG-UK) Consortium | Aminu S. Jahun, Yasmin Chaudhry, Grant Hall, Iliana Georgana, Myra Hosmillo, Martin D. Curran, Malte Pinckert, Surendra Parmar, Ian Goodfellow |
| EPI_ISL_682105, EPI_ISL_682108, EPI_ISL_682111, EPI_ISL_682124, EPI_ISL_682125, EPI_ISL_682129, EPI_ISL_682133, EPI_ISL_682161, EPI_ISL_682162, EPI_ISL_682163, EPI_ISL_682164, EPI_ISL_682165, EPI_ISL_682166, EPI_ISL_682167, EPI_ISL_682168, EPI_ISL_682169, EPI_ISL_682170, EPI_ISL_682171, EPI_ISL_682172, EPI_ISL_682173, EPI_ISL_682174, EPI_ISL_682175, EPI_ISL_682176, EPI_ISL_682177, EPI_ISL_682178, EPI_ISL_682179, EPI_ISL_682180, EPI_ISL_682181, EPI_ISL_682182, EPI_ISL_682183, EPI_ISL_682184, EPI_ISL_682185, EPI_ISL_682186, EPI_ISL_682187, EPI_ISL_682188, EPI_ISL_682189, EPI_ISL_682190, EPI_ISL_682191, EPI_ISL_682192, EPI_ISL_682193, EPI_ISL_682194, EPI_ISL_682195, EPI_ISL_682196, EPI_ISL_682197, EPI_ISL_682198, EPI_ISL_682200, EPI_ISL_682201, EPI_ISL_682202, EPI_ISL_682203, EPI_ISL_682204, EPI_ISL_682205, EPI_ISL_682206, EPI_ISL_682207, EPI_ISL_682208, EPI_ISL_682209, EPI_ISL_682210, EPI_ISL_682211, EPI_ISL_682212, EPI_ISL_682213, EPI_ISL_682214, EPI_ISL_682215, EPI_ISL_682216, EPI_ISL_682217, EPI_ISL_682218, EPI_ISL_682219, EPI_ISL_682220, EPI_ISL_682221, EPI_ISL_682222, EPI_ISL_682223, EPI_ISL_682224, EPI_ISL_682225, EPI_ISL_682226, EPI_ISL_682227, EPI_ISL_682228, EPI_ISL_682229, EPI_ISL_682230, EPI_ISL_682231, EPI_ISL_682232, EPI_ISL_682233, EPI_ISL_682234 | | | |
| see above | University of Michigan Clinical Microbiology Laboratory | Lauring Lab, University of Michigan, Department of Microbiology and Immunology | Valesano |
| EPI_ISL_682967, EPI_ISL_682968, EPI_ISL_682969, EPI_ISL_682970, EPI_ISL_682971, EPI_ISL_682972, EPI_ISL_682973, EPI_ISL_682974, EPI_ISL_682975, EPI_ISL_682976, EPI_ISL_682977, EPI_ISL_682978, EPI_ISL_682979, EPI_ISL_682980, EPI_ISL_682981, EPI_ISL_682982, EPI_ISL_683038, EPI_ISL_683039, EPI_ISL_683040, EPI_ISL_683041, EPI_ISL_683042, EPI_ISL_683043, EPI_ISL_683044, EPI_ISL_683045, EPI_ISL_683046, EPI_ISL_683047, EPI_ISL_683048, EPI_ISL_683049, EPI_ISL_683050, EPI_ISL_683051, EPI_ISL_683052, EPI_ISL_683053, EPI_ISL_683054, EPI_ISL_683055, EPI_ISL_683056, EPI_ISL_683057, EPI_ISL_683058, EPI_ISL_683059, EPI_ISL_683060, EPI_ISL_683061, EPI_ISL_683062, EPI_ISL_683063, EPI_ISL_683064, EPI_ISL_683065, EPI_ISL_683066, EPI_ISL_683067, EPI_ISL_683068, EPI_ISL_683069, EPI_ISL_683070, EPI_ISL_683071, EPI_ISL_683072, EPI_ISL_683073, EPI_ISL_683074, EPI_ISL_683075, EPI_ISL_683076, EPI_ISL_683077, EPI_ISL_683078, EPI_ISL_683079, EPI_ISL_683080, EPI_ISL_683081, EPI_ISL_683082, EPI_ISL_683083, EPI_ISL_683084, EPI_ISL_683085, EPI_ISL_683086 | | | |
| see above | Department of Virus and Microbiological Special Diagnostics, Statens Serum Institut, Copenhagen, Denmark | Albertsen Lab, Department of Chemistry and Bioscience, Aalborg University, Denmark | Danish Covid-19 Genome Consortium |
| EPI_ISL_683334, EPI_ISL_683402 | CNR Virus des Infections Respiratoires - France SUD | CNR Virus des Infections Respiratoires - France SUD | Antonin Bal, Gregory Destras, Gwendolyne Burfin, Quentin Semanas, Martine Valette, Bruno Lina, Laurence Josset |
| EPI_ISL_683611, EPI_ISL_683612, EPI_ISL_683613, EPI_ISL_683614, EPI_ISL_683615, EPI_ISL_683616, EPI_ISL_683617, EPI_ISL_683618, EPI_ISL_683647, EPI_ISL_683651, EPI_ISL_683656 | | | |
| see above | Servicio de Microbiología, Laboratori Clínic Metropolitana Nord. Hospital Universitari Germans Trias i Pujol. Institut d'Investigació en Ciències de la Salut Germans Trias i Pujol (IGTP) | SeqCOVID-SPAIN consortium/IBV(CSIC) | Elisa Martró, Antoni E. Bordoy, Anna Not, Adrián Antuori, Anabel Fernández, Nona Romaní, Verónica Saludes, Cristina Casañ and SeqCOVID-SPAIN consortium |
| EPI_ISL_683698, EPI_ISL_683699, EPI_ISL_683700, EPI_ISL_683701, EPI_ISL_683709, EPI_ISL_683711 | Essentia Health-St. Mary's Medical Center | Minnesota Department of Health, Public Health Laboratory | Alexandra Lorentz, Jacob Garfin, Matt Plumb, and Xiong Wang |
| EPI_ISL_683735, EPI_ISL_683736, EPI_ISL_683737, EPI_ISL_683738, EPI_ISL_683739, EPI_ISL_683740, EPI_ISL_683741, EPI_ISL_683742, EPI_ISL_683743 | Minnesota Department of Health, Public Health Laboratory | Minnesota Department of Health, Public Health Laboratory | Alexandra Lorentz, Jacob Garfin, Matt Plumb, and Xiong Wang |
| EPI_ISL_683773 | DOHMH Corona | New York City Public Health Laboratory | Jade Wang, et al. |
| EPI_ISL_683774 | DOHMH PHL | New York City Public Health Laboratory | Jade Wang, et al. |
| EPI_ISL_683775, EPI_ISL_683776 | DOHMH Jamaica | New York City Public Health Laboratory | Jade Wang, et al. |
| EPI_ISL_683777 | DOHMH Corona | New York City Public Health Laboratory | Jade Wang, et al. |
| EPI_ISL_683778 | DOHMH Morrisania | New York City Public Health Laboratory | Jade Wang, et al. |
| EPI_ISL_683859 | DOHMH Central Harlem | New York City Public Health Laboratory | Jade Wang, et al. |
| EPI_ISL_686541, EPI_ISL_686542, EPI_ISL_686543 | Respiratory Virus Unit, Microbiology Services Colindale, Public Health England | COVID-19 Genomics UK (COG-UK) Consortium | PHE Covid Sequencing Team |
| EPI_ISL_692802, EPI_ISL_692803, EPI_ISL_692804, EPI_ISL_692805, EPI_ISL_692808 | Massachusetts State Public Health Laboratory | Massachusetts State Public Health Laboratory | Andrew Lang, Timelia Fink, Glen Gallagher, Sandra Smole |

| | | | |
|---|---|---|---|
| EPI_ISL_693661, EPI_ISL_693665, EPI_ISL_693666, EPI_ISL_693669, EPI_ISL_693672, EPI_ISL_693675, EPI_ISL_693677, EPI_ISL_693678, EPI_ISL_693679, EPI_ISL_693682, EPI_ISL_693684, EPI_ISL_693685, EPI_ISL_693686 | | | |
| see above | The National Institute of Public Health | State Veterinary Institute Prague | Nagy,A;Jirincova,H;Trnka,D;Vecerova,J |
| EPI_ISL_693767, EPI_ISL_693769 | General practitioner | National Reference Center for Viruses of Respiratory Infections, Institut Pasteur, Paris | Marion Barbet, Sylvie Behillil, Méline Bizard, Angela Brisebarre, Camille Capel, Etienne Simon-Lorière, Vincent Enouf, Maud Vanpeene, Sylvie van der Werf |
| EPI_ISL_693775, EPI_ISL_693776, EPI_ISL_693777, EPI_ISL_693778, EPI_ISL_693779, EPI_ISL_693780, EPI_ISL_693781, EPI_ISL_693782, EPI_ISL_693783, EPI_ISL_693784, EPI_ISL_693785, EPI_ISL_693786, EPI_ISL_693787, EPI_ISL_693788, EPI_ISL_693789, EPI_ISL_693790, EPI_ISL_693791, EPI_ISL_693792, EPI_ISL_693793, EPI_ISL_693794, EPI_ISL_693795, EPI_ISL_693796, EPI_ISL_693797, EPI_ISL_693798, EPI_ISL_693799, EPI_ISL_693800, EPI_ISL_693801, EPI_ISL_693802, EPI_ISL_693803, EPI_ISL_693804, EPI_ISL_693805, EPI_ISL_693806, EPI_ISL_693807, EPI_ISL_693808, EPI_ISL_693809, EPI_ISL_693810, EPI_ISL_693811, EPI_ISL_693812, EPI_ISL_693813, EPI_ISL_693814, EPI_ISL_693815, EPI_ISL_693816, EPI_ISL_693817, EPI_ISL_693818, EPI_ISL_693819, EPI_ISL_693820, EPI_ISL_693821, EPI_ISL_693822, EPI_ISL_693823, EPI_ISL_693824, EPI_ISL_693825 | | | |
| see above | Viollier AG | Department of Biosystems Science and Engineering, ETH Zürich | Christian Beisel, Sarah Nadeau, Chaoran Chen, Ivan Topolsky, Pedro Ferreira, Philipp Jablonski, Susana Posada-Céspedes, Tobias Schär, Ina Nissen, Natascha Santacroce, Elodie Burcklen, Christiane Beckmann, Maurice Redondo, Olivier Kobel, Christoph Noppen, Sophie Seidel, Noemie Santamaria de Souza, Niko Beerenwinkel, Tanja Stadler |
| EPI_ISL_700431 | Thembalethu CDC wc THC | NHLS/UCT | Houriiyah Tegally, Arash Iranzadeh, Deelan Doolabh, Lynn Tyers, Bruna Galvao, Innocent Mudau, Marvin Hsiao, Kruger Marais, Diana Hardie, Stephen Korsman, Carolyn Williamson |
| EPI_ISL_700439 | Pacaltsdorp Clinic wc PAC | NHLS/UCT | Houriiyah Tegally, Arash Iranzadeh, Deelan Doolabh, Lynn Tyers, Bruna Galvao, Innocent Mudau, Marvin Hsiao, Kruger Marais, Diana Hardie, Stephen Korsman, Carolyn Williamson |
| EPI_ISL_700458 | Hornlee Clinic wc HLC | NHLS/UCT | Houriiyah Tegally, Arash Iranzadeh, Deelan Doolabh, Lynn Tyers, Bruna Galvao, Innocent Mudau, Marvin Hsiao, Kruger Marais, Diana Hardie, Stephen Korsman, Carolyn Williamson |
| EPI_ISL_700467, EPI_ISL_700496 | Conville CDC wc CVC | NHLS/UCT | Houriiyah Tegally, Arash Iranzadeh, Deelan Doolabh, Lynn Tyers, Bruna Galvao, Innocent Mudau, Marvin Hsiao, Kruger Marais, Diana Hardie, Stephen Korsman, Carolyn Williamson |
| EPI_ISL_700500 | Plettenberg Bay Clinic wc PLC | NHLS/UCT | Houriiyah Tegally, Arash Iranzadeh, Deelan Doolabh, Lynn Tyers, Bruna Galvao, Innocent Mudau, Marvin Hsiao, Kruger Marais, Diana Hardie, Stephen Korsman, Carolyn Williamson |
| EPI_ISL_700548 | Alma CDC wc AHC | NHLS/UCT | Arash Iranzadeh, Deelan Doolabh, Lynn Tyers, Bruna Galvao, Innocent Mudau, Marvin Hsiao, Kruger Marais, Diana Hardie, Stephen Korsman, Carolyn Williamson |
| EPI_ISL_700554 | Conville CDC wc CVC | NHLS/UCT | Houriiyah Tegally, Arash Iranzadeh, Deelan Doolabh, Lynn Tyers, Bruna Galvao, Innocent Mudau, Marvin Hsiao, Kruger Marais, Diana Hardie, Stephen Korsman, Carolyn Williamson |
| EPI_ISL_702569, EPI_ISL_702667, EPI_ISL_702779 | Liverpool Clinical Laboratories | COVID-19 Genomics UK (COG-UK) Consortium | Sam Haldenby, Anita Lucaci, Steve Paterson, Julian Hiscox, Alistair Darby, M Almsaud, A Alrezaihi, Muhannad Alruwaili, Stuart D Armstrong, Jones Benjamin, Eleanor G Bentley, Anu Chawla, Jordan J Clark, Angela Cowell, Richard Eccles, Isabel García-Dorival, Matthew Gemmell, Alessandro Gerada, PKF Gilmore, Richard Gregory, Ximeng Han, Catherine Hartley, Margaret Hughes, Miren Iturriza-Gomara, James Johnson, L Luu, Jenifer Manson, Charlotte Nelson, Elaine O'Toole, Cassie Olateju, Rebekah Penrice-Randal , Lucille Rainbow, N.P Randle, Trevor Ian Robinson, Parul Sharma, Ghada T Shawli, James P Stewart, Neil Swainston, Ecaterina Vamos, Joanne Watts, Mark Whitehead |
| EPI_ISL_702784 | Northumbria University / South Tees Hospitals NHS Foundation Trust / North Cumbria Integrated Care NHS Foundation Trust / North Tees and Hartlepool NHS Foundation Trust / Newcastle Hospitals NHS Foundation Trust | COVID-19 Genomics UK (COG-UK) Consortium | Darren L Smith,Andrew Nelson,Matthew Bashton,Greg R Young,Joshua Loh,John Allan,Mohammad A Tariq,Giles S Holt,Gary Black,Wen C Yew,Lynn Dover,Paul Baker,Steve Liggett,Sarah Essex,Jane Greenaway,Debra Padgett,Clive Graham,Garren Scott,Edward Barton,Emma Swindells,Brendan Payne,Jennifer Collins,Yusri Taha,Gary Eltringham |
| EPI_ISL_702792, EPI_ISL_702877 | Liverpool Clinical Laboratories | COVID-19 Genomics UK (COG-UK) Consortium | Sam Haldenby, Anita Lucaci, Steve Paterson, Julian Hiscox, Alistair Darby, M Almsaud, A Alrezaihi, Muhannad Alruwaili, Stuart D Armstrong, Jones Benjamin, Eleanor G Bentley, Anu Chawla, Jordan J Clark, Angela Cowell, Richard Eccles, Isabel García-Dorival, Matthew Gemmell, Alessandro Gerada, PKF Gilmore, Richard Gregory, Ximeng Han, Catherine Hartley, Margaret Hughes, Miren Iturriza-Gomara, James Johnson, L Luu, Jenifer Manson, Charlotte Nelson, Elaine O'Toole, Cassie Olateju, Rebekah Penrice-Randal , Lucille Rainbow, N.P Randle, Trevor Ian Robinson, Parul Sharma, Ghada T Shawli, James P Stewart, Neil Swainston, Ecaterina Vamos, Joanne Watts, Mark Whitehead |
| EPI_ISL_702937 | Virology Department, Royal Infirmary of Edinburgh, NHS Lothian / School of Biological Sciences, University of Edinburgh / Institute of Genetics and Molecular Medicine, University of Edinburgh | COVID-19 Genomics UK (COG-UK) Consortium | McHugh M, Dewar R, Rooke S, Gallagher M, Balcaza C, O'Toole Á, Scher E, Hill V, McCrone JT, Colquhoun R, Yu X, Jackson B, Rambaut A, Williams TC, Templeton K |
| EPI_ISL_702953 | Liverpool Clinical Laboratories | COVID-19 Genomics UK (COG-UK) Consortium | Sam Haldenby, Anita Lucaci, Steve Paterson, Julian Hiscox, Alistair Darby, M Almsaud, A Alrezaihi, Muhannad Alruwaili, Stuart D Armstrong, Jones Benjamin, Eleanor G Bentley, Anu Chawla, Jordan J Clark, Angela Cowell, Richard Eccles, Isabel García-Dorival, Matthew Gemmell, Alessandro Gerada, PKF Gilmore, Richard Gregory, Ximeng Han, Catherine Hartley, Margaret Hughes, Miren Iturriza-Gomara, James Johnson, L Luu, Jenifer Manson, Charlotte Nelson, Elaine O'Toole, Cassie Olateju, Rebekah Penrice-Randal , Lucille Rainbow, N.P Randle, Trevor Ian Robinson, Parul Sharma, Ghada T Shawli, James P Stewart, Neil Swainston, Ecaterina Vamos, Joanne Watts, Mark Whitehead |
| EPI_ISL_703048 | Virology Department, Sheffield Teaching Hospitals NHS Foundation Trust/Department of Infection, Immunity and Cardiovascular Disease, The Medical School, University of Sheffield | COVID-19 Genomics UK (COG-UK) Consortium | Thushan de Silva, Matthew Parker, Nikki Smith, Adri Angyal, Rebecca Brown, Luke Green, Rachel Tucker, Paul Parsons, Danielle Groves, Katie Johnson, Laura Carrilero, Alex Keeley, Dave Partridge, Matthew Wyles, Benjamin Lindsey, Mehmet Yavuz, Mohammad Raza, Cariad Evans |
| EPI_ISL_703085, EPI_ISL_703881, EPI_ISL_704057, EPI_ISL_704129 | Liverpool Clinical Laboratories | COVID-19 Genomics UK (COG-UK) Consortium | Sam Haldenby, Anita Lucaci, Steve Paterson, Julian Hiscox, Alistair Darby, M Almsaud, A Alrezaihi, Muhannad Alruwaili, Stuart D Armstrong, Jones Benjamin, Eleanor G Bentley, Anu Chawla, Jordan J Clark, Angela Cowell, Richard Eccles, Isabel García-Dorival, Matthew Gemmell, Alessandro Gerada, PKF Gilmore, Richard Gregory, Ximeng Han, Catherine Hartley, Margaret Hughes, Miren Iturriza-Gomara, James Johnson, L Luu, Jenifer Manson, Charlotte Nelson, Elaine O'Toole, Cassie Olateju, Rebekah Penrice-Randal , Lucille Rainbow, N.P Randle, Trevor Ian Robinson, Parul Sharma, Ghada T Shawli, James P Stewart, Neil Swainston, Ecaterina Vamos, Joanne Watts, Mark Whitehead |
| EPI_ISL_704676, EPI_ISL_704679, EPI_ISL_704692, EPI_ISL_704695, EPI_ISL_704706, EPI_ISL_704815, EPI_ISL_704836, EPI_ISL_704879, EPI_ISL_704885 | Oxford Viromics, NDM, University of Oxford; Oxford University Hospitals; Basingstoke and North Hampshire Hospital | COVID-19 Genomics UK (COG-UK) Consortium | Tanya Golubchik, David Bonsall, George Macintyre, Amy Trebes, Mariateresa de Cesare, Catrin Moore, Alex Mobbs, Anita Justice, Robert Shaw, Monique Andersson, Timothy Peto, Emma Wise, Nathan Moore, Jessica Lynch, Nick Cortes, Matilde Mori, Stephen Kidd, David Buck, John Todd, Christophe Fraser |
| EPI_ISL_705224 | Northumbria University / South Tees Hospitals NHS Foundation Trust / North Cumbria Integrated Care NHS Foundation Trust / North Tees and Hartlepool NHS Foundation Trust / Newcastle Hospitals NHS Foundation Trust | COVID-19 Genomics UK (COG-UK) Consortium | Darren L Smith,Andrew Nelson,Matthew Bashton,Greg R Young,Joshua Loh,John Allan,Mohammad A Tariq,Giles S Holt,Gary Black,Wen C Yew,Lynn Dover,Paul Baker,Steve Liggett,Sarah Essex,Jane Greenaway,Debra Padgett,Clive Graham,Garren Scott,Edward Barton,Emma Swindells,Brendan Payne,Jennifer Collins,Yusri Taha,Gary Eltringham |
| EPI_ISL_705236 | University College London, Great Ormond Street Hospital for Children NHS Foundation Trust, Imperial College Healthcare NHS Trust | COVID-19 Genomics UK (COG-UK) Consortium | Sergi Castellano, Rachel Williams, Mark Kristiansen, Paola Resende Silva, Sunando Roy, Tony Brooks, Helena Tutill, Paola Niola, Patricia Dyal, Charlotte Williams, Leysa Forrest, Yasmin Panchbhaya, Jacqueline Findlay, Samuel Weeks, Julianne Brown, Kathryn Harris, Paul Randell, James Price, Alison Holmes, Judith Breuer |
| EPI_ISL_705259 | Liverpool Clinical Laboratories | COVID-19 Genomics UK (COG-UK) Consortium | Sam Haldenby, Anita Lucaci, Steve Paterson, Julian Hiscox, Alistair Darby, M Almsaud, A Alrezaihi, Muhannad Alruwaili, Stuart D Armstrong, Jones Benjamin, Eleanor G Bentley, Anu Chawla, Jordan J Clark, Angela Cowell, Richard Eccles, Isabel García-Dorival, Matthew Gemmell, Alessandro Gerada, PKF Gilmore, Richard Gregory, Ximeng Han, Catherine Hartley, Margaret Hughes, Miren Iturriza-Gomara, James Johnson, L Luu, Jenifer Manson, Charlotte Nelson, Elaine O'Toole, Cassie Olateju, Rebekah Penrice-Randal , Lucille Rainbow, N.P Randle, Trevor Ian Robinson, Parul Sharma, Ghada T Shawli, James P Stewart, Neil Swainston, Ecaterina Vamos, Joanne Watts, Mark Whitehead |
| EPI_ISL_705280, EPI_ISL_705281 | Northumbria University / South Tees Hospitals NHS Foundation Trust / North Cumbria Integrated Care NHS Foundation Trust / North Tees and Hartlepool NHS | COVID-19 Genomics UK (COG-UK) Consortium | Darren L Smith,Andrew Nelson,Matthew Bashton,Greg R Young,Joshua Loh,John Allan,Mohammad A Tariq,Giles S Holt,Gary Black,Wen C Yew,Lynn Dover,Paul Baker,Steve Liggett,Sarah Essex,Jane Greenaway,Debra Padgett,Clive Graham,Garren Scott,Edward Barton,Emma Swindells,Brendan Payne,Jennifer Collins,Yusri Taha,Gary Eltringham |

| | | | |
|---|---|---|---|
| | Foundation Trust / Newcastle Hospitals NHS Foundation Trust | | |
| EPI_ISL_705284, EPI_ISL_705314, EPI_ISL_705322 | Liverpool Clinical Laboratories | COVID-19 Genomics UK (COG-UK) Consortium | Sam Haldenby, Anita Lucaci, Steve Paterson, Julian Hiscox, Alistair Darby, M Almsaud, A Alrezaihi, Muhannad Alruwaili, Stuart D Armstrong, Jones Benjamin, Eleanor G Bentley, Anu Chawla, Jordan J Clark, Angela Cowell, Richard Eccles, Isabel García-Dorival, Matthew Gemmell, Alessandro Gerada, PKF Gilmore, Richard Gregory, Ximeng Han, Catherine Hartley, Margaret Hughes, Miren Iturriza-Gomara, James Johnson, L Luu, Jenifer Manson, Charlotte Nelson, Elaine O'Toole, Cassie Olateju, Rebekah Penrice-Randal , Lucille Rainbow, N.P Randle, Trevor Ian Robinson, Parul Sharma, Ghada T Shawli, James P Stewart, Neil Swainston, Ecaterina Vamos, Joanne Watts, Mark Whitehead |
| EPI_ISL_705338 | University College London, Great Ormond Street Hospital for Children NHS Foundation Trust, Imperial College Healthcare NHS Trust | COVID-19 Genomics UK (COG-UK) Consortium | Sergi Castellano, Rachel Williams, Mark Kristiansen, Paola Resende Silva, Sunando Roy, Tony Brooks, Helena Tutill, Paola Niola, Patricia Dyal, Charlotte Williams, Leysa Forrest, Yasmin Panchbhaya, Jacqueline Findlay, Samuel Weeks, Julianne Brown, Kathryn Harris, Paul Randell, James Price, Alison Holmes, Judith Breuer |
| EPI_ISL_705340, EPI_ISL_705360, EPI_ISL_705362 | Liverpool Clinical Laboratories | COVID-19 Genomics UK (COG-UK) Consortium | Sam Haldenby, Anita Lucaci, Steve Paterson, Julian Hiscox, Alistair Darby, M Almsaud, A Alrezaihi, Muhannad Alruwaili, Stuart D Armstrong, Jones Benjamin, Eleanor G Bentley, Anu Chawla, Jordan J Clark, Angela Cowell, Richard Eccles, Isabel García-Dorival, Matthew Gemmell, Alessandro Gerada, PKF Gilmore, Richard Gregory, Ximeng Han, Catherine Hartley, Margaret Hughes, Miren Iturriza-Gomara, James Johnson, L Luu, Jenifer Manson, Charlotte Nelson, Elaine O'Toole, Cassie Olateju, Rebekah Penrice-Randal , Lucille Rainbow, N.P Randle, Trevor Ian Robinson, Parul Sharma, Ghada T Shawli, James P Stewart, Neil Swainston, Ecaterina Vamos, Joanne Watts, Mark Whitehead |
| EPI_ISL_705395 | University College London, Great Ormond Street Hospital for Children NHS Foundation Trust, Imperial College Healthcare NHS Trust | COVID-19 Genomics UK (COG-UK) Consortium | Sergi Castellano, Rachel Williams, Mark Kristiansen, Paola Resende Silva, Sunando Roy, Tony Brooks, Helena Tutill, Paola Niola, Patricia Dyal, Charlotte Williams, Leysa Forrest, Yasmin Panchbhaya, Jacqueline Findlay, Samuel Weeks, Julianne Brown, Kathryn Harris, Paul Randell, James Price, Alison Holmes, Judith Breuer |
| EPI_ISL_705407, EPI_ISL_705411 | Liverpool Clinical Laboratories | COVID-19 Genomics UK (COG-UK) Consortium | Sam Haldenby, Anita Lucaci, Steve Paterson, Julian Hiscox, Alistair Darby, M Almsaud, A Alrezaihi, Muhannad Alruwaili, Stuart D Armstrong, Jones Benjamin, Eleanor G Bentley, Anu Chawla, Jordan J Clark, Angela Cowell, Richard Eccles, Isabel García-Dorival, Matthew Gemmell, Alessandro Gerada, PKF Gilmore, Richard Gregory, Ximeng Han, Catherine Hartley, Margaret Hughes, Miren Iturriza-Gomara, James Johnson, L Luu, Jenifer Manson, Charlotte Nelson, Elaine O'Toole, Cassie Olateju, Rebekah Penrice-Randal , Lucille Rainbow, N.P Randle, Trevor Ian Robinson, Parul Sharma, Ghada T Shawli, James P Stewart, Neil Swainston, Ecaterina Vamos, Joanne Watts, Mark Whitehead |
| EPI_ISL_705429, EPI_ISL_705430, EPI_ISL_705453, EPI_ISL_705454, EPI_ISL_705456, EPI_ISL_705467, EPI_ISL_705468, EPI_ISL_705470, EPI_ISL_705474, EPI_ISL_705481, EPI_ISL_705497 | | | |
| see above | Oxford Viromics, NDM, University of Oxford; Oxford University Hospitals; Basingstoke and North Hampshire Hospital | COVID-19 Genomics UK (COG-UK) Consortium | Tanya Golubchik, David Bonsall, George Macintyre, Amy Trebes, Mariateresa de Cesare, Catrin Moore, Alex Mobbs, Anita Justice, Robert Shaw, Monique Andersson, Timothy Peto, Emma Wise, Nathan Moore, Jessica Lynch, Nick Cortes, Matilde Mori, Stephen Kidd, David Buck, John Todd, Christophe Fraser |
| EPI_ISL_705772, EPI_ISL_705799 | Virology Department, Royal Infirmary of Edinburgh, NHS Lothian / School of Biological Sciences, University of Edinburgh / Institute of Genetics and Molecular Medicine, University of Edinburgh | COVID-19 Genomics UK (COG-UK) Consortium | McHugh M, Dewar R, Rooke S, Gallagher M, Balcaza C, O'Toole Á, Scher E, Hill V, McCrone JT, Colquhoun R, Yu X, Jackson B, Rambaut A, Williams TC, Templeton K |
| EPI_ISL_706025, EPI_ISL_706026, EPI_ISL_706027, EPI_ISL_706032, EPI_ISL_706033, EPI_ISL_706034, EPI_ISL_706037, EPI_ISL_706038, EPI_ISL_706058, EPI_ISL_706059, EPI_ISL_706060, EPI_ISL_706063, EPI_ISL_706065, EPI_ISL_706066, EPI_ISL_706068, EPI_ISL_706069, EPI_ISL_706070, EPI_ISL_706072, EPI_ISL_706073, EPI_ISL_706081, EPI_ISL_706085, EPI_ISL_706087, EPI_ISL_706088, EPI_ISL_706089, EPI_ISL_706090, EPI_ISL_706091, EPI_ISL_706093, EPI_ISL_706095, EPI_ISL_706096, EPI_ISL_706097, EPI_ISL_706098, EPI_ISL_706101, EPI_ISL_706102, EPI_ISL_706104, EPI_ISL_706105, EPI_ISL_706106, EPI_ISL_706110, EPI_ISL_706134, EPI_ISL_706136, EPI_ISL_706144, EPI_ISL_706145, EPI_ISL_706147, EPI_ISL_706149, EPI_ISL_706150, EPI_ISL_706156, EPI_ISL_706158, EPI_ISL_706160, EPI_ISL_706163, EPI_ISL_706166, EPI_ISL_706168, EPI_ISL_706169, EPI_ISL_706170, EPI_ISL_706185, EPI_ISL_706201, EPI_ISL_706202, EPI_ISL_706203, EPI_ISL_706204, EPI_ISL_706205, EPI_ISL_706206, EPI_ISL_706207, EPI_ISL_706208, EPI_ISL_706209, EPI_ISL_706210, EPI_ISL_706211, EPI_ISL_706212, EPI_ISL_706213, EPI_ISL_706214, EPI_ISL_706216, EPI_ISL_706218, EPI_ISL_706222, EPI_ISL_706223, EPI_ISL_706229, EPI_ISL_706230, EPI_ISL_706239, EPI_ISL_706240, EPI_ISL_706241, EPI_ISL_706243, EPI_ISL_706244, EPI_ISL_706247, EPI_ISL_706269, EPI_ISL_706270, EPI_ISL_706271, EPI_ISL_706272, EPI_ISL_706273, EPI_ISL_706274, EPI_ISL_706275 | | | |
| see above | Oxford Viromics, NDM, University of Oxford; Oxford University Hospitals; Basingstoke and North Hampshire Hospital | COVID-19 Genomics UK (COG-UK) Consortium | Tanya Golubchik, David Bonsall, George Macintyre, Amy Trebes, Mariateresa de Cesare, Catrin Moore, Alex Mobbs, Anita Justice, Robert Shaw, Monique Andersson, Timothy Peto, Emma Wise, Nathan Moore, Jessica Lynch, Nick Cortes, Matilde Mori, Stephen Kidd, David Buck, John Todd, Christophe Fraser |
| EPI_ISL_706354 | Northumbria University / South Tees Hospitals NHS Foundation Trust / North Cumbria Integrated Care NHS Foundation Trust / North Tees and Hartlepool NHS Foundation Trust / Newcastle Hospitals NHS Foundation Trust | COVID-19 Genomics UK (COG-UK) Consortium | Darren L Smith,Andrew Nelson,Matthew Bashton,Greg R Young,Joshua Loh,John Allan,Mohammad A Tariq,Giles S Holt,Gary Black,Wen C Yew,Lynn Dover,Paul Baker,Steve Liggett,Sarah Essex,Jane Greenaway,Debra Padgett,Clive Graham,Garren Scott,Edward Barton,Emma Swindells,Brendan Payne,Jennifer Collins,Yusri Taha,Gary Eltringham |
| EPI_ISL_706442, EPI_ISL_706483, EPI_ISL_706659 | Wales Specialist Virology Centre Sequencing lab: Pathogen Genomics Unit | COVID-19 Genomics UK (COG-UK) Consortium | Catherine Moore, Johnathan Evans, Laura Gifford, Malorie Perry, Simon Cottrell, Angela Marchbank, Alec Birchley, Alexander Adams, Amy Gaskin, Bree Gatica-Wilcox, Jason Coombes, Joel Southgate, Lauren Gilbert, Lee Graham, Nicole Pacchiarini, Sara Kumziene-Summerhayes, Sarah Taylor, Sophie Jones, Sara Rey, Matthew Bull, Joanne Watkins, Sally Corden, Tom Connor |
| EPI_ISL_707808, EPI_ISL_707813, EPI_ISL_707819, EPI_ISL_707824, EPI_ISL_707825, EPI_ISL_707826 | University of Michigan Clinical Microbiology Laboratory | Lauring Lab, University of Michigan, Department of Microbiology and Immunology | Valesano |
| EPI_ISL_707894 | Area of Virology, Serology and Virology Division (SAViD), New South Wales Health Pathology Randwick | Virology Research Laboratory; Area of Virology, Serology and Virology Division (SAViD), New South Wales Health Pathology Randwick | Foster, C.; Au, J.; Ruiz Silva, M.; Deveson, I.; Bull, R.; Van Hal, S.; Rawlinson, W. |
| EPI_ISL_708040 | Norwegian Institute of Public Health, Department of Virology | Norwegian Institute of Public Health, Department of Virology | Kathrine Stene-Johansen, Kamilla Heddeland Instefjord, Hilde Elshaug, Marie Paulsen Madsen, Rasmus Riis Kopperud, Hilde Vollan, Karoline Bragstad, Olav Hungnes |
| EPI_ISL_708060, EPI_ISL_708061, EPI_ISL_708062, EPI_ISL_708063, EPI_ISL_708064 | Unilabs Laboratory Medicine | Norwegian Institute of Public Health, Department of Virology | Kathrine Stene-Johansen, Kamilla Heddeland Instefjord, Hilde Elshaug, Marie Paulsen Madsen, Rasmus Riis Kopperud, Hilde Vollan, Karoline Bragstad, Olav Hungnes |
| EPI_ISL_708065, EPI_ISL_708066, EPI_ISL_708067, EPI_ISL_708068 | University Hospital of Northern Norway, Department for Microbiology and Infectious Disease Control | Norwegian Institute of Public Health, Department of Virology | Kathrine Stene-Johansen, Kamilla Heddeland Instefjord, Hilde Elshaug, Marie Paulsen Madsen, Rasmus Riis Kopperud, Hilde Vollan, Karoline Bragstad, Olav Hungnes |
| EPI_ISL_708078 | Foerde Hospital, Department of Microbiology | Norwegian Institute of Public Health, Department of Virology | Kathrine Stene-Johansen, Kamilla Heddeland Instefjord, Hilde Elshaug, Marie Paulsen Madsen, Rasmus Riis Kopperud, Hilde Vollan, Karoline Bragstad, Olav Hungnes |
| EPI_ISL_708080 | University Hospital of Northern Norway, Department for Microbiology and Infectious Disease Control | Norwegian Institute of Public Health, Department of Virology | Kathrine Stene-Johansen, Kamilla Heddeland Instefjord, Hilde Elshaug, Marie Paulsen Madsen, Rasmus Riis Kopperud, Hilde Vollan, Karoline Bragstad, Olav Hungnes |
| EPI_ISL_708081 | Dept. of Medical Microbiology, Stavanger University Hospital, Helse Stavanger HF | Norwegian Institute of Public Health, Department of Virology | Kathrine Stene-Johansen, Kamilla Heddeland Instefjord, Hilde Elshaug, Marie Paulsen Madsen, Rasmus Riis Kopperud, Hilde Vollan, Karoline Bragstad, Olav Hungnes |
| EPI_ISL_708086, EPI_ISL_708087, EPI_ISL_708088, EPI_ISL_708089, EPI_ISL_708090, EPI_ISL_708092, EPI_ISL_708094 | University Hospital of Northern Norway, Department for Microbiology and Infectious Disease Control | Norwegian Institute of Public Health, Department of Virology | Kathrine Stene-Johansen, Kamilla Heddeland Instefjord, Hilde Elshaug, Marie Paulsen Madsen, Rasmus Riis Kopperud, Hilde Vollan, Karoline Bragstad, Olav Hungnes |
| EPI_ISL_708096 | Oslo University Hospital, Department of Medical Microbiology | Norwegian Institute of Public Health, Department of Virology | Kathrine Stene-Johansen, Kamilla Heddeland Instefjord, Hilde Elshaug, Marie Paulsen Madsen, Rasmus Riis Kopperud, Hilde Vollan, Karoline Bragstad, Olav Hungnes |
| EPI_ISL_708097, EPI_ISL_708101, EPI_ISL_708104, EPI_ISL_708105 | Nordland Hospital - Bodo, Laboratory Department, Molecular Biology Unit | Norwegian Institute of Public Health, Department of Virology | Kathrine Stene-Johansen, Kamilla Heddeland Instefjord, Hilde Elshaug, Marie Paulsen Madsen, Rasmus Riis Kopperud, Hilde Vollan, Karoline Bragstad, Olav Hungnes |
| EPI_ISL_708110, EPI_ISL_708111, EPI_ISL_708112 | Ostfold Hospital Trust - Kalnes, Centre for Laboratory Medicine, Section for gene technology and infection serology | Norwegian Institute of Public Health, Department of Virology | Kathrine Stene-Johansen, Kamilla Heddeland Instefjord, Hilde Elshaug, Marie Paulsen Madsen, Rasmus Riis Kopperud, Hilde Vollan, Karoline Bragstad, Olav Hungnes |
| EPI_ISL_708319, EPI_ISL_708347 | Michigan Department of Health and Human Services, Bureau of Laboratories | Michigan Department of Health and Human Services, Bureau of Laboratories | Blankenship HM, Riner D, Soehnlen MK |

| | | | |
|---|---|---|---|
| EPI_ISL_708440, EPI_ISL_708444 | Delaware Public Health Lab | Delaware Public Health Lab | Gregory Hovan |
| EPI_ISL_708462, EPI_ISL_708463 | Minnesota Department of Health, Public Health Laboratory | Minnesota Department of Health, Public Health Laboratory | Alexandra Lorentz, Jacob Garfin, Matt Plumb, and Xiong Wang |
| EPI_ISL_708484 | Essentia Health-St. Mary's Medical Center | Minnesota Department of Health, Public Health Laboratory | Alexandra Lorentz, Jacob Garfin, Matt Plumb, and Xiong Wang |
| EPI_ISL_708532, EPI_ISL_708533, EPI_ISL_708536, EPI_ISL_708542, EPI_ISL_708543, EPI_ISL_708547, EPI_ISL_708549, EPI_ISL_708553, EPI_ISL_708554, EPI_ISL_708556, EPI_ISL_708557, EPI_ISL_708559, EPI_ISL_708561, EPI_ISL_708566, EPI_ISL_708567, EPI_ISL_708568, EPI_ISL_708569, EPI_ISL_708570, EPI_ISL_708573, EPI_ISL_708576, EPI_ISL_708579, EPI_ISL_708581, EPI_ISL_708586, EPI_ISL_708588, EPI_ISL_708589, EPI_ISL_708590, EPI_ISL_708591, EPI_ISL_708592, EPI_ISL_708594, EPI_ISL_708596, EPI_ISL_708597, EPI_ISL_708598, EPI_ISL_708599, EPI_ISL_708600, EPI_ISL_708604, EPI_ISL_708605 | | | |
| see above | Michigan Department of Health and Human Services, Bureau of Laboratories | Michigan Department of Health and Human Services, Bureau of Laboratories | Blankenship HM, Riner D, Soehnlen MK |
| EPI_ISL_710037, EPI_ISL_710038 | University of Michigan Clinical Microbiology Laboratory | Lauring Lab, University of Michigan, Department of Microbiology and Immunology | Valesano |
| EPI_ISL_710121 | South Eastern Area Laboratory Services (SEALS) | CIDM-PH et al. | CIDM-PH et al. |
| EPI_ISL_710592, EPI_ISL_710593 | MLF Care AB | The Public Health Agency of Sweden | Department of Microbiology, The Public Health Agency of Sweden |
| EPI_ISL_710601, EPI_ISL_710602 | Klinisk mikrobiologi | The Public Health Agency of Sweden | Department of Microbiology, The Public Health Agency of Sweden |
| EPI_ISL_710903, EPI_ISL_710904, EPI_ISL_710905, EPI_ISL_710906 | Lighthouse Lab in Glasgow | Wellcome Sanger Institute for the COVID-19 Genomics UK (COG-UK) Consortium | Harper VanSteenhouse, Yumi Kasai, David Gray, Carol Clugston, Anna Dominiczak and Alex Alderton, Roberto Amato, Sonia Goncalves, Ewan Harrison, David K. Jackson, Ian Johnston, Dominic Kwiatkowski, Cordelia Langford, John Sillitoe on behalf of the Wellcome Sanger Institute COVID-19 Surveillance Team |
| EPI_ISL_710916, EPI_ISL_710924, EPI_ISL_710930 | Lighthouse Lab in Alderley Park | Wellcome Sanger Institute for the COVID-19 Genomics UK (COG-UK) Consortium | Jacquelyn Wynn, Mairead Hyland, The Lighthouse Lab in Alderley Park and Alex Alderton, Roberto Amato, Sonia Goncalves, Ewan Harrison, David K. Jackson, Ian Johnston, Dominic Kwiatkowski, Cordelia Langford, John Sillitoe on behalf of the Wellcome Sanger Institute COVID-19 Surveillance Team |
| EPI_ISL_710995, EPI_ISL_710996, EPI_ISL_710997, EPI_ISL_710998, EPI_ISL_710999, EPI_ISL_711000, EPI_ISL_711001, EPI_ISL_711002, EPI_ISL_711003, EPI_ISL_711004, EPI_ISL_711005, EPI_ISL_711006, EPI_ISL_711007 | | | |
| see above | Lighthouse Lab in Glasgow | Wellcome Sanger Institute for the COVID-19 Genomics UK (COG-UK) Consortium | Harper VanSteenhouse, Yumi Kasai, David Gray, Carol Clugston, Anna Dominiczak and Alex Alderton, Roberto Amato, Sonia Goncalves, Ewan Harrison, David K. Jackson, Ian Johnston, Dominic Kwiatkowski, Cordelia Langford, John Sillitoe on behalf of the Wellcome Sanger Institute COVID-19 Surveillance Team |
| EPI_ISL_711023, EPI_ISL_711024 | Lighthouse Lab in Cambridge | Wellcome Sanger Institute for the COVID-19 Genomics UK (COG-UK) Consortium | Rob Howes, The Lighthouse Lab in Cambridge and Alex Alderton, Roberto Amato, Sonia Goncalves, Ewan Harrison, David K. Jackson, Ian Johnston, Dominic Kwiatkowski, Cordelia Langford, John Sillitoe on behalf of the Wellcome Sanger Institute COVID-19 Surveillance Team |
| EPI_ISL_712087, EPI_ISL_712088, EPI_ISL_712095 | Port Elizabeth Provincial Hospital, National Health Laboratory Services, Eastern Cape, South Africa | National Institute for Communicable Diseases of the National Health Laboratory Service | Mohale T, Ntuli N, Mahlangu B, Allam M, Ismail A, Bhiman JN |
| EPI_ISL_714107, EPI_ISL_714227, EPI_ISL_714252, EPI_ISL_714269, EPI_ISL_714377, EPI_ISL_714476, EPI_ISL_714923, EPI_ISL_714924, EPI_ISL_714925, EPI_ISL_714926, EPI_ISL_714927 | | | |
| see above | Department of Virus and Microbiological Special Diagnostics, Statens Serum Institut, Copenhagen, Denmark | Albertsen Lab, Department of Chemistry and Bioscience, Aalborg University, Denmark | Danish Covid-19 Genome Consortium |
| EPI_ISL_717808, EPI_ISL_717825, EPI_ISL_717826, EPI_ISL_717827, EPI_ISL_717890, EPI_ISL_717891, EPI_ISL_717892, EPI_ISL_717942, EPI_ISL_717943, EPI_ISL_717944, EPI_ISL_717945, EPI_ISL_717946, EPI_ISL_717947, EPI_ISL_717948, EPI_ISL_717949, EPI_ISL_717950 | | | |
| see above | Laboratorio de Virologia Molecular / UFRJ | Bioinformatics Laboratory / LNCC | Carolina M Voloch, Ronaldo da Silva F Jr, Luiz G P de Almeida, Cynthia C Cardoso, Otavio Bustrolini, Alexandra L Gerber, Ana Paula de C Guimarães, Diana Mariani, Andréa Cony Cavalcanti, Claudia dos Santos Rodrigues, Terezinha M P P Castiñeira, Amilcar Tanuri, Ana Tereza R de Vasconcelos |
| EPI_ISL_717995, EPI_ISL_717996 | Lab voor klinische biologie | Onderzoeksgroep Virologie | Laurens Lambrechts, Nick Vereecke, Marthe Pauwels, Bruno Verhasselt, Linos Vandekerckhove, Hans Nauwynck, Sebastiaan Theuns |
| EPI_ISL_718188 | Ministry of Health Hospitals | Institute of Health and Community Medicine | David Perera, Ooi Mong How, Chua Hock Hin, Tonnii Sia Loong Loong, Wong Jyn Shan, Wong Kiing Aik, Chan Chia Jui |
| EPI_ISL_718191 | Borneo Medical Centre | Institute of Health and Community Medicine | David Perera, Ooi Mong How, Chua Hock Hin, Tonnii Sia Loong Loong, Wong Jyn Shan, Wong Kiing Aik, Chan Chia Jui |
| EPI_ISL_718248 | Hospital | National Reference Center for Viruses of Respiratory Infections, Institut Pasteur, Paris | Marion Barbet, Sylvie Behillil, Méline Bizard, Angela Brisebarre, Camille Capel, Etienne Simon-Lorière, Vincent Enouf, Maud Vanpeene, Sylvie van der Werf, Gisèle Lagathu |
| EPI_ISL_718262 | National Institute for Infectious Diseases, INMI, "L. Spallanzani" IRCCS | National Institute for Infectious Diseases, INMI, "L. Spallanzani" IRCCS | C.E.M Gruber, F Messina, M Rueca, B Bartolini, E Giombini, MR Capobianchi, A Di Caro |
| EPI_ISL_718264 | National Institute for Infectious Diseases, INMI, "L. Spallanzani" IRCCS | National Institute for Infectious Diseases, INMI, "L. Spallanzani" IRCCS | F Messina, M Rueca, B Bartolini, C.E.M Gruber, E Giombini, MR Capobianchi, A Di Caro |
| EPI_ISL_718265 | National Institute for Infectious Diseases, INMI, "L. Spallanzani" IRCCS | National Institute for Infectious Diseases, INMI, "L. Spallanzani" IRCCS | B Bartolini, M Rueca, C.E.M Gruber, F Messina, E Giombini, A Di Caro, MR Capobianchi |
| EPI_ISL_721534 | Lighthouse Lab in Glasgow | Wellcome Sanger Institute for the COVID-19 Genomics UK (COG-UK) Consortium | Harper VanSteenhouse, Yumi Kasai, David Gray, Carol Clugston, Anna Dominiczak and Alex Alderton, Roberto Amato, Sonia Goncalves, Ewan Harrison, David K. Jackson, Ian Johnston, Dominic Kwiatkowski, Cordelia Langford, John Sillitoe on behalf of the Wellcome Sanger Institute COVID-19 Surveillance Team |
| EPI_ISL_722275, EPI_ISL_722282, EPI_ISL_722295 | Dutch COVID-19 response team | Erasmus Medical Center | Bas Oude Munnink, Reina Sikkema, David Nieuwenhuijse, Irina Chestakova, Anne van der Linden, Marjan Boter, Emmanuelle Munger, Corine GeurtsvanKessel, Annemiek van der Eijk, Richard Molenkamp, Marion Koopmans, on behalf of the Dutch national COVID-19 response team. |
| EPI_ISL_722320, EPI_ISL_722448, EPI_ISL_722449, EPI_ISL_722450, EPI_ISL_722451 | Dutch COVID-19 response team | Erasmus Medical Center | OH consortium |
| EPI_ISL_722679, EPI_ISL_722680, EPI_ISL_722681, EPI_ISL_722698, EPI_ISL_722699, EPI_ISL_722700, EPI_ISL_722701, EPI_ISL_722702, EPI_ISL_722703, EPI_ISL_722704, EPI_ISL_722705, EPI_ISL_722706, EPI_ISL_722707, EPI_ISL_722708, EPI_ISL_722709, EPI_ISL_722710, EPI_ISL_722850 | | | |
| see above | Dutch COVID-19 response team | Erasmus Medical Center | Bas Oude Munnink, Reina Sikkema, David Nieuwenhuijse, Irina Chestakova, Anne van der Linden, Marjan Boter, Emmanuelle Munger, Corine GeurtsvanKessel, Annemiek van der Eijk, Richard Molenkamp, Marion Koopmans, on behalf of the Dutch national COVID-19 response team. |
| EPI_ISL_723141 | Mayo Clinic & Mayo Clinic Laboratories | Minnesota Department of Health, Public Health Laboratory | Alexandra Lorentz, Jacob Garfin, Matt Plumb, and Xiong Wang |
| EPI_ISL_723496, EPI_ISL_723497, EPI_ISL_723498 | Virginia Division of Consolidated Laboratory Services (DCLS) | Virginia Division of Consolidated Laboratory Services (DCLS) | Virginia DCLS |
| EPI_ISL_723525, EPI_ISL_723526, EPI_ISL_723539, EPI_ISL_723547, EPI_ISL_723556 | Centre for Enzyme Innovation, University of Portsmouth / Translational Research Laboratory, Portsmouth Hospitals NHS Trust | COVID-19 Genomics UK (COG-UK) Consortium | Angela Beckett,Yann Bourgeois,Garry Scarlett,Sharon Glaysher,Scott Elliott,Kelly Bicknell,Robert Impey,Allyson Lloyd,Sarah Wyllie,Ethan Butcher,Anoop Chauhan,Samuel Robson |
| EPI_ISL_724620, EPI_ISL_724629, EPI_ISL_724660, EPI_ISL_724662, EPI_ISL_724664, EPI_ISL_724675, EPI_ISL_724683, EPI_ISL_724684, EPI_ISL_724708, EPI_ISL_724716, EPI_ISL_724740 | | | |
| see above | University College London, Great Ormond Street Hospital for Children NHS Foundation Trust, Imperial College Healthcare NHS Trust | COVID-19 Genomics UK (COG-UK) Consortium | Sergi Castellano, Rachel Williams, Mark Kristiansen, Paola Resende Silva, Sunando Roy, Tony Brooks, Helena Tutill, Paola Niola, Patricia Dyal, Charlotte Williams, Leysa Forrest, Yasmin Panchbhaya, Jacqueline Findlay, Samuel Weeks, Julianne Brown, Kathryn Harris, Paul Randell, James Price, Alison Holmes, Judith Breuer |
| EPI_ISL_724963, EPI_ISL_724980, EPI_ISL_724981, EPI_ISL_724982 | Northumbria University / South Tees Hospitals NHS Foundation Trust / North Cumbria Integrated Care NHS Foundation Trust / North Tees and Hartlepool NHS Foundation Trust / Newcastle Hospitals NHS Foundation Trust | COVID-19 Genomics UK (COG-UK) Consortium | Darren L Smith,Andrew Nelson,Matthew Bashton,Greg R Young,Joshua Loh,John Allan,Mohammad A Tariq,Giles S Holt,Gary Black,Wen C Yew,Lynn Dover,Paul Baker,Steve Liggett,Sarah Essex,Jane Greenaway,Debra Padgett,Clive Graham,Garren Scott,Edward Barton,Emma Swindells,Brendan Payne,Jennifer Collins,Yusri Taha,Gary Eltringham |
| EPI_ISL_725271 | Quadram Institute Bioscience | COVID-19 Genomics UK (COG-UK) Consortium | Dave J. Baker, Gemma L. Kay, Alp Aydin, Thanh Le-Viet, Steven Rudder, Ana P. Tedim, Anastasia Kolyva, Maria Diaz, Leonardo de Oliveira Martins, Nabil-Fareed Alikhan, Lizzie Meadows, Rachael Stanley, Ngozi Elumogo, Muhammed Yasir, Nicholas M. Thomson, Alexander J Trotter, Rachel Gilroy, |

| | | | |
|---|---|---|---|
| | | | Samuel Bloomfield, Claire Stuart, Andrew Bell, Reenesh Prakash, Samir Dervisevic, Alison E. Mather, John Wain, Mark Webber, Andrew J. Page, Justin O'Grady |
| EPI_ISL_727750 | Centre for Enzyme Innovation, University of Portsmouth / Translational Research Laboratory, Portsmouth Hospitals NHS Trust | COVID-19 Genomics UK (COG-UK) Consortium | Angela Beckett,Yann Bourgeois,Garry Scarlett,Sharon Glaysher,Scott Elliott,Kelly Bicknell,Robert Impey,Allyson Lloyd,Sarah Wyllie,Ethan Butcher,Anoop Chauhan,Samuel Robson |
| EPI_ISL_728134, EPI_ISL_728135, EPI_ISL_728136, EPI_ISL_728137, EPI_ISL_728138, EPI_ISL_728139, EPI_ISL_728140 | University of Wisconsin-Madison AIDS Vaccine Research Laboratories | University of Wisconsin-Madison AIDS Vaccine Research Laboratories | Gage Moreno, Katarina Braun, et al. AIDS Vaccine Research Laboratories |
| EPI_ISL_728169, EPI_ISL_728170 | Institute for Medical Research, Infectious Disease Research Centre, National Institutes of Health, Ministry of Health Malaysia | Institute for Medical Research, Infectious Disease Research Centre, National Institutes of Health, Ministry of Health Malaysia | Suppiah J, Kamel K, Mohd-Zawawi Z, Thayan R |
| EPI_ISL_728773, EPI_ISL_728777, EPI_ISL_728781, EPI_ISL_728785, EPI_ISL_728788, EPI_ISL_728793, EPI_ISL_728804, EPI_ISL_728807, EPI_ISL_728810, EPI_ISL_728816, EPI_ISL_728821, EPI_ISL_728826, EPI_ISL_728830, EPI_ISL_728836, EPI_ISL_728840, EPI_ISL_728844, EPI_ISL_728855, EPI_ISL_728860, EPI_ISL_728861, EPI_ISL_728877, EPI_ISL_728915, EPI_ISL_728934, EPI_ISL_728935, EPI_ISL_728941, EPI_ISL_728946, EPI_ISL_728970, EPI_ISL_728971, EPI_ISL_728985, EPI_ISL_729192, EPI_ISL_729193, EPI_ISL_729194, EPI_ISL_729195, EPI_ISL_729196, EPI_ISL_729197, EPI_ISL_729198, EPI_ISL_729199, EPI_ISL_729200, EPI_ISL_729204, EPI_ISL_729205, EPI_ISL_729206, EPI_ISL_729207, EPI_ISL_729208, EPI_ISL_729209, EPI_ISL_729210, EPI_ISL_729256 | | | |
| see above | Viollier AG | Department of Biosystems Science and Engineering, ETH Zürich | Chaoran Chen, Sarah Nadeau, Catharine Aquino, Ivan Topolsky, Pedro Ferreira, Philipp Jablonski, Susana Posada-Céspedes, Andreia Cabral de Gouvea, Maria Domenica Moccia, Simon Grüter, Timothy Sykes, Lennart Opitz, Ralph Schlapbach, Christiane Beckmann, Maurice Redondo, Olivier Kobel, Christoph Noppen, Sophie Seidel, Noemie Santamaria de Souza, Niko Beerenwinkel, Tanja Stadler |
| EPI_ISL_729341, EPI_ISL_729349, EPI_ISL_729369, EPI_ISL_729401, EPI_ISL_729414, EPI_ISL_729427, EPI_ISL_729428, EPI_ISL_729429, EPI_ISL_729430, EPI_ISL_729444, EPI_ISL_729462, EPI_ISL_729471, EPI_ISL_729620, EPI_ISL_729622, EPI_ISL_729623, EPI_ISL_729657, EPI_ISL_729660, EPI_ISL_729664 | | | |
| see above | A. Krumbholz, Labor Dr. Krause und Kollegen MVZ GmbH, Kiel | Charité Universitätsmedizin Berlin, Institut für Virologie | Victor M Corman, Barbara Mühlemann, Jörn Beheim-Schwarzbach, Talitha Veith, Julia Schneider, Terry Jones, Christian Drosten |
| EPI_ISL_729767, EPI_ISL_729768 | Yale Pathology Lab | Grubaugh Lab - Yale School of Public Health | Joseph Fauver, Tara Alpert, Anderson Brito, Annie Watkins, Anne Wyllie, Chantal Vogels, Mary Petrone, Chaney Kalinich, Isabel Ott, Arnau Casanovas, Catherine Muenker, Adam Moore, Alice Lu, Maria Tokuyama, Patrick Wong, Peiwen Lu, Saad Omer, Richard Martinello, Allison Nelson, Shelli Farhadian, Akiko Iwasaki, Charlese Dela Cruz, Albert Ko, Nathan Grubaugh |
| EPI_ISL_730055, EPI_ISL_730058, EPI_ISL_730060 | Connecticut Department of Health | Grubaugh Lab - Yale School of Public Health | Joseph Fauver, Tara Alpert, Anderson Brito, Annie Watkins, Anne Wyllie, Chantal Vogels, Mary Petrone, Chaney Kalinich, Isabel Ott, Arnau Casanovas, Catherine Muenker, Adam Moore, Alice Lu, Maria Tokuyama, Patrick Wong, Peiwen Lu, Saad Omer, Richard Martinello, Allison Nelson, Shelli Farhadian, Akiko Iwasaki, Charlese Dela Cruz, Albert Ko, Nathan Grubaugh |
| EPI_ISL_730168, EPI_ISL_730177, EPI_ISL_730186, EPI_ISL_730188, EPI_ISL_730190, EPI_ISL_730191, EPI_ISL_730194, EPI_ISL_730196 | San Diego County Public Health Laboratory | Andersen lab at Scripps Research | SEARCH Alliance San Diego with Tracy Basler, Jovan Shephard, Brett Austin |
| EPI_ISL_730517, EPI_ISL_730523, EPI_ISL_730528, EPI_ISL_730532, EPI_ISL_730538, EPI_ISL_730543, EPI_ISL_730544, EPI_ISL_730548, EPI_ISL_730549, EPI_ISL_730554, EPI_ISL_730555, EPI_ISL_730560, EPI_ISL_730562 | | | |
| see above | Biolab Diagnostic Laboratories | Andersen lab at Scripps Research | Issa Abu-Dayyeh, Ahmad Tibi, Lama Hussein, Lina Mohammad, Zein Naber, Amid Abdelnour with SEARCH Alliance San Diego |
| EPI_ISL_731911 | Instituto Nacional de Saude (INSA) | Instituto Nacional de Saude (INSA) | Borges et al |
| EPI_ISL_732374, EPI_ISL_732385, EPI_ISL_732456 | National Virus Reference Laboratory | National Virus Reference Laboratory | Michael Carr, Gabriel Gonzalez, Jonathan Dean, Daniel Hare, Cillian F De Gascun |
| EPI_ISL_732677, EPI_ISL_732678 | CNR Virus des Infections Respiratoires - France SUD | CNR Virus des Infections Respiratoires - France SUD | Antonin Bal, Gregory Destras, Claudia Gonzalez, Gwendolyne Burfin, Quentin Semanas, Martine Valette, Bruno Lina, Laurence Josset |
| EPI_ISL_732792 | Centro de Investigación Biomédica de La Rioja - Hospital San Pedro Logroño | SeqCOVID-SPAIN consortium/IBV(CSIC) | María de Toro, José Manuel Azcona Gutiérrez, María Pilar Bea Escudero, Miriam Blasco Alberdi and SeqCOVID-SPAIN consortium |
| EPI_ISL_733483, EPI_ISL_733484 | Omsk Research Institute of Natural Focal Infections | WHO National Influenza Centre Russian Federation | Andrey Komissarov, Artem Fadeev, Anna Ivanova, Kseniya Komissarova, Dmitry Bazhenov, Daria Danilenko, Ekaterina Gradoboeva, Elena Poleshchuk, Aleksei Vasilenko, Valery Yakimenko, Ksenia Safina, Elena Nabieva, Georgii Bazykin, Dmitry Lioznov |
| EPI_ISL_734207, EPI_ISL_734208, EPI_ISL_734209, EPI_ISL_734210, EPI_ISL_734211, EPI_ISL_734212, EPI_ISL_734214, EPI_ISL_734215, EPI_ISL_734219, EPI_ISL_734220, EPI_ISL_734221, EPI_ISL_734222, EPI_ISL_734223, EPI_ISL_734278 | | | |
| see above | Virginia Division of Consolidated Laboratory Services (DCLS) | Virginia Division of Consolidated Laboratory Services (DCLS) | Virginia DCLS |
| EPI_ISL_734414 | Wadsworth Center, New York State Department.of Health | Wadsworth Center, New York State Department.of Health | Kirsten St. George, Daryl M. Lamson, Alexis Russel, Jonathan Plitnick, Navjot Singh, John Kelly, Sara Griesemer, Erasmus Schneider, Erica Lasek-Nesselquist |
| EPI_ISL_734471, EPI_ISL_734472, EPI_ISL_734473, EPI_ISL_734474 | Masonic Medical Research Institute | Wadsworth Center, New York State Department.of Health | Kirsten St. George, Nathan Tucker, Ryan D. Pfeiffer, Daryl M. Lamson, Alexis Russel, Jonathan Plitnick, Navjot Singh, John Kelly, Sara Griesemer, Erasmus Schneider, Erica Lasek-Nesselquist |
| EPI_ISL_734483, EPI_ISL_735215, EPI_ISL_735216, EPI_ISL_735217, EPI_ISL_735218, EPI_ISL_735219, EPI_ISL_735220 | UZ Leuven, National Reference Laboratory for Coronaviruses, Laboratory Medicine, Leuven, Belgium | KU Leuven, Rega Institute, Clinical and Epidemiological Virology | Tony Wawina-Bokalanga, Joan Marti-Carerras, Bert Vanmechelen, Piet Maes |
| EPI_ISL_737014, EPI_ISL_737015 | The National Institute of Public Health | State Veterinary Institute Prague | Nagy,A;Jirincova,H;Trnka,D;Vecerova,J |
| EPI_ISL_737140 | Michigan Department of Health and Human Services, Bureau of Laboratories | Michigan Department of Health and Human Services, Bureau of Laboratories | Blankenship HM, Riner D, Soehnlen MK |
| EPI_ISL_738947, EPI_ISL_738953, EPI_ISL_739433, EPI_ISL_739442, EPI_ISL_739530 | Humboldt County Public Health Laboratory | Chan-Zuckerberg Biohub | CZB Cliahub Consortium |
| EPI_ISL_739699, EPI_ISL_739701, EPI_ISL_739755, EPI_ISL_739809, EPI_ISL_739810, EPI_ISL_739816, EPI_ISL_739817, EPI_ISL_739829, EPI_ISL_739843, EPI_ISL_739844, EPI_ISL_739851, EPI_ISL_739892, EPI_ISL_739905, EPI_ISL_739916, EPI_ISL_739933, EPI_ISL_739963, EPI_ISL_739998, EPI_ISL_740007, EPI_ISL_740047, EPI_ISL_740123, EPI_ISL_740190, EPI_ISL_740231, EPI_ISL_740270, EPI_ISL_740295, EPI_ISL_740302, EPI_ISL_740332, EPI_ISL_740348, EPI_ISL_740364, EPI_ISL_740372, EPI_ISL_740434, EPI_ISL_740436, EPI_ISL_740450, EPI_ISL_740499, EPI_ISL_740504, EPI_ISL_740506 | | | |
| see above | Laboratoire national de santé, Microbiology, Virology | Laboratoire national de santé, Microbiology, Microbial Genomics Platform | Anke Wienecke-Baldacchino, Catherine Ragimbeau,Jessica Tapp, Fatu Djabi, Lise Pignon, Raoul Salmon, Tamir Abdelrahman |
| EPI_ISL_741367, EPI_ISL_741368, EPI_ISL_741370, EPI_ISL_741373, EPI_ISL_741384, EPI_ISL_741385, EPI_ISL_741386, EPI_ISL_741387, EPI_ISL_741388, EPI_ISL_741390, EPI_ISL_741393, EPI_ISL_741394, EPI_ISL_741404, EPI_ISL_741405, EPI_ISL_741406, EPI_ISL_741409, EPI_ISL_741411, EPI_ISL_741412, EPI_ISL_741413, EPI_ISL_741415, EPI_ISL_741418 | | | |
| see above | Oxford Viromics, NDM, University of Oxford; Oxford University Hospitals; Basingstoke and North Hampshire Hospital | COVID-19 Genomics UK (COG-UK) Consortium | Tanya Golubchik, David Bonsall, George Macintyre, Amy Trebes, Mariateresa de Cesare, Catrin Moore, Alex Mobbs, Anita Justice, Robert Shaw, Monique Andersson, Timothy Peto, Emma Wise, Nathan Moore, Jessica Lynch, Nick Cortes, Matilde Mori, Stephen Kidd, David Buck, John Todd, Christophe Fraser |
| EPI_ISL_741609 | Queens Medical Centre, Clinical Microbiology Department / DeepSeq Nottingham | COVID-19 Genomics UK (COG-UK) Consortium | Gemma Clark, Wendy Smith, Manjinder Khakh, Vicki M Fleming, Michelle M Lister, Hannah Howson-Wells, Jonathan Ball, Patrick McClure, Joseph Chappell, Theocharis Tsoleridis, Nadine Holmes, Matthew Carlisle, Christopher Moore, Fei Sang, Johnny Debebe, Victoria Wright, Matthew Loose |
| EPI_ISL_744168, EPI_ISL_744172, EPI_ISL_744194, EPI_ISL_744217, EPI_ISL_744218, EPI_ISL_744258, EPI_ISL_744347, EPI_ISL_744359, EPI_ISL_744389, EPI_ISL_744419, EPI_ISL_744428, EPI_ISL_744440, EPI_ISL_744443, EPI_ISL_744460, EPI_ISL_744501, EPI_ISL_744534, EPI_ISL_744536, EPI_ISL_744565, EPI_ISL_744567, EPI_ISL_744572, EPI_ISL_744575, EPI_ISL_744581, EPI_ISL_744588, EPI_ISL_744624, EPI_ISL_744659, EPI_ISL_744680, EPI_ISL_744684, EPI_ISL_744686, EPI_ISL_744712, EPI_ISL_744732, EPI_ISL_744742, EPI_ISL_744788, EPI_ISL_744808, EPI_ISL_744829, EPI_ISL_744877, EPI_ISL_744907, EPI_ISL_744947 | | | |
| see above | Laboratoire national de santé, Microbiology, Virology | Laboratoire national de santé, Microbiology, Microbial Genomics Platform | Anke Wienecke-Baldacchino, Catherine Ragimbeau,Jessica Tapp, Fatu Djabi, Lise Pignon, Raoul Salmon, Tamir Abdelrahman |

| | | | |
|---|---|---|---|
| EPI_ISL_745313 | CHU Clermont-Ferrand | CNR Virus des Infections Respiratoires - France SUD | Antonin Bal, Gregory Destras, Gwendolyne Burfin, Hadrien Règue, Quentin Semanas, Martine Valette, Bruno Lina, Christine Archimbaud, Amélie Brebion, Hélène Chabrolles, Martine Chambon, Audrey Mirand, Christel Regagnon, Maxime Bisseux, Patricia Combes, Cécile Henquell, Laurence Josset |
| EPI_ISL_745441, EPI_ISL_745442, EPI_ISL_745443 | DOHMH Morrisania | New York City Public Health Laboratory | Jade Wang, et al. |
| EPI_ISL_745444 | DOHMH Jamaica | New York City Public Health Laboratory | Jade Wang, et al. |
| EPI_ISL_747093, EPI_ISL_747100, EPI_ISL_747101, EPI_ISL_747117, EPI_ISL_747120, EPI_ISL_747122, EPI_ISL_747123, EPI_ISL_747124, EPI_ISL_747125, EPI_ISL_747126, EPI_ISL_747136, EPI_ISL_747137, EPI_ISL_747138, EPI_ISL_747139, EPI_ISL_747140, EPI_ISL_747141, EPI_ISL_747142, EPI_ISL_747143 | | | |
| see above | Respiratory Viruses Branch, Centers for Disease Control and Prevention | Respiratory Viruses Branch, Centers for Disease Control and Prevention | Queen,K., Li,Y., Tao,Y., Uehara,A., Montmayeur,A., Paden,C.R., Cook,P.W., Marine,R., Sheth,M., Wang,H., Lee,J., Tong,S. |
| EPI_ISL_748122, EPI_ISL_748123 | Department of Virus and Microbiological Special Diagnostics, Statens Serum Institut, Copenhagen, Denmark | Albertsen Lab, Department of Chemistry and Bioscience, Aalborg University, Denmark | Danish Covid-19 Genome Consortium |
| EPI_ISL_751450, EPI_ISL_751451, EPI_ISL_751452, EPI_ISL_751453, EPI_ISL_751498 | CHU Purpan - Laboratoire de Virologie - Institut Fédératif de Biologie | CHU Purpan - Laboratoire de Virologie - Institut Fédératif de Biologie | Latour J., Ranger N., Dubois M., Carcenac R., Harter A., Boyer P., Tremeaux P., Izopet J. |
| EPI_ISL_751545 | NYSDOH Wadsworth Center, Virology Lab | Genomics and Discovery, Respiratory Viruses Branch, Division of Viral Diseases, Centers for Disease Control and Prevention | Krista Queen, Yan Li, Ying Tao, Jing Zhang, Anna Uehara, Anna Montmayeur, Clinton R. Paden, Peter W. Cook,Rachel Marine, Mili Sheth, Haibin Wang, Justin Lee, Suxiang Tong |
| EPI_ISL_751572 | KS Health and Environmental Laboratories | Genomics and Discovery, Respiratory Viruses Branch, Division of Viral Diseases, Centers for Disease Control and Prevention | Krista Queen, Yan Li, Ying Tao, Jing Zhang, Anna Uehara, Anna Montmayeur, Clinton R. Paden, Peter W. Cook,Rachel Marine, Mili Sheth, Haibin Wang, Justin Lee, Suxiang Tong |
| EPI_ISL_751584, EPI_ISL_751587 | NH Dept. of Health and Human Services Public Health Labs | Genomics and Discovery, Respiratory Viruses Branch, Division of Viral Diseases, Centers for Disease Control and Prevention | Krista Queen, Yan Li, Ying Tao, Jing Zhang, Anna Uehara, Anna Montmayeur, Clinton R. Paden, Peter W. Cook,Rachel Marine, Mili Sheth, Haibin Wang, Justin Lee, Suxiang Tong |
| EPI_ISL_751677 | KS Health and Environmental Laboratories | Genomics and Discovery, Respiratory Viruses Branch, Division of Viral Diseases, Centers for Disease Control and Prevention | Krista Queen, Yan Li, Ying Tao, Jing Zhang, Anna Uehara, Anna Montmayeur, Clinton R. Paden, Peter W. Cook,Rachel Marine, Mili Sheth, Haibin Wang, Justin Lee, Suxiang Tong |
| EPI_ISL_751678 | NH Dept. of Health and Human Services Public Health Labs | Genomics and Discovery, Respiratory Viruses Branch, Division of Viral Diseases, Centers for Disease Control and Prevention | Krista Queen, Yan Li, Ying Tao, Jing Zhang, Anna Uehara, Anna Montmayeur, Clinton R. Paden, Peter W. Cook,Rachel Marine, Mili Sheth, Haibin Wang, Justin Lee, Suxiang Tong |
| EPI_ISL_751681 | NE Public Health Laboratory | Genomics and Discovery, Respiratory Viruses Branch, Division of Viral Diseases, Centers for Disease Control and Prevention | Krista Queen, Yan Li, Ying Tao, Jing Zhang, Anna Uehara, Anna Montmayeur, Clinton R. Paden, Peter W. Cook,Rachel Marine, Mili Sheth, Haibin Wang, Justin Lee, Suxiang Tong |
| EPI_ISL_751707 | KS Health and Environmental Laboratories | Genomics and Discovery, Respiratory Viruses Branch, Division of Viral Diseases, Centers for Disease Control and Prevention | Krista Queen, Yan Li, Ying Tao, Jing Zhang, Anna Uehara, Anna Montmayeur, Clinton R. Paden, Peter W. Cook,Rachel Marine, Mili Sheth, Haibin Wang, Justin Lee, Suxiang Tong |
| EPI_ISL_751712 | NH Dept. of Health and Human Services Public Health Labs | Genomics and Discovery, Respiratory Viruses Branch, Division of Viral Diseases, Centers for Disease Control and Prevention | Krista Queen, Yan Li, Ying Tao, Jing Zhang, Anna Uehara, Anna Montmayeur, Clinton R. Paden, Peter W. Cook,Rachel Marine, Mili Sheth, Haibin Wang, Justin Lee, Suxiang Tong |
| EPI_ISL_751724, EPI_ISL_751730 | KS Health and Environmental Laboratories | Genomics and Discovery, Respiratory Viruses Branch, Division of Viral Diseases, Centers for Disease Control and Prevention | Krista Queen, Yan Li, Ying Tao, Jing Zhang, Anna Uehara, Anna Montmayeur, Clinton R. Paden, Peter W. Cook,Rachel Marine, Mili Sheth, Haibin Wang, Justin Lee, Suxiang Tong |
| EPI_ISL_753055, EPI_ISL_753056, EPI_ISL_753057, EPI_ISL_753058, EPI_ISL_753059, EPI_ISL_753060, EPI_ISL_753061, EPI_ISL_753062, EPI_ISL_753121, EPI_ISL_753122 | State Laboratories Division, Hawaii State Department of Health | State Laboratories Division, Hawaii State Department of Health | Pamela O'Brien, Sabrina Diemert, Drew Kuwazaki, Razvan Sultana, Edward Desmond |
| EPI_ISL_753262, EPI_ISL_753263, EPI_ISL_753264, EPI_ISL_753265, EPI_ISL_753266, EPI_ISL_753267, EPI_ISL_753268, EPI_ISL_753269, EPI_ISL_753270, EPI_ISL_753271, EPI_ISL_753272, EPI_ISL_753276, EPI_ISL_753285 | | | |
| see above | Clinical virology Laboratory, Children's Hospital Los Angeles | Center for Personalized Medicine, Children's Hospital Los Angeles | Gai et al |
| EPI_ISL_754400, EPI_ISL_754510, EPI_ISL_754511, EPI_ISL_754512 | Wadsworth Center, New York State Department.of Health | Wadsworth Center, New York State Department.of Health | Kirsten St. George, Daryl M. Lamson, Alexis Russel, Matthew Shudt, Melissa A Leisner, Jonathan Plitnick, Navjot Singh, John Kelly, Sara Griesemer, Erasmus Schneider, Erica Lasek-Nesselquist |
| EPI_ISL_756276 | State Laboratories Division, Hawaii State Department of Health | State Laboratories Division, Hawaii State Department of Health | Pamela O'Brien, Sabrina Diemert, Drew Kuwazaki, Razvan Sultana, Edward Desmond |
| EPI_ISL_757700 | Department of Virus and Microbiological Special Diagnostics, Statens Serum Institut, Copenhagen, Denmark | Albertsen Lab, Department of Chemistry and Bioscience, Aalborg University, Denmark | Danish Covid-19 Genome Consortium |
| EPI_ISL_760148, EPI_ISL_760149, EPI_ISL_760153, EPI_ISL_760174, EPI_ISL_760215 | Division of Emerging Infectious Diseases, Bureau of Infectious Diseases Diagnosis Control, Korea Disease Control and Prevention Agency | Division of Emerging Infectious Diseases, Bureau of Infectious Diseases Diagnosis Control, Korea Disease Control and Prevention Agency | Ae Kyung Park, Il-Hwan Kim, Heui Man Kim, Jeong-Min Kim, Namjoo Lee, Chaeyoung Lee, Sang Hee Woo, Eun-Jin Kim |
| EPI_ISL_764356 | Virology Department, Royal Infirmary of Edinburgh, NHS Lothian / School of Biological Sciences, University of Edinburgh / Institute of Genetics and Molecular Medicine, University of Edinburgh | COVID-19 Genomics UK (COG-UK) Consortium | McHugh M, Dewar R, Rooke S, Gallagher M, Balcaza C, O'Toole Á, Scher E, Hill V, McCrone JT, Colquhoun R, Yu X, Jackson B, Rambaut A, Williams TC, Templeton K |
| EPI_ISL_764432, EPI_ISL_764435 | Regional Virus Laboratory, Belfast Health and Social Care Trust | COVID-19 Genomics UK (COG-UK) Consortium | Conall McCaughey, James McKenna, Tanya Curran, Susan Feeney, Alison Watt, Ciara Cox, Mairead Connor, Zoltan Molnar, David Simpson, Derek Fairley |
| EPI_ISL_765786, EPI_ISL_765787, EPI_ISL_765788, EPI_ISL_765789, EPI_ISL_765790, EPI_ISL_765791 | Massachusetts General Hospital | Infectious Disease Program, Broad Institute of Harvard and MIT | Lemieux,J.E., Siddle,K.J., Shaw,B., Adams,G., Pierce,V., Turbett,S., Anahtar,M., Branda,J., Slater,D., Harris,J., Lin,A.E., Gladden-Young,A., Lagerborg,K., Rudy,M., DeRuff,K., Carter,A., Normandin,E., Bauer,M., Reilly,S., Tomkins-Tinch,C., Loreth,C., Chaluvadi,S., Neumann,A., Cusick,C., Chapman,S.B., Gnirke,A., Flowers,K., Cerrato,F., Birren,B.W., Gallagher,G., Smole,S., Park,D.J., MacInnis,B.L., Ryan,E., LaRocque,R., Rosenberg,E. and Sabeti,P.C. |
| EPI_ISL_766623, EPI_ISL_766624 | Klinisk mikrobiologi | The Public Health Agency of Sweden | Department of Microbiology, The Public Health Agency of Sweden |
| EPI_ISL_768726 | Jena University Hospital, Institute for Infectious Diseases and Infection Control | Institute of infectious medicine & hospital hygiene, CaSe-Group | Spott, Riccardo; Marquet, Mike; Pletz, Matthias W.; Brandt, Christian |
| EPI_ISL_768784, EPI_ISL_768785, EPI_ISL_768787, EPI_ISL_768788, EPI_ISL_768833 | AIID | Irish Coronavirus Sequencing Consortium - National Virus Reference Laboratory | Michael Carr, Gabriel Gonzalez, Alejandro Abner Garcia Leon, Patrick Mallon |
| EPI_ISL_769997 | Area De Salud Catedral Noreste | Inciensa, Instituto Costarricense de Investigación y | Francisco Duarte, Hebleen Porras, Claudio Soto-Garita, Estela Cordero, Adriana Godínez, Melany Calderón & Mariel López |

| | | | |
|---|---|---|---|
| | | Enseñanza en Nutrición y Salud | |
| EPI_ISL_770778, EPI_ISL_770779 | Minnesota Department of Health, Public Health Laboratory | Minnesota Department of Health, Public Health Laboratory | Alexandra Lorentz, Jacob Garfin, Matt Plumb, and Xiong Wang |
| EPI_ISL_775295 | Nordland Hospital - Bodo, Laboratory Department, Molecular Biology Unit | Norwegian Institute of Public Health, Department of Virology | Kathrine Stene-Johansen, Kamilla Heddeland Instefjord, Hilde Elshaug, Atiya R Ali,Marie Paulsen Madsen, Rasmus Riis Kopperud, Hilde Vollan, Karoline Bragstad, Olav Hungnes |
| EPI_ISL_775328, EPI_ISL_775329 | Dept. of Medical Microbiology, Stavanger University Hospital, Helse Stavanger HF | Norwegian Institute of Public Health, Department of Virology | Kathrine Stene-Johansen, Kamilla Heddeland Instefjord, Hilde Elshaug, Atiya R Ali,Marie Paulsen Madsen, Rasmus Riis Kopperud, Hilde Vollan, Karoline Bragstad, Olav Hungnes |
| EPI_ISL_775455 | University Hospital of Northern Norway, Department for Microbiology and Infectious Disease Control | Norwegian Institute of Public Health, Department of Virology | Kathrine Stene-Johansen, Kamilla Heddeland Instefjord, Hilde Elshaug, Atiya R Ali,Marie Paulsen Madsen, Rasmus Riis Kopperud, Hilde Vollan, Karoline Bragstad, Olav Hungnes |
| EPI_ISL_775486 | Nordland Hospital - Bodo, Laboratory Department, Molecular Biology Unit | Norwegian Institute of Public Health, Department of Virology | Kathrine Stene-Johansen, Kamilla Heddeland Instefjord, Hilde Elshaug, Atiya R Ali,Marie Paulsen Madsen, Rasmus Riis Kopperud, Hilde Vollan, Karoline Bragstad, Olav Hungnes |
| EPI_ISL_775504 | Vestfold Hospital, Toensberg Department of Microbiology | Norwegian Institute of Public Health, Department of Virology | Kathrine Stene-Johansen, Kamilla Heddeland Instefjord, Hilde Elshaug, Atiya R Ali,Marie Paulsen Madsen, Rasmus Riis Kopperud, Hilde Vollan, Karoline Bragstad, Olav Hungnes |
| EPI_ISL_779315 | Utah Public Health Laboratory, Utah Public Health Laboratory Infectious Disease submission group | Utah Public Health Laboratory, Utah Public Health Laboratory Infectious Disease submission group | Gallagher,T., Young,E.L., Oakeson,K.F. |
| EPI_ISL_780048 | Hospital General Universitario Gregorio Marañón | SeqCOVID-SPAIN consortium/IBV(CSIC) | Darío García de Viedma, Laura Pérez-Lago, Marta Herranz, Jon Sicilia, Julia Suárez, Pilar Catalán, Patricia Muñoz and SeqCOVID-SPAIN consortium |
| EPI_ISL_780407 | Bermuda Government Molecular Diagnostics Laboratory (MDL) | Respiratory Virus Unit, National Infection Service, Public Health England | PHE Covid Sequencing Team, Dr Carika Weldon (Bermuda), Dr Ayoola Oyinloye (Bermuda) |
| EPI_ISL_786001, EPI_ISL_786002, EPI_ISL_786003, EPI_ISL_786004, EPI_ISL_786005, EPI_ISL_786006, EPI_ISL_786012, EPI_ISL_786013, EPI_ISL_786014, EPI_ISL_786015, EPI_ISL_786016, EPI_ISL_786017, EPI_ISL_786018, EPI_ISL_786024, EPI_ISL_786025, EPI_ISL_786026, EPI_ISL_786027, EPI_ISL_786028, EPI_ISL_786029, EPI_ISL_786030, EPI_ISL_786036, EPI_ISL_786037, EPI_ISL_786038, EPI_ISL_786039, EPI_ISL_786040, EPI_ISL_786041, EPI_ISL_786042, EPI_ISL_786048, EPI_ISL_786049, EPI_ISL_786050, EPI_ISL_786051, EPI_ISL_786052, EPI_ISL_786053, EPI_ISL_786054, EPI_ISL_786060, EPI_ISL_786061, EPI_ISL_786062, EPI_ISL_786063, EPI_ISL_786064, EPI_ISL_786065, EPI_ISL_786066, EPI_ISL_786072, EPI_ISL_786073, EPI_ISL_786074, EPI_ISL_786075, EPI_ISL_786076, EPI_ISL_786077, EPI_ISL_786078, EPI_ISL_786084, EPI_ISL_786085, EPI_ISL_786086, EPI_ISL_786087, EPI_ISL_786088, EPI_ISL_786089, EPI_ISL_786090 | | | |
| see above | Houston Methodist Hospital | Houston Methodist Hospital | S. Wesley Long, Randall J. Olsen, Paul A. Christensen, David W. Bernard, James J. Davis, Maulik Shukla, Marcus Nguyen, Matthew Ojeda Saavedra, Prasanti Yerramilli, Layne Pruitt, Sishir Subedi, Heather Hendrickson, and James M. Musser |
| EPI_ISL_790568 | Florida Bureau of Public Health Laboratories | Florida Bureau of Public Health Laboratories | Sarah Schmedes, Jason Blanton |
| EPI_ISL_791336, EPI_ISL_791355, EPI_ISL_791361, EPI_ISL_791362, EPI_ISL_791378 | Johns Hopkins Hospital Department of Pathology | Johns Hopkins Hospital Department of Pathology | C. Paul Morris, Chun Huai Luo, Heba H. Mostafa |
| EPI_ISL_792017 | Plateforme COVID IDF | National Reference Center for Viruses of Respiratory Infections, Institut Pasteur, Paris | Marion Barbet, Sylvie Behillil, Méline Bizard, Angela Brisebarre, Camille Capel, Etienne Simon-Lorière, Vincent Enouf, Maud Vanpeene, Sylvie van der Werf,Jacques Fourgeaud |
| EPI_ISL_792627 | LACEN-PB | Laboratory of Respiratory Viruses and Measles, Oswaldo Cruz Institute, FIOCRUZ | Paola Resende, Luciana Appolinario, Fernando Motta, Anna Carolina Paixao, Ana Carolina Mendonca, João Felipe Bezerra, Romero Henrique Teixeira de Vasconcelos, Dalane Loudal Florentino Teixeira, Thiago Franco de Oliveira Carneiro, Marilda Siqueira |
| EPI_ISL_792646 | LACEN-PR | Laboratory of Respiratory Viruses and Measles, Oswaldo Cruz Institute, FIOCRUZ | Paola Resende, Luciana Appolinario, Fernando Motta, Anna Carolina Paixao, Ana Carolina Mendonca, Maria do Carmo Debur, Irina Nastassja Riediger, Marilda Siqueira |
| EPI_ISL_794719 | PathWest Laboratory Medicine WA | PathWest Laboratory Medicine WA Microbial Surveillance Unit | PathWest Laboratory Medicine WA Microbial Surveillance Unit |
| EPI_ISL_794828 | ULSS 7 Pedemontana - Distretto 2 | Istituto Zooprofilattico Sperimentale delle Venezie | Adelaide Milani, Alessia Schivo, Annalisa Salviato, Erika Giorgia Quaranta, Ambra Pastori, Bianca Zecchin, Alice Fusaro, Isabella Monne, Calogero Terregino, Antonia Ricci |
| EPI_ISL_796649 | Akershus University Hospital, Department for Microbiology and Infectious Disease Control | Norwegian Institute of Public Health, Department of Virology | Kathrine Stene-Johansen, Kamilla Heddeland Instefjord, Hilde Elshaug, Atiya R Ali,Marie Paulsen Madsen, Rasmus Riis Kopperud, Hilde Vollan, Karoline Bragstad, Olav Hungnes |
| EPI_ISL_801395 | Laboratorio de Ecologia de Doencas Transmissiveis na Amazonia, Instituto Leonidas e Maria Deane - Fiocruz Amazonia | Laboratorio de Ecologia de Doencas Transmissiveis na Amazonia, Instituto Leonidas e Maria Deane - Fiocruz Amazonia | Valdinete Nascimento, Victor Souza, André Corado, Fernanda Nascimento, George Silva, Ágatha Costa, Debora Duarte, Luciana Gonçalves, Maria Júlia Brandão, Michele Jesus, Felipe Naveca |
| EPI_ISL_801482, EPI_ISL_801489 | Dutch COVID-19 response team | Erasmus Medical Center | Bas Oude Munnink, Reina Sikkema, David Nieuwenhuijse, Irina Chestakova, Anne van der Linden, Marjan Boter, Emmanuelle Munger, Corine GeurtsvanKessel, Annemiek van der Eijk, Richard Molenkamp, Marion Koopmans, on behalf of the Dutch national COVID-19 response team. |
| EPI_ISL_802502, EPI_ISL_802504 | Connecticut Department of Health | Grubaugh Lab - Yale School of Public Health | Tara Alpert, Joseph Fauver, Anderson Brito, Mallery Breban, Anne Wyllie, Chantal Vogels, Mary Petrone, Chaney Kalinich, Isabel Ott, Nathan Grubaugh |
| EPI_ISL_802544 | Dutch COVID-19 response team | Erasmus Medical Center | Bas Oude Munnink, Reina Sikkema, David Nieuwenhuijse, Irina Chestakova, Anne van der Linden, Marjan Boter, Emmanuelle Munger, Corine GeurtsvanKessel, Annemiek van der Eijk, Richard Molenkamp, Marion Koopmans, on behalf of the Dutch national COVID-19 response team. |
| EPI_ISL_803428, EPI_ISL_803429, EPI_ISL_803431, EPI_ISL_803432, EPI_ISL_803433, EPI_ISL_803434, EPI_ISL_803435, EPI_ISL_803436, EPI_ISL_803479, EPI_ISL_803480, EPI_ISL_803481, EPI_ISL_803482, EPI_ISL_803483, EPI_ISL_803484, EPI_ISL_803485 | | | |
| see above | Wisconsin State Laboratory of Hygiene Communicable Disease Division | Wisconsin State Laboratory of Hygiene Communicable Disease Division | Kelsey R. Florek, Abigail C. Shockey |
| EPI_ISL_804570, EPI_ISL_804571, EPI_ISL_804572, EPI_ISL_804573 | MEPHI, Aix Marseille University | MEPHI, Aix Marseille University | Anthony LEVASSEUR |
| EPI_ISL_812143, EPI_ISL_812144, EPI_ISL_812182, EPI_ISL_812183, EPI_ISL_812184, EPI_ISL_812185, EPI_ISL_812186, EPI_ISL_812187, EPI_ISL_812188, EPI_ISL_812189, EPI_ISL_812190, EPI_ISL_812191, EPI_ISL_812192, EPI_ISL_812193, EPI_ISL_812194 | | | |
| see above | GA Department of Public Health Laboratory | Pathogen Discovery, Respiratory Viruses Branch, Division of Viral Diseases, Centers for Disease Control and Prevention | Yan Li, Ying Tao, Anna Montmayeur, Jing Zhang, Brian Lynch, Krista Queen, Anna Uehara, Rachel Marine, Peter Cook, Clinton R. Paden, Haibin Wang, Suxiang Tong |
| EPI_ISL_812997, EPI_ISL_812998, EPI_ISL_812999, EPI_ISL_813000, EPI_ISL_813001, EPI_ISL_813002, EPI_ISL_813003, EPI_ISL_813004, EPI_ISL_813005, EPI_ISL_813006 | University of Birmingham | COVID-19 Genomics UK (COG-UK) Consortium | Institute of Microbiology, University of Birmingham: Claire McMurray, Joanne Stockton, Samuel Nicholls, Radoslaw Poplawski, Will Rowe, Josh Quick, Nicholas Loman. University of Birmingham Testing Laboratory: Celina M Whalley, Andrew Bosworth, Charlotte Poxon, Kasun Wanigasooriya, Oliver Pickles, Mike Kidd, Alex Richter, Andrew D Beggs PHE Heartlands Lab: Husam Osman, Andrew Bosworth. Queen Elizabeth Hospital: Anna Casey |
| EPI_ISL_814685, EPI_ISL_814686 | Virology Department, Royal Infirmary of Edinburgh, NHS Lothian / School of Biological Sciences, University of Edinburgh / Institute of Genetics and Molecular Medicine, University of Edinburgh | COVID-19 Genomics UK (COG-UK) Consortium | McHugh M, Dewar R, Rooke S, Gallagher M, Balcaza C, O'Toole Á, Scher E, Hill V, McCrone JT, Colquhoun R, Yu X, Jackson B, Rambaut A, Williams TC, Templeton K |
| EPI_ISL_815479, EPI_ISL_815556, EPI_ISL_815604, EPI_ISL_816128, EPI_ISL_816129, EPI_ISL_816130, EPI_ISL_816131 | Department of Virus and Microbiological Special Diagnostics, Statens Serum Institut, Copenhagen, Denmark | Albertsen Lab, Department of Chemistry and Bioscience, Aalborg University, Denmark | Danish Covid-19 Genome Consortium |
| EPI_ISL_819456 | Northumbria University / South Tees Hospitals NHS Foundation Trust / North Cumbria Integrated Care NHS Foundation Trust / North Tees and Hartlepool NHS Foundation Trust / Newcastle Hospitals NHS Foundation Trust | COVID-19 Genomics UK (COG-UK) Consortium | Darren L Smith,Andrew Nelson,Matthew Bashton,Greg R Young,Joshua Loh,John Allan,Mohammad A Tariq,Giles S Holt,Gary Black,Wen C Yew,Lynn Dover,Paul Baker,Steve Liggett,Sarah Essex,Jane Greenaway,Debra Padgett,Clive Graham,Garren Scott,Edward Barton,Emma Swindells,Brendan Payne,Jennifer Collins,Yusri Taha,Gary Eltringham |

| | | | |
|---|---|---|---|
| EPI_ISL_825337, EPI_ISL_825349 | Hospital Universitari Vall d'Hebron - Vall d'Hebron Institut de Rercerca | Hospital Universitari Vall d'Hebron | Cristina Andrés, Maria Piñana, Josep F Abril, Damir Garcia-Cehic, Ariadna Rando, Juliana Esperalba, Maria Gema Codina, Carla Castillo, Maria Carmen Martin, Tomàs Pumarola, Josep Quer, Andrés Antón |
| EPI_ISL_825549, EPI_ISL_825593, EPI_ISL_825594 | Respiratory Virus Unit, National Infection Service, Public Health England | COVID-19 Genomics UK (COG-UK) Consortium | PHE Covid Sequencing Team |
| EPI_ISL_826944, EPI_ISL_826945, EPI_ISL_826946, EPI_ISL_826947, EPI_ISL_826948, EPI_ISL_826950, EPI_ISL_826951, EPI_ISL_826954, EPI_ISL_827211, EPI_ISL_827344, EPI_ISL_827508, EPI_ISL_827509, EPI_ISL_827512, EPI_ISL_827581, EPI_ISL_827832, EPI_ISL_827833, EPI_ISL_827834, EPI_ISL_827836, EPI_ISL_827837, EPI_ISL_827838, EPI_ISL_827839, EPI_ISL_827840, EPI_ISL_827901, EPI_ISL_827941, EPI_ISL_827944, EPI_ISL_827946, EPI_ISL_827947, EPI_ISL_827988, EPI_ISL_828024, EPI_ISL_828150, EPI_ISL_828274, EPI_ISL_828288 | | | |
| see above | deCODE genetics | deCODE genetics | Daniel F Gudbjartsson; Agnar Helgason; Hakon Jonsson; Olafur T Magnusson; Pall Melsted; Gudmundur L Norddahl; Jona Saemundsdottir; Asgeir Sigurdsson; Patrick Sulem; Arna B Agustsdottir; Hannes Eggertsson; Berglind Eiriksdottir; Run Fridriksdottir; Elisabet E Gardarsdottir; Gudmundur Georgsson; Olafia S Gretarsdottir; Kjartan R Gudmundsson; Thora R Gunnarsdottir; Arnaldur Gylfason; Hilma Holm; Brynjar O Jensson; Aslaug Jonasdottir; Kamilla S Josefsdottir; Thordur Kristjansson; Droplaug N Magnusdottir; Solvi Rognvaldsson; Louise le Roux; Gudrun Sigmundsdottir; Gardar Sveinbjornsson; Kristin E Sveinsdottir; Maney Sveinsdottir; Emil A Thorarensen; Bjarni Thorbjornsson; Gisli Masson; Ingileif Jonsdottir; Alma Moller; Thorolfur Gudnason; Karl G Kristinsson; Unnur Thorsteinsdottir; Kari Stefansson |
| EPI_ISL_828554, EPI_ISL_828635 | The National University Hospital of Iceland | deCODE genetics | Daniel F Gudbjartsson; Agnar Helgason; Hakon Jonsson; Olafur T Magnusson; Pall Melsted; Gudmundur L Norddahl; Jona Saemundsdottir; Asgeir Sigurdsson; Patrick Sulem; Arna B Agustsdottir; Hannes Eggertsson; Berglind Eiriksdottir; Run Fridriksdottir; Elisabet E Gardarsdottir; Gudmundur Georgsson; Olafia S Gretarsdottir; Kjartan R Gudmundsson; Thora R Gunnarsdottir; Arnaldur Gylfason; Hilma Holm; Brynjar O Jensson; Aslaug Jonasdottir; Kamilla S Josefsdottir; Thordur Kristjansson; Droplaug N Magnusdottir; Solvi Rognvaldsson; Louise le Roux; Gudrun Sigmundsdottir; Gardar Sveinbjornsson; Kristin E Sveinsdottir; Maney Sveinsdottir; Emil A Thorarensen; Bjarni Thorbjornsson; Gisli Masson; Ingileif Jonsdottir; Alma Moller; Thorolfur Gudnason; Karl G Kristinsson; Unnur Thorsteinsdottir; Kari Stefansson |
| EPI_ISL_828888, EPI_ISL_828899, EPI_ISL_828901, EPI_ISL_828903, EPI_ISL_829222, EPI_ISL_829404, EPI_ISL_829681, EPI_ISL_829717, EPI_ISL_829738, EPI_ISL_829781, EPI_ISL_829783 | | | |
| see above | deCODE genetics | deCODE genetics | Daniel F Gudbjartsson; Agnar Helgason; Hakon Jonsson; Olafur T Magnusson; Pall Melsted; Gudmundur L Norddahl; Jona Saemundsdottir; Asgeir Sigurdsson; Patrick Sulem; Arna B Agustsdottir; Hannes Eggertsson; Berglind Eiriksdottir; Run Fridriksdottir; Elisabet E Gardarsdottir; Gudmundur Georgsson; Olafia S Gretarsdottir; Kjartan R Gudmundsson; Thora R Gunnarsdottir; Arnaldur Gylfason; Hilma Holm; Brynjar O Jensson; Aslaug Jonasdottir; Kamilla S Josefsdottir; Thordur Kristjansson; Droplaug N Magnusdottir; Solvi Rognvaldsson; Louise le Roux; Gudrun Sigmundsdottir; Gardar Sveinbjornsson; Kristin E Sveinsdottir; Maney Sveinsdottir; Emil A Thorarensen; Bjarni Thorbjornsson; Gisli Masson; Ingileif Jonsdottir; Alma Moller; Thorolfur Gudnason; Karl G Kristinsson; Unnur Thorsteinsdottir; Kari Stefansson |
| EPI_ISL_829793 | The National University Hospital of Iceland | deCODE genetics | Daniel F Gudbjartsson; Agnar Helgason; Hakon Jonsson; Olafur T Magnusson; Pall Melsted; Gudmundur L Norddahl; Jona Saemundsdottir; Asgeir Sigurdsson; Patrick Sulem; Arna B Agustsdottir; Hannes Eggertsson; Berglind Eiriksdottir; Run Fridriksdottir; Elisabet E Gardarsdottir; Gudmundur Georgsson; Olafia S Gretarsdottir; Kjartan R Gudmundsson; Thora R Gunnarsdottir; Arnaldur Gylfason; Hilma Holm; Brynjar O Jensson; Aslaug Jonasdottir; Kamilla S Josefsdottir; Thordur Kristjansson; Droplaug N Magnusdottir; Solvi Rognvaldsson; Louise le Roux; Gudrun Sigmundsdottir; Gardar Sveinbjornsson; Kristin E Sveinsdottir; Maney Sveinsdottir; Emil A Thorarensen; Bjarni Thorbjornsson; Gisli Masson; Ingileif Jonsdottir; Alma Moller; Thorolfur Gudnason; Karl G Kristinsson; Unnur Thorsteinsdottir; Kari Stefansson |
| EPI_ISL_829828, EPI_ISL_829829, EPI_ISL_829835, EPI_ISL_829932, EPI_ISL_829950, EPI_ISL_829962, EPI_ISL_829978, EPI_ISL_829987, EPI_ISL_830070 | deCODE genetics | deCODE genetics | Daniel F Gudbjartsson; Agnar Helgason; Hakon Jonsson; Olafur T Magnusson; Pall Melsted; Gudmundur L Norddahl; Jona Saemundsdottir; Asgeir Sigurdsson; Patrick Sulem; Arna B Agustsdottir; Hannes Eggertsson; Berglind Eiriksdottir; Run Fridriksdottir; Elisabet E Gardarsdottir; Gudmundur Georgsson; Olafia S Gretarsdottir; Kjartan R Gudmundsson; Thora R Gunnarsdottir; Arnaldur Gylfason; Hilma Holm; Brynjar O Jensson; Aslaug Jonasdottir; Kamilla S Josefsdottir; Thordur Kristjansson; Droplaug N Magnusdottir; Solvi Rognvaldsson; Louise le Roux; Gudrun Sigmundsdottir; Gardar Sveinbjornsson; Kristin E Sveinsdottir; Maney Sveinsdottir; Emil A Thorarensen; Bjarni Thorbjornsson; Gisli Masson; Ingileif Jonsdottir; Alma Moller; Thorolfur Gudnason; Karl G Kristinsson; Unnur Thorsteinsdottir; Kari Stefansson |
| EPI_ISL_830264 | The National University Hospital of Iceland | deCODE genetics | Daniel F Gudbjartsson; Agnar Helgason; Hakon Jonsson; Olafur T Magnusson; Pall Melsted; Gudmundur L Norddahl; Jona Saemundsdottir; Asgeir Sigurdsson; Patrick Sulem; Arna B Agustsdottir; Hannes Eggertsson; Berglind Eiriksdottir; Run Fridriksdottir; Elisabet E Gardarsdottir; Gudmundur Georgsson; Olafia S Gretarsdottir; Kjartan R Gudmundsson; Thora R Gunnarsdottir; Arnaldur Gylfason; Hilma Holm; Brynjar O Jensson; Aslaug Jonasdottir; Kamilla S Josefsdottir; Thordur Kristjansson; Droplaug N Magnusdottir; Solvi Rognvaldsson; Louise le Roux; Gudrun Sigmundsdottir; Gardar Sveinbjornsson; Kristin E Sveinsdottir; Maney Sveinsdottir; Emil A Thorarensen; Bjarni Thorbjornsson; Gisli Masson; Ingileif Jonsdottir; Alma Moller; Thorolfur Gudnason; Karl G Kristinsson; Unnur Thorsteinsdottir; Kari Stefansson |
| EPI_ISL_831222, EPI_ISL_831223 | Hospital Universitario La Paz (Madrid) | SeqCOVID-SPAIN consortium/IBV(CSIC) | María Rodríguez-Tejedor, Elias Dahdouh, Fernando Lázaro-Perona, Jesús Mingorance and SeqCOVID-SPAIN consortium |
| EPI_ISL_831784, EPI_ISL_831786, EPI_ISL_831788, EPI_ISL_831790, EPI_ISL_831792, EPI_ISL_831793, EPI_ISL_831795, EPI_ISL_831813 | United States Air Force School of Aerospace Medicine | United States Air Force School of Aerospace Medicine | Anthony Fries, Jennifer Meyer, William Gruner, Amanda Javorina, Sarah Purves, Clarise Starr, Elizabeth Macias |
| EPI_ISL_832665, EPI_ISL_832666, EPI_ISL_832667, EPI_ISL_832668, EPI_ISL_832669, EPI_ISL_832670, EPI_ISL_832671, EPI_ISL_832672, EPI_ISL_832673, EPI_ISL_832674, EPI_ISL_832685, EPI_ISL_832686, EPI_ISL_832693 | | | |
| see above | OHSU Lab Services Molecular Microbiology Lab | Oregon SARS-CoV-2 Genome Sequencing Center | Brendan L. O'Connell, Ruth V. Nichols, Sally Grindstaff, Alec J. Hirsch, Donna Hansel, Guang Fan, Daniel N. Streblow, William B. Messer, Andrew C. Adey, Benjamin N. Bimber, Brian J. O'Roak |
| EPI_ISL_833461, EPI_ISL_833469, EPI_ISL_833473 | CHU Purpan - Laboratoire de Virologie - Institut Fédératif de Biologie | CHU Purpan - Laboratoire de Virologie - Institut Fédératif de Biologie | Latour J., Ranger N., Dubois M., Carcenac R., Harter A., Boyer P., Tremeaux P., Izopet J. |
| EPI_ISL_838313 | Virology Department, Royal Infirmary of Edinburgh, NHS Lothian / School of Biological Sciences, University of Edinburgh / Institute of Genetics and Molecular Medicine, University of Edinburgh | COVID-19 Genomics UK (COG-UK) Consortium | McHugh M, Dewar R, Rooke S, Gallagher M, Balcaza C, O'Toole Á, Scher E, Hill V, McCrone JT, Colquhoun R, Yu X, Jackson B, Rambaut A, Williams TC, Templeton K |
| EPI_ISL_840202 | Oxford Viromics, NDM, University of Oxford; Oxford University Hospitals; Basingstoke and North Hampshire Hospital | COVID-19 Genomics UK (COG-UK) Consortium | Tanya Golubchik, David Bonsall, George Macintyre, Amy Trebes, Mariateresa de Cesare, Catrin Moore, Alex Mobbs, Anita Justice, Robert Shaw, Monique Andersson, Timothy Peto, Emma Wise, Nathan Moore, Jessica Lynch, Nick Cortes, Matilde Mori, Stephen Kidd, David Buck, John Todd, Christophe Fraser |
| EPI_ISL_842353 | Virology Department, Sheffield Teaching Hospitals NHS Foundation Trust/Department of Infection, Immunity and Cardiovascular Disease, The Medical School, University of Sheffield | COVID-19 Genomics UK (COG-UK) Consortium | Thushan de Silva, Matthew Parker, Nikki Smith, Adri Angyal, Rebecca Brown, Luke Green, Rachel Tucker, Paul Parsons, Danielle Groves, Katie Johnson, Laura Carrilero, Alex Keeley, Dave Partridge, Matthew Wyles, Benjamin Lindsey, Mehmet Yavuz, Mohammad Raza, Cariad Evans |
| EPI_ISL_842939 | Barts Health NHS Trust | COVID-19 Genomics UK (COG-UK) Consortium | CUTINO-MOGUEL, Maria-Teresa; HARRINGTON, David; OWOYEMI, Dola; SHYLINI, Raghavendran; BROAD, Claire; KELE, Beatrix |
| EPI_ISL_844542 | Department of Virus and Microbiological Special Diagnostics, Statens Serum Institut, Copenhagen, Denmark | Albertsen Lab, Department of Chemistry and Bioscience, Aalborg University, Denmark | Danish Covid-19 Genome Consortium |
| EPI_ISL_845755 | Emory Molecular Diagnostics Laboratory, Emory Healthcare | Piantadosi Lab, Emory Department of Pathology | Ahmed Babiker, Anne Piantadosi |
| EPI_ISL_847704, EPI_ISL_847729 | California Department of Public Health | Chiu Laboratory, University of California, San Francisco | Charles Chiu, Xianding (Wayne) Deng, Candace Wang, Brian Bushnell, Scot Federman, Jill Hacker, Debra Wadford |
| EPI_ISL_847831 | Yale New Haven Hospital | Grubaugh Lab - Yale School of Public Health | Tara Alpert, Joseph Fauver, Anderson Brito, Mallery Breban, Anne Wyllie, Chantal Vogels, Mary Petrone, Chaney Kalinich, Isabel Ott, Nathan Grubaugh |
| EPI_ISL_848382, EPI_ISL_848383, EPI_ISL_848384 | Illinois Department of Public Health | Gagnon Lab, Southern Illinois University | Keith Gagnon |
| EPI_ISL_848557 | Evandro Chagas Institute | Evandro Chagas Institute | Santos, M.C.; Silva, A.M.; Junior, W.D.C.; Barbagelata, L.S.; Ferreira, J.A.; Sousa, E.M.A.; da Silva, P.S.; Pinheiro, K.C.; L.C.; Sousa Junior, E.C. |
| EPI_ISL_848871, EPI_ISL_848872, EPI_ISL_848873 | Florida Bureau of Public Health Laboratories | Florida Bureau of Public Health Laboratories | Sarah Schmedes, Jason Blanton |

| | | | |
|---|---|---|---|
| EPI_ISL_849440, EPI_ISL_849500, EPI_ISL_849501, EPI_ISL_849502, EPI_ISL_849564 | Seattle Flu Study | Seattle Flu Study | Deborah A. Nickerson, Chris D. Frazar, Jover Lee, Benjamin Pelle, Matthew Richardson, Amanda Adler, Elisabeth Brandstetter, Peter D. Han, Kairsten Fay, Misja Ilcisin, Kirsten Lacombe, Thomas R. Sibley, Melissa Truong, Caitlin R. Wolf, Michael Boeckh, Janet A. Englund, Michael Famulare, Barry R. Lutz, Mark J. Rieder, Lea M. Starita, Matthew Thompson, Jay Shendure, Trevor Bedford, Helen Y. Chu |
| EPI_ISL_849944 | UC Davis- Department of Pathology and Laboratory Medicine | Chan-Zuckerberg Biohub | CZB Cliahub Consortium |
| EPI_ISL_852595, EPI_ISL_852596 | Max von Pettenkofer Institute, Virology, National Reference Center for Retroviruses, LMU München | Laboratory for Functional Genome Analysis, Dept. Genomics, Gene Center of the LMU Munich | Max Muenchhoff, Stefan Krebs, Alexander Graf, Oliver Keppler, Helmut Blum |
| EPI_ISL_853325 | UPMC Clinical Microbiology Laboratory | Microbial Genome Sequencing Center; Microbial Genomic Epidemiology Laboratory | Mustapha M. Mustapha, Jane W. Marsh, Dan Snyder, Marissa P. Griffith, Stephanie L. Mitchell, Vatsala R. Srinivasa, Kady D. Waggle, Chinelo Ezeonwuku, Vaughn S. Cooper, Lee H. Harrison |
| EPI_ISL_853752 | Institute for Water Quality and Resource Management, Technical University Vienna | Bergthaler laboratory, CeMM Research Center for Molecular Medicine of the Austrian Academy of Sciences | Lukas Endler, Alexandra Popa, Benedikt Agerer, Jakob-Wendelin Genger, Alexander Lercher, Anna Schedl, Thomas Penz, Michael Schuster, Jan Laine, Martin Senekowitsch, Christoph Bock, Andreas Bergthaler |
| EPI_ISL_853941 | Department of Microbiology, University Innsbruck | Bergthaler laboratory, CeMM Research Center for Molecular Medicine of the Austrian Academy of Sciences | Lukas Endler, Alexandra Popa, Benedikt Agerer, Jakob-Wendelin Genger, Alexander Lercher, Anna Schedl, Thomas Penz, Michael Schuster, Jan Laine, Martin Senekowitsch, Christoph Bock, Andreas Bergthaler |
| EPI_ISL_853975, EPI_ISL_853984, EPI_ISL_853985, EPI_ISL_853994, EPI_ISL_853995, EPI_ISL_854003, EPI_ISL_854004, EPI_ISL_854010, EPI_ISL_854017, EPI_ISL_854030, EPI_ISL_854031, EPI_ISL_854040, EPI_ISL_854228, EPI_ISL_854272, EPI_ISL_854280 | | | |
| see above | Austrian Agency for Health and Food Safety (AGES) | Bergthaler laboratory, CeMM Research Center for Molecular Medicine of the Austrian Academy of Sciences | Lukas Endler, Alexandra Popa, Benedikt Agerer, Jakob-Wendelin Genger, Alexander Lercher, Anna Schedl, Thomas Penz, Michael Schuster, Jan Laine, Martin Senekowitsch, Christoph Bock, Andreas Bergthaler |
| EPI_ISL_855525, EPI_ISL_855537, EPI_ISL_855545 | KEMRI-Wellcome Trust Research Programme/KEMRI-CGMR-C Kilifi | KEMRI-Wellcome Trust Research Programme/KEMRI-CGMR-C Kilifi | Githinji et al |
| EPI_ISL_856797, EPI_ISL_856798, EPI_ISL_856799 | Servicio Virosis Respiratorias-Departamento Virología-INEI | Instituto Nacional Enfermedades Infecciosas C.G.Malbran | Baumeister E., Avaro M., Benedetti E., Russo M., Dattero ME, Pontoriero A., Cisterna D., Molina V., Perandones C., Tuduri E., Lorenzo F., Poklepovich T., Campos J. |
| EPI_ISL_860737, EPI_ISL_860738, EPI_ISL_860741, EPI_ISL_860742, EPI_ISL_860777, EPI_ISL_860778, EPI_ISL_860779, EPI_ISL_860780, EPI_ISL_860781 | Swiss National Reference Centre for Influenza | Swiss National Reference Centre for Influenza | Ana Rita Gonçalves Cabecinhas, Samuel Cordey, Florian Laubscher,Christoph Grünig, Laurent Kaiser |
| EPI_ISL_860805, EPI_ISL_860806, EPI_ISL_860812 | WHO/Minsk | Charité Universitätsmedizin Berlin, Institut für Virologie | Victor M Corman, Barbara Mühlemann, Jörn Beheim-Schwarzbach, Talitha Veith, Julia Tesch, Tobias Bleicker, Julia Schneider, Shmialiova Natallia, Sivets Natallia, Terry Jones, Christian Drosten |
| EPI_ISL_861672 | Day Hospital de Ermelino Matarazzo | Instituto Adolfo Lutz, Interdiciplinary Procedures Center, Strategic Laboratory | Claudio Tavares Sacchi, Claudia Regina Gonçalves, Erica Valessa Ramos Gomes, Karoline Rodrigues Campos |
| EPI_ISL_861723 | Department of Medical Microbiology, Hospital Pengajar Universiti Putra Malaysia | Malaysia Genome Institute | Mohd Noor Mat Isa, Syafinaz Amin-Nordin, Irni Suhayu Sapian, Hui-Yee Chee, Yusuf Muhammad Noor, Nurhezreen Md Iqbal, Enizza Kasim, Siti Noraini Othman, Mohd Faizal Abu Bakar, Shamsidar Sopie, Azrin Ahmad, Narcisse Joseph, Muhammad MI, Avisha Richards, Nor Zahrin Hasran, Nor Azfa Johari |
| EPI_ISL_861804, EPI_ISL_861854 | Hospital General Universitario Gregorio Marañón | SeqCOVID-SPAIN consortium/IBV(CSIC) | Darío García de Viedma, Laura Pérez-Lago, Pedro J Sola-Campoy, Sergio Buenestado-Serrano, Marta Herranz, Victor Manuel de la Cueva, Julia Suárez, Pilar Catalán, Patricia Muñoz and SeqCOVID-SPAIN consortium |
| EPI_ISL_865164, EPI_ISL_865165, EPI_ISL_865166, EPI_ISL_865167, EPI_ISL_865168, EPI_ISL_865170 | Liverpool Clinical Laboratories | COVID-19 Genomics UK (COG-UK) Consortium | Sam Haldenby, Anita Lucaci, Steve Paterson, Julian Hiscox, Alistair Darby, M Almsaud, A Alrezaihi, Muhannad Alruwaili, Stuart D Armstrong, Jones Benjamin, Eleanor G Bentley, Anu Chawla, Jordan J Clark, Angela Cowell, Richard Eccles, Isabel García-Dorival, Matthew Gemmell, Alessandro Gerada, PKF Gilmore, Richard Gregory, Ximeng Han, Catherine Hartley, Margaret Hughes, Miren Iturriza-Gomara, James Johnson, L Luu, Jenifer Manson, Charlotte Nelson, Elaine O'Toole, Cassie Olateju, Rebekah Penrice-Randal , Lucille Rainbow, N.P Randle, Trevor Ian Robinson, Parul Sharma, Ghada T Shawli, James P Stewart, Neil Swainston, Ecaterina Vamos, Joanne Watts, Mark Whitehead |
| EPI_ISL_867963 | Centre for Enzyme Innovation, University of Portsmouth / Translational Research Laboratory, Portsmouth Hospitals NHS Trust | COVID-19 Genomics UK (COG-UK) Consortium | Angela Beckett,Yann Bourgeois,Garry Scarlett,Sharon Glaysher,Scott Elliott,Kelly Bicknell,Robert Impey,Allyson Lloyd,Sarah Wyllie,Ethan Butcher,Anoop Chauhan,Samuel Robson |
| EPI_ISL_871787 | Department of Virus and Microbiological Special Diagnostics, Statens Serum Institut, Copenhagen, Denmark | Aalborg University | Danish Covid-19 Genome Consortium |
| EPI_ISL_872682, EPI_ISL_872683 | Rhode Island Department of Health | Infectious Disease Program, Broad Institute of Harvard and MIT | Lemieux,J.E., Siddle,K.J., Huard,R., King,E., Azevedo,K., Miller,A., Adams,G., Gladden-Young,A., Lagerborg,K., Rudy,M., DeRuff,K., Carter,A., Normandin,E., Bauer,M., Reilly,S., Tomkins-Tinch,C., Loreth,C., Chaluvadi,S., Birren,B.W., Gallagher,G., Smole,S., Park,D.J., MacInnis,B.L., and Sabeti,P.C. |
| EPI_ISL_872746, EPI_ISL_872747, EPI_ISL_872748 | Colorado Mesa University | Infectious Disease Program, Broad Institute of Harvard and MIT | Lemieux,J.E., Siddle,K.J., Marshall,J., ONeill,M., Bronson,A., Adams,G., Gladden-Young,A., Lagerborg,K., Rudy,M., DeRuff,K., Carter,A., Normandin,E., Bauer,M., Reilly,S., Tomkins-Tinch,C., Loreth,C., Chaluvadi,S., Birren,B.W., Gallagher,G., Smole,S., Park,D.J., MacInnis,B.L., and Sabeti,P.C. |
| EPI_ISL_875874, EPI_ISL_875877, EPI_ISL_875879, EPI_ISL_875882, EPI_ISL_875883, EPI_ISL_875884, EPI_ISL_875886, EPI_ISL_875890, EPI_ISL_875891, EPI_ISL_875892, EPI_ISL_875893, EPI_ISL_875894, EPI_ISL_875895, EPI_ISL_875899, EPI_ISL_875901, EPI_ISL_875906, EPI_ISL_875907, EPI_ISL_875908, EPI_ISL_875909, EPI_ISL_875910, EPI_ISL_875911, EPI_ISL_875912, EPI_ISL_875915, EPI_ISL_875921, EPI_ISL_875922, EPI_ISL_875924, EPI_ISL_875934, EPI_ISL_875938, EPI_ISL_875939, EPI_ISL_875945, EPI_ISL_875946, EPI_ISL_875949, EPI_ISL_875950, EPI_ISL_875952, EPI_ISL_875953, EPI_ISL_875958, EPI_ISL_876428, EPI_ISL_876429, EPI_ISL_876514, EPI_ISL_876515, EPI_ISL_876516, EPI_ISL_876517 | | | |
| see above | Eurofins Diatherix | Hudsonalpha Genome Sequencing Center | Jane Grimwood, Melissa Williams, Lori H. Handley, Joshua Stough, Leslie Malone, Stefan Brzezinski, Ada Stewart, Teresa Jones, Jenell Webber, John Lovell, Jennifer Cart, and Jeremy Schmutz |
| EPI_ISL_876848, EPI_ISL_876849 | Quest Diagnostics | Quest Diagnostics | Rosenthal,S.H., Gerasimova,A., Kagan,R.M., Anderson, B., Hua, M., Liu Y., Bernstein, L.E., Livingston, K.E., Perez, A., Shalhout, D.F., Shlyakhter, I.A., Owen, R., Tanpaiboon, P., Lacbawan, F. |
| EPI_ISL_882696, EPI_ISL_882733, EPI_ISL_882744, EPI_ISL_882745, EPI_ISL_882751, EPI_ISL_882753, EPI_ISL_882758, EPI_ISL_882759 | 1.AO Universitaria 'S. Giovanni di Dio e Ruggi D'Aragona, Scuola Medica Salernitana' Hospital / 2.UOC di Virologia e Microbiologia, Università della Campania 'L. Vanvitelli' / 3.AO Universitaria 'Federico II' Napoli Hospital / 4.AORN 'San Giuseppe Moscati' Avellino Hospital / 5.AO 'San Pio - presidio G. Rummo' Benevento Hospital / 6.AO 'Sant'Anna e San Sebastiano' Caserta Hospital / 7.PO 'Maria Santissima Addolorata' Eboli Hospital / 8.Biogem Istituto di Ricerche Genetiche | 1. Genome Research Center for Health (CRGS) / 2. Laboratory of Molecular Medicine and Genomics(LMMGe) / 3. Center for Research in Pure and Applied Mathematics (CRMPA) | Giorgio Giurato, Francesca Rizzo, Alessandro Weisz, Gianluigi Franci, Giovanni Nassa, Pasquale Pagliano, Roberta Tarallo, Elena Alexandrova, Ylenia D'Agostino, Carlo Ferravante, Jessica Lamberti, Viola Melone, Domenico Memoli, Valeria Mirici Cappa, Domenico Palumbo, Giovanni Pecoraro, Assunta Sellitto, Oriana Strianese, Ilaria Terenzi, Giuseppe Fenza, Aniello Gentile, Antonello Saccomanno, Sonia Amabile, Teresa Rocco, Annamaria Salvati, Emilia Vaccaro, Massimiliano Galdiero, Michele Cennamo, Giuseppe Portella, Maria Grazia Foti, Mariarosaria Ingino, Maria Landi, Maurizio Fumi, Vincenzo Rocco, Rita Greco, Vittoria Letizia, Arnolfo Petruzziello, Maddalena Schioppa, Gregorio Goffredi, Francesca Marciano, Michele Caraglia, Alessia Cossu, Marianna Scrima |
| EPI_ISL_882961 | Laboratorium Diagnostyi Mikrobiologicznej z Pracowni Prtka Grulicy SPSzW im. Jana Boego w Lublinie | National Institute of Public Health - National Institute of Hygiene | Wokowicz Tomasz, Zacharczuk Katarzyna, Gawor Jan |
| EPI_ISL_884400 | Infectious Diseases, Quest Diagnostics | Infectious Diseases, Quest Diagnostics | Rosenthal,S.H., Gerasimova,A., Kagan,R.M., Anderson,B., Bernstein,L.E., Livingston,K.E., Hua,M., Liu,Y., Shalhout,D.F., Owen,R., Lacbawan,F. |
| EPI_ISL_887159 | Massachusetts General Hospital | Infectious Disease Program, Broad Institute of Harvard and MIT | Lemieux,J.E., Siddle,K.J., Shaw,B., Adams,G., Pierce,V., Turbett,S., Anahtar,M., Branda,J., Slater,D., Harris,J., Lin,A.E., Gladden-Young,A., Lagerborg,K., Rudy,M., DeRuff,K., Carter,A., Normandin,E., Bauer,M., Reilly,S., Tomkins-Tinch,C., Loreth,C., Chaluvadi,S., Neumann,A., Cusick,C., Chapman,S.B., Gnirke,A., Flowers,K., Cerrato,F., Birren,B.W., Gallagher,G., Smole,S., Park,D.J., MacInnis,B.L., Ryan,E., LaRocque,R., Rosenberg,E. and Sabeti,P.C. |
| EPI_ISL_888835, EPI_ISL_888887 | Michigan Department of Health and Human Services, Bureau of Laboratories | Michigan Department of Health and Human Services, Bureau of Laboratories | Blankenship HM, Riner D, Soehnlen MK |
| EPI_ISL_891190 | DPH, Massachusetts State Public Health Lab | DPH, Massachusetts State Public Health Lab | Lang,A.S., Fink,T., Gallagher,G.R., Smole,S.C. |
| EPI_ISL_900430 | MEPHI, Aix Marseille University | MEPHI, Aix Marseille University | Anthony LEVASSEUR |

| | | | |
|---|---|---|---|
| EPI_ISL_903977 | NYSDOH Wadsworth Center, Virology Lab | Genomics and Discovery, Respiratory Viruses Branch, Division of Viral Diseases, Centers for Disease Control and Prevention | Krista Queen, Yan Li, Ying Tao, Jing Zhang, Anna Uehara, Anna Montmayeur, Clinton R. Paden, Peter W. Cook, Rachel Marine, Mili Sheth, Jasmine Padilla, Sarah Nobles, Mark Burroughs, Lori Rowe, Haibin Wang, Ben L. Rambo-Martin, Dhwani Batra, Justin Lee, Suxiang Tong |
| EPI_ISL_912473, EPI_ISL_912510, EPI_ISL_912531 | NHLS Universitas Academic | UFS Virology | PA Bester, MM Nyaga, P Nthiga, MT Mogotsi, D Goedhals, T de Oliveira |
| EPI_ISL_913097 | Center for Virology | Center for Virology | Jeremy V. Camp, Irene Goerzer, Monika Redlberger-Fritz, Stephan W. Aberle |
| EPI_ISL_924421, EPI_ISL_924429 | Virology Department, Sheffield Teaching Hospitals NHS Foundation Trust/Department of Infection, Immunity and Cardiovascular Disease, The Medical School, University of Sheffield | COVID-19 Genomics UK (COG-UK) Consortium | Thushan de Silva, Matthew Parker, Nikki Smith, Adri Angyal, Rebecca Brown, Luke Green, Rachel Tucker, Paul Parsons, Danielle Groves, Katie Johnson, Laura Carrilero, Alex Keeley, Dave Partridge, Matthew Wyles, Benjamin Lindsey, Mehmet Yavuz, Mohammad Raza, Cariad Evans |
| EPI_ISL_925093, EPI_ISL_925096, EPI_ISL_925103, EPI_ISL_925106, EPI_ISL_925115, EPI_ISL_925118, EPI_ISL_925125, EPI_ISL_925128 | 1.AO Universitaria 'S. Giovanni di Dio e Ruggi D'Aragona, Scuola Medica Salernitana' Hospital / 2.UOC di Virologia e Microbiologia, Università della Campania 'L. Vanvitelli' / 3.AO Universitaria 'Federico II' Napoli Hospital / 4.AORN 'San Giuseppe Moscati' Avellino Hospital / 5.AO 'San Pio - presidio G. Rummo' Benevento Hospital / 6.AO 'Sant'Anna e San Sebastiano' Caserta Hospital / 7.PO 'Maria Santissima Addolorata' Eboli Hospital / 8.Biogem Istituto di Ricerche Genetiche | 1. Genome Research Center for Health (CRGS) / 2. Laboratory of Molecular Medicine and Genomics(LMMGe) / 3. Center for Research in Pure and Applied Mathematics (CRMPA) | Giorgio Giurato, Francesca Rizzo, Alessandro Weisz, Gianluigi Franci, Giovanni Nassa, Pasquale Pagliano, Roberta Tarallo, Elena Alexandrova, Ylenia D'Agostino, Carlo Ferravante, Jessica Lamberti, Viola Melone, Domenico Memoli, Valeria Mirici Cappa, Domenico Palumbo, Giovanni Pecoraro, Assunta Sellitto, Oriana Strianese, Ilaria Terenzi, Giuseppe Fenza, Aniello Gentile, Antonello Saccomanno, Sonia Amabile, Teresa Rocco, Annamaria Salvati, Emilia Vaccaro, Massimiliano Galdiero, Michele Cennamo, Giuseppe Portella, Maria Grazia Foti, Mariarosaria Ingino, Maria Landi, Maurizio Fumi, Vincenzo Rocco, Rita Greco, Vittoria Letizia, Arnolfo Petruzziello, Maddalena Schioppa, Gregorio Goffredi, Francesca Marciano, Michele Caraglia, Alessia Cossu, Marianna Scrima |
| EPI_ISL_925142, EPI_ISL_925143, EPI_ISL_925144, EPI_ISL_925145, EPI_ISL_925146, EPI_ISL_925148, EPI_ISL_925149, EPI_ISL_925150 | Virginia DCLS | Virginia DCLS | Virginia DCLS |
| EPI_ISL_925463, EPI_ISL_925464 | Department of Clinical Microbiology | GIGA Medical Genomics | Keith Durkin, Maria Artesi, Sébastien Bontems, Raphaël Boreux, Bouchra Boujemla, Cécile Meex, Pierrette Melin, Marie-Pierre Hayette, Vincent Bours |
| EPI_ISL_925901, EPI_ISL_925902, EPI_ISL_925903, EPI_ISL_925904, EPI_ISL_925905, EPI_ISL_925906, EPI_ISL_925907 | Nucleic Acid Testing, National Reference Laboratory | GIGA Medical Genomics | Yvan Butera, Keith Durkin, Maria Artesi, Bouchra Boujemla, Robert Rutayisire, Patrick Tuyisenge, Esperence Umumararungu, Sébastien Bontems, Marie-Pierre Hayette, Nathalie Renotte, Swaibu Gatare, Jacob Souopgui, Sabin Nsanzimana, Vincent Bours, Léon Mutesa |
| EPI_ISL_926075, EPI_ISL_926386, EPI_ISL_926465, EPI_ISL_926794, EPI_ISL_926833, EPI_ISL_926955, EPI_ISL_927091, EPI_ISL_927398, EPI_ISL_927621, EPI_ISL_927887, EPI_ISL_928043, EPI_ISL_928105, EPI_ISL_928305, EPI_ISL_928412, EPI_ISL_928550, EPI_ISL_928588, EPI_ISL_928629, EPI_ISL_928682, EPI_ISL_929197, EPI_ISL_929214, EPI_ISL_929229, EPI_ISL_930075, EPI_ISL_930222 | | | |
| see above | Department of Virus and Microbiological Special Diagnostics, Statens Serum Institut, Copenhagen, Denmark | Aalborg University | Danish Covid-19 Genome Consortium |
| EPI_ISL_931072, EPI_ISL_931073 | University Hospital Basel, Clinical Virology | University Hospital Basel, Clinical Bacteriology | Tim Roloff, Madlen Stange, Helena MB Seth-Smith, Alfredo Mari, Karoline Leuzinger, Julia Bielicki, Manuel Battegay, Hans Hirsch, Adrian Egli |
| EPI_ISL_933660 | General Hospital - Veles | Research Center for Genetic Engineering and Biotechnology "Georgi D. Efremov" , Macedonian Academy of Sciences and Arts | RCGEB - MASA |
| EPI_ISL_933713 | General Hospital - Prilep | Research Center for Genetic Engineering and Biotechnology "Georgi D. Efremov" , Macedonian Academy of Sciences and Arts | RCGEB - MASA |
| EPI_ISL_933717 | General Hospital - Strumica | Research Center for Genetic Engineering and Biotechnology "Georgi D. Efremov" , Macedonian Academy of Sciences and Arts | RCGEB - MASA |
| EPI_ISL_933720 | General Hospital - Tetovo | Research Center for Genetic Engineering and Biotechnology "Georgi D. Efremov" , Macedonian Academy of Sciences and Arts | RCGEB - MASA |
| EPI_ISL_933722 | Gerontology Institute - Skopje | Research Center for Genetic Engineering and Biotechnology "Georgi D. Efremov" , Macedonian Academy of Sciences and Arts | RCGEB - MASA |
| EPI_ISL_933726 | General Hospital - Strumica | Research Center for Genetic Engineering and Biotechnology "Georgi D. Efremov" , Macedonian Academy of Sciences and Arts | RCGEB - MASA |
| EPI_ISL_936601, EPI_ISL_936602, EPI_ISL_936603, EPI_ISL_936604, EPI_ISL_936605 | Northwestern Memorial Hospital | Ozer Lab | Ramon Lorenzo-Redondo, Lacy M. Simons, Chad J. Achenbach, Lawrence J. Jennings, Michael G. Ison, Judd F. Hultquist, Egon A. Ozer |
| EPI_ISL_937024, EPI_ISL_937049 | Quest Diagnostics | Quest Diagnostics | Rosenthal,S.H., Gerasimova,A., Kagan,R.M., Anderson, B., Livingston, K.E., Hua, M., Liu Y., Shalhout, D.F., Owen, R., Lacbawan, F. |
| EPI_ISL_940138 | National Institute for Communicable Diseases,National Health Laboratory Services, Gauteng, South Africa | National Institute for Communicable Diseases of the National Health Laboratory Service | Amoako DG, Mohale T, Ntuli N, Mahlangu B, Allam M, Ismail A, Bhiman JN |
| EPI_ISL_940155, EPI_ISL_940156 | NHLS Universitas Academic | UFS Virology | PA Bester, MM Nyaga, P Nthiga, MT Mogotsi, D Goedhals, T de Oliveira |
| EPI_ISL_940173, EPI_ISL_940175, EPI_ISL_940176, EPI_ISL_940177, EPI_ISL_940178, EPI_ISL_940179, EPI_ISL_940192 | Hôpital Bichat Claude Bernard, Laboratoire de Virologie | IAME UMR1137 Inserm, Université de Paris, Hôpital Bichat | Antoine Bridier-Nahmias, Amélie Recoing, Quentin Le Hingrat, Lena Daniel, Siham Hamri, Gilles Collin, Alexandre Storto, Mélanie Bertine, Charlotte Charpentier, Nadhira Houhou-Fidouh, Diane Descamps, Benoit Visseaux |
| EPI_ISL_940872 | Vaccines and Infectious Diseases Analytics Research Unit (VIDA) | KRISP, KZN Research Innovation and Sequencing Platform | Baillie Vicky, du Plessis Jeanine, Giandhari Jennifer, Pillay Sureshnee, Naidoo Yeshnee, Tegally Houriiyah, de Oliveira Tulio, Madhi Shabir |
| EPI_ISL_954190 | 1.AO Universitaria 'S. Giovanni di Dio e Ruggi D'Aragona, Scuola Medica Salernitana' Hospital / 2.UOC di Virologia e Microbiologia, Università della Campania 'L. Vanvitelli' / 3.AO Universitaria 'Federico II' Napoli Hospital / 4.AORN 'San Giuseppe Moscati' Avellino Hospital / 5.AO 'San Pio - presidio G. Rummo' Benevento Hospital / 6.AO 'Sant'Anna e San Sebastiano' Caserta Hospital / 7.PO 'Maria Santissima Addolorata' Eboli Hospital / 8.Biogem Istituto di Ricerche Genetiche | 1. Genome Research Center for Health (CRGS) / 2. Laboratory of Molecular Medicine and Genomics(LMMGe) / 3. Center for Research in Pure and Applied Mathematics (CRMPA) | Giorgio Giurato, Francesca Rizzo, Alessandro Weisz, Gianluigi Franci, Giovanni Nassa, Pasquale Pagliano, Roberta Tarallo, Elena Alexandrova, Ylenia D'Agostino, Carlo Ferravante, Jessica Lamberti, Viola Melone, Domenico Memoli, Valeria Mirici Cappa, Domenico Palumbo, Giovanni Pecoraro, Assunta Sellitto, Oriana Strianese, Ilaria Terenzi, Giuseppe Fenza, Aniello Gentile, Antonello Saccomanno, Sonia Amabile, Teresa Rocco, Annamaria Salvati, Emilia Vaccaro, Massimiliano Galdiero, Michele Cennamo, Giuseppe Portella, Maria Grazia Foti, Mariarosaria Ingino, Maria Landi, Maurizio Fumi, Vincenzo Rocco, Rita Greco, Vittoria Letizia, Arnolfo Petruzziello, Maddalena Schioppa, Gregorio Goffredi, Francesca Marciano, Michele Caraglia, Alessia Cossu, Marianna Scrima, Edmondo Adorisio, Morena D'Avenia, Michela Iacobellis, Rosanna Piluscio, Giorgio Dirani, Vittorio Sambri, Simona Semprini, Silvia Zanoli, Francesco Curcio, Stefania Marzinotto, Andreina Baj, Fausto Sessa. |
| EPI_ISL_955689 | Orange County Public Health Lab | Chan-Zuckerberg Biohub | CZB Cliahub Consortium |
| EPI_ISL_960120 | Knysna Hospital wc KNY | National Health Laboratory Service/UCT | Arash Iranzadeh, Deelan Doolabh, Lynn Tyers, Bruna Galvao, Innocent Mudau, Marvin Hsiao, Kruger Marais, Diana Hardie, Stephen Korsman, Carolyn |

| | | | |
|---|---|---|---|
| | | | Williamson |
| EPI_ISL_960121 | Mitchells Plain Hospital wc MPH | National Health Laboratory Service/UCT | Arash Iranzadeh, Deelan Doolabh, Lynn Tyers, Bruna Galvao, Innocent Mudau, Marvin Hsiao, Kruger Marais, Diana Hardie, Stephen Korsman, Carolyn Williamson |
| EPI_ISL_960122 | Conville CDC wc CVC | National Health Laboratory Service/UCT | Arash Iranzadeh, Deelan Doolabh, Lynn Tyers, Bruna Galvao, Innocent Mudau, Marvin Hsiao, Kruger Marais, Diana Hardie, Stephen Korsman, Carolyn Williamson |
| EPI_ISL_960123 | Kwanokuthula CDC wc KWA | National Health Laboratory Service/UCT | Arash Iranzadeh, Deelan Doolabh, Lynn Tyers, Bruna Galvao, Innocent Mudau, Marvin Hsiao, Kruger Marais, Diana Hardie, Stephen Korsman, Carolyn Williamson |
| EPI_ISL_960124 | Heideveld Emergency Centre | National Health Laboratory Service/UCT | Arash Iranzadeh, Deelan Doolabh, Lynn Tyers, Bruna Galvao, Innocent Mudau, Marvin Hsiao, Kruger Marais, Diana Hardie, Stephen Korsman, Carolyn Williamson |
| EPI_ISL_961914, EPI_ISL_961958, EPI_ISL_962101, EPI_ISL_962122, EPI_ISL_962177 | Illinois Department of Public Health | Gagnon Lab, Southern Illinois University | Keith Gagnon |
| EPI_ISL_965362, EPI_ISL_965377, EPI_ISL_965378, EPI_ISL_965379, EPI_ISL_965380 | University of Liège COVID-19 testing center | GIGA Medical Genomics | Keith Durkin, Maria Artesi, Bouchra Boujemla, Emmanuel André, Marc Van Ranst, Fabrice Bureau, Laurent Gillet, Wouter Coppieters, Vincent Bours |
| EPI_ISL_965623 | Dutch COVID-19 response team | Medical Microbiology, Maastricht University Medical Centre | Jozef Dingemans*, Brian van der Veer*, Erik Beuken, Carmen Reumkens, Lieke van Alphen, Christian Hoebe, Paul Savelkoul |
| EPI_ISL_968905, EPI_ISL_968907, EPI_ISL_968908, EPI_ISL_968909 | KEMRI-Wellcome Trust Research Programme/KEMRI-CGMR-C Kilifi | KEMRI-Wellcome Trust Research Programme/KEMRI-CGMR-C Kilifi | Githinji et al |
| EPI_ISL_974825, EPI_ISL_974826, EPI_ISL_974827, EPI_ISL_974828, EPI_ISL_974829, EPI_ISL_974830, EPI_ISL_974831, EPI_ISL_974832, EPI_ISL_974833, EPI_ISL_974834, EPI_ISL_974835, EPI_ISL_974836, EPI_ISL_974837, EPI_ISL_974838, EPI_ISL_974839, EPI_ISL_974840, EPI_ISL_974841, EPI_ISL_974842, EPI_ISL_974843, EPI_ISL_974844, EPI_ISL_974845, EPI_ISL_974846, EPI_ISL_974847, EPI_ISL_974848, EPI_ISL_974849, EPI_ISL_974850, EPI_ISL_974851, EPI_ISL_974852, EPI_ISL_974854, EPI_ISL_974855, EPI_ISL_974856, EPI_ISL_974857, EPI_ISL_974858, EPI_ISL_974859, EPI_ISL_974860, EPI_ISL_974861, EPI_ISL_974862, EPI_ISL_974863, EPI_ISL_974864, EPI_ISL_974865, EPI_ISL_974866, EPI_ISL_974867, EPI_ISL_974868, EPI_ISL_974869, EPI_ISL_974870, EPI_ISL_974871, EPI_ISL_974872, EPI_ISL_974873, EPI_ISL_974874, EPI_ISL_974875, EPI_ISL_974876, EPI_ISL_974877, EPI_ISL_974878, EPI_ISL_974879, EPI_ISL_974880, EPI_ISL_974881, EPI_ISL_974882, EPI_ISL_974883, EPI_ISL_974884, EPI_ISL_974885, EPI_ISL_974886, EPI_ISL_974887, EPI_ISL_974888, EPI_ISL_974889 | | | |
| see above | BCCDC Public Health Laboratory | BCCDC Public Health Laboratory | Prystajecky Natalie, Linda Hoang, Dan Fornika, John Tyson, Shannon Russell, Kim Macdonald, Kimia Kamelian, Ana Pacagnella, Corrinne Ng, Loretta Janz, Robert Azana Terry Snutch, Mel Krajden |
| EPI_ISL_977589 | Department of Medical Microbiology, Hospital Pengajar Universiti Putra Malaysia | Malaysia Genome Institute | Mohd Noor Mat Isa, Syafinaz Amin-Nordin, Irni Suhayu Sapian, Hui-Yee Chee, Yusuf Muhammad Noor, Nurhezreen Md Iqbal, Enizza Kasim, Siti Noraini Othman, Mohd Faizal Abu Bakar, Shamsidar Sopie, Azrin Ahmad, Narcisse Joseph, Muhammad MI, Avisha Richards, Nor Zahrin Hasran, Nor Azfa Johari |
