## Supplementary material for "Detection and characterization of the SARS-CoV-2 lineage B.1.526 in New York": Supp. Table 3 GISAID Acknowledgment part 2: gisaid_hcov-19_acknowledgement_table_2021_02_13_00-7.pdf

We gratefully acknowledge the following Authors from the Originating laboratories responsible for obtaining the specimens, as well as the Submitting laboratories where the genome data were generated and shared via GISAID, on which this research is based.

All Submitters of data may be contacted directly via www.gisaid.org

Authors are sorted alphabetically.

| Accession ID | Originating Laboratory | Submitting Laboratory | Authors |
|---|---|---|---|
| EPI_ISL_635199 | Bamrasnaradura Infectious Disease Institute | Molecular HIV and Emerging Infectious Diseases Laboratory | Sumonmal UTTAYAMAKUL |
| EPI_ISL_636734, EPI_ISL_636735 | Respiratory Virus Unit, Microbiology Services Colindale, Public Health England | Respiratory Virus Unit, Microbiology Services Colindale, Public Health England | PHE Covid Sequencing Team |
| EPI_ISL_637096 | LabTests | Institute of Environmental Science and Research (ESR) | Xiaoyun Ren, Matt Storey, Nikki Freed, Muhammad Faisal, Jing Wang, Hermes Perez, Anja Werno, Antje van der Linden, Arlo Upton, Chris Mansell, David Hammer, Dragana Drinkovic, Gary McAuliffe, Hana Sofia Andersson, James Ussher, Jill Sherwood, Josh Freeman, Julia Howard, Juliet Elvy, Mary DeAlmeida, Matt Blakiston, Matthew Rogers, Max Bloomfield, Michael Addidle, Michelle Balm, Sally Roberts, Sarah Jefferies, Sharmini Muttaiyah, Susan Morpeth, Susan Taylor, Timothy Blackmore, Vani Sathyendran, Veronica Playle, Virginia Hope, Erasmus Smit, Lauren Jelly, Olin Silander, Joep de Ligt |
| EPI_ISL_637298 | Queens Medical Centre, Clinical Microbiology Department / DeepSeq Nottingham | COVID-19 Genomics UK (COG-UK) Consortium | Gemma Clark, Wendy Smith, Manjinder Khakh, Vicki M Fleming, Michelle M Lister, Hannah Howson-Wells, Jonathan Ball, Patrick McClure, Joseph Chappell, Theocharis Tsoleridis, Nadine Holmes, Matthew Carlisle, Christopher Moore, Fei Sang, Johnny Debebe, Victoria Wright, Matthew Loose |
| EPI_ISL_637382, EPI_ISL_637383 | Northumbria University / South Tees Hospitals NHS Foundation Trust / North Cumbria Integrated Care NHS Foundation Trust / North Tees and Hartlepool NHS Foundation Trust / Newcastle Hospitals NHS Foundation Trust | COVID-19 Genomics UK (COG-UK) Consortium | Darren L Smith,Andrew Nelson,Matthew Bashton,Greg R Young,Joshua Loh,John Allan,Mohammad A Tariq,Giles S Holt,Gary Black,Wen C Yew,Lynn Dover,Paul Baker,Steve Liggett,Sarah Essex,Jane Greenaway,Debra Padgett,Clive Graham,Garren Scott,Edward Barton,Emma Swindells,Brendan Payne,Jennifer Collins,Yusri Taha,Gary Eltringham |
| EPI_ISL_637800 | Queens Medical Centre, Clinical Microbiology Department / DeepSeq Nottingham | COVID-19 Genomics UK (COG-UK) Consortium | Gemma Clark, Wendy Smith, Manjinder Khakh, Vicki M Fleming, Michelle M Lister, Hannah Howson-Wells, Jonathan Ball, Patrick McClure, Joseph Chappell, Theocharis Tsoleridis, Nadine Holmes, Matthew Carlisle, Christopher Moore, Fei Sang, Johnny Debebe, Victoria Wright, Matthew Loose |
| EPI_ISL_637988 | Northumbria University / South Tees Hospitals NHS Foundation Trust / North Cumbria Integrated Care NHS Foundation Trust / North Tees and Hartlepool NHS Foundation Trust / Newcastle Hospitals NHS Foundation Trust | COVID-19 Genomics UK (COG-UK) Consortium | Darren L Smith,Andrew Nelson,Matthew Bashton,Greg R Young,Joshua Loh,John Allan,Mohammad A Tariq,Giles S Holt,Gary Black,Wen C Yew,Lynn Dover,Paul Baker,Steve Liggett,Sarah Essex,Jane Greenaway,Debra Padgett,Clive Graham,Garren Scott,Edward Barton,Emma Swindells,Brendan Payne,Jennifer Collins,Yusri Taha,Gary Eltringham |
| EPI_ISL_638609, EPI_ISL_638610, EPI_ISL_638611, EPI_ISL_638612, EPI_ISL_638613 | Queens Medical Centre, Clinical Microbiology Department / DeepSeq Nottingham | COVID-19 Genomics UK (COG-UK) Consortium | Gemma Clark, Wendy Smith, Manjinder Khakh, Vicki M Fleming, Michelle M Lister, Hannah Howson-Wells, Jonathan Ball, Patrick McClure, Joseph Chappell, Theocharis Tsoleridis, Nadine Holmes, Matthew Carlisle, Christopher Moore, Fei Sang, Johnny Debebe, Victoria Wright, Matthew Loose |
| EPI_ISL_645120, EPI_ISL_645122, EPI_ISL_645123, EPI_ISL_645125, EPI_ISL_645126, EPI_ISL_645127, EPI_ISL_645130 | National Public Health Laboratory, National Centre for Infectious Diseases | National Public Health Laboratory, National Centre for Infectious Diseases | Tze Minn Mak, Sophie Octavia, Zhenyang Zhou, Lin Cui, Raymond Tzer Pin Lin |
| EPI_ISL_649122 | Middlemore Hospital | Institute of Environmental Science and Research (ESR) | Xiaoyun Ren, Matt Storey, Nikki Freed, Muhammad Faisal, Jing Wang, Hermes Perez, Anja Werno, Antje van der Linden, Arlo Upton, Chris Mansell, David Hammer, Dragana Drinkovic, Gary McAuliffe, Hana Sofia Andersson, James Ussher, Jill Sherwood, Josh Freeman, Julia Howard, Juliet Elvy, Mary DeAlmeida, Matt Blakiston, Matthew Rogers, Max Bloomfield, Michael Addidle, Michelle Balm, Sally Roberts, Sarah Jefferies, Sharmini Muttaiyah, Susan Morpeth, Susan Taylor, Timothy Blackmore, Vani Sathyendran, Veronica Playle, Virginia Hope, Erasmus Smit, Lauren Jelly, Olin Silander, Joep de Ligt |
| EPI_ISL_649123 | Wellington SCL (WN) | Institute of Environmental Science and Research (ESR) | Xiaoyun Ren, Matt Storey, Nikki Freed, Muhammad Faisal, Jing Wang, Hermes Perez, Anja Werno, Antje van der Linden, Arlo Upton, Chris Mansell, David Hammer, Dragana Drinkovic, Gary McAuliffe, Hana Sofia Andersson, James Ussher, Jill Sherwood, Josh Freeman, Julia Howard, Juliet Elvy, Mary DeAlmeida, Matt Blakiston, Matthew Rogers, Max Bloomfield, Michael Addidle, Michelle Balm, Sally Roberts, Sarah Jefferies, Sharmini Muttaiyah, Susan Morpeth, Susan Taylor, Timothy Blackmore, Vani Sathyendran, Veronica Playle, Virginia Hope, Erasmus Smit, Lauren Jelly, Olin Silander, Joep de Ligt |
| EPI_ISL_650119, EPI_ISL_650126 | University of Birmingham | COVID-19 Genomics UK (COG-UK) Consortium | Institute of Microbiology, University of Birmingham: Claire McMurray, Joanne Stockton, Samuel Nicholls, Radoslaw Poplawski, Will Rowe, Josh Quick, Nicholas Loman. University of Birmingham Testing Laboratory: Celina M Whalley, Andrew Bosworth, Charlotte Poxon, Kasun Wanigasooriya, Oliver Pickles, Mike Kidd, Alex Richter, Andrew D Beggs PHE Heartlands Lab: Husam Osman, Andrew Bosworth. Queen Elizabeth Hospital: Anna Casey |
| EPI_ISL_650134 | Virology Department, Sheffield Teaching Hospitals NHS Foundation Trust/Department of Infection, Immunity and Cardiovascular Disease, The Medical School, University of Sheffield | COVID-19 Genomics UK (COG-UK) Consortium | Thushan de Silva, Matthew Parker, Nikki Smith, Adri Angyal, Rebecca Brown, Luke Green, Rachel Tucker, Paul Parsons, Danielle Groves, Katie Johnson, Laura Carrilero, Alex Keeley, Dave Partridge, Matthew Wyles, Benjamin Lindsey, Mehmet Yavuz, Mohammad Raza, Cariad Evans |
| EPI_ISL_650188, EPI_ISL_650211 | Quadram Institute Bioscience | COVID-19 Genomics UK (COG-UK) Consortium | Dave J. Baker, Gemma L. Kay, Alp Aydin, Thanh Le-Viet, Steven Rudder, Ana P. Tedim, Anastasia Kolyva, Maria Diaz, Leonardo de Oliveira Martins, Nabil-Fareed Alikhan, Lizzie Meadows, Rachael Stanley, Ngozi Elumogo, Muhammed Yasir, Nicholas M. Thomson, Alexander J Trotter, Rachel Gilroy, Samuel Bloomfield, Claire Stuart, Andrew Bell, Reenesh Prakash, Samir Dervisevic, Alison E. Mather, John Wain, Mark Webber, Andrew J. Page, Justin O'Grady |
| EPI_ISL_650212, EPI_ISL_650216 | University of Birmingham | COVID-19 Genomics UK (COG-UK) Consortium | Institute of Microbiology, University of Birmingham: Claire McMurray, Joanne Stockton, Samuel Nicholls, Radoslaw Poplawski, Will Rowe, Josh Quick, Nicholas Loman. University of Birmingham Testing Laboratory: Celina M Whalley, Andrew Bosworth, Charlotte Poxon, Kasun Wanigasooriya, Oliver Pickles, Mike Kidd, Alex Richter, Andrew D Beggs PHE Heartlands Lab: Husam Osman, Andrew Bosworth. Queen Elizabeth Hospital: Anna Casey |
| EPI_ISL_650227, EPI_ISL_650230, EPI_ISL_650294 | Quadram Institute Bioscience | COVID-19 Genomics UK (COG-UK) Consortium | Dave J. Baker, Gemma L. Kay, Alp Aydin, Thanh Le-Viet, Steven Rudder, Ana P. Tedim, Anastasia Kolyva, Maria Diaz, Leonardo de Oliveira Martins, Nabil-Fareed Alikhan, Lizzie Meadows, Rachael Stanley, Ngozi Elumogo, Muhammed Yasir, Nicholas M. Thomson, Alexander J Trotter, Rachel Gilroy, Samuel Bloomfield, Claire Stuart, Andrew Bell, Reenesh Prakash, Samir Dervisevic, Alison E. Mather, John Wain, Mark Webber, Andrew J. Page, Justin O'Grady |
| EPI_ISL_650295 | Queens Medical Centre, Clinical Microbiology Department / DeepSeq Nottingham | COVID-19 Genomics UK (COG-UK) Consortium | Gemma Clark, Wendy Smith, Manjinder Khakh, Vicki M Fleming, Michelle M Lister, Hannah Howson-Wells, Jonathan Ball, Patrick McClure, Joseph Chappell, Theocharis Tsoleridis, Nadine Holmes, Matthew Carlisle, Christopher Moore, Fei Sang, Johnny Debebe, Victoria Wright, Matthew Loose |
| EPI_ISL_650327, EPI_ISL_650332 | Quadram Institute Bioscience | COVID-19 Genomics UK (COG-UK) Consortium | Dave J. Baker, Gemma L. Kay, Alp Aydin, Thanh Le-Viet, Steven Rudder, Ana P. Tedim, Anastasia Kolyva, Maria Diaz, Leonardo de Oliveira Martins, Nabil-Fareed Alikhan, Lizzie Meadows, Rachael Stanley, Ngozi Elumogo, Muhammed Yasir, Nicholas M. Thomson, Alexander J Trotter, Rachel Gilroy, Samuel Bloomfield, Claire Stuart, Andrew Bell, Reenesh Prakash, Samir Dervisevic, Alison E. Mather, John Wain, Mark Webber, Andrew J. Page, Justin O'Grady |
| EPI_ISL_650350, EPI_ISL_650373 | University of Birmingham | COVID-19 Genomics UK (COG-UK) Consortium | Institute of Microbiology, University of Birmingham: Claire McMurray, Joanne Stockton, Samuel Nicholls, Radoslaw Poplawski, Will Rowe, Josh Quick, Nicholas Loman. University of Birmingham Testing Laboratory: Celina M Whalley, Andrew Bosworth, Charlotte Poxon, Kasun Wanigasooriya, Oliver Pickles, Mike Kidd, Alex Richter, Andrew D Beggs PHE Heartlands Lab: Husam Osman, Andrew Bosworth. Queen Elizabeth Hospital: Anna Casey |
| EPI_ISL_650421, EPI_ISL_650429, EPI_ISL_650457, EPI_ISL_650458, EPI_ISL_650461, EPI_ISL_650465, EPI_ISL_650466, EPI_ISL_650467, EPI_ISL_650474, EPI_ISL_650475, EPI_ISL_650485, EPI_ISL_650489, EPI_ISL_650555 | | | |
| see above | Quadram Institute Bioscience | COVID-19 Genomics UK (COG-UK) Consortium | Dave J. Baker, Gemma L. Kay, Alp Aydin, Thanh Le-Viet, Steven Rudder, Ana P. Tedim, Anastasia Kolyva, Maria Diaz, Leonardo de Oliveira Martins, |

| | | | |
|---|---|---|---|
| | | | Nabil-Fareed Alikhan, Lizzie Meadows, Rachael Stanley, Ngozi Elumogo, Muhammed Yasir, Nicholas M. Thomson, Alexander J Trotter, Rachel Gilroy, Samuel Bloomfield, Claire Stuart, Andrew Bell, Reenesh Prakash, Samir Dervisevic, Alison E. Mather, John Wain, Mark Webber, Andrew J. Page, Justin O'Grady |
| EPI_ISL_650582, EPI_ISL_650648 | University of Birmingham | COVID-19 Genomics UK (COG-UK) Consortium | Institute of Microbiology, University of Birmingham: Claire McMurray, Joanne Stockton, Samuel Nicholls, Radoslaw Poplawski, Will Rowe, Josh Quick, Nicholas Loman. University of Birmingham Testing Laboratory: Celina M Whalley, Andrew Bosworth, Charlotte Poxon, Kasun Wanigasooriya, Oliver Pickles, Mike Kidd, Alex Richter, Andrew D Beggs PHE Heartlands Lab: Husam Osman, Andrew Bosworth. Queen Elizabeth Hospital: Anna Casey |
| EPI_ISL_650685, EPI_ISL_650686 | Quadram Institute Bioscience | COVID-19 Genomics UK (COG-UK) Consortium | Dave J. Baker, Gemma L. Kay, Alp Aydin, Thanh Le-Viet, Steven Rudder, Ana P. Tedim, Anastasia Kolyva, Maria Diaz, Leonardo de Oliveira Martins, Nabil-Fareed Alikhan, Lizzie Meadows, Rachael Stanley, Ngozi Elumogo, Muhammed Yasir, Nicholas M. Thomson, Alexander J Trotter, Rachel Gilroy, Samuel Bloomfield, Claire Stuart, Andrew Bell, Reenesh Prakash, Samir Dervisevic, Alison E. Mather, John Wain, Mark Webber, Andrew J. Page, Justin O'Grady |
| EPI_ISL_650717 | University of Birmingham | COVID-19 Genomics UK (COG-UK) Consortium | Institute of Microbiology, University of Birmingham: Claire McMurray, Joanne Stockton, Samuel Nicholls, Radoslaw Poplawski, Will Rowe, Josh Quick, Nicholas Loman. University of Birmingham Testing Laboratory: Celina M Whalley, Andrew Bosworth, Charlotte Poxon, Kasun Wanigasooriya, Oliver Pickles, Mike Kidd, Alex Richter, Andrew D Beggs PHE Heartlands Lab: Husam Osman, Andrew Bosworth. Queen Elizabeth Hospital: Anna Casey |
| EPI_ISL_650719, EPI_ISL_650720, EPI_ISL_650722 | Quadram Institute Bioscience | COVID-19 Genomics UK (COG-UK) Consortium | Dave J. Baker, Gemma L. Kay, Alp Aydin, Thanh Le-Viet, Steven Rudder, Ana P. Tedim, Anastasia Kolyva, Maria Diaz, Leonardo de Oliveira Martins, Nabil-Fareed Alikhan, Lizzie Meadows, Rachael Stanley, Ngozi Elumogo, Muhammed Yasir, Nicholas M. Thomson, Alexander J Trotter, Rachel Gilroy, Samuel Bloomfield, Claire Stuart, Andrew Bell, Reenesh Prakash, Samir Dervisevic, Alison E. Mather, John Wain, Mark Webber, Andrew J. Page, Justin O'Grady |
| EPI_ISL_650738 | Queens Medical Centre, Clinical Microbiology Department / DeepSeq Nottingham | COVID-19 Genomics UK (COG-UK) Consortium | Gemma Clark, Wendy Smith, Manjinder Khakh, Vicki M Fleming, Michelle M Lister, Hannah Howson-Wells, Jonathan Ball, Patrick McClure, Joseph Chappell, Theocharis Tsoleridis, Nadine Holmes, Matthew Carlisle, Christopher Moore, Fei Sang, Johnny Debebe, Victoria Wright, Matthew Loose |
| EPI_ISL_650760, EPI_ISL_650825 | University of Birmingham | COVID-19 Genomics UK (COG-UK) Consortium | Institute of Microbiology, University of Birmingham: Claire McMurray, Joanne Stockton, Samuel Nicholls, Radoslaw Poplawski, Will Rowe, Josh Quick, Nicholas Loman. University of Birmingham Testing Laboratory: Celina M Whalley, Andrew Bosworth, Charlotte Poxon, Kasun Wanigasooriya, Oliver Pickles, Mike Kidd, Alex Richter, Andrew D Beggs PHE Heartlands Lab: Husam Osman, Andrew Bosworth. Queen Elizabeth Hospital: Anna Casey |
| EPI_ISL_650880 | Quadram Institute Bioscience | COVID-19 Genomics UK (COG-UK) Consortium | Dave J. Baker, Gemma L. Kay, Alp Aydin, Thanh Le-Viet, Steven Rudder, Ana P. Tedim, Anastasia Kolyva, Maria Diaz, Leonardo de Oliveira Martins, Nabil-Fareed Alikhan, Lizzie Meadows, Rachael Stanley, Ngozi Elumogo, Muhammed Yasir, Nicholas M. Thomson, Alexander J Trotter, Rachel Gilroy, Samuel Bloomfield, Claire Stuart, Andrew Bell, Reenesh Prakash, Samir Dervisevic, Alison E. Mather, John Wain, Mark Webber, Andrew J. Page, Justin O'Grady |
| EPI_ISL_651000 | University of Birmingham | COVID-19 Genomics UK (COG-UK) Consortium | Institute of Microbiology, University of Birmingham: Claire McMurray, Joanne Stockton, Samuel Nicholls, Radoslaw Poplawski, Will Rowe, Josh Quick, Nicholas Loman. University of Birmingham Testing Laboratory: Celina M Whalley, Andrew Bosworth, Charlotte Poxon, Kasun Wanigasooriya, Oliver Pickles, Mike Kidd, Alex Richter, Andrew D Beggs PHE Heartlands Lab: Husam Osman, Andrew Bosworth. Queen Elizabeth Hospital: Anna Casey |
| EPI_ISL_651011 | West of Scotland Specialist Virology Centre, NHSGGC / MRC-University of Glasgow Centre for Virus Research | COVID-19 Genomics UK (COG-UK) Consortium | Ana da Silva Filipe, Natasha Johnson, Kathy Smollett, Daniel Mair, Stephen Carmichael, Alice Broos, Lily Tong, Jenna Nichols, Kyriaki Nomikou; Sarah McDonald; Richard Orton, Joseph Hughes, Sreenu Vattipally, David L Robertson; Alasdair MacLean, Rory Gunson; Sharif Shaaban, Matthew Holden; Rachel Blacow, Guy Mollett, Kathy Li, James Shepherd, Antonia Ho, Emma Thomson |
| EPI_ISL_651077 | Virology Department, Sheffield Teaching Hospitals NHS Foundation Trust/Department of Infection, Immunity and Cardiovascular Disease, The Medical School, University of Sheffield | COVID-19 Genomics UK (COG-UK) Consortium | Thushan de Silva, Matthew Parker, Nikki Smith, Adri Angyal, Rebecca Brown, Luke Green, Rachel Tucker, Paul Parsons, Danielle Groves, Katie Johnson, Laura Carrilero, Alex Keeley, Dave Partridge, Matthew Wyles, Benjamin Lindsey, Mehmet Yavuz, Mohammad Raza, Cariad Evans |
| EPI_ISL_651090, EPI_ISL_651091 | Quadram Institute Bioscience | COVID-19 Genomics UK (COG-UK) Consortium | Dave J. Baker, Gemma L. Kay, Alp Aydin, Thanh Le-Viet, Steven Rudder, Ana P. Tedim, Anastasia Kolyva, Maria Diaz, Leonardo de Oliveira Martins, Nabil-Fareed Alikhan, Lizzie Meadows, Rachael Stanley, Ngozi Elumogo, Muhammed Yasir, Nicholas M. Thomson, Alexander J Trotter, Rachel Gilroy, Samuel Bloomfield, Claire Stuart, Andrew Bell, Reenesh Prakash, Samir Dervisevic, Alison E. Mather, John Wain, Mark Webber, Andrew J. Page, Justin O'Grady |
| EPI_ISL_651098 | University of Birmingham | COVID-19 Genomics UK (COG-UK) Consortium | Institute of Microbiology, University of Birmingham: Claire McMurray, Joanne Stockton, Samuel Nicholls, Radoslaw Poplawski, Will Rowe, Josh Quick, Nicholas Loman. University of Birmingham Testing Laboratory: Celina M Whalley, Andrew Bosworth, Charlotte Poxon, Kasun Wanigasooriya, Oliver Pickles, Mike Kidd, Alex Richter, Andrew D Beggs PHE Heartlands Lab: Husam Osman, Andrew Bosworth. Queen Elizabeth Hospital: Anna Casey |
| EPI_ISL_651139 | Quadram Institute Bioscience | COVID-19 Genomics UK (COG-UK) Consortium | Dave J. Baker, Gemma L. Kay, Alp Aydin, Thanh Le-Viet, Steven Rudder, Ana P. Tedim, Anastasia Kolyva, Maria Diaz, Leonardo de Oliveira Martins, Nabil-Fareed Alikhan, Lizzie Meadows, Rachael Stanley, Ngozi Elumogo, Muhammed Yasir, Nicholas M. Thomson, Alexander J Trotter, Rachel Gilroy, Samuel Bloomfield, Claire Stuart, Andrew Bell, Reenesh Prakash, Samir Dervisevic, Alison E. Mather, John Wain, Mark Webber, Andrew J. Page, Justin O'Grady |
| EPI_ISL_651144 | University of Birmingham | COVID-19 Genomics UK (COG-UK) Consortium | Institute of Microbiology, University of Birmingham: Claire McMurray, Joanne Stockton, Samuel Nicholls, Radoslaw Poplawski, Will Rowe, Josh Quick, Nicholas Loman. University of Birmingham Testing Laboratory: Celina M Whalley, Andrew Bosworth, Charlotte Poxon, Kasun Wanigasooriya, Oliver Pickles, Mike Kidd, Alex Richter, Andrew D Beggs PHE Heartlands Lab: Husam Osman, Andrew Bosworth. Queen Elizabeth Hospital: Anna Casey |
| EPI_ISL_651157 | Quadram Institute Bioscience | COVID-19 Genomics UK (COG-UK) Consortium | Dave J. Baker, Gemma L. Kay, Alp Aydin, Thanh Le-Viet, Steven Rudder, Ana P. Tedim, Anastasia Kolyva, Maria Diaz, Leonardo de Oliveira Martins, Nabil-Fareed Alikhan, Lizzie Meadows, Rachael Stanley, Ngozi Elumogo, Muhammed Yasir, Nicholas M. Thomson, Alexander J Trotter, Rachel Gilroy, Samuel Bloomfield, Claire Stuart, Andrew Bell, Reenesh Prakash, Samir Dervisevic, Alison E. Mather, John Wain, Mark Webber, Andrew J. Page, Justin O'Grady |
| EPI_ISL_651174, EPI_ISL_651195 | University of Birmingham | COVID-19 Genomics UK (COG-UK) Consortium | Institute of Microbiology, University of Birmingham: Claire McMurray, Joanne Stockton, Samuel Nicholls, Radoslaw Poplawski, Will Rowe, Josh Quick, Nicholas Loman. University of Birmingham Testing Laboratory: Celina M Whalley, Andrew Bosworth, Charlotte Poxon, Kasun Wanigasooriya, Oliver Pickles, Mike Kidd, Alex Richter, Andrew D Beggs PHE Heartlands Lab: Husam Osman, Andrew Bosworth. Queen Elizabeth Hospital: Anna Casey |
| EPI_ISL_651203, EPI_ISL_651204, EPI_ISL_651210, EPI_ISL_651220, EPI_ISL_651254 | Quadram Institute Bioscience | COVID-19 Genomics UK (COG-UK) Consortium | Dave J. Baker, Gemma L. Kay, Alp Aydin, Thanh Le-Viet, Steven Rudder, Ana P. Tedim, Anastasia Kolyva, Maria Diaz, Leonardo de Oliveira Martins, Nabil-Fareed Alikhan, Lizzie Meadows, Rachael Stanley, Ngozi Elumogo, Muhammed Yasir, Nicholas M. Thomson, Alexander J Trotter, Rachel Gilroy, Samuel Bloomfield, Claire Stuart, Andrew Bell, Reenesh Prakash, Samir Dervisevic, Alison E. Mather, John Wain, Mark Webber, Andrew J. Page, Justin O'Grady |
| EPI_ISL_651266, EPI_ISL_651342, EPI_ISL_651353 | University of Birmingham | COVID-19 Genomics UK (COG-UK) Consortium | Institute of Microbiology, University of Birmingham: Claire McMurray, Joanne Stockton, Samuel Nicholls, Radoslaw Poplawski, Will Rowe, Josh Quick, Nicholas Loman. University of Birmingham Testing Laboratory: Celina M Whalley, Andrew Bosworth, Charlotte Poxon, Kasun Wanigasooriya, Oliver Pickles, Mike Kidd, Alex Richter, Andrew D Beggs PHE Heartlands Lab: Husam Osman, Andrew Bosworth. Queen Elizabeth Hospital: Anna Casey |
| EPI_ISL_651371 | Quadram Institute Bioscience | COVID-19 Genomics UK (COG-UK) Consortium | Dave J. Baker, Gemma L. Kay, Alp Aydin, Thanh Le-Viet, Steven Rudder, Ana P. Tedim, Anastasia Kolyva, Maria Diaz, Leonardo de Oliveira Martins, Nabil-Fareed Alikhan, Lizzie Meadows, Rachael Stanley, Ngozi Elumogo, Muhammed Yasir, Nicholas M. Thomson, Alexander J Trotter, Rachel Gilroy, Samuel Bloomfield, Claire Stuart, Andrew Bell, Reenesh Prakash, Samir Dervisevic, Alison E. Mather, John Wain, Mark Webber, Andrew J. Page, Justin O'Grady |
| EPI_ISL_651390 | University of Birmingham | COVID-19 Genomics UK (COG-UK) Consortium | Institute of Microbiology, University of Birmingham: Claire McMurray, Joanne Stockton, Samuel Nicholls, Radoslaw Poplawski, Will Rowe, Josh Quick, Nicholas Loman. University of Birmingham Testing Laboratory: Celina M Whalley, Andrew Bosworth, Charlotte Poxon, Kasun Wanigasooriya, Oliver Pickles, Mike Kidd, Alex Richter, Andrew D Beggs PHE Heartlands Lab: Husam Osman, Andrew Bosworth. Queen Elizabeth Hospital: Anna Casey |
| EPI_ISL_651392 | Quadram Institute Bioscience | COVID-19 Genomics UK (COG-UK) Consortium | Dave J. Baker, Gemma L. Kay, Alp Aydin, Thanh Le-Viet, Steven Rudder, Ana P. Tedim, Anastasia Kolyva, Maria Diaz, Leonardo de Oliveira Martins, Nabil-Fareed Alikhan, Lizzie Meadows, Rachael Stanley, Ngozi Elumogo, Muhammed Yasir, Nicholas M. Thomson, Alexander J Trotter, Rachel Gilroy, Samuel Bloomfield, Claire Stuart, Andrew Bell, Reenesh Prakash, Samir Dervisevic, Alison E. Mather, John Wain, Mark Webber, Andrew J. Page, Justin O'Grady |
| EPI_ISL_651422 | University of Birmingham | COVID-19 Genomics UK (COG-UK) Consortium | Institute of Microbiology, University of Birmingham: Claire McMurray, Joanne Stockton, Samuel Nicholls, Radoslaw Poplawski, Will Rowe, Josh Quick, |

| | | | |
|---|---|---|---|
| | | | Nicholas Loman. University of Birmingham Testing Laboratory: Celina M Whalley, Andrew Bosworth, Charlotte Poxon, Kasun Wanigasooriya, Oliver Pickles, Mike Kidd, Alex Richter, Andrew D Beggs PHE Heartlands Lab: Husam Osman, Andrew Bosworth. Queen Elizabeth Hospital: Anna Casey |
| EPI_ISL_651496 | West of Scotland Specialist Virology Centre, NHSGGC / MRC-University of Glasgow Centre for Virus Research | COVID-19 Genomics UK (COG-UK) Consortium | Ana da Silva Filipe, Natasha Johnson, Kathy Smollett, Daniel Mair, Stephen Carmichael, Alice Broos, Lily Tong, Jenna Nichols, Kyriaki Nomikou; Sarah McDonald; Richard Orton, Joseph Hughes, Sreenu Vattipally, David L Robertson; Alasdair MacLean, Rory Gunson; Sharif Shaaban, Matthew Holden; Rachel Blacow, Guy Mollett, Kathy Li, James Shepherd, Antonia Ho, Emma Thomson |
| EPI_ISL_651633, EPI_ISL_651635, EPI_ISL_651636, EPI_ISL_651637, EPI_ISL_651638, EPI_ISL_651639, EPI_ISL_651640, EPI_ISL_651641, EPI_ISL_651642, EPI_ISL_651643, EPI_ISL_651644, EPI_ISL_651645, EPI_ISL_651646, EPI_ISL_651647, EPI_ISL_651648, EPI_ISL_651649, EPI_ISL_651650, EPI_ISL_651651, EPI_ISL_651652 | | | |
| see above | University of Birmingham | COVID-19 Genomics UK (COG-UK) Consortium | Institute of Microbiology, University of Birmingham: Claire McMurray, Joanne Stockton, Samuel Nicholls, Radoslaw Poplawski, Will Rowe, Josh Quick, Nicholas Loman. University of Birmingham Testing Laboratory: Celina M Whalley, Andrew Bosworth, Charlotte Poxon, Kasun Wanigasooriya, Oliver Pickles, Mike Kidd, Alex Richter, Andrew D Beggs PHE Heartlands Lab: Husam Osman, Andrew Bosworth. Queen Elizabeth Hospital: Anna Casey |
| EPI_ISL_651888, EPI_ISL_651895, EPI_ISL_651898, EPI_ISL_651902, EPI_ISL_651903, EPI_ISL_651908, EPI_ISL_651909, EPI_ISL_651912, EPI_ISL_651916, EPI_ISL_651919, EPI_ISL_651920, EPI_ISL_651921, EPI_ISL_651923, EPI_ISL_651925, EPI_ISL_651927, EPI_ISL_651929, EPI_ISL_651932, EPI_ISL_651934, EPI_ISL_651941, EPI_ISL_651945, EPI_ISL_651948, EPI_ISL_651949, EPI_ISL_651954 | | | |
| see above | West of Scotland Specialist Virology Centre, NHSGGC / MRC-University of Glasgow Centre for Virus Research | COVID-19 Genomics UK (COG-UK) Consortium | Ana da Silva Filipe, Natasha Johnson, Kathy Smollett, Daniel Mair, Stephen Carmichael, Alice Broos, Lily Tong, Jenna Nichols, Kyriaki Nomikou; Sarah McDonald; Richard Orton, Joseph Hughes, Sreenu Vattipally, David L Robertson; Alasdair MacLean, Rory Gunson; Sharif Shaaban, Matthew Holden; Rachel Blacow, Guy Mollett, Kathy Li, James Shepherd, Antonia Ho, Emma Thomson |
| EPI_ISL_652351, EPI_ISL_652357, EPI_ISL_652360 | Quadram Institute Bioscience | COVID-19 Genomics UK (COG-UK) Consortium | Dave J. Baker, Gemma L. Kay, Alp Aydin, Thanh Le-Viet, Steven Rudder, Ana P. Tedim, Anastasia Kolyva, Maria Diaz, Leonardo de Oliveira Martins, Nabil-Fareed Alikhan, Lizzie Meadows, Rachael Stanley, Ngozi Elumogo, Muhammed Yasir, Nicholas M. Thomson, Alexander J Trotter, Rachel Gilroy, Samuel Bloomfield, Claire Stuart, Andrew Bell, Reenesh Prakash, Samir Dervisevic, Alison E. Mather, John Wain, Mark Webber, Andrew J. Page, Justin O'Grady |
| EPI_ISL_652365, EPI_ISL_652366, EPI_ISL_652367, EPI_ISL_652368, EPI_ISL_652369, EPI_ISL_652370, EPI_ISL_652371, EPI_ISL_652372 | Queens Medical Centre, Clinical Microbiology Department / DeepSeq Nottingham | COVID-19 Genomics UK (COG-UK) Consortium | Gemma Clark, Wendy Smith, Manjinder Khakh, Vicki M Fleming, Michelle M Lister, Hannah Howson-Wells, Jonathan Ball, Patrick McClure, Joseph Chappell, Theocharis Tsoleridis, Nadine Holmes, Matthew Carlisle, Christopher Moore, Fei Sang, Johnny Debebe, Victoria Wright, Matthew Loose |
| EPI_ISL_653070, EPI_ISL_653076, EPI_ISL_653078, EPI_ISL_653079, EPI_ISL_653080, EPI_ISL_653089 | Virology Department, Sheffield Teaching Hospitals NHS Foundation Trust/Department of Infection, Immunity and Cardiovascular Disease, The Medical School, University of Sheffield | COVID-19 Genomics UK (COG-UK) Consortium | Thushan de Silva, Matthew Parker, Nikki Smith, Adri Angyal, Rebecca Brown, Luke Green, Rachel Tucker, Paul Parsons, Danielle Groves, Katie Johnson, Laura Carrilero, Alex Keeley, Dave Partridge, Matthew Wyles, Benjamin Lindsey, Mehmet Yavuz, Mohammad Raza, Cariad Evans |
| EPI_ISL_654811, EPI_ISL_654813, EPI_ISL_654814, EPI_ISL_654815, EPI_ISL_654817 | National Public Health Laboratory, National Centre for Infectious Diseases | National Public Health Laboratory, National Centre for Infectious Diseases | Tze Minn Mak, Sophie Octavia, Zhenyang Zhou, Lin Cui, Raymond Tzer Pin Lin |
| EPI_ISL_654959, EPI_ISL_654960, EPI_ISL_654961, EPI_ISL_654962, EPI_ISL_654963, EPI_ISL_654964, EPI_ISL_654965, EPI_ISL_654966, EPI_ISL_654967, EPI_ISL_654968, EPI_ISL_654969, EPI_ISL_654970, EPI_ISL_654971, EPI_ISL_654972, EPI_ISL_654973, EPI_ISL_654974, EPI_ISL_654975, EPI_ISL_654976, EPI_ISL_654977, EPI_ISL_654978, EPI_ISL_654979, EPI_ISL_654980, EPI_ISL_654981, EPI_ISL_654982, EPI_ISL_654983, EPI_ISL_654984, EPI_ISL_654985, EPI_ISL_654986, EPI_ISL_654987, EPI_ISL_654988, EPI_ISL_654989, EPI_ISL_654990, EPI_ISL_654991, EPI_ISL_654992, EPI_ISL_654993, EPI_ISL_654994, EPI_ISL_654995, EPI_ISL_654996, EPI_ISL_654997, EPI_ISL_654998, EPI_ISL_654999, EPI_ISL_655000, EPI_ISL_655001, EPI_ISL_655002, EPI_ISL_655003, EPI_ISL_655004, EPI_ISL_655005, EPI_ISL_655006, EPI_ISL_655007, EPI_ISL_655008, EPI_ISL_655009, EPI_ISL_655010, EPI_ISL_655011, EPI_ISL_655012, EPI_ISL_655013, EPI_ISL_655014, EPI_ISL_655015, EPI_ISL_655016, EPI_ISL_655017, EPI_ISL_655018, EPI_ISL_655019, EPI_ISL_655020, EPI_ISL_655021, EPI_ISL_655022, EPI_ISL_655023, EPI_ISL_655024, EPI_ISL_655025, EPI_ISL_655026, EPI_ISL_655027, EPI_ISL_655028, EPI_ISL_655029, EPI_ISL_655030, EPI_ISL_655031, EPI_ISL_655032, EPI_ISL_655033, EPI_ISL_655034, EPI_ISL_655035, EPI_ISL_655036, EPI_ISL_655037, EPI_ISL_655038, EPI_ISL_655039, EPI_ISL_655040, EPI_ISL_655041, EPI_ISL_655042, EPI_ISL_655043, EPI_ISL_655044, EPI_ISL_655045, EPI_ISL_655046, EPI_ISL_655047, EPI_ISL_655048, EPI_ISL_655049, EPI_ISL_655050, EPI_ISL_655051, EPI_ISL_655052, EPI_ISL_655053, EPI_ISL_655054, EPI_ISL_655055, EPI_ISL_655056, EPI_ISL_655057, EPI_ISL_655058, EPI_ISL_655059, EPI_ISL_655060, EPI_ISL_655061, EPI_ISL_655062, EPI_ISL_655063, EPI_ISL_655064, EPI_ISL_655065, EPI_ISL_655066, EPI_ISL_655067, EPI_ISL_655068, EPI_ISL_655069, EPI_ISL_655070, EPI_ISL_655071, EPI_ISL_655072, EPI_ISL_655073, EPI_ISL_655074, EPI_ISL_655075, EPI_ISL_655076, EPI_ISL_655077, EPI_ISL_655078, EPI_ISL_655079, EPI_ISL_655080, EPI_ISL_655081, EPI_ISL_655082, EPI_ISL_655083, EPI_ISL_655084, EPI_ISL_655085, EPI_ISL_655086, EPI_ISL_655087, EPI_ISL_655088, EPI_ISL_655089, EPI_ISL_655090, EPI_ISL_655091, EPI_ISL_655092, EPI_ISL_655093, EPI_ISL_655094, EPI_ISL_655095, EPI_ISL_655096, EPI_ISL_655097, EPI_ISL_655098, EPI_ISL_655099, EPI_ISL_655100, EPI_ISL_655101, EPI_ISL_655102, EPI_ISL_655103, EPI_ISL_655104, EPI_ISL_655105, EPI_ISL_655106, EPI_ISL_655107, EPI_ISL_655108, EPI_ISL_655109, EPI_ISL_655110, EPI_ISL_655111, EPI_ISL_655112, EPI_ISL_655113, EPI_ISL_655114, EPI_ISL_655115, EPI_ISL_655116, EPI_ISL_655117, EPI_ISL_655118, EPI_ISL_655119, EPI_ISL_655120, EPI_ISL_655121, EPI_ISL_655122, EPI_ISL_655123, EPI_ISL_655124, EPI_ISL_655125, EPI_ISL_655126, EPI_ISL_655127, EPI_ISL_655128, EPI_ISL_655129, EPI_ISL_655130, EPI_ISL_655131, EPI_ISL_655132, EPI_ISL_655133, EPI_ISL_655134, EPI_ISL_655135, EPI_ISL_655136, EPI_ISL_655137, EPI_ISL_655138, EPI_ISL_655139, EPI_ISL_655140, EPI_ISL_655141, EPI_ISL_655142, EPI_ISL_655143, EPI_ISL_655144, EPI_ISL_655145, EPI_ISL_655146, EPI_ISL_655147, EPI_ISL_655148, EPI_ISL_655149, EPI_ISL_655150, EPI_ISL_655151, EPI_ISL_655152, EPI_ISL_655153, EPI_ISL_655154, EPI_ISL_655155, EPI_ISL_655156, EPI_ISL_655157, EPI_ISL_655158, EPI_ISL_655159, EPI_ISL_655160, EPI_ISL_655161, EPI_ISL_655162, EPI_ISL_655163, EPI_ISL_655164, EPI_ISL_655165, EPI_ISL_655166, EPI_ISL_655167, EPI_ISL_655168, EPI_ISL_655169, EPI_ISL_655170, EPI_ISL_655171, EPI_ISL_655172, EPI_ISL_655173, EPI_ISL_655174, EPI_ISL_655175, EPI_ISL_655176, EPI_ISL_655177, EPI_ISL_655178, EPI_ISL_655179, EPI_ISL_655180, EPI_ISL_655181, EPI_ISL_655182, EPI_ISL_655183, EPI_ISL_655184, EPI_ISL_655185, EPI_ISL_655186, EPI_ISL_655187, EPI_ISL_655188, EPI_ISL_655189, EPI_ISL_655190, EPI_ISL_655191, EPI_ISL_655192, EPI_ISL_655193, EPI_ISL_655194, EPI_ISL_655195, EPI_ISL_655196, EPI_ISL_655197, EPI_ISL_655198, EPI_ISL_655199, EPI_ISL_655200, EPI_ISL_655201, EPI_ISL_655202, EPI_ISL_655203, EPI_ISL_655204, EPI_ISL_655205, EPI_ISL_655206, EPI_ISL_655207, EPI_ISL_655208, EPI_ISL_655209, EPI_ISL_655210, EPI_ISL_655211, EPI_ISL_655212, EPI_ISL_655213, EPI_ISL_655214, EPI_ISL_655215, EPI_ISL_655216, EPI_ISL_655217, EPI_ISL_655218, EPI_ISL_655219, EPI_ISL_655220, EPI_ISL_655221, EPI_ISL_655222, EPI_ISL_655223, EPI_ISL_655224, EPI_ISL_655225, EPI_ISL_655226, EPI_ISL_655227, EPI_ISL_655228, EPI_ISL_655229, EPI_ISL_655230, EPI_ISL_655231, EPI_ISL_655232, EPI_ISL_655233, EPI_ISL_655234, EPI_ISL_655235, EPI_ISL_655236, EPI_ISL_655237, EPI_ISL_655238, EPI_ISL_655239, EPI_ISL_655240, EPI_ISL_655241, EPI_ISL_655242, EPI_ISL_655243, EPI_ISL_655244, EPI_ISL_655245, EPI_ISL_655246, EPI_ISL_655247, EPI_ISL_655248, EPI_ISL_655249, EPI_ISL_655250, EPI_ISL_655251, EPI_ISL_655252, EPI_ISL_655253, EPI_ISL_655254, EPI_ISL_655255, EPI_ISL_655256, EPI_ISL_655257, EPI_ISL_655258, EPI_ISL_655259, EPI_ISL_655260, EPI_ISL_655261, EPI_ISL_655262, EPI_ISL_655263, EPI_ISL_655264, EPI_ISL_655265, EPI_ISL_655266, EPI_ISL_655267, EPI_ISL_655268, EPI_ISL_655269, EPI_ISL_655270, EPI_ISL_655271, EPI_ISL_655272 | | | |
| see above | Lighthouse Lab in Glasgow | Wellcome Sanger Institute for the COVID-19 Genomics UK (COG-UK) Consortium | Harper VanSteenhouse, Yumi Kasai, David Gray, Carol Clugston, Anna Dominiczak and Alex Alderton, Roberto Amato, Sonia Goncalves, Ewan Harrison, David K. Jackson, Ian Johnston, Dominic Kwiatkowski, Cordelia Langford, John Sillitoe on behalf of the Wellcome Sanger Institute COVID-19 Surveillance Team |
| EPI_ISL_655273, EPI_ISL_655277, EPI_ISL_655281, EPI_ISL_655282, EPI_ISL_655283, EPI_ISL_655285, EPI_ISL_655286, EPI_ISL_655287, EPI_ISL_655294, EPI_ISL_655296, EPI_ISL_655298, EPI_ISL_655309, EPI_ISL_655312, EPI_ISL_655313, EPI_ISL_655314, EPI_ISL_655317, EPI_ISL_655318, EPI_ISL_655320, EPI_ISL_655322, EPI_ISL_655323, EPI_ISL_655327, EPI_ISL_655328, EPI_ISL_655329, EPI_ISL_655334, EPI_ISL_655335, EPI_ISL_655336, EPI_ISL_655337, EPI_ISL_655338, EPI_ISL_655343, EPI_ISL_655344, EPI_ISL_655346, EPI_ISL_655347, EPI_ISL_655349, EPI_ISL_655350, EPI_ISL_655351, EPI_ISL_655352, EPI_ISL_655354, EPI_ISL_655356, EPI_ISL_655357, EPI_ISL_655359, EPI_ISL_655363, EPI_ISL_655367, EPI_ISL_655368, EPI_ISL_655370, EPI_ISL_655371, EPI_ISL_655372, EPI_ISL_655373, EPI_ISL_655377, EPI_ISL_655378, EPI_ISL_655380, EPI_ISL_655381, EPI_ISL_655383, EPI_ISL_655386, EPI_ISL_655390, EPI_ISL_655392, EPI_ISL_655395, EPI_ISL_655396, EPI_ISL_655397, EPI_ISL_655399, EPI_ISL_655402, EPI_ISL_655406, EPI_ISL_655408, EPI_ISL_655409, EPI_ISL_655412, EPI_ISL_655416, EPI_ISL_655417, EPI_ISL_655420, EPI_ISL_655421, EPI_ISL_655425, EPI_ISL_655432, EPI_ISL_655436, EPI_ISL_655437, EPI_ISL_655440, EPI_ISL_655442, EPI_ISL_655443, EPI_ISL_655445, EPI_ISL_655446, EPI_ISL_655448, EPI_ISL_655449, EPI_ISL_655455, EPI_ISL_655458, EPI_ISL_655461, EPI_ISL_655463, EPI_ISL_655464, EPI_ISL_655465, EPI_ISL_655467, EPI_ISL_655468, EPI_ISL_655469, EPI_ISL_655473, EPI_ISL_655475, EPI_ISL_655477, EPI_ISL_655479, EPI_ISL_655481, EPI_ISL_655487, EPI_ISL_655489, EPI_ISL_655491, EPI_ISL_655493, EPI_ISL_655495, EPI_ISL_655496, EPI_ISL_655497, EPI_ISL_655501, EPI_ISL_655507, EPI_ISL_655510, EPI_ISL_655512, EPI_ISL_655513, EPI_ISL_655519, EPI_ISL_655520, EPI_ISL_655521, EPI_ISL_655523, EPI_ISL_655524, EPI_ISL_655530, EPI_ISL_655532, EPI_ISL_655533, EPI_ISL_655535, EPI_ISL_655536, EPI_ISL_655537, EPI_ISL_655540, EPI_ISL_655545, EPI_ISL_655549, EPI_ISL_655551, EPI_ISL_655553, EPI_ISL_655555, EPI_ISL_655556, EPI_ISL_655557, EPI_ISL_655560, EPI_ISL_655561, EPI_ISL_655562, EPI_ISL_655564, EPI_ISL_655568, EPI_ISL_655569, EPI_ISL_655570, EPI_ISL_655574, EPI_ISL_655575, EPI_ISL_655582, EPI_ISL_655583, EPI_ISL_655586, EPI_ISL_655587, EPI_ISL_655589, EPI_ISL_655591, EPI_ISL_655594, EPI_ISL_655595, EPI_ISL_655596, EPI_ISL_655601, EPI_ISL_655603, EPI_ISL_655605, EPI_ISL_655607, EPI_ISL_655608, EPI_ISL_655610, EPI_ISL_655612, EPI_ISL_655613, EPI_ISL_655614, EPI_ISL_655616, EPI_ISL_655619, EPI_ISL_655620, EPI_ISL_655622, EPI_ISL_655623, EPI_ISL_655624, EPI_ISL_655625, EPI_ISL_655627, EPI_ISL_655631, EPI_ISL_655639, EPI_ISL_655640, EPI_ISL_655641, EPI_ISL_655642, EPI_ISL_655643, EPI_ISL_655644, EPI_ISL_655646, EPI_ISL_655647, EPI_ISL_655648, EPI_ISL_655649, EPI_ISL_655650 | | | |
| see above | Lighthouse Lab in Alderley Park | Wellcome Sanger Institute for the COVID-19 Genomics UK (COG-UK) Consortium | Jacquelyn Wynn, Mairead Hyland, The Lighthouse Lab in Alderley Park and Alex Alderton, Roberto Amato, Sonia Goncalves, Ewan Harrison, David K. Jackson, Ian Johnston, Dominic Kwiatkowski, Cordelia Langford, John Sillitoe on behalf of the Wellcome Sanger Institute COVID-19 Surveillance Team |
| EPI_ISL_655651 | Lighthouse Lab in Glasgow | Wellcome Sanger Institute for the COVID-19 Genomics UK (COG-UK) Consortium | Harper VanSteenhouse, Yumi Kasai, David Gray, Carol Clugston, Anna Dominiczak and Alex Alderton, Roberto Amato, Sonia Goncalves, Ewan Harrison, David K. Jackson, Ian Johnston, Dominic Kwiatkowski, Cordelia Langford, John Sillitoe on behalf of the Wellcome Sanger Institute COVID-19 Surveillance Team |
| EPI_ISL_655652, EPI_ISL_655655, EPI_ISL_655656, EPI_ISL_655657, EPI_ISL_655658, EPI_ISL_655659 | Lighthouse Lab in Alderley Park | Wellcome Sanger Institute for the COVID-19 Genomics UK (COG-UK) Consortium | Jacquelyn Wynn, Mairead Hyland, The Lighthouse Lab in Alderley Park and Alex Alderton, Roberto Amato, Sonia Goncalves, Ewan Harrison, David K. Jackson, Ian Johnston, Dominic Kwiatkowski, Cordelia Langford, John Sillitoe on behalf of the Wellcome Sanger Institute COVID-19 Surveillance Team |
| EPI_ISL_655660, EPI_ISL_655662 | Lighthouse Lab in Glasgow | Wellcome Sanger Institute for the COVID-19 Genomics UK (COG-UK) Consortium | Harper VanSteenhouse, Yumi Kasai, David Gray, Carol Clugston, Anna Dominiczak and Alex Alderton, Roberto Amato, Sonia Goncalves, Ewan Harrison, David K. Jackson, Ian Johnston, Dominic Kwiatkowski, Cordelia Langford, John Sillitoe on behalf of the Wellcome Sanger Institute COVID-19 Surveillance Team |
| EPI_ISL_655664, EPI_ISL_655666, EPI_ISL_655667, EPI_ISL_655668, EPI_ISL_655669, EPI_ISL_655671 | Lighthouse Lab in Alderley Park | Wellcome Sanger Institute for the COVID-19 Genomics UK (COG-UK) Consortium | Jacquelyn Wynn, Mairead Hyland, The Lighthouse Lab in Alderley Park and Alex Alderton, Roberto Amato, Sonia Goncalves, Ewan Harrison, David K. Jackson, Ian Johnston, Dominic Kwiatkowski, Cordelia Langford, John Sillitoe on behalf of the Wellcome Sanger Institute COVID-19 Surveillance Team |
| EPI_ISL_655672 | Lighthouse Lab in Glasgow | Wellcome Sanger Institute for the COVID-19 Genomics UK (COG-UK) Consortium | Harper VanSteenhouse, Yumi Kasai, David Gray, Carol Clugston, Anna Dominiczak and Alex Alderton, Roberto Amato, Sonia Goncalves, Ewan Harrison, David K. Jackson, Ian Johnston, Dominic Kwiatkowski, Cordelia Langford, John Sillitoe on behalf of the Wellcome Sanger Institute COVID-19 Surveillance Team |

| | | | |
|---|---|---|---|
| EPI_ISL_655675, EPI_ISL_655676 | Lighthouse Lab in Alderley Park | Wellcome Sanger Institute for the COVID-19 Genomics UK (COG-UK) Consortium | Jacquelyn Wynn, Mairead Hyland, The Lighthouse Lab in Alderley Park and Alex Alderton, Roberto Amato, Sonia Goncalves, Ewan Harrison, David K. Jackson, Ian Johnston, Dominic Kwiatkowski, Cordelia Langford, John Sillitoe on behalf of the Wellcome Sanger Institute COVID-19 Surveillance Team |
| EPI_ISL_655677 | Lighthouse Lab in Glasgow | Wellcome Sanger Institute for the COVID-19 Genomics UK (COG-UK) Consortium | Harper VanSteenhouse, Yumi Kasai, David Gray, Carol Clugston, Anna Dominiczak and Alex Alderton, Roberto Amato, Sonia Goncalves, Ewan Harrison, David K. Jackson, Ian Johnston, Dominic Kwiatkowski, Cordelia Langford, John Sillitoe on behalf of the Wellcome Sanger Institute COVID-19 Surveillance Team |
| EPI_ISL_655678, EPI_ISL_655679, EPI_ISL_655680, EPI_ISL_655681, EPI_ISL_655683, EPI_ISL_655684, EPI_ISL_655685, EPI_ISL_655687, EPI_ISL_655688, EPI_ISL_655689, EPI_ISL_655690, EPI_ISL_655691, EPI_ISL_655692, EPI_ISL_655693 | | | |
| see above | Lighthouse Lab in Alderley Park | Wellcome Sanger Institute for the COVID-19 Genomics UK (COG-UK) Consortium | Jacquelyn Wynn, Mairead Hyland, The Lighthouse Lab in Alderley Park and Alex Alderton, Roberto Amato, Sonia Goncalves, Ewan Harrison, David K. Jackson, Ian Johnston, Dominic Kwiatkowski, Cordelia Langford, John Sillitoe on behalf of the Wellcome Sanger Institute COVID-19 Surveillance Team |
| EPI_ISL_655694, EPI_ISL_655695 | Lighthouse Lab in Glasgow | Wellcome Sanger Institute for the COVID-19 Genomics UK (COG-UK) Consortium | Harper VanSteenhouse, Yumi Kasai, David Gray, Carol Clugston, Anna Dominiczak and Alex Alderton, Roberto Amato, Sonia Goncalves, Ewan Harrison, David K. Jackson, Ian Johnston, Dominic Kwiatkowski, Cordelia Langford, John Sillitoe on behalf of the Wellcome Sanger Institute COVID-19 Surveillance Team |
| EPI_ISL_655696 | Lighthouse Lab in Alderley Park | Wellcome Sanger Institute for the COVID-19 Genomics UK (COG-UK) Consortium | Jacquelyn Wynn, Mairead Hyland, The Lighthouse Lab in Alderley Park and Alex Alderton, Roberto Amato, Sonia Goncalves, Ewan Harrison, David K. Jackson, Ian Johnston, Dominic Kwiatkowski, Cordelia Langford, John Sillitoe on behalf of the Wellcome Sanger Institute COVID-19 Surveillance Team |
| EPI_ISL_655697 | Lighthouse Lab in Glasgow | Wellcome Sanger Institute for the COVID-19 Genomics UK (COG-UK) Consortium | Harper VanSteenhouse, Yumi Kasai, David Gray, Carol Clugston, Anna Dominiczak and Alex Alderton, Roberto Amato, Sonia Goncalves, Ewan Harrison, David K. Jackson, Ian Johnston, Dominic Kwiatkowski, Cordelia Langford, John Sillitoe on behalf of the Wellcome Sanger Institute COVID-19 Surveillance Team |
| EPI_ISL_655698 | Lighthouse Lab in Alderley Park | Wellcome Sanger Institute for the COVID-19 Genomics UK (COG-UK) Consortium | Jacquelyn Wynn, Mairead Hyland, The Lighthouse Lab in Alderley Park and Alex Alderton, Roberto Amato, Sonia Goncalves, Ewan Harrison, David K. Jackson, Ian Johnston, Dominic Kwiatkowski, Cordelia Langford, John Sillitoe on behalf of the Wellcome Sanger Institute COVID-19 Surveillance Team |
| EPI_ISL_655699 | Lighthouse Lab in Glasgow | Wellcome Sanger Institute for the COVID-19 Genomics UK (COG-UK) Consortium | Harper VanSteenhouse, Yumi Kasai, David Gray, Carol Clugston, Anna Dominiczak and Alex Alderton, Roberto Amato, Sonia Goncalves, Ewan Harrison, David K. Jackson, Ian Johnston, Dominic Kwiatkowski, Cordelia Langford, John Sillitoe on behalf of the Wellcome Sanger Institute COVID-19 Surveillance Team |
| EPI_ISL_655700, EPI_ISL_655702, EPI_ISL_655703, EPI_ISL_655706, EPI_ISL_655707, EPI_ISL_655710 | Lighthouse Lab in Alderley Park | Wellcome Sanger Institute for the COVID-19 Genomics UK (COG-UK) Consortium | Jacquelyn Wynn, Mairead Hyland, The Lighthouse Lab in Alderley Park and Alex Alderton, Roberto Amato, Sonia Goncalves, Ewan Harrison, David K. Jackson, Ian Johnston, Dominic Kwiatkowski, Cordelia Langford, John Sillitoe on behalf of the Wellcome Sanger Institute COVID-19 Surveillance Team |
| EPI_ISL_655711 | Lighthouse Lab in Glasgow | Wellcome Sanger Institute for the COVID-19 Genomics UK (COG-UK) Consortium | Harper VanSteenhouse, Yumi Kasai, David Gray, Carol Clugston, Anna Dominiczak and Alex Alderton, Roberto Amato, Sonia Goncalves, Ewan Harrison, David K. Jackson, Ian Johnston, Dominic Kwiatkowski, Cordelia Langford, John Sillitoe on behalf of the Wellcome Sanger Institute COVID-19 Surveillance Team |
| EPI_ISL_655712, EPI_ISL_655713, EPI_ISL_655717, EPI_ISL_655718 | Lighthouse Lab in Alderley Park | Wellcome Sanger Institute for the COVID-19 Genomics UK (COG-UK) Consortium | Jacquelyn Wynn, Mairead Hyland, The Lighthouse Lab in Alderley Park and Alex Alderton, Roberto Amato, Sonia Goncalves, Ewan Harrison, David K. Jackson, Ian Johnston, Dominic Kwiatkowski, Cordelia Langford, John Sillitoe on behalf of the Wellcome Sanger Institute COVID-19 Surveillance Team |
| EPI_ISL_655719 | Lighthouse Lab in Glasgow | Wellcome Sanger Institute for the COVID-19 Genomics UK (COG-UK) Consortium | Harper VanSteenhouse, Yumi Kasai, David Gray, Carol Clugston, Anna Dominiczak and Alex Alderton, Roberto Amato, Sonia Goncalves, Ewan Harrison, David K. Jackson, Ian Johnston, Dominic Kwiatkowski, Cordelia Langford, John Sillitoe on behalf of the Wellcome Sanger Institute COVID-19 Surveillance Team |
| EPI_ISL_655721, EPI_ISL_655722 | Lighthouse Lab in Alderley Park | Wellcome Sanger Institute for the COVID-19 Genomics UK (COG-UK) Consortium | Jacquelyn Wynn, Mairead Hyland, The Lighthouse Lab in Alderley Park and Alex Alderton, Roberto Amato, Sonia Goncalves, Ewan Harrison, David K. Jackson, Ian Johnston, Dominic Kwiatkowski, Cordelia Langford, John Sillitoe on behalf of the Wellcome Sanger Institute COVID-19 Surveillance Team |
| EPI_ISL_655723 | Lighthouse Lab in Glasgow | Wellcome Sanger Institute for the COVID-19 Genomics UK (COG-UK) Consortium | Harper VanSteenhouse, Yumi Kasai, David Gray, Carol Clugston, Anna Dominiczak and Alex Alderton, Roberto Amato, Sonia Goncalves, Ewan Harrison, David K. Jackson, Ian Johnston, Dominic Kwiatkowski, Cordelia Langford, John Sillitoe on behalf of the Wellcome Sanger Institute COVID-19 Surveillance Team |
| EPI_ISL_655724, EPI_ISL_655726, EPI_ISL_655728, EPI_ISL_655731, EPI_ISL_655732, EPI_ISL_655733, EPI_ISL_655734 | Lighthouse Lab in Alderley Park | Wellcome Sanger Institute for the COVID-19 Genomics UK (COG-UK) Consortium | Jacquelyn Wynn, Mairead Hyland, The Lighthouse Lab in Alderley Park and Alex Alderton, Roberto Amato, Sonia Goncalves, Ewan Harrison, David K. Jackson, Ian Johnston, Dominic Kwiatkowski, Cordelia Langford, John Sillitoe on behalf of the Wellcome Sanger Institute COVID-19 Surveillance Team |
| EPI_ISL_655735 | Lighthouse Lab in Glasgow | Wellcome Sanger Institute for the COVID-19 Genomics UK (COG-UK) Consortium | Harper VanSteenhouse, Yumi Kasai, David Gray, Carol Clugston, Anna Dominiczak and Alex Alderton, Roberto Amato, Sonia Goncalves, Ewan Harrison, David K. Jackson, Ian Johnston, Dominic Kwiatkowski, Cordelia Langford, John Sillitoe on behalf of the Wellcome Sanger Institute COVID-19 Surveillance Team |
| EPI_ISL_655736 | Lighthouse Lab in Alderley Park | Wellcome Sanger Institute for the COVID-19 Genomics UK (COG-UK) Consortium | Jacquelyn Wynn, Mairead Hyland, The Lighthouse Lab in Alderley Park and Alex Alderton, Roberto Amato, Sonia Goncalves, Ewan Harrison, David K. Jackson, Ian Johnston, Dominic Kwiatkowski, Cordelia Langford, John Sillitoe on behalf of the Wellcome Sanger Institute COVID-19 Surveillance Team |
| EPI_ISL_655737, EPI_ISL_655739 | Lighthouse Lab in Glasgow | Wellcome Sanger Institute for the COVID-19 Genomics UK (COG-UK) Consortium | Harper VanSteenhouse, Yumi Kasai, David Gray, Carol Clugston, Anna Dominiczak and Alex Alderton, Roberto Amato, Sonia Goncalves, Ewan Harrison, David K. Jackson, Ian Johnston, Dominic Kwiatkowski, Cordelia Langford, John Sillitoe on behalf of the Wellcome Sanger Institute COVID-19 Surveillance Team |
| EPI_ISL_655740, EPI_ISL_655741, EPI_ISL_655743 | Lighthouse Lab in Alderley Park | Wellcome Sanger Institute for the COVID-19 Genomics UK (COG-UK) Consortium | Jacquelyn Wynn, Mairead Hyland, The Lighthouse Lab in Alderley Park and Alex Alderton, Roberto Amato, Sonia Goncalves, Ewan Harrison, David K. Jackson, Ian Johnston, Dominic Kwiatkowski, Cordelia Langford, John Sillitoe on behalf of the Wellcome Sanger Institute COVID-19 Surveillance Team |
| EPI_ISL_655744, EPI_ISL_655745 | Lighthouse Lab in Glasgow | Wellcome Sanger Institute for the COVID-19 Genomics UK (COG-UK) Consortium | Harper VanSteenhouse, Yumi Kasai, David Gray, Carol Clugston, Anna Dominiczak and Alex Alderton, Roberto Amato, Sonia Goncalves, Ewan Harrison, David K. Jackson, Ian Johnston, Dominic Kwiatkowski, Cordelia Langford, John Sillitoe on behalf of the Wellcome Sanger Institute COVID-19 Surveillance Team |
| EPI_ISL_655746, EPI_ISL_655747, EPI_ISL_655749, EPI_ISL_655751, EPI_ISL_655752, EPI_ISL_655753, EPI_ISL_655754 | Lighthouse Lab in Alderley Park | Wellcome Sanger Institute for the COVID-19 Genomics UK (COG-UK) Consortium | Jacquelyn Wynn, Mairead Hyland, The Lighthouse Lab in Alderley Park and Alex Alderton, Roberto Amato, Sonia Goncalves, Ewan Harrison, David K. Jackson, Ian Johnston, Dominic Kwiatkowski, Cordelia Langford, John Sillitoe on behalf of the Wellcome Sanger Institute COVID-19 Surveillance Team |
| EPI_ISL_655756 | Lighthouse Lab in Glasgow | Wellcome Sanger Institute for the COVID-19 Genomics UK (COG-UK) Consortium | Harper VanSteenhouse, Yumi Kasai, David Gray, Carol Clugston, Anna Dominiczak and Alex Alderton, Roberto Amato, Sonia Goncalves, Ewan Harrison, David K. Jackson, Ian Johnston, Dominic Kwiatkowski, Cordelia Langford, John Sillitoe on behalf of the Wellcome Sanger Institute COVID-19 Surveillance Team |
| EPI_ISL_655757, EPI_ISL_655758, EPI_ISL_655759, EPI_ISL_655761 | Lighthouse Lab in Alderley Park | Wellcome Sanger Institute for the COVID-19 Genomics UK (COG-UK) Consortium | Jacquelyn Wynn, Mairead Hyland, The Lighthouse Lab in Alderley Park and Alex Alderton, Roberto Amato, Sonia Goncalves, Ewan Harrison, David K. Jackson, Ian Johnston, Dominic Kwiatkowski, Cordelia Langford, John Sillitoe on behalf of the Wellcome Sanger Institute COVID-19 Surveillance Team |
| EPI_ISL_655762 | Lighthouse Lab in Glasgow | Wellcome Sanger Institute for the COVID-19 Genomics UK (COG-UK) Consortium | Harper VanSteenhouse, Yumi Kasai, David Gray, Carol Clugston, Anna Dominiczak and Alex Alderton, Roberto Amato, Sonia Goncalves, Ewan Harrison, David K. Jackson, Ian Johnston, Dominic Kwiatkowski, Cordelia Langford, John Sillitoe on behalf of the Wellcome Sanger Institute COVID-19 Surveillance Team |
| EPI_ISL_655763, EPI_ISL_655765, EPI_ISL_655766 | Lighthouse Lab in Alderley Park | Wellcome Sanger Institute for the COVID-19 Genomics UK (COG-UK) Consortium | Jacquelyn Wynn, Mairead Hyland, The Lighthouse Lab in Alderley Park and Alex Alderton, Roberto Amato, Sonia Goncalves, Ewan Harrison, David K. Jackson, Ian Johnston, Dominic Kwiatkowski, Cordelia Langford, John Sillitoe on behalf of the Wellcome Sanger Institute COVID-19 Surveillance Team |
| EPI_ISL_655768, EPI_ISL_655769, EPI_ISL_655770 | Lighthouse Lab in Glasgow | Wellcome Sanger Institute for the COVID-19 Genomics UK (COG-UK) Consortium | Harper VanSteenhouse, Yumi Kasai, David Gray, Carol Clugston, Anna Dominiczak and Alex Alderton, Roberto Amato, Sonia Goncalves, Ewan Harrison, David K. Jackson, Ian Johnston, Dominic Kwiatkowski, Cordelia Langford, John Sillitoe on behalf of the Wellcome Sanger Institute COVID-19 Surveillance Team |
| EPI_ISL_655771 | Lighthouse Lab in Alderley Park | Wellcome Sanger Institute for the COVID-19 Genomics UK (COG-UK) Consortium | Jacquelyn Wynn, Mairead Hyland, The Lighthouse Lab in Alderley Park and Alex Alderton, Roberto Amato, Sonia Goncalves, Ewan Harrison, David K. Jackson, Ian Johnston, Dominic Kwiatkowski, Cordelia Langford, John Sillitoe on behalf of the Wellcome Sanger Institute COVID-19 Surveillance Team |

| | | | |
|---|---|---|---|
| EPI_ISL_655773 | Lighthouse Lab in Glasgow | Wellcome Sanger Institute for the COVID-19 Genomics UK (COG-UK) Consortium | Harper VanSteenhouse, Yumi Kasai, David Gray, Carol Clugston, Anna Dominiczak and Alex Alderton, Roberto Amato, Sonia Goncalves, Ewan Harrison, David K. Jackson, Ian Johnston, Dominic Kwiatkowski, Cordelia Langford, John Sillitoe on behalf of the Wellcome Sanger Institute COVID-19 Surveillance Team |
| EPI_ISL_655774, EPI_ISL_655775, EPI_ISL_655776, EPI_ISL_655777, EPI_ISL_655778 | Lighthouse Lab in Alderley Park | Wellcome Sanger Institute for the COVID-19 Genomics UK (COG-UK) Consortium | Jacquelyn Wynn, Mairead Hyland, The Lighthouse Lab in Alderley Park and Alex Alderton, Roberto Amato, Sonia Goncalves, Ewan Harrison, David K. Jackson, Ian Johnston, Dominic Kwiatkowski, Cordelia Langford, John Sillitoe on behalf of the Wellcome Sanger Institute COVID-19 Surveillance Team |
| EPI_ISL_655779, EPI_ISL_655780 | Lighthouse Lab in Glasgow | Wellcome Sanger Institute for the COVID-19 Genomics UK (COG-UK) Consortium | Harper VanSteenhouse, Yumi Kasai, David Gray, Carol Clugston, Anna Dominiczak and Alex Alderton, Roberto Amato, Sonia Goncalves, Ewan Harrison, David K. Jackson, Ian Johnston, Dominic Kwiatkowski, Cordelia Langford, John Sillitoe on behalf of the Wellcome Sanger Institute COVID-19 Surveillance Team |
| EPI_ISL_655782 | Lighthouse Lab in Alderley Park | Wellcome Sanger Institute for the COVID-19 Genomics UK (COG-UK) Consortium | Jacquelyn Wynn, Mairead Hyland, The Lighthouse Lab in Alderley Park and Alex Alderton, Roberto Amato, Sonia Goncalves, Ewan Harrison, David K. Jackson, Ian Johnston, Dominic Kwiatkowski, Cordelia Langford, John Sillitoe on behalf of the Wellcome Sanger Institute COVID-19 Surveillance Team |
| EPI_ISL_655783 | Lighthouse Lab in Glasgow | Wellcome Sanger Institute for the COVID-19 Genomics UK (COG-UK) Consortium | Harper VanSteenhouse, Yumi Kasai, David Gray, Carol Clugston, Anna Dominiczak and Alex Alderton, Roberto Amato, Sonia Goncalves, Ewan Harrison, David K. Jackson, Ian Johnston, Dominic Kwiatkowski, Cordelia Langford, John Sillitoe on behalf of the Wellcome Sanger Institute COVID-19 Surveillance Team |
| EPI_ISL_655784, EPI_ISL_655785, EPI_ISL_655786, EPI_ISL_655787 | Lighthouse Lab in Alderley Park | Wellcome Sanger Institute for the COVID-19 Genomics UK (COG-UK) Consortium | Jacquelyn Wynn, Mairead Hyland, The Lighthouse Lab in Alderley Park and Alex Alderton, Roberto Amato, Sonia Goncalves, Ewan Harrison, David K. Jackson, Ian Johnston, Dominic Kwiatkowski, Cordelia Langford, John Sillitoe on behalf of the Wellcome Sanger Institute COVID-19 Surveillance Team |
| EPI_ISL_655788 | Lighthouse Lab in Glasgow | Wellcome Sanger Institute for the COVID-19 Genomics UK (COG-UK) Consortium | Harper VanSteenhouse, Yumi Kasai, David Gray, Carol Clugston, Anna Dominiczak and Alex Alderton, Roberto Amato, Sonia Goncalves, Ewan Harrison, David K. Jackson, Ian Johnston, Dominic Kwiatkowski, Cordelia Langford, John Sillitoe on behalf of the Wellcome Sanger Institute COVID-19 Surveillance Team |
| EPI_ISL_655789, EPI_ISL_655790, EPI_ISL_655791, EPI_ISL_655793, EPI_ISL_655794, EPI_ISL_655796 | Lighthouse Lab in Alderley Park | Wellcome Sanger Institute for the COVID-19 Genomics UK (COG-UK) Consortium | Jacquelyn Wynn, Mairead Hyland, The Lighthouse Lab in Alderley Park and Alex Alderton, Roberto Amato, Sonia Goncalves, Ewan Harrison, David K. Jackson, Ian Johnston, Dominic Kwiatkowski, Cordelia Langford, John Sillitoe on behalf of the Wellcome Sanger Institute COVID-19 Surveillance Team |
| EPI_ISL_655798 | Lighthouse Lab in Glasgow | Wellcome Sanger Institute for the COVID-19 Genomics UK (COG-UK) Consortium | Harper VanSteenhouse, Yumi Kasai, David Gray, Carol Clugston, Anna Dominiczak and Alex Alderton, Roberto Amato, Sonia Goncalves, Ewan Harrison, David K. Jackson, Ian Johnston, Dominic Kwiatkowski, Cordelia Langford, John Sillitoe on behalf of the Wellcome Sanger Institute COVID-19 Surveillance Team |
| EPI_ISL_655799, EPI_ISL_655800, EPI_ISL_655801, EPI_ISL_655802 | Lighthouse Lab in Alderley Park | Wellcome Sanger Institute for the COVID-19 Genomics UK (COG-UK) Consortium | Jacquelyn Wynn, Mairead Hyland, The Lighthouse Lab in Alderley Park and Alex Alderton, Roberto Amato, Sonia Goncalves, Ewan Harrison, David K. Jackson, Ian Johnston, Dominic Kwiatkowski, Cordelia Langford, John Sillitoe on behalf of the Wellcome Sanger Institute COVID-19 Surveillance Team |
| EPI_ISL_655804 | Lighthouse Lab in Glasgow | Wellcome Sanger Institute for the COVID-19 Genomics UK (COG-UK) Consortium | Harper VanSteenhouse, Yumi Kasai, David Gray, Carol Clugston, Anna Dominiczak and Alex Alderton, Roberto Amato, Sonia Goncalves, Ewan Harrison, David K. Jackson, Ian Johnston, Dominic Kwiatkowski, Cordelia Langford, John Sillitoe on behalf of the Wellcome Sanger Institute COVID-19 Surveillance Team |
| EPI_ISL_655805 | Lighthouse Lab in Alderley Park | Wellcome Sanger Institute for the COVID-19 Genomics UK (COG-UK) Consortium | Jacquelyn Wynn, Mairead Hyland, The Lighthouse Lab in Alderley Park and Alex Alderton, Roberto Amato, Sonia Goncalves, Ewan Harrison, David K. Jackson, Ian Johnston, Dominic Kwiatkowski, Cordelia Langford, John Sillitoe on behalf of the Wellcome Sanger Institute COVID-19 Surveillance Team |
| EPI_ISL_655806, EPI_ISL_655807, EPI_ISL_655809 | Lighthouse Lab in Glasgow | Wellcome Sanger Institute for the COVID-19 Genomics UK (COG-UK) Consortium | Harper VanSteenhouse, Yumi Kasai, David Gray, Carol Clugston, Anna Dominiczak and Alex Alderton, Roberto Amato, Sonia Goncalves, Ewan Harrison, David K. Jackson, Ian Johnston, Dominic Kwiatkowski, Cordelia Langford, John Sillitoe on behalf of the Wellcome Sanger Institute COVID-19 Surveillance Team |
| EPI_ISL_655811, EPI_ISL_655815 | Lighthouse Lab in Alderley Park | Wellcome Sanger Institute for the COVID-19 Genomics UK (COG-UK) Consortium | Jacquelyn Wynn, Mairead Hyland, The Lighthouse Lab in Alderley Park and Alex Alderton, Roberto Amato, Sonia Goncalves, Ewan Harrison, David K. Jackson, Ian Johnston, Dominic Kwiatkowski, Cordelia Langford, John Sillitoe on behalf of the Wellcome Sanger Institute COVID-19 Surveillance Team |
| EPI_ISL_655817, EPI_ISL_655819 | Lighthouse Lab in Glasgow | Wellcome Sanger Institute for the COVID-19 Genomics UK (COG-UK) Consortium | Harper VanSteenhouse, Yumi Kasai, David Gray, Carol Clugston, Anna Dominiczak and Alex Alderton, Roberto Amato, Sonia Goncalves, Ewan Harrison, David K. Jackson, Ian Johnston, Dominic Kwiatkowski, Cordelia Langford, John Sillitoe on behalf of the Wellcome Sanger Institute COVID-19 Surveillance Team |
| EPI_ISL_655821 | Lighthouse Lab in Alderley Park | Wellcome Sanger Institute for the COVID-19 Genomics UK (COG-UK) Consortium | Jacquelyn Wynn, Mairead Hyland, The Lighthouse Lab in Alderley Park and Alex Alderton, Roberto Amato, Sonia Goncalves, Ewan Harrison, David K. Jackson, Ian Johnston, Dominic Kwiatkowski, Cordelia Langford, John Sillitoe on behalf of the Wellcome Sanger Institute COVID-19 Surveillance Team |
| EPI_ISL_655822 | Lighthouse Lab in Glasgow | Wellcome Sanger Institute for the COVID-19 Genomics UK (COG-UK) Consortium | Harper VanSteenhouse, Yumi Kasai, David Gray, Carol Clugston, Anna Dominiczak and Alex Alderton, Roberto Amato, Sonia Goncalves, Ewan Harrison, David K. Jackson, Ian Johnston, Dominic Kwiatkowski, Cordelia Langford, John Sillitoe on behalf of the Wellcome Sanger Institute COVID-19 Surveillance Team |
| EPI_ISL_655823, EPI_ISL_655824 | Lighthouse Lab in Alderley Park | Wellcome Sanger Institute for the COVID-19 Genomics UK (COG-UK) Consortium | Jacquelyn Wynn, Mairead Hyland, The Lighthouse Lab in Alderley Park and Alex Alderton, Roberto Amato, Sonia Goncalves, Ewan Harrison, David K. Jackson, Ian Johnston, Dominic Kwiatkowski, Cordelia Langford, John Sillitoe on behalf of the Wellcome Sanger Institute COVID-19 Surveillance Team |
| EPI_ISL_655828 | Lighthouse Lab in Glasgow | Wellcome Sanger Institute for the COVID-19 Genomics UK (COG-UK) Consortium | Harper VanSteenhouse, Yumi Kasai, David Gray, Carol Clugston, Anna Dominiczak and Alex Alderton, Roberto Amato, Sonia Goncalves, Ewan Harrison, David K. Jackson, Ian Johnston, Dominic Kwiatkowski, Cordelia Langford, John Sillitoe on behalf of the Wellcome Sanger Institute COVID-19 Surveillance Team |
| EPI_ISL_655830 | Lighthouse Lab in Alderley Park | Wellcome Sanger Institute for the COVID-19 Genomics UK (COG-UK) Consortium | Jacquelyn Wynn, Mairead Hyland, The Lighthouse Lab in Alderley Park and Alex Alderton, Roberto Amato, Sonia Goncalves, Ewan Harrison, David K. Jackson, Ian Johnston, Dominic Kwiatkowski, Cordelia Langford, John Sillitoe on behalf of the Wellcome Sanger Institute COVID-19 Surveillance Team |
| EPI_ISL_655832 | Lighthouse Lab in Glasgow | Wellcome Sanger Institute for the COVID-19 Genomics UK (COG-UK) Consortium | Harper VanSteenhouse, Yumi Kasai, David Gray, Carol Clugston, Anna Dominiczak and Alex Alderton, Roberto Amato, Sonia Goncalves, Ewan Harrison, David K. Jackson, Ian Johnston, Dominic Kwiatkowski, Cordelia Langford, John Sillitoe on behalf of the Wellcome Sanger Institute COVID-19 Surveillance Team |
| EPI_ISL_655833, EPI_ISL_655835, EPI_ISL_655836, EPI_ISL_655837, EPI_ISL_655838, EPI_ISL_655839, EPI_ISL_655840 | Lighthouse Lab in Alderley Park | Wellcome Sanger Institute for the COVID-19 Genomics UK (COG-UK) Consortium | Jacquelyn Wynn, Mairead Hyland, The Lighthouse Lab in Alderley Park and Alex Alderton, Roberto Amato, Sonia Goncalves, Ewan Harrison, David K. Jackson, Ian Johnston, Dominic Kwiatkowski, Cordelia Langford, John Sillitoe on behalf of the Wellcome Sanger Institute COVID-19 Surveillance Team |
| EPI_ISL_655841 | Lighthouse Lab in Glasgow | Wellcome Sanger Institute for the COVID-19 Genomics UK (COG-UK) Consortium | Harper VanSteenhouse, Yumi Kasai, David Gray, Carol Clugston, Anna Dominiczak and Alex Alderton, Roberto Amato, Sonia Goncalves, Ewan Harrison, David K. Jackson, Ian Johnston, Dominic Kwiatkowski, Cordelia Langford, John Sillitoe on behalf of the Wellcome Sanger Institute COVID-19 Surveillance Team |
| EPI_ISL_655842, EPI_ISL_655843, EPI_ISL_655845 | Lighthouse Lab in Alderley Park | Wellcome Sanger Institute for the COVID-19 Genomics UK (COG-UK) Consortium | Jacquelyn Wynn, Mairead Hyland, The Lighthouse Lab in Alderley Park and Alex Alderton, Roberto Amato, Sonia Goncalves, Ewan Harrison, David K. Jackson, Ian Johnston, Dominic Kwiatkowski, Cordelia Langford, John Sillitoe on behalf of the Wellcome Sanger Institute COVID-19 Surveillance Team |
| EPI_ISL_655847 | Lighthouse Lab in Glasgow | Wellcome Sanger Institute for the COVID-19 Genomics UK (COG-UK) Consortium | Harper VanSteenhouse, Yumi Kasai, David Gray, Carol Clugston, Anna Dominiczak and Alex Alderton, Roberto Amato, Sonia Goncalves, Ewan Harrison, David K. Jackson, Ian Johnston, Dominic Kwiatkowski, Cordelia Langford, John Sillitoe on behalf of the Wellcome Sanger Institute COVID-19 Surveillance Team |
| EPI_ISL_655848 | Lighthouse Lab in Alderley Park | Wellcome Sanger Institute for the COVID-19 Genomics UK (COG-UK) Consortium | Jacquelyn Wynn, Mairead Hyland, The Lighthouse Lab in Alderley Park and Alex Alderton, Roberto Amato, Sonia Goncalves, Ewan Harrison, David K. Jackson, Ian Johnston, Dominic Kwiatkowski, Cordelia Langford, John Sillitoe on behalf of the Wellcome Sanger Institute COVID-19 Surveillance Team |
| EPI_ISL_655850 | Lighthouse Lab in Glasgow | Wellcome Sanger Institute for the COVID-19 Genomics UK (COG-UK) Consortium | Harper VanSteenhouse, Yumi Kasai, David Gray, Carol Clugston, Anna Dominiczak and Alex Alderton, Roberto Amato, Sonia Goncalves, Ewan Harrison, David K. Jackson, Ian Johnston, Dominic Kwiatkowski, Cordelia Langford, John Sillitoe on behalf of the Wellcome Sanger Institute COVID-19 Surveillance Team |
| EPI_ISL_655852, EPI_ISL_655853 | Lighthouse Lab in Alderley Park | Wellcome Sanger Institute for the COVID-19 Genomics UK (COG-UK) Consortium | Jacquelyn Wynn, Mairead Hyland, The Lighthouse Lab in Alderley Park and Alex Alderton, Roberto Amato, Sonia Goncalves, Ewan Harrison, David K. Jackson, Ian Johnston, Dominic Kwiatkowski, Cordelia Langford, John Sillitoe on behalf of the Wellcome Sanger Institute COVID-19 Surveillance Team |

| | | | |
|---|---|---|---|
| EPI_ISL_655854 | Lighthouse Lab in Glasgow | Wellcome Sanger Institute for the COVID-19 Genomics UK (COG-UK) Consortium | Harper VanSteenhouse, Yumi Kasai, David Gray, Carol Clugston, Anna Dominiczak and Alex Alderton, Roberto Amato, Sonia Goncalves, Ewan Harrison, David K. Jackson, Ian Johnston, Dominic Kwiatkowski, Cordelia Langford, John Sillitoe on behalf of the Wellcome Sanger Institute COVID-19 Surveillance Team |
| EPI_ISL_655855, EPI_ISL_655856, EPI_ISL_655857, EPI_ISL_655859 | Lighthouse Lab in Alderley Park | Wellcome Sanger Institute for the COVID-19 Genomics UK (COG-UK) Consortium | Jacquelyn Wynn, Mairead Hyland, The Lighthouse Lab in Alderley Park and Alex Alderton, Roberto Amato, Sonia Goncalves, Ewan Harrison, David K. Jackson, Ian Johnston, Dominic Kwiatkowski, Cordelia Langford, John Sillitoe on behalf of the Wellcome Sanger Institute COVID-19 Surveillance Team |
| EPI_ISL_655861 | Lighthouse Lab in Glasgow | Wellcome Sanger Institute for the COVID-19 Genomics UK (COG-UK) Consortium | Harper VanSteenhouse, Yumi Kasai, David Gray, Carol Clugston, Anna Dominiczak and Alex Alderton, Roberto Amato, Sonia Goncalves, Ewan Harrison, David K. Jackson, Ian Johnston, Dominic Kwiatkowski, Cordelia Langford, John Sillitoe on behalf of the Wellcome Sanger Institute COVID-19 Surveillance Team |
| EPI_ISL_655863 | Lighthouse Lab in Alderley Park | Wellcome Sanger Institute for the COVID-19 Genomics UK (COG-UK) Consortium | Jacquelyn Wynn, Mairead Hyland, The Lighthouse Lab in Alderley Park and Alex Alderton, Roberto Amato, Sonia Goncalves, Ewan Harrison, David K. Jackson, Ian Johnston, Dominic Kwiatkowski, Cordelia Langford, John Sillitoe on behalf of the Wellcome Sanger Institute COVID-19 Surveillance Team |
| EPI_ISL_655864, EPI_ISL_655865 | Lighthouse Lab in Glasgow | Wellcome Sanger Institute for the COVID-19 Genomics UK (COG-UK) Consortium | Harper VanSteenhouse, Yumi Kasai, David Gray, Carol Clugston, Anna Dominiczak and Alex Alderton, Roberto Amato, Sonia Goncalves, Ewan Harrison, David K. Jackson, Ian Johnston, Dominic Kwiatkowski, Cordelia Langford, John Sillitoe on behalf of the Wellcome Sanger Institute COVID-19 Surveillance Team |
| EPI_ISL_655867 | Lighthouse Lab in Alderley Park | Wellcome Sanger Institute for the COVID-19 Genomics UK (COG-UK) Consortium | Jacquelyn Wynn, Mairead Hyland, The Lighthouse Lab in Alderley Park and Alex Alderton, Roberto Amato, Sonia Goncalves, Ewan Harrison, David K. Jackson, Ian Johnston, Dominic Kwiatkowski, Cordelia Langford, John Sillitoe on behalf of the Wellcome Sanger Institute COVID-19 Surveillance Team |
| EPI_ISL_655868 | Lighthouse Lab in Glasgow | Wellcome Sanger Institute for the COVID-19 Genomics UK (COG-UK) Consortium | Harper VanSteenhouse, Yumi Kasai, David Gray, Carol Clugston, Anna Dominiczak and Alex Alderton, Roberto Amato, Sonia Goncalves, Ewan Harrison, David K. Jackson, Ian Johnston, Dominic Kwiatkowski, Cordelia Langford, John Sillitoe on behalf of the Wellcome Sanger Institute COVID-19 Surveillance Team |
| EPI_ISL_655870, EPI_ISL_655871, EPI_ISL_655872 | Lighthouse Lab in Alderley Park | Wellcome Sanger Institute for the COVID-19 Genomics UK (COG-UK) Consortium | Jacquelyn Wynn, Mairead Hyland, The Lighthouse Lab in Alderley Park and Alex Alderton, Roberto Amato, Sonia Goncalves, Ewan Harrison, David K. Jackson, Ian Johnston, Dominic Kwiatkowski, Cordelia Langford, John Sillitoe on behalf of the Wellcome Sanger Institute COVID-19 Surveillance Team |
| EPI_ISL_655873, EPI_ISL_655874 | Lighthouse Lab in Glasgow | Wellcome Sanger Institute for the COVID-19 Genomics UK (COG-UK) Consortium | Harper VanSteenhouse, Yumi Kasai, David Gray, Carol Clugston, Anna Dominiczak and Alex Alderton, Roberto Amato, Sonia Goncalves, Ewan Harrison, David K. Jackson, Ian Johnston, Dominic Kwiatkowski, Cordelia Langford, John Sillitoe on behalf of the Wellcome Sanger Institute COVID-19 Surveillance Team |
| EPI_ISL_655877, EPI_ISL_655878 | Lighthouse Lab in Alderley Park | Wellcome Sanger Institute for the COVID-19 Genomics UK (COG-UK) Consortium | Jacquelyn Wynn, Mairead Hyland, The Lighthouse Lab in Alderley Park and Alex Alderton, Roberto Amato, Sonia Goncalves, Ewan Harrison, David K. Jackson, Ian Johnston, Dominic Kwiatkowski, Cordelia Langford, John Sillitoe on behalf of the Wellcome Sanger Institute COVID-19 Surveillance Team |
| EPI_ISL_655879 | Lighthouse Lab in Glasgow | Wellcome Sanger Institute for the COVID-19 Genomics UK (COG-UK) Consortium | Harper VanSteenhouse, Yumi Kasai, David Gray, Carol Clugston, Anna Dominiczak and Alex Alderton, Roberto Amato, Sonia Goncalves, Ewan Harrison, David K. Jackson, Ian Johnston, Dominic Kwiatkowski, Cordelia Langford, John Sillitoe on behalf of the Wellcome Sanger Institute COVID-19 Surveillance Team |
| EPI_ISL_655881, EPI_ISL_655882 | Lighthouse Lab in Alderley Park | Wellcome Sanger Institute for the COVID-19 Genomics UK (COG-UK) Consortium | Jacquelyn Wynn, Mairead Hyland, The Lighthouse Lab in Alderley Park and Alex Alderton, Roberto Amato, Sonia Goncalves, Ewan Harrison, David K. Jackson, Ian Johnston, Dominic Kwiatkowski, Cordelia Langford, John Sillitoe on behalf of the Wellcome Sanger Institute COVID-19 Surveillance Team |
| EPI_ISL_655883 | Lighthouse Lab in Glasgow | Wellcome Sanger Institute for the COVID-19 Genomics UK (COG-UK) Consortium | Harper VanSteenhouse, Yumi Kasai, David Gray, Carol Clugston, Anna Dominiczak and Alex Alderton, Roberto Amato, Sonia Goncalves, Ewan Harrison, David K. Jackson, Ian Johnston, Dominic Kwiatkowski, Cordelia Langford, John Sillitoe on behalf of the Wellcome Sanger Institute COVID-19 Surveillance Team |
| EPI_ISL_655885, EPI_ISL_655888, EPI_ISL_655890 | Lighthouse Lab in Alderley Park | Wellcome Sanger Institute for the COVID-19 Genomics UK (COG-UK) Consortium | Jacquelyn Wynn, Mairead Hyland, The Lighthouse Lab in Alderley Park and Alex Alderton, Roberto Amato, Sonia Goncalves, Ewan Harrison, David K. Jackson, Ian Johnston, Dominic Kwiatkowski, Cordelia Langford, John Sillitoe on behalf of the Wellcome Sanger Institute COVID-19 Surveillance Team |
| EPI_ISL_655891 | Lighthouse Lab in Glasgow | Wellcome Sanger Institute for the COVID-19 Genomics UK (COG-UK) Consortium | Harper VanSteenhouse, Yumi Kasai, David Gray, Carol Clugston, Anna Dominiczak and Alex Alderton, Roberto Amato, Sonia Goncalves, Ewan Harrison, David K. Jackson, Ian Johnston, Dominic Kwiatkowski, Cordelia Langford, John Sillitoe on behalf of the Wellcome Sanger Institute COVID-19 Surveillance Team |
| EPI_ISL_655892, EPI_ISL_655894, EPI_ISL_655895, EPI_ISL_655897, EPI_ISL_655898, EPI_ISL_655899, EPI_ISL_655900, EPI_ISL_655901, EPI_ISL_655903, EPI_ISL_655904, EPI_ISL_655905, EPI_ISL_655906 | | | |
| see above | Lighthouse Lab in Alderley Park | Wellcome Sanger Institute for the COVID-19 Genomics UK (COG-UK) Consortium | Jacquelyn Wynn, Mairead Hyland, The Lighthouse Lab in Alderley Park and Alex Alderton, Roberto Amato, Sonia Goncalves, Ewan Harrison, David K. Jackson, Ian Johnston, Dominic Kwiatkowski, Cordelia Langford, John Sillitoe on behalf of the Wellcome Sanger Institute COVID-19 Surveillance Team |
| EPI_ISL_655907 | Lighthouse Lab in Glasgow | Wellcome Sanger Institute for the COVID-19 Genomics UK (COG-UK) Consortium | Harper VanSteenhouse, Yumi Kasai, David Gray, Carol Clugston, Anna Dominiczak and Alex Alderton, Roberto Amato, Sonia Goncalves, Ewan Harrison, David K. Jackson, Ian Johnston, Dominic Kwiatkowski, Cordelia Langford, John Sillitoe on behalf of the Wellcome Sanger Institute COVID-19 Surveillance Team |
| EPI_ISL_655908, EPI_ISL_655909, EPI_ISL_655910 | Lighthouse Lab in Alderley Park | Wellcome Sanger Institute for the COVID-19 Genomics UK (COG-UK) Consortium | Jacquelyn Wynn, Mairead Hyland, The Lighthouse Lab in Alderley Park and Alex Alderton, Roberto Amato, Sonia Goncalves, Ewan Harrison, David K. Jackson, Ian Johnston, Dominic Kwiatkowski, Cordelia Langford, John Sillitoe on behalf of the Wellcome Sanger Institute COVID-19 Surveillance Team |
| EPI_ISL_655911, EPI_ISL_655912 | Lighthouse Lab in Glasgow | Wellcome Sanger Institute for the COVID-19 Genomics UK (COG-UK) Consortium | Harper VanSteenhouse, Yumi Kasai, David Gray, Carol Clugston, Anna Dominiczak and Alex Alderton, Roberto Amato, Sonia Goncalves, Ewan Harrison, David K. Jackson, Ian Johnston, Dominic Kwiatkowski, Cordelia Langford, John Sillitoe on behalf of the Wellcome Sanger Institute COVID-19 Surveillance Team |
| EPI_ISL_655914 | Lighthouse Lab in Alderley Park | Wellcome Sanger Institute for the COVID-19 Genomics UK (COG-UK) Consortium | Jacquelyn Wynn, Mairead Hyland, The Lighthouse Lab in Alderley Park and Alex Alderton, Roberto Amato, Sonia Goncalves, Ewan Harrison, David K. Jackson, Ian Johnston, Dominic Kwiatkowski, Cordelia Langford, John Sillitoe on behalf of the Wellcome Sanger Institute COVID-19 Surveillance Team |
| EPI_ISL_655918 | Lighthouse Lab in Glasgow | Wellcome Sanger Institute for the COVID-19 Genomics UK (COG-UK) Consortium | Harper VanSteenhouse, Yumi Kasai, David Gray, Carol Clugston, Anna Dominiczak and Alex Alderton, Roberto Amato, Sonia Goncalves, Ewan Harrison, David K. Jackson, Ian Johnston, Dominic Kwiatkowski, Cordelia Langford, John Sillitoe on behalf of the Wellcome Sanger Institute COVID-19 Surveillance Team |
| EPI_ISL_655919, EPI_ISL_655923 | Lighthouse Lab in Alderley Park | Wellcome Sanger Institute for the COVID-19 Genomics UK (COG-UK) Consortium | Jacquelyn Wynn, Mairead Hyland, The Lighthouse Lab in Alderley Park and Alex Alderton, Roberto Amato, Sonia Goncalves, Ewan Harrison, David K. Jackson, Ian Johnston, Dominic Kwiatkowski, Cordelia Langford, John Sillitoe on behalf of the Wellcome Sanger Institute COVID-19 Surveillance Team |
| EPI_ISL_655924 | Lighthouse Lab in Glasgow | Wellcome Sanger Institute for the COVID-19 Genomics UK (COG-UK) Consortium | Harper VanSteenhouse, Yumi Kasai, David Gray, Carol Clugston, Anna Dominiczak and Alex Alderton, Roberto Amato, Sonia Goncalves, Ewan Harrison, David K. Jackson, Ian Johnston, Dominic Kwiatkowski, Cordelia Langford, John Sillitoe on behalf of the Wellcome Sanger Institute COVID-19 Surveillance Team |
| EPI_ISL_655925, EPI_ISL_655926, EPI_ISL_655927 | Lighthouse Lab in Alderley Park | Wellcome Sanger Institute for the COVID-19 Genomics UK (COG-UK) Consortium | Jacquelyn Wynn, Mairead Hyland, The Lighthouse Lab in Alderley Park and Alex Alderton, Roberto Amato, Sonia Goncalves, Ewan Harrison, David K. Jackson, Ian Johnston, Dominic Kwiatkowski, Cordelia Langford, John Sillitoe on behalf of the Wellcome Sanger Institute COVID-19 Surveillance Team |
| EPI_ISL_655929 | Lighthouse Lab in Glasgow | Wellcome Sanger Institute for the COVID-19 Genomics UK (COG-UK) Consortium | Harper VanSteenhouse, Yumi Kasai, David Gray, Carol Clugston, Anna Dominiczak and Alex Alderton, Roberto Amato, Sonia Goncalves, Ewan Harrison, David K. Jackson, Ian Johnston, Dominic Kwiatkowski, Cordelia Langford, John Sillitoe on behalf of the Wellcome Sanger Institute COVID-19 Surveillance Team |
| EPI_ISL_655930, EPI_ISL_655931, EPI_ISL_655932 | Lighthouse Lab in Alderley Park | Wellcome Sanger Institute for the COVID-19 Genomics UK (COG-UK) Consortium | Jacquelyn Wynn, Mairead Hyland, The Lighthouse Lab in Alderley Park and Alex Alderton, Roberto Amato, Sonia Goncalves, Ewan Harrison, David K. Jackson, Ian Johnston, Dominic Kwiatkowski, Cordelia Langford, John Sillitoe on behalf of the Wellcome Sanger Institute COVID-19 Surveillance Team |
| EPI_ISL_655933 | Lighthouse Lab in Glasgow | Wellcome Sanger Institute for the COVID-19 Genomics UK (COG-UK) Consortium | Harper VanSteenhouse, Yumi Kasai, David Gray, Carol Clugston, Anna Dominiczak and Alex Alderton, Roberto Amato, Sonia Goncalves, Ewan Harrison, David K. Jackson, Ian Johnston, Dominic Kwiatkowski, Cordelia Langford, John Sillitoe on behalf of the Wellcome Sanger Institute COVID-19 Surveillance Team |
| EPI_ISL_655935, EPI_ISL_655936, EPI_ISL_655937, EPI_ISL_655940, EPI_ISL_655943, EPI_ISL_655944, EPI_ISL_655946 | Lighthouse Lab in Alderley Park | Wellcome Sanger Institute for the COVID-19 Genomics UK (COG-UK) Consortium | Jacquelyn Wynn, Mairead Hyland, The Lighthouse Lab in Alderley Park and Alex Alderton, Roberto Amato, Sonia Goncalves, Ewan Harrison, David K. Jackson, Ian Johnston, Dominic Kwiatkowski, Cordelia Langford, John Sillitoe on behalf of the Wellcome Sanger Institute COVID-19 Surveillance Team |

| | | | |
|---|---|---|---|
| EPI_ISL_655947, EPI_ISL_655948 | Lighthouse Lab in Glasgow | Wellcome Sanger Institute for the COVID-19 Genomics UK (COG-UK) Consortium | Harper VanSteenhouse, Yumi Kasai, David Gray, Carol Clugston, Anna Dominiczak and Alex Alderton, Roberto Amato, Sonia Goncalves, Ewan Harrison, David K. Jackson, Ian Johnston, Dominic Kwiatkowski, Cordelia Langford, John Sillitoe on behalf of the Wellcome Sanger Institute COVID-19 Surveillance Team |
| EPI_ISL_655949, EPI_ISL_655951, EPI_ISL_655952, EPI_ISL_655954 | Lighthouse Lab in Alderley Park | Wellcome Sanger Institute for the COVID-19 Genomics UK (COG-UK) Consortium | Jacquelyn Wynn, Mairead Hyland, The Lighthouse Lab in Alderley Park and Alex Alderton, Roberto Amato, Sonia Goncalves, Ewan Harrison, David K. Jackson, Ian Johnston, Dominic Kwiatkowski, Cordelia Langford, John Sillitoe on behalf of the Wellcome Sanger Institute COVID-19 Surveillance Team |
| EPI_ISL_655957, EPI_ISL_655959 | Lighthouse Lab in Glasgow | Wellcome Sanger Institute for the COVID-19 Genomics UK (COG-UK) Consortium | Harper VanSteenhouse, Yumi Kasai, David Gray, Carol Clugston, Anna Dominiczak and Alex Alderton, Roberto Amato, Sonia Goncalves, Ewan Harrison, David K. Jackson, Ian Johnston, Dominic Kwiatkowski, Cordelia Langford, John Sillitoe on behalf of the Wellcome Sanger Institute COVID-19 Surveillance Team |
| EPI_ISL_655961, EPI_ISL_655963 | Lighthouse Lab in Alderley Park | Wellcome Sanger Institute for the COVID-19 Genomics UK (COG-UK) Consortium | Jacquelyn Wynn, Mairead Hyland, The Lighthouse Lab in Alderley Park and Alex Alderton, Roberto Amato, Sonia Goncalves, Ewan Harrison, David K. Jackson, Ian Johnston, Dominic Kwiatkowski, Cordelia Langford, John Sillitoe on behalf of the Wellcome Sanger Institute COVID-19 Surveillance Team |
| EPI_ISL_655967 | Lighthouse Lab in Glasgow | Wellcome Sanger Institute for the COVID-19 Genomics UK (COG-UK) Consortium | Harper VanSteenhouse, Yumi Kasai, David Gray, Carol Clugston, Anna Dominiczak and Alex Alderton, Roberto Amato, Sonia Goncalves, Ewan Harrison, David K. Jackson, Ian Johnston, Dominic Kwiatkowski, Cordelia Langford, John Sillitoe on behalf of the Wellcome Sanger Institute COVID-19 Surveillance Team |
| EPI_ISL_655968, EPI_ISL_655970 | Lighthouse Lab in Alderley Park | Wellcome Sanger Institute for the COVID-19 Genomics UK (COG-UK) Consortium | Jacquelyn Wynn, Mairead Hyland, The Lighthouse Lab in Alderley Park and Alex Alderton, Roberto Amato, Sonia Goncalves, Ewan Harrison, David K. Jackson, Ian Johnston, Dominic Kwiatkowski, Cordelia Langford, John Sillitoe on behalf of the Wellcome Sanger Institute COVID-19 Surveillance Team |
| EPI_ISL_655971 | Lighthouse Lab in Glasgow | Wellcome Sanger Institute for the COVID-19 Genomics UK (COG-UK) Consortium | Harper VanSteenhouse, Yumi Kasai, David Gray, Carol Clugston, Anna Dominiczak and Alex Alderton, Roberto Amato, Sonia Goncalves, Ewan Harrison, David K. Jackson, Ian Johnston, Dominic Kwiatkowski, Cordelia Langford, John Sillitoe on behalf of the Wellcome Sanger Institute COVID-19 Surveillance Team |
| EPI_ISL_655972 | Lighthouse Lab in Alderley Park | Wellcome Sanger Institute for the COVID-19 Genomics UK (COG-UK) Consortium | Jacquelyn Wynn, Mairead Hyland, The Lighthouse Lab in Alderley Park and Alex Alderton, Roberto Amato, Sonia Goncalves, Ewan Harrison, David K. Jackson, Ian Johnston, Dominic Kwiatkowski, Cordelia Langford, John Sillitoe on behalf of the Wellcome Sanger Institute COVID-19 Surveillance Team |
| EPI_ISL_655973 | Lighthouse Lab in Glasgow | Wellcome Sanger Institute for the COVID-19 Genomics UK (COG-UK) Consortium | Harper VanSteenhouse, Yumi Kasai, David Gray, Carol Clugston, Anna Dominiczak and Alex Alderton, Roberto Amato, Sonia Goncalves, Ewan Harrison, David K. Jackson, Ian Johnston, Dominic Kwiatkowski, Cordelia Langford, John Sillitoe on behalf of the Wellcome Sanger Institute COVID-19 Surveillance Team |
| EPI_ISL_655974 | Lighthouse Lab in Alderley Park | Wellcome Sanger Institute for the COVID-19 Genomics UK (COG-UK) Consortium | Jacquelyn Wynn, Mairead Hyland, The Lighthouse Lab in Alderley Park and Alex Alderton, Roberto Amato, Sonia Goncalves, Ewan Harrison, David K. Jackson, Ian Johnston, Dominic Kwiatkowski, Cordelia Langford, John Sillitoe on behalf of the Wellcome Sanger Institute COVID-19 Surveillance Team |
| EPI_ISL_655975 | Lighthouse Lab in Glasgow | Wellcome Sanger Institute for the COVID-19 Genomics UK (COG-UK) Consortium | Harper VanSteenhouse, Yumi Kasai, David Gray, Carol Clugston, Anna Dominiczak and Alex Alderton, Roberto Amato, Sonia Goncalves, Ewan Harrison, David K. Jackson, Ian Johnston, Dominic Kwiatkowski, Cordelia Langford, John Sillitoe on behalf of the Wellcome Sanger Institute COVID-19 Surveillance Team |
| EPI_ISL_656345, EPI_ISL_656346, EPI_ISL_656348, EPI_ISL_656349, EPI_ISL_656350, EPI_ISL_656351, EPI_ISL_656352, EPI_ISL_656353, EPI_ISL_656355, EPI_ISL_656359, EPI_ISL_656364, EPI_ISL_656368, EPI_ISL_656369, EPI_ISL_656370, EPI_ISL_656371, EPI_ISL_656374, EPI_ISL_656375, EPI_ISL_656376, EPI_ISL_656379, EPI_ISL_656382, EPI_ISL_656383, EPI_ISL_656384, EPI_ISL_656389, EPI_ISL_656390, EPI_ISL_656392, EPI_ISL_656394, EPI_ISL_656395, EPI_ISL_656396, EPI_ISL_656398, EPI_ISL_656399, EPI_ISL_656400, EPI_ISL_656401, EPI_ISL_656402, EPI_ISL_656404, EPI_ISL_656405, EPI_ISL_656406, EPI_ISL_656410, EPI_ISL_656411, EPI_ISL_656414, EPI_ISL_656415, EPI_ISL_656420, EPI_ISL_656421, EPI_ISL_656423, EPI_ISL_656425, EPI_ISL_656426, EPI_ISL_656429, EPI_ISL_656430, EPI_ISL_656432, EPI_ISL_656433, EPI_ISL_656435, EPI_ISL_656439, EPI_ISL_656440, EPI_ISL_656445, EPI_ISL_656446, EPI_ISL_656447, EPI_ISL_656452, EPI_ISL_656453, EPI_ISL_656459, EPI_ISL_656463, EPI_ISL_656465, EPI_ISL_656466, EPI_ISL_656468, EPI_ISL_656469, EPI_ISL_656470, EPI_ISL_656472, EPI_ISL_656473, EPI_ISL_656474, EPI_ISL_656475, EPI_ISL_656477, EPI_ISL_656478, EPI_ISL_656481, EPI_ISL_656482, EPI_ISL_656483, EPI_ISL_656485, EPI_ISL_656486, EPI_ISL_656487, EPI_ISL_656488, EPI_ISL_656491, EPI_ISL_656493, EPI_ISL_656495, EPI_ISL_656496, EPI_ISL_656499, EPI_ISL_656501, EPI_ISL_656504, EPI_ISL_656506, EPI_ISL_656508, EPI_ISL_656510, EPI_ISL_656513, EPI_ISL_656514, EPI_ISL_656515, EPI_ISL_656516, EPI_ISL_656518, EPI_ISL_656520, EPI_ISL_656521, EPI_ISL_656522, EPI_ISL_656523, EPI_ISL_656524, EPI_ISL_656526, EPI_ISL_656529, EPI_ISL_656533, EPI_ISL_656538, EPI_ISL_656539, EPI_ISL_656541, EPI_ISL_656544, EPI_ISL_656549, EPI_ISL_656552, EPI_ISL_656553, EPI_ISL_656554, EPI_ISL_656555, EPI_ISL_656556, EPI_ISL_656561, EPI_ISL_656564, EPI_ISL_656565, EPI_ISL_656569, EPI_ISL_656573, EPI_ISL_656574, EPI_ISL_656577, EPI_ISL_656578, EPI_ISL_656581, EPI_ISL_656582, EPI_ISL_656584, EPI_ISL_656587, EPI_ISL_656589, EPI_ISL_656590, EPI_ISL_656591, EPI_ISL_656592, EPI_ISL_656595, EPI_ISL_656597, EPI_ISL_656599, EPI_ISL_656601, EPI_ISL_656602, EPI_ISL_656603, EPI_ISL_656605, EPI_ISL_656606, EPI_ISL_656607, EPI_ISL_656608, EPI_ISL_656609, EPI_ISL_656610, EPI_ISL_656613, EPI_ISL_656615, EPI_ISL_656616, EPI_ISL_656620, EPI_ISL_656623, EPI_ISL_656625, EPI_ISL_656629, EPI_ISL_656634, EPI_ISL_656635, EPI_ISL_656636, EPI_ISL_656639, EPI_ISL_656640, EPI_ISL_656641, EPI_ISL_656642, EPI_ISL_656644, EPI_ISL_656646, EPI_ISL_656647, EPI_ISL_656649, EPI_ISL_656651, EPI_ISL_656654, EPI_ISL_656655, EPI_ISL_656660, EPI_ISL_656668, EPI_ISL_656671, EPI_ISL_656672, EPI_ISL_656675, EPI_ISL_656676, EPI_ISL_656679, EPI_ISL_656680 | | | |
| see above | Lighthouse Lab in Alderley Park | Wellcome Sanger Institute for the COVID-19 Genomics UK (COG-UK) Consortium | Jacquelyn Wynn, Mairead Hyland, The Lighthouse Lab in Alderley Park and Alex Alderton, Roberto Amato, Sonia Goncalves, Ewan Harrison, David K. Jackson, Ian Johnston, Dominic Kwiatkowski, Cordelia Langford, John Sillitoe on behalf of the Wellcome Sanger Institute COVID-19 Surveillance Team |
| EPI_ISL_656686, EPI_ISL_656697, EPI_ISL_656698, EPI_ISL_656709, EPI_ISL_656710, EPI_ISL_656721, EPI_ISL_656748, EPI_ISL_656751, EPI_ISL_656754, EPI_ISL_656756, EPI_ISL_656761, EPI_ISL_656765, EPI_ISL_656768, EPI_ISL_656770, EPI_ISL_656773, EPI_ISL_656780, EPI_ISL_656782, EPI_ISL_656793, EPI_ISL_656794, EPI_ISL_656802, EPI_ISL_656813, EPI_ISL_656839, EPI_ISL_656852, EPI_ISL_656878, EPI_ISL_656907, EPI_ISL_656914, EPI_ISL_656937, EPI_ISL_656943, EPI_ISL_656944, EPI_ISL_656989, EPI_ISL_656993, EPI_ISL_656998, EPI_ISL_657009, EPI_ISL_657018, EPI_ISL_657356, EPI_ISL_657357, EPI_ISL_657358, EPI_ISL_657362, EPI_ISL_657363, EPI_ISL_657366, EPI_ISL_657368, EPI_ISL_657376, EPI_ISL_657378, EPI_ISL_657380, EPI_ISL_657383, EPI_ISL_657387, EPI_ISL_657389, EPI_ISL_657395, EPI_ISL_657397, EPI_ISL_657399, EPI_ISL_657402, EPI_ISL_657403, EPI_ISL_657422, EPI_ISL_657427, EPI_ISL_657431, EPI_ISL_657432, EPI_ISL_657433, EPI_ISL_657437, EPI_ISL_657444, EPI_ISL_657445, EPI_ISL_657447, EPI_ISL_657448, EPI_ISL_657449, EPI_ISL_657454, EPI_ISL_657470, EPI_ISL_657471, EPI_ISL_657474, EPI_ISL_657475, EPI_ISL_657476, EPI_ISL_657477, EPI_ISL_657489, EPI_ISL_657498, EPI_ISL_657499, EPI_ISL_657501, EPI_ISL_657504, EPI_ISL_657506, EPI_ISL_657507, EPI_ISL_657508, EPI_ISL_657511, EPI_ISL_657514, EPI_ISL_657516, EPI_ISL_657517, EPI_ISL_657521, EPI_ISL_657522, EPI_ISL_657525, EPI_ISL_657528, EPI_ISL_657529, EPI_ISL_657530, EPI_ISL_657534, EPI_ISL_657538, EPI_ISL_657540, EPI_ISL_657541, EPI_ISL_657545, EPI_ISL_657546, EPI_ISL_657548, EPI_ISL_657549, EPI_ISL_657552, EPI_ISL_657559, EPI_ISL_657562, EPI_ISL_657565, EPI_ISL_657566, EPI_ISL_657569, EPI_ISL_657572, EPI_ISL_657573, EPI_ISL_657576, EPI_ISL_657580, EPI_ISL_657582, EPI_ISL_657586, EPI_ISL_657592, EPI_ISL_657595, EPI_ISL_657598, EPI_ISL_657600, EPI_ISL_657603, EPI_ISL_657612, EPI_ISL_657613, EPI_ISL_657618, EPI_ISL_657619, EPI_ISL_657620, EPI_ISL_657621, EPI_ISL_657623, EPI_ISL_657626, EPI_ISL_657628, EPI_ISL_657630, EPI_ISL_657634, EPI_ISL_657641, EPI_ISL_657645, EPI_ISL_657649, EPI_ISL_657653, EPI_ISL_657654, EPI_ISL_657655, EPI_ISL_657656, EPI_ISL_657658, EPI_ISL_657660, EPI_ISL_657668, EPI_ISL_657669, EPI_ISL_657670, EPI_ISL_657673, EPI_ISL_657674, EPI_ISL_657675, EPI_ISL_657677, EPI_ISL_657679, EPI_ISL_657684, EPI_ISL_657685, EPI_ISL_657689, EPI_ISL_657690, EPI_ISL_657691, EPI_ISL_657698, EPI_ISL_657699, EPI_ISL_657701, EPI_ISL_657704, EPI_ISL_657707, EPI_ISL_657708, EPI_ISL_657750, EPI_ISL_657766, EPI_ISL_657833, EPI_ISL_657882, EPI_ISL_657883, EPI_ISL_657907, EPI_ISL_657952, EPI_ISL_657994, EPI_ISL_658011, EPI_ISL_658032, EPI_ISL_658033, EPI_ISL_658045, EPI_ISL_658064, EPI_ISL_658076, EPI_ISL_658099, EPI_ISL_658122, EPI_ISL_658133, EPI_ISL_658142, EPI_ISL_658144, EPI_ISL_658148, EPI_ISL_658153, EPI_ISL_658162, EPI_ISL_658178, EPI_ISL_658196, EPI_ISL_658202, EPI_ISL_658203, EPI_ISL_658206, EPI_ISL_658207, EPI_ISL_658220, EPI_ISL_658228, EPI_ISL_658246, EPI_ISL_658258, EPI_ISL_658263, EPI_ISL_658270, EPI_ISL_658274, EPI_ISL_658307, EPI_ISL_658314, EPI_ISL_658331, EPI_ISL_658343, EPI_ISL_658355, EPI_ISL_658356, EPI_ISL_658361, EPI_ISL_658382, EPI_ISL_658398 | | | |
| see above | Lighthouse Lab in Cambridge | Wellcome Sanger Institute for the COVID-19 Genomics UK (COG-UK) Consortium | Rob Howes, The Lighthouse Lab in Cambridge and Alex Alderton, Roberto Amato, Sonia Goncalves, Ewan Harrison, David K. Jackson, Ian Johnston, Dominic Kwiatkowski, Cordelia Langford, John Sillitoe on behalf of the Wellcome Sanger Institute COVID-19 Surveillance Team |
| EPI_ISL_660221, EPI_ISL_660222, EPI_ISL_660233 | KRISP, KZN Research Innovation and Sequencing Platform | KRISP, KZN Research Innovation and Sequencing Platform | Giandhari J, Pillay S, Lessells R, Mdlalose K, York D, Khan S, Tegally H, Wilkinson E, de Oliveira T |
| EPI_ISL_660608, EPI_ISL_660610, EPI_ISL_660612 | NHLS-IALCH | KRISP, KZN Research Innovation and Sequencing Platform | Giandhari J, Pillay S, Lessells R, Mdlalose K, York D, Khan S, Tegally H, Wilkinson E, de Oliveira T |
| EPI_ISL_661139, EPI_ISL_661140, EPI_ISL_661141, EPI_ISL_661142, EPI_ISL_661143, EPI_ISL_661144, EPI_ISL_661145, EPI_ISL_661146, EPI_ISL_661147, EPI_ISL_661148, EPI_ISL_661149, EPI_ISL_661150, EPI_ISL_661151, EPI_ISL_661152, EPI_ISL_661153, EPI_ISL_661154, EPI_ISL_661155, EPI_ISL_661156, EPI_ISL_661157, EPI_ISL_661158, EPI_ISL_661159, EPI_ISL_661160, EPI_ISL_661161, EPI_ISL_661162, EPI_ISL_661163, EPI_ISL_661164, EPI_ISL_661165, EPI_ISL_661166, EPI_ISL_661167, EPI_ISL_661168, EPI_ISL_661169, EPI_ISL_661170, EPI_ISL_661171, EPI_ISL_661172 | | | |
| see above | Gundersen Molecular Diagnostics Laboratory | Kabara Cancer Research Institute | Craig S. Richmond, Paraic A. Kenny |
| EPI_ISL_661255 | LabPLUS | Institute of Environmental Science and Research (ESR) | Xiaoyun Ren, Matt Storey, Nikki Freed, Muhammad Faisal, Jing Wang, Hermes Perez, Anja Werno, Antje van der Linden, Arlo Upton, Chris Mansell, David Hammer, Dragana Drinkovic, Gary McAuliffe, Hana Sofia Andersson, James Ussher, Jill Sherwood, Josh Freeman, Julia Howard, Juliet Elvy, Mary DeAlmeida, Matt Blakiston, Matthew Rogers, Max Bloomfield, Michael Addidle, Michelle Balm, Sally Roberts, Sarah Jefferies, Sharmini Muttaiyah, Susan Morpeth, Susan Taylor, Timothy Blackmore, Vani Sathyendran, Veronica Playle, Virginia Hope, Erasmus Smit, Lauren Jelly, Olin Silander, Joep de Ligt |
| EPI_ISL_661276, EPI_ISL_661277, EPI_ISL_661302 | Klinisk mikrobiologi | The Public Health Agency of Sweden | Department of Microbiology, The Public Health Agency of Sweden |
| EPI_ISL_661314, EPI_ISL_661315, EPI_ISL_661316, EPI_ISL_661317, EPI_ISL_661318, EPI_ISL_661319, EPI_ISL_661320, EPI_ISL_661321, EPI_ISL_661322, EPI_ISL_661323, EPI_ISL_661324, EPI_ISL_661325, EPI_ISL_661326, EPI_ISL_661327, EPI_ISL_661328, EPI_ISL_661329, EPI_ISL_661330, EPI_ISL_661331, EPI_ISL_661332, EPI_ISL_661333, EPI_ISL_661334, EPI_ISL_661335, EPI_ISL_661336, EPI_ISL_661337, EPI_ISL_661338, EPI_ISL_661339, EPI_ISL_661340, EPI_ISL_661341, EPI_ISL_661342, EPI_ISL_661343, EPI_ISL_661344, EPI_ISL_661345, EPI_ISL_661346, EPI_ISL_661347, EPI_ISL_661348, EPI_ISL_661349, EPI_ISL_661350, EPI_ISL_661351, EPI_ISL_661352, EPI_ISL_661353, EPI_ISL_661354, EPI_ISL_661355, EPI_ISL_661356, EPI_ISL_661357, EPI_ISL_661358, EPI_ISL_661359, EPI_ISL_661360, EPI_ISL_661361, EPI_ISL_661362, EPI_ISL_661363, EPI_ISL_661364, EPI_ISL_661365, EPI_ISL_661366, EPI_ISL_661367, EPI_ISL_661368, EPI_ISL_661369, EPI_ISL_661370, EPI_ISL_661371, EPI_ISL_661372, EPI_ISL_661373, EPI_ISL_661374, EPI_ISL_661375, EPI_ISL_661376, EPI_ISL_661377, EPI_ISL_661378, EPI_ISL_661379, EPI_ISL_661380, EPI_ISL_661381, EPI_ISL_661382, EPI_ISL_661383, EPI_ISL_661384, EPI_ISL_661385, EPI_ISL_661386, EPI_ISL_661387, EPI_ISL_661388, EPI_ISL_661389, EPI_ISL_661390, EPI_ISL_661391, EPI_ISL_661392, EPI_ISL_661393, EPI_ISL_661394, EPI_ISL_661395, EPI_ISL_661396, EPI_ISL_661397, EPI_ISL_661398, EPI_ISL_661399, EPI_ISL_661400, EPI_ISL_661401, EPI_ISL_661402, EPI_ISL_661403, EPI_ISL_661404, EPI_ISL_661405, EPI_ISL_661406, EPI_ISL_661407, EPI_ISL_661408, EPI_ISL_661409, EPI_ISL_661410, EPI_ISL_661411, EPI_ISL_661412, EPI_ISL_661413, EPI_ISL_661414, EPI_ISL_661415, EPI_ISL_661416, EPI_ISL_661417, EPI_ISL_661418, EPI_ISL_661419, EPI_ISL_661420, EPI_ISL_661421, EPI_ISL_661422, EPI_ISL_661423, EPI_ISL_661424, EPI_ISL_661425, EPI_ISL_661426, EPI_ISL_661427, EPI_ISL_661428, EPI_ISL_661429, EPI_ISL_661430, EPI_ISL_661431, EPI_ISL_661432, EPI_ISL_661433, EPI_ISL_661434, EPI_ISL_661435, EPI_ISL_661436, EPI_ISL_661437, EPI_ISL_661438, EPI_ISL_661439, EPI_ISL_661440, EPI_ISL_661441, EPI_ISL_661442, EPI_ISL_661443, EPI_ISL_661444, EPI_ISL_661445, EPI_ISL_661446, EPI_ISL_661447, EPI_ISL_661448, EPI_ISL_661449, EPI_ISL_661450, EPI_ISL_661451, EPI_ISL_661452, EPI_ISL_661453, EPI_ISL_661454, EPI_ISL_661455, EPI_ISL_661456, EPI_ISL_661457, EPI_ISL_661458, EPI_ISL_661459, EPI_ISL_661460, EPI_ISL_661461, EPI_ISL_661462, EPI_ISL_661463, EPI_ISL_661464, EPI_ISL_661465, EPI_ISL_661466, EPI_ISL_661467, EPI_ISL_661468, EPI_ISL_661469, EPI_ISL_661470, EPI_ISL_661471, EPI_ISL_661472, EPI_ISL_661473, EPI_ISL_661474, EPI_ISL_661475, EPI_ISL_661476, EPI_ISL_661477, EPI_ISL_661478, EPI_ISL_661479, EPI_ISL_661480, EPI_ISL_661481, EPI_ISL_661482, EPI_ISL_661483, EPI_ISL_661484, EPI_ISL_661485, EPI_ISL_661486, EPI_ISL_661487, EPI_ISL_661488, EPI_ISL_661489, EPI_ISL_661490, EPI_ISL_661491, EPI_ISL_661492, EPI_ISL_661493, EPI_ISL_661494, EPI_ISL_661495, EPI_ISL_661496, EPI_ISL_661497, EPI_ISL_661498, EPI_ISL_661499, EPI_ISL_661500, EPI_ISL_661501, EPI_ISL_661502, EPI_ISL_661503, EPI_ISL_661504, EPI_ISL_661505, EPI_ISL_661506, EPI_ISL_661507, EPI_ISL_661508, EPI_ISL_661509, EPI_ISL_661510, EPI_ISL_661511, | | | |

| | | | |
|---|---|---|---|
| EPI_ISL_661512, EPI_ISL_661513, EPI_ISL_661514, EPI_ISL_661515, EPI_ISL_661517, EPI_ISL_661518, EPI_ISL_661519, EPI_ISL_661520, EPI_ISL_661521, EPI_ISL_661522, EPI_ISL_661523, EPI_ISL_661524, EPI_ISL_661525, EPI_ISL_661526, EPI_ISL_661527, EPI_ISL_661528, EPI_ISL_661529, EPI_ISL_661530, EPI_ISL_661531, EPI_ISL_661532, EPI_ISL_661533, EPI_ISL_661534, EPI_ISL_661535, EPI_ISL_661536, EPI_ISL_661537, EPI_ISL_661538, EPI_ISL_661539, EPI_ISL_661540, EPI_ISL_661541, EPI_ISL_661542, EPI_ISL_661543, EPI_ISL_661544, EPI_ISL_661545, EPI_ISL_661546, EPI_ISL_661547, EPI_ISL_661548, EPI_ISL_661549, EPI_ISL_661550, EPI_ISL_661551, EPI_ISL_661552, EPI_ISL_661553, EPI_ISL_661554, EPI_ISL_661555, EPI_ISL_661556, EPI_ISL_661557, EPI_ISL_661558, EPI_ISL_661559, EPI_ISL_661560, EPI_ISL_661561, EPI_ISL_661562, EPI_ISL_661563, EPI_ISL_661564, EPI_ISL_661565, EPI_ISL_661566, EPI_ISL_661567, EPI_ISL_661568, EPI_ISL_661569, EPI_ISL_661570, EPI_ISL_661571, EPI_ISL_661572, EPI_ISL_661573, EPI_ISL_661574, EPI_ISL_661575, EPI_ISL_661576, EPI_ISL_661577, EPI_ISL_661578, EPI_ISL_661579, EPI_ISL_661580, EPI_ISL_661581, EPI_ISL_661582, EPI_ISL_661583, EPI_ISL_661584, EPI_ISL_661585, EPI_ISL_661586, EPI_ISL_661587, EPI_ISL_661588, EPI_ISL_661589, EPI_ISL_661590, EPI_ISL_661591, EPI_ISL_661592, EPI_ISL_661593, EPI_ISL_661594, EPI_ISL_661595, EPI_ISL_661596, EPI_ISL_661597, EPI_ISL_661598, EPI_ISL_661599, EPI_ISL_661600, EPI_ISL_661601, EPI_ISL_661602, EPI_ISL_661603, EPI_ISL_661604, EPI_ISL_661605, EPI_ISL_661606, EPI_ISL_661607, EPI_ISL_661608, EPI_ISL_661609, EPI_ISL_661610, EPI_ISL_661611, EPI_ISL_661612, EPI_ISL_661613, EPI_ISL_661614, EPI_ISL_661615, EPI_ISL_661616, EPI_ISL_661617, EPI_ISL_661618, EPI_ISL_661620, EPI_ISL_661621, EPI_ISL_661622, EPI_ISL_661623, EPI_ISL_661624, EPI_ISL_661625, EPI_ISL_661626, EPI_ISL_661627, EPI_ISL_661628, EPI_ISL_661629, EPI_ISL_661630, EPI_ISL_661631 | | | |
| see above | Lighthouse Lab in Glasgow | Wellcome Sanger Institute for the COVID-19 Genomics UK (COG-UK) Consortium | Harper VanSteenhouse, Yumi Kasai, David Gray, Carol Clugston, Anna Dominiczak and Alex Alderton, Roberto Amato, Sonia Goncalves, Ewan Harrison, David K. Jackson, Ian Johnston, Dominic Kwiatkowski, Cordelia Langford, John Sillitoe on behalf of the Wellcome Sanger Institute COVID-19 Surveillance Team |
| EPI_ISL_661640, EPI_ISL_661647, EPI_ISL_661651, EPI_ISL_661652, EPI_ISL_661654, EPI_ISL_661664, EPI_ISL_661675, EPI_ISL_661684, EPI_ISL_661687, EPI_ISL_661691, EPI_ISL_661695, EPI_ISL_661696, EPI_ISL_661710, EPI_ISL_661711, EPI_ISL_661712, EPI_ISL_661713, EPI_ISL_661717, EPI_ISL_661721, EPI_ISL_661729, EPI_ISL_661732, EPI_ISL_661734, EPI_ISL_661741, EPI_ISL_661742, EPI_ISL_661743, EPI_ISL_661747, EPI_ISL_661749, EPI_ISL_661751, EPI_ISL_661752, EPI_ISL_661761, EPI_ISL_661778, EPI_ISL_661781, EPI_ISL_661789, EPI_ISL_661791, EPI_ISL_661797, EPI_ISL_661799, EPI_ISL_661801, EPI_ISL_661805, EPI_ISL_661811, EPI_ISL_661815, EPI_ISL_661817, EPI_ISL_661830, EPI_ISL_661833, EPI_ISL_661834, EPI_ISL_661844, EPI_ISL_661845, EPI_ISL_661863, EPI_ISL_661869, EPI_ISL_661874, EPI_ISL_661876, EPI_ISL_661881, EPI_ISL_661883, EPI_ISL_661886, EPI_ISL_661887, EPI_ISL_661895, EPI_ISL_661897, EPI_ISL_661900, EPI_ISL_661902, EPI_ISL_661903, EPI_ISL_661917, EPI_ISL_661918, EPI_ISL_661926, EPI_ISL_661930, EPI_ISL_661941, EPI_ISL_661947, EPI_ISL_661953, EPI_ISL_661954 | | | |
| see above | Lighthouse Lab in Cambridge | Wellcome Sanger Institute for the COVID-19 Genomics UK (COG-UK) Consortium | Rob Howes, The Lighthouse Lab in Cambridge and Alex Alderton, Roberto Amato, Sonia Goncalves, Ewan Harrison, David K. Jackson, Ian Johnston, Dominic Kwiatkowski, Cordelia Langford, John Sillitoe on behalf of the Wellcome Sanger Institute COVID-19 Surveillance Team |
| EPI_ISL_661957, EPI_ISL_661959, EPI_ISL_661960, EPI_ISL_661963, EPI_ISL_661967, EPI_ISL_661969, EPI_ISL_661971, EPI_ISL_661972, EPI_ISL_661973, EPI_ISL_661975, EPI_ISL_661976, EPI_ISL_661979, EPI_ISL_661981, EPI_ISL_661984, EPI_ISL_661987, EPI_ISL_661993, EPI_ISL_661994, EPI_ISL_661996, EPI_ISL_661997, EPI_ISL_662003, EPI_ISL_662005, EPI_ISL_662007, EPI_ISL_662009, EPI_ISL_662011, EPI_ISL_662012, EPI_ISL_662016, EPI_ISL_662021, EPI_ISL_662023, EPI_ISL_662024, EPI_ISL_662026, EPI_ISL_662027, EPI_ISL_662028, EPI_ISL_662030, EPI_ISL_662033, EPI_ISL_662035, EPI_ISL_662037, EPI_ISL_662039, EPI_ISL_662040, EPI_ISL_662041, EPI_ISL_662042, EPI_ISL_662044, EPI_ISL_662052, EPI_ISL_662054, EPI_ISL_662059, EPI_ISL_662064, EPI_ISL_662066, EPI_ISL_662069, EPI_ISL_662072, EPI_ISL_662073, EPI_ISL_662075, EPI_ISL_662076, EPI_ISL_662077, EPI_ISL_662082, EPI_ISL_662083, EPI_ISL_662084, EPI_ISL_662085, EPI_ISL_662087, EPI_ISL_662088, EPI_ISL_662089, EPI_ISL_662090, EPI_ISL_662091, EPI_ISL_662092, EPI_ISL_662094, EPI_ISL_662095, EPI_ISL_662096, EPI_ISL_662102, EPI_ISL_662103, EPI_ISL_662104, EPI_ISL_662109, EPI_ISL_662110, EPI_ISL_662111, EPI_ISL_662112, EPI_ISL_662113, EPI_ISL_662117, EPI_ISL_662119, EPI_ISL_662121, EPI_ISL_662125, EPI_ISL_662126, EPI_ISL_662127, EPI_ISL_662128, EPI_ISL_662130, EPI_ISL_662134, EPI_ISL_662136, EPI_ISL_662139, EPI_ISL_662142, EPI_ISL_662144, EPI_ISL_662147, EPI_ISL_662149, EPI_ISL_662152, EPI_ISL_662153, EPI_ISL_662154, EPI_ISL_662155, EPI_ISL_662157, EPI_ISL_662158, EPI_ISL_662161, EPI_ISL_662168, EPI_ISL_662171, EPI_ISL_662172, EPI_ISL_662180, EPI_ISL_662185, EPI_ISL_662188, EPI_ISL_662189, EPI_ISL_662190, EPI_ISL_662193, EPI_ISL_662195, EPI_ISL_662197, EPI_ISL_662199, EPI_ISL_662201, EPI_ISL_662204, EPI_ISL_662205, EPI_ISL_662210, EPI_ISL_662211, EPI_ISL_662213, EPI_ISL_662216, EPI_ISL_662220, EPI_ISL_662221, EPI_ISL_662222, EPI_ISL_662224, EPI_ISL_662225, EPI_ISL_662226, EPI_ISL_662230, EPI_ISL_662232, EPI_ISL_662241, EPI_ISL_662242, EPI_ISL_662243, EPI_ISL_662246, EPI_ISL_662250, EPI_ISL_662251, EPI_ISL_662253, EPI_ISL_662256, EPI_ISL_662257, EPI_ISL_662259, EPI_ISL_662261, EPI_ISL_662262, EPI_ISL_662264, EPI_ISL_662268, EPI_ISL_662271, EPI_ISL_662277, EPI_ISL_662278, EPI_ISL_662280, EPI_ISL_662282, EPI_ISL_662283, EPI_ISL_662284, EPI_ISL_662287, EPI_ISL_662288, EPI_ISL_662289, EPI_ISL_662291, EPI_ISL_662293, EPI_ISL_662294, EPI_ISL_662295, EPI_ISL_662297, EPI_ISL_662298, EPI_ISL_662299, EPI_ISL_662301, EPI_ISL_662302, EPI_ISL_662303, EPI_ISL_662304, EPI_ISL_662306, EPI_ISL_662307, EPI_ISL_662310, EPI_ISL_662311, EPI_ISL_662313, EPI_ISL_662314, EPI_ISL_662315, EPI_ISL_662316, EPI_ISL_662317, EPI_ISL_662318, EPI_ISL_662319, EPI_ISL_662321, EPI_ISL_662322, EPI_ISL_662323, EPI_ISL_662324, EPI_ISL_662325, EPI_ISL_662326, EPI_ISL_662328, EPI_ISL_662329, EPI_ISL_662330, EPI_ISL_662331, EPI_ISL_662332, EPI_ISL_662333, EPI_ISL_662335, EPI_ISL_662336, EPI_ISL_662337, EPI_ISL_662338, EPI_ISL_662339, EPI_ISL_662340, EPI_ISL_662341, EPI_ISL_662345, EPI_ISL_662346, EPI_ISL_662347, EPI_ISL_662348, EPI_ISL_662349, EPI_ISL_662350, EPI_ISL_662351, EPI_ISL_662353, EPI_ISL_662355, EPI_ISL_662357, EPI_ISL_662360, EPI_ISL_662361, EPI_ISL_662362, EPI_ISL_662363, EPI_ISL_662364, EPI_ISL_662366, EPI_ISL_662367, EPI_ISL_662368, EPI_ISL_662370, EPI_ISL_662371, EPI_ISL_662372, EPI_ISL_662376, EPI_ISL_662377, EPI_ISL_662378, EPI_ISL_662379, EPI_ISL_662380, EPI_ISL_662383, EPI_ISL_662384, EPI_ISL_662386, EPI_ISL_662387, EPI_ISL_662388, EPI_ISL_662392, EPI_ISL_662395, EPI_ISL_662396, EPI_ISL_662397, EPI_ISL_662398, EPI_ISL_662399, EPI_ISL_662400, EPI_ISL_662401, EPI_ISL_662402, EPI_ISL_662404, EPI_ISL_662406, EPI_ISL_662407, EPI_ISL_662408, EPI_ISL_662409, EPI_ISL_662413, EPI_ISL_662414, EPI_ISL_662416, EPI_ISL_662417, EPI_ISL_662420, EPI_ISL_662421, EPI_ISL_662422, EPI_ISL_662423, EPI_ISL_662424, EPI_ISL_662425, EPI_ISL_662426, EPI_ISL_662428, EPI_ISL_662430, EPI_ISL_662432, EPI_ISL_662434, EPI_ISL_662435, EPI_ISL_662436, EPI_ISL_662437, EPI_ISL_662438, EPI_ISL_662439, EPI_ISL_662440, EPI_ISL_662441, EPI_ISL_662442, EPI_ISL_662443, EPI_ISL_662445, EPI_ISL_662447, EPI_ISL_662448, EPI_ISL_662449, EPI_ISL_662450, EPI_ISL_662451, EPI_ISL_662453, EPI_ISL_662454, EPI_ISL_662455, EPI_ISL_662456, EPI_ISL_662458, EPI_ISL_662459, EPI_ISL_662460, EPI_ISL_662461, EPI_ISL_662464, EPI_ISL_662465, EPI_ISL_662466, EPI_ISL_662467, EPI_ISL_662469, EPI_ISL_662470, EPI_ISL_662471, EPI_ISL_662473, EPI_ISL_662475, EPI_ISL_662476, EPI_ISL_662479, EPI_ISL_662481, EPI_ISL_662482, EPI_ISL_662483, EPI_ISL_662484, EPI_ISL_662485, EPI_ISL_662490, EPI_ISL_662491, EPI_ISL_662493, EPI_ISL_662494, EPI_ISL_662495, EPI_ISL_662497, EPI_ISL_662498, EPI_ISL_662500, EPI_ISL_662501, EPI_ISL_662503, EPI_ISL_662504, EPI_ISL_662505, EPI_ISL_662506, EPI_ISL_662509, EPI_ISL_662510, EPI_ISL_662511, EPI_ISL_662512, EPI_ISL_662513, EPI_ISL_662515, EPI_ISL_662516, EPI_ISL_662517, EPI_ISL_662519, EPI_ISL_662521, EPI_ISL_662522, EPI_ISL_662523, EPI_ISL_662524, EPI_ISL_662525, EPI_ISL_662527, EPI_ISL_662528, EPI_ISL_662529, EPI_ISL_662530, EPI_ISL_662531, EPI_ISL_662532, EPI_ISL_662533, EPI_ISL_662534, EPI_ISL_662536, EPI_ISL_662537, EPI_ISL_662538, EPI_ISL_662539, EPI_ISL_662540, EPI_ISL_662541, EPI_ISL_662542, EPI_ISL_662543, EPI_ISL_662544, EPI_ISL_662545, EPI_ISL_662546, EPI_ISL_662547, EPI_ISL_662548, EPI_ISL_662550, EPI_ISL_662551, EPI_ISL_662552, EPI_ISL_662554, EPI_ISL_662555, EPI_ISL_662556, EPI_ISL_662557, EPI_ISL_662559, EPI_ISL_662560, EPI_ISL_662561, EPI_ISL_662562, EPI_ISL_662563, EPI_ISL_662565, EPI_ISL_662566, EPI_ISL_662567, EPI_ISL_662568, EPI_ISL_662569, EPI_ISL_662570, EPI_ISL_662571, EPI_ISL_662572, EPI_ISL_662573, EPI_ISL_662574, EPI_ISL_662577, EPI_ISL_662582, EPI_ISL_662584, EPI_ISL_662585, EPI_ISL_662586, EPI_ISL_662587, EPI_ISL_662589, EPI_ISL_662590, EPI_ISL_662591, EPI_ISL_662592, EPI_ISL_662594, EPI_ISL_662595, EPI_ISL_662596, EPI_ISL_662598, EPI_ISL_662599, EPI_ISL_662600, EPI_ISL_662601, EPI_ISL_662602, EPI_ISL_662603, EPI_ISL_662606, EPI_ISL_662607, EPI_ISL_662608, EPI_ISL_662611, EPI_ISL_662612, EPI_ISL_662613, EPI_ISL_662615, EPI_ISL_662618, EPI_ISL_662619, EPI_ISL_662620 | | | |
| see above | Lighthouse Lab in Glasgow | Wellcome Sanger Institute for the COVID-19 Genomics UK (COG-UK) Consortium | Harper VanSteenhouse, Yumi Kasai, David Gray, Carol Clugston, Anna Dominiczak and Alex Alderton, Roberto Amato, Sonia Goncalves, Ewan Harrison, David K. Jackson, Ian Johnston, Dominic Kwiatkowski, Cordelia Langford, John Sillitoe on behalf of the Wellcome Sanger Institute COVID-19 Surveillance Team |
| EPI_ISL_662623, EPI_ISL_662624 | Lighthouse Lab in Alderley Park | Wellcome Sanger Institute for the COVID-19 Genomics UK (COG-UK) Consortium | Jacquelyn Wynn, Mairead Hyland, The Lighthouse Lab in Alderley Park and Alex Alderton, Roberto Amato, Sonia Goncalves, Ewan Harrison, David K. Jackson, Ian Johnston, Dominic Kwiatkowski, Cordelia Langford, John Sillitoe on behalf of the Wellcome Sanger Institute COVID-19 Surveillance Team |
| EPI_ISL_662627 | Lighthouse Lab in Glasgow | Wellcome Sanger Institute for the COVID-19 Genomics UK (COG-UK) Consortium | Harper VanSteenhouse, Yumi Kasai, David Gray, Carol Clugston, Anna Dominiczak and Alex Alderton, Roberto Amato, Sonia Goncalves, Ewan Harrison, David K. Jackson, Ian Johnston, Dominic Kwiatkowski, Cordelia Langford, John Sillitoe on behalf of the Wellcome Sanger Institute COVID-19 Surveillance Team |
| EPI_ISL_662630 | Lighthouse Lab in Alderley Park | Wellcome Sanger Institute for the COVID-19 Genomics UK (COG-UK) Consortium | Jacquelyn Wynn, Mairead Hyland, The Lighthouse Lab in Alderley Park and Alex Alderton, Roberto Amato, Sonia Goncalves, Ewan Harrison, David K. Jackson, Ian Johnston, Dominic Kwiatkowski, Cordelia Langford, John Sillitoe on behalf of the Wellcome Sanger Institute COVID-19 Surveillance Team |
| EPI_ISL_664049, EPI_ISL_664050, EPI_ISL_664053, EPI_ISL_664054, EPI_ISL_664055, EPI_ISL_664056, EPI_ISL_664057, EPI_ISL_664058, EPI_ISL_664060, EPI_ISL_664061, EPI_ISL_664062, EPI_ISL_664063, EPI_ISL_664064, EPI_ISL_664065, EPI_ISL_664066, EPI_ISL_664067, EPI_ISL_664068, EPI_ISL_664069, EPI_ISL_664070, EPI_ISL_664071, EPI_ISL_664072, EPI_ISL_664073, EPI_ISL_664074, EPI_ISL_664075, EPI_ISL_664076, EPI_ISL_664077, EPI_ISL_664078, EPI_ISL_664079, EPI_ISL_664080, EPI_ISL_664081, EPI_ISL_664082, EPI_ISL_664083, EPI_ISL_664084, EPI_ISL_664085, EPI_ISL_664086, EPI_ISL_664087, EPI_ISL_664088, EPI_ISL_664093, EPI_ISL_664094, EPI_ISL_664095, EPI_ISL_664096, EPI_ISL_664098, EPI_ISL_664099, EPI_ISL_664100 | | | |
| see above | Respiratory Virus Unit, Microbiology Services Colindale, Public Health England | COVID-19 Genomics UK (COG-UK) Consortium | PHE Covid Sequencing Team |
| EPI_ISL_664109 | Department of Pathology, University of Cambridge | COVID-19 Genomics UK (COG-UK) Consortium | Aminu S. Jahun, Yasmin Chaudhry, Grant Hall, Iliana Georgana, Myra Hosmillo, Martin D. Curran, Malte Pinckert, Surendra Parmar, Ian Goodfellow |
| EPI_ISL_664115 | University of Exeter | COVID-19 Genomics UK (COG-UK) Consortium | Ben Temperton,Aaron Jeffries,Michelle Michelsen,Joanna Warwick-Dugdale,Audrey Farbos,Robyn Manley,Stephen Michell,Jane Masoli |
| EPI_ISL_664126 | Department of Pathology, University of Cambridge | COVID-19 Genomics UK (COG-UK) Consortium | Aminu S. Jahun, Yasmin Chaudhry, Grant Hall, Iliana Georgana, Myra Hosmillo, Martin D. Curran, Malte Pinckert, Surendra Parmar, Ian Goodfellow |
| EPI_ISL_664136 | University of Exeter | COVID-19 Genomics UK (COG-UK) Consortium | Ben Temperton,Aaron Jeffries,Michelle Michelsen,Joanna Warwick-Dugdale,Audrey Farbos,Robyn Manley,Stephen Michell,Jane Masoli |
| EPI_ISL_664143 | Department of Pathology, University of Cambridge | COVID-19 Genomics UK (COG-UK) Consortium | Aminu S. Jahun, Yasmin Chaudhry, Grant Hall, Iliana Georgana, Myra Hosmillo, Martin D. Curran, Malte Pinckert, Surendra Parmar, Ian Goodfellow |
| EPI_ISL_664144, EPI_ISL_664145 | University of Exeter | COVID-19 Genomics UK (COG-UK) Consortium | Ben Temperton,Aaron Jeffries,Michelle Michelsen,Joanna Warwick-Dugdale,Audrey Farbos,Robyn Manley,Stephen Michell,Jane Masoli |
| EPI_ISL_664152 | Department of Pathology, University of Cambridge | COVID-19 Genomics UK (COG-UK) Consortium | Aminu S. Jahun, Yasmin Chaudhry, Grant Hall, Iliana Georgana, Myra Hosmillo, Martin D. Curran, Malte Pinckert, Surendra Parmar, Ian Goodfellow |
| EPI_ISL_664153 | University of Exeter | COVID-19 Genomics UK (COG-UK) Consortium | Ben Temperton,Aaron Jeffries,Michelle Michelsen,Joanna Warwick-Dugdale,Audrey Farbos,Robyn Manley,Stephen Michell,Jane Masoli |
| EPI_ISL_664157, EPI_ISL_664160, EPI_ISL_664161 | Department of Pathology, University of Cambridge | COVID-19 Genomics UK (COG-UK) Consortium | Aminu S. Jahun, Yasmin Chaudhry, Grant Hall, Iliana Georgana, Myra Hosmillo, Martin D. Curran, Malte Pinckert, Surendra Parmar, Ian Goodfellow |
| EPI_ISL_664173 | University of Exeter | COVID-19 Genomics UK (COG-UK) Consortium | Ben Temperton,Aaron Jeffries,Michelle Michelsen,Joanna Warwick-Dugdale,Audrey Farbos,Robyn Manley,Stephen Michell,Jane Masoli |
| EPI_ISL_664174, EPI_ISL_664177, EPI_ISL_664178, EPI_ISL_664180, EPI_ISL_664181, EPI_ISL_664182, EPI_ISL_664185 | Department of Pathology, University of Cambridge | COVID-19 Genomics UK (COG-UK) Consortium | Aminu S. Jahun, Yasmin Chaudhry, Grant Hall, Iliana Georgana, Myra Hosmillo, Martin D. Curran, Malte Pinckert, Surendra Parmar, Ian Goodfellow |
| EPI_ISL_664193, EPI_ISL_664194, EPI_ISL_664210, EPI_ISL_664211 | University of Exeter | COVID-19 Genomics UK (COG-UK) Consortium | Ben Temperton,Aaron Jeffries,Michelle Michelsen,Joanna Warwick-Dugdale,Audrey Farbos,Robyn Manley,Stephen Michell,Jane Masoli |
| EPI_ISL_664213 | Department of Pathology, University of Cambridge | COVID-19 Genomics UK (COG-UK) Consortium | Aminu S. Jahun, Yasmin Chaudhry, Grant Hall, Iliana Georgana, Myra Hosmillo, Martin D. Curran, Malte Pinckert, Surendra Parmar, Ian Goodfellow |
| EPI_ISL_664214, EPI_ISL_664216 | University of Exeter | COVID-19 Genomics UK (COG-UK) Consortium | Ben Temperton,Aaron Jeffries,Michelle Michelsen,Joanna Warwick-Dugdale,Audrey Farbos,Robyn Manley,Stephen Michell,Jane Masoli |
| EPI_ISL_664221, EPI_ISL_664237 | Department of Pathology, University of Cambridge | COVID-19 Genomics UK (COG-UK) Consortium | Aminu S. Jahun, Yasmin Chaudhry, Grant Hall, Iliana Georgana, Myra Hosmillo, Martin D. Curran, Malte Pinckert, Surendra Parmar, Ian Goodfellow |
| EPI_ISL_664239, EPI_ISL_664254, EPI_ISL_664256, EPI_ISL_664258, EPI_ISL_664259, EPI_ISL_664260, EPI_ISL_664262, EPI_ISL_664264, EPI_ISL_664266, EPI_ISL_664267, EPI_ISL_664268, EPI_ISL_664269, EPI_ISL_664270, EPI_ISL_664271, EPI_ISL_664272, EPI_ISL_664273, EPI_ISL_664274, EPI_ISL_664275, | | | |

| | | | |
|---|---|---|---|
| EPI_ISL_664279, EPI_ISL_664285 | | | |
| see above | University of Exeter | COVID-19 Genomics UK (COG-UK) Consortium | Ben Temperton,Aaron Jeffries,Michelle Michelsen,Joanna Warwick-Dugdale,Audrey Farbos,Robyn Manley,Stephen Michell,Jane Masoli |
| EPI_ISL_664286 | Department of Pathology, University of Cambridge | COVID-19 Genomics UK (COG-UK) Consortium | Aminu S. Jahun, Yasmin Chaudhry, Grant Hall, Iliana Georgana, Myra Hosmillo, Martin D. Curran, Malte Pinckert, Surendra Parmar, Ian Goodfellow |
| EPI_ISL_664296, EPI_ISL_664297 | University of Exeter | COVID-19 Genomics UK (COG-UK) Consortium | Ben Temperton,Aaron Jeffries,Michelle Michelsen,Joanna Warwick-Dugdale,Audrey Farbos,Robyn Manley,Stephen Michell,Jane Masoli |
| EPI_ISL_664306, EPI_ISL_664307, EPI_ISL_664308, EPI_ISL_664312, EPI_ISL_664313, EPI_ISL_664314, EPI_ISL_664322, EPI_ISL_664323, EPI_ISL_664324 | Department of Pathology, University of Cambridge | COVID-19 Genomics UK (COG-UK) Consortium | Aminu S. Jahun, Yasmin Chaudhry, Grant Hall, Iliana Georgana, Myra Hosmillo, Martin D. Curran, Malte Pinckert, Surendra Parmar, Ian Goodfellow |
| EPI_ISL_664341 | Liverpool Clinical Laboratories | COVID-19 Genomics UK (COG-UK) Consortium | Sam Haldenby, Anita Lucaci, Steve Paterson, Julian Hiscox, Alistair Darby, M Almsaud, A Alrezaihi, Muhannad Alruwaili, Stuart D Armstrong, Jones Benjamin, Eleanor G Bentley, Anu Chawla, Jordan J Clark, Angela Cowell, Richard Eccles, Isabel García-Dorival, Matthew Gemmell, Alessandro Gerada, PKF Gilmore, Richard Gregory, Ximeng Han, Catherine Hartley, Margaret Hughes, Miren Iturriza-Gomara, James Johnson, L Luu, Jenifer Manson, Charlotte Nelson, Elaine O'Toole, Cassie Olateju, Rebekah Penrice-Randal , Lucille Rainbow, N.P Randle, Trevor Ian Robinson, Parul Sharma, Ghada T Shawli, James P Stewart, Neil Swainston, Ecaterina Vamos, Joanne Watts, Mark Whitehead |
| EPI_ISL_664401, EPI_ISL_664408, EPI_ISL_664409, EPI_ISL_664410, EPI_ISL_664411, EPI_ISL_664412, EPI_ISL_664413, EPI_ISL_664414, EPI_ISL_664415, EPI_ISL_664416, EPI_ISL_664417, EPI_ISL_664418, EPI_ISL_664419, EPI_ISL_664420 | | | |
| see above | University of Exeter | COVID-19 Genomics UK (COG-UK) Consortium | Ben Temperton,Aaron Jeffries,Michelle Michelsen,Joanna Warwick-Dugdale,Audrey Farbos,Robyn Manley,Stephen Michell,Jane Masoli |
| EPI_ISL_664421, EPI_ISL_664428, EPI_ISL_664432, EPI_ISL_664433, EPI_ISL_664434 | Department of Pathology, University of Cambridge | COVID-19 Genomics UK (COG-UK) Consortium | Aminu S. Jahun, Yasmin Chaudhry, Grant Hall, Iliana Georgana, Myra Hosmillo, Martin D. Curran, Malte Pinckert, Surendra Parmar, Ian Goodfellow |
| EPI_ISL_664436 | Liverpool Clinical Laboratories | COVID-19 Genomics UK (COG-UK) Consortium | Sam Haldenby, Anita Lucaci, Steve Paterson, Julian Hiscox, Alistair Darby, M Almsaud, A Alrezaihi, Muhannad Alruwaili, Stuart D Armstrong, Jones Benjamin, Eleanor G Bentley, Anu Chawla, Jordan J Clark, Angela Cowell, Richard Eccles, Isabel García-Dorival, Matthew Gemmell, Alessandro Gerada, PKF Gilmore, Richard Gregory, Ximeng Han, Catherine Hartley, Margaret Hughes, Miren Iturriza-Gomara, James Johnson, L Luu, Jenifer Manson, Charlotte Nelson, Elaine O'Toole, Cassie Olateju, Rebekah Penrice-Randal , Lucille Rainbow, N.P Randle, Trevor Ian Robinson, Parul Sharma, Ghada T Shawli, James P Stewart, Neil Swainston, Ecaterina Vamos, Joanne Watts, Mark Whitehead |
| EPI_ISL_664459 | University of Exeter | COVID-19 Genomics UK (COG-UK) Consortium | Ben Temperton,Aaron Jeffries,Michelle Michelsen,Joanna Warwick-Dugdale,Audrey Farbos,Robyn Manley,Stephen Michell,Jane Masoli |
| EPI_ISL_664481 | Liverpool Clinical Laboratories | COVID-19 Genomics UK (COG-UK) Consortium | Sam Haldenby, Anita Lucaci, Steve Paterson, Julian Hiscox, Alistair Darby, M Almsaud, A Alrezaihi, Muhannad Alruwaili, Stuart D Armstrong, Jones Benjamin, Eleanor G Bentley, Anu Chawla, Jordan J Clark, Angela Cowell, Richard Eccles, Isabel García-Dorival, Matthew Gemmell, Alessandro Gerada, PKF Gilmore, Richard Gregory, Ximeng Han, Catherine Hartley, Margaret Hughes, Miren Iturriza-Gomara, James Johnson, L Luu, Jenifer Manson, Charlotte Nelson, Elaine O'Toole, Cassie Olateju, Rebekah Penrice-Randal , Lucille Rainbow, N.P Randle, Trevor Ian Robinson, Parul Sharma, Ghada T Shawli, James P Stewart, Neil Swainston, Ecaterina Vamos, Joanne Watts, Mark Whitehead |
| EPI_ISL_664509 | University of Exeter | COVID-19 Genomics UK (COG-UK) Consortium | Ben Temperton,Aaron Jeffries,Michelle Michelsen,Joanna Warwick-Dugdale,Audrey Farbos,Robyn Manley,Stephen Michell,Jane Masoli |
| EPI_ISL_664514, EPI_ISL_664518, EPI_ISL_664528, EPI_ISL_664534 | Department of Pathology, University of Cambridge | COVID-19 Genomics UK (COG-UK) Consortium | Aminu S. Jahun, Yasmin Chaudhry, Grant Hall, Iliana Georgana, Myra Hosmillo, Martin D. Curran, Malte Pinckert, Surendra Parmar, Ian Goodfellow |
| EPI_ISL_664539 | Liverpool Clinical Laboratories | COVID-19 Genomics UK (COG-UK) Consortium | Sam Haldenby, Anita Lucaci, Steve Paterson, Julian Hiscox, Alistair Darby, M Almsaud, A Alrezaihi, Muhannad Alruwaili, Stuart D Armstrong, Jones Benjamin, Eleanor G Bentley, Anu Chawla, Jordan J Clark, Angela Cowell, Richard Eccles, Isabel García-Dorival, Matthew Gemmell, Alessandro Gerada, PKF Gilmore, Richard Gregory, Ximeng Han, Catherine Hartley, Margaret Hughes, Miren Iturriza-Gomara, James Johnson, L Luu, Jenifer Manson, Charlotte Nelson, Elaine O'Toole, Cassie Olateju, Rebekah Penrice-Randal , Lucille Rainbow, N.P Randle, Trevor Ian Robinson, Parul Sharma, Ghada T Shawli, James P Stewart, Neil Swainston, Ecaterina Vamos, Joanne Watts, Mark Whitehead |
| EPI_ISL_664545 | University of Exeter | COVID-19 Genomics UK (COG-UK) Consortium | Ben Temperton,Aaron Jeffries,Michelle Michelsen,Joanna Warwick-Dugdale,Audrey Farbos,Robyn Manley,Stephen Michell,Jane Masoli |
| EPI_ISL_664550, EPI_ISL_664551, EPI_ISL_664580, EPI_ISL_664581, EPI_ISL_664584, EPI_ISL_664585, EPI_ISL_664586, EPI_ISL_664587 | Department of Pathology, University of Cambridge | COVID-19 Genomics UK (COG-UK) Consortium | Aminu S. Jahun, Yasmin Chaudhry, Grant Hall, Iliana Georgana, Myra Hosmillo, Martin D. Curran, Malte Pinckert, Surendra Parmar, Ian Goodfellow |
| EPI_ISL_664595, EPI_ISL_664596, EPI_ISL_664597 | University of Exeter | COVID-19 Genomics UK (COG-UK) Consortium | Ben Temperton,Aaron Jeffries,Michelle Michelsen,Joanna Warwick-Dugdale,Audrey Farbos,Robyn Manley,Stephen Michell,Jane Masoli |
| EPI_ISL_664598 | Department of Pathology, University of Cambridge | COVID-19 Genomics UK (COG-UK) Consortium | Aminu S. Jahun, Yasmin Chaudhry, Grant Hall, Iliana Georgana, Myra Hosmillo, Martin D. Curran, Malte Pinckert, Surendra Parmar, Ian Goodfellow |
| EPI_ISL_664608, EPI_ISL_664611 | University of Exeter | COVID-19 Genomics UK (COG-UK) Consortium | Ben Temperton,Aaron Jeffries,Michelle Michelsen,Joanna Warwick-Dugdale,Audrey Farbos,Robyn Manley,Stephen Michell,Jane Masoli |
| EPI_ISL_664615 | Liverpool Clinical Laboratories | COVID-19 Genomics UK (COG-UK) Consortium | Sam Haldenby, Anita Lucaci, Steve Paterson, Julian Hiscox, Alistair Darby, M Almsaud, A Alrezaihi, Muhannad Alruwaili, Stuart D Armstrong, Jones Benjamin, Eleanor G Bentley, Anu Chawla, Jordan J Clark, Angela Cowell, Richard Eccles, Isabel García-Dorival, Matthew Gemmell, Alessandro Gerada, PKF Gilmore, Richard Gregory, Ximeng Han, Catherine Hartley, Margaret Hughes, Miren Iturriza-Gomara, James Johnson, L Luu, Jenifer Manson, Charlotte Nelson, Elaine O'Toole, Cassie Olateju, Rebekah Penrice-Randal , Lucille Rainbow, N.P Randle, Trevor Ian Robinson, Parul Sharma, Ghada T Shawli, James P Stewart, Neil Swainston, Ecaterina Vamos, Joanne Watts, Mark Whitehead |
| EPI_ISL_664622 | Department of Pathology, University of Cambridge | COVID-19 Genomics UK (COG-UK) Consortium | Aminu S. Jahun, Yasmin Chaudhry, Grant Hall, Iliana Georgana, Myra Hosmillo, Martin D. Curran, Malte Pinckert, Surendra Parmar, Ian Goodfellow |
| EPI_ISL_664624 | Liverpool Clinical Laboratories | COVID-19 Genomics UK (COG-UK) Consortium | Sam Haldenby, Anita Lucaci, Steve Paterson, Julian Hiscox, Alistair Darby, M Almsaud, A Alrezaihi, Muhannad Alruwaili, Stuart D Armstrong, Jones Benjamin, Eleanor G Bentley, Anu Chawla, Jordan J Clark, Angela Cowell, Richard Eccles, Isabel García-Dorival, Matthew Gemmell, Alessandro Gerada, PKF Gilmore, Richard Gregory, Ximeng Han, Catherine Hartley, Margaret Hughes, Miren Iturriza-Gomara, James Johnson, L Luu, Jenifer Manson, Charlotte Nelson, Elaine O'Toole, Cassie Olateju, Rebekah Penrice-Randal , Lucille Rainbow, N.P Randle, Trevor Ian Robinson, Parul Sharma, Ghada T Shawli, James P Stewart, Neil Swainston, Ecaterina Vamos, Joanne Watts, Mark Whitehead |
| EPI_ISL_664625 | Department of Pathology, University of Cambridge | COVID-19 Genomics UK (COG-UK) Consortium | Aminu S. Jahun, Yasmin Chaudhry, Grant Hall, Iliana Georgana, Myra Hosmillo, Martin D. Curran, Malte Pinckert, Surendra Parmar, Ian Goodfellow |
| EPI_ISL_664626 | University of Exeter | COVID-19 Genomics UK (COG-UK) Consortium | Ben Temperton,Aaron Jeffries,Michelle Michelsen,Joanna Warwick-Dugdale,Audrey Farbos,Robyn Manley,Stephen Michell,Jane Masoli |
| EPI_ISL_664627 | Department of Pathology, University of Cambridge | COVID-19 Genomics UK (COG-UK) Consortium | Aminu S. Jahun, Yasmin Chaudhry, Grant Hall, Iliana Georgana, Myra Hosmillo, Martin D. Curran, Malte Pinckert, Surendra Parmar, Ian Goodfellow |
| EPI_ISL_664640 | Liverpool Clinical Laboratories | COVID-19 Genomics UK (COG-UK) Consortium | Sam Haldenby, Anita Lucaci, Steve Paterson, Julian Hiscox, Alistair Darby, M Almsaud, A Alrezaihi, Muhannad Alruwaili, Stuart D Armstrong, Jones Benjamin, Eleanor G Bentley, Anu Chawla, Jordan J Clark, Angela Cowell, Richard Eccles, Isabel García-Dorival, Matthew Gemmell, Alessandro Gerada, PKF Gilmore, Richard Gregory, Ximeng Han, Catherine Hartley, Margaret Hughes, Miren Iturriza-Gomara, James Johnson, L Luu, Jenifer Manson, Charlotte Nelson, Elaine O'Toole, Cassie Olateju, Rebekah Penrice-Randal , Lucille Rainbow, N.P Randle, Trevor Ian Robinson, Parul Sharma, Ghada T Shawli, James P Stewart, Neil Swainston, Ecaterina Vamos, Joanne Watts, Mark Whitehead |
| EPI_ISL_664647 | Department of Pathology, University of Cambridge | COVID-19 Genomics UK (COG-UK) Consortium | Aminu S. Jahun, Yasmin Chaudhry, Grant Hall, Iliana Georgana, Myra Hosmillo, Martin D. Curran, Malte Pinckert, Surendra Parmar, Ian Goodfellow |
| EPI_ISL_664648 | Liverpool Clinical Laboratories | COVID-19 Genomics UK (COG-UK) Consortium | Sam Haldenby, Anita Lucaci, Steve Paterson, Julian Hiscox, Alistair Darby, M Almsaud, A Alrezaihi, Muhannad Alruwaili, Stuart D Armstrong, Jones Benjamin, Eleanor G Bentley, Anu Chawla, Jordan J Clark, Angela Cowell, Richard Eccles, Isabel García-Dorival, Matthew Gemmell, Alessandro Gerada, PKF Gilmore, Richard Gregory, Ximeng Han, Catherine Hartley, Margaret Hughes, Miren Iturriza-Gomara, James Johnson, L Luu, Jenifer Manson, Charlotte Nelson, Elaine O'Toole, Cassie Olateju, Rebekah Penrice-Randal , Lucille Rainbow, N.P Randle, Trevor Ian Robinson, Parul Sharma, Ghada T Shawli, James P Stewart, Neil Swainston, Ecaterina Vamos, Joanne Watts, Mark Whitehead |
| EPI_ISL_664660, EPI_ISL_664662, | Department of Pathology, University of Cambridge | COVID-19 Genomics UK (COG-UK) Consortium | Aminu S. Jahun, Yasmin Chaudhry, Grant Hall, Iliana Georgana, Myra Hosmillo, Martin D. Curran, Malte Pinckert, Surendra Parmar, Ian Goodfellow |

| | | | |
|---|---|---|---|
| EPI_ISL_664666, EPI_ISL_664680 | | | |
| EPI_ISL_664685 | Liverpool Clinical Laboratories | COVID-19 Genomics UK (COG-UK) Consortium | Sam Haldenby, Anita Lucaci, Steve Paterson, Julian Hiscox, Alistair Darby, M Almsaud, A Alrezaihi, Muhannad Alruwaili, Stuart D Armstrong, Jones Benjamin, Eleanor G Bentley, Anu Chawla, Jordan J Clark, Angela Cowell, Richard Eccles, Isabel García-Dorival, Matthew Gemmell, Alessandro Gerada, PKF Gilmore, Richard Gregory, Ximeng Han, Catherine Hartley, Margaret Hughes, Miren Iturriza-Gomara, James Johnson, L Luu, Jenifer Manson, Charlotte Nelson, Elaine O'Toole, Cassie Olateju, Rebekah Penrice-Randal , Lucille Rainbow, N.P Randle, Trevor Ian Robinson, Parul Sharma, Ghada T Shawli, James P Stewart, Neil Swainston, Ecaterina Vamos, Joanne Watts, Mark Whitehead |
| EPI_ISL_664687 | University of Exeter | COVID-19 Genomics UK (COG-UK) Consortium | Ben Temperton,Aaron Jeffries,Michelle Michelsen,Joanna Warwick-Dugdale,Audrey Farbos,Robyn Manley,Stephen Michell,Jane Masoli |
| EPI_ISL_664697 | Department of Pathology, University of Cambridge | COVID-19 Genomics UK (COG-UK) Consortium | Aminu S. Jahun, Yasmin Chaudhry, Grant Hall, Iliana Georgana, Myra Hosmillo, Martin D. Curran, Malte Pinckert, Surendra Parmar, Ian Goodfellow |
| EPI_ISL_664700 | Liverpool Clinical Laboratories | COVID-19 Genomics UK (COG-UK) Consortium | Sam Haldenby, Anita Lucaci, Steve Paterson, Julian Hiscox, Alistair Darby, M Almsaud, A Alrezaihi, Muhannad Alruwaili, Stuart D Armstrong, Jones Benjamin, Eleanor G Bentley, Anu Chawla, Jordan J Clark, Angela Cowell, Richard Eccles, Isabel García-Dorival, Matthew Gemmell, Alessandro Gerada, PKF Gilmore, Richard Gregory, Ximeng Han, Catherine Hartley, Margaret Hughes, Miren Iturriza-Gomara, James Johnson, L Luu, Jenifer Manson, Charlotte Nelson, Elaine O'Toole, Cassie Olateju, Rebekah Penrice-Randal , Lucille Rainbow, N.P Randle, Trevor Ian Robinson, Parul Sharma, Ghada T Shawli, James P Stewart, Neil Swainston, Ecaterina Vamos, Joanne Watts, Mark Whitehead |
| EPI_ISL_664719 | Department of Pathology, University of Cambridge | COVID-19 Genomics UK (COG-UK) Consortium | Aminu S. Jahun, Yasmin Chaudhry, Grant Hall, Iliana Georgana, Myra Hosmillo, Martin D. Curran, Malte Pinckert, Surendra Parmar, Ian Goodfellow |
| EPI_ISL_664729 | Liverpool Clinical Laboratories | COVID-19 Genomics UK (COG-UK) Consortium | Sam Haldenby, Anita Lucaci, Steve Paterson, Julian Hiscox, Alistair Darby, M Almsaud, A Alrezaihi, Muhannad Alruwaili, Stuart D Armstrong, Jones Benjamin, Eleanor G Bentley, Anu Chawla, Jordan J Clark, Angela Cowell, Richard Eccles, Isabel García-Dorival, Matthew Gemmell, Alessandro Gerada, PKF Gilmore, Richard Gregory, Ximeng Han, Catherine Hartley, Margaret Hughes, Miren Iturriza-Gomara, James Johnson, L Luu, Jenifer Manson, Charlotte Nelson, Elaine O'Toole, Cassie Olateju, Rebekah Penrice-Randal , Lucille Rainbow, N.P Randle, Trevor Ian Robinson, Parul Sharma, Ghada T Shawli, James P Stewart, Neil Swainston, Ecaterina Vamos, Joanne Watts, Mark Whitehead |
| EPI_ISL_664801, EPI_ISL_664802, EPI_ISL_664803, EPI_ISL_664804, EPI_ISL_664805, EPI_ISL_664806, EPI_ISL_664807, EPI_ISL_664808, EPI_ISL_664809, EPI_ISL_664810, EPI_ISL_664811, EPI_ISL_664812, EPI_ISL_664813, EPI_ISL_664814, EPI_ISL_664815, EPI_ISL_664816, EPI_ISL_664817, EPI_ISL_664818, EPI_ISL_664819, EPI_ISL_664820, EPI_ISL_664821, EPI_ISL_664822, EPI_ISL_664823, EPI_ISL_664824, EPI_ISL_664825, EPI_ISL_664826, EPI_ISL_664827, EPI_ISL_664828, EPI_ISL_664829, EPI_ISL_664830, EPI_ISL_664831, EPI_ISL_664832, EPI_ISL_664834, EPI_ISL_664959, EPI_ISL_664960, EPI_ISL_664961, EPI_ISL_664962, EPI_ISL_664964, EPI_ISL_664965, EPI_ISL_664966, EPI_ISL_664967, EPI_ISL_664968, EPI_ISL_664971, EPI_ISL_664972, EPI_ISL_664973, EPI_ISL_664974, EPI_ISL_664975, EPI_ISL_664976, EPI_ISL_664977, EPI_ISL_664978, EPI_ISL_664979, EPI_ISL_664989, EPI_ISL_664990, EPI_ISL_664992, EPI_ISL_664993, EPI_ISL_664999, EPI_ISL_665000, EPI_ISL_665001, EPI_ISL_665002, EPI_ISL_665003 | | | |
| see above | Department of Pathology, University of Cambridge | COVID-19 Genomics UK (COG-UK) Consortium | Aminu S. Jahun, Yasmin Chaudhry, Grant Hall, Iliana Georgana, Myra Hosmillo, Martin D. Curran, Malte Pinckert, Surendra Parmar, Ian Goodfellow |
| EPI_ISL_665070, EPI_ISL_665071, EPI_ISL_665080, EPI_ISL_665083, EPI_ISL_665084, EPI_ISL_665091, EPI_ISL_665092, EPI_ISL_665093, EPI_ISL_665094, EPI_ISL_665095, EPI_ISL_665096, EPI_ISL_665097, EPI_ISL_665098, EPI_ISL_665099, EPI_ISL_665100, EPI_ISL_665101, EPI_ISL_665102, EPI_ISL_665103, EPI_ISL_665104, EPI_ISL_665105, EPI_ISL_665106, EPI_ISL_665107, EPI_ISL_665108, EPI_ISL_665109, EPI_ISL_665110, EPI_ISL_665111 | | | |
| see above | University of Exeter | COVID-19 Genomics UK (COG-UK) Consortium | Ben Temperton,Aaron Jeffries,Michelle Michelsen,Joanna Warwick-Dugdale,Audrey Farbos,Robyn Manley,Stephen Michell,Jane Masoli |
| EPI_ISL_665113 | Liverpool Clinical Laboratories | COVID-19 Genomics UK (COG-UK) Consortium | Sam Haldenby, Anita Lucaci, Steve Paterson, Julian Hiscox, Alistair Darby, M Almsaud, A Alrezaihi, Muhannad Alruwaili, Stuart D Armstrong, Jones Benjamin, Eleanor G Bentley, Anu Chawla, Jordan J Clark, Angela Cowell, Richard Eccles, Isabel García-Dorival, Matthew Gemmell, Alessandro Gerada, PKF Gilmore, Richard Gregory, Ximeng Han, Catherine Hartley, Margaret Hughes, Miren Iturriza-Gomara, James Johnson, L Luu, Jenifer Manson, Charlotte Nelson, Elaine O'Toole, Cassie Olateju, Rebekah Penrice-Randal , Lucille Rainbow, N.P Randle, Trevor Ian Robinson, Parul Sharma, Ghada T Shawli, James P Stewart, Neil Swainston, Ecaterina Vamos, Joanne Watts, Mark Whitehead |
| EPI_ISL_665266 | University College London, Great Ormond Street Hospital for Children NHS Foundation Trust, Imperial College Healthcare NHS Trust | COVID-19 Genomics UK (COG-UK) Consortium | Sergi Castellano, Rachel Williams, Mark Kristiansen, Paola Resende Silva, Sunando Roy, Tony Brooks, Helena Tutill, Paola Niola, Patricia Dyal, Charlotte Williams, Leysa Forrest, Yasmin Panchbhaya, Jacqueline Findlay, Samuel Weeks, Julianne Brown, Kathryn Harris, Paul Randell, James Price, Alison Holmes, Judith Breuer |
| EPI_ISL_665283 | West of Scotland Specialist Virology Centre, NHSGGC / MRC-University of Glasgow Centre for Virus Research | COVID-19 Genomics UK (COG-UK) Consortium | Ana da Silva Filipe, Natasha Johnson, Kathy Smollett, Daniel Mair, Stephen Carmichael, Alice Broos, Lily Tong, Jenna Nichols, Kyriaki Nomikou; Sarah McDonald; Richard Orton, Joseph Hughes, Sreenu Vattipally, David L Robertson; Alasdair MacLean, Rory Gunson; Sharif Shaaban, Matthew Holden; Rachel Blacow, Guy Mollett, Kathy Li, James Shepherd, Antonia Ho, Emma Thomson |
| EPI_ISL_665289 | Wales Specialist Virology Centre Sequencing lab: Pathogen Genomics Unit | COVID-19 Genomics UK (COG-UK) Consortium | Catherine Moore, Johnathan Evans, Laura Gifford, Malorie Perry, Simon Cottrell, Angela Marchbank, Alec Birchley, Alexander Adams, Amy Gaskin, Bree Gatica-Wilcox, Jason Coombes, Joel Southgate, Lauren Gilbert, Lee Graham, Nicole Pacchiarini, Sara Kumziene-Summerhayes, Sarah Taylor, Sophie Jones, Sara Rey, Matthew Bull, Joanne Watkins, Sally Corden, Tom Connor |
| EPI_ISL_665297, EPI_ISL_665299 | University College London Hospital | COVID-19 Genomics UK (COG-UK) Consortium | Judith Heaney, Matthew Byott, Catherine Houlihan, Dan Frampton, Stuart Kirk, Moira Spyer and Eleni Nastouli |
| EPI_ISL_665339 | University College London, Great Ormond Street Hospital for Children NHS Foundation Trust, Imperial College Healthcare NHS Trust | COVID-19 Genomics UK (COG-UK) Consortium | Sergi Castellano, Rachel Williams, Mark Kristiansen, Paola Resende Silva, Sunando Roy, Tony Brooks, Helena Tutill, Paola Niola, Patricia Dyal, Charlotte Williams, Leysa Forrest, Yasmin Panchbhaya, Jacqueline Findlay, Samuel Weeks, Julianne Brown, Kathryn Harris, Paul Randell, James Price, Alison Holmes, Judith Breuer |
| EPI_ISL_665348 | Centre for Enzyme Innovation, University of Portsmouth / Translational Research Laboratory, Portsmouth Hospitals NHS Trust | COVID-19 Genomics UK (COG-UK) Consortium | Angela Beckett,Yann Bourgeois,Garry Scarlett,Sharon Glaysher,Scott Elliott,Kelly Bicknell,Robert Impey,Allyson Lloyd,Sarah Wyllie,Ethan Butcher,Anoop Chauhan,Samuel Robson |
| EPI_ISL_665352 | University College London Hospital | COVID-19 Genomics UK (COG-UK) Consortium | Judith Heaney, Matthew Byott, Catherine Houlihan, Dan Frampton, Stuart Kirk, Moira Spyer and Eleni Nastouli |
| EPI_ISL_665413 | University College London, Great Ormond Street Hospital for Children NHS Foundation Trust, Imperial College Healthcare NHS Trust | COVID-19 Genomics UK (COG-UK) Consortium | Sergi Castellano, Rachel Williams, Mark Kristiansen, Paola Resende Silva, Sunando Roy, Tony Brooks, Helena Tutill, Paola Niola, Patricia Dyal, Charlotte Williams, Leysa Forrest, Yasmin Panchbhaya, Jacqueline Findlay, Samuel Weeks, Julianne Brown, Kathryn Harris, Paul Randell, James Price, Alison Holmes, Judith Breuer |
| EPI_ISL_665431 | Quadram Institute Bioscience | COVID-19 Genomics UK (COG-UK) Consortium | Dave J. Baker, Gemma L. Kay, Alp Aydin, Thanh Le-Viet, Steven Rudder, Ana P. Tedim, Anastasia Kolyva, Maria Diaz, Leonardo de Oliveira Martins, Nabil-Fareed Alikhan, Lizzie Meadows, Rachael Stanley, Ngozi Elumogo, Muhammed Yasir, Nicholas M. Thomson, Alexander J Trotter, Rachel Gilroy, Samuel Bloomfield, Claire Stuart, Andrew Bell, Reenesh Prakash, Samir Dervisevic, Alison E. Mather, John Wain, Mark Webber, Andrew J. Page, Justin O'Grady |
| EPI_ISL_665441 | Queens Medical Centre, Clinical Microbiology Department / DeepSeq Nottingham | COVID-19 Genomics UK (COG-UK) Consortium | Gemma Clark, Wendy Smith, Manjinder Khakh, Vicki M Fleming, Michelle M Lister, Hannah Howson-Wells, Jonathan Ball, Patrick McClure, Joseph Chappell, Theocharis Tsoleridis, Nadine Holmes, Matthew Carlisle, Christopher Moore, Fei Sang, Johnny Debebe, Victoria Wright, Matthew Loose |
| EPI_ISL_665462 | Wales Specialist Virology Centre Sequencing lab: Pathogen Genomics Unit | COVID-19 Genomics UK (COG-UK) Consortium | Catherine Moore, Johnathan Evans, Laura Gifford, Malorie Perry, Simon Cottrell, Angela Marchbank, Alec Birchley, Alexander Adams, Amy Gaskin, Bree Gatica-Wilcox, Jason Coombes, Joel Southgate, Lauren Gilbert, Lee Graham, Nicole Pacchiarini, Sara Kumziene-Summerhayes, Sarah Taylor, Sophie Jones, Sara Rey, Matthew Bull, Joanne Watkins, Sally Corden, Tom Connor |
| EPI_ISL_665466, EPI_ISL_665517 | University College London Hospital | COVID-19 Genomics UK (COG-UK) Consortium | Judith Heaney, Matthew Byott, Catherine Houlihan, Dan Frampton, Stuart Kirk, Moira Spyer and Eleni Nastouli |
| EPI_ISL_665522, EPI_ISL_665523, EPI_ISL_665529, EPI_ISL_665535 | West of Scotland Specialist Virology Centre, NHSGGC / MRC-University of Glasgow Centre for Virus Research | COVID-19 Genomics UK (COG-UK) Consortium | Ana da Silva Filipe, Natasha Johnson, Kathy Smollett, Daniel Mair, Stephen Carmichael, Alice Broos, Lily Tong, Jenna Nichols, Kyriaki Nomikou; Sarah McDonald; Richard Orton, Joseph Hughes, Sreenu Vattipally, David L Robertson; Alasdair MacLean, Rory Gunson; Sharif Shaaban, Matthew Holden; Rachel Blacow, Guy Mollett, Kathy Li, James Shepherd, Antonia Ho, Emma Thomson |
| EPI_ISL_665539 | Wales Specialist Virology Centre Sequencing lab: Pathogen Genomics Unit | COVID-19 Genomics UK (COG-UK) Consortium | Catherine Moore, Johnathan Evans, Laura Gifford, Malorie Perry, Simon Cottrell, Angela Marchbank, Alec Birchley, Alexander Adams, Amy Gaskin, Bree Gatica-Wilcox, Jason Coombes, Joel Southgate, Lauren Gilbert, Lee Graham, Nicole Pacchiarini, Sara Kumziene-Summerhayes, Sarah Taylor, Sophie Jones, Sara Rey, Matthew Bull, Joanne Watkins, Sally Corden, Tom Connor |
| EPI_ISL_665541, EPI_ISL_665564, EPI_ISL_665565, EPI_ISL_665566 | West of Scotland Specialist Virology Centre, NHSGGC / MRC-University of Glasgow Centre for Virus Research | COVID-19 Genomics UK (COG-UK) Consortium | Ana da Silva Filipe, Natasha Johnson, Kathy Smollett, Daniel Mair, Stephen Carmichael, Alice Broos, Lily Tong, Jenna Nichols, Kyriaki Nomikou; Sarah McDonald; Richard Orton, Joseph Hughes, Sreenu Vattipally, David L Robertson; Alasdair MacLean, Rory Gunson; Sharif Shaaban, Matthew Holden; Rachel Blacow, Guy Mollett, Kathy Li, James Shepherd, Antonia Ho, Emma Thomson |
| EPI_ISL_665572 | University College London Hospital | COVID-19 Genomics UK (COG-UK) Consortium | Judith Heaney, Matthew Byott, Catherine Houlihan, Dan Frampton, Stuart Kirk, Moira Spyer and Eleni Nastouli |
| EPI_ISL_665578 | West of Scotland Specialist Virology Centre, NHSGGC / | COVID-19 Genomics UK (COG-UK) Consortium | Ana da Silva Filipe, Natasha Johnson, Kathy Smollett, Daniel Mair, Stephen Carmichael, Alice Broos, Lily Tong, Jenna Nichols, Kyriaki Nomikou; Sarah |

| | | | |
|---|---|---|---|
| | MRC-University of Glasgow Centre for Virus Research | | McDonald; Richard Orton, Joseph Hughes, Sreenu Vattipally, David L Robertson; Alasdair MacLean, Rory Gunson; Sharif Shaaban, Matthew Holden; Rachel Blacow, Guy Mollett, Kathy Li, James Shepherd, Antonia Ho, Emma Thomson |
| EPI_ISL_665634, EPI_ISL_665635 | University College London Hospital | COVID-19 Genomics UK (COG-UK) Consortium | Judith Heaney, Matthew Byott, Catherine Houlihan, Dan Frampton, Stuart Kirk, Moira Spyer and Eleni Nastouli |
| EPI_ISL_665640 | West of Scotland Specialist Virology Centre, NHSGGC / MRC-University of Glasgow Centre for Virus Research | COVID-19 Genomics UK (COG-UK) Consortium | Ana da Silva Filipe, Natasha Johnson, Kathy Smollett, Daniel Mair, Stephen Carmichael, Alice Broos, Lily Tong, Jenna Nichols, Kyriaki Nomikou; Sarah McDonald; Richard Orton, Joseph Hughes, Sreenu Vattipally, David L Robertson; Alasdair MacLean, Rory Gunson; Sharif Shaaban, Matthew Holden; Rachel Blacow, Guy Mollett, Kathy Li, James Shepherd, Antonia Ho, Emma Thomson |
| EPI_ISL_665644 | University College London Hospital | COVID-19 Genomics UK (COG-UK) Consortium | Judith Heaney, Matthew Byott, Catherine Houlihan, Dan Frampton, Stuart Kirk, Moira Spyer and Eleni Nastouli |
| EPI_ISL_665650 | Queens Medical Centre, Clinical Microbiology Department / DeepSeq Nottingham | COVID-19 Genomics UK (COG-UK) Consortium | Gemma Clark, Wendy Smith, Manjinder Khakh, Vicki M Fleming, Michelle M Lister, Hannah Howson-Wells, Jonathan Ball, Patrick McClure, Joseph Chappell, Theocharis Tsoleridis, Nadine Holmes, Matthew Carlisle, Christopher Moore, Fei Sang, Johnny Debebe, Victoria Wright, Matthew Loose |
| EPI_ISL_665670 | Wales Specialist Virology Centre Sequencing lab: Pathogen Genomics Unit | COVID-19 Genomics UK (COG-UK) Consortium | Catherine Moore, Johnathan Evans, Laura Gifford, Malorie Perry, Simon Cottrell, Angela Marchbank, Alec Birchley, Alexander Adams, Amy Gaskin, Bree Gatica-Wilcox, Jason Coombes, Joel Southgate, Lauren Gilbert, Lee Graham, Nicole Pacchiarini, Sara Kumziene-Summerhayes, Sarah Taylor, Sophie Jones, Sara Rey, Matthew Bull, Joanne Watkins, Sally Corden, Tom Connor |
| EPI_ISL_665759 | University College London Hospital | COVID-19 Genomics UK (COG-UK) Consortium | Judith Heaney, Matthew Byott, Catherine Houlihan, Dan Frampton, Stuart Kirk, Moira Spyer and Eleni Nastouli |
| EPI_ISL_665777, EPI_ISL_665778, EPI_ISL_665779 | Wales Specialist Virology Centre Sequencing lab: Pathogen Genomics Unit | COVID-19 Genomics UK (COG-UK) Consortium | Catherine Moore, Johnathan Evans, Laura Gifford, Malorie Perry, Simon Cottrell, Angela Marchbank, Alec Birchley, Alexander Adams, Amy Gaskin, Bree Gatica-Wilcox, Jason Coombes, Joel Southgate, Lauren Gilbert, Lee Graham, Nicole Pacchiarini, Sara Kumziene-Summerhayes, Sarah Taylor, Sophie Jones, Sara Rey, Matthew Bull, Joanne Watkins, Sally Corden, Tom Connor |
| EPI_ISL_665792, EPI_ISL_665804 | West of Scotland Specialist Virology Centre, NHSGGC / MRC-University of Glasgow Centre for Virus Research | COVID-19 Genomics UK (COG-UK) Consortium | Ana da Silva Filipe, Natasha Johnson, Kathy Smollett, Daniel Mair, Stephen Carmichael, Alice Broos, Lily Tong, Jenna Nichols, Kyriaki Nomikou; Sarah McDonald; Richard Orton, Joseph Hughes, Sreenu Vattipally, David L Robertson; Alasdair MacLean, Rory Gunson; Sharif Shaaban, Matthew Holden; Rachel Blacow, Guy Mollett, Kathy Li, James Shepherd, Antonia Ho, Emma Thomson |
| EPI_ISL_665813, EPI_ISL_665814, EPI_ISL_665815, EPI_ISL_665823, EPI_ISL_665827, EPI_ISL_665828, EPI_ISL_665829, EPI_ISL_665830 | University College London Hospital | COVID-19 Genomics UK (COG-UK) Consortium | Judith Heaney, Matthew Byott, Catherine Houlihan, Dan Frampton, Stuart Kirk, Moira Spyer and Eleni Nastouli |
| EPI_ISL_665899 | Queens Medical Centre, Clinical Microbiology Department / DeepSeq Nottingham | COVID-19 Genomics UK (COG-UK) Consortium | Gemma Clark, Wendy Smith, Manjinder Khakh, Vicki M Fleming, Michelle M Lister, Hannah Howson-Wells, Jonathan Ball, Patrick McClure, Joseph Chappell, Theocharis Tsoleridis, Nadine Holmes, Matthew Carlisle, Christopher Moore, Fei Sang, Johnny Debebe, Victoria Wright, Matthew Loose |
| EPI_ISL_666023, EPI_ISL_666024, EPI_ISL_666025, EPI_ISL_666026, EPI_ISL_666027, EPI_ISL_666028 | West of Scotland Specialist Virology Centre, NHSGGC / MRC-University of Glasgow Centre for Virus Research | COVID-19 Genomics UK (COG-UK) Consortium | Ana da Silva Filipe, Natasha Johnson, Kathy Smollett, Daniel Mair, Stephen Carmichael, Alice Broos, Lily Tong, Jenna Nichols, Kyriaki Nomikou; Sarah McDonald; Richard Orton, Joseph Hughes, Sreenu Vattipally, David L Robertson; Alasdair MacLean, Rory Gunson; Sharif Shaaban, Matthew Holden; Rachel Blacow, Guy Mollett, Kathy Li, James Shepherd, Antonia Ho, Emma Thomson |
| EPI_ISL_666083 | Virology Department, Royal Infirmary of Edinburgh, NHS Lothian / School of Biological Sciences, University of Edinburgh / Institute of Genetics and Molecular Medicine, University of Edinburgh | COVID-19 Genomics UK (COG-UK) Consortium | McHugh M, Dewar R, Rooke S, Gallagher M, Balcaza C, O'Toole Á, Scher E, Hill V, McCrone JT, Colquhoun R, Yu X, Jackson B, Rambaut A, Williams TC, Templeton K |
| EPI_ISL_666130, EPI_ISL_666300, EPI_ISL_666375, EPI_ISL_666390, EPI_ISL_666404, EPI_ISL_666433, EPI_ISL_666438, EPI_ISL_666455, EPI_ISL_666469, EPI_ISL_666508, EPI_ISL_666521, EPI_ISL_666522, EPI_ISL_666524, EPI_ISL_666542, EPI_ISL_666560, EPI_ISL_666566, EPI_ISL_666571, EPI_ISL_666576, EPI_ISL_666578, EPI_ISL_666590 | | | |
| see above | Wales Specialist Virology Centre Sequencing lab: Pathogen Genomics Unit | COVID-19 Genomics UK (COG-UK) Consortium | Catherine Moore, Johnathan Evans, Laura Gifford, Malorie Perry, Simon Cottrell, Angela Marchbank, Alec Birchley, Alexander Adams, Amy Gaskin, Bree Gatica-Wilcox, Jason Coombes, Joel Southgate, Lauren Gilbert, Lee Graham, Nicole Pacchiarini, Sara Kumziene-Summerhayes, Sarah Taylor, Sophie Jones, Sara Rey, Matthew Bull, Joanne Watkins, Sally Corden, Tom Connor |
| EPI_ISL_666739, EPI_ISL_666767 | Respiratory Virus Unit, Microbiology Services Colindale, Public Health England | COVID-19 Genomics UK (COG-UK) Consortium | PHE Covid Sequencing Team |
| EPI_ISL_666799, EPI_ISL_666800, EPI_ISL_666801, EPI_ISL_666802, EPI_ISL_666803, EPI_ISL_666804, EPI_ISL_666805, EPI_ISL_666806, EPI_ISL_666807, EPI_ISL_666808, EPI_ISL_666809, EPI_ISL_666810, EPI_ISL_666811, EPI_ISL_666812, EPI_ISL_666813 | | | |
| see above | Maryland Public Health Laboratory | Maryland Public Health Laboratory | Maryland Department of Health Laboratories Administration |
| EPI_ISL_666887, EPI_ISL_666888, EPI_ISL_666889 | Department of Clinical Microbiology | GIGA Medical Genomics | Keith Durkin, Maria Artesi, Justine Defêche, Gilles Darcis, Michel Moutschen, Sébastien Bontems, Raphaël Boreux, Bouchra Boujemla, Cécile Meex, Pierrette Melin, Marie-Pierre Hayette, Vincent Bours |
| EPI_ISL_667054, EPI_ISL_667055, EPI_ISL_667064, EPI_ISL_667065, EPI_ISL_667066, EPI_ISL_667070, EPI_ISL_667071, EPI_ISL_667072, EPI_ISL_667073 | San Diego County Public Health Laboratory | Andersen lab at Scripps Research | SEARCH Alliance San Diego with Tracy Basler, Jovan Shephard, Brett Austin |
| EPI_ISL_667538, EPI_ISL_667544, EPI_ISL_667545, EPI_ISL_667546, EPI_ISL_667547 | OHSU Lab Services Molecular Microbiology Lab | Oregon SARS-CoV-2 Genome Sequencing Center | Brendan L. O'Connell, Ruth V. Nichols, Sally Grindstaff, Alec J. Hirsch, Donna Hansel, Guang Fan, Daniel N. Streblow, William B. Messer, Andrew C. Adey, Benjamin N. Bimber, Brian J. O'Roak |
| EPI_ISL_667804 | South Eastern Area Laboratory Services (SEALS) | NSW Health Pathology - Institute of Clinical Pathology and Medical Research; Westmead Hospital; University of Sydney | CIDM-PH et al. |
| EPI_ISL_667829, EPI_ISL_667861, EPI_ISL_667865, EPI_ISL_667870, EPI_ISL_667902, EPI_ISL_667906, EPI_ISL_667933, EPI_ISL_667955, EPI_ISL_667961, EPI_ISL_667988, EPI_ISL_667991, EPI_ISL_668027, EPI_ISL_668071, EPI_ISL_668072, EPI_ISL_668080 | | | |
| see above | Lighthouse Lab in Cambridge | Wellcome Sanger Institute for the COVID-19 Genomics UK (COG-UK) Consortium | Rob Howes, The Lighthouse Lab in Cambridge and Alex Alderton, Roberto Amato, Sonia Goncalves, Ewan Harrison, David K. Jackson, Ian Johnston, Dominic Kwiatkowski, Cordelia Langford, John Sillitoe on behalf of the Wellcome Sanger Institute COVID-19 Surveillance Team |
| EPI_ISL_668348 | Lighthouse Lab in Glasgow | Wellcome Sanger Institute for the COVID-19 Genomics UK (COG-UK) Consortium | Harper VanSteenhouse, Yumi Kasai, David Gray, Carol Clugston, Anna Dominiczak and Alex Alderton, Roberto Amato, Sonia Goncalves, Ewan Harrison, David K. Jackson, Ian Johnston, Dominic Kwiatkowski, Cordelia Langford, John Sillitoe on behalf of the Wellcome Sanger Institute COVID-19 Surveillance Team (http://www.sanger.ac.uk/covid-team) |
| EPI_ISL_668452 | Centre for Dengue Research, Department of Immunology and Molecular Medicine | Centre for Dengue Research, Department of Immunology and Molecular Medicine | Chandima Jeewandara, Deshni Jayathilaka, Dinuka Ariyaratne, Diyanath Ranasinghe, Laksiri Gomes, Gathsaurie Neelika Malavige |
| EPI_ISL_668458, EPI_ISL_668459, EPI_ISL_668460, EPI_ISL_668461, EPI_ISL_668462, EPI_ISL_668463, EPI_ISL_668464, EPI_ISL_668466, EPI_ISL_668469, EPI_ISL_668470, EPI_ISL_668471, EPI_ISL_668472, EPI_ISL_668473, EPI_ISL_668474, EPI_ISL_668475, EPI_ISL_668476, EPI_ISL_668477, EPI_ISL_668478, EPI_ISL_668479, EPI_ISL_668480, EPI_ISL_668481, EPI_ISL_668482, EPI_ISL_668483, EPI_ISL_668484, EPI_ISL_668485, EPI_ISL_668486, EPI_ISL_668487, EPI_ISL_668488, EPI_ISL_668489, EPI_ISL_668490, EPI_ISL_668491, EPI_ISL_668492, EPI_ISL_668493, EPI_ISL_668494, EPI_ISL_668495, EPI_ISL_668496, EPI_ISL_668497, EPI_ISL_668498, EPI_ISL_668499, EPI_ISL_668500, EPI_ISL_668503, EPI_ISL_668504, EPI_ISL_668505, EPI_ISL_668506, EPI_ISL_668507, EPI_ISL_668508, EPI_ISL_668509, EPI_ISL_668511, EPI_ISL_668512, EPI_ISL_668513, EPI_ISL_668514, EPI_ISL_668515, EPI_ISL_668516, EPI_ISL_668520, EPI_ISL_668528, EPI_ISL_668529, EPI_ISL_668531, EPI_ISL_668532, EPI_ISL_668533, EPI_ISL_668538, EPI_ISL_668539, EPI_ISL_668540, EPI_ISL_668541, EPI_ISL_668542, EPI_ISL_668543, EPI_ISL_668545, EPI_ISL_668547, EPI_ISL_668548, EPI_ISL_668549, EPI_ISL_668550, EPI_ISL_668551, EPI_ISL_668552, EPI_ISL_668553, EPI_ISL_668554, EPI_ISL_668555, EPI_ISL_668556, EPI_ISL_668557, EPI_ISL_668560, EPI_ISL_668564, EPI_ISL_668565, EPI_ISL_668566, EPI_ISL_668567, EPI_ISL_668568, EPI_ISL_668569, EPI_ISL_668570, EPI_ISL_668571, EPI_ISL_668573, EPI_ISL_668574, EPI_ISL_668575, EPI_ISL_668576, EPI_ISL_668577, EPI_ISL_668578, EPI_ISL_668582, EPI_ISL_668583, EPI_ISL_668584, EPI_ISL_668585, EPI_ISL_668586, EPI_ISL_668587, EPI_ISL_668588, EPI_ISL_668589, EPI_ISL_668590, EPI_ISL_668591, EPI_ISL_668593, EPI_ISL_668594, EPI_ISL_668598, EPI_ISL_668599, EPI_ISL_668600, EPI_ISL_668602, EPI_ISL_668603, EPI_ISL_668604, EPI_ISL_668605, EPI_ISL_668607, EPI_ISL_668608, EPI_ISL_668611, EPI_ISL_668613, EPI_ISL_668614, EPI_ISL_668616, EPI_ISL_668617, EPI_ISL_668618, EPI_ISL_668619, EPI_ISL_668620, EPI_ISL_668621, EPI_ISL_668622, EPI_ISL_668623, EPI_ISL_668624, EPI_ISL_668625, EPI_ISL_668626, EPI_ISL_668627, EPI_ISL_668628, EPI_ISL_668629, EPI_ISL_668630, EPI_ISL_668631, EPI_ISL_668632, EPI_ISL_668634, EPI_ISL_668637, EPI_ISL_668638, EPI_ISL_668639, EPI_ISL_668640, EPI_ISL_668641, EPI_ISL_668642, EPI_ISL_668644, EPI_ISL_668645, EPI_ISL_668646, EPI_ISL_668647, EPI_ISL_668648, EPI_ISL_668649, EPI_ISL_668650, EPI_ISL_668651, EPI_ISL_668654, EPI_ISL_668655, EPI_ISL_668656, EPI_ISL_668657, EPI_ISL_668658, EPI_ISL_668659, EPI_ISL_668660, EPI_ISL_668661, EPI_ISL_668662, EPI_ISL_668675, EPI_ISL_668676, EPI_ISL_668677, EPI_ISL_668678, EPI_ISL_668679, EPI_ISL_668680, EPI_ISL_668681, EPI_ISL_668682, EPI_ISL_668683, EPI_ISL_668684, EPI_ISL_668685, EPI_ISL_668686, EPI_ISL_668687, EPI_ISL_668688, EPI_ISL_668689, EPI_ISL_668690, EPI_ISL_668691, EPI_ISL_668692, EPI_ISL_668693, EPI_ISL_668694, EPI_ISL_668695, EPI_ISL_668696, EPI_ISL_668697, EPI_ISL_668698, EPI_ISL_668699, EPI_ISL_668700, EPI_ISL_668701, EPI_ISL_668702, EPI_ISL_668703, EPI_ISL_668704, EPI_ISL_668705, EPI_ISL_668706, EPI_ISL_668707, EPI_ISL_668708, EPI_ISL_668709, EPI_ISL_668710, EPI_ISL_668711, EPI_ISL_668712, EPI_ISL_668713, EPI_ISL_668714, EPI_ISL_668715, EPI_ISL_668716, EPI_ISL_668717, EPI_ISL_668720, EPI_ISL_668721, EPI_ISL_668722, EPI_ISL_668723, EPI_ISL_668724, EPI_ISL_668725, EPI_ISL_668726, EPI_ISL_668729, EPI_ISL_668731, EPI_ISL_668732, EPI_ISL_668733, EPI_ISL_668734, EPI_ISL_668736, EPI_ISL_668737, EPI_ISL_668739, EPI_ISL_668740, EPI_ISL_668741, EPI_ISL_668742, EPI_ISL_668745, EPI_ISL_668746, EPI_ISL_668747, EPI_ISL_668748, EPI_ISL_668749, EPI_ISL_668750, EPI_ISL_668751, EPI_ISL_668752, EPI_ISL_668753, EPI_ISL_668754, EPI_ISL_668755, EPI_ISL_668756, EPI_ISL_668775, EPI_ISL_668776, EPI_ISL_668777, EPI_ISL_668778, | | | |

| | | | |
|---|---|---|---|
| EPI_ISL_668779, EPI_ISL_668780, EPI_ISL_668781, EPI_ISL_668782, EPI_ISL_668783, EPI_ISL_668784, EPI_ISL_668785, EPI_ISL_668786, EPI_ISL_668787, EPI_ISL_668788, EPI_ISL_668789, EPI_ISL_668790, EPI_ISL_668791, EPI_ISL_668792, EPI_ISL_668793, EPI_ISL_668794, EPI_ISL_668795, EPI_ISL_668796, EPI_ISL_668797, EPI_ISL_668798, EPI_ISL_668799, EPI_ISL_668800, EPI_ISL_668801, EPI_ISL_668802, EPI_ISL_668803, EPI_ISL_668804, EPI_ISL_668805, EPI_ISL_668806, EPI_ISL_668807, EPI_ISL_668808, EPI_ISL_668809, EPI_ISL_668810, EPI_ISL_668811, EPI_ISL_668812, EPI_ISL_668813, EPI_ISL_668814, EPI_ISL_668815, EPI_ISL_668816, EPI_ISL_668817, EPI_ISL_668818, EPI_ISL_668819, EPI_ISL_668820, EPI_ISL_668821, EPI_ISL_668822, EPI_ISL_668823, EPI_ISL_668824, EPI_ISL_668825, EPI_ISL_668826, EPI_ISL_668827, EPI_ISL_668828, EPI_ISL_668831, EPI_ISL_668832, EPI_ISL_668833, EPI_ISL_668834, EPI_ISL_668835, EPI_ISL_668836, EPI_ISL_668837, EPI_ISL_668838, EPI_ISL_668841, EPI_ISL_668842, EPI_ISL_668843, EPI_ISL_668844, EPI_ISL_668845, EPI_ISL_668846, EPI_ISL_668847, EPI_ISL_668848, EPI_ISL_668849, EPI_ISL_668851, EPI_ISL_668852, EPI_ISL_668957, EPI_ISL_668958, EPI_ISL_668959, EPI_ISL_668960, EPI_ISL_668961, EPI_ISL_668962, EPI_ISL_668963, EPI_ISL_668964, EPI_ISL_668965, EPI_ISL_668966, EPI_ISL_668967, EPI_ISL_668968, EPI_ISL_668969, EPI_ISL_668970, EPI_ISL_668971, EPI_ISL_668972, EPI_ISL_668973, EPI_ISL_668974, EPI_ISL_668975, EPI_ISL_668976, EPI_ISL_668977, EPI_ISL_668978, EPI_ISL_668979, EPI_ISL_668980, EPI_ISL_668981, EPI_ISL_668982, EPI_ISL_668983, EPI_ISL_668984, EPI_ISL_668985, EPI_ISL_668986, EPI_ISL_668987, EPI_ISL_668988, EPI_ISL_668989, EPI_ISL_668990, EPI_ISL_668991, EPI_ISL_668992, EPI_ISL_668993, EPI_ISL_668994, EPI_ISL_668995, EPI_ISL_668996, EPI_ISL_668997, EPI_ISL_668998, EPI_ISL_668999, EPI_ISL_669000, EPI_ISL_669001, EPI_ISL_669002, EPI_ISL_669003, EPI_ISL_669004, EPI_ISL_669005, EPI_ISL_669006, EPI_ISL_669007, EPI_ISL_669008, EPI_ISL_669009, EPI_ISL_669010, EPI_ISL_669011, EPI_ISL_669012, EPI_ISL_669013, EPI_ISL_669014, EPI_ISL_669015, EPI_ISL_669016, EPI_ISL_669017, EPI_ISL_669018, EPI_ISL_669019, EPI_ISL_669020, EPI_ISL_669021, EPI_ISL_669022, EPI_ISL_669023, EPI_ISL_669024, EPI_ISL_669025, EPI_ISL_669026, EPI_ISL_669027, EPI_ISL_669028, EPI_ISL_669029, EPI_ISL_669030, EPI_ISL_669031, EPI_ISL_669032, EPI_ISL_669033, EPI_ISL_669034, EPI_ISL_669035, EPI_ISL_669036, EPI_ISL_669037, EPI_ISL_669038, EPI_ISL_669039, EPI_ISL_669040, EPI_ISL_669041, EPI_ISL_669042, EPI_ISL_669043, EPI_ISL_669044, EPI_ISL_669045, EPI_ISL_669046, EPI_ISL_669047, EPI_ISL_669048, EPI_ISL_669049, EPI_ISL_669050, EPI_ISL_669051, EPI_ISL_669052, EPI_ISL_669053, EPI_ISL_669054, EPI_ISL_669055, EPI_ISL_669056, EPI_ISL_669057, EPI_ISL_669058, EPI_ISL_669059, EPI_ISL_669060, EPI_ISL_669061, EPI_ISL_669062, EPI_ISL_669063, EPI_ISL_669064, EPI_ISL_669065, EPI_ISL_669066, EPI_ISL_669067, EPI_ISL_669068, EPI_ISL_669069, EPI_ISL_669070, EPI_ISL_669071, EPI_ISL_669072, EPI_ISL_669073, EPI_ISL_669074, EPI_ISL_669075, EPI_ISL_669076, EPI_ISL_669077, EPI_ISL_669078, EPI_ISL_669079, EPI_ISL_669080, EPI_ISL_669081, EPI_ISL_669082, EPI_ISL_669083, EPI_ISL_669084, EPI_ISL_669085, EPI_ISL_669086, EPI_ISL_669087, EPI_ISL_669088, EPI_ISL_669089, EPI_ISL_669090, EPI_ISL_669091, EPI_ISL_669092, EPI_ISL_669093, EPI_ISL_669094, EPI_ISL_669095, EPI_ISL_669096, EPI_ISL_669097, EPI_ISL_669098, EPI_ISL_669099, EPI_ISL_669100, EPI_ISL_669101, EPI_ISL_669102, EPI_ISL_669103, EPI_ISL_669104, EPI_ISL_669105, EPI_ISL_669106, EPI_ISL_669107, EPI_ISL_669108, EPI_ISL_669109, EPI_ISL_669110, EPI_ISL_669111, EPI_ISL_669112, EPI_ISL_669113, EPI_ISL_669114, EPI_ISL_669115, EPI_ISL_669116, EPI_ISL_669117, EPI_ISL_669118, EPI_ISL_669119, EPI_ISL_669120, EPI_ISL_669121, EPI_ISL_669122, EPI_ISL_669123, EPI_ISL_669124, EPI_ISL_669125, EPI_ISL_669126, EPI_ISL_669127, EPI_ISL_669128, EPI_ISL_669129, EPI_ISL_669130, EPI_ISL_669131, EPI_ISL_669132, EPI_ISL_669133, EPI_ISL_669134, EPI_ISL_669135, EPI_ISL_669136, EPI_ISL_669137, EPI_ISL_669138, EPI_ISL_669139, EPI_ISL_669140, EPI_ISL_669141, EPI_ISL_669142, EPI_ISL_669143, EPI_ISL_669144, EPI_ISL_669145, EPI_ISL_669146, EPI_ISL_669147, EPI_ISL_669148, EPI_ISL_669149, EPI_ISL_669150, EPI_ISL_669151, EPI_ISL_669152, EPI_ISL_669153, EPI_ISL_669154, EPI_ISL_669155, EPI_ISL_669156, EPI_ISL_669157, EPI_ISL_669158, EPI_ISL_669159, EPI_ISL_669160, EPI_ISL_669161, EPI_ISL_669162, EPI_ISL_669163, EPI_ISL_669164, EPI_ISL_669165, EPI_ISL_669166, EPI_ISL_669167, EPI_ISL_669168, EPI_ISL_669169, EPI_ISL_669170, EPI_ISL_669171, EPI_ISL_669172, EPI_ISL_669173, EPI_ISL_669174, EPI_ISL_669175, EPI_ISL_669176, EPI_ISL_669177, EPI_ISL_669178, EPI_ISL_669179, EPI_ISL_669180, EPI_ISL_669181, EPI_ISL_669182, EPI_ISL_669183, EPI_ISL_669184, EPI_ISL_669185, EPI_ISL_669186, EPI_ISL_669187, EPI_ISL_669188, EPI_ISL_669189, EPI_ISL_669190, EPI_ISL_669191, EPI_ISL_669192, EPI_ISL_669193, EPI_ISL_669194, EPI_ISL_669195, EPI_ISL_669196, EPI_ISL_669197, EPI_ISL_669198, EPI_ISL_669199, EPI_ISL_669200, EPI_ISL_669201, EPI_ISL_669202, EPI_ISL_669203, EPI_ISL_669204, EPI_ISL_669205, EPI_ISL_669206, EPI_ISL_669207, EPI_ISL_669208, EPI_ISL_669209, EPI_ISL_669210, EPI_ISL_669211, EPI_ISL_669212, EPI_ISL_669213, EPI_ISL_669214, EPI_ISL_669215, EPI_ISL_669216, EPI_ISL_669217, EPI_ISL_669218, EPI_ISL_669219, EPI_ISL_669220, EPI_ISL_669221, EPI_ISL_669222, EPI_ISL_669223, EPI_ISL_669224, EPI_ISL_669225, EPI_ISL_669226, EPI_ISL_669227, EPI_ISL_669228, EPI_ISL_669229, EPI_ISL_669230, EPI_ISL_669231, EPI_ISL_669232, EPI_ISL_669233, EPI_ISL_669234, EPI_ISL_669235, EPI_ISL_669236, EPI_ISL_669237, EPI_ISL_669238, EPI_ISL_669239, EPI_ISL_669240, EPI_ISL_669241, EPI_ISL_669242, EPI_ISL_669243, EPI_ISL_669244, EPI_ISL_669245, EPI_ISL_669246, EPI_ISL_669247, EPI_ISL_669248, EPI_ISL_669249, EPI_ISL_669250, EPI_ISL_669251, EPI_ISL_669252, EPI_ISL_669253, EPI_ISL_669254, EPI_ISL_669255, EPI_ISL_669256, EPI_ISL_669257, EPI_ISL_669258, EPI_ISL_669259, EPI_ISL_669260, EPI_ISL_669261, EPI_ISL_669262, EPI_ISL_669263, EPI_ISL_669264, EPI_ISL_669265, EPI_ISL_669266, EPI_ISL_669267, EPI_ISL_669268, EPI_ISL_669269, EPI_ISL_669270, EPI_ISL_669271, EPI_ISL_669272, EPI_ISL_669273, EPI_ISL_669274, EPI_ISL_669275, EPI_ISL_669276, EPI_ISL_669277, EPI_ISL_669278, EPI_ISL_669279, EPI_ISL_669280, EPI_ISL_669281, EPI_ISL_669282, EPI_ISL_669283, EPI_ISL_669284, EPI_ISL_669285, EPI_ISL_669286, EPI_ISL_669287, EPI_ISL_669288, EPI_ISL_669289, EPI_ISL_669290, EPI_ISL_669291, EPI_ISL_669292, EPI_ISL_669293, EPI_ISL_669294, EPI_ISL_669295, EPI_ISL_669296, EPI_ISL_669297, EPI_ISL_669298, EPI_ISL_669299, EPI_ISL_669300, EPI_ISL_669301, EPI_ISL_669302, EPI_ISL_669303, EPI_ISL_669304, EPI_ISL_669305, EPI_ISL_669306, EPI_ISL_669307, EPI_ISL_669308, EPI_ISL_669309, EPI_ISL_669310, EPI_ISL_669311, EPI_ISL_669312, EPI_ISL_669313, EPI_ISL_669314, EPI_ISL_669315, EPI_ISL_669316, EPI_ISL_669317, EPI_ISL_669318, EPI_ISL_669319, EPI_ISL_669320, EPI_ISL_669321, EPI_ISL_669322, EPI_ISL_669323, EPI_ISL_669324, EPI_ISL_669325, EPI_ISL_669326, EPI_ISL_669327, EPI_ISL_669328, EPI_ISL_669329, EPI_ISL_669330, EPI_ISL_669331, EPI_ISL_669332, EPI_ISL_669333, EPI_ISL_669334, EPI_ISL_669335, EPI_ISL_669336, EPI_ISL_669337, EPI_ISL_669338, EPI_ISL_669339, EPI_ISL_669340, EPI_ISL_669341, EPI_ISL_669342, EPI_ISL_669343, EPI_ISL_669344, EPI_ISL_669345, EPI_ISL_669346, EPI_ISL_669347, EPI_ISL_669348, EPI_ISL_669349, EPI_ISL_669350, EPI_ISL_669351, EPI_ISL_669352, EPI_ISL_669353, EPI_ISL_669354, EPI_ISL_669355, EPI_ISL_669356, EPI_ISL_669357, EPI_ISL_669358, EPI_ISL_669359, EPI_ISL_669360, EPI_ISL_669361, EPI_ISL_669362, EPI_ISL_669363, EPI_ISL_669364, EPI_ISL_669365, EPI_ISL_669366, EPI_ISL_669367, EPI_ISL_669368, EPI_ISL_669369, EPI_ISL_669370, EPI_ISL_669371, EPI_ISL_669372, EPI_ISL_669373, EPI_ISL_669374, EPI_ISL_669375, EPI_ISL_669376, EPI_ISL_669377, EPI_ISL_669378, EPI_ISL_669379, EPI_ISL_669380, EPI_ISL_669381, EPI_ISL_669382, EPI_ISL_669383, EPI_ISL_669384, EPI_ISL_669385, EPI_ISL_669386, EPI_ISL_669387, EPI_ISL_669388, EPI_ISL_669389, EPI_ISL_669390, EPI_ISL_669391, EPI_ISL_669392, EPI_ISL_669393, EPI_ISL_669394, EPI_ISL_669395, EPI_ISL_669396, EPI_ISL_669397, EPI_ISL_669398, EPI_ISL_669399, EPI_ISL_669400, EPI_ISL_669401, EPI_ISL_669402, EPI_ISL_669403, EPI_ISL_669404, EPI_ISL_669405, EPI_ISL_669406, EPI_ISL_669407, EPI_ISL_669408, EPI_ISL_669409, EPI_ISL_669410, EPI_ISL_669411, EPI_ISL_669412, EPI_ISL_669413, EPI_ISL_669414, EPI_ISL_669415, EPI_ISL_669416, EPI_ISL_669417, EPI_ISL_669418, EPI_ISL_669419, EPI_ISL_669420, EPI_ISL_669421, EPI_ISL_669422, EPI_ISL_669423, EPI_ISL_669424, EPI_ISL_669425, EPI_ISL_669426, EPI_ISL_669427, EPI_ISL_669428, EPI_ISL_669429, EPI_ISL_669430, EPI_ISL_669431, EPI_ISL_669432, EPI_ISL_669433, EPI_ISL_669434, EPI_ISL_669435, EPI_ISL_669436, EPI_ISL_669437, EPI_ISL_669438, EPI_ISL_669439, EPI_ISL_669440, EPI_ISL_669441, EPI_ISL_669442, EPI_ISL_669443, EPI_ISL_669444, EPI_ISL_669445, EPI_ISL_669446, EPI_ISL_669447, EPI_ISL_669448, EPI_ISL_669449, EPI_ISL_669450, EPI_ISL_669451, EPI_ISL_669452, EPI_ISL_669453, EPI_ISL_669454, EPI_ISL_669455, EPI_ISL_669456, EPI_ISL_669457, EPI_ISL_669459, EPI_ISL_669463, EPI_ISL_669465, EPI_ISL_669486, EPI_ISL_669495, EPI_ISL_669503, EPI_ISL_669504, EPI_ISL_669521, EPI_ISL_669525, EPI_ISL_669537, EPI_ISL_669548, EPI_ISL_669564, EPI_ISL_669583, EPI_ISL_669612, EPI_ISL_669613, EPI_ISL_669636, EPI_ISL_669637, EPI_ISL_669638, EPI_ISL_669639, EPI_ISL_669640, EPI_ISL_669641, EPI_ISL_669642, EPI_ISL_669643, EPI_ISL_669731, EPI_ISL_669733, EPI_ISL_669734, EPI_ISL_669735, EPI_ISL_669757, EPI_ISL_669760, EPI_ISL_669773, EPI_ISL_669774, EPI_ISL_669785, EPI_ISL_669786, EPI_ISL_669787, EPI_ISL_669817, EPI_ISL_669833, EPI_ISL_669834, EPI_ISL_669835, EPI_ISL_669836, EPI_ISL_669837, EPI_ISL_669838, EPI_ISL_669839, EPI_ISL_669840, EPI_ISL_669841, EPI_ISL_669842, EPI_ISL_669843, EPI_ISL_669844, EPI_ISL_669845, EPI_ISL_669846, EPI_ISL_669847, EPI_ISL_669848, EPI_ISL_669849, EPI_ISL_669850, EPI_ISL_669851, EPI_ISL_669852, EPI_ISL_669853, EPI_ISL_669854, EPI_ISL_669855, EPI_ISL_669856, EPI_ISL_669857, EPI_ISL_669858, EPI_ISL_669859, EPI_ISL_669860, EPI_ISL_669861, EPI_ISL_669862, EPI_ISL_669863, EPI_ISL_669864, EPI_ISL_669865, EPI_ISL_669866, EPI_ISL_669867, EPI_ISL_669868, EPI_ISL_669869, EPI_ISL_669870, EPI_ISL_669871, EPI_ISL_669872, EPI_ISL_669873, EPI_ISL_669874, EPI_ISL_669875, EPI_ISL_669876, EPI_ISL_669877, EPI_ISL_669878, EPI_ISL_669879, EPI_ISL_669880, EPI_ISL_669881 | | | |
| see above | Department of Virus and Microbiological Special Diagnostics, Statens Serum Institut, Copenhagen, Denmark | Albertsen Lab, Department of Chemistry and Bioscience, Aalborg University, Denmark | Danish Covid-19 Genome Consortium |
| EPI_ISL_673310, EPI_ISL_673385, EPI_ISL_673441, EPI_ISL_673447, EPI_ISL_673851 | Lighthouse Lab in Cambridge | Wellcome Sanger Institute for the COVID-19 Genomics UK (COG-UK) Consortium | Rob Howes, The Lighthouse Lab in Cambridge and Alex Alderton, Roberto Amato, Sonia Goncalves, Ewan Harrison, David K. Jackson, Ian Johnston, Dominic Kwiatkowski, Cordelia Langford, John Sillitoe on behalf of the Wellcome Sanger Institute COVID-19 Surveillance Team |
| EPI_ISL_673995 | Lighthouse Lab in Milton Keynes | Wellcome Sanger Institute for the COVID-19 Genomics UK (COG-UK) Consortium | The Lighthouse Lab in Milton Keynes and Alex Alderton, Roberto Amato, Sonia Goncalves, Ewan Harrison, David K. Jackson, Ian Johnston, Dominic Kwiatkowski, Cordelia Langford, John Sillitoe on behalf of the Wellcome Sanger Institute COVID-19 Surveillance Team |
| EPI_ISL_674223, EPI_ISL_674418 | Lighthouse Lab in Alderley Park | Wellcome Sanger Institute for the COVID-19 Genomics UK (COG-UK) Consortium | Jacquelyn Wynn, Mairead Hyland, The Lighthouse Lab in Alderley Park and Alex Alderton, Roberto Amato, Sonia Goncalves, Ewan Harrison, David K. Jackson, Ian Johnston, Dominic Kwiatkowski, Cordelia Langford, John Sillitoe on behalf of the Wellcome Sanger Institute COVID-19 Surveillance Team |
| EPI_ISL_674832 | Lighthouse Lab in Milton Keynes | Wellcome Sanger Institute for the COVID-19 Genomics UK (COG-UK) Consortium | The Lighthouse Lab in Milton Keynes and Alex Alderton, Roberto Amato, Sonia Goncalves, Ewan Harrison, David K. Jackson, Ian Johnston, Dominic Kwiatkowski, Cordelia Langford, John Sillitoe on behalf of the Wellcome Sanger Institute COVID-19 Surveillance Team |
| EPI_ISL_675720, EPI_ISL_675721, EPI_ISL_675746, EPI_ISL_675855, EPI_ISL_675925 | Lighthouse Lab in Alderley Park | Wellcome Sanger Institute for the COVID-19 Genomics UK (COG-UK) Consortium | Jacquelyn Wynn, Mairead Hyland, The Lighthouse Lab in Alderley Park and Alex Alderton, Roberto Amato, Sonia Goncalves, Ewan Harrison, David K. Jackson, Ian Johnston, Dominic Kwiatkowski, Cordelia Langford, John Sillitoe on behalf of the Wellcome Sanger Institute COVID-19 Surveillance Team |
| EPI_ISL_676313, EPI_ISL_676314, EPI_ISL_676315, EPI_ISL_676316, EPI_ISL_676317, EPI_ISL_676318, EPI_ISL_676319, EPI_ISL_676320, EPI_ISL_676321, EPI_ISL_676322, EPI_ISL_676323, EPI_ISL_676324, EPI_ISL_676325, EPI_ISL_676326, EPI_ISL_676327, EPI_ISL_676328, EPI_ISL_676329, EPI_ISL_676330, EPI_ISL_676331, EPI_ISL_676332, EPI_ISL_676333, EPI_ISL_676334, EPI_ISL_676335, EPI_ISL_676336, EPI_ISL_676337, EPI_ISL_676338, EPI_ISL_676339, EPI_ISL_676340, EPI_ISL_676341, EPI_ISL_676342, EPI_ISL_676343, EPI_ISL_676344, EPI_ISL_676345, EPI_ISL_676346, EPI_ISL_676347, EPI_ISL_676348, EPI_ISL_676349, EPI_ISL_676350, EPI_ISL_676351, EPI_ISL_676352, EPI_ISL_676359, EPI_ISL_676360, EPI_ISL_676361, EPI_ISL_676362, EPI_ISL_676363, EPI_ISL_676364, EPI_ISL_676365, EPI_ISL_676366, EPI_ISL_676367, EPI_ISL_676368, EPI_ISL_676369, EPI_ISL_676370, EPI_ISL_676371, EPI_ISL_676372, EPI_ISL_676373, EPI_ISL_676374, EPI_ISL_676375, EPI_ISL_676376, EPI_ISL_676377, EPI_ISL_676378, EPI_ISL_676379, EPI_ISL_676380, EPI_ISL_676381, EPI_ISL_676382, EPI_ISL_676384, EPI_ISL_676385, EPI_ISL_676386, EPI_ISL_676387, EPI_ISL_676388, EPI_ISL_676389, EPI_ISL_676391, EPI_ISL_676392, EPI_ISL_676393, EPI_ISL_676394 | | | |
| see above | Lighthouse Lab in Glasgow | Wellcome Sanger Institute for the COVID-19 Genomics UK (COG-UK) Consortium | Harper VanSteenhouse, Yumi Kasai, David Gray, Carol Clugston, Anna Dominiczak and Alex Alderton, Roberto Amato, Sonia Goncalves, Ewan Harrison, David K. Jackson, Ian Johnston, Dominic Kwiatkowski, Cordelia Langford, John Sillitoe on behalf of the Wellcome Sanger Institute COVID-19 Surveillance Team |
| EPI_ISL_676505 | Klinisk mikrobiologi | The Public Health Agency of Sweden | Department of Microbiology, The Public Health Agency of Sweden |
| EPI_ISL_677203, EPI_ISL_677204, EPI_ISL_677205, EPI_ISL_677206, EPI_ISL_677207, EPI_ISL_677208, EPI_ISL_677209, EPI_ISL_677210 | Virginia Division of Consolidated Laboratory Services | Virginia Division of Consolidated Laboratory Services | Virginia DCLS |
| EPI_ISL_677507, EPI_ISL_677508 | University of Wisconsin-Madison AIDS Vaccine Research Laboratories | University of Wisconsin-Madison AIDS Vaccine Research Laboratories | Gage Moreno, Katarina Braun, et al. AIDS Vaccine Research Laboratories |
| EPI_ISL_678574, EPI_ISL_678625, EPI_ISL_678627, EPI_ISL_678630, EPI_ISL_678632, EPI_ISL_678633, EPI_ISL_678634, EPI_ISL_678636, EPI_ISL_678642 | NHLS-IALCH | KRISP, KZN Research Innovation and Sequencing Platform | Giandhari J, Pillay S, Lessells R, ChimukangaraB, Mdlalose K, York D, Khan S, Tegally H, Wilkinson E, de Oliveira T |
| EPI_ISL_678705 | Respiratory Virus Unit, Microbiology Services Colindale, Public Health England | COVID-19 Genomics UK (COG-UK) Consortium | PHE Covid Sequencing Team |
| EPI_ISL_678832, EPI_ISL_678833, EPI_ISL_678834, EPI_ISL_678835, EPI_ISL_678836, EPI_ISL_678837, EPI_ISL_678838, EPI_ISL_678839, EPI_ISL_678840, EPI_ISL_678841, EPI_ISL_678842, EPI_ISL_678843, EPI_ISL_678844, EPI_ISL_678845, EPI_ISL_678846, EPI_ISL_678847, EPI_ISL_678848, EPI_ISL_678849, EPI_ISL_678850, EPI_ISL_678851, EPI_ISL_678852, EPI_ISL_678853, EPI_ISL_678854, EPI_ISL_678855, EPI_ISL_678856, EPI_ISL_678857, EPI_ISL_678858, EPI_ISL_678859, EPI_ISL_678860, EPI_ISL_678861, EPI_ISL_678862, EPI_ISL_678863, EPI_ISL_678864, EPI_ISL_678865, EPI_ISL_678866, EPI_ISL_678867, EPI_ISL_678868, EPI_ISL_678869, EPI_ISL_678870, EPI_ISL_678871, EPI_ISL_678872, EPI_ISL_678873, EPI_ISL_678874, EPI_ISL_678875, EPI_ISL_678876, EPI_ISL_678877, EPI_ISL_678878, EPI_ISL_678879, EPI_ISL_678880, EPI_ISL_678881, EPI_ISL_678882, EPI_ISL_678883, EPI_ISL_678884, EPI_ISL_678885, EPI_ISL_678886, EPI_ISL_678887, EPI_ISL_678888, EPI_ISL_678889, EPI_ISL_678890, EPI_ISL_678891, EPI_ISL_678892, EPI_ISL_678893, EPI_ISL_678894, EPI_ISL_678895, EPI_ISL_678896, EPI_ISL_678897, EPI_ISL_678898, EPI_ISL_678899, EPI_ISL_678900, EPI_ISL_678901, EPI_ISL_678902, EPI_ISL_678903, EPI_ISL_678904, EPI_ISL_678905, EPI_ISL_678906, EPI_ISL_678907, EPI_ISL_678908, EPI_ISL_678909, EPI_ISL_678910, EPI_ISL_678911, EPI_ISL_678912, EPI_ISL_678913, EPI_ISL_678914, EPI_ISL_678915 | | | |
| see above | Department of Pathology, University of Cambridge | COVID-19 Genomics UK (COG-UK) Consortium | Aminu S. Jahun, Yasmin Chaudhry, Grant Hall, Iliana Georgana, Myra Hosmillo, Martin D. Curran, Malte Pinckert, Surendra Parmar, Ian Goodfellow |

| | | | |
|---|---|---|---|
| EPI_ISL_679008, EPI_ISL_679009, EPI_ISL_679010, EPI_ISL_679011, EPI_ISL_679012, EPI_ISL_679013, EPI_ISL_679014, EPI_ISL_679015, EPI_ISL_679016, EPI_ISL_679017, EPI_ISL_679018, EPI_ISL_679019, EPI_ISL_679020, EPI_ISL_679021, EPI_ISL_679022, EPI_ISL_679023, EPI_ISL_679024, EPI_ISL_679025, EPI_ISL_679026, EPI_ISL_679027, EPI_ISL_679028, EPI_ISL_679029, EPI_ISL_679030, EPI_ISL_679031, EPI_ISL_679032, EPI_ISL_679033, EPI_ISL_679034, EPI_ISL_679035, EPI_ISL_679036, EPI_ISL_679037, EPI_ISL_679038, EPI_ISL_679039, EPI_ISL_679040, EPI_ISL_679041, EPI_ISL_679042, EPI_ISL_679043, EPI_ISL_679044, EPI_ISL_679045, EPI_ISL_679046, EPI_ISL_679047, EPI_ISL_679048, EPI_ISL_679049, EPI_ISL_679050, EPI_ISL_679053, EPI_ISL_679054, EPI_ISL_679055, EPI_ISL_679056, EPI_ISL_679057, EPI_ISL_679058, EPI_ISL_679059, EPI_ISL_679060, EPI_ISL_679061, EPI_ISL_679062, EPI_ISL_679063, EPI_ISL_679064, EPI_ISL_679065, EPI_ISL_679066, EPI_ISL_679067, EPI_ISL_679068, EPI_ISL_679069, EPI_ISL_679070, EPI_ISL_679071, EPI_ISL_679072, EPI_ISL_679073, EPI_ISL_679074, EPI_ISL_679075, EPI_ISL_679076, EPI_ISL_679077, EPI_ISL_679078, EPI_ISL_679079, EPI_ISL_679080, EPI_ISL_679081, EPI_ISL_679082, EPI_ISL_679083, EPI_ISL_679084, EPI_ISL_679085, EPI_ISL_679086, EPI_ISL_679087, EPI_ISL_679088, EPI_ISL_679089, EPI_ISL_679090, EPI_ISL_679091, EPI_ISL_679092, EPI_ISL_679093, EPI_ISL_679094, EPI_ISL_679095, EPI_ISL_679096, EPI_ISL_679097, EPI_ISL_679098, EPI_ISL_679099, EPI_ISL_679100, EPI_ISL_679101, EPI_ISL_679102, EPI_ISL_679103, EPI_ISL_679129 | | | |
| see above | University of Birmingham | COVID-19 Genomics UK (COG-UK) Consortium | Institute of Microbiology, University of Birmingham: Claire McMurray, Joanne Stockton, Samuel Nicholls, Radoslaw Poplawski, Will Rowe, Josh Quick, Nicholas Loman. University of Birmingham Testing Laboratory: Celina M Whalley, Andrew Bosworth, Charlotte Poxon, Kasun Wanigasooriya, Oliver Pickles, Mike Kidd, Alex Richter, Andrew D Beggs PHE Heartlands Lab: Husam Osman, Andrew Bosworth. Queen Elizabeth Hospital: Anna Casey |
| EPI_ISL_679154, EPI_ISL_679155, EPI_ISL_679158 | Department of Pathology, University of Cambridge | COVID-19 Genomics UK (COG-UK) Consortium | Aminu S. Jahun, Yasmin Chaudhry, Grant Hall, Iliana Georgana, Myra Hosmillo, Martin D. Curran, Malte Pinckert, Surendra Parmar, Ian Goodfellow |
| EPI_ISL_679381, EPI_ISL_679382, EPI_ISL_679383, EPI_ISL_679384, EPI_ISL_679385, EPI_ISL_679386, EPI_ISL_679387, EPI_ISL_679388 | Liverpool Clinical Laboratories | COVID-19 Genomics UK (COG-UK) Consortium | Sam Haldenby, Anita Lucaci, Steve Paterson, Julian Hiscox, Alistair Darby, M Almsaud, A Alrezaihi, Muhannad Alruwaili, Stuart D Armstrong, Jones Benjamin, Eleanor G Bentley, Anu Chawla, Jordan J Clark, Angela Cowell, Richard Eccles, Isabel García-Dorival, Matthew Gemmell, Alessandro Gerada, PKF Gilmore, Richard Gregory, Ximeng Han, Catherine Hartley, Margaret Hughes, Miren Iturriza-Gomara, James Johnson, L Luu, Jenifer Manson, Charlotte Nelson, Elaine O'Toole, Cassie Olateju, Rebekah Penrice-Randal , Lucille Rainbow, N.P Randle, Trevor Ian Robinson, Parul Sharma, Ghada T Shawli, James P Stewart, Neil Swainston, Ecaterina Vamos, Joanne Watts, Mark Whitehead |
| EPI_ISL_679478, EPI_ISL_679480, EPI_ISL_679481, EPI_ISL_679482, EPI_ISL_679484, EPI_ISL_679485, EPI_ISL_679486, EPI_ISL_679487, EPI_ISL_679488, EPI_ISL_679489, EPI_ISL_679490, EPI_ISL_679491, EPI_ISL_679493, EPI_ISL_679494, EPI_ISL_679495, EPI_ISL_679496, EPI_ISL_679497, EPI_ISL_679498, EPI_ISL_679499, EPI_ISL_679500, EPI_ISL_679502, EPI_ISL_679505, EPI_ISL_679621, EPI_ISL_679622, EPI_ISL_679623 | | | |
| see above | University College London, Great Ormond Street Hospital for Children NHS Foundation Trust, Imperial College Healthcare NHS Trust | COVID-19 Genomics UK (COG-UK) Consortium | Sergi Castellano, Rachel Williams, Mark Kristiansen, Paola Resende Silva, Sunando Roy, Tony Brooks, Helena Tutill, Paola Niola, Patricia Dyal, Charlotte Williams, Leysa Forrest, Yasmin Panchbhaya, Jacqueline Findlay, Samuel Weeks, Julianne Brown, Kathryn Harris, Paul Randell, James Price, Alison Holmes, Judith Breuer |
| EPI_ISL_680101, EPI_ISL_680117, EPI_ISL_680120, EPI_ISL_680122, EPI_ISL_680123, EPI_ISL_680126, EPI_ISL_680127, EPI_ISL_680129, EPI_ISL_680130, EPI_ISL_680133, EPI_ISL_680139, EPI_ISL_680145, EPI_ISL_680149, EPI_ISL_680159, EPI_ISL_680160, EPI_ISL_680163, EPI_ISL_680165, EPI_ISL_680166, EPI_ISL_680169, EPI_ISL_680172, EPI_ISL_680176, EPI_ISL_680177, EPI_ISL_680179, EPI_ISL_680182, EPI_ISL_680184, EPI_ISL_680188, EPI_ISL_680189, EPI_ISL_680195, EPI_ISL_680197 | | | |
| see above | Virology Department, Sheffield Teaching Hospitals NHS Foundation Trust/Department of Infection, Immunity and Cardiovascular Disease, The Medical School, University of Sheffield | COVID-19 Genomics UK (COG-UK) Consortium | Thushan de Silva, Matthew Parker, Nikki Smith, Adri Angyal, Rebecca Brown, Luke Green, Rachel Tucker, Paul Parsons, Danielle Groves, Katie Johnson, Laura Carrilero, Alex Keeley, Dave Partridge, Matthew Wyles, Benjamin Lindsey, Mehmet Yavuz, Mohammad Raza, Cariad Evans |
| EPI_ISL_680446, EPI_ISL_680447, EPI_ISL_680448 | West of Scotland Specialist Virology Centre, NHSGGC / MRC-University of Glasgow Centre for Virus Research | COVID-19 Genomics UK (COG-UK) Consortium | Ana da Silva Filipe, Natasha Johnson, Kathy Smollett, Daniel Mair, Stephen Carmichael, Alice Broos, Lily Tong, Jenna Nichols, Kyriaki Nomikou; Sarah McDonald; Richard Orton, Joseph Hughes, Sreenu Vattipally, David L Robertson; Alasdair MacLean, Rory Gunson; Sharif Shaaban, Matthew Holden; Rachel Blacow, Guy Mollett, Kathy Li, James Shepherd, Antonia Ho, Emma Thomson |
| EPI_ISL_680510, EPI_ISL_680538, EPI_ISL_680546, EPI_ISL_680547 | Virology Department, Royal Infirmary of Edinburgh, NHS Lothian / School of Biological Sciences, University of Edinburgh / Institute of Genetics and Molecular Medicine, University of Edinburgh | COVID-19 Genomics UK (COG-UK) Consortium | McHugh M, Dewar R, Rooke S, Gallagher M, Balcaza C, O'Toole Á, Scher E, Hill V, McCrone JT, Colquhoun R, Yu X, Jackson B, Rambaut A, Williams TC, Templeton K |
| EPI_ISL_680586, EPI_ISL_680606, EPI_ISL_680637, EPI_ISL_680638, EPI_ISL_680639, EPI_ISL_680671, EPI_ISL_680672, EPI_ISL_680677, EPI_ISL_680683, EPI_ISL_680690, EPI_ISL_680692, EPI_ISL_680693, EPI_ISL_680694, EPI_ISL_680695, EPI_ISL_680696, EPI_ISL_680697, EPI_ISL_680698, EPI_ISL_680701, EPI_ISL_680713, EPI_ISL_680742, EPI_ISL_680754, EPI_ISL_680761, EPI_ISL_680772, EPI_ISL_680777, EPI_ISL_680778, EPI_ISL_680779, EPI_ISL_680780, EPI_ISL_680781, EPI_ISL_680782, EPI_ISL_680785, EPI_ISL_680786, EPI_ISL_680787, EPI_ISL_680788, EPI_ISL_680789, EPI_ISL_680790, EPI_ISL_680791, EPI_ISL_680792, EPI_ISL_680793, EPI_ISL_680794, EPI_ISL_680795, EPI_ISL_680796, EPI_ISL_680797, EPI_ISL_680798, EPI_ISL_680799, EPI_ISL_680800, EPI_ISL_680801, EPI_ISL_680802, EPI_ISL_680803, EPI_ISL_680804, EPI_ISL_680805, EPI_ISL_680806, EPI_ISL_680807, EPI_ISL_680808, EPI_ISL_680809, EPI_ISL_680810, EPI_ISL_680811, EPI_ISL_680812, EPI_ISL_680813, EPI_ISL_680814, EPI_ISL_680815, EPI_ISL_680816, EPI_ISL_680817, EPI_ISL_680818, EPI_ISL_680819, EPI_ISL_680822, EPI_ISL_680823, EPI_ISL_680824, EPI_ISL_680825, EPI_ISL_680826, EPI_ISL_680827, EPI_ISL_680828, EPI_ISL_680829, EPI_ISL_680830, EPI_ISL_680831, EPI_ISL_680832, EPI_ISL_680833, EPI_ISL_680834, EPI_ISL_680835, EPI_ISL_680836, EPI_ISL_680837, EPI_ISL_680838, EPI_ISL_680840, EPI_ISL_680841, EPI_ISL_680842, EPI_ISL_680843, EPI_ISL_680844, EPI_ISL_680845, EPI_ISL_680846, EPI_ISL_680847, EPI_ISL_680852, EPI_ISL_680853, EPI_ISL_680854, EPI_ISL_680855, EPI_ISL_680857, EPI_ISL_680953, EPI_ISL_680954, EPI_ISL_680957, EPI_ISL_680958, EPI_ISL_680970, EPI_ISL_680971, EPI_ISL_680972, EPI_ISL_680973, EPI_ISL_680974, EPI_ISL_680975, EPI_ISL_680976, EPI_ISL_680977, EPI_ISL_680978, EPI_ISL_680979, EPI_ISL_680980, EPI_ISL_680981, EPI_ISL_680982, EPI_ISL_680983, EPI_ISL_680984, EPI_ISL_680985, EPI_ISL_680986, EPI_ISL_680987, EPI_ISL_680988, EPI_ISL_680989, EPI_ISL_680990, EPI_ISL_680991, EPI_ISL_680992, EPI_ISL_680993, EPI_ISL_680994, EPI_ISL_680995, EPI_ISL_680996, EPI_ISL_680997, EPI_ISL_680998, EPI_ISL_680999, EPI_ISL_681000, EPI_ISL_681001, EPI_ISL_681002, EPI_ISL_681003, EPI_ISL_681004, EPI_ISL_681005, EPI_ISL_681006, EPI_ISL_681007, EPI_ISL_681008, EPI_ISL_681009, EPI_ISL_681010, EPI_ISL_681011, EPI_ISL_681012, EPI_ISL_681018, EPI_ISL_681021, EPI_ISL_681022, EPI_ISL_681023, EPI_ISL_681024, EPI_ISL_681025, EPI_ISL_681026, EPI_ISL_681027, EPI_ISL_681044, EPI_ISL_681048, EPI_ISL_681050, EPI_ISL_681054, EPI_ISL_681055, EPI_ISL_681056, EPI_ISL_681065, EPI_ISL_681068, EPI_ISL_681069, EPI_ISL_681070, EPI_ISL_681071, EPI_ISL_681073, EPI_ISL_681074, EPI_ISL_681077, EPI_ISL_681078, EPI_ISL_681079, EPI_ISL_681080, EPI_ISL_681081, EPI_ISL_681082, EPI_ISL_681083, EPI_ISL_681084, EPI_ISL_681085, EPI_ISL_681086, EPI_ISL_681087, EPI_ISL_681088, EPI_ISL_681089, EPI_ISL_681090, EPI_ISL_681091, EPI_ISL_681092, EPI_ISL_681093, EPI_ISL_681094, EPI_ISL_681095, EPI_ISL_681096, EPI_ISL_681097, EPI_ISL_681098, EPI_ISL_681099, EPI_ISL_681100, EPI_ISL_681101, EPI_ISL_681102, EPI_ISL_681103, EPI_ISL_681104, EPI_ISL_681105, EPI_ISL_681106, EPI_ISL_681107, EPI_ISL_681108, EPI_ISL_681109, EPI_ISL_681110, EPI_ISL_681111, EPI_ISL_681112, EPI_ISL_681113, EPI_ISL_681114, EPI_ISL_681115, EPI_ISL_681116, EPI_ISL_681117, EPI_ISL_681118, EPI_ISL_681119, EPI_ISL_681120, EPI_ISL_681121, EPI_ISL_681122, EPI_ISL_681123, EPI_ISL_681124, EPI_ISL_681125, EPI_ISL_681126, EPI_ISL_681127, EPI_ISL_681128, EPI_ISL_681129, EPI_ISL_681130, EPI_ISL_681131, EPI_ISL_681132, EPI_ISL_681133, EPI_ISL_681134, EPI_ISL_681135, EPI_ISL_681136, EPI_ISL_681137, EPI_ISL_681138, EPI_ISL_681139, EPI_ISL_681140, EPI_ISL_681141, EPI_ISL_681142, EPI_ISL_681143, EPI_ISL_681144, EPI_ISL_681145, EPI_ISL_681146, EPI_ISL_681147, EPI_ISL_681148, EPI_ISL_681149, EPI_ISL_681150, EPI_ISL_681151, EPI_ISL_681152, EPI_ISL_681153, EPI_ISL_681154, EPI_ISL_681155, EPI_ISL_681156, EPI_ISL_681157, EPI_ISL_681158, EPI_ISL_681159, EPI_ISL_681160, EPI_ISL_681161, EPI_ISL_681162, EPI_ISL_681163, EPI_ISL_681164, EPI_ISL_681165, EPI_ISL_681166, EPI_ISL_681167, EPI_ISL_681168, EPI_ISL_681169, EPI_ISL_681170, EPI_ISL_681171, EPI_ISL_681172, EPI_ISL_681173, EPI_ISL_681174, EPI_ISL_681175, EPI_ISL_681177, EPI_ISL_681178, EPI_ISL_681179, EPI_ISL_681180, EPI_ISL_681181, EPI_ISL_681182, EPI_ISL_681183, EPI_ISL_681184, EPI_ISL_681185, EPI_ISL_681186, EPI_ISL_681187, EPI_ISL_681188, EPI_ISL_681189, EPI_ISL_681190, EPI_ISL_681191, EPI_ISL_681192, EPI_ISL_681193, EPI_ISL_681194, EPI_ISL_681195, EPI_ISL_681196, EPI_ISL_681197, EPI_ISL_681198, EPI_ISL_681199, EPI_ISL_681200, EPI_ISL_681201, EPI_ISL_681202, EPI_ISL_681203, EPI_ISL_681204, EPI_ISL_681205, EPI_ISL_681206, EPI_ISL_681207, EPI_ISL_681208, EPI_ISL_681209, EPI_ISL_681210, EPI_ISL_681211, EPI_ISL_681212, EPI_ISL_681213, EPI_ISL_681214, EPI_ISL_681215, EPI_ISL_681216, EPI_ISL_681217, EPI_ISL_681218, EPI_ISL_681219, EPI_ISL_681220, EPI_ISL_681221, EPI_ISL_681222, EPI_ISL_681223, EPI_ISL_681224, EPI_ISL_681225, EPI_ISL_681226, EPI_ISL_681227, EPI_ISL_681228, EPI_ISL_681229, EPI_ISL_681230, EPI_ISL_681231, EPI_ISL_681232, EPI_ISL_681233, EPI_ISL_681234, EPI_ISL_681235, EPI_ISL_681236, EPI_ISL_681237, EPI_ISL_681238, EPI_ISL_681239, EPI_ISL_681240, EPI_ISL_681241, EPI_ISL_681242, EPI_ISL_681243, EPI_ISL_681244, EPI_ISL_681245, EPI_ISL_681247, EPI_ISL_681248, EPI_ISL_681249, EPI_ISL_681250, EPI_ISL_681251, EPI_ISL_681252, EPI_ISL_681253, EPI_ISL_681254, EPI_ISL_681255 | | | |
| see above | Wales Specialist Virology Centre Sequencing lab: Pathogen Genomics Unit | COVID-19 Genomics UK (COG-UK) Consortium | Catherine Moore, Johnathan Evans, Laura Gifford, Malorie Perry, Simon Cottrell, Angela Marchbank, Alec Birchley, Alexander Adams, Amy Gaskin, Bree Gatica-Wilcox, Jason Coombes, Joel Southgate, Lauren Gilbert, Lee Graham, Nicole Pacchiarini, Sara Kumziene-Summerhayes, Sarah Taylor, Sophie Jones, Sara Rey, Matthew Bull, Joanne Watkins, Sally Corden, Tom Connor |
| EPI_ISL_681302, EPI_ISL_681303, EPI_ISL_681318, EPI_ISL_681319 | Communicable Disease Laboratory, Public Health Directorate | Communicable Disease Laboratory, Public Health Directorate | Alwasti,H., Altaif,Z., AlHujairi,Z., AlAbbas,Z. |
| EPI_ISL_682060, EPI_ISL_682061, EPI_ISL_682064, EPI_ISL_682065, EPI_ISL_682066, EPI_ISL_682067, EPI_ISL_682068, EPI_ISL_682070, EPI_ISL_682071, EPI_ISL_682072, EPI_ISL_682073, EPI_ISL_682075, EPI_ISL_682076, EPI_ISL_682077, EPI_ISL_682078 | | | |
| see above | Maryland Public Health Laboratory | Maryland Public Health Laboratory | Maryland Department of Health Laboratories Administration |
| EPI_ISL_682317 | Communicable Disease Laboratory, Public Health Directorate | Communicable Disease Laboratory, Public Health Directorate | Alwasti,H., Altaif,Z., AlHujairi,Z., AlAbbas,Z. |
| EPI_ISL_682930, EPI_ISL_682931, EPI_ISL_682932, EPI_ISL_682933, EPI_ISL_682934, EPI_ISL_682935, EPI_ISL_682936, EPI_ISL_682937, EPI_ISL_682938, EPI_ISL_682939, EPI_ISL_682940, EPI_ISL_682941, EPI_ISL_682942, EPI_ISL_682943, EPI_ISL_682944, EPI_ISL_682945, EPI_ISL_682946, EPI_ISL_682947, EPI_ISL_682948, EPI_ISL_682949, EPI_ISL_682950, EPI_ISL_682951, EPI_ISL_682952, EPI_ISL_682953, EPI_ISL_682954, EPI_ISL_682955, EPI_ISL_682956, EPI_ISL_682957, EPI_ISL_682958, EPI_ISL_682959, EPI_ISL_682960, EPI_ISL_682961, EPI_ISL_682962, EPI_ISL_682963, EPI_ISL_682964, EPI_ISL_682965, EPI_ISL_682966, EPI_ISL_683111, EPI_ISL_683112, EPI_ISL_683113, EPI_ISL_683114, EPI_ISL_683115, EPI_ISL_683116, EPI_ISL_683117, EPI_ISL_683118, EPI_ISL_683119, EPI_ISL_683120, EPI_ISL_683121, EPI_ISL_683122, EPI_ISL_683123, EPI_ISL_683124, EPI_ISL_683125, EPI_ISL_683126, EPI_ISL_683127, EPI_ISL_683128, EPI_ISL_683129, EPI_ISL_683130, EPI_ISL_683131, EPI_ISL_683132, EPI_ISL_683133, EPI_ISL_683134, EPI_ISL_683135, EPI_ISL_683136, EPI_ISL_683137, EPI_ISL_683138, EPI_ISL_683139, EPI_ISL_683140, EPI_ISL_683141, EPI_ISL_683142, EPI_ISL_683143, EPI_ISL_683144, EPI_ISL_683145, EPI_ISL_683146, EPI_ISL_683147, EPI_ISL_683148, EPI_ISL_683149, EPI_ISL_683150, EPI_ISL_683151, EPI_ISL_683152, EPI_ISL_683153, EPI_ISL_683154, EPI_ISL_683155, EPI_ISL_683156, EPI_ISL_683157, EPI_ISL_683158, EPI_ISL_683159, EPI_ISL_683160 | | | |
| see above | Department of Virus and Microbiological Special Diagnostics, Statens Serum Institut, Copenhagen, Denmark | Albertsen Lab, Department of Chemistry and Bioscience, Aalborg University, Denmark | Danish Covid-19 Genome Consortium |
| EPI_ISL_683343, EPI_ISL_683344, EPI_ISL_683345, EPI_ISL_683346, EPI_ISL_683347, EPI_ISL_683348, EPI_ISL_683349, EPI_ISL_683350, EPI_ISL_683351 | CNR Virus des Infections Respiratoires - France SUD | CNR Virus des Infections Respiratoires - France SUD | Antonin Bal, Gregory Destras, Gwendolyne Burfin, Quentin Semanas, Martine Valette, Bruno Lina, Laurence Josset |
| EPI_ISL_683481, EPI_ISL_683516, EPI_ISL_683517, EPI_ISL_683518, EPI_ISL_683523 | Respiratory Virus Unit, Microbiology Services Colindale, Public Health England | COVID-19 Genomics UK (COG-UK) Consortium | PHE Covid Sequencing Team |
| EPI_ISL_683598, EPI_ISL_683606, | Servicio de Microbiología, Laboratori Clínic Metropolitana | SeqCOVID-SPAIN consortium/IBV(CSIC) | Elisa Martró, Antoni E. Bordoy, Anna Not, Adrián Antuori, Anabel Fernández, Nona Romaní, Verónica Saludes, Cristina Casañ and SeqCOVID-SPAIN |

| | | | |
|---|---|---|---|
| EPI_ISL_683607, EPI_ISL_683621, EPI_ISL_683625, EPI_ISL_683627, EPI_ISL_683632 | Nord. Hospital Universitari Germans Trias i Pujol. Institut d'Investigació en Ciències de la Salut Germans Trias i Pujol (IGTP) | | consortium |
| EPI_ISL_683730, EPI_ISL_683731, EPI_ISL_683732, EPI_ISL_683733 | Minnesota Department of Health, Public Health Laboratory | Minnesota Department of Health, Public Health Laboratory | Alexandra Lorentz, Jacob Garfin, Matt Plumb, and Xiong Wang |
| EPI_ISL_683836 | DOHMH PHL | New York City Public Health Laboratory | Jade Wang, et al. |
| EPI_ISL_683841 | DOHMH Corona | New York City Public Health Laboratory | Jade Wang, et al. |
| EPI_ISL_683845 | DOHMH Morrisania | New York City Public Health Laboratory | Jade Wang, et al. |
| EPI_ISL_683846 | DOHMH PHL | New York City Public Health Laboratory | Jade Wang, et al. |
| EPI_ISL_683857 | DOHMH Corona | New York City Public Health Laboratory | Jade Wang, et al. |
| EPI_ISL_683861, EPI_ISL_683862 | DOHMH Jamaica | New York City Public Health Laboratory | Jade Wang, et al. |
| EPI_ISL_683863 | DOHMH Morrisania | New York City Public Health Laboratory | Jade Wang, et al. |
| EPI_ISL_683864 | DOHMH Jamaica | New York City Public Health Laboratory | Jade Wang, et al. |
| EPI_ISL_683865 | DOHMH Chelsea | New York City Public Health Laboratory | Jade Wang, et al. |
| EPI_ISL_683867, EPI_ISL_683870, EPI_ISL_683873 | DOHMH Corona | New York City Public Health Laboratory | Jade Wang, et al. |
| EPI_ISL_683874 | DOHMH Jamaica | New York City Public Health Laboratory | Jade Wang, et al. |
| EPI_ISL_683876 | DOHMH Corona | New York City Public Health Laboratory | Jade Wang, et al. |
| EPI_ISL_683878 | DOHMH Jamaica | New York City Public Health Laboratory | Jade Wang, et al. |
| EPI_ISL_683879, EPI_ISL_683880 | DOHMH Morrisania | New York City Public Health Laboratory | Jade Wang, et al. |
| EPI_ISL_683896, EPI_ISL_683897, EPI_ISL_683898, EPI_ISL_683899, EPI_ISL_683900 | DOHMH Corona | New York City Public Health Laboratory | Jade Wang, et al. |
| EPI_ISL_683901 | DOHMH Jamaica | New York City Public Health Laboratory | Jade Wang, et al. |
| EPI_ISL_683902 | DOHMH Riverside | New York City Public Health Laboratory | Jade Wang, et al. |
| EPI_ISL_683903 | DOHMH Jamaica | New York City Public Health Laboratory | Jade Wang, et al. |
| EPI_ISL_683904 | DOHMH Crown Heights | New York City Public Health Laboratory | Jade Wang, et al. |
| EPI_ISL_683905 | DOHMH Morrisania | New York City Public Health Laboratory | Jade Wang, et al. |
| EPI_ISL_683906 | DOHMH Fort Greene | New York City Public Health Laboratory | Jade Wang, et al. |
| EPI_ISL_683907 | DOHMH Riverside | New York City Public Health Laboratory | Jade Wang, et al. |
| EPI_ISL_683908, EPI_ISL_683909 | DOHMH Corona | New York City Public Health Laboratory | Jade Wang, et al. |
| EPI_ISL_683910 | DOHMH Crown Heights | New York City Public Health Laboratory | Jade Wang, et al. |
| EPI_ISL_683911, EPI_ISL_683912 | DOHMH Corona | New York City Public Health Laboratory | Jade Wang, et al. |
| EPI_ISL_683913 | DOHMH Fort Greene | New York City Public Health Laboratory | Jade Wang, et al. |
| EPI_ISL_683914, EPI_ISL_683915, EPI_ISL_683916 | DOHMH Jamaica | New York City Public Health Laboratory | Jade Wang, et al. |
| EPI_ISL_683917, EPI_ISL_683918, EPI_ISL_683919 | DOHMH Morrisania | New York City Public Health Laboratory | Jade Wang, et al. |
| EPI_ISL_683952, EPI_ISL_683953 | DOHMH Corona | New York City Public Health Laboratory | Jade Wang, et al. |
| EPI_ISL_683954 | DOHMH PHL | New York City Public Health Laboratory | Jade Wang, et al. |
| EPI_ISL_683955 | DOHMH Riverside | New York City Public Health Laboratory | Jade Wang, et al. |
| EPI_ISL_683956 | DOHMH Jamaica | New York City Public Health Laboratory | Jade Wang, et al. |
| EPI_ISL_683957 | DOHMH Chelsea | New York City Public Health Laboratory | Jade Wang, et al. |
| EPI_ISL_683958 | DOHMH Jamaica | New York City Public Health Laboratory | Jade Wang, et al. |
| EPI_ISL_683959 | DOHMH Morrisania | New York City Public Health Laboratory | Jade Wang, et al. |
| EPI_ISL_683960 | DOHMH Corona | New York City Public Health Laboratory | Jade Wang, et al. |
| EPI_ISL_683961 | DOHMH Chelsea | New York City Public Health Laboratory | Jade Wang, et al. |
| EPI_ISL_683962 | DOHMH Central Harlem | New York City Public Health Laboratory | Jade Wang, et al. |
| EPI_ISL_683963 | DOHMH Fort Greene | New York City Public Health Laboratory | Jade Wang, et al. |
| EPI_ISL_683964 | DOHMH Central Harlem | New York City Public Health Laboratory | Jade Wang, et al. |
| EPI_ISL_683965, EPI_ISL_683966 | DOHMH Morrisania | New York City Public Health Laboratory | Jade Wang, et al. |
| EPI_ISL_686108, EPI_ISL_686560, EPI_ISL_686561, EPI_ISL_686562, EPI_ISL_686563, EPI_ISL_686564, EPI_ISL_686565, EPI_ISL_686566, EPI_ISL_686567, EPI_ISL_686568, EPI_ISL_686569, EPI_ISL_686570, EPI_ISL_686571, EPI_ISL_686572, EPI_ISL_686573, EPI_ISL_686574, EPI_ISL_686575, EPI_ISL_686576, EPI_ISL_686577, EPI_ISL_686578, EPI_ISL_686579 | | | |
| see above | Respiratory Virus Unit, Microbiology Services Colindale, Public Health England | COVID-19 Genomics UK (COG-UK) Consortium | PHE Covid Sequencing Team |
| EPI_ISL_692786, EPI_ISL_692787, EPI_ISL_692788, EPI_ISL_692794, EPI_ISL_692796, EPI_ISL_692798 | Massachusetts State Public Health Laboratory | Massachusetts State Public Health Laboratory | Andrew Lang, Timelia Fink, Glen Gallagher, Sandra Smole |
| EPI_ISL_693283, EPI_ISL_693284 | unknown | Public Health Virology Laboratory, Forensic and Scientific Services (PHV-FSS) | Son Nguyen et al. |
| EPI_ISL_693302 | Department of Laboratory Medicine, National Taiwan University Hospital | Microbial Genomics Core Lab, National Taiwan University Centers of Genomic and Precision Medicine | Shiou-Hwei Yeh, You-Yu Lin, Ya-Yun Lai, Chiao-Ling Li, Shan-Chwen Chang, Pei-Jer Chen, Sui-Yuan Chang |
| EPI_ISL_693393, EPI_ISL_693400, EPI_ISL_693409, EPI_ISL_693410, EPI_ISL_693411, EPI_ISL_693414, EPI_ISL_693415, EPI_ISL_693419, EPI_ISL_693420, EPI_ISL_693422, EPI_ISL_693433, EPI_ISL_693485 | | | |

| | | | |
|---|---|---|---|
| see above | Respiratory Virus Unit, Microbiology Services Colindale, Public Health England | COVID-19 Genomics UK (COG-UK) Consortium | PHE Covid Sequencing Team |
| EPI_ISL_693771, EPI_ISL_693772 | General practitioner | National Reference Center for Viruses of Respiratory Infections, Institut Pasteur, Paris | Marion Barbet, Sylvie Behillil, Méline Bizard, Angela Brisebarre, Camille Capel, Etienne Simon-Lorière, Vincent Enouf, Maud Vanpeene, Sylvie van der Werf |
| EPI_ISL_693875, EPI_ISL_693876, EPI_ISL_693877, EPI_ISL_693878, EPI_ISL_693879, EPI_ISL_693880, EPI_ISL_693881, EPI_ISL_693882, EPI_ISL_693883, EPI_ISL_693884, EPI_ISL_693885, EPI_ISL_693886, EPI_ISL_693887, EPI_ISL_693888, EPI_ISL_693889, EPI_ISL_693890, EPI_ISL_693891, EPI_ISL_693892, EPI_ISL_693893, EPI_ISL_693894, EPI_ISL_693942, EPI_ISL_693943 | | | |
| see above | Viollier AG | Department of Biosystems Science and Engineering, ETH Zürich | Christian Beisel, Sarah Nadeau, Chaoran Chen, Ivan Topolsky, Pedro Ferreira, Philipp Jablonski, Susana Posada-Céspedes, Tobias Schär, Ina Nissen, Natascha Santacroce, Elodie Burcklen, Christiane Beckmann, Maurice Redondo, Olivier Kobel, Christoph Noppen, Sophie Seidel, Noemie Santamaria de Souza, Niko Beerenwinkel, Tanja Stadler |
| EPI_ISL_699653 | South Eastern Area Laboratory Services (SEALS) | NSW Health Pathology - Institute of Clinical Pathology and Medical Research; Westmead Hospital; University of Sydney | CIDM-PH et al. |
| EPI_ISL_702474 | Department of Pathology, University of Cambridge | COVID-19 Genomics UK (COG-UK) Consortium | Aminu S. Jahun, Yasmin Chaudhry, Grant Hall, Iliana Georgana, Myra Hosmillo, Martin D. Curran, Malte Pinckert, Surendra Parmar, Ian Goodfellow |
| EPI_ISL_702498 | Liverpool Clinical Laboratories | COVID-19 Genomics UK (COG-UK) Consortium | Sam Haldenby, Anita Lucaci, Steve Paterson, Julian Hiscox, Alistair Darby, M Almsaud, A Alrezaihi, Muhannad Alruwaili, Stuart D Armstrong, Jones Benjamin, Eleanor G Bentley, Anu Chawla, Jordan J Clark, Angela Cowell, Richard Eccles, Isabel García-Dorival, Matthew Gemmell, Alessandro Gerada, PKF Gilmore, Richard Gregory, Ximeng Han, Catherine Hartley, Margaret Hughes, Miren Iturriza-Gomara, James Johnson, L Luu, Jenifer Manson, Charlotte Nelson, Elaine O'Toole, Cassie Olateju, Rebekah Penrice-Randal , Lucille Rainbow, N.P Randle, Trevor Ian Robinson, Parul Sharma, Ghada T Shawli, James P Stewart, Neil Swainston, Ecaterina Vamos, Joanne Watts, Mark Whitehead |
| EPI_ISL_702528 | Virology Department, Royal Infirmary of Edinburgh, NHS Lothian / School of Biological Sciences, University of Edinburgh / Institute of Genetics and Molecular Medicine, University of Edinburgh | COVID-19 Genomics UK (COG-UK) Consortium | McHugh M, Dewar R, Rooke S, Gallagher M, Balcaza C, O'Toole Á, Scher E, Hill V, McCrone JT, Colquhoun R, Yu X, Jackson B, Rambaut A, Williams TC, Templeton K |
| EPI_ISL_702536, EPI_ISL_702582, EPI_ISL_702587 | Liverpool Clinical Laboratories | COVID-19 Genomics UK (COG-UK) Consortium | Sam Haldenby, Anita Lucaci, Steve Paterson, Julian Hiscox, Alistair Darby, M Almsaud, A Alrezaihi, Muhannad Alruwaili, Stuart D Armstrong, Jones Benjamin, Eleanor G Bentley, Anu Chawla, Jordan J Clark, Angela Cowell, Richard Eccles, Isabel García-Dorival, Matthew Gemmell, Alessandro Gerada, PKF Gilmore, Richard Gregory, Ximeng Han, Catherine Hartley, Margaret Hughes, Miren Iturriza-Gomara, James Johnson, L Luu, Jenifer Manson, Charlotte Nelson, Elaine O'Toole, Cassie Olateju, Rebekah Penrice-Randal , Lucille Rainbow, N.P Randle, Trevor Ian Robinson, Parul Sharma, Ghada T Shawli, James P Stewart, Neil Swainston, Ecaterina Vamos, Joanne Watts, Mark Whitehead |
| EPI_ISL_702594 | Virology Department, Royal Infirmary of Edinburgh, NHS Lothian / School of Biological Sciences, University of Edinburgh / Institute of Genetics and Molecular Medicine, University of Edinburgh | COVID-19 Genomics UK (COG-UK) Consortium | McHugh M, Dewar R, Rooke S, Gallagher M, Balcaza C, O'Toole Á, Scher E, Hill V, McCrone JT, Colquhoun R, Yu X, Jackson B, Rambaut A, Williams TC, Templeton K |
| EPI_ISL_702597, EPI_ISL_702647, EPI_ISL_702656, EPI_ISL_702664, EPI_ISL_702687, EPI_ISL_702710, EPI_ISL_702721 | Liverpool Clinical Laboratories | COVID-19 Genomics UK (COG-UK) Consortium | Sam Haldenby, Anita Lucaci, Steve Paterson, Julian Hiscox, Alistair Darby, M Almsaud, A Alrezaihi, Muhannad Alruwaili, Stuart D Armstrong, Jones Benjamin, Eleanor G Bentley, Anu Chawla, Jordan J Clark, Angela Cowell, Richard Eccles, Isabel García-Dorival, Matthew Gemmell, Alessandro Gerada, PKF Gilmore, Richard Gregory, Ximeng Han, Catherine Hartley, Margaret Hughes, Miren Iturriza-Gomara, James Johnson, L Luu, Jenifer Manson, Charlotte Nelson, Elaine O'Toole, Cassie Olateju, Rebekah Penrice-Randal , Lucille Rainbow, N.P Randle, Trevor Ian Robinson, Parul Sharma, Ghada T Shawli, James P Stewart, Neil Swainston, Ecaterina Vamos, Joanne Watts, Mark Whitehead |
| EPI_ISL_702723 | Department of Pathology, University of Cambridge | COVID-19 Genomics UK (COG-UK) Consortium | Aminu S. Jahun, Yasmin Chaudhry, Grant Hall, Iliana Georgana, Myra Hosmillo, Martin D. Curran, Malte Pinckert, Surendra Parmar, Ian Goodfellow |
| EPI_ISL_702726, EPI_ISL_702739, EPI_ISL_702753, EPI_ISL_702756 | Liverpool Clinical Laboratories | COVID-19 Genomics UK (COG-UK) Consortium | Sam Haldenby, Anita Lucaci, Steve Paterson, Julian Hiscox, Alistair Darby, M Almsaud, A Alrezaihi, Muhannad Alruwaili, Stuart D Armstrong, Jones Benjamin, Eleanor G Bentley, Anu Chawla, Jordan J Clark, Angela Cowell, Richard Eccles, Isabel García-Dorival, Matthew Gemmell, Alessandro Gerada, PKF Gilmore, Richard Gregory, Ximeng Han, Catherine Hartley, Margaret Hughes, Miren Iturriza-Gomara, James Johnson, L Luu, Jenifer Manson, Charlotte Nelson, Elaine O'Toole, Cassie Olateju, Rebekah Penrice-Randal , Lucille Rainbow, N.P Randle, Trevor Ian Robinson, Parul Sharma, Ghada T Shawli, James P Stewart, Neil Swainston, Ecaterina Vamos, Joanne Watts, Mark Whitehead |
| EPI_ISL_702782, EPI_ISL_702787 | Department of Pathology, University of Cambridge | COVID-19 Genomics UK (COG-UK) Consortium | Aminu S. Jahun, Yasmin Chaudhry, Grant Hall, Iliana Georgana, Myra Hosmillo, Martin D. Curran, Malte Pinckert, Surendra Parmar, Ian Goodfellow |
| EPI_ISL_702828 | Liverpool Clinical Laboratories | COVID-19 Genomics UK (COG-UK) Consortium | Sam Haldenby, Anita Lucaci, Steve Paterson, Julian Hiscox, Alistair Darby, M Almsaud, A Alrezaihi, Muhannad Alruwaili, Stuart D Armstrong, Jones Benjamin, Eleanor G Bentley, Anu Chawla, Jordan J Clark, Angela Cowell, Richard Eccles, Isabel García-Dorival, Matthew Gemmell, Alessandro Gerada, PKF Gilmore, Richard Gregory, Ximeng Han, Catherine Hartley, Margaret Hughes, Miren Iturriza-Gomara, James Johnson, L Luu, Jenifer Manson, Charlotte Nelson, Elaine O'Toole, Cassie Olateju, Rebekah Penrice-Randal , Lucille Rainbow, N.P Randle, Trevor Ian Robinson, Parul Sharma, Ghada T Shawli, James P Stewart, Neil Swainston, Ecaterina Vamos, Joanne Watts, Mark Whitehead |
| EPI_ISL_702834 | Department of Pathology, University of Cambridge | COVID-19 Genomics UK (COG-UK) Consortium | Aminu S. Jahun, Yasmin Chaudhry, Grant Hall, Iliana Georgana, Myra Hosmillo, Martin D. Curran, Malte Pinckert, Surendra Parmar, Ian Goodfellow |
| EPI_ISL_702835 | Northumbria University / South Tees Hospitals NHS Foundation Trust / North Cumbria Integrated Care NHS Foundation Trust / North Tees and Hartlepool NHS Foundation Trust / Newcastle Hospitals NHS Foundation Trust | COVID-19 Genomics UK (COG-UK) Consortium | Darren L Smith,Andrew Nelson,Matthew Bashton,Greg R Young,Joshua Loh,John Allan,Mohammad A Tariq,Giles S Holt,Gary Black,Wen C Yew,Lynn Dover,Paul Baker,Steve Liggett,Sarah Essex,Jane Greenaway,Debra Padgett,Clive Graham,Garren Scott,Edward Barton,Emma Swindells,Brendan Payne,Jennifer Collins,Yusri Taha,Gary Eltringham |
| EPI_ISL_702847 | Department of Pathology, University of Cambridge | COVID-19 Genomics UK (COG-UK) Consortium | Aminu S. Jahun, Yasmin Chaudhry, Grant Hall, Iliana Georgana, Myra Hosmillo, Martin D. Curran, Malte Pinckert, Surendra Parmar, Ian Goodfellow |
| EPI_ISL_702854 | Liverpool Clinical Laboratories | COVID-19 Genomics UK (COG-UK) Consortium | Sam Haldenby, Anita Lucaci, Steve Paterson, Julian Hiscox, Alistair Darby, M Almsaud, A Alrezaihi, Muhannad Alruwaili, Stuart D Armstrong, Jones Benjamin, Eleanor G Bentley, Anu Chawla, Jordan J Clark, Angela Cowell, Richard Eccles, Isabel García-Dorival, Matthew Gemmell, Alessandro Gerada, PKF Gilmore, Richard Gregory, Ximeng Han, Catherine Hartley, Margaret Hughes, Miren Iturriza-Gomara, James Johnson, L Luu, Jenifer Manson, Charlotte Nelson, Elaine O'Toole, Cassie Olateju, Rebekah Penrice-Randal , Lucille Rainbow, N.P Randle, Trevor Ian Robinson, Parul Sharma, Ghada T Shawli, James P Stewart, Neil Swainston, Ecaterina Vamos, Joanne Watts, Mark Whitehead |
| EPI_ISL_702860 | Department of Pathology, University of Cambridge | COVID-19 Genomics UK (COG-UK) Consortium | Aminu S. Jahun, Yasmin Chaudhry, Grant Hall, Iliana Georgana, Myra Hosmillo, Martin D. Curran, Malte Pinckert, Surendra Parmar, Ian Goodfellow |
| EPI_ISL_702865, EPI_ISL_702869, EPI_ISL_702888, EPI_ISL_702900, EPI_ISL_702913, EPI_ISL_702920, EPI_ISL_702922, EPI_ISL_702966 | Liverpool Clinical Laboratories | COVID-19 Genomics UK (COG-UK) Consortium | Sam Haldenby, Anita Lucaci, Steve Paterson, Julian Hiscox, Alistair Darby, M Almsaud, A Alrezaihi, Muhannad Alruwaili, Stuart D Armstrong, Jones Benjamin, Eleanor G Bentley, Anu Chawla, Jordan J Clark, Angela Cowell, Richard Eccles, Isabel García-Dorival, Matthew Gemmell, Alessandro Gerada, PKF Gilmore, Richard Gregory, Ximeng Han, Catherine Hartley, Margaret Hughes, Miren Iturriza-Gomara, James Johnson, L Luu, Jenifer Manson, Charlotte Nelson, Elaine O'Toole, Cassie Olateju, Rebekah Penrice-Randal , Lucille Rainbow, N.P Randle, Trevor Ian Robinson, Parul Sharma, Ghada T Shawli, James P Stewart, Neil Swainston, Ecaterina Vamos, Joanne Watts, Mark Whitehead |
| EPI_ISL_702971, EPI_ISL_702977 | Department of Pathology, University of Cambridge | COVID-19 Genomics UK (COG-UK) Consortium | Aminu S. Jahun, Yasmin Chaudhry, Grant Hall, Iliana Georgana, Myra Hosmillo, Martin D. Curran, Malte Pinckert, Surendra Parmar, Ian Goodfellow |
| EPI_ISL_703005 | Northumbria University / South Tees Hospitals NHS Foundation Trust / North Cumbria Integrated Care NHS Foundation Trust / North Tees and Hartlepool NHS Foundation Trust / Newcastle Hospitals NHS Foundation Trust | COVID-19 Genomics UK (COG-UK) Consortium | Darren L Smith,Andrew Nelson,Matthew Bashton,Greg R Young,Joshua Loh,John Allan,Mohammad A Tariq,Giles S Holt,Gary Black,Wen C Yew,Lynn Dover,Paul Baker,Steve Liggett,Sarah Essex,Jane Greenaway,Debra Padgett,Clive Graham,Garren Scott,Edward Barton,Emma Swindells,Brendan Payne,Jennifer Collins,Yusri Taha,Gary Eltringham |
| EPI_ISL_703046, EPI_ISL_703059 | Liverpool Clinical Laboratories | COVID-19 Genomics UK (COG-UK) Consortium | Sam Haldenby, Anita Lucaci, Steve Paterson, Julian Hiscox, Alistair Darby, M Almsaud, A Alrezaihi, Muhannad Alruwaili, Stuart D Armstrong, Jones Benjamin, Eleanor G Bentley, Anu Chawla, Jordan J Clark, Angela Cowell, Richard Eccles, Isabel García-Dorival, Matthew Gemmell, Alessandro Gerada, PKF Gilmore, Richard Gregory, Ximeng Han, Catherine Hartley, Margaret Hughes, Miren Iturriza-Gomara, James Johnson, L Luu, Jenifer Manson, Charlotte Nelson, Elaine O'Toole, Cassie Olateju, Rebekah Penrice-Randal , Lucille Rainbow, N.P Randle, Trevor Ian Robinson, Parul Sharma, Ghada T Shawli, James P Stewart, Neil Swainston, Ecaterina Vamos, Joanne Watts, Mark Whitehead |

| | | | |
|---|---|---|---|
| EPI_ISL_703093, EPI_ISL_703115 | Department of Pathology, University of Cambridge | COVID-19 Genomics UK (COG-UK) Consortium | Aminu S. Jahun, Yasmin Chaudhry, Grant Hall, Iliana Georgana, Myra Hosmillo, Martin D. Curran, Malte Pinckert, Surendra Parmar, Ian Goodfellow |
| EPI_ISL_703121, EPI_ISL_703150, EPI_ISL_703158, EPI_ISL_703225, EPI_ISL_703230, EPI_ISL_703256 | Liverpool Clinical Laboratories | COVID-19 Genomics UK (COG-UK) Consortium | Sam Haldenby, Anita Lucaci, Steve Paterson, Julian Hiscox, Alistair Darby, M Almsaud, A Alrezaihi, Muhannad Alruwaili, Stuart D Armstrong, Jones Benjamin, Eleanor G Bentley, Anu Chawla, Jordan J Clark, Angela Cowell, Richard Eccles, Isabel García-Dorival, Matthew Gemmell, Alessandro Gerada, PKF Gilmore, Richard Gregory, Ximeng Han, Catherine Hartley, Margaret Hughes, Miren Iturriza-Gomara, James Johnson, L Luu, Jenifer Manson, Charlotte Nelson, Elaine O'Toole, Cassie Olateju, Rebekah Penrice-Randal , Lucille Rainbow, N.P Randle, Trevor Ian Robinson, Parul Sharma, Ghada T Shawli, James P Stewart, Neil Swainston, Ecaterina Vamos, Joanne Watts, Mark Whitehead |
| EPI_ISL_703287 | West of Scotland Specialist Virology Centre, NHSGGC / MRC-University of Glasgow Centre for Virus Research | COVID-19 Genomics UK (COG-UK) Consortium | Ana da Silva Filipe, Natasha Johnson, Kathy Smollett, Daniel Mair, Stephen Carmichael, Alice Broos, Lily Tong, Jenna Nichols, Kyriaki Nomikou; Sarah McDonald; Richard Orton, Joseph Hughes, Sreenu Vattipally, David L Robertson; Alasdair MacLean, Rory Gunson; Sharif Shaaban, Matthew Holden; Rachel Blacow, Guy Mollett, Kathy Li, James Shepherd, Antonia Ho, Emma Thomson |
| EPI_ISL_703308, EPI_ISL_703316 | Liverpool Clinical Laboratories | COVID-19 Genomics UK (COG-UK) Consortium | Sam Haldenby, Anita Lucaci, Steve Paterson, Julian Hiscox, Alistair Darby, M Almsaud, A Alrezaihi, Muhannad Alruwaili, Stuart D Armstrong, Jones Benjamin, Eleanor G Bentley, Anu Chawla, Jordan J Clark, Angela Cowell, Richard Eccles, Isabel García-Dorival, Matthew Gemmell, Alessandro Gerada, PKF Gilmore, Richard Gregory, Ximeng Han, Catherine Hartley, Margaret Hughes, Miren Iturriza-Gomara, James Johnson, L Luu, Jenifer Manson, Charlotte Nelson, Elaine O'Toole, Cassie Olateju, Rebekah Penrice-Randal , Lucille Rainbow, N.P Randle, Trevor Ian Robinson, Parul Sharma, Ghada T Shawli, James P Stewart, Neil Swainston, Ecaterina Vamos, Joanne Watts, Mark Whitehead |
| EPI_ISL_703432, EPI_ISL_703435, EPI_ISL_703438, EPI_ISL_703440, EPI_ISL_703459, EPI_ISL_703462, EPI_ISL_703465, EPI_ISL_703468, EPI_ISL_703471, EPI_ISL_703473, EPI_ISL_703513, EPI_ISL_703516, EPI_ISL_703521 | | | |
| see above | Department of Pathology, University of Cambridge | COVID-19 Genomics UK (COG-UK) Consortium | Aminu S. Jahun, Yasmin Chaudhry, Grant Hall, Iliana Georgana, Myra Hosmillo, Martin D. Curran, Malte Pinckert, Surendra Parmar, Ian Goodfellow |
| EPI_ISL_703533, EPI_ISL_703535 | Northumbria University / South Tees Hospitals NHS Foundation Trust / North Cumbria Integrated Care NHS Foundation Trust / North Tees and Hartlepool NHS Foundation Trust / Newcastle Hospitals NHS Foundation Trust | COVID-19 Genomics UK (COG-UK) Consortium | Darren L Smith,Andrew Nelson,Matthew Bashton,Greg R Young,Joshua Loh,John Allan,Mohammad A Tariq,Giles S Holt,Gary Black,Wen C Yew,Lynn Dover,Paul Baker,Steve Liggett,Sarah Essex,Jane Greenaway,Debra Padgett,Clive Graham,Garren Scott,Edward Barton,Emma Swindells,Brendan Payne,Jennifer Collins,Yusri Taha,Gary Eltringham |
| EPI_ISL_703596, EPI_ISL_703652 | Liverpool Clinical Laboratories | COVID-19 Genomics UK (COG-UK) Consortium | Sam Haldenby, Anita Lucaci, Steve Paterson, Julian Hiscox, Alistair Darby, M Almsaud, A Alrezaihi, Muhannad Alruwaili, Stuart D Armstrong, Jones Benjamin, Eleanor G Bentley, Anu Chawla, Jordan J Clark, Angela Cowell, Richard Eccles, Isabel García-Dorival, Matthew Gemmell, Alessandro Gerada, PKF Gilmore, Richard Gregory, Ximeng Han, Catherine Hartley, Margaret Hughes, Miren Iturriza-Gomara, James Johnson, L Luu, Jenifer Manson, Charlotte Nelson, Elaine O'Toole, Cassie Olateju, Rebekah Penrice-Randal , Lucille Rainbow, N.P Randle, Trevor Ian Robinson, Parul Sharma, Ghada T Shawli, James P Stewart, Neil Swainston, Ecaterina Vamos, Joanne Watts, Mark Whitehead |
| EPI_ISL_703701 | Northumbria University / South Tees Hospitals NHS Foundation Trust / North Cumbria Integrated Care NHS Foundation Trust / North Tees and Hartlepool NHS Foundation Trust / Newcastle Hospitals NHS Foundation Trust | COVID-19 Genomics UK (COG-UK) Consortium | Darren L Smith,Andrew Nelson,Matthew Bashton,Greg R Young,Joshua Loh,John Allan,Mohammad A Tariq,Giles S Holt,Gary Black,Wen C Yew,Lynn Dover,Paul Baker,Steve Liggett,Sarah Essex,Jane Greenaway,Debra Padgett,Clive Graham,Garren Scott,Edward Barton,Emma Swindells,Brendan Payne,Jennifer Collins,Yusri Taha,Gary Eltringham |
| EPI_ISL_703712 | Liverpool Clinical Laboratories | COVID-19 Genomics UK (COG-UK) Consortium | Sam Haldenby, Anita Lucaci, Steve Paterson, Julian Hiscox, Alistair Darby, M Almsaud, A Alrezaihi, Muhannad Alruwaili, Stuart D Armstrong, Jones Benjamin, Eleanor G Bentley, Anu Chawla, Jordan J Clark, Angela Cowell, Richard Eccles, Isabel García-Dorival, Matthew Gemmell, Alessandro Gerada, PKF Gilmore, Richard Gregory, Ximeng Han, Catherine Hartley, Margaret Hughes, Miren Iturriza-Gomara, James Johnson, L Luu, Jenifer Manson, Charlotte Nelson, Elaine O'Toole, Cassie Olateju, Rebekah Penrice-Randal , Lucille Rainbow, N.P Randle, Trevor Ian Robinson, Parul Sharma, Ghada T Shawli, James P Stewart, Neil Swainston, Ecaterina Vamos, Joanne Watts, Mark Whitehead |
| EPI_ISL_703715 | Department of Pathology, University of Cambridge | COVID-19 Genomics UK (COG-UK) Consortium | Aminu S. Jahun, Yasmin Chaudhry, Grant Hall, Iliana Georgana, Myra Hosmillo, Martin D. Curran, Malte Pinckert, Surendra Parmar, Ian Goodfellow |
| EPI_ISL_703718 | West of Scotland Specialist Virology Centre, NHSGGC / MRC-University of Glasgow Centre for Virus Research | COVID-19 Genomics UK (COG-UK) Consortium | Ana da Silva Filipe, Natasha Johnson, Kathy Smollett, Daniel Mair, Stephen Carmichael, Alice Broos, Lily Tong, Jenna Nichols, Kyriaki Nomikou; Sarah McDonald; Richard Orton, Joseph Hughes, Sreenu Vattipally, David L Robertson; Alasdair MacLean, Rory Gunson; Sharif Shaaban, Matthew Holden; Rachel Blacow, Guy Mollett, Kathy Li, James Shepherd, Antonia Ho, Emma Thomson |
| EPI_ISL_703725, EPI_ISL_703731 | Liverpool Clinical Laboratories | COVID-19 Genomics UK (COG-UK) Consortium | Sam Haldenby, Anita Lucaci, Steve Paterson, Julian Hiscox, Alistair Darby, M Almsaud, A Alrezaihi, Muhannad Alruwaili, Stuart D Armstrong, Jones Benjamin, Eleanor G Bentley, Anu Chawla, Jordan J Clark, Angela Cowell, Richard Eccles, Isabel García-Dorival, Matthew Gemmell, Alessandro Gerada, PKF Gilmore, Richard Gregory, Ximeng Han, Catherine Hartley, Margaret Hughes, Miren Iturriza-Gomara, James Johnson, L Luu, Jenifer Manson, Charlotte Nelson, Elaine O'Toole, Cassie Olateju, Rebekah Penrice-Randal , Lucille Rainbow, N.P Randle, Trevor Ian Robinson, Parul Sharma, Ghada T Shawli, James P Stewart, Neil Swainston, Ecaterina Vamos, Joanne Watts, Mark Whitehead |
| EPI_ISL_703745 | University College London, Great Ormond Street Hospital for Children NHS Foundation Trust, Imperial College Healthcare NHS Trust | COVID-19 Genomics UK (COG-UK) Consortium | Sergi Castellano, Rachel Williams, Mark Kristiansen, Paola Resende Silva, Sunando Roy, Tony Brooks, Helena Tutill, Paola Niola, Patricia Dyal, Charlotte Williams, Leysa Forrest, Yasmin Panchbhaya, Jacqueline Findlay, Samuel Weeks, Julianne Brown, Kathryn Harris, Paul Randell, James Price, Alison Holmes, Judith Breuer |
| EPI_ISL_703888, EPI_ISL_703935 | Liverpool Clinical Laboratories | COVID-19 Genomics UK (COG-UK) Consortium | Sam Haldenby, Anita Lucaci, Steve Paterson, Julian Hiscox, Alistair Darby, M Almsaud, A Alrezaihi, Muhannad Alruwaili, Stuart D Armstrong, Jones Benjamin, Eleanor G Bentley, Anu Chawla, Jordan J Clark, Angela Cowell, Richard Eccles, Isabel García-Dorival, Matthew Gemmell, Alessandro Gerada, PKF Gilmore, Richard Gregory, Ximeng Han, Catherine Hartley, Margaret Hughes, Miren Iturriza-Gomara, James Johnson, L Luu, Jenifer Manson, Charlotte Nelson, Elaine O'Toole, Cassie Olateju, Rebekah Penrice-Randal , Lucille Rainbow, N.P Randle, Trevor Ian Robinson, Parul Sharma, Ghada T Shawli, James P Stewart, Neil Swainston, Ecaterina Vamos, Joanne Watts, Mark Whitehead |
| EPI_ISL_703950 | Virology Department, Royal Infirmary of Edinburgh, NHS Lothian / School of Biological Sciences, University of Edinburgh / Institute of Genetics and Molecular Medicine, University of Edinburgh | COVID-19 Genomics UK (COG-UK) Consortium | McHugh M, Dewar R, Rooke S, Gallagher M, Balcaza C, O'Toole Á, Scher E, Hill V, McCrone JT, Colquhoun R, Yu X, Jackson B, Rambaut A, Williams TC, Templeton K |
| EPI_ISL_703990, EPI_ISL_704012, EPI_ISL_704020, EPI_ISL_704028, EPI_ISL_704068, EPI_ISL_704079, EPI_ISL_704097 | Liverpool Clinical Laboratories | COVID-19 Genomics UK (COG-UK) Consortium | Sam Haldenby, Anita Lucaci, Steve Paterson, Julian Hiscox, Alistair Darby, M Almsaud, A Alrezaihi, Muhannad Alruwaili, Stuart D Armstrong, Jones Benjamin, Eleanor G Bentley, Anu Chawla, Jordan J Clark, Angela Cowell, Richard Eccles, Isabel García-Dorival, Matthew Gemmell, Alessandro Gerada, PKF Gilmore, Richard Gregory, Ximeng Han, Catherine Hartley, Margaret Hughes, Miren Iturriza-Gomara, James Johnson, L Luu, Jenifer Manson, Charlotte Nelson, Elaine O'Toole, Cassie Olateju, Rebekah Penrice-Randal , Lucille Rainbow, N.P Randle, Trevor Ian Robinson, Parul Sharma, Ghada T Shawli, James P Stewart, Neil Swainston, Ecaterina Vamos, Joanne Watts, Mark Whitehead |
| EPI_ISL_704187 | Virology Department, Royal Infirmary of Edinburgh, NHS Lothian / School of Biological Sciences, University of Edinburgh / Institute of Genetics and Molecular Medicine, University of Edinburgh | COVID-19 Genomics UK (COG-UK) Consortium | McHugh M, Dewar R, Rooke S, Gallagher M, Balcaza C, O'Toole Á, Scher E, Hill V, McCrone JT, Colquhoun R, Yu X, Jackson B, Rambaut A, Williams TC, Templeton K |
| EPI_ISL_704206 | West of Scotland Specialist Virology Centre, NHSGGC / MRC-University of Glasgow Centre for Virus Research | COVID-19 Genomics UK (COG-UK) Consortium | Ana da Silva Filipe, Natasha Johnson, Kathy Smollett, Daniel Mair, Stephen Carmichael, Alice Broos, Lily Tong, Jenna Nichols, Kyriaki Nomikou; Sarah McDonald; Richard Orton, Joseph Hughes, Sreenu Vattipally, David L Robertson; Alasdair MacLean, Rory Gunson; Sharif Shaaban, Matthew Holden; Rachel Blacow, Guy Mollett, Kathy Li, James Shepherd, Antonia Ho, Emma Thomson |
| EPI_ISL_704219, EPI_ISL_704222 | Department of Pathology, University of Cambridge | COVID-19 Genomics UK (COG-UK) Consortium | Aminu S. Jahun, Yasmin Chaudhry, Grant Hall, Iliana Georgana, Myra Hosmillo, Martin D. Curran, Malte Pinckert, Surendra Parmar, Ian Goodfellow |
| EPI_ISL_704231 | Liverpool Clinical Laboratories | COVID-19 Genomics UK (COG-UK) Consortium | Sam Haldenby, Anita Lucaci, Steve Paterson, Julian Hiscox, Alistair Darby, M Almsaud, A Alrezaihi, Muhannad Alruwaili, Stuart D Armstrong, Jones Benjamin, Eleanor G Bentley, Anu Chawla, Jordan J Clark, Angela Cowell, Richard Eccles, Isabel García-Dorival, Matthew Gemmell, Alessandro Gerada, PKF Gilmore, Richard Gregory, Ximeng Han, Catherine Hartley, Margaret Hughes, Miren Iturriza-Gomara, James Johnson, L Luu, Jenifer Manson, Charlotte Nelson, Elaine O'Toole, Cassie Olateju, Rebekah Penrice-Randal , Lucille Rainbow, N.P Randle, Trevor Ian Robinson, Parul Sharma, Ghada T Shawli, James P Stewart, Neil Swainston, Ecaterina Vamos, Joanne Watts, Mark Whitehead |
| EPI_ISL_704273, EPI_ISL_704280, EPI_ISL_704283, EPI_ISL_704286 | Department of Pathology, University of Cambridge | COVID-19 Genomics UK (COG-UK) Consortium | Aminu S. Jahun, Yasmin Chaudhry, Grant Hall, Iliana Georgana, Myra Hosmillo, Martin D. Curran, Malte Pinckert, Surendra Parmar, Ian Goodfellow |

| | | | |
|---|---|---|---|
| EPI_ISL_704315 | Queens Medical Centre, Clinical Microbiology Department / DeepSeq Nottingham | COVID-19 Genomics UK (COG-UK) Consortium | Gemma Clark, Wendy Smith, Manjinder Khakh, Vicki M Fleming, Michelle M Lister, Hannah Howson-Wells, Jonathan Ball, Patrick McClure, Joseph Chappell, Theocharis Tsoleridis, Nadine Holmes, Matthew Carlisle, Christopher Moore, Fei Sang, Johnny Debebe, Victoria Wright, Matthew Loose |
| EPI_ISL_704389, EPI_ISL_704392, EPI_ISL_704395, EPI_ISL_704427, EPI_ISL_704430, EPI_ISL_704433, EPI_ISL_704435, EPI_ISL_704438, EPI_ISL_704443, EPI_ISL_704446, EPI_ISL_704449, EPI_ISL_704452, EPI_ISL_704454, EPI_ISL_704457, EPI_ISL_704460, EPI_ISL_704462, EPI_ISL_704465, EPI_ISL_704467, EPI_ISL_704957 | | | |
| see above | Department of Pathology, University of Cambridge | COVID-19 Genomics UK (COG-UK) Consortium | Aminu S. Jahun, Yasmin Chaudhry, Grant Hall, Iliana Georgana, Myra Hosmillo, Martin D. Curran, Malte Pinckert, Surendra Parmar, Ian Goodfellow |
| EPI_ISL_704970, EPI_ISL_704986, EPI_ISL_705013, EPI_ISL_705040 | Liverpool Clinical Laboratories | COVID-19 Genomics UK (COG-UK) Consortium | Sam Haldenby, Anita Lucaci, Steve Paterson, Julian Hiscox, Alistair Darby, M Almsaud, A Alrezaihi, Muhannad Alruwaili, Stuart D Armstrong, Jones Benjamin, Eleanor G Bentley, Anu Chawla, Jordan J Clark, Angela Cowell, Richard Eccles, Isabel García-Dorival, Matthew Gemmell, Alessandro Gerada, PKF Gilmore, Richard Gregory, Ximeng Han, Catherine Hartley, Margaret Hughes, Miren Iturriza-Gomara, James Johnson, L Luu, Jenifer Manson, Charlotte Nelson, Elaine O'Toole, Cassie Olateju, Rebekah Penrice-Randal , Lucille Rainbow, N.P Randle, Trevor Ian Robinson, Parul Sharma, Ghada T Shawli, James P Stewart, Neil Swainston, Ecaterina Vamos, Joanne Watts, Mark Whitehead |
| EPI_ISL_705070 | Wales Specialist Virology Centre Sequencing lab: Pathogen Genomics Unit | COVID-19 Genomics UK (COG-UK) Consortium | Catherine Moore, Johnathan Evans, Laura Gifford, Malorie Perry, Simon Cottrell, Angela Marchbank, Alec Birchley, Alexander Adams, Amy Gaskin, Bree Gatica-Wilcox, Jason Coombes, Joel Southgate, Lauren Gilbert, Lee Graham, Nicole Pacchiarini, Sara Kumziene-Summerhayes, Sarah Taylor, Sophie Jones, Sara Rey, Matthew Bull, Joanne Watkins, Sally Corden, Tom Connor |
| EPI_ISL_705190 | Liverpool Clinical Laboratories | COVID-19 Genomics UK (COG-UK) Consortium | Sam Haldenby, Anita Lucaci, Steve Paterson, Julian Hiscox, Alistair Darby, M Almsaud, A Alrezaihi, Muhannad Alruwaili, Stuart D Armstrong, Jones Benjamin, Eleanor G Bentley, Anu Chawla, Jordan J Clark, Angela Cowell, Richard Eccles, Isabel García-Dorival, Matthew Gemmell, Alessandro Gerada, PKF Gilmore, Richard Gregory, Ximeng Han, Catherine Hartley, Margaret Hughes, Miren Iturriza-Gomara, James Johnson, L Luu, Jenifer Manson, Charlotte Nelson, Elaine O'Toole, Cassie Olateju, Rebekah Penrice-Randal , Lucille Rainbow, N.P Randle, Trevor Ian Robinson, Parul Sharma, Ghada T Shawli, James P Stewart, Neil Swainston, Ecaterina Vamos, Joanne Watts, Mark Whitehead |
| EPI_ISL_705212 | Northumbria University / South Tees Hospitals NHS Foundation Trust / North Cumbria Integrated Care NHS Foundation Trust / North Tees and Hartlepool NHS Foundation Trust / Newcastle Hospitals NHS Foundation Trust | COVID-19 Genomics UK (COG-UK) Consortium | Darren L Smith,Andrew Nelson,Matthew Bashton,Greg R Young,Joshua Loh,John Allan,Mohammad A Tariq,Giles S Holt,Gary Black,Wen C Yew,Lynn Dover,Paul Baker,Steve Liggett,Sarah Essex,Jane Greenaway,Debra Padgett,Clive Graham,Garren Scott,Edward Barton,Emma Swindells,Brendan Payne,Jennifer Collins,Yusri Taha,Gary Eltringham |
| EPI_ISL_705238, EPI_ISL_705260, EPI_ISL_705264, EPI_ISL_705275 | Liverpool Clinical Laboratories | COVID-19 Genomics UK (COG-UK) Consortium | Sam Haldenby, Anita Lucaci, Steve Paterson, Julian Hiscox, Alistair Darby, M Almsaud, A Alrezaihi, Muhannad Alruwaili, Stuart D Armstrong, Jones Benjamin, Eleanor G Bentley, Anu Chawla, Jordan J Clark, Angela Cowell, Richard Eccles, Isabel García-Dorival, Matthew Gemmell, Alessandro Gerada, PKF Gilmore, Richard Gregory, Ximeng Han, Catherine Hartley, Margaret Hughes, Miren Iturriza-Gomara, James Johnson, L Luu, Jenifer Manson, Charlotte Nelson, Elaine O'Toole, Cassie Olateju, Rebekah Penrice-Randal , Lucille Rainbow, N.P Randle, Trevor Ian Robinson, Parul Sharma, Ghada T Shawli, James P Stewart, Neil Swainston, Ecaterina Vamos, Joanne Watts, Mark Whitehead |
| EPI_ISL_705290 | Department of Pathology, University of Cambridge | COVID-19 Genomics UK (COG-UK) Consortium | Aminu S. Jahun, Yasmin Chaudhry, Grant Hall, Iliana Georgana, Myra Hosmillo, Martin D. Curran, Malte Pinckert, Surendra Parmar, Ian Goodfellow |
| EPI_ISL_705312, EPI_ISL_705320, EPI_ISL_705330 | Liverpool Clinical Laboratories | COVID-19 Genomics UK (COG-UK) Consortium | Sam Haldenby, Anita Lucaci, Steve Paterson, Julian Hiscox, Alistair Darby, M Almsaud, A Alrezaihi, Muhannad Alruwaili, Stuart D Armstrong, Jones Benjamin, Eleanor G Bentley, Anu Chawla, Jordan J Clark, Angela Cowell, Richard Eccles, Isabel García-Dorival, Matthew Gemmell, Alessandro Gerada, PKF Gilmore, Richard Gregory, Ximeng Han, Catherine Hartley, Margaret Hughes, Miren Iturriza-Gomara, James Johnson, L Luu, Jenifer Manson, Charlotte Nelson, Elaine O'Toole, Cassie Olateju, Rebekah Penrice-Randal , Lucille Rainbow, N.P Randle, Trevor Ian Robinson, Parul Sharma, Ghada T Shawli, James P Stewart, Neil Swainston, Ecaterina Vamos, Joanne Watts, Mark Whitehead |
| EPI_ISL_705333, EPI_ISL_705346 | Department of Pathology, University of Cambridge | COVID-19 Genomics UK (COG-UK) Consortium | Aminu S. Jahun, Yasmin Chaudhry, Grant Hall, Iliana Georgana, Myra Hosmillo, Martin D. Curran, Malte Pinckert, Surendra Parmar, Ian Goodfellow |
| EPI_ISL_705359, EPI_ISL_705363 | Liverpool Clinical Laboratories | COVID-19 Genomics UK (COG-UK) Consortium | Sam Haldenby, Anita Lucaci, Steve Paterson, Julian Hiscox, Alistair Darby, M Almsaud, A Alrezaihi, Muhannad Alruwaili, Stuart D Armstrong, Jones Benjamin, Eleanor G Bentley, Anu Chawla, Jordan J Clark, Angela Cowell, Richard Eccles, Isabel García-Dorival, Matthew Gemmell, Alessandro Gerada, PKF Gilmore, Richard Gregory, Ximeng Han, Catherine Hartley, Margaret Hughes, Miren Iturriza-Gomara, James Johnson, L Luu, Jenifer Manson, Charlotte Nelson, Elaine O'Toole, Cassie Olateju, Rebekah Penrice-Randal , Lucille Rainbow, N.P Randle, Trevor Ian Robinson, Parul Sharma, Ghada T Shawli, James P Stewart, Neil Swainston, Ecaterina Vamos, Joanne Watts, Mark Whitehead |
| EPI_ISL_705365, EPI_ISL_705382 | Department of Pathology, University of Cambridge | COVID-19 Genomics UK (COG-UK) Consortium | Aminu S. Jahun, Yasmin Chaudhry, Grant Hall, Iliana Georgana, Myra Hosmillo, Martin D. Curran, Malte Pinckert, Surendra Parmar, Ian Goodfellow |
| EPI_ISL_705406, EPI_ISL_705409 | Liverpool Clinical Laboratories | COVID-19 Genomics UK (COG-UK) Consortium | Sam Haldenby, Anita Lucaci, Steve Paterson, Julian Hiscox, Alistair Darby, M Almsaud, A Alrezaihi, Muhannad Alruwaili, Stuart D Armstrong, Jones Benjamin, Eleanor G Bentley, Anu Chawla, Jordan J Clark, Angela Cowell, Richard Eccles, Isabel García-Dorival, Matthew Gemmell, Alessandro Gerada, PKF Gilmore, Richard Gregory, Ximeng Han, Catherine Hartley, Margaret Hughes, Miren Iturriza-Gomara, James Johnson, L Luu, Jenifer Manson, Charlotte Nelson, Elaine O'Toole, Cassie Olateju, Rebekah Penrice-Randal , Lucille Rainbow, N.P Randle, Trevor Ian Robinson, Parul Sharma, Ghada T Shawli, James P Stewart, Neil Swainston, Ecaterina Vamos, Joanne Watts, Mark Whitehead |
| EPI_ISL_705418, EPI_ISL_705419 | Wales Specialist Virology Centre Sequencing lab: Pathogen Genomics Unit | COVID-19 Genomics UK (COG-UK) Consortium | Catherine Moore, Johnathan Evans, Laura Gifford, Malorie Perry, Simon Cottrell, Angela Marchbank, Alec Birchley, Alexander Adams, Amy Gaskin, Bree Gatica-Wilcox, Jason Coombes, Joel Southgate, Lauren Gilbert, Lee Graham, Nicole Pacchiarini, Sara Kumziene-Summerhayes, Sarah Taylor, Sophie Jones, Sara Rey, Matthew Bull, Joanne Watkins, Sally Corden, Tom Connor |
| EPI_ISL_705423, EPI_ISL_705424, EPI_ISL_705449 | Liverpool Clinical Laboratories | COVID-19 Genomics UK (COG-UK) Consortium | Sam Haldenby, Anita Lucaci, Steve Paterson, Julian Hiscox, Alistair Darby, M Almsaud, A Alrezaihi, Muhannad Alruwaili, Stuart D Armstrong, Jones Benjamin, Eleanor G Bentley, Anu Chawla, Jordan J Clark, Angela Cowell, Richard Eccles, Isabel García-Dorival, Matthew Gemmell, Alessandro Gerada, PKF Gilmore, Richard Gregory, Ximeng Han, Catherine Hartley, Margaret Hughes, Miren Iturriza-Gomara, James Johnson, L Luu, Jenifer Manson, Charlotte Nelson, Elaine O'Toole, Cassie Olateju, Rebekah Penrice-Randal , Lucille Rainbow, N.P Randle, Trevor Ian Robinson, Parul Sharma, Ghada T Shawli, James P Stewart, Neil Swainston, Ecaterina Vamos, Joanne Watts, Mark Whitehead |
| EPI_ISL_705460 | Wales Specialist Virology Centre Sequencing lab: Pathogen Genomics Unit | COVID-19 Genomics UK (COG-UK) Consortium | Catherine Moore, Johnathan Evans, Laura Gifford, Malorie Perry, Simon Cottrell, Angela Marchbank, Alec Birchley, Alexander Adams, Amy Gaskin, Bree Gatica-Wilcox, Jason Coombes, Joel Southgate, Lauren Gilbert, Lee Graham, Nicole Pacchiarini, Sara Kumziene-Summerhayes, Sarah Taylor, Sophie Jones, Sara Rey, Matthew Bull, Joanne Watkins, Sally Corden, Tom Connor |
| EPI_ISL_705525 | Liverpool Clinical Laboratories | COVID-19 Genomics UK (COG-UK) Consortium | Sam Haldenby, Anita Lucaci, Steve Paterson, Julian Hiscox, Alistair Darby, M Almsaud, A Alrezaihi, Muhannad Alruwaili, Stuart D Armstrong, Jones Benjamin, Eleanor G Bentley, Anu Chawla, Jordan J Clark, Angela Cowell, Richard Eccles, Isabel García-Dorival, Matthew Gemmell, Alessandro Gerada, PKF Gilmore, Richard Gregory, Ximeng Han, Catherine Hartley, Margaret Hughes, Miren Iturriza-Gomara, James Johnson, L Luu, Jenifer Manson, Charlotte Nelson, Elaine O'Toole, Cassie Olateju, Rebekah Penrice-Randal , Lucille Rainbow, N.P Randle, Trevor Ian Robinson, Parul Sharma, Ghada T Shawli, James P Stewart, Neil Swainston, Ecaterina Vamos, Joanne Watts, Mark Whitehead |
| EPI_ISL_705527 | Wales Specialist Virology Centre Sequencing lab: Pathogen Genomics Unit | COVID-19 Genomics UK (COG-UK) Consortium | Catherine Moore, Johnathan Evans, Laura Gifford, Malorie Perry, Simon Cottrell, Angela Marchbank, Alec Birchley, Alexander Adams, Amy Gaskin, Bree Gatica-Wilcox, Jason Coombes, Joel Southgate, Lauren Gilbert, Lee Graham, Nicole Pacchiarini, Sara Kumziene-Summerhayes, Sarah Taylor, Sophie Jones, Sara Rey, Matthew Bull, Joanne Watkins, Sally Corden, Tom Connor |
| EPI_ISL_705530, EPI_ISL_705531, EPI_ISL_705532, EPI_ISL_705533, EPI_ISL_705534, EPI_ISL_705535 | Department of Pathology, University of Cambridge | COVID-19 Genomics UK (COG-UK) Consortium | Aminu S. Jahun, Yasmin Chaudhry, Grant Hall, Iliana Georgana, Myra Hosmillo, Martin D. Curran, Malte Pinckert, Surendra Parmar, Ian Goodfellow |
| EPI_ISL_705636, EPI_ISL_705637 | West of Scotland Specialist Virology Centre, NHSGGC / MRC-University of Glasgow Centre for Virus Research | COVID-19 Genomics UK (COG-UK) Consortium | Ana da Silva Filipe, Natasha Johnson, Kathy Smollett, Daniel Mair, Stephen Carmichael, Alice Broos, Lily Tong, Jenna Nichols, Kyriaki Nomikou; Sarah McDonald; Richard Orton, Joseph Hughes, Sreenu Vattipally, David L Robertson; Alasdair MacLean, Rory Gunson; Sharif Shaaban, Matthew Holden; Rachel Blacow, Guy Mollett, Kathy Li, James Shepherd, Antonia Ho, Emma Thomson |
| EPI_ISL_705764, EPI_ISL_705765, EPI_ISL_705766, EPI_ISL_705767, EPI_ISL_705770, EPI_ISL_705773 | Virology Department, Royal Infirmary of Edinburgh, NHS Lothian / School of Biological Sciences, University of Edinburgh / Institute of Genetics and Molecular Medicine, University of Edinburgh | COVID-19 Genomics UK (COG-UK) Consortium | McHugh M, Dewar R, Rooke S, Gallagher M, Balcaza C, O'Toole Á, Scher E, Hill V, McCrone JT, Colquhoun R, Yu X, Jackson B, Rambaut A, Williams TC, Templeton K |
| EPI_ISL_705806, EPI_ISL_705807, EPI_ISL_705808, EPI_ISL_705865, EPI_ISL_705866, EPI_ISL_705867, EPI_ISL_705868, EPI_ISL_705870, EPI_ISL_705872, EPI_ISL_705876, EPI_ISL_705878, EPI_ISL_705895, EPI_ISL_705896, EPI_ISL_705897, EPI_ISL_705898, EPI_ISL_705899, EPI_ISL_705900, EPI_ISL_705901, | | | |

| | | | |
|---|---|---|---|
| EPI_ISL_705902, EPI_ISL_705903, EPI_ISL_705904, EPI_ISL_705905, EPI_ISL_705906, EPI_ISL_705907, EPI_ISL_705908, EPI_ISL_705909, EPI_ISL_705910, EPI_ISL_705911, EPI_ISL_705912, EPI_ISL_705913, EPI_ISL_705914, EPI_ISL_705915, EPI_ISL_705916, EPI_ISL_705917, EPI_ISL_705918, EPI_ISL_705920 | | | |
| see above | Liverpool Clinical Laboratories | COVID-19 Genomics UK (COG-UK) Consortium | Sam Haldenby, Anita Lucaci, Steve Paterson, Julian Hiscox, Alistair Darby, M Almsaud, A Alrezaihi, Muhannad Alruwaili, Stuart D Armstrong, Jones Benjamin, Eleanor G Bentley, Anu Chawla, Jordan J Clark, Angela Cowell, Richard Eccles, Isabel García-Dorival, Matthew Gemmell, Alessandro Gerada, PKF Gilmore, Richard Gregory, Ximeng Han, Catherine Hartley, Margaret Hughes, Miren Iturriza-Gomara, James Johnson, L Luu, Jenifer Manson, Charlotte Nelson, Elaine O'Toole, Cassie Olateju, Rebekah Penrice-Randal , Lucille Rainbow, N.P Randle, Trevor Ian Robinson, Parul Sharma, Ghada T Shawli, James P Stewart, Neil Swainston, Ecaterina Vamos, Joanne Watts, Mark Whitehead |
| EPI_ISL_706128 | Oxford Viromics, NDM, University of Oxford; Oxford University Hospitals; Basingstoke and North Hampshire Hospital | COVID-19 Genomics UK (COG-UK) Consortium | Tanya Golubchik, David Bonsall, George Macintyre, Amy Trebes, Mariateresa de Cesare, Catrin Moore, Alex Mobbs, Anita Justice, Robert Shaw, Monique Andersson, Timothy Peto, Emma Wise, Nathan Moore, Jessica Lynch, Nick Cortes, Matilde Mori, Stephen Kidd, David Buck, John Todd, Christophe Fraser |
| EPI_ISL_706277, EPI_ISL_706278, EPI_ISL_706286, EPI_ISL_706289, EPI_ISL_706292, EPI_ISL_706293, EPI_ISL_706296 | Northumbria University / South Tees Hospitals NHS Foundation Trust / North Cumbria Integrated Care NHS Foundation Trust / North Tees and Hartlepool NHS Foundation Trust / Newcastle Hospitals NHS Foundation Trust | COVID-19 Genomics UK (COG-UK) Consortium | Darren L Smith,Andrew Nelson,Matthew Bashton,Greg R Young,Joshua Loh,John Allan,Mohammad A Tariq,Giles S Holt,Gary Black,Wen C Yew,Lynn Dover,Paul Baker,Steve Liggett,Sarah Essex,Jane Greenaway,Debra Padgett,Clive Graham,Garren Scott,Edward Barton,Emma Swindells,Brendan Payne,Jennifer Collins,Yusri Taha,Gary Eltringham |
| EPI_ISL_706493, EPI_ISL_706495, EPI_ISL_706496, EPI_ISL_706589, EPI_ISL_706590, EPI_ISL_706591, EPI_ISL_706594, EPI_ISL_706595, EPI_ISL_706604, EPI_ISL_706605, EPI_ISL_706606, EPI_ISL_706615, EPI_ISL_706616, EPI_ISL_706617, EPI_ISL_706618, EPI_ISL_706663, EPI_ISL_706664, EPI_ISL_706665, EPI_ISL_706675, EPI_ISL_706676, EPI_ISL_706677, EPI_ISL_706678, EPI_ISL_706687, EPI_ISL_706688, EPI_ISL_706715, EPI_ISL_706716, EPI_ISL_706717, EPI_ISL_706718, EPI_ISL_706719, EPI_ISL_706720, EPI_ISL_706721, EPI_ISL_706722, EPI_ISL_706723, EPI_ISL_706724, EPI_ISL_706725, EPI_ISL_706726, EPI_ISL_706727, EPI_ISL_706728, EPI_ISL_706729, EPI_ISL_706730, EPI_ISL_706731, EPI_ISL_706732, EPI_ISL_706733, EPI_ISL_706734, EPI_ISL_706735, EPI_ISL_706736, EPI_ISL_706737, EPI_ISL_706738, EPI_ISL_706739, EPI_ISL_706740, EPI_ISL_706741, EPI_ISL_706742, EPI_ISL_706743, EPI_ISL_706744, EPI_ISL_706745, EPI_ISL_706746, EPI_ISL_706747, EPI_ISL_706748, EPI_ISL_706749, EPI_ISL_706750, EPI_ISL_706751, EPI_ISL_706752, EPI_ISL_706753, EPI_ISL_706754, EPI_ISL_706755, EPI_ISL_706756, EPI_ISL_706757, EPI_ISL_706758, EPI_ISL_706759, EPI_ISL_706760, EPI_ISL_706761, EPI_ISL_706762, EPI_ISL_706763 | | | |
| see above | Wales Specialist Virology Centre Sequencing lab: Pathogen Genomics Unit | COVID-19 Genomics UK (COG-UK) Consortium | Catherine Moore, Johnathan Evans, Laura Gifford, Malorie Perry, Simon Cottrell, Angela Marchbank, Alec Birchley, Alexander Adams, Amy Gaskin, Bree Gatica-Wilcox, Jason Coombes, Joel Southgate, Lauren Gilbert, Lee Graham, Nicole Pacchiarini, Sara Kumziene-Summerhayes, Sarah Taylor, Sophie Jones, Sara Rey, Matthew Bull, Joanne Watkins, Sally Corden, Tom Connor |
| EPI_ISL_708119 | Oslo University Hospital, Department of Medical Microbiology | Norwegian Institute of Public Health, Department of Virology | Kathrine Stene-Johansen, Kamilla Heddeland Instefjord, Hilde Elshaug, Marie Paulsen Madsen, Rasmus Riis Kopperud, Hilde Vollan, Karoline Bragstad, Olav Hungnes |
| EPI_ISL_708122, EPI_ISL_708123, EPI_ISL_708124, EPI_ISL_708125 | Hospital of Southern Norway - Kristiansand, Department of Medical Microbiology | Norwegian Institute of Public Health, Department of Virology | Kathrine Stene-Johansen, Kamilla Heddeland Instefjord, Hilde Elshaug, Marie Paulsen Madsen, Rasmus Riis Kopperud, Hilde Vollan, Karoline Bragstad, Olav Hungnes |
| EPI_ISL_708126, EPI_ISL_708127, EPI_ISL_708128, EPI_ISL_708129, EPI_ISL_708130 | Furst Medical Laboratory | Norwegian Institute of Public Health, Department of Virology | Kathrine Stene-Johansen, Kamilla Heddeland Instefjord, Hilde Elshaug, Marie Paulsen Madsen, Rasmus Riis Kopperud, Hilde Vollan, Karoline Bragstad, Olav Hungnes |
| EPI_ISL_708132, EPI_ISL_708133, EPI_ISL_708134, EPI_ISL_708135 | Dept. of Medical Microbiology, Stavanger University Hospital, Helse Stavanger HF | Norwegian Institute of Public Health, Department of Virology | Kathrine Stene-Johansen, Kamilla Heddeland Instefjord, Hilde Elshaug, Marie Paulsen Madsen, Rasmus Riis Kopperud, Hilde Vollan, Karoline Bragstad, Olav Hungnes |
| EPI_ISL_708143, EPI_ISL_708144, EPI_ISL_708146 | Department of Medical Microbiology, St. Olavs hospital | Norwegian Institute of Public Health, Department of Virology | Kathrine Stene-Johansen, Kamilla Heddeland Instefjord, Hilde Elshaug, Marie Paulsen Madsen, Rasmus Riis Kopperud, Hilde Vollan, Karoline Bragstad, Olav Hungnes |
| EPI_ISL_708147, EPI_ISL_708148, EPI_ISL_708149, EPI_ISL_708150 | Unilabs Laboratory Medicine | Norwegian Institute of Public Health, Department of Virology | Kathrine Stene-Johansen, Kamilla Heddeland Instefjord, Hilde Elshaug, Marie Paulsen Madsen, Rasmus Riis Kopperud, Hilde Vollan, Karoline Bragstad, Olav Hungnes |
| EPI_ISL_708176, EPI_ISL_708177, EPI_ISL_708178 | Foerde Hospital, Department of Microbiology | Norwegian Institute of Public Health, Department of Virology | Kathrine Stene-Johansen, Kamilla Heddeland Instefjord, Hilde Elshaug, Marie Paulsen Madsen, Rasmus Riis Kopperud, Hilde Vollan, Karoline Bragstad, Olav Hungnes |
| EPI_ISL_708234, EPI_ISL_708258, EPI_ISL_708274, EPI_ISL_708275, EPI_ISL_708276, EPI_ISL_708278, EPI_ISL_708279, EPI_ISL_708280, EPI_ISL_708281, EPI_ISL_708282, EPI_ISL_708284, EPI_ISL_708285, EPI_ISL_708286, EPI_ISL_708287, EPI_ISL_708288, EPI_ISL_708289, EPI_ISL_708290, EPI_ISL_708291, EPI_ISL_708292, EPI_ISL_708293, EPI_ISL_708298, EPI_ISL_708299, EPI_ISL_708300, EPI_ISL_708301, EPI_ISL_708302, EPI_ISL_708303, EPI_ISL_708305, EPI_ISL_708316, EPI_ISL_708321, EPI_ISL_708322, EPI_ISL_708325, EPI_ISL_708326, EPI_ISL_708330, EPI_ISL_708331, EPI_ISL_708335, EPI_ISL_708337, EPI_ISL_708338, EPI_ISL_708346, EPI_ISL_708348, EPI_ISL_708350, EPI_ISL_708352, EPI_ISL_708355, EPI_ISL_708356, EPI_ISL_708357, EPI_ISL_708360, EPI_ISL_708363, EPI_ISL_708364, EPI_ISL_708371, EPI_ISL_708372, EPI_ISL_708373, EPI_ISL_708375, EPI_ISL_708377 | | | |
| see above | Michigan Department of Health and Human Services, Bureau of Laboratories | Michigan Department of Health and Human Services, Bureau of Laboratories | Blankenship HM, Riner D, Soehnlen MK |
| EPI_ISL_708479, EPI_ISL_708480, EPI_ISL_708482 | Minnesota Department of Health, Public Health Laboratory | Minnesota Department of Health, Public Health Laboratory | Alexandra Lorentz, Jacob Garfin, Matt Plumb, and Xiong Wang |
| EPI_ISL_708540, EPI_ISL_708546 | Michigan Department of Health and Human Services, Bureau of Laboratories | Michigan Department of Health and Human Services, Bureau of Laboratories | Blankenship HM, Riner D, Soehnlen MK |
| EPI_ISL_708808 | Regional medical sciences center 6 chonburi | National Institute of Health, Department of Medical Sciences, Ministry of Public Health, Thailand | Pilailuk Okada; Siripaporn Phuygun; Thanutsapa Thanadachakul; Sittiporn Parnmen; Pakorn Piromtong; Warawan Wongboot; Sunthareeya Waicharoen; Malinee Chittaganpitch |
| EPI_ISL_708810 | Regional medical sciences center 2 Phitsanulok | National Institute of Health, Department of Medical Sciences, Ministry of Public Health, Thailand | Pilailuk Okada; Siripaporn Phuygun; Thanutsapa Thanadachakul; Sittiporn Parnmen; Pakorn Piromtong; Warawan Wongboot; Sunthareeya Waicharoen; Malinee Chittaganpitch |
| EPI_ISL_709886, EPI_ISL_709893 | Lighthouse Lab in Milton Keynes | Wellcome Sanger Institute for the COVID-19 Genomics UK (COG-UK) Consortium | The Lighthouse Lab in Milton Keynes and Alex Alderton, Roberto Amato, Sonia Goncalves, Ewan Harrison, David K. Jackson, Ian Johnston, Dominic Kwiatkowski, Cordelia Langford, John Sillitoe on behalf of the Wellcome Sanger Institute COVID-19 Surveillance Team |
| EPI_ISL_710090, EPI_ISL_710091, EPI_ISL_710092, EPI_ISL_710093, EPI_ISL_710095 | Maryland Public Health Laboratory | Maryland Public Health Laboratory | Maryland Department of Health Laboratories Administration |
| EPI_ISL_710862, EPI_ISL_710863, EPI_ISL_710864 | Lighthouse Lab in Glasgow | Wellcome Sanger Institute for the COVID-19 Genomics UK (COG-UK) Consortium | Harper VanSteenhouse, Yumi Kasai, David Gray, Carol Clugston, Anna Dominiczak and Alex Alderton, Roberto Amato, Sonia Goncalves, Ewan Harrison, David K. Jackson, Ian Johnston, Dominic Kwiatkowski, Cordelia Langford, John Sillitoe on behalf of the Wellcome Sanger Institute COVID-19 Surveillance Team |
| EPI_ISL_710865 | Lighthouse Lab in Alderley Park | Wellcome Sanger Institute for the COVID-19 Genomics UK (COG-UK) Consortium | Jacquelyn Wynn, Mairead Hyland, The Lighthouse Lab in Alderley Park and Alex Alderton, Roberto Amato, Sonia Goncalves, Ewan Harrison, David K. Jackson, Ian Johnston, Dominic Kwiatkowski, Cordelia Langford, John Sillitoe on behalf of the Wellcome Sanger Institute COVID-19 Surveillance Team |
| EPI_ISL_710867, EPI_ISL_710868 | Lighthouse Lab in Glasgow | Wellcome Sanger Institute for the COVID-19 Genomics UK (COG-UK) Consortium | Harper VanSteenhouse, Yumi Kasai, David Gray, Carol Clugston, Anna Dominiczak and Alex Alderton, Roberto Amato, Sonia Goncalves, Ewan Harrison, David K. Jackson, Ian Johnston, Dominic Kwiatkowski, Cordelia Langford, John Sillitoe on behalf of the Wellcome Sanger Institute COVID-19 Surveillance Team |
| EPI_ISL_710869 | Lighthouse Lab in Alderley Park | Wellcome Sanger Institute for the COVID-19 Genomics UK (COG-UK) Consortium | Jacquelyn Wynn, Mairead Hyland, The Lighthouse Lab in Alderley Park and Alex Alderton, Roberto Amato, Sonia Goncalves, Ewan Harrison, David K. Jackson, Ian Johnston, Dominic Kwiatkowski, Cordelia Langford, John Sillitoe on behalf of the Wellcome Sanger Institute COVID-19 Surveillance Team |
| EPI_ISL_710872 | Lighthouse Lab in Glasgow | Wellcome Sanger Institute for the COVID-19 Genomics UK (COG-UK) Consortium | Harper VanSteenhouse, Yumi Kasai, David Gray, Carol Clugston, Anna Dominiczak and Alex Alderton, Roberto Amato, Sonia Goncalves, Ewan Harrison, David K. Jackson, Ian Johnston, Dominic Kwiatkowski, Cordelia Langford, John Sillitoe on behalf of the Wellcome Sanger Institute COVID-19 Surveillance Team |
| EPI_ISL_714174, EPI_ISL_714210, EPI_ISL_714275, EPI_ISL_714282, EPI_ISL_714363, EPI_ISL_714411, EPI_ISL_714412, EPI_ISL_714475, EPI_ISL_714904, EPI_ISL_714905, EPI_ISL_714906, EPI_ISL_714907, EPI_ISL_714908, EPI_ISL_714909, EPI_ISL_714910, EPI_ISL_714911, EPI_ISL_714912, EPI_ISL_714913, EPI_ISL_714914, EPI_ISL_714915, EPI_ISL_714916, EPI_ISL_714917, EPI_ISL_714918, EPI_ISL_714919, EPI_ISL_714920, EPI_ISL_714921, EPI_ISL_714922 | | | |
| see above | Department of Virus and Microbiological Special Diagnostics, Statens Serum Institut, Copenhagen, Denmark | Albertsen Lab, Department of Chemistry and Bioscience, Aalborg University, Denmark | Danish Covid-19 Genome Consortium |
| EPI_ISL_717586 | Lab voor klinische biologie | Onderzoeksgroep Virologie | Nick Vereecke, Laurens Lambrechts, Marthe Pauwels, Bruno Verhasselt, Linos Vandekerckhove, Hans Nauwynck, Sebastiaan Theuns |

| | | | |
|---|---|---|---|
| EPI_ISL_717590, EPI_ISL_717632 | Lab voor klinische biologie | Onderzoeksgroep Virologie | Laurens Lambrechts, Nick Vereecke, Marthe Pauwels, Bruno Verhasselt, Linos Vandekerckhove, Hans Nauwynck, Sebastiaan Theuns |
| EPI_ISL_717633, EPI_ISL_717983 | Lab voor klinische biologie | Onderzoeksgroep Virologie | Nick Vereecke, Laurens Lambrechts, Marthe Pauwels, Bruno Verhasselt, Linos Vandekerckhove, Hans Nauwynck, Sebastiaan Theuns |
| EPI_ISL_718000 | Lab voor klinische biologie | Onderzoeksgroep Virologie | Laurens Lambrechts, Nick Vereecke, Marthe Pauwels, Bruno Verhasselt, Linos Vandekerckhove, Hans Nauwynck, Sebastiaan Theuns |
| EPI_ISL_718004, EPI_ISL_718005 | Lab voor klinische biologie | Onderzoeksgroep Virologie | Nick Vereecke, Laurens Lambrechts, Marthe Pauwels, Bruno Verhasselt, Linos Vandekerckhove, Hans Nauwynck, Sebastiaan Theuns |
| EPI_ISL_718084, EPI_ISL_718085, EPI_ISL_718086, EPI_ISL_718087, EPI_ISL_718088, EPI_ISL_718089, EPI_ISL_718090, EPI_ISL_718091, EPI_ISL_718092, EPI_ISL_718093, EPI_ISL_718094, EPI_ISL_718095 | | | |
| see above | ZOTZ KLIMAS MVZ Düsseldorf-Centrum GbR ÜBAG für Labormedizin, Genetik, Zytologie, Pathologie | Center of Medical Microbiology, Virology, and Hospital Hygiene, University of Duesseldorf | Maximilian Damagnez, Alexander Dilthey, Ashley-Jane Duplessis, Patrick Finzer, Katrin Hoffmann, Torsten Houwaart, Lisanna Hülse, Malte Kohns Vasconcelos, Marek Korencak, Nadine Lübke, Jessica Nicolai, Klaus Pfeffer, Daniel Strelow, Jörg Timm, Andreas Walker, Tobias Wienemann, Rainer Zotz |
| EPI_ISL_718180, EPI_ISL_718183, EPI_ISL_718184, EPI_ISL_718189, EPI_ISL_718199 | Ministry of Health Hospitals | Institute of Health and Community Medicine | David Perera, Ooi Mong How, Chua Hock Hin, Tonnii Sia Loong Loong, Wong Jyn Shan, Wong Kiing Aik, Chan Chia Jui |
| EPI_ISL_718235 | Hospital | National Reference Center for Viruses of Respiratory Infections, Institut Pasteur, Paris | Marion Barbet, Sylvie Behillil, Méline Bizard, Angela Brisebarre, Camille Capel, Etienne Simon-Lorière, Vincent Enouf, Maud Vanpeene, Sylvie van der Werf, Gisèle Lagathu |
| EPI_ISL_722202, EPI_ISL_722204, EPI_ISL_722205, EPI_ISL_722207 | University Hospital Zurich | Institute of Medical Virology, University of Zurich | Stefan Schmutz, Verena Kufner, Maryam Zaheri, Gabriela Ziltener, Aline Wolfensberger, Thomas Scheier, Jürg Böni, Michael Huber, Alexandra Trkola |
| EPI_ISL_722242, EPI_ISL_722243, EPI_ISL_722244, EPI_ISL_722258, EPI_ISL_722259, EPI_ISL_722262, EPI_ISL_722263 | Servicio de Microbiología, Hospital Miguel Servet, Zaragoza | SeqCOVID-SPAIN consortium/IBV(CSIC) | Antonio Rezusta López, Alexander Tristancho Baró, Ana Milagro, Yolanda Gracia Grataloup, Nieves Martínez Cameo and SeqCOVID-SPAIN consortium |
| EPI_ISL_722280, EPI_ISL_722291, EPI_ISL_722292, EPI_ISL_722304, EPI_ISL_722312, EPI_ISL_722323, EPI_ISL_722379, EPI_ISL_722401, EPI_ISL_722426, EPI_ISL_722427, EPI_ISL_722428, EPI_ISL_722453, EPI_ISL_722608, EPI_ISL_722609, EPI_ISL_722610, EPI_ISL_722611, EPI_ISL_722612, EPI_ISL_722613, EPI_ISL_722614, EPI_ISL_722615, EPI_ISL_722616, EPI_ISL_722617, EPI_ISL_722618, EPI_ISL_722750, EPI_ISL_722751, EPI_ISL_722752, EPI_ISL_722753, EPI_ISL_722754, EPI_ISL_722755, EPI_ISL_722756, EPI_ISL_722757 | | | |
| see above | Dutch COVID-19 response team | Erasmus Medical Center | Bas Oude Munnink, Reina Sikkema, David Nieuwenhuijse, Irina Chestakova, Anne van der Linden, Marjan Boter, Emmanuelle Munger, Corine GeurtsvanKessel, Annemiek van der Eijk, Richard Molenkamp, Marion Koopmans, on behalf of the Dutch national COVID-19 response team. |
| EPI_ISL_723119 | Minnesota Department of Health, Public Health Laboratory | Minnesota Department of Health, Public Health Laboratory | Alexandra Lorentz, Jacob Garfin, Matt Plumb, and Xiong Wang |
| EPI_ISL_723144 | Mayo Clinic & Mayo Clinic Laboratories | Minnesota Department of Health, Public Health Laboratory | Alexandra Lorentz, Jacob Garfin, Matt Plumb, and Xiong Wang |
| EPI_ISL_723164, EPI_ISL_723166, EPI_ISL_723178, EPI_ISL_723180, EPI_ISL_723219, EPI_ISL_723223, EPI_ISL_723224, EPI_ISL_723232, EPI_ISL_723238, EPI_ISL_723258, EPI_ISL_723271, EPI_ISL_723275, EPI_ISL_723276, EPI_ISL_723283, EPI_ISL_723298, EPI_ISL_723299, EPI_ISL_723300, EPI_ISL_723303, EPI_ISL_723326, EPI_ISL_723427, EPI_ISL_723452, EPI_ISL_723453 | | | |
| see above | Dutch COVID-19 response team | National Institute for Public Health and the Environment (RIVM) | Adam Meijer, Harry Vennema, Jeroen Cremer, Sharon van den Brink, Bas van der Veer, AnneMarie van den Brandt, Florian Zwagemaker, Dennis Schmitz, Chantal Reusken, on behalf of the national COVID-19 response team |
| EPI_ISL_723558, EPI_ISL_723559, EPI_ISL_723560, EPI_ISL_723561, EPI_ISL_723562, EPI_ISL_723563, EPI_ISL_723564, EPI_ISL_723565, EPI_ISL_723566, EPI_ISL_723567, EPI_ISL_723568, EPI_ISL_723569, EPI_ISL_723570, EPI_ISL_723571, EPI_ISL_723572, EPI_ISL_723573, EPI_ISL_723574, EPI_ISL_723575, EPI_ISL_723576, EPI_ISL_723577, EPI_ISL_723578, EPI_ISL_723579, EPI_ISL_723580, EPI_ISL_723581, EPI_ISL_723582, EPI_ISL_723584, EPI_ISL_723585, EPI_ISL_723586, EPI_ISL_723587, EPI_ISL_723588, EPI_ISL_723589, EPI_ISL_723590, EPI_ISL_723591, EPI_ISL_723593, EPI_ISL_723595, EPI_ISL_723601, EPI_ISL_723603, EPI_ISL_723604, EPI_ISL_723605, EPI_ISL_723606, EPI_ISL_723607, EPI_ISL_723608, EPI_ISL_723609, EPI_ISL_723610, EPI_ISL_723611, EPI_ISL_723612, EPI_ISL_723613, EPI_ISL_723614, EPI_ISL_723615, EPI_ISL_723616, EPI_ISL_723617, EPI_ISL_723618, EPI_ISL_723619, EPI_ISL_723620, EPI_ISL_723621, EPI_ISL_723622, EPI_ISL_723623, EPI_ISL_723624, EPI_ISL_723625, EPI_ISL_723626, EPI_ISL_723627 | | | |
| see above | Northumbria University / South Tees Hospitals NHS Foundation Trust / North Cumbria Integrated Care NHS Foundation Trust / North Tees and Hartlepool NHS Foundation Trust / Newcastle Hospitals NHS Foundation Trust | COVID-19 Genomics UK (COG-UK) Consortium | Darren L Smith,Andrew Nelson,Matthew Bashton,Greg R Young,Joshua Loh,John Allan,Mohammad A Tariq,Giles S Holt,Gary Black,Wen C Yew,Lynn Dover,Paul Baker,Steve Liggett,Sarah Essex,Jane Greenaway,Debra Padgett,Clive Graham,Garren Scott,Edward Barton,Emma Swindells,Brendan Payne,Jennifer Collins,Yusri Taha,Gary Eltringham |
| EPI_ISL_723635, EPI_ISL_723636, EPI_ISL_723637, EPI_ISL_723638, EPI_ISL_723639, EPI_ISL_723640, EPI_ISL_723641, EPI_ISL_723642, EPI_ISL_723643, EPI_ISL_723644, EPI_ISL_723645, EPI_ISL_723647, EPI_ISL_723659, EPI_ISL_723660, EPI_ISL_723661, EPI_ISL_723662, EPI_ISL_723663, EPI_ISL_723664, EPI_ISL_723665, EPI_ISL_723729, EPI_ISL_723964, EPI_ISL_723967, EPI_ISL_723982, EPI_ISL_724016, EPI_ISL_724021, EPI_ISL_724022, EPI_ISL_724023, EPI_ISL_724024, EPI_ISL_724025, EPI_ISL_724043 | | | |
| see above | Oxford Viromics, NDM, University of Oxford; Oxford University Hospitals; Basingstoke and North Hampshire Hospital | COVID-19 Genomics UK (COG-UK) Consortium | Tanya Golubchik, David Bonsall, George Macintyre, Amy Trebes, Mariateresa de Cesare, Catrin Moore, Alex Mobbs, Anita Justice, Robert Shaw, Monique Andersson, Timothy Peto, Emma Wise, Nathan Moore, Jessica Lynch, Nick Cortes, Matilde Mori, Stephen Kidd, David Buck, John Todd, Christophe Fraser |
| EPI_ISL_724611, EPI_ISL_724630, EPI_ISL_724636, EPI_ISL_724639, EPI_ISL_724642, EPI_ISL_724648, EPI_ISL_724649, EPI_ISL_724650, EPI_ISL_724651, EPI_ISL_724652, EPI_ISL_724661, EPI_ISL_724665, EPI_ISL_724686, EPI_ISL_724689, EPI_ISL_724691, EPI_ISL_724695, EPI_ISL_724713, EPI_ISL_724723, EPI_ISL_724734, EPI_ISL_724735, EPI_ISL_724736, EPI_ISL_724738, EPI_ISL_724739, EPI_ISL_724744, EPI_ISL_724752, EPI_ISL_724754, EPI_ISL_724756, EPI_ISL_724758, EPI_ISL_724777 | | | |
| see above | University College London, Great Ormond Street Hospital for Children NHS Foundation Trust, Imperial College Healthcare NHS Trust | COVID-19 Genomics UK (COG-UK) Consortium | Sergi Castellano, Rachel Williams, Mark Kristiansen, Paola Resende Silva, Sunando Roy, Tony Brooks, Helena Tutill, Paola Niola, Patricia Dyal, Charlotte Williams, Leysa Forrest, Yasmin Panchbhaya, Jacqueline Findlay, Samuel Weeks, Julianne Brown, Kathryn Harris, Paul Randell, James Price, Alison Holmes, Judith Breuer |
| EPI_ISL_724971, EPI_ISL_724972, EPI_ISL_724995, EPI_ISL_724996, EPI_ISL_724997, EPI_ISL_724998, EPI_ISL_724999, EPI_ISL_725000, EPI_ISL_725001 | Northumbria University / South Tees Hospitals NHS Foundation Trust / North Cumbria Integrated Care NHS Foundation Trust / North Tees and Hartlepool NHS Foundation Trust / Newcastle Hospitals NHS Foundation Trust | COVID-19 Genomics UK (COG-UK) Consortium | Darren L Smith,Andrew Nelson,Matthew Bashton,Greg R Young,Joshua Loh,John Allan,Mohammad A Tariq,Giles S Holt,Gary Black,Wen C Yew,Lynn Dover,Paul Baker,Steve Liggett,Sarah Essex,Jane Greenaway,Debra Padgett,Clive Graham,Garren Scott,Edward Barton,Emma Swindells,Brendan Payne,Jennifer Collins,Yusri Taha,Gary Eltringham |
| EPI_ISL_725113, EPI_ISL_725116, EPI_ISL_725117, EPI_ISL_725121, EPI_ISL_725128, EPI_ISL_725130, EPI_ISL_725136, EPI_ISL_725166, EPI_ISL_725167, EPI_ISL_725185, EPI_ISL_725190, EPI_ISL_725193, EPI_ISL_725282, EPI_ISL_725345 | | | |
| see above | Quadram Institute Bioscience | COVID-19 Genomics UK (COG-UK) Consortium | Dave J. Baker, Gemma L. Kay, Alp Aydin, Thanh Le-Viet, Steven Rudder, Ana P. Tedim, Anastasia Kolyva, Maria Diaz, Leonardo de Oliveira Martins, Nabil-Fareed Alikhan, Lizzie Meadows, Rachael Stanley, Ngozi Elumogo, Muhammed Yasir, Nicholas M. Thomson, Alexander J Trotter, Rachel Gilroy, Samuel Bloomfield, Claire Stuart, Andrew Bell, Reenesh Prakash, Samir Dervisevic, Alison E. Mather, John Wain, Mark Webber, Andrew J. Page, Justin O'Grady |
| EPI_ISL_725986, EPI_ISL_725987, EPI_ISL_725988, EPI_ISL_725989, EPI_ISL_726033, EPI_ISL_726035, EPI_ISL_726036, EPI_ISL_726037, EPI_ISL_726038, EPI_ISL_726039, EPI_ISL_726040, EPI_ISL_726041, EPI_ISL_726042, EPI_ISL_726044 | | | |
| see above | Wales Specialist Virology Centre Sequencing lab: Pathogen Genomics Unit | COVID-19 Genomics UK (COG-UK) Consortium | Catherine Moore, Johnathan Evans, Laura Gifford, Malorie Perry, Simon Cottrell, Angela Marchbank, Alec Birchley, Alexander Adams, Amy Gaskin, Bree Gatica-Wilcox, Jason Coombes, Joel Southgate, Lauren Gilbert, Lee Graham, Nicole Pacchiarini, Sara Kumziene-Summerhayes, Sarah Taylor, Sophie Jones, Sara Rey, Matthew Bull, Joanne Watkins, Sally Corden, Tom Connor |
| EPI_ISL_727698, EPI_ISL_727699, EPI_ISL_727700, EPI_ISL_727701, EPI_ISL_727735, EPI_ISL_727736, EPI_ISL_727743, EPI_ISL_727744, EPI_ISL_727745, EPI_ISL_727747, EPI_ISL_727748 | | | |
| see above | Centre for Enzyme Innovation, University of Portsmouth / Translational Research Laboratory, Portsmouth Hospitals NHS Trust | COVID-19 Genomics UK (COG-UK) Consortium | Angela Beckett,Yann Bourgeois,Garry Scarlett,Sharon Glaysher,Scott Elliott,Kelly Bicknell,Robert Impey,Allyson Lloyd,Sarah Wyllie,Ethan Butcher,Anoop Chauhan,Samuel Robson |
| EPI_ISL_727997 | Virology Department, Sheffield Teaching Hospitals NHS Foundation Trust/Department of Infection, Immunity and Cardiovascular Disease, The Medical School, University of Sheffield | COVID-19 Genomics UK (COG-UK) Consortium | Thushan de Silva, Matthew Parker, Nikki Smith, Adri Angyal, Rebecca Brown, Luke Green, Rachel Tucker, Paul Parsons, Danielle Groves, Katie Johnson, Laura Carrilero, Alex Keeley, Dave Partridge, Matthew Wyles, Benjamin Lindsey, Mehmet Yavuz, Mohammad Raza, Cariad Evans |
| EPI_ISL_728009, EPI_ISL_728043, EPI_ISL_728075, EPI_ISL_728078 | University of Wisconsin-Madison AIDS Vaccine Research Laboratories | University of Wisconsin-Madison AIDS Vaccine Research Laboratories | Gage Moreno, Katarina Braun, et al. AIDS Vaccine Research Laboratories |
| EPI_ISL_728212, EPI_ISL_728241, | Institute for Medical Research, Infectious Disease Research | Institute for Medical Research, Infectious Disease Research | Suppiah J, Kamel K, Mohd-Zawawi Z, Thayan R |

| | | | |
|---|---|---|---|
| EPI_ISL_728242, EPI_ISL_728243, EPI_ISL_728244, EPI_ISL_728245, EPI_ISL_728246 | Centre, National Institutes of Health, Ministry of Health Malaysia | Centre, National Institutes of Health, Ministry of Health Malaysia | |
| EPI_ISL_728445, EPI_ISL_728446, EPI_ISL_728447, EPI_ISL_728448, EPI_ISL_728449, EPI_ISL_728450, EPI_ISL_728451, EPI_ISL_728452, EPI_ISL_728453, EPI_ISL_728454, EPI_ISL_728455, EPI_ISL_728456, EPI_ISL_728476, EPI_ISL_728477, EPI_ISL_728478, EPI_ISL_728479, EPI_ISL_728480, EPI_ISL_728481, EPI_ISL_728482, EPI_ISL_728483, EPI_ISL_728484, EPI_ISL_728485, EPI_ISL_728486, EPI_ISL_728487, EPI_ISL_728488, EPI_ISL_728489, EPI_ISL_728490, EPI_ISL_728491, EPI_ISL_728492, EPI_ISL_728493, EPI_ISL_728494, EPI_ISL_728495, EPI_ISL_728496, EPI_ISL_728497, EPI_ISL_728498, EPI_ISL_728499, EPI_ISL_728500, EPI_ISL_728501, EPI_ISL_728502, EPI_ISL_728503, EPI_ISL_728504, EPI_ISL_728505, EPI_ISL_728506, EPI_ISL_728507, EPI_ISL_728508, EPI_ISL_728509, EPI_ISL_728510, EPI_ISL_728511, EPI_ISL_728512, EPI_ISL_728513, EPI_ISL_728514, EPI_ISL_728515 | | | |
| see above | University of Michigan Clinical Microbiology Laboratory | Lauring Lab, University of Michigan, Department of Microbiology and Immunology | Valesano |
| EPI_ISL_728552 | University of Wisconsin-Madison AIDS Vaccine Research Laboratories | University of Wisconsin-Madison AIDS Vaccine Research Laboratories | Gage Moreno, Katarina Braun, et al. AIDS Vaccine Research Laboratories |
| EPI_ISL_728728 | Dutch COVID-19 response team | National Institute for Public Health and the Environment (RIVM) | Adam Meijer, Harry Vennema, Jeroen Cremer, Sharon van den Brink, Bas van der Veer, AnneMarie van den Brandt, Florian Zwagemaker, Dennis Schmitz, Chantal Reusken, on behalf of the national COVID-19 response team |
| EPI_ISL_728778, EPI_ISL_728782, EPI_ISL_728800, EPI_ISL_728802, EPI_ISL_728824, EPI_ISL_728831, EPI_ISL_728833, EPI_ISL_728858, EPI_ISL_728859, EPI_ISL_728862, EPI_ISL_728866, EPI_ISL_728868, EPI_ISL_728879, EPI_ISL_728880, EPI_ISL_728884, EPI_ISL_728893, EPI_ISL_728938, EPI_ISL_728940, EPI_ISL_728950, EPI_ISL_728960, EPI_ISL_728981, EPI_ISL_728987, EPI_ISL_728989, EPI_ISL_728993, EPI_ISL_729055, EPI_ISL_729238, EPI_ISL_729239, EPI_ISL_729240, EPI_ISL_729241, EPI_ISL_729242, EPI_ISL_729243, EPI_ISL_729244, EPI_ISL_729245, EPI_ISL_729246, EPI_ISL_729247, EPI_ISL_729248, EPI_ISL_729249, EPI_ISL_729250, EPI_ISL_729251, EPI_ISL_729252, EPI_ISL_729253, EPI_ISL_729254, EPI_ISL_729255 | | | |
| see above | Viollier AG | Department of Biosystems Science and Engineering, ETH Zürich | Chaoran Chen, Sarah Nadeau, Catharine Aquino, Ivan Topolsky, Pedro Ferreira, Philipp Jablonski, Susana Posada-Céspedes, Andreia Cabral de Gouvea, Maria Domenica Moccia, Simon Grüter, Timothy Sykes, Lennart Opitz, Ralph Schlapbach, Christiane Beckmann, Maurice Redondo, Olivier Kobel, Christoph Noppen, Sophie Seidel, Noemie Santamaria de Souza, Niko Beerenwinkel, Tanja Stadler |
| EPI_ISL_730049, EPI_ISL_730050 | Yale COVID-19 Biorepository | Grubaugh Lab - Yale School of Public Health | Joseph Fauver, Tara Alpert, Anderson Brito, Annie Watkins, Anne Wyllie, Chantal Vogels, Mary Petrone, Chaney Kalinich, Isabel Ott, Arnau Casanovas, Catherine Muenker, Adam Moore, Alice Lu, Maria Tokuyama, Patrick Wong, Peiwen Lu, Saad Omer, Richard Martinello, Allison Nelson, Shelli Farhadian, Akiko Iwasaki, Charlese Dela Cruz, Albert Ko, Nathan Grubaugh |
| EPI_ISL_730086, EPI_ISL_730166, EPI_ISL_730169, EPI_ISL_730182, EPI_ISL_730195, EPI_ISL_730304, EPI_ISL_730308, EPI_ISL_730329, EPI_ISL_730338, EPI_ISL_730342, EPI_ISL_730346 | | | |
| see above | San Diego County Public Health Laboratory | Andersen lab at Scripps Research | SEARCH Alliance San Diego with Tracy Basler, Jovan Shephard, Brett Austin |
| EPI_ISL_730564, EPI_ISL_730565 | University of Michigan Clinical Microbiology Laboratory | Lauring Lab, University of Michigan, Department of Microbiology and Immunology | Valesano |
| EPI_ISL_730640, EPI_ISL_730641, EPI_ISL_730642, EPI_ISL_730643, EPI_ISL_730644, EPI_ISL_730645, EPI_ISL_730646, EPI_ISL_730647, EPI_ISL_730648, EPI_ISL_730649, EPI_ISL_730650, EPI_ISL_730651 | | | |
| see above | CNR Virus des Infections Respiratoires - France SUD | CNR Virus des Infections Respiratoires - France SUD | Antonin Bal, Gregory Destras, Claudia Gonzalez, Gwendolyne Burfin, Quentin Semanas, Martine Valette, Bruno Lina, Laurence Josset |
| EPI_ISL_731902 | Instituto Nacional de Saude (INSA) | Instituto Nacional de Saude (INSA) | Borges et al |
| EPI_ISL_731903, EPI_ISL_731912 | Instituto Nacional de Saude (INSA) and Instituto Gulbenkian de Ciencia (IGC) | Instituto Nacional de Saude (INSA) and Instituto Gulbenkian de Ciencia (IGC) | Borges et al |
| EPI_ISL_731913, EPI_ISL_731916, EPI_ISL_731922, EPI_ISL_731923 | Instituto Nacional de Saude (INSA) | Instituto Nacional de Saude (INSA) | Borges et al |
| EPI_ISL_731926 | Instituto Nacional de Saude (INSA) and Instituto Gulbenkian de Ciencia (IGC) | Instituto Nacional de Saude (INSA) and Instituto Gulbenkian de Ciencia (IGC) | Borges et al |
| EPI_ISL_731931, EPI_ISL_731937, EPI_ISL_731938, EPI_ISL_731939, EPI_ISL_731949, EPI_ISL_731950, EPI_ISL_731958, EPI_ISL_731959, EPI_ISL_731960, EPI_ISL_731961, EPI_ISL_731962, EPI_ISL_731963, EPI_ISL_731977, EPI_ISL_731978, EPI_ISL_731979, EPI_ISL_731980, EPI_ISL_731981, EPI_ISL_731982, EPI_ISL_731983, EPI_ISL_731984, EPI_ISL_731985, EPI_ISL_731986, EPI_ISL_731987, EPI_ISL_731988, EPI_ISL_731989, EPI_ISL_731990, EPI_ISL_731991, EPI_ISL_732000, EPI_ISL_732001, EPI_ISL_732002, EPI_ISL_732003, EPI_ISL_732004, EPI_ISL_732005, EPI_ISL_732006, EPI_ISL_732007, EPI_ISL_732025, EPI_ISL_732026, EPI_ISL_732027, EPI_ISL_732028, EPI_ISL_732029, EPI_ISL_732030, EPI_ISL_732031, EPI_ISL_732032, EPI_ISL_732033, EPI_ISL_732034, EPI_ISL_732035, EPI_ISL_732053, EPI_ISL_732054, EPI_ISL_732055, EPI_ISL_732056, EPI_ISL_732057, EPI_ISL_732058, EPI_ISL_732059, EPI_ISL_732060, EPI_ISL_732061, EPI_ISL_732062, EPI_ISL_732063, EPI_ISL_732064, EPI_ISL_732065, EPI_ISL_732066, EPI_ISL_732067, EPI_ISL_732068, EPI_ISL_732081, EPI_ISL_732082, EPI_ISL_732083, EPI_ISL_732084, EPI_ISL_732087, EPI_ISL_732088, EPI_ISL_732090, EPI_ISL_732091, EPI_ISL_732092, EPI_ISL_732093, EPI_ISL_732098, EPI_ISL_732108, EPI_ISL_732109, EPI_ISL_732110 | | | |
| see above | Instituto Nacional de Saude (INSA) | Instituto Nacional de Saude (INSA) | Borges et al |
| EPI_ISL_732118, EPI_ISL_732119, EPI_ISL_732120, EPI_ISL_732148, EPI_ISL_732149, EPI_ISL_732150, EPI_ISL_732151, EPI_ISL_732152, EPI_ISL_732153, EPI_ISL_732154, EPI_ISL_732155, EPI_ISL_732167, EPI_ISL_732168, EPI_ISL_732169, EPI_ISL_732170, EPI_ISL_732171, EPI_ISL_732172, EPI_ISL_732173, EPI_ISL_732174, EPI_ISL_732196, EPI_ISL_732197, EPI_ISL_732198, EPI_ISL_732199, EPI_ISL_732200, EPI_ISL_732201, EPI_ISL_732202, EPI_ISL_732203, EPI_ISL_732204, EPI_ISL_732205, EPI_ISL_732210, EPI_ISL_732211, EPI_ISL_732225, EPI_ISL_732246, EPI_ISL_732247, EPI_ISL_732248, EPI_ISL_732249, EPI_ISL_732250, EPI_ISL_732251, EPI_ISL_732252, EPI_ISL_732253, EPI_ISL_732254, EPI_ISL_732255, EPI_ISL_732256, EPI_ISL_732257, EPI_ISL_732267, EPI_ISL_732268, EPI_ISL_732269, EPI_ISL_732270, EPI_ISL_732271, EPI_ISL_732272, EPI_ISL_732273, EPI_ISL_732274, EPI_ISL_732275, EPI_ISL_732276, EPI_ISL_732277, EPI_ISL_732278, EPI_ISL_732279, EPI_ISL_732280, EPI_ISL_732281, EPI_ISL_732282, EPI_ISL_732283 | | | |
| see above | Instituto Nacional de Saude (INSA) and Instituto Gulbenkian de Ciencia (IGC) | Instituto Nacional de Saude (INSA) and Instituto Gulbenkian de Ciencia (IGC) | Borges et al |
| EPI_ISL_732378, EPI_ISL_732393, EPI_ISL_732395, EPI_ISL_732410, EPI_ISL_732411, EPI_ISL_732412, EPI_ISL_732413, EPI_ISL_732414, EPI_ISL_732415, EPI_ISL_732416, EPI_ISL_732417, EPI_ISL_732418, EPI_ISL_732419, EPI_ISL_732420, EPI_ISL_732421, EPI_ISL_732422, EPI_ISL_732424, EPI_ISL_732429, EPI_ISL_732430, EPI_ISL_732431, EPI_ISL_732434, EPI_ISL_732436, EPI_ISL_732437, EPI_ISL_732482 | | | |
| see above | National Virus Reference Laboratory | National Virus Reference Laboratory | Michael Carr, Gabriel Gonzalez, Jonathan Dean, Daniel Hare, Cillian F De Gascun |
| EPI_ISL_732683 | CNR Virus des Infections Respiratoires - France SUD | CNR Virus des Infections Respiratoires - France SUD | Antonin Bal, Gregory Destras, Claudia Gonzalez, Gwendolyne Burfin, Quentin Semanas, Martine Valette, Bruno Lina, Laurence Josset |
| EPI_ISL_732755, EPI_ISL_732772, EPI_ISL_732788, EPI_ISL_732789, EPI_ISL_732798 | Centro de Investigación Biomédica de La Rioja - Hospital San Pedro Logroño | SeqCOVID-SPAIN consortium/IBV(CSIC) | María de Toro, José Manuel Azcona Gutiérrez, María Pilar Bea Escudero, Miriam Blasco Alberdi and SeqCOVID-SPAIN consortium |
| EPI_ISL_733241 | WHO National Influenza Centre Russian Federation | WHO National Influenza Centre Russian Federation | Andrey Komissarov, Artem Fadeev, Anna Ivanova, Kseniya Komissarova, Dmitry Bazhenov, Daria Danilenko, Ksenia Safina, Elena Nabieva, Georgii Bazykin, Dmitry Lioznov |
| EPI_ISL_733301, EPI_ISL_733302, EPI_ISL_733303, EPI_ISL_733304, EPI_ISL_733305, EPI_ISL_733306, EPI_ISL_733307, EPI_ISL_733308, EPI_ISL_733309, EPI_ISL_733310, EPI_ISL_733314, EPI_ISL_733315, EPI_ISL_733316, EPI_ISL_733317, EPI_ISL_733326, EPI_ISL_733327, EPI_ISL_733328, EPI_ISL_733329, EPI_ISL_733331, EPI_ISL_733332, EPI_ISL_733334, EPI_ISL_733337, EPI_ISL_733338, EPI_ISL_733348, EPI_ISL_733349, EPI_ISL_733362, EPI_ISL_733367, EPI_ISL_733370, EPI_ISL_733371, EPI_ISL_733372, EPI_ISL_733374, EPI_ISL_733377, EPI_ISL_733383, EPI_ISL_733384, EPI_ISL_733385, EPI_ISL_733386, EPI_ISL_733387, EPI_ISL_733389, EPI_ISL_733390 | | | |
| see above | HELIX LLC | WHO National Influenza Centre Russian Federation | Andrey Komissarov, Artem Fadeev, Anna Ivanova, Kseniya Komissarova, Dmitry Bazhenov, Daria Danilenko, Ksenia Safina, Elena Nabieva, Georgii Bazykin, Dmitry Lioznov |
| EPI_ISL_733486, EPI_ISL_733487, EPI_ISL_733488 | Omsk Research Institute of Natural Focal Infections | WHO National Influenza Centre Russian Federation | Andrey Komissarov, Artem Fadeev, Anna Ivanova, Kseniya Komissarova, Dmitry Bazhenov, Daria Danilenko, Ekaterina Gradoboeva, Elena Poleshchuk, Aleksei Vasilenko, Valery Yakimenko, Ksenia Safina, Elena Nabieva, Georgii Bazykin, Dmitry Lioznov |
| EPI_ISL_733571 | Queen Mary Hospital | Hong Kong Department of Health | Alan K.L. Tsang, Peter C.W. Yip, Edman T.K. Lam, Rickjason C.W. Chan, Dominic N.C. Tsang |
| EPI_ISL_734279 | Virginia Division of Consolidated Laboratory Services (DCLS) | Virginia Division of Consolidated Laboratory Services (DCLS) | Virginia DCLS |
| EPI_ISL_734288, EPI_ISL_734291, EPI_ISL_734292, EPI_ISL_734295, EPI_ISL_734427, EPI_ISL_734428, EPI_ISL_734429, EPI_ISL_734430, EPI_ISL_734431, EPI_ISL_734432, EPI_ISL_734433, EPI_ISL_734434, EPI_ISL_734435, EPI_ISL_734436, EPI_ISL_734437, EPI_ISL_734438, EPI_ISL_734439, EPI_ISL_734440, EPI_ISL_734441, EPI_ISL_734442, EPI_ISL_734443, EPI_ISL_734444, EPI_ISL_734445, EPI_ISL_734446 | | | |
| see above | Wadsworth Center, New York State Department.of Health | Wadsworth Center, New York State Department.of Health | Kirsten St. George, Daryl M. Lamson, Alexis Russel, Jonathan Plitnick, Navjot Singh, John Kelly, Sara Griesemer, Erasmus Schneider, Erica Lasek-Nesselquist |
| EPI_ISL_737020, EPI_ISL_737021, EPI_ISL_737024, EPI_ISL_737027 | The National Institute of Public Health | State Veterinary Institute Prague | Nagy,A;Jirincova,H;Trnka,D;Vecerova,J |
| EPI_ISL_737104, EPI_ISL_737106, EPI_ISL_737110, EPI_ISL_737119, EPI_ISL_737122, EPI_ISL_737126, EPI_ISL_737130, EPI_ISL_737133, EPI_ISL_737136, EPI_ISL_737137, EPI_ISL_737138, EPI_ISL_737139, EPI_ISL_737158, EPI_ISL_737171, EPI_ISL_737172, EPI_ISL_737173, EPI_ISL_737184, EPI_ISL_737186, | | | |

| | | | |
|---|---|---|---|
| EPI_ISL_737187, EPI_ISL_737189, EPI_ISL_737190, EPI_ISL_737191, EPI_ISL_737192, EPI_ISL_737195, EPI_ISL_737271, EPI_ISL_737277, EPI_ISL_737278, EPI_ISL_737296, EPI_ISL_737298 | | | |
| see above | Michigan Department of Health and Human Services, Bureau of Laboratories | Michigan Department of Health and Human Services, Bureau of Laboratories | Blankenship HM, Riner D, Soehnlen MK |
| EPI_ISL_737839 | Viollier AG | Department of Biosystems Science and Engineering, ETH Zürich | Chaoran Chen, Sarah Nadeau, Ivan Topolsky, Emmanouil Dermitzakis, Keith Harshman, Ioannis Xenarios, Henri Pegeot, Lorenzo Cerutti, Deborah Penet, Philipp Jablonski, Lara Fuhrmann, David Dreifuss, Katharina Jahn, Christiane Beckmann, Maurice Redondo, Olivier Kobel, Christoph Noppen, Sophie Seidel, Noemie Santamaria de Souza, Niko Beerenwinkel, Tanja Stadler |
| EPI_ISL_738107 | Instituto Nacional de Saude (INSA) | Instituto Nacional de Saude (INSA) | Borges et al |
| EPI_ISL_738500, EPI_ISL_738517, EPI_ISL_738526, EPI_ISL_738536, EPI_ISL_738560 | Santa Clara County Public Health Laboratory | Chan-Zuckerberg Biohub | CZB Cliahub Consortium |
| EPI_ISL_738571 | Orange County Public Health Lab | Chan-Zuckerberg Biohub | CZB Cliahub Consortium |
| EPI_ISL_738600 | Humboldt County Public Health Laboratory | Chan-Zuckerberg Biohub | CZB Cliahub Consortium |
| EPI_ISL_738644, EPI_ISL_738672, EPI_ISL_738706 | Santa Clara County Public Health Laboratory | Chan-Zuckerberg Biohub | CZB Cliahub Consortium |
| EPI_ISL_738757 | Orange County Public Health Lab | Chan-Zuckerberg Biohub | CZB Cliahub Consortium |
| EPI_ISL_738759 | Humboldt County Public Health Laboratory | Chan-Zuckerberg Biohub | CZB Cliahub Consortium |
| EPI_ISL_738765 | Santa Clara County Public Health Laboratory | Chan-Zuckerberg Biohub | CZB Cliahub Consortium |
| EPI_ISL_738779, EPI_ISL_738795, EPI_ISL_738810, EPI_ISL_738827, EPI_ISL_738852 | Humboldt County Public Health Laboratory | Chan-Zuckerberg Biohub | CZB Cliahub Consortium |
| EPI_ISL_738882, EPI_ISL_738888 | Orange County Public Health Lab | Chan-Zuckerberg Biohub | CZB Cliahub Consortium |
| EPI_ISL_738912 | Humboldt County Public Health Laboratory | Chan-Zuckerberg Biohub | CZB Cliahub Consortium |
| EPI_ISL_738930, EPI_ISL_738946 | Orange County Public Health Lab | Chan-Zuckerberg Biohub | CZB Cliahub Consortium |
| EPI_ISL_738972 | Santa Clara County Public Health Laboratory | Chan-Zuckerberg Biohub | CZB Cliahub Consortium |
| EPI_ISL_738985 | Humboldt County Public Health Laboratory | Chan-Zuckerberg Biohub | CZB Cliahub Consortium |
| EPI_ISL_739010 | Santa Clara County Public Health Laboratory | Chan-Zuckerberg Biohub | CZB Cliahub Consortium |
| EPI_ISL_739042 | Orange County Public Health Lab | Chan-Zuckerberg Biohub | CZB Cliahub Consortium |
| EPI_ISL_739080 | Santa Clara County Public Health Laboratory | Chan-Zuckerberg Biohub | CZB Cliahub Consortium |
| EPI_ISL_739108 | Orange County Public Health Lab | Chan-Zuckerberg Biohub | CZB Cliahub Consortium |
| EPI_ISL_739122, EPI_ISL_739130 | Santa Clara County Public Health Laboratory | Chan-Zuckerberg Biohub | CZB Cliahub Consortium |
| EPI_ISL_739143, EPI_ISL_739151, EPI_ISL_739156 | Orange County Public Health Lab | Chan-Zuckerberg Biohub | CZB Cliahub Consortium |
| EPI_ISL_739167, EPI_ISL_739168, EPI_ISL_739184 | Santa Clara County Public Health Laboratory | Chan-Zuckerberg Biohub | CZB Cliahub Consortium |
| EPI_ISL_739185, EPI_ISL_739239, EPI_ISL_739245, EPI_ISL_739291 | Orange County Public Health Lab | Chan-Zuckerberg Biohub | CZB Cliahub Consortium |
| EPI_ISL_739330, EPI_ISL_739367 | Humboldt County Public Health Laboratory | Chan-Zuckerberg Biohub | CZB Cliahub Consortium |
| EPI_ISL_739384 | Orange County Public Health Lab | Chan-Zuckerberg Biohub | CZB Cliahub Consortium |
| EPI_ISL_739411 | Humboldt County Public Health Laboratory | Chan-Zuckerberg Biohub | CZB Cliahub Consortium |
| EPI_ISL_739420, EPI_ISL_739423 | Orange County Public Health Lab | Chan-Zuckerberg Biohub | CZB Cliahub Consortium |
| EPI_ISL_739451 | Santa Clara County Public Health Laboratory | Chan-Zuckerberg Biohub | CZB Cliahub Consortium |
| EPI_ISL_739477, EPI_ISL_739494 | Orange County Public Health Lab | Chan-Zuckerberg Biohub | CZB Cliahub Consortium |
| EPI_ISL_739527 | Humboldt County Public Health Laboratory | Chan-Zuckerberg Biohub | CZB Cliahub Consortium |
| EPI_ISL_739536 | Santa Clara County Public Health Laboratory | Chan-Zuckerberg Biohub | CZB Cliahub Consortium |
| EPI_ISL_739553 | Humboldt County Public Health Laboratory | Chan-Zuckerberg Biohub | CZB Cliahub Consortium |
| EPI_ISL_739561 | Santa Clara County Public Health Laboratory | Chan-Zuckerberg Biohub | CZB Cliahub Consortium |
| EPI_ISL_739603 | San Francisco Public Health Laboratory | Chan-Zuckerberg Biohub | CZB Cliahub Consortium |
| EPI_ISL_739620, EPI_ISL_739637, EPI_ISL_739658 | Orange County Public Health Lab | Chan-Zuckerberg Biohub | CZB Cliahub Consortium |
| EPI_ISL_739673 | Instituto Nacional de Salud, Bogotá, Colombia | Instituto Nacional de Salud, Bogotá, Colombia | Katherine Laiton-Donato, Diego A. Álvarez-Díaz, Carlos Franco-Muñoz, Mauricio Pacheco-Montealegre, Jonathan Reales, Diego Andrés Prada, Sheryl Corchuelo, Magdalena Weisner, Martha Lucia Ospina Martinez, Marcela Mercado-Reyes |
| EPI_ISL_739693, EPI_ISL_739709, EPI_ISL_739737, EPI_ISL_739746, EPI_ISL_739758, EPI_ISL_739796, EPI_ISL_739799, EPI_ISL_739812, EPI_ISL_739813, EPI_ISL_739839, EPI_ISL_739869, EPI_ISL_739878, EPI_ISL_739903, EPI_ISL_739931, EPI_ISL_739940, EPI_ISL_739973, EPI_ISL_739975, EPI_ISL_739988, EPI_ISL_740016, EPI_ISL_740024, EPI_ISL_740073, EPI_ISL_740087, EPI_ISL_740104, EPI_ISL_740121, EPI_ISL_740133, EPI_ISL_740178, EPI_ISL_740184, EPI_ISL_740185, EPI_ISL_740234, EPI_ISL_740277, EPI_ISL_740321, EPI_ISL_740338, EPI_ISL_740353, EPI_ISL_740394, EPI_ISL_740425, EPI_ISL_740431, EPI_ISL_740481, EPI_ISL_740495, EPI_ISL_740515 | | | |
| see above | Laboratoire national de santé, Microbiology, Virology | Laboratoire national de santé, Microbiology, Microbial Genomics Platform | Anke Wienecke-Baldacchino, Catherine Ragimbeau,Jessica Tapp, Fatu Djabi, Lise Pignon, Raoul Salmon, Tamir Abdelrahman |
| EPI_ISL_741032, EPI_ISL_741033, EPI_ISL_741054, EPI_ISL_741055, EPI_ISL_741085, EPI_ISL_741105, EPI_ISL_741107, EPI_ISL_741108, EPI_ISL_741109, EPI_ISL_741110, EPI_ISL_741150 | | | |
| see above | Oxford Viromics, NDM, University of Oxford; Oxford University Hospitals; Basingstoke and North Hampshire Hospital | COVID-19 Genomics UK (COG-UK) Consortium | Tanya Golubchik, David Bonsall, George Macintyre, Amy Trebes, Mariateresa de Cesare, Catrin Moore, Alex Mobbs, Anita Justice, Robert Shaw, Monique Andersson, Timothy Peto, Emma Wise, Nathan Moore, Jessica Lynch, Nick Cortes, Matilde Mori, Stephen Kidd, David Buck, John Todd, Christophe Fraser |
| EPI_ISL_741552 | Quadram Institute Bioscience | COVID-19 Genomics UK (COG-UK) Consortium | Dave J. Baker, Gemma L. Kay, Alp Aydin, Thanh Le-Viet, Steven Rudder, Ana P. Tedim, Anastasia Kolyva, Maria Diaz, Leonardo de Oliveira Martins, Nabil-Fareed Alikhan, Lizzie Meadows, Rachael Stanley, Ngozi Elumogo, Muhammed Yasir, Nicholas M. Thomson, Alexander J Trotter, Rachel Gilroy, Samuel Bloomfield, Claire Stuart, Andrew Bell, Reenesh Prakash, Samir Dervisevic, Alison E. Mather, John Wain, Mark Webber, Andrew J. Page, Justin O'Grady |
| EPI_ISL_741724, EPI_ISL_741725, EPI_ISL_741726, EPI_ISL_741727, | Centre for Enzyme Innovation, University of Portsmouth / Translational Research Laboratory, Portsmouth Hospitals | COVID-19 Genomics UK (COG-UK) Consortium | Angela Beckett,Yann Bourgeois,Garry Scarlett,Sharon Glaysher,Scott Elliott,Kelly Bicknell,Robert Impey,Allyson Lloyd,Sarah Wyllie,Ethan Butcher,Anoop Chauhan,Samuel Robson |

| | | | |
|---|---|---|---|
| EPI_ISL_741745, EPI_ISL_741753, EPI_ISL_741755 | NHS Trust | | |
| EPI_ISL_742118, EPI_ISL_742119, EPI_ISL_742120, EPI_ISL_742121, EPI_ISL_742122, EPI_ISL_742126, EPI_ISL_742127, EPI_ISL_742128, EPI_ISL_742160, EPI_ISL_742161, EPI_ISL_742162, EPI_ISL_742163, EPI_ISL_742164, EPI_ISL_742165, EPI_ISL_742166, EPI_ISL_742167, EPI_ISL_742168, EPI_ISL_742169, EPI_ISL_742170, EPI_ISL_742171, EPI_ISL_742172, EPI_ISL_742173 | | | |
| see above | Regional Virus Laboratory, Belfast Health and Social Care Trust | COVID-19 Genomics UK (COG-UK) Consortium | Conall McCaughey, James McKenna, Tanya Curran, Susan Feeney, Alison Watt, Ciara Cox, Mairead Connor, Zoltan Molnar, David Simpson, Derek Fairley |
| EPI_ISL_742202, EPI_ISL_742245, EPI_ISL_742246 | Virology Department, Royal Infirmary of Edinburgh, NHS Lothian / School of Biological Sciences, University of Edinburgh / Institute of Genetics and Molecular Medicine, University of Edinburgh | COVID-19 Genomics UK (COG-UK) Consortium | McHugh M, Dewar R, Rooke S, Gallagher M, Balcaza C, O'Toole Á, Scher E, Hill V, McCrone JT, Colquhoun R, Yu X, Jackson B, Rambaut A, Williams TC, Templeton K |
| EPI_ISL_742281, EPI_ISL_742282, EPI_ISL_742318, EPI_ISL_742319, EPI_ISL_742320, EPI_ISL_742321, EPI_ISL_742323, EPI_ISL_742324 | Wales Specialist Virology Centre Sequencing lab: Pathogen Genomics Unit | COVID-19 Genomics UK (COG-UK) Consortium | Catherine Moore, Johnathan Evans, Laura Gifford, Malorie Perry, Simon Cottrell, Angela Marchbank, Alec Birchley, Alexander Adams, Amy Gaskin, Bree Gatica-Wilcox, Jason Coombes, Joel Southgate, Lauren Gilbert, Lee Graham, Nicole Pacchiarini, Sara Kumziene-Summerhayes, Sarah Taylor, Sophie Jones, Sara Rey, Matthew Bull, Joanne Watkins, Sally Corden, Tom Connor |
| EPI_ISL_744144, EPI_ISL_744160, EPI_ISL_744184, EPI_ISL_744222, EPI_ISL_744259, EPI_ISL_744265, EPI_ISL_744266, EPI_ISL_744274, EPI_ISL_744302, EPI_ISL_744327, EPI_ISL_744350, EPI_ISL_744365, EPI_ISL_744370, EPI_ISL_744387, EPI_ISL_744425, EPI_ISL_744446, EPI_ISL_744448, EPI_ISL_744470, EPI_ISL_744488, EPI_ISL_744517, EPI_ISL_744520, EPI_ISL_744541, EPI_ISL_744549, EPI_ISL_744571, EPI_ISL_744618, EPI_ISL_744622, EPI_ISL_744636, EPI_ISL_744645, EPI_ISL_744654, EPI_ISL_744678, EPI_ISL_744767, EPI_ISL_744772, EPI_ISL_744821, EPI_ISL_744823, EPI_ISL_744831, EPI_ISL_744841, EPI_ISL_744883, EPI_ISL_744885, EPI_ISL_744887, EPI_ISL_744891, EPI_ISL_744904, EPI_ISL_744909, EPI_ISL_744914, EPI_ISL_744918, EPI_ISL_744920, EPI_ISL_744943, EPI_ISL_744981, EPI_ISL_744985, EPI_ISL_744987 | | | |
| see above | Laboratoire national de santé, Microbiology, Virology | Laboratoire national de santé, Microbiology, Microbial Genomics Platform | Anke Wienecke-Baldacchino, Catherine Ragimbeau,Jessica Tapp, Fatu Djabi, Lise Pignon, Raoul Salmon, Tamir Abdelrahman |
| EPI_ISL_745229, EPI_ISL_745230, EPI_ISL_745231, EPI_ISL_745232, EPI_ISL_745244, EPI_ISL_745245, EPI_ISL_745246, EPI_ISL_745247, EPI_ISL_745248, EPI_ISL_745249, EPI_ISL_745250, EPI_ISL_745251, EPI_ISL_745252, EPI_ISL_745253, EPI_ISL_745254, EPI_ISL_745255, EPI_ISL_745256, EPI_ISL_745257, EPI_ISL_745258 | | | |
| see above | Molecular diagnostic laboratory of Federal Budget Institution of Science "Central Research Institute of Epidemiology" of The Federal Service on Customers' Rights Protection and Human Well-being Surveillance | Group of Genomics and Postgenomic Technologies of Central Research Institute of Epidemilology | Samoilov AE, Kaptelova VV, Korneenko EV, Dudorova AV, Saenko SS, Speranskaya AS, Tivanova EV, Shipulina OY, Akimkin VG |
| EPI_ISL_746964 | Utah Public Health Laboratory | Utah Public Health Laboratory | Erin Young, Kelly Oakeson, Tara Gallagher |
| EPI_ISL_747051, EPI_ISL_747054, EPI_ISL_747056, EPI_ISL_747058, EPI_ISL_747103, EPI_ISL_747104, EPI_ISL_747105, EPI_ISL_747106, EPI_ISL_747110, EPI_ISL_747111, EPI_ISL_747153, EPI_ISL_747154, EPI_ISL_747173, EPI_ISL_747174, EPI_ISL_747175, EPI_ISL_747176, EPI_ISL_747177, EPI_ISL_747178 | | | |
| see above | Respiratory Viruses Branch, Centers for Disease Control and Prevention | Respiratory Viruses Branch, Centers for Disease Control and Prevention | Queen,K., Li,Y., Tao,Y., Uehara,A., Montmayeur,A., Paden,C.R., Cook,P.W., Marine,R., Sheth,M., Wang,H., Lee,J., Tong,S. |
| EPI_ISL_747385, EPI_ISL_747396, EPI_ISL_747407, EPI_ISL_747455 | Division of Emerging Infectious Diseases, Bureau of Infectious Diseases Diagnosis Control, Korea Disease Control and Prevention Agency | Division of Emerging Infectious Diseases, Bureau of Infectious Diseases Diagnosis Control, Korea Disease Control and Prevention Agency | Ae Kyung Park, Il-Hwan Kim, Heui Man Kim, Jeong-Min Kim, Namjoo Lee, Chaeyoung Lee, Sang Hee Woo, Eun-Jin Kim |
| EPI_ISL_747466, EPI_ISL_747467, EPI_ISL_747469, EPI_ISL_747484 | Ospedale Mater Salutis | Istituto Zooprofilattico Sperimentale delle Venezie | Adelaide Milani, Alessia Schivo, Annalisa Salviato, Erika Giorgia Quaranta, Ambra Pastori, Bianca Zecchin, Alice Fusaro, Isabella Monne, Calogero Terregino, Antonia Ricci |
| EPI_ISL_747600, EPI_ISL_747647, EPI_ISL_748119, EPI_ISL_748120, EPI_ISL_748121 | Department of Virus and Microbiological Special Diagnostics, Statens Serum Institut, Copenhagen, Denmark | Albertsen Lab, Department of Chemistry and Bioscience, Aalborg University, Denmark | Danish Covid-19 Genome Consortium |
| EPI_ISL_751449, EPI_ISL_751491, EPI_ISL_751492, EPI_ISL_751493, EPI_ISL_751494 | CHU Purpan - Laboratoire de Virologie - Institut Fédératif de Biologie | CHU Purpan - Laboratoire de Virologie - Institut Fédératif de Biologie | Latour J., Ranger N., Dubois M., Carcenac R., Harter A., Boyer P., Tremeaux P., Izopet J. |
| EPI_ISL_751554 | CT-Dr. Katherine A. Kelley State Public Health Lab | Genomics and Discovery, Respiratory Viruses Branch, Division of Viral Diseases, Centers for Disease Control and Prevention | Krista Queen, Yan Li, Ying Tao, Jing Zhang, Anna Uehara, Anna Montmayeur, Clinton R. Paden, Peter W. Cook,Rachel Marine, Mili Sheth, Haibin Wang, Justin Lee, Suxiang Tong |
| EPI_ISL_751573 | MN PHL Division, Minnesota Department of Health | Genomics and Discovery, Respiratory Viruses Branch, Division of Viral Diseases, Centers for Disease Control and Prevention | Krista Queen, Yan Li, Ying Tao, Jing Zhang, Anna Uehara, Anna Montmayeur, Clinton R. Paden, Peter W. Cook,Rachel Marine, Mili Sheth, Haibin Wang, Justin Lee, Suxiang Tong |
| EPI_ISL_751586, EPI_ISL_751609 | OH Department of Health Laboratory | Genomics and Discovery, Respiratory Viruses Branch, Division of Viral Diseases, Centers for Disease Control and Prevention | Krista Queen, Yan Li, Ying Tao, Jing Zhang, Anna Uehara, Anna Montmayeur, Clinton R. Paden, Peter W. Cook,Rachel Marine, Mili Sheth, Haibin Wang, Justin Lee, Suxiang Tong |
| EPI_ISL_751635 | MO State Public Health Laboratory | Genomics and Discovery, Respiratory Viruses Branch, Division of Viral Diseases, Centers for Disease Control and Prevention | Krista Queen, Yan Li, Ying Tao, Jing Zhang, Anna Uehara, Anna Montmayeur, Clinton R. Paden, Peter W. Cook,Rachel Marine, Mili Sheth, Haibin Wang, Justin Lee, Suxiang Tong |
| EPI_ISL_751653, EPI_ISL_751654 | MN PHL Division, Minnesota Department of Health | Genomics and Discovery, Respiratory Viruses Branch, Division of Viral Diseases, Centers for Disease Control and Prevention | Krista Queen, Yan Li, Ying Tao, Jing Zhang, Anna Uehara, Anna Montmayeur, Clinton R. Paden, Peter W. Cook,Rachel Marine, Mili Sheth, Haibin Wang, Justin Lee, Suxiang Tong |
| EPI_ISL_751657 | MO State Public Health Laboratory | Genomics and Discovery, Respiratory Viruses Branch, Division of Viral Diseases, Centers for Disease Control and Prevention | Krista Queen, Yan Li, Ying Tao, Jing Zhang, Anna Uehara, Anna Montmayeur, Clinton R. Paden, Peter W. Cook,Rachel Marine, Mili Sheth, Haibin Wang, Justin Lee, Suxiang Tong |
| EPI_ISL_751658 | NYSDOH Wadsworth Center, Virology Lab | Genomics and Discovery, Respiratory Viruses Branch, Division of Viral Diseases, Centers for Disease Control and Prevention | Krista Queen, Yan Li, Ying Tao, Jing Zhang, Anna Uehara, Anna Montmayeur, Clinton R. Paden, Peter W. Cook,Rachel Marine, Mili Sheth, Haibin Wang, Justin Lee, Suxiang Tong |
| EPI_ISL_751685, EPI_ISL_751686, EPI_ISL_751690 | VA-Division of Consolidated Laboratory Services | Genomics and Discovery, Respiratory Viruses Branch, Division of Viral Diseases, Centers for Disease Control and Prevention | Krista Queen, Yan Li, Ying Tao, Jing Zhang, Anna Uehara, Anna Montmayeur, Clinton R. Paden, Peter W. Cook,Rachel Marine, Mili Sheth, Haibin Wang, Justin Lee, Suxiang Tong |
| EPI_ISL_751719 | ID Bureau of Laboratories | Genomics and Discovery, Respiratory Viruses Branch, Division of Viral Diseases, Centers for Disease Control and Prevention | Krista Queen, Yan Li, Ying Tao, Jing Zhang, Anna Uehara, Anna Montmayeur, Clinton R. Paden, Peter W. Cook,Rachel Marine, Mili Sheth, Haibin Wang, Justin Lee, Suxiang Tong |
| EPI_ISL_751745, EPI_ISL_751747 | VA-Division of Consolidated Laboratory Services | Genomics and Discovery, Respiratory Viruses Branch, Division of Viral Diseases, Centers for Disease Control and Prevention | Krista Queen, Yan Li, Ying Tao, Jing Zhang, Anna Uehara, Anna Montmayeur, Clinton R. Paden, Peter W. Cook,Rachel Marine, Mili Sheth, Haibin Wang, Justin Lee, Suxiang Tong |
| EPI_ISL_751774 | MO State Public Health Laboratory | Genomics and Discovery, Respiratory Viruses Branch, Division of Viral Diseases, Centers for Disease Control and Prevention | Krista Queen, Yan Li, Ying Tao, Jing Zhang, Anna Uehara, Anna Montmayeur, Clinton R. Paden, Peter W. Cook,Rachel Marine, Mili Sheth, Haibin Wang, Justin Lee, Suxiang Tong |
| EPI_ISL_751777 | OH Department of Health Laboratory | Genomics and Discovery, Respiratory Viruses Branch, | Krista Queen, Yan Li, Ying Tao, Jing Zhang, Anna Uehara, Anna Montmayeur, Clinton R. Paden, Peter W. Cook,Rachel Marine, Mili Sheth, Haibin Wang, |

| | | | |
|---|---|---|---|
| | | Division of Viral Diseases, Centers for Disease Control and Prevention | Justin Lee, Suxiang Tong |
| EPI_ISL_751791 | AL Dept. of Public Health Bureau of Clinical Laboratories | Genomics and Discovery, Respiratory Viruses Branch, Division of Viral Diseases, Centers for Disease Control and Prevention | Krista Queen, Yan Li, Ying Tao, Jing Zhang, Anna Uehara, Anna Montmayeur, Clinton R. Paden, Peter W. Cook,Rachel Marine, Mili Sheth, Haibin Wang, Justin Lee, Suxiang Tong |
| EPI_ISL_753150 | State Laboratories Division, Hawaii State Department of Health | State Laboratories Division, Hawaii State Department of Health | Pamela O'Brien, Sabrina Diemert, Drew Kuwazaki, Razvan Sultana, Edward Desmond |
| EPI_ISL_753291, EPI_ISL_753292, EPI_ISL_753293, EPI_ISL_753294, EPI_ISL_753295, EPI_ISL_753296, EPI_ISL_753297, EPI_ISL_753298, EPI_ISL_753299, EPI_ISL_753300, EPI_ISL_753301, EPI_ISL_753302, EPI_ISL_753303, EPI_ISL_753304, EPI_ISL_753327, EPI_ISL_753328, EPI_ISL_753329 | | | |
| see above | Clinical virology Laboratory, Children's Hospital Los Angeles | Center for Personalized Medicine, Children's Hospital Los Angeles | Gai et al |
| EPI_ISL_753735, EPI_ISL_753832, EPI_ISL_753916 | Charité Universitätsmedizin Berlin, Institut für Virologie/Labor Berlin | Charité Universitätsmedizin Berlin, Institut für Virologie | Victor M Corman, Jörn Beheim-Schwarzbach, Barbara Mühlemann, Julia Schneider, Talitha Veith, Terry Jones, Christian Drosten |
| EPI_ISL_754078 | National Public Health Laboratory, National Centre for Infectious Diseases | National Public Health Laboratory, National Centre for Infectious Diseases | Tze Minn Mak, Sophie Octavia, Zhenyang Zhou, Lin Cui, Raymond Tzer Pin Lin |
| EPI_ISL_754139 | CHU Purpan - Laboratoire de Virologie - Institut Fédératif de Biologie | CHU Purpan - Laboratoire de Virologie - Institut Fédératif de Biologie | Latour J., Ranger N., Dubois M., Carcenac R., Harter A., Boyer P., Tremeaux P., Izopet J. |
| EPI_ISL_754196, EPI_ISL_754197, EPI_ISL_754198, EPI_ISL_754199, EPI_ISL_754200, EPI_ISL_754201, EPI_ISL_754202, EPI_ISL_754203, EPI_ISL_754204, EPI_ISL_754205, EPI_ISL_754206, EPI_ISL_754207, EPI_ISL_754208, EPI_ISL_754209, EPI_ISL_754210, EPI_ISL_754211, EPI_ISL_754212, EPI_ISL_754213, EPI_ISL_754214 | | | |
| see above | Molecular diagnostic laboratory of Federal Budget Institution of Science "Central Research Institute of Epidemiology" of The Federal Service on Customers' Rights Protection and Human Well-being Surveillance | Group of Genomics and Postgenomic Technologies of Central Research Institute of Epidemilology | Samoilov AE, Kaptelova VV, Korneenko EV, Saenko SS, Dudorova AV, Speranskaya AS, Tivanova EV, Shipulina OY, Akimkin VG |
| EPI_ISL_754401, EPI_ISL_754525, EPI_ISL_754541, EPI_ISL_754545, EPI_ISL_754575, EPI_ISL_754576 | Wadsworth Center, New York State Department.of Health | Wadsworth Center, New York State Department.of Health | Kirsten St. George, Daryl M. Lamson, Alexis Russel, Matthew Shudt, Melissa A Leisner, Jonathan Plitnick, Navjot Singh, John Kelly, Sara Griesemer, Erasmus Schneider, Erica Lasek-Nesselquist |
| EPI_ISL_755761, EPI_ISL_755762, EPI_ISL_755763 | Toronto Invasive Bacterial Diseases Network | McMaster University | Allison McGeer, Patryk Aftanas, Hooman Derakhshani, Angel Li, Kuganya Nirmalarajah, Emily Panousis, Ahmed Draia, Jalees Nasir, Michael Surette, Samira Mubareka, Andrew G. McArthur |
| EPI_ISL_760163, EPI_ISL_760164, EPI_ISL_760165, EPI_ISL_760191, EPI_ISL_760192, EPI_ISL_760195, EPI_ISL_760198, EPI_ISL_760206, EPI_ISL_760210, EPI_ISL_760211, EPI_ISL_760212, EPI_ISL_760213, EPI_ISL_760224 | | | |
| see above | Division of Emerging Infectious Diseases, Bureau of Infectious Diseases Diagnosis Control, Korea Disease Control and Prevention Agency | Division of Emerging Infectious Diseases, Bureau of Infectious Diseases Diagnosis Control, Korea Disease Control and Prevention Agency | Ae Kyung Park, Il-Hwan Kim, Heui Man Kim, Jeong-Min Kim, Namjoo Lee, Chaeyoung Lee, Sang Hee Woo, Eun-Jin Kim |
| EPI_ISL_762997 | Respiratory Virus Unit, National Infection Service, Public Health England | COVID-19 Genomics UK (COG-UK) Consortium | PHE Covid Sequencing Team |
| EPI_ISL_763110, EPI_ISL_763135, EPI_ISL_763136, EPI_ISL_763188, EPI_ISL_763233, EPI_ISL_763307 | Dutch COVID-19 response team | Erasmus Medical Center | Bas Oude Munnink, Reina Sikkema, David Nieuwenhuijse, Irina Chestakova, Anne van der Linden, Marjan Boter, Emmanuelle Munger, Corine GeurtsvanKessel, Annemiek van der Eijk, Richard Molenkamp, Marion Koopmans, on behalf of the Dutch national COVID-19 response team. |
| EPI_ISL_763373, EPI_ISL_763642, EPI_ISL_764431, EPI_ISL_764433, EPI_ISL_764440 | Regional Virus Laboratory, Belfast Health and Social Care Trust | COVID-19 Genomics UK (COG-UK) Consortium | Conall McCaughey, James McKenna, Tanya Curran, Susan Feeney, Alison Watt, Ciara Cox, Mairead Connor, Zoltan Molnar, David Simpson, Derek Fairley |
| EPI_ISL_765212 | Instituto Nacional de Saude (INSA) | Instituto Nacional de Saude (INSA) | Borges et al |
| EPI_ISL_765216, EPI_ISL_765218 | Instituto Nacional de Saude (INSA) and Instituto Gulbenkian de Ciencia (IGC) | Instituto Nacional de Saude (INSA) and Instituto Gulbenkian de Ciencia (IGC) | Borges et al |
| EPI_ISL_765574, EPI_ISL_765575, EPI_ISL_765576, EPI_ISL_765577, EPI_ISL_765578, EPI_ISL_765579, EPI_ISL_765580, EPI_ISL_765581, EPI_ISL_765582 | Colorado Mesa University | Infectious Disease Program, Broad Institute of Harvard and MIT | Lemieux,J.E., Siddle,K.J., Shaw,B., Adams,G., Pierce,V., Turbett,S., Anahtar,M., Branda,J., Slater,D., Harris,J., Lin,A.E., Gladden-Young,A., Lagerborg,K., Rudy,M., DeRuff,K., Carter,A., Normandin,E., Bauer,M., Reilly,S., Tomkins-Tinch,C., Loreth,C., Chaluvadi,S., Neumann,A., Cusick,C., Chapman,S.B., Gnirke,A., Flowers,K., Cerrato,F., Birren,B.W., Gallagher,G., Smole,S., Park,D.J., MacInnis,B.L., Ryan,E., LaRocque,R., Rosenberg,E. and Sabeti,P.C. |
| EPI_ISL_765687, EPI_ISL_765806, EPI_ISL_765807, EPI_ISL_765808, EPI_ISL_765809, EPI_ISL_765810, EPI_ISL_765811, EPI_ISL_765812, EPI_ISL_765813, EPI_ISL_765814, EPI_ISL_765815, EPI_ISL_765816, EPI_ISL_765817, EPI_ISL_765825, EPI_ISL_765826, EPI_ISL_765827, EPI_ISL_765828, EPI_ISL_765829, EPI_ISL_765830 | | | |
| see above | Massachusetts General Hospital | Infectious Disease Program, Broad Institute of Harvard and MIT | Lemieux,J.E., Siddle,K.J., Shaw,B., Adams,G., Pierce,V., Turbett,S., Anahtar,M., Branda,J., Slater,D., Harris,J., Lin,A.E., Gladden-Young,A., Lagerborg,K., Rudy,M., DeRuff,K., Carter,A., Normandin,E., Bauer,M., Reilly,S., Tomkins-Tinch,C., Loreth,C., Chaluvadi,S., Neumann,A., Cusick,C., Chapman,S.B., Gnirke,A., Flowers,K., Cerrato,F., Birren,B.W., Gallagher,G., Smole,S., Park,D.J., MacInnis,B.L., Ryan,E., LaRocque,R., Rosenberg,E. and Sabeti,P.C. |
| EPI_ISL_766101, EPI_ISL_766263, EPI_ISL_766264, EPI_ISL_766265, EPI_ISL_766266 | Respiratory Virus Unit, National Infection Service, Public Health England | COVID-19 Genomics UK (COG-UK) Consortium | PHE Covid Sequencing Team |
| EPI_ISL_766977, EPI_ISL_766998 | Delaware Public Health Laboratory | Delaware Public Health Laboratory | Gregory Hovan |
| EPI_ISL_767357 | Michigan Department of Health and Human Services, Bureau of Laboratories | Michigan Department of Health and Human Services, Bureau of Laboratories | Blankenship HM, Riner D, Soehnlen MK |
| EPI_ISL_768795, EPI_ISL_768796, EPI_ISL_768797 | AIID | Irish Coronavirus Sequencing Consortium - National Virus Reference Laboratory | Michael Carr, Gabriel Gonzalez, Alejandro Abner Garcia Leon, Patrick Mallon |
| EPI_ISL_770811, EPI_ISL_770813 | Essentia Health-St. Mary's Medical Center | Minnesota Department of Health, Public Health Laboratory | Alexandra Lorentz, Jacob Garfin, Matt Plumb, and Xiong Wang |
| EPI_ISL_771185, EPI_ISL_771246 | Colorado Department of Public Health and Environment | Colorado Department of Puplic Health and Environment | Laura Bankers, Molly C. Hetherington-Rauth, Diana Ir, Shannon Ely, Shannon R. Matzinger, Sarah Elizabeth Totten, Emily A. Travanty |
| EPI_ISL_775302 | Nordland Hospital - Bodo, Laboratory Department, Molecular Biology Unit | Norwegian Institute of Public Health, Department of Virology | Kathrine Stene-Johansen, Kamilla Heddeland Instefjord, Hilde Elshaug, Atiya R Ali,Marie Paulsen Madsen, Rasmus Riis Kopperud, Hilde Vollan, Karoline Bragstad, Olav Hungnes |
| EPI_ISL_775304 | Furst Medical Laboratory | Norwegian Institute of Public Health, Department of Virology | Kathrine Stene-Johansen, Kamilla Heddeland Instefjord, Hilde Elshaug, Atiya R Ali,Marie Paulsen Madsen, Rasmus Riis Kopperud, Hilde Vollan, Karoline Bragstad, Olav Hungnes |
| EPI_ISL_775333 | Foerde Hospital, Department of Microbiology | Norwegian Institute of Public Health, Department of Virology | Kathrine Stene-Johansen, Kamilla Heddeland Instefjord, Hilde Elshaug, Atiya R Ali,Marie Paulsen Madsen, Rasmus Riis Kopperud, Hilde Vollan, Karoline Bragstad, Olav Hungnes |
| EPI_ISL_775341, EPI_ISL_775342, EPI_ISL_775345 | Ostfold Hospital Trust - Kalnes, Centre for Laboratory Medicine, Section for gene technology and infection serology | Norwegian Institute of Public Health, Department of Virology | Kathrine Stene-Johansen, Kamilla Heddeland Instefjord, Hilde Elshaug, Atiya R Ali,Marie Paulsen Madsen, Rasmus Riis Kopperud, Hilde Vollan, Karoline Bragstad, Olav Hungnes |
| EPI_ISL_775500 | Vestfold Hospital, Toensberg Department of Microbiology | Norwegian Institute of Public Health, Department of Virology | Kathrine Stene-Johansen, Kamilla Heddeland Instefjord, Hilde Elshaug, Atiya R Ali,Marie Paulsen Madsen, Rasmus Riis Kopperud, Hilde Vollan, Karoline Bragstad, Olav Hungnes |

| | | | |
|---|---|---|---|
| EPI_ISL_775509, EPI_ISL_775512 | Norwegian Institute of Public Health, Department of Virology | Norwegian Institute of Public Health, Department of Virology | Kathrine Stene-Johansen, Kamilla Heddeland Instefjord, Hilde Elshaug, Atiya R Ali,Marie Paulsen Madsen, Rasmus Riis Kopperud, Hilde Vollan, Karoline Bragstad, Olav Hungnes |
| EPI_ISL_779405, EPI_ISL_779406 | Royal Darwin Hospital Pathology | MDU-PHL | Meumann, E., Caly L., Seemann T., Sait, M.L., Druce J., Sherry, N.L. |
| EPI_ISL_780064 | Hospital General Universitario Gregorio Marañón | SeqCOVID-SPAIN consortium/IBV(CSIC) | Darío García de Viedma, Laura Pérez-Lago, Marta Herranz, Jon Sicilia, Julia Suárez, Pilar Catalán, Patricia Muñoz and SeqCOVID-SPAIN consortium |
| EPI_ISL_780382, EPI_ISL_780384, EPI_ISL_780392, EPI_ISL_780393, EPI_ISL_780394 | Bermuda Government Molecular Diagnostics Laboratory (MDL) | Respiratory Virus Unit, National Infection Service, Public Health England | PHE Covid Sequencing Team, Dr Carika Weldon (Bermuda), Dr Ayoola Oyinloye (Bermuda) |
| EPI_ISL_783367, EPI_ISL_783368, EPI_ISL_783370, EPI_ISL_783374, EPI_ISL_783376, EPI_ISL_783377, EPI_ISL_783379, EPI_ISL_783383, EPI_ISL_783394, EPI_ISL_783398, EPI_ISL_783399, EPI_ISL_783400, EPI_ISL_783408, EPI_ISL_783410, EPI_ISL_783412, EPI_ISL_783413, EPI_ISL_783423, EPI_ISL_783425, EPI_ISL_783426, EPI_ISL_783427, EPI_ISL_783428, EPI_ISL_783429, EPI_ISL_783433, EPI_ISL_783445, EPI_ISL_783449, EPI_ISL_783454, EPI_ISL_783457, EPI_ISL_783462, EPI_ISL_783464, EPI_ISL_783465, EPI_ISL_783475, EPI_ISL_783478, EPI_ISL_783490, EPI_ISL_783501, EPI_ISL_783515, EPI_ISL_783516, EPI_ISL_783517, EPI_ISL_783518, EPI_ISL_783522, EPI_ISL_783525, EPI_ISL_783526, EPI_ISL_783529, EPI_ISL_783534 | | | |
| see above | Lighthouse Lab in Cambridge | Wellcome Sanger Institute for the COVID-19 Genomics UK (COG-UK) Consortium | Rob Howes, The Lighthouse Lab in Cambridge and Alex Alderton, Roberto Amato, Sonia Goncalves, Ewan Harrison, David K. Jackson, Ian Johnston, Dominic Kwiatkowski, Cordelia Langford, John Sillitoe on behalf of the Wellcome Sanger Institute COVID-19 Surveillance Team |
| EPI_ISL_784421, EPI_ISL_784467, EPI_ISL_784485, EPI_ISL_784493, EPI_ISL_784494, EPI_ISL_784512, EPI_ISL_784517, EPI_ISL_784518, EPI_ISL_784529, EPI_ISL_784533, EPI_ISL_784538, EPI_ISL_784539, EPI_ISL_784541, EPI_ISL_784544, EPI_ISL_784555, EPI_ISL_784561, EPI_ISL_784565, EPI_ISL_784567, EPI_ISL_784569, EPI_ISL_784574, EPI_ISL_784640, EPI_ISL_784651, EPI_ISL_784663, EPI_ISL_784664 | | | |
| see above | Houston Methodist Hospital | Houston Methodist Hospital | S. Wesley Long, Randall J. Olsen, Paul A. Christensen, David W. Bernard, James J. Davis, Maulik Shukla, Marcus Nguyen, Matthew Ojeda Saavedra, Prasanti Yerramilli, Layne Pruitt, Sishir Subedi, Heather Hendrickson, and James M. Musser |
| EPI_ISL_790569 | Florida Bureau of Public Health Laboratories | Florida Bureau of Public Health Laboratories | Sarah Schmedes, Jason Blanton |
| EPI_ISL_791348, EPI_ISL_791363, EPI_ISL_791365, EPI_ISL_791371, EPI_ISL_791372, EPI_ISL_791375, EPI_ISL_791376 | Johns Hopkins Hospital Department of Pathology | Johns Hopkins Hospital Department of Pathology | C. Paul Morris, Chun Huai Luo, Heba H. Mostafa |
| EPI_ISL_791980 | RSUD Blambangan Banyuwangi | National Institute of Health Research and Development | Ikawati,HD;Subangkit;Pawestri,HA;Nugraha,AA;Puspa,KD;Noor,RI;Pangesti,KNA;Soekarso,T;Puspandari,N;Setiawaty,V |
| EPI_ISL_791985 | RS Santa Maria Pekanbaru | National Institute of Health Research and Development | Puspa,KD;Subangkit;Pawestri,HA;Ikawati,HD;Nugraha,AA;Fridayenti;Pangesti,KNA;Soekarso,T;Puspandari,N;Setiawaty,V |
| EPI_ISL_792631, EPI_ISL_792632, EPI_ISL_792633, EPI_ISL_792634, EPI_ISL_792635, EPI_ISL_792636, EPI_ISL_792637, EPI_ISL_792638 | LACEN-PB | Laboratory of Respiratory Viruses and Measles, Oswaldo Cruz Institute, FIOCRUZ | Paola Resende, Luciana Appolinario, Fernando Motta, Anna Carolina Paixao, Ana Carolina Mendonca, João Felipe Bezerra, Romero Henrique Teixeira de Vasconcelos, Dalane Loudal Florentino Teixeira, Thiago Franco de Oliveira Carneiro, Marilda Siqueira |
| EPI_ISL_792647, EPI_ISL_792653 | LACEN-PR | Laboratory of Respiratory Viruses and Measles, Oswaldo Cruz Institute, FIOCRUZ | Paola Resende, Luciana Appolinario, Fernando Motta, Anna Carolina Paixao, Ana Carolina Mendonca, Maria do Carmo Debur, Irina Nastassja Riediger, Marilda Siqueira |
| EPI_ISL_792684, EPI_ISL_792685, EPI_ISL_792689 | The National Institute of Public Health | State Veterinary Institute Prague | Nagy,A;Jirincova,H;Trnka,D;Vecerova,J |
| EPI_ISL_792839 | Department of Virus and Microbiological Special Diagnostics, Statens Serum Institut, Copenhagen, Denmark | Albertsen Lab, Department of Chemistry and Bioscience, Aalborg University, Denmark | Danish Covid-19 Genome Consortium |
| EPI_ISL_794686, EPI_ISL_794711 | PathWest Laboratory Medicine WA | PathWest Laboratory Medicine WA Microbial Surveillance Unit | PathWest Laboratory Medicine WA Microbial Surveillance Unit |
| EPI_ISL_796763 | Instituto Nacional de Saude (INSA) | Instituto Nacional de Saude (INSA) | Borges et al |
| EPI_ISL_802726, EPI_ISL_802727, EPI_ISL_802728, EPI_ISL_802729, EPI_ISL_802730, EPI_ISL_802733, EPI_ISL_802734, EPI_ISL_802735, EPI_ISL_802736, EPI_ISL_802737, EPI_ISL_802749 | | | |
| see above | Hospital Clínic de Barcelona | Instituto de Salud Carlos III | Iglesias-Caballero, M. Molinero Calamita, M. González-Esguevillas, M. Camarero, S. Pozo, F. Casas, I. Jiménez, P. Jiménez, M. Zaballos, A. Monzón, S. Varona, S. Juliá, M. Cuesta, I, M.A Marcos. |
| EPI_ISL_802862, EPI_ISL_802863 | Vilnius University Hospital Santaros Klinikos, Vilnius University | Institute of Biotechnology, Life Sciences Center, Vilnius University | Emilija Vasiliunaite, Milda Norkiene, Albertas Timinskas, Alma Gedvilaite, Aurelija Zvirbliene, Daniel Naumovas, Laimonas Griskevicius |
| EPI_ISL_803099, EPI_ISL_803103 | Respiratory Viruses Branch, Centers for Disease Control and Prevention | Respiratory Viruses Branch, Centers for Disease Control and Prevention | Queen,K., Li,Y., Tao,Y., Uehara,A., Montmayeur,A., Paden,C.R., Cook,P.W., Marine,R., Sheth,M., Wang,H., Lee,J., Tong,S. |
| EPI_ISL_804258 | Respiratory Virus Unit, National Infection Service, Public Health England | COVID-19 Genomics UK (COG-UK) Consortium | PHE Covid Sequencing Team |
| EPI_ISL_804577, EPI_ISL_804578, EPI_ISL_804579 | MEPHI, Aix Marseille University | MEPHI, Aix Marseille University | Anthony LEVASSEUR |
| EPI_ISL_804620, EPI_ISL_804648, EPI_ISL_804714, EPI_ISL_804718, EPI_ISL_804722, EPI_ISL_804727, EPI_ISL_804729, EPI_ISL_804733, EPI_ISL_804734, EPI_ISL_804742, EPI_ISL_804753 | | | |
| see above | Michigan Department of Health and Human Services, Bureau of Laboratories | Michigan Department of Health and Human Services, Bureau of Laboratories | Blankenship HM, Riner D, Soehnlen MK |
| EPI_ISL_806990, EPI_ISL_806992, EPI_ISL_806993, EPI_ISL_806994, EPI_ISL_806995, EPI_ISL_806996, EPI_ISL_806997, EPI_ISL_806999, EPI_ISL_807000, EPI_ISL_807001, EPI_ISL_807002, EPI_ISL_807003, EPI_ISL_807005, EPI_ISL_807007, EPI_ISL_807016, EPI_ISL_807017, EPI_ISL_807019, EPI_ISL_807020 | | | |
| see above | Washington State Department of Health | Seattle Flu Study | Deborah A. Nickerson, Chris D. Frazar, Jover Lee, Benjamin Pelle, Matthew Richardson, Amanda Adler, Elisabeth Brandstetter, Peter D. Han, Kairsten Fay, Misja Ilcisin, Kirsten Lacombe, Thomas R. Sibley, Melissa Truong, Caitlin R. Wolf, Romesh Gautom, Geoff Melly, Brian Hiatt, Philip Dykema, Scott Lindquist, Michael Boeckh, Janet A. Englund, Michael Famulare, Barry R. Lutz, Mark J. Rieder, Lea M. Starita, Matthew Thompson, Helen Y. Chu, Jay Shendure, Trevor Bedford |
| EPI_ISL_813019, EPI_ISL_813020, EPI_ISL_813021, EPI_ISL_813022, EPI_ISL_813024, EPI_ISL_813029 | University of Birmingham | COVID-19 Genomics UK (COG-UK) Consortium | Institute of Microbiology, University of Birmingham: Claire McMurray, Joanne Stockton, Samuel Nicholls, Radoslaw Poplawski, Will Rowe, Josh Quick, Nicholas Loman. University of Birmingham Testing Laboratory: Celina M Whalley, Andrew Bosworth, Charlotte Poxon, Kasun Wanigasooriya, Oliver Pickles, Mike Kidd, Alex Richter, Andrew D Beggs PHE Heartlands Lab: Husam Osman, Andrew Bosworth. Queen Elizabeth Hospital: Anna Casey |
| EPI_ISL_814333, EPI_ISL_814569, EPI_ISL_814570 | Virology Department, Royal Infirmary of Edinburgh, NHS Lothian / School of Biological Sciences, University of Edinburgh / Institute of Genetics and Molecular Medicine, University of Edinburgh | COVID-19 Genomics UK (COG-UK) Consortium | McHugh M, Dewar R, Rooke S, Gallagher M, Balcaza C, O'Toole Á, Scher E, Hill V, McCrone JT, Colquhoun R, Yu X, Jackson B, Rambaut A, Williams TC, Templeton K |
| EPI_ISL_815473, EPI_ISL_815482, EPI_ISL_815486, EPI_ISL_815487, EPI_ISL_815601, EPI_ISL_815602, EPI_ISL_815603, EPI_ISL_815641, EPI_ISL_815642, EPI_ISL_815643, EPI_ISL_815644, EPI_ISL_815645, EPI_ISL_816111, EPI_ISL_816112, EPI_ISL_816113, EPI_ISL_816114, EPI_ISL_816115, EPI_ISL_816116, EPI_ISL_816117, EPI_ISL_816118, EPI_ISL_816119, EPI_ISL_816120, EPI_ISL_816121, EPI_ISL_816122, EPI_ISL_816123, EPI_ISL_816124, EPI_ISL_816125, EPI_ISL_816126, EPI_ISL_816127 | | | |
| see above | Department of Virus and Microbiological Special Diagnostics, Statens Serum Institut, Copenhagen, Denmark | Albertsen Lab, Department of Chemistry and Bioscience, Aalborg University, Denmark | Danish Covid-19 Genome Consortium |
| EPI_ISL_816645 | Virology Department, Sheffield Teaching Hospitals NHS Foundation Trust/Department of Infection, Immunity and Cardiovascular Disease, The Medical School, University of Sheffield | COVID-19 Genomics UK (COG-UK) Consortium | Thushan de Silva, Matthew Parker, Nikki Smith, Adri Angyal, Rebecca Brown, Luke Green, Rachel Tucker, Paul Parsons, Danielle Groves, Katie Johnson, Laura Carrilero, Alex Keeley, Dave Partridge, Matthew Wyles, Benjamin Lindsey, Mehmet Yavuz, Mohammad Raza, Cariad Evans |
| EPI_ISL_819461 | Northumbria University / South Tees Hospitals NHS | COVID-19 Genomics UK (COG-UK) Consortium | Darren L Smith,Andrew Nelson,Matthew Bashton,Greg R Young,Joshua Loh,John Allan,Mohammad A Tariq,Giles S Holt,Gary Black,Wen C Yew,Lynn |

| | | | |
|---|---|---|---|
| | Foundation Trust / North Cumbria Integrated Care NHS Foundation Trust / North Tees and Hartlepool NHS Foundation Trust / Newcastle Hospitals NHS Foundation Trust | | Dover,Paul Baker,Steve Liggett,Sarah Essex,Jane Greenaway,Debra Padgett,Clive Graham,Garren Scott,Edward Barton,Emma Swindells,Brendan Payne,Jennifer Collins,Yusri Taha,Gary Eltringham |
| EPI_ISL_824062 | Dutch COVID-19 response team | National Institute for Public Health and the Environment (RIVM) | Adam Meijer, Harry Vennema, Jeroen Cremer, Sharon van den Brink, Bas van der Veer, AnneMarie van den Brandt, Florian Zwagemaker, Dennis Schmitz, Chantal Reusken, on behalf of the national COVID-19 response team |
| EPI_ISL_824390, EPI_ISL_824391 | California Department of Public Health | California Department of Public Health | CDPH IDLB COVIDNet |
| EPI_ISL_824787, EPI_ISL_824788, EPI_ISL_824789 | Arizona State Public Health Laboratory | Arizona State Public Health Laboratory | Trung Huynh, Jessica Escobar, Katherine Fullerton, Nobuko Fukushima, Stacy White, Linda Getsinger, Victor Waddell |
| EPI_ISL_825037 | B.J. Medical College and Civil hospital, Ahmedabad | Gujarat Biotechnology Research Centre | Pranay Shah, Janvi Raval, Zarna Patel, Nitin Savaliya, Dinesh Kumar, Zuber Saiyed, Labdhi Pandya, Afzal Ansari, Nikha Trivedi, Apurvasinh Puvar, Ramesh Pandit, Kamlesh J Upadhyay, Nilima Shah, Sanjay Kapadia, Dipa Kinariwala, Chaitanya Joshi, Madhvi Joshi |
| EPI_ISL_825038 | B.J. Medical College and Civil hospital, Ahmedabad | Gujarat Biotechnology Research Centre | Zuber Saiyed, Labdhi Pandya, Afzal Ansari, Nikha Trivedi, Apurvasinh Puvar, Ramesh Pandit, Janvi Raval, Zarna Patel, Nitin Savaliya, Dinesh Kumar, Pranay Shah, Kamlesh J Upadhyay, Sanjay Kapadia, Dipa Kinariwala, Chaitanya Joshi, Madhvi Joshi |
| EPI_ISL_825039 | B.J. Medical College and Civil hospital, Ahmedabad | Gujarat Biotechnology Research Centre | Labdhi Pandya, Afzal Ansari, Nikha Trivedi, Apurvasinh Puvar, Ramesh Pandit, Janvi Raval, Zarna Patel, Nitin Savaliya, Dinesh Kumar, Zuber Saiyed, Pranay Shah, Kamlesh J Upadhyay, Sanjay Kapadia, Dipa Kinariwala, Chaitanya Joshi, Madhvi Joshi |
| EPI_ISL_825040 | B.J. Medical College and Civil hospital, Ahmedabad | Gujarat Biotechnology Research Centre | Afzal Ansari, Nikha Trivedi, Apurvasinh Puvar, Ramesh Pandit, Janvi Raval, Zarna Patel, Nitin Savaliya, Dinesh Kumar, Zuber Saiyed, Labdhi Pandya, Pranay Shah, Kamlesh J Upadhyay, Sanjay Kapadia, Dipa Kinariwala, Chaitanya Joshi, Madhvi Joshi |
| EPI_ISL_825041 | B.J. Medical College and Civil hospital, Ahmedabad | Gujarat Biotechnology Research Centre | Nikha Trivedi, Apurvasinh Puvar, Ramesh Pandit, Janvi Raval, Zarna Patel, Nitin Savaliya, Dinesh Kumar, Zuber Saiyed, Labdhi Pandya, Afzal Ansari, Pranay Shah, Kamlesh J Upadhyay, Sanjay Kapadia, Dipa Kinariwala, Chaitanya Joshi, Madhvi Joshi |
| EPI_ISL_825355, EPI_ISL_825356 | Hospital Universitari Vall d'Hebron - Vall d'Hebron Institut de Rercerca | Hospital Universitari Vall d'Hebron | Cristina Andrés, Maria Piñana, Josep F Abril, Damir Garcia-Cehic, Ariadna Rando, Juliana Esperalba, Maria Gema Codina, Carla Castillo, Maria Carmen Martin, Tomàs Pumarola, Josep Quer, Andrés Antón |
| EPI_ISL_826320, EPI_ISL_826321, EPI_ISL_826322, EPI_ISL_826324, EPI_ISL_826325, EPI_ISL_826326, EPI_ISL_826327, EPI_ISL_826328, EPI_ISL_826329, EPI_ISL_826330, EPI_ISL_826332, EPI_ISL_826333, EPI_ISL_826334, EPI_ISL_826335, EPI_ISL_826336, EPI_ISL_826337, EPI_ISL_826338, EPI_ISL_826341, EPI_ISL_826342, EPI_ISL_826343, EPI_ISL_826344, EPI_ISL_826345, EPI_ISL_826346, EPI_ISL_826347, EPI_ISL_826348, EPI_ISL_826350, EPI_ISL_826351, EPI_ISL_826352, EPI_ISL_826353, EPI_ISL_826355, EPI_ISL_826356, EPI_ISL_826357, EPI_ISL_826358, EPI_ISL_826359, EPI_ISL_826360, EPI_ISL_826361, EPI_ISL_826362, EPI_ISL_826363, EPI_ISL_826364, EPI_ISL_826365, EPI_ISL_826366, EPI_ISL_826367, EPI_ISL_826368, EPI_ISL_826369, EPI_ISL_826370, EPI_ISL_826371, EPI_ISL_826372, EPI_ISL_826373, EPI_ISL_826374, EPI_ISL_826375, EPI_ISL_826376, EPI_ISL_826377, EPI_ISL_826378, EPI_ISL_826379, EPI_ISL_826380, EPI_ISL_826381, EPI_ISL_826382, EPI_ISL_826383, EPI_ISL_826384, EPI_ISL_826385, EPI_ISL_826386, EPI_ISL_826387, EPI_ISL_826388, EPI_ISL_826389, EPI_ISL_826390, EPI_ISL_826391, EPI_ISL_826392, EPI_ISL_826393, EPI_ISL_826394, EPI_ISL_826395, EPI_ISL_826396, EPI_ISL_826397, EPI_ISL_826398, EPI_ISL_826399, EPI_ISL_826400, EPI_ISL_826401, EPI_ISL_826402, EPI_ISL_826403, EPI_ISL_826404, EPI_ISL_826405, EPI_ISL_826406, EPI_ISL_826407, EPI_ISL_826408, EPI_ISL_826409, EPI_ISL_826410, EPI_ISL_826411, EPI_ISL_826412, EPI_ISL_826413, EPI_ISL_826414, EPI_ISL_826415, EPI_ISL_826416, EPI_ISL_826417, EPI_ISL_826418, EPI_ISL_826419, EPI_ISL_826420, EPI_ISL_826421, EPI_ISL_826422, EPI_ISL_826423, EPI_ISL_826424, EPI_ISL_826425, EPI_ISL_826426, EPI_ISL_826427, EPI_ISL_826428, EPI_ISL_826429, EPI_ISL_826430, EPI_ISL_826431, EPI_ISL_826442, EPI_ISL_826447, EPI_ISL_826450, EPI_ISL_826453, EPI_ISL_826454, EPI_ISL_826455, EPI_ISL_826456 | | | |
| see above | University of Michigan Clinical Microbiology Laboratory | Lauring Lab, University of Michigan, Department of Microbiology and Immunology | Valesano |
| EPI_ISL_826711, EPI_ISL_826712, EPI_ISL_826931, EPI_ISL_826933 | deCODE genetics | deCODE genetics | Daniel F Gudbjartsson; Agnar Helgason; Hakon Jonsson; Olafur T Magnusson; Pall Melsted; Gudmundur L Norddahl; Jona Saemundsdottir; Asgeir Sigurdsson; Patrick Sulem; Arna B Agustsdottir; Hannes Eggertsson; Berglind Eiriksdottir; Run Fridriksdottir; Elisabet E Gardarsdottir; Gudmundur Georgsson; Olafia S Gretarsdottir; Kjartan R Gudmundsson; Thora R Gunnarsdottir; Arnaldur Gylfason; Hilma Holm; Brynjar O Jensson; Aslaug Jonasdottir; Kamilla S Josefsdottir; Thordur Kristjansson; Droplaug N Magnusdottir; Solvi Rognvaldsson; Louise le Roux; Gudrun Sigmundsdottir; Gardar Sveinbjornsson; Kristin E Sveinsdottir; Maney Sveinsdottir; Emil A Thorarensen; Bjarni Thorbjornsson; Gisli Masson; Ingileif Jonsdottir; Alma Moller; Thorolfur Gudnason; Karl G Kristinsson; Unnur Thorsteinsdottir; Kari Stefansson |
| EPI_ISL_827438, EPI_ISL_827439, EPI_ISL_827441, EPI_ISL_827443 | University of Michigan Clinical Microbiology Laboratory | Lauring Lab, University of Michigan, Department of Microbiology and Immunology | Valesano |
| EPI_ISL_827547, EPI_ISL_827624, EPI_ISL_827721, EPI_ISL_827734, EPI_ISL_828182, EPI_ISL_828183, EPI_ISL_828185, EPI_ISL_828186, EPI_ISL_828187, EPI_ISL_828386 | deCODE genetics | deCODE genetics | Daniel F Gudbjartsson; Agnar Helgason; Hakon Jonsson; Olafur T Magnusson; Pall Melsted; Gudmundur L Norddahl; Jona Saemundsdottir; Asgeir Sigurdsson; Patrick Sulem; Arna B Agustsdottir; Hannes Eggertsson; Berglind Eiriksdottir; Run Fridriksdottir; Elisabet E Gardarsdottir; Gudmundur Georgsson; Olafia S Gretarsdottir; Kjartan R Gudmundsson; Thora R Gunnarsdottir; Arnaldur Gylfason; Hilma Holm; Brynjar O Jensson; Aslaug Jonasdottir; Kamilla S Josefsdottir; Thordur Kristjansson; Droplaug N Magnusdottir; Solvi Rognvaldsson; Louise le Roux; Gudrun Sigmundsdottir; Gardar Sveinbjornsson; Kristin E Sveinsdottir; Maney Sveinsdottir; Emil A Thorarensen; Bjarni Thorbjornsson; Gisli Masson; Ingileif Jonsdottir; Alma Moller; Thorolfur Gudnason; Karl G Kristinsson; Unnur Thorsteinsdottir; Kari Stefansson |
| EPI_ISL_828399, EPI_ISL_828400, EPI_ISL_828401, EPI_ISL_828402 | The National University Hospital of Iceland | deCODE genetics | Daniel F Gudbjartsson; Agnar Helgason; Hakon Jonsson; Olafur T Magnusson; Pall Melsted; Gudmundur L Norddahl; Jona Saemundsdottir; Asgeir Sigurdsson; Patrick Sulem; Arna B Agustsdottir; Hannes Eggertsson; Berglind Eiriksdottir; Run Fridriksdottir; Elisabet E Gardarsdottir; Gudmundur Georgsson; Olafia S Gretarsdottir; Kjartan R Gudmundsson; Thora R Gunnarsdottir; Arnaldur Gylfason; Hilma Holm; Brynjar O Jensson; Aslaug Jonasdottir; Kamilla S Josefsdottir; Thordur Kristjansson; Droplaug N Magnusdottir; Solvi Rognvaldsson; Louise le Roux; Gudrun Sigmundsdottir; Gardar Sveinbjornsson; Kristin E Sveinsdottir; Maney Sveinsdottir; Emil A Thorarensen; Bjarni Thorbjornsson; Gisli Masson; Ingileif Jonsdottir; Alma Moller; Thorolfur Gudnason; Karl G Kristinsson; Unnur Thorsteinsdottir; Kari Stefansson |
| EPI_ISL_828403, EPI_ISL_828404 | deCODE genetics | deCODE genetics | Daniel F Gudbjartsson; Agnar Helgason; Hakon Jonsson; Olafur T Magnusson; Pall Melsted; Gudmundur L Norddahl; Jona Saemundsdottir; Asgeir Sigurdsson; Patrick Sulem; Arna B Agustsdottir; Hannes Eggertsson; Berglind Eiriksdottir; Run Fridriksdottir; Elisabet E Gardarsdottir; Gudmundur Georgsson; Olafia S Gretarsdottir; Kjartan R Gudmundsson; Thora R Gunnarsdottir; Arnaldur Gylfason; Hilma Holm; Brynjar O Jensson; Aslaug Jonasdottir; Kamilla S Josefsdottir; Thordur Kristjansson; Droplaug N Magnusdottir; Solvi Rognvaldsson; Louise le Roux; Gudrun Sigmundsdottir; Gardar Sveinbjornsson; Kristin E Sveinsdottir; Maney Sveinsdottir; Emil A Thorarensen; Bjarni Thorbjornsson; Gisli Masson; Ingileif Jonsdottir; Alma Moller; Thorolfur Gudnason; Karl G Kristinsson; Unnur Thorsteinsdottir; Kari Stefansson |
| EPI_ISL_828405, EPI_ISL_828406, EPI_ISL_828408 | The National University Hospital of Iceland | deCODE genetics | Daniel F Gudbjartsson; Agnar Helgason; Hakon Jonsson; Olafur T Magnusson; Pall Melsted; Gudmundur L Norddahl; Jona Saemundsdottir; Asgeir Sigurdsson; Patrick Sulem; Arna B Agustsdottir; Hannes Eggertsson; Berglind Eiriksdottir; Run Fridriksdottir; Elisabet E Gardarsdottir; Gudmundur Georgsson; Olafia S Gretarsdottir; Kjartan R Gudmundsson; Thora R Gunnarsdottir; Arnaldur Gylfason; Hilma Holm; Brynjar O Jensson; Aslaug Jonasdottir; Kamilla S Josefsdottir; Thordur Kristjansson; Droplaug N Magnusdottir; Solvi Rognvaldsson; Louise le Roux; Gudrun Sigmundsdottir; Gardar Sveinbjornsson; Kristin E Sveinsdottir; Maney Sveinsdottir; Emil A Thorarensen; Bjarni Thorbjornsson; Gisli Masson; Ingileif Jonsdottir; Alma Moller; Thorolfur Gudnason; Karl G Kristinsson; Unnur Thorsteinsdottir; Kari Stefansson |
| EPI_ISL_828451, EPI_ISL_828452, EPI_ISL_828453, EPI_ISL_828454, EPI_ISL_828733, EPI_ISL_828798, EPI_ISL_828800, EPI_ISL_828923, EPI_ISL_829175, EPI_ISL_829186, EPI_ISL_829280, EPI_ISL_829284, EPI_ISL_829285 | | | |
| see above | deCODE genetics | deCODE genetics | Daniel F Gudbjartsson; Agnar Helgason; Hakon Jonsson; Olafur T Magnusson; Pall Melsted; Gudmundur L Norddahl; Jona Saemundsdottir; Asgeir Sigurdsson; Patrick Sulem; Arna B Agustsdottir; Hannes Eggertsson; Berglind Eiriksdottir; Run Fridriksdottir; Elisabet E Gardarsdottir; Gudmundur Georgsson; Olafia S Gretarsdottir; Kjartan R Gudmundsson; Thora R Gunnarsdottir; Arnaldur Gylfason; Hilma Holm; Brynjar O Jensson; Aslaug Jonasdottir; Kamilla S Josefsdottir; Thordur Kristjansson; Droplaug N Magnusdottir; Solvi Rognvaldsson; Louise le Roux; Gudrun Sigmundsdottir; Gardar Sveinbjornsson; Kristin E Sveinsdottir; Maney Sveinsdottir; Emil A Thorarensen; Bjarni Thorbjornsson; Gisli Masson; Ingileif Jonsdottir; Alma Moller; Thorolfur Gudnason; Karl G Kristinsson; Unnur Thorsteinsdottir; Kari Stefansson |
| EPI_ISL_829286 | The National University Hospital of Iceland | deCODE genetics | Daniel F Gudbjartsson; Agnar Helgason; Hakon Jonsson; Olafur T Magnusson; Pall Melsted; Gudmundur L Norddahl; Jona Saemundsdottir; Asgeir Sigurdsson; Patrick Sulem; Arna B Agustsdottir; Hannes Eggertsson; Berglind Eiriksdottir; Run Fridriksdottir; Elisabet E Gardarsdottir; Gudmundur Georgsson; Olafia S Gretarsdottir; Kjartan R Gudmundsson; Thora R Gunnarsdottir; Arnaldur Gylfason; Hilma Holm; Brynjar O Jensson; Aslaug Jonasdottir; Kamilla S Josefsdottir; Thordur Kristjansson; Droplaug N Magnusdottir; Solvi Rognvaldsson; Louise le Roux; Gudrun Sigmundsdottir; Gardar Sveinbjornsson; Kristin E Sveinsdottir; Maney Sveinsdottir; Emil A Thorarensen; Bjarni Thorbjornsson; Gisli Masson; Ingileif Jonsdottir; Alma Moller; Thorolfur Gudnason; Karl G Kristinsson; Unnur Thorsteinsdottir; Kari Stefansson |
| EPI_ISL_829288, EPI_ISL_829289, EPI_ISL_829313, EPI_ISL_829428, EPI_ISL_829432, EPI_ISL_829448, EPI_ISL_829450, EPI_ISL_829451, EPI_ISL_829965, EPI_ISL_830051, EPI_ISL_830053, EPI_ISL_830162, EPI_ISL_830383, EPI_ISL_830384, EPI_ISL_830385, EPI_ISL_830386, EPI_ISL_830388, EPI_ISL_830392 | | | |

| | | | |
|---|---|---|---|
| see above | deCODE genetics | deCODE genetics | Daniel F Gudbjartsson; Agnar Helgason; Hakon Jonsson; Olafur T Magnusson; Pall Melsted; Gudmundur L Norddahl; Jona Saemundsdottir; Asgeir Sigurdsson; Patrick Sulem; Arna B Agustsdottir; Hannes Eggertsson; Berglind Eiriksdottir; Run Fridriksdottir; Elisabet E Gardarsdottir; Gudmundur Georgsson; Olafia S Gretarsdottir; Kjartan R Gudmundsson; Thora R Gunnarsdottir; Arnaldur Gylfason; Hilma Holm; Brynjar O Jensson; Aslaug Jonasdottir; Kamilla S Josefsdottir; Thordur Kristjansson; Droplaug N Magnusdottir; Solvi Rognvaldsson; Louise le Roux; Gudrun Sigmundsdottir; Gardar Sveinbjornsson; Kristin E Sveinsdottir; Maney Sveinsdottir; Emil A Thorarensen; Bjarni Thorbjornsson; Gisli Masson; Ingileif Jonsdottir; Alma Moller; Thorolfur Gudnason; Karl G Kristinsson; Unnur Thorsteinsdottir; Kari Stefansson |
| EPI_ISL_831221, EPI_ISL_831236, EPI_ISL_831237, EPI_ISL_831238, EPI_ISL_831239, EPI_ISL_831240 | Hospital Universitario La Paz (Madrid) | SeqCOVID-SPAIN consortium/IBV(CSIC) | María Rodríguez-Tejedor, Elias Dahdouh, Fernando Lázaro-Perona, Jesús Mingorance and SeqCOVID-SPAIN consortium |
| EPI_ISL_831650 | Institute for Infectious Diseases, University of Bern, Switzerland | Institute for Infectious Diseases, University of Bern, Switzerland | Michel C Koch, Christian Baumann, Miguel A Terrazos Miani, Cora Sägesser, Pascal Bittel, Stephen L Leib, Peter Keller, Franziska Suter-Riniker, Alban Ramette |
| EPI_ISL_831719, EPI_ISL_831738, EPI_ISL_831746, EPI_ISL_831761, EPI_ISL_831797, EPI_ISL_831799, EPI_ISL_831800, EPI_ISL_831801, EPI_ISL_831817, EPI_ISL_831818, EPI_ISL_831819, EPI_ISL_831820 | | | |
| see above | United States Air Force School of Aerospace Medicine | United States Air Force School of Aerospace Medicine | Anthony Fries, Jennifer Meyer, William Gruner, Amanda Javorina, Sarah Purves, Clarise Starr, Elizabeth Macias |
| EPI_ISL_832726, EPI_ISL_832727, EPI_ISL_832728, EPI_ISL_832729, EPI_ISL_832730, EPI_ISL_832731, EPI_ISL_832733, EPI_ISL_832734, EPI_ISL_832735, EPI_ISL_832736, EPI_ISL_832737, EPI_ISL_832738, EPI_ISL_832739, EPI_ISL_832740, EPI_ISL_832741, EPI_ISL_832742, EPI_ISL_832743, EPI_ISL_832747, EPI_ISL_832748, EPI_ISL_832749, EPI_ISL_832750 | | | |
| see above | OHSU Lab Services Molecular Microbiology Lab | Oregon SARS-CoV-2 Genome Sequencing Center | Brendan L. O'Connell, Ruth V. Nichols, Sally Grindstaff, Alec J. Hirsch, Donna Hansel, Guang Fan, Daniel N. Streblow, William B. Messer, Andrew C. Adey, Benjamin N. Bimber, Brian J. O'Roak |
| EPI_ISL_833470, EPI_ISL_833477 | CHU Purpan - Laboratoire de Virologie - Institut Fédératif de Biologie | CHU Purpan - Laboratoire de Virologie - Institut Fédératif de Biologie | Latour J., Ranger N., Dubois M., Carcenac R., Harter A., Boyer P., Tremeaux P., Izopet J. |
| EPI_ISL_840234, EPI_ISL_840235, EPI_ISL_840236, EPI_ISL_840237, EPI_ISL_840238, EPI_ISL_840241, EPI_ISL_840244 | Oxford Viromics, NDM, University of Oxford; Oxford University Hospitals; Basingstoke and North Hampshire Hospital | COVID-19 Genomics UK (COG-UK) Consortium | Tanya Golubchik, David Bonsall, George Macintyre, Amy Trebes, Mariateresa de Cesare, Catrin Moore, Alex Mobbs, Anita Justice, Robert Shaw, Monique Andersson, Timothy Peto, Emma Wise, Nathan Moore, Jessica Lynch, Nick Cortes, Matilde Mori, Stephen Kidd, David Buck, John Todd, Christophe Fraser |
| EPI_ISL_842870, EPI_ISL_842878, EPI_ISL_842879, EPI_ISL_842880, EPI_ISL_842881, EPI_ISL_842882, EPI_ISL_842883, EPI_ISL_842884, EPI_ISL_842885, EPI_ISL_842886, EPI_ISL_842888, EPI_ISL_842889, EPI_ISL_842890, EPI_ISL_842891, EPI_ISL_842892, EPI_ISL_842893, EPI_ISL_842894, EPI_ISL_842895, EPI_ISL_842896, EPI_ISL_842897, EPI_ISL_842898, EPI_ISL_842902, EPI_ISL_842904, EPI_ISL_842905, EPI_ISL_842906, EPI_ISL_842907, EPI_ISL_842908, EPI_ISL_842909, EPI_ISL_842919, EPI_ISL_842920, EPI_ISL_842921, EPI_ISL_842922, EPI_ISL_842932 | | | |
| see above | Barts Health NHS Trust | COVID-19 Genomics UK (COG-UK) Consortium | CUTINO-MOGUEL, Maria-Teresa; HARRINGTON, David; OWOYEMI, Dola; SHYLINI, Raghavendran; BROAD, Claire; KELE, Beatrix |
| EPI_ISL_844254, EPI_ISL_844262, EPI_ISL_844541 | Department of Virus and Microbiological Special Diagnostics, Statens Serum Institut, Copenhagen, Denmark | Albertsen Lab, Department of Chemistry and Bioscience, Aalborg University, Denmark | Danish Covid-19 Genome Consortium |
| EPI_ISL_846705, EPI_ISL_846706, EPI_ISL_846708, EPI_ISL_846709 | University of Michigan Clinical Microbiology Laboratory | Lauring Lab, University of Michigan, Department of Microbiology and Immunology | Valesano |
| EPI_ISL_847703 | California Department of Public Health | Chiu Laboratory, University of California, San Francisco | Charles Chiu, Xianding (Wayne) Deng, Candace Wang, Brian Bushnell, Scot Federman, Jill Hacker, Debra Wadford |
| EPI_ISL_847827 | COVID-19 National Reference Laboratory | COVID-19 National Reference Laboratory | Tahmineh Jalali, Mohammad Hassan Pouriayevali, Zahra Ahmadi, Marzieh Sadjadi, Mahsa Tavakoli, Zahra Fereydouni, Setareh Kashanian, Sanam Azad-Manjiri, Tahereh Mohammadi, Zabihollah Shoja, Parastoo Yekta, Farideh Niknam, Hessam Nemati, Ahmad Ghasemi, Sahar Khakifirouz, Sepideh Gerdooei, Maryam Rostamtabar,Sana Eybpoosh, Mohammad Mehdi Mortazavipour, Mohamad Sadegh Shams Nosrati, Zeynab VeisiZadeh, Amitis Ramezani, Kayhan Azadmanesh, Mostafa Salehi-Vaziri |
| EPI_ISL_848452 | Illinois Department of Public Health | Gagnon Lab, Southern Illinois University | Keith Gagnon |
| EPI_ISL_849612 | Seattle Flu Study | Seattle Flu Study | Deborah A. Nickerson, Chris D. Frazar, Jover Lee, Benjamin Pelle, Matthew Richardson, Amanda Adler, Elisabeth Brandstetter, Peter D. Han, Kairsten Fay, Misja Ilcisin, Kirsten Lacombe, Thomas R. Sibley, Melissa Truong, Caitlin R. Wolf, Michael Boeckh, Janet A. Englund, Michael Famulare, Barry R. Lutz, Mark J. Rieder, Lea M. Starita, Matthew Thompson, Jay Shendure, Trevor Bedford, Helen Y. Chu |
| EPI_ISL_853313, EPI_ISL_853317, EPI_ISL_853326 | UPMC Clinical Microbiology Laboratory | Microbial Genome Sequencing Center; Microbial Genomic Epidemiology Laboratory | Mustapha M. Mustapha, Jane W. Marsh, Dan Snyder, Marissa P. Griffith, Stephanie L. Mitchell, Vatsala R. Srinivasa, Kady D. Waggle, Chinelo Ezeonwuku, Vaughn S. Cooper, Lee H. Harrison |
| EPI_ISL_853754, EPI_ISL_853769, EPI_ISL_853770, EPI_ISL_853771, EPI_ISL_853772, EPI_ISL_853773, EPI_ISL_853774, EPI_ISL_853798, EPI_ISL_853799, EPI_ISL_853805, EPI_ISL_853874, EPI_ISL_853875, EPI_ISL_853876, EPI_ISL_853917, EPI_ISL_853918, EPI_ISL_853919, EPI_ISL_853920, EPI_ISL_853921, EPI_ISL_853922, EPI_ISL_853923, EPI_ISL_854217, EPI_ISL_854255, EPI_ISL_854256, EPI_ISL_854257, EPI_ISL_854259, EPI_ISL_854260, EPI_ISL_854300, EPI_ISL_854302, EPI_ISL_854303 | | | |
| see above | Pharmgenetix GmbH | Bergthaler laboratory, CeMM Research Center for Molecular Medicine of the Austrian Academy of Sciences | Lukas Endler, Alexandra Popa, Benedikt Agerer, Jakob-Wendelin Genger, Alexander Lercher, Anna Schedl, Thomas Penz, Michael Schuster, Jan Laine, Martin Senekowitsch, Christoph Bock, Andreas Bergthaler |
| EPI_ISL_854443, EPI_ISL_854444, EPI_ISL_854445 | SARATOGA HOSPITAL LABORATORY | Wadsworth Center, New York State Department of Health | Kirsten St. George, Daryl M. Lamson, Alexis Russel, Matthew Shudt, Melissa A Leisner, Jonathan Plitnick, Navjot Singh, John Kelly, Erasmus Schneider, Erica Lasek-Nesselquist |
| EPI_ISL_855496, EPI_ISL_855524 | KEMRI-Wellcome Trust Research Programme/KEMRI-CGMR-C Kilifi | KEMRI-Wellcome Trust Research Programme/KEMRI-CGMR-C Kilifi | Githinji et al |
| EPI_ISL_857492, EPI_ISL_857493 | Swiss National Reference Centre for Influenza | Swiss National Reference Centre for Influenza | Ana Rita Goncalves,Samuel Cordey, Laurent Kaiser, Lorenzo Cerutti, Henri Pegeot, Melyssa Elies, Keith Harshman, Ioannis Xenarios, Emmanouil Dermitzakis |
| EPI_ISL_857517 | Swiss National Reference Centre for Influenza | Swiss National Reference Centre for Influenza | Tim Roloff, Ana Rita Gonçalves, Madlen Stange, Helena MB Seth-Smith, Alfredo Mari, Karoline Leuzinger, Julia Bielicki, Manuel Battegay, Hans Hirsch, Laurent Kaiser, Adrian Egli |
| EPI_ISL_860137, EPI_ISL_860148, EPI_ISL_860157 | Keio University School of Medicine | Keio University School of Medicine | Kenjiro Kosaki, Yuka Iwasaki, Hirotsugu Ishizu, Haruhiko Siomi, Kodai Abe |
| EPI_ISL_860809 | WHO/Minsk | Charité Universitätsmedizin Berlin, Institut für Virologie | Victor M Corman, Barbara Mühlemann, Jörn Beheim-Schwarzbach, Talitha Veith, Julia Tesch, Tobias Bleicker, Julia Schneider, Shmialiova Natallia, Sivets Natallia, Terry Jones, Christian Drosten |
| EPI_ISL_861853 | Hospital General Universitario Gregorio Marañón | SeqCOVID-SPAIN consortium/IBV(CSIC) | Darío García de Viedma, Laura Pérez-Lago, Pedro J Sola-Campoy, Sergio Buenestado-Serrano, Marta Herranz, Victor Manuel de la Cueva, Julia Suárez, Pilar Catalán, Patricia Muñoz and SeqCOVID-SPAIN consortium |
| EPI_ISL_861877, EPI_ISL_861892, EPI_ISL_861899 | LATE - Laboratório de Técnicas Especiais - Hospital Israelita Albert Einstein | LATE - Laboratório de Técnicas Especiais - Hospital Israelita Albert Einstein | Deyvid Amgarten, Fernanda de Mello Malta, Raquel Riyuzo, Ana Paula Moreira Salles, Pedro Henrique Sebe Rodrigues, João Renato Rebello Pinho |
| EPI_ISL_862075 | National Influenza Center, Virology Department | National Influenza Center | V Salimi,NZ Shafiei Jandaghi, J Yavarian, A Nejati, K Sadeghi, N Ghavvami,F Ajaminejad and T Mokhtari Azad |
| EPI_ISL_864572 | Institute of Medical Microbiology and Hospital Hygiene | Institute of Medical Microbiology and Hospital Hygiene | Prof. Dr. Achim Kaasch, Aljoscha Tersteegen |
| EPI_ISL_865162 | Virology Department, Royal Infirmary of Edinburgh, NHS Lothian / School of Biological Sciences, University of Edinburgh / Institute of Genetics and Molecular Medicine, University of Edinburgh | COVID-19 Genomics UK (COG-UK) Consortium | McHugh M, Dewar R, Rooke S, Gallagher M, Balcaza C, O'Toole Á, Scher E, Hill V, McCrone JT, Colquhoun R, Yu X, Jackson B, Rambaut A, Williams TC, Templeton K |
| EPI_ISL_865171 | Liverpool Clinical Laboratories | COVID-19 Genomics UK (COG-UK) Consortium | Sam Haldenby, Anita Lucaci, Steve Paterson, Julian Hiscox, Alistair Darby, M Almsaud, A Alrezaihi, Muhannad Alruwaili, Stuart D Armstrong, Jones Benjamin, Eleanor G Bentley, Anu Chawla, Jordan J Clark, Angela Cowell, Richard Eccles, Isabel García-Dorival, Matthew Gemmell, Alessandro Gerada, PKF Gilmore, Richard Gregory, Ximeng Han, Catherine Hartley, Margaret Hughes, Miren Iturriza-Gomara, James Johnson, L Luu, Jenifer Manson, Charlotte Nelson, Elaine O'Toole, Cassie Olateju, Rebekah Penrice-Randal , Lucille Rainbow, N.P Randle, Trevor Ian Robinson, Parul Sharma, Ghada T Shawli, James P Stewart, Neil Swainston, Ecaterina Vamos, Joanne Watts, Mark Whitehead |

| | | | |
|---|---|---|---|
| EPI_ISL_865684 | University College London, Great Ormond Street Hospital for Children NHS Foundation Trust, Imperial College Healthcare NHS Trust | COVID-19 Genomics UK (COG-UK) Consortium | Sergi Castellano, Rachel Williams, Mark Kristiansen, Paola Resende Silva, Sunando Roy, Tony Brooks, Helena Tutill, Paola Niola, Patricia Dyal, Charlotte Williams, Leysa Forrest, Yasmin Panchbhaya, Jacqueline Findlay, Samuel Weeks, Julianne Brown, Kathryn Harris, Paul Randell, James Price, Alison Holmes, Judith Breuer |
| EPI_ISL_866048 | University College London Hospital | COVID-19 Genomics UK (COG-UK) Consortium | Judith Heaney, Matthew Byott, Catherine Houlihan, Dan Frampton, Stuart Kirk, Moira Spyer and Eleni Nastouli |
| EPI_ISL_866901, EPI_ISL_866902, EPI_ISL_866903 | Queens Medical Centre, Clinical Microbiology Department / DeepSeq Nottingham | COVID-19 Genomics UK (COG-UK) Consortium | Gemma Clark, Wendy Smith, Manjinder Khakh, Vicki M Fleming, Michelle M Lister, Hannah Howson-Wells, Jonathan Ball, Patrick McClure, Joseph Chappell, Theocharis Tsoleridis, Nadine Holmes, Matthew Carlisle, Christopher Moore, Fei Sang, Johnny Debebe, Victoria Wright, Matthew Loose |
| EPI_ISL_871786 | Department of Virus and Microbiological Special Diagnostics, Statens Serum Institut, Copenhagen, Denmark | Aalborg University | Danish Covid-19 Genome Consortium |
| EPI_ISL_872605 | Nigeria Centre for Disease Control (NCDC) | African Centre of Excellence for Genomics of Infectious Diseases (ACEGID), Redeemer's University | Oluniyi P.E. et al |
| EPI_ISL_872689 | Rhode Island Department of Health | Infectious Disease Program, Broad Institute of Harvard and MIT | Lemieux,J.E., Siddle,K.J., Huard,R., King,E., Azevedo,K., Miller,A., Adams,G., Gladden-Young,A., Lagerborg,K., Rudy,M., DeRuff,K., Carter,A., Normandin,E., Bauer,M., Reilly,S., Tomkins-Tinch,C., Loreth,C., Chaluvadi,S., Birren,B.W., Gallagher,G., Smole,S., Park,D.J., MacInnis,B.L., and Sabeti,P.C. |
| EPI_ISL_872757, EPI_ISL_872758, EPI_ISL_872759, EPI_ISL_872760, EPI_ISL_872761, EPI_ISL_872762, EPI_ISL_872763, EPI_ISL_872764, EPI_ISL_872765, EPI_ISL_872766, EPI_ISL_872767, EPI_ISL_872768, EPI_ISL_872769, EPI_ISL_872770, EPI_ISL_872771, EPI_ISL_872772, EPI_ISL_872773, EPI_ISL_872774, EPI_ISL_872775, EPI_ISL_872776, EPI_ISL_872777 | | | |
| see above | Colorado Mesa University | Infectious Disease Program, Broad Institute of Harvard and MIT | Lemieux,J.E., Siddle,K.J., Marshall,J., ONeill,M., Bronson,A., Adams,G., Gladden-Young,A., Lagerborg,K., Rudy,M., DeRuff,K., Carter,A., Normandin,E., Bauer,M., Reilly,S., Tomkins-Tinch,C., Loreth,C., Chaluvadi,S., Birren,B.W., Gallagher,G., Smole,S., Park,D.J., MacInnis,B.L., and Sabeti,P.C. |
| EPI_ISL_873148, EPI_ISL_873149, EPI_ISL_873150 | University of Michigan Clinical Microbiology Laboratory | Lauring Lab, University of Michigan, Department of Microbiology and Immunology | Valesano |
| EPI_ISL_875531 | Institute of Virology, Biomedical Research Center of the Slovak Academy of Sciences, Bratislava | Faculty of Natural Sciences, Comenius University, Bratislava | Viktória abanová, Kristína Boršová, Broa Brejová, Viktória Hodorová, Sabina Fumaová Havlíková, Juraj Kopáek, Martina Liková, ubomíra Lukáiková, Martina Neboháová, Monika Sláviková, Tomáš Vina, Jozef Nosek, Boris Klempa |
| EPI_ISL_875535 | Institute of Virology, Biomedical Research Center of the Slovak Academy of Sciences, Bratislava | Faculty of Natural Sciences, Comenius University, Bratislava | Broa Brejová, Viktória abanová, Kristína Boršová, Viktória Hodorová, Sabina Fumaová Havlíková, Juraj Kopáek, Martina Liková, ubomíra Lukáiková, Martina Neboháová, Monika Sláviková, Tomáš Vina, Jozef Nosek, Boris Klempa |
| EPI_ISL_875536 | Institute of Virology, Biomedical Research Center of the Slovak Academy of Sciences, Bratislava | Faculty of Natural Sciences, Comenius University, Bratislava | Viktória abanová, Kristína Boršová, Broa Brejová, Viktória Hodorová, Sabina Fumaová Havlíková, Juraj Kopáek, Martina Liková, ubomíra Lukáiková, Martina Neboháová, Monika Sláviková, Tomáš Vina, Jozef Nosek, Boris Klempa |
| EPI_ISL_875537 | Institute of Virology, Biomedical Research Center of the Slovak Academy of Sciences, Bratislava | Faculty of Natural Sciences, Comenius University, Bratislava | Kristína Boršová, Viktória abanová, Broa Brejová, Viktória Hodorová, Sabina Fumaová Havlíková, Juraj Kopáek, Martina Liková, ubomíra Lukáiková, Martina Neboháová, Monika Sláviková, Tomáš Vina, Boris Klempa, Jozef Nosek |
| EPI_ISL_875552 | ULSS 2 Treviso | Istituto Zooprofilattico Sperimentale delle Venezie | Adelaide Milani, Alessia Schivo, Annalisa Salviato, Erika Giorgia Quaranta, Ambra Pastori, Bianca Zecchin, Alice Fusaro, Isabella Monne, Calogero Terregino, Antonia Ricci |
| EPI_ISL_876331, EPI_ISL_876332, EPI_ISL_876333, EPI_ISL_876335, EPI_ISL_876336, EPI_ISL_876337, EPI_ISL_876338, EPI_ISL_876339, EPI_ISL_876340, EPI_ISL_876341, EPI_ISL_876342, EPI_ISL_876343, EPI_ISL_876344, EPI_ISL_876345, EPI_ISL_876346, EPI_ISL_876347, EPI_ISL_876348, EPI_ISL_876349, EPI_ISL_876350, EPI_ISL_876351, EPI_ISL_876352, EPI_ISL_876353, EPI_ISL_876354, EPI_ISL_876355, EPI_ISL_876356, EPI_ISL_876357, EPI_ISL_876358, EPI_ISL_876359, EPI_ISL_876360, EPI_ISL_876361, EPI_ISL_876362, EPI_ISL_876363, EPI_ISL_876364, EPI_ISL_876365, EPI_ISL_876366, EPI_ISL_876367, EPI_ISL_876368, EPI_ISL_876369, EPI_ISL_876370, EPI_ISL_876371, EPI_ISL_876372, EPI_ISL_876373, EPI_ISL_876374, EPI_ISL_876375, EPI_ISL_876376, EPI_ISL_876377, EPI_ISL_876378, EPI_ISL_876379, EPI_ISL_876380, EPI_ISL_876381, EPI_ISL_876382, EPI_ISL_876383, EPI_ISL_876384, EPI_ISL_876385, EPI_ISL_876386, EPI_ISL_876387, EPI_ISL_876388, EPI_ISL_876389, EPI_ISL_876390, EPI_ISL_876391, EPI_ISL_876392, EPI_ISL_876393, EPI_ISL_876394, EPI_ISL_876395, EPI_ISL_876396, EPI_ISL_876397, EPI_ISL_876398, EPI_ISL_876399, EPI_ISL_876400, EPI_ISL_876401, EPI_ISL_876402, EPI_ISL_876403, EPI_ISL_876404 | | | |
| see above | Eurofins Diatherix | Hudsonalpha Genome Sequencing Center | Jane Grimwood, Melissa Williams, Lori H. Handley, Joshua Stough, Leslie Malone, Stefan Brzezinski, Ada Stewart, Teresa Jones, Jenell Webber, John Lovell, Jennifer Cart, and Jeremy Schmutz |
| EPI_ISL_876525 | Florida Bureau of Public Health Laboratories | Florida Bureau of Public Health Laboratories | Sarah Schmedes, Jason Blanton |
| EPI_ISL_876832, EPI_ISL_876833 | Quest Diagnostics | Quest Diagnostics | Rosenthal,S.H., Gerasimova,A., Kagan,R.M., Anderson, B., Hua, M., Liu Y., Bernstein, L.E., Livingston, K.E., Perez, A., Shalhout, D.F., Shlyakhter, I.A., Owen, R., Tanpaiboon, P., Lacbawan, F. |
| EPI_ISL_877237, EPI_ISL_877238, EPI_ISL_877239, EPI_ISL_877240, EPI_ISL_877241, EPI_ISL_877242, EPI_ISL_877243, EPI_ISL_877244, EPI_ISL_877245, EPI_ISL_877246, EPI_ISL_877247, EPI_ISL_877248, EPI_ISL_877249, EPI_ISL_877250, EPI_ISL_877251, EPI_ISL_877252, EPI_ISL_877253, EPI_ISL_877254, EPI_ISL_877255, EPI_ISL_877256, EPI_ISL_877257, EPI_ISL_877258, EPI_ISL_877259, EPI_ISL_877260, EPI_ISL_877261, EPI_ISL_877262, EPI_ISL_877263, EPI_ISL_877264, EPI_ISL_877265, EPI_ISL_877266, EPI_ISL_877267, EPI_ISL_877268, EPI_ISL_877269, EPI_ISL_877270, EPI_ISL_877271, EPI_ISL_877272, EPI_ISL_877273, EPI_ISL_877274, EPI_ISL_877275, EPI_ISL_877276, EPI_ISL_877277, EPI_ISL_877278, EPI_ISL_877279, EPI_ISL_877280, EPI_ISL_877281, EPI_ISL_877282, EPI_ISL_877283, EPI_ISL_877284, EPI_ISL_877285, EPI_ISL_877286, EPI_ISL_877287, EPI_ISL_877288, EPI_ISL_877289, EPI_ISL_877290, EPI_ISL_877291, EPI_ISL_877292, EPI_ISL_877293, EPI_ISL_877294, EPI_ISL_877295, EPI_ISL_877296, EPI_ISL_877297, EPI_ISL_877298, EPI_ISL_877299, EPI_ISL_877300, EPI_ISL_877301, EPI_ISL_877302, EPI_ISL_877303, EPI_ISL_877304, EPI_ISL_877305, EPI_ISL_877306, EPI_ISL_877307, EPI_ISL_877308, EPI_ISL_877309, EPI_ISL_877310, EPI_ISL_877311, EPI_ISL_877312, EPI_ISL_877313, EPI_ISL_877314, EPI_ISL_877315, EPI_ISL_877316, EPI_ISL_877317, EPI_ISL_877318, EPI_ISL_877319, EPI_ISL_877320, EPI_ISL_877321, EPI_ISL_877322, EPI_ISL_877323, EPI_ISL_877324, EPI_ISL_877325, EPI_ISL_877326, EPI_ISL_877327, EPI_ISL_877328 | | | |
| see above | Eurofins Diatherix | Hudsonalpha Genome Sequencing Center | Jane Grimwood, Melissa Williams, Lori H. Handley, Joshua Stough, Leslie Malone, Stefan Brzezinski, Ada Stewart, Teresa Jones, Jenell Webber, John Lovell, Jennifer Cart, and Jeremy Schmutz |
| EPI_ISL_877436 | National laboratory of health, environment and food Celje | Institute of Microbiology and Immunology, Faculty of Medicine, University of Ljubljana | Samo Zakotnik, Tomaž Mark Zorec, Matic Brvar, Miša Korva, Mario Poljak, Tatjana Avši - Županc |
| EPI_ISL_884326, EPI_ISL_884372, EPI_ISL_884384, EPI_ISL_884422, EPI_ISL_884423 | Infectious Diseases, Quest Diagnostics | Infectious Diseases, Quest Diagnostics | Rosenthal,S.H., Gerasimova,A., Kagan,R.M., Anderson,B., Bernstein,L.E., Livingston,K.E., Hua,M., Liu,Y., Shalhout,D.F., Owen,R., Lacbawan,F. |
| EPI_ISL_887162, EPI_ISL_887163 | Massachusetts General Hospital | Infectious Disease Program, Broad Institute of Harvard and MIT | Lemieux,J.E., Siddle,K.J., Shaw,B., Adams,G., Pierce,V., Turbett,S., Anahtar,M., Branda,J., Slater,D., Harris,J., Lin,A.E., Gladden-Young,A., Lagerborg,K., Rudy,M., DeRuff,K., Carter,A., Normandin,E., Bauer,M., Reilly,S., Tomkins-Tinch,C., Loreth,C., Chaluvadi,S., Neumann,A., Cusick,C., Chapman,S.B., Gnirke,A., Flowers,K., Cerrato,F., Birren,B.W., Gallagher,G., Smole,S., Park,D.J., MacInnis,B.L., Ryan,E., LaRocque,R., Rosenberg,E. and Sabeti,P.C. |
| EPI_ISL_888985 | RS Hermina Serpong | Eijkman Institute for Molecular Biology, Ministry of Research and Technology/National Agency for Research and Innovation | Edison Johar, Frilasita A Yudhaputri, Hidayat Trimarsanto, Iskandar Adnan, Lydia V. Panggalo, Sukma Oktavianthi, Willy Agustine, Safarina G Malik, Khin Saw Myint, Amin Soebandrio |
| EPI_ISL_889009 | RS Mitra Keluarga Bintaro | Eijkman Institute for Molecular Biology, Ministry of Research and Technology/National Agency for Research and Innovation | Edison Johar, Frilasita A Yudhaputri, Hidayat Trimarsanto, Iskandar Adnan, Lydia V. Panggalo, Sukma Oktavianthi, Willy Agustine, Safarina G Malik, Khin Saw Myint, Amin Soebandrio |
| EPI_ISL_891192 | DPH, Massachusetts State Public Health Lab | DPH, Massachusetts State Public Health Lab | Lang,A.S., Fink,T., Gallagher,G.R., Smole,S.C. |
| EPI_ISL_896377 | University of Medicine and Pharmacy of Craiova | "Stefan cel Mare" University Metagenomics Lab | Lobiuc Andrei, Gheorghita Roxana |
| EPI_ISL_900054 | Houston Health Dept. | Houston Health Dept. | Ryker Penn, Pamela Brown |
| EPI_ISL_900205 | MEPHI, Aix Marseille University | MEPHI, Aix Marseille University | Anthony LEVASSEUR |
| EPI_ISL_903631 | WA State Department of Health | Genomics and Discovery, Respiratory Viruses Branch, Division of Viral Diseases, Centers for Disease Control and Prevention | Krista Queen, Yan Li, Ying Tao, Jing Zhang, Anna Uehara, Anna Montmayeur, Clinton R. Paden, Peter W. Cook, Rachel Marine, Mili Sheth, Jasmine Padilla, Sarah Nobles, Mark Burroughs, Lori Rowe, Haibin Wang, Ben L. Rambo-Martin, Dhwani Batra, Justin Lee, Suxiang Tong |
| EPI_ISL_903918 | AZ SPHL, Arizona Department of Health Services | Genomics and Discovery, Respiratory Viruses Branch, Division of Viral Diseases, Centers for Disease Control and Prevention | Krista Queen, Yan Li, Ying Tao, Jing Zhang, Anna Uehara, Anna Montmayeur, Clinton R. Paden, Peter W. Cook, Rachel Marine, Mili Sheth, Jasmine Padilla, Sarah Nobles, Mark Burroughs, Lori Rowe, Haibin Wang, Ben L. Rambo-Martin, Dhwani Batra, Justin Lee, Suxiang Tong |
| EPI_ISL_903965, EPI_ISL_903968 | WA State Department of Health | Genomics and Discovery, Respiratory Viruses Branch, Division of Viral Diseases, Centers for Disease Control and | Krista Queen, Yan Li, Ying Tao, Jing Zhang, Anna Uehara, Anna Montmayeur, Clinton R. Paden, Peter W. Cook, Rachel Marine, Mili Sheth, Jasmine Padilla, Sarah Nobles, Mark Burroughs, Lori Rowe, Haibin Wang, Ben L. Rambo-Martin, Dhwani Batra, Justin Lee, Suxiang Tong |

| | | | |
|---|---|---|---|
| | | Prevention | |
| EPI_ISL_904010 | Veterinary Specialized Institute Kraljevo | Veterinary Specialized Institute "Kraljevo", Serbia | Vidanovic,D., Tesovic,B., Knezevic,A., Jovanovic,T., Jankovic,M., Sekler,M., Banovic Djeri,B., Petrovic,T., Volkening,J., Afonso,C. |
| EPI_ISL_904465, EPI_ISL_904466, EPI_ISL_904467 | Dutch COVID-19 response team | Erasmus Medical Center | Bas Oude Munnink, Reina Sikkema, David Nieuwenhuijse, Irina Chestakova, Anne van der Linden, Marjan Boter, Emmanuelle Munger, Corine GeurtsvanKessel, Annemiek van der Eijk, Richard Molenkamp, Marion Koopmans, on behalf of the Dutch national COVID-19 response team. |
| EPI_ISL_906262 | University of Wisconsin-Madison AIDS Vaccine Research Laboratories | University of Wisconsin-Madison AIDS Vaccine Research Laboratories | Gage Moreno, Katarina Braun, et al. AIDS Vaccine Research Laboratories |
| EPI_ISL_911236, EPI_ISL_911237 | Laboratoire national de sante, Microbiology, Virology | Laboratoire national de sante, Microbiology, Microbial Genomics Platform | Anke Wienecke-Baldacchino, Catherine Ragimbeau,Jessica Tapp, Fatu Djabi, Lise Pignon, Raoul Salmon, Tamir Abdelrahman |
| EPI_ISL_911685, EPI_ISL_911701 | Alaska State Virology Laboratory | Alaska State Virology Laboratory | Stephanie DeRonde, Lisa Smith, Ph.D., Jack Chen, Ph.D. |
| EPI_ISL_912464, EPI_ISL_912470, EPI_ISL_912502, EPI_ISL_912525, EPI_ISL_912535 | NHLS Universitas Academic | UFS Virology | PA Bester, MM Nyaga, P Nthiga, MT Mogotsi, D Goedhals, T de Oliveira |
| EPI_ISL_913063 | Labor Mustafa Wien | Center for Virology | Jeremy V. Camp, Irene Goerzer, Monika Redlberger-Fritz, Stephan W. Aberle |
| EPI_ISL_915408, EPI_ISL_915409 | Keio University School of Medicine | Keio University School of Medicine | Kenjiro Kosaki, Yuka Iwasaki, Hirotsugu Ishizu, Haruhiko Siomi, Kodai Abe |
| EPI_ISL_918178, EPI_ISL_918210, EPI_ISL_918212, EPI_ISL_918217, EPI_ISL_918222 | Innovative Genomics Institute, UC Berkeley | Innovative Genomics Institute, UC Berkeley | Stacia Wyman, Haridha Shivram, Phil Frankino, Liana Lareau, Shana McDevitt, Justin Choi |
| EPI_ISL_918373, EPI_ISL_918374, EPI_ISL_918375 | Virology Unit, Institut Pasteur du Cambodge | Virology Unit, Institut Pasteur du Cambodge | Sokhoun Yann, Ly Sovann, Kraing Sidonn, Yi Sengdoeurn, Chin Savuth, Chau Darapheak, Etienne Simon-Loriere, Veasna Duong, Erik A Karlsson |
| EPI_ISL_918517 | LACEN - Laboratório Central de Saúde Pública do Para | Evandro Chagas Institute | Santos, M.C.; Silva, A.M.; Junior, W.D.C.; Barbagelata, L.S.; Ferreira, J.A.; Sousa, E.M.A.; da Silva, P.S.; Pinheiro, K.C.; L.C.; Sousa Junior, E.C. |
| EPI_ISL_918531, EPI_ISL_918533 | LACEN - Laboratório Central de Saúde Pública do Amazonas | Evandro Chagas Institute | Santos, M.C.; Silva, A.M.; Junior, W.D.C.; Barbagelata, L.S.; Ferreira, J.A.; Sousa, E.M.A.; da Silva, P.S.; Pinheiro, K.C.; L.C.; Sousa Junior, E.C. |
| EPI_ISL_919254 | West of Scotland Specialist Virology Centre, NHSGGC / MRC-University of Glasgow Centre for Virus Research | COVID-19 Genomics UK (COG-UK) Consortium | Ana da Silva Filipe, Natasha Johnson, Kathy Smollett, Daniel Mair, Stephen Carmichael, Alice Broos, Lily Tong, Jenna Nichols, Kyriaki Nomikou; Sarah McDonald; Richard Orton, Joseph Hughes, Sreenu Vattipally, David L Robertson; Alasdair MacLean, Rory Gunson; Sharif Shaaban, Matthew Holden; Rachel Blacow, Guy Mollett, Kathy Li, James Shepherd, Antonia Ho, Emma Thomson |
| EPI_ISL_920209, EPI_ISL_920213, EPI_ISL_920529, EPI_ISL_920530 | University College London Hospital | COVID-19 Genomics UK (COG-UK) Consortium | Judith Heaney, Matthew Byott, Catherine Houlihan, Dan Frampton, Stuart Kirk, Moira Spyer and Eleni Nastouli |
| EPI_ISL_923213, EPI_ISL_923216, EPI_ISL_923226 | Centre for Enzyme Innovation, University of Portsmouth / Translational Research Laboratory, Portsmouth Hospitals NHS Trust | COVID-19 Genomics UK (COG-UK) Consortium | Angela Beckett,Salman Goudarzi,Christopher Fearn,Kate Cook,Katie Loveson,Sharon Glaysher,Scott Elliott,Samuel Robson |
| EPI_ISL_924427 | Virology Department, Sheffield Teaching Hospitals NHS Foundation Trust/Department of Infection, Immunity and Cardiovascular Disease, The Medical School, University of Sheffield | COVID-19 Genomics UK (COG-UK) Consortium | Thushan de Silva, Matthew Parker, Nikki Smith, Adri Angyal, Rebecca Brown, Luke Green, Rachel Tucker, Paul Parsons, Danielle Groves, Katie Johnson, Laura Carrilero, Alex Keeley, Dave Partridge, Matthew Wyles, Benjamin Lindsey, Mehmet Yavuz, Mohammad Raza, Cariad Evans |
| EPI_ISL_925174, EPI_ISL_925175, EPI_ISL_925176, EPI_ISL_925177 | Virginia DCLS | Virginia DCLS | Virginia DCLS |
| EPI_ISL_925493, EPI_ISL_925494 | Department of Clinical Microbiology | GIGA Medical Genomics | Keith Durkin, Maria Artesi, Sébastien Bontems, Raphaël Boreux, Bouchra Boujemla, Cécile Meex, Pierrette Melin, Marie-Pierre Hayette, Vincent Bours |
| EPI_ISL_925846 | LACEN - Laboratório Central de Saúde Pública do Amazonas | Evandro Chagas Institute Virology | Santos, M.C.; Silva, A.M.; Junior, W.D.C.; Barbagelata, L.S.; Ferreira, J.A.; Sousa, E.M.A.; da Silva, P.S.; Pinheiro, K.C.; L.C.; Sousa Junior, E.C. |
| EPI_ISL_926030, EPI_ISL_926043, EPI_ISL_926048, EPI_ISL_926073, EPI_ISL_926083, EPI_ISL_926134, EPI_ISL_926270, EPI_ISL_926350, EPI_ISL_926407, EPI_ISL_926451, EPI_ISL_926538, EPI_ISL_926684, EPI_ISL_926865, EPI_ISL_927168, EPI_ISL_927270, EPI_ISL_927293, EPI_ISL_927566, EPI_ISL_927958, EPI_ISL_927959, EPI_ISL_928071, EPI_ISL_928113, EPI_ISL_928443, EPI_ISL_928820, EPI_ISL_928940, EPI_ISL_928985, EPI_ISL_929030, EPI_ISL_929034, EPI_ISL_929356, EPI_ISL_929531, EPI_ISL_929603, EPI_ISL_929612, EPI_ISL_929624, EPI_ISL_929689, EPI_ISL_929770, EPI_ISL_930027, EPI_ISL_930059, EPI_ISL_930341 | | | |
| see above | Department of Virus and Microbiological Special Diagnostics, Statens Serum Institut, Copenhagen, Denmark | Aalborg University | Danish Covid-19 Genome Consortium |
| EPI_ISL_931081, EPI_ISL_931238, EPI_ISL_931342 | University Hospital Basel, Clinical Virology | University Hospital Basel, Clinical Bacteriology | Tim Roloff, Madlen Stange, Helena MB Seth-Smith, Alfredo Mari, Karoline Leuzinger, Julia Bielicki, Manuel Battegay, Hans Hirsch, Adrian Egli |
| EPI_ISL_933803 | General Hospital - Ohrid | Research Center for Genetic Engineering and Biotechnology "Georgi D. Efremov" , Macedonian Academy of Sciences and Arts | RCGEB - MASA |
| EPI_ISL_933804 | Institute for Lung Diseases in Children - Skopje | Research Center for Genetic Engineering and Biotechnology "Georgi D. Efremov" , Macedonian Academy of Sciences and Arts | RCGEB - MASA |
| EPI_ISL_934092 | General Hospital - Kumanovo | Research Center for Genetic Engineering and Biotechnology "Georgi D. Efremov" , Macedonian Academy of Sciences and Arts | RCGEB - MASA |
| EPI_ISL_934314 | General Hospital - Bitola | Research Center for Genetic Engineering and Biotechnology "Georgi D. Efremov" , Macedonian Academy of Sciences and Arts | RCGEB - MASA |
| EPI_ISL_934413 | General Hospital - Shtip | Research Center for Genetic Engineering and Biotechnology "Georgi D. Efremov" , Macedonian Academy of Sciences and Arts | RCGEB - MASA |
| EPI_ISL_934415 | General Hospital - Veles | Research Center for Genetic Engineering and Biotechnology "Georgi D. Efremov" , Macedonian Academy of Sciences and Arts | RCGEB - MASA |
| EPI_ISL_934540, EPI_ISL_934541, EPI_ISL_934543, EPI_ISL_934545, EPI_ISL_934546, EPI_ISL_934547 | Institut für Virologie am Department für Hygiene, Mikrobiologie und Public Health | Bergthaler laboratory, CeMM Research Center for Molecular Medicine of the Austrian Academy of Sciences | Lukas Endler, Anna Schedl, Thomas Penz, Benedikt Agerer, Maelle Le Moing, Michael Schuster, Bekir Erguner, Jan Laine, Martin Senekowitsch, Christoph Bock, Andreas Bergthaler |
| EPI_ISL_935892, EPI_ISL_935954, EPI_ISL_935955, EPI_ISL_935956, EPI_ISL_935958, EPI_ISL_935959, EPI_ISL_935960, EPI_ISL_935964, EPI_ISL_935965, EPI_ISL_935966, EPI_ISL_935969 | | | |
| see above | Cadham Provincial laboratory | National Microbiology Laboratory (NML) | Anna Majer, Shari Tyson, Grace Seo, Philip Mabon, Elsie Grudeski, Rhiannon Huzarewich, Russell Mandes, Anneliese Landgraff, Jennifer Tanner, Natalie Knox, Morag Graham, Gary Van Domselaar, Paul Van Caeseele, Jared Bullard, David Alexander, Kerry Dust, Nathalie Bastien, Yan Li, Timothy Booth, Darian Hole, Madison Chapel, Kirsten Biggar, CanCOGeN's metadata curation team, Public Health Agency of Canada CanCOGeN team |
| EPI_ISL_936488 | Hospital Queen Elizabeth | Institute for Medical Research, Infectious Disease Research Centre, National Institutes of Health, Ministry of Health Malaysia | Suppiah J, Kamel K, Azizan MA, Thayan R |

| | | | |
|---|---|---|---|
| EPI_ISL_936610, EPI_ISL_936611, EPI_ISL_936614 | Northwestern Memorial Hospital | Ozer Lab | Ramon Lorenzo-Redondo, Lacy M. Simons, Chad J. Achenbach, Lawrence J. Jennings, Michael G. Ison, Judd F. Hultquist, Egon A. Ozer |
| EPI_ISL_937058 | Quest Diagnostics | Quest Diagnostics | Rosenthal,S.H., Gerasimova,A., Kagan,R.M., Anderson, B., Livingston, K.E., Hua, M., Liu Y., Shalhout, D.F., Owen, R., Lacbawan, F. |
| EPI_ISL_940140 | National Institute for Communicable Diseases,National Health Laboratory Services, Gauteng, South Africa | National Institute for Communicable Diseases of the National Health Laboratory Service | Amoako DG, Mohale T, Ntuli N, Mahlangu B, Allam M, Ismail A, Bhiman JN |
| EPI_ISL_940147, EPI_ISL_940151 | NHLS Universitas Academic | UFS Virology | PA Bester, MM Nyaga, P Nthiga, MT Mogotsi, D Goedhals, T de Oliveira |
| EPI_ISL_940171, EPI_ISL_940172, EPI_ISL_940339, EPI_ISL_940341, EPI_ISL_940342, EPI_ISL_940345, EPI_ISL_940346, EPI_ISL_940348 | Hôpital Bichat Claude Bernard, Laboratoire de Virologie | IAME UMR1137 Inserm, Université de Paris, Hôpital Bichat | Antoine Bridier-Nahmias, Amélie Recoing, Quentin Le Hingrat, Lena Daniel, Siham Hamri, Gilles Collin, Alexandre Storto, Mélanie Bertine, Charlotte Charpentier, Nadhira Houhou-Fidouh, Diane Descamps, Benoit Visseaux |
| EPI_ISL_941203 | Servicio de Microbiología. Hospital Clínico Universitario de Valencia | SeqCOVID-SPAIN consortium/IBV(CSIC) | David Navarro Ortega, Eliseo Albert Vicent, Ignacio Torres and SeqCOVID-SPAIN consortium |
| EPI_ISL_942378, EPI_ISL_942379, EPI_ISL_942380, EPI_ISL_942381, EPI_ISL_942382, EPI_ISL_942383, EPI_ISL_942384, EPI_ISL_942385, EPI_ISL_942386, EPI_ISL_942387, EPI_ISL_942388, EPI_ISL_942389, EPI_ISL_942390, EPI_ISL_942392, EPI_ISL_942393, EPI_ISL_942394, EPI_ISL_942395, EPI_ISL_942396, EPI_ISL_942397, EPI_ISL_942398, EPI_ISL_942399, EPI_ISL_942400 | | | |
| see above | Virginia DCLS | Virginia DCLS | Virginia DCLS |
| EPI_ISL_942408, EPI_ISL_942409, EPI_ISL_942410, EPI_ISL_942411, EPI_ISL_942412, EPI_ISL_942413, EPI_ISL_942414 | Gundersen Molecular Diagnostics Laboratory | Kabara Cancer Research Institute | Craig S. Richmond, Paraic A. Kenny |
| EPI_ISL_942415, EPI_ISL_942416, EPI_ISL_942417, EPI_ISL_942418 | Gundersen Clinical Microbiology Laboratory | Kabara Cancer Research Institute | Craig S. Richmond, Paraic A. Kenny |
| EPI_ISL_942419, EPI_ISL_942420, EPI_ISL_942421, EPI_ISL_942422, EPI_ISL_942940 | Gundersen Molecular Diagnostics Laboratory | Kabara Cancer Research Institute | Craig S. Richmond, Paraic A. Kenny |
| EPI_ISL_947249, EPI_ISL_947252 | RS Bina Husada Cibinong | Eijkman Institute for Molecular Biology, Ministry of Research and Technology/National Agency for Research and Innovation | Edison Johar, Frilasita A Yudhaputri, Hidayat Trimarsanto, Iskandar Adnan, Lydia V. Panggalo, Sukma Oktavianthi, Willy Agustine, Safarina G Malik, Khin Saw Myint, Amin Soebandrio |
| EPI_ISL_947255 | RSU Meilia | Eijkman Institute for Molecular Biology, Ministry of Research and Technology/National Agency for Research and Innovation | Edison Johar, Frilasita A Yudhaputri, Hidayat Trimarsanto, Iskandar Adnan, Lydia V. Panggalo, Sukma Oktavianthi, Willy Agustine, Safarina G Malik, Khin Saw Myint, Amin Soebandrio |
| EPI_ISL_947257 | RS Bina Husada Cibinong | Eijkman Institute for Molecular Biology, Ministry of Research and Technology/National Agency for Research and Innovation | Edison Johar, Frilasita A Yudhaputri, Hidayat Trimarsanto, Iskandar Adnan, Lydia V. Panggalo, Sukma Oktavianthi, Willy Agustine, Safarina G Malik, Khin Saw Myint, Amin Soebandrio |
| EPI_ISL_947260 | RSU Bunda Mulia | Eijkman Institute for Molecular Biology, Ministry of Research and Technology/National Agency for Research and Innovation | Iskandar Adnan, Lydia V. Panggalo, Sukma Oktavianthi, Willy Agustine, Edison Johar, Hidayat Trimarsanto, Frilasita A Yudhaputri, Safarina G Malik, Khin Saw Myint, Amin Soebandrio |
| EPI_ISL_949327, EPI_ISL_949328, EPI_ISL_949329, EPI_ISL_949330, EPI_ISL_949331, EPI_ISL_949332, EPI_ISL_949333, EPI_ISL_949334, EPI_ISL_949337, EPI_ISL_949338, EPI_ISL_949339, EPI_ISL_949340, EPI_ISL_949341, EPI_ISL_949342 | | | |
| see above | University of Birmingham | COVID-19 Genomics UK (COG-UK) Consortium | Institute of Microbiology, University of Birmingham: Claire McMurray, Joanne Stockton, Samuel Nicholls, Radoslaw Poplawski, Will Rowe, Josh Quick, Nicholas Loman. University of Birmingham Testing Laboratory: Celina M Whalley, Andrew Bosworth, Charlotte Poxon, Kasun Wanigasooriya, Oliver Pickles, Mike Kidd, Alex Richter, Andrew D Beggs PHE Heartlands Lab: Husam Osman, Andrew Bosworth. Queen Elizabeth Hospital: Anna Casey |
| EPI_ISL_950637 | Queens Medical Centre, Clinical Microbiology Department / DeepSeq Nottingham | COVID-19 Genomics UK (COG-UK) Consortium | Gemma Clark, Wendy Smith, Manjinder Khakh, Vicki M Fleming, Michelle M Lister, Hannah Howson-Wells, Jonathan Ball, Patrick McClure, Joseph Chappell, Theocharis Tsoleridis, Nadine Holmes, Matthew Carlisle, Christopher Moore, Fei Sang, Johnny Debebe, Victoria Wright, Matthew Loose |
| EPI_ISL_952403 | Centre for Enzyme Innovation, University of Portsmouth / Translational Research Laboratory, Portsmouth Hospitals NHS Trust | COVID-19 Genomics UK (COG-UK) Consortium | Angela Beckett,Salman Goudarzi,Christopher Fearn,Kate Cook,Katie Loveson,Sharon Glaysher,Scott Elliott,Samuel Robson |
| EPI_ISL_955676, EPI_ISL_955677, EPI_ISL_955680, EPI_ISL_955682, EPI_ISL_955686, EPI_ISL_955687, EPI_ISL_955688, EPI_ISL_955690 | Orange County Public Health Lab | Chan-Zuckerberg Biohub | CZB Cliahub Consortium |
| EPI_ISL_955975 | Division of Emerging Infectious Diseases, Bureau of Infectious Diseases Diagnosis Control, Korea Disease Control and Prevention Agency | Division of Emerging Infectious Diseases, Bureau of Infectious Diseases Diagnosis Control, Korea Disease Control and Prevention Agency | Ae Kyung Park, Il-Hwan Kim, Heui Man Kim, Jeong-Min Kim, Namjoo Lee, Chae Young Lee, Sang Hee Woo, Eun-Jin Kim |
| EPI_ISL_956321 | Laboratory Medicine | Department of Laboratory Medicine, Lin-Kou Chang Gung Memorial Hospital, Taoyuan, Taiwan | Kuo-Chien Tsao, Yu-Nong Gong, Shu-Li Yang, Yi-Chun Liu, Chung-Guei Huang, Mei-Jen Hsiao, Po-Wei Huang, Cheng-Ta Yang, Cheng-Hsun Chiu, Peng-Nien Huang, Kuo-Ming Lee, Guang-Wu Chen, Shin-Ru Shih |
| EPI_ISL_959295, EPI_ISL_959296, EPI_ISL_959297, EPI_ISL_959298, EPI_ISL_959299, EPI_ISL_959300, EPI_ISL_959301, EPI_ISL_959304, EPI_ISL_959305 | Centre de Recerca en Salut Animal (IRTA/CRESA) | IrsiCaixa - Can Ruti CovidSeq | Marc Noguera-Julian, Mariona Parera, Maria Casadellà, Pilar Armengol, Francesc Catala-Moll, Roger Paredes, Bonaventura Clotet J. Segalés, M. Puig, J. Rodon, C. Avila-Nieto, J. Carrillo, G. Cantero, M.T. Terrón, S. Cruz, N. Izquierdo-Useros, E. Vidal, J. Blanco, B. Clotet, J. Vergara-Alert |
| EPI_ISL_960347, EPI_ISL_960356, EPI_ISL_960383 | University of Wisconsin-Madison AIDS Vaccine Research Laboratories | University of Wisconsin-Madison AIDS Vaccine Research Laboratories | Gage Moreno, Katarina Braun, et al. AIDS Vaccine Research Laboratories |
| EPI_ISL_961994 | Illinois Department of Public Health | Gagnon Lab, Southern Illinois University | Keith Gagnon |
| EPI_ISL_965357, EPI_ISL_965358, EPI_ISL_965383, EPI_ISL_965384, EPI_ISL_965386, EPI_ISL_965459, EPI_ISL_965465 | University of Liège COVID-19 testing center | GIGA Medical Genomics | Keith Durkin, Maria Artesi, Bouchra Boujemla, Emmanuel André, Marc Van Ranst, Fabrice Bureau, Laurent Gillet, Wouter Coppieters, Vincent Bours |
| EPI_ISL_965537, EPI_ISL_965539, EPI_ISL_965542, EPI_ISL_965552, EPI_ISL_965604, EPI_ISL_965622, EPI_ISL_965645, EPI_ISL_965678, EPI_ISL_965710, EPI_ISL_965784 | Dutch COVID-19 response team | Medical Microbiology, Maastricht University Medical Centre | Jozef Dingemans*, Brian van der Veer*, Erik Beuken, Carmen Reumkens, Lieke van Alphen, Christian Hoebe, Paul Savelkoul |
| EPI_ISL_968190, EPI_ISL_968191, EPI_ISL_968192, EPI_ISL_968193 | Arizona State Public Health Laboratory | Arizona State Public Health Laboratory | Trung Huynh, Jessica Escobar, Katherine Fullerton, Nobuko Fukushima, Stacy White, Linda Getsinger, Victor Waddell |

| | | | |
|---|---|---|---|
| EPI_ISL_968914, EPI_ISL_968916, EPI_ISL_968919, EPI_ISL_968929, EPI_ISL_968985, EPI_ISL_968986, EPI_ISL_968987, EPI_ISL_968988, EPI_ISL_968992 | KEMRI-Wellcome Trust Research Programme/KEMRI-CGMR-C Kilifi | KEMRI-Wellcome Trust Research Programme/KEMRI-CGMR-C Kilifi | Githinji et al |
| EPI_ISL_973207 | Department of Virus and Microbiological Special Diagnostics, Statens Serum Institut, Copenhagen, Denmark | Aalborg University | Danish Covid-19 Genome Consortium |
| EPI_ISL_975156, EPI_ISL_975157, EPI_ISL_975158, EPI_ISL_975159, EPI_ISL_975160, EPI_ISL_975161, EPI_ISL_975162, EPI_ISL_975163, EPI_ISL_975164, EPI_ISL_975165, EPI_ISL_975166, EPI_ISL_975167, EPI_ISL_975168, EPI_ISL_975169, EPI_ISL_975170, EPI_ISL_975171, EPI_ISL_975172, EPI_ISL_975173, EPI_ISL_975174, EPI_ISL_975175, EPI_ISL_975176, EPI_ISL_975177, EPI_ISL_975178, EPI_ISL_975179, EPI_ISL_975180, EPI_ISL_975181, EPI_ISL_975182, EPI_ISL_975183, EPI_ISL_975184, EPI_ISL_975185, EPI_ISL_975186, EPI_ISL_975187, EPI_ISL_975188, EPI_ISL_975189, EPI_ISL_975190, EPI_ISL_975191, EPI_ISL_975192, EPI_ISL_975193, EPI_ISL_975194, EPI_ISL_975195, EPI_ISL_975196, EPI_ISL_975197, EPI_ISL_975198, EPI_ISL_975199, EPI_ISL_975200, EPI_ISL_975201, EPI_ISL_975202, EPI_ISL_975203, EPI_ISL_975204, EPI_ISL_975205, EPI_ISL_975206, EPI_ISL_975207, EPI_ISL_975208, EPI_ISL_975209, EPI_ISL_975210, EPI_ISL_975211, EPI_ISL_975212, EPI_ISL_975213, EPI_ISL_975214, EPI_ISL_975215, EPI_ISL_975216, EPI_ISL_975217, EPI_ISL_975218, EPI_ISL_975219, EPI_ISL_975220, EPI_ISL_975221, EPI_ISL_975222, EPI_ISL_975223, EPI_ISL_975224, EPI_ISL_975225, EPI_ISL_975226, EPI_ISL_975227, EPI_ISL_975228, EPI_ISL_975229, EPI_ISL_975230, EPI_ISL_975231, EPI_ISL_975232, EPI_ISL_975233, EPI_ISL_975234, EPI_ISL_975235, EPI_ISL_975236, EPI_ISL_975237, EPI_ISL_975238, EPI_ISL_975239, EPI_ISL_975240, EPI_ISL_975241, EPI_ISL_975242, EPI_ISL_975243, EPI_ISL_975244, EPI_ISL_975245, EPI_ISL_975246, EPI_ISL_975247, EPI_ISL_975248, EPI_ISL_975249, EPI_ISL_975250, EPI_ISL_975251, EPI_ISL_975252, EPI_ISL_975253, EPI_ISL_975254, EPI_ISL_975255, EPI_ISL_975256, EPI_ISL_975257, EPI_ISL_975258, EPI_ISL_975259, EPI_ISL_975260, EPI_ISL_975261, EPI_ISL_975262, EPI_ISL_975263, EPI_ISL_975264, EPI_ISL_975265, EPI_ISL_975266, EPI_ISL_975267, EPI_ISL_975268, EPI_ISL_975269, EPI_ISL_975270, EPI_ISL_975271, EPI_ISL_975272, EPI_ISL_975273, EPI_ISL_975274, EPI_ISL_975275, EPI_ISL_975276, EPI_ISL_975277, EPI_ISL_975278, EPI_ISL_975279, EPI_ISL_975280, EPI_ISL_975281, EPI_ISL_975282, EPI_ISL_975283, EPI_ISL_975284, EPI_ISL_975285, EPI_ISL_975286, EPI_ISL_975287, EPI_ISL_975288, EPI_ISL_975289, EPI_ISL_975290, EPI_ISL_975291, EPI_ISL_975292, EPI_ISL_975293, EPI_ISL_975294, EPI_ISL_975295, EPI_ISL_975296, EPI_ISL_975297, EPI_ISL_975298, EPI_ISL_975299, EPI_ISL_975300, EPI_ISL_975301 | | | |
| see above | BCCDC Public Health Laboratory | BCCDC Public Health Laboratory | Prystajecky Natalie, Linda Hoang, Dan Fornika, John Tyson, Shannon Russell, Kim Macdonald, Kimia Kamelian, Ana Pacagnella, Corrinne Ng, Loretta Janz, Robert Azana Terry Snutch, Mel Krajden |
| EPI_ISL_977504, EPI_ISL_977508 | NCSLPH | NCSLPH | Chase K, Miller MC, Greene S, Glover W |
| EPI_ISL_977892, EPI_ISL_977893, EPI_ISL_977894, EPI_ISL_977895, EPI_ISL_977896, EPI_ISL_977897, EPI_ISL_977899, EPI_ISL_977900, EPI_ISL_978970, EPI_ISL_978971 | Chiu Laboratory, University of California, San Francisco | Chiu Laboratory, University of California, San Francisco | Charles Chiu, Xianding (Wayne) Deng, Candace Wang, Venice Servellita, Jill Hacker, Debra Wadford |
| EPI_ISL_982512, EPI_ISL_982513, EPI_ISL_982514 | Kentucky State Public Health Lab | Kentucky State Public Health Lab | Stephanie Lunn, Karim George, Joshua Tobias, William Grooms, Vaneet Arora, Matthew Johnson, Rachel Zinner, Rhonda Lucas |
